# Molecular evolution and structural analyses of the spike glycoprotein from Brazilian SARS-CoV-2 genomes: the impact of the fixation of selected mutations

**DOI:** 10.1101/2021.07.16.452571

**Authors:** Patrícia Aline Gröhs Ferrareze, Ricardo Ariel Zimerman, Vinícius Bonetti Franceschi, Gabriel Dickin Caldana, Paulo Augusto Netz, Claudia Elizabeth Thompson

## Abstract

The COVID-19 pandemic caused by *Severe acute respiratory syndrome coronavirus 2* (SARS-CoV-2) has reached by July 2021 almost 200 million cases and more than 4 million deaths worldwide since its beginning in late 2019, leading to enhanced concern in the scientific community and the general population. One of the most important pieces of this host-pathogen interaction is the spike protein, which binds to the human Angiotensin-converting enzyme 2 (hACE2) cell receptor, mediates the membrane fusion and is the major target of neutralizing antibodies against SARS-CoV-2. The multiple amino acid substitutions observed in this region, specially in the Receptor Binding Domain (RBD), mainly after almost one year of its emergence (late 2020), have enhanced the hACE2 binding affinity and led to several modifications in the mechanisms of SARS-CoV-2 pathogenesis, improving the viral fitness and/or promoting immune evasion, with potential impact in the vaccine development. In this way, the present work aimed to evaluate the effect of positively selected mutations fixed in the Brazilian SARS-CoV-2 lineages and to check for mutational evidence of coevolution. Additionally, we evaluated the impact of selected mutations identified in some of the VOC and VOI lineages (C.37, B.1.1.7, P.1, and P.2) of Brazilian samples on the structural stability of the spike protein, as well as their possible association with more aggressive infection profiles by estimating the binding affinity in the RBD-hACE2 complex. We identified 48 sites under selective pressure in Brazilian spike sequences, 17 of them with the strongest evidence by the HyPhy tests, including VOC related mutation sites 138, 142, 222, 262, 484, 681, and 845, among others. The coevolutionary analysis identified a number of 28 coevolving sites that were found not to be conditionally independent, such as the couple E484K - N501Y from P.1 and B.1.351 lineages. Finally, the molecular dynamics and free energy estimates showed the structural stabilizing effect and the higher impact of E484K for the improvement of the binding affinity between the spike RBD and the hACE2 in P.1 and P.2 lineages, as well as the stabilizing and destabilizing effects for the positively selected sites.

## INTRODUCTION

The *severe acute respiratory syndrome coronavirus 2* (SARS-CoV-2) is the etiological agent of the COVID-19 pandemic. The virus is composed of an enveloped positive-sense single-stranded RNA genome, which encodes 16 non-structural proteins (nsp1-16), four structural proteins (Spike, Membrane, Envelope, and Nucleocapsid), and other accessory proteins (ORFs 3a, 6, 7a, 7b, 8, 10) (Fehr and Perlman, 2015; Liu et al., 2014). The spike (S) glycoprotein is necessary for the viral binding to ACE2 host cell receptor, it is common to multiple coronaviruses (*e. g.*, MERS-CoV and SARS-CoV) (Fehr and Perlman, 2015), and is the major target of neutralizing antibodies against SARS-CoV-2 (Walls et al., 2020; Yuan et al., 2020). Spike has a homotrimeric form where each monomer (protomer) is composed by a S1 subunit that mediates the receptor binding and an alpha-helix rich S2 subunit that promotes the subsequent membrane fusion (Hoffmann et al., 2020).

Since the emergence of the first SARS-CoV-2 in the early Wuhan epidemic, several mutations have been identified, both occurring isolated or as a signature of a broader mutational complex. Although S protein represents only 13% of the viral genome, substitution at this site has been overrepresented suggesting immunodominance of this protein, specially at its Receptor Binding Domain (RBD) (Greaney et al., 2020; Weisblum et al., 2020). These mutations can enhance receptor binding, either directly or allosterically, alter viral fitness or promote immune evasion, ensuing occasional reinfection and possible impacting the efficacy of developed vaccines (Greaney et al., 2020; Harvey et al., 2021; Korber et al., 2020; Weisblum et al., 2020).

The S protein mutation D614G (aspartic acid to glycine substitution at amino acid position 614) has been progressively dominant worldwide since March 2020, playing an essential role in infectivity and augmented viral load (Groves et al., 2021; C. B. Jackson et al., 2021; Korber et al., 2020; Yurkovetskiy et al., 2020). This substitution may have emerged independently and promptly became dominant in several viral strains, outcompeting D614 harboring viruses, even where they originally appeared. D614G abolishes a hydrogen bond between this position and a threonine present in S2 of the neighbour protomer. The consequent conformational changes acts allosterically favouring the maintenance of the Receptor Binding Domain (RBD) in an activated “up” or “open” position, which exhibits its ACE2 binding site (Receptor Binding Motif) for longer periods. This allows a more efficient binding to ACE2 (Groves et al., 2021), with consequent higher replication and viral loads. However, G614 mutants are similarly (or even more) susceptible to immune neutralization as the original D614 variant (Plante et al., 2020; Weissman et al., 2021). Notably, this enhanced epitope exposure of “up” RBD may be a “weakness”, since RBD is poor in predicted O and N-linked glycan sites required for immune shielding. In theory, this undesirable effect could be compensated by accumulation of other RBD mutations, leading to increased infectiousness, and rendering D614G unnecessary. Specifically, the “HMN 19B variant”, recently described in France, is a direct descendent from the earlier 19B that have predominated before D614G harboring lineages took over (Fourati et al., 2021).

The codon 501 in RBD has been considered another major mutational hotspot. Substitutions at this position have been associated with increased binding affinity to ACE2 (Gu et al., 2020; Starr et al., 2020). The non-synonymous mutation from an asparagine to a tyrosine at position 501 (N501Y) has first appeared in samples from Wales and other parts of the world. However, its actual emergence and dissemination have been related to coevolution with different mutations. The deletion of H69-V70 in the N-terminal-domain, co-occurring in B.1.1.7 lineage, has been particularly important for the N501Y emergence (Meng et al., 2021). However, N501Y has increasingly been demonstrated in coevolution with many different mutational signatures, for instance, in the Variants of Concern (VOCs) B.1.351 and P.1, as well as in the more recent HMN 19B aforementioned. Data suggests that some N501Y harboring viruses could be 50% more transmissible and up to 61% more lethal, mainly due to major changes in the electrostatic interaction between Spike and ACE2 receptors (Washington et al., 2021; Davies et al., 2021). This would be related to the substitution of a relatively small asparagine (N) for a larger aromatic tyrosine (Y), allowing for an extra contact site with ACE2 (Ali et al., 2021; Gu et al., 2020; Tang et al., 2021). Interesantly, N501Y increases host diversity allowing direct infection of non-humanized mice (Rathnasinghe et al., 2021).

The E484K mutation in RBD has been drowning progressive attention. It is an amino acid replacement of glutamic acid to lysine at amino acid position 484, which could significantly alter the complementarity of antibodies to the RBD region leading to immune evasion (Baum et al., 2020; Greaney et al., 2020; Weisblum et al., 2020). In fact, both in a Deep Mutational Scanning (DMS) experiment and in a selective pressure study using 19 types of mAbs, different substitutions at this position (E484K/Q/P/A/D/G) were consistently demonstrated as being the most critical for decreasing antibody neutralization titers (Greaney et al., 2020; Harvey et al., 2021). Since this mutation has been arising independently in multiple lineages (Ferrareze et al., 2021), at the beginning it may represent a common evolutionary solution for viral maintenance. E484K emerged in a few selected lineages where it co-exists with other signature substitutions (e.g. in B.1.351 and in P1), similarly to the phenomenon previously described for N501Y. Importantly, the combination of E484K and N501Y seems to induce more conformational changes than the N501Y mutant alone, potentially altering antibodies binding to this region and resulting in the immune evasion phenomena (Nelson et al., 2021).

The emergence of three independently evolving lineages (B.1.1.7, B.1.351, and P.1) (Faria et al., 2021; Rambaut et al., 2020; Tegally et al., 2021), all characterized by a constellation of mutations (including the aforementioned del 69-70, K417N/T, E484K, and N501Y convergent mutations) in RBD, has prompted concerns about the evolutionary forces leading to the SARS-CoV-2 adaptation. The presence of these sets of consistently present substitutions are highly suggestive of coevolutionary mutational processes (Martin et al., 2021). In addition to these convergent mutations, other substitutions of these VOCs are also associated with positive selection. For example, there is evidence that eight out of the 10 lineage-defining mutations in the spike protein of the P.1 lineage are under diversifying positive selection (Faria et al., 2021). It is possible to detect selection along prespecified lineages that affect certain subsets of codons in a protein-coding gene. Sites 484 and 501 (located in the RBD) definitively show a pattern of nucleotide diversification that can be linked with positive selection (Tegally et al., 2021). On the other hand, alternative evolutionary processes that could have led to this plethora of substitutions have been rarely described thus far. Recombination, a recurrent and well documented process of the molecular evolution of coronaviruses, has been described between B.1.1.7 and other lineages (Jackson et al., 2021), but their real role in SARS-CoV-2 evolution remains elusive.

The present work aimed to evaluate the effect of positively selected mutations fixed in the Brazilian SARS-CoV-2 lineages and to check for mutational coevolution evidence. Additionally, we evaluated the impact of some mutations identified in some VOC and VOI lineages (C.37, B.1.1.7, P.1 and P.2) of Brazilian samples on the structural stability of the spike protein, as well as their possible association with more aggressive infection profiles by the binding affinity estimation in the RBD-hACE2 complex.

## METHODOLOGY

### SPIKE SEQUENCE ANALYSES

#### Genome sequence retrieving

After the exclusion of incomplete or low coverage (Ns > 5%) sequences, 11,078, Brazilian genomes were recovered from the GISAID database between January 01^st^, 2020 and June 06^th^, 2021 (submission date up to June 06^th^, 2021).

#### Spike phylogenetic analysis

The multiple sequence alignment with NC_045512.2 as reference was performed by the MAFFT v.7 web server (Katoh et al., 2019) with default parameters and 1PAM / κ=2’ scoring matrix for closely related DNA sequences. For the spike sequence analysis, the region between the positions 21,562 and 25,384 (spike regions in the NC_045512.2 reference genome) were selected from the multiple sequence alignment previously performed. The deletion of sequence duplicates (identical spike sequences) kept 2,901 unique spike sequences. The clade assignment and variant calling was performed by Nextclade (https://clades.nextstrain.org/). The phylogenetic analysis was started by the inference of the best evolutionary model by ModelTest-NG (Darriba et al., 2020), which identified GTR+R4 in all selection strategies. The phylogenetic tree reconstruction was performed by the Maximum Likelihood method in the IQ-TREE program (Nguyen et al., 2015), using 1,000 replicates of ultrafast bootstrap (Hoang et al., 2018) and a Shimodaira-Hasegawa-like approximate likelihood ratio test (SH-aLRT) with 1,000 replicates (Guindon et al., 2010), 2,000 iterations and the optimization of the UFBoot trees by NNI on bootstrap alignment. The tree visualization was performed by FigTree software (http://tree.bio.ed.ac.uk/software/figtree/).

#### Spike selection and coevolutionary analyses

The multiple sequence alignment of the 2,901 unique spike sequences and the phylogenetic tree previously built were used as input in the HyPhy program. To perform different selection tests, the methods: (i) Fast Unconstrained Bayesian AppRoximation (FUBAR) (Murrell et al., 2013), (ii) Fixed Effects Likelihood (FEL) (Kosakovsky Pond and Frost, 2005), and (iii) Single-Likelihood Ancestor Counting (SLAC) (Kosakovsky Pond and Frost, 2005) were evaluated. The analysis of coevolution across sites in the spike sequences was performed by BGM (Bayesian Graphical Model) (Poon et al., 2007), a tool for detecting coevolutionary interactions between amino acid positions in a protein by a Markov Chain Monte Carlo (MCMC) method.

### SPIKE STRUCTURAL ANALYSES

#### Spike structural stability

The analysis of the spike structural stability by changes in free energies upon mutation (ΔΔG kcal/mol) was performed using PDB ID 6XR8 (chain A) as the structure of the full-length prefusion conformation of the SARS-CoV-2 spike protein (Cai et al., 2020) using five methodologies: the web server DynaMut (Rodrigues et al., 2018), FoldX Suite v5.0 (Schymkowitz et al., 2005), and the web servers iMutant3.0 (Capriotti et al., 2005), MAESTRO (Laimer et al., 2016), and PremPS (Chen et al., 2020).

##### DynaMut

The web-server DynaMut was selected to perform the analyses of vibrational entropy and total energy using the PDB ID 6XR8 (chain A). The DynaMut implements a Normal Mode Analysis (NMA) through Bio3D (Grant et al., 2006) and ENCoM (Frappier and Najmanovich, 2014) approaches, providing rapid and simplified access to insightful analyses about protein motions. Moreover, DynaMut also enables rapid analysis of the impact of mutations on dynamics and stability of proteins resulting from vibrational entropy changes.

##### FoldX

For the energy estimation, the analysis and correction of structural problems resulting from crystallography in PDB ID 6XR8 were performed using the RepairPDB command and additional parameters --*water=crystal*, --*pH=7.4* and --*temperature=309.65*. Posteriorly, the mutagenesis process was applied for all the positively selected sites with the BuildModel function (additional parameters: --*water=crystal*, --*pH=7.4*, --*temperature=309.65* and --*numberOfRuns=5*). The average values for each ΔΔG (based on the 5 runs) were classified in seven categories according to the reported Foldx accuracy (0.46 kcal/mol): highly stabilizing mutations (ΔΔG < −1.84 kcal/mol), stabilizing mutations (−1.84 kcal/mol ≤ ΔΔG < −0.92 kcal/mol), slightly stabilizing mutations (−0.92 kcal/mol ≤ ΔΔG < −0.46 kcal/mol), neutral mutations (−0.46 kcal/mol < ΔΔG ≤ +0.46 kcal/mol), slightly destabilizing mutations (+0.46 kcal/mol < ΔΔG ≤ +0.92 kcal/mol), destabilizing mutations (+0.92 kcal/mol < ΔΔG ≤ +1.84 kcal/mol), and highly destabilizing mutations (ΔΔG > +1.84 kcal/mol) (Studer et al., 2014).

##### iMutant3

The iMutant web server was selected to estimate the free energy changes by a support vector machine (SVM)-based tool. This system predicted the sign of the protein stability change upon mutation and as a regression estimator predicted the related ΔΔG values in the physiological pH (7.4) and temperature (36.5 ℃).

##### MAESTRO

The Multi AgEnt STability pRedictiOn web server was used to estimate the changes in unfolding free energy upon point mutation through a machine learning system.

##### PremPS

The estimation of the unfolding Gibbs free energies was performed by a random forest regression scoring function using the ProTherm database for parameterization. Negative values indicate stabilization by the decrease of the free energies.

#### Spike RBD comparative homology modelling

To perform an accurate estimation of the binding free energy associated with mutations in the spike protein belonging to different lineages, spike RBD - ACE2 protein complexes were modelled for the reference SARS-CoV-2 spike (YP_009724390.1) and lineages C.37, B.1.1.7, P.1, P.2, and P.2+452. The PDB file 6M0J was selected as the best template (2.45 Å and 194 amino acids) to the modeling using the MODELLER pipeline (Webb and Sali, 2016). Five models were generated for each sequence. The thirty resulting structures were evaluated in relation to stereochemical parameters and structural quality by the programs PROCHECK (Laskowski et al., 1993) and VERIFY3D (Eisenberg et al., 1997), available on SAVES v6.0 web server (https://saves.mbi.ucla.edu/). The analysis of the Ramachandran plot statistics, G-factors and residue properties (PROCHECK), as well as the evaluation of the 3D-1D score using VERIFY3D, allowed the selection of the best modelled RBD structures to be subsequently used for the energy calculations. The generation of the spike RBD - ACE2 complexes for each different lineage was performed by the PyMOL software using the structural alignment of the RBD models with the 6M0J template, which was the source of the ACE2 structural coordinates.

#### Molecular dynamics and binding free energy estimation

The structures of the fragments comprising residues 333 until 526 of the wild-type (reference) spike protein, as well as of the variants P.1 (K417T, E484K, N501Y), P.2 (E484K), P.2+452 (E484K, L452V), C.37 (L452Q, F490S), and B.1.1.7 (N501Y) complexed with the human ACE2 protein (residues 19 until 615) in the pdb format were used as input for classical molecular dynamics simulations using GROMACS (Abraham et al., 2015) with the AMBER03 force field (Duan et al., 2003).

The spike-ACE2 complexes were simulated in cubic boxes with periodic boundary conditions, solvated with TIP3P water molecules (Jorgensen et al., 1983) and with sodium and chloride ions corresponding to physiological concentration. The van der Waals interactions were calculated using a cutoff radius of 1.2 nm and the electrostatic interactions were calculated with the Particle Mesh Ewald (PME) method (Darden et al., 1993). All systems were initially energetically minimized using conjugate gradients and steepest descent algorithms. After minimization, they were submitted to a 500 ps where the coordinates of both proteins were restrained, allowing the solvent and ions to relax without disturbing the geometry of the complex. After the position restrained simulation, a thermalization phase consisting of a sequence of three unrestricted molecular dynamics simulations with 5 ns each in temperatures of 200 K, 240 K, and 280 K was carried out. The production phase consisted of 200 ns long simulation runs, in the NPT ensemble, using a Nosé-Hoover thermostat (Hoover, 1985; Nosé, 1984) and a Parrinello-Rahman barostat (Parrinello and Rahman, 1981).

After simulations, the trajectories were visually analyzed using VMD (Humphrey et al., 1996) and GROMACS tools to quantify the structural and thermodynamic stability of the complexes, the number of hydrogen bonds and the interface area. The interface area was computed taking SASA (Solvent Accessible Surface Area) (Eisenhaber et al., 1995) of ACE2 and spike minus SASA of the complex and dividing the result by two (because the contact was counted twice).

#### Spike RBD-ACE2 binding free energy calculation

The spike-ACE2 binding free energy was calculated using Molecular Mechanics Poisson-Boltzmann Surface Area (MM/PBSA) binding free energy calculations (Baker et al., 2001; Homeyer and Gohlke, 2012), employing sets of 200 configurations (1ns spaced snapshots obtained from the molecular dynamics trajectories) for each system. The calculations were carried out using the program g_mmpbsa (Kumari et al., 2014), which is compatible with GROMACS, using as settings a gridspace of 0.5 A, salt concentration of 0.150 M, solute dielectric constant of 2, and estimating the nonpolar solvation energy using the solvent accessible surface area (SASA). The contributions of the residues to the binding free energy were calculated with the same program.

## RESULTS

### Spike protein phylogenetic analyses

The spike phylogenetic analysis of 2,901 unique sequences showed the formation of a large monophyletic group containing two main subclades: one formed by P.1 and P.1.2 sequences and another formed by the B.1.1.7. B.1.1.28 and P.2 sequences were located at the basis of the P.1 subclade, while some B.1.1.33/N.9 sequences are basal to the B.1.1.7 subclade. Interesting to note that B.1.1.7 formed one monophyletic group (Figure 1A). Based on spike sequences, the P.2, B.1.1.28 and B.1.1.33 lineages did not form individual clades. Finally, it was possible to observe the formation of a large basal clade with B.1.1.28 and B.1.1.33 sequences, with the B lineage (among others) as its ancestor. Despite P.2 being considered as derivative from B.1.1.28 by the inclusion of the E484K mutation (D614G+E484K+V1176F), the presence of this substitution along the tree (as well D614G and V1176F) indicated that the formation of all the small clades is probably due to the high variable combinations of mutations found in these lineages.

**Figure 1.**
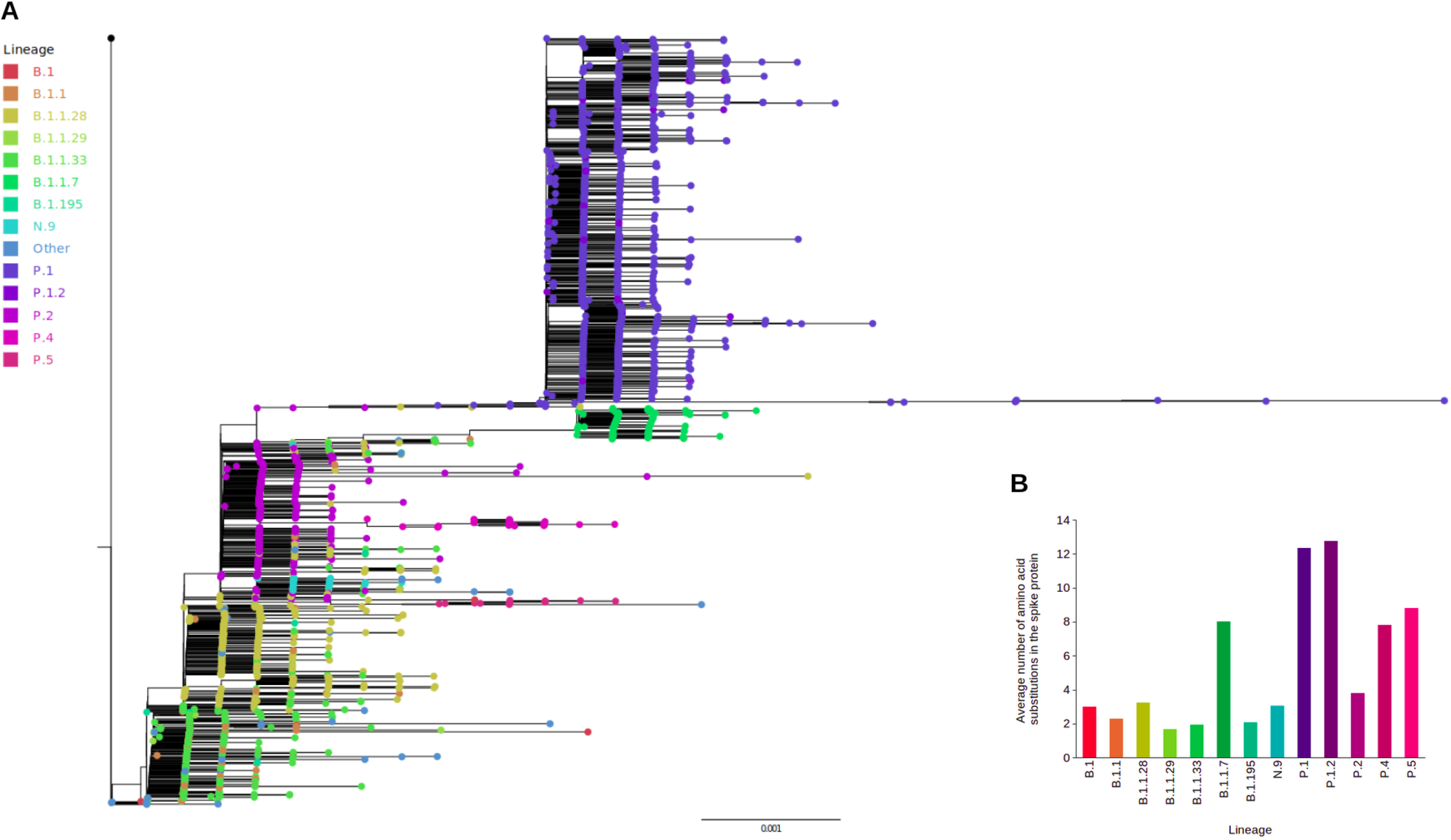
Spike phylogenetic and genomic analyses. (A) Phylogenetic tree of 2,901 unique Brazilian spike nucleotide sequences available until June 06, 2021. Node tips are colored by PANGO lineages represented by ≥ 10 genomes. “Other” defines lineages representing < 10 genomes. The tree is rooted using the reference spike nucleotide sequence (NC_045512.2). (B) Average number of amino acid substitution events in the spike protein by lineage. The average values were calculated based on the spike set of 2,901 brazilian sequences between January, 2020 and June 06, 2021. This spike set represents the nucleotide sequence variability for 11,054 brazilian genomes.

The genetic analysis of these 2,901 unique Brazilian spike sequences showed that 575 amino acid sites presented missense substitutions in different lineages, with a mutation rate of 7.79 amino acid substitutions per spike sequence/genome, since January 2020. With a range between 0 and 16 amino acid substitutions and a different average count in each lineage (Figure 1B), the spike sequence set was mostly represented by lineages such as P.1 (45.71%), P.2 (14.34%), B.1.1.28 (13.82%), B.1.1.33 (10.62%), and B.1,1.7 (4.03%), from 53 identified lineages. The evaluation of the mutation rate before and after the P.1 first occurrence (October 2020) indicated that the spike sequences showed an estimated amino acid substitution of 2.12 events per genome between January and September, 2020. However, from October 2020 up to June 06 2021, this rate was increased to 8.99 amino acid mutations for each spike sequence.

### Spike selection analyses

The selection analysis performed with HyPhy for site-to-site tests evaluated the presence of diversifying and purifying selection in the spike protein using a set of 2,901 sequences, which represents the genetic variability for 11,054 genomes (24 sequences were excluded due to the presence of >5% of ambiguous sites or truncating insertions in the final alignment). The FUBAR method identified 20 sites under adaptive pressure along the phylogeny (Table 1 and Supplementary File 2). Among these, the mutation in site 484 is represented by three missense substitutions: (i) E484K, which was found in 71.36% of all genomes, belonging to 11 different lineages, (ii) E484Q, present in 0.22% of the sequences, including the B.1, B.1.1.28 and P.5 lineages, and (iii) a new substitution found in only one genome, E484D. Other sites related to known mutations from VOC lineages such as P.1, B.1.1.7, B.1.351, and B.1.617.2 were identified as potentially fixed substitutions. In this group, there are sites with the mutations L5F (0.95% of the spike sequences), A67S/V/P (0.12/0.08/0.01%), G75V (0.11%), D138Y/H (55.04/0.11%), G142D (0.03%), A222V/S/P (0.49/0.05/0.01%), A262S/D (0.50/0.04%), K417T/N (49.11/0.05%), N501Y (61.12%), T572I (0.34%), P681H/L/R (3.03/0.40/0.03%), A688V/S/G (0.19/0.04/0.03%), A845S/V (1.27/0.04%), V1176F (82.12%), among others.

**Table 1.**
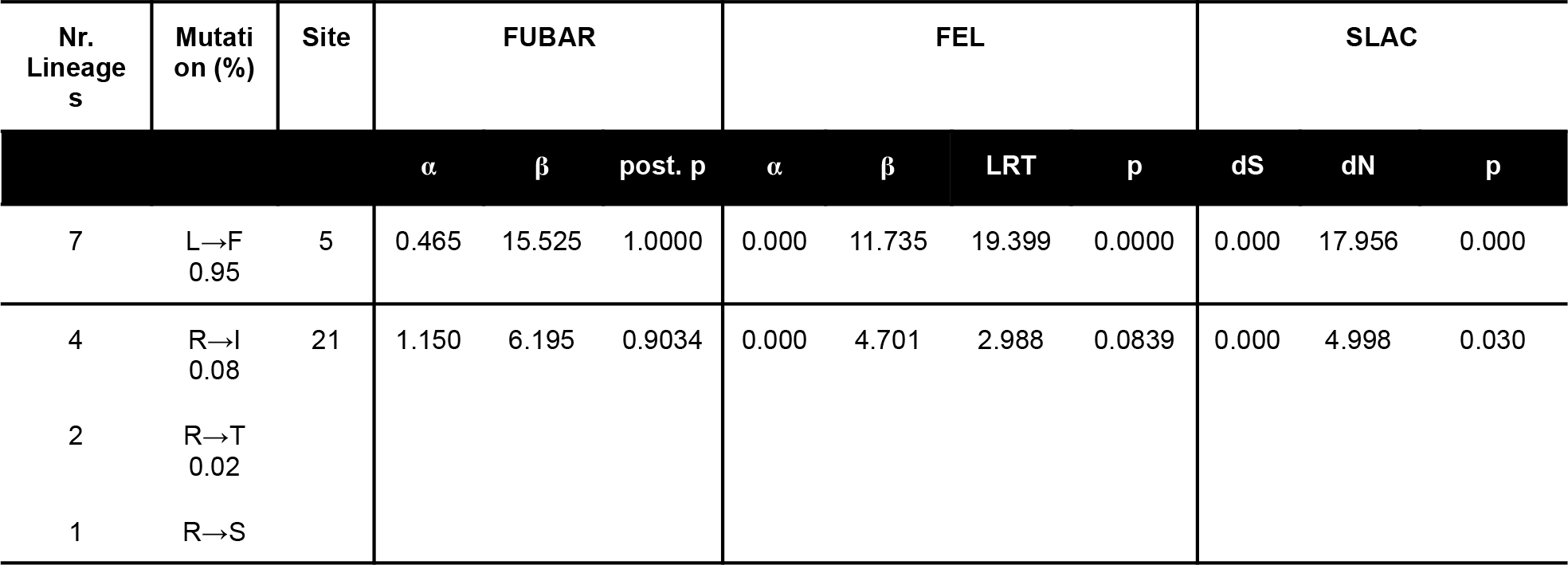
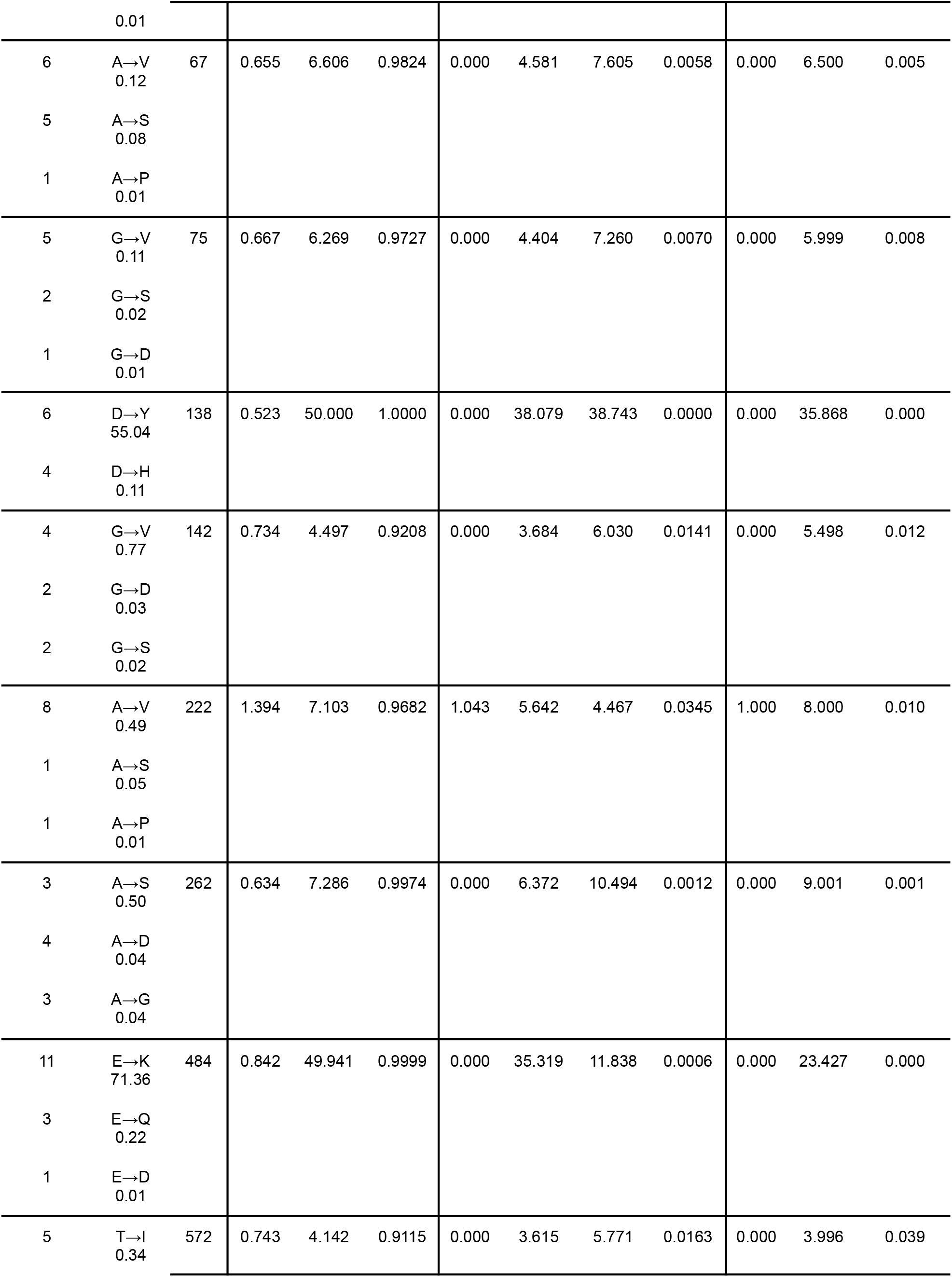
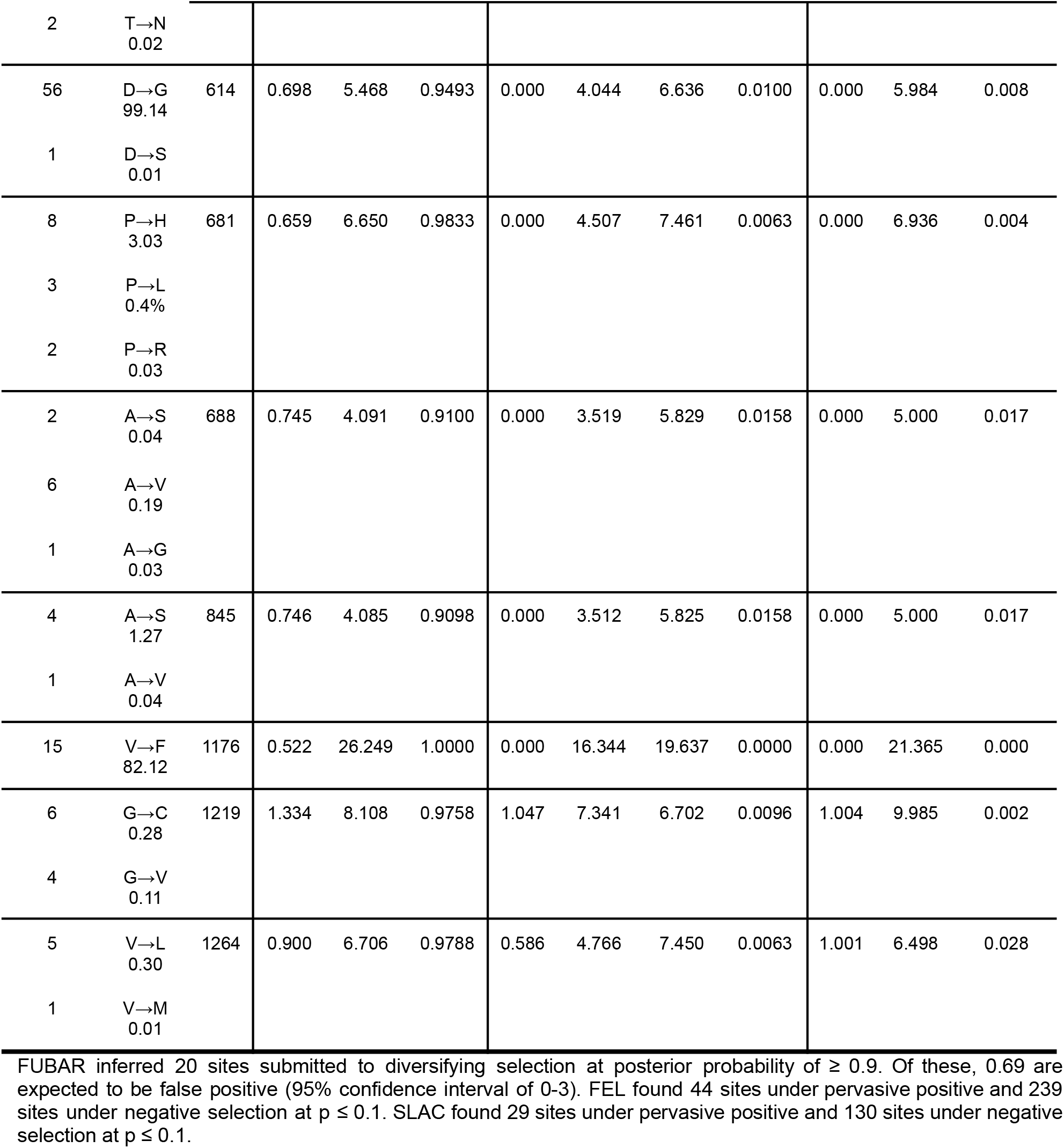
Positively selected sites detected by all tested HyPhy evolutionary methods (FUBAR, FEL and SLAC) for pervasive site-level selection and its frequencies (%) in the genome set (n=11,078).

The FEL method was mainly tested to detect negatively selected sites, since more powerful methods such as FUBAR do not evaluate purifying selection. With the initial assumption that the selective pressure for each site is constant along the entire phylogeny, FEL indicated sites that are evolving under positive and negative selective pressures in the spike protein sequence. As result, 239 sites were marked as targets of purifying selection (Supplementary File 3) along the phylogeny, while other 44 sites were suggested to be under adaptive selection. Interestingly, all positively selected sites indicated by the FUBAR method were found by FEL (Table 1 and Figure 2A), including those with low frequency in the analysed genomes. Some of them were indeed related to the VOC lineages. Among the sites identified only by the FEL test are the residues 29, 49, 63, 76, 78, 367, 483, 553, 585, 689, 747, 1,027, 1,084, 1,124, 1,133, and 1,167 (Supplementary File 2).

**Figure 2.**
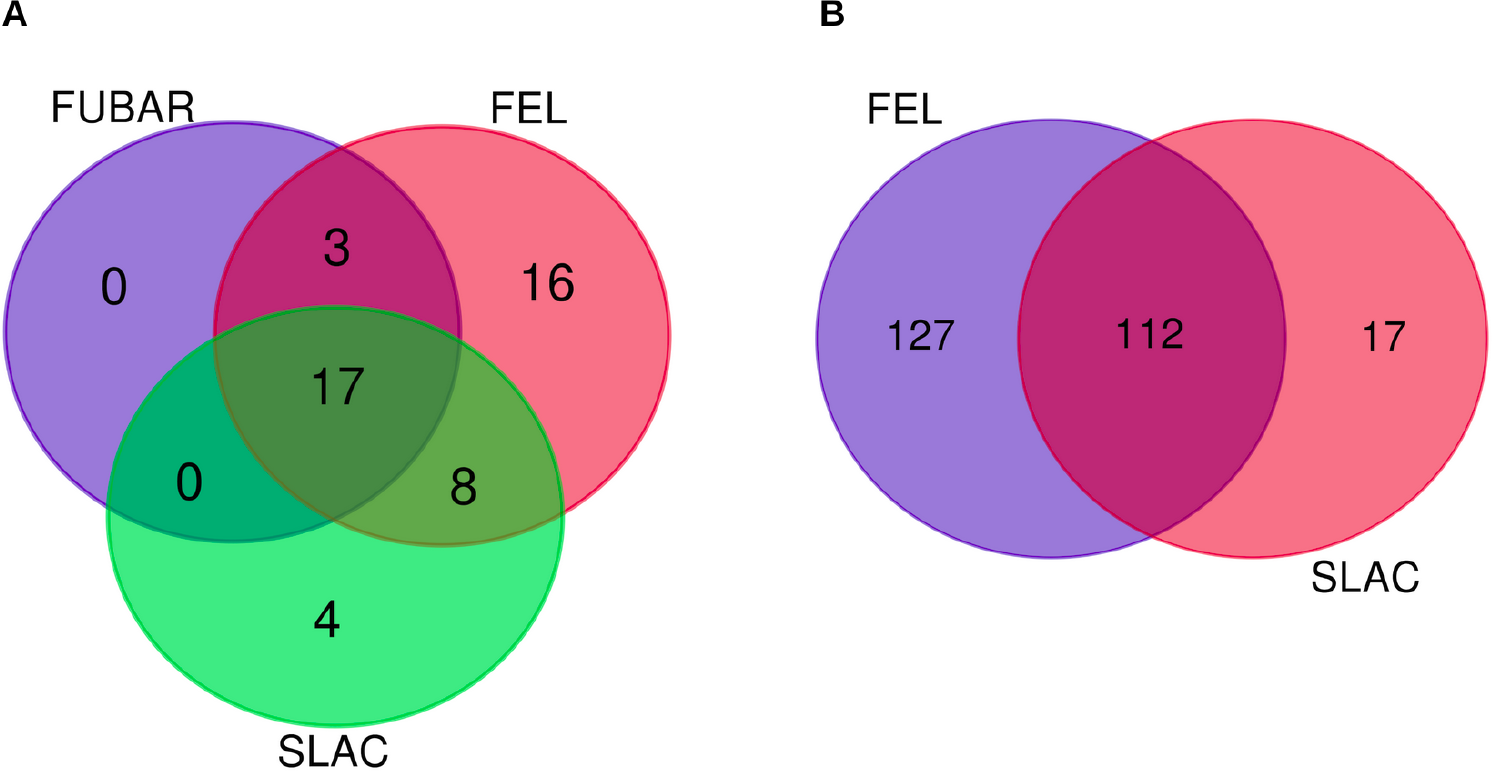
Number of predicted sites under adaptive and purifying selection by each HyPhy method. (A) Positively selected sites shared by FUBAR, FEL, and SLAC methods. (B) Sites under purifying selection detected by FEL and SLAC.

Finally, using a modified version of the Suzuki-Gojobori counting algorithm, the SLAC method assumes that the selective pressure for each site is constant. Despite being the weakest method of this group (Kosakovsky Pond and Frost, 2005), the analysis of pervasive site-selection with SLAC indicated the presence of 130 sites under negative selection, of which 112 were previously found by FEL (Supplementary File 3 and Figure 2B). In relation to the 29 positively selected sites, 17 of them were previously identified by the FUBAR and FEL methods (Table 1), with eight also found by FEL (12, 26, 98, 570, 653, 684, 846, and 1,078) and four only identified by SLAC (27, 701, 769, and 1,228). Considering the consensus sites identified by the three selection methods, the residues 5, 21, 67, 75, 138, 142, 222, 262, 484, 572, 614, 681, 688, 845, 1,176, 1,219, and 1,264 presented the strongest evidences for positive selection pressure on the spike protein (Table 1 and Figure 2A). Following are those recognized by two methods such as sites 12, 26, 98, 417, 501, 570, 653, 684, 1,078, and 1,260.

These missense substitutions were identified in up to 15 different lineages (except for mutations in site 614). This is the case of V1176F, which arose as a B.1.1.28 lineage-defining mutation and is now spread along the phylogenetic tree. The arithmetical average calculation (without site 614) showed an occurrence of 3.22 lineages per each amino acid substitution type, while N501Y was found in 10 lineages, N501T was described in only four. Despite the occurrence of different substitutions in the same site for different lineages, generating an average of 2.19 amino acid mutation possibilities for each mutated site, some lineages were predominantly found. From the 37 identified lineages (excluding those from site 614), the variability of P.1 genomes comprised 45 of the 48 sites under adaptive selection considering all predicted sites by FUBAR, FEL, and SLAC, in some cases, covering multiple substitutions on the same site (e.g.: A67S/V/P, P681H/L/R). For the sites predicted by the three methods (n = 17), the lineages P.2 and B.1.1.28 were identified in the analysis of all events (Supplementary File 2).

### Spike coevolutionary analysis

In order to evaluate if the spike protein sites could be under a coevolutionary process, we performed a HyPhy BGM analysis. The Bayesian Graphical Model (BGM) method is a tool for detecting coevolutionary interactions between amino acid positions in a protein. This method works by detecting pairs of positions with correlated mutations in protein multiple sequence alignments. It also performs a statistical analysis of the distribution of mutations in the branches of the tree. The coevolutionary analysis detected mutational correlations between 62 pairs of sites (Supplementary File 4), of which 28 achieved a P ≥ 0.8 (Table 2). Of these, sites 262, 484, 501, and 681 were identified under diversifying selection and are suggested to coevolve in a conditional manner. Specifically, sites 484 and 501 seem to coevolve. In fact, the E484K + N501Y combination is found in three VOCs: B.1.351, P.1, P.1.1 and P.1.2, besides others such as the Variants of Interest (VOIs) as P.2. For the B.1.1.7 lineage, although the E484K mutation is not lineage-defining, it can be present (with N501Y) in some variants.

**Table 2.**
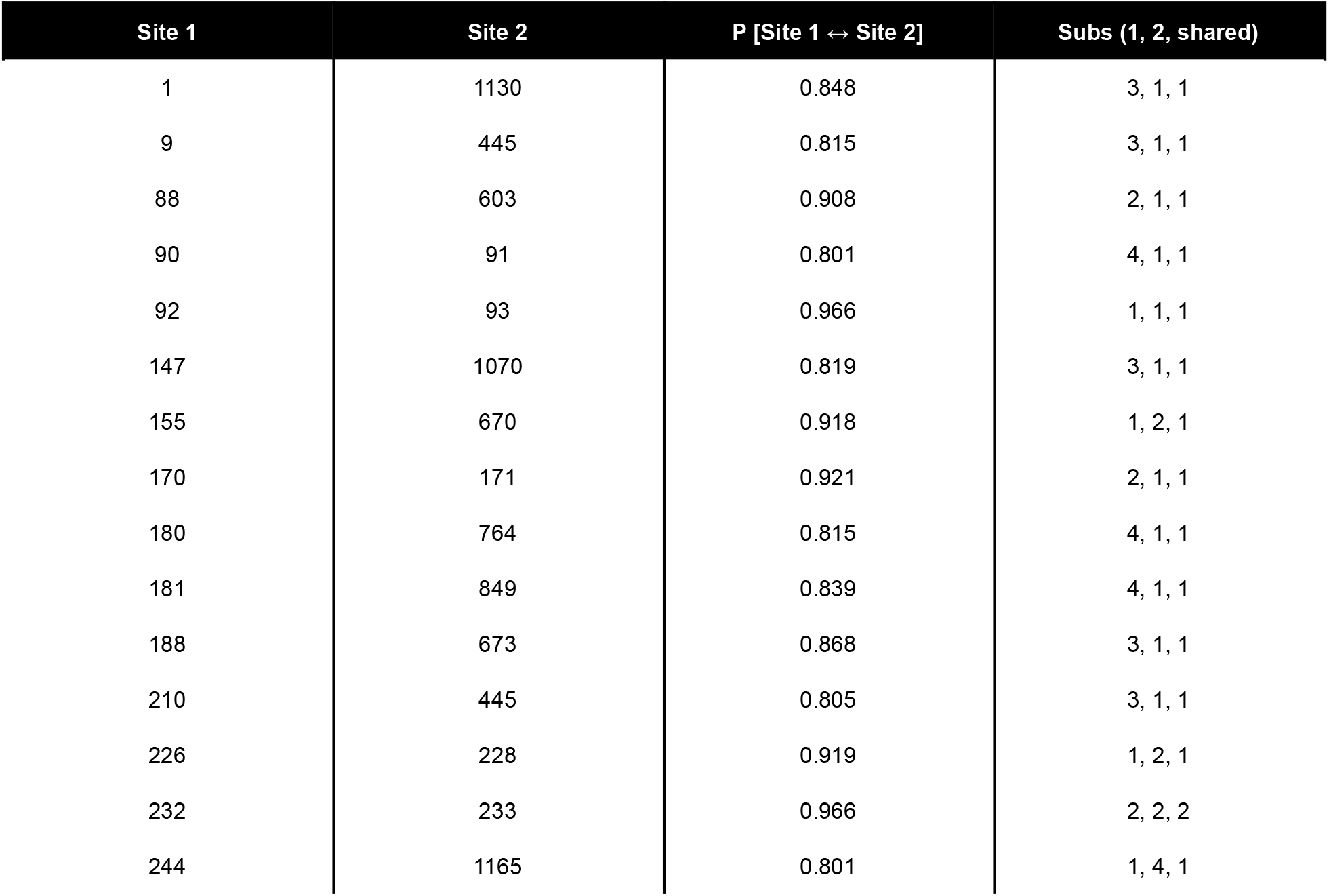
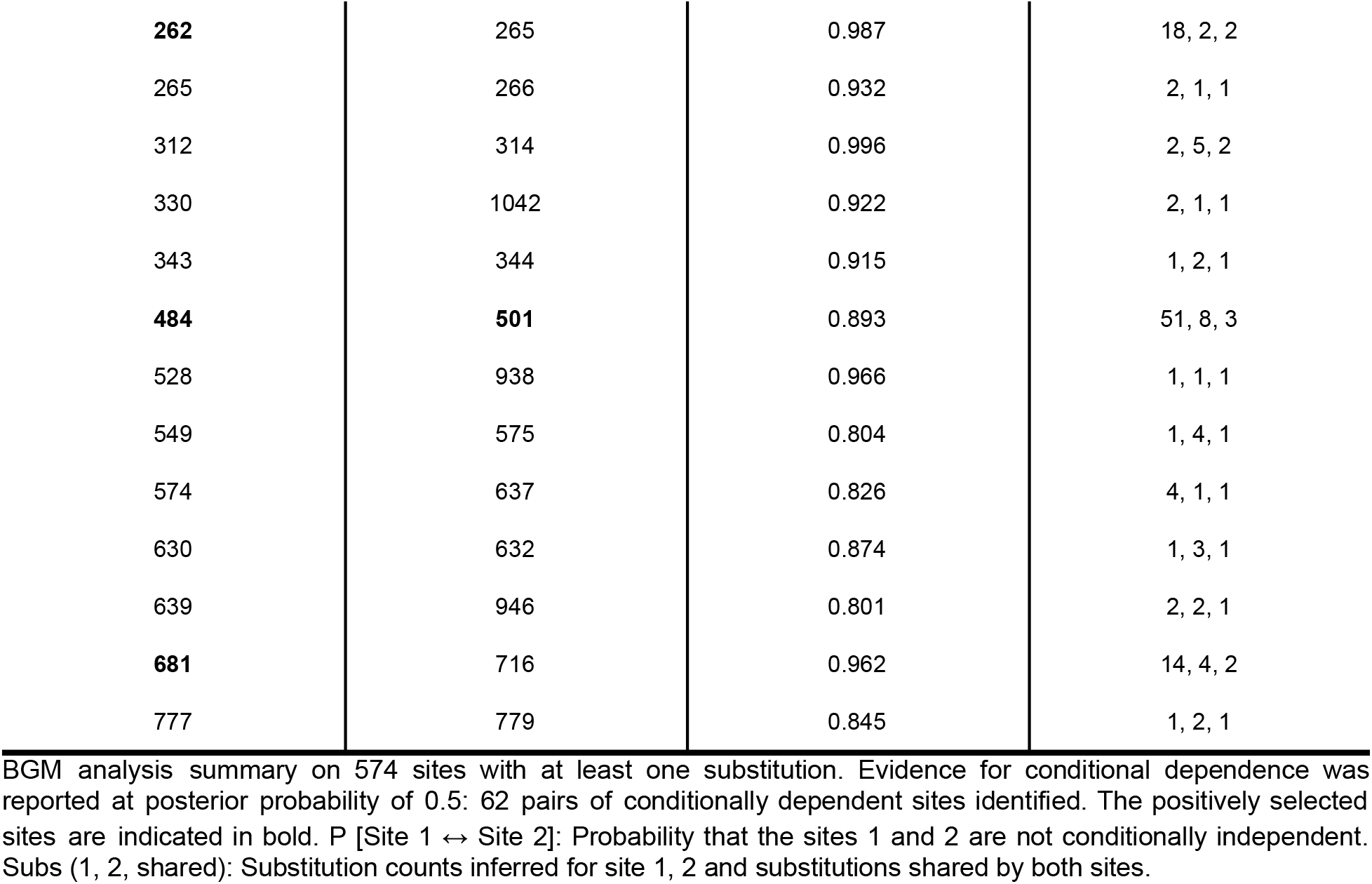
Coevolutionary analysis: Bayesian Graphical Model (BGM) inference to substitution histories at individual sites with P ≥ 0.8.

### Spike structural analyses

To analyze the impact of the spike sites under adaptive selection on the protein stability and host-pathogen interaction, two main categories of structural analyses were applied related to: (i) the folding/unfolding free energies and the vibrational entropy: a reference spike structure PDB ID 6XR8 (relative to the NC045512.2 genome) was used to estimate the energy changes caused by these positively selected single mutations in the prefusion conformation of the spike protein and (ii) protein molecular dynamics: SARS-CoV-2 spike RBD models belonging to different viral lineages in a complex with the human ACE2 were simulated using molecular dynamics and submitted to estimation of binding free energies to provide a better understanding of the effect of those sites under positive selection (single and combined) in the viral fitness.

### Spike structural stability

The evaluation of the structural stability of the spike protein considered the results of five different methodologies in order to find a majority consensus. The application of DynaMut, FoldX, iMutant, MAESTRO, and PremPS provided the estimate of the unfolding and total free energies, as well as the vibrational entropy of the mutated structures. Despite some sites presenting opposite trends in relation to the stabilizing/destabilizing impact, which denotes the specific differences in energy calculations among these algorithms (Figure 3A), a majority consensus result (found by 3 or more tests) was obtained (Figure 3B and Table 3). Sites without resolution in the crystallographic structure, located at N/C-terminal regions, as those in the furin cleavage site (among others), were not analyzed for the energy estimation.

**Figure 3.**
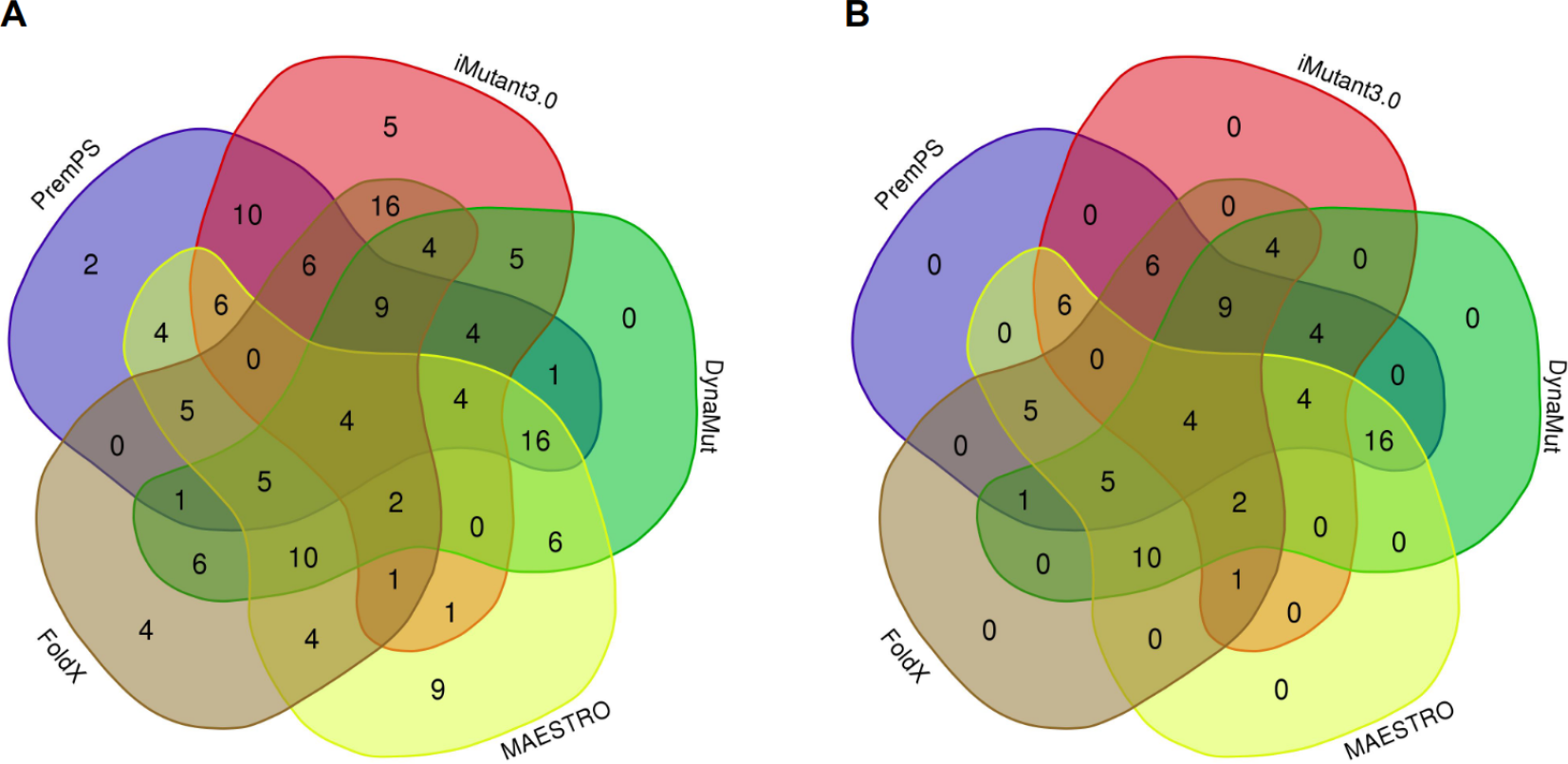
Results for the effect of mutations on the spike protein using different methods. **(A)** Distribution of the number of mutations and their stabilizing/destabilizing effect for each tested method considering 77 mutations in 35 positively selected sites. **(B)** Analysis of all 77 mutations in 35 positively selected sites, considering only the majority consensus, i.e. stabilizing/destabilizing effect appearing in three or more methods.

**Table 3.**
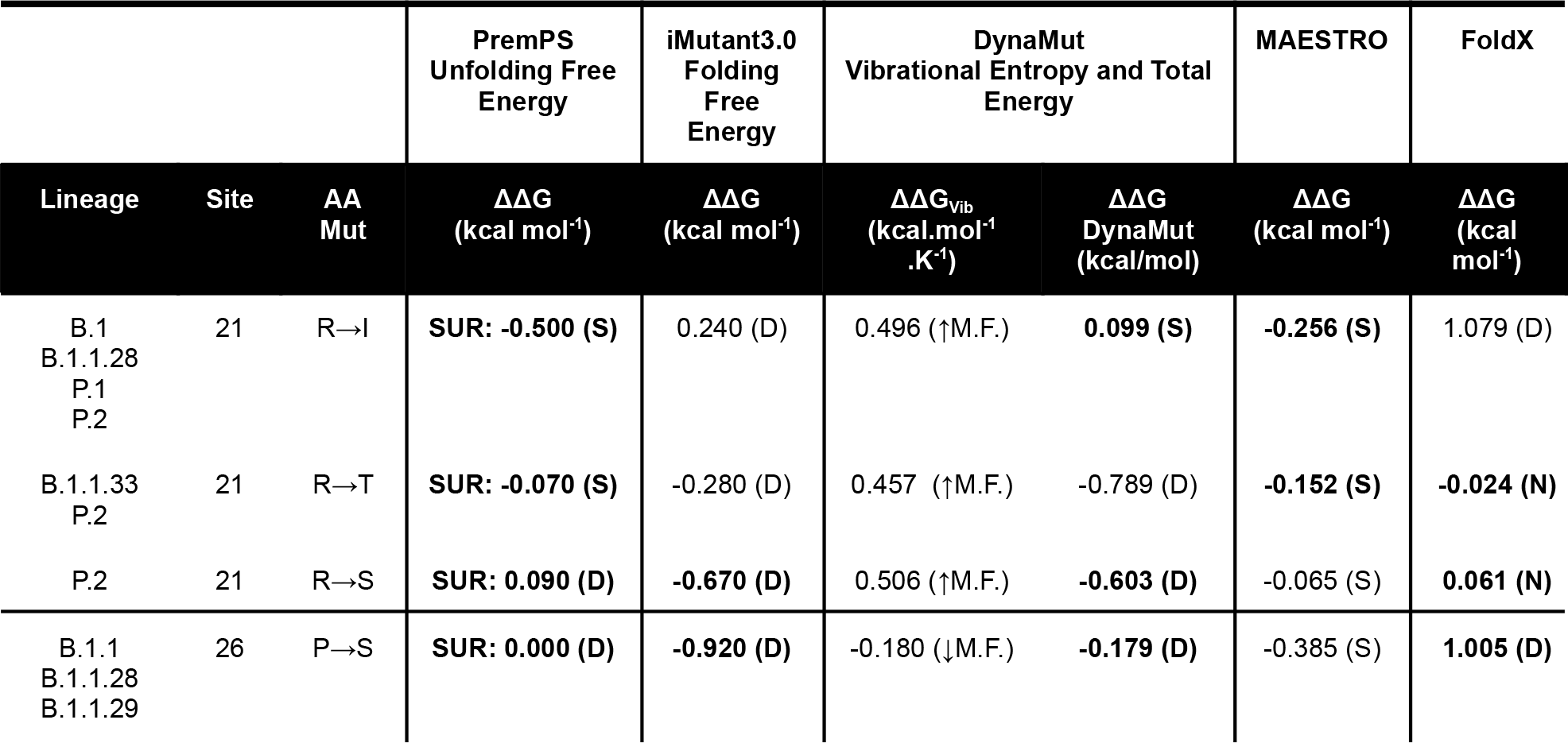
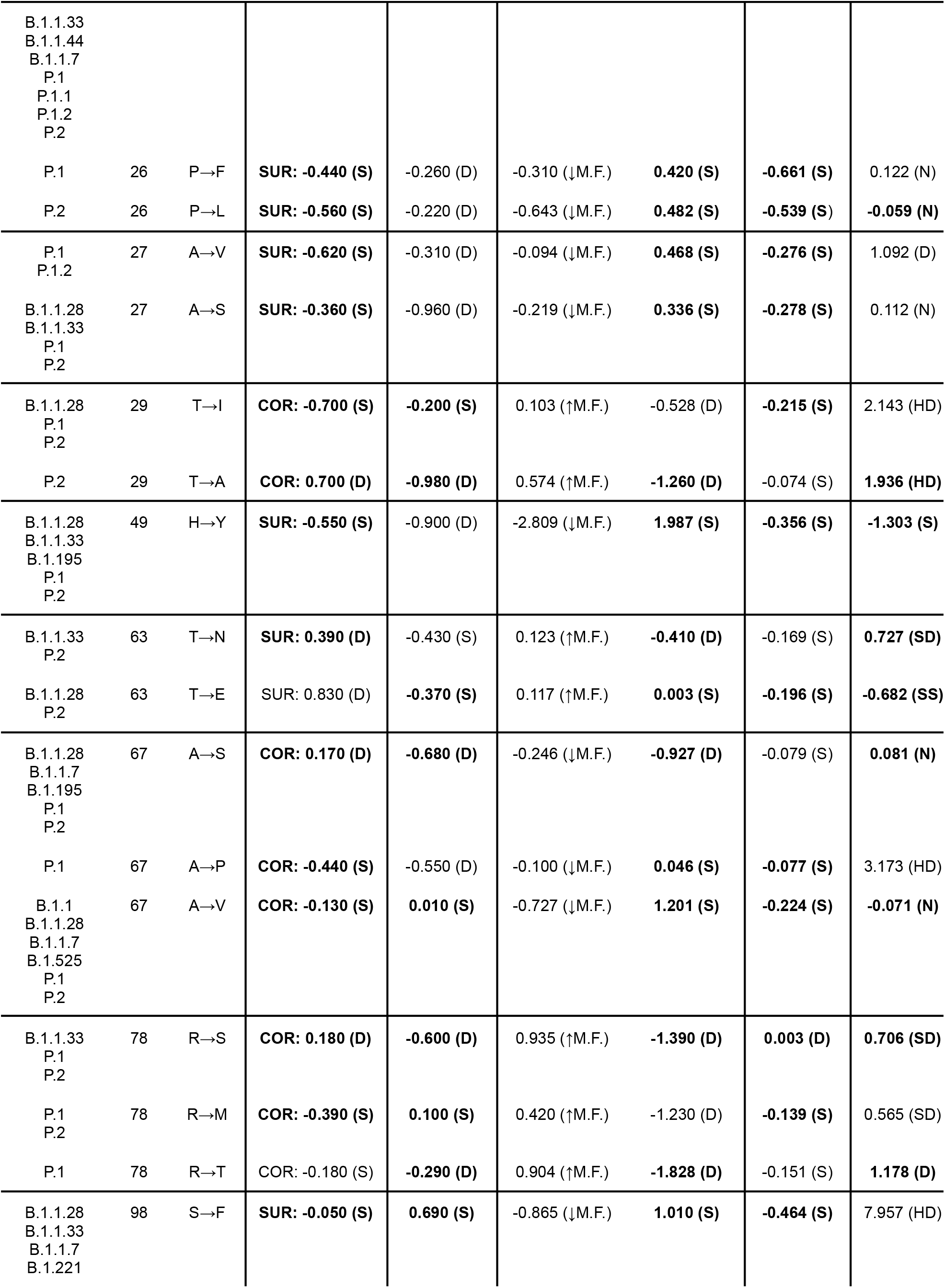
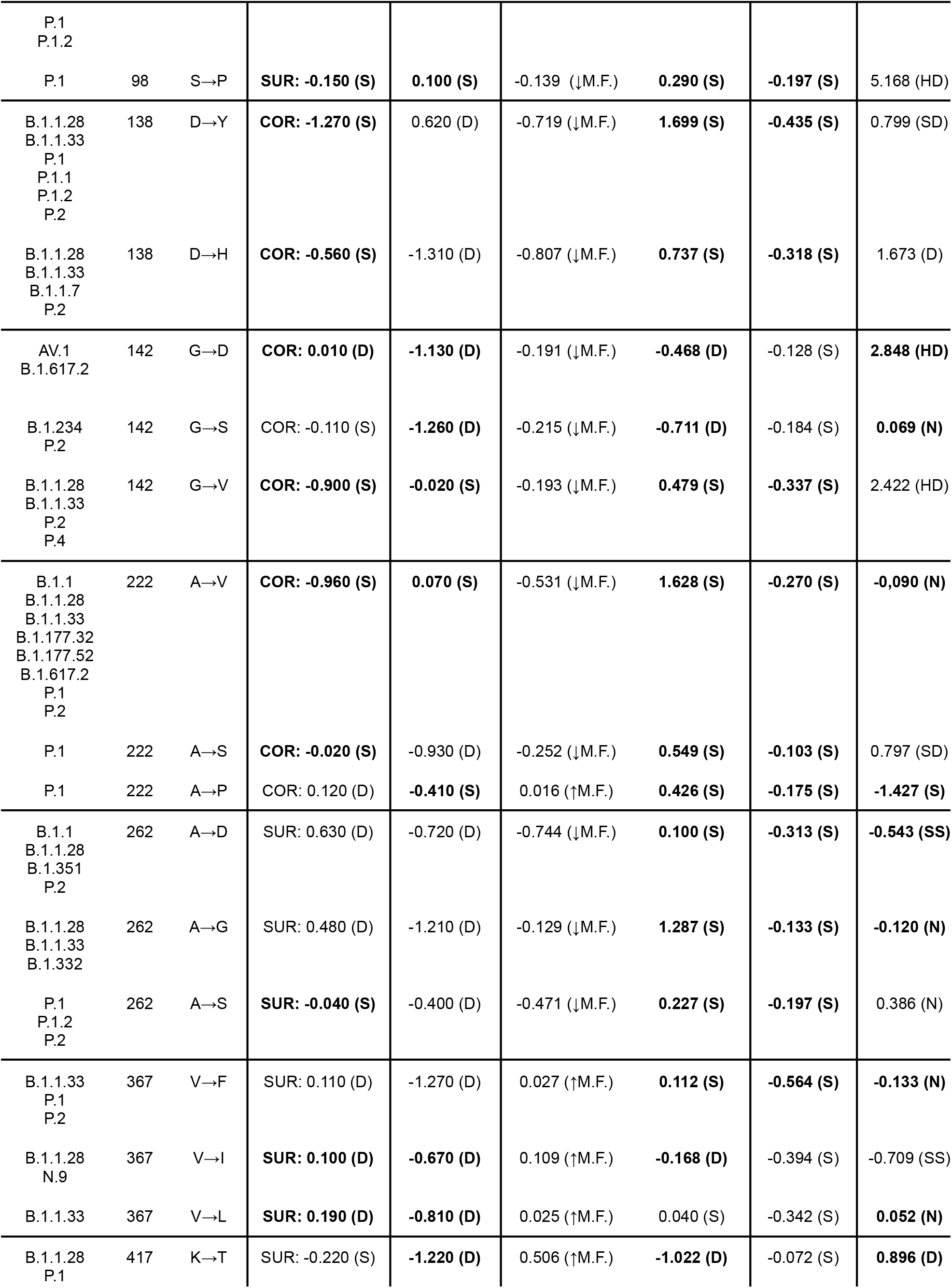
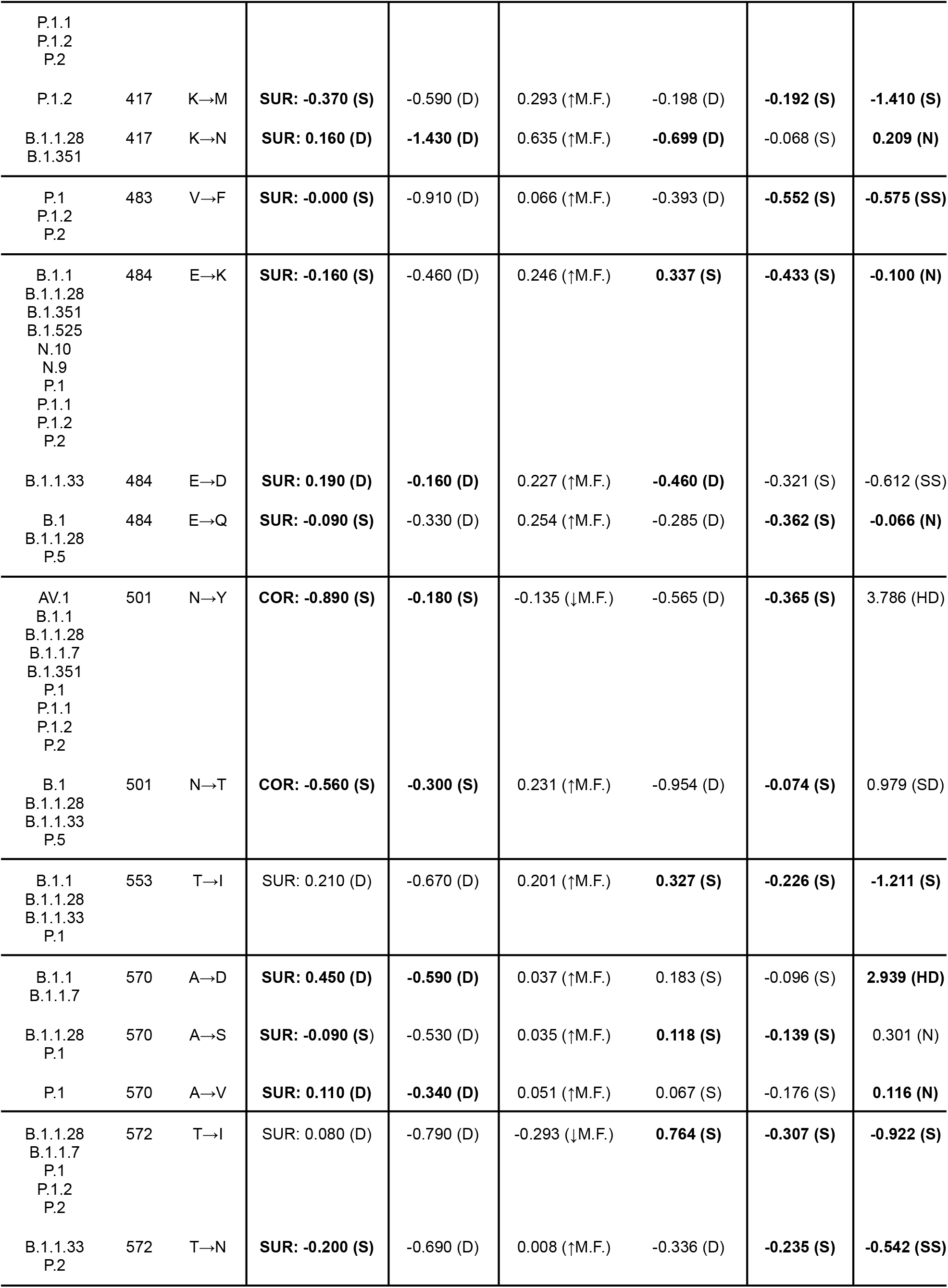
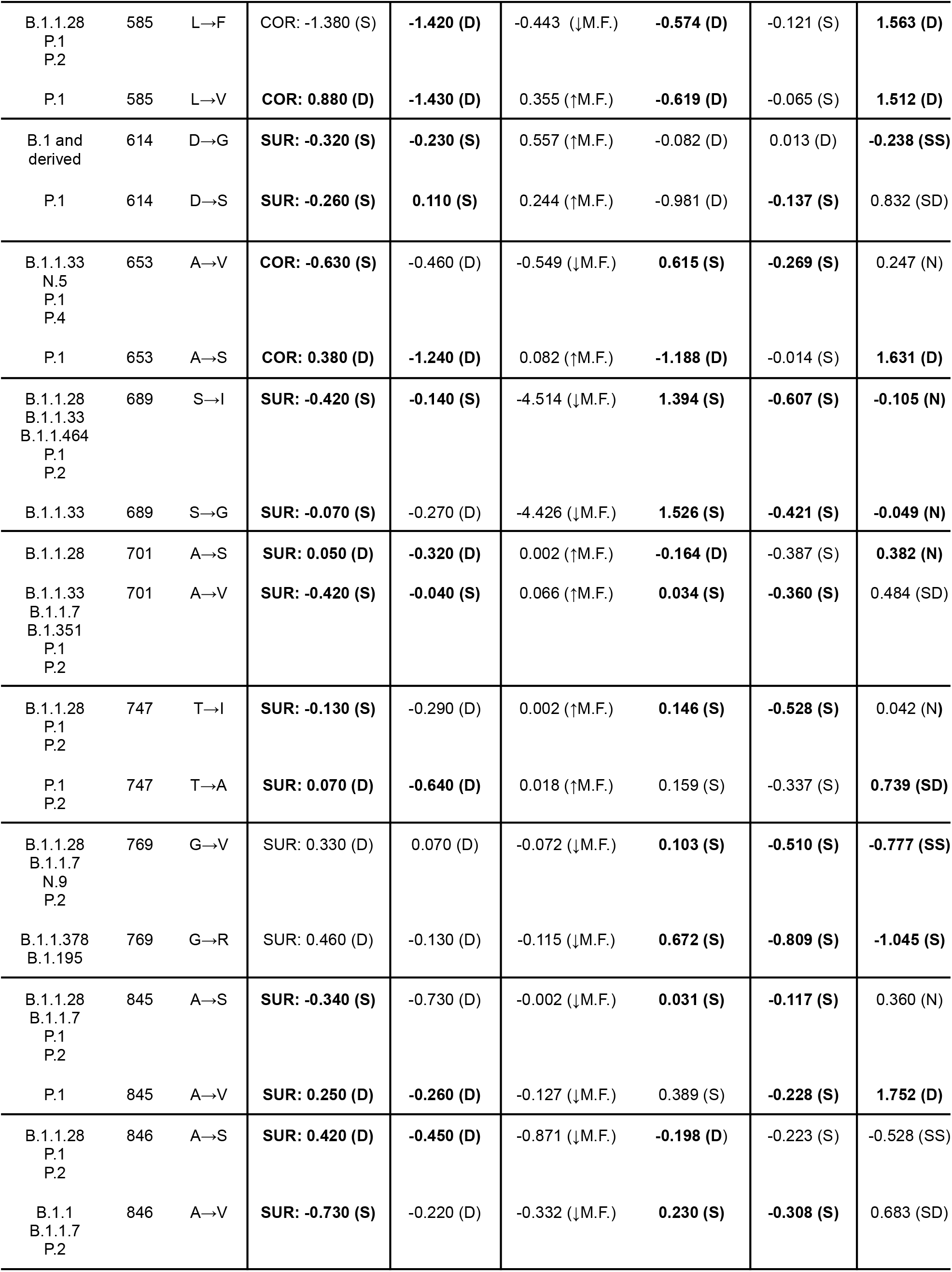
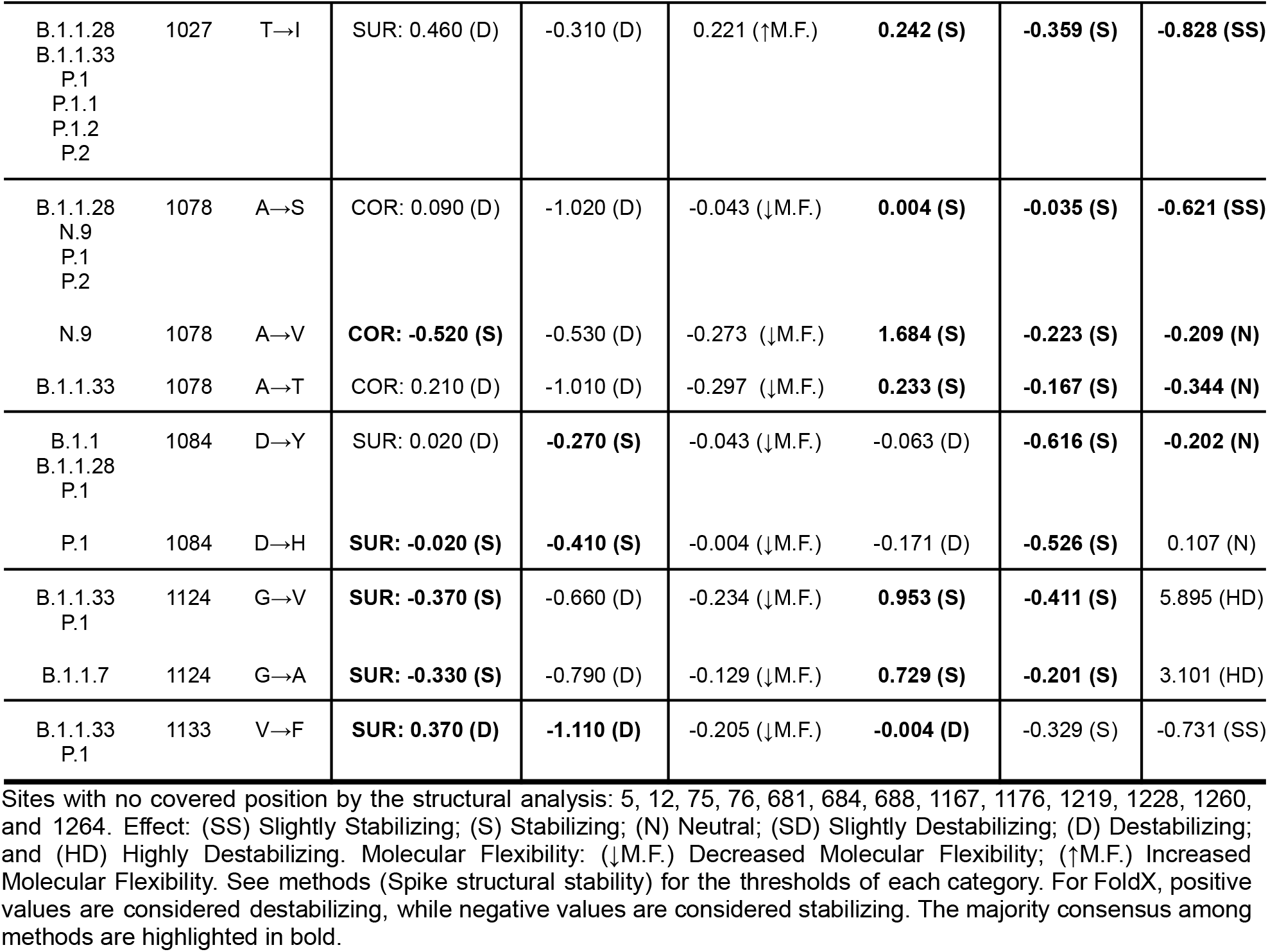
Energy estimation for positively selected mutations.

The PremPS algorithm was used to calculate the changes in unfolding Gibbs free energy generated by each single mutation in the full-lenght prefusion conformation of the spike protein. As observed in Table 3, important mutations for the VOC lineages B.1.1.351, B.1.1.7 and P.1, such as E484K and N501Y, seem to stabilize the protein structure by the decrease of the unfolding free energy change (−0.16 and −0.89 kcal/mol, respectively). Substitutions as those from residues 138 and 585 presented the lowest ΔΔG values, which may suggest a higher stabilizing effect of the mutations D138Y and L585F in lineages as P.1 and P.2, among others. However, for site 585, a destabilizing effect was found by three methods (iMutant, DynaMut and FoldX).

The analysis of the impact of the positively selected mutations on protein dynamics and stability was performed with the DynaMut web-server. The higher vibrational entropy (ΔΔG_Vib_ ENCoM) found in sites 21 (R→S), 78 (R→S/T) and 417 (K→T/N) indicated an increased molecular flexibility (>0.5 kcal/mol) due to the loss of molecular interactions. Moreover, the consensus results correlated with the predicted destabilizing impact. In other sites, such as 49, the conformational molecular rigidity caused by a reduced vibrational entropy (−2.809 kcal/mol) follows the stabilizing effect predicted by DynaMut, MAESTRO, and FoldX. Especially, site 689 achieved the lowest ΔΔG_vib_, with an estimated reduction in the molecular flexibility of −4.514 kcal/mol in the serine to isoleucine substitution. Although the substitution of a negatively charged glutamic acid by a positively charged lysine increased the vibrational entropy in the E484K-mutated protein, slightly decreasing the molecular rigidity (ΔΔG_vib_ = +0.246 kcal/mol), a potential stabilizing effect of this mutation was predicted by PremPs, Dynamut, MAESTRO, and FoldX analyses.

The iMutant3.0 web server was used to evaluate the changes in the thermodynamic stability of folded proteins. Using the physiological pH (7.4) and temperature (36.5 °C) as parameters, the SVM algorithm provided the estimation of the free energy change (ΔΔG) between the wild-type and the mutated 6XR8 spike structures. According to the analysis of the thermodynamic properties, the mutations marked as positively selected seem to destabilize the protein structure. However, the majority consensus evaluation did not confirm this trend, suggesting 51 substitution events (from 33 sites) as stabilizing mutations.

Finally, the FoldX pipeline repaired the 6XR8 PDB file by the correction of residues with bad torsion angles or van der Waals clashes and performed the energy minimization testing different rotamer combinations. As result of the mutagenesis and total energy estimation, the mutations in sites T29A, G142D, and A570D were considered as highly destabilizing, increasing the total energy in 1.936, 2.848 and 2.939 kcal/mol, respectively. Despite some energy differences being categorized as neutral, the increase or decrease of the math signal suggests the potential effect of these mutations on the structural stability of spike, in the tested conditions. The comparison between the mutational effect predicted by the five methods and the majority consensus result are presented in Figure 4. The evaluation of all existing substitutions for each positively selected site by the majority consensus indicated the prevalence of stabilizing mutations, which was observed for the important VOC and VOI mutated sites. The analysis of the positively selected sites according to the prevalent substitution per site obtained a similar profile, with 25.71% of the mutated sites generating a destabilizing effect in the spike prefusion structure (Figure 5).

**Figure 4.**
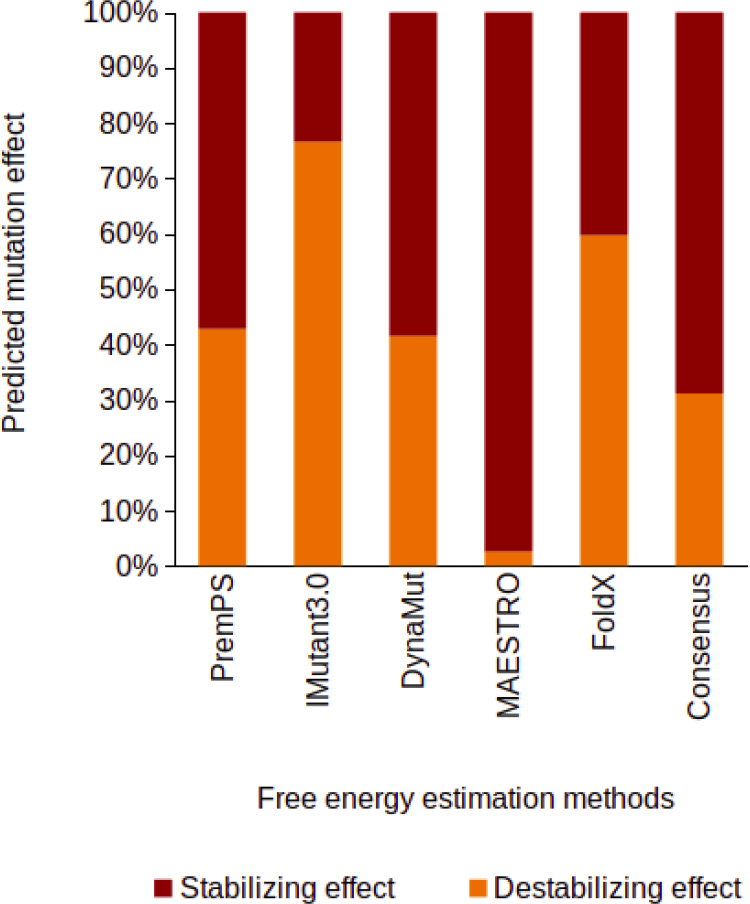
Predicted mutation effect on the positively selected sites according to the tested methods (PremPS, iMutant3.0, DynaMut, MAESTRO, and FoldX) and the majority consensus results (energy trend estimated by three or more methods).

**Figure 5.**
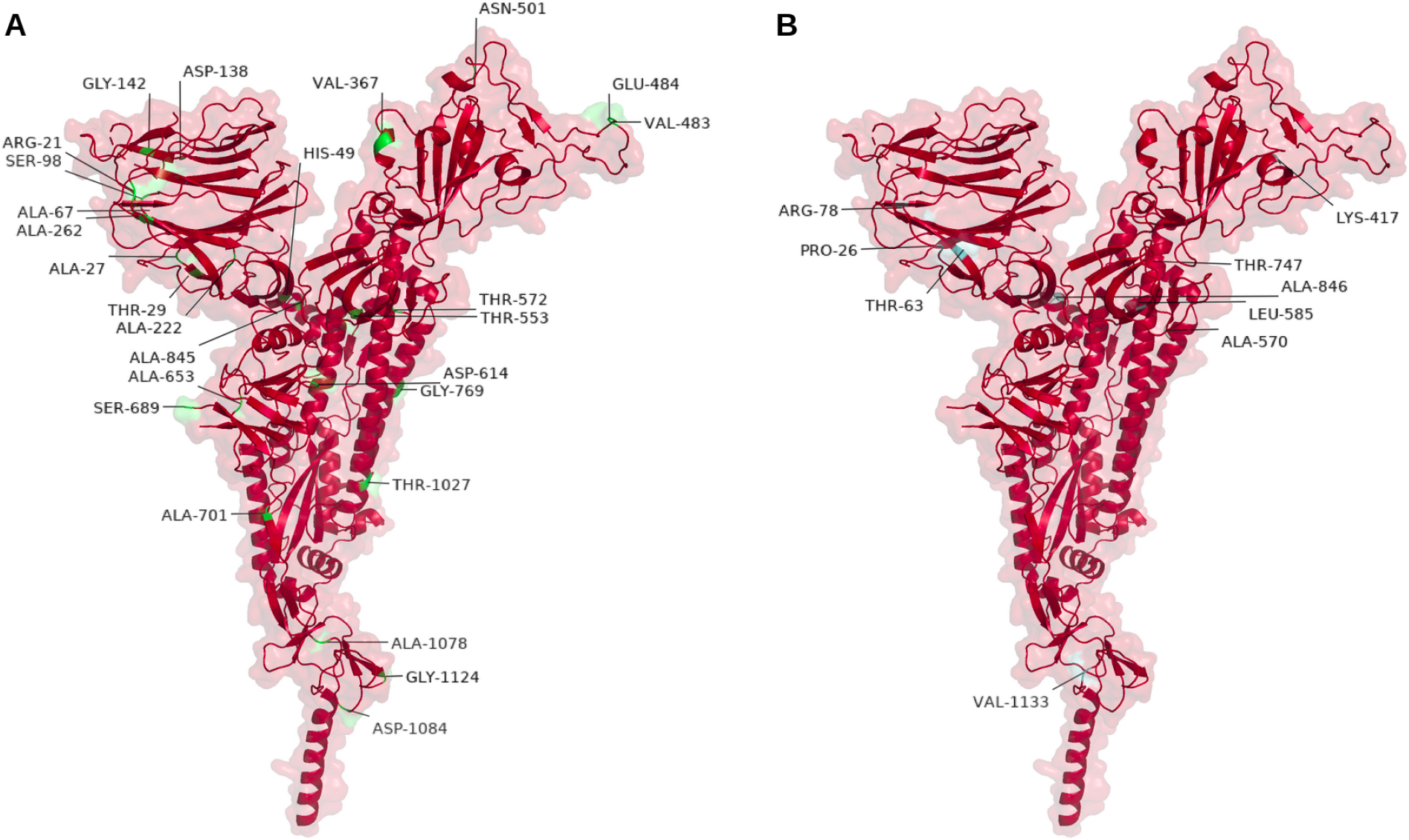
Identification of the stabilizing and destabilizing mutations in the prefusion spike protein structure (6XR8, chain A). (A) Stabilizing mutations (highlighted in green and labeled) related to the positively selected sites, according to the consensus majority analysis. (B) Destabilizing mutations (highlighted in cyan and labeled) related to the positively selected sites, according to the majority consensus analysis. For visualization purposes, sites with multiple possible mutations were represented by the prevalent one and its respective effect (stabilizing/destabilizing).

### Spike RBD-ACE2 structural modelling

In total, 30 theoretical models with 194 amino acid length for the spike RBD - ACE2 protein complexes were obtained using PDB ID 6M0J as template, being five models generated for each lineage (reference, P.1, P.2, C.37, B.1.1.7, and P.2+452). The model quality was evaluated by analysis of DOPE score and through PROCHECK/VERIFY3D parameters (Supplementary File 5). The models covered the region between the amino acid positions 333 and 526 of the reference spike protein sequence (relative to the genome NC_045512.2) and from five SARS-CoV-2 lineages: C.37 (L452Q + F490S), B.1.1.7 (N501Y), P.1 (K417T + E484K + N501Y), P.2 (E484K), and P.2+452 (E484K + L452V) (Figure 6). The final selected models for the reference as well as for the lineages C.37, B.1.1.7, P.1, P.2, and P.2+452 had 88.7 up to 89.9% of the residues in favoured regions and 10.1 up to 11.3% of the residues in allowed regions, which indicated the good model quality. There were no residues in generously allowed or disallowed regions (Figure S1). The overall G-Factor values ranged between −0.06 and −0.12. For all selected models, the compatibility of the atomic model with the amino acid sequence was defined with 100% of the residues achieving an average 3D-1D score ≥ 0.2 in the VERIFY3D analysis.

**Figure 6.**
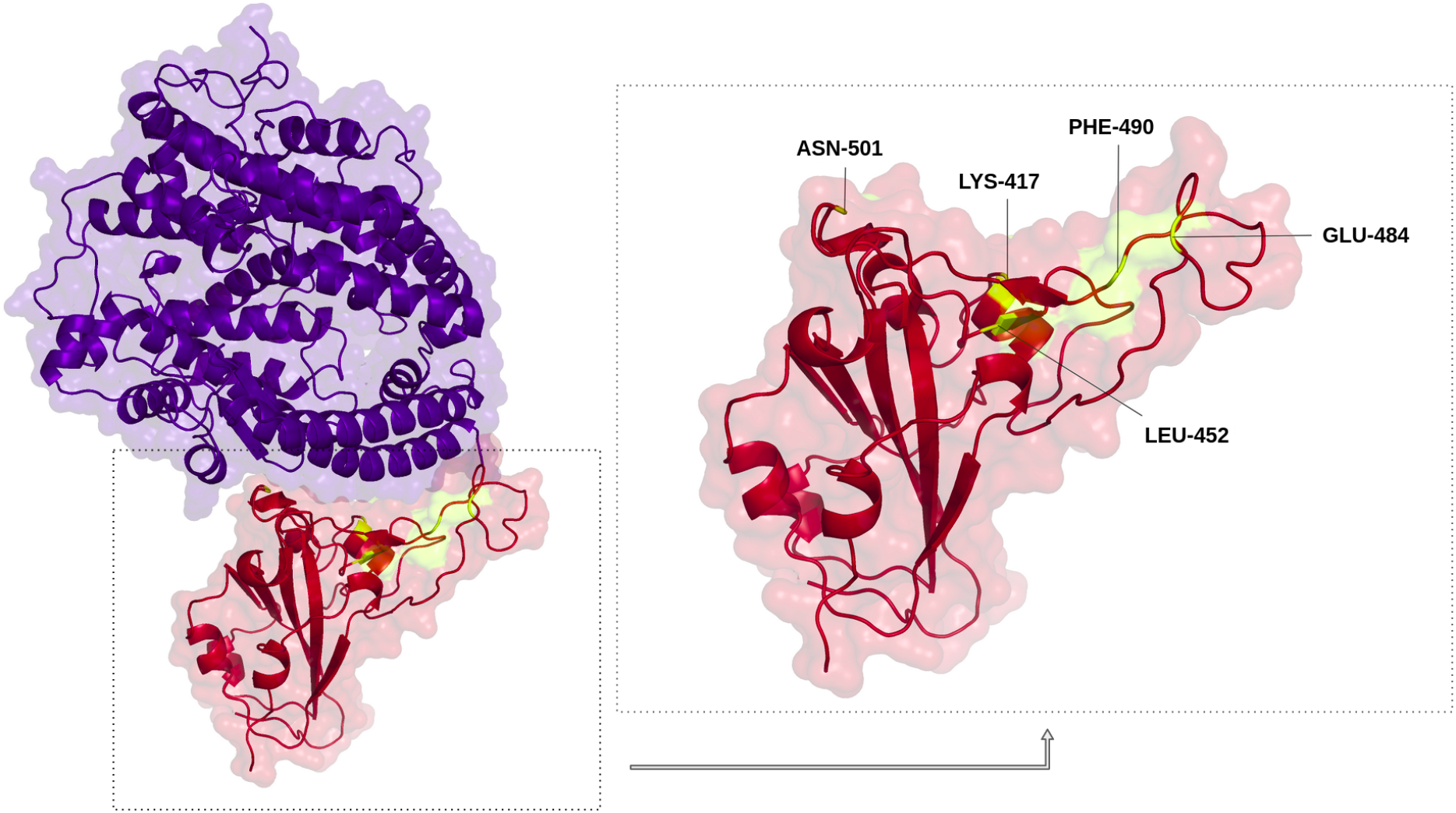
Spike Receptor Binding Domain (RBD) from NC_045512.2 reference genome complexed with human Angiotensin Converting Enzyme 2 (hACE2). The ACE2 structure is colored in blue. The localization of the mutated residues for the modelled RBD structures from lineages C.37 (L452, F490), B.1.1.7 (N501), P.1 (K417, E484, N501), P.2 (E484), and P.2+452 (L452, E484) are highlighted in yellow and labeled.

### Molecular dynamics and binding free energy estimation

The analysis of the RMSD, center of mass and minimum distances indicated that all systems were structurally stable, in the considered time window, with overall conservation of the arrangement of the complex, as can also be seen by visual inspection of the snapshots (Figures 7 and 8). The lowest changes were found in the reference and lineages P.1 and C.37, meaning higher structural stability. Some structural changes in spike were found in the P.2 and B.1.1.7 lineages, whereas for P.2+452 and B.1.1.7 the structural changes were also noticeable in relation to the ACE2 molecule. For all systems, the spike-hACE2 interaction was characterized by the presence of many close contacts, which is reflected in a substantial interface area, and many hydrogen bonds (Figure 9 and Table 4).

**Figure 7.**
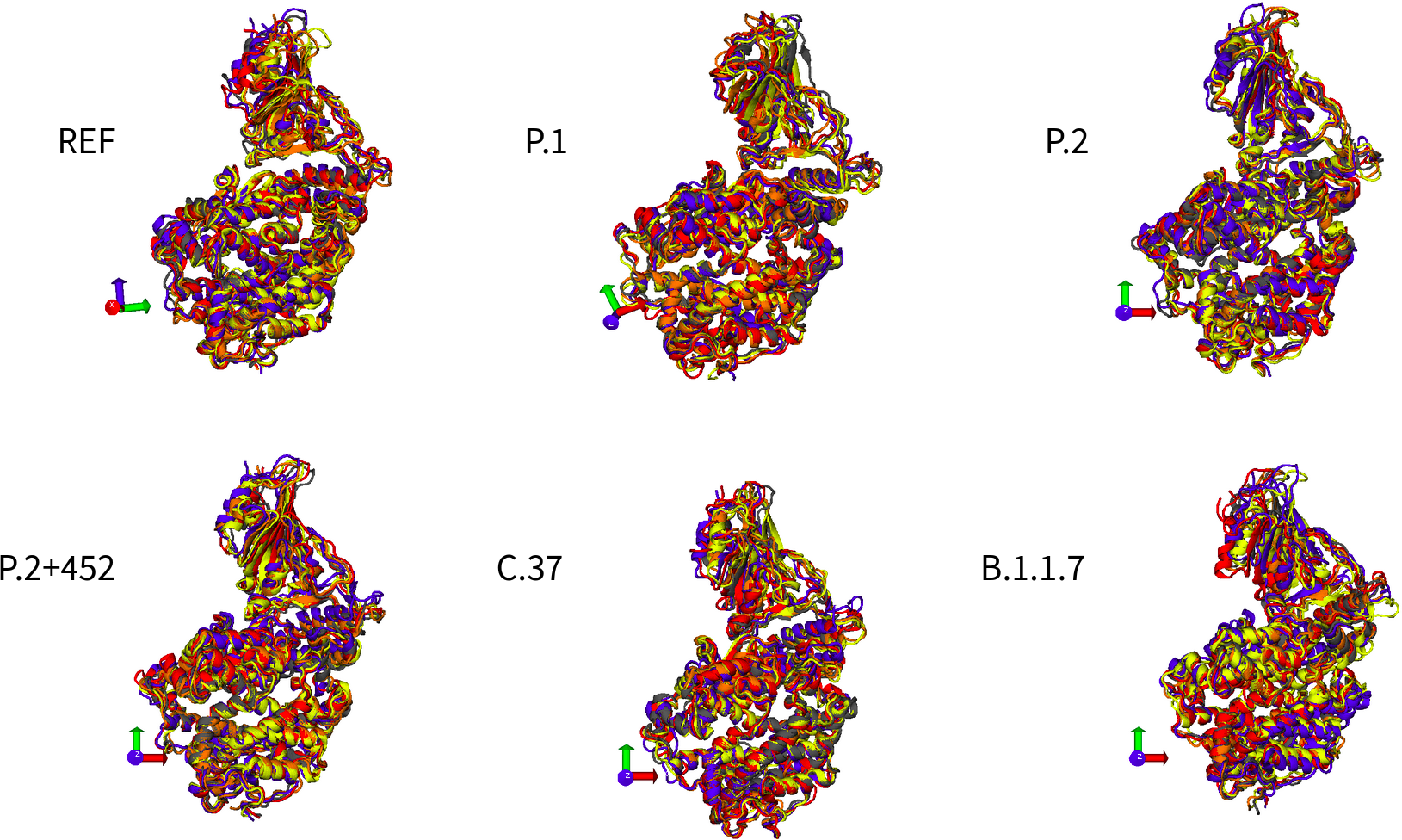
Snapshots from molecular dynamics (MD) simulations in 0, 50, 100, 150, and 200 ns for the spike RBD - ACE2 protein complexes related to the reference (REF) and lineages P.1, P.2, P.2+452, C.37, and B.1.1.7.

**Figure 8.**
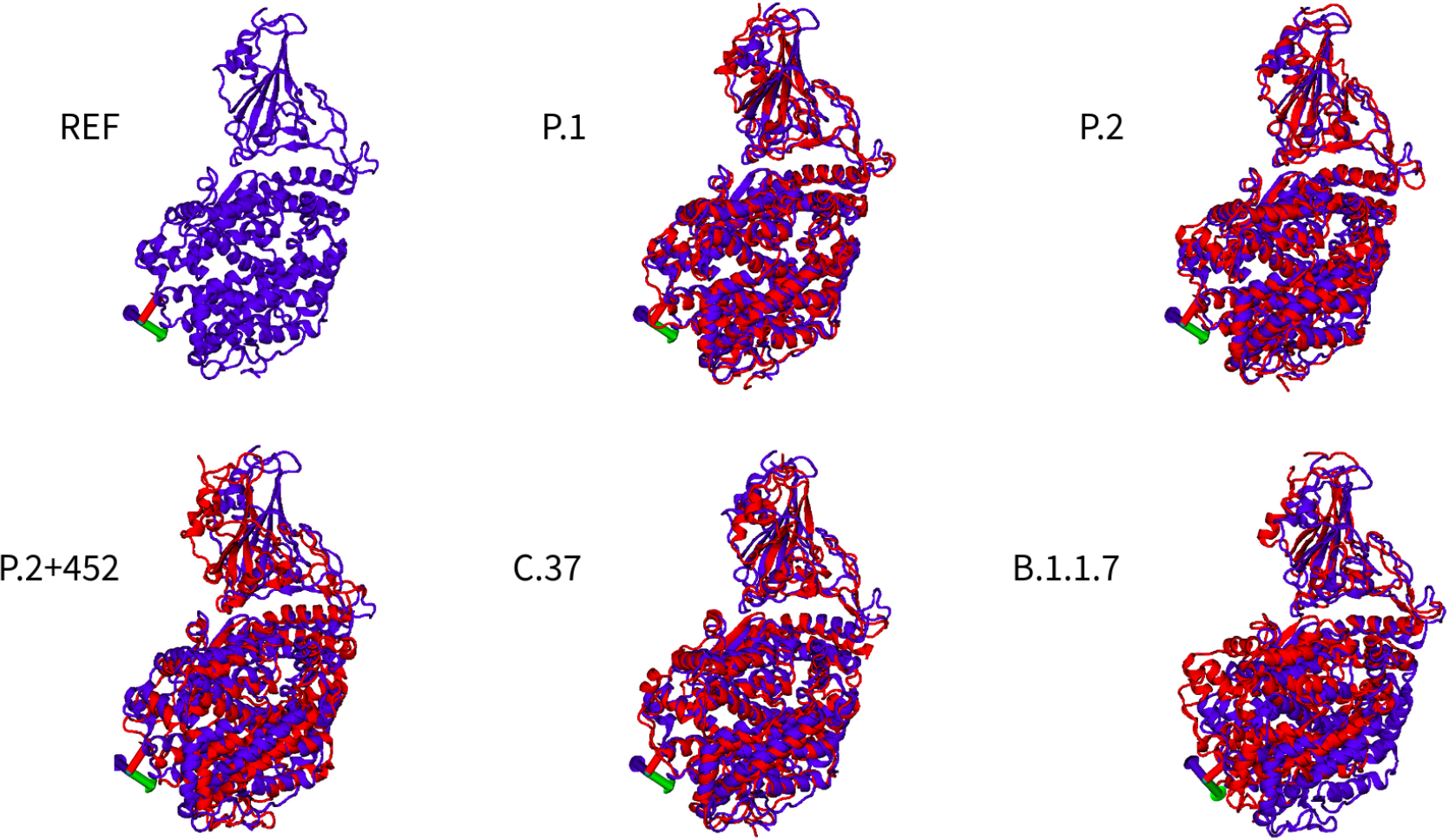
Snapshots from molecular dynamics (MD) simulations - the final configuration fitted in relation to the reference (REF) - for the spike RBD - ACE2 protein complexes belonging to lineages P.1, P.2, P.2+452, C.37, and B.1.1.7.

**Figure 9.**
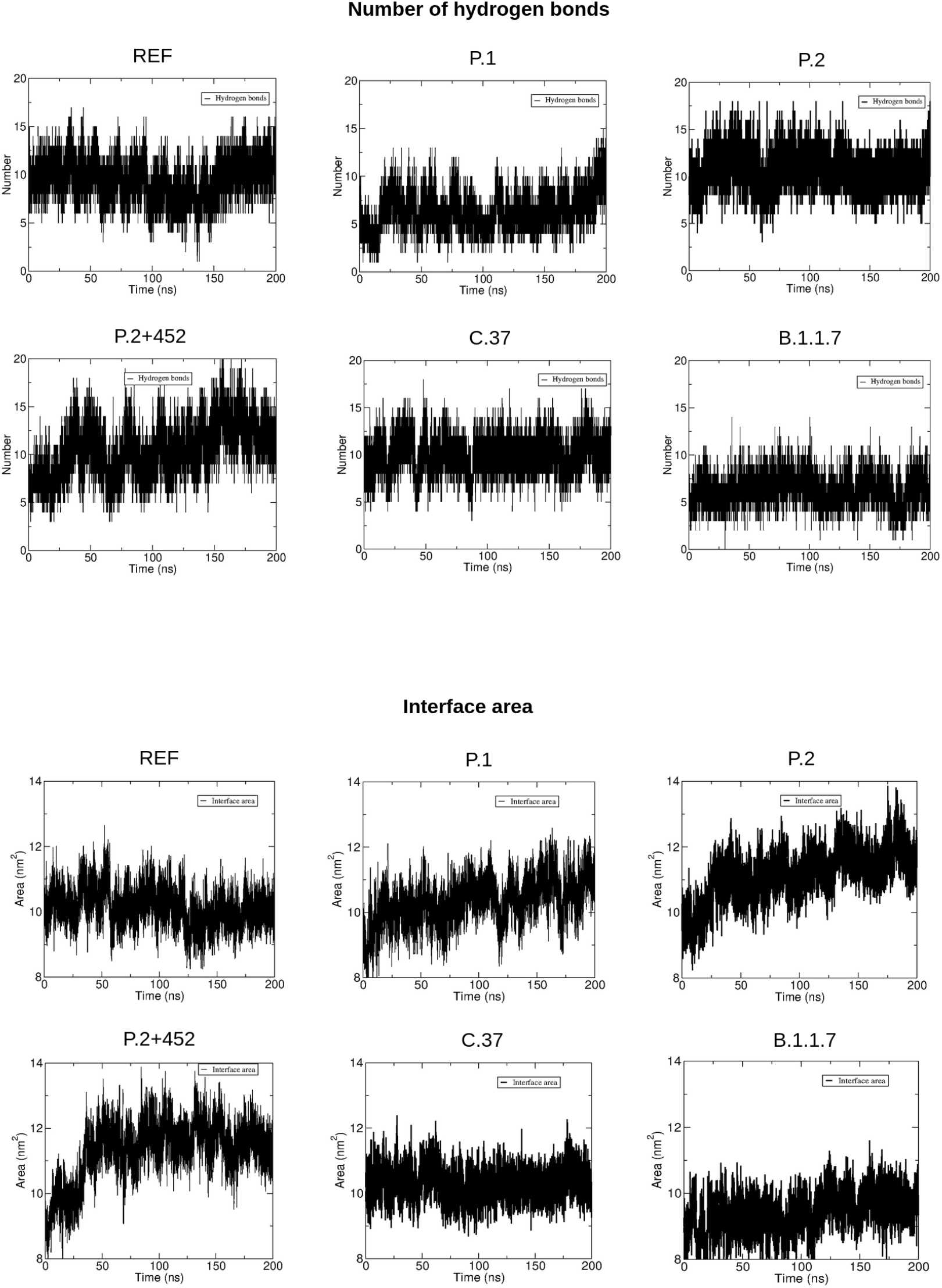
Number of hydrogen bonds and interface area for the spike RBD - ACE2 protein complexes related to the reference (REF) and lineages P.1, P.2, P.2+452, C.37, and B.1.1.7.

**Table 4.**
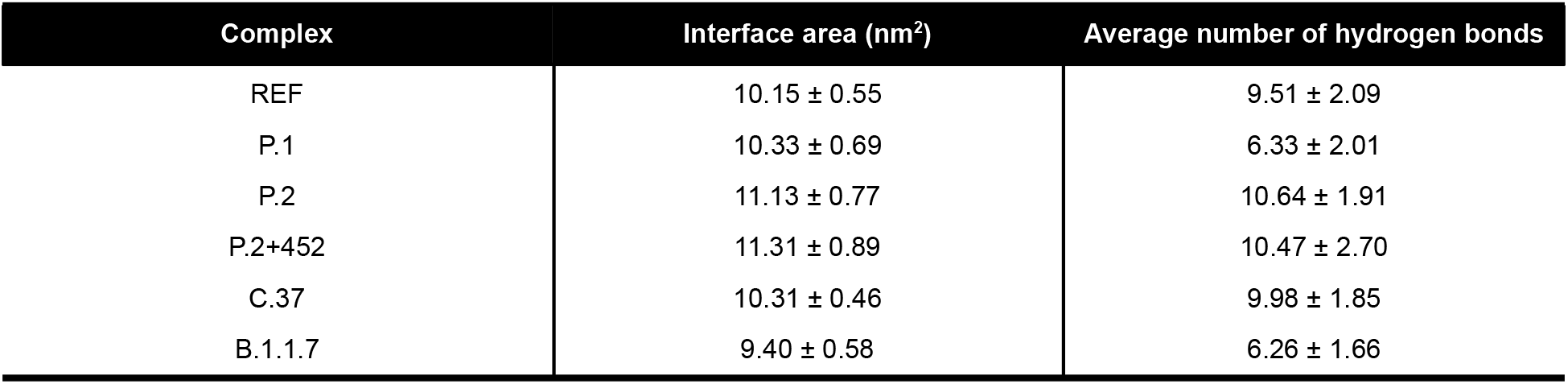
Interface area of the complexes (the half of the sum of solvent-accessible areas of spike and ACE2 minus solvent-accessible area of the complex) and number of hydrogen bonds, calculated as average over the MD simulations.

Despite the distance plots with similar results, indicating that the relative position between the spike and ACE2 is not altered during the simulation time, the lineages P.2+452, C.37 and B.1.1.7 showed a small alteration in the distance, with a slight approximation between spike and ACE2 (Figure S2). In the reference system, there were on average 9.51 ± 2.09 hydrogen bonds and an interface area 10.15 ± 0.55 nm^2^. The systems P.2, P.2+452, and C.37 displayed a higher number of hydrogen bonds and interface area than the reference, whereas the lineage P.1 showed larger interface area but less hydrogen bonds and the lineage B.1.1.7 showed smaller interface area and number of hydrogen bonds. About the number and temporal evolution of the hydrogen bonds, the order followed: P.2 > P.2+452 > C.37 > reference > P.1 > B.1.1.7. The intensity of the hydrogen bonds in the lineages P.2 and P.2+452 is reflected in the interaction intensity with ACE2. As for the contact area between spike and ACE2, proportional to the intensity of the van der Waals interactions (also reflected in the interaction intensity), the order followed with a slight change: P.2+452 > P.2 > P.1 > C.37 > reference > B.1.1.7 (Figure 9 and Table 5).

**Table 5.** Binding free energies and their components, calculated using MM/PBSA, for the interaction between ACE2 and the lineages a) REF, b) P.1, c) P.2, d) P.2+452, e) C.37, and f) B.1.1.7.

| Complex | Van der Waals Energy (kJ/mol) | Electrostatic Energy (kJ/mol) | Polar Solvation Energy (kJ/mol) | SASA Energy (kJ/mol) | Binding Energy (kJ/mol) |
| --- | --- | --- | --- | --- | --- |
| REF | -406.772 +/- 2.081 | -1405.564 +/- 7.144 | 724.141 +/- 10.657 | -45.611 +/- 0.286 | **-1134.283 +/- 10.350** |
| P.1 | -417.644 +/- 2.970 | -1627.348 +/- 7.132 | 533.145 +/- 11.232 | -45.812 +/- 0.302 | **-1558.114 +/- 10.124** |
| P.2 | -448.576 +/- 3.682 | -2348.699 +/- 7.801 | 811.214 +/- 11.373 | -49.739 +/- 0.325 | **-2035.611 +/- 10.218** |
| P.2+452 | -437.065 +/- 3.066 | -2494.514 +/- 8.456 | 911.602 +/- 11.865 | -50.055 +/- 0.351 | **-2069.977 +/- 9.690** |
| C.37 | -417.849 +/- 2.285 | -1412.224 +/- 8.546 | 740.556 +/- 12.203 | -46.253 +/- 0.257 | **-1135.494 +/- 9.501** |
| B.1.1.7 | -374.295 +/- 2.474 | -1295.292 +/- 5.377 | 571.656 +/- 10.855 | -41.636 +/- 0.274 | **-1140.138 +/- 11.183** |

In all cases, the binding free energy of the mutant spikes was found to be more negative (stronger binding) than the native one and the order is roughly similar to the intensity of interface area or hydrogen bonds: P.2+452 > P.2 > P.1 > B.1.1.7 > C.37 > reference. Specifically, for P.2+452 and P.2, the binding intensity was significantly stronger than the reference, while for B.1.1.7, C.37 and reference the interaction intensity was almost the same (Table 5).

In relation to the hotspots for the RBD-ACE2 interaction, the residue distances for the wild-type (reference) and the mutated sites were calculated (Supplementary Figures S3-S7). The spike K417 interacts with hACE2 residues T27, D30, K31, and H34 in the wild type, while the mutation for T417 in P.1 maintains the same contacts. In the P.2 lineage, K417 is affected by the E484K mutation, changing the residue interactions to the hACE2 D30, N33, H34, and P389, which is the same profile identified in B.1.1.7, despite the higher distance variation during the simulation time. The impact of the E484K + L452V combination in P.2+452 changes the residue interaction of K417 to D30, H34, Q388, and P389. For the C.37 lineage, the effect of L452Q and F490S modified the residue interaction of K417 to T27, D30, H34, and P389. The residue N501 in the wild type RBD-ACE2 complex interacts with L351, G352, K353, G354, D355, and Y41. This same pattern is kept by P.1, with a slight reduction in the Y501-K353 distance. N501 of P.2, P.2+452 and C.37 lineages presented the same ACE2 interactions. However, C.37 showed a slight increase in the distance value for F356 and D355 in the middle and end of the simulation time. Finally, Y501 in B.1.1.7 showed the highest variation, creating a new interaction with hACE2 residue H34.

In relation to the magnitude of interaction, all three top-ranked lineages bear the same E484K mutation, which yielded a particularly strong binding, remarkably stronger than in the native case. Indeed, the analysis of the contribution of the residues to the free energy of binding showed that the mutation of the negatively charged residue GLU 484 to the positive charged residue LYS 484 yielded a substantial negative free energy contribution to the stabilization of the complex. The mutation in the residue 501 (ASN to TYR, both polar non charged residues) presented in the lineages P.1 and B.1.1.7 can also contribute to the stabilization of the complex, but much less than E484K (Table 6).

**Table 6.** Contribution of mutations to the Binding Free Energy changes (kJ/mol).

| REF |  | P.1 |  | P.2 |  | P.2+452 |  | C.37 |  | B.1.1.7 |  |
| --- | --- | --- | --- | --- | --- | --- | --- | --- | --- | --- | --- |
| Lys417 | -236.4 | **Thr417** | **-4.6** | Lys417 | -239.9 | Lys417 | -236.8 | Lys417 | -241.3 | Lys417 | -239.8 |
| Leu452 | -1.6 | Leu452 | -2.0 | Leu452 | -1.6 | **Val452** | **0** | **Gln452** | **-3.1** | Leu452 | -1.8 |
| Glu484 | 249.5 | **Lys484** | **-211.4** | **Lys484** | **-218.0** | **Lys484** | -234.5 | Glu484 | 228.4 | Glu484 | 224.3 |
| Phe490 | 0.4 | Phe490 | 1.2 | Phe490 | 3.4 | Phe490 | 0.4 | **Ser490** | **3.2** | Phe490 | 1.3 |
| Asn501 | -12.0 | **Tyr501** | **-15.7** | Asn501 | -12.5 | Asn501 | -10.7 | Asn501 | -12.6 | **Tyr501** | -15.5 |

## DISCUSSION

The spike protein is one of the most rapidly evolving regions in the SARS-CoV-2 genome, with slightly different evolution rates according to the clades, emerging in the beginning and middle 2020 (Pereson et al., 2020). The emergence of the P.1 lineage, at the end 2020, with a possibly increased rate of the molecular evolution, represents an important change in the SARS-CoV-2 evolutionary history (Faria et al., 2021). That is the main reason why lineage assignments should be carefully evaluated, since a considerable number of sequences generally described as P.2 or B.1.1.28 (e.g.) presents a mutation set consistent with the P.1 lineage and are grouped in the same monophyletic clade as well. A systematic and detailed phylogenomic analysis should be conducted in order to better understand and classify new SARS-CoV-2 genomes. The prevalence of P.1 containing mutations in the set of sites indicated to be under adaptive selective pressure may suggest the improvement of viral fitness in this lineage. In fact, a high number of mutations were found in P.1 variants, besides the 10 lineage-defining mutations (https://outbreak.info/situation-reports?pango=P.1). Interestingly, eight of them were already found to be positively selected in the study of Faria and colleagues (2021), using a small set of P.1 and P.2 sequences. However, our analyses included 2,901 unique spike sequences from several lineages and identified pervasive adaptive selection signatures in sites 26, 138, 417, 484, 501, and 1,027, whose mutations are found occurring in more than nine Brazilian lineages. Therefore, the possible change of a neutral genetic drift to a strong selective pressure may be related to the P.1 mutational advantage facing the population-level immunity, according to Van Egeren et al. (2021). In this regard, it is interesting to note that P.1 has emerged in a region with up to 75% demonstrated previous seroprevalence of SARS-CoV-2. Similarly, lineage-defining mutated sites from B.1.1.7 (570 and 681) and C.37 lineages (75 and 76) were also identified in the positive selection tests, strongly suggesting that similar phenomena of selective pressure have been the crucial in the emergence of others lineages as well.

According to Shindyalov and colleagues (1994), mutations in certain positions are fixed in a correlated manner along the evolution. This may be the result of structural or functional constraints imposed for the maintenance of the protein integrity and stability. Although close chain neighbours have higher probabilities to mutate in a conditional way, even distant structural positions can follow this pattern through a number of different mechanisms. Coevolution may be the result of compensatory, fitness recovery interactions. Alternatively, mutations that enhance infectivity, such as N501Y, may increase the chance of further viral evolution. Recently, for example, the acquisition of type I interferon resistance, a seemingly key factor for pathogenesis in SARS-CoV, has been found in the newer S harboring mutant lineages (Guo et al., 2021; Kim and Shin, 2021). A third potential cause of viral coevolution through mutation linkage could be viral immune escape. The H69-V70 deletion, for instance, present in NTD from the B.1.1.7 lineage, abolishes an important immunogenic loop by allowing the inward movement of the amino acids 71-75 from that domain. In this work, the analysis of coevolution with the Bayesian Graphical Model showed that 28 pairs of sites (including the couple 484 - 501) can be statistically related with a posterior probability ≥ 0.8, indicating that these sites are probably not conditionally independent. The co-occurrence of E484K and N501Y is an interesting fact, since N501Y emerged independently in P.1 and B.1.351 lineages (Winger and Caspari, 2021), as well E484K (Ferrareze et al., 2021).

We have demonstrated that, individually, some mutations may stabilize or destabilize the protein structure, since their occurrence triggers different effects on energy balances and potentially affects viral fitness. As shown by the result of our majority consensus analysis and demonstrated by Pucci and Rooman (2021), mutations such as A222V (B.1.617.2), N501Y (B.1.1.7, P.1, B.1.351), and D614G generate a stabilizing effect in the spike structure. The D614G, specifically, reduces furin cleavage, lowering the risk of premature S1 shedding (Winger and Caspari, 2021). About the set of P.1 lineage-defining mutations found in the RBD region, the prevalent E484K substitution, located in the loop region outside the direct hACE2 interface, also stabilizes the spike protein structure, reducing the unfolding free energy changes. This effect was also predicted for E484Q, identified in Brazilian SARS-CoV-2 viruses from B.1, B.1.1.28, and P.5 lineages. However, E484D (B.1.1.33), represented by a single sequence in the genome set (and 44 sequences worldwide https://outbreak.info/situation-reports?pango&muts=S%3AE484D) seems to destabilize the spike structure. The analysis of the multiple substitutions potentially fixed by sites under adaptive selection with the strongest evidence (i.e., those 17 sites identified by FUBAR, FEL, and SLAC) showed that sites such as 138 (P.1, P.1.1, P.1.2, P.2), 222 (B.1.617.2, P.1, P.2), 262 (P.1, P.1.2, P.2, B.1.351), 572 (B.1.1.7, P.1, P.1.2, P.2), 614, and 845 (B.1.1.7, P.1, P.2) presented a concordant stabilizing trend for all the evaluated substitutions. However, the high frequency observed for the mutations D138Y (55.04%) and D614G (99.14%) can not be extended to any other site substitutions. In fact, considering 17 sites under positive selection identified by the three HyPhy methods, only six had substitution frequencies above 1% in the whole genome set (138, 484, 614, 681, 845, and 1,176).

The association of N501Y with D614G (B.1.1.7, P.1, B.1.351) improves the thermostability as well as increases the binding affinity with hACE2 (Winger and Caspari, 2021). Several studies (Chen et al., 2021; Hark Gan et al., 2021) have demonstrated the increased binding affinity upon the amino acid changes E484K+N501Y+K417T (P.1), superposing the K417T effect that decreases the binding affinity. The threonine substitution at site 417 avoids the salt bridge formation with D30 in hACE2. Consequently, there is expected diminution in the binding affinity for the mutated structures (Winger and Caspari, 2021). As previously elucidated, the binding free energy between RBD and ACE2 reflects the infectivity (Qu et al., 2005). Previous analysis of the substitutions L452R, F490S, and N501Y presented relatively high binding free energy changes suggesting that these sites may generate more infectious lineages (Wang et al., 2021). Our findings corroborates the improved binding pattern found in P.1 models, showing the greater contribution of E484K mutation to the binding free energy reduction and the RBD-hACE complex stabilization in P.1 and P.2 lineages, especially in the presence of a mutated L452V site. Despite L452Q (C.37) suggesting similar properties to L452R (B.1.617.2), increasing viral infectivity (Acevedo et al., 2021), nothing is known about the valine substitution and its effect on P.2 viruses. However, a significant interaction effect with E484K was observed in this study, also with indication of structural changes in the ACE2 molecule (as for B.1.1.7 model). Interestingly, the escape from neutralizing antibodies may be related to mutants with increased ACE2 binding affinities (Van Egeren et al., 2021). As shown by our molecular dynamic simulations, the comparison of the combined binding energies for the evaluated lineages indicates a major reduction in the binding free energy of those containing E484K, with only a small difference between the reference and lineages such as C.37 and B.1.1.7.

In our data, mutations such as P26S (B.1.1.7, P.1, P.1.1, P.1.2, P.2), T63N (P.2), R78S/T (P.1, P.2), K417T/N (P.1, P.1.1, P.1.2, P.2, B.1.351), A570D/V (B.1.1.7, P.1), L585F/V (P.1, P.2), T747A (P.1, P.2), A846S (P.1, P.2), and V1133F (P.1), among others, in positively selected sites, seem to impact the spike prefusion structure with a destabilizing effect. Except for P26S and K417T, which are found in 58.45% and 49.11% of the spike sequences from the genome set (n=11,078), respectively, the remaining mutations are observed in very low frequencies.

The selection of theoretically “destabilizing” or “stabilizing” mutations still remains to be completely elucidated. One possible explanation for fixation of such mutations would be the occurrence of “copy choice” recombinations capable of selecting very quickly for a set of mutations that have coevolved for the aforementioned reasons. The other possibility would be rapid evolution mediated by lineages enhanced with greater replicative fitness, as a consequence of key RBD mutations and/or other allosteric interactions (*e.g.*, D614G). The higher viral turnover would account for a tremendous selective pressure and fast viral selection, provided the viruses find an appropriate environment for transmissions. In this regard, it is interesting to note that both B.1.351 and P.1 appeared to have emerged in a very intense selective pressure scenario. In some Amazonas State cities in Brazil, where the P.1 lineage has emerged, up to 75% exposure were found for previous first-wave lineage. Similarly, B.1.351 has emerged in South Africa in a scenario with up to 30% of the population being previously exposed to other lineages. Taken together this evidence supports the role of consecutive accumulation of mutations rather than recombinations in order to explain the emergence of multiple lineages with specific sets of mutations, characteristic of the VOC viruses. Finally, it is possible that an immunosuppressed host would be critical for selecting these viruses, since the first occurrence of important, but intrinsically “destabilizing” substitution, would prevent further evolution if the virus encounters a robust immune response. In other hand, an immunocompromised patient would allow the progressively mutated virus to evolve, eventually acquiring fitness recovery and/or escape mutants permitting transmission to normal hosts. Although this hypothesis is merely speculative at this time, previous works involving multiple genotyping of a single chronically immunosuppressed patient (Choi et al., 2020; Kemp et al., 2021) suggested that this could be a way of rapidly selecting virus harboring mutations that, on an individual basis, are “destabilizing”.

## Availability of data and materials

Full tables acknowledging the authors and corresponding labs submitting sequencing data used in this study can be found in Supplementary File S1. Additional information used and/or analysed during the current study are available from the corresponding author on reasonable request.

## Author’s contributions

**Patrícia A. G. Ferrareze:** Conceptualization, Methodology, Software, Validation, Formal analysis, Investigation, Data Curation, Visualization, Writing - Original Draft, Writing - Review & Editing. **Ricardo A. Zimerman:** Formal analysis, Investigation, Writing - Original Draft, Writing - Review & Editing. **Vinicius B. Franceschi:** Investigation, Visualization, Writing - Original Draft, Writing - Review & Editing. **Gabriel D. Caldana:** Writing - Original Draft, Writing - Review & Editing. **Paulo A. Netz:** Methodology, Software, Formal analysis, Investigation, Resources, Visualization, Writing - Original Draft, Writing - Review & Editing. **Claudia E. Thompson:** Conceptualization, Methodology, Software, Formal analysis, Investigation, Resources, Writing - Original Draft, Writing - Review & Editing, Supervision, Project administration. All authors have read and approved the manuscript.

## Acknowledgements

We thank Amanda M. Mayer for suggestions in the introduction section. We also thank the administrators of the GISAID database and research groups across the world for supporting the rapid and transparent sharing of genomic data during the COVID-19 pandemic.

---

**Figure S1.**
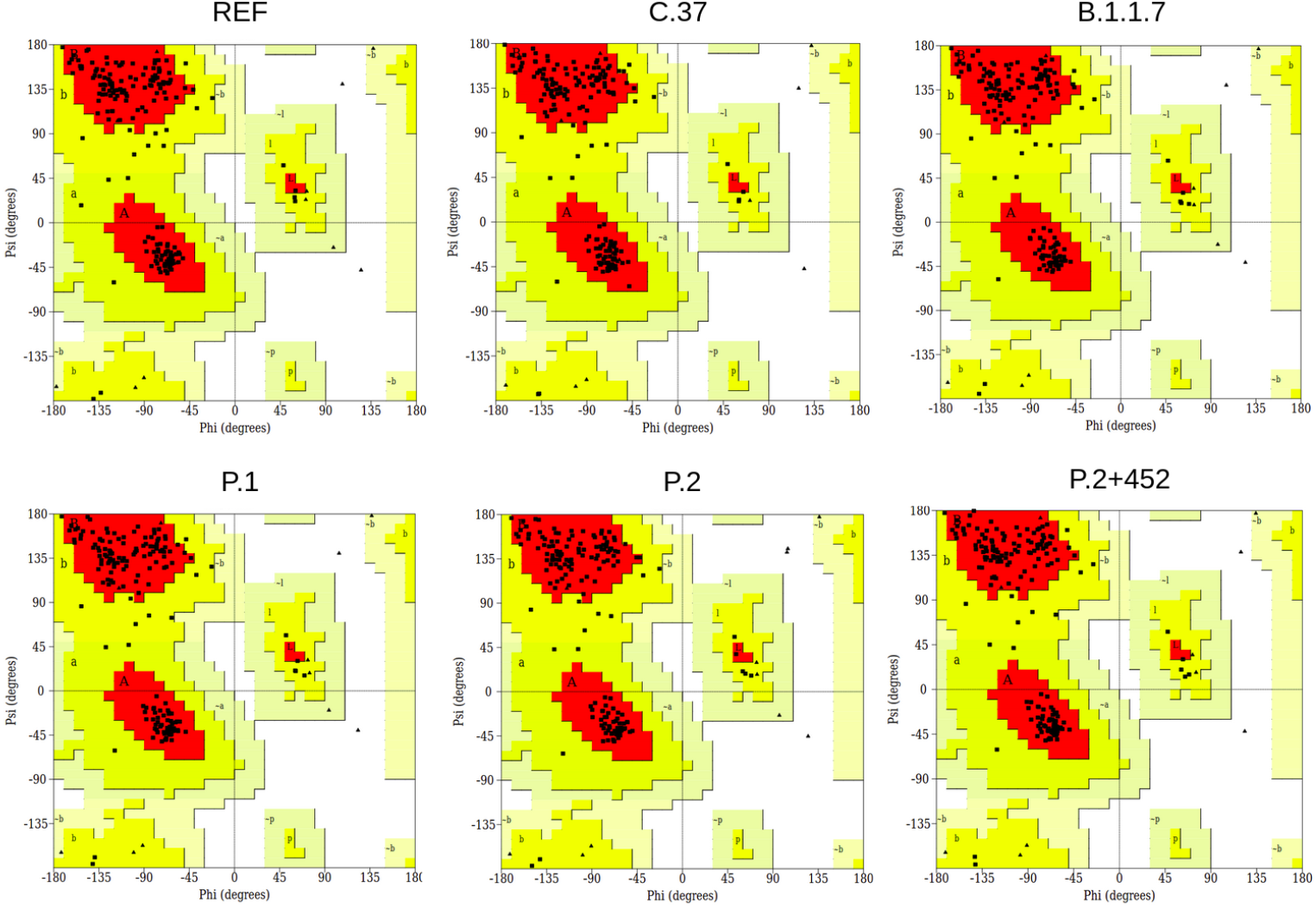
Ramachandran plots indicating the quality of the modeled structures of the spike RBD - hACE2 protein complexes for the reference and lineages P.1, P.2, C.37, B.1.1.7, and P.2+452.

**Figure S2.**
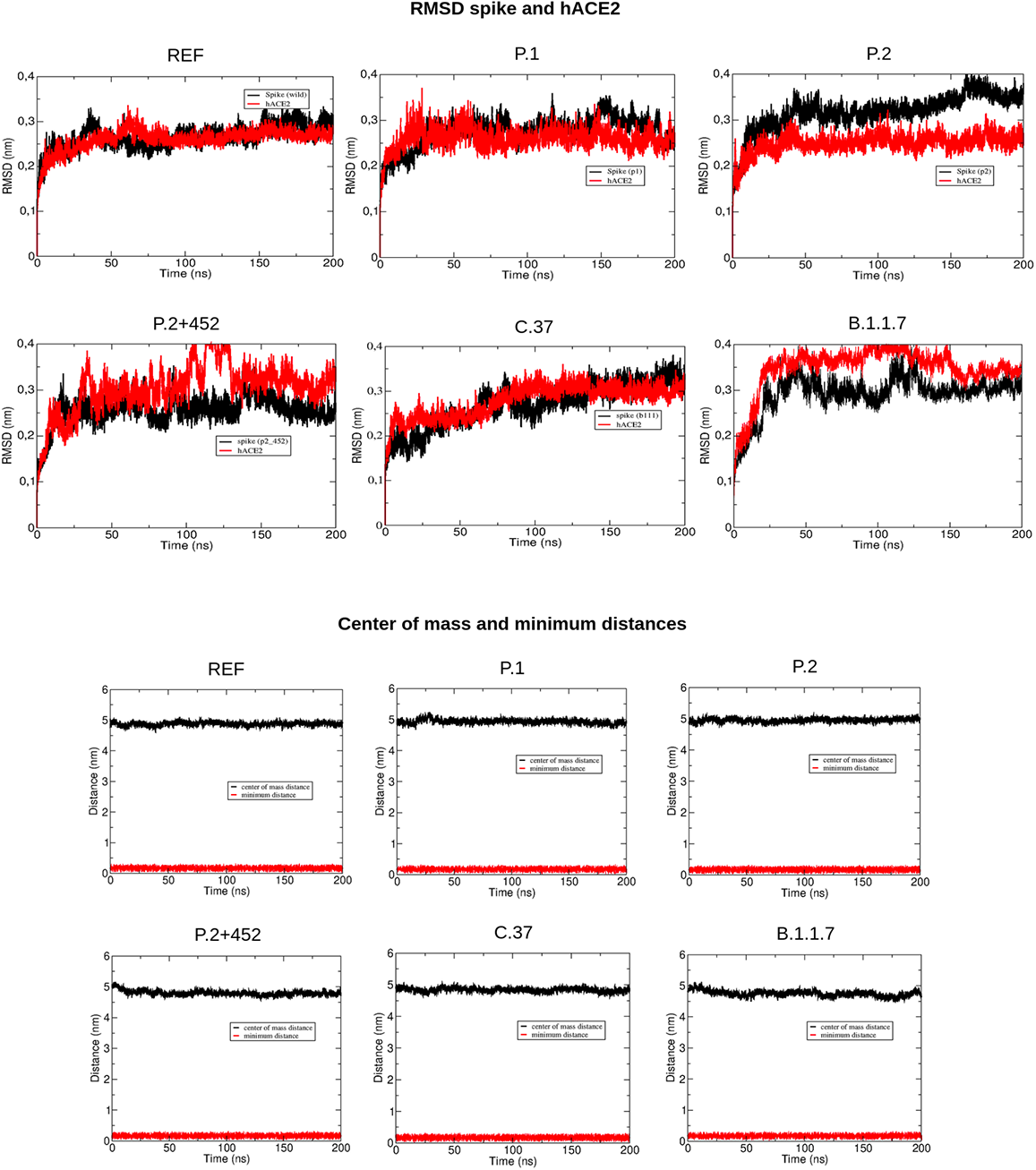
Root-mean-square deviation (RMSD), center of mass and minimum distances for the spike RBD - ACE2 protein complexes related to the reference (REF) and lineages P.1, P.2, P.2+452, C.37, and B.1.1.7.

**Figure S3.**
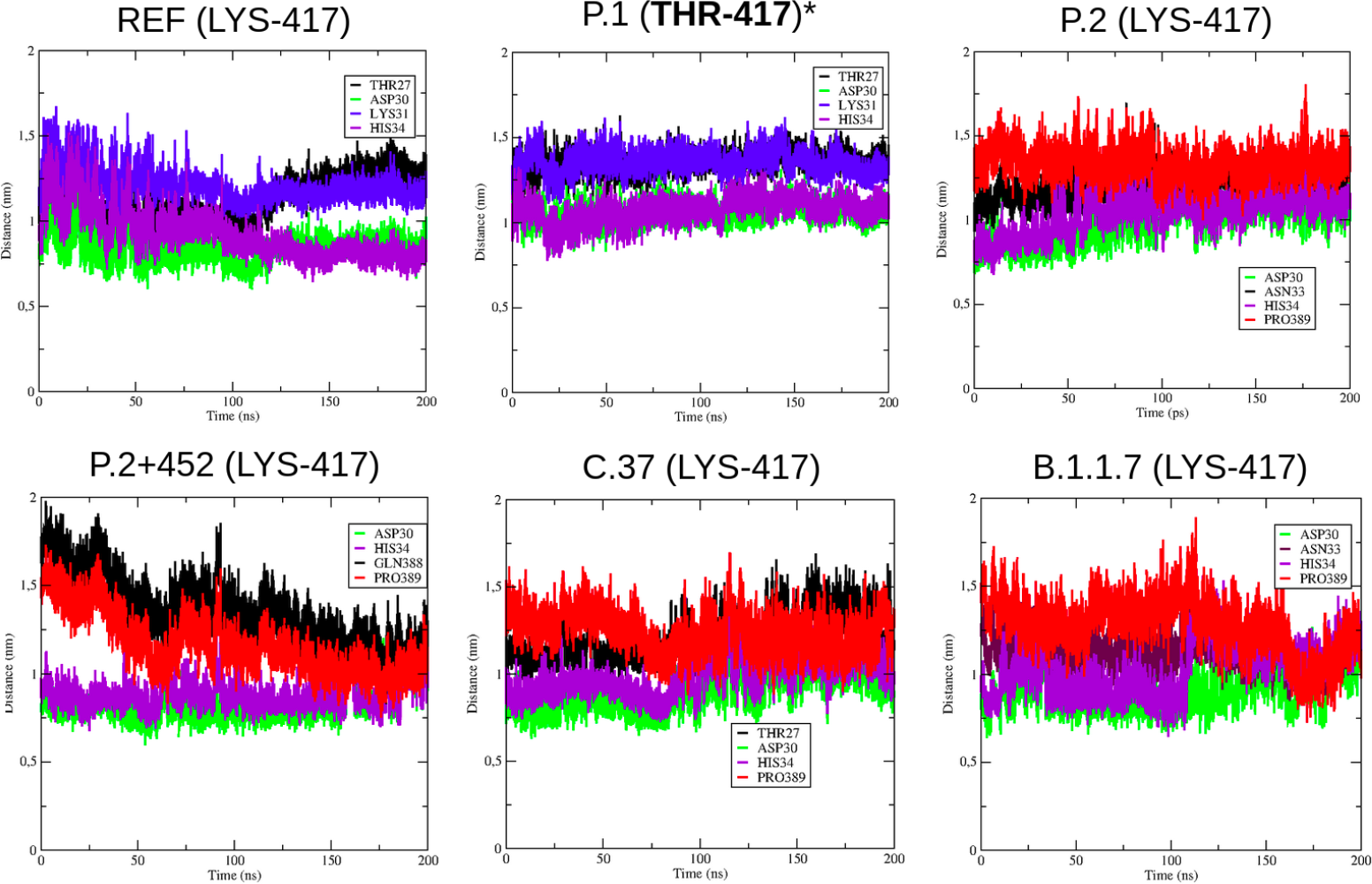
Structural distance measurements to the residue 417 in the reference RBD-ACE2 complex and for the lineages C.37, B.1.1.7, P.1, P.2, and P.2+452.

**Figure S4.**
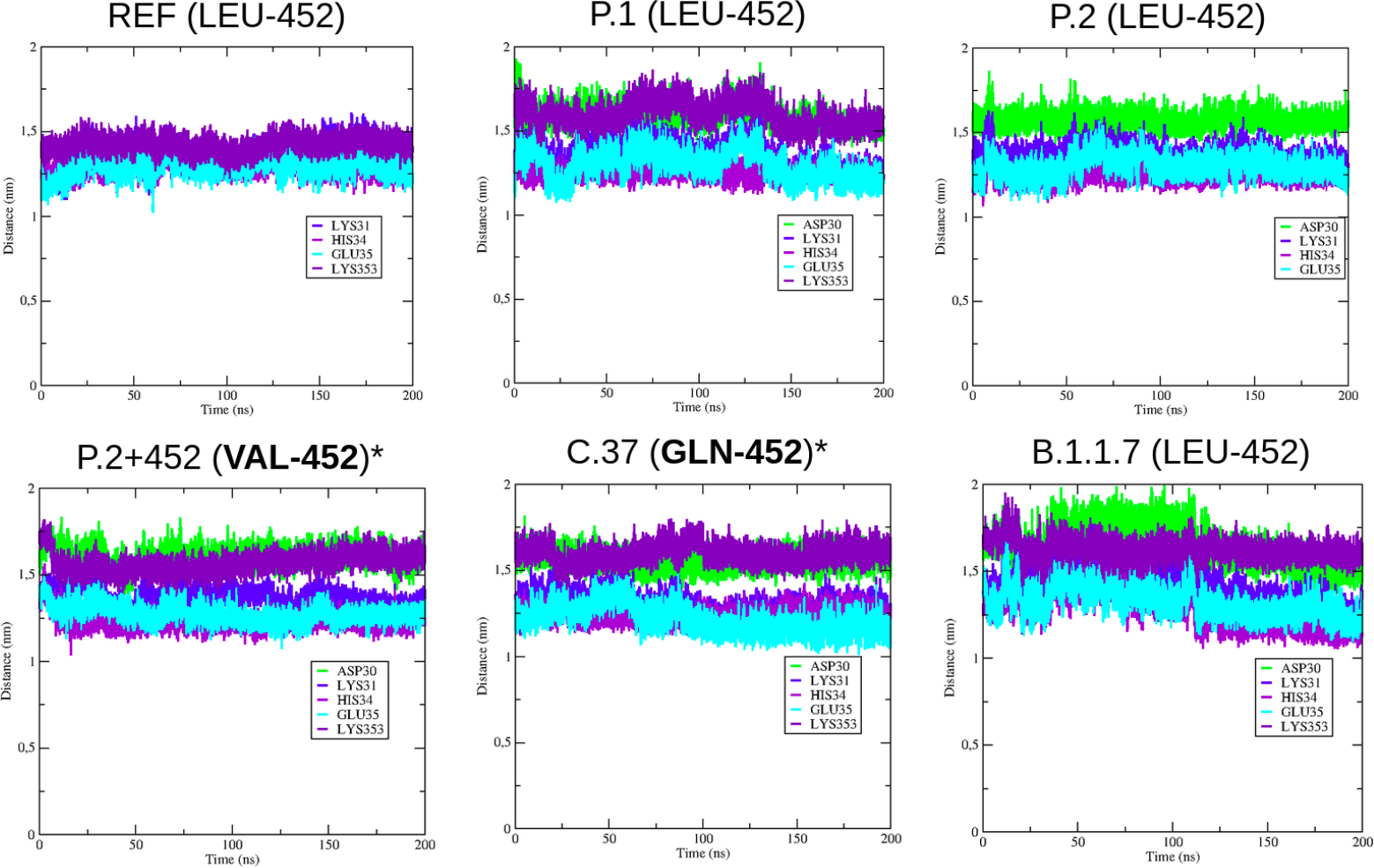
Structural distance measurements to the residue 452 in the reference RBD-ACE2 complex and for the lineages C.37, B.1.1.7, P.1, P.2, and P.2+452.

**Figure S5.**
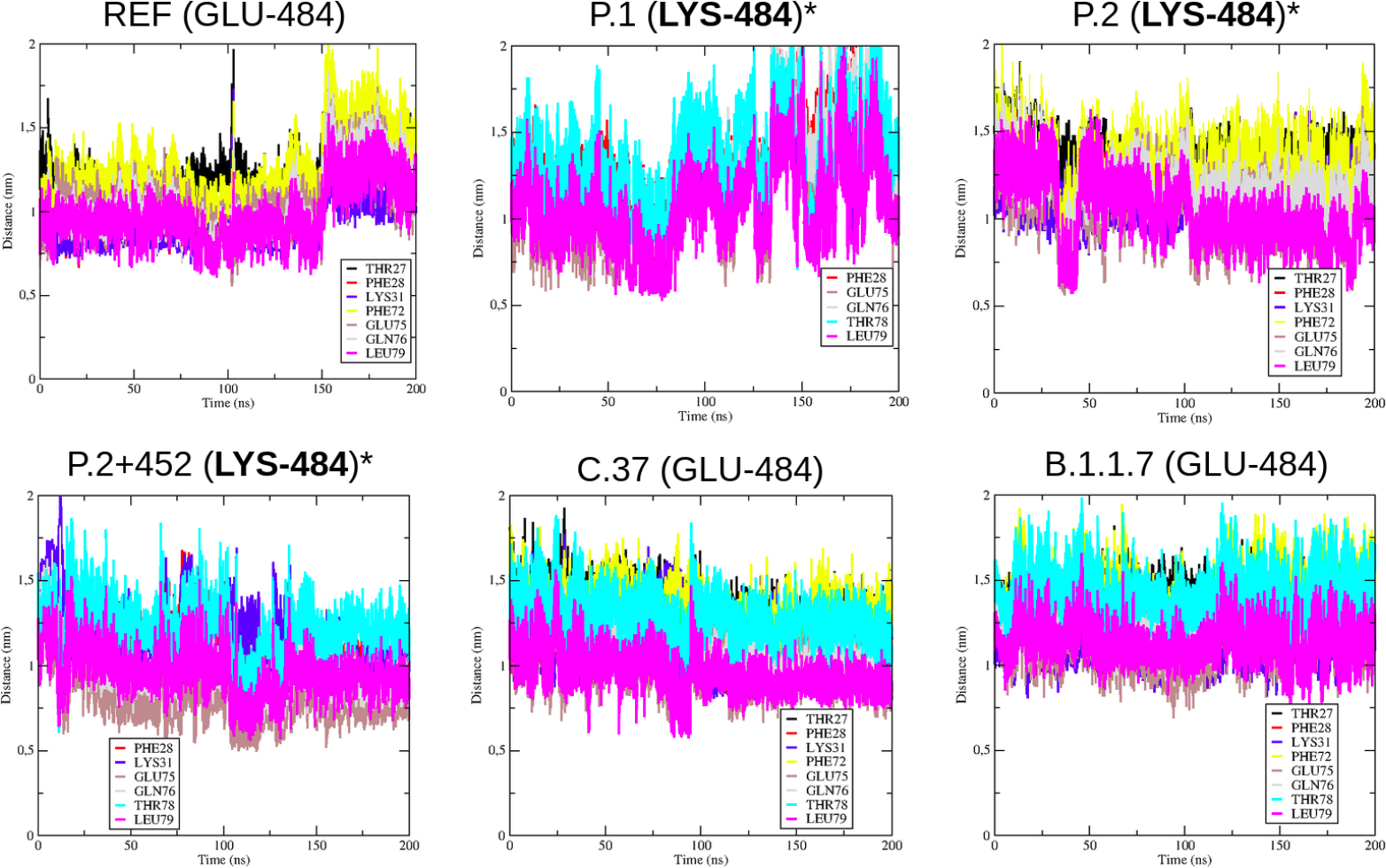
Structural distance measurements to the residue 484 in the reference RBD-ACE2 complex and for the lineages C.37, B.1.1.7, P.1, P.2, and P.2+452.

**Figure S6.**
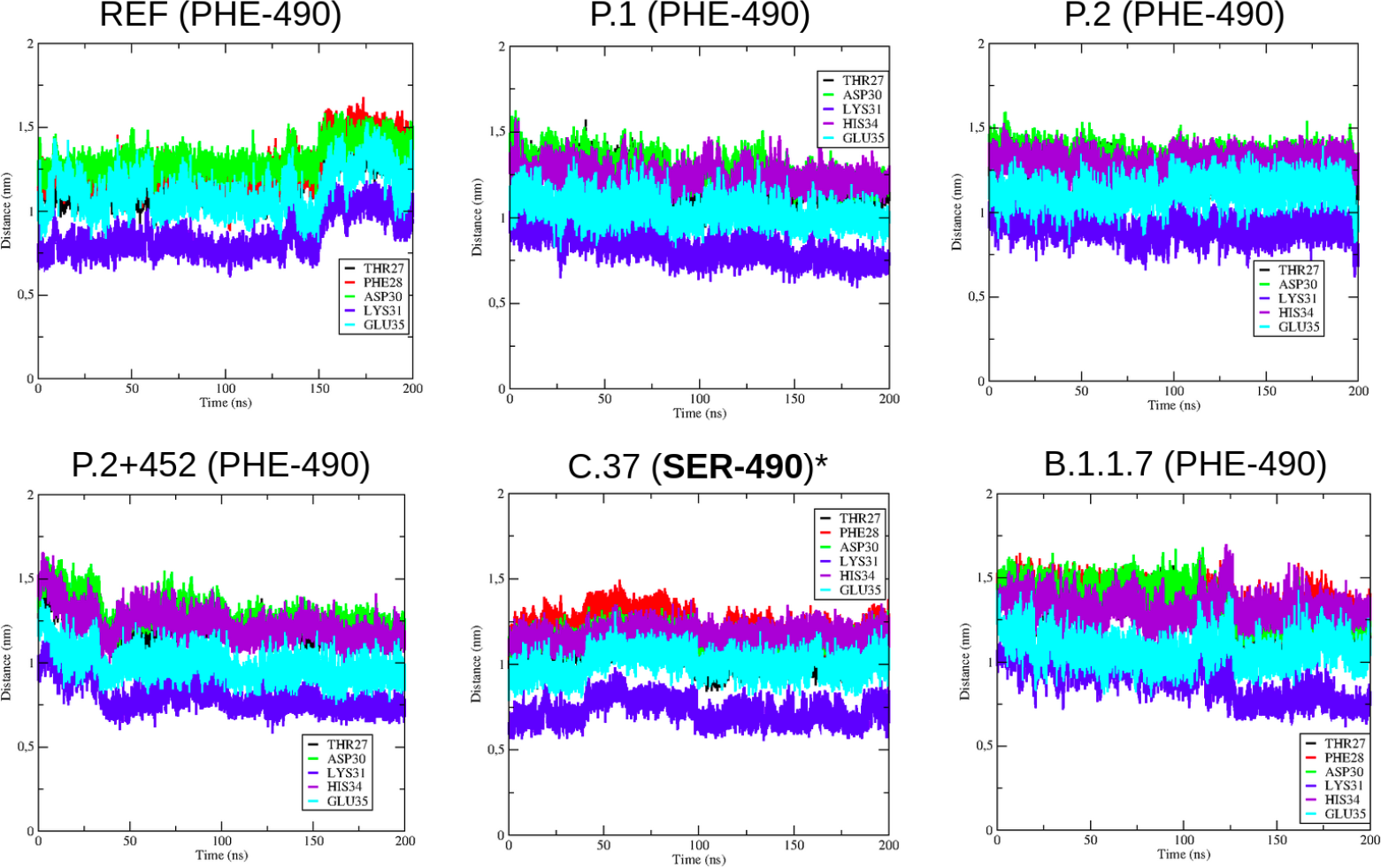
Structural distance measurements to the residue 490 in the reference RBD-ACE2 complex and for the lineages C.37, B.1.1.7, P.1, P.2, and P.2+452.

**Figure S7.**
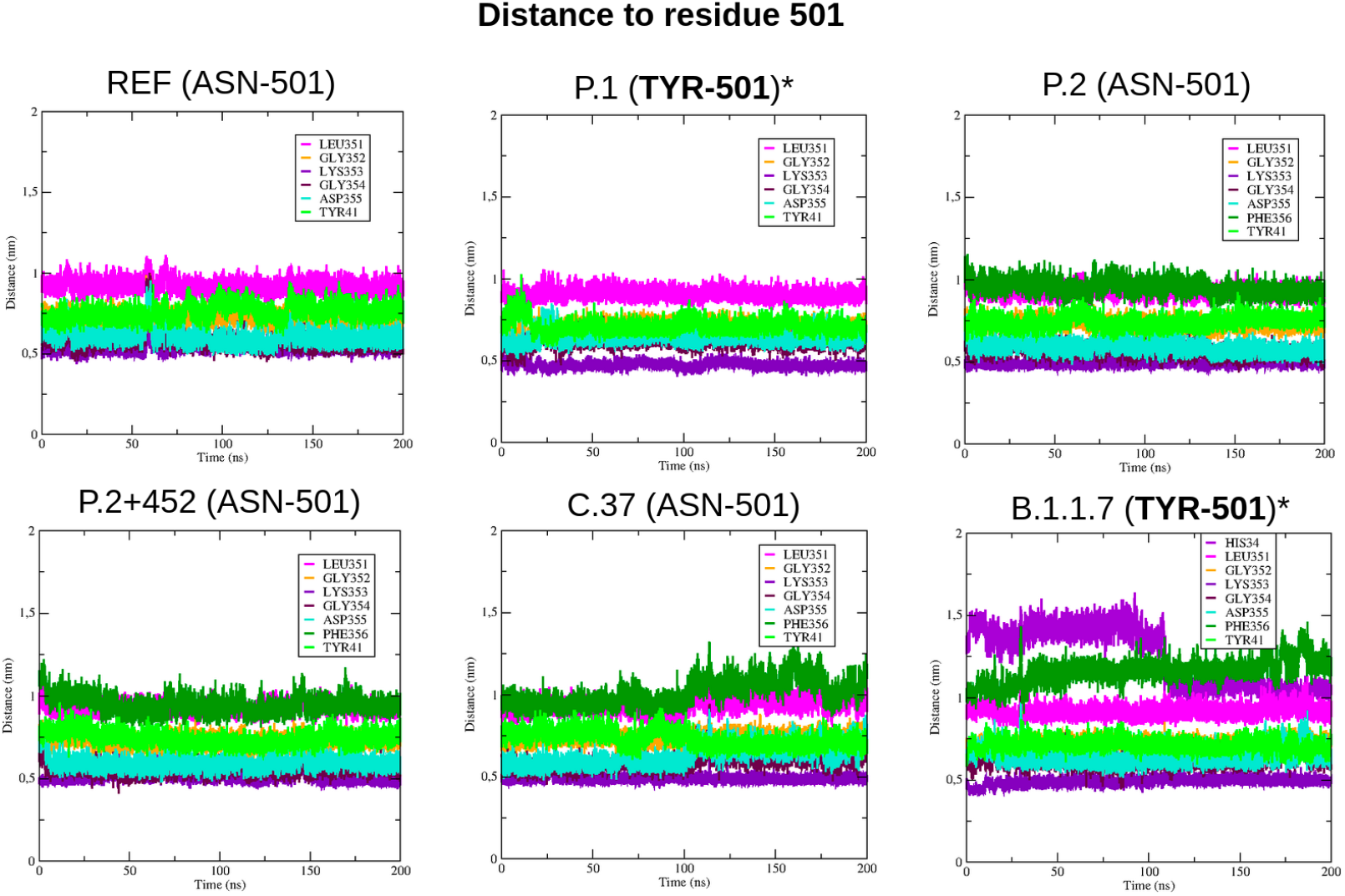
Structural distance measurements to the residue 501 in the reference RBD-ACE2 complex and for the lineages C.37, B.1.1.7, P.1, P.2, and P.2+452.

## Supplementary files

**Supplementary File 1.** Acknowledgement list of the 11,078 GISAID genomes used in this study.

**Supplementary File 2.** Complete table of positively selected sites identified by FUBAR, FEL, and SLAC methods.

**Supplementary File 3.** Table of sites under purifying selection according to the FEL and SLAC methods.

**Supplementary File 4.** Complete table of sites not conditionally independent identified by the Bayesian Graphical Model (BGM).

**Supplementary File 5.** Table of evaluated parameters from PROCHECK/VERIFY3D for the spike RBD models selection.

