## Supplementary File 1 for "Molecular evolution and structural analyses of the spike glycoprotein from Brazilian SARS-CoV-2 genomes: the impact of the fixation of selected mutations"

We gratefully acknowledge the following Authors from the Originating laboratories responsible for obtaining the specimens, as well as the Submitting laboratories where the genome data were generated and shared via GISAID, on which this research is based.

All Submitters of data may be contacted directly via www.gisaid.org

Authors are sorted alphabetically.

| Accession ID | Originating Laboratory | Submitting Laboratory | Authors |
|---|---|---|---|
| EPI_ISL_1000670, EPI_ISL_1000671, EPI_ISL_1000673, EPI_ISL_1000675, EPI_ISL_1000677 | Instituto de Biotecnologia - UNESP-Botucatu-SP | Instituto de Biotecnologia - UNESP-Botucatu-SP | Leila Sabrina Ullmann; Fábio Sossai Possebon, Camila Dantas Malossi, Paula Rahal, Paulo Inacio da Costa, João Pessoa Araújo Jr. |
| EPI_ISL_1023841 | Centro de Desenvolvimento Tecnológico em Saúde - CDTS | Centro de Desenvolvimento Tecnológico em Saúde - CDTS | Souza,T.M., Fintelman-Rodrigues,N., De Paula,A.D., Saraiva,F.B., Ferreira,M.A. and Sacramento,C.Q. |
| EPI_ISL_1034304, EPI_ISL_1034305, EPI_ISL_1034306 | Laboratorio de Ecologia de Doencas Transmissiveis na Amazonia, Instituto Leonidas e Maria Deane - Fiocruz Amazonia | Laboratorio de Ecologia de Doencas Transmissiveis na Amazonia, Instituto Leonidas e Maria Deane - Fiocruz Amazonia | Valdinete Nascimento, Victor Souza, André Corado, Fernanda Nascimento, George Silva, Ágatha Costa, Debora Duarte, Karina Pessoa, Matilde Mejía, Luciana Gonçalves, Maria Júlia Brandão, Michele Jesus, Felipe Naveca on behalf of the Fiocruz COVID-19 Genomic Surveillance Network |
| EPI_ISL_1039691, EPI_ISL_1039692, EPI_ISL_1039693, EPI_ISL_1039694, EPI_ISL_1039695 | LACEN do Estado de Goias | Instituto Adolfo Lutz, Interdiciplinary Procedures Center, Strategic Laboratory | Claudio Tavares Sacchi, Claudia Regina Gonçalves, Erica Valessa Ramos Gomes, Karoline Rodrigues Campos |
| EPI_ISL_1039696 | Instituto Adolfo Lutz - Regional de Presidente Prudente | Instituto Adolfo Lutz, Interdiciplinary Procedures Center, Strategic Laboratory | Claudio Tavares Sacchi, Claudia Regina Gonçalves, Erica Valessa Ramos Gomes, Karoline Rodrigues Campos |
| EPI_ISL_1039697 | Instituto Adolfo Lutz Central | Instituto Adolfo Lutz, Interdiciplinary Procedures Center, Strategic Laboratory | Claudio Tavares Sacchi, Claudia Regina Gonçalves, Erica Valessa Ramos Gomes, Karoline Rodrigues Campos |
| EPI_ISL_1039698 | Lab Loc - Itapecerica da Serra | Instituto Adolfo Lutz, Interdiciplinary Procedures Center, Strategic Laboratory | Claudio Tavares Sacchi, Claudia Regina Gonçalves, Erica Valessa Ramos Gomes, Karoline Rodrigues Campos |
| EPI_ISL_1039699 | Instituto Adolfo Lutz - Regional de Taubate | Instituto Adolfo Lutz, Interdiciplinary Procedures Center, Strategic Laboratory | Claudio Tavares Sacchi, Claudia Regina Gonçalves, Erica Valessa Ramos Gomes, Karoline Rodrigues Campos |
| EPI_ISL_1039700, EPI_ISL_1039701 | Instituto Adolfo Lutz Central | Instituto Adolfo Lutz, Interdiciplinary Procedures Center, Strategic Laboratory | Claudio Tavares Sacchi, Claudia Regina Gonçalves, Erica Valessa Ramos Gomes, Karoline Rodrigues Campos |
| EPI_ISL_1039702 | Instituto Adolfo Lutz - Regional de Aracatuba | Instituto Adolfo Lutz, Interdiciplinary Procedures Center, Strategic Laboratory | Claudio Tavares Sacchi, Claudia Regina Gonçalves, Erica Valessa Ramos Gomes, Karoline Rodrigues Campos |
| EPI_ISL_1039703 | Instituto Adolfo Lutz - Regional de Taubate | Instituto Adolfo Lutz, Interdiciplinary Procedures Center, Strategic Laboratory | Claudio Tavares Sacchi, Claudia Regina Gonçalves, Erica Valessa Ramos Gomes, Karoline Rodrigues Campos |
| EPI_ISL_1039704 | Lab Loc - Itapecerica da Serra | Instituto Adolfo Lutz, Interdiciplinary Procedures Center, Strategic Laboratory | Claudio Tavares Sacchi, Claudia Regina Gonçalves, Erica Valessa Ramos Gomes, Karoline Rodrigues Campos |
| EPI_ISL_1039705, EPI_ISL_1039706, EPI_ISL_1039707, EPI_ISL_1039708, EPI_ISL_1039709, EPI_ISL_1039710 | Instituto Adolfo Lutz Central | Instituto Adolfo Lutz, Interdiciplinary Procedures Center, Strategic Laboratory | Claudio Tavares Sacchi, Claudia Regina Gonçalves, Erica Valessa Ramos Gomes, Karoline Rodrigues Campos |
| EPI_ISL_1040823 | Secretaria Municipal de Saude de Piracaia | Instituto Adolfo Lutz, Interdiciplinary Procedures Center, Strategic Laboratory | Claudio Tavares Sacchi, Claudia Regina Gonçalves, Erica Valessa Ramos Gomes, Karoline Rodrigues Campos |
| EPI_ISL_1040824, EPI_ISL_1040825, EPI_ISL_1040826, EPI_ISL_1040827, EPI_ISL_1040828, EPI_ISL_1040829, EPI_ISL_1040830, EPI_ISL_1040831, EPI_ISL_1040832, EPI_ISL_1040833, EPI_ISL_1040834, EPI_ISL_1040835, EPI_ISL_1040836, EPI_ISL_1040837, EPI_ISL_1040838, EPI_ISL_1040839, EPI_ISL_1040840, EPI_ISL_1040841, EPI_ISL_1040842, EPI_ISL_1040843, EPI_ISL_1040844, EPI_ISL_1040845, EPI_ISL_1040846, EPI_ISL_1040847, EPI_ISL_1040848, EPI_ISL_1040849, EPI_ISL_1040850, EPI_ISL_1040851 | | | |
| see above | LACEN do Mato Grosso do Sul | Instituto Adolfo Lutz, Interdiciplinary Procedures Center, Strategic Laboratory | Claudio Tavares Sacchi, Claudia Regina Gonçalves, Erica Valessa Ramos Gomes, Karoline Rodrigues Campos |
| EPI_ISL_1041509 | LACEN do Estado de Goias | Instituto Adolfo Lutz, Interdiciplinary Procedures Center, Strategic Laboratory | Claudio Tavares Sacchi, Claudia Regina Gonçalves, Erica Valessa Ramos Gomes, Karoline Rodrigues Campos |
| EPI_ISL_1061031, EPI_ISL_1061032 | CDL Laboratorio Santos e Vidal LTDA. | Instituto de Medicina Tropical de Sao Paulo | Brazil-UK Centre for Arbovirus Discovery Diagnosis Genomics and Epidemiology (CADDE) Genomic Network - Instituto de Medicina Tropical |
| EPI_ISL_1063789 | Evandro Chagas Institute | Evandro Chagas Institute Virology | Santos, M.C.; Silva, A.M.; Junior, W.D.C.; Barbagelata, L.S.; Ferreira, J.A.; Sousa, E.M.A.; da Silva, P.S.; Pinheiro, K.C.; L.C.; Sousa Junior, E.C. |
| EPI_ISL_1067728 | Center for Biotechnology and Cell Therapy, São Rafael Hospital, Salvador, Brazil | Central Public Health Laboratory - LACEN -Bahia, Salvador, Brazil | Stephane Tosta, Luciana Oliveira, Vanessa Nardy,Patrícia Cajado,Marcela Gómez, Breno Dominguez, Jaqueline Gomes, Vagner Fonseca,Marta Giovanetti,Luiz Alcantara, Felicidade Pereira, Arabela Leal |
| EPI_ISL_1067729, EPI_ISL_1067730, EPI_ISL_1067731 | Central Public Health Laboratory - LACEN -Bahia, Salvador, Brazil | Central Public Health Laboratory - LACEN -Bahia, Salvador, Brazil | Stephane Tosta, Luciana Oliveira, Vanessa Nardy,Patrícia Cajado,Marcela Gómez, Breno Dominguez, Jaqueline Gomes, Vagner Fonseca,Marta Giovanetti,Luiz Alcantara, Felicidade Pereira, Arabela Leal |
| EPI_ISL_1067732 | Center for Biotechnology and Cell Therapy, São Rafael Hospital, Salvador, Brazil | Central Public Health Laboratory - LACEN -Bahia, Salvador, Brazil | Stephane Tosta, Luciana Oliveira, Vanessa Nardy,Patrícia Cajado,Marcela Gómez, Breno Dominguez, Jaqueline Gomes, Vagner Fonseca,Marta Giovanetti,Luiz Alcantara, Felicidade Pereira, Arabela Leal |
| EPI_ISL_1067733, EPI_ISL_1067734, EPI_ISL_1067735 | Central Public Health Laboratory - LACEN -Bahia, Salvador, Brazil | Central Public Health Laboratory - LACEN -Bahia, Salvador, Brazil | Stephane Tosta, Luciana Oliveira, Vanessa Nardy,Patrícia Cajado,Marcela Gómez, Breno Dominguez, Jaqueline Gomes, Vagner Fonseca,Marta Giovanetti,Luiz Alcantara, Felicidade Pereira, Arabela Leal |
| EPI_ISL_1067736 | Center for Biotechnology and Cell Therapy, São Rafael Hospital, Salvador, Brazil | Central Public Health Laboratory - LACEN -Bahia, Salvador, Brazil | Stephane Tosta, Luciana Oliveira, Vanessa Nardy,Patrícia Cajado,Marcela Gómez, Breno Dominguez, Jaqueline Gomes, Vagner Fonseca,Marta Giovanetti,Luiz Alcantara, Felicidade Pereira, Arabela Leal |
| EPI_ISL_1067737, EPI_ISL_1067738 | Central Public Health Laboratory - LACEN -Bahia, Salvador, Brazil | Central Public Health Laboratory - LACEN -Bahia, Salvador, Brazil | Stephane Tosta, Luciana Oliveira, Vanessa Nardy,Patrícia Cajado,Marcela Gómez, Breno Dominguez, Jaqueline Gomes, Vagner Fonseca,Marta Giovanetti,Luiz Alcantara, Felicidade Pereira, Arabela Leal |
| EPI_ISL_1068078, EPI_ISL_1068079, EPI_ISL_1068080, EPI_ISL_1068081, EPI_ISL_1068082, EPI_ISL_1068083, EPI_ISL_1068084, EPI_ISL_1068085, EPI_ISL_1068086, EPI_ISL_1068087, EPI_ISL_1068088, EPI_ISL_1068089, EPI_ISL_1068090, EPI_ISL_1068091, EPI_ISL_1068092, EPI_ISL_1068093, EPI_ISL_1068094, EPI_ISL_1068095, EPI_ISL_1068096, EPI_ISL_1068097, EPI_ISL_1068098, EPI_ISL_1068099, EPI_ISL_1068100, EPI_ISL_1068101, EPI_ISL_1068102, EPI_ISL_1068103, EPI_ISL_1068104, EPI_ISL_1068105, EPI_ISL_1068106, EPI_ISL_1068107, EPI_ISL_1068108, EPI_ISL_1068109, EPI_ISL_1068110, EPI_ISL_1068111, EPI_ISL_1068112, EPI_ISL_1068113, EPI_ISL_1068114, EPI_ISL_1068115, EPI_ISL_1068117, EPI_ISL_1068118, EPI_ISL_1068119, EPI_ISL_1068120, EPI_ISL_1068121, EPI_ISL_1068122, EPI_ISL_1068123, EPI_ISL_1068124, EPI_ISL_1068125, EPI_ISL_1068126, EPI_ISL_1068127, EPI_ISL_1068128, EPI_ISL_1068129, EPI_ISL_1068130, EPI_ISL_1068131, EPI_ISL_1068132, EPI_ISL_1068133, EPI_ISL_1068134, EPI_ISL_1068135, EPI_ISL_1068136, EPI_ISL_1068137, EPI_ISL_1068138, EPI_ISL_1068139, EPI_ISL_1068140, EPI_ISL_1068141, EPI_ISL_1068142, EPI_ISL_1068143, EPI_ISL_1068144, EPI_ISL_1068145, EPI_ISL_1068146, EPI_ISL_1068147, EPI_ISL_1068148, EPI_ISL_1068149, EPI_ISL_1068150, EPI_ISL_1068151, EPI_ISL_1068152, EPI_ISL_1068153, EPI_ISL_1068154, EPI_ISL_1068155, EPI_ISL_1068156, EPI_ISL_1068157, EPI_ISL_1068158, EPI_ISL_1068159, EPI_ISL_1068160, EPI_ISL_1068161, EPI_ISL_1068162, EPI_ISL_1068163, EPI_ISL_1068164, EPI_ISL_1068165, EPI_ISL_1068166, EPI_ISL_1068167, EPI_ISL_1068168, EPI_ISL_1068169, EPI_ISL_1068170, EPI_ISL_1068171, EPI_ISL_1068172, EPI_ISL_1068173, EPI_ISL_1068174, EPI_ISL_1068175, EPI_ISL_1068176, EPI_ISL_1068177, EPI_ISL_1068178, EPI_ISL_1068179, EPI_ISL_1068180, EPI_ISL_1068181, EPI_ISL_1068182, EPI_ISL_1068183, EPI_ISL_1068184, EPI_ISL_1068185, EPI_ISL_1068186, EPI_ISL_1068187, EPI_ISL_1068188, EPI_ISL_1068189, EPI_ISL_1068190, EPI_ISL_1068191, EPI_ISL_1068192, EPI_ISL_1068193, EPI_ISL_1068194, EPI_ISL_1068195, EPI_ISL_1068196, EPI_ISL_1068197, EPI_ISL_1068198, EPI_ISL_1068201, EPI_ISL_1068203, EPI_ISL_1068204, EPI_ISL_1068205, EPI_ISL_1068206, EPI_ISL_1068207, EPI_ISL_1068208, EPI_ISL_1068209, EPI_ISL_1068210, EPI_ISL_1068211, EPI_ISL_1068212, EPI_ISL_1068213, EPI_ISL_1068214, EPI_ISL_1068215, EPI_ISL_1068216, EPI_ISL_1068217, EPI_ISL_1068218, EPI_ISL_1068219, EPI_ISL_1068221, EPI_ISL_1068222, EPI_ISL_1068223, EPI_ISL_1068224, EPI_ISL_1068225, EPI_ISL_1068226, EPI_ISL_1068227, EPI_ISL_1068228, EPI_ISL_1068229, EPI_ISL_1068230, EPI_ISL_1068231, EPI_ISL_1068232, EPI_ISL_1068233, EPI_ISL_1068234, EPI_ISL_1068235, EPI_ISL_1068236, EPI_ISL_1068237, EPI_ISL_1068238, EPI_ISL_1068239, EPI_ISL_1068240, EPI_ISL_1068241, EPI_ISL_1068242, EPI_ISL_1068243, EPI_ISL_1068244, EPI_ISL_1068245, EPI_ISL_1068246, EPI_ISL_1068247, EPI_ISL_1068248, EPI_ISL_1068249, EPI_ISL_1068250, EPI_ISL_1068251, EPI_ISL_1068252, | | | |

| | | | |
|---|---|---|---|
| EPI_ISL_1068253, EPI_ISL_1068254, EPI_ISL_1068255, EPI_ISL_1068256, EPI_ISL_1068257, EPI_ISL_1068258, EPI_ISL_1068259, EPI_ISL_1068260, EPI_ISL_1068261, EPI_ISL_1068262, EPI_ISL_1068263, EPI_ISL_1068264, EPI_ISL_1068265, EPI_ISL_1068266, EPI_ISL_1068268, EPI_ISL_1068269, EPI_ISL_1068270, EPI_ISL_1068271, EPI_ISL_1068272, EPI_ISL_1068273, EPI_ISL_1068274, EPI_ISL_1068275, EPI_ISL_1068276, EPI_ISL_1068277, EPI_ISL_1068278, EPI_ISL_1068279, EPI_ISL_1068280, EPI_ISL_1068281, EPI_ISL_1068282, EPI_ISL_1068283, EPI_ISL_1068284, EPI_ISL_1068285, EPI_ISL_1068286, EPI_ISL_1068287, EPI_ISL_1068288, EPI_ISL_1068289, EPI_ISL_1068290, EPI_ISL_1068291, EPI_ISL_1068292 | | | |
| see above | Laboratorio de Ecologia de Doencas Transmissiveis na Amazonia, Instituto Leonidas e Maria Deane - Fiocruz Amazonia | Laboratorio de Ecologia de Doencas Transmissiveis na Amazonia, Instituto Leonidas e Maria Deane - Fiocruz Amazonia | Valdinete Nascimento, Victor Souza, André Corado, Fernanda Nascimento, George Silva, Ágatha Costa, Debora Duarte, Karina Pessoa, Matilde Mejía, Luciana Gonçalves, Maria Júlia Brandão, Michele Jesus, Felipe Naveca on behalf of the Fiocruz COVID-19 Genomic Surveillance Network |
| EPI_ISL_1068315, EPI_ISL_1068316, EPI_ISL_1068317, EPI_ISL_1068318, EPI_ISL_1068319, EPI_ISL_1068320, EPI_ISL_1068321, EPI_ISL_1068322, EPI_ISL_1068323, EPI_ISL_1068363, EPI_ISL_1068364, EPI_ISL_1068365, EPI_ISL_1068366, EPI_ISL_1068367, EPI_ISL_1068368, EPI_ISL_1068369, EPI_ISL_1068370, EPI_ISL_1068371, EPI_ISL_1068372, EPI_ISL_1068373, EPI_ISL_1068374, EPI_ISL_1068375, EPI_ISL_1068376, EPI_ISL_1068377, EPI_ISL_1068378, EPI_ISL_1068379, EPI_ISL_1068380, EPI_ISL_1068382, EPI_ISL_1068383, EPI_ISL_1068384, EPI_ISL_1068385, EPI_ISL_1068386, EPI_ISL_1068387, EPI_ISL_1068389, EPI_ISL_1068390, EPI_ISL_1068392, EPI_ISL_1068393, EPI_ISL_1068394 | | | |
| see above | Central Public Health Laboratory - LACEN -Bahia, Salvador, Brazil | Central Public Health Laboratory - LACEN -Bahia, Salvador, Brazil | Stephane Tosta, Luciana Oliveira, Vanessa Nardy,Patrícia Cajado,Marcela Gómez, Breno Dominguez, Jaqueline Gomes, Vagner Fonseca,Marta Giovanetti,Luiz Alcantara, Felicidade Pereira, Arabela Leal |
| EPI_ISL_1078981, EPI_ISL_1078983, EPI_ISL_1078984, EPI_ISL_1078986, EPI_ISL_1078987, EPI_ISL_1078988, EPI_ISL_1078989, EPI_ISL_1078991, EPI_ISL_1078992, EPI_ISL_1078993, EPI_ISL_1078995, EPI_ISL_1078996, EPI_ISL_1078997, EPI_ISL_1078999, EPI_ISL_1079000, EPI_ISL_1079002, EPI_ISL_1079003, EPI_ISL_1079004, EPI_ISL_1079006, EPI_ISL_1079007, EPI_ISL_1079008, EPI_ISL_1079158, EPI_ISL_1079159, EPI_ISL_1079161, EPI_ISL_1079162, EPI_ISL_1079163, EPI_ISL_1079165, EPI_ISL_1079166 | | | |
| see above | IAL Regional de Bauru | Instituto Adolfo Lutz, Interdiciplinary Procedures Center, Strategic Laboratory | Claudio Tavares Sacchi, Claudia Regina Gonçalves, Erica Valessa Ramos Gomes, Karoline Rodrigues Campos |
| EPI_ISL_1086034 | Diagnosticos da America - DASA | Instituto Adolfo Lutz, Interdiciplinary Procedures Center, Strategic Laboratory | Claudio Tavares Sacchi, Claudia Regina Gonçalves, Erica Valessa Ramos Gomes, Karoline Rodrigues Campos, Caio Vinicius Dias Lopes |
| EPI_ISL_1086035, EPI_ISL_1086036 | IAL Regional de Bauru | Instituto Adolfo Lutz, Interdiciplinary Procedures Center, Strategic Laboratory | Claudio Tavares Sacchi, Claudia Regina Gonçalves, Erica Valessa Ramos Gomes, Karoline Rodrigues Campos, Caio Vinicius Dias Lopes |
| EPI_ISL_1086037, EPI_ISL_1086038, EPI_ISL_1086039, EPI_ISL_1086040, EPI_ISL_1086041, EPI_ISL_1086042, EPI_ISL_1086043 | Diagnosticos da America - DASA | Instituto Adolfo Lutz, Interdiciplinary Procedures Center, Strategic Laboratory | Claudio Tavares Sacchi, Claudia Regina Gonçalves, Erica Valessa Ramos Gomes, Karoline Rodrigues Campos, Caio Vinicius Dias Lopes |
| EPI_ISL_1086044, EPI_ISL_1086045, EPI_ISL_1086046, EPI_ISL_1086047, EPI_ISL_1086048, EPI_ISL_1086049, EPI_ISL_1086050, EPI_ISL_1086051, EPI_ISL_1086052, EPI_ISL_1086053, EPI_ISL_1086054, EPI_ISL_1086055, EPI_ISL_1086056, EPI_ISL_1086057 | | | |
| see above | IAL Regional de Bauru | Instituto Adolfo Lutz, Interdiciplinary Procedures Center, Strategic Laboratory | Claudio Tavares Sacchi, Claudia Regina Gonçalves, Erica Valessa Ramos Gomes, Karoline Rodrigues Campos, Caio Vinicius Dias Lopes |
| EPI_ISL_1086373, EPI_ISL_1086374 | LACEN - Laboratório Central de Saúde Pública do Maranhao | Evandro Chagas Institute | Santos, M.C.; Silva, A.M.; Junior, W.D.C.; Barbagelata, L.S.; Ferreira, J.A.; Sousa, E.M.A.; da Silva, P.S.; Pinheiro, K.C.; L.C.; Sousa Junior, E.C. |
| EPI_ISL_1086375, EPI_ISL_1086376 | LACEN - Laboratório Central de Saúde Pública do Rio Grande do Norte | Evandro Chagas Institute | Santos, M.C.; Silva, A.M.; Junior, W.D.C.; Barbagelata, L.S.; Ferreira, J.A.; Sousa, E.M.A.; da Silva, P.S.; Pinheiro, K.C.; L.C.; Sousa Junior, E.C. |
| EPI_ISL_1086377 | LACEN - Laboratório Central de Saúde Pública do Paraiba | Evandro Chagas Institute | Santos, M.C.; Silva, A.M.; Junior, W.D.C.; Barbagelata, L.S.; Ferreira, J.A.; Sousa, E.M.A.; da Silva, P.S.; Pinheiro, K.C.; L.C.; Sousa Junior, E.C. |
| EPI_ISL_1092360 | IAL Regional de Bauru | Instituto Adolfo Lutz, Interdiciplinary Procedures Center, Strategic Laboratory | Claudio Tavares Sacchi, Claudia Regina Gonçalves, Erica Valessa Ramos Gomes, Karoline Rodrigues Campos |
| EPI_ISL_1092725 | Diagnosticos da America - DASA | Instituto Adolfo Lutz, Interdiciplinary Procedures Center, Strategic Laboratory | Claudio Tavares Sacchi, Claudia Regina Gonçalves, Erica Valessa Ramos Gomes, Karoline Rodrigues Campos |
| EPI_ISL_1095913 | IAL Regional de Bauru | Instituto Adolfo Lutz, Interdiciplinary Procedures Center, Strategic Laboratory | Claudio Tavares Sacchi, Claudia Regina Gonçalves, Erica Valessa Ramos Gomes, Karoline Rodrigues Campos |
| EPI_ISL_1096120 | IAL Regional de Bauru | Instituto Adolfo Lutz, Interdiciplinary Procedures Center, Strategic Laboratory | Claudio Tavares Sacchi, Claudia Regina Gonçalves, Erica Valessa Ramos Gomes, Karoline Rodrigues Campos, Caio Vinicius Dias Lopes |
| EPI_ISL_1096121 | Diagnosticos da America - DASA | Instituto Adolfo Lutz, Interdiciplinary Procedures Center, Strategic Laboratory | Claudio Tavares Sacchi, Claudia Regina Gonçalves, Erica Valessa Ramos Gomes, Karoline Rodrigues Campos, Caio Vinicius Dias Lopes |
| EPI_ISL_1096122, EPI_ISL_1096123, EPI_ISL_1096124, EPI_ISL_1096125, EPI_ISL_1096126, EPI_ISL_1096127, EPI_ISL_1096128, EPI_ISL_1096129, EPI_ISL_1096130, EPI_ISL_1096131, EPI_ISL_1096132, EPI_ISL_1096133, EPI_ISL_1096134 | | | |
| see above | IAL Regional de Bauru | Instituto Adolfo Lutz, Interdiciplinary Procedures Center, Strategic Laboratory | Claudio Tavares Sacchi, Claudia Regina Gonçalves, Erica Valessa Ramos Gomes, Karoline Rodrigues Campos, Caio Vinicius Dias Lopes |
| EPI_ISL_1096135 | Diagnosticos da America - DASA | Instituto Adolfo Lutz, Interdiciplinary Procedures Center, Strategic Laboratory | Claudio Tavares Sacchi, Claudia Regina Gonçalves, Erica Valessa Ramos Gomes, Karoline Rodrigues Campos, Caio Vinicius Dias Lopes |
| EPI_ISL_1096136 | IAL Regional de Bauru | Instituto Adolfo Lutz, Interdiciplinary Procedures Center, Strategic Laboratory | Claudio Tavares Sacchi, Claudia Regina Gonçalves, Erica Valessa Ramos Gomes, Karoline Rodrigues Campos, Caio Vinicius Dias Lopes |
| EPI_ISL_1109632 | Instituto de Biotecnologia - UNESP-Botucatu-SP | Instituto de Biotecnologia - UNESP-Botucatu-SP | Leila Sabrina Ullmann; Fábio Sossai Possebon; Cecília Artico Banho; Cíntia Bittar; Guilherme Campos; Helena Lage Ferreira; Jorge A. Petroli Marchesi; Lívia Sacchetto; Maisa C. Pereira Parra; Marília Moraes; Maurício L. Nogueira; Paula Rahal; Paulo Inacio da Costa; João Pessoa Araújo Jr. |
| EPI_ISL_1117379, EPI_ISL_1117380, EPI_ISL_1117381, EPI_ISL_1117382, EPI_ISL_1117383, EPI_ISL_1117384, EPI_ISL_1117385, EPI_ISL_1117386, EPI_ISL_1117387, EPI_ISL_1117388, EPI_ISL_1117389, EPI_ISL_1117390, EPI_ISL_1117391, EPI_ISL_1117392, EPI_ISL_1117393, EPI_ISL_1117394, EPI_ISL_1117395, EPI_ISL_1117396, EPI_ISL_1117397, EPI_ISL_1117398, EPI_ISL_1117399, EPI_ISL_1117400, EPI_ISL_1117401, EPI_ISL_1117402, EPI_ISL_1117403, EPI_ISL_1117404, EPI_ISL_1117405, EPI_ISL_1117406, EPI_ISL_1117407, EPI_ISL_1117408, EPI_ISL_1117409, EPI_ISL_1117410, EPI_ISL_1117411, EPI_ISL_1117412, EPI_ISL_1117413, EPI_ISL_1117414, EPI_ISL_1117415, EPI_ISL_1117416, EPI_ISL_1117417, EPI_ISL_1117418, EPI_ISL_1117419, EPI_ISL_1117420, EPI_ISL_1117421, EPI_ISL_1117422, EPI_ISL_1117423, EPI_ISL_1117424, EPI_ISL_1117425, EPI_ISL_1117426, EPI_ISL_1117427, EPI_ISL_1117428, EPI_ISL_1117429, EPI_ISL_1117430, EPI_ISL_1117431, EPI_ISL_1117432, EPI_ISL_1117433, EPI_ISL_1117434, EPI_ISL_1117435, EPI_ISL_1117436, EPI_ISL_1117437, EPI_ISL_1117438, EPI_ISL_1117439, EPI_ISL_1117440, EPI_ISL_1117441, EPI_ISL_1117442, EPI_ISL_1117443, EPI_ISL_1117444, EPI_ISL_1117445, EPI_ISL_1117446, EPI_ISL_1117447, EPI_ISL_1117448, EPI_ISL_1117449 | | | |
| see above | Nucleo de Pesquisa em Inovacao Terapeutica - UFPE | LABBE, Federal University of Pernambuco | Wilson Jose da Silva Junior, Marcos da Silveira Regueira Neto, Heidi Lacerda Alves da Cruz, Bruno Sampaio, Reginaldo Goncalves de Lima Neto, Maira Galdino da Rocha Pitta, Michelly Cristiny Pereira, Marco Katzenberger, Valdir de Queiroz Balbino |
| EPI_ISL_1121307 | Hospital E Antonio Policarpo de Oliveira | Instituto Adolfo Lutz, Interdiciplinary Procedures Center, Strategic Laboratory | Claudio Tavares Sacchi, Claudia Regina Gonçalves, Erica Valessa Ramos Gomes, Karoline Rodrigues Campos, Caio Vinicius Dias Lopes |
| EPI_ISL_1121308 | Hospital e Pronto Socorro Portinari | Instituto Adolfo Lutz, Interdiciplinary Procedures Center, Strategic Laboratory | Claudio Tavares Sacchi, Claudia Regina Gonçalves, Erica Valessa Ramos Gomes, Karoline Rodrigues Campos, Caio Vinicius Dias Lopes |
| EPI_ISL_1121309 | UBS Jose Francisco Rezende | Instituto Adolfo Lutz, Interdiciplinary Procedures Center, Strategic Laboratory | Claudio Tavares Sacchi, Claudia Regina Gonçalves, Erica Valessa Ramos Gomes, Karoline Rodrigues Campos, Caio Vinicius Dias Lopes |
| EPI_ISL_1121310 | IAL Regional de Bauru | Instituto Adolfo Lutz, Interdiciplinary Procedures Center, Strategic Laboratory | Claudio Tavares Sacchi, Claudia Regina Gonçalves, Erica Valessa Ramos Gomes, Karoline Rodrigues Campos, Caio Vinicius Dias Lopes |
| EPI_ISL_1121311 | Centro de Saude II Dr. Jose Paione Mococa | Instituto Adolfo Lutz, Interdiciplinary Procedures Center, Strategic Laboratory | Claudio Tavares Sacchi, Claudia Regina Gonçalves, Erica Valessa Ramos Gomes, Karoline Rodrigues Campos, Caio Vinicius Dias Lopes |
| EPI_ISL_1121312, EPI_ISL_1121313, EPI_ISL_1121314, EPI_ISL_1121315 | IAL Regional de Bauru | Instituto Adolfo Lutz, Interdiciplinary Procedures Center, Strategic Laboratory | Claudio Tavares Sacchi, Claudia Regina Gonçalves, Erica Valessa Ramos Gomes, Karoline Rodrigues Campos, Caio Vinicius Dias Lopes |
| EPI_ISL_1121316 | LACEN do Rio Grande do Sul | Instituto Adolfo Lutz, Interdiciplinary Procedures Center, Strategic Laboratory | Claudio Tavares Sacchi, Claudia Regina Gonçalves, Erica Valessa Ramos Gomes, Karoline Rodrigues Campos, Caio Vinicius Dias Lopes |
| EPI_ISL_1121317 | IAL Regional de Bauru | Instituto Adolfo Lutz, Interdiciplinary Procedures Center, | Claudio Tavares Sacchi, Claudia Regina Gonçalves, Erica Valessa Ramos Gomes, Karoline Rodrigues Campos, Caio Vinicius Dias Lopes |

| | | | |
|---|---|---|---|
| | | Strategic Laboratory | |
| EPI_ISL_1121318, EPI_ISL_1121319 | Hospital de Campanha COVID 19 Caieiras | Instituto Adolfo Lutz, Interdiciplinary Procedures Center, Strategic Laboratory | Claudio Tavares Sacchi, Claudia Regina Gonçalves, Erica Valessa Ramos Gomes, Karoline Rodrigues Campos, Caio Vinicius Dias Lopes |
| EPI_ISL_1121320, EPI_ISL_1121321 | IAL Regional de Bauru | Instituto Adolfo Lutz, Interdiciplinary Procedures Center, Strategic Laboratory | Claudio Tavares Sacchi, Claudia Regina Gonçalves, Erica Valessa Ramos Gomes, Karoline Rodrigues Campos, Caio Vinicius Dias Lopes |
| EPI_ISL_1121322 | Santa Casa de Santa Isabel | Instituto Adolfo Lutz, Interdiciplinary Procedures Center, Strategic Laboratory | Claudio Tavares Sacchi, Claudia Regina Gonçalves, Erica Valessa Ramos Gomes, Karoline Rodrigues Campos, Caio Vinicius Dias Lopes |
| EPI_ISL_1121323, EPI_ISL_1121324 | Complexo Hospitalar Padre Bentode Guarulhos | Instituto Adolfo Lutz, Interdiciplinary Procedures Center, Strategic Laboratory | Claudio Tavares Sacchi, Claudia Regina Gonçalves, Erica Valessa Ramos Gomes, Karoline Rodrigues Campos, Caio Vinicius Dias Lopes |
| EPI_ISL_1121325 | IAL Regional de Santos | Instituto Adolfo Lutz, Interdiciplinary Procedures Center, Strategic Laboratory | Claudio Tavares Sacchi, Claudia Regina Gonçalves, Erica Valessa Ramos Gomes, Karoline Rodrigues Campos, Caio Vinicius Dias Lopes |
| EPI_ISL_1121326 | IAL Regional de Bauru | Instituto Adolfo Lutz, Interdiciplinary Procedures Center, Strategic Laboratory | Claudio Tavares Sacchi, Claudia Regina Gonçalves, Erica Valessa Ramos Gomes, Karoline Rodrigues Campos, Caio Vinicius Dias Lopes |
| EPI_ISL_1121327, EPI_ISL_1121328, EPI_ISL_1121329, EPI_ISL_1121330 | LACEN do Mato Grosso do Sul | Instituto Adolfo Lutz, Interdiciplinary Procedures Center, Strategic Laboratory | Claudio Tavares Sacchi, Claudia Regina Gonçalves, Erica Valessa Ramos Gomes, Karoline Rodrigues Campos, Caio Vinicius Dias Lopes |
| EPI_ISL_1123372 | UPA I Santa Isabel | Instituto Adolfo Lutz, Interdiciplinary Procedures Center, Strategic Laboratory | Claudio Tavares Sacchi, Claudia Regina Gonçalves, Erica Valessa Ramos Gomes, Karoline Rodrigues Campos, Caio Vinicius Dias Lopes |
| EPI_ISL_1123373 | Grupo Tecnico de Vigilancia Sanitaria e Epidemiologica | Instituto Adolfo Lutz, Interdiciplinary Procedures Center, Strategic Laboratory | Claudio Tavares Sacchi, Claudia Regina Gonçalves, Erica Valessa Ramos Gomes, Karoline Rodrigues Campos, Caio Vinicius Dias Lopes |
| EPI_ISL_1123374, EPI_ISL_1123375 | IAL Regional de Santos | Instituto Adolfo Lutz, Interdiciplinary Procedures Center, Strategic Laboratory | Claudio Tavares Sacchi, Claudia Regina Gonçalves, Erica Valessa Ramos Gomes, Karoline Rodrigues Campos, Caio Vinicius Dias Lopes |
| EPI_ISL_1133120, EPI_ISL_1133121, EPI_ISL_1133122, EPI_ISL_1133123, EPI_ISL_1133124, EPI_ISL_1133125, EPI_ISL_1133126, EPI_ISL_1133127, EPI_ISL_1133128, EPI_ISL_1133129, EPI_ISL_1133130, EPI_ISL_1133131, EPI_ISL_1133132, EPI_ISL_1133133, EPI_ISL_1133134, EPI_ISL_1133135, EPI_ISL_1133136, EPI_ISL_1133137, EPI_ISL_1133138, EPI_ISL_1133139, EPI_ISL_1133140, EPI_ISL_1133141, EPI_ISL_1133142, EPI_ISL_1133143, EPI_ISL_1133144 | | | |
| see above | LABCOVID_HCPA | LABRESIS_HCPA | Martins AF, Wink PL, Volpato F, Rosset C, de Paris F, Monteiro F, Barth AL |
| EPI_ISL_1133256, EPI_ISL_1133258, EPI_ISL_1133259, EPI_ISL_1133260, EPI_ISL_1133261, EPI_ISL_1133262, EPI_ISL_1133263, EPI_ISL_1133265, EPI_ISL_1133266, EPI_ISL_1133267, EPI_ISL_1133268, EPI_ISL_1133269, EPI_ISL_1133272, EPI_ISL_1133273, EPI_ISL_1133274, EPI_ISL_1133276, EPI_ISL_1133277, EPI_ISL_1133278 | | | |
| see above | Laboratório Hermes Pardini | Laboratório de Biologia Integrativa, Instituto de Ciências Biológicas, Universidade Federal de Minas Gerais | Filipe Romero Rebello Moreira, Diego Menezes Bonfim, Danielle Alves Gomes Zauli, Joice do Prado Silva, Aline Brito de Lima, Frederico Scott Varella Malta, Alessandro Clayton de Souza Ferreira, Victor Cavalcanti Pardini, Daniel Costa Queiroz, Rafael Marques de Souza, Victor Emmanuel Viana Geddes, Walyson Coelho Costa, Wagner Carlos Santos Magalhaes, Rennan Garcias Moreira, Carolina Moreira Voloch, Renan Pedra de Souza, Renato Santana Aguiar. |
| EPI_ISL_1139052, EPI_ISL_1139053, EPI_ISL_1139054, EPI_ISL_1139055, EPI_ISL_1139056, EPI_ISL_1139057, EPI_ISL_1139058, EPI_ISL_1139059, EPI_ISL_1139060, EPI_ISL_1139061, EPI_ISL_1139062, EPI_ISL_1139063, EPI_ISL_1139064, EPI_ISL_1139065, EPI_ISL_1139066, EPI_ISL_1139067, EPI_ISL_1139068 | | | |
| see above | LACEN do Mato Grosso do Sul | Instituto Adolfo Lutz, Interdiciplinary Procedures Center, Strategic Laboratory | Claudio Tavares Sacchi, Claudia Regina Gonçalves, Erica Valessa Ramos Gomes, Karoline Rodrigues Campos, Caio Vinicius Dias Lopes |
| EPI_ISL_1139069 | SMS Aruja | Instituto Adolfo Lutz, Interdiciplinary Procedures Center, Strategic Laboratory | Claudio Tavares Sacchi, Claudia Regina Gonçalves, Erica Valessa Ramos Gomes, Karoline Rodrigues Campos, Caio Vinicius Dias Lopes |
| EPI_ISL_1139070 | IAL Regional de Ribeirao Preto | Instituto Adolfo Lutz, Interdiciplinary Procedures Center, Strategic Laboratory | Claudio Tavares Sacchi, Claudia Regina Gonçalves, Erica Valessa Ramos Gomes, Karoline Rodrigues Campos, Caio Vinicius Dias Lopes |
| EPI_ISL_1139071, EPI_ISL_1139072, EPI_ISL_1139073, EPI_ISL_1139074 | Instituto Adolfo Lutz Central | Instituto Adolfo Lutz, Interdiciplinary Procedures Center, Strategic Laboratory | Claudio Tavares Sacchi, Claudia Regina Gonçalves, Erica Valessa Ramos Gomes, Karoline Rodrigues Campos, Caio Vinicius Dias Lopes |
| EPI_ISL_1139075, EPI_ISL_1139077 | Lab Loc - Itapecerica da Serra | Instituto Adolfo Lutz, Interdiciplinary Procedures Center, Strategic Laboratory | Claudio Tavares Sacchi, Claudia Regina Gonçalves, Erica Valessa Ramos Gomes, Karoline Rodrigues Campos, Caio Vinicius Dias Lopes |
| EPI_ISL_1161401, EPI_ISL_1161402, EPI_ISL_1161403, EPI_ISL_1161404, EPI_ISL_1161405, EPI_ISL_1161406, EPI_ISL_1161407, EPI_ISL_1161408, EPI_ISL_1161409, EPI_ISL_1161410, EPI_ISL_1161412, EPI_ISL_1161413, EPI_ISL_1161414, EPI_ISL_1161415, EPI_ISL_1163530, EPI_ISL_1163693, EPI_ISL_1163694, EPI_ISL_1163695, EPI_ISL_1163696, EPI_ISL_1163697, EPI_ISL_1163698, EPI_ISL_1163699, EPI_ISL_1163700, EPI_ISL_1163701, EPI_ISL_1163702, EPI_ISL_1163703, EPI_ISL_1163704, EPI_ISL_1163705, EPI_ISL_1163706, EPI_ISL_1163707, EPI_ISL_1163708, EPI_ISL_1163709, EPI_ISL_1163710, EPI_ISL_1163711, EPI_ISL_1163712, EPI_ISL_1163713, EPI_ISL_1163714, EPI_ISL_1163715, EPI_ISL_1163716 | | | |
| see above | LABCOVID_HCPA | LABRESIS_HCPA | Martins AF, Wink PL, Volpato F, Rosset C, de Paris F, Monteiro F, Zavascki AP, Barth AL |
| EPI_ISL_1163736 | LABCOVID_HCPA | LABCOVID_HCPA | Martins AF, Wink PL, Volpato F, Rosset C, de Paris F, Monteiro F, Zavascki AP, Barth AL |
| EPI_ISL_1163737 | LABCOVID_HCPA | LABRESIS_HCPA | Martins AF, Wink PL, Volpato F, Rosset C, de Paris F, Monteiro F, Zavascki AP, Barth AL |
| EPI_ISL_1164970, EPI_ISL_1164971 | LACEN - Laboratório Central de Saúde Pública do Ceará | Evandro Chagas Institute | Santos, M.C.; Silva, A.M.; Junior, W.D.C.; Barbagelata, L.S.; Ferreira, J.A.; Sousa, E.M.A.; da Silva, P.S.; Pinheiro, K.C.; L.C.; Sousa Junior, E.C. |
| EPI_ISL_1164972 | LACEN - Laboratório Central de Saúde Pública do Pará | Evandro Chagas Institute | Santos, M.C.; Silva, A.M.; Junior, W.D.C.; Barbagelata, L.S.; Ferreira, J.A.; Sousa, E.M.A.; da Silva, P.S.; Pinheiro, K.C.; L.C.; Sousa Junior, E.C. |
| EPI_ISL_1164973 | LACEN - Laboratório Central de Saúde Pública do Ceará | Evandro Chagas Institute | Santos, M.C.; Silva, A.M.; Junior, W.D.C.; Barbagelata, L.S.; Ferreira, J.A.; Sousa, E.M.A.; da Silva, P.S.; Pinheiro, K.C.; L.C.; Sousa Junior, E.C. |
| EPI_ISL_1164974, EPI_ISL_1164975 | LACEN - Laboratório Central de Saúde Pública do Pará | Evandro Chagas Institute | Santos, M.C.; Silva, A.M.; Junior, W.D.C.; Barbagelata, L.S.; Ferreira, J.A.; Sousa, E.M.A.; da Silva, P.S.; Pinheiro, K.C.; L.C.; Sousa Junior, E.C. |
| EPI_ISL_1164976 | LACEN - Laboratório Central de Saúde Pública do Amapá | Evandro Chagas Institute | Santos, M.C.; Silva, A.M.; Junior, W.D.C.; Barbagelata, L.S.; Ferreira, J.A.; Sousa, E.M.A.; da Silva, P.S.; Pinheiro, K.C.; L.C.; Sousa Junior, E.C. |
| EPI_ISL_1164978 | LACEN - Laboratório Central de Saúde Pública do Pará | Evandro Chagas Institute | Santos, M.C.; Silva, A.M.; Junior, W.D.C.; Barbagelata, L.S.; Ferreira, J.A.; Sousa, E.M.A.; da Silva, P.S.; Pinheiro, K.C.; L.C.; Sousa Junior, E.C. |
| EPI_ISL_1164979 | LACEN - Laboratório Central de Saúde Pública do Maranhao | Evandro Chagas Institute | Santos, M.C.; Silva, A.M.; Junior, W.D.C.; Barbagelata, L.S.; Ferreira, J.A.; Sousa, E.M.A.; da Silva, P.S.; Pinheiro, K.C.; L.C.; Sousa Junior, E.C. |
| EPI_ISL_1164980 | LACEN - Laboratório Central de Saúde Pública do Ceará | Evandro Chagas Institute | Santos, M.C.; Silva, A.M.; Junior, W.D.C.; Barbagelata, L.S.; Ferreira, J.A.; Sousa, E.M.A.; da Silva, P.S.; Pinheiro, K.C.; L.C.; Sousa Junior, E.C. |
| EPI_ISL_1164981, EPI_ISL_1164982 | LACEN - Laboratório Central de Saúde Pública do Amapá | Evandro Chagas Institute | Santos, M.C.; Silva, A.M.; Junior, W.D.C.; Barbagelata, L.S.; Ferreira, J.A.; Sousa, E.M.A.; da Silva, P.S.; Pinheiro, K.C.; L.C.; Sousa Junior, E.C. |
| EPI_ISL_1164983 | LACEN - Laboratório Central de Saúde Pública do Pará | Evandro Chagas Institute | Santos, M.C.; Silva, A.M.; Junior, W.D.C.; Barbagelata, L.S.; Ferreira, J.A.; Sousa, E.M.A.; da Silva, P.S.; Pinheiro, K.C.; L.C.; Sousa Junior, E.C. |
| EPI_ISL_1164984, EPI_ISL_1164985 | LACEN - Laboratório Central de Saúde Pública do Amapá | Evandro Chagas Institute | Santos, M.C.; Silva, A.M.; Junior, W.D.C.; Barbagelata, L.S.; Ferreira, J.A.; Sousa, E.M.A.; da Silva, P.S.; Pinheiro, K.C.; L.C.; Sousa Junior, E.C. |
| EPI_ISL_1164986 | LACEN - Laboratório Central de Saúde Pública do Ceará | Evandro Chagas Institute | Santos, M.C.; Silva, A.M.; Junior, W.D.C.; Barbagelata, L.S.; Ferreira, J.A.; Sousa, E.M.A.; da Silva, P.S.; Pinheiro, K.C.; L.C.; Sousa Junior, E.C. |
| EPI_ISL_1164987, EPI_ISL_1164988 | LACEN - Laboratório Central de Saúde Pública do Rio Grande do Norte | Evandro Chagas Institute | Santos, M.C.; Silva, A.M.; Junior, W.D.C.; Barbagelata, L.S.; Ferreira, J.A.; Sousa, E.M.A.; da Silva, P.S.; Pinheiro, K.C.; L.C.; Sousa Junior, E.C. |
| EPI_ISL_1164989, EPI_ISL_1164990, EPI_ISL_1164991, EPI_ISL_1164992 | LACEN - Laboratório Central de Saúde Pública do Paraiba | Evandro Chagas Institute | Santos, M.C.; Silva, A.M.; Junior, W.D.C.; Barbagelata, L.S.; Ferreira, J.A.; Sousa, E.M.A.; da Silva, P.S.; Pinheiro, K.C.; L.C.; Sousa Junior, E.C. |
| EPI_ISL_1164993 | LACEN - Laboratório Central de Saúde Pública do Ceará | Evandro Chagas Institute | Santos, M.C.; Silva, A.M.; Junior, W.D.C.; Barbagelata, L.S.; Ferreira, J.A.; Sousa, E.M.A.; da Silva, P.S.; Pinheiro, K.C.; L.C.; Sousa Junior, E.C. |
| EPI_ISL_1164994 | LACEN - Laboratório Central de Saúde Pública do Paraiba | Evandro Chagas Institute | Santos, M.C.; Silva, A.M.; Junior, W.D.C.; Barbagelata, L.S.; Ferreira, J.A.; Sousa, E.M.A.; da Silva, P.S.; Pinheiro, K.C.; L.C.; Sousa Junior, E.C. |
| EPI_ISL_1164995 | LACEN - Laboratório Central de Saúde Pública do Ceará | Evandro Chagas Institute | Santos, M.C.; Silva, A.M.; Junior, W.D.C.; Barbagelata, L.S.; Ferreira, J.A.; Sousa, E.M.A.; da Silva, P.S.; Pinheiro, K.C.; L.C.; Sousa Junior, E.C. |
| EPI_ISL_1164996 | LACEN - Laboratório Central de Saúde Pública do Maranhao | Evandro Chagas Institute | Santos, M.C.; Silva, A.M.; Junior, W.D.C.; Barbagelata, L.S.; Ferreira, J.A.; Sousa, E.M.A.; da Silva, P.S.; Pinheiro, K.C.; L.C.; Sousa Junior, E.C. |

| | | | |
|---|---|---|---|
| EPI_ISL_1164999 | LACEN - Laboratório Central de Saúde Pública do Paraiba | Evandro Chagas Institute | Santos, M.C.; Silva, A.M.; Junior, W.D.C.; Barbagelata, L.S.; Ferreira, J.A.; Sousa, E.M.A.; da Silva, P.S.; Pinheiro, K.C.; L.C.; Sousa Junior, E.C. |
| EPI_ISL_1166615 | LACEN - Laboratório Central de Saúde Pública do Rio Grande do Norte | Evandro Chagas Institute Virology | Santos, M.C.; Silva, A.M.; Junior, W.D.C.; Barbagelata, L.S.; Ferreira, J.A.; Sousa, E.M.A.; da Silva, P.S.; Pinheiro, K.C.; L.C.; Sousa Junior, E.C. |
| EPI_ISL_1171619 | IAL Regional de Presidente Prudente | Instituto Adolfo Lutz, Interdiciplinary Procedures Center, Strategic Laboratory | Claudio Tavares Sacchi, Claudia Regina Gonçalves, Erica Valessa Ramos Gomes, Karoline Rodrigues Campos |
| EPI_ISL_1171620 | Instituto Adolfo Lutz Central | Instituto Adolfo Lutz, Interdiciplinary Procedures Center, Strategic Laboratory | Claudio Tavares Sacchi, Claudia Regina Gonçalves, Erica Valessa Ramos Gomes, Karoline Rodrigues Campos, Caio Vinicius Dias Lopes |
| EPI_ISL_1171621 | LACEN do Mato Grosso do Sul | Instituto Adolfo Lutz, Interdiciplinary Procedures Center, Strategic Laboratory | Claudio Tavares Sacchi, Claudia Regina Gonçalves, Erica Valessa Ramos Gomes, Karoline Rodrigues Campos, Caio Vinicius Dias Lopes |
| EPI_ISL_1171622, EPI_ISL_1171623, EPI_ISL_1171624, EPI_ISL_1171625, EPI_ISL_1171626, EPI_ISL_1171627, EPI_ISL_1171628, EPI_ISL_1171629, EPI_ISL_1171630, EPI_ISL_1171631, EPI_ISL_1171632, EPI_ISL_1171633, EPI_ISL_1171634, EPI_ISL_1171635, EPI_ISL_1171636, EPI_ISL_1171637, EPI_ISL_1171638, EPI_ISL_1171639, EPI_ISL_1171640 | | | |
| see above | IAL Regional de Santos | Instituto Adolfo Lutz, Interdiciplinary Procedures Center, Strategic Laboratory | Claudio Tavares Sacchi, Claudia Regina Gonçalves, Erica Valessa Ramos Gomes, Karoline Rodrigues Campos, Caio Vinicius Dias Lopes |
| EPI_ISL_1171641, EPI_ISL_1171642, EPI_ISL_1171643, EPI_ISL_1171644, EPI_ISL_1171645, EPI_ISL_1171646, EPI_ISL_1171647 | IAL Regional de Marilia | Instituto Adolfo Lutz, Interdiciplinary Procedures Center, Strategic Laboratory | Claudio Tavares Sacchi, Claudia Regina Gonçalves, Erica Valessa Ramos Gomes, Karoline Rodrigues Campos, Caio Vinicius Dias Lopes |
| EPI_ISL_1171648, EPI_ISL_1171649, EPI_ISL_1171650 | Centro de Saude Dr. Jose Paione em Mococa | Instituto Adolfo Lutz, Interdiciplinary Procedures Center, Strategic Laboratory | Claudio Tavares Sacchi, Claudia Regina Gonçalves, Erica Valessa Ramos Gomes, Karoline Rodrigues Campos, Caio Vinicius Dias Lopes |
| EPI_ISL_1171651, EPI_ISL_1171652, EPI_ISL_1171653, EPI_ISL_1171654, EPI_ISL_1171655, EPI_ISL_1171656, EPI_ISL_1171657, EPI_ISL_1171658, EPI_ISL_1171659, EPI_ISL_1171660, EPI_ISL_1171661, EPI_ISL_1171662, EPI_ISL_1171663, EPI_ISL_1171664, EPI_ISL_1171665, EPI_ISL_1171666, EPI_ISL_1171667, EPI_ISL_1171668, EPI_ISL_1171669, EPI_ISL_1171670, EPI_ISL_1171671, EPI_ISL_1171672, EPI_ISL_1171673, EPI_ISL_1171674 | | | |
| see above | IAL Regional de Presidente Prudente | Instituto Adolfo Lutz, Interdiciplinary Procedures Center, Strategic Laboratory | Claudio Tavares Sacchi, Claudia Regina Gonçalves, Erica Valessa Ramos Gomes, Karoline Rodrigues Campos, Caio Vinicius Dias Lopes |
| EPI_ISL_1172014 | HC_FMUSP | Laboratório de Parasitologia Médica - Instituto de Medicina Tropical - Universidade de São Paulo | Brazil-UK Centre for Arbovirus Discovery Diagnosis Genomics and Epidemiology (CADDE) Genomic Network - Instituto de Medicina Tropical |
| EPI_ISL_1181356 | Oswaldo Cruz Foundation, FIOCRUZ - Ceara (Fiocruz-CE) | Laboratory of Respiratory Viruses and Measles, Oswaldo Cruz Institute, FIOCRUZ | Paola Resende, Luciana Appolinario, Fernando Motta, Anna Carolina Paixao, Ana Carolina Mendonca, Alice Sampaio Rocha, Renata Serrano Lopes, Thais de Oliveira Costa, Joaquim César do Nascimento Sousa Júnior, Fabio Miyajima, Marilda Siqueira on behalf of the Fiocruz COVID-19 Genomic Surveillance Network |
| EPI_ISL_1181360, EPI_ISL_1181361 | Laboratory of Respiratory Viruses and Measles, Oswaldo Cruz Institute, FIOCRUZ | Laboratory of Respiratory Viruses and Measles, Oswaldo Cruz Institute, FIOCRUZ | Paola Resende, Luciana Appolinario, Fernando Motta, Anna Carolina Paixao, Ana Carolina Mendonca, Alice Sampaio Rocha, Renata Serrano Lopes, Marilda Siqueira on behalf of the Fiocruz COVID-19 Genomic Surveillance Network |
| EPI_ISL_1181362 | Instituto Aggeu Magalhães - Oswaldo Cruz Foundation, FIOCRUZ (FIOCRUZ-PE) | Laboratory of Respiratory Viruses and Measles, Oswaldo Cruz Institute, FIOCRUZ | Paola Resende, Luciana Appolinario, Fernando Motta, Anna Carolina Paixao, Ana Carolina Mendonca, Alice Sampaio Rocha, Renata Serrano Lopes, Gabriel Wallau, Marilda Siqueira on behalf of the Fiocruz COVID-19 Genomic Surveillance Network |
| EPI_ISL_1181363 | Laboratory of Respiratory Viruses and Measles, Oswaldo Cruz Institute, FIOCRUZ | Laboratory of Respiratory Viruses and Measles, Oswaldo Cruz Institute, FIOCRUZ | Paola Resende, Luciana Appolinario, Fernando Motta, Anna Carolina Paixao, Ana Carolina Mendonca, Alice Sampaio Rocha, Renata Serrano Lopes, Marilda Siqueira on behalf of the Fiocruz COVID-19 Genomic Surveillance Network |
| EPI_ISL_1181365 | Laboratorio Central de Saude Publica do Estado do Alagoas (LACEN-AL) | Laboratory of Respiratory Viruses and Measles, Oswaldo Cruz Institute, FIOCRUZ | Paola Resende, Luciana Appolinario, Fernando Motta, Anna Carolina Paixao, Ana Carolina Mendonca, Alice Sampaio Rocha, Renata Serrano Lopes, Anderson Brandao Leite, Marilda Siqueira on behalf of the Fiocruz COVID-19 Genomic Surveillance Network |
| EPI_ISL_1181367, EPI_ISL_1181369 | Laboratory of Respiratory Viruses and Measles, Oswaldo Cruz Institute, FIOCRUZ | Laboratory of Respiratory Viruses and Measles, Oswaldo Cruz Institute, FIOCRUZ | Paola Resende, Luciana Appolinario, Fernando Motta, Anna Carolina Paixao, Ana Carolina Mendonca, Alice Sampaio Rocha, Renata Serrano Lopes, Marilda Siqueira on behalf of the Fiocruz COVID-19 Genomic Surveillance Network |
| EPI_ISL_1181379, EPI_ISL_1181380 | Gonçalo Moniz Institute, FIOCRUZ, Bahia | Laboratory of Respiratory Viruses and Measles, Oswaldo Cruz Institute, FIOCRUZ | Paola Resende, Luciana Appolinario, Fernando Motta, Anna Carolina Paixao, Ana Carolina Mendonca, Alice Sampaio Rocha, Renata Serrano Lopes, Tiago Graf, Ricardo Khouri, Marilda Siqueira on behalf of the Fiocruz COVID-19 Genomic Surveillance Network |
| EPI_ISL_1181381, EPI_ISL_1181382, EPI_ISL_1181383, EPI_ISL_1181384, EPI_ISL_1181385, EPI_ISL_1181386, EPI_ISL_1181387, EPI_ISL_1181388, EPI_ISL_1181389 | Laboratory of Respiratory Viruses and Measles, Oswaldo Cruz Institute, FIOCRUZ | Laboratory of Respiratory Viruses and Measles, Oswaldo Cruz Institute, FIOCRUZ | Paola Resende, Luciana Appolinario, Fernando Motta, Anna Carolina Paixao, Ana Carolina Mendonca, Alice Sampaio Rocha, Renata Serrano Lopes, Marilda Siqueira on behalf of the Fiocruz COVID-19 Genomic Surveillance Network |
| EPI_ISL_1181390 | Laboratorio Central de Saude Publica do Estado do Parana (LACEN-PR) | Laboratory of Respiratory Viruses and Measles, Oswaldo Cruz Institute, FIOCRUZ | Paola Resende, Luciana Appolinario, Fernando Motta, Anna Carolina Paixao, Ana Carolina Mendonca, Alice Sampaio Rocha, Renata Serrano Lopes, Maria do Carmo Debur, Irina Nastassja Riediger, Marilda Siqueira on behalf of the Fiocruz COVID-19 Genomic Surveillance Network |
| EPI_ISL_1181391 | Laboratory of Respiratory Viruses and Measles, Oswaldo Cruz Institute, FIOCRUZ | Laboratory of Respiratory Viruses and Measles, Oswaldo Cruz Institute, FIOCRUZ | Paola Resende, Luciana Appolinario, Fernando Motta, Anna Carolina Paixao, Ana Carolina Mendonca, Alice Sampaio Rocha, Renata Serrano Lopes, Marilda Siqueira on behalf of the Fiocruz COVID-19 Genomic Surveillance Network |
| EPI_ISL_1181392 | Laboratorio Central de Saude Publica do Estado do Parana (LACEN-PR) | Laboratory of Respiratory Viruses and Measles, Oswaldo Cruz Institute, FIOCRUZ | Paola Resende, Luciana Appolinario, Fernando Motta, Anna Carolina Paixao, Ana Carolina Mendonca, Alice Sampaio Rocha, Renata Serrano Lopes, Maria do Carmo Debur, Irina Nastassja Riediger, Marilda Siqueira on behalf of the Fiocruz COVID-19 Genomic Surveillance Network |
| EPI_ISL_1181393 | Laboratory of Respiratory Viruses and Measles, Oswaldo Cruz Institute, FIOCRUZ | Laboratory of Respiratory Viruses and Measles, Oswaldo Cruz Institute, FIOCRUZ | Paola Resende, Luciana Appolinario, Fernando Motta, Anna Carolina Paixao, Ana Carolina Mendonca, Alice Sampaio Rocha, Renata Serrano Lopes, Marilda Siqueira on behalf of the Fiocruz COVID-19 Genomic Surveillance Network |
| EPI_ISL_1181394 | Laboratorio Central de Saude Publica do Estado do Parana (LACEN-PR) | Laboratory of Respiratory Viruses and Measles, Oswaldo Cruz Institute, FIOCRUZ | Paola Resende, Luciana Appolinario, Fernando Motta, Anna Carolina Paixao, Ana Carolina Mendonca, Alice Sampaio Rocha, Renata Serrano Lopes, Maria do Carmo Debur, Irina Nastassja Riediger, Marilda Siqueira on behalf of the Fiocruz COVID-19 Genomic Surveillance Network |
| EPI_ISL_1181395 | Laboratorio Central de Saude Publica do Estado do Espitito Santo (LACEN-ES) | Laboratory of Respiratory Viruses and Measles, Oswaldo Cruz Institute, FIOCRUZ | Paola Resende, Luciana Appolinario, Fernando Motta, Anna Carolina Paixao, Ana Carolina Mendonca, Alice Sampaio Rocha, Renata Serrano Lopes, Rodrigo Ribeiro Rodrigues, Marilda Siqueira on behalf of the Fiocruz COVID-19 Genomic Surveillance Network |
| EPI_ISL_1181396 | Laboratorio Central de Saude Publica do Estado do Alagoas (LACEN-AL) | Laboratory of Respiratory Viruses and Measles, Oswaldo Cruz Institute, FIOCRUZ | Paola Resende, Luciana Appolinario, Fernando Motta, Anna Carolina Paixao, Ana Carolina Mendonca, Alice Sampaio Rocha, Renata Serrano Lopes, Anderson Brandao Leite, Marilda Siqueira on behalf of the Fiocruz COVID-19 Genomic Surveillance Network |
| EPI_ISL_1181397 | Laboratorio Central de Saude Publica do Estado do Espitito Santo (LACEN-ES) | Laboratory of Respiratory Viruses and Measles, Oswaldo Cruz Institute, FIOCRUZ | Paola Resende, Luciana Appolinario, Fernando Motta, Anna Carolina Paixao, Ana Carolina Mendonca, Alice Sampaio Rocha, Renata Serrano Lopes, Rodrigo Ribeiro Rodrigues, Marilda Siqueira on behalf of the Fiocruz COVID-19 Genomic Surveillance Network |
| EPI_ISL_1181398 | Laboratory of Respiratory Viruses and Measles, Oswaldo Cruz Institute, FIOCRUZ | Laboratory of Respiratory Viruses and Measles, Oswaldo Cruz Institute, FIOCRUZ | Paola Resende, Luciana Appolinario, Fernando Motta, Anna Carolina Paixao, Ana Carolina Mendonca, Alice Sampaio Rocha, Renata Serrano Lopes, Marilda Siqueira on behalf of the Fiocruz COVID-19 Genomic Surveillance Network |
| EPI_ISL_1181399 | Laboratorio Central de Saude Publica do Estado de Minas Gerais (LACEN-MG) | Laboratory of Respiratory Viruses and Measles, Oswaldo Cruz Institute, FIOCRUZ | Paola Resende, Luciana Appolinario, Fernando Motta, Anna Carolina Paixao, Ana Carolina Mendonca, Alice Sampaio Rocha, Renata Serrano Lopes, Felipe Iani, Marilda Siqueira on behalf of the Fiocruz COVID-19 Genomic Surveillance Network |
| EPI_ISL_1181400, EPI_ISL_1181401 | Laboratorio Central de Saude Publica do Estado Maranhao (LACEN-MA) | Laboratory of Respiratory Viruses and Measles, Oswaldo Cruz Institute, FIOCRUZ | Paola Resende, Luciana Appolinario, Fernando Motta, Anna Carolina Paixao, Ana Carolina Mendonca, Alice Sampaio Rocha, Renata Serrano Lopes, Lidio Gonçalves Lima Neto, Marilda Siqueira on behalf of the Fiocruz COVID-19 Genomic Surveillance Network |
| EPI_ISL_1181402 | Laboratory of Respiratory Viruses and Measles, Oswaldo Cruz Institute, FIOCRUZ | Laboratory of Respiratory Viruses and Measles, Oswaldo Cruz Institute, FIOCRUZ | Paola Resende, Luciana Appolinario, Fernando Motta, Anna Carolina Paixao, Ana Carolina Mendonca, Alice Sampaio Rocha, Renata Serrano Lopes, Marilda Siqueira on behalf of the Fiocruz COVID-19 Genomic Surveillance Network |
| EPI_ISL_1181403, EPI_ISL_1181404 | Laboratorio Central de Saude Publica do Estado de Santa Catarina (LACEN-SC) | Laboratory of Respiratory Viruses and Measles, Oswaldo Cruz Institute, FIOCRUZ | Paola Resende, Luciana Appolinario, Fernando Motta, Anna Carolina Paixao, Ana Carolina Mendonca, Alice Sampaio Rocha, Renata Serrano Lopes, Darcita Buerger Rovaris, Sandra Bianchini Fernandes, Marilda Siqueira on behalf of the Fiocruz COVID-19 Genomic Surveillance Network |

| | | | |
|---|---|---|---|
| EPI_ISL_1181405, EPI_ISL_1181406, EPI_ISL_1181407 | Laboratorio Central de Saude Publica do Estado de Sergipe (LACEN-SE) | Laboratory of Respiratory Viruses and Measles, Oswaldo Cruz Institute, FIOCRUZ | Paola Resende, Luciana Appolinario, Fernando Motta, Anna Carolina Paixao, Ana Carolina Mendonca, Alice Sampaio Rocha, Renata Serrano Lopes, Cliomar Alves dos Santos, Marilda Siqueira on behalf of the Fiocruz COVID-19 Genomic Surveillance Network |
| EPI_ISL_1181408 | Gonçalo Moniz Institute, FIOCRUZ, Bahia | Laboratory of Respiratory Viruses and Measles, Oswaldo Cruz Institute, FIOCRUZ | Paola Resende, Luciana Appolinario, Fernando Motta, Anna Carolina Paixao, Ana Carolina Mendonca, Alice Sampaio Rocha, Renata Serrano Lopes, Tiago Graf, Ricardo Khouri, Marilda Siqueira on behalf of the Fiocruz COVID-19 Genomic Surveillance Network |
| EPI_ISL_1181409 | Laboratorio Central de Saude Publica do Estado de Sergipe (LACEN-SE) | Laboratory of Respiratory Viruses and Measles, Oswaldo Cruz Institute, FIOCRUZ | Paola Resende, Luciana Appolinario, Fernando Motta, Anna Carolina Paixao, Ana Carolina Mendonca, Alice Sampaio Rocha, Renata Serrano Lopes, Cliomar Alves dos Santos, Marilda Siqueira on behalf of the Fiocruz COVID-19 Genomic Surveillance Network |
| EPI_ISL_1181410 | Laboratory of Respiratory Viruses and Measles, Oswaldo Cruz Institute, FIOCRUZ | Laboratory of Respiratory Viruses and Measles, Oswaldo Cruz Institute, FIOCRUZ | Paola Resende, Luciana Appolinario, Fernando Motta, Anna Carolina Paixao, Ana Carolina Mendonca, Alice Sampaio Rocha, Renata Serrano Lopes, Marilda Siqueira on behalf of the Fiocruz COVID-19 Genomic Surveillance Network |
| EPI_ISL_1181411 | Laboratorio Central de Saude Publica do Estado do Rio Grande do Sul (LACEN-RS) | Laboratory of Respiratory Viruses and Measles, Oswaldo Cruz Institute, FIOCRUZ | Paola Resende, Luciana Appolinario, Fernando Motta, Anna Carolina Paixao, Ana Carolina Mendonca, Alice Sampaio Rocha, Renata Serrano Lopes, Tatiana Schaffer Gregianini, Marilda Siqueira on behalf of the Fiocruz COVID-19 Genomic Surveillance Network |
| EPI_ISL_1181412 | Laboratorio Central de Saude Publica do Estado de Santa Catarina (LACEN-SC) | Laboratory of Respiratory Viruses and Measles, Oswaldo Cruz Institute, FIOCRUZ | Paola Resende, Luciana Appolinario, Fernando Motta, Anna Carolina Paixao, Ana Carolina Mendonca, Alice Sampaio Rocha, Renata Serrano Lopes, Darcita Buerger Rovaris, Sandra Bianchini Fernandes, Marilda Siqueira on behalf of the Fiocruz COVID-19 Genomic Surveillance Network |
| EPI_ISL_1181413, EPI_ISL_1181414 | Laboratorio Central de Saude Publica do Estado Maranhao (LACEN-MA) | Laboratory of Respiratory Viruses and Measles, Oswaldo Cruz Institute, FIOCRUZ | Paola Resende, Luciana Appolinario, Fernando Motta, Anna Carolina Paixao, Ana Carolina Mendonca, Alice Sampaio Rocha, Renata Serrano Lopes, Lidio Gonçalves Lima Neto, Marilda Siqueira on behalf of the Fiocruz COVID-19 Genomic Surveillance Network |
| EPI_ISL_1181415 | Laboratory of Respiratory Viruses and Measles, Oswaldo Cruz Institute, FIOCRUZ | Laboratory of Respiratory Viruses and Measles, Oswaldo Cruz Institute, FIOCRUZ | Paola Resende, Luciana Appolinario, Fernando Motta, Anna Carolina Paixao, Ana Carolina Mendonca, Alice Sampaio Rocha, Renata Serrano Lopes, Marilda Siqueira on behalf of the Fiocruz COVID-19 Genomic Surveillance Network |
| EPI_ISL_1181416 | Laboratorio Central de Saude Publica do Estado da Paraiba (LACEN-PB) | Laboratory of Respiratory Viruses and Measles, Oswaldo Cruz Institute, FIOCRUZ | Paola Resende, Luciana Appolinario, Fernando Motta, Anna Carolina Paixao, Ana Carolina Mendonca, Alice Sampaio Rocha, Renata Serrano Lopes, Joao Felipe Bezerra, Dalane Loudal Florentino Teixeira, Marilda Siqueira on behalf of the Fiocruz COVID-19 Genomic Surveillance Network |
| EPI_ISL_1181418 | Laboratorio Central de Saude Publica do Estado do Espitito Santo (LACEN-ES) | Laboratory of Respiratory Viruses and Measles, Oswaldo Cruz Institute, FIOCRUZ | Paola Resende, Luciana Appolinario, Fernando Motta, Anna Carolina Paixao, Ana Carolina Mendonca, Alice Sampaio Rocha, Renata Serrano Lopes, Rodrigo Ribeiro Rodrigues, Marilda Siqueira on behalf of the Fiocruz COVID-19 Genomic Surveillance Network |
| EPI_ISL_1181419 | Gonçalo Moniz Institute, FIOCRUZ, Bahia | Laboratory of Respiratory Viruses and Measles, Oswaldo Cruz Institute, FIOCRUZ | Paola Resende, Luciana Appolinario, Fernando Motta, Anna Carolina Paixao, Ana Carolina Mendonca, Alice Sampaio Rocha, Renata Serrano Lopes, Tiago Graf, Ricardo Khouri, Marilda Siqueira on behalf of the Fiocruz COVID-19 Genomic Surveillance Network |
| EPI_ISL_1181420, EPI_ISL_1181421 | Laboratorio Central de Saude Publica do Estado do Espitito Santo (LACEN-ES) | Laboratory of Respiratory Viruses and Measles, Oswaldo Cruz Institute, FIOCRUZ | Paola Resende, Luciana Appolinario, Fernando Motta, Anna Carolina Paixao, Ana Carolina Mendonca, Alice Sampaio Rocha, Renata Serrano Lopes, Rodrigo Ribeiro Rodrigues, Marilda Siqueira on behalf of the Fiocruz COVID-19 Genomic Surveillance Network |
| EPI_ISL_1181422 | Laboratorio Central de Saude Publica do Estado da Paraiba (LACEN-PB) | Laboratory of Respiratory Viruses and Measles, Oswaldo Cruz Institute, FIOCRUZ | Paola Resende, Luciana Appolinario, Fernando Motta, Anna Carolina Paixao, Ana Carolina Mendonca, Alice Sampaio Rocha, Renata Serrano Lopes, Joao Felipe Bezerra, Dalane Loudal Florentino Teixeira, Marilda Siqueira on behalf of the Fiocruz COVID-19 Genomic Surveillance Network |
| EPI_ISL_1181423 | Laboratorio Central de Saude Publica do Estado do Espitito Santo (LACEN-ES) | Laboratory of Respiratory Viruses and Measles, Oswaldo Cruz Institute, FIOCRUZ | Paola Resende, Luciana Appolinario, Fernando Motta, Anna Carolina Paixao, Ana Carolina Mendonca, Alice Sampaio Rocha, Renata Serrano Lopes, Rodrigo Ribeiro Rodrigues, Marilda Siqueira on behalf of the Fiocruz COVID-19 Genomic Surveillance Network |
| EPI_ISL_1181424 | Gonçalo Moniz Institute, FIOCRUZ, Bahia | Laboratory of Respiratory Viruses and Measles, Oswaldo Cruz Institute, FIOCRUZ | Paola Resende, Luciana Appolinario, Fernando Motta, Anna Carolina Paixao, Ana Carolina Mendonca, Alice Sampaio Rocha, Renata Serrano Lopes, Tiago Graf, Ricardo Khouri, Marilda Siqueira on behalf of the Fiocruz COVID-19 Genomic Surveillance Network |
| EPI_ISL_1181425 | Laboratorio Central de Saude Publica do Estado do Rio de Janeiro (LACEN-RJ) | Laboratory of Respiratory Viruses and Measles, Oswaldo Cruz Institute, FIOCRUZ | Paola Resende, Luciana Appolinario, Fernando Motta, Anna Carolina Paixao, Ana Carolina Mendonca, Alice Sampaio Rocha, Renata Serrano Lopes, Andrea Cony Cavalcanti, Marilda Siqueira on behalf of the Fiocruz COVID-19 Genomic Surveillance Network |
| EPI_ISL_1181426, EPI_ISL_1181427, EPI_ISL_1181428 | Laboratorio Central de Saude Publica do Estado de Minas Gerais (LACEN-MG) | Laboratory of Respiratory Viruses and Measles, Oswaldo Cruz Institute, FIOCRUZ | Paola Resende, Luciana Appolinario, Fernando Motta, Anna Carolina Paixao, Ana Carolina Mendonca, Alice Sampaio Rocha, Renata Serrano Lopes, Felipe Iani, Marilda Siqueira on behalf of the Fiocruz COVID-19 Genomic Surveillance Network |
| EPI_ISL_1181429 | Gonçalo Moniz Institute, FIOCRUZ, Bahia | Laboratory of Respiratory Viruses and Measles, Oswaldo Cruz Institute, FIOCRUZ | Paola Resende, Luciana Appolinario, Fernando Motta, Anna Carolina Paixao, Ana Carolina Mendonca, Alice Sampaio Rocha, Renata Serrano Lopes, Tiago Graf, Ricardo Khouri, Marilda Siqueira on behalf of the Fiocruz COVID-19 Genomic Surveillance Network |
| EPI_ISL_1181430 | Laboratory of Respiratory Viruses and Measles, Oswaldo Cruz Institute, FIOCRUZ | Laboratory of Respiratory Viruses and Measles, Oswaldo Cruz Institute, FIOCRUZ | Paola Resende, Luciana Appolinario, Fernando Motta, Anna Carolina Paixao, Ana Carolina Mendonca, Alice Sampaio Rocha, Renata Serrano Lopes, Marilda Siqueira on behalf of the Fiocruz COVID-19 Genomic Surveillance Network |
| EPI_ISL_1181431 | Laboratorio Central de Saude Publica do Estado do Rio de Janeiro (LACEN-RJ) | Laboratory of Respiratory Viruses and Measles, Oswaldo Cruz Institute, FIOCRUZ | Paola Resende, Luciana Appolinario, Fernando Motta, Anna Carolina Paixao, Ana Carolina Mendonca, Alice Sampaio Rocha, Renata Serrano Lopes, Andrea Cony Cavalcanti, Marilda Siqueira on behalf of the Fiocruz COVID-19 Genomic Surveillance Network |
| EPI_ISL_1181432 | Laboratorio Central de Saude Publica do Estado do Espitito Santo (LACEN-ES) | Laboratory of Respiratory Viruses and Measles, Oswaldo Cruz Institute, FIOCRUZ | Paola Resende, Luciana Appolinario, Fernando Motta, Anna Carolina Paixao, Ana Carolina Mendonca, Alice Sampaio Rocha, Renata Serrano Lopes, Rodrigo Ribeiro Rodrigues, Marilda Siqueira on behalf of the Fiocruz COVID-19 Genomic Surveillance Network |
| EPI_ISL_1181433, EPI_ISL_1181434 | Laboratorio Central de Saude Publica do Estado do Rio Grande do Sul (LACEN-RS) | Laboratory of Respiratory Viruses and Measles, Oswaldo Cruz Institute, FIOCRUZ | Paola Resende, Luciana Appolinario, Fernando Motta, Anna Carolina Paixao, Ana Carolina Mendonca, Alice Sampaio Rocha, Renata Serrano Lopes, Tatiana Schaffer Gregianini, Marilda Siqueira on behalf of the Fiocruz COVID-19 Genomic Surveillance Network |
| EPI_ISL_1181435, EPI_ISL_1181436 | Federal University of Southern Bahia (UFSB - Universidade Federal do Sul da Bahia) | Laboratory of Respiratory Viruses and Measles, Oswaldo Cruz Institute, FIOCRUZ | Paola Resende, Luciana Appolinario, Fernando Motta, Anna Carolina Paixao, Ana Carolina Mendonca, Alice Sampaio Rocha, Renata Serrano Lopes, Thiago Mafra, Marilda Siqueira on behalf of the Fiocruz COVID-19 Genomic Surveillance Network |
| EPI_ISL_1181437, EPI_ISL_1181438, EPI_ISL_1181439 | Laboratory of Respiratory Viruses and Measles, Oswaldo Cruz Institute, FIOCRUZ | Laboratory of Respiratory Viruses and Measles, Oswaldo Cruz Institute, FIOCRUZ | Paola Resende, Luciana Appolinario, Fernando Motta, Anna Carolina Paixao, Ana Carolina Mendonca, Alice Sampaio Rocha, Renata Serrano Lopes, Marilda Siqueira on behalf of the Fiocruz COVID-19 Genomic Surveillance Network |
| EPI_ISL_1181440, EPI_ISL_1181441 | Gonçalo Moniz Institute, FIOCRUZ, Bahia | Laboratory of Respiratory Viruses and Measles, Oswaldo Cruz Institute, FIOCRUZ | Paola Resende, Luciana Appolinario, Fernando Motta, Anna Carolina Paixao, Ana Carolina Mendonca, Alice Sampaio Rocha, Renata Serrano Lopes, Tiago Graf, Ricardo Khouri, Marilda Siqueira on behalf of the Fiocruz COVID-19 Genomic Surveillance Network |
| EPI_ISL_1181442 | Laboratory of Respiratory Viruses and Measles, Oswaldo Cruz Institute, FIOCRUZ | Laboratory of Respiratory Viruses and Measles, Oswaldo Cruz Institute, FIOCRUZ | Paola Resende, Luciana Appolinario, Fernando Motta, Anna Carolina Paixao, Ana Carolina Mendonca, Alice Sampaio Rocha, Renata Serrano Lopes, Marilda Siqueira on behalf of the Fiocruz COVID-19 Genomic Surveillance Network |
| EPI_ISL_1181443 | Federal University of Southern Bahia (UFSB - Universidade Federal do Sul da Bahia) | Laboratory of Respiratory Viruses and Measles, Oswaldo Cruz Institute, FIOCRUZ | Paola Resende, Luciana Appolinario, Fernando Motta, Anna Carolina Paixao, Ana Carolina Mendonca, Alice Sampaio Rocha, Renata Serrano Lopes, Thiago Mafra, Marilda Siqueira on behalf of the Fiocruz COVID-19 Genomic Surveillance Network |
| EPI_ISL_1181444, EPI_ISL_1181445 | Gonçalo Moniz Institute, FIOCRUZ, Bahia | Laboratory of Respiratory Viruses and Measles, Oswaldo Cruz Institute, FIOCRUZ | Paola Resende, Luciana Appolinario, Fernando Motta, Anna Carolina Paixao, Ana Carolina Mendonca, Alice Sampaio Rocha, Renata Serrano Lopes, Tiago Graf, Ricardo Khouri, Marilda Siqueira on behalf of the Fiocruz COVID-19 Genomic Surveillance Network |
| EPI_ISL_1181446, EPI_ISL_1181447, EPI_ISL_1181448 | Laboratory of Respiratory Viruses and Measles, Oswaldo Cruz Institute, FIOCRUZ | Laboratory of Respiratory Viruses and Measles, Oswaldo Cruz Institute, FIOCRUZ | Paola Resende, Luciana Appolinario, Fernando Motta, Anna Carolina Paixao, Ana Carolina Mendonca, Alice Sampaio Rocha, Renata Serrano Lopes, Marilda Siqueira on behalf of the Fiocruz COVID-19 Genomic Surveillance Network |
| EPI_ISL_1181449 | Gonçalo Moniz Institute, FIOCRUZ, Bahia | Laboratory of Respiratory Viruses and Measles, Oswaldo Cruz Institute, FIOCRUZ | Paola Resende, Luciana Appolinario, Fernando Motta, Anna Carolina Paixao, Ana Carolina Mendonca, Alice Sampaio Rocha, Renata Serrano Lopes, Tiago Graf, Ricardo Khouri, Marilda Siqueira on behalf of the Fiocruz COVID-19 Genomic Surveillance Network |
| EPI_ISL_1181450 | Federal University of Southern Bahia (UFSB - Universidade Federal do Sul da Bahia) | Laboratory of Respiratory Viruses and Measles, Oswaldo Cruz Institute, FIOCRUZ | Paola Resende, Luciana Appolinario, Fernando Motta, Anna Carolina Paixao, Ana Carolina Mendonca, Alice Sampaio Rocha, Renata Serrano Lopes, Thiago Mafra, Marilda Siqueira on behalf of the Fiocruz COVID-19 Genomic Surveillance Network |
| EPI_ISL_1181451 | Laboratorio Central de Saude Publica do Estado do Rio Grande do Sul (LACEN-RS) | Laboratory of Respiratory Viruses and Measles, Oswaldo Cruz Institute, FIOCRUZ | Paola Resende, Luciana Appolinario, Fernando Motta, Anna Carolina Paixao, Ana Carolina Mendonca, Alice Sampaio Rocha, Renata Serrano Lopes, Tatiana Schaffer Gregianini, Marilda Siqueira on behalf of the Fiocruz COVID-19 Genomic Surveillance Network |
| EPI_ISL_1181452 | Laboratorio de Imunologia de Transplantes de Goias LTDA (HLAGYN) | Laboratory of Respiratory Viruses and Measles, Oswaldo Cruz Institute, FIOCRUZ | Paola Resende, Luciana Appolinario, Fernando Motta, Anna Carolina Paixao, Ana Carolina Mendonca, Alice Sampaio Rocha, Renata Serrano Lopes , Alessandro Leonardo Alvares Magalhaes, Fernando Antonio Vinhal dos Santos, Erika Lopes Rocha Batista, Marilda Siqueira on behalf of the Fiocruz COVID-19 Genomic Surveillance Network |
| EPI_ISL_1181453 | Laboratorio Central de Saude Publica do Estado de Santa Catarina (LACEN-SC) | Laboratory of Respiratory Viruses and Measles, Oswaldo Cruz Institute, FIOCRUZ | Paola Resende, Luciana Appolinario, Fernando Motta, Anna Carolina Paixao, Ana Carolina Mendonca, Alice Sampaio Rocha, Renata Serrano Lopes, Darcita Buerger Rovaris, Sandra Bianchini Fernandes, Marilda Siqueira on behalf of the Fiocruz COVID-19 Genomic Surveillance Network |

EPI_ISL_1181454, EPI_ISL_1181455, EPI_ISL_1181456, EPI_ISL_1181457, EPI_ISL_1181458, EPI_ISL_1181459, EPI_ISL_1181460, EPI_ISL_1181461, EPI_ISL_1181462, EPI_ISL_1181463, EPI_ISL_1181464, EPI_ISL_1181465, EPI_ISL_1181466, EPI_ISL_1181467

| | | | |
|---|---|---|---|
| see above | Laboratory of Respiratory Viruses and Measles, Oswaldo Cruz Institute, FIOCRUZ | Laboratory of Respiratory Viruses and Measles, Oswaldo Cruz Institute, FIOCRUZ | Paola Resende, Luciana Appolinario, Fernando Motta, Anna Carolina Paixao, Ana Carolina Mendonca, Alice Sampaio Rocha, Renata Serrano Lopes, Marilda Siqueira on behalf of the Fiocruz COVID-19 Genomic Surveillance Network |
| EPI_ISL_1181468 | Laboratorio Central de Saude Publica do Estado do Rio Grande do Sul (LACEN-RS) | Laboratory of Respiratory Viruses and Measles, Oswaldo Cruz Institute, FIOCRUZ | Paola Resende, Luciana Appolinario, Fernando Motta, Anna Carolina Paixao, Ana Carolina Mendonca, Alice Sampaio Rocha, Renata Serrano Lopes, Tatiana Schaffer Gregianini, Marilda Siqueira on behalf of the Fiocruz COVID-19 Genomic Surveillance Network |
| EPI_ISL_1181469 | Federal University of Mato Grosso (UFMT) | Laboratory of Respiratory Viruses and Measles, Oswaldo Cruz Institute, FIOCRUZ | Paola Resende, Luciana Appolinario, Fernando Motta, Anna Carolina Paixao, Ana Carolina Mendonca, Alice Sampaio Rocha, Renata Serrano Lopes, Renata Dezengrini, Marilda Siqueira on behalf of the Fiocruz COVID-19 Genomic Surveillance Network |
| EPI_ISL_1181470 | Laboratory of Respiratory Viruses and Measles, Oswaldo Cruz Institute, FIOCRUZ | Laboratory of Respiratory Viruses and Measles, Oswaldo Cruz Institute, FIOCRUZ | Paola Resende, Luciana Appolinario, Fernando Motta, Anna Carolina Paixao, Ana Carolina Mendonca, Alice Sampaio Rocha, Renata Serrano Lopes, Marilda Siqueira on behalf of the Fiocruz COVID-19 Genomic Surveillance Network |
| EPI_ISL_1181471, EPI_ISL_1181472, EPI_ISL_1181473 | Laboratorio de Imunologia de Transplantes de Goias LTDA (HLAGYN) | Laboratory of Respiratory Viruses and Measles, Oswaldo Cruz Institute, FIOCRUZ | Paola Resende, Luciana Appolinario, Fernando Motta, Anna Carolina Paixao, Ana Carolina Mendonca, Alice Sampaio Rocha, Renata Serrano Lopes , Alessandro Leonardo Alvares Magalhaes, Fernando Antonio Vinhal dos Santos, Erika Lopes Rocha Batista, Marilda Siqueira on behalf of the Fiocruz COVID-19 Genomic Surveillance Network |
| EPI_ISL_1181474 | Laboratorio Central de Saude Publica do Estado do Parana (LACEN-PR) | Laboratory of Respiratory Viruses and Measles, Oswaldo Cruz Institute, FIOCRUZ | Paola Resende, Luciana Appolinario, Fernando Motta, Anna Carolina Paixao, Ana Carolina Mendonca, Alice Sampaio Rocha, Renata Serrano Lopes, Maria do Carmo Debur, Irina Nastassja Riediger, Marilda Siqueira on behalf of the Fiocruz COVID-19 Genomic Surveillance Network |
| EPI_ISL_1181475, EPI_ISL_1181476, EPI_ISL_1181477 | Laboratory of Respiratory Viruses and Measles, Oswaldo Cruz Institute, FIOCRUZ | Laboratory of Respiratory Viruses and Measles, Oswaldo Cruz Institute, FIOCRUZ | Paola Resende, Luciana Appolinario, Fernando Motta, Anna Carolina Paixao, Ana Carolina Mendonca, Alice Sampaio Rocha, Renata Serrano Lopes, Marilda Siqueira on behalf of the Fiocruz COVID-19 Genomic Surveillance Network |
| EPI_ISL_1181478, EPI_ISL_1181479, EPI_ISL_1181480 | Laboratorio de Imunologia de Transplantes de Goias LTDA (HLAGYN) | Laboratory of Respiratory Viruses and Measles, Oswaldo Cruz Institute, FIOCRUZ | Paola Resende, Luciana Appolinario, Fernando Motta, Anna Carolina Paixao, Ana Carolina Mendonca, Alice Sampaio Rocha, Renata Serrano Lopes , Alessandro Leonardo Alvares Magalhaes, Fernando Antonio Vinhal dos Santos, Erika Lopes Rocha Batista, Marilda Siqueira on behalf of the Fiocruz COVID-19 Genomic Surveillance Network |
| EPI_ISL_1181481, EPI_ISL_1181482, EPI_ISL_1181483 | Laboratory of Respiratory Viruses and Measles, Oswaldo Cruz Institute, FIOCRUZ | Laboratory of Respiratory Viruses and Measles, Oswaldo Cruz Institute, FIOCRUZ | Paola Resende, Luciana Appolinario, Fernando Motta, Anna Carolina Paixao, Ana Carolina Mendonca, Alice Sampaio Rocha, Renata Serrano Lopes, Marilda Siqueira on behalf of the Fiocruz COVID-19 Genomic Surveillance Network |
| EPI_ISL_1181484, EPI_ISL_1181485 | Gonçalo Moniz Institute, FIOCRUZ, Bahia | Laboratory of Respiratory Viruses and Measles, Oswaldo Cruz Institute, FIOCRUZ | Paola Resende, Luciana Appolinario, Fernando Motta, Anna Carolina Paixao, Ana Carolina Mendonca, Alice Sampaio Rocha, Renata Serrano Lopes, Tiago Graf, Ricardo Khouri, Marilda Siqueira on behalf of the Fiocruz COVID-19 Genomic Surveillance Network |
| EPI_ISL_1181486 | Laboratory of Respiratory Viruses and Measles, Oswaldo Cruz Institute, FIOCRUZ | Laboratory of Respiratory Viruses and Measles, Oswaldo Cruz Institute, FIOCRUZ | Paola Resende, Luciana Appolinario, Fernando Motta, Anna Carolina Paixao, Ana Carolina Mendonca, Alice Sampaio Rocha, Renata Serrano Lopes, Marilda Siqueira on behalf of the Fiocruz COVID-19 Genomic Surveillance Network |
| EPI_ISL_1181487 | Laboratorio de Imunologia de Transplantes de Goias LTDA (HLAGYN) | Laboratory of Respiratory Viruses and Measles, Oswaldo Cruz Institute, FIOCRUZ | Paola Resende, Luciana Appolinario, Fernando Motta, Anna Carolina Paixao, Ana Carolina Mendonca, Alice Sampaio Rocha, Renata Serrano Lopes , Alessandro Leonardo Alvares Magalhaes, Fernando Antonio Vinhal dos Santos, Erika Lopes Rocha Batista, Marilda Siqueira on behalf of the Fiocruz COVID-19 Genomic Surveillance Network |
| EPI_ISL_1181488, EPI_ISL_1181489, EPI_ISL_1181490, EPI_ISL_1181491 | Laboratory of Respiratory Viruses and Measles, Oswaldo Cruz Institute, FIOCRUZ | Laboratory of Respiratory Viruses and Measles, Oswaldo Cruz Institute, FIOCRUZ | Paola Resende, Luciana Appolinario, Fernando Motta, Anna Carolina Paixao, Ana Carolina Mendonca, Alice Sampaio Rocha, Renata Serrano Lopes, Marilda Siqueira on behalf of the Fiocruz COVID-19 Genomic Surveillance Network |
| EPI_ISL_1181492 | Laboratorio de Imunologia de Transplantes de Goias LTDA (HLAGYN) | Laboratory of Respiratory Viruses and Measles, Oswaldo Cruz Institute, FIOCRUZ | Paola Resende, Luciana Appolinario, Fernando Motta, Anna Carolina Paixao, Ana Carolina Mendonca, Alice Sampaio Rocha, Renata Serrano Lopes , Alessandro Leonardo Alvares Magalhaes, Fernando Antonio Vinhal dos Santos, Erika Lopes Rocha Batista, Marilda Siqueira on behalf of the Fiocruz COVID-19 Genomic Surveillance Network |
| EPI_ISL_1181493, EPI_ISL_1181494, EPI_ISL_1181495, EPI_ISL_1181496, EPI_ISL_1181497 | Laboratory of Respiratory Viruses and Measles, Oswaldo Cruz Institute, FIOCRUZ | Laboratory of Respiratory Viruses and Measles, Oswaldo Cruz Institute, FIOCRUZ | Paola Resende, Luciana Appolinario, Fernando Motta, Anna Carolina Paixao, Ana Carolina Mendonca, Alice Sampaio Rocha, Renata Serrano Lopes, Marilda Siqueira on behalf of the Fiocruz COVID-19 Genomic Surveillance Network |
| EPI_ISL_1181498 | Laboratorio Central de Saude Publica do Estado do Parana (LACEN-PR) | Laboratory of Respiratory Viruses and Measles, Oswaldo Cruz Institute, FIOCRUZ | Paola Resende, Luciana Appolinario, Fernando Motta, Anna Carolina Paixao, Ana Carolina Mendonca, Alice Sampaio Rocha, Renata Serrano Lopes, Maria do Carmo Debur, Irina Nastassja Riediger, Marilda Siqueira on behalf of the Fiocruz COVID-19 Genomic Surveillance Network |
| EPI_ISL_1181499, EPI_ISL_1181500, EPI_ISL_1181501, EPI_ISL_1181502, EPI_ISL_1181503, EPI_ISL_1181504, EPI_ISL_1181505, EPI_ISL_1181506, EPI_ISL_1181507, EPI_ISL_1181508, EPI_ISL_1181509, EPI_ISL_1181510, EPI_ISL_1181511, EPI_ISL_1181512, EPI_ISL_1181513, EPI_ISL_1181514, EPI_ISL_1181515 | | | |
| see above | Laboratory of Respiratory Viruses and Measles, Oswaldo Cruz Institute, FIOCRUZ | Laboratory of Respiratory Viruses and Measles, Oswaldo Cruz Institute, FIOCRUZ | Paola Resende, Luciana Appolinario, Fernando Motta, Anna Carolina Paixao, Ana Carolina Mendonca, Alice Sampaio Rocha, Renata Serrano Lopes, Marilda Siqueira on behalf of the Fiocruz COVID-19 Genomic Surveillance Network |
| EPI_ISL_1181516 | Laboratorio de Imunologia de Transplantes de Goias LTDA (HLAGYN) | Laboratory of Respiratory Viruses and Measles, Oswaldo Cruz Institute, FIOCRUZ | Paola Resende, Luciana Appolinario, Fernando Motta, Anna Carolina Paixao, Ana Carolina Mendonca, Alice Sampaio Rocha, Renata Serrano Lopes , Alessandro Leonardo Alvares Magalhaes, Fernando Antonio Vinhal dos Santos, Erika Lopes Rocha Batista, Marilda Siqueira on behalf of the Fiocruz COVID-19 Genomic Surveillance Network |
| EPI_ISL_1181517 | Laboratorio Central de Saude Publica do Estado do Parana (LACEN-PR) | Laboratory of Respiratory Viruses and Measles, Oswaldo Cruz Institute, FIOCRUZ | Paola Resende, Luciana Appolinario, Fernando Motta, Anna Carolina Paixao, Ana Carolina Mendonca, Alice Sampaio Rocha, Renata Serrano Lopes, Maria do Carmo Debur, Irina Nastassja Riediger, Marilda Siqueira on behalf of the Fiocruz COVID-19 Genomic Surveillance Network |
| EPI_ISL_1181518, EPI_ISL_1181519 | Laboratory of Respiratory Viruses and Measles, Oswaldo Cruz Institute, FIOCRUZ | Laboratory of Respiratory Viruses and Measles, Oswaldo Cruz Institute, FIOCRUZ | Paola Resende, Luciana Appolinario, Fernando Motta, Anna Carolina Paixao, Ana Carolina Mendonca, Alice Sampaio Rocha, Renata Serrano Lopes, Marilda Siqueira on behalf of the Fiocruz COVID-19 Genomic Surveillance Network |
| EPI_ISL_1181520 | Oswaldo Cruz Foundation, FIOCRUZ - Ceara (Fiocruz-CE) | Laboratory of Respiratory Viruses and Measles, Oswaldo Cruz Institute, FIOCRUZ | Paola Resende, Luciana Appolinario, Fernando Motta, Anna Carolina Paixao, Ana Carolina Mendonca, Alice Sampaio Rocha, Renata Serrano Lopes, Thais de Oliveira Costa, Joaquim César do Nascimento Sousa Júnior, Fabio Miyajima, Marilda Siqueira on behalf of the Fiocruz COVID-19 Genomic Surveillance Network |
| EPI_ISL_1181521 | Laboratory of Respiratory Viruses and Measles, Oswaldo Cruz Institute, FIOCRUZ | Laboratory of Respiratory Viruses and Measles, Oswaldo Cruz Institute, FIOCRUZ | Paola Resende, Luciana Appolinario, Fernando Motta, Anna Carolina Paixao, Ana Carolina Mendonca, Alice Sampaio Rocha, Renata Serrano Lopes, Marilda Siqueira on behalf of the Fiocruz COVID-19 Genomic Surveillance Network |
| EPI_ISL_1181522 | Oswaldo Cruz Foundation, FIOCRUZ - Ceara (Fiocruz-CE) | Laboratory of Respiratory Viruses and Measles, Oswaldo Cruz Institute, FIOCRUZ | Paola Resende, Luciana Appolinario, Fernando Motta, Anna Carolina Paixao, Ana Carolina Mendonca, Alice Sampaio Rocha, Renata Serrano Lopes, Thais de Oliveira Costa, Joaquim César do Nascimento Sousa Júnior, Fabio Miyajima, Marilda Siqueira on behalf of the Fiocruz COVID-19 Genomic Surveillance Network |
| EPI_ISL_1181523 | Laboratory of Respiratory Viruses and Measles, Oswaldo Cruz Institute, FIOCRUZ | Laboratory of Respiratory Viruses and Measles, Oswaldo Cruz Institute, FIOCRUZ | Paola Resende, Luciana Appolinario, Fernando Motta, Anna Carolina Paixao, Ana Carolina Mendonca, Alice Sampaio Rocha, Renata Serrano Lopes, Marilda Siqueira on behalf of the Fiocruz COVID-19 Genomic Surveillance Network |
| EPI_ISL_1181524 | Laboratorio Central de Saude Publica do Estado do Rio Grande do Sul (LACEN-RS) | Laboratory of Respiratory Viruses and Measles, Oswaldo Cruz Institute, FIOCRUZ | Paola Resende, Luciana Appolinario, Fernando Motta, Anna Carolina Paixao, Ana Carolina Mendonca, Alice Sampaio Rocha, Renata Serrano Lopes, Tatiana Schaffer Gregianini, Marilda Siqueira on behalf of the Fiocruz COVID-19 Genomic Surveillance Network |
| EPI_ISL_1181525 | Laboratorio de Imunologia de Transplantes de Goias LTDA (HLAGYN) | Laboratory of Respiratory Viruses and Measles, Oswaldo Cruz Institute, FIOCRUZ | Paola Resende, Luciana Appolinario, Fernando Motta, Anna Carolina Paixao, Ana Carolina Mendonca, Alice Sampaio Rocha, Renata Serrano Lopes , Alessandro Leonardo Alvares Magalhaes, Fernando Antonio Vinhal dos Santos, Erika Lopes Rocha Batista, Marilda Siqueira on behalf of the Fiocruz COVID-19 Genomic Surveillance Network |
| EPI_ISL_1181526, EPI_ISL_1181527, EPI_ISL_1181528 | Laboratorio Central de Saude Publica do Estado do Parana (LACEN-PR) | Laboratory of Respiratory Viruses and Measles, Oswaldo Cruz Institute, FIOCRUZ | Paola Resende, Luciana Appolinario, Fernando Motta, Anna Carolina Paixao, Ana Carolina Mendonca, Alice Sampaio Rocha, Renata Serrano Lopes, Maria do Carmo Debur, Irina Nastassja Riediger, Marilda Siqueira on behalf of the Fiocruz COVID-19 Genomic Surveillance Network |
| EPI_ISL_1181529 | Laboratory of Respiratory Viruses and Measles, Oswaldo Cruz Institute, FIOCRUZ | Laboratory of Respiratory Viruses and Measles, Oswaldo Cruz Institute, FIOCRUZ | Paola Resende, Luciana Appolinario, Fernando Motta, Anna Carolina Paixao, Ana Carolina Mendonca, Alice Sampaio Rocha, Renata Serrano Lopes, Marilda Siqueira on behalf of the Fiocruz COVID-19 Genomic Surveillance Network |
| EPI_ISL_1181530 | Laboratorio Central de Saude Publica do Estado do Parana (LACEN-PR) | Laboratory of Respiratory Viruses and Measles, Oswaldo Cruz Institute, FIOCRUZ | Paola Resende, Luciana Appolinario, Fernando Motta, Anna Carolina Paixao, Ana Carolina Mendonca, Alice Sampaio Rocha, Renata Serrano Lopes, Maria do Carmo Debur, Irina Nastassja Riediger, Marilda Siqueira on behalf of the Fiocruz COVID-19 Genomic Surveillance Network |
| EPI_ISL_1181531 | Laboratory of Respiratory Viruses and Measles, Oswaldo Cruz Institute, FIOCRUZ | Laboratory of Respiratory Viruses and Measles, Oswaldo Cruz Institute, FIOCRUZ | Paola Resende, Luciana Appolinario, Fernando Motta, Anna Carolina Paixao, Ana Carolina Mendonca, Alice Sampaio Rocha, Renata Serrano Lopes, Marilda Siqueira on behalf of the Fiocruz COVID-19 Genomic Surveillance Network |
| EPI_ISL_1181532 | Laboratorio de Imunologia de Transplantes de Goias LTDA (HLAGYN) | Laboratory of Respiratory Viruses and Measles, Oswaldo Cruz Institute, FIOCRUZ | Paola Resende, Luciana Appolinario, Fernando Motta, Anna Carolina Paixao, Ana Carolina Mendonca, Alice Sampaio Rocha, Renata Serrano Lopes , Alessandro Leonardo Alvares Magalhaes, Fernando Antonio Vinhal dos Santos, Erika Lopes Rocha Batista, Marilda Siqueira on behalf of the Fiocruz |

| | | | |
|---|---|---|---|
| | | | COVID-19 Genomic Surveillance Network |
| EPI_ISL_1181533, EPI_ISL_1181534 | Laboratorio Central de Saude Publica do Estado do Parana (LACEN-PR) | Laboratory of Respiratory Viruses and Measles, Oswaldo Cruz Institute, FIOCRUZ | Paola Resende, Luciana Appolinario, Fernando Motta, Anna Carolina Paixao, Ana Carolina Mendonca, Alice Sampaio Rocha, Renata Serrano Lopes, Maria do Carmo Debur, Irina Nastassja Riediger, Marilda Siqueira on behalf of the Fiocruz COVID-19 Genomic Surveillance Network |
| EPI_ISL_1181535 | Laboratorio de Imunologia de Transplantes de Goias LTDA (HLAGYN) | Laboratory of Respiratory Viruses and Measles, Oswaldo Cruz Institute, FIOCRUZ | Paola Resende, Luciana Appolinario, Fernando Motta, Anna Carolina Paixao, Ana Carolina Mendonca, Alice Sampaio Rocha, Renata Serrano Lopes , Alessandro Leonardo Alvares Magalhaes, Fernando Antonio Vinhal dos Santos, Erika Lopes Rocha Batista, Marilda Siqueira on behalf of the Fiocruz COVID-19 Genomic Surveillance Network |
| EPI_ISL_1181536, EPI_ISL_1181537, EPI_ISL_1181538, EPI_ISL_1181539, EPI_ISL_1181540, EPI_ISL_1181541, EPI_ISL_1181542, EPI_ISL_1181543, EPI_ISL_1181544, EPI_ISL_1181545 | Laboratorio Central de Saude Publica do Estado do Parana (LACEN-PR) | Laboratory of Respiratory Viruses and Measles, Oswaldo Cruz Institute, FIOCRUZ | Paola Resende, Luciana Appolinario, Fernando Motta, Anna Carolina Paixao, Ana Carolina Mendonca, Alice Sampaio Rocha, Renata Serrano Lopes, Maria do Carmo Debur, Irina Nastassja Riediger, Marilda Siqueira on behalf of the Fiocruz COVID-19 Genomic Surveillance Network |
| EPI_ISL_1181546, EPI_ISL_1181547, EPI_ISL_1181548, EPI_ISL_1181549, EPI_ISL_1181550 | Laboratorio de Imunologia de Transplantes de Goias LTDA (HLAGYN) | Laboratory of Respiratory Viruses and Measles, Oswaldo Cruz Institute, FIOCRUZ | Paola Resende, Luciana Appolinario, Fernando Motta, Anna Carolina Paixao, Ana Carolina Mendonca, Alice Sampaio Rocha, Renata Serrano Lopes , Alessandro Leonardo Alvares Magalhaes, Fernando Antonio Vinhal dos Santos, Erika Lopes Rocha Batista, Marilda Siqueira on behalf of the Fiocruz COVID-19 Genomic Surveillance Network |
| EPI_ISL_1181551, EPI_ISL_1181552, EPI_ISL_1181553, EPI_ISL_1181554, EPI_ISL_1181555, EPI_ISL_1181556, EPI_ISL_1181557, EPI_ISL_1181558, EPI_ISL_1181559 | Laboratorio Central de Saude Publica do Estado do Parana (LACEN-PR) | Laboratory of Respiratory Viruses and Measles, Oswaldo Cruz Institute, FIOCRUZ | Paola Resende, Luciana Appolinario, Fernando Motta, Anna Carolina Paixao, Ana Carolina Mendonca, Alice Sampaio Rocha, Renata Serrano Lopes, Maria do Carmo Debur, Irina Nastassja Riediger, Marilda Siqueira on behalf of the Fiocruz COVID-19 Genomic Surveillance Network |
| EPI_ISL_1181560, EPI_ISL_1181561, EPI_ISL_1181562 | Laboratory of Respiratory Viruses and Measles, Oswaldo Cruz Institute, FIOCRUZ | Laboratory of Respiratory Viruses and Measles, Oswaldo Cruz Institute, FIOCRUZ | Paola Resende, Luciana Appolinario, Fernando Motta, Anna Carolina Paixao, Ana Carolina Mendonca, Alice Sampaio Rocha, Renata Serrano Lopes, Marilda Siqueira on behalf of the Fiocruz COVID-19 Genomic Surveillance Network |
| EPI_ISL_1181563, EPI_ISL_1181564, EPI_ISL_1181565, EPI_ISL_1181566 | Laboratorio Central de Saude Publica do Estado do Parana (LACEN-PR) | Laboratory of Respiratory Viruses and Measles, Oswaldo Cruz Institute, FIOCRUZ | Paola Resende, Luciana Appolinario, Fernando Motta, Anna Carolina Paixao, Ana Carolina Mendonca, Alice Sampaio Rocha, Renata Serrano Lopes, Maria do Carmo Debur, Irina Nastassja Riediger, Marilda Siqueira on behalf of the Fiocruz COVID-19 Genomic Surveillance Network |
| EPI_ISL_1181567 | Laboratorio Central de Saude Publica do Estado do Rio Grande do Sul (LACEN-RS) | Laboratory of Respiratory Viruses and Measles, Oswaldo Cruz Institute, FIOCRUZ | Paola Resende, Luciana Appolinario, Fernando Motta, Anna Carolina Paixao, Ana Carolina Mendonca, Alice Sampaio Rocha, Renata Serrano Lopes, Tatiana Schaffer Gregianini, Marilda Siqueira on behalf of the Fiocruz COVID-19 Genomic Surveillance Network |
| EPI_ISL_1181568 | Laboratorio Central de Saude Publica do Estado do Parana (LACEN-PR) | Laboratory of Respiratory Viruses and Measles, Oswaldo Cruz Institute, FIOCRUZ | Paola Resende, Luciana Appolinario, Fernando Motta, Anna Carolina Paixao, Ana Carolina Mendonca, Alice Sampaio Rocha, Renata Serrano Lopes, Maria do Carmo Debur, Irina Nastassja Riediger, Marilda Siqueira on behalf of the Fiocruz COVID-19 Genomic Surveillance Network |
| EPI_ISL_1181569 | Laboratorio de Imunologia de Transplantes de Goias LTDA (HLAGYN) | Laboratory of Respiratory Viruses and Measles, Oswaldo Cruz Institute, FIOCRUZ | Paola Resende, Luciana Appolinario, Fernando Motta, Anna Carolina Paixao, Ana Carolina Mendonca, Alice Sampaio Rocha, Renata Serrano Lopes , Alessandro Leonardo Alvares Magalhaes, Fernando Antonio Vinhal dos Santos, Erika Lopes Rocha Batista, Marilda Siqueira on behalf of the Fiocruz COVID-19 Genomic Surveillance Network |
| EPI_ISL_1181570 | Laboratorio Central de Saude Publica do Estado do Parana (LACEN-PR) | Laboratory of Respiratory Viruses and Measles, Oswaldo Cruz Institute, FIOCRUZ | Paola Resende, Luciana Appolinario, Fernando Motta, Anna Carolina Paixao, Ana Carolina Mendonca, Alice Sampaio Rocha, Renata Serrano Lopes, Maria do Carmo Debur, Irina Nastassja Riediger, Marilda Siqueira on behalf of the Fiocruz COVID-19 Genomic Surveillance Network |
| EPI_ISL_1181571, EPI_ISL_1181572, EPI_ISL_1181573, EPI_ISL_1181574 | Laboratory of Respiratory Viruses and Measles, Oswaldo Cruz Institute, FIOCRUZ | Laboratory of Respiratory Viruses and Measles, Oswaldo Cruz Institute, FIOCRUZ | Paola Resende, Luciana Appolinario, Fernando Motta, Anna Carolina Paixao, Ana Carolina Mendonca, Alice Sampaio Rocha, Renata Serrano Lopes, Marilda Siqueira on behalf of the Fiocruz COVID-19 Genomic Surveillance Network |
| EPI_ISL_1181575 | Oswaldo Cruz Foundation, FIOCRUZ - Ceara (Fiocruz-CE) | Laboratory of Respiratory Viruses and Measles, Oswaldo Cruz Institute, FIOCRUZ | Paola Resende, Luciana Appolinario, Fernando Motta, Anna Carolina Paixao, Ana Carolina Mendonca, Alice Sampaio Rocha, Renata Serrano Lopes, Thais de Oliveira Costa, Joaquim César do Nascimento Sousa Júnior, Fabio Miyajima, Marilda Siqueira on behalf of the Fiocruz COVID-19 Genomic Surveillance Network |
| EPI_ISL_1181576 | Laboratorio Central de Saude Publica do Estado do Parana (LACEN-PR) | Laboratory of Respiratory Viruses and Measles, Oswaldo Cruz Institute, FIOCRUZ | Paola Resende, Luciana Appolinario, Fernando Motta, Anna Carolina Paixao, Ana Carolina Mendonca, Alice Sampaio Rocha, Renata Serrano Lopes, Maria do Carmo Debur, Irina Nastassja Riediger, Marilda Siqueira on behalf of the Fiocruz COVID-19 Genomic Surveillance Network |
| EPI_ISL_1181577 | Oswaldo Cruz Foundation, FIOCRUZ - Ceara (Fiocruz-CE) | Laboratory of Respiratory Viruses and Measles, Oswaldo Cruz Institute, FIOCRUZ | Paola Resende, Luciana Appolinario, Fernando Motta, Anna Carolina Paixao, Ana Carolina Mendonca, Alice Sampaio Rocha, Renata Serrano Lopes, Thais de Oliveira Costa, Joaquim César do Nascimento Sousa Júnior, Fabio Miyajima, Marilda Siqueira on behalf of the Fiocruz COVID-19 Genomic Surveillance Network |
| EPI_ISL_1181578, EPI_ISL_1181579 | Laboratory of Respiratory Viruses and Measles, Oswaldo Cruz Institute, FIOCRUZ | Laboratory of Respiratory Viruses and Measles, Oswaldo Cruz Institute, FIOCRUZ | Paola Resende, Luciana Appolinario, Fernando Motta, Anna Carolina Paixao, Ana Carolina Mendonca, Alice Sampaio Rocha, Renata Serrano Lopes, Marilda Siqueira on behalf of the Fiocruz COVID-19 Genomic Surveillance Network |
| EPI_ISL_1181580, EPI_ISL_1181581 | Laboratorio Central de Saude Publica do Estado do Parana (LACEN-PR) | Laboratory of Respiratory Viruses and Measles, Oswaldo Cruz Institute, FIOCRUZ | Paola Resende, Luciana Appolinario, Fernando Motta, Anna Carolina Paixao, Ana Carolina Mendonca, Alice Sampaio Rocha, Renata Serrano Lopes, Maria do Carmo Debur, Irina Nastassja Riediger, Marilda Siqueira on behalf of the Fiocruz COVID-19 Genomic Surveillance Network |
| EPI_ISL_1181582, EPI_ISL_1181583 | Laboratorio Central de Saude Publica do Estado do Espitito Santo (LACEN-ES) | Laboratory of Respiratory Viruses and Measles, Oswaldo Cruz Institute, FIOCRUZ | Paola Resende, Luciana Appolinario, Fernando Motta, Anna Carolina Paixao, Ana Carolina Mendonca, Alice Sampaio Rocha, Renata Serrano Lopes, Rodrigo Ribeiro Rodrigues, Marilda Siqueira on behalf of the Fiocruz COVID-19 Genomic Surveillance Network |
| EPI_ISL_1181584 | Laboratorio Central de Saude Publica do Estado do Parana (LACEN-PR) | Laboratory of Respiratory Viruses and Measles, Oswaldo Cruz Institute, FIOCRUZ | Paola Resende, Luciana Appolinario, Fernando Motta, Anna Carolina Paixao, Ana Carolina Mendonca, Alice Sampaio Rocha, Renata Serrano Lopes, Maria do Carmo Debur, Irina Nastassja Riediger, Marilda Siqueira on behalf of the Fiocruz COVID-19 Genomic Surveillance Network |
| EPI_ISL_1181585 | Laboratorio Central de Saude Publica do Estado do Espitito Santo (LACEN-ES) | Laboratory of Respiratory Viruses and Measles, Oswaldo Cruz Institute, FIOCRUZ | Paola Resende, Luciana Appolinario, Fernando Motta, Anna Carolina Paixao, Ana Carolina Mendonca, Alice Sampaio Rocha, Renata Serrano Lopes, Rodrigo Ribeiro Rodrigues, Marilda Siqueira on behalf of the Fiocruz COVID-19 Genomic Surveillance Network |
| EPI_ISL_1181586 | Oswaldo Cruz Foundation, FIOCRUZ - Ceara (Fiocruz-CE) | Laboratory of Respiratory Viruses and Measles, Oswaldo Cruz Institute, FIOCRUZ | Paola Resende, Luciana Appolinario, Fernando Motta, Anna Carolina Paixao, Ana Carolina Mendonca, Alice Sampaio Rocha, Renata Serrano Lopes, Thais de Oliveira Costa, Joaquim César do Nascimento Sousa Júnior, Fabio Miyajima, Marilda Siqueira on behalf of the Fiocruz COVID-19 Genomic Surveillance Network |
| EPI_ISL_1181587 | Laboratory of Respiratory Viruses and Measles, Oswaldo Cruz Institute, FIOCRUZ | Laboratory of Respiratory Viruses and Measles, Oswaldo Cruz Institute, FIOCRUZ | Paola Resende, Luciana Appolinario, Fernando Motta, Anna Carolina Paixao, Ana Carolina Mendonca, Alice Sampaio Rocha, Renata Serrano Lopes, Marilda Siqueira on behalf of the Fiocruz COVID-19 Genomic Surveillance Network |
| EPI_ISL_1181588 | Oswaldo Cruz Foundation, FIOCRUZ - Ceara (Fiocruz-CE) | Laboratory of Respiratory Viruses and Measles, Oswaldo Cruz Institute, FIOCRUZ | Paola Resende, Luciana Appolinario, Fernando Motta, Anna Carolina Paixao, Ana Carolina Mendonca, Alice Sampaio Rocha, Renata Serrano Lopes, Thais de Oliveira Costa, Joaquim César do Nascimento Sousa Júnior, Fabio Miyajima, Marilda Siqueira on behalf of the Fiocruz COVID-19 Genomic Surveillance Network |
| EPI_ISL_1181589 | Laboratory of Respiratory Viruses and Measles, Oswaldo Cruz Institute, FIOCRUZ | Laboratory of Respiratory Viruses and Measles, Oswaldo Cruz Institute, FIOCRUZ | Paola Resende, Luciana Appolinario, Fernando Motta, Anna Carolina Paixao, Ana Carolina Mendonca, Alice Sampaio Rocha, Renata Serrano Lopes, Marilda Siqueira on behalf of the Fiocruz COVID-19 Genomic Surveillance Network |
| EPI_ISL_1181590 | Laboratorio Central de Saude Publica do Estado de Sergipe (LACEN-SE) | Laboratory of Respiratory Viruses and Measles, Oswaldo Cruz Institute, FIOCRUZ | Paola Resende, Luciana Appolinario, Fernando Motta, Anna Carolina Paixao, Ana Carolina Mendonca, Alice Sampaio Rocha, Renata Serrano Lopes, Cliomar Alves dos Santos, Marilda Siqueira on behalf of the Fiocruz COVID-19 Genomic Surveillance Network |
| EPI_ISL_1181591 | Oswaldo Cruz Foundation, FIOCRUZ - Ceara (Fiocruz-CE) | Laboratory of Respiratory Viruses and Measles, Oswaldo Cruz Institute, FIOCRUZ | Paola Resende, Luciana Appolinario, Fernando Motta, Anna Carolina Paixao, Ana Carolina Mendonca, Alice Sampaio Rocha, Renata Serrano Lopes, Thais de Oliveira Costa, Joaquim César do Nascimento Sousa Júnior, Fabio Miyajima, Marilda Siqueira on behalf of the Fiocruz COVID-19 Genomic Surveillance Network |
| EPI_ISL_1181592 | Laboratorio Central de Saude Publica do Estado do Alagoas (LACEN-AL) | Laboratory of Respiratory Viruses and Measles, Oswaldo Cruz Institute, FIOCRUZ | Paola Resende, Luciana Appolinario, Fernando Motta, Anna Carolina Paixao, Ana Carolina Mendonca, Alice Sampaio Rocha, Renata Serrano Lopes, Anderson Brandao Leite, Marilda Siqueira on behalf of the Fiocruz COVID-19 Genomic Surveillance Network |
| EPI_ISL_1181594 | Laboratory of Respiratory Viruses and Measles, Oswaldo Cruz Institute, FIOCRUZ | Laboratory of Respiratory Viruses and Measles, Oswaldo Cruz Institute, FIOCRUZ | Paola Resende, Luciana Appolinario, Fernando Motta, Anna Carolina Paixao, Ana Carolina Mendonca, Alice Sampaio Rocha, Renata Serrano Lopes, Marilda Siqueira on behalf of the Fiocruz COVID-19 Genomic Surveillance Network |
| EPI_ISL_1181595 | Laboratorio Central de Saude Publica do Estado de Sergipe (LACEN-SE) | Laboratory of Respiratory Viruses and Measles, Oswaldo Cruz Institute, FIOCRUZ | Paola Resende, Luciana Appolinario, Fernando Motta, Anna Carolina Paixao, Ana Carolina Mendonca, Alice Sampaio Rocha, Renata Serrano Lopes, Cliomar Alves dos Santos, Marilda Siqueira on behalf of the Fiocruz COVID-19 Genomic Surveillance Network |

| | | | |
|---|---|---|---|
| EPI_ISL_1181596 | Oswaldo Cruz Foundation, FIOCRUZ - Ceara (Fiocruz-CE) | Laboratory of Respiratory Viruses and Measles, Oswaldo Cruz Institute, FIOCRUZ | Paola Resende, Luciana Appolinario, Fernando Motta, Anna Carolina Paixao, Ana Carolina Mendonca, Alice Sampaio Rocha, Renata Serrano Lopes, Thais de Oliveira Costa, Joaquim César do Nascimento Sousa Júnior, Fabio Miyajima, Marilda Siqueira on behalf of the Fiocruz COVID-19 Genomic Surveillance Network |
| EPI_ISL_1181597 | Laboratory of Respiratory Viruses and Measles, Oswaldo Cruz Institute, FIOCRUZ | Laboratory of Respiratory Viruses and Measles, Oswaldo Cruz Institute, FIOCRUZ | Paola Resende, Luciana Appolinario, Fernando Motta, Anna Carolina Paixao, Ana Carolina Mendonca, Alice Sampaio Rocha, Renata Serrano Lopes, Marilda Siqueira on behalf of the Fiocruz COVID-19 Genomic Surveillance Network |
| EPI_ISL_1181598 | Oswaldo Cruz Foundation, FIOCRUZ - Ceara (Fiocruz-CE) | Laboratory of Respiratory Viruses and Measles, Oswaldo Cruz Institute, FIOCRUZ | Paola Resende, Luciana Appolinario, Fernando Motta, Anna Carolina Paixao, Ana Carolina Mendonca, Alice Sampaio Rocha, Renata Serrano Lopes, Thais de Oliveira Costa, Joaquim César do Nascimento Sousa Júnior, Fabio Miyajima, Marilda Siqueira on behalf of the Fiocruz COVID-19 Genomic Surveillance Network |
| EPI_ISL_1181599 | Laboratory of Respiratory Viruses and Measles, Oswaldo Cruz Institute, FIOCRUZ | Laboratory of Respiratory Viruses and Measles, Oswaldo Cruz Institute, FIOCRUZ | Paola Resende, Luciana Appolinario, Fernando Motta, Anna Carolina Paixao, Ana Carolina Mendonca, Alice Sampaio Rocha, Renata Serrano Lopes, Marilda Siqueira on behalf of the Fiocruz COVID-19 Genomic Surveillance Network |
| EPI_ISL_1181600, EPI_ISL_1181601, EPI_ISL_1181602, EPI_ISL_1181603 | Laboratorio Central de Saude Publica do Estado do Rio de Janeiro (LACEN-RJ) | Laboratory of Respiratory Viruses and Measles, Oswaldo Cruz Institute, FIOCRUZ | Paola Resende, Luciana Appolinario, Fernando Motta, Anna Carolina Paixao, Ana Carolina Mendonca, Alice Sampaio Rocha, Renata Serrano Lopes, Andrea Cony Cavalcanti, Marilda Siqueira on behalf of the Fiocruz COVID-19 Genomic Surveillance Network |
| EPI_ISL_1181604 | Oswaldo Cruz Foundation, FIOCRUZ - Ceara (Fiocruz-CE) | Laboratory of Respiratory Viruses and Measles, Oswaldo Cruz Institute, FIOCRUZ | Paola Resende, Luciana Appolinario, Fernando Motta, Anna Carolina Paixao, Ana Carolina Mendonca, Alice Sampaio Rocha, Renata Serrano Lopes, Thais de Oliveira Costa, Joaquim César do Nascimento Sousa Júnior, Fabio Miyajima, Marilda Siqueira on behalf of the Fiocruz COVID-19 Genomic Surveillance Network |
| EPI_ISL_1181605 | Laboratorio Central de Saude Publica do Estado do Parana (LACEN-PR) | Laboratory of Respiratory Viruses and Measles, Oswaldo Cruz Institute, FIOCRUZ | Paola Resende, Luciana Appolinario, Fernando Motta, Anna Carolina Paixao, Ana Carolina Mendonca, Alice Sampaio Rocha, Renata Serrano Lopes, Maria do Carmo Debur, Irina Nastassja Riediger, Marilda Siqueira on behalf of the Fiocruz COVID-19 Genomic Surveillance Network |
| EPI_ISL_1181606, EPI_ISL_1181607 | Laboratorio Central de Saude Publica do Estado do Rio de Janeiro (LACEN-RJ) | Laboratory of Respiratory Viruses and Measles, Oswaldo Cruz Institute, FIOCRUZ | Paola Resende, Luciana Appolinario, Fernando Motta, Anna Carolina Paixao, Ana Carolina Mendonca, Alice Sampaio Rocha, Renata Serrano Lopes, Andrea Cony Cavalcanti, Marilda Siqueira on behalf of the Fiocruz COVID-19 Genomic Surveillance Network |
| EPI_ISL_1181608 | Laboratory of Respiratory Viruses and Measles, Oswaldo Cruz Institute, FIOCRUZ | Laboratory of Respiratory Viruses and Measles, Oswaldo Cruz Institute, FIOCRUZ | Paola Resende, Luciana Appolinario, Fernando Motta, Anna Carolina Paixao, Ana Carolina Mendonca, Alice Sampaio Rocha, Renata Serrano Lopes, Marilda Siqueira on behalf of the Fiocruz COVID-19 Genomic Surveillance Network |
| EPI_ISL_1181609, EPI_ISL_1181610 | Laboratorio Central de Saude Publica do Estado do Parana (LACEN-PR) | Laboratory of Respiratory Viruses and Measles, Oswaldo Cruz Institute, FIOCRUZ | Paola Resende, Luciana Appolinario, Fernando Motta, Anna Carolina Paixao, Ana Carolina Mendonca, Alice Sampaio Rocha, Renata Serrano Lopes, Maria do Carmo Debur, Irina Nastassja Riediger, Marilda Siqueira on behalf of the Fiocruz COVID-19 Genomic Surveillance Network |
| EPI_ISL_1181611, EPI_ISL_1181612, EPI_ISL_1181613 | Laboratorio Central de Saude Publica do Estado do Rio de Janeiro (LACEN-RJ) | Laboratory of Respiratory Viruses and Measles, Oswaldo Cruz Institute, FIOCRUZ | Paola Resende, Luciana Appolinario, Fernando Motta, Anna Carolina Paixao, Ana Carolina Mendonca, Alice Sampaio Rocha, Renata Serrano Lopes, Andrea Cony Cavalcanti, Marilda Siqueira on behalf of the Fiocruz COVID-19 Genomic Surveillance Network |
| EPI_ISL_1181614, EPI_ISL_1181615, EPI_ISL_1181616, EPI_ISL_1181617, EPI_ISL_1181618, EPI_ISL_1181619 | Laboratorio Central de Saude Publica do Estado do Parana (LACEN-PR) | Laboratory of Respiratory Viruses and Measles, Oswaldo Cruz Institute, FIOCRUZ | Paola Resende, Luciana Appolinario, Fernando Motta, Anna Carolina Paixao, Ana Carolina Mendonca, Alice Sampaio Rocha, Renata Serrano Lopes, Maria do Carmo Debur, Irina Nastassja Riediger, Marilda Siqueira on behalf of the Fiocruz COVID-19 Genomic Surveillance Network |
| EPI_ISL_1181620 | Laboratorio Central de Saude Publica do Estado do Rio Grande do Sul (LACEN-RS) | Laboratory of Respiratory Viruses and Measles, Oswaldo Cruz Institute, FIOCRUZ | Paola Resende, Luciana Appolinario, Fernando Motta, Anna Carolina Paixao, Ana Carolina Mendonca, Alice Sampaio Rocha, Renata Serrano Lopes, Tatiana Schaffer Gregianini, Marilda Siqueira on behalf of the Fiocruz COVID-19 Genomic Surveillance Network |
| EPI_ISL_1181621, EPI_ISL_1181622 | Oswaldo Cruz Foundation, FIOCRUZ - Ceara (Fiocruz-CE) | Laboratory of Respiratory Viruses and Measles, Oswaldo Cruz Institute, FIOCRUZ | Paola Resende, Luciana Appolinario, Fernando Motta, Anna Carolina Paixao, Ana Carolina Mendonca, Alice Sampaio Rocha, Renata Serrano Lopes, Thais de Oliveira Costa, Joaquim César do Nascimento Sousa Júnior, Fabio Miyajima, Marilda Siqueira on behalf of the Fiocruz COVID-19 Genomic Surveillance Network |
| EPI_ISL_1181623 | Laboratorio Central de Saude Publica do Estado do Parana (LACEN-PR) | Laboratory of Respiratory Viruses and Measles, Oswaldo Cruz Institute, FIOCRUZ | Paola Resende, Luciana Appolinario, Fernando Motta, Anna Carolina Paixao, Ana Carolina Mendonca, Alice Sampaio Rocha, Renata Serrano Lopes, Maria do Carmo Debur, Irina Nastassja Riediger, Marilda Siqueira on behalf of the Fiocruz COVID-19 Genomic Surveillance Network |
| EPI_ISL_1181624 | Laboratorio Central de Saude Publica do Estado do Rio de Janeiro (LACEN-RJ) | Laboratory of Respiratory Viruses and Measles, Oswaldo Cruz Institute, FIOCRUZ | Paola Resende, Luciana Appolinario, Fernando Motta, Anna Carolina Paixao, Ana Carolina Mendonca, Alice Sampaio Rocha, Renata Serrano Lopes, Andrea Cony Cavalcanti, Marilda Siqueira on behalf of the Fiocruz COVID-19 Genomic Surveillance Network |
| EPI_ISL_1182543, EPI_ISL_1182544, EPI_ISL_1182545 | Fundação Ezequiel Dias (FUNED) | Coordenação Geral de Laboratórios de Saúde Pública (CGLAB/DAEVS/SVS/MS) | Vagner Fonseca, et al. |
| EPI_ISL_1182546 | Laboratório Central do Estado do Paraná | Coordenação Geral de Laboratórios de Saúde Pública (CGLAB/DAEVS/SVS/MS) | Vagner Fonseca, et al. |
| EPI_ISL_1182547 | Laboratório Central de Saúde Pública do Rio Grande do Sul | Coordenação Geral de Laboratórios de Saúde Pública (CGLAB/DAEVS/SVS/MS) | Vagner Fonseca, et al. |
| EPI_ISL_1182548, EPI_ISL_1182549, EPI_ISL_1182550, EPI_ISL_1182551 | Fundação Ezequiel Dias (FUNED) | Coordenação Geral de Laboratórios de Saúde Pública (CGLAB/DAEVS/SVS/MS) | Vagner Fonseca, et al. |
| EPI_ISL_1182552, EPI_ISL_1182553, EPI_ISL_1182554 | Laboratório Central do Estado do Rio de Janeiro | Coordenação Geral de Laboratórios de Saúde Pública (CGLAB/DAEVS/SVS/MS) | Vagner Fonseca, et al. |
| EPI_ISL_1182555 | Fundação Ezequiel Dias (FUNED) | Coordenação Geral de Laboratórios de Saúde Pública (CGLAB/DAEVS/SVS/MS) | Vagner Fonseca, et al. |
| EPI_ISL_1182556, EPI_ISL_1182557, EPI_ISL_1182558 | Laboratório Central do Estado do Rio de Janeiro | Coordenação Geral de Laboratórios de Saúde Pública (CGLAB/DAEVS/SVS/MS) | Vagner Fonseca, et al. |
| EPI_ISL_1182559, EPI_ISL_1182560, EPI_ISL_1182561 | Fundação Ezequiel Dias (FUNED) | Coordenação Geral de Laboratórios de Saúde Pública (CGLAB/DAEVS/SVS/MS) | Vagner Fonseca, et al. |
| EPI_ISL_1182562, EPI_ISL_1182563 | Laboratório Central do Estado do Paraná | Coordenação Geral de Laboratórios de Saúde Pública (CGLAB/DAEVS/SVS/MS) | Vagner Fonseca, et al. |
| EPI_ISL_1182564 | Fundação Ezequiel Dias (FUNED) | Coordenação Geral de Laboratórios de Saúde Pública (CGLAB/DAEVS/SVS/MS) | Vagner Fonseca, et al. |
| EPI_ISL_1182565 | Laboratório Central do Estado do Paraná | Coordenação Geral de Laboratórios de Saúde Pública (CGLAB/DAEVS/SVS/MS) | Vagner Fonseca, et al. |
| EPI_ISL_1182566 | Fundação Ezequiel Dias (FUNED) | Coordenação Geral de Laboratórios de Saúde Pública (CGLAB/DAEVS/SVS/MS) | Vagner Fonseca, et al. |
| EPI_ISL_1182567, EPI_ISL_1182568 | Laboratório Central do Estado do Paraná | Coordenação Geral de Laboratórios de Saúde Pública (CGLAB/DAEVS/SVS/MS) | Vagner Fonseca, et al. |
| EPI_ISL_1182569, EPI_ISL_1182570 | Fundação Ezequiel Dias (FUNED) | Coordenação Geral de Laboratórios de Saúde Pública (CGLAB/DAEVS/SVS/MS) | Vagner Fonseca, et al. |
| EPI_ISL_1182571, EPI_ISL_1182572 | Laboratório Central do Estado do Paraná | Coordenação Geral de Laboratórios de Saúde Pública (CGLAB/DAEVS/SVS/MS) | Vagner Fonseca, et al. |
| EPI_ISL_1182573, EPI_ISL_1182574 | Fundação Ezequiel Dias (FUNED) | Coordenação Geral de Laboratórios de Saúde Pública (CGLAB/DAEVS/SVS/MS) | Vagner Fonseca, et al. |
| EPI_ISL_1182575, EPI_ISL_1182576 | Laboratório Central do Estado do Paraná | Coordenação Geral de Laboratórios de Saúde Pública (CGLAB/DAEVS/SVS/MS) | Vagner Fonseca, et al. |

| | | | |
|---|---|---|---|
| EPI_ISL_1182577, EPI_ISL_1182578, EPI_ISL_1182579 | Fundação Ezequiel Dias (FUNED) | Coordenação Geral de Laboratórios de Saúde Pública (CGLAB/DAEVS/SVS/MS) | Vagner Fonseca, et al. |
| EPI_ISL_1182580, EPI_ISL_1182581, EPI_ISL_1182582, EPI_ISL_1182583, EPI_ISL_1182584 | Laboratório Central do Estado do Paraná | Coordenação Geral de Laboratórios de Saúde Pública (CGLAB/DAEVS/SVS/MS) | Vagner Fonseca, et al. |
| EPI_ISL_1182585, EPI_ISL_1182586, EPI_ISL_1182587 | Fundação Ezequiel Dias (FUNED) | Coordenação Geral de Laboratórios de Saúde Pública (CGLAB/DAEVS/SVS/MS) | Vagner Fonseca, et al. |
| EPI_ISL_1182588, EPI_ISL_1182589 | Laboratório Central do Estado do Paraná | Coordenação Geral de Laboratórios de Saúde Pública (CGLAB/DAEVS/SVS/MS) | Vagner Fonseca, et al. |
| EPI_ISL_1182590, EPI_ISL_1182591 | Fundação Ezequiel Dias (FUNED) | Coordenação Geral de Laboratórios de Saúde Pública (CGLAB/DAEVS/SVS/MS) | Vagner Fonseca, et al. |
| EPI_ISL_1182592 | Laboratório Central do Estado do Paraná | Coordenação Geral de Laboratórios de Saúde Pública (CGLAB/DAEVS/SVS/MS) | Vagner Fonseca, et al. |
| EPI_ISL_1182593 | Fundação Ezequiel Dias (FUNED) | Coordenação Geral de Laboratórios de Saúde Pública (CGLAB/DAEVS/SVS/MS) | Vagner Fonseca, et al. |
| EPI_ISL_1182594, EPI_ISL_1182595, EPI_ISL_1182596 | Laboratório Central do Estado do Paraná | Coordenação Geral de Laboratórios de Saúde Pública (CGLAB/DAEVS/SVS/MS) | Vagner Fonseca, et al. |
| EPI_ISL_1182597 | Laboratório Central de Saúde Pública do Rio Grande do Sul | Coordenação Geral de Laboratórios de Saúde Pública (CGLAB/DAEVS/SVS/MS) | Vagner Fonseca, et al. |
| EPI_ISL_1182598 | Fundação Ezequiel Dias (FUNED) | Coordenação Geral de Laboratórios de Saúde Pública (CGLAB/DAEVS/SVS/MS) | Vagner Fonseca, et al. |
| EPI_ISL_1182599 | Laboratório Central de Saúde Pública do Rio Grande do Sul | Coordenação Geral de Laboratórios de Saúde Pública (CGLAB/DAEVS/SVS/MS) | Vagner Fonseca, et al. |
| EPI_ISL_1182600, EPI_ISL_1182601, EPI_ISL_1182602 | Fundação Ezequiel Dias (FUNED) | Coordenação Geral de Laboratórios de Saúde Pública (CGLAB/DAEVS/SVS/MS) | Vagner Fonseca, et al. |
| EPI_ISL_1182603, EPI_ISL_1182604 | Laboratório Central do Estado do Paraná | Coordenação Geral de Laboratórios de Saúde Pública (CGLAB/DAEVS/SVS/MS) | Vagner Fonseca, et al. |
| EPI_ISL_1182605, EPI_ISL_1182606 | Laboratório Central de Saúde Pública do Rio Grande do Sul | Coordenação Geral de Laboratórios de Saúde Pública (CGLAB/DAEVS/SVS/MS) | Vagner Fonseca, et al. |
| EPI_ISL_1182607 | Fundação Ezequiel Dias (FUNED) | Coordenação Geral de Laboratórios de Saúde Pública (CGLAB/DAEVS/SVS/MS) | Vagner Fonseca, et al. |
| EPI_ISL_1182608 | Laboratório Central de Saúde Pública do Rio Grande do Sul | Coordenação Geral de Laboratórios de Saúde Pública (CGLAB/DAEVS/SVS/MS) | Vagner Fonseca, et al. |
| EPI_ISL_1182609 | Fundação Ezequiel Dias (FUNED) | Coordenação Geral de Laboratórios de Saúde Pública (CGLAB/DAEVS/SVS/MS) | Vagner Fonseca, et al. |
| EPI_ISL_1182610 | Laboratório Central de Saúde Pública do Rio Grande do Sul | Coordenação Geral de Laboratórios de Saúde Pública (CGLAB/DAEVS/SVS/MS) | Vagner Fonseca, et al. |
| EPI_ISL_1182611, EPI_ISL_1182612 | Fundação Ezequiel Dias (FUNED) | Coordenação Geral de Laboratórios de Saúde Pública (CGLAB/DAEVS/SVS/MS) | Vagner Fonseca, et al. |
| EPI_ISL_1182613, EPI_ISL_1182614 | Laboratório Central do Estado do Paraná | Coordenação Geral de Laboratórios de Saúde Pública (CGLAB/DAEVS/SVS/MS) | Vagner Fonseca, et al. |
| EPI_ISL_1182615 | Fundação Ezequiel Dias (FUNED) | Coordenação Geral de Laboratórios de Saúde Pública (CGLAB/DAEVS/SVS/MS) | Vagner Fonseca, et al. |
| EPI_ISL_1182616 | Laboratório Central de Saúde Pública do Rio Grande do Sul | Coordenação Geral de Laboratórios de Saúde Pública (CGLAB/DAEVS/SVS/MS) | Vagner Fonseca, et al. |
| EPI_ISL_1182617 | Laboratório Central do Estado do Paraná | Coordenação Geral de Laboratórios de Saúde Pública (CGLAB/DAEVS/SVS/MS) | Vagner Fonseca, et al. |
| EPI_ISL_1182618 | Fundação Ezequiel Dias (FUNED) | Coordenação Geral de Laboratórios de Saúde Pública (CGLAB/DAEVS/SVS/MS) | Vagner Fonseca, et al. |
| EPI_ISL_1182619, EPI_ISL_1182620, EPI_ISL_1182621 | Laboratório Central de Saúde Pública do Rio Grande do Sul | Coordenação Geral de Laboratórios de Saúde Pública (CGLAB/DAEVS/SVS/MS) | Vagner Fonseca, et al. |
| EPI_ISL_1182622 | Laboratório Central do Estado do Paraná | Coordenação Geral de Laboratórios de Saúde Pública (CGLAB/DAEVS/SVS/MS) | Vagner Fonseca, et al. |
| EPI_ISL_1182623 | Laboratório Central de Saúde Pública do Rio Grande do Sul | Coordenação Geral de Laboratórios de Saúde Pública (CGLAB/DAEVS/SVS/MS) | Vagner Fonseca, et al. |
| EPI_ISL_1182624 | Laboratório Central do Estado do Paraná | Coordenação Geral de Laboratórios de Saúde Pública (CGLAB/DAEVS/SVS/MS) | Vagner Fonseca, et al. |
| EPI_ISL_1182625 | Fundação Ezequiel Dias (FUNED) | Coordenação Geral de Laboratórios de Saúde Pública (CGLAB/DAEVS/SVS/MS) | Vagner Fonseca, et al. |
| EPI_ISL_1195275, EPI_ISL_1195276 | CENTRO DE REFERENCIA EM SINDROMES GRIPAIS | Epiclin | Fernando Hayashi Sant'Anna, Ana Paula Muterle, Janira Prichula, Juliana Comerlato, Carolina Comerlato, Eliana Márcia Da Ros Wendland |
| EPI_ISL_1195277 | Unidade de Atendimento DST AIDS TB e Han | Epiclin | Fernando Hayashi Sant'Anna, Ana Paula Muterle, Janira Prichula, Juliana Comerlato, Carolina Comerlato, Eliana Márcia Da Ros Wendland |
| EPI_ISL_1195278 | DIRETORIA DE VIGILANCIA EM SAUDE | Epiclin | Fernando Hayashi Sant'Anna, Ana Paula Muterle, Janira Prichula, Juliana Comerlato, Carolina Comerlato, Eliana Márcia Da Ros Wendland |
| EPI_ISL_1195279 | CENTRO DE REFERENCIA EM SINDROMES GRIPAIS | Epiclin | Fernando Hayashi Sant'Anna, Ana Paula Muterle, Janira Prichula, Juliana Comerlato, Carolina Comerlato, Eliana Márcia Da Ros Wendland |
| EPI_ISL_1195280 | SECRETARIA MUNICIPAL DE SAUDE DE ARARICA | Epiclin | Fernando Hayashi Sant'Anna, Ana Paula Muterle, Janira Prichula, Juliana Comerlato, Carolina Comerlato, Eliana Márcia Da Ros Wendland |
| EPI_ISL_1195281 | FUNDACAO DE SAUDE PUBLICA SAO CAMILO DE ESTEIO | Epiclin | Fernando Hayashi Sant'Anna, Ana Paula Muterle, Janira Prichula, Juliana Comerlato, Carolina Comerlato, Eliana Márcia Da Ros Wendland |
| EPI_ISL_1195282 | UNIDADE SANITARIA DE IGREJINHA | Epiclin | Fernando Hayashi Sant'Anna, Ana Paula Muterle, Janira Prichula, Juliana Comerlato, Carolina Comerlato, Eliana Márcia Da Ros Wendland |
| EPI_ISL_1195283 | SECRETARIA MUNICIPAL DE SAUDE DE TRES COROAS | Epiclin | Fernando Hayashi Sant'Anna, Ana Paula Muterle, Janira Prichula, Juliana Comerlato, Carolina Comerlato, Eliana Márcia Da Ros Wendland |
| EPI_ISL_1195284 | Centro de Especialidades Triunfo | Epiclin | Fernando Hayashi Sant'Anna, Ana Paula Muterle, Janira Prichula, Juliana Comerlato, Carolina Comerlato, Eliana Márcia Da Ros Wendland |
| EPI_ISL_1195285, EPI_ISL_1195286 | SECRETARIA MUNICIPAL DE SAUDE DE TRES COROAS | Epiclin | Fernando Hayashi Sant'Anna, Ana Paula Muterle, Janira Prichula, Juliana Comerlato, Carolina Comerlato, Eliana Márcia Da Ros Wendland |

| | | | |
|---|---|---|---|
| EPI_ISL_1195287 | SECRETARIA MUNICIPAL DE SAUDE DE SAO LEOPOLDO | Epiclin | Fernando Hayashi Sant'Anna, Ana Paula Muterle, Janira Prichula, Juliana Comerlato, Carolina Comerlato, Eliana Márcia Da Ros Wendland |
| EPI_ISL_1195288 | DIRETORIA DE VIGILANCIA EM SAUDE | Epiclin | Fernando Hayashi Sant'Anna, Ana Paula Muterle, Janira Prichula, Juliana Comerlato, Carolina Comerlato, Eliana Márcia Da Ros Wendland |
| EPI_ISL_1195289 | SECRETARIA MUNICIPAL DE SAUDE DE SAO LEOPOLDO | Epiclin | Fernando Hayashi Sant'Anna, Ana Paula Muterle, Janira Prichula, Juliana Comerlato, Carolina Comerlato, Eliana Márcia Da Ros Wendland |
| EPI_ISL_1195290 | SECRETARIA MUNICIPAL DE SAUDE DE TRES COROAS | Epiclin | Fernando Hayashi Sant'Anna, Ana Paula Muterle, Janira Prichula, Juliana Comerlato, Carolina Comerlato, Eliana Márcia Da Ros Wendland |
| EPI_ISL_1195291 | DIRETORIA DE VIGILANCIA EM SAUDE | Epiclin | Fernando Hayashi Sant'Anna, Ana Paula Muterle, Janira Prichula, Juliana Comerlato, Carolina Comerlato, Eliana Márcia Da Ros Wendland |
| EPI_ISL_1195292 | SECRETARIA MUNICIPAL DE SAUDE DE TRES COROAS | Epiclin | Fernando Hayashi Sant'Anna, Ana Paula Muterle, Janira Prichula, Juliana Comerlato, Carolina Comerlato, Eliana Márcia Da Ros Wendland |
| EPI_ISL_1195293 | SECRETARIA MUNICIPAL DE SAUDE DE TAQUARA | Epiclin | Fernando Hayashi Sant'Anna, Ana Paula Muterle, Janira Prichula, Juliana Comerlato, Carolina Comerlato, Eliana Márcia Da Ros Wendland |
| EPI_ISL_1196282, EPI_ISL_1196283, EPI_ISL_1196284 | LACEN do Mato Grosso do Sul | Instituto Adolfo Lutz, Interdiciplinary Procedures Center, Strategic Laboratory | Claudio Tavares Sacchi, Claudia Regina Gonçalves, Erica Valessa Ramos Gomes, Karoline Rodrigues Campos, Caio Vinicius Dias Lopes |
| EPI_ISL_1196285, EPI_ISL_1196286, EPI_ISL_1196287, EPI_ISL_1196288, EPI_ISL_1196289, EPI_ISL_1196290, EPI_ISL_1196291, EPI_ISL_1196292, EPI_ISL_1196293, EPI_ISL_1196294 | LACEN do Distrito Federal | Instituto Adolfo Lutz, Interdiciplinary Procedures Center, Strategic Laboratory | Claudio Tavares Sacchi, Claudia Regina Gonçalves, Erica Valessa Ramos Gomes, Karoline Rodrigues Campos, Caio Vinicius Dias Lopes |
| EPI_ISL_1196295 | UBS Otacilio Firmino Lopes | Instituto Adolfo Lutz, Interdiciplinary Procedures Center, Strategic Laboratory | Claudio Tavares Sacchi, Claudia Regina Gonçalves, Erica Valessa Ramos Gomes, Karoline Rodrigues Campos, Caio Vinicius Dias Lopes |
| EPI_ISL_1196296 | Centro de Saude II Dr. Jose Paione Mococa | Instituto Adolfo Lutz, Interdiciplinary Procedures Center, Strategic Laboratory | Claudio Tavares Sacchi, Claudia Regina Gonçalves, Erica Valessa Ramos Gomes, Karoline Rodrigues Campos, Caio Vinicius Dias Lopes |
| EPI_ISL_1196297, EPI_ISL_1196298, EPI_ISL_1196299, EPI_ISL_1196300, EPI_ISL_1196301, EPI_ISL_1196302 | IAL Regional de Marilia | Instituto Adolfo Lutz, Interdiciplinary Procedures Center, Strategic Laboratory | Claudio Tavares Sacchi, Claudia Regina Gonçalves, Erica Valessa Ramos Gomes, Karoline Rodrigues Campos, Caio Vinicius Dias Lopes |
| EPI_ISL_1201883 | LACEN do Mato Grosso do Sul | Instituto Adolfo Lutz, Interdiciplinary Procedures Center, Strategic Laboratory | Claudio Tavares Sacchi, Claudia Regina Gonçalves, Erica Valessa Ramos Gomes, Karoline Rodrigues Campos, Caio Vinicius Dias Lopes |
| EPI_ISL_1201884, EPI_ISL_1201885, EPI_ISL_1201886, EPI_ISL_1201887, EPI_ISL_1201888 | Aeroporto Internacional de Guarulhos | Instituto Adolfo Lutz, Interdiciplinary Procedures Center, Strategic Laboratory | Claudio Tavares Sacchi, Claudia Regina Gonçalves, Erica Valessa Ramos Gomes, Karoline Rodrigues Campos, Caio Vinicius Dias Lopes |
| EPI_ISL_1201889, EPI_ISL_1201890, EPI_ISL_1201891, EPI_ISL_1201892 | IAL Regional de Marilia | Instituto Adolfo Lutz, Interdiciplinary Procedures Center, Strategic Laboratory | Claudio Tavares Sacchi, Claudia Regina Gonçalves, Erica Valessa Ramos Gomes, Karoline Rodrigues Campos, Caio Vinicius Dias Lopes |
| EPI_ISL_1201893 | IAL Regional de Sorocaba | Instituto Adolfo Lutz, Interdiciplinary Procedures Center, Strategic Laboratory | Claudio Tavares Sacchi, Claudia Regina Gonçalves, Erica Valessa Ramos Gomes, Karoline Rodrigues Campos, Caio Vinicius Dias Lopes |
| EPI_ISL_1201894, EPI_ISL_1201895, EPI_ISL_1201896 | Diagnosticos da America - DASA | Instituto Adolfo Lutz, Interdiciplinary Procedures Center, Strategic Laboratory | Claudio Tavares Sacchi, Claudia Regina Gonçalves, Erica Valessa Ramos Gomes, Karoline Rodrigues Campos, Caio Vinicius Dias Lopes |
| EPI_ISL_1213160, EPI_ISL_1213161 | LBM/UFPB | Bioinformatics Laboratory / LNCC | Alessandra P Lamarca, Luiz G P de Almeida, Ronaldo da Silva Francisco Jr, Lucymara Fassarella Agnez Lima, Kátia Castanho Scortecci, Vinícius Pietta Perez, Otavio J. Brustolini, Eduardo Sérgio Soares Sousa, Danielle Angst Secco, Angela Maria Guimarães Santos, George Rego Albuquerque, Ana Paula Melo Mariano, Bianca Mendes Maciel, Alexandra L Gerber, Ana Paula de C Guimarães, Paulo Ricardo Nascimento, Francisco Paulo Freire Neto, Sandra Rocha Gadelha, Luís Cristóvão Porto, Eloiza Helena Campana, Selma Maria Bezerra Jeronimo, Ana Tereza R Vasconcelos |
| EPI_ISL_1213163 | Laboratório HLA/UERJ | Bioinformatics Laboratory / LNCC | Alessandra P Lamarca, Luiz G P de Almeida, Ronaldo da Silva Francisco Jr, Lucymara Fassarella Agnez Lima, Kátia Castanho Scortecci, Vinícius Pietta Perez, Otavio J. Brustolini, Eduardo Sérgio Soares Sousa, Danielle Angst Secco, Angela Maria Guimarães Santos, George Rego Albuquerque, Ana Paula Melo Mariano, Bianca Mendes Maciel, Alexandra L Gerber, Ana Paula de C Guimarães, Paulo Ricardo Nascimento, Francisco Paulo Freire Neto, Sandra Rocha Gadelha, Luís Cristóvão Porto, Eloiza Helena Campana, Selma Maria Bezerra Jeronimo, Ana Tereza R Vasconcelos |
| EPI_ISL_1213168, EPI_ISL_1213170 | LAFEM/UESC | Bioinformatics Laboratory / LNCC | Alessandra P Lamarca, Luiz G P de Almeida, Ronaldo da Silva Francisco Jr, Lucymara Fassarella Agnez Lima, Kátia Castanho Scortecci, Vinícius Pietta Perez, Otavio J. Brustolini, Eduardo Sérgio Soares Sousa, Danielle Angst Secco, Angela Maria Guimarães Santos, George Rego Albuquerque, Ana Paula Melo Mariano, Bianca Mendes Maciel, Alexandra L Gerber, Ana Paula de C Guimarães, Paulo Ricardo Nascimento, Francisco Paulo Freire Neto, Sandra Rocha Gadelha, Luís Cristóvão Porto, Eloiza Helena Campana, Selma Maria Bezerra Jeronimo, Ana Tereza R Vasconcelos |
| EPI_ISL_1213171 | Laboratório HLA/UERJ | Bioinformatics Laboratory / LNCC | Alessandra P Lamarca, Luiz G P de Almeida, Ronaldo da Silva Francisco Jr, Lucymara Fassarella Agnez Lima, Kátia Castanho Scortecci, Vinícius Pietta Perez, Otavio J. Brustolini, Eduardo Sérgio Soares Sousa, Danielle Angst Secco, Angela Maria Guimarães Santos, George Rego Albuquerque, Ana Paula Melo Mariano, Bianca Mendes Maciel, Alexandra L Gerber, Ana Paula de C Guimarães, Paulo Ricardo Nascimento, Francisco Paulo Freire Neto, Sandra Rocha Gadelha, Luís Cristóvão Porto, Eloiza Helena Campana, Selma Maria Bezerra Jeronimo, Ana Tereza R Vasconcelos |
| EPI_ISL_1213173, EPI_ISL_1213175, EPI_ISL_1213177, EPI_ISL_1213178, EPI_ISL_1213180, EPI_ISL_1213182, EPI_ISL_1213183, EPI_ISL_1213185, EPI_ISL_1213187 | IMT-UFRN/RN | Bioinformatics Laboratory / LNCC | Alessandra P Lamarca, Luiz G P de Almeida, Ronaldo da Silva Francisco Jr, Lucymara Fassarella Agnez Lima, Kátia Castanho Scortecci, Vinícius Pietta Perez, Otavio J. Brustolini, Eduardo Sérgio Soares Sousa, Danielle Angst Secco, Angela Maria Guimarães Santos, George Rego Albuquerque, Ana Paula Melo Mariano, Bianca Mendes Maciel, Alexandra L Gerber, Ana Paula de C Guimarães, Paulo Ricardo Nascimento, Francisco Paulo Freire Neto, Sandra Rocha Gadelha, Luís Cristóvão Porto, Eloiza Helena Campana, Selma Maria Bezerra Jeronimo, Ana Tereza R Vasconcelos |
| EPI_ISL_1213189, EPI_ISL_1213190 | LBM/UFPB | Bioinformatics Laboratory / LNCC | Alessandra P Lamarca, Luiz G P de Almeida, Ronaldo da Silva Francisco Jr, Lucymara Fassarella Agnez Lima, Kátia Castanho Scortecci, Vinícius Pietta Perez, Otavio J. Brustolini, Eduardo Sérgio Soares Sousa, Danielle Angst Secco, Angela Maria Guimarães Santos, George Rego Albuquerque, Ana Paula Melo Mariano, Bianca Mendes Maciel, Alexandra L Gerber, Ana Paula de C Guimarães, Paulo Ricardo Nascimento, Francisco Paulo Freire Neto, Sandra Rocha Gadelha, Luís Cristóvão Porto, Eloiza Helena Campana, Selma Maria Bezerra Jeronimo, Ana Tereza R Vasconcelos |
| EPI_ISL_1213192, EPI_ISL_1213194, EPI_ISL_1213196, EPI_ISL_1213197, EPI_ISL_1213199 | IMT-UFRN/RN | Bioinformatics Laboratory / LNCC | Alessandra P Lamarca, Luiz G P de Almeida, Ronaldo da Silva Francisco Jr, Lucymara Fassarella Agnez Lima, Kátia Castanho Scortecci, Vinícius Pietta Perez, Otavio J. Brustolini, Eduardo Sérgio Soares Sousa, Danielle Angst Secco, Angela Maria Guimarães Santos, George Rego Albuquerque, Ana Paula Melo Mariano, Bianca Mendes Maciel, Alexandra L Gerber, Ana Paula de C Guimarães, Paulo Ricardo Nascimento, Francisco Paulo Freire Neto, Sandra Rocha Gadelha, Luís Cristóvão Porto, Eloiza Helena Campana, Selma Maria Bezerra Jeronimo, Ana Tereza R Vasconcelos |
| EPI_ISL_1213201, EPI_ISL_1213202, EPI_ISL_1213204, EPI_ISL_1213206, EPI_ISL_1213207, EPI_ISL_1213213 | LBM/UFPB | Bioinformatics Laboratory / LNCC | Alessandra P Lamarca, Luiz G P de Almeida, Ronaldo da Silva Francisco Jr, Lucymara Fassarella Agnez Lima, Kátia Castanho Scortecci, Vinícius Pietta Perez, Otavio J. Brustolini, Eduardo Sérgio Soares Sousa, Danielle Angst Secco, Angela Maria Guimarães Santos, George Rego Albuquerque, Ana Paula Melo Mariano, Bianca Mendes Maciel, Alexandra L Gerber, Ana Paula de C Guimarães, Paulo Ricardo Nascimento, Francisco Paulo Freire Neto, Sandra Rocha Gadelha, Luís Cristóvão Porto, Eloiza Helena Campana, Selma Maria Bezerra Jeronimo, Ana Tereza R Vasconcelos |
| EPI_ISL_1213218, EPI_ISL_1213220, EPI_ISL_1213222, EPI_ISL_1213224, EPI_ISL_1213226, EPI_ISL_1213227, EPI_ISL_1213229, EPI_ISL_1213230, EPI_ISL_1213232, EPI_ISL_1213234, EPI_ISL_1213236, EPI_ISL_1213237, EPI_ISL_1213239, EPI_ISL_1213241, EPI_ISL_1213242, EPI_ISL_1213244, EPI_ISL_1213245 | | | |
| see above | LAFEM/UESC | Bioinformatics Laboratory / LNCC | Alessandra P Lamarca, Luiz G P de Almeida, Ronaldo da Silva Francisco Jr, Lucymara Fassarella Agnez Lima, Kátia Castanho Scortecci, Vinícius Pietta Perez, Otavio J. Brustolini, Eduardo Sérgio Soares Sousa, Danielle Angst Secco, Angela Maria Guimarães Santos, George Rego Albuquerque, Ana Paula Melo Mariano, Bianca Mendes Maciel, Alexandra L Gerber, Ana Paula de C Guimarães, Paulo Ricardo Nascimento, Francisco Paulo Freire Neto, Sandra Rocha Gadelha, Luís Cristóvão Porto, Eloiza Helena Campana, Selma Maria Bezerra Jeronimo, Ana Tereza R Vasconcelos |
| EPI_ISL_1213248, EPI_ISL_1213249 | Laboratório HLA/UERJ | Bioinformatics Laboratory / LNCC | Alessandra P Lamarca, Luiz G P de Almeida, Ronaldo da Silva Francisco Jr, Lucymara Fassarella Agnez Lima, Kátia Castanho Scortecci, Vinícius Pietta Perez, Otavio J. Brustolini, Eduardo Sérgio Soares Sousa, Danielle Angst Secco, Angela Maria Guimarães Santos, George Rego Albuquerque, Ana Paula Melo Mariano, Bianca Mendes Maciel, Alexandra L Gerber, Ana Paula de C Guimarães, Paulo Ricardo Nascimento, Francisco Paulo Freire Neto, Sandra Rocha Gadelha, Luís Cristóvão Porto, Eloiza Helena Campana, Selma Maria Bezerra Jeronimo, Ana Tereza R Vasconcelos |

| | | | |
|---|---|---|---|
| EPI_ISL_1213251, EPI_ISL_1213252, EPI_ISL_1213254, EPI_ISL_1213256, EPI_ISL_1213258, EPI_ISL_1213259, EPI_ISL_1213261, EPI_ISL_1213263, EPI_ISL_1213265, EPI_ISL_1213267, EPI_ISL_1213269 | | | |
| see above | LAFEM/UESC | Bioinformatics Laboratory / LNCC | Alessandra P Lamarca, Luiz G P de Almeida, Ronaldo da Silva Francisco Jr, Lucymara Fassarella Agnez Lima, Kátia Castanho Scortecci, Vinícius Pietta Perez, Otavio J. Brustolini, Eduardo Sérgio Soares Sousa, Danielle Angst Secco, Angela Maria Guimarães Santos, George Rego Albuquerque, Ana Paula Melo Mariano, Bianca Mendes Maciel, Alexandra L Gerber, Ana Paula de C Guimarães, Paulo Ricardo Nascimento, Francisco Paulo Freire Neto, Sandra Rocha Gadelha, Luís Cristóvão Porto, Eloiza Helena Campana, Selma Maria Bezerra Jeronimo, Ana Tereza R Vasconcelos |
| EPI_ISL_1213270, EPI_ISL_1213272 | Laboratório HLA/UERJ | Bioinformatics Laboratory / LNCC | Alessandra P Lamarca, Luiz G P de Almeida, Ronaldo da Silva Francisco Jr, Lucymara Fassarella Agnez Lima, Kátia Castanho Scortecci, Vinícius Pietta Perez, Otavio J. Brustolini, Eduardo Sérgio Soares Sousa, Danielle Angst Secco, Angela Maria Guimarães Santos, George Rego Albuquerque, Ana Paula Melo Mariano, Bianca Mendes Maciel, Alexandra L Gerber, Ana Paula de C Guimarães, Paulo Ricardo Nascimento, Francisco Paulo Freire Neto, Sandra Rocha Gadelha, Luís Cristóvão Porto, Eloiza Helena Campana, Selma Maria Bezerra Jeronimo, Ana Tereza R Vasconcelos |
| EPI_ISL_1213274 | LAFEM/UESC | Bioinformatics Laboratory / LNCC | Alessandra P Lamarca, Luiz G P de Almeida, Ronaldo da Silva Francisco Jr, Lucymara Fassarella Agnez Lima, Kátia Castanho Scortecci, Vinícius Pietta Perez, Otavio J. Brustolini, Eduardo Sérgio Soares Sousa, Danielle Angst Secco, Angela Maria Guimarães Santos, George Rego Albuquerque, Ana Paula Melo Mariano, Bianca Mendes Maciel, Alexandra L Gerber, Ana Paula de C Guimarães, Paulo Ricardo Nascimento, Francisco Paulo Freire Neto, Sandra Rocha Gadelha, Luís Cristóvão Porto, Eloiza Helena Campana, Selma Maria Bezerra Jeronimo, Ana Tereza R Vasconcelos |
| EPI_ISL_1213275, EPI_ISL_1213277, EPI_ISL_1213279, EPI_ISL_1213281, EPI_ISL_1213282, EPI_ISL_1213284, EPI_ISL_1213286, EPI_ISL_1213288, EPI_ISL_1213289, EPI_ISL_1213291, EPI_ISL_1213293 | | | |
| see above | IMT-UFRN/RN | Bioinformatics Laboratory / LNCC | Alessandra P Lamarca, Luiz G P de Almeida, Ronaldo da Silva Francisco Jr, Lucymara Fassarella Agnez Lima, Kátia Castanho Scortecci, Vinícius Pietta Perez, Otavio J. Brustolini, Eduardo Sérgio Soares Sousa, Danielle Angst Secco, Angela Maria Guimarães Santos, George Rego Albuquerque, Ana Paula Melo Mariano, Bianca Mendes Maciel, Alexandra L Gerber, Ana Paula de C Guimarães, Paulo Ricardo Nascimento, Francisco Paulo Freire Neto, Sandra Rocha Gadelha, Luís Cristóvão Porto, Eloiza Helena Campana, Selma Maria Bezerra Jeronimo, Ana Tereza R Vasconcelos |
| EPI_ISL_1213294, EPI_ISL_1213297, EPI_ISL_1213298, EPI_ISL_1213300, EPI_ISL_1213302, EPI_ISL_1213303, EPI_ISL_1213305 | LAFEM/UESC | Bioinformatics Laboratory / LNCC | Alessandra P Lamarca, Luiz G P de Almeida, Ronaldo da Silva Francisco Jr, Lucymara Fassarella Agnez Lima, Kátia Castanho Scortecci, Vinícius Pietta Perez, Otavio J. Brustolini, Eduardo Sérgio Soares Sousa, Danielle Angst Secco, Angela Maria Guimarães Santos, George Rego Albuquerque, Ana Paula Melo Mariano, Bianca Mendes Maciel, Alexandra L Gerber, Ana Paula de C Guimarães, Paulo Ricardo Nascimento, Francisco Paulo Freire Neto, Sandra Rocha Gadelha, Luís Cristóvão Porto, Eloiza Helena Campana, Selma Maria Bezerra Jeronimo, Ana Tereza R Vasconcelos |
| EPI_ISL_1213307, EPI_ISL_1213309, EPI_ISL_1213310, EPI_ISL_1213312, EPI_ISL_1213314 | Laboratório HLA/UERJ | Bioinformatics Laboratory / LNCC | Alessandra P Lamarca, Luiz G P de Almeida, Ronaldo da Silva Francisco Jr, Lucymara Fassarella Agnez Lima, Kátia Castanho Scortecci, Vinícius Pietta Perez, Otavio J. Brustolini, Eduardo Sérgio Soares Sousa, Danielle Angst Secco, Angela Maria Guimarães Santos, George Rego Albuquerque, Ana Paula Melo Mariano, Bianca Mendes Maciel, Alexandra L Gerber, Ana Paula de C Guimarães, Paulo Ricardo Nascimento, Francisco Paulo Freire Neto, Sandra Rocha Gadelha, Luís Cristóvão Porto, Eloiza Helena Campana, Selma Maria Bezerra Jeronimo, Ana Tereza R Vasconcelos |
| EPI_ISL_1213315, EPI_ISL_1213317, EPI_ISL_1213319, EPI_ISL_1213320, EPI_ISL_1213322 | IMT-UFRN/RN | Bioinformatics Laboratory / LNCC | Alessandra P Lamarca, Luiz G P de Almeida, Ronaldo da Silva Francisco Jr, Lucymara Fassarella Agnez Lima, Kátia Castanho Scortecci, Vinícius Pietta Perez, Otavio J. Brustolini, Eduardo Sérgio Soares Sousa, Danielle Angst Secco, Angela Maria Guimarães Santos, George Rego Albuquerque, Ana Paula Melo Mariano, Bianca Mendes Maciel, Alexandra L Gerber, Ana Paula de C Guimarães, Paulo Ricardo Nascimento, Francisco Paulo Freire Neto, Sandra Rocha Gadelha, Luís Cristóvão Porto, Eloiza Helena Campana, Selma Maria Bezerra Jeronimo, Ana Tereza R Vasconcelos |
| EPI_ISL_1213324, EPI_ISL_1213326, EPI_ISL_1213328 | LAFEM/UESC | Bioinformatics Laboratory / LNCC | Alessandra P Lamarca, Luiz G P de Almeida, Ronaldo da Silva Francisco Jr, Lucymara Fassarella Agnez Lima, Kátia Castanho Scortecci, Vinícius Pietta Perez, Otavio J. Brustolini, Eduardo Sérgio Soares Sousa, Danielle Angst Secco, Angela Maria Guimarães Santos, George Rego Albuquerque, Ana Paula Melo Mariano, Bianca Mendes Maciel, Alexandra L Gerber, Ana Paula de C Guimarães, Paulo Ricardo Nascimento, Francisco Paulo Freire Neto, Sandra Rocha Gadelha, Luís Cristóvão Porto, Eloiza Helena Campana, Selma Maria Bezerra Jeronimo, Ana Tereza R Vasconcelos |
| EPI_ISL_1213329 | IMT-UFRN/RN | Bioinformatics Laboratory / LNCC | Alessandra P Lamarca, Luiz G P de Almeida, Ronaldo da Silva Francisco Jr, Lucymara Fassarella Agnez Lima, Kátia Castanho Scortecci, Vinícius Pietta Perez, Otavio J. Brustolini, Eduardo Sérgio Soares Sousa, Danielle Angst Secco, Angela Maria Guimarães Santos, George Rego Albuquerque, Ana Paula Melo Mariano, Bianca Mendes Maciel, Alexandra L Gerber, Ana Paula de C Guimarães, Paulo Ricardo Nascimento, Francisco Paulo Freire Neto, Sandra Rocha Gadelha, Luís Cristóvão Porto, Eloiza Helena Campana, Selma Maria Bezerra Jeronimo, Ana Tereza R Vasconcelos |
| EPI_ISL_1213331, EPI_ISL_1213333, EPI_ISL_1213335 | LAFEM/UESC | Bioinformatics Laboratory / LNCC | Alessandra P Lamarca, Luiz G P de Almeida, Ronaldo da Silva Francisco Jr, Lucymara Fassarella Agnez Lima, Kátia Castanho Scortecci, Vinícius Pietta Perez, Otavio J. Brustolini, Eduardo Sérgio Soares Sousa, Danielle Angst Secco, Angela Maria Guimarães Santos, George Rego Albuquerque, Ana Paula Melo Mariano, Bianca Mendes Maciel, Alexandra L Gerber, Ana Paula de C Guimarães, Paulo Ricardo Nascimento, Francisco Paulo Freire Neto, Sandra Rocha Gadelha, Luís Cristóvão Porto, Eloiza Helena Campana, Selma Maria Bezerra Jeronimo, Ana Tereza R Vasconcelos |
| EPI_ISL_1213336 | LBM/UFPB | Bioinformatics Laboratory / LNCC | Alessandra P Lamarca, Luiz G P de Almeida, Ronaldo da Silva Francisco Jr, Lucymara Fassarella Agnez Lima, Kátia Castanho Scortecci, Vinícius Pietta Perez, Otavio J. Brustolini, Eduardo Sérgio Soares Sousa, Danielle Angst Secco, Angela Maria Guimarães Santos, George Rego Albuquerque, Ana Paula Melo Mariano, Bianca Mendes Maciel, Alexandra L Gerber, Ana Paula de C Guimarães, Paulo Ricardo Nascimento, Francisco Paulo Freire Neto, Sandra Rocha Gadelha, Luís Cristóvão Porto, Eloiza Helena Campana, Selma Maria Bezerra Jeronimo, Ana Tereza R Vasconcelos |
| EPI_ISL_1213339, EPI_ISL_1213340 | Laboratório HLA/UERJ | Bioinformatics Laboratory / LNCC | Alessandra P Lamarca, Luiz G P de Almeida, Ronaldo da Silva Francisco Jr, Lucymara Fassarella Agnez Lima, Kátia Castanho Scortecci, Vinícius Pietta Perez, Otavio J. Brustolini, Eduardo Sérgio Soares Sousa, Danielle Angst Secco, Angela Maria Guimarães Santos, George Rego Albuquerque, Ana Paula Melo Mariano, Bianca Mendes Maciel, Alexandra L Gerber, Ana Paula de C Guimarães, Paulo Ricardo Nascimento, Francisco Paulo Freire Neto, Sandra Rocha Gadelha, Luís Cristóvão Porto, Eloiza Helena Campana, Selma Maria Bezerra Jeronimo, Ana Tereza R Vasconcelos |
| EPI_ISL_1213341, EPI_ISL_1213342 | LAFEM/UESC | Bioinformatics Laboratory / LNCC | Alessandra P Lamarca, Luiz G P de Almeida, Ronaldo da Silva Francisco Jr, Lucymara Fassarella Agnez Lima, Kátia Castanho Scortecci, Vinícius Pietta Perez, Otavio J. Brustolini, Eduardo Sérgio Soares Sousa, Danielle Angst Secco, Angela Maria Guimarães Santos, George Rego Albuquerque, Ana Paula Melo Mariano, Bianca Mendes Maciel, Alexandra L Gerber, Ana Paula de C Guimarães, Paulo Ricardo Nascimento, Francisco Paulo Freire Neto, Sandra Rocha Gadelha, Luís Cristóvão Porto, Eloiza Helena Campana, Selma Maria Bezerra Jeronimo, Ana Tereza R Vasconcelos |
| EPI_ISL_1213343, EPI_ISL_1213344 | LBM/UFPB | Bioinformatics Laboratory / LNCC | Alessandra P Lamarca, Luiz G P de Almeida, Ronaldo da Silva Francisco Jr, Lucymara Fassarella Agnez Lima, Kátia Castanho Scortecci, Vinícius Pietta Perez, Otavio J. Brustolini, Eduardo Sérgio Soares Sousa, Danielle Angst Secco, Angela Maria Guimarães Santos, George Rego Albuquerque, Ana Paula Melo Mariano, Bianca Mendes Maciel, Alexandra L Gerber, Ana Paula de C Guimarães, Paulo Ricardo Nascimento, Francisco Paulo Freire Neto, Sandra Rocha Gadelha, Luís Cristóvão Porto, Eloiza Helena Campana, Selma Maria Bezerra Jeronimo, Ana Tereza R Vasconcelos |
| EPI_ISL_1213345, EPI_ISL_1213346, EPI_ISL_1213348, EPI_ISL_1213350, EPI_ISL_1213352, EPI_ISL_1213353, EPI_ISL_1213355, EPI_ISL_1213357, EPI_ISL_1213358, EPI_ISL_1213360, EPI_ISL_1213362, EPI_ISL_1213364 | | | |
| see above | IMT-UFRN/RN | Bioinformatics Laboratory / LNCC | Alessandra P Lamarca, Luiz G P de Almeida, Ronaldo da Silva Francisco Jr, Lucymara Fassarella Agnez Lima, Kátia Castanho Scortecci, Vinícius Pietta Perez, Otavio J. Brustolini, Eduardo Sérgio Soares Sousa, Danielle Angst Secco, Angela Maria Guimarães Santos, George Rego Albuquerque, Ana Paula Melo Mariano, Bianca Mendes Maciel, Alexandra L Gerber, Ana Paula de C Guimarães, Paulo Ricardo Nascimento, Francisco Paulo Freire Neto, Sandra Rocha Gadelha, Luís Cristóvão Porto, Eloiza Helena Campana, Selma Maria Bezerra Jeronimo, Ana Tereza R Vasconcelos |
| EPI_ISL_1213365 | LAFEM/UESC | Bioinformatics Laboratory / LNCC | Alessandra P Lamarca, Luiz G P de Almeida, Ronaldo da Silva Francisco Jr, Lucymara Fassarella Agnez Lima, Kátia Castanho Scortecci, Vinícius Pietta Perez, Otavio J. Brustolini, Eduardo Sérgio Soares Sousa, Danielle Angst Secco, Angela Maria Guimarães Santos, George Rego Albuquerque, Ana Paula Melo Mariano, Bianca Mendes Maciel, Alexandra L Gerber, Ana Paula de C Guimarães, Paulo Ricardo Nascimento, Francisco Paulo Freire Neto, Sandra Rocha Gadelha, Luís Cristóvão Porto, Eloiza Helena Campana, Selma Maria Bezerra Jeronimo, Ana Tereza R Vasconcelos |
| EPI_ISL_1213367, EPI_ISL_1213369, EPI_ISL_1213370 | LBM/UFPB | Bioinformatics Laboratory / LNCC | Alessandra P Lamarca, Luiz G P de Almeida, Ronaldo da Silva Francisco Jr, Lucymara Fassarella Agnez Lima, Kátia Castanho Scortecci, Vinícius Pietta Perez, Otavio J. Brustolini, Eduardo Sérgio Soares Sousa, Danielle Angst Secco, Angela Maria Guimarães Santos, George Rego Albuquerque, Ana Paula Melo Mariano, Bianca Mendes Maciel, Alexandra L Gerber, Ana Paula de C Guimarães, Paulo Ricardo Nascimento, Francisco Paulo Freire Neto, Sandra Rocha Gadelha, Luís Cristóvão Porto, Eloiza Helena Campana, Selma Maria Bezerra Jeronimo, Ana Tereza R Vasconcelos |
| EPI_ISL_1213372, EPI_ISL_1213374, EPI_ISL_1213376, EPI_ISL_1213378, EPI_ISL_1213379, EPI_ISL_1213381, EPI_ISL_1213382 | Laboratório HLA/UERJ | Bioinformatics Laboratory / LNCC | Alessandra P Lamarca, Luiz G P de Almeida, Ronaldo da Silva Francisco Jr, Lucymara Fassarella Agnez Lima, Kátia Castanho Scortecci, Vinícius Pietta Perez, Otavio J. Brustolini, Eduardo Sérgio Soares Sousa, Danielle Angst Secco, Angela Maria Guimarães Santos, George Rego Albuquerque, Ana Paula Melo Mariano, Bianca Mendes Maciel, Alexandra L Gerber, Ana Paula de C Guimarães, Paulo Ricardo Nascimento, Francisco Paulo Freire Neto, Sandra Rocha Gadelha, Luís Cristóvão Porto, Eloiza Helena Campana, Selma Maria Bezerra Jeronimo, Ana Tereza R Vasconcelos |
| EPI_ISL_1213384, EPI_ISL_1213386 | LAFEM/UESC | Bioinformatics Laboratory / LNCC | Alessandra P Lamarca, Luiz G P de Almeida, Ronaldo da Silva Francisco Jr, Lucymara Fassarella Agnez Lima, Kátia Castanho Scortecci, Vinícius Pietta Perez, Otavio J. Brustolini, Eduardo Sérgio Soares Sousa, Danielle Angst Secco, Angela Maria Guimarães Santos, George Rego Albuquerque, Ana Paula |

| | | | |
|---|---|---|---|
| | | | Melo Mariano, Bianca Mendes Maciel, Alexandra L Gerber, Ana Paula de C Guimarães, Paulo Ricardo Nascimento, Francisco Paulo Freire Neto, Sandra Rocha Gadelha, Luís Cristóvão Porto, Eloiza Helena Campana, Selma Maria Bezerra Jeronimo, Ana Tereza R Vasconcelos |
| EPI_ISL_1213388 | LBM/UFPB | Bioinformatics Laboratory / LNCC | Alessandra P Lamarca, Luiz G P de Almeida, Ronaldo da Silva Francisco Jr, Lucymara Fassarella Agnez Lima, Kátia Castanho Scortecci, Vinícius Pietta Perez, Otavio J. Brustolini, Eduardo Sérgio Soares Sousa, Danielle Angst Secco, Angela Maria Guimarães Santos, George Rego Albuquerque, Ana Paula Melo Mariano, Bianca Mendes Maciel, Alexandra L Gerber, Ana Paula de C Guimarães, Paulo Ricardo Nascimento, Francisco Paulo Freire Neto, Sandra Rocha Gadelha, Luís Cristóvão Porto, Eloiza Helena Campana, Selma Maria Bezerra Jeronimo, Ana Tereza R Vasconcelos |
| EPI_ISL_1213390 | LAFEM/UESC | Bioinformatics Laboratory / LNCC | Alessandra P Lamarca, Luiz G P de Almeida, Ronaldo da Silva Francisco Jr, Lucymara Fassarella Agnez Lima, Kátia Castanho Scortecci, Vinícius Pietta Perez, Otavio J. Brustolini, Eduardo Sérgio Soares Sousa, Danielle Angst Secco, Angela Maria Guimarães Santos, George Rego Albuquerque, Ana Paula Melo Mariano, Bianca Mendes Maciel, Alexandra L Gerber, Ana Paula de C Guimarães, Paulo Ricardo Nascimento, Francisco Paulo Freire Neto, Sandra Rocha Gadelha, Luís Cristóvão Porto, Eloiza Helena Campana, Selma Maria Bezerra Jeronimo, Ana Tereza R Vasconcelos |
| EPI_ISL_1213392 | LBM/UFPB | Bioinformatics Laboratory / LNCC | Alessandra P Lamarca, Luiz G P de Almeida, Ronaldo da Silva Francisco Jr, Lucymara Fassarella Agnez Lima, Kátia Castanho Scortecci, Vinícius Pietta Perez, Otavio J. Brustolini, Eduardo Sérgio Soares Sousa, Danielle Angst Secco, Angela Maria Guimarães Santos, George Rego Albuquerque, Ana Paula Melo Mariano, Bianca Mendes Maciel, Alexandra L Gerber, Ana Paula de C Guimarães, Paulo Ricardo Nascimento, Francisco Paulo Freire Neto, Sandra Rocha Gadelha, Luís Cristóvão Porto, Eloiza Helena Campana, Selma Maria Bezerra Jeronimo, Ana Tereza R Vasconcelos |
| EPI_ISL_1213393, EPI_ISL_1213395, EPI_ISL_1213397 | Laboratório HLA/UERJ | Bioinformatics Laboratory / LNCC | Alessandra P Lamarca, Luiz G P de Almeida, Ronaldo da Silva Francisco Jr, Lucymara Fassarella Agnez Lima, Kátia Castanho Scortecci, Vinícius Pietta Perez, Otavio J. Brustolini, Eduardo Sérgio Soares Sousa, Danielle Angst Secco, Angela Maria Guimarães Santos, George Rego Albuquerque, Ana Paula Melo Mariano, Bianca Mendes Maciel, Alexandra L Gerber, Ana Paula de C Guimarães, Paulo Ricardo Nascimento, Francisco Paulo Freire Neto, Sandra Rocha Gadelha, Luís Cristóvão Porto, Eloiza Helena Campana, Selma Maria Bezerra Jeronimo, Ana Tereza R Vasconcelos |
| EPI_ISL_1213399, EPI_ISL_1213401, EPI_ISL_1213402 | LAFEM/UESC | Bioinformatics Laboratory / LNCC | Alessandra P Lamarca, Luiz G P de Almeida, Ronaldo da Silva Francisco Jr, Lucymara Fassarella Agnez Lima, Kátia Castanho Scortecci, Vinícius Pietta Perez, Otavio J. Brustolini, Eduardo Sérgio Soares Sousa, Danielle Angst Secco, Angela Maria Guimarães Santos, George Rego Albuquerque, Ana Paula Melo Mariano, Bianca Mendes Maciel, Alexandra L Gerber, Ana Paula de C Guimarães, Paulo Ricardo Nascimento, Francisco Paulo Freire Neto, Sandra Rocha Gadelha, Luís Cristóvão Porto, Eloiza Helena Campana, Selma Maria Bezerra Jeronimo, Ana Tereza R Vasconcelos |
| EPI_ISL_1213404, EPI_ISL_1213406 | LBM/UFPB | Bioinformatics Laboratory / LNCC | Alessandra P Lamarca, Luiz G P de Almeida, Ronaldo da Silva Francisco Jr, Lucymara Fassarella Agnez Lima, Kátia Castanho Scortecci, Vinícius Pietta Perez, Otavio J. Brustolini, Eduardo Sérgio Soares Sousa, Danielle Angst Secco, Angela Maria Guimarães Santos, George Rego Albuquerque, Ana Paula Melo Mariano, Bianca Mendes Maciel, Alexandra L Gerber, Ana Paula de C Guimarães, Paulo Ricardo Nascimento, Francisco Paulo Freire Neto, Sandra Rocha Gadelha, Luís Cristóvão Porto, Eloiza Helena Campana, Selma Maria Bezerra Jeronimo, Ana Tereza R Vasconcelos |
| EPI_ISL_1213408, EPI_ISL_1213410 | Laboratório HLA/UERJ | Bioinformatics Laboratory / LNCC | Alessandra P Lamarca, Luiz G P de Almeida, Ronaldo da Silva Francisco Jr, Lucymara Fassarella Agnez Lima, Kátia Castanho Scortecci, Vinícius Pietta Perez, Otavio J. Brustolini, Eduardo Sérgio Soares Sousa, Danielle Angst Secco, Angela Maria Guimarães Santos, George Rego Albuquerque, Ana Paula Melo Mariano, Bianca Mendes Maciel, Alexandra L Gerber, Ana Paula de C Guimarães, Paulo Ricardo Nascimento, Francisco Paulo Freire Neto, Sandra Rocha Gadelha, Luís Cristóvão Porto, Eloiza Helena Campana, Selma Maria Bezerra Jeronimo, Ana Tereza R Vasconcelos |
| EPI_ISL_1213411, EPI_ISL_1213413, EPI_ISL_1213415 | LAFEM/UESC | Bioinformatics Laboratory / LNCC | Alessandra P Lamarca, Luiz G P de Almeida, Ronaldo da Silva Francisco Jr, Lucymara Fassarella Agnez Lima, Kátia Castanho Scortecci, Vinícius Pietta Perez, Otavio J. Brustolini, Eduardo Sérgio Soares Sousa, Danielle Angst Secco, Angela Maria Guimarães Santos, George Rego Albuquerque, Ana Paula Melo Mariano, Bianca Mendes Maciel, Alexandra L Gerber, Ana Paula de C Guimarães, Paulo Ricardo Nascimento, Francisco Paulo Freire Neto, Sandra Rocha Gadelha, Luís Cristóvão Porto, Eloiza Helena Campana, Selma Maria Bezerra Jeronimo, Ana Tereza R Vasconcelos |
| EPI_ISL_1213417, EPI_ISL_1213418, EPI_ISL_1213420, EPI_ISL_1213422, EPI_ISL_1213424, EPI_ISL_1213425 | Laboratório HLA/UERJ | Bioinformatics Laboratory / LNCC | Alessandra P Lamarca, Luiz G P de Almeida, Ronaldo da Silva Francisco Jr, Lucymara Fassarella Agnez Lima, Kátia Castanho Scortecci, Vinícius Pietta Perez, Otavio J. Brustolini, Eduardo Sérgio Soares Sousa, Danielle Angst Secco, Angela Maria Guimarães Santos, George Rego Albuquerque, Ana Paula Melo Mariano, Bianca Mendes Maciel, Alexandra L Gerber, Ana Paula de C Guimarães, Paulo Ricardo Nascimento, Francisco Paulo Freire Neto, Sandra Rocha Gadelha, Luís Cristóvão Porto, Eloiza Helena Campana, Selma Maria Bezerra Jeronimo, Ana Tereza R Vasconcelos |
| EPI_ISL_1213427, EPI_ISL_1213429, EPI_ISL_1213431, EPI_ISL_1213433, EPI_ISL_1213434, EPI_ISL_1213436, EPI_ISL_1213438, EPI_ISL_1213439, EPI_ISL_1213441, EPI_ISL_1213443, EPI_ISL_1213444, EPI_ISL_1213446, EPI_ISL_1213448, EPI_ISL_1213449, EPI_ISL_1213451, EPI_ISL_1213453, EPI_ISL_1213454, EPI_ISL_1213456, EPI_ISL_1213458, EPI_ISL_1213459, EPI_ISL_1213461, EPI_ISL_1213463 | | | |
| see above | LBM/UFPB | Bioinformatics Laboratory / LNCC | Alessandra P Lamarca, Luiz G P de Almeida, Ronaldo da Silva Francisco Jr, Lucymara Fassarella Agnez Lima, Kátia Castanho Scortecci, Vinícius Pietta Perez, Otavio J. Brustolini, Eduardo Sérgio Soares Sousa, Danielle Angst Secco, Angela Maria Guimarães Santos, George Rego Albuquerque, Ana Paula Melo Mariano, Bianca Mendes Maciel, Alexandra L Gerber, Ana Paula de C Guimarães, Paulo Ricardo Nascimento, Francisco Paulo Freire Neto, Sandra Rocha Gadelha, Luís Cristóvão Porto, Eloiza Helena Campana, Selma Maria Bezerra Jeronimo, Ana Tereza R Vasconcelos |
| EPI_ISL_1219021 | Hospital Aviccena | Instituto Adolfo Lutz, Interdiciplinary Procedures Center, Strategic Laboratory | Claudio Tavares Sacchi, Claudia Regina Gonçalves, Erica Valessa Ramos Gomes, Karoline Rodrigues Campos, Caio Vinicius Dias Lopes |
| EPI_ISL_1219022, EPI_ISL_1219023, EPI_ISL_1219024, EPI_ISL_1219025, EPI_ISL_1219026 | IAL Regional de Sorocaba | Instituto Adolfo Lutz, Interdiciplinary Procedures Center, Strategic Laboratory | Claudio Tavares Sacchi, Claudia Regina Gonçalves, Erica Valessa Ramos Gomes, Karoline Rodrigues Campos, Caio Vinicius Dias Lopes |
| EPI_ISL_1219027 | IAL Regional de Presidente Prudente | Instituto Adolfo Lutz, Interdiciplinary Procedures Center, Strategic Laboratory | Claudio Tavares Sacchi, Claudia Regina Gonçalves, Erica Valessa Ramos Gomes, Karoline Rodrigues Campos, Caio Vinicius Dias Lopes |
| EPI_ISL_1219028, EPI_ISL_1219029, EPI_ISL_1219030, EPI_ISL_1219031, EPI_ISL_1219032, EPI_ISL_1219033, EPI_ISL_1219034, EPI_ISL_1219035, EPI_ISL_1219036 | Aeroporto Internacional de Guarulhos | Instituto Adolfo Lutz, Interdiciplinary Procedures Center, Strategic Laboratory | Claudio Tavares Sacchi, Claudia Regina Gonçalves, Erica Valessa Ramos Gomes, Karoline Rodrigues Campos, Caio Vinicius Dias Lopes |
| EPI_ISL_1219037 | IAL Regional de Presidente Prudente | Instituto Adolfo Lutz, Interdiciplinary Procedures Center, Strategic Laboratory | Claudio Tavares Sacchi, Claudia Regina Gonçalves, Erica Valessa Ramos Gomes, Karoline Rodrigues Campos, Caio Vinicius Dias Lopes |
| EPI_ISL_1219132, EPI_ISL_1219133 | Laboratorio Central de Saude Publica do Estado do Parana (LACEN-PR) | Laboratory of Respiratory Viruses and Measles, Oswaldo Cruz Institute, FIOCRUZ | Paola Resende, Luciana Appolinario, Fernando Motta, Anna Carolina Paixao, Ana Carolina Mendonca, Alice Sampaio Rocha, Renata Serrano Lopes, Maria do Carmo Debur, Irina Nastassja Riediger, Marilda Siqueira on behalf of the Fiocruz COVID-19 Genomic Surveillance Network |
| EPI_ISL_1219134, EPI_ISL_1219135 | Laboratorio Central de Saude Publica do Estado do Alagoas (LACEN-AL) | Laboratory of Respiratory Viruses and Measles, Oswaldo Cruz Institute, FIOCRUZ | Paola Resende, Luciana Appolinario, Fernando Motta, Anna Carolina Paixao, Ana Carolina Mendonca, Alice Sampaio Rocha, Renata Serrano Lopes, Anderson Brandao Leite, Marilda Siqueira on behalf of the Fiocruz COVID-19 Genomic Surveillance Network |
| EPI_ISL_1219136 | Gonçalo Moniz Institute, FIOCRUZ, Bahia | Laboratory of Respiratory Viruses and Measles, Oswaldo Cruz Institute, FIOCRUZ | Paola Resende, Luciana Appolinario, Fernando Motta, Anna Carolina Paixao, Ana Carolina Mendonca, Alice Sampaio Rocha, Renata Serrano Lopes, Tiago Graf, Ricardo Khouri, Marilda Siqueira on behalf of the Fiocruz COVID-19 Genomic Surveillance Network |
| EPI_ISL_1219137 | Laboratorio Central de Saude Publica do Estado de Minas Gerais (LACEN-MG) | Laboratory of Respiratory Viruses and Measles, Oswaldo Cruz Institute, FIOCRUZ | Paola Resende, Luciana Appolinario, Fernando Motta, Anna Carolina Paixao, Ana Carolina Mendonca, Alice Sampaio Rocha, Renata Serrano Lopes, Felipe Iani, Marilda Siqueira on behalf of the Fiocruz COVID-19 Genomic Surveillance Network |
| EPI_ISL_1239012 | Laboratório Central de Saúde Pública do Estado de Pernambuco (LACEN-PE) | WallauLab, Aggeu Magalhaes Institute | Marcelo Henrique dos Santos Paiva, Duschinka Ribeiro Duarte Guedes, Cássia Docena, Matheus Filgueira Bezerra, Filipe Zimmer Dezordi, Laís Ceschini Machado, Larissa Krokovsky, Elisama Helvecio, Alexandre Freitas da Silva, Antonio Mauro Rezende, Sinval Pinto Brandão Filho, Constância Flávia Junqueira Ayres, Gabriel Luz Wallau on behalf of the Fiocruz COVID-19 Genomic Surveillance Network |
| EPI_ISL_1239111, EPI_ISL_1239112, EPI_ISL_1239113 | Laboratório Central de Saúde Pública Noel Nutels | Coordenação Geral de Laboratórios de Saúde Pública (CGLAB) | Vagner Fonseca et al, |
| EPI_ISL_1239114, EPI_ISL_1239115, EPI_ISL_1239116, EPI_ISL_1239117, EPI_ISL_1239118, EPI_ISL_1239119, EPI_ISL_1239120, EPI_ISL_1239121, EPI_ISL_1239122, EPI_ISL_1239123 | Laboratório Central de Saúde Pública do Espírito Santo | Coordenação Geral de Laboratórios de Saúde Pública (CGLAB) | Vagner Fonseca et al, |

| | | | |
|---|---|---|---|
| EPI_ISL_1239124 | Fundação Ezequiel Dias | Coordenação Geral de Laboratórios de Saúde Pública (CGLAB) | Vagner Fonseca et al, |
| EPI_ISL_1239125, EPI_ISL_1239126, EPI_ISL_1239127, EPI_ISL_1239128, EPI_ISL_1239129, EPI_ISL_1239130, EPI_ISL_1239131, EPI_ISL_1239132, EPI_ISL_1239133, EPI_ISL_1239134, EPI_ISL_1239135, EPI_ISL_1239136 | | | |
| see above | Laboratório Central de Saúde Pública do Espírito Santo | Coordenação Geral de Laboratórios de Saúde Pública (CGLAB) | Vagner Fonseca et al, |
| EPI_ISL_1239137, EPI_ISL_1239138, EPI_ISL_1239139 | Fundação Ezequiel Dias | Coordenação Geral de Laboratórios de Saúde Pública (CGLAB) | Vagner Fonseca et al, |
| EPI_ISL_1240639, EPI_ISL_1240640, EPI_ISL_1240641 | Laboratório Central de Saúde Pública do Espírito Santo | Coordenação Geral de Laboratórios de Saúde Pública (CGLAB) | Vagner Fonseca et al. |
| EPI_ISL_1240642 | Fundação Ezequiel Dias | Coordenação Geral de Laboratórios de Saúde Pública (CGLAB) | Vagner Fonseca et al. |
| EPI_ISL_1251222, EPI_ISL_1251223, EPI_ISL_1251224, EPI_ISL_1251225 | Raimundo Reginaldo de Almeida | MOA Lab | Vanessa C. Nicolete, Priscila T. Rodrigues, Marcelo U. Ferreira |
| EPI_ISL_1261122, EPI_ISL_1261123 | Laboratorio de Ecologia de Doencas Transmissiveis na Amazonia, Instituto Leonidas e Maria Deane - Fiocruz Amazonia | Laboratorio de Ecologia de Doencas Transmissiveis na Amazonia, Instituto Leonidas e Maria Deane - Fiocruz Amazonia | Valdinete Nascimento, Victor Souza, André Corado, Fernanda Nascimento, George Silva, Ágatha Costa, Debora Duarte, Karina Pessoa, Matilde Mejía, Luciana Gonçalves, Maria Júlia Brandão, Michele Jesus, Felipe Naveca |
| EPI_ISL_1261683 | LACEN - Laboratório Central de Saúde Pública do Amazonas | Evandro Chagas Institute | Santos, M.C.; Silva, A.M.; Junior, W.D.C.; Barbagelata, L.S.; Ferreira, J.A.; Sousa, E.M.A.; da Silva, P.S.; Pinheiro, K.C.; L.C.; Sousa Junior, E.C. |
| EPI_ISL_1261684 | LACEN - Laboratório Central de Saúde Pública do Ceará | Evandro Chagas Institute | Santos, M.C.; Silva, A.M.; Junior, W.D.C.; Barbagelata, L.S.; Ferreira, J.A.; Sousa, E.M.A.; da Silva, P.S.; Pinheiro, K.C.; L.C.; Sousa Junior, E.C. |
| EPI_ISL_1261685 | LACEN - Laboratório Central de Saúde Pública do Amazonas | Evandro Chagas Institute | Santos, M.C.; Silva, A.M.; Junior, W.D.C.; Barbagelata, L.S.; Ferreira, J.A.; Sousa, E.M.A.; da Silva, P.S.; Pinheiro, K.C.; L.C.; Sousa Junior, E.C. |
| EPI_ISL_1261686 | LACEN - Laboratório Central de Saúde Pública do Amapá | Evandro Chagas Institute | Santos, M.C.; Silva, A.M.; Junior, W.D.C.; Barbagelata, L.S.; Ferreira, J.A.; Sousa, E.M.A.; da Silva, P.S.; Pinheiro, K.C.; L.C.; Sousa Junior, E.C. |
| EPI_ISL_1261687 | LACEN - Laboratório Central de Saúde Pública de Roraima | Evandro Chagas Institute | Santos, M.C.; Silva, A.M.; Junior, W.D.C.; Barbagelata, L.S.; Ferreira, J.A.; Sousa, E.M.A.; da Silva, P.S.; Pinheiro, K.C.; L.C.; Sousa Junior, E.C. |
| EPI_ISL_1261688, EPI_ISL_1261689 | LACEN - Laboratório Central de Saúde Pública do Amapá | Evandro Chagas Institute | Santos, M.C.; Silva, A.M.; Junior, W.D.C.; Barbagelata, L.S.; Ferreira, J.A.; Sousa, E.M.A.; da Silva, P.S.; Pinheiro, K.C.; L.C.; Sousa Junior, E.C. |
| EPI_ISL_1261690 | LACEN - Laboratório Central de Saúde Pública do Amazonas | Evandro Chagas Institute | Santos, M.C.; Silva, A.M.; Junior, W.D.C.; Barbagelata, L.S.; Ferreira, J.A.; Sousa, E.M.A.; da Silva, P.S.; Pinheiro, K.C.; L.C.; Sousa Junior, E.C. |
| EPI_ISL_1261691 | Laboratório Paulo C. Azevedo | Evandro Chagas Institute | Santos, M.C.; Silva, A.M.; Junior, W.D.C.; Barbagelata, L.S.; Ferreira, J.A.; Sousa, E.M.A.; da Silva, P.S.; Pinheiro, K.C.; L.C.; Sousa Junior, E.C. |
| EPI_ISL_1261692 | LACEN - Laboratório Central de Saúde Pública do Amapá | Evandro Chagas Institute | Santos, M.C.; Silva, A.M.; Junior, W.D.C.; Barbagelata, L.S.; Ferreira, J.A.; Sousa, E.M.A.; da Silva, P.S.; Pinheiro, K.C.; L.C.; Sousa Junior, E.C. |
| EPI_ISL_1261693 | LACEN - Laboratório Central de Saúde Pública do Ceará | Evandro Chagas Institute | Santos, M.C.; Silva, A.M.; Junior, W.D.C.; Barbagelata, L.S.; Ferreira, J.A.; Sousa, E.M.A.; da Silva, P.S.; Pinheiro, K.C.; L.C.; Sousa Junior, E.C. |
| EPI_ISL_1261694 | LACEN - Laboratório Central de Saúde Pública do Amazonas | Evandro Chagas Institute | Santos, M.C.; Silva, A.M.; Junior, W.D.C.; Barbagelata, L.S.; Ferreira, J.A.; Sousa, E.M.A.; da Silva, P.S.; Pinheiro, K.C.; L.C.; Sousa Junior, E.C. |
| EPI_ISL_1261696 | LACEN - Laboratório Central de Saúde Pública do Amapá | Evandro Chagas Institute | Santos, M.C.; Silva, A.M.; Junior, W.D.C.; Barbagelata, L.S.; Ferreira, J.A.; Sousa, E.M.A.; da Silva, P.S.; Pinheiro, K.C.; L.C.; Sousa Junior, E.C. |
| EPI_ISL_1261697 | LACEN - Laboratório Central de Saúde Pública do Ceará | Evandro Chagas Institute | Santos, M.C.; Silva, A.M.; Junior, W.D.C.; Barbagelata, L.S.; Ferreira, J.A.; Sousa, E.M.A.; da Silva, P.S.; Pinheiro, K.C.; L.C.; Sousa Junior, E.C. |
| EPI_ISL_1261698, EPI_ISL_1261699 | LACEN - Laboratório Central de Saúde Pública de Pernambuco | Evandro Chagas Institute | Santos, M.C.; Silva, A.M.; Junior, W.D.C.; Barbagelata, L.S.; Ferreira, J.A.; Sousa, E.M.A.; da Silva, P.S.; Pinheiro, K.C.; L.C.; Sousa Junior, E.C. |
| EPI_ISL_1261700 | LACEN - Laboratório Central de Saúde Pública do Maranhao | Evandro Chagas Institute | Santos, M.C.; Silva, A.M.; Junior, W.D.C.; Barbagelata, L.S.; Ferreira, J.A.; Sousa, E.M.A.; da Silva, P.S.; Pinheiro, K.C.; L.C.; Sousa Junior, E.C. |
| EPI_ISL_1272236 | Universidade Federal do Norte do Tocantins (UFNT) | Laboratório de Bioinformática e Biotecnologia (Labinftec/UFT) | Ueric José Borges de Souza, Fabrício Souza Campos, Raíssa Nunes dos Santos, José Carlos Ribeiro Júnior, Rogério Fernandes Carvalho, Monike da Silva Oliveira, Bergmann Morais Ribeiro, Fernando Lucas Melo |
| EPI_ISL_1289960 | Laboratory of Virology, Ribeirão Preto General Hospital, Ribeirão Preto Medical School, University of São Paulo | Laboratory of Oncology, Blood Center of Ribeirão Preto, Ribeirão Preto School of Medicine, University of São Paulo | CAMPOS, MR; SANTOS, A.L.P.; YAMAMOTO, A.Y.; COLLI, L.M.; FONSECA, B.A.L.; BELLISSIMO-RODRIGUES, F. |
| EPI_ISL_1293051 | LACEN do Distrito Federal | Instituto Adolfo Lutz, Interdiciplinary Procedures Center, Strategic Laboratory | Claudio Tavares Sacchi, Claudia Regina Gonçalves, Erica Valessa Ramos Gomes, Karoline Rodrigues Campos, Caio Vinicius Dias Lopes |
| EPI_ISL_1293052, EPI_ISL_1293053, EPI_ISL_1293054, EPI_ISL_1293055 | LACEN de Rondonia | Instituto Adolfo Lutz, Interdiciplinary Procedures Center, Strategic Laboratory | Claudio Tavares Sacchi, Claudia Regina Gonçalves, Erica Valessa Ramos Gomes, Karoline Rodrigues Campos, Caio Vinicius Dias Lopes |
| EPI_ISL_1293056, EPI_ISL_1293057, EPI_ISL_1293058, EPI_ISL_1293059, EPI_ISL_1293060, EPI_ISL_1293061, EPI_ISL_1293062, EPI_ISL_1293063, EPI_ISL_1293064, EPI_ISL_1293065, EPI_ISL_1293067, EPI_ISL_1293068, EPI_ISL_1293069, EPI_ISL_1293070, EPI_ISL_1293071, EPI_ISL_1293072, EPI_ISL_1293073, EPI_ISL_1293074, EPI_ISL_1293075, EPI_ISL_1293076, EPI_ISL_1293077, EPI_ISL_1293078, EPI_ISL_1293079, EPI_ISL_1293080, EPI_ISL_1293081 | | | |
| see above | IAL Regional de Sorocaba | Instituto Adolfo Lutz, Interdiciplinary Procedures Center, Strategic Laboratory | Claudio Tavares Sacchi, Claudia Regina Gonçalves, Erica Valessa Ramos Gomes, Karoline Rodrigues Campos, Caio Vinicius Dias Lopes |
| EPI_ISL_1303499, EPI_ISL_1303500, EPI_ISL_1303501, EPI_ISL_1303502, EPI_ISL_1303503, EPI_ISL_1303504, EPI_ISL_1303505 | LACEN de Rondonia | Instituto Adolfo Lutz, Interdiciplinary Procedures Center, Strategic Laboratory | Claudio Tavares Sacchi, Claudia Regina Gonçalves, Erica Valessa Ramos Gomes, Karoline Rodrigues Campos, Caio Vinicius Dias Lopes |
| EPI_ISL_1303506, EPI_ISL_1303507, EPI_ISL_1303508 | LACEN do Distrito Federal | Instituto Adolfo Lutz, Interdiciplinary Procedures Center, Strategic Laboratory | Claudio Tavares Sacchi, Claudia Regina Gonçalves, Erica Valessa Ramos Gomes, Karoline Rodrigues Campos, Caio Vinicius Dias Lopes |
| EPI_ISL_1303509 | LACEN do Estado de Tocantins | Instituto Adolfo Lutz, Interdiciplinary Procedures Center, Strategic Laboratory | Claudio Tavares Sacchi, Claudia Regina Gonçalves, Erica Valessa Ramos Gomes, Karoline Rodrigues Campos, Caio Vinicius Dias Lopes |
| EPI_ISL_1303510, EPI_ISL_1303511, EPI_ISL_1303512, EPI_ISL_1303513, EPI_ISL_1303514, EPI_ISL_1303515, EPI_ISL_1303516, EPI_ISL_1303517 | LACEN do Estado de Goias | Instituto Adolfo Lutz, Interdiciplinary Procedures Center, Strategic Laboratory | Claudio Tavares Sacchi, Claudia Regina Gonçalves, Erica Valessa Ramos Gomes, Karoline Rodrigues Campos, Caio Vinicius Dias Lopes |
| EPI_ISL_1303518, EPI_ISL_1303519, EPI_ISL_1303520, EPI_ISL_1303521, EPI_ISL_1303522, EPI_ISL_1303523, EPI_ISL_1303524, EPI_ISL_1303525, EPI_ISL_1303526, EPI_ISL_1303527, EPI_ISL_1303528, EPI_ISL_1303529, EPI_ISL_1303530, EPI_ISL_1303531, EPI_ISL_1303532, EPI_ISL_1303533, EPI_ISL_1303534 | | | |
| see above | IAL Regional de São Jose do Rio Preto | Instituto Adolfo Lutz, Interdiciplinary Procedures Center, Strategic Laboratory | Claudio Tavares Sacchi, Claudia Regina Gonçalves, Erica Valessa Ramos Gomes, Karoline Rodrigues Campos, Caio Vinicius Dias Lopes |
| EPI_ISL_1303535, EPI_ISL_1303536 | Hospital Heliopolis | Instituto Adolfo Lutz, Interdiciplinary Procedures Center, Strategic Laboratory | Claudio Tavares Sacchi, Claudia Regina Gonçalves, Erica Valessa Ramos Gomes, Karoline Rodrigues Campos, Caio Vinicius Dias Lopes |
| EPI_ISL_1303537 | Hospital Estadual de Vila Alpina | Instituto Adolfo Lutz, Interdiciplinary Procedures Center, Strategic Laboratory | Claudio Tavares Sacchi, Claudia Regina Gonçalves, Erica Valessa Ramos Gomes, Karoline Rodrigues Campos, Caio Vinicius Dias Lopes |
| EPI_ISL_1303538, EPI_ISL_1303539 | Hospital Presidente | Instituto Adolfo Lutz, Interdiciplinary Procedures Center, Strategic Laboratory | Claudio Tavares Sacchi, Claudia Regina Gonçalves, Erica Valessa Ramos Gomes, Karoline Rodrigues Campos, Caio Vinicius Dias Lopes |
| EPI_ISL_1303540, EPI_ISL_1303541 | Hospital Municipal Cidade Tiradentes Carmen Prudente | Instituto Adolfo Lutz, Interdiciplinary Procedures Center, Strategic Laboratory | Claudio Tavares Sacchi, Claudia Regina Gonçalves, Erica Valessa Ramos Gomes, Karoline Rodrigues Campos, Caio Vinicius Dias Lopes |
| EPI_ISL_1303542, EPI_ISL_1303543 | Hospital Estadual de Campanha Barradas | Instituto Adolfo Lutz, Interdiciplinary Procedures Center, | Claudio Tavares Sacchi, Claudia Regina Gonçalves, Erica Valessa Ramos Gomes, Karoline Rodrigues Campos, Caio Vinicius Dias Lopes |

| | | | |
|---|---|---|---|
| | | Strategic Laboratory | |
| EPI_ISL_1303544, EPI_ISL_1303545 | Hospital Municipal Cidade Tiradentes Carmen Prudente | Instituto Adolfo Lutz, Interdiciplinary Procedures Center, Strategic Laboratory | Claudio Tavares Sacchi, Claudia Regina Gonçalves, Erica Valessa Ramos Gomes, Karoline Rodrigues Campos, Caio Vinicius Dias Lopes |
| EPI_ISL_1303546, EPI_ISL_1303547, EPI_ISL_1303548, EPI_ISL_1303549 | UPA Vila Santa Catarina | Instituto Adolfo Lutz, Interdiciplinary Procedures Center, Strategic Laboratory | Claudio Tavares Sacchi, Claudia Regina Gonçalves, Erica Valessa Ramos Gomes, Karoline Rodrigues Campos, Caio Vinicius Dias Lopes |
| EPI_ISL_1324137, EPI_ISL_1324139, EPI_ISL_1324140, EPI_ISL_1324142, EPI_ISL_1324145, EPI_ISL_1324147, EPI_ISL_1324149 | UW Virology Lab | UW Virology Lab | Pavitra Roychoudhury, Hong Xie, Lasata Shrestha, Shah Mohamed Bakhash, Michelle Lin, Margaret Mills, Noah Baker, Sean Ellis, Saraswathi Sathees, Meei-Li Huang, Keith R Jerome, Alexander Greninger |
| EPI_ISL_1358285 | Centro de Treinamento e Referencia DST AIDS | Instituto Adolfo Lutz, Interdiciplinary Procedures Center, Strategic Laboratory | Claudio Tavares Sacchi, Claudia Regina Gonçalves, Erica Valessa Ramos Gomes, Karoline Rodrigues Campos, Caio Vinicius Dias Lopes |
| EPI_ISL_1358286 | Hospital Municipal Josanias Castanha Braga | Instituto Adolfo Lutz, Interdiciplinary Procedures Center, Strategic Laboratory | Claudio Tavares Sacchi, Claudia Regina Gonçalves, Erica Valessa Ramos Gomes, Karoline Rodrigues Campos, Caio Vinicius Dias Lopes |
| EPI_ISL_1358287 | Hospital Nipo Brasileiro | Instituto Adolfo Lutz, Interdiciplinary Procedures Center, Strategic Laboratory | Claudio Tavares Sacchi, Claudia Regina Gonçalves, Erica Valessa Ramos Gomes, Karoline Rodrigues Campos, Caio Vinicius Dias Lopes |
| EPI_ISL_1358288, EPI_ISL_1358289, EPI_ISL_1358290 | IAL Regional de Aracatuba | Instituto Adolfo Lutz, Interdiciplinary Procedures Center, Strategic Laboratory | Claudio Tavares Sacchi, Claudia Regina Gonçalves, Erica Valessa Ramos Gomes, Karoline Rodrigues Campos, Caio Vinicius Dias Lopes |
| EPI_ISL_1358291, EPI_ISL_1358292, EPI_ISL_1358293, EPI_ISL_1358294, EPI_ISL_1358295, EPI_ISL_1358296, EPI_ISL_1358297, EPI_ISL_1358298, EPI_ISL_1358299 | IAL Regional de Santo Andre | Instituto Adolfo Lutz, Interdiciplinary Procedures Center, Strategic Laboratory | Claudio Tavares Sacchi, Claudia Regina Gonçalves, Erica Valessa Ramos Gomes, Karoline Rodrigues Campos, Caio Vinicius Dias Lopes |
| EPI_ISL_1358300, EPI_ISL_1358301, EPI_ISL_1358302, EPI_ISL_1358303 | Lacen de Tocantins | Instituto Adolfo Lutz, Interdiciplinary Procedures Center, Strategic Laboratory | Claudio Tavares Sacchi, Claudia Regina Gonçalves, Erica Valessa Ramos Gomes, Karoline Rodrigues Campos, Caio Vinicius Dias Lopes |
| EPI_ISL_1358304, EPI_ISL_1358305, EPI_ISL_1358306, EPI_ISL_1358307, EPI_ISL_1358308, EPI_ISL_1358309, EPI_ISL_1358310, EPI_ISL_1358311, EPI_ISL_1358312, EPI_ISL_1358313, EPI_ISL_1358314, EPI_ISL_1358315, EPI_ISL_1358316, EPI_ISL_1358317 | | | |
| see above | LACEN do Mato Grosso do Sul | Instituto Adolfo Lutz, Interdiciplinary Procedures Center, Strategic Laboratory | Claudio Tavares Sacchi, Claudia Regina Gonçalves, Erica Valessa Ramos Gomes, Karoline Rodrigues Campos, Caio Vinicius Dias Lopes |
| EPI_ISL_1358318, EPI_ISL_1358319, EPI_ISL_1358320, EPI_ISL_1358321, EPI_ISL_1358322 | UPA Vila Santa Catarina | Instituto Adolfo Lutz, Interdiciplinary Procedures Center, Strategic Laboratory | Claudio Tavares Sacchi, Claudia Regina Gonçalves, Erica Valessa Ramos Gomes, Karoline Rodrigues Campos, Caio Vinicius Dias Lopes |
| EPI_ISL_1365747 | Associação Fundo de Incentivo a Pesquisa | Associação Fundo de Incentivo a Pesquisa | Priscila Farias Tempaku, Juliana Nogueira Martins Rodrigues, Erika Rodrigues de Oliveira, Soraya Sgambatti de Andrade, Debora Ribeiro Ramadan, Sergio Tufik |
| EPI_ISL_1381043, EPI_ISL_1381044, EPI_ISL_1381045, EPI_ISL_1381046, EPI_ISL_1381047, EPI_ISL_1381048, EPI_ISL_1381049, EPI_ISL_1381050, EPI_ISL_1381051, EPI_ISL_1381052, EPI_ISL_1381053, EPI_ISL_1381054, EPI_ISL_1381055, EPI_ISL_1381056, EPI_ISL_1381057, EPI_ISL_1381058, EPI_ISL_1381059, EPI_ISL_1381060, EPI_ISL_1381061, EPI_ISL_1381062, EPI_ISL_1381063, EPI_ISL_1381064, EPI_ISL_1381065 | | | |
| see above | IAL Regional de Santo Andre | Instituto Adolfo Lutz, Interdiciplinary Procedures Center, Strategic Laboratory | Claudio Tavares Sacchi, Claudia Regina Gonçalves, Erica Valessa Ramos Gomes, Karoline Rodrigues Campos, Caio Vinicius Dias Lopes |
| EPI_ISL_1381066, EPI_ISL_1381067 | LACEN do Mato Grosso do Sul | Instituto Adolfo Lutz, Interdiciplinary Procedures Center, Strategic Laboratory | Claudio Tavares Sacchi, Claudia Regina Gonçalves, Erica Valessa Ramos Gomes, Karoline Rodrigues Campos, Caio Vinicius Dias Lopes |
| EPI_ISL_1381068 | Conjunto Hospitalar do Mandaqui de Sao Paulo | Instituto Adolfo Lutz, Interdiciplinary Procedures Center, Strategic Laboratory | Claudio Tavares Sacchi, Claudia Regina Gonçalves, Erica Valessa Ramos Gomes, Karoline Rodrigues Campos, Caio Vinicius Dias Lopes |
| EPI_ISL_1381069 | Hospital Heliopolis | Instituto Adolfo Lutz, Interdiciplinary Procedures Center, Strategic Laboratory | Claudio Tavares Sacchi, Claudia Regina Gonçalves, Erica Valessa Ramos Gomes, Karoline Rodrigues Campos, Caio Vinicius Dias Lopes |
| EPI_ISL_1381070, EPI_ISL_1381071 | Hospital Municipal Cidade Tiradentes Carmem Prudente | Instituto Adolfo Lutz, Interdiciplinary Procedures Center, Strategic Laboratory | Claudio Tavares Sacchi, Claudia Regina Gonçalves, Erica Valessa Ramos Gomes, Karoline Rodrigues Campos, Caio Vinicius Dias Lopes |
| EPI_ISL_1402429, EPI_ISL_1402430 | Laboratory of Respiratory Viruses and Measles, Oswaldo Cruz Institute, FIOCRUZ | Laboratory of Respiratory Viruses and Measles, Oswaldo Cruz Institute, FIOCRUZ | Paola Resende, Alex Pauvolid-Correa, Mia Ferreira Araujo, Ana Beatriz Machado Lima, Luciana Appolinario, Fernando Motta, Anna Carolina Paixao, Ana Carolina Mendonca, Alice Sampaio Rocha, Renata Serrano Lopes, Marilda Siqueira on behalf of the Fiocruz COVID-19 Genomic Surveillance Network |
| EPI_ISL_1402431 | Laboratory of Respiratory Viruses and Measles, Oswaldo Cruz Institute, FIOCRUZ | Laboratory of Respiratory Viruses and Measles, Oswaldo Cruz Institute, FIOCRUZ | Paola Resende, Felipe Naveca, Alex Pauvolid-Correa, Mia Ferreira Araujo, Ana Beatriz Machado Lima, Luciana Appolinario, Fernando Motta, Anna Carolina Paixao, Ana Carolina Mendonca, Alice Sampaio Rocha, Renata Serrano Lopes, Marilda Siqueira on behalf of the Fiocruz COVID-19 Genomic Surveillance Network |
| EPI_ISL_1443196, EPI_ISL_1443197, EPI_ISL_1443198 | Hospital Aliança | Hospital São Rafael - IDOR | Isadora Cristina de Siqueira, Aquiles Assunção Camelier, Elves A.P. Maciel, Margarida Celia L. C. Neves, Carolina Kymie Vasques Nonaka, Karoline Almeida Félix de Sousa, Victor Costa Araujo, Yasmin Santos Freitas Macêdo, Aurea Angelica Paste, Bruno Solano de Freitas Souza, Tiago Gräf |
| EPI_ISL_1445065 | UNIDADE MISTA DE TAIACU | Instituto Butantan / Mendelics | Dimas Tadeu Covas, Sandra Coccuzzo Sampaio, Maria Carolina Elias, José Salvatore Leister Patané, Vincent Louis Viala, Antonio Jorge Martins, Ricardo Haddad, Claudia Renata dos Santos Barros, Elaine Cristina Marqueze, Raul Machado Neto, Debora Botequio Moretti, Bibiana Santos, João Paulo Kitajima, Erika Freitas, David Schlesinger, Simone Kashima, Evandra Strazza Rodrigues, Svetoslav Nanev Slavov, Elaine Vieira dos Santos, Rafael dos Santos Bezerra, Luiz Carlos Junior de Alcantara, Marta Giovanetti, Vagner Fonseca, Flavia Aburjaile, Rodrigo Tocantins Calado. |
| EPI_ISL_1445070 | USF ARCADAS | Instituto Butantan / Mendelics | Dimas Tadeu Covas, Sandra Coccuzzo Sampaio, Maria Carolina Elias, José Salvatore Leister Patané, Vincent Louis Viala, Antonio Jorge Martins, Ricardo Haddad, Claudia Renata dos Santos Barros, Elaine Cristina Marqueze, Raul Machado Neto, Debora Botequio Moretti, Bibiana Santos, João Paulo Kitajima, Erika Freitas, David Schlesinger, Simone Kashima, Evandra Strazza Rodrigues, Svetoslav Nanev Slavov, Elaine Vieira dos Santos, Rafael dos Santos Bezerra, Luiz Carlos Junior de Alcantara, Marta Giovanetti, Vagner Fonseca, Flavia Aburjaile, Rodrigo Tocantins Calado. |
| EPI_ISL_1445071 | USF TRES PONTES | Instituto Butantan / Mendelics | Dimas Tadeu Covas, Sandra Coccuzzo Sampaio, Maria Carolina Elias, José Salvatore Leister Patané, Vincent Louis Viala, Antonio Jorge Martins, Ricardo Haddad, Claudia Renata dos Santos Barros, Elaine Cristina Marqueze, Raul Machado Neto, Debora Botequio Moretti, Bibiana Santos, João Paulo Kitajima, Erika Freitas, David Schlesinger, Simone Kashima, Evandra Strazza Rodrigues, Svetoslav Nanev Slavov, Elaine Vieira dos Santos, Rafael dos Santos Bezerra, Luiz Carlos Junior de Alcantara, Marta Giovanetti, Vagner Fonseca, Flavia Aburjaile, Rodrigo Tocantins Calado. |
| EPI_ISL_1445072 | USF JARDIM SAO DIMAS | Instituto Butantan / Mendelics | Dimas Tadeu Covas, Sandra Coccuzzo Sampaio, Maria Carolina Elias, José Salvatore Leister Patané, Vincent Louis Viala, Antonio Jorge Martins, Ricardo Haddad, Claudia Renata dos Santos Barros, Elaine Cristina Marqueze, Raul Machado Neto, Debora Botequio Moretti, Bibiana Santos, João Paulo Kitajima, Erika Freitas, David Schlesinger, Simone Kashima, Evandra Strazza Rodrigues, Svetoslav Nanev Slavov, Elaine Vieira dos Santos, Rafael dos Santos Bezerra, Luiz Carlos Junior de Alcantara, Marta Giovanetti, Vagner Fonseca, Flavia Aburjaile, Rodrigo Tocantins Calado. |
| EPI_ISL_1445078, EPI_ISL_1445079 | CENTRO DE SAUDE II DR GABRIEL MESQUITA VARGEM GDE DO SUL | Instituto Butantan / Mendelics | Dimas Tadeu Covas, Sandra Coccuzzo Sampaio, Maria Carolina Elias, José Salvatore Leister Patané, Vincent Louis Viala, Antonio Jorge Martins, Ricardo Haddad, Claudia Renata dos Santos Barros, Elaine Cristina Marqueze, Raul Machado Neto, Debora Botequio Moretti, Bibiana Santos, João Paulo Kitajima, Erika Freitas, David Schlesinger, Simone Kashima, Evandra Strazza Rodrigues, Svetoslav Nanev Slavov, Elaine Vieira dos Santos, Rafael dos Santos Bezerra, Luiz Carlos Junior de Alcantara, Marta Giovanetti, Vagner Fonseca, Flavia Aburjaile, Rodrigo Tocantins Calado. |
| EPI_ISL_1445082 | SECRETARIA MUNICIPAL DE SAUDE SOROCABA | Instituto Butantan / Mendelics | Dimas Tadeu Covas, Sandra Coccuzzo Sampaio, Maria Carolina Elias, José Salvatore Leister Patané, Vincent Louis Viala, Antonio Jorge Martins, Ricardo Haddad, Claudia Renata dos Santos Barros, Elaine Cristina Marqueze, Raul Machado Neto, Debora Botequio Moretti, Bibiana Santos, João Paulo |

| | | | |
|---|---|---|---|
| | | | Kitajima, Erika Freitas, David Schlesinger, Simone Kashima, Evandra Strazza Rodrigues, Svetoslav Nanev Slavov, Elaine Vieira dos Santos, Rafael dos Santos Bezerra, Luiz Carlos Junior de Alcantara, Marta Giovanetti, Vagner Fonseca, Flavia Aburjaile, Rodrigo Tocantins Calado. |
| EPI_ISL_1445084 | SMS SECRETARIA MUNICIPAL DE SAUDE DE BOITUVA | Instituto Butantan / Mendelics | Dimas Tadeu Covas, Sandra Coccuzzo Sampaio, Maria Carolina Elias, José Salvatore Leister Patané, Vincent Louis Viala, Antonio Jorge Martins, Ricardo Haddad, Claudia Renata dos Santos Barros, Elaine Cristina Marqueze, Raul Machado Neto, Debora Botequio Moretti, Bibiana Santos, João Paulo Kitajima, Erika Freitas, David Schlesinger, Simone Kashima, Evandra Strazza Rodrigues, Svetoslav Nanev Slavov, Elaine Vieira dos Santos, Rafael dos Santos Bezerra, Luiz Carlos Junior de Alcantara, Marta Giovanetti, Vagner Fonseca, Flavia Aburjaile, Rodrigo Tocantins Calado. |
| EPI_ISL_1445088 | SECRETARIA MUNICIPAL DE SAUDE SOROCABA | Instituto Butantan / Mendelics | Dimas Tadeu Covas, Sandra Coccuzzo Sampaio, Maria Carolina Elias, José Salvatore Leister Patané, Vincent Louis Viala, Antonio Jorge Martins, Ricardo Haddad, Claudia Renata dos Santos Barros, Elaine Cristina Marqueze, Raul Machado Neto, Debora Botequio Moretti, Bibiana Santos, João Paulo Kitajima, Erika Freitas, David Schlesinger, Simone Kashima, Evandra Strazza Rodrigues, Svetoslav Nanev Slavov, Elaine Vieira dos Santos, Rafael dos Santos Bezerra, Luiz Carlos Junior de Alcantara, Marta Giovanetti, Vagner Fonseca, Flavia Aburjaile, Rodrigo Tocantins Calado. |
| EPI_ISL_1445089 | CENTRO DE SAUDE II MAIRINQUE MAIRINQUE | Instituto Butantan / Mendelics | Dimas Tadeu Covas, Sandra Coccuzzo Sampaio, Maria Carolina Elias, José Salvatore Leister Patané, Vincent Louis Viala, Antonio Jorge Martins, Ricardo Haddad, Claudia Renata dos Santos Barros, Elaine Cristina Marqueze, Raul Machado Neto, Debora Botequio Moretti, Bibiana Santos, João Paulo Kitajima, Erika Freitas, David Schlesinger, Simone Kashima, Evandra Strazza Rodrigues, Svetoslav Nanev Slavov, Elaine Vieira dos Santos, Rafael dos Santos Bezerra, Luiz Carlos Junior de Alcantara, Marta Giovanetti, Vagner Fonseca, Flavia Aburjaile, Rodrigo Tocantins Calado. |
| EPI_ISL_1445090 | POLICLINICA COVID 19 ITAPETININGA | Instituto Butantan / Mendelics | Dimas Tadeu Covas, Sandra Coccuzzo Sampaio, Maria Carolina Elias, José Salvatore Leister Patané, Vincent Louis Viala, Antonio Jorge Martins, Ricardo Haddad, Claudia Renata dos Santos Barros, Elaine Cristina Marqueze, Raul Machado Neto, Debora Botequio Moretti, Bibiana Santos, João Paulo Kitajima, Erika Freitas, David Schlesinger, Simone Kashima, Evandra Strazza Rodrigues, Svetoslav Nanev Slavov, Elaine Vieira dos Santos, Rafael dos Santos Bezerra, Luiz Carlos Junior de Alcantara, Marta Giovanetti, Vagner Fonseca, Flavia Aburjaile, Rodrigo Tocantins Calado. |
| EPI_ISL_1445091, EPI_ISL_1445092, EPI_ISL_1445093, EPI_ISL_1445094, EPI_ISL_1445095 | SECRETARIA MUNICIPAL DE SAUDE DE PIRACAIA | Instituto Butantan / Mendelics | Dimas Tadeu Covas, Sandra Coccuzzo Sampaio, Maria Carolina Elias, José Salvatore Leister Patané, Vincent Louis Viala, Antonio Jorge Martins, Ricardo Haddad, Claudia Renata dos Santos Barros, Elaine Cristina Marqueze, Raul Machado Neto, Debora Botequio Moretti, Bibiana Santos, João Paulo Kitajima, Erika Freitas, David Schlesinger, Simone Kashima, Evandra Strazza Rodrigues, Svetoslav Nanev Slavov, Elaine Vieira dos Santos, Rafael dos Santos Bezerra, Luiz Carlos Junior de Alcantara, Marta Giovanetti, Vagner Fonseca, Flavia Aburjaile, Rodrigo Tocantins Calado. |
| EPI_ISL_1445096, EPI_ISL_1445097, EPI_ISL_1445098, EPI_ISL_1445099, EPI_ISL_1445100 | PRONTO ATENDIMENTO VILA PADRE ANCHIETA | Instituto Butantan / Mendelics | Dimas Tadeu Covas, Sandra Coccuzzo Sampaio, Maria Carolina Elias, José Salvatore Leister Patané, Vincent Louis Viala, Antonio Jorge Martins, Ricardo Haddad, Claudia Renata dos Santos Barros, Elaine Cristina Marqueze, Raul Machado Neto, Debora Botequio Moretti, Bibiana Santos, João Paulo Kitajima, Erika Freitas, David Schlesinger, Simone Kashima, Evandra Strazza Rodrigues, Svetoslav Nanev Slavov, Elaine Vieira dos Santos, Rafael dos Santos Bezerra, Luiz Carlos Junior de Alcantara, Marta Giovanetti, Vagner Fonseca, Flavia Aburjaile, Rodrigo Tocantins Calado. |
| EPI_ISL_1445101 | SECRETARIA MUNICIPAL DE SAUDE DE PIRACAIA | Instituto Butantan / Mendelics | Dimas Tadeu Covas, Sandra Coccuzzo Sampaio, Maria Carolina Elias, José Salvatore Leister Patané, Vincent Louis Viala, Antonio Jorge Martins, Ricardo Haddad, Claudia Renata dos Santos Barros, Elaine Cristina Marqueze, Raul Machado Neto, Debora Botequio Moretti, Bibiana Santos, João Paulo Kitajima, Erika Freitas, David Schlesinger, Simone Kashima, Evandra Strazza Rodrigues, Svetoslav Nanev Slavov, Elaine Vieira dos Santos, Rafael dos Santos Bezerra, Luiz Carlos Junior de Alcantara, Marta Giovanetti, Vagner Fonseca, Flavia Aburjaile, Rodrigo Tocantins Calado. |
| EPI_ISL_1445102, EPI_ISL_1445103, EPI_ISL_1445104, EPI_ISL_1445105, EPI_ISL_1445106, EPI_ISL_1445107, EPI_ISL_1445108 | PRONTO ATENDIMENTO VILA PADRE ANCHIETA | Instituto Butantan / Mendelics | Dimas Tadeu Covas, Sandra Coccuzzo Sampaio, Maria Carolina Elias, José Salvatore Leister Patané, Vincent Louis Viala, Antonio Jorge Martins, Ricardo Haddad, Claudia Renata dos Santos Barros, Elaine Cristina Marqueze, Raul Machado Neto, Debora Botequio Moretti, Bibiana Santos, João Paulo Kitajima, Erika Freitas, David Schlesinger, Simone Kashima, Evandra Strazza Rodrigues, Svetoslav Nanev Slavov, Elaine Vieira dos Santos, Rafael dos Santos Bezerra, Luiz Carlos Junior de Alcantara, Marta Giovanetti, Vagner Fonseca, Flavia Aburjaile, Rodrigo Tocantins Calado. |
| EPI_ISL_1445109, EPI_ISL_1445110 | COMPLEXO HOSPITALAR OURO VERDE DE CAMPINAS | Instituto Butantan / Mendelics | Dimas Tadeu Covas, Sandra Coccuzzo Sampaio, Maria Carolina Elias, José Salvatore Leister Patané, Vincent Louis Viala, Antonio Jorge Martins, Ricardo Haddad, Claudia Renata dos Santos Barros, Elaine Cristina Marqueze, Raul Machado Neto, Debora Botequio Moretti, Bibiana Santos, João Paulo Kitajima, Erika Freitas, David Schlesinger, Simone Kashima, Evandra Strazza Rodrigues, Svetoslav Nanev Slavov, Elaine Vieira dos Santos, Rafael dos Santos Bezerra, Luiz Carlos Junior de Alcantara, Marta Giovanetti, Vagner Fonseca, Flavia Aburjaile, Rodrigo Tocantins Calado. |
| EPI_ISL_1445111, EPI_ISL_1445112 | PRONTO ATENDIMENTO VILA PADRE ANCHIETA | Instituto Butantan / Mendelics | Dimas Tadeu Covas, Sandra Coccuzzo Sampaio, Maria Carolina Elias, José Salvatore Leister Patané, Vincent Louis Viala, Antonio Jorge Martins, Ricardo Haddad, Claudia Renata dos Santos Barros, Elaine Cristina Marqueze, Raul Machado Neto, Debora Botequio Moretti, Bibiana Santos, João Paulo Kitajima, Erika Freitas, David Schlesinger, Simone Kashima, Evandra Strazza Rodrigues, Svetoslav Nanev Slavov, Elaine Vieira dos Santos, Rafael dos Santos Bezerra, Luiz Carlos Junior de Alcantara, Marta Giovanetti, Vagner Fonseca, Flavia Aburjaile, Rodrigo Tocantins Calado. |
| EPI_ISL_1445113 | UNIDADE RESPIRATORIA NOVA HORTOLANDIA | Instituto Butantan / Mendelics | Dimas Tadeu Covas, Sandra Coccuzzo Sampaio, Maria Carolina Elias, José Salvatore Leister Patané, Vincent Louis Viala, Antonio Jorge Martins, Ricardo Haddad, Claudia Renata dos Santos Barros, Elaine Cristina Marqueze, Raul Machado Neto, Debora Botequio Moretti, Bibiana Santos, João Paulo Kitajima, Erika Freitas, David Schlesinger, Simone Kashima, Evandra Strazza Rodrigues, Svetoslav Nanev Slavov, Elaine Vieira dos Santos, Rafael dos Santos Bezerra, Luiz Carlos Junior de Alcantara, Marta Giovanetti, Vagner Fonseca, Flavia Aburjaile, Rodrigo Tocantins Calado. |
| EPI_ISL_1445114, EPI_ISL_1445115, EPI_ISL_1445116, EPI_ISL_1445117, EPI_ISL_1445118 | COMPLEXO HOSPITALAR OURO VERDE DE CAMPINAS | Instituto Butantan / Mendelics | Dimas Tadeu Covas, Sandra Coccuzzo Sampaio, Maria Carolina Elias, José Salvatore Leister Patané, Vincent Louis Viala, Antonio Jorge Martins, Ricardo Haddad, Claudia Renata dos Santos Barros, Elaine Cristina Marqueze, Raul Machado Neto, Debora Botequio Moretti, Bibiana Santos, João Paulo Kitajima, Erika Freitas, David Schlesinger, Simone Kashima, Evandra Strazza Rodrigues, Svetoslav Nanev Slavov, Elaine Vieira dos Santos, Rafael dos Santos Bezerra, Luiz Carlos Junior de Alcantara, Marta Giovanetti, Vagner Fonseca, Flavia Aburjaile, Rodrigo Tocantins Calado. |
| EPI_ISL_1445119 | UNIDADE RESPIRATORIA NOVA HORTOLANDIA | Instituto Butantan / Mendelics | Dimas Tadeu Covas, Sandra Coccuzzo Sampaio, Maria Carolina Elias, José Salvatore Leister Patané, Vincent Louis Viala, Antonio Jorge Martins, Ricardo Haddad, Claudia Renata dos Santos Barros, Elaine Cristina Marqueze, Raul Machado Neto, Debora Botequio Moretti, Bibiana Santos, João Paulo Kitajima, Erika Freitas, David Schlesinger, Simone Kashima, Evandra Strazza Rodrigues, Svetoslav Nanev Slavov, Elaine Vieira dos Santos, Rafael dos Santos Bezerra, Luiz Carlos Junior de Alcantara, Marta Giovanetti, Vagner Fonseca, Flavia Aburjaile, Rodrigo Tocantins Calado. |
| EPI_ISL_1445120, EPI_ISL_1445121, EPI_ISL_1445122, EPI_ISL_1445123, EPI_ISL_1445124, EPI_ISL_1445125, EPI_ISL_1445126 | UNIDADE DE PRONTO ATENDIMENTO UPA DRA ANA OLIVIA BENTIVOGLIO | Instituto Butantan / Mendelics | Dimas Tadeu Covas, Sandra Coccuzzo Sampaio, Maria Carolina Elias, José Salvatore Leister Patané, Vincent Louis Viala, Antonio Jorge Martins, Ricardo Haddad, Claudia Renata dos Santos Barros, Elaine Cristina Marqueze, Raul Machado Neto, Debora Botequio Moretti, Bibiana Santos, João Paulo Kitajima, Erika Freitas, David Schlesinger, Simone Kashima, Evandra Strazza Rodrigues, Svetoslav Nanev Slavov, Elaine Vieira dos Santos, Rafael dos Santos Bezerra, Luiz Carlos Junior de Alcantara, Marta Giovanetti, Vagner Fonseca, Flavia Aburjaile, Rodrigo Tocantins Calado. |
| EPI_ISL_1445127, EPI_ISL_1445128, EPI_ISL_1445129 | UNIDADE RESPIRATORIA NOVA HORTOLANDIA | Instituto Butantan / Mendelics | Dimas Tadeu Covas, Sandra Coccuzzo Sampaio, Maria Carolina Elias, José Salvatore Leister Patané, Vincent Louis Viala, Antonio Jorge Martins, Ricardo Haddad, Claudia Renata dos Santos Barros, Elaine Cristina Marqueze, Raul Machado Neto, Debora Botequio Moretti, Bibiana Santos, João Paulo Kitajima, Erika Freitas, David Schlesinger, Simone Kashima, Evandra Strazza Rodrigues, Svetoslav Nanev Slavov, Elaine Vieira dos Santos, Rafael dos Santos Bezerra, Luiz Carlos Junior de Alcantara, Marta Giovanetti, Vagner Fonseca, Flavia Aburjaile, Rodrigo Tocantins Calado. |
| EPI_ISL_1445130, EPI_ISL_1445131, EPI_ISL_1445132 | PRONTO ATENDIMENTO DA PONTE DE SAO JOAO | Instituto Butantan / Mendelics | Dimas Tadeu Covas, Sandra Coccuzzo Sampaio, Maria Carolina Elias, José Salvatore Leister Patané, Vincent Louis Viala, Antonio Jorge Martins, Ricardo Haddad, Claudia Renata dos Santos Barros, Elaine Cristina Marqueze, Raul Machado Neto, Debora Botequio Moretti, Bibiana Santos, João Paulo Kitajima, Erika Freitas, David Schlesinger, Simone Kashima, Evandra Strazza Rodrigues, Svetoslav Nanev Slavov, Elaine Vieira dos Santos, Rafael dos Santos Bezerra, Luiz Carlos Junior de Alcantara, Marta Giovanetti, Vagner Fonseca, Flavia Aburjaile, Rodrigo Tocantins Calado. |
| EPI_ISL_1445133, EPI_ISL_1445134, EPI_ISL_1445135 | SECRETARIA MUNICIPAL DE SAUDE SOROCABA | Instituto Butantan / Mendelics | Dimas Tadeu Covas, Sandra Coccuzzo Sampaio, Maria Carolina Elias, José Salvatore Leister Patané, Vincent Louis Viala, Antonio Jorge Martins, Ricardo Haddad, Claudia Renata dos Santos Barros, Elaine Cristina Marqueze, Raul Machado Neto, Debora Botequio Moretti, Bibiana Santos, João Paulo Kitajima, Erika Freitas, David Schlesinger, Simone Kashima, Evandra Strazza Rodrigues, Svetoslav Nanev Slavov, Elaine Vieira dos Santos, Rafael dos Santos Bezerra, Luiz Carlos Junior de Alcantara, Marta Giovanetti, Vagner Fonseca, Flavia Aburjaile, Rodrigo Tocantins Calado. |
| EPI_ISL_1445136 | SMS IPERO | Instituto Butantan / Mendelics | Dimas Tadeu Covas, Sandra Coccuzzo Sampaio, Maria Carolina Elias, José Salvatore Leister Patané, Vincent Louis Viala, Antonio Jorge Martins, Ricardo Haddad, Claudia Renata dos Santos Barros, Elaine Cristina Marqueze, Raul Machado Neto, Debora Botequio Moretti, Bibiana Santos, João Paulo Kitajima, Erika Freitas, David Schlesinger, Simone Kashima, Evandra Strazza Rodrigues, Svetoslav Nanev Slavov, Elaine Vieira dos Santos, Rafael dos Santos Bezerra, Luiz Carlos Junior de Alcantara, Marta Giovanetti, Vagner Fonseca, Flavia Aburjaile, Rodrigo Tocantins Calado. |
| EPI_ISL_1445137, EPI_ISL_1445138, EPI_ISL_1445139 | SECRETARIA MUNICIPAL DE SAUDE SOROCABA | Instituto Butantan / Mendelics | Dimas Tadeu Covas, Sandra Coccuzzo Sampaio, Maria Carolina Elias, José Salvatore Leister Patané, Vincent Louis Viala, Antonio Jorge Martins, Ricardo Haddad, Claudia Renata dos Santos Barros, Elaine Cristina Marqueze, Raul Machado Neto, Debora Botequio Moretti, Bibiana Santos, João Paulo Kitajima, Erika Freitas, David Schlesinger, Simone Kashima, Evandra Strazza Rodrigues, Svetoslav Nanev Slavov, Elaine Vieira dos Santos, Rafael dos Santos Bezerra, Luiz Carlos Junior de Alcantara, Marta Giovanetti, Vagner Fonseca, Flavia Aburjaile, Rodrigo Tocantins Calado. |

| | | | |
|---|---|---|---|
| EPI_ISL_1445140 | UNIDADE MISTA DE SAUDE CAPELA DO ALTO | Instituto Butantan / Mendelics | Dimas Tadeu Covas, Sandra Coccuzzo Sampaio, Maria Carolina Elias, José Salvatore Leister Patané, Vincent Louis Viala, Antonio Jorge Martins, Ricardo Haddad, Claudia Renata dos Santos Barros, Elaine Cristina Marqueze, Raul Machado Neto, Debora Botequio Moretti, Bibiana Santos, João Paulo Kitajima, Erika Freitas, David Schlesinger, Simone Kashima, Evandra Strazza Rodrigues, Svetoslav Nanev Slavov, Elaine Vieira dos Santos, Rafael dos Santos Bezerra, Luiz Carlos Junior de Alcantara, Marta Giovanetti, Vagner Fonseca, Flavia Aburjaile, Rodrigo Tocantins Calado. |
| EPI_ISL_1445142, EPI_ISL_1445143, EPI_ISL_1445144, EPI_ISL_1445145, EPI_ISL_1445146 | CS II EGIDIO BRUNHARA MORRO AGUDO | Instituto Butantan / Mendelics | Dimas Tadeu Covas, Sandra Coccuzzo Sampaio, Maria Carolina Elias, José Salvatore Leister Patané, Vincent Louis Viala, Antonio Jorge Martins, Ricardo Haddad, Claudia Renata dos Santos Barros, Elaine Cristina Marqueze, Raul Machado Neto, Debora Botequio Moretti, Bibiana Santos, João Paulo Kitajima, Erika Freitas, David Schlesinger, Simone Kashima, Evandra Strazza Rodrigues, Svetoslav Nanev Slavov, Elaine Vieira dos Santos, Rafael dos Santos Bezerra, Luiz Carlos Junior de Alcantara, Marta Giovanetti, Vagner Fonseca, Flavia Aburjaile, Rodrigo Tocantins Calado. |
| EPI_ISL_1445147, EPI_ISL_1445148, EPI_ISL_1445149, EPI_ISL_1445150, EPI_ISL_1445151, EPI_ISL_1445152 | CENTRO DE REFERENCIA DO IDOSO DR HUMBERTO MENDES DE CARVALHO | Instituto Butantan / Mendelics | Dimas Tadeu Covas, Sandra Coccuzzo Sampaio, Maria Carolina Elias, José Salvatore Leister Patané, Vincent Louis Viala, Antonio Jorge Martins, Ricardo Haddad, Claudia Renata dos Santos Barros, Elaine Cristina Marqueze, Raul Machado Neto, Debora Botequio Moretti, Bibiana Santos, João Paulo Kitajima, Erika Freitas, David Schlesinger, Simone Kashima, Evandra Strazza Rodrigues, Svetoslav Nanev Slavov, Elaine Vieira dos Santos, Rafael dos Santos Bezerra, Luiz Carlos Junior de Alcantara, Marta Giovanetti, Vagner Fonseca, Flavia Aburjaile, Rodrigo Tocantins Calado. |
| EPI_ISL_1445153, EPI_ISL_1445154 | SAE SERVICO DE ATENDIMENTO ESPECIALIZADO | Instituto Butantan / Mendelics | Dimas Tadeu Covas, Sandra Coccuzzo Sampaio, Maria Carolina Elias, José Salvatore Leister Patané, Vincent Louis Viala, Antonio Jorge Martins, Ricardo Haddad, Claudia Renata dos Santos Barros, Elaine Cristina Marqueze, Raul Machado Neto, Debora Botequio Moretti, Bibiana Santos, João Paulo Kitajima, Erika Freitas, David Schlesinger, Simone Kashima, Evandra Strazza Rodrigues, Svetoslav Nanev Slavov, Elaine Vieira dos Santos, Rafael dos Santos Bezerra, Luiz Carlos Junior de Alcantara, Marta Giovanetti, Vagner Fonseca, Flavia Aburjaile, Rodrigo Tocantins Calado. |
| EPI_ISL_1445155 | LABORATORIO DE FRANCA | Instituto Butantan / Mendelics | Dimas Tadeu Covas, Sandra Coccuzzo Sampaio, Maria Carolina Elias, José Salvatore Leister Patané, Vincent Louis Viala, Antonio Jorge Martins, Ricardo Haddad, Claudia Renata dos Santos Barros, Elaine Cristina Marqueze, Raul Machado Neto, Debora Botequio Moretti, Bibiana Santos, João Paulo Kitajima, Erika Freitas, David Schlesinger, Simone Kashima, Evandra Strazza Rodrigues, Svetoslav Nanev Slavov, Elaine Vieira dos Santos, Rafael dos Santos Bezerra, Luiz Carlos Junior de Alcantara, Marta Giovanetti, Vagner Fonseca, Flavia Aburjaile, Rodrigo Tocantins Calado. |
| EPI_ISL_1445156 | SAE SERVICO DE ATENDIMENTO ESPECIALIZADO | Instituto Butantan / Mendelics | Dimas Tadeu Covas, Sandra Coccuzzo Sampaio, Maria Carolina Elias, José Salvatore Leister Patané, Vincent Louis Viala, Antonio Jorge Martins, Ricardo Haddad, Claudia Renata dos Santos Barros, Elaine Cristina Marqueze, Raul Machado Neto, Debora Botequio Moretti, Bibiana Santos, João Paulo Kitajima, Erika Freitas, David Schlesinger, Simone Kashima, Evandra Strazza Rodrigues, Svetoslav Nanev Slavov, Elaine Vieira dos Santos, Rafael dos Santos Bezerra, Luiz Carlos Junior de Alcantara, Marta Giovanetti, Vagner Fonseca, Flavia Aburjaile, Rodrigo Tocantins Calado. |
| EPI_ISL_1445157, EPI_ISL_1445158, EPI_ISL_1445159, EPI_ISL_1445160, EPI_ISL_1445161 | SECAO CENTRO DE DIAGNOSTICO SECEDI | Instituto Butantan / Mendelics | Dimas Tadeu Covas, Sandra Coccuzzo Sampaio, Maria Carolina Elias, José Salvatore Leister Patané, Vincent Louis Viala, Antonio Jorge Martins, Ricardo Haddad, Claudia Renata dos Santos Barros, Elaine Cristina Marqueze, Raul Machado Neto, Debora Botequio Moretti, Bibiana Santos, João Paulo Kitajima, Erika Freitas, David Schlesinger, Simone Kashima, Evandra Strazza Rodrigues, Svetoslav Nanev Slavov, Elaine Vieira dos Santos, Rafael dos Santos Bezerra, Luiz Carlos Junior de Alcantara, Marta Giovanetti, Vagner Fonseca, Flavia Aburjaile, Rodrigo Tocantins Calado. |
| EPI_ISL_1445162, EPI_ISL_1445163, EPI_ISL_1445164, EPI_ISL_1445165, EPI_ISL_1445166, EPI_ISL_1445167 | SERV DE VIG SANITARIA EPIDEMIO E CTRL DE ZOONOZES GUARUJA | Instituto Butantan / Mendelics | Dimas Tadeu Covas, Sandra Coccuzzo Sampaio, Maria Carolina Elias, José Salvatore Leister Patané, Vincent Louis Viala, Antonio Jorge Martins, Ricardo Haddad, Claudia Renata dos Santos Barros, Elaine Cristina Marqueze, Raul Machado Neto, Debora Botequio Moretti, Bibiana Santos, João Paulo Kitajima, Erika Freitas, David Schlesinger, Simone Kashima, Evandra Strazza Rodrigues, Svetoslav Nanev Slavov, Elaine Vieira dos Santos, Rafael dos Santos Bezerra, Luiz Carlos Junior de Alcantara, Marta Giovanetti, Vagner Fonseca, Flavia Aburjaile, Rodrigo Tocantins Calado. |
| EPI_ISL_1445168, EPI_ISL_1445169, EPI_ISL_1445170 | SECAO CENTRO DE DIAGNOSTICO SECEDI | Instituto Butantan / Mendelics | Dimas Tadeu Covas, Sandra Coccuzzo Sampaio, Maria Carolina Elias, José Salvatore Leister Patané, Vincent Louis Viala, Antonio Jorge Martins, Ricardo Haddad, Claudia Renata dos Santos Barros, Elaine Cristina Marqueze, Raul Machado Neto, Debora Botequio Moretti, Bibiana Santos, João Paulo Kitajima, Erika Freitas, David Schlesinger, Simone Kashima, Evandra Strazza Rodrigues, Svetoslav Nanev Slavov, Elaine Vieira dos Santos, Rafael dos Santos Bezerra, Luiz Carlos Junior de Alcantara, Marta Giovanetti, Vagner Fonseca, Flavia Aburjaile, Rodrigo Tocantins Calado. |
| EPI_ISL_1445171 | SECRETARIA MUNICIPAL DE SAUDE SOROCABA | Instituto Butantan / Mendelics | Dimas Tadeu Covas, Sandra Coccuzzo Sampaio, Maria Carolina Elias, José Salvatore Leister Patané, Vincent Louis Viala, Antonio Jorge Martins, Ricardo Haddad, Claudia Renata dos Santos Barros, Elaine Cristina Marqueze, Raul Machado Neto, Debora Botequio Moretti, Bibiana Santos, João Paulo Kitajima, Erika Freitas, David Schlesinger, Simone Kashima, Evandra Strazza Rodrigues, Svetoslav Nanev Slavov, Elaine Vieira dos Santos, Rafael dos Santos Bezerra, Luiz Carlos Junior de Alcantara, Marta Giovanetti, Vagner Fonseca, Flavia Aburjaile, Rodrigo Tocantins Calado. |
| EPI_ISL_1445174 | PRONTO SOCORRO MUNICIPAL DE TATUI | Instituto Butantan / Mendelics | Dimas Tadeu Covas, Sandra Coccuzzo Sampaio, Maria Carolina Elias, José Salvatore Leister Patané, Vincent Louis Viala, Antonio Jorge Martins, Ricardo Haddad, Claudia Renata dos Santos Barros, Elaine Cristina Marqueze, Raul Machado Neto, Debora Botequio Moretti, Bibiana Santos, João Paulo Kitajima, Erika Freitas, David Schlesinger, Simone Kashima, Evandra Strazza Rodrigues, Svetoslav Nanev Slavov, Elaine Vieira dos Santos, Rafael dos Santos Bezerra, Luiz Carlos Junior de Alcantara, Marta Giovanetti, Vagner Fonseca, Flavia Aburjaile, Rodrigo Tocantins Calado. |
| EPI_ISL_1445175 | DEPARTAMENTO DE SAUDE COLETIVA | Instituto Butantan / Mendelics | Dimas Tadeu Covas, Sandra Coccuzzo Sampaio, Maria Carolina Elias, José Salvatore Leister Patané, Vincent Louis Viala, Antonio Jorge Martins, Ricardo Haddad, Claudia Renata dos Santos Barros, Elaine Cristina Marqueze, Raul Machado Neto, Debora Botequio Moretti, Bibiana Santos, João Paulo Kitajima, Erika Freitas, David Schlesinger, Simone Kashima, Evandra Strazza Rodrigues, Svetoslav Nanev Slavov, Elaine Vieira dos Santos, Rafael dos Santos Bezerra, Luiz Carlos Junior de Alcantara, Marta Giovanetti, Vagner Fonseca, Flavia Aburjaile, Rodrigo Tocantins Calado. |
| EPI_ISL_1445176 | SECRETARIA MUNICIPAL DE SAUDE SOROCABA | Instituto Butantan / Mendelics | Dimas Tadeu Covas, Sandra Coccuzzo Sampaio, Maria Carolina Elias, José Salvatore Leister Patané, Vincent Louis Viala, Antonio Jorge Martins, Ricardo Haddad, Claudia Renata dos Santos Barros, Elaine Cristina Marqueze, Raul Machado Neto, Debora Botequio Moretti, Bibiana Santos, João Paulo Kitajima, Erika Freitas, David Schlesinger, Simone Kashima, Evandra Strazza Rodrigues, Svetoslav Nanev Slavov, Elaine Vieira dos Santos, Rafael dos Santos Bezerra, Luiz Carlos Junior de Alcantara, Marta Giovanetti, Vagner Fonseca, Flavia Aburjaile, Rodrigo Tocantins Calado. |
| EPI_ISL_1445177 | SAUDE COLETIVA CAPAO BONITO | Instituto Butantan / Mendelics | Dimas Tadeu Covas, Sandra Coccuzzo Sampaio, Maria Carolina Elias, José Salvatore Leister Patané, Vincent Louis Viala, Antonio Jorge Martins, Ricardo Haddad, Claudia Renata dos Santos Barros, Elaine Cristina Marqueze, Raul Machado Neto, Debora Botequio Moretti, Bibiana Santos, João Paulo Kitajima, Erika Freitas, David Schlesinger, Simone Kashima, Evandra Strazza Rodrigues, Svetoslav Nanev Slavov, Elaine Vieira dos Santos, Rafael dos Santos Bezerra, Luiz Carlos Junior de Alcantara, Marta Giovanetti, Vagner Fonseca, Flavia Aburjaile, Rodrigo Tocantins Calado. |
| EPI_ISL_1445178, EPI_ISL_1445180 | SECRETARIA MUNICIPAL DE SAUDE SOROCABA | Instituto Butantan / Mendelics | Dimas Tadeu Covas, Sandra Coccuzzo Sampaio, Maria Carolina Elias, José Salvatore Leister Patané, Vincent Louis Viala, Antonio Jorge Martins, Ricardo Haddad, Claudia Renata dos Santos Barros, Elaine Cristina Marqueze, Raul Machado Neto, Debora Botequio Moretti, Bibiana Santos, João Paulo Kitajima, Erika Freitas, David Schlesinger, Simone Kashima, Evandra Strazza Rodrigues, Svetoslav Nanev Slavov, Elaine Vieira dos Santos, Rafael dos Santos Bezerra, Luiz Carlos Junior de Alcantara, Marta Giovanetti, Vagner Fonseca, Flavia Aburjaile, Rodrigo Tocantins Calado. |
| EPI_ISL_1445181 | HOSPITAL REGIONAL DE ITAPETININGA | Instituto Butantan / Mendelics | Dimas Tadeu Covas, Sandra Coccuzzo Sampaio, Maria Carolina Elias, José Salvatore Leister Patané, Vincent Louis Viala, Antonio Jorge Martins, Ricardo Haddad, Claudia Renata dos Santos Barros, Elaine Cristina Marqueze, Raul Machado Neto, Debora Botequio Moretti, Bibiana Santos, João Paulo Kitajima, Erika Freitas, David Schlesinger, Simone Kashima, Evandra Strazza Rodrigues, Svetoslav Nanev Slavov, Elaine Vieira dos Santos, Rafael dos Santos Bezerra, Luiz Carlos Junior de Alcantara, Marta Giovanetti, Vagner Fonseca, Flavia Aburjaile, Rodrigo Tocantins Calado. |
| EPI_ISL_1445182 | SECRETARIA MUNICIPAL DE SAUDE SOROCABA | Instituto Butantan / Mendelics | Dimas Tadeu Covas, Sandra Coccuzzo Sampaio, Maria Carolina Elias, José Salvatore Leister Patané, Vincent Louis Viala, Antonio Jorge Martins, Ricardo Haddad, Claudia Renata dos Santos Barros, Elaine Cristina Marqueze, Raul Machado Neto, Debora Botequio Moretti, Bibiana Santos, João Paulo Kitajima, Erika Freitas, David Schlesinger, Simone Kashima, Evandra Strazza Rodrigues, Svetoslav Nanev Slavov, Elaine Vieira dos Santos, Rafael dos Santos Bezerra, Luiz Carlos Junior de Alcantara, Marta Giovanetti, Vagner Fonseca, Flavia Aburjaile, Rodrigo Tocantins Calado. |
| EPI_ISL_1445183, EPI_ISL_1445184 | UNIDADE BASICA DE SAUDE DA FAMILIA BILAC | Instituto Butantan / Mendelics | Dimas Tadeu Covas, Sandra Coccuzzo Sampaio, Maria Carolina Elias, José Salvatore Leister Patané, Vincent Louis Viala, Antonio Jorge Martins, Ricardo Haddad, Claudia Renata dos Santos Barros, Elaine Cristina Marqueze, Raul Machado Neto, Debora Botequio Moretti, Bibiana Santos, João Paulo Kitajima, Erika Freitas, David Schlesinger, Simone Kashima, Evandra Strazza Rodrigues, Svetoslav Nanev Slavov, Elaine Vieira dos Santos, Rafael dos Santos Bezerra, Luiz Carlos Junior de Alcantara, Marta Giovanetti, Vagner Fonseca, Flavia Aburjaile, Rodrigo Tocantins Calado. |
| EPI_ISL_1445185 | PRONTO ATENDIMENTO SAO JOSE | Instituto Butantan / Mendelics | Dimas Tadeu Covas, Sandra Coccuzzo Sampaio, Maria Carolina Elias, José Salvatore Leister Patané, Vincent Louis Viala, Antonio Jorge Martins, Ricardo Haddad, Claudia Renata dos Santos Barros, Elaine Cristina Marqueze, Raul Machado Neto, Debora Botequio Moretti, Bibiana Santos, João Paulo Kitajima, Erika Freitas, David Schlesinger, Simone Kashima, Evandra Strazza Rodrigues, Svetoslav Nanev Slavov, Elaine Vieira dos Santos, Rafael dos Santos Bezerra, Luiz Carlos Junior de Alcantara, Marta Giovanetti, Vagner Fonseca, Flavia Aburjaile, Rodrigo Tocantins Calado. |
| EPI_ISL_1445186 | PRONTO ATENDIMENTO VILA PADRE ANCHIETA | Instituto Butantan / Mendelics | Dimas Tadeu Covas, Sandra Coccuzzo Sampaio, Maria Carolina Elias, José Salvatore Leister Patané, Vincent Louis Viala, Antonio Jorge Martins, Ricardo Haddad, Claudia Renata dos Santos Barros, Elaine Cristina Marqueze, Raul Machado Neto, Debora Botequio Moretti, Bibiana Santos, João Paulo |

| | | | |
|---|---|---|---|
| | | | Kitajima, Erika Freitas, David Schlesinger, Simone Kashima, Evandra Strazza Rodrigues, Svetoslav Nanev Slavov, Elaine Vieira dos Santos, Rafael dos Santos Bezerra, Luiz Carlos Junior de Alcantara, Marta Giovanetti, Vagner Fonseca, Flavia Aburjaile, Rodrigo Tocantins Calado. |
| EPI_ISL_1445187 | PRONTO ATENDIMENTO SAO JOSE | Instituto Butantan / Mendelics | Dimas Tadeu Covas, Sandra Coccuzzo Sampaio, Maria Carolina Elias, José Salvatore Leister Patané, Vincent Louis Viala, Antonio Jorge Martins, Ricardo Haddad, Claudia Renata dos Santos Barros, Elaine Cristina Marqueze, Raul Machado Neto, Debora Botequio Moretti, Bibiana Santos, João Paulo Kitajima, Erika Freitas, David Schlesinger, Simone Kashima, Evandra Strazza Rodrigues, Svetoslav Nanev Slavov, Elaine Vieira dos Santos, Rafael dos Santos Bezerra, Luiz Carlos Junior de Alcantara, Marta Giovanetti, Vagner Fonseca, Flavia Aburjaile, Rodrigo Tocantins Calado. |
| EPI_ISL_1445188, EPI_ISL_1445189, EPI_ISL_1445190, EPI_ISL_1445191, EPI_ISL_1445192 | PRONTO ATENDIMENTO VILA PADRE ANCHIETA | Instituto Butantan / Mendelics | Dimas Tadeu Covas, Sandra Coccuzzo Sampaio, Maria Carolina Elias, José Salvatore Leister Patané, Vincent Louis Viala, Antonio Jorge Martins, Ricardo Haddad, Claudia Renata dos Santos Barros, Elaine Cristina Marqueze, Raul Machado Neto, Debora Botequio Moretti, Bibiana Santos, João Paulo Kitajima, Erika Freitas, David Schlesinger, Simone Kashima, Evandra Strazza Rodrigues, Svetoslav Nanev Slavov, Elaine Vieira dos Santos, Rafael dos Santos Bezerra, Luiz Carlos Junior de Alcantara, Marta Giovanetti, Vagner Fonseca, Flavia Aburjaile, Rodrigo Tocantins Calado. |
| EPI_ISL_1445193, EPI_ISL_1445194 | PRONTO ATENDIMENTO SAO JOSE | Instituto Butantan / Mendelics | Dimas Tadeu Covas, Sandra Coccuzzo Sampaio, Maria Carolina Elias, José Salvatore Leister Patané, Vincent Louis Viala, Antonio Jorge Martins, Ricardo Haddad, Claudia Renata dos Santos Barros, Elaine Cristina Marqueze, Raul Machado Neto, Debora Botequio Moretti, Bibiana Santos, João Paulo Kitajima, Erika Freitas, David Schlesinger, Simone Kashima, Evandra Strazza Rodrigues, Svetoslav Nanev Slavov, Elaine Vieira dos Santos, Rafael dos Santos Bezerra, Luiz Carlos Junior de Alcantara, Marta Giovanetti, Vagner Fonseca, Flavia Aburjaile, Rodrigo Tocantins Calado. |
| EPI_ISL_1445195 | CENTRO DE SAUDE DE BORA | Instituto Butantan / Mendelics | Dimas Tadeu Covas, Sandra Coccuzzo Sampaio, Maria Carolina Elias, José Salvatore Leister Patané, Vincent Louis Viala, Antonio Jorge Martins, Ricardo Haddad, Claudia Renata dos Santos Barros, Elaine Cristina Marqueze, Raul Machado Neto, Debora Botequio Moretti, Bibiana Santos, João Paulo Kitajima, Erika Freitas, David Schlesinger, Simone Kashima, Evandra Strazza Rodrigues, Svetoslav Nanev Slavov, Elaine Vieira dos Santos, Rafael dos Santos Bezerra, Luiz Carlos Junior de Alcantara, Marta Giovanetti, Vagner Fonseca, Flavia Aburjaile, Rodrigo Tocantins Calado. |
| EPI_ISL_1445196 | UBS SAO JUDAS | Instituto Butantan / Mendelics | Dimas Tadeu Covas, Sandra Coccuzzo Sampaio, Maria Carolina Elias, José Salvatore Leister Patané, Vincent Louis Viala, Antonio Jorge Martins, Ricardo Haddad, Claudia Renata dos Santos Barros, Elaine Cristina Marqueze, Raul Machado Neto, Debora Botequio Moretti, Bibiana Santos, João Paulo Kitajima, Erika Freitas, David Schlesinger, Simone Kashima, Evandra Strazza Rodrigues, Svetoslav Nanev Slavov, Elaine Vieira dos Santos, Rafael dos Santos Bezerra, Luiz Carlos Junior de Alcantara, Marta Giovanetti, Vagner Fonseca, Flavia Aburjaile, Rodrigo Tocantins Calado. |
| EPI_ISL_1445197 | CENTRO DE SAUDE DE JULIO MESQUITA | Instituto Butantan / Mendelics | Dimas Tadeu Covas, Sandra Coccuzzo Sampaio, Maria Carolina Elias, José Salvatore Leister Patané, Vincent Louis Viala, Antonio Jorge Martins, Ricardo Haddad, Claudia Renata dos Santos Barros, Elaine Cristina Marqueze, Raul Machado Neto, Debora Botequio Moretti, Bibiana Santos, João Paulo Kitajima, Erika Freitas, David Schlesinger, Simone Kashima, Evandra Strazza Rodrigues, Svetoslav Nanev Slavov, Elaine Vieira dos Santos, Rafael dos Santos Bezerra, Luiz Carlos Junior de Alcantara, Marta Giovanetti, Vagner Fonseca, Flavia Aburjaile, Rodrigo Tocantins Calado. |
| EPI_ISL_1445198 | UNIDADE MISTA DE TAIACU | Instituto Butantan / Mendelics | Dimas Tadeu Covas, Sandra Coccuzzo Sampaio, Maria Carolina Elias, José Salvatore Leister Patané, Vincent Louis Viala, Antonio Jorge Martins, Ricardo Haddad, Claudia Renata dos Santos Barros, Elaine Cristina Marqueze, Raul Machado Neto, Debora Botequio Moretti, Bibiana Santos, João Paulo Kitajima, Erika Freitas, David Schlesinger, Simone Kashima, Evandra Strazza Rodrigues, Svetoslav Nanev Slavov, Elaine Vieira dos Santos, Rafael dos Santos Bezerra, Luiz Carlos Junior de Alcantara, Marta Giovanetti, Vagner Fonseca, Flavia Aburjaile, Rodrigo Tocantins Calado. |
| EPI_ISL_1445199 | CENTRO DE SAUDE III SALES OLIVEIRA | Instituto Butantan / Mendelics | Dimas Tadeu Covas, Sandra Coccuzzo Sampaio, Maria Carolina Elias, José Salvatore Leister Patané, Vincent Louis Viala, Antonio Jorge Martins, Ricardo Haddad, Claudia Renata dos Santos Barros, Elaine Cristina Marqueze, Raul Machado Neto, Debora Botequio Moretti, Bibiana Santos, João Paulo Kitajima, Erika Freitas, David Schlesinger, Simone Kashima, Evandra Strazza Rodrigues, Svetoslav Nanev Slavov, Elaine Vieira dos Santos, Rafael dos Santos Bezerra, Luiz Carlos Junior de Alcantara, Marta Giovanetti, Vagner Fonseca, Flavia Aburjaile, Rodrigo Tocantins Calado. |
| EPI_ISL_1445200 | HOSPITAL E MATERNIDADE NOSSA SENHORA DA AJUDA | Instituto Butantan / Mendelics | Dimas Tadeu Covas, Sandra Coccuzzo Sampaio, Maria Carolina Elias, José Salvatore Leister Patané, Vincent Louis Viala, Antonio Jorge Martins, Ricardo Haddad, Claudia Renata dos Santos Barros, Elaine Cristina Marqueze, Raul Machado Neto, Debora Botequio Moretti, Bibiana Santos, João Paulo Kitajima, Erika Freitas, David Schlesinger, Simone Kashima, Evandra Strazza Rodrigues, Svetoslav Nanev Slavov, Elaine Vieira dos Santos, Rafael dos Santos Bezerra, Luiz Carlos Junior de Alcantara, Marta Giovanetti, Vagner Fonseca, Flavia Aburjaile, Rodrigo Tocantins Calado. |
| EPI_ISL_1445201 | USAFA TUDE BASTOS | Instituto Butantan / Mendelics | Dimas Tadeu Covas, Sandra Coccuzzo Sampaio, Maria Carolina Elias, José Salvatore Leister Patané, Vincent Louis Viala, Antonio Jorge Martins, Ricardo Haddad, Claudia Renata dos Santos Barros, Elaine Cristina Marqueze, Raul Machado Neto, Debora Botequio Moretti, Bibiana Santos, João Paulo Kitajima, Erika Freitas, David Schlesinger, Simone Kashima, Evandra Strazza Rodrigues, Svetoslav Nanev Slavov, Elaine Vieira dos Santos, Rafael dos Santos Bezerra, Luiz Carlos Junior de Alcantara, Marta Giovanetti, Vagner Fonseca, Flavia Aburjaile, Rodrigo Tocantins Calado. |
| EPI_ISL_1445202 | USAFA MARACANA | Instituto Butantan / Mendelics | Dimas Tadeu Covas, Sandra Coccuzzo Sampaio, Maria Carolina Elias, José Salvatore Leister Patané, Vincent Louis Viala, Antonio Jorge Martins, Ricardo Haddad, Claudia Renata dos Santos Barros, Elaine Cristina Marqueze, Raul Machado Neto, Debora Botequio Moretti, Bibiana Santos, João Paulo Kitajima, Erika Freitas, David Schlesinger, Simone Kashima, Evandra Strazza Rodrigues, Svetoslav Nanev Slavov, Elaine Vieira dos Santos, Rafael dos Santos Bezerra, Luiz Carlos Junior de Alcantara, Marta Giovanetti, Vagner Fonseca, Flavia Aburjaile, Rodrigo Tocantins Calado. |
| EPI_ISL_1445203 | USAFA GUILHERMINA | Instituto Butantan / Mendelics | Dimas Tadeu Covas, Sandra Coccuzzo Sampaio, Maria Carolina Elias, José Salvatore Leister Patané, Vincent Louis Viala, Antonio Jorge Martins, Ricardo Haddad, Claudia Renata dos Santos Barros, Elaine Cristina Marqueze, Raul Machado Neto, Debora Botequio Moretti, Bibiana Santos, João Paulo Kitajima, Erika Freitas, David Schlesinger, Simone Kashima, Evandra Strazza Rodrigues, Svetoslav Nanev Slavov, Elaine Vieira dos Santos, Rafael dos Santos Bezerra, Luiz Carlos Junior de Alcantara, Marta Giovanetti, Vagner Fonseca, Flavia Aburjaile, Rodrigo Tocantins Calado. |
| EPI_ISL_1445204 | USAFA MARACANA | Instituto Butantan / Mendelics | Dimas Tadeu Covas, Sandra Coccuzzo Sampaio, Maria Carolina Elias, José Salvatore Leister Patané, Vincent Louis Viala, Antonio Jorge Martins, Ricardo Haddad, Claudia Renata dos Santos Barros, Elaine Cristina Marqueze, Raul Machado Neto, Debora Botequio Moretti, Bibiana Santos, João Paulo Kitajima, Erika Freitas, David Schlesinger, Simone Kashima, Evandra Strazza Rodrigues, Svetoslav Nanev Slavov, Elaine Vieira dos Santos, Rafael dos Santos Bezerra, Luiz Carlos Junior de Alcantara, Marta Giovanetti, Vagner Fonseca, Flavia Aburjaile, Rodrigo Tocantins Calado. |
| EPI_ISL_1445205 | UBS FRANCISCA LIMA DE LIRA | Instituto Butantan / Mendelics | Dimas Tadeu Covas, Sandra Coccuzzo Sampaio, Maria Carolina Elias, José Salvatore Leister Patané, Vincent Louis Viala, Antonio Jorge Martins, Ricardo Haddad, Claudia Renata dos Santos Barros, Elaine Cristina Marqueze, Raul Machado Neto, Debora Botequio Moretti, Bibiana Santos, João Paulo Kitajima, Erika Freitas, David Schlesinger, Simone Kashima, Evandra Strazza Rodrigues, Svetoslav Nanev Slavov, Elaine Vieira dos Santos, Rafael dos Santos Bezerra, Luiz Carlos Junior de Alcantara, Marta Giovanetti, Vagner Fonseca, Flavia Aburjaile, Rodrigo Tocantins Calado. |
| EPI_ISL_1445206 | UBS DARCY ALVES E ROBALINHO | Instituto Butantan / Mendelics | Dimas Tadeu Covas, Sandra Coccuzzo Sampaio, Maria Carolina Elias, José Salvatore Leister Patané, Vincent Louis Viala, Antonio Jorge Martins, Ricardo Haddad, Claudia Renata dos Santos Barros, Elaine Cristina Marqueze, Raul Machado Neto, Debora Botequio Moretti, Bibiana Santos, João Paulo Kitajima, Erika Freitas, David Schlesinger, Simone Kashima, Evandra Strazza Rodrigues, Svetoslav Nanev Slavov, Elaine Vieira dos Santos, Rafael dos Santos Bezerra, Luiz Carlos Junior de Alcantara, Marta Giovanetti, Vagner Fonseca, Flavia Aburjaile, Rodrigo Tocantins Calado. |
| EPI_ISL_1445207 | PPA MARCIO VALDEVINO BATISTA | Instituto Butantan / Mendelics | Dimas Tadeu Covas, Sandra Coccuzzo Sampaio, Maria Carolina Elias, José Salvatore Leister Patané, Vincent Louis Viala, Antonio Jorge Martins, Ricardo Haddad, Claudia Renata dos Santos Barros, Elaine Cristina Marqueze, Raul Machado Neto, Debora Botequio Moretti, Bibiana Santos, João Paulo Kitajima, Erika Freitas, David Schlesinger, Simone Kashima, Evandra Strazza Rodrigues, Svetoslav Nanev Slavov, Elaine Vieira dos Santos, Rafael dos Santos Bezerra, Luiz Carlos Junior de Alcantara, Marta Giovanetti, Vagner Fonseca, Flavia Aburjaile, Rodrigo Tocantins Calado. |
| EPI_ISL_1445209 | UBS OTACILIO FIRMINO LOPES | Instituto Butantan / Mendelics | Dimas Tadeu Covas, Sandra Coccuzzo Sampaio, Maria Carolina Elias, José Salvatore Leister Patané, Vincent Louis Viala, Antonio Jorge Martins, Ricardo Haddad, Claudia Renata dos Santos Barros, Elaine Cristina Marqueze, Raul Machado Neto, Debora Botequio Moretti, Bibiana Santos, João Paulo Kitajima, Erika Freitas, David Schlesinger, Simone Kashima, Evandra Strazza Rodrigues, Svetoslav Nanev Slavov, Elaine Vieira dos Santos, Rafael dos Santos Bezerra, Luiz Carlos Junior de Alcantara, Marta Giovanetti, Vagner Fonseca, Flavia Aburjaile, Rodrigo Tocantins Calado. |
| EPI_ISL_1445210, EPI_ISL_1445211 | UBS JOSE SABINO FERREIRA | Instituto Butantan / Mendelics | Dimas Tadeu Covas, Sandra Coccuzzo Sampaio, Maria Carolina Elias, José Salvatore Leister Patané, Vincent Louis Viala, Antonio Jorge Martins, Ricardo Haddad, Claudia Renata dos Santos Barros, Elaine Cristina Marqueze, Raul Machado Neto, Debora Botequio Moretti, Bibiana Santos, João Paulo Kitajima, Erika Freitas, David Schlesinger, Simone Kashima, Evandra Strazza Rodrigues, Svetoslav Nanev Slavov, Elaine Vieira dos Santos, Rafael dos Santos Bezerra, Luiz Carlos Junior de Alcantara, Marta Giovanetti, Vagner Fonseca, Flavia Aburjaile, Rodrigo Tocantins Calado. |
| EPI_ISL_1445212 | UBS IRMA AGUEDA MARIA JAIME | Instituto Butantan / Mendelics | Dimas Tadeu Covas, Sandra Coccuzzo Sampaio, Maria Carolina Elias, José Salvatore Leister Patané, Vincent Louis Viala, Antonio Jorge Martins, Ricardo Haddad, Claudia Renata dos Santos Barros, Elaine Cristina Marqueze, Raul Machado Neto, Debora Botequio Moretti, Bibiana Santos, João Paulo Kitajima, Erika Freitas, David Schlesinger, Simone Kashima, Evandra Strazza Rodrigues, Svetoslav Nanev Slavov, Elaine Vieira dos Santos, Rafael dos Santos Bezerra, Luiz Carlos Junior de Alcantara, Marta Giovanetti, Vagner Fonseca, Flavia Aburjaile, Rodrigo Tocantins Calado. |

| | | | |
|---|---|---|---|
| EPI_ISL_1445213 | UBS OTACILIO FIRMINO LOPES | Instituto Butantan / Mendelics | Dimas Tadeu Covas, Sandra Coccuzzo Sampaio, Maria Carolina Elias, José Salvatore Leister Patané, Vincent Louis Viala, Antonio Jorge Martins, Ricardo Haddad, Claudia Renata dos Santos Barros, Elaine Cristina Marqueze, Raul Machado Neto, Debora Botequio Moretti, Bibiana Santos, João Paulo Kitajima, Erika Freitas, David Schlesinger, Simone Kashima, Evandra Strazza Rodrigues, Svetoslav Nanev Slavov, Elaine Vieira dos Santos, Rafael dos Santos Bezerra, Luiz Carlos Junior de Alcantara, Marta Giovanetti, Vagner Fonseca, Flavia Aburjaile, Rodrigo Tocantins Calado. |
| EPI_ISL_1445214 | UBS JOSE HILARIO DOS SANTOS | Instituto Butantan / Mendelics | Dimas Tadeu Covas, Sandra Coccuzzo Sampaio, Maria Carolina Elias, José Salvatore Leister Patané, Vincent Louis Viala, Antonio Jorge Martins, Ricardo Haddad, Claudia Renata dos Santos Barros, Elaine Cristina Marqueze, Raul Machado Neto, Debora Botequio Moretti, Bibiana Santos, João Paulo Kitajima, Erika Freitas, David Schlesinger, Simone Kashima, Evandra Strazza Rodrigues, Svetoslav Nanev Slavov, Elaine Vieira dos Santos, Rafael dos Santos Bezerra, Luiz Carlos Junior de Alcantara, Marta Giovanetti, Vagner Fonseca, Flavia Aburjaile, Rodrigo Tocantins Calado. |
| EPI_ISL_1445215, EPI_ISL_1445216 | UBS HELENA MARREY | Instituto Butantan / Mendelics | Dimas Tadeu Covas, Sandra Coccuzzo Sampaio, Maria Carolina Elias, José Salvatore Leister Patané, Vincent Louis Viala, Antonio Jorge Martins, Ricardo Haddad, Claudia Renata dos Santos Barros, Elaine Cristina Marqueze, Raul Machado Neto, Debora Botequio Moretti, Bibiana Santos, João Paulo Kitajima, Erika Freitas, David Schlesinger, Simone Kashima, Evandra Strazza Rodrigues, Svetoslav Nanev Slavov, Elaine Vieira dos Santos, Rafael dos Santos Bezerra, Luiz Carlos Junior de Alcantara, Marta Giovanetti, Vagner Fonseca, Flavia Aburjaile, Rodrigo Tocantins Calado. |
| EPI_ISL_1445217 | UBS EMILIA COSME CERQUEIRA | Instituto Butantan / Mendelics | Dimas Tadeu Covas, Sandra Coccuzzo Sampaio, Maria Carolina Elias, José Salvatore Leister Patané, Vincent Louis Viala, Antonio Jorge Martins, Ricardo Haddad, Claudia Renata dos Santos Barros, Elaine Cristina Marqueze, Raul Machado Neto, Debora Botequio Moretti, Bibiana Santos, João Paulo Kitajima, Erika Freitas, David Schlesinger, Simone Kashima, Evandra Strazza Rodrigues, Svetoslav Nanev Slavov, Elaine Vieira dos Santos, Rafael dos Santos Bezerra, Luiz Carlos Junior de Alcantara, Marta Giovanetti, Vagner Fonseca, Flavia Aburjaile, Rodrigo Tocantins Calado. |
| EPI_ISL_1445218 | UBS SYLVIO JOAO L DE LUCIA | Instituto Butantan / Mendelics | Dimas Tadeu Covas, Sandra Coccuzzo Sampaio, Maria Carolina Elias, José Salvatore Leister Patané, Vincent Louis Viala, Antonio Jorge Martins, Ricardo Haddad, Claudia Renata dos Santos Barros, Elaine Cristina Marqueze, Raul Machado Neto, Debora Botequio Moretti, Bibiana Santos, João Paulo Kitajima, Erika Freitas, David Schlesinger, Simone Kashima, Evandra Strazza Rodrigues, Svetoslav Nanev Slavov, Elaine Vieira dos Santos, Rafael dos Santos Bezerra, Luiz Carlos Junior de Alcantara, Marta Giovanetti, Vagner Fonseca, Flavia Aburjaile, Rodrigo Tocantins Calado. |
| EPI_ISL_1445219, EPI_ISL_1445220 | UBS EMILIA COSME CERQUEIRA | Instituto Butantan / Mendelics | Dimas Tadeu Covas, Sandra Coccuzzo Sampaio, Maria Carolina Elias, José Salvatore Leister Patané, Vincent Louis Viala, Antonio Jorge Martins, Ricardo Haddad, Claudia Renata dos Santos Barros, Elaine Cristina Marqueze, Raul Machado Neto, Debora Botequio Moretti, Bibiana Santos, João Paulo Kitajima, Erika Freitas, David Schlesinger, Simone Kashima, Evandra Strazza Rodrigues, Svetoslav Nanev Slavov, Elaine Vieira dos Santos, Rafael dos Santos Bezerra, Luiz Carlos Junior de Alcantara, Marta Giovanetti, Vagner Fonseca, Flavia Aburjaile, Rodrigo Tocantins Calado. |
| EPI_ISL_1445221, EPI_ISL_1445222, EPI_ISL_1445223 | UBS HELENA MARREY | Instituto Butantan / Mendelics | Dimas Tadeu Covas, Sandra Coccuzzo Sampaio, Maria Carolina Elias, José Salvatore Leister Patané, Vincent Louis Viala, Antonio Jorge Martins, Ricardo Haddad, Claudia Renata dos Santos Barros, Elaine Cristina Marqueze, Raul Machado Neto, Debora Botequio Moretti, Bibiana Santos, João Paulo Kitajima, Erika Freitas, David Schlesinger, Simone Kashima, Evandra Strazza Rodrigues, Svetoslav Nanev Slavov, Elaine Vieira dos Santos, Rafael dos Santos Bezerra, Luiz Carlos Junior de Alcantara, Marta Giovanetti, Vagner Fonseca, Flavia Aburjaile, Rodrigo Tocantins Calado. |
| EPI_ISL_1445224 | PPA FENELON GUEDES PEREIRA | Instituto Butantan / Mendelics | Dimas Tadeu Covas, Sandra Coccuzzo Sampaio, Maria Carolina Elias, José Salvatore Leister Patané, Vincent Louis Viala, Antonio Jorge Martins, Ricardo Haddad, Claudia Renata dos Santos Barros, Elaine Cristina Marqueze, Raul Machado Neto, Debora Botequio Moretti, Bibiana Santos, João Paulo Kitajima, Erika Freitas, David Schlesinger, Simone Kashima, Evandra Strazza Rodrigues, Svetoslav Nanev Slavov, Elaine Vieira dos Santos, Rafael dos Santos Bezerra, Luiz Carlos Junior de Alcantara, Marta Giovanetti, Vagner Fonseca, Flavia Aburjaile, Rodrigo Tocantins Calado. |
| EPI_ISL_1445225 | PRONTO SOCORRO DR ANTONIO FLAVIO FRANCA | Instituto Butantan / Mendelics | Dimas Tadeu Covas, Sandra Coccuzzo Sampaio, Maria Carolina Elias, José Salvatore Leister Patané, Vincent Louis Viala, Antonio Jorge Martins, Ricardo Haddad, Claudia Renata dos Santos Barros, Elaine Cristina Marqueze, Raul Machado Neto, Debora Botequio Moretti, Bibiana Santos, João Paulo Kitajima, Erika Freitas, David Schlesinger, Simone Kashima, Evandra Strazza Rodrigues, Svetoslav Nanev Slavov, Elaine Vieira dos Santos, Rafael dos Santos Bezerra, Luiz Carlos Junior de Alcantara, Marta Giovanetti, Vagner Fonseca, Flavia Aburjaile, Rodrigo Tocantins Calado. |
| EPI_ISL_1445226 | UBS SYLVIO JOAO L DE LUCIA | Instituto Butantan / Mendelics | Dimas Tadeu Covas, Sandra Coccuzzo Sampaio, Maria Carolina Elias, José Salvatore Leister Patané, Vincent Louis Viala, Antonio Jorge Martins, Ricardo Haddad, Claudia Renata dos Santos Barros, Elaine Cristina Marqueze, Raul Machado Neto, Debora Botequio Moretti, Bibiana Santos, João Paulo Kitajima, Erika Freitas, David Schlesinger, Simone Kashima, Evandra Strazza Rodrigues, Svetoslav Nanev Slavov, Elaine Vieira dos Santos, Rafael dos Santos Bezerra, Luiz Carlos Junior de Alcantara, Marta Giovanetti, Vagner Fonseca, Flavia Aburjaile, Rodrigo Tocantins Calado. |
| EPI_ISL_1445227 | UBS EMILIA COSME CERQUEIRA | Instituto Butantan / Mendelics | Dimas Tadeu Covas, Sandra Coccuzzo Sampaio, Maria Carolina Elias, José Salvatore Leister Patané, Vincent Louis Viala, Antonio Jorge Martins, Ricardo Haddad, Claudia Renata dos Santos Barros, Elaine Cristina Marqueze, Raul Machado Neto, Debora Botequio Moretti, Bibiana Santos, João Paulo Kitajima, Erika Freitas, David Schlesinger, Simone Kashima, Evandra Strazza Rodrigues, Svetoslav Nanev Slavov, Elaine Vieira dos Santos, Rafael dos Santos Bezerra, Luiz Carlos Junior de Alcantara, Marta Giovanetti, Vagner Fonseca, Flavia Aburjaile, Rodrigo Tocantins Calado. |
| EPI_ISL_1445228 | UBS IRMA AGUEDA MARIA JAIME | Instituto Butantan / Mendelics | Dimas Tadeu Covas, Sandra Coccuzzo Sampaio, Maria Carolina Elias, José Salvatore Leister Patané, Vincent Louis Viala, Antonio Jorge Martins, Ricardo Haddad, Claudia Renata dos Santos Barros, Elaine Cristina Marqueze, Raul Machado Neto, Debora Botequio Moretti, Bibiana Santos, João Paulo Kitajima, Erika Freitas, David Schlesinger, Simone Kashima, Evandra Strazza Rodrigues, Svetoslav Nanev Slavov, Elaine Vieira dos Santos, Rafael dos Santos Bezerra, Luiz Carlos Junior de Alcantara, Marta Giovanetti, Vagner Fonseca, Flavia Aburjaile, Rodrigo Tocantins Calado. |
| EPI_ISL_1445229, EPI_ISL_1445230, EPI_ISL_1445231, EPI_ISL_1445232, EPI_ISL_1445233, EPI_ISL_1445234, EPI_ISL_1445235, EPI_ISL_1445236, EPI_ISL_1445237 | VIGILANCIA EPIDEMIOLOGICA | Instituto Butantan / Mendelics | Dimas Tadeu Covas, Sandra Coccuzzo Sampaio, Maria Carolina Elias, José Salvatore Leister Patané, Vincent Louis Viala, Antonio Jorge Martins, Ricardo Haddad, Claudia Renata dos Santos Barros, Elaine Cristina Marqueze, Raul Machado Neto, Debora Botequio Moretti, Bibiana Santos, João Paulo Kitajima, Erika Freitas, David Schlesinger, Simone Kashima, Evandra Strazza Rodrigues, Svetoslav Nanev Slavov, Elaine Vieira dos Santos, Rafael dos Santos Bezerra, Luiz Carlos Junior de Alcantara, Marta Giovanetti, Vagner Fonseca, Flavia Aburjaile, Rodrigo Tocantins Calado. |
| EPI_ISL_1445238, EPI_ISL_1445239, EPI_ISL_1445240, EPI_ISL_1445241, EPI_ISL_1445242, EPI_ISL_1445243, EPI_ISL_1445244, EPI_ISL_1445245, EPI_ISL_1445246, EPI_ISL_1445247, EPI_ISL_1445248 | | | |
| see above | SECAO CENTRO DE DIAGNOSTICO SECEDI | Instituto Butantan / Mendelics | Dimas Tadeu Covas, Sandra Coccuzzo Sampaio, Maria Carolina Elias, José Salvatore Leister Patané, Vincent Louis Viala, Antonio Jorge Martins, Ricardo Haddad, Claudia Renata dos Santos Barros, Elaine Cristina Marqueze, Raul Machado Neto, Debora Botequio Moretti, Bibiana Santos, João Paulo Kitajima, Erika Freitas, David Schlesinger, Simone Kashima, Evandra Strazza Rodrigues, Svetoslav Nanev Slavov, Elaine Vieira dos Santos, Rafael dos Santos Bezerra, Luiz Carlos Junior de Alcantara, Marta Giovanetti, Vagner Fonseca, Flavia Aburjaile, Rodrigo Tocantins Calado. |
| EPI_ISL_1445249, EPI_ISL_1445250, EPI_ISL_1445251 | VIGILANCIA EPIDEMIOLOGICA | Instituto Butantan / Mendelics | Dimas Tadeu Covas, Sandra Coccuzzo Sampaio, Maria Carolina Elias, José Salvatore Leister Patané, Vincent Louis Viala, Antonio Jorge Martins, Ricardo Haddad, Claudia Renata dos Santos Barros, Elaine Cristina Marqueze, Raul Machado Neto, Debora Botequio Moretti, Bibiana Santos, João Paulo Kitajima, Erika Freitas, David Schlesinger, Simone Kashima, Evandra Strazza Rodrigues, Svetoslav Nanev Slavov, Elaine Vieira dos Santos, Rafael dos Santos Bezerra, Luiz Carlos Junior de Alcantara, Marta Giovanetti, Vagner Fonseca, Flavia Aburjaile, Rodrigo Tocantins Calado. |
| EPI_ISL_1445252, EPI_ISL_1445253 | SECAO CENTRO DE DIAGNOSTICO SECEDI | Instituto Butantan / Mendelics | Dimas Tadeu Covas, Sandra Coccuzzo Sampaio, Maria Carolina Elias, José Salvatore Leister Patané, Vincent Louis Viala, Antonio Jorge Martins, Ricardo Haddad, Claudia Renata dos Santos Barros, Elaine Cristina Marqueze, Raul Machado Neto, Debora Botequio Moretti, Bibiana Santos, João Paulo Kitajima, Erika Freitas, David Schlesinger, Simone Kashima, Evandra Strazza Rodrigues, Svetoslav Nanev Slavov, Elaine Vieira dos Santos, Rafael dos Santos Bezerra, Luiz Carlos Junior de Alcantara, Marta Giovanetti, Vagner Fonseca, Flavia Aburjaile, Rodrigo Tocantins Calado. |
| EPI_ISL_1445254, EPI_ISL_1445255, EPI_ISL_1445256, EPI_ISL_1445257, EPI_ISL_1445258, EPI_ISL_1445259, EPI_ISL_1445260, EPI_ISL_1445261, EPI_ISL_1445263 | VIGILANCIA EPIDEMIOLOGICA | Instituto Butantan / Mendelics | Dimas Tadeu Covas, Sandra Coccuzzo Sampaio, Maria Carolina Elias, José Salvatore Leister Patané, Vincent Louis Viala, Antonio Jorge Martins, Ricardo Haddad, Claudia Renata dos Santos Barros, Elaine Cristina Marqueze, Raul Machado Neto, Debora Botequio Moretti, Bibiana Santos, João Paulo Kitajima, Erika Freitas, David Schlesinger, Simone Kashima, Evandra Strazza Rodrigues, Svetoslav Nanev Slavov, Elaine Vieira dos Santos, Rafael dos Santos Bezerra, Luiz Carlos Junior de Alcantara, Marta Giovanetti, Vagner Fonseca, Flavia Aburjaile, Rodrigo Tocantins Calado. |
| EPI_ISL_1445264 | AMBULATORIO MEDICO DE ESPECIALIDADES DE PERUIBE | Instituto Butantan / Mendelics | Dimas Tadeu Covas, Sandra Coccuzzo Sampaio, Maria Carolina Elias, José Salvatore Leister Patané, Vincent Louis Viala, Antonio Jorge Martins, Ricardo Haddad, Claudia Renata dos Santos Barros, Elaine Cristina Marqueze, Raul Machado Neto, Debora Botequio Moretti, Bibiana Santos, João Paulo Kitajima, Erika Freitas, David Schlesinger, Simone Kashima, Evandra Strazza Rodrigues, Svetoslav Nanev Slavov, Elaine Vieira dos Santos, Rafael dos Santos Bezerra, Luiz Carlos Junior de Alcantara, Marta Giovanetti, Vagner Fonseca, Flavia Aburjaile, Rodrigo Tocantins Calado. |
| EPI_ISL_1445265 | VIGILANCIA EPIDEMIOLOGICA | Instituto Butantan / Mendelics | Dimas Tadeu Covas, Sandra Coccuzzo Sampaio, Maria Carolina Elias, José Salvatore Leister Patané, Vincent Louis Viala, Antonio Jorge Martins, Ricardo Haddad, Claudia Renata dos Santos Barros, Elaine Cristina Marqueze, Raul Machado Neto, Debora Botequio Moretti, Bibiana Santos, João Paulo Kitajima, Erika Freitas, David Schlesinger, Simone Kashima, Evandra Strazza Rodrigues, Svetoslav Nanev Slavov, Elaine Vieira dos Santos, Rafael dos |

| | | | |
|---|---|---|---|
| | | | Santos Bezerra, Luiz Carlos Junior de Alcantara, Marta Giovanetti, Vagner Fonseca, Flavia Aburjaile, Rodrigo Tocantins Calado. |
| EPI_ISL_1445266, EPI_ISL_1445267 | AMBULATORIO MEDICO DE ESPECIALIDADES DE PERUIBE | Instituto Butantan / Mendelics | Dimas Tadeu Covas, Sandra Coccuzzo Sampaio, Maria Carolina Elias, José Salvatore Leister Patané, Vincent Louis Viala, Antonio Jorge Martins, Ricardo Haddad, Claudia Renata dos Santos Barros, Elaine Cristina Marqueze, Raul Machado Neto, Debora Botequio Moretti, Bibiana Santos, João Paulo Kitajima, Erika Freitas, David Schlesinger, Simone Kashima, Evandra Strazza Rodrigues, Svetoslav Nanev Slavov, Elaine Vieira dos Santos, Rafael dos Santos Bezerra, Luiz Carlos Junior de Alcantara, Marta Giovanetti, Vagner Fonseca, Flavia Aburjaile, Rodrigo Tocantins Calado. |
| EPI_ISL_1445268 | VIGILANCIA EPIDEMIOLOGICA | Instituto Butantan / Mendelics | Dimas Tadeu Covas, Sandra Coccuzzo Sampaio, Maria Carolina Elias, José Salvatore Leister Patané, Vincent Louis Viala, Antonio Jorge Martins, Ricardo Haddad, Claudia Renata dos Santos Barros, Elaine Cristina Marqueze, Raul Machado Neto, Debora Botequio Moretti, Bibiana Santos, João Paulo Kitajima, Erika Freitas, David Schlesinger, Simone Kashima, Evandra Strazza Rodrigues, Svetoslav Nanev Slavov, Elaine Vieira dos Santos, Rafael dos Santos Bezerra, Luiz Carlos Junior de Alcantara, Marta Giovanetti, Vagner Fonseca, Flavia Aburjaile, Rodrigo Tocantins Calado. |
| EPI_ISL_1445269 | SERV DE VIG SANITARIA EPIDEMIO E CTRL DE ZOONOZES GUARUJA | Instituto Butantan / Mendelics | Dimas Tadeu Covas, Sandra Coccuzzo Sampaio, Maria Carolina Elias, José Salvatore Leister Patané, Vincent Louis Viala, Antonio Jorge Martins, Ricardo Haddad, Claudia Renata dos Santos Barros, Elaine Cristina Marqueze, Raul Machado Neto, Debora Botequio Moretti, Bibiana Santos, João Paulo Kitajima, Erika Freitas, David Schlesinger, Simone Kashima, Evandra Strazza Rodrigues, Svetoslav Nanev Slavov, Elaine Vieira dos Santos, Rafael dos Santos Bezerra, Luiz Carlos Junior de Alcantara, Marta Giovanetti, Vagner Fonseca, Flavia Aburjaile, Rodrigo Tocantins Calado. |
| EPI_ISL_1445270, EPI_ISL_1445271 | VIGILANCIA EPIDEMIOLOGICA | Instituto Butantan / Mendelics | Dimas Tadeu Covas, Sandra Coccuzzo Sampaio, Maria Carolina Elias, José Salvatore Leister Patané, Vincent Louis Viala, Antonio Jorge Martins, Ricardo Haddad, Claudia Renata dos Santos Barros, Elaine Cristina Marqueze, Raul Machado Neto, Debora Botequio Moretti, Bibiana Santos, João Paulo Kitajima, Erika Freitas, David Schlesinger, Simone Kashima, Evandra Strazza Rodrigues, Svetoslav Nanev Slavov, Elaine Vieira dos Santos, Rafael dos Santos Bezerra, Luiz Carlos Junior de Alcantara, Marta Giovanetti, Vagner Fonseca, Flavia Aburjaile, Rodrigo Tocantins Calado. |
| EPI_ISL_1445272 | SERV DE VIG SANITARIA EPIDEMIO E CTRL DE ZOONOZES GUARUJA | Instituto Butantan / Mendelics | Dimas Tadeu Covas, Sandra Coccuzzo Sampaio, Maria Carolina Elias, José Salvatore Leister Patané, Vincent Louis Viala, Antonio Jorge Martins, Ricardo Haddad, Claudia Renata dos Santos Barros, Elaine Cristina Marqueze, Raul Machado Neto, Debora Botequio Moretti, Bibiana Santos, João Paulo Kitajima, Erika Freitas, David Schlesinger, Simone Kashima, Evandra Strazza Rodrigues, Svetoslav Nanev Slavov, Elaine Vieira dos Santos, Rafael dos Santos Bezerra, Luiz Carlos Junior de Alcantara, Marta Giovanetti, Vagner Fonseca, Flavia Aburjaile, Rodrigo Tocantins Calado. |
| EPI_ISL_1445273, EPI_ISL_1445274 | AMBULATORIO MEDICO DE ESPECIALIDADES DE PERUIBE | Instituto Butantan / Mendelics | Dimas Tadeu Covas, Sandra Coccuzzo Sampaio, Maria Carolina Elias, José Salvatore Leister Patané, Vincent Louis Viala, Antonio Jorge Martins, Ricardo Haddad, Claudia Renata dos Santos Barros, Elaine Cristina Marqueze, Raul Machado Neto, Debora Botequio Moretti, Bibiana Santos, João Paulo Kitajima, Erika Freitas, David Schlesinger, Simone Kashima, Evandra Strazza Rodrigues, Svetoslav Nanev Slavov, Elaine Vieira dos Santos, Rafael dos Santos Bezerra, Luiz Carlos Junior de Alcantara, Marta Giovanetti, Vagner Fonseca, Flavia Aburjaile, Rodrigo Tocantins Calado. |
| EPI_ISL_1446194 | LATE - Laboratório de Técnicas Especiais - Hospital Israelita Albert Einstein | LATE - Laboratório de Técnicas Especiais - Hospital Israelita Albert Einstein | Deyvid Amgarten, Fernanda de Mello Malta, Raquel Riyuzo, Ana Paula Moreira Salles, Pedro Henrique Sebe Rodrigues, João Renato Rebello Pinho |
| EPI_ISL_1464626, EPI_ISL_1464627, EPI_ISL_1464628, EPI_ISL_1464629, EPI_ISL_1464630, EPI_ISL_1464631, EPI_ISL_1464632, EPI_ISL_1464633, EPI_ISL_1464634, EPI_ISL_1464635, EPI_ISL_1464636, EPI_ISL_1464637, EPI_ISL_1464638, EPI_ISL_1464639, EPI_ISL_1464640, EPI_ISL_1464641, EPI_ISL_1464642, EPI_ISL_1464643, EPI_ISL_1464644, EPI_ISL_1464645, EPI_ISL_1464646, EPI_ISL_1464647, EPI_ISL_1464648, EPI_ISL_1464649, EPI_ISL_1464650, EPI_ISL_1464651, EPI_ISL_1464652, EPI_ISL_1464653, EPI_ISL_1464654, EPI_ISL_1464655, EPI_ISL_1464656, EPI_ISL_1464657, EPI_ISL_1464658, EPI_ISL_1464659, EPI_ISL_1464660, EPI_ISL_1464661, EPI_ISL_1464662, EPI_ISL_1464663, EPI_ISL_1464664, EPI_ISL_1464665, EPI_ISL_1464666, EPI_ISL_1464667, EPI_ISL_1464668, EPI_ISL_1464669, EPI_ISL_1464670, EPI_ISL_1464671, EPI_ISL_1464672, EPI_ISL_1464673, EPI_ISL_1464674, EPI_ISL_1464675, EPI_ISL_1464676, EPI_ISL_1464677 | | | |
| see above | Laboratório de Virologia - UNIFESP | Laboratory of Respiratory Viruses and Measles, Oswaldo Cruz Institute, FIOCRUZ | Paola Resende, Nancy Belei, Luciana Appolinario, Fernando Motta, Anna Carolina Paixao, Ana Carolina Mendonca, Alice Sampaio Rocha, Renata Serrano Lopes, Marilda Siqueira on behalf of the Fiocruz COVID-19 Genomic Surveillance Network |
| EPI_ISL_1465186, EPI_ISL_1465188, EPI_ISL_1465189, EPI_ISL_1465191, EPI_ISL_1465192, EPI_ISL_1465194, EPI_ISL_1465195, EPI_ISL_1465196, EPI_ISL_1465198, EPI_ISL_1465199, EPI_ISL_1465201, EPI_ISL_1465202, EPI_ISL_1465203, EPI_ISL_1465205, EPI_ISL_1465206, EPI_ISL_1465208, EPI_ISL_1465209, EPI_ISL_1465210, EPI_ISL_1465212, EPI_ISL_1465213, EPI_ISL_1465215, EPI_ISL_1465216, EPI_ISL_1465217, EPI_ISL_1465219, EPI_ISL_1465220, EPI_ISL_1465221, EPI_ISL_1465222, EPI_ISL_1465224, EPI_ISL_1465225, EPI_ISL_1465246, EPI_ISL_1465248, EPI_ISL_1465249, EPI_ISL_1465250, EPI_ISL_1465252, EPI_ISL_1465253, EPI_ISL_1465254, EPI_ISL_1465255, EPI_ISL_1465257, EPI_ISL_1465258, EPI_ISL_1465259, EPI_ISL_1465261, EPI_ISL_1465262, EPI_ISL_1465264, EPI_ISL_1465265, EPI_ISL_1465267, EPI_ISL_1465268, EPI_ISL_1465270, EPI_ISL_1465271, EPI_ISL_1465273, EPI_ISL_1465274, EPI_ISL_1465275, EPI_ISL_1465277, EPI_ISL_1465278 | | | |
| see above | Laboratorio Central de Saude Publica do Estado do Maranhao (LACEN-MA) | Laboratory of Respiratory Viruses and Measles, Oswaldo Cruz Institute, FIOCRUZ | Paola Resende, Luciana Appolinario, Fernando Motta, Anna Carolina Paixao, Ana Carolina Mendonca, Alice Sampaio Rocha, Renata Serrano Lopes, Lidio Gonçalves Lima Neto, Marilda Siqueira on behalf of the Fiocruz COVID-19 Genomic Surveillance Network |
| EPI_ISL_1468411 | Centro de Saude II Matao | Instituto Adolfo Lutz, Interdiciplinary Procedures Center, Strategic Laboratory | Claudio Tavares Sacchi, Claudia Regina Gonçalves, Erica Valessa Ramos Gomes, Karoline Rodrigues Campos, Caio Vinicius Dias Lopes |
| EPI_ISL_1468412 | Hospital Estadual de Mirandopolis | Instituto Adolfo Lutz, Interdiciplinary Procedures Center, Strategic Laboratory | Claudio Tavares Sacchi, Claudia Regina Gonçalves, Erica Valessa Ramos Gomes, Karoline Rodrigues Campos, Caio Vinicius Dias Lopes |
| EPI_ISL_1468413, EPI_ISL_1468414, EPI_ISL_1468415 | LACEN do Estado de Goias | Instituto Adolfo Lutz, Interdiciplinary Procedures Center, Strategic Laboratory | Claudio Tavares Sacchi, Claudia Regina Gonçalves, Erica Valessa Ramos Gomes, Karoline Rodrigues Campos, Caio Vinicius Dias Lopes |
| EPI_ISL_1468416 | Secretaria Municipal de Saude de Andradina | Instituto Adolfo Lutz, Interdiciplinary Procedures Center, Strategic Laboratory | Claudio Tavares Sacchi, Claudia Regina Gonçalves, Erica Valessa Ramos Gomes, Karoline Rodrigues Campos, Caio Vinicius Dias Lopes |
| EPI_ISL_1468417 | Santa Casa de Aracatuba Hospital Sagrado Coracao de Jesus | Instituto Adolfo Lutz, Interdiciplinary Procedures Center, Strategic Laboratory | Claudio Tavares Sacchi, Claudia Regina Gonçalves, Erica Valessa Ramos Gomes, Karoline Rodrigues Campos, Caio Vinicius Dias Lopes |
| EPI_ISL_1468418 | Secretaria Municipal de Saude de Birigui | Instituto Adolfo Lutz, Interdiciplinary Procedures Center, Strategic Laboratory | Claudio Tavares Sacchi, Claudia Regina Gonçalves, Erica Valessa Ramos Gomes, Karoline Rodrigues Campos, Caio Vinicius Dias Lopes |
| EPI_ISL_1468419 | Santa Casa de Aracatuba Hospital Sagrado Coracao de Jesus | Instituto Adolfo Lutz, Interdiciplinary Procedures Center, Strategic Laboratory | Claudio Tavares Sacchi, Claudia Regina Gonçalves, Erica Valessa Ramos Gomes, Karoline Rodrigues Campos, Caio Vinicius Dias Lopes |
| EPI_ISL_1468420, EPI_ISL_1468421 | Santa Casa de Birigui | Instituto Adolfo Lutz, Interdiciplinary Procedures Center, Strategic Laboratory | Claudio Tavares Sacchi, Claudia Regina Gonçalves, Erica Valessa Ramos Gomes, Karoline Rodrigues Campos, Caio Vinicius Dias Lopes |
| EPI_ISL_1468422, EPI_ISL_1468423 | Penitenciaria Compacta de Avanhandava | Instituto Adolfo Lutz, Interdiciplinary Procedures Center, Strategic Laboratory | Claudio Tavares Sacchi, Claudia Regina Gonçalves, Erica Valessa Ramos Gomes, Karoline Rodrigues Campos, Caio Vinicius Dias Lopes |
| EPI_ISL_1468424 | Santa Casa de Aracatuba Hospital Sagrado Coracao de Jesus | Instituto Adolfo Lutz, Interdiciplinary Procedures Center, Strategic Laboratory | Claudio Tavares Sacchi, Claudia Regina Gonçalves, Erica Valessa Ramos Gomes, Karoline Rodrigues Campos, Caio Vinicius Dias Lopes |
| EPI_ISL_1468425 | Santa Casa de Birigui | Instituto Adolfo Lutz, Interdiciplinary Procedures Center, Strategic Laboratory | Claudio Tavares Sacchi, Claudia Regina Gonçalves, Erica Valessa Ramos Gomes, Karoline Rodrigues Campos, Caio Vinicius Dias Lopes |
| EPI_ISL_1468426, EPI_ISL_1468427 | Centro de Saude II Matao | Instituto Adolfo Lutz, Interdiciplinary Procedures Center, Strategic Laboratory | Claudio Tavares Sacchi, Claudia Regina Gonçalves, Erica Valessa Ramos Gomes, Karoline Rodrigues Campos, Caio Vinicius Dias Lopes |
| EPI_ISL_1468428 | SAE Servico de Atendimento Especializado | Instituto Adolfo Lutz, Interdiciplinary Procedures Center, Strategic Laboratory | Claudio Tavares Sacchi, Claudia Regina Gonçalves, Erica Valessa Ramos Gomes, Karoline Rodrigues Campos, Caio Vinicius Dias Lopes |
| EPI_ISL_1468429, EPI_ISL_1468430 | Centro de Saude II Matao | Instituto Adolfo Lutz, Interdiciplinary Procedures Center, Strategic Laboratory | Claudio Tavares Sacchi, Claudia Regina Gonçalves, Erica Valessa Ramos Gomes, Karoline Rodrigues Campos, Caio Vinicius Dias Lopes |
| EPI_ISL_1468431 | LACEN do Estado de Goias | Instituto Adolfo Lutz, Interdiciplinary Procedures Center, Strategic Laboratory | Claudio Tavares Sacchi, Claudia Regina Gonçalves, Erica Valessa Ramos Gomes, Karoline Rodrigues Campos, Caio Vinicius Dias Lopes |
| EPI_ISL_1468432, EPI_ISL_1468433, EPI_ISL_1468434, EPI_ISL_1468435, EPI_ISL_1468436, EPI_ISL_1468437, EPI_ISL_1468438, EPI_ISL_1468439, EPI_ISL_1468440, EPI_ISL_1468441 | LACEN do Mato Grosso do Sul | Instituto Adolfo Lutz, Interdiciplinary Procedures Center, Strategic Laboratory | Claudio Tavares Sacchi, Claudia Regina Gonçalves, Erica Valessa Ramos Gomes, Karoline Rodrigues Campos, Caio Vinicius Dias Lopes |

| | | | |
|---|---|---|---|
| EPI_ISL_1468442 | Santa Casa de Birigui | Instituto Adolfo Lutz, Interdiciplinary Procedures Center, Strategic Laboratory | Claudio Tavares Sacchi, Claudia Regina Gonçalves, Erica Valessa Ramos Gomes, Karoline Rodrigues Campos, Caio Vinicius Dias Lopes |
| EPI_ISL_1468443 | Hospital Estadual de Mirandopolis | Instituto Adolfo Lutz, Interdiciplinary Procedures Center, Strategic Laboratory | Claudio Tavares Sacchi, Claudia Regina Gonçalves, Erica Valessa Ramos Gomes, Karoline Rodrigues Campos, Caio Vinicius Dias Lopes |
| EPI_ISL_1468444 | Secretaria Municipal de Saude de Birigui | Instituto Adolfo Lutz, Interdiciplinary Procedures Center, Strategic Laboratory | Claudio Tavares Sacchi, Claudia Regina Gonçalves, Erica Valessa Ramos Gomes, Karoline Rodrigues Campos, Caio Vinicius Dias Lopes |
| EPI_ISL_1468445 | Santa Casa de Misericordia de Pereira Barreto | Instituto Adolfo Lutz, Interdiciplinary Procedures Center, Strategic Laboratory | Claudio Tavares Sacchi, Claudia Regina Gonçalves, Erica Valessa Ramos Gomes, Karoline Rodrigues Campos, Caio Vinicius Dias Lopes |
| EPI_ISL_1468446 | Secretaria Municipal de Saude de Birigui | Instituto Adolfo Lutz, Interdiciplinary Procedures Center, Strategic Laboratory | Claudio Tavares Sacchi, Claudia Regina Gonçalves, Erica Valessa Ramos Gomes, Karoline Rodrigues Campos, Caio Vinicius Dias Lopes |
| EPI_ISL_1468447 | Secretaria Municipal de Saude de Andradina | Instituto Adolfo Lutz, Interdiciplinary Procedures Center, Strategic Laboratory | Claudio Tavares Sacchi, Claudia Regina Gonçalves, Erica Valessa Ramos Gomes, Karoline Rodrigues Campos, Caio Vinicius Dias Lopes |
| EPI_ISL_1468448 | Santa Casa de Birigui | Instituto Adolfo Lutz, Interdiciplinary Procedures Center, Strategic Laboratory | Claudio Tavares Sacchi, Claudia Regina Gonçalves, Erica Valessa Ramos Gomes, Karoline Rodrigues Campos, Caio Vinicius Dias Lopes |
| EPI_ISL_1468449, EPI_ISL_1468450 | Santa Casa de Aracatuba Hospital Sagrado Coracao de Jesus | Instituto Adolfo Lutz, Interdiciplinary Procedures Center, Strategic Laboratory | Claudio Tavares Sacchi, Claudia Regina Gonçalves, Erica Valessa Ramos Gomes, Karoline Rodrigues Campos, Caio Vinicius Dias Lopes |
| EPI_ISL_1468451 | UBS IV Guararapes | Instituto Adolfo Lutz, Interdiciplinary Procedures Center, Strategic Laboratory | Claudio Tavares Sacchi, Claudia Regina Gonçalves, Erica Valessa Ramos Gomes, Karoline Rodrigues Campos, Caio Vinicius Dias Lopes |
| EPI_ISL_1468452 | Centro de Atendimento COVID | Instituto Adolfo Lutz, Interdiciplinary Procedures Center, Strategic Laboratory | Claudio Tavares Sacchi, Claudia Regina Gonçalves, Erica Valessa Ramos Gomes, Karoline Rodrigues Campos, Caio Vinicius Dias Lopes |
| EPI_ISL_1468453 | Santa Casa de Misericordia de Pereira Barreto | Instituto Adolfo Lutz, Interdiciplinary Procedures Center, Strategic Laboratory | Claudio Tavares Sacchi, Claudia Regina Gonçalves, Erica Valessa Ramos Gomes, Karoline Rodrigues Campos, Caio Vinicius Dias Lopes |
| EPI_ISL_1468454 | Santa Casa de Andradina | Instituto Adolfo Lutz, Interdiciplinary Procedures Center, Strategic Laboratory | Claudio Tavares Sacchi, Claudia Regina Gonçalves, Erica Valessa Ramos Gomes, Karoline Rodrigues Campos, Caio Vinicius Dias Lopes |
| EPI_ISL_1468455 | UBS 02 Jardim Toselar Birigui | Instituto Adolfo Lutz, Interdiciplinary Procedures Center, Strategic Laboratory | Claudio Tavares Sacchi, Claudia Regina Gonçalves, Erica Valessa Ramos Gomes, Karoline Rodrigues Campos, Caio Vinicius Dias Lopes |
| EPI_ISL_1468456, EPI_ISL_1468457 | UBS Dr Alfredo Dantas de Souza Umuarama | Instituto Adolfo Lutz, Interdiciplinary Procedures Center, Strategic Laboratory | Claudio Tavares Sacchi, Claudia Regina Gonçalves, Erica Valessa Ramos Gomes, Karoline Rodrigues Campos, Caio Vinicius Dias Lopes |
| EPI_ISL_1468458, EPI_ISL_1468459 | Secretaria Municipal de Saude de Valparaiso SP | Instituto Adolfo Lutz, Interdiciplinary Procedures Center, Strategic Laboratory | Claudio Tavares Sacchi, Claudia Regina Gonçalves, Erica Valessa Ramos Gomes, Karoline Rodrigues Campos, Caio Vinicius Dias Lopes |
| EPI_ISL_1468460 | Santa Casa de Birigui | Instituto Adolfo Lutz, Interdiciplinary Procedures Center, Strategic Laboratory | Claudio Tavares Sacchi, Claudia Regina Gonçalves, Erica Valessa Ramos Gomes, Karoline Rodrigues Campos, Caio Vinicius Dias Lopes |
| EPI_ISL_1468461 | Santa Casa de Aracatuba Hospital Sagrado Coracao de Jesus | Instituto Adolfo Lutz, Interdiciplinary Procedures Center, Strategic Laboratory | Claudio Tavares Sacchi, Claudia Regina Gonçalves, Erica Valessa Ramos Gomes, Karoline Rodrigues Campos, Caio Vinicius Dias Lopes |
| EPI_ISL_1468462 | Servico Especial de Saude de Araraquara Sesa Araraquara | Instituto Adolfo Lutz, Interdiciplinary Procedures Center, Strategic Laboratory | Claudio Tavares Sacchi, Claudia Regina Gonçalves, Erica Valessa Ramos Gomes, Karoline Rodrigues Campos, Caio Vinicius Dias Lopes |
| EPI_ISL_1468463 | Centro de Saude II Matao | Instituto Adolfo Lutz, Interdiciplinary Procedures Center, Strategic Laboratory | Claudio Tavares Sacchi, Claudia Regina Gonçalves, Erica Valessa Ramos Gomes, Karoline Rodrigues Campos, Caio Vinicius Dias Lopes |
| EPI_ISL_1468464, EPI_ISL_1468465 | Sae servico de Atendimento Especializado | Instituto Adolfo Lutz, Interdiciplinary Procedures Center, Strategic Laboratory | Claudio Tavares Sacchi, Claudia Regina Gonçalves, Erica Valessa Ramos Gomes, Karoline Rodrigues Campos, Caio Vinicius Dias Lopes |
| EPI_ISL_1468466 | Santa Casa de Sao Carlos | Instituto Adolfo Lutz, Interdiciplinary Procedures Center, Strategic Laboratory | Claudio Tavares Sacchi, Claudia Regina Gonçalves, Erica Valessa Ramos Gomes, Karoline Rodrigues Campos, Caio Vinicius Dias Lopes |
| EPI_ISL_1468467 | Secretaria Municipal de Saude Descalvado | Instituto Adolfo Lutz, Interdiciplinary Procedures Center, Strategic Laboratory | Claudio Tavares Sacchi, Claudia Regina Gonçalves, Erica Valessa Ramos Gomes, Karoline Rodrigues Campos, Caio Vinicius Dias Lopes |
| EPI_ISL_1468468 | Secretaria municipal de saude de Itapolis | Instituto Adolfo Lutz, Interdiciplinary Procedures Center, Strategic Laboratory | Claudio Tavares Sacchi, Claudia Regina Gonçalves, Erica Valessa Ramos Gomes, Karoline Rodrigues Campos, Caio Vinicius Dias Lopes |
| EPI_ISL_1468469 | Secretaria Municipal de Saude Porto Ferreira | Instituto Adolfo Lutz, Interdiciplinary Procedures Center, Strategic Laboratory | Claudio Tavares Sacchi, Claudia Regina Gonçalves, Erica Valessa Ramos Gomes, Karoline Rodrigues Campos, Caio Vinicius Dias Lopes |
| EPI_ISL_1468470, EPI_ISL_1468471 | Secretaria Municipal de Saude Descalvado | Instituto Adolfo Lutz, Interdiciplinary Procedures Center, Strategic Laboratory | Claudio Tavares Sacchi, Claudia Regina Gonçalves, Erica Valessa Ramos Gomes, Karoline Rodrigues Campos, Caio Vinicius Dias Lopes |
| EPI_ISL_1468472, EPI_ISL_1468473 | Centro de Saude II Matao | Instituto Adolfo Lutz, Interdiciplinary Procedures Center, Strategic Laboratory | Claudio Tavares Sacchi, Claudia Regina Gonçalves, Erica Valessa Ramos Gomes, Karoline Rodrigues Campos, Caio Vinicius Dias Lopes |
| EPI_ISL_1468474 | Sae servico de Atendimento Especializado | Instituto Adolfo Lutz, Interdiciplinary Procedures Center, Strategic Laboratory | Claudio Tavares Sacchi, Claudia Regina Gonçalves, Erica Valessa Ramos Gomes, Karoline Rodrigues Campos, Caio Vinicius Dias Lopes |
| EPI_ISL_1469549 | FUNDACAO HOSPITALAR DE SAPUCAIA DO SUL | Epiclin | Fernando Hayashi Sant'Anna, Ana Paula Muterle, Janira Prichula, Juliana Comerlato, Carolina Comerlato, Eliana Márcia Da Ros Wendland |
| EPI_ISL_1469550 | Vigilância em Saúde de Sapucaia do Sul | Epiclin | Fernando Hayashi Sant'Anna, Ana Paula Muterle, Janira Prichula, Juliana Comerlato, Carolina Comerlato, Eliana Márcia Da Ros Wendland |
| EPI_ISL_1469552 | Unidade Sanitária de Igrejinha | Epiclin | Fernando Hayashi Sant'Anna, Ana Paula Muterle, Janira Prichula, Juliana Comerlato, Carolina Comerlato, Eliana Márcia Da Ros Wendland |
| EPI_ISL_1469554 | Fundação de Saúde Pública São Camilo de Esteio | Epiclin | Fernando Hayashi Sant'Anna, Ana Paula Muterle, Janira Prichula, Juliana Comerlato, Carolina Comerlato, Eliana Márcia Da Ros Wendland |
| EPI_ISL_1469555 | FUNDACAO DE SAUDE PUBLICA DE NOVO HAMBURGO FSNH | Epiclin | Fernando Hayashi Sant'Anna, Ana Paula Muterle, Janira Prichula, Juliana Comerlato, Carolina Comerlato, Eliana Márcia Da Ros Wendland |
| EPI_ISL_1469557 | Centro de Serviços Especializados Santa Rita | Epiclin | Fernando Hayashi Sant'Anna, Ana Paula Muterle, Janira Prichula, Juliana Comerlato, Carolina Comerlato, Eliana Márcia Da Ros Wendland |
| EPI_ISL_1469558 | DIRETORIA DE VIGILANCIA EM SAUDE | Epiclin | Fernando Hayashi Sant'Anna, Ana Paula Muterle, Janira Prichula, Juliana Comerlato, Carolina Comerlato, Eliana Márcia Da Ros Wendland |
| EPI_ISL_1469559 | Secretaria Municipal de Saúde de Três Coroas | Epiclin | Fernando Hayashi Sant'Anna, Ana Paula Muterle, Janira Prichula, Juliana Comerlato, Carolina Comerlato, Eliana Márcia Da Ros Wendland |
| EPI_ISL_1469560 | Diretoria de Vigilância em Saúde | Epiclin | Fernando Hayashi Sant'Anna, Ana Paula Muterle, Janira Prichula, Juliana Comerlato, Carolina Comerlato, Eliana Márcia Da Ros Wendland |
| EPI_ISL_1469561 | Fundação de Saúde Pública São Camilo de Esteio | Epiclin | Fernando Hayashi Sant'Anna, Ana Paula Muterle, Janira Prichula, Juliana Comerlato, Carolina Comerlato, Eliana Márcia Da Ros Wendland |
| EPI_ISL_1469562 | Hospital Dia e Pronto Atendimento | Epiclin | Fernando Hayashi Sant'Anna, Ana Paula Muterle, Janira Prichula, Juliana Comerlato, Carolina Comerlato, Eliana Márcia Da Ros Wendland |
| EPI_ISL_1469564 | Pronto Atendimento Campo Bom | Epiclin | Fernando Hayashi Sant'Anna, Ana Paula Muterle, Janira Prichula, Juliana Comerlato, Carolina Comerlato, Eliana Márcia Da Ros Wendland |
| EPI_ISL_1469566 | DIRETORIA DE VIGILANCIA EM SAUDE | Epiclin | Fernando Hayashi Sant'Anna, Ana Paula Muterle, Janira Prichula, Juliana Comerlato, Carolina Comerlato, Eliana Márcia Da Ros Wendland |

| | | | |
|---|---|---|---|
| EPI_ISL_1469567 | Unidade Sanitária de Igrejinha | Epiclin | Fernando Hayashi Sant'Anna, Ana Paula Muterle, Janira Prichula, Juliana Comerlato, Carolina Comerlato, Eliana Márcia Da Ros Wendland |
| EPI_ISL_1469568 | SECRETARIA MUNICIPAL DE SAUDE DE SAO LEOPOLDO | Epiclin | Fernando Hayashi Sant'Anna, Ana Paula Muterle, Janira Prichula, Juliana Comerlato, Carolina Comerlato, Eliana Márcia Da Ros Wendland |
| EPI_ISL_1469569 | Fundação de Saúde Pública São Camilo de Esteio | Epiclin | Fernando Hayashi Sant'Anna, Ana Paula Muterle, Janira Prichula, Juliana Comerlato, Carolina Comerlato, Eliana Márcia Da Ros Wendland |
| EPI_ISL_1469570 | Hospital Universitário | Epiclin | Fernando Hayashi Sant'Anna, Ana Paula Muterle, Janira Prichula, Juliana Comerlato, Carolina Comerlato, Eliana Márcia Da Ros Wendland |
| EPI_ISL_1469571 | Unidade de Pronto Atendimento de Sapucaia do Sul | Epiclin | Fernando Hayashi Sant'Anna, Ana Paula Muterle, Janira Prichula, Juliana Comerlato, Carolina Comerlato, Eliana Márcia Da Ros Wendland |
| EPI_ISL_1469572 | FUNDACAO DE SAUDE PUBLICA DE NOVO HAMBURGO FSNH | Epiclin | Fernando Hayashi Sant'Anna, Ana Paula Muterle, Janira Prichula, Juliana Comerlato, Carolina Comerlato, Eliana Márcia Da Ros Wendland |
| EPI_ISL_1469573 | VIGILANCIA EM SAUDE NH | Epiclin | Fernando Hayashi Sant'Anna, Ana Paula Muterle, Janira Prichula, Juliana Comerlato, Carolina Comerlato, Eliana Márcia Da Ros Wendland |
| EPI_ISL_1469575 | HOSPITAL SAO FRANCISCO DE ASSIS | Epiclin | Fernando Hayashi Sant'Anna, Ana Paula Muterle, Janira Prichula, Juliana Comerlato, Carolina Comerlato, Eliana Márcia Da Ros Wendland |
| EPI_ISL_1469576 | Unidade Sanitária de Igrejinha | Epiclin | Fernando Hayashi Sant'Anna, Ana Paula Muterle, Janira Prichula, Juliana Comerlato, Carolina Comerlato, Eliana Márcia Da Ros Wendland |
| EPI_ISL_1469577 | Unidade de Pronto Atendimento de Sapucaia do Sul | Epiclin | Fernando Hayashi Sant'Anna, Ana Paula Muterle, Janira Prichula, Juliana Comerlato, Carolina Comerlato, Eliana Márcia Da Ros Wendland |
| EPI_ISL_1469578 | HOSPITAL SAO FRANCISCO DE ASSIS | Epiclin | Fernando Hayashi Sant'Anna, Ana Paula Muterle, Janira Prichula, Juliana Comerlato, Carolina Comerlato, Eliana Márcia Da Ros Wendland |
| EPI_ISL_1469579 | DIRETORIA DE VIGILANCIA EM SAUDE | Epiclin | Fernando Hayashi Sant'Anna, Ana Paula Muterle, Janira Prichula, Juliana Comerlato, Carolina Comerlato, Eliana Márcia Da Ros Wendland |
| EPI_ISL_1469580 | Unidade de Pronto Atendimento de Sapucaia do Sul | Epiclin | Fernando Hayashi Sant'Anna, Ana Paula Muterle, Janira Prichula, Juliana Comerlato, Carolina Comerlato, Eliana Márcia Da Ros Wendland |
| EPI_ISL_1469581 | Hospital Dia e Pronto Atendimento | Epiclin | Fernando Hayashi Sant'Anna, Ana Paula Muterle, Janira Prichula, Juliana Comerlato, Carolina Comerlato, Eliana Márcia Da Ros Wendland |
| EPI_ISL_1469582 | Unidade de Pronto Atendimento de Sapucaia do Sul | Epiclin | Fernando Hayashi Sant'Anna, Ana Paula Muterle, Janira Prichula, Juliana Comerlato, Carolina Comerlato, Eliana Márcia Da Ros Wendland |
| EPI_ISL_1469583 | UNIDADE SANITARIA DE IGREJINHA | Epiclin | Fernando Hayashi Sant'Anna, Ana Paula Muterle, Janira Prichula, Juliana Comerlato, Carolina Comerlato, Eliana Márcia Da Ros Wendland |
| EPI_ISL_1469584 | CENTRO MUNICIPAL DE SAUDE DE ROLANTE | Epiclin | Fernando Hayashi Sant'Anna, Ana Paula Muterle, Janira Prichula, Juliana Comerlato, Carolina Comerlato, Eliana Márcia Da Ros Wendland |
| EPI_ISL_1469585 | Fundação Hospitalar de Sapucaia do Sul | Epiclin | Fernando Hayashi Sant'Anna, Ana Paula Muterle, Janira Prichula, Juliana Comerlato, Carolina Comerlato, Eliana Márcia Da Ros Wendland |
| EPI_ISL_1469586 | FUNDACAO HOSPITALAR SAO JOSE | Epiclin | Fernando Hayashi Sant'Anna, Ana Paula Muterle, Janira Prichula, Juliana Comerlato, Carolina Comerlato, Eliana Márcia Da Ros Wendland |
| EPI_ISL_1469587 | Hospital Sapiranga | Epiclin | Fernando Hayashi Sant'Anna, Ana Paula Muterle, Janira Prichula, Juliana Comerlato, Carolina Comerlato, Eliana Márcia Da Ros Wendland |
| EPI_ISL_1469588 | Secretaria Municipal de Saúde de Taquara | Epiclin | Fernando Hayashi Sant'Anna, Ana Paula Muterle, Janira Prichula, Juliana Comerlato, Carolina Comerlato, Eliana Márcia Da Ros Wendland |
| EPI_ISL_1469589 | UNIDADE DE PRONTO ATENDIMENTO DE SAPUCAIA DO SUL UPA | Epiclin | Fernando Hayashi Sant'Anna, Ana Paula Muterle, Janira Prichula, Juliana Comerlato, Carolina Comerlato, Eliana Márcia Da Ros Wendland |
| EPI_ISL_1469590 | DIRETORIA DE VIGILANCIA EM SAUDE | Epiclin | Fernando Hayashi Sant'Anna, Ana Paula Muterle, Janira Prichula, Juliana Comerlato, Carolina Comerlato, Eliana Márcia Da Ros Wendland |
| EPI_ISL_1469591 | Hospital Nossa Senhora das Graças | Epiclin | Fernando Hayashi Sant'Anna, Ana Paula Muterle, Janira Prichula, Juliana Comerlato, Carolina Comerlato, Eliana Márcia Da Ros Wendland |
| EPI_ISL_1469592 | Diretoria de Vigilância em Saúde | Epiclin | Fernando Hayashi Sant'Anna, Ana Paula Muterle, Janira Prichula, Juliana Comerlato, Carolina Comerlato, Eliana Márcia Da Ros Wendland |
| EPI_ISL_1469593 | Hospital Sapiranga | Epiclin | Fernando Hayashi Sant'Anna, Ana Paula Muterle, Janira Prichula, Juliana Comerlato, Carolina Comerlato, Eliana Márcia Da Ros Wendland |
| EPI_ISL_1469594 | DIRETORIA DE VIGILANCIA EM SAUDE | Epiclin | Fernando Hayashi Sant'Anna, Ana Paula Muterle, Janira Prichula, Juliana Comerlato, Carolina Comerlato, Eliana Márcia Da Ros Wendland |
| EPI_ISL_1469596 | CENTRO DE REFERENCIA EM SINDROMES GRIPAIS | Epiclin | Fernando Hayashi Sant'Anna, Ana Paula Muterle, Janira Prichula, Juliana Comerlato, Carolina Comerlato, Eliana Márcia Da Ros Wendland |
| EPI_ISL_1469597 | Fundação de Saúde Pública de Novo Hamburgo | Epiclin | Fernando Hayashi Sant'Anna, Ana Paula Muterle, Janira Prichula, Juliana Comerlato, Carolina Comerlato, Eliana Márcia Da Ros Wendland |
| EPI_ISL_1469598 | Diretoria de Vigilância em Saúde | Epiclin | Fernando Hayashi Sant'Anna, Ana Paula Muterle, Janira Prichula, Juliana Comerlato, Carolina Comerlato, Eliana Márcia Da Ros Wendland |
| EPI_ISL_1469600 | Hospital de Portão | Epiclin | Fernando Hayashi Sant'Anna, Ana Paula Muterle, Janira Prichula, Juliana Comerlato, Carolina Comerlato, Eliana Márcia Da Ros Wendland |
| EPI_ISL_1469601 | SECRETARIA MUNICIPAL DE SAUDE DE TAQUARA | Epiclin | Fernando Hayashi Sant'Anna, Ana Paula Muterle, Janira Prichula, Juliana Comerlato, Carolina Comerlato, Eliana Márcia Da Ros Wendland |
| EPI_ISL_1469602 | Pronto Atendimento Campo Bom | Epiclin | Fernando Hayashi Sant'Anna, Ana Paula Muterle, Janira Prichula, Juliana Comerlato, Carolina Comerlato, Eliana Márcia Da Ros Wendland |
| EPI_ISL_1469603 | Centro de Referência em Síndromes Gripais | Epiclin | Fernando Hayashi Sant'Anna, Ana Paula Muterle, Janira Prichula, Juliana Comerlato, Carolina Comerlato, Eliana Márcia Da Ros Wendland |
| EPI_ISL_1469604 | COORDENADORIA GERAL DE VIGILANCIA EM SAUDE | Epiclin | Fernando Hayashi Sant'Anna, Ana Paula Muterle, Janira Prichula, Juliana Comerlato, Carolina Comerlato, Eliana Márcia Da Ros Wendland |
| EPI_ISL_1469606 | Fundação Hospitalar de Sapucaia do Sul | Epiclin | Fernando Hayashi Sant'Anna, Ana Paula Muterle, Janira Prichula, Juliana Comerlato, Carolina Comerlato, Eliana Márcia Da Ros Wendland |
| EPI_ISL_1469607 | Secretaria Municipal de Saúde de Três Coroas | Epiclin | Fernando Hayashi Sant'Anna, Ana Paula Muterle, Janira Prichula, Juliana Comerlato, Carolina Comerlato, Eliana Márcia Da Ros Wendland |
| EPI_ISL_1469608 | COORDENADORIA GERAL DE VIGILANCIA EM SAUDE | Epiclin | Fernando Hayashi Sant'Anna, Ana Paula Muterle, Janira Prichula, Juliana Comerlato, Carolina Comerlato, Eliana Márcia Da Ros Wendland |
| EPI_ISL_1469609, EPI_ISL_1469610 | Diretoria de Vigilância em Saúde | Epiclin | Fernando Hayashi Sant'Anna, Ana Paula Muterle, Janira Prichula, Juliana Comerlato, Carolina Comerlato, Eliana Márcia Da Ros Wendland |
| EPI_ISL_1469611 | DIRETORIA DE VIGILANCIA EM SAUDE | Epiclin | Fernando Hayashi Sant'Anna, Ana Paula Muterle, Janira Prichula, Juliana Comerlato, Carolina Comerlato, Eliana Márcia Da Ros Wendland |
| EPI_ISL_1469612 | FUNDACAO DE SAUDE PUBLICA DE NOVO HAMBURGO FSNH | Epiclin | Fernando Hayashi Sant'Anna, Ana Paula Muterle, Janira Prichula, Juliana Comerlato, Carolina Comerlato, Eliana Márcia Da Ros Wendland |
| EPI_ISL_1469613 | Fundação de Saúde Pública de Novo Hamburgo | Epiclin | Fernando Hayashi Sant'Anna, Ana Paula Muterle, Janira Prichula, Juliana Comerlato, Carolina Comerlato, Eliana Márcia Da Ros Wendland |
| EPI_ISL_1469614 | Unidade Sanitária de Igrejinha | Epiclin | Fernando Hayashi Sant'Anna, Ana Paula Muterle, Janira Prichula, Juliana Comerlato, Carolina Comerlato, Eliana Márcia Da Ros Wendland |
| EPI_ISL_1469615 | Pronto Atendimento Campo Bom | Epiclin | Fernando Hayashi Sant'Anna, Ana Paula Muterle, Janira Prichula, Juliana Comerlato, Carolina Comerlato, Eliana Márcia Da Ros Wendland |
| EPI_ISL_1469616 | Diretoria de Vigilância em Saúde | Epiclin | Fernando Hayashi Sant'Anna, Ana Paula Muterle, Janira Prichula, Juliana Comerlato, Carolina Comerlato, Eliana Márcia Da Ros Wendland |
| EPI_ISL_1469617, EPI_ISL_1469619 | Fundação Hospitalar de Sapucaia do Sul | Epiclin | Fernando Hayashi Sant'Anna, Ana Paula Muterle, Janira Prichula, Juliana Comerlato, Carolina Comerlato, Eliana Márcia Da Ros Wendland |
| EPI_ISL_1469620 | FUNDACAO DE SAUDE PUBLICA DE NOVO HAMBURGO FSNH | Epiclin | Fernando Hayashi Sant'Anna, Ana Paula Muterle, Janira Prichula, Juliana Comerlato, Carolina Comerlato, Eliana Márcia Da Ros Wendland |
| EPI_ISL_1469621 | Pronto Atendimento Campo Bom | Epiclin | Fernando Hayashi Sant'Anna, Ana Paula Muterle, Janira Prichula, Juliana Comerlato, Carolina Comerlato, Eliana Márcia Da Ros Wendland |
| EPI_ISL_1469623 | Diretoria de Vigilância em Saúde | Epiclin | Fernando Hayashi Sant'Anna, Ana Paula Muterle, Janira Prichula, Juliana Comerlato, Carolina Comerlato, Eliana Márcia Da Ros Wendland |
| EPI_ISL_1469624 | HOSPITAL MUNICIPAL GETULIO VARGAS | Epiclin | Fernando Hayashi Sant'Anna, Ana Paula Muterle, Janira Prichula, Juliana Comerlato, Carolina Comerlato, Eliana Márcia Da Ros Wendland |
| EPI_ISL_1469625 | Diretoria de Vigilância em Saúde | Epiclin | Fernando Hayashi Sant'Anna, Ana Paula Muterle, Janira Prichula, Juliana Comerlato, Carolina Comerlato, Eliana Márcia Da Ros Wendland |
| EPI_ISL_1469626 | Fundação de Saúde Pública de Novo Hamburgo | Epiclin | Fernando Hayashi Sant'Anna, Ana Paula Muterle, Janira Prichula, Juliana Comerlato, Carolina Comerlato, Eliana Márcia Da Ros Wendland |
| EPI_ISL_1469627 | Diretoria de Vigilância em Saúde | Epiclin | Fernando Hayashi Sant'Anna, Ana Paula Muterle, Janira Prichula, Juliana Comerlato, Carolina Comerlato, Eliana Márcia Da Ros Wendland |
| EPI_ISL_1469628 | CENTRO DE SERVICOS ESPECIALIZADOS SANTA RITA DE CASSIA | Epiclin | Fernando Hayashi Sant'Anna, Ana Paula Muterle, Janira Prichula, Juliana Comerlato, Carolina Comerlato, Eliana Márcia Da Ros Wendland |
| EPI_ISL_1469629 | CENTRO MUNICIPAL DE SAUDE DE ROLANTE | Epiclin | Fernando Hayashi Sant'Anna, Ana Paula Muterle, Janira Prichula, Juliana Comerlato, Carolina Comerlato, Eliana Márcia Da Ros Wendland |

| | | | |
|---|---|---|---|
| EPI_ISL_1469630 | FUNDACAO DE SAUDE PUBLICA DE NOVO HAMBURGO FSNH | Epiclin | Fernando Hayashi Sant'Anna, Ana Paula Muterle, Janira Prichula, Juliana Comerlato, Carolina Comerlato, Eliana Márcia Da Ros Wendland |
| EPI_ISL_1469631 | HOSPITAL MUNICIPAL GETULIO VARGAS | Epiclin | Fernando Hayashi Sant'Anna, Ana Paula Muterle, Janira Prichula, Juliana Comerlato, Carolina Comerlato, Eliana Márcia Da Ros Wendland |
| EPI_ISL_1469633 | DIRETORIA DE VIGILANCIA EM SAUDE | Epiclin | Fernando Hayashi Sant'Anna, Ana Paula Muterle, Janira Prichula, Juliana Comerlato, Carolina Comerlato, Eliana Márcia Da Ros Wendland |
| EPI_ISL_1469634 | Diretoria de Vigilância em Saúde | Epiclin | Fernando Hayashi Sant'Anna, Ana Paula Muterle, Janira Prichula, Juliana Comerlato, Carolina Comerlato, Eliana Márcia Da Ros Wendland |
| EPI_ISL_1469635, EPI_ISL_1469636 | COORDENADORIA GERAL DE VIGILANCIA EM SAUDE | Epiclin | Fernando Hayashi Sant'Anna, Ana Paula Muterle, Janira Prichula, Juliana Comerlato, Carolina Comerlato, Eliana Márcia Da Ros Wendland |
| EPI_ISL_1469637, EPI_ISL_1469638 | DIRETORIA DE VIGILANCIA EM SAUDE | Epiclin | Fernando Hayashi Sant'Anna, Ana Paula Muterle, Janira Prichula, Juliana Comerlato, Carolina Comerlato, Eliana Márcia Da Ros Wendland |
| EPI_ISL_1469639 | SECRETARIA MUNICIPAL DE SAUDE DE SAO LEOPOLDO | Epiclin | Fernando Hayashi Sant'Anna, Ana Paula Muterle, Janira Prichula, Juliana Comerlato, Carolina Comerlato, Eliana Márcia Da Ros Wendland |
| EPI_ISL_1469640 | Fundação de Saúde Pública São Camilo de Esteio | Epiclin | Fernando Hayashi Sant'Anna, Ana Paula Muterle, Janira Prichula, Juliana Comerlato, Carolina Comerlato, Eliana Márcia Da Ros Wendland |
| EPI_ISL_1469641 | COORDENADORIA GERAL DE VIGILANCIA EM SAUDE | Epiclin | Fernando Hayashi Sant'Anna, Ana Paula Muterle, Janira Prichula, Juliana Comerlato, Carolina Comerlato, Eliana Márcia Da Ros Wendland |
| EPI_ISL_1469642 | CENTRO DE REFERENCIA EM SINDROMES GRIPAIS | Epiclin | Fernando Hayashi Sant'Anna, Ana Paula Muterle, Janira Prichula, Juliana Comerlato, Carolina Comerlato, Eliana Márcia Da Ros Wendland |
| EPI_ISL_1469643 | HOSPITAL SAO FRANCISCO DE ASSIS | Epiclin | Fernando Hayashi Sant'Anna, Ana Paula Muterle, Janira Prichula, Juliana Comerlato, Carolina Comerlato, Eliana Márcia Da Ros Wendland |
| EPI_ISL_1469644 | Diretoria de Vigilância em Saúde | Epiclin | Fernando Hayashi Sant'Anna, Ana Paula Muterle, Janira Prichula, Juliana Comerlato, Carolina Comerlato, Eliana Márcia Da Ros Wendland |
| EPI_ISL_1469645 | SECRETARIA MUNICIPAL DE SAUDE DE TRES COROAS | Epiclin | Fernando Hayashi Sant'Anna, Ana Paula Muterle, Janira Prichula, Juliana Comerlato, Carolina Comerlato, Eliana Márcia Da Ros Wendland |
| EPI_ISL_1469646 | UNIDADE DE PRONTO ATENDIMENTO DE SAPUCAIA DO SUL UPA | Epiclin | Fernando Hayashi Sant'Anna, Ana Paula Muterle, Janira Prichula, Juliana Comerlato, Carolina Comerlato, Eliana Márcia Da Ros Wendland |
| EPI_ISL_1469647 | Secretaria Municipal de Saúde de Taquara | Epiclin | Fernando Hayashi Sant'Anna, Ana Paula Muterle, Janira Prichula, Juliana Comerlato, Carolina Comerlato, Eliana Márcia Da Ros Wendland |
| EPI_ISL_1469648 | HOSPITAL SAO FRANCISCO DE ASSIS | Epiclin | Fernando Hayashi Sant'Anna, Ana Paula Muterle, Janira Prichula, Juliana Comerlato, Carolina Comerlato, Eliana Márcia Da Ros Wendland |
| EPI_ISL_1469649 | Unidade de Atendimento DST AIDS TB e Han | Epiclin | Fernando Hayashi Sant'Anna, Ana Paula Muterle, Janira Prichula, Juliana Comerlato, Carolina Comerlato, Eliana Márcia Da Ros Wendland |
| EPI_ISL_1469651 | FUNDACAO DE SAUDE PUBLICA DE NOVO HAMBURGO FSNH | Epiclin | Fernando Hayashi Sant'Anna, Ana Paula Muterle, Janira Prichula, Juliana Comerlato, Carolina Comerlato, Eliana Márcia Da Ros Wendland |
| EPI_ISL_1469652 | DIRETORIA DE VIGILANCIA EM SAUDE | Epiclin | Fernando Hayashi Sant'Anna, Ana Paula Muterle, Janira Prichula, Juliana Comerlato, Carolina Comerlato, Eliana Márcia Da Ros Wendland |
| EPI_ISL_1469653 | Fundação de Saúde Pública de Novo Hamburgo | Epiclin | Fernando Hayashi Sant'Anna, Ana Paula Muterle, Janira Prichula, Juliana Comerlato, Carolina Comerlato, Eliana Márcia Da Ros Wendland |
| EPI_ISL_1469656 | Diretoria de Vigilância em Saúde | Epiclin | Fernando Hayashi Sant'Anna, Ana Paula Muterle, Janira Prichula, Juliana Comerlato, Carolina Comerlato, Eliana Márcia Da Ros Wendland |
| EPI_ISL_1469657 | Pronto Atendimento Campo Bom | Epiclin | Fernando Hayashi Sant'Anna, Ana Paula Muterle, Janira Prichula, Juliana Comerlato, Carolina Comerlato, Eliana Márcia Da Ros Wendland |
| EPI_ISL_1469658 | CENTRO DE ESPECIALIDADES TRIUNFO | Epiclin | Fernando Hayashi Sant'Anna, Ana Paula Muterle, Janira Prichula, Juliana Comerlato, Carolina Comerlato, Eliana Márcia Da Ros Wendland |
| EPI_ISL_1469659 | DIRETORIA DE VIGILANCIA EM SAUDE | Epiclin | Fernando Hayashi Sant'Anna, Ana Paula Muterle, Janira Prichula, Juliana Comerlato, Carolina Comerlato, Eliana Márcia Da Ros Wendland |
| EPI_ISL_1469660 | UNIDADE BASICA DE SAUDE DE RIOZINHO | Epiclin | Fernando Hayashi Sant'Anna, Ana Paula Muterle, Janira Prichula, Juliana Comerlato, Carolina Comerlato, Eliana Márcia Da Ros Wendland |
| EPI_ISL_1469661 | FUNDACAO DE SAUDE PUBLICA DE NOVO HAMBURGO FSNH | Epiclin | Fernando Hayashi Sant'Anna, Ana Paula Muterle, Janira Prichula, Juliana Comerlato, Carolina Comerlato, Eliana Márcia Da Ros Wendland |
| EPI_ISL_1469662 | CENTRO DE SAUDE DR BRUNO CASSEL | Epiclin | Fernando Hayashi Sant'Anna, Ana Paula Muterle, Janira Prichula, Juliana Comerlato, Carolina Comerlato, Eliana Márcia Da Ros Wendland |
| EPI_ISL_1469663 | DIRETORIA DE VIGILANCIA EM SAUDE | Epiclin | Fernando Hayashi Sant'Anna, Ana Paula Muterle, Janira Prichula, Juliana Comerlato, Carolina Comerlato, Eliana Márcia Da Ros Wendland |
| EPI_ISL_1469664 | Unidade de Atendimento DST AIDS TB e Han | Epiclin | Fernando Hayashi Sant'Anna, Ana Paula Muterle, Janira Prichula, Juliana Comerlato, Carolina Comerlato, Eliana Márcia Da Ros Wendland |
| EPI_ISL_1469665 | FUNDACAO DE SAUDE PUBLICA SAO CAMILO DE ESTEIO | Epiclin | Fernando Hayashi Sant'Anna, Ana Paula Muterle, Janira Prichula, Juliana Comerlato, Carolina Comerlato, Eliana Márcia Da Ros Wendland |
| EPI_ISL_1469666 | Secretaria Municipal de Saúde de Montenegro | Epiclin | Fernando Hayashi Sant'Anna, Ana Paula Muterle, Janira Prichula, Juliana Comerlato, Carolina Comerlato, Eliana Márcia Da Ros Wendland |
| EPI_ISL_1469667 | Centro de Referência em Síndromes Gripais | Epiclin | Fernando Hayashi Sant'Anna, Ana Paula Muterle, Janira Prichula, Juliana Comerlato, Carolina Comerlato, Eliana Márcia Da Ros Wendland |
| EPI_ISL_1469668 | Hospital Unimed Vale do Caí | Epiclin | Fernando Hayashi Sant'Anna, Ana Paula Muterle, Janira Prichula, Juliana Comerlato, Carolina Comerlato, Eliana Márcia Da Ros Wendland |
| EPI_ISL_1469669 | Diretoria de Vigilância em Saúde | Epiclin | Fernando Hayashi Sant'Anna, Ana Paula Muterle, Janira Prichula, Juliana Comerlato, Carolina Comerlato, Eliana Márcia Da Ros Wendland |
| EPI_ISL_1469670 | Hospital São Francisco de Assis | Epiclin | Fernando Hayashi Sant'Anna, Ana Paula Muterle, Janira Prichula, Juliana Comerlato, Carolina Comerlato, Eliana Márcia Da Ros Wendland |
| EPI_ISL_1469672 | Diretoria de Vigilância em Saúde | Epiclin | Fernando Hayashi Sant'Anna, Ana Paula Muterle, Janira Prichula, Juliana Comerlato, Carolina Comerlato, Eliana Márcia Da Ros Wendland |
| EPI_ISL_1469673 | Fundação de Saúde Pública de Novo Hamburgo | Epiclin | Fernando Hayashi Sant'Anna, Ana Paula Muterle, Janira Prichula, Juliana Comerlato, Carolina Comerlato, Eliana Márcia Da Ros Wendland |
| EPI_ISL_1469674, EPI_ISL_1469675 | DIRETORIA DE VIGILANCIA EM SAUDE | Epiclin | Fernando Hayashi Sant'Anna, Ana Paula Muterle, Janira Prichula, Juliana Comerlato, Carolina Comerlato, Eliana Márcia Da Ros Wendland |
| EPI_ISL_1469676 | HOSPITAL SAO FRANCISCO DE ASSIS | Epiclin | Fernando Hayashi Sant'Anna, Ana Paula Muterle, Janira Prichula, Juliana Comerlato, Carolina Comerlato, Eliana Márcia Da Ros Wendland |
| EPI_ISL_1469677 | DIRETORIA DE VIGILANCIA EM SAUDE | Epiclin | Fernando Hayashi Sant'Anna, Ana Paula Muterle, Janira Prichula, Juliana Comerlato, Carolina Comerlato, Eliana Márcia Da Ros Wendland |
| EPI_ISL_1469678 | SECRETARIA MUNICIPAL DE SAUDE DE TRES COROAS | Epiclin | Fernando Hayashi Sant'Anna, Ana Paula Muterle, Janira Prichula, Juliana Comerlato, Carolina Comerlato, Eliana Márcia Da Ros Wendland |
| EPI_ISL_1469679 | DIRETORIA DE VIGILANCIA EM SAUDE | Epiclin | Fernando Hayashi Sant'Anna, Ana Paula Muterle, Janira Prichula, Juliana Comerlato, Carolina Comerlato, Eliana Márcia Da Ros Wendland |
| EPI_ISL_1469680 | UNIDADE DE ATENDIMENTO DST AIDS TB E HAN | Epiclin | Fernando Hayashi Sant'Anna, Ana Paula Muterle, Janira Prichula, Juliana Comerlato, Carolina Comerlato, Eliana Márcia Da Ros Wendland |
| EPI_ISL_1469681 | DIRETORIA DE VIGILANCIA EM SAUDE | Epiclin | Fernando Hayashi Sant'Anna, Ana Paula Muterle, Janira Prichula, Juliana Comerlato, Carolina Comerlato, Eliana Márcia Da Ros Wendland |
| EPI_ISL_1469682 | CENTRO DE SERVICOS ESPECIALIZADOS SANTA RITA DE CASSIA | Epiclin | Fernando Hayashi Sant'Anna, Ana Paula Muterle, Janira Prichula, Juliana Comerlato, Carolina Comerlato, Eliana Márcia Da Ros Wendland |
| EPI_ISL_1469683 | SECRETARIA MUNICIPAL DE SAUDE DE TAQUARA | Epiclin | Fernando Hayashi Sant'Anna, Ana Paula Muterle, Janira Prichula, Juliana Comerlato, Carolina Comerlato, Eliana Márcia Da Ros Wendland |
| EPI_ISL_1469684 | DIRETORIA DE VIGILANCIA EM SAUDE | Epiclin | Fernando Hayashi Sant'Anna, Ana Paula Muterle, Janira Prichula, Juliana Comerlato, Carolina Comerlato, Eliana Márcia Da Ros Wendland |
| EPI_ISL_1469685 | HOSPITAL SAO FRANCISCO DE ASSIS | Epiclin | Fernando Hayashi Sant'Anna, Ana Paula Muterle, Janira Prichula, Juliana Comerlato, Carolina Comerlato, Eliana Márcia Da Ros Wendland |
| EPI_ISL_1469686 | DIRETORIA DE VIGILANCIA EM SAUDE | Epiclin | Fernando Hayashi Sant'Anna, Ana Paula Muterle, Janira Prichula, Juliana Comerlato, Carolina Comerlato, Eliana Márcia Da Ros Wendland |
| EPI_ISL_1469687 | SECRETARIA MUNICIPAL DE SAUDE DE TAQUARA | Epiclin | Fernando Hayashi Sant'Anna, Ana Paula Muterle, Janira Prichula, Juliana Comerlato, Carolina Comerlato, Eliana Márcia Da Ros Wendland |
| EPI_ISL_1469688 | DIRETORIA DE VIGILANCIA EM SAUDE | Epiclin | Fernando Hayashi Sant'Anna, Ana Paula Muterle, Janira Prichula, Juliana Comerlato, Carolina Comerlato, Eliana Márcia Da Ros Wendland |
| EPI_ISL_1469689 | FUNDACAO DE SAUDE PUBLICA SAO CAMILO DE ESTEIO | Epiclin | Fernando Hayashi Sant'Anna, Ana Paula Muterle, Janira Prichula, Juliana Comerlato, Carolina Comerlato, Eliana Márcia Da Ros Wendland |
| EPI_ISL_1469692 | DIRETORIA DE VIGILANCIA EM SAUDE | Epiclin | Fernando Hayashi Sant'Anna, Ana Paula Muterle, Janira Prichula, Juliana Comerlato, Carolina Comerlato, Eliana Márcia Da Ros Wendland |
| EPI_ISL_1469693 | FUNDACAO DE SAUDE PUBLICA SAO CAMILO DE ESTEIO | Epiclin | Fernando Hayashi Sant'Anna, Ana Paula Muterle, Janira Prichula, Juliana Comerlato, Carolina Comerlato, Eliana Márcia Da Ros Wendland |
| EPI_ISL_1469694 | UNIDADE BASICA DE SAUDE PARECI NOVO | Epiclin | Fernando Hayashi Sant'Anna, Ana Paula Muterle, Janira Prichula, Juliana Comerlato, Carolina Comerlato, Eliana Márcia Da Ros Wendland |

| | | | |
|---|---|---|---|
| EPI_ISL_1469695 | Unidade de Pronto Atendimento de Sapucaia do Sul | Epiclin | Fernando Hayashi Sant'Anna, Ana Paula Muterle, Janira Prichula, Juliana Comerlato, Carolina Comerlato, Eliana Márcia Da Ros Wendland |
| EPI_ISL_1469696 | Vigilância em Saúde de Sapucaia do Sul | Epiclin | Fernando Hayashi Sant'Anna, Ana Paula Muterle, Janira Prichula, Juliana Comerlato, Carolina Comerlato, Eliana Márcia Da Ros Wendland |
| EPI_ISL_1469697 | Hospital Regina | Epiclin | Fernando Hayashi Sant'Anna, Ana Paula Muterle, Janira Prichula, Juliana Comerlato, Carolina Comerlato, Eliana Márcia Da Ros Wendland |
| EPI_ISL_1469698 | CENTRO DE SERVICOS ESPECIALIZADOS SANTA RITA DE CASSIA | Epiclin | Fernando Hayashi Sant'Anna, Ana Paula Muterle, Janira Prichula, Juliana Comerlato, Carolina Comerlato, Eliana Márcia Da Ros Wendland |
| EPI_ISL_1469699 | HOSPITAL MUNICIPAL GETULIO VARGAS | Epiclin | Fernando Hayashi Sant'Anna, Ana Paula Muterle, Janira Prichula, Juliana Comerlato, Carolina Comerlato, Eliana Márcia Da Ros Wendland |
| EPI_ISL_1469703 | Unidade de Pronto Atendimento de Sapucaia do Sul | Epiclin | Fernando Hayashi Sant'Anna, Ana Paula Muterle, Janira Prichula, Juliana Comerlato, Carolina Comerlato, Eliana Márcia Da Ros Wendland |
| EPI_ISL_1469704 | FUNDACAO DE SAUDE PUBLICA SAO CAMILO DE ESTEIO | Epiclin | Fernando Hayashi Sant'Anna, Ana Paula Muterle, Janira Prichula, Juliana Comerlato, Carolina Comerlato, Eliana Márcia Da Ros Wendland |
| EPI_ISL_1469705 | HOSPITAL DE CAMPO BOM DR LAURO REUS | Epiclin | Fernando Hayashi Sant'Anna, Ana Paula Muterle, Janira Prichula, Juliana Comerlato, Carolina Comerlato, Eliana Márcia Da Ros Wendland |
| EPI_ISL_1469706 | CENTRO DE SAUDE DR BRUNO CASSEL | Epiclin | Fernando Hayashi Sant'Anna, Ana Paula Muterle, Janira Prichula, Juliana Comerlato, Carolina Comerlato, Eliana Márcia Da Ros Wendland |
| EPI_ISL_1469707 | Hospital Bom Jesus | Epiclin | Fernando Hayashi Sant'Anna, Ana Paula Muterle, Janira Prichula, Juliana Comerlato, Carolina Comerlato, Eliana Márcia Da Ros Wendland |
| EPI_ISL_1469708 | Diretoria de Vigilância em Saúde | Epiclin | Fernando Hayashi Sant'Anna, Ana Paula Muterle, Janira Prichula, Juliana Comerlato, Carolina Comerlato, Eliana Márcia Da Ros Wendland |
| EPI_ISL_1469709 | SECRETARIA MUNICIPAL DE SAUDE DE TAQUARA | Epiclin | Fernando Hayashi Sant'Anna, Ana Paula Muterle, Janira Prichula, Juliana Comerlato, Carolina Comerlato, Eliana Márcia Da Ros Wendland |
| EPI_ISL_1469710 | DIRETORIA DE VIGILANCIA EM SAUDE | Epiclin | Fernando Hayashi Sant'Anna, Ana Paula Muterle, Janira Prichula, Juliana Comerlato, Carolina Comerlato, Eliana Márcia Da Ros Wendland |
| EPI_ISL_1469711 | Hospital Dia e Pronto Atendimento | Epiclin | Fernando Hayashi Sant'Anna, Ana Paula Muterle, Janira Prichula, Juliana Comerlato, Carolina Comerlato, Eliana Márcia Da Ros Wendland |
| EPI_ISL_1469712 | Pronto Atendimento Campo Bom | Epiclin | Fernando Hayashi Sant'Anna, Ana Paula Muterle, Janira Prichula, Juliana Comerlato, Carolina Comerlato, Eliana Márcia Da Ros Wendland |
| EPI_ISL_1469713 | Fundação Hospitalar de Sapucaia do Sul | Epiclin | Fernando Hayashi Sant'Anna, Ana Paula Muterle, Janira Prichula, Juliana Comerlato, Carolina Comerlato, Eliana Márcia Da Ros Wendland |
| EPI_ISL_1469714 | Unidade de Atendimento DST AIDS TB e Han | Epiclin | Fernando Hayashi Sant'Anna, Ana Paula Muterle, Janira Prichula, Juliana Comerlato, Carolina Comerlato, Eliana Márcia Da Ros Wendland |
| EPI_ISL_1469715 | COORDENADORIA GERAL DE VIGILANCIA EM SAUDE | Epiclin | Fernando Hayashi Sant'Anna, Ana Paula Muterle, Janira Prichula, Juliana Comerlato, Carolina Comerlato, Eliana Márcia Da Ros Wendland |
| EPI_ISL_1469719 | UNIDADE DE PRONTO ATENDIMENTO DE SAPUCAIA DO SUL UPA | Epiclin | Fernando Hayashi Sant'Anna, Ana Paula Muterle, Janira Prichula, Juliana Comerlato, Carolina Comerlato, Eliana Márcia Da Ros Wendland |
| EPI_ISL_1469720 | UNIDADE SANITARIA DE IGREJINHA | Epiclin | Fernando Hayashi Sant'Anna, Ana Paula Muterle, Janira Prichula, Juliana Comerlato, Carolina Comerlato, Eliana Márcia Da Ros Wendland |
| EPI_ISL_1469721 | Pronto Atendimento Cruzeiro do Sul | Epiclin | Fernando Hayashi Sant'Anna, Ana Paula Muterle, Janira Prichula, Juliana Comerlato, Carolina Comerlato, Eliana Márcia Da Ros Wendland |
| EPI_ISL_1469722 | UNIDADE SANITARIA DE IGREJINHA | Epiclin | Fernando Hayashi Sant'Anna, Ana Paula Muterle, Janira Prichula, Juliana Comerlato, Carolina Comerlato, Eliana Márcia Da Ros Wendland |
| EPI_ISL_1469723 | FUNDACAO DE SAUDE PUBLICA SAO CAMILO DE ESTEIO | Epiclin | Fernando Hayashi Sant'Anna, Ana Paula Muterle, Janira Prichula, Juliana Comerlato, Carolina Comerlato, Eliana Márcia Da Ros Wendland |
| EPI_ISL_1469724 | Fundação Hospitalar de Sapucaia do Sul | Epiclin | Fernando Hayashi Sant'Anna, Ana Paula Muterle, Janira Prichula, Juliana Comerlato, Carolina Comerlato, Eliana Márcia Da Ros Wendland |
| EPI_ISL_1469725 | DIRETORIA DE VIGILANCIA EM SAUDE | Epiclin | Fernando Hayashi Sant'Anna, Ana Paula Muterle, Janira Prichula, Juliana Comerlato, Carolina Comerlato, Eliana Márcia Da Ros Wendland |
| EPI_ISL_1469726 | HOSPITAL SAPIRANGA | Epiclin | Fernando Hayashi Sant'Anna, Ana Paula Muterle, Janira Prichula, Juliana Comerlato, Carolina Comerlato, Eliana Márcia Da Ros Wendland |
| EPI_ISL_1469727 | Secretaria Municipal de Saúde de São Leopoldo | Epiclin | Fernando Hayashi Sant'Anna, Ana Paula Muterle, Janira Prichula, Juliana Comerlato, Carolina Comerlato, Eliana Márcia Da Ros Wendland |
| EPI_ISL_1469729 | Centro de Especialidades Triunfo | Epiclin | Fernando Hayashi Sant'Anna, Ana Paula Muterle, Janira Prichula, Juliana Comerlato, Carolina Comerlato, Eliana Márcia Da Ros Wendland |
| EPI_ISL_1469730 | CENTRO DE REFERENCIA EM SINDROMES GRIPAIS | Epiclin | Fernando Hayashi Sant'Anna, Ana Paula Muterle, Janira Prichula, Juliana Comerlato, Carolina Comerlato, Eliana Márcia Da Ros Wendland |
| EPI_ISL_1469731 | DIRETORIA DE VIGILANCIA EM SAUDE | Epiclin | Fernando Hayashi Sant'Anna, Ana Paula Muterle, Janira Prichula, Juliana Comerlato, Carolina Comerlato, Eliana Márcia Da Ros Wendland |
| EPI_ISL_1469732 | CENTRO DE REFERENCIA EM SINDROMES GRIPAIS | Epiclin | Fernando Hayashi Sant'Anna, Ana Paula Muterle, Janira Prichula, Juliana Comerlato, Carolina Comerlato, Eliana Márcia Da Ros Wendland |
| EPI_ISL_1469733 | UNIDADE DE PRONTO ATENDIMENTO DE SAPUCAIA DO SUL UPA | Epiclin | Fernando Hayashi Sant'Anna, Ana Paula Muterle, Janira Prichula, Juliana Comerlato, Carolina Comerlato, Eliana Márcia Da Ros Wendland |
| EPI_ISL_1469734 | Hospital Dia e Pronto Atendimento | Epiclin | Fernando Hayashi Sant'Anna, Ana Paula Muterle, Janira Prichula, Juliana Comerlato, Carolina Comerlato, Eliana Márcia Da Ros Wendland |
| EPI_ISL_1469736 | SECRETARIA MUNICIPAL DE SAUDE DE SAO LEOPOLDO | Epiclin | Fernando Hayashi Sant'Anna, Ana Paula Muterle, Janira Prichula, Juliana Comerlato, Carolina Comerlato, Eliana Márcia Da Ros Wendland |
| EPI_ISL_1469737 | CENTRO DE REFERENCIA EM SINDROMES GRIPAIS | Epiclin | Fernando Hayashi Sant'Anna, Ana Paula Muterle, Janira Prichula, Juliana Comerlato, Carolina Comerlato, Eliana Márcia Da Ros Wendland |
| EPI_ISL_1469740 | SECRETARIA MUNICIPAL DE SAUDE DE TRES COROAS | Epiclin | Fernando Hayashi Sant'Anna, Ana Paula Muterle, Janira Prichula, Juliana Comerlato, Carolina Comerlato, Eliana Márcia Da Ros Wendland |
| EPI_ISL_1469741 | Centro de Referência em Síndromes Gripais | Epiclin | Fernando Hayashi Sant'Anna, Ana Paula Muterle, Janira Prichula, Juliana Comerlato, Carolina Comerlato, Eliana Márcia Da Ros Wendland |
| EPI_ISL_1469742 | CENTRO DE REFERENCIA EM SINDROMES GRIPAIS | Epiclin | Fernando Hayashi Sant'Anna, Ana Paula Muterle, Janira Prichula, Juliana Comerlato, Carolina Comerlato, Eliana Márcia Da Ros Wendland |
| EPI_ISL_1469743 | DIRETORIA DE VIGILANCIA EM SAUDE | Epiclin | Fernando Hayashi Sant'Anna, Ana Paula Muterle, Janira Prichula, Juliana Comerlato, Carolina Comerlato, Eliana Márcia Da Ros Wendland |
| EPI_ISL_1469744 | HOSPITAL SAO FRANCISCO DE ASSIS | Epiclin | Fernando Hayashi Sant'Anna, Ana Paula Muterle, Janira Prichula, Juliana Comerlato, Carolina Comerlato, Eliana Márcia Da Ros Wendland |
| EPI_ISL_1469746 | HOSPITAL MONTENEGRO | Epiclin | Fernando Hayashi Sant'Anna, Ana Paula Muterle, Janira Prichula, Juliana Comerlato, Carolina Comerlato, Eliana Márcia Da Ros Wendland |
| EPI_ISL_1469747 | Unidade de Pronto Atendimento de Sapucaia do Sul | Epiclin | Fernando Hayashi Sant'Anna, Ana Paula Muterle, Janira Prichula, Juliana Comerlato, Carolina Comerlato, Eliana Márcia Da Ros Wendland |
| EPI_ISL_1469748 | Diretoria de Vigilância em Saúde | Epiclin | Fernando Hayashi Sant'Anna, Ana Paula Muterle, Janira Prichula, Juliana Comerlato, Carolina Comerlato, Eliana Márcia Da Ros Wendland |
| EPI_ISL_1469749 | Centro de Referência em Síndromes Gripais | Epiclin | Fernando Hayashi Sant'Anna, Ana Paula Muterle, Janira Prichula, Juliana Comerlato, Carolina Comerlato, Eliana Márcia Da Ros Wendland |
| EPI_ISL_1469751 | UNIDADE BASICA DE SAUDE DE RIOZINHO | Epiclin | Fernando Hayashi Sant'Anna, Ana Paula Muterle, Janira Prichula, Juliana Comerlato, Carolina Comerlato, Eliana Márcia Da Ros Wendland |
| EPI_ISL_1469752 | Fundação Hospitalar de Sapucaia do Sul | Epiclin | Fernando Hayashi Sant'Anna, Ana Paula Muterle, Janira Prichula, Juliana Comerlato, Carolina Comerlato, Eliana Márcia Da Ros Wendland |
| EPI_ISL_1469753 | Diretoria de Vigilância em Saúde | Epiclin | Fernando Hayashi Sant'Anna, Ana Paula Muterle, Janira Prichula, Juliana Comerlato, Carolina Comerlato, Eliana Márcia Da Ros Wendland |
| EPI_ISL_1469754 | Hospital Municipal Getúlio Vargas | Epiclin | Fernando Hayashi Sant'Anna, Ana Paula Muterle, Janira Prichula, Juliana Comerlato, Carolina Comerlato, Eliana Márcia Da Ros Wendland |
| EPI_ISL_1469756 | Diretoria de Vigilância em Saúde | Epiclin | Fernando Hayashi Sant'Anna, Ana Paula Muterle, Janira Prichula, Juliana Comerlato, Carolina Comerlato, Eliana Márcia Da Ros Wendland |
| EPI_ISL_1469757 | Secretaria Municipal de Saúde de São Leopoldo | Epiclin | Fernando Hayashi Sant'Anna, Ana Paula Muterle, Janira Prichula, Juliana Comerlato, Carolina Comerlato, Eliana Márcia Da Ros Wendland |
| EPI_ISL_1469759 | HOSPITAL MUNICIPAL GETULIO VARGAS | Epiclin | Fernando Hayashi Sant'Anna, Ana Paula Muterle, Janira Prichula, Juliana Comerlato, Carolina Comerlato, Eliana Márcia Da Ros Wendland |
| EPI_ISL_1469760 | DIRETORIA DE VIGILANCIA EM SAUDE | Epiclin | Fernando Hayashi Sant'Anna, Ana Paula Muterle, Janira Prichula, Juliana Comerlato, Carolina Comerlato, Eliana Márcia Da Ros Wendland |
| EPI_ISL_1469761 | Diretoria de Vigilância em Saúde | Epiclin | Fernando Hayashi Sant'Anna, Ana Paula Muterle, Janira Prichula, Juliana Comerlato, Carolina Comerlato, Eliana Márcia Da Ros Wendland |
| EPI_ISL_1469762 | Hospital Nossa Senhora das Graças | Epiclin | Fernando Hayashi Sant'Anna, Ana Paula Muterle, Janira Prichula, Juliana Comerlato, Carolina Comerlato, Eliana Márcia Da Ros Wendland |
| EPI_ISL_1469763 | Unidade Básica de Saúde de Riozinho | Epiclin | Fernando Hayashi Sant'Anna, Ana Paula Muterle, Janira Prichula, Juliana Comerlato, Carolina Comerlato, Eliana Márcia Da Ros Wendland |
| EPI_ISL_1469766 | Coordenadoria Geral de Vigilância em Saúde · Vigilância em Saúde | Epiclin | Fernando Hayashi Sant'Anna, Ana Paula Muterle, Janira Prichula, Juliana Comerlato, Carolina Comerlato, Eliana Márcia Da Ros Wendland |

| | | | |
|---|---|---|---|
| EPI_ISL_1469767 | Fundação de Saúde Pública de Novo Hamburgo | Epiclin | Fernando Hayashi Sant'Anna, Ana Paula Muterle, Janira Prichula, Juliana Comerlato, Carolina Comerlato, Eliana Márcia Da Ros Wendland |
| EPI_ISL_1469771 | Hospital de Portão | Epiclin | Fernando Hayashi Sant'Anna, Ana Paula Muterle, Janira Prichula, Juliana Comerlato, Carolina Comerlato, Eliana Márcia Da Ros Wendland |
| EPI_ISL_1469773 | Hospital Universitário | Epiclin | Fernando Hayashi Sant'Anna, Ana Paula Muterle, Janira Prichula, Juliana Comerlato, Carolina Comerlato, Eliana Márcia Da Ros Wendland |
| EPI_ISL_1469774 | Hospital Universitário de Canoas | Epiclin | Fernando Hayashi Sant'Anna, Ana Paula Muterle, Janira Prichula, Juliana Comerlato, Carolina Comerlato, Eliana Márcia Da Ros Wendland |
| EPI_ISL_1469777 | Centro de Referência em Síndromes Gripais | Epiclin | Fernando Hayashi Sant'Anna, Ana Paula Muterle, Janira Prichula, Juliana Comerlato, Carolina Comerlato, Eliana Márcia Da Ros Wendland |
| EPI_ISL_1469778 | Hospital Municipal Getúlio Vargas | Epiclin | Fernando Hayashi Sant'Anna, Ana Paula Muterle, Janira Prichula, Juliana Comerlato, Carolina Comerlato, Eliana Márcia Da Ros Wendland |
| EPI_ISL_1469779 | Centro de Especialidades Triunfo | Epiclin | Fernando Hayashi Sant'Anna, Ana Paula Muterle, Janira Prichula, Juliana Comerlato, Carolina Comerlato, Eliana Márcia Da Ros Wendland |
| EPI_ISL_1469781 | DIRETORIA DE VIGILANCIA EM SAUDE | Epiclin | Fernando Hayashi Sant'Anna, Ana Paula Muterle, Janira Prichula, Juliana Comerlato, Carolina Comerlato, Eliana Márcia Da Ros Wendland |
| EPI_ISL_1469782 | COORDENADORIA GERAL DE VIGILANCIA EM SAUDE | Epiclin | Fernando Hayashi Sant'Anna, Ana Paula Muterle, Janira Prichula, Juliana Comerlato, Carolina Comerlato, Eliana Márcia Da Ros Wendland |
| EPI_ISL_1469783 | Diretoria de Vigilância em Saúde | Epiclin | Fernando Hayashi Sant'Anna, Ana Paula Muterle, Janira Prichula, Juliana Comerlato, Carolina Comerlato, Eliana Márcia Da Ros Wendland |
| EPI_ISL_1469785 | FUNDACAO DE SAUDE PUBLICA DE NOVO HAMBURGO FSNH | Epiclin | Fernando Hayashi Sant'Anna, Ana Paula Muterle, Janira Prichula, Juliana Comerlato, Carolina Comerlato, Eliana Márcia Da Ros Wendland |
| EPI_ISL_1469786 | Diretoria de Vigilância em Saúde | Epiclin | Fernando Hayashi Sant'Anna, Ana Paula Muterle, Janira Prichula, Juliana Comerlato, Carolina Comerlato, Eliana Márcia Da Ros Wendland |
| EPI_ISL_1469789 | Hospital São Francisco de Assis | Epiclin | Fernando Hayashi Sant'Anna, Ana Paula Muterle, Janira Prichula, Juliana Comerlato, Carolina Comerlato, Eliana Márcia Da Ros Wendland |
| EPI_ISL_1469791 | DIRETORIA DE VIGILANCIA EM SAUDE | Epiclin | Fernando Hayashi Sant'Anna, Ana Paula Muterle, Janira Prichula, Juliana Comerlato, Carolina Comerlato, Eliana Márcia Da Ros Wendland |
| EPI_ISL_1469792 | Diretoria de Vigilância em Saúde | Epiclin | Fernando Hayashi Sant'Anna, Ana Paula Muterle, Janira Prichula, Juliana Comerlato, Carolina Comerlato, Eliana Márcia Da Ros Wendland |
| EPI_ISL_1469794, EPI_ISL_1469795 | Hospital Dia e Pronto Atendimento | Epiclin | Fernando Hayashi Sant'Anna, Ana Paula Muterle, Janira Prichula, Juliana Comerlato, Carolina Comerlato, Eliana Márcia Da Ros Wendland |
| EPI_ISL_1469796, EPI_ISL_1469797 | Diretoria de Vigilância em Saúde | Epiclin | Fernando Hayashi Sant'Anna, Ana Paula Muterle, Janira Prichula, Juliana Comerlato, Carolina Comerlato, Eliana Márcia Da Ros Wendland |
| EPI_ISL_1469798 | DIRETORIA DE VIGILANCIA EM SAUDE | Epiclin | Fernando Hayashi Sant'Anna, Ana Paula Muterle, Janira Prichula, Juliana Comerlato, Carolina Comerlato, Eliana Márcia Da Ros Wendland |
| EPI_ISL_1469799 | Hospital Regina | Epiclin | Fernando Hayashi Sant'Anna, Ana Paula Muterle, Janira Prichula, Juliana Comerlato, Carolina Comerlato, Eliana Márcia Da Ros Wendland |
| EPI_ISL_1469802 | Diretoria de Vigilância em Saúde | Epiclin | Fernando Hayashi Sant'Anna, Ana Paula Muterle, Janira Prichula, Juliana Comerlato, Carolina Comerlato, Eliana Márcia Da Ros Wendland |
| EPI_ISL_1469803 | Secretaria Municipal de Saúde de Três Coroas | Epiclin | Fernando Hayashi Sant'Anna, Ana Paula Muterle, Janira Prichula, Juliana Comerlato, Carolina Comerlato, Eliana Márcia Da Ros Wendland |
| EPI_ISL_1469806 | Centro de Referência em Síndromes Gripais | Epiclin | Fernando Hayashi Sant'Anna, Ana Paula Muterle, Janira Prichula, Juliana Comerlato, Carolina Comerlato, Eliana Márcia Da Ros Wendland |
| EPI_ISL_1469807 | Hospital São Francisco de Assis | Epiclin | Fernando Hayashi Sant'Anna, Ana Paula Muterle, Janira Prichula, Juliana Comerlato, Carolina Comerlato, Eliana Márcia Da Ros Wendland |
| EPI_ISL_1469808 | Diretoria de Vigilância em Saúde | Epiclin | Fernando Hayashi Sant'Anna, Ana Paula Muterle, Janira Prichula, Juliana Comerlato, Carolina Comerlato, Eliana Márcia Da Ros Wendland |
| EPI_ISL_1469809 | Fundação Hospitalar de Sapucaia do Sul | Epiclin | Fernando Hayashi Sant'Anna, Ana Paula Muterle, Janira Prichula, Juliana Comerlato, Carolina Comerlato, Eliana Márcia Da Ros Wendland |
| EPI_ISL_1469816, EPI_ISL_1469819 | Diretoria de Vigilância em Saúde | Epiclin | Fernando Hayashi Sant'Anna, Ana Paula Muterle, Janira Prichula, Juliana Comerlato, Carolina Comerlato, Eliana Márcia Da Ros Wendland |
| EPI_ISL_1469821 | Centro de Saúde Bruno Cassel | Epiclin | Fernando Hayashi Sant'Anna, Ana Paula Muterle, Janira Prichula, Juliana Comerlato, Carolina Comerlato, Eliana Márcia Da Ros Wendland |
| EPI_ISL_1469823 | Diretoria de Vigilância em Saúde | Epiclin | Fernando Hayashi Sant'Anna, Ana Paula Muterle, Janira Prichula, Juliana Comerlato, Carolina Comerlato, Eliana Márcia Da Ros Wendland |
| EPI_ISL_1469824 | Hospital Regina | Epiclin | Fernando Hayashi Sant'Anna, Ana Paula Muterle, Janira Prichula, Juliana Comerlato, Carolina Comerlato, Eliana Márcia Da Ros Wendland |
| EPI_ISL_1469825 | Pronto Atendimento Campo Bom | Epiclin | Fernando Hayashi Sant'Anna, Ana Paula Muterle, Janira Prichula, Juliana Comerlato, Carolina Comerlato, Eliana Márcia Da Ros Wendland |
| EPI_ISL_1469826 | Fundação de Saúde Pública de Novo Hamburgo | Epiclin | Fernando Hayashi Sant'Anna, Ana Paula Muterle, Janira Prichula, Juliana Comerlato, Carolina Comerlato, Eliana Márcia Da Ros Wendland |
| EPI_ISL_1469827 | Hospital de Portão | Epiclin | Fernando Hayashi Sant'Anna, Ana Paula Muterle, Janira Prichula, Juliana Comerlato, Carolina Comerlato, Eliana Márcia Da Ros Wendland |
| EPI_ISL_1469828 | Fundação de Saúde Pública de Novo Hamburgo | Epiclin | Fernando Hayashi Sant'Anna, Ana Paula Muterle, Janira Prichula, Juliana Comerlato, Carolina Comerlato, Eliana Márcia Da Ros Wendland |
| EPI_ISL_1469831 | Fundação Hospitalar de Sapucaia do Sul | Epiclin | Fernando Hayashi Sant'Anna, Ana Paula Muterle, Janira Prichula, Juliana Comerlato, Carolina Comerlato, Eliana Márcia Da Ros Wendland |
| EPI_ISL_1469833 | Hospital Nossa Senhora das Graças | Epiclin | Fernando Hayashi Sant'Anna, Ana Paula Muterle, Janira Prichula, Juliana Comerlato, Carolina Comerlato, Eliana Márcia Da Ros Wendland |
| EPI_ISL_1469834 | HOSPITAL SAO FRANCISCO DE ASSIS | Epiclin | Fernando Hayashi Sant'Anna, Ana Paula Muterle, Janira Prichula, Juliana Comerlato, Carolina Comerlato, Eliana Márcia Da Ros Wendland |
| EPI_ISL_1469835 | Diretoria de Vigilância em Saúde | Epiclin | Fernando Hayashi Sant'Anna, Ana Paula Muterle, Janira Prichula, Juliana Comerlato, Carolina Comerlato, Eliana Márcia Da Ros Wendland |
| EPI_ISL_1469838 | Secretaria Municipal de Saúde de Três Coroas | Epiclin | Fernando Hayashi Sant'Anna, Ana Paula Muterle, Janira Prichula, Juliana Comerlato, Carolina Comerlato, Eliana Márcia Da Ros Wendland |
| EPI_ISL_1469840, EPI_ISL_1469843 | Hospital Sapiranga | Epiclin | Fernando Hayashi Sant'Anna, Ana Paula Muterle, Janira Prichula, Juliana Comerlato, Carolina Comerlato, Eliana Márcia Da Ros Wendland |
| EPI_ISL_1469844 | Hospital Universitário de Canoas | Epiclin | Fernando Hayashi Sant'Anna, Ana Paula Muterle, Janira Prichula, Juliana Comerlato, Carolina Comerlato, Eliana Márcia Da Ros Wendland |
| EPI_ISL_1469845 | Fundação Hospitalar de Sapucaia do Sul | Epiclin | Fernando Hayashi Sant'Anna, Ana Paula Muterle, Janira Prichula, Juliana Comerlato, Carolina Comerlato, Eliana Márcia Da Ros Wendland |
| EPI_ISL_1469848 | Diretoria de Vigilância em Saúde | Epiclin | Fernando Hayashi Sant'Anna, Ana Paula Muterle, Janira Prichula, Juliana Comerlato, Carolina Comerlato, Eliana Márcia Da Ros Wendland |
| EPI_ISL_1469849 | Hospital Universitário de Canoas | Epiclin | Fernando Hayashi Sant'Anna, Ana Paula Muterle, Janira Prichula, Juliana Comerlato, Carolina Comerlato, Eliana Márcia Da Ros Wendland |
| EPI_ISL_1469850 | Fundação de Saúde Pública São Camilo de Esteio | Epiclin | Fernando Hayashi Sant'Anna, Ana Paula Muterle, Janira Prichula, Juliana Comerlato, Carolina Comerlato, Eliana Márcia Da Ros Wendland |
| EPI_ISL_1479119 | DIRETORIA DE VIGILANCIA EM SAUDE | Epiclin | Fernando Hayashi Sant'Anna, Ana Paula Muterle, Janira Prichula, Juliana Comerlato, Carolina Comerlato, Eliana Márcia Da Ros Wendland |
| EPI_ISL_1479120 | Diretoria de Vigilância em Saúde | Epiclin | Fernando Hayashi Sant'Anna, Ana Paula Muterle, Janira Prichula, Juliana Comerlato, Carolina Comerlato, Eliana Márcia Da Ros Wendland |
| EPI_ISL_1479121 | DIRETORIA DE VIGILANCIA EM SAUDE | Epiclin | Fernando Hayashi Sant'Anna, Ana Paula Muterle, Janira Prichula, Juliana Comerlato, Carolina Comerlato, Eliana Márcia Da Ros Wendland |
| EPI_ISL_1479123 | Fundação de Saúde Pública de Novo Hamburgo | Epiclin | Fernando Hayashi Sant'Anna, Ana Paula Muterle, Janira Prichula, Juliana Comerlato, Carolina Comerlato, Eliana Márcia Da Ros Wendland |
| EPI_ISL_1479124 | UNIDADE DE SAUDE NOVA HARTZ | Epiclin | Fernando Hayashi Sant'Anna, Ana Paula Muterle, Janira Prichula, Juliana Comerlato, Carolina Comerlato, Eliana Márcia Da Ros Wendland |
| EPI_ISL_1479125 | CENTRO DE REFERENCIA EM SINDROMES GRIPAIS | Epiclin | Fernando Hayashi Sant'Anna, Ana Paula Muterle, Janira Prichula, Juliana Comerlato, Carolina Comerlato, Eliana Márcia Da Ros Wendland |
| EPI_ISL_1479126 | UNIDADE DE ATENDIMENTO DST AIDS TB E HAN | Epiclin | Fernando Hayashi Sant'Anna, Ana Paula Muterle, Janira Prichula, Juliana Comerlato, Carolina Comerlato, Eliana Márcia Da Ros Wendland |
| EPI_ISL_1479127 | Hospital Regina | Epiclin | Fernando Hayashi Sant'Anna, Ana Paula Muterle, Janira Prichula, Juliana Comerlato, Carolina Comerlato, Eliana Márcia Da Ros Wendland |
| EPI_ISL_1479128 | HOSPITAL SAO FRANCISCO DE ASSIS | Epiclin | Fernando Hayashi Sant'Anna, Ana Paula Muterle, Janira Prichula, Juliana Comerlato, Carolina Comerlato, Eliana Márcia Da Ros Wendland |
| EPI_ISL_1479129 | Pronto Atendimento Campo Bom | Epiclin | Fernando Hayashi Sant'Anna, Ana Paula Muterle, Janira Prichula, Juliana Comerlato, Carolina Comerlato, Eliana Márcia Da Ros Wendland |
| EPI_ISL_1479131, EPI_ISL_1479132 | Fundação de Saúde Pública de Novo Hamburgo | Epiclin | Fernando Hayashi Sant'Anna, Ana Paula Muterle, Janira Prichula, Juliana Comerlato, Carolina Comerlato, Eliana Márcia Da Ros Wendland |
| EPI_ISL_1493572 | Centro de Saude II Dr Alcides Facundo Arroyo | Instituto Adolfo Lutz, Interdiciplinary Procedures Center, Strategic Laboratory | Claudio Tavares Sacchi, Claudia Regina Gonçalves, Erica Valessa Ramos Gomes, Karoline Rodrigues Campos, Caio Vinicius Dias Lopes |
| EPI_ISL_1493573, EPI_ISL_1493574, EPI_ISL_1493575, EPI_ISL_1493576, | LACEN do Estado de Goias | Instituto Adolfo Lutz, Interdiciplinary Procedures Center, Strategic Laboratory | Claudio Tavares Sacchi, Claudia Regina Gonçalves, Erica Valessa Ramos Gomes, Karoline Rodrigues Campos, Caio Vinicius Dias Lopes |

| | | | |
|---|---|---|---|
| EPI_ISL_1493577 | | | |
| EPI_ISL_1493578, EPI_ISL_1493579 | LACEN do Estado de Rondonia | Instituto Adolfo Lutz, Interdiciplinary Procedures Center, Strategic Laboratory | Claudio Tavares Sacchi, Claudia Regina Gonçalves, Erica Valessa Ramos Gomes, Karoline Rodrigues Campos, Caio Vinicius Dias Lopes |
| EPI_ISL_1493580 | CS II Dr Miguel Vitaliano Orlandia | Instituto Adolfo Lutz, Interdiciplinary Procedures Center, Strategic Laboratory | Claudio Tavares Sacchi, Claudia Regina Gonçalves, Erica Valessa Ramos Gomes, Karoline Rodrigues Campos, Caio Vinicius Dias Lopes |
| EPI_ISL_1493581 | Sae servico de Atendimento Especializado | Instituto Adolfo Lutz, Interdiciplinary Procedures Center, Strategic Laboratory | Claudio Tavares Sacchi, Claudia Regina Gonçalves, Erica Valessa Ramos Gomes, Karoline Rodrigues Campos, Caio Vinicius Dias Lopes |
| EPI_ISL_1493582 | CS II Dr Miguel Vitaliano Orlandia | Instituto Adolfo Lutz, Interdiciplinary Procedures Center, Strategic Laboratory | Claudio Tavares Sacchi, Claudia Regina Gonçalves, Erica Valessa Ramos Gomes, Karoline Rodrigues Campos, Caio Vinicius Dias Lopes |
| EPI_ISL_1493584 | LACEN do Estado de Rondonia | Instituto Adolfo Lutz, Interdiciplinary Procedures Center, Strategic Laboratory | Claudio Tavares Sacchi, Claudia Regina Gonçalves, Erica Valessa Ramos Gomes, Karoline Rodrigues Campos, Caio Vinicius Dias Lopes |
| EPI_ISL_1493585 | Unidade de Saude Dr Phebo de Oliveira Roge Ferreira | Instituto Adolfo Lutz, Interdiciplinary Procedures Center, Strategic Laboratory | Claudio Tavares Sacchi, Claudia Regina Gonçalves, Erica Valessa Ramos Gomes, Karoline Rodrigues Campos, Caio Vinicius Dias Lopes |
| EPI_ISL_1493586 | Hospital Sao Marcos da Samamorro Agudo | Instituto Adolfo Lutz, Interdiciplinary Procedures Center, Strategic Laboratory | Claudio Tavares Sacchi, Claudia Regina Gonçalves, Erica Valessa Ramos Gomes, Karoline Rodrigues Campos, Caio Vinicius Dias Lopes |
| EPI_ISL_1493587, EPI_ISL_1493588 | CS III de Patrocinio Paulista | Instituto Adolfo Lutz, Interdiciplinary Procedures Center, Strategic Laboratory | Claudio Tavares Sacchi, Claudia Regina Gonçalves, Erica Valessa Ramos Gomes, Karoline Rodrigues Campos, Caio Vinicius Dias Lopes |
| EPI_ISL_1493589 | CS II Dr Jose Ferreira Telles | Instituto Adolfo Lutz, Interdiciplinary Procedures Center, Strategic Laboratory | Claudio Tavares Sacchi, Claudia Regina Gonçalves, Erica Valessa Ramos Gomes, Karoline Rodrigues Campos, Caio Vinicius Dias Lopes |
| EPI_ISL_1493590 | Hospital Sao Marcos da Samamorro Agudo | Instituto Adolfo Lutz, Interdiciplinary Procedures Center, Strategic Laboratory | Claudio Tavares Sacchi, Claudia Regina Gonçalves, Erica Valessa Ramos Gomes, Karoline Rodrigues Campos, Caio Vinicius Dias Lopes |
| EPI_ISL_1493591 | Centro de Saude II Dr Alcides Facundo Arroyo | Instituto Adolfo Lutz, Interdiciplinary Procedures Center, Strategic Laboratory | Claudio Tavares Sacchi, Claudia Regina Gonçalves, Erica Valessa Ramos Gomes, Karoline Rodrigues Campos, Caio Vinicius Dias Lopes |
| EPI_ISL_1493592 | Santa Casa de Guaira | Instituto Adolfo Lutz, Interdiciplinary Procedures Center, Strategic Laboratory | Claudio Tavares Sacchi, Claudia Regina Gonçalves, Erica Valessa Ramos Gomes, Karoline Rodrigues Campos, Caio Vinicius Dias Lopes |
| EPI_ISL_1493593 | CS II Dr Jahyr de Paula Ribeiro Guara | Instituto Adolfo Lutz, Interdiciplinary Procedures Center, Strategic Laboratory | Claudio Tavares Sacchi, Claudia Regina Gonçalves, Erica Valessa Ramos Gomes, Karoline Rodrigues Campos, Caio Vinicius Dias Lopes |
| EPI_ISL_1493594 | Unidade de Saude Dr Phebo de Oliveira Roge Ferreira | Instituto Adolfo Lutz, Interdiciplinary Procedures Center, Strategic Laboratory | Claudio Tavares Sacchi, Claudia Regina Gonçalves, Erica Valessa Ramos Gomes, Karoline Rodrigues Campos, Caio Vinicius Dias Lopes |
| EPI_ISL_1493595, EPI_ISL_1493596, EPI_ISL_1493597, EPI_ISL_1493598, EPI_ISL_1493599, EPI_ISL_1493600 | LACEN do Estado de Rondonia | Instituto Adolfo Lutz, Interdiciplinary Procedures Center, Strategic Laboratory | Claudio Tavares Sacchi, Claudia Regina Gonçalves, Erica Valessa Ramos Gomes, Karoline Rodrigues Campos, Caio Vinicius Dias Lopes |
| EPI_ISL_1494923 | CS II Dr Jose Ferreira Telles | Instituto Adolfo Lutz, Interdiciplinary Procedures Center, Strategic Laboratory | Claudio Tavares Sacchi, Claudia Regina Gonçalves, Erica Valessa Ramos Gomes, Karoline Rodrigues Campos, Caio Vinicius Dias Lopes |
| EPI_ISL_1494924 | LACEN do Estado de Rondonia | Instituto Adolfo Lutz, Interdiciplinary Procedures Center, Strategic Laboratory | Claudio Tavares Sacchi, Claudia Regina Gonçalves, Erica Valessa Ramos Gomes, Karoline Rodrigues Campos, Caio Vinicius Dias Lopes |
| EPI_ISL_1494960, EPI_ISL_1494963, EPI_ISL_1494964, EPI_ISL_1494965, EPI_ISL_1494966, EPI_ISL_1494967, EPI_ISL_1494968, EPI_ISL_1494969, EPI_ISL_1494970, EPI_ISL_1494971, EPI_ISL_1494975, EPI_ISL_1494976, EPI_ISL_1494977, EPI_ISL_1494979, EPI_ISL_1494980, EPI_ISL_1494981, EPI_ISL_1494983, EPI_ISL_1494985, EPI_ISL_1494986, EPI_ISL_1494988, EPI_ISL_1494989, EPI_ISL_1494990, EPI_ISL_1494991, EPI_ISL_1494992, EPI_ISL_1494994, EPI_ISL_1494995, EPI_ISL_1494996, EPI_ISL_1494997, EPI_ISL_1494998, EPI_ISL_1494999, EPI_ISL_1495002, EPI_ISL_1495004, EPI_ISL_1495005, EPI_ISL_1495006, EPI_ISL_1495008, EPI_ISL_1495009, EPI_ISL_1495010, EPI_ISL_1495011, EPI_ISL_1495013, EPI_ISL_1495014, EPI_ISL_1495015, EPI_ISL_1495016, EPI_ISL_1495017, EPI_ISL_1495018, EPI_ISL_1495019, EPI_ISL_1495020, EPI_ISL_1495021, EPI_ISL_1495022, EPI_ISL_1495023, EPI_ISL_1495024, EPI_ISL_1495025, EPI_ISL_1495026, EPI_ISL_1495029, EPI_ISL_1495030, EPI_ISL_1495031, EPI_ISL_1495034, EPI_ISL_1495035, EPI_ISL_1495038, EPI_ISL_1495041, EPI_ISL_1495042, EPI_ISL_1497548 | | | |
| see above | Laboratório de Biologia Integrativa | Laboratório de Biologia Integrativa | Filipe Romero Rebello Moreira, Diego Menezes Bonfim, Victor Emmanuel Viana Geddes, Danielle Alves Gomes Zauli, Joice do Prado Silva, Aline Brito de Lima, Frederico Scott Varella Malta, Alessandro Clayton de Souza Ferreira, Victor Cavalcanti Pardini, Daniel Costa Queiroz, Rafael Marques de Souza, Lucyene Miguita Luiz, Paula Luize Camargos Fonseca, Rennan Garcias Moreira, Nuno Rodrigues Faria, Carolina Moreira Voloch, Renan Pedra de Souza, Renato Santana Aguiar |
| EPI_ISL_1498263 | Associação Fundo de Incentivo à Pesquisa (AFIP) | Associação Fundo de Incentivo à Pesquisa (AFIP) | Priscila Farias Tempaku, Juliana Nogueira Martins Rodrigues, Erika Rodrigues de Oliveira, Debora R. Ramadan, Soraya Sgambatti de Andrade, Sergio Tufik. |
| EPI_ISL_1498380 | Associação Fundo de Incentivo à Pesquisa (AFIP). | Associação Fundo de Incentivo à Pesquisa (AFIP). | Priscila Farias Tempaku, Juliana Nogueira Martins Rodrigues, Erika Rodrigues de Oliveira, Debora R. Ramadan, Soraya Sgambatti de Andrade, Sergio Tufik. |
| EPI_ISL_1498585, EPI_ISL_1498821, EPI_ISL_1498823, EPI_ISL_1498824, EPI_ISL_1498825, EPI_ISL_1498916 | Associação Fundo de Incentivo à Pesquisa (AFIP) | Associação Fundo de Incentivo à Pesquisa (AFIP) | Priscila Farias Tempaku, Juliana Nogueira Martins Rodrigues, Erika Rodrigues de Oliveira, Debora R. Ramadan, Soraya Sgambatti de Andrade, Sergio Tufik. |
| EPI_ISL_1498917 | Hospital Estadual de Mirandopolis | Instituto Adolfo Lutz, Interdiciplinary Procedures Center, Strategic Laboratory | Claudio Tavares Sacchi, Claudia Regina Gonçalves, Erica Valessa Ramos Gomes, Karoline Rodrigues Campos, Caio Vinicius Dias Lopes |
| EPI_ISL_1498919, EPI_ISL_1499020, EPI_ISL_1499105, EPI_ISL_1499114, EPI_ISL_1499201, EPI_ISL_1499202, EPI_ISL_1499203, EPI_ISL_1499205, EPI_ISL_1499297, EPI_ISL_1499298, EPI_ISL_1499299, EPI_ISL_1499300 | | | |
| see above | Associação Fundo de Incentivo à Pesquisa (AFIP) | Associação Fundo de Incentivo à Pesquisa (AFIP) | Priscila Farias Tempaku, Juliana Nogueira Martins Rodrigues, Erika Rodrigues de Oliveira, Debora R. Ramadan, Soraya Sgambatti de Andrade, Sergio Tufik. |
| EPI_ISL_1509639 | UBS de Auriflama | Instituto Adolfo Lutz, Interdiciplinary Procedures Center, Strategic Laboratory | Claudio Tavares Sacchi, Claudia Regina Gonçalves, Erica Valessa Ramos Gomes, Karoline Rodrigues Campos, Caio Vinicius Dias Lopes, Leonardo Jose Tadeu de Araujo |
| EPI_ISL_1509720 | UBS de Auriflama | Instituto Adolfo Lutz, Interdiciplinary Procedures Center, Strategic Laboratory | Claudio Tavares Sacchi, Claudia Regina Gonçalves, Erica Valessa Ramos Gomes, Karoline Rodrigues Campos, Caio Vinicius Dias Lopes |
| EPI_ISL_1511641 | Laboratorio de Patologia Clinica - UNICAMP | Laboratorio de Estudos de Virus Emergentes | Mariene R. Amorim, William M. Souza, Antonio C. G. Carlos Jr, Daniel A. Toledo-Teixeira, Karina Bispo-dos-Santos, Camila L. Simeoni, Pierina L. Parise, Aline Vieira, Julia Forato, Ingra M. Claro, Luciana S. Mofatto, Natalia S. Brunetti, Emerson S.S. França, Gisele A. Pedroso, Barbara F. N. Carvalho, Tania R. Zaccariotto, Kamila C. S. Krywacz, André S. Vieira, Marcelo A. Mori, Alessandro S. Farias, Maria H. P. Pavan, Luís Felipe Bachur, Luís G. O. Cardoso, Fernando R. Spilki, Ester C. Sabino, Nuno R. Faria, Magnun N. N. Santos, Rodrigo Angerami, Patricia A. F. Leme, Angelica Schreiber, Maria L. Moretti, Fabiana Granja, José Luiz Proenca-Modena |
| EPI_ISL_1511643 | Genomic and molecular Biology Group, A.C.Camargo Cancer Center | Laboratory of Bioinformatics and Computational Biology, A.C.Camargo Cancer Center | Giovana Torrezan, Dirce Carraro, Israel Tojal, Alexandre Defelicibus |
| EPI_ISL_1520107, EPI_ISL_1520108, EPI_ISL_1520109 | LACEN do Estado de Rondonia | Instituto Adolfo Lutz, Interdiciplinary Procedures Center, Strategic Laboratory | Claudio Tavares Sacchi, Claudia Regina Gonçalves, Erica Valessa Ramos Gomes, Karoline Rodrigues Campos, Caio Vinicius Dias Lopes |
| EPI_ISL_1520110, EPI_ISL_1520111, EPI_ISL_1520112 | Hospital Municipal Reynaldo Guerra Cajati | Instituto Adolfo Lutz, Interdiciplinary Procedures Center, Strategic Laboratory | Claudio Tavares Sacchi, Claudia Regina Gonçalves, Erica Valessa Ramos Gomes, Karoline Rodrigues Campos, Caio Vinicius Dias Lopes |
| EPI_ISL_1520113 | Hospital Santo Antonio de Juquia Juquia | Instituto Adolfo Lutz, Interdiciplinary Procedures Center, | Claudio Tavares Sacchi, Claudia Regina Gonçalves, Erica Valessa Ramos Gomes, Karoline Rodrigues Campos, Caio Vinicius Dias Lopes |

| | | | |
|---|---|---|---|
| | | Strategic Laboratory | |
| EPI_ISL_1520114 | UBS III de Pariquera Acu Pariquera Acu | Instituto Adolfo Lutz, Interdiciplinary Procedures Center, Strategic Laboratory | Claudio Tavares Sacchi, Claudia Regina Gonçalves, Erica Valessa Ramos Gomes, Karoline Rodrigues Campos, Caio Vinicius Dias Lopes |
| EPI_ISL_1520115, EPI_ISL_1520116 | Unidade de Pronto Atendimento UPA | Instituto Adolfo Lutz, Interdiciplinary Procedures Center, Strategic Laboratory | Claudio Tavares Sacchi, Claudia Regina Gonçalves, Erica Valessa Ramos Gomes, Karoline Rodrigues Campos, Caio Vinicius Dias Lopes |
| EPI_ISL_1520117, EPI_ISL_1520118, EPI_ISL_1520119, EPI_ISL_1520120, EPI_ISL_1520121, EPI_ISL_1520122, EPI_ISL_1520123, EPI_ISL_1520124, EPI_ISL_1520125, EPI_ISL_1520126, EPI_ISL_1520127, EPI_ISL_1520128 | | | |
| see above | Centro de Saude II Dr Jose de Felipe Espito Santo do Pinhal SP | Instituto Adolfo Lutz, Interdiciplinary Procedures Center, Strategic Laboratory | Claudio Tavares Sacchi, Claudia Regina Gonçalves, Erica Valessa Ramos Gomes, Karoline Rodrigues Campos, Caio Vinicius Dias Lopes |
| EPI_ISL_1520129, EPI_ISL_1520130, EPI_ISL_1520131, EPI_ISL_1520132, EPI_ISL_1520133, EPI_ISL_1520134, EPI_ISL_1520135, EPI_ISL_1520136, EPI_ISL_1520137 | Centro de Saude II Dr Jose Paione Mococa | Instituto Adolfo Lutz, Interdiciplinary Procedures Center, Strategic Laboratory | Claudio Tavares Sacchi, Claudia Regina Gonçalves, Erica Valessa Ramos Gomes, Karoline Rodrigues Campos, Caio Vinicius Dias Lopes |
| EPI_ISL_1533609 | Laboratorio de Ecologia de Doencas Transmissiveis na Amazonia, Instituto Leonidas e Maria Deane - Fiocruz Amazonia | Laboratorio de Ecologia de Doencas Transmissiveis na Amazonia, Instituto Leonidas e Maria Deane - Fiocruz Amazonia | Valdinete Nascimento, Victor Souza, André Corado, Fernanda Nascimento, George Silva, Ágatha Costa, Debora Duarte, Karina Pessoa, Matilde Mejía, Luciana Gonçalves, Maria Júlia Brandão, Michele Jesus, Felipe Naveca |
| EPI_ISL_1533689, EPI_ISL_1533690, EPI_ISL_1533691 | Centro de Saude II Dr. Jose Paione Mococa | Instituto Adolfo Lutz, Interdiciplinary Procedures Center, Strategic Laboratory | Claudio Tavares Sacchi, Claudia Regina Gonçalves, Erica Valessa Ramos Gomes, Karoline Rodrigues Campos, Caio Vinicius Dias Lopes, Leonardo Jose Tadeu de Araujo |
| EPI_ISL_1533692 | Santa Casa de Sao Paulo | Instituto Adolfo Lutz, Interdiciplinary Procedures Center, Strategic Laboratory | Claudio Tavares Sacchi, Claudia Regina Gonçalves, Erica Valessa Ramos Gomes, Karoline Rodrigues Campos, Caio Vinicius Dias Lopes, Leonardo Jose Tadeu de Araujo |
| EPI_ISL_1533693 | Secretaria Municipal de Saude de Ubatuba | Instituto Adolfo Lutz, Interdiciplinary Procedures Center, Strategic Laboratory | Claudio Tavares Sacchi, Claudia Regina Gonçalves, Erica Valessa Ramos Gomes, Karoline Rodrigues Campos, Caio Vinicius Dias Lopes, Leonardo Jose Tadeu de Araujo |
| EPI_ISL_1533694 | Santa Casa de Birigui | Instituto Adolfo Lutz, Interdiciplinary Procedures Center, Strategic Laboratory | Claudio Tavares Sacchi, Claudia Regina Gonçalves, Erica Valessa Ramos Gomes, Karoline Rodrigues Campos, Caio Vinicius Dias Lopes, Leonardo Jose Tadeu de Araujo |
| EPI_ISL_1533695 | Hospital Municipal Dr Mario Gatti | Instituto Adolfo Lutz, Interdiciplinary Procedures Center, Strategic Laboratory | Claudio Tavares Sacchi, Claudia Regina Gonçalves, Erica Valessa Ramos Gomes, Karoline Rodrigues Campos, Caio Vinicius Dias Lopes, Leonardo Jose Tadeu de Araujo |
| EPI_ISL_1533696 | Hospital Municipal de Pedreira | Instituto Adolfo Lutz, Interdiciplinary Procedures Center, Strategic Laboratory | Claudio Tavares Sacchi, Claudia Regina Gonçalves, Erica Valessa Ramos Gomes, Karoline Rodrigues Campos, Caio Vinicius Dias Lopes, Leonardo Jose Tadeu de Araujo |
| EPI_ISL_1533697 | Hospital Estadual de Vila Alpina | Instituto Adolfo Lutz, Interdiciplinary Procedures Center, Strategic Laboratory | Claudio Tavares Sacchi, Claudia Regina Gonçalves, Erica Valessa Ramos Gomes, Karoline Rodrigues Campos, Caio Vinicius Dias Lopes, Leonardo Jose Tadeu de Araujo |
| EPI_ISL_1533698 | Secretaria Municipal de Saude de Birigui | Instituto Adolfo Lutz, Interdiciplinary Procedures Center, Strategic Laboratory | Claudio Tavares Sacchi, Claudia Regina Gonçalves, Erica Valessa Ramos Gomes, Karoline Rodrigues Campos, Caio Vinicius Dias Lopes, Leonardo Jose Tadeu de Araujo |
| EPI_ISL_1533699 | Hospital das Clinicas Luzia de Pinho Melo | Instituto Adolfo Lutz, Interdiciplinary Procedures Center, Strategic Laboratory | Claudio Tavares Sacchi, Claudia Regina Gonçalves, Erica Valessa Ramos Gomes, Karoline Rodrigues Campos, Caio Vinicius Dias Lopes, Leonardo Jose Tadeu de Araujo |
| EPI_ISL_1533700 | Hospital Estadual de Sapopemba Sao Paulo | Instituto Adolfo Lutz, Interdiciplinary Procedures Center, Strategic Laboratory | Claudio Tavares Sacchi, Claudia Regina Gonçalves, Erica Valessa Ramos Gomes, Karoline Rodrigues Campos, Caio Vinicius Dias Lopes, Leonardo Jose Tadeu de Araujo |
| EPI_ISL_1533701 | Hospital Estadual de Vila Alpina Org Social Seconci Sao Paulo | Instituto Adolfo Lutz, Interdiciplinary Procedures Center, Strategic Laboratory | Claudio Tavares Sacchi, Claudia Regina Gonçalves, Erica Valessa Ramos Gomes, Karoline Rodrigues Campos, Caio Vinicius Dias Lopes, Leonardo Jose Tadeu de Araujo |
| EPI_ISL_1533702 | Hospital Universitario da USP Sao Paulo | Instituto Adolfo Lutz, Interdiciplinary Procedures Center, Strategic Laboratory | Claudio Tavares Sacchi, Claudia Regina Gonçalves, Erica Valessa Ramos Gomes, Karoline Rodrigues Campos, Caio Vinicius Dias Lopes, Leonardo Jose Tadeu de Araujo |
| EPI_ISL_1533703 | Pronto Atendimento Sao Jose | Instituto Adolfo Lutz, Interdiciplinary Procedures Center, Strategic Laboratory | Claudio Tavares Sacchi, Claudia Regina Gonçalves, Erica Valessa Ramos Gomes, Karoline Rodrigues Campos, Caio Vinicius Dias Lopes, Leonardo Jose Tadeu de Araujo |
| EPI_ISL_1533704 | Santa Casa de Atibaia Pro Saude | Instituto Adolfo Lutz, Interdiciplinary Procedures Center, Strategic Laboratory | Claudio Tavares Sacchi, Claudia Regina Gonçalves, Erica Valessa Ramos Gomes, Karoline Rodrigues Campos, Caio Vinicius Dias Lopes, Leonardo Jose Tadeu de Araujo |
| EPI_ISL_1533705 | Vigilancia em Saude | Instituto Adolfo Lutz, Interdiciplinary Procedures Center, Strategic Laboratory | Claudio Tavares Sacchi, Claudia Regina Gonçalves, Erica Valessa Ramos Gomes, Karoline Rodrigues Campos, Caio Vinicius Dias Lopes, Leonardo Jose Tadeu de Araujo |
| EPI_ISL_1533706 | PS Mun Santana Lauro Ribas Braga | Instituto Adolfo Lutz, Interdiciplinary Procedures Center, Strategic Laboratory | Claudio Tavares Sacchi, Claudia Regina Gonçalves, Erica Valessa Ramos Gomes, Karoline Rodrigues Campos, Caio Vinicius Dias Lopes, Leonardo Jose Tadeu de Araujo |
| EPI_ISL_1533707 | Hosp Mun Planalto Waldomiro de Paula | Instituto Adolfo Lutz, Interdiciplinary Procedures Center, Strategic Laboratory | Claudio Tavares Sacchi, Claudia Regina Gonçalves, Erica Valessa Ramos Gomes, Karoline Rodrigues Campos, Caio Vinicius Dias Lopes, Leonardo Jose Tadeu de Araujo |
| EPI_ISL_1533708 | Hospital Municipal Dr Mario Gatti Campinas | Instituto Adolfo Lutz, Interdiciplinary Procedures Center, Strategic Laboratory | Claudio Tavares Sacchi, Claudia Regina Gonçalves, Erica Valessa Ramos Gomes, Karoline Rodrigues Campos, Caio Vinicius Dias Lopes, Leonardo Jose Tadeu de Araujo |
| EPI_ISL_1533709, EPI_ISL_1533710 | Hospital Estadual de Vila Alpina Org Social Seconci Sao Paulo | Instituto Adolfo Lutz, Interdiciplinary Procedures Center, Strategic Laboratory | Claudio Tavares Sacchi, Claudia Regina Gonçalves, Erica Valessa Ramos Gomes, Karoline Rodrigues Campos, Caio Vinicius Dias Lopes, Leonardo Jose Tadeu de Araujo |
| EPI_ISL_1533711 | Hospital Santa Clara | Instituto Adolfo Lutz, Interdiciplinary Procedures Center, Strategic Laboratory | Claudio Tavares Sacchi, Claudia Regina Gonçalves, Erica Valessa Ramos Gomes, Karoline Rodrigues Campos, Caio Vinicius Dias Lopes, Leonardo Jose Tadeu de Araujo |
| EPI_ISL_1533712 | Hospital Universitario da USP Sao Paulo | Instituto Adolfo Lutz, Interdiciplinary Procedures Center, Strategic Laboratory | Claudio Tavares Sacchi, Claudia Regina Gonçalves, Erica Valessa Ramos Gomes, Karoline Rodrigues Campos, Caio Vinicius Dias Lopes, Leonardo Jose Tadeu de Araujo |
| EPI_ISL_1533713 | Unidade de Pronto Atendimento Jd Amanda | Instituto Adolfo Lutz, Interdiciplinary Procedures Center, Strategic Laboratory | Claudio Tavares Sacchi, Claudia Regina Gonçalves, Erica Valessa Ramos Gomes, Karoline Rodrigues Campos, Caio Vinicius Dias Lopes, Leonardo Jose Tadeu de Araujo |
| EPI_ISL_1533714 | Pronto Socorro Dr Osmar Mesquita | Instituto Adolfo Lutz, Interdiciplinary Procedures Center, Strategic Laboratory | Claudio Tavares Sacchi, Claudia Regina Gonçalves, Erica Valessa Ramos Gomes, Karoline Rodrigues Campos, Caio Vinicius Dias Lopes, Leonardo Jose Tadeu de Araujo |
| EPI_ISL_1533715 | Santa Casa de Aracatuba Hospital Sagrado Coracao de Jesus | Instituto Adolfo Lutz, Interdiciplinary Procedures Center, Strategic Laboratory | Claudio Tavares Sacchi, Claudia Regina Gonçalves, Erica Valessa Ramos Gomes, Karoline Rodrigues Campos, Caio Vinicius Dias Lopes, Leonardo Jose Tadeu de Araujo |
| EPI_ISL_1533716 | Vigilancia Em Saude | Instituto Adolfo Lutz, Interdiciplinary Procedures Center, Strategic Laboratory | Claudio Tavares Sacchi, Claudia Regina Gonçalves, Erica Valessa Ramos Gomes, Karoline Rodrigues Campos, Caio Vinicius Dias Lopes, Leonardo Jose Tadeu de Araujo |
| EPI_ISL_1533717 | Unidade de Pronto Atendimento Jd Amanda | Instituto Adolfo Lutz, Interdiciplinary Procedures Center, Strategic Laboratory | Claudio Tavares Sacchi, Claudia Regina Gonçalves, Erica Valessa Ramos Gomes, Karoline Rodrigues Campos, Caio Vinicius Dias Lopes, Leonardo Jose Tadeu de Araujo |
| EPI_ISL_1533718 | Servico de Verificacao de Obitos Svo Guarulhos | Instituto Adolfo Lutz, Interdiciplinary Procedures Center, Strategic Laboratory | Claudio Tavares Sacchi, Claudia Regina Gonçalves, Erica Valessa Ramos Gomes, Karoline Rodrigues Campos, Caio Vinicius Dias Lopes, Leonardo Jose Tadeu de Araujo |
| EPI_ISL_1533719 | Policlinica Munic da Est Turistica de Holambra | Instituto Adolfo Lutz, Interdiciplinary Procedures Center, | Claudio Tavares Sacchi, Claudia Regina Gonçalves, Erica Valessa Ramos Gomes, Karoline Rodrigues Campos, Caio Vinicius Dias Lopes, Leonardo Jose |

| | | | |
|---|---|---|---|
| | | Strategic Laboratory | Tadeu de Araujo |
| EPI_ISL_1533720 | Santa Casa de Penapolis | Instituto Adolfo Lutz, Interdiciplinary Procedures Center, Strategic Laboratory | Claudio Tavares Sacchi, Claudia Regina Gonçalves, Erica Valessa Ramos Gomes, Karoline Rodrigues Campos, Caio Vinicius Dias Lopes, Leonardo Jose Tadeu de Araujo |
| EPI_ISL_1533721 | Santa Casa de Aracatuba Hospital Sagrado Coracao De Jesus | Instituto Adolfo Lutz, Interdiciplinary Procedures Center, Strategic Laboratory | Claudio Tavares Sacchi, Claudia Regina Gonçalves, Erica Valessa Ramos Gomes, Karoline Rodrigues Campos, Caio Vinicius Dias Lopes, Leonardo Jose Tadeu de Araujo |
| EPI_ISL_1533722 | Secretaria Municipal de Saude De Piracaia | Instituto Adolfo Lutz, Interdiciplinary Procedures Center, Strategic Laboratory | Claudio Tavares Sacchi, Claudia Regina Gonçalves, Erica Valessa Ramos Gomes, Karoline Rodrigues Campos, Caio Vinicius Dias Lopes, Leonardo Jose Tadeu de Araujo |
| EPI_ISL_1533723 | Hospital Geral de Itaquaquecetuba | Instituto Adolfo Lutz, Interdiciplinary Procedures Center, Strategic Laboratory | Claudio Tavares Sacchi, Claudia Regina Gonçalves, Erica Valessa Ramos Gomes, Karoline Rodrigues Campos, Caio Vinicius Dias Lopes, Leonardo Jose Tadeu de Araujo |
| EPI_ISL_1533724 | Diretoria Municipal de Saude | Instituto Adolfo Lutz, Interdiciplinary Procedures Center, Strategic Laboratory | Claudio Tavares Sacchi, Claudia Regina Gonçalves, Erica Valessa Ramos Gomes, Karoline Rodrigues Campos, Caio Vinicius Dias Lopes, Leonardo Jose Tadeu de Araujo |
| EPI_ISL_1533725 | Hospital Geral de Guarulhos | Instituto Adolfo Lutz, Interdiciplinary Procedures Center, Strategic Laboratory | Claudio Tavares Sacchi, Claudia Regina Gonçalves, Erica Valessa Ramos Gomes, Karoline Rodrigues Campos, Caio Vinicius Dias Lopes, Leonardo Jose Tadeu de Araujo |
| EPI_ISL_1533726 | Hospital Estadual de Campanha Covid 19 Barradas | Instituto Adolfo Lutz, Interdiciplinary Procedures Center, Strategic Laboratory | Claudio Tavares Sacchi, Claudia Regina Gonçalves, Erica Valessa Ramos Gomes, Karoline Rodrigues Campos, Caio Vinicius Dias Lopes, Leonardo Jose Tadeu de Araujo |
| EPI_ISL_1533727 | Hospital Universitario da USP Sao Paulo | Instituto Adolfo Lutz, Interdiciplinary Procedures Center, Strategic Laboratory | Claudio Tavares Sacchi, Claudia Regina Gonçalves, Erica Valessa Ramos Gomes, Karoline Rodrigues Campos, Caio Vinicius Dias Lopes, Leonardo Jose Tadeu de Araujo |
| EPI_ISL_1533977 | Laboratorio de Virologia Molecular / UFRJ | Laboratory of Respiratory Viruses and Measles, Oswaldo Cruz Institute, FIOCRUZ | Paola Resende, Carolina M Voloch, Luciana Appolinario, Fernando Motta, Anna Carolina Paixao, Ana Carolina Mendonca, Alice Sampaio Rocha, Renata Serrano Lopes, Amilcar Tanuri, Marilda Siqueira on behalf of the Fiocruz COVID-19 Genomic Surveillance Network |
| EPI_ISL_1533978 | Laboratorio Central de Saude Publica do Estado do Parana (LACEN-PR) | Laboratory of Respiratory Viruses and Measles, Oswaldo Cruz Institute, FIOCRUZ | Paola Resende, Luciana Appolinario, Fernando Motta, Anna Carolina Paixao, Ana Carolina Mendonca, Alice Sampaio Rocha, Renata Serrano Lopes, Maria do Carmo Debur, Irina Nastassja Riediger, Marilda Siqueira on behalf of the Fiocruz COVID-19 Genomic Surveillance Network |
| EPI_ISL_1533979 | Laboratorio Central de Saude Publica do Estado do Rio Grande do Sul (LACEN-RS) | Laboratory of Respiratory Viruses and Measles, Oswaldo Cruz Institute, FIOCRUZ | Paola Resende, Luciana Appolinario, Fernando Motta, Anna Carolina Paixao, Ana Carolina Mendonca, Alice Sampaio Rocha, Renata Serrano Lopes, Richard Salvato, Tatiana Schaffer Gregianini, Marilda Siqueira on behalf of the Fiocruz COVID-19 Genomic Surveillance Network |
| EPI_ISL_1533980, EPI_ISL_1533983 | Laboratory of Respiratory Viruses and Measles, Oswaldo Cruz Institute, FIOCRUZ | Laboratory of Respiratory Viruses and Measles, Oswaldo Cruz Institute, FIOCRUZ | Paola Resende, Luciana Appolinario, Fernando Motta, Anna Carolina Paixao, Ana Carolina Mendonca, Alice Sampaio Rocha, Renata Serrano Lopes, Marilda Siqueira on behalf of the Fiocruz COVID-19 Genomic Surveillance Network |
| EPI_ISL_1533990 | Laboratorio Central de Saude Publica do Estado de Santa Catarina (LACEN-SC) | Laboratory of Respiratory Viruses and Measles, Oswaldo Cruz Institute, FIOCRUZ | Paola Resende, Luciana Appolinario, Fernando Motta, Anna Carolina Paixao, Ana Carolina Mendonca, Alice Sampaio Rocha, Renata Serrano Lopes, Darcita Buerger Rovaris, Sandra Bianchini Fernandes, Marilda Siqueira on behalf of the Fiocruz COVID-19 Genomic Surveillance Network |
| EPI_ISL_1533991 | Laboratorio de Virologia Molecular / UFRJ | Laboratory of Respiratory Viruses and Measles, Oswaldo Cruz Institute, FIOCRUZ | Paola Resende, Carolina M Voloch, Luciana Appolinario, Fernando Motta, Anna Carolina Paixao, Ana Carolina Mendonca, Alice Sampaio Rocha, Renata Serrano Lopes, Amilcar Tanuri, Marilda Siqueira on behalf of the Fiocruz COVID-19 Genomic Surveillance Network |
| EPI_ISL_1533992, EPI_ISL_1533993, EPI_ISL_1533994 | Laboratorio Central de Saude Publica do Estado de Santa Catarina (LACEN-SC) | Laboratory of Respiratory Viruses and Measles, Oswaldo Cruz Institute, FIOCRUZ | Paola Resende, Luciana Appolinario, Fernando Motta, Anna Carolina Paixao, Ana Carolina Mendonca, Alice Sampaio Rocha, Renata Serrano Lopes, Darcita Buerger Rovaris, Sandra Bianchini Fernandes, Marilda Siqueira on behalf of the Fiocruz COVID-19 Genomic Surveillance Network |
| EPI_ISL_1533995 | Laboratorio de Virologia Molecular / UFRJ | Laboratory of Respiratory Viruses and Measles, Oswaldo Cruz Institute, FIOCRUZ | Paola Resende, Carolina M Voloch, Luciana Appolinario, Fernando Motta, Anna Carolina Paixao, Ana Carolina Mendonca, Alice Sampaio Rocha, Renata Serrano Lopes, Amilcar Tanuri, Marilda Siqueira on behalf of the Fiocruz COVID-19 Genomic Surveillance Network |
| EPI_ISL_1533996, EPI_ISL_1533997 | Laboratorio Central de Saude Publica do Estado de Santa Catarina (LACEN-SC) | Laboratory of Respiratory Viruses and Measles, Oswaldo Cruz Institute, FIOCRUZ | Paola Resende, Luciana Appolinario, Fernando Motta, Anna Carolina Paixao, Ana Carolina Mendonca, Alice Sampaio Rocha, Renata Serrano Lopes, Darcita Buerger Rovaris, Sandra Bianchini Fernandes, Marilda Siqueira on behalf of the Fiocruz COVID-19 Genomic Surveillance Network |
| EPI_ISL_1533998 | Laboratorio de Virologia Molecular / UFRJ | Laboratory of Respiratory Viruses and Measles, Oswaldo Cruz Institute, FIOCRUZ | Paola Resende, Carolina M Voloch, Luciana Appolinario, Fernando Motta, Anna Carolina Paixao, Ana Carolina Mendonca, Alice Sampaio Rocha, Renata Serrano Lopes, Amilcar Tanuri, Marilda Siqueira on behalf of the Fiocruz COVID-19 Genomic Surveillance Network |
| EPI_ISL_1533999, EPI_ISL_1534000, EPI_ISL_1534001 | Laboratorio Central de Saude Publica do Estado do Parana (LACEN-PR) | Laboratory of Respiratory Viruses and Measles, Oswaldo Cruz Institute, FIOCRUZ | Paola Resende, Luciana Appolinario, Fernando Motta, Anna Carolina Paixao, Ana Carolina Mendonca, Alice Sampaio Rocha, Renata Serrano Lopes, Maria do Carmo Debur, Irina Nastassja Riediger, Marilda Siqueira on behalf of the Fiocruz COVID-19 Genomic Surveillance Network |
| EPI_ISL_1534002, EPI_ISL_1534003 | Laboratorio Central de Saude Publica do Estado de Santa Catarina (LACEN-SC) | Laboratory of Respiratory Viruses and Measles, Oswaldo Cruz Institute, FIOCRUZ | Paola Resende, Luciana Appolinario, Fernando Motta, Anna Carolina Paixao, Ana Carolina Mendonca, Alice Sampaio Rocha, Renata Serrano Lopes, Darcita Buerger Rovaris, Sandra Bianchini Fernandes, Marilda Siqueira on behalf of the Fiocruz COVID-19 Genomic Surveillance Network |
| EPI_ISL_1534004 | Laboratorio Central de Saude Publica do Estado de Sergipe (LACEN-SE) | Laboratory of Respiratory Viruses and Measles, Oswaldo Cruz Institute, FIOCRUZ | Paola Resende, Luciana Appolinario, Fernando Motta, Anna Carolina Paixao, Ana Carolina Mendonca, Alice Sampaio Rocha, Renata Serrano Lopes, Cliomar Alves dos Santos, Marilda Siqueira on behalf of the Fiocruz COVID-19 Genomic Surveillance Network |
| EPI_ISL_1534005 | Laboratory of Respiratory Viruses and Measles, Oswaldo Cruz Institute, FIOCRUZ | Laboratory of Respiratory Viruses and Measles, Oswaldo Cruz Institute, FIOCRUZ | Paola Resende, Luciana Appolinario, Fernando Motta, Anna Carolina Paixao, Ana Carolina Mendonca, Alice Sampaio Rocha, Renata Serrano Lopes, Marilda Siqueira on behalf of the Fiocruz COVID-19 Genomic Surveillance Network |
| EPI_ISL_1534006, EPI_ISL_1534007, EPI_ISL_1534008, EPI_ISL_1534009, EPI_ISL_1534010 | Laboratorio Central de Saude Publica do Estado de Santa Catarina (LACEN-SC) | Laboratory of Respiratory Viruses and Measles, Oswaldo Cruz Institute, FIOCRUZ | Paola Resende, Luciana Appolinario, Fernando Motta, Anna Carolina Paixao, Ana Carolina Mendonca, Alice Sampaio Rocha, Renata Serrano Lopes, Darcita Buerger Rovaris, Sandra Bianchini Fernandes, Marilda Siqueira on behalf of the Fiocruz COVID-19 Genomic Surveillance Network |
| EPI_ISL_1534011 | Laboratory of Respiratory Viruses and Measles, Oswaldo Cruz Institute, FIOCRUZ | Laboratory of Respiratory Viruses and Measles, Oswaldo Cruz Institute, FIOCRUZ | Paola Resende, Luciana Appolinario, Fernando Motta, Anna Carolina Paixao, Ana Carolina Mendonca, Alice Sampaio Rocha, Renata Serrano Lopes, Marilda Siqueira on behalf of the Fiocruz COVID-19 Genomic Surveillance Network |
| EPI_ISL_1534012 | Laboratorio Central de Saude Publica do Estado do Rio de Janeiro (LACEN-RJ) | Laboratory of Respiratory Viruses and Measles, Oswaldo Cruz Institute, FIOCRUZ | Paola Resende, Luciana Appolinario, Fernando Motta, Anna Carolina Paixao, Ana Carolina Mendonca, Alice Sampaio Rocha, Renata Serrano Lopes, Andrea Cony Cavalcanti, Marilda Siqueira on behalf of the Fiocruz COVID-19 Genomic Surveillance Network |
| EPI_ISL_1534013 | Laboratorio Central de Saude Publica do Estado do Rio Grande do Sul (LACEN-RS) | Laboratory of Respiratory Viruses and Measles, Oswaldo Cruz Institute, FIOCRUZ | Paola Resende, Luciana Appolinario, Fernando Motta, Anna Carolina Paixao, Ana Carolina Mendonca, Alice Sampaio Rocha, Renata Serrano Lopes, Richard Salvato, Tatiana Schaffer Gregianini, Marilda Siqueira on behalf of the Fiocruz COVID-19 Genomic Surveillance Network |
| EPI_ISL_1534014, EPI_ISL_1534015, EPI_ISL_1534016 | Laboratory of Respiratory Viruses and Measles, Oswaldo Cruz Institute, FIOCRUZ | Laboratory of Respiratory Viruses and Measles, Oswaldo Cruz Institute, FIOCRUZ | Paola Resende, Luciana Appolinario, Fernando Motta, Anna Carolina Paixao, Ana Carolina Mendonca, Alice Sampaio Rocha, Renata Serrano Lopes, Marilda Siqueira on behalf of the Fiocruz COVID-19 Genomic Surveillance Network |
| EPI_ISL_1550388 | Instituto Oswaldo Cruz | Laboratorio de Virologia, Faculdade de Medicina, Universidade Federal de Mato Grosso, campus Cuiabá | María de Fátima Ferreira, Renata Dezengrini Slhessarenko, Janeth Aracely Ramirez Pavon, Luciano Nakazato, Valeria Dutra, Rosane Christiane Hahn, Paola Cristina Resende |
| EPI_ISL_1550878 | Instituto Oswaldo Cruz | Laboratorio de Virologia, Faculdade de Medicina, Universidade Federal de Mato Grosso, campus Cuiabá | Maria de Fatima Ferrerira, Renata Dezengrini Slhessarenko, Janeth Aracely Ramirez Pavon, Luciano Nakazato, Valeria Dutra , Rosane Hahn |
| EPI_ISL_1578453 | Laboratório de Biologia Molecular do Hospital das Clínicas da Faculdade de Medicina de Botucatu/SP | Laboratórios Genômica Funcional (FCA/UNESP) e Biologia Molecular (FMB-HC/UNESP) - Rede de Vigilância Genômica (Vigenômica)/UNESP | Patrícia Akemi Assato; Felipe Allan da Silva da Costa; Bianca Cechetto Carlos; Flavia Hebner Barbosa Trovão; Guilherme Targino Valente; Rejane Maria Tommasini Grotto; Jayme A. Souza-Neto. |
| EPI_ISL_1578740, EPI_ISL_1579258, EPI_ISL_1580269, EPI_ISL_1580502, EPI_ISL_1580504 | Laboratório de Biologia Molecular do Hospital das Clínicas da Faculdade de Medicina de Botucatu/SP | Laboratórios de Genômica Funcional (FCA/UNESP) e Biologia Molecular (FMB-HC/UNESP) - Rede de Vigilância Genômica (Vigenômica)/UNESP | Patrícia Akemi Assato; Felipe Allan da Silva da Costa; Bianca Cechetto Carlos; Flavia Hebner Barbosa Trovão; Guilherme Targino Valente; Rejane Maria Tommasini Grotto; Jayme A. Souza-Neto. |
| EPI_ISL_1583641, EPI_ISL_1583642, EPI_ISL_1583643, EPI_ISL_1583644, EPI_ISL_1583645, EPI_ISL_1583646, EPI_ISL_1583647, EPI_ISL_1583648, EPI_ISL_1583649, EPI_ISL_1583650, EPI_ISL_1583651, EPI_ISL_1583652, EPI_ISL_1583653, EPI_ISL_1583654, EPI_ISL_1583655, EPI_ISL_1583656, EPI_ISL_1583657, EPI_ISL_1583658, EPI_ISL_1583659, EPI_ISL_1583660, EPI_ISL_1583661, EPI_ISL_1583662, EPI_ISL_1583663, EPI_ISL_1583664, EPI_ISL_1583665, EPI_ISL_1583666, EPI_ISL_1583667, EPI_ISL_1583668, EPI_ISL_1583669, EPI_ISL_1583670, EPI_ISL_1583671, EPI_ISL_1583672, EPI_ISL_1583673, EPI_ISL_1583674 | | | |
| see above | Central Public Health Laboratory - LACEN -Bahia, Salvador, Brazil | Central Public Health Laboratory - LACEN -Bahia, Salvador, Brazil | Stephane Tosta, Luciana Oliveira, Vanessa Nardy,Patrícia Cajado,Marcela Gómez, Breno Dominguez, Jaqueline Gomes, Vagner Fonseca,Marta Giovanetti,Luiz Alcantara, Felicidade Pereira, Arabela Leal |
| EPI_ISL_1583675 | DNA Laboratory | Central Public Health Laboratory - LACEN -Bahia, Salvador, | Stephane Tosta, Luciana Oliveira, Vanessa Nardy,Patrícia Cajado,Marcela Gómez, Breno Dominguez, Jaqueline Gomes, Vagner Fonseca,Marta |

| | | | |
|---|---|---|---|
| | | Brazil | Giovanetti,Luiz Alcantara, Felicidade Pereira, Arabela Leal |
| EPI_ISL_1583676, EPI_ISL_1583677, EPI_ISL_1583678, EPI_ISL_1583679, EPI_ISL_1583680, EPI_ISL_1583681, EPI_ISL_1583682, EPI_ISL_1583683, EPI_ISL_1583686, EPI_ISL_1583689, EPI_ISL_1583691, EPI_ISL_1583694, EPI_ISL_1583697, EPI_ISL_1583700, EPI_ISL_1583703, EPI_ISL_1583705, EPI_ISL_1583708, EPI_ISL_1583710, EPI_ISL_1583714, EPI_ISL_1583719, EPI_ISL_1583722, EPI_ISL_1583725, EPI_ISL_1583727, EPI_ISL_1583730, EPI_ISL_1583733 | | | |
| see above | Central Public Health Laboratory - LACEN -Bahia, Salvador, Brazil | Central Public Health Laboratory - LACEN -Bahia, Salvador, Brazil | Stephane Tosta, Luciana Oliveira, Vanessa Nardy,Patrícia Cajado,Marcela Gómez, Breno Dominguez, Jaqueline Gomes, Vagner Fonseca,Marta Giovanetti,Luiz Alcantara, Felicidade Pereira, Arabela Leal |
| EPI_ISL_1592230 | Laboratório Hermes Pardini | Laboratório de Biologia Integrativa, Instituto de Ciências Biológicas, Universidade Federal de Minas Gerais | Filipe Romero Rebello Moreira, Diego Menezes Bonfim, Danielle Alves Gomes Zauli, Joice do Prado Silva, Aline Brito de Lima, Frederico Scott Varella Malta, Alessandro Clayton de Souza Ferreira, Victor Cavalcanti Pardini, Daniel Costa Queiroz, Rafael Marques de Souza, Victor Emmanuel Viana Geddes, Walyson Coelho Costa, Wagner Carlos Santos Magalhaes, Rennan Garcias Moreira, Carolina Moreira Voloch, Renan Pedra de Souza, Renato Santana Aguiar. |
| EPI_ISL_1608161, EPI_ISL_1608162, EPI_ISL_1608163, EPI_ISL_1608164, EPI_ISL_1608165, EPI_ISL_1608166, EPI_ISL_1608167, EPI_ISL_1608168, EPI_ISL_1608169, EPI_ISL_1608170, EPI_ISL_1608171 | | | |
| see above | Hospital São Rafael - IDOR | Hospital São Rafael - IDOR | Carolina Kymie Vasques Nonaka, Tiago Grãf, Camila Araújo de Lorenzo Barcia, Vanessa Ferreira Costa, Janderson Lopes de Oliveira, Rogério da Hora Passos, Clarissa Araújo Gurgel Rocha, Iasmin Nogueira Bastos, Maria Clara Brito de Santana, Ian Marinho Santos, Karoline Almeida Felix de Sousa, Isadora Cristina de Siqueira, Thamires Gomes Lopes Weber, Ana Verena Almeida Mendes, Bruno Solano de Freitas Souza |
| EPI_ISL_1620638 | Hospital Municipal Dr. Ignacio Proença de Gouvea | Instituto Butantan / Mendelics | Alexander Roberto Precioso, Dimas Tadeu Covas, Sandra Coccuzzo Sampaio, Maria Carolina Elias, José Salvatore Leister Patané, Vincent Louis Viala, Antonio Jorge Martins, Ricardo Haddad, Claudia Renata dos Santos Barros, Elaine Cristina Marqueze, Raul Machado Neto, Debora Botequio Moretti, Bibiana Santos, João Paulo Kitajima, Erika Freitas, David Schlesinger, Simone Kashima, Evandra Strazza Rodrigues, Svetoslav Nanev Slavov, Elaine Vieira dos Santos, Rafael dos Santos Bezerra, Luiz Carlos Junior de Alcantara, Marta Giovanetti, Vagner Fonseca, Flavia Aburjaile, Rodrigo Tocantins Calado. |
| EPI_ISL_1620639 | Fleury | Instituto Butantan / Mendelics | Alexander Roberto Precioso, Dimas Tadeu Covas, Sandra Coccuzzo Sampaio, Maria Carolina Elias, José Salvatore Leister Patané, Vincent Louis Viala, Antonio Jorge Martins, Ricardo Haddad, Claudia Renata dos Santos Barros, Elaine Cristina Marqueze, Raul Machado Neto, Debora Botequio Moretti, Bibiana Santos, João Paulo Kitajima, Erika Freitas, David Schlesinger, Simone Kashima, Evandra Strazza Rodrigues, Svetoslav Nanev Slavov, Elaine Vieira dos Santos, Rafael dos Santos Bezerra, Luiz Carlos Junior de Alcantara, Marta Giovanetti, Vagner Fonseca, Flavia Aburjaile, Rodrigo Tocantins Calado. |
| EPI_ISL_1625972 | Ama J Angela | Instituto Adolfo Lutz, Interdiciplinary Procedures Center, Strategic Laboratory | Claudio Tavares Sacchi, Claudia Regina Gonçalves, Erica Valessa Ramos Gomes, Karoline Rodrigues Campos, Caio Vinicius Dias Lopes, Leonardo Jose Tadeu de Araujo, Katia Correa de Oliveira Santos |
| EPI_ISL_1625973, EPI_ISL_1625974 | Santa Casa de Aracatuba Hospital Sagrado Coracao de Jesus | Instituto Adolfo Lutz, Interdiciplinary Procedures Center, Strategic Laboratory | Claudio Tavares Sacchi, Claudia Regina Gonçalves, Erica Valessa Ramos Gomes, Karoline Rodrigues Campos, Caio Vinicius Dias Lopes, Leonardo Jose Tadeu de Araujo, Katia Correa de Oliveira Santos |
| EPI_ISL_1625975 | CS II Dr Antonio Vicoso Moreira De Rezende Sumare | Instituto Adolfo Lutz, Interdiciplinary Procedures Center, Strategic Laboratory | Claudio Tavares Sacchi, Claudia Regina Gonçalves, Erica Valessa Ramos Gomes, Karoline Rodrigues Campos, Caio Vinicius Dias Lopes, Leonardo Jose Tadeu de Araujo, Katia Correa de Oliveira Santos |
| EPI_ISL_1625976 | Centro de Saude III de Divinolandia | Instituto Adolfo Lutz, Interdiciplinary Procedures Center, Strategic Laboratory | Claudio Tavares Sacchi, Claudia Regina Gonçalves, Erica Valessa Ramos Gomes, Karoline Rodrigues Campos, Caio Vinicius Dias Lopes, Leonardo Jose Tadeu de Araujo, Katia Correa de Oliveira Santos |
| EPI_ISL_1625977, EPI_ISL_1625978, EPI_ISL_1625979, EPI_ISL_1625980, EPI_ISL_1625981, EPI_ISL_1625982, EPI_ISL_1625983, EPI_ISL_1625984, EPI_ISL_1625985, EPI_ISL_1625986, EPI_ISL_1625987, EPI_ISL_1625988, EPI_ISL_1625989, EPI_ISL_1625990, EPI_ISL_1625991, EPI_ISL_1625992, EPI_ISL_1625993, EPI_ISL_1625994, EPI_ISL_1625995, EPI_ISL_1625996, EPI_ISL_1625997, EPI_ISL_1625998, EPI_ISL_1625999, EPI_ISL_1626000, EPI_ISL_1626001, EPI_ISL_1626002, EPI_ISL_1626003, EPI_ISL_1626004, EPI_ISL_1626005, EPI_ISL_1626006, EPI_ISL_1626007, EPI_ISL_1626008, EPI_ISL_1626009, EPI_ISL_1626606 | | | |
| see above | Instituto Adolfo Lutz - Regional de Rio Claro | Instituto Adolfo Lutz, Interdiciplinary Procedures Center, Strategic Laboratory | Claudio Tavares Sacchi, Claudia Regina Gonçalves, Erica Valessa Ramos Gomes, Karoline Rodrigues Campos, Caio Vinicius Dias Lopes, Leonardo Jose Tadeu de Araujo, Katia Correa de Oliveira Santos |
| EPI_ISL_1628344 | Hospital Geral de Pedreira | Instituto Adolfo Lutz, Interdiciplinary Procedures Center, Strategic Laboratory | Claudio Tavares Sacchi, Claudia Regina Gonçalves, Erica Valessa Ramos Gomes, Karoline Rodrigues Campos, Caio Vinicius Dias Lopes, Leonardo Jose Tadeu de Araujo, Katia Correa de Oliveira Santos |
| EPI_ISL_1628345, EPI_ISL_1628346 | Instituto Adolfo Lutz - Regional de Rio Claro | Instituto Adolfo Lutz, Interdiciplinary Procedures Center, Strategic Laboratory | Claudio Tavares Sacchi, Claudia Regina Gonçalves, Erica Valessa Ramos Gomes, Karoline Rodrigues Campos, Caio Vinicius Dias Lopes, Leonardo Jose Tadeu de Araujo, Katia Correa de Oliveira Santos |
| EPI_ISL_1628347, EPI_ISL_1628348, EPI_ISL_1628349, EPI_ISL_1628350, EPI_ISL_1628351, EPI_ISL_1628352, EPI_ISL_1628353, EPI_ISL_1628354, EPI_ISL_1628355, EPI_ISL_1628356, EPI_ISL_1628357, EPI_ISL_1628358, EPI_ISL_1628359, EPI_ISL_1628360, EPI_ISL_1628361, EPI_ISL_1628362 | | | |
| see above | Instituto Adolfo Lutz Central | Instituto Adolfo Lutz, Interdiciplinary Procedures Center, Strategic Laboratory | Claudio Tavares Sacchi, Claudia Regina Gonçalves, Erica Valessa Ramos Gomes, Karoline Rodrigues Campos, Caio Vinicius Dias Lopes, Leonardo Jose Tadeu de Araujo, Katia Correa de Oliveira Santos |
| EPI_ISL_1628363, EPI_ISL_1628364, EPI_ISL_1628365 | LACEN do Estado de Goias | Instituto Adolfo Lutz, Interdiciplinary Procedures Center, Strategic Laboratory | Claudio Tavares Sacchi, Claudia Regina Gonçalves, Erica Valessa Ramos Gomes, Karoline Rodrigues Campos, Caio Vinicius Dias Lopes, Leonardo Jose Tadeu de Araujo, Katia Correa de Oliveira Santos |
| EPI_ISL_1628366 | CS III de Patrocinio Paulista | Instituto Adolfo Lutz, Interdiciplinary Procedures Center, Strategic Laboratory | Claudio Tavares Sacchi, Claudia Regina Gonçalves, Erica Valessa Ramos Gomes, Karoline Rodrigues Campos, Caio Vinicius Dias Lopes, Leonardo Jose Tadeu de Araujo, Katia Correa de Oliveira Santos |
| EPI_ISL_1628367 | Santa Casa de Guaira | Instituto Adolfo Lutz, Interdiciplinary Procedures Center, Strategic Laboratory | Claudio Tavares Sacchi, Claudia Regina Gonçalves, Erica Valessa Ramos Gomes, Karoline Rodrigues Campos, Caio Vinicius Dias Lopes, Leonardo Jose Tadeu de Araujo, Katia Correa de Oliveira Santos |
| EPI_ISL_1628368 | Centro Medico Social Comunitario Januario Teodoro de Souza | Instituto Adolfo Lutz, Interdiciplinary Procedures Center, Strategic Laboratory | Claudio Tavares Sacchi, Claudia Regina Gonçalves, Erica Valessa Ramos Gomes, Karoline Rodrigues Campos, Caio Vinicius Dias Lopes, Leonardo Jose Tadeu de Araujo, Katia Correa de Oliveira Santos |
| EPI_ISL_1628369 | UPA Dr Luis Atilio Losi Viana Ribeirao Preto | Instituto Adolfo Lutz, Interdiciplinary Procedures Center, Strategic Laboratory | Claudio Tavares Sacchi, Claudia Regina Gonçalves, Erica Valessa Ramos Gomes, Karoline Rodrigues Campos, Caio Vinicius Dias Lopes, Leonardo Jose Tadeu de Araujo, Katia Correa de Oliveira Santos |
| EPI_ISL_1628370 | Sae Servico De Atendimento Especializado | Instituto Adolfo Lutz, Interdiciplinary Procedures Center, Strategic Laboratory | Claudio Tavares Sacchi, Claudia Regina Gonçalves, Erica Valessa Ramos Gomes, Karoline Rodrigues Campos, Caio Vinicius Dias Lopes, Leonardo Jose Tadeu de Araujo, Katia Correa de Oliveira Santos |
| EPI_ISL_1628371 | Centro De Saude II Ibitinga | Instituto Adolfo Lutz, Interdiciplinary Procedures Center, Strategic Laboratory | Claudio Tavares Sacchi, Claudia Regina Gonçalves, Erica Valessa Ramos Gomes, Karoline Rodrigues Campos, Caio Vinicius Dias Lopes, Leonardo Jose Tadeu de Araujo, Katia Correa de Oliveira Santos |
| EPI_ISL_1628372 | Hospital Sao Marcos da Sama Morro Agudo | Instituto Adolfo Lutz, Interdiciplinary Procedures Center, Strategic Laboratory | Claudio Tavares Sacchi, Claudia Regina Gonçalves, Erica Valessa Ramos Gomes, Karoline Rodrigues Campos, Caio Vinicius Dias Lopes, Leonardo Jose Tadeu de Araujo, Katia Correa de Oliveira Santos |
| EPI_ISL_1628373 | UPA De Bebedouro | Instituto Adolfo Lutz, Interdiciplinary Procedures Center, Strategic Laboratory | Claudio Tavares Sacchi, Claudia Regina Gonçalves, Erica Valessa Ramos Gomes, Karoline Rodrigues Campos, Caio Vinicius Dias Lopes, Leonardo Jose Tadeu de Araujo, Katia Correa de Oliveira Santos |
| EPI_ISL_1628374 | Santa Casa De Cravinhos | Instituto Adolfo Lutz, Interdiciplinary Procedures Center, Strategic Laboratory | Claudio Tavares Sacchi, Claudia Regina Gonçalves, Erica Valessa Ramos Gomes, Karoline Rodrigues Campos, Caio Vinicius Dias Lopes, Leonardo Jose Tadeu de Araujo, Katia Correa de Oliveira Santos |
| EPI_ISL_1628375, EPI_ISL_1628376 | Secretaria Municipal de Saude de Guariba | Instituto Adolfo Lutz, Interdiciplinary Procedures Center, Strategic Laboratory | Claudio Tavares Sacchi, Claudia Regina Gonçalves, Erica Valessa Ramos Gomes, Karoline Rodrigues Campos, Caio Vinicius Dias Lopes, Leonardo Jose Tadeu de Araujo, Katia Correa de Oliveira Santos |
| EPI_ISL_1628377 | Centro De Saude II Ibitinga | Instituto Adolfo Lutz, Interdiciplinary Procedures Center, Strategic Laboratory | Claudio Tavares Sacchi, Claudia Regina Gonçalves, Erica Valessa Ramos Gomes, Karoline Rodrigues Campos, Caio Vinicius Dias Lopes, Leonardo Jose Tadeu de Araujo, Katia Correa de Oliveira Santos |
| EPI_ISL_1628378 | Vigilancia Epidemiologica Jardinopolis | Instituto Adolfo Lutz, Interdiciplinary Procedures Center, Strategic Laboratory | Claudio Tavares Sacchi, Claudia Regina Gonçalves, Erica Valessa Ramos Gomes, Karoline Rodrigues Campos, Caio Vinicius Dias Lopes, Leonardo Jose Tadeu de Araujo, Katia Correa de Oliveira Santos |
| EPI_ISL_1628379 | UPA Dr Luis Atilio Losi Viana Ribeirao Preto | Instituto Adolfo Lutz, Interdiciplinary Procedures Center, Strategic Laboratory | Claudio Tavares Sacchi, Claudia Regina Gonçalves, Erica Valessa Ramos Gomes, Karoline Rodrigues Campos, Caio Vinicius Dias Lopes, Leonardo Jose Tadeu de Araujo, Katia Correa de Oliveira Santos |
| EPI_ISL_1628380 | Santa Casa De Pitangueiras | Instituto Adolfo Lutz, Interdiciplinary Procedures Center, Strategic Laboratory | Claudio Tavares Sacchi, Claudia Regina Gonçalves, Erica Valessa Ramos Gomes, Karoline Rodrigues Campos, Caio Vinicius Dias Lopes, Leonardo Jose Tadeu de Araujo, Katia Correa de Oliveira Santos |

| | | | |
|---|---|---|---|
| EPI_ISL_1629809 | Laboratório de Microbiologia Molecular - Universidade FEEVALE | Molecular Microbiology Laboratory | Alana Witt Hansen, Fágner Henrique Heldt, Fernando Rosado Spilki, Flávio Silveira, Juliana Schons Gulart, Juliane Deise Fleck, Mariana Soares da Silva, Meriane Demoliner, Matheus Nunes Weber, Paula Rodrigues de Almeida, Michele Filippi. |
| EPI_ISL_1630158 | Laboratório de Microbiologia Molecular - Universidade FEEVALE | Laboratório de Microbiologia Molecular - Universidade FEEVALE | Alana Witt Hansen, Fágner Henrique Heldt, Fernando Rosado Spilki, Flávio Silveira, Juliana Schons Gularte, Juliane Deise Fleck, Mariana Soares da Silva, Meriane Demoliner, Matheus Nunes Weber, Paula Rodrigues de Almeida, Micheli Filippi. |
| EPI_ISL_1652237 | Fleury | Instituto Butantan / Mendelics | Alexander Roberto Precioso, Dimas Tadeu Covas, Sandra Coccuzzo Sampaio, Maria Carolina Elias, José Salvatore Leister Patané, Vincent Louis Viala, Antonio Jorge Martins, Ricardo Haddad, Claudia Renata dos Santos Barros, Elaine Cristina Marqueze, Raul Machado Neto, Debora Botequio Moretti, Bibiana Santos, João Paulo Kitajima, Erika Freitas, David Schlesinger, Simone Kashima, Evandra Strazza Rodrigues, Svetoslav Nanev Slavov, Elaine Vieira dos Santos, Rafael dos Santos Bezerra, Luiz Carlos Junior de Alcantara, Marta Giovanetti, Vagner Fonseca, Flavia Aburjaile, Rodrigo Tocantins Calado. |
| EPI_ISL_1661250, EPI_ISL_1661251, EPI_ISL_1661252 | Laboratorio de Ecologia de Doencas Transmissiveis na Amazonia, Instituto Leonidas e Maria Deane - Fiocruz Amazonia | Laboratorio de Ecologia de Doencas Transmissiveis na Amazonia, Instituto Leonidas e Maria Deane - Fiocruz Amazonia | Valdinete Nascimento, Victor Souza, André Corado, Fernanda Nascimento, George Silva, Ágatha Costa, Debora Duarte, Karina Pessoa, Matilde Mejía, Luciana Gonçalves, Maria Júlia Brandão, Michele Jesus, Felipe Naveca |
| EPI_ISL_1664113, EPI_ISL_1664114, EPI_ISL_1664115, EPI_ISL_1664116, EPI_ISL_1664117, EPI_ISL_1664118, EPI_ISL_1664119, EPI_ISL_1664120, EPI_ISL_1664121, EPI_ISL_1664122, EPI_ISL_1664123, EPI_ISL_1664124, EPI_ISL_1664125, EPI_ISL_1664126, EPI_ISL_1664127, EPI_ISL_1664128, EPI_ISL_1664129, EPI_ISL_1664130, EPI_ISL_1664131, EPI_ISL_1664132, EPI_ISL_1664133, EPI_ISL_1664134, EPI_ISL_1664135, EPI_ISL_1664136, EPI_ISL_1664137, EPI_ISL_1664138, EPI_ISL_1664139, EPI_ISL_1664140, EPI_ISL_1664141, EPI_ISL_1664142, EPI_ISL_1664143, EPI_ISL_1664144, EPI_ISL_1664145, EPI_ISL_1664146, EPI_ISL_1664147, EPI_ISL_1664148, EPI_ISL_1664149, EPI_ISL_1664150, EPI_ISL_1664151, EPI_ISL_1664152, EPI_ISL_1664153, EPI_ISL_1664154, EPI_ISL_1664155, EPI_ISL_1664156, EPI_ISL_1664157, EPI_ISL_1664158, EPI_ISL_1664159, EPI_ISL_1664160, EPI_ISL_1664161, EPI_ISL_1664162, EPI_ISL_1664163, EPI_ISL_1664164, EPI_ISL_1664165, EPI_ISL_1664166, EPI_ISL_1664167, EPI_ISL_1664168, EPI_ISL_1664169, EPI_ISL_1664170, EPI_ISL_1664171, EPI_ISL_1664172, EPI_ISL_1664173, EPI_ISL_1664174, EPI_ISL_1664175, EPI_ISL_1664176, EPI_ISL_1664177, EPI_ISL_1664178, EPI_ISL_1664179, EPI_ISL_1664180, EPI_ISL_1664181, EPI_ISL_1664182, EPI_ISL_1664183, EPI_ISL_1664184, EPI_ISL_1664185, EPI_ISL_1664186, EPI_ISL_1664187, EPI_ISL_1664188, EPI_ISL_1664189, EPI_ISL_1664190, EPI_ISL_1664191, EPI_ISL_1664192, EPI_ISL_1664193, EPI_ISL_1664194, EPI_ISL_1664195, EPI_ISL_1664196, EPI_ISL_1664197, EPI_ISL_1664198, EPI_ISL_1664199, EPI_ISL_1664200, EPI_ISL_1664201, EPI_ISL_1664202 | | | |
| see above | Laboratorio Central Noel Nutels | Bioinformatics Laboratory / LNCC | Luiz G P de Almeida, Alessandra P Lamarca, Ronaldo da Silva F Jr, Liliane Cavalcante, Alexandra L Gerber, Ana Paula de C Guimarães, Douglas Terra Machado, Cassia Alves, Diana Mariani, Thais Felix Cruz, Mario Sergio Ribeiro, Silvia Carvalho, Flávio Dias da Silva, Marcio Henrique de Oliveira Garcia, Leandro Magalhães de Souza, Cristiane Gomes da Silva, Caio Luiz Pereira Ribeiro, Andréa Cony Cavalcanti, Claudia Maria Braga de Mello, Amilcar Tanuri, Ana Tereza R Vasconcelos |
| EPI_ISL_1678584, EPI_ISL_1690551, EPI_ISL_1694627, EPI_ISL_1694628, EPI_ISL_1694631, EPI_ISL_1694632, EPI_ISL_1694634, EPI_ISL_1695909 | Instituto Estadual do Cérebro Paulo Niemayer (IECPN) | Laboratório de Imunofarmacologia | Thiago Moreno Lopes Souza |
| EPI_ISL_1701311 | Laboratório de diagnóstico molecular da COVID-19 - Bahia, Santo Antônio de Jesus, Brazil | Laboratório Baculovírus | Josilene R Pinheiro, Jaime H Amorim, Aline Belmok, Fernando L Melo, Bergmann M Ribeiro |
| EPI_ISL_1707688 | UBDS Dr Marco Antonio Sahao Vila Virginia | Instituto Adolfo Lutz, Interdiciplinary Procedures Center, Strategic Laboratory | Claudio Tavares Sacchi, Claudia Regina Gonçalves, Erica Valessa Ramos Gomes, Karoline Rodrigues Campos, Caio Vinicius Dias Lopes, Leonardo Jose Tadeu de Araujo, Katia Correa de Oliveira Santos |
| EPI_ISL_1707689, EPI_ISL_1707690, EPI_ISL_1707691, EPI_ISL_1707692 | Instituto Adolfo Lutz - Regional de Taubate | Instituto Adolfo Lutz, Interdiciplinary Procedures Center, Strategic Laboratory | Claudio Tavares Sacchi, Claudia Regina Gonçalves, Erica Valessa Ramos Gomes, Karoline Rodrigues Campos, Caio Vinicius Dias Lopes, Leonardo Jose Tadeu de Araujo, Katia Correa de Oliveira Santos |
| EPI_ISL_1708318 | Fundação Hospitalar de Hematologia e Hemoterapia do Amazonas | Laboratório de Estudos de Vírus Emergentes | William M. Souza; Mariene R. Amorim, Renata Sesti-Costa, Lais D. Coimbra, Daniel A. Toledo-Teixeira, Pierina L. Parise, Priscilla P. Barbosa, Karina Bispo-dos-Santos, Luciana S. Mofatto, Camila L. Simeoni, Natalia S. Brunetti, Ingra M. Claro, Adriana S. S. Duarte, Thais M. Coletti, Audrey B. Zangirolami, Carolina Costa-Lima, Arilson Bernardo S. P. Gomes, Lucas I. Buscaratti, Flavia C. Sales, Vitor A. Costa, Lucas A.M. Franco, Darlan S. Candido, Oliver G. Pybus, Jaqueline G. de Jesus, Camila A.M. Silva, Mariana S. Ramundo, Giulia M. Ferreira, Mariana C. Pinho, Leandro M. Souza, Esmenia C. Rocha, Pamela S. Andrade, Myuki A.E. Crispim, Grazielle C. Maktura, Erika R. Manuli, Magnun N.N. Santos, Cecilia C. Camilo, Rodrigo N. Angerami, Maria L. Moretti, Fernando R. Spilki, Clarice W. Arns, Marcelo Addas-Carvalho, Bruno D. Benites, Marcelo A.S. Mori, Nelson Gaburo, Christopher Dye, Chieh-Hsi Wu, Henrique Marques-Souza, Rafael E. Marques, Alessandro S. Farias, Michael S. Diamond, Nuno R. Faria, Ester C. Sabino, Fabia |
| EPI_ISL_1715134 | Centro de Saude II Ibitinga | Instituto Adolfo Lutz, Interdiciplinary Procedures Center, Strategic Laboratory | Claudio Tavares Sacchi, Claudia Regina Gonçalves, Erica Valessa Ramos Gomes, Karoline Rodrigues Campos, Caio Vinicius Dias Lopes, Leonardo Jose Tadeu de Araujo, Katia Correa de Oliveira Santos |
| EPI_ISL_1715135, EPI_ISL_1715136 | CS II Dr Jose Ferreira Telles | Instituto Adolfo Lutz, Interdiciplinary Procedures Center, Strategic Laboratory | Claudio Tavares Sacchi, Claudia Regina Gonçalves, Erica Valessa Ramos Gomes, Karoline Rodrigues Campos, Caio Vinicius Dias Lopes, Leonardo Jose Tadeu de Araujo, Katia Correa de Oliveira Santos |
| EPI_ISL_1715137 | Secretaria Municipal De Saude | Instituto Adolfo Lutz, Interdiciplinary Procedures Center, Strategic Laboratory | Claudio Tavares Sacchi, Claudia Regina Gonçalves, Erica Valessa Ramos Gomes, Karoline Rodrigues Campos, Caio Vinicius Dias Lopes, Leonardo Jose Tadeu de Araujo, Katia Correa de Oliveira Santos |
| EPI_ISL_1715138 | Santa Casa De Cravinhos | Instituto Adolfo Lutz, Interdiciplinary Procedures Center, Strategic Laboratory | Claudio Tavares Sacchi, Claudia Regina Gonçalves, Erica Valessa Ramos Gomes, Karoline Rodrigues Campos, Caio Vinicius Dias Lopes, Leonardo Jose Tadeu de Araujo, Katia Correa de Oliveira Santos |
| EPI_ISL_1715139 | Sae Servico De Atendimento Especializado | Instituto Adolfo Lutz, Interdiciplinary Procedures Center, Strategic Laboratory | Claudio Tavares Sacchi, Claudia Regina Gonçalves, Erica Valessa Ramos Gomes, Karoline Rodrigues Campos, Caio Vinicius Dias Lopes, Leonardo Jose Tadeu de Araujo, Katia Correa de Oliveira Santos |
| EPI_ISL_1715140 | UPA Dr Luis Atilio Losi Viana Ribeirao Preto | Instituto Adolfo Lutz, Interdiciplinary Procedures Center, Strategic Laboratory | Claudio Tavares Sacchi, Claudia Regina Gonçalves, Erica Valessa Ramos Gomes, Karoline Rodrigues Campos, Caio Vinicius Dias Lopes, Leonardo Jose Tadeu de Araujo, Katia Correa de Oliveira Santos |
| EPI_ISL_1715141 | Centro De Saude II Ibitinga | Instituto Adolfo Lutz, Interdiciplinary Procedures Center, Strategic Laboratory | Claudio Tavares Sacchi, Claudia Regina Gonçalves, Erica Valessa Ramos Gomes, Karoline Rodrigues Campos, Caio Vinicius Dias Lopes, Leonardo Jose Tadeu de Araujo, Katia Correa de Oliveira Santos |
| EPI_ISL_1715142 | CS III de Patrocinio Paulista | Instituto Adolfo Lutz, Interdiciplinary Procedures Center, Strategic Laboratory | Claudio Tavares Sacchi, Claudia Regina Gonçalves, Erica Valessa Ramos Gomes, Karoline Rodrigues Campos, Caio Vinicius Dias Lopes, Leonardo Jose Tadeu de Araujo, Katia Correa de Oliveira Santos |
| EPI_ISL_1715143 | Secretaria Municipal de Saude De Guariba | Instituto Adolfo Lutz, Interdiciplinary Procedures Center, Strategic Laboratory | Claudio Tavares Sacchi, Claudia Regina Gonçalves, Erica Valessa Ramos Gomes, Karoline Rodrigues Campos, Caio Vinicius Dias Lopes, Leonardo Jose Tadeu de Araujo, Katia Correa de Oliveira Santos |
| EPI_ISL_1715144 | UBDS Dr Joao Baptista Quartim Central | Instituto Adolfo Lutz, Interdiciplinary Procedures Center, Strategic Laboratory | Claudio Tavares Sacchi, Claudia Regina Gonçalves, Erica Valessa Ramos Gomes, Karoline Rodrigues Campos, Caio Vinicius Dias Lopes, Leonardo Jose Tadeu de Araujo, Katia Correa de Oliveira Santos |
| EPI_ISL_1715145 | UPA de Bebedouro | Instituto Adolfo Lutz, Interdiciplinary Procedures Center, Strategic Laboratory | Claudio Tavares Sacchi, Claudia Regina Gonçalves, Erica Valessa Ramos Gomes, Karoline Rodrigues Campos, Caio Vinicius Dias Lopes, Leonardo Jose Tadeu de Araujo, Katia Correa de Oliveira Santos |
| EPI_ISL_1715146, EPI_ISL_1715147 | Instituto Adolfo Lutz - Regional de Aracatuba | Instituto Adolfo Lutz, Interdiciplinary Procedures Center, Strategic Laboratory | Claudio Tavares Sacchi, Claudia Regina Gonçalves, Erica Valessa Ramos Gomes, Karoline Rodrigues Campos, Caio Vinicius Dias Lopes, Leonardo Jose Tadeu de Araujo, Katia Correa de Oliveira Santos |
| EPI_ISL_1715148 | Instituto Adolfo Lutz - Regional de Ribeirao Preto | Instituto Adolfo Lutz, Interdiciplinary Procedures Center, Strategic Laboratory | Claudio Tavares Sacchi, Claudia Regina Gonçalves, Erica Valessa Ramos Gomes, Karoline Rodrigues Campos, Caio Vinicius Dias Lopes, Leonardo Jose Tadeu de Araujo, Katia Correa de Oliveira Santos |
| EPI_ISL_1715149, EPI_ISL_1715150, EPI_ISL_1715151, EPI_ISL_1715152, EPI_ISL_1715153, EPI_ISL_1715154, EPI_ISL_1715155, EPI_ISL_1715156, EPI_ISL_1715157, EPI_ISL_1715158, EPI_ISL_1715159 | | | |
| see above | Instituto Adolfo Lutz Central | Instituto Adolfo Lutz, Interdiciplinary Procedures Center, Strategic Laboratory | Claudio Tavares Sacchi, Claudia Regina Gonçalves, Erica Valessa Ramos Gomes, Karoline Rodrigues Campos, Caio Vinicius Dias Lopes, Leonardo Jose Tadeu de Araujo, Katia Correa de Oliveira Santos |
| EPI_ISL_1716407, EPI_ISL_1716408, EPI_ISL_1716441, EPI_ISL_1716442, EPI_ISL_1716443, EPI_ISL_1716445, EPI_ISL_1716447, EPI_ISL_1716448, EPI_ISL_1716453, EPI_ISL_1716455 | Laboratorio Saude | LGBio (Laboratorio de Genetica & Biodiversidade) | Mariana Pires de Campos Telles, Daniela de Melo e Silva, Elisangela de Paula Silveira Lacerda, Renata de Oliveira Dias, Rhewter Nunes, Cintia Pelegrineti Targueta de Azevedo Brito, Ramilla dos Santos Braga, Thais Guimarães Castro, Thays Millena Alves Pedroso, Amanda Alves de Melo, Francyelli Mello Andrade |

| | | | |
|---|---|---|---|
| EPI_ISL_1716494 | LACEN (Laboratorio de Saude Publica Dr. Giovanni Cysneiros) | LGBio (Laboratorio de Genetica & Biodiversidade) | Mariana Pires de Campos Telles, Daniela de Melo e Silva, Elisangela de Paula Silveira Lacerda, Renata de Oliveira Dias, Rhewter Nunes, Cintia Pelegrineti Targueta de Azevedo Brito, Ramilla dos Santos Braga, Thais Guimarães Castro, Thays Millena Alves Pedroso, Amanda Alves de Melo, Aparecido Divino da Cruz, Luiz Augusto Pereira, Thais Cidália Vieira Gigonzac, Marc Alexandre Duarte Gigonzac, Alex Honda Bernardes, Francyelli Mello Andrade |
| EPI_ISL_1716877 | SECRETARIA MUNICIPAL DE SAUDE SOROCABA | Instituto Butantan / Mendelics | Instituto Butantan: Alexander Roberto Precioso, Dimas Tadeu Covas, Sandra Coccuzzo Sampaio, Maria Carolina Elias, José Salvatore Leister Patané, Vincent Louis Viala, Antonio Jorge Martins, Ricardo Haddad, Claudia Renata dos Santos Barros, Elaine Cristina Marqueze, Raul Machado Neto, Debora Botequio Moretti. Centro de Genômica Funcional da ESALQ: Luiz Lehmann Coutinho, Ricardo Augusto Brassaloti, Raquel de Lello Rocha Campos Cassano. NGS Soluções Genômicas: Pilar Drummond Sampaio Corrêa Mariani. FZEA-USP Pirassununga: Mirele Daiana Poleti, Jessika Cristina Chagas Lesbon, Elisangela Chicaroni Mattos, Heidge Fukumasu. USP-Botucatu: Rejane Maria Tommasini Grotto, Jayme A. Souza-Neto, Guilherme Targino Valente, Patricia Akemi Assato, Felipe Allan da Silva da Costa, Bianca Cechetto Carlos. Mendelics: Bibiana Santos, João Paulo Kitajima, Erika Freitas, David Schlesinger. Hemocentro Ribeirão Preto: Simone Kashima, Evandra Strazza Rodrigues, Svetoslav Nanev Slavov, Elaine Vieira dos Santos, Rafael dos Santos Bezerra, Luiz Carlos Junior de Alcantara, Marta Giovanetti, Vagner Fonseca, Flavia Aburjaile, Rodrigo Tocantins Calado. |
| EPI_ISL_1716878 | VIGILANCIA EPIDEMIOLOGICA JARDINOPOLIS SP | Instituto Butantan / ESALQ-USP | Instituto Butantan: Alexander Roberto Precioso, Dimas Tadeu Covas, Sandra Coccuzzo Sampaio, Maria Carolina Elias, José Salvatore Leister Patané, Vincent Louis Viala, Antonio Jorge Martins, Ricardo Haddad, Claudia Renata dos Santos Barros, Elaine Cristina Marqueze, Raul Machado Neto, Debora Botequio Moretti. Centro de Genômica Funcional da ESALQ: Luiz Lehmann Coutinho, Ricardo Augusto Brassaloti, Raquel de Lello Rocha Campos Cassano. NGS Soluções Genômicas: Pilar Drummond Sampaio Corrêa Mariani. FZEA-USP Pirassununga: Mirele Daiana Poleti, Jessika Cristina Chagas Lesbon, Elisangela Chicaroni Mattos, Heidge Fukumasu. USP-Botucatu: Rejane Maria Tommasini Grotto, Jayme A. Souza-Neto, Guilherme Targino Valente, Patricia Akemi Assato, Felipe Allan da Silva da Costa, Bianca Cechetto Carlos. Mendelics: Bibiana Santos, João Paulo Kitajima, Erika Freitas, David Schlesinger. Hemocentro Ribeirão Preto: Simone Kashima, Evandra Strazza Rodrigues, Svetoslav Nanev Slavov, Elaine Vieira dos Santos, Rafael dos Santos Bezerra, Luiz Carlos Junior de Alcantara, Marta Giovanetti, Vagner Fonseca, Flavia Aburjaile, Rodrigo Tocantins Calado. |
| EPI_ISL_1716879 | CENTRO MEDICO PMESP | Instituto Butantan / ESALQ-USP | Instituto Butantan: Alexander Roberto Precioso, Dimas Tadeu Covas, Sandra Coccuzzo Sampaio, Maria Carolina Elias, José Salvatore Leister Patané, Vincent Louis Viala, Antonio Jorge Martins, Ricardo Haddad, Claudia Renata dos Santos Barros, Elaine Cristina Marqueze, Raul Machado Neto, Debora Botequio Moretti. Centro de Genômica Funcional da ESALQ: Luiz Lehmann Coutinho, Ricardo Augusto Brassaloti, Raquel de Lello Rocha Campos Cassano. NGS Soluções Genômicas: Pilar Drummond Sampaio Corrêa Mariani. FZEA-USP Pirassununga: Mirele Daiana Poleti, Jessika Cristina Chagas Lesbon, Elisangela Chicaroni Mattos, Heidge Fukumasu. USP-Botucatu: Rejane Maria Tommasini Grotto, Jayme A. Souza-Neto, Guilherme Targino Valente, Patricia Akemi Assato, Felipe Allan da Silva da Costa, Bianca Cechetto Carlos. Mendelics: Bibiana Santos, João Paulo Kitajima, Erika Freitas, David Schlesinger. Hemocentro Ribeirão Preto: Simone Kashima, Evandra Strazza Rodrigues, Svetoslav Nanev Slavov, Elaine Vieira dos Santos, Rafael dos Santos Bezerra, Luiz Carlos Junior de Alcantara, Marta Giovanetti, Vagner Fonseca, Flavia Aburjaile, Rodrigo Tocantins Calado. |
| EPI_ISL_1731574 | UBDS Dr Marco Antonio Sahao Vila Virginia | Instituto Adolfo Lutz, Interdiciplinary Procedures Center, Strategic Laboratory | Claudio Tavares Sacchi, Claudia Regina Gonçalves, Erica Valessa Ramos Gomes, Karoline Rodrigues Campos, Caio Vinicius Dias Lopes, Leonardo Jose Tadeu de Araujo, Katia Correa de Oliveira Santos |
| EPI_ISL_1731575 | Casa de Caridade Sao Vicente de Paulo Cajuru | Instituto Adolfo Lutz, Interdiciplinary Procedures Center, Strategic Laboratory | Claudio Tavares Sacchi, Claudia Regina Gonçalves, Erica Valessa Ramos Gomes, Karoline Rodrigues Campos, Caio Vinicius Dias Lopes, Leonardo Jose Tadeu de Araujo, Katia Correa de Oliveira Santos |
| EPI_ISL_1731576 | UPA Dr Luis Atilio Losi Viana Ribeirao Preto | Instituto Adolfo Lutz, Interdiciplinary Procedures Center, Strategic Laboratory | Claudio Tavares Sacchi, Claudia Regina Gonçalves, Erica Valessa Ramos Gomes, Karoline Rodrigues Campos, Caio Vinicius Dias Lopes, Leonardo Jose Tadeu de Araujo, Katia Correa de Oliveira Santos |
| EPI_ISL_1731577 | Hospital Municipal Dr Mario Gatti Campinas | Instituto Adolfo Lutz, Interdiciplinary Procedures Center, Strategic Laboratory | Claudio Tavares Sacchi, Claudia Regina Gonçalves, Erica Valessa Ramos Gomes, Karoline Rodrigues Campos, Caio Vinicius Dias Lopes, Leonardo Jose Tadeu de Araujo, Katia Correa de Oliveira Santos |
| EPI_ISL_1731578, EPI_ISL_1731579, EPI_ISL_1731580 | Instituto Adolfo Lutz - Regional de Aracatuba | Instituto Adolfo Lutz, Interdiciplinary Procedures Center, Strategic Laboratory | Claudio Tavares Sacchi, Claudia Regina Gonçalves, Erica Valessa Ramos Gomes, Karoline Rodrigues Campos, Caio Vinicius Dias Lopes, Leonardo Jose Tadeu de Araujo, Katia Correa de Oliveira Santos |
| EPI_ISL_1731581, EPI_ISL_1731582 | Instituto Adolfo Lutz - Regional de Ribeirao Preto | Instituto Adolfo Lutz, Interdiciplinary Procedures Center, Strategic Laboratory | Claudio Tavares Sacchi, Claudia Regina Gonçalves, Erica Valessa Ramos Gomes, Karoline Rodrigues Campos, Caio Vinicius Dias Lopes, Leonardo Jose Tadeu de Araujo, Katia Correa de Oliveira Santos |
| EPI_ISL_1731583, EPI_ISL_1731584, EPI_ISL_1731585, EPI_ISL_1731586, EPI_ISL_1731587, EPI_ISL_1731588, EPI_ISL_1731589, EPI_ISL_1731590, EPI_ISL_1731591, EPI_ISL_1731592, EPI_ISL_1731593, EPI_ISL_1731594, EPI_ISL_1731595, EPI_ISL_1731596, EPI_ISL_1731597, EPI_ISL_1731598, EPI_ISL_1731599, EPI_ISL_1731600, EPI_ISL_1731601, EPI_ISL_1731602, EPI_ISL_1731603, EPI_ISL_1731604, EPI_ISL_1731605, EPI_ISL_1731606, EPI_ISL_1731607, EPI_ISL_1731608, EPI_ISL_1731609, EPI_ISL_1731610, EPI_ISL_1731611, EPI_ISL_1731612 | | | |
| see above | Instituto Adolfo Lutz Central | Instituto Adolfo Lutz, Interdiciplinary Procedures Center, Strategic Laboratory | Claudio Tavares Sacchi, Claudia Regina Gonçalves, Erica Valessa Ramos Gomes, Karoline Rodrigues Campos, Caio Vinicius Dias Lopes, Leonardo Jose Tadeu de Araujo, Katia Correa de Oliveira Santos |
| EPI_ISL_1732275 | SECAO CENTRO DE DIAGNOSTICO SECEDI | Instituto Butantan / Mendelics | Instituto Butantan: Alexander Roberto Precioso, Dimas Tadeu Covas, Sandra Coccuzzo Sampaio, Maria Carolina Elias, José Salvatore Leister Patané, Vincent Louis Viala, Antonio Jorge Martins, Ricardo Haddad, Claudia Renata dos Santos Barros, Elaine Cristina Marqueze, Raul Machado Neto, Debora Botequio Moretti. Centro de Genômica Funcional da ESALQ: Luiz Lehmann Coutinho, Ricardo Augusto Brassaloti, Raquel de Lello Rocha Campos Cassano. NGS Soluções Genômicas: Pilar Drummond Sampaio Corrêa Mariani. FZEA-USP Pirassununga: Mirele Daiana Poleti, Jessika Cristina Chagas Lesbon, Elisangela Chicaroni Mattos, Heidge Fukumasu. USP-Botucatu: Rejane Maria Tommasini Grotto, Jayme A. Souza-Neto, Guilherme Targino Valente, Patricia Akemi Assato, Felipe Allan da Silva da Costa, Bianca Cechetto Carlos. Mendelics: Bibiana Santos, João Paulo Kitajima, Erika Freitas, David Schlesinger. Hemocentro Ribeirão Preto: Simone Kashima, Evandra Strazza Rodrigues, Svetoslav Nanev Slavov, Elaine Vieira dos Santos, Rafael dos Santos Bezerra, Luiz Carlos Junior de Alcantara, Marta Giovanetti, Vagner Fonseca, Flavia Aburjaile, Rodrigo Tocantins Calado. |
| EPI_ISL_1734841, EPI_ISL_1734843, EPI_ISL_1734844, EPI_ISL_1734846, EPI_ISL_1734847, EPI_ISL_1734849, EPI_ISL_1734850, EPI_ISL_1734852, EPI_ISL_1734853, EPI_ISL_1734855, EPI_ISL_1734857, EPI_ISL_1734858, EPI_ISL_1734860, EPI_ISL_1734861, EPI_ISL_1734863, EPI_ISL_1734864, EPI_ISL_1734866, EPI_ISL_1734868, EPI_ISL_1734869, EPI_ISL_1734871, EPI_ISL_1734872, EPI_ISL_1734874, EPI_ISL_1734875, EPI_ISL_1734877, EPI_ISL_1734878, EPI_ISL_1734880, EPI_ISL_1734881, EPI_ISL_1734883, EPI_ISL_1734884 | | | |
| see above | Instituto de Biotecnologia - UNESP-Botucatu-SP | Instituto de Biotecnologia - UNESP-Botucatu-SP | Fábio Sossai Possebon; Leila Sabrina Ullmann; Cecília Artico Banho; Cíntia Bittar; Guilherme Campos; Helena Lage Ferreira; Jorge A. Petroli Marchesi; Lívia Sacchetto; Maisa C. Pereira Parra; Marília Moraes; Maurício L. Nogueira; Paula Rahal; Paulo Inacio da Costa; João Pessoa Araújo Jr. |
| EPI_ISL_1752637 | UBDS Dr Marco Antonio Sahao Vila Virginia | Instituto Adolfo Lutz, Interdiciplinary Procedures Center, Strategic Laboratory | Claudio Tavares Sacchi, Claudia Regina Gonçalves, Erica Valessa Ramos Gomes, Karoline Rodrigues Campos, Caio Vinicius Dias Lopes, Leonardo Jose Tadeu de Araujo, Katia Correa de Oliveira Santos |
| EPI_ISL_1752638 | Casa de Caridade Sao Vicente de Paulo Cajuru | Instituto Adolfo Lutz, Interdiciplinary Procedures Center, Strategic Laboratory | Claudio Tavares Sacchi, Claudia Regina Gonçalves, Erica Valessa Ramos Gomes, Karoline Rodrigues Campos, Caio Vinicius Dias Lopes, Leonardo Jose Tadeu de Araujo, Katia Correa de Oliveira Santos |
| EPI_ISL_1752639 | UPA Dr Luis Atilio Losi Viana Ribeirao Preto | Instituto Adolfo Lutz, Interdiciplinary Procedures Center, Strategic Laboratory | Claudio Tavares Sacchi, Claudia Regina Gonçalves, Erica Valessa Ramos Gomes, Karoline Rodrigues Campos, Caio Vinicius Dias Lopes, Leonardo Jose Tadeu de Araujo, Katia Correa de Oliveira Santos |
| EPI_ISL_1752640 | Hospital Municipal Dr Mario Gatti Campinas | Instituto Adolfo Lutz, Interdiciplinary Procedures Center, Strategic Laboratory | Claudio Tavares Sacchi, Claudia Regina Gonçalves, Erica Valessa Ramos Gomes, Karoline Rodrigues Campos, Caio Vinicius Dias Lopes, Leonardo Jose Tadeu de Araujo, Katia Correa de Oliveira Santos |
| EPI_ISL_1752641, EPI_ISL_1752642, EPI_ISL_1752643 | Instituto Adolfo Lutz - Regional de Aracatuba | Instituto Adolfo Lutz, Interdiciplinary Procedures Center, Strategic Laboratory | Claudio Tavares Sacchi, Claudia Regina Gonçalves, Erica Valessa Ramos Gomes, Karoline Rodrigues Campos, Caio Vinicius Dias Lopes, Leonardo Jose Tadeu de Araujo, Katia Correa de Oliveira Santos |
| EPI_ISL_1752644, EPI_ISL_1752645 | Instituto Adolfo Lutz - Regional de Ribeirao Preto | Instituto Adolfo Lutz, Interdiciplinary Procedures Center, Strategic Laboratory | Claudio Tavares Sacchi, Claudia Regina Gonçalves, Erica Valessa Ramos Gomes, Karoline Rodrigues Campos, Caio Vinicius Dias Lopes, Leonardo Jose Tadeu de Araujo, Katia Correa de Oliveira Santos |
| EPI_ISL_1752646, EPI_ISL_1752647, EPI_ISL_1752648, EPI_ISL_1752649, EPI_ISL_1752650, EPI_ISL_1752651, EPI_ISL_1752652, EPI_ISL_1752653, EPI_ISL_1752654, EPI_ISL_1752655, EPI_ISL_1752656, EPI_ISL_1752657, EPI_ISL_1752658, EPI_ISL_1752659, EPI_ISL_1752660, EPI_ISL_1752661, EPI_ISL_1752662, EPI_ISL_1752663, EPI_ISL_1752664, EPI_ISL_1752665, EPI_ISL_1752666, EPI_ISL_1752667, EPI_ISL_1752668 | | | |
| see above | Instituto Adolfo Lutz Central | Instituto Adolfo Lutz, Interdiciplinary Procedures Center, Strategic Laboratory | Claudio Tavares Sacchi, Claudia Regina Gonçalves, Erica Valessa Ramos Gomes, Karoline Rodrigues Campos, Caio Vinicius Dias Lopes, Leonardo Jose Tadeu de Araujo, Katia Correa de Oliveira Santos |
| EPI_ISL_1754186 | Laboratório de Pesquisa em Virologia, FAMERP, SJRP | Laboratório de Pesquisa em Virologia, FAMERP, SJRP | Cecília Artico Banho; Fábio Sossai Possebon; Leila Sabrina Ullmann; Cíntia Bittar; Guilherme Campos; Helena Lage Ferreira; Jorge A. Petroli Marchesi; Lívia Sacchetto; Maisa C. Pereira Parra; Marília Moraes; Paula Rahal; Paulo Inacio da Costa; João Pessoa Araújo Jr.; Maurício L. Nogueira |
| EPI_ISL_1785610, EPI_ISL_1785611, | Laboratório de Pesquisa em Virologia, FAMERP, SJRP | Laboratório de Pesquisa em Virologia, FAMERP, SJRP | Fábio Sossai Possebon; Leila Sabrina Ullmann; Cecília Artico Banho; Cíntia Bittar; Guilherme Campos; Helena Lage Ferreira; Jorge A. Petroli Marchesi; |

| | | | |
|---|---|---|---|
| EPI_ISL_1785612 | | | Lívia Sacchetto; Maisa C. Pereira Parra; Marília Moraes; Maurício L. Nogueira; Paula Rahal; Paulo Inacio da Costa; João Pessoa Araújo Jr. |
| EPI_ISL_1786560, EPI_ISL_1786561, EPI_ISL_1786562, EPI_ISL_1786563, EPI_ISL_1786564, EPI_ISL_1786565, EPI_ISL_1786566, EPI_ISL_1786567, EPI_ISL_1786568, EPI_ISL_1786569 | Laboratorio de Virologia Clinica do HCFMRP-USP | Hocentro de Ribeirao Preto - Laboratório de Biologia Molecular | Svetoslav Nanev Slavov, Marta Giovanetti, Rafael dos Santos Bezerra, Vagner Fonseca, Elaine Vieira Santos, Evandra Strazza Rodrigues, Talita Adelino, Joilson Xavier, Josiane Serrano Borges, Mariane Evaristo, Glauco de Carvalho Pereira, Aparecida Yulie Yamamoto, Diego Villa Clé, Rodrigo Tocantins Calado, Dimas Tadeu Covas, Luiz Carlos Junior Alcantara, Simone Kashima |
| EPI_ISL_1795063 | UNIDADE DE VIGILANCIA EM SAUDE | Instituto Butantan / ESALQ-Piracicaba | Instituto Butantan: Alexander Roberto Precioso, Dimas Tadeu Covas, Sandra Coccuzzo Sampaio, Maria Carolina Elias, José Salvatore Leister Patané, Vincent Louis Viala, Antonio Jorge Martins, Ricardo Haddad, Claudia Renata dos Santos Barros, Elaine Cristina Marqueze, Raul Machado Neto, Debora Botequio Moretti. Centro de Genômica Funcional da ESALQ: Luiz Lehmann Coutinho, Ricardo Augusto Brassaloti, Raquel de Lello Rocha Campos Cassano. NGS Soluções Genômicas: Pilar Drummond Sampaio Corrêa Mariani. FZEA-USP Pirassununga: Mirele Daiana Poleti, Jessika Cristina Chagas Lesbon, Elisangela Chicaroni Mattos, Heidge Fukumasu. USP-Botucatu: Rejane Maria Tommasini Grotto, Jayme A. Souza-Neto, Guilherme Targino Valente, Patricia Akemi Assato, Felipe Allan da Silva da Costa, Bianca Cechetto Carlos. Mendelics: Bibiana Santos, João Paulo Kitajima, Erika Freitas, David Schlesinger. Hemocentro Ribeirão Preto: Simone Kashima, Evandra Strazza Rodrigues, Svetoslav Nanev Slavov, Elaine Vieira dos Santos, Rafael dos Santos Bezerra, Luiz Carlos Junior de Alcantara, Marta Giovanetti, Vagner Fonseca, Flavia Aburjaile, Rodrigo Tocantins Calado. |
| EPI_ISL_1795064 | USF SALERNO | Instituto Butantan / ESALQ-Piracicaba | Instituto Butantan: Alexander Roberto Precioso, Dimas Tadeu Covas, Sandra Coccuzzo Sampaio, Maria Carolina Elias, José Salvatore Leister Patané, Vincent Louis Viala, Antonio Jorge Martins, Ricardo Haddad, Claudia Renata dos Santos Barros, Elaine Cristina Marqueze, Raul Machado Neto, Debora Botequio Moretti. Centro de Genômica Funcional da ESALQ: Luiz Lehmann Coutinho, Ricardo Augusto Brassaloti, Raquel de Lello Rocha Campos Cassano. NGS Soluções Genômicas: Pilar Drummond Sampaio Corrêa Mariani. FZEA-USP Pirassununga: Mirele Daiana Poleti, Jessika Cristina Chagas Lesbon, Elisangela Chicaroni Mattos, Heidge Fukumasu. USP-Botucatu: Rejane Maria Tommasini Grotto, Jayme A. Souza-Neto, Guilherme Targino Valente, Patricia Akemi Assato, Felipe Allan da Silva da Costa, Bianca Cechetto Carlos. Mendelics: Bibiana Santos, João Paulo Kitajima, Erika Freitas, David Schlesinger. Hemocentro Ribeirão Preto: Simone Kashima, Evandra Strazza Rodrigues, Svetoslav Nanev Slavov, Elaine Vieira dos Santos, Rafael dos Santos Bezerra, Luiz Carlos Junior de Alcantara, Marta Giovanetti, Vagner Fonseca, Flavia Aburjaile, Rodrigo Tocantins Calado. |
| EPI_ISL_1795065 | UNIDADE DE SAUDE DA FAMILIA JOSE ADALBERTO LELLIS GARCIA | Instituto Butantan / ESALQ-Piracicaba | Instituto Butantan: Alexander Roberto Precioso, Dimas Tadeu Covas, Sandra Coccuzzo Sampaio, Maria Carolina Elias, José Salvatore Leister Patané, Vincent Louis Viala, Antonio Jorge Martins, Ricardo Haddad, Claudia Renata dos Santos Barros, Elaine Cristina Marqueze, Raul Machado Neto, Debora Botequio Moretti. Centro de Genômica Funcional da ESALQ: Luiz Lehmann Coutinho, Ricardo Augusto Brassaloti, Raquel de Lello Rocha Campos Cassano. NGS Soluções Genômicas: Pilar Drummond Sampaio Corrêa Mariani. FZEA-USP Pirassununga: Mirele Daiana Poleti, Jessika Cristina Chagas Lesbon, Elisangela Chicaroni Mattos, Heidge Fukumasu. USP-Botucatu: Rejane Maria Tommasini Grotto, Jayme A. Souza-Neto, Guilherme Targino Valente, Patricia Akemi Assato, Felipe Allan da Silva da Costa, Bianca Cechetto Carlos. Mendelics: Bibiana Santos, João Paulo Kitajima, Erika Freitas, David Schlesinger. Hemocentro Ribeirão Preto: Simone Kashima, Evandra Strazza Rodrigues, Svetoslav Nanev Slavov, Elaine Vieira dos Santos, Rafael dos Santos Bezerra, Luiz Carlos Junior de Alcantara, Marta Giovanetti, Vagner Fonseca, Flavia Aburjaile, Rodrigo Tocantins Calado. |
| EPI_ISL_1795066, EPI_ISL_1795067, EPI_ISL_1795068, EPI_ISL_1795069, EPI_ISL_1795070, EPI_ISL_1795071 | LABORATORIO DE FRANCA | Instituto Butantan / ESALQ-Piracicaba | Instituto Butantan: Alexander Roberto Precioso, Dimas Tadeu Covas, Sandra Coccuzzo Sampaio, Maria Carolina Elias, José Salvatore Leister Patané, Vincent Louis Viala, Antonio Jorge Martins, Ricardo Haddad, Claudia Renata dos Santos Barros, Elaine Cristina Marqueze, Raul Machado Neto, Debora Botequio Moretti. Centro de Genômica Funcional da ESALQ: Luiz Lehmann Coutinho, Ricardo Augusto Brassaloti, Raquel de Lello Rocha Campos Cassano. NGS Soluções Genômicas: Pilar Drummond Sampaio Corrêa Mariani. FZEA-USP Pirassununga: Mirele Daiana Poleti, Jessika Cristina Chagas Lesbon, Elisangela Chicaroni Mattos, Heidge Fukumasu. USP-Botucatu: Rejane Maria Tommasini Grotto, Jayme A. Souza-Neto, Guilherme Targino Valente, Patricia Akemi Assato, Felipe Allan da Silva da Costa, Bianca Cechetto Carlos. Mendelics: Bibiana Santos, João Paulo Kitajima, Erika Freitas, David Schlesinger. Hemocentro Ribeirão Preto: Simone Kashima, Evandra Strazza Rodrigues, Svetoslav Nanev Slavov, Elaine Vieira dos Santos, Rafael dos Santos Bezerra, Luiz Carlos Junior de Alcantara, Marta Giovanetti, Vagner Fonseca, Flavia Aburjaile, Rodrigo Tocantins Calado. |
| EPI_ISL_1795072, EPI_ISL_1795073, EPI_ISL_1795074 | LABORATORIO DR PAULO EMILIO DALESSANDRO PINDAMONHANGABA | Instituto Butantan / ESALQ-Piracicaba | Instituto Butantan: Alexander Roberto Precioso, Dimas Tadeu Covas, Sandra Coccuzzo Sampaio, Maria Carolina Elias, José Salvatore Leister Patané, Vincent Louis Viala, Antonio Jorge Martins, Ricardo Haddad, Claudia Renata dos Santos Barros, Elaine Cristina Marqueze, Raul Machado Neto, Debora Botequio Moretti. Centro de Genômica Funcional da ESALQ: Luiz Lehmann Coutinho, Ricardo Augusto Brassaloti, Raquel de Lello Rocha Campos Cassano. NGS Soluções Genômicas: Pilar Drummond Sampaio Corrêa Mariani. FZEA-USP Pirassununga: Mirele Daiana Poleti, Jessika Cristina Chagas Lesbon, Elisangela Chicaroni Mattos, Heidge Fukumasu. USP-Botucatu: Rejane Maria Tommasini Grotto, Jayme A. Souza-Neto, Guilherme Targino Valente, Patricia Akemi Assato, Felipe Allan da Silva da Costa, Bianca Cechetto Carlos. Mendelics: Bibiana Santos, João Paulo Kitajima, Erika Freitas, David Schlesinger. Hemocentro Ribeirão Preto: Simone Kashima, Evandra Strazza Rodrigues, Svetoslav Nanev Slavov, Elaine Vieira dos Santos, Rafael dos Santos Bezerra, Luiz Carlos Junior de Alcantara, Marta Giovanetti, Vagner Fonseca, Flavia Aburjaile, Rodrigo Tocantins Calado. |
| EPI_ISL_1795075, EPI_ISL_1795076 | NUCLEO SAUDE III N HABITACIONAL PRESIDENTE GEISEL BAURU | Instituto Butantan / ESALQ-Piracicaba | Instituto Butantan: Alexander Roberto Precioso, Dimas Tadeu Covas, Sandra Coccuzzo Sampaio, Maria Carolina Elias, José Salvatore Leister Patané, Vincent Louis Viala, Antonio Jorge Martins, Ricardo Haddad, Claudia Renata dos Santos Barros, Elaine Cristina Marqueze, Raul Machado Neto, Debora Botequio Moretti. Centro de Genômica Funcional da ESALQ: Luiz Lehmann Coutinho, Ricardo Augusto Brassaloti, Raquel de Lello Rocha Campos Cassano. NGS Soluções Genômicas: Pilar Drummond Sampaio Corrêa Mariani. FZEA-USP Pirassununga: Mirele Daiana Poleti, Jessika Cristina Chagas Lesbon, Elisangela Chicaroni Mattos, Heidge Fukumasu. USP-Botucatu: Rejane Maria Tommasini Grotto, Jayme A. Souza-Neto, Guilherme Targino Valente, Patricia Akemi Assato, Felipe Allan da Silva da Costa, Bianca Cechetto Carlos. Mendelics: Bibiana Santos, João Paulo Kitajima, Erika Freitas, David Schlesinger. Hemocentro Ribeirão Preto: Simone Kashima, Evandra Strazza Rodrigues, Svetoslav Nanev Slavov, Elaine Vieira dos Santos, Rafael dos Santos Bezerra, Luiz Carlos Junior de Alcantara, Marta Giovanetti, Vagner Fonseca, Flavia Aburjaile, Rodrigo Tocantins Calado. |
| EPI_ISL_1795077 | MARIA TEREZINHA BOGNAR CHECHETO | Instituto Butantan / ESALQ-Piracicaba | Instituto Butantan: Alexander Roberto Precioso, Dimas Tadeu Covas, Sandra Coccuzzo Sampaio, Maria Carolina Elias, José Salvatore Leister Patané, Vincent Louis Viala, Antonio Jorge Martins, Ricardo Haddad, Claudia Renata dos Santos Barros, Elaine Cristina Marqueze, Raul Machado Neto, Debora Botequio Moretti. Centro de Genômica Funcional da ESALQ: Luiz Lehmann Coutinho, Ricardo Augusto Brassaloti, Raquel de Lello Rocha Campos Cassano. NGS Soluções Genômicas: Pilar Drummond Sampaio Corrêa Mariani. FZEA-USP Pirassununga: Mirele Daiana Poleti, Jessika Cristina Chagas Lesbon, Elisangela Chicaroni Mattos, Heidge Fukumasu. USP-Botucatu: Rejane Maria Tommasini Grotto, Jayme A. Souza-Neto, Guilherme Targino Valente, Patricia Akemi Assato, Felipe Allan da Silva da Costa, Bianca Cechetto Carlos. Mendelics: Bibiana Santos, João Paulo Kitajima, Erika Freitas, David Schlesinger. Hemocentro Ribeirão Preto: Simone Kashima, Evandra Strazza Rodrigues, Svetoslav Nanev Slavov, Elaine Vieira dos Santos, Rafael dos Santos Bezerra, Luiz Carlos Junior de Alcantara, Marta Giovanetti, Vagner Fonseca, Flavia Aburjaile, Rodrigo Tocantins Calado. |
| EPI_ISL_1795078 | LABORATORIO DE FRANCA | Instituto Butantan / ESALQ-Piracicaba | Instituto Butantan: Alexander Roberto Precioso, Dimas Tadeu Covas, Sandra Coccuzzo Sampaio, Maria Carolina Elias, José Salvatore Leister Patané, Vincent Louis Viala, Antonio Jorge Martins, Ricardo Haddad, Claudia Renata dos Santos Barros, Elaine Cristina Marqueze, Raul Machado Neto, Debora Botequio Moretti. Centro de Genômica Funcional da ESALQ: Luiz Lehmann Coutinho, Ricardo Augusto Brassaloti, Raquel de Lello Rocha Campos Cassano. NGS Soluções Genômicas: Pilar Drummond Sampaio Corrêa Mariani. FZEA-USP Pirassununga: Mirele Daiana Poleti, Jessika Cristina Chagas Lesbon, Elisangela Chicaroni Mattos, Heidge Fukumasu. USP-Botucatu: Rejane Maria Tommasini Grotto, Jayme A. Souza-Neto, Guilherme Targino Valente, Patricia Akemi Assato, Felipe Allan da Silva da Costa, Bianca Cechetto Carlos. Mendelics: Bibiana Santos, João Paulo Kitajima, Erika Freitas, David Schlesinger. Hemocentro Ribeirão Preto: Simone Kashima, Evandra Strazza Rodrigues, Svetoslav Nanev Slavov, Elaine Vieira dos Santos, Rafael dos Santos Bezerra, Luiz Carlos Junior de Alcantara, Marta Giovanetti, Vagner Fonseca, Flavia Aburjaile, Rodrigo Tocantins Calado. |
| EPI_ISL_1795079 | CS DE PARANAPUA | Instituto Butantan / ESALQ-Piracicaba | Instituto Butantan: Alexander Roberto Precioso, Dimas Tadeu Covas, Sandra Coccuzzo Sampaio, Maria Carolina Elias, José Salvatore Leister Patané, Vincent Louis Viala, Antonio Jorge Martins, Ricardo Haddad, Claudia Renata dos Santos Barros, Elaine Cristina Marqueze, Raul Machado Neto, Debora Botequio Moretti. Centro de Genômica Funcional da ESALQ: Luiz Lehmann Coutinho, Ricardo Augusto Brassaloti, Raquel de Lello Rocha Campos Cassano. NGS Soluções Genômicas: Pilar Drummond Sampaio Corrêa Mariani. FZEA-USP Pirassununga: Mirele Daiana Poleti, Jessika Cristina Chagas Lesbon, Elisangela Chicaroni Mattos, Heidge Fukumasu. USP-Botucatu: Rejane Maria Tommasini Grotto, Jayme A. Souza-Neto, Guilherme Targino Valente, Patricia Akemi Assato, Felipe Allan da Silva da Costa, Bianca Cechetto Carlos. Mendelics: Bibiana Santos, João Paulo Kitajima, Erika Freitas, David Schlesinger. Hemocentro Ribeirão Preto: Simone Kashima, Evandra Strazza Rodrigues, Svetoslav Nanev Slavov, Elaine Vieira dos Santos, Rafael dos Santos Bezerra, Luiz Carlos Junior de Alcantara, Marta Giovanetti, Vagner Fonseca, Flavia Aburjaile, Rodrigo Tocantins Calado. |
| EPI_ISL_1795080 | CENTRO MEDICO PMESP | Instituto Butantan / ESALQ-Piracicaba | Instituto Butantan: Alexander Roberto Precioso, Dimas Tadeu Covas, Sandra Coccuzzo Sampaio, Maria Carolina Elias, José Salvatore Leister Patané, Vincent Louis Viala, Antonio Jorge Martins, Ricardo Haddad, Claudia Renata dos Santos Barros, Elaine Cristina Marqueze, Raul Machado Neto, Debora Botequio Moretti. Centro de Genômica Funcional da ESALQ: Luiz Lehmann Coutinho, Ricardo Augusto Brassaloti, Raquel de Lello Rocha Campos Cassano. NGS Soluções Genômicas: Pilar Drummond Sampaio Corrêa Mariani. FZEA-USP Pirassununga: Mirele Daiana Poleti, Jessika Cristina Chagas |

| | | | |
|---|---|---|---|
| | | | Lesbon, Elisangela Chicaroni Mattos, Heidge Fukumasu. USP-Botucatu: Rejane Maria Tommasini Grotto, Jayme A. Souza-Neto, Guilherme Targino Valente, Patricia Akemi Assato, Felipe Allan da Silva da Costa, Bianca Cechetto Carlos. Mendelics: Bibiana Santos, João Paulo Kitajima, Erika Freitas, David Schlesinger. Hemocentro Ribeirão Preto: Simone Kashima, Evandra Strazza Rodrigues, Svetoslav Nanev Slavov, Elaine Vieira dos Santos, Rafael dos Santos Bezerra, Luiz Carlos Junior de Alcantara, Marta Giovanetti, Vagner Fonseca, Flavia Aburjaile, Rodrigo Tocantins Calado. |
| EPI_ISL_1795081 | CENTRO DE SAUDE III AFFONSO LUZZI SANTA CRUZ DAS PALMEIRAS | Instituto Butantan / ESALQ-Piracicaba | Instituto Butantan: Alexander Roberto Precioso, Dimas Tadeu Covas, Sandra Coccuzzo Sampaio, Maria Carolina Elias, José Salvatore Leister Patané, Vincent Louis Viala, Antonio Jorge Martins, Ricardo Haddad, Claudia Renata dos Santos Barros, Elaine Cristina Marqueze, Raul Machado Neto, Debora Botequio Moretti. Centro de Genômica Funcional da ESALQ: Luiz Lehmann Coutinho, Ricardo Augusto Brassaloti, Raquel de Lello Rocha Campos Cassano. NGS Soluções Genômicas: Pilar Drummond Sampaio Corrêa Mariani. FZEA-USP Pirassununga: Mirele Daiana Poleti, Jessika Cristina Chagas Lesbon, Elisangela Chicaroni Mattos, Heidge Fukumasu. USP-Botucatu: Rejane Maria Tommasini Grotto, Jayme A. Souza-Neto, Guilherme Targino Valente, Patricia Akemi Assato, Felipe Allan da Silva da Costa, Bianca Cechetto Carlos. Mendelics: Bibiana Santos, João Paulo Kitajima, Erika Freitas, David Schlesinger. Hemocentro Ribeirão Preto: Simone Kashima, Evandra Strazza Rodrigues, Svetoslav Nanev Slavov, Elaine Vieira dos Santos, Rafael dos Santos Bezerra, Luiz Carlos Junior de Alcantara, Marta Giovanetti, Vagner Fonseca, Flavia Aburjaile, Rodrigo Tocantins Calado. |
| EPI_ISL_1795082, EPI_ISL_1795083 | POLICLINICA HORTOLANDIA | Instituto Butantan / ESALQ-Piracicaba | Instituto Butantan: Alexander Roberto Precioso, Dimas Tadeu Covas, Sandra Coccuzzo Sampaio, Maria Carolina Elias, José Salvatore Leister Patané, Vincent Louis Viala, Antonio Jorge Martins, Ricardo Haddad, Claudia Renata dos Santos Barros, Elaine Cristina Marqueze, Raul Machado Neto, Debora Botequio Moretti. Centro de Genômica Funcional da ESALQ: Luiz Lehmann Coutinho, Ricardo Augusto Brassaloti, Raquel de Lello Rocha Campos Cassano. NGS Soluções Genômicas: Pilar Drummond Sampaio Corrêa Mariani. FZEA-USP Pirassununga: Mirele Daiana Poleti, Jessika Cristina Chagas Lesbon, Elisangela Chicaroni Mattos, Heidge Fukumasu. USP-Botucatu: Rejane Maria Tommasini Grotto, Jayme A. Souza-Neto, Guilherme Targino Valente, Patricia Akemi Assato, Felipe Allan da Silva da Costa, Bianca Cechetto Carlos. Mendelics: Bibiana Santos, João Paulo Kitajima, Erika Freitas, David Schlesinger. Hemocentro Ribeirão Preto: Simone Kashima, Evandra Strazza Rodrigues, Svetoslav Nanev Slavov, Elaine Vieira dos Santos, Rafael dos Santos Bezerra, Luiz Carlos Junior de Alcantara, Marta Giovanetti, Vagner Fonseca, Flavia Aburjaile, Rodrigo Tocantins Calado. |
| EPI_ISL_1795084, EPI_ISL_1795085, EPI_ISL_1795086 | PRONTO SOCORRO MUNICIPAL TAMBAU | Instituto Butantan / ESALQ-Piracicaba | Instituto Butantan: Alexander Roberto Precioso, Dimas Tadeu Covas, Sandra Coccuzzo Sampaio, Maria Carolina Elias, José Salvatore Leister Patané, Vincent Louis Viala, Antonio Jorge Martins, Ricardo Haddad, Claudia Renata dos Santos Barros, Elaine Cristina Marqueze, Raul Machado Neto, Debora Botequio Moretti. Centro de Genômica Funcional da ESALQ: Luiz Lehmann Coutinho, Ricardo Augusto Brassaloti, Raquel de Lello Rocha Campos Cassano. NGS Soluções Genômicas: Pilar Drummond Sampaio Corrêa Mariani. FZEA-USP Pirassununga: Mirele Daiana Poleti, Jessika Cristina Chagas Lesbon, Elisangela Chicaroni Mattos, Heidge Fukumasu. USP-Botucatu: Rejane Maria Tommasini Grotto, Jayme A. Souza-Neto, Guilherme Targino Valente, Patricia Akemi Assato, Felipe Allan da Silva da Costa, Bianca Cechetto Carlos. Mendelics: Bibiana Santos, João Paulo Kitajima, Erika Freitas, David Schlesinger. Hemocentro Ribeirão Preto: Simone Kashima, Evandra Strazza Rodrigues, Svetoslav Nanev Slavov, Elaine Vieira dos Santos, Rafael dos Santos Bezerra, Luiz Carlos Junior de Alcantara, Marta Giovanetti, Vagner Fonseca, Flavia Aburjaile, Rodrigo Tocantins Calado. |
| EPI_ISL_1795087, EPI_ISL_1795088, EPI_ISL_1795089, EPI_ISL_1795090 | CENTRO DE SAUDE III AFFONSO LUZZI SANTA CRUZ DAS PALMEIRAS | Instituto Butantan / ESALQ-Piracicaba | Instituto Butantan: Alexander Roberto Precioso, Dimas Tadeu Covas, Sandra Coccuzzo Sampaio, Maria Carolina Elias, José Salvatore Leister Patané, Vincent Louis Viala, Antonio Jorge Martins, Ricardo Haddad, Claudia Renata dos Santos Barros, Elaine Cristina Marqueze, Raul Machado Neto, Debora Botequio Moretti. Centro de Genômica Funcional da ESALQ: Luiz Lehmann Coutinho, Ricardo Augusto Brassaloti, Raquel de Lello Rocha Campos Cassano. NGS Soluções Genômicas: Pilar Drummond Sampaio Corrêa Mariani. FZEA-USP Pirassununga: Mirele Daiana Poleti, Jessika Cristina Chagas Lesbon, Elisangela Chicaroni Mattos, Heidge Fukumasu. USP-Botucatu: Rejane Maria Tommasini Grotto, Jayme A. Souza-Neto, Guilherme Targino Valente, Patricia Akemi Assato, Felipe Allan da Silva da Costa, Bianca Cechetto Carlos. Mendelics: Bibiana Santos, João Paulo Kitajima, Erika Freitas, David Schlesinger. Hemocentro Ribeirão Preto: Simone Kashima, Evandra Strazza Rodrigues, Svetoslav Nanev Slavov, Elaine Vieira dos Santos, Rafael dos Santos Bezerra, Luiz Carlos Junior de Alcantara, Marta Giovanetti, Vagner Fonseca, Flavia Aburjaile, Rodrigo Tocantins Calado. |
| EPI_ISL_1795091, EPI_ISL_1795092 | PRONTO SOCORRO MUNICIPAL TAMBAU | Instituto Butantan / ESALQ-Piracicaba | Instituto Butantan: Alexander Roberto Precioso, Dimas Tadeu Covas, Sandra Coccuzzo Sampaio, Maria Carolina Elias, José Salvatore Leister Patané, Vincent Louis Viala, Antonio Jorge Martins, Ricardo Haddad, Claudia Renata dos Santos Barros, Elaine Cristina Marqueze, Raul Machado Neto, Debora Botequio Moretti. Centro de Genômica Funcional da ESALQ: Luiz Lehmann Coutinho, Ricardo Augusto Brassaloti, Raquel de Lello Rocha Campos Cassano. NGS Soluções Genômicas: Pilar Drummond Sampaio Corrêa Mariani. FZEA-USP Pirassununga: Mirele Daiana Poleti, Jessika Cristina Chagas Lesbon, Elisangela Chicaroni Mattos, Heidge Fukumasu. USP-Botucatu: Rejane Maria Tommasini Grotto, Jayme A. Souza-Neto, Guilherme Targino Valente, Patricia Akemi Assato, Felipe Allan da Silva da Costa, Bianca Cechetto Carlos. Mendelics: Bibiana Santos, João Paulo Kitajima, Erika Freitas, David Schlesinger. Hemocentro Ribeirão Preto: Simone Kashima, Evandra Strazza Rodrigues, Svetoslav Nanev Slavov, Elaine Vieira dos Santos, Rafael dos Santos Bezerra, Luiz Carlos Junior de Alcantara, Marta Giovanetti, Vagner Fonseca, Flavia Aburjaile, Rodrigo Tocantins Calado. |
| EPI_ISL_1795098, EPI_ISL_1795099 | ESALQ | Instituto Butantan / ESALQ-Piracicaba | Instituto Butantan: Alexander Roberto Precioso, Dimas Tadeu Covas, Sandra Coccuzzo Sampaio, Maria Carolina Elias, José Salvatore Leister Patané, Vincent Louis Viala, Antonio Jorge Martins, Ricardo Haddad, Claudia Renata dos Santos Barros, Elaine Cristina Marqueze, Raul Machado Neto, Debora Botequio Moretti. Centro de Genômica Funcional da ESALQ: Luiz Lehmann Coutinho, Ricardo Augusto Brassaloti, Raquel de Lello Rocha Campos Cassano. NGS Soluções Genômicas: Pilar Drummond Sampaio Corrêa Mariani. FZEA-USP Pirassununga: Mirele Daiana Poleti, Jessika Cristina Chagas Lesbon, Elisangela Chicaroni Mattos, Heidge Fukumasu. USP-Botucatu: Rejane Maria Tommasini Grotto, Jayme A. Souza-Neto, Guilherme Targino Valente, Patricia Akemi Assato, Felipe Allan da Silva da Costa, Bianca Cechetto Carlos. Mendelics: Bibiana Santos, João Paulo Kitajima, Erika Freitas, David Schlesinger. Hemocentro Ribeirão Preto: Simone Kashima, Evandra Strazza Rodrigues, Svetoslav Nanev Slavov, Elaine Vieira dos Santos, Rafael dos Santos Bezerra, Luiz Carlos Junior de Alcantara, Marta Giovanetti, Vagner Fonseca, Flavia Aburjaile, Rodrigo Tocantins Calado. |
| EPI_ISL_1795100 | LABORATORIO MUNICIPAL DE PIRACICABA | Instituto Butantan / ESALQ-Piracicaba | Instituto Butantan: Alexander Roberto Precioso, Dimas Tadeu Covas, Sandra Coccuzzo Sampaio, Maria Carolina Elias, José Salvatore Leister Patané, Vincent Louis Viala, Antonio Jorge Martins, Ricardo Haddad, Claudia Renata dos Santos Barros, Elaine Cristina Marqueze, Raul Machado Neto, Debora Botequio Moretti. Centro de Genômica Funcional da ESALQ: Luiz Lehmann Coutinho, Ricardo Augusto Brassaloti, Raquel de Lello Rocha Campos Cassano. NGS Soluções Genômicas: Pilar Drummond Sampaio Corrêa Mariani. FZEA-USP Pirassununga: Mirele Daiana Poleti, Jessika Cristina Chagas Lesbon, Elisangela Chicaroni Mattos, Heidge Fukumasu. USP-Botucatu: Rejane Maria Tommasini Grotto, Jayme A. Souza-Neto, Guilherme Targino Valente, Patricia Akemi Assato, Felipe Allan da Silva da Costa, Bianca Cechetto Carlos. Mendelics: Bibiana Santos, João Paulo Kitajima, Erika Freitas, David Schlesinger. Hemocentro Ribeirão Preto: Simone Kashima, Evandra Strazza Rodrigues, Svetoslav Nanev Slavov, Elaine Vieira dos Santos, Rafael dos Santos Bezerra, Luiz Carlos Junior de Alcantara, Marta Giovanetti, Vagner Fonseca, Flavia Aburjaile, Rodrigo Tocantins Calado. |
| EPI_ISL_1795101 | HOSPITAL DOS FORNCEDORES | Instituto Butantan / ESALQ-Piracicaba | Instituto Butantan: Alexander Roberto Precioso, Dimas Tadeu Covas, Sandra Coccuzzo Sampaio, Maria Carolina Elias, José Salvatore Leister Patané, Vincent Louis Viala, Antonio Jorge Martins, Ricardo Haddad, Claudia Renata dos Santos Barros, Elaine Cristina Marqueze, Raul Machado Neto, Debora Botequio Moretti. Centro de Genômica Funcional da ESALQ: Luiz Lehmann Coutinho, Ricardo Augusto Brassaloti, Raquel de Lello Rocha Campos Cassano. NGS Soluções Genômicas: Pilar Drummond Sampaio Corrêa Mariani. FZEA-USP Pirassununga: Mirele Daiana Poleti, Jessika Cristina Chagas Lesbon, Elisangela Chicaroni Mattos, Heidge Fukumasu. USP-Botucatu: Rejane Maria Tommasini Grotto, Jayme A. Souza-Neto, Guilherme Targino Valente, Patricia Akemi Assato, Felipe Allan da Silva da Costa, Bianca Cechetto Carlos. Mendelics: Bibiana Santos, João Paulo Kitajima, Erika Freitas, David Schlesinger. Hemocentro Ribeirão Preto: Simone Kashima, Evandra Strazza Rodrigues, Svetoslav Nanev Slavov, Elaine Vieira dos Santos, Rafael dos Santos Bezerra, Luiz Carlos Junior de Alcantara, Marta Giovanetti, Vagner Fonseca, Flavia Aburjaile, Rodrigo Tocantins Calado. |
| EPI_ISL_1795102 | LABORATORIO MUNICIPAL DE PIRACICABA | Instituto Butantan / ESALQ-Piracicaba | Instituto Butantan: Alexander Roberto Precioso, Dimas Tadeu Covas, Sandra Coccuzzo Sampaio, Maria Carolina Elias, José Salvatore Leister Patané, Vincent Louis Viala, Antonio Jorge Martins, Ricardo Haddad, Claudia Renata dos Santos Barros, Elaine Cristina Marqueze, Raul Machado Neto, Debora Botequio Moretti. Centro de Genômica Funcional da ESALQ: Luiz Lehmann Coutinho, Ricardo Augusto Brassaloti, Raquel de Lello Rocha Campos Cassano. NGS Soluções Genômicas: Pilar Drummond Sampaio Corrêa Mariani. FZEA-USP Pirassununga: Mirele Daiana Poleti, Jessika Cristina Chagas Lesbon, Elisangela Chicaroni Mattos, Heidge Fukumasu. USP-Botucatu: Rejane Maria Tommasini Grotto, Jayme A. Souza-Neto, Guilherme Targino Valente, Patricia Akemi Assato, Felipe Allan da Silva da Costa, Bianca Cechetto Carlos. Mendelics: Bibiana Santos, João Paulo Kitajima, Erika Freitas, David Schlesinger. Hemocentro Ribeirão Preto: Simone Kashima, Evandra Strazza Rodrigues, Svetoslav Nanev Slavov, Elaine Vieira dos Santos, Rafael dos Santos Bezerra, Luiz Carlos Junior de Alcantara, Marta Giovanetti, Vagner Fonseca, Flavia Aburjaile, Rodrigo Tocantins Calado. |
| EPI_ISL_1795103 | HOSPITAL DOS FORNCEDORES | Instituto Butantan / ESALQ-Piracicaba | Instituto Butantan: Alexander Roberto Precioso, Dimas Tadeu Covas, Sandra Coccuzzo Sampaio, Maria Carolina Elias, José Salvatore Leister Patané, Vincent Louis Viala, Antonio Jorge Martins, Ricardo Haddad, Claudia Renata dos Santos Barros, Elaine Cristina Marqueze, Raul Machado Neto, Debora Botequio Moretti. Centro de Genômica Funcional da ESALQ: Luiz Lehmann Coutinho, Ricardo Augusto Brassaloti, Raquel de Lello Rocha Campos Cassano. NGS Soluções Genômicas: Pilar Drummond Sampaio Corrêa Mariani. FZEA-USP Pirassununga: Mirele Daiana Poleti, Jessika Cristina Chagas Lesbon, Elisangela Chicaroni Mattos, Heidge Fukumasu. USP-Botucatu: Rejane Maria Tommasini Grotto, Jayme A. Souza-Neto, Guilherme Targino Valente, Patricia Akemi Assato, Felipe Allan da Silva da Costa, Bianca Cechetto Carlos. Mendelics: Bibiana Santos, João Paulo Kitajima, Erika Freitas, |

| | | | |
|---|---|---|---|
| | | | David Schlesinger. Hemocentro Ribeirão Preto: Simone Kashima, Evandra Strazza Rodrigues, Svetoslav Nanev Slavov, Elaine Vieira dos Santos, Rafael dos Santos Bezerra, Luiz Carlos Junior de Alcantara, Marta Giovanetti, Vagner Fonseca, Flavia Aburjaile, Rodrigo Tocantins Calado. |
| EPI_ISL_1795104, EPI_ISL_1795105 | LABORATORIO MUNICIPAL DE PIRACICABA | Instituto Butantan / ESALQ-Piracicaba | Instituto Butantan: Alexander Roberto Precioso, Dimas Tadeu Covas, Sandra Coccuzzo Sampaio, Maria Carolina Elias, José Salvatore Leister Patané, Vincent Louis Viala, Antonio Jorge Martins, Ricardo Haddad, Claudia Renata dos Santos Barros, Elaine Cristina Marqueze, Raul Machado Neto, Debora Botequio Moretti. Centro de Genômica Funcional da ESALQ: Luiz Lehmann Coutinho, Ricardo Augusto Brassaloti, Raquel de Lello Rocha Campos Cassano. NGS Soluções Genômicas: Pilar Drummond Sampaio Corrêa Mariani. FZEA-USP Pirassununga: Mirele Daiana Poleti, Jessika Cristina Chagas Lesbon, Elisangela Chicaroni Mattos, Heidge Fukumasu. USP-Botucatu: Rejane Maria Tommasini Grotto, Jayme A. Souza-Neto, Guilherme Targino Valente, Patricia Akemi Assato, Felipe Allan da Silva da Costa, Bianca Cechetto Carlos. Mendelics: Bibiana Santos, João Paulo Kitajima, Erika Freitas, David Schlesinger. Hemocentro Ribeirão Preto: Simone Kashima, Evandra Strazza Rodrigues, Svetoslav Nanev Slavov, Elaine Vieira dos Santos, Rafael dos Santos Bezerra, Luiz Carlos Junior de Alcantara, Marta Giovanetti, Vagner Fonseca, Flavia Aburjaile, Rodrigo Tocantins Calado. |
| EPI_ISL_1795106 | HOSPITAL DOS FORNCEDORES | Instituto Butantan / ESALQ-Piracicaba | Instituto Butantan: Alexander Roberto Precioso, Dimas Tadeu Covas, Sandra Coccuzzo Sampaio, Maria Carolina Elias, José Salvatore Leister Patané, Vincent Louis Viala, Antonio Jorge Martins, Ricardo Haddad, Claudia Renata dos Santos Barros, Elaine Cristina Marqueze, Raul Machado Neto, Debora Botequio Moretti. Centro de Genômica Funcional da ESALQ: Luiz Lehmann Coutinho, Ricardo Augusto Brassaloti, Raquel de Lello Rocha Campos Cassano. NGS Soluções Genômicas: Pilar Drummond Sampaio Corrêa Mariani. FZEA-USP Pirassununga: Mirele Daiana Poleti, Jessika Cristina Chagas Lesbon, Elisangela Chicaroni Mattos, Heidge Fukumasu. USP-Botucatu: Rejane Maria Tommasini Grotto, Jayme A. Souza-Neto, Guilherme Targino Valente, Patricia Akemi Assato, Felipe Allan da Silva da Costa, Bianca Cechetto Carlos. Mendelics: Bibiana Santos, João Paulo Kitajima, Erika Freitas, David Schlesinger. Hemocentro Ribeirão Preto: Simone Kashima, Evandra Strazza Rodrigues, Svetoslav Nanev Slavov, Elaine Vieira dos Santos, Rafael dos Santos Bezerra, Luiz Carlos Junior de Alcantara, Marta Giovanetti, Vagner Fonseca, Flavia Aburjaile, Rodrigo Tocantins Calado. |
| EPI_ISL_1795107 | COORDENADORIA MUNICIPAL DE SAUDE DE IRACEMAPOLIS | Instituto Butantan / ESALQ-Piracicaba | Instituto Butantan: Alexander Roberto Precioso, Dimas Tadeu Covas, Sandra Coccuzzo Sampaio, Maria Carolina Elias, José Salvatore Leister Patané, Vincent Louis Viala, Antonio Jorge Martins, Ricardo Haddad, Claudia Renata dos Santos Barros, Elaine Cristina Marqueze, Raul Machado Neto, Debora Botequio Moretti. Centro de Genômica Funcional da ESALQ: Luiz Lehmann Coutinho, Ricardo Augusto Brassaloti, Raquel de Lello Rocha Campos Cassano. NGS Soluções Genômicas: Pilar Drummond Sampaio Corrêa Mariani. FZEA-USP Pirassununga: Mirele Daiana Poleti, Jessika Cristina Chagas Lesbon, Elisangela Chicaroni Mattos, Heidge Fukumasu. USP-Botucatu: Rejane Maria Tommasini Grotto, Jayme A. Souza-Neto, Guilherme Targino Valente, Patricia Akemi Assato, Felipe Allan da Silva da Costa, Bianca Cechetto Carlos. Mendelics: Bibiana Santos, João Paulo Kitajima, Erika Freitas, David Schlesinger. Hemocentro Ribeirão Preto: Simone Kashima, Evandra Strazza Rodrigues, Svetoslav Nanev Slavov, Elaine Vieira dos Santos, Rafael dos Santos Bezerra, Luiz Carlos Junior de Alcantara, Marta Giovanetti, Vagner Fonseca, Flavia Aburjaile, Rodrigo Tocantins Calado. |
| EPI_ISL_1795108 | SECRETARIA DE SAUDE DE SAO PEDRO | Instituto Butantan / ESALQ-Piracicaba | Instituto Butantan: Alexander Roberto Precioso, Dimas Tadeu Covas, Sandra Coccuzzo Sampaio, Maria Carolina Elias, José Salvatore Leister Patané, Vincent Louis Viala, Antonio Jorge Martins, Ricardo Haddad, Claudia Renata dos Santos Barros, Elaine Cristina Marqueze, Raul Machado Neto, Debora Botequio Moretti. Centro de Genômica Funcional da ESALQ: Luiz Lehmann Coutinho, Ricardo Augusto Brassaloti, Raquel de Lello Rocha Campos Cassano. NGS Soluções Genômicas: Pilar Drummond Sampaio Corrêa Mariani. FZEA-USP Pirassununga: Mirele Daiana Poleti, Jessika Cristina Chagas Lesbon, Elisangela Chicaroni Mattos, Heidge Fukumasu. USP-Botucatu: Rejane Maria Tommasini Grotto, Jayme A. Souza-Neto, Guilherme Targino Valente, Patricia Akemi Assato, Felipe Allan da Silva da Costa, Bianca Cechetto Carlos. Mendelics: Bibiana Santos, João Paulo Kitajima, Erika Freitas, David Schlesinger. Hemocentro Ribeirão Preto: Simone Kashima, Evandra Strazza Rodrigues, Svetoslav Nanev Slavov, Elaine Vieira dos Santos, Rafael dos Santos Bezerra, Luiz Carlos Junior de Alcantara, Marta Giovanetti, Vagner Fonseca, Flavia Aburjaile, Rodrigo Tocantins Calado. |
| EPI_ISL_1795109 | CENTRO DE ESPECIALIDDS MEDICAS IRMA LEOPOLDINA PIRASSUNUNGA | Instituto Butantan / ESALQ-Piracicaba | Instituto Butantan: Alexander Roberto Precioso, Dimas Tadeu Covas, Sandra Coccuzzo Sampaio, Maria Carolina Elias, José Salvatore Leister Patané, Vincent Louis Viala, Antonio Jorge Martins, Ricardo Haddad, Claudia Renata dos Santos Barros, Elaine Cristina Marqueze, Raul Machado Neto, Debora Botequio Moretti. Centro de Genômica Funcional da ESALQ: Luiz Lehmann Coutinho, Ricardo Augusto Brassaloti, Raquel de Lello Rocha Campos Cassano. NGS Soluções Genômicas: Pilar Drummond Sampaio Corrêa Mariani. FZEA-USP Pirassununga: Mirele Daiana Poleti, Jessika Cristina Chagas Lesbon, Elisangela Chicaroni Mattos, Heidge Fukumasu. USP-Botucatu: Rejane Maria Tommasini Grotto, Jayme A. Souza-Neto, Guilherme Targino Valente, Patricia Akemi Assato, Felipe Allan da Silva da Costa, Bianca Cechetto Carlos. Mendelics: Bibiana Santos, João Paulo Kitajima, Erika Freitas, David Schlesinger. Hemocentro Ribeirão Preto: Simone Kashima, Evandra Strazza Rodrigues, Svetoslav Nanev Slavov, Elaine Vieira dos Santos, Rafael dos Santos Bezerra, Luiz Carlos Junior de Alcantara, Marta Giovanetti, Vagner Fonseca, Flavia Aburjaile, Rodrigo Tocantins Calado. |
| EPI_ISL_1795110 | UNIDADE ESF DR LUIZ SPINA | Instituto Butantan / ESALQ-Piracicaba | Instituto Butantan: Alexander Roberto Precioso, Dimas Tadeu Covas, Sandra Coccuzzo Sampaio, Maria Carolina Elias, José Salvatore Leister Patané, Vincent Louis Viala, Antonio Jorge Martins, Ricardo Haddad, Claudia Renata dos Santos Barros, Elaine Cristina Marqueze, Raul Machado Neto, Debora Botequio Moretti. Centro de Genômica Funcional da ESALQ: Luiz Lehmann Coutinho, Ricardo Augusto Brassaloti, Raquel de Lello Rocha Campos Cassano. NGS Soluções Genômicas: Pilar Drummond Sampaio Corrêa Mariani. FZEA-USP Pirassununga: Mirele Daiana Poleti, Jessika Cristina Chagas Lesbon, Elisangela Chicaroni Mattos, Heidge Fukumasu. USP-Botucatu: Rejane Maria Tommasini Grotto, Jayme A. Souza-Neto, Guilherme Targino Valente, Patricia Akemi Assato, Felipe Allan da Silva da Costa, Bianca Cechetto Carlos. Mendelics: Bibiana Santos, João Paulo Kitajima, Erika Freitas, David Schlesinger. Hemocentro Ribeirão Preto: Simone Kashima, Evandra Strazza Rodrigues, Svetoslav Nanev Slavov, Elaine Vieira dos Santos, Rafael dos Santos Bezerra, Luiz Carlos Junior de Alcantara, Marta Giovanetti, Vagner Fonseca, Flavia Aburjaile, Rodrigo Tocantins Calado. |
| EPI_ISL_1795111, EPI_ISL_1795112 | SECRETARIA MUNICIPAL DA SAUDE DE GUARIBA | Instituto Butantan / ESALQ-Piracicaba | Instituto Butantan: Alexander Roberto Precioso, Dimas Tadeu Covas, Sandra Coccuzzo Sampaio, Maria Carolina Elias, José Salvatore Leister Patané, Vincent Louis Viala, Antonio Jorge Martins, Ricardo Haddad, Claudia Renata dos Santos Barros, Elaine Cristina Marqueze, Raul Machado Neto, Debora Botequio Moretti. Centro de Genômica Funcional da ESALQ: Luiz Lehmann Coutinho, Ricardo Augusto Brassaloti, Raquel de Lello Rocha Campos Cassano. NGS Soluções Genômicas: Pilar Drummond Sampaio Corrêa Mariani. FZEA-USP Pirassununga: Mirele Daiana Poleti, Jessika Cristina Chagas Lesbon, Elisangela Chicaroni Mattos, Heidge Fukumasu. USP-Botucatu: Rejane Maria Tommasini Grotto, Jayme A. Souza-Neto, Guilherme Targino Valente, Patricia Akemi Assato, Felipe Allan da Silva da Costa, Bianca Cechetto Carlos. Mendelics: Bibiana Santos, João Paulo Kitajima, Erika Freitas, David Schlesinger. Hemocentro Ribeirão Preto: Simone Kashima, Evandra Strazza Rodrigues, Svetoslav Nanev Slavov, Elaine Vieira dos Santos, Rafael dos Santos Bezerra, Luiz Carlos Junior de Alcantara, Marta Giovanetti, Vagner Fonseca, Flavia Aburjaile, Rodrigo Tocantins Calado. |
| EPI_ISL_1795113 | AMBULATORIO DE ATENDIMENTO DE DST DE GUARIBA | Instituto Butantan / ESALQ-Piracicaba | Instituto Butantan: Alexander Roberto Precioso, Dimas Tadeu Covas, Sandra Coccuzzo Sampaio, Maria Carolina Elias, José Salvatore Leister Patané, Vincent Louis Viala, Antonio Jorge Martins, Ricardo Haddad, Claudia Renata dos Santos Barros, Elaine Cristina Marqueze, Raul Machado Neto, Debora Botequio Moretti. Centro de Genômica Funcional da ESALQ: Luiz Lehmann Coutinho, Ricardo Augusto Brassaloti, Raquel de Lello Rocha Campos Cassano. NGS Soluções Genômicas: Pilar Drummond Sampaio Corrêa Mariani. FZEA-USP Pirassununga: Mirele Daiana Poleti, Jessika Cristina Chagas Lesbon, Elisangela Chicaroni Mattos, Heidge Fukumasu. USP-Botucatu: Rejane Maria Tommasini Grotto, Jayme A. Souza-Neto, Guilherme Targino Valente, Patricia Akemi Assato, Felipe Allan da Silva da Costa, Bianca Cechetto Carlos. Mendelics: Bibiana Santos, João Paulo Kitajima, Erika Freitas, David Schlesinger. Hemocentro Ribeirão Preto: Simone Kashima, Evandra Strazza Rodrigues, Svetoslav Nanev Slavov, Elaine Vieira dos Santos, Rafael dos Santos Bezerra, Luiz Carlos Junior de Alcantara, Marta Giovanetti, Vagner Fonseca, Flavia Aburjaile, Rodrigo Tocantins Calado. |
| EPI_ISL_1795114, EPI_ISL_1795115 | CENTRO DE SAUDE III BORBOREMA | Instituto Butantan / ESALQ-Piracicaba | Instituto Butantan: Alexander Roberto Precioso, Dimas Tadeu Covas, Sandra Coccuzzo Sampaio, Maria Carolina Elias, José Salvatore Leister Patané, Vincent Louis Viala, Antonio Jorge Martins, Ricardo Haddad, Claudia Renata dos Santos Barros, Elaine Cristina Marqueze, Raul Machado Neto, Debora Botequio Moretti. Centro de Genômica Funcional da ESALQ: Luiz Lehmann Coutinho, Ricardo Augusto Brassaloti, Raquel de Lello Rocha Campos Cassano. NGS Soluções Genômicas: Pilar Drummond Sampaio Corrêa Mariani. FZEA-USP Pirassununga: Mirele Daiana Poleti, Jessika Cristina Chagas Lesbon, Elisangela Chicaroni Mattos, Heidge Fukumasu. USP-Botucatu: Rejane Maria Tommasini Grotto, Jayme A. Souza-Neto, Guilherme Targino Valente, Patricia Akemi Assato, Felipe Allan da Silva da Costa, Bianca Cechetto Carlos. Mendelics: Bibiana Santos, João Paulo Kitajima, Erika Freitas, David Schlesinger. Hemocentro Ribeirão Preto: Simone Kashima, Evandra Strazza Rodrigues, Svetoslav Nanev Slavov, Elaine Vieira dos Santos, Rafael dos Santos Bezerra, Luiz Carlos Junior de Alcantara, Marta Giovanetti, Vagner Fonseca, Flavia Aburjaile, Rodrigo Tocantins Calado. |
| EPI_ISL_1795116, EPI_ISL_1795117, EPI_ISL_1795118, EPI_ISL_1795119 | UNIDADE MISTA DE LUIZ ANTONIO | Instituto Butantan / ESALQ-Piracicaba | Instituto Butantan: Alexander Roberto Precioso, Dimas Tadeu Covas, Sandra Coccuzzo Sampaio, Maria Carolina Elias, José Salvatore Leister Patané, Vincent Louis Viala, Antonio Jorge Martins, Ricardo Haddad, Claudia Renata dos Santos Barros, Elaine Cristina Marqueze, Raul Machado Neto, Debora Botequio Moretti. Centro de Genômica Funcional da ESALQ: Luiz Lehmann Coutinho, Ricardo Augusto Brassaloti, Raquel de Lello Rocha Campos Cassano. NGS Soluções Genômicas: Pilar Drummond Sampaio Corrêa Mariani. FZEA-USP Pirassununga: Mirele Daiana Poleti, Jessika Cristina Chagas Lesbon, Elisangela Chicaroni Mattos, Heidge Fukumasu. USP-Botucatu: Rejane Maria Tommasini Grotto, Jayme A. Souza-Neto, Guilherme Targino Valente, Patricia Akemi Assato, Felipe Allan da Silva da Costa, Bianca Cechetto Carlos. Mendelics: Bibiana Santos, João Paulo Kitajima, Erika Freitas, David Schlesinger. Hemocentro Ribeirão Preto: Simone Kashima, Evandra Strazza Rodrigues, Svetoslav Nanev Slavov, Elaine Vieira dos Santos, Rafael dos Santos Bezerra, Luiz Carlos Junior de Alcantara, Marta Giovanetti, Vagner Fonseca, Flavia Aburjaile, Rodrigo Tocantins Calado. |

| | | | |
|---|---|---|---|
| EPI_ISL_1795120, EPI_ISL_1795121 | CENTRO DE SAUDE DE CAJOBI | Instituto Butantan / ESALQ-Piracicaba | Instituto Butantan: Alexander Roberto Precioso, Dimas Tadeu Covas, Sandra Coccuzzo Sampaio, Maria Carolina Elias, José Salvatore Leister Patané, Vincent Louis Viala, Antonio Jorge Martins, Ricardo Haddad, Claudia Renata dos Santos Barros, Elaine Cristina Marqueze, Raul Machado Neto, Debora Botequio Moretti. Centro de Genômica Funcional da ESALQ: Luiz Lehmann Coutinho, Ricardo Augusto Brassaloti, Raquel de Lello Rocha Campos Cassano. NGS Soluções Genômicas: Pilar Drummond Sampaio Corrêa Mariani. FZEA-USP Pirassununga: Mirele Daiana Poleti, Jessika Cristina Chagas Lesbon, Elisangela Chicaroni Mattos, Heidge Fukumasu. USP-Botucatu: Rejane Maria Tommasini Grotto, Jayme A. Souza-Neto, Guilherme Targino Valente, Patricia Akemi Assato, Felipe Allan da Silva da Costa, Bianca Cechetto Carlos. Mendelics: Bibiana Santos, João Paulo Kitajima, Erika Freitas, David Schlesinger. Hemocentro Ribeirão Preto: Simone Kashima, Evandra Strazza Rodrigues, Svetoslav Nanev Slavov, Elaine Vieira dos Santos, Rafael dos Santos Bezerra, Luiz Carlos Junior de Alcantara, Marta Giovanetti, Vagner Fonseca, Flavia Aburjaile, Rodrigo Tocantins Calado. |
| EPI_ISL_1795122, EPI_ISL_1795123, EPI_ISL_1795124 | UNIDADE DE VIGILANCIA EM SAUDE | Instituto Butantan / ESALQ-Piracicaba | Instituto Butantan: Alexander Roberto Precioso, Dimas Tadeu Covas, Sandra Coccuzzo Sampaio, Maria Carolina Elias, José Salvatore Leister Patané, Vincent Louis Viala, Antonio Jorge Martins, Ricardo Haddad, Claudia Renata dos Santos Barros, Elaine Cristina Marqueze, Raul Machado Neto, Debora Botequio Moretti. Centro de Genômica Funcional da ESALQ: Luiz Lehmann Coutinho, Ricardo Augusto Brassaloti, Raquel de Lello Rocha Campos Cassano. NGS Soluções Genômicas: Pilar Drummond Sampaio Corrêa Mariani. FZEA-USP Pirassununga: Mirele Daiana Poleti, Jessika Cristina Chagas Lesbon, Elisangela Chicaroni Mattos, Heidge Fukumasu. USP-Botucatu: Rejane Maria Tommasini Grotto, Jayme A. Souza-Neto, Guilherme Targino Valente, Patricia Akemi Assato, Felipe Allan da Silva da Costa, Bianca Cechetto Carlos. Mendelics: Bibiana Santos, João Paulo Kitajima, Erika Freitas, David Schlesinger. Hemocentro Ribeirão Preto: Simone Kashima, Evandra Strazza Rodrigues, Svetoslav Nanev Slavov, Elaine Vieira dos Santos, Rafael dos Santos Bezerra, Luiz Carlos Junior de Alcantara, Marta Giovanetti, Vagner Fonseca, Flavia Aburjaile, Rodrigo Tocantins Calado. |
| EPI_ISL_1795125 | CENTRO DE SAUDE III CANDIDO RODRIGUES | Instituto Butantan / ESALQ-Piracicaba | Instituto Butantan: Alexander Roberto Precioso, Dimas Tadeu Covas, Sandra Coccuzzo Sampaio, Maria Carolina Elias, José Salvatore Leister Patané, Vincent Louis Viala, Antonio Jorge Martins, Ricardo Haddad, Claudia Renata dos Santos Barros, Elaine Cristina Marqueze, Raul Machado Neto, Debora Botequio Moretti. Centro de Genômica Funcional da ESALQ: Luiz Lehmann Coutinho, Ricardo Augusto Brassaloti, Raquel de Lello Rocha Campos Cassano. NGS Soluções Genômicas: Pilar Drummond Sampaio Corrêa Mariani. FZEA-USP Pirassununga: Mirele Daiana Poleti, Jessika Cristina Chagas Lesbon, Elisangela Chicaroni Mattos, Heidge Fukumasu. USP-Botucatu: Rejane Maria Tommasini Grotto, Jayme A. Souza-Neto, Guilherme Targino Valente, Patricia Akemi Assato, Felipe Allan da Silva da Costa, Bianca Cechetto Carlos. Mendelics: Bibiana Santos, João Paulo Kitajima, Erika Freitas, David Schlesinger. Hemocentro Ribeirão Preto: Simone Kashima, Evandra Strazza Rodrigues, Svetoslav Nanev Slavov, Elaine Vieira dos Santos, Rafael dos Santos Bezerra, Luiz Carlos Junior de Alcantara, Marta Giovanetti, Vagner Fonseca, Flavia Aburjaile, Rodrigo Tocantins Calado. |
| EPI_ISL_1795126 | CENTRO DE SAUDE III SALES OLIVEIRA | Instituto Butantan / ESALQ-Piracicaba | Instituto Butantan: Alexander Roberto Precioso, Dimas Tadeu Covas, Sandra Coccuzzo Sampaio, Maria Carolina Elias, José Salvatore Leister Patané, Vincent Louis Viala, Antonio Jorge Martins, Ricardo Haddad, Claudia Renata dos Santos Barros, Elaine Cristina Marqueze, Raul Machado Neto, Debora Botequio Moretti. Centro de Genômica Funcional da ESALQ: Luiz Lehmann Coutinho, Ricardo Augusto Brassaloti, Raquel de Lello Rocha Campos Cassano. NGS Soluções Genômicas: Pilar Drummond Sampaio Corrêa Mariani. FZEA-USP Pirassununga: Mirele Daiana Poleti, Jessika Cristina Chagas Lesbon, Elisangela Chicaroni Mattos, Heidge Fukumasu. USP-Botucatu: Rejane Maria Tommasini Grotto, Jayme A. Souza-Neto, Guilherme Targino Valente, Patricia Akemi Assato, Felipe Allan da Silva da Costa, Bianca Cechetto Carlos. Mendelics: Bibiana Santos, João Paulo Kitajima, Erika Freitas, David Schlesinger. Hemocentro Ribeirão Preto: Simone Kashima, Evandra Strazza Rodrigues, Svetoslav Nanev Slavov, Elaine Vieira dos Santos, Rafael dos Santos Bezerra, Luiz Carlos Junior de Alcantara, Marta Giovanetti, Vagner Fonseca, Flavia Aburjaile, Rodrigo Tocantins Calado. |
| EPI_ISL_1795127 | CENTRO DE SAUDE III CANDIDO RODRIGUES | Instituto Butantan / ESALQ-Piracicaba | Instituto Butantan: Alexander Roberto Precioso, Dimas Tadeu Covas, Sandra Coccuzzo Sampaio, Maria Carolina Elias, José Salvatore Leister Patané, Vincent Louis Viala, Antonio Jorge Martins, Ricardo Haddad, Claudia Renata dos Santos Barros, Elaine Cristina Marqueze, Raul Machado Neto, Debora Botequio Moretti. Centro de Genômica Funcional da ESALQ: Luiz Lehmann Coutinho, Ricardo Augusto Brassaloti, Raquel de Lello Rocha Campos Cassano. NGS Soluções Genômicas: Pilar Drummond Sampaio Corrêa Mariani. FZEA-USP Pirassununga: Mirele Daiana Poleti, Jessika Cristina Chagas Lesbon, Elisangela Chicaroni Mattos, Heidge Fukumasu. USP-Botucatu: Rejane Maria Tommasini Grotto, Jayme A. Souza-Neto, Guilherme Targino Valente, Patricia Akemi Assato, Felipe Allan da Silva da Costa, Bianca Cechetto Carlos. Mendelics: Bibiana Santos, João Paulo Kitajima, Erika Freitas, David Schlesinger. Hemocentro Ribeirão Preto: Simone Kashima, Evandra Strazza Rodrigues, Svetoslav Nanev Slavov, Elaine Vieira dos Santos, Rafael dos Santos Bezerra, Luiz Carlos Junior de Alcantara, Marta Giovanetti, Vagner Fonseca, Flavia Aburjaile, Rodrigo Tocantins Calado. |
| EPI_ISL_1795128, EPI_ISL_1795129, EPI_ISL_1795130, EPI_ISL_1795131, EPI_ISL_1795132, EPI_ISL_1795133, EPI_ISL_1795134, EPI_ISL_1795135 | UNIDADE DE VIGILANCIA EM SAUDE | Instituto Butantan / ESALQ-Piracicaba | Instituto Butantan: Alexander Roberto Precioso, Dimas Tadeu Covas, Sandra Coccuzzo Sampaio, Maria Carolina Elias, José Salvatore Leister Patané, Vincent Louis Viala, Antonio Jorge Martins, Ricardo Haddad, Claudia Renata dos Santos Barros, Elaine Cristina Marqueze, Raul Machado Neto, Debora Botequio Moretti. Centro de Genômica Funcional da ESALQ: Luiz Lehmann Coutinho, Ricardo Augusto Brassaloti, Raquel de Lello Rocha Campos Cassano. NGS Soluções Genômicas: Pilar Drummond Sampaio Corrêa Mariani. FZEA-USP Pirassununga: Mirele Daiana Poleti, Jessika Cristina Chagas Lesbon, Elisangela Chicaroni Mattos, Heidge Fukumasu. USP-Botucatu: Rejane Maria Tommasini Grotto, Jayme A. Souza-Neto, Guilherme Targino Valente, Patricia Akemi Assato, Felipe Allan da Silva da Costa, Bianca Cechetto Carlos. Mendelics: Bibiana Santos, João Paulo Kitajima, Erika Freitas, David Schlesinger. Hemocentro Ribeirão Preto: Simone Kashima, Evandra Strazza Rodrigues, Svetoslav Nanev Slavov, Elaine Vieira dos Santos, Rafael dos Santos Bezerra, Luiz Carlos Junior de Alcantara, Marta Giovanetti, Vagner Fonseca, Flavia Aburjaile, Rodrigo Tocantins Calado. |
| EPI_ISL_1795136, EPI_ISL_1795137, EPI_ISL_1795138, EPI_ISL_1795139, EPI_ISL_1795140 | UNIDADE MISTA DE VISTA ALEGRE DO ALTO VISTA ALEGRE DO ALTO | Instituto Butantan / ESALQ-Piracicaba | Instituto Butantan: Alexander Roberto Precioso, Dimas Tadeu Covas, Sandra Coccuzzo Sampaio, Maria Carolina Elias, José Salvatore Leister Patané, Vincent Louis Viala, Antonio Jorge Martins, Ricardo Haddad, Claudia Renata dos Santos Barros, Elaine Cristina Marqueze, Raul Machado Neto, Debora Botequio Moretti. Centro de Genômica Funcional da ESALQ: Luiz Lehmann Coutinho, Ricardo Augusto Brassaloti, Raquel de Lello Rocha Campos Cassano. NGS Soluções Genômicas: Pilar Drummond Sampaio Corrêa Mariani. FZEA-USP Pirassununga: Mirele Daiana Poleti, Jessika Cristina Chagas Lesbon, Elisangela Chicaroni Mattos, Heidge Fukumasu. USP-Botucatu: Rejane Maria Tommasini Grotto, Jayme A. Souza-Neto, Guilherme Targino Valente, Patricia Akemi Assato, Felipe Allan da Silva da Costa, Bianca Cechetto Carlos. Mendelics: Bibiana Santos, João Paulo Kitajima, Erika Freitas, David Schlesinger. Hemocentro Ribeirão Preto: Simone Kashima, Evandra Strazza Rodrigues, Svetoslav Nanev Slavov, Elaine Vieira dos Santos, Rafael dos Santos Bezerra, Luiz Carlos Junior de Alcantara, Marta Giovanetti, Vagner Fonseca, Flavia Aburjaile, Rodrigo Tocantins Calado. |
| EPI_ISL_1795141, EPI_ISL_1795142, EPI_ISL_1795143, EPI_ISL_1795144, EPI_ISL_1795145, EPI_ISL_1795146, EPI_ISL_1795147, EPI_ISL_1795148 | POLICLINICA HORTOLANDIA | Instituto Butantan / ESALQ-Piracicaba | Instituto Butantan: Alexander Roberto Precioso, Dimas Tadeu Covas, Sandra Coccuzzo Sampaio, Maria Carolina Elias, José Salvatore Leister Patané, Vincent Louis Viala, Antonio Jorge Martins, Ricardo Haddad, Claudia Renata dos Santos Barros, Elaine Cristina Marqueze, Raul Machado Neto, Debora Botequio Moretti. Centro de Genômica Funcional da ESALQ: Luiz Lehmann Coutinho, Ricardo Augusto Brassaloti, Raquel de Lello Rocha Campos Cassano. NGS Soluções Genômicas: Pilar Drummond Sampaio Corrêa Mariani. FZEA-USP Pirassununga: Mirele Daiana Poleti, Jessika Cristina Chagas Lesbon, Elisangela Chicaroni Mattos, Heidge Fukumasu. USP-Botucatu: Rejane Maria Tommasini Grotto, Jayme A. Souza-Neto, Guilherme Targino Valente, Patricia Akemi Assato, Felipe Allan da Silva da Costa, Bianca Cechetto Carlos. Mendelics: Bibiana Santos, João Paulo Kitajima, Erika Freitas, David Schlesinger. Hemocentro Ribeirão Preto: Simone Kashima, Evandra Strazza Rodrigues, Svetoslav Nanev Slavov, Elaine Vieira dos Santos, Rafael dos Santos Bezerra, Luiz Carlos Junior de Alcantara, Marta Giovanetti, Vagner Fonseca, Flavia Aburjaile, Rodrigo Tocantins Calado. |
| EPI_ISL_1795149, EPI_ISL_1795150 | UNIDADE MISTA DE VISTA ALEGRE DO ALTO VISTA ALEGRE DO ALTO | Instituto Butantan / ESALQ-Piracicaba | Instituto Butantan: Alexander Roberto Precioso, Dimas Tadeu Covas, Sandra Coccuzzo Sampaio, Maria Carolina Elias, José Salvatore Leister Patané, Vincent Louis Viala, Antonio Jorge Martins, Ricardo Haddad, Claudia Renata dos Santos Barros, Elaine Cristina Marqueze, Raul Machado Neto, Debora Botequio Moretti. Centro de Genômica Funcional da ESALQ: Luiz Lehmann Coutinho, Ricardo Augusto Brassaloti, Raquel de Lello Rocha Campos Cassano. NGS Soluções Genômicas: Pilar Drummond Sampaio Corrêa Mariani. FZEA-USP Pirassununga: Mirele Daiana Poleti, Jessika Cristina Chagas Lesbon, Elisangela Chicaroni Mattos, Heidge Fukumasu. USP-Botucatu: Rejane Maria Tommasini Grotto, Jayme A. Souza-Neto, Guilherme Targino Valente, Patricia Akemi Assato, Felipe Allan da Silva da Costa, Bianca Cechetto Carlos. Mendelics: Bibiana Santos, João Paulo Kitajima, Erika Freitas, David Schlesinger. Hemocentro Ribeirão Preto: Simone Kashima, Evandra Strazza Rodrigues, Svetoslav Nanev Slavov, Elaine Vieira dos Santos, Rafael dos Santos Bezerra, Luiz Carlos Junior de Alcantara, Marta Giovanetti, Vagner Fonseca, Flavia Aburjaile, Rodrigo Tocantins Calado. |
| EPI_ISL_1795151 | CENTRO DE SAUDE DE CAJOBI | Instituto Butantan / ESALQ-Piracicaba | Instituto Butantan: Alexander Roberto Precioso, Dimas Tadeu Covas, Sandra Coccuzzo Sampaio, Maria Carolina Elias, José Salvatore Leister Patané, Vincent Louis Viala, Antonio Jorge Martins, Ricardo Haddad, Claudia Renata dos Santos Barros, Elaine Cristina Marqueze, Raul Machado Neto, Debora Botequio Moretti. Centro de Genômica Funcional da ESALQ: Luiz Lehmann Coutinho, Ricardo Augusto Brassaloti, Raquel de Lello Rocha Campos Cassano. NGS Soluções Genômicas: Pilar Drummond Sampaio Corrêa Mariani. FZEA-USP Pirassununga: Mirele Daiana Poleti, Jessika Cristina Chagas Lesbon, Elisangela Chicaroni Mattos, Heidge Fukumasu. USP-Botucatu: Rejane Maria Tommasini Grotto, Jayme A. Souza-Neto, Guilherme Targino Valente, Patricia Akemi Assato, Felipe Allan da Silva da Costa, Bianca Cechetto Carlos. Mendelics: Bibiana Santos, João Paulo Kitajima, Erika Freitas, David Schlesinger. Hemocentro Ribeirão Preto: Simone Kashima, Evandra Strazza Rodrigues, Svetoslav Nanev Slavov, Elaine Vieira dos Santos, Rafael dos Santos Bezerra, Luiz Carlos Junior de Alcantara, Marta Giovanetti, Vagner Fonseca, Flavia Aburjaile, Rodrigo Tocantins Calado. |
| EPI_ISL_1795152, EPI_ISL_1795153, EPI_ISL_1795154, EPI_ISL_1795155, | POLICLINICA HORTOLANDIA | Instituto Butantan / ESALQ-Piracicaba | Instituto Butantan: Alexander Roberto Precioso, Dimas Tadeu Covas, Sandra Coccuzzo Sampaio, Maria Carolina Elias, José Salvatore Leister Patané, Vincent Louis Viala, Antonio Jorge Martins, Ricardo Haddad, Claudia Renata dos Santos Barros, Elaine Cristina Marqueze, Raul Machado Neto, Debora |

| | | | |
|---|---|---|---|
| EPI_ISL_1795156, EPI_ISL_1795157, EPI_ISL_1795158, EPI_ISL_1795159 | | | Botequio Moretti. Centro de Genômica Funcional da ESALQ: Luiz Lehmann Coutinho, Ricardo Augusto Brassaloti, Raquel de Lello Rocha Campos Cassano. NGS Soluções Genômicas: Pilar Drummond Sampaio Corrêa Mariani. FZEA-USP Pirassununga: Mirele Daiana Poleti, Jessika Cristina Chagas Lesbon, Elisangela Chicaroni Mattos, Heidge Fukumasu. USP-Botucatu: Rejane Maria Tommasini Grotto, Jayme A. Souza-Neto, Guilherme Targino Valente, Patricia Akemi Assato, Felipe Allan da Silva da Costa, Bianca Cechetto Carlos. Mendelics: Bibiana Santos, João Paulo Kitajima, Erika Freitas, David Schlesinger. Hemocentro Ribeirão Preto: Simone Kashima, Evandra Strazza Rodrigues, Svetoslav Nanev Slavov, Elaine Vieira dos Santos, Rafael dos Santos Bezerra, Luiz Carlos Junior de Alcantara, Marta Giovanetti, Vagner Fonseca, Flavia Aburjaile, Rodrigo Tocantins Calado. |
| EPI_ISL_1795160 | UBS JARDIM ITAMARATY | Instituto Butantan / ESALQ-Piracicaba | Instituto Butantan: Alexander Roberto Precioso, Dimas Tadeu Covas, Sandra Coccuzzo Sampaio, Maria Carolina Elias, José Salvatore Leister Patané, Vincent Louis Viala, Antonio Jorge Martins, Ricardo Haddad, Claudia Renata dos Santos Barros, Elaine Cristina Marqueze, Raul Machado Neto, Debora Botequio Moretti. Centro de Genômica Funcional da ESALQ: Luiz Lehmann Coutinho, Ricardo Augusto Brassaloti, Raquel de Lello Rocha Campos Cassano. NGS Soluções Genômicas: Pilar Drummond Sampaio Corrêa Mariani. FZEA-USP Pirassununga: Mirele Daiana Poleti, Jessika Cristina Chagas Lesbon, Elisangela Chicaroni Mattos, Heidge Fukumasu. USP-Botucatu: Rejane Maria Tommasini Grotto, Jayme A. Souza-Neto, Guilherme Targino Valente, Patricia Akemi Assato, Felipe Allan da Silva da Costa, Bianca Cechetto Carlos. Mendelics: Bibiana Santos, João Paulo Kitajima, Erika Freitas, David Schlesinger. Hemocentro Ribeirão Preto: Simone Kashima, Evandra Strazza Rodrigues, Svetoslav Nanev Slavov, Elaine Vieira dos Santos, Rafael dos Santos Bezerra, Luiz Carlos Junior de Alcantara, Marta Giovanetti, Vagner Fonseca, Flavia Aburjaile, Rodrigo Tocantins Calado. |
| EPI_ISL_1795161 | CS III VILA ODILON | Instituto Butantan / ESALQ-Piracicaba | Instituto Butantan: Alexander Roberto Precioso, Dimas Tadeu Covas, Sandra Coccuzzo Sampaio, Maria Carolina Elias, José Salvatore Leister Patané, Vincent Louis Viala, Antonio Jorge Martins, Ricardo Haddad, Claudia Renata dos Santos Barros, Elaine Cristina Marqueze, Raul Machado Neto, Debora Botequio Moretti. Centro de Genômica Funcional da ESALQ: Luiz Lehmann Coutinho, Ricardo Augusto Brassaloti, Raquel de Lello Rocha Campos Cassano. NGS Soluções Genômicas: Pilar Drummond Sampaio Corrêa Mariani. FZEA-USP Pirassununga: Mirele Daiana Poleti, Jessika Cristina Chagas Lesbon, Elisangela Chicaroni Mattos, Heidge Fukumasu. USP-Botucatu: Rejane Maria Tommasini Grotto, Jayme A. Souza-Neto, Guilherme Targino Valente, Patricia Akemi Assato, Felipe Allan da Silva da Costa, Bianca Cechetto Carlos. Mendelics: Bibiana Santos, João Paulo Kitajima, Erika Freitas, David Schlesinger. Hemocentro Ribeirão Preto: Simone Kashima, Evandra Strazza Rodrigues, Svetoslav Nanev Slavov, Elaine Vieira dos Santos, Rafael dos Santos Bezerra, Luiz Carlos Junior de Alcantara, Marta Giovanetti, Vagner Fonseca, Flavia Aburjaile, Rodrigo Tocantins Calado. |
| EPI_ISL_1795162, EPI_ISL_1795163, EPI_ISL_1795164 | CENTRO DE SAUDE DR PLINIO ALBERS DE GALIA | Instituto Butantan / ESALQ-Piracicaba | Instituto Butantan: Alexander Roberto Precioso, Dimas Tadeu Covas, Sandra Coccuzzo Sampaio, Maria Carolina Elias, José Salvatore Leister Patané, Vincent Louis Viala, Antonio Jorge Martins, Ricardo Haddad, Claudia Renata dos Santos Barros, Elaine Cristina Marqueze, Raul Machado Neto, Debora Botequio Moretti. Centro de Genômica Funcional da ESALQ: Luiz Lehmann Coutinho, Ricardo Augusto Brassaloti, Raquel de Lello Rocha Campos Cassano. NGS Soluções Genômicas: Pilar Drummond Sampaio Corrêa Mariani. FZEA-USP Pirassununga: Mirele Daiana Poleti, Jessika Cristina Chagas Lesbon, Elisangela Chicaroni Mattos, Heidge Fukumasu. USP-Botucatu: Rejane Maria Tommasini Grotto, Jayme A. Souza-Neto, Guilherme Targino Valente, Patricia Akemi Assato, Felipe Allan da Silva da Costa, Bianca Cechetto Carlos. Mendelics: Bibiana Santos, João Paulo Kitajima, Erika Freitas, David Schlesinger. Hemocentro Ribeirão Preto: Simone Kashima, Evandra Strazza Rodrigues, Svetoslav Nanev Slavov, Elaine Vieira dos Santos, Rafael dos Santos Bezerra, Luiz Carlos Junior de Alcantara, Marta Giovanetti, Vagner Fonseca, Flavia Aburjaile, Rodrigo Tocantins Calado. |
| EPI_ISL_1795165 | CS DE ALVARO CARVALHO | Instituto Butantan / ESALQ-Piracicaba | Instituto Butantan: Alexander Roberto Precioso, Dimas Tadeu Covas, Sandra Coccuzzo Sampaio, Maria Carolina Elias, José Salvatore Leister Patané, Vincent Louis Viala, Antonio Jorge Martins, Ricardo Haddad, Claudia Renata dos Santos Barros, Elaine Cristina Marqueze, Raul Machado Neto, Debora Botequio Moretti. Centro de Genômica Funcional da ESALQ: Luiz Lehmann Coutinho, Ricardo Augusto Brassaloti, Raquel de Lello Rocha Campos Cassano. NGS Soluções Genômicas: Pilar Drummond Sampaio Corrêa Mariani. FZEA-USP Pirassununga: Mirele Daiana Poleti, Jessika Cristina Chagas Lesbon, Elisangela Chicaroni Mattos, Heidge Fukumasu. USP-Botucatu: Rejane Maria Tommasini Grotto, Jayme A. Souza-Neto, Guilherme Targino Valente, Patricia Akemi Assato, Felipe Allan da Silva da Costa, Bianca Cechetto Carlos. Mendelics: Bibiana Santos, João Paulo Kitajima, Erika Freitas, David Schlesinger. Hemocentro Ribeirão Preto: Simone Kashima, Evandra Strazza Rodrigues, Svetoslav Nanev Slavov, Elaine Vieira dos Santos, Rafael dos Santos Bezerra, Luiz Carlos Junior de Alcantara, Marta Giovanetti, Vagner Fonseca, Flavia Aburjaile, Rodrigo Tocantins Calado. |
| EPI_ISL_1795166 | CENTRO DE SAUDE DR PLINIO ALBERS DE GALIA | Instituto Butantan / ESALQ-Piracicaba | Instituto Butantan: Alexander Roberto Precioso, Dimas Tadeu Covas, Sandra Coccuzzo Sampaio, Maria Carolina Elias, José Salvatore Leister Patané, Vincent Louis Viala, Antonio Jorge Martins, Ricardo Haddad, Claudia Renata dos Santos Barros, Elaine Cristina Marqueze, Raul Machado Neto, Debora Botequio Moretti. Centro de Genômica Funcional da ESALQ: Luiz Lehmann Coutinho, Ricardo Augusto Brassaloti, Raquel de Lello Rocha Campos Cassano. NGS Soluções Genômicas: Pilar Drummond Sampaio Corrêa Mariani. FZEA-USP Pirassununga: Mirele Daiana Poleti, Jessika Cristina Chagas Lesbon, Elisangela Chicaroni Mattos, Heidge Fukumasu. USP-Botucatu: Rejane Maria Tommasini Grotto, Jayme A. Souza-Neto, Guilherme Targino Valente, Patricia Akemi Assato, Felipe Allan da Silva da Costa, Bianca Cechetto Carlos. Mendelics: Bibiana Santos, João Paulo Kitajima, Erika Freitas, David Schlesinger. Hemocentro Ribeirão Preto: Simone Kashima, Evandra Strazza Rodrigues, Svetoslav Nanev Slavov, Elaine Vieira dos Santos, Rafael dos Santos Bezerra, Luiz Carlos Junior de Alcantara, Marta Giovanetti, Vagner Fonseca, Flavia Aburjaile, Rodrigo Tocantins Calado. |
| EPI_ISL_1795167, EPI_ISL_1795168, EPI_ISL_1795169 | CENTRO DE SAUDE DE FLORINEA | Instituto Butantan / ESALQ-Piracicaba | Instituto Butantan: Alexander Roberto Precioso, Dimas Tadeu Covas, Sandra Coccuzzo Sampaio, Maria Carolina Elias, José Salvatore Leister Patané, Vincent Louis Viala, Antonio Jorge Martins, Ricardo Haddad, Claudia Renata dos Santos Barros, Elaine Cristina Marqueze, Raul Machado Neto, Debora Botequio Moretti. Centro de Genômica Funcional da ESALQ: Luiz Lehmann Coutinho, Ricardo Augusto Brassaloti, Raquel de Lello Rocha Campos Cassano. NGS Soluções Genômicas: Pilar Drummond Sampaio Corrêa Mariani. FZEA-USP Pirassununga: Mirele Daiana Poleti, Jessika Cristina Chagas Lesbon, Elisangela Chicaroni Mattos, Heidge Fukumasu. USP-Botucatu: Rejane Maria Tommasini Grotto, Jayme A. Souza-Neto, Guilherme Targino Valente, Patricia Akemi Assato, Felipe Allan da Silva da Costa, Bianca Cechetto Carlos. Mendelics: Bibiana Santos, João Paulo Kitajima, Erika Freitas, David Schlesinger. Hemocentro Ribeirão Preto: Simone Kashima, Evandra Strazza Rodrigues, Svetoslav Nanev Slavov, Elaine Vieira dos Santos, Rafael dos Santos Bezerra, Luiz Carlos Junior de Alcantara, Marta Giovanetti, Vagner Fonseca, Flavia Aburjaile, Rodrigo Tocantins Calado. |
| EPI_ISL_1795170 | CENTRO DE SAUDE DE MARACAI | Instituto Butantan / ESALQ-Piracicaba | Instituto Butantan: Alexander Roberto Precioso, Dimas Tadeu Covas, Sandra Coccuzzo Sampaio, Maria Carolina Elias, José Salvatore Leister Patané, Vincent Louis Viala, Antonio Jorge Martins, Ricardo Haddad, Claudia Renata dos Santos Barros, Elaine Cristina Marqueze, Raul Machado Neto, Debora Botequio Moretti. Centro de Genômica Funcional da ESALQ: Luiz Lehmann Coutinho, Ricardo Augusto Brassaloti, Raquel de Lello Rocha Campos Cassano. NGS Soluções Genômicas: Pilar Drummond Sampaio Corrêa Mariani. FZEA-USP Pirassununga: Mirele Daiana Poleti, Jessika Cristina Chagas Lesbon, Elisangela Chicaroni Mattos, Heidge Fukumasu. USP-Botucatu: Rejane Maria Tommasini Grotto, Jayme A. Souza-Neto, Guilherme Targino Valente, Patricia Akemi Assato, Felipe Allan da Silva da Costa, Bianca Cechetto Carlos. Mendelics: Bibiana Santos, João Paulo Kitajima, Erika Freitas, David Schlesinger. Hemocentro Ribeirão Preto: Simone Kashima, Evandra Strazza Rodrigues, Svetoslav Nanev Slavov, Elaine Vieira dos Santos, Rafael dos Santos Bezerra, Luiz Carlos Junior de Alcantara, Marta Giovanetti, Vagner Fonseca, Flavia Aburjaile, Rodrigo Tocantins Calado. |
| EPI_ISL_1795171 | CS DE OSCAR BRESSANE PSF | Instituto Butantan / ESALQ-Piracicaba | Instituto Butantan: Alexander Roberto Precioso, Dimas Tadeu Covas, Sandra Coccuzzo Sampaio, Maria Carolina Elias, José Salvatore Leister Patané, Vincent Louis Viala, Antonio Jorge Martins, Ricardo Haddad, Claudia Renata dos Santos Barros, Elaine Cristina Marqueze, Raul Machado Neto, Debora Botequio Moretti. Centro de Genômica Funcional da ESALQ: Luiz Lehmann Coutinho, Ricardo Augusto Brassaloti, Raquel de Lello Rocha Campos Cassano. NGS Soluções Genômicas: Pilar Drummond Sampaio Corrêa Mariani. FZEA-USP Pirassununga: Mirele Daiana Poleti, Jessika Cristina Chagas Lesbon, Elisangela Chicaroni Mattos, Heidge Fukumasu. USP-Botucatu: Rejane Maria Tommasini Grotto, Jayme A. Souza-Neto, Guilherme Targino Valente, Patricia Akemi Assato, Felipe Allan da Silva da Costa, Bianca Cechetto Carlos. Mendelics: Bibiana Santos, João Paulo Kitajima, Erika Freitas, David Schlesinger. Hemocentro Ribeirão Preto: Simone Kashima, Evandra Strazza Rodrigues, Svetoslav Nanev Slavov, Elaine Vieira dos Santos, Rafael dos Santos Bezerra, Luiz Carlos Junior de Alcantara, Marta Giovanetti, Vagner Fonseca, Flavia Aburjaile, Rodrigo Tocantins Calado. |
| EPI_ISL_1795172 | CENTRO DE SAUDE DE MARACAI | Instituto Butantan / ESALQ-Piracicaba | Instituto Butantan: Alexander Roberto Precioso, Dimas Tadeu Covas, Sandra Coccuzzo Sampaio, Maria Carolina Elias, José Salvatore Leister Patané, Vincent Louis Viala, Antonio Jorge Martins, Ricardo Haddad, Claudia Renata dos Santos Barros, Elaine Cristina Marqueze, Raul Machado Neto, Debora Botequio Moretti. Centro de Genômica Funcional da ESALQ: Luiz Lehmann Coutinho, Ricardo Augusto Brassaloti, Raquel de Lello Rocha Campos Cassano. NGS Soluções Genômicas: Pilar Drummond Sampaio Corrêa Mariani. FZEA-USP Pirassununga: Mirele Daiana Poleti, Jessika Cristina Chagas Lesbon, Elisangela Chicaroni Mattos, Heidge Fukumasu. USP-Botucatu: Rejane Maria Tommasini Grotto, Jayme A. Souza-Neto, Guilherme Targino Valente, Patricia Akemi Assato, Felipe Allan da Silva da Costa, Bianca Cechetto Carlos. Mendelics: Bibiana Santos, João Paulo Kitajima, Erika Freitas, David Schlesinger. Hemocentro Ribeirão Preto: Simone Kashima, Evandra Strazza Rodrigues, Svetoslav Nanev Slavov, Elaine Vieira dos Santos, Rafael dos Santos Bezerra, Luiz Carlos Junior de Alcantara, Marta Giovanetti, Vagner Fonseca, Flavia Aburjaile, Rodrigo Tocantins Calado. |
| EPI_ISL_1795173 | CS DE OSCAR BRESSANE PSF | Instituto Butantan / ESALQ-Piracicaba | Instituto Butantan: Alexander Roberto Precioso, Dimas Tadeu Covas, Sandra Coccuzzo Sampaio, Maria Carolina Elias, José Salvatore Leister Patané, Vincent Louis Viala, Antonio Jorge Martins, Ricardo Haddad, Claudia Renata dos Santos Barros, Elaine Cristina Marqueze, Raul Machado Neto, Debora Botequio Moretti. Centro de Genômica Funcional da ESALQ: Luiz Lehmann Coutinho, Ricardo Augusto Brassaloti, Raquel de Lello Rocha Campos Cassano. NGS Soluções Genômicas: Pilar Drummond Sampaio Corrêa Mariani. FZEA-USP Pirassununga: Mirele Daiana Poleti, Jessika Cristina Chagas |

| | | | |
|---|---|---|---|
| | | | Lesbon, Elisangela Chicaroni Mattos, Heidge Fukumasu. USP-Botucatu: Rejane Maria Tommasini Grotto, Jayme A. Souza-Neto, Guilherme Targino Valente, Patricia Akemi Assato, Felipe Allan da Silva da Costa, Bianca Cechetto Carlos. Mendelics: Bibiana Santos, João Paulo Kitajima, Erika Freitas, David Schlesinger. Hemocentro Ribeirão Preto: Simone Kashima, Evandra Strazza Rodrigues, Svetoslav Nanev Slavov, Elaine Vieira dos Santos, Rafael dos Santos Bezerra, Luiz Carlos Junior de Alcantara, Marta Giovanetti, Vagner Fonseca, Flavia Aburjaile, Rodrigo Tocantins Calado. |
| EPI_ISL_1795174, EPI_ISL_1795175, EPI_ISL_1795176 | USF SALERNO | Instituto Butantan / ESALQ-Piracicaba | Instituto Butantan: Alexander Roberto Precioso, Dimas Tadeu Covas, Sandra Coccuzzo Sampaio, Maria Carolina Elias, José Salvatore Leister Patané, Vincent Louis Viala, Antonio Jorge Martins, Ricardo Haddad, Claudia Renata dos Santos Barros, Elaine Cristina Marqueze, Raul Machado Neto, Debora Botequio Moretti. Centro de Genômica Funcional da ESALQ: Luiz Lehmann Coutinho, Ricardo Augusto Brassaloti, Raquel de Lello Rocha Campos Cassano. NGS Soluções Genômicas: Pilar Drummond Sampaio Corrêa Mariani. FZEA-USP Pirassununga: Mirele Daiana Poleti, Jessika Cristina Chagas Lesbon, Elisangela Chicaroni Mattos, Heidge Fukumasu. USP-Botucatu: Rejane Maria Tommasini Grotto, Jayme A. Souza-Neto, Guilherme Targino Valente, Patricia Akemi Assato, Felipe Allan da Silva da Costa, Bianca Cechetto Carlos. Mendelics: Bibiana Santos, João Paulo Kitajima, Erika Freitas, David Schlesinger. Hemocentro Ribeirão Preto: Simone Kashima, Evandra Strazza Rodrigues, Svetoslav Nanev Slavov, Elaine Vieira dos Santos, Rafael dos Santos Bezerra, Luiz Carlos Junior de Alcantara, Marta Giovanetti, Vagner Fonseca, Flavia Aburjaile, Rodrigo Tocantins Calado. |
| EPI_ISL_1795177, EPI_ISL_1795178, EPI_ISL_1795179, EPI_ISL_1795180, EPI_ISL_1795181 | POLICLINICA HORTOLANDIA | Instituto Butantan / ESALQ-Piracicaba | Instituto Butantan: Alexander Roberto Precioso, Dimas Tadeu Covas, Sandra Coccuzzo Sampaio, Maria Carolina Elias, José Salvatore Leister Patané, Vincent Louis Viala, Antonio Jorge Martins, Ricardo Haddad, Claudia Renata dos Santos Barros, Elaine Cristina Marqueze, Raul Machado Neto, Debora Botequio Moretti. Centro de Genômica Funcional da ESALQ: Luiz Lehmann Coutinho, Ricardo Augusto Brassaloti, Raquel de Lello Rocha Campos Cassano. NGS Soluções Genômicas: Pilar Drummond Sampaio Corrêa Mariani. FZEA-USP Pirassununga: Mirele Daiana Poleti, Jessika Cristina Chagas Lesbon, Elisangela Chicaroni Mattos, Heidge Fukumasu. USP-Botucatu: Rejane Maria Tommasini Grotto, Jayme A. Souza-Neto, Guilherme Targino Valente, Patricia Akemi Assato, Felipe Allan da Silva da Costa, Bianca Cechetto Carlos. Mendelics: Bibiana Santos, João Paulo Kitajima, Erika Freitas, David Schlesinger. Hemocentro Ribeirão Preto: Simone Kashima, Evandra Strazza Rodrigues, Svetoslav Nanev Slavov, Elaine Vieira dos Santos, Rafael dos Santos Bezerra, Luiz Carlos Junior de Alcantara, Marta Giovanetti, Vagner Fonseca, Flavia Aburjaile, Rodrigo Tocantins Calado. |
| EPI_ISL_1795182 | DEPARTAMENTO DE SAUDE MUNICIPAL SOCORRO SP | Instituto Butantan / ESALQ-Piracicaba | Instituto Butantan: Alexander Roberto Precioso, Dimas Tadeu Covas, Sandra Coccuzzo Sampaio, Maria Carolina Elias, José Salvatore Leister Patané, Vincent Louis Viala, Antonio Jorge Martins, Ricardo Haddad, Claudia Renata dos Santos Barros, Elaine Cristina Marqueze, Raul Machado Neto, Debora Botequio Moretti. Centro de Genômica Funcional da ESALQ: Luiz Lehmann Coutinho, Ricardo Augusto Brassaloti, Raquel de Lello Rocha Campos Cassano. NGS Soluções Genômicas: Pilar Drummond Sampaio Corrêa Mariani. FZEA-USP Pirassununga: Mirele Daiana Poleti, Jessika Cristina Chagas Lesbon, Elisangela Chicaroni Mattos, Heidge Fukumasu. USP-Botucatu: Rejane Maria Tommasini Grotto, Jayme A. Souza-Neto, Guilherme Targino Valente, Patricia Akemi Assato, Felipe Allan da Silva da Costa, Bianca Cechetto Carlos. Mendelics: Bibiana Santos, João Paulo Kitajima, Erika Freitas, David Schlesinger. Hemocentro Ribeirão Preto: Simone Kashima, Evandra Strazza Rodrigues, Svetoslav Nanev Slavov, Elaine Vieira dos Santos, Rafael dos Santos Bezerra, Luiz Carlos Junior de Alcantara, Marta Giovanetti, Vagner Fonseca, Flavia Aburjaile, Rodrigo Tocantins Calado. |
| EPI_ISL_1795183, EPI_ISL_1795184, EPI_ISL_1795185, EPI_ISL_1795186 | POLICLINICA HORTOLANDIA | Instituto Butantan / ESALQ-Piracicaba | Instituto Butantan: Alexander Roberto Precioso, Dimas Tadeu Covas, Sandra Coccuzzo Sampaio, Maria Carolina Elias, José Salvatore Leister Patané, Vincent Louis Viala, Antonio Jorge Martins, Ricardo Haddad, Claudia Renata dos Santos Barros, Elaine Cristina Marqueze, Raul Machado Neto, Debora Botequio Moretti. Centro de Genômica Funcional da ESALQ: Luiz Lehmann Coutinho, Ricardo Augusto Brassaloti, Raquel de Lello Rocha Campos Cassano. NGS Soluções Genômicas: Pilar Drummond Sampaio Corrêa Mariani. FZEA-USP Pirassununga: Mirele Daiana Poleti, Jessika Cristina Chagas Lesbon, Elisangela Chicaroni Mattos, Heidge Fukumasu. USP-Botucatu: Rejane Maria Tommasini Grotto, Jayme A. Souza-Neto, Guilherme Targino Valente, Patricia Akemi Assato, Felipe Allan da Silva da Costa, Bianca Cechetto Carlos. Mendelics: Bibiana Santos, João Paulo Kitajima, Erika Freitas, David Schlesinger. Hemocentro Ribeirão Preto: Simone Kashima, Evandra Strazza Rodrigues, Svetoslav Nanev Slavov, Elaine Vieira dos Santos, Rafael dos Santos Bezerra, Luiz Carlos Junior de Alcantara, Marta Giovanetti, Vagner Fonseca, Flavia Aburjaile, Rodrigo Tocantins Calado. |
| EPI_ISL_1795187 | USF SALERNO | Instituto Butantan / ESALQ-Piracicaba | Instituto Butantan: Alexander Roberto Precioso, Dimas Tadeu Covas, Sandra Coccuzzo Sampaio, Maria Carolina Elias, José Salvatore Leister Patané, Vincent Louis Viala, Antonio Jorge Martins, Ricardo Haddad, Claudia Renata dos Santos Barros, Elaine Cristina Marqueze, Raul Machado Neto, Debora Botequio Moretti. Centro de Genômica Funcional da ESALQ: Luiz Lehmann Coutinho, Ricardo Augusto Brassaloti, Raquel de Lello Rocha Campos Cassano. NGS Soluções Genômicas: Pilar Drummond Sampaio Corrêa Mariani. FZEA-USP Pirassununga: Mirele Daiana Poleti, Jessika Cristina Chagas Lesbon, Elisangela Chicaroni Mattos, Heidge Fukumasu. USP-Botucatu: Rejane Maria Tommasini Grotto, Jayme A. Souza-Neto, Guilherme Targino Valente, Patricia Akemi Assato, Felipe Allan da Silva da Costa, Bianca Cechetto Carlos. Mendelics: Bibiana Santos, João Paulo Kitajima, Erika Freitas, David Schlesinger. Hemocentro Ribeirão Preto: Simone Kashima, Evandra Strazza Rodrigues, Svetoslav Nanev Slavov, Elaine Vieira dos Santos, Rafael dos Santos Bezerra, Luiz Carlos Junior de Alcantara, Marta Giovanetti, Vagner Fonseca, Flavia Aburjaile, Rodrigo Tocantins Calado. |
| EPI_ISL_1795188 | POLICLINICA HORTOLANDIA | Instituto Butantan / ESALQ-Piracicaba | Instituto Butantan: Alexander Roberto Precioso, Dimas Tadeu Covas, Sandra Coccuzzo Sampaio, Maria Carolina Elias, José Salvatore Leister Patané, Vincent Louis Viala, Antonio Jorge Martins, Ricardo Haddad, Claudia Renata dos Santos Barros, Elaine Cristina Marqueze, Raul Machado Neto, Debora Botequio Moretti. Centro de Genômica Funcional da ESALQ: Luiz Lehmann Coutinho, Ricardo Augusto Brassaloti, Raquel de Lello Rocha Campos Cassano. NGS Soluções Genômicas: Pilar Drummond Sampaio Corrêa Mariani. FZEA-USP Pirassununga: Mirele Daiana Poleti, Jessika Cristina Chagas Lesbon, Elisangela Chicaroni Mattos, Heidge Fukumasu. USP-Botucatu: Rejane Maria Tommasini Grotto, Jayme A. Souza-Neto, Guilherme Targino Valente, Patricia Akemi Assato, Felipe Allan da Silva da Costa, Bianca Cechetto Carlos. Mendelics: Bibiana Santos, João Paulo Kitajima, Erika Freitas, David Schlesinger. Hemocentro Ribeirão Preto: Simone Kashima, Evandra Strazza Rodrigues, Svetoslav Nanev Slavov, Elaine Vieira dos Santos, Rafael dos Santos Bezerra, Luiz Carlos Junior de Alcantara, Marta Giovanetti, Vagner Fonseca, Flavia Aburjaile, Rodrigo Tocantins Calado. |
| EPI_ISL_1795189 | USF SALERNO | Instituto Butantan / ESALQ-Piracicaba | Instituto Butantan: Alexander Roberto Precioso, Dimas Tadeu Covas, Sandra Coccuzzo Sampaio, Maria Carolina Elias, José Salvatore Leister Patané, Vincent Louis Viala, Antonio Jorge Martins, Ricardo Haddad, Claudia Renata dos Santos Barros, Elaine Cristina Marqueze, Raul Machado Neto, Debora Botequio Moretti. Centro de Genômica Funcional da ESALQ: Luiz Lehmann Coutinho, Ricardo Augusto Brassaloti, Raquel de Lello Rocha Campos Cassano. NGS Soluções Genômicas: Pilar Drummond Sampaio Corrêa Mariani. FZEA-USP Pirassununga: Mirele Daiana Poleti, Jessika Cristina Chagas Lesbon, Elisangela Chicaroni Mattos, Heidge Fukumasu. USP-Botucatu: Rejane Maria Tommasini Grotto, Jayme A. Souza-Neto, Guilherme Targino Valente, Patricia Akemi Assato, Felipe Allan da Silva da Costa, Bianca Cechetto Carlos. Mendelics: Bibiana Santos, João Paulo Kitajima, Erika Freitas, David Schlesinger. Hemocentro Ribeirão Preto: Simone Kashima, Evandra Strazza Rodrigues, Svetoslav Nanev Slavov, Elaine Vieira dos Santos, Rafael dos Santos Bezerra, Luiz Carlos Junior de Alcantara, Marta Giovanetti, Vagner Fonseca, Flavia Aburjaile, Rodrigo Tocantins Calado. |
| EPI_ISL_1795190, EPI_ISL_1795191 | POLICLINICA HORTOLANDIA | Instituto Butantan / ESALQ-Piracicaba | Instituto Butantan: Alexander Roberto Precioso, Dimas Tadeu Covas, Sandra Coccuzzo Sampaio, Maria Carolina Elias, José Salvatore Leister Patané, Vincent Louis Viala, Antonio Jorge Martins, Ricardo Haddad, Claudia Renata dos Santos Barros, Elaine Cristina Marqueze, Raul Machado Neto, Debora Botequio Moretti. Centro de Genômica Funcional da ESALQ: Luiz Lehmann Coutinho, Ricardo Augusto Brassaloti, Raquel de Lello Rocha Campos Cassano. NGS Soluções Genômicas: Pilar Drummond Sampaio Corrêa Mariani. FZEA-USP Pirassununga: Mirele Daiana Poleti, Jessika Cristina Chagas Lesbon, Elisangela Chicaroni Mattos, Heidge Fukumasu. USP-Botucatu: Rejane Maria Tommasini Grotto, Jayme A. Souza-Neto, Guilherme Targino Valente, Patricia Akemi Assato, Felipe Allan da Silva da Costa, Bianca Cechetto Carlos. Mendelics: Bibiana Santos, João Paulo Kitajima, Erika Freitas, David Schlesinger. Hemocentro Ribeirão Preto: Simone Kashima, Evandra Strazza Rodrigues, Svetoslav Nanev Slavov, Elaine Vieira dos Santos, Rafael dos Santos Bezerra, Luiz Carlos Junior de Alcantara, Marta Giovanetti, Vagner Fonseca, Flavia Aburjaile, Rodrigo Tocantins Calado. |
| EPI_ISL_1795192, EPI_ISL_1795193 | USF SALERNO | Instituto Butantan / ESALQ-Piracicaba | Instituto Butantan: Alexander Roberto Precioso, Dimas Tadeu Covas, Sandra Coccuzzo Sampaio, Maria Carolina Elias, José Salvatore Leister Patané, Vincent Louis Viala, Antonio Jorge Martins, Ricardo Haddad, Claudia Renata dos Santos Barros, Elaine Cristina Marqueze, Raul Machado Neto, Debora Botequio Moretti. Centro de Genômica Funcional da ESALQ: Luiz Lehmann Coutinho, Ricardo Augusto Brassaloti, Raquel de Lello Rocha Campos Cassano. NGS Soluções Genômicas: Pilar Drummond Sampaio Corrêa Mariani. FZEA-USP Pirassununga: Mirele Daiana Poleti, Jessika Cristina Chagas Lesbon, Elisangela Chicaroni Mattos, Heidge Fukumasu. USP-Botucatu: Rejane Maria Tommasini Grotto, Jayme A. Souza-Neto, Guilherme Targino Valente, Patricia Akemi Assato, Felipe Allan da Silva da Costa, Bianca Cechetto Carlos. Mendelics: Bibiana Santos, João Paulo Kitajima, Erika Freitas, David Schlesinger. Hemocentro Ribeirão Preto: Simone Kashima, Evandra Strazza Rodrigues, Svetoslav Nanev Slavov, Elaine Vieira dos Santos, Rafael dos Santos Bezerra, Luiz Carlos Junior de Alcantara, Marta Giovanetti, Vagner Fonseca, Flavia Aburjaile, Rodrigo Tocantins Calado. |
| EPI_ISL_1795194, EPI_ISL_1795195, EPI_ISL_1795196, EPI_ISL_1795197, EPI_ISL_1795198, EPI_ISL_1795199, EPI_ISL_1795200, EPI_ISL_1795201, EPI_ISL_1795202, EPI_ISL_1795203, EPI_ISL_1795204, EPI_ISL_1795205, EPI_ISL_1795206, EPI_ISL_1795207 | | | |
| see above | AMBULATORIO DE ESPECIALIDADE V E MOGI MIRIM | Instituto Butantan / ESALQ-Piracicaba | Instituto Butantan: Alexander Roberto Precioso, Dimas Tadeu Covas, Sandra Coccuzzo Sampaio, Maria Carolina Elias, José Salvatore Leister Patané, Vincent Louis Viala, Antonio Jorge Martins, Ricardo Haddad, Claudia Renata dos Santos Barros, Elaine Cristina Marqueze, Raul Machado Neto, Debora Botequio Moretti. Centro de Genômica Funcional da ESALQ: Luiz Lehmann Coutinho, Ricardo Augusto Brassaloti, Raquel de Lello Rocha Campos Cassano. NGS Soluções Genômicas: Pilar Drummond Sampaio Corrêa Mariani. FZEA-USP Pirassununga: Mirele Daiana Poleti, Jessika Cristina Chagas Lesbon, Elisangela Chicaroni Mattos, Heidge Fukumasu. USP-Botucatu: Rejane Maria Tommasini Grotto, Jayme A. Souza-Neto, Guilherme Targino |

| | | | |
|---|---|---|---|
| | | | Valente, Patricia Akemi Assato, Felipe Allan da Silva da Costa, Bianca Cechetto Carlos. Mendelics: Bibiana Santos, João Paulo Kitajima, Erika Freitas, David Schlesinger. Hemocentro Ribeirão Preto: Simone Kashima, Evandra Strazza Rodrigues, Svetoslav Nanev Slavov, Elaine Vieira dos Santos, Rafael dos Santos Bezerra, Luiz Carlos Junior de Alcantara, Marta Giovanetti, Vagner Fonseca, Flavia Aburjaile, Rodrigo Tocantins Calado. |
| EPI_ISL_1795208, EPI_ISL_1795209 | PRONTO SOCORRO MUNICIPAL TAMBAU | Instituto Butantan / ESALQ-Piracicaba | Instituto Butantan: Alexander Roberto Precioso, Dimas Tadeu Covas, Sandra Coccuzzo Sampaio, Maria Carolina Elias, José Salvatore Leister Patané, Vincent Louis Viala, Antonio Jorge Martins, Ricardo Haddad, Claudia Renata dos Santos Barros, Elaine Cristina Marqueze, Raul Machado Neto, Debora Botequio Moretti. Centro de Genômica Funcional da ESALQ: Luiz Lehmann Coutinho, Ricardo Augusto Brassaloti, Raquel de Lello Rocha Campos Cassano. NGS Soluções Genômicas: Pilar Drummond Sampaio Corrêa Mariani. FZEA-USP Pirassununga: Mirele Daiana Poleti, Jessika Cristina Chagas Lesbon, Elisangela Chicaroni Mattos, Heidge Fukumasu. USP-Botucatu: Rejane Maria Tommasini Grotto, Jayme A. Souza-Neto, Guilherme Targino Valente, Patricia Akemi Assato, Felipe Allan da Silva da Costa, Bianca Cechetto Carlos. Mendelics: Bibiana Santos, João Paulo Kitajima, Erika Freitas, David Schlesinger. Hemocentro Ribeirão Preto: Simone Kashima, Evandra Strazza Rodrigues, Svetoslav Nanev Slavov, Elaine Vieira dos Santos, Rafael dos Santos Bezerra, Luiz Carlos Junior de Alcantara, Marta Giovanetti, Vagner Fonseca, Flavia Aburjaile, Rodrigo Tocantins Calado. |
| EPI_ISL_1795210 | AMBULATORIO DE ESPECIALIDADE V E MOGI MIRIM | Instituto Butantan / ESALQ-Piracicaba | Instituto Butantan: Alexander Roberto Precioso, Dimas Tadeu Covas, Sandra Coccuzzo Sampaio, Maria Carolina Elias, José Salvatore Leister Patané, Vincent Louis Viala, Antonio Jorge Martins, Ricardo Haddad, Claudia Renata dos Santos Barros, Elaine Cristina Marqueze, Raul Machado Neto, Debora Botequio Moretti. Centro de Genômica Funcional da ESALQ: Luiz Lehmann Coutinho, Ricardo Augusto Brassaloti, Raquel de Lello Rocha Campos Cassano. NGS Soluções Genômicas: Pilar Drummond Sampaio Corrêa Mariani. FZEA-USP Pirassununga: Mirele Daiana Poleti, Jessika Cristina Chagas Lesbon, Elisangela Chicaroni Mattos, Heidge Fukumasu. USP-Botucatu: Rejane Maria Tommasini Grotto, Jayme A. Souza-Neto, Guilherme Targino Valente, Patricia Akemi Assato, Felipe Allan da Silva da Costa, Bianca Cechetto Carlos. Mendelics: Bibiana Santos, João Paulo Kitajima, Erika Freitas, David Schlesinger. Hemocentro Ribeirão Preto: Simone Kashima, Evandra Strazza Rodrigues, Svetoslav Nanev Slavov, Elaine Vieira dos Santos, Rafael dos Santos Bezerra, Luiz Carlos Junior de Alcantara, Marta Giovanetti, Vagner Fonseca, Flavia Aburjaile, Rodrigo Tocantins Calado. |
| EPI_ISL_1795211 | HOSPITAL MUNICIPAL DR MARIO GATTI CAMPINAS | Instituto Butantan / ESALQ-Piracicaba | Instituto Butantan: Alexander Roberto Precioso, Dimas Tadeu Covas, Sandra Coccuzzo Sampaio, Maria Carolina Elias, José Salvatore Leister Patané, Vincent Louis Viala, Antonio Jorge Martins, Ricardo Haddad, Claudia Renata dos Santos Barros, Elaine Cristina Marqueze, Raul Machado Neto, Debora Botequio Moretti. Centro de Genômica Funcional da ESALQ: Luiz Lehmann Coutinho, Ricardo Augusto Brassaloti, Raquel de Lello Rocha Campos Cassano. NGS Soluções Genômicas: Pilar Drummond Sampaio Corrêa Mariani. FZEA-USP Pirassununga: Mirele Daiana Poleti, Jessika Cristina Chagas Lesbon, Elisangela Chicaroni Mattos, Heidge Fukumasu. USP-Botucatu: Rejane Maria Tommasini Grotto, Jayme A. Souza-Neto, Guilherme Targino Valente, Patricia Akemi Assato, Felipe Allan da Silva da Costa, Bianca Cechetto Carlos. Mendelics: Bibiana Santos, João Paulo Kitajima, Erika Freitas, David Schlesinger. Hemocentro Ribeirão Preto: Simone Kashima, Evandra Strazza Rodrigues, Svetoslav Nanev Slavov, Elaine Vieira dos Santos, Rafael dos Santos Bezerra, Luiz Carlos Junior de Alcantara, Marta Giovanetti, Vagner Fonseca, Flavia Aburjaile, Rodrigo Tocantins Calado. |
| EPI_ISL_1795212 | CENTRO DE SAUDE II DR ALCIDES FACUNDO ARROYO | Instituto Butantan / ESALQ-Piracicaba | Instituto Butantan: Alexander Roberto Precioso, Dimas Tadeu Covas, Sandra Coccuzzo Sampaio, Maria Carolina Elias, José Salvatore Leister Patané, Vincent Louis Viala, Antonio Jorge Martins, Ricardo Haddad, Claudia Renata dos Santos Barros, Elaine Cristina Marqueze, Raul Machado Neto, Debora Botequio Moretti. Centro de Genômica Funcional da ESALQ: Luiz Lehmann Coutinho, Ricardo Augusto Brassaloti, Raquel de Lello Rocha Campos Cassano. NGS Soluções Genômicas: Pilar Drummond Sampaio Corrêa Mariani. FZEA-USP Pirassununga: Mirele Daiana Poleti, Jessika Cristina Chagas Lesbon, Elisangela Chicaroni Mattos, Heidge Fukumasu. USP-Botucatu: Rejane Maria Tommasini Grotto, Jayme A. Souza-Neto, Guilherme Targino Valente, Patricia Akemi Assato, Felipe Allan da Silva da Costa, Bianca Cechetto Carlos. Mendelics: Bibiana Santos, João Paulo Kitajima, Erika Freitas, David Schlesinger. Hemocentro Ribeirão Preto: Simone Kashima, Evandra Strazza Rodrigues, Svetoslav Nanev Slavov, Elaine Vieira dos Santos, Rafael dos Santos Bezerra, Luiz Carlos Junior de Alcantara, Marta Giovanetti, Vagner Fonseca, Flavia Aburjaile, Rodrigo Tocantins Calado. |
| EPI_ISL_1795213, EPI_ISL_1795214, EPI_ISL_1795215, EPI_ISL_1795216 | PRONTO SOCORRO MUNICIPAL DE SEVERINIA | Instituto Butantan / ESALQ-Piracicaba | Instituto Butantan: Alexander Roberto Precioso, Dimas Tadeu Covas, Sandra Coccuzzo Sampaio, Maria Carolina Elias, José Salvatore Leister Patané, Vincent Louis Viala, Antonio Jorge Martins, Ricardo Haddad, Claudia Renata dos Santos Barros, Elaine Cristina Marqueze, Raul Machado Neto, Debora Botequio Moretti. Centro de Genômica Funcional da ESALQ: Luiz Lehmann Coutinho, Ricardo Augusto Brassaloti, Raquel de Lello Rocha Campos Cassano. NGS Soluções Genômicas: Pilar Drummond Sampaio Corrêa Mariani. FZEA-USP Pirassununga: Mirele Daiana Poleti, Jessika Cristina Chagas Lesbon, Elisangela Chicaroni Mattos, Heidge Fukumasu. USP-Botucatu: Rejane Maria Tommasini Grotto, Jayme A. Souza-Neto, Guilherme Targino Valente, Patricia Akemi Assato, Felipe Allan da Silva da Costa, Bianca Cechetto Carlos. Mendelics: Bibiana Santos, João Paulo Kitajima, Erika Freitas, David Schlesinger. Hemocentro Ribeirão Preto: Simone Kashima, Evandra Strazza Rodrigues, Svetoslav Nanev Slavov, Elaine Vieira dos Santos, Rafael dos Santos Bezerra, Luiz Carlos Junior de Alcantara, Marta Giovanetti, Vagner Fonseca, Flavia Aburjaile, Rodrigo Tocantins Calado. |
| EPI_ISL_1795217 | VIGILANCIA EPIDEMIOLOGICA JARDINOPOLIS SP | Instituto Butantan / ESALQ-Piracicaba | Instituto Butantan: Alexander Roberto Precioso, Dimas Tadeu Covas, Sandra Coccuzzo Sampaio, Maria Carolina Elias, José Salvatore Leister Patané, Vincent Louis Viala, Antonio Jorge Martins, Ricardo Haddad, Claudia Renata dos Santos Barros, Elaine Cristina Marqueze, Raul Machado Neto, Debora Botequio Moretti. Centro de Genômica Funcional da ESALQ: Luiz Lehmann Coutinho, Ricardo Augusto Brassaloti, Raquel de Lello Rocha Campos Cassano. NGS Soluções Genômicas: Pilar Drummond Sampaio Corrêa Mariani. FZEA-USP Pirassununga: Mirele Daiana Poleti, Jessika Cristina Chagas Lesbon, Elisangela Chicaroni Mattos, Heidge Fukumasu. USP-Botucatu: Rejane Maria Tommasini Grotto, Jayme A. Souza-Neto, Guilherme Targino Valente, Patricia Akemi Assato, Felipe Allan da Silva da Costa, Bianca Cechetto Carlos. Mendelics: Bibiana Santos, João Paulo Kitajima, Erika Freitas, David Schlesinger. Hemocentro Ribeirão Preto: Simone Kashima, Evandra Strazza Rodrigues, Svetoslav Nanev Slavov, Elaine Vieira dos Santos, Rafael dos Santos Bezerra, Luiz Carlos Junior de Alcantara, Marta Giovanetti, Vagner Fonseca, Flavia Aburjaile, Rodrigo Tocantins Calado. |
| EPI_ISL_1795218 | VIGILANCIA EPIDIMIOLOGICA DE ARACARIGUAMA | Instituto Butantan / ESALQ-Piracicaba | Instituto Butantan: Alexander Roberto Precioso, Dimas Tadeu Covas, Sandra Coccuzzo Sampaio, Maria Carolina Elias, José Salvatore Leister Patané, Vincent Louis Viala, Antonio Jorge Martins, Ricardo Haddad, Claudia Renata dos Santos Barros, Elaine Cristina Marqueze, Raul Machado Neto, Debora Botequio Moretti. Centro de Genômica Funcional da ESALQ: Luiz Lehmann Coutinho, Ricardo Augusto Brassaloti, Raquel de Lello Rocha Campos Cassano. NGS Soluções Genômicas: Pilar Drummond Sampaio Corrêa Mariani. FZEA-USP Pirassununga: Mirele Daiana Poleti, Jessika Cristina Chagas Lesbon, Elisangela Chicaroni Mattos, Heidge Fukumasu. USP-Botucatu: Rejane Maria Tommasini Grotto, Jayme A. Souza-Neto, Guilherme Targino Valente, Patricia Akemi Assato, Felipe Allan da Silva da Costa, Bianca Cechetto Carlos. Mendelics: Bibiana Santos, João Paulo Kitajima, Erika Freitas, David Schlesinger. Hemocentro Ribeirão Preto: Simone Kashima, Evandra Strazza Rodrigues, Svetoslav Nanev Slavov, Elaine Vieira dos Santos, Rafael dos Santos Bezerra, Luiz Carlos Junior de Alcantara, Marta Giovanetti, Vagner Fonseca, Flavia Aburjaile, Rodrigo Tocantins Calado. |
| EPI_ISL_1795219 | PRONTO ATENDIMENTO UNIDADE SAUDE ADALBERTO ROCHA GUAREI | Instituto Butantan / ESALQ-Piracicaba | Instituto Butantan: Alexander Roberto Precioso, Dimas Tadeu Covas, Sandra Coccuzzo Sampaio, Maria Carolina Elias, José Salvatore Leister Patané, Vincent Louis Viala, Antonio Jorge Martins, Ricardo Haddad, Claudia Renata dos Santos Barros, Elaine Cristina Marqueze, Raul Machado Neto, Debora Botequio Moretti. Centro de Genômica Funcional da ESALQ: Luiz Lehmann Coutinho, Ricardo Augusto Brassaloti, Raquel de Lello Rocha Campos Cassano. NGS Soluções Genômicas: Pilar Drummond Sampaio Corrêa Mariani. FZEA-USP Pirassununga: Mirele Daiana Poleti, Jessika Cristina Chagas Lesbon, Elisangela Chicaroni Mattos, Heidge Fukumasu. USP-Botucatu: Rejane Maria Tommasini Grotto, Jayme A. Souza-Neto, Guilherme Targino Valente, Patricia Akemi Assato, Felipe Allan da Silva da Costa, Bianca Cechetto Carlos. Mendelics: Bibiana Santos, João Paulo Kitajima, Erika Freitas, David Schlesinger. Hemocentro Ribeirão Preto: Simone Kashima, Evandra Strazza Rodrigues, Svetoslav Nanev Slavov, Elaine Vieira dos Santos, Rafael dos Santos Bezerra, Luiz Carlos Junior de Alcantara, Marta Giovanetti, Vagner Fonseca, Flavia Aburjaile, Rodrigo Tocantins Calado. |
| EPI_ISL_1795220 | UNIDADE SENTINELA COVID19 | Instituto Butantan / ESALQ-Piracicaba | Instituto Butantan: Alexander Roberto Precioso, Dimas Tadeu Covas, Sandra Coccuzzo Sampaio, Maria Carolina Elias, José Salvatore Leister Patané, Vincent Louis Viala, Antonio Jorge Martins, Ricardo Haddad, Claudia Renata dos Santos Barros, Elaine Cristina Marqueze, Raul Machado Neto, Debora Botequio Moretti. Centro de Genômica Funcional da ESALQ: Luiz Lehmann Coutinho, Ricardo Augusto Brassaloti, Raquel de Lello Rocha Campos Cassano. NGS Soluções Genômicas: Pilar Drummond Sampaio Corrêa Mariani. FZEA-USP Pirassununga: Mirele Daiana Poleti, Jessika Cristina Chagas Lesbon, Elisangela Chicaroni Mattos, Heidge Fukumasu. USP-Botucatu: Rejane Maria Tommasini Grotto, Jayme A. Souza-Neto, Guilherme Targino Valente, Patricia Akemi Assato, Felipe Allan da Silva da Costa, Bianca Cechetto Carlos. Mendelics: Bibiana Santos, João Paulo Kitajima, Erika Freitas, David Schlesinger. Hemocentro Ribeirão Preto: Simone Kashima, Evandra Strazza Rodrigues, Svetoslav Nanev Slavov, Elaine Vieira dos Santos, Rafael dos Santos Bezerra, Luiz Carlos Junior de Alcantara, Marta Giovanetti, Vagner Fonseca, Flavia Aburjaile, Rodrigo Tocantins Calado. |
| EPI_ISL_1795221 | HOSPITAL REGIONAL DE ITAPETININGA | Instituto Butantan / ESALQ-Piracicaba | Instituto Butantan: Alexander Roberto Precioso, Dimas Tadeu Covas, Sandra Coccuzzo Sampaio, Maria Carolina Elias, José Salvatore Leister Patané, Vincent Louis Viala, Antonio Jorge Martins, Ricardo Haddad, Claudia Renata dos Santos Barros, Elaine Cristina Marqueze, Raul Machado Neto, Debora Botequio Moretti. Centro de Genômica Funcional da ESALQ: Luiz Lehmann Coutinho, Ricardo Augusto Brassaloti, Raquel de Lello Rocha Campos Cassano. NGS Soluções Genômicas: Pilar Drummond Sampaio Corrêa Mariani. FZEA-USP Pirassununga: Mirele Daiana Poleti, Jessika Cristina Chagas Lesbon, Elisangela Chicaroni Mattos, Heidge Fukumasu. USP-Botucatu: Rejane Maria Tommasini Grotto, Jayme A. Souza-Neto, Guilherme Targino |

| | | | |
|---|---|---|---|
| | | | Valente, Patricia Akemi Assato, Felipe Allan da Silva da Costa, Bianca Cechetto Carlos. Mendelics: Bibiana Santos, João Paulo Kitajima, Erika Freitas, David Schlesinger. Hemocentro Ribeirão Preto: Simone Kashima, Evandra Strazza Rodrigues, Svetoslav Nanev Slavov, Elaine Vieira dos Santos, Rafael dos Santos Bezerra, Luiz Carlos Junior de Alcantara, Marta Giovanetti, Vagner Fonseca, Flavia Aburjaile, Rodrigo Tocantins Calado. |
| EPI_ISL_1795222 | UNIDADE MISTA ARACOIABA DA SERRA | Instituto Butantan / ESALQ-Piracicaba | Instituto Butantan: Alexander Roberto Precioso, Dimas Tadeu Covas, Sandra Coccuzzo Sampaio, Maria Carolina Elias, José Salvatore Leister Patané, Vincent Louis Viala, Antonio Jorge Martins, Ricardo Haddad, Claudia Renata dos Santos Barros, Elaine Cristina Marqueze, Raul Machado Neto, Debora Botequio Moretti. Centro de Genômica Funcional da ESALQ: Luiz Lehmann Coutinho, Ricardo Augusto Brassaloti, Raquel de Lello Rocha Campos Cassano. NGS Soluções Genômicas: Pilar Drummond Sampaio Corrêa Mariani. FZEA-USP Pirassununga: Mirele Daiana Poleti, Jessika Cristina Chagas Lesbon, Elisangela Chicaroni Mattos, Heidge Fukumasu. USP-Botucatu: Rejane Maria Tommasini Grotto, Jayme A. Souza-Neto, Guilherme Targino Valente, Patricia Akemi Assato, Felipe Allan da Silva da Costa, Bianca Cechetto Carlos. Mendelics: Bibiana Santos, João Paulo Kitajima, Erika Freitas, David Schlesinger. Hemocentro Ribeirão Preto: Simone Kashima, Evandra Strazza Rodrigues, Svetoslav Nanev Slavov, Elaine Vieira dos Santos, Rafael dos Santos Bezerra, Luiz Carlos Junior de Alcantara, Marta Giovanetti, Vagner Fonseca, Flavia Aburjaile, Rodrigo Tocantins Calado. |
| EPI_ISL_1795223 | HOSPITAL REGIONAL DE ITAPETININGA | Instituto Butantan / ESALQ-Piracicaba | Instituto Butantan: Alexander Roberto Precioso, Dimas Tadeu Covas, Sandra Coccuzzo Sampaio, Maria Carolina Elias, José Salvatore Leister Patané, Vincent Louis Viala, Antonio Jorge Martins, Ricardo Haddad, Claudia Renata dos Santos Barros, Elaine Cristina Marqueze, Raul Machado Neto, Debora Botequio Moretti. Centro de Genômica Funcional da ESALQ: Luiz Lehmann Coutinho, Ricardo Augusto Brassaloti, Raquel de Lello Rocha Campos Cassano. NGS Soluções Genômicas: Pilar Drummond Sampaio Corrêa Mariani. FZEA-USP Pirassununga: Mirele Daiana Poleti, Jessika Cristina Chagas Lesbon, Elisangela Chicaroni Mattos, Heidge Fukumasu. USP-Botucatu: Rejane Maria Tommasini Grotto, Jayme A. Souza-Neto, Guilherme Targino Valente, Patricia Akemi Assato, Felipe Allan da Silva da Costa, Bianca Cechetto Carlos. Mendelics: Bibiana Santos, João Paulo Kitajima, Erika Freitas, David Schlesinger. Hemocentro Ribeirão Preto: Simone Kashima, Evandra Strazza Rodrigues, Svetoslav Nanev Slavov, Elaine Vieira dos Santos, Rafael dos Santos Bezerra, Luiz Carlos Junior de Alcantara, Marta Giovanetti, Vagner Fonseca, Flavia Aburjaile, Rodrigo Tocantins Calado. |
| EPI_ISL_1795224 | UNIDADE BASICA DE SAUDE DR APOLONIO MORAES E SOUZA | Instituto Butantan / ESALQ-Piracicaba | Instituto Butantan: Alexander Roberto Precioso, Dimas Tadeu Covas, Sandra Coccuzzo Sampaio, Maria Carolina Elias, José Salvatore Leister Patané, Vincent Louis Viala, Antonio Jorge Martins, Ricardo Haddad, Claudia Renata dos Santos Barros, Elaine Cristina Marqueze, Raul Machado Neto, Debora Botequio Moretti. Centro de Genômica Funcional da ESALQ: Luiz Lehmann Coutinho, Ricardo Augusto Brassaloti, Raquel de Lello Rocha Campos Cassano. NGS Soluções Genômicas: Pilar Drummond Sampaio Corrêa Mariani. FZEA-USP Pirassununga: Mirele Daiana Poleti, Jessika Cristina Chagas Lesbon, Elisangela Chicaroni Mattos, Heidge Fukumasu. USP-Botucatu: Rejane Maria Tommasini Grotto, Jayme A. Souza-Neto, Guilherme Targino Valente, Patricia Akemi Assato, Felipe Allan da Silva da Costa, Bianca Cechetto Carlos. Mendelics: Bibiana Santos, João Paulo Kitajima, Erika Freitas, David Schlesinger. Hemocentro Ribeirão Preto: Simone Kashima, Evandra Strazza Rodrigues, Svetoslav Nanev Slavov, Elaine Vieira dos Santos, Rafael dos Santos Bezerra, Luiz Carlos Junior de Alcantara, Marta Giovanetti, Vagner Fonseca, Flavia Aburjaile, Rodrigo Tocantins Calado. |
| EPI_ISL_1795225 | UNIDADE BASICA DE SAUDE DR WILSON HAYEK SAIHG | Instituto Butantan / ESALQ-Piracicaba | Instituto Butantan: Alexander Roberto Precioso, Dimas Tadeu Covas, Sandra Coccuzzo Sampaio, Maria Carolina Elias, José Salvatore Leister Patané, Vincent Louis Viala, Antonio Jorge Martins, Ricardo Haddad, Claudia Renata dos Santos Barros, Elaine Cristina Marqueze, Raul Machado Neto, Debora Botequio Moretti. Centro de Genômica Funcional da ESALQ: Luiz Lehmann Coutinho, Ricardo Augusto Brassaloti, Raquel de Lello Rocha Campos Cassano. NGS Soluções Genômicas: Pilar Drummond Sampaio Corrêa Mariani. FZEA-USP Pirassununga: Mirele Daiana Poleti, Jessika Cristina Chagas Lesbon, Elisangela Chicaroni Mattos, Heidge Fukumasu. USP-Botucatu: Rejane Maria Tommasini Grotto, Jayme A. Souza-Neto, Guilherme Targino Valente, Patricia Akemi Assato, Felipe Allan da Silva da Costa, Bianca Cechetto Carlos. Mendelics: Bibiana Santos, João Paulo Kitajima, Erika Freitas, David Schlesinger. Hemocentro Ribeirão Preto: Simone Kashima, Evandra Strazza Rodrigues, Svetoslav Nanev Slavov, Elaine Vieira dos Santos, Rafael dos Santos Bezerra, Luiz Carlos Junior de Alcantara, Marta Giovanetti, Vagner Fonseca, Flavia Aburjaile, Rodrigo Tocantins Calado. |
| EPI_ISL_1795226 | UNIDADE BASICA DR JOSE PARASSU CARVALHO | Instituto Butantan / ESALQ-Piracicaba | Instituto Butantan: Alexander Roberto Precioso, Dimas Tadeu Covas, Sandra Coccuzzo Sampaio, Maria Carolina Elias, José Salvatore Leister Patané, Vincent Louis Viala, Antonio Jorge Martins, Ricardo Haddad, Claudia Renata dos Santos Barros, Elaine Cristina Marqueze, Raul Machado Neto, Debora Botequio Moretti. Centro de Genômica Funcional da ESALQ: Luiz Lehmann Coutinho, Ricardo Augusto Brassaloti, Raquel de Lello Rocha Campos Cassano. NGS Soluções Genômicas: Pilar Drummond Sampaio Corrêa Mariani. FZEA-USP Pirassununga: Mirele Daiana Poleti, Jessika Cristina Chagas Lesbon, Elisangela Chicaroni Mattos, Heidge Fukumasu. USP-Botucatu: Rejane Maria Tommasini Grotto, Jayme A. Souza-Neto, Guilherme Targino Valente, Patricia Akemi Assato, Felipe Allan da Silva da Costa, Bianca Cechetto Carlos. Mendelics: Bibiana Santos, João Paulo Kitajima, Erika Freitas, David Schlesinger. Hemocentro Ribeirão Preto: Simone Kashima, Evandra Strazza Rodrigues, Svetoslav Nanev Slavov, Elaine Vieira dos Santos, Rafael dos Santos Bezerra, Luiz Carlos Junior de Alcantara, Marta Giovanetti, Vagner Fonseca, Flavia Aburjaile, Rodrigo Tocantins Calado. |
| EPI_ISL_1795227 | UNIDADE BASICA DE SAUDE DR LOTFALLAH MIZIARA | Instituto Butantan / ESALQ-Piracicaba | Instituto Butantan: Alexander Roberto Precioso, Dimas Tadeu Covas, Sandra Coccuzzo Sampaio, Maria Carolina Elias, José Salvatore Leister Patané, Vincent Louis Viala, Antonio Jorge Martins, Ricardo Haddad, Claudia Renata dos Santos Barros, Elaine Cristina Marqueze, Raul Machado Neto, Debora Botequio Moretti. Centro de Genômica Funcional da ESALQ: Luiz Lehmann Coutinho, Ricardo Augusto Brassaloti, Raquel de Lello Rocha Campos Cassano. NGS Soluções Genômicas: Pilar Drummond Sampaio Corrêa Mariani. FZEA-USP Pirassununga: Mirele Daiana Poleti, Jessika Cristina Chagas Lesbon, Elisangela Chicaroni Mattos, Heidge Fukumasu. USP-Botucatu: Rejane Maria Tommasini Grotto, Jayme A. Souza-Neto, Guilherme Targino Valente, Patricia Akemi Assato, Felipe Allan da Silva da Costa, Bianca Cechetto Carlos. Mendelics: Bibiana Santos, João Paulo Kitajima, Erika Freitas, David Schlesinger. Hemocentro Ribeirão Preto: Simone Kashima, Evandra Strazza Rodrigues, Svetoslav Nanev Slavov, Elaine Vieira dos Santos, Rafael dos Santos Bezerra, Luiz Carlos Junior de Alcantara, Marta Giovanetti, Vagner Fonseca, Flavia Aburjaile, Rodrigo Tocantins Calado. |
| EPI_ISL_1795228, EPI_ISL_1795229 | CENTRO DE REFERENCIA DO IDOSO DR HUMBERTO MENDES DE CARVALHO | Instituto Butantan / ESALQ-Piracicaba | Instituto Butantan: Alexander Roberto Precioso, Dimas Tadeu Covas, Sandra Coccuzzo Sampaio, Maria Carolina Elias, José Salvatore Leister Patané, Vincent Louis Viala, Antonio Jorge Martins, Ricardo Haddad, Claudia Renata dos Santos Barros, Elaine Cristina Marqueze, Raul Machado Neto, Debora Botequio Moretti. Centro de Genômica Funcional da ESALQ: Luiz Lehmann Coutinho, Ricardo Augusto Brassaloti, Raquel de Lello Rocha Campos Cassano. NGS Soluções Genômicas: Pilar Drummond Sampaio Corrêa Mariani. FZEA-USP Pirassununga: Mirele Daiana Poleti, Jessika Cristina Chagas Lesbon, Elisangela Chicaroni Mattos, Heidge Fukumasu. USP-Botucatu: Rejane Maria Tommasini Grotto, Jayme A. Souza-Neto, Guilherme Targino Valente, Patricia Akemi Assato, Felipe Allan da Silva da Costa, Bianca Cechetto Carlos. Mendelics: Bibiana Santos, João Paulo Kitajima, Erika Freitas, David Schlesinger. Hemocentro Ribeirão Preto: Simone Kashima, Evandra Strazza Rodrigues, Svetoslav Nanev Slavov, Elaine Vieira dos Santos, Rafael dos Santos Bezerra, Luiz Carlos Junior de Alcantara, Marta Giovanetti, Vagner Fonseca, Flavia Aburjaile, Rodrigo Tocantins Calado. |
| EPI_ISL_1795230, EPI_ISL_1795231 | UNIDADE DE SAUDE DA FAMILIA JOSE ADALBERTO LELLIS GARCIA | Instituto Butantan / ESALQ-Piracicaba | Instituto Butantan: Alexander Roberto Precioso, Dimas Tadeu Covas, Sandra Coccuzzo Sampaio, Maria Carolina Elias, José Salvatore Leister Patané, Vincent Louis Viala, Antonio Jorge Martins, Ricardo Haddad, Claudia Renata dos Santos Barros, Elaine Cristina Marqueze, Raul Machado Neto, Debora Botequio Moretti. Centro de Genômica Funcional da ESALQ: Luiz Lehmann Coutinho, Ricardo Augusto Brassaloti, Raquel de Lello Rocha Campos Cassano. NGS Soluções Genômicas: Pilar Drummond Sampaio Corrêa Mariani. FZEA-USP Pirassununga: Mirele Daiana Poleti, Jessika Cristina Chagas Lesbon, Elisangela Chicaroni Mattos, Heidge Fukumasu. USP-Botucatu: Rejane Maria Tommasini Grotto, Jayme A. Souza-Neto, Guilherme Targino Valente, Patricia Akemi Assato, Felipe Allan da Silva da Costa, Bianca Cechetto Carlos. Mendelics: Bibiana Santos, João Paulo Kitajima, Erika Freitas, David Schlesinger. Hemocentro Ribeirão Preto: Simone Kashima, Evandra Strazza Rodrigues, Svetoslav Nanev Slavov, Elaine Vieira dos Santos, Rafael dos Santos Bezerra, Luiz Carlos Junior de Alcantara, Marta Giovanetti, Vagner Fonseca, Flavia Aburjaile, Rodrigo Tocantins Calado. |
| EPI_ISL_1795232, EPI_ISL_1795233 | CS DE URUPES | Instituto Butantan / ESALQ-Piracicaba | Instituto Butantan: Alexander Roberto Precioso, Dimas Tadeu Covas, Sandra Coccuzzo Sampaio, Maria Carolina Elias, José Salvatore Leister Patané, Vincent Louis Viala, Antonio Jorge Martins, Ricardo Haddad, Claudia Renata dos Santos Barros, Elaine Cristina Marqueze, Raul Machado Neto, Debora Botequio Moretti. Centro de Genômica Funcional da ESALQ: Luiz Lehmann Coutinho, Ricardo Augusto Brassaloti, Raquel de Lello Rocha Campos Cassano. NGS Soluções Genômicas: Pilar Drummond Sampaio Corrêa Mariani. FZEA-USP Pirassununga: Mirele Daiana Poleti, Jessika Cristina Chagas Lesbon, Elisangela Chicaroni Mattos, Heidge Fukumasu. USP-Botucatu: Rejane Maria Tommasini Grotto, Jayme A. Souza-Neto, Guilherme Targino Valente, Patricia Akemi Assato, Felipe Allan da Silva da Costa, Bianca Cechetto Carlos. Mendelics: Bibiana Santos, João Paulo Kitajima, Erika Freitas, David Schlesinger. Hemocentro Ribeirão Preto: Simone Kashima, Evandra Strazza Rodrigues, Svetoslav Nanev Slavov, Elaine Vieira dos Santos, Rafael dos Santos Bezerra, Luiz Carlos Junior de Alcantara, Marta Giovanetti, Vagner Fonseca, Flavia Aburjaile, Rodrigo Tocantins Calado. |
| EPI_ISL_1795234, EPI_ISL_1795235, EPI_ISL_1795236, EPI_ISL_1795237, EPI_ISL_1795238 | CS DE BALSAMO | Instituto Butantan / ESALQ-Piracicaba | Instituto Butantan: Alexander Roberto Precioso, Dimas Tadeu Covas, Sandra Coccuzzo Sampaio, Maria Carolina Elias, José Salvatore Leister Patané, Vincent Louis Viala, Antonio Jorge Martins, Ricardo Haddad, Claudia Renata dos Santos Barros, Elaine Cristina Marqueze, Raul Machado Neto, Debora Botequio Moretti. Centro de Genômica Funcional da ESALQ: Luiz Lehmann Coutinho, Ricardo Augusto Brassaloti, Raquel de Lello Rocha Campos Cassano. NGS Soluções Genômicas: Pilar Drummond Sampaio Corrêa Mariani. FZEA-USP Pirassununga: Mirele Daiana Poleti, Jessika Cristina Chagas Lesbon, Elisangela Chicaroni Mattos, Heidge Fukumasu. USP-Botucatu: Rejane Maria Tommasini Grotto, Jayme A. Souza-Neto, Guilherme Targino |

| | | | |
|---|---|---|---|
| | | | Valente, Patricia Akemi Assato, Felipe Allan da Silva da Costa, Bianca Cechetto Carlos. Mendelics: Bibiana Santos, João Paulo Kitajima, Erika Freitas, David Schlesinger. Hemocentro Ribeirão Preto: Simone Kashima, Evandra Strazza Rodrigues, Svetoslav Nanev Slavov, Elaine Vieira dos Santos, Rafael dos Santos Bezerra, Luiz Carlos Junior de Alcantara, Marta Giovanetti, Vagner Fonseca, Flavia Aburjaile, Rodrigo Tocantins Calado. |
| EPI_ISL_1795239 | CS DE URUPES | Instituto Butantan / ESALQ-Piracicaba | Instituto Butantan: Alexander Roberto Precioso, Dimas Tadeu Covas, Sandra Coccuzzo Sampaio, Maria Carolina Elias, José Salvatore Leister Patané, Vincent Louis Viala, Antonio Jorge Martins, Ricardo Haddad, Claudia Renata dos Santos Barros, Elaine Cristina Marqueze, Raul Machado Neto, Debora Botequio Moretti. Centro de Genômica Funcional da ESALQ: Luiz Lehmann Coutinho, Ricardo Augusto Brassaloti, Raquel de Lello Rocha Campos Cassano. NGS Soluções Genômicas: Pilar Drummond Sampaio Corrêa Mariani. FZEA-USP Pirassununga: Mirele Daiana Poleti, Jessika Cristina Chagas Lesbon, Elisangela Chicaroni Mattos, Heidge Fukumasu. USP-Botucatu: Rejane Maria Tommasini Grotto, Jayme A. Souza-Neto, Guilherme Targino Valente, Patricia Akemi Assato, Felipe Allan da Silva da Costa, Bianca Cechetto Carlos. Mendelics: Bibiana Santos, João Paulo Kitajima, Erika Freitas, David Schlesinger. Hemocentro Ribeirão Preto: Simone Kashima, Evandra Strazza Rodrigues, Svetoslav Nanev Slavov, Elaine Vieira dos Santos, Rafael dos Santos Bezerra, Luiz Carlos Junior de Alcantara, Marta Giovanetti, Vagner Fonseca, Flavia Aburjaile, Rodrigo Tocantins Calado. |
| EPI_ISL_1795240 | CS DE SEBASTIANOPOLIS DO SUL | Instituto Butantan / ESALQ-Piracicaba | Instituto Butantan: Alexander Roberto Precioso, Dimas Tadeu Covas, Sandra Coccuzzo Sampaio, Maria Carolina Elias, José Salvatore Leister Patané, Vincent Louis Viala, Antonio Jorge Martins, Ricardo Haddad, Claudia Renata dos Santos Barros, Elaine Cristina Marqueze, Raul Machado Neto, Debora Botequio Moretti. Centro de Genômica Funcional da ESALQ: Luiz Lehmann Coutinho, Ricardo Augusto Brassaloti, Raquel de Lello Rocha Campos Cassano. NGS Soluções Genômicas: Pilar Drummond Sampaio Corrêa Mariani. FZEA-USP Pirassununga: Mirele Daiana Poleti, Jessika Cristina Chagas Lesbon, Elisangela Chicaroni Mattos, Heidge Fukumasu. USP-Botucatu: Rejane Maria Tommasini Grotto, Jayme A. Souza-Neto, Guilherme Targino Valente, Patricia Akemi Assato, Felipe Allan da Silva da Costa, Bianca Cechetto Carlos. Mendelics: Bibiana Santos, João Paulo Kitajima, Erika Freitas, David Schlesinger. Hemocentro Ribeirão Preto: Simone Kashima, Evandra Strazza Rodrigues, Svetoslav Nanev Slavov, Elaine Vieira dos Santos, Rafael dos Santos Bezerra, Luiz Carlos Junior de Alcantara, Marta Giovanetti, Vagner Fonseca, Flavia Aburjaile, Rodrigo Tocantins Calado. |
| EPI_ISL_1795241 | CS DE BALSAMO | Instituto Butantan / ESALQ-Piracicaba | Instituto Butantan: Alexander Roberto Precioso, Dimas Tadeu Covas, Sandra Coccuzzo Sampaio, Maria Carolina Elias, José Salvatore Leister Patané, Vincent Louis Viala, Antonio Jorge Martins, Ricardo Haddad, Claudia Renata dos Santos Barros, Elaine Cristina Marqueze, Raul Machado Neto, Debora Botequio Moretti. Centro de Genômica Funcional da ESALQ: Luiz Lehmann Coutinho, Ricardo Augusto Brassaloti, Raquel de Lello Rocha Campos Cassano. NGS Soluções Genômicas: Pilar Drummond Sampaio Corrêa Mariani. FZEA-USP Pirassununga: Mirele Daiana Poleti, Jessika Cristina Chagas Lesbon, Elisangela Chicaroni Mattos, Heidge Fukumasu. USP-Botucatu: Rejane Maria Tommasini Grotto, Jayme A. Souza-Neto, Guilherme Targino Valente, Patricia Akemi Assato, Felipe Allan da Silva da Costa, Bianca Cechetto Carlos. Mendelics: Bibiana Santos, João Paulo Kitajima, Erika Freitas, David Schlesinger. Hemocentro Ribeirão Preto: Simone Kashima, Evandra Strazza Rodrigues, Svetoslav Nanev Slavov, Elaine Vieira dos Santos, Rafael dos Santos Bezerra, Luiz Carlos Junior de Alcantara, Marta Giovanetti, Vagner Fonseca, Flavia Aburjaile, Rodrigo Tocantins Calado. |
| EPI_ISL_1795242, EPI_ISL_1795243, EPI_ISL_1795244 | CS DE URUPES | Instituto Butantan / ESALQ-Piracicaba | Instituto Butantan: Alexander Roberto Precioso, Dimas Tadeu Covas, Sandra Coccuzzo Sampaio, Maria Carolina Elias, José Salvatore Leister Patané, Vincent Louis Viala, Antonio Jorge Martins, Ricardo Haddad, Claudia Renata dos Santos Barros, Elaine Cristina Marqueze, Raul Machado Neto, Debora Botequio Moretti. Centro de Genômica Funcional da ESALQ: Luiz Lehmann Coutinho, Ricardo Augusto Brassaloti, Raquel de Lello Rocha Campos Cassano. NGS Soluções Genômicas: Pilar Drummond Sampaio Corrêa Mariani. FZEA-USP Pirassununga: Mirele Daiana Poleti, Jessika Cristina Chagas Lesbon, Elisangela Chicaroni Mattos, Heidge Fukumasu. USP-Botucatu: Rejane Maria Tommasini Grotto, Jayme A. Souza-Neto, Guilherme Targino Valente, Patricia Akemi Assato, Felipe Allan da Silva da Costa, Bianca Cechetto Carlos. Mendelics: Bibiana Santos, João Paulo Kitajima, Erika Freitas, David Schlesinger. Hemocentro Ribeirão Preto: Simone Kashima, Evandra Strazza Rodrigues, Svetoslav Nanev Slavov, Elaine Vieira dos Santos, Rafael dos Santos Bezerra, Luiz Carlos Junior de Alcantara, Marta Giovanetti, Vagner Fonseca, Flavia Aburjaile, Rodrigo Tocantins Calado. |
| EPI_ISL_1795245, EPI_ISL_1795246, EPI_ISL_1795247 | EMERGENCIA RESPIRATORIA DE NOVA GRANADA | Instituto Butantan / ESALQ-Piracicaba | Instituto Butantan: Alexander Roberto Precioso, Dimas Tadeu Covas, Sandra Coccuzzo Sampaio, Maria Carolina Elias, José Salvatore Leister Patané, Vincent Louis Viala, Antonio Jorge Martins, Ricardo Haddad, Claudia Renata dos Santos Barros, Elaine Cristina Marqueze, Raul Machado Neto, Debora Botequio Moretti. Centro de Genômica Funcional da ESALQ: Luiz Lehmann Coutinho, Ricardo Augusto Brassaloti, Raquel de Lello Rocha Campos Cassano. NGS Soluções Genômicas: Pilar Drummond Sampaio Corrêa Mariani. FZEA-USP Pirassununga: Mirele Daiana Poleti, Jessika Cristina Chagas Lesbon, Elisangela Chicaroni Mattos, Heidge Fukumasu. USP-Botucatu: Rejane Maria Tommasini Grotto, Jayme A. Souza-Neto, Guilherme Targino Valente, Patricia Akemi Assato, Felipe Allan da Silva da Costa, Bianca Cechetto Carlos. Mendelics: Bibiana Santos, João Paulo Kitajima, Erika Freitas, David Schlesinger. Hemocentro Ribeirão Preto: Simone Kashima, Evandra Strazza Rodrigues, Svetoslav Nanev Slavov, Elaine Vieira dos Santos, Rafael dos Santos Bezerra, Luiz Carlos Junior de Alcantara, Marta Giovanetti, Vagner Fonseca, Flavia Aburjaile, Rodrigo Tocantins Calado. |
| EPI_ISL_1795248, EPI_ISL_1795249, EPI_ISL_1795250 | ESF NOVA TANABI II | Instituto Butantan / ESALQ-Piracicaba | Instituto Butantan: Alexander Roberto Precioso, Dimas Tadeu Covas, Sandra Coccuzzo Sampaio, Maria Carolina Elias, José Salvatore Leister Patané, Vincent Louis Viala, Antonio Jorge Martins, Ricardo Haddad, Claudia Renata dos Santos Barros, Elaine Cristina Marqueze, Raul Machado Neto, Debora Botequio Moretti. Centro de Genômica Funcional da ESALQ: Luiz Lehmann Coutinho, Ricardo Augusto Brassaloti, Raquel de Lello Rocha Campos Cassano. NGS Soluções Genômicas: Pilar Drummond Sampaio Corrêa Mariani. FZEA-USP Pirassununga: Mirele Daiana Poleti, Jessika Cristina Chagas Lesbon, Elisangela Chicaroni Mattos, Heidge Fukumasu. USP-Botucatu: Rejane Maria Tommasini Grotto, Jayme A. Souza-Neto, Guilherme Targino Valente, Patricia Akemi Assato, Felipe Allan da Silva da Costa, Bianca Cechetto Carlos. Mendelics: Bibiana Santos, João Paulo Kitajima, Erika Freitas, David Schlesinger. Hemocentro Ribeirão Preto: Simone Kashima, Evandra Strazza Rodrigues, Svetoslav Nanev Slavov, Elaine Vieira dos Santos, Rafael dos Santos Bezerra, Luiz Carlos Junior de Alcantara, Marta Giovanetti, Vagner Fonseca, Flavia Aburjaile, Rodrigo Tocantins Calado. |
| EPI_ISL_1795251, EPI_ISL_1795252, EPI_ISL_1795253, EPI_ISL_1795254, EPI_ISL_1795255, EPI_ISL_1795256, EPI_ISL_1795257 | UBS II DE TANABI MILTON MARTINS PERCHES | Instituto Butantan / ESALQ-Piracicaba | Instituto Butantan: Alexander Roberto Precioso, Dimas Tadeu Covas, Sandra Coccuzzo Sampaio, Maria Carolina Elias, José Salvatore Leister Patané, Vincent Louis Viala, Antonio Jorge Martins, Ricardo Haddad, Claudia Renata dos Santos Barros, Elaine Cristina Marqueze, Raul Machado Neto, Debora Botequio Moretti. Centro de Genômica Funcional da ESALQ: Luiz Lehmann Coutinho, Ricardo Augusto Brassaloti, Raquel de Lello Rocha Campos Cassano. NGS Soluções Genômicas: Pilar Drummond Sampaio Corrêa Mariani. FZEA-USP Pirassununga: Mirele Daiana Poleti, Jessika Cristina Chagas Lesbon, Elisangela Chicaroni Mattos, Heidge Fukumasu. USP-Botucatu: Rejane Maria Tommasini Grotto, Jayme A. Souza-Neto, Guilherme Targino Valente, Patricia Akemi Assato, Felipe Allan da Silva da Costa, Bianca Cechetto Carlos. Mendelics: Bibiana Santos, João Paulo Kitajima, Erika Freitas, David Schlesinger. Hemocentro Ribeirão Preto: Simone Kashima, Evandra Strazza Rodrigues, Svetoslav Nanev Slavov, Elaine Vieira dos Santos, Rafael dos Santos Bezerra, Luiz Carlos Junior de Alcantara, Marta Giovanetti, Vagner Fonseca, Flavia Aburjaile, Rodrigo Tocantins Calado. |
| EPI_ISL_1795258 | CS DE PLANALTO | Instituto Butantan / ESALQ-Piracicaba | Instituto Butantan: Alexander Roberto Precioso, Dimas Tadeu Covas, Sandra Coccuzzo Sampaio, Maria Carolina Elias, José Salvatore Leister Patané, Vincent Louis Viala, Antonio Jorge Martins, Ricardo Haddad, Claudia Renata dos Santos Barros, Elaine Cristina Marqueze, Raul Machado Neto, Debora Botequio Moretti. Centro de Genômica Funcional da ESALQ: Luiz Lehmann Coutinho, Ricardo Augusto Brassaloti, Raquel de Lello Rocha Campos Cassano. NGS Soluções Genômicas: Pilar Drummond Sampaio Corrêa Mariani. FZEA-USP Pirassununga: Mirele Daiana Poleti, Jessika Cristina Chagas Lesbon, Elisangela Chicaroni Mattos, Heidge Fukumasu. USP-Botucatu: Rejane Maria Tommasini Grotto, Jayme A. Souza-Neto, Guilherme Targino Valente, Patricia Akemi Assato, Felipe Allan da Silva da Costa, Bianca Cechetto Carlos. Mendelics: Bibiana Santos, João Paulo Kitajima, Erika Freitas, David Schlesinger. Hemocentro Ribeirão Preto: Simone Kashima, Evandra Strazza Rodrigues, Svetoslav Nanev Slavov, Elaine Vieira dos Santos, Rafael dos Santos Bezerra, Luiz Carlos Junior de Alcantara, Marta Giovanetti, Vagner Fonseca, Flavia Aburjaile, Rodrigo Tocantins Calado. |
| EPI_ISL_1795259, EPI_ISL_1795260, EPI_ISL_1795261, EPI_ISL_1795262, EPI_ISL_1795263 | CS DE ICEM | Instituto Butantan / ESALQ-Piracicaba | Instituto Butantan: Alexander Roberto Precioso, Dimas Tadeu Covas, Sandra Coccuzzo Sampaio, Maria Carolina Elias, José Salvatore Leister Patané, Vincent Louis Viala, Antonio Jorge Martins, Ricardo Haddad, Claudia Renata dos Santos Barros, Elaine Cristina Marqueze, Raul Machado Neto, Debora Botequio Moretti. Centro de Genômica Funcional da ESALQ: Luiz Lehmann Coutinho, Ricardo Augusto Brassaloti, Raquel de Lello Rocha Campos Cassano. NGS Soluções Genômicas: Pilar Drummond Sampaio Corrêa Mariani. FZEA-USP Pirassununga: Mirele Daiana Poleti, Jessika Cristina Chagas Lesbon, Elisangela Chicaroni Mattos, Heidge Fukumasu. USP-Botucatu: Rejane Maria Tommasini Grotto, Jayme A. Souza-Neto, Guilherme Targino Valente, Patricia Akemi Assato, Felipe Allan da Silva da Costa, Bianca Cechetto Carlos. Mendelics: Bibiana Santos, João Paulo Kitajima, Erika Freitas, David Schlesinger. Hemocentro Ribeirão Preto: Simone Kashima, Evandra Strazza Rodrigues, Svetoslav Nanev Slavov, Elaine Vieira dos Santos, Rafael dos Santos Bezerra, Luiz Carlos Junior de Alcantara, Marta Giovanetti, Vagner Fonseca, Flavia Aburjaile, Rodrigo Tocantins Calado. |
| EPI_ISL_1795264 | CS DE PARANAPUA | Instituto Butantan / ESALQ-Piracicaba | Instituto Butantan: Alexander Roberto Precioso, Dimas Tadeu Covas, Sandra Coccuzzo Sampaio, Maria Carolina Elias, José Salvatore Leister Patané, Vincent Louis Viala, Antonio Jorge Martins, Ricardo Haddad, Claudia Renata dos Santos Barros, Elaine Cristina Marqueze, Raul Machado Neto, Debora Botequio Moretti. Centro de Genômica Funcional da ESALQ: Luiz Lehmann Coutinho, Ricardo Augusto Brassaloti, Raquel de Lello Rocha Campos Cassano. NGS Soluções Genômicas: Pilar Drummond Sampaio Corrêa Mariani. FZEA-USP Pirassununga: Mirele Daiana Poleti, Jessika Cristina Chagas Lesbon, Elisangela Chicaroni Mattos, Heidge Fukumasu. USP-Botucatu: Rejane Maria Tommasini Grotto, Jayme A. Souza-Neto, Guilherme Targino |

| | | | |
|---|---|---|---|
| | | | Valente, Patricia Akemi Assato, Felipe Allan da Silva da Costa, Bianca Cechetto Carlos. Mendelics: Bibiana Santos, João Paulo Kitajima, Erika Freitas, David Schlesinger. Hemocentro Ribeirão Preto: Simone Kashima, Evandra Strazza Rodrigues, Svetoslav Nanev Slavov, Elaine Vieira dos Santos, Rafael dos Santos Bezerra, Luiz Carlos Junior de Alcantara, Marta Giovanetti, Vagner Fonseca, Flavia Aburjaile, Rodrigo Tocantins Calado. |
| EPI_ISL_1795265 | CS DE ICEM | Instituto Butantan / ESALQ-Piracicaba | Instituto Butantan: Alexander Roberto Precioso, Dimas Tadeu Covas, Sandra Coccuzzo Sampaio, Maria Carolina Elias, José Salvatore Leister Patané, Vincent Louis Viala, Antonio Jorge Martins, Ricardo Haddad, Claudia Renata dos Santos Barros, Elaine Cristina Marqueze, Raul Machado Neto, Debora Botequio Moretti. Centro de Genômica Funcional da ESALQ: Luiz Lehmann Coutinho, Ricardo Augusto Brassaloti, Raquel de Lello Rocha Campos Cassano. NGS Soluções Genômicas: Pilar Drummond Sampaio Corrêa Mariani. FZEA-USP Pirassununga: Mirele Daiana Poleti, Jessika Cristina Chagas Lesbon, Elisangela Chicaroni Mattos, Heidge Fukumasu. USP-Botucatu: Rejane Maria Tommasini Grotto, Jayme A. Souza-Neto, Guilherme Targino Valente, Patricia Akemi Assato, Felipe Allan da Silva da Costa, Bianca Cechetto Carlos. Mendelics: Bibiana Santos, João Paulo Kitajima, Erika Freitas, David Schlesinger. Hemocentro Ribeirão Preto: Simone Kashima, Evandra Strazza Rodrigues, Svetoslav Nanev Slavov, Elaine Vieira dos Santos, Rafael dos Santos Bezerra, Luiz Carlos Junior de Alcantara, Marta Giovanetti, Vagner Fonseca, Flavia Aburjaile, Rodrigo Tocantins Calado. |
| EPI_ISL_1795266, EPI_ISL_1795267, EPI_ISL_1795268 | UBS LUIS FACHIN IPIGUA | Instituto Butantan / ESALQ-Piracicaba | Instituto Butantan: Alexander Roberto Precioso, Dimas Tadeu Covas, Sandra Coccuzzo Sampaio, Maria Carolina Elias, José Salvatore Leister Patané, Vincent Louis Viala, Antonio Jorge Martins, Ricardo Haddad, Claudia Renata dos Santos Barros, Elaine Cristina Marqueze, Raul Machado Neto, Debora Botequio Moretti. Centro de Genômica Funcional da ESALQ: Luiz Lehmann Coutinho, Ricardo Augusto Brassaloti, Raquel de Lello Rocha Campos Cassano. NGS Soluções Genômicas: Pilar Drummond Sampaio Corrêa Mariani. FZEA-USP Pirassununga: Mirele Daiana Poleti, Jessika Cristina Chagas Lesbon, Elisangela Chicaroni Mattos, Heidge Fukumasu. USP-Botucatu: Rejane Maria Tommasini Grotto, Jayme A. Souza-Neto, Guilherme Targino Valente, Patricia Akemi Assato, Felipe Allan da Silva da Costa, Bianca Cechetto Carlos. Mendelics: Bibiana Santos, João Paulo Kitajima, Erika Freitas, David Schlesinger. Hemocentro Ribeirão Preto: Simone Kashima, Evandra Strazza Rodrigues, Svetoslav Nanev Slavov, Elaine Vieira dos Santos, Rafael dos Santos Bezerra, Luiz Carlos Junior de Alcantara, Marta Giovanetti, Vagner Fonseca, Flavia Aburjaile, Rodrigo Tocantins Calado. |
| EPI_ISL_1795269, EPI_ISL_1795270 | UBS TEREZA GALLO UCHOA | Instituto Butantan / ESALQ-Piracicaba | Instituto Butantan: Alexander Roberto Precioso, Dimas Tadeu Covas, Sandra Coccuzzo Sampaio, Maria Carolina Elias, José Salvatore Leister Patané, Vincent Louis Viala, Antonio Jorge Martins, Ricardo Haddad, Claudia Renata dos Santos Barros, Elaine Cristina Marqueze, Raul Machado Neto, Debora Botequio Moretti. Centro de Genômica Funcional da ESALQ: Luiz Lehmann Coutinho, Ricardo Augusto Brassaloti, Raquel de Lello Rocha Campos Cassano. NGS Soluções Genômicas: Pilar Drummond Sampaio Corrêa Mariani. FZEA-USP Pirassununga: Mirele Daiana Poleti, Jessika Cristina Chagas Lesbon, Elisangela Chicaroni Mattos, Heidge Fukumasu. USP-Botucatu: Rejane Maria Tommasini Grotto, Jayme A. Souza-Neto, Guilherme Targino Valente, Patricia Akemi Assato, Felipe Allan da Silva da Costa, Bianca Cechetto Carlos. Mendelics: Bibiana Santos, João Paulo Kitajima, Erika Freitas, David Schlesinger. Hemocentro Ribeirão Preto: Simone Kashima, Evandra Strazza Rodrigues, Svetoslav Nanev Slavov, Elaine Vieira dos Santos, Rafael dos Santos Bezerra, Luiz Carlos Junior de Alcantara, Marta Giovanetti, Vagner Fonseca, Flavia Aburjaile, Rodrigo Tocantins Calado. |
| EPI_ISL_1795271 | CS DE PARANAPUA | Instituto Butantan / ESALQ-Piracicaba | Instituto Butantan: Alexander Roberto Precioso, Dimas Tadeu Covas, Sandra Coccuzzo Sampaio, Maria Carolina Elias, José Salvatore Leister Patané, Vincent Louis Viala, Antonio Jorge Martins, Ricardo Haddad, Claudia Renata dos Santos Barros, Elaine Cristina Marqueze, Raul Machado Neto, Debora Botequio Moretti. Centro de Genômica Funcional da ESALQ: Luiz Lehmann Coutinho, Ricardo Augusto Brassaloti, Raquel de Lello Rocha Campos Cassano. NGS Soluções Genômicas: Pilar Drummond Sampaio Corrêa Mariani. FZEA-USP Pirassununga: Mirele Daiana Poleti, Jessika Cristina Chagas Lesbon, Elisangela Chicaroni Mattos, Heidge Fukumasu. USP-Botucatu: Rejane Maria Tommasini Grotto, Jayme A. Souza-Neto, Guilherme Targino Valente, Patricia Akemi Assato, Felipe Allan da Silva da Costa, Bianca Cechetto Carlos. Mendelics: Bibiana Santos, João Paulo Kitajima, Erika Freitas, David Schlesinger. Hemocentro Ribeirão Preto: Simone Kashima, Evandra Strazza Rodrigues, Svetoslav Nanev Slavov, Elaine Vieira dos Santos, Rafael dos Santos Bezerra, Luiz Carlos Junior de Alcantara, Marta Giovanetti, Vagner Fonseca, Flavia Aburjaile, Rodrigo Tocantins Calado. |
| EPI_ISL_1795272, EPI_ISL_1795273 | UBS TEREZA GALLO UCHOA | Instituto Butantan / ESALQ-Piracicaba | Instituto Butantan: Alexander Roberto Precioso, Dimas Tadeu Covas, Sandra Coccuzzo Sampaio, Maria Carolina Elias, José Salvatore Leister Patané, Vincent Louis Viala, Antonio Jorge Martins, Ricardo Haddad, Claudia Renata dos Santos Barros, Elaine Cristina Marqueze, Raul Machado Neto, Debora Botequio Moretti. Centro de Genômica Funcional da ESALQ: Luiz Lehmann Coutinho, Ricardo Augusto Brassaloti, Raquel de Lello Rocha Campos Cassano. NGS Soluções Genômicas: Pilar Drummond Sampaio Corrêa Mariani. FZEA-USP Pirassununga: Mirele Daiana Poleti, Jessika Cristina Chagas Lesbon, Elisangela Chicaroni Mattos, Heidge Fukumasu. USP-Botucatu: Rejane Maria Tommasini Grotto, Jayme A. Souza-Neto, Guilherme Targino Valente, Patricia Akemi Assato, Felipe Allan da Silva da Costa, Bianca Cechetto Carlos. Mendelics: Bibiana Santos, João Paulo Kitajima, Erika Freitas, David Schlesinger. Hemocentro Ribeirão Preto: Simone Kashima, Evandra Strazza Rodrigues, Svetoslav Nanev Slavov, Elaine Vieira dos Santos, Rafael dos Santos Bezerra, Luiz Carlos Junior de Alcantara, Marta Giovanetti, Vagner Fonseca, Flavia Aburjaile, Rodrigo Tocantins Calado. |
| EPI_ISL_1795274, EPI_ISL_1795275 | CENTRO MEDICO PMESP | Instituto Butantan / ESALQ-Piracicaba | Instituto Butantan: Alexander Roberto Precioso, Dimas Tadeu Covas, Sandra Coccuzzo Sampaio, Maria Carolina Elias, José Salvatore Leister Patané, Vincent Louis Viala, Antonio Jorge Martins, Ricardo Haddad, Claudia Renata dos Santos Barros, Elaine Cristina Marqueze, Raul Machado Neto, Debora Botequio Moretti. Centro de Genômica Funcional da ESALQ: Luiz Lehmann Coutinho, Ricardo Augusto Brassaloti, Raquel de Lello Rocha Campos Cassano. NGS Soluções Genômicas: Pilar Drummond Sampaio Corrêa Mariani. FZEA-USP Pirassununga: Mirele Daiana Poleti, Jessika Cristina Chagas Lesbon, Elisangela Chicaroni Mattos, Heidge Fukumasu. USP-Botucatu: Rejane Maria Tommasini Grotto, Jayme A. Souza-Neto, Guilherme Targino Valente, Patricia Akemi Assato, Felipe Allan da Silva da Costa, Bianca Cechetto Carlos. Mendelics: Bibiana Santos, João Paulo Kitajima, Erika Freitas, David Schlesinger. Hemocentro Ribeirão Preto: Simone Kashima, Evandra Strazza Rodrigues, Svetoslav Nanev Slavov, Elaine Vieira dos Santos, Rafael dos Santos Bezerra, Luiz Carlos Junior de Alcantara, Marta Giovanetti, Vagner Fonseca, Flavia Aburjaile, Rodrigo Tocantins Calado. |
| EPI_ISL_1795276, EPI_ISL_1795277, EPI_ISL_1795278, EPI_ISL_1795279, EPI_ISL_1795280, EPI_ISL_1795281, EPI_ISL_1795282, EPI_ISL_1795283, EPI_ISL_1795284, EPI_ISL_1795285, EPI_ISL_1795286 | | | |
| see above | CS DE PALESTINA | Instituto Butantan / ESALQ-Piracicaba | Instituto Butantan: Alexander Roberto Precioso, Dimas Tadeu Covas, Sandra Coccuzzo Sampaio, Maria Carolina Elias, José Salvatore Leister Patané, Vincent Louis Viala, Antonio Jorge Martins, Ricardo Haddad, Claudia Renata dos Santos Barros, Elaine Cristina Marqueze, Raul Machado Neto, Debora Botequio Moretti. Centro de Genômica Funcional da ESALQ: Luiz Lehmann Coutinho, Ricardo Augusto Brassaloti, Raquel de Lello Rocha Campos Cassano. NGS Soluções Genômicas: Pilar Drummond Sampaio Corrêa Mariani. FZEA-USP Pirassununga: Mirele Daiana Poleti, Jessika Cristina Chagas Lesbon, Elisangela Chicaroni Mattos, Heidge Fukumasu. USP-Botucatu: Rejane Maria Tommasini Grotto, Jayme A. Souza-Neto, Guilherme Targino Valente, Patricia Akemi Assato, Felipe Allan da Silva da Costa, Bianca Cechetto Carlos. Mendelics: Bibiana Santos, João Paulo Kitajima, Erika Freitas, David Schlesinger. Hemocentro Ribeirão Preto: Simone Kashima, Evandra Strazza Rodrigues, Svetoslav Nanev Slavov, Elaine Vieira dos Santos, Rafael dos Santos Bezerra, Luiz Carlos Junior de Alcantara, Marta Giovanetti, Vagner Fonseca, Flavia Aburjaile, Rodrigo Tocantins Calado. |
| EPI_ISL_1795287 | UBS ALCIMINIO DE ASSIS LOURENCO BADY BASSITT | Instituto Butantan / ESALQ-Piracicaba | Instituto Butantan: Alexander Roberto Precioso, Dimas Tadeu Covas, Sandra Coccuzzo Sampaio, Maria Carolina Elias, José Salvatore Leister Patané, Vincent Louis Viala, Antonio Jorge Martins, Ricardo Haddad, Claudia Renata dos Santos Barros, Elaine Cristina Marqueze, Raul Machado Neto, Debora Botequio Moretti. Centro de Genômica Funcional da ESALQ: Luiz Lehmann Coutinho, Ricardo Augusto Brassaloti, Raquel de Lello Rocha Campos Cassano. NGS Soluções Genômicas: Pilar Drummond Sampaio Corrêa Mariani. FZEA-USP Pirassununga: Mirele Daiana Poleti, Jessika Cristina Chagas Lesbon, Elisangela Chicaroni Mattos, Heidge Fukumasu. USP-Botucatu: Rejane Maria Tommasini Grotto, Jayme A. Souza-Neto, Guilherme Targino Valente, Patricia Akemi Assato, Felipe Allan da Silva da Costa, Bianca Cechetto Carlos. Mendelics: Bibiana Santos, João Paulo Kitajima, Erika Freitas, David Schlesinger. Hemocentro Ribeirão Preto: Simone Kashima, Evandra Strazza Rodrigues, Svetoslav Nanev Slavov, Elaine Vieira dos Santos, Rafael dos Santos Bezerra, Luiz Carlos Junior de Alcantara, Marta Giovanetti, Vagner Fonseca, Flavia Aburjaile, Rodrigo Tocantins Calado. |
| EPI_ISL_1795288 | CS DE NIPOA | Instituto Butantan / ESALQ-Piracicaba | Instituto Butantan: Alexander Roberto Precioso, Dimas Tadeu Covas, Sandra Coccuzzo Sampaio, Maria Carolina Elias, José Salvatore Leister Patané, Vincent Louis Viala, Antonio Jorge Martins, Ricardo Haddad, Claudia Renata dos Santos Barros, Elaine Cristina Marqueze, Raul Machado Neto, Debora Botequio Moretti. Centro de Genômica Funcional da ESALQ: Luiz Lehmann Coutinho, Ricardo Augusto Brassaloti, Raquel de Lello Rocha Campos Cassano. NGS Soluções Genômicas: Pilar Drummond Sampaio Corrêa Mariani. FZEA-USP Pirassununga: Mirele Daiana Poleti, Jessika Cristina Chagas Lesbon, Elisangela Chicaroni Mattos, Heidge Fukumasu. USP-Botucatu: Rejane Maria Tommasini Grotto, Jayme A. Souza-Neto, Guilherme Targino Valente, Patricia Akemi Assato, Felipe Allan da Silva da Costa, Bianca Cechetto Carlos. Mendelics: Bibiana Santos, João Paulo Kitajima, Erika Freitas, David Schlesinger. Hemocentro Ribeirão Preto: Simone Kashima, Evandra Strazza Rodrigues, Svetoslav Nanev Slavov, Elaine Vieira dos Santos, Rafael dos Santos Bezerra, Luiz Carlos Junior de Alcantara, Marta Giovanetti, Vagner Fonseca, Flavia Aburjaile, Rodrigo Tocantins Calado. |
| EPI_ISL_1795289 | UBS ALCIMINIO DE ASSIS LOURENCO BADY BASSITT | Instituto Butantan / ESALQ-Piracicaba | Instituto Butantan: Alexander Roberto Precioso, Dimas Tadeu Covas, Sandra Coccuzzo Sampaio, Maria Carolina Elias, José Salvatore Leister Patané, Vincent Louis Viala, Antonio Jorge Martins, Ricardo Haddad, Claudia Renata dos Santos Barros, Elaine Cristina Marqueze, Raul Machado Neto, Debora Botequio Moretti. Centro de Genômica Funcional da ESALQ: Luiz Lehmann Coutinho, Ricardo Augusto Brassaloti, Raquel de Lello Rocha Campos Cassano. NGS Soluções Genômicas: Pilar Drummond Sampaio Corrêa Mariani. FZEA-USP Pirassununga: Mirele Daiana Poleti, Jessika Cristina Chagas Lesbon, Elisangela Chicaroni Mattos, Heidge Fukumasu. USP-Botucatu: Rejane Maria Tommasini Grotto, Jayme A. Souza-Neto, Guilherme Targino Valente, Patricia Akemi Assato, Felipe Allan da Silva da Costa, Bianca Cechetto Carlos. Mendelics: Bibiana Santos, João Paulo Kitajima, Erika Freitas, |

| | | | |
|---|---|---|---|
| | | | David Schlesinger. Hemocentro Ribeirão Preto: Simone Kashima, Evandra Strazza Rodrigues, Svetoslav Nanev Slavov, Elaine Vieira dos Santos, Rafael dos Santos Bezerra, Luiz Carlos Junior de Alcantara, Marta Giovanetti, Vagner Fonseca, Flavia Aburjaile, Rodrigo Tocantins Calado. |
| EPI_ISL_1795290 | USF EUCALIPTOS | Instituto Butantan / ESALQ-Piracicaba | Instituto Butantan: Alexander Roberto Precioso, Dimas Tadeu Covas, Sandra Coccuzzo Sampaio, Maria Carolina Elias, José Salvatore Leister Patané, Vincent Louis Viala, Antonio Jorge Martins, Ricardo Haddad, Claudia Renata dos Santos Barros, Elaine Cristina Marqueze, Raul Machado Neto, Debora Botequio Moretti. Centro de Genômica Funcional da ESALQ: Luiz Lehmann Coutinho, Ricardo Augusto Brassaloti, Raquel de Lello Rocha Campos Cassano. NGS Soluções Genômicas: Pilar Drummond Sampaio Corrêa Mariani. FZEA-USP Pirassununga: Mirele Daiana Poleti, Jessika Cristina Chagas Lesbon, Elisangela Chicaroni Mattos, Heidge Fukumasu. USP-Botucatu: Rejane Maria Tommasini Grotto, Jayme A. Souza-Neto, Guilherme Targino Valente, Patricia Akemi Assato, Felipe Allan da Silva da Costa, Bianca Cechetto Carlos. Mendelics: Bibiana Santos, João Paulo Kitajima, Erika Freitas, David Schlesinger. Hemocentro Ribeirão Preto: Simone Kashima, Evandra Strazza Rodrigues, Svetoslav Nanev Slavov, Elaine Vieira dos Santos, Rafael dos Santos Bezerra, Luiz Carlos Junior de Alcantara, Marta Giovanetti, Vagner Fonseca, Flavia Aburjaile, Rodrigo Tocantins Calado. |
| EPI_ISL_1795291, EPI_ISL_1795292 | AMBULATORIO DE ESPECIALIDADE V E MOGI MIRIM | Instituto Butantan / ESALQ-Piracicaba | Instituto Butantan: Alexander Roberto Precioso, Dimas Tadeu Covas, Sandra Coccuzzo Sampaio, Maria Carolina Elias, José Salvatore Leister Patané, Vincent Louis Viala, Antonio Jorge Martins, Ricardo Haddad, Claudia Renata dos Santos Barros, Elaine Cristina Marqueze, Raul Machado Neto, Debora Botequio Moretti. Centro de Genômica Funcional da ESALQ: Luiz Lehmann Coutinho, Ricardo Augusto Brassaloti, Raquel de Lello Rocha Campos Cassano. NGS Soluções Genômicas: Pilar Drummond Sampaio Corrêa Mariani. FZEA-USP Pirassununga: Mirele Daiana Poleti, Jessika Cristina Chagas Lesbon, Elisangela Chicaroni Mattos, Heidge Fukumasu. USP-Botucatu: Rejane Maria Tommasini Grotto, Jayme A. Souza-Neto, Guilherme Targino Valente, Patricia Akemi Assato, Felipe Allan da Silva da Costa, Bianca Cechetto Carlos. Mendelics: Bibiana Santos, João Paulo Kitajima, Erika Freitas, David Schlesinger. Hemocentro Ribeirão Preto: Simone Kashima, Evandra Strazza Rodrigues, Svetoslav Nanev Slavov, Elaine Vieira dos Santos, Rafael dos Santos Bezerra, Luiz Carlos Junior de Alcantara, Marta Giovanetti, Vagner Fonseca, Flavia Aburjaile, Rodrigo Tocantins Calado. |
| EPI_ISL_1795293, EPI_ISL_1795294 | UNIDADE DE SAUDE DE ITOBI ALCIBIADES PIRES | Instituto Butantan / ESALQ-Piracicaba | Instituto Butantan: Alexander Roberto Precioso, Dimas Tadeu Covas, Sandra Coccuzzo Sampaio, Maria Carolina Elias, José Salvatore Leister Patané, Vincent Louis Viala, Antonio Jorge Martins, Ricardo Haddad, Claudia Renata dos Santos Barros, Elaine Cristina Marqueze, Raul Machado Neto, Debora Botequio Moretti. Centro de Genômica Funcional da ESALQ: Luiz Lehmann Coutinho, Ricardo Augusto Brassaloti, Raquel de Lello Rocha Campos Cassano. NGS Soluções Genômicas: Pilar Drummond Sampaio Corrêa Mariani. FZEA-USP Pirassununga: Mirele Daiana Poleti, Jessika Cristina Chagas Lesbon, Elisangela Chicaroni Mattos, Heidge Fukumasu. USP-Botucatu: Rejane Maria Tommasini Grotto, Jayme A. Souza-Neto, Guilherme Targino Valente, Patricia Akemi Assato, Felipe Allan da Silva da Costa, Bianca Cechetto Carlos. Mendelics: Bibiana Santos, João Paulo Kitajima, Erika Freitas, David Schlesinger. Hemocentro Ribeirão Preto: Simone Kashima, Evandra Strazza Rodrigues, Svetoslav Nanev Slavov, Elaine Vieira dos Santos, Rafael dos Santos Bezerra, Luiz Carlos Junior de Alcantara, Marta Giovanetti, Vagner Fonseca, Flavia Aburjaile, Rodrigo Tocantins Calado. |
| EPI_ISL_1795295 | USF GUACUANO | Instituto Butantan / ESALQ-Piracicaba | Instituto Butantan: Alexander Roberto Precioso, Dimas Tadeu Covas, Sandra Coccuzzo Sampaio, Maria Carolina Elias, José Salvatore Leister Patané, Vincent Louis Viala, Antonio Jorge Martins, Ricardo Haddad, Claudia Renata dos Santos Barros, Elaine Cristina Marqueze, Raul Machado Neto, Debora Botequio Moretti. Centro de Genômica Funcional da ESALQ: Luiz Lehmann Coutinho, Ricardo Augusto Brassaloti, Raquel de Lello Rocha Campos Cassano. NGS Soluções Genômicas: Pilar Drummond Sampaio Corrêa Mariani. FZEA-USP Pirassununga: Mirele Daiana Poleti, Jessika Cristina Chagas Lesbon, Elisangela Chicaroni Mattos, Heidge Fukumasu. USP-Botucatu: Rejane Maria Tommasini Grotto, Jayme A. Souza-Neto, Guilherme Targino Valente, Patricia Akemi Assato, Felipe Allan da Silva da Costa, Bianca Cechetto Carlos. Mendelics: Bibiana Santos, João Paulo Kitajima, Erika Freitas, David Schlesinger. Hemocentro Ribeirão Preto: Simone Kashima, Evandra Strazza Rodrigues, Svetoslav Nanev Slavov, Elaine Vieira dos Santos, Rafael dos Santos Bezerra, Luiz Carlos Junior de Alcantara, Marta Giovanetti, Vagner Fonseca, Flavia Aburjaile, Rodrigo Tocantins Calado. |
| EPI_ISL_1795296, EPI_ISL_1795297 | CENTRO DE SAUDE III AFFONSO LUZZI SANTA CRUZ DAS PALMEIRAS | Instituto Butantan / ESALQ-Piracicaba | Instituto Butantan: Alexander Roberto Precioso, Dimas Tadeu Covas, Sandra Coccuzzo Sampaio, Maria Carolina Elias, José Salvatore Leister Patané, Vincent Louis Viala, Antonio Jorge Martins, Ricardo Haddad, Claudia Renata dos Santos Barros, Elaine Cristina Marqueze, Raul Machado Neto, Debora Botequio Moretti. Centro de Genômica Funcional da ESALQ: Luiz Lehmann Coutinho, Ricardo Augusto Brassaloti, Raquel de Lello Rocha Campos Cassano. NGS Soluções Genômicas: Pilar Drummond Sampaio Corrêa Mariani. FZEA-USP Pirassununga: Mirele Daiana Poleti, Jessika Cristina Chagas Lesbon, Elisangela Chicaroni Mattos, Heidge Fukumasu. USP-Botucatu: Rejane Maria Tommasini Grotto, Jayme A. Souza-Neto, Guilherme Targino Valente, Patricia Akemi Assato, Felipe Allan da Silva da Costa, Bianca Cechetto Carlos. Mendelics: Bibiana Santos, João Paulo Kitajima, Erika Freitas, David Schlesinger. Hemocentro Ribeirão Preto: Simone Kashima, Evandra Strazza Rodrigues, Svetoslav Nanev Slavov, Elaine Vieira dos Santos, Rafael dos Santos Bezerra, Luiz Carlos Junior de Alcantara, Marta Giovanetti, Vagner Fonseca, Flavia Aburjaile, Rodrigo Tocantins Calado. |
| EPI_ISL_1795298 | UNIDADE DE SAUDE DE ITOBI ALCIBIADES PIRES | Instituto Butantan / ESALQ-Piracicaba | Instituto Butantan: Alexander Roberto Precioso, Dimas Tadeu Covas, Sandra Coccuzzo Sampaio, Maria Carolina Elias, José Salvatore Leister Patané, Vincent Louis Viala, Antonio Jorge Martins, Ricardo Haddad, Claudia Renata dos Santos Barros, Elaine Cristina Marqueze, Raul Machado Neto, Debora Botequio Moretti. Centro de Genômica Funcional da ESALQ: Luiz Lehmann Coutinho, Ricardo Augusto Brassaloti, Raquel de Lello Rocha Campos Cassano. NGS Soluções Genômicas: Pilar Drummond Sampaio Corrêa Mariani. FZEA-USP Pirassununga: Mirele Daiana Poleti, Jessika Cristina Chagas Lesbon, Elisangela Chicaroni Mattos, Heidge Fukumasu. USP-Botucatu: Rejane Maria Tommasini Grotto, Jayme A. Souza-Neto, Guilherme Targino Valente, Patricia Akemi Assato, Felipe Allan da Silva da Costa, Bianca Cechetto Carlos. Mendelics: Bibiana Santos, João Paulo Kitajima, Erika Freitas, David Schlesinger. Hemocentro Ribeirão Preto: Simone Kashima, Evandra Strazza Rodrigues, Svetoslav Nanev Slavov, Elaine Vieira dos Santos, Rafael dos Santos Bezerra, Luiz Carlos Junior de Alcantara, Marta Giovanetti, Vagner Fonseca, Flavia Aburjaile, Rodrigo Tocantins Calado. |
| EPI_ISL_1795299 | CENTRO DE SAUDE III AFFONSO LUZZI SANTA CRUZ DAS PALMEIRAS | Instituto Butantan / ESALQ-Piracicaba | Instituto Butantan: Alexander Roberto Precioso, Dimas Tadeu Covas, Sandra Coccuzzo Sampaio, Maria Carolina Elias, José Salvatore Leister Patané, Vincent Louis Viala, Antonio Jorge Martins, Ricardo Haddad, Claudia Renata dos Santos Barros, Elaine Cristina Marqueze, Raul Machado Neto, Debora Botequio Moretti. Centro de Genômica Funcional da ESALQ: Luiz Lehmann Coutinho, Ricardo Augusto Brassaloti, Raquel de Lello Rocha Campos Cassano. NGS Soluções Genômicas: Pilar Drummond Sampaio Corrêa Mariani. FZEA-USP Pirassununga: Mirele Daiana Poleti, Jessika Cristina Chagas Lesbon, Elisangela Chicaroni Mattos, Heidge Fukumasu. USP-Botucatu: Rejane Maria Tommasini Grotto, Jayme A. Souza-Neto, Guilherme Targino Valente, Patricia Akemi Assato, Felipe Allan da Silva da Costa, Bianca Cechetto Carlos. Mendelics: Bibiana Santos, João Paulo Kitajima, Erika Freitas, David Schlesinger. Hemocentro Ribeirão Preto: Simone Kashima, Evandra Strazza Rodrigues, Svetoslav Nanev Slavov, Elaine Vieira dos Santos, Rafael dos Santos Bezerra, Luiz Carlos Junior de Alcantara, Marta Giovanetti, Vagner Fonseca, Flavia Aburjaile, Rodrigo Tocantins Calado. |
| EPI_ISL_1795300, EPI_ISL_1795301, EPI_ISL_1795302 | PRONTO SOCORRO MUNICIPAL TAMBAU | Instituto Butantan / ESALQ-Piracicaba | Instituto Butantan: Alexander Roberto Precioso, Dimas Tadeu Covas, Sandra Coccuzzo Sampaio, Maria Carolina Elias, José Salvatore Leister Patané, Vincent Louis Viala, Antonio Jorge Martins, Ricardo Haddad, Claudia Renata dos Santos Barros, Elaine Cristina Marqueze, Raul Machado Neto, Debora Botequio Moretti. Centro de Genômica Funcional da ESALQ: Luiz Lehmann Coutinho, Ricardo Augusto Brassaloti, Raquel de Lello Rocha Campos Cassano. NGS Soluções Genômicas: Pilar Drummond Sampaio Corrêa Mariani. FZEA-USP Pirassununga: Mirele Daiana Poleti, Jessika Cristina Chagas Lesbon, Elisangela Chicaroni Mattos, Heidge Fukumasu. USP-Botucatu: Rejane Maria Tommasini Grotto, Jayme A. Souza-Neto, Guilherme Targino Valente, Patricia Akemi Assato, Felipe Allan da Silva da Costa, Bianca Cechetto Carlos. Mendelics: Bibiana Santos, João Paulo Kitajima, Erika Freitas, David Schlesinger. Hemocentro Ribeirão Preto: Simone Kashima, Evandra Strazza Rodrigues, Svetoslav Nanev Slavov, Elaine Vieira dos Santos, Rafael dos Santos Bezerra, Luiz Carlos Junior de Alcantara, Marta Giovanetti, Vagner Fonseca, Flavia Aburjaile, Rodrigo Tocantins Calado. |
| EPI_ISL_1795303 | LABORATORIO DE FRANCA | Instituto Butantan / ESALQ-Piracicaba | Instituto Butantan: Alexander Roberto Precioso, Dimas Tadeu Covas, Sandra Coccuzzo Sampaio, Maria Carolina Elias, José Salvatore Leister Patané, Vincent Louis Viala, Antonio Jorge Martins, Ricardo Haddad, Claudia Renata dos Santos Barros, Elaine Cristina Marqueze, Raul Machado Neto, Debora Botequio Moretti. Centro de Genômica Funcional da ESALQ: Luiz Lehmann Coutinho, Ricardo Augusto Brassaloti, Raquel de Lello Rocha Campos Cassano. NGS Soluções Genômicas: Pilar Drummond Sampaio Corrêa Mariani. FZEA-USP Pirassununga: Mirele Daiana Poleti, Jessika Cristina Chagas Lesbon, Elisangela Chicaroni Mattos, Heidge Fukumasu. USP-Botucatu: Rejane Maria Tommasini Grotto, Jayme A. Souza-Neto, Guilherme Targino Valente, Patricia Akemi Assato, Felipe Allan da Silva da Costa, Bianca Cechetto Carlos. Mendelics: Bibiana Santos, João Paulo Kitajima, Erika Freitas, David Schlesinger. Hemocentro Ribeirão Preto: Simone Kashima, Evandra Strazza Rodrigues, Svetoslav Nanev Slavov, Elaine Vieira dos Santos, Rafael dos Santos Bezerra, Luiz Carlos Junior de Alcantara, Marta Giovanetti, Vagner Fonseca, Flavia Aburjaile, Rodrigo Tocantins Calado. |
| EPI_ISL_1795304 | PRONTO SOCORRO MUNICIPAL TAMBAU | Instituto Butantan / ESALQ-Piracicaba | Instituto Butantan: Alexander Roberto Precioso, Dimas Tadeu Covas, Sandra Coccuzzo Sampaio, Maria Carolina Elias, José Salvatore Leister Patané, Vincent Louis Viala, Antonio Jorge Martins, Ricardo Haddad, Claudia Renata dos Santos Barros, Elaine Cristina Marqueze, Raul Machado Neto, Debora Botequio Moretti. Centro de Genômica Funcional da ESALQ: Luiz Lehmann Coutinho, Ricardo Augusto Brassaloti, Raquel de Lello Rocha Campos Cassano. NGS Soluções Genômicas: Pilar Drummond Sampaio Corrêa Mariani. FZEA-USP Pirassununga: Mirele Daiana Poleti, Jessika Cristina Chagas Lesbon, Elisangela Chicaroni Mattos, Heidge Fukumasu. USP-Botucatu: Rejane Maria Tommasini Grotto, Jayme A. Souza-Neto, Guilherme Targino Valente, Patricia Akemi Assato, Felipe Allan da Silva da Costa, Bianca Cechetto Carlos. Mendelics: Bibiana Santos, João Paulo Kitajima, Erika Freitas, David Schlesinger. Hemocentro Ribeirão Preto: Simone Kashima, Evandra Strazza Rodrigues, Svetoslav Nanev Slavov, Elaine Vieira dos Santos, Rafael dos Santos Bezerra, Luiz Carlos Junior de Alcantara, Marta Giovanetti, Vagner Fonseca, Flavia Aburjaile, Rodrigo Tocantins Calado. |

| | | | |
|---|---|---|---|
| EPI_ISL_1795305 | UNIDADE DA SAUDE DO ADULTO CASA BRANCA PREFEITURA | Instituto Butantan / ESALQ-Piracicaba | Instituto Butantan: Alexander Roberto Precioso, Dimas Tadeu Covas, Sandra Coccuzzo Sampaio, Maria Carolina Elias, José Salvatore Leister Patané, Vincent Louis Viala, Antonio Jorge Martins, Ricardo Haddad, Claudia Renata dos Santos Barros, Elaine Cristina Marqueze, Raul Machado Neto, Debora Botequio Moretti. Centro de Genômica Funcional da ESALQ: Luiz Lehmann Coutinho, Ricardo Augusto Brassaloti, Raquel de Lello Rocha Campos Cassano. NGS Soluções Genômicas: Pilar Drummond Sampaio Corrêa Mariani. FZEA-USP Pirassununga: Mirele Daiana Poleti, Jessika Cristina Chagas Lesbon, Elisangela Chicaroni Mattos, Heidge Fukumasu. USP-Botucatu: Rejane Maria Tommasini Grotto, Jayme A. Souza-Neto, Guilherme Targino Valente, Patricia Akemi Assato, Felipe Allan da Silva da Costa, Bianca Cechetto Carlos. Mendelics: Bibiana Santos, João Paulo Kitajima, Erika Freitas, David Schlesinger. Hemocentro Ribeirão Preto: Simone Kashima, Evandra Strazza Rodrigues, Svetoslav Nanev Slavov, Elaine Vieira dos Santos, Rafael dos Santos Bezerra, Luiz Carlos Junior de Alcantara, Marta Giovanetti, Vagner Fonseca, Flavia Aburjaile, Rodrigo Tocantins Calado. |
| EPI_ISL_1795306, EPI_ISL_1795307, EPI_ISL_1795308, EPI_ISL_1795309, EPI_ISL_1795310, EPI_ISL_1795311, EPI_ISL_1795312, EPI_ISL_1795313, EPI_ISL_1795314, EPI_ISL_1795315, EPI_ISL_1795316, EPI_ISL_1795317, EPI_ISL_1795318, EPI_ISL_1795319, EPI_ISL_1795320, EPI_ISL_1795321, EPI_ISL_1795322, EPI_ISL_1795323 | | | |
| see above | LABORATORIO DE FRANCA | Instituto Butantan / ESALQ-Piracicaba | Instituto Butantan: Alexander Roberto Precioso, Dimas Tadeu Covas, Sandra Coccuzzo Sampaio, Maria Carolina Elias, José Salvatore Leister Patané, Vincent Louis Viala, Antonio Jorge Martins, Ricardo Haddad, Claudia Renata dos Santos Barros, Elaine Cristina Marqueze, Raul Machado Neto, Debora Botequio Moretti. Centro de Genômica Funcional da ESALQ: Luiz Lehmann Coutinho, Ricardo Augusto Brassaloti, Raquel de Lello Rocha Campos Cassano. NGS Soluções Genômicas: Pilar Drummond Sampaio Corrêa Mariani. FZEA-USP Pirassununga: Mirele Daiana Poleti, Jessika Cristina Chagas Lesbon, Elisangela Chicaroni Mattos, Heidge Fukumasu. USP-Botucatu: Rejane Maria Tommasini Grotto, Jayme A. Souza-Neto, Guilherme Targino Valente, Patricia Akemi Assato, Felipe Allan da Silva da Costa, Bianca Cechetto Carlos. Mendelics: Bibiana Santos, João Paulo Kitajima, Erika Freitas, David Schlesinger. Hemocentro Ribeirão Preto: Simone Kashima, Evandra Strazza Rodrigues, Svetoslav Nanev Slavov, Elaine Vieira dos Santos, Rafael dos Santos Bezerra, Luiz Carlos Junior de Alcantara, Marta Giovanetti, Vagner Fonseca, Flavia Aburjaile, Rodrigo Tocantins Calado. |
| EPI_ISL_1795324 | UBS II DE NARANDIBA | Instituto Butantan / ESALQ-Piracicaba | Instituto Butantan: Alexander Roberto Precioso, Dimas Tadeu Covas, Sandra Coccuzzo Sampaio, Maria Carolina Elias, José Salvatore Leister Patané, Vincent Louis Viala, Antonio Jorge Martins, Ricardo Haddad, Claudia Renata dos Santos Barros, Elaine Cristina Marqueze, Raul Machado Neto, Debora Botequio Moretti. Centro de Genômica Funcional da ESALQ: Luiz Lehmann Coutinho, Ricardo Augusto Brassaloti, Raquel de Lello Rocha Campos Cassano. NGS Soluções Genômicas: Pilar Drummond Sampaio Corrêa Mariani. FZEA-USP Pirassununga: Mirele Daiana Poleti, Jessika Cristina Chagas Lesbon, Elisangela Chicaroni Mattos, Heidge Fukumasu. USP-Botucatu: Rejane Maria Tommasini Grotto, Jayme A. Souza-Neto, Guilherme Targino Valente, Patricia Akemi Assato, Felipe Allan da Silva da Costa, Bianca Cechetto Carlos. Mendelics: Bibiana Santos, João Paulo Kitajima, Erika Freitas, David Schlesinger. Hemocentro Ribeirão Preto: Simone Kashima, Evandra Strazza Rodrigues, Svetoslav Nanev Slavov, Elaine Vieira dos Santos, Rafael dos Santos Bezerra, Luiz Carlos Junior de Alcantara, Marta Giovanetti, Vagner Fonseca, Flavia Aburjaile, Rodrigo Tocantins Calado. |
| EPI_ISL_1795325 | CENTRO DE ESPECIALIDADES DE PRIMAVERA | Instituto Butantan / ESALQ-Piracicaba | Instituto Butantan: Alexander Roberto Precioso, Dimas Tadeu Covas, Sandra Coccuzzo Sampaio, Maria Carolina Elias, José Salvatore Leister Patané, Vincent Louis Viala, Antonio Jorge Martins, Ricardo Haddad, Claudia Renata dos Santos Barros, Elaine Cristina Marqueze, Raul Machado Neto, Debora Botequio Moretti. Centro de Genômica Funcional da ESALQ: Luiz Lehmann Coutinho, Ricardo Augusto Brassaloti, Raquel de Lello Rocha Campos Cassano. NGS Soluções Genômicas: Pilar Drummond Sampaio Corrêa Mariani. FZEA-USP Pirassununga: Mirele Daiana Poleti, Jessika Cristina Chagas Lesbon, Elisangela Chicaroni Mattos, Heidge Fukumasu. USP-Botucatu: Rejane Maria Tommasini Grotto, Jayme A. Souza-Neto, Guilherme Targino Valente, Patricia Akemi Assato, Felipe Allan da Silva da Costa, Bianca Cechetto Carlos. Mendelics: Bibiana Santos, João Paulo Kitajima, Erika Freitas, David Schlesinger. Hemocentro Ribeirão Preto: Simone Kashima, Evandra Strazza Rodrigues, Svetoslav Nanev Slavov, Elaine Vieira dos Santos, Rafael dos Santos Bezerra, Luiz Carlos Junior de Alcantara, Marta Giovanetti, Vagner Fonseca, Flavia Aburjaile, Rodrigo Tocantins Calado. |
| EPI_ISL_1795326 | UBS II DE REGENTE FEIJO | Instituto Butantan / ESALQ-Piracicaba | Instituto Butantan: Alexander Roberto Precioso, Dimas Tadeu Covas, Sandra Coccuzzo Sampaio, Maria Carolina Elias, José Salvatore Leister Patané, Vincent Louis Viala, Antonio Jorge Martins, Ricardo Haddad, Claudia Renata dos Santos Barros, Elaine Cristina Marqueze, Raul Machado Neto, Debora Botequio Moretti. Centro de Genômica Funcional da ESALQ: Luiz Lehmann Coutinho, Ricardo Augusto Brassaloti, Raquel de Lello Rocha Campos Cassano. NGS Soluções Genômicas: Pilar Drummond Sampaio Corrêa Mariani. FZEA-USP Pirassununga: Mirele Daiana Poleti, Jessika Cristina Chagas Lesbon, Elisangela Chicaroni Mattos, Heidge Fukumasu. USP-Botucatu: Rejane Maria Tommasini Grotto, Jayme A. Souza-Neto, Guilherme Targino Valente, Patricia Akemi Assato, Felipe Allan da Silva da Costa, Bianca Cechetto Carlos. Mendelics: Bibiana Santos, João Paulo Kitajima, Erika Freitas, David Schlesinger. Hemocentro Ribeirão Preto: Simone Kashima, Evandra Strazza Rodrigues, Svetoslav Nanev Slavov, Elaine Vieira dos Santos, Rafael dos Santos Bezerra, Luiz Carlos Junior de Alcantara, Marta Giovanetti, Vagner Fonseca, Flavia Aburjaile, Rodrigo Tocantins Calado. |
| EPI_ISL_1795327 | CENTRO DE SAUDE II JUNQUEIROPOLIS | Instituto Butantan / ESALQ-Piracicaba | Instituto Butantan: Alexander Roberto Precioso, Dimas Tadeu Covas, Sandra Coccuzzo Sampaio, Maria Carolina Elias, José Salvatore Leister Patané, Vincent Louis Viala, Antonio Jorge Martins, Ricardo Haddad, Claudia Renata dos Santos Barros, Elaine Cristina Marqueze, Raul Machado Neto, Debora Botequio Moretti. Centro de Genômica Funcional da ESALQ: Luiz Lehmann Coutinho, Ricardo Augusto Brassaloti, Raquel de Lello Rocha Campos Cassano. NGS Soluções Genômicas: Pilar Drummond Sampaio Corrêa Mariani. FZEA-USP Pirassununga: Mirele Daiana Poleti, Jessika Cristina Chagas Lesbon, Elisangela Chicaroni Mattos, Heidge Fukumasu. USP-Botucatu: Rejane Maria Tommasini Grotto, Jayme A. Souza-Neto, Guilherme Targino Valente, Patricia Akemi Assato, Felipe Allan da Silva da Costa, Bianca Cechetto Carlos. Mendelics: Bibiana Santos, João Paulo Kitajima, Erika Freitas, David Schlesinger. Hemocentro Ribeirão Preto: Simone Kashima, Evandra Strazza Rodrigues, Svetoslav Nanev Slavov, Elaine Vieira dos Santos, Rafael dos Santos Bezerra, Luiz Carlos Junior de Alcantara, Marta Giovanetti, Vagner Fonseca, Flavia Aburjaile, Rodrigo Tocantins Calado. |
| EPI_ISL_1795328 | UBS II DR EXPEDITO SHIZUO KUROCE | Instituto Butantan / ESALQ-Piracicaba | Instituto Butantan: Alexander Roberto Precioso, Dimas Tadeu Covas, Sandra Coccuzzo Sampaio, Maria Carolina Elias, José Salvatore Leister Patané, Vincent Louis Viala, Antonio Jorge Martins, Ricardo Haddad, Claudia Renata dos Santos Barros, Elaine Cristina Marqueze, Raul Machado Neto, Debora Botequio Moretti. Centro de Genômica Funcional da ESALQ: Luiz Lehmann Coutinho, Ricardo Augusto Brassaloti, Raquel de Lello Rocha Campos Cassano. NGS Soluções Genômicas: Pilar Drummond Sampaio Corrêa Mariani. FZEA-USP Pirassununga: Mirele Daiana Poleti, Jessika Cristina Chagas Lesbon, Elisangela Chicaroni Mattos, Heidge Fukumasu. USP-Botucatu: Rejane Maria Tommasini Grotto, Jayme A. Souza-Neto, Guilherme Targino Valente, Patricia Akemi Assato, Felipe Allan da Silva da Costa, Bianca Cechetto Carlos. Mendelics: Bibiana Santos, João Paulo Kitajima, Erika Freitas, David Schlesinger. Hemocentro Ribeirão Preto: Simone Kashima, Evandra Strazza Rodrigues, Svetoslav Nanev Slavov, Elaine Vieira dos Santos, Rafael dos Santos Bezerra, Luiz Carlos Junior de Alcantara, Marta Giovanetti, Vagner Fonseca, Flavia Aburjaile, Rodrigo Tocantins Calado. |
| EPI_ISL_1795329 | CENTRO DE ESPECIALIDADES DE PRIMAVERA | Instituto Butantan / ESALQ-Piracicaba | Instituto Butantan: Alexander Roberto Precioso, Dimas Tadeu Covas, Sandra Coccuzzo Sampaio, Maria Carolina Elias, José Salvatore Leister Patané, Vincent Louis Viala, Antonio Jorge Martins, Ricardo Haddad, Claudia Renata dos Santos Barros, Elaine Cristina Marqueze, Raul Machado Neto, Debora Botequio Moretti. Centro de Genômica Funcional da ESALQ: Luiz Lehmann Coutinho, Ricardo Augusto Brassaloti, Raquel de Lello Rocha Campos Cassano. NGS Soluções Genômicas: Pilar Drummond Sampaio Corrêa Mariani. FZEA-USP Pirassununga: Mirele Daiana Poleti, Jessika Cristina Chagas Lesbon, Elisangela Chicaroni Mattos, Heidge Fukumasu. USP-Botucatu: Rejane Maria Tommasini Grotto, Jayme A. Souza-Neto, Guilherme Targino Valente, Patricia Akemi Assato, Felipe Allan da Silva da Costa, Bianca Cechetto Carlos. Mendelics: Bibiana Santos, João Paulo Kitajima, Erika Freitas, David Schlesinger. Hemocentro Ribeirão Preto: Simone Kashima, Evandra Strazza Rodrigues, Svetoslav Nanev Slavov, Elaine Vieira dos Santos, Rafael dos Santos Bezerra, Luiz Carlos Junior de Alcantara, Marta Giovanetti, Vagner Fonseca, Flavia Aburjaile, Rodrigo Tocantins Calado. |
| EPI_ISL_1795330 | UBS II DR EXPEDITO SHIZUO KUROCE | Instituto Butantan / ESALQ-Piracicaba | Instituto Butantan: Alexander Roberto Precioso, Dimas Tadeu Covas, Sandra Coccuzzo Sampaio, Maria Carolina Elias, José Salvatore Leister Patané, Vincent Louis Viala, Antonio Jorge Martins, Ricardo Haddad, Claudia Renata dos Santos Barros, Elaine Cristina Marqueze, Raul Machado Neto, Debora Botequio Moretti. Centro de Genômica Funcional da ESALQ: Luiz Lehmann Coutinho, Ricardo Augusto Brassaloti, Raquel de Lello Rocha Campos Cassano. NGS Soluções Genômicas: Pilar Drummond Sampaio Corrêa Mariani. FZEA-USP Pirassununga: Mirele Daiana Poleti, Jessika Cristina Chagas Lesbon, Elisangela Chicaroni Mattos, Heidge Fukumasu. USP-Botucatu: Rejane Maria Tommasini Grotto, Jayme A. Souza-Neto, Guilherme Targino Valente, Patricia Akemi Assato, Felipe Allan da Silva da Costa, Bianca Cechetto Carlos. Mendelics: Bibiana Santos, João Paulo Kitajima, Erika Freitas, David Schlesinger. Hemocentro Ribeirão Preto: Simone Kashima, Evandra Strazza Rodrigues, Svetoslav Nanev Slavov, Elaine Vieira dos Santos, Rafael dos Santos Bezerra, Luiz Carlos Junior de Alcantara, Marta Giovanetti, Vagner Fonseca, Flavia Aburjaile, Rodrigo Tocantins Calado. |
| EPI_ISL_1795331 | CENTRO DE SAUDE II JUNQUEIROPOLIS | Instituto Butantan / ESALQ-Piracicaba | Instituto Butantan: Alexander Roberto Precioso, Dimas Tadeu Covas, Sandra Coccuzzo Sampaio, Maria Carolina Elias, José Salvatore Leister Patané, Vincent Louis Viala, Antonio Jorge Martins, Ricardo Haddad, Claudia Renata dos Santos Barros, Elaine Cristina Marqueze, Raul Machado Neto, Debora Botequio Moretti. Centro de Genômica Funcional da ESALQ: Luiz Lehmann Coutinho, Ricardo Augusto Brassaloti, Raquel de Lello Rocha Campos Cassano. NGS Soluções Genômicas: Pilar Drummond Sampaio Corrêa Mariani. FZEA-USP Pirassununga: Mirele Daiana Poleti, Jessika Cristina Chagas Lesbon, Elisangela Chicaroni Mattos, Heidge Fukumasu. USP-Botucatu: Rejane Maria Tommasini Grotto, Jayme A. Souza-Neto, Guilherme Targino Valente, Patricia Akemi Assato, Felipe Allan da Silva da Costa, Bianca Cechetto Carlos. Mendelics: Bibiana Santos, João Paulo Kitajima, Erika Freitas, David Schlesinger. Hemocentro Ribeirão Preto: Simone Kashima, Evandra Strazza Rodrigues, Svetoslav Nanev Slavov, Elaine Vieira dos Santos, Rafael dos Santos Bezerra, Luiz Carlos Junior de Alcantara, Marta Giovanetti, Vagner Fonseca, Flavia Aburjaile, Rodrigo Tocantins Calado. |

| | | | |
|---|---|---|---|
| EPI_ISL_1795332 | UBS III DE RANCHARIA | Instituto Butantan / ESALQ-Piracicaba | Instituto Butantan: Alexander Roberto Precioso, Dimas Tadeu Covas, Sandra Coccuzzo Sampaio, Maria Carolina Elias, José Salvatore Leister Patané, Vincent Louis Viala, Antonio Jorge Martins, Ricardo Haddad, Claudia Renata dos Santos Barros, Elaine Cristina Marqueze, Raul Machado Neto, Debora Botequio Moretti. Centro de Genômica Funcional da ESALQ: Luiz Lehmann Coutinho, Ricardo Augusto Brassaloti, Raquel de Lello Rocha Campos Cassano. NGS Soluções Genômicas: Pilar Drummond Sampaio Corrêa Mariani. FZEA-USP Pirassununga: Mirele Daiana Poleti, Jessika Cristina Chagas Lesbon, Elisangela Chicaroni Mattos, Heidge Fukumasu. USP-Botucatu: Rejane Maria Tommasini Grotto, Jayme A. Souza-Neto, Guilherme Targino Valente, Patricia Akemi Assato, Felipe Allan da Silva da Costa, Bianca Cechetto Carlos. Mendelics: Bibiana Santos, João Paulo Kitajima, Erika Freitas, David Schlesinger. Hemocentro Ribeirão Preto: Simone Kashima, Evandra Strazza Rodrigues, Svetoslav Nanev Slavov, Elaine Vieira dos Santos, Rafael dos Santos Bezerra, Luiz Carlos Junior de Alcantara, Marta Giovanetti, Vagner Fonseca, Flavia Aburjaile, Rodrigo Tocantins Calado. |
| EPI_ISL_1795333 | CENTRO DE ESPECIALIDADES DE PRIMAVERA | Instituto Butantan / ESALQ-Piracicaba | Instituto Butantan: Alexander Roberto Precioso, Dimas Tadeu Covas, Sandra Coccuzzo Sampaio, Maria Carolina Elias, José Salvatore Leister Patané, Vincent Louis Viala, Antonio Jorge Martins, Ricardo Haddad, Claudia Renata dos Santos Barros, Elaine Cristina Marqueze, Raul Machado Neto, Debora Botequio Moretti. Centro de Genômica Funcional da ESALQ: Luiz Lehmann Coutinho, Ricardo Augusto Brassaloti, Raquel de Lello Rocha Campos Cassano. NGS Soluções Genômicas: Pilar Drummond Sampaio Corrêa Mariani. FZEA-USP Pirassununga: Mirele Daiana Poleti, Jessika Cristina Chagas Lesbon, Elisangela Chicaroni Mattos, Heidge Fukumasu. USP-Botucatu: Rejane Maria Tommasini Grotto, Jayme A. Souza-Neto, Guilherme Targino Valente, Patricia Akemi Assato, Felipe Allan da Silva da Costa, Bianca Cechetto Carlos. Mendelics: Bibiana Santos, João Paulo Kitajima, Erika Freitas, David Schlesinger. Hemocentro Ribeirão Preto: Simone Kashima, Evandra Strazza Rodrigues, Svetoslav Nanev Slavov, Elaine Vieira dos Santos, Rafael dos Santos Bezerra, Luiz Carlos Junior de Alcantara, Marta Giovanetti, Vagner Fonseca, Flavia Aburjaile, Rodrigo Tocantins Calado. |
| EPI_ISL_1795334 | UBS MARCIA CRISTIANE DA SILVA DE OURO VERDE | Instituto Butantan / ESALQ-Piracicaba | Instituto Butantan: Alexander Roberto Precioso, Dimas Tadeu Covas, Sandra Coccuzzo Sampaio, Maria Carolina Elias, José Salvatore Leister Patané, Vincent Louis Viala, Antonio Jorge Martins, Ricardo Haddad, Claudia Renata dos Santos Barros, Elaine Cristina Marqueze, Raul Machado Neto, Debora Botequio Moretti. Centro de Genômica Funcional da ESALQ: Luiz Lehmann Coutinho, Ricardo Augusto Brassaloti, Raquel de Lello Rocha Campos Cassano. NGS Soluções Genômicas: Pilar Drummond Sampaio Corrêa Mariani. FZEA-USP Pirassununga: Mirele Daiana Poleti, Jessika Cristina Chagas Lesbon, Elisangela Chicaroni Mattos, Heidge Fukumasu. USP-Botucatu: Rejane Maria Tommasini Grotto, Jayme A. Souza-Neto, Guilherme Targino Valente, Patricia Akemi Assato, Felipe Allan da Silva da Costa, Bianca Cechetto Carlos. Mendelics: Bibiana Santos, João Paulo Kitajima, Erika Freitas, David Schlesinger. Hemocentro Ribeirão Preto: Simone Kashima, Evandra Strazza Rodrigues, Svetoslav Nanev Slavov, Elaine Vieira dos Santos, Rafael dos Santos Bezerra, Luiz Carlos Junior de Alcantara, Marta Giovanetti, Vagner Fonseca, Flavia Aburjaile, Rodrigo Tocantins Calado. |
| EPI_ISL_1795335 | UBS III DE RANCHARIA | Instituto Butantan / ESALQ-Piracicaba | Instituto Butantan: Alexander Roberto Precioso, Dimas Tadeu Covas, Sandra Coccuzzo Sampaio, Maria Carolina Elias, José Salvatore Leister Patané, Vincent Louis Viala, Antonio Jorge Martins, Ricardo Haddad, Claudia Renata dos Santos Barros, Elaine Cristina Marqueze, Raul Machado Neto, Debora Botequio Moretti. Centro de Genômica Funcional da ESALQ: Luiz Lehmann Coutinho, Ricardo Augusto Brassaloti, Raquel de Lello Rocha Campos Cassano. NGS Soluções Genômicas: Pilar Drummond Sampaio Corrêa Mariani. FZEA-USP Pirassununga: Mirele Daiana Poleti, Jessika Cristina Chagas Lesbon, Elisangela Chicaroni Mattos, Heidge Fukumasu. USP-Botucatu: Rejane Maria Tommasini Grotto, Jayme A. Souza-Neto, Guilherme Targino Valente, Patricia Akemi Assato, Felipe Allan da Silva da Costa, Bianca Cechetto Carlos. Mendelics: Bibiana Santos, João Paulo Kitajima, Erika Freitas, David Schlesinger. Hemocentro Ribeirão Preto: Simone Kashima, Evandra Strazza Rodrigues, Svetoslav Nanev Slavov, Elaine Vieira dos Santos, Rafael dos Santos Bezerra, Luiz Carlos Junior de Alcantara, Marta Giovanetti, Vagner Fonseca, Flavia Aburjaile, Rodrigo Tocantins Calado. |
| EPI_ISL_1795336 | PAS JOAO ANTONIO DO NASCIMENTO | Instituto Butantan / ESALQ-Piracicaba | Instituto Butantan: Alexander Roberto Precioso, Dimas Tadeu Covas, Sandra Coccuzzo Sampaio, Maria Carolina Elias, José Salvatore Leister Patané, Vincent Louis Viala, Antonio Jorge Martins, Ricardo Haddad, Claudia Renata dos Santos Barros, Elaine Cristina Marqueze, Raul Machado Neto, Debora Botequio Moretti. Centro de Genômica Funcional da ESALQ: Luiz Lehmann Coutinho, Ricardo Augusto Brassaloti, Raquel de Lello Rocha Campos Cassano. NGS Soluções Genômicas: Pilar Drummond Sampaio Corrêa Mariani. FZEA-USP Pirassununga: Mirele Daiana Poleti, Jessika Cristina Chagas Lesbon, Elisangela Chicaroni Mattos, Heidge Fukumasu. USP-Botucatu: Rejane Maria Tommasini Grotto, Jayme A. Souza-Neto, Guilherme Targino Valente, Patricia Akemi Assato, Felipe Allan da Silva da Costa, Bianca Cechetto Carlos. Mendelics: Bibiana Santos, João Paulo Kitajima, Erika Freitas, David Schlesinger. Hemocentro Ribeirão Preto: Simone Kashima, Evandra Strazza Rodrigues, Svetoslav Nanev Slavov, Elaine Vieira dos Santos, Rafael dos Santos Bezerra, Luiz Carlos Junior de Alcantara, Marta Giovanetti, Vagner Fonseca, Flavia Aburjaile, Rodrigo Tocantins Calado. |
| EPI_ISL_1795337, EPI_ISL_1795338 | HOSPITAL MUNICIPAL REYNALDO GUERRA CAJATI | Instituto Butantan / ESALQ-Piracicaba | Instituto Butantan: Alexander Roberto Precioso, Dimas Tadeu Covas, Sandra Coccuzzo Sampaio, Maria Carolina Elias, José Salvatore Leister Patané, Vincent Louis Viala, Antonio Jorge Martins, Ricardo Haddad, Claudia Renata dos Santos Barros, Elaine Cristina Marqueze, Raul Machado Neto, Debora Botequio Moretti. Centro de Genômica Funcional da ESALQ: Luiz Lehmann Coutinho, Ricardo Augusto Brassaloti, Raquel de Lello Rocha Campos Cassano. NGS Soluções Genômicas: Pilar Drummond Sampaio Corrêa Mariani. FZEA-USP Pirassununga: Mirele Daiana Poleti, Jessika Cristina Chagas Lesbon, Elisangela Chicaroni Mattos, Heidge Fukumasu. USP-Botucatu: Rejane Maria Tommasini Grotto, Jayme A. Souza-Neto, Guilherme Targino Valente, Patricia Akemi Assato, Felipe Allan da Silva da Costa, Bianca Cechetto Carlos. Mendelics: Bibiana Santos, João Paulo Kitajima, Erika Freitas, David Schlesinger. Hemocentro Ribeirão Preto: Simone Kashima, Evandra Strazza Rodrigues, Svetoslav Nanev Slavov, Elaine Vieira dos Santos, Rafael dos Santos Bezerra, Luiz Carlos Junior de Alcantara, Marta Giovanetti, Vagner Fonseca, Flavia Aburjaile, Rodrigo Tocantins Calado. |
| EPI_ISL_1795339, EPI_ISL_1795340 | VIGILANCIA EM SAUDE | Instituto Butantan / ESALQ-Piracicaba | Instituto Butantan: Alexander Roberto Precioso, Dimas Tadeu Covas, Sandra Coccuzzo Sampaio, Maria Carolina Elias, José Salvatore Leister Patané, Vincent Louis Viala, Antonio Jorge Martins, Ricardo Haddad, Claudia Renata dos Santos Barros, Elaine Cristina Marqueze, Raul Machado Neto, Debora Botequio Moretti. Centro de Genômica Funcional da ESALQ: Luiz Lehmann Coutinho, Ricardo Augusto Brassaloti, Raquel de Lello Rocha Campos Cassano. NGS Soluções Genômicas: Pilar Drummond Sampaio Corrêa Mariani. FZEA-USP Pirassununga: Mirele Daiana Poleti, Jessika Cristina Chagas Lesbon, Elisangela Chicaroni Mattos, Heidge Fukumasu. USP-Botucatu: Rejane Maria Tommasini Grotto, Jayme A. Souza-Neto, Guilherme Targino Valente, Patricia Akemi Assato, Felipe Allan da Silva da Costa, Bianca Cechetto Carlos. Mendelics: Bibiana Santos, João Paulo Kitajima, Erika Freitas, David Schlesinger. Hemocentro Ribeirão Preto: Simone Kashima, Evandra Strazza Rodrigues, Svetoslav Nanev Slavov, Elaine Vieira dos Santos, Rafael dos Santos Bezerra, Luiz Carlos Junior de Alcantara, Marta Giovanetti, Vagner Fonseca, Flavia Aburjaile, Rodrigo Tocantins Calado. |
| EPI_ISL_1795341 | HOSPITAL MUNICIPAL REYNALDO GUERRA CAJATI | Instituto Butantan / ESALQ-Piracicaba | Instituto Butantan: Alexander Roberto Precioso, Dimas Tadeu Covas, Sandra Coccuzzo Sampaio, Maria Carolina Elias, José Salvatore Leister Patané, Vincent Louis Viala, Antonio Jorge Martins, Ricardo Haddad, Claudia Renata dos Santos Barros, Elaine Cristina Marqueze, Raul Machado Neto, Debora Botequio Moretti. Centro de Genômica Funcional da ESALQ: Luiz Lehmann Coutinho, Ricardo Augusto Brassaloti, Raquel de Lello Rocha Campos Cassano. NGS Soluções Genômicas: Pilar Drummond Sampaio Corrêa Mariani. FZEA-USP Pirassununga: Mirele Daiana Poleti, Jessika Cristina Chagas Lesbon, Elisangela Chicaroni Mattos, Heidge Fukumasu. USP-Botucatu: Rejane Maria Tommasini Grotto, Jayme A. Souza-Neto, Guilherme Targino Valente, Patricia Akemi Assato, Felipe Allan da Silva da Costa, Bianca Cechetto Carlos. Mendelics: Bibiana Santos, João Paulo Kitajima, Erika Freitas, David Schlesinger. Hemocentro Ribeirão Preto: Simone Kashima, Evandra Strazza Rodrigues, Svetoslav Nanev Slavov, Elaine Vieira dos Santos, Rafael dos Santos Bezerra, Luiz Carlos Junior de Alcantara, Marta Giovanetti, Vagner Fonseca, Flavia Aburjaile, Rodrigo Tocantins Calado. |
| EPI_ISL_1795342, EPI_ISL_1795343 | VIGILANCIA EM SAUDE | Instituto Butantan / ESALQ-Piracicaba | Instituto Butantan: Alexander Roberto Precioso, Dimas Tadeu Covas, Sandra Coccuzzo Sampaio, Maria Carolina Elias, José Salvatore Leister Patané, Vincent Louis Viala, Antonio Jorge Martins, Ricardo Haddad, Claudia Renata dos Santos Barros, Elaine Cristina Marqueze, Raul Machado Neto, Debora Botequio Moretti. Centro de Genômica Funcional da ESALQ: Luiz Lehmann Coutinho, Ricardo Augusto Brassaloti, Raquel de Lello Rocha Campos Cassano. NGS Soluções Genômicas: Pilar Drummond Sampaio Corrêa Mariani. FZEA-USP Pirassununga: Mirele Daiana Poleti, Jessika Cristina Chagas Lesbon, Elisangela Chicaroni Mattos, Heidge Fukumasu. USP-Botucatu: Rejane Maria Tommasini Grotto, Jayme A. Souza-Neto, Guilherme Targino Valente, Patricia Akemi Assato, Felipe Allan da Silva da Costa, Bianca Cechetto Carlos. Mendelics: Bibiana Santos, João Paulo Kitajima, Erika Freitas, David Schlesinger. Hemocentro Ribeirão Preto: Simone Kashima, Evandra Strazza Rodrigues, Svetoslav Nanev Slavov, Elaine Vieira dos Santos, Rafael dos Santos Bezerra, Luiz Carlos Junior de Alcantara, Marta Giovanetti, Vagner Fonseca, Flavia Aburjaile, Rodrigo Tocantins Calado. |
| EPI_ISL_1795344 | HOSPITAL E MATERNIDADE NOSSA SENHORA DA AJUDA | Instituto Butantan / ESALQ-Piracicaba | Instituto Butantan: Alexander Roberto Precioso, Dimas Tadeu Covas, Sandra Coccuzzo Sampaio, Maria Carolina Elias, José Salvatore Leister Patané, Vincent Louis Viala, Antonio Jorge Martins, Ricardo Haddad, Claudia Renata dos Santos Barros, Elaine Cristina Marqueze, Raul Machado Neto, Debora Botequio Moretti. Centro de Genômica Funcional da ESALQ: Luiz Lehmann Coutinho, Ricardo Augusto Brassaloti, Raquel de Lello Rocha Campos Cassano. NGS Soluções Genômicas: Pilar Drummond Sampaio Corrêa Mariani. FZEA-USP Pirassununga: Mirele Daiana Poleti, Jessika Cristina Chagas Lesbon, Elisangela Chicaroni Mattos, Heidge Fukumasu. USP-Botucatu: Rejane Maria Tommasini Grotto, Jayme A. Souza-Neto, Guilherme Targino Valente, Patricia Akemi Assato, Felipe Allan da Silva da Costa, Bianca Cechetto Carlos. Mendelics: Bibiana Santos, João Paulo Kitajima, Erika Freitas, David Schlesinger. Hemocentro Ribeirão Preto: Simone Kashima, Evandra Strazza Rodrigues, Svetoslav Nanev Slavov, Elaine Vieira dos Santos, Rafael dos Santos Bezerra, Luiz Carlos Junior de Alcantara, Marta Giovanetti, Vagner Fonseca, Flavia Aburjaile, Rodrigo Tocantins Calado. |
| EPI_ISL_1795345 | HOSPITAL DE CAMPANHA COVID 19 MUNICIPIO DE TAUBATE | Instituto Butantan / ESALQ-Piracicaba | Instituto Butantan: Alexander Roberto Precioso, Dimas Tadeu Covas, Sandra Coccuzzo Sampaio, Maria Carolina Elias, José Salvatore Leister Patané, Vincent Louis Viala, Antonio Jorge Martins, Ricardo Haddad, Claudia Renata dos Santos Barros, Elaine Cristina Marqueze, Raul Machado Neto, Debora |

| | | | |
|---|---|---|---|
| | | | Botequio Moretti. Centro de Genômica Funcional da ESALQ: Luiz Lehmann Coutinho, Ricardo Augusto Brassaloti, Raquel de Lello Rocha Campos Cassano. NGS Soluções Genômicas: Pilar Drummond Sampaio Corrêa Mariani. FZEA-USP Pirassununga: Mirele Daiana Poleti, Jessika Cristina Chagas Lesbon, Elisangela Chicaroni Mattos, Heidge Fukumasu. USP-Botucatu: Rejane Maria Tommasini Grotto, Jayme A. Souza-Neto, Guilherme Targino Valente, Patricia Akemi Assato, Felipe Allan da Silva da Costa, Bianca Cechetto Carlos. Mendelics: Bibiana Santos, João Paulo Kitajima, Erika Freitas, David Schlesinger. Hemocentro Ribeirão Preto: Simone Kashima, Evandra Strazza Rodrigues, Svetoslav Nanev Slavov, Elaine Vieira dos Santos, Rafael dos Santos Bezerra, Luiz Carlos Junior de Alcantara, Marta Giovanetti, Vagner Fonseca, Flavia Aburjaile, Rodrigo Tocantins Calado. |
| EPI_ISL_1795346, EPI_ISL_1795347, EPI_ISL_1795348, EPI_ISL_1795349, EPI_ISL_1795350, EPI_ISL_1795351, EPI_ISL_1795352, EPI_ISL_1795353, EPI_ISL_1795354, EPI_ISL_1795355, EPI_ISL_1795356, EPI_ISL_1795357, EPI_ISL_1795358, EPI_ISL_1795359, EPI_ISL_1795360 | | | |
| see above | LABORATORIO DR PAULO EMILIO DALESSANDRO PINDAMONHANGABA | Instituto Butantan / ESALQ-Piracicaba | Instituto Butantan: Alexander Roberto Precioso, Dimas Tadeu Covas, Sandra Coccuzzo Sampaio, Maria Carolina Elias, José Salvatore Leister Patané, Vincent Louis Viala, Antonio Jorge Martins, Ricardo Haddad, Claudia Renata dos Santos Barros, Elaine Cristina Marqueze, Raul Machado Neto, Debora Botequio Moretti. Centro de Genômica Funcional da ESALQ: Luiz Lehmann Coutinho, Ricardo Augusto Brassaloti, Raquel de Lello Rocha Campos Cassano. NGS Soluções Genômicas: Pilar Drummond Sampaio Corrêa Mariani. FZEA-USP Pirassununga: Mirele Daiana Poleti, Jessika Cristina Chagas Lesbon, Elisangela Chicaroni Mattos, Heidge Fukumasu. USP-Botucatu: Rejane Maria Tommasini Grotto, Jayme A. Souza-Neto, Guilherme Targino Valente, Patricia Akemi Assato, Felipe Allan da Silva da Costa, Bianca Cechetto Carlos. Mendelics: Bibiana Santos, João Paulo Kitajima, Erika Freitas, David Schlesinger. Hemocentro Ribeirão Preto: Simone Kashima, Evandra Strazza Rodrigues, Svetoslav Nanev Slavov, Elaine Vieira dos Santos, Rafael dos Santos Bezerra, Luiz Carlos Junior de Alcantara, Marta Giovanetti, Vagner Fonseca, Flavia Aburjaile, Rodrigo Tocantins Calado. |
| EPI_ISL_1795361 | SANTA CASA DE MISERICORDIA DE UBATUBA | Instituto Butantan / ESALQ-Piracicaba | Instituto Butantan: Alexander Roberto Precioso, Dimas Tadeu Covas, Sandra Coccuzzo Sampaio, Maria Carolina Elias, José Salvatore Leister Patané, Vincent Louis Viala, Antonio Jorge Martins, Ricardo Haddad, Claudia Renata dos Santos Barros, Elaine Cristina Marqueze, Raul Machado Neto, Debora Botequio Moretti. Centro de Genômica Funcional da ESALQ: Luiz Lehmann Coutinho, Ricardo Augusto Brassaloti, Raquel de Lello Rocha Campos Cassano. NGS Soluções Genômicas: Pilar Drummond Sampaio Corrêa Mariani. FZEA-USP Pirassununga: Mirele Daiana Poleti, Jessika Cristina Chagas Lesbon, Elisangela Chicaroni Mattos, Heidge Fukumasu. USP-Botucatu: Rejane Maria Tommasini Grotto, Jayme A. Souza-Neto, Guilherme Targino Valente, Patricia Akemi Assato, Felipe Allan da Silva da Costa, Bianca Cechetto Carlos. Mendelics: Bibiana Santos, João Paulo Kitajima, Erika Freitas, David Schlesinger. Hemocentro Ribeirão Preto: Simone Kashima, Evandra Strazza Rodrigues, Svetoslav Nanev Slavov, Elaine Vieira dos Santos, Rafael dos Santos Bezerra, Luiz Carlos Junior de Alcantara, Marta Giovanetti, Vagner Fonseca, Flavia Aburjaile, Rodrigo Tocantins Calado. |
| EPI_ISL_1795362, EPI_ISL_1795363 | UNIDADE BASICA DE SAUDE DE PIQUETE | Instituto Butantan / ESALQ-Piracicaba | Instituto Butantan: Alexander Roberto Precioso, Dimas Tadeu Covas, Sandra Coccuzzo Sampaio, Maria Carolina Elias, José Salvatore Leister Patané, Vincent Louis Viala, Antonio Jorge Martins, Ricardo Haddad, Claudia Renata dos Santos Barros, Elaine Cristina Marqueze, Raul Machado Neto, Debora Botequio Moretti. Centro de Genômica Funcional da ESALQ: Luiz Lehmann Coutinho, Ricardo Augusto Brassaloti, Raquel de Lello Rocha Campos Cassano. NGS Soluções Genômicas: Pilar Drummond Sampaio Corrêa Mariani. FZEA-USP Pirassununga: Mirele Daiana Poleti, Jessika Cristina Chagas Lesbon, Elisangela Chicaroni Mattos, Heidge Fukumasu. USP-Botucatu: Rejane Maria Tommasini Grotto, Jayme A. Souza-Neto, Guilherme Targino Valente, Patricia Akemi Assato, Felipe Allan da Silva da Costa, Bianca Cechetto Carlos. Mendelics: Bibiana Santos, João Paulo Kitajima, Erika Freitas, David Schlesinger. Hemocentro Ribeirão Preto: Simone Kashima, Evandra Strazza Rodrigues, Svetoslav Nanev Slavov, Elaine Vieira dos Santos, Rafael dos Santos Bezerra, Luiz Carlos Junior de Alcantara, Marta Giovanetti, Vagner Fonseca, Flavia Aburjaile, Rodrigo Tocantins Calado. |
| EPI_ISL_1795364 | SECRETARIA DE SAUDE | Instituto Butantan / ESALQ-Piracicaba | Instituto Butantan: Alexander Roberto Precioso, Dimas Tadeu Covas, Sandra Coccuzzo Sampaio, Maria Carolina Elias, José Salvatore Leister Patané, Vincent Louis Viala, Antonio Jorge Martins, Ricardo Haddad, Claudia Renata dos Santos Barros, Elaine Cristina Marqueze, Raul Machado Neto, Debora Botequio Moretti. Centro de Genômica Funcional da ESALQ: Luiz Lehmann Coutinho, Ricardo Augusto Brassaloti, Raquel de Lello Rocha Campos Cassano. NGS Soluções Genômicas: Pilar Drummond Sampaio Corrêa Mariani. FZEA-USP Pirassununga: Mirele Daiana Poleti, Jessika Cristina Chagas Lesbon, Elisangela Chicaroni Mattos, Heidge Fukumasu. USP-Botucatu: Rejane Maria Tommasini Grotto, Jayme A. Souza-Neto, Guilherme Targino Valente, Patricia Akemi Assato, Felipe Allan da Silva da Costa, Bianca Cechetto Carlos. Mendelics: Bibiana Santos, João Paulo Kitajima, Erika Freitas, David Schlesinger. Hemocentro Ribeirão Preto: Simone Kashima, Evandra Strazza Rodrigues, Svetoslav Nanev Slavov, Elaine Vieira dos Santos, Rafael dos Santos Bezerra, Luiz Carlos Junior de Alcantara, Marta Giovanetti, Vagner Fonseca, Flavia Aburjaile, Rodrigo Tocantins Calado. |
| EPI_ISL_1795365, EPI_ISL_1795366 | LABORATORIO DR PAULO EMILIO DALESSANDRO PINDAMONHANGABA | Instituto Butantan / ESALQ-Piracicaba | Instituto Butantan: Alexander Roberto Precioso, Dimas Tadeu Covas, Sandra Coccuzzo Sampaio, Maria Carolina Elias, José Salvatore Leister Patané, Vincent Louis Viala, Antonio Jorge Martins, Ricardo Haddad, Claudia Renata dos Santos Barros, Elaine Cristina Marqueze, Raul Machado Neto, Debora Botequio Moretti. Centro de Genômica Funcional da ESALQ: Luiz Lehmann Coutinho, Ricardo Augusto Brassaloti, Raquel de Lello Rocha Campos Cassano. NGS Soluções Genômicas: Pilar Drummond Sampaio Corrêa Mariani. FZEA-USP Pirassununga: Mirele Daiana Poleti, Jessika Cristina Chagas Lesbon, Elisangela Chicaroni Mattos, Heidge Fukumasu. USP-Botucatu: Rejane Maria Tommasini Grotto, Jayme A. Souza-Neto, Guilherme Targino Valente, Patricia Akemi Assato, Felipe Allan da Silva da Costa, Bianca Cechetto Carlos. Mendelics: Bibiana Santos, João Paulo Kitajima, Erika Freitas, David Schlesinger. Hemocentro Ribeirão Preto: Simone Kashima, Evandra Strazza Rodrigues, Svetoslav Nanev Slavov, Elaine Vieira dos Santos, Rafael dos Santos Bezerra, Luiz Carlos Junior de Alcantara, Marta Giovanetti, Vagner Fonseca, Flavia Aburjaile, Rodrigo Tocantins Calado. |
| EPI_ISL_1795367 | AMBULATORIO MEDICO MUNICIPAL DE AGUDOS | Instituto Butantan / ESALQ-Piracicaba | Instituto Butantan: Alexander Roberto Precioso, Dimas Tadeu Covas, Sandra Coccuzzo Sampaio, Maria Carolina Elias, José Salvatore Leister Patané, Vincent Louis Viala, Antonio Jorge Martins, Ricardo Haddad, Claudia Renata dos Santos Barros, Elaine Cristina Marqueze, Raul Machado Neto, Debora Botequio Moretti. Centro de Genômica Funcional da ESALQ: Luiz Lehmann Coutinho, Ricardo Augusto Brassaloti, Raquel de Lello Rocha Campos Cassano. NGS Soluções Genômicas: Pilar Drummond Sampaio Corrêa Mariani. FZEA-USP Pirassununga: Mirele Daiana Poleti, Jessika Cristina Chagas Lesbon, Elisangela Chicaroni Mattos, Heidge Fukumasu. USP-Botucatu: Rejane Maria Tommasini Grotto, Jayme A. Souza-Neto, Guilherme Targino Valente, Patricia Akemi Assato, Felipe Allan da Silva da Costa, Bianca Cechetto Carlos. Mendelics: Bibiana Santos, João Paulo Kitajima, Erika Freitas, David Schlesinger. Hemocentro Ribeirão Preto: Simone Kashima, Evandra Strazza Rodrigues, Svetoslav Nanev Slavov, Elaine Vieira dos Santos, Rafael dos Santos Bezerra, Luiz Carlos Junior de Alcantara, Marta Giovanetti, Vagner Fonseca, Flavia Aburjaile, Rodrigo Tocantins Calado. |
| EPI_ISL_1795368 | CENTRO INTEGRADO DE SAUDE | Instituto Butantan / ESALQ-Piracicaba | Instituto Butantan: Alexander Roberto Precioso, Dimas Tadeu Covas, Sandra Coccuzzo Sampaio, Maria Carolina Elias, José Salvatore Leister Patané, Vincent Louis Viala, Antonio Jorge Martins, Ricardo Haddad, Claudia Renata dos Santos Barros, Elaine Cristina Marqueze, Raul Machado Neto, Debora Botequio Moretti. Centro de Genômica Funcional da ESALQ: Luiz Lehmann Coutinho, Ricardo Augusto Brassaloti, Raquel de Lello Rocha Campos Cassano. NGS Soluções Genômicas: Pilar Drummond Sampaio Corrêa Mariani. FZEA-USP Pirassununga: Mirele Daiana Poleti, Jessika Cristina Chagas Lesbon, Elisangela Chicaroni Mattos, Heidge Fukumasu. USP-Botucatu: Rejane Maria Tommasini Grotto, Jayme A. Souza-Neto, Guilherme Targino Valente, Patricia Akemi Assato, Felipe Allan da Silva da Costa, Bianca Cechetto Carlos. Mendelics: Bibiana Santos, João Paulo Kitajima, Erika Freitas, David Schlesinger. Hemocentro Ribeirão Preto: Simone Kashima, Evandra Strazza Rodrigues, Svetoslav Nanev Slavov, Elaine Vieira dos Santos, Rafael dos Santos Bezerra, Luiz Carlos Junior de Alcantara, Marta Giovanetti, Vagner Fonseca, Flavia Aburjaile, Rodrigo Tocantins Calado. |
| EPI_ISL_1795369 | AMBULATORIO MEDICO MUNICIPAL DE AGUDOS | Instituto Butantan / ESALQ-Piracicaba | Instituto Butantan: Alexander Roberto Precioso, Dimas Tadeu Covas, Sandra Coccuzzo Sampaio, Maria Carolina Elias, José Salvatore Leister Patané, Vincent Louis Viala, Antonio Jorge Martins, Ricardo Haddad, Claudia Renata dos Santos Barros, Elaine Cristina Marqueze, Raul Machado Neto, Debora Botequio Moretti. Centro de Genômica Funcional da ESALQ: Luiz Lehmann Coutinho, Ricardo Augusto Brassaloti, Raquel de Lello Rocha Campos Cassano. NGS Soluções Genômicas: Pilar Drummond Sampaio Corrêa Mariani. FZEA-USP Pirassununga: Mirele Daiana Poleti, Jessika Cristina Chagas Lesbon, Elisangela Chicaroni Mattos, Heidge Fukumasu. USP-Botucatu: Rejane Maria Tommasini Grotto, Jayme A. Souza-Neto, Guilherme Targino Valente, Patricia Akemi Assato, Felipe Allan da Silva da Costa, Bianca Cechetto Carlos. Mendelics: Bibiana Santos, João Paulo Kitajima, Erika Freitas, David Schlesinger. Hemocentro Ribeirão Preto: Simone Kashima, Evandra Strazza Rodrigues, Svetoslav Nanev Slavov, Elaine Vieira dos Santos, Rafael dos Santos Bezerra, Luiz Carlos Junior de Alcantara, Marta Giovanetti, Vagner Fonseca, Flavia Aburjaile, Rodrigo Tocantins Calado. |
| EPI_ISL_1795370 | CENTRO INTEGRADO DE SAUDE | Instituto Butantan / ESALQ-Piracicaba | Instituto Butantan: Alexander Roberto Precioso, Dimas Tadeu Covas, Sandra Coccuzzo Sampaio, Maria Carolina Elias, José Salvatore Leister Patané, Vincent Louis Viala, Antonio Jorge Martins, Ricardo Haddad, Claudia Renata dos Santos Barros, Elaine Cristina Marqueze, Raul Machado Neto, Debora Botequio Moretti. Centro de Genômica Funcional da ESALQ: Luiz Lehmann Coutinho, Ricardo Augusto Brassaloti, Raquel de Lello Rocha Campos Cassano. NGS Soluções Genômicas: Pilar Drummond Sampaio Corrêa Mariani. FZEA-USP Pirassununga: Mirele Daiana Poleti, Jessika Cristina Chagas Lesbon, Elisangela Chicaroni Mattos, Heidge Fukumasu. USP-Botucatu: Rejane Maria Tommasini Grotto, Jayme A. Souza-Neto, Guilherme Targino Valente, Patricia Akemi Assato, Felipe Allan da Silva da Costa, Bianca Cechetto Carlos. Mendelics: Bibiana Santos, João Paulo Kitajima, Erika Freitas, David Schlesinger. Hemocentro Ribeirão Preto: Simone Kashima, Evandra Strazza Rodrigues, Svetoslav Nanev Slavov, Elaine Vieira dos Santos, Rafael dos Santos Bezerra, Luiz Carlos Junior de Alcantara, Marta Giovanetti, Vagner Fonseca, Flavia Aburjaile, Rodrigo Tocantins Calado. |
| EPI_ISL_1795371, EPI_ISL_1795372 | HOSPITAL SANTA THEREZINHA BROTAS | Instituto Butantan / ESALQ-Piracicaba | Instituto Butantan: Alexander Roberto Precioso, Dimas Tadeu Covas, Sandra Coccuzzo Sampaio, Maria Carolina Elias, José Salvatore Leister Patané, Vincent Louis Viala, Antonio Jorge Martins, Ricardo Haddad, Claudia Renata dos Santos Barros, Elaine Cristina Marqueze, Raul Machado Neto, Debora Botequio Moretti. Centro de Genômica Funcional da ESALQ: Luiz Lehmann Coutinho, Ricardo Augusto Brassaloti, Raquel de Lello Rocha Campos |

| | | | |
|---|---|---|---|
| | | | Cassano. NGS Soluções Genômicas: Pilar Drummond Sampaio Corrêa Mariani. FZEA-USP Pirassununga: Mirele Daiana Poleti, Jessika Cristina Chagas Lesbon, Elisangela Chicaroni Mattos, Heidge Fukumasu. USP-Botucatu: Rejane Maria Tommasini Grotto, Jayme A. Souza-Neto, Guilherme Targino Valente, Patricia Akemi Assato, Felipe Allan da Silva da Costa, Bianca Cechetto Carlos. Mendelics: Bibiana Santos, João Paulo Kitajima, Erika Freitas, David Schlesinger. Hemocentro Ribeirão Preto: Simone Kashima, Evandra Strazza Rodrigues, Svetoslav Nanev Slavov, Elaine Vieira dos Santos, Rafael dos Santos Bezerra, Luiz Carlos Junior de Alcantara, Marta Giovanetti, Vagner Fonseca, Flavia Aburjaile, Rodrigo Tocantins Calado. |
| EPI_ISL_1795373 | SECRETARIA MUNICIPAL DE SAUDE DE PEDERNEIRAS | Instituto Butantan / ESALQ-Piracicaba | Instituto Butantan: Alexander Roberto Precioso, Dimas Tadeu Covas, Sandra Coccuzzo Sampaio, Maria Carolina Elias, José Salvatore Leister Patané, Vincent Louis Viala, Antonio Jorge Martins, Ricardo Haddad, Claudia Renata dos Santos Barros, Elaine Cristina Marqueze, Raul Machado Neto, Debora Botequio Moretti. Centro de Genômica Funcional da ESALQ: Luiz Lehmann Coutinho, Ricardo Augusto Brassaloti, Raquel de Lello Rocha Campos Cassano. NGS Soluções Genômicas: Pilar Drummond Sampaio Corrêa Mariani. FZEA-USP Pirassununga: Mirele Daiana Poleti, Jessika Cristina Chagas Lesbon, Elisangela Chicaroni Mattos, Heidge Fukumasu. USP-Botucatu: Rejane Maria Tommasini Grotto, Jayme A. Souza-Neto, Guilherme Targino Valente, Patricia Akemi Assato, Felipe Allan da Silva da Costa, Bianca Cechetto Carlos. Mendelics: Bibiana Santos, João Paulo Kitajima, Erika Freitas, David Schlesinger. Hemocentro Ribeirão Preto: Simone Kashima, Evandra Strazza Rodrigues, Svetoslav Nanev Slavov, Elaine Vieira dos Santos, Rafael dos Santos Bezerra, Luiz Carlos Junior de Alcantara, Marta Giovanetti, Vagner Fonseca, Flavia Aburjaile, Rodrigo Tocantins Calado. |
| EPI_ISL_1795374, EPI_ISL_1795375 | SANTA CASA DE DOIS CORREGOS | Instituto Butantan / ESALQ-Piracicaba | Instituto Butantan: Alexander Roberto Precioso, Dimas Tadeu Covas, Sandra Coccuzzo Sampaio, Maria Carolina Elias, José Salvatore Leister Patané, Vincent Louis Viala, Antonio Jorge Martins, Ricardo Haddad, Claudia Renata dos Santos Barros, Elaine Cristina Marqueze, Raul Machado Neto, Debora Botequio Moretti. Centro de Genômica Funcional da ESALQ: Luiz Lehmann Coutinho, Ricardo Augusto Brassaloti, Raquel de Lello Rocha Campos Cassano. NGS Soluções Genômicas: Pilar Drummond Sampaio Corrêa Mariani. FZEA-USP Pirassununga: Mirele Daiana Poleti, Jessika Cristina Chagas Lesbon, Elisangela Chicaroni Mattos, Heidge Fukumasu. USP-Botucatu: Rejane Maria Tommasini Grotto, Jayme A. Souza-Neto, Guilherme Targino Valente, Patricia Akemi Assato, Felipe Allan da Silva da Costa, Bianca Cechetto Carlos. Mendelics: Bibiana Santos, João Paulo Kitajima, Erika Freitas, David Schlesinger. Hemocentro Ribeirão Preto: Simone Kashima, Evandra Strazza Rodrigues, Svetoslav Nanev Slavov, Elaine Vieira dos Santos, Rafael dos Santos Bezerra, Luiz Carlos Junior de Alcantara, Marta Giovanetti, Vagner Fonseca, Flavia Aburjaile, Rodrigo Tocantins Calado. |
| EPI_ISL_1795376, EPI_ISL_1795377, EPI_ISL_1795378 | ESF MINEIROS DO TIETE | Instituto Butantan / ESALQ-Piracicaba | Instituto Butantan: Alexander Roberto Precioso, Dimas Tadeu Covas, Sandra Coccuzzo Sampaio, Maria Carolina Elias, José Salvatore Leister Patané, Vincent Louis Viala, Antonio Jorge Martins, Ricardo Haddad, Claudia Renata dos Santos Barros, Elaine Cristina Marqueze, Raul Machado Neto, Debora Botequio Moretti. Centro de Genômica Funcional da ESALQ: Luiz Lehmann Coutinho, Ricardo Augusto Brassaloti, Raquel de Lello Rocha Campos Cassano. NGS Soluções Genômicas: Pilar Drummond Sampaio Corrêa Mariani. FZEA-USP Pirassununga: Mirele Daiana Poleti, Jessika Cristina Chagas Lesbon, Elisangela Chicaroni Mattos, Heidge Fukumasu. USP-Botucatu: Rejane Maria Tommasini Grotto, Jayme A. Souza-Neto, Guilherme Targino Valente, Patricia Akemi Assato, Felipe Allan da Silva da Costa, Bianca Cechetto Carlos. Mendelics: Bibiana Santos, João Paulo Kitajima, Erika Freitas, David Schlesinger. Hemocentro Ribeirão Preto: Simone Kashima, Evandra Strazza Rodrigues, Svetoslav Nanev Slavov, Elaine Vieira dos Santos, Rafael dos Santos Bezerra, Luiz Carlos Junior de Alcantara, Marta Giovanetti, Vagner Fonseca, Flavia Aburjaile, Rodrigo Tocantins Calado. |
| EPI_ISL_1795379 | NUCLEO DE SAUDE VILA FALCAO DE BAURU | Instituto Butantan / ESALQ-Piracicaba | Instituto Butantan: Alexander Roberto Precioso, Dimas Tadeu Covas, Sandra Coccuzzo Sampaio, Maria Carolina Elias, José Salvatore Leister Patané, Vincent Louis Viala, Antonio Jorge Martins, Ricardo Haddad, Claudia Renata dos Santos Barros, Elaine Cristina Marqueze, Raul Machado Neto, Debora Botequio Moretti. Centro de Genômica Funcional da ESALQ: Luiz Lehmann Coutinho, Ricardo Augusto Brassaloti, Raquel de Lello Rocha Campos Cassano. NGS Soluções Genômicas: Pilar Drummond Sampaio Corrêa Mariani. FZEA-USP Pirassununga: Mirele Daiana Poleti, Jessika Cristina Chagas Lesbon, Elisangela Chicaroni Mattos, Heidge Fukumasu. USP-Botucatu: Rejane Maria Tommasini Grotto, Jayme A. Souza-Neto, Guilherme Targino Valente, Patricia Akemi Assato, Felipe Allan da Silva da Costa, Bianca Cechetto Carlos. Mendelics: Bibiana Santos, João Paulo Kitajima, Erika Freitas, David Schlesinger. Hemocentro Ribeirão Preto: Simone Kashima, Evandra Strazza Rodrigues, Svetoslav Nanev Slavov, Elaine Vieira dos Santos, Rafael dos Santos Bezerra, Luiz Carlos Junior de Alcantara, Marta Giovanetti, Vagner Fonseca, Flavia Aburjaile, Rodrigo Tocantins Calado. |
| EPI_ISL_1795380 | HOSPITAL MATERNIDADE SAO JOSE ITAPUI | Instituto Butantan / ESALQ-Piracicaba | Instituto Butantan: Alexander Roberto Precioso, Dimas Tadeu Covas, Sandra Coccuzzo Sampaio, Maria Carolina Elias, José Salvatore Leister Patané, Vincent Louis Viala, Antonio Jorge Martins, Ricardo Haddad, Claudia Renata dos Santos Barros, Elaine Cristina Marqueze, Raul Machado Neto, Debora Botequio Moretti. Centro de Genômica Funcional da ESALQ: Luiz Lehmann Coutinho, Ricardo Augusto Brassaloti, Raquel de Lello Rocha Campos Cassano. NGS Soluções Genômicas: Pilar Drummond Sampaio Corrêa Mariani. FZEA-USP Pirassununga: Mirele Daiana Poleti, Jessika Cristina Chagas Lesbon, Elisangela Chicaroni Mattos, Heidge Fukumasu. USP-Botucatu: Rejane Maria Tommasini Grotto, Jayme A. Souza-Neto, Guilherme Targino Valente, Patricia Akemi Assato, Felipe Allan da Silva da Costa, Bianca Cechetto Carlos. Mendelics: Bibiana Santos, João Paulo Kitajima, Erika Freitas, David Schlesinger. Hemocentro Ribeirão Preto: Simone Kashima, Evandra Strazza Rodrigues, Svetoslav Nanev Slavov, Elaine Vieira dos Santos, Rafael dos Santos Bezerra, Luiz Carlos Junior de Alcantara, Marta Giovanetti, Vagner Fonseca, Flavia Aburjaile, Rodrigo Tocantins Calado. |
| EPI_ISL_1795381 | SECRETARIA MUNICIPAL DE SAUDE DE MACATUBA | Instituto Butantan / ESALQ-Piracicaba | Instituto Butantan: Alexander Roberto Precioso, Dimas Tadeu Covas, Sandra Coccuzzo Sampaio, Maria Carolina Elias, José Salvatore Leister Patané, Vincent Louis Viala, Antonio Jorge Martins, Ricardo Haddad, Claudia Renata dos Santos Barros, Elaine Cristina Marqueze, Raul Machado Neto, Debora Botequio Moretti. Centro de Genômica Funcional da ESALQ: Luiz Lehmann Coutinho, Ricardo Augusto Brassaloti, Raquel de Lello Rocha Campos Cassano. NGS Soluções Genômicas: Pilar Drummond Sampaio Corrêa Mariani. FZEA-USP Pirassununga: Mirele Daiana Poleti, Jessika Cristina Chagas Lesbon, Elisangela Chicaroni Mattos, Heidge Fukumasu. USP-Botucatu: Rejane Maria Tommasini Grotto, Jayme A. Souza-Neto, Guilherme Targino Valente, Patricia Akemi Assato, Felipe Allan da Silva da Costa, Bianca Cechetto Carlos. Mendelics: Bibiana Santos, João Paulo Kitajima, Erika Freitas, David Schlesinger. Hemocentro Ribeirão Preto: Simone Kashima, Evandra Strazza Rodrigues, Svetoslav Nanev Slavov, Elaine Vieira dos Santos, Rafael dos Santos Bezerra, Luiz Carlos Junior de Alcantara, Marta Giovanetti, Vagner Fonseca, Flavia Aburjaile, Rodrigo Tocantins Calado. |
| EPI_ISL_1795382, EPI_ISL_1795383 | LABORATORIO DR PAULO EMILIO DALESSANDRO PINDAMONHANGABA | Instituto Butantan / ESALQ-Piracicaba | Instituto Butantan: Alexander Roberto Precioso, Dimas Tadeu Covas, Sandra Coccuzzo Sampaio, Maria Carolina Elias, José Salvatore Leister Patané, Vincent Louis Viala, Antonio Jorge Martins, Ricardo Haddad, Claudia Renata dos Santos Barros, Elaine Cristina Marqueze, Raul Machado Neto, Debora Botequio Moretti. Centro de Genômica Funcional da ESALQ: Luiz Lehmann Coutinho, Ricardo Augusto Brassaloti, Raquel de Lello Rocha Campos Cassano. NGS Soluções Genômicas: Pilar Drummond Sampaio Corrêa Mariani. FZEA-USP Pirassununga: Mirele Daiana Poleti, Jessika Cristina Chagas Lesbon, Elisangela Chicaroni Mattos, Heidge Fukumasu. USP-Botucatu: Rejane Maria Tommasini Grotto, Jayme A. Souza-Neto, Guilherme Targino Valente, Patricia Akemi Assato, Felipe Allan da Silva da Costa, Bianca Cechetto Carlos. Mendelics: Bibiana Santos, João Paulo Kitajima, Erika Freitas, David Schlesinger. Hemocentro Ribeirão Preto: Simone Kashima, Evandra Strazza Rodrigues, Svetoslav Nanev Slavov, Elaine Vieira dos Santos, Rafael dos Santos Bezerra, Luiz Carlos Junior de Alcantara, Marta Giovanetti, Vagner Fonseca, Flavia Aburjaile, Rodrigo Tocantins Calado. |
| EPI_ISL_1795384 | USF SALERNO | Instituto Butantan / ESALQ-Piracicaba | Instituto Butantan: Alexander Roberto Precioso, Dimas Tadeu Covas, Sandra Coccuzzo Sampaio, Maria Carolina Elias, José Salvatore Leister Patané, Vincent Louis Viala, Antonio Jorge Martins, Ricardo Haddad, Claudia Renata dos Santos Barros, Elaine Cristina Marqueze, Raul Machado Neto, Debora Botequio Moretti. Centro de Genômica Funcional da ESALQ: Luiz Lehmann Coutinho, Ricardo Augusto Brassaloti, Raquel de Lello Rocha Campos Cassano. NGS Soluções Genômicas: Pilar Drummond Sampaio Corrêa Mariani. FZEA-USP Pirassununga: Mirele Daiana Poleti, Jessika Cristina Chagas Lesbon, Elisangela Chicaroni Mattos, Heidge Fukumasu. USP-Botucatu: Rejane Maria Tommasini Grotto, Jayme A. Souza-Neto, Guilherme Targino Valente, Patricia Akemi Assato, Felipe Allan da Silva da Costa, Bianca Cechetto Carlos. Mendelics: Bibiana Santos, João Paulo Kitajima, Erika Freitas, David Schlesinger. Hemocentro Ribeirão Preto: Simone Kashima, Evandra Strazza Rodrigues, Svetoslav Nanev Slavov, Elaine Vieira dos Santos, Rafael dos Santos Bezerra, Luiz Carlos Junior de Alcantara, Marta Giovanetti, Vagner Fonseca, Flavia Aburjaile, Rodrigo Tocantins Calado. |
| EPI_ISL_1795385, EPI_ISL_1795386 | UNIDADE DE VIGILANCIA EM SAUDE | Instituto Butantan / ESALQ-Piracicaba | Instituto Butantan: Alexander Roberto Precioso, Dimas Tadeu Covas, Sandra Coccuzzo Sampaio, Maria Carolina Elias, José Salvatore Leister Patané, Vincent Louis Viala, Antonio Jorge Martins, Ricardo Haddad, Claudia Renata dos Santos Barros, Elaine Cristina Marqueze, Raul Machado Neto, Debora Botequio Moretti. Centro de Genômica Funcional da ESALQ: Luiz Lehmann Coutinho, Ricardo Augusto Brassaloti, Raquel de Lello Rocha Campos Cassano. NGS Soluções Genômicas: Pilar Drummond Sampaio Corrêa Mariani. FZEA-USP Pirassununga: Mirele Daiana Poleti, Jessika Cristina Chagas Lesbon, Elisangela Chicaroni Mattos, Heidge Fukumasu. USP-Botucatu: Rejane Maria Tommasini Grotto, Jayme A. Souza-Neto, Guilherme Targino Valente, Patricia Akemi Assato, Felipe Allan da Silva da Costa, Bianca Cechetto Carlos. Mendelics: Bibiana Santos, João Paulo Kitajima, Erika Freitas, David Schlesinger. Hemocentro Ribeirão Preto: Simone Kashima, Evandra Strazza Rodrigues, Svetoslav Nanev Slavov, Elaine Vieira dos Santos, Rafael dos Santos Bezerra, Luiz Carlos Junior de Alcantara, Marta Giovanetti, Vagner Fonseca, Flavia Aburjaile, Rodrigo Tocantins Calado. |
| EPI_ISL_1795387, EPI_ISL_1795388 | POLICLINICA HORTOLANDIA | Instituto Butantan / ESALQ-Piracicaba | Instituto Butantan: Alexander Roberto Precioso, Dimas Tadeu Covas, Sandra Coccuzzo Sampaio, Maria Carolina Elias, José Salvatore Leister Patané, Vincent Louis Viala, Antonio Jorge Martins, Ricardo Haddad, Claudia Renata dos Santos Barros, Elaine Cristina Marqueze, Raul Machado Neto, Debora Botequio Moretti. Centro de Genômica Funcional da ESALQ: Luiz Lehmann Coutinho, Ricardo Augusto Brassaloti, Raquel de Lello Rocha Campos Cassano. NGS Soluções Genômicas: Pilar Drummond Sampaio Corrêa Mariani. FZEA-USP Pirassununga: Mirele Daiana Poleti, Jessika Cristina Chagas Lesbon, Elisangela Chicaroni Mattos, Heidge Fukumasu. USP-Botucatu: Rejane Maria Tommasini Grotto, Jayme A. Souza-Neto, Guilherme Targino |

| | | | |
|---|---|---|---|
| | | | Valente, Patricia Akemi Assato, Felipe Allan da Silva da Costa, Bianca Cechetto Carlos. Mendelics: Bibiana Santos, João Paulo Kitajima, Erika Freitas, David Schlesinger. Hemocentro Ribeirão Preto: Simone Kashima, Evandra Strazza Rodrigues, Svetoslav Nanev Slavov, Elaine Vieira dos Santos, Rafael dos Santos Bezerra, Luiz Carlos Junior de Alcantara, Marta Giovanetti, Vagner Fonseca, Flavia Aburjaile, Rodrigo Tocantins Calado. |
| EPI_ISL_1795389 | CS DE BALSAMO | Instituto Butantan / ESALQ-Piracicaba | Instituto Butantan: Alexander Roberto Precioso, Dimas Tadeu Covas, Sandra Coccuzzo Sampaio, Maria Carolina Elias, José Salvatore Leister Patané, Vincent Louis Viala, Antonio Jorge Martins, Ricardo Haddad, Claudia Renata dos Santos Barros, Elaine Cristina Marqueze, Raul Machado Neto, Debora Botequio Moretti. Centro de Genômica Funcional da ESALQ: Luiz Lehmann Coutinho, Ricardo Augusto Brassaloti, Raquel de Lello Rocha Campos Cassano. NGS Soluções Genômicas: Pilar Drummond Sampaio Corrêa Mariani. FZEA-USP Pirassununga: Mirele Daiana Poleti, Jessika Cristina Chagas Lesbon, Elisangela Chicaroni Mattos, Heidge Fukumasu. USP-Botucatu: Rejane Maria Tommasini Grotto, Jayme A. Souza-Neto, Guilherme Targino Valente, Patricia Akemi Assato, Felipe Allan da Silva da Costa, Bianca Cechetto Carlos. Mendelics: Bibiana Santos, João Paulo Kitajima, Erika Freitas, David Schlesinger. Hemocentro Ribeirão Preto: Simone Kashima, Evandra Strazza Rodrigues, Svetoslav Nanev Slavov, Elaine Vieira dos Santos, Rafael dos Santos Bezerra, Luiz Carlos Junior de Alcantara, Marta Giovanetti, Vagner Fonseca, Flavia Aburjaile, Rodrigo Tocantins Calado. |
| EPI_ISL_1795390 | CS DE OSCAR BRESSANE PSF | Instituto Butantan / ESALQ-Piracicaba | Instituto Butantan: Alexander Roberto Precioso, Dimas Tadeu Covas, Sandra Coccuzzo Sampaio, Maria Carolina Elias, José Salvatore Leister Patané, Vincent Louis Viala, Antonio Jorge Martins, Ricardo Haddad, Claudia Renata dos Santos Barros, Elaine Cristina Marqueze, Raul Machado Neto, Debora Botequio Moretti. Centro de Genômica Funcional da ESALQ: Luiz Lehmann Coutinho, Ricardo Augusto Brassaloti, Raquel de Lello Rocha Campos Cassano. NGS Soluções Genômicas: Pilar Drummond Sampaio Corrêa Mariani. FZEA-USP Pirassununga: Mirele Daiana Poleti, Jessika Cristina Chagas Lesbon, Elisangela Chicaroni Mattos, Heidge Fukumasu. USP-Botucatu: Rejane Maria Tommasini Grotto, Jayme A. Souza-Neto, Guilherme Targino Valente, Patricia Akemi Assato, Felipe Allan da Silva da Costa, Bianca Cechetto Carlos. Mendelics: Bibiana Santos, João Paulo Kitajima, Erika Freitas, David Schlesinger. Hemocentro Ribeirão Preto: Simone Kashima, Evandra Strazza Rodrigues, Svetoslav Nanev Slavov, Elaine Vieira dos Santos, Rafael dos Santos Bezerra, Luiz Carlos Junior de Alcantara, Marta Giovanetti, Vagner Fonseca, Flavia Aburjaile, Rodrigo Tocantins Calado. |
| EPI_ISL_1795391 | USF ROSA CRUZ | Instituto Butantan / ESALQ-Piracicaba | Instituto Butantan: Alexander Roberto Precioso, Dimas Tadeu Covas, Sandra Coccuzzo Sampaio, Maria Carolina Elias, José Salvatore Leister Patané, Vincent Louis Viala, Antonio Jorge Martins, Ricardo Haddad, Claudia Renata dos Santos Barros, Elaine Cristina Marqueze, Raul Machado Neto, Debora Botequio Moretti. Centro de Genômica Funcional da ESALQ: Luiz Lehmann Coutinho, Ricardo Augusto Brassaloti, Raquel de Lello Rocha Campos Cassano. NGS Soluções Genômicas: Pilar Drummond Sampaio Corrêa Mariani. FZEA-USP Pirassununga: Mirele Daiana Poleti, Jessika Cristina Chagas Lesbon, Elisangela Chicaroni Mattos, Heidge Fukumasu. USP-Botucatu: Rejane Maria Tommasini Grotto, Jayme A. Souza-Neto, Guilherme Targino Valente, Patricia Akemi Assato, Felipe Allan da Silva da Costa, Bianca Cechetto Carlos. Mendelics: Bibiana Santos, João Paulo Kitajima, Erika Freitas, David Schlesinger. Hemocentro Ribeirão Preto: Simone Kashima, Evandra Strazza Rodrigues, Svetoslav Nanev Slavov, Elaine Vieira dos Santos, Rafael dos Santos Bezerra, Luiz Carlos Junior de Alcantara, Marta Giovanetti, Vagner Fonseca, Flavia Aburjaile, Rodrigo Tocantins Calado. |
| EPI_ISL_1795392 | POLICLINICA HORTOLANDIA | Instituto Butantan / ESALQ-Piracicaba | Instituto Butantan: Alexander Roberto Precioso, Dimas Tadeu Covas, Sandra Coccuzzo Sampaio, Maria Carolina Elias, José Salvatore Leister Patané, Vincent Louis Viala, Antonio Jorge Martins, Ricardo Haddad, Claudia Renata dos Santos Barros, Elaine Cristina Marqueze, Raul Machado Neto, Debora Botequio Moretti. Centro de Genômica Funcional da ESALQ: Luiz Lehmann Coutinho, Ricardo Augusto Brassaloti, Raquel de Lello Rocha Campos Cassano. NGS Soluções Genômicas: Pilar Drummond Sampaio Corrêa Mariani. FZEA-USP Pirassununga: Mirele Daiana Poleti, Jessika Cristina Chagas Lesbon, Elisangela Chicaroni Mattos, Heidge Fukumasu. USP-Botucatu: Rejane Maria Tommasini Grotto, Jayme A. Souza-Neto, Guilherme Targino Valente, Patricia Akemi Assato, Felipe Allan da Silva da Costa, Bianca Cechetto Carlos. Mendelics: Bibiana Santos, João Paulo Kitajima, Erika Freitas, David Schlesinger. Hemocentro Ribeirão Preto: Simone Kashima, Evandra Strazza Rodrigues, Svetoslav Nanev Slavov, Elaine Vieira dos Santos, Rafael dos Santos Bezerra, Luiz Carlos Junior de Alcantara, Marta Giovanetti, Vagner Fonseca, Flavia Aburjaile, Rodrigo Tocantins Calado. |
| EPI_ISL_1795393, EPI_ISL_1795394 | LABORATORIO DE FRANCA | Instituto Butantan / ESALQ-Piracicaba | Instituto Butantan: Alexander Roberto Precioso, Dimas Tadeu Covas, Sandra Coccuzzo Sampaio, Maria Carolina Elias, José Salvatore Leister Patané, Vincent Louis Viala, Antonio Jorge Martins, Ricardo Haddad, Claudia Renata dos Santos Barros, Elaine Cristina Marqueze, Raul Machado Neto, Debora Botequio Moretti. Centro de Genômica Funcional da ESALQ: Luiz Lehmann Coutinho, Ricardo Augusto Brassaloti, Raquel de Lello Rocha Campos Cassano. NGS Soluções Genômicas: Pilar Drummond Sampaio Corrêa Mariani. FZEA-USP Pirassununga: Mirele Daiana Poleti, Jessika Cristina Chagas Lesbon, Elisangela Chicaroni Mattos, Heidge Fukumasu. USP-Botucatu: Rejane Maria Tommasini Grotto, Jayme A. Souza-Neto, Guilherme Targino Valente, Patricia Akemi Assato, Felipe Allan da Silva da Costa, Bianca Cechetto Carlos. Mendelics: Bibiana Santos, João Paulo Kitajima, Erika Freitas, David Schlesinger. Hemocentro Ribeirão Preto: Simone Kashima, Evandra Strazza Rodrigues, Svetoslav Nanev Slavov, Elaine Vieira dos Santos, Rafael dos Santos Bezerra, Luiz Carlos Junior de Alcantara, Marta Giovanetti, Vagner Fonseca, Flavia Aburjaile, Rodrigo Tocantins Calado. |
| EPI_ISL_1795395, EPI_ISL_1795396, EPI_ISL_1795397 | UBS III DE RANCHARIA | Instituto Butantan / ESALQ-Piracicaba | Instituto Butantan: Alexander Roberto Precioso, Dimas Tadeu Covas, Sandra Coccuzzo Sampaio, Maria Carolina Elias, José Salvatore Leister Patané, Vincent Louis Viala, Antonio Jorge Martins, Ricardo Haddad, Claudia Renata dos Santos Barros, Elaine Cristina Marqueze, Raul Machado Neto, Debora Botequio Moretti. Centro de Genômica Funcional da ESALQ: Luiz Lehmann Coutinho, Ricardo Augusto Brassaloti, Raquel de Lello Rocha Campos Cassano. NGS Soluções Genômicas: Pilar Drummond Sampaio Corrêa Mariani. FZEA-USP Pirassununga: Mirele Daiana Poleti, Jessika Cristina Chagas Lesbon, Elisangela Chicaroni Mattos, Heidge Fukumasu. USP-Botucatu: Rejane Maria Tommasini Grotto, Jayme A. Souza-Neto, Guilherme Targino Valente, Patricia Akemi Assato, Felipe Allan da Silva da Costa, Bianca Cechetto Carlos. Mendelics: Bibiana Santos, João Paulo Kitajima, Erika Freitas, David Schlesinger. Hemocentro Ribeirão Preto: Simone Kashima, Evandra Strazza Rodrigues, Svetoslav Nanev Slavov, Elaine Vieira dos Santos, Rafael dos Santos Bezerra, Luiz Carlos Junior de Alcantara, Marta Giovanetti, Vagner Fonseca, Flavia Aburjaile, Rodrigo Tocantins Calado. |
| EPI_ISL_1795398 | SMS SECRETARIA MUNICIPAL DE SAUDE DE BOITUVA | Instituto Butantan / ESALQ-Piracicaba | Instituto Butantan: Alexander Roberto Precioso, Dimas Tadeu Covas, Sandra Coccuzzo Sampaio, Maria Carolina Elias, José Salvatore Leister Patané, Vincent Louis Viala, Antonio Jorge Martins, Ricardo Haddad, Claudia Renata dos Santos Barros, Elaine Cristina Marqueze, Raul Machado Neto, Debora Botequio Moretti. Centro de Genômica Funcional da ESALQ: Luiz Lehmann Coutinho, Ricardo Augusto Brassaloti, Raquel de Lello Rocha Campos Cassano. NGS Soluções Genômicas: Pilar Drummond Sampaio Corrêa Mariani. FZEA-USP Pirassununga: Mirele Daiana Poleti, Jessika Cristina Chagas Lesbon, Elisangela Chicaroni Mattos, Heidge Fukumasu. USP-Botucatu: Rejane Maria Tommasini Grotto, Jayme A. Souza-Neto, Guilherme Targino Valente, Patricia Akemi Assato, Felipe Allan da Silva da Costa, Bianca Cechetto Carlos. Mendelics: Bibiana Santos, João Paulo Kitajima, Erika Freitas, David Schlesinger. Hemocentro Ribeirão Preto: Simone Kashima, Evandra Strazza Rodrigues, Svetoslav Nanev Slavov, Elaine Vieira dos Santos, Rafael dos Santos Bezerra, Luiz Carlos Junior de Alcantara, Marta Giovanetti, Vagner Fonseca, Flavia Aburjaile, Rodrigo Tocantins Calado. |
| EPI_ISL_1795399 | POLICLINICA HORTOLANDIA | Instituto Butantan / ESALQ-Piracicaba | Instituto Butantan: Alexander Roberto Precioso, Dimas Tadeu Covas, Sandra Coccuzzo Sampaio, Maria Carolina Elias, José Salvatore Leister Patané, Vincent Louis Viala, Antonio Jorge Martins, Ricardo Haddad, Claudia Renata dos Santos Barros, Elaine Cristina Marqueze, Raul Machado Neto, Debora Botequio Moretti. Centro de Genômica Funcional da ESALQ: Luiz Lehmann Coutinho, Ricardo Augusto Brassaloti, Raquel de Lello Rocha Campos Cassano. NGS Soluções Genômicas: Pilar Drummond Sampaio Corrêa Mariani. FZEA-USP Pirassununga: Mirele Daiana Poleti, Jessika Cristina Chagas Lesbon, Elisangela Chicaroni Mattos, Heidge Fukumasu. USP-Botucatu: Rejane Maria Tommasini Grotto, Jayme A. Souza-Neto, Guilherme Targino Valente, Patricia Akemi Assato, Felipe Allan da Silva da Costa, Bianca Cechetto Carlos. Mendelics: Bibiana Santos, João Paulo Kitajima, Erika Freitas, David Schlesinger. Hemocentro Ribeirão Preto: Simone Kashima, Evandra Strazza Rodrigues, Svetoslav Nanev Slavov, Elaine Vieira dos Santos, Rafael dos Santos Bezerra, Luiz Carlos Junior de Alcantara, Marta Giovanetti, Vagner Fonseca, Flavia Aburjaile, Rodrigo Tocantins Calado. |
| EPI_ISL_1795400, EPI_ISL_1795401 | LABORATORIO MUNICIPAL DE ANALISES CLINICAS DE RIO CLARO | Instituto Butantan / ESALQ-Piracicaba | Instituto Butantan: Alexander Roberto Precioso, Dimas Tadeu Covas, Sandra Coccuzzo Sampaio, Maria Carolina Elias, José Salvatore Leister Patané, Vincent Louis Viala, Antonio Jorge Martins, Ricardo Haddad, Claudia Renata dos Santos Barros, Elaine Cristina Marqueze, Raul Machado Neto, Debora Botequio Moretti. Centro de Genômica Funcional da ESALQ: Luiz Lehmann Coutinho, Ricardo Augusto Brassaloti, Raquel de Lello Rocha Campos Cassano. NGS Soluções Genômicas: Pilar Drummond Sampaio Corrêa Mariani. FZEA-USP Pirassununga: Mirele Daiana Poleti, Jessika Cristina Chagas Lesbon, Elisangela Chicaroni Mattos, Heidge Fukumasu. USP-Botucatu: Rejane Maria Tommasini Grotto, Jayme A. Souza-Neto, Guilherme Targino Valente, Patricia Akemi Assato, Felipe Allan da Silva da Costa, Bianca Cechetto Carlos. Mendelics: Bibiana Santos, João Paulo Kitajima, Erika Freitas, David Schlesinger. Hemocentro Ribeirão Preto: Simone Kashima, Evandra Strazza Rodrigues, Svetoslav Nanev Slavov, Elaine Vieira dos Santos, Rafael dos Santos Bezerra, Luiz Carlos Junior de Alcantara, Marta Giovanetti, Vagner Fonseca, Flavia Aburjaile, Rodrigo Tocantins Calado. |
| EPI_ISL_1795402 | SECRETARIA MUNICIPAL DE SAUDE DE CORDEIROPOLIS | Instituto Butantan / ESALQ-Piracicaba | Instituto Butantan: Alexander Roberto Precioso, Dimas Tadeu Covas, Sandra Coccuzzo Sampaio, Maria Carolina Elias, José Salvatore Leister Patané, Vincent Louis Viala, Antonio Jorge Martins, Ricardo Haddad, Claudia Renata dos Santos Barros, Elaine Cristina Marqueze, Raul Machado Neto, Debora Botequio Moretti. Centro de Genômica Funcional da ESALQ: Luiz Lehmann Coutinho, Ricardo Augusto Brassaloti, Raquel de Lello Rocha Campos Cassano. NGS Soluções Genômicas: Pilar Drummond Sampaio Corrêa Mariani. FZEA-USP Pirassununga: Mirele Daiana Poleti, Jessika Cristina Chagas Lesbon, Elisangela Chicaroni Mattos, Heidge Fukumasu. USP-Botucatu: Rejane Maria Tommasini Grotto, Jayme A. Souza-Neto, Guilherme Targino |

| | | | |
|---|---|---|---|
| | | | Valente, Patricia Akemi Assato, Felipe Allan da Silva da Costa, Bianca Cechetto Carlos. Mendelics: Bibiana Santos, João Paulo Kitajima, Erika Freitas, David Schlesinger. Hemocentro Ribeirão Preto: Simone Kashima, Evandra Strazza Rodrigues, Svetoslav Nanev Slavov, Elaine Vieira dos Santos, Rafael dos Santos Bezerra, Luiz Carlos Junior de Alcantara, Marta Giovanetti, Vagner Fonseca, Flavia Aburjaile, Rodrigo Tocantins Calado. |
| EPI_ISL_1795403 | UNIDADE MISTA DE LUIZ ANTONIO | Instituto Butantan / ESALQ-Piracicaba | Instituto Butantan: Alexander Roberto Precioso, Dimas Tadeu Covas, Sandra Coccuzzo Sampaio, Maria Carolina Elias, José Salvatore Leister Patané, Vincent Louis Viala, Antonio Jorge Martins, Ricardo Haddad, Claudia Renata dos Santos Barros, Elaine Cristina Marqueze, Raul Machado Neto, Debora Botequio Moretti. Centro de Genômica Funcional da ESALQ: Luiz Lehmann Coutinho, Ricardo Augusto Brassaloti, Raquel de Lello Rocha Campos Cassano. NGS Soluções Genômicas: Pilar Drummond Sampaio Corrêa Mariani. FZEA-USP Pirassununga: Mirele Daiana Poleti, Jessika Cristina Chagas Lesbon, Elisangela Chicaroni Mattos, Heidge Fukumasu. USP-Botucatu: Rejane Maria Tommasini Grotto, Jayme A. Souza-Neto, Guilherme Targino Valente, Patricia Akemi Assato, Felipe Allan da Silva da Costa, Bianca Cechetto Carlos. Mendelics: Bibiana Santos, João Paulo Kitajima, Erika Freitas, David Schlesinger. Hemocentro Ribeirão Preto: Simone Kashima, Evandra Strazza Rodrigues, Svetoslav Nanev Slavov, Elaine Vieira dos Santos, Rafael dos Santos Bezerra, Luiz Carlos Junior de Alcantara, Marta Giovanetti, Vagner Fonseca, Flavia Aburjaile, Rodrigo Tocantins Calado. |
| EPI_ISL_1795404 | POLICLINICA HORTOLANDIA | Instituto Butantan / ESALQ-Piracicaba | Instituto Butantan: Alexander Roberto Precioso, Dimas Tadeu Covas, Sandra Coccuzzo Sampaio, Maria Carolina Elias, José Salvatore Leister Patané, Vincent Louis Viala, Antonio Jorge Martins, Ricardo Haddad, Claudia Renata dos Santos Barros, Elaine Cristina Marqueze, Raul Machado Neto, Debora Botequio Moretti. Centro de Genômica Funcional da ESALQ: Luiz Lehmann Coutinho, Ricardo Augusto Brassaloti, Raquel de Lello Rocha Campos Cassano. NGS Soluções Genômicas: Pilar Drummond Sampaio Corrêa Mariani. FZEA-USP Pirassununga: Mirele Daiana Poleti, Jessika Cristina Chagas Lesbon, Elisangela Chicaroni Mattos, Heidge Fukumasu. USP-Botucatu: Rejane Maria Tommasini Grotto, Jayme A. Souza-Neto, Guilherme Targino Valente, Patricia Akemi Assato, Felipe Allan da Silva da Costa, Bianca Cechetto Carlos. Mendelics: Bibiana Santos, João Paulo Kitajima, Erika Freitas, David Schlesinger. Hemocentro Ribeirão Preto: Simone Kashima, Evandra Strazza Rodrigues, Svetoslav Nanev Slavov, Elaine Vieira dos Santos, Rafael dos Santos Bezerra, Luiz Carlos Junior de Alcantara, Marta Giovanetti, Vagner Fonseca, Flavia Aburjaile, Rodrigo Tocantins Calado. |
| EPI_ISL_1795405 | UBS CAIC OURINHOS | Instituto Butantan / ESALQ-Piracicaba | Instituto Butantan: Alexander Roberto Precioso, Dimas Tadeu Covas, Sandra Coccuzzo Sampaio, Maria Carolina Elias, José Salvatore Leister Patané, Vincent Louis Viala, Antonio Jorge Martins, Ricardo Haddad, Claudia Renata dos Santos Barros, Elaine Cristina Marqueze, Raul Machado Neto, Debora Botequio Moretti. Centro de Genômica Funcional da ESALQ: Luiz Lehmann Coutinho, Ricardo Augusto Brassaloti, Raquel de Lello Rocha Campos Cassano. NGS Soluções Genômicas: Pilar Drummond Sampaio Corrêa Mariani. FZEA-USP Pirassununga: Mirele Daiana Poleti, Jessika Cristina Chagas Lesbon, Elisangela Chicaroni Mattos, Heidge Fukumasu. USP-Botucatu: Rejane Maria Tommasini Grotto, Jayme A. Souza-Neto, Guilherme Targino Valente, Patricia Akemi Assato, Felipe Allan da Silva da Costa, Bianca Cechetto Carlos. Mendelics: Bibiana Santos, João Paulo Kitajima, Erika Freitas, David Schlesinger. Hemocentro Ribeirão Preto: Simone Kashima, Evandra Strazza Rodrigues, Svetoslav Nanev Slavov, Elaine Vieira dos Santos, Rafael dos Santos Bezerra, Luiz Carlos Junior de Alcantara, Marta Giovanetti, Vagner Fonseca, Flavia Aburjaile, Rodrigo Tocantins Calado. |
| EPI_ISL_1795406 | CS DE OSCAR BRESSANE PSF | Instituto Butantan / ESALQ-Piracicaba | Instituto Butantan: Alexander Roberto Precioso, Dimas Tadeu Covas, Sandra Coccuzzo Sampaio, Maria Carolina Elias, José Salvatore Leister Patané, Vincent Louis Viala, Antonio Jorge Martins, Ricardo Haddad, Claudia Renata dos Santos Barros, Elaine Cristina Marqueze, Raul Machado Neto, Debora Botequio Moretti. Centro de Genômica Funcional da ESALQ: Luiz Lehmann Coutinho, Ricardo Augusto Brassaloti, Raquel de Lello Rocha Campos Cassano. NGS Soluções Genômicas: Pilar Drummond Sampaio Corrêa Mariani. FZEA-USP Pirassununga: Mirele Daiana Poleti, Jessika Cristina Chagas Lesbon, Elisangela Chicaroni Mattos, Heidge Fukumasu. USP-Botucatu: Rejane Maria Tommasini Grotto, Jayme A. Souza-Neto, Guilherme Targino Valente, Patricia Akemi Assato, Felipe Allan da Silva da Costa, Bianca Cechetto Carlos. Mendelics: Bibiana Santos, João Paulo Kitajima, Erika Freitas, David Schlesinger. Hemocentro Ribeirão Preto: Simone Kashima, Evandra Strazza Rodrigues, Svetoslav Nanev Slavov, Elaine Vieira dos Santos, Rafael dos Santos Bezerra, Luiz Carlos Junior de Alcantara, Marta Giovanetti, Vagner Fonseca, Flavia Aburjaile, Rodrigo Tocantins Calado. |
| EPI_ISL_1795407 | CENTRO DE SAUDE DE RIBEIRAO DO SUL | Instituto Butantan / ESALQ-Piracicaba | Instituto Butantan: Alexander Roberto Precioso, Dimas Tadeu Covas, Sandra Coccuzzo Sampaio, Maria Carolina Elias, José Salvatore Leister Patané, Vincent Louis Viala, Antonio Jorge Martins, Ricardo Haddad, Claudia Renata dos Santos Barros, Elaine Cristina Marqueze, Raul Machado Neto, Debora Botequio Moretti. Centro de Genômica Funcional da ESALQ: Luiz Lehmann Coutinho, Ricardo Augusto Brassaloti, Raquel de Lello Rocha Campos Cassano. NGS Soluções Genômicas: Pilar Drummond Sampaio Corrêa Mariani. FZEA-USP Pirassununga: Mirele Daiana Poleti, Jessika Cristina Chagas Lesbon, Elisangela Chicaroni Mattos, Heidge Fukumasu. USP-Botucatu: Rejane Maria Tommasini Grotto, Jayme A. Souza-Neto, Guilherme Targino Valente, Patricia Akemi Assato, Felipe Allan da Silva da Costa, Bianca Cechetto Carlos. Mendelics: Bibiana Santos, João Paulo Kitajima, Erika Freitas, David Schlesinger. Hemocentro Ribeirão Preto: Simone Kashima, Evandra Strazza Rodrigues, Svetoslav Nanev Slavov, Elaine Vieira dos Santos, Rafael dos Santos Bezerra, Luiz Carlos Junior de Alcantara, Marta Giovanetti, Vagner Fonseca, Flavia Aburjaile, Rodrigo Tocantins Calado. |
| EPI_ISL_1795408 | CS DE OSCAR BRESSANE PSF | Instituto Butantan / ESALQ-Piracicaba | Instituto Butantan: Alexander Roberto Precioso, Dimas Tadeu Covas, Sandra Coccuzzo Sampaio, Maria Carolina Elias, José Salvatore Leister Patané, Vincent Louis Viala, Antonio Jorge Martins, Ricardo Haddad, Claudia Renata dos Santos Barros, Elaine Cristina Marqueze, Raul Machado Neto, Debora Botequio Moretti. Centro de Genômica Funcional da ESALQ: Luiz Lehmann Coutinho, Ricardo Augusto Brassaloti, Raquel de Lello Rocha Campos Cassano. NGS Soluções Genômicas: Pilar Drummond Sampaio Corrêa Mariani. FZEA-USP Pirassununga: Mirele Daiana Poleti, Jessika Cristina Chagas Lesbon, Elisangela Chicaroni Mattos, Heidge Fukumasu. USP-Botucatu: Rejane Maria Tommasini Grotto, Jayme A. Souza-Neto, Guilherme Targino Valente, Patricia Akemi Assato, Felipe Allan da Silva da Costa, Bianca Cechetto Carlos. Mendelics: Bibiana Santos, João Paulo Kitajima, Erika Freitas, David Schlesinger. Hemocentro Ribeirão Preto: Simone Kashima, Evandra Strazza Rodrigues, Svetoslav Nanev Slavov, Elaine Vieira dos Santos, Rafael dos Santos Bezerra, Luiz Carlos Junior de Alcantara, Marta Giovanetti, Vagner Fonseca, Flavia Aburjaile, Rodrigo Tocantins Calado. |
| EPI_ISL_1795409, EPI_ISL_1795410 | CENTRO DE SAUDE DE MARACAI | Instituto Butantan / ESALQ-Piracicaba | Instituto Butantan: Alexander Roberto Precioso, Dimas Tadeu Covas, Sandra Coccuzzo Sampaio, Maria Carolina Elias, José Salvatore Leister Patané, Vincent Louis Viala, Antonio Jorge Martins, Ricardo Haddad, Claudia Renata dos Santos Barros, Elaine Cristina Marqueze, Raul Machado Neto, Debora Botequio Moretti. Centro de Genômica Funcional da ESALQ: Luiz Lehmann Coutinho, Ricardo Augusto Brassaloti, Raquel de Lello Rocha Campos Cassano. NGS Soluções Genômicas: Pilar Drummond Sampaio Corrêa Mariani. FZEA-USP Pirassununga: Mirele Daiana Poleti, Jessika Cristina Chagas Lesbon, Elisangela Chicaroni Mattos, Heidge Fukumasu. USP-Botucatu: Rejane Maria Tommasini Grotto, Jayme A. Souza-Neto, Guilherme Targino Valente, Patricia Akemi Assato, Felipe Allan da Silva da Costa, Bianca Cechetto Carlos. Mendelics: Bibiana Santos, João Paulo Kitajima, Erika Freitas, David Schlesinger. Hemocentro Ribeirão Preto: Simone Kashima, Evandra Strazza Rodrigues, Svetoslav Nanev Slavov, Elaine Vieira dos Santos, Rafael dos Santos Bezerra, Luiz Carlos Junior de Alcantara, Marta Giovanetti, Vagner Fonseca, Flavia Aburjaile, Rodrigo Tocantins Calado. |
| EPI_ISL_1795411 | POLICLINICA HORTOLANDIA | Instituto Butantan / ESALQ-Piracicaba | Instituto Butantan: Alexander Roberto Precioso, Dimas Tadeu Covas, Sandra Coccuzzo Sampaio, Maria Carolina Elias, José Salvatore Leister Patané, Vincent Louis Viala, Antonio Jorge Martins, Ricardo Haddad, Claudia Renata dos Santos Barros, Elaine Cristina Marqueze, Raul Machado Neto, Debora Botequio Moretti. Centro de Genômica Funcional da ESALQ: Luiz Lehmann Coutinho, Ricardo Augusto Brassaloti, Raquel de Lello Rocha Campos Cassano. NGS Soluções Genômicas: Pilar Drummond Sampaio Corrêa Mariani. FZEA-USP Pirassununga: Mirele Daiana Poleti, Jessika Cristina Chagas Lesbon, Elisangela Chicaroni Mattos, Heidge Fukumasu. USP-Botucatu: Rejane Maria Tommasini Grotto, Jayme A. Souza-Neto, Guilherme Targino Valente, Patricia Akemi Assato, Felipe Allan da Silva da Costa, Bianca Cechetto Carlos. Mendelics: Bibiana Santos, João Paulo Kitajima, Erika Freitas, David Schlesinger. Hemocentro Ribeirão Preto: Simone Kashima, Evandra Strazza Rodrigues, Svetoslav Nanev Slavov, Elaine Vieira dos Santos, Rafael dos Santos Bezerra, Luiz Carlos Junior de Alcantara, Marta Giovanetti, Vagner Fonseca, Flavia Aburjaile, Rodrigo Tocantins Calado. |
| EPI_ISL_1795412 | USF SALERNO | Instituto Butantan / ESALQ-Piracicaba | Instituto Butantan: Alexander Roberto Precioso, Dimas Tadeu Covas, Sandra Coccuzzo Sampaio, Maria Carolina Elias, José Salvatore Leister Patané, Vincent Louis Viala, Antonio Jorge Martins, Ricardo Haddad, Claudia Renata dos Santos Barros, Elaine Cristina Marqueze, Raul Machado Neto, Debora Botequio Moretti. Centro de Genômica Funcional da ESALQ: Luiz Lehmann Coutinho, Ricardo Augusto Brassaloti, Raquel de Lello Rocha Campos Cassano. NGS Soluções Genômicas: Pilar Drummond Sampaio Corrêa Mariani. FZEA-USP Pirassununga: Mirele Daiana Poleti, Jessika Cristina Chagas Lesbon, Elisangela Chicaroni Mattos, Heidge Fukumasu. USP-Botucatu: Rejane Maria Tommasini Grotto, Jayme A. Souza-Neto, Guilherme Targino Valente, Patricia Akemi Assato, Felipe Allan da Silva da Costa, Bianca Cechetto Carlos. Mendelics: Bibiana Santos, João Paulo Kitajima, Erika Freitas, David Schlesinger. Hemocentro Ribeirão Preto: Simone Kashima, Evandra Strazza Rodrigues, Svetoslav Nanev Slavov, Elaine Vieira dos Santos, Rafael dos Santos Bezerra, Luiz Carlos Junior de Alcantara, Marta Giovanetti, Vagner Fonseca, Flavia Aburjaile, Rodrigo Tocantins Calado. |
| EPI_ISL_1795413 | PRONTO SOCORRO MUNICIPAL TAMBAU | Instituto Butantan / ESALQ-Piracicaba | Instituto Butantan: Alexander Roberto Precioso, Dimas Tadeu Covas, Sandra Coccuzzo Sampaio, Maria Carolina Elias, José Salvatore Leister Patané, Vincent Louis Viala, Antonio Jorge Martins, Ricardo Haddad, Claudia Renata dos Santos Barros, Elaine Cristina Marqueze, Raul Machado Neto, Debora Botequio Moretti. Centro de Genômica Funcional da ESALQ: Luiz Lehmann Coutinho, Ricardo Augusto Brassaloti, Raquel de Lello Rocha Campos Cassano. NGS Soluções Genômicas: Pilar Drummond Sampaio Corrêa Mariani. FZEA-USP Pirassununga: Mirele Daiana Poleti, Jessika Cristina Chagas Lesbon, Elisangela Chicaroni Mattos, Heidge Fukumasu. USP-Botucatu: Rejane Maria Tommasini Grotto, Jayme A. Souza-Neto, Guilherme Targino |

| | | | |
|---|---|---|---|
| | | | Valente, Patricia Akemi Assato, Felipe Allan da Silva da Costa, Bianca Cechetto Carlos. Mendelics: Bibiana Santos, João Paulo Kitajima, Erika Freitas, David Schlesinger. Hemocentro Ribeirão Preto: Simone Kashima, Evandra Strazza Rodrigues, Svetoslav Nanev Slavov, Elaine Vieira dos Santos, Rafael dos Santos Bezerra, Luiz Carlos Junior de Alcantara, Marta Giovanetti, Vagner Fonseca, Flavia Aburjaile, Rodrigo Tocantins Calado. |
| EPI_ISL_1795414 | HOSPITAL MUNICIPAL DE IBIUNA IBIUNA SP | Instituto Butantan / ESALQ-Piracicaba | Instituto Butantan: Alexander Roberto Precioso, Dimas Tadeu Covas, Sandra Coccuzzo Sampaio, Maria Carolina Elias, José Salvatore Leister Patané, Vincent Louis Viala, Antonio Jorge Martins, Ricardo Haddad, Claudia Renata dos Santos Barros, Elaine Cristina Marqueze, Raul Machado Neto, Debora Botequio Moretti. Centro de Genômica Funcional da ESALQ: Luiz Lehmann Coutinho, Ricardo Augusto Brassaloti, Raquel de Lello Rocha Campos Cassano. NGS Soluções Genômicas: Pilar Drummond Sampaio Corrêa Mariani. FZEA-USP Pirassununga: Mirele Daiana Poleti, Jessika Cristina Chagas Lesbon, Elisangela Chicaroni Mattos, Heidge Fukumasu. USP-Botucatu: Rejane Maria Tommasini Grotto, Jayme A. Souza-Neto, Guilherme Targino Valente, Patricia Akemi Assato, Felipe Allan da Silva da Costa, Bianca Cechetto Carlos. Mendelics: Bibiana Santos, João Paulo Kitajima, Erika Freitas, David Schlesinger. Hemocentro Ribeirão Preto: Simone Kashima, Evandra Strazza Rodrigues, Svetoslav Nanev Slavov, Elaine Vieira dos Santos, Rafael dos Santos Bezerra, Luiz Carlos Junior de Alcantara, Marta Giovanetti, Vagner Fonseca, Flavia Aburjaile, Rodrigo Tocantins Calado. |
| EPI_ISL_1795415 | UBS DE SANTA SALETE | Instituto Butantan / ESALQ-Piracicaba | Instituto Butantan: Alexander Roberto Precioso, Dimas Tadeu Covas, Sandra Coccuzzo Sampaio, Maria Carolina Elias, José Salvatore Leister Patané, Vincent Louis Viala, Antonio Jorge Martins, Ricardo Haddad, Claudia Renata dos Santos Barros, Elaine Cristina Marqueze, Raul Machado Neto, Debora Botequio Moretti. Centro de Genômica Funcional da ESALQ: Luiz Lehmann Coutinho, Ricardo Augusto Brassaloti, Raquel de Lello Rocha Campos Cassano. NGS Soluções Genômicas: Pilar Drummond Sampaio Corrêa Mariani. FZEA-USP Pirassununga: Mirele Daiana Poleti, Jessika Cristina Chagas Lesbon, Elisangela Chicaroni Mattos, Heidge Fukumasu. USP-Botucatu: Rejane Maria Tommasini Grotto, Jayme A. Souza-Neto, Guilherme Targino Valente, Patricia Akemi Assato, Felipe Allan da Silva da Costa, Bianca Cechetto Carlos. Mendelics: Bibiana Santos, João Paulo Kitajima, Erika Freitas, David Schlesinger. Hemocentro Ribeirão Preto: Simone Kashima, Evandra Strazza Rodrigues, Svetoslav Nanev Slavov, Elaine Vieira dos Santos, Rafael dos Santos Bezerra, Luiz Carlos Junior de Alcantara, Marta Giovanetti, Vagner Fonseca, Flavia Aburjaile, Rodrigo Tocantins Calado. |
| EPI_ISL_1795416 | CS DE MARAPOAMA | Instituto Butantan / ESALQ-Piracicaba | Instituto Butantan: Alexander Roberto Precioso, Dimas Tadeu Covas, Sandra Coccuzzo Sampaio, Maria Carolina Elias, José Salvatore Leister Patané, Vincent Louis Viala, Antonio Jorge Martins, Ricardo Haddad, Claudia Renata dos Santos Barros, Elaine Cristina Marqueze, Raul Machado Neto, Debora Botequio Moretti. Centro de Genômica Funcional da ESALQ: Luiz Lehmann Coutinho, Ricardo Augusto Brassaloti, Raquel de Lello Rocha Campos Cassano. NGS Soluções Genômicas: Pilar Drummond Sampaio Corrêa Mariani. FZEA-USP Pirassununga: Mirele Daiana Poleti, Jessika Cristina Chagas Lesbon, Elisangela Chicaroni Mattos, Heidge Fukumasu. USP-Botucatu: Rejane Maria Tommasini Grotto, Jayme A. Souza-Neto, Guilherme Targino Valente, Patricia Akemi Assato, Felipe Allan da Silva da Costa, Bianca Cechetto Carlos. Mendelics: Bibiana Santos, João Paulo Kitajima, Erika Freitas, David Schlesinger. Hemocentro Ribeirão Preto: Simone Kashima, Evandra Strazza Rodrigues, Svetoslav Nanev Slavov, Elaine Vieira dos Santos, Rafael dos Santos Bezerra, Luiz Carlos Junior de Alcantara, Marta Giovanetti, Vagner Fonseca, Flavia Aburjaile, Rodrigo Tocantins Calado. |
| EPI_ISL_1795417 | AMBULATORIO DE ESPECIALIDADE V E MOGI MIRIM | Instituto Butantan / ESALQ-Piracicaba | Instituto Butantan: Alexander Roberto Precioso, Dimas Tadeu Covas, Sandra Coccuzzo Sampaio, Maria Carolina Elias, José Salvatore Leister Patané, Vincent Louis Viala, Antonio Jorge Martins, Ricardo Haddad, Claudia Renata dos Santos Barros, Elaine Cristina Marqueze, Raul Machado Neto, Debora Botequio Moretti. Centro de Genômica Funcional da ESALQ: Luiz Lehmann Coutinho, Ricardo Augusto Brassaloti, Raquel de Lello Rocha Campos Cassano. NGS Soluções Genômicas: Pilar Drummond Sampaio Corrêa Mariani. FZEA-USP Pirassununga: Mirele Daiana Poleti, Jessika Cristina Chagas Lesbon, Elisangela Chicaroni Mattos, Heidge Fukumasu. USP-Botucatu: Rejane Maria Tommasini Grotto, Jayme A. Souza-Neto, Guilherme Targino Valente, Patricia Akemi Assato, Felipe Allan da Silva da Costa, Bianca Cechetto Carlos. Mendelics: Bibiana Santos, João Paulo Kitajima, Erika Freitas, David Schlesinger. Hemocentro Ribeirão Preto: Simone Kashima, Evandra Strazza Rodrigues, Svetoslav Nanev Slavov, Elaine Vieira dos Santos, Rafael dos Santos Bezerra, Luiz Carlos Junior de Alcantara, Marta Giovanetti, Vagner Fonseca, Flavia Aburjaile, Rodrigo Tocantins Calado. |
| EPI_ISL_1795418, EPI_ISL_1795419, EPI_ISL_1795420, EPI_ISL_1795421, EPI_ISL_1795422, EPI_ISL_1795423 | LABORATORIO DE FRANCA | Instituto Butantan / ESALQ-Piracicaba | Instituto Butantan: Alexander Roberto Precioso, Dimas Tadeu Covas, Sandra Coccuzzo Sampaio, Maria Carolina Elias, José Salvatore Leister Patané, Vincent Louis Viala, Antonio Jorge Martins, Ricardo Haddad, Claudia Renata dos Santos Barros, Elaine Cristina Marqueze, Raul Machado Neto, Debora Botequio Moretti. Centro de Genômica Funcional da ESALQ: Luiz Lehmann Coutinho, Ricardo Augusto Brassaloti, Raquel de Lello Rocha Campos Cassano. NGS Soluções Genômicas: Pilar Drummond Sampaio Corrêa Mariani. FZEA-USP Pirassununga: Mirele Daiana Poleti, Jessika Cristina Chagas Lesbon, Elisangela Chicaroni Mattos, Heidge Fukumasu. USP-Botucatu: Rejane Maria Tommasini Grotto, Jayme A. Souza-Neto, Guilherme Targino Valente, Patricia Akemi Assato, Felipe Allan da Silva da Costa, Bianca Cechetto Carlos. Mendelics: Bibiana Santos, João Paulo Kitajima, Erika Freitas, David Schlesinger. Hemocentro Ribeirão Preto: Simone Kashima, Evandra Strazza Rodrigues, Svetoslav Nanev Slavov, Elaine Vieira dos Santos, Rafael dos Santos Bezerra, Luiz Carlos Junior de Alcantara, Marta Giovanetti, Vagner Fonseca, Flavia Aburjaile, Rodrigo Tocantins Calado. |
| EPI_ISL_1795424 | UBS II DE REGENTE FEIJO | Instituto Butantan / ESALQ-Piracicaba | Instituto Butantan: Alexander Roberto Precioso, Dimas Tadeu Covas, Sandra Coccuzzo Sampaio, Maria Carolina Elias, José Salvatore Leister Patané, Vincent Louis Viala, Antonio Jorge Martins, Ricardo Haddad, Claudia Renata dos Santos Barros, Elaine Cristina Marqueze, Raul Machado Neto, Debora Botequio Moretti. Centro de Genômica Funcional da ESALQ: Luiz Lehmann Coutinho, Ricardo Augusto Brassaloti, Raquel de Lello Rocha Campos Cassano. NGS Soluções Genômicas: Pilar Drummond Sampaio Corrêa Mariani. FZEA-USP Pirassununga: Mirele Daiana Poleti, Jessika Cristina Chagas Lesbon, Elisangela Chicaroni Mattos, Heidge Fukumasu. USP-Botucatu: Rejane Maria Tommasini Grotto, Jayme A. Souza-Neto, Guilherme Targino Valente, Patricia Akemi Assato, Felipe Allan da Silva da Costa, Bianca Cechetto Carlos. Mendelics: Bibiana Santos, João Paulo Kitajima, Erika Freitas, David Schlesinger. Hemocentro Ribeirão Preto: Simone Kashima, Evandra Strazza Rodrigues, Svetoslav Nanev Slavov, Elaine Vieira dos Santos, Rafael dos Santos Bezerra, Luiz Carlos Junior de Alcantara, Marta Giovanetti, Vagner Fonseca, Flavia Aburjaile, Rodrigo Tocantins Calado. |
| EPI_ISL_1795425 | UBS III DE RANCHARIA | Instituto Butantan / ESALQ-Piracicaba | Instituto Butantan: Alexander Roberto Precioso, Dimas Tadeu Covas, Sandra Coccuzzo Sampaio, Maria Carolina Elias, José Salvatore Leister Patané, Vincent Louis Viala, Antonio Jorge Martins, Ricardo Haddad, Claudia Renata dos Santos Barros, Elaine Cristina Marqueze, Raul Machado Neto, Debora Botequio Moretti. Centro de Genômica Funcional da ESALQ: Luiz Lehmann Coutinho, Ricardo Augusto Brassaloti, Raquel de Lello Rocha Campos Cassano. NGS Soluções Genômicas: Pilar Drummond Sampaio Corrêa Mariani. FZEA-USP Pirassununga: Mirele Daiana Poleti, Jessika Cristina Chagas Lesbon, Elisangela Chicaroni Mattos, Heidge Fukumasu. USP-Botucatu: Rejane Maria Tommasini Grotto, Jayme A. Souza-Neto, Guilherme Targino Valente, Patricia Akemi Assato, Felipe Allan da Silva da Costa, Bianca Cechetto Carlos. Mendelics: Bibiana Santos, João Paulo Kitajima, Erika Freitas, David Schlesinger. Hemocentro Ribeirão Preto: Simone Kashima, Evandra Strazza Rodrigues, Svetoslav Nanev Slavov, Elaine Vieira dos Santos, Rafael dos Santos Bezerra, Luiz Carlos Junior de Alcantara, Marta Giovanetti, Vagner Fonseca, Flavia Aburjaile, Rodrigo Tocantins Calado. |
| EPI_ISL_1795426 | PRONTO SOCORRO MUNICIPAL DE PARAIBUNA | Instituto Butantan / ESALQ-Piracicaba | Instituto Butantan: Alexander Roberto Precioso, Dimas Tadeu Covas, Sandra Coccuzzo Sampaio, Maria Carolina Elias, José Salvatore Leister Patané, Vincent Louis Viala, Antonio Jorge Martins, Ricardo Haddad, Claudia Renata dos Santos Barros, Elaine Cristina Marqueze, Raul Machado Neto, Debora Botequio Moretti. Centro de Genômica Funcional da ESALQ: Luiz Lehmann Coutinho, Ricardo Augusto Brassaloti, Raquel de Lello Rocha Campos Cassano. NGS Soluções Genômicas: Pilar Drummond Sampaio Corrêa Mariani. FZEA-USP Pirassununga: Mirele Daiana Poleti, Jessika Cristina Chagas Lesbon, Elisangela Chicaroni Mattos, Heidge Fukumasu. USP-Botucatu: Rejane Maria Tommasini Grotto, Jayme A. Souza-Neto, Guilherme Targino Valente, Patricia Akemi Assato, Felipe Allan da Silva da Costa, Bianca Cechetto Carlos. Mendelics: Bibiana Santos, João Paulo Kitajima, Erika Freitas, David Schlesinger. Hemocentro Ribeirão Preto: Simone Kashima, Evandra Strazza Rodrigues, Svetoslav Nanev Slavov, Elaine Vieira dos Santos, Rafael dos Santos Bezerra, Luiz Carlos Junior de Alcantara, Marta Giovanetti, Vagner Fonseca, Flavia Aburjaile, Rodrigo Tocantins Calado. |
| EPI_ISL_1795427 | CENTRO DE SAUDE DE NATIVIDADE DA SERRA | Instituto Butantan / ESALQ-Piracicaba | Instituto Butantan: Alexander Roberto Precioso, Dimas Tadeu Covas, Sandra Coccuzzo Sampaio, Maria Carolina Elias, José Salvatore Leister Patané, Vincent Louis Viala, Antonio Jorge Martins, Ricardo Haddad, Claudia Renata dos Santos Barros, Elaine Cristina Marqueze, Raul Machado Neto, Debora Botequio Moretti. Centro de Genômica Funcional da ESALQ: Luiz Lehmann Coutinho, Ricardo Augusto Brassaloti, Raquel de Lello Rocha Campos Cassano. NGS Soluções Genômicas: Pilar Drummond Sampaio Corrêa Mariani. FZEA-USP Pirassununga: Mirele Daiana Poleti, Jessika Cristina Chagas Lesbon, Elisangela Chicaroni Mattos, Heidge Fukumasu. USP-Botucatu: Rejane Maria Tommasini Grotto, Jayme A. Souza-Neto, Guilherme Targino Valente, Patricia Akemi Assato, Felipe Allan da Silva da Costa, Bianca Cechetto Carlos. Mendelics: Bibiana Santos, João Paulo Kitajima, Erika Freitas, David Schlesinger. Hemocentro Ribeirão Preto: Simone Kashima, Evandra Strazza Rodrigues, Svetoslav Nanev Slavov, Elaine Vieira dos Santos, Rafael dos Santos Bezerra, Luiz Carlos Junior de Alcantara, Marta Giovanetti, Vagner Fonseca, Flavia Aburjaile, Rodrigo Tocantins Calado. |
| EPI_ISL_1795428 | LABORATORIO DR PAULO EMILIO DALESSANDRO PINDAMONHANGABA | Instituto Butantan / ESALQ-Piracicaba | Instituto Butantan: Alexander Roberto Precioso, Dimas Tadeu Covas, Sandra Coccuzzo Sampaio, Maria Carolina Elias, José Salvatore Leister Patané, Vincent Louis Viala, Antonio Jorge Martins, Ricardo Haddad, Claudia Renata dos Santos Barros, Elaine Cristina Marqueze, Raul Machado Neto, Debora Botequio Moretti. Centro de Genômica Funcional da ESALQ: Luiz Lehmann Coutinho, Ricardo Augusto Brassaloti, Raquel de Lello Rocha Campos Cassano. NGS Soluções Genômicas: Pilar Drummond Sampaio Corrêa Mariani. FZEA-USP Pirassununga: Mirele Daiana Poleti, Jessika Cristina Chagas Lesbon, Elisangela Chicaroni Mattos, Heidge Fukumasu. USP-Botucatu: Rejane Maria Tommasini Grotto, Jayme A. Souza-Neto, Guilherme Targino |

| | | | |
|---|---|---|---|
| | | | Valente, Patricia Akemi Assato, Felipe Allan da Silva da Costa, Bianca Cechetto Carlos. Mendelics: Bibiana Santos, João Paulo Kitajima, Erika Freitas, David Schlesinger. Hemocentro Ribeirão Preto: Simone Kashima, Evandra Strazza Rodrigues, Svetoslav Nanev Slavov, Elaine Vieira dos Santos, Rafael dos Santos Bezerra, Luiz Carlos Junior de Alcantara, Marta Giovanetti, Vagner Fonseca, Flavia Aburjaile, Rodrigo Tocantins Calado. |
| EPI_ISL_1795429 | CENTRO INTEGRADO DE SAUDE | Instituto Butantan / ESALQ-Piracicaba | Instituto Butantan: Alexander Roberto Precioso, Dimas Tadeu Covas, Sandra Coccuzzo Sampaio, Maria Carolina Elias, José Salvatore Leister Patané, Vincent Louis Viala, Antonio Jorge Martins, Ricardo Haddad, Claudia Renata dos Santos Barros, Elaine Cristina Marqueze, Raul Machado Neto, Debora Botequio Moretti. Centro de Genômica Funcional da ESALQ: Luiz Lehmann Coutinho, Ricardo Augusto Brassaloti, Raquel de Lello Rocha Campos Cassano. NGS Soluções Genômicas: Pilar Drummond Sampaio Corrêa Mariani. FZEA-USP Pirassununga: Mirele Daiana Poleti, Jessika Cristina Chagas Lesbon, Elisangela Chicaroni Mattos, Heidge Fukumasu. USP-Botucatu: Rejane Maria Tommasini Grotto, Jayme A. Souza-Neto, Guilherme Targino Valente, Patricia Akemi Assato, Felipe Allan da Silva da Costa, Bianca Cechetto Carlos. Mendelics: Bibiana Santos, João Paulo Kitajima, Erika Freitas, David Schlesinger. Hemocentro Ribeirão Preto: Simone Kashima, Evandra Strazza Rodrigues, Svetoslav Nanev Slavov, Elaine Vieira dos Santos, Rafael dos Santos Bezerra, Luiz Carlos Junior de Alcantara, Marta Giovanetti, Vagner Fonseca, Flavia Aburjaile, Rodrigo Tocantins Calado. |
| EPI_ISL_1799498 | Laboratório de Microbiologia Molecular - Universidade FEEVALE | Molecular Microbiology Laboratory | Alana Witt Hansen, Fágner Henrique Heldt, Fernando Rosado Spilki, Flávio Silveira, Juliana Schons Gularte, Juliane Deise Fleck, Mariana Soares da Silva, Meriane Demoliner, Matheus Nunes Weber, Paula Rodrigues de Almeida, Micheli Filippi. |
| EPI_ISL_1799499, EPI_ISL_1799501, EPI_ISL_1799502, EPI_ISL_1799505, EPI_ISL_1799507 | Laboratório de Microbiologia Molecular - Universidade FEEVALE | Molecular Microbiology Laboratory | Alana Witt Hansen, Fágner Henrique Heldt, Fernando Rosado Spilki, Flávio Silveira, Juliana Schons Gulart, Juliane Deise Fleck, Mariana Soares da Silva, Meriane Demoliner, Matheus Nunes Weber, Paula Rodrigues de Almeida, Michele Filippi. |
| EPI_ISL_1821204 | Santa Casa de Cravinhos | Instituto Adolfo Lutz, Interdiciplinary Procedures Center, Strategic Laboratory | Claudio Tavares Sacchi, Claudia Regina Gonçalves, Erica Valessa Ramos Gomes, Karoline Rodrigues Campos, Caio Vinicius Dias Lopes, Leonardo Jose Tadeu de Araujo |
| EPI_ISL_1821205 | UPA Dr Luis Atilio Losi Viana Ribeirao Preto | Instituto Adolfo Lutz, Interdiciplinary Procedures Center, Strategic Laboratory | Claudio Tavares Sacchi, Claudia Regina Gonçalves, Erica Valessa Ramos Gomes, Karoline Rodrigues Campos, Caio Vinicius Dias Lopes, Leonardo Jose Tadeu de Araujo |
| EPI_ISL_1821206 | Sae Servico de Atendimento Especializado | Instituto Adolfo Lutz, Interdiciplinary Procedures Center, Strategic Laboratory | Claudio Tavares Sacchi, Claudia Regina Gonçalves, Erica Valessa Ramos Gomes, Karoline Rodrigues Campos, Caio Vinicius Dias Lopes, Leonardo Jose Tadeu de Araujo |
| EPI_ISL_1821207 | Secretaria Municipal de Saúde de Lins | Instituto Adolfo Lutz, Interdiciplinary Procedures Center, Strategic Laboratory | Claudio Tavares Sacchi, Claudia Regina Gonçalves, Erica Valessa Ramos Gomes, Karoline Rodrigues Campos, Caio Vinicius Dias Lopes, Leonardo Jose Tadeu de Araujo |
| EPI_ISL_1821208 | Hospital de Base de Bauru Hospital de Base Setima Regiao | Instituto Adolfo Lutz, Interdiciplinary Procedures Center, Strategic Laboratory | Claudio Tavares Sacchi, Claudia Regina Gonçalves, Erica Valessa Ramos Gomes, Karoline Rodrigues Campos, Caio Vinicius Dias Lopes, Leonardo Jose Tadeu de Araujo |
| EPI_ISL_1821209 | CS de Paulo de Faria | Instituto Adolfo Lutz, Interdiciplinary Procedures Center, Strategic Laboratory | Claudio Tavares Sacchi, Claudia Regina Gonçalves, Erica Valessa Ramos Gomes, Karoline Rodrigues Campos, Caio Vinicius Dias Lopes, Leonardo Jose Tadeu de Araujo |
| EPI_ISL_1821210 | Hospital Universitario de Marilia | Instituto Adolfo Lutz, Interdiciplinary Procedures Center, Strategic Laboratory | Claudio Tavares Sacchi, Claudia Regina Gonçalves, Erica Valessa Ramos Gomes, Karoline Rodrigues Campos, Caio Vinicius Dias Lopes, Leonardo Jose Tadeu de Araujo |
| EPI_ISL_1821211, EPI_ISL_1821212, EPI_ISL_1821213 | Diagnóstico da America S/A | Instituto Adolfo Lutz, Interdiciplinary Procedures Center, Strategic Laboratory | Claudio Tavares Sacchi, Claudia Regina Gonçalves, Erica Valessa Ramos Gomes, Karoline Rodrigues Campos, Caio Vinicius Dias Lopes, Leonardo Jose Tadeu de Araujo |
| EPI_ISL_1821214, EPI_ISL_1821215, EPI_ISL_1821216, EPI_ISL_1821217, EPI_ISL_1821218, EPI_ISL_1821219, EPI_ISL_1821220, EPI_ISL_1821221, EPI_ISL_1821222, EPI_ISL_1821223, EPI_ISL_1821224 | | | |
| see above | Instituto Adolfo Lutz - Regional de Aracatuba | Instituto Adolfo Lutz, Interdiciplinary Procedures Center, Strategic Laboratory | Claudio Tavares Sacchi, Claudia Regina Gonçalves, Erica Valessa Ramos Gomes, Karoline Rodrigues Campos, Caio Vinicius Dias Lopes, Leonardo Jose Tadeu de Araujo |
| EPI_ISL_1821225, EPI_ISL_1821226, EPI_ISL_1821227, EPI_ISL_1821228, EPI_ISL_1821229, EPI_ISL_1821230, EPI_ISL_1821231, EPI_ISL_1821232, EPI_ISL_1821233, EPI_ISL_1821234, EPI_ISL_1821235, EPI_ISL_1821236, EPI_ISL_1821237, EPI_ISL_1821238, EPI_ISL_1821239, EPI_ISL_1821240, EPI_ISL_1821241, EPI_ISL_1821242, EPI_ISL_1821243, EPI_ISL_1821244, EPI_ISL_1821245 | | | |
| see above | Instituto Adolfo Lutz - Regional de Marilia | Instituto Adolfo Lutz, Interdiciplinary Procedures Center, Strategic Laboratory | Claudio Tavares Sacchi, Claudia Regina Gonçalves, Erica Valessa Ramos Gomes, Karoline Rodrigues Campos, Caio Vinicius Dias Lopes, Leonardo Jose Tadeu de Araujo |
| EPI_ISL_1821246, EPI_ISL_1821247, EPI_ISL_1821248, EPI_ISL_1821249, EPI_ISL_1821250, EPI_ISL_1821251, EPI_ISL_1821252, EPI_ISL_1821253, EPI_ISL_1821254, EPI_ISL_1821255, EPI_ISL_1821256, EPI_ISL_1821257, EPI_ISL_1821258, EPI_ISL_1821259, EPI_ISL_1821260, EPI_ISL_1821261, EPI_ISL_1821262, EPI_ISL_1821263, EPI_ISL_1821264, EPI_ISL_1821265 | | | |
| see above | Instituto Adolfo Lutz - Regional de Ribeirao Preto | Instituto Adolfo Lutz, Interdiciplinary Procedures Center, Strategic Laboratory | Claudio Tavares Sacchi, Claudia Regina Gonçalves, Erica Valessa Ramos Gomes, Karoline Rodrigues Campos, Caio Vinicius Dias Lopes, Leonardo Jose Tadeu de Araujo |
| EPI_ISL_1821266, EPI_ISL_1821267, EPI_ISL_1821268, EPI_ISL_1821269, EPI_ISL_1821270, EPI_ISL_1821271, EPI_ISL_1821272, EPI_ISL_1821273 | Instituto Adolfo Lutz Central | Instituto Adolfo Lutz, Interdiciplinary Procedures Center, Strategic Laboratory | Claudio Tavares Sacchi, Claudia Regina Gonçalves, Erica Valessa Ramos Gomes, Karoline Rodrigues Campos, Caio Vinicius Dias Lopes, Leonardo Jose Tadeu de Araujo |
| EPI_ISL_412964 | Hospital Israelita Albert Einstein | Instituto Adolfo Lutz Interdisciplinary Procedures Center Strategic Laboratory | Jaqueline Goes de Jesus, Claudio Tavares Sacchi, Daniela Bernardes Borges da Silva, Ingra Morales Claro, Flávia Cristina da Silva Sales, Claudia Regina Gonçalves, Joshua Quick, Maria do Carmo, Sampaio Tavares Timenetsky, Nicholas James Loman, Andrew Rambaut, Ester Cerdeira Sabino, Nuno Rodrigues Faria |
| EPI_ISL_413016 | Hospital Israelita Albert Einstein | Instituto Adolfo Lutz, Interdisciplinary Procedures Center, Strategic Laboratory | Jaqueline Goes de Jesus, Claudio Tavares Sacchi, Fabiana Cristina Pereira dos Santos, Ingra Morales Claro, Flávia Cristina da Silva Sales, Claudia Regina Gonçalves, Joshua Quick, Maria do Carmo Sampaio Tavares Timenetsky, Nicholas James Loman, Andrew Rambaut, Ester Cerdeira Sabino, Nuno Rodrigues Faria |
| EPI_ISL_414014 | Hospital Israelita Albert Einstein | Instituto Adolfo Lutz, Interdiciplinary Procedures Center, Strategic Laboratory | Claudio Tavares Sacchi, Claudia Regina Gonçalves, Katia Correia dos Santos, Carlos Henrique Camargo, Maria do Carmo Sampaito Tavares Timenetsky, Terezinha Maria de Paiva, Ester Cerdeira Sabino |
| EPI_ISL_414015 | Hospital São Joaquim Beneficencia Portuguesa | Instituto Adolfo Lutz, Interdiciplinary Procedures Center, Strategic Laboratory | Claudio Tavares Sacchi, Claudia Regina Gonçalves, SimoneGuadagnucci Morillo, Carlos Henrique Camargo, Maria do Carmo Sampaito Tavares Timenetsky, Fabiana Cristina Pereira dos Santos Terezinha Maria de Paiva, Ester Cerdeira Sabino |
| EPI_ISL_414016 | Hospital São Joaquim Beneficencia Portuguesa | Instituto Adolfo Lutz, Interdiciplinary Procedures Center, Strategic Laboratory | Claudio Tavares Sacchi, Claudia Regina Gonçalves, Audrey Cilli, Carlos Henrique Camargo, Maria do Carmo Sampaito Tavares Timenetsky, Daniela Bernardes Borges da Silva, Terezinha Maria de Paiva, Ester Cerdeira Sabino |
| EPI_ISL_414017 | Hospital São Joaquim Beneficencia Portuguesa | Instituto Adolfo Lutz, Interdiciplinary Procedures Center, Strategic Laboratory | Claudio Tavares Sacchi, Claudia Regina Gonçalves, Fabiana Cristina Pereira dos Santos, Carlos Henrique Camargo, Maria do Carmo Sampaito Tavares Timenetsky, Daniela Bernardes Borges da Silva, Terezinha Maria de Paiva, Ester Cerdeira Sabino |
| EPI_ISL_414045 | Laboratório Central de Saúde Pública Noel Nutels (LACEN-RJ) | Laboratory of Respiratory Viruses and Measles, Oswaldo Cruz Institute, FIOCRUZ | Paola Resende, Alisson Fabri, Joilson Xavier, Sunando Roy, Fernando Motta, Aline Mattos, Milene Miranda, Cristiana Garcia, Braulia Caetano, Maria Ogrzewalska, Jonathan Lopes, Luciana Appolinario, Maria Nóbrega, Marilda SiqueiraFiocruz COVID-19 Genomic Surveillance Network |
| EPI_ISL_415105 | Laboratório Central de Saúde Pública Professor Gonçalo Moniz (LACEN-BA) | Laboratory of Respiratory Viruses and Measles, Oswaldo Cruz Institute, FIOCRUZ | Paola Resende, Allison Fabri, Joilson Xavier, Sunando Roy, Fernando Motta, Aline Mattos, Milene Miranda, Cristiana Garcia, Braulia Caetano, Maria Ogrzewalska, Jonathan Lopes, Luciana Appolinario, Maria Nóbrega, Marilda Siqueira Fiocruz COVID-19 Genomic Surveillance Network |
| EPI_ISL_415128 | Laboratório Central de Saúde Pública do Estado do Espírito Santo (LACEN-ES) | Laboratory of Respiratory Viruses and Measles, Oswaldo Cruz Institute, FIOCRUZ | Paola Resende, Allison Fabri, Joilson Xavier, Sunando Roy, Fernando Motta, Aline Mattos, Milene Miranda, Cristiana Garcia, Braulia Caetano, Maria Ogrzewalska, Jonathan Lopes, Luciana Appolinario, Maria Nóbrega, Marilda Siqueira on behalf of the Fiocruz COVID-19 Genomic Surveillance Network |
| EPI_ISL_416028 | National Influenza Center - Instituto Adolfo Lutz | Instituto Adolfo Lutz, Interdiciplinary Procedures Center, Strategic Laboratory | Claudio Tavares Sacchi, Claudia Regina Gonçalves, Carlos Henrique Camargo, Fabiana Cristina Pereira dos Santos, Daniela Bernardes Borges da Silva, Simone Guadagnucci Morillo, Adriano Abbud, Adriana Bugno, Maria do Carmo Sampaio Tavares Timenetsky, Terezinha Maria de Paiva |
| EPI_ISL_416029 | Laboiratório Fleury | Instituto Adolfo Lutz, Interdiciplinary Procedures Center, Strategic Laboratory | Claudio Tavares Sacchi, Claudia Regina Gonçalves, Carlos Henrique Camargo, Fabiana Cristina Pereira dos Santos, Daniela Bernardes Borges da Silva, Simone Guadagnucci Morillo, Adriano Abbud, Adriana Bugno, Maria do Carmo Sampaio Tavares Timenetsky, Terezinha Maria de Paiva |
| EPI_ISL_416031, EPI_ISL_416032 | National Influenza Center - Instituto Adolfo Lutz | Instituto Adolfo Lutz, Interdiciplinary Procedures Center, Strategic Laboratory | Claudio Tavares Sacchi, Claudia Regina Gonçalves, Carlos Henrique Camargo, Fabiana Cristina Pereira dos Santos, Daniela Bernardes Borges da Silva, Simone Guadagnucci Morillo, Adriano Abbud, Adriana Bugno, Maria do Carmo Sampaio Tavares Timenetsky, Terezinha Maria de Paiva |

| | | | |
|---|---|---|---|
| EPI_ISL_416033, EPI_ISL_416034 | Hospital Israelita Albert Einstein | Instituto Adolfo Lutz, Interdiciplinary Procedures Center, Strategic Laboratory | Claudio Tavares Sacchi, Claudia Regina Gonçalves, Carlos Henrique Camargo, Erica Valessa Ramos Gomes, Fabiana Cristina Pereira dos Santos, Daniela Bernardes Borges da Silva, Simone Guadagnucci Morillo, Adriano Abbud, Adriana Bugno, Maria do Carmo Sampaio Tavares Timenetsky, Terezinha Maria de Paiva |
| EPI_ISL_416035, EPI_ISL_416036 | National Influenza Center - Instituto Adolfo Lutz | Instituto Adolfo Lutz, Interdiciplinary Procedures Center, Strategic Laboratory | Claudio Tavares Sacchi, Claudia Regina Gonçalves, Carlos Henrique Camargo, Erica Valessa Ramos Gomes, Fabiana Cristina Pereira dos Santos, Daniela Bernardes Borges da Silva, Simone Guadagnucci Morillo, Adriano Abbud, Adriana Bugno, Maria do Carmo Sampaio Tavares Timenetsky, Terezinha Maria de Paiva |
| EPI_ISL_417034 | Laboratorio de Ecologia de Doencas Transmissiveis na Amazonia, Instituto Leonidas e Maria Deane - Fiocruz Amazonia | Laboratorio de Ecologia de Doencas Transmissiveis na Amazonia, Instituto Leonidas e Maria Deane - Fiocruz Amazonia | Valdinete Nascimento, André Corado, Fernanda Nascimento, Ágatha Costa, Debora Duarte, Luciana Gonçalves, Michele Jesus, Sérgio Luz, Felipe Naveca on behalf of the Fiocruz COVID-19 Genomic Surveillance Network |
| EPI_ISL_426580 | Instituto Sabin | Laboratory of Virology | Fernando L Melo,Gustavo Barra, Ticiane H Santa-Rita, Pedro G Mesquita, Ikaro A Andrade, Tatsuya Nagata, Bergmann M Ribeiro |
| EPI_ISL_427292 | Laboratório Central de Saúde Pública do Estado de Alagoas (LACEN-AL) | Laboratory of Respiratory Viruses and Measles, Oswaldo Cruz Institute, FIOCRUZ | Paola Resende, Fernando Motta, Luciana Appolinario, Sunando Roy, Aline Mattos, Milene Miranda, Cristiana Garcia, Braulia Caetano, Maria Ogrzewalska, Priscila Born, Jonathan Lopes, Marilda Siqueira on behalf of the Fiocruz COVID-19 Genomic Surveillance Network |
| EPI_ISL_427293 | Laboratório Central de Saúde Pública Professor Gonçalo Moniz (LACEN-BA) | Laboratory of Respiratory Viruses and Measles, Oswaldo Cruz Institute, FIOCRUZ | Paola Resende, Fernando Motta, Luciana Appolinario, Sunando Roy, Aline Mattos, Milene Miranda, Cristiana Garcia, Braulia Caetano, Maria Ogrzewalska, Priscila Born, Jonathan Lopes, Marilda Siqueira on behalf of the Fiocruz COVID-19 Genomic Surveillance Network |
| EPI_ISL_427294, EPI_ISL_427295, EPI_ISL_427296, EPI_ISL_427297, EPI_ISL_427298, EPI_ISL_427299, EPI_ISL_427300, EPI_ISL_427301, EPI_ISL_427302, EPI_ISL_427303, EPI_ISL_427304 | | | |
| see above | Laboratory of Respiratory Viruses and Measles, Oswaldo Cruz Institute, FIOCRUZ | Laboratory of Respiratory Viruses and Measles, Oswaldo Cruz Institute, FIOCRUZ | Paola Resende, Fernando Motta, Luciana Appolinario, Sunando Roy, Aline Mattos, Milene Miranda, Cristiana Garcia, Braulia Caetano, Maria Ogrzewalska, Priscila Born, Jonathan Lopes, Marilda Siqueira on behalf of the Fiocruz COVID-19 Genomic Surveillance Network |
| EPI_ISL_427305, EPI_ISL_427306 | Laboratório Central de Saúde Pública do Estado de Santa Catarina (LACEN-SC) | Laboratory of Respiratory Viruses and Measles, Oswaldo Cruz Institute, FIOCRUZ | Paola Resende, Fernando Motta, Luciana Appolinario, Sunando Roy, Aline Mattos, Milene Miranda, Cristiana Garcia, Braulia Caetano, Maria Ogrzewalska, Priscila Born, Jonathan Lopes, Marilda Siqueira on behalf of the Fiocruz COVID-19 Genomic Surveillance Network |
| EPI_ISL_429667, EPI_ISL_429669, EPI_ISL_429671, EPI_ISL_429674, EPI_ISL_429676, EPI_ISL_429681, EPI_ISL_429687, EPI_ISL_429688, EPI_ISL_429689, EPI_ISL_429695, EPI_ISL_429702 | | | |
| see above | Central Public Health Laboratory/Octávio Magalhães Institute (IOM) from the Ezequiel Dias Foundation (FUNED) | Instituto Octávio Magalhães / Fundação Ezequiel Dias (IOM/Funed) | Talita Adelino, Joilson Xavier, Marta Giovanetti, Vagner Fonseca, Marcos Vinícius Silva, Luiz Carlos Junior Alcantara, Marluce Aparecida Assunção Oliveira |
| EPI_ISL_450506 | Clinical Laboratory, Hospital Israelita Albert Einstein | Clinical Laboratory, Hospital Israelita Albert Einstein | Malta,F., Amgarten,D., de Oliveira,D.B.L., Araujo,D.B., Machado,R.R.G., Santana,R.A.F., Mangueira,C.L.P., Durigon,E.L. and Pinho,J.R.R. |
| EPI_ISL_450873 | Evandro Chagas Institute | Evandro Chagas Institute | Santos, M.C.; Silva, A.M.; Junior, W.D.C.; Barbagelata, L.S.; Ferreira, J.A.; Sousa, E.M.A.; da Silva, P.S.; Martins, L.C.; Sousa Junior, E.C.; Viana, G.M.R |
| EPI_ISL_450874 | Evandro Chagas Institute | Evandro Chagas Institute | Santos, M.C.; Silva, A.M.; Junior, W.D.C.; Barbagelata, L.S.; Ferreira, J.A.; Sousa, E.M.A.; da Silva, P.S.; Martins, L.C.;Sousa Junior, E.C.;Viana, G.M.R |
| EPI_ISL_456071, EPI_ISL_456072, EPI_ISL_456073, EPI_ISL_456074, EPI_ISL_456075 | Laboratory of Respiratory Viruses and Measles, Oswaldo Cruz Institute, FIOCRUZ | Laboratory of Respiratory Viruses and Measles, Oswaldo Cruz Institute, FIOCRUZ | Paola Resende, Luciana Appolinario, Fernando Motta, Aline Mattos, Milene Miranda, Cristiana Garcia, Braulia Caetano, Maria Ogrzewalska, Jonathan Lopes, Marilda Siqueira on behalf of the Fiocruz COVID-19 Genomic Surveillance Network |
| EPI_ISL_456076, EPI_ISL_456077 | Laboratório Central de Saúde Pública Noel Nutels (LACEN-RJ) | Laboratory of Respiratory Viruses and Measles, Oswaldo Cruz Institute, FIOCRUZ | Paola Resende, Luciana Appolinario, Fernando Motta, Aline Mattos, Milene Miranda, Cristiana Garcia, Braulia Caetano, Maria Ogrzewalska, Jonathan Lopes, Marilda Siqueira on behalf of the Fiocruz COVID-19 Genomic Surveillance Network |
| EPI_ISL_456079, EPI_ISL_456080, EPI_ISL_456081 | Laboratory of Respiratory Viruses and Measles, Oswaldo Cruz Institute, FIOCRUZ | Laboratory of Respiratory Viruses and Measles, Oswaldo Cruz Institute, FIOCRUZ | Paola Resende, Luciana Appolinario, Fernando Motta, Aline Mattos, Milene Miranda, Cristiana Garcia, Braulia Caetano, Maria Ogrzewalska, Jonathan Lopes, Marilda Siqueira on behalf of the Fiocruz COVID-19 Genomic Surveillance Network |
| EPI_ISL_456082, EPI_ISL_456083 | Laboratório Central de Saúde Pública Noel Nutels (LACEN-RJ) | Laboratory of Respiratory Viruses and Measles, Oswaldo Cruz Institute, FIOCRUZ | Paola Resende, Luciana Appolinario, Fernando Motta, Aline Mattos, Milene Miranda, Cristiana Garcia, Braulia Caetano, Maria Ogrzewalska, Jonathan Lopes, Marilda Siqueira on behalf of the Fiocruz COVID-19 Genomic Surveillance Network |
| EPI_ISL_456084, EPI_ISL_456085, EPI_ISL_456086, EPI_ISL_456087 | Laboratory of Respiratory Viruses and Measles, Oswaldo Cruz Institute, FIOCRUZ | Laboratory of Respiratory Viruses and Measles, Oswaldo Cruz Institute, FIOCRUZ | Paola Resende, Luciana Appolinario, Fernando Motta, Aline Mattos, Milene Miranda, Cristiana Garcia, Braulia Caetano, Maria Ogrzewalska, Jonathan Lopes, Marilda Siqueira on behalf of the Fiocruz COVID-19 Genomic Surveillance Network |
| EPI_ISL_456088 | Laboratório Central de Saúde Pública Noel Nutels (LACEN-RJ) | Laboratory of Respiratory Viruses and Measles, Oswaldo Cruz Institute, FIOCRUZ | Paola Resende, Luciana Appolinario, Fernando Motta, Aline Mattos, Milene Miranda, Cristiana Garcia, Braulia Caetano, Maria Ogrzewalska, Jonathan Lopes, Marilda Siqueira on behalf of the Fiocruz COVID-19 Genomic Surveillance Network |
| EPI_ISL_456089, EPI_ISL_456090, EPI_ISL_456091, EPI_ISL_456092, EPI_ISL_456093, EPI_ISL_456094, EPI_ISL_456095, EPI_ISL_456096, EPI_ISL_456097, EPI_ISL_456098, EPI_ISL_456099, EPI_ISL_456100, EPI_ISL_456101, EPI_ISL_456102, EPI_ISL_456103, EPI_ISL_456104, EPI_ISL_456105, EPI_ISL_456106 | | | |
| see above | Laboratory of Respiratory Viruses and Measles, Oswaldo Cruz Institute, FIOCRUZ | Laboratory of Respiratory Viruses and Measles, Oswaldo Cruz Institute, FIOCRUZ | Paola Resende, Luciana Appolinario, Fernando Motta, Aline Mattos, Milene Miranda, Cristiana Garcia, Braulia Caetano, Maria Ogrzewalska, Jonathan Lopes, Marilda Siqueira on behalf of the Fiocruz COVID-19 Genomic Surveillance Network |
| EPI_ISL_458138, EPI_ISL_458139, EPI_ISL_458140, EPI_ISL_458141, EPI_ISL_458142, EPI_ISL_458143, EPI_ISL_458144, EPI_ISL_458145, EPI_ISL_458146, EPI_ISL_458147, EPI_ISL_458148, EPI_ISL_458149 | | | |
| see above | Evandro Chagas Institute | Evandro Chagas Institute | Santos, M.C.; Silva, A.M.; Junior, W.D.C.; Barbagelata, L.S.; Ferreira, J.A.; Sousa, E.M.A.; da Silva, P.S.; Resque, H.R; Martins, L.C.; Sousa Junior, E.C.;Viana, G.M.R |
| EPI_ISL_467344, EPI_ISL_467345, EPI_ISL_467346, EPI_ISL_467347, EPI_ISL_467348, EPI_ISL_467349, EPI_ISL_467350, EPI_ISL_467351, EPI_ISL_467352, EPI_ISL_467353, EPI_ISL_467354, EPI_ISL_467355, EPI_ISL_467356, EPI_ISL_467357, EPI_ISL_467358, EPI_ISL_467359, EPI_ISL_467360, EPI_ISL_467362, EPI_ISL_467363, EPI_ISL_467364, EPI_ISL_467365, EPI_ISL_467366, EPI_ISL_467367, EPI_ISL_467368, EPI_ISL_467369, EPI_ISL_467370, EPI_ISL_467371 | | | |
| see above | Laboratory of Respiratory Viruses and Measles, Oswaldo Cruz Institute, FIOCRUZ | Laboratory of Respiratory Viruses and Measles, Oswaldo Cruz Institute, FIOCRUZ | Paola Resende, Luciana Appolinario, Fernando Motta, Anna Carolina Paixão, Ana Carolina Mendonça, Aline Mattos, Milene Miranda, Cristiana Garcia, Braulia Caetano, Maria Ogrzewalska, Jonathan Lopes, Marilda Siqueira on behalf of the Fiocruz COVID-19 Genomic Surveillance Network |
| EPI_ISL_468305, EPI_ISL_468307 | Centro de Vigilancia a Saude de Diadema | Instituto Adolfo Lutz, Interdiciplinary Procedures Center, Strategic Laboratory | Claudio Tavares Sacchi, Claudia Regina Gonçalves, Erica Valessa Ramos Gomes |
| EPI_ISL_468308 | Hospital Municipal do Tatuape Carmino Caricchio | Instituto Adolfo Lutz, Interdiciplinary Procedures Center, Strategic Laboratory | Claudio Tavares Sacchi, Claudia Regina Gonçalves, Erica Valessa Ramos Gomes |
| EPI_ISL_468310 | Hospital Sao Paulo de Ensino da UNIFESP | Instituto Adolfo Lutz, Interdiciplinary Procedures Center, Strategic Laboratory | Claudio Tavares Sacchi, Claudia Regina Gonçalves, Erica Valessa Ramos Gomes |
| EPI_ISL_468311, EPI_ISL_468312 | Hospital Municipal Dr Ignacio Proenca de Gouvea | Instituto Adolfo Lutz, Interdiciplinary Procedures Center, Strategic Laboratory | Claudio Tavares Sacchi, Claudia Regina Gonçalves, Erica Valessa Ramos Gomes |
| EPI_ISL_468313 | Vigilancia Epidemiologica de São Bernardo do Campo | Instituto Adolfo Lutz, Interdiciplinary Procedures Center, Strategic Laboratory | Claudio Tavares Sacchi, Claudia Regina Gonçalves, Erica Valessa Ramos Gomes |
| EPI_ISL_468314 | CTA Centro de Testagem e Aconselhamento | Instituto Adolfo Lutz, Interdiciplinary Procedures Center, Strategic Laboratory | Claudio Tavares Sacchi, Claudia Regina Gonçalves, Erica Valessa Ramos Gomes |
| EPI_ISL_468315 | Hospital Municipal do Tatuape Carmino Caricchio | Instituto Adolfo Lutz, Interdiciplinary Procedures Center, Strategic Laboratory | Claudio Tavares Sacchi, Claudia Regina Gonçalves, Erica Valessa Ramos Gomes |
| EPI_ISL_468316 | UPA Vila Assis | Instituto Adolfo Lutz, Interdiciplinary Procedures Center, Strategic Laboratory | Claudio Tavares Sacchi, Claudia Regina Gonçalves, Erica Valessa Ramos Gomes |
| EPI_ISL_468318 | Hospital Universitario da USP | Instituto Adolfo Lutz, Interdiciplinary Procedures Center, Strategic Laboratory | Claudio Tavares Sacchi, Claudia Regina Gonçalves, Erica Valessa Ramos Gomes |
| EPI_ISL_468319 | Vigilancia Epidemiologica de São Bernardo do Campo | Instituto Adolfo Lutz, Interdiciplinary Procedures Center, Strategic Laboratory | Claudio Tavares Sacchi, Claudia Regina Gonçalves, Erica Valessa Ramos Gomes |
| EPI_ISL_468320 | Secretaria Municipal de Saude de Hortolandia | Instituto Adolfo Lutz, Interdiciplinary Procedures Center, | Claudio Tavares Sacchi, Claudia Regina Gonçalves, Erica Valessa Ramos Gomes |

| | | | |
|---|---|---|---|
| | | Strategic Laboratory | |
| EPI_ISL_468321 | Hospital Universitario da USP | Instituto Adolfo Lutz, Interdiciplinary Procedures Center, Strategic Laboratory | Claudio Tavares Sacchi, Claudia Regina Gonçalves, Erica Valessa Ramos Gomes |
| EPI_ISL_471539 | Hospital Universitario da USP Sao Paulo | Instituto Adolfo Lutz, Interdiciplinary Procedures Center, Strategic Laboratory | Claudio Tavares Sacchi, Claudia Regina Gonçalves, Erica Valessa Ramos Gomes |
| EPI_ISL_471541 | Hospital Geral Santa Marcelina | Instituto Adolfo Lutz, Interdiciplinary Procedures Center, Strategic Laboratory | Claudio Tavares Sacchi, Claudia Regina Gonçalves, Erica Valessa Ramos Gomes |
| EPI_ISL_471542 | Secretaria de Saude de Mogi das Cruzes | Instituto Adolfo Lutz, Interdiciplinary Procedures Center, Strategic Laboratory | Claudio Tavares Sacchi, Claudia Regina Gonçalves, Erica Valessa Ramos Gomes |
| EPI_ISL_471543 | Centro de Saude I Tacito Leite de Carvalho e Silva | Instituto Adolfo Lutz, Interdiciplinary Procedures Center, Strategic Laboratory | Claudio Tavares Sacchi, Claudia Regina Gonçalves, Erica Valessa Ramos Gomes |
| EPI_ISL_471545 | Hospital Sao Paulo de Ensino da Unifesp | Instituto Adolfo Lutz, Interdiciplinary Procedures Center, Strategic Laboratory | Claudio Tavares Sacchi, Claudia Regina Gonçalves, Erica Valessa Ramos Gomes |
| EPI_ISL_471546 | AMA DR Jose Soares Hungria | Instituto Adolfo Lutz, Interdiciplinary Procedures Center, Strategic Laboratory | Claudio Tavares Sacchi, Claudia Regina Gonçalves, Erica Valessa Ramos Gomes |
| EPI_ISL_471548 | Hospital do Servidor Público Estadual Francisco Morato de Oliveira | Instituto Adolfo Lutz, Interdiciplinary Procedures Center, Strategic Laboratory | Claudio Tavares Sacchi, Claudia Regina Gonçalves, Erica Valessa Ramos Gomes |
| EPI_ISL_471549 | Hospital Municipal Carmen Prudente | Instituto Adolfo Lutz, Interdiciplinary Procedures Center, Strategic Laboratory | Claudio Tavares Sacchi, Claudia Regina Gonçalves, Erica Valessa Ramos Gomes |
| EPI_ISL_471551 | Hospital Sao Paulo de Ensino da Unifesp | Instituto Adolfo Lutz, Interdiciplinary Procedures Center, Strategic Laboratory | Claudio Tavares Sacchi, Claudia Regina Gonçalves, Erica Valessa Ramos Gomes |
| EPI_ISL_471552 | Hospital Sancta Maggiore | Instituto Adolfo Lutz, Interdiciplinary Procedures Center, Strategic Laboratory | Claudio Tavares Sacchi, Claudia Regina Gonçalves, Erica Valessa Ramos Gomes |
| EPI_ISL_471554 | Hospital Bosque da Saúde | Instituto Adolfo Lutz, Interdiciplinary Procedures Center, Strategic Laboratory | Claudio Tavares Sacchi, Claudia Regina Gonçalves, Erica Valessa Ramos Gomes |
| EPI_ISL_471556 | Pronto Socorro Jose Ibrahin | Instituto Adolfo Lutz, Interdiciplinary Procedures Center, Strategic Laboratory | Claudio Tavares Sacchi, Claudia Regina Gonçalves, Erica Valessa Ramos Gomes |
| EPI_ISL_471562, EPI_ISL_471581, EPI_ISL_471582 | Hosp. Municipal Prof. Dr. Alípio Corrêa Netto | Instituto Adolfo Lutz, Interdiciplinary Procedures Center, Strategic Laboratory | Claudio Tavares Sacchi, Claudia Regina Gonçalves, Erica Valessa Ramos Gomes |
| EPI_ISL_471647 | Hospital Municipal de Barueri Dr. Francisco Moran | Instituto Adolfo Lutz, Interdiciplinary Procedures Center, Strategic Laboratory | Claudio Tavares Sacchi, Claudia Regina Gonçalves, Erica Valessa Ramos Gomes |
| EPI_ISL_471648 | UBS e Pronto Socorro Jd. Jacira | Instituto Adolfo Lutz, Interdiciplinary Procedures Center, Strategic Laboratory | Claudio Tavares Sacchi, Claudia Regina Gonçalves, Erica Valessa Ramos Gomes |
| EPI_ISL_476221 | Laboratory Fleury | Instituto de Medicina Tropical da Univesidade de São Paulo | Samples: Celso Granato; Sequencing: Ingra Morales Claro, Jaqueline Goes de Jesus, Erika Regina Manuli, Flavia Cristina da Silva Sales, Thais de Moura Coletti, Camila Alves Maia da Silva, Mariana Severo Ramundo, Giula Magalhaes Ferreira, Darlan da Silva Candido, Julien Theze, Nuno Faria, Ester Sabino |
| EPI_ISL_476282, EPI_ISL_476288, EPI_ISL_476289, EPI_ISL_476297 | DB Diagnósticos do Brasil | Instituto de Medicina Tropical da Univesidade de São Paulo | Samples: Nelson Gaburo Jr; Sequencing: Ingra Morales Claro, Jaqueline Goes de Jesus, Erika Regina Manuli, Flavia Cristina da Silva Sales, Thais de Moura Coletti, Camila Alves Maia da Silva, Mariana Severo Ramundo, Giula Magalhaes Ferreira, Darlan da Silva Candido, Julien Theze, Nuno Faria, Ester Sabino |
| EPI_ISL_476341 | Laboratório de Patologia Clínica - UNICAMP | Laboratório de Estudos de Vírus Emergentes - UNICAMP | José Luiz Proença-Modena, Magnun Nueldo Nunes dos Santos, Angelica Schreiber, Julia Forato,Camila Simeoni, Marcilio Jorge Fumagalli, Mariene Ribeiro Amorim, Darlan da Silva Candido, Nuno Rodrigues Faria, Julien Theze, Luiz Gonzaga,Jaqueline Goes Jesus e William Marciel de Souza |
| EPI_ISL_476373 | Hospital da Clínicas da Faculdade de Medicina da Universidade de São Paulo | Instituto de Medicina Tropical da Univesidade de São Paulo | Samples: Ingra Morales Claro, Erika Regina Manuli, Cecilia Salete Alencar, Carolina S. Lazar, Silvia F. Costa; Sequencing: Ingra Morales Claro, Jaqueline Goes de Jesus, Erika Regina Manuli, Flavia Cristina da Silva Sales, Thais de Moura Coletti, Camila Alves Maia da Silva, Mariana Severo Ramundo, Giula Magalhaes Ferreira, Darlan da Silva Candido, Julien Theze, Nuno Faria, Ester Sabino |
| EPI_ISL_476395, EPI_ISL_476398 | Laboratório de Patologia Clínica - UNICAMP | Laboratório de Estudos de Vírus Emergentes - UNICAMP | José Luiz Proença-Modena, Magnun Nueldo Nunes dos Santos, Angelica Schreiber, Julia Forato,Camila Simeoni, Marcilio Jorge Fumagalli, Mariene Ribeiro Amorim, Darlan da Silva Candido, Nuno Rodrigues Faria, Julien Theze, Luiz Gonzaga,Jaqueline Goes Jesus e William Marciel de Souza |
| EPI_ISL_476435, EPI_ISL_476439, EPI_ISL_476445, EPI_ISL_476446, EPI_ISL_476469, EPI_ISL_476490 | Hospital da Clínicas da Faculdade de Medicina da Universidade de São Paulo | Instituto de Medicina Tropical da Univesidade de São Paulo | Samples: Ingra Morales Claro, Erika Regina Manuli, Cecilia Salete Alencar, Carolina S. Lazar, Silvia F. Costa; Sequencing: Ingra Morales Claro, Jaqueline Goes de Jesus, Erika Regina Manuli, Flavia Cristina da Silva Sales, Thais de Moura Coletti, Camila Alves Maia da Silva, Mariana Severo Ramundo, Giula Magalhaes Ferreira, Darlan da Silva Candido, Julien Theze, Nuno Faria, Ester Sabino |
| EPI_ISL_483065 | Centro de Desenvolvimento Tecnologico em Saude, Fundacao Oswaldo Cruz | Centro de Desenvolvimento Tecnologico em Saude, Fundacao Oswaldo Cruz | Souza,T.M., Fintelman-Rodrigues,N., De Paula,A.D., Tschoeke,D., Barroso,S.P., Gregorio,M.L., Oliveira,J.S., Saraiva,F.B., Ferreira,M.A., Sacramento,C.Q. |
| EPI_ISL_486427 | unknown | Clinical Laboratory, Hospital Israelita Albert Einstein | Amgarteb,D., Malta,F., Guedes,R.L., Santana,R.A., de Menezes,F.G., Mangueira,C.L. and Pinho,J.R. |
| EPI_ISL_486429 | unknown | Clinical Laboratory, Hospital Israelita Albert Einstein | Malta,F., Amgarten,D., Guedes,R.L., Santana,R.A., de Menezes,F.G., Mangueira,C.L. and Pinho,J.R. |
| EPI_ISL_492032 | Instituto de Biologia do Exército | Laboratório Metabolismo Macromolecular FirminoTorres de Castro, Instituto de Biofísica Carlos Chagas Filho, Universidade Federal do Rio de Janeiro | Bianca Catarina Azevedo Cabral, Aline Rosa Vianna de Souza , Marcos Dornelas-Ribeiro, Tatiana LS Nogueira, Nádia Vaez Gonçalves da Cruz, Caleb GM Santos, Elizabeth Valentin, Marcio da Costa Cipitelli, Virginia Sara Grancieri do Amaral, Rodrigo Soares de Moura Neto, Clarissa Damaso, Rosane Silva |
| EPI_ISL_492033 | Instituto de Biologia do Exército | Laboratório Metabolismo Macromolecular FirminoTorres de Castro, Instituto de Biofísica Carlos Chagas Filho, Universidade Federal do Rio de Janeiro | Bianca Catarina Azevedo Cabral, Aline Rosa Vianna de Souza, Caleb GM Santos, Marcos Dornelas-Ribeiro, Tatiana LS Nogueira, Nádia Vaez Gonçalves da Cruz, Elizabeth Valentin, Marcio da Costa Cipitelli, Virginia Sara Grancieri do Amaral, Rodrigo Soares de Moura Neto, Clarissa Damaso, Rosane Silva |
| EPI_ISL_492034 | Instituto de Biologia do Exército | Laboratório Metabolismo Macromolecular FirminoTorres de Castro, Instituto de Biofísica Carlos Chagas Filho, Universidade Federal do Rio de Janeiro | Bianca Catarina Azevedo Cabral, Aline Rosa Vianna de Souza, Nádia Vaez Gonçalves da Cruz, Caleb GM Santos, Marcos Dornelas-Ribeiro, Tatiana LS Nogueira, Elizabeth Valentin, Marcio da Costa Cipitelli, Virginia Sara Grancieri do Amaral, Rodrigo Soares de Moura Neto, Clarissa Damaso, Rosane Silva |
| EPI_ISL_492035 | Instituto de Biologia do Exército | Laboratório Metabolismo Macromolecular FirminoTorres de Castro, Instituto de Biofísica Carlos Chagas Filho, Universidade Federal do Rio de Janeiro | Bianca Catarina Azevedo Cabral, Aline Rosa Vianna de Souza, Tatiana LS Nogueira, Nádia Vaez Gonçalves da Cruz, Caleb GM Santos, Marcos Dornelas-Ribeiro, Elizabeth Valentin, Marcio da Costa Cipitelli, Virginia Sara Grancieri do Amaral, Rodrigo Soares de Moura Neto, Clarissa Damaso, Rosane Silva |
| EPI_ISL_492036 | Instituto de Biologia do Exército | Laboratório Metabolismo Macromolecular FirminoTorres de Castro, Instituto de Biofísica Carlos Chagas Filho, Universidade Federal do Rio de Janeiro | Bianca Catarina Azevedo Cabral, Aline Rosa Vianna de Souza , Marcos Dornelas-Ribeiro, Tatiana LS Nogueira, Nádia Vaez Gonçalves da Cruz, Caleb GM Santos, Elizabeth Valentin, Marcio da Costa Cipitelli, Virginia Sara Grancieri do Amaral, Rodrigo Soares de Moura Neto, Clarissa Damaso, Rosane Silva |
| EPI_ISL_492037 | Instituto de Biologia do Exército | Laboratório Metabolismo Macromolecular FirminoTorres de Castro, Instituto de Biofísica Carlos Chagas Filho, Universidade Federal do Rio de Janeiro | Bianca Catarina Azevedo Cabral, Aline Rosa Vianna de Souza, Caleb GM Santos, Marcos Dornelas-Ribeiro, Tatiana LS Nogueira, Nádia Vaez Gonçalves da Cruz, Elizabeth Valentin, Marcio da Costa Cipitelli, Virginia Sara Grancieri do Amaral, Rodrigo Soares de Moura Neto, Clarissa Damaso, Rosane Silva |
| EPI_ISL_492038 | Instituto de Biologia do Exército | Laboratório Metabolismo Macromolecular FirminoTorres de | Bianca Catarina Azevedo Cabral, Aline Rosa Vianna de Souza, Nádia Vaez Gonçalves da Cruz, Caleb GM Santos, Marcos Dornelas-Ribeiro, Tatiana LS |

| | | | |
|---|---|---|---|
| | | Castro, Instituto de Biofísica Carlos Chagas Filho, Universidade Federal do Rio de Janeiro | Nogueira, Elizabeth Valentin, Marcio da Costa Cipitelli, Virginia Sara Grancieri do Amaral, Rodrigo Soares de Moura Neto, Clarissa Damaso, Rosane Silva |
| EPI_ISL_492039 | Instituto de Biologia do Exército | Laboratório Metabolismo Macromolecular FirminoTorres de Castro, Instituto de Biofísica Carlos Chagas Filho, Universidade Federal do Rio de Janeiro | Bianca Catarina Azevedo Cabral, Aline Rosa Vianna de Souza, Tatiana LS Nogueira, Nádia Vaez Gonçalves da Cruz, Caleb GM Santos, Marcos Dornelas-Ribeiro, Elizabeth Valentin, Marcio da Costa Cipitelli, Virginia Sara Grancieri do Amaral, Rodrigo Soares de Moura Neto, Clarissa Damaso, Rosane Silva |
| EPI_ISL_492040 | Instituto de Biologia do Exército | Laboratório Metabolismo Macromolecular FirminoTorres de Castro, Instituto de Biofísica Carlos Chagas Filho, Universidade Federal do Rio de Janeiro | Bianca Catarina Azevedo Cabral, Aline Rosa Vianna de Souza , Marcos Dornelas-Ribeiro, Tatiana LS Nogueira, Nádia Vaez Gonçalves da Cruz, Caleb GM Santos, Elizabeth Valentin, Marcio da Costa Cipitelli, Virginia Sara Grancieri do Amaral, Rodrigo Soares de Moura Neto, Clarissa Damaso, Rosane Silva |
| EPI_ISL_492041 | Instituto de Biologia do Exército | Laboratório Metabolismo Macromolecular FirminoTorres de Castro, Instituto de Biofísica Carlos Chagas Filho, Universidade Federal do Rio de Janeiro | Bianca Catarina Azevedo Cabral, Aline Rosa Vianna de Souza, Caleb GM Santos, Marcos Dornelas-Ribeiro, Tatiana LS Nogueira, Nádia Vaez Gonçalves da Cruz, Elizabeth Valentin, Marcio da Costa Cipitelli, Virginia Sara Grancieri do Amaral, Rodrigo Soares de Moura Neto, Clarissa Damaso, Rosane Silva |
| EPI_ISL_492042 | Instituto de Biologia do Exército | Laboratório Metabolismo Macromolecular FirminoTorres de Castro, Instituto de Biofísica Carlos Chagas Filho, Universidade Federal do Rio de Janeiro | Bianca Catarina Azevedo Cabral, Aline Rosa Vianna de Souza, Nádia Vaez Gonçalves da Cruz, Caleb GM Santos, Marcos Dornelas-Ribeiro, Tatiana LS Nogueira, Elizabeth Valentin, Marcio da Costa Cipitelli, Virginia Sara Grancieri do Amaral, Rodrigo Soares de Moura Neto, Clarissa Damaso, Rosane Silva |
| EPI_ISL_492043 | Instituto de Biologia do Exército | Laboratório Metabolismo Macromolecular FirminoTorres de Castro, Instituto de Biofísica Carlos Chagas Filho, Universidade Federal do Rio de Janeiro | Bianca Catarina Azevedo Cabral, Aline Rosa Vianna de Souza, Tatiana LS Nogueira, Nádia Vaez Gonçalves da Cruz, Caleb GM Santos, Marcos Dornelas-Ribeiro, Elizabeth Valentin, Marcio da Costa Cipitelli, Virginia Sara Grancieri do Amaral, Rodrigo Soares de Moura Neto, Clarissa Damaso, Rosane Silva |
| EPI_ISL_492044 | Instituto de Biologia do Exército | Laboratório Metabolismo Macromolecular FirminoTorres de Castro, Instituto de Biofísica Carlos Chagas Filho, Universidade Federal do Rio de Janeiro | Bianca Catarina Azevedo Cabral, Aline Rosa Vianna de Souza , Marcos Dornelas-Ribeiro, Tatiana LS Nogueira, Nádia Vaez Gonçalves da Cruz, Caleb GM Santos, Elizabeth Valentin, Marcio da Costa Cipitelli, Virginia Sara Grancieri do Amaral, Rodrigo Soares de Moura Neto, Clarissa Damaso, Rosane Silva |
| EPI_ISL_492045 | Instituto de Biologia do Exército | Laboratório Metabolismo Macromolecular FirminoTorres de Castro, Instituto de Biofísica Carlos Chagas Filho, Universidade Federal do Rio de Janeiro | Bianca Catarina Azevedo Cabral, Aline Rosa Vianna de Souza, Caleb GM Santos, Marcos Dornelas-Ribeiro, Tatiana LS Nogueira, Nádia Vaez Gonçalves da Cruz, Elizabeth Valentin, Marcio da Costa Cipitelli, Virginia Sara Grancieri do Amaral, Rodrigo Soares de Moura Neto, Clarissa Damaso, Rosane Silva |
| EPI_ISL_492046 | Instituto de Biologia do Exército | Laboratório Metabolismo Macromolecular FirminoTorres de Castro, Instituto de Biofísica Carlos Chagas Filho, Universidade Federal do Rio de Janeiro | Bianca Catarina Azevedo Cabral, Aline Rosa Vianna de Souza, Nádia Vaez Gonçalves da Cruz, Caleb GM Santos, Marcos Dornelas-Ribeiro, Tatiana LS Nogueira, Elizabeth Valentin, Marcio da Costa Cipitelli, Virginia Sara Grancieri do Amaral, Rodrigo Soares de Moura Neto, Clarissa Damaso, Rosane Silva |
| EPI_ISL_492047 | Instituto de Biologia do Exército | Laboratório Metabolismo Macromolecular FirminoTorres de Castro, Instituto de Biofísica Carlos Chagas Filho, Universidade Federal do Rio de Janeiro | Bianca Catarina Azevedo Cabral, Aline Rosa Vianna de Souza, Tatiana LS Nogueira, Nádia Vaez Gonçalves da Cruz, Caleb GM Santos, Marcos Dornelas-Ribeiro, Elizabeth Valentin, Marcio da Costa Cipitelli, Virginia Sara Grancieri do Amaral, Rodrigo Soares de Moura Neto, Clarissa Damaso, Rosane Silva |
| EPI_ISL_492048 | Instituto de Biologia do Exército | Laboratório Metabolismo Macromolecular FirminoTorres de Castro, Instituto de Biofísica Carlos Chagas Filho, Universidade Federal do Rio de Janeiro | Bianca Catarina Azevedo Cabral, Aline Rosa Vianna de Souza , Marcos Dornelas-Ribeiro, Tatiana LS Nogueira, Nádia Vaez Gonçalves da Cruz, Caleb GM Santos, Elizabeth Valentin, Marcio da Costa Cipitelli, Virginia Sara Grancieri do Amaral, Rodrigo Soares de Moura Neto, Clarissa Damaso, Rosane Silva |
| EPI_ISL_500460, EPI_ISL_500461, EPI_ISL_500462, EPI_ISL_500463, EPI_ISL_500471, EPI_ISL_500473, EPI_ISL_500475, EPI_ISL_500477, EPI_ISL_500480, EPI_ISL_500481, EPI_ISL_500482, EPI_ISL_500483, EPI_ISL_500484, EPI_ISL_500485, EPI_ISL_500486, EPI_ISL_500872 | | | |
| see above | Laboratório Central de Saúde Pública do Estado de Pernambuco (LACEN-PE) | WallauLab, Aggeu Magalhaes Institute | Marcelo Henrique Santos Paiva, Duschinka Ribeiro Duarte Guedes, Cássia Docena, Matheus Filgueira Bezerra, Filipe Zimmer Dezordi, Laís Ceschini Machado, Larissa Krokovsky, Elisama Helvecio, Alexandre Freitas da Silva, Luydson Richardson Silva Vasconcelos, Antonio Mauro Rezende, Severino Jefferson Ribeiro da Silva, Kamila Gaudêncio da Silva Sales, Bruna Santos Lima Figueiredo de Sá, Derciliano Lopes da Cruz, Claudio Eduardo Cavalcanti, Armando de Menezes Neto, Caroline Targino Alves da Silva, Renata Pessôa Germano Mendes, Maria Almerice Lopes da Silva, Tiago Gräf, Paola Cristina Resende, Gonzalo Bello, Michelle da Silva Barros, Wheverton Ricardo Correia do Nascimento, Rodrigo Moraes Loyo Arcoverde, Luciane Caroline Albuquerque Bezerra, Sinval Pinto Brandão Filho, Constância Flávia Junqueira Ayres, Gabriel Luz Wallau on behalf of the Fiocruz COVID-19 Genomic Surveillance Network |
| EPI_ISL_502779 | LACEN/PE | LABBE, Federal University of Pernambuco | WILSON JOSE DA SILVA JUNIOR, HEIDI LACERDA ALVES DA CRUZ, MARCOS DA SILVEIRA REGUEIRA NETO, BRUNO SAMPAIO, SERGIO DE SA LEITAO PAIVA JUNIOR, ZILDENE DE SOUSA SILVEIRA, MAIRA GALDINO DA ROCHA PITTA, MICHELLY CRISTINY PEREIRA, MARCOS ANTONIO DE MORAIS JUNIOR, ANTONIO CARLOS DE FREITAS, VALDIR DE QUEIROZ BALBINO. |
| EPI_ISL_502875 | LACEN/PE | LABBE, Federal University of Pernambuco | WILSON JOSE DA SILVA JUNIOR, HEIDI LACERDA ALVES DA CRUZ, MARCOS DA SILVEIRA REGUEIRA NETO, BRUNO SAMPAIO, SERGIO DE SA LEITAO PAIVA JUNIOR, ZILDENE DE SOUSA SILVEIRA, MAIRA GALDINO DA ROCHA PITTA, MICHELLY CRISTINY PEREIRA, REGINALDO GONCALVES DE LIMA NETO, MARCOS ANTONIO DE MORAIS JUNIOR, ANTONIO CARLOS DE FREITAS, VALDIR DE QUEIROZ BALBINO. |
| EPI_ISL_509430, EPI_ISL_509431, EPI_ISL_509432, EPI_ISL_509433, EPI_ISL_509434, EPI_ISL_509435 | Centro de Desenvolvimento Tecnologico em Saude, Fundacao Oswaldo Cruz | Centro de Desenvolvimento Tecnologico em Saude, Fundacao Oswaldo Cruz | Souza,T.M., Fintelman-Rodrigues,N., De Paula,A.D., Saraiva,F.B., Ferreira,M.A., Sacramento,C.Q., Medeiros,M.A. |
| EPI_ISL_510535 | Molecular Virology, Instituto Carlos Chagas / Fiocruz Paraná | Universidade Federal do Parana (UFPR) | Suzukawa,A., Tscha,M., Zanluca,C., Raboni,S., Duarte dos Santos,C. |
| EPI_ISL_510536 | Centro de Desenvolvimento Tecnologico em Saude, Fundacao Oswaldo Cruz | Centro de Desenvolvimento Tecnologico em Saude, Fundacao Oswaldo Cruz | Souza,T.M., Fintelman-Rodrigues,N., De Paula,A.D., Saraiva,F.B., Ferreira,M.A., Sacramento,C.Q. and Medeiros,M.A. |
| EPI_ISL_510541 | Centro de Desenvolvimento Tecnologico em Saude, Fundacao Oswaldo Cruz | Centro de Desenvolvimento Tecnologico em Saude, Fundacao Oswaldo Cruz | Souza,T.M., Fintelman-Rodrigues,N., De Paula,A.D., Saraiva,F.B., Ferreira,M.A., Sacramento,C.Q. and Medeiros,M.A. |
| EPI_ISL_513513, EPI_ISL_513514, EPI_ISL_513515, EPI_ISL_513516, EPI_ISL_513518, EPI_ISL_513519, EPI_ISL_513520, EPI_ISL_513521, EPI_ISL_513522, EPI_ISL_513524, EPI_ISL_513525, EPI_ISL_513526, EPI_ISL_513527, EPI_ISL_513529, EPI_ISL_513530, EPI_ISL_513531, EPI_ISL_513532, EPI_ISL_513533, EPI_ISL_513534, EPI_ISL_513535, EPI_ISL_513536, EPI_ISL_513537, EPI_ISL_513538, EPI_ISL_513539, EPI_ISL_513540, EPI_ISL_513541, EPI_ISL_513542, EPI_ISL_513543, EPI_ISL_513544, EPI_ISL_513545, EPI_ISL_513546, EPI_ISL_513547, EPI_ISL_513548, EPI_ISL_513549, EPI_ISL_513550, EPI_ISL_513551, EPI_ISL_513553, EPI_ISL_513556, EPI_ISL_513557, EPI_ISL_513558, EPI_ISL_513559, EPI_ISL_513560, EPI_ISL_513561, EPI_ISL_513562, EPI_ISL_513563, EPI_ISL_513564, EPI_ISL_513567, EPI_ISL_513568, EPI_ISL_513571, EPI_ISL_513572, EPI_ISL_513573, EPI_ISL_513574, EPI_ISL_513575, EPI_ISL_513576, EPI_ISL_513577, EPI_ISL_513578, EPI_ISL_513579, EPI_ISL_513580, EPI_ISL_513581, EPI_ISL_513582, EPI_ISL_513583 | | | |
| see above | Programa de Oncovirologia, Instituto Nacional de Câncer | Programa de Oncovirologia, Instituto Nacional de Câncer | Juliana D. Siqueira, Livia R. Goes, Brunna M. Alves, Claudia Cicala,James Arthos, João P.B. Viola, Andreia C. de Melo, Marcelo A. Soares |
| EPI_ISL_514131 | Rondônia Central Public Health Laboratory (LACEN/RO), vinculated to State Health Secretariat of Rondônia (SESAU/RO) | Molecular Virology Laboratory of Oswaldo Cruz Foundation of Rondônia | Luan Felipo Botelho-Souza, Felipe Souza Nogueira-Lima, Tárcio Peixoto Roca, Alcione de Oliveira dos Santos, Felipe Gomes Naveca, Adriana Cristina Salvador Maia, Cicileia Correia da Silva, Aline Linhares Ferreira de Melo Mendonça, Celina Aparecida Bertoni Lugtenburg, Camila Flávia Gomes Azzi, Juliana Loca Furtado, Suelen Cavalcante, Rita de Cássia Pontello Rampazzo, Caio Henrique Nemeth Santos, Alice Paula Di Sabatino Guimarães, Jansen Fernandes de Medeiros, Fernando Rodrigues Máximo, Juan Miguel Vilallobos-Salcedo and Deusilene Souza Vieira1 |
| EPI_ISL_514132 | Rondônia Central Public Health Laboratory (LACEN/RO), vinculated to State Health Secretariat of Rondônia (SESAU/RO) | Molecular Virology Laboratory of Oswaldo Cruz Foundation of Rondônia | Luan Felipo Botelho-Souza, Felipe Souza Nogueira-Lima, Tárcio Peixoto Roca, Alcione de Oliveira dos Santos, Felipe Gomes Naveca, Adriana Cristina Salvador Maia, Cicileia Correia da Silva, Aline Linhares Ferreira de Melo Mendonça, Celina Aparecida Bertoni Lugtenburg, Camila Flávia Gomes Azzi, Juliana Loca Furtado, Suelen Cavalcante, Rita de Cássia Pontello Rampazzo, Caio Henrique Nemeth Santos, Alice Paula Di Sabatino Guimarães, Jansen Fernandes de Medeiros, Fernando Rodrigues Máximo, Juan Miguel Vilallobos-Salcedo and Deusilene Souza Vieira. |
| EPI_ISL_514133, EPI_ISL_514134, EPI_ISL_514135, EPI_ISL_514136, EPI_ISL_514137, EPI_ISL_514138 | Rondônia Central Public Health Laboratory (LACEN/RO), vinculated to State Health Secretariat of Rondônia (SESAU/RO) | Molecular Virology Laboratory of Oswaldo Cruz Foundation of Rondônia | Luan Felipo Botelho-Souza, Felipe Souza Nogueira-Lima, Tárcio Peixoto Roca, Alcione de Oliveira dos Santos, Felipe Gomes Naveca, Adriana Cristina Salvador Maia, Cicileia Correia da Silva, Aline Linhares Ferreira de Melo Mendonça, Celina Aparecida Bertoni Lugtenburg, Camila Flávia Gomes Azzi, Juliana Loca Furtado, Suelen Cavalcante, Rita de Cássia Pontello Rampazzo, Caio Henrique Nemeth Santos, Alice Paula Di Sabatino Guimarães, Jansen Fernandes de Medeiros, Fernando Rodrigues Máximo, Juan Miguel Vilallobos-Salcedo and Deusilene Souza Vieira |
| EPI_ISL_515520 | Hospital Municipal do Tatuape Carmino Caricchio | Instituto Adolfo Lutz, Interdiciplinary Procedures Center, Strategic Laboratory | Claudio Tavares Sacchi, Claudia Regina Gonçalves, Erica Valessa Ramos Gomes |
| EPI_ISL_515521 | Hospital Municipal Dr Waldemar Tebaldi | Instituto Adolfo Lutz, Interdiciplinary Procedures Center, Strategic Laboratory | Claudio Tavares Sacchi, Claudia Regina Gonçalves, Erica Valessa Ramos Gomes |

| | | | |
|---|---|---|---|
| EPI_ISL_515522 | UPA 24HS de Itatiba | Instituto Adolfo Lutz, Interdiciplinary Procedures Center, Strategic Laboratory | Claudio Tavares Sacchi, Claudia Regina Gonçalves, Erica Valessa Ramos Gomes |
| EPI_ISL_515523, EPI_ISL_515524 | PS Municipal Dr Lauro Ribas Braga | Instituto Adolfo Lutz, Interdiciplinary Procedures Center, Strategic Laboratory | Claudio Tavares Sacchi, Claudia Regina Gonçalves, Erica Valessa Ramos Gomes |
| EPI_ISL_515525 | National Influenza Center - Instituto Adolfo Lutz | Instituto Adolfo Lutz, Interdiciplinary Procedures Center, Strategic Laboratory | Claudio Tavares Sacchi, Claudia Regina Gonçalves, Erica Valessa Ramos Gomes |
| EPI_ISL_515526 | Hospital Municipal do Tatuape Carmino Caricchio | Instituto Adolfo Lutz, Interdiciplinary Procedures Center, Strategic Laboratory | Claudio Tavares Sacchi, Claudia Regina Gonçalves, Erica Valessa Ramos Gomes |
| EPI_ISL_515527 | Hospital Santa Clara | Instituto Adolfo Lutz, Interdiciplinary Procedures Center, Strategic Laboratory | Claudio Tavares Sacchi, Claudia Regina Gonçalves, Erica Valessa Ramos Gomes |
| EPI_ISL_515528 | Hospital Sao Paulo de Ensino da Unifesp | Instituto Adolfo Lutz, Interdiciplinary Procedures Center, Strategic Laboratory | Claudio Tavares Sacchi, Claudia Regina Gonçalves, Erica Valessa Ramos Gomes |
| EPI_ISL_515529 | Pronto Socorro Municipal Julio Tupy | Instituto Adolfo Lutz, Interdiciplinary Procedures Center, Strategic Laboratory | Claudio Tavares Sacchi, Claudia Regina Gonçalves, Erica Valessa Ramos Gomes |
| EPI_ISL_515541 | Hospital Montemagno | Instituto Adolfo Lutz, Interdiciplinary Procedures Center, Strategic Laboratory | Claudio Tavares Sacchi, Claudia Regina Gonçalves, Erica Valessa Ramos Gomes |
| EPI_ISL_515542 | Vigilância Epidemiológica de Leme | Instituto Adolfo Lutz, Interdiciplinary Procedures Center, Strategic Laboratory | Claudio Tavares Sacchi, Claudia Regina Gonçalves, Erica Valessa Ramos Gomes |
| EPI_ISL_515543 | Serviço de Vigilância Sanitária e Epidemiológica | Instituto Adolfo Lutz, Interdiciplinary Procedures Center, Strategic Laboratory | Claudio Tavares Sacchi, Claudia Regina Gonçalves, Erica Valessa Ramos Gomes |
| EPI_ISL_515544 | Ama Dr Jose Soares Hungria | Instituto Adolfo Lutz, Interdiciplinary Procedures Center, Strategic Laboratory | Claudio Tavares Sacchi, Claudia Regina Gonçalves, Erica Valessa Ramos Gomes |
| EPI_ISL_515545 | Hospital Sao Paulo de Ensino da Unifesp | Instituto Adolfo Lutz, Interdiciplinary Procedures Center, Strategic Laboratory | Claudio Tavares Sacchi, Claudia Regina Gonçalves, Erica Valessa Ramos Gomes |
| EPI_ISL_515546 | Hospital Municipal do Tatuape Carmino Caricchio | Instituto Adolfo Lutz, Interdiciplinary Procedures Center, Strategic Laboratory | Claudio Tavares Sacchi, Claudia Regina Gonçalves, Erica Valessa Ramos Gomes |
| EPI_ISL_515547 | Centro Medico da Policia Militar do Estado de Sao Paulo | Instituto Adolfo Lutz, Interdiciplinary Procedures Center, Strategic Laboratory | Claudio Tavares Sacchi, Claudia Regina Gonçalves, Erica Valessa Ramos Gomes |
| EPI_ISL_515548 | Hospital Municipal Dr. Jose Soares Hungria | Instituto Adolfo Lutz, Interdiciplinary Procedures Center, Strategic Laboratory | Claudio Tavares Sacchi, Claudia Regina Gonçalves, Erica Valessa Ramos Gomes |
| EPI_ISL_515550 | UPA Vila Santa Catarina | Instituto Adolfo Lutz, Interdiciplinary Procedures Center, Strategic Laboratory | Claudio Tavares Sacchi, Claudia Regina Gonçalves, Erica Valessa Ramos Gomes |
| EPI_ISL_515551, EPI_ISL_515552 | Hospital Municipal do Tatuape Carmino Caricchio | Instituto Adolfo Lutz, Interdiciplinary Procedures Center, Strategic Laboratory | Claudio Tavares Sacchi, Claudia Regina Gonçalves, Erica Valessa Ramos Gomes |
| EPI_ISL_515553 | Hospital Municipal Dr. Ignacio Proença de Gouvea | Instituto Adolfo Lutz, Interdiciplinary Procedures Center, Strategic Laboratory | Claudio Tavares Sacchi, Claudia Regina Gonçalves, Erica Valessa Ramos Gomes |
| EPI_ISL_515554 | Pronto Socorro Municipal de Perus | Instituto Adolfo Lutz, Interdiciplinary Procedures Center, Strategic Laboratory | Claudio Tavares Sacchi, Claudia Regina Gonçalves, Erica Valessa Ramos Gomes |
| EPI_ISL_515555 | Hospital Geral de Vila Nova Cachoeirinha | Instituto Adolfo Lutz, Interdiciplinary Procedures Center, Strategic Laboratory | Claudio Tavares Sacchi, Claudia Regina Gonçalves, Erica Valessa Ramos Gomes |
| EPI_ISL_515557 | Hospital Municipal Dr. Moysés Deutsch | Instituto Adolfo Lutz, Interdiciplinary Procedures Center, Strategic Laboratory | Claudio Tavares Sacchi, Claudia Regina Gonçalves, Erica Valessa Ramos Gomes |
| EPI_ISL_515559, EPI_ISL_515560 | Hospital Sao Paulo de Ensino da Unifesp | Instituto Adolfo Lutz, Interdiciplinary Procedures Center, Strategic Laboratory | Claudio Tavares Sacchi, Claudia Regina Gonçalves, Erica Valessa Ramos Gomes |
| EPI_ISL_515561 | Hospital Montemagno | Instituto Adolfo Lutz, Interdiciplinary Procedures Center, Strategic Laboratory | Claudio Tavares Sacchi, Claudia Regina Gonçalves, Erica Valessa Ramos Gomes |
| EPI_ISL_515562 | Hospital Municipal Doutor Alexandre Zaio | Instituto Adolfo Lutz, Interdiciplinary Procedures Center, Strategic Laboratory | Claudio Tavares Sacchi, Claudia Regina Gonçalves, Erica Valessa Ramos Gomes |
| EPI_ISL_515563 | Hospital Municipal Dr. Jose Soares Hungria | Instituto Adolfo Lutz, Interdiciplinary Procedures Center, Strategic Laboratory | Claudio Tavares Sacchi, Claudia Regina Gonçalves, Erica Valessa Ramos Gomes |
| EPI_ISL_515564 | Hosp. Municipal Prof. Dr. Alípio Corrêa Netto | Instituto Adolfo Lutz, Interdiciplinary Procedures Center, Strategic Laboratory | Claudio Tavares Sacchi, Claudia Regina Gonçalves, Erica Valessa Ramos Gomes |
| EPI_ISL_515565 | Hospital do Servidor Público Estadual Francisco Morato de Oliveira | Instituto Adolfo Lutz, Interdiciplinary Procedures Center, Strategic Laboratory | Claudio Tavares Sacchi, Claudia Regina Gonçalves, Erica Valessa Ramos Gomes |
| EPI_ISL_515566 | PS Municipal Dr Lauro Ribas Braga | Instituto Adolfo Lutz, Interdiciplinary Procedures Center, Strategic Laboratory | Claudio Tavares Sacchi, Claudia Regina Gonçalves, Erica Valessa Ramos Gomes |
| EPI_ISL_523955 | Hospital Municipal do Tatuape Carmino Caricchio | Instituto Adolfo Lutz, Interdiciplinary Procedures Center, Strategic Laboratory | Claudio Tavares Sacchi, Claudia Regina Gonçalves, Erica Valessa Ramos Gomes |
| EPI_ISL_523956 | Hospital Regional de Assis | Instituto Adolfo Lutz, Interdiciplinary Procedures Center, Strategic Laboratory | Claudio Tavares Sacchi, Claudia Regina Gonçalves, Erica Valessa Ramos Gomes |
| EPI_ISL_523957 | Hospital Itamaraty | Instituto Adolfo Lutz, Interdiciplinary Procedures Center, Strategic Laboratory | Claudio Tavares Sacchi, Claudia Regina Gonçalves, Erica Valessa Ramos Gomes |
| EPI_ISL_523958, EPI_ISL_523959 | Pronto Socorro Municipal de Perus | Instituto Adolfo Lutz, Interdiciplinary Procedures Center, Strategic Laboratory | Claudio Tavares Sacchi, Claudia Regina Gonçalves, Erica Valessa Ramos Gomes |
| EPI_ISL_523961 | Pronto Socorro Municipal 21 de Junho | Instituto Adolfo Lutz, Interdiciplinary Procedures Center, Strategic Laboratory | Claudio Tavares Sacchi, Claudia Regina Gonçalves, Erica Valessa Ramos Gomes |
| EPI_ISL_523963 | UBS Vila Silvia | Instituto Adolfo Lutz, Interdiciplinary Procedures Center, Strategic Laboratory | Claudio Tavares Sacchi, Claudia Regina Gonçalves, Erica Valessa Ramos Gomes |
| EPI_ISL_523965 | Hospital do Servidor Público Estadual Francisco Morato de Oliveira | Instituto Adolfo Lutz, Interdiciplinary Procedures Center, Strategic Laboratory | Claudio Tavares Sacchi, Claudia Regina Gonçalves, Erica Valessa Ramos Gomes |

| | | | |
|---|---|---|---|
| EPI_ISL_523967 | Hospital Sancta Maggiore | Instituto Adolfo Lutz, Interdiciplinary Procedures Center, Strategic Laboratory | Claudio Tavares Sacchi, Claudia Regina Gonçalves, Erica Valessa Ramos Gomes |
| EPI_ISL_523969 | Hospital Sao Paulo de Ensino da Unifesp | Instituto Adolfo Lutz, Interdiciplinary Procedures Center, Strategic Laboratory | Claudio Tavares Sacchi, Claudia Regina Gonçalves, Erica Valessa Ramos Gomes |
| EPI_ISL_523970 | Conjunto Hospitalar do Mandaqui | Instituto Adolfo Lutz, Interdiciplinary Procedures Center, Strategic Laboratory | Claudio Tavares Sacchi, Claudia Regina Gonçalves, Erica Valessa Ramos Gomes |
| EPI_ISL_523971 | Hospital Geral Santa Marcelina | Instituto Adolfo Lutz, Interdiciplinary Procedures Center, Strategic Laboratory | Claudio Tavares Sacchi, Claudia Regina Gonçalves, Erica Valessa Ramos Gomes |
| EPI_ISL_523972 | Hospital do Servidor Público Estadual Francisco Morato de Oliveira | Instituto Adolfo Lutz, Interdiciplinary Procedures Center, Strategic Laboratory | Claudio Tavares Sacchi, Claudia Regina Gonçalves, Erica Valessa Ramos Gomes |
| EPI_ISL_523973 | PS Municipal Dona Maria Antonieta Ferreira de Barros | Instituto Adolfo Lutz, Interdiciplinary Procedures Center, Strategic Laboratory | Claudio Tavares Sacchi, Claudia Regina Gonçalves, Erica Valessa Ramos Gomes |
| EPI_ISL_523974 | Hospital Municipal do Tatuape Carmino Caricchio | Instituto Adolfo Lutz, Interdiciplinary Procedures Center, Strategic Laboratory | Claudio Tavares Sacchi, Claudia Regina Gonçalves, Erica Valessa Ramos Gomes |
| EPI_ISL_523975 | UPA Tito Lopes | Instituto Adolfo Lutz, Interdiciplinary Procedures Center, Strategic Laboratory | Claudio Tavares Sacchi, Claudia Regina Gonçalves, Erica Valessa Ramos Gomes |
| EPI_ISL_523976 | Hospital Municipal do Tatuape Carmino Caricchio | Instituto Adolfo Lutz, Interdiciplinary Procedures Center, Strategic Laboratory | Claudio Tavares Sacchi, Claudia Regina Gonçalves, Erica Valessa Ramos Gomes |
| EPI_ISL_523977 | Hosp. Municipal Prof. Dr. Alípio Corrêa Netto | Instituto Adolfo Lutz, Interdiciplinary Procedures Center, Strategic Laboratory | Claudio Tavares Sacchi, Claudia Regina Gonçalves, Erica Valessa Ramos Gomes |
| EPI_ISL_523978 | Hospital do Servidor Público Estadual Francisco Morato de Oliveira | Instituto Adolfo Lutz, Interdiciplinary Procedures Center, Strategic Laboratory | Claudio Tavares Sacchi, Claudia Regina Gonçalves, Erica Valessa Ramos Gomes |
| EPI_ISL_523980 | UPA Tito Lopes | Instituto Adolfo Lutz, Interdiciplinary Procedures Center, Strategic Laboratory | Claudio Tavares Sacchi, Claudia Regina Gonçalves, Erica Valessa Ramos Gomes |
| EPI_ISL_523981 | Hospital Sao Paulo de Ensino da Unifesp | Instituto Adolfo Lutz, Interdiciplinary Procedures Center, Strategic Laboratory | Claudio Tavares Sacchi, Claudia Regina Gonçalves, Erica Valessa Ramos Gomes |
| EPI_ISL_523982 | Hospital do Servidor Público Estadual Francisco Morato de Oliveira | Instituto Adolfo Lutz, Interdiciplinary Procedures Center, Strategic Laboratory | Claudio Tavares Sacchi, Claudia Regina Gonçalves, Erica Valessa Ramos Gomes |
| EPI_ISL_523983 | UPA Campo Limpo | Instituto Adolfo Lutz, Interdiciplinary Procedures Center, Strategic Laboratory | Claudio Tavares Sacchi, Claudia Regina Gonçalves, Erica Valessa Ramos Gomes |
| EPI_ISL_523984 | Ama Dr Jose Soares Hungria | Instituto Adolfo Lutz, Interdiciplinary Procedures Center, Strategic Laboratory | Claudio Tavares Sacchi, Claudia Regina Gonçalves, Erica Valessa Ramos Gomes |
| EPI_ISL_523985 | Hospital Municipal Dr. Benedicto Montenegro | Instituto Adolfo Lutz, Interdiciplinary Procedures Center, Strategic Laboratory | Claudio Tavares Sacchi, Claudia Regina Gonçalves, Erica Valessa Ramos Gomes |
| EPI_ISL_523986 | Ama Dr Jose Soares Hungria | Instituto Adolfo Lutz, Interdiciplinary Procedures Center, Strategic Laboratory | Claudio Tavares Sacchi, Claudia Regina Gonçalves, Erica Valessa Ramos Gomes |
| EPI_ISL_523988 | Hospital Sao Paulo de Ensino da Unifesp | Instituto Adolfo Lutz, Interdiciplinary Procedures Center, Strategic Laboratory | Claudio Tavares Sacchi, Claudia Regina Gonçalves, Erica Valessa Ramos Gomes |
| EPI_ISL_523989 | AMA Jardim Joamar | Instituto Adolfo Lutz, Interdiciplinary Procedures Center, Strategic Laboratory | Claudio Tavares Sacchi, Claudia Regina Gonçalves, Erica Valessa Ramos Gomes |
| EPI_ISL_523990 | AMA Jardim Peri | Instituto Adolfo Lutz, Interdiciplinary Procedures Center, Strategic Laboratory | Claudio Tavares Sacchi, Claudia Regina Gonçalves, Erica Valessa Ramos Gomes |
| EPI_ISL_523991, EPI_ISL_523992 | Hospital Municipal Carmen Prudente | Instituto Adolfo Lutz, Interdiciplinary Procedures Center, Strategic Laboratory | Claudio Tavares Sacchi, Claudia Regina Gonçalves, Erica Valessa Ramos Gomes |
| EPI_ISL_524462 | Hospital Metropolitano | Instituto Adolfo Lutz, Interdiciplinary Procedures Center, Strategic Laboratory | Claudio Tavares Sacchi, Claudia Regina Gonçalves, Erica Valessa Ramos Gomes |
| EPI_ISL_524463 | Hospital Regional de Cotia | Instituto Adolfo Lutz, Interdiciplinary Procedures Center, Strategic Laboratory | Claudio Tavares Sacchi, Claudia Regina Gonçalves, Erica Valessa Ramos Gomes |
| EPI_ISL_524464 | Santa Casa de Santa Isabel | Instituto Adolfo Lutz, Interdiciplinary Procedures Center, Strategic Laboratory | Claudio Tavares Sacchi, Claudia Regina Gonçalves, Erica Valessa Ramos Gomes |
| EPI_ISL_524465 | PS Municipal Dr. Caetano Virgílio Neto | Instituto Adolfo Lutz, Interdiciplinary Procedures Center, Strategic Laboratory | Claudio Tavares Sacchi, Claudia Regina Gonçalves, Erica Valessa Ramos Gomes |
| EPI_ISL_524466 | PS Municipal Dr Lauro Ribas Braga | Instituto Adolfo Lutz, Interdiciplinary Procedures Center, Strategic Laboratory | Claudio Tavares Sacchi, Claudia Regina Gonçalves, Erica Valessa Ramos Gomes |
| EPI_ISL_524467 | Hospital Municipal Dr. Moysés Deutsch | Instituto Adolfo Lutz, Interdiciplinary Procedures Center, Strategic Laboratory | Claudio Tavares Sacchi, Claudia Regina Gonçalves, Erica Valessa Ramos Gomes |
| EPI_ISL_524468 | Hospital Municipal Vereador Jose Storopolli | Instituto Adolfo Lutz, Interdiciplinary Procedures Center, Strategic Laboratory | Claudio Tavares Sacchi, Claudia Regina Gonçalves, Erica Valessa Ramos Gomes |
| EPI_ISL_524469 | Santa Casa de Misericordia de Sao Paulo | Instituto Adolfo Lutz, Interdiciplinary Procedures Center, Strategic Laboratory | Claudio Tavares Sacchi, Claudia Regina Gonçalves, Erica Valessa Ramos Gomes |
| EPI_ISL_524470 | Hospital do Servidor Público Estadual Francisco Morato de Oliveira | Instituto Adolfo Lutz, Interdiciplinary Procedures Center, Strategic Laboratory | Claudio Tavares Sacchi, Claudia Regina Gonçalves, Erica Valessa Ramos Gomes |
| EPI_ISL_524783, EPI_ISL_524784, EPI_ISL_524785, EPI_ISL_524786, EPI_ISL_524787, EPI_ISL_524788, EPI_ISL_524789, EPI_ISL_524790, EPI_ISL_524791, EPI_ISL_524792, EPI_ISL_524793, EPI_ISL_524794, EPI_ISL_524795, EPI_ISL_524796, EPI_ISL_524797, EPI_ISL_524798, EPI_ISL_524799, EPI_ISL_524800, EPI_ISL_524801 | | | |
| see above | Evandro Chagas Institute | Evandro Chagas Institute | Santos, M.C.; Silva, A.M.; Junior, W.D.C.; Barbagelata, L.S.; Ferreira, J.A.; Sousa, E.M.A.; da Silva, P.S.; Resque, H.R; Martins, L.C.; Sousa Junior, E.C.;Viana, G.M.R |
| EPI_ISL_527856 | Hospital Municipal Prof. Waldomiro de Paula | Instituto Adolfo Lutz, Interdiciplinary Procedures Center, Strategic Laboratory | Claudio Tavares Sacchi, Claudia Regina Gonçalves, Erica Valessa Ramos Gomes |
| EPI_ISL_527857 | Hospital Regional Vale do Ribeira | Instituto Adolfo Lutz, Interdiciplinary Procedures Center, Strategic Laboratory | Claudio Tavares Sacchi, Claudia Regina Gonçalves, Erica Valessa Ramos Gomes |

| | | | |
|---|---|---|---|
| EPI_ISL_527858 | Pronto Atendimento Sancta Maggiore Jardim Paulista | Instituto Adolfo Lutz, Interdiciplinary Procedures Center, Strategic Laboratory | Claudio Tavares Sacchi, Claudia Regina Gonçalves, Erica Valessa Ramos Gomes |
| EPI_ISL_527859 | Hospital Municipal Vereador Jose Storopolli | Instituto Adolfo Lutz, Interdiciplinary Procedures Center, Strategic Laboratory | Claudio Tavares Sacchi, Claudia Regina Gonçalves, Erica Valessa Ramos Gomes |
| EPI_ISL_527860 | Hospital Municipal de Parelheiros Josanias Castanha Braga | Instituto Adolfo Lutz, Interdiciplinary Procedures Center, Strategic Laboratory | Claudio Tavares Sacchi, Claudia Regina Gonçalves, Erica Valessa Ramos Gomes |
| EPI_ISL_527861 | Hospital e Maternidade Celso Pierro | Instituto Adolfo Lutz, Interdiciplinary Procedures Center, Strategic Laboratory | Av. Dr. Arnaldo, 355 - Brazil, Cerqueira Cesar, São Paulo - SP, 01246-1301 |
| EPI_ISL_527862 | Hospital Municipal de Urgência | Instituto Adolfo Lutz, Interdiciplinary Procedures Center, Strategic Laboratory | Claudio Tavares Sacchi, Claudia Regina Gonçalves, Erica Valessa Ramos Gomes |
| EPI_ISL_527863 | Hospital Municipal do Tatuape Carmino Caricchio | Instituto Adolfo Lutz, Interdiciplinary Procedures Center, Strategic Laboratory | Claudio Tavares Sacchi, Claudia Regina Gonçalves, Erica Valessa Ramos Gomes |
| EPI_ISL_527864 | Hospital e Pronto Socorro Comunitário Vila Iolanda | Instituto Adolfo Lutz, Interdiciplinary Procedures Center, Strategic Laboratory | Claudio Tavares Sacchi, Claudia Regina Gonçalves, Erica Valessa Ramos Gomes |
| EPI_ISL_527865 | Hospital e Maternidade São Cristóvão | Instituto Adolfo Lutz, Interdiciplinary Procedures Center, Strategic Laboratory | Claudio Tavares Sacchi, Claudia Regina Gonçalves, Erica Valessa Ramos Gomes |
| EPI_ISL_527866 | PS Municipal Dr Lauro Ribas Braga | Instituto Adolfo Lutz, Interdiciplinary Procedures Center, Strategic Laboratory | Av. Dr. Arnaldo, 355 - Brazil, Cerqueira Cesar, São Paulo - SP, 01246-1301 |
| EPI_ISL_527867 | Pronto Socorro Municipal - Balneario São José | Instituto Adolfo Lutz, Interdiciplinary Procedures Center, Strategic Laboratory | Claudio Tavares Sacchi, Claudia Regina Gonçalves, Erica Valessa Ramos Gomes |
| EPI_ISL_527868 | Hospital e Maternidade do Braz | Instituto Adolfo Lutz, Interdiciplinary Procedures Center, Strategic Laboratory | Claudio Tavares Sacchi, Claudia Regina Gonçalves, Erica Valessa Ramos Gomes |
| EPI_ISL_527869 | Hospital Municipal Carmen Prudente | Instituto Adolfo Lutz, Interdiciplinary Procedures Center, Strategic Laboratory | Claudio Tavares Sacchi, Claudia Regina Gonçalves, Erica Valessa Ramos Gomes |
| EPI_ISL_527870 | Hospital Municipal Mário Gatti | Instituto Adolfo Lutz, Interdiciplinary Procedures Center, Strategic Laboratory | Claudio Tavares Sacchi, Claudia Regina Gonçalves, Erica Valessa Ramos Gomes |
| EPI_ISL_528539 | LVM/UFRJ | LNCC | Gustavo M. Romário M. de Souza; Bruno B. Bezerra; Lucio A. Caldas; Fabio Limonte; Elena Cobos; Sharton V. A. Coelho; Luiz Almeida; Luiza Higga; Isadora A. Correa; Diana Marianni; Luciana B. Arruda; Marcelo Bozza; Orlando Ferreira; Wanderley de Souza; Ana Teresa R. Vasconcelos; Terezinha M. Castineiras; Amilcar Tanuri; Luciana J. Costa. |
| EPI_ISL_528637, EPI_ISL_528638 | LVM/UFRJ | Bioinformatics Laboratory / LNCC | Gustavo D. P. Silva; M. Romário M. de Souza; Bruno B. Bezerra; Lucio A. Caldas; Fabio Limonte; Elena Cobos; Sharton V. A. Coelho; Luiz Almeida; Luiza Higga; Isadora A. Correa; Diana Marianni; Luciana B. Arruda; Marcelo Bozza; Orlando Ferreira; Wanderley de Souza; Ana Teresa R. Vasconcelos; Terezinha M. Castineiras; Amilcar Tanuri; Luciana J. Costa |
| EPI_ISL_529139 | Centro de Desenvolvimento Tecnologico em Saude, Fundacao Oswaldo Cruz | Centro de Desenvolvimento Tecnologico em Saude, Fundacao Oswaldo Cruz | Souza,T.M., Fintelman-Rodrigues,N., De Paula,A.D., Saraiva,F.B., Ferreira,M.A., Sacramento,C.Q., Medeiros,M.A. |
| EPI_ISL_529140 | Centro de Desenvolvimento Tecnologico em Saude, Fundacao Oswaldo Cruz | Centro de Desenvolvimento Tecnologico em Saude, Fundacao Oswaldo Cruz | Souza,T.M., Fintelman-Rodrigues,N., De Paula,A.D., Saraiva,F.B., Ferreira,M.A., Sacramento,C.Q., Medeiros,M.A. |
| EPI_ISL_534311 | UPA III 26 de Agosto | Instituto Adolfo Lutz, Interdiciplinary Procedures Center, Strategic Laboratory | Claudio Tavares Sacchi, Claudia Regina Gonçalves, Erica Valessa Ramos Gomes |
| EPI_ISL_534312 | Distrito Sanitario Sul | Instituto Adolfo Lutz, Interdiciplinary Procedures Center, Strategic Laboratory | Claudio Tavares Sacchi, Claudia Regina Gonçalves, Erica Valessa Ramos Gomes |
| EPI_ISL_534313 | Hospital da Sta Casa de Sto Amaro | Instituto Adolfo Lutz, Interdiciplinary Procedures Center, Strategic Laboratory | Claudio Tavares Sacchi, Claudia Regina Gonçalves, Erica Valessa Ramos Gomes |
| EPI_ISL_534314 | Hospital Universitario da USP de SP | Instituto Adolfo Lutz, Interdiciplinary Procedures Center, Strategic Laboratory | Claudio Tavares Sacchi, Claudia Regina Gonçalves, Erica Valessa Ramos Gomes |
| EPI_ISL_534315 | Serviço de Verificação de Óbitos SVO Guaruulhos | Instituto Adolfo Lutz, Interdiciplinary Procedures Center, Strategic Laboratory | Claudio Tavares Sacchi, Claudia Regina Gonçalves, Erica Valessa Ramos Gomes |
| EPI_ISL_534316 | OS Mun Santana Lauro Ribas Braga | Instituto Adolfo Lutz, Interdiciplinary Procedures Center, Strategic Laboratory | Claudio Tavares Sacchi, Claudia Regina Gonçalves, Erica Valessa Ramos Gomes |
| EPI_ISL_534317 | Hospital Geral de Itapevi | Instituto Adolfo Lutz, Interdiciplinary Procedures Center, Strategic Laboratory | Claudio Tavares Sacchi, Claudia Regina Gonçalves, Erica Valessa Ramos Gomes |
| EPI_ISL_534318 | Hospital Municipal Antonio Giglio | Instituto Adolfo Lutz, Interdiciplinary Procedures Center, Strategic Laboratory | Claudio Tavares Sacchi, Claudia Regina Gonçalves, Erica Valessa Ramos Gomes |
| EPI_ISL_534319, EPI_ISL_534320 | Hospital do Serv Pub ESTAFCO Morato de Oliveira | Instituto Adolfo Lutz, Interdiciplinary Procedures Center, Strategic Laboratory | Claudio Tavares Sacchi, Claudia Regina Gonçalves, Erica Valessa Ramos Gomes |
| EPI_ISL_534321 | PS e Maternidade Nair Fonseca Leitao Arantes | Instituto Adolfo Lutz, Interdiciplinary Procedures Center, Strategic Laboratory | Claudio Tavares Sacchi, Claudia Regina Gonçalves, Erica Valessa Ramos Gomes |
| EPI_ISL_534322 | PS Mun Julio Tupy | Instituto Adolfo Lutz, Interdiciplinary Procedures Center, Strategic Laboratory | Claudio Tavares Sacchi, Claudia Regina Gonçalves, Erica Valessa Ramos Gomes |
| EPI_ISL_534323 | Hospital e Pronto Socorro Comunitario Vila Yolanda | Instituto Adolfo Lutz, Interdiciplinary Procedures Center, Strategic Laboratory | Claudio Tavares Sacchi, Claudia Regina Gonçalves, Erica Valessa Ramos Gomes |
| EPI_ISL_534324 | Hospital Mun Ver Jose Storopolli | Instituto Adolfo Lutz, Interdiciplinary Procedures Center, Strategic Laboratory | Claudio Tavares Sacchi, Claudia Regina Gonçalves, Erica Valessa Ramos Gomes |
| EPI_ISL_534325 | Unidade de Vigilancia em Saude de Guarulhos | Instituto Adolfo Lutz, Interdiciplinary Procedures Center, Strategic Laboratory | Claudio Tavares Sacchi, Claudia Regina Gonçalves, Erica Valessa Ramos Gomes |
| EPI_ISL_534326 | Notre Dame Intermedica Saude AS | Instituto Adolfo Lutz, Interdiciplinary Procedures Center, Strategic Laboratory | Claudio Tavares Sacchi, Claudia Regina Gonçalves, Erica Valessa Ramos Gomes |
| EPI_ISL_541340, EPI_ISL_541341, EPI_ISL_541342, EPI_ISL_541343, EPI_ISL_541344, EPI_ISL_541345, EPI_ISL_541346 | Laboratório Central de Saúde Pública do Estado do Paraná (LACEN-PR) | Laboratory of Respiratory Viruses and Measles, Oswaldo Cruz Institute, FIOCRUZ | Paola Resende, Luciana Appolinario, Fernando Motta, Anna Carolina Paixão, Ana Carolina Mendonça, Jonathan Lopes, Irina Riediger, Maria do Carmo Debur, Marilda Siqueira on behalf of the Fiocruz COVID-19 Genomic Surveillance Network |
| EPI_ISL_541347, EPI_ISL_541348, | Laboratory of Respiratory Viruses and Measles, Oswaldo Cruz | Laboratory of Respiratory Viruses and Measles, Oswaldo Cruz | Paola Resende, Luciana Appolinario, Fernando Motta, Anna Carolina Paixão, Ana Carolina Mendonça, Jonathan Lopes, Marilda Siqueira on behalf of the |

| | | | |
|---|---|---|---|
| EPI_ISL_541349, EPI_ISL_541350, EPI_ISL_541351 | Institute, FIOCRUZ | Institute, FIOCRUZ | Fiocruz COVID-19 Genomic Surveillance Network |
| EPI_ISL_541352 | Laboratory of Respiratory Viruses and Measles, Oswaldo Cruz Institute, FIOCRUZ | Laboratory of Respiratory Viruses and Measles, Oswaldo Cruz Institute, FIOCRUZ | Paola Resende, Roxana Loayza, Cinthia Avila, Luciana Appolinario, Fernando Motta, Anna Carolina Paixao, Ana Carolina Mendonca, Marilda Siqueira on behalf of the Fiocruz COVID-19 Genomic Surveillance Network |
| EPI_ISL_541353, EPI_ISL_541354, EPI_ISL_541355 | Laboratory of Respiratory Viruses and Measles, Oswaldo Cruz Institute, FIOCRUZ | Laboratory of Respiratory Viruses and Measles, Oswaldo Cruz Institute, FIOCRUZ | Paola Resende, Luciana Appolinario, Fernando Motta, Anna Carolina Paixão, Ana Carolina Mendonça, Jonathan Lopes, Marilda Siqueira on behalf of the Fiocruz COVID-19 Genomic Surveillance Network |
| EPI_ISL_541356 | Laboratory of Respiratory Viruses and Measles, Oswaldo Cruz Institute, FIOCRUZ | Laboratory of Respiratory Viruses and Measles, Oswaldo Cruz Institute, FIOCRUZ | Paola Resende, Roxana Loayza, Cinthia Avila, Luciana Appolinario, Fernando Motta, Anna Carolina Paixao, Ana Carolina Mendonca, Marilda Siqueira on behalf of the Fiocruz COVID-19 Genomic Surveillance Network |
| EPI_ISL_541357, EPI_ISL_541358 | Laboratory of Respiratory Viruses and Measles, Oswaldo Cruz Institute, FIOCRUZ | Laboratory of Respiratory Viruses and Measles, Oswaldo Cruz Institute, FIOCRUZ | Paola Resende, Luciana Appolinario, Fernando Motta, Anna Carolina Paixão, Ana Carolina Mendonça, Jonathan Lopes, Marilda Siqueira on behalf of the Fiocruz COVID-19 Genomic Surveillance Network |
| EPI_ISL_541359, EPI_ISL_541360, EPI_ISL_541361 | Laboratory of Respiratory Viruses and Measles, Oswaldo Cruz Institute, FIOCRUZ | Laboratory of Respiratory Viruses and Measles, Oswaldo Cruz Institute, FIOCRUZ | Paola Resende, Roxana Loayza, Cinthia Avila, Luciana Appolinario, Fernando Motta, Anna Carolina Paixao, Ana Carolina Mendonca, Marilda Siqueira on behalf of the Fiocruz COVID-19 Genomic Surveillance Network |
| EPI_ISL_541362, EPI_ISL_541363 | Laboratory of Respiratory Viruses and Measles, Oswaldo Cruz Institute, FIOCRUZ | Laboratory of Respiratory Viruses and Measles, Oswaldo Cruz Institute, FIOCRUZ | Paola Resende, Luciana Appolinario, Fernando Motta, Anna Carolina Paixão, Ana Carolina Mendonça, Jonathan Lopes, Marilda Siqueira on behalf of the Fiocruz COVID-19 Genomic Surveillance Network |
| EPI_ISL_541364, EPI_ISL_541365, EPI_ISL_541366 | Laboratory of Respiratory Viruses and Measles, Oswaldo Cruz Institute, FIOCRUZ | Laboratory of Respiratory Viruses and Measles, Oswaldo Cruz Institute, FIOCRUZ | Paola Resende, Roxana Loayza, Cinthia Avila, Luciana Appolinario, Fernando Motta, Anna Carolina Paixao, Ana Carolina Mendonca, Marilda Siqueira on behalf of the Fiocruz COVID-19 Genomic Surveillance Network |
| EPI_ISL_541367 | Laboratory of Respiratory Viruses and Measles, Oswaldo Cruz Institute, FIOCRUZ | Laboratory of Respiratory Viruses and Measles, Oswaldo Cruz Institute, FIOCRUZ | Paola Resende, Luciana Appolinario, Fernando Motta, Anna Carolina Paixão, Ana Carolina Mendonça, Jonathan Lopes, Marilda Siqueira on behalf of the Fiocruz COVID-19 Genomic Surveillance Network |
| EPI_ISL_541368, EPI_ISL_541369 | Laboratory of Respiratory Viruses and Measles, Oswaldo Cruz Institute, FIOCRUZ | Laboratory of Respiratory Viruses and Measles, Oswaldo Cruz Institute, FIOCRUZ | Paola Resende, Roxana Loayza, Cinthia Avila, Luciana Appolinario, Fernando Motta, Anna Carolina Paixao, Ana Carolina Mendonca, Marilda Siqueira on behalf of the Fiocruz COVID-19 Genomic Surveillance Network |
| EPI_ISL_541370, EPI_ISL_541371 | Laboratório Central de Saúde Pública do Estado de Santa Catarina (LACEN-SC) | Laboratory of Respiratory Viruses and Measles, Oswaldo Cruz Institute, FIOCRUZ | Paola Resende, Luciana Appolinario, Fernando Motta, Anna Carolina Paixão, Ana Carolina Mendonça, Jonathan Lopes, Sandra Bianchini, Marilda Siqueira on behalf of the Fiocruz COVID-19 Genomic Surveillance Network |
| EPI_ISL_541372, EPI_ISL_541373, EPI_ISL_541374, EPI_ISL_541375, EPI_ISL_541376, EPI_ISL_541377, EPI_ISL_541378, EPI_ISL_541379, EPI_ISL_541380, EPI_ISL_541381, EPI_ISL_541382, EPI_ISL_541383, EPI_ISL_541384, EPI_ISL_541385, EPI_ISL_541386, EPI_ISL_541387, EPI_ISL_541388, EPI_ISL_541389, EPI_ISL_541390, EPI_ISL_541391, EPI_ISL_541393, EPI_ISL_541394, EPI_ISL_541395, EPI_ISL_541396 | | | |
| see above | Laboratório Central de Saúde Pública do Estado de Sergipe (LACEN-SE) | Laboratory of Respiratory Viruses and Measles, Oswaldo Cruz Institute, FIOCRUZ | Paola Resende, Luciana Appolinario, Fernando Motta, Anna Carolina Paixão, Ana Carolina Mendonça, Jonathan Lopes, Clioma Santos, Marilda Siqueira on behalf of the Fiocruz COVID-19 Genomic Surveillance Network |
| EPI_ISL_547570 | PS Municipal Dr Augusto Gomes de Mattos | Instituto Adolfo Lutz, Interdiciplinary Procedures Center, Strategic Laboratory | Claudio Tavares Sacchi, Claudia Regina Gonçalves, Erica Valessa Ramos Gomes, Karoline Rodrigues Campos |
| EPI_ISL_547571 | Hospital Municipal Antônio Giglio | Instituto Adolfo Lutz, Interdiciplinary Procedures Center, Strategic Laboratory | Claudio Tavares Sacchi, Claudia Regina Gonçalves, Erica Valessa Ramos Gomes, Karoline Rodrigues Campos |
| EPI_ISL_547573 | Vigilância em Saúde de Cajamar | Instituto Adolfo Lutz, Interdiciplinary Procedures Center, Strategic Laboratory | Claudio Tavares Sacchi, Claudia Regina Gonçalves, Erica Valessa Ramos Gomes, Karoline Rodrigues Campos |
| EPI_ISL_547574 | Hospital Universitario da USP | Instituto Adolfo Lutz, Interdiciplinary Procedures Center, Strategic Laboratory | Claudio Tavares Sacchi, Claudia Regina Gonçalves, Erica Valessa Ramos Gomes, Karoline Rodrigues Campos |
| EPI_ISL_547575 | SVO Jundiaí | Instituto Adolfo Lutz, Interdiciplinary Procedures Center, Strategic Laboratory | Claudio Tavares Sacchi, Claudia Regina Gonçalves, Erica Valessa Ramos Gomes, Karoline Rodrigues Campos |
| EPI_ISL_547576 | Secretaria Municipal de Saúde | Instituto Adolfo Lutz, Interdiciplinary Procedures Center, Strategic Laboratory | Claudio Tavares Sacchi, Claudia Regina Gonçalves, Erica Valessa Ramos Gomes, Karoline Rodrigues Campos |
| EPI_ISL_547577 | Hospital e Maternidade Nossa Senhora das Graças | Instituto Adolfo Lutz, Interdiciplinary Procedures Center, Strategic Laboratory | Claudio Tavares Sacchi, Claudia Regina Gonçalves, Erica Valessa Ramos Gomes, Karoline Rodrigues Campos |
| EPI_ISL_547578 | Hospital Doutor Domingos Leonardo Cerávolo | Instituto Adolfo Lutz, Interdiciplinary Procedures Center, Strategic Laboratory | Claudio Tavares Sacchi, Claudia Regina Gonçalves, Erica Valessa Ramos Gomes, Karoline Rodrigues Campos |
| EPI_ISL_547579 | Santa Casa de Misericórdia de Araçatuba | Instituto Adolfo Lutz, Interdiciplinary Procedures Center, Strategic Laboratory | Claudio Tavares Sacchi, Claudia Regina Gonçalves, Erica Valessa Ramos Gomes, Karoline Rodrigues Campos |
| EPI_ISL_547580 | Santa Casa da Misericórdia de Presidente Prudente | Instituto Adolfo Lutz, Interdiciplinary Procedures Center, Strategic Laboratory | Claudio Tavares Sacchi, Claudia Regina Gonçalves, Erica Valessa Ramos Gomes, Karoline Rodrigues Campos |
| EPI_ISL_572334, EPI_ISL_572335, EPI_ISL_572351, EPI_ISL_572353, EPI_ISL_572366, EPI_ISL_572371, EPI_ISL_572386 | Laboratório Central de Saúde Pública do Estado de Pernambuco (LACEN-PE) | WallauLab, Aggeu Magalhaes Institute | Marcelo Henrique Santos Paiva, Duschinka Ribeiro Duarte Guedes, Cássia Docena, Matheus Filgueira Bezerra, Filipe Zimmer Dezordi, Laís Ceschini Machado, Larissa Krokovsky, Elisama Helvecio, Alexandre Freitas da Silva, Luydson Richardson Silva Vasconcelos, Antonio Mauro Rezende, Severino Jefferson Ribeiro da Silva, Kamila Gaudêncio da Silva Sales, Bruna Santos Lima Figueiredo de Sá, Derciliano Lopes da Cruz, Claudio Eduardo Cavalcanti, Armando de Menezes Neto, Caroline Targino Alves da Silva, Renata Pessôa Germano Mendes, Maria Almerice Lopes da Silva, Tiago Gräf, Paola Cristina Resende, Gonzalo Bello, Michelle da Silva Barros, Wheverton Ricardo Correia do Nascimento, Rodrigo Moraes Loyo Arcoverde, Luciane Caroline Albuquerque Bezerra, Sinval Pinto Brandão Filho, Constância Flávia Junqueira Ayres, Gabriel Luz Wallau on behalf of the Fiocruz COVID-19 Genomic Surveillance Network |
| EPI_ISL_574577 | Hospital Municipal Dr. Ignacio Proença de Gouvea | Instituto Adolfo Lutz, Interdiciplinary Procedures Center, Strategic Laboratory | Claudio Tavares Sacchi, Claudia Regina Gonçalves, Erica Valessa Ramos Gomes, Karoline Rodrigues Campos |
| EPI_ISL_574578 | Hospital Municipal Mário Gatti | Instituto Adolfo Lutz, Interdiciplinary Procedures Center, Strategic Laboratory | Claudio Tavares Sacchi, Claudia Regina Gonçalves, Erica Valessa Ramos Gomes, Karoline Rodrigues Campos |
| EPI_ISL_574579 | Hospital Municipal Dr. Ignacio Proença de Gouvea | Instituto Adolfo Lutz, Interdiciplinary Procedures Center, Strategic Laboratory | Claudio Tavares Sacchi, Claudia Regina Gonçalves, Erica Valessa Ramos Gomes, Karoline Rodrigues Campos |
| EPI_ISL_574580 | Hospital Cidade Tiradentes Carmen Prudente | Instituto Adolfo Lutz, Interdiciplinary Procedures Center, Strategic Laboratory | Claudio Tavares Sacchi, Claudia Regina Gonçalves, Erica Valessa Ramos Gomes, Karoline Rodrigues Campos |
| EPI_ISL_574582 | Hospital Municipal Dr. Jose Soares Hungria | Instituto Adolfo Lutz, Interdiciplinary Procedures Center, Strategic Laboratory | Claudio Tavares Sacchi, Claudia Regina Gonçalves, Erica Valessa Ramos Gomes, Karoline Rodrigues Campos |
| EPI_ISL_574583 | Secretaria Municipal de Saude de Jandira | Instituto Adolfo Lutz, Interdiciplinary Procedures Center, Strategic Laboratory | Claudio Tavares Sacchi, Claudia Regina Gonçalves, Erica Valessa Ramos Gomes, Karoline Rodrigues Campos |
| EPI_ISL_574588 | Hospital Estadual Sumare | Instituto Adolfo Lutz, Interdiciplinary Procedures Center, Strategic Laboratory | Claudio Tavares Sacchi, Claudia Regina Gonçalves, Erica Valessa Ramos Gomes, Karoline Rodrigues Campos |
| EPI_ISL_574589 | Hospital Municipal Dr. Jose Soares Hungria | Instituto Adolfo Lutz, Interdiciplinary Procedures Center, Strategic Laboratory | Claudio Tavares Sacchi, Claudia Regina Gonçalves, Erica Valessa Ramos Gomes, Karoline Rodrigues Campos |
| EPI_ISL_574590 | Unidade de Printo Atendimento UPA I Santa Isabel | Instituto Adolfo Lutz, Interdiciplinary Procedures Center, Strategic Laboratory | Claudio Tavares Sacchi, Claudia Regina Gonçalves, Erica Valessa Ramos Gomes, Karoline Rodrigues Campos |

| | | | |
|---|---|---|---|
| EPI_ISL_574591, EPI_ISL_574592 | Hospital Domingos Leonardo Ceravolo Presidente Prudente | Instituto Adolfo Lutz, Interdiciplinary Procedures Center, Strategic Laboratory | Claudio Tavares Sacchi, Claudia Regina Gonçalves, Erica Valessa Ramos Gomes, Karoline Rodrigues Campos |
| EPI_ISL_574593 | CS II Dr. Antonio Vicoso Moreira de Rezende Sumare | Instituto Adolfo Lutz, Interdiciplinary Procedures Center, Strategic Laboratory | Claudio Tavares Sacchi, Claudia Regina Gonçalves, Erica Valessa Ramos Gomes, Karoline Rodrigues Campos |
| EPI_ISL_574594 | Hospital Escola da Universidade de Taubate | Instituto Adolfo Lutz, Interdiciplinary Procedures Center, Strategic Laboratory | Claudio Tavares Sacchi, Claudia Regina Gonçalves, Erica Valessa Ramos Gomes, Karoline Rodrigues Campos |
| EPI_ISL_574595 | Hospital Geral de Vila Penteado Dr. Jose Pamgella | Instituto Adolfo Lutz, Interdiciplinary Procedures Center, Strategic Laboratory | Claudio Tavares Sacchi, Claudia Regina Gonçalves, Erica Valessa Ramos Gomes, Karoline Rodrigues Campos |
| EPI_ISL_574596 | CS II Dr. Antonio Vicoso Moreira de Rezende Sumare | Instituto Adolfo Lutz, Interdiciplinary Procedures Center, Strategic Laboratory | Claudio Tavares Sacchi, Claudia Regina Gonçalves, Erica Valessa Ramos Gomes, Karoline Rodrigues Campos |
| EPI_ISL_574597 | Secretaria Municipal de Saude de Jarinu | Instituto Adolfo Lutz, Interdiciplinary Procedures Center, Strategic Laboratory | Claudio Tavares Sacchi, Claudia Regina Gonçalves, Erica Valessa Ramos Gomes, Karoline Rodrigues Campos |
| EPI_ISL_574598 | Servico de Verificacao de Obito SVO | Instituto Adolfo Lutz, Interdiciplinary Procedures Center, Strategic Laboratory | Claudio Tavares Sacchi, Claudia Regina Gonçalves, Erica Valessa Ramos Gomes, Karoline Rodrigues Campos |
| EPI_ISL_583490 | Hospital Estadual Sumare | Instituto Adolfo Lutz, Interdiciplinary Procedures Center, Strategic Laboratory | Claudio Tavares Sacchi, Claudia Regina Gonçalves, Erica Valessa Ramos Gomes, Karoline Rodrigues Campos |
| EPI_ISL_583491 | Centro de Saude Esf IV Zona Rual Domingos de SJ Rio Pardo | Instituto Adolfo Lutz, Interdiciplinary Procedures Center, Strategic Laboratory | Claudio Tavares Sacchi, Claudia Regina Gonçalves, Erica Valessa Ramos Gomes, Karoline Rodrigues Campos |
| EPI_ISL_583492 | Santa Casa Anna Cintra | Instituto Adolfo Lutz, Interdiciplinary Procedures Center, Strategic Laboratory | Claudio Tavares Sacchi, Claudia Regina Gonçalves, Erica Valessa Ramos Gomes, Karoline Rodrigues Campos |
| EPI_ISL_583493 | Vigilância em Saúde de Cajamar | Instituto Adolfo Lutz, Interdiciplinary Procedures Center, Strategic Laboratory | Claudio Tavares Sacchi, Claudia Regina Gonçalves, Erica Valessa Ramos Gomes, Karoline Rodrigues Campos |
| EPI_ISL_583494 | CS II Dr. Antonio Vicoso Moreira de Rezende Sumare | Instituto Adolfo Lutz, Interdiciplinary Procedures Center, Strategic Laboratory | Claudio Tavares Sacchi, Claudia Regina Gonçalves, Erica Valessa Ramos Gomes, Karoline Rodrigues Campos |
| EPI_ISL_583495 | Serviço de Verificação de Óbitos SVO Guaruulhos | Instituto Adolfo Lutz, Interdiciplinary Procedures Center, Strategic Laboratory | Claudio Tavares Sacchi, Claudia Regina Gonçalves, Erica Valessa Ramos Gomes, Karoline Rodrigues Campos |
| EPI_ISL_583496 | UPA Jandira | Instituto Adolfo Lutz, Interdiciplinary Procedures Center, Strategic Laboratory | Claudio Tavares Sacchi, Claudia Regina Gonçalves, Erica Valessa Ramos Gomes, Karoline Rodrigues Campos |
| EPI_ISL_583497 | Complexo Hospitalar Ouro Verde de Campinas | Instituto Adolfo Lutz, Interdiciplinary Procedures Center, Strategic Laboratory | Claudio Tavares Sacchi, Claudia Regina Gonçalves, Erica Valessa Ramos Gomes, Karoline Rodrigues Campos |
| EPI_ISL_583498 | Hospital Municipal Dr. Waldemar Tebaldi | Instituto Adolfo Lutz, Interdiciplinary Procedures Center, Strategic Laboratory | Claudio Tavares Sacchi, Claudia Regina Gonçalves, Erica Valessa Ramos Gomes, Karoline Rodrigues Campos |
| EPI_ISL_583499 | Distrito Sanitario Sul Campinas | Instituto Adolfo Lutz, Interdiciplinary Procedures Center, Strategic Laboratory | Claudio Tavares Sacchi, Claudia Regina Gonçalves, Erica Valessa Ramos Gomes, Karoline Rodrigues Campos |
| EPI_ISL_583500 | Centro de Saude I Tacito Leite de Carvalho e Silva | Instituto Adolfo Lutz, Interdiciplinary Procedures Center, Strategic Laboratory | Claudio Tavares Sacchi, Claudia Regina Gonçalves, Erica Valessa Ramos Gomes, Karoline Rodrigues Campos |
| EPI_ISL_583501 | Hospital Estadual de CampanhaCOVID 19 Barradas | Instituto Adolfo Lutz, Interdiciplinary Procedures Center, Strategic Laboratory | Claudio Tavares Sacchi, Claudia Regina Gonçalves, Erica Valessa Ramos Gomes, Karoline Rodrigues Campos |
| EPI_ISL_583502 | Serv de Vig Sanitaria Epidemio e CTRL de Zoonoses Guaruja | Instituto Adolfo Lutz, Interdiciplinary Procedures Center, Strategic Laboratory | Claudio Tavares Sacchi, Claudia Regina Gonçalves, Erica Valessa Ramos Gomes, Karoline Rodrigues Campos |
| EPI_ISL_583503 | CTA Centro de Testagem e Aconselhamento | Instituto Adolfo Lutz, Interdiciplinary Procedures Center, Strategic Laboratory | Claudio Tavares Sacchi, Claudia Regina Gonçalves, Erica Valessa Ramos Gomes, Karoline Rodrigues Campos |
| EPI_ISL_583504, EPI_ISL_583505 | Casa de Saude Stella Maris | Instituto Adolfo Lutz, Interdiciplinary Procedures Center, Strategic Laboratory | Claudio Tavares Sacchi, Claudia Regina Gonçalves, Erica Valessa Ramos Gomes, Karoline Rodrigues Campos |
| EPI_ISL_603021 | Pronto Socorro Dr. Conrado Cesarino Nuvolini | Instituto Adolfo Lutz, Interdiciplinary Procedures Center, Strategic Laboratory | Claudio Tavares Sacchi, Claudia Regina Gonçalves, Erica Valessa Ramos Gomes, Karoline Rodrigues Campos |
| EPI_ISL_603022 | Departamento de Vigilância à Saúde | Instituto Adolfo Lutz, Interdiciplinary Procedures Center, Strategic Laboratory | Claudio Tavares Sacchi, Claudia Regina Gonçalves, Erica Valessa Ramos Gomes, Karoline Rodrigues Campos |
| EPI_ISL_603023 | Vigilância em Saúde Visa Sul | Instituto Adolfo Lutz, Interdiciplinary Procedures Center, Strategic Laboratory | Claudio Tavares Sacchi, Claudia Regina Gonçalves, Erica Valessa Ramos Gomes, Karoline Rodrigues Campos |
| EPI_ISL_603024 | Santa Casa de Misericordia de Araçatuba | Instituto Adolfo Lutz, Interdiciplinary Procedures Center, Strategic Laboratory | Claudio Tavares Sacchi, Claudia Regina Gonçalves, Erica Valessa Ramos Gomes, Karoline Rodrigues Campos |
| EPI_ISL_603025 | UPA Central de Caraguatatuba | Instituto Adolfo Lutz, Interdiciplinary Procedures Center, Strategic Laboratory | Claudio Tavares Sacchi, Claudia Regina Gonçalves, Erica Valessa Ramos Gomes, Karoline Rodrigues Campos |
| EPI_ISL_603026 | Santa Casa da Misericórdia de Presidente Prudente | Instituto Adolfo Lutz, Interdiciplinary Procedures Center, Strategic Laboratory | Claudio Tavares Sacchi, Claudia Regina Gonçalves, Erica Valessa Ramos Gomes, Karoline Rodrigues Campos |
| EPI_ISL_603027 | Santa Casa de Misericordia de Araçatuba | Instituto Adolfo Lutz, Interdiciplinary Procedures Center, Strategic Laboratory | Claudio Tavares Sacchi, Claudia Regina Gonçalves, Erica Valessa Ramos Gomes, Karoline Rodrigues Campos |
| EPI_ISL_603028 | Hospital Municipal Santa Ana | Instituto Adolfo Lutz, Interdiciplinary Procedures Center, Strategic Laboratory | Claudio Tavares Sacchi, Claudia Regina Gonçalves, Erica Valessa Ramos Gomes, Karoline Rodrigues Campos |
| EPI_ISL_603029 | Hospital Municipal Mário Gatti | Instituto Adolfo Lutz, Interdiciplinary Procedures Center, Strategic Laboratory | Claudio Tavares Sacchi, Claudia Regina Gonçalves, Erica Valessa Ramos Gomes, Karoline Rodrigues Campos |
| EPI_ISL_603030 | Hospital Domingos Leonardo Ceravolo Presidente Prudente | Instituto Adolfo Lutz, Interdiciplinary Procedures Center, Strategic Laboratory | Claudio Tavares Sacchi, Claudia Regina Gonçalves, Erica Valessa Ramos Gomes, Karoline Rodrigues Campos |
| EPI_ISL_603031 | Santa Casa de Presidente Epitácio | Instituto Adolfo Lutz, Interdiciplinary Procedures Center, Strategic Laboratory | Claudio Tavares Sacchi, Claudia Regina Gonçalves, Erica Valessa Ramos Gomes, Karoline Rodrigues Campos |
| EPI_ISL_603032 | Santa Casa da Misericórdia de Presidente Prudente | Instituto Adolfo Lutz, Interdiciplinary Procedures Center, Strategic Laboratory | Claudio Tavares Sacchi, Claudia Regina Gonçalves, Erica Valessa Ramos Gomes, Karoline Rodrigues Campos |
| EPI_ISL_603033 | Vigilancia Epidemiologica de São Bernardo do Campo | Instituto Adolfo Lutz, Interdiciplinary Procedures Center, Strategic Laboratory | Claudio Tavares Sacchi, Claudia Regina Gonçalves, Erica Valessa Ramos Gomes, Karoline Rodrigues Campos |

| | | | |
|---|---|---|---|
| EPI_ISL_603034 | Departamento de Vigilância à Saúde | Instituto Adolfo Lutz, Interdiciplinary Procedures Center, Strategic Laboratory | Claudio Tavares Sacchi, Claudia Regina Gonçalves, Erica Valessa Ramos Gomes, Karoline Rodrigues Campos |
| EPI_ISL_603035 | Secretaria Municipal de Saúde | Instituto Adolfo Lutz, Interdiciplinary Procedures Center, Strategic Laboratory | Claudio Tavares Sacchi, Claudia Regina Gonçalves, Erica Valessa Ramos Gomes, Karoline Rodrigues Campos |
| EPI_ISL_603036 | Hospital Santa Ana | Instituto Adolfo Lutz, Interdiciplinary Procedures Center, Strategic Laboratory | Claudio Tavares Sacchi, Claudia Regina Gonçalves, Erica Valessa Ramos Gomes, Karoline Rodrigues Campos |
| EPI_ISL_603037 | Hospital Geral de Pedreira | Instituto Adolfo Lutz, Interdiciplinary Procedures Center, Strategic Laboratory | Claudio Tavares Sacchi, Claudia Regina Gonçalves, Erica Valessa Ramos Gomes, Karoline Rodrigues Campos |
| EPI_ISL_603038 | Santa Casa de Misericordia de Araçatuba | Instituto Adolfo Lutz, Interdiciplinary Procedures Center, Strategic Laboratory | Claudio Tavares Sacchi, Claudia Regina Gonçalves, Erica Valessa Ramos Gomes, Karoline Rodrigues Campos |
| EPI_ISL_603039 | Hospital Municipal Mário Gatti | Instituto Adolfo Lutz, Interdiciplinary Procedures Center, Strategic Laboratory | Claudio Tavares Sacchi, Claudia Regina Gonçalves, Erica Valessa Ramos Gomes, Karoline Rodrigues Campos |
| EPI_ISL_613563, EPI_ISL_613564, EPI_ISL_613707, EPI_ISL_613708 | Laboratory of Molecular Biology, Blood Center of Ribeirão Preto | Laboratory of Molecular Biology, Blood Center of Ribeirão Preto, Faculty of Medicine of Ribeirão Preto, University of São Paulo | Svetoslav N Slavov, Marta Giovanetti, Vagner Fonseca, Elaine V Santos, Evandra S Rodrigues, Talita Adelino, Joilson Xavier, Glauco de Carvalho Pereira, Aparecida Y Yamamoto, Diego Villa Clé, Rodrigo T Calado; Dimas T Covas, Luiz CJ Alcantara, Simone Kashima |
| EPI_ISL_613709, EPI_ISL_613951 | Laboratory of Molecular Biology, Blood Center of Ribeirão Preto, Faculty of Medicine of Ribeirão Preto, University of São Paulo | Laboratory of Molecular Biology, Blood Center of Ribeirão Preto, Faculty of Medicine of Ribeirão Preto, University of São Paulo | Svetoslav N Slavov, Marta Giovanetti, Vagner Fonseca, Elaine V Santos, Evandra S Rodrigues, Talita Adelino, Joilson Xavier, Glauco de Carvalho Pereira, Aparecida Y Yamamoto, Diego Villa Clé, Rodrigo T Calado; Dimas T Covas, Luiz CJ Alcantara, Simone Kashima |
| EPI_ISL_623108, EPI_ISL_623110, EPI_ISL_623112, EPI_ISL_623114, EPI_ISL_623116, EPI_ISL_623118, EPI_ISL_623119, EPI_ISL_623120, EPI_ISL_623124, EPI_ISL_623126, EPI_ISL_623129, EPI_ISL_623130, EPI_ISL_623132, EPI_ISL_623134, EPI_ISL_623136, EPI_ISL_623138, EPI_ISL_623140, EPI_ISL_623142, EPI_ISL_623143, EPI_ISL_623144, EPI_ISL_623145, EPI_ISL_623146, EPI_ISL_623147, EPI_ISL_623148, EPI_ISL_623149, EPI_ISL_623152, EPI_ISL_623154, EPI_ISL_623157, EPI_ISL_623158, EPI_ISL_623160, EPI_ISL_623161, EPI_ISL_623162, EPI_ISL_623164, EPI_ISL_623165, EPI_ISL_623166, EPI_ISL_623168 | | | |
| see above | Laboratorio de Virologia Molecular / UFRJ | Bioinformatics Laboratory / LNCC | Carolina M Voloch, Ronaldo S Francisco Jr, Luiz G P de Almeida, Otavio J. Brustolini, Cynthia C Cardoso, Alexandra L Gerber, Ana Paula de C Guimarães, Diana Mariani, Covid19-UFRJ Workgroup, Luís Cristóvão Pôrto, Renato S Aguiar, Terezinha M P P Castiñeiras, Orlando C. Ferreira, Amilcar Tanuri, Ana Tereza R de Vasconcelos |
| EPI_ISL_636737, EPI_ISL_636834, EPI_ISL_636835, EPI_ISL_636837 | Laboratório de Imunofarmacologia - Instituto Oswaldo Cruz | Laboratório de Imunofarmacologia - Instituto Oswaldo Cruz | Souza,T.M., Fintelman-Rodrigues,N., De Paula,A.D., Saraiva,F.B., Ferreira,M.A. and Sacramento,C.Q. |
| EPI_ISL_672705, EPI_ISL_672711, EPI_ISL_672719, EPI_ISL_672720, EPI_ISL_672748 | Institute of Tropical Medicine at the University of São Paulo (IMT-USP) | Laboratório de Parasitologia Médica - Instituto de Medicina Tropical - Universidade de São Paulo | Brazil-UK Centre for Arbovirus Discovery Diagnosis Genomics and Epidemiology (CADDE) Genomic Network - Instituto de Medicina Tropical |
| EPI_ISL_693195 | Hospital e Pronto Socorro Portinari | Instituto Adolfo Lutz, Interdiciplinary Procedures Center, Strategic Laboratory | Claudio Tavares Sacchi, Claudia Regina Gonçalves, Erica Valessa Ramos Gomes, Karoline Rodrigues Campos |
| EPI_ISL_693196 | Hospital Santa Clara | Instituto Adolfo Lutz, Interdiciplinary Procedures Center, Strategic Laboratory | Claudio Tavares Sacchi, Claudia Regina Gonçalves, Erica Valessa Ramos Gomes, Karoline Rodrigues Campos |
| EPI_ISL_693198 | Santa Casa de Misericordia de Sao Paulo - Hospital Central | Instituto Adolfo Lutz, Interdiciplinary Procedures Center, Strategic Laboratory | Claudio Tavares Sacchi, Claudia Regina Gonçalves, Erica Valessa Ramos Gomes, Karoline Rodrigues Campos |
| EPI_ISL_693199 | Hospital do Servidor Publico Estadual Francisco Morato de Oliveira | Instituto Adolfo Lutz, Interdiciplinary Procedures Center, Strategic Laboratory | Claudio Tavares Sacchi, Claudia Regina Gonçalves, Erica Valessa Ramos Gomes, Karoline Rodrigues Campos |
| EPI_ISL_693200 | Hospital e Maternidade Mairipora | Instituto Adolfo Lutz, Interdiciplinary Procedures Center, Strategic Laboratory | Claudio Tavares Sacchi, Claudia Regina Gonçalves, Erica Valessa Ramos Gomes, Karoline Rodrigues Campos |
| EPI_ISL_693201 | Hospital Sao Paulo de Ensino da Unifesp | Instituto Adolfo Lutz, Interdiciplinary Procedures Center, Strategic Laboratory | Claudio Tavares Sacchi, Claudia Regina Gonçalves, Erica Valessa Ramos Gomes, Karoline Rodrigues Campos |
| EPI_ISL_693202 | Pronto Socorro Municipal Prof. Joao Catarin Mezomo | Instituto Adolfo Lutz, Interdiciplinary Procedures Center, Strategic Laboratory | Claudio Tavares Sacchi, Claudia Regina Gonçalves, Erica Valessa Ramos Gomes, Karoline Rodrigues Campos |
| EPI_ISL_693203 | Hospital Municipal Doutor Arthur Ribeiro de Saboya | Instituto Adolfo Lutz, Interdiciplinary Procedures Center, Strategic Laboratory | Claudio Tavares Sacchi, Claudia Regina Gonçalves, Erica Valessa Ramos Gomes, Karoline Rodrigues Campos |
| EPI_ISL_693204 | Pronto Socorro Dr. Conrado Cesarino Nuvolini | Instituto Adolfo Lutz, Interdiciplinary Procedures Center, Strategic Laboratory | Claudio Tavares Sacchi, Claudia Regina Gonçalves, Erica Valessa Ramos Gomes, Karoline Rodrigues Campos |
| EPI_ISL_693205 | Hospital de Campanha Covid-19 Assis | Instituto Adolfo Lutz, Interdiciplinary Procedures Center, Strategic Laboratory | Claudio Tavares Sacchi, Claudia Regina Gonçalves, Erica Valessa Ramos Gomes, Karoline Rodrigues Campos |
| EPI_ISL_693206 | Hospital Municipal Mario Gatti | Instituto Adolfo Lutz, Interdiciplinary Procedures Center, Strategic Laboratory | Claudio Tavares Sacchi, Claudia Regina Gonçalves, Erica Valessa Ramos Gomes, Karoline Rodrigues Campos |
| EPI_ISL_693207 | Cs II Doutor Antonio Vicoso Moreira de Rezende | Instituto Adolfo Lutz, Interdiciplinary Procedures Center, Strategic Laboratory | Claudio Tavares Sacchi, Claudia Regina Gonçalves, Erica Valessa Ramos Gomes, Karoline Rodrigues Campos |
| EPI_ISL_693208, EPI_ISL_693209 | Hospital Municipal Antonio Giglio | Instituto Adolfo Lutz, Interdiciplinary Procedures Center, Strategic Laboratory | Claudio Tavares Sacchi, Claudia Regina Gonçalves, Erica Valessa Ramos Gomes, Karoline Rodrigues Campos |
| EPI_ISL_693210 | Pronto-Socorro Dr. Osmar Mesquita | Instituto Adolfo Lutz, Interdiciplinary Procedures Center, Strategic Laboratory | Claudio Tavares Sacchi, Claudia Regina Gonçalves, Erica Valessa Ramos Gomes, Karoline Rodrigues Campos |
| EPI_ISL_693211 | Santa Casa de Misericordia e Maternidade | Instituto Adolfo Lutz, Interdiciplinary Procedures Center, Strategic Laboratory | Claudio Tavares Sacchi, Claudia Regina Gonçalves, Erica Valessa Ramos Gomes, Karoline Rodrigues Campos |
| EPI_ISL_693212 | Santa Casa de Misericordia de Braganca Paulista | Instituto Adolfo Lutz, Interdiciplinary Procedures Center, Strategic Laboratory | Claudio Tavares Sacchi, Claudia Regina Gonçalves, Erica Valessa Ramos Gomes, Karoline Rodrigues Campos |
| EPI_ISL_693213 | Hospital E Maternidade Municipal Governador Mario Covas | Instituto Adolfo Lutz, Interdiciplinary Procedures Center, Strategic Laboratory | Claudio Tavares Sacchi, Claudia Regina Gonçalves, Erica Valessa Ramos Gomes, Karoline Rodrigues Campos |
| EPI_ISL_693214 | Unidade de Pronto Atendimento Central de Caraguatatuba | Instituto Adolfo Lutz, Interdiciplinary Procedures Center, Strategic Laboratory | Claudio Tavares Sacchi, Claudia Regina Gonçalves, Erica Valessa Ramos Gomes, Karoline Rodrigues Campos |
| EPI_ISL_693215 | Secretaria Municipal de Saúde de Iracemapolis | Instituto Adolfo Lutz, Interdiciplinary Procedures Center, Strategic Laboratory | Claudio Tavares Sacchi, Claudia Regina Gonçalves, Erica Valessa Ramos Gomes, Karoline Rodrigues Campos |
| EPI_ISL_693216, EPI_ISL_693217 | Unidade de Vigilância Epiudemiológica de Araras | Instituto Adolfo Lutz, Interdiciplinary Procedures Center, Strategic Laboratory | Claudio Tavares Sacchi, Claudia Regina Gonçalves, Erica Valessa Ramos Gomes, Karoline Rodrigues Campos |
| EPI_ISL_693218 | Hospital Domingos Leonardo Ceravolo Presidente Prudente | Instituto Adolfo Lutz, Interdiciplinary Procedures Center, Strategic Laboratory | Claudio Tavares Sacchi, Claudia Regina Gonçalves, Erica Valessa Ramos Gomes, Karoline Rodrigues Campos |
| EPI_ISL_693219 | Santa Casa da Misericórdia de Presidente Prudente | Instituto Adolfo Lutz, Interdiciplinary Procedures Center, | Claudio Tavares Sacchi, Claudia Regina Gonçalves, Erica Valessa Ramos Gomes, Karoline Rodrigues Campos |

| | | | |
|---|---|---|---|
| | | Strategic Laboratory | |
| EPI_ISL_693220 | Laboratório Municipal de Piracicaba | Instituto Adolfo Lutz, Interdiciplinary Procedures Center, Strategic Laboratory | Claudio Tavares Sacchi, Claudia Regina Gonçalves, Erica Valessa Ramos Gomes, Karoline Rodrigues Campos |
| EPI_ISL_693221, EPI_ISL_693222 | Secretaria Municipal de Saúde de Birigui | Instituto Adolfo Lutz, Interdiciplinary Procedures Center, Strategic Laboratory | Claudio Tavares Sacchi, Claudia Regina Gonçalves, Erica Valessa Ramos Gomes, Karoline Rodrigues Campos |
| EPI_ISL_693223, EPI_ISL_693224 | Laboratório Municipal de Piracicaba | Instituto Adolfo Lutz, Interdiciplinary Procedures Center, Strategic Laboratory | Claudio Tavares Sacchi, Claudia Regina Gonçalves, Erica Valessa Ramos Gomes, Karoline Rodrigues Campos |
| EPI_ISL_693225 | Ubs Vila Rosa - Olímpia Gomes De Almeida | Instituto Adolfo Lutz, Interdiciplinary Procedures Center, Strategic Laboratory | Claudio Tavares Sacchi, Claudia Regina Gonçalves, Erica Valessa Ramos Gomes, Karoline Rodrigues Campos |
| EPI_ISL_693226 | Unidade de Pronto Atendimento Sao José | Instituto Adolfo Lutz, Interdiciplinary Procedures Center, Strategic Laboratory | Claudio Tavares Sacchi, Claudia Regina Gonçalves, Erica Valessa Ramos Gomes, Karoline Rodrigues Campos |
| EPI_ISL_693227 | UBS Vila Marchi | Instituto Adolfo Lutz, Interdiciplinary Procedures Center, Strategic Laboratory | Claudio Tavares Sacchi, Claudia Regina Gonçalves, Erica Valessa Ramos Gomes, Karoline Rodrigues Campos |
| EPI_ISL_693228 | Secretaria Municipal de Sorocaba | Instituto Adolfo Lutz, Interdiciplinary Procedures Center, Strategic Laboratory | Claudio Tavares Sacchi, Claudia Regina Gonçalves, Erica Valessa Ramos Gomes, Karoline Rodrigues Campos |
| EPI_ISL_693229 | Hospital 8 de Maio | Instituto Adolfo Lutz, Interdiciplinary Procedures Center, Strategic Laboratory | Claudio Tavares Sacchi, Claudia Regina Gonçalves, Erica Valessa Ramos Gomes, Karoline Rodrigues Campos |
| EPI_ISL_693230 | Hospital e Pronto Socorro Portinari | Instituto Adolfo Lutz, Interdiciplinary Procedures Center, Strategic Laboratory | Claudio Tavares Sacchi, Claudia Regina Gonçalves, Erica Valessa Ramos Gomes, Karoline Rodrigues Campos |
| EPI_ISL_693231 | Pronto Socorro Municipal de Santa Branca | Instituto Adolfo Lutz, Interdiciplinary Procedures Center, Strategic Laboratory | Claudio Tavares Sacchi, Claudia Regina Gonçalves, Erica Valessa Ramos Gomes, Karoline Rodrigues Campos |
| EPI_ISL_693232 | Hospital e Pronto Socorro Portinari | Instituto Adolfo Lutz, Interdiciplinary Procedures Center, Strategic Laboratory | Claudio Tavares Sacchi, Claudia Regina Gonçalves, Erica Valessa Ramos Gomes, Karoline Rodrigues Campos |
| EPI_ISL_693233 | Hospital Santa Cruz | Instituto Adolfo Lutz, Interdiciplinary Procedures Center, Strategic Laboratory | Claudio Tavares Sacchi, Claudia Regina Gonçalves, Erica Valessa Ramos Gomes, Karoline Rodrigues Campos |
| EPI_ISL_693234 | Upa Vereador Jose Da Rocha Goncalves | Instituto Adolfo Lutz, Interdiciplinary Procedures Center, Strategic Laboratory | Claudio Tavares Sacchi, Claudia Regina Gonçalves, Erica Valessa Ramos Gomes, Karoline Rodrigues Campos |
| EPI_ISL_693235 | Casmi Centro Atendimento Saude da Mulher e Infancia | Instituto Adolfo Lutz, Interdiciplinary Procedures Center, Strategic Laboratory | Claudio Tavares Sacchi, Claudia Regina Gonçalves, Erica Valessa Ramos Gomes, Karoline Rodrigues Campos |
| EPI_ISL_693236 | Hospital Santa Marcelina Sao Paulo | Instituto Adolfo Lutz, Interdiciplinary Procedures Center, Strategic Laboratory | Claudio Tavares Sacchi, Claudia Regina Gonçalves, Erica Valessa Ramos Gomes, Karoline Rodrigues Campos |
| EPI_ISL_693237 | UPA Santa Isabel | Instituto Adolfo Lutz, Interdiciplinary Procedures Center, Strategic Laboratory | Claudio Tavares Sacchi, Claudia Regina Gonçalves, Erica Valessa Ramos Gomes, Karoline Rodrigues Campos |
| EPI_ISL_693238, EPI_ISL_693239 | Secao Centro de Diagnostico Secedi | Instituto Adolfo Lutz, Interdiciplinary Procedures Center, Strategic Laboratory | Claudio Tavares Sacchi, Claudia Regina Gonçalves, Erica Valessa Ramos Gomes, Karoline Rodrigues Campos |
| EPI_ISL_693240 | Centro de Vigilância a Saude de Diadema | Instituto Adolfo Lutz, Interdiciplinary Procedures Center, Strategic Laboratory | Claudio Tavares Sacchi, Claudia Regina Gonçalves, Erica Valessa Ramos Gomes, Karoline Rodrigues Campos |
| EPI_ISL_693241 | Hospital e Maternidade Sao Lucas | Instituto Adolfo Lutz, Interdiciplinary Procedures Center, Strategic Laboratory | Claudio Tavares Sacchi, Claudia Regina Gonçalves, Erica Valessa Ramos Gomes, Karoline Rodrigues Campos |
| EPI_ISL_693242 | Centro de Vigilância a Saude de Diadema | Instituto Adolfo Lutz, Interdiciplinary Procedures Center, Strategic Laboratory | Claudio Tavares Sacchi, Claudia Regina Gonçalves, Erica Valessa Ramos Gomes, Karoline Rodrigues Campos |
| EPI_ISL_693243 | Laboratório Municipal de Piracicaba | Instituto Adolfo Lutz, Interdiciplinary Procedures Center, Strategic Laboratory | Claudio Tavares Sacchi, Claudia Regina Gonçalves, Erica Valessa Ramos Gomes, Karoline Rodrigues Campos |
| EPI_ISL_693244 | Centro Médico da Polícia Militar do Estado de Sao Paulo | Instituto Adolfo Lutz, Interdiciplinary Procedures Center, Strategic Laboratory | Claudio Tavares Sacchi, Claudia Regina Gonçalves, Erica Valessa Ramos Gomes, Karoline Rodrigues Campos |
| EPI_ISL_693245 | UPA Santa Isabel | Instituto Adolfo Lutz, Interdiciplinary Procedures Center, Strategic Laboratory | Claudio Tavares Sacchi, Claudia Regina Gonçalves, Erica Valessa Ramos Gomes, Karoline Rodrigues Campos |
| EPI_ISL_693246 | Laboratorio Municipal de Rio Grande da Serra | Instituto Adolfo Lutz, Interdiciplinary Procedures Center, Strategic Laboratory | Claudio Tavares Sacchi, Claudia Regina Gonçalves, Erica Valessa Ramos Gomes, Karoline Rodrigues Campos |
| EPI_ISL_693247 | Secao Centro de Diagnostico Secedi | Instituto Adolfo Lutz, Interdiciplinary Procedures Center, Strategic Laboratory | Claudio Tavares Sacchi, Claudia Regina Gonçalves, Erica Valessa Ramos Gomes, Karoline Rodrigues Campos |
| EPI_ISL_693248 | Centro Municipal de Epidemiologia e Imunizações | Instituto Adolfo Lutz, Interdiciplinary Procedures Center, Strategic Laboratory | Claudio Tavares Sacchi, Claudia Regina Gonçalves, Erica Valessa Ramos Gomes, Karoline Rodrigues Campos |
| EPI_ISL_708529 | Secretária Municipal de Saude de Fernandópolis | Instituto Adolfo Lutz, Interdiciplinary Procedures Center, Strategic Laboratory | Claudio Tavares Sacchi, Claudia Regina Gonçalves, Erica Valessa Ramos Gomes, Carlos Henrique Camargo, Karoline Rodrigues Campos, Fernanda Modesto Tolentino Binhardi, Maricelia Navarro Pinheiro Flores, Marcia Maria Costa Nunes Soares, Janaina Other Martins Montanha |
| EPI_ISL_708530 | Secretaria Municipal de Saude de Fernandópolis | Instituto Adolfo Lutz, Interdiciplinary Procedures Center, Strategic Laboratory | Claudio Tavares Sacchi, Claudia Regina Gonçalves, Erica Valessa Ramos Gomes, Carlos Henrique Camargo, Karoline Rodrigues Campos, Fernanda Modesto Tolentino Binhardi, Maricelia Navarro Pinheiro Flores, Marcia Maria Costa Nunes Soares, Janaina Other Martins Montanha |
| EPI_ISL_717806, EPI_ISL_717807, EPI_ISL_717808 | Laboratorio de Virologia Molecular / UFRJ | Bioinformatics Laboratory / LNCC | Carolina M Voloch, Ronaldo da Silva F Jr, Luiz G P de Almeida, Cynthia C Cardoso, Otavio Bustrolini, Alexandra L Gerber, Ana Paula de C Guimarães, Diana Mariani, Andréa Cony Cavalcanti, Claudia dos Santos Rodrigues, Terezinha M P P Castiñeira, Amilcar Tanuri, Ana Tereza R de Vasconcelos |
| EPI_ISL_717809 | LACEN Dr. Francisco Rimolo Neto | Bioinformatics Laboratory / LNCC | Carolina M Voloch, Ronaldo da Silva F Jr, Luiz G P de Almeida, Cynthia C Cardoso, Otavio Bustrolini, Alexandra L Gerber, Ana Paula de C Guimarães, Diana Mariani, Andréa Cony Cavalcanti, Claudia dos Santos Rodrigues, Terezinha M P P Castiñeira, Amilcar Tanuri, Ana Tereza R de Vasconcelos |
| EPI_ISL_717810, EPI_ISL_717811, EPI_ISL_717812, EPI_ISL_717813, EPI_ISL_717814, EPI_ISL_717815, EPI_ISL_717816, EPI_ISL_717817, EPI_ISL_717818, EPI_ISL_717819, EPI_ISL_717820, EPI_ISL_717821, EPI_ISL_717822, EPI_ISL_717823, EPI_ISL_717824, EPI_ISL_717825, EPI_ISL_717826, EPI_ISL_717827, EPI_ISL_717828, EPI_ISL_717829, EPI_ISL_717830, EPI_ISL_717831 | | | |
| see above | Laboratorio de Virologia Molecular / UFRJ | Bioinformatics Laboratory / LNCC | Carolina M Voloch, Ronaldo da Silva F Jr, Luiz G P de Almeida, Cynthia C Cardoso, Otavio Bustrolini, Alexandra L Gerber, Ana Paula de C Guimarães, Diana Mariani, Andréa Cony Cavalcanti, Claudia dos Santos Rodrigues, Terezinha M P P Castiñeira, Amilcar Tanuri, Ana Tereza R de Vasconcelos |
| EPI_ISL_717832, EPI_ISL_717833, EPI_ISL_717834, EPI_ISL_717835, EPI_ISL_717836 | LACEN Dr. Francisco Rimolo Neto | Bioinformatics Laboratory / LNCC | Carolina M Voloch, Ronaldo da Silva F Jr, Luiz G P de Almeida, Cynthia C Cardoso, Otavio Bustrolini, Alexandra L Gerber, Ana Paula de C Guimarães, Diana Mariani, Andréa Cony Cavalcanti, Claudia dos Santos Rodrigues, Terezinha M P P Castiñeira, Amilcar Tanuri, Ana Tereza R de Vasconcelos |
| EPI_ISL_717837, EPI_ISL_717838, EPI_ISL_717839, EPI_ISL_717840 | Laboratorio de Virologia Molecular / UFRJ | Bioinformatics Laboratory / LNCC | Carolina M Voloch, Ronaldo da Silva F Jr, Luiz G P de Almeida, Cynthia C Cardoso, Otavio Bustrolini, Alexandra L Gerber, Ana Paula de C Guimarães, Diana Mariani, Andréa Cony Cavalcanti, Claudia dos Santos Rodrigues, Terezinha M P P Castiñeira, Amilcar Tanuri, Ana Tereza R de Vasconcelos |

| | | | |
|---|---|---|---|
| EPI_ISL_717841 | LACEN Dr. Francisco Rimolo Neto | Bioinformatics Laboratory / LNCC | Carolina M Voloch, Ronaldo da Silva F Jr, Luiz G P de Almeida, Cynthia C Cardoso, Otavio Bustrolini, Alexandra L Gerber, Ana Paula de C Guimarães, Diana Mariani, Andréa Cony Cavalcanti, Claudia dos Santos Rodrigues, Terezinha M P P Castiñeira, Amilcar Tanuri, Ana Tereza R de Vasconcelos |
| EPI_ISL_717842, EPI_ISL_717843, EPI_ISL_717844, EPI_ISL_717845, EPI_ISL_717846, EPI_ISL_717847, EPI_ISL_717848, EPI_ISL_717849, EPI_ISL_717850, EPI_ISL_717851, EPI_ISL_717852, EPI_ISL_717853, EPI_ISL_717854, EPI_ISL_717855, EPI_ISL_717856, EPI_ISL_717857, EPI_ISL_717858, EPI_ISL_717859, EPI_ISL_717860, EPI_ISL_717861, EPI_ISL_717862, EPI_ISL_717863, EPI_ISL_717864, EPI_ISL_717865, EPI_ISL_717866, EPI_ISL_717867, EPI_ISL_717868, EPI_ISL_717869, EPI_ISL_717870, EPI_ISL_717871, EPI_ISL_717872, EPI_ISL_717873, EPI_ISL_717874, EPI_ISL_717875, EPI_ISL_717876, EPI_ISL_717877, EPI_ISL_717878, EPI_ISL_717879, EPI_ISL_717880, EPI_ISL_717881, EPI_ISL_717882, EPI_ISL_717883, EPI_ISL_717884, EPI_ISL_717885, EPI_ISL_717886, EPI_ISL_717887, EPI_ISL_717888, EPI_ISL_717889, EPI_ISL_717890, EPI_ISL_717891, EPI_ISL_717892, EPI_ISL_717893, EPI_ISL_717894, EPI_ISL_717895, EPI_ISL_717896, EPI_ISL_717897, EPI_ISL_717898 | | | |
| see above | Laboratorio de Virologia Molecular / UFRJ | Bioinformatics Laboratory / LNCC | Carolina M Voloch, Ronaldo da Silva F Jr, Luiz G P de Almeida, Cynthia C Cardoso, Otavio Bustrolini, Alexandra L Gerber, Ana Paula de C Guimarães, Diana Mariani, Andréa Cony Cavalcanti, Claudia dos Santos Rodrigues, Terezinha M P P Castiñeira, Amilcar Tanuri, Ana Tereza R de Vasconcelos |
| EPI_ISL_717899, EPI_ISL_717900, EPI_ISL_717901, EPI_ISL_717902, EPI_ISL_717903, EPI_ISL_717904, EPI_ISL_717905, EPI_ISL_717906, EPI_ISL_717907, EPI_ISL_717908, EPI_ISL_717909 | | | |
| see above | LACEN RJ - Noel Nutels | Bioinformatics Laboratory / LNCC | Carolina M Voloch, Ronaldo da Silva F Jr, Luiz G P de Almeida, Cynthia C Cardoso, Otavio Bustrolini, Alexandra L Gerber, Ana Paula de C Guimarães, Diana Mariani, Andréa Cony Cavalcanti, Claudia dos Santos Rodrigues, Terezinha M P P Castiñeira, Amilcar Tanuri, Ana Tereza R de Vasconcelos |
| EPI_ISL_717910, EPI_ISL_717911, EPI_ISL_717912, EPI_ISL_717913, EPI_ISL_717914, EPI_ISL_717915, EPI_ISL_717916, EPI_ISL_717917, EPI_ISL_717918, EPI_ISL_717919 | LACEN Dr. Francisco Rimolo Neto | Bioinformatics Laboratory / LNCC | Carolina M Voloch, Ronaldo da Silva F Jr, Luiz G P de Almeida, Cynthia C Cardoso, Otavio Bustrolini, Alexandra L Gerber, Ana Paula de C Guimarães, Diana Mariani, Andréa Cony Cavalcanti, Claudia dos Santos Rodrigues, Terezinha M P P Castiñeira, Amilcar Tanuri, Ana Tereza R de Vasconcelos |
| EPI_ISL_717920, EPI_ISL_717921, EPI_ISL_717922, EPI_ISL_717923, EPI_ISL_717924, EPI_ISL_717925, EPI_ISL_717926, EPI_ISL_717927, EPI_ISL_717928, EPI_ISL_717929, EPI_ISL_717930, EPI_ISL_717931, EPI_ISL_717932, EPI_ISL_717933, EPI_ISL_717934, EPI_ISL_717935, EPI_ISL_717936, EPI_ISL_717937, EPI_ISL_717938, EPI_ISL_717939, EPI_ISL_717940, EPI_ISL_717941, EPI_ISL_717942, EPI_ISL_717943, EPI_ISL_717944, EPI_ISL_717945, EPI_ISL_717946, EPI_ISL_717947, EPI_ISL_717948, EPI_ISL_717949, EPI_ISL_717950, EPI_ISL_717951, EPI_ISL_717952, EPI_ISL_717953, EPI_ISL_717954, EPI_ISL_717955, EPI_ISL_717956, EPI_ISL_717957 | | | |
| see above | Laboratorio de Virologia Molecular / UFRJ | Bioinformatics Laboratory / LNCC | Carolina M Voloch, Ronaldo da Silva F Jr, Luiz G P de Almeida, Cynthia C Cardoso, Otavio Bustrolini, Alexandra L Gerber, Ana Paula de C Guimarães, Diana Mariani, Andréa Cony Cavalcanti, Claudia dos Santos Rodrigues, Terezinha M P P Castiñeira, Amilcar Tanuri, Ana Tereza R de Vasconcelos |
| EPI_ISL_717958 | LACEN Dr. Francisco Rimolo Neto | Bioinformatics Laboratory / LNCC | Carolina M Voloch, Ronaldo da Silva F Jr, Luiz G P de Almeida, Cynthia C Cardoso, Otavio Bustrolini, Alexandra L Gerber, Ana Paula de C Guimarães, Diana Mariani, Andréa Cony Cavalcanti, Claudia dos Santos Rodrigues, Terezinha M P P Castiñeira, Amilcar Tanuri, Ana Tereza R de Vasconcelos |
| EPI_ISL_717959, EPI_ISL_717960, EPI_ISL_717961 | Laboratorio de Virologia Molecular / UFRJ | Bioinformatics Laboratory / LNCC | Carolina M Voloch, Ronaldo da Silva F Jr, Luiz G P de Almeida, Cynthia C Cardoso, Otavio Bustrolini, Alexandra L Gerber, Ana Paula de C Guimarães, Diana Mariani, Andréa Cony Cavalcanti, Claudia dos Santos Rodrigues, Terezinha M P P Castiñeira, Amilcar Tanuri, Ana Tereza R de Vasconcelos |
| EPI_ISL_717962 | LACEN RJ - Noel Nutels | Bioinformatics Laboratory / LNCC | Carolina M Voloch, Ronaldo da Silva F Jr, Luiz G P de Almeida, Cynthia C Cardoso, Otavio Bustrolini, Alexandra L Gerber, Ana Paula de C Guimarães, Diana Mariani, Andréa Cony Cavalcanti, Claudia dos Santos Rodrigues, Terezinha M P P Castiñeira, Amilcar Tanuri, Ana Tereza R de Vasconcelos |
| EPI_ISL_717963, EPI_ISL_717964 | LACEN Dr. Francisco Rimolo Neto | Bioinformatics Laboratory / LNCC | Carolina M Voloch, Ronaldo da Silva F Jr, Luiz G P de Almeida, Cynthia C Cardoso, Otavio Bustrolini, Alexandra L Gerber, Ana Paula de C Guimarães, Diana Mariani, Andréa Cony Cavalcanti, Claudia dos Santos Rodrigues, Terezinha M P P Castiñeira, Amilcar Tanuri, Ana Tereza R de Vasconcelos |
| EPI_ISL_722129 | Hospital das Clínicas Universidade de São Paulo Medical School | Laboratório de Parasitologia Médica - Instituto de Medicina Tropical - Universidade de São Paulo | Brazil-UK Centre for Arbovirus Discovery Diagnosis Genomics and Epidemiology (CADDE) Genomic Network - Instituto de Medicina Tropical |
| EPI_ISL_729794, EPI_ISL_729797, EPI_ISL_729798, EPI_ISL_729800, EPI_ISL_729801, EPI_ISL_729802, EPI_ISL_729803, EPI_ISL_729804, EPI_ISL_729805, EPI_ISL_729806, EPI_ISL_729807, EPI_ISL_729808, EPI_ISL_729809, EPI_ISL_729810, EPI_ISL_729811, EPI_ISL_729812, EPI_ISL_729813, EPI_ISL_729814, EPI_ISL_729815, EPI_ISL_729816, EPI_ISL_729817, EPI_ISL_729818, EPI_ISL_729819, EPI_ISL_729820, EPI_ISL_729821, EPI_ISL_729822, EPI_ISL_729823, EPI_ISL_729824, EPI_ISL_729825, EPI_ISL_729826, EPI_ISL_729827, EPI_ISL_729828, EPI_ISL_729829, EPI_ISL_729830, EPI_ISL_729831, EPI_ISL_729832, EPI_ISL_729833, EPI_ISL_729834, EPI_ISL_729835, EPI_ISL_729836, EPI_ISL_729837, EPI_ISL_729838, EPI_ISL_729839, EPI_ISL_729840, EPI_ISL_729841, EPI_ISL_729842, EPI_ISL_729843, EPI_ISL_729844, EPI_ISL_729845, EPI_ISL_729846, EPI_ISL_729847, EPI_ISL_729848, EPI_ISL_729849, EPI_ISL_729850, EPI_ISL_729851, EPI_ISL_729852, EPI_ISL_729853, EPI_ISL_729854, EPI_ISL_729855, EPI_ISL_729856, EPI_ISL_729857, EPI_ISL_729858, EPI_ISL_729859, EPI_ISL_729860, EPI_ISL_729861 | | | |
| see above | Laboratório Central de Saúde Pública do Estado do Rio Grande do Sul (LACEN-RS) | Laboratory of Respiratory Viruses and Measles, Oswaldo Cruz Institute, FIOCRUZ | Paola Resende, Luciana Appolinario, Fernando Motta, Anna Carolina Paixão, Ana Carolina Mendonça, Tatiana Schaffer Gregianini, Marilda Tereza Mar da Rosa, Marilda Siqueira on behalf of the Fiocruz COVID-19 Genomic Surveillance Network |
| EPI_ISL_735396 | Hospital de Camplanha COVID 19 SER | Instituto Adolfo Lutz, Interdiciplinary Procedures Center, Strategic Laboratory | Claudio Tavares Sacchi, Claudia Regina Gonçalves, Erica Valessa Ramos Gomes, Karoline Rodrigues Campos |
| EPI_ISL_735397 | Unidade Respiratória Nova Hortolandia | Instituto Adolfo Lutz, Interdiciplinary Procedures Center, Strategic Laboratory | Claudio Tavares Sacchi, Claudia Regina Gonçalves, Erica Valessa Ramos Gomes, Karoline Rodrigues Campos |
| EPI_ISL_735398 | Laboratorio Fleury | Instituto Adolfo Lutz, Interdiciplinary Procedures Center, Strategic Laboratory | Claudio Tavares Sacchi, Claudia Regina Gonçalves, Erica Valessa Ramos Gomes, Karoline Rodrigues Campos |
| EPI_ISL_735399 | Hospital Municiplal Dr Ignacio de gouvea | Instituto Adolfo Lutz, Interdiciplinary Procedures Center, Strategic Laboratory | Claudio Tavares Sacchi, Claudia Regina Gonçalves, Erica Valessa Ramos Gomes, Karoline Rodrigues Campos |
| EPI_ISL_735400 | Instituto Adolfo Lutz - Regional de Santos | Instituto Adolfo Lutz, Interdiciplinary Procedures Center, Strategic Laboratory | Claudio Tavares Sacchi, Claudia Regina Gonçalves, Erica Valessa Ramos Gomes, Karoline Rodrigues Campos |
| EPI_ISL_735401, EPI_ISL_735402, EPI_ISL_735403, EPI_ISL_735404 | Instituto Adolfo Lutz - Regional de Rio Claro | Instituto Adolfo Lutz, Interdiciplinary Procedures Center, Strategic Laboratory | Claudio Tavares Sacchi, Claudia Regina Gonçalves, Erica Valessa Ramos Gomes, Karoline Rodrigues Campos |
| EPI_ISL_735405 | Secretaria Minucipal de Saude de Birigui | Instituto Adolfo Lutz, Interdiciplinary Procedures Center, Strategic Laboratory | Claudio Tavares Sacchi, Claudia Regina Gonçalves, Erica Valessa Ramos Gomes, Karoline Rodrigues Campos |
| EPI_ISL_735406 | Unidade de Pronto Atendimento UPA I Sta Isabel | Instituto Adolfo Lutz, Interdiciplinary Procedures Center, Strategic Laboratory | Claudio Tavares Sacchi, Claudia Regina Gonçalves, Erica Valessa Ramos Gomes, Karoline Rodrigues Campos |
| EPI_ISL_735407 | Santa Casa de Marilia | Instituto Adolfo Lutz, Interdiciplinary Procedures Center, Strategic Laboratory | Claudio Tavares Sacchi, Claudia Regina Gonçalves, Erica Valessa Ramos Gomes, Karoline Rodrigues Campos |
| EPI_ISL_735408 | COVID 19 Centro de Combate ao Coronavirus CCC Jandira | Instituto Adolfo Lutz, Interdiciplinary Procedures Center, Strategic Laboratory | Claudio Tavares Sacchi, Claudia Regina Gonçalves, Erica Valessa Ramos Gomes, Karoline Rodrigues Campos |
| EPI_ISL_735409 | Unidade de Pronto Atendimento Carlos Lourenco | Instituto Adolfo Lutz, Interdiciplinary Procedures Center, Strategic Laboratory | Claudio Tavares Sacchi, Claudia Regina Gonçalves, Erica Valessa Ramos Gomes, Karoline Rodrigues Campos |
| EPI_ISL_735410 | Instituto Adolfo Lutz - Regional de Rio Claro | Instituto Adolfo Lutz, Interdiciplinary Procedures Center, Strategic Laboratory | Claudio Tavares Sacchi, Claudia Regina Gonçalves, Erica Valessa Ramos Gomes, Karoline Rodrigues Campos |
| EPI_ISL_735411 | Centro de Vigilancia a Saude de Diadema | Instituto Adolfo Lutz, Interdiciplinary Procedures Center, Strategic Laboratory | Claudio Tavares Sacchi, Claudia Regina Gonçalves, Erica Valessa Ramos Gomes, Karoline Rodrigues Campos |
| EPI_ISL_735412 | Hospital e Pronto Socorro Portinari | Instituto Adolfo Lutz, Interdiciplinary Procedures Center, Strategic Laboratory | Claudio Tavares Sacchi, Claudia Regina Gonçalves, Erica Valessa Ramos Gomes, Karoline Rodrigues Campos |
| EPI_ISL_735413 | Militello Centro de Diagnosticos e Biopesquisa Clinica | Instituto Adolfo Lutz, Interdiciplinary Procedures Center, Strategic Laboratory | Claudio Tavares Sacchi, Claudia Regina Gonçalves, Erica Valessa Ramos Gomes, Karoline Rodrigues Campos |
| EPI_ISL_735414, EPI_ISL_735415 | Unidade de Pronto Atendimento de Agenor de Campos | Instituto Adolfo Lutz, Interdiciplinary Procedures Center, Strategic Laboratory | Claudio Tavares Sacchi, Claudia Regina Gonçalves, Erica Valessa Ramos Gomes, Karoline Rodrigues Campos |
| EPI_ISL_735416 | Centro de Saude II Dr Jose Paione Mococa | Instituto Adolfo Lutz, Interdiciplinary Procedures Center, Strategic Laboratory | Claudio Tavares Sacchi, Claudia Regina Gonçalves, Erica Valessa Ramos Gomes, Karoline Rodrigues Campos |
| EPI_ISL_735417 | Unidade de Pronto Atendimento de Agenor de Campos | Instituto Adolfo Lutz, Interdiciplinary Procedures Center, | Claudio Tavares Sacchi, Claudia Regina Gonçalves, Erica Valessa Ramos Gomes, Karoline Rodrigues Campos |

| | | | |
|---|---|---|---|
| | | Strategic Laboratory | |
| EPI_ISL_735418 | Hospital Regional do Vale do Paraiba | Instituto Adolfo Lutz, Interdiciplinary Procedures Center, Strategic Laboratory | Claudio Tavares Sacchi, Claudia Regina Gonçalves, Erica Valessa Ramos Gomes, Karoline Rodrigues Campos |
| EPI_ISL_735419 | UBS Alvarenga | Instituto Adolfo Lutz, Interdiciplinary Procedures Center, Strategic Laboratory | Claudio Tavares Sacchi, Claudia Regina Gonçalves, Erica Valessa Ramos Gomes, Karoline Rodrigues Campos |
| EPI_ISL_735420 | UBS Riacho Grande | Instituto Adolfo Lutz, Interdiciplinary Procedures Center, Strategic Laboratory | Claudio Tavares Sacchi, Claudia Regina Gonçalves, Erica Valessa Ramos Gomes, Karoline Rodrigues Campos |
| EPI_ISL_735421 | UBS Sta Terezinha | Instituto Adolfo Lutz, Interdiciplinary Procedures Center, Strategic Laboratory | Claudio Tavares Sacchi, Claudia Regina Gonçalves, Erica Valessa Ramos Gomes, Karoline Rodrigues Campos |
| EPI_ISL_735422 | UBS Dematchi | Instituto Adolfo Lutz, Interdiciplinary Procedures Center, Strategic Laboratory | Claudio Tavares Sacchi, Claudia Regina Gonçalves, Erica Valessa Ramos Gomes, Karoline Rodrigues Campos |
| EPI_ISL_735423, EPI_ISL_735424 | Centro de Vigilancia a Saude de Diadema | Instituto Adolfo Lutz, Interdiciplinary Procedures Center, Strategic Laboratory | Claudio Tavares Sacchi, Claudia Regina Gonçalves, Erica Valessa Ramos Gomes, Karoline Rodrigues Campos |
| EPI_ISL_735425 | Hospital e Maternidade Sao Lucas | Instituto Adolfo Lutz, Interdiciplinary Procedures Center, Strategic Laboratory | Claudio Tavares Sacchi, Claudia Regina Gonçalves, Erica Valessa Ramos Gomes, Karoline Rodrigues Campos |
| EPI_ISL_735426 | Centro de Vigilancia a Saude de Diadema | Instituto Adolfo Lutz, Interdiciplinary Procedures Center, Strategic Laboratory | Claudio Tavares Sacchi, Claudia Regina Gonçalves, Erica Valessa Ramos Gomes, Karoline Rodrigues Campos |
| EPI_ISL_735427 | Instituto Adolfo Lutz - Regional de Santos | Instituto Adolfo Lutz, Interdiciplinary Procedures Center, Strategic Laboratory | Claudio Tavares Sacchi, Claudia Regina Gonçalves, Erica Valessa Ramos Gomes, Karoline Rodrigues Campos |
| EPI_ISL_735428, EPI_ISL_735429 | Hospital Nipo Brasileiro | Instituto Adolfo Lutz, Interdiciplinary Procedures Center, Strategic Laboratory | Claudio Tavares Sacchi, Claudia Regina Gonçalves, Erica Valessa Ramos Gomes, Karoline Rodrigues Campos |
| EPI_ISL_735430 | Instituto Adolfo Lutz - Regional de Santos | Instituto Adolfo Lutz, Interdiciplinary Procedures Center, Strategic Laboratory | Claudio Tavares Sacchi, Claudia Regina Gonçalves, Erica Valessa Ramos Gomes, Karoline Rodrigues Campos |
| EPI_ISL_735431, EPI_ISL_735432 | Hospital Nipo Brasileiro | Instituto Adolfo Lutz, Interdiciplinary Procedures Center, Strategic Laboratory | Claudio Tavares Sacchi, Claudia Regina Gonçalves, Erica Valessa Ramos Gomes, Karoline Rodrigues Campos |
| EPI_ISL_735433 | Posto de Atendimento Saude Cidade Pasc Cajati | Instituto Adolfo Lutz, Interdiciplinary Procedures Center, Strategic Laboratory | Claudio Tavares Sacchi, Claudia Regina Gonçalves, Erica Valessa Ramos Gomes, Karoline Rodrigues Campos |
| EPI_ISL_755640 | Instituto Adolfo Lutz - Central | Instituto Adolfo Lutz, Interdiciplinary Procedures Center, Strategic Laboratory | Claudio Tavares Sacchi, Claudia Regina Gonçalves, Erica Valessa Ramos Gomes, Karoline Rodrigues Campos |
| EPI_ISL_755641 | Instituto Adolfo Lutz - Regional de Santo Andre | Instituto Adolfo Lutz, Interdiciplinary Procedures Center, Strategic Laboratory | Claudio Tavares Sacchi, Claudia Regina Gonçalves, Erica Valessa Ramos Gomes, Karoline Rodrigues Campos |
| EPI_ISL_755642, EPI_ISL_755643 | Instituto Adolfo Lutz - Central | Instituto Adolfo Lutz, Interdiciplinary Procedures Center, Strategic Laboratory | Claudio Tavares Sacchi, Claudia Regina Gonçalves, Erica Valessa Ramos Gomes, Karoline Rodrigues Campos |
| EPI_ISL_755644, EPI_ISL_755645 | Lab LOC - Itapecerica da Serra | Instituto Adolfo Lutz, Interdiciplinary Procedures Center, Strategic Laboratory | Claudio Tavares Sacchi, Claudia Regina Gonçalves, Erica Valessa Ramos Gomes, Karoline Rodrigues Campos |
| EPI_ISL_755646, EPI_ISL_755647 | Instituto Adolfo Lutz - Regional de Santo Andre | Instituto Adolfo Lutz, Interdiciplinary Procedures Center, Strategic Laboratory | Claudio Tavares Sacchi, Claudia Regina Gonçalves, Erica Valessa Ramos Gomes, Karoline Rodrigues Campos |
| EPI_ISL_755648 | Instituto Adolfo Lutz - Regional de Taubate | Instituto Adolfo Lutz, Interdiciplinary Procedures Center, Strategic Laboratory | Claudio Tavares Sacchi, Claudia Regina Gonçalves, Erica Valessa Ramos Gomes, Karoline Rodrigues Campos |
| EPI_ISL_755649 | Instituto Adolfo Lutz - Regional de Santo Andre | Instituto Adolfo Lutz, Interdiciplinary Procedures Center, Strategic Laboratory | Claudio Tavares Sacchi, Claudia Regina Gonçalves, Erica Valessa Ramos Gomes, Karoline Rodrigues Campos |
| EPI_ISL_755650 | Instituto Adolfo Lutz - Regional de Taubate | Instituto Adolfo Lutz, Interdiciplinary Procedures Center, Strategic Laboratory | Claudio Tavares Sacchi, Claudia Regina Gonçalves, Erica Valessa Ramos Gomes, Karoline Rodrigues Campos |
| EPI_ISL_755651 | Instituto Adolfo Lutz - Central | Instituto Adolfo Lutz, Interdiciplinary Procedures Center, Strategic Laboratory | Claudio Tavares Sacchi, Claudia Regina Gonçalves, Erica Valessa Ramos Gomes, Karoline Rodrigues Campos |
| EPI_ISL_755652 | Lab LOC - Itapecerica da Serra | Instituto Adolfo Lutz, Interdiciplinary Procedures Center, Strategic Laboratory | Claudio Tavares Sacchi, Claudia Regina Gonçalves, Erica Valessa Ramos Gomes, Karoline Rodrigues Campos |
| EPI_ISL_755653, EPI_ISL_755654 | Instituto Adolfo Lutz - Central | Instituto Adolfo Lutz, Interdiciplinary Procedures Center, Strategic Laboratory | Claudio Tavares Sacchi, Claudia Regina Gonçalves, Erica Valessa Ramos Gomes, Karoline Rodrigues Campos |
| EPI_ISL_755655 | Instituto Adolfo Lutz - Regional de Campinas | Instituto Adolfo Lutz, Interdiciplinary Procedures Center, Strategic Laboratory | Claudio Tavares Sacchi, Claudia Regina Gonçalves, Erica Valessa Ramos Gomes, Karoline Rodrigues Campos |
| EPI_ISL_756293, EPI_ISL_756294 | Center for Biotechnology and Cell Therapy, São Rafael Hospital, Salvador, Brazil | Center for Biotechnology and Cell Therapy, São Rafael Hospital, Salvador, Brazil | Carolina Kymie Vasques Nonaka, Marília Miranda Franco, Tiago Gräf, Ana Verena Almeida Mendes, Renato Santana de Aguiar, Marta Giovanetti, Bruno Solano de Freitas Souza |
| EPI_ISL_763074, EPI_ISL_763075 | Diagnosticos da America - DASA | Instituto Adolfo Lutz, Interdiciplinary Procedures Center, Strategic Laboratory | Claudio Tavares Sacchi, Claudia Regina Gonçalves, Erica Valessa Ramos Gomes, Karoline Rodrigues Campos |
| EPI_ISL_770551, EPI_ISL_770552, EPI_ISL_770553, EPI_ISL_770554, EPI_ISL_770555, EPI_ISL_770556, EPI_ISL_770557, EPI_ISL_770558, EPI_ISL_770559, EPI_ISL_770560, EPI_ISL_770561, EPI_ISL_770562, EPI_ISL_770563, EPI_ISL_770564, EPI_ISL_770565, EPI_ISL_770566, EPI_ISL_770567, EPI_ISL_770568, EPI_ISL_770569, EPI_ISL_770570, EPI_ISL_770571, EPI_ISL_770572, EPI_ISL_770573, EPI_ISL_770574, EPI_ISL_770575, EPI_ISL_770576, EPI_ISL_770577, EPI_ISL_770578, EPI_ISL_770579, EPI_ISL_770580, EPI_ISL_770581, EPI_ISL_770582, EPI_ISL_770583, EPI_ISL_770584, EPI_ISL_770585, EPI_ISL_770586, EPI_ISL_770587, EPI_ISL_770588, EPI_ISL_770589, EPI_ISL_770590, EPI_ISL_770591, EPI_ISL_770592, EPI_ISL_770593, EPI_ISL_770594, EPI_ISL_770595, EPI_ISL_770596, EPI_ISL_770597, EPI_ISL_770598, EPI_ISL_770599, EPI_ISL_770600, EPI_ISL_770601, EPI_ISL_770602, EPI_ISL_770603, EPI_ISL_770604, EPI_ISL_770605, EPI_ISL_770606, EPI_ISL_770607, EPI_ISL_770608, EPI_ISL_770609, EPI_ISL_770610, EPI_ISL_770611, EPI_ISL_770612, EPI_ISL_770613, EPI_ISL_770614, EPI_ISL_770615, EPI_ISL_770616, EPI_ISL_770617, EPI_ISL_770618, EPI_ISL_770619, EPI_ISL_770620, EPI_ISL_770621, EPI_ISL_770622, EPI_ISL_770623, EPI_ISL_770624, EPI_ISL_770625, EPI_ISL_770626, EPI_ISL_770627, EPI_ISL_770628, EPI_ISL_770629, EPI_ISL_770630 | | | |
| see above | Laboratório de Microbiologia Molecular - Universidade FEEVALE | Bioinformatics Laboratory / LNCC | Felipe Benites, Fernando Rosado Spilki, Alana Witt Hansen, Juliane Deise Fleck, Juliana Schons, Meriane Demoliner, Ana Karolina Eisen Antunes, Fagner Henrique Heldt, Larissa Mallmann, Bruna Hermann, Ana Luiza Ziulkoski, Vyctoria Goes, Karoline Schallenberger, Matheus Nunes Weber, Paula Rodrigues de Almeida, Alessandra Pavan Lamarca da Silva, Ronaldo da Silva F Jr , Luiz G P de Almeida, Alexandra L Gerber , Ana Paula de C Guimarães,Ana Tereza R de Vasconcelos |
| EPI_ISL_776750, EPI_ISL_776751, EPI_ISL_776752, EPI_ISL_776753, EPI_ISL_776754, EPI_ISL_776755, EPI_ISL_776756 | Instituto Adolfo Lutz - Central | Instituto Adolfo Lutz, Interdiciplinary Procedures Center, Strategic Laboratory | Claudio Tavares Sacchi, Claudia Regina Gonçalves, Erica Valessa Ramos Gomes, Karoline Rodrigues Campos |
| EPI_ISL_776757, EPI_ISL_776758 | Instituto Adolfo Lutz - Regional de Marilia | Instituto Adolfo Lutz, Interdiciplinary Procedures Center, Strategic Laboratory | Claudio Tavares Sacchi, Claudia Regina Gonçalves, Erica Valessa Ramos Gomes, Karoline Rodrigues Campos |
| EPI_ISL_776759, EPI_ISL_776760, EPI_ISL_776761, EPI_ISL_776762, | Instituto Adolfo Lutz - Central | Instituto Adolfo Lutz, Interdiciplinary Procedures Center, Strategic Laboratory | Claudio Tavares Sacchi, Claudia Regina Gonçalves, Erica Valessa Ramos Gomes, Karoline Rodrigues Campos |

| | | | |
|---|---|---|---|
| EPI_ISL_776763 | | | |
| EPI_ISL_776764, EPI_ISL_776765, EPI_ISL_776766 | Instituto Adolfo Lutz - Regional de Santo Andre | Instituto Adolfo Lutz, Interdiciplinary Procedures Center, Strategic Laboratory | Claudio Tavares Sacchi, Claudia Regina Gonçalves, Erica Valessa Ramos Gomes, Karoline Rodrigues Campos |
| EPI_ISL_776767 | Instituto Adolfo Lutz - Regional de Marilia | Instituto Adolfo Lutz, Interdiciplinary Procedures Center, Strategic Laboratory | Claudio Tavares Sacchi, Claudia Regina Gonçalves, Erica Valessa Ramos Gomes, Karoline Rodrigues Campos |
| EPI_ISL_776768 | Instituto Adolfo Lutz - Regional de Aracatuba | Instituto Adolfo Lutz, Interdiciplinary Procedures Center, Strategic Laboratory | Claudio Tavares Sacchi, Claudia Regina Gonçalves, Erica Valessa Ramos Gomes, Karoline Rodrigues Campos |
| EPI_ISL_776769 | Instituto Adolfo Lutz - Regional de Santo Andre | Instituto Adolfo Lutz, Interdiciplinary Procedures Center, Strategic Laboratory | Claudio Tavares Sacchi, Claudia Regina Gonçalves, Erica Valessa Ramos Gomes, Karoline Rodrigues Campos |
| EPI_ISL_779155, EPI_ISL_779156, EPI_ISL_779157, EPI_ISL_779158, EPI_ISL_779159, EPI_ISL_779160, EPI_ISL_779161, EPI_ISL_779162, EPI_ISL_779163, EPI_ISL_779164, EPI_ISL_779165, EPI_ISL_779166, EPI_ISL_779167, EPI_ISL_779168 | | | |
| see above | Laboratório de Microbiologia Molecular - Universidade FEEVALE | Bioinformatics Laboratory / LNCC | Felipe Benites, Fernando Rosado Spilki, Alana Witt Hansen, Juliane Deise Fleck, Juliana Schons, Meriane Demoliner, Ana Karolina Eisen Antunes, Fagner Henrique Heldt, Larissa Mallmann, Bruna Hermann, Ana Luiza Ziulkoski, Vyctoria Goes, Karoline Schallenberger, Matheus Nunes Weber, Paula Rodrigues de Almeida, Alessandra Pavan Lamarca da Silva, Ronaldo da Silva F Jr , Luiz G P de Almeida, Alexandra L Gerber , Ana Paula de C Guimarães,Ana Tereza R de Vasconcelos |
| EPI_ISL_792101, EPI_ISL_792102 | Instituto Adolfo Lutz - Central | Instituto Adolfo Lutz, Interdiciplinary Procedures Center, Strategic Laboratory | Claudio Tavares Sacchi, Claudia Regina Gonçalves, Erica Valessa Ramos Gomes, Karoline Rodrigues Campos |
| EPI_ISL_792103 | Instituto Adolfo Lutz - Regional de Santo Andre | Instituto Adolfo Lutz, Interdiciplinary Procedures Center, Strategic Laboratory | Claudio Tavares Sacchi, Claudia Regina Gonçalves, Erica Valessa Ramos Gomes, Karoline Rodrigues Campos |
| EPI_ISL_792104, EPI_ISL_792105, EPI_ISL_792106, EPI_ISL_792107, EPI_ISL_792108, EPI_ISL_792109, EPI_ISL_792110, EPI_ISL_792111, EPI_ISL_792112, EPI_ISL_792113, EPI_ISL_792114 | | | |
| see above | Instituto Adolfo Lutz - Central | Instituto Adolfo Lutz, Interdiciplinary Procedures Center, Strategic Laboratory | Claudio Tavares Sacchi, Claudia Regina Gonçalves, Erica Valessa Ramos Gomes, Karoline Rodrigues Campos |
| EPI_ISL_792115, EPI_ISL_792116 | Instituto Adolfo Lutz - Regional de Taubate | Instituto Adolfo Lutz, Interdiciplinary Procedures Center, Strategic Laboratory | Claudio Tavares Sacchi, Claudia Regina Gonçalves, Erica Valessa Ramos Gomes, Karoline Rodrigues Campos |
| EPI_ISL_792560 | Laboratorio de Ecologia de Doencas Transmissiveis na Amazonia, Instituto Leonidas e Maria Deane - Fiocruz Amazonia | Laboratorio de Ecologia de Doencas Transmissiveis na Amazonia, Instituto Leonidas e Maria Deane - Fiocruz Amazonia | Valdinete Nascimento, Victor Souza, André Corado, Fernanda Nascimento, George Silva, Ágatha Costa, Karina Pessoa, Debora Duarte, Luciana Gonçalves, Maria Júlia Brandão, Michele Jesus, Felipe Naveca on behalf of the Fiocruz COVID-19 Genomic Surveillance Network |
| EPI_ISL_792561, EPI_ISL_792562, EPI_ISL_792563, EPI_ISL_792564, EPI_ISL_792565, EPI_ISL_792566, EPI_ISL_792567, EPI_ISL_792568, EPI_ISL_792569, EPI_ISL_792570, EPI_ISL_792571, EPI_ISL_792572, EPI_ISL_792573, EPI_ISL_792574, EPI_ISL_792575, EPI_ISL_792576, EPI_ISL_792577, EPI_ISL_792578, EPI_ISL_792579, EPI_ISL_792580, EPI_ISL_792581, EPI_ISL_792582, EPI_ISL_792583, EPI_ISL_792584, EPI_ISL_792585, EPI_ISL_792586, EPI_ISL_792587, EPI_ISL_792588, EPI_ISL_792589, EPI_ISL_792590, EPI_ISL_792591, EPI_ISL_792592, EPI_ISL_792593, EPI_ISL_792594, EPI_ISL_792595, EPI_ISL_792596, EPI_ISL_792597, EPI_ISL_792598, EPI_ISL_792599, EPI_ISL_792600, EPI_ISL_792601, EPI_ISL_792602, EPI_ISL_792603, EPI_ISL_792604, EPI_ISL_792605, EPI_ISL_792606, EPI_ISL_792607, EPI_ISL_792608, EPI_ISL_792609, EPI_ISL_792610, EPI_ISL_792611, EPI_ISL_792612, EPI_ISL_792613, EPI_ISL_792614, EPI_ISL_792615, EPI_ISL_792616, EPI_ISL_792617, EPI_ISL_792618, EPI_ISL_792619, EPI_ISL_792620, EPI_ISL_792621, EPI_ISL_792622, EPI_ISL_792623, EPI_ISL_792624, EPI_ISL_792625, EPI_ISL_792626, EPI_ISL_792627, EPI_ISL_792628, EPI_ISL_792629, EPI_ISL_792630, EPI_ISL_792631, EPI_ISL_792632, EPI_ISL_792633, EPI_ISL_792634, EPI_ISL_792635, EPI_ISL_792636, EPI_ISL_792637, EPI_ISL_792638 | | | |
| see above | Laboratório Central de Saúde Pública do Estado da Paraíba (LACEN-PB) | Laboratory of Respiratory Viruses and Measles, Oswaldo Cruz Institute, FIOCRUZ | Paola Resende, Luciana Appolinario, Fernando Motta, Anna Carolina Paixao, Ana Carolina Mendonca, João Felipe Bezerra, Romero Henrique Teixeira de Vasconcelos, Dalane Loudal Florentino Teixeira, Thiago Franco de Oliveira Carneiro, Marilda Siqueira on behalf of the Fiocruz COVID-19 Genomic Surveillance Network |
| EPI_ISL_792639, EPI_ISL_792640, EPI_ISL_792641, EPI_ISL_792643, EPI_ISL_792644 | Laboratório Central de Saúde Pública do Estado de Alagoas (LACEN-AL) | Laboratory of Respiratory Viruses and Measles, Oswaldo Cruz Institute, FIOCRUZ | Paola Resende, Luciana Appolinario, Fernando Motta, Anna Carolina Paixao, Ana Carolina Mendonca, Anderson Brandao Leite, Marilda Siqueira on behalf of the Fiocruz COVID-19 Genomic Surveillance Network |
| EPI_ISL_792645, EPI_ISL_792646, EPI_ISL_792647, EPI_ISL_792648, EPI_ISL_792649, EPI_ISL_792650, EPI_ISL_792651, EPI_ISL_792652, EPI_ISL_792653, EPI_ISL_792654 | Laboratório Central de Saúde Pública do Estado do Paraná (LACEN-PR) | Laboratory of Respiratory Viruses and Measles, Oswaldo Cruz Institute, FIOCRUZ | Paola Resende, Luciana Appolinario, Fernando Motta, Anna Carolina Paixao, Ana Carolina Mendonca, Maria do Carmo Debur, Irina Nastassja Riediger, Marilda Siqueira on behalf of the Fiocruz COVID-19 Genomic Surveillance Network |
| EPI_ISL_801386, EPI_ISL_801387, EPI_ISL_801388, EPI_ISL_801389, EPI_ISL_801390, EPI_ISL_801391, EPI_ISL_801392, EPI_ISL_801393, EPI_ISL_801394, EPI_ISL_801395, EPI_ISL_801396 | | | |
| see above | Laboratorio de Ecologia de Doencas Transmissiveis na Amazonia, Instituto Leonidas e Maria Deane - Fiocruz Amazonia | Laboratorio de Ecologia de Doencas Transmissiveis na Amazonia, Instituto Leonidas e Maria Deane - Fiocruz Amazonia | Valdinete Nascimento, Victor Souza, André Corado, Fernanda Nascimento, George Silva, Ágatha Costa, Debora Duarte, Luciana Gonçalves, Maria Júlia Brandão, Michele Jesus, Felipe Naveca on behalf of the Fiocruz COVID-19 Genomic Surveillance Network |
| EPI_ISL_801397, EPI_ISL_801398, EPI_ISL_801399, EPI_ISL_801400, EPI_ISL_801401, EPI_ISL_801402, EPI_ISL_801403 | Laboratório Central de Saúde Pública do Estado do Amazonas (LACEN-AM) | Laboratorio de Ecologia de Doencas Transmissiveis na Amazonia, Instituto Leonidas e Maria Deane - Fiocruz Amazonia | Valdinete Nascimento, Victor Souza, André Corado, Fernanda Nascimento, George Silva, Ágatha Costa, Debora Duarte, Luciana Gonçalves, Maria Júlia Brandão, Michele Jesus, Felipe Naveca on behalf of the Fiocruz COVID-19 Genomic Surveillance Network |
| EPI_ISL_811148, EPI_ISL_811149 | Laboratorio de Ecologia de Doencas Transmissiveis na Amazonia, Instituto Leonidas e Maria Deane - Fiocruz Amazonia | Laboratorio de Ecologia de Doencas Transmissiveis na Amazonia, Instituto Leonidas e Maria Deane - Fiocruz Amazonia | Valdinete Nascimento, Victor Souza, André Corado, Fernanda Nascimento, George Silva, Ágatha Costa, Debora Duarte, Luciana Gonçalves, Matilde Mejía, Karina Pessoa, Maria Júlia Brandão, Michele Jesus, Felipe Naveca on behalf of the Fiocruz COVID-19 Genomic Surveillance Network |
| EPI_ISL_831474, EPI_ISL_831645, EPI_ISL_831646, EPI_ISL_831660, EPI_ISL_831678, EPI_ISL_831681, EPI_ISL_831683, EPI_ISL_831685, EPI_ISL_831688, EPI_ISL_831689, EPI_ISL_831892, EPI_ISL_831898, EPI_ISL_831913, EPI_ISL_831938, EPI_ISL_831939, EPI_ISL_831940, EPI_ISL_832009, EPI_ISL_832010, EPI_ISL_832011, EPI_ISL_832012, EPI_ISL_832013 | | | |
| see above | Laboratório de Microbiologia Molecular - Universidade FEEVALE | Universidade Federal de Ciências da Saúde de Porto Alegre | Vinícius Bonetti Franceschi, Amanda de Menezes Mayer, Gabriel Dickin Caldana, Carla Andretta Moreira Neves, Patrícia Aline Gröhs Ferrareze, Gabriela Bettella Cybis, Ricardo Ariel Zimerman, Lívia Kmetzsch, Fernando Rosado Spilki, Claudia Elizabeth Thompson |
| EPI_ISL_833131, EPI_ISL_833132, EPI_ISL_833133, EPI_ISL_833134, EPI_ISL_833135, EPI_ISL_833136, EPI_ISL_833137, EPI_ISL_833138, EPI_ISL_833139, EPI_ISL_833140 | Laboratorio de Ecologia de Doencas Transmissiveis na Amazonia, Instituto Leonidas e Maria Deane - Fiocruz Amazonia | Laboratorio de Ecologia de Doencas Transmissiveis na Amazonia, Instituto Leonidas e Maria Deane - Fiocruz Amazonia | Valdinete Nascimento, Victor Souza, André Corado, Fernanda Nascimento, George Silva, Ágatha Costa, Debora Duarte, Karina Pessoa, Matilde Mejía, Luciana Gonçalves, Maria Júlia Brandão, Michele Jesus, Felipe Naveca on behalf of the Fiocruz COVID-19 Genomic Surveillance Network |
| EPI_ISL_833152, EPI_ISL_833153, EPI_ISL_833154, EPI_ISL_833155 | Instituto Adolfo Lutz - Central | Instituto Adolfo Lutz, Interdiciplinary Procedures Center, Strategic Laboratory | Claudio Tavares Sacchi, Claudia Regina Gonçalves, Erica Valessa Ramos Gomes, Karoline Rodrigues Campos |
| EPI_ISL_833156 | Instituto Adolfo Lutz - Regional de Sorocaba | Instituto Adolfo Lutz, Interdiciplinary Procedures Center, Strategic Laboratory | Claudio Tavares Sacchi, Claudia Regina Gonçalves, Erica Valessa Ramos Gomes, Karoline Rodrigues Campos |
| EPI_ISL_833157, EPI_ISL_833158 | Instituto Adolfo Lutz - Regional de Santo Andre | Instituto Adolfo Lutz, Interdiciplinary Procedures Center, Strategic Laboratory | Claudio Tavares Sacchi, Claudia Regina Gonçalves, Erica Valessa Ramos Gomes, Karoline Rodrigues Campos |
| EPI_ISL_833159 | Instituto Adolfo Lutz - Central | Instituto Adolfo Lutz, Interdiciplinary Procedures Center, Strategic Laboratory | Claudio Tavares Sacchi, Claudia Regina Gonçalves, Erica Valessa Ramos Gomes, Karoline Rodrigues Campos |
| EPI_ISL_833160 | Instituto Adolfo Lutz - Regional de Santo Andre | Instituto Adolfo Lutz, Interdiciplinary Procedures Center, Strategic Laboratory | Claudio Tavares Sacchi, Claudia Regina Gonçalves, Erica Valessa Ramos Gomes, Karoline Rodrigues Campos |
| EPI_ISL_833161 | Instituto Adolfo Lutz - Central | Instituto Adolfo Lutz, Interdiciplinary Procedures Center, | Claudio Tavares Sacchi, Claudia Regina Gonçalves, Erica Valessa Ramos Gomes, Karoline Rodrigues Campos |

| | | | |
|---|---|---|---|
| | | Strategic Laboratory | |
| EPI_ISL_833162 | Lab LOC - Itapecerica da Serra | Instituto Adolfo Lutz, Interdiciplinary Procedures Center, Strategic Laboratory | Claudio Tavares Sacchi, Claudia Regina Gonçalves, Erica Valessa Ramos Gomes, Karoline Rodrigues Campos |
| EPI_ISL_833163 | Instituto Adolfo Lutz - Regional de Marilia | Instituto Adolfo Lutz, Interdiciplinary Procedures Center, Strategic Laboratory | Claudio Tavares Sacchi, Claudia Regina Gonçalves, Erica Valessa Ramos Gomes, Karoline Rodrigues Campos |
| EPI_ISL_833164 | Secretaria Municipal de Saude de Santa Barbara d'oeste | Instituto Adolfo Lutz, Interdiciplinary Procedures Center, Strategic Laboratory | Claudio Tavares Sacchi, Claudia Regina Gonçalves, Erica Valessa Ramos Gomes, Karoline Rodrigues Campos |
| EPI_ISL_833165, EPI_ISL_833166 | Hospital Samaritano | Instituto Adolfo Lutz, Interdiciplinary Procedures Center, Strategic Laboratory | Claudio Tavares Sacchi, Claudia Regina Gonçalves, Erica Valessa Ramos Gomes, Karoline Rodrigues Campos |
| EPI_ISL_833167, EPI_ISL_833168, EPI_ISL_833169, EPI_ISL_833170, EPI_ISL_833171, EPI_ISL_833172, EPI_ISL_833173, EPI_ISL_833174, EPI_ISL_833175, EPI_ISL_833176 | DB Diagnosticos do Brasil | Instituto Adolfo Lutz, Interdiciplinary Procedures Center, Strategic Laboratory | Claudio Tavares Sacchi, Claudia Regina Gonçalves, Erica Valessa Ramos Gomes, Karoline Rodrigues Campos |
| EPI_ISL_836143 | Hospital de Campanha COVID-19 de Mairipora | Instituto Adolfo Lutz, Interdiciplinary Procedures Center, Strategic Laboratory | Claudio Tavares Sacchi, Claudia Regina Gonçalves, Erica Valessa Ramos Gomes, Karoline Rodrigues Campos |
| EPI_ISL_836977 | Hospital Municiplal Dr. Jose de Carvalho Florence | Instituto Adolfo Lutz, Interdiciplinary Procedures Center, Strategic Laboratory | Claudio Tavares Sacchi, Claudia Regina Gonçalves, Erica Valessa Ramos Gomes, Karoline Rodrigues Campos |
| EPI_ISL_836978 | Irmandade da Santa Casa de Misericordia de Lorena | Instituto Adolfo Lutz, Interdiciplinary Procedures Center, Strategic Laboratory | Claudio Tavares Sacchi, Claudia Regina Gonçalves, Erica Valessa Ramos Gomes, Karoline Rodrigues Campos |
| EPI_ISL_837053 | UBS Darcy Alves e Robalinho | Instituto Adolfo Lutz, Interdiciplinary Procedures Center, Strategic Laboratory | Claudio Tavares Sacchi, Claudia Regina Gonçalves, Erica Valessa Ramos Gomes, Karoline Rodrigues Campos |
| EPI_ISL_837054 | UBS Jose Sabino Ferreira | Instituto Adolfo Lutz, Interdiciplinary Procedures Center, Strategic Laboratory | Claudio Tavares Sacchi, Claudia Regina Gonçalves, Erica Valessa Ramos Gomes, Karoline Rodrigues Campos |
| EPI_ISL_848555, EPI_ISL_848556, EPI_ISL_848557, EPI_ISL_848561, EPI_ISL_848562, EPI_ISL_848563, EPI_ISL_848564, EPI_ISL_848565, EPI_ISL_848566, EPI_ISL_848567, EPI_ISL_848568, EPI_ISL_848569, EPI_ISL_848570, EPI_ISL_848571, EPI_ISL_848572, EPI_ISL_848573, EPI_ISL_848574, EPI_ISL_848575, EPI_ISL_848576, EPI_ISL_848577, EPI_ISL_848578, EPI_ISL_848579, EPI_ISL_848580, EPI_ISL_848581, EPI_ISL_848582, EPI_ISL_848583, EPI_ISL_848584, EPI_ISL_848585, EPI_ISL_848586, EPI_ISL_848587, EPI_ISL_848588, EPI_ISL_848589, EPI_ISL_848590, EPI_ISL_848591, EPI_ISL_848592, EPI_ISL_848593, EPI_ISL_848594, EPI_ISL_848595, EPI_ISL_848596, EPI_ISL_848597, EPI_ISL_848598, EPI_ISL_848599, EPI_ISL_848600, EPI_ISL_848601, EPI_ISL_848602, EPI_ISL_848606, EPI_ISL_848607, EPI_ISL_848608, EPI_ISL_848609, EPI_ISL_848610, EPI_ISL_848611, EPI_ISL_848612, EPI_ISL_848613, EPI_ISL_848614, EPI_ISL_848615, EPI_ISL_848616, EPI_ISL_848617, EPI_ISL_848618, EPI_ISL_848619, EPI_ISL_848620, EPI_ISL_848621, EPI_ISL_848622, EPI_ISL_848623, EPI_ISL_848624, EPI_ISL_848625, EPI_ISL_848626, EPI_ISL_848627, EPI_ISL_848628, EPI_ISL_848629 | | | |
| see above | Evandro Chagas Institute | Evandro Chagas Institute | Santos, M.C.; Silva, A.M.; Junior, W.D.C.; Barbagelata, L.S.; Ferreira, J.A.; Sousa, E.M.A.; da Silva, P.S.; Pinheiro, K.C.; L.C.; Sousa Junior, E.C. |
| EPI_ISL_860317, EPI_ISL_860633 | Instituto Adolfo Lutz - Regional de Sorocaba | Instituto Adolfo Lutz, Interdiciplinary Procedures Center, Strategic Laboratory | Claudio Tavares Sacchi, Claudia Regina Gonçalves, Erica Valessa Ramos Gomes, Karoline Rodrigues Campos |
| EPI_ISL_861242 | Instituto de Biotecnologia - UNESP-Botucatu-SP | Instituto de Biotecnologia - UNESP-Botucatu-SP | Leila Sabrina Ullmann; Fábio Sossai Possebon, Camila Dantas Malossi, Paula Rahal, Paulo Inacio da Costa, João Pessoa Araújo Jr. |
| EPI_ISL_861625, EPI_ISL_861626, EPI_ISL_861627 | Instituto Adolfo Lutz - Central | Instituto Adolfo Lutz, Interdiciplinary Procedures Center, Strategic Laboratory | Claudio Tavares Sacchi, Claudia Regina Gonçalves, Erica Valessa Ramos Gomes, Karoline Rodrigues Campos |
| EPI_ISL_861628 | Laboratorio Municipal de Guarulhos | Instituto Adolfo Lutz, Interdiciplinary Procedures Center, Strategic Laboratory | Claudio Tavares Sacchi, Claudia Regina Gonçalves, Erica Valessa Ramos Gomes, Karoline Rodrigues Campos |
| EPI_ISL_861629, EPI_ISL_861630, EPI_ISL_861631, EPI_ISL_861632, EPI_ISL_861633, EPI_ISL_861634 | Instituto Adolfo Lutz - Central | Instituto Adolfo Lutz, Interdiciplinary Procedures Center, Strategic Laboratory | Claudio Tavares Sacchi, Claudia Regina Gonçalves, Erica Valessa Ramos Gomes, Karoline Rodrigues Campos |
| EPI_ISL_861635 | Hospital e Maternidade Madre Theodora | Instituto Adolfo Lutz, Interdiciplinary Procedures Center, Strategic Laboratory | Claudio Tavares Sacchi, Claudia Regina Gonçalves, Erica Valessa Ramos Gomes, Karoline Rodrigues Campos |
| EPI_ISL_861636 | Hospital Geral de Sao Mateus São Paulo | Instituto Adolfo Lutz, Interdiciplinary Procedures Center, Strategic Laboratory | Claudio Tavares Sacchi, Claudia Regina Gonçalves, Erica Valessa Ramos Gomes, Karoline Rodrigues Campos |
| EPI_ISL_861637 | Hospital Nipo Brasileiro | Instituto Adolfo Lutz, Interdiciplinary Procedures Center, Strategic Laboratory | Claudio Tavares Sacchi, Claudia Regina Gonçalves, Erica Valessa Ramos Gomes, Karoline Rodrigues Campos |
| EPI_ISL_861638 | Hospital Municipal Dr. Moyses Deutsch | Instituto Adolfo Lutz, Interdiciplinary Procedures Center, Strategic Laboratory | Claudio Tavares Sacchi, Claudia Regina Gonçalves, Erica Valessa Ramos Gomes, Karoline Rodrigues Campos |
| EPI_ISL_861639 | Hospital Sao Paulo de Ensino da Unifesp | Instituto Adolfo Lutz, Interdiciplinary Procedures Center, Strategic Laboratory | Claudio Tavares Sacchi, Claudia Regina Gonçalves, Erica Valessa Ramos Gomes, Karoline Rodrigues Campos |
| EPI_ISL_861640, EPI_ISL_861641 | Hospital Municipal Dr. Moyses Deutsch | Instituto Adolfo Lutz, Interdiciplinary Procedures Center, Strategic Laboratory | Claudio Tavares Sacchi, Claudia Regina Gonçalves, Erica Valessa Ramos Gomes, Karoline Rodrigues Campos |
| EPI_ISL_861642, EPI_ISL_861643 | Instituto Adolfo Lutz - Central | Instituto Adolfo Lutz, Interdiciplinary Procedures Center, Strategic Laboratory | Claudio Tavares Sacchi, Claudia Regina Gonçalves, Erica Valessa Ramos Gomes, Karoline Rodrigues Campos |
| EPI_ISL_861644 | Hospital Santa Virginia | Instituto Adolfo Lutz, Interdiciplinary Procedures Center, Strategic Laboratory | Claudio Tavares Sacchi, Claudia Regina Gonçalves, Erica Valessa Ramos Gomes, Karoline Rodrigues Campos |
| EPI_ISL_861645 | Hospital e Pronto Socorro Comunitario Vila Iolanda | Instituto Adolfo Lutz, Interdiciplinary Procedures Center, Strategic Laboratory | Claudio Tavares Sacchi, Claudia Regina Gonçalves, Erica Valessa Ramos Gomes, Karoline Rodrigues Campos |
| EPI_ISL_861646, EPI_ISL_861647 | Hospital Santa Marcelina Sao Paulo | Instituto Adolfo Lutz, Interdiciplinary Procedures Center, Strategic Laboratory | Claudio Tavares Sacchi, Claudia Regina Gonçalves, Erica Valessa Ramos Gomes, Karoline Rodrigues Campos |
| EPI_ISL_861648 | Hospital e Pronto Socorro Portinari | Instituto Adolfo Lutz, Interdiciplinary Procedures Center, Strategic Laboratory | Claudio Tavares Sacchi, Claudia Regina Gonçalves, Erica Valessa Ramos Gomes, Karoline Rodrigues Campos |
| EPI_ISL_861649 | Hospital Renascença Campinas | Instituto Adolfo Lutz, Interdiciplinary Procedures Center, Strategic Laboratory | Claudio Tavares Sacchi, Claudia Regina Gonçalves, Erica Valessa Ramos Gomes, Karoline Rodrigues Campos |
| EPI_ISL_861650 | Hospital Santa Marcelina Sao Paulo | Instituto Adolfo Lutz, Interdiciplinary Procedures Center, Strategic Laboratory | Claudio Tavares Sacchi, Claudia Regina Gonçalves, Erica Valessa Ramos Gomes, Karoline Rodrigues Campos |
| EPI_ISL_861651 | AMA Jardim Brasil | Instituto Adolfo Lutz, Interdiciplinary Procedures Center, Strategic Laboratory | Claudio Tavares Sacchi, Claudia Regina Gonçalves, Erica Valessa Ramos Gomes, Karoline Rodrigues Campos |
| EPI_ISL_861652 | AMA Wamberto Dias da Costa | Instituto Adolfo Lutz, Interdiciplinary Procedures Center, Strategic Laboratory | Claudio Tavares Sacchi, Claudia Regina Gonçalves, Erica Valessa Ramos Gomes, Karoline Rodrigues Campos |
| EPI_ISL_861653 | Hospital Santa Virgínia | Instituto Adolfo Lutz, Interdiciplinary Procedures Center, Strategic Laboratory | Claudio Tavares Sacchi, Claudia Regina Gonçalves, Erica Valessa Ramos Gomes, Karoline Rodrigues Campos |

| | | | |
|---|---|---|---|
| EPI_ISL_861654, EPI_ISL_861655 | Hospital Santa Marcelina Sao Paulo | Instituto Adolfo Lutz, Interdiciplinary Procedures Center, Strategic Laboratory | Claudio Tavares Sacchi, Claudia Regina Gonçalves, Erica Valessa Ramos Gomes, Karoline Rodrigues Campos |
| EPI_ISL_861656 | UPA de Jandira | Instituto Adolfo Lutz, Interdiciplinary Procedures Center, Strategic Laboratory | Claudio Tavares Sacchi, Claudia Regina Gonçalves, Erica Valessa Ramos Gomes, Karoline Rodrigues Campos |
| EPI_ISL_861657 | Hospital e Maternidade Sino Brasileiro | Instituto Adolfo Lutz, Interdiciplinary Procedures Center, Strategic Laboratory | Claudio Tavares Sacchi, Claudia Regina Gonçalves, Erica Valessa Ramos Gomes, Karoline Rodrigues Campos |
| EPI_ISL_861658 | Hospital Municipal Antônio Giglio | Instituto Adolfo Lutz, Interdiciplinary Procedures Center, Strategic Laboratory | Claudio Tavares Sacchi, Claudia Regina Gonçalves, Erica Valessa Ramos Gomes, Karoline Rodrigues Campos |
| EPI_ISL_861659, EPI_ISL_861660, EPI_ISL_861661 | PS e Maternidade Nair Fonseca Leitao Arantes | Instituto Adolfo Lutz, Interdiciplinary Procedures Center, Strategic Laboratory | Claudio Tavares Sacchi, Claudia Regina Gonçalves, Erica Valessa Ramos Gomes, Karoline Rodrigues Campos |
| EPI_ISL_861662 | CS I Tacito Leite de Carvalho e Silva | Instituto Adolfo Lutz, Interdiciplinary Procedures Center, Strategic Laboratory | Claudio Tavares Sacchi, Claudia Regina Gonçalves, Erica Valessa Ramos Gomes, Karoline Rodrigues Campos |
| EPI_ISL_861663 | Instituto Adolfo Lutz - Central | Instituto Adolfo Lutz, Interdiciplinary Procedures Center, Strategic Laboratory | Claudio Tavares Sacchi, Claudia Regina Gonçalves, Erica Valessa Ramos Gomes, Karoline Rodrigues Campos |
| EPI_ISL_861664 | Instituto Adolfo Lutz - Regional de Campinas | Instituto Adolfo Lutz, Interdiciplinary Procedures Center, Strategic Laboratory | Claudio Tavares Sacchi, Claudia Regina Gonçalves, Erica Valessa Ramos Gomes, Karoline Rodrigues Campos |
| EPI_ISL_861665 | Instituto Adolfo Lutz - Regional de Taubate | Instituto Adolfo Lutz, Interdiciplinary Procedures Center, Strategic Laboratory | Claudio Tavares Sacchi, Claudia Regina Gonçalves, Erica Valessa Ramos Gomes, Karoline Rodrigues Campos |
| EPI_ISL_861666 | PSF Dr. Antonio Pires de Almeida | Instituto Adolfo Lutz, Interdiciplinary Procedures Center, Strategic Laboratory | Claudio Tavares Sacchi, Claudia Regina Gonçalves, Erica Valessa Ramos Gomes, Karoline Rodrigues Campos |
| EPI_ISL_861667 | Instituto Adolfo Lutz - Regional de Rio Claro | Instituto Adolfo Lutz, Interdiciplinary Procedures Center, Strategic Laboratory | Claudio Tavares Sacchi, Claudia Regina Gonçalves, Erica Valessa Ramos Gomes, Karoline Rodrigues Campos |
| EPI_ISL_861668 | Centro de Triagem Covid19 | Instituto Adolfo Lutz, Interdiciplinary Procedures Center, Strategic Laboratory | Claudio Tavares Sacchi, Claudia Regina Gonçalves, Erica Valessa Ramos Gomes, Karoline Rodrigues Campos |
| EPI_ISL_861669 | Lab LOC - Itapecerica da Serra | Instituto Adolfo Lutz, Interdiciplinary Procedures Center, Strategic Laboratory | Claudio Tavares Sacchi, Claudia Regina Gonçalves, Erica Valessa Ramos Gomes, Karoline Rodrigues Campos |
| EPI_ISL_861670 | Instituto Adolfo Lutz - Regional de Taubate | Instituto Adolfo Lutz, Interdiciplinary Procedures Center, Strategic Laboratory | Claudio Tavares Sacchi, Claudia Regina Gonçalves, Erica Valessa Ramos Gomes, Karoline Rodrigues Campos |
| EPI_ISL_861671 | Hospital Municipal Prefeito Waldemar Costa Filho | Instituto Adolfo Lutz, Interdiciplinary Procedures Center, Strategic Laboratory | Claudio Tavares Sacchi, Claudia Regina Gonçalves, Erica Valessa Ramos Gomes, Karoline Rodrigues Campos |
| EPI_ISL_861672 | Day Hospital de Ermelino Matarazzo | Instituto Adolfo Lutz, Interdiciplinary Procedures Center, Strategic Laboratory | Claudio Tavares Sacchi, Claudia Regina Gonçalves, Erica Valessa Ramos Gomes, Karoline Rodrigues Campos |
| EPI_ISL_861673 | PA Novo Osasco | Instituto Adolfo Lutz, Interdiciplinary Procedures Center, Strategic Laboratory | Claudio Tavares Sacchi, Claudia Regina Gonçalves, Erica Valessa Ramos Gomes, Karoline Rodrigues Campos |
| EPI_ISL_861674, EPI_ISL_861675 | UPA Central de Caraguatatuba | Instituto Adolfo Lutz, Interdiciplinary Procedures Center, Strategic Laboratory | Claudio Tavares Sacchi, Claudia Regina Gonçalves, Erica Valessa Ramos Gomes, Karoline Rodrigues Campos |
| EPI_ISL_861676 | UPA Vila Santa Catarina | Instituto Adolfo Lutz, Interdiciplinary Procedures Center, Strategic Laboratory | Claudio Tavares Sacchi, Claudia Regina Gonçalves, Erica Valessa Ramos Gomes, Karoline Rodrigues Campos |
| EPI_ISL_861677 | Instituto Adolfo Lutz - Central | Instituto Adolfo Lutz, Interdiciplinary Procedures Center, Strategic Laboratory | Claudio Tavares Sacchi, Claudia Regina Gonçalves, Erica Valessa Ramos Gomes, Karoline Rodrigues Campos |
| EPI_ISL_861678 | Laboratorio Municipal de Guarulhos | Instituto Adolfo Lutz, Interdiciplinary Procedures Center, Strategic Laboratory | Claudio Tavares Sacchi, Claudia Regina Gonçalves, Erica Valessa Ramos Gomes, Karoline Rodrigues Campos |
| EPI_ISL_861679 | Instituto Adolfo Lutz - Regional de Taubate | Instituto Adolfo Lutz, Interdiciplinary Procedures Center, Strategic Laboratory | Claudio Tavares Sacchi, Claudia Regina Gonçalves, Erica Valessa Ramos Gomes, Karoline Rodrigues Campos |
| EPI_ISL_861680 | Hospital e Pronto Socorro Portinari | Instituto Adolfo Lutz, Interdiciplinary Procedures Center, Strategic Laboratory | Claudio Tavares Sacchi, Claudia Regina Gonçalves, Erica Valessa Ramos Gomes, Karoline Rodrigues Campos |
| EPI_ISL_861682 | UPA Vila Santa Catarina | Instituto Adolfo Lutz, Interdiciplinary Procedures Center, Strategic Laboratory | Claudio Tavares Sacchi, Claudia Regina Gonçalves, Erica Valessa Ramos Gomes, Karoline Rodrigues Campos |
| EPI_ISL_861683 | Complexo Hospitalar Padre Bento de Guarulhos | Instituto Adolfo Lutz, Interdiciplinary Procedures Center, Strategic Laboratory | Claudio Tavares Sacchi, Claudia Regina Gonçalves, Erica Valessa Ramos Gomes, Karoline Rodrigues Campos |
| EPI_ISL_861684, EPI_ISL_861685 | Hospital Pronto Socorro Itaquera | Instituto Adolfo Lutz, Interdiciplinary Procedures Center, Strategic Laboratory | Claudio Tavares Sacchi, Claudia Regina Gonçalves, Erica Valessa Ramos Gomes, Karoline Rodrigues Campos |
| EPI_ISL_861867, EPI_ISL_861868, EPI_ISL_861869, EPI_ISL_861870, EPI_ISL_861871, EPI_ISL_861872, EPI_ISL_861873, EPI_ISL_861874, EPI_ISL_861875, EPI_ISL_861876, EPI_ISL_861877, EPI_ISL_861878, EPI_ISL_861879, EPI_ISL_861880, EPI_ISL_861881, EPI_ISL_861882, EPI_ISL_861883, EPI_ISL_861884, EPI_ISL_861885, EPI_ISL_861886, EPI_ISL_861887, EPI_ISL_861888 | | | |
| see above | LATE - Laboratório de Técnicas Especiais - Hospital Israelita Albert Einstein | LATE - Laboratório de Técnicas Especiais - Hospital Israelita Albert Einstein | Deyvid Amgarten, Fernanda de Mello Malta, Raquel Riyuzo, Ana Paula Moreira Salles, Pedro Henrique Sebe Rodrigues, João Renato Rebello Pinho |
| EPI_ISL_861889 | Genomika Einstein | LATE - Laboratório de Técnicas Especiais - Hospital Israelita Albert Einstein | Deyvid Amgarten, Fernanda de Mello Malta, Raquel Riyuzo, Ana Paula Moreira Salles, Pedro Henrique Sebe Rodrigues, João Bosco Oliveira Filho, João Renato Rebello Pinho |
| EPI_ISL_861890, EPI_ISL_861891, EPI_ISL_861892, EPI_ISL_861893, EPI_ISL_861894, EPI_ISL_861895, EPI_ISL_861896, EPI_ISL_861897, EPI_ISL_861898, EPI_ISL_861899, EPI_ISL_861900, EPI_ISL_861901, EPI_ISL_861902, EPI_ISL_861903, EPI_ISL_861904, EPI_ISL_861905, EPI_ISL_861906, EPI_ISL_861907, EPI_ISL_861908, EPI_ISL_861909, EPI_ISL_861910 | | | |
| see above | LATE - Laboratório de Técnicas Especiais - Hospital Israelita Albert Einstein | LATE - Laboratório de Técnicas Especiais - Hospital Israelita Albert Einstein | Deyvid Amgarten, Fernanda de Mello Malta, Raquel Riyuzo, Ana Paula Moreira Salles, Pedro Henrique Sebe Rodrigues, João Renato Rebello Pinho |
| EPI_ISL_861911 | Genomika Einstein | LATE - Laboratório de Técnicas Especiais - Hospital Israelita Albert Einstein | Deyvid Amgarten, Fernanda de Mello Malta, Raquel Riyuzo, Ana Paula Moreira Salles, Pedro Henrique Sebe Rodrigues, João Bosco Oliveira Filho, João Renato Rebello Pinho |
| EPI_ISL_861912, EPI_ISL_861913 | LATE - Laboratório de Técnicas Especiais - Hospital Israelita Albert Einstein | LATE - Laboratório de Técnicas Especiais - Hospital Israelita Albert Einstein | Deyvid Amgarten, Fernanda de Mello Malta, Raquel Riyuzo, Ana Paula Moreira Salles, Pedro Henrique Sebe Rodrigues, João Renato Rebello Pinho |
| EPI_ISL_861914 | Genomika Einstein | LATE - Laboratório de Técnicas Especiais - Hospital Israelita Albert Einstein | Deyvid Amgarten, Fernanda de Mello Malta, Raquel Riyuzo, Ana Paula Moreira Salles, Pedro Henrique Sebe Rodrigues, João Bosco Oliveira Filho, João Renato Rebello Pinho |
| EPI_ISL_861915 | LATE - Laboratório de Técnicas Especiais - Hospital Israelita Albert Einstein | LATE - Laboratório de Técnicas Especiais - Hospital Israelita Albert Einstein | Deyvid Amgarten, Fernanda de Mello Malta, Raquel Riyuzo, Ana Paula Moreira Salles, Pedro Henrique Sebe Rodrigues, João Renato Rebello Pinho |

| | | | |
|---|---|---|---|
| EPI_ISL_872191, EPI_ISL_872192 | Conjunto Hospitalar do Mandaqui de Sao Paulo | Instituto Adolfo Lutz, Interdiciplinary Procedures Center, Strategic Laboratory | Claudio Tavares Sacchi, Claudia Regina Gonçalves, Erica Valessa Ramos Gomes, Karoline Rodrigues Campos, Katia Correa de Oliveira Santos, Ana Lucia de Carvalho Avelino, Fabiana Cristina Pereira dos Santos |
| EPI_ISL_875540, EPI_ISL_875541, EPI_ISL_875542, EPI_ISL_875543, EPI_ISL_875544, EPI_ISL_875545, EPI_ISL_875546, EPI_ISL_875547, EPI_ISL_875548, EPI_ISL_875549, EPI_ISL_875550 | | | |
| see above | Instituto de Biotecnologia - UNESP-Botucatu-SP | Instituto de Biotecnologia - UNESP-Botucatu-SP | Leila Sabrina Ullmann; Fábio Sossai Possebon, Camila Dantas Malossi, Paula Rahal, Paulo Inacio da Costa, João Pessoa Araújo Jr. |
| EPI_ISL_875688 | National Influenza Center - Instituto Adolfo Lutz | Instituto Adolfo Lutz, Interdiciplinary Procedures Center, Strategic Laboratory | Claudio Tavares Sacchi, Claudia Regina Gonçalves, Erica Valessa Ramos Gomes, Karoline Rodrigues Campos, Katia Correa de Oliveira Santos, Ana Lucia de Carvalho Avelino, Clovis Roberto Abe Constantino |
| EPI_ISL_875689 | Hospital do Servidor Publico | Instituto Adolfo Lutz, Interdiciplinary Procedures Center, Strategic Laboratory | Claudio Tavares Sacchi, Claudia Regina Gonçalves, Erica Valessa Ramos Gomes, Karoline Rodrigues Campos |
| EPI_ISL_882657 | LACEN do Estado do Piaui, Dr. Costa Alvarenga | Instituto Adolfo Lutz, Interdiciplinary Procedures Center, Strategic Laboratory | Claudio Tavares Sacchi, Claudia Regina Gonçalves, Erica Valessa Ramos Gomes, Karoline Rodrigues Campos |
| EPI_ISL_882658 | Secretaria Municipal de Saude | Instituto Adolfo Lutz, Interdiciplinary Procedures Center, Strategic Laboratory | Claudio Tavares Sacchi, Claudia Regina Gonçalves, Erica Valessa Ramos Gomes, Karoline Rodrigues Campos |
| EPI_ISL_882659 | Centro de Triagem Covid19 | Instituto Adolfo Lutz, Interdiciplinary Procedures Center, Strategic Laboratory | Claudio Tavares Sacchi, Claudia Regina Gonçalves, Erica Valessa Ramos Gomes, Karoline Rodrigues Campos |
| EPI_ISL_882660 | Hospital Municipal Prefeito Waldemar Costa Filho | Instituto Adolfo Lutz, Interdiciplinary Procedures Center, Strategic Laboratory | Claudio Tavares Sacchi, Claudia Regina Gonçalves, Erica Valessa Ramos Gomes, Karoline Rodrigues Campos |
| EPI_ISL_882661, EPI_ISL_882662 | Hospital de Santa Barbara de Goias | Instituto Adolfo Lutz, Interdiciplinary Procedures Center, Strategic Laboratory | Claudio Tavares Sacchi, Claudia Regina Gonçalves, Erica Valessa Ramos Gomes, Karoline Rodrigues Campos |
| EPI_ISL_882663, EPI_ISL_882664 | LACEN do Distrito Federal | Instituto Adolfo Lutz, Interdiciplinary Procedures Center, Strategic Laboratory | Claudio Tavares Sacchi, Claudia Regina Gonçalves, Erica Valessa Ramos Gomes, Karoline Rodrigues Campos |
| EPI_ISL_882665 | Unidade de Pronto Atendimento Dra Zilda Arns | Instituto Adolfo Lutz, Interdiciplinary Procedures Center, Strategic Laboratory | Claudio Tavares Sacchi, Claudia Regina Gonçalves, Erica Valessa Ramos Gomes, Karoline Rodrigues Campos |
| EPI_ISL_882666 | UMS de Juquitiba | Instituto Adolfo Lutz, Interdiciplinary Procedures Center, Strategic Laboratory | Claudio Tavares Sacchi, Claudia Regina Gonçalves, Erica Valessa Ramos Gomes, Karoline Rodrigues Campos |
| EPI_ISL_882667 | Hospital de Clinicas Caieiras | Instituto Adolfo Lutz, Interdiciplinary Procedures Center, Strategic Laboratory | Claudio Tavares Sacchi, Claudia Regina Gonçalves, Erica Valessa Ramos Gomes, Karoline Rodrigues Campos |
| EPI_ISL_882668 | UMS de Juquitiba | Instituto Adolfo Lutz, Interdiciplinary Procedures Center, Strategic Laboratory | Claudio Tavares Sacchi, Claudia Regina Gonçalves, Erica Valessa Ramos Gomes, Karoline Rodrigues Campos |
| EPI_ISL_882669 | UPA Vila Santa Catarina | Instituto Adolfo Lutz, Interdiciplinary Procedures Center, Strategic Laboratory | Claudio Tavares Sacchi, Claudia Regina Gonçalves, Erica Valessa Ramos Gomes, Karoline Rodrigues Campos |
| EPI_ISL_882670 | Hospital Samaritano Paulista | Instituto Adolfo Lutz, Interdiciplinary Procedures Center, Strategic Laboratory | Claudio Tavares Sacchi, Claudia Regina Gonçalves, Erica Valessa Ramos Gomes, Karoline Rodrigues Campos |
| EPI_ISL_882671, EPI_ISL_882672, EPI_ISL_882673 | Hospital Municipal Dr. Guido Guida | Instituto Adolfo Lutz, Interdiciplinary Procedures Center, Strategic Laboratory | Claudio Tavares Sacchi, Claudia Regina Gonçalves, Erica Valessa Ramos Gomes, Karoline Rodrigues Campos |
| EPI_ISL_888671, EPI_ISL_888672 | Instituto de Biotecnologia - UNESP-Botucatu-SP | Instituto de Biotecnologia - UNESP-Botucatu-SP | Leila Sabrina Ullmann; Fábio Sossai Possebon, Camila Dantas Malossi, Paula Rahal, Paulo Inacio da Costa, João Pessoa Araújo Jr. |
| EPI_ISL_904120, EPI_ISL_904121 | LACEN - Laboratório Central de Saúde Pública do Pará | Evandro Chagas Institute | Santos, M.C.; Silva, A.M.; Junior, W.D.C.; Barbagelata, L.S.; Ferreira, J.A.; Sousa, E.M.A.; da Silva, P.S.; Pinheiro, K.C.; L.C.; Sousa Junior, E.C. |
| EPI_ISL_906065 | Day Hospital de Ermelino Matarazzo | Instituto Adolfo Lutz, Interdiciplinary Procedures Center, Strategic Laboratory | Claudio Tavares Sacchi, Claudia Regina Gonçalves, Erica Valessa Ramos Gomes, Karoline Rodrigues Campos |
| EPI_ISL_906066 | Hospital Nipo Brasileiro | Instituto Adolfo Lutz, Interdiciplinary Procedures Center, Strategic Laboratory | Claudio Tavares Sacchi, Claudia Regina Gonçalves, Erica Valessa Ramos Gomes, Karoline Rodrigues Campos |
| EPI_ISL_906067 | PS e Maternidade Nair Fonseca Leitao Arantes | Instituto Adolfo Lutz, Interdiciplinary Procedures Center, Strategic Laboratory | Claudio Tavares Sacchi, Claudia Regina Gonçalves, Erica Valessa Ramos Gomes, Karoline Rodrigues Campos |
| EPI_ISL_906068, EPI_ISL_906069 | Instituto Adolfo Lutz - Regional de Campinas | Instituto Adolfo Lutz, Interdiciplinary Procedures Center, Strategic Laboratory | Claudio Tavares Sacchi, Claudia Regina Gonçalves, Erica Valessa Ramos Gomes, Karoline Rodrigues Campos |
| EPI_ISL_906070 | UPA Dr. Akira Tada | Instituto Adolfo Lutz, Interdiciplinary Procedures Center, Strategic Laboratory | Claudio Tavares Sacchi, Claudia Regina Gonçalves, Erica Valessa Ramos Gomes, Karoline Rodrigues Campos |
| EPI_ISL_906071 | LACEN-PI DR. Costa Alvarenga | Instituto Adolfo Lutz, Interdiciplinary Procedures Center, Strategic Laboratory | Claudio Tavares Sacchi, Claudia Regina Gonçalves, Erica Valessa Ramos Gomes, Karoline Rodrigues Campos |
| EPI_ISL_906072 | UPA Dr. Akira Tada | Instituto Adolfo Lutz, Interdiciplinary Procedures Center, Strategic Laboratory | Claudio Tavares Sacchi, Claudia Regina Gonçalves, Erica Valessa Ramos Gomes, Karoline Rodrigues Campos |
| EPI_ISL_906073 | Hospital Vila Lobos | Instituto Adolfo Lutz, Interdiciplinary Procedures Center, Strategic Laboratory | Claudio Tavares Sacchi, Claudia Regina Gonçalves, Erica Valessa Ramos Gomes, Karoline Rodrigues Campos |
| EPI_ISL_906074 | Searom Laboratorio Diagnostico | Instituto Adolfo Lutz, Interdiciplinary Procedures Center, Strategic Laboratory | Claudio Tavares Sacchi, Claudia Regina Gonçalves, Erica Valessa Ramos Gomes, Karoline Rodrigues Campos |
| EPI_ISL_906075 | Hospital Geral de Vila Penteado Dr Jose Pangella Sao Paulo | Instituto Adolfo Lutz, Interdiciplinary Procedures Center, Strategic Laboratory | Claudio Tavares Sacchi, Claudia Regina Gonçalves, Erica Valessa Ramos Gomes, Karoline Rodrigues Campos |
| EPI_ISL_906076, EPI_ISL_906077 | Hospital Sao Luiz Sao Caetano | Instituto Adolfo Lutz, Interdiciplinary Procedures Center, Strategic Laboratory | Claudio Tavares Sacchi, Claudia Regina Gonçalves, Erica Valessa Ramos Gomes, Karoline Rodrigues Campos |
| EPI_ISL_906078 | Hospital Carlos Chagas | Instituto Adolfo Lutz, Interdiciplinary Procedures Center, Strategic Laboratory | Claudio Tavares Sacchi, Claudia Regina Gonçalves, Erica Valessa Ramos Gomes, Karoline Rodrigues Campos |
| EPI_ISL_906079 | Hospital Paulistano Paulista | Instituto Adolfo Lutz, Interdiciplinary Procedures Center, Strategic Laboratory | Claudio Tavares Sacchi, Claudia Regina Gonçalves, Erica Valessa Ramos Gomes, Karoline Rodrigues Campos |
| EPI_ISL_906080, EPI_ISL_906081 | Hospital Beneficiencia Portuguesa | Instituto Adolfo Lutz, Interdiciplinary Procedures Center, Strategic Laboratory | Claudio Tavares Sacchi, Claudia Regina Gonçalves, Erica Valessa Ramos Gomes, Karoline Rodrigues Campos |
| EPI_ISL_918499, EPI_ISL_918500, EPI_ISL_918501, EPI_ISL_918502, EPI_ISL_918503, EPI_ISL_918504, EPI_ISL_918505, EPI_ISL_918506, EPI_ISL_918507, EPI_ISL_918508, EPI_ISL_918509, EPI_ISL_918510, EPI_ISL_918511, EPI_ISL_918512 | | | |
| see above | LACEN - Laboratório Central de Saúde Pública do Amazonas | Evandro Chagas Institute | Santos, M.C.; Silva, A.M.; Junior, W.D.C.; Barbagelata, L.S.; Ferreira, J.A.; Sousa, E.M.A.; da Silva, P.S.; Pinheiro, K.C.; L.C.; Sousa Junior, E.C. |
| EPI_ISL_918513 | LACEN - Laboratório Central de Saúde Pública do Roraima | Evandro Chagas Institute | Santos, M.C.; Silva, A.M.; Junior, W.D.C.; Barbagelata, L.S.; Ferreira, J.A.; Sousa, E.M.A.; da Silva, P.S.; Pinheiro, K.C.; L.C.; Sousa Junior, E.C. |
| EPI_ISL_918514 | Evandro Chagas Institute | Evandro Chagas Institute | Santos, M.C.; Silva, A.M.; Junior, W.D.C.; Barbagelata, L.S.; Ferreira, J.A.; Sousa, E.M.A.; da Silva, P.S.; Pinheiro, K.C.; L.C.; Sousa Junior, E.C. |
| EPI_ISL_918515, EPI_ISL_918516, | LACEN - Laboratório Central de Saúde Pública do Para | Evandro Chagas Institute | Santos, M.C.; Silva, A.M.; Junior, W.D.C.; Barbagelata, L.S.; Ferreira, J.A.; Sousa, E.M.A.; da Silva, P.S.; Pinheiro, K.C.; L.C.; Sousa Junior, E.C. |

| | | | |
|---|---|---|---|
| EPI_ISL_918517 | | | |
| EPI_ISL_918518, EPI_ISL_918519, EPI_ISL_918520, EPI_ISL_918521 | Evandro Chagas Institute | Evandro Chagas Institute | Santos, M.C.; Silva, A.M.; Junior, W.D.C.; Barbagelata, L.S.; Ferreira, J.A.; Sousa, E.M.A.; da Silva, P.S.; Pinheiro, K.C.; L.C.; Sousa Junior, E.C. |
| EPI_ISL_918522, EPI_ISL_918523, EPI_ISL_918524, EPI_ISL_918525, EPI_ISL_918526, EPI_ISL_918527, EPI_ISL_918528, EPI_ISL_918529, EPI_ISL_918530 | LACEN - Laboratório Central de Saúde Pública do Para | Evandro Chagas Institute | Santos, M.C.; Silva, A.M.; Junior, W.D.C.; Barbagelata, L.S.; Ferreira, J.A.; Sousa, E.M.A.; da Silva, P.S.; Pinheiro, K.C.; L.C.; Sousa Junior, E.C. |
| EPI_ISL_918531, EPI_ISL_918532, EPI_ISL_918533, EPI_ISL_918534, EPI_ISL_918535 | LACEN - Laboratório Central de Saúde Pública do Amazonas | Evandro Chagas Institute | Santos, M.C.; Silva, A.M.; Junior, W.D.C.; Barbagelata, L.S.; Ferreira, J.A.; Sousa, E.M.A.; da Silva, P.S.; Pinheiro, K.C.; L.C.; Sousa Junior, E.C. |
| EPI_ISL_918536, EPI_ISL_918537, EPI_ISL_918538, EPI_ISL_918539, EPI_ISL_918540, EPI_ISL_918541, EPI_ISL_918542, EPI_ISL_918543, EPI_ISL_918544 | LACEN - Laboratório Central de Saúde Pública do Ceara | Evandro Chagas Institute | Santos, M.C.; Silva, A.M.; Junior, W.D.C.; Barbagelata, L.S.; Ferreira, J.A.; Sousa, E.M.A.; da Silva, P.S.; Pinheiro, K.C.; L.C.; Sousa Junior, E.C. |
| EPI_ISL_918545, EPI_ISL_918546, EPI_ISL_918547, EPI_ISL_918548, EPI_ISL_918549, EPI_ISL_918550 | LACEN - Laboratório Central de Saúde Pública do Para | Evandro Chagas Institute | Santos, M.C.; Silva, A.M.; Junior, W.D.C.; Barbagelata, L.S.; Ferreira, J.A.; Sousa, E.M.A.; da Silva, P.S.; Pinheiro, K.C.; L.C.; Sousa Junior, E.C. |
| EPI_ISL_918551 | LACEN - Laboratório Central de Saúde Pública do Amapa | Evandro Chagas Institute | Santos, M.C.; Silva, A.M.; Junior, W.D.C.; Barbagelata, L.S.; Ferreira, J.A.; Sousa, E.M.A.; da Silva, P.S.; Pinheiro, K.C.; L.C.; Sousa Junior, E.C. |
| EPI_ISL_918552 | LACEN - Laboratório Central de Saúde Pública do Para | Evandro Chagas Institute | Santos, M.C.; Silva, A.M.; Junior, W.D.C.; Barbagelata, L.S.; Ferreira, J.A.; Sousa, E.M.A.; da Silva, P.S.; Pinheiro, K.C.; L.C.; Sousa Junior, E.C. |
| EPI_ISL_918553, EPI_ISL_918554, EPI_ISL_918555, EPI_ISL_918556, EPI_ISL_918557, EPI_ISL_918558, EPI_ISL_918559, EPI_ISL_918560, EPI_ISL_918561 | LACEN - Laboratório Central de Saúde Pública do Amapa | Evandro Chagas Institute | Santos, M.C.; Silva, A.M.; Junior, W.D.C.; Barbagelata, L.S.; Ferreira, J.A.; Sousa, E.M.A.; da Silva, P.S.; Pinheiro, K.C.; L.C.; Sousa Junior, E.C. |
| EPI_ISL_925846, EPI_ISL_925916, EPI_ISL_926446 | LACEN - Laboratório Central de Saúde Pública do Amazonas | Evandro Chagas Institute Virology | Santos, M.C.; Silva, A.M.; Junior, W.D.C.; Barbagelata, L.S.; Ferreira, J.A.; Sousa, E.M.A.; da Silva, P.S.; Pinheiro, K.C.; L.C.; Sousa Junior, E.C. |
| EPI_ISL_930855, EPI_ISL_930856, EPI_ISL_930857, EPI_ISL_930858 | Central Laboratory of Public Health of Rio Grande do Sul (Lacen-RS) | State Center for Health Surveillance of the Health Department of the State of Rio Grande do Sul (CEVS/SES-RS) | Barcellos R, Campos A, Dornelles C, Godinho F, Gonzalez A, Gregianini T, Molina C, Salvato R, Schaurich A, |
| EPI_ISL_940608 | Laboratório Sao Lucas | Instituto Adolfo Lutz, Interdiciplinary Procedures Center, Strategic Laboratory | Claudio Tavares Sacchi, Claudia Regina Gonçalves, Erica Valessa Ramos Gomes, Karoline Rodrigues Campos |
| EPI_ISL_940609 | Hospital e Maternidade Celso Pierro | Instituto Adolfo Lutz, Interdiciplinary Procedures Center, Strategic Laboratory | Claudio Tavares Sacchi, Claudia Regina Gonçalves, Erica Valessa Ramos Gomes, Karoline Rodrigues Campos |
| EPI_ISL_940610, EPI_ISL_940611 | Hospital Paulistano Paulista | Instituto Adolfo Lutz, Interdiciplinary Procedures Center, Strategic Laboratory | Claudio Tavares Sacchi, Claudia Regina Gonçalves, Erica Valessa Ramos Gomes, Karoline Rodrigues Campos |
| EPI_ISL_940612 | Hospital de Campanha para Enfrentamento do Coronavirus - Goiania | Instituto Adolfo Lutz, Interdiciplinary Procedures Center, Strategic Laboratory | Claudio Tavares Sacchi, Claudia Regina Gonçalves, Erica Valessa Ramos Gomes, Karoline Rodrigues Campos |
| EPI_ISL_940613, EPI_ISL_940614, EPI_ISL_940615, EPI_ISL_940616, EPI_ISL_940617, EPI_ISL_940618 | LACEN-PI DR. Costa Alvarenga | Instituto Adolfo Lutz, Interdiciplinary Procedures Center, Strategic Laboratory | Claudio Tavares Sacchi, Claudia Regina Gonçalves, Erica Valessa Ramos Gomes, Karoline Rodrigues Campos |
| EPI_ISL_940619, EPI_ISL_940620, EPI_ISL_940621, EPI_ISL_940622, EPI_ISL_940623, EPI_ISL_940624, EPI_ISL_940625 | Hospital Sao Joaquim - Beneficiencia Portuguesa | Instituto Adolfo Lutz, Interdiciplinary Procedures Center, Strategic Laboratory | Claudio Tavares Sacchi, Claudia Regina Gonçalves, Erica Valessa Ramos Gomes, Karoline Rodrigues Campos |
| EPI_ISL_940626, EPI_ISL_940627 | Hospital Central Sao Caetano do Sul | Instituto Adolfo Lutz, Interdiciplinary Procedures Center, Strategic Laboratory | Claudio Tavares Sacchi, Claudia Regina Gonçalves, Erica Valessa Ramos Gomes, Karoline Rodrigues Campos |
| EPI_ISL_940628 | Unidade Mista de Iguape | Instituto Adolfo Lutz, Interdiciplinary Procedures Center, Strategic Laboratory | Claudio Tavares Sacchi, Claudia Regina Gonçalves, Erica Valessa Ramos Gomes, Karoline Rodrigues Campos |
| EPI_ISL_940629 | Hospital Municipal Josanias Castanha Braga | Instituto Adolfo Lutz, Interdiciplinary Procedures Center, Strategic Laboratory | Claudio Tavares Sacchi, Claudia Regina Gonçalves, Erica Valessa Ramos Gomes, Karoline Rodrigues Campos |
| EPI_ISL_940630 | Hospital Geral de Sao Paulo | Instituto Adolfo Lutz, Interdiciplinary Procedures Center, Strategic Laboratory | Claudio Tavares Sacchi, Claudia Regina Gonçalves, Erica Valessa Ramos Gomes, Karoline Rodrigues Campos |
| EPI_ISL_942375, EPI_ISL_942897, EPI_ISL_942898, EPI_ISL_942930, EPI_ISL_942931 | Central Laboratory of Public Health of Rio Grande do Sul (Lacen-RS) | State Center for Health Surveillance of the Health Department of the State of Rio Grande do Sul (CEVS/SES-RS) | Barcellos R, Campos A, Crescente L, Da Silva A, Dornelles C, Fonseca V, Garay L, Godinho F, Gonzalez A, Gregianini T, Molina C, Salvato R, Schaurich A |
| EPI_ISL_943574, EPI_ISL_943575, EPI_ISL_943576, EPI_ISL_943577, EPI_ISL_943580, EPI_ISL_943581, EPI_ISL_943582, EPI_ISL_943584, EPI_ISL_943586, EPI_ISL_943587, EPI_ISL_943588, EPI_ISL_943589, EPI_ISL_943590, EPI_ISL_943591, EPI_ISL_943592, EPI_ISL_943593, EPI_ISL_943595, EPI_ISL_943596, EPI_ISL_943606, EPI_ISL_943607, EPI_ISL_943608, EPI_ISL_943609, EPI_ISL_943610 | | | |
| see above | Central Laboratory of Public Health of Rio Grande do Sul (Lacen-RS) | State Center for Health Surveillance of the Health Department of the State of Rio Grande do Sul (CEVS/SES-RS) | Aline Campos, Amanda da Silva, Anelise Schaurich, Claudia Dornelles, Cynthia Molina, Fernanda Godinho, Lara Crescente, Leticia Garay, Regina Barcellos, Richard Salvato, Tatiana Gregianini, Vagner Fonseca |
| EPI_ISL_943967, EPI_ISL_943968, EPI_ISL_943969, EPI_ISL_943970, EPI_ISL_943971 | Hospital Geral de Sao Paulo | Instituto Adolfo Lutz, Interdiciplinary Procedures Center, Strategic Laboratory | Claudio Tavares Sacchi, Claudia Regina Gonçalves, Erica Valessa Ramos Gomes, Karoline Rodrigues Campos |
| EPI_ISL_943973, EPI_ISL_943974, EPI_ISL_943975, EPI_ISL_943976, EPI_ISL_943977, EPI_ISL_943978, EPI_ISL_943979, EPI_ISL_943980, EPI_ISL_943981, EPI_ISL_943982, EPI_ISL_943983, EPI_ISL_943985, EPI_ISL_943986, EPI_ISL_943987 | | | |
| see above | LACEN do Estado de Tocantins | Instituto Adolfo Lutz, Interdiciplinary Procedures Center, Strategic Laboratory | Claudio Tavares Sacchi, Claudia Regina Gonçalves, Erica Valessa Ramos Gomes, Karoline Rodrigues Campos |
| EPI_ISL_943988, EPI_ISL_943989, EPI_ISL_943990 | LACEN do Estado de Goias | Instituto Adolfo Lutz, Interdiciplinary Procedures Center, Strategic Laboratory | Claudio Tavares Sacchi, Claudia Regina Gonçalves, Erica Valessa Ramos Gomes, Karoline Rodrigues Campos |
| EPI_ISL_943991 | LACEN do Estado de Tocantins | Instituto Adolfo Lutz, Interdiciplinary Procedures Center, Strategic Laboratory | Claudio Tavares Sacchi, Claudia Regina Gonçalves, Erica Valessa Ramos Gomes, Karoline Rodrigues Campos |
| EPI_ISL_977471 | Instituto Adolfo Lutz - Regional de Presidente Prudente | Instituto Adolfo Lutz, Interdiciplinary Procedures Center, Strategic Laboratory | Claudio Tavares Sacchi, Claudia Regina Gonçalves, Erica Valessa Ramos Gomes, Karoline Rodrigues Campos |
| EPI_ISL_977472, EPI_ISL_977473, | Instituto Adolfo Lutz Central | Instituto Adolfo Lutz, Interdiciplinary Procedures Center, | Claudio Tavares Sacchi, Claudia Regina Gonçalves, Erica Valessa Ramos Gomes, Karoline Rodrigues Campos |

| | | | |
|---|---|---|---|
| EPI_ISL_977474 | | Strategic Laboratory | |
| EPI_ISL_977475 | Instituto Adolfo Lutz - Regional de Presidente Prudente | Instituto Adolfo Lutz, Interdiciplinary Procedures Center, Strategic Laboratory | Claudio Tavares Sacchi, Claudia Regina Gonçalves, Erica Valessa Ramos Gomes, Karoline Rodrigues Campos |
| EPI_ISL_977476, EPI_ISL_977477 | Instituto Adolfo Lutz Central | Instituto Adolfo Lutz, Interdiciplinary Procedures Center, Strategic Laboratory | Claudio Tavares Sacchi, Claudia Regina Gonçalves, Erica Valessa Ramos Gomes, Karoline Rodrigues Campos |
| EPI_ISL_977478 | Instituto Adolfo Lutz - Regional de Presidente Prudente | Instituto Adolfo Lutz, Interdiciplinary Procedures Center, Strategic Laboratory | Claudio Tavares Sacchi, Claudia Regina Gonçalves, Erica Valessa Ramos Gomes, Karoline Rodrigues Campos |
| EPI_ISL_977479 | Lab Loc - Itapecerica da Serra | Instituto Adolfo Lutz, Interdiciplinary Procedures Center, Strategic Laboratory | Claudio Tavares Sacchi, Claudia Regina Gonçalves, Erica Valessa Ramos Gomes, Karoline Rodrigues Campos |
| EPI_ISL_977480, EPI_ISL_977481 | Instituto Adolfo Lutz - Regional de Presidente Prudente | Instituto Adolfo Lutz, Interdiciplinary Procedures Center, Strategic Laboratory | Claudio Tavares Sacchi, Claudia Regina Gonçalves, Erica Valessa Ramos Gomes, Karoline Rodrigues Campos |
| EPI_ISL_977482 | Instituto Adolfo Lutz - Regional de Aracatuba | Instituto Adolfo Lutz, Interdiciplinary Procedures Center, Strategic Laboratory | Claudio Tavares Sacchi, Claudia Regina Gonçalves, Erica Valessa Ramos Gomes, Karoline Rodrigues Campos |
| EPI_ISL_977483, EPI_ISL_977484 | Instituto Adolfo Lutz Central | Instituto Adolfo Lutz, Interdiciplinary Procedures Center, Strategic Laboratory | Claudio Tavares Sacchi, Claudia Regina Gonçalves, Erica Valessa Ramos Gomes, Karoline Rodrigues Campos |
| EPI_ISL_977485 | Instituto Adolfo Lutz - Regional de Presidente Prudente | Instituto Adolfo Lutz, Interdiciplinary Procedures Center, Strategic Laboratory | Claudio Tavares Sacchi, Claudia Regina Gonçalves, Erica Valessa Ramos Gomes, Karoline Rodrigues Campos |
| EPI_ISL_977486 | Instituto Adolfo Lutz - Regional de Santo Andre | Instituto Adolfo Lutz, Interdiciplinary Procedures Center, Strategic Laboratory | Claudio Tavares Sacchi, Claudia Regina Gonçalves, Erica Valessa Ramos Gomes, Karoline Rodrigues Campos |
| EPI_ISL_977487 | Instituto Adolfo Lutz Central | Instituto Adolfo Lutz, Interdiciplinary Procedures Center, Strategic Laboratory | Claudio Tavares Sacchi, Claudia Regina Gonçalves, Erica Valessa Ramos Gomes, Karoline Rodrigues Campos |
| EPI_ISL_977488 | Instituto Adolfo Lutz - Regional de Presidente Prudente | Instituto Adolfo Lutz, Interdiciplinary Procedures Center, Strategic Laboratory | Claudio Tavares Sacchi, Claudia Regina Gonçalves, Erica Valessa Ramos Gomes, Karoline Rodrigues Campos |
| EPI_ISL_977489 | UPA Dr. Akira Tada | Instituto Adolfo Lutz, Interdiciplinary Procedures Center, Strategic Laboratory | Claudio Tavares Sacchi, Claudia Regina Gonçalves, Erica Valessa Ramos Gomes, Karoline Rodrigues Campos |
| EPI_ISL_977490 | Hospital Municipal Guido Guida | Instituto Adolfo Lutz, Interdiciplinary Procedures Center, Strategic Laboratory | Claudio Tavares Sacchi, Claudia Regina Gonçalves, Erica Valessa Ramos Gomes, Karoline Rodrigues Campos |
| EPI_ISL_977500 | LATE - Laboratório de Técnicas Especiais - Hospital Israelita Albert Einstein | LATE - Laboratório de Técnicas Especiais - Hospital Israelita Albert Einstein | Deyvid Amgarten, Fernanda de Mello Malta, Raquel Riyuzo, Ana Paula Moreira Salles, Pedro Henrique Sebe Rodrigues, João Renato Rebello Pinho |
| EPI_ISL_978495, EPI_ISL_978496, EPI_ISL_978497, EPI_ISL_978498, EPI_ISL_978499, EPI_ISL_978500, EPI_ISL_978502, EPI_ISL_978503, EPI_ISL_978504, EPI_ISL_978505, EPI_ISL_978506, EPI_ISL_978507, EPI_ISL_978508, EPI_ISL_978509, EPI_ISL_978510, EPI_ISL_978511, EPI_ISL_978512, EPI_ISL_978513, EPI_ISL_978514, EPI_ISL_978515, EPI_ISL_978516, EPI_ISL_978517, EPI_ISL_978518, EPI_ISL_978519, EPI_ISL_978520, EPI_ISL_978521, EPI_ISL_978522, EPI_ISL_978523, EPI_ISL_978524, EPI_ISL_978525, EPI_ISL_978526, EPI_ISL_978527, EPI_ISL_978528, EPI_ISL_978529, EPI_ISL_978530, EPI_ISL_978531, EPI_ISL_978532, EPI_ISL_978533 | | | |
| see above | Central Public Health Laboratory - LACEN -Bahia, Salvador, Brazil | Central Public Health Laboratory - LACEN -Bahia, Salvador, Brazil | Stephane Tosta, Luciana Oliveira, Vanessa Nardy,Patrícia Cajado,Marcela Gómez, Breno Dominguez, Jaqueline Gomes, Vagner Fonseca,Marta Giovanetti,Luiz Alcantara, Felicidade Pereira, Arabela Leal |
| EPI_ISL_981383, EPI_ISL_981385, EPI_ISL_981387 | IAL Regional de Bauru | Instituto Adolfo Lutz, Interdiciplinary Procedures Center, Strategic Laboratory | Claudio Tavares Sacchi, Claudia Regina Gonçalves, Erica Valessa Ramos Gomes, Karoline Rodrigues Campos |
| EPI_ISL_983863, EPI_ISL_983864, EPI_ISL_983865, EPI_ISL_983867, EPI_ISL_983868 | Central Laboratory of Public Health of Rio Grande do Sul (Lacen-RS) | State Center for Health Surveillance of the Health Department of the State of Rio Grande do Sul (CEVS/SES-RS) | Aline Campos, Cynthia Molina, Lara Crescente, Leticia Garay, Ludmila Fiorenzano Baethgen, Richard Salvato, Tatiana Gregianini |
| EPI_ISL_984242 | Instituto Adolfo Lutz Central | Instituto Adolfo Lutz, Interdiciplinary Procedures Center, Strategic Laboratory | Claudio Tavares Sacchi, Claudia Regina Gonçalves, Erica Valessa Ramos Gomes, Karoline Rodrigues Campos |
| EPI_ISL_984243, EPI_ISL_984244 | Instituto Adolfo Lutz - Regional de Marilia | Instituto Adolfo Lutz, Interdiciplinary Procedures Center, Strategic Laboratory | Claudio Tavares Sacchi, Claudia Regina Gonçalves, Erica Valessa Ramos Gomes, Karoline Rodrigues Campos |
| EPI_ISL_984246 | Instituto Adolfo Lutz Central | Instituto Adolfo Lutz, Interdiciplinary Procedures Center, Strategic Laboratory | Claudio Tavares Sacchi, Claudia Regina Gonçalves, Erica Valessa Ramos Gomes, Karoline Rodrigues Campos |
| EPI_ISL_984247, EPI_ISL_984248, EPI_ISL_984249, EPI_ISL_984250, EPI_ISL_984251, EPI_ISL_984252, EPI_ISL_984253, EPI_ISL_984254, EPI_ISL_984255, EPI_ISL_984256, EPI_ISL_984257, EPI_ISL_984258, EPI_ISL_984259, EPI_ISL_984260, EPI_ISL_984261, EPI_ISL_984262 | | | |
| see above | IAL Regional de Marilia | Instituto Adolfo Lutz, Interdiciplinary Procedures Center, Strategic Laboratory | Claudio Tavares Sacchi, Claudia Regina Gonçalves, Erica Valessa Ramos Gomes, Karoline Rodrigues Campos |
| EPI_ISL_984263 | IAL Regional de Bauru | Instituto Adolfo Lutz, Interdiciplinary Procedures Center, Strategic Laboratory | Claudio Tavares Sacchi, Claudia Regina Gonçalves, Erica Valessa Ramos Gomes, Karoline Rodrigues Campos |
| EPI_ISL_984619, EPI_ISL_984620, EPI_ISL_984621 | Central Laboratory of Public Health of Rio Grande do Sul (Lacen-RS) | State Center for Health Surveillance of the Health Department of the State of Rio Grande do Sul (CEVS/SES-RS) | Aline Campos, Cynthia Molina, Lara Crescente, Leticia Garay, Ludmila Fiorenzano Baethgen, Richard Salvato, Tatiana Gregianini |
| EPI_ISL_985170 | Instituto Adolfo Lutz - Regional de Presidente Prudente | Instituto Adolfo Lutz, Interdiciplinary Procedures Center, Strategic Laboratory | Claudio Tavares Sacchi, Claudia Regina Gonçalves, Erica Valessa Ramos Gomes, Karoline Rodrigues Campos |
| EPI_ISL_985171, EPI_ISL_985172, EPI_ISL_985173, EPI_ISL_985174 | Instituto Adolfo Lutz - Regional de Taubate | Instituto Adolfo Lutz, Interdiciplinary Procedures Center, Strategic Laboratory | Claudio Tavares Sacchi, Claudia Regina Gonçalves, Erica Valessa Ramos Gomes, Karoline Rodrigues Campos |
| EPI_ISL_985175, EPI_ISL_985176, EPI_ISL_985177 | Instituto Adolfo Lutz Central | Instituto Adolfo Lutz, Interdiciplinary Procedures Center, Strategic Laboratory | Claudio Tavares Sacchi, Claudia Regina Gonçalves, Erica Valessa Ramos Gomes, Karoline Rodrigues Campos |
| EPI_ISL_985178 | Lab Loc - Itapecerica da Serra | Instituto Adolfo Lutz, Interdiciplinary Procedures Center, Strategic Laboratory | Claudio Tavares Sacchi, Claudia Regina Gonçalves, Erica Valessa Ramos Gomes, Karoline Rodrigues Campos |
| EPI_ISL_985303, EPI_ISL_985304, EPI_ISL_985305, EPI_ISL_985306, EPI_ISL_985307, EPI_ISL_985308, EPI_ISL_985309, EPI_ISL_985310, EPI_ISL_985311, EPI_ISL_985312, EPI_ISL_985313, EPI_ISL_985314, EPI_ISL_985315, EPI_ISL_985316, EPI_ISL_985317 | | | |
| see above | LACEN do Estado de Goias | Instituto Adolfo Lutz, Interdiciplinary Procedures Center, Strategic Laboratory | Claudio Tavares Sacchi, Claudia Regina Gonçalves, Erica Valessa Ramos Gomes, Karoline Rodrigues Campos |
| EPI_ISL_985318, EPI_ISL_985319 | LACEN de Santa Catarina | Instituto Adolfo Lutz, Interdiciplinary Procedures Center, Strategic Laboratory | Claudio Tavares Sacchi, Claudia Regina Gonçalves, Erica Valessa Ramos Gomes, Karoline Rodrigues Campos |

We gratefully acknowledge the following Authors from the Originating laboratories responsible for obtaining the specimens, as well as the Submitting laboratories where the genome data were generated and shared via GISAID, on which this research is based.

All Submitters of data may be contacted directly via www.gisaid.org

Authors are sorted alphabetically.

| Accession ID | Originating Laboratory | Submitting Laboratory | Authors |
| --- | --- | --- | --- |
| EPI_ISL_1858267, EPI_ISL_1858269, EPI_ISL_1858271, EPI_ISL_1858273, EPI_ISL_1858275, EPI_ISL_1858277, EPI_ISL_1858279, EPI_ISL_1858281, EPI_ISL_1858282, EPI_ISL_1858284, EPI_ISL_1858286, EPI_ISL_1858288, EPI_ISL_1858290, EPI_ISL_1858292, EPI_ISL_1858294, EPI_ISL_1858295, EPI_ISL_1858297, EPI_ISL_1858299, EPI_ISL_1858301, EPI_ISL_1858303, EPI_ISL_1858304, EPI_ISL_1858306, EPI_ISL_1858308, EPI_ISL_1858310, EPI_ISL_1858312, EPI_ISL_1858314, EPI_ISL_1858315, EPI_ISL_1858317, EPI_ISL_1858319, EPI_ISL_1858321, EPI_ISL_1858323, EPI_ISL_1858325, EPI_ISL_1858327, EPI_ISL_1858329, EPI_ISL_1858331, EPI_ISL_1858333, EPI_ISL_1858334, EPI_ISL_1858336, EPI_ISL_1858338, EPI_ISL_1858340, EPI_ISL_1858342, EPI_ISL_1858344, EPI_ISL_1858346, EPI_ISL_1858348, EPI_ISL_1858350, EPI_ISL_1858352, EPI_ISL_1858354, EPI_ISL_1858356, EPI_ISL_1858357, EPI_ISL_1858359, EPI_ISL_1858361, EPI_ISL_1858363, EPI_ISL_1858365, EPI_ISL_1858367, EPI_ISL_1858369, EPI_ISL_1858371, EPI_ISL_1858373, EPI_ISL_1858375, EPI_ISL_1858377, EPI_ISL_1858379, EPI_ISL_1858380, EPI_ISL_1858383, EPI_ISL_1858384, EPI_ISL_1858387, EPI_ISL_1858389, EPI_ISL_1858391, EPI_ISL_1858392, EPI_ISL_1858395, EPI_ISL_1858397, EPI_ISL_1858399, EPI_ISL_1858401, EPI_ISL_1858403, EPI_ISL_1858409, EPI_ISL_1858411, EPI_ISL_1858414, EPI_ISL_1858416, EPI_ISL_1858418, EPI_ISL_1858420, EPI_ISL_1858422, EPI_ISL_1858423, EPI_ISL_1858426, EPI_ISL_1858428, EPI_ISL_1858430, EPI_ISL_1858432, EPI_ISL_1858434, EPI_ISL_1858436, EPI_ISL_1858438, EPI_ISL_1858440, EPI_ISL_1858442, EPI_ISL_1858444, EPI_ISL_1858446, EPI_ISL_1858449, EPI_ISL_1858451, EPI_ISL_1858453, EPI_ISL_1858454, EPI_ISL_1858456, EPI_ISL_1858458, EPI_ISL_1858460, EPI_ISL_1858462, EPI_ISL_1858464, EPI_ISL_1858466, EPI_ISL_1858468, EPI_ISL_1858470, EPI_ISL_1858473, EPI_ISL_1858475, EPI_ISL_1858477, EPI_ISL_1858479, EPI_ISL_1858481, EPI_ISL_1858483, EPI_ISL_1858486, EPI_ISL_1858488, EPI_ISL_1858490, EPI_ISL_1858492, EPI_ISL_1858494, EPI_ISL_1858496, EPI_ISL_1858498, EPI_ISL_1858500, EPI_ISL_1858502, EPI_ISL_1858504, EPI_ISL_1858506, EPI_ISL_1858508, EPI_ISL_1858512, EPI_ISL_1858514, EPI_ISL_1858516, EPI_ISL_1858518, EPI_ISL_1858520, EPI_ISL_1858522, EPI_ISL_1858526, EPI_ISL_1858528, EPI_ISL_1858530, EPI_ISL_1858532, EPI_ISL_1858536, EPI_ISL_1858538, EPI_ISL_1858540, EPI_ISL_1858542, EPI_ISL_1858544, EPI_ISL_1858546, EPI_ISL_1858548, EPI_ISL_1858549, EPI_ISL_1858551, EPI_ISL_1858554, EPI_ISL_1858555, EPI_ISL_1858557, EPI_ISL_1858559, EPI_ISL_1858561, EPI_ISL_1858563, EPI_ISL_1858565, EPI_ISL_1858567, EPI_ISL_1858571, EPI_ISL_1858572, EPI_ISL_1858575, EPI_ISL_1858577, EPI_ISL_1858578, EPI_ISL_1858580, EPI_ISL_1858582, EPI_ISL_1858584, EPI_ISL_1858586, EPI_ISL_1858588, EPI_ISL_1858590, EPI_ISL_1858592, EPI_ISL_1858594, EPI_ISL_1858596, EPI_ISL_1858598, EPI_ISL_1858599, EPI_ISL_1858603, EPI_ISL_1858606, EPI_ISL_1858607, EPI_ISL_1858610, EPI_ISL_1858611, EPI_ISL_1858613, EPI_ISL_1858615, EPI_ISL_1858617, EPI_ISL_1858619, EPI_ISL_1858621, EPI_ISL_1858623, EPI_ISL_1858625, EPI_ISL_1858627, EPI_ISL_1858629, EPI_ISL_1858631, EPI_ISL_1858633, EPI_ISL_1858636, EPI_ISL_1858637, EPI_ISL_1858639, EPI_ISL_1858641, EPI_ISL_1858643, EPI_ISL_1858645, EPI_ISL_1858648, EPI_ISL_1858650, EPI_ISL_1858651, EPI_ISL_1858655, EPI_ISL_1858656, EPI_ISL_1858658, EPI_ISL_1858660, EPI_ISL_1858662, EPI_ISL_1858664, EPI_ISL_1858666, EPI_ISL_1858668, EPI_ISL_1858670, EPI_ISL_1858672, EPI_ISL_1858674, EPI_ISL_1858675, EPI_ISL_1858677, EPI_ISL_1858680, EPI_ISL_1858682, EPI_ISL_1858684, EPI_ISL_1858686, EPI_ISL_1858688, EPI_ISL_1858691, EPI_ISL_1858693, EPI_ISL_1858694, EPI_ISL_1858697, EPI_ISL_1858701, EPI_ISL_1858703, EPI_ISL_1858705, EPI_ISL_1858707, EPI_ISL_1858709, EPI_ISL_1858711, EPI_ISL_1858713, EPI_ISL_1858715, EPI_ISL_1858717, EPI_ISL_1858719, EPI_ISL_1858721, EPI_ISL_1858722, EPI_ISL_1858724, EPI_ISL_1858726, EPI_ISL_1858728, EPI_ISL_1858730, EPI_ISL_1858732, EPI_ISL_1858734, EPI_ISL_1858736, EPI_ISL_1858738, EPI_ISL_1858740, EPI_ISL_1858741, EPI_ISL_1858743, EPI_ISL_1858745, EPI_ISL_1858747, EPI_ISL_1858751, EPI_ISL_1858752, EPI_ISL_1858754, EPI_ISL_1858756 | | | |
| see above | Laboratorio Central Noel Nutels | Bioinformatics Laboratory / LNCC | Luiz G P de Almeida, Alessandra P Lamarca, Ronaldo da Silva F Jr, Liliane Cavalcante, Alexandra L Gerber, Ana Paula de C Guimarães, Douglas Terra Machado, Cassia Alves, Diana Mariani, Thais Felix Cruz, Mario Sergio Ribeiro, Silvia Carvalho, Flávio Dias da Silva, Marcio Henrique de Oliveira Garcia, Leandro Magalhães de Souza, Cristiane Gomes da Silva, Caio Luiz Pereira Ribeiro, Andréa Cony Cavalcanti, Claudia Maria Braga de Mello, Amilcar Tanuri, Ana Tereza R Vasconcelos |
| EPI_ISL_1858758, EPI_ISL_1858761, EPI_ISL_1858763, EPI_ISL_1858765, EPI_ISL_1858767, EPI_ISL_1858771, EPI_ISL_1858775, EPI_ISL_1858777, EPI_ISL_1858779, EPI_ISL_1858781, EPI_ISL_1858783, EPI_ISL_1858786, EPI_ISL_1858788, EPI_ISL_1858790, EPI_ISL_1858792, EPI_ISL_1858794, EPI_ISL_1858796, EPI_ISL_1858798, EPI_ISL_1858800, EPI_ISL_1858802, EPI_ISL_1858803, EPI_ISL_1858805, EPI_ISL_1858807, EPI_ISL_1858809, EPI_ISL_1858811, EPI_ISL_1858814, EPI_ISL_1858816, EPI_ISL_1858818, EPI_ISL_1858819, EPI_ISL_1858822, EPI_ISL_1858824, EPI_ISL_1858826, EPI_ISL_1858828, EPI_ISL_1858830, EPI_ISL_1858832, EPI_ISL_1858834, EPI_ISL_1858836, EPI_ISL_1858838, EPI_ISL_1858840, EPI_ISL_1858842, EPI_ISL_1858843, EPI_ISL_1858846, EPI_ISL_1858847, EPI_ISL_1858850, EPI_ISL_1858852, EPI_ISL_1858853, EPI_ISL_1858856, EPI_ISL_1858858, EPI_ISL_1858860, EPI_ISL_1858862, EPI_ISL_1858864, EPI_ISL_1858866, EPI_ISL_1858868, EPI_ISL_1858870, EPI_ISL_1858872, EPI_ISL_1858878, EPI_ISL_1858880, EPI_ISL_1858882, EPI_ISL_1858884, EPI_ISL_1858886, EPI_ISL_1858888, EPI_ISL_1858890, EPI_ISL_1858892, EPI_ISL_1858894, EPI_ISL_1858896, EPI_ISL_1858898, EPI_ISL_1858900, EPI_ISL_1858902, EPI_ISL_1858904, EPI_ISL_1858906, EPI_ISL_1858908, EPI_ISL_1858910, EPI_ISL_1858912, EPI_ISL_1858914, EPI_ISL_1858916, EPI_ISL_1858918, EPI_ISL_1858920, EPI_ISL_1858922, EPI_ISL_1858924, EPI_ISL_1858926, EPI_ISL_1858928, EPI_ISL_1858930, EPI_ISL_1858933, EPI_ISL_1858934, EPI_ISL_1858936, EPI_ISL_1858938, EPI_ISL_1858940, EPI_ISL_1858942, EPI_ISL_1858944, EPI_ISL_1858946, EPI_ISL_1858948, EPI_ISL_1858950, EPI_ISL_1858952, EPI_ISL_1858954, EPI_ISL_1858957, EPI_ISL_1858960, EPI_ISL_1858963, EPI_ISL_1858964, EPI_ISL_1858966, EPI_ISL_1858968, EPI_ISL_1858969, EPI_ISL_1858970, EPI_ISL_1858971, EPI_ISL_1858973, EPI_ISL_1858975, EPI_ISL_1858977, EPI_ISL_1858979, EPI_ISL_1858981, EPI_ISL_1858984, EPI_ISL_1858986, EPI_ISL_1858988, EPI_ISL_1858990, EPI_ISL_1858992, EPI_ISL_1858994, EPI_ISL_1858996, EPI_ISL_1858998, EPI_ISL_1859000, EPI_ISL_1859002, EPI_ISL_1859005, EPI_ISL_1859007, EPI_ISL_1859008 | | | |
| see above | Unidade de apoio ao diagnóstico da COVID - UNADIG | Bioinformatics Laboratory / LNCC | Luiz G P de Almeida, Alessandra P Lamarca, Ronaldo da Silva F Jr, Liliane Cavalcante, Alexandra L Gerber, Ana Paula de C Guimarães, Douglas Terra Machado, Cassia Alves, Diana Mariani, Thais Felix Cruz, Mario Sergio Ribeiro, Silvia Carvalho, Flávio Dias da Silva, Marcio Henrique de Oliveira Garcia, Leandro Magalhães de Souza, Cristiane Gomes da Silva, Caio Luiz Pereira Ribeiro, Andréa Cony Cavalcanti, Claudia Maria Braga de Mello, Amilcar Tanuri, Ana Tereza R Vasconcelos |
| EPI_ISL_1911958, EPI_ISL_1911959, EPI_ISL_1911960, EPI_ISL_1911961, EPI_ISL_1911962, EPI_ISL_1911964 | Instituto de Biotecnologia - UNESP-Botucatu-SP | Instituto de Biotecnologia - UNESP-Botucatu-SP | Fábio Sossai Possebon; Leila Sabrina Ullmann; Cecília Artico Banho; Cíntia Bittar; Guilherme Campos; Helena Lage Ferreira; Jorge A. Petroli Marchesi; Lívia Sacchetto; Maisa C. Pereira Parra; Marília Moraes; Maurício L. Nogueira; Paula Rahal; Paulo Inacio da Costa; João Pessoa Araújo Jr. |
| EPI_ISL_1966059, EPI_ISL_1966060, EPI_ISL_1966061, EPI_ISL_1966062, EPI_ISL_1966063, EPI_ISL_1966064, EPI_ISL_1966065, EPI_ISL_1966066, EPI_ISL_1966067, EPI_ISL_1966068, EPI_ISL_1966069 | | | |
| see above | VIGILANCIA EPIDEMIOLOGICA | Instituto Butantan / Mendelics | Instituto Butantan: Dimas Tadeu Covas, Sandra Coccuzzo Sampaio, Maria Carolina Elias, José Salvatore Leister Patané, Vincent Louis Viala, Antonio Jorge Martins, Ricardo Haddad, Claudia Renata dos Santos Barros, Elaine Cristina Marqueze, Raul Machado Neto, Debora Botequio Moretti, Jardelina de Souza Todao Bernardino, Loyze Paola Oliveira de Lima, Luiz Aurelio de Campos Crispin. Centro de Genômica Funcional da ESALQ: Luiz Lehmann Coutinho, Ricardo Augusto Brassaloti, Raquel de Lello Rocha Campos Cassano. NGS Soluções Genômicas: Pilar Drummond Sampaio Corrêa Mariani. FZEA-USP Pirassununga: Mirele Daiana Poleti, Jessika Cristina Chagas Lesbon, Elisangela Chicaroni Mattos, Heidge Fukumasu. USP-Botucatu: Rejane Maria Tommasini Grotto, Jayme A. Souza-Neto, Guilherme Targino Valente, Patricia Akemi Assato, Felipe Allan da Silva da Costa, Bianca Cechetto Carlos. Mendelics: Bibiana Santos, João Paulo Kitajima, Erika Freitas, David Schlesinger. Hemocentro Ribeirão Preto: Simone Kashima, Evandra Strazza Rodrigues, Svetoslav Nanev Slavov, Elaine Vieira dos Santos, Rafael dos Santos Bezerra, Luiz Carlos Junior de Alcantara, Marta Giovanetti, Vagner Fonseca, Flavia Aburjaile, Rodrigo Tocantins Calado. FAMERP-SJRP: Cecília Artico Banho, Lívia Sacchetto, Fábio Sossai Possebon, Leila Sabrina Ullmann, Cíntia Bittar, Guilherme Campos, Helena Lage Ferreira, Jorge A. Petroli Marchesi, Maisa C. Pereira Parra, Marília Moraes, Paula Rahal, Paulo Inacio da Costa, João Pessoa Araújo Jr., Maurício Lacerda Nogueira. Prefeitura de Sao Paulo: Melissa Palmieri. |
| EPI_ISL_1966070, EPI_ISL_1966071, EPI_ISL_1966072, EPI_ISL_1966073, EPI_ISL_1966074, EPI_ISL_1966075, EPI_ISL_1966076, EPI_ISL_1966077, EPI_ISL_1966078, EPI_ISL_1966079 | SECAO CENTRO DE DIAGNOSTICO SECEDI | Instituto Butantan / Mendelics | Instituto Butantan: Dimas Tadeu Covas, Sandra Coccuzzo Sampaio, Maria Carolina Elias, José Salvatore Leister Patané, Vincent Louis Viala, Antonio Jorge Martins, Ricardo Haddad, Claudia Renata dos Santos Barros, Elaine Cristina Marqueze, Raul Machado Neto, Debora Botequio Moretti, Jardelina de Souza Todao Bernardino, Loyze Paola Oliveira de Lima, Luiz Aurelio de Campos Crispin. Centro de Genômica Funcional da ESALQ: Luiz Lehmann Coutinho, Ricardo Augusto Brassaloti, Raquel de Lello Rocha Campos Cassano. NGS Soluções Genômicas: Pilar Drummond Sampaio Corrêa Mariani. FZEA-USP Pirassununga: Mirele Daiana Poleti, Jessika Cristina Chagas Lesbon, Elisangela Chicaroni Mattos, Heidge Fukumasu. USP-Botucatu: Rejane Maria Tommasini Grotto, Jayme A. Souza-Neto, Guilherme Targino Valente, Patricia Akemi Assato, Felipe Allan da Silva da Costa, Bianca Cechetto Carlos. Mendelics: Bibiana Santos, João Paulo Kitajima, Erika Freitas, David Schlesinger. Hemocentro Ribeirão Preto: Simone Kashima, Evandra Strazza Rodrigues, Svetoslav Nanev Slavov, Elaine Vieira dos Santos, Rafael dos Santos Bezerra, Luiz Carlos Junior de Alcantara, Marta Giovanetti, Vagner Fonseca, Flavia Aburjaile, Rodrigo Tocantins Calado. FAMERP-SJRP: Cecília Artico Banho, Lívia Sacchetto, Fábio Sossai Possebon, Leila Sabrina Ullmann, Cíntia Bittar, Guilherme Campos, Helena Lage Ferreira, Jorge A. Petroli Marchesi, Maisa C. Pereira Parra, Marília Moraes, Paula Rahal, Paulo Inacio da Costa, João Pessoa Araújo Jr., Maurício Lacerda Nogueira. Prefeitura de Sao Paulo: Melissa Palmieri. |
| EPI_ISL_1966080, EPI_ISL_1966081, EPI_ISL_1966082, EPI_ISL_1966083, EPI_ISL_1966084, EPI_ISL_1966085, EPI_ISL_1966086, EPI_ISL_1966087, EPI_ISL_1966088, EPI_ISL_1966089, EPI_ISL_1966090, EPI_ISL_1966091, EPI_ISL_1966092 | | | |
| see above | VIGILANCIA EPIDEMIOLOGICA | Instituto Butantan / Mendelics | Instituto Butantan: Dimas Tadeu Covas, Sandra Coccuzzo Sampaio, Maria Carolina Elias, José Salvatore Leister Patané, Vincent Louis Viala, Antonio Jorge Martins, Ricardo Haddad, Claudia Renata dos Santos Barros, Elaine Cristina Marqueze, Raul Machado Neto, Debora Botequio Moretti, Jardelina de Souza Todao Bernardino, Loyze Paola Oliveira de Lima, Luiz Aurelio de Campos Crispin. Centro de Genômica Funcional da ESALQ: Luiz Lehmann Coutinho, Ricardo Augusto Brassaloti, Raquel de Lello Rocha Campos Cassano. NGS Soluções Genômicas: Pilar Drummond Sampaio Corrêa Mariani. FZEA-USP Pirassununga: Mirele Daiana Poleti, Jessika Cristina Chagas Lesbon, Elisangela Chicaroni Mattos, Heidge Fukumasu. USP-Botucatu: Rejane Maria Tommasini Grotto, Jayme A. Souza-Neto, Guilherme Targino Valente, Patricia Akemi Assato, Felipe Allan da Silva da Costa, Bianca Cechetto Carlos. Mendelics: Bibiana Santos, João Paulo Kitajima, Erika Freitas, David Schlesinger. Hemocentro Ribeirão Preto: Simone Kashima, Evandra Strazza Rodrigues, Svetoslav Nanev Slavov, Elaine Vieira dos Santos, Rafael dos Santos Bezerra, Luiz Carlos Junior de Alcantara, Marta Giovanetti, Vagner Fonseca, Flavia Aburjaile, Rodrigo Tocantins Calado. FAMERP-SJRP: Cecília Artico Banho, Lívia Sacchetto, Fábio Sossai Possebon, Leila Sabrina Ullmann, Cíntia Bittar, Guilherme Campos, Helena Lage Ferreira, Jorge A. Petroli Marchesi, Maisa C. Pereira Parra, Marília Moraes, Paula Rahal, Paulo Inacio da Costa, João Pessoa Araújo Jr., Maurício Lacerda Nogueira. Prefeitura de Sao Paulo: Melissa Palmieri. |
| EPI_ISL_1966095, EPI_ISL_1966096, | AMBULATORIO MEDICO DE ESPECIALIDADES DE | Instituto Butantan / Mendelics | Instituto Butantan: Dimas Tadeu Covas, Sandra Coccuzzo Sampaio, Maria Carolina Elias, José Salvatore Leister Patané, Vincent Louis Viala, Antonio |

| | | | |
|---|---|---|---|
| EPI_ISL_1966097, EPI_ISL_1966098 | PERUIBE | | Jorge Martins, Ricardo Haddad, Claudia Renata dos Santos Barros, Elaine Cristina Marqueze, Raul Machado Neto, Debora Botequio Moretti, Jardelina de Souza Todao Bernardino, Loyze Paola Oliveira de Lima, Luiz Aurelio de Campos Crispin. Centro de Genômica Funcional da ESALQ: Luiz Lehmann Coutinho, Ricardo Augusto Brassaloti, Raquel de Lello Rocha Campos Cassano. NGS Soluções Genômicas: Pilar Drummond Sampaio Corrêa Mariani. FZEA-USP Pirassununga: Mirele Daiana Poleti, Jessika Cristina Chagas Lesbon, Elisangela Chicaroni Mattos, Heidge Fukumasu. USP-Botucatu: Rejane Maria Tommasini Grotto, Jayme A. Souza-Neto, Guilherme Targino Valente, Patricia Akemi Assato, Felipe Allan da Silva da Costa, Bianca Cechetto Carlos. Mendelics: Bibiana Santos, João Paulo Kitajima, Erika Freitas, David Schlesinger. Hemocentro Ribeirão Preto: Simone Kashima, Evandra Strazza Rodrigues, Svetoslav Nanev Slavov, Elaine Vieira dos Santos, Rafael dos Santos Bezerra, Luiz Carlos Junior de Alcantara, Marta Giovanetti, Vagner Fonseca, Flavia Aburjaile, Rodrigo Tocantins Calado. FAMERP-SJRP: Cecília Artico Banho, Lívia Sacchetto, Fábio Sossai Possebon, Leila Sabrina Ullmann, Cíntia Bittar, Guilherme Campos, Helena Lage Ferreira, Jorge A. Petroli Marchesi, Maisa C. Pereira Parra, Marília Moraes, Paula Rahal, Paulo Inacio da Costa, João Pessoa Araújo Jr., Maurício Lacerda Nogueira. Prefeitura de Sao Paulo: Melissa Palmieri. |
| EPI_ISL_1966099, EPI_ISL_1966101 | SECAO CENTRO DE DIAGNOSTICO SECEDI | Instituto Butantan / Mendelics | Instituto Butantan: Dimas Tadeu Covas, Sandra Coccuzzo Sampaio, Maria Carolina Elias, José Salvatore Leister Patané, Vincent Louis Viala, Antonio Jorge Martins, Ricardo Haddad, Claudia Renata dos Santos Barros, Elaine Cristina Marqueze, Raul Machado Neto, Debora Botequio Moretti, Jardelina de Souza Todao Bernardino, Loyze Paola Oliveira de Lima, Luiz Aurelio de Campos Crispin. Centro de Genômica Funcional da ESALQ: Luiz Lehmann Coutinho, Ricardo Augusto Brassaloti, Raquel de Lello Rocha Campos Cassano. NGS Soluções Genômicas: Pilar Drummond Sampaio Corrêa Mariani. FZEA-USP Pirassununga: Mirele Daiana Poleti, Jessika Cristina Chagas Lesbon, Elisangela Chicaroni Mattos, Heidge Fukumasu. USP-Botucatu: Rejane Maria Tommasini Grotto, Jayme A. Souza-Neto, Guilherme Targino Valente, Patricia Akemi Assato, Felipe Allan da Silva da Costa, Bianca Cechetto Carlos. Mendelics: Bibiana Santos, João Paulo Kitajima, Erika Freitas, David Schlesinger. Hemocentro Ribeirão Preto: Simone Kashima, Evandra Strazza Rodrigues, Svetoslav Nanev Slavov, Elaine Vieira dos Santos, Rafael dos Santos Bezerra, Luiz Carlos Junior de Alcantara, Marta Giovanetti, Vagner Fonseca, Flavia Aburjaile, Rodrigo Tocantins Calado. FAMERP-SJRP: Cecília Artico Banho, Lívia Sacchetto, Fábio Sossai Possebon, Leila Sabrina Ullmann, Cíntia Bittar, Guilherme Campos, Helena Lage Ferreira, Jorge A. Petroli Marchesi, Maisa C. Pereira Parra, Marília Moraes, Paula Rahal, Paulo Inacio da Costa, João Pessoa Araújo Jr., Maurício Lacerda Nogueira. Prefeitura de Sao Paulo: Melissa Palmieri. |
| EPI_ISL_1966102 | USF TRES PONTES | Instituto Butantan / Mendelics | Instituto Butantan: Dimas Tadeu Covas, Sandra Coccuzzo Sampaio, Maria Carolina Elias, José Salvatore Leister Patané, Vincent Louis Viala, Antonio Jorge Martins, Ricardo Haddad, Claudia Renata dos Santos Barros, Elaine Cristina Marqueze, Raul Machado Neto, Debora Botequio Moretti, Jardelina de Souza Todao Bernardino, Loyze Paola Oliveira de Lima, Luiz Aurelio de Campos Crispin. Centro de Genômica Funcional da ESALQ: Luiz Lehmann Coutinho, Ricardo Augusto Brassaloti, Raquel de Lello Rocha Campos Cassano. NGS Soluções Genômicas: Pilar Drummond Sampaio Corrêa Mariani. FZEA-USP Pirassununga: Mirele Daiana Poleti, Jessika Cristina Chagas Lesbon, Elisangela Chicaroni Mattos, Heidge Fukumasu. USP-Botucatu: Rejane Maria Tommasini Grotto, Jayme A. Souza-Neto, Guilherme Targino Valente, Patricia Akemi Assato, Felipe Allan da Silva da Costa, Bianca Cechetto Carlos. Mendelics: Bibiana Santos, João Paulo Kitajima, Erika Freitas, David Schlesinger. Hemocentro Ribeirão Preto: Simone Kashima, Evandra Strazza Rodrigues, Svetoslav Nanev Slavov, Elaine Vieira dos Santos, Rafael dos Santos Bezerra, Luiz Carlos Junior de Alcantara, Marta Giovanetti, Vagner Fonseca, Flavia Aburjaile, Rodrigo Tocantins Calado. FAMERP-SJRP: Cecília Artico Banho, Lívia Sacchetto, Fábio Sossai Possebon, Leila Sabrina Ullmann, Cíntia Bittar, Guilherme Campos, Helena Lage Ferreira, Jorge A. Petroli Marchesi, Maisa C. Pereira Parra, Marília Moraes, Paula Rahal, Paulo Inacio da Costa, João Pessoa Araújo Jr., Maurício Lacerda Nogueira. Prefeitura de Sao Paulo: Melissa Palmieri. |
| EPI_ISL_1966103 | SMS SECRETARIA MUNICIPAL DE SAUDE DE BOITUVA | Instituto Butantan / Mendelics | Instituto Butantan: Dimas Tadeu Covas, Sandra Coccuzzo Sampaio, Maria Carolina Elias, José Salvatore Leister Patané, Vincent Louis Viala, Antonio Jorge Martins, Ricardo Haddad, Claudia Renata dos Santos Barros, Elaine Cristina Marqueze, Raul Machado Neto, Debora Botequio Moretti, Jardelina de Souza Todao Bernardino, Loyze Paola Oliveira de Lima, Luiz Aurelio de Campos Crispin. Centro de Genômica Funcional da ESALQ: Luiz Lehmann Coutinho, Ricardo Augusto Brassaloti, Raquel de Lello Rocha Campos Cassano. NGS Soluções Genômicas: Pilar Drummond Sampaio Corrêa Mariani. FZEA-USP Pirassununga: Mirele Daiana Poleti, Jessika Cristina Chagas Lesbon, Elisangela Chicaroni Mattos, Heidge Fukumasu. USP-Botucatu: Rejane Maria Tommasini Grotto, Jayme A. Souza-Neto, Guilherme Targino Valente, Patricia Akemi Assato, Felipe Allan da Silva da Costa, Bianca Cechetto Carlos. Mendelics: Bibiana Santos, João Paulo Kitajima, Erika Freitas, David Schlesinger. Hemocentro Ribeirão Preto: Simone Kashima, Evandra Strazza Rodrigues, Svetoslav Nanev Slavov, Elaine Vieira dos Santos, Rafael dos Santos Bezerra, Luiz Carlos Junior de Alcantara, Marta Giovanetti, Vagner Fonseca, Flavia Aburjaile, Rodrigo Tocantins Calado. FAMERP-SJRP: Cecília Artico Banho, Lívia Sacchetto, Fábio Sossai Possebon, Leila Sabrina Ullmann, Cíntia Bittar, Guilherme Campos, Helena Lage Ferreira, Jorge A. Petroli Marchesi, Maisa C. Pereira Parra, Marília Moraes, Paula Rahal, Paulo Inacio da Costa, João Pessoa Araújo Jr., Maurício Lacerda Nogueira. Prefeitura de Sao Paulo: Melissa Palmieri. |
| EPI_ISL_1966104 | POLICLINICA COVID 19 ITAPETININGA | Instituto Butantan / Mendelics | Instituto Butantan: Dimas Tadeu Covas, Sandra Coccuzzo Sampaio, Maria Carolina Elias, José Salvatore Leister Patané, Vincent Louis Viala, Antonio Jorge Martins, Ricardo Haddad, Claudia Renata dos Santos Barros, Elaine Cristina Marqueze, Raul Machado Neto, Debora Botequio Moretti, Jardelina de Souza Todao Bernardino, Loyze Paola Oliveira de Lima, Luiz Aurelio de Campos Crispin. Centro de Genômica Funcional da ESALQ: Luiz Lehmann Coutinho, Ricardo Augusto Brassaloti, Raquel de Lello Rocha Campos Cassano. NGS Soluções Genômicas: Pilar Drummond Sampaio Corrêa Mariani. FZEA-USP Pirassununga: Mirele Daiana Poleti, Jessika Cristina Chagas Lesbon, Elisangela Chicaroni Mattos, Heidge Fukumasu. USP-Botucatu: Rejane Maria Tommasini Grotto, Jayme A. Souza-Neto, Guilherme Targino Valente, Patricia Akemi Assato, Felipe Allan da Silva da Costa, Bianca Cechetto Carlos. Mendelics: Bibiana Santos, João Paulo Kitajima, Erika Freitas, David Schlesinger. Hemocentro Ribeirão Preto: Simone Kashima, Evandra Strazza Rodrigues, Svetoslav Nanev Slavov, Elaine Vieira dos Santos, Rafael dos Santos Bezerra, Luiz Carlos Junior de Alcantara, Marta Giovanetti, Vagner Fonseca, Flavia Aburjaile, Rodrigo Tocantins Calado. FAMERP-SJRP: Cecília Artico Banho, Lívia Sacchetto, Fábio Sossai Possebon, Leila Sabrina Ullmann, Cíntia Bittar, Guilherme Campos, Helena Lage Ferreira, Jorge A. Petroli Marchesi, Maisa C. Pereira Parra, Marília Moraes, Paula Rahal, Paulo Inacio da Costa, João Pessoa Araújo Jr., Maurício Lacerda Nogueira. Prefeitura de Sao Paulo: Melissa Palmieri. |
| EPI_ISL_1966107 | SECRETARIA MUNICIPAL DE SAUDE SOROCABA | Instituto Butantan / Mendelics | Instituto Butantan: Dimas Tadeu Covas, Sandra Coccuzzo Sampaio, Maria Carolina Elias, José Salvatore Leister Patané, Vincent Louis Viala, Antonio Jorge Martins, Ricardo Haddad, Claudia Renata dos Santos Barros, Elaine Cristina Marqueze, Raul Machado Neto, Debora Botequio Moretti, Jardelina de Souza Todao Bernardino, Loyze Paola Oliveira de Lima, Luiz Aurelio de Campos Crispin. Centro de Genômica Funcional da ESALQ: Luiz Lehmann Coutinho, Ricardo Augusto Brassaloti, Raquel de Lello Rocha Campos Cassano. NGS Soluções Genômicas: Pilar Drummond Sampaio Corrêa Mariani. FZEA-USP Pirassununga: Mirele Daiana Poleti, Jessika Cristina Chagas Lesbon, Elisangela Chicaroni Mattos, Heidge Fukumasu. USP-Botucatu: Rejane Maria Tommasini Grotto, Jayme A. Souza-Neto, Guilherme Targino Valente, Patricia Akemi Assato, Felipe Allan da Silva da Costa, Bianca Cechetto Carlos. Mendelics: Bibiana Santos, João Paulo Kitajima, Erika Freitas, David Schlesinger. Hemocentro Ribeirão Preto: Simone Kashima, Evandra Strazza Rodrigues, Svetoslav Nanev Slavov, Elaine Vieira dos Santos, Rafael dos Santos Bezerra, Luiz Carlos Junior de Alcantara, Marta Giovanetti, Vagner Fonseca, Flavia Aburjaile, Rodrigo Tocantins Calado. FAMERP-SJRP: Cecília Artico Banho, Lívia Sacchetto, Fábio Sossai Possebon, Leila Sabrina Ullmann, Cíntia Bittar, Guilherme Campos, Helena Lage Ferreira, Jorge A. Petroli Marchesi, Maisa C. Pereira Parra, Marília Moraes, Paula Rahal, Paulo Inacio da Costa, João Pessoa Araújo Jr., Maurício Lacerda Nogueira. Prefeitura de Sao Paulo: Melissa Palmieri. |
| EPI_ISL_1966109, EPI_ISL_1966110 | CENTRO DE SAUDE II DR GABRIEL MESQUITA VARGEM GDE DO SUL | Instituto Butantan / Mendelics | Instituto Butantan: Dimas Tadeu Covas, Sandra Coccuzzo Sampaio, Maria Carolina Elias, José Salvatore Leister Patané, Vincent Louis Viala, Antonio Jorge Martins, Ricardo Haddad, Claudia Renata dos Santos Barros, Elaine Cristina Marqueze, Raul Machado Neto, Debora Botequio Moretti, Jardelina de Souza Todao Bernardino, Loyze Paola Oliveira de Lima, Luiz Aurelio de Campos Crispin. Centro de Genômica Funcional da ESALQ: Luiz Lehmann Coutinho, Ricardo Augusto Brassaloti, Raquel de Lello Rocha Campos Cassano. NGS Soluções Genômicas: Pilar Drummond Sampaio Corrêa Mariani. FZEA-USP Pirassununga: Mirele Daiana Poleti, Jessika Cristina Chagas Lesbon, Elisangela Chicaroni Mattos, Heidge Fukumasu. USP-Botucatu: Rejane Maria Tommasini Grotto, Jayme A. Souza-Neto, Guilherme Targino Valente, Patricia Akemi Assato, Felipe Allan da Silva da Costa, Bianca Cechetto Carlos. Mendelics: Bibiana Santos, João Paulo Kitajima, Erika Freitas, David Schlesinger. Hemocentro Ribeirão Preto: Simone Kashima, Evandra Strazza Rodrigues, Svetoslav Nanev Slavov, Elaine Vieira dos Santos, Rafael dos Santos Bezerra, Luiz Carlos Junior de Alcantara, Marta Giovanetti, Vagner Fonseca, Flavia Aburjaile, Rodrigo Tocantins Calado. FAMERP-SJRP: Cecília Artico Banho, Lívia Sacchetto, Fábio Sossai Possebon, Leila Sabrina Ullmann, Cíntia Bittar, Guilherme Campos, Helena Lage Ferreira, Jorge A. Petroli Marchesi, Maisa C. Pereira Parra, Marília Moraes, Paula Rahal, Paulo Inacio da Costa, João Pessoa Araújo Jr., Maurício Lacerda Nogueira. Prefeitura de Sao Paulo: Melissa Palmieri. |
| EPI_ISL_1966112 | SAUDE COLETIVA CAPAO BONITO | Instituto Butantan / Mendelics | Instituto Butantan: Dimas Tadeu Covas, Sandra Coccuzzo Sampaio, Maria Carolina Elias, José Salvatore Leister Patané, Vincent Louis Viala, Antonio Jorge Martins, Ricardo Haddad, Claudia Renata dos Santos Barros, Elaine Cristina Marqueze, Raul Machado Neto, Debora Botequio Moretti, Jardelina de Souza Todao Bernardino, Loyze Paola Oliveira de Lima, Luiz Aurelio de Campos Crispin. Centro de Genômica Funcional da ESALQ: Luiz Lehmann Coutinho, Ricardo Augusto Brassaloti, Raquel de Lello Rocha Campos Cassano. NGS Soluções Genômicas: Pilar Drummond Sampaio Corrêa Mariani. FZEA-USP Pirassununga: Mirele Daiana Poleti, Jessika Cristina Chagas Lesbon, Elisangela Chicaroni Mattos, Heidge Fukumasu. USP-Botucatu: Rejane Maria Tommasini Grotto, Jayme A. Souza-Neto, Guilherme Targino Valente, Patricia Akemi Assato, Felipe Allan da Silva da Costa, Bianca Cechetto Carlos. Mendelics: Bibiana Santos, João Paulo Kitajima, Erika Freitas, David Schlesinger. Hemocentro Ribeirão Preto: Simone Kashima, Evandra Strazza Rodrigues, Svetoslav Nanev Slavov, Elaine Vieira dos Santos, Rafael dos Santos Bezerra, Luiz Carlos Junior de Alcantara, Marta Giovanetti, Vagner |

| | | | |
|---|---|---|---|
| | | | Fonseca, Flavia Aburjaile, Rodrigo Tocantins Calado. FAMERP-SJRP: Cecília Artico Banho, Lívia Sacchetto, Fábio Sossai Possebon, Leila Sabrina Ullmann, Cíntia Bittar, Guilherme Campos, Helena Lage Ferreira, Jorge A. Petroli Marchesi, Maisa C. Pereira Parra, Marília Moraes, Paula Rahal, Paulo Inacio da Costa, João Pessoa Araújo Jr., Maurício Lacerda Nogueira. Prefeitura de Sao Paulo: Melissa Palmieri. |
| EPI_ISL_1966113, EPI_ISL_1966114, EPI_ISL_1966115, EPI_ISL_1966116, EPI_ISL_1966117 | CS II EGIDIO BRUNHARA MORRO AGUDO | Instituto Butantan / Mendelics | Instituto Butantan: Dimas Tadeu Covas, Sandra Coccuzzo Sampaio, Maria Carolina Elias, José Salvatore Leister Patané, Vincent Louis Viala, Antonio Jorge Martins, Ricardo Haddad, Claudia Renata dos Santos Barros, Elaine Cristina Marqueze, Raul Machado Neto, Debora Botequio Moretti, Jardelina de Souza Todao Bernardino, Loyze Paola Oliveira de Lima, Luiz Aurelio de Campos Crispin. Centro de Genômica Funcional da ESALQ: Luiz Lehmann Coutinho, Ricardo Augusto Brassaloti, Raquel de Lello Rocha Campos Cassano. NGS Soluções Genômicas: Pilar Drummond Sampaio Corrêa Mariani. FZEA-USP Pirassununga: Mirele Daiana Poleti, Jessika Cristina Chagas Lesbon, Elisangela Chicaroni Mattos, Heidge Fukumasu. USP-Botucatu: Rejane Maria Tommasini Grotto, Jayme A. Souza-Neto, Guilherme Targino Valente, Patricia Akemi Assato, Felipe Allan da Silva da Costa, Bianca Cechetto Carlos. Mendelics: Bibiana Santos, João Paulo Kitajima, Erika Freitas, David Schlesinger. Hemocentro Ribeirão Preto: Simone Kashima, Evandra Strazza Rodrigues, Svetoslav Nanev Slavov, Elaine Vieira dos Santos, Rafael dos Santos Bezerra, Luiz Carlos Junior de Alcantara, Marta Giovanetti, Vagner Fonseca, Flavia Aburjaile, Rodrigo Tocantins Calado. FAMERP-SJRP: Cecília Artico Banho, Lívia Sacchetto, Fábio Sossai Possebon, Leila Sabrina Ullmann, Cíntia Bittar, Guilherme Campos, Helena Lage Ferreira, Jorge A. Petroli Marchesi, Maisa C. Pereira Parra, Marília Moraes, Paula Rahal, Paulo Inacio da Costa, João Pessoa Araújo Jr., Maurício Lacerda Nogueira. Prefeitura de Sao Paulo: Melissa Palmieri. |
| EPI_ISL_1966118, EPI_ISL_1966119, EPI_ISL_1966120, EPI_ISL_1966121, EPI_ISL_1966122, EPI_ISL_1966123 | CENTRO DE REFERENCIA DO IDOSO DR HUMBERTO MENDES DE CARVALHO | Instituto Butantan / Mendelics | Instituto Butantan: Dimas Tadeu Covas, Sandra Coccuzzo Sampaio, Maria Carolina Elias, José Salvatore Leister Patané, Vincent Louis Viala, Antonio Jorge Martins, Ricardo Haddad, Claudia Renata dos Santos Barros, Elaine Cristina Marqueze, Raul Machado Neto, Debora Botequio Moretti, Jardelina de Souza Todao Bernardino, Loyze Paola Oliveira de Lima, Luiz Aurelio de Campos Crispin. Centro de Genômica Funcional da ESALQ: Luiz Lehmann Coutinho, Ricardo Augusto Brassaloti, Raquel de Lello Rocha Campos Cassano. NGS Soluções Genômicas: Pilar Drummond Sampaio Corrêa Mariani. FZEA-USP Pirassununga: Mirele Daiana Poleti, Jessika Cristina Chagas Lesbon, Elisangela Chicaroni Mattos, Heidge Fukumasu. USP-Botucatu: Rejane Maria Tommasini Grotto, Jayme A. Souza-Neto, Guilherme Targino Valente, Patricia Akemi Assato, Felipe Allan da Silva da Costa, Bianca Cechetto Carlos. Mendelics: Bibiana Santos, João Paulo Kitajima, Erika Freitas, David Schlesinger. Hemocentro Ribeirão Preto: Simone Kashima, Evandra Strazza Rodrigues, Svetoslav Nanev Slavov, Elaine Vieira dos Santos, Rafael dos Santos Bezerra, Luiz Carlos Junior de Alcantara, Marta Giovanetti, Vagner Fonseca, Flavia Aburjaile, Rodrigo Tocantins Calado. FAMERP-SJRP: Cecília Artico Banho, Lívia Sacchetto, Fábio Sossai Possebon, Leila Sabrina Ullmann, Cíntia Bittar, Guilherme Campos, Helena Lage Ferreira, Jorge A. Petroli Marchesi, Maisa C. Pereira Parra, Marília Moraes, Paula Rahal, Paulo Inacio da Costa, João Pessoa Araújo Jr., Maurício Lacerda Nogueira. Prefeitura de Sao Paulo: Melissa Palmieri. |
| EPI_ISL_1966124, EPI_ISL_1966125 | SECRETARIA MUNICIPAL DE SAUDE SOROCABA | Instituto Butantan / Mendelics | Instituto Butantan: Dimas Tadeu Covas, Sandra Coccuzzo Sampaio, Maria Carolina Elias, José Salvatore Leister Patané, Vincent Louis Viala, Antonio Jorge Martins, Ricardo Haddad, Claudia Renata dos Santos Barros, Elaine Cristina Marqueze, Raul Machado Neto, Debora Botequio Moretti, Jardelina de Souza Todao Bernardino, Loyze Paola Oliveira de Lima, Luiz Aurelio de Campos Crispin. Centro de Genômica Funcional da ESALQ: Luiz Lehmann Coutinho, Ricardo Augusto Brassaloti, Raquel de Lello Rocha Campos Cassano. NGS Soluções Genômicas: Pilar Drummond Sampaio Corrêa Mariani. FZEA-USP Pirassununga: Mirele Daiana Poleti, Jessika Cristina Chagas Lesbon, Elisangela Chicaroni Mattos, Heidge Fukumasu. USP-Botucatu: Rejane Maria Tommasini Grotto, Jayme A. Souza-Neto, Guilherme Targino Valente, Patricia Akemi Assato, Felipe Allan da Silva da Costa, Bianca Cechetto Carlos. Mendelics: Bibiana Santos, João Paulo Kitajima, Erika Freitas, David Schlesinger. Hemocentro Ribeirão Preto: Simone Kashima, Evandra Strazza Rodrigues, Svetoslav Nanev Slavov, Elaine Vieira dos Santos, Rafael dos Santos Bezerra, Luiz Carlos Junior de Alcantara, Marta Giovanetti, Vagner Fonseca, Flavia Aburjaile, Rodrigo Tocantins Calado. FAMERP-SJRP: Cecília Artico Banho, Lívia Sacchetto, Fábio Sossai Possebon, Leila Sabrina Ullmann, Cíntia Bittar, Guilherme Campos, Helena Lage Ferreira, Jorge A. Petroli Marchesi, Maisa C. Pereira Parra, Marília Moraes, Paula Rahal, Paulo Inacio da Costa, João Pessoa Araújo Jr., Maurício Lacerda Nogueira. Prefeitura de Sao Paulo: Melissa Palmieri. |
| EPI_ISL_1966126, EPI_ISL_1966127, EPI_ISL_1966128 | SAE SERVICO DE ATENDIMENTO ESPECIALIZADO | Instituto Butantan / Mendelics | Instituto Butantan: Dimas Tadeu Covas, Sandra Coccuzzo Sampaio, Maria Carolina Elias, José Salvatore Leister Patané, Vincent Louis Viala, Antonio Jorge Martins, Ricardo Haddad, Claudia Renata dos Santos Barros, Elaine Cristina Marqueze, Raul Machado Neto, Debora Botequio Moretti, Jardelina de Souza Todao Bernardino, Loyze Paola Oliveira de Lima, Luiz Aurelio de Campos Crispin. Centro de Genômica Funcional da ESALQ: Luiz Lehmann Coutinho, Ricardo Augusto Brassaloti, Raquel de Lello Rocha Campos Cassano. NGS Soluções Genômicas: Pilar Drummond Sampaio Corrêa Mariani. FZEA-USP Pirassununga: Mirele Daiana Poleti, Jessika Cristina Chagas Lesbon, Elisangela Chicaroni Mattos, Heidge Fukumasu. USP-Botucatu: Rejane Maria Tommasini Grotto, Jayme A. Souza-Neto, Guilherme Targino Valente, Patricia Akemi Assato, Felipe Allan da Silva da Costa, Bianca Cechetto Carlos. Mendelics: Bibiana Santos, João Paulo Kitajima, Erika Freitas, David Schlesinger. Hemocentro Ribeirão Preto: Simone Kashima, Evandra Strazza Rodrigues, Svetoslav Nanev Slavov, Elaine Vieira dos Santos, Rafael dos Santos Bezerra, Luiz Carlos Junior de Alcantara, Marta Giovanetti, Vagner Fonseca, Flavia Aburjaile, Rodrigo Tocantins Calado. FAMERP-SJRP: Cecília Artico Banho, Lívia Sacchetto, Fábio Sossai Possebon, Leila Sabrina Ullmann, Cíntia Bittar, Guilherme Campos, Helena Lage Ferreira, Jorge A. Petroli Marchesi, Maisa C. Pereira Parra, Marília Moraes, Paula Rahal, Paulo Inacio da Costa, João Pessoa Araújo Jr., Maurício Lacerda Nogueira. Prefeitura de Sao Paulo: Melissa Palmieri. |
| EPI_ISL_1966129 | PRONTO SOCORRO MUNICIPAL DE TATUI | Instituto Butantan / Mendelics | Instituto Butantan: Dimas Tadeu Covas, Sandra Coccuzzo Sampaio, Maria Carolina Elias, José Salvatore Leister Patané, Vincent Louis Viala, Antonio Jorge Martins, Ricardo Haddad, Claudia Renata dos Santos Barros, Elaine Cristina Marqueze, Raul Machado Neto, Debora Botequio Moretti, Jardelina de Souza Todao Bernardino, Loyze Paola Oliveira de Lima, Luiz Aurelio de Campos Crispin. Centro de Genômica Funcional da ESALQ: Luiz Lehmann Coutinho, Ricardo Augusto Brassaloti, Raquel de Lello Rocha Campos Cassano. NGS Soluções Genômicas: Pilar Drummond Sampaio Corrêa Mariani. FZEA-USP Pirassununga: Mirele Daiana Poleti, Jessika Cristina Chagas Lesbon, Elisangela Chicaroni Mattos, Heidge Fukumasu. USP-Botucatu: Rejane Maria Tommasini Grotto, Jayme A. Souza-Neto, Guilherme Targino Valente, Patricia Akemi Assato, Felipe Allan da Silva da Costa, Bianca Cechetto Carlos. Mendelics: Bibiana Santos, João Paulo Kitajima, Erika Freitas, David Schlesinger. Hemocentro Ribeirão Preto: Simone Kashima, Evandra Strazza Rodrigues, Svetoslav Nanev Slavov, Elaine Vieira dos Santos, Rafael dos Santos Bezerra, Luiz Carlos Junior de Alcantara, Marta Giovanetti, Vagner Fonseca, Flavia Aburjaile, Rodrigo Tocantins Calado. FAMERP-SJRP: Cecília Artico Banho, Lívia Sacchetto, Fábio Sossai Possebon, Leila Sabrina Ullmann, Cíntia Bittar, Guilherme Campos, Helena Lage Ferreira, Jorge A. Petroli Marchesi, Maisa C. Pereira Parra, Marília Moraes, Paula Rahal, Paulo Inacio da Costa, João Pessoa Araújo Jr., Maurício Lacerda Nogueira. Prefeitura de Sao Paulo: Melissa Palmieri. |
| EPI_ISL_1966131 | CENTRO DE SAUDE II MAIRINQUE MAIRINQUE | Instituto Butantan / Mendelics | Instituto Butantan: Dimas Tadeu Covas, Sandra Coccuzzo Sampaio, Maria Carolina Elias, José Salvatore Leister Patané, Vincent Louis Viala, Antonio Jorge Martins, Ricardo Haddad, Claudia Renata dos Santos Barros, Elaine Cristina Marqueze, Raul Machado Neto, Debora Botequio Moretti, Jardelina de Souza Todao Bernardino, Loyze Paola Oliveira de Lima, Luiz Aurelio de Campos Crispin. Centro de Genômica Funcional da ESALQ: Luiz Lehmann Coutinho, Ricardo Augusto Brassaloti, Raquel de Lello Rocha Campos Cassano. NGS Soluções Genômicas: Pilar Drummond Sampaio Corrêa Mariani. FZEA-USP Pirassununga: Mirele Daiana Poleti, Jessika Cristina Chagas Lesbon, Elisangela Chicaroni Mattos, Heidge Fukumasu. USP-Botucatu: Rejane Maria Tommasini Grotto, Jayme A. Souza-Neto, Guilherme Targino Valente, Patricia Akemi Assato, Felipe Allan da Silva da Costa, Bianca Cechetto Carlos. Mendelics: Bibiana Santos, João Paulo Kitajima, Erika Freitas, David Schlesinger. Hemocentro Ribeirão Preto: Simone Kashima, Evandra Strazza Rodrigues, Svetoslav Nanev Slavov, Elaine Vieira dos Santos, Rafael dos Santos Bezerra, Luiz Carlos Junior de Alcantara, Marta Giovanetti, Vagner Fonseca, Flavia Aburjaile, Rodrigo Tocantins Calado. FAMERP-SJRP: Cecília Artico Banho, Lívia Sacchetto, Fábio Sossai Possebon, Leila Sabrina Ullmann, Cíntia Bittar, Guilherme Campos, Helena Lage Ferreira, Jorge A. Petroli Marchesi, Maisa C. Pereira Parra, Marília Moraes, Paula Rahal, Paulo Inacio da Costa, João Pessoa Araújo Jr., Maurício Lacerda Nogueira. Prefeitura de Sao Paulo: Melissa Palmieri. |
| EPI_ISL_1966132 | UNIDADE MISTA DE TAIACU | Instituto Butantan / Mendelics | Instituto Butantan: Dimas Tadeu Covas, Sandra Coccuzzo Sampaio, Maria Carolina Elias, José Salvatore Leister Patané, Vincent Louis Viala, Antonio Jorge Martins, Ricardo Haddad, Claudia Renata dos Santos Barros, Elaine Cristina Marqueze, Raul Machado Neto, Debora Botequio Moretti, Jardelina de Souza Todao Bernardino, Loyze Paola Oliveira de Lima, Luiz Aurelio de Campos Crispin. Centro de Genômica Funcional da ESALQ: Luiz Lehmann Coutinho, Ricardo Augusto Brassaloti, Raquel de Lello Rocha Campos Cassano. NGS Soluções Genômicas: Pilar Drummond Sampaio Corrêa Mariani. FZEA-USP Pirassununga: Mirele Daiana Poleti, Jessika Cristina Chagas Lesbon, Elisangela Chicaroni Mattos, Heidge Fukumasu. USP-Botucatu: Rejane Maria Tommasini Grotto, Jayme A. Souza-Neto, Guilherme Targino Valente, Patricia Akemi Assato, Felipe Allan da Silva da Costa, Bianca Cechetto Carlos. Mendelics: Bibiana Santos, João Paulo Kitajima, Erika Freitas, David Schlesinger. Hemocentro Ribeirão Preto: Simone Kashima, Evandra Strazza Rodrigues, Svetoslav Nanev Slavov, Elaine Vieira dos Santos, Rafael dos Santos Bezerra, Luiz Carlos Junior de Alcantara, Marta Giovanetti, Vagner Fonseca, Flavia Aburjaile, Rodrigo Tocantins Calado. FAMERP-SJRP: Cecília Artico Banho, Lívia Sacchetto, Fábio Sossai Possebon, Leila Sabrina Ullmann, Cíntia Bittar, Guilherme Campos, Helena Lage Ferreira, Jorge A. Petroli Marchesi, Maisa C. Pereira Parra, Marília Moraes, Paula Rahal, Paulo Inacio da Costa, João Pessoa Araújo Jr., Maurício Lacerda Nogueira. Prefeitura de Sao Paulo: Melissa Palmieri. |
| EPI_ISL_1966133, EPI_ISL_1966134, EPI_ISL_1966135, EPI_ISL_1966136, EPI_ISL_1966137, EPI_ISL_1966138, EPI_ISL_1966139, EPI_ISL_1966140, EPI_ISL_1966141, EPI_ISL_1966142, EPI_ISL_1966143, EPI_ISL_1966144 | | | |
| see above | PRONTO ATENDIMENTO VILA PADRE ANCHIETA | Instituto Butantan / Mendelics | Instituto Butantan: Dimas Tadeu Covas, Sandra Coccuzzo Sampaio, Maria Carolina Elias, José Salvatore Leister Patané, Vincent Louis Viala, Antonio Jorge Martins, Ricardo Haddad, Claudia Renata dos Santos Barros, Elaine Cristina Marqueze, Raul Machado Neto, Debora Botequio Moretti, Jardelina de |

| | | | |
|---|---|---|---|
| | | | Souza Todao Bernardino, Loyze Paola Oliveira de Lima, Luiz Aurelio de Campos Crispin. Centro de Genômica Funcional da ESALQ: Luiz Lehmann Coutinho, Ricardo Augusto Brassaloti, Raquel de Lello Rocha Campos Cassano. NGS Soluções Genômicas: Pilar Drummond Sampaio Corrêa Mariani. FZEA-USP Pirassununga: Mirele Daiana Poleti, Jessika Cristina Chagas Lesbon, Elisangela Chicaroni Mattos, Heidge Fukumasu. USP-Botucatu: Rejane Maria Tommasini Grotto, Jayme A. Souza-Neto, Guilherme Targino Valente, Patricia Akemi Assato, Felipe Allan da Silva da Costa, Bianca Cechetto Carlos. Mendelics: Bibiana Santos, João Paulo Kitajima, Erika Freitas, David Schlesinger. Hemocentro Ribeirão Preto: Simone Kashima, Evandra Strazza Rodrigues, Svetoslav Nanev Slavov, Elaine Vieira dos Santos, Rafael dos Santos Bezerra, Luiz Carlos Junior de Alcantara, Marta Giovanetti, Vagner Fonseca, Flavia Aburjaile, Rodrigo Tocantins Calado. FAMERP-SJRP: Cecília Artico Banho, Lívia Sacchetto, Fábio Sossai Possebon, Leila Sabrina Ullmann, Cíntia Bittar, Guilherme Campos, Helena Lage Ferreira, Jorge A. Petroli Marchesi, Maisa C. Pereira Parra, Marília Moraes, Paula Rahal, Paulo Inacio da Costa, João Pessoa Araújo Jr., Maurício Lacerda Nogueira. Prefeitura de Sao Paulo: Melissa Palmieri. |
| EPI_ISL_1966145, EPI_ISL_1966146 | COMPLEXO HOSPITALAR OURO VERDE DE CAMPINAS | Instituto Butantan / Mendelics | Instituto Butantan: Dimas Tadeu Covas, Sandra Coccuzzo Sampaio, Maria Carolina Elias, José Salvatore Leister Patané, Vincent Louis Viala, Antonio Jorge Martins, Ricardo Haddad, Claudia Renata dos Santos Barros, Elaine Cristina Marqueze, Raul Machado Neto, Debora Botequio Moretti, Jardelina de Souza Todao Bernardino, Loyze Paola Oliveira de Lima, Luiz Aurelio de Campos Crispin. Centro de Genômica Funcional da ESALQ: Luiz Lehmann Coutinho, Ricardo Augusto Brassaloti, Raquel de Lello Rocha Campos Cassano. NGS Soluções Genômicas: Pilar Drummond Sampaio Corrêa Mariani. FZEA-USP Pirassununga: Mirele Daiana Poleti, Jessika Cristina Chagas Lesbon, Elisangela Chicaroni Mattos, Heidge Fukumasu. USP-Botucatu: Rejane Maria Tommasini Grotto, Jayme A. Souza-Neto, Guilherme Targino Valente, Patricia Akemi Assato, Felipe Allan da Silva da Costa, Bianca Cechetto Carlos. Mendelics: Bibiana Santos, João Paulo Kitajima, Erika Freitas, David Schlesinger. Hemocentro Ribeirão Preto: Simone Kashima, Evandra Strazza Rodrigues, Svetoslav Nanev Slavov, Elaine Vieira dos Santos, Rafael dos Santos Bezerra, Luiz Carlos Junior de Alcantara, Marta Giovanetti, Vagner Fonseca, Flavia Aburjaile, Rodrigo Tocantins Calado. FAMERP-SJRP: Cecília Artico Banho, Lívia Sacchetto, Fábio Sossai Possebon, Leila Sabrina Ullmann, Cíntia Bittar, Guilherme Campos, Helena Lage Ferreira, Jorge A. Petroli Marchesi, Maisa C. Pereira Parra, Marília Moraes, Paula Rahal, Paulo Inacio da Costa, João Pessoa Araújo Jr., Maurício Lacerda Nogueira. Prefeitura de Sao Paulo: Melissa Palmieri. |
| EPI_ISL_1966147 | HOSPITAL REGIONAL DE ITAPETININGA | Instituto Butantan / Mendelics | Instituto Butantan: Dimas Tadeu Covas, Sandra Coccuzzo Sampaio, Maria Carolina Elias, José Salvatore Leister Patané, Vincent Louis Viala, Antonio Jorge Martins, Ricardo Haddad, Claudia Renata dos Santos Barros, Elaine Cristina Marqueze, Raul Machado Neto, Debora Botequio Moretti, Jardelina de Souza Todao Bernardino, Loyze Paola Oliveira de Lima, Luiz Aurelio de Campos Crispin. Centro de Genômica Funcional da ESALQ: Luiz Lehmann Coutinho, Ricardo Augusto Brassaloti, Raquel de Lello Rocha Campos Cassano. NGS Soluções Genômicas: Pilar Drummond Sampaio Corrêa Mariani. FZEA-USP Pirassununga: Mirele Daiana Poleti, Jessika Cristina Chagas Lesbon, Elisangela Chicaroni Mattos, Heidge Fukumasu. USP-Botucatu: Rejane Maria Tommasini Grotto, Jayme A. Souza-Neto, Guilherme Targino Valente, Patricia Akemi Assato, Felipe Allan da Silva da Costa, Bianca Cechetto Carlos. Mendelics: Bibiana Santos, João Paulo Kitajima, Erika Freitas, David Schlesinger. Hemocentro Ribeirão Preto: Simone Kashima, Evandra Strazza Rodrigues, Svetoslav Nanev Slavov, Elaine Vieira dos Santos, Rafael dos Santos Bezerra, Luiz Carlos Junior de Alcantara, Marta Giovanetti, Vagner Fonseca, Flavia Aburjaile, Rodrigo Tocantins Calado. FAMERP-SJRP: Cecília Artico Banho, Lívia Sacchetto, Fábio Sossai Possebon, Leila Sabrina Ullmann, Cíntia Bittar, Guilherme Campos, Helena Lage Ferreira, Jorge A. Petroli Marchesi, Maisa C. Pereira Parra, Marília Moraes, Paula Rahal, Paulo Inacio da Costa, João Pessoa Araújo Jr., Maurício Lacerda Nogueira. Prefeitura de Sao Paulo: Melissa Palmieri. |
| EPI_ISL_1966148, EPI_ISL_1966149, EPI_ISL_1966150, EPI_ISL_1966151, EPI_ISL_1966152, EPI_ISL_1966153 | SECRETARIA MUNICIPAL DE SAUDE DE PIRACAIA | Instituto Butantan / Mendelics | Instituto Butantan: Dimas Tadeu Covas, Sandra Coccuzzo Sampaio, Maria Carolina Elias, José Salvatore Leister Patané, Vincent Louis Viala, Antonio Jorge Martins, Ricardo Haddad, Claudia Renata dos Santos Barros, Elaine Cristina Marqueze, Raul Machado Neto, Debora Botequio Moretti, Jardelina de Souza Todao Bernardino, Loyze Paola Oliveira de Lima, Luiz Aurelio de Campos Crispin. Centro de Genômica Funcional da ESALQ: Luiz Lehmann Coutinho, Ricardo Augusto Brassaloti, Raquel de Lello Rocha Campos Cassano. NGS Soluções Genômicas: Pilar Drummond Sampaio Corrêa Mariani. FZEA-USP Pirassununga: Mirele Daiana Poleti, Jessika Cristina Chagas Lesbon, Elisangela Chicaroni Mattos, Heidge Fukumasu. USP-Botucatu: Rejane Maria Tommasini Grotto, Jayme A. Souza-Neto, Guilherme Targino Valente, Patricia Akemi Assato, Felipe Allan da Silva da Costa, Bianca Cechetto Carlos. Mendelics: Bibiana Santos, João Paulo Kitajima, Erika Freitas, David Schlesinger. Hemocentro Ribeirão Preto: Simone Kashima, Evandra Strazza Rodrigues, Svetoslav Nanev Slavov, Elaine Vieira dos Santos, Rafael dos Santos Bezerra, Luiz Carlos Junior de Alcantara, Marta Giovanetti, Vagner Fonseca, Flavia Aburjaile, Rodrigo Tocantins Calado. FAMERP-SJRP: Cecília Artico Banho, Lívia Sacchetto, Fábio Sossai Possebon, Leila Sabrina Ullmann, Cíntia Bittar, Guilherme Campos, Helena Lage Ferreira, Jorge A. Petroli Marchesi, Maisa C. Pereira Parra, Marília Moraes, Paula Rahal, Paulo Inacio da Costa, João Pessoa Araújo Jr., Maurício Lacerda Nogueira. Prefeitura de Sao Paulo: Melissa Palmieri. |
| EPI_ISL_1966154, EPI_ISL_1966155 | SECRETARIA MUNICIPAL DE SAUDE SOROCABA | Instituto Butantan / Mendelics | Instituto Butantan: Dimas Tadeu Covas, Sandra Coccuzzo Sampaio, Maria Carolina Elias, José Salvatore Leister Patané, Vincent Louis Viala, Antonio Jorge Martins, Ricardo Haddad, Claudia Renata dos Santos Barros, Elaine Cristina Marqueze, Raul Machado Neto, Debora Botequio Moretti, Jardelina de Souza Todao Bernardino, Loyze Paola Oliveira de Lima, Luiz Aurelio de Campos Crispin. Centro de Genômica Funcional da ESALQ: Luiz Lehmann Coutinho, Ricardo Augusto Brassaloti, Raquel de Lello Rocha Campos Cassano. NGS Soluções Genômicas: Pilar Drummond Sampaio Corrêa Mariani. FZEA-USP Pirassununga: Mirele Daiana Poleti, Jessika Cristina Chagas Lesbon, Elisangela Chicaroni Mattos, Heidge Fukumasu. USP-Botucatu: Rejane Maria Tommasini Grotto, Jayme A. Souza-Neto, Guilherme Targino Valente, Patricia Akemi Assato, Felipe Allan da Silva da Costa, Bianca Cechetto Carlos. Mendelics: Bibiana Santos, João Paulo Kitajima, Erika Freitas, David Schlesinger. Hemocentro Ribeirão Preto: Simone Kashima, Evandra Strazza Rodrigues, Svetoslav Nanev Slavov, Elaine Vieira dos Santos, Rafael dos Santos Bezerra, Luiz Carlos Junior de Alcantara, Marta Giovanetti, Vagner Fonseca, Flavia Aburjaile, Rodrigo Tocantins Calado. FAMERP-SJRP: Cecília Artico Banho, Lívia Sacchetto, Fábio Sossai Possebon, Leila Sabrina Ullmann, Cíntia Bittar, Guilherme Campos, Helena Lage Ferreira, Jorge A. Petroli Marchesi, Maisa C. Pereira Parra, Marília Moraes, Paula Rahal, Paulo Inacio da Costa, João Pessoa Araújo Jr., Maurício Lacerda Nogueira. Prefeitura de Sao Paulo: Melissa Palmieri. |
| EPI_ISL_1966156 | PRONTO ATENDIMENTO VILA PADRE ANCHIETA | Instituto Butantan / Mendelics | Instituto Butantan: Dimas Tadeu Covas, Sandra Coccuzzo Sampaio, Maria Carolina Elias, José Salvatore Leister Patané, Vincent Louis Viala, Antonio Jorge Martins, Ricardo Haddad, Claudia Renata dos Santos Barros, Elaine Cristina Marqueze, Raul Machado Neto, Debora Botequio Moretti, Jardelina de Souza Todao Bernardino, Loyze Paola Oliveira de Lima, Luiz Aurelio de Campos Crispin. Centro de Genômica Funcional da ESALQ: Luiz Lehmann Coutinho, Ricardo Augusto Brassaloti, Raquel de Lello Rocha Campos Cassano. NGS Soluções Genômicas: Pilar Drummond Sampaio Corrêa Mariani. FZEA-USP Pirassununga: Mirele Daiana Poleti, Jessika Cristina Chagas Lesbon, Elisangela Chicaroni Mattos, Heidge Fukumasu. USP-Botucatu: Rejane Maria Tommasini Grotto, Jayme A. Souza-Neto, Guilherme Targino Valente, Patricia Akemi Assato, Felipe Allan da Silva da Costa, Bianca Cechetto Carlos. Mendelics: Bibiana Santos, João Paulo Kitajima, Erika Freitas, David Schlesinger. Hemocentro Ribeirão Preto: Simone Kashima, Evandra Strazza Rodrigues, Svetoslav Nanev Slavov, Elaine Vieira dos Santos, Rafael dos Santos Bezerra, Luiz Carlos Junior de Alcantara, Marta Giovanetti, Vagner Fonseca, Flavia Aburjaile, Rodrigo Tocantins Calado. FAMERP-SJRP: Cecília Artico Banho, Lívia Sacchetto, Fábio Sossai Possebon, Leila Sabrina Ullmann, Cíntia Bittar, Guilherme Campos, Helena Lage Ferreira, Jorge A. Petroli Marchesi, Maisa C. Pereira Parra, Marília Moraes, Paula Rahal, Paulo Inacio da Costa, João Pessoa Araújo Jr., Maurício Lacerda Nogueira. Prefeitura de Sao Paulo: Melissa Palmieri. |
| EPI_ISL_1966157, EPI_ISL_1966158, EPI_ISL_1966159, EPI_ISL_1966160, EPI_ISL_1966161 | COMPLEXO HOSPITALAR OURO VERDE DE CAMPINAS | Instituto Butantan / Mendelics | Instituto Butantan: Dimas Tadeu Covas, Sandra Coccuzzo Sampaio, Maria Carolina Elias, José Salvatore Leister Patané, Vincent Louis Viala, Antonio Jorge Martins, Ricardo Haddad, Claudia Renata dos Santos Barros, Elaine Cristina Marqueze, Raul Machado Neto, Debora Botequio Moretti, Jardelina de Souza Todao Bernardino, Loyze Paola Oliveira de Lima, Luiz Aurelio de Campos Crispin. Centro de Genômica Funcional da ESALQ: Luiz Lehmann Coutinho, Ricardo Augusto Brassaloti, Raquel de Lello Rocha Campos Cassano. NGS Soluções Genômicas: Pilar Drummond Sampaio Corrêa Mariani. FZEA-USP Pirassununga: Mirele Daiana Poleti, Jessika Cristina Chagas Lesbon, Elisangela Chicaroni Mattos, Heidge Fukumasu. USP-Botucatu: Rejane Maria Tommasini Grotto, Jayme A. Souza-Neto, Guilherme Targino Valente, Patricia Akemi Assato, Felipe Allan da Silva da Costa, Bianca Cechetto Carlos. Mendelics: Bibiana Santos, João Paulo Kitajima, Erika Freitas, David Schlesinger. Hemocentro Ribeirão Preto: Simone Kashima, Evandra Strazza Rodrigues, Svetoslav Nanev Slavov, Elaine Vieira dos Santos, Rafael dos Santos Bezerra, Luiz Carlos Junior de Alcantara, Marta Giovanetti, Vagner Fonseca, Flavia Aburjaile, Rodrigo Tocantins Calado. FAMERP-SJRP: Cecília Artico Banho, Lívia Sacchetto, Fábio Sossai Possebon, Leila Sabrina Ullmann, Cíntia Bittar, Guilherme Campos, Helena Lage Ferreira, Jorge A. Petroli Marchesi, Maisa C. Pereira Parra, Marília Moraes, Paula Rahal, Paulo Inacio da Costa, João Pessoa Araújo Jr., Maurício Lacerda Nogueira. Prefeitura de Sao Paulo: Melissa Palmieri. |
| EPI_ISL_1966162, EPI_ISL_1966163, EPI_ISL_1966164, EPI_ISL_1966165, EPI_ISL_1966166 | UNIDADE RESPIRATORIA NOVA HORTOLANDIA | Instituto Butantan / Mendelics | Instituto Butantan: Dimas Tadeu Covas, Sandra Coccuzzo Sampaio, Maria Carolina Elias, José Salvatore Leister Patané, Vincent Louis Viala, Antonio Jorge Martins, Ricardo Haddad, Claudia Renata dos Santos Barros, Elaine Cristina Marqueze, Raul Machado Neto, Debora Botequio Moretti, Jardelina de Souza Todao Bernardino, Loyze Paola Oliveira de Lima, Luiz Aurelio de Campos Crispin. Centro de Genômica Funcional da ESALQ: Luiz Lehmann Coutinho, Ricardo Augusto Brassaloti, Raquel de Lello Rocha Campos Cassano. NGS Soluções Genômicas: Pilar Drummond Sampaio Corrêa Mariani. FZEA-USP Pirassununga: Mirele Daiana Poleti, Jessika Cristina Chagas Lesbon, Elisangela Chicaroni Mattos, Heidge Fukumasu. USP-Botucatu: Rejane Maria Tommasini Grotto, Jayme A. Souza-Neto, Guilherme Targino Valente, Patricia Akemi Assato, Felipe Allan da Silva da Costa, Bianca Cechetto Carlos. Mendelics: Bibiana Santos, João Paulo Kitajima, Erika Freitas, David Schlesinger. Hemocentro Ribeirão Preto: Simone Kashima, Evandra Strazza Rodrigues, Svetoslav Nanev Slavov, Elaine Vieira dos Santos, Rafael dos Santos Bezerra, Luiz Carlos Junior de Alcantara, Marta Giovanetti, Vagner Fonseca, Flavia Aburjaile, Rodrigo Tocantins Calado. FAMERP-SJRP: Cecília Artico Banho, Lívia Sacchetto, Fábio Sossai Possebon, Leila Sabrina |

| | | | |
|---|---|---|---|
| | | | Ullmann, Cíntia Bittar, Guilherme Campos, Helena Lage Ferreira, Jorge A. Petroli Marchesi, Maisa C. Pereira Parra, Marília Moraes, Paula Rahal, Paulo Inacio da Costa, João Pessoa Araújo Jr., Maurício Lacerda Nogueira. Prefeitura de Sao Paulo: Melissa Palmieri. |
| EPI_ISL_1966167, EPI_ISL_1966168, EPI_ISL_1966169, EPI_ISL_1966170, EPI_ISL_1966171, EPI_ISL_1966172, EPI_ISL_1966173 | UNIDADE DE PRONTO ATENDIMENTO UPA DRA ANA OLIVIA BENTIVOGLIO | Instituto Butantan / Mendelics | Instituto Butantan: Dimas Tadeu Covas, Sandra Coccuzzo Sampaio, Maria Carolina Elias, José Salvatore Leister Patané, Vincent Louis Viala, Antonio Jorge Martins, Ricardo Haddad, Claudia Renata dos Santos Barros, Elaine Cristina Marqueze, Raul Machado Neto, Debora Botequio Moretti, Jardelina de Souza Todao Bernardino, Loyze Paola Oliveira de Lima, Luiz Aurelio de Campos Crispin. Centro de Genômica Funcional da ESALQ: Luiz Lehmann Coutinho, Ricardo Augusto Brassaloti, Raquel de Lello Rocha Campos Cassano. NGS Soluções Genômicas: Pilar Drummond Sampaio Corrêa Mariani. FZEA-USP Pirassununga: Mirele Daiana Poleti, Jessika Cristina Chagas Lesbon, Elisangela Chicaroni Mattos, Heidge Fukumasu. USP-Botucatu: Rejane Maria Tommasini Grotto, Jayme A. Souza-Neto, Guilherme Targino Valente, Patricia Akemi Assato, Felipe Allan da Silva da Costa, Bianca Cechetto Carlos. Mendelics: Bibiana Santos, João Paulo Kitajima, Erika Freitas, David Schlesinger. Hemocentro Ribeirão Preto: Simone Kashima, Evandra Strazza Rodrigues, Svetoslav Nanev Slavov, Elaine Vieira dos Santos, Rafael dos Santos Bezerra, Luiz Carlos Junior de Alcantara, Marta Giovanetti, Vagner Fonseca, Flavia Aburjaile, Rodrigo Tocantins Calado. FAMERP-SJRP: Cecília Artico Banho, Lívia Sacchetto, Fábio Sossai Possebon, Leila Sabrina Ullmann, Cíntia Bittar, Guilherme Campos, Helena Lage Ferreira, Jorge A. Petroli Marchesi, Maisa C. Pereira Parra, Marília Moraes, Paula Rahal, Paulo Inacio da Costa, João Pessoa Araújo Jr., Maurício Lacerda Nogueira. Prefeitura de Sao Paulo: Melissa Palmieri. |
| EPI_ISL_1966174, EPI_ISL_1966175, EPI_ISL_1966176 | PRONTO ATENDIMENTO DA PONTE DE SAO JOAO | Instituto Butantan / Mendelics | Instituto Butantan: Dimas Tadeu Covas, Sandra Coccuzzo Sampaio, Maria Carolina Elias, José Salvatore Leister Patané, Vincent Louis Viala, Antonio Jorge Martins, Ricardo Haddad, Claudia Renata dos Santos Barros, Elaine Cristina Marqueze, Raul Machado Neto, Debora Botequio Moretti, Jardelina de Souza Todao Bernardino, Loyze Paola Oliveira de Lima, Luiz Aurelio de Campos Crispin. Centro de Genômica Funcional da ESALQ: Luiz Lehmann Coutinho, Ricardo Augusto Brassaloti, Raquel de Lello Rocha Campos Cassano. NGS Soluções Genômicas: Pilar Drummond Sampaio Corrêa Mariani. FZEA-USP Pirassununga: Mirele Daiana Poleti, Jessika Cristina Chagas Lesbon, Elisangela Chicaroni Mattos, Heidge Fukumasu. USP-Botucatu: Rejane Maria Tommasini Grotto, Jayme A. Souza-Neto, Guilherme Targino Valente, Patricia Akemi Assato, Felipe Allan da Silva da Costa, Bianca Cechetto Carlos. Mendelics: Bibiana Santos, João Paulo Kitajima, Erika Freitas, David Schlesinger. Hemocentro Ribeirão Preto: Simone Kashima, Evandra Strazza Rodrigues, Svetoslav Nanev Slavov, Elaine Vieira dos Santos, Rafael dos Santos Bezerra, Luiz Carlos Junior de Alcantara, Marta Giovanetti, Vagner Fonseca, Flavia Aburjaile, Rodrigo Tocantins Calado. FAMERP-SJRP: Cecília Artico Banho, Lívia Sacchetto, Fábio Sossai Possebon, Leila Sabrina Ullmann, Cíntia Bittar, Guilherme Campos, Helena Lage Ferreira, Jorge A. Petroli Marchesi, Maisa C. Pereira Parra, Marília Moraes, Paula Rahal, Paulo Inacio da Costa, João Pessoa Araújo Jr., Maurício Lacerda Nogueira. Prefeitura de Sao Paulo: Melissa Palmieri. |
| EPI_ISL_1966177, EPI_ISL_1966178 | SECRETARIA MUNICIPAL DE SAUDE SOROCABA | Instituto Butantan / Mendelics | Instituto Butantan: Dimas Tadeu Covas, Sandra Coccuzzo Sampaio, Maria Carolina Elias, José Salvatore Leister Patané, Vincent Louis Viala, Antonio Jorge Martins, Ricardo Haddad, Claudia Renata dos Santos Barros, Elaine Cristina Marqueze, Raul Machado Neto, Debora Botequio Moretti, Jardelina de Souza Todao Bernardino, Loyze Paola Oliveira de Lima, Luiz Aurelio de Campos Crispin. Centro de Genômica Funcional da ESALQ: Luiz Lehmann Coutinho, Ricardo Augusto Brassaloti, Raquel de Lello Rocha Campos Cassano. NGS Soluções Genômicas: Pilar Drummond Sampaio Corrêa Mariani. FZEA-USP Pirassununga: Mirele Daiana Poleti, Jessika Cristina Chagas Lesbon, Elisangela Chicaroni Mattos, Heidge Fukumasu. USP-Botucatu: Rejane Maria Tommasini Grotto, Jayme A. Souza-Neto, Guilherme Targino Valente, Patricia Akemi Assato, Felipe Allan da Silva da Costa, Bianca Cechetto Carlos. Mendelics: Bibiana Santos, João Paulo Kitajima, Erika Freitas, David Schlesinger. Hemocentro Ribeirão Preto: Simone Kashima, Evandra Strazza Rodrigues, Svetoslav Nanev Slavov, Elaine Vieira dos Santos, Rafael dos Santos Bezerra, Luiz Carlos Junior de Alcantara, Marta Giovanetti, Vagner Fonseca, Flavia Aburjaile, Rodrigo Tocantins Calado. FAMERP-SJRP: Cecília Artico Banho, Lívia Sacchetto, Fábio Sossai Possebon, Leila Sabrina Ullmann, Cíntia Bittar, Guilherme Campos, Helena Lage Ferreira, Jorge A. Petroli Marchesi, Maisa C. Pereira Parra, Marília Moraes, Paula Rahal, Paulo Inacio da Costa, João Pessoa Araújo Jr., Maurício Lacerda Nogueira. Prefeitura de Sao Paulo: Melissa Palmieri. |
| EPI_ISL_1966179 | PRONTO ATENDIMENTO VILA PADRE ANCHIETA | Instituto Butantan / Mendelics | Instituto Butantan: Dimas Tadeu Covas, Sandra Coccuzzo Sampaio, Maria Carolina Elias, José Salvatore Leister Patané, Vincent Louis Viala, Antonio Jorge Martins, Ricardo Haddad, Claudia Renata dos Santos Barros, Elaine Cristina Marqueze, Raul Machado Neto, Debora Botequio Moretti, Jardelina de Souza Todao Bernardino, Loyze Paola Oliveira de Lima, Luiz Aurelio de Campos Crispin. Centro de Genômica Funcional da ESALQ: Luiz Lehmann Coutinho, Ricardo Augusto Brassaloti, Raquel de Lello Rocha Campos Cassano. NGS Soluções Genômicas: Pilar Drummond Sampaio Corrêa Mariani. FZEA-USP Pirassununga: Mirele Daiana Poleti, Jessika Cristina Chagas Lesbon, Elisangela Chicaroni Mattos, Heidge Fukumasu. USP-Botucatu: Rejane Maria Tommasini Grotto, Jayme A. Souza-Neto, Guilherme Targino Valente, Patricia Akemi Assato, Felipe Allan da Silva da Costa, Bianca Cechetto Carlos. Mendelics: Bibiana Santos, João Paulo Kitajima, Erika Freitas, David Schlesinger. Hemocentro Ribeirão Preto: Simone Kashima, Evandra Strazza Rodrigues, Svetoslav Nanev Slavov, Elaine Vieira dos Santos, Rafael dos Santos Bezerra, Luiz Carlos Junior de Alcantara, Marta Giovanetti, Vagner Fonseca, Flavia Aburjaile, Rodrigo Tocantins Calado. FAMERP-SJRP: Cecília Artico Banho, Lívia Sacchetto, Fábio Sossai Possebon, Leila Sabrina Ullmann, Cíntia Bittar, Guilherme Campos, Helena Lage Ferreira, Jorge A. Petroli Marchesi, Maisa C. Pereira Parra, Marília Moraes, Paula Rahal, Paulo Inacio da Costa, João Pessoa Araújo Jr., Maurício Lacerda Nogueira. Prefeitura de Sao Paulo: Melissa Palmieri. |
| EPI_ISL_1966180 | UNIDADE MISTA DE SAUDE CAPELA DO ALTO | Instituto Butantan / Mendelics | Instituto Butantan: Dimas Tadeu Covas, Sandra Coccuzzo Sampaio, Maria Carolina Elias, José Salvatore Leister Patané, Vincent Louis Viala, Antonio Jorge Martins, Ricardo Haddad, Claudia Renata dos Santos Barros, Elaine Cristina Marqueze, Raul Machado Neto, Debora Botequio Moretti, Jardelina de Souza Todao Bernardino, Loyze Paola Oliveira de Lima, Luiz Aurelio de Campos Crispin. Centro de Genômica Funcional da ESALQ: Luiz Lehmann Coutinho, Ricardo Augusto Brassaloti, Raquel de Lello Rocha Campos Cassano. NGS Soluções Genômicas: Pilar Drummond Sampaio Corrêa Mariani. FZEA-USP Pirassununga: Mirele Daiana Poleti, Jessika Cristina Chagas Lesbon, Elisangela Chicaroni Mattos, Heidge Fukumasu. USP-Botucatu: Rejane Maria Tommasini Grotto, Jayme A. Souza-Neto, Guilherme Targino Valente, Patricia Akemi Assato, Felipe Allan da Silva da Costa, Bianca Cechetto Carlos. Mendelics: Bibiana Santos, João Paulo Kitajima, Erika Freitas, David Schlesinger. Hemocentro Ribeirão Preto: Simone Kashima, Evandra Strazza Rodrigues, Svetoslav Nanev Slavov, Elaine Vieira dos Santos, Rafael dos Santos Bezerra, Luiz Carlos Junior de Alcantara, Marta Giovanetti, Vagner Fonseca, Flavia Aburjaile, Rodrigo Tocantins Calado. FAMERP-SJRP: Cecília Artico Banho, Lívia Sacchetto, Fábio Sossai Possebon, Leila Sabrina Ullmann, Cíntia Bittar, Guilherme Campos, Helena Lage Ferreira, Jorge A. Petroli Marchesi, Maisa C. Pereira Parra, Marília Moraes, Paula Rahal, Paulo Inacio da Costa, João Pessoa Araújo Jr., Maurício Lacerda Nogueira. Prefeitura de Sao Paulo: Melissa Palmieri. |
| EPI_ISL_1966181, EPI_ISL_1966182, EPI_ISL_1966183, EPI_ISL_1966184, EPI_ISL_1966185, EPI_ISL_1966186 | SERV DE VIG SANITARIA EPIDEMIO E CTRL DE ZOONOZES GUARUJA | Instituto Butantan / Mendelics | Instituto Butantan: Dimas Tadeu Covas, Sandra Coccuzzo Sampaio, Maria Carolina Elias, José Salvatore Leister Patané, Vincent Louis Viala, Antonio Jorge Martins, Ricardo Haddad, Claudia Renata dos Santos Barros, Elaine Cristina Marqueze, Raul Machado Neto, Debora Botequio Moretti, Jardelina de Souza Todao Bernardino, Loyze Paola Oliveira de Lima, Luiz Aurelio de Campos Crispin. Centro de Genômica Funcional da ESALQ: Luiz Lehmann Coutinho, Ricardo Augusto Brassaloti, Raquel de Lello Rocha Campos Cassano. NGS Soluções Genômicas: Pilar Drummond Sampaio Corrêa Mariani. FZEA-USP Pirassununga: Mirele Daiana Poleti, Jessika Cristina Chagas Lesbon, Elisangela Chicaroni Mattos, Heidge Fukumasu. USP-Botucatu: Rejane Maria Tommasini Grotto, Jayme A. Souza-Neto, Guilherme Targino Valente, Patricia Akemi Assato, Felipe Allan da Silva da Costa, Bianca Cechetto Carlos. Mendelics: Bibiana Santos, João Paulo Kitajima, Erika Freitas, David Schlesinger. Hemocentro Ribeirão Preto: Simone Kashima, Evandra Strazza Rodrigues, Svetoslav Nanev Slavov, Elaine Vieira dos Santos, Rafael dos Santos Bezerra, Luiz Carlos Junior de Alcantara, Marta Giovanetti, Vagner Fonseca, Flavia Aburjaile, Rodrigo Tocantins Calado. FAMERP-SJRP: Cecília Artico Banho, Lívia Sacchetto, Fábio Sossai Possebon, Leila Sabrina Ullmann, Cíntia Bittar, Guilherme Campos, Helena Lage Ferreira, Jorge A. Petroli Marchesi, Maisa C. Pereira Parra, Marília Moraes, Paula Rahal, Paulo Inacio da Costa, João Pessoa Araújo Jr., Maurício Lacerda Nogueira. Prefeitura de Sao Paulo: Melissa Palmieri. |
| EPI_ISL_1966187 | SMS IPERO | Instituto Butantan / Mendelics | Instituto Butantan: Dimas Tadeu Covas, Sandra Coccuzzo Sampaio, Maria Carolina Elias, José Salvatore Leister Patané, Vincent Louis Viala, Antonio Jorge Martins, Ricardo Haddad, Claudia Renata dos Santos Barros, Elaine Cristina Marqueze, Raul Machado Neto, Debora Botequio Moretti, Jardelina de Souza Todao Bernardino, Loyze Paola Oliveira de Lima, Luiz Aurelio de Campos Crispin. Centro de Genômica Funcional da ESALQ: Luiz Lehmann Coutinho, Ricardo Augusto Brassaloti, Raquel de Lello Rocha Campos Cassano. NGS Soluções Genômicas: Pilar Drummond Sampaio Corrêa Mariani. FZEA-USP Pirassununga: Mirele Daiana Poleti, Jessika Cristina Chagas Lesbon, Elisangela Chicaroni Mattos, Heidge Fukumasu. USP-Botucatu: Rejane Maria Tommasini Grotto, Jayme A. Souza-Neto, Guilherme Targino Valente, Patricia Akemi Assato, Felipe Allan da Silva da Costa, Bianca Cechetto Carlos. Mendelics: Bibiana Santos, João Paulo Kitajima, Erika Freitas, David Schlesinger. Hemocentro Ribeirão Preto: Simone Kashima, Evandra Strazza Rodrigues, Svetoslav Nanev Slavov, Elaine Vieira dos Santos, Rafael dos Santos Bezerra, Luiz Carlos Junior de Alcantara, Marta Giovanetti, Vagner Fonseca, Flavia Aburjaile, Rodrigo Tocantins Calado. FAMERP-SJRP: Cecília Artico Banho, Lívia Sacchetto, Fábio Sossai Possebon, Leila Sabrina Ullmann, Cíntia Bittar, Guilherme Campos, Helena Lage Ferreira, Jorge A. Petroli Marchesi, Maisa C. Pereira Parra, Marília Moraes, Paula Rahal, Paulo Inacio da Costa, João Pessoa Araújo Jr., Maurício Lacerda Nogueira. Prefeitura de Sao Paulo: Melissa Palmieri. |
| EPI_ISL_1966188 | UBS SALTO DE SAO JOSE | Instituto Butantan / Mendelics | Instituto Butantan: Dimas Tadeu Covas, Sandra Coccuzzo Sampaio, Maria Carolina Elias, José Salvatore Leister Patané, Vincent Louis Viala, Antonio Jorge Martins, Ricardo Haddad, Claudia Renata dos Santos Barros, Elaine Cristina Marqueze, Raul Machado Neto, Debora Botequio Moretti, Jardelina de Souza Todao Bernardino, Loyze Paola Oliveira de Lima, Luiz Aurelio de Campos Crispin. Centro de Genômica Funcional da ESALQ: Luiz Lehmann Coutinho, Ricardo Augusto Brassaloti, Raquel de Lello Rocha Campos Cassano. NGS Soluções Genômicas: Pilar Drummond Sampaio Corrêa Mariani. |

| | | | |
|---|---|---|---|
| | | | FZEA-USP Pirassununga: Mirele Daiana Poleti, Jessika Cristina Chagas Lesbon, Elisangela Chicaroni Mattos, Heidge Fukumasu. USP-Botucatu: Rejane Maria Tommasini Grotto, Jayme A. Souza-Neto, Guilherme Targino Valente, Patricia Akemi Assato, Felipe Allan da Silva da Costa, Bianca Cechetto Carlos. Mendelics: Bibiana Santos, João Paulo Kitajima, Erika Freitas, David Schlesinger. Hemocentro Ribeirão Preto: Simone Kashima, Evandra Strazza Rodrigues, Svetoslav Nanev Slavov, Elaine Vieira dos Santos, Rafael dos Santos Bezerra, Luiz Carlos Junior de Alcantara, Marta Giovanetti, Vagner Fonseca, Flavia Aburjaile, Rodrigo Tocantins Calado. FAMERP-SJRP: Cecília Artico Banho, Lívia Sacchetto, Fábio Sossai Possebon, Leila Sabrina Ullmann, Cíntia Bittar, Guilherme Campos, Helena Lage Ferreira, Jorge A. Petroli Marchesi, Maisa C. Pereira Parra, Marília Moraes, Paula Rahal, Paulo Inacio da Costa, João Pessoa Araújo Jr., Maurício Lacerda Nogueira. Prefeitura de Sao Paulo: Melissa Palmieri. |
| EPI_ISL_1966189, EPI_ISL_1966190, EPI_ISL_1966191, EPI_ISL_1966192, EPI_ISL_1966193, EPI_ISL_1966194, EPI_ISL_1966195, EPI_ISL_1966196 | SECAO CENTRO DE DIAGNOSTICO SECEDI | Instituto Butantan / Mendelics | Instituto Butantan: Dimas Tadeu Covas, Sandra Coccuzzo Sampaio, Maria Carolina Elias, José Salvatore Leister Patané, Vincent Louis Viala, Antonio Jorge Martins, Ricardo Haddad, Claudia Renata dos Santos Barros, Elaine Cristina Marqueze, Raul Machado Neto, Debora Botequio Moretti, Jardelina de Souza Todao Bernardino, Loyze Paola Oliveira de Lima, Luiz Aurelio de Campos Crispin. Centro de Genômica Funcional da ESALQ: Luiz Lehmann Coutinho, Ricardo Augusto Brassaloti, Raquel de Lello Rocha Campos Cassano. NGS Soluções Genômicas: Pilar Drummond Sampaio Corrêa Mariani. FZEA-USP Pirassununga: Mirele Daiana Poleti, Jessika Cristina Chagas Lesbon, Elisangela Chicaroni Mattos, Heidge Fukumasu. USP-Botucatu: Rejane Maria Tommasini Grotto, Jayme A. Souza-Neto, Guilherme Targino Valente, Patricia Akemi Assato, Felipe Allan da Silva da Costa, Bianca Cechetto Carlos. Mendelics: Bibiana Santos, João Paulo Kitajima, Erika Freitas, David Schlesinger. Hemocentro Ribeirão Preto: Simone Kashima, Evandra Strazza Rodrigues, Svetoslav Nanev Slavov, Elaine Vieira dos Santos, Rafael dos Santos Bezerra, Luiz Carlos Junior de Alcantara, Marta Giovanetti, Vagner Fonseca, Flavia Aburjaile, Rodrigo Tocantins Calado. FAMERP-SJRP: Cecília Artico Banho, Lívia Sacchetto, Fábio Sossai Possebon, Leila Sabrina Ullmann, Cíntia Bittar, Guilherme Campos, Helena Lage Ferreira, Jorge A. Petroli Marchesi, Maisa C. Pereira Parra, Marília Moraes, Paula Rahal, Paulo Inacio da Costa, João Pessoa Araújo Jr., Maurício Lacerda Nogueira. Prefeitura de Sao Paulo: Melissa Palmieri. |
| EPI_ISL_1966197, EPI_ISL_1966198, EPI_ISL_1966199, EPI_ISL_1966200 | SECRETARIA MUNICIPAL DE SAUDE SOROCABA | Instituto Butantan / Mendelics | Instituto Butantan: Dimas Tadeu Covas, Sandra Coccuzzo Sampaio, Maria Carolina Elias, José Salvatore Leister Patané, Vincent Louis Viala, Antonio Jorge Martins, Ricardo Haddad, Claudia Renata dos Santos Barros, Elaine Cristina Marqueze, Raul Machado Neto, Debora Botequio Moretti, Jardelina de Souza Todao Bernardino, Loyze Paola Oliveira de Lima, Luiz Aurelio de Campos Crispin. Centro de Genômica Funcional da ESALQ: Luiz Lehmann Coutinho, Ricardo Augusto Brassaloti, Raquel de Lello Rocha Campos Cassano. NGS Soluções Genômicas: Pilar Drummond Sampaio Corrêa Mariani. FZEA-USP Pirassununga: Mirele Daiana Poleti, Jessika Cristina Chagas Lesbon, Elisangela Chicaroni Mattos, Heidge Fukumasu. USP-Botucatu: Rejane Maria Tommasini Grotto, Jayme A. Souza-Neto, Guilherme Targino Valente, Patricia Akemi Assato, Felipe Allan da Silva da Costa, Bianca Cechetto Carlos. Mendelics: Bibiana Santos, João Paulo Kitajima, Erika Freitas, David Schlesinger. Hemocentro Ribeirão Preto: Simone Kashima, Evandra Strazza Rodrigues, Svetoslav Nanev Slavov, Elaine Vieira dos Santos, Rafael dos Santos Bezerra, Luiz Carlos Junior de Alcantara, Marta Giovanetti, Vagner Fonseca, Flavia Aburjaile, Rodrigo Tocantins Calado. FAMERP-SJRP: Cecília Artico Banho, Lívia Sacchetto, Fábio Sossai Possebon, Leila Sabrina Ullmann, Cíntia Bittar, Guilherme Campos, Helena Lage Ferreira, Jorge A. Petroli Marchesi, Maisa C. Pereira Parra, Marília Moraes, Paula Rahal, Paulo Inacio da Costa, João Pessoa Araújo Jr., Maurício Lacerda Nogueira. Prefeitura de Sao Paulo: Melissa Palmieri. |
| EPI_ISL_1966201 | UNIDADE BASICA DE SAUDE DA FAMILIA BILAC | Instituto Butantan / Mendelics | Instituto Butantan: Dimas Tadeu Covas, Sandra Coccuzzo Sampaio, Maria Carolina Elias, José Salvatore Leister Patané, Vincent Louis Viala, Antonio Jorge Martins, Ricardo Haddad, Claudia Renata dos Santos Barros, Elaine Cristina Marqueze, Raul Machado Neto, Debora Botequio Moretti, Jardelina de Souza Todao Bernardino, Loyze Paola Oliveira de Lima, Luiz Aurelio de Campos Crispin. Centro de Genômica Funcional da ESALQ: Luiz Lehmann Coutinho, Ricardo Augusto Brassaloti, Raquel de Lello Rocha Campos Cassano. NGS Soluções Genômicas: Pilar Drummond Sampaio Corrêa Mariani. FZEA-USP Pirassununga: Mirele Daiana Poleti, Jessika Cristina Chagas Lesbon, Elisangela Chicaroni Mattos, Heidge Fukumasu. USP-Botucatu: Rejane Maria Tommasini Grotto, Jayme A. Souza-Neto, Guilherme Targino Valente, Patricia Akemi Assato, Felipe Allan da Silva da Costa, Bianca Cechetto Carlos. Mendelics: Bibiana Santos, João Paulo Kitajima, Erika Freitas, David Schlesinger. Hemocentro Ribeirão Preto: Simone Kashima, Evandra Strazza Rodrigues, Svetoslav Nanev Slavov, Elaine Vieira dos Santos, Rafael dos Santos Bezerra, Luiz Carlos Junior de Alcantara, Marta Giovanetti, Vagner Fonseca, Flavia Aburjaile, Rodrigo Tocantins Calado. FAMERP-SJRP: Cecília Artico Banho, Lívia Sacchetto, Fábio Sossai Possebon, Leila Sabrina Ullmann, Cíntia Bittar, Guilherme Campos, Helena Lage Ferreira, Jorge A. Petroli Marchesi, Maisa C. Pereira Parra, Marília Moraes, Paula Rahal, Paulo Inacio da Costa, João Pessoa Araújo Jr., Maurício Lacerda Nogueira. Prefeitura de Sao Paulo: Melissa Palmieri. |
| EPI_ISL_1966202 | UBS JOSE SABINO FERREIRA | Instituto Butantan / Mendelics | Instituto Butantan: Dimas Tadeu Covas, Sandra Coccuzzo Sampaio, Maria Carolina Elias, José Salvatore Leister Patané, Vincent Louis Viala, Antonio Jorge Martins, Ricardo Haddad, Claudia Renata dos Santos Barros, Elaine Cristina Marqueze, Raul Machado Neto, Debora Botequio Moretti, Jardelina de Souza Todao Bernardino, Loyze Paola Oliveira de Lima, Luiz Aurelio de Campos Crispin. Centro de Genômica Funcional da ESALQ: Luiz Lehmann Coutinho, Ricardo Augusto Brassaloti, Raquel de Lello Rocha Campos Cassano. NGS Soluções Genômicas: Pilar Drummond Sampaio Corrêa Mariani. FZEA-USP Pirassununga: Mirele Daiana Poleti, Jessika Cristina Chagas Lesbon, Elisangela Chicaroni Mattos, Heidge Fukumasu. USP-Botucatu: Rejane Maria Tommasini Grotto, Jayme A. Souza-Neto, Guilherme Targino Valente, Patricia Akemi Assato, Felipe Allan da Silva da Costa, Bianca Cechetto Carlos. Mendelics: Bibiana Santos, João Paulo Kitajima, Erika Freitas, David Schlesinger. Hemocentro Ribeirão Preto: Simone Kashima, Evandra Strazza Rodrigues, Svetoslav Nanev Slavov, Elaine Vieira dos Santos, Rafael dos Santos Bezerra, Luiz Carlos Junior de Alcantara, Marta Giovanetti, Vagner Fonseca, Flavia Aburjaile, Rodrigo Tocantins Calado. FAMERP-SJRP: Cecília Artico Banho, Lívia Sacchetto, Fábio Sossai Possebon, Leila Sabrina Ullmann, Cíntia Bittar, Guilherme Campos, Helena Lage Ferreira, Jorge A. Petroli Marchesi, Maisa C. Pereira Parra, Marília Moraes, Paula Rahal, Paulo Inacio da Costa, João Pessoa Araújo Jr., Maurício Lacerda Nogueira. Prefeitura de Sao Paulo: Melissa Palmieri. |
| EPI_ISL_1966203 | UBS IRMA AGUEDA MARIA JAIME | Instituto Butantan / Mendelics | Instituto Butantan: Dimas Tadeu Covas, Sandra Coccuzzo Sampaio, Maria Carolina Elias, José Salvatore Leister Patané, Vincent Louis Viala, Antonio Jorge Martins, Ricardo Haddad, Claudia Renata dos Santos Barros, Elaine Cristina Marqueze, Raul Machado Neto, Debora Botequio Moretti, Jardelina de Souza Todao Bernardino, Loyze Paola Oliveira de Lima, Luiz Aurelio de Campos Crispin. Centro de Genômica Funcional da ESALQ: Luiz Lehmann Coutinho, Ricardo Augusto Brassaloti, Raquel de Lello Rocha Campos Cassano. NGS Soluções Genômicas: Pilar Drummond Sampaio Corrêa Mariani. FZEA-USP Pirassununga: Mirele Daiana Poleti, Jessika Cristina Chagas Lesbon, Elisangela Chicaroni Mattos, Heidge Fukumasu. USP-Botucatu: Rejane Maria Tommasini Grotto, Jayme A. Souza-Neto, Guilherme Targino Valente, Patricia Akemi Assato, Felipe Allan da Silva da Costa, Bianca Cechetto Carlos. Mendelics: Bibiana Santos, João Paulo Kitajima, Erika Freitas, David Schlesinger. Hemocentro Ribeirão Preto: Simone Kashima, Evandra Strazza Rodrigues, Svetoslav Nanev Slavov, Elaine Vieira dos Santos, Rafael dos Santos Bezerra, Luiz Carlos Junior de Alcantara, Marta Giovanetti, Vagner Fonseca, Flavia Aburjaile, Rodrigo Tocantins Calado. FAMERP-SJRP: Cecília Artico Banho, Lívia Sacchetto, Fábio Sossai Possebon, Leila Sabrina Ullmann, Cíntia Bittar, Guilherme Campos, Helena Lage Ferreira, Jorge A. Petroli Marchesi, Maisa C. Pereira Parra, Marília Moraes, Paula Rahal, Paulo Inacio da Costa, João Pessoa Araújo Jr., Maurício Lacerda Nogueira. Prefeitura de Sao Paulo: Melissa Palmieri. |
| EPI_ISL_1966204 | UBS JOSE HILARIO DOS SANTOS | Instituto Butantan / Mendelics | Instituto Butantan: Dimas Tadeu Covas, Sandra Coccuzzo Sampaio, Maria Carolina Elias, José Salvatore Leister Patané, Vincent Louis Viala, Antonio Jorge Martins, Ricardo Haddad, Claudia Renata dos Santos Barros, Elaine Cristina Marqueze, Raul Machado Neto, Debora Botequio Moretti, Jardelina de Souza Todao Bernardino, Loyze Paola Oliveira de Lima, Luiz Aurelio de Campos Crispin. Centro de Genômica Funcional da ESALQ: Luiz Lehmann Coutinho, Ricardo Augusto Brassaloti, Raquel de Lello Rocha Campos Cassano. NGS Soluções Genômicas: Pilar Drummond Sampaio Corrêa Mariani. FZEA-USP Pirassununga: Mirele Daiana Poleti, Jessika Cristina Chagas Lesbon, Elisangela Chicaroni Mattos, Heidge Fukumasu. USP-Botucatu: Rejane Maria Tommasini Grotto, Jayme A. Souza-Neto, Guilherme Targino Valente, Patricia Akemi Assato, Felipe Allan da Silva da Costa, Bianca Cechetto Carlos. Mendelics: Bibiana Santos, João Paulo Kitajima, Erika Freitas, David Schlesinger. Hemocentro Ribeirão Preto: Simone Kashima, Evandra Strazza Rodrigues, Svetoslav Nanev Slavov, Elaine Vieira dos Santos, Rafael dos Santos Bezerra, Luiz Carlos Junior de Alcantara, Marta Giovanetti, Vagner Fonseca, Flavia Aburjaile, Rodrigo Tocantins Calado. FAMERP-SJRP: Cecília Artico Banho, Lívia Sacchetto, Fábio Sossai Possebon, Leila Sabrina Ullmann, Cíntia Bittar, Guilherme Campos, Helena Lage Ferreira, Jorge A. Petroli Marchesi, Maisa C. Pereira Parra, Marília Moraes, Paula Rahal, Paulo Inacio da Costa, João Pessoa Araújo Jr., Maurício Lacerda Nogueira. Prefeitura de Sao Paulo: Melissa Palmieri. |
| EPI_ISL_1966206 | UBS OTACILIO FIRMINO LOPES | Instituto Butantan / Mendelics | Instituto Butantan: Dimas Tadeu Covas, Sandra Coccuzzo Sampaio, Maria Carolina Elias, José Salvatore Leister Patané, Vincent Louis Viala, Antonio Jorge Martins, Ricardo Haddad, Claudia Renata dos Santos Barros, Elaine Cristina Marqueze, Raul Machado Neto, Debora Botequio Moretti, Jardelina de Souza Todao Bernardino, Loyze Paola Oliveira de Lima, Luiz Aurelio de Campos Crispin. Centro de Genômica Funcional da ESALQ: Luiz Lehmann Coutinho, Ricardo Augusto Brassaloti, Raquel de Lello Rocha Campos Cassano. NGS Soluções Genômicas: Pilar Drummond Sampaio Corrêa Mariani. FZEA-USP Pirassununga: Mirele Daiana Poleti, Jessika Cristina Chagas Lesbon, Elisangela Chicaroni Mattos, Heidge Fukumasu. USP-Botucatu: Rejane Maria Tommasini Grotto, Jayme A. Souza-Neto, Guilherme Targino Valente, Patricia Akemi Assato, Felipe Allan da Silva da Costa, Bianca Cechetto Carlos. Mendelics: Bibiana Santos, João Paulo Kitajima, Erika Freitas, David Schlesinger. Hemocentro Ribeirão Preto: Simone Kashima, Evandra Strazza Rodrigues, Svetoslav Nanev Slavov, Elaine Vieira dos Santos, Rafael dos Santos Bezerra, Luiz Carlos Junior de Alcantara, Marta Giovanetti, Vagner Fonseca, Flavia Aburjaile, Rodrigo Tocantins Calado. FAMERP-SJRP: Cecília Artico Banho, Lívia Sacchetto, Fábio Sossai Possebon, Leila Sabrina Ullmann, Cíntia Bittar, Guilherme Campos, Helena Lage Ferreira, Jorge A. Petroli Marchesi, Maisa C. Pereira Parra, Marília Moraes, Paula Rahal, Paulo Inacio da Costa, João Pessoa Araújo Jr., Maurício Lacerda Nogueira. Prefeitura de Sao Paulo: Melissa Palmieri. |

| | | | |
|---|---|---|---|
| EPI_ISL_1966207 | UBS FRANCISCA LIMA DE LIRA | Instituto Butantan / Mendelics | Instituto Butantan: Dimas Tadeu Covas, Sandra Coccuzzo Sampaio, Maria Carolina Elias, José Salvatore Leister Patané, Vincent Louis Viala, Antonio Jorge Martins, Ricardo Haddad, Claudia Renata dos Santos Barros, Elaine Cristina Marqueze, Raul Machado Neto, Debora Botequio Moretti, Jardelina de Souza Todao Bernardino, Loyze Paola Oliveira de Lima, Luiz Aurelio de Campos Crispin. Centro de Genômica Funcional da ESALQ: Luiz Lehmann Coutinho, Ricardo Augusto Brassaloti, Raquel de Lello Rocha Campos Cassano. NGS Soluções Genômicas: Pilar Drummond Sampaio Corrêa Mariani. FZEA-USP Pirassununga: Mirele Daiana Poleti, Jessika Cristina Chagas Lesbon, Elisangela Chicaroni Mattos, Heidge Fukumasu. USP-Botucatu: Rejane Maria Tommasini Grotto, Jayme A. Souza-Neto, Guilherme Targino Valente, Patricia Akemi Assato, Felipe Allan da Silva da Costa, Bianca Cechetto Carlos. Mendelics: Bibiana Santos, João Paulo Kitajima, Erika Freitas, David Schlesinger. Hemocentro Ribeirão Preto: Simone Kashima, Evandra Strazza Rodrigues, Svetoslav Nanev Slavov, Elaine Vieira dos Santos, Rafael dos Santos Bezerra, Luiz Carlos Junior de Alcantara, Marta Giovanetti, Vagner Fonseca, Flavia Aburjaile, Rodrigo Tocantins Calado. FAMERP-SJRP: Cecília Artico Banho, Lívia Sacchetto, Fábio Sossai Possebon, Leila Sabrina Ullmann, Cíntia Bittar, Guilherme Campos, Helena Lage Ferreira, Jorge A. Petroli Marchesi, Maisa C. Pereira Parra, Marília Moraes, Paula Rahal, Paulo Inacio da Costa, João Pessoa Araújo Jr., Maurício Lacerda Nogueira. Prefeitura de Sao Paulo: Melissa Palmieri. |
| EPI_ISL_1966208 | UBS OTACILIO FIRMINO LOPES | Instituto Butantan / Mendelics | Instituto Butantan: Dimas Tadeu Covas, Sandra Coccuzzo Sampaio, Maria Carolina Elias, José Salvatore Leister Patané, Vincent Louis Viala, Antonio Jorge Martins, Ricardo Haddad, Claudia Renata dos Santos Barros, Elaine Cristina Marqueze, Raul Machado Neto, Debora Botequio Moretti, Jardelina de Souza Todao Bernardino, Loyze Paola Oliveira de Lima, Luiz Aurelio de Campos Crispin. Centro de Genômica Funcional da ESALQ: Luiz Lehmann Coutinho, Ricardo Augusto Brassaloti, Raquel de Lello Rocha Campos Cassano. NGS Soluções Genômicas: Pilar Drummond Sampaio Corrêa Mariani. FZEA-USP Pirassununga: Mirele Daiana Poleti, Jessika Cristina Chagas Lesbon, Elisangela Chicaroni Mattos, Heidge Fukumasu. USP-Botucatu: Rejane Maria Tommasini Grotto, Jayme A. Souza-Neto, Guilherme Targino Valente, Patricia Akemi Assato, Felipe Allan da Silva da Costa, Bianca Cechetto Carlos. Mendelics: Bibiana Santos, João Paulo Kitajima, Erika Freitas, David Schlesinger. Hemocentro Ribeirão Preto: Simone Kashima, Evandra Strazza Rodrigues, Svetoslav Nanev Slavov, Elaine Vieira dos Santos, Rafael dos Santos Bezerra, Luiz Carlos Junior de Alcantara, Marta Giovanetti, Vagner Fonseca, Flavia Aburjaile, Rodrigo Tocantins Calado. FAMERP-SJRP: Cecília Artico Banho, Lívia Sacchetto, Fábio Sossai Possebon, Leila Sabrina Ullmann, Cíntia Bittar, Guilherme Campos, Helena Lage Ferreira, Jorge A. Petroli Marchesi, Maisa C. Pereira Parra, Marília Moraes, Paula Rahal, Paulo Inacio da Costa, João Pessoa Araújo Jr., Maurício Lacerda Nogueira. Prefeitura de Sao Paulo: Melissa Palmieri. |
| EPI_ISL_1966209, EPI_ISL_1966210 | UBS HELENA MARREY | Instituto Butantan / Mendelics | Instituto Butantan: Dimas Tadeu Covas, Sandra Coccuzzo Sampaio, Maria Carolina Elias, José Salvatore Leister Patané, Vincent Louis Viala, Antonio Jorge Martins, Ricardo Haddad, Claudia Renata dos Santos Barros, Elaine Cristina Marqueze, Raul Machado Neto, Debora Botequio Moretti, Jardelina de Souza Todao Bernardino, Loyze Paola Oliveira de Lima, Luiz Aurelio de Campos Crispin. Centro de Genômica Funcional da ESALQ: Luiz Lehmann Coutinho, Ricardo Augusto Brassaloti, Raquel de Lello Rocha Campos Cassano. NGS Soluções Genômicas: Pilar Drummond Sampaio Corrêa Mariani. FZEA-USP Pirassununga: Mirele Daiana Poleti, Jessika Cristina Chagas Lesbon, Elisangela Chicaroni Mattos, Heidge Fukumasu. USP-Botucatu: Rejane Maria Tommasini Grotto, Jayme A. Souza-Neto, Guilherme Targino Valente, Patricia Akemi Assato, Felipe Allan da Silva da Costa, Bianca Cechetto Carlos. Mendelics: Bibiana Santos, João Paulo Kitajima, Erika Freitas, David Schlesinger. Hemocentro Ribeirão Preto: Simone Kashima, Evandra Strazza Rodrigues, Svetoslav Nanev Slavov, Elaine Vieira dos Santos, Rafael dos Santos Bezerra, Luiz Carlos Junior de Alcantara, Marta Giovanetti, Vagner Fonseca, Flavia Aburjaile, Rodrigo Tocantins Calado. FAMERP-SJRP: Cecília Artico Banho, Lívia Sacchetto, Fábio Sossai Possebon, Leila Sabrina Ullmann, Cíntia Bittar, Guilherme Campos, Helena Lage Ferreira, Jorge A. Petroli Marchesi, Maisa C. Pereira Parra, Marília Moraes, Paula Rahal, Paulo Inacio da Costa, João Pessoa Araújo Jr., Maurício Lacerda Nogueira. Prefeitura de Sao Paulo: Melissa Palmieri. |
| EPI_ISL_1966211 | UBS EMILIA COSME CERQUEIRA | Instituto Butantan / Mendelics | Instituto Butantan: Dimas Tadeu Covas, Sandra Coccuzzo Sampaio, Maria Carolina Elias, José Salvatore Leister Patané, Vincent Louis Viala, Antonio Jorge Martins, Ricardo Haddad, Claudia Renata dos Santos Barros, Elaine Cristina Marqueze, Raul Machado Neto, Debora Botequio Moretti, Jardelina de Souza Todao Bernardino, Loyze Paola Oliveira de Lima, Luiz Aurelio de Campos Crispin. Centro de Genômica Funcional da ESALQ: Luiz Lehmann Coutinho, Ricardo Augusto Brassaloti, Raquel de Lello Rocha Campos Cassano. NGS Soluções Genômicas: Pilar Drummond Sampaio Corrêa Mariani. FZEA-USP Pirassununga: Mirele Daiana Poleti, Jessika Cristina Chagas Lesbon, Elisangela Chicaroni Mattos, Heidge Fukumasu. USP-Botucatu: Rejane Maria Tommasini Grotto, Jayme A. Souza-Neto, Guilherme Targino Valente, Patricia Akemi Assato, Felipe Allan da Silva da Costa, Bianca Cechetto Carlos. Mendelics: Bibiana Santos, João Paulo Kitajima, Erika Freitas, David Schlesinger. Hemocentro Ribeirão Preto: Simone Kashima, Evandra Strazza Rodrigues, Svetoslav Nanev Slavov, Elaine Vieira dos Santos, Rafael dos Santos Bezerra, Luiz Carlos Junior de Alcantara, Marta Giovanetti, Vagner Fonseca, Flavia Aburjaile, Rodrigo Tocantins Calado. FAMERP-SJRP: Cecília Artico Banho, Lívia Sacchetto, Fábio Sossai Possebon, Leila Sabrina Ullmann, Cíntia Bittar, Guilherme Campos, Helena Lage Ferreira, Jorge A. Petroli Marchesi, Maisa C. Pereira Parra, Marília Moraes, Paula Rahal, Paulo Inacio da Costa, João Pessoa Araújo Jr., Maurício Lacerda Nogueira. Prefeitura de Sao Paulo: Melissa Palmieri. |
| EPI_ISL_1966212 | UBS SYLVIO JOAO L DE LUCIA | Instituto Butantan / Mendelics | Instituto Butantan: Dimas Tadeu Covas, Sandra Coccuzzo Sampaio, Maria Carolina Elias, José Salvatore Leister Patané, Vincent Louis Viala, Antonio Jorge Martins, Ricardo Haddad, Claudia Renata dos Santos Barros, Elaine Cristina Marqueze, Raul Machado Neto, Debora Botequio Moretti, Jardelina de Souza Todao Bernardino, Loyze Paola Oliveira de Lima, Luiz Aurelio de Campos Crispin. Centro de Genômica Funcional da ESALQ: Luiz Lehmann Coutinho, Ricardo Augusto Brassaloti, Raquel de Lello Rocha Campos Cassano. NGS Soluções Genômicas: Pilar Drummond Sampaio Corrêa Mariani. FZEA-USP Pirassununga: Mirele Daiana Poleti, Jessika Cristina Chagas Lesbon, Elisangela Chicaroni Mattos, Heidge Fukumasu. USP-Botucatu: Rejane Maria Tommasini Grotto, Jayme A. Souza-Neto, Guilherme Targino Valente, Patricia Akemi Assato, Felipe Allan da Silva da Costa, Bianca Cechetto Carlos. Mendelics: Bibiana Santos, João Paulo Kitajima, Erika Freitas, David Schlesinger. Hemocentro Ribeirão Preto: Simone Kashima, Evandra Strazza Rodrigues, Svetoslav Nanev Slavov, Elaine Vieira dos Santos, Rafael dos Santos Bezerra, Luiz Carlos Junior de Alcantara, Marta Giovanetti, Vagner Fonseca, Flavia Aburjaile, Rodrigo Tocantins Calado. FAMERP-SJRP: Cecília Artico Banho, Lívia Sacchetto, Fábio Sossai Possebon, Leila Sabrina Ullmann, Cíntia Bittar, Guilherme Campos, Helena Lage Ferreira, Jorge A. Petroli Marchesi, Maisa C. Pereira Parra, Marília Moraes, Paula Rahal, Paulo Inacio da Costa, João Pessoa Araújo Jr., Maurício Lacerda Nogueira. Prefeitura de Sao Paulo: Melissa Palmieri. |
| EPI_ISL_1966213, EPI_ISL_1966214 | UBS EMILIA COSME CERQUEIRA | Instituto Butantan / Mendelics | Instituto Butantan: Dimas Tadeu Covas, Sandra Coccuzzo Sampaio, Maria Carolina Elias, José Salvatore Leister Patané, Vincent Louis Viala, Antonio Jorge Martins, Ricardo Haddad, Claudia Renata dos Santos Barros, Elaine Cristina Marqueze, Raul Machado Neto, Debora Botequio Moretti, Jardelina de Souza Todao Bernardino, Loyze Paola Oliveira de Lima, Luiz Aurelio de Campos Crispin. Centro de Genômica Funcional da ESALQ: Luiz Lehmann Coutinho, Ricardo Augusto Brassaloti, Raquel de Lello Rocha Campos Cassano. NGS Soluções Genômicas: Pilar Drummond Sampaio Corrêa Mariani. FZEA-USP Pirassununga: Mirele Daiana Poleti, Jessika Cristina Chagas Lesbon, Elisangela Chicaroni Mattos, Heidge Fukumasu. USP-Botucatu: Rejane Maria Tommasini Grotto, Jayme A. Souza-Neto, Guilherme Targino Valente, Patricia Akemi Assato, Felipe Allan da Silva da Costa, Bianca Cechetto Carlos. Mendelics: Bibiana Santos, João Paulo Kitajima, Erika Freitas, David Schlesinger. Hemocentro Ribeirão Preto: Simone Kashima, Evandra Strazza Rodrigues, Svetoslav Nanev Slavov, Elaine Vieira dos Santos, Rafael dos Santos Bezerra, Luiz Carlos Junior de Alcantara, Marta Giovanetti, Vagner Fonseca, Flavia Aburjaile, Rodrigo Tocantins Calado. FAMERP-SJRP: Cecília Artico Banho, Lívia Sacchetto, Fábio Sossai Possebon, Leila Sabrina Ullmann, Cíntia Bittar, Guilherme Campos, Helena Lage Ferreira, Jorge A. Petroli Marchesi, Maisa C. Pereira Parra, Marília Moraes, Paula Rahal, Paulo Inacio da Costa, João Pessoa Araújo Jr., Maurício Lacerda Nogueira. Prefeitura de Sao Paulo: Melissa Palmieri. |
| EPI_ISL_1966215, EPI_ISL_1966216, EPI_ISL_1966217 | UBS HELENA MARREY | Instituto Butantan / Mendelics | Instituto Butantan: Dimas Tadeu Covas, Sandra Coccuzzo Sampaio, Maria Carolina Elias, José Salvatore Leister Patané, Vincent Louis Viala, Antonio Jorge Martins, Ricardo Haddad, Claudia Renata dos Santos Barros, Elaine Cristina Marqueze, Raul Machado Neto, Debora Botequio Moretti, Jardelina de Souza Todao Bernardino, Loyze Paola Oliveira de Lima, Luiz Aurelio de Campos Crispin. Centro de Genômica Funcional da ESALQ: Luiz Lehmann Coutinho, Ricardo Augusto Brassaloti, Raquel de Lello Rocha Campos Cassano. NGS Soluções Genômicas: Pilar Drummond Sampaio Corrêa Mariani. FZEA-USP Pirassununga: Mirele Daiana Poleti, Jessika Cristina Chagas Lesbon, Elisangela Chicaroni Mattos, Heidge Fukumasu. USP-Botucatu: Rejane Maria Tommasini Grotto, Jayme A. Souza-Neto, Guilherme Targino Valente, Patricia Akemi Assato, Felipe Allan da Silva da Costa, Bianca Cechetto Carlos. Mendelics: Bibiana Santos, João Paulo Kitajima, Erika Freitas, David Schlesinger. Hemocentro Ribeirão Preto: Simone Kashima, Evandra Strazza Rodrigues, Svetoslav Nanev Slavov, Elaine Vieira dos Santos, Rafael dos Santos Bezerra, Luiz Carlos Junior de Alcantara, Marta Giovanetti, Vagner Fonseca, Flavia Aburjaile, Rodrigo Tocantins Calado. FAMERP-SJRP: Cecília Artico Banho, Lívia Sacchetto, Fábio Sossai Possebon, Leila Sabrina Ullmann, Cíntia Bittar, Guilherme Campos, Helena Lage Ferreira, Jorge A. Petroli Marchesi, Maisa C. Pereira Parra, Marília Moraes, Paula Rahal, Paulo Inacio da Costa, João Pessoa Araújo Jr., Maurício Lacerda Nogueira. Prefeitura de Sao Paulo: Melissa Palmieri. |
| EPI_ISL_1966218 | PPA FENELON GUEDES PEREIRA | Instituto Butantan / Mendelics | Instituto Butantan: Dimas Tadeu Covas, Sandra Coccuzzo Sampaio, Maria Carolina Elias, José Salvatore Leister Patané, Vincent Louis Viala, Antonio Jorge Martins, Ricardo Haddad, Claudia Renata dos Santos Barros, Elaine Cristina Marqueze, Raul Machado Neto, Debora Botequio Moretti, Jardelina de Souza Todao Bernardino, Loyze Paola Oliveira de Lima, Luiz Aurelio de Campos Crispin. Centro de Genômica Funcional da ESALQ: Luiz Lehmann Coutinho, Ricardo Augusto Brassaloti, Raquel de Lello Rocha Campos Cassano. NGS Soluções Genômicas: Pilar Drummond Sampaio Corrêa Mariani. FZEA-USP Pirassununga: Mirele Daiana Poleti, Jessika Cristina Chagas Lesbon, Elisangela Chicaroni Mattos, Heidge Fukumasu. USP-Botucatu: Rejane Maria Tommasini Grotto, Jayme A. Souza-Neto, Guilherme Targino Valente, Patricia Akemi Assato, Felipe Allan da Silva da Costa, Bianca Cechetto Carlos. Mendelics: Bibiana Santos, João Paulo Kitajima, Erika Freitas, David Schlesinger. Hemocentro Ribeirão Preto: Simone Kashima, Evandra Strazza |

| | | | |
|---|---|---|---|
| | | | Rodrigues, Svetoslav Nanev Slavov, Elaine Vieira dos Santos, Rafael dos Santos Bezerra, Luiz Carlos Junior de Alcantara, Marta Giovanetti, Vagner Fonseca, Flavia Aburjaile, Rodrigo Tocantins Calado. FAMERP-SJRP: Cecília Artico Banho, Lívia Sacchetto, Fábio Sossai Possebon, Leila Sabrina Ullmann, Cíntia Bittar, Guilherme Campos, Helena Lage Ferreira, Jorge A. Petroli Marchesi, Maisa C. Pereira Parra, Marília Moraes, Paula Rahal, Paulo Inacio da Costa, João Pessoa Araújo Jr., Maurício Lacerda Nogueira. Prefeitura de Sao Paulo: Melissa Palmieri. |
| EPI_ISL_1966219 | CENTRO DE SAUDE DE BORA | Instituto Butantan / Mendelics | Instituto Butantan: Dimas Tadeu Covas, Sandra Coccuzzo Sampaio, Maria Carolina Elias, José Salvatore Leister Patané, Vincent Louis Viala, Antonio Jorge Martins, Ricardo Haddad, Claudia Renata dos Santos Barros, Elaine Cristina Marqueze, Raul Machado Neto, Debora Botequio Moretti, Jardelina de Souza Todao Bernardino, Loyze Paola Oliveira de Lima, Luiz Aurelio de Campos Crispin. Centro de Genômica Funcional da ESALQ: Luiz Lehmann Coutinho, Ricardo Augusto Brassaloti, Raquel de Lello Rocha Campos Cassano. NGS Soluções Genômicas: Pilar Drummond Sampaio Corrêa Mariani. FZEA-USP Pirassununga: Mirele Daiana Poleti, Jessika Cristina Chagas Lesbon, Elisangela Chicaroni Mattos, Heidge Fukumasu. USP-Botucatu: Rejane Maria Tommasini Grotto, Jayme A. Souza-Neto, Guilherme Targino Valente, Patricia Akemi Assato, Felipe Allan da Silva da Costa, Bianca Cechetto Carlos. Mendelics: Bibiana Santos, João Paulo Kitajima, Erika Freitas, David Schlesinger. Hemocentro Ribeirão Preto: Simone Kashima, Evandra Strazza Rodrigues, Svetoslav Nanev Slavov, Elaine Vieira dos Santos, Rafael dos Santos Bezerra, Luiz Carlos Junior de Alcantara, Marta Giovanetti, Vagner Fonseca, Flavia Aburjaile, Rodrigo Tocantins Calado. FAMERP-SJRP: Cecília Artico Banho, Lívia Sacchetto, Fábio Sossai Possebon, Leila Sabrina Ullmann, Cíntia Bittar, Guilherme Campos, Helena Lage Ferreira, Jorge A. Petroli Marchesi, Maisa C. Pereira Parra, Marília Moraes, Paula Rahal, Paulo Inacio da Costa, João Pessoa Araújo Jr., Maurício Lacerda Nogueira. Prefeitura de Sao Paulo: Melissa Palmieri. |
| EPI_ISL_1966220 | PRONTO ATENDIMENTO SAO JOSE | Instituto Butantan / Mendelics | Instituto Butantan: Dimas Tadeu Covas, Sandra Coccuzzo Sampaio, Maria Carolina Elias, José Salvatore Leister Patané, Vincent Louis Viala, Antonio Jorge Martins, Ricardo Haddad, Claudia Renata dos Santos Barros, Elaine Cristina Marqueze, Raul Machado Neto, Debora Botequio Moretti, Jardelina de Souza Todao Bernardino, Loyze Paola Oliveira de Lima, Luiz Aurelio de Campos Crispin. Centro de Genômica Funcional da ESALQ: Luiz Lehmann Coutinho, Ricardo Augusto Brassaloti, Raquel de Lello Rocha Campos Cassano. NGS Soluções Genômicas: Pilar Drummond Sampaio Corrêa Mariani. FZEA-USP Pirassununga: Mirele Daiana Poleti, Jessika Cristina Chagas Lesbon, Elisangela Chicaroni Mattos, Heidge Fukumasu. USP-Botucatu: Rejane Maria Tommasini Grotto, Jayme A. Souza-Neto, Guilherme Targino Valente, Patricia Akemi Assato, Felipe Allan da Silva da Costa, Bianca Cechetto Carlos. Mendelics: Bibiana Santos, João Paulo Kitajima, Erika Freitas, David Schlesinger. Hemocentro Ribeirão Preto: Simone Kashima, Evandra Strazza Rodrigues, Svetoslav Nanev Slavov, Elaine Vieira dos Santos, Rafael dos Santos Bezerra, Luiz Carlos Junior de Alcantara, Marta Giovanetti, Vagner Fonseca, Flavia Aburjaile, Rodrigo Tocantins Calado. FAMERP-SJRP: Cecília Artico Banho, Lívia Sacchetto, Fábio Sossai Possebon, Leila Sabrina Ullmann, Cíntia Bittar, Guilherme Campos, Helena Lage Ferreira, Jorge A. Petroli Marchesi, Maisa C. Pereira Parra, Marília Moraes, Paula Rahal, Paulo Inacio da Costa, João Pessoa Araújo Jr., Maurício Lacerda Nogueira. Prefeitura de Sao Paulo: Melissa Palmieri. |
| EPI_ISL_1966221 | PRONTO ATENDIMENTO VILA PADRE ANCHIETA | Instituto Butantan / Mendelics | Instituto Butantan: Dimas Tadeu Covas, Sandra Coccuzzo Sampaio, Maria Carolina Elias, José Salvatore Leister Patané, Vincent Louis Viala, Antonio Jorge Martins, Ricardo Haddad, Claudia Renata dos Santos Barros, Elaine Cristina Marqueze, Raul Machado Neto, Debora Botequio Moretti, Jardelina de Souza Todao Bernardino, Loyze Paola Oliveira de Lima, Luiz Aurelio de Campos Crispin. Centro de Genômica Funcional da ESALQ: Luiz Lehmann Coutinho, Ricardo Augusto Brassaloti, Raquel de Lello Rocha Campos Cassano. NGS Soluções Genômicas: Pilar Drummond Sampaio Corrêa Mariani. FZEA-USP Pirassununga: Mirele Daiana Poleti, Jessika Cristina Chagas Lesbon, Elisangela Chicaroni Mattos, Heidge Fukumasu. USP-Botucatu: Rejane Maria Tommasini Grotto, Jayme A. Souza-Neto, Guilherme Targino Valente, Patricia Akemi Assato, Felipe Allan da Silva da Costa, Bianca Cechetto Carlos. Mendelics: Bibiana Santos, João Paulo Kitajima, Erika Freitas, David Schlesinger. Hemocentro Ribeirão Preto: Simone Kashima, Evandra Strazza Rodrigues, Svetoslav Nanev Slavov, Elaine Vieira dos Santos, Rafael dos Santos Bezerra, Luiz Carlos Junior de Alcantara, Marta Giovanetti, Vagner Fonseca, Flavia Aburjaile, Rodrigo Tocantins Calado. FAMERP-SJRP: Cecília Artico Banho, Lívia Sacchetto, Fábio Sossai Possebon, Leila Sabrina Ullmann, Cíntia Bittar, Guilherme Campos, Helena Lage Ferreira, Jorge A. Petroli Marchesi, Maisa C. Pereira Parra, Marília Moraes, Paula Rahal, Paulo Inacio da Costa, João Pessoa Araújo Jr., Maurício Lacerda Nogueira. Prefeitura de Sao Paulo: Melissa Palmieri. |
| EPI_ISL_1966222 | PRONTO ATENDIMENTO SAO JOSE | Instituto Butantan / Mendelics | Instituto Butantan: Dimas Tadeu Covas, Sandra Coccuzzo Sampaio, Maria Carolina Elias, José Salvatore Leister Patané, Vincent Louis Viala, Antonio Jorge Martins, Ricardo Haddad, Claudia Renata dos Santos Barros, Elaine Cristina Marqueze, Raul Machado Neto, Debora Botequio Moretti, Jardelina de Souza Todao Bernardino, Loyze Paola Oliveira de Lima, Luiz Aurelio de Campos Crispin. Centro de Genômica Funcional da ESALQ: Luiz Lehmann Coutinho, Ricardo Augusto Brassaloti, Raquel de Lello Rocha Campos Cassano. NGS Soluções Genômicas: Pilar Drummond Sampaio Corrêa Mariani. FZEA-USP Pirassununga: Mirele Daiana Poleti, Jessika Cristina Chagas Lesbon, Elisangela Chicaroni Mattos, Heidge Fukumasu. USP-Botucatu: Rejane Maria Tommasini Grotto, Jayme A. Souza-Neto, Guilherme Targino Valente, Patricia Akemi Assato, Felipe Allan da Silva da Costa, Bianca Cechetto Carlos. Mendelics: Bibiana Santos, João Paulo Kitajima, Erika Freitas, David Schlesinger. Hemocentro Ribeirão Preto: Simone Kashima, Evandra Strazza Rodrigues, Svetoslav Nanev Slavov, Elaine Vieira dos Santos, Rafael dos Santos Bezerra, Luiz Carlos Junior de Alcantara, Marta Giovanetti, Vagner Fonseca, Flavia Aburjaile, Rodrigo Tocantins Calado. FAMERP-SJRP: Cecília Artico Banho, Lívia Sacchetto, Fábio Sossai Possebon, Leila Sabrina Ullmann, Cíntia Bittar, Guilherme Campos, Helena Lage Ferreira, Jorge A. Petroli Marchesi, Maisa C. Pereira Parra, Marília Moraes, Paula Rahal, Paulo Inacio da Costa, João Pessoa Araújo Jr., Maurício Lacerda Nogueira. Prefeitura de Sao Paulo: Melissa Palmieri. |
| EPI_ISL_1966223, EPI_ISL_1966224, EPI_ISL_1966225 | PRONTO ATENDIMENTO VILA PADRE ANCHIETA | Instituto Butantan / Mendelics | Instituto Butantan: Dimas Tadeu Covas, Sandra Coccuzzo Sampaio, Maria Carolina Elias, José Salvatore Leister Patané, Vincent Louis Viala, Antonio Jorge Martins, Ricardo Haddad, Claudia Renata dos Santos Barros, Elaine Cristina Marqueze, Raul Machado Neto, Debora Botequio Moretti, Jardelina de Souza Todao Bernardino, Loyze Paola Oliveira de Lima, Luiz Aurelio de Campos Crispin. Centro de Genômica Funcional da ESALQ: Luiz Lehmann Coutinho, Ricardo Augusto Brassaloti, Raquel de Lello Rocha Campos Cassano. NGS Soluções Genômicas: Pilar Drummond Sampaio Corrêa Mariani. FZEA-USP Pirassununga: Mirele Daiana Poleti, Jessika Cristina Chagas Lesbon, Elisangela Chicaroni Mattos, Heidge Fukumasu. USP-Botucatu: Rejane Maria Tommasini Grotto, Jayme A. Souza-Neto, Guilherme Targino Valente, Patricia Akemi Assato, Felipe Allan da Silva da Costa, Bianca Cechetto Carlos. Mendelics: Bibiana Santos, João Paulo Kitajima, Erika Freitas, David Schlesinger. Hemocentro Ribeirão Preto: Simone Kashima, Evandra Strazza Rodrigues, Svetoslav Nanev Slavov, Elaine Vieira dos Santos, Rafael dos Santos Bezerra, Luiz Carlos Junior de Alcantara, Marta Giovanetti, Vagner Fonseca, Flavia Aburjaile, Rodrigo Tocantins Calado. FAMERP-SJRP: Cecília Artico Banho, Lívia Sacchetto, Fábio Sossai Possebon, Leila Sabrina Ullmann, Cíntia Bittar, Guilherme Campos, Helena Lage Ferreira, Jorge A. Petroli Marchesi, Maisa C. Pereira Parra, Marília Moraes, Paula Rahal, Paulo Inacio da Costa, João Pessoa Araújo Jr., Maurício Lacerda Nogueira. Prefeitura de Sao Paulo: Melissa Palmieri. |
| EPI_ISL_1966226 | PRONTO ATENDIMENTO SAO JOSE | Instituto Butantan / Mendelics | Instituto Butantan: Dimas Tadeu Covas, Sandra Coccuzzo Sampaio, Maria Carolina Elias, José Salvatore Leister Patané, Vincent Louis Viala, Antonio Jorge Martins, Ricardo Haddad, Claudia Renata dos Santos Barros, Elaine Cristina Marqueze, Raul Machado Neto, Debora Botequio Moretti, Jardelina de Souza Todao Bernardino, Loyze Paola Oliveira de Lima, Luiz Aurelio de Campos Crispin. Centro de Genômica Funcional da ESALQ: Luiz Lehmann Coutinho, Ricardo Augusto Brassaloti, Raquel de Lello Rocha Campos Cassano. NGS Soluções Genômicas: Pilar Drummond Sampaio Corrêa Mariani. FZEA-USP Pirassununga: Mirele Daiana Poleti, Jessika Cristina Chagas Lesbon, Elisangela Chicaroni Mattos, Heidge Fukumasu. USP-Botucatu: Rejane Maria Tommasini Grotto, Jayme A. Souza-Neto, Guilherme Targino Valente, Patricia Akemi Assato, Felipe Allan da Silva da Costa, Bianca Cechetto Carlos. Mendelics: Bibiana Santos, João Paulo Kitajima, Erika Freitas, David Schlesinger. Hemocentro Ribeirão Preto: Simone Kashima, Evandra Strazza Rodrigues, Svetoslav Nanev Slavov, Elaine Vieira dos Santos, Rafael dos Santos Bezerra, Luiz Carlos Junior de Alcantara, Marta Giovanetti, Vagner Fonseca, Flavia Aburjaile, Rodrigo Tocantins Calado. FAMERP-SJRP: Cecília Artico Banho, Lívia Sacchetto, Fábio Sossai Possebon, Leila Sabrina Ullmann, Cíntia Bittar, Guilherme Campos, Helena Lage Ferreira, Jorge A. Petroli Marchesi, Maisa C. Pereira Parra, Marília Moraes, Paula Rahal, Paulo Inacio da Costa, João Pessoa Araújo Jr., Maurício Lacerda Nogueira. Prefeitura de Sao Paulo: Melissa Palmieri. |
| EPI_ISL_1966227 | UBS SAO JUDAS | Instituto Butantan / Mendelics | Instituto Butantan: Dimas Tadeu Covas, Sandra Coccuzzo Sampaio, Maria Carolina Elias, José Salvatore Leister Patané, Vincent Louis Viala, Antonio Jorge Martins, Ricardo Haddad, Claudia Renata dos Santos Barros, Elaine Cristina Marqueze, Raul Machado Neto, Debora Botequio Moretti, Jardelina de Souza Todao Bernardino, Loyze Paola Oliveira de Lima, Luiz Aurelio de Campos Crispin. Centro de Genômica Funcional da ESALQ: Luiz Lehmann Coutinho, Ricardo Augusto Brassaloti, Raquel de Lello Rocha Campos Cassano. NGS Soluções Genômicas: Pilar Drummond Sampaio Corrêa Mariani. FZEA-USP Pirassununga: Mirele Daiana Poleti, Jessika Cristina Chagas Lesbon, Elisangela Chicaroni Mattos, Heidge Fukumasu. USP-Botucatu: Rejane Maria Tommasini Grotto, Jayme A. Souza-Neto, Guilherme Targino Valente, Patricia Akemi Assato, Felipe Allan da Silva da Costa, Bianca Cechetto Carlos. Mendelics: Bibiana Santos, João Paulo Kitajima, Erika Freitas, David Schlesinger. Hemocentro Ribeirão Preto: Simone Kashima, Evandra Strazza Rodrigues, Svetoslav Nanev Slavov, Elaine Vieira dos Santos, Rafael dos Santos Bezerra, Luiz Carlos Junior de Alcantara, Marta Giovanetti, Vagner Fonseca, Flavia Aburjaile, Rodrigo Tocantins Calado. FAMERP-SJRP: Cecília Artico Banho, Lívia Sacchetto, Fábio Sossai Possebon, Leila Sabrina Ullmann, Cíntia Bittar, Guilherme Campos, Helena Lage Ferreira, Jorge A. Petroli Marchesi, Maisa C. Pereira Parra, Marília Moraes, Paula Rahal, Paulo Inacio da Costa, João Pessoa Araújo Jr., Maurício Lacerda Nogueira. Prefeitura de Sao Paulo: Melissa Palmieri. |
| EPI_ISL_1966228 | CENTRO DE SAUDE DE JULIO MESQUITA | Instituto Butantan / Mendelics | Instituto Butantan: Dimas Tadeu Covas, Sandra Coccuzzo Sampaio, Maria Carolina Elias, José Salvatore Leister Patané, Vincent Louis Viala, Antonio Jorge Martins, Ricardo Haddad, Claudia Renata dos Santos Barros, Elaine Cristina Marqueze, Raul Machado Neto, Debora Botequio Moretti, Jardelina de |

| | | | |
|---|---|---|---|
| | | | Souza Todao Bernardino, Loyze Paola Oliveira de Lima, Luiz Aurelio de Campos Crispin. Centro de Genômica Funcional da ESALQ: Luiz Lehmann Coutinho, Ricardo Augusto Brassaloti, Raquel de Lello Rocha Campos Cassano. NGS Soluções Genômicas: Pilar Drummond Sampaio Corrêa Mariani. FZEA-USP Pirassununga: Mirele Daiana Poleti, Jessika Cristina Chagas Lesbon, Elisangela Chicaroni Mattos, Heidge Fukumasu. USP-Botucatu: Rejane Maria Tommasini Grotto, Jayme A. Souza-Neto, Guilherme Targino Valente, Patricia Akemi Assato, Felipe Allan da Silva da Costa, Bianca Cechetto Carlos. Mendelics: Bibiana Santos, João Paulo Kitajima, Erika Freitas, David Schlesinger. Hemocentro Ribeirão Preto: Simone Kashima, Evandra Strazza Rodrigues, Svetoslav Nanev Slavov, Elaine Vieira dos Santos, Rafael dos Santos Bezerra, Luiz Carlos Junior de Alcantara, Marta Giovanetti, Vagner Fonseca, Flavia Aburjaile, Rodrigo Tocantins Calado. FAMERP-SJRP: Cecília Artico Banho, Lívia Sacchetto, Fábio Sossai Possebon, Leila Sabrina Ullmann, Cíntia Bittar, Guilherme Campos, Helena Lage Ferreira, Jorge A. Petroli Marchesi, Maisa C. Pereira Parra, Marília Moraes, Paula Rahal, Paulo Inacio da Costa, João Pessoa Araújo Jr., Maurício Lacerda Nogueira. Prefeitura de Sao Paulo: Melissa Palmieri. |
| EPI_ISL_1966229 | UBS SYLVIO JOAO L DE LUCIA | Instituto Butantan / Mendelics | Instituto Butantan: Dimas Tadeu Covas, Sandra Coccuzzo Sampaio, Maria Carolina Elias, José Salvatore Leister Patané, Vincent Louis Viala, Antonio Jorge Martins, Ricardo Haddad, Claudia Renata dos Santos Barros, Elaine Cristina Marqueze, Raul Machado Neto, Debora Botequio Moretti, Jardelina de Souza Todao Bernardino, Loyze Paola Oliveira de Lima, Luiz Aurelio de Campos Crispin. Centro de Genômica Funcional da ESALQ: Luiz Lehmann Coutinho, Ricardo Augusto Brassaloti, Raquel de Lello Rocha Campos Cassano. NGS Soluções Genômicas: Pilar Drummond Sampaio Corrêa Mariani. FZEA-USP Pirassununga: Mirele Daiana Poleti, Jessika Cristina Chagas Lesbon, Elisangela Chicaroni Mattos, Heidge Fukumasu. USP-Botucatu: Rejane Maria Tommasini Grotto, Jayme A. Souza-Neto, Guilherme Targino Valente, Patricia Akemi Assato, Felipe Allan da Silva da Costa, Bianca Cechetto Carlos. Mendelics: Bibiana Santos, João Paulo Kitajima, Erika Freitas, David Schlesinger. Hemocentro Ribeirão Preto: Simone Kashima, Evandra Strazza Rodrigues, Svetoslav Nanev Slavov, Elaine Vieira dos Santos, Rafael dos Santos Bezerra, Luiz Carlos Junior de Alcantara, Marta Giovanetti, Vagner Fonseca, Flavia Aburjaile, Rodrigo Tocantins Calado. FAMERP-SJRP: Cecília Artico Banho, Lívia Sacchetto, Fábio Sossai Possebon, Leila Sabrina Ullmann, Cíntia Bittar, Guilherme Campos, Helena Lage Ferreira, Jorge A. Petroli Marchesi, Maisa C. Pereira Parra, Marília Moraes, Paula Rahal, Paulo Inacio da Costa, João Pessoa Araújo Jr., Maurício Lacerda Nogueira. Prefeitura de Sao Paulo: Melissa Palmieri. |
| EPI_ISL_1966230 | UBS EMILIA COSME CERQUEIRA | Instituto Butantan / Mendelics | Instituto Butantan: Dimas Tadeu Covas, Sandra Coccuzzo Sampaio, Maria Carolina Elias, José Salvatore Leister Patané, Vincent Louis Viala, Antonio Jorge Martins, Ricardo Haddad, Claudia Renata dos Santos Barros, Elaine Cristina Marqueze, Raul Machado Neto, Debora Botequio Moretti, Jardelina de Souza Todao Bernardino, Loyze Paola Oliveira de Lima, Luiz Aurelio de Campos Crispin. Centro de Genômica Funcional da ESALQ: Luiz Lehmann Coutinho, Ricardo Augusto Brassaloti, Raquel de Lello Rocha Campos Cassano. NGS Soluções Genômicas: Pilar Drummond Sampaio Corrêa Mariani. FZEA-USP Pirassununga: Mirele Daiana Poleti, Jessika Cristina Chagas Lesbon, Elisangela Chicaroni Mattos, Heidge Fukumasu. USP-Botucatu: Rejane Maria Tommasini Grotto, Jayme A. Souza-Neto, Guilherme Targino Valente, Patricia Akemi Assato, Felipe Allan da Silva da Costa, Bianca Cechetto Carlos. Mendelics: Bibiana Santos, João Paulo Kitajima, Erika Freitas, David Schlesinger. Hemocentro Ribeirão Preto: Simone Kashima, Evandra Strazza Rodrigues, Svetoslav Nanev Slavov, Elaine Vieira dos Santos, Rafael dos Santos Bezerra, Luiz Carlos Junior de Alcantara, Marta Giovanetti, Vagner Fonseca, Flavia Aburjaile, Rodrigo Tocantins Calado. FAMERP-SJRP: Cecília Artico Banho, Lívia Sacchetto, Fábio Sossai Possebon, Leila Sabrina Ullmann, Cíntia Bittar, Guilherme Campos, Helena Lage Ferreira, Jorge A. Petroli Marchesi, Maisa C. Pereira Parra, Marília Moraes, Paula Rahal, Paulo Inacio da Costa, João Pessoa Araújo Jr., Maurício Lacerda Nogueira. Prefeitura de Sao Paulo: Melissa Palmieri. |
| EPI_ISL_1966231 | UBS IRMA AGUEDA MARIA JAIME | Instituto Butantan / Mendelics | Instituto Butantan: Dimas Tadeu Covas, Sandra Coccuzzo Sampaio, Maria Carolina Elias, José Salvatore Leister Patané, Vincent Louis Viala, Antonio Jorge Martins, Ricardo Haddad, Claudia Renata dos Santos Barros, Elaine Cristina Marqueze, Raul Machado Neto, Debora Botequio Moretti, Jardelina de Souza Todao Bernardino, Loyze Paola Oliveira de Lima, Luiz Aurelio de Campos Crispin. Centro de Genômica Funcional da ESALQ: Luiz Lehmann Coutinho, Ricardo Augusto Brassaloti, Raquel de Lello Rocha Campos Cassano. NGS Soluções Genômicas: Pilar Drummond Sampaio Corrêa Mariani. FZEA-USP Pirassununga: Mirele Daiana Poleti, Jessika Cristina Chagas Lesbon, Elisangela Chicaroni Mattos, Heidge Fukumasu. USP-Botucatu: Rejane Maria Tommasini Grotto, Jayme A. Souza-Neto, Guilherme Targino Valente, Patricia Akemi Assato, Felipe Allan da Silva da Costa, Bianca Cechetto Carlos. Mendelics: Bibiana Santos, João Paulo Kitajima, Erika Freitas, David Schlesinger. Hemocentro Ribeirão Preto: Simone Kashima, Evandra Strazza Rodrigues, Svetoslav Nanev Slavov, Elaine Vieira dos Santos, Rafael dos Santos Bezerra, Luiz Carlos Junior de Alcantara, Marta Giovanetti, Vagner Fonseca, Flavia Aburjaile, Rodrigo Tocantins Calado. FAMERP-SJRP: Cecília Artico Banho, Lívia Sacchetto, Fábio Sossai Possebon, Leila Sabrina Ullmann, Cíntia Bittar, Guilherme Campos, Helena Lage Ferreira, Jorge A. Petroli Marchesi, Maisa C. Pereira Parra, Marília Moraes, Paula Rahal, Paulo Inacio da Costa, João Pessoa Araújo Jr., Maurício Lacerda Nogueira. Prefeitura de Sao Paulo: Melissa Palmieri. |
| EPI_ISL_1966232 | USAFA GUILHERMINA | Instituto Butantan / Mendelics | Instituto Butantan: Dimas Tadeu Covas, Sandra Coccuzzo Sampaio, Maria Carolina Elias, José Salvatore Leister Patané, Vincent Louis Viala, Antonio Jorge Martins, Ricardo Haddad, Claudia Renata dos Santos Barros, Elaine Cristina Marqueze, Raul Machado Neto, Debora Botequio Moretti, Jardelina de Souza Todao Bernardino, Loyze Paola Oliveira de Lima, Luiz Aurelio de Campos Crispin. Centro de Genômica Funcional da ESALQ: Luiz Lehmann Coutinho, Ricardo Augusto Brassaloti, Raquel de Lello Rocha Campos Cassano. NGS Soluções Genômicas: Pilar Drummond Sampaio Corrêa Mariani. FZEA-USP Pirassununga: Mirele Daiana Poleti, Jessika Cristina Chagas Lesbon, Elisangela Chicaroni Mattos, Heidge Fukumasu. USP-Botucatu: Rejane Maria Tommasini Grotto, Jayme A. Souza-Neto, Guilherme Targino Valente, Patricia Akemi Assato, Felipe Allan da Silva da Costa, Bianca Cechetto Carlos. Mendelics: Bibiana Santos, João Paulo Kitajima, Erika Freitas, David Schlesinger. Hemocentro Ribeirão Preto: Simone Kashima, Evandra Strazza Rodrigues, Svetoslav Nanev Slavov, Elaine Vieira dos Santos, Rafael dos Santos Bezerra, Luiz Carlos Junior de Alcantara, Marta Giovanetti, Vagner Fonseca, Flavia Aburjaile, Rodrigo Tocantins Calado. FAMERP-SJRP: Cecília Artico Banho, Lívia Sacchetto, Fábio Sossai Possebon, Leila Sabrina Ullmann, Cíntia Bittar, Guilherme Campos, Helena Lage Ferreira, Jorge A. Petroli Marchesi, Maisa C. Pereira Parra, Marília Moraes, Paula Rahal, Paulo Inacio da Costa, João Pessoa Araújo Jr., Maurício Lacerda Nogueira. Prefeitura de Sao Paulo: Melissa Palmieri. |
| EPI_ISL_1966233 | USAFA MARACANA | Instituto Butantan / Mendelics | Instituto Butantan: Dimas Tadeu Covas, Sandra Coccuzzo Sampaio, Maria Carolina Elias, José Salvatore Leister Patané, Vincent Louis Viala, Antonio Jorge Martins, Ricardo Haddad, Claudia Renata dos Santos Barros, Elaine Cristina Marqueze, Raul Machado Neto, Debora Botequio Moretti, Jardelina de Souza Todao Bernardino, Loyze Paola Oliveira de Lima, Luiz Aurelio de Campos Crispin. Centro de Genômica Funcional da ESALQ: Luiz Lehmann Coutinho, Ricardo Augusto Brassaloti, Raquel de Lello Rocha Campos Cassano. NGS Soluções Genômicas: Pilar Drummond Sampaio Corrêa Mariani. FZEA-USP Pirassununga: Mirele Daiana Poleti, Jessika Cristina Chagas Lesbon, Elisangela Chicaroni Mattos, Heidge Fukumasu. USP-Botucatu: Rejane Maria Tommasini Grotto, Jayme A. Souza-Neto, Guilherme Targino Valente, Patricia Akemi Assato, Felipe Allan da Silva da Costa, Bianca Cechetto Carlos. Mendelics: Bibiana Santos, João Paulo Kitajima, Erika Freitas, David Schlesinger. Hemocentro Ribeirão Preto: Simone Kashima, Evandra Strazza Rodrigues, Svetoslav Nanev Slavov, Elaine Vieira dos Santos, Rafael dos Santos Bezerra, Luiz Carlos Junior de Alcantara, Marta Giovanetti, Vagner Fonseca, Flavia Aburjaile, Rodrigo Tocantins Calado. FAMERP-SJRP: Cecília Artico Banho, Lívia Sacchetto, Fábio Sossai Possebon, Leila Sabrina Ullmann, Cíntia Bittar, Guilherme Campos, Helena Lage Ferreira, Jorge A. Petroli Marchesi, Maisa C. Pereira Parra, Marília Moraes, Paula Rahal, Paulo Inacio da Costa, João Pessoa Araújo Jr., Maurício Lacerda Nogueira. Prefeitura de Sao Paulo: Melissa Palmieri. |
| EPI_ISL_1966234 | USAFA TUDE BASTOS | Instituto Butantan / Mendelics | Instituto Butantan: Dimas Tadeu Covas, Sandra Coccuzzo Sampaio, Maria Carolina Elias, José Salvatore Leister Patané, Vincent Louis Viala, Antonio Jorge Martins, Ricardo Haddad, Claudia Renata dos Santos Barros, Elaine Cristina Marqueze, Raul Machado Neto, Debora Botequio Moretti, Jardelina de Souza Todao Bernardino, Loyze Paola Oliveira de Lima, Luiz Aurelio de Campos Crispin. Centro de Genômica Funcional da ESALQ: Luiz Lehmann Coutinho, Ricardo Augusto Brassaloti, Raquel de Lello Rocha Campos Cassano. NGS Soluções Genômicas: Pilar Drummond Sampaio Corrêa Mariani. FZEA-USP Pirassununga: Mirele Daiana Poleti, Jessika Cristina Chagas Lesbon, Elisangela Chicaroni Mattos, Heidge Fukumasu. USP-Botucatu: Rejane Maria Tommasini Grotto, Jayme A. Souza-Neto, Guilherme Targino Valente, Patricia Akemi Assato, Felipe Allan da Silva da Costa, Bianca Cechetto Carlos. Mendelics: Bibiana Santos, João Paulo Kitajima, Erika Freitas, David Schlesinger. Hemocentro Ribeirão Preto: Simone Kashima, Evandra Strazza Rodrigues, Svetoslav Nanev Slavov, Elaine Vieira dos Santos, Rafael dos Santos Bezerra, Luiz Carlos Junior de Alcantara, Marta Giovanetti, Vagner Fonseca, Flavia Aburjaile, Rodrigo Tocantins Calado. FAMERP-SJRP: Cecília Artico Banho, Lívia Sacchetto, Fábio Sossai Possebon, Leila Sabrina Ullmann, Cíntia Bittar, Guilherme Campos, Helena Lage Ferreira, Jorge A. Petroli Marchesi, Maisa C. Pereira Parra, Marília Moraes, Paula Rahal, Paulo Inacio da Costa, João Pessoa Araújo Jr., Maurício Lacerda Nogueira. Prefeitura de Sao Paulo: Melissa Palmieri. |
| EPI_ISL_1966235 | CENTRO DE SAUDE III SALES OLIVEIRA | Instituto Butantan / Mendelics | Instituto Butantan: Dimas Tadeu Covas, Sandra Coccuzzo Sampaio, Maria Carolina Elias, José Salvatore Leister Patané, Vincent Louis Viala, Antonio Jorge Martins, Ricardo Haddad, Claudia Renata dos Santos Barros, Elaine Cristina Marqueze, Raul Machado Neto, Debora Botequio Moretti, Jardelina de Souza Todao Bernardino, Loyze Paola Oliveira de Lima, Luiz Aurelio de Campos Crispin. Centro de Genômica Funcional da ESALQ: Luiz Lehmann Coutinho, Ricardo Augusto Brassaloti, Raquel de Lello Rocha Campos Cassano. NGS Soluções Genômicas: Pilar Drummond Sampaio Corrêa Mariani. FZEA-USP Pirassununga: Mirele Daiana Poleti, Jessika Cristina Chagas Lesbon, Elisangela Chicaroni Mattos, Heidge Fukumasu. USP-Botucatu: Rejane Maria Tommasini Grotto, Jayme A. Souza-Neto, Guilherme Targino Valente, Patricia Akemi Assato, Felipe Allan da Silva da Costa, Bianca Cechetto Carlos. Mendelics: Bibiana Santos, João Paulo Kitajima, Erika Freitas, David Schlesinger. Hemocentro Ribeirão Preto: Simone Kashima, Evandra Strazza Rodrigues, Svetoslav Nanev Slavov, Elaine Vieira dos Santos, Rafael dos Santos Bezerra, Luiz Carlos Junior de Alcantara, Marta Giovanetti, Vagner Fonseca, Flavia Aburjaile, Rodrigo Tocantins Calado. FAMERP-SJRP: Cecília Artico Banho, Lívia Sacchetto, Fábio Sossai Possebon, Leila Sabrina |

| | | | |
|---|---|---|---|
| | | | Ullmann, Cíntia Bittar, Guilherme Campos, Helena Lage Ferreira, Jorge A. Petroli Marchesi, Maisa C. Pereira Parra, Marília Moraes, Paula Rahal, Paulo Inacio da Costa, João Pessoa Araújo Jr., Maurício Lacerda Nogueira. Prefeitura de Sao Paulo: Melissa Palmieri. |
| EPI_ISL_1966236 | UNIDADE MISTA DE TAIACU | Instituto Butantan / Mendelics | Instituto Butantan: Dimas Tadeu Covas, Sandra Coccuzzo Sampaio, Maria Carolina Elias, José Salvatore Leister Patané, Vincent Louis Viala, Antonio Jorge Martins, Ricardo Haddad, Claudia Renata dos Santos Barros, Elaine Cristina Marqueze, Raul Machado Neto, Debora Botequio Moretti, Jardelina de Souza Todao Bernardino, Loyze Paola Oliveira de Lima, Luiz Aurelio de Campos Crispin. Centro de Genômica Funcional da ESALQ: Luiz Lehmann Coutinho, Ricardo Augusto Brassaloti, Raquel de Lello Rocha Campos Cassano. NGS Soluções Genômicas: Pilar Drummond Sampaio Corrêa Mariani. FZEA-USP Pirassununga: Mirele Daiana Poleti, Jessika Cristina Chagas Lesbon, Elisangela Chicaroni Mattos, Heidge Fukumasu. USP-Botucatu: Rejane Maria Tommasini Grotto, Jayme A. Souza-Neto, Guilherme Targino Valente, Patricia Akemi Assato, Felipe Allan da Silva da Costa, Bianca Cechetto Carlos. Mendelics: Bibiana Santos, João Paulo Kitajima, Erika Freitas, David Schlesinger. Hemocentro Ribeirão Preto: Simone Kashima, Evandra Strazza Rodrigues, Svetoslav Nanev Slavov, Elaine Vieira dos Santos, Rafael dos Santos Bezerra, Luiz Carlos Junior de Alcantara, Marta Giovanetti, Vagner Fonseca, Flavia Aburjaile, Rodrigo Tocantins Calado. FAMERP-SJRP: Cecília Artico Banho, Lívia Sacchetto, Fábio Sossai Possebon, Leila Sabrina Ullmann, Cíntia Bittar, Guilherme Campos, Helena Lage Ferreira, Jorge A. Petroli Marchesi, Maisa C. Pereira Parra, Marília Moraes, Paula Rahal, Paulo Inacio da Costa, João Pessoa Araújo Jr., Maurício Lacerda Nogueira. Prefeitura de Sao Paulo: Melissa Palmieri. |
| EPI_ISL_1966237 | UPA DE BEBEDOURO | Instituto Butantan / Mendelics | Instituto Butantan: Dimas Tadeu Covas, Sandra Coccuzzo Sampaio, Maria Carolina Elias, José Salvatore Leister Patané, Vincent Louis Viala, Antonio Jorge Martins, Ricardo Haddad, Claudia Renata dos Santos Barros, Elaine Cristina Marqueze, Raul Machado Neto, Debora Botequio Moretti, Jardelina de Souza Todao Bernardino, Loyze Paola Oliveira de Lima, Luiz Aurelio de Campos Crispin. Centro de Genômica Funcional da ESALQ: Luiz Lehmann Coutinho, Ricardo Augusto Brassaloti, Raquel de Lello Rocha Campos Cassano. NGS Soluções Genômicas: Pilar Drummond Sampaio Corrêa Mariani. FZEA-USP Pirassununga: Mirele Daiana Poleti, Jessika Cristina Chagas Lesbon, Elisangela Chicaroni Mattos, Heidge Fukumasu. USP-Botucatu: Rejane Maria Tommasini Grotto, Jayme A. Souza-Neto, Guilherme Targino Valente, Patricia Akemi Assato, Felipe Allan da Silva da Costa, Bianca Cechetto Carlos. Mendelics: Bibiana Santos, João Paulo Kitajima, Erika Freitas, David Schlesinger. Hemocentro Ribeirão Preto: Simone Kashima, Evandra Strazza Rodrigues, Svetoslav Nanev Slavov, Elaine Vieira dos Santos, Rafael dos Santos Bezerra, Luiz Carlos Junior de Alcantara, Marta Giovanetti, Vagner Fonseca, Flavia Aburjaile, Rodrigo Tocantins Calado. FAMERP-SJRP: Cecília Artico Banho, Lívia Sacchetto, Fábio Sossai Possebon, Leila Sabrina Ullmann, Cíntia Bittar, Guilherme Campos, Helena Lage Ferreira, Jorge A. Petroli Marchesi, Maisa C. Pereira Parra, Marília Moraes, Paula Rahal, Paulo Inacio da Costa, João Pessoa Araújo Jr., Maurício Lacerda Nogueira. Prefeitura de Sao Paulo: Melissa Palmieri. |
| EPI_ISL_1966238 | CS II DR SEBASTIAO RIBEIRO DO VALLE | Instituto Butantan / Mendelics | Instituto Butantan: Dimas Tadeu Covas, Sandra Coccuzzo Sampaio, Maria Carolina Elias, José Salvatore Leister Patané, Vincent Louis Viala, Antonio Jorge Martins, Ricardo Haddad, Claudia Renata dos Santos Barros, Elaine Cristina Marqueze, Raul Machado Neto, Debora Botequio Moretti, Jardelina de Souza Todao Bernardino, Loyze Paola Oliveira de Lima, Luiz Aurelio de Campos Crispin. Centro de Genômica Funcional da ESALQ: Luiz Lehmann Coutinho, Ricardo Augusto Brassaloti, Raquel de Lello Rocha Campos Cassano. NGS Soluções Genômicas: Pilar Drummond Sampaio Corrêa Mariani. FZEA-USP Pirassununga: Mirele Daiana Poleti, Jessika Cristina Chagas Lesbon, Elisangela Chicaroni Mattos, Heidge Fukumasu. USP-Botucatu: Rejane Maria Tommasini Grotto, Jayme A. Souza-Neto, Guilherme Targino Valente, Patricia Akemi Assato, Felipe Allan da Silva da Costa, Bianca Cechetto Carlos. Mendelics: Bibiana Santos, João Paulo Kitajima, Erika Freitas, David Schlesinger. Hemocentro Ribeirão Preto: Simone Kashima, Evandra Strazza Rodrigues, Svetoslav Nanev Slavov, Elaine Vieira dos Santos, Rafael dos Santos Bezerra, Luiz Carlos Junior de Alcantara, Marta Giovanetti, Vagner Fonseca, Flavia Aburjaile, Rodrigo Tocantins Calado. FAMERP-SJRP: Cecília Artico Banho, Lívia Sacchetto, Fábio Sossai Possebon, Leila Sabrina Ullmann, Cíntia Bittar, Guilherme Campos, Helena Lage Ferreira, Jorge A. Petroli Marchesi, Maisa C. Pereira Parra, Marília Moraes, Paula Rahal, Paulo Inacio da Costa, João Pessoa Araújo Jr., Maurício Lacerda Nogueira. Prefeitura de Sao Paulo: Melissa Palmieri. |
| EPI_ISL_1966239, EPI_ISL_1966240, EPI_ISL_1966241 | CENTRO DE SAUDE DR MARIO DIAS DE AGUIAR CAPIVARI | Instituto Butantan / Mendelics | Instituto Butantan: Dimas Tadeu Covas, Sandra Coccuzzo Sampaio, Maria Carolina Elias, José Salvatore Leister Patané, Vincent Louis Viala, Antonio Jorge Martins, Ricardo Haddad, Claudia Renata dos Santos Barros, Elaine Cristina Marqueze, Raul Machado Neto, Debora Botequio Moretti, Jardelina de Souza Todao Bernardino, Loyze Paola Oliveira de Lima, Luiz Aurelio de Campos Crispin. Centro de Genômica Funcional da ESALQ: Luiz Lehmann Coutinho, Ricardo Augusto Brassaloti, Raquel de Lello Rocha Campos Cassano. NGS Soluções Genômicas: Pilar Drummond Sampaio Corrêa Mariani. FZEA-USP Pirassununga: Mirele Daiana Poleti, Jessika Cristina Chagas Lesbon, Elisangela Chicaroni Mattos, Heidge Fukumasu. USP-Botucatu: Rejane Maria Tommasini Grotto, Jayme A. Souza-Neto, Guilherme Targino Valente, Patricia Akemi Assato, Felipe Allan da Silva da Costa, Bianca Cechetto Carlos. Mendelics: Bibiana Santos, João Paulo Kitajima, Erika Freitas, David Schlesinger. Hemocentro Ribeirão Preto: Simone Kashima, Evandra Strazza Rodrigues, Svetoslav Nanev Slavov, Elaine Vieira dos Santos, Rafael dos Santos Bezerra, Luiz Carlos Junior de Alcantara, Marta Giovanetti, Vagner Fonseca, Flavia Aburjaile, Rodrigo Tocantins Calado. FAMERP-SJRP: Cecília Artico Banho, Lívia Sacchetto, Fábio Sossai Possebon, Leila Sabrina Ullmann, Cíntia Bittar, Guilherme Campos, Helena Lage Ferreira, Jorge A. Petroli Marchesi, Maisa C. Pereira Parra, Marília Moraes, Paula Rahal, Paulo Inacio da Costa, João Pessoa Araújo Jr., Maurício Lacerda Nogueira. Prefeitura de Sao Paulo: Melissa Palmieri. |
| EPI_ISL_1966242 | SANTA CASA DE MIS E MATERNIDADE N SRA DA CONCEICAO CUNHA | Instituto Butantan / Mendelics | Instituto Butantan: Dimas Tadeu Covas, Sandra Coccuzzo Sampaio, Maria Carolina Elias, José Salvatore Leister Patané, Vincent Louis Viala, Antonio Jorge Martins, Ricardo Haddad, Claudia Renata dos Santos Barros, Elaine Cristina Marqueze, Raul Machado Neto, Debora Botequio Moretti, Jardelina de Souza Todao Bernardino, Loyze Paola Oliveira de Lima, Luiz Aurelio de Campos Crispin. Centro de Genômica Funcional da ESALQ: Luiz Lehmann Coutinho, Ricardo Augusto Brassaloti, Raquel de Lello Rocha Campos Cassano. NGS Soluções Genômicas: Pilar Drummond Sampaio Corrêa Mariani. FZEA-USP Pirassununga: Mirele Daiana Poleti, Jessika Cristina Chagas Lesbon, Elisangela Chicaroni Mattos, Heidge Fukumasu. USP-Botucatu: Rejane Maria Tommasini Grotto, Jayme A. Souza-Neto, Guilherme Targino Valente, Patricia Akemi Assato, Felipe Allan da Silva da Costa, Bianca Cechetto Carlos. Mendelics: Bibiana Santos, João Paulo Kitajima, Erika Freitas, David Schlesinger. Hemocentro Ribeirão Preto: Simone Kashima, Evandra Strazza Rodrigues, Svetoslav Nanev Slavov, Elaine Vieira dos Santos, Rafael dos Santos Bezerra, Luiz Carlos Junior de Alcantara, Marta Giovanetti, Vagner Fonseca, Flavia Aburjaile, Rodrigo Tocantins Calado. FAMERP-SJRP: Cecília Artico Banho, Lívia Sacchetto, Fábio Sossai Possebon, Leila Sabrina Ullmann, Cíntia Bittar, Guilherme Campos, Helena Lage Ferreira, Jorge A. Petroli Marchesi, Maisa C. Pereira Parra, Marília Moraes, Paula Rahal, Paulo Inacio da Costa, João Pessoa Araújo Jr., Maurício Lacerda Nogueira. Prefeitura de Sao Paulo: Melissa Palmieri. |
| EPI_ISL_1966243 | UNIDADE MISTA DE SAUDE DE DUMONT | Instituto Butantan / Mendelics | Instituto Butantan: Dimas Tadeu Covas, Sandra Coccuzzo Sampaio, Maria Carolina Elias, José Salvatore Leister Patané, Vincent Louis Viala, Antonio Jorge Martins, Ricardo Haddad, Claudia Renata dos Santos Barros, Elaine Cristina Marqueze, Raul Machado Neto, Debora Botequio Moretti, Jardelina de Souza Todao Bernardino, Loyze Paola Oliveira de Lima, Luiz Aurelio de Campos Crispin. Centro de Genômica Funcional da ESALQ: Luiz Lehmann Coutinho, Ricardo Augusto Brassaloti, Raquel de Lello Rocha Campos Cassano. NGS Soluções Genômicas: Pilar Drummond Sampaio Corrêa Mariani. FZEA-USP Pirassununga: Mirele Daiana Poleti, Jessika Cristina Chagas Lesbon, Elisangela Chicaroni Mattos, Heidge Fukumasu. USP-Botucatu: Rejane Maria Tommasini Grotto, Jayme A. Souza-Neto, Guilherme Targino Valente, Patricia Akemi Assato, Felipe Allan da Silva da Costa, Bianca Cechetto Carlos. Mendelics: Bibiana Santos, João Paulo Kitajima, Erika Freitas, David Schlesinger. Hemocentro Ribeirão Preto: Simone Kashima, Evandra Strazza Rodrigues, Svetoslav Nanev Slavov, Elaine Vieira dos Santos, Rafael dos Santos Bezerra, Luiz Carlos Junior de Alcantara, Marta Giovanetti, Vagner Fonseca, Flavia Aburjaile, Rodrigo Tocantins Calado. FAMERP-SJRP: Cecília Artico Banho, Lívia Sacchetto, Fábio Sossai Possebon, Leila Sabrina Ullmann, Cíntia Bittar, Guilherme Campos, Helena Lage Ferreira, Jorge A. Petroli Marchesi, Maisa C. Pereira Parra, Marília Moraes, Paula Rahal, Paulo Inacio da Costa, João Pessoa Araújo Jr., Maurício Lacerda Nogueira. Prefeitura de Sao Paulo: Melissa Palmieri. |
| EPI_ISL_1966244 | POSTO MEDICO NATAL DIEGUES DE ESTIVA GERBI | Instituto Butantan / Mendelics | Instituto Butantan: Dimas Tadeu Covas, Sandra Coccuzzo Sampaio, Maria Carolina Elias, José Salvatore Leister Patané, Vincent Louis Viala, Antonio Jorge Martins, Ricardo Haddad, Claudia Renata dos Santos Barros, Elaine Cristina Marqueze, Raul Machado Neto, Debora Botequio Moretti, Jardelina de Souza Todao Bernardino, Loyze Paola Oliveira de Lima, Luiz Aurelio de Campos Crispin. Centro de Genômica Funcional da ESALQ: Luiz Lehmann Coutinho, Ricardo Augusto Brassaloti, Raquel de Lello Rocha Campos Cassano. NGS Soluções Genômicas: Pilar Drummond Sampaio Corrêa Mariani. FZEA-USP Pirassununga: Mirele Daiana Poleti, Jessika Cristina Chagas Lesbon, Elisangela Chicaroni Mattos, Heidge Fukumasu. USP-Botucatu: Rejane Maria Tommasini Grotto, Jayme A. Souza-Neto, Guilherme Targino Valente, Patricia Akemi Assato, Felipe Allan da Silva da Costa, Bianca Cechetto Carlos. Mendelics: Bibiana Santos, João Paulo Kitajima, Erika Freitas, David Schlesinger. Hemocentro Ribeirão Preto: Simone Kashima, Evandra Strazza Rodrigues, Svetoslav Nanev Slavov, Elaine Vieira dos Santos, Rafael dos Santos Bezerra, Luiz Carlos Junior de Alcantara, Marta Giovanetti, Vagner Fonseca, Flavia Aburjaile, Rodrigo Tocantins Calado. FAMERP-SJRP: Cecília Artico Banho, Lívia Sacchetto, Fábio Sossai Possebon, Leila Sabrina Ullmann, Cíntia Bittar, Guilherme Campos, Helena Lage Ferreira, Jorge A. Petroli Marchesi, Maisa C. Pereira Parra, Marília Moraes, Paula Rahal, Paulo Inacio da Costa, João Pessoa Araújo Jr., Maurício Lacerda Nogueira. Prefeitura de Sao Paulo: Melissa Palmieri. |
| EPI_ISL_1966245 | UNIDADE BASICA DE SAUDE DR ORESTES DE MOURA PINTO | Instituto Butantan / Mendelics | Instituto Butantan: Dimas Tadeu Covas, Sandra Coccuzzo Sampaio, Maria Carolina Elias, José Salvatore Leister Patané, Vincent Louis Viala, Antonio Jorge Martins, Ricardo Haddad, Claudia Renata dos Santos Barros, Elaine Cristina Marqueze, Raul Machado Neto, Debora Botequio Moretti, Jardelina de Souza Todao Bernardino, Loyze Paola Oliveira de Lima, Luiz Aurelio de Campos Crispin. Centro de Genômica Funcional da ESALQ: Luiz Lehmann Coutinho, Ricardo Augusto Brassaloti, Raquel de Lello Rocha Campos Cassano. NGS Soluções Genômicas: Pilar Drummond Sampaio Corrêa Mariani. |

| | | | |
|---|---|---|---|
| | | | FZEA-USP Pirassununga: Mirele Daiana Poleti, Jessika Cristina Chagas Lesbon, Elisangela Chicaroni Mattos, Heidge Fukumasu. USP-Botucatu: Rejane Maria Tommasini Grotto, Jayme A. Souza-Neto, Guilherme Targino Valente, Patricia Akemi Assato, Felipe Allan da Silva da Costa, Bianca Cechetto Carlos. Mendelics: Bibiana Santos, João Paulo Kitajima, Erika Freitas, David Schlesinger. Hemocentro Ribeirão Preto: Simone Kashima, Evandra Strazza Rodrigues, Svetoslav Nanev Slavov, Elaine Vieira dos Santos, Rafael dos Santos Bezerra, Luiz Carlos Junior de Alcantara, Marta Giovanetti, Vagner Fonseca, Flavia Aburjaile, Rodrigo Tocantins Calado. FAMERP-SJRP: Cecília Artico Banho, Lívia Sacchetto, Fábio Sossai Possebon, Leila Sabrina Ullmann, Cíntia Bittar, Guilherme Campos, Helena Lage Ferreira, Jorge A. Petroli Marchesi, Maisa C. Pereira Parra, Marília Moraes, Paula Rahal, Paulo Inacio da Costa, João Pessoa Araújo Jr., Maurício Lacerda Nogueira. Prefeitura de Sao Paulo: Melissa Palmieri. |
| EPI_ISL_1966246 | UNIDADE MISTA DE IGUAPE | Instituto Butantan / Mendelics | Instituto Butantan: Dimas Tadeu Covas, Sandra Coccuzzo Sampaio, Maria Carolina Elias, José Salvatore Leister Patané, Vincent Louis Viala, Antonio Jorge Martins, Ricardo Haddad, Claudia Renata dos Santos Barros, Elaine Cristina Marqueze, Raul Machado Neto, Debora Botequio Moretti, Jardelina de Souza Todao Bernardino, Loyze Paola Oliveira de Lima, Luiz Aurelio de Campos Crispin. Centro de Genômica Funcional da ESALQ: Luiz Lehmann Coutinho, Ricardo Augusto Brassaloti, Raquel de Lello Rocha Campos Cassano. NGS Soluções Genômicas: Pilar Drummond Sampaio Corrêa Mariani. FZEA-USP Pirassununga: Mirele Daiana Poleti, Jessika Cristina Chagas Lesbon, Elisangela Chicaroni Mattos, Heidge Fukumasu. USP-Botucatu: Rejane Maria Tommasini Grotto, Jayme A. Souza-Neto, Guilherme Targino Valente, Patricia Akemi Assato, Felipe Allan da Silva da Costa, Bianca Cechetto Carlos. Mendelics: Bibiana Santos, João Paulo Kitajima, Erika Freitas, David Schlesinger. Hemocentro Ribeirão Preto: Simone Kashima, Evandra Strazza Rodrigues, Svetoslav Nanev Slavov, Elaine Vieira dos Santos, Rafael dos Santos Bezerra, Luiz Carlos Junior de Alcantara, Marta Giovanetti, Vagner Fonseca, Flavia Aburjaile, Rodrigo Tocantins Calado. FAMERP-SJRP: Cecília Artico Banho, Lívia Sacchetto, Fábio Sossai Possebon, Leila Sabrina Ullmann, Cíntia Bittar, Guilherme Campos, Helena Lage Ferreira, Jorge A. Petroli Marchesi, Maisa C. Pereira Parra, Marília Moraes, Paula Rahal, Paulo Inacio da Costa, João Pessoa Araújo Jr., Maurício Lacerda Nogueira. Prefeitura de Sao Paulo: Melissa Palmieri. |
| EPI_ISL_1966247 | SECRETARIA DE SAUDE DO MUNICIPIO DE IPUA | Instituto Butantan / Mendelics | Instituto Butantan: Dimas Tadeu Covas, Sandra Coccuzzo Sampaio, Maria Carolina Elias, José Salvatore Leister Patané, Vincent Louis Viala, Antonio Jorge Martins, Ricardo Haddad, Claudia Renata dos Santos Barros, Elaine Cristina Marqueze, Raul Machado Neto, Debora Botequio Moretti, Jardelina de Souza Todao Bernardino, Loyze Paola Oliveira de Lima, Luiz Aurelio de Campos Crispin. Centro de Genômica Funcional da ESALQ: Luiz Lehmann Coutinho, Ricardo Augusto Brassaloti, Raquel de Lello Rocha Campos Cassano. NGS Soluções Genômicas: Pilar Drummond Sampaio Corrêa Mariani. FZEA-USP Pirassununga: Mirele Daiana Poleti, Jessika Cristina Chagas Lesbon, Elisangela Chicaroni Mattos, Heidge Fukumasu. USP-Botucatu: Rejane Maria Tommasini Grotto, Jayme A. Souza-Neto, Guilherme Targino Valente, Patricia Akemi Assato, Felipe Allan da Silva da Costa, Bianca Cechetto Carlos. Mendelics: Bibiana Santos, João Paulo Kitajima, Erika Freitas, David Schlesinger. Hemocentro Ribeirão Preto: Simone Kashima, Evandra Strazza Rodrigues, Svetoslav Nanev Slavov, Elaine Vieira dos Santos, Rafael dos Santos Bezerra, Luiz Carlos Junior de Alcantara, Marta Giovanetti, Vagner Fonseca, Flavia Aburjaile, Rodrigo Tocantins Calado. FAMERP-SJRP: Cecília Artico Banho, Lívia Sacchetto, Fábio Sossai Possebon, Leila Sabrina Ullmann, Cíntia Bittar, Guilherme Campos, Helena Lage Ferreira, Jorge A. Petroli Marchesi, Maisa C. Pereira Parra, Marília Moraes, Paula Rahal, Paulo Inacio da Costa, João Pessoa Araújo Jr., Maurício Lacerda Nogueira. Prefeitura de Sao Paulo: Melissa Palmieri. |
| EPI_ISL_1966248, EPI_ISL_1966249 | UNIDADE DE PRONTO ATENDIMENTO DE JALES | Instituto Butantan / Mendelics | Instituto Butantan: Dimas Tadeu Covas, Sandra Coccuzzo Sampaio, Maria Carolina Elias, José Salvatore Leister Patané, Vincent Louis Viala, Antonio Jorge Martins, Ricardo Haddad, Claudia Renata dos Santos Barros, Elaine Cristina Marqueze, Raul Machado Neto, Debora Botequio Moretti, Jardelina de Souza Todao Bernardino, Loyze Paola Oliveira de Lima, Luiz Aurelio de Campos Crispin. Centro de Genômica Funcional da ESALQ: Luiz Lehmann Coutinho, Ricardo Augusto Brassaloti, Raquel de Lello Rocha Campos Cassano. NGS Soluções Genômicas: Pilar Drummond Sampaio Corrêa Mariani. FZEA-USP Pirassununga: Mirele Daiana Poleti, Jessika Cristina Chagas Lesbon, Elisangela Chicaroni Mattos, Heidge Fukumasu. USP-Botucatu: Rejane Maria Tommasini Grotto, Jayme A. Souza-Neto, Guilherme Targino Valente, Patricia Akemi Assato, Felipe Allan da Silva da Costa, Bianca Cechetto Carlos. Mendelics: Bibiana Santos, João Paulo Kitajima, Erika Freitas, David Schlesinger. Hemocentro Ribeirão Preto: Simone Kashima, Evandra Strazza Rodrigues, Svetoslav Nanev Slavov, Elaine Vieira dos Santos, Rafael dos Santos Bezerra, Luiz Carlos Junior de Alcantara, Marta Giovanetti, Vagner Fonseca, Flavia Aburjaile, Rodrigo Tocantins Calado. FAMERP-SJRP: Cecília Artico Banho, Lívia Sacchetto, Fábio Sossai Possebon, Leila Sabrina Ullmann, Cíntia Bittar, Guilherme Campos, Helena Lage Ferreira, Jorge A. Petroli Marchesi, Maisa C. Pereira Parra, Marília Moraes, Paula Rahal, Paulo Inacio da Costa, João Pessoa Araújo Jr., Maurício Lacerda Nogueira. Prefeitura de Sao Paulo: Melissa Palmieri. |
| EPI_ISL_1966250 | UNIDADE SAUDE DA FAMILIA DRA ZILDA ARNS NEUMANN JALES | Instituto Butantan / Mendelics | Instituto Butantan: Dimas Tadeu Covas, Sandra Coccuzzo Sampaio, Maria Carolina Elias, José Salvatore Leister Patané, Vincent Louis Viala, Antonio Jorge Martins, Ricardo Haddad, Claudia Renata dos Santos Barros, Elaine Cristina Marqueze, Raul Machado Neto, Debora Botequio Moretti, Jardelina de Souza Todao Bernardino, Loyze Paola Oliveira de Lima, Luiz Aurelio de Campos Crispin. Centro de Genômica Funcional da ESALQ: Luiz Lehmann Coutinho, Ricardo Augusto Brassaloti, Raquel de Lello Rocha Campos Cassano. NGS Soluções Genômicas: Pilar Drummond Sampaio Corrêa Mariani. FZEA-USP Pirassununga: Mirele Daiana Poleti, Jessika Cristina Chagas Lesbon, Elisangela Chicaroni Mattos, Heidge Fukumasu. USP-Botucatu: Rejane Maria Tommasini Grotto, Jayme A. Souza-Neto, Guilherme Targino Valente, Patricia Akemi Assato, Felipe Allan da Silva da Costa, Bianca Cechetto Carlos. Mendelics: Bibiana Santos, João Paulo Kitajima, Erika Freitas, David Schlesinger. Hemocentro Ribeirão Preto: Simone Kashima, Evandra Strazza Rodrigues, Svetoslav Nanev Slavov, Elaine Vieira dos Santos, Rafael dos Santos Bezerra, Luiz Carlos Junior de Alcantara, Marta Giovanetti, Vagner Fonseca, Flavia Aburjaile, Rodrigo Tocantins Calado. FAMERP-SJRP: Cecília Artico Banho, Lívia Sacchetto, Fábio Sossai Possebon, Leila Sabrina Ullmann, Cíntia Bittar, Guilherme Campos, Helena Lage Ferreira, Jorge A. Petroli Marchesi, Maisa C. Pereira Parra, Marília Moraes, Paula Rahal, Paulo Inacio da Costa, João Pessoa Araújo Jr., Maurício Lacerda Nogueira. Prefeitura de Sao Paulo: Melissa Palmieri. |
| EPI_ISL_1966251 | ESF DR LUIS ERNESTO SANDI MORI JALES | Instituto Butantan / Mendelics | Instituto Butantan: Dimas Tadeu Covas, Sandra Coccuzzo Sampaio, Maria Carolina Elias, José Salvatore Leister Patané, Vincent Louis Viala, Antonio Jorge Martins, Ricardo Haddad, Claudia Renata dos Santos Barros, Elaine Cristina Marqueze, Raul Machado Neto, Debora Botequio Moretti, Jardelina de Souza Todao Bernardino, Loyze Paola Oliveira de Lima, Luiz Aurelio de Campos Crispin. Centro de Genômica Funcional da ESALQ: Luiz Lehmann Coutinho, Ricardo Augusto Brassaloti, Raquel de Lello Rocha Campos Cassano. NGS Soluções Genômicas: Pilar Drummond Sampaio Corrêa Mariani. FZEA-USP Pirassununga: Mirele Daiana Poleti, Jessika Cristina Chagas Lesbon, Elisangela Chicaroni Mattos, Heidge Fukumasu. USP-Botucatu: Rejane Maria Tommasini Grotto, Jayme A. Souza-Neto, Guilherme Targino Valente, Patricia Akemi Assato, Felipe Allan da Silva da Costa, Bianca Cechetto Carlos. Mendelics: Bibiana Santos, João Paulo Kitajima, Erika Freitas, David Schlesinger. Hemocentro Ribeirão Preto: Simone Kashima, Evandra Strazza Rodrigues, Svetoslav Nanev Slavov, Elaine Vieira dos Santos, Rafael dos Santos Bezerra, Luiz Carlos Junior de Alcantara, Marta Giovanetti, Vagner Fonseca, Flavia Aburjaile, Rodrigo Tocantins Calado. FAMERP-SJRP: Cecília Artico Banho, Lívia Sacchetto, Fábio Sossai Possebon, Leila Sabrina Ullmann, Cíntia Bittar, Guilherme Campos, Helena Lage Ferreira, Jorge A. Petroli Marchesi, Maisa C. Pereira Parra, Marília Moraes, Paula Rahal, Paulo Inacio da Costa, João Pessoa Araújo Jr., Maurício Lacerda Nogueira. Prefeitura de Sao Paulo: Melissa Palmieri. |
| EPI_ISL_1966252, EPI_ISL_1966253, EPI_ISL_1966254, EPI_ISL_1966255 | UBS DR MOHANNA ADAS | Instituto Butantan / Mendelics | Instituto Butantan: Dimas Tadeu Covas, Sandra Coccuzzo Sampaio, Maria Carolina Elias, José Salvatore Leister Patané, Vincent Louis Viala, Antonio Jorge Martins, Ricardo Haddad, Claudia Renata dos Santos Barros, Elaine Cristina Marqueze, Raul Machado Neto, Debora Botequio Moretti, Jardelina de Souza Todao Bernardino, Loyze Paola Oliveira de Lima, Luiz Aurelio de Campos Crispin. Centro de Genômica Funcional da ESALQ: Luiz Lehmann Coutinho, Ricardo Augusto Brassaloti, Raquel de Lello Rocha Campos Cassano. NGS Soluções Genômicas: Pilar Drummond Sampaio Corrêa Mariani. FZEA-USP Pirassununga: Mirele Daiana Poleti, Jessika Cristina Chagas Lesbon, Elisangela Chicaroni Mattos, Heidge Fukumasu. USP-Botucatu: Rejane Maria Tommasini Grotto, Jayme A. Souza-Neto, Guilherme Targino Valente, Patricia Akemi Assato, Felipe Allan da Silva da Costa, Bianca Cechetto Carlos. Mendelics: Bibiana Santos, João Paulo Kitajima, Erika Freitas, David Schlesinger. Hemocentro Ribeirão Preto: Simone Kashima, Evandra Strazza Rodrigues, Svetoslav Nanev Slavov, Elaine Vieira dos Santos, Rafael dos Santos Bezerra, Luiz Carlos Junior de Alcantara, Marta Giovanetti, Vagner Fonseca, Flavia Aburjaile, Rodrigo Tocantins Calado. FAMERP-SJRP: Cecília Artico Banho, Lívia Sacchetto, Fábio Sossai Possebon, Leila Sabrina Ullmann, Cíntia Bittar, Guilherme Campos, Helena Lage Ferreira, Jorge A. Petroli Marchesi, Maisa C. Pereira Parra, Marília Moraes, Paula Rahal, Paulo Inacio da Costa, João Pessoa Araújo Jr., Maurício Lacerda Nogueira. Prefeitura de Sao Paulo: Melissa Palmieri. |
| EPI_ISL_1966256 | CENTRO INTEGRADO DE SAUDE | Instituto Butantan / Mendelics | Instituto Butantan: Dimas Tadeu Covas, Sandra Coccuzzo Sampaio, Maria Carolina Elias, José Salvatore Leister Patané, Vincent Louis Viala, Antonio Jorge Martins, Ricardo Haddad, Claudia Renata dos Santos Barros, Elaine Cristina Marqueze, Raul Machado Neto, Debora Botequio Moretti, Jardelina de Souza Todao Bernardino, Loyze Paola Oliveira de Lima, Luiz Aurelio de Campos Crispin. Centro de Genômica Funcional da ESALQ: Luiz Lehmann Coutinho, Ricardo Augusto Brassaloti, Raquel de Lello Rocha Campos Cassano. NGS Soluções Genômicas: Pilar Drummond Sampaio Corrêa Mariani. FZEA-USP Pirassununga: Mirele Daiana Poleti, Jessika Cristina Chagas Lesbon, Elisangela Chicaroni Mattos, Heidge Fukumasu. USP-Botucatu: Rejane Maria Tommasini Grotto, Jayme A. Souza-Neto, Guilherme Targino Valente, Patricia Akemi Assato, Felipe Allan da Silva da Costa, Bianca Cechetto Carlos. Mendelics: Bibiana Santos, João Paulo Kitajima, Erika Freitas, David Schlesinger. Hemocentro Ribeirão Preto: Simone Kashima, Evandra Strazza Rodrigues, Svetoslav Nanev Slavov, Elaine Vieira dos Santos, Rafael dos Santos Bezerra, Luiz Carlos Junior de Alcantara, Marta Giovanetti, Vagner Fonseca, Flavia Aburjaile, Rodrigo Tocantins Calado. FAMERP-SJRP: Cecília Artico Banho, Lívia Sacchetto, Fábio Sossai Possebon, Leila Sabrina Ullmann, Cíntia Bittar, Guilherme Campos, Helena Lage Ferreira, Jorge A. Petroli Marchesi, Maisa C. Pereira Parra, Marília Moraes, Paula Rahal, Paulo Inacio da Costa, João Pessoa Araújo Jr., Maurício Lacerda Nogueira. Prefeitura de Sao Paulo: Melissa Palmieri. |

| | | | |
|---|---|---|---|
| EPI_ISL_1966257, EPI_ISL_1966258 | CS DE RUBINEIA | Instituto Butantan / Mendelics | Instituto Butantan: Dimas Tadeu Covas, Sandra Coccuzzo Sampaio, Maria Carolina Elias, José Salvatore Leister Patané, Vincent Louis Viala, Antonio Jorge Martins, Ricardo Haddad, Claudia Renata dos Santos Barros, Elaine Cristina Marqueze, Raul Machado Neto, Debora Botequio Moretti, Jardelina de Souza Todao Bernardino, Loyze Paola Oliveira de Lima, Luiz Aurelio de Campos Crispin. Centro de Genômica Funcional da ESALQ: Luiz Lehmann Coutinho, Ricardo Augusto Brassaloti, Raquel de Lello Rocha Campos Cassano. NGS Soluções Genômicas: Pilar Drummond Sampaio Corrêa Mariani. FZEA-USP Pirassununga: Mirele Daiana Poleti, Jessika Cristina Chagas Lesbon, Elisangela Chicaroni Mattos, Heidge Fukumasu. USP-Botucatu: Rejane Maria Tommasini Grotto, Jayme A. Souza-Neto, Guilherme Targino Valente, Patricia Akemi Assato, Felipe Allan da Silva da Costa, Bianca Cechetto Carlos. Mendelics: Bibiana Santos, João Paulo Kitajima, Erika Freitas, David Schlesinger. Hemocentro Ribeirão Preto: Simone Kashima, Evandra Strazza Rodrigues, Svetoslav Nanev Slavov, Elaine Vieira dos Santos, Rafael dos Santos Bezerra, Luiz Carlos Junior de Alcantara, Marta Giovanetti, Vagner Fonseca, Flavia Aburjaile, Rodrigo Tocantins Calado. FAMERP-SJRP: Cecília Artico Banho, Lívia Sacchetto, Fábio Sossai Possebon, Leila Sabrina Ullmann, Cíntia Bittar, Guilherme Campos, Helena Lage Ferreira, Jorge A. Petroli Marchesi, Maisa C. Pereira Parra, Marília Moraes, Paula Rahal, Paulo Inacio da Costa, João Pessoa Araújo Jr., Maurício Lacerda Nogueira. Prefeitura de Sao Paulo: Melissa Palmieri. |
| EPI_ISL_1966259 | SANTA CASA DE MISERICORDIA DE UBATUBA | Instituto Butantan / Mendelics | Instituto Butantan: Dimas Tadeu Covas, Sandra Coccuzzo Sampaio, Maria Carolina Elias, José Salvatore Leister Patané, Vincent Louis Viala, Antonio Jorge Martins, Ricardo Haddad, Claudia Renata dos Santos Barros, Elaine Cristina Marqueze, Raul Machado Neto, Debora Botequio Moretti, Jardelina de Souza Todao Bernardino, Loyze Paola Oliveira de Lima, Luiz Aurelio de Campos Crispin. Centro de Genômica Funcional da ESALQ: Luiz Lehmann Coutinho, Ricardo Augusto Brassaloti, Raquel de Lello Rocha Campos Cassano. NGS Soluções Genômicas: Pilar Drummond Sampaio Corrêa Mariani. FZEA-USP Pirassununga: Mirele Daiana Poleti, Jessika Cristina Chagas Lesbon, Elisangela Chicaroni Mattos, Heidge Fukumasu. USP-Botucatu: Rejane Maria Tommasini Grotto, Jayme A. Souza-Neto, Guilherme Targino Valente, Patricia Akemi Assato, Felipe Allan da Silva da Costa, Bianca Cechetto Carlos. Mendelics: Bibiana Santos, João Paulo Kitajima, Erika Freitas, David Schlesinger. Hemocentro Ribeirão Preto: Simone Kashima, Evandra Strazza Rodrigues, Svetoslav Nanev Slavov, Elaine Vieira dos Santos, Rafael dos Santos Bezerra, Luiz Carlos Junior de Alcantara, Marta Giovanetti, Vagner Fonseca, Flavia Aburjaile, Rodrigo Tocantins Calado. FAMERP-SJRP: Cecília Artico Banho, Lívia Sacchetto, Fábio Sossai Possebon, Leila Sabrina Ullmann, Cíntia Bittar, Guilherme Campos, Helena Lage Ferreira, Jorge A. Petroli Marchesi, Maisa C. Pereira Parra, Marília Moraes, Paula Rahal, Paulo Inacio da Costa, João Pessoa Araújo Jr., Maurício Lacerda Nogueira. Prefeitura de Sao Paulo: Melissa Palmieri. |
| EPI_ISL_1966260, EPI_ISL_1966261, EPI_ISL_1966262, EPI_ISL_1966263, EPI_ISL_1966264 | SMS SECRETARIA MUNICIPAL DE SAUDE DE BOITUVA | Instituto Butantan / Mendelics | Instituto Butantan: Dimas Tadeu Covas, Sandra Coccuzzo Sampaio, Maria Carolina Elias, José Salvatore Leister Patané, Vincent Louis Viala, Antonio Jorge Martins, Ricardo Haddad, Claudia Renata dos Santos Barros, Elaine Cristina Marqueze, Raul Machado Neto, Debora Botequio Moretti, Jardelina de Souza Todao Bernardino, Loyze Paola Oliveira de Lima, Luiz Aurelio de Campos Crispin. Centro de Genômica Funcional da ESALQ: Luiz Lehmann Coutinho, Ricardo Augusto Brassaloti, Raquel de Lello Rocha Campos Cassano. NGS Soluções Genômicas: Pilar Drummond Sampaio Corrêa Mariani. FZEA-USP Pirassununga: Mirele Daiana Poleti, Jessika Cristina Chagas Lesbon, Elisangela Chicaroni Mattos, Heidge Fukumasu. USP-Botucatu: Rejane Maria Tommasini Grotto, Jayme A. Souza-Neto, Guilherme Targino Valente, Patricia Akemi Assato, Felipe Allan da Silva da Costa, Bianca Cechetto Carlos. Mendelics: Bibiana Santos, João Paulo Kitajima, Erika Freitas, David Schlesinger. Hemocentro Ribeirão Preto: Simone Kashima, Evandra Strazza Rodrigues, Svetoslav Nanev Slavov, Elaine Vieira dos Santos, Rafael dos Santos Bezerra, Luiz Carlos Junior de Alcantara, Marta Giovanetti, Vagner Fonseca, Flavia Aburjaile, Rodrigo Tocantins Calado. FAMERP-SJRP: Cecília Artico Banho, Lívia Sacchetto, Fábio Sossai Possebon, Leila Sabrina Ullmann, Cíntia Bittar, Guilherme Campos, Helena Lage Ferreira, Jorge A. Petroli Marchesi, Maisa C. Pereira Parra, Marília Moraes, Paula Rahal, Paulo Inacio da Costa, João Pessoa Araújo Jr., Maurício Lacerda Nogueira. Prefeitura de Sao Paulo: Melissa Palmieri. |
| EPI_ISL_1966265 | UBS II DE NARANDIBA | Instituto Butantan / Mendelics | Instituto Butantan: Dimas Tadeu Covas, Sandra Coccuzzo Sampaio, Maria Carolina Elias, José Salvatore Leister Patané, Vincent Louis Viala, Antonio Jorge Martins, Ricardo Haddad, Claudia Renata dos Santos Barros, Elaine Cristina Marqueze, Raul Machado Neto, Debora Botequio Moretti, Jardelina de Souza Todao Bernardino, Loyze Paola Oliveira de Lima, Luiz Aurelio de Campos Crispin. Centro de Genômica Funcional da ESALQ: Luiz Lehmann Coutinho, Ricardo Augusto Brassaloti, Raquel de Lello Rocha Campos Cassano. NGS Soluções Genômicas: Pilar Drummond Sampaio Corrêa Mariani. FZEA-USP Pirassununga: Mirele Daiana Poleti, Jessika Cristina Chagas Lesbon, Elisangela Chicaroni Mattos, Heidge Fukumasu. USP-Botucatu: Rejane Maria Tommasini Grotto, Jayme A. Souza-Neto, Guilherme Targino Valente, Patricia Akemi Assato, Felipe Allan da Silva da Costa, Bianca Cechetto Carlos. Mendelics: Bibiana Santos, João Paulo Kitajima, Erika Freitas, David Schlesinger. Hemocentro Ribeirão Preto: Simone Kashima, Evandra Strazza Rodrigues, Svetoslav Nanev Slavov, Elaine Vieira dos Santos, Rafael dos Santos Bezerra, Luiz Carlos Junior de Alcantara, Marta Giovanetti, Vagner Fonseca, Flavia Aburjaile, Rodrigo Tocantins Calado. FAMERP-SJRP: Cecília Artico Banho, Lívia Sacchetto, Fábio Sossai Possebon, Leila Sabrina Ullmann, Cíntia Bittar, Guilherme Campos, Helena Lage Ferreira, Jorge A. Petroli Marchesi, Maisa C. Pereira Parra, Marília Moraes, Paula Rahal, Paulo Inacio da Costa, João Pessoa Araújo Jr., Maurício Lacerda Nogueira. Prefeitura de Sao Paulo: Melissa Palmieri. |
| EPI_ISL_1966266 | UBS IRMA AGUEDA MARIA JAIME | Instituto Butantan / Mendelics | Instituto Butantan: Dimas Tadeu Covas, Sandra Coccuzzo Sampaio, Maria Carolina Elias, José Salvatore Leister Patané, Vincent Louis Viala, Antonio Jorge Martins, Ricardo Haddad, Claudia Renata dos Santos Barros, Elaine Cristina Marqueze, Raul Machado Neto, Debora Botequio Moretti, Jardelina de Souza Todao Bernardino, Loyze Paola Oliveira de Lima, Luiz Aurelio de Campos Crispin. Centro de Genômica Funcional da ESALQ: Luiz Lehmann Coutinho, Ricardo Augusto Brassaloti, Raquel de Lello Rocha Campos Cassano. NGS Soluções Genômicas: Pilar Drummond Sampaio Corrêa Mariani. FZEA-USP Pirassununga: Mirele Daiana Poleti, Jessika Cristina Chagas Lesbon, Elisangela Chicaroni Mattos, Heidge Fukumasu. USP-Botucatu: Rejane Maria Tommasini Grotto, Jayme A. Souza-Neto, Guilherme Targino Valente, Patricia Akemi Assato, Felipe Allan da Silva da Costa, Bianca Cechetto Carlos. Mendelics: Bibiana Santos, João Paulo Kitajima, Erika Freitas, David Schlesinger. Hemocentro Ribeirão Preto: Simone Kashima, Evandra Strazza Rodrigues, Svetoslav Nanev Slavov, Elaine Vieira dos Santos, Rafael dos Santos Bezerra, Luiz Carlos Junior de Alcantara, Marta Giovanetti, Vagner Fonseca, Flavia Aburjaile, Rodrigo Tocantins Calado. FAMERP-SJRP: Cecília Artico Banho, Lívia Sacchetto, Fábio Sossai Possebon, Leila Sabrina Ullmann, Cíntia Bittar, Guilherme Campos, Helena Lage Ferreira, Jorge A. Petroli Marchesi, Maisa C. Pereira Parra, Marília Moraes, Paula Rahal, Paulo Inacio da Costa, João Pessoa Araújo Jr., Maurício Lacerda Nogueira. Prefeitura de Sao Paulo: Melissa Palmieri. |
| EPI_ISL_1966267, EPI_ISL_1966268, EPI_ISL_1966269 | VIGILANCIA EM SAUDE SAE VIGILANCIA EPIDEMIOLOGICA | Instituto Butantan / Mendelics | Instituto Butantan: Dimas Tadeu Covas, Sandra Coccuzzo Sampaio, Maria Carolina Elias, José Salvatore Leister Patané, Vincent Louis Viala, Antonio Jorge Martins, Ricardo Haddad, Claudia Renata dos Santos Barros, Elaine Cristina Marqueze, Raul Machado Neto, Debora Botequio Moretti, Jardelina de Souza Todao Bernardino, Loyze Paola Oliveira de Lima, Luiz Aurelio de Campos Crispin. Centro de Genômica Funcional da ESALQ: Luiz Lehmann Coutinho, Ricardo Augusto Brassaloti, Raquel de Lello Rocha Campos Cassano. NGS Soluções Genômicas: Pilar Drummond Sampaio Corrêa Mariani. FZEA-USP Pirassununga: Mirele Daiana Poleti, Jessika Cristina Chagas Lesbon, Elisangela Chicaroni Mattos, Heidge Fukumasu. USP-Botucatu: Rejane Maria Tommasini Grotto, Jayme A. Souza-Neto, Guilherme Targino Valente, Patricia Akemi Assato, Felipe Allan da Silva da Costa, Bianca Cechetto Carlos. Mendelics: Bibiana Santos, João Paulo Kitajima, Erika Freitas, David Schlesinger. Hemocentro Ribeirão Preto: Simone Kashima, Evandra Strazza Rodrigues, Svetoslav Nanev Slavov, Elaine Vieira dos Santos, Rafael dos Santos Bezerra, Luiz Carlos Junior de Alcantara, Marta Giovanetti, Vagner Fonseca, Flavia Aburjaile, Rodrigo Tocantins Calado. FAMERP-SJRP: Cecília Artico Banho, Lívia Sacchetto, Fábio Sossai Possebon, Leila Sabrina Ullmann, Cíntia Bittar, Guilherme Campos, Helena Lage Ferreira, Jorge A. Petroli Marchesi, Maisa C. Pereira Parra, Marília Moraes, Paula Rahal, Paulo Inacio da Costa, João Pessoa Araújo Jr., Maurício Lacerda Nogueira. Prefeitura de Sao Paulo: Melissa Palmieri. |
| EPI_ISL_1966270, EPI_ISL_1966271, EPI_ISL_1966272, EPI_ISL_1966273, EPI_ISL_1966274, EPI_ISL_1966275, EPI_ISL_1966276, EPI_ISL_1966277 | SECRETARIA MUNICIPAL DE SAUDE DE ANDRADINA | Instituto Butantan / Mendelics | Instituto Butantan: Dimas Tadeu Covas, Sandra Coccuzzo Sampaio, Maria Carolina Elias, José Salvatore Leister Patané, Vincent Louis Viala, Antonio Jorge Martins, Ricardo Haddad, Claudia Renata dos Santos Barros, Elaine Cristina Marqueze, Raul Machado Neto, Debora Botequio Moretti, Jardelina de Souza Todao Bernardino, Loyze Paola Oliveira de Lima, Luiz Aurelio de Campos Crispin. Centro de Genômica Funcional da ESALQ: Luiz Lehmann Coutinho, Ricardo Augusto Brassaloti, Raquel de Lello Rocha Campos Cassano. NGS Soluções Genômicas: Pilar Drummond Sampaio Corrêa Mariani. FZEA-USP Pirassununga: Mirele Daiana Poleti, Jessika Cristina Chagas Lesbon, Elisangela Chicaroni Mattos, Heidge Fukumasu. USP-Botucatu: Rejane Maria Tommasini Grotto, Jayme A. Souza-Neto, Guilherme Targino Valente, Patricia Akemi Assato, Felipe Allan da Silva da Costa, Bianca Cechetto Carlos. Mendelics: Bibiana Santos, João Paulo Kitajima, Erika Freitas, David Schlesinger. Hemocentro Ribeirão Preto: Simone Kashima, Evandra Strazza Rodrigues, Svetoslav Nanev Slavov, Elaine Vieira dos Santos, Rafael dos Santos Bezerra, Luiz Carlos Junior de Alcantara, Marta Giovanetti, Vagner Fonseca, Flavia Aburjaile, Rodrigo Tocantins Calado. FAMERP-SJRP: Cecília Artico Banho, Lívia Sacchetto, Fábio Sossai Possebon, Leila Sabrina Ullmann, Cíntia Bittar, Guilherme Campos, Helena Lage Ferreira, Jorge A. Petroli Marchesi, Maisa C. Pereira Parra, Marília Moraes, Paula Rahal, Paulo Inacio da Costa, João Pessoa Araújo Jr., Maurício Lacerda Nogueira. Prefeitura de Sao Paulo: Melissa Palmieri. |
| EPI_ISL_1966278, EPI_ISL_1966279, EPI_ISL_1966280, EPI_ISL_1966281, EPI_ISL_1966282, EPI_ISL_1966283, EPI_ISL_1966284, EPI_ISL_1966285, EPI_ISL_1966286 | UBS DR ALFREDO DANTAS DE SOUZA UMUARAMA | Instituto Butantan / Mendelics | Instituto Butantan: Dimas Tadeu Covas, Sandra Coccuzzo Sampaio, Maria Carolina Elias, José Salvatore Leister Patané, Vincent Louis Viala, Antonio Jorge Martins, Ricardo Haddad, Claudia Renata dos Santos Barros, Elaine Cristina Marqueze, Raul Machado Neto, Debora Botequio Moretti, Jardelina de Souza Todao Bernardino, Loyze Paola Oliveira de Lima, Luiz Aurelio de Campos Crispin. Centro de Genômica Funcional da ESALQ: Luiz Lehmann Coutinho, Ricardo Augusto Brassaloti, Raquel de Lello Rocha Campos Cassano. NGS Soluções Genômicas: Pilar Drummond Sampaio Corrêa Mariani. FZEA-USP Pirassununga: Mirele Daiana Poleti, Jessika Cristina Chagas Lesbon, Elisangela Chicaroni Mattos, Heidge Fukumasu. USP-Botucatu: Rejane Maria Tommasini Grotto, Jayme A. Souza-Neto, Guilherme Targino Valente, Patricia Akemi Assato, Felipe Allan da Silva da Costa, Bianca Cechetto Carlos. Mendelics: Bibiana Santos, João Paulo Kitajima, Erika Freitas, David Schlesinger. Hemocentro Ribeirão Preto: Simone Kashima, Evandra Strazza |

| | | | |
|---|---|---|---|
| | | | Rodrigues, Svetoslav Nanev Slavov, Elaine Vieira dos Santos, Rafael dos Santos Bezerra, Luiz Carlos Junior de Alcantara, Marta Giovanetti, Vagner Fonseca, Flavia Aburjaile, Rodrigo Tocantins Calado. FAMERP-SJRP: Cecília Artico Banho, Lívia Sacchetto, Fábio Sossai Possebon, Leila Sabrina Ullmann, Cíntia Bittar, Guilherme Campos, Helena Lage Ferreira, Jorge A. Petroli Marchesi, Maisa C. Pereira Parra, Marília Moraes, Paula Rahal, Paulo Inacio da Costa, João Pessoa Araújo Jr., Maurício Lacerda Nogueira. Prefeitura de Sao Paulo: Melissa Palmieri. |
| EPI_ISL_1966287 | UBS DE AURIFLAMA | Instituto Butantan / Mendelics | Instituto Butantan: Dimas Tadeu Covas, Sandra Coccuzzo Sampaio, Maria Carolina Elias, José Salvatore Leister Patané, Vincent Louis Viala, Antonio Jorge Martins, Ricardo Haddad, Claudia Renata dos Santos Barros, Elaine Cristina Marqueze, Raul Machado Neto, Debora Botequio Moretti, Jardelina de Souza Todao Bernardino, Loyze Paola Oliveira de Lima, Luiz Aurelio de Campos Crispin. Centro de Genômica Funcional da ESALQ: Luiz Lehmann Coutinho, Ricardo Augusto Brassaloti, Raquel de Lello Rocha Campos Cassano. NGS Soluções Genômicas: Pilar Drummond Sampaio Corrêa Mariani. FZEA-USP Pirassununga: Mirele Daiana Poleti, Jessika Cristina Chagas Lesbon, Elisangela Chicaroni Mattos, Heidge Fukumasu. USP-Botucatu: Rejane Maria Tommasini Grotto, Jayme A. Souza-Neto, Guilherme Targino Valente, Patricia Akemi Assato, Felipe Allan da Silva da Costa, Bianca Cechetto Carlos. Mendelics: Bibiana Santos, João Paulo Kitajima, Erika Freitas, David Schlesinger. Hemocentro Ribeirão Preto: Simone Kashima, Evandra Strazza Rodrigues, Svetoslav Nanev Slavov, Elaine Vieira dos Santos, Rafael dos Santos Bezerra, Luiz Carlos Junior de Alcantara, Marta Giovanetti, Vagner Fonseca, Flavia Aburjaile, Rodrigo Tocantins Calado. FAMERP-SJRP: Cecília Artico Banho, Lívia Sacchetto, Fábio Sossai Possebon, Leila Sabrina Ullmann, Cíntia Bittar, Guilherme Campos, Helena Lage Ferreira, Jorge A. Petroli Marchesi, Maisa C. Pereira Parra, Marília Moraes, Paula Rahal, Paulo Inacio da Costa, João Pessoa Araújo Jr., Maurício Lacerda Nogueira. Prefeitura de Sao Paulo: Melissa Palmieri. |
| EPI_ISL_1966288 | UNIDADE BASICA DE SAUDE DR BARTOLOMEU MARAGLIANO VENERE | Instituto Butantan / Mendelics | Instituto Butantan: Dimas Tadeu Covas, Sandra Coccuzzo Sampaio, Maria Carolina Elias, José Salvatore Leister Patané, Vincent Louis Viala, Antonio Jorge Martins, Ricardo Haddad, Claudia Renata dos Santos Barros, Elaine Cristina Marqueze, Raul Machado Neto, Debora Botequio Moretti, Jardelina de Souza Todao Bernardino, Loyze Paola Oliveira de Lima, Luiz Aurelio de Campos Crispin. Centro de Genômica Funcional da ESALQ: Luiz Lehmann Coutinho, Ricardo Augusto Brassaloti, Raquel de Lello Rocha Campos Cassano. NGS Soluções Genômicas: Pilar Drummond Sampaio Corrêa Mariani. FZEA-USP Pirassununga: Mirele Daiana Poleti, Jessika Cristina Chagas Lesbon, Elisangela Chicaroni Mattos, Heidge Fukumasu. USP-Botucatu: Rejane Maria Tommasini Grotto, Jayme A. Souza-Neto, Guilherme Targino Valente, Patricia Akemi Assato, Felipe Allan da Silva da Costa, Bianca Cechetto Carlos. Mendelics: Bibiana Santos, João Paulo Kitajima, Erika Freitas, David Schlesinger. Hemocentro Ribeirão Preto: Simone Kashima, Evandra Strazza Rodrigues, Svetoslav Nanev Slavov, Elaine Vieira dos Santos, Rafael dos Santos Bezerra, Luiz Carlos Junior de Alcantara, Marta Giovanetti, Vagner Fonseca, Flavia Aburjaile, Rodrigo Tocantins Calado. FAMERP-SJRP: Cecília Artico Banho, Lívia Sacchetto, Fábio Sossai Possebon, Leila Sabrina Ullmann, Cíntia Bittar, Guilherme Campos, Helena Lage Ferreira, Jorge A. Petroli Marchesi, Maisa C. Pereira Parra, Marília Moraes, Paula Rahal, Paulo Inacio da Costa, João Pessoa Araújo Jr., Maurício Lacerda Nogueira. Prefeitura de Sao Paulo: Melissa Palmieri. |
| EPI_ISL_1966289 | UBS DR ALLY ALAHMAR DE BARRETOS | Instituto Butantan / Mendelics | Instituto Butantan: Dimas Tadeu Covas, Sandra Coccuzzo Sampaio, Maria Carolina Elias, José Salvatore Leister Patané, Vincent Louis Viala, Antonio Jorge Martins, Ricardo Haddad, Claudia Renata dos Santos Barros, Elaine Cristina Marqueze, Raul Machado Neto, Debora Botequio Moretti, Jardelina de Souza Todao Bernardino, Loyze Paola Oliveira de Lima, Luiz Aurelio de Campos Crispin. Centro de Genômica Funcional da ESALQ: Luiz Lehmann Coutinho, Ricardo Augusto Brassaloti, Raquel de Lello Rocha Campos Cassano. NGS Soluções Genômicas: Pilar Drummond Sampaio Corrêa Mariani. FZEA-USP Pirassununga: Mirele Daiana Poleti, Jessika Cristina Chagas Lesbon, Elisangela Chicaroni Mattos, Heidge Fukumasu. USP-Botucatu: Rejane Maria Tommasini Grotto, Jayme A. Souza-Neto, Guilherme Targino Valente, Patricia Akemi Assato, Felipe Allan da Silva da Costa, Bianca Cechetto Carlos. Mendelics: Bibiana Santos, João Paulo Kitajima, Erika Freitas, David Schlesinger. Hemocentro Ribeirão Preto: Simone Kashima, Evandra Strazza Rodrigues, Svetoslav Nanev Slavov, Elaine Vieira dos Santos, Rafael dos Santos Bezerra, Luiz Carlos Junior de Alcantara, Marta Giovanetti, Vagner Fonseca, Flavia Aburjaile, Rodrigo Tocantins Calado. FAMERP-SJRP: Cecília Artico Banho, Lívia Sacchetto, Fábio Sossai Possebon, Leila Sabrina Ullmann, Cíntia Bittar, Guilherme Campos, Helena Lage Ferreira, Jorge A. Petroli Marchesi, Maisa C. Pereira Parra, Marília Moraes, Paula Rahal, Paulo Inacio da Costa, João Pessoa Araújo Jr., Maurício Lacerda Nogueira. Prefeitura de Sao Paulo: Melissa Palmieri. |
| EPI_ISL_1966290, EPI_ISL_1966291 | UNIDADE BASICA DE SAUDE DA FAMILIA BILAC | Instituto Butantan / Mendelics | Instituto Butantan: Dimas Tadeu Covas, Sandra Coccuzzo Sampaio, Maria Carolina Elias, José Salvatore Leister Patané, Vincent Louis Viala, Antonio Jorge Martins, Ricardo Haddad, Claudia Renata dos Santos Barros, Elaine Cristina Marqueze, Raul Machado Neto, Debora Botequio Moretti, Jardelina de Souza Todao Bernardino, Loyze Paola Oliveira de Lima, Luiz Aurelio de Campos Crispin. Centro de Genômica Funcional da ESALQ: Luiz Lehmann Coutinho, Ricardo Augusto Brassaloti, Raquel de Lello Rocha Campos Cassano. NGS Soluções Genômicas: Pilar Drummond Sampaio Corrêa Mariani. FZEA-USP Pirassununga: Mirele Daiana Poleti, Jessika Cristina Chagas Lesbon, Elisangela Chicaroni Mattos, Heidge Fukumasu. USP-Botucatu: Rejane Maria Tommasini Grotto, Jayme A. Souza-Neto, Guilherme Targino Valente, Patricia Akemi Assato, Felipe Allan da Silva da Costa, Bianca Cechetto Carlos. Mendelics: Bibiana Santos, João Paulo Kitajima, Erika Freitas, David Schlesinger. Hemocentro Ribeirão Preto: Simone Kashima, Evandra Strazza Rodrigues, Svetoslav Nanev Slavov, Elaine Vieira dos Santos, Rafael dos Santos Bezerra, Luiz Carlos Junior de Alcantara, Marta Giovanetti, Vagner Fonseca, Flavia Aburjaile, Rodrigo Tocantins Calado. FAMERP-SJRP: Cecília Artico Banho, Lívia Sacchetto, Fábio Sossai Possebon, Leila Sabrina Ullmann, Cíntia Bittar, Guilherme Campos, Helena Lage Ferreira, Jorge A. Petroli Marchesi, Maisa C. Pereira Parra, Marília Moraes, Paula Rahal, Paulo Inacio da Costa, João Pessoa Araújo Jr., Maurício Lacerda Nogueira. Prefeitura de Sao Paulo: Melissa Palmieri. |
| EPI_ISL_1966292 | CENTRO DE SAUDE DE FLORINEA | Instituto Butantan / Mendelics | Instituto Butantan: Dimas Tadeu Covas, Sandra Coccuzzo Sampaio, Maria Carolina Elias, José Salvatore Leister Patané, Vincent Louis Viala, Antonio Jorge Martins, Ricardo Haddad, Claudia Renata dos Santos Barros, Elaine Cristina Marqueze, Raul Machado Neto, Debora Botequio Moretti, Jardelina de Souza Todao Bernardino, Loyze Paola Oliveira de Lima, Luiz Aurelio de Campos Crispin. Centro de Genômica Funcional da ESALQ: Luiz Lehmann Coutinho, Ricardo Augusto Brassaloti, Raquel de Lello Rocha Campos Cassano. NGS Soluções Genômicas: Pilar Drummond Sampaio Corrêa Mariani. FZEA-USP Pirassununga: Mirele Daiana Poleti, Jessika Cristina Chagas Lesbon, Elisangela Chicaroni Mattos, Heidge Fukumasu. USP-Botucatu: Rejane Maria Tommasini Grotto, Jayme A. Souza-Neto, Guilherme Targino Valente, Patricia Akemi Assato, Felipe Allan da Silva da Costa, Bianca Cechetto Carlos. Mendelics: Bibiana Santos, João Paulo Kitajima, Erika Freitas, David Schlesinger. Hemocentro Ribeirão Preto: Simone Kashima, Evandra Strazza Rodrigues, Svetoslav Nanev Slavov, Elaine Vieira dos Santos, Rafael dos Santos Bezerra, Luiz Carlos Junior de Alcantara, Marta Giovanetti, Vagner Fonseca, Flavia Aburjaile, Rodrigo Tocantins Calado. FAMERP-SJRP: Cecília Artico Banho, Lívia Sacchetto, Fábio Sossai Possebon, Leila Sabrina Ullmann, Cíntia Bittar, Guilherme Campos, Helena Lage Ferreira, Jorge A. Petroli Marchesi, Maisa C. Pereira Parra, Marília Moraes, Paula Rahal, Paulo Inacio da Costa, João Pessoa Araújo Jr., Maurício Lacerda Nogueira. Prefeitura de Sao Paulo: Melissa Palmieri. |
| EPI_ISL_1966293 | CENTRO DE SAUDE III GLICERIO GIMAIEL ELIAS GIMAIEL | Instituto Butantan / Mendelics | Instituto Butantan: Dimas Tadeu Covas, Sandra Coccuzzo Sampaio, Maria Carolina Elias, José Salvatore Leister Patané, Vincent Louis Viala, Antonio Jorge Martins, Ricardo Haddad, Claudia Renata dos Santos Barros, Elaine Cristina Marqueze, Raul Machado Neto, Debora Botequio Moretti, Jardelina de Souza Todao Bernardino, Loyze Paola Oliveira de Lima, Luiz Aurelio de Campos Crispin. Centro de Genômica Funcional da ESALQ: Luiz Lehmann Coutinho, Ricardo Augusto Brassaloti, Raquel de Lello Rocha Campos Cassano. NGS Soluções Genômicas: Pilar Drummond Sampaio Corrêa Mariani. FZEA-USP Pirassununga: Mirele Daiana Poleti, Jessika Cristina Chagas Lesbon, Elisangela Chicaroni Mattos, Heidge Fukumasu. USP-Botucatu: Rejane Maria Tommasini Grotto, Jayme A. Souza-Neto, Guilherme Targino Valente, Patricia Akemi Assato, Felipe Allan da Silva da Costa, Bianca Cechetto Carlos. Mendelics: Bibiana Santos, João Paulo Kitajima, Erika Freitas, David Schlesinger. Hemocentro Ribeirão Preto: Simone Kashima, Evandra Strazza Rodrigues, Svetoslav Nanev Slavov, Elaine Vieira dos Santos, Rafael dos Santos Bezerra, Luiz Carlos Junior de Alcantara, Marta Giovanetti, Vagner Fonseca, Flavia Aburjaile, Rodrigo Tocantins Calado. FAMERP-SJRP: Cecília Artico Banho, Lívia Sacchetto, Fábio Sossai Possebon, Leila Sabrina Ullmann, Cíntia Bittar, Guilherme Campos, Helena Lage Ferreira, Jorge A. Petroli Marchesi, Maisa C. Pereira Parra, Marília Moraes, Paula Rahal, Paulo Inacio da Costa, João Pessoa Araújo Jr., Maurício Lacerda Nogueira. Prefeitura de Sao Paulo: Melissa Palmieri. |
| EPI_ISL_1966294, EPI_ISL_1966295, EPI_ISL_1966296, EPI_ISL_1966297, EPI_ISL_1966298, EPI_ISL_1966299, EPI_ISL_1966300, EPI_ISL_1966301 | VIGILANCIA EPIDEMIOLOGICA DE IBATE | Instituto Butantan / Mendelics | Instituto Butantan: Dimas Tadeu Covas, Sandra Coccuzzo Sampaio, Maria Carolina Elias, José Salvatore Leister Patané, Vincent Louis Viala, Antonio Jorge Martins, Ricardo Haddad, Claudia Renata dos Santos Barros, Elaine Cristina Marqueze, Raul Machado Neto, Debora Botequio Moretti, Jardelina de Souza Todao Bernardino, Loyze Paola Oliveira de Lima, Luiz Aurelio de Campos Crispin. Centro de Genômica Funcional da ESALQ: Luiz Lehmann Coutinho, Ricardo Augusto Brassaloti, Raquel de Lello Rocha Campos Cassano. NGS Soluções Genômicas: Pilar Drummond Sampaio Corrêa Mariani. FZEA-USP Pirassununga: Mirele Daiana Poleti, Jessika Cristina Chagas Lesbon, Elisangela Chicaroni Mattos, Heidge Fukumasu. USP-Botucatu: Rejane Maria Tommasini Grotto, Jayme A. Souza-Neto, Guilherme Targino Valente, Patricia Akemi Assato, Felipe Allan da Silva da Costa, Bianca Cechetto Carlos. Mendelics: Bibiana Santos, João Paulo Kitajima, Erika Freitas, David Schlesinger. Hemocentro Ribeirão Preto: Simone Kashima, Evandra Strazza Rodrigues, Svetoslav Nanev Slavov, Elaine Vieira dos Santos, Rafael dos Santos Bezerra, Luiz Carlos Junior de Alcantara, Marta Giovanetti, Vagner Fonseca, Flavia Aburjaile, Rodrigo Tocantins Calado. FAMERP-SJRP: Cecília Artico Banho, Lívia Sacchetto, Fábio Sossai Possebon, Leila Sabrina Ullmann, Cíntia Bittar, Guilherme Campos, Helena Lage Ferreira, Jorge A. Petroli Marchesi, Maisa C. Pereira Parra, Marília Moraes, Paula Rahal, Paulo Inacio da Costa, João Pessoa Araújo Jr., Maurício Lacerda Nogueira. Prefeitura de Sao Paulo: Melissa Palmieri. |
| EPI_ISL_1966302, EPI_ISL_1966303, EPI_ISL_1966304, EPI_ISL_1966305 | PRONTO SOCORRO DE AGENOR DE CAMPOS MONGAGUA | Instituto Butantan / Mendelics | Instituto Butantan: Dimas Tadeu Covas, Sandra Coccuzzo Sampaio, Maria Carolina Elias, José Salvatore Leister Patané, Vincent Louis Viala, Antonio Jorge Martins, Ricardo Haddad, Claudia Renata dos Santos Barros, Elaine Cristina Marqueze, Raul Machado Neto, Debora Botequio Moretti, Jardelina de |

| | | | |
|---|---|---|---|
| | | | Souza Todao Bernardino, Loyze Paola Oliveira de Lima, Luiz Aurelio de Campos Crispin. Centro de Genômica Funcional da ESALQ: Luiz Lehmann Coutinho, Ricardo Augusto Brassaloti, Raquel de Lello Rocha Campos Cassano. NGS Soluções Genômicas: Pilar Drummond Sampaio Corrêa Mariani. FZEA-USP Pirassununga: Mirele Daiana Poleti, Jessika Cristina Chagas Lesbon, Elisangela Chicaroni Mattos, Heidge Fukumasu. USP-Botucatu: Rejane Maria Tommasini Grotto, Jayme A. Souza-Neto, Guilherme Targino Valente, Patricia Akemi Assato, Felipe Allan da Silva da Costa, Bianca Cechetto Carlos. Mendelics: Bibiana Santos, João Paulo Kitajima, Erika Freitas, David Schlesinger. Hemocentro Ribeirão Preto: Simone Kashima, Evandra Strazza Rodrigues, Svetoslav Nanev Slavov, Elaine Vieira dos Santos, Rafael dos Santos Bezerra, Luiz Carlos Junior de Alcantara, Marta Giovanetti, Vagner Fonseca, Flavia Aburjaile, Rodrigo Tocantins Calado. FAMERP-SJRP: Cecília Artico Banho, Lívia Sacchetto, Fábio Sossai Possebon, Leila Sabrina Ullmann, Cíntia Bittar, Guilherme Campos, Helena Lage Ferreira, Jorge A. Petroli Marchesi, Maisa C. Pereira Parra, Marília Moraes, Paula Rahal, Paulo Inacio da Costa, João Pessoa Araújo Jr., Maurício Lacerda Nogueira. Prefeitura de Sao Paulo: Melissa Palmieri. |
| EPI_ISL_1966306, EPI_ISL_1966307 | UNIDADE DE PRONTO ATENDIMENTO DE AGENOR DE CAMPOS | Instituto Butantan / Mendelics | Instituto Butantan: Dimas Tadeu Covas, Sandra Coccuzzo Sampaio, Maria Carolina Elias, José Salvatore Leister Patané, Vincent Louis Viala, Antonio Jorge Martins, Ricardo Haddad, Claudia Renata dos Santos Barros, Elaine Cristina Marqueze, Raul Machado Neto, Debora Botequio Moretti, Jardelina de Souza Todao Bernardino, Loyze Paola Oliveira de Lima, Luiz Aurelio de Campos Crispin. Centro de Genômica Funcional da ESALQ: Luiz Lehmann Coutinho, Ricardo Augusto Brassaloti, Raquel de Lello Rocha Campos Cassano. NGS Soluções Genômicas: Pilar Drummond Sampaio Corrêa Mariani. FZEA-USP Pirassununga: Mirele Daiana Poleti, Jessika Cristina Chagas Lesbon, Elisangela Chicaroni Mattos, Heidge Fukumasu. USP-Botucatu: Rejane Maria Tommasini Grotto, Jayme A. Souza-Neto, Guilherme Targino Valente, Patricia Akemi Assato, Felipe Allan da Silva da Costa, Bianca Cechetto Carlos. Mendelics: Bibiana Santos, João Paulo Kitajima, Erika Freitas, David Schlesinger. Hemocentro Ribeirão Preto: Simone Kashima, Evandra Strazza Rodrigues, Svetoslav Nanev Slavov, Elaine Vieira dos Santos, Rafael dos Santos Bezerra, Luiz Carlos Junior de Alcantara, Marta Giovanetti, Vagner Fonseca, Flavia Aburjaile, Rodrigo Tocantins Calado. FAMERP-SJRP: Cecília Artico Banho, Lívia Sacchetto, Fábio Sossai Possebon, Leila Sabrina Ullmann, Cíntia Bittar, Guilherme Campos, Helena Lage Ferreira, Jorge A. Petroli Marchesi, Maisa C. Pereira Parra, Marília Moraes, Paula Rahal, Paulo Inacio da Costa, João Pessoa Araújo Jr., Maurício Lacerda Nogueira. Prefeitura de Sao Paulo: Melissa Palmieri. |
| EPI_ISL_1966308 | PRONTO SOCORRO DE AGENOR DE CAMPOS MONGAGUA | Instituto Butantan / Mendelics | Instituto Butantan: Dimas Tadeu Covas, Sandra Coccuzzo Sampaio, Maria Carolina Elias, José Salvatore Leister Patané, Vincent Louis Viala, Antonio Jorge Martins, Ricardo Haddad, Claudia Renata dos Santos Barros, Elaine Cristina Marqueze, Raul Machado Neto, Debora Botequio Moretti, Jardelina de Souza Todao Bernardino, Loyze Paola Oliveira de Lima, Luiz Aurelio de Campos Crispin. Centro de Genômica Funcional da ESALQ: Luiz Lehmann Coutinho, Ricardo Augusto Brassaloti, Raquel de Lello Rocha Campos Cassano. NGS Soluções Genômicas: Pilar Drummond Sampaio Corrêa Mariani. FZEA-USP Pirassununga: Mirele Daiana Poleti, Jessika Cristina Chagas Lesbon, Elisangela Chicaroni Mattos, Heidge Fukumasu. USP-Botucatu: Rejane Maria Tommasini Grotto, Jayme A. Souza-Neto, Guilherme Targino Valente, Patricia Akemi Assato, Felipe Allan da Silva da Costa, Bianca Cechetto Carlos. Mendelics: Bibiana Santos, João Paulo Kitajima, Erika Freitas, David Schlesinger. Hemocentro Ribeirão Preto: Simone Kashima, Evandra Strazza Rodrigues, Svetoslav Nanev Slavov, Elaine Vieira dos Santos, Rafael dos Santos Bezerra, Luiz Carlos Junior de Alcantara, Marta Giovanetti, Vagner Fonseca, Flavia Aburjaile, Rodrigo Tocantins Calado. FAMERP-SJRP: Cecília Artico Banho, Lívia Sacchetto, Fábio Sossai Possebon, Leila Sabrina Ullmann, Cíntia Bittar, Guilherme Campos, Helena Lage Ferreira, Jorge A. Petroli Marchesi, Maisa C. Pereira Parra, Marília Moraes, Paula Rahal, Paulo Inacio da Costa, João Pessoa Araújo Jr., Maurício Lacerda Nogueira. Prefeitura de Sao Paulo: Melissa Palmieri. |
| EPI_ISL_1966309, EPI_ISL_1966310 | UNIDADE DE PRONTO ATENDIMENTO DE AGENOR DE CAMPOS | Instituto Butantan / Mendelics | Instituto Butantan: Dimas Tadeu Covas, Sandra Coccuzzo Sampaio, Maria Carolina Elias, José Salvatore Leister Patané, Vincent Louis Viala, Antonio Jorge Martins, Ricardo Haddad, Claudia Renata dos Santos Barros, Elaine Cristina Marqueze, Raul Machado Neto, Debora Botequio Moretti, Jardelina de Souza Todao Bernardino, Loyze Paola Oliveira de Lima, Luiz Aurelio de Campos Crispin. Centro de Genômica Funcional da ESALQ: Luiz Lehmann Coutinho, Ricardo Augusto Brassaloti, Raquel de Lello Rocha Campos Cassano. NGS Soluções Genômicas: Pilar Drummond Sampaio Corrêa Mariani. FZEA-USP Pirassununga: Mirele Daiana Poleti, Jessika Cristina Chagas Lesbon, Elisangela Chicaroni Mattos, Heidge Fukumasu. USP-Botucatu: Rejane Maria Tommasini Grotto, Jayme A. Souza-Neto, Guilherme Targino Valente, Patricia Akemi Assato, Felipe Allan da Silva da Costa, Bianca Cechetto Carlos. Mendelics: Bibiana Santos, João Paulo Kitajima, Erika Freitas, David Schlesinger. Hemocentro Ribeirão Preto: Simone Kashima, Evandra Strazza Rodrigues, Svetoslav Nanev Slavov, Elaine Vieira dos Santos, Rafael dos Santos Bezerra, Luiz Carlos Junior de Alcantara, Marta Giovanetti, Vagner Fonseca, Flavia Aburjaile, Rodrigo Tocantins Calado. FAMERP-SJRP: Cecília Artico Banho, Lívia Sacchetto, Fábio Sossai Possebon, Leila Sabrina Ullmann, Cíntia Bittar, Guilherme Campos, Helena Lage Ferreira, Jorge A. Petroli Marchesi, Maisa C. Pereira Parra, Marília Moraes, Paula Rahal, Paulo Inacio da Costa, João Pessoa Araújo Jr., Maurício Lacerda Nogueira. Prefeitura de Sao Paulo: Melissa Palmieri. |
| EPI_ISL_1966311, EPI_ISL_1966312, EPI_ISL_1966313, EPI_ISL_1966314, EPI_ISL_1966315 | PRONTO SOCORRO DE AGENOR DE CAMPOS MONGAGUA | Instituto Butantan / Mendelics | Instituto Butantan: Dimas Tadeu Covas, Sandra Coccuzzo Sampaio, Maria Carolina Elias, José Salvatore Leister Patané, Vincent Louis Viala, Antonio Jorge Martins, Ricardo Haddad, Claudia Renata dos Santos Barros, Elaine Cristina Marqueze, Raul Machado Neto, Debora Botequio Moretti, Jardelina de Souza Todao Bernardino, Loyze Paola Oliveira de Lima, Luiz Aurelio de Campos Crispin. Centro de Genômica Funcional da ESALQ: Luiz Lehmann Coutinho, Ricardo Augusto Brassaloti, Raquel de Lello Rocha Campos Cassano. NGS Soluções Genômicas: Pilar Drummond Sampaio Corrêa Mariani. FZEA-USP Pirassununga: Mirele Daiana Poleti, Jessika Cristina Chagas Lesbon, Elisangela Chicaroni Mattos, Heidge Fukumasu. USP-Botucatu: Rejane Maria Tommasini Grotto, Jayme A. Souza-Neto, Guilherme Targino Valente, Patricia Akemi Assato, Felipe Allan da Silva da Costa, Bianca Cechetto Carlos. Mendelics: Bibiana Santos, João Paulo Kitajima, Erika Freitas, David Schlesinger. Hemocentro Ribeirão Preto: Simone Kashima, Evandra Strazza Rodrigues, Svetoslav Nanev Slavov, Elaine Vieira dos Santos, Rafael dos Santos Bezerra, Luiz Carlos Junior de Alcantara, Marta Giovanetti, Vagner Fonseca, Flavia Aburjaile, Rodrigo Tocantins Calado. FAMERP-SJRP: Cecília Artico Banho, Lívia Sacchetto, Fábio Sossai Possebon, Leila Sabrina Ullmann, Cíntia Bittar, Guilherme Campos, Helena Lage Ferreira, Jorge A. Petroli Marchesi, Maisa C. Pereira Parra, Marília Moraes, Paula Rahal, Paulo Inacio da Costa, João Pessoa Araújo Jr., Maurício Lacerda Nogueira. Prefeitura de Sao Paulo: Melissa Palmieri. |
| EPI_ISL_1966316 | UNIDADE DE PRONTO ATENDIMENTO DE AGENOR DE CAMPOS | Instituto Butantan / Mendelics | Instituto Butantan: Dimas Tadeu Covas, Sandra Coccuzzo Sampaio, Maria Carolina Elias, José Salvatore Leister Patané, Vincent Louis Viala, Antonio Jorge Martins, Ricardo Haddad, Claudia Renata dos Santos Barros, Elaine Cristina Marqueze, Raul Machado Neto, Debora Botequio Moretti, Jardelina de Souza Todao Bernardino, Loyze Paola Oliveira de Lima, Luiz Aurelio de Campos Crispin. Centro de Genômica Funcional da ESALQ: Luiz Lehmann Coutinho, Ricardo Augusto Brassaloti, Raquel de Lello Rocha Campos Cassano. NGS Soluções Genômicas: Pilar Drummond Sampaio Corrêa Mariani. FZEA-USP Pirassununga: Mirele Daiana Poleti, Jessika Cristina Chagas Lesbon, Elisangela Chicaroni Mattos, Heidge Fukumasu. USP-Botucatu: Rejane Maria Tommasini Grotto, Jayme A. Souza-Neto, Guilherme Targino Valente, Patricia Akemi Assato, Felipe Allan da Silva da Costa, Bianca Cechetto Carlos. Mendelics: Bibiana Santos, João Paulo Kitajima, Erika Freitas, David Schlesinger. Hemocentro Ribeirão Preto: Simone Kashima, Evandra Strazza Rodrigues, Svetoslav Nanev Slavov, Elaine Vieira dos Santos, Rafael dos Santos Bezerra, Luiz Carlos Junior de Alcantara, Marta Giovanetti, Vagner Fonseca, Flavia Aburjaile, Rodrigo Tocantins Calado. FAMERP-SJRP: Cecília Artico Banho, Lívia Sacchetto, Fábio Sossai Possebon, Leila Sabrina Ullmann, Cíntia Bittar, Guilherme Campos, Helena Lage Ferreira, Jorge A. Petroli Marchesi, Maisa C. Pereira Parra, Marília Moraes, Paula Rahal, Paulo Inacio da Costa, João Pessoa Araújo Jr., Maurício Lacerda Nogueira. Prefeitura de Sao Paulo: Melissa Palmieri. |
| EPI_ISL_1966317, EPI_ISL_1966318 | PRONTO SOCORRO DE AGENOR DE CAMPOS MONGAGUA | Instituto Butantan / Mendelics | Instituto Butantan: Dimas Tadeu Covas, Sandra Coccuzzo Sampaio, Maria Carolina Elias, José Salvatore Leister Patané, Vincent Louis Viala, Antonio Jorge Martins, Ricardo Haddad, Claudia Renata dos Santos Barros, Elaine Cristina Marqueze, Raul Machado Neto, Debora Botequio Moretti, Jardelina de Souza Todao Bernardino, Loyze Paola Oliveira de Lima, Luiz Aurelio de Campos Crispin. Centro de Genômica Funcional da ESALQ: Luiz Lehmann Coutinho, Ricardo Augusto Brassaloti, Raquel de Lello Rocha Campos Cassano. NGS Soluções Genômicas: Pilar Drummond Sampaio Corrêa Mariani. FZEA-USP Pirassununga: Mirele Daiana Poleti, Jessika Cristina Chagas Lesbon, Elisangela Chicaroni Mattos, Heidge Fukumasu. USP-Botucatu: Rejane Maria Tommasini Grotto, Jayme A. Souza-Neto, Guilherme Targino Valente, Patricia Akemi Assato, Felipe Allan da Silva da Costa, Bianca Cechetto Carlos. Mendelics: Bibiana Santos, João Paulo Kitajima, Erika Freitas, David Schlesinger. Hemocentro Ribeirão Preto: Simone Kashima, Evandra Strazza Rodrigues, Svetoslav Nanev Slavov, Elaine Vieira dos Santos, Rafael dos Santos Bezerra, Luiz Carlos Junior de Alcantara, Marta Giovanetti, Vagner Fonseca, Flavia Aburjaile, Rodrigo Tocantins Calado. FAMERP-SJRP: Cecília Artico Banho, Lívia Sacchetto, Fábio Sossai Possebon, Leila Sabrina Ullmann, Cíntia Bittar, Guilherme Campos, Helena Lage Ferreira, Jorge A. Petroli Marchesi, Maisa C. Pereira Parra, Marília Moraes, Paula Rahal, Paulo Inacio da Costa, João Pessoa Araújo Jr., Maurício Lacerda Nogueira. Prefeitura de Sao Paulo: Melissa Palmieri. |
| EPI_ISL_1966319 | UNIDADE DE PRONTO ATENDIMENTO DE AGENOR DE CAMPOS | Instituto Butantan / Mendelics | Instituto Butantan: Dimas Tadeu Covas, Sandra Coccuzzo Sampaio, Maria Carolina Elias, José Salvatore Leister Patané, Vincent Louis Viala, Antonio Jorge Martins, Ricardo Haddad, Claudia Renata dos Santos Barros, Elaine Cristina Marqueze, Raul Machado Neto, Debora Botequio Moretti, Jardelina de Souza Todao Bernardino, Loyze Paola Oliveira de Lima, Luiz Aurelio de Campos Crispin. Centro de Genômica Funcional da ESALQ: Luiz Lehmann Coutinho, Ricardo Augusto Brassaloti, Raquel de Lello Rocha Campos Cassano. NGS Soluções Genômicas: Pilar Drummond Sampaio Corrêa Mariani. FZEA-USP Pirassununga: Mirele Daiana Poleti, Jessika Cristina Chagas Lesbon, Elisangela Chicaroni Mattos, Heidge Fukumasu. USP-Botucatu: Rejane Maria Tommasini Grotto, Jayme A. Souza-Neto, Guilherme Targino Valente, Patricia Akemi Assato, Felipe Allan da Silva da Costa, Bianca Cechetto Carlos. Mendelics: Bibiana Santos, João Paulo Kitajima, Erika Freitas, David Schlesinger. Hemocentro Ribeirão Preto: Simone Kashima, Evandra Strazza Rodrigues, Svetoslav Nanev Slavov, Elaine Vieira dos Santos, Rafael dos Santos Bezerra, Luiz Carlos Junior de Alcantara, Marta Giovanetti, Vagner Fonseca, Flavia Aburjaile, Rodrigo Tocantins Calado. FAMERP-SJRP: Cecília Artico Banho, Lívia Sacchetto, Fábio Sossai Possebon, Leila Sabrina |

| | | | |
|---|---|---|---|
| | | | Ullmann, Cíntia Bittar, Guilherme Campos, Helena Lage Ferreira, Jorge A. Petroli Marchesi, Maisa C. Pereira Parra, Marília Moraes, Paula Rahal, Paulo Inacio da Costa, João Pessoa Araújo Jr., Maurício Lacerda Nogueira. Prefeitura de Sao Paulo: Melissa Palmieri. |
| EPI_ISL_1966321, EPI_ISL_1966322 | UPA SILVINA | Instituto Butantan / Mendelics | Instituto Butantan: Dimas Tadeu Covas, Sandra Coccuzzo Sampaio, Maria Carolina Elias, José Salvatore Leister Patané, Vincent Louis Viala, Antonio Jorge Martins, Ricardo Haddad, Claudia Renata dos Santos Barros, Elaine Cristina Marqueze, Raul Machado Neto, Debora Botequio Moretti, Jardelina de Souza Todao Bernardino, Loyze Paola Oliveira de Lima, Luiz Aurelio de Campos Crispin. Centro de Genômica Funcional da ESALQ: Luiz Lehmann Coutinho, Ricardo Augusto Brassaloti, Raquel de Lello Rocha Campos Cassano. NGS Soluções Genômicas: Pilar Drummond Sampaio Corrêa Mariani. FZEA-USP Pirassununga: Mirele Daiana Poleti, Jessika Cristina Chagas Lesbon, Elisangela Chicaroni Mattos, Heidge Fukumasu. USP-Botucatu: Rejane Maria Tommasini Grotto, Jayme A. Souza-Neto, Guilherme Targino Valente, Patricia Akemi Assato, Felipe Allan da Silva da Costa, Bianca Cechetto Carlos. Mendelics: Bibiana Santos, João Paulo Kitajima, Erika Freitas, David Schlesinger. Hemocentro Ribeirão Preto: Simone Kashima, Evandra Strazza Rodrigues, Svetoslav Nanev Slavov, Elaine Vieira dos Santos, Rafael dos Santos Bezerra, Luiz Carlos Junior de Alcantara, Marta Giovanetti, Vagner Fonseca, Flavia Aburjaile, Rodrigo Tocantins Calado. FAMERP-SJRP: Cecília Artico Banho, Lívia Sacchetto, Fábio Sossai Possebon, Leila Sabrina Ullmann, Cíntia Bittar, Guilherme Campos, Helena Lage Ferreira, Jorge A. Petroli Marchesi, Maisa C. Pereira Parra, Marília Moraes, Paula Rahal, Paulo Inacio da Costa, João Pessoa Araújo Jr., Maurício Lacerda Nogueira. Prefeitura de Sao Paulo: Melissa Palmieri. |
| EPI_ISL_1966323, EPI_ISL_1966324, EPI_ISL_1966325, EPI_ISL_1966326, EPI_ISL_1966327, EPI_ISL_1966328, EPI_ISL_1966329, EPI_ISL_1966330, EPI_ISL_1966331, EPI_ISL_1966332, EPI_ISL_1966333, EPI_ISL_1966334, EPI_ISL_1966335, EPI_ISL_1966336, EPI_ISL_1966337, EPI_ISL_1966338, EPI_ISL_1966339, EPI_ISL_1966340 | | | |
| see above | Ambulatório IB | Instituto Butantan / Mendelics | Instituto Butantan: Dimas Tadeu Covas, Sandra Coccuzzo Sampaio, Maria Carolina Elias, José Salvatore Leister Patané, Vincent Louis Viala, Antonio Jorge Martins, Ricardo Haddad, Claudia Renata dos Santos Barros, Elaine Cristina Marqueze, Raul Machado Neto, Debora Botequio Moretti, Jardelina de Souza Todao Bernardino, Loyze Paola Oliveira de Lima, Luiz Aurelio de Campos Crispin. Centro de Genômica Funcional da ESALQ: Luiz Lehmann Coutinho, Ricardo Augusto Brassaloti, Raquel de Lello Rocha Campos Cassano. NGS Soluções Genômicas: Pilar Drummond Sampaio Corrêa Mariani. FZEA-USP Pirassununga: Mirele Daiana Poleti, Jessika Cristina Chagas Lesbon, Elisangela Chicaroni Mattos, Heidge Fukumasu. USP-Botucatu: Rejane Maria Tommasini Grotto, Jayme A. Souza-Neto, Guilherme Targino Valente, Patricia Akemi Assato, Felipe Allan da Silva da Costa, Bianca Cechetto Carlos. Mendelics: Bibiana Santos, João Paulo Kitajima, Erika Freitas, David Schlesinger. Hemocentro Ribeirão Preto: Simone Kashima, Evandra Strazza Rodrigues, Svetoslav Nanev Slavov, Elaine Vieira dos Santos, Rafael dos Santos Bezerra, Luiz Carlos Junior de Alcantara, Marta Giovanetti, Vagner Fonseca, Flavia Aburjaile, Rodrigo Tocantins Calado. FAMERP-SJRP: Cecília Artico Banho, Lívia Sacchetto, Fábio Sossai Possebon, Leila Sabrina Ullmann, Cíntia Bittar, Guilherme Campos, Helena Lage Ferreira, Jorge A. Petroli Marchesi, Maisa C. Pereira Parra, Marília Moraes, Paula Rahal, Paulo Inacio da Costa, João Pessoa Araújo Jr., Maurício Lacerda Nogueira. Prefeitura de Sao Paulo: Melissa Palmieri. |
| EPI_ISL_1966342 | UPA 24 HORAS CENTRO | Instituto Butantan / Mendelics | Instituto Butantan: Dimas Tadeu Covas, Sandra Coccuzzo Sampaio, Maria Carolina Elias, José Salvatore Leister Patané, Vincent Louis Viala, Antonio Jorge Martins, Ricardo Haddad, Claudia Renata dos Santos Barros, Elaine Cristina Marqueze, Raul Machado Neto, Debora Botequio Moretti, Jardelina de Souza Todao Bernardino, Loyze Paola Oliveira de Lima, Luiz Aurelio de Campos Crispin. Centro de Genômica Funcional da ESALQ: Luiz Lehmann Coutinho, Ricardo Augusto Brassaloti, Raquel de Lello Rocha Campos Cassano. NGS Soluções Genômicas: Pilar Drummond Sampaio Corrêa Mariani. FZEA-USP Pirassununga: Mirele Daiana Poleti, Jessika Cristina Chagas Lesbon, Elisangela Chicaroni Mattos, Heidge Fukumasu. USP-Botucatu: Rejane Maria Tommasini Grotto, Jayme A. Souza-Neto, Guilherme Targino Valente, Patricia Akemi Assato, Felipe Allan da Silva da Costa, Bianca Cechetto Carlos. Mendelics: Bibiana Santos, João Paulo Kitajima, Erika Freitas, David Schlesinger. Hemocentro Ribeirão Preto: Simone Kashima, Evandra Strazza Rodrigues, Svetoslav Nanev Slavov, Elaine Vieira dos Santos, Rafael dos Santos Bezerra, Luiz Carlos Junior de Alcantara, Marta Giovanetti, Vagner Fonseca, Flavia Aburjaile, Rodrigo Tocantins Calado. FAMERP-SJRP: Cecília Artico Banho, Lívia Sacchetto, Fábio Sossai Possebon, Leila Sabrina Ullmann, Cíntia Bittar, Guilherme Campos, Helena Lage Ferreira, Jorge A. Petroli Marchesi, Maisa C. Pereira Parra, Marília Moraes, Paula Rahal, Paulo Inacio da Costa, João Pessoa Araújo Jr., Maurício Lacerda Nogueira. Prefeitura de Sao Paulo: Melissa Palmieri. |
| EPI_ISL_1966343, EPI_ISL_1966344, EPI_ISL_1966345 | CENTRO DE SAUDE III TABATINGA | Instituto Butantan / Mendelics | Instituto Butantan: Dimas Tadeu Covas, Sandra Coccuzzo Sampaio, Maria Carolina Elias, José Salvatore Leister Patané, Vincent Louis Viala, Antonio Jorge Martins, Ricardo Haddad, Claudia Renata dos Santos Barros, Elaine Cristina Marqueze, Raul Machado Neto, Debora Botequio Moretti, Jardelina de Souza Todao Bernardino, Loyze Paola Oliveira de Lima, Luiz Aurelio de Campos Crispin. Centro de Genômica Funcional da ESALQ: Luiz Lehmann Coutinho, Ricardo Augusto Brassaloti, Raquel de Lello Rocha Campos Cassano. NGS Soluções Genômicas: Pilar Drummond Sampaio Corrêa Mariani. FZEA-USP Pirassununga: Mirele Daiana Poleti, Jessika Cristina Chagas Lesbon, Elisangela Chicaroni Mattos, Heidge Fukumasu. USP-Botucatu: Rejane Maria Tommasini Grotto, Jayme A. Souza-Neto, Guilherme Targino Valente, Patricia Akemi Assato, Felipe Allan da Silva da Costa, Bianca Cechetto Carlos. Mendelics: Bibiana Santos, João Paulo Kitajima, Erika Freitas, David Schlesinger. Hemocentro Ribeirão Preto: Simone Kashima, Evandra Strazza Rodrigues, Svetoslav Nanev Slavov, Elaine Vieira dos Santos, Rafael dos Santos Bezerra, Luiz Carlos Junior de Alcantara, Marta Giovanetti, Vagner Fonseca, Flavia Aburjaile, Rodrigo Tocantins Calado. FAMERP-SJRP: Cecília Artico Banho, Lívia Sacchetto, Fábio Sossai Possebon, Leila Sabrina Ullmann, Cíntia Bittar, Guilherme Campos, Helena Lage Ferreira, Jorge A. Petroli Marchesi, Maisa C. Pereira Parra, Marília Moraes, Paula Rahal, Paulo Inacio da Costa, João Pessoa Araújo Jr., Maurício Lacerda Nogueira. Prefeitura de Sao Paulo: Melissa Palmieri. |
| EPI_ISL_1966346, EPI_ISL_1966347, EPI_ISL_1966348, EPI_ISL_1966349 | SECRETARIA MUNICIPAL DE SAUDE DE VALPARAISO SP | Instituto Butantan / Mendelics | Instituto Butantan: Dimas Tadeu Covas, Sandra Coccuzzo Sampaio, Maria Carolina Elias, José Salvatore Leister Patané, Vincent Louis Viala, Antonio Jorge Martins, Ricardo Haddad, Claudia Renata dos Santos Barros, Elaine Cristina Marqueze, Raul Machado Neto, Debora Botequio Moretti, Jardelina de Souza Todao Bernardino, Loyze Paola Oliveira de Lima, Luiz Aurelio de Campos Crispin. Centro de Genômica Funcional da ESALQ: Luiz Lehmann Coutinho, Ricardo Augusto Brassaloti, Raquel de Lello Rocha Campos Cassano. NGS Soluções Genômicas: Pilar Drummond Sampaio Corrêa Mariani. FZEA-USP Pirassununga: Mirele Daiana Poleti, Jessika Cristina Chagas Lesbon, Elisangela Chicaroni Mattos, Heidge Fukumasu. USP-Botucatu: Rejane Maria Tommasini Grotto, Jayme A. Souza-Neto, Guilherme Targino Valente, Patricia Akemi Assato, Felipe Allan da Silva da Costa, Bianca Cechetto Carlos. Mendelics: Bibiana Santos, João Paulo Kitajima, Erika Freitas, David Schlesinger. Hemocentro Ribeirão Preto: Simone Kashima, Evandra Strazza Rodrigues, Svetoslav Nanev Slavov, Elaine Vieira dos Santos, Rafael dos Santos Bezerra, Luiz Carlos Junior de Alcantara, Marta Giovanetti, Vagner Fonseca, Flavia Aburjaile, Rodrigo Tocantins Calado. FAMERP-SJRP: Cecília Artico Banho, Lívia Sacchetto, Fábio Sossai Possebon, Leila Sabrina Ullmann, Cíntia Bittar, Guilherme Campos, Helena Lage Ferreira, Jorge A. Petroli Marchesi, Maisa C. Pereira Parra, Marília Moraes, Paula Rahal, Paulo Inacio da Costa, João Pessoa Araújo Jr., Maurício Lacerda Nogueira. Prefeitura de Sao Paulo: Melissa Palmieri. |
| EPI_ISL_1966350 | CENTRO DE SAUDE DR RENATO DE CARVALHO RIBEIRO ANGATUBA | Instituto Butantan / Mendelics | Instituto Butantan: Dimas Tadeu Covas, Sandra Coccuzzo Sampaio, Maria Carolina Elias, José Salvatore Leister Patané, Vincent Louis Viala, Antonio Jorge Martins, Ricardo Haddad, Claudia Renata dos Santos Barros, Elaine Cristina Marqueze, Raul Machado Neto, Debora Botequio Moretti, Jardelina de Souza Todao Bernardino, Loyze Paola Oliveira de Lima, Luiz Aurelio de Campos Crispin. Centro de Genômica Funcional da ESALQ: Luiz Lehmann Coutinho, Ricardo Augusto Brassaloti, Raquel de Lello Rocha Campos Cassano. NGS Soluções Genômicas: Pilar Drummond Sampaio Corrêa Mariani. FZEA-USP Pirassununga: Mirele Daiana Poleti, Jessika Cristina Chagas Lesbon, Elisangela Chicaroni Mattos, Heidge Fukumasu. USP-Botucatu: Rejane Maria Tommasini Grotto, Jayme A. Souza-Neto, Guilherme Targino Valente, Patricia Akemi Assato, Felipe Allan da Silva da Costa, Bianca Cechetto Carlos. Mendelics: Bibiana Santos, João Paulo Kitajima, Erika Freitas, David Schlesinger. Hemocentro Ribeirão Preto: Simone Kashima, Evandra Strazza Rodrigues, Svetoslav Nanev Slavov, Elaine Vieira dos Santos, Rafael dos Santos Bezerra, Luiz Carlos Junior de Alcantara, Marta Giovanetti, Vagner Fonseca, Flavia Aburjaile, Rodrigo Tocantins Calado. FAMERP-SJRP: Cecília Artico Banho, Lívia Sacchetto, Fábio Sossai Possebon, Leila Sabrina Ullmann, Cíntia Bittar, Guilherme Campos, Helena Lage Ferreira, Jorge A. Petroli Marchesi, Maisa C. Pereira Parra, Marília Moraes, Paula Rahal, Paulo Inacio da Costa, João Pessoa Araújo Jr., Maurício Lacerda Nogueira. Prefeitura de Sao Paulo: Melissa Palmieri. |
| EPI_ISL_1966351 | ESF VILA GLORIA I | Instituto Butantan / Mendelics | Instituto Butantan: Dimas Tadeu Covas, Sandra Coccuzzo Sampaio, Maria Carolina Elias, José Salvatore Leister Patané, Vincent Louis Viala, Antonio Jorge Martins, Ricardo Haddad, Claudia Renata dos Santos Barros, Elaine Cristina Marqueze, Raul Machado Neto, Debora Botequio Moretti, Jardelina de Souza Todao Bernardino, Loyze Paola Oliveira de Lima, Luiz Aurelio de Campos Crispin. Centro de Genômica Funcional da ESALQ: Luiz Lehmann Coutinho, Ricardo Augusto Brassaloti, Raquel de Lello Rocha Campos Cassano. NGS Soluções Genômicas: Pilar Drummond Sampaio Corrêa Mariani. FZEA-USP Pirassununga: Mirele Daiana Poleti, Jessika Cristina Chagas Lesbon, Elisangela Chicaroni Mattos, Heidge Fukumasu. USP-Botucatu: Rejane Maria Tommasini Grotto, Jayme A. Souza-Neto, Guilherme Targino Valente, Patricia Akemi Assato, Felipe Allan da Silva da Costa, Bianca Cechetto Carlos. Mendelics: Bibiana Santos, João Paulo Kitajima, Erika Freitas, David Schlesinger. Hemocentro Ribeirão Preto: Simone Kashima, Evandra Strazza Rodrigues, Svetoslav Nanev Slavov, Elaine Vieira dos Santos, Rafael dos Santos Bezerra, Luiz Carlos Junior de Alcantara, Marta Giovanetti, Vagner Fonseca, Flavia Aburjaile, Rodrigo Tocantins Calado. FAMERP-SJRP: Cecília Artico Banho, Lívia Sacchetto, Fábio Sossai Possebon, Leila Sabrina Ullmann, Cíntia Bittar, Guilherme Campos, Helena Lage Ferreira, Jorge A. Petroli Marchesi, Maisa C. Pereira Parra, Marília Moraes, Paula Rahal, Paulo Inacio da Costa, João Pessoa Araújo Jr., Maurício Lacerda Nogueira. Prefeitura de Sao Paulo: Melissa Palmieri. |
| EPI_ISL_1966352, EPI_ISL_1966353, EPI_ISL_1966354, EPI_ISL_1966355, | UNIDADE REFERENCIAL SUDOESTE | Instituto Butantan / Mendelics | Instituto Butantan: Dimas Tadeu Covas, Sandra Coccuzzo Sampaio, Maria Carolina Elias, José Salvatore Leister Patané, Vincent Louis Viala, Antonio Jorge Martins, Ricardo Haddad, Claudia Renata dos Santos Barros, Elaine Cristina Marqueze, Raul Machado Neto, Debora Botequio Moretti, Jardelina de |

| | | | |
|---|---|---|---|
| EPI_ISL_1966356 | | | Souza Todao Bernardino, Loyze Paola Oliveira de Lima, Luiz Aurelio de Campos Crispin. Centro de Genômica Funcional da ESALQ: Luiz Lehmann Coutinho, Ricardo Augusto Brassaloti, Raquel de Lello Rocha Campos Cassano. NGS Soluções Genômicas: Pilar Drummond Sampaio Corrêa Mariani. FZEA-USP Pirassununga: Mirele Daiana Poleti, Jessika Cristina Chagas Lesbon, Elisangela Chicaroni Mattos, Heidge Fukumasu. USP-Botucatu: Rejane Maria Tommasini Grotto, Jayme A. Souza-Neto, Guilherme Targino Valente, Patricia Akemi Assato, Felipe Allan da Silva da Costa, Bianca Cechetto Carlos. Mendelics: Bibiana Santos, João Paulo Kitajima, Erika Freitas, David Schlesinger. Hemocentro Ribeirão Preto: Simone Kashima, Evandra Strazza Rodrigues, Svetoslav Nanev Slavov, Elaine Vieira dos Santos, Rafael dos Santos Bezerra, Luiz Carlos Junior de Alcantara, Marta Giovanetti, Vagner Fonseca, Flavia Aburjaile, Rodrigo Tocantins Calado. FAMERP-SJRP: Cecília Artico Banho, Lívia Sacchetto, Fábio Sossai Possebon, Leila Sabrina Ullmann, Cíntia Bittar, Guilherme Campos, Helena Lage Ferreira, Jorge A. Petroli Marchesi, Maisa C. Pereira Parra, Marília Moraes, Paula Rahal, Paulo Inacio da Costa, João Pessoa Araújo Jr., Maurício Lacerda Nogueira. Prefeitura de Sao Paulo: Melissa Palmieri. |
| EPI_ISL_1966357, EPI_ISL_1966358, EPI_ISL_1966359 | SMS SECRETARIA MUNICIPAL DE SAUDE DE BOITUVA | Instituto Butantan / Mendelics | Instituto Butantan: Dimas Tadeu Covas, Sandra Coccuzzo Sampaio, Maria Carolina Elias, José Salvatore Leister Patané, Vincent Louis Viala, Antonio Jorge Martins, Ricardo Haddad, Claudia Renata dos Santos Barros, Elaine Cristina Marqueze, Raul Machado Neto, Debora Botequio Moretti, Jardelina de Souza Todao Bernardino, Loyze Paola Oliveira de Lima, Luiz Aurelio de Campos Crispin. Centro de Genômica Funcional da ESALQ: Luiz Lehmann Coutinho, Ricardo Augusto Brassaloti, Raquel de Lello Rocha Campos Cassano. NGS Soluções Genômicas: Pilar Drummond Sampaio Corrêa Mariani. FZEA-USP Pirassununga: Mirele Daiana Poleti, Jessika Cristina Chagas Lesbon, Elisangela Chicaroni Mattos, Heidge Fukumasu. USP-Botucatu: Rejane Maria Tommasini Grotto, Jayme A. Souza-Neto, Guilherme Targino Valente, Patricia Akemi Assato, Felipe Allan da Silva da Costa, Bianca Cechetto Carlos. Mendelics: Bibiana Santos, João Paulo Kitajima, Erika Freitas, David Schlesinger. Hemocentro Ribeirão Preto: Simone Kashima, Evandra Strazza Rodrigues, Svetoslav Nanev Slavov, Elaine Vieira dos Santos, Rafael dos Santos Bezerra, Luiz Carlos Junior de Alcantara, Marta Giovanetti, Vagner Fonseca, Flavia Aburjaile, Rodrigo Tocantins Calado. FAMERP-SJRP: Cecília Artico Banho, Lívia Sacchetto, Fábio Sossai Possebon, Leila Sabrina Ullmann, Cíntia Bittar, Guilherme Campos, Helena Lage Ferreira, Jorge A. Petroli Marchesi, Maisa C. Pereira Parra, Marília Moraes, Paula Rahal, Paulo Inacio da Costa, João Pessoa Araújo Jr., Maurício Lacerda Nogueira. Prefeitura de Sao Paulo: Melissa Palmieri. |
| EPI_ISL_1966360 | SECRETARIA DE SAUDE CABREUVA | Instituto Butantan / Mendelics | Instituto Butantan: Dimas Tadeu Covas, Sandra Coccuzzo Sampaio, Maria Carolina Elias, José Salvatore Leister Patané, Vincent Louis Viala, Antonio Jorge Martins, Ricardo Haddad, Claudia Renata dos Santos Barros, Elaine Cristina Marqueze, Raul Machado Neto, Debora Botequio Moretti, Jardelina de Souza Todao Bernardino, Loyze Paola Oliveira de Lima, Luiz Aurelio de Campos Crispin. Centro de Genômica Funcional da ESALQ: Luiz Lehmann Coutinho, Ricardo Augusto Brassaloti, Raquel de Lello Rocha Campos Cassano. NGS Soluções Genômicas: Pilar Drummond Sampaio Corrêa Mariani. FZEA-USP Pirassununga: Mirele Daiana Poleti, Jessika Cristina Chagas Lesbon, Elisangela Chicaroni Mattos, Heidge Fukumasu. USP-Botucatu: Rejane Maria Tommasini Grotto, Jayme A. Souza-Neto, Guilherme Targino Valente, Patricia Akemi Assato, Felipe Allan da Silva da Costa, Bianca Cechetto Carlos. Mendelics: Bibiana Santos, João Paulo Kitajima, Erika Freitas, David Schlesinger. Hemocentro Ribeirão Preto: Simone Kashima, Evandra Strazza Rodrigues, Svetoslav Nanev Slavov, Elaine Vieira dos Santos, Rafael dos Santos Bezerra, Luiz Carlos Junior de Alcantara, Marta Giovanetti, Vagner Fonseca, Flavia Aburjaile, Rodrigo Tocantins Calado. FAMERP-SJRP: Cecília Artico Banho, Lívia Sacchetto, Fábio Sossai Possebon, Leila Sabrina Ullmann, Cíntia Bittar, Guilherme Campos, Helena Lage Ferreira, Jorge A. Petroli Marchesi, Maisa C. Pereira Parra, Marília Moraes, Paula Rahal, Paulo Inacio da Costa, João Pessoa Araújo Jr., Maurício Lacerda Nogueira. Prefeitura de Sao Paulo: Melissa Palmieri. |
| EPI_ISL_1966361, EPI_ISL_1966363 | PRONTO ATENDIMENTO SAO JOSE | Instituto Butantan / Mendelics | Instituto Butantan: Dimas Tadeu Covas, Sandra Coccuzzo Sampaio, Maria Carolina Elias, José Salvatore Leister Patané, Vincent Louis Viala, Antonio Jorge Martins, Ricardo Haddad, Claudia Renata dos Santos Barros, Elaine Cristina Marqueze, Raul Machado Neto, Debora Botequio Moretti, Jardelina de Souza Todao Bernardino, Loyze Paola Oliveira de Lima, Luiz Aurelio de Campos Crispin. Centro de Genômica Funcional da ESALQ: Luiz Lehmann Coutinho, Ricardo Augusto Brassaloti, Raquel de Lello Rocha Campos Cassano. NGS Soluções Genômicas: Pilar Drummond Sampaio Corrêa Mariani. FZEA-USP Pirassununga: Mirele Daiana Poleti, Jessika Cristina Chagas Lesbon, Elisangela Chicaroni Mattos, Heidge Fukumasu. USP-Botucatu: Rejane Maria Tommasini Grotto, Jayme A. Souza-Neto, Guilherme Targino Valente, Patricia Akemi Assato, Felipe Allan da Silva da Costa, Bianca Cechetto Carlos. Mendelics: Bibiana Santos, João Paulo Kitajima, Erika Freitas, David Schlesinger. Hemocentro Ribeirão Preto: Simone Kashima, Evandra Strazza Rodrigues, Svetoslav Nanev Slavov, Elaine Vieira dos Santos, Rafael dos Santos Bezerra, Luiz Carlos Junior de Alcantara, Marta Giovanetti, Vagner Fonseca, Flavia Aburjaile, Rodrigo Tocantins Calado. FAMERP-SJRP: Cecília Artico Banho, Lívia Sacchetto, Fábio Sossai Possebon, Leila Sabrina Ullmann, Cíntia Bittar, Guilherme Campos, Helena Lage Ferreira, Jorge A. Petroli Marchesi, Maisa C. Pereira Parra, Marília Moraes, Paula Rahal, Paulo Inacio da Costa, João Pessoa Araújo Jr., Maurício Lacerda Nogueira. Prefeitura de Sao Paulo: Melissa Palmieri. |
| EPI_ISL_1966364 | HOSPITAL MUNICIPAL DR MARIO GATTI CAMPINAS | Instituto Butantan / Mendelics | Instituto Butantan: Dimas Tadeu Covas, Sandra Coccuzzo Sampaio, Maria Carolina Elias, José Salvatore Leister Patané, Vincent Louis Viala, Antonio Jorge Martins, Ricardo Haddad, Claudia Renata dos Santos Barros, Elaine Cristina Marqueze, Raul Machado Neto, Debora Botequio Moretti, Jardelina de Souza Todao Bernardino, Loyze Paola Oliveira de Lima, Luiz Aurelio de Campos Crispin. Centro de Genômica Funcional da ESALQ: Luiz Lehmann Coutinho, Ricardo Augusto Brassaloti, Raquel de Lello Rocha Campos Cassano. NGS Soluções Genômicas: Pilar Drummond Sampaio Corrêa Mariani. FZEA-USP Pirassununga: Mirele Daiana Poleti, Jessika Cristina Chagas Lesbon, Elisangela Chicaroni Mattos, Heidge Fukumasu. USP-Botucatu: Rejane Maria Tommasini Grotto, Jayme A. Souza-Neto, Guilherme Targino Valente, Patricia Akemi Assato, Felipe Allan da Silva da Costa, Bianca Cechetto Carlos. Mendelics: Bibiana Santos, João Paulo Kitajima, Erika Freitas, David Schlesinger. Hemocentro Ribeirão Preto: Simone Kashima, Evandra Strazza Rodrigues, Svetoslav Nanev Slavov, Elaine Vieira dos Santos, Rafael dos Santos Bezerra, Luiz Carlos Junior de Alcantara, Marta Giovanetti, Vagner Fonseca, Flavia Aburjaile, Rodrigo Tocantins Calado. FAMERP-SJRP: Cecília Artico Banho, Lívia Sacchetto, Fábio Sossai Possebon, Leila Sabrina Ullmann, Cíntia Bittar, Guilherme Campos, Helena Lage Ferreira, Jorge A. Petroli Marchesi, Maisa C. Pereira Parra, Marília Moraes, Paula Rahal, Paulo Inacio da Costa, João Pessoa Araújo Jr., Maurício Lacerda Nogueira. Prefeitura de Sao Paulo: Melissa Palmieri. |
| EPI_ISL_1966365, EPI_ISL_1966366 | CENTRO DE SAUDE DE CANITAR | Instituto Butantan / Mendelics | Instituto Butantan: Dimas Tadeu Covas, Sandra Coccuzzo Sampaio, Maria Carolina Elias, José Salvatore Leister Patané, Vincent Louis Viala, Antonio Jorge Martins, Ricardo Haddad, Claudia Renata dos Santos Barros, Elaine Cristina Marqueze, Raul Machado Neto, Debora Botequio Moretti, Jardelina de Souza Todao Bernardino, Loyze Paola Oliveira de Lima, Luiz Aurelio de Campos Crispin. Centro de Genômica Funcional da ESALQ: Luiz Lehmann Coutinho, Ricardo Augusto Brassaloti, Raquel de Lello Rocha Campos Cassano. NGS Soluções Genômicas: Pilar Drummond Sampaio Corrêa Mariani. FZEA-USP Pirassununga: Mirele Daiana Poleti, Jessika Cristina Chagas Lesbon, Elisangela Chicaroni Mattos, Heidge Fukumasu. USP-Botucatu: Rejane Maria Tommasini Grotto, Jayme A. Souza-Neto, Guilherme Targino Valente, Patricia Akemi Assato, Felipe Allan da Silva da Costa, Bianca Cechetto Carlos. Mendelics: Bibiana Santos, João Paulo Kitajima, Erika Freitas, David Schlesinger. Hemocentro Ribeirão Preto: Simone Kashima, Evandra Strazza Rodrigues, Svetoslav Nanev Slavov, Elaine Vieira dos Santos, Rafael dos Santos Bezerra, Luiz Carlos Junior de Alcantara, Marta Giovanetti, Vagner Fonseca, Flavia Aburjaile, Rodrigo Tocantins Calado. FAMERP-SJRP: Cecília Artico Banho, Lívia Sacchetto, Fábio Sossai Possebon, Leila Sabrina Ullmann, Cíntia Bittar, Guilherme Campos, Helena Lage Ferreira, Jorge A. Petroli Marchesi, Maisa C. Pereira Parra, Marília Moraes, Paula Rahal, Paulo Inacio da Costa, João Pessoa Araújo Jr., Maurício Lacerda Nogueira. Prefeitura de Sao Paulo: Melissa Palmieri. |
| EPI_ISL_1966367, EPI_ISL_1966368 | CENTRO DE SAUDE DE CHAVANTES | Instituto Butantan / Mendelics | Instituto Butantan: Dimas Tadeu Covas, Sandra Coccuzzo Sampaio, Maria Carolina Elias, José Salvatore Leister Patané, Vincent Louis Viala, Antonio Jorge Martins, Ricardo Haddad, Claudia Renata dos Santos Barros, Elaine Cristina Marqueze, Raul Machado Neto, Debora Botequio Moretti, Jardelina de Souza Todao Bernardino, Loyze Paola Oliveira de Lima, Luiz Aurelio de Campos Crispin. Centro de Genômica Funcional da ESALQ: Luiz Lehmann Coutinho, Ricardo Augusto Brassaloti, Raquel de Lello Rocha Campos Cassano. NGS Soluções Genômicas: Pilar Drummond Sampaio Corrêa Mariani. FZEA-USP Pirassununga: Mirele Daiana Poleti, Jessika Cristina Chagas Lesbon, Elisangela Chicaroni Mattos, Heidge Fukumasu. USP-Botucatu: Rejane Maria Tommasini Grotto, Jayme A. Souza-Neto, Guilherme Targino Valente, Patricia Akemi Assato, Felipe Allan da Silva da Costa, Bianca Cechetto Carlos. Mendelics: Bibiana Santos, João Paulo Kitajima, Erika Freitas, David Schlesinger. Hemocentro Ribeirão Preto: Simone Kashima, Evandra Strazza Rodrigues, Svetoslav Nanev Slavov, Elaine Vieira dos Santos, Rafael dos Santos Bezerra, Luiz Carlos Junior de Alcantara, Marta Giovanetti, Vagner Fonseca, Flavia Aburjaile, Rodrigo Tocantins Calado. FAMERP-SJRP: Cecília Artico Banho, Lívia Sacchetto, Fábio Sossai Possebon, Leila Sabrina Ullmann, Cíntia Bittar, Guilherme Campos, Helena Lage Ferreira, Jorge A. Petroli Marchesi, Maisa C. Pereira Parra, Marília Moraes, Paula Rahal, Paulo Inacio da Costa, João Pessoa Araújo Jr., Maurício Lacerda Nogueira. Prefeitura de Sao Paulo: Melissa Palmieri. |
| EPI_ISL_1966369, EPI_ISL_1966370, EPI_ISL_1966371, EPI_ISL_1966372, EPI_ISL_1966373 | UPA DR FRANCO DA ROCHA | Instituto Butantan / Mendelics | Instituto Butantan: Dimas Tadeu Covas, Sandra Coccuzzo Sampaio, Maria Carolina Elias, José Salvatore Leister Patané, Vincent Louis Viala, Antonio Jorge Martins, Ricardo Haddad, Claudia Renata dos Santos Barros, Elaine Cristina Marqueze, Raul Machado Neto, Debora Botequio Moretti, Jardelina de Souza Todao Bernardino, Loyze Paola Oliveira de Lima, Luiz Aurelio de Campos Crispin. Centro de Genômica Funcional da ESALQ: Luiz Lehmann Coutinho, Ricardo Augusto Brassaloti, Raquel de Lello Rocha Campos Cassano. NGS Soluções Genômicas: Pilar Drummond Sampaio Corrêa Mariani. FZEA-USP Pirassununga: Mirele Daiana Poleti, Jessika Cristina Chagas Lesbon, Elisangela Chicaroni Mattos, Heidge Fukumasu. USP-Botucatu: Rejane Maria Tommasini Grotto, Jayme A. Souza-Neto, Guilherme Targino Valente, Patricia Akemi Assato, Felipe Allan da Silva da Costa, Bianca Cechetto Carlos. Mendelics: Bibiana Santos, João Paulo Kitajima, Erika Freitas, David Schlesinger. Hemocentro Ribeirão Preto: Simone Kashima, Evandra Strazza Rodrigues, Svetoslav Nanev Slavov, Elaine Vieira dos Santos, Rafael dos Santos Bezerra, Luiz Carlos Junior de Alcantara, Marta Giovanetti, Vagner Fonseca, Flavia Aburjaile, Rodrigo Tocantins Calado. FAMERP-SJRP: Cecília Artico Banho, Lívia Sacchetto, Fábio Sossai Possebon, Leila Sabrina |

| | | | |
|---|---|---|---|
| | | | Ullmann, Cíntia Bittar, Guilherme Campos, Helena Lage Ferreira, Jorge A. Petroli Marchesi, Maisa C. Pereira Parra, Marília Moraes, Paula Rahal, Paulo Inacio da Costa, João Pessoa Araújo Jr., Maurício Lacerda Nogueira. Prefeitura de Sao Paulo: Melissa Palmieri. |
| EPI_ISL_1966374 | USF II DE HERCULANDIA | Instituto Butantan / Mendelics | Instituto Butantan: Dimas Tadeu Covas, Sandra Coccuzzo Sampaio, Maria Carolina Elias, José Salvatore Leister Patané, Vincent Louis Viala, Antonio Jorge Martins, Ricardo Haddad, Claudia Renata dos Santos Barros, Elaine Cristina Marqueze, Raul Machado Neto, Debora Botequio Moretti, Jardelina de Souza Todao Bernardino, Loyze Paola Oliveira de Lima, Luiz Aurelio de Campos Crispin. Centro de Genômica Funcional da ESALQ: Luiz Lehmann Coutinho, Ricardo Augusto Brassaloti, Raquel de Lello Rocha Campos Cassano. NGS Soluções Genômicas: Pilar Drummond Sampaio Corrêa Mariani. FZEA-USP Pirassununga: Mirele Daiana Poleti, Jessika Cristina Chagas Lesbon, Elisangela Chicaroni Mattos, Heidge Fukumasu. USP-Botucatu: Rejane Maria Tommasini Grotto, Jayme A. Souza-Neto, Guilherme Targino Valente, Patricia Akemi Assato, Felipe Allan da Silva da Costa, Bianca Cechetto Carlos. Mendelics: Bibiana Santos, João Paulo Kitajima, Erika Freitas, David Schlesinger. Hemocentro Ribeirão Preto: Simone Kashima, Evandra Strazza Rodrigues, Svetoslav Nanev Slavov, Elaine Vieira dos Santos, Rafael dos Santos Bezerra, Luiz Carlos Junior de Alcantara, Marta Giovanetti, Vagner Fonseca, Flavia Aburjaile, Rodrigo Tocantins Calado. FAMERP-SJRP: Cecília Artico Banho, Lívia Sacchetto, Fábio Sossai Possebon, Leila Sabrina Ullmann, Cíntia Bittar, Guilherme Campos, Helena Lage Ferreira, Jorge A. Petroli Marchesi, Maisa C. Pereira Parra, Marília Moraes, Paula Rahal, Paulo Inacio da Costa, João Pessoa Araújo Jr., Maurício Lacerda Nogueira. Prefeitura de Sao Paulo: Melissa Palmieri. |
| EPI_ISL_1966375, EPI_ISL_1966376 | UNIDADE RESPIRATORIA NOVA HORTOLANDIA | Instituto Butantan / Mendelics | Instituto Butantan: Dimas Tadeu Covas, Sandra Coccuzzo Sampaio, Maria Carolina Elias, José Salvatore Leister Patané, Vincent Louis Viala, Antonio Jorge Martins, Ricardo Haddad, Claudia Renata dos Santos Barros, Elaine Cristina Marqueze, Raul Machado Neto, Debora Botequio Moretti, Jardelina de Souza Todao Bernardino, Loyze Paola Oliveira de Lima, Luiz Aurelio de Campos Crispin. Centro de Genômica Funcional da ESALQ: Luiz Lehmann Coutinho, Ricardo Augusto Brassaloti, Raquel de Lello Rocha Campos Cassano. NGS Soluções Genômicas: Pilar Drummond Sampaio Corrêa Mariani. FZEA-USP Pirassununga: Mirele Daiana Poleti, Jessika Cristina Chagas Lesbon, Elisangela Chicaroni Mattos, Heidge Fukumasu. USP-Botucatu: Rejane Maria Tommasini Grotto, Jayme A. Souza-Neto, Guilherme Targino Valente, Patricia Akemi Assato, Felipe Allan da Silva da Costa, Bianca Cechetto Carlos. Mendelics: Bibiana Santos, João Paulo Kitajima, Erika Freitas, David Schlesinger. Hemocentro Ribeirão Preto: Simone Kashima, Evandra Strazza Rodrigues, Svetoslav Nanev Slavov, Elaine Vieira dos Santos, Rafael dos Santos Bezerra, Luiz Carlos Junior de Alcantara, Marta Giovanetti, Vagner Fonseca, Flavia Aburjaile, Rodrigo Tocantins Calado. FAMERP-SJRP: Cecília Artico Banho, Lívia Sacchetto, Fábio Sossai Possebon, Leila Sabrina Ullmann, Cíntia Bittar, Guilherme Campos, Helena Lage Ferreira, Jorge A. Petroli Marchesi, Maisa C. Pereira Parra, Marília Moraes, Paula Rahal, Paulo Inacio da Costa, João Pessoa Araújo Jr., Maurício Lacerda Nogueira. Prefeitura de Sao Paulo: Melissa Palmieri. |
| EPI_ISL_1966377, EPI_ISL_1966378 | SMS IPERO | Instituto Butantan / Mendelics | Instituto Butantan: Dimas Tadeu Covas, Sandra Coccuzzo Sampaio, Maria Carolina Elias, José Salvatore Leister Patané, Vincent Louis Viala, Antonio Jorge Martins, Ricardo Haddad, Claudia Renata dos Santos Barros, Elaine Cristina Marqueze, Raul Machado Neto, Debora Botequio Moretti, Jardelina de Souza Todao Bernardino, Loyze Paola Oliveira de Lima, Luiz Aurelio de Campos Crispin. Centro de Genômica Funcional da ESALQ: Luiz Lehmann Coutinho, Ricardo Augusto Brassaloti, Raquel de Lello Rocha Campos Cassano. NGS Soluções Genômicas: Pilar Drummond Sampaio Corrêa Mariani. FZEA-USP Pirassununga: Mirele Daiana Poleti, Jessika Cristina Chagas Lesbon, Elisangela Chicaroni Mattos, Heidge Fukumasu. USP-Botucatu: Rejane Maria Tommasini Grotto, Jayme A. Souza-Neto, Guilherme Targino Valente, Patricia Akemi Assato, Felipe Allan da Silva da Costa, Bianca Cechetto Carlos. Mendelics: Bibiana Santos, João Paulo Kitajima, Erika Freitas, David Schlesinger. Hemocentro Ribeirão Preto: Simone Kashima, Evandra Strazza Rodrigues, Svetoslav Nanev Slavov, Elaine Vieira dos Santos, Rafael dos Santos Bezerra, Luiz Carlos Junior de Alcantara, Marta Giovanetti, Vagner Fonseca, Flavia Aburjaile, Rodrigo Tocantins Calado. FAMERP-SJRP: Cecília Artico Banho, Lívia Sacchetto, Fábio Sossai Possebon, Leila Sabrina Ullmann, Cíntia Bittar, Guilherme Campos, Helena Lage Ferreira, Jorge A. Petroli Marchesi, Maisa C. Pereira Parra, Marília Moraes, Paula Rahal, Paulo Inacio da Costa, João Pessoa Araújo Jr., Maurício Lacerda Nogueira. Prefeitura de Sao Paulo: Melissa Palmieri. |
| EPI_ISL_1966380 | HOSPITAL REGIONAL DE ITAPETININGA | Instituto Butantan / Mendelics | Instituto Butantan: Dimas Tadeu Covas, Sandra Coccuzzo Sampaio, Maria Carolina Elias, José Salvatore Leister Patané, Vincent Louis Viala, Antonio Jorge Martins, Ricardo Haddad, Claudia Renata dos Santos Barros, Elaine Cristina Marqueze, Raul Machado Neto, Debora Botequio Moretti, Jardelina de Souza Todao Bernardino, Loyze Paola Oliveira de Lima, Luiz Aurelio de Campos Crispin. Centro de Genômica Funcional da ESALQ: Luiz Lehmann Coutinho, Ricardo Augusto Brassaloti, Raquel de Lello Rocha Campos Cassano. NGS Soluções Genômicas: Pilar Drummond Sampaio Corrêa Mariani. FZEA-USP Pirassununga: Mirele Daiana Poleti, Jessika Cristina Chagas Lesbon, Elisangela Chicaroni Mattos, Heidge Fukumasu. USP-Botucatu: Rejane Maria Tommasini Grotto, Jayme A. Souza-Neto, Guilherme Targino Valente, Patricia Akemi Assato, Felipe Allan da Silva da Costa, Bianca Cechetto Carlos. Mendelics: Bibiana Santos, João Paulo Kitajima, Erika Freitas, David Schlesinger. Hemocentro Ribeirão Preto: Simone Kashima, Evandra Strazza Rodrigues, Svetoslav Nanev Slavov, Elaine Vieira dos Santos, Rafael dos Santos Bezerra, Luiz Carlos Junior de Alcantara, Marta Giovanetti, Vagner Fonseca, Flavia Aburjaile, Rodrigo Tocantins Calado. FAMERP-SJRP: Cecília Artico Banho, Lívia Sacchetto, Fábio Sossai Possebon, Leila Sabrina Ullmann, Cíntia Bittar, Guilherme Campos, Helena Lage Ferreira, Jorge A. Petroli Marchesi, Maisa C. Pereira Parra, Marília Moraes, Paula Rahal, Paulo Inacio da Costa, João Pessoa Araújo Jr., Maurício Lacerda Nogueira. Prefeitura de Sao Paulo: Melissa Palmieri. |
| EPI_ISL_1966381, EPI_ISL_1966382, EPI_ISL_1966383, EPI_ISL_1966384 | POLICLINICA COVID 19 ITAPETININGA | Instituto Butantan / Mendelics | Instituto Butantan: Dimas Tadeu Covas, Sandra Coccuzzo Sampaio, Maria Carolina Elias, José Salvatore Leister Patané, Vincent Louis Viala, Antonio Jorge Martins, Ricardo Haddad, Claudia Renata dos Santos Barros, Elaine Cristina Marqueze, Raul Machado Neto, Debora Botequio Moretti, Jardelina de Souza Todao Bernardino, Loyze Paola Oliveira de Lima, Luiz Aurelio de Campos Crispin. Centro de Genômica Funcional da ESALQ: Luiz Lehmann Coutinho, Ricardo Augusto Brassaloti, Raquel de Lello Rocha Campos Cassano. NGS Soluções Genômicas: Pilar Drummond Sampaio Corrêa Mariani. FZEA-USP Pirassununga: Mirele Daiana Poleti, Jessika Cristina Chagas Lesbon, Elisangela Chicaroni Mattos, Heidge Fukumasu. USP-Botucatu: Rejane Maria Tommasini Grotto, Jayme A. Souza-Neto, Guilherme Targino Valente, Patricia Akemi Assato, Felipe Allan da Silva da Costa, Bianca Cechetto Carlos. Mendelics: Bibiana Santos, João Paulo Kitajima, Erika Freitas, David Schlesinger. Hemocentro Ribeirão Preto: Simone Kashima, Evandra Strazza Rodrigues, Svetoslav Nanev Slavov, Elaine Vieira dos Santos, Rafael dos Santos Bezerra, Luiz Carlos Junior de Alcantara, Marta Giovanetti, Vagner Fonseca, Flavia Aburjaile, Rodrigo Tocantins Calado. FAMERP-SJRP: Cecília Artico Banho, Lívia Sacchetto, Fábio Sossai Possebon, Leila Sabrina Ullmann, Cíntia Bittar, Guilherme Campos, Helena Lage Ferreira, Jorge A. Petroli Marchesi, Maisa C. Pereira Parra, Marília Moraes, Paula Rahal, Paulo Inacio da Costa, João Pessoa Araújo Jr., Maurício Lacerda Nogueira. Prefeitura de Sao Paulo: Melissa Palmieri. |
| EPI_ISL_1966385, EPI_ISL_1966386, EPI_ISL_1966387, EPI_ISL_1966388 | UNIDADE DE PRONTO ATENDIMENTO UPA DRA ANA OLIVIA BENTIVOGLIO | Instituto Butantan / Mendelics | Instituto Butantan: Dimas Tadeu Covas, Sandra Coccuzzo Sampaio, Maria Carolina Elias, José Salvatore Leister Patané, Vincent Louis Viala, Antonio Jorge Martins, Ricardo Haddad, Claudia Renata dos Santos Barros, Elaine Cristina Marqueze, Raul Machado Neto, Debora Botequio Moretti, Jardelina de Souza Todao Bernardino, Loyze Paola Oliveira de Lima, Luiz Aurelio de Campos Crispin. Centro de Genômica Funcional da ESALQ: Luiz Lehmann Coutinho, Ricardo Augusto Brassaloti, Raquel de Lello Rocha Campos Cassano. NGS Soluções Genômicas: Pilar Drummond Sampaio Corrêa Mariani. FZEA-USP Pirassununga: Mirele Daiana Poleti, Jessika Cristina Chagas Lesbon, Elisangela Chicaroni Mattos, Heidge Fukumasu. USP-Botucatu: Rejane Maria Tommasini Grotto, Jayme A. Souza-Neto, Guilherme Targino Valente, Patricia Akemi Assato, Felipe Allan da Silva da Costa, Bianca Cechetto Carlos. Mendelics: Bibiana Santos, João Paulo Kitajima, Erika Freitas, David Schlesinger. Hemocentro Ribeirão Preto: Simone Kashima, Evandra Strazza Rodrigues, Svetoslav Nanev Slavov, Elaine Vieira dos Santos, Rafael dos Santos Bezerra, Luiz Carlos Junior de Alcantara, Marta Giovanetti, Vagner Fonseca, Flavia Aburjaile, Rodrigo Tocantins Calado. FAMERP-SJRP: Cecília Artico Banho, Lívia Sacchetto, Fábio Sossai Possebon, Leila Sabrina Ullmann, Cíntia Bittar, Guilherme Campos, Helena Lage Ferreira, Jorge A. Petroli Marchesi, Maisa C. Pereira Parra, Marília Moraes, Paula Rahal, Paulo Inacio da Costa, João Pessoa Araújo Jr., Maurício Lacerda Nogueira. Prefeitura de Sao Paulo: Melissa Palmieri. |
| EPI_ISL_1966389, EPI_ISL_1966390, EPI_ISL_1966391, EPI_ISL_1966392 | SECRETARIA MUNICIPAL DA SAUDE DE JOANOPOLIS | Instituto Butantan / Mendelics | Instituto Butantan: Dimas Tadeu Covas, Sandra Coccuzzo Sampaio, Maria Carolina Elias, José Salvatore Leister Patané, Vincent Louis Viala, Antonio Jorge Martins, Ricardo Haddad, Claudia Renata dos Santos Barros, Elaine Cristina Marqueze, Raul Machado Neto, Debora Botequio Moretti, Jardelina de Souza Todao Bernardino, Loyze Paola Oliveira de Lima, Luiz Aurelio de Campos Crispin. Centro de Genômica Funcional da ESALQ: Luiz Lehmann Coutinho, Ricardo Augusto Brassaloti, Raquel de Lello Rocha Campos Cassano. NGS Soluções Genômicas: Pilar Drummond Sampaio Corrêa Mariani. FZEA-USP Pirassununga: Mirele Daiana Poleti, Jessika Cristina Chagas Lesbon, Elisangela Chicaroni Mattos, Heidge Fukumasu. USP-Botucatu: Rejane Maria Tommasini Grotto, Jayme A. Souza-Neto, Guilherme Targino Valente, Patricia Akemi Assato, Felipe Allan da Silva da Costa, Bianca Cechetto Carlos. Mendelics: Bibiana Santos, João Paulo Kitajima, Erika Freitas, David Schlesinger. Hemocentro Ribeirão Preto: Simone Kashima, Evandra Strazza Rodrigues, Svetoslav Nanev Slavov, Elaine Vieira dos Santos, Rafael dos Santos Bezerra, Luiz Carlos Junior de Alcantara, Marta Giovanetti, Vagner Fonseca, Flavia Aburjaile, Rodrigo Tocantins Calado. FAMERP-SJRP: Cecília Artico Banho, Lívia Sacchetto, Fábio Sossai Possebon, Leila Sabrina Ullmann, Cíntia Bittar, Guilherme Campos, Helena Lage Ferreira, Jorge A. Petroli Marchesi, Maisa C. Pereira Parra, Marília Moraes, Paula Rahal, Paulo Inacio da Costa, João Pessoa Araújo Jr., Maurício Lacerda Nogueira. Prefeitura de Sao Paulo: Melissa Palmieri. |
| EPI_ISL_1966393, EPI_ISL_1966394 | ADECIO POMPEU CARVALHO | Instituto Butantan / Mendelics | Instituto Butantan: Dimas Tadeu Covas, Sandra Coccuzzo Sampaio, Maria Carolina Elias, José Salvatore Leister Patané, Vincent Louis Viala, Antonio Jorge Martins, Ricardo Haddad, Claudia Renata dos Santos Barros, Elaine Cristina Marqueze, Raul Machado Neto, Debora Botequio Moretti, Jardelina de Souza Todao Bernardino, Loyze Paola Oliveira de Lima, Luiz Aurelio de Campos Crispin. Centro de Genômica Funcional da ESALQ: Luiz Lehmann Coutinho, Ricardo Augusto Brassaloti, Raquel de Lello Rocha Campos Cassano. NGS Soluções Genômicas: Pilar Drummond Sampaio Corrêa Mariani. |

| | | | |
|---|---|---|---|
| | | | FZEA-USP Pirassununga: Mirele Daiana Poleti, Jessika Cristina Chagas Lesbon, Elisangela Chicaroni Mattos, Heidge Fukumasu. USP-Botucatu: Rejane Maria Tommasini Grotto, Jayme A. Souza-Neto, Guilherme Targino Valente, Patricia Akemi Assato, Felipe Allan da Silva da Costa, Bianca Cechetto Carlos. Mendelics: Bibiana Santos, João Paulo Kitajima, Erika Freitas, David Schlesinger. Hemocentro Ribeirão Preto: Simone Kashima, Evandra Strazza Rodrigues, Svetoslav Nanev Slavov, Elaine Vieira dos Santos, Rafael dos Santos Bezerra, Luiz Carlos Junior de Alcantara, Marta Giovanetti, Vagner Fonseca, Flavia Aburjaile, Rodrigo Tocantins Calado. FAMERP-SJRP: Cecília Artico Banho, Lívia Sacchetto, Fábio Sossai Possebon, Leila Sabrina Ullmann, Cíntia Bittar, Guilherme Campos, Helena Lage Ferreira, Jorge A. Petroli Marchesi, Maisa C. Pereira Parra, Marília Moraes, Paula Rahal, Paulo Inacio da Costa, João Pessoa Araújo Jr., Maurício Lacerda Nogueira. Prefeitura de Sao Paulo: Melissa Palmieri. |
| EPI_ISL_1966395 | SAE CTA SERV ATEND ESPEC TESTAGEM E ACONSELHAMENTO | Instituto Butantan / Mendelics | Instituto Butantan: Dimas Tadeu Covas, Sandra Coccuzzo Sampaio, Maria Carolina Elias, José Salvatore Leister Patané, Vincent Louis Viala, Antonio Jorge Martins, Ricardo Haddad, Claudia Renata dos Santos Barros, Elaine Cristina Marqueze, Raul Machado Neto, Debora Botequio Moretti, Jardelina de Souza Todao Bernardino, Loyze Paola Oliveira de Lima, Luiz Aurelio de Campos Crispin. Centro de Genômica Funcional da ESALQ: Luiz Lehmann Coutinho, Ricardo Augusto Brassaloti, Raquel de Lello Rocha Campos Cassano. NGS Soluções Genômicas: Pilar Drummond Sampaio Corrêa Mariani. FZEA-USP Pirassununga: Mirele Daiana Poleti, Jessika Cristina Chagas Lesbon, Elisangela Chicaroni Mattos, Heidge Fukumasu. USP-Botucatu: Rejane Maria Tommasini Grotto, Jayme A. Souza-Neto, Guilherme Targino Valente, Patricia Akemi Assato, Felipe Allan da Silva da Costa, Bianca Cechetto Carlos. Mendelics: Bibiana Santos, João Paulo Kitajima, Erika Freitas, David Schlesinger. Hemocentro Ribeirão Preto: Simone Kashima, Evandra Strazza Rodrigues, Svetoslav Nanev Slavov, Elaine Vieira dos Santos, Rafael dos Santos Bezerra, Luiz Carlos Junior de Alcantara, Marta Giovanetti, Vagner Fonseca, Flavia Aburjaile, Rodrigo Tocantins Calado. FAMERP-SJRP: Cecília Artico Banho, Lívia Sacchetto, Fábio Sossai Possebon, Leila Sabrina Ullmann, Cíntia Bittar, Guilherme Campos, Helena Lage Ferreira, Jorge A. Petroli Marchesi, Maisa C. Pereira Parra, Marília Moraes, Paula Rahal, Paulo Inacio da Costa, João Pessoa Araújo Jr., Maurício Lacerda Nogueira. Prefeitura de Sao Paulo: Melissa Palmieri. |
| EPI_ISL_1966396, EPI_ISL_1966397 | PSF DR ANTONIO PIRES DE ALMEIDA PORTO FELIZ | Instituto Butantan / Mendelics | Instituto Butantan: Dimas Tadeu Covas, Sandra Coccuzzo Sampaio, Maria Carolina Elias, José Salvatore Leister Patané, Vincent Louis Viala, Antonio Jorge Martins, Ricardo Haddad, Claudia Renata dos Santos Barros, Elaine Cristina Marqueze, Raul Machado Neto, Debora Botequio Moretti, Jardelina de Souza Todao Bernardino, Loyze Paola Oliveira de Lima, Luiz Aurelio de Campos Crispin. Centro de Genômica Funcional da ESALQ: Luiz Lehmann Coutinho, Ricardo Augusto Brassaloti, Raquel de Lello Rocha Campos Cassano. NGS Soluções Genômicas: Pilar Drummond Sampaio Corrêa Mariani. FZEA-USP Pirassununga: Mirele Daiana Poleti, Jessika Cristina Chagas Lesbon, Elisangela Chicaroni Mattos, Heidge Fukumasu. USP-Botucatu: Rejane Maria Tommasini Grotto, Jayme A. Souza-Neto, Guilherme Targino Valente, Patricia Akemi Assato, Felipe Allan da Silva da Costa, Bianca Cechetto Carlos. Mendelics: Bibiana Santos, João Paulo Kitajima, Erika Freitas, David Schlesinger. Hemocentro Ribeirão Preto: Simone Kashima, Evandra Strazza Rodrigues, Svetoslav Nanev Slavov, Elaine Vieira dos Santos, Rafael dos Santos Bezerra, Luiz Carlos Junior de Alcantara, Marta Giovanetti, Vagner Fonseca, Flavia Aburjaile, Rodrigo Tocantins Calado. FAMERP-SJRP: Cecília Artico Banho, Lívia Sacchetto, Fábio Sossai Possebon, Leila Sabrina Ullmann, Cíntia Bittar, Guilherme Campos, Helena Lage Ferreira, Jorge A. Petroli Marchesi, Maisa C. Pereira Parra, Marília Moraes, Paula Rahal, Paulo Inacio da Costa, João Pessoa Araújo Jr., Maurício Lacerda Nogueira. Prefeitura de Sao Paulo: Melissa Palmieri. |
| EPI_ISL_1966398 | CENTRO DE SAUDE DE SALMOURAO | Instituto Butantan / Mendelics | Instituto Butantan: Dimas Tadeu Covas, Sandra Coccuzzo Sampaio, Maria Carolina Elias, José Salvatore Leister Patané, Vincent Louis Viala, Antonio Jorge Martins, Ricardo Haddad, Claudia Renata dos Santos Barros, Elaine Cristina Marqueze, Raul Machado Neto, Debora Botequio Moretti, Jardelina de Souza Todao Bernardino, Loyze Paola Oliveira de Lima, Luiz Aurelio de Campos Crispin. Centro de Genômica Funcional da ESALQ: Luiz Lehmann Coutinho, Ricardo Augusto Brassaloti, Raquel de Lello Rocha Campos Cassano. NGS Soluções Genômicas: Pilar Drummond Sampaio Corrêa Mariani. FZEA-USP Pirassununga: Mirele Daiana Poleti, Jessika Cristina Chagas Lesbon, Elisangela Chicaroni Mattos, Heidge Fukumasu. USP-Botucatu: Rejane Maria Tommasini Grotto, Jayme A. Souza-Neto, Guilherme Targino Valente, Patricia Akemi Assato, Felipe Allan da Silva da Costa, Bianca Cechetto Carlos. Mendelics: Bibiana Santos, João Paulo Kitajima, Erika Freitas, David Schlesinger. Hemocentro Ribeirão Preto: Simone Kashima, Evandra Strazza Rodrigues, Svetoslav Nanev Slavov, Elaine Vieira dos Santos, Rafael dos Santos Bezerra, Luiz Carlos Junior de Alcantara, Marta Giovanetti, Vagner Fonseca, Flavia Aburjaile, Rodrigo Tocantins Calado. FAMERP-SJRP: Cecília Artico Banho, Lívia Sacchetto, Fábio Sossai Possebon, Leila Sabrina Ullmann, Cíntia Bittar, Guilherme Campos, Helena Lage Ferreira, Jorge A. Petroli Marchesi, Maisa C. Pereira Parra, Marília Moraes, Paula Rahal, Paulo Inacio da Costa, João Pessoa Araújo Jr., Maurício Lacerda Nogueira. Prefeitura de Sao Paulo: Melissa Palmieri. |
| EPI_ISL_1966399 | CENTRO DE SAUDE II | Instituto Butantan / Mendelics | Instituto Butantan: Dimas Tadeu Covas, Sandra Coccuzzo Sampaio, Maria Carolina Elias, José Salvatore Leister Patané, Vincent Louis Viala, Antonio Jorge Martins, Ricardo Haddad, Claudia Renata dos Santos Barros, Elaine Cristina Marqueze, Raul Machado Neto, Debora Botequio Moretti, Jardelina de Souza Todao Bernardino, Loyze Paola Oliveira de Lima, Luiz Aurelio de Campos Crispin. Centro de Genômica Funcional da ESALQ: Luiz Lehmann Coutinho, Ricardo Augusto Brassaloti, Raquel de Lello Rocha Campos Cassano. NGS Soluções Genômicas: Pilar Drummond Sampaio Corrêa Mariani. FZEA-USP Pirassununga: Mirele Daiana Poleti, Jessika Cristina Chagas Lesbon, Elisangela Chicaroni Mattos, Heidge Fukumasu. USP-Botucatu: Rejane Maria Tommasini Grotto, Jayme A. Souza-Neto, Guilherme Targino Valente, Patricia Akemi Assato, Felipe Allan da Silva da Costa, Bianca Cechetto Carlos. Mendelics: Bibiana Santos, João Paulo Kitajima, Erika Freitas, David Schlesinger. Hemocentro Ribeirão Preto: Simone Kashima, Evandra Strazza Rodrigues, Svetoslav Nanev Slavov, Elaine Vieira dos Santos, Rafael dos Santos Bezerra, Luiz Carlos Junior de Alcantara, Marta Giovanetti, Vagner Fonseca, Flavia Aburjaile, Rodrigo Tocantins Calado. FAMERP-SJRP: Cecília Artico Banho, Lívia Sacchetto, Fábio Sossai Possebon, Leila Sabrina Ullmann, Cíntia Bittar, Guilherme Campos, Helena Lage Ferreira, Jorge A. Petroli Marchesi, Maisa C. Pereira Parra, Marília Moraes, Paula Rahal, Paulo Inacio da Costa, João Pessoa Araújo Jr., Maurício Lacerda Nogueira. Prefeitura de Sao Paulo: Melissa Palmieri. |
| EPI_ISL_1966400 | CLINICA SALTO SAUDE SANTA CRUZ | Instituto Butantan / Mendelics | Instituto Butantan: Dimas Tadeu Covas, Sandra Coccuzzo Sampaio, Maria Carolina Elias, José Salvatore Leister Patané, Vincent Louis Viala, Antonio Jorge Martins, Ricardo Haddad, Claudia Renata dos Santos Barros, Elaine Cristina Marqueze, Raul Machado Neto, Debora Botequio Moretti, Jardelina de Souza Todao Bernardino, Loyze Paola Oliveira de Lima, Luiz Aurelio de Campos Crispin. Centro de Genômica Funcional da ESALQ: Luiz Lehmann Coutinho, Ricardo Augusto Brassaloti, Raquel de Lello Rocha Campos Cassano. NGS Soluções Genômicas: Pilar Drummond Sampaio Corrêa Mariani. FZEA-USP Pirassununga: Mirele Daiana Poleti, Jessika Cristina Chagas Lesbon, Elisangela Chicaroni Mattos, Heidge Fukumasu. USP-Botucatu: Rejane Maria Tommasini Grotto, Jayme A. Souza-Neto, Guilherme Targino Valente, Patricia Akemi Assato, Felipe Allan da Silva da Costa, Bianca Cechetto Carlos. Mendelics: Bibiana Santos, João Paulo Kitajima, Erika Freitas, David Schlesinger. Hemocentro Ribeirão Preto: Simone Kashima, Evandra Strazza Rodrigues, Svetoslav Nanev Slavov, Elaine Vieira dos Santos, Rafael dos Santos Bezerra, Luiz Carlos Junior de Alcantara, Marta Giovanetti, Vagner Fonseca, Flavia Aburjaile, Rodrigo Tocantins Calado. FAMERP-SJRP: Cecília Artico Banho, Lívia Sacchetto, Fábio Sossai Possebon, Leila Sabrina Ullmann, Cíntia Bittar, Guilherme Campos, Helena Lage Ferreira, Jorge A. Petroli Marchesi, Maisa C. Pereira Parra, Marília Moraes, Paula Rahal, Paulo Inacio da Costa, João Pessoa Araújo Jr., Maurício Lacerda Nogueira. Prefeitura de Sao Paulo: Melissa Palmieri. |
| EPI_ISL_1966401 | HOSP E MATERNIDADE MUNICIPAL N SRA MONTE SERRAT | Instituto Butantan / Mendelics | Instituto Butantan: Dimas Tadeu Covas, Sandra Coccuzzo Sampaio, Maria Carolina Elias, José Salvatore Leister Patané, Vincent Louis Viala, Antonio Jorge Martins, Ricardo Haddad, Claudia Renata dos Santos Barros, Elaine Cristina Marqueze, Raul Machado Neto, Debora Botequio Moretti, Jardelina de Souza Todao Bernardino, Loyze Paola Oliveira de Lima, Luiz Aurelio de Campos Crispin. Centro de Genômica Funcional da ESALQ: Luiz Lehmann Coutinho, Ricardo Augusto Brassaloti, Raquel de Lello Rocha Campos Cassano. NGS Soluções Genômicas: Pilar Drummond Sampaio Corrêa Mariani. FZEA-USP Pirassununga: Mirele Daiana Poleti, Jessika Cristina Chagas Lesbon, Elisangela Chicaroni Mattos, Heidge Fukumasu. USP-Botucatu: Rejane Maria Tommasini Grotto, Jayme A. Souza-Neto, Guilherme Targino Valente, Patricia Akemi Assato, Felipe Allan da Silva da Costa, Bianca Cechetto Carlos. Mendelics: Bibiana Santos, João Paulo Kitajima, Erika Freitas, David Schlesinger. Hemocentro Ribeirão Preto: Simone Kashima, Evandra Strazza Rodrigues, Svetoslav Nanev Slavov, Elaine Vieira dos Santos, Rafael dos Santos Bezerra, Luiz Carlos Junior de Alcantara, Marta Giovanetti, Vagner Fonseca, Flavia Aburjaile, Rodrigo Tocantins Calado. FAMERP-SJRP: Cecília Artico Banho, Lívia Sacchetto, Fábio Sossai Possebon, Leila Sabrina Ullmann, Cíntia Bittar, Guilherme Campos, Helena Lage Ferreira, Jorge A. Petroli Marchesi, Maisa C. Pereira Parra, Marília Moraes, Paula Rahal, Paulo Inacio da Costa, João Pessoa Araújo Jr., Maurício Lacerda Nogueira. Prefeitura de Sao Paulo: Melissa Palmieri. |
| EPI_ISL_1966402, EPI_ISL_1966403, EPI_ISL_1966404, EPI_ISL_1966405 | UNIDADE DE PRONTO ATENDIMENTO UPA I SANTA ISABEL | Instituto Butantan / Mendelics | Instituto Butantan: Dimas Tadeu Covas, Sandra Coccuzzo Sampaio, Maria Carolina Elias, José Salvatore Leister Patané, Vincent Louis Viala, Antonio Jorge Martins, Ricardo Haddad, Claudia Renata dos Santos Barros, Elaine Cristina Marqueze, Raul Machado Neto, Debora Botequio Moretti, Jardelina de Souza Todao Bernardino, Loyze Paola Oliveira de Lima, Luiz Aurelio de Campos Crispin. Centro de Genômica Funcional da ESALQ: Luiz Lehmann Coutinho, Ricardo Augusto Brassaloti, Raquel de Lello Rocha Campos Cassano. NGS Soluções Genômicas: Pilar Drummond Sampaio Corrêa Mariani. FZEA-USP Pirassununga: Mirele Daiana Poleti, Jessika Cristina Chagas Lesbon, Elisangela Chicaroni Mattos, Heidge Fukumasu. USP-Botucatu: Rejane Maria Tommasini Grotto, Jayme A. Souza-Neto, Guilherme Targino Valente, Patricia Akemi Assato, Felipe Allan da Silva da Costa, Bianca Cechetto Carlos. Mendelics: Bibiana Santos, João Paulo Kitajima, Erika Freitas, David Schlesinger. Hemocentro Ribeirão Preto: Simone Kashima, Evandra Strazza Rodrigues, Svetoslav Nanev Slavov, Elaine Vieira dos Santos, Rafael dos Santos Bezerra, Luiz Carlos Junior de Alcantara, Marta Giovanetti, Vagner Fonseca, Flavia Aburjaile, Rodrigo Tocantins Calado. FAMERP-SJRP: Cecília Artico Banho, Lívia Sacchetto, Fábio Sossai Possebon, Leila Sabrina Ullmann, Cíntia Bittar, Guilherme Campos, Helena Lage Ferreira, Jorge A. Petroli Marchesi, Maisa C. Pereira Parra, Marília Moraes, Paula Rahal, Paulo Inacio da Costa, João Pessoa Araújo Jr., Maurício Lacerda Nogueira. Prefeitura de Sao Paulo: Melissa Palmieri. |

| | | | |
|---|---|---|---|
| EPI_ISL_1966406 | HOSPITAL GERAL DE VILA NOVA CACHOEIRINHA SAO PAULO | Instituto Butantan / Mendelics | Instituto Butantan: Dimas Tadeu Covas, Sandra Coccuzzo Sampaio, Maria Carolina Elias, José Salvatore Leister Patané, Vincent Louis Viala, Antonio Jorge Martins, Ricardo Haddad, Claudia Renata dos Santos Barros, Elaine Cristina Marqueze, Raul Machado Neto, Debora Botequio Moretti, Jardelina de Souza Todao Bernardino, Loyze Paola Oliveira de Lima, Luiz Aurelio de Campos Crispin. Centro de Genômica Funcional da ESALQ: Luiz Lehmann Coutinho, Ricardo Augusto Brassaloti, Raquel de Lello Rocha Campos Cassano. NGS Soluções Genômicas: Pilar Drummond Sampaio Corrêa Mariani. FZEA-USP Pirassununga: Mirele Daiana Poleti, Jessika Cristina Chagas Lesbon, Elisangela Chicaroni Mattos, Heidge Fukumasu. USP-Botucatu: Rejane Maria Tommasini Grotto, Jayme A. Souza-Neto, Guilherme Targino Valente, Patricia Akemi Assato, Felipe Allan da Silva da Costa, Bianca Cechetto Carlos. Mendelics: Bibiana Santos, João Paulo Kitajima, Erika Freitas, David Schlesinger. Hemocentro Ribeirão Preto: Simone Kashima, Evandra Strazza Rodrigues, Svetoslav Nanev Slavov, Elaine Vieira dos Santos, Rafael dos Santos Bezerra, Luiz Carlos Junior de Alcantara, Marta Giovanetti, Vagner Fonseca, Flavia Aburjaile, Rodrigo Tocantins Calado. FAMERP-SJRP: Cecília Artico Banho, Lívia Sacchetto, Fábio Sossai Possebon, Leila Sabrina Ullmann, Cíntia Bittar, Guilherme Campos, Helena Lage Ferreira, Jorge A. Petroli Marchesi, Maisa C. Pereira Parra, Marília Moraes, Paula Rahal, Paulo Inacio da Costa, João Pessoa Araújo Jr., Maurício Lacerda Nogueira. Prefeitura de Sao Paulo: Melissa Palmieri. |
| EPI_ISL_1966407 | CENTRO MEDICO PMESP | Instituto Butantan / Mendelics | Instituto Butantan: Dimas Tadeu Covas, Sandra Coccuzzo Sampaio, Maria Carolina Elias, José Salvatore Leister Patané, Vincent Louis Viala, Antonio Jorge Martins, Ricardo Haddad, Claudia Renata dos Santos Barros, Elaine Cristina Marqueze, Raul Machado Neto, Debora Botequio Moretti, Jardelina de Souza Todao Bernardino, Loyze Paola Oliveira de Lima, Luiz Aurelio de Campos Crispin. Centro de Genômica Funcional da ESALQ: Luiz Lehmann Coutinho, Ricardo Augusto Brassaloti, Raquel de Lello Rocha Campos Cassano. NGS Soluções Genômicas: Pilar Drummond Sampaio Corrêa Mariani. FZEA-USP Pirassununga: Mirele Daiana Poleti, Jessika Cristina Chagas Lesbon, Elisangela Chicaroni Mattos, Heidge Fukumasu. USP-Botucatu: Rejane Maria Tommasini Grotto, Jayme A. Souza-Neto, Guilherme Targino Valente, Patricia Akemi Assato, Felipe Allan da Silva da Costa, Bianca Cechetto Carlos. Mendelics: Bibiana Santos, João Paulo Kitajima, Erika Freitas, David Schlesinger. Hemocentro Ribeirão Preto: Simone Kashima, Evandra Strazza Rodrigues, Svetoslav Nanev Slavov, Elaine Vieira dos Santos, Rafael dos Santos Bezerra, Luiz Carlos Junior de Alcantara, Marta Giovanetti, Vagner Fonseca, Flavia Aburjaile, Rodrigo Tocantins Calado. FAMERP-SJRP: Cecília Artico Banho, Lívia Sacchetto, Fábio Sossai Possebon, Leila Sabrina Ullmann, Cíntia Bittar, Guilherme Campos, Helena Lage Ferreira, Jorge A. Petroli Marchesi, Maisa C. Pereira Parra, Marília Moraes, Paula Rahal, Paulo Inacio da Costa, João Pessoa Araújo Jr., Maurício Lacerda Nogueira. Prefeitura de Sao Paulo: Melissa Palmieri. |
| EPI_ISL_1966408 | HOSP STA VIRGINIA | Instituto Butantan / Mendelics | Instituto Butantan: Dimas Tadeu Covas, Sandra Coccuzzo Sampaio, Maria Carolina Elias, José Salvatore Leister Patané, Vincent Louis Viala, Antonio Jorge Martins, Ricardo Haddad, Claudia Renata dos Santos Barros, Elaine Cristina Marqueze, Raul Machado Neto, Debora Botequio Moretti, Jardelina de Souza Todao Bernardino, Loyze Paola Oliveira de Lima, Luiz Aurelio de Campos Crispin. Centro de Genômica Funcional da ESALQ: Luiz Lehmann Coutinho, Ricardo Augusto Brassaloti, Raquel de Lello Rocha Campos Cassano. NGS Soluções Genômicas: Pilar Drummond Sampaio Corrêa Mariani. FZEA-USP Pirassununga: Mirele Daiana Poleti, Jessika Cristina Chagas Lesbon, Elisangela Chicaroni Mattos, Heidge Fukumasu. USP-Botucatu: Rejane Maria Tommasini Grotto, Jayme A. Souza-Neto, Guilherme Targino Valente, Patricia Akemi Assato, Felipe Allan da Silva da Costa, Bianca Cechetto Carlos. Mendelics: Bibiana Santos, João Paulo Kitajima, Erika Freitas, David Schlesinger. Hemocentro Ribeirão Preto: Simone Kashima, Evandra Strazza Rodrigues, Svetoslav Nanev Slavov, Elaine Vieira dos Santos, Rafael dos Santos Bezerra, Luiz Carlos Junior de Alcantara, Marta Giovanetti, Vagner Fonseca, Flavia Aburjaile, Rodrigo Tocantins Calado. FAMERP-SJRP: Cecília Artico Banho, Lívia Sacchetto, Fábio Sossai Possebon, Leila Sabrina Ullmann, Cíntia Bittar, Guilherme Campos, Helena Lage Ferreira, Jorge A. Petroli Marchesi, Maisa C. Pereira Parra, Marília Moraes, Paula Rahal, Paulo Inacio da Costa, João Pessoa Araújo Jr., Maurício Lacerda Nogueira. Prefeitura de Sao Paulo: Melissa Palmieri. |
| EPI_ISL_1966409 | HOSPITAL GERAL DE VILA NOVA CACHOEIRINHA SAO PAULO | Instituto Butantan / Mendelics | Instituto Butantan: Dimas Tadeu Covas, Sandra Coccuzzo Sampaio, Maria Carolina Elias, José Salvatore Leister Patané, Vincent Louis Viala, Antonio Jorge Martins, Ricardo Haddad, Claudia Renata dos Santos Barros, Elaine Cristina Marqueze, Raul Machado Neto, Debora Botequio Moretti, Jardelina de Souza Todao Bernardino, Loyze Paola Oliveira de Lima, Luiz Aurelio de Campos Crispin. Centro de Genômica Funcional da ESALQ: Luiz Lehmann Coutinho, Ricardo Augusto Brassaloti, Raquel de Lello Rocha Campos Cassano. NGS Soluções Genômicas: Pilar Drummond Sampaio Corrêa Mariani. FZEA-USP Pirassununga: Mirele Daiana Poleti, Jessika Cristina Chagas Lesbon, Elisangela Chicaroni Mattos, Heidge Fukumasu. USP-Botucatu: Rejane Maria Tommasini Grotto, Jayme A. Souza-Neto, Guilherme Targino Valente, Patricia Akemi Assato, Felipe Allan da Silva da Costa, Bianca Cechetto Carlos. Mendelics: Bibiana Santos, João Paulo Kitajima, Erika Freitas, David Schlesinger. Hemocentro Ribeirão Preto: Simone Kashima, Evandra Strazza Rodrigues, Svetoslav Nanev Slavov, Elaine Vieira dos Santos, Rafael dos Santos Bezerra, Luiz Carlos Junior de Alcantara, Marta Giovanetti, Vagner Fonseca, Flavia Aburjaile, Rodrigo Tocantins Calado. FAMERP-SJRP: Cecília Artico Banho, Lívia Sacchetto, Fábio Sossai Possebon, Leila Sabrina Ullmann, Cíntia Bittar, Guilherme Campos, Helena Lage Ferreira, Jorge A. Petroli Marchesi, Maisa C. Pereira Parra, Marília Moraes, Paula Rahal, Paulo Inacio da Costa, João Pessoa Araújo Jr., Maurício Lacerda Nogueira. Prefeitura de Sao Paulo: Melissa Palmieri. |
| EPI_ISL_1966411 | CENTRO MEDICO PMESP | Instituto Butantan / Mendelics | Instituto Butantan: Dimas Tadeu Covas, Sandra Coccuzzo Sampaio, Maria Carolina Elias, José Salvatore Leister Patané, Vincent Louis Viala, Antonio Jorge Martins, Ricardo Haddad, Claudia Renata dos Santos Barros, Elaine Cristina Marqueze, Raul Machado Neto, Debora Botequio Moretti, Jardelina de Souza Todao Bernardino, Loyze Paola Oliveira de Lima, Luiz Aurelio de Campos Crispin. Centro de Genômica Funcional da ESALQ: Luiz Lehmann Coutinho, Ricardo Augusto Brassaloti, Raquel de Lello Rocha Campos Cassano. NGS Soluções Genômicas: Pilar Drummond Sampaio Corrêa Mariani. FZEA-USP Pirassununga: Mirele Daiana Poleti, Jessika Cristina Chagas Lesbon, Elisangela Chicaroni Mattos, Heidge Fukumasu. USP-Botucatu: Rejane Maria Tommasini Grotto, Jayme A. Souza-Neto, Guilherme Targino Valente, Patricia Akemi Assato, Felipe Allan da Silva da Costa, Bianca Cechetto Carlos. Mendelics: Bibiana Santos, João Paulo Kitajima, Erika Freitas, David Schlesinger. Hemocentro Ribeirão Preto: Simone Kashima, Evandra Strazza Rodrigues, Svetoslav Nanev Slavov, Elaine Vieira dos Santos, Rafael dos Santos Bezerra, Luiz Carlos Junior de Alcantara, Marta Giovanetti, Vagner Fonseca, Flavia Aburjaile, Rodrigo Tocantins Calado. FAMERP-SJRP: Cecília Artico Banho, Lívia Sacchetto, Fábio Sossai Possebon, Leila Sabrina Ullmann, Cíntia Bittar, Guilherme Campos, Helena Lage Ferreira, Jorge A. Petroli Marchesi, Maisa C. Pereira Parra, Marília Moraes, Paula Rahal, Paulo Inacio da Costa, João Pessoa Araújo Jr., Maurício Lacerda Nogueira. Prefeitura de Sao Paulo: Melissa Palmieri. |
| EPI_ISL_1966412 | CENTRO DE SAUDE DE SPEDRO DO TURVO | Instituto Butantan / Mendelics | Instituto Butantan: Dimas Tadeu Covas, Sandra Coccuzzo Sampaio, Maria Carolina Elias, José Salvatore Leister Patané, Vincent Louis Viala, Antonio Jorge Martins, Ricardo Haddad, Claudia Renata dos Santos Barros, Elaine Cristina Marqueze, Raul Machado Neto, Debora Botequio Moretti, Jardelina de Souza Todao Bernardino, Loyze Paola Oliveira de Lima, Luiz Aurelio de Campos Crispin. Centro de Genômica Funcional da ESALQ: Luiz Lehmann Coutinho, Ricardo Augusto Brassaloti, Raquel de Lello Rocha Campos Cassano. NGS Soluções Genômicas: Pilar Drummond Sampaio Corrêa Mariani. FZEA-USP Pirassununga: Mirele Daiana Poleti, Jessika Cristina Chagas Lesbon, Elisangela Chicaroni Mattos, Heidge Fukumasu. USP-Botucatu: Rejane Maria Tommasini Grotto, Jayme A. Souza-Neto, Guilherme Targino Valente, Patricia Akemi Assato, Felipe Allan da Silva da Costa, Bianca Cechetto Carlos. Mendelics: Bibiana Santos, João Paulo Kitajima, Erika Freitas, David Schlesinger. Hemocentro Ribeirão Preto: Simone Kashima, Evandra Strazza Rodrigues, Svetoslav Nanev Slavov, Elaine Vieira dos Santos, Rafael dos Santos Bezerra, Luiz Carlos Junior de Alcantara, Marta Giovanetti, Vagner Fonseca, Flavia Aburjaile, Rodrigo Tocantins Calado. FAMERP-SJRP: Cecília Artico Banho, Lívia Sacchetto, Fábio Sossai Possebon, Leila Sabrina Ullmann, Cíntia Bittar, Guilherme Campos, Helena Lage Ferreira, Jorge A. Petroli Marchesi, Maisa C. Pereira Parra, Marília Moraes, Paula Rahal, Paulo Inacio da Costa, João Pessoa Araújo Jr., Maurício Lacerda Nogueira. Prefeitura de Sao Paulo: Melissa Palmieri. |
| EPI_ISL_1966413, EPI_ISL_1966414, EPI_ISL_1966415, EPI_ISL_1966416, EPI_ISL_1966417, EPI_ISL_1966418, EPI_ISL_1966419, EPI_ISL_1966420, EPI_ISL_1966421, EPI_ISL_1966422, EPI_ISL_1966423, EPI_ISL_1966424, EPI_ISL_1966425, EPI_ISL_1966426, EPI_ISL_1966427, EPI_ISL_1966428, EPI_ISL_1966429, EPI_ISL_1966430, EPI_ISL_1966431, EPI_ISL_1966432, EPI_ISL_1966433 | | | |
| see above | PSF PARQUE EMILIA | Instituto Butantan / Mendelics | Instituto Butantan: Dimas Tadeu Covas, Sandra Coccuzzo Sampaio, Maria Carolina Elias, José Salvatore Leister Patané, Vincent Louis Viala, Antonio Jorge Martins, Ricardo Haddad, Claudia Renata dos Santos Barros, Elaine Cristina Marqueze, Raul Machado Neto, Debora Botequio Moretti, Jardelina de Souza Todao Bernardino, Loyze Paola Oliveira de Lima, Luiz Aurelio de Campos Crispin. Centro de Genômica Funcional da ESALQ: Luiz Lehmann Coutinho, Ricardo Augusto Brassaloti, Raquel de Lello Rocha Campos Cassano. NGS Soluções Genômicas: Pilar Drummond Sampaio Corrêa Mariani. FZEA-USP Pirassununga: Mirele Daiana Poleti, Jessika Cristina Chagas Lesbon, Elisangela Chicaroni Mattos, Heidge Fukumasu. USP-Botucatu: Rejane Maria Tommasini Grotto, Jayme A. Souza-Neto, Guilherme Targino Valente, Patricia Akemi Assato, Felipe Allan da Silva da Costa, Bianca Cechetto Carlos. Mendelics: Bibiana Santos, João Paulo Kitajima, Erika Freitas, David Schlesinger. Hemocentro Ribeirão Preto: Simone Kashima, Evandra Strazza Rodrigues, Svetoslav Nanev Slavov, Elaine Vieira dos Santos, Rafael dos Santos Bezerra, Luiz Carlos Junior de Alcantara, Marta Giovanetti, Vagner Fonseca, Flavia Aburjaile, Rodrigo Tocantins Calado. FAMERP-SJRP: Cecília Artico Banho, Lívia Sacchetto, Fábio Sossai Possebon, Leila Sabrina Ullmann, Cíntia Bittar, Guilherme Campos, Helena Lage Ferreira, Jorge A. Petroli Marchesi, Maisa C. Pereira Parra, Marília Moraes, Paula Rahal, Paulo Inacio da Costa, João Pessoa Araújo Jr., Maurício Lacerda Nogueira. Prefeitura de Sao Paulo: Melissa Palmieri. |
| EPI_ISL_1966434, EPI_ISL_1966435, EPI_ISL_1966436, EPI_ISL_1966437, EPI_ISL_1966438 | UPA DR AKIRA TADA | Instituto Butantan / Mendelics | Instituto Butantan: Dimas Tadeu Covas, Sandra Coccuzzo Sampaio, Maria Carolina Elias, José Salvatore Leister Patané, Vincent Louis Viala, Antonio Jorge Martins, Ricardo Haddad, Claudia Renata dos Santos Barros, Elaine Cristina Marqueze, Raul Machado Neto, Debora Botequio Moretti, Jardelina de Souza Todao Bernardino, Loyze Paola Oliveira de Lima, Luiz Aurelio de Campos Crispin. Centro de Genômica Funcional da ESALQ: Luiz Lehmann Coutinho, Ricardo Augusto Brassaloti, Raquel de Lello Rocha Campos Cassano. NGS Soluções Genômicas: Pilar Drummond Sampaio Corrêa Mariani. |

| | | | |
|---|---|---|---|
| | | | FZEA-USP Pirassununga: Mirele Daiana Poleti, Jessika Cristina Chagas Lesbon, Elisangela Chicaroni Mattos, Heidge Fukumasu. USP-Botucatu: Rejane Maria Tommasini Grotto, Jayme A. Souza-Neto, Guilherme Targino Valente, Patricia Akemi Assato, Felipe Allan da Silva da Costa, Bianca Cechetto Carlos. Mendelics: Bibiana Santos, João Paulo Kitajima, Erika Freitas, David Schlesinger. Hemocentro Ribeirão Preto: Simone Kashima, Evandra Strazza Rodrigues, Svetoslav Nanev Slavov, Elaine Vieira dos Santos, Rafael dos Santos Bezerra, Luiz Carlos Junior de Alcantara, Marta Giovanetti, Vagner Fonseca, Flavia Aburjaile, Rodrigo Tocantins Calado. FAMERP-SJRP: Cecília Artico Banho, Lívia Sacchetto, Fábio Sossai Possebon, Leila Sabrina Ullmann, Cíntia Bittar, Guilherme Campos, Helena Lage Ferreira, Jorge A. Petroli Marchesi, Maisa C. Pereira Parra, Marília Moraes, Paula Rahal, Paulo Inacio da Costa, João Pessoa Araújo Jr., Maurício Lacerda Nogueira. Prefeitura de Sao Paulo: Melissa Palmieri. |
| EPI_ISL_1966439 | DEPARTAMENTO DE SAUDE COLETIVA | Instituto Butantan / Mendelics | Instituto Butantan: Dimas Tadeu Covas, Sandra Coccuzzo Sampaio, Maria Carolina Elias, José Salvatore Leister Patané, Vincent Louis Viala, Antonio Jorge Martins, Ricardo Haddad, Claudia Renata dos Santos Barros, Elaine Cristina Marqueze, Raul Machado Neto, Debora Botequio Moretti, Jardelina de Souza Todao Bernardino, Loyze Paola Oliveira de Lima, Luiz Aurelio de Campos Crispin. Centro de Genômica Funcional da ESALQ: Luiz Lehmann Coutinho, Ricardo Augusto Brassaloti, Raquel de Lello Rocha Campos Cassano. NGS Soluções Genômicas: Pilar Drummond Sampaio Corrêa Mariani. FZEA-USP Pirassununga: Mirele Daiana Poleti, Jessika Cristina Chagas Lesbon, Elisangela Chicaroni Mattos, Heidge Fukumasu. USP-Botucatu: Rejane Maria Tommasini Grotto, Jayme A. Souza-Neto, Guilherme Targino Valente, Patricia Akemi Assato, Felipe Allan da Silva da Costa, Bianca Cechetto Carlos. Mendelics: Bibiana Santos, João Paulo Kitajima, Erika Freitas, David Schlesinger. Hemocentro Ribeirão Preto: Simone Kashima, Evandra Strazza Rodrigues, Svetoslav Nanev Slavov, Elaine Vieira dos Santos, Rafael dos Santos Bezerra, Luiz Carlos Junior de Alcantara, Marta Giovanetti, Vagner Fonseca, Flavia Aburjaile, Rodrigo Tocantins Calado. FAMERP-SJRP: Cecília Artico Banho, Lívia Sacchetto, Fábio Sossai Possebon, Leila Sabrina Ullmann, Cíntia Bittar, Guilherme Campos, Helena Lage Ferreira, Jorge A. Petroli Marchesi, Maisa C. Pereira Parra, Marília Moraes, Paula Rahal, Paulo Inacio da Costa, João Pessoa Araújo Jr., Maurício Lacerda Nogueira. Prefeitura de Sao Paulo: Melissa Palmieri. |
| EPI_ISL_1966440, EPI_ISL_1966441, EPI_ISL_1966442, EPI_ISL_1966443 | UBS DR ALFREDO DANTAS DE SOUZA UMUARAMA | Instituto Butantan / Mendelics | Instituto Butantan: Dimas Tadeu Covas, Sandra Coccuzzo Sampaio, Maria Carolina Elias, José Salvatore Leister Patané, Vincent Louis Viala, Antonio Jorge Martins, Ricardo Haddad, Claudia Renata dos Santos Barros, Elaine Cristina Marqueze, Raul Machado Neto, Debora Botequio Moretti, Jardelina de Souza Todao Bernardino, Loyze Paola Oliveira de Lima, Luiz Aurelio de Campos Crispin. Centro de Genômica Funcional da ESALQ: Luiz Lehmann Coutinho, Ricardo Augusto Brassaloti, Raquel de Lello Rocha Campos Cassano. NGS Soluções Genômicas: Pilar Drummond Sampaio Corrêa Mariani. FZEA-USP Pirassununga: Mirele Daiana Poleti, Jessika Cristina Chagas Lesbon, Elisangela Chicaroni Mattos, Heidge Fukumasu. USP-Botucatu: Rejane Maria Tommasini Grotto, Jayme A. Souza-Neto, Guilherme Targino Valente, Patricia Akemi Assato, Felipe Allan da Silva da Costa, Bianca Cechetto Carlos. Mendelics: Bibiana Santos, João Paulo Kitajima, Erika Freitas, David Schlesinger. Hemocentro Ribeirão Preto: Simone Kashima, Evandra Strazza Rodrigues, Svetoslav Nanev Slavov, Elaine Vieira dos Santos, Rafael dos Santos Bezerra, Luiz Carlos Junior de Alcantara, Marta Giovanetti, Vagner Fonseca, Flavia Aburjaile, Rodrigo Tocantins Calado. FAMERP-SJRP: Cecília Artico Banho, Lívia Sacchetto, Fábio Sossai Possebon, Leila Sabrina Ullmann, Cíntia Bittar, Guilherme Campos, Helena Lage Ferreira, Jorge A. Petroli Marchesi, Maisa C. Pereira Parra, Marília Moraes, Paula Rahal, Paulo Inacio da Costa, João Pessoa Araújo Jr., Maurício Lacerda Nogueira. Prefeitura de Sao Paulo: Melissa Palmieri. |
| EPI_ISL_1966444 | UBS DE AURIFLAMA | Instituto Butantan / Mendelics | Instituto Butantan: Dimas Tadeu Covas, Sandra Coccuzzo Sampaio, Maria Carolina Elias, José Salvatore Leister Patané, Vincent Louis Viala, Antonio Jorge Martins, Ricardo Haddad, Claudia Renata dos Santos Barros, Elaine Cristina Marqueze, Raul Machado Neto, Debora Botequio Moretti, Jardelina de Souza Todao Bernardino, Loyze Paola Oliveira de Lima, Luiz Aurelio de Campos Crispin. Centro de Genômica Funcional da ESALQ: Luiz Lehmann Coutinho, Ricardo Augusto Brassaloti, Raquel de Lello Rocha Campos Cassano. NGS Soluções Genômicas: Pilar Drummond Sampaio Corrêa Mariani. FZEA-USP Pirassununga: Mirele Daiana Poleti, Jessika Cristina Chagas Lesbon, Elisangela Chicaroni Mattos, Heidge Fukumasu. USP-Botucatu: Rejane Maria Tommasini Grotto, Jayme A. Souza-Neto, Guilherme Targino Valente, Patricia Akemi Assato, Felipe Allan da Silva da Costa, Bianca Cechetto Carlos. Mendelics: Bibiana Santos, João Paulo Kitajima, Erika Freitas, David Schlesinger. Hemocentro Ribeirão Preto: Simone Kashima, Evandra Strazza Rodrigues, Svetoslav Nanev Slavov, Elaine Vieira dos Santos, Rafael dos Santos Bezerra, Luiz Carlos Junior de Alcantara, Marta Giovanetti, Vagner Fonseca, Flavia Aburjaile, Rodrigo Tocantins Calado. FAMERP-SJRP: Cecília Artico Banho, Lívia Sacchetto, Fábio Sossai Possebon, Leila Sabrina Ullmann, Cíntia Bittar, Guilherme Campos, Helena Lage Ferreira, Jorge A. Petroli Marchesi, Maisa C. Pereira Parra, Marília Moraes, Paula Rahal, Paulo Inacio da Costa, João Pessoa Araújo Jr., Maurício Lacerda Nogueira. Prefeitura de Sao Paulo: Melissa Palmieri. |
| EPI_ISL_1966445 | UBS DR ALLY ALAHMAR DE BARRETOS | Instituto Butantan / Mendelics | Instituto Butantan: Dimas Tadeu Covas, Sandra Coccuzzo Sampaio, Maria Carolina Elias, José Salvatore Leister Patané, Vincent Louis Viala, Antonio Jorge Martins, Ricardo Haddad, Claudia Renata dos Santos Barros, Elaine Cristina Marqueze, Raul Machado Neto, Debora Botequio Moretti, Jardelina de Souza Todao Bernardino, Loyze Paola Oliveira de Lima, Luiz Aurelio de Campos Crispin. Centro de Genômica Funcional da ESALQ: Luiz Lehmann Coutinho, Ricardo Augusto Brassaloti, Raquel de Lello Rocha Campos Cassano. NGS Soluções Genômicas: Pilar Drummond Sampaio Corrêa Mariani. FZEA-USP Pirassununga: Mirele Daiana Poleti, Jessika Cristina Chagas Lesbon, Elisangela Chicaroni Mattos, Heidge Fukumasu. USP-Botucatu: Rejane Maria Tommasini Grotto, Jayme A. Souza-Neto, Guilherme Targino Valente, Patricia Akemi Assato, Felipe Allan da Silva da Costa, Bianca Cechetto Carlos. Mendelics: Bibiana Santos, João Paulo Kitajima, Erika Freitas, David Schlesinger. Hemocentro Ribeirão Preto: Simone Kashima, Evandra Strazza Rodrigues, Svetoslav Nanev Slavov, Elaine Vieira dos Santos, Rafael dos Santos Bezerra, Luiz Carlos Junior de Alcantara, Marta Giovanetti, Vagner Fonseca, Flavia Aburjaile, Rodrigo Tocantins Calado. FAMERP-SJRP: Cecília Artico Banho, Lívia Sacchetto, Fábio Sossai Possebon, Leila Sabrina Ullmann, Cíntia Bittar, Guilherme Campos, Helena Lage Ferreira, Jorge A. Petroli Marchesi, Maisa C. Pereira Parra, Marília Moraes, Paula Rahal, Paulo Inacio da Costa, João Pessoa Araújo Jr., Maurício Lacerda Nogueira. Prefeitura de Sao Paulo: Melissa Palmieri. |
| EPI_ISL_1966446, EPI_ISL_1966447, EPI_ISL_1966448, EPI_ISL_1966449, EPI_ISL_1966450, EPI_ISL_1966451, EPI_ISL_1966452, EPI_ISL_1966453, EPI_ISL_1966454, EPI_ISL_1966455, EPI_ISL_1966456, EPI_ISL_1966457, EPI_ISL_1966458, EPI_ISL_1966459 | | | |
| see above | NUCLEO DE SAUDE VILA FALCAO DE BAURU | Instituto Butantan / Mendelics | Instituto Butantan: Dimas Tadeu Covas, Sandra Coccuzzo Sampaio, Maria Carolina Elias, José Salvatore Leister Patané, Vincent Louis Viala, Antonio Jorge Martins, Ricardo Haddad, Claudia Renata dos Santos Barros, Elaine Cristina Marqueze, Raul Machado Neto, Debora Botequio Moretti, Jardelina de Souza Todao Bernardino, Loyze Paola Oliveira de Lima, Luiz Aurelio de Campos Crispin. Centro de Genômica Funcional da ESALQ: Luiz Lehmann Coutinho, Ricardo Augusto Brassaloti, Raquel de Lello Rocha Campos Cassano. NGS Soluções Genômicas: Pilar Drummond Sampaio Corrêa Mariani. FZEA-USP Pirassununga: Mirele Daiana Poleti, Jessika Cristina Chagas Lesbon, Elisangela Chicaroni Mattos, Heidge Fukumasu. USP-Botucatu: Rejane Maria Tommasini Grotto, Jayme A. Souza-Neto, Guilherme Targino Valente, Patricia Akemi Assato, Felipe Allan da Silva da Costa, Bianca Cechetto Carlos. Mendelics: Bibiana Santos, João Paulo Kitajima, Erika Freitas, David Schlesinger. Hemocentro Ribeirão Preto: Simone Kashima, Evandra Strazza Rodrigues, Svetoslav Nanev Slavov, Elaine Vieira dos Santos, Rafael dos Santos Bezerra, Luiz Carlos Junior de Alcantara, Marta Giovanetti, Vagner Fonseca, Flavia Aburjaile, Rodrigo Tocantins Calado. FAMERP-SJRP: Cecília Artico Banho, Lívia Sacchetto, Fábio Sossai Possebon, Leila Sabrina Ullmann, Cíntia Bittar, Guilherme Campos, Helena Lage Ferreira, Jorge A. Petroli Marchesi, Maisa C. Pereira Parra, Marília Moraes, Paula Rahal, Paulo Inacio da Costa, João Pessoa Araújo Jr., Maurício Lacerda Nogueira. Prefeitura de Sao Paulo: Melissa Palmieri. |
| EPI_ISL_1966460, EPI_ISL_1966461, EPI_ISL_1966462 | UBS IV GUARARAPES | Instituto Butantan / Mendelics | Instituto Butantan: Dimas Tadeu Covas, Sandra Coccuzzo Sampaio, Maria Carolina Elias, José Salvatore Leister Patané, Vincent Louis Viala, Antonio Jorge Martins, Ricardo Haddad, Claudia Renata dos Santos Barros, Elaine Cristina Marqueze, Raul Machado Neto, Debora Botequio Moretti, Jardelina de Souza Todao Bernardino, Loyze Paola Oliveira de Lima, Luiz Aurelio de Campos Crispin. Centro de Genômica Funcional da ESALQ: Luiz Lehmann Coutinho, Ricardo Augusto Brassaloti, Raquel de Lello Rocha Campos Cassano. NGS Soluções Genômicas: Pilar Drummond Sampaio Corrêa Mariani. FZEA-USP Pirassununga: Mirele Daiana Poleti, Jessika Cristina Chagas Lesbon, Elisangela Chicaroni Mattos, Heidge Fukumasu. USP-Botucatu: Rejane Maria Tommasini Grotto, Jayme A. Souza-Neto, Guilherme Targino Valente, Patricia Akemi Assato, Felipe Allan da Silva da Costa, Bianca Cechetto Carlos. Mendelics: Bibiana Santos, João Paulo Kitajima, Erika Freitas, David Schlesinger. Hemocentro Ribeirão Preto: Simone Kashima, Evandra Strazza Rodrigues, Svetoslav Nanev Slavov, Elaine Vieira dos Santos, Rafael dos Santos Bezerra, Luiz Carlos Junior de Alcantara, Marta Giovanetti, Vagner Fonseca, Flavia Aburjaile, Rodrigo Tocantins Calado. FAMERP-SJRP: Cecília Artico Banho, Lívia Sacchetto, Fábio Sossai Possebon, Leila Sabrina Ullmann, Cíntia Bittar, Guilherme Campos, Helena Lage Ferreira, Jorge A. Petroli Marchesi, Maisa C. Pereira Parra, Marília Moraes, Paula Rahal, Paulo Inacio da Costa, João Pessoa Araújo Jr., Maurício Lacerda Nogueira. Prefeitura de Sao Paulo: Melissa Palmieri. |
| EPI_ISL_1966463 | CENTRO DE SAUDE II IBITINGA | Instituto Butantan / Mendelics | Instituto Butantan: Dimas Tadeu Covas, Sandra Coccuzzo Sampaio, Maria Carolina Elias, José Salvatore Leister Patané, Vincent Louis Viala, Antonio Jorge Martins, Ricardo Haddad, Claudia Renata dos Santos Barros, Elaine Cristina Marqueze, Raul Machado Neto, Debora Botequio Moretti, Jardelina de Souza Todao Bernardino, Loyze Paola Oliveira de Lima, Luiz Aurelio de Campos Crispin. Centro de Genômica Funcional da ESALQ: Luiz Lehmann Coutinho, Ricardo Augusto Brassaloti, Raquel de Lello Rocha Campos Cassano. NGS Soluções Genômicas: Pilar Drummond Sampaio Corrêa Mariani. FZEA-USP Pirassununga: Mirele Daiana Poleti, Jessika Cristina Chagas Lesbon, Elisangela Chicaroni Mattos, Heidge Fukumasu. USP-Botucatu: Rejane Maria Tommasini Grotto, Jayme A. Souza-Neto, Guilherme Targino Valente, Patricia Akemi Assato, Felipe Allan da Silva da Costa, Bianca Cechetto Carlos. Mendelics: Bibiana Santos, João Paulo Kitajima, Erika Freitas, David Schlesinger. Hemocentro Ribeirão Preto: Simone Kashima, Evandra Strazza Rodrigues, Svetoslav Nanev Slavov, Elaine Vieira dos Santos, Rafael dos Santos Bezerra, Luiz Carlos Junior de Alcantara, Marta Giovanetti, Vagner Fonseca, Flavia Aburjaile, Rodrigo Tocantins Calado. FAMERP-SJRP: Cecília Artico Banho, Lívia Sacchetto, Fábio Sossai Possebon, Leila Sabrina |

| | | | |
|---|---|---|---|
| | | | Ullmann, Cíntia Bittar, Guilherme Campos, Helena Lage Ferreira, Jorge A. Petroli Marchesi, Maisa C. Pereira Parra, Marília Moraes, Paula Rahal, Paulo Inacio da Costa, João Pessoa Araújo Jr., Maurício Lacerda Nogueira. Prefeitura de Sao Paulo: Melissa Palmieri. |
| EPI_ISL_1966464, EPI_ISL_1966465 | POLICLINICA COVID 19 ITAPETININGA | Instituto Butantan / Mendelics | Instituto Butantan: Dimas Tadeu Covas, Sandra Coccuzzo Sampaio, Maria Carolina Elias, José Salvatore Leister Patané, Vincent Louis Viala, Antonio Jorge Martins, Ricardo Haddad, Claudia Renata dos Santos Barros, Elaine Cristina Marqueze, Raul Machado Neto, Debora Botequio Moretti, Jardelina de Souza Todao Bernardino, Loyze Paola Oliveira de Lima, Luiz Aurelio de Campos Crispin. Centro de Genômica Funcional da ESALQ: Luiz Lehmann Coutinho, Ricardo Augusto Brassaloti, Raquel de Lello Rocha Campos Cassano. NGS Soluções Genômicas: Pilar Drummond Sampaio Corrêa Mariani. FZEA-USP Pirassununga: Mirele Daiana Poleti, Jessika Cristina Chagas Lesbon, Elisangela Chicaroni Mattos, Heidge Fukumasu. USP-Botucatu: Rejane Maria Tommasini Grotto, Jayme A. Souza-Neto, Guilherme Targino Valente, Patricia Akemi Assato, Felipe Allan da Silva da Costa, Bianca Cechetto Carlos. Mendelics: Bibiana Santos, João Paulo Kitajima, Erika Freitas, David Schlesinger. Hemocentro Ribeirão Preto: Simone Kashima, Evandra Strazza Rodrigues, Svetoslav Nanev Slavov, Elaine Vieira dos Santos, Rafael dos Santos Bezerra, Luiz Carlos Junior de Alcantara, Marta Giovanetti, Vagner Fonseca, Flavia Aburjaile, Rodrigo Tocantins Calado. FAMERP-SJRP: Cecília Artico Banho, Lívia Sacchetto, Fábio Sossai Possebon, Leila Sabrina Ullmann, Cíntia Bittar, Guilherme Campos, Helena Lage Ferreira, Jorge A. Petroli Marchesi, Maisa C. Pereira Parra, Marília Moraes, Paula Rahal, Paulo Inacio da Costa, João Pessoa Araújo Jr., Maurício Lacerda Nogueira. Prefeitura de Sao Paulo: Melissa Palmieri. |
| EPI_ISL_1966466 | HOSPITAL REGIONAL DE ITAPETININGA | Instituto Butantan / Mendelics | Instituto Butantan: Dimas Tadeu Covas, Sandra Coccuzzo Sampaio, Maria Carolina Elias, José Salvatore Leister Patané, Vincent Louis Viala, Antonio Jorge Martins, Ricardo Haddad, Claudia Renata dos Santos Barros, Elaine Cristina Marqueze, Raul Machado Neto, Debora Botequio Moretti, Jardelina de Souza Todao Bernardino, Loyze Paola Oliveira de Lima, Luiz Aurelio de Campos Crispin. Centro de Genômica Funcional da ESALQ: Luiz Lehmann Coutinho, Ricardo Augusto Brassaloti, Raquel de Lello Rocha Campos Cassano. NGS Soluções Genômicas: Pilar Drummond Sampaio Corrêa Mariani. FZEA-USP Pirassununga: Mirele Daiana Poleti, Jessika Cristina Chagas Lesbon, Elisangela Chicaroni Mattos, Heidge Fukumasu. USP-Botucatu: Rejane Maria Tommasini Grotto, Jayme A. Souza-Neto, Guilherme Targino Valente, Patricia Akemi Assato, Felipe Allan da Silva da Costa, Bianca Cechetto Carlos. Mendelics: Bibiana Santos, João Paulo Kitajima, Erika Freitas, David Schlesinger. Hemocentro Ribeirão Preto: Simone Kashima, Evandra Strazza Rodrigues, Svetoslav Nanev Slavov, Elaine Vieira dos Santos, Rafael dos Santos Bezerra, Luiz Carlos Junior de Alcantara, Marta Giovanetti, Vagner Fonseca, Flavia Aburjaile, Rodrigo Tocantins Calado. FAMERP-SJRP: Cecília Artico Banho, Lívia Sacchetto, Fábio Sossai Possebon, Leila Sabrina Ullmann, Cíntia Bittar, Guilherme Campos, Helena Lage Ferreira, Jorge A. Petroli Marchesi, Maisa C. Pereira Parra, Marília Moraes, Paula Rahal, Paulo Inacio da Costa, João Pessoa Araújo Jr., Maurício Lacerda Nogueira. Prefeitura de Sao Paulo: Melissa Palmieri. |
| EPI_ISL_1966467 | POLICLINICA COVID 19 ITAPETININGA | Instituto Butantan / Mendelics | Instituto Butantan: Dimas Tadeu Covas, Sandra Coccuzzo Sampaio, Maria Carolina Elias, José Salvatore Leister Patané, Vincent Louis Viala, Antonio Jorge Martins, Ricardo Haddad, Claudia Renata dos Santos Barros, Elaine Cristina Marqueze, Raul Machado Neto, Debora Botequio Moretti, Jardelina de Souza Todao Bernardino, Loyze Paola Oliveira de Lima, Luiz Aurelio de Campos Crispin. Centro de Genômica Funcional da ESALQ: Luiz Lehmann Coutinho, Ricardo Augusto Brassaloti, Raquel de Lello Rocha Campos Cassano. NGS Soluções Genômicas: Pilar Drummond Sampaio Corrêa Mariani. FZEA-USP Pirassununga: Mirele Daiana Poleti, Jessika Cristina Chagas Lesbon, Elisangela Chicaroni Mattos, Heidge Fukumasu. USP-Botucatu: Rejane Maria Tommasini Grotto, Jayme A. Souza-Neto, Guilherme Targino Valente, Patricia Akemi Assato, Felipe Allan da Silva da Costa, Bianca Cechetto Carlos. Mendelics: Bibiana Santos, João Paulo Kitajima, Erika Freitas, David Schlesinger. Hemocentro Ribeirão Preto: Simone Kashima, Evandra Strazza Rodrigues, Svetoslav Nanev Slavov, Elaine Vieira dos Santos, Rafael dos Santos Bezerra, Luiz Carlos Junior de Alcantara, Marta Giovanetti, Vagner Fonseca, Flavia Aburjaile, Rodrigo Tocantins Calado. FAMERP-SJRP: Cecília Artico Banho, Lívia Sacchetto, Fábio Sossai Possebon, Leila Sabrina Ullmann, Cíntia Bittar, Guilherme Campos, Helena Lage Ferreira, Jorge A. Petroli Marchesi, Maisa C. Pereira Parra, Marília Moraes, Paula Rahal, Paulo Inacio da Costa, João Pessoa Araújo Jr., Maurício Lacerda Nogueira. Prefeitura de Sao Paulo: Melissa Palmieri. |
| EPI_ISL_1966468 | SECRETARIA MUNICIPAL DE SAUDE | Instituto Butantan / Mendelics | Instituto Butantan: Dimas Tadeu Covas, Sandra Coccuzzo Sampaio, Maria Carolina Elias, José Salvatore Leister Patané, Vincent Louis Viala, Antonio Jorge Martins, Ricardo Haddad, Claudia Renata dos Santos Barros, Elaine Cristina Marqueze, Raul Machado Neto, Debora Botequio Moretti, Jardelina de Souza Todao Bernardino, Loyze Paola Oliveira de Lima, Luiz Aurelio de Campos Crispin. Centro de Genômica Funcional da ESALQ: Luiz Lehmann Coutinho, Ricardo Augusto Brassaloti, Raquel de Lello Rocha Campos Cassano. NGS Soluções Genômicas: Pilar Drummond Sampaio Corrêa Mariani. FZEA-USP Pirassununga: Mirele Daiana Poleti, Jessika Cristina Chagas Lesbon, Elisangela Chicaroni Mattos, Heidge Fukumasu. USP-Botucatu: Rejane Maria Tommasini Grotto, Jayme A. Souza-Neto, Guilherme Targino Valente, Patricia Akemi Assato, Felipe Allan da Silva da Costa, Bianca Cechetto Carlos. Mendelics: Bibiana Santos, João Paulo Kitajima, Erika Freitas, David Schlesinger. Hemocentro Ribeirão Preto: Simone Kashima, Evandra Strazza Rodrigues, Svetoslav Nanev Slavov, Elaine Vieira dos Santos, Rafael dos Santos Bezerra, Luiz Carlos Junior de Alcantara, Marta Giovanetti, Vagner Fonseca, Flavia Aburjaile, Rodrigo Tocantins Calado. FAMERP-SJRP: Cecília Artico Banho, Lívia Sacchetto, Fábio Sossai Possebon, Leila Sabrina Ullmann, Cíntia Bittar, Guilherme Campos, Helena Lage Ferreira, Jorge A. Petroli Marchesi, Maisa C. Pereira Parra, Marília Moraes, Paula Rahal, Paulo Inacio da Costa, João Pessoa Araújo Jr., Maurício Lacerda Nogueira. Prefeitura de Sao Paulo: Melissa Palmieri. |
| EPI_ISL_1966469, EPI_ISL_1966470 | UBS II DA FAMILIA SEGISFREDO MAMEDE PINTO LOURDES | Instituto Butantan / Mendelics | Instituto Butantan: Dimas Tadeu Covas, Sandra Coccuzzo Sampaio, Maria Carolina Elias, José Salvatore Leister Patané, Vincent Louis Viala, Antonio Jorge Martins, Ricardo Haddad, Claudia Renata dos Santos Barros, Elaine Cristina Marqueze, Raul Machado Neto, Debora Botequio Moretti, Jardelina de Souza Todao Bernardino, Loyze Paola Oliveira de Lima, Luiz Aurelio de Campos Crispin. Centro de Genômica Funcional da ESALQ: Luiz Lehmann Coutinho, Ricardo Augusto Brassaloti, Raquel de Lello Rocha Campos Cassano. NGS Soluções Genômicas: Pilar Drummond Sampaio Corrêa Mariani. FZEA-USP Pirassununga: Mirele Daiana Poleti, Jessika Cristina Chagas Lesbon, Elisangela Chicaroni Mattos, Heidge Fukumasu. USP-Botucatu: Rejane Maria Tommasini Grotto, Jayme A. Souza-Neto, Guilherme Targino Valente, Patricia Akemi Assato, Felipe Allan da Silva da Costa, Bianca Cechetto Carlos. Mendelics: Bibiana Santos, João Paulo Kitajima, Erika Freitas, David Schlesinger. Hemocentro Ribeirão Preto: Simone Kashima, Evandra Strazza Rodrigues, Svetoslav Nanev Slavov, Elaine Vieira dos Santos, Rafael dos Santos Bezerra, Luiz Carlos Junior de Alcantara, Marta Giovanetti, Vagner Fonseca, Flavia Aburjaile, Rodrigo Tocantins Calado. FAMERP-SJRP: Cecília Artico Banho, Lívia Sacchetto, Fábio Sossai Possebon, Leila Sabrina Ullmann, Cíntia Bittar, Guilherme Campos, Helena Lage Ferreira, Jorge A. Petroli Marchesi, Maisa C. Pereira Parra, Marília Moraes, Paula Rahal, Paulo Inacio da Costa, João Pessoa Araújo Jr., Maurício Lacerda Nogueira. Prefeitura de Sao Paulo: Melissa Palmieri. |
| EPI_ISL_1966473 | CENTRO DE SAUDE II DR ALCIDES FACUNDO ARROYO | Instituto Butantan / Mendelics | Instituto Butantan: Dimas Tadeu Covas, Sandra Coccuzzo Sampaio, Maria Carolina Elias, José Salvatore Leister Patané, Vincent Louis Viala, Antonio Jorge Martins, Ricardo Haddad, Claudia Renata dos Santos Barros, Elaine Cristina Marqueze, Raul Machado Neto, Debora Botequio Moretti, Jardelina de Souza Todao Bernardino, Loyze Paola Oliveira de Lima, Luiz Aurelio de Campos Crispin. Centro de Genômica Funcional da ESALQ: Luiz Lehmann Coutinho, Ricardo Augusto Brassaloti, Raquel de Lello Rocha Campos Cassano. NGS Soluções Genômicas: Pilar Drummond Sampaio Corrêa Mariani. FZEA-USP Pirassununga: Mirele Daiana Poleti, Jessika Cristina Chagas Lesbon, Elisangela Chicaroni Mattos, Heidge Fukumasu. USP-Botucatu: Rejane Maria Tommasini Grotto, Jayme A. Souza-Neto, Guilherme Targino Valente, Patricia Akemi Assato, Felipe Allan da Silva da Costa, Bianca Cechetto Carlos. Mendelics: Bibiana Santos, João Paulo Kitajima, Erika Freitas, David Schlesinger. Hemocentro Ribeirão Preto: Simone Kashima, Evandra Strazza Rodrigues, Svetoslav Nanev Slavov, Elaine Vieira dos Santos, Rafael dos Santos Bezerra, Luiz Carlos Junior de Alcantara, Marta Giovanetti, Vagner Fonseca, Flavia Aburjaile, Rodrigo Tocantins Calado. FAMERP-SJRP: Cecília Artico Banho, Lívia Sacchetto, Fábio Sossai Possebon, Leila Sabrina Ullmann, Cíntia Bittar, Guilherme Campos, Helena Lage Ferreira, Jorge A. Petroli Marchesi, Maisa C. Pereira Parra, Marília Moraes, Paula Rahal, Paulo Inacio da Costa, João Pessoa Araújo Jr., Maurício Lacerda Nogueira. Prefeitura de Sao Paulo: Melissa Palmieri. |
| EPI_ISL_1966474 | PAS SILVIA COVAS | Instituto Butantan / Mendelics | Instituto Butantan: Dimas Tadeu Covas, Sandra Coccuzzo Sampaio, Maria Carolina Elias, José Salvatore Leister Patané, Vincent Louis Viala, Antonio Jorge Martins, Ricardo Haddad, Claudia Renata dos Santos Barros, Elaine Cristina Marqueze, Raul Machado Neto, Debora Botequio Moretti, Jardelina de Souza Todao Bernardino, Loyze Paola Oliveira de Lima, Luiz Aurelio de Campos Crispin. Centro de Genômica Funcional da ESALQ: Luiz Lehmann Coutinho, Ricardo Augusto Brassaloti, Raquel de Lello Rocha Campos Cassano. NGS Soluções Genômicas: Pilar Drummond Sampaio Corrêa Mariani. FZEA-USP Pirassununga: Mirele Daiana Poleti, Jessika Cristina Chagas Lesbon, Elisangela Chicaroni Mattos, Heidge Fukumasu. USP-Botucatu: Rejane Maria Tommasini Grotto, Jayme A. Souza-Neto, Guilherme Targino Valente, Patricia Akemi Assato, Felipe Allan da Silva da Costa, Bianca Cechetto Carlos. Mendelics: Bibiana Santos, João Paulo Kitajima, Erika Freitas, David Schlesinger. Hemocentro Ribeirão Preto: Simone Kashima, Evandra Strazza Rodrigues, Svetoslav Nanev Slavov, Elaine Vieira dos Santos, Rafael dos Santos Bezerra, Luiz Carlos Junior de Alcantara, Marta Giovanetti, Vagner Fonseca, Flavia Aburjaile, Rodrigo Tocantins Calado. FAMERP-SJRP: Cecília Artico Banho, Lívia Sacchetto, Fábio Sossai Possebon, Leila Sabrina Ullmann, Cíntia Bittar, Guilherme Campos, Helena Lage Ferreira, Jorge A. Petroli Marchesi, Maisa C. Pereira Parra, Marília Moraes, Paula Rahal, Paulo Inacio da Costa, João Pessoa Araújo Jr., Maurício Lacerda Nogueira. Prefeitura de Sao Paulo: Melissa Palmieri. |
| EPI_ISL_1966475 | PAS CIDADE JARDIM | Instituto Butantan / Mendelics | Instituto Butantan: Dimas Tadeu Covas, Sandra Coccuzzo Sampaio, Maria Carolina Elias, José Salvatore Leister Patané, Vincent Louis Viala, Antonio Jorge Martins, Ricardo Haddad, Claudia Renata dos Santos Barros, Elaine Cristina Marqueze, Raul Machado Neto, Debora Botequio Moretti, Jardelina de Souza Todao Bernardino, Loyze Paola Oliveira de Lima, Luiz Aurelio de Campos Crispin. Centro de Genômica Funcional da ESALQ: Luiz Lehmann Coutinho, Ricardo Augusto Brassaloti, Raquel de Lello Rocha Campos Cassano. NGS Soluções Genômicas: Pilar Drummond Sampaio Corrêa Mariani. |

| | | | |
|---|---|---|---|
| | | | FZEA-USP Pirassununga: Mirele Daiana Poleti, Jessika Cristina Chagas Lesbon, Elisangela Chicaroni Mattos, Heidge Fukumasu. USP-Botucatu: Rejane Maria Tommasini Grotto, Jayme A. Souza-Neto, Guilherme Targino Valente, Patricia Akemi Assato, Felipe Allan da Silva da Costa, Bianca Cechetto Carlos. Mendelics: Bibiana Santos, João Paulo Kitajima, Erika Freitas, David Schlesinger. Hemocentro Ribeirão Preto: Simone Kashima, Evandra Strazza Rodrigues, Svetoslav Nanev Slavov, Elaine Vieira dos Santos, Rafael dos Santos Bezerra, Luiz Carlos Junior de Alcantara, Marta Giovanetti, Vagner Fonseca, Flavia Aburjaile, Rodrigo Tocantins Calado. FAMERP-SJRP: Cecília Artico Banho, Lívia Sacchetto, Fábio Sossai Possebon, Leila Sabrina Ullmann, Cíntia Bittar, Guilherme Campos, Helena Lage Ferreira, Jorge A. Petroli Marchesi, Maisa C. Pereira Parra, Marília Moraes, Paula Rahal, Paulo Inacio da Costa, João Pessoa Araújo Jr., Maurício Lacerda Nogueira. Prefeitura de Sao Paulo: Melissa Palmieri. |
| EPI_ISL_1966476 | SECRETARIA DE SAUDE PUBLICA DE PRAIA GRANDE | Instituto Butantan / Mendelics | Instituto Butantan: Dimas Tadeu Covas, Sandra Coccuzzo Sampaio, Maria Carolina Elias, José Salvatore Leister Patané, Vincent Louis Viala, Antonio Jorge Martins, Ricardo Haddad, Claudia Renata dos Santos Barros, Elaine Cristina Marqueze, Raul Machado Neto, Debora Botequio Moretti, Jardelina de Souza Todao Bernardino, Loyze Paola Oliveira de Lima, Luiz Aurelio de Campos Crispin. Centro de Genômica Funcional da ESALQ: Luiz Lehmann Coutinho, Ricardo Augusto Brassaloti, Raquel de Lello Rocha Campos Cassano. NGS Soluções Genômicas: Pilar Drummond Sampaio Corrêa Mariani. FZEA-USP Pirassununga: Mirele Daiana Poleti, Jessika Cristina Chagas Lesbon, Elisangela Chicaroni Mattos, Heidge Fukumasu. USP-Botucatu: Rejane Maria Tommasini Grotto, Jayme A. Souza-Neto, Guilherme Targino Valente, Patricia Akemi Assato, Felipe Allan da Silva da Costa, Bianca Cechetto Carlos. Mendelics: Bibiana Santos, João Paulo Kitajima, Erika Freitas, David Schlesinger. Hemocentro Ribeirão Preto: Simone Kashima, Evandra Strazza Rodrigues, Svetoslav Nanev Slavov, Elaine Vieira dos Santos, Rafael dos Santos Bezerra, Luiz Carlos Junior de Alcantara, Marta Giovanetti, Vagner Fonseca, Flavia Aburjaile, Rodrigo Tocantins Calado. FAMERP-SJRP: Cecília Artico Banho, Lívia Sacchetto, Fábio Sossai Possebon, Leila Sabrina Ullmann, Cíntia Bittar, Guilherme Campos, Helena Lage Ferreira, Jorge A. Petroli Marchesi, Maisa C. Pereira Parra, Marília Moraes, Paula Rahal, Paulo Inacio da Costa, João Pessoa Araújo Jr., Maurício Lacerda Nogueira. Prefeitura de Sao Paulo: Melissa Palmieri. |
| EPI_ISL_1966477, EPI_ISL_1966478, EPI_ISL_1966479 | CENTRO DE SAUDE III RINCAO | Instituto Butantan / Mendelics | Instituto Butantan: Dimas Tadeu Covas, Sandra Coccuzzo Sampaio, Maria Carolina Elias, José Salvatore Leister Patané, Vincent Louis Viala, Antonio Jorge Martins, Ricardo Haddad, Claudia Renata dos Santos Barros, Elaine Cristina Marqueze, Raul Machado Neto, Debora Botequio Moretti, Jardelina de Souza Todao Bernardino, Loyze Paola Oliveira de Lima, Luiz Aurelio de Campos Crispin. Centro de Genômica Funcional da ESALQ: Luiz Lehmann Coutinho, Ricardo Augusto Brassaloti, Raquel de Lello Rocha Campos Cassano. NGS Soluções Genômicas: Pilar Drummond Sampaio Corrêa Mariani. FZEA-USP Pirassununga: Mirele Daiana Poleti, Jessika Cristina Chagas Lesbon, Elisangela Chicaroni Mattos, Heidge Fukumasu. USP-Botucatu: Rejane Maria Tommasini Grotto, Jayme A. Souza-Neto, Guilherme Targino Valente, Patricia Akemi Assato, Felipe Allan da Silva da Costa, Bianca Cechetto Carlos. Mendelics: Bibiana Santos, João Paulo Kitajima, Erika Freitas, David Schlesinger. Hemocentro Ribeirão Preto: Simone Kashima, Evandra Strazza Rodrigues, Svetoslav Nanev Slavov, Elaine Vieira dos Santos, Rafael dos Santos Bezerra, Luiz Carlos Junior de Alcantara, Marta Giovanetti, Vagner Fonseca, Flavia Aburjaile, Rodrigo Tocantins Calado. FAMERP-SJRP: Cecília Artico Banho, Lívia Sacchetto, Fábio Sossai Possebon, Leila Sabrina Ullmann, Cíntia Bittar, Guilherme Campos, Helena Lage Ferreira, Jorge A. Petroli Marchesi, Maisa C. Pereira Parra, Marília Moraes, Paula Rahal, Paulo Inacio da Costa, João Pessoa Araújo Jr., Maurício Lacerda Nogueira. Prefeitura de Sao Paulo: Melissa Palmieri. |
| EPI_ISL_1966480 | CS III SANTOPOLIS DO AGUAPEI | Instituto Butantan / Mendelics | Instituto Butantan: Dimas Tadeu Covas, Sandra Coccuzzo Sampaio, Maria Carolina Elias, José Salvatore Leister Patané, Vincent Louis Viala, Antonio Jorge Martins, Ricardo Haddad, Claudia Renata dos Santos Barros, Elaine Cristina Marqueze, Raul Machado Neto, Debora Botequio Moretti, Jardelina de Souza Todao Bernardino, Loyze Paola Oliveira de Lima, Luiz Aurelio de Campos Crispin. Centro de Genômica Funcional da ESALQ: Luiz Lehmann Coutinho, Ricardo Augusto Brassaloti, Raquel de Lello Rocha Campos Cassano. NGS Soluções Genômicas: Pilar Drummond Sampaio Corrêa Mariani. FZEA-USP Pirassununga: Mirele Daiana Poleti, Jessika Cristina Chagas Lesbon, Elisangela Chicaroni Mattos, Heidge Fukumasu. USP-Botucatu: Rejane Maria Tommasini Grotto, Jayme A. Souza-Neto, Guilherme Targino Valente, Patricia Akemi Assato, Felipe Allan da Silva da Costa, Bianca Cechetto Carlos. Mendelics: Bibiana Santos, João Paulo Kitajima, Erika Freitas, David Schlesinger. Hemocentro Ribeirão Preto: Simone Kashima, Evandra Strazza Rodrigues, Svetoslav Nanev Slavov, Elaine Vieira dos Santos, Rafael dos Santos Bezerra, Luiz Carlos Junior de Alcantara, Marta Giovanetti, Vagner Fonseca, Flavia Aburjaile, Rodrigo Tocantins Calado. FAMERP-SJRP: Cecília Artico Banho, Lívia Sacchetto, Fábio Sossai Possebon, Leila Sabrina Ullmann, Cíntia Bittar, Guilherme Campos, Helena Lage Ferreira, Jorge A. Petroli Marchesi, Maisa C. Pereira Parra, Marília Moraes, Paula Rahal, Paulo Inacio da Costa, João Pessoa Araújo Jr., Maurício Lacerda Nogueira. Prefeitura de Sao Paulo: Melissa Palmieri. |
| EPI_ISL_1966481, EPI_ISL_1966482, EPI_ISL_1966483, EPI_ISL_1966484, EPI_ISL_1966485, EPI_ISL_1966486, EPI_ISL_1966487, EPI_ISL_1966488, EPI_ISL_1966489, EPI_ISL_1966490, EPI_ISL_1966491, EPI_ISL_1966492, EPI_ISL_1966493, EPI_ISL_1966494, EPI_ISL_1966495, EPI_ISL_1966496 | | | |
| see above | SECAO CENTRO DE DIAGNOSTICO SECEDI | Instituto Butantan / Mendelics | Instituto Butantan: Dimas Tadeu Covas, Sandra Coccuzzo Sampaio, Maria Carolina Elias, José Salvatore Leister Patané, Vincent Louis Viala, Antonio Jorge Martins, Ricardo Haddad, Claudia Renata dos Santos Barros, Elaine Cristina Marqueze, Raul Machado Neto, Debora Botequio Moretti, Jardelina de Souza Todao Bernardino, Loyze Paola Oliveira de Lima, Luiz Aurelio de Campos Crispin. Centro de Genômica Funcional da ESALQ: Luiz Lehmann Coutinho, Ricardo Augusto Brassaloti, Raquel de Lello Rocha Campos Cassano. NGS Soluções Genômicas: Pilar Drummond Sampaio Corrêa Mariani. FZEA-USP Pirassununga: Mirele Daiana Poleti, Jessika Cristina Chagas Lesbon, Elisangela Chicaroni Mattos, Heidge Fukumasu. USP-Botucatu: Rejane Maria Tommasini Grotto, Jayme A. Souza-Neto, Guilherme Targino Valente, Patricia Akemi Assato, Felipe Allan da Silva da Costa, Bianca Cechetto Carlos. Mendelics: Bibiana Santos, João Paulo Kitajima, Erika Freitas, David Schlesinger. Hemocentro Ribeirão Preto: Simone Kashima, Evandra Strazza Rodrigues, Svetoslav Nanev Slavov, Elaine Vieira dos Santos, Rafael dos Santos Bezerra, Luiz Carlos Junior de Alcantara, Marta Giovanetti, Vagner Fonseca, Flavia Aburjaile, Rodrigo Tocantins Calado. FAMERP-SJRP: Cecília Artico Banho, Lívia Sacchetto, Fábio Sossai Possebon, Leila Sabrina Ullmann, Cíntia Bittar, Guilherme Campos, Helena Lage Ferreira, Jorge A. Petroli Marchesi, Maisa C. Pereira Parra, Marília Moraes, Paula Rahal, Paulo Inacio da Costa, João Pessoa Araújo Jr., Maurício Lacerda Nogueira. Prefeitura de Sao Paulo: Melissa Palmieri. |
| EPI_ISL_1966497, EPI_ISL_1966498, EPI_ISL_1966499, EPI_ISL_1966500, EPI_ISL_1966501, EPI_ISL_1966502, EPI_ISL_1966503, EPI_ISL_1966504, EPI_ISL_1966505, EPI_ISL_1966506 | Ambulatório IB | Instituto Butantan / Mendelics | Instituto Butantan: Dimas Tadeu Covas, Sandra Coccuzzo Sampaio, Maria Carolina Elias, José Salvatore Leister Patané, Vincent Louis Viala, Antonio Jorge Martins, Ricardo Haddad, Claudia Renata dos Santos Barros, Elaine Cristina Marqueze, Raul Machado Neto, Debora Botequio Moretti, Jardelina de Souza Todao Bernardino, Loyze Paola Oliveira de Lima, Luiz Aurelio de Campos Crispin. Centro de Genômica Funcional da ESALQ: Luiz Lehmann Coutinho, Ricardo Augusto Brassaloti, Raquel de Lello Rocha Campos Cassano. NGS Soluções Genômicas: Pilar Drummond Sampaio Corrêa Mariani. FZEA-USP Pirassununga: Mirele Daiana Poleti, Jessika Cristina Chagas Lesbon, Elisangela Chicaroni Mattos, Heidge Fukumasu. USP-Botucatu: Rejane Maria Tommasini Grotto, Jayme A. Souza-Neto, Guilherme Targino Valente, Patricia Akemi Assato, Felipe Allan da Silva da Costa, Bianca Cechetto Carlos. Mendelics: Bibiana Santos, João Paulo Kitajima, Erika Freitas, David Schlesinger. Hemocentro Ribeirão Preto: Simone Kashima, Evandra Strazza Rodrigues, Svetoslav Nanev Slavov, Elaine Vieira dos Santos, Rafael dos Santos Bezerra, Luiz Carlos Junior de Alcantara, Marta Giovanetti, Vagner Fonseca, Flavia Aburjaile, Rodrigo Tocantins Calado. FAMERP-SJRP: Cecília Artico Banho, Lívia Sacchetto, Fábio Sossai Possebon, Leila Sabrina Ullmann, Cíntia Bittar, Guilherme Campos, Helena Lage Ferreira, Jorge A. Petroli Marchesi, Maisa C. Pereira Parra, Marília Moraes, Paula Rahal, Paulo Inacio da Costa, João Pessoa Araújo Jr., Maurício Lacerda Nogueira. Prefeitura de Sao Paulo: Melissa Palmieri. |
| EPI_ISL_1966507 | CENTRO DE SAUDE SAO ROQUE DR JOSE CARVALHO BRITO | Instituto Butantan / Mendelics | Instituto Butantan: Dimas Tadeu Covas, Sandra Coccuzzo Sampaio, Maria Carolina Elias, José Salvatore Leister Patané, Vincent Louis Viala, Antonio Jorge Martins, Ricardo Haddad, Claudia Renata dos Santos Barros, Elaine Cristina Marqueze, Raul Machado Neto, Debora Botequio Moretti, Jardelina de Souza Todao Bernardino, Loyze Paola Oliveira de Lima, Luiz Aurelio de Campos Crispin. Centro de Genômica Funcional da ESALQ: Luiz Lehmann Coutinho, Ricardo Augusto Brassaloti, Raquel de Lello Rocha Campos Cassano. NGS Soluções Genômicas: Pilar Drummond Sampaio Corrêa Mariani. FZEA-USP Pirassununga: Mirele Daiana Poleti, Jessika Cristina Chagas Lesbon, Elisangela Chicaroni Mattos, Heidge Fukumasu. USP-Botucatu: Rejane Maria Tommasini Grotto, Jayme A. Souza-Neto, Guilherme Targino Valente, Patricia Akemi Assato, Felipe Allan da Silva da Costa, Bianca Cechetto Carlos. Mendelics: Bibiana Santos, João Paulo Kitajima, Erika Freitas, David Schlesinger. Hemocentro Ribeirão Preto: Simone Kashima, Evandra Strazza Rodrigues, Svetoslav Nanev Slavov, Elaine Vieira dos Santos, Rafael dos Santos Bezerra, Luiz Carlos Junior de Alcantara, Marta Giovanetti, Vagner Fonseca, Flavia Aburjaile, Rodrigo Tocantins Calado. FAMERP-SJRP: Cecília Artico Banho, Lívia Sacchetto, Fábio Sossai Possebon, Leila Sabrina Ullmann, Cíntia Bittar, Guilherme Campos, Helena Lage Ferreira, Jorge A. Petroli Marchesi, Maisa C. Pereira Parra, Marília Moraes, Paula Rahal, Paulo Inacio da Costa, João Pessoa Araújo Jr., Maurício Lacerda Nogueira. Prefeitura de Sao Paulo: Melissa Palmieri. |
| EPI_ISL_1966508, EPI_ISL_1966509 | PRONTO SOCORRO MUNICIPAL DE SEVERINIA | Instituto Butantan / Mendelics | Instituto Butantan: Dimas Tadeu Covas, Sandra Coccuzzo Sampaio, Maria Carolina Elias, José Salvatore Leister Patané, Vincent Louis Viala, Antonio Jorge Martins, Ricardo Haddad, Claudia Renata dos Santos Barros, Elaine Cristina Marqueze, Raul Machado Neto, Debora Botequio Moretti, Jardelina de Souza Todao Bernardino, Loyze Paola Oliveira de Lima, Luiz Aurelio de Campos Crispin. Centro de Genômica Funcional da ESALQ: Luiz Lehmann Coutinho, Ricardo Augusto Brassaloti, Raquel de Lello Rocha Campos Cassano. NGS Soluções Genômicas: Pilar Drummond Sampaio Corrêa Mariani. FZEA-USP Pirassununga: Mirele Daiana Poleti, Jessika Cristina Chagas Lesbon, Elisangela Chicaroni Mattos, Heidge Fukumasu. USP-Botucatu: Rejane Maria Tommasini Grotto, Jayme A. Souza-Neto, Guilherme Targino Valente, Patricia Akemi Assato, Felipe Allan da Silva da Costa, Bianca Cechetto Carlos. Mendelics: Bibiana Santos, João Paulo Kitajima, Erika Freitas, David Schlesinger. Hemocentro Ribeirão Preto: Simone Kashima, Evandra Strazza Rodrigues, Svetoslav Nanev Slavov, Elaine Vieira dos Santos, Rafael dos Santos Bezerra, Luiz Carlos Junior de Alcantara, Marta Giovanetti, Vagner Fonseca, Flavia Aburjaile, Rodrigo Tocantins Calado. FAMERP-SJRP: Cecília Artico Banho, Lívia Sacchetto, Fábio Sossai Possebon, Leila Sabrina |

| | | | |
|---|---|---|---|
| | | | Ullmann, Cíntia Bittar, Guilherme Campos, Helena Lage Ferreira, Jorge A. Petroli Marchesi, Maisa C. Pereira Parra, Marília Moraes, Paula Rahal, Paulo Inacio da Costa, João Pessoa Araújo Jr., Maurício Lacerda Nogueira. Prefeitura de Sao Paulo: Melissa Palmieri. |
| EPI_ISL_1966510 | UBS DE SUD MENNUCCI | Instituto Butantan / Mendelics | Instituto Butantan: Dimas Tadeu Covas, Sandra Coccuzzo Sampaio, Maria Carolina Elias, José Salvatore Leister Patané, Vincent Louis Viala, Antonio Jorge Martins, Ricardo Haddad, Claudia Renata dos Santos Barros, Elaine Cristina Marqueze, Raul Machado Neto, Debora Botequio Moretti, Jardelina de Souza Todao Bernardino, Loyze Paola Oliveira de Lima, Luiz Aurelio de Campos Crispin. Centro de Genômica Funcional da ESALQ: Luiz Lehmann Coutinho, Ricardo Augusto Brassaloti, Raquel de Lello Rocha Campos Cassano. NGS Soluções Genômicas: Pilar Drummond Sampaio Corrêa Mariani. FZEA-USP Pirassununga: Mirele Daiana Poleti, Jessika Cristina Chagas Lesbon, Elisangela Chicaroni Mattos, Heidge Fukumasu. USP-Botucatu: Rejane Maria Tommasini Grotto, Jayme A. Souza-Neto, Guilherme Targino Valente, Patricia Akemi Assato, Felipe Allan da Silva da Costa, Bianca Cechetto Carlos. Mendelics: Bibiana Santos, João Paulo Kitajima, Erika Freitas, David Schlesinger. Hemocentro Ribeirão Preto: Simone Kashima, Evandra Strazza Rodrigues, Svetoslav Nanev Slavov, Elaine Vieira dos Santos, Rafael dos Santos Bezerra, Luiz Carlos Junior de Alcantara, Marta Giovanetti, Vagner Fonseca, Flavia Aburjaile, Rodrigo Tocantins Calado. FAMERP-SJRP: Cecília Artico Banho, Lívia Sacchetto, Fábio Sossai Possebon, Leila Sabrina Ullmann, Cíntia Bittar, Guilherme Campos, Helena Lage Ferreira, Jorge A. Petroli Marchesi, Maisa C. Pereira Parra, Marília Moraes, Paula Rahal, Paulo Inacio da Costa, João Pessoa Araújo Jr., Maurício Lacerda Nogueira. Prefeitura de Sao Paulo: Melissa Palmieri. |
| EPI_ISL_1966511 | UBS 2 DE SUZANAPOLIS | Instituto Butantan / Mendelics | Instituto Butantan: Dimas Tadeu Covas, Sandra Coccuzzo Sampaio, Maria Carolina Elias, José Salvatore Leister Patané, Vincent Louis Viala, Antonio Jorge Martins, Ricardo Haddad, Claudia Renata dos Santos Barros, Elaine Cristina Marqueze, Raul Machado Neto, Debora Botequio Moretti, Jardelina de Souza Todao Bernardino, Loyze Paola Oliveira de Lima, Luiz Aurelio de Campos Crispin. Centro de Genômica Funcional da ESALQ: Luiz Lehmann Coutinho, Ricardo Augusto Brassaloti, Raquel de Lello Rocha Campos Cassano. NGS Soluções Genômicas: Pilar Drummond Sampaio Corrêa Mariani. FZEA-USP Pirassununga: Mirele Daiana Poleti, Jessika Cristina Chagas Lesbon, Elisangela Chicaroni Mattos, Heidge Fukumasu. USP-Botucatu: Rejane Maria Tommasini Grotto, Jayme A. Souza-Neto, Guilherme Targino Valente, Patricia Akemi Assato, Felipe Allan da Silva da Costa, Bianca Cechetto Carlos. Mendelics: Bibiana Santos, João Paulo Kitajima, Erika Freitas, David Schlesinger. Hemocentro Ribeirão Preto: Simone Kashima, Evandra Strazza Rodrigues, Svetoslav Nanev Slavov, Elaine Vieira dos Santos, Rafael dos Santos Bezerra, Luiz Carlos Junior de Alcantara, Marta Giovanetti, Vagner Fonseca, Flavia Aburjaile, Rodrigo Tocantins Calado. FAMERP-SJRP: Cecília Artico Banho, Lívia Sacchetto, Fábio Sossai Possebon, Leila Sabrina Ullmann, Cíntia Bittar, Guilherme Campos, Helena Lage Ferreira, Jorge A. Petroli Marchesi, Maisa C. Pereira Parra, Marília Moraes, Paula Rahal, Paulo Inacio da Costa, João Pessoa Araújo Jr., Maurício Lacerda Nogueira. Prefeitura de Sao Paulo: Melissa Palmieri. |
| EPI_ISL_1966512 | UNIDADE MISTA DE TAIACU | Instituto Butantan / Mendelics | Dimas Tadeu Covas, Antonio Jorge Martins, Claudia Renata dos Santos Barros, David Schlesinger, Debora Botequio Moretti, Elaine Cristina Marqueze, Elaine Vieira Santos, Evandra Strazza Rodrigues, Heidge Fukumasu, Jayme Augusto de Souza-Neto, José Salvatore Leister Patané, Luiz Alcantara, Luiz Lehmann Coutinho, Maria Carolina Elias, Maurício Lacerda Nogueira, Rafael dos Santos Bezerra, Raul Machado Neto, Rejane Maria Tommasini Grotto, Ricardo Haddad, Sandra Coccuzzo Sampaio Vessoni, Simone Kashima, Svetoslav Nanev Slavov, Vincent Louis Viala |
| EPI_ISL_1966513, EPI_ISL_1966514, EPI_ISL_1966515, EPI_ISL_1966516 | UNIDADE BASICA DE SAUDE DE TAQUARAL CAETANO PITELLI | Instituto Butantan / Mendelics | Instituto Butantan: Dimas Tadeu Covas, Sandra Coccuzzo Sampaio, Maria Carolina Elias, José Salvatore Leister Patané, Vincent Louis Viala, Antonio Jorge Martins, Ricardo Haddad, Claudia Renata dos Santos Barros, Elaine Cristina Marqueze, Raul Machado Neto, Debora Botequio Moretti, Jardelina de Souza Todao Bernardino, Loyze Paola Oliveira de Lima, Luiz Aurelio de Campos Crispin. Centro de Genômica Funcional da ESALQ: Luiz Lehmann Coutinho, Ricardo Augusto Brassaloti, Raquel de Lello Rocha Campos Cassano. NGS Soluções Genômicas: Pilar Drummond Sampaio Corrêa Mariani. FZEA-USP Pirassununga: Mirele Daiana Poleti, Jessika Cristina Chagas Lesbon, Elisangela Chicaroni Mattos, Heidge Fukumasu. USP-Botucatu: Rejane Maria Tommasini Grotto, Jayme A. Souza-Neto, Guilherme Targino Valente, Patricia Akemi Assato, Felipe Allan da Silva da Costa, Bianca Cechetto Carlos. Mendelics: Bibiana Santos, João Paulo Kitajima, Erika Freitas, David Schlesinger. Hemocentro Ribeirão Preto: Simone Kashima, Evandra Strazza Rodrigues, Svetoslav Nanev Slavov, Elaine Vieira dos Santos, Rafael dos Santos Bezerra, Luiz Carlos Junior de Alcantara, Marta Giovanetti, Vagner Fonseca, Flavia Aburjaile, Rodrigo Tocantins Calado. FAMERP-SJRP: Cecília Artico Banho, Lívia Sacchetto, Fábio Sossai Possebon, Leila Sabrina Ullmann, Cíntia Bittar, Guilherme Campos, Helena Lage Ferreira, Jorge A. Petroli Marchesi, Maisa C. Pereira Parra, Marília Moraes, Paula Rahal, Paulo Inacio da Costa, João Pessoa Araújo Jr., Maurício Lacerda Nogueira. Prefeitura de Sao Paulo: Melissa Palmieri. |
| EPI_ISL_1966517, EPI_ISL_1966518, EPI_ISL_1966519, EPI_ISL_1966520 | SAE SERVICO DE ATENDIMENTO ESPECIALIZADO | Instituto Butantan / Mendelics | Instituto Butantan: Dimas Tadeu Covas, Sandra Coccuzzo Sampaio, Maria Carolina Elias, José Salvatore Leister Patané, Vincent Louis Viala, Antonio Jorge Martins, Ricardo Haddad, Claudia Renata dos Santos Barros, Elaine Cristina Marqueze, Raul Machado Neto, Debora Botequio Moretti, Jardelina de Souza Todao Bernardino, Loyze Paola Oliveira de Lima, Luiz Aurelio de Campos Crispin. Centro de Genômica Funcional da ESALQ: Luiz Lehmann Coutinho, Ricardo Augusto Brassaloti, Raquel de Lello Rocha Campos Cassano. NGS Soluções Genômicas: Pilar Drummond Sampaio Corrêa Mariani. FZEA-USP Pirassununga: Mirele Daiana Poleti, Jessika Cristina Chagas Lesbon, Elisangela Chicaroni Mattos, Heidge Fukumasu. USP-Botucatu: Rejane Maria Tommasini Grotto, Jayme A. Souza-Neto, Guilherme Targino Valente, Patricia Akemi Assato, Felipe Allan da Silva da Costa, Bianca Cechetto Carlos. Mendelics: Bibiana Santos, João Paulo Kitajima, Erika Freitas, David Schlesinger. Hemocentro Ribeirão Preto: Simone Kashima, Evandra Strazza Rodrigues, Svetoslav Nanev Slavov, Elaine Vieira dos Santos, Rafael dos Santos Bezerra, Luiz Carlos Junior de Alcantara, Marta Giovanetti, Vagner Fonseca, Flavia Aburjaile, Rodrigo Tocantins Calado. FAMERP-SJRP: Cecília Artico Banho, Lívia Sacchetto, Fábio Sossai Possebon, Leila Sabrina Ullmann, Cíntia Bittar, Guilherme Campos, Helena Lage Ferreira, Jorge A. Petroli Marchesi, Maisa C. Pereira Parra, Marília Moraes, Paula Rahal, Paulo Inacio da Costa, João Pessoa Araújo Jr., Maurício Lacerda Nogueira. Prefeitura de Sao Paulo: Melissa Palmieri. |
| EPI_ISL_1966521 | UNIDADE MISTA DE VISTA ALEGRE DO ALTO VISTA ALEGRE DO ALTO | Instituto Butantan / Mendelics | Instituto Butantan: Dimas Tadeu Covas, Sandra Coccuzzo Sampaio, Maria Carolina Elias, José Salvatore Leister Patané, Vincent Louis Viala, Antonio Jorge Martins, Ricardo Haddad, Claudia Renata dos Santos Barros, Elaine Cristina Marqueze, Raul Machado Neto, Debora Botequio Moretti, Jardelina de Souza Todao Bernardino, Loyze Paola Oliveira de Lima, Luiz Aurelio de Campos Crispin. Centro de Genômica Funcional da ESALQ: Luiz Lehmann Coutinho, Ricardo Augusto Brassaloti, Raquel de Lello Rocha Campos Cassano. NGS Soluções Genômicas: Pilar Drummond Sampaio Corrêa Mariani. FZEA-USP Pirassununga: Mirele Daiana Poleti, Jessika Cristina Chagas Lesbon, Elisangela Chicaroni Mattos, Heidge Fukumasu. USP-Botucatu: Rejane Maria Tommasini Grotto, Jayme A. Souza-Neto, Guilherme Targino Valente, Patricia Akemi Assato, Felipe Allan da Silva da Costa, Bianca Cechetto Carlos. Mendelics: Bibiana Santos, João Paulo Kitajima, Erika Freitas, David Schlesinger. Hemocentro Ribeirão Preto: Simone Kashima, Evandra Strazza Rodrigues, Svetoslav Nanev Slavov, Elaine Vieira dos Santos, Rafael dos Santos Bezerra, Luiz Carlos Junior de Alcantara, Marta Giovanetti, Vagner Fonseca, Flavia Aburjaile, Rodrigo Tocantins Calado. FAMERP-SJRP: Cecília Artico Banho, Lívia Sacchetto, Fábio Sossai Possebon, Leila Sabrina Ullmann, Cíntia Bittar, Guilherme Campos, Helena Lage Ferreira, Jorge A. Petroli Marchesi, Maisa C. Pereira Parra, Marília Moraes, Paula Rahal, Paulo Inacio da Costa, João Pessoa Araújo Jr., Maurício Lacerda Nogueira. Prefeitura de Sao Paulo: Melissa Palmieri. |
| EPI_ISL_1966522 | CENTRO DE SAUDE DE ADAMANTINA | Instituto Butantan / FZEA-USP (Pirassununga) | Instituto Butantan: Dimas Tadeu Covas, Sandra Coccuzzo Sampaio, Maria Carolina Elias, José Salvatore Leister Patané, Vincent Louis Viala, Antonio Jorge Martins, Ricardo Haddad, Claudia Renata dos Santos Barros, Elaine Cristina Marqueze, Raul Machado Neto, Debora Botequio Moretti, Jardelina de Souza Todao Bernardino, Loyze Paola Oliveira de Lima, Luiz Aurelio de Campos Crispin. Centro de Genômica Funcional da ESALQ: Luiz Lehmann Coutinho, Ricardo Augusto Brassaloti, Raquel de Lello Rocha Campos Cassano. NGS Soluções Genômicas: Pilar Drummond Sampaio Corrêa Mariani. FZEA-USP Pirassununga: Mirele Daiana Poleti, Jessika Cristina Chagas Lesbon, Elisangela Chicaroni Mattos, Heidge Fukumasu. USP-Botucatu: Rejane Maria Tommasini Grotto, Jayme A. Souza-Neto, Guilherme Targino Valente, Patricia Akemi Assato, Felipe Allan da Silva da Costa, Bianca Cechetto Carlos. Mendelics: Bibiana Santos, João Paulo Kitajima, Erika Freitas, David Schlesinger. Hemocentro Ribeirão Preto: Simone Kashima, Evandra Strazza Rodrigues, Svetoslav Nanev Slavov, Elaine Vieira dos Santos, Rafael dos Santos Bezerra, Luiz Carlos Junior de Alcantara, Marta Giovanetti, Vagner Fonseca, Flavia Aburjaile, Rodrigo Tocantins Calado. FAMERP-SJRP: Cecília Artico Banho, Lívia Sacchetto, Fábio Sossai Possebon, Leila Sabrina Ullmann, Cíntia Bittar, Guilherme Campos, Helena Lage Ferreira, Jorge A. Petroli Marchesi, Maisa C. Pereira Parra, Marília Moraes, Paula Rahal, Paulo Inacio da Costa, João Pessoa Araújo Jr., Maurício Lacerda Nogueira. Prefeitura de Sao Paulo: Melissa Palmieri. |
| EPI_ISL_1966524 | CS DE BALSAMO | Instituto Butantan / FZEA-USP (Pirassununga) | Instituto Butantan: Dimas Tadeu Covas, Sandra Coccuzzo Sampaio, Maria Carolina Elias, José Salvatore Leister Patané, Vincent Louis Viala, Antonio Jorge Martins, Ricardo Haddad, Claudia Renata dos Santos Barros, Elaine Cristina Marqueze, Raul Machado Neto, Debora Botequio Moretti, Jardelina de Souza Todao Bernardino, Loyze Paola Oliveira de Lima, Luiz Aurelio de Campos Crispin. Centro de Genômica Funcional da ESALQ: Luiz Lehmann Coutinho, Ricardo Augusto Brassaloti, Raquel de Lello Rocha Campos Cassano. NGS Soluções Genômicas: Pilar Drummond Sampaio Corrêa Mariani. FZEA-USP Pirassununga: Mirele Daiana Poleti, Jessika Cristina Chagas Lesbon, Elisangela Chicaroni Mattos, Heidge Fukumasu. USP-Botucatu: Rejane Maria Tommasini Grotto, Jayme A. Souza-Neto, Guilherme Targino Valente, Patricia Akemi Assato, Felipe Allan da Silva da Costa, Bianca Cechetto Carlos. Mendelics: Bibiana Santos, João Paulo Kitajima, Erika Freitas, David Schlesinger. Hemocentro Ribeirão Preto: Simone Kashima, Evandra Strazza Rodrigues, Svetoslav Nanev Slavov, Elaine Vieira dos Santos, Rafael dos Santos Bezerra, Luiz Carlos Junior de Alcantara, Marta Giovanetti, Vagner Fonseca, Flavia Aburjaile, Rodrigo Tocantins Calado. FAMERP-SJRP: Cecília Artico Banho, Lívia Sacchetto, Fábio Sossai Possebon, Leila Sabrina Ullmann, Cíntia Bittar, Guilherme Campos, Helena Lage Ferreira, Jorge A. Petroli Marchesi, Maisa C. Pereira Parra, Marília Moraes, Paula Rahal, Paulo Inacio da Costa, João Pessoa Araújo Jr., Maurício Lacerda Nogueira. Prefeitura de Sao Paulo: Melissa Palmieri. |

| | | | |
|---|---|---|---|
| EPI_ISL_1966525 | VIGILANCIA EM SAUDE | Instituto Butantan / FZEA-USP (Pirassununga) | Instituto Butantan: Dimas Tadeu Covas, Sandra Coccuzzo Sampaio, Maria Carolina Elias, José Salvatore Leister Patané, Vincent Louis Viala, Antonio Jorge Martins, Ricardo Haddad, Claudia Renata dos Santos Barros, Elaine Cristina Marqueze, Raul Machado Neto, Debora Botequio Moretti, Jardelina de Souza Todao Bernardino, Loyze Paola Oliveira de Lima, Luiz Aurelio de Campos Crispin. Centro de Genômica Funcional da ESALQ: Luiz Lehmann Coutinho, Ricardo Augusto Brassaloti, Raquel de Lello Rocha Campos Cassano. NGS Soluções Genômicas: Pilar Drummond Sampaio Corrêa Mariani. FZEA-USP Pirassununga: Mirele Daiana Poleti, Jessika Cristina Chagas Lesbon, Elisangela Chicaroni Mattos, Heidge Fukumasu. USP-Botucatu: Rejane Maria Tommasini Grotto, Jayme A. Souza-Neto, Guilherme Targino Valente, Patricia Akemi Assato, Felipe Allan da Silva da Costa, Bianca Cechetto Carlos. Mendelics: Bibiana Santos, João Paulo Kitajima, Erika Freitas, David Schlesinger. Hemocentro Ribeirão Preto: Simone Kashima, Evandra Strazza Rodrigues, Svetoslav Nanev Slavov, Elaine Vieira dos Santos, Rafael dos Santos Bezerra, Luiz Carlos Junior de Alcantara, Marta Giovanetti, Vagner Fonseca, Flavia Aburjaile, Rodrigo Tocantins Calado. FAMERP-SJRP: Cecília Artico Banho, Lívia Sacchetto, Fábio Sossai Possebon, Leila Sabrina Ullmann, Cíntia Bittar, Guilherme Campos, Helena Lage Ferreira, Jorge A. Petroli Marchesi, Maisa C. Pereira Parra, Marília Moraes, Paula Rahal, Paulo Inacio da Costa, João Pessoa Araújo Jr., Maurício Lacerda Nogueira. Prefeitura de Sao Paulo: Melissa Palmieri. |
| EPI_ISL_1966526, EPI_ISL_1966527, EPI_ISL_1966528, EPI_ISL_1966529 | USF PAULISTA FERNANDOPOLIS ANTONIO PIVATO | Instituto Butantan / FZEA-USP (Pirassununga) | Instituto Butantan: Dimas Tadeu Covas, Sandra Coccuzzo Sampaio, Maria Carolina Elias, José Salvatore Leister Patané, Vincent Louis Viala, Antonio Jorge Martins, Ricardo Haddad, Claudia Renata dos Santos Barros, Elaine Cristina Marqueze, Raul Machado Neto, Debora Botequio Moretti, Jardelina de Souza Todao Bernardino, Loyze Paola Oliveira de Lima, Luiz Aurelio de Campos Crispin. Centro de Genômica Funcional da ESALQ: Luiz Lehmann Coutinho, Ricardo Augusto Brassaloti, Raquel de Lello Rocha Campos Cassano. NGS Soluções Genômicas: Pilar Drummond Sampaio Corrêa Mariani. FZEA-USP Pirassununga: Mirele Daiana Poleti, Jessika Cristina Chagas Lesbon, Elisangela Chicaroni Mattos, Heidge Fukumasu. USP-Botucatu: Rejane Maria Tommasini Grotto, Jayme A. Souza-Neto, Guilherme Targino Valente, Patricia Akemi Assato, Felipe Allan da Silva da Costa, Bianca Cechetto Carlos. Mendelics: Bibiana Santos, João Paulo Kitajima, Erika Freitas, David Schlesinger. Hemocentro Ribeirão Preto: Simone Kashima, Evandra Strazza Rodrigues, Svetoslav Nanev Slavov, Elaine Vieira dos Santos, Rafael dos Santos Bezerra, Luiz Carlos Junior de Alcantara, Marta Giovanetti, Vagner Fonseca, Flavia Aburjaile, Rodrigo Tocantins Calado. FAMERP-SJRP: Cecília Artico Banho, Lívia Sacchetto, Fábio Sossai Possebon, Leila Sabrina Ullmann, Cíntia Bittar, Guilherme Campos, Helena Lage Ferreira, Jorge A. Petroli Marchesi, Maisa C. Pereira Parra, Marília Moraes, Paula Rahal, Paulo Inacio da Costa, João Pessoa Araújo Jr., Maurício Lacerda Nogueira. Prefeitura de Sao Paulo: Melissa Palmieri. |
| EPI_ISL_1966530 | HOSPITAL MUNICIPAL DR TABAJARA RAMOS | Instituto Butantan / FZEA-USP (Pirassununga) | Instituto Butantan: Dimas Tadeu Covas, Sandra Coccuzzo Sampaio, Maria Carolina Elias, José Salvatore Leister Patané, Vincent Louis Viala, Antonio Jorge Martins, Ricardo Haddad, Claudia Renata dos Santos Barros, Elaine Cristina Marqueze, Raul Machado Neto, Debora Botequio Moretti, Jardelina de Souza Todao Bernardino, Loyze Paola Oliveira de Lima, Luiz Aurelio de Campos Crispin. Centro de Genômica Funcional da ESALQ: Luiz Lehmann Coutinho, Ricardo Augusto Brassaloti, Raquel de Lello Rocha Campos Cassano. NGS Soluções Genômicas: Pilar Drummond Sampaio Corrêa Mariani. FZEA-USP Pirassununga: Mirele Daiana Poleti, Jessika Cristina Chagas Lesbon, Elisangela Chicaroni Mattos, Heidge Fukumasu. USP-Botucatu: Rejane Maria Tommasini Grotto, Jayme A. Souza-Neto, Guilherme Targino Valente, Patricia Akemi Assato, Felipe Allan da Silva da Costa, Bianca Cechetto Carlos. Mendelics: Bibiana Santos, João Paulo Kitajima, Erika Freitas, David Schlesinger. Hemocentro Ribeirão Preto: Simone Kashima, Evandra Strazza Rodrigues, Svetoslav Nanev Slavov, Elaine Vieira dos Santos, Rafael dos Santos Bezerra, Luiz Carlos Junior de Alcantara, Marta Giovanetti, Vagner Fonseca, Flavia Aburjaile, Rodrigo Tocantins Calado. FAMERP-SJRP: Cecília Artico Banho, Lívia Sacchetto, Fábio Sossai Possebon, Leila Sabrina Ullmann, Cíntia Bittar, Guilherme Campos, Helena Lage Ferreira, Jorge A. Petroli Marchesi, Maisa C. Pereira Parra, Marília Moraes, Paula Rahal, Paulo Inacio da Costa, João Pessoa Araújo Jr., Maurício Lacerda Nogueira. Prefeitura de Sao Paulo: Melissa Palmieri. |
| EPI_ISL_1966531, EPI_ISL_1966533 | EMERGENCIA RESPIRATORIA DE NOVA GRANADA | Instituto Butantan / FZEA-USP (Pirassununga) | Instituto Butantan: Dimas Tadeu Covas, Sandra Coccuzzo Sampaio, Maria Carolina Elias, José Salvatore Leister Patané, Vincent Louis Viala, Antonio Jorge Martins, Ricardo Haddad, Claudia Renata dos Santos Barros, Elaine Cristina Marqueze, Raul Machado Neto, Debora Botequio Moretti, Jardelina de Souza Todao Bernardino, Loyze Paola Oliveira de Lima, Luiz Aurelio de Campos Crispin. Centro de Genômica Funcional da ESALQ: Luiz Lehmann Coutinho, Ricardo Augusto Brassaloti, Raquel de Lello Rocha Campos Cassano. NGS Soluções Genômicas: Pilar Drummond Sampaio Corrêa Mariani. FZEA-USP Pirassununga: Mirele Daiana Poleti, Jessika Cristina Chagas Lesbon, Elisangela Chicaroni Mattos, Heidge Fukumasu. USP-Botucatu: Rejane Maria Tommasini Grotto, Jayme A. Souza-Neto, Guilherme Targino Valente, Patricia Akemi Assato, Felipe Allan da Silva da Costa, Bianca Cechetto Carlos. Mendelics: Bibiana Santos, João Paulo Kitajima, Erika Freitas, David Schlesinger. Hemocentro Ribeirão Preto: Simone Kashima, Evandra Strazza Rodrigues, Svetoslav Nanev Slavov, Elaine Vieira dos Santos, Rafael dos Santos Bezerra, Luiz Carlos Junior de Alcantara, Marta Giovanetti, Vagner Fonseca, Flavia Aburjaile, Rodrigo Tocantins Calado. FAMERP-SJRP: Cecília Artico Banho, Lívia Sacchetto, Fábio Sossai Possebon, Leila Sabrina Ullmann, Cíntia Bittar, Guilherme Campos, Helena Lage Ferreira, Jorge A. Petroli Marchesi, Maisa C. Pereira Parra, Marília Moraes, Paula Rahal, Paulo Inacio da Costa, João Pessoa Araújo Jr., Maurício Lacerda Nogueira. Prefeitura de Sao Paulo: Melissa Palmieri. |
| EPI_ISL_1966534 | UBS DR HELIO MIGLIARI | Instituto Butantan / FZEA-USP (Pirassununga) | Instituto Butantan: Dimas Tadeu Covas, Sandra Coccuzzo Sampaio, Maria Carolina Elias, José Salvatore Leister Patané, Vincent Louis Viala, Antonio Jorge Martins, Ricardo Haddad, Claudia Renata dos Santos Barros, Elaine Cristina Marqueze, Raul Machado Neto, Debora Botequio Moretti, Jardelina de Souza Todao Bernardino, Loyze Paola Oliveira de Lima, Luiz Aurelio de Campos Crispin. Centro de Genômica Funcional da ESALQ: Luiz Lehmann Coutinho, Ricardo Augusto Brassaloti, Raquel de Lello Rocha Campos Cassano. NGS Soluções Genômicas: Pilar Drummond Sampaio Corrêa Mariani. FZEA-USP Pirassununga: Mirele Daiana Poleti, Jessika Cristina Chagas Lesbon, Elisangela Chicaroni Mattos, Heidge Fukumasu. USP-Botucatu: Rejane Maria Tommasini Grotto, Jayme A. Souza-Neto, Guilherme Targino Valente, Patricia Akemi Assato, Felipe Allan da Silva da Costa, Bianca Cechetto Carlos. Mendelics: Bibiana Santos, João Paulo Kitajima, Erika Freitas, David Schlesinger. Hemocentro Ribeirão Preto: Simone Kashima, Evandra Strazza Rodrigues, Svetoslav Nanev Slavov, Elaine Vieira dos Santos, Rafael dos Santos Bezerra, Luiz Carlos Junior de Alcantara, Marta Giovanetti, Vagner Fonseca, Flavia Aburjaile, Rodrigo Tocantins Calado. FAMERP-SJRP: Cecília Artico Banho, Lívia Sacchetto, Fábio Sossai Possebon, Leila Sabrina Ullmann, Cíntia Bittar, Guilherme Campos, Helena Lage Ferreira, Jorge A. Petroli Marchesi, Maisa C. Pereira Parra, Marília Moraes, Paula Rahal, Paulo Inacio da Costa, João Pessoa Araújo Jr., Maurício Lacerda Nogueira. Prefeitura de Sao Paulo: Melissa Palmieri. |
| EPI_ISL_1966535 | UBS REGIAO OESTE DE OURINHOS | Instituto Butantan / FZEA-USP (Pirassununga) | Instituto Butantan: Dimas Tadeu Covas, Sandra Coccuzzo Sampaio, Maria Carolina Elias, José Salvatore Leister Patané, Vincent Louis Viala, Antonio Jorge Martins, Ricardo Haddad, Claudia Renata dos Santos Barros, Elaine Cristina Marqueze, Raul Machado Neto, Debora Botequio Moretti, Jardelina de Souza Todao Bernardino, Loyze Paola Oliveira de Lima, Luiz Aurelio de Campos Crispin. Centro de Genômica Funcional da ESALQ: Luiz Lehmann Coutinho, Ricardo Augusto Brassaloti, Raquel de Lello Rocha Campos Cassano. NGS Soluções Genômicas: Pilar Drummond Sampaio Corrêa Mariani. FZEA-USP Pirassununga: Mirele Daiana Poleti, Jessika Cristina Chagas Lesbon, Elisangela Chicaroni Mattos, Heidge Fukumasu. USP-Botucatu: Rejane Maria Tommasini Grotto, Jayme A. Souza-Neto, Guilherme Targino Valente, Patricia Akemi Assato, Felipe Allan da Silva da Costa, Bianca Cechetto Carlos. Mendelics: Bibiana Santos, João Paulo Kitajima, Erika Freitas, David Schlesinger. Hemocentro Ribeirão Preto: Simone Kashima, Evandra Strazza Rodrigues, Svetoslav Nanev Slavov, Elaine Vieira dos Santos, Rafael dos Santos Bezerra, Luiz Carlos Junior de Alcantara, Marta Giovanetti, Vagner Fonseca, Flavia Aburjaile, Rodrigo Tocantins Calado. FAMERP-SJRP: Cecília Artico Banho, Lívia Sacchetto, Fábio Sossai Possebon, Leila Sabrina Ullmann, Cíntia Bittar, Guilherme Campos, Helena Lage Ferreira, Jorge A. Petroli Marchesi, Maisa C. Pereira Parra, Marília Moraes, Paula Rahal, Paulo Inacio da Costa, João Pessoa Araújo Jr., Maurício Lacerda Nogueira. Prefeitura de Sao Paulo: Melissa Palmieri. |
| EPI_ISL_1966536, EPI_ISL_1966537, EPI_ISL_1966538, EPI_ISL_1966539, EPI_ISL_1966540 | CENTRO DE SAUDE I TACITO LEITE DE CARVALHO E SILVA | Instituto Butantan / FZEA-USP (Pirassununga) | Instituto Butantan: Dimas Tadeu Covas, Sandra Coccuzzo Sampaio, Maria Carolina Elias, José Salvatore Leister Patané, Vincent Louis Viala, Antonio Jorge Martins, Ricardo Haddad, Claudia Renata dos Santos Barros, Elaine Cristina Marqueze, Raul Machado Neto, Debora Botequio Moretti, Jardelina de Souza Todao Bernardino, Loyze Paola Oliveira de Lima, Luiz Aurelio de Campos Crispin. Centro de Genômica Funcional da ESALQ: Luiz Lehmann Coutinho, Ricardo Augusto Brassaloti, Raquel de Lello Rocha Campos Cassano. NGS Soluções Genômicas: Pilar Drummond Sampaio Corrêa Mariani. FZEA-USP Pirassununga: Mirele Daiana Poleti, Jessika Cristina Chagas Lesbon, Elisangela Chicaroni Mattos, Heidge Fukumasu. USP-Botucatu: Rejane Maria Tommasini Grotto, Jayme A. Souza-Neto, Guilherme Targino Valente, Patricia Akemi Assato, Felipe Allan da Silva da Costa, Bianca Cechetto Carlos. Mendelics: Bibiana Santos, João Paulo Kitajima, Erika Freitas, David Schlesinger. Hemocentro Ribeirão Preto: Simone Kashima, Evandra Strazza Rodrigues, Svetoslav Nanev Slavov, Elaine Vieira dos Santos, Rafael dos Santos Bezerra, Luiz Carlos Junior de Alcantara, Marta Giovanetti, Vagner Fonseca, Flavia Aburjaile, Rodrigo Tocantins Calado. FAMERP-SJRP: Cecília Artico Banho, Lívia Sacchetto, Fábio Sossai Possebon, Leila Sabrina Ullmann, Cíntia Bittar, Guilherme Campos, Helena Lage Ferreira, Jorge A. Petroli Marchesi, Maisa C. Pereira Parra, Marília Moraes, Paula Rahal, Paulo Inacio da Costa, João Pessoa Araújo Jr., Maurício Lacerda Nogueira. Prefeitura de Sao Paulo: Melissa Palmieri. |
| EPI_ISL_1966541 | CENTRO DE SAUDE III AFFONSO LUZZI SANTA CRUZ DAS PALMEIRAS | Instituto Butantan / FZEA-USP (Pirassununga) | Instituto Butantan: Dimas Tadeu Covas, Sandra Coccuzzo Sampaio, Maria Carolina Elias, José Salvatore Leister Patané, Vincent Louis Viala, Antonio Jorge Martins, Ricardo Haddad, Claudia Renata dos Santos Barros, Elaine Cristina Marqueze, Raul Machado Neto, Debora Botequio Moretti, Jardelina de Souza Todao Bernardino, Loyze Paola Oliveira de Lima, Luiz Aurelio de Campos Crispin. Centro de Genômica Funcional da ESALQ: Luiz Lehmann Coutinho, Ricardo Augusto Brassaloti, Raquel de Lello Rocha Campos Cassano. NGS Soluções Genômicas: Pilar Drummond Sampaio Corrêa Mariani. FZEA-USP Pirassununga: Mirele Daiana Poleti, Jessika Cristina Chagas Lesbon, Elisangela Chicaroni Mattos, Heidge Fukumasu. USP-Botucatu: Rejane Maria Tommasini Grotto, Jayme A. Souza-Neto, Guilherme Targino Valente, Patricia Akemi Assato, Felipe Allan da Silva da Costa, Bianca Cechetto Carlos. Mendelics: Bibiana Santos, João Paulo Kitajima, Erika Freitas, David Schlesinger. Hemocentro Ribeirão Preto: Simone Kashima, Evandra Strazza |

| | | | |
|---|---|---|---|
| | | | Rodrigues, Svetoslav Nanev Slavov, Elaine Vieira dos Santos, Rafael dos Santos Bezerra, Luiz Carlos Junior de Alcantara, Marta Giovanetti, Vagner Fonseca, Flavia Aburjaile, Rodrigo Tocantins Calado. FAMERP-SJRP: Cecília Artico Banho, Lívia Sacchetto, Fábio Sossai Possebon, Leila Sabrina Ullmann, Cíntia Bittar, Guilherme Campos, Helena Lage Ferreira, Jorge A. Petroli Marchesi, Maisa C. Pereira Parra, Marília Moraes, Paula Rahal, Paulo Inacio da Costa, João Pessoa Araújo Jr., Maurício Lacerda Nogueira. Prefeitura de Sao Paulo: Melissa Palmieri. |
| EPI_ISL_1966542 | USF DR APARECIDO RODRIGUES MOUCO SCRPARDO | Instituto Butantan / FZEA-USP (Pirassununga) | Instituto Butantan: Dimas Tadeu Covas, Sandra Coccuzzo Sampaio, Maria Carolina Elias, José Salvatore Leister Patané, Vincent Louis Viala, Antonio Jorge Martins, Ricardo Haddad, Claudia Renata dos Santos Barros, Elaine Cristina Marqueze, Raul Machado Neto, Debora Botequio Moretti, Jardelina de Souza Todao Bernardino, Loyze Paola Oliveira de Lima, Luiz Aurelio de Campos Crispin. Centro de Genômica Funcional da ESALQ: Luiz Lehmann Coutinho, Ricardo Augusto Brassaloti, Raquel de Lello Rocha Campos Cassano. NGS Soluções Genômicas: Pilar Drummond Sampaio Corrêa Mariani. FZEA-USP Pirassununga: Mirele Daiana Poleti, Jessika Cristina Chagas Lesbon, Elisangela Chicaroni Mattos, Heidge Fukumasu. USP-Botucatu: Rejane Maria Tommasini Grotto, Jayme A. Souza-Neto, Guilherme Targino Valente, Patricia Akemi Assato, Felipe Allan da Silva da Costa, Bianca Cechetto Carlos. Mendelics: Bibiana Santos, João Paulo Kitajima, Erika Freitas, David Schlesinger. Hemocentro Ribeirão Preto: Simone Kashima, Evandra Strazza Rodrigues, Svetoslav Nanev Slavov, Elaine Vieira dos Santos, Rafael dos Santos Bezerra, Luiz Carlos Junior de Alcantara, Marta Giovanetti, Vagner Fonseca, Flavia Aburjaile, Rodrigo Tocantins Calado. FAMERP-SJRP: Cecília Artico Banho, Lívia Sacchetto, Fábio Sossai Possebon, Leila Sabrina Ullmann, Cíntia Bittar, Guilherme Campos, Helena Lage Ferreira, Jorge A. Petroli Marchesi, Maisa C. Pereira Parra, Marília Moraes, Paula Rahal, Paulo Inacio da Costa, João Pessoa Araújo Jr., Maurício Lacerda Nogueira. Prefeitura de Sao Paulo: Melissa Palmieri. |
| EPI_ISL_1966543, EPI_ISL_1966544 | UBS DR ABELARDO PINHEIRO GUIMARAES SCRPARDO | Instituto Butantan / FZEA-USP (Pirassununga) | Instituto Butantan: Dimas Tadeu Covas, Sandra Coccuzzo Sampaio, Maria Carolina Elias, José Salvatore Leister Patané, Vincent Louis Viala, Antonio Jorge Martins, Ricardo Haddad, Claudia Renata dos Santos Barros, Elaine Cristina Marqueze, Raul Machado Neto, Debora Botequio Moretti, Jardelina de Souza Todao Bernardino, Loyze Paola Oliveira de Lima, Luiz Aurelio de Campos Crispin. Centro de Genômica Funcional da ESALQ: Luiz Lehmann Coutinho, Ricardo Augusto Brassaloti, Raquel de Lello Rocha Campos Cassano. NGS Soluções Genômicas: Pilar Drummond Sampaio Corrêa Mariani. FZEA-USP Pirassununga: Mirele Daiana Poleti, Jessika Cristina Chagas Lesbon, Elisangela Chicaroni Mattos, Heidge Fukumasu. USP-Botucatu: Rejane Maria Tommasini Grotto, Jayme A. Souza-Neto, Guilherme Targino Valente, Patricia Akemi Assato, Felipe Allan da Silva da Costa, Bianca Cechetto Carlos. Mendelics: Bibiana Santos, João Paulo Kitajima, Erika Freitas, David Schlesinger. Hemocentro Ribeirão Preto: Simone Kashima, Evandra Strazza Rodrigues, Svetoslav Nanev Slavov, Elaine Vieira dos Santos, Rafael dos Santos Bezerra, Luiz Carlos Junior de Alcantara, Marta Giovanetti, Vagner Fonseca, Flavia Aburjaile, Rodrigo Tocantins Calado. FAMERP-SJRP: Cecília Artico Banho, Lívia Sacchetto, Fábio Sossai Possebon, Leila Sabrina Ullmann, Cíntia Bittar, Guilherme Campos, Helena Lage Ferreira, Jorge A. Petroli Marchesi, Maisa C. Pereira Parra, Marília Moraes, Paula Rahal, Paulo Inacio da Costa, João Pessoa Araújo Jr., Maurício Lacerda Nogueira. Prefeitura de Sao Paulo: Melissa Palmieri. |
| EPI_ISL_1966545, EPI_ISL_1966546 | UBS DR WALDOMIRO FERREIRA NEVES SCRPARDO | Instituto Butantan / FZEA-USP (Pirassununga) | Instituto Butantan: Dimas Tadeu Covas, Sandra Coccuzzo Sampaio, Maria Carolina Elias, José Salvatore Leister Patané, Vincent Louis Viala, Antonio Jorge Martins, Ricardo Haddad, Claudia Renata dos Santos Barros, Elaine Cristina Marqueze, Raul Machado Neto, Debora Botequio Moretti, Jardelina de Souza Todao Bernardino, Loyze Paola Oliveira de Lima, Luiz Aurelio de Campos Crispin. Centro de Genômica Funcional da ESALQ: Luiz Lehmann Coutinho, Ricardo Augusto Brassaloti, Raquel de Lello Rocha Campos Cassano. NGS Soluções Genômicas: Pilar Drummond Sampaio Corrêa Mariani. FZEA-USP Pirassununga: Mirele Daiana Poleti, Jessika Cristina Chagas Lesbon, Elisangela Chicaroni Mattos, Heidge Fukumasu. USP-Botucatu: Rejane Maria Tommasini Grotto, Jayme A. Souza-Neto, Guilherme Targino Valente, Patricia Akemi Assato, Felipe Allan da Silva da Costa, Bianca Cechetto Carlos. Mendelics: Bibiana Santos, João Paulo Kitajima, Erika Freitas, David Schlesinger. Hemocentro Ribeirão Preto: Simone Kashima, Evandra Strazza Rodrigues, Svetoslav Nanev Slavov, Elaine Vieira dos Santos, Rafael dos Santos Bezerra, Luiz Carlos Junior de Alcantara, Marta Giovanetti, Vagner Fonseca, Flavia Aburjaile, Rodrigo Tocantins Calado. FAMERP-SJRP: Cecília Artico Banho, Lívia Sacchetto, Fábio Sossai Possebon, Leila Sabrina Ullmann, Cíntia Bittar, Guilherme Campos, Helena Lage Ferreira, Jorge A. Petroli Marchesi, Maisa C. Pereira Parra, Marília Moraes, Paula Rahal, Paulo Inacio da Costa, João Pessoa Araújo Jr., Maurício Lacerda Nogueira. Prefeitura de Sao Paulo: Melissa Palmieri. |
| EPI_ISL_1966547 | CS DE SANTA RITA DOESTE | Instituto Butantan / FZEA-USP (Pirassununga) | Instituto Butantan: Dimas Tadeu Covas, Sandra Coccuzzo Sampaio, Maria Carolina Elias, José Salvatore Leister Patané, Vincent Louis Viala, Antonio Jorge Martins, Ricardo Haddad, Claudia Renata dos Santos Barros, Elaine Cristina Marqueze, Raul Machado Neto, Debora Botequio Moretti, Jardelina de Souza Todao Bernardino, Loyze Paola Oliveira de Lima, Luiz Aurelio de Campos Crispin. Centro de Genômica Funcional da ESALQ: Luiz Lehmann Coutinho, Ricardo Augusto Brassaloti, Raquel de Lello Rocha Campos Cassano. NGS Soluções Genômicas: Pilar Drummond Sampaio Corrêa Mariani. FZEA-USP Pirassununga: Mirele Daiana Poleti, Jessika Cristina Chagas Lesbon, Elisangela Chicaroni Mattos, Heidge Fukumasu. USP-Botucatu: Rejane Maria Tommasini Grotto, Jayme A. Souza-Neto, Guilherme Targino Valente, Patricia Akemi Assato, Felipe Allan da Silva da Costa, Bianca Cechetto Carlos. Mendelics: Bibiana Santos, João Paulo Kitajima, Erika Freitas, David Schlesinger. Hemocentro Ribeirão Preto: Simone Kashima, Evandra Strazza Rodrigues, Svetoslav Nanev Slavov, Elaine Vieira dos Santos, Rafael dos Santos Bezerra, Luiz Carlos Junior de Alcantara, Marta Giovanetti, Vagner Fonseca, Flavia Aburjaile, Rodrigo Tocantins Calado. FAMERP-SJRP: Cecília Artico Banho, Lívia Sacchetto, Fábio Sossai Possebon, Leila Sabrina Ullmann, Cíntia Bittar, Guilherme Campos, Helena Lage Ferreira, Jorge A. Petroli Marchesi, Maisa C. Pereira Parra, Marília Moraes, Paula Rahal, Paulo Inacio da Costa, João Pessoa Araújo Jr., Maurício Lacerda Nogueira. Prefeitura de Sao Paulo: Melissa Palmieri. |
| EPI_ISL_1966548 | UBS II DOUTOR ORLANDO BERTOLLI | Instituto Butantan / FZEA-USP (Pirassununga) | Instituto Butantan: Dimas Tadeu Covas, Sandra Coccuzzo Sampaio, Maria Carolina Elias, José Salvatore Leister Patané, Vincent Louis Viala, Antonio Jorge Martins, Ricardo Haddad, Claudia Renata dos Santos Barros, Elaine Cristina Marqueze, Raul Machado Neto, Debora Botequio Moretti, Jardelina de Souza Todao Bernardino, Loyze Paola Oliveira de Lima, Luiz Aurelio de Campos Crispin. Centro de Genômica Funcional da ESALQ: Luiz Lehmann Coutinho, Ricardo Augusto Brassaloti, Raquel de Lello Rocha Campos Cassano. NGS Soluções Genômicas: Pilar Drummond Sampaio Corrêa Mariani. FZEA-USP Pirassununga: Mirele Daiana Poleti, Jessika Cristina Chagas Lesbon, Elisangela Chicaroni Mattos, Heidge Fukumasu. USP-Botucatu: Rejane Maria Tommasini Grotto, Jayme A. Souza-Neto, Guilherme Targino Valente, Patricia Akemi Assato, Felipe Allan da Silva da Costa, Bianca Cechetto Carlos. Mendelics: Bibiana Santos, João Paulo Kitajima, Erika Freitas, David Schlesinger. Hemocentro Ribeirão Preto: Simone Kashima, Evandra Strazza Rodrigues, Svetoslav Nanev Slavov, Elaine Vieira dos Santos, Rafael dos Santos Bezerra, Luiz Carlos Junior de Alcantara, Marta Giovanetti, Vagner Fonseca, Flavia Aburjaile, Rodrigo Tocantins Calado. FAMERP-SJRP: Cecília Artico Banho, Lívia Sacchetto, Fábio Sossai Possebon, Leila Sabrina Ullmann, Cíntia Bittar, Guilherme Campos, Helena Lage Ferreira, Jorge A. Petroli Marchesi, Maisa C. Pereira Parra, Marília Moraes, Paula Rahal, Paulo Inacio da Costa, João Pessoa Araújo Jr., Maurício Lacerda Nogueira. Prefeitura de Sao Paulo: Melissa Palmieri. |
| EPI_ISL_1966549, EPI_ISL_1966550 | UNIDADE DE VIGILANCIA EM SAUDE | Instituto Butantan / FZEA-USP (Pirassununga) | Instituto Butantan: Dimas Tadeu Covas, Sandra Coccuzzo Sampaio, Maria Carolina Elias, José Salvatore Leister Patané, Vincent Louis Viala, Antonio Jorge Martins, Ricardo Haddad, Claudia Renata dos Santos Barros, Elaine Cristina Marqueze, Raul Machado Neto, Debora Botequio Moretti, Jardelina de Souza Todao Bernardino, Loyze Paola Oliveira de Lima, Luiz Aurelio de Campos Crispin. Centro de Genômica Funcional da ESALQ: Luiz Lehmann Coutinho, Ricardo Augusto Brassaloti, Raquel de Lello Rocha Campos Cassano. NGS Soluções Genômicas: Pilar Drummond Sampaio Corrêa Mariani. FZEA-USP Pirassununga: Mirele Daiana Poleti, Jessika Cristina Chagas Lesbon, Elisangela Chicaroni Mattos, Heidge Fukumasu. USP-Botucatu: Rejane Maria Tommasini Grotto, Jayme A. Souza-Neto, Guilherme Targino Valente, Patricia Akemi Assato, Felipe Allan da Silva da Costa, Bianca Cechetto Carlos. Mendelics: Bibiana Santos, João Paulo Kitajima, Erika Freitas, David Schlesinger. Hemocentro Ribeirão Preto: Simone Kashima, Evandra Strazza Rodrigues, Svetoslav Nanev Slavov, Elaine Vieira dos Santos, Rafael dos Santos Bezerra, Luiz Carlos Junior de Alcantara, Marta Giovanetti, Vagner Fonseca, Flavia Aburjaile, Rodrigo Tocantins Calado. FAMERP-SJRP: Cecília Artico Banho, Lívia Sacchetto, Fábio Sossai Possebon, Leila Sabrina Ullmann, Cíntia Bittar, Guilherme Campos, Helena Lage Ferreira, Jorge A. Petroli Marchesi, Maisa C. Pereira Parra, Marília Moraes, Paula Rahal, Paulo Inacio da Costa, João Pessoa Araújo Jr., Maurício Lacerda Nogueira. Prefeitura de Sao Paulo: Melissa Palmieri. |
| EPI_ISL_1966552, EPI_ISL_1966553 | UBS II DE TANABI MILTON MARTINS PERCHES | Instituto Butantan / FZEA-USP (Pirassununga) | Instituto Butantan: Dimas Tadeu Covas, Sandra Coccuzzo Sampaio, Maria Carolina Elias, José Salvatore Leister Patané, Vincent Louis Viala, Antonio Jorge Martins, Ricardo Haddad, Claudia Renata dos Santos Barros, Elaine Cristina Marqueze, Raul Machado Neto, Debora Botequio Moretti, Jardelina de Souza Todao Bernardino, Loyze Paola Oliveira de Lima, Luiz Aurelio de Campos Crispin. Centro de Genômica Funcional da ESALQ: Luiz Lehmann Coutinho, Ricardo Augusto Brassaloti, Raquel de Lello Rocha Campos Cassano. NGS Soluções Genômicas: Pilar Drummond Sampaio Corrêa Mariani. FZEA-USP Pirassununga: Mirele Daiana Poleti, Jessika Cristina Chagas Lesbon, Elisangela Chicaroni Mattos, Heidge Fukumasu. USP-Botucatu: Rejane Maria Tommasini Grotto, Jayme A. Souza-Neto, Guilherme Targino Valente, Patricia Akemi Assato, Felipe Allan da Silva da Costa, Bianca Cechetto Carlos. Mendelics: Bibiana Santos, João Paulo Kitajima, Erika Freitas, David Schlesinger. Hemocentro Ribeirão Preto: Simone Kashima, Evandra Strazza Rodrigues, Svetoslav Nanev Slavov, Elaine Vieira dos Santos, Rafael dos Santos Bezerra, Luiz Carlos Junior de Alcantara, Marta Giovanetti, Vagner Fonseca, Flavia Aburjaile, Rodrigo Tocantins Calado. FAMERP-SJRP: Cecília Artico Banho, Lívia Sacchetto, Fábio Sossai Possebon, Leila Sabrina Ullmann, Cíntia Bittar, Guilherme Campos, Helena Lage Ferreira, Jorge A. Petroli Marchesi, Maisa C. Pereira Parra, Marília Moraes, Paula Rahal, Paulo Inacio da Costa, João Pessoa Araújo Jr., Maurício Lacerda Nogueira. Prefeitura de Sao Paulo: Melissa Palmieri. |
| EPI_ISL_1966554 | CENTRO DE SAUDE II SIZENANDO NABUCO TAPIRATIBA | Instituto Butantan / FZEA-USP (Pirassununga) | Instituto Butantan: Dimas Tadeu Covas, Sandra Coccuzzo Sampaio, Maria Carolina Elias, José Salvatore Leister Patané, Vincent Louis Viala, Antonio Jorge Martins, Ricardo Haddad, Claudia Renata dos Santos Barros, Elaine Cristina Marqueze, Raul Machado Neto, Debora Botequio Moretti, Jardelina de |

| | | | |
|---|---|---|---|
| | | | Souza Todao Bernardino, Loyze Paola Oliveira de Lima, Luiz Aurelio de Campos Crispin. Centro de Genômica Funcional da ESALQ: Luiz Lehmann Coutinho, Ricardo Augusto Brassaloti, Raquel de Lello Rocha Campos Cassano. NGS Soluções Genômicas: Pilar Drummond Sampaio Corrêa Mariani. FZEA-USP Pirassununga: Mirele Daiana Poleti, Jessika Cristina Chagas Lesbon, Elisangela Chicaroni Mattos, Heidge Fukumasu. USP-Botucatu: Rejane Maria Tommasini Grotto, Jayme A. Souza-Neto, Guilherme Targino Valente, Patricia Akemi Assato, Felipe Allan da Silva da Costa, Bianca Cechetto Carlos. Mendelics: Bibiana Santos, João Paulo Kitajima, Erika Freitas, David Schlesinger. Hemocentro Ribeirão Preto: Simone Kashima, Evandra Strazza Rodrigues, Svetoslav Nanev Slavov, Elaine Vieira dos Santos, Rafael dos Santos Bezerra, Luiz Carlos Junior de Alcantara, Marta Giovanetti, Vagner Fonseca, Flavia Aburjaile, Rodrigo Tocantins Calado. FAMERP-SJRP: Cecília Artico Banho, Lívia Sacchetto, Fábio Sossai Possebon, Leila Sabrina Ullmann, Cíntia Bittar, Guilherme Campos, Helena Lage Ferreira, Jorge A. Petroli Marchesi, Maisa C. Pereira Parra, Marília Moraes, Paula Rahal, Paulo Inacio da Costa, João Pessoa Araújo Jr., Maurício Lacerda Nogueira. Prefeitura de Sao Paulo: Melissa Palmieri. |
| EPI_ISL_1966555 | UNIDADE BASICA DE SAUDE II DE TARABAI | Instituto Butantan / FZEA-USP (Pirassununga) | Instituto Butantan: Dimas Tadeu Covas, Sandra Coccuzzo Sampaio, Maria Carolina Elias, José Salvatore Leister Patané, Vincent Louis Viala, Antonio Jorge Martins, Ricardo Haddad, Claudia Renata dos Santos Barros, Elaine Cristina Marqueze, Raul Machado Neto, Debora Botequio Moretti, Jardelina de Souza Todao Bernardino, Loyze Paola Oliveira de Lima, Luiz Aurelio de Campos Crispin. Centro de Genômica Funcional da ESALQ: Luiz Lehmann Coutinho, Ricardo Augusto Brassaloti, Raquel de Lello Rocha Campos Cassano. NGS Soluções Genômicas: Pilar Drummond Sampaio Corrêa Mariani. FZEA-USP Pirassununga: Mirele Daiana Poleti, Jessika Cristina Chagas Lesbon, Elisangela Chicaroni Mattos, Heidge Fukumasu. USP-Botucatu: Rejane Maria Tommasini Grotto, Jayme A. Souza-Neto, Guilherme Targino Valente, Patricia Akemi Assato, Felipe Allan da Silva da Costa, Bianca Cechetto Carlos. Mendelics: Bibiana Santos, João Paulo Kitajima, Erika Freitas, David Schlesinger. Hemocentro Ribeirão Preto: Simone Kashima, Evandra Strazza Rodrigues, Svetoslav Nanev Slavov, Elaine Vieira dos Santos, Rafael dos Santos Bezerra, Luiz Carlos Junior de Alcantara, Marta Giovanetti, Vagner Fonseca, Flavia Aburjaile, Rodrigo Tocantins Calado. FAMERP-SJRP: Cecília Artico Banho, Lívia Sacchetto, Fábio Sossai Possebon, Leila Sabrina Ullmann, Cíntia Bittar, Guilherme Campos, Helena Lage Ferreira, Jorge A. Petroli Marchesi, Maisa C. Pereira Parra, Marília Moraes, Paula Rahal, Paulo Inacio da Costa, João Pessoa Araújo Jr., Maurício Lacerda Nogueira. Prefeitura de Sao Paulo: Melissa Palmieri. |
| EPI_ISL_1966556, EPI_ISL_1966557, EPI_ISL_1966558, EPI_ISL_1966559, EPI_ISL_1966561 | VIGILANCIA EM SAUDE SAE VIGILANCIA EPIDEMIOLOGICA | Instituto Butantan / Mendelics | Instituto Butantan: Dimas Tadeu Covas, Sandra Coccuzzo Sampaio, Maria Carolina Elias, José Salvatore Leister Patané, Vincent Louis Viala, Antonio Jorge Martins, Ricardo Haddad, Claudia Renata dos Santos Barros, Elaine Cristina Marqueze, Raul Machado Neto, Debora Botequio Moretti, Jardelina de Souza Todao Bernardino, Loyze Paola Oliveira de Lima, Luiz Aurelio de Campos Crispin. Centro de Genômica Funcional da ESALQ: Luiz Lehmann Coutinho, Ricardo Augusto Brassaloti, Raquel de Lello Rocha Campos Cassano. NGS Soluções Genômicas: Pilar Drummond Sampaio Corrêa Mariani. FZEA-USP Pirassununga: Mirele Daiana Poleti, Jessika Cristina Chagas Lesbon, Elisangela Chicaroni Mattos, Heidge Fukumasu. USP-Botucatu: Rejane Maria Tommasini Grotto, Jayme A. Souza-Neto, Guilherme Targino Valente, Patricia Akemi Assato, Felipe Allan da Silva da Costa, Bianca Cechetto Carlos. Mendelics: Bibiana Santos, João Paulo Kitajima, Erika Freitas, David Schlesinger. Hemocentro Ribeirão Preto: Simone Kashima, Evandra Strazza Rodrigues, Svetoslav Nanev Slavov, Elaine Vieira dos Santos, Rafael dos Santos Bezerra, Luiz Carlos Junior de Alcantara, Marta Giovanetti, Vagner Fonseca, Flavia Aburjaile, Rodrigo Tocantins Calado. FAMERP-SJRP: Cecília Artico Banho, Lívia Sacchetto, Fábio Sossai Possebon, Leila Sabrina Ullmann, Cíntia Bittar, Guilherme Campos, Helena Lage Ferreira, Jorge A. Petroli Marchesi, Maisa C. Pereira Parra, Marília Moraes, Paula Rahal, Paulo Inacio da Costa, João Pessoa Araújo Jr., Maurício Lacerda Nogueira. Prefeitura de Sao Paulo: Melissa Palmieri. |
| EPI_ISL_1966562, EPI_ISL_1966563, EPI_ISL_1966564 | NUCLEO DE SAUDE VILA FALCAO DE BAURU | Instituto Butantan / Mendelics | Instituto Butantan: Dimas Tadeu Covas, Sandra Coccuzzo Sampaio, Maria Carolina Elias, José Salvatore Leister Patané, Vincent Louis Viala, Antonio Jorge Martins, Ricardo Haddad, Claudia Renata dos Santos Barros, Elaine Cristina Marqueze, Raul Machado Neto, Debora Botequio Moretti, Jardelina de Souza Todao Bernardino, Loyze Paola Oliveira de Lima, Luiz Aurelio de Campos Crispin. Centro de Genômica Funcional da ESALQ: Luiz Lehmann Coutinho, Ricardo Augusto Brassaloti, Raquel de Lello Rocha Campos Cassano. NGS Soluções Genômicas: Pilar Drummond Sampaio Corrêa Mariani. FZEA-USP Pirassununga: Mirele Daiana Poleti, Jessika Cristina Chagas Lesbon, Elisangela Chicaroni Mattos, Heidge Fukumasu. USP-Botucatu: Rejane Maria Tommasini Grotto, Jayme A. Souza-Neto, Guilherme Targino Valente, Patricia Akemi Assato, Felipe Allan da Silva da Costa, Bianca Cechetto Carlos. Mendelics: Bibiana Santos, João Paulo Kitajima, Erika Freitas, David Schlesinger. Hemocentro Ribeirão Preto: Simone Kashima, Evandra Strazza Rodrigues, Svetoslav Nanev Slavov, Elaine Vieira dos Santos, Rafael dos Santos Bezerra, Luiz Carlos Junior de Alcantara, Marta Giovanetti, Vagner Fonseca, Flavia Aburjaile, Rodrigo Tocantins Calado. FAMERP-SJRP: Cecília Artico Banho, Lívia Sacchetto, Fábio Sossai Possebon, Leila Sabrina Ullmann, Cíntia Bittar, Guilherme Campos, Helena Lage Ferreira, Jorge A. Petroli Marchesi, Maisa C. Pereira Parra, Marília Moraes, Paula Rahal, Paulo Inacio da Costa, João Pessoa Araújo Jr., Maurício Lacerda Nogueira. Prefeitura de Sao Paulo: Melissa Palmieri. |
| EPI_ISL_1966565 | UPA BELA VISTA | Instituto Butantan / Mendelics | Instituto Butantan: Dimas Tadeu Covas, Sandra Coccuzzo Sampaio, Maria Carolina Elias, José Salvatore Leister Patané, Vincent Louis Viala, Antonio Jorge Martins, Ricardo Haddad, Claudia Renata dos Santos Barros, Elaine Cristina Marqueze, Raul Machado Neto, Debora Botequio Moretti, Jardelina de Souza Todao Bernardino, Loyze Paola Oliveira de Lima, Luiz Aurelio de Campos Crispin. Centro de Genômica Funcional da ESALQ: Luiz Lehmann Coutinho, Ricardo Augusto Brassaloti, Raquel de Lello Rocha Campos Cassano. NGS Soluções Genômicas: Pilar Drummond Sampaio Corrêa Mariani. FZEA-USP Pirassununga: Mirele Daiana Poleti, Jessika Cristina Chagas Lesbon, Elisangela Chicaroni Mattos, Heidge Fukumasu. USP-Botucatu: Rejane Maria Tommasini Grotto, Jayme A. Souza-Neto, Guilherme Targino Valente, Patricia Akemi Assato, Felipe Allan da Silva da Costa, Bianca Cechetto Carlos. Mendelics: Bibiana Santos, João Paulo Kitajima, Erika Freitas, David Schlesinger. Hemocentro Ribeirão Preto: Simone Kashima, Evandra Strazza Rodrigues, Svetoslav Nanev Slavov, Elaine Vieira dos Santos, Rafael dos Santos Bezerra, Luiz Carlos Junior de Alcantara, Marta Giovanetti, Vagner Fonseca, Flavia Aburjaile, Rodrigo Tocantins Calado. FAMERP-SJRP: Cecília Artico Banho, Lívia Sacchetto, Fábio Sossai Possebon, Leila Sabrina Ullmann, Cíntia Bittar, Guilherme Campos, Helena Lage Ferreira, Jorge A. Petroli Marchesi, Maisa C. Pereira Parra, Marília Moraes, Paula Rahal, Paulo Inacio da Costa, João Pessoa Araújo Jr., Maurício Lacerda Nogueira. Prefeitura de Sao Paulo: Melissa Palmieri. |
| EPI_ISL_1966566, EPI_ISL_1966567, EPI_ISL_1966568 | NUCLEO DE SAUDE VILA FALCAO DE BAURU | Instituto Butantan / Mendelics | Instituto Butantan: Dimas Tadeu Covas, Sandra Coccuzzo Sampaio, Maria Carolina Elias, José Salvatore Leister Patané, Vincent Louis Viala, Antonio Jorge Martins, Ricardo Haddad, Claudia Renata dos Santos Barros, Elaine Cristina Marqueze, Raul Machado Neto, Debora Botequio Moretti, Jardelina de Souza Todao Bernardino, Loyze Paola Oliveira de Lima, Luiz Aurelio de Campos Crispin. Centro de Genômica Funcional da ESALQ: Luiz Lehmann Coutinho, Ricardo Augusto Brassaloti, Raquel de Lello Rocha Campos Cassano. NGS Soluções Genômicas: Pilar Drummond Sampaio Corrêa Mariani. FZEA-USP Pirassununga: Mirele Daiana Poleti, Jessika Cristina Chagas Lesbon, Elisangela Chicaroni Mattos, Heidge Fukumasu. USP-Botucatu: Rejane Maria Tommasini Grotto, Jayme A. Souza-Neto, Guilherme Targino Valente, Patricia Akemi Assato, Felipe Allan da Silva da Costa, Bianca Cechetto Carlos. Mendelics: Bibiana Santos, João Paulo Kitajima, Erika Freitas, David Schlesinger. Hemocentro Ribeirão Preto: Simone Kashima, Evandra Strazza Rodrigues, Svetoslav Nanev Slavov, Elaine Vieira dos Santos, Rafael dos Santos Bezerra, Luiz Carlos Junior de Alcantara, Marta Giovanetti, Vagner Fonseca, Flavia Aburjaile, Rodrigo Tocantins Calado. FAMERP-SJRP: Cecília Artico Banho, Lívia Sacchetto, Fábio Sossai Possebon, Leila Sabrina Ullmann, Cíntia Bittar, Guilherme Campos, Helena Lage Ferreira, Jorge A. Petroli Marchesi, Maisa C. Pereira Parra, Marília Moraes, Paula Rahal, Paulo Inacio da Costa, João Pessoa Araújo Jr., Maurício Lacerda Nogueira. Prefeitura de Sao Paulo: Melissa Palmieri. |
| EPI_ISL_1966569 | UPA BELA VISTA | Instituto Butantan / Mendelics | Instituto Butantan: Dimas Tadeu Covas, Sandra Coccuzzo Sampaio, Maria Carolina Elias, José Salvatore Leister Patané, Vincent Louis Viala, Antonio Jorge Martins, Ricardo Haddad, Claudia Renata dos Santos Barros, Elaine Cristina Marqueze, Raul Machado Neto, Debora Botequio Moretti, Jardelina de Souza Todao Bernardino, Loyze Paola Oliveira de Lima, Luiz Aurelio de Campos Crispin. Centro de Genômica Funcional da ESALQ: Luiz Lehmann Coutinho, Ricardo Augusto Brassaloti, Raquel de Lello Rocha Campos Cassano. NGS Soluções Genômicas: Pilar Drummond Sampaio Corrêa Mariani. FZEA-USP Pirassununga: Mirele Daiana Poleti, Jessika Cristina Chagas Lesbon, Elisangela Chicaroni Mattos, Heidge Fukumasu. USP-Botucatu: Rejane Maria Tommasini Grotto, Jayme A. Souza-Neto, Guilherme Targino Valente, Patricia Akemi Assato, Felipe Allan da Silva da Costa, Bianca Cechetto Carlos. Mendelics: Bibiana Santos, João Paulo Kitajima, Erika Freitas, David Schlesinger. Hemocentro Ribeirão Preto: Simone Kashima, Evandra Strazza Rodrigues, Svetoslav Nanev Slavov, Elaine Vieira dos Santos, Rafael dos Santos Bezerra, Luiz Carlos Junior de Alcantara, Marta Giovanetti, Vagner Fonseca, Flavia Aburjaile, Rodrigo Tocantins Calado. FAMERP-SJRP: Cecília Artico Banho, Lívia Sacchetto, Fábio Sossai Possebon, Leila Sabrina Ullmann, Cíntia Bittar, Guilherme Campos, Helena Lage Ferreira, Jorge A. Petroli Marchesi, Maisa C. Pereira Parra, Marília Moraes, Paula Rahal, Paulo Inacio da Costa, João Pessoa Araújo Jr., Maurício Lacerda Nogueira. Prefeitura de Sao Paulo: Melissa Palmieri. |
| EPI_ISL_1966570, EPI_ISL_1966571, EPI_ISL_1966572 | NUCLEO DE SAUDE VILA FALCAO DE BAURU | Instituto Butantan / Mendelics | Instituto Butantan: Dimas Tadeu Covas, Sandra Coccuzzo Sampaio, Maria Carolina Elias, José Salvatore Leister Patané, Vincent Louis Viala, Antonio Jorge Martins, Ricardo Haddad, Claudia Renata dos Santos Barros, Elaine Cristina Marqueze, Raul Machado Neto, Debora Botequio Moretti, Jardelina de Souza Todao Bernardino, Loyze Paola Oliveira de Lima, Luiz Aurelio de Campos Crispin. Centro de Genômica Funcional da ESALQ: Luiz Lehmann Coutinho, Ricardo Augusto Brassaloti, Raquel de Lello Rocha Campos Cassano. NGS Soluções Genômicas: Pilar Drummond Sampaio Corrêa Mariani. FZEA-USP Pirassununga: Mirele Daiana Poleti, Jessika Cristina Chagas Lesbon, Elisangela Chicaroni Mattos, Heidge Fukumasu. USP-Botucatu: Rejane Maria Tommasini Grotto, Jayme A. Souza-Neto, Guilherme Targino Valente, Patricia Akemi Assato, Felipe Allan da Silva da Costa, Bianca Cechetto Carlos. Mendelics: Bibiana Santos, João Paulo Kitajima, Erika Freitas, David Schlesinger. Hemocentro Ribeirão Preto: Simone Kashima, Evandra Strazza Rodrigues, Svetoslav Nanev Slavov, Elaine Vieira dos Santos, Rafael dos Santos Bezerra, Luiz Carlos Junior de Alcantara, Marta Giovanetti, Vagner Fonseca, Flavia Aburjaile, Rodrigo Tocantins Calado. FAMERP-SJRP: Cecília Artico Banho, Lívia Sacchetto, Fábio Sossai Possebon, Leila Sabrina |

| | | | |
|---|---|---|---|
| | | | Ullmann, Cíntia Bittar, Guilherme Campos, Helena Lage Ferreira, Jorge A. Petroli Marchesi, Maisa C. Pereira Parra, Marília Moraes, Paula Rahal, Paulo Inacio da Costa, João Pessoa Araújo Jr., Maurício Lacerda Nogueira. Prefeitura de Sao Paulo: Melissa Palmieri. |
| EPI_ISL_1966573 | UPA BELA VISTA | Instituto Butantan / Mendelics | Instituto Butantan: Dimas Tadeu Covas, Sandra Coccuzzo Sampaio, Maria Carolina Elias, José Salvatore Leister Patané, Vincent Louis Viala, Antonio Jorge Martins, Ricardo Haddad, Claudia Renata dos Santos Barros, Elaine Cristina Marqueze, Raul Machado Neto, Debora Botequio Moretti, Jardelina de Souza Todao Bernardino, Loyze Paola Oliveira de Lima, Luiz Aurelio de Campos Crispin. Centro de Genômica Funcional da ESALQ: Luiz Lehmann Coutinho, Ricardo Augusto Brassaloti, Raquel de Lello Rocha Campos Cassano. NGS Soluções Genômicas: Pilar Drummond Sampaio Corrêa Mariani. FZEA-USP Pirassununga: Mirele Daiana Poleti, Jessika Cristina Chagas Lesbon, Elisangela Chicaroni Mattos, Heidge Fukumasu. USP-Botucatu: Rejane Maria Tommasini Grotto, Jayme A. Souza-Neto, Guilherme Targino Valente, Patricia Akemi Assato, Felipe Allan da Silva da Costa, Bianca Cechetto Carlos. Mendelics: Bibiana Santos, João Paulo Kitajima, Erika Freitas, David Schlesinger. Hemocentro Ribeirão Preto: Simone Kashima, Evandra Strazza Rodrigues, Svetoslav Nanev Slavov, Elaine Vieira dos Santos, Rafael dos Santos Bezerra, Luiz Carlos Junior de Alcantara, Marta Giovanetti, Vagner Fonseca, Flavia Aburjaile, Rodrigo Tocantins Calado. FAMERP-SJRP: Cecília Artico Banho, Lívia Sacchetto, Fábio Sossai Possebon, Leila Sabrina Ullmann, Cíntia Bittar, Guilherme Campos, Helena Lage Ferreira, Jorge A. Petroli Marchesi, Maisa C. Pereira Parra, Marília Moraes, Paula Rahal, Paulo Inacio da Costa, João Pessoa Araújo Jr., Maurício Lacerda Nogueira. Prefeitura de Sao Paulo: Melissa Palmieri. |
| EPI_ISL_1966574, EPI_ISL_1966575, EPI_ISL_1966576, EPI_ISL_1966577, EPI_ISL_1966578, EPI_ISL_1966579, EPI_ISL_1966580, EPI_ISL_1966581, EPI_ISL_1966582, EPI_ISL_1966583, EPI_ISL_1966584, EPI_ISL_1966585, EPI_ISL_1966586, EPI_ISL_1966587, EPI_ISL_1966588, EPI_ISL_1966589, EPI_ISL_1966590, EPI_ISL_1966591, EPI_ISL_1966592 | | | |
| see above | NUCLEO DE SAUDE VILA FALCAO DE BAURU | Instituto Butantan / Mendelics | Instituto Butantan: Dimas Tadeu Covas, Sandra Coccuzzo Sampaio, Maria Carolina Elias, José Salvatore Leister Patané, Vincent Louis Viala, Antonio Jorge Martins, Ricardo Haddad, Claudia Renata dos Santos Barros, Elaine Cristina Marqueze, Raul Machado Neto, Debora Botequio Moretti, Jardelina de Souza Todao Bernardino, Loyze Paola Oliveira de Lima, Luiz Aurelio de Campos Crispin. Centro de Genômica Funcional da ESALQ: Luiz Lehmann Coutinho, Ricardo Augusto Brassaloti, Raquel de Lello Rocha Campos Cassano. NGS Soluções Genômicas: Pilar Drummond Sampaio Corrêa Mariani. FZEA-USP Pirassununga: Mirele Daiana Poleti, Jessika Cristina Chagas Lesbon, Elisangela Chicaroni Mattos, Heidge Fukumasu. USP-Botucatu: Rejane Maria Tommasini Grotto, Jayme A. Souza-Neto, Guilherme Targino Valente, Patricia Akemi Assato, Felipe Allan da Silva da Costa, Bianca Cechetto Carlos. Mendelics: Bibiana Santos, João Paulo Kitajima, Erika Freitas, David Schlesinger. Hemocentro Ribeirão Preto: Simone Kashima, Evandra Strazza Rodrigues, Svetoslav Nanev Slavov, Elaine Vieira dos Santos, Rafael dos Santos Bezerra, Luiz Carlos Junior de Alcantara, Marta Giovanetti, Vagner Fonseca, Flavia Aburjaile, Rodrigo Tocantins Calado. FAMERP-SJRP: Cecília Artico Banho, Lívia Sacchetto, Fábio Sossai Possebon, Leila Sabrina Ullmann, Cíntia Bittar, Guilherme Campos, Helena Lage Ferreira, Jorge A. Petroli Marchesi, Maisa C. Pereira Parra, Marília Moraes, Paula Rahal, Paulo Inacio da Costa, João Pessoa Araújo Jr., Maurício Lacerda Nogueira. Prefeitura de Sao Paulo: Melissa Palmieri. |
| EPI_ISL_1966593, EPI_ISL_1966594, EPI_ISL_1966595, EPI_ISL_1966596, EPI_ISL_1966597, EPI_ISL_1966598, EPI_ISL_1966599, EPI_ISL_1966600, EPI_ISL_1966601 | UPA DE BEBEDOURO | Instituto Butantan / Mendelics | Instituto Butantan: Dimas Tadeu Covas, Sandra Coccuzzo Sampaio, Maria Carolina Elias, José Salvatore Leister Patané, Vincent Louis Viala, Antonio Jorge Martins, Ricardo Haddad, Claudia Renata dos Santos Barros, Elaine Cristina Marqueze, Raul Machado Neto, Debora Botequio Moretti, Jardelina de Souza Todao Bernardino, Loyze Paola Oliveira de Lima, Luiz Aurelio de Campos Crispin. Centro de Genômica Funcional da ESALQ: Luiz Lehmann Coutinho, Ricardo Augusto Brassaloti, Raquel de Lello Rocha Campos Cassano. NGS Soluções Genômicas: Pilar Drummond Sampaio Corrêa Mariani. FZEA-USP Pirassununga: Mirele Daiana Poleti, Jessika Cristina Chagas Lesbon, Elisangela Chicaroni Mattos, Heidge Fukumasu. USP-Botucatu: Rejane Maria Tommasini Grotto, Jayme A. Souza-Neto, Guilherme Targino Valente, Patricia Akemi Assato, Felipe Allan da Silva da Costa, Bianca Cechetto Carlos. Mendelics: Bibiana Santos, João Paulo Kitajima, Erika Freitas, David Schlesinger. Hemocentro Ribeirão Preto: Simone Kashima, Evandra Strazza Rodrigues, Svetoslav Nanev Slavov, Elaine Vieira dos Santos, Rafael dos Santos Bezerra, Luiz Carlos Junior de Alcantara, Marta Giovanetti, Vagner Fonseca, Flavia Aburjaile, Rodrigo Tocantins Calado. FAMERP-SJRP: Cecília Artico Banho, Lívia Sacchetto, Fábio Sossai Possebon, Leila Sabrina Ullmann, Cíntia Bittar, Guilherme Campos, Helena Lage Ferreira, Jorge A. Petroli Marchesi, Maisa C. Pereira Parra, Marília Moraes, Paula Rahal, Paulo Inacio da Costa, João Pessoa Araújo Jr., Maurício Lacerda Nogueira. Prefeitura de Sao Paulo: Melissa Palmieri. |
| EPI_ISL_1966602, EPI_ISL_1966603, EPI_ISL_1966604, EPI_ISL_1966605, EPI_ISL_1966606 | UNIDADE RESPIRATORIA NOVA HORTOLANDIA | Instituto Butantan / Mendelics | Instituto Butantan: Dimas Tadeu Covas, Sandra Coccuzzo Sampaio, Maria Carolina Elias, José Salvatore Leister Patané, Vincent Louis Viala, Antonio Jorge Martins, Ricardo Haddad, Claudia Renata dos Santos Barros, Elaine Cristina Marqueze, Raul Machado Neto, Debora Botequio Moretti, Jardelina de Souza Todao Bernardino, Loyze Paola Oliveira de Lima, Luiz Aurelio de Campos Crispin. Centro de Genômica Funcional da ESALQ: Luiz Lehmann Coutinho, Ricardo Augusto Brassaloti, Raquel de Lello Rocha Campos Cassano. NGS Soluções Genômicas: Pilar Drummond Sampaio Corrêa Mariani. FZEA-USP Pirassununga: Mirele Daiana Poleti, Jessika Cristina Chagas Lesbon, Elisangela Chicaroni Mattos, Heidge Fukumasu. USP-Botucatu: Rejane Maria Tommasini Grotto, Jayme A. Souza-Neto, Guilherme Targino Valente, Patricia Akemi Assato, Felipe Allan da Silva da Costa, Bianca Cechetto Carlos. Mendelics: Bibiana Santos, João Paulo Kitajima, Erika Freitas, David Schlesinger. Hemocentro Ribeirão Preto: Simone Kashima, Evandra Strazza Rodrigues, Svetoslav Nanev Slavov, Elaine Vieira dos Santos, Rafael dos Santos Bezerra, Luiz Carlos Junior de Alcantara, Marta Giovanetti, Vagner Fonseca, Flavia Aburjaile, Rodrigo Tocantins Calado. FAMERP-SJRP: Cecília Artico Banho, Lívia Sacchetto, Fábio Sossai Possebon, Leila Sabrina Ullmann, Cíntia Bittar, Guilherme Campos, Helena Lage Ferreira, Jorge A. Petroli Marchesi, Maisa C. Pereira Parra, Marília Moraes, Paula Rahal, Paulo Inacio da Costa, João Pessoa Araújo Jr., Maurício Lacerda Nogueira. Prefeitura de Sao Paulo: Melissa Palmieri. |
| EPI_ISL_1966607 | VIGILANCIA EPIDEMIOLOGICA DE IBATE | Instituto Butantan / Mendelics | Instituto Butantan: Dimas Tadeu Covas, Sandra Coccuzzo Sampaio, Maria Carolina Elias, José Salvatore Leister Patané, Vincent Louis Viala, Antonio Jorge Martins, Ricardo Haddad, Claudia Renata dos Santos Barros, Elaine Cristina Marqueze, Raul Machado Neto, Debora Botequio Moretti, Jardelina de Souza Todao Bernardino, Loyze Paola Oliveira de Lima, Luiz Aurelio de Campos Crispin. Centro de Genômica Funcional da ESALQ: Luiz Lehmann Coutinho, Ricardo Augusto Brassaloti, Raquel de Lello Rocha Campos Cassano. NGS Soluções Genômicas: Pilar Drummond Sampaio Corrêa Mariani. FZEA-USP Pirassununga: Mirele Daiana Poleti, Jessika Cristina Chagas Lesbon, Elisangela Chicaroni Mattos, Heidge Fukumasu. USP-Botucatu: Rejane Maria Tommasini Grotto, Jayme A. Souza-Neto, Guilherme Targino Valente, Patricia Akemi Assato, Felipe Allan da Silva da Costa, Bianca Cechetto Carlos. Mendelics: Bibiana Santos, João Paulo Kitajima, Erika Freitas, David Schlesinger. Hemocentro Ribeirão Preto: Simone Kashima, Evandra Strazza Rodrigues, Svetoslav Nanev Slavov, Elaine Vieira dos Santos, Rafael dos Santos Bezerra, Luiz Carlos Junior de Alcantara, Marta Giovanetti, Vagner Fonseca, Flavia Aburjaile, Rodrigo Tocantins Calado. FAMERP-SJRP: Cecília Artico Banho, Lívia Sacchetto, Fábio Sossai Possebon, Leila Sabrina Ullmann, Cíntia Bittar, Guilherme Campos, Helena Lage Ferreira, Jorge A. Petroli Marchesi, Maisa C. Pereira Parra, Marília Moraes, Paula Rahal, Paulo Inacio da Costa, João Pessoa Araújo Jr., Maurício Lacerda Nogueira. Prefeitura de Sao Paulo: Melissa Palmieri. |
| EPI_ISL_1966608, EPI_ISL_1966609, EPI_ISL_1966610, EPI_ISL_1966611 | LABORATORIO MUNICIPAL DE ANALISES CLINICAS DE ITANHAEM | Instituto Butantan / Mendelics | Instituto Butantan: Dimas Tadeu Covas, Sandra Coccuzzo Sampaio, Maria Carolina Elias, José Salvatore Leister Patané, Vincent Louis Viala, Antonio Jorge Martins, Ricardo Haddad, Claudia Renata dos Santos Barros, Elaine Cristina Marqueze, Raul Machado Neto, Debora Botequio Moretti, Jardelina de Souza Todao Bernardino, Loyze Paola Oliveira de Lima, Luiz Aurelio de Campos Crispin. Centro de Genômica Funcional da ESALQ: Luiz Lehmann Coutinho, Ricardo Augusto Brassaloti, Raquel de Lello Rocha Campos Cassano. NGS Soluções Genômicas: Pilar Drummond Sampaio Corrêa Mariani. FZEA-USP Pirassununga: Mirele Daiana Poleti, Jessika Cristina Chagas Lesbon, Elisangela Chicaroni Mattos, Heidge Fukumasu. USP-Botucatu: Rejane Maria Tommasini Grotto, Jayme A. Souza-Neto, Guilherme Targino Valente, Patricia Akemi Assato, Felipe Allan da Silva da Costa, Bianca Cechetto Carlos. Mendelics: Bibiana Santos, João Paulo Kitajima, Erika Freitas, David Schlesinger. Hemocentro Ribeirão Preto: Simone Kashima, Evandra Strazza Rodrigues, Svetoslav Nanev Slavov, Elaine Vieira dos Santos, Rafael dos Santos Bezerra, Luiz Carlos Junior de Alcantara, Marta Giovanetti, Vagner Fonseca, Flavia Aburjaile, Rodrigo Tocantins Calado. FAMERP-SJRP: Cecília Artico Banho, Lívia Sacchetto, Fábio Sossai Possebon, Leila Sabrina Ullmann, Cíntia Bittar, Guilherme Campos, Helena Lage Ferreira, Jorge A. Petroli Marchesi, Maisa C. Pereira Parra, Marília Moraes, Paula Rahal, Paulo Inacio da Costa, João Pessoa Araújo Jr., Maurício Lacerda Nogueira. Prefeitura de Sao Paulo: Melissa Palmieri. |
| EPI_ISL_1966612, EPI_ISL_1966613, EPI_ISL_1966614, EPI_ISL_1966615 | UNIDADE DE PRONTO ATENDIMENTO UPA DRA ANA OLIVIA BENTIVOGLIO | Instituto Butantan / Mendelics | Instituto Butantan: Dimas Tadeu Covas, Sandra Coccuzzo Sampaio, Maria Carolina Elias, José Salvatore Leister Patané, Vincent Louis Viala, Antonio Jorge Martins, Ricardo Haddad, Claudia Renata dos Santos Barros, Elaine Cristina Marqueze, Raul Machado Neto, Debora Botequio Moretti, Jardelina de Souza Todao Bernardino, Loyze Paola Oliveira de Lima, Luiz Aurelio de Campos Crispin. Centro de Genômica Funcional da ESALQ: Luiz Lehmann Coutinho, Ricardo Augusto Brassaloti, Raquel de Lello Rocha Campos Cassano. NGS Soluções Genômicas: Pilar Drummond Sampaio Corrêa Mariani. FZEA-USP Pirassununga: Mirele Daiana Poleti, Jessika Cristina Chagas Lesbon, Elisangela Chicaroni Mattos, Heidge Fukumasu. USP-Botucatu: Rejane Maria Tommasini Grotto, Jayme A. Souza-Neto, Guilherme Targino Valente, Patricia Akemi Assato, Felipe Allan da Silva da Costa, Bianca Cechetto Carlos. Mendelics: Bibiana Santos, João Paulo Kitajima, Erika Freitas, David Schlesinger. Hemocentro Ribeirão Preto: Simone Kashima, Evandra Strazza Rodrigues, Svetoslav Nanev Slavov, Elaine Vieira dos Santos, Rafael dos Santos Bezerra, Luiz Carlos Junior de Alcantara, Marta Giovanetti, Vagner Fonseca, Flavia Aburjaile, Rodrigo Tocantins Calado. FAMERP-SJRP: Cecília Artico Banho, Lívia Sacchetto, Fábio Sossai Possebon, Leila Sabrina Ullmann, Cíntia Bittar, Guilherme Campos, Helena Lage Ferreira, Jorge A. Petroli Marchesi, Maisa C. Pereira Parra, Marília Moraes, Paula Rahal, Paulo Inacio da Costa, João Pessoa Araújo Jr., Maurício Lacerda Nogueira. Prefeitura de Sao Paulo: Melissa Palmieri. |
| EPI_ISL_1966616, EPI_ISL_1966617, EPI_ISL_1966618, EPI_ISL_1966619, | POLICLINICA HORTOLANDIA | Instituto Butantan / Mendelics | Instituto Butantan: Dimas Tadeu Covas, Sandra Coccuzzo Sampaio, Maria Carolina Elias, José Salvatore Leister Patané, Vincent Louis Viala, Antonio Jorge Martins, Ricardo Haddad, Claudia Renata dos Santos Barros, Elaine Cristina Marqueze, Raul Machado Neto, Debora Botequio Moretti, Jardelina de |

| | | | |
|---|---|---|---|
| EPI_ISL_1966620, EPI_ISL_1966621, EPI_ISL_1966622, EPI_ISL_1966623, EPI_ISL_1966624, EPI_ISL_1966625 | | | Souza Todao Bernardino, Loyze Paola Oliveira de Lima, Luiz Aurelio de Campos Crispin. Centro de Genômica Funcional da ESALQ: Luiz Lehmann Coutinho, Ricardo Augusto Brassaloti, Raquel de Lello Rocha Campos Cassano. NGS Soluções Genômicas: Pilar Drummond Sampaio Corrêa Mariani. FZEA-USP Pirassununga: Mirele Daiana Poleti, Jessika Cristina Chagas Lesbon, Elisangela Chicaroni Mattos, Heidge Fukumasu. USP-Botucatu: Rejane Maria Tommasini Grotto, Jayme A. Souza-Neto, Guilherme Targino Valente, Patricia Akemi Assato, Felipe Allan da Silva da Costa, Bianca Cechetto Carlos. Mendelics: Bibiana Santos, João Paulo Kitajima, Erika Freitas, David Schlesinger. Hemocentro Ribeirão Preto: Simone Kashima, Evandra Strazza Rodrigues, Svetoslav Nanev Slavov, Elaine Vieira dos Santos, Rafael dos Santos Bezerra, Luiz Carlos Junior de Alcantara, Marta Giovanetti, Vagner Fonseca, Flavia Aburjaile, Rodrigo Tocantins Calado. FAMERP-SJRP: Cecília Artico Banho, Lívia Sacchetto, Fábio Sossai Possebon, Leila Sabrina Ullmann, Cíntia Bittar, Guilherme Campos, Helena Lage Ferreira, Jorge A. Petroli Marchesi, Maisa C. Pereira Parra, Marília Moraes, Paula Rahal, Paulo Inacio da Costa, João Pessoa Araújo Jr., Maurício Lacerda Nogueira. Prefeitura de Sao Paulo: Melissa Palmieri. |
| EPI_ISL_1966626 | USF MARGARIDA JANETE FERRARI GANZAROLI | Instituto Butantan / Mendelics | Instituto Butantan: Dimas Tadeu Covas, Sandra Coccuzzo Sampaio, Maria Carolina Elias, José Salvatore Leister Patané, Vincent Louis Viala, Antonio Jorge Martins, Ricardo Haddad, Claudia Renata dos Santos Barros, Elaine Cristina Marqueze, Raul Machado Neto, Debora Botequio Moretti, Jardelina de Souza Todao Bernardino, Loyze Paola Oliveira de Lima, Luiz Aurelio de Campos Crispin. Centro de Genômica Funcional da ESALQ: Luiz Lehmann Coutinho, Ricardo Augusto Brassaloti, Raquel de Lello Rocha Campos Cassano. NGS Soluções Genômicas: Pilar Drummond Sampaio Corrêa Mariani. FZEA-USP Pirassununga: Mirele Daiana Poleti, Jessika Cristina Chagas Lesbon, Elisangela Chicaroni Mattos, Heidge Fukumasu. USP-Botucatu: Rejane Maria Tommasini Grotto, Jayme A. Souza-Neto, Guilherme Targino Valente, Patricia Akemi Assato, Felipe Allan da Silva da Costa, Bianca Cechetto Carlos. Mendelics: Bibiana Santos, João Paulo Kitajima, Erika Freitas, David Schlesinger. Hemocentro Ribeirão Preto: Simone Kashima, Evandra Strazza Rodrigues, Svetoslav Nanev Slavov, Elaine Vieira dos Santos, Rafael dos Santos Bezerra, Luiz Carlos Junior de Alcantara, Marta Giovanetti, Vagner Fonseca, Flavia Aburjaile, Rodrigo Tocantins Calado. FAMERP-SJRP: Cecília Artico Banho, Lívia Sacchetto, Fábio Sossai Possebon, Leila Sabrina Ullmann, Cíntia Bittar, Guilherme Campos, Helena Lage Ferreira, Jorge A. Petroli Marchesi, Maisa C. Pereira Parra, Marília Moraes, Paula Rahal, Paulo Inacio da Costa, João Pessoa Araújo Jr., Maurício Lacerda Nogueira. Prefeitura de Sao Paulo: Melissa Palmieri. |
| EPI_ISL_1966627, EPI_ISL_1966629, EPI_ISL_1966630, EPI_ISL_1966631, EPI_ISL_1966632, EPI_ISL_1966633 | SECAO CENTRO DE DIAGNOSTICO SECEDI | Instituto Butantan / Mendelics | Instituto Butantan: Dimas Tadeu Covas, Sandra Coccuzzo Sampaio, Maria Carolina Elias, José Salvatore Leister Patané, Vincent Louis Viala, Antonio Jorge Martins, Ricardo Haddad, Claudia Renata dos Santos Barros, Elaine Cristina Marqueze, Raul Machado Neto, Debora Botequio Moretti, Jardelina de Souza Todao Bernardino, Loyze Paola Oliveira de Lima, Luiz Aurelio de Campos Crispin. Centro de Genômica Funcional da ESALQ: Luiz Lehmann Coutinho, Ricardo Augusto Brassaloti, Raquel de Lello Rocha Campos Cassano. NGS Soluções Genômicas: Pilar Drummond Sampaio Corrêa Mariani. FZEA-USP Pirassununga: Mirele Daiana Poleti, Jessika Cristina Chagas Lesbon, Elisangela Chicaroni Mattos, Heidge Fukumasu. USP-Botucatu: Rejane Maria Tommasini Grotto, Jayme A. Souza-Neto, Guilherme Targino Valente, Patricia Akemi Assato, Felipe Allan da Silva da Costa, Bianca Cechetto Carlos. Mendelics: Bibiana Santos, João Paulo Kitajima, Erika Freitas, David Schlesinger. Hemocentro Ribeirão Preto: Simone Kashima, Evandra Strazza Rodrigues, Svetoslav Nanev Slavov, Elaine Vieira dos Santos, Rafael dos Santos Bezerra, Luiz Carlos Junior de Alcantara, Marta Giovanetti, Vagner Fonseca, Flavia Aburjaile, Rodrigo Tocantins Calado. FAMERP-SJRP: Cecília Artico Banho, Lívia Sacchetto, Fábio Sossai Possebon, Leila Sabrina Ullmann, Cíntia Bittar, Guilherme Campos, Helena Lage Ferreira, Jorge A. Petroli Marchesi, Maisa C. Pereira Parra, Marília Moraes, Paula Rahal, Paulo Inacio da Costa, João Pessoa Araújo Jr., Maurício Lacerda Nogueira. Prefeitura de Sao Paulo: Melissa Palmieri. |
| EPI_ISL_1966634, EPI_ISL_1966635, EPI_ISL_1966636, EPI_ISL_1966637, EPI_ISL_1966638, EPI_ISL_1966639 | CENTRO DE SAUDE III TABATINGA | Instituto Butantan / Mendelics | Instituto Butantan: Dimas Tadeu Covas, Sandra Coccuzzo Sampaio, Maria Carolina Elias, José Salvatore Leister Patané, Vincent Louis Viala, Antonio Jorge Martins, Ricardo Haddad, Claudia Renata dos Santos Barros, Elaine Cristina Marqueze, Raul Machado Neto, Debora Botequio Moretti, Jardelina de Souza Todao Bernardino, Loyze Paola Oliveira de Lima, Luiz Aurelio de Campos Crispin. Centro de Genômica Funcional da ESALQ: Luiz Lehmann Coutinho, Ricardo Augusto Brassaloti, Raquel de Lello Rocha Campos Cassano. NGS Soluções Genômicas: Pilar Drummond Sampaio Corrêa Mariani. FZEA-USP Pirassununga: Mirele Daiana Poleti, Jessika Cristina Chagas Lesbon, Elisangela Chicaroni Mattos, Heidge Fukumasu. USP-Botucatu: Rejane Maria Tommasini Grotto, Jayme A. Souza-Neto, Guilherme Targino Valente, Patricia Akemi Assato, Felipe Allan da Silva da Costa, Bianca Cechetto Carlos. Mendelics: Bibiana Santos, João Paulo Kitajima, Erika Freitas, David Schlesinger. Hemocentro Ribeirão Preto: Simone Kashima, Evandra Strazza Rodrigues, Svetoslav Nanev Slavov, Elaine Vieira dos Santos, Rafael dos Santos Bezerra, Luiz Carlos Junior de Alcantara, Marta Giovanetti, Vagner Fonseca, Flavia Aburjaile, Rodrigo Tocantins Calado. FAMERP-SJRP: Cecília Artico Banho, Lívia Sacchetto, Fábio Sossai Possebon, Leila Sabrina Ullmann, Cíntia Bittar, Guilherme Campos, Helena Lage Ferreira, Jorge A. Petroli Marchesi, Maisa C. Pereira Parra, Marília Moraes, Paula Rahal, Paulo Inacio da Costa, João Pessoa Araújo Jr., Maurício Lacerda Nogueira. Prefeitura de Sao Paulo: Melissa Palmieri. |
| EPI_ISL_1966640, EPI_ISL_1966641, EPI_ISL_1966642, EPI_ISL_1966643 | HOSPITAL MUNICIPAL SANTO ANTONIO | Instituto Butantan / Mendelics | Instituto Butantan: Dimas Tadeu Covas, Sandra Coccuzzo Sampaio, Maria Carolina Elias, José Salvatore Leister Patané, Vincent Louis Viala, Antonio Jorge Martins, Ricardo Haddad, Claudia Renata dos Santos Barros, Elaine Cristina Marqueze, Raul Machado Neto, Debora Botequio Moretti, Jardelina de Souza Todao Bernardino, Loyze Paola Oliveira de Lima, Luiz Aurelio de Campos Crispin. Centro de Genômica Funcional da ESALQ: Luiz Lehmann Coutinho, Ricardo Augusto Brassaloti, Raquel de Lello Rocha Campos Cassano. NGS Soluções Genômicas: Pilar Drummond Sampaio Corrêa Mariani. FZEA-USP Pirassununga: Mirele Daiana Poleti, Jessika Cristina Chagas Lesbon, Elisangela Chicaroni Mattos, Heidge Fukumasu. USP-Botucatu: Rejane Maria Tommasini Grotto, Jayme A. Souza-Neto, Guilherme Targino Valente, Patricia Akemi Assato, Felipe Allan da Silva da Costa, Bianca Cechetto Carlos. Mendelics: Bibiana Santos, João Paulo Kitajima, Erika Freitas, David Schlesinger. Hemocentro Ribeirão Preto: Simone Kashima, Evandra Strazza Rodrigues, Svetoslav Nanev Slavov, Elaine Vieira dos Santos, Rafael dos Santos Bezerra, Luiz Carlos Junior de Alcantara, Marta Giovanetti, Vagner Fonseca, Flavia Aburjaile, Rodrigo Tocantins Calado. FAMERP-SJRP: Cecília Artico Banho, Lívia Sacchetto, Fábio Sossai Possebon, Leila Sabrina Ullmann, Cíntia Bittar, Guilherme Campos, Helena Lage Ferreira, Jorge A. Petroli Marchesi, Maisa C. Pereira Parra, Marília Moraes, Paula Rahal, Paulo Inacio da Costa, João Pessoa Araújo Jr., Maurício Lacerda Nogueira. Prefeitura de Sao Paulo: Melissa Palmieri. |
| EPI_ISL_1966644, EPI_ISL_1966645 | HOSPITAL MUNICIPAL DR WALDEMAR TEBALDI | Instituto Butantan / Mendelics | Instituto Butantan: Dimas Tadeu Covas, Sandra Coccuzzo Sampaio, Maria Carolina Elias, José Salvatore Leister Patané, Vincent Louis Viala, Antonio Jorge Martins, Ricardo Haddad, Claudia Renata dos Santos Barros, Elaine Cristina Marqueze, Raul Machado Neto, Debora Botequio Moretti, Jardelina de Souza Todao Bernardino, Loyze Paola Oliveira de Lima, Luiz Aurelio de Campos Crispin. Centro de Genômica Funcional da ESALQ: Luiz Lehmann Coutinho, Ricardo Augusto Brassaloti, Raquel de Lello Rocha Campos Cassano. NGS Soluções Genômicas: Pilar Drummond Sampaio Corrêa Mariani. FZEA-USP Pirassununga: Mirele Daiana Poleti, Jessika Cristina Chagas Lesbon, Elisangela Chicaroni Mattos, Heidge Fukumasu. USP-Botucatu: Rejane Maria Tommasini Grotto, Jayme A. Souza-Neto, Guilherme Targino Valente, Patricia Akemi Assato, Felipe Allan da Silva da Costa, Bianca Cechetto Carlos. Mendelics: Bibiana Santos, João Paulo Kitajima, Erika Freitas, David Schlesinger. Hemocentro Ribeirão Preto: Simone Kashima, Evandra Strazza Rodrigues, Svetoslav Nanev Slavov, Elaine Vieira dos Santos, Rafael dos Santos Bezerra, Luiz Carlos Junior de Alcantara, Marta Giovanetti, Vagner Fonseca, Flavia Aburjaile, Rodrigo Tocantins Calado. FAMERP-SJRP: Cecília Artico Banho, Lívia Sacchetto, Fábio Sossai Possebon, Leila Sabrina Ullmann, Cíntia Bittar, Guilherme Campos, Helena Lage Ferreira, Jorge A. Petroli Marchesi, Maisa C. Pereira Parra, Marília Moraes, Paula Rahal, Paulo Inacio da Costa, João Pessoa Araújo Jr., Maurício Lacerda Nogueira. Prefeitura de Sao Paulo: Melissa Palmieri. |
| EPI_ISL_1966646 | HOSPITAL MUNICIPAL DR WALDEMAR TEBALDI | Instituto Butantan / Mendelics | Dimas Tadeu Covas, Antonio Jorge Martins, Claudia Renata dos Santos Barros, David Schlesinger, Debora Botequio Moretti, Elaine Cristina Marqueze, Elaine Vieira Santos, Evandra Strazza Rodrigues, Heidge Fukumasu, Jayme Augusto de Souza-Neto, José Salvatore Leister Patané, Luiz Alcantara, Luiz Lehmann Coutinho, Maria Carolina Elias, Maurício Lacerda Nogueira, Rafael dos Santos Bezerra, Raul Machado Neto, Rejane Maria Tommasini Grotto, Ricardo Haddad, Sandra Coccuzzo Sampaio Vessoni, Simone Kashima, Svetoslav Nanev Slavov, Vincent Louis Viala |
| EPI_ISL_1966647, EPI_ISL_1966648, EPI_ISL_1966649 | UNIDADE MISTA DE BOM JESUS DOS PERDOES | Instituto Butantan / Mendelics | Instituto Butantan: Dimas Tadeu Covas, Sandra Coccuzzo Sampaio, Maria Carolina Elias, José Salvatore Leister Patané, Vincent Louis Viala, Antonio Jorge Martins, Ricardo Haddad, Claudia Renata dos Santos Barros, Elaine Cristina Marqueze, Raul Machado Neto, Debora Botequio Moretti, Jardelina de Souza Todao Bernardino, Loyze Paola Oliveira de Lima, Luiz Aurelio de Campos Crispin. Centro de Genômica Funcional da ESALQ: Luiz Lehmann Coutinho, Ricardo Augusto Brassaloti, Raquel de Lello Rocha Campos Cassano. NGS Soluções Genômicas: Pilar Drummond Sampaio Corrêa Mariani. FZEA-USP Pirassununga: Mirele Daiana Poleti, Jessika Cristina Chagas Lesbon, Elisangela Chicaroni Mattos, Heidge Fukumasu. USP-Botucatu: Rejane Maria Tommasini Grotto, Jayme A. Souza-Neto, Guilherme Targino Valente, Patricia Akemi Assato, Felipe Allan da Silva da Costa, Bianca Cechetto Carlos. Mendelics: Bibiana Santos, João Paulo Kitajima, Erika Freitas, David Schlesinger. Hemocentro Ribeirão Preto: Simone Kashima, Evandra Strazza Rodrigues, Svetoslav Nanev Slavov, Elaine Vieira dos Santos, Rafael dos Santos Bezerra, Luiz Carlos Junior de Alcantara, Marta Giovanetti, Vagner Fonseca, Flavia Aburjaile, Rodrigo Tocantins Calado. FAMERP-SJRP: Cecília Artico Banho, Lívia Sacchetto, Fábio Sossai Possebon, Leila Sabrina Ullmann, Cíntia Bittar, Guilherme Campos, Helena Lage Ferreira, Jorge A. Petroli Marchesi, Maisa C. Pereira Parra, Marília Moraes, Paula Rahal, Paulo Inacio da Costa, João Pessoa Araújo Jr., Maurício Lacerda Nogueira. Prefeitura de Sao Paulo: Melissa Palmieri. |
| EPI_ISL_1966650, EPI_ISL_1966651, EPI_ISL_1966652, EPI_ISL_1966653, | UPA UNIDADE DE PRONTO ATENDIMENTO 24 HORAS BOM JESUS | Instituto Butantan / Mendelics | Instituto Butantan: Dimas Tadeu Covas, Sandra Coccuzzo Sampaio, Maria Carolina Elias, José Salvatore Leister Patané, Vincent Louis Viala, Antonio Jorge Martins, Ricardo Haddad, Claudia Renata dos Santos Barros, Elaine Cristina Marqueze, Raul Machado Neto, Debora Botequio Moretti, Jardelina de Souza Todao Bernardino, Loyze Paola Oliveira de Lima, Luiz Aurelio de Campos Crispin. Centro de Genômica Funcional da ESALQ: Luiz Lehmann Coutinho, Ricardo Augusto Brassaloti, Raquel de Lello Rocha Campos Cassano. NGS Soluções Genômicas: Pilar Drummond Sampaio Corrêa Mariani. |

| | | | |
|---|---|---|---|
| EPI_ISL_1966654, EPI_ISL_1966655, EPI_ISL_1966656, EPI_ISL_1966657, EPI_ISL_1966658 | | | FZEA-USP Pirassununga: Mirele Daiana Poleti, Jessika Cristina Chagas Lesbon, Elisangela Chicaroni Mattos, Heidge Fukumasu. USP-Botucatu: Rejane Maria Tommasini Grotto, Jayme A. Souza-Neto, Guilherme Targino Valente, Patricia Akemi Assato, Felipe Allan da Silva da Costa, Bianca Cechetto Carlos. Mendelics: Bibiana Santos, João Paulo Kitajima, Erika Freitas, David Schlesinger. Hemocentro Ribeirão Preto: Simone Kashima, Evandra Strazza Rodrigues, Svetoslav Nanev Slavov, Elaine Vieira dos Santos, Rafael dos Santos Bezerra, Luiz Carlos Junior de Alcantara, Marta Giovanetti, Vagner Fonseca, Flavia Aburjaile, Rodrigo Tocantins Calado. FAMERP-SJRP: Cecília Artico Banho, Lívia Sacchetto, Fábio Sossai Possebon, Leila Sabrina Ullmann, Cíntia Bittar, Guilherme Campos, Helena Lage Ferreira, Jorge A. Petroli Marchesi, Maisa C. Pereira Parra, Marília Moraes, Paula Rahal, Paulo Inacio da Costa, João Pessoa Araújo Jr., Maurício Lacerda Nogueira. Prefeitura de Sao Paulo: Melissa Palmieri. |
| EPI_ISL_1966659 | UPA UNIDADE DE PRONTO ATENDIMENTO 24 HORAS BOM JESUS | Instituto Butantan / Mendelics | Dimas Tadeu Covas, Antonio Jorge Martins, Claudia Renata dos Santos Barros, David Schlesinger, Debora Botequio Moretti, Elaine Cristina Marqueze, Elaine Vieira Santos, Evandra Strazza Rodrigues, Heidge Fukumasu, Jayme Augusto de Souza-Neto, José Salvatore Leister Patané, Luiz Alcantara, Luiz Lehmann Coutinho, Maria Carolina Elias, Maurício Lacerda Nogueira, Rafael dos Santos Bezerra, Raul Machado Neto, Rejane Maria Tommasini Grotto, Ricardo Haddad, Sandra Coccuzzo Sampaio Vessoni, Simone Kashima, Svetoslav Nanev Slavov, Vincent Louis Viala |
| EPI_ISL_1966660, EPI_ISL_1966661, EPI_ISL_1966662, EPI_ISL_1966663, EPI_ISL_1966665, EPI_ISL_1966666, EPI_ISL_1966667, EPI_ISL_1966668, EPI_ISL_1966669 | UPA UNIDADE DE PRONTO ATENDIMENTO 24 HORAS BOM JESUS | Instituto Butantan / Mendelics | Instituto Butantan: Dimas Tadeu Covas, Sandra Coccuzzo Sampaio, Maria Carolina Elias, José Salvatore Leister Patané, Vincent Louis Viala, Antonio Jorge Martins, Ricardo Haddad, Claudia Renata dos Santos Barros, Elaine Cristina Marqueze, Raul Machado Neto, Debora Botequio Moretti, Jardelina de Souza Todao Bernardino, Loyze Paola Oliveira de Lima, Luiz Aurelio de Campos Crispin. Centro de Genômica Funcional da ESALQ: Luiz Lehmann Coutinho, Ricardo Augusto Brassaloti, Raquel de Lello Rocha Campos Cassano. NGS Soluções Genômicas: Pilar Drummond Sampaio Corrêa Mariani. FZEA-USP Pirassununga: Mirele Daiana Poleti, Jessika Cristina Chagas Lesbon, Elisangela Chicaroni Mattos, Heidge Fukumasu. USP-Botucatu: Rejane Maria Tommasini Grotto, Jayme A. Souza-Neto, Guilherme Targino Valente, Patricia Akemi Assato, Felipe Allan da Silva da Costa, Bianca Cechetto Carlos. Mendelics: Bibiana Santos, João Paulo Kitajima, Erika Freitas, David Schlesinger. Hemocentro Ribeirão Preto: Simone Kashima, Evandra Strazza Rodrigues, Svetoslav Nanev Slavov, Elaine Vieira dos Santos, Rafael dos Santos Bezerra, Luiz Carlos Junior de Alcantara, Marta Giovanetti, Vagner Fonseca, Flavia Aburjaile, Rodrigo Tocantins Calado. FAMERP-SJRP: Cecília Artico Banho, Lívia Sacchetto, Fábio Sossai Possebon, Leila Sabrina Ullmann, Cíntia Bittar, Guilherme Campos, Helena Lage Ferreira, Jorge A. Petroli Marchesi, Maisa C. Pereira Parra, Marília Moraes, Paula Rahal, Paulo Inacio da Costa, João Pessoa Araújo Jr., Maurício Lacerda Nogueira. Prefeitura de Sao Paulo: Melissa Palmieri. |
| EPI_ISL_1966670, EPI_ISL_1966671, EPI_ISL_1966672, EPI_ISL_1966673 | HOSPITAL MUNICIPAL REYNALDO GUERRA CAJATI | Instituto Butantan / Mendelics | Instituto Butantan: Dimas Tadeu Covas, Sandra Coccuzzo Sampaio, Maria Carolina Elias, José Salvatore Leister Patané, Vincent Louis Viala, Antonio Jorge Martins, Ricardo Haddad, Claudia Renata dos Santos Barros, Elaine Cristina Marqueze, Raul Machado Neto, Debora Botequio Moretti, Jardelina de Souza Todao Bernardino, Loyze Paola Oliveira de Lima, Luiz Aurelio de Campos Crispin. Centro de Genômica Funcional da ESALQ: Luiz Lehmann Coutinho, Ricardo Augusto Brassaloti, Raquel de Lello Rocha Campos Cassano. NGS Soluções Genômicas: Pilar Drummond Sampaio Corrêa Mariani. FZEA-USP Pirassununga: Mirele Daiana Poleti, Jessika Cristina Chagas Lesbon, Elisangela Chicaroni Mattos, Heidge Fukumasu. USP-Botucatu: Rejane Maria Tommasini Grotto, Jayme A. Souza-Neto, Guilherme Targino Valente, Patricia Akemi Assato, Felipe Allan da Silva da Costa, Bianca Cechetto Carlos. Mendelics: Bibiana Santos, João Paulo Kitajima, Erika Freitas, David Schlesinger. Hemocentro Ribeirão Preto: Simone Kashima, Evandra Strazza Rodrigues, Svetoslav Nanev Slavov, Elaine Vieira dos Santos, Rafael dos Santos Bezerra, Luiz Carlos Junior de Alcantara, Marta Giovanetti, Vagner Fonseca, Flavia Aburjaile, Rodrigo Tocantins Calado. FAMERP-SJRP: Cecília Artico Banho, Lívia Sacchetto, Fábio Sossai Possebon, Leila Sabrina Ullmann, Cíntia Bittar, Guilherme Campos, Helena Lage Ferreira, Jorge A. Petroli Marchesi, Maisa C. Pereira Parra, Marília Moraes, Paula Rahal, Paulo Inacio da Costa, João Pessoa Araújo Jr., Maurício Lacerda Nogueira. Prefeitura de Sao Paulo: Melissa Palmieri. |
| EPI_ISL_1966674 | ESF NOVA ALEXANDRIA | Instituto Butantan / Mendelics | Instituto Butantan: Dimas Tadeu Covas, Sandra Coccuzzo Sampaio, Maria Carolina Elias, José Salvatore Leister Patané, Vincent Louis Viala, Antonio Jorge Martins, Ricardo Haddad, Claudia Renata dos Santos Barros, Elaine Cristina Marqueze, Raul Machado Neto, Debora Botequio Moretti, Jardelina de Souza Todao Bernardino, Loyze Paola Oliveira de Lima, Luiz Aurelio de Campos Crispin. Centro de Genômica Funcional da ESALQ: Luiz Lehmann Coutinho, Ricardo Augusto Brassaloti, Raquel de Lello Rocha Campos Cassano. NGS Soluções Genômicas: Pilar Drummond Sampaio Corrêa Mariani. FZEA-USP Pirassununga: Mirele Daiana Poleti, Jessika Cristina Chagas Lesbon, Elisangela Chicaroni Mattos, Heidge Fukumasu. USP-Botucatu: Rejane Maria Tommasini Grotto, Jayme A. Souza-Neto, Guilherme Targino Valente, Patricia Akemi Assato, Felipe Allan da Silva da Costa, Bianca Cechetto Carlos. Mendelics: Bibiana Santos, João Paulo Kitajima, Erika Freitas, David Schlesinger. Hemocentro Ribeirão Preto: Simone Kashima, Evandra Strazza Rodrigues, Svetoslav Nanev Slavov, Elaine Vieira dos Santos, Rafael dos Santos Bezerra, Luiz Carlos Junior de Alcantara, Marta Giovanetti, Vagner Fonseca, Flavia Aburjaile, Rodrigo Tocantins Calado. FAMERP-SJRP: Cecília Artico Banho, Lívia Sacchetto, Fábio Sossai Possebon, Leila Sabrina Ullmann, Cíntia Bittar, Guilherme Campos, Helena Lage Ferreira, Jorge A. Petroli Marchesi, Maisa C. Pereira Parra, Marília Moraes, Paula Rahal, Paulo Inacio da Costa, João Pessoa Araújo Jr., Maurício Lacerda Nogueira. Prefeitura de Sao Paulo: Melissa Palmieri. |
| EPI_ISL_1966675, EPI_ISL_1966676, EPI_ISL_1966677, EPI_ISL_1966678, EPI_ISL_1966679, EPI_ISL_1966680 | LABORATORIO DE FRANCA | Instituto Butantan / Mendelics | Instituto Butantan: Dimas Tadeu Covas, Sandra Coccuzzo Sampaio, Maria Carolina Elias, José Salvatore Leister Patané, Vincent Louis Viala, Antonio Jorge Martins, Ricardo Haddad, Claudia Renata dos Santos Barros, Elaine Cristina Marqueze, Raul Machado Neto, Debora Botequio Moretti, Jardelina de Souza Todao Bernardino, Loyze Paola Oliveira de Lima, Luiz Aurelio de Campos Crispin. Centro de Genômica Funcional da ESALQ: Luiz Lehmann Coutinho, Ricardo Augusto Brassaloti, Raquel de Lello Rocha Campos Cassano. NGS Soluções Genômicas: Pilar Drummond Sampaio Corrêa Mariani. FZEA-USP Pirassununga: Mirele Daiana Poleti, Jessika Cristina Chagas Lesbon, Elisangela Chicaroni Mattos, Heidge Fukumasu. USP-Botucatu: Rejane Maria Tommasini Grotto, Jayme A. Souza-Neto, Guilherme Targino Valente, Patricia Akemi Assato, Felipe Allan da Silva da Costa, Bianca Cechetto Carlos. Mendelics: Bibiana Santos, João Paulo Kitajima, Erika Freitas, David Schlesinger. Hemocentro Ribeirão Preto: Simone Kashima, Evandra Strazza Rodrigues, Svetoslav Nanev Slavov, Elaine Vieira dos Santos, Rafael dos Santos Bezerra, Luiz Carlos Junior de Alcantara, Marta Giovanetti, Vagner Fonseca, Flavia Aburjaile, Rodrigo Tocantins Calado. FAMERP-SJRP: Cecília Artico Banho, Lívia Sacchetto, Fábio Sossai Possebon, Leila Sabrina Ullmann, Cíntia Bittar, Guilherme Campos, Helena Lage Ferreira, Jorge A. Petroli Marchesi, Maisa C. Pereira Parra, Marília Moraes, Paula Rahal, Paulo Inacio da Costa, João Pessoa Araújo Jr., Maurício Lacerda Nogueira. Prefeitura de Sao Paulo: Melissa Palmieri. |
| EPI_ISL_1966681, EPI_ISL_1966682 | UNIDADE MISTA DE IGUAPE | Instituto Butantan / Mendelics | Instituto Butantan: Dimas Tadeu Covas, Sandra Coccuzzo Sampaio, Maria Carolina Elias, José Salvatore Leister Patané, Vincent Louis Viala, Antonio Jorge Martins, Ricardo Haddad, Claudia Renata dos Santos Barros, Elaine Cristina Marqueze, Raul Machado Neto, Debora Botequio Moretti, Jardelina de Souza Todao Bernardino, Loyze Paola Oliveira de Lima, Luiz Aurelio de Campos Crispin. Centro de Genômica Funcional da ESALQ: Luiz Lehmann Coutinho, Ricardo Augusto Brassaloti, Raquel de Lello Rocha Campos Cassano. NGS Soluções Genômicas: Pilar Drummond Sampaio Corrêa Mariani. FZEA-USP Pirassununga: Mirele Daiana Poleti, Jessika Cristina Chagas Lesbon, Elisangela Chicaroni Mattos, Heidge Fukumasu. USP-Botucatu: Rejane Maria Tommasini Grotto, Jayme A. Souza-Neto, Guilherme Targino Valente, Patricia Akemi Assato, Felipe Allan da Silva da Costa, Bianca Cechetto Carlos. Mendelics: Bibiana Santos, João Paulo Kitajima, Erika Freitas, David Schlesinger. Hemocentro Ribeirão Preto: Simone Kashima, Evandra Strazza Rodrigues, Svetoslav Nanev Slavov, Elaine Vieira dos Santos, Rafael dos Santos Bezerra, Luiz Carlos Junior de Alcantara, Marta Giovanetti, Vagner Fonseca, Flavia Aburjaile, Rodrigo Tocantins Calado. FAMERP-SJRP: Cecília Artico Banho, Lívia Sacchetto, Fábio Sossai Possebon, Leila Sabrina Ullmann, Cíntia Bittar, Guilherme Campos, Helena Lage Ferreira, Jorge A. Petroli Marchesi, Maisa C. Pereira Parra, Marília Moraes, Paula Rahal, Paulo Inacio da Costa, João Pessoa Araújo Jr., Maurício Lacerda Nogueira. Prefeitura de Sao Paulo: Melissa Palmieri. |
| EPI_ISL_1966683, EPI_ISL_1966685, EPI_ISL_1966686, EPI_ISL_1966687, EPI_ISL_1966688, EPI_ISL_1966689, EPI_ISL_1966690 | LABORATORIO MUNICIPAL DE ANALISES CLINICAS DE ITANHAEM | Instituto Butantan / Mendelics | Instituto Butantan: Dimas Tadeu Covas, Sandra Coccuzzo Sampaio, Maria Carolina Elias, José Salvatore Leister Patané, Vincent Louis Viala, Antonio Jorge Martins, Ricardo Haddad, Claudia Renata dos Santos Barros, Elaine Cristina Marqueze, Raul Machado Neto, Debora Botequio Moretti, Jardelina de Souza Todao Bernardino, Loyze Paola Oliveira de Lima, Luiz Aurelio de Campos Crispin. Centro de Genômica Funcional da ESALQ: Luiz Lehmann Coutinho, Ricardo Augusto Brassaloti, Raquel de Lello Rocha Campos Cassano. NGS Soluções Genômicas: Pilar Drummond Sampaio Corrêa Mariani. FZEA-USP Pirassununga: Mirele Daiana Poleti, Jessika Cristina Chagas Lesbon, Elisangela Chicaroni Mattos, Heidge Fukumasu. USP-Botucatu: Rejane Maria Tommasini Grotto, Jayme A. Souza-Neto, Guilherme Targino Valente, Patricia Akemi Assato, Felipe Allan da Silva da Costa, Bianca Cechetto Carlos. Mendelics: Bibiana Santos, João Paulo Kitajima, Erika Freitas, David Schlesinger. Hemocentro Ribeirão Preto: Simone Kashima, Evandra Strazza Rodrigues, Svetoslav Nanev Slavov, Elaine Vieira dos Santos, Rafael dos Santos Bezerra, Luiz Carlos Junior de Alcantara, Marta Giovanetti, Vagner Fonseca, Flavia Aburjaile, Rodrigo Tocantins Calado. FAMERP-SJRP: Cecília Artico Banho, Lívia Sacchetto, Fábio Sossai Possebon, Leila Sabrina Ullmann, Cíntia Bittar, Guilherme Campos, Helena Lage Ferreira, Jorge A. Petroli Marchesi, Maisa C. Pereira Parra, Marília Moraes, Paula Rahal, Paulo Inacio da Costa, João Pessoa Araújo Jr., Maurício Lacerda Nogueira. Prefeitura de Sao Paulo: Melissa Palmieri. |
| EPI_ISL_1966691, EPI_ISL_1966692 | PRONTO ATENDIMENTO MUNICIPAL DE JACUPIRANGA | Instituto Butantan / Mendelics | Instituto Butantan: Dimas Tadeu Covas, Sandra Coccuzzo Sampaio, Maria Carolina Elias, José Salvatore Leister Patané, Vincent Louis Viala, Antonio Jorge Martins, Ricardo Haddad, Claudia Renata dos Santos Barros, Elaine Cristina Marqueze, Raul Machado Neto, Debora Botequio Moretti, Jardelina de Souza Todao Bernardino, Loyze Paola Oliveira de Lima, Luiz Aurelio de Campos Crispin. Centro de Genômica Funcional da ESALQ: Luiz Lehmann Coutinho, Ricardo Augusto Brassaloti, Raquel de Lello Rocha Campos Cassano. NGS Soluções Genômicas: Pilar Drummond Sampaio Corrêa Mariani. FZEA-USP Pirassununga: Mirele Daiana Poleti, Jessika Cristina Chagas Lesbon, Elisangela Chicaroni Mattos, Heidge Fukumasu. USP-Botucatu: Rejane Maria Tommasini Grotto, Jayme A. Souza-Neto, Guilherme Targino Valente, Patricia Akemi Assato, Felipe Allan da Silva da Costa, Bianca Cechetto |

| | | | |
|---|---|---|---|
| | | | Carlos. Mendelics: Bibiana Santos, João Paulo Kitajima, Erika Freitas, David Schlesinger. Hemocentro Ribeirão Preto: Simone Kashima, Evandra Strazza Rodrigues, Svetoslav Nanev Slavov, Elaine Vieira dos Santos, Rafael dos Santos Bezerra, Luiz Carlos Junior de Alcantara, Marta Giovanetti, Vagner Fonseca, Flavia Aburjaile, Rodrigo Tocantins Calado. FAMERP-SJRP: Cecília Artico Banho, Lívia Sacchetto, Fábio Sossai Possebon, Leila Sabrina Ullmann, Cíntia Bittar, Guilherme Campos, Helena Lage Ferreira, Jorge A. Petroli Marchesi, Maisa C. Pereira Parra, Marília Moraes, Paula Rahal, Paulo Inacio da Costa, João Pessoa Araújo Jr., Maurício Lacerda Nogueira. Prefeitura de Sao Paulo: Melissa Palmieri. |
| EPI_ISL_1966693 | SECRETARIA MUNICIPAL DE SAUDE DE JARINU | Instituto Butantan / Mendelics | Instituto Butantan: Dimas Tadeu Covas, Sandra Coccuzzo Sampaio, Maria Carolina Elias, José Salvatore Leister Patané, Vincent Louis Viala, Antonio Jorge Martins, Ricardo Haddad, Claudia Renata dos Santos Barros, Elaine Cristina Marqueze, Raul Machado Neto, Debora Botequio Moretti, Jardelina de Souza Todao Bernardino, Loyze Paola Oliveira de Lima, Luiz Aurelio de Campos Crispin. Centro de Genômica Funcional da ESALQ: Luiz Lehmann Coutinho, Ricardo Augusto Brassaloti, Raquel de Lello Rocha Campos Cassano. NGS Soluções Genômicas: Pilar Drummond Sampaio Corrêa Mariani. FZEA-USP Pirassununga: Mirele Daiana Poleti, Jessika Cristina Chagas Lesbon, Elisangela Chicaroni Mattos, Heidge Fukumasu. USP-Botucatu: Rejane Maria Tommasini Grotto, Jayme A. Souza-Neto, Guilherme Targino Valente, Patricia Akemi Assato, Felipe Allan da Silva da Costa, Bianca Cechetto Carlos. Mendelics: Bibiana Santos, João Paulo Kitajima, Erika Freitas, David Schlesinger. Hemocentro Ribeirão Preto: Simone Kashima, Evandra Strazza Rodrigues, Svetoslav Nanev Slavov, Elaine Vieira dos Santos, Rafael dos Santos Bezerra, Luiz Carlos Junior de Alcantara, Marta Giovanetti, Vagner Fonseca, Flavia Aburjaile, Rodrigo Tocantins Calado. FAMERP-SJRP: Cecília Artico Banho, Lívia Sacchetto, Fábio Sossai Possebon, Leila Sabrina Ullmann, Cíntia Bittar, Guilherme Campos, Helena Lage Ferreira, Jorge A. Petroli Marchesi, Maisa C. Pereira Parra, Marília Moraes, Paula Rahal, Paulo Inacio da Costa, João Pessoa Araújo Jr., Maurício Lacerda Nogueira. Prefeitura de Sao Paulo: Melissa Palmieri. |
| EPI_ISL_1966694 | HOSPITAL SANTO ANTONIO DE JUQUIA JUQUIA | Instituto Butantan / Mendelics | Instituto Butantan: Dimas Tadeu Covas, Sandra Coccuzzo Sampaio, Maria Carolina Elias, José Salvatore Leister Patané, Vincent Louis Viala, Antonio Jorge Martins, Ricardo Haddad, Claudia Renata dos Santos Barros, Elaine Cristina Marqueze, Raul Machado Neto, Debora Botequio Moretti, Jardelina de Souza Todao Bernardino, Loyze Paola Oliveira de Lima, Luiz Aurelio de Campos Crispin. Centro de Genômica Funcional da ESALQ: Luiz Lehmann Coutinho, Ricardo Augusto Brassaloti, Raquel de Lello Rocha Campos Cassano. NGS Soluções Genômicas: Pilar Drummond Sampaio Corrêa Mariani. FZEA-USP Pirassununga: Mirele Daiana Poleti, Jessika Cristina Chagas Lesbon, Elisangela Chicaroni Mattos, Heidge Fukumasu. USP-Botucatu: Rejane Maria Tommasini Grotto, Jayme A. Souza-Neto, Guilherme Targino Valente, Patricia Akemi Assato, Felipe Allan da Silva da Costa, Bianca Cechetto Carlos. Mendelics: Bibiana Santos, João Paulo Kitajima, Erika Freitas, David Schlesinger. Hemocentro Ribeirão Preto: Simone Kashima, Evandra Strazza Rodrigues, Svetoslav Nanev Slavov, Elaine Vieira dos Santos, Rafael dos Santos Bezerra, Luiz Carlos Junior de Alcantara, Marta Giovanetti, Vagner Fonseca, Flavia Aburjaile, Rodrigo Tocantins Calado. FAMERP-SJRP: Cecília Artico Banho, Lívia Sacchetto, Fábio Sossai Possebon, Leila Sabrina Ullmann, Cíntia Bittar, Guilherme Campos, Helena Lage Ferreira, Jorge A. Petroli Marchesi, Maisa C. Pereira Parra, Marília Moraes, Paula Rahal, Paulo Inacio da Costa, João Pessoa Araújo Jr., Maurício Lacerda Nogueira. Prefeitura de Sao Paulo: Melissa Palmieri. |
| EPI_ISL_1966695 | CS IOLANDA FLEURY DE CAMPOS GUEDES NUPORANGA | Instituto Butantan / Mendelics | Instituto Butantan: Dimas Tadeu Covas, Sandra Coccuzzo Sampaio, Maria Carolina Elias, José Salvatore Leister Patané, Vincent Louis Viala, Antonio Jorge Martins, Ricardo Haddad, Claudia Renata dos Santos Barros, Elaine Cristina Marqueze, Raul Machado Neto, Debora Botequio Moretti, Jardelina de Souza Todao Bernardino, Loyze Paola Oliveira de Lima, Luiz Aurelio de Campos Crispin. Centro de Genômica Funcional da ESALQ: Luiz Lehmann Coutinho, Ricardo Augusto Brassaloti, Raquel de Lello Rocha Campos Cassano. NGS Soluções Genômicas: Pilar Drummond Sampaio Corrêa Mariani. FZEA-USP Pirassununga: Mirele Daiana Poleti, Jessika Cristina Chagas Lesbon, Elisangela Chicaroni Mattos, Heidge Fukumasu. USP-Botucatu: Rejane Maria Tommasini Grotto, Jayme A. Souza-Neto, Guilherme Targino Valente, Patricia Akemi Assato, Felipe Allan da Silva da Costa, Bianca Cechetto Carlos. Mendelics: Bibiana Santos, João Paulo Kitajima, Erika Freitas, David Schlesinger. Hemocentro Ribeirão Preto: Simone Kashima, Evandra Strazza Rodrigues, Svetoslav Nanev Slavov, Elaine Vieira dos Santos, Rafael dos Santos Bezerra, Luiz Carlos Junior de Alcantara, Marta Giovanetti, Vagner Fonseca, Flavia Aburjaile, Rodrigo Tocantins Calado. FAMERP-SJRP: Cecília Artico Banho, Lívia Sacchetto, Fábio Sossai Possebon, Leila Sabrina Ullmann, Cíntia Bittar, Guilherme Campos, Helena Lage Ferreira, Jorge A. Petroli Marchesi, Maisa C. Pereira Parra, Marília Moraes, Paula Rahal, Paulo Inacio da Costa, João Pessoa Araújo Jr., Maurício Lacerda Nogueira. Prefeitura de Sao Paulo: Melissa Palmieri. |
| EPI_ISL_1966696 | UBS III DE PARIQUERA ACU PARIQUERA ACU | Instituto Butantan / Mendelics | Instituto Butantan: Dimas Tadeu Covas, Sandra Coccuzzo Sampaio, Maria Carolina Elias, José Salvatore Leister Patané, Vincent Louis Viala, Antonio Jorge Martins, Ricardo Haddad, Claudia Renata dos Santos Barros, Elaine Cristina Marqueze, Raul Machado Neto, Debora Botequio Moretti, Jardelina de Souza Todao Bernardino, Loyze Paola Oliveira de Lima, Luiz Aurelio de Campos Crispin. Centro de Genômica Funcional da ESALQ: Luiz Lehmann Coutinho, Ricardo Augusto Brassaloti, Raquel de Lello Rocha Campos Cassano. NGS Soluções Genômicas: Pilar Drummond Sampaio Corrêa Mariani. FZEA-USP Pirassununga: Mirele Daiana Poleti, Jessika Cristina Chagas Lesbon, Elisangela Chicaroni Mattos, Heidge Fukumasu. USP-Botucatu: Rejane Maria Tommasini Grotto, Jayme A. Souza-Neto, Guilherme Targino Valente, Patricia Akemi Assato, Felipe Allan da Silva da Costa, Bianca Cechetto Carlos. Mendelics: Bibiana Santos, João Paulo Kitajima, Erika Freitas, David Schlesinger. Hemocentro Ribeirão Preto: Simone Kashima, Evandra Strazza Rodrigues, Svetoslav Nanev Slavov, Elaine Vieira dos Santos, Rafael dos Santos Bezerra, Luiz Carlos Junior de Alcantara, Marta Giovanetti, Vagner Fonseca, Flavia Aburjaile, Rodrigo Tocantins Calado. FAMERP-SJRP: Cecília Artico Banho, Lívia Sacchetto, Fábio Sossai Possebon, Leila Sabrina Ullmann, Cíntia Bittar, Guilherme Campos, Helena Lage Ferreira, Jorge A. Petroli Marchesi, Maisa C. Pereira Parra, Marília Moraes, Paula Rahal, Paulo Inacio da Costa, João Pessoa Araújo Jr., Maurício Lacerda Nogueira. Prefeitura de Sao Paulo: Melissa Palmieri. |
| EPI_ISL_1966697, EPI_ISL_1966698, EPI_ISL_1966699, EPI_ISL_1966700, EPI_ISL_1966701, EPI_ISL_1966702, EPI_ISL_1966703 | AMBULATORIO MEDICO DE ESPECIALIDADES DE PERUIBE | Instituto Butantan / Mendelics | Instituto Butantan: Dimas Tadeu Covas, Sandra Coccuzzo Sampaio, Maria Carolina Elias, José Salvatore Leister Patané, Vincent Louis Viala, Antonio Jorge Martins, Ricardo Haddad, Claudia Renata dos Santos Barros, Elaine Cristina Marqueze, Raul Machado Neto, Debora Botequio Moretti, Jardelina de Souza Todao Bernardino, Loyze Paola Oliveira de Lima, Luiz Aurelio de Campos Crispin. Centro de Genômica Funcional da ESALQ: Luiz Lehmann Coutinho, Ricardo Augusto Brassaloti, Raquel de Lello Rocha Campos Cassano. NGS Soluções Genômicas: Pilar Drummond Sampaio Corrêa Mariani. FZEA-USP Pirassununga: Mirele Daiana Poleti, Jessika Cristina Chagas Lesbon, Elisangela Chicaroni Mattos, Heidge Fukumasu. USP-Botucatu: Rejane Maria Tommasini Grotto, Jayme A. Souza-Neto, Guilherme Targino Valente, Patricia Akemi Assato, Felipe Allan da Silva da Costa, Bianca Cechetto Carlos. Mendelics: Bibiana Santos, João Paulo Kitajima, Erika Freitas, David Schlesinger. Hemocentro Ribeirão Preto: Simone Kashima, Evandra Strazza Rodrigues, Svetoslav Nanev Slavov, Elaine Vieira dos Santos, Rafael dos Santos Bezerra, Luiz Carlos Junior de Alcantara, Marta Giovanetti, Vagner Fonseca, Flavia Aburjaile, Rodrigo Tocantins Calado. FAMERP-SJRP: Cecília Artico Banho, Lívia Sacchetto, Fábio Sossai Possebon, Leila Sabrina Ullmann, Cíntia Bittar, Guilherme Campos, Helena Lage Ferreira, Jorge A. Petroli Marchesi, Maisa C. Pereira Parra, Marília Moraes, Paula Rahal, Paulo Inacio da Costa, João Pessoa Araújo Jr., Maurício Lacerda Nogueira. Prefeitura de Sao Paulo: Melissa Palmieri. |
| EPI_ISL_1966704 | USAFA REAL | Instituto Butantan / Mendelics | Instituto Butantan: Dimas Tadeu Covas, Sandra Coccuzzo Sampaio, Maria Carolina Elias, José Salvatore Leister Patané, Vincent Louis Viala, Antonio Jorge Martins, Ricardo Haddad, Claudia Renata dos Santos Barros, Elaine Cristina Marqueze, Raul Machado Neto, Debora Botequio Moretti, Jardelina de Souza Todao Bernardino, Loyze Paola Oliveira de Lima, Luiz Aurelio de Campos Crispin. Centro de Genômica Funcional da ESALQ: Luiz Lehmann Coutinho, Ricardo Augusto Brassaloti, Raquel de Lello Rocha Campos Cassano. NGS Soluções Genômicas: Pilar Drummond Sampaio Corrêa Mariani. FZEA-USP Pirassununga: Mirele Daiana Poleti, Jessika Cristina Chagas Lesbon, Elisangela Chicaroni Mattos, Heidge Fukumasu. USP-Botucatu: Rejane Maria Tommasini Grotto, Jayme A. Souza-Neto, Guilherme Targino Valente, Patricia Akemi Assato, Felipe Allan da Silva da Costa, Bianca Cechetto Carlos. Mendelics: Bibiana Santos, João Paulo Kitajima, Erika Freitas, David Schlesinger. Hemocentro Ribeirão Preto: Simone Kashima, Evandra Strazza Rodrigues, Svetoslav Nanev Slavov, Elaine Vieira dos Santos, Rafael dos Santos Bezerra, Luiz Carlos Junior de Alcantara, Marta Giovanetti, Vagner Fonseca, Flavia Aburjaile, Rodrigo Tocantins Calado. FAMERP-SJRP: Cecília Artico Banho, Lívia Sacchetto, Fábio Sossai Possebon, Leila Sabrina Ullmann, Cíntia Bittar, Guilherme Campos, Helena Lage Ferreira, Jorge A. Petroli Marchesi, Maisa C. Pereira Parra, Marília Moraes, Paula Rahal, Paulo Inacio da Costa, João Pessoa Araújo Jr., Maurício Lacerda Nogueira. Prefeitura de Sao Paulo: Melissa Palmieri. |
| EPI_ISL_1966705 | USAFA ALOHA | Instituto Butantan / Mendelics | Instituto Butantan: Dimas Tadeu Covas, Sandra Coccuzzo Sampaio, Maria Carolina Elias, José Salvatore Leister Patané, Vincent Louis Viala, Antonio Jorge Martins, Ricardo Haddad, Claudia Renata dos Santos Barros, Elaine Cristina Marqueze, Raul Machado Neto, Debora Botequio Moretti, Jardelina de Souza Todao Bernardino, Loyze Paola Oliveira de Lima, Luiz Aurelio de Campos Crispin. Centro de Genômica Funcional da ESALQ: Luiz Lehmann Coutinho, Ricardo Augusto Brassaloti, Raquel de Lello Rocha Campos Cassano. NGS Soluções Genômicas: Pilar Drummond Sampaio Corrêa Mariani. FZEA-USP Pirassununga: Mirele Daiana Poleti, Jessika Cristina Chagas Lesbon, Elisangela Chicaroni Mattos, Heidge Fukumasu. USP-Botucatu: Rejane Maria Tommasini Grotto, Jayme A. Souza-Neto, Guilherme Targino Valente, Patricia Akemi Assato, Felipe Allan da Silva da Costa, Bianca Cechetto Carlos. Mendelics: Bibiana Santos, João Paulo Kitajima, Erika Freitas, David Schlesinger. Hemocentro Ribeirão Preto: Simone Kashima, Evandra Strazza Rodrigues, Svetoslav Nanev Slavov, Elaine Vieira dos Santos, Rafael dos Santos Bezerra, Luiz Carlos Junior de Alcantara, Marta Giovanetti, Vagner Fonseca, Flavia Aburjaile, Rodrigo Tocantins Calado. FAMERP-SJRP: Cecília Artico Banho, Lívia Sacchetto, Fábio Sossai Possebon, Leila Sabrina Ullmann, Cíntia Bittar, Guilherme Campos, Helena Lage Ferreira, Jorge A. Petroli Marchesi, Maisa C. Pereira Parra, Marília Moraes, Paula Rahal, Paulo Inacio da Costa, João Pessoa Araújo Jr., Maurício Lacerda Nogueira. Prefeitura de Sao Paulo: Melissa Palmieri. |
| EPI_ISL_1966706 | AMBULATORIO TUPI | Instituto Butantan / Mendelics | Instituto Butantan: Dimas Tadeu Covas, Sandra Coccuzzo Sampaio, Maria Carolina Elias, José Salvatore Leister Patané, Vincent Louis Viala, Antonio |

| | | | |
|---|---|---|---|
| | | | Jorge Martins, Ricardo Haddad, Claudia Renata dos Santos Barros, Elaine Cristina Marqueze, Raul Machado Neto, Debora Botequio Moretti, Jardelina de Souza Todao Bernardino, Loyze Paola Oliveira de Lima, Luiz Aurelio de Campos Crispin. Centro de Genômica Funcional da ESALQ: Luiz Lehmann Coutinho, Ricardo Augusto Brassaloti, Raquel de Lello Rocha Campos Cassano. NGS Soluções Genômicas: Pilar Drummond Sampaio Corrêa Mariani. FZEA-USP Pirassununga: Mirele Daiana Poleti, Jessika Cristina Chagas Lesbon, Elisangela Chicaroni Mattos, Heidge Fukumasu. USP-Botucatu: Rejane Maria Tommasini Grotto, Jayme A. Souza-Neto, Guilherme Targino Valente, Patricia Akemi Assato, Felipe Allan da Silva da Costa, Bianca Cechetto Carlos. Mendelics: Bibiana Santos, João Paulo Kitajima, Erika Freitas, David Schlesinger. Hemocentro Ribeirão Preto: Simone Kashima, Evandra Strazza Rodrigues, Svetoslav Nanev Slavov, Elaine Vieira dos Santos, Rafael dos Santos Bezerra, Luiz Carlos Junior de Alcantara, Marta Giovanetti, Vagner Fonseca, Flavia Aburjaile, Rodrigo Tocantins Calado. FAMERP-SJRP: Cecília Artico Banho, Lívia Sacchetto, Fábio Sossai Possebon, Leila Sabrina Ullmann, Cíntia Bittar, Guilherme Campos, Helena Lage Ferreira, Jorge A. Petroli Marchesi, Maisa C. Pereira Parra, Marília Moraes, Paula Rahal, Paulo Inacio da Costa, João Pessoa Araújo Jr., Maurício Lacerda Nogueira. Prefeitura de Sao Paulo: Melissa Palmieri. |
| EPI_ISL_1966707 | USAFA GUARAMAR | Instituto Butantan / Mendelics | Instituto Butantan: Dimas Tadeu Covas, Sandra Coccuzzo Sampaio, Maria Carolina Elias, José Salvatore Leister Patané, Vincent Louis Viala, Antonio Jorge Martins, Ricardo Haddad, Claudia Renata dos Santos Barros, Elaine Cristina Marqueze, Raul Machado Neto, Debora Botequio Moretti, Jardelina de Souza Todao Bernardino, Loyze Paola Oliveira de Lima, Luiz Aurelio de Campos Crispin. Centro de Genômica Funcional da ESALQ: Luiz Lehmann Coutinho, Ricardo Augusto Brassaloti, Raquel de Lello Rocha Campos Cassano. NGS Soluções Genômicas: Pilar Drummond Sampaio Corrêa Mariani. FZEA-USP Pirassununga: Mirele Daiana Poleti, Jessika Cristina Chagas Lesbon, Elisangela Chicaroni Mattos, Heidge Fukumasu. USP-Botucatu: Rejane Maria Tommasini Grotto, Jayme A. Souza-Neto, Guilherme Targino Valente, Patricia Akemi Assato, Felipe Allan da Silva da Costa, Bianca Cechetto Carlos. Mendelics: Bibiana Santos, João Paulo Kitajima, Erika Freitas, David Schlesinger. Hemocentro Ribeirão Preto: Simone Kashima, Evandra Strazza Rodrigues, Svetoslav Nanev Slavov, Elaine Vieira dos Santos, Rafael dos Santos Bezerra, Luiz Carlos Junior de Alcantara, Marta Giovanetti, Vagner Fonseca, Flavia Aburjaile, Rodrigo Tocantins Calado. FAMERP-SJRP: Cecília Artico Banho, Lívia Sacchetto, Fábio Sossai Possebon, Leila Sabrina Ullmann, Cíntia Bittar, Guilherme Campos, Helena Lage Ferreira, Jorge A. Petroli Marchesi, Maisa C. Pereira Parra, Marília Moraes, Paula Rahal, Paulo Inacio da Costa, João Pessoa Araújo Jr., Maurício Lacerda Nogueira. Prefeitura de Sao Paulo: Melissa Palmieri. |
| EPI_ISL_1966708 | UNIDADE DE PRONTO ATENDIMENTO UPA | Instituto Butantan / Mendelics | Instituto Butantan: Dimas Tadeu Covas, Sandra Coccuzzo Sampaio, Maria Carolina Elias, José Salvatore Leister Patané, Vincent Louis Viala, Antonio Jorge Martins, Ricardo Haddad, Claudia Renata dos Santos Barros, Elaine Cristina Marqueze, Raul Machado Neto, Debora Botequio Moretti, Jardelina de Souza Todao Bernardino, Loyze Paola Oliveira de Lima, Luiz Aurelio de Campos Crispin. Centro de Genômica Funcional da ESALQ: Luiz Lehmann Coutinho, Ricardo Augusto Brassaloti, Raquel de Lello Rocha Campos Cassano. NGS Soluções Genômicas: Pilar Drummond Sampaio Corrêa Mariani. FZEA-USP Pirassununga: Mirele Daiana Poleti, Jessika Cristina Chagas Lesbon, Elisangela Chicaroni Mattos, Heidge Fukumasu. USP-Botucatu: Rejane Maria Tommasini Grotto, Jayme A. Souza-Neto, Guilherme Targino Valente, Patricia Akemi Assato, Felipe Allan da Silva da Costa, Bianca Cechetto Carlos. Mendelics: Bibiana Santos, João Paulo Kitajima, Erika Freitas, David Schlesinger. Hemocentro Ribeirão Preto: Simone Kashima, Evandra Strazza Rodrigues, Svetoslav Nanev Slavov, Elaine Vieira dos Santos, Rafael dos Santos Bezerra, Luiz Carlos Junior de Alcantara, Marta Giovanetti, Vagner Fonseca, Flavia Aburjaile, Rodrigo Tocantins Calado. FAMERP-SJRP: Cecília Artico Banho, Lívia Sacchetto, Fábio Sossai Possebon, Leila Sabrina Ullmann, Cíntia Bittar, Guilherme Campos, Helena Lage Ferreira, Jorge A. Petroli Marchesi, Maisa C. Pereira Parra, Marília Moraes, Paula Rahal, Paulo Inacio da Costa, João Pessoa Araújo Jr., Maurício Lacerda Nogueira. Prefeitura de Sao Paulo: Melissa Palmieri. |
| EPI_ISL_1966710, EPI_ISL_1966711, EPI_ISL_1966712, EPI_ISL_1966713, EPI_ISL_1966714 | SECRETARIA MUNCIPAL DE SAUDE DE SANTA BARBARA D OESTE | Instituto Butantan / Mendelics | Instituto Butantan: Dimas Tadeu Covas, Sandra Coccuzzo Sampaio, Maria Carolina Elias, José Salvatore Leister Patané, Vincent Louis Viala, Antonio Jorge Martins, Ricardo Haddad, Claudia Renata dos Santos Barros, Elaine Cristina Marqueze, Raul Machado Neto, Debora Botequio Moretti, Jardelina de Souza Todao Bernardino, Loyze Paola Oliveira de Lima, Luiz Aurelio de Campos Crispin. Centro de Genômica Funcional da ESALQ: Luiz Lehmann Coutinho, Ricardo Augusto Brassaloti, Raquel de Lello Rocha Campos Cassano. NGS Soluções Genômicas: Pilar Drummond Sampaio Corrêa Mariani. FZEA-USP Pirassununga: Mirele Daiana Poleti, Jessika Cristina Chagas Lesbon, Elisangela Chicaroni Mattos, Heidge Fukumasu. USP-Botucatu: Rejane Maria Tommasini Grotto, Jayme A. Souza-Neto, Guilherme Targino Valente, Patricia Akemi Assato, Felipe Allan da Silva da Costa, Bianca Cechetto Carlos. Mendelics: Bibiana Santos, João Paulo Kitajima, Erika Freitas, David Schlesinger. Hemocentro Ribeirão Preto: Simone Kashima, Evandra Strazza Rodrigues, Svetoslav Nanev Slavov, Elaine Vieira dos Santos, Rafael dos Santos Bezerra, Luiz Carlos Junior de Alcantara, Marta Giovanetti, Vagner Fonseca, Flavia Aburjaile, Rodrigo Tocantins Calado. FAMERP-SJRP: Cecília Artico Banho, Lívia Sacchetto, Fábio Sossai Possebon, Leila Sabrina Ullmann, Cíntia Bittar, Guilherme Campos, Helena Lage Ferreira, Jorge A. Petroli Marchesi, Maisa C. Pereira Parra, Marília Moraes, Paula Rahal, Paulo Inacio da Costa, João Pessoa Araújo Jr., Maurício Lacerda Nogueira. Prefeitura de Sao Paulo: Melissa Palmieri. |
| EPI_ISL_1966716, EPI_ISL_1966717, EPI_ISL_1966718, EPI_ISL_1966719 | SECAO CENTRO DE DIAGNOSTICO SECEDI | Instituto Butantan / Mendelics | Instituto Butantan: Dimas Tadeu Covas, Sandra Coccuzzo Sampaio, Maria Carolina Elias, José Salvatore Leister Patané, Vincent Louis Viala, Antonio Jorge Martins, Ricardo Haddad, Claudia Renata dos Santos Barros, Elaine Cristina Marqueze, Raul Machado Neto, Debora Botequio Moretti, Jardelina de Souza Todao Bernardino, Loyze Paola Oliveira de Lima, Luiz Aurelio de Campos Crispin. Centro de Genômica Funcional da ESALQ: Luiz Lehmann Coutinho, Ricardo Augusto Brassaloti, Raquel de Lello Rocha Campos Cassano. NGS Soluções Genômicas: Pilar Drummond Sampaio Corrêa Mariani. FZEA-USP Pirassununga: Mirele Daiana Poleti, Jessika Cristina Chagas Lesbon, Elisangela Chicaroni Mattos, Heidge Fukumasu. USP-Botucatu: Rejane Maria Tommasini Grotto, Jayme A. Souza-Neto, Guilherme Targino Valente, Patricia Akemi Assato, Felipe Allan da Silva da Costa, Bianca Cechetto Carlos. Mendelics: Bibiana Santos, João Paulo Kitajima, Erika Freitas, David Schlesinger. Hemocentro Ribeirão Preto: Simone Kashima, Evandra Strazza Rodrigues, Svetoslav Nanev Slavov, Elaine Vieira dos Santos, Rafael dos Santos Bezerra, Luiz Carlos Junior de Alcantara, Marta Giovanetti, Vagner Fonseca, Flavia Aburjaile, Rodrigo Tocantins Calado. FAMERP-SJRP: Cecília Artico Banho, Lívia Sacchetto, Fábio Sossai Possebon, Leila Sabrina Ullmann, Cíntia Bittar, Guilherme Campos, Helena Lage Ferreira, Jorge A. Petroli Marchesi, Maisa C. Pereira Parra, Marília Moraes, Paula Rahal, Paulo Inacio da Costa, João Pessoa Araújo Jr., Maurício Lacerda Nogueira. Prefeitura de Sao Paulo: Melissa Palmieri. |
| EPI_ISL_1966720 | UNIDADE DE VIGILANCIA EM SAUDE | Instituto Butantan / Mendelics | Instituto Butantan: Dimas Tadeu Covas, Sandra Coccuzzo Sampaio, Maria Carolina Elias, José Salvatore Leister Patané, Vincent Louis Viala, Antonio Jorge Martins, Ricardo Haddad, Claudia Renata dos Santos Barros, Elaine Cristina Marqueze, Raul Machado Neto, Debora Botequio Moretti, Jardelina de Souza Todao Bernardino, Loyze Paola Oliveira de Lima, Luiz Aurelio de Campos Crispin. Centro de Genômica Funcional da ESALQ: Luiz Lehmann Coutinho, Ricardo Augusto Brassaloti, Raquel de Lello Rocha Campos Cassano. NGS Soluções Genômicas: Pilar Drummond Sampaio Corrêa Mariani. FZEA-USP Pirassununga: Mirele Daiana Poleti, Jessika Cristina Chagas Lesbon, Elisangela Chicaroni Mattos, Heidge Fukumasu. USP-Botucatu: Rejane Maria Tommasini Grotto, Jayme A. Souza-Neto, Guilherme Targino Valente, Patricia Akemi Assato, Felipe Allan da Silva da Costa, Bianca Cechetto Carlos. Mendelics: Bibiana Santos, João Paulo Kitajima, Erika Freitas, David Schlesinger. Hemocentro Ribeirão Preto: Simone Kashima, Evandra Strazza Rodrigues, Svetoslav Nanev Slavov, Elaine Vieira dos Santos, Rafael dos Santos Bezerra, Luiz Carlos Junior de Alcantara, Marta Giovanetti, Vagner Fonseca, Flavia Aburjaile, Rodrigo Tocantins Calado. FAMERP-SJRP: Cecília Artico Banho, Lívia Sacchetto, Fábio Sossai Possebon, Leila Sabrina Ullmann, Cíntia Bittar, Guilherme Campos, Helena Lage Ferreira, Jorge A. Petroli Marchesi, Maisa C. Pereira Parra, Marília Moraes, Paula Rahal, Paulo Inacio da Costa, João Pessoa Araújo Jr., Maurício Lacerda Nogueira. Prefeitura de Sao Paulo: Melissa Palmieri. |
| EPI_ISL_1966721, EPI_ISL_1966722, EPI_ISL_1966723 | UNIDADE DE PRONTO ATENDIMENTO DE VARZEA PAULISTA UPA II | Instituto Butantan / Mendelics | Instituto Butantan: Dimas Tadeu Covas, Sandra Coccuzzo Sampaio, Maria Carolina Elias, José Salvatore Leister Patané, Vincent Louis Viala, Antonio Jorge Martins, Ricardo Haddad, Claudia Renata dos Santos Barros, Elaine Cristina Marqueze, Raul Machado Neto, Debora Botequio Moretti, Jardelina de Souza Todao Bernardino, Loyze Paola Oliveira de Lima, Luiz Aurelio de Campos Crispin. Centro de Genômica Funcional da ESALQ: Luiz Lehmann Coutinho, Ricardo Augusto Brassaloti, Raquel de Lello Rocha Campos Cassano. NGS Soluções Genômicas: Pilar Drummond Sampaio Corrêa Mariani. FZEA-USP Pirassununga: Mirele Daiana Poleti, Jessika Cristina Chagas Lesbon, Elisangela Chicaroni Mattos, Heidge Fukumasu. USP-Botucatu: Rejane Maria Tommasini Grotto, Jayme A. Souza-Neto, Guilherme Targino Valente, Patricia Akemi Assato, Felipe Allan da Silva da Costa, Bianca Cechetto Carlos. Mendelics: Bibiana Santos, João Paulo Kitajima, Erika Freitas, David Schlesinger. Hemocentro Ribeirão Preto: Simone Kashima, Evandra Strazza Rodrigues, Svetoslav Nanev Slavov, Elaine Vieira dos Santos, Rafael dos Santos Bezerra, Luiz Carlos Junior de Alcantara, Marta Giovanetti, Vagner Fonseca, Flavia Aburjaile, Rodrigo Tocantins Calado. FAMERP-SJRP: Cecília Artico Banho, Lívia Sacchetto, Fábio Sossai Possebon, Leila Sabrina Ullmann, Cíntia Bittar, Guilherme Campos, Helena Lage Ferreira, Jorge A. Petroli Marchesi, Maisa C. Pereira Parra, Marília Moraes, Paula Rahal, Paulo Inacio da Costa, João Pessoa Araújo Jr., Maurício Lacerda Nogueira. Prefeitura de Sao Paulo: Melissa Palmieri. |
| EPI_ISL_1966724, EPI_ISL_1966725, EPI_ISL_1966726 | GRUPO TECNICO DE VIGILANCIA SANITARIA E EPIDEMIOLOGICA | Instituto Butantan | Instituto Butantan: Dimas Tadeu Covas, Sandra Coccuzzo Sampaio, Maria Carolina Elias, José Salvatore Leister Patané, Vincent Louis Viala, Antonio Jorge Martins, Ricardo Haddad, Claudia Renata dos Santos Barros, Elaine Cristina Marqueze, Raul Machado Neto, Debora Botequio Moretti, Jardelina de Souza Todao Bernardino, Loyze Paola Oliveira de Lima, Luiz Aurelio de Campos Crispin. Centro de Genômica Funcional da ESALQ: Luiz Lehmann Coutinho, Ricardo Augusto Brassaloti, Raquel de Lello Rocha Campos Cassano. NGS Soluções Genômicas: Pilar Drummond Sampaio Corrêa Mariani. FZEA-USP Pirassununga: Mirele Daiana Poleti, Jessika Cristina Chagas Lesbon, Elisangela Chicaroni Mattos, Heidge Fukumasu. USP-Botucatu: Rejane Maria Tommasini Grotto, Jayme A. Souza-Neto, Guilherme Targino Valente, Patricia Akemi Assato, Felipe Allan da Silva da Costa, Bianca Cechetto Carlos. Mendelics: Bibiana Santos, João Paulo Kitajima, Erika Freitas, David Schlesinger. Hemocentro Ribeirão Preto: Simone Kashima, Evandra Strazza Rodrigues, Svetoslav Nanev Slavov, Elaine Vieira dos Santos, Rafael dos Santos Bezerra, Luiz Carlos Junior de Alcantara, Marta Giovanetti, Vagner |

| | | | |
|---|---|---|---|
| | | | Fonseca, Flavia Aburjaile, Rodrigo Tocantins Calado. FAMERP-SJRP: Cecília Artico Banho, Lívia Sacchetto, Fábio Sossai Possebon, Leila Sabrina Ullmann, Cíntia Bittar, Guilherme Campos, Helena Lage Ferreira, Jorge A. Petroli Marchesi, Maisa C. Pereira Parra, Marília Moraes, Paula Rahal, Paulo Inacio da Costa, João Pessoa Araújo Jr., Maurício Lacerda Nogueira. Prefeitura de Sao Paulo: Melissa Palmieri. |
| EPI_ISL_1966727 | PPA MARCIO VALDEVINO BATISTA | Instituto Butantan | Instituto Butantan: Dimas Tadeu Covas, Sandra Coccuzzo Sampaio, Maria Carolina Elias, José Salvatore Leister Patané, Vincent Louis Viala, Antonio Jorge Martins, Ricardo Haddad, Claudia Renata dos Santos Barros, Elaine Cristina Marqueze, Raul Machado Neto, Debora Botequio Moretti, Jardelina de Souza Todao Bernardino, Loyze Paola Oliveira de Lima, Luiz Aurelio de Campos Crispin. Centro de Genômica Funcional da ESALQ: Luiz Lehmann Coutinho, Ricardo Augusto Brassaloti, Raquel de Lello Rocha Campos Cassano. NGS Soluções Genômicas: Pilar Drummond Sampaio Corrêa Mariani. FZEA-USP Pirassununga: Mirele Daiana Poleti, Jessika Cristina Chagas Lesbon, Elisangela Chicaroni Mattos, Heidge Fukumasu. USP-Botucatu: Rejane Maria Tommasini Grotto, Jayme A. Souza-Neto, Guilherme Targino Valente, Patricia Akemi Assato, Felipe Allan da Silva da Costa, Bianca Cechetto Carlos. Mendelics: Bibiana Santos, João Paulo Kitajima, Erika Freitas, David Schlesinger. Hemocentro Ribeirão Preto: Simone Kashima, Evandra Strazza Rodrigues, Svetoslav Nanev Slavov, Elaine Vieira dos Santos, Rafael dos Santos Bezerra, Luiz Carlos Junior de Alcantara, Marta Giovanetti, Vagner Fonseca, Flavia Aburjaile, Rodrigo Tocantins Calado. FAMERP-SJRP: Cecília Artico Banho, Lívia Sacchetto, Fábio Sossai Possebon, Leila Sabrina Ullmann, Cíntia Bittar, Guilherme Campos, Helena Lage Ferreira, Jorge A. Petroli Marchesi, Maisa C. Pereira Parra, Marília Moraes, Paula Rahal, Paulo Inacio da Costa, João Pessoa Araújo Jr., Maurício Lacerda Nogueira. Prefeitura de Sao Paulo: Melissa Palmieri. |
| EPI_ISL_1966728 | UBS JOSE SABINO FERREIRA | Instituto Butantan | Instituto Butantan: Dimas Tadeu Covas, Sandra Coccuzzo Sampaio, Maria Carolina Elias, José Salvatore Leister Patané, Vincent Louis Viala, Antonio Jorge Martins, Ricardo Haddad, Claudia Renata dos Santos Barros, Elaine Cristina Marqueze, Raul Machado Neto, Debora Botequio Moretti, Jardelina de Souza Todao Bernardino, Loyze Paola Oliveira de Lima, Luiz Aurelio de Campos Crispin. Centro de Genômica Funcional da ESALQ: Luiz Lehmann Coutinho, Ricardo Augusto Brassaloti, Raquel de Lello Rocha Campos Cassano. NGS Soluções Genômicas: Pilar Drummond Sampaio Corrêa Mariani. FZEA-USP Pirassununga: Mirele Daiana Poleti, Jessika Cristina Chagas Lesbon, Elisangela Chicaroni Mattos, Heidge Fukumasu. USP-Botucatu: Rejane Maria Tommasini Grotto, Jayme A. Souza-Neto, Guilherme Targino Valente, Patricia Akemi Assato, Felipe Allan da Silva da Costa, Bianca Cechetto Carlos. Mendelics: Bibiana Santos, João Paulo Kitajima, Erika Freitas, David Schlesinger. Hemocentro Ribeirão Preto: Simone Kashima, Evandra Strazza Rodrigues, Svetoslav Nanev Slavov, Elaine Vieira dos Santos, Rafael dos Santos Bezerra, Luiz Carlos Junior de Alcantara, Marta Giovanetti, Vagner Fonseca, Flavia Aburjaile, Rodrigo Tocantins Calado. FAMERP-SJRP: Cecília Artico Banho, Lívia Sacchetto, Fábio Sossai Possebon, Leila Sabrina Ullmann, Cíntia Bittar, Guilherme Campos, Helena Lage Ferreira, Jorge A. Petroli Marchesi, Maisa C. Pereira Parra, Marília Moraes, Paula Rahal, Paulo Inacio da Costa, João Pessoa Araújo Jr., Maurício Lacerda Nogueira. Prefeitura de Sao Paulo: Melissa Palmieri. |
| EPI_ISL_1966729 | UPA JOSE CAMPOS BARRETO ZEQUINHA BARRETO | Instituto Butantan | Instituto Butantan: Dimas Tadeu Covas, Sandra Coccuzzo Sampaio, Maria Carolina Elias, José Salvatore Leister Patané, Vincent Louis Viala, Antonio Jorge Martins, Ricardo Haddad, Claudia Renata dos Santos Barros, Elaine Cristina Marqueze, Raul Machado Neto, Debora Botequio Moretti, Jardelina de Souza Todao Bernardino, Loyze Paola Oliveira de Lima, Luiz Aurelio de Campos Crispin. Centro de Genômica Funcional da ESALQ: Luiz Lehmann Coutinho, Ricardo Augusto Brassaloti, Raquel de Lello Rocha Campos Cassano. NGS Soluções Genômicas: Pilar Drummond Sampaio Corrêa Mariani. FZEA-USP Pirassununga: Mirele Daiana Poleti, Jessika Cristina Chagas Lesbon, Elisangela Chicaroni Mattos, Heidge Fukumasu. USP-Botucatu: Rejane Maria Tommasini Grotto, Jayme A. Souza-Neto, Guilherme Targino Valente, Patricia Akemi Assato, Felipe Allan da Silva da Costa, Bianca Cechetto Carlos. Mendelics: Bibiana Santos, João Paulo Kitajima, Erika Freitas, David Schlesinger. Hemocentro Ribeirão Preto: Simone Kashima, Evandra Strazza Rodrigues, Svetoslav Nanev Slavov, Elaine Vieira dos Santos, Rafael dos Santos Bezerra, Luiz Carlos Junior de Alcantara, Marta Giovanetti, Vagner Fonseca, Flavia Aburjaile, Rodrigo Tocantins Calado. FAMERP-SJRP: Cecília Artico Banho, Lívia Sacchetto, Fábio Sossai Possebon, Leila Sabrina Ullmann, Cíntia Bittar, Guilherme Campos, Helena Lage Ferreira, Jorge A. Petroli Marchesi, Maisa C. Pereira Parra, Marília Moraes, Paula Rahal, Paulo Inacio da Costa, João Pessoa Araújo Jr., Maurício Lacerda Nogueira. Prefeitura de Sao Paulo: Melissa Palmieri. |
| EPI_ISL_1966730 | UBS JOSE SABINO FERREIRA | Instituto Butantan | Instituto Butantan: Dimas Tadeu Covas, Sandra Coccuzzo Sampaio, Maria Carolina Elias, José Salvatore Leister Patané, Vincent Louis Viala, Antonio Jorge Martins, Ricardo Haddad, Claudia Renata dos Santos Barros, Elaine Cristina Marqueze, Raul Machado Neto, Debora Botequio Moretti, Jardelina de Souza Todao Bernardino, Loyze Paola Oliveira de Lima, Luiz Aurelio de Campos Crispin. Centro de Genômica Funcional da ESALQ: Luiz Lehmann Coutinho, Ricardo Augusto Brassaloti, Raquel de Lello Rocha Campos Cassano. NGS Soluções Genômicas: Pilar Drummond Sampaio Corrêa Mariani. FZEA-USP Pirassununga: Mirele Daiana Poleti, Jessika Cristina Chagas Lesbon, Elisangela Chicaroni Mattos, Heidge Fukumasu. USP-Botucatu: Rejane Maria Tommasini Grotto, Jayme A. Souza-Neto, Guilherme Targino Valente, Patricia Akemi Assato, Felipe Allan da Silva da Costa, Bianca Cechetto Carlos. Mendelics: Bibiana Santos, João Paulo Kitajima, Erika Freitas, David Schlesinger. Hemocentro Ribeirão Preto: Simone Kashima, Evandra Strazza Rodrigues, Svetoslav Nanev Slavov, Elaine Vieira dos Santos, Rafael dos Santos Bezerra, Luiz Carlos Junior de Alcantara, Marta Giovanetti, Vagner Fonseca, Flavia Aburjaile, Rodrigo Tocantins Calado. FAMERP-SJRP: Cecília Artico Banho, Lívia Sacchetto, Fábio Sossai Possebon, Leila Sabrina Ullmann, Cíntia Bittar, Guilherme Campos, Helena Lage Ferreira, Jorge A. Petroli Marchesi, Maisa C. Pereira Parra, Marília Moraes, Paula Rahal, Paulo Inacio da Costa, João Pessoa Araújo Jr., Maurício Lacerda Nogueira. Prefeitura de Sao Paulo: Melissa Palmieri. |
| EPI_ISL_1966731, EPI_ISL_1966732, EPI_ISL_1966733, EPI_ISL_1966734 | UBS RAIMUNDA CAVALCANTE DE SOUZA | Instituto Butantan | Instituto Butantan: Dimas Tadeu Covas, Sandra Coccuzzo Sampaio, Maria Carolina Elias, José Salvatore Leister Patané, Vincent Louis Viala, Antonio Jorge Martins, Ricardo Haddad, Claudia Renata dos Santos Barros, Elaine Cristina Marqueze, Raul Machado Neto, Debora Botequio Moretti, Jardelina de Souza Todao Bernardino, Loyze Paola Oliveira de Lima, Luiz Aurelio de Campos Crispin. Centro de Genômica Funcional da ESALQ: Luiz Lehmann Coutinho, Ricardo Augusto Brassaloti, Raquel de Lello Rocha Campos Cassano. NGS Soluções Genômicas: Pilar Drummond Sampaio Corrêa Mariani. FZEA-USP Pirassununga: Mirele Daiana Poleti, Jessika Cristina Chagas Lesbon, Elisangela Chicaroni Mattos, Heidge Fukumasu. USP-Botucatu: Rejane Maria Tommasini Grotto, Jayme A. Souza-Neto, Guilherme Targino Valente, Patricia Akemi Assato, Felipe Allan da Silva da Costa, Bianca Cechetto Carlos. Mendelics: Bibiana Santos, João Paulo Kitajima, Erika Freitas, David Schlesinger. Hemocentro Ribeirão Preto: Simone Kashima, Evandra Strazza Rodrigues, Svetoslav Nanev Slavov, Elaine Vieira dos Santos, Rafael dos Santos Bezerra, Luiz Carlos Junior de Alcantara, Marta Giovanetti, Vagner Fonseca, Flavia Aburjaile, Rodrigo Tocantins Calado. FAMERP-SJRP: Cecília Artico Banho, Lívia Sacchetto, Fábio Sossai Possebon, Leila Sabrina Ullmann, Cíntia Bittar, Guilherme Campos, Helena Lage Ferreira, Jorge A. Petroli Marchesi, Maisa C. Pereira Parra, Marília Moraes, Paula Rahal, Paulo Inacio da Costa, João Pessoa Araújo Jr., Maurício Lacerda Nogueira. Prefeitura de Sao Paulo: Melissa Palmieri. |
| EPI_ISL_1966735, EPI_ISL_1966736 | PRONTO SOCORRO PESTANA P S ANDRE SACCO | Instituto Butantan | Instituto Butantan: Dimas Tadeu Covas, Sandra Coccuzzo Sampaio, Maria Carolina Elias, José Salvatore Leister Patané, Vincent Louis Viala, Antonio Jorge Martins, Ricardo Haddad, Claudia Renata dos Santos Barros, Elaine Cristina Marqueze, Raul Machado Neto, Debora Botequio Moretti, Jardelina de Souza Todao Bernardino, Loyze Paola Oliveira de Lima, Luiz Aurelio de Campos Crispin. Centro de Genômica Funcional da ESALQ: Luiz Lehmann Coutinho, Ricardo Augusto Brassaloti, Raquel de Lello Rocha Campos Cassano. NGS Soluções Genômicas: Pilar Drummond Sampaio Corrêa Mariani. FZEA-USP Pirassununga: Mirele Daiana Poleti, Jessika Cristina Chagas Lesbon, Elisangela Chicaroni Mattos, Heidge Fukumasu. USP-Botucatu: Rejane Maria Tommasini Grotto, Jayme A. Souza-Neto, Guilherme Targino Valente, Patricia Akemi Assato, Felipe Allan da Silva da Costa, Bianca Cechetto Carlos. Mendelics: Bibiana Santos, João Paulo Kitajima, Erika Freitas, David Schlesinger. Hemocentro Ribeirão Preto: Simone Kashima, Evandra Strazza Rodrigues, Svetoslav Nanev Slavov, Elaine Vieira dos Santos, Rafael dos Santos Bezerra, Luiz Carlos Junior de Alcantara, Marta Giovanetti, Vagner Fonseca, Flavia Aburjaile, Rodrigo Tocantins Calado. FAMERP-SJRP: Cecília Artico Banho, Lívia Sacchetto, Fábio Sossai Possebon, Leila Sabrina Ullmann, Cíntia Bittar, Guilherme Campos, Helena Lage Ferreira, Jorge A. Petroli Marchesi, Maisa C. Pereira Parra, Marília Moraes, Paula Rahal, Paulo Inacio da Costa, João Pessoa Araújo Jr., Maurício Lacerda Nogueira. Prefeitura de Sao Paulo: Melissa Palmieri. |
| EPI_ISL_1966737 | HOSP MUN JOSANIAS CASTANHA BRAGA | Instituto Butantan | Instituto Butantan: Dimas Tadeu Covas, Sandra Coccuzzo Sampaio, Maria Carolina Elias, José Salvatore Leister Patané, Vincent Louis Viala, Antonio Jorge Martins, Ricardo Haddad, Claudia Renata dos Santos Barros, Elaine Cristina Marqueze, Raul Machado Neto, Debora Botequio Moretti, Jardelina de Souza Todao Bernardino, Loyze Paola Oliveira de Lima, Luiz Aurelio de Campos Crispin. Centro de Genômica Funcional da ESALQ: Luiz Lehmann Coutinho, Ricardo Augusto Brassaloti, Raquel de Lello Rocha Campos Cassano. NGS Soluções Genômicas: Pilar Drummond Sampaio Corrêa Mariani. FZEA-USP Pirassununga: Mirele Daiana Poleti, Jessika Cristina Chagas Lesbon, Elisangela Chicaroni Mattos, Heidge Fukumasu. USP-Botucatu: Rejane Maria Tommasini Grotto, Jayme A. Souza-Neto, Guilherme Targino Valente, Patricia Akemi Assato, Felipe Allan da Silva da Costa, Bianca Cechetto Carlos. Mendelics: Bibiana Santos, João Paulo Kitajima, Erika Freitas, David Schlesinger. Hemocentro Ribeirão Preto: Simone Kashima, Evandra Strazza Rodrigues, Svetoslav Nanev Slavov, Elaine Vieira dos Santos, Rafael dos Santos Bezerra, Luiz Carlos Junior de Alcantara, Marta Giovanetti, Vagner Fonseca, Flavia Aburjaile, Rodrigo Tocantins Calado. FAMERP-SJRP: Cecília Artico Banho, Lívia Sacchetto, Fábio Sossai Possebon, Leila Sabrina Ullmann, Cíntia Bittar, Guilherme Campos, Helena Lage Ferreira, Jorge A. Petroli Marchesi, Maisa C. Pereira Parra, Marília Moraes, Paula Rahal, Paulo Inacio da Costa, João Pessoa Araújo Jr., Maurício Lacerda Nogueira. Prefeitura de Sao Paulo: Melissa Palmieri. |
| EPI_ISL_1966738 | HOSPITAL GERAL DE VILA NOVA CACHOEIRINHA SAO PAULO | Instituto Butantan | Instituto Butantan: Dimas Tadeu Covas, Sandra Coccuzzo Sampaio, Maria Carolina Elias, José Salvatore Leister Patané, Vincent Louis Viala, Antonio Jorge Martins, Ricardo Haddad, Claudia Renata dos Santos Barros, Elaine Cristina Marqueze, Raul Machado Neto, Debora Botequio Moretti, Jardelina de Souza Todao Bernardino, Loyze Paola Oliveira de Lima, Luiz Aurelio de Campos Crispin. Centro de Genômica Funcional da ESALQ: Luiz Lehmann |

| | | | |
|---|---|---|---|
| | | | Coutinho, Ricardo Augusto Brassaloti, Raquel de Lello Rocha Campos Cassano. NGS Soluções Genômicas: Pilar Drummond Sampaio Corrêa Mariani. FZEA-USP Pirassununga: Mirele Daiana Poleti, Jessika Cristina Chagas Lesbon, Elisangela Chicaroni Mattos, Heidge Fukumasu. USP-Botucatu: Rejane Maria Tommasini Grotto, Jayme A. Souza-Neto, Guilherme Targino Valente, Patricia Akemi Assato, Felipe Allan da Silva da Costa, Bianca Cechetto Carlos. Mendelics: Bibiana Santos, João Paulo Kitajima, Erika Freitas, David Schlesinger. Hemocentro Ribeirão Preto: Simone Kashima, Evandra Strazza Rodrigues, Svetoslav Nanev Slavov, Elaine Vieira dos Santos, Rafael dos Santos Bezerra, Luiz Carlos Junior de Alcantara, Marta Giovanetti, Vagner Fonseca, Flavia Aburjaile, Rodrigo Tocantins Calado. FAMERP-SJRP: Cecília Artico Banho, Lívia Sacchetto, Fábio Sossai Possebon, Leila Sabrina Ullmann, Cíntia Bittar, Guilherme Campos, Helena Lage Ferreira, Jorge A. Petroli Marchesi, Maisa C. Pereira Parra, Marília Moraes, Paula Rahal, Paulo Inacio da Costa, João Pessoa Araújo Jr., Maurício Lacerda Nogueira. Prefeitura de Sao Paulo: Melissa Palmieri. |
| EPI_ISL_1966739, EPI_ISL_1966740, EPI_ISL_1966741 | CS DE AMERICO DE CAMPOS | Instituto Butantan / Mendelics | Instituto Butantan: Dimas Tadeu Covas, Sandra Coccuzzo Sampaio, Maria Carolina Elias, José Salvatore Leister Patané, Vincent Louis Viala, Antonio Jorge Martins, Ricardo Haddad, Claudia Renata dos Santos Barros, Elaine Cristina Marqueze, Raul Machado Neto, Debora Botequio Moretti, Jardelina de Souza Todao Bernardino, Loyze Paola Oliveira de Lima, Luiz Aurelio de Campos Crispin. Centro de Genômica Funcional da ESALQ: Luiz Lehmann Coutinho, Ricardo Augusto Brassaloti, Raquel de Lello Rocha Campos Cassano. NGS Soluções Genômicas: Pilar Drummond Sampaio Corrêa Mariani. FZEA-USP Pirassununga: Mirele Daiana Poleti, Jessika Cristina Chagas Lesbon, Elisangela Chicaroni Mattos, Heidge Fukumasu. USP-Botucatu: Rejane Maria Tommasini Grotto, Jayme A. Souza-Neto, Guilherme Targino Valente, Patricia Akemi Assato, Felipe Allan da Silva da Costa, Bianca Cechetto Carlos. Mendelics: Bibiana Santos, João Paulo Kitajima, Erika Freitas, David Schlesinger. Hemocentro Ribeirão Preto: Simone Kashima, Evandra Strazza Rodrigues, Svetoslav Nanev Slavov, Elaine Vieira dos Santos, Rafael dos Santos Bezerra, Luiz Carlos Junior de Alcantara, Marta Giovanetti, Vagner Fonseca, Flavia Aburjaile, Rodrigo Tocantins Calado. FAMERP-SJRP: Cecília Artico Banho, Lívia Sacchetto, Fábio Sossai Possebon, Leila Sabrina Ullmann, Cíntia Bittar, Guilherme Campos, Helena Lage Ferreira, Jorge A. Petroli Marchesi, Maisa C. Pereira Parra, Marília Moraes, Paula Rahal, Paulo Inacio da Costa, João Pessoa Araújo Jr., Maurício Lacerda Nogueira. Prefeitura de Sao Paulo: Melissa Palmieri. |
| EPI_ISL_1966742, EPI_ISL_1966743, EPI_ISL_1966744 | CENTRO DE SAUDE DR RENATO DE CARVALHO RIBEIRO ANGATUBA | Instituto Butantan / Mendelics | Instituto Butantan: Dimas Tadeu Covas, Sandra Coccuzzo Sampaio, Maria Carolina Elias, José Salvatore Leister Patané, Vincent Louis Viala, Antonio Jorge Martins, Ricardo Haddad, Claudia Renata dos Santos Barros, Elaine Cristina Marqueze, Raul Machado Neto, Debora Botequio Moretti, Jardelina de Souza Todao Bernardino, Loyze Paola Oliveira de Lima, Luiz Aurelio de Campos Crispin. Centro de Genômica Funcional da ESALQ: Luiz Lehmann Coutinho, Ricardo Augusto Brassaloti, Raquel de Lello Rocha Campos Cassano. NGS Soluções Genômicas: Pilar Drummond Sampaio Corrêa Mariani. FZEA-USP Pirassununga: Mirele Daiana Poleti, Jessika Cristina Chagas Lesbon, Elisangela Chicaroni Mattos, Heidge Fukumasu. USP-Botucatu: Rejane Maria Tommasini Grotto, Jayme A. Souza-Neto, Guilherme Targino Valente, Patricia Akemi Assato, Felipe Allan da Silva da Costa, Bianca Cechetto Carlos. Mendelics: Bibiana Santos, João Paulo Kitajima, Erika Freitas, David Schlesinger. Hemocentro Ribeirão Preto: Simone Kashima, Evandra Strazza Rodrigues, Svetoslav Nanev Slavov, Elaine Vieira dos Santos, Rafael dos Santos Bezerra, Luiz Carlos Junior de Alcantara, Marta Giovanetti, Vagner Fonseca, Flavia Aburjaile, Rodrigo Tocantins Calado. FAMERP-SJRP: Cecília Artico Banho, Lívia Sacchetto, Fábio Sossai Possebon, Leila Sabrina Ullmann, Cíntia Bittar, Guilherme Campos, Helena Lage Ferreira, Jorge A. Petroli Marchesi, Maisa C. Pereira Parra, Marília Moraes, Paula Rahal, Paulo Inacio da Costa, João Pessoa Araújo Jr., Maurício Lacerda Nogueira. Prefeitura de Sao Paulo: Melissa Palmieri. |
| EPI_ISL_1966745 | VIGILANCIA EPIDIMIOLOGICA DE ARACARIGUAMA | Instituto Butantan / Mendelics | Instituto Butantan: Dimas Tadeu Covas, Sandra Coccuzzo Sampaio, Maria Carolina Elias, José Salvatore Leister Patané, Vincent Louis Viala, Antonio Jorge Martins, Ricardo Haddad, Claudia Renata dos Santos Barros, Elaine Cristina Marqueze, Raul Machado Neto, Debora Botequio Moretti, Jardelina de Souza Todao Bernardino, Loyze Paola Oliveira de Lima, Luiz Aurelio de Campos Crispin. Centro de Genômica Funcional da ESALQ: Luiz Lehmann Coutinho, Ricardo Augusto Brassaloti, Raquel de Lello Rocha Campos Cassano. NGS Soluções Genômicas: Pilar Drummond Sampaio Corrêa Mariani. FZEA-USP Pirassununga: Mirele Daiana Poleti, Jessika Cristina Chagas Lesbon, Elisangela Chicaroni Mattos, Heidge Fukumasu. USP-Botucatu: Rejane Maria Tommasini Grotto, Jayme A. Souza-Neto, Guilherme Targino Valente, Patricia Akemi Assato, Felipe Allan da Silva da Costa, Bianca Cechetto Carlos. Mendelics: Bibiana Santos, João Paulo Kitajima, Erika Freitas, David Schlesinger. Hemocentro Ribeirão Preto: Simone Kashima, Evandra Strazza Rodrigues, Svetoslav Nanev Slavov, Elaine Vieira dos Santos, Rafael dos Santos Bezerra, Luiz Carlos Junior de Alcantara, Marta Giovanetti, Vagner Fonseca, Flavia Aburjaile, Rodrigo Tocantins Calado. FAMERP-SJRP: Cecília Artico Banho, Lívia Sacchetto, Fábio Sossai Possebon, Leila Sabrina Ullmann, Cíntia Bittar, Guilherme Campos, Helena Lage Ferreira, Jorge A. Petroli Marchesi, Maisa C. Pereira Parra, Marília Moraes, Paula Rahal, Paulo Inacio da Costa, João Pessoa Araújo Jr., Maurício Lacerda Nogueira. Prefeitura de Sao Paulo: Melissa Palmieri. |
| EPI_ISL_1966746, EPI_ISL_1966747, EPI_ISL_1966748 | SMS SECRETARIA MUNICIPAL DE SAUDE DE BOITUVA | Instituto Butantan / Mendelics | Instituto Butantan: Dimas Tadeu Covas, Sandra Coccuzzo Sampaio, Maria Carolina Elias, José Salvatore Leister Patané, Vincent Louis Viala, Antonio Jorge Martins, Ricardo Haddad, Claudia Renata dos Santos Barros, Elaine Cristina Marqueze, Raul Machado Neto, Debora Botequio Moretti, Jardelina de Souza Todao Bernardino, Loyze Paola Oliveira de Lima, Luiz Aurelio de Campos Crispin. Centro de Genômica Funcional da ESALQ: Luiz Lehmann Coutinho, Ricardo Augusto Brassaloti, Raquel de Lello Rocha Campos Cassano. NGS Soluções Genômicas: Pilar Drummond Sampaio Corrêa Mariani. FZEA-USP Pirassununga: Mirele Daiana Poleti, Jessika Cristina Chagas Lesbon, Elisangela Chicaroni Mattos, Heidge Fukumasu. USP-Botucatu: Rejane Maria Tommasini Grotto, Jayme A. Souza-Neto, Guilherme Targino Valente, Patricia Akemi Assato, Felipe Allan da Silva da Costa, Bianca Cechetto Carlos. Mendelics: Bibiana Santos, João Paulo Kitajima, Erika Freitas, David Schlesinger. Hemocentro Ribeirão Preto: Simone Kashima, Evandra Strazza Rodrigues, Svetoslav Nanev Slavov, Elaine Vieira dos Santos, Rafael dos Santos Bezerra, Luiz Carlos Junior de Alcantara, Marta Giovanetti, Vagner Fonseca, Flavia Aburjaile, Rodrigo Tocantins Calado. FAMERP-SJRP: Cecília Artico Banho, Lívia Sacchetto, Fábio Sossai Possebon, Leila Sabrina Ullmann, Cíntia Bittar, Guilherme Campos, Helena Lage Ferreira, Jorge A. Petroli Marchesi, Maisa C. Pereira Parra, Marília Moraes, Paula Rahal, Paulo Inacio da Costa, João Pessoa Araújo Jr., Maurício Lacerda Nogueira. Prefeitura de Sao Paulo: Melissa Palmieri. |
| EPI_ISL_1966749, EPI_ISL_1966750 | CENTRO DE SAUDE III BORBOREMA | Instituto Butantan / Mendelics | Instituto Butantan: Dimas Tadeu Covas, Sandra Coccuzzo Sampaio, Maria Carolina Elias, José Salvatore Leister Patané, Vincent Louis Viala, Antonio Jorge Martins, Ricardo Haddad, Claudia Renata dos Santos Barros, Elaine Cristina Marqueze, Raul Machado Neto, Debora Botequio Moretti, Jardelina de Souza Todao Bernardino, Loyze Paola Oliveira de Lima, Luiz Aurelio de Campos Crispin. Centro de Genômica Funcional da ESALQ: Luiz Lehmann Coutinho, Ricardo Augusto Brassaloti, Raquel de Lello Rocha Campos Cassano. NGS Soluções Genômicas: Pilar Drummond Sampaio Corrêa Mariani. FZEA-USP Pirassununga: Mirele Daiana Poleti, Jessika Cristina Chagas Lesbon, Elisangela Chicaroni Mattos, Heidge Fukumasu. USP-Botucatu: Rejane Maria Tommasini Grotto, Jayme A. Souza-Neto, Guilherme Targino Valente, Patricia Akemi Assato, Felipe Allan da Silva da Costa, Bianca Cechetto Carlos. Mendelics: Bibiana Santos, João Paulo Kitajima, Erika Freitas, David Schlesinger. Hemocentro Ribeirão Preto: Simone Kashima, Evandra Strazza Rodrigues, Svetoslav Nanev Slavov, Elaine Vieira dos Santos, Rafael dos Santos Bezerra, Luiz Carlos Junior de Alcantara, Marta Giovanetti, Vagner Fonseca, Flavia Aburjaile, Rodrigo Tocantins Calado. FAMERP-SJRP: Cecília Artico Banho, Lívia Sacchetto, Fábio Sossai Possebon, Leila Sabrina Ullmann, Cíntia Bittar, Guilherme Campos, Helena Lage Ferreira, Jorge A. Petroli Marchesi, Maisa C. Pereira Parra, Marília Moraes, Paula Rahal, Paulo Inacio da Costa, João Pessoa Araújo Jr., Maurício Lacerda Nogueira. Prefeitura de Sao Paulo: Melissa Palmieri. |
| EPI_ISL_1966751 | PRONTO ATENDIMENTO MUNICIPAL ITALO SANTUCCI | Instituto Butantan / Mendelics | Instituto Butantan: Dimas Tadeu Covas, Sandra Coccuzzo Sampaio, Maria Carolina Elias, José Salvatore Leister Patané, Vincent Louis Viala, Antonio Jorge Martins, Ricardo Haddad, Claudia Renata dos Santos Barros, Elaine Cristina Marqueze, Raul Machado Neto, Debora Botequio Moretti, Jardelina de Souza Todao Bernardino, Loyze Paola Oliveira de Lima, Luiz Aurelio de Campos Crispin. Centro de Genômica Funcional da ESALQ: Luiz Lehmann Coutinho, Ricardo Augusto Brassaloti, Raquel de Lello Rocha Campos Cassano. NGS Soluções Genômicas: Pilar Drummond Sampaio Corrêa Mariani. FZEA-USP Pirassununga: Mirele Daiana Poleti, Jessika Cristina Chagas Lesbon, Elisangela Chicaroni Mattos, Heidge Fukumasu. USP-Botucatu: Rejane Maria Tommasini Grotto, Jayme A. Souza-Neto, Guilherme Targino Valente, Patricia Akemi Assato, Felipe Allan da Silva da Costa, Bianca Cechetto Carlos. Mendelics: Bibiana Santos, João Paulo Kitajima, Erika Freitas, David Schlesinger. Hemocentro Ribeirão Preto: Simone Kashima, Evandra Strazza Rodrigues, Svetoslav Nanev Slavov, Elaine Vieira dos Santos, Rafael dos Santos Bezerra, Luiz Carlos Junior de Alcantara, Marta Giovanetti, Vagner Fonseca, Flavia Aburjaile, Rodrigo Tocantins Calado. FAMERP-SJRP: Cecília Artico Banho, Lívia Sacchetto, Fábio Sossai Possebon, Leila Sabrina Ullmann, Cíntia Bittar, Guilherme Campos, Helena Lage Ferreira, Jorge A. Petroli Marchesi, Maisa C. Pereira Parra, Marília Moraes, Paula Rahal, Paulo Inacio da Costa, João Pessoa Araújo Jr., Maurício Lacerda Nogueira. Prefeitura de Sao Paulo: Melissa Palmieri. |
| EPI_ISL_1966752, EPI_ISL_1966753, EPI_ISL_1966754 | UNIDADE SENTINELA COVID19 | Instituto Butantan / Mendelics | Instituto Butantan: Dimas Tadeu Covas, Sandra Coccuzzo Sampaio, Maria Carolina Elias, José Salvatore Leister Patané, Vincent Louis Viala, Antonio Jorge Martins, Ricardo Haddad, Claudia Renata dos Santos Barros, Elaine Cristina Marqueze, Raul Machado Neto, Debora Botequio Moretti, Jardelina de Souza Todao Bernardino, Loyze Paola Oliveira de Lima, Luiz Aurelio de Campos Crispin. Centro de Genômica Funcional da ESALQ: Luiz Lehmann Coutinho, Ricardo Augusto Brassaloti, Raquel de Lello Rocha Campos Cassano. NGS Soluções Genômicas: Pilar Drummond Sampaio Corrêa Mariani. FZEA-USP Pirassununga: Mirele Daiana Poleti, Jessika Cristina Chagas Lesbon, Elisangela Chicaroni Mattos, Heidge Fukumasu. USP-Botucatu: Rejane Maria Tommasini Grotto, Jayme A. Souza-Neto, Guilherme Targino Valente, Patricia Akemi Assato, Felipe Allan da Silva da Costa, Bianca Cechetto Carlos. Mendelics: Bibiana Santos, João Paulo Kitajima, Erika Freitas, David Schlesinger. Hemocentro Ribeirão Preto: Simone Kashima, Evandra Strazza Rodrigues, Svetoslav Nanev Slavov, Elaine Vieira dos Santos, Rafael dos Santos Bezerra, Luiz Carlos Junior de Alcantara, Marta Giovanetti, Vagner |

| | | | |
|---|---|---|---|
| | | | Fonseca, Flavia Aburjaile, Rodrigo Tocantins Calado. FAMERP-SJRP: Cecília Artico Banho, Lívia Sacchetto, Fábio Sossai Possebon, Leila Sabrina Ullmann, Cíntia Bittar, Guilherme Campos, Helena Lage Ferreira, Jorge A. Petroli Marchesi, Maisa C. Pereira Parra, Marília Moraes, Paula Rahal, Paulo Inacio da Costa, João Pessoa Araújo Jr., Maurício Lacerda Nogueira. Prefeitura de Sao Paulo: Melissa Palmieri. |
| EPI_ISL_1966755 | UNIDADE DE SAUDE DA FAMILIA I COLOMBIA | Instituto Butantan / Mendelics | Instituto Butantan: Dimas Tadeu Covas, Sandra Coccuzzo Sampaio, Maria Carolina Elias, José Salvatore Leister Patané, Vincent Louis Viala, Antonio Jorge Martins, Ricardo Haddad, Claudia Renata dos Santos Barros, Elaine Cristina Marqueze, Raul Machado Neto, Debora Botequio Moretti, Jardelina de Souza Todao Bernardino, Loyze Paola Oliveira de Lima, Luiz Aurelio de Campos Crispin. Centro de Genômica Funcional da ESALQ: Luiz Lehmann Coutinho, Ricardo Augusto Brassaloti, Raquel de Lello Rocha Campos Cassano. NGS Soluções Genômicas: Pilar Drummond Sampaio Corrêa Mariani. FZEA-USP Pirassununga: Mirele Daiana Poleti, Jessika Cristina Chagas Lesbon, Elisangela Chicaroni Mattos, Heidge Fukumasu. USP-Botucatu: Rejane Maria Tommasini Grotto, Jayme A. Souza-Neto, Guilherme Targino Valente, Patricia Akemi Assato, Felipe Allan da Silva da Costa, Bianca Cechetto Carlos. Mendelics: Bibiana Santos, João Paulo Kitajima, Erika Freitas, David Schlesinger. Hemocentro Ribeirão Preto: Simone Kashima, Evandra Strazza Rodrigues, Svetoslav Nanev Slavov, Elaine Vieira dos Santos, Rafael dos Santos Bezerra, Luiz Carlos Junior de Alcantara, Marta Giovanetti, Vagner Fonseca, Flavia Aburjaile, Rodrigo Tocantins Calado. FAMERP-SJRP: Cecília Artico Banho, Lívia Sacchetto, Fábio Sossai Possebon, Leila Sabrina Ullmann, Cíntia Bittar, Guilherme Campos, Helena Lage Ferreira, Jorge A. Petroli Marchesi, Maisa C. Pereira Parra, Marília Moraes, Paula Rahal, Paulo Inacio da Costa, João Pessoa Araújo Jr., Maurício Lacerda Nogueira. Prefeitura de Sao Paulo: Melissa Palmieri. |
| EPI_ISL_1966756, EPI_ISL_1966757, EPI_ISL_1966758, EPI_ISL_1966759, EPI_ISL_1966760 | LABORATORIO DE FRANCA | Instituto Butantan / Mendelics | Instituto Butantan: Dimas Tadeu Covas, Sandra Coccuzzo Sampaio, Maria Carolina Elias, José Salvatore Leister Patané, Vincent Louis Viala, Antonio Jorge Martins, Ricardo Haddad, Claudia Renata dos Santos Barros, Elaine Cristina Marqueze, Raul Machado Neto, Debora Botequio Moretti, Jardelina de Souza Todao Bernardino, Loyze Paola Oliveira de Lima, Luiz Aurelio de Campos Crispin. Centro de Genômica Funcional da ESALQ: Luiz Lehmann Coutinho, Ricardo Augusto Brassaloti, Raquel de Lello Rocha Campos Cassano. NGS Soluções Genômicas: Pilar Drummond Sampaio Corrêa Mariani. FZEA-USP Pirassununga: Mirele Daiana Poleti, Jessika Cristina Chagas Lesbon, Elisangela Chicaroni Mattos, Heidge Fukumasu. USP-Botucatu: Rejane Maria Tommasini Grotto, Jayme A. Souza-Neto, Guilherme Targino Valente, Patricia Akemi Assato, Felipe Allan da Silva da Costa, Bianca Cechetto Carlos. Mendelics: Bibiana Santos, João Paulo Kitajima, Erika Freitas, David Schlesinger. Hemocentro Ribeirão Preto: Simone Kashima, Evandra Strazza Rodrigues, Svetoslav Nanev Slavov, Elaine Vieira dos Santos, Rafael dos Santos Bezerra, Luiz Carlos Junior de Alcantara, Marta Giovanetti, Vagner Fonseca, Flavia Aburjaile, Rodrigo Tocantins Calado. FAMERP-SJRP: Cecília Artico Banho, Lívia Sacchetto, Fábio Sossai Possebon, Leila Sabrina Ullmann, Cíntia Bittar, Guilherme Campos, Helena Lage Ferreira, Jorge A. Petroli Marchesi, Maisa C. Pereira Parra, Marília Moraes, Paula Rahal, Paulo Inacio da Costa, João Pessoa Araújo Jr., Maurício Lacerda Nogueira. Prefeitura de Sao Paulo: Melissa Palmieri. |
| EPI_ISL_1966763 | HOSPITAL MUNICIPAL DE IBIUNA IBIUNA SP | Instituto Butantan / Mendelics | Instituto Butantan: Dimas Tadeu Covas, Sandra Coccuzzo Sampaio, Maria Carolina Elias, José Salvatore Leister Patané, Vincent Louis Viala, Antonio Jorge Martins, Ricardo Haddad, Claudia Renata dos Santos Barros, Elaine Cristina Marqueze, Raul Machado Neto, Debora Botequio Moretti, Jardelina de Souza Todao Bernardino, Loyze Paola Oliveira de Lima, Luiz Aurelio de Campos Crispin. Centro de Genômica Funcional da ESALQ: Luiz Lehmann Coutinho, Ricardo Augusto Brassaloti, Raquel de Lello Rocha Campos Cassano. NGS Soluções Genômicas: Pilar Drummond Sampaio Corrêa Mariani. FZEA-USP Pirassununga: Mirele Daiana Poleti, Jessika Cristina Chagas Lesbon, Elisangela Chicaroni Mattos, Heidge Fukumasu. USP-Botucatu: Rejane Maria Tommasini Grotto, Jayme A. Souza-Neto, Guilherme Targino Valente, Patricia Akemi Assato, Felipe Allan da Silva da Costa, Bianca Cechetto Carlos. Mendelics: Bibiana Santos, João Paulo Kitajima, Erika Freitas, David Schlesinger. Hemocentro Ribeirão Preto: Simone Kashima, Evandra Strazza Rodrigues, Svetoslav Nanev Slavov, Elaine Vieira dos Santos, Rafael dos Santos Bezerra, Luiz Carlos Junior de Alcantara, Marta Giovanetti, Vagner Fonseca, Flavia Aburjaile, Rodrigo Tocantins Calado. FAMERP-SJRP: Cecília Artico Banho, Lívia Sacchetto, Fábio Sossai Possebon, Leila Sabrina Ullmann, Cíntia Bittar, Guilherme Campos, Helena Lage Ferreira, Jorge A. Petroli Marchesi, Maisa C. Pereira Parra, Marília Moraes, Paula Rahal, Paulo Inacio da Costa, João Pessoa Araújo Jr., Maurício Lacerda Nogueira. Prefeitura de Sao Paulo: Melissa Palmieri. |
| EPI_ISL_1966764, EPI_ISL_1966765 | CS DE ICEM | Instituto Butantan / Mendelics | Instituto Butantan: Dimas Tadeu Covas, Sandra Coccuzzo Sampaio, Maria Carolina Elias, José Salvatore Leister Patané, Vincent Louis Viala, Antonio Jorge Martins, Ricardo Haddad, Claudia Renata dos Santos Barros, Elaine Cristina Marqueze, Raul Machado Neto, Debora Botequio Moretti, Jardelina de Souza Todao Bernardino, Loyze Paola Oliveira de Lima, Luiz Aurelio de Campos Crispin. Centro de Genômica Funcional da ESALQ: Luiz Lehmann Coutinho, Ricardo Augusto Brassaloti, Raquel de Lello Rocha Campos Cassano. NGS Soluções Genômicas: Pilar Drummond Sampaio Corrêa Mariani. FZEA-USP Pirassununga: Mirele Daiana Poleti, Jessika Cristina Chagas Lesbon, Elisangela Chicaroni Mattos, Heidge Fukumasu. USP-Botucatu: Rejane Maria Tommasini Grotto, Jayme A. Souza-Neto, Guilherme Targino Valente, Patricia Akemi Assato, Felipe Allan da Silva da Costa, Bianca Cechetto Carlos. Mendelics: Bibiana Santos, João Paulo Kitajima, Erika Freitas, David Schlesinger. Hemocentro Ribeirão Preto: Simone Kashima, Evandra Strazza Rodrigues, Svetoslav Nanev Slavov, Elaine Vieira dos Santos, Rafael dos Santos Bezerra, Luiz Carlos Junior de Alcantara, Marta Giovanetti, Vagner Fonseca, Flavia Aburjaile, Rodrigo Tocantins Calado. FAMERP-SJRP: Cecília Artico Banho, Lívia Sacchetto, Fábio Sossai Possebon, Leila Sabrina Ullmann, Cíntia Bittar, Guilherme Campos, Helena Lage Ferreira, Jorge A. Petroli Marchesi, Maisa C. Pereira Parra, Marília Moraes, Paula Rahal, Paulo Inacio da Costa, João Pessoa Araújo Jr., Maurício Lacerda Nogueira. Prefeitura de Sao Paulo: Melissa Palmieri. |
| EPI_ISL_1966767, EPI_ISL_1966768, EPI_ISL_1966770 | POLICLINICA COVID 19 ITAPETININGA | Instituto Butantan / Mendelics | Instituto Butantan: Dimas Tadeu Covas, Sandra Coccuzzo Sampaio, Maria Carolina Elias, José Salvatore Leister Patané, Vincent Louis Viala, Antonio Jorge Martins, Ricardo Haddad, Claudia Renata dos Santos Barros, Elaine Cristina Marqueze, Raul Machado Neto, Debora Botequio Moretti, Jardelina de Souza Todao Bernardino, Loyze Paola Oliveira de Lima, Luiz Aurelio de Campos Crispin. Centro de Genômica Funcional da ESALQ: Luiz Lehmann Coutinho, Ricardo Augusto Brassaloti, Raquel de Lello Rocha Campos Cassano. NGS Soluções Genômicas: Pilar Drummond Sampaio Corrêa Mariani. FZEA-USP Pirassununga: Mirele Daiana Poleti, Jessika Cristina Chagas Lesbon, Elisangela Chicaroni Mattos, Heidge Fukumasu. USP-Botucatu: Rejane Maria Tommasini Grotto, Jayme A. Souza-Neto, Guilherme Targino Valente, Patricia Akemi Assato, Felipe Allan da Silva da Costa, Bianca Cechetto Carlos. Mendelics: Bibiana Santos, João Paulo Kitajima, Erika Freitas, David Schlesinger. Hemocentro Ribeirão Preto: Simone Kashima, Evandra Strazza Rodrigues, Svetoslav Nanev Slavov, Elaine Vieira dos Santos, Rafael dos Santos Bezerra, Luiz Carlos Junior de Alcantara, Marta Giovanetti, Vagner Fonseca, Flavia Aburjaile, Rodrigo Tocantins Calado. FAMERP-SJRP: Cecília Artico Banho, Lívia Sacchetto, Fábio Sossai Possebon, Leila Sabrina Ullmann, Cíntia Bittar, Guilherme Campos, Helena Lage Ferreira, Jorge A. Petroli Marchesi, Maisa C. Pereira Parra, Marília Moraes, Paula Rahal, Paulo Inacio da Costa, João Pessoa Araújo Jr., Maurício Lacerda Nogueira. Prefeitura de Sao Paulo: Melissa Palmieri. |
| EPI_ISL_1966771 | SECRETARIA MUNICIPAL DA SAUDE DE ITAPEVA SP | Instituto Butantan / Mendelics | Instituto Butantan: Dimas Tadeu Covas, Sandra Coccuzzo Sampaio, Maria Carolina Elias, José Salvatore Leister Patané, Vincent Louis Viala, Antonio Jorge Martins, Ricardo Haddad, Claudia Renata dos Santos Barros, Elaine Cristina Marqueze, Raul Machado Neto, Debora Botequio Moretti, Jardelina de Souza Todao Bernardino, Loyze Paola Oliveira de Lima, Luiz Aurelio de Campos Crispin. Centro de Genômica Funcional da ESALQ: Luiz Lehmann Coutinho, Ricardo Augusto Brassaloti, Raquel de Lello Rocha Campos Cassano. NGS Soluções Genômicas: Pilar Drummond Sampaio Corrêa Mariani. FZEA-USP Pirassununga: Mirele Daiana Poleti, Jessika Cristina Chagas Lesbon, Elisangela Chicaroni Mattos, Heidge Fukumasu. USP-Botucatu: Rejane Maria Tommasini Grotto, Jayme A. Souza-Neto, Guilherme Targino Valente, Patricia Akemi Assato, Felipe Allan da Silva da Costa, Bianca Cechetto Carlos. Mendelics: Bibiana Santos, João Paulo Kitajima, Erika Freitas, David Schlesinger. Hemocentro Ribeirão Preto: Simone Kashima, Evandra Strazza Rodrigues, Svetoslav Nanev Slavov, Elaine Vieira dos Santos, Rafael dos Santos Bezerra, Luiz Carlos Junior de Alcantara, Marta Giovanetti, Vagner Fonseca, Flavia Aburjaile, Rodrigo Tocantins Calado. FAMERP-SJRP: Cecília Artico Banho, Lívia Sacchetto, Fábio Sossai Possebon, Leila Sabrina Ullmann, Cíntia Bittar, Guilherme Campos, Helena Lage Ferreira, Jorge A. Petroli Marchesi, Maisa C. Pereira Parra, Marília Moraes, Paula Rahal, Paulo Inacio da Costa, João Pessoa Araújo Jr., Maurício Lacerda Nogueira. Prefeitura de Sao Paulo: Melissa Palmieri. |
| EPI_ISL_1966772 | SECRETARIA MUNICIPAL DA SAUDE DE ITARARE | Instituto Butantan / Mendelics | Instituto Butantan: Dimas Tadeu Covas, Sandra Coccuzzo Sampaio, Maria Carolina Elias, José Salvatore Leister Patané, Vincent Louis Viala, Antonio Jorge Martins, Ricardo Haddad, Claudia Renata dos Santos Barros, Elaine Cristina Marqueze, Raul Machado Neto, Debora Botequio Moretti, Jardelina de Souza Todao Bernardino, Loyze Paola Oliveira de Lima, Luiz Aurelio de Campos Crispin. Centro de Genômica Funcional da ESALQ: Luiz Lehmann Coutinho, Ricardo Augusto Brassaloti, Raquel de Lello Rocha Campos Cassano. NGS Soluções Genômicas: Pilar Drummond Sampaio Corrêa Mariani. FZEA-USP Pirassununga: Mirele Daiana Poleti, Jessika Cristina Chagas Lesbon, Elisangela Chicaroni Mattos, Heidge Fukumasu. USP-Botucatu: Rejane Maria Tommasini Grotto, Jayme A. Souza-Neto, Guilherme Targino Valente, Patricia Akemi Assato, Felipe Allan da Silva da Costa, Bianca Cechetto Carlos. Mendelics: Bibiana Santos, João Paulo Kitajima, Erika Freitas, David Schlesinger. Hemocentro Ribeirão Preto: Simone Kashima, Evandra Strazza Rodrigues, Svetoslav Nanev Slavov, Elaine Vieira dos Santos, Rafael dos Santos Bezerra, Luiz Carlos Junior de Alcantara, Marta Giovanetti, Vagner Fonseca, Flavia Aburjaile, Rodrigo Tocantins Calado. FAMERP-SJRP: Cecília Artico Banho, Lívia Sacchetto, Fábio Sossai Possebon, Leila Sabrina Ullmann, Cíntia Bittar, Guilherme Campos, Helena Lage Ferreira, Jorge A. Petroli Marchesi, Maisa C. Pereira Parra, Marília Moraes, Paula Rahal, Paulo Inacio da Costa, João Pessoa Araújo Jr., Maurício Lacerda Nogueira. Prefeitura de Sao Paulo: Melissa Palmieri. |
| EPI_ISL_1966774, EPI_ISL_1966775 | VIGILANCIA EPIDEMIOLOGICA JARDINOPOLIS SP | Instituto Butantan / Mendelics | Instituto Butantan: Dimas Tadeu Covas, Sandra Coccuzzo Sampaio, Maria Carolina Elias, José Salvatore Leister Patané, Vincent Louis Viala, Antonio Jorge Martins, Ricardo Haddad, Claudia Renata dos Santos Barros, Elaine Cristina Marqueze, Raul Machado Neto, Debora Botequio Moretti, Jardelina de Souza Todao Bernardino, Loyze Paola Oliveira de Lima, Luiz Aurelio de Campos Crispin. Centro de Genômica Funcional da ESALQ: Luiz Lehmann |

| | | | |
|---|---|---|---|
| | | | Coutinho, Ricardo Augusto Brassaloti, Raquel de Lello Rocha Campos Cassano. NGS Soluções Genômicas: Pilar Drummond Sampaio Corrêa Mariani. FZEA-USP Pirassununga: Mirele Daiana Poleti, Jessika Cristina Chagas Lesbon, Elisangela Chicaroni Mattos, Heidge Fukumasu. USP-Botucatu: Rejane Maria Tommasini Grotto, Jayme A. Souza-Neto, Guilherme Targino Valente, Patricia Akemi Assato, Felipe Allan da Silva da Costa, Bianca Cechetto Carlos. Mendelics: Bibiana Santos, João Paulo Kitajima, Erika Freitas, David Schlesinger. Hemocentro Ribeirão Preto: Simone Kashima, Evandra Strazza Rodrigues, Svetoslav Nanev Slavov, Elaine Vieira dos Santos, Rafael dos Santos Bezerra, Luiz Carlos Junior de Alcantara, Marta Giovanetti, Vagner Fonseca, Flavia Aburjaile, Rodrigo Tocantins Calado. FAMERP-SJRP: Cecília Artico Banho, Lívia Sacchetto, Fábio Sossai Possebon, Leila Sabrina Ullmann, Cíntia Bittar, Guilherme Campos, Helena Lage Ferreira, Jorge A. Petroli Marchesi, Maisa C. Pereira Parra, Marília Moraes, Paula Rahal, Paulo Inacio da Costa, João Pessoa Araújo Jr., Maurício Lacerda Nogueira. Prefeitura de Sao Paulo: Melissa Palmieri. |
| EPI_ISL_1966776 | AMB MEDICO DR FRANCISCO FELIX DE MENDONCA SAUDADE | Instituto Butantan / Mendelics | Instituto Butantan: Dimas Tadeu Covas, Sandra Coccuzzo Sampaio, Maria Carolina Elias, José Salvatore Leister Patané, Vincent Louis Viala, Antonio Jorge Martins, Ricardo Haddad, Claudia Renata dos Santos Barros, Elaine Cristina Marqueze, Raul Machado Neto, Debora Botequio Moretti, Jardelina de Souza Todao Bernardino, Loyze Paola Oliveira de Lima, Luiz Aurelio de Campos Crispin. Centro de Genômica Funcional da ESALQ: Luiz Lehmann Coutinho, Ricardo Augusto Brassaloti, Raquel de Lello Rocha Campos Cassano. NGS Soluções Genômicas: Pilar Drummond Sampaio Corrêa Mariani. FZEA-USP Pirassununga: Mirele Daiana Poleti, Jessika Cristina Chagas Lesbon, Elisangela Chicaroni Mattos, Heidge Fukumasu. USP-Botucatu: Rejane Maria Tommasini Grotto, Jayme A. Souza-Neto, Guilherme Targino Valente, Patricia Akemi Assato, Felipe Allan da Silva da Costa, Bianca Cechetto Carlos. Mendelics: Bibiana Santos, João Paulo Kitajima, Erika Freitas, David Schlesinger. Hemocentro Ribeirão Preto: Simone Kashima, Evandra Strazza Rodrigues, Svetoslav Nanev Slavov, Elaine Vieira dos Santos, Rafael dos Santos Bezerra, Luiz Carlos Junior de Alcantara, Marta Giovanetti, Vagner Fonseca, Flavia Aburjaile, Rodrigo Tocantins Calado. FAMERP-SJRP: Cecília Artico Banho, Lívia Sacchetto, Fábio Sossai Possebon, Leila Sabrina Ullmann, Cíntia Bittar, Guilherme Campos, Helena Lage Ferreira, Jorge A. Petroli Marchesi, Maisa C. Pereira Parra, Marília Moraes, Paula Rahal, Paulo Inacio da Costa, João Pessoa Araújo Jr., Maurício Lacerda Nogueira. Prefeitura de Sao Paulo: Melissa Palmieri. |
| EPI_ISL_1966777, EPI_ISL_1966778, EPI_ISL_1966779, EPI_ISL_1966780, EPI_ISL_1966781 | AMBULATORIO DR JOAO LANIA CENTRO JOSE BONIFACIO | Instituto Butantan / Mendelics | Instituto Butantan: Dimas Tadeu Covas, Sandra Coccuzzo Sampaio, Maria Carolina Elias, José Salvatore Leister Patané, Vincent Louis Viala, Antonio Jorge Martins, Ricardo Haddad, Claudia Renata dos Santos Barros, Elaine Cristina Marqueze, Raul Machado Neto, Debora Botequio Moretti, Jardelina de Souza Todao Bernardino, Loyze Paola Oliveira de Lima, Luiz Aurelio de Campos Crispin. Centro de Genômica Funcional da ESALQ: Luiz Lehmann Coutinho, Ricardo Augusto Brassaloti, Raquel de Lello Rocha Campos Cassano. NGS Soluções Genômicas: Pilar Drummond Sampaio Corrêa Mariani. FZEA-USP Pirassununga: Mirele Daiana Poleti, Jessika Cristina Chagas Lesbon, Elisangela Chicaroni Mattos, Heidge Fukumasu. USP-Botucatu: Rejane Maria Tommasini Grotto, Jayme A. Souza-Neto, Guilherme Targino Valente, Patricia Akemi Assato, Felipe Allan da Silva da Costa, Bianca Cechetto Carlos. Mendelics: Bibiana Santos, João Paulo Kitajima, Erika Freitas, David Schlesinger. Hemocentro Ribeirão Preto: Simone Kashima, Evandra Strazza Rodrigues, Svetoslav Nanev Slavov, Elaine Vieira dos Santos, Rafael dos Santos Bezerra, Luiz Carlos Junior de Alcantara, Marta Giovanetti, Vagner Fonseca, Flavia Aburjaile, Rodrigo Tocantins Calado. FAMERP-SJRP: Cecília Artico Banho, Lívia Sacchetto, Fábio Sossai Possebon, Leila Sabrina Ullmann, Cíntia Bittar, Guilherme Campos, Helena Lage Ferreira, Jorge A. Petroli Marchesi, Maisa C. Pereira Parra, Marília Moraes, Paula Rahal, Paulo Inacio da Costa, João Pessoa Araújo Jr., Maurício Lacerda Nogueira. Prefeitura de Sao Paulo: Melissa Palmieri. |
| EPI_ISL_1966782, EPI_ISL_1966783 | UBS DE MENDONCA | Instituto Butantan / Mendelics | Instituto Butantan: Dimas Tadeu Covas, Sandra Coccuzzo Sampaio, Maria Carolina Elias, José Salvatore Leister Patané, Vincent Louis Viala, Antonio Jorge Martins, Ricardo Haddad, Claudia Renata dos Santos Barros, Elaine Cristina Marqueze, Raul Machado Neto, Debora Botequio Moretti, Jardelina de Souza Todao Bernardino, Loyze Paola Oliveira de Lima, Luiz Aurelio de Campos Crispin. Centro de Genômica Funcional da ESALQ: Luiz Lehmann Coutinho, Ricardo Augusto Brassaloti, Raquel de Lello Rocha Campos Cassano. NGS Soluções Genômicas: Pilar Drummond Sampaio Corrêa Mariani. FZEA-USP Pirassununga: Mirele Daiana Poleti, Jessika Cristina Chagas Lesbon, Elisangela Chicaroni Mattos, Heidge Fukumasu. USP-Botucatu: Rejane Maria Tommasini Grotto, Jayme A. Souza-Neto, Guilherme Targino Valente, Patricia Akemi Assato, Felipe Allan da Silva da Costa, Bianca Cechetto Carlos. Mendelics: Bibiana Santos, João Paulo Kitajima, Erika Freitas, David Schlesinger. Hemocentro Ribeirão Preto: Simone Kashima, Evandra Strazza Rodrigues, Svetoslav Nanev Slavov, Elaine Vieira dos Santos, Rafael dos Santos Bezerra, Luiz Carlos Junior de Alcantara, Marta Giovanetti, Vagner Fonseca, Flavia Aburjaile, Rodrigo Tocantins Calado. FAMERP-SJRP: Cecília Artico Banho, Lívia Sacchetto, Fábio Sossai Possebon, Leila Sabrina Ullmann, Cíntia Bittar, Guilherme Campos, Helena Lage Ferreira, Jorge A. Petroli Marchesi, Maisa C. Pereira Parra, Marília Moraes, Paula Rahal, Paulo Inacio da Costa, João Pessoa Araújo Jr., Maurício Lacerda Nogueira. Prefeitura de Sao Paulo: Melissa Palmieri. |
| EPI_ISL_1966784, EPI_ISL_1966785 | CS DE MIRASSOLANDIA | Instituto Butantan / Mendelics | Instituto Butantan: Dimas Tadeu Covas, Sandra Coccuzzo Sampaio, Maria Carolina Elias, José Salvatore Leister Patané, Vincent Louis Viala, Antonio Jorge Martins, Ricardo Haddad, Claudia Renata dos Santos Barros, Elaine Cristina Marqueze, Raul Machado Neto, Debora Botequio Moretti, Jardelina de Souza Todao Bernardino, Loyze Paola Oliveira de Lima, Luiz Aurelio de Campos Crispin. Centro de Genômica Funcional da ESALQ: Luiz Lehmann Coutinho, Ricardo Augusto Brassaloti, Raquel de Lello Rocha Campos Cassano. NGS Soluções Genômicas: Pilar Drummond Sampaio Corrêa Mariani. FZEA-USP Pirassununga: Mirele Daiana Poleti, Jessika Cristina Chagas Lesbon, Elisangela Chicaroni Mattos, Heidge Fukumasu. USP-Botucatu: Rejane Maria Tommasini Grotto, Jayme A. Souza-Neto, Guilherme Targino Valente, Patricia Akemi Assato, Felipe Allan da Silva da Costa, Bianca Cechetto Carlos. Mendelics: Bibiana Santos, João Paulo Kitajima, Erika Freitas, David Schlesinger. Hemocentro Ribeirão Preto: Simone Kashima, Evandra Strazza Rodrigues, Svetoslav Nanev Slavov, Elaine Vieira dos Santos, Rafael dos Santos Bezerra, Luiz Carlos Junior de Alcantara, Marta Giovanetti, Vagner Fonseca, Flavia Aburjaile, Rodrigo Tocantins Calado. FAMERP-SJRP: Cecília Artico Banho, Lívia Sacchetto, Fábio Sossai Possebon, Leila Sabrina Ullmann, Cíntia Bittar, Guilherme Campos, Helena Lage Ferreira, Jorge A. Petroli Marchesi, Maisa C. Pereira Parra, Marília Moraes, Paula Rahal, Paulo Inacio da Costa, João Pessoa Araújo Jr., Maurício Lacerda Nogueira. Prefeitura de Sao Paulo: Melissa Palmieri. |
| EPI_ISL_1966786 | PSF SETOR LESTE NHANDEARA | Instituto Butantan / Mendelics | Instituto Butantan: Dimas Tadeu Covas, Sandra Coccuzzo Sampaio, Maria Carolina Elias, José Salvatore Leister Patané, Vincent Louis Viala, Antonio Jorge Martins, Ricardo Haddad, Claudia Renata dos Santos Barros, Elaine Cristina Marqueze, Raul Machado Neto, Debora Botequio Moretti, Jardelina de Souza Todao Bernardino, Loyze Paola Oliveira de Lima, Luiz Aurelio de Campos Crispin. Centro de Genômica Funcional da ESALQ: Luiz Lehmann Coutinho, Ricardo Augusto Brassaloti, Raquel de Lello Rocha Campos Cassano. NGS Soluções Genômicas: Pilar Drummond Sampaio Corrêa Mariani. FZEA-USP Pirassununga: Mirele Daiana Poleti, Jessika Cristina Chagas Lesbon, Elisangela Chicaroni Mattos, Heidge Fukumasu. USP-Botucatu: Rejane Maria Tommasini Grotto, Jayme A. Souza-Neto, Guilherme Targino Valente, Patricia Akemi Assato, Felipe Allan da Silva da Costa, Bianca Cechetto Carlos. Mendelics: Bibiana Santos, João Paulo Kitajima, Erika Freitas, David Schlesinger. Hemocentro Ribeirão Preto: Simone Kashima, Evandra Strazza Rodrigues, Svetoslav Nanev Slavov, Elaine Vieira dos Santos, Rafael dos Santos Bezerra, Luiz Carlos Junior de Alcantara, Marta Giovanetti, Vagner Fonseca, Flavia Aburjaile, Rodrigo Tocantins Calado. FAMERP-SJRP: Cecília Artico Banho, Lívia Sacchetto, Fábio Sossai Possebon, Leila Sabrina Ullmann, Cíntia Bittar, Guilherme Campos, Helena Lage Ferreira, Jorge A. Petroli Marchesi, Maisa C. Pereira Parra, Marília Moraes, Paula Rahal, Paulo Inacio da Costa, João Pessoa Araújo Jr., Maurício Lacerda Nogueira. Prefeitura de Sao Paulo: Melissa Palmieri. |
| EPI_ISL_1966787, EPI_ISL_1966788, EPI_ISL_1966789, EPI_ISL_1966790, EPI_ISL_1966791 | CS DE NIPOA | Instituto Butantan / Mendelics | Instituto Butantan: Dimas Tadeu Covas, Sandra Coccuzzo Sampaio, Maria Carolina Elias, José Salvatore Leister Patané, Vincent Louis Viala, Antonio Jorge Martins, Ricardo Haddad, Claudia Renata dos Santos Barros, Elaine Cristina Marqueze, Raul Machado Neto, Debora Botequio Moretti, Jardelina de Souza Todao Bernardino, Loyze Paola Oliveira de Lima, Luiz Aurelio de Campos Crispin. Centro de Genômica Funcional da ESALQ: Luiz Lehmann Coutinho, Ricardo Augusto Brassaloti, Raquel de Lello Rocha Campos Cassano. NGS Soluções Genômicas: Pilar Drummond Sampaio Corrêa Mariani. FZEA-USP Pirassununga: Mirele Daiana Poleti, Jessika Cristina Chagas Lesbon, Elisangela Chicaroni Mattos, Heidge Fukumasu. USP-Botucatu: Rejane Maria Tommasini Grotto, Jayme A. Souza-Neto, Guilherme Targino Valente, Patricia Akemi Assato, Felipe Allan da Silva da Costa, Bianca Cechetto Carlos. Mendelics: Bibiana Santos, João Paulo Kitajima, Erika Freitas, David Schlesinger. Hemocentro Ribeirão Preto: Simone Kashima, Evandra Strazza Rodrigues, Svetoslav Nanev Slavov, Elaine Vieira dos Santos, Rafael dos Santos Bezerra, Luiz Carlos Junior de Alcantara, Marta Giovanetti, Vagner Fonseca, Flavia Aburjaile, Rodrigo Tocantins Calado. FAMERP-SJRP: Cecília Artico Banho, Lívia Sacchetto, Fábio Sossai Possebon, Leila Sabrina Ullmann, Cíntia Bittar, Guilherme Campos, Helena Lage Ferreira, Jorge A. Petroli Marchesi, Maisa C. Pereira Parra, Marília Moraes, Paula Rahal, Paulo Inacio da Costa, João Pessoa Araújo Jr., Maurício Lacerda Nogueira. Prefeitura de Sao Paulo: Melissa Palmieri. |
| EPI_ISL_1966792, EPI_ISL_1966793, EPI_ISL_1966794 | CENTRO DE SAUDE DE OSVALDO CRUZ | Instituto Butantan / Mendelics | Instituto Butantan: Dimas Tadeu Covas, Sandra Coccuzzo Sampaio, Maria Carolina Elias, José Salvatore Leister Patané, Vincent Louis Viala, Antonio Jorge Martins, Ricardo Haddad, Claudia Renata dos Santos Barros, Elaine Cristina Marqueze, Raul Machado Neto, Debora Botequio Moretti, Jardelina de Souza Todao Bernardino, Loyze Paola Oliveira de Lima, Luiz Aurelio de Campos Crispin. Centro de Genômica Funcional da ESALQ: Luiz Lehmann Coutinho, Ricardo Augusto Brassaloti, Raquel de Lello Rocha Campos Cassano. NGS Soluções Genômicas: Pilar Drummond Sampaio Corrêa Mariani. FZEA-USP Pirassununga: Mirele Daiana Poleti, Jessika Cristina Chagas Lesbon, Elisangela Chicaroni Mattos, Heidge Fukumasu. USP-Botucatu: Rejane Maria Tommasini Grotto, Jayme A. Souza-Neto, Guilherme Targino Valente, Patricia Akemi Assato, Felipe Allan da Silva da Costa, Bianca Cechetto Carlos. Mendelics: Bibiana Santos, João Paulo Kitajima, Erika Freitas, David Schlesinger. Hemocentro Ribeirão Preto: Simone Kashima, Evandra Strazza Rodrigues, Svetoslav Nanev Slavov, Elaine Vieira dos Santos, Rafael dos Santos Bezerra, Luiz Carlos Junior de Alcantara, Marta Giovanetti, Vagner |

| | | | |
|---|---|---|---|
| | | | Fonseca, Flavia Aburjaile, Rodrigo Tocantins Calado. FAMERP-SJRP: Cecília Artico Banho, Lívia Sacchetto, Fábio Sossai Possebon, Leila Sabrina Ullmann, Cíntia Bittar, Guilherme Campos, Helena Lage Ferreira, Jorge A. Petroli Marchesi, Maisa C. Pereira Parra, Marília Moraes, Paula Rahal, Paulo Inacio da Costa, João Pessoa Araújo Jr., Maurício Lacerda Nogueira. Prefeitura de Sao Paulo: Melissa Palmieri. |
| EPI_ISL_1966795 | CS DE PALESTINA | Instituto Butantan / Mendelics | Instituto Butantan: Dimas Tadeu Covas, Sandra Coccuzzo Sampaio, Maria Carolina Elias, José Salvatore Leister Patané, Vincent Louis Viala, Antonio Jorge Martins, Ricardo Haddad, Claudia Renata dos Santos Barros, Elaine Cristina Marqueze, Raul Machado Neto, Debora Botequio Moretti, Jardelina de Souza Todao Bernardino, Loyze Paola Oliveira de Lima, Luiz Aurelio de Campos Crispin. Centro de Genômica Funcional da ESALQ: Luiz Lehmann Coutinho, Ricardo Augusto Brassaloti, Raquel de Lello Rocha Campos Cassano. NGS Soluções Genômicas: Pilar Drummond Sampaio Corrêa Mariani. FZEA-USP Pirassununga: Mirele Daiana Poleti, Jessika Cristina Chagas Lesbon, Elisangela Chicaroni Mattos, Heidge Fukumasu. USP-Botucatu: Rejane Maria Tommasini Grotto, Jayme A. Souza-Neto, Guilherme Targino Valente, Patricia Akemi Assato, Felipe Allan da Silva da Costa, Bianca Cechetto Carlos. Mendelics: Bibiana Santos, João Paulo Kitajima, Erika Freitas, David Schlesinger. Hemocentro Ribeirão Preto: Simone Kashima, Evandra Strazza Rodrigues, Svetoslav Nanev Slavov, Elaine Vieira dos Santos, Rafael dos Santos Bezerra, Luiz Carlos Junior de Alcantara, Marta Giovanetti, Vagner Fonseca, Flavia Aburjaile, Rodrigo Tocantins Calado. FAMERP-SJRP: Cecília Artico Banho, Lívia Sacchetto, Fábio Sossai Possebon, Leila Sabrina Ullmann, Cíntia Bittar, Guilherme Campos, Helena Lage Ferreira, Jorge A. Petroli Marchesi, Maisa C. Pereira Parra, Marília Moraes, Paula Rahal, Paulo Inacio da Costa, João Pessoa Araújo Jr., Maurício Lacerda Nogueira. Prefeitura de Sao Paulo: Melissa Palmieri. |
| EPI_ISL_1966796 | CENTRO DE SAUDE TEREZINHA DE MORAES ARSILLA | Instituto Butantan / Mendelics | Instituto Butantan: Dimas Tadeu Covas, Sandra Coccuzzo Sampaio, Maria Carolina Elias, José Salvatore Leister Patané, Vincent Louis Viala, Antonio Jorge Martins, Ricardo Haddad, Claudia Renata dos Santos Barros, Elaine Cristina Marqueze, Raul Machado Neto, Debora Botequio Moretti, Jardelina de Souza Todao Bernardino, Loyze Paola Oliveira de Lima, Luiz Aurelio de Campos Crispin. Centro de Genômica Funcional da ESALQ: Luiz Lehmann Coutinho, Ricardo Augusto Brassaloti, Raquel de Lello Rocha Campos Cassano. NGS Soluções Genômicas: Pilar Drummond Sampaio Corrêa Mariani. FZEA-USP Pirassununga: Mirele Daiana Poleti, Jessika Cristina Chagas Lesbon, Elisangela Chicaroni Mattos, Heidge Fukumasu. USP-Botucatu: Rejane Maria Tommasini Grotto, Jayme A. Souza-Neto, Guilherme Targino Valente, Patricia Akemi Assato, Felipe Allan da Silva da Costa, Bianca Cechetto Carlos. Mendelics: Bibiana Santos, João Paulo Kitajima, Erika Freitas, David Schlesinger. Hemocentro Ribeirão Preto: Simone Kashima, Evandra Strazza Rodrigues, Svetoslav Nanev Slavov, Elaine Vieira dos Santos, Rafael dos Santos Bezerra, Luiz Carlos Junior de Alcantara, Marta Giovanetti, Vagner Fonseca, Flavia Aburjaile, Rodrigo Tocantins Calado. FAMERP-SJRP: Cecília Artico Banho, Lívia Sacchetto, Fábio Sossai Possebon, Leila Sabrina Ullmann, Cíntia Bittar, Guilherme Campos, Helena Lage Ferreira, Jorge A. Petroli Marchesi, Maisa C. Pereira Parra, Marília Moraes, Paula Rahal, Paulo Inacio da Costa, João Pessoa Araújo Jr., Maurício Lacerda Nogueira. Prefeitura de Sao Paulo: Melissa Palmieri. |
| EPI_ISL_1966798, EPI_ISL_1966799 | PSF DR ANTONIO PIRES DE ALMEIDA PORTO FELIZ | Instituto Butantan / Mendelics | Instituto Butantan: Dimas Tadeu Covas, Sandra Coccuzzo Sampaio, Maria Carolina Elias, José Salvatore Leister Patané, Vincent Louis Viala, Antonio Jorge Martins, Ricardo Haddad, Claudia Renata dos Santos Barros, Elaine Cristina Marqueze, Raul Machado Neto, Debora Botequio Moretti, Jardelina de Souza Todao Bernardino, Loyze Paola Oliveira de Lima, Luiz Aurelio de Campos Crispin. Centro de Genômica Funcional da ESALQ: Luiz Lehmann Coutinho, Ricardo Augusto Brassaloti, Raquel de Lello Rocha Campos Cassano. NGS Soluções Genômicas: Pilar Drummond Sampaio Corrêa Mariani. FZEA-USP Pirassununga: Mirele Daiana Poleti, Jessika Cristina Chagas Lesbon, Elisangela Chicaroni Mattos, Heidge Fukumasu. USP-Botucatu: Rejane Maria Tommasini Grotto, Jayme A. Souza-Neto, Guilherme Targino Valente, Patricia Akemi Assato, Felipe Allan da Silva da Costa, Bianca Cechetto Carlos. Mendelics: Bibiana Santos, João Paulo Kitajima, Erika Freitas, David Schlesinger. Hemocentro Ribeirão Preto: Simone Kashima, Evandra Strazza Rodrigues, Svetoslav Nanev Slavov, Elaine Vieira dos Santos, Rafael dos Santos Bezerra, Luiz Carlos Junior de Alcantara, Marta Giovanetti, Vagner Fonseca, Flavia Aburjaile, Rodrigo Tocantins Calado. FAMERP-SJRP: Cecília Artico Banho, Lívia Sacchetto, Fábio Sossai Possebon, Leila Sabrina Ullmann, Cíntia Bittar, Guilherme Campos, Helena Lage Ferreira, Jorge A. Petroli Marchesi, Maisa C. Pereira Parra, Marília Moraes, Paula Rahal, Paulo Inacio da Costa, João Pessoa Araújo Jr., Maurício Lacerda Nogueira. Prefeitura de Sao Paulo: Melissa Palmieri. |
| EPI_ISL_1966800 | UNIDADE DE PRONTO ATENDIMENTO UPA | Instituto Butantan / Mendelics | Instituto Butantan: Dimas Tadeu Covas, Sandra Coccuzzo Sampaio, Maria Carolina Elias, José Salvatore Leister Patané, Vincent Louis Viala, Antonio Jorge Martins, Ricardo Haddad, Claudia Renata dos Santos Barros, Elaine Cristina Marqueze, Raul Machado Neto, Debora Botequio Moretti, Jardelina de Souza Todao Bernardino, Loyze Paola Oliveira de Lima, Luiz Aurelio de Campos Crispin. Centro de Genômica Funcional da ESALQ: Luiz Lehmann Coutinho, Ricardo Augusto Brassaloti, Raquel de Lello Rocha Campos Cassano. NGS Soluções Genômicas: Pilar Drummond Sampaio Corrêa Mariani. FZEA-USP Pirassununga: Mirele Daiana Poleti, Jessika Cristina Chagas Lesbon, Elisangela Chicaroni Mattos, Heidge Fukumasu. USP-Botucatu: Rejane Maria Tommasini Grotto, Jayme A. Souza-Neto, Guilherme Targino Valente, Patricia Akemi Assato, Felipe Allan da Silva da Costa, Bianca Cechetto Carlos. Mendelics: Bibiana Santos, João Paulo Kitajima, Erika Freitas, David Schlesinger. Hemocentro Ribeirão Preto: Simone Kashima, Evandra Strazza Rodrigues, Svetoslav Nanev Slavov, Elaine Vieira dos Santos, Rafael dos Santos Bezerra, Luiz Carlos Junior de Alcantara, Marta Giovanetti, Vagner Fonseca, Flavia Aburjaile, Rodrigo Tocantins Calado. FAMERP-SJRP: Cecília Artico Banho, Lívia Sacchetto, Fábio Sossai Possebon, Leila Sabrina Ullmann, Cíntia Bittar, Guilherme Campos, Helena Lage Ferreira, Jorge A. Petroli Marchesi, Maisa C. Pereira Parra, Marília Moraes, Paula Rahal, Paulo Inacio da Costa, João Pessoa Araújo Jr., Maurício Lacerda Nogueira. Prefeitura de Sao Paulo: Melissa Palmieri. |
| EPI_ISL_1966801, EPI_ISL_1966802, EPI_ISL_1966803, EPI_ISL_1966804, EPI_ISL_1966805, EPI_ISL_1966806 | CENTRO DE SAUDE DE RIBEIRAO DO SUL | Instituto Butantan / Mendelics | Instituto Butantan: Dimas Tadeu Covas, Sandra Coccuzzo Sampaio, Maria Carolina Elias, José Salvatore Leister Patané, Vincent Louis Viala, Antonio Jorge Martins, Ricardo Haddad, Claudia Renata dos Santos Barros, Elaine Cristina Marqueze, Raul Machado Neto, Debora Botequio Moretti, Jardelina de Souza Todao Bernardino, Loyze Paola Oliveira de Lima, Luiz Aurelio de Campos Crispin. Centro de Genômica Funcional da ESALQ: Luiz Lehmann Coutinho, Ricardo Augusto Brassaloti, Raquel de Lello Rocha Campos Cassano. NGS Soluções Genômicas: Pilar Drummond Sampaio Corrêa Mariani. FZEA-USP Pirassununga: Mirele Daiana Poleti, Jessika Cristina Chagas Lesbon, Elisangela Chicaroni Mattos, Heidge Fukumasu. USP-Botucatu: Rejane Maria Tommasini Grotto, Jayme A. Souza-Neto, Guilherme Targino Valente, Patricia Akemi Assato, Felipe Allan da Silva da Costa, Bianca Cechetto Carlos. Mendelics: Bibiana Santos, João Paulo Kitajima, Erika Freitas, David Schlesinger. Hemocentro Ribeirão Preto: Simone Kashima, Evandra Strazza Rodrigues, Svetoslav Nanev Slavov, Elaine Vieira dos Santos, Rafael dos Santos Bezerra, Luiz Carlos Junior de Alcantara, Marta Giovanetti, Vagner Fonseca, Flavia Aburjaile, Rodrigo Tocantins Calado. FAMERP-SJRP: Cecília Artico Banho, Lívia Sacchetto, Fábio Sossai Possebon, Leila Sabrina Ullmann, Cíntia Bittar, Guilherme Campos, Helena Lage Ferreira, Jorge A. Petroli Marchesi, Maisa C. Pereira Parra, Marília Moraes, Paula Rahal, Paulo Inacio da Costa, João Pessoa Araújo Jr., Maurício Lacerda Nogueira. Prefeitura de Sao Paulo: Melissa Palmieri. |
| EPI_ISL_1966807 | UBS DR WALDOMIRO FERREIRA NEVES SCRPARDO | Instituto Butantan / Mendelics | Instituto Butantan: Dimas Tadeu Covas, Sandra Coccuzzo Sampaio, Maria Carolina Elias, José Salvatore Leister Patané, Vincent Louis Viala, Antonio Jorge Martins, Ricardo Haddad, Claudia Renata dos Santos Barros, Elaine Cristina Marqueze, Raul Machado Neto, Debora Botequio Moretti, Jardelina de Souza Todao Bernardino, Loyze Paola Oliveira de Lima, Luiz Aurelio de Campos Crispin. Centro de Genômica Funcional da ESALQ: Luiz Lehmann Coutinho, Ricardo Augusto Brassaloti, Raquel de Lello Rocha Campos Cassano. NGS Soluções Genômicas: Pilar Drummond Sampaio Corrêa Mariani. FZEA-USP Pirassununga: Mirele Daiana Poleti, Jessika Cristina Chagas Lesbon, Elisangela Chicaroni Mattos, Heidge Fukumasu. USP-Botucatu: Rejane Maria Tommasini Grotto, Jayme A. Souza-Neto, Guilherme Targino Valente, Patricia Akemi Assato, Felipe Allan da Silva da Costa, Bianca Cechetto Carlos. Mendelics: Bibiana Santos, João Paulo Kitajima, Erika Freitas, David Schlesinger. Hemocentro Ribeirão Preto: Simone Kashima, Evandra Strazza Rodrigues, Svetoslav Nanev Slavov, Elaine Vieira dos Santos, Rafael dos Santos Bezerra, Luiz Carlos Junior de Alcantara, Marta Giovanetti, Vagner Fonseca, Flavia Aburjaile, Rodrigo Tocantins Calado. FAMERP-SJRP: Cecília Artico Banho, Lívia Sacchetto, Fábio Sossai Possebon, Leila Sabrina Ullmann, Cíntia Bittar, Guilherme Campos, Helena Lage Ferreira, Jorge A. Petroli Marchesi, Maisa C. Pereira Parra, Marília Moraes, Paula Rahal, Paulo Inacio da Costa, João Pessoa Araújo Jr., Maurício Lacerda Nogueira. Prefeitura de Sao Paulo: Melissa Palmieri. |
| EPI_ISL_1966808 | CS II DR JOSE CARQUEIJO SCRPARDO | Instituto Butantan / Mendelics | Instituto Butantan: Dimas Tadeu Covas, Sandra Coccuzzo Sampaio, Maria Carolina Elias, José Salvatore Leister Patané, Vincent Louis Viala, Antonio Jorge Martins, Ricardo Haddad, Claudia Renata dos Santos Barros, Elaine Cristina Marqueze, Raul Machado Neto, Debora Botequio Moretti, Jardelina de Souza Todao Bernardino, Loyze Paola Oliveira de Lima, Luiz Aurelio de Campos Crispin. Centro de Genômica Funcional da ESALQ: Luiz Lehmann Coutinho, Ricardo Augusto Brassaloti, Raquel de Lello Rocha Campos Cassano. NGS Soluções Genômicas: Pilar Drummond Sampaio Corrêa Mariani. FZEA-USP Pirassununga: Mirele Daiana Poleti, Jessika Cristina Chagas Lesbon, Elisangela Chicaroni Mattos, Heidge Fukumasu. USP-Botucatu: Rejane Maria Tommasini Grotto, Jayme A. Souza-Neto, Guilherme Targino Valente, Patricia Akemi Assato, Felipe Allan da Silva da Costa, Bianca Cechetto Carlos. Mendelics: Bibiana Santos, João Paulo Kitajima, Erika Freitas, David Schlesinger. Hemocentro Ribeirão Preto: Simone Kashima, Evandra Strazza Rodrigues, Svetoslav Nanev Slavov, Elaine Vieira dos Santos, Rafael dos Santos Bezerra, Luiz Carlos Junior de Alcantara, Marta Giovanetti, Vagner Fonseca, Flavia Aburjaile, Rodrigo Tocantins Calado. FAMERP-SJRP: Cecília Artico Banho, Lívia Sacchetto, Fábio Sossai Possebon, Leila Sabrina Ullmann, Cíntia Bittar, Guilherme Campos, Helena Lage Ferreira, Jorge A. Petroli Marchesi, Maisa C. Pereira Parra, Marília Moraes, Paula Rahal, Paulo Inacio da Costa, João Pessoa Araújo Jr., Maurício Lacerda Nogueira. Prefeitura de Sao Paulo: Melissa Palmieri. |
| EPI_ISL_1966809 | UBS DR ABELARDO PINHEIRO GUIMARAES SCRPARDO | Instituto Butantan / Mendelics | Instituto Butantan: Dimas Tadeu Covas, Sandra Coccuzzo Sampaio, Maria Carolina Elias, José Salvatore Leister Patané, Vincent Louis Viala, Antonio Jorge Martins, Ricardo Haddad, Claudia Renata dos Santos Barros, Elaine Cristina Marqueze, Raul Machado Neto, Debora Botequio Moretti, Jardelina de Souza Todao Bernardino, Loyze Paola Oliveira de Lima, Luiz Aurelio de Campos Crispin. Centro de Genômica Funcional da ESALQ: Luiz Lehmann |

| | | | |
|---|---|---|---|
| | | | Coutinho, Ricardo Augusto Brassaloti, Raquel de Lello Rocha Campos Cassano. NGS Soluções Genômicas: Pilar Drummond Sampaio Corrêa Mariani. FZEA-USP Pirassununga: Mirele Daiana Poleti, Jessika Cristina Chagas Lesbon, Elisangela Chicaroni Mattos, Heidge Fukumasu. USP-Botucatu: Rejane Maria Tommasini Grotto, Jayme A. Souza-Neto, Guilherme Targino Valente, Patricia Akemi Assato, Felipe Allan da Silva da Costa, Bianca Cechetto Carlos. Mendelics: Bibiana Santos, João Paulo Kitajima, Erika Freitas, David Schlesinger. Hemocentro Ribeirão Preto: Simone Kashima, Evandra Strazza Rodrigues, Svetoslav Nanev Slavov, Elaine Vieira dos Santos, Rafael dos Santos Bezerra, Luiz Carlos Junior de Alcantara, Marta Giovanetti, Vagner Fonseca, Flavia Aburjaile, Rodrigo Tocantins Calado. FAMERP-SJRP: Cecília Artico Banho, Lívia Sacchetto, Fábio Sossai Possebon, Leila Sabrina Ullmann, Cíntia Bittar, Guilherme Campos, Helena Lage Ferreira, Jorge A. Petroli Marchesi, Maisa C. Pereira Parra, Marília Moraes, Paula Rahal, Paulo Inacio da Costa, João Pessoa Araújo Jr., Maurício Lacerda Nogueira. Prefeitura de Sao Paulo: Melissa Palmieri. |
| EPI_ISL_1966810 | USF PARQUE DAS NACOES SCRPARDO | Instituto Butantan / Mendelics | Instituto Butantan: Dimas Tadeu Covas, Sandra Coccuzzo Sampaio, Maria Carolina Elias, José Salvatore Leister Patané, Vincent Louis Viala, Antonio Jorge Martins, Ricardo Haddad, Claudia Renata dos Santos Barros, Elaine Cristina Marqueze, Raul Machado Neto, Debora Botequio Moretti, Jardelina de Souza Todao Bernardino, Loyze Paola Oliveira de Lima, Luiz Aurelio de Campos Crispin. Centro de Genômica Funcional da ESALQ: Luiz Lehmann Coutinho, Ricardo Augusto Brassaloti, Raquel de Lello Rocha Campos Cassano. NGS Soluções Genômicas: Pilar Drummond Sampaio Corrêa Mariani. FZEA-USP Pirassununga: Mirele Daiana Poleti, Jessika Cristina Chagas Lesbon, Elisangela Chicaroni Mattos, Heidge Fukumasu. USP-Botucatu: Rejane Maria Tommasini Grotto, Jayme A. Souza-Neto, Guilherme Targino Valente, Patricia Akemi Assato, Felipe Allan da Silva da Costa, Bianca Cechetto Carlos. Mendelics: Bibiana Santos, João Paulo Kitajima, Erika Freitas, David Schlesinger. Hemocentro Ribeirão Preto: Simone Kashima, Evandra Strazza Rodrigues, Svetoslav Nanev Slavov, Elaine Vieira dos Santos, Rafael dos Santos Bezerra, Luiz Carlos Junior de Alcantara, Marta Giovanetti, Vagner Fonseca, Flavia Aburjaile, Rodrigo Tocantins Calado. FAMERP-SJRP: Cecília Artico Banho, Lívia Sacchetto, Fábio Sossai Possebon, Leila Sabrina Ullmann, Cíntia Bittar, Guilherme Campos, Helena Lage Ferreira, Jorge A. Petroli Marchesi, Maisa C. Pereira Parra, Marília Moraes, Paula Rahal, Paulo Inacio da Costa, João Pessoa Araújo Jr., Maurício Lacerda Nogueira. Prefeitura de Sao Paulo: Melissa Palmieri. |
| EPI_ISL_1966811 | UBS DR ABELARDO PINHEIRO GUIMARAES SCRPARDO | Instituto Butantan / Mendelics | Instituto Butantan: Dimas Tadeu Covas, Sandra Coccuzzo Sampaio, Maria Carolina Elias, José Salvatore Leister Patané, Vincent Louis Viala, Antonio Jorge Martins, Ricardo Haddad, Claudia Renata dos Santos Barros, Elaine Cristina Marqueze, Raul Machado Neto, Debora Botequio Moretti, Jardelina de Souza Todao Bernardino, Loyze Paola Oliveira de Lima, Luiz Aurelio de Campos Crispin. Centro de Genômica Funcional da ESALQ: Luiz Lehmann Coutinho, Ricardo Augusto Brassaloti, Raquel de Lello Rocha Campos Cassano. NGS Soluções Genômicas: Pilar Drummond Sampaio Corrêa Mariani. FZEA-USP Pirassununga: Mirele Daiana Poleti, Jessika Cristina Chagas Lesbon, Elisangela Chicaroni Mattos, Heidge Fukumasu. USP-Botucatu: Rejane Maria Tommasini Grotto, Jayme A. Souza-Neto, Guilherme Targino Valente, Patricia Akemi Assato, Felipe Allan da Silva da Costa, Bianca Cechetto Carlos. Mendelics: Bibiana Santos, João Paulo Kitajima, Erika Freitas, David Schlesinger. Hemocentro Ribeirão Preto: Simone Kashima, Evandra Strazza Rodrigues, Svetoslav Nanev Slavov, Elaine Vieira dos Santos, Rafael dos Santos Bezerra, Luiz Carlos Junior de Alcantara, Marta Giovanetti, Vagner Fonseca, Flavia Aburjaile, Rodrigo Tocantins Calado. FAMERP-SJRP: Cecília Artico Banho, Lívia Sacchetto, Fábio Sossai Possebon, Leila Sabrina Ullmann, Cíntia Bittar, Guilherme Campos, Helena Lage Ferreira, Jorge A. Petroli Marchesi, Maisa C. Pereira Parra, Marília Moraes, Paula Rahal, Paulo Inacio da Costa, João Pessoa Araújo Jr., Maurício Lacerda Nogueira. Prefeitura de Sao Paulo: Melissa Palmieri. |
| EPI_ISL_1966812 | USF JARDIM SAO JOAO SCRPARDO | Instituto Butantan / Mendelics | Instituto Butantan: Dimas Tadeu Covas, Sandra Coccuzzo Sampaio, Maria Carolina Elias, José Salvatore Leister Patané, Vincent Louis Viala, Antonio Jorge Martins, Ricardo Haddad, Claudia Renata dos Santos Barros, Elaine Cristina Marqueze, Raul Machado Neto, Debora Botequio Moretti, Jardelina de Souza Todao Bernardino, Loyze Paola Oliveira de Lima, Luiz Aurelio de Campos Crispin. Centro de Genômica Funcional da ESALQ: Luiz Lehmann Coutinho, Ricardo Augusto Brassaloti, Raquel de Lello Rocha Campos Cassano. NGS Soluções Genômicas: Pilar Drummond Sampaio Corrêa Mariani. FZEA-USP Pirassununga: Mirele Daiana Poleti, Jessika Cristina Chagas Lesbon, Elisangela Chicaroni Mattos, Heidge Fukumasu. USP-Botucatu: Rejane Maria Tommasini Grotto, Jayme A. Souza-Neto, Guilherme Targino Valente, Patricia Akemi Assato, Felipe Allan da Silva da Costa, Bianca Cechetto Carlos. Mendelics: Bibiana Santos, João Paulo Kitajima, Erika Freitas, David Schlesinger. Hemocentro Ribeirão Preto: Simone Kashima, Evandra Strazza Rodrigues, Svetoslav Nanev Slavov, Elaine Vieira dos Santos, Rafael dos Santos Bezerra, Luiz Carlos Junior de Alcantara, Marta Giovanetti, Vagner Fonseca, Flavia Aburjaile, Rodrigo Tocantins Calado. FAMERP-SJRP: Cecília Artico Banho, Lívia Sacchetto, Fábio Sossai Possebon, Leila Sabrina Ullmann, Cíntia Bittar, Guilherme Campos, Helena Lage Ferreira, Jorge A. Petroli Marchesi, Maisa C. Pereira Parra, Marília Moraes, Paula Rahal, Paulo Inacio da Costa, João Pessoa Araújo Jr., Maurício Lacerda Nogueira. Prefeitura de Sao Paulo: Melissa Palmieri. |
| EPI_ISL_1966813 | UBS DR WALDOMIRO FERREIRA NEVES SCRPARDO | Instituto Butantan / Mendelics | Instituto Butantan: Dimas Tadeu Covas, Sandra Coccuzzo Sampaio, Maria Carolina Elias, José Salvatore Leister Patané, Vincent Louis Viala, Antonio Jorge Martins, Ricardo Haddad, Claudia Renata dos Santos Barros, Elaine Cristina Marqueze, Raul Machado Neto, Debora Botequio Moretti, Jardelina de Souza Todao Bernardino, Loyze Paola Oliveira de Lima, Luiz Aurelio de Campos Crispin. Centro de Genômica Funcional da ESALQ: Luiz Lehmann Coutinho, Ricardo Augusto Brassaloti, Raquel de Lello Rocha Campos Cassano. NGS Soluções Genômicas: Pilar Drummond Sampaio Corrêa Mariani. FZEA-USP Pirassununga: Mirele Daiana Poleti, Jessika Cristina Chagas Lesbon, Elisangela Chicaroni Mattos, Heidge Fukumasu. USP-Botucatu: Rejane Maria Tommasini Grotto, Jayme A. Souza-Neto, Guilherme Targino Valente, Patricia Akemi Assato, Felipe Allan da Silva da Costa, Bianca Cechetto Carlos. Mendelics: Bibiana Santos, João Paulo Kitajima, Erika Freitas, David Schlesinger. Hemocentro Ribeirão Preto: Simone Kashima, Evandra Strazza Rodrigues, Svetoslav Nanev Slavov, Elaine Vieira dos Santos, Rafael dos Santos Bezerra, Luiz Carlos Junior de Alcantara, Marta Giovanetti, Vagner Fonseca, Flavia Aburjaile, Rodrigo Tocantins Calado. FAMERP-SJRP: Cecília Artico Banho, Lívia Sacchetto, Fábio Sossai Possebon, Leila Sabrina Ullmann, Cíntia Bittar, Guilherme Campos, Helena Lage Ferreira, Jorge A. Petroli Marchesi, Maisa C. Pereira Parra, Marília Moraes, Paula Rahal, Paulo Inacio da Costa, João Pessoa Araújo Jr., Maurício Lacerda Nogueira. Prefeitura de Sao Paulo: Melissa Palmieri. |
| EPI_ISL_1966814, EPI_ISL_1966815 | CENTRO DE SAUDE II SAO MIGUEL ARCANJO | Instituto Butantan / Mendelics | Instituto Butantan: Dimas Tadeu Covas, Sandra Coccuzzo Sampaio, Maria Carolina Elias, José Salvatore Leister Patané, Vincent Louis Viala, Antonio Jorge Martins, Ricardo Haddad, Claudia Renata dos Santos Barros, Elaine Cristina Marqueze, Raul Machado Neto, Debora Botequio Moretti, Jardelina de Souza Todao Bernardino, Loyze Paola Oliveira de Lima, Luiz Aurelio de Campos Crispin. Centro de Genômica Funcional da ESALQ: Luiz Lehmann Coutinho, Ricardo Augusto Brassaloti, Raquel de Lello Rocha Campos Cassano. NGS Soluções Genômicas: Pilar Drummond Sampaio Corrêa Mariani. FZEA-USP Pirassununga: Mirele Daiana Poleti, Jessika Cristina Chagas Lesbon, Elisangela Chicaroni Mattos, Heidge Fukumasu. USP-Botucatu: Rejane Maria Tommasini Grotto, Jayme A. Souza-Neto, Guilherme Targino Valente, Patricia Akemi Assato, Felipe Allan da Silva da Costa, Bianca Cechetto Carlos. Mendelics: Bibiana Santos, João Paulo Kitajima, Erika Freitas, David Schlesinger. Hemocentro Ribeirão Preto: Simone Kashima, Evandra Strazza Rodrigues, Svetoslav Nanev Slavov, Elaine Vieira dos Santos, Rafael dos Santos Bezerra, Luiz Carlos Junior de Alcantara, Marta Giovanetti, Vagner Fonseca, Flavia Aburjaile, Rodrigo Tocantins Calado. FAMERP-SJRP: Cecília Artico Banho, Lívia Sacchetto, Fábio Sossai Possebon, Leila Sabrina Ullmann, Cíntia Bittar, Guilherme Campos, Helena Lage Ferreira, Jorge A. Petroli Marchesi, Maisa C. Pereira Parra, Marília Moraes, Paula Rahal, Paulo Inacio da Costa, João Pessoa Araújo Jr., Maurício Lacerda Nogueira. Prefeitura de Sao Paulo: Melissa Palmieri. |
| EPI_ISL_1966816 | AMBULATORIO COVID 19 | Instituto Butantan / Mendelics | Instituto Butantan: Dimas Tadeu Covas, Sandra Coccuzzo Sampaio, Maria Carolina Elias, José Salvatore Leister Patané, Vincent Louis Viala, Antonio Jorge Martins, Ricardo Haddad, Claudia Renata dos Santos Barros, Elaine Cristina Marqueze, Raul Machado Neto, Debora Botequio Moretti, Jardelina de Souza Todao Bernardino, Loyze Paola Oliveira de Lima, Luiz Aurelio de Campos Crispin. Centro de Genômica Funcional da ESALQ: Luiz Lehmann Coutinho, Ricardo Augusto Brassaloti, Raquel de Lello Rocha Campos Cassano. NGS Soluções Genômicas: Pilar Drummond Sampaio Corrêa Mariani. FZEA-USP Pirassununga: Mirele Daiana Poleti, Jessika Cristina Chagas Lesbon, Elisangela Chicaroni Mattos, Heidge Fukumasu. USP-Botucatu: Rejane Maria Tommasini Grotto, Jayme A. Souza-Neto, Guilherme Targino Valente, Patricia Akemi Assato, Felipe Allan da Silva da Costa, Bianca Cechetto Carlos. Mendelics: Bibiana Santos, João Paulo Kitajima, Erika Freitas, David Schlesinger. Hemocentro Ribeirão Preto: Simone Kashima, Evandra Strazza Rodrigues, Svetoslav Nanev Slavov, Elaine Vieira dos Santos, Rafael dos Santos Bezerra, Luiz Carlos Junior de Alcantara, Marta Giovanetti, Vagner Fonseca, Flavia Aburjaile, Rodrigo Tocantins Calado. FAMERP-SJRP: Cecília Artico Banho, Lívia Sacchetto, Fábio Sossai Possebon, Leila Sabrina Ullmann, Cíntia Bittar, Guilherme Campos, Helena Lage Ferreira, Jorge A. Petroli Marchesi, Maisa C. Pereira Parra, Marília Moraes, Paula Rahal, Paulo Inacio da Costa, João Pessoa Araújo Jr., Maurício Lacerda Nogueira. Prefeitura de Sao Paulo: Melissa Palmieri. |
| EPI_ISL_1966817 | UNIDADE BASICA DE SAUDE DE TARUMA | Instituto Butantan / Mendelics | Instituto Butantan: Dimas Tadeu Covas, Sandra Coccuzzo Sampaio, Maria Carolina Elias, José Salvatore Leister Patané, Vincent Louis Viala, Antonio Jorge Martins, Ricardo Haddad, Claudia Renata dos Santos Barros, Elaine Cristina Marqueze, Raul Machado Neto, Debora Botequio Moretti, Jardelina de Souza Todao Bernardino, Loyze Paola Oliveira de Lima, Luiz Aurelio de Campos Crispin. Centro de Genômica Funcional da ESALQ: Luiz Lehmann Coutinho, Ricardo Augusto Brassaloti, Raquel de Lello Rocha Campos Cassano. NGS Soluções Genômicas: Pilar Drummond Sampaio Corrêa Mariani. FZEA-USP Pirassununga: Mirele Daiana Poleti, Jessika Cristina Chagas Lesbon, Elisangela Chicaroni Mattos, Heidge Fukumasu. USP-Botucatu: Rejane Maria Tommasini Grotto, Jayme A. Souza-Neto, Guilherme Targino Valente, Patricia Akemi Assato, Felipe Allan da Silva da Costa, Bianca Cechetto Carlos. Mendelics: Bibiana Santos, João Paulo Kitajima, Erika Freitas, David Schlesinger. Hemocentro Ribeirão Preto: Simone Kashima, Evandra Strazza Rodrigues, Svetoslav Nanev Slavov, Elaine Vieira dos Santos, Rafael dos Santos Bezerra, Luiz Carlos Junior de Alcantara, Marta Giovanetti, Vagner |

| | | | |
|---|---|---|---|
| | | | Fonseca, Flavia Aburjaile, Rodrigo Tocantins Calado. FAMERP-SJRP: Cecília Artico Banho, Lívia Sacchetto, Fábio Sossai Possebon, Leila Sabrina Ullmann, Cíntia Bittar, Guilherme Campos, Helena Lage Ferreira, Jorge A. Petroli Marchesi, Maisa C. Pereira Parra, Marília Moraes, Paula Rahal, Paulo Inacio da Costa, João Pessoa Araújo Jr., Maurício Lacerda Nogueira. Prefeitura de Sao Paulo: Melissa Palmieri. |
| EPI_ISL_1966818, EPI_ISL_1966820, EPI_ISL_1966821 | CENTRO DE SAUDE II TIETE DR RUI SILVEIRA MELO | Instituto Butantan / Mendelics | Instituto Butantan: Dimas Tadeu Covas, Sandra Coccuzzo Sampaio, Maria Carolina Elias, José Salvatore Leister Patané, Vincent Louis Viala, Antonio Jorge Martins, Ricardo Haddad, Claudia Renata dos Santos Barros, Elaine Cristina Marqueze, Raul Machado Neto, Debora Botequio Moretti, Jardelina de Souza Todao Bernardino, Loyze Paola Oliveira de Lima, Luiz Aurelio de Campos Crispin. Centro de Genômica Funcional da ESALQ: Luiz Lehmann Coutinho, Ricardo Augusto Brassaloti, Raquel de Lello Rocha Campos Cassano. NGS Soluções Genômicas: Pilar Drummond Sampaio Corrêa Mariani. FZEA-USP Pirassununga: Mirele Daiana Poleti, Jessika Cristina Chagas Lesbon, Elisangela Chicaroni Mattos, Heidge Fukumasu. USP-Botucatu: Rejane Maria Tommasini Grotto, Jayme A. Souza-Neto, Guilherme Targino Valente, Patricia Akemi Assato, Felipe Allan da Silva da Costa, Bianca Cechetto Carlos. Mendelics: Bibiana Santos, João Paulo Kitajima, Erika Freitas, David Schlesinger. Hemocentro Ribeirão Preto: Simone Kashima, Evandra Strazza Rodrigues, Svetoslav Nanev Slavov, Elaine Vieira dos Santos, Rafael dos Santos Bezerra, Luiz Carlos Junior de Alcantara, Marta Giovanetti, Vagner Fonseca, Flavia Aburjaile, Rodrigo Tocantins Calado. FAMERP-SJRP: Cecília Artico Banho, Lívia Sacchetto, Fábio Sossai Possebon, Leila Sabrina Ullmann, Cíntia Bittar, Guilherme Campos, Helena Lage Ferreira, Jorge A. Petroli Marchesi, Maisa C. Pereira Parra, Marília Moraes, Paula Rahal, Paulo Inacio da Costa, João Pessoa Araújo Jr., Maurício Lacerda Nogueira. Prefeitura de Sao Paulo: Melissa Palmieri. |
| EPI_ISL_1966822 | UNIDADE DE SAUDE DA FAMILIA DE UBIRAJARA | Instituto Butantan / Mendelics | Instituto Butantan: Dimas Tadeu Covas, Sandra Coccuzzo Sampaio, Maria Carolina Elias, José Salvatore Leister Patané, Vincent Louis Viala, Antonio Jorge Martins, Ricardo Haddad, Claudia Renata dos Santos Barros, Elaine Cristina Marqueze, Raul Machado Neto, Debora Botequio Moretti, Jardelina de Souza Todao Bernardino, Loyze Paola Oliveira de Lima, Luiz Aurelio de Campos Crispin. Centro de Genômica Funcional da ESALQ: Luiz Lehmann Coutinho, Ricardo Augusto Brassaloti, Raquel de Lello Rocha Campos Cassano. NGS Soluções Genômicas: Pilar Drummond Sampaio Corrêa Mariani. FZEA-USP Pirassununga: Mirele Daiana Poleti, Jessika Cristina Chagas Lesbon, Elisangela Chicaroni Mattos, Heidge Fukumasu. USP-Botucatu: Rejane Maria Tommasini Grotto, Jayme A. Souza-Neto, Guilherme Targino Valente, Patricia Akemi Assato, Felipe Allan da Silva da Costa, Bianca Cechetto Carlos. Mendelics: Bibiana Santos, João Paulo Kitajima, Erika Freitas, David Schlesinger. Hemocentro Ribeirão Preto: Simone Kashima, Evandra Strazza Rodrigues, Svetoslav Nanev Slavov, Elaine Vieira dos Santos, Rafael dos Santos Bezerra, Luiz Carlos Junior de Alcantara, Marta Giovanetti, Vagner Fonseca, Flavia Aburjaile, Rodrigo Tocantins Calado. FAMERP-SJRP: Cecília Artico Banho, Lívia Sacchetto, Fábio Sossai Possebon, Leila Sabrina Ullmann, Cíntia Bittar, Guilherme Campos, Helena Lage Ferreira, Jorge A. Petroli Marchesi, Maisa C. Pereira Parra, Marília Moraes, Paula Rahal, Paulo Inacio da Costa, João Pessoa Araújo Jr., Maurício Lacerda Nogueira. Prefeitura de Sao Paulo: Melissa Palmieri. |
| EPI_ISL_1966823 | PRONTO ATENDIMENTO MUNICIPAL ALUMINIO | Instituto Butantan / ESALQ-USP (Piracicaba) | Instituto Butantan: Dimas Tadeu Covas, Sandra Coccuzzo Sampaio, Maria Carolina Elias, José Salvatore Leister Patané, Vincent Louis Viala, Antonio Jorge Martins, Ricardo Haddad, Claudia Renata dos Santos Barros, Elaine Cristina Marqueze, Raul Machado Neto, Debora Botequio Moretti, Jardelina de Souza Todao Bernardino, Loyze Paola Oliveira de Lima, Luiz Aurelio de Campos Crispin. Centro de Genômica Funcional da ESALQ: Luiz Lehmann Coutinho, Ricardo Augusto Brassaloti, Raquel de Lello Rocha Campos Cassano. NGS Soluções Genômicas: Pilar Drummond Sampaio Corrêa Mariani. FZEA-USP Pirassununga: Mirele Daiana Poleti, Jessika Cristina Chagas Lesbon, Elisangela Chicaroni Mattos, Heidge Fukumasu. USP-Botucatu: Rejane Maria Tommasini Grotto, Jayme A. Souza-Neto, Guilherme Targino Valente, Patricia Akemi Assato, Felipe Allan da Silva da Costa, Bianca Cechetto Carlos. Mendelics: Bibiana Santos, João Paulo Kitajima, Erika Freitas, David Schlesinger. Hemocentro Ribeirão Preto: Simone Kashima, Evandra Strazza Rodrigues, Svetoslav Nanev Slavov, Elaine Vieira dos Santos, Rafael dos Santos Bezerra, Luiz Carlos Junior de Alcantara, Marta Giovanetti, Vagner Fonseca, Flavia Aburjaile, Rodrigo Tocantins Calado. FAMERP-SJRP: Cecília Artico Banho, Lívia Sacchetto, Fábio Sossai Possebon, Leila Sabrina Ullmann, Cíntia Bittar, Guilherme Campos, Helena Lage Ferreira, Jorge A. Petroli Marchesi, Maisa C. Pereira Parra, Marília Moraes, Paula Rahal, Paulo Inacio da Costa, João Pessoa Araújo Jr., Maurício Lacerda Nogueira. Prefeitura de Sao Paulo: Melissa Palmieri. |
| EPI_ISL_1966824, EPI_ISL_1966825, EPI_ISL_1966826, EPI_ISL_1966827, EPI_ISL_1966828, EPI_ISL_1966829 | CENTRO DE SAUDE DR RENATO DE CARVALHO RIBEIRO ANGATUBA | Instituto Butantan / ESALQ-USP (Piracicaba) | Instituto Butantan: Dimas Tadeu Covas, Sandra Coccuzzo Sampaio, Maria Carolina Elias, José Salvatore Leister Patané, Vincent Louis Viala, Antonio Jorge Martins, Ricardo Haddad, Claudia Renata dos Santos Barros, Elaine Cristina Marqueze, Raul Machado Neto, Debora Botequio Moretti, Jardelina de Souza Todao Bernardino, Loyze Paola Oliveira de Lima, Luiz Aurelio de Campos Crispin. Centro de Genômica Funcional da ESALQ: Luiz Lehmann Coutinho, Ricardo Augusto Brassaloti, Raquel de Lello Rocha Campos Cassano. NGS Soluções Genômicas: Pilar Drummond Sampaio Corrêa Mariani. FZEA-USP Pirassununga: Mirele Daiana Poleti, Jessika Cristina Chagas Lesbon, Elisangela Chicaroni Mattos, Heidge Fukumasu. USP-Botucatu: Rejane Maria Tommasini Grotto, Jayme A. Souza-Neto, Guilherme Targino Valente, Patricia Akemi Assato, Felipe Allan da Silva da Costa, Bianca Cechetto Carlos. Mendelics: Bibiana Santos, João Paulo Kitajima, Erika Freitas, David Schlesinger. Hemocentro Ribeirão Preto: Simone Kashima, Evandra Strazza Rodrigues, Svetoslav Nanev Slavov, Elaine Vieira dos Santos, Rafael dos Santos Bezerra, Luiz Carlos Junior de Alcantara, Marta Giovanetti, Vagner Fonseca, Flavia Aburjaile, Rodrigo Tocantins Calado. FAMERP-SJRP: Cecília Artico Banho, Lívia Sacchetto, Fábio Sossai Possebon, Leila Sabrina Ullmann, Cíntia Bittar, Guilherme Campos, Helena Lage Ferreira, Jorge A. Petroli Marchesi, Maisa C. Pereira Parra, Marília Moraes, Paula Rahal, Paulo Inacio da Costa, João Pessoa Araújo Jr., Maurício Lacerda Nogueira. Prefeitura de Sao Paulo: Melissa Palmieri. |
| EPI_ISL_1966830, EPI_ISL_1966831 | VIGILANCIA EPIDIMIOLOGICA DE ARACARIGUAMA | Instituto Butantan / ESALQ-USP (Piracicaba) | Instituto Butantan: Dimas Tadeu Covas, Sandra Coccuzzo Sampaio, Maria Carolina Elias, José Salvatore Leister Patané, Vincent Louis Viala, Antonio Jorge Martins, Ricardo Haddad, Claudia Renata dos Santos Barros, Elaine Cristina Marqueze, Raul Machado Neto, Debora Botequio Moretti, Jardelina de Souza Todao Bernardino, Loyze Paola Oliveira de Lima, Luiz Aurelio de Campos Crispin. Centro de Genômica Funcional da ESALQ: Luiz Lehmann Coutinho, Ricardo Augusto Brassaloti, Raquel de Lello Rocha Campos Cassano. NGS Soluções Genômicas: Pilar Drummond Sampaio Corrêa Mariani. FZEA-USP Pirassununga: Mirele Daiana Poleti, Jessika Cristina Chagas Lesbon, Elisangela Chicaroni Mattos, Heidge Fukumasu. USP-Botucatu: Rejane Maria Tommasini Grotto, Jayme A. Souza-Neto, Guilherme Targino Valente, Patricia Akemi Assato, Felipe Allan da Silva da Costa, Bianca Cechetto Carlos. Mendelics: Bibiana Santos, João Paulo Kitajima, Erika Freitas, David Schlesinger. Hemocentro Ribeirão Preto: Simone Kashima, Evandra Strazza Rodrigues, Svetoslav Nanev Slavov, Elaine Vieira dos Santos, Rafael dos Santos Bezerra, Luiz Carlos Junior de Alcantara, Marta Giovanetti, Vagner Fonseca, Flavia Aburjaile, Rodrigo Tocantins Calado. FAMERP-SJRP: Cecília Artico Banho, Lívia Sacchetto, Fábio Sossai Possebon, Leila Sabrina Ullmann, Cíntia Bittar, Guilherme Campos, Helena Lage Ferreira, Jorge A. Petroli Marchesi, Maisa C. Pereira Parra, Marília Moraes, Paula Rahal, Paulo Inacio da Costa, João Pessoa Araújo Jr., Maurício Lacerda Nogueira. Prefeitura de Sao Paulo: Melissa Palmieri. |
| EPI_ISL_1966832 | ESTRATEGIA DE SAUDE DA FAMILIA DE ARCO IRIS | Instituto Butantan / ESALQ-USP (Piracicaba) | Instituto Butantan: Dimas Tadeu Covas, Sandra Coccuzzo Sampaio, Maria Carolina Elias, José Salvatore Leister Patané, Vincent Louis Viala, Antonio Jorge Martins, Ricardo Haddad, Claudia Renata dos Santos Barros, Elaine Cristina Marqueze, Raul Machado Neto, Debora Botequio Moretti, Jardelina de Souza Todao Bernardino, Loyze Paola Oliveira de Lima, Luiz Aurelio de Campos Crispin. Centro de Genômica Funcional da ESALQ: Luiz Lehmann Coutinho, Ricardo Augusto Brassaloti, Raquel de Lello Rocha Campos Cassano. NGS Soluções Genômicas: Pilar Drummond Sampaio Corrêa Mariani. FZEA-USP Pirassununga: Mirele Daiana Poleti, Jessika Cristina Chagas Lesbon, Elisangela Chicaroni Mattos, Heidge Fukumasu. USP-Botucatu: Rejane Maria Tommasini Grotto, Jayme A. Souza-Neto, Guilherme Targino Valente, Patricia Akemi Assato, Felipe Allan da Silva da Costa, Bianca Cechetto Carlos. Mendelics: Bibiana Santos, João Paulo Kitajima, Erika Freitas, David Schlesinger. Hemocentro Ribeirão Preto: Simone Kashima, Evandra Strazza Rodrigues, Svetoslav Nanev Slavov, Elaine Vieira dos Santos, Rafael dos Santos Bezerra, Luiz Carlos Junior de Alcantara, Marta Giovanetti, Vagner Fonseca, Flavia Aburjaile, Rodrigo Tocantins Calado. FAMERP-SJRP: Cecília Artico Banho, Lívia Sacchetto, Fábio Sossai Possebon, Leila Sabrina Ullmann, Cíntia Bittar, Guilherme Campos, Helena Lage Ferreira, Jorge A. Petroli Marchesi, Maisa C. Pereira Parra, Marília Moraes, Paula Rahal, Paulo Inacio da Costa, João Pessoa Araújo Jr., Maurício Lacerda Nogueira. Prefeitura de Sao Paulo: Melissa Palmieri. |
| EPI_ISL_1966833 | CENTRO DE SAUDE I AVARE | Instituto Butantan / ESALQ-USP (Piracicaba) | Instituto Butantan: Dimas Tadeu Covas, Sandra Coccuzzo Sampaio, Maria Carolina Elias, José Salvatore Leister Patané, Vincent Louis Viala, Antonio Jorge Martins, Ricardo Haddad, Claudia Renata dos Santos Barros, Elaine Cristina Marqueze, Raul Machado Neto, Debora Botequio Moretti, Jardelina de Souza Todao Bernardino, Loyze Paola Oliveira de Lima, Luiz Aurelio de Campos Crispin. Centro de Genômica Funcional da ESALQ: Luiz Lehmann Coutinho, Ricardo Augusto Brassaloti, Raquel de Lello Rocha Campos Cassano. NGS Soluções Genômicas: Pilar Drummond Sampaio Corrêa Mariani. FZEA-USP Pirassununga: Mirele Daiana Poleti, Jessika Cristina Chagas Lesbon, Elisangela Chicaroni Mattos, Heidge Fukumasu. USP-Botucatu: Rejane Maria Tommasini Grotto, Jayme A. Souza-Neto, Guilherme Targino Valente, Patricia Akemi Assato, Felipe Allan da Silva da Costa, Bianca Cechetto Carlos. Mendelics: Bibiana Santos, João Paulo Kitajima, Erika Freitas, David Schlesinger. Hemocentro Ribeirão Preto: Simone Kashima, Evandra Strazza Rodrigues, Svetoslav Nanev Slavov, Elaine Vieira dos Santos, Rafael dos Santos Bezerra, Luiz Carlos Junior de Alcantara, Marta Giovanetti, Vagner Fonseca, Flavia Aburjaile, Rodrigo Tocantins Calado. FAMERP-SJRP: Cecília Artico Banho, Lívia Sacchetto, Fábio Sossai Possebon, Leila Sabrina Ullmann, Cíntia Bittar, Guilherme Campos, Helena Lage Ferreira, Jorge A. Petroli Marchesi, Maisa C. Pereira Parra, Marília Moraes, Paula Rahal, Paulo Inacio da Costa, João Pessoa Araújo Jr., Maurício Lacerda Nogueira. Prefeitura de Sao Paulo: Melissa Palmieri. |
| EPI_ISL_1966834 | CS DE BALSAMO | Instituto Butantan / ESALQ-USP (Piracicaba) | Instituto Butantan: Dimas Tadeu Covas, Sandra Coccuzzo Sampaio, Maria Carolina Elias, José Salvatore Leister Patané, Vincent Louis Viala, Antonio Jorge Martins, Ricardo Haddad, Claudia Renata dos Santos Barros, Elaine Cristina Marqueze, Raul Machado Neto, Debora Botequio Moretti, Jardelina de Souza Todao Bernardino, Loyze Paola Oliveira de Lima, Luiz Aurelio de Campos Crispin. Centro de Genômica Funcional da ESALQ: Luiz Lehmann |

| | | | |
|---|---|---|---|
| | | | Coutinho, Ricardo Augusto Brassaloti, Raquel de Lello Rocha Campos Cassano. NGS Soluções Genômicas: Pilar Drummond Sampaio Corrêa Mariani. FZEA-USP Pirassununga: Mirele Daiana Poleti, Jessika Cristina Chagas Lesbon, Elisangela Chicaroni Mattos, Heidge Fukumasu. USP-Botucatu: Rejane Maria Tommasini Grotto, Jayme A. Souza-Neto, Guilherme Targino Valente, Patricia Akemi Assato, Felipe Allan da Silva da Costa, Bianca Cechetto Carlos. Mendelics: Bibiana Santos, João Paulo Kitajima, Erika Freitas, David Schlesinger. Hemocentro Ribeirão Preto: Simone Kashima, Evandra Strazza Rodrigues, Svetoslav Nanev Slavov, Elaine Vieira dos Santos, Rafael dos Santos Bezerra, Luiz Carlos Junior de Alcantara, Marta Giovanetti, Vagner Fonseca, Flavia Aburjaile, Rodrigo Tocantins Calado. FAMERP-SJRP: Cecília Artico Banho, Lívia Sacchetto, Fábio Sossai Possebon, Leila Sabrina Ullmann, Cíntia Bittar, Guilherme Campos, Helena Lage Ferreira, Jorge A. Petroli Marchesi, Maisa C. Pereira Parra, Marília Moraes, Paula Rahal, Paulo Inacio da Costa, João Pessoa Araújo Jr., Maurício Lacerda Nogueira. Prefeitura de Sao Paulo: Melissa Palmieri. |
| EPI_ISL_1966835, EPI_ISL_1966836, EPI_ISL_1966837, EPI_ISL_1966838, EPI_ISL_1966839 | GRUPO TECNICO DE VIGILANCIA SANITARIA E EPIDEMIOLOGICA | Instituto Butantan / ESALQ-USP (Piracicaba) | Instituto Butantan: Dimas Tadeu Covas, Sandra Coccuzzo Sampaio, Maria Carolina Elias, José Salvatore Leister Patané, Vincent Louis Viala, Antonio Jorge Martins, Ricardo Haddad, Claudia Renata dos Santos Barros, Elaine Cristina Marqueze, Raul Machado Neto, Debora Botequio Moretti, Jardelina de Souza Todao Bernardino, Loyze Paola Oliveira de Lima, Luiz Aurelio de Campos Crispin. Centro de Genômica Funcional da ESALQ: Luiz Lehmann Coutinho, Ricardo Augusto Brassaloti, Raquel de Lello Rocha Campos Cassano. NGS Soluções Genômicas: Pilar Drummond Sampaio Corrêa Mariani. FZEA-USP Pirassununga: Mirele Daiana Poleti, Jessika Cristina Chagas Lesbon, Elisangela Chicaroni Mattos, Heidge Fukumasu. USP-Botucatu: Rejane Maria Tommasini Grotto, Jayme A. Souza-Neto, Guilherme Targino Valente, Patricia Akemi Assato, Felipe Allan da Silva da Costa, Bianca Cechetto Carlos. Mendelics: Bibiana Santos, João Paulo Kitajima, Erika Freitas, David Schlesinger. Hemocentro Ribeirão Preto: Simone Kashima, Evandra Strazza Rodrigues, Svetoslav Nanev Slavov, Elaine Vieira dos Santos, Rafael dos Santos Bezerra, Luiz Carlos Junior de Alcantara, Marta Giovanetti, Vagner Fonseca, Flavia Aburjaile, Rodrigo Tocantins Calado. FAMERP-SJRP: Cecília Artico Banho, Lívia Sacchetto, Fábio Sossai Possebon, Leila Sabrina Ullmann, Cíntia Bittar, Guilherme Campos, Helena Lage Ferreira, Jorge A. Petroli Marchesi, Maisa C. Pereira Parra, Marília Moraes, Paula Rahal, Paulo Inacio da Costa, João Pessoa Araújo Jr., Maurício Lacerda Nogueira. Prefeitura de Sao Paulo: Melissa Palmieri. |
| EPI_ISL_1966840, EPI_ISL_1966841, EPI_ISL_1966842, EPI_ISL_1966843, EPI_ISL_1966844 | SMS SECRETARIA MUNICIPAL DE SAUDE DE BOITUVA | Instituto Butantan / ESALQ-USP (Piracicaba) | Instituto Butantan: Dimas Tadeu Covas, Sandra Coccuzzo Sampaio, Maria Carolina Elias, José Salvatore Leister Patané, Vincent Louis Viala, Antonio Jorge Martins, Ricardo Haddad, Claudia Renata dos Santos Barros, Elaine Cristina Marqueze, Raul Machado Neto, Debora Botequio Moretti, Jardelina de Souza Todao Bernardino, Loyze Paola Oliveira de Lima, Luiz Aurelio de Campos Crispin. Centro de Genômica Funcional da ESALQ: Luiz Lehmann Coutinho, Ricardo Augusto Brassaloti, Raquel de Lello Rocha Campos Cassano. NGS Soluções Genômicas: Pilar Drummond Sampaio Corrêa Mariani. FZEA-USP Pirassununga: Mirele Daiana Poleti, Jessika Cristina Chagas Lesbon, Elisangela Chicaroni Mattos, Heidge Fukumasu. USP-Botucatu: Rejane Maria Tommasini Grotto, Jayme A. Souza-Neto, Guilherme Targino Valente, Patricia Akemi Assato, Felipe Allan da Silva da Costa, Bianca Cechetto Carlos. Mendelics: Bibiana Santos, João Paulo Kitajima, Erika Freitas, David Schlesinger. Hemocentro Ribeirão Preto: Simone Kashima, Evandra Strazza Rodrigues, Svetoslav Nanev Slavov, Elaine Vieira dos Santos, Rafael dos Santos Bezerra, Luiz Carlos Junior de Alcantara, Marta Giovanetti, Vagner Fonseca, Flavia Aburjaile, Rodrigo Tocantins Calado. FAMERP-SJRP: Cecília Artico Banho, Lívia Sacchetto, Fábio Sossai Possebon, Leila Sabrina Ullmann, Cíntia Bittar, Guilherme Campos, Helena Lage Ferreira, Jorge A. Petroli Marchesi, Maisa C. Pereira Parra, Marília Moraes, Paula Rahal, Paulo Inacio da Costa, João Pessoa Araújo Jr., Maurício Lacerda Nogueira. Prefeitura de Sao Paulo: Melissa Palmieri. |
| EPI_ISL_1966845 | PRONTO ATENDIMENTO MUNICIPAL ITALO SANTUCCI | Instituto Butantan / ESALQ-USP (Piracicaba) | Instituto Butantan: Dimas Tadeu Covas, Sandra Coccuzzo Sampaio, Maria Carolina Elias, José Salvatore Leister Patané, Vincent Louis Viala, Antonio Jorge Martins, Ricardo Haddad, Claudia Renata dos Santos Barros, Elaine Cristina Marqueze, Raul Machado Neto, Debora Botequio Moretti, Jardelina de Souza Todao Bernardino, Loyze Paola Oliveira de Lima, Luiz Aurelio de Campos Crispin. Centro de Genômica Funcional da ESALQ: Luiz Lehmann Coutinho, Ricardo Augusto Brassaloti, Raquel de Lello Rocha Campos Cassano. NGS Soluções Genômicas: Pilar Drummond Sampaio Corrêa Mariani. FZEA-USP Pirassununga: Mirele Daiana Poleti, Jessika Cristina Chagas Lesbon, Elisangela Chicaroni Mattos, Heidge Fukumasu. USP-Botucatu: Rejane Maria Tommasini Grotto, Jayme A. Souza-Neto, Guilherme Targino Valente, Patricia Akemi Assato, Felipe Allan da Silva da Costa, Bianca Cechetto Carlos. Mendelics: Bibiana Santos, João Paulo Kitajima, Erika Freitas, David Schlesinger. Hemocentro Ribeirão Preto: Simone Kashima, Evandra Strazza Rodrigues, Svetoslav Nanev Slavov, Elaine Vieira dos Santos, Rafael dos Santos Bezerra, Luiz Carlos Junior de Alcantara, Marta Giovanetti, Vagner Fonseca, Flavia Aburjaile, Rodrigo Tocantins Calado. FAMERP-SJRP: Cecília Artico Banho, Lívia Sacchetto, Fábio Sossai Possebon, Leila Sabrina Ullmann, Cíntia Bittar, Guilherme Campos, Helena Lage Ferreira, Jorge A. Petroli Marchesi, Maisa C. Pereira Parra, Marília Moraes, Paula Rahal, Paulo Inacio da Costa, João Pessoa Araújo Jr., Maurício Lacerda Nogueira. Prefeitura de Sao Paulo: Melissa Palmieri. |
| EPI_ISL_1966846 | HOSPITAL MUNICIPAL REYNALDO GUERRA CAJATI | Instituto Butantan / ESALQ-USP (Piracicaba) | Instituto Butantan: Dimas Tadeu Covas, Sandra Coccuzzo Sampaio, Maria Carolina Elias, José Salvatore Leister Patané, Vincent Louis Viala, Antonio Jorge Martins, Ricardo Haddad, Claudia Renata dos Santos Barros, Elaine Cristina Marqueze, Raul Machado Neto, Debora Botequio Moretti, Jardelina de Souza Todao Bernardino, Loyze Paola Oliveira de Lima, Luiz Aurelio de Campos Crispin. Centro de Genômica Funcional da ESALQ: Luiz Lehmann Coutinho, Ricardo Augusto Brassaloti, Raquel de Lello Rocha Campos Cassano. NGS Soluções Genômicas: Pilar Drummond Sampaio Corrêa Mariani. FZEA-USP Pirassununga: Mirele Daiana Poleti, Jessika Cristina Chagas Lesbon, Elisangela Chicaroni Mattos, Heidge Fukumasu. USP-Botucatu: Rejane Maria Tommasini Grotto, Jayme A. Souza-Neto, Guilherme Targino Valente, Patricia Akemi Assato, Felipe Allan da Silva da Costa, Bianca Cechetto Carlos. Mendelics: Bibiana Santos, João Paulo Kitajima, Erika Freitas, David Schlesinger. Hemocentro Ribeirão Preto: Simone Kashima, Evandra Strazza Rodrigues, Svetoslav Nanev Slavov, Elaine Vieira dos Santos, Rafael dos Santos Bezerra, Luiz Carlos Junior de Alcantara, Marta Giovanetti, Vagner Fonseca, Flavia Aburjaile, Rodrigo Tocantins Calado. FAMERP-SJRP: Cecília Artico Banho, Lívia Sacchetto, Fábio Sossai Possebon, Leila Sabrina Ullmann, Cíntia Bittar, Guilherme Campos, Helena Lage Ferreira, Jorge A. Petroli Marchesi, Maisa C. Pereira Parra, Marília Moraes, Paula Rahal, Paulo Inacio da Costa, João Pessoa Araújo Jr., Maurício Lacerda Nogueira. Prefeitura de Sao Paulo: Melissa Palmieri. |
| EPI_ISL_1966847, EPI_ISL_1966848, EPI_ISL_1966849, EPI_ISL_1966850, EPI_ISL_1966851, EPI_ISL_1966852 | UBS SAO JUDAS | Instituto Butantan / ESALQ-USP (Piracicaba) | Instituto Butantan: Dimas Tadeu Covas, Sandra Coccuzzo Sampaio, Maria Carolina Elias, José Salvatore Leister Patané, Vincent Louis Viala, Antonio Jorge Martins, Ricardo Haddad, Claudia Renata dos Santos Barros, Elaine Cristina Marqueze, Raul Machado Neto, Debora Botequio Moretti, Jardelina de Souza Todao Bernardino, Loyze Paola Oliveira de Lima, Luiz Aurelio de Campos Crispin. Centro de Genômica Funcional da ESALQ: Luiz Lehmann Coutinho, Ricardo Augusto Brassaloti, Raquel de Lello Rocha Campos Cassano. NGS Soluções Genômicas: Pilar Drummond Sampaio Corrêa Mariani. FZEA-USP Pirassununga: Mirele Daiana Poleti, Jessika Cristina Chagas Lesbon, Elisangela Chicaroni Mattos, Heidge Fukumasu. USP-Botucatu: Rejane Maria Tommasini Grotto, Jayme A. Souza-Neto, Guilherme Targino Valente, Patricia Akemi Assato, Felipe Allan da Silva da Costa, Bianca Cechetto Carlos. Mendelics: Bibiana Santos, João Paulo Kitajima, Erika Freitas, David Schlesinger. Hemocentro Ribeirão Preto: Simone Kashima, Evandra Strazza Rodrigues, Svetoslav Nanev Slavov, Elaine Vieira dos Santos, Rafael dos Santos Bezerra, Luiz Carlos Junior de Alcantara, Marta Giovanetti, Vagner Fonseca, Flavia Aburjaile, Rodrigo Tocantins Calado. FAMERP-SJRP: Cecília Artico Banho, Lívia Sacchetto, Fábio Sossai Possebon, Leila Sabrina Ullmann, Cíntia Bittar, Guilherme Campos, Helena Lage Ferreira, Jorge A. Petroli Marchesi, Maisa C. Pereira Parra, Marília Moraes, Paula Rahal, Paulo Inacio da Costa, João Pessoa Araújo Jr., Maurício Lacerda Nogueira. Prefeitura de Sao Paulo: Melissa Palmieri. |
| EPI_ISL_1966853 | ESF ROSA ROSSI MAIA | Instituto Butantan / ESALQ-USP (Piracicaba) | Instituto Butantan: Dimas Tadeu Covas, Sandra Coccuzzo Sampaio, Maria Carolina Elias, José Salvatore Leister Patané, Vincent Louis Viala, Antonio Jorge Martins, Ricardo Haddad, Claudia Renata dos Santos Barros, Elaine Cristina Marqueze, Raul Machado Neto, Debora Botequio Moretti, Jardelina de Souza Todao Bernardino, Loyze Paola Oliveira de Lima, Luiz Aurelio de Campos Crispin. Centro de Genômica Funcional da ESALQ: Luiz Lehmann Coutinho, Ricardo Augusto Brassaloti, Raquel de Lello Rocha Campos Cassano. NGS Soluções Genômicas: Pilar Drummond Sampaio Corrêa Mariani. FZEA-USP Pirassununga: Mirele Daiana Poleti, Jessika Cristina Chagas Lesbon, Elisangela Chicaroni Mattos, Heidge Fukumasu. USP-Botucatu: Rejane Maria Tommasini Grotto, Jayme A. Souza-Neto, Guilherme Targino Valente, Patricia Akemi Assato, Felipe Allan da Silva da Costa, Bianca Cechetto Carlos. Mendelics: Bibiana Santos, João Paulo Kitajima, Erika Freitas, David Schlesinger. Hemocentro Ribeirão Preto: Simone Kashima, Evandra Strazza Rodrigues, Svetoslav Nanev Slavov, Elaine Vieira dos Santos, Rafael dos Santos Bezerra, Luiz Carlos Junior de Alcantara, Marta Giovanetti, Vagner Fonseca, Flavia Aburjaile, Rodrigo Tocantins Calado. FAMERP-SJRP: Cecília Artico Banho, Lívia Sacchetto, Fábio Sossai Possebon, Leila Sabrina Ullmann, Cíntia Bittar, Guilherme Campos, Helena Lage Ferreira, Jorge A. Petroli Marchesi, Maisa C. Pereira Parra, Marília Moraes, Paula Rahal, Paulo Inacio da Costa, João Pessoa Araújo Jr., Maurício Lacerda Nogueira. Prefeitura de Sao Paulo: Melissa Palmieri. |
| EPI_ISL_1966854, EPI_ISL_1966857 | UNIDADE SENTINELA COVID19 | Instituto Butantan / ESALQ-USP (Piracicaba) | Instituto Butantan: Dimas Tadeu Covas, Sandra Coccuzzo Sampaio, Maria Carolina Elias, José Salvatore Leister Patané, Vincent Louis Viala, Antonio Jorge Martins, Ricardo Haddad, Claudia Renata dos Santos Barros, Elaine Cristina Marqueze, Raul Machado Neto, Debora Botequio Moretti, Jardelina de Souza Todao Bernardino, Loyze Paola Oliveira de Lima, Luiz Aurelio de Campos Crispin. Centro de Genômica Funcional da ESALQ: Luiz Lehmann Coutinho, Ricardo Augusto Brassaloti, Raquel de Lello Rocha Campos Cassano. NGS Soluções Genômicas: Pilar Drummond Sampaio Corrêa Mariani. FZEA-USP Pirassununga: Mirele Daiana Poleti, Jessika Cristina Chagas Lesbon, Elisangela Chicaroni Mattos, Heidge Fukumasu. USP-Botucatu: Rejane Maria Tommasini Grotto, Jayme A. Souza-Neto, Guilherme Targino Valente, Patricia Akemi Assato, Felipe Allan da Silva da Costa, Bianca Cechetto Carlos. Mendelics: Bibiana Santos, João Paulo Kitajima, Erika Freitas, David Schlesinger. Hemocentro Ribeirão Preto: Simone Kashima, Evandra Strazza Rodrigues, Svetoslav Nanev Slavov, Elaine Vieira dos Santos, Rafael dos Santos Bezerra, Luiz Carlos Junior de Alcantara, Marta Giovanetti, Vagner |

| | | | |
|---|---|---|---|
| | | | Fonseca, Flavia Aburjaile, Rodrigo Tocantins Calado. FAMERP-SJRP: Cecília Artico Banho, Lívia Sacchetto, Fábio Sossai Possebon, Leila Sabrina Ullmann, Cíntia Bittar, Guilherme Campos, Helena Lage Ferreira, Jorge A. Petroli Marchesi, Maisa C. Pereira Parra, Marília Moraes, Paula Rahal, Paulo Inacio da Costa, João Pessoa Araújo Jr., Maurício Lacerda Nogueira. Prefeitura de Sao Paulo: Melissa Palmieri. |
| EPI_ISL_1966858, EPI_ISL_1966859, EPI_ISL_1966860 | UNIDADE SENTINELA VILA APARECIDA | Instituto Butantan / ESALQ-USP (Piracicaba) | Instituto Butantan: Dimas Tadeu Covas, Sandra Coccuzzo Sampaio, Maria Carolina Elias, José Salvatore Leister Patané, Vincent Louis Viala, Antonio Jorge Martins, Ricardo Haddad, Claudia Renata dos Santos Barros, Elaine Cristina Marqueze, Raul Machado Neto, Debora Botequio Moretti, Jardelina de Souza Todao Bernardino, Loyze Paola Oliveira de Lima, Luiz Aurelio de Campos Crispin. Centro de Genômica Funcional da ESALQ: Luiz Lehmann Coutinho, Ricardo Augusto Brassaloti, Raquel de Lello Rocha Campos Cassano. NGS Soluções Genômicas: Pilar Drummond Sampaio Corrêa Mariani. FZEA-USP Pirassununga: Mirele Daiana Poleti, Jessika Cristina Chagas Lesbon, Elisangela Chicaroni Mattos, Heidge Fukumasu. USP-Botucatu: Rejane Maria Tommasini Grotto, Jayme A. Souza-Neto, Guilherme Targino Valente, Patricia Akemi Assato, Felipe Allan da Silva da Costa, Bianca Cechetto Carlos. Mendelics: Bibiana Santos, João Paulo Kitajima, Erika Freitas, David Schlesinger. Hemocentro Ribeirão Preto: Simone Kashima, Evandra Strazza Rodrigues, Svetoslav Nanev Slavov, Elaine Vieira dos Santos, Rafael dos Santos Bezerra, Luiz Carlos Junior de Alcantara, Marta Giovanetti, Vagner Fonseca, Flavia Aburjaile, Rodrigo Tocantins Calado. FAMERP-SJRP: Cecília Artico Banho, Lívia Sacchetto, Fábio Sossai Possebon, Leila Sabrina Ullmann, Cíntia Bittar, Guilherme Campos, Helena Lage Ferreira, Jorge A. Petroli Marchesi, Maisa C. Pereira Parra, Marília Moraes, Paula Rahal, Paulo Inacio da Costa, João Pessoa Araújo Jr., Maurício Lacerda Nogueira. Prefeitura de Sao Paulo: Melissa Palmieri. |
| EPI_ISL_1966861, EPI_ISL_1966862 | UNIDADE SENTINELA COVID19 | Instituto Butantan / ESALQ-USP (Piracicaba) | Instituto Butantan: Dimas Tadeu Covas, Sandra Coccuzzo Sampaio, Maria Carolina Elias, José Salvatore Leister Patané, Vincent Louis Viala, Antonio Jorge Martins, Ricardo Haddad, Claudia Renata dos Santos Barros, Elaine Cristina Marqueze, Raul Machado Neto, Debora Botequio Moretti, Jardelina de Souza Todao Bernardino, Loyze Paola Oliveira de Lima, Luiz Aurelio de Campos Crispin. Centro de Genômica Funcional da ESALQ: Luiz Lehmann Coutinho, Ricardo Augusto Brassaloti, Raquel de Lello Rocha Campos Cassano. NGS Soluções Genômicas: Pilar Drummond Sampaio Corrêa Mariani. FZEA-USP Pirassununga: Mirele Daiana Poleti, Jessika Cristina Chagas Lesbon, Elisangela Chicaroni Mattos, Heidge Fukumasu. USP-Botucatu: Rejane Maria Tommasini Grotto, Jayme A. Souza-Neto, Guilherme Targino Valente, Patricia Akemi Assato, Felipe Allan da Silva da Costa, Bianca Cechetto Carlos. Mendelics: Bibiana Santos, João Paulo Kitajima, Erika Freitas, David Schlesinger. Hemocentro Ribeirão Preto: Simone Kashima, Evandra Strazza Rodrigues, Svetoslav Nanev Slavov, Elaine Vieira dos Santos, Rafael dos Santos Bezerra, Luiz Carlos Junior de Alcantara, Marta Giovanetti, Vagner Fonseca, Flavia Aburjaile, Rodrigo Tocantins Calado. FAMERP-SJRP: Cecília Artico Banho, Lívia Sacchetto, Fábio Sossai Possebon, Leila Sabrina Ullmann, Cíntia Bittar, Guilherme Campos, Helena Lage Ferreira, Jorge A. Petroli Marchesi, Maisa C. Pereira Parra, Marília Moraes, Paula Rahal, Paulo Inacio da Costa, João Pessoa Araújo Jr., Maurício Lacerda Nogueira. Prefeitura de Sao Paulo: Melissa Palmieri. |
| EPI_ISL_1966863, EPI_ISL_1966864, EPI_ISL_1966865, EPI_ISL_1966866, EPI_ISL_1966867 | UNIDADE MISTA DE SAUDE CAPELA DO ALTO | Instituto Butantan / ESALQ-USP (Piracicaba) | Instituto Butantan: Dimas Tadeu Covas, Sandra Coccuzzo Sampaio, Maria Carolina Elias, José Salvatore Leister Patané, Vincent Louis Viala, Antonio Jorge Martins, Ricardo Haddad, Claudia Renata dos Santos Barros, Elaine Cristina Marqueze, Raul Machado Neto, Debora Botequio Moretti, Jardelina de Souza Todao Bernardino, Loyze Paola Oliveira de Lima, Luiz Aurelio de Campos Crispin. Centro de Genômica Funcional da ESALQ: Luiz Lehmann Coutinho, Ricardo Augusto Brassaloti, Raquel de Lello Rocha Campos Cassano. NGS Soluções Genômicas: Pilar Drummond Sampaio Corrêa Mariani. FZEA-USP Pirassununga: Mirele Daiana Poleti, Jessika Cristina Chagas Lesbon, Elisangela Chicaroni Mattos, Heidge Fukumasu. USP-Botucatu: Rejane Maria Tommasini Grotto, Jayme A. Souza-Neto, Guilherme Targino Valente, Patricia Akemi Assato, Felipe Allan da Silva da Costa, Bianca Cechetto Carlos. Mendelics: Bibiana Santos, João Paulo Kitajima, Erika Freitas, David Schlesinger. Hemocentro Ribeirão Preto: Simone Kashima, Evandra Strazza Rodrigues, Svetoslav Nanev Slavov, Elaine Vieira dos Santos, Rafael dos Santos Bezerra, Luiz Carlos Junior de Alcantara, Marta Giovanetti, Vagner Fonseca, Flavia Aburjaile, Rodrigo Tocantins Calado. FAMERP-SJRP: Cecília Artico Banho, Lívia Sacchetto, Fábio Sossai Possebon, Leila Sabrina Ullmann, Cíntia Bittar, Guilherme Campos, Helena Lage Ferreira, Jorge A. Petroli Marchesi, Maisa C. Pereira Parra, Marília Moraes, Paula Rahal, Paulo Inacio da Costa, João Pessoa Araújo Jr., Maurício Lacerda Nogueira. Prefeitura de Sao Paulo: Melissa Palmieri. |
| EPI_ISL_1966868, EPI_ISL_1966869 | CS DE CATIGUA | Instituto Butantan / ESALQ-USP (Piracicaba) | Instituto Butantan: Dimas Tadeu Covas, Sandra Coccuzzo Sampaio, Maria Carolina Elias, José Salvatore Leister Patané, Vincent Louis Viala, Antonio Jorge Martins, Ricardo Haddad, Claudia Renata dos Santos Barros, Elaine Cristina Marqueze, Raul Machado Neto, Debora Botequio Moretti, Jardelina de Souza Todao Bernardino, Loyze Paola Oliveira de Lima, Luiz Aurelio de Campos Crispin. Centro de Genômica Funcional da ESALQ: Luiz Lehmann Coutinho, Ricardo Augusto Brassaloti, Raquel de Lello Rocha Campos Cassano. NGS Soluções Genômicas: Pilar Drummond Sampaio Corrêa Mariani. FZEA-USP Pirassununga: Mirele Daiana Poleti, Jessika Cristina Chagas Lesbon, Elisangela Chicaroni Mattos, Heidge Fukumasu. USP-Botucatu: Rejane Maria Tommasini Grotto, Jayme A. Souza-Neto, Guilherme Targino Valente, Patricia Akemi Assato, Felipe Allan da Silva da Costa, Bianca Cechetto Carlos. Mendelics: Bibiana Santos, João Paulo Kitajima, Erika Freitas, David Schlesinger. Hemocentro Ribeirão Preto: Simone Kashima, Evandra Strazza Rodrigues, Svetoslav Nanev Slavov, Elaine Vieira dos Santos, Rafael dos Santos Bezerra, Luiz Carlos Junior de Alcantara, Marta Giovanetti, Vagner Fonseca, Flavia Aburjaile, Rodrigo Tocantins Calado. FAMERP-SJRP: Cecília Artico Banho, Lívia Sacchetto, Fábio Sossai Possebon, Leila Sabrina Ullmann, Cíntia Bittar, Guilherme Campos, Helena Lage Ferreira, Jorge A. Petroli Marchesi, Maisa C. Pereira Parra, Marília Moraes, Paula Rahal, Paulo Inacio da Costa, João Pessoa Araújo Jr., Maurício Lacerda Nogueira. Prefeitura de Sao Paulo: Melissa Palmieri. |
| EPI_ISL_1966870, EPI_ISL_1966871, EPI_ISL_1966872, EPI_ISL_1966873, EPI_ISL_1966874, EPI_ISL_1966875 | UBS ATALAIA DR MARIO AGUIAR FILHO | Instituto Butantan / ESALQ-USP (Piracicaba) | Instituto Butantan: Dimas Tadeu Covas, Sandra Coccuzzo Sampaio, Maria Carolina Elias, José Salvatore Leister Patané, Vincent Louis Viala, Antonio Jorge Martins, Ricardo Haddad, Claudia Renata dos Santos Barros, Elaine Cristina Marqueze, Raul Machado Neto, Debora Botequio Moretti, Jardelina de Souza Todao Bernardino, Loyze Paola Oliveira de Lima, Luiz Aurelio de Campos Crispin. Centro de Genômica Funcional da ESALQ: Luiz Lehmann Coutinho, Ricardo Augusto Brassaloti, Raquel de Lello Rocha Campos Cassano. NGS Soluções Genômicas: Pilar Drummond Sampaio Corrêa Mariani. FZEA-USP Pirassununga: Mirele Daiana Poleti, Jessika Cristina Chagas Lesbon, Elisangela Chicaroni Mattos, Heidge Fukumasu. USP-Botucatu: Rejane Maria Tommasini Grotto, Jayme A. Souza-Neto, Guilherme Targino Valente, Patricia Akemi Assato, Felipe Allan da Silva da Costa, Bianca Cechetto Carlos. Mendelics: Bibiana Santos, João Paulo Kitajima, Erika Freitas, David Schlesinger. Hemocentro Ribeirão Preto: Simone Kashima, Evandra Strazza Rodrigues, Svetoslav Nanev Slavov, Elaine Vieira dos Santos, Rafael dos Santos Bezerra, Luiz Carlos Junior de Alcantara, Marta Giovanetti, Vagner Fonseca, Flavia Aburjaile, Rodrigo Tocantins Calado. FAMERP-SJRP: Cecília Artico Banho, Lívia Sacchetto, Fábio Sossai Possebon, Leila Sabrina Ullmann, Cíntia Bittar, Guilherme Campos, Helena Lage Ferreira, Jorge A. Petroli Marchesi, Maisa C. Pereira Parra, Marília Moraes, Paula Rahal, Paulo Inacio da Costa, João Pessoa Araújo Jr., Maurício Lacerda Nogueira. Prefeitura de Sao Paulo: Melissa Palmieri. |
| EPI_ISL_1966876 | PRONTO SOCORRO CAUCAIA DO ALTO | Instituto Butantan / ESALQ-USP (Piracicaba) | Instituto Butantan: Dimas Tadeu Covas, Sandra Coccuzzo Sampaio, Maria Carolina Elias, José Salvatore Leister Patané, Vincent Louis Viala, Antonio Jorge Martins, Ricardo Haddad, Claudia Renata dos Santos Barros, Elaine Cristina Marqueze, Raul Machado Neto, Debora Botequio Moretti, Jardelina de Souza Todao Bernardino, Loyze Paola Oliveira de Lima, Luiz Aurelio de Campos Crispin. Centro de Genômica Funcional da ESALQ: Luiz Lehmann Coutinho, Ricardo Augusto Brassaloti, Raquel de Lello Rocha Campos Cassano. NGS Soluções Genômicas: Pilar Drummond Sampaio Corrêa Mariani. FZEA-USP Pirassununga: Mirele Daiana Poleti, Jessika Cristina Chagas Lesbon, Elisangela Chicaroni Mattos, Heidge Fukumasu. USP-Botucatu: Rejane Maria Tommasini Grotto, Jayme A. Souza-Neto, Guilherme Targino Valente, Patricia Akemi Assato, Felipe Allan da Silva da Costa, Bianca Cechetto Carlos. Mendelics: Bibiana Santos, João Paulo Kitajima, Erika Freitas, David Schlesinger. Hemocentro Ribeirão Preto: Simone Kashima, Evandra Strazza Rodrigues, Svetoslav Nanev Slavov, Elaine Vieira dos Santos, Rafael dos Santos Bezerra, Luiz Carlos Junior de Alcantara, Marta Giovanetti, Vagner Fonseca, Flavia Aburjaile, Rodrigo Tocantins Calado. FAMERP-SJRP: Cecília Artico Banho, Lívia Sacchetto, Fábio Sossai Possebon, Leila Sabrina Ullmann, Cíntia Bittar, Guilherme Campos, Helena Lage Ferreira, Jorge A. Petroli Marchesi, Maisa C. Pereira Parra, Marília Moraes, Paula Rahal, Paulo Inacio da Costa, João Pessoa Araújo Jr., Maurício Lacerda Nogueira. Prefeitura de Sao Paulo: Melissa Palmieri. |
| EPI_ISL_1966877, EPI_ISL_1966878, EPI_ISL_1966879, EPI_ISL_1966880, EPI_ISL_1966881, EPI_ISL_1966882, EPI_ISL_1966883, EPI_ISL_1966884, EPI_ISL_1966885, EPI_ISL_1966886, EPI_ISL_1966887, EPI_ISL_1966888 | | | |
| see above | UBS ATALAIA DR MARIO AGUIAR FILHO | Instituto Butantan / ESALQ-USP (Piracicaba) | Instituto Butantan: Dimas Tadeu Covas, Sandra Coccuzzo Sampaio, Maria Carolina Elias, José Salvatore Leister Patané, Vincent Louis Viala, Antonio Jorge Martins, Ricardo Haddad, Claudia Renata dos Santos Barros, Elaine Cristina Marqueze, Raul Machado Neto, Debora Botequio Moretti, Jardelina de Souza Todao Bernardino, Loyze Paola Oliveira de Lima, Luiz Aurelio de Campos Crispin. Centro de Genômica Funcional da ESALQ: Luiz Lehmann Coutinho, Ricardo Augusto Brassaloti, Raquel de Lello Rocha Campos Cassano. NGS Soluções Genômicas: Pilar Drummond Sampaio Corrêa Mariani. FZEA-USP Pirassununga: Mirele Daiana Poleti, Jessika Cristina Chagas Lesbon, Elisangela Chicaroni Mattos, Heidge Fukumasu. USP-Botucatu: Rejane Maria Tommasini Grotto, Jayme A. Souza-Neto, Guilherme Targino Valente, Patricia Akemi Assato, Felipe Allan da Silva da Costa, Bianca Cechetto Carlos. Mendelics: Bibiana Santos, João Paulo Kitajima, Erika Freitas, David Schlesinger. Hemocentro Ribeirão Preto: Simone Kashima, Evandra Strazza Rodrigues, Svetoslav Nanev Slavov, Elaine Vieira dos Santos, Rafael dos Santos Bezerra, Luiz Carlos Junior de Alcantara, Marta Giovanetti, Vagner Fonseca, Flavia Aburjaile, Rodrigo Tocantins Calado. FAMERP-SJRP: Cecília Artico Banho, Lívia Sacchetto, Fábio Sossai Possebon, Leila Sabrina Ullmann, Cíntia Bittar, Guilherme Campos, Helena Lage Ferreira, Jorge A. Petroli Marchesi, Maisa C. Pereira Parra, Marília Moraes, Paula Rahal, Paulo Inacio da Costa, João Pessoa Araújo Jr., Maurício Lacerda Nogueira. Prefeitura de Sao Paulo: Melissa Palmieri. |
| EPI_ISL_1966889 | UPA ATALAIA DRA RITA DE CASSIA SORIO | Instituto Butantan / ESALQ-USP (Piracicaba) | Instituto Butantan: Dimas Tadeu Covas, Sandra Coccuzzo Sampaio, Maria Carolina Elias, José Salvatore Leister Patané, Vincent Louis Viala, Antonio Jorge Martins, Ricardo Haddad, Claudia Renata dos Santos Barros, Elaine Cristina Marqueze, Raul Machado Neto, Debora Botequio Moretti, Jardelina de |

| | | | |
|---|---|---|---|
| | | | Souza Todao Bernardino, Loyze Paola Oliveira de Lima, Luiz Aurelio de Campos Crispin. Centro de Genômica Funcional da ESALQ: Luiz Lehmann Coutinho, Ricardo Augusto Brassaloti, Raquel de Lello Rocha Campos Cassano. NGS Soluções Genômicas: Pilar Drummond Sampaio Corrêa Mariani. FZEA-USP Pirassununga: Mirele Daiana Poleti, Jessika Cristina Chagas Lesbon, Elisangela Chicaroni Mattos, Heidge Fukumasu. USP-Botucatu: Rejane Maria Tommasini Grotto, Jayme A. Souza-Neto, Guilherme Targino Valente, Patricia Akemi Assato, Felipe Allan da Silva da Costa, Bianca Cechetto Carlos. Mendelics: Bibiana Santos, João Paulo Kitajima, Erika Freitas, David Schlesinger. Hemocentro Ribeirão Preto: Simone Kashima, Evandra Strazza Rodrigues, Svetoslav Nanev Slavov, Elaine Vieira dos Santos, Rafael dos Santos Bezerra, Luiz Carlos Junior de Alcantara, Marta Giovanetti, Vagner Fonseca, Flavia Aburjaile, Rodrigo Tocantins Calado. FAMERP-SJRP: Cecília Artico Banho, Lívia Sacchetto, Fábio Sossai Possebon, Leila Sabrina Ullmann, Cíntia Bittar, Guilherme Campos, Helena Lage Ferreira, Jorge A. Petroli Marchesi, Maisa C. Pereira Parra, Marília Moraes, Paula Rahal, Paulo Inacio da Costa, João Pessoa Araújo Jr., Maurício Lacerda Nogueira. Prefeitura de Sao Paulo: Melissa Palmieri. |
| EPI_ISL_1966890 | UBS ATALAIA DR MARIO AGUIAR FILHO | Instituto Butantan / ESALQ-USP (Piracicaba) | Instituto Butantan: Dimas Tadeu Covas, Sandra Coccuzzo Sampaio, Maria Carolina Elias, José Salvatore Leister Patané, Vincent Louis Viala, Antonio Jorge Martins, Ricardo Haddad, Claudia Renata dos Santos Barros, Elaine Cristina Marqueze, Raul Machado Neto, Debora Botequio Moretti, Jardelina de Souza Todao Bernardino, Loyze Paola Oliveira de Lima, Luiz Aurelio de Campos Crispin. Centro de Genômica Funcional da ESALQ: Luiz Lehmann Coutinho, Ricardo Augusto Brassaloti, Raquel de Lello Rocha Campos Cassano. NGS Soluções Genômicas: Pilar Drummond Sampaio Corrêa Mariani. FZEA-USP Pirassununga: Mirele Daiana Poleti, Jessika Cristina Chagas Lesbon, Elisangela Chicaroni Mattos, Heidge Fukumasu. USP-Botucatu: Rejane Maria Tommasini Grotto, Jayme A. Souza-Neto, Guilherme Targino Valente, Patricia Akemi Assato, Felipe Allan da Silva da Costa, Bianca Cechetto Carlos. Mendelics: Bibiana Santos, João Paulo Kitajima, Erika Freitas, David Schlesinger. Hemocentro Ribeirão Preto: Simone Kashima, Evandra Strazza Rodrigues, Svetoslav Nanev Slavov, Elaine Vieira dos Santos, Rafael dos Santos Bezerra, Luiz Carlos Junior de Alcantara, Marta Giovanetti, Vagner Fonseca, Flavia Aburjaile, Rodrigo Tocantins Calado. FAMERP-SJRP: Cecília Artico Banho, Lívia Sacchetto, Fábio Sossai Possebon, Leila Sabrina Ullmann, Cíntia Bittar, Guilherme Campos, Helena Lage Ferreira, Jorge A. Petroli Marchesi, Maisa C. Pereira Parra, Marília Moraes, Paula Rahal, Paulo Inacio da Costa, João Pessoa Araújo Jr., Maurício Lacerda Nogueira. Prefeitura de Sao Paulo: Melissa Palmieri. |
| EPI_ISL_1966891, EPI_ISL_1966892, EPI_ISL_1966893, EPI_ISL_1966894 | UPA ATALAIA DRA RITA DE CASSIA SORIO | Instituto Butantan / ESALQ-USP (Piracicaba) | Instituto Butantan: Dimas Tadeu Covas, Sandra Coccuzzo Sampaio, Maria Carolina Elias, José Salvatore Leister Patané, Vincent Louis Viala, Antonio Jorge Martins, Ricardo Haddad, Claudia Renata dos Santos Barros, Elaine Cristina Marqueze, Raul Machado Neto, Debora Botequio Moretti, Jardelina de Souza Todao Bernardino, Loyze Paola Oliveira de Lima, Luiz Aurelio de Campos Crispin. Centro de Genômica Funcional da ESALQ: Luiz Lehmann Coutinho, Ricardo Augusto Brassaloti, Raquel de Lello Rocha Campos Cassano. NGS Soluções Genômicas: Pilar Drummond Sampaio Corrêa Mariani. FZEA-USP Pirassununga: Mirele Daiana Poleti, Jessika Cristina Chagas Lesbon, Elisangela Chicaroni Mattos, Heidge Fukumasu. USP-Botucatu: Rejane Maria Tommasini Grotto, Jayme A. Souza-Neto, Guilherme Targino Valente, Patricia Akemi Assato, Felipe Allan da Silva da Costa, Bianca Cechetto Carlos. Mendelics: Bibiana Santos, João Paulo Kitajima, Erika Freitas, David Schlesinger. Hemocentro Ribeirão Preto: Simone Kashima, Evandra Strazza Rodrigues, Svetoslav Nanev Slavov, Elaine Vieira dos Santos, Rafael dos Santos Bezerra, Luiz Carlos Junior de Alcantara, Marta Giovanetti, Vagner Fonseca, Flavia Aburjaile, Rodrigo Tocantins Calado. FAMERP-SJRP: Cecília Artico Banho, Lívia Sacchetto, Fábio Sossai Possebon, Leila Sabrina Ullmann, Cíntia Bittar, Guilherme Campos, Helena Lage Ferreira, Jorge A. Petroli Marchesi, Maisa C. Pereira Parra, Marília Moraes, Paula Rahal, Paulo Inacio da Costa, João Pessoa Araújo Jr., Maurício Lacerda Nogueira. Prefeitura de Sao Paulo: Melissa Palmieri. |
| EPI_ISL_1966895, EPI_ISL_1966896, EPI_ISL_1966897, EPI_ISL_1966898, EPI_ISL_1966899, EPI_ISL_1966900, EPI_ISL_1966901, EPI_ISL_1966902, EPI_ISL_1966903, EPI_ISL_1966904 | CENTRO DE APOIO EPIDEMIOLOGICO | Instituto Butantan / ESALQ-USP (Piracicaba) | Instituto Butantan: Dimas Tadeu Covas, Sandra Coccuzzo Sampaio, Maria Carolina Elias, José Salvatore Leister Patané, Vincent Louis Viala, Antonio Jorge Martins, Ricardo Haddad, Claudia Renata dos Santos Barros, Elaine Cristina Marqueze, Raul Machado Neto, Debora Botequio Moretti, Jardelina de Souza Todao Bernardino, Loyze Paola Oliveira de Lima, Luiz Aurelio de Campos Crispin. Centro de Genômica Funcional da ESALQ: Luiz Lehmann Coutinho, Ricardo Augusto Brassaloti, Raquel de Lello Rocha Campos Cassano. NGS Soluções Genômicas: Pilar Drummond Sampaio Corrêa Mariani. FZEA-USP Pirassununga: Mirele Daiana Poleti, Jessika Cristina Chagas Lesbon, Elisangela Chicaroni Mattos, Heidge Fukumasu. USP-Botucatu: Rejane Maria Tommasini Grotto, Jayme A. Souza-Neto, Guilherme Targino Valente, Patricia Akemi Assato, Felipe Allan da Silva da Costa, Bianca Cechetto Carlos. Mendelics: Bibiana Santos, João Paulo Kitajima, Erika Freitas, David Schlesinger. Hemocentro Ribeirão Preto: Simone Kashima, Evandra Strazza Rodrigues, Svetoslav Nanev Slavov, Elaine Vieira dos Santos, Rafael dos Santos Bezerra, Luiz Carlos Junior de Alcantara, Marta Giovanetti, Vagner Fonseca, Flavia Aburjaile, Rodrigo Tocantins Calado. FAMERP-SJRP: Cecília Artico Banho, Lívia Sacchetto, Fábio Sossai Possebon, Leila Sabrina Ullmann, Cíntia Bittar, Guilherme Campos, Helena Lage Ferreira, Jorge A. Petroli Marchesi, Maisa C. Pereira Parra, Marília Moraes, Paula Rahal, Paulo Inacio da Costa, João Pessoa Araújo Jr., Maurício Lacerda Nogueira. Prefeitura de Sao Paulo: Melissa Palmieri. |
| EPI_ISL_1966905, EPI_ISL_1966906 | LABORATORIO DE FRANCA | Instituto Butantan / ESALQ-USP (Piracicaba) | Instituto Butantan: Dimas Tadeu Covas, Sandra Coccuzzo Sampaio, Maria Carolina Elias, José Salvatore Leister Patané, Vincent Louis Viala, Antonio Jorge Martins, Ricardo Haddad, Claudia Renata dos Santos Barros, Elaine Cristina Marqueze, Raul Machado Neto, Debora Botequio Moretti, Jardelina de Souza Todao Bernardino, Loyze Paola Oliveira de Lima, Luiz Aurelio de Campos Crispin. Centro de Genômica Funcional da ESALQ: Luiz Lehmann Coutinho, Ricardo Augusto Brassaloti, Raquel de Lello Rocha Campos Cassano. NGS Soluções Genômicas: Pilar Drummond Sampaio Corrêa Mariani. FZEA-USP Pirassununga: Mirele Daiana Poleti, Jessika Cristina Chagas Lesbon, Elisangela Chicaroni Mattos, Heidge Fukumasu. USP-Botucatu: Rejane Maria Tommasini Grotto, Jayme A. Souza-Neto, Guilherme Targino Valente, Patricia Akemi Assato, Felipe Allan da Silva da Costa, Bianca Cechetto Carlos. Mendelics: Bibiana Santos, João Paulo Kitajima, Erika Freitas, David Schlesinger. Hemocentro Ribeirão Preto: Simone Kashima, Evandra Strazza Rodrigues, Svetoslav Nanev Slavov, Elaine Vieira dos Santos, Rafael dos Santos Bezerra, Luiz Carlos Junior de Alcantara, Marta Giovanetti, Vagner Fonseca, Flavia Aburjaile, Rodrigo Tocantins Calado. FAMERP-SJRP: Cecília Artico Banho, Lívia Sacchetto, Fábio Sossai Possebon, Leila Sabrina Ullmann, Cíntia Bittar, Guilherme Campos, Helena Lage Ferreira, Jorge A. Petroli Marchesi, Maisa C. Pereira Parra, Marília Moraes, Paula Rahal, Paulo Inacio da Costa, João Pessoa Araújo Jr., Maurício Lacerda Nogueira. Prefeitura de Sao Paulo: Melissa Palmieri. |
| EPI_ISL_1966907, EPI_ISL_1966908 | USF I DE GUAIMBE | Instituto Butantan / ESALQ-USP (Piracicaba) | Instituto Butantan: Dimas Tadeu Covas, Sandra Coccuzzo Sampaio, Maria Carolina Elias, José Salvatore Leister Patané, Vincent Louis Viala, Antonio Jorge Martins, Ricardo Haddad, Claudia Renata dos Santos Barros, Elaine Cristina Marqueze, Raul Machado Neto, Debora Botequio Moretti, Jardelina de Souza Todao Bernardino, Loyze Paola Oliveira de Lima, Luiz Aurelio de Campos Crispin. Centro de Genômica Funcional da ESALQ: Luiz Lehmann Coutinho, Ricardo Augusto Brassaloti, Raquel de Lello Rocha Campos Cassano. NGS Soluções Genômicas: Pilar Drummond Sampaio Corrêa Mariani. FZEA-USP Pirassununga: Mirele Daiana Poleti, Jessika Cristina Chagas Lesbon, Elisangela Chicaroni Mattos, Heidge Fukumasu. USP-Botucatu: Rejane Maria Tommasini Grotto, Jayme A. Souza-Neto, Guilherme Targino Valente, Patricia Akemi Assato, Felipe Allan da Silva da Costa, Bianca Cechetto Carlos. Mendelics: Bibiana Santos, João Paulo Kitajima, Erika Freitas, David Schlesinger. Hemocentro Ribeirão Preto: Simone Kashima, Evandra Strazza Rodrigues, Svetoslav Nanev Slavov, Elaine Vieira dos Santos, Rafael dos Santos Bezerra, Luiz Carlos Junior de Alcantara, Marta Giovanetti, Vagner Fonseca, Flavia Aburjaile, Rodrigo Tocantins Calado. FAMERP-SJRP: Cecília Artico Banho, Lívia Sacchetto, Fábio Sossai Possebon, Leila Sabrina Ullmann, Cíntia Bittar, Guilherme Campos, Helena Lage Ferreira, Jorge A. Petroli Marchesi, Maisa C. Pereira Parra, Marília Moraes, Paula Rahal, Paulo Inacio da Costa, João Pessoa Araújo Jr., Maurício Lacerda Nogueira. Prefeitura de Sao Paulo: Melissa Palmieri. |
| EPI_ISL_1966909 | CS II DR JAHYR DE PAULA RIBEIRO GUARA | Instituto Butantan / ESALQ-USP (Piracicaba) | Instituto Butantan: Dimas Tadeu Covas, Sandra Coccuzzo Sampaio, Maria Carolina Elias, José Salvatore Leister Patané, Vincent Louis Viala, Antonio Jorge Martins, Ricardo Haddad, Claudia Renata dos Santos Barros, Elaine Cristina Marqueze, Raul Machado Neto, Debora Botequio Moretti, Jardelina de Souza Todao Bernardino, Loyze Paola Oliveira de Lima, Luiz Aurelio de Campos Crispin. Centro de Genômica Funcional da ESALQ: Luiz Lehmann Coutinho, Ricardo Augusto Brassaloti, Raquel de Lello Rocha Campos Cassano. NGS Soluções Genômicas: Pilar Drummond Sampaio Corrêa Mariani. FZEA-USP Pirassununga: Mirele Daiana Poleti, Jessika Cristina Chagas Lesbon, Elisangela Chicaroni Mattos, Heidge Fukumasu. USP-Botucatu: Rejane Maria Tommasini Grotto, Jayme A. Souza-Neto, Guilherme Targino Valente, Patricia Akemi Assato, Felipe Allan da Silva da Costa, Bianca Cechetto Carlos. Mendelics: Bibiana Santos, João Paulo Kitajima, Erika Freitas, David Schlesinger. Hemocentro Ribeirão Preto: Simone Kashima, Evandra Strazza Rodrigues, Svetoslav Nanev Slavov, Elaine Vieira dos Santos, Rafael dos Santos Bezerra, Luiz Carlos Junior de Alcantara, Marta Giovanetti, Vagner Fonseca, Flavia Aburjaile, Rodrigo Tocantins Calado. FAMERP-SJRP: Cecília Artico Banho, Lívia Sacchetto, Fábio Sossai Possebon, Leila Sabrina Ullmann, Cíntia Bittar, Guilherme Campos, Helena Lage Ferreira, Jorge A. Petroli Marchesi, Maisa C. Pereira Parra, Marília Moraes, Paula Rahal, Paulo Inacio da Costa, João Pessoa Araújo Jr., Maurício Lacerda Nogueira. Prefeitura de Sao Paulo: Melissa Palmieri. |
| EPI_ISL_1966910, EPI_ISL_1966911 | PR S DA FAMILIA UNIDADE DE SAUDE ADALBERTO ROCHA GUAREI | Instituto Butantan / ESALQ-USP (Piracicaba) | Instituto Butantan: Dimas Tadeu Covas, Sandra Coccuzzo Sampaio, Maria Carolina Elias, José Salvatore Leister Patané, Vincent Louis Viala, Antonio Jorge Martins, Ricardo Haddad, Claudia Renata dos Santos Barros, Elaine Cristina Marqueze, Raul Machado Neto, Debora Botequio Moretti, Jardelina de Souza Todao Bernardino, Loyze Paola Oliveira de Lima, Luiz Aurelio de Campos Crispin. Centro de Genômica Funcional da ESALQ: Luiz Lehmann Coutinho, Ricardo Augusto Brassaloti, Raquel de Lello Rocha Campos Cassano. NGS Soluções Genômicas: Pilar Drummond Sampaio Corrêa Mariani. FZEA-USP Pirassununga: Mirele Daiana Poleti, Jessika Cristina Chagas Lesbon, Elisangela Chicaroni Mattos, Heidge Fukumasu. USP-Botucatu: Rejane Maria Tommasini Grotto, Jayme A. Souza-Neto, Guilherme Targino Valente, Patricia Akemi Assato, Felipe Allan da Silva da Costa, Bianca Cechetto Carlos. Mendelics: Bibiana Santos, João Paulo Kitajima, Erika Freitas, David Schlesinger. Hemocentro Ribeirão Preto: Simone Kashima, Evandra Strazza Rodrigues, Svetoslav Nanev Slavov, Elaine Vieira dos Santos, Rafael dos Santos Bezerra, Luiz Carlos Junior de Alcantara, Marta Giovanetti, Vagner Fonseca, Flavia Aburjaile, Rodrigo Tocantins Calado. FAMERP-SJRP: Cecília Artico Banho, Lívia Sacchetto, Fábio Sossai Possebon, Leila Sabrina |

| | | | |
|---|---|---|---|
| | | | Ullmann, Cíntia Bittar, Guilherme Campos, Helena Lage Ferreira, Jorge A. Petroli Marchesi, Maisa C. Pereira Parra, Marília Moraes, Paula Rahal, Paulo Inacio da Costa, João Pessoa Araújo Jr., Maurício Lacerda Nogueira. Prefeitura de Sao Paulo: Melissa Palmieri. |
| EPI_ISL_1966912 | CENTRO DE SAUDE II DR ARCY BANDEIRA IBIUNA SP | Instituto Butantan / ESALQ-USP (Piracicaba) | Instituto Butantan: Dimas Tadeu Covas, Sandra Coccuzzo Sampaio, Maria Carolina Elias, José Salvatore Leister Patané, Vincent Louis Viala, Antonio Jorge Martins, Ricardo Haddad, Claudia Renata dos Santos Barros, Elaine Cristina Marqueze, Raul Machado Neto, Debora Botequio Moretti, Jardelina de Souza Todao Bernardino, Loyze Paola Oliveira de Lima, Luiz Aurelio de Campos Crispin. Centro de Genômica Funcional da ESALQ: Luiz Lehmann Coutinho, Ricardo Augusto Brassaloti, Raquel de Lello Rocha Campos Cassano. NGS Soluções Genômicas: Pilar Drummond Sampaio Corrêa Mariani. FZEA-USP Pirassununga: Mirele Daiana Poleti, Jessika Cristina Chagas Lesbon, Elisangela Chicaroni Mattos, Heidge Fukumasu. USP-Botucatu: Rejane Maria Tommasini Grotto, Jayme A. Souza-Neto, Guilherme Targino Valente, Patricia Akemi Assato, Felipe Allan da Silva da Costa, Bianca Cechetto Carlos. Mendelics: Bibiana Santos, João Paulo Kitajima, Erika Freitas, David Schlesinger. Hemocentro Ribeirão Preto: Simone Kashima, Evandra Strazza Rodrigues, Svetoslav Nanev Slavov, Elaine Vieira dos Santos, Rafael dos Santos Bezerra, Luiz Carlos Junior de Alcantara, Marta Giovanetti, Vagner Fonseca, Flavia Aburjaile, Rodrigo Tocantins Calado. FAMERP-SJRP: Cecília Artico Banho, Lívia Sacchetto, Fábio Sossai Possebon, Leila Sabrina Ullmann, Cíntia Bittar, Guilherme Campos, Helena Lage Ferreira, Jorge A. Petroli Marchesi, Maisa C. Pereira Parra, Marília Moraes, Paula Rahal, Paulo Inacio da Costa, João Pessoa Araújo Jr., Maurício Lacerda Nogueira. Prefeitura de Sao Paulo: Melissa Palmieri. |
| EPI_ISL_1966913, EPI_ISL_1966914 | HOSPITAL MUNICIPAL DE IBIUNA IBIUNA SP | Instituto Butantan / ESALQ-USP (Piracicaba) | Instituto Butantan: Dimas Tadeu Covas, Sandra Coccuzzo Sampaio, Maria Carolina Elias, José Salvatore Leister Patané, Vincent Louis Viala, Antonio Jorge Martins, Ricardo Haddad, Claudia Renata dos Santos Barros, Elaine Cristina Marqueze, Raul Machado Neto, Debora Botequio Moretti, Jardelina de Souza Todao Bernardino, Loyze Paola Oliveira de Lima, Luiz Aurelio de Campos Crispin. Centro de Genômica Funcional da ESALQ: Luiz Lehmann Coutinho, Ricardo Augusto Brassaloti, Raquel de Lello Rocha Campos Cassano. NGS Soluções Genômicas: Pilar Drummond Sampaio Corrêa Mariani. FZEA-USP Pirassununga: Mirele Daiana Poleti, Jessika Cristina Chagas Lesbon, Elisangela Chicaroni Mattos, Heidge Fukumasu. USP-Botucatu: Rejane Maria Tommasini Grotto, Jayme A. Souza-Neto, Guilherme Targino Valente, Patricia Akemi Assato, Felipe Allan da Silva da Costa, Bianca Cechetto Carlos. Mendelics: Bibiana Santos, João Paulo Kitajima, Erika Freitas, David Schlesinger. Hemocentro Ribeirão Preto: Simone Kashima, Evandra Strazza Rodrigues, Svetoslav Nanev Slavov, Elaine Vieira dos Santos, Rafael dos Santos Bezerra, Luiz Carlos Junior de Alcantara, Marta Giovanetti, Vagner Fonseca, Flavia Aburjaile, Rodrigo Tocantins Calado. FAMERP-SJRP: Cecília Artico Banho, Lívia Sacchetto, Fábio Sossai Possebon, Leila Sabrina Ullmann, Cíntia Bittar, Guilherme Campos, Helena Lage Ferreira, Jorge A. Petroli Marchesi, Maisa C. Pereira Parra, Marília Moraes, Paula Rahal, Paulo Inacio da Costa, João Pessoa Araújo Jr., Maurício Lacerda Nogueira. Prefeitura de Sao Paulo: Melissa Palmieri. |
| EPI_ISL_1966915 | UNIDADE MISTA DE IGUAPE | Instituto Butantan / ESALQ-USP (Piracicaba) | Instituto Butantan: Dimas Tadeu Covas, Sandra Coccuzzo Sampaio, Maria Carolina Elias, José Salvatore Leister Patané, Vincent Louis Viala, Antonio Jorge Martins, Ricardo Haddad, Claudia Renata dos Santos Barros, Elaine Cristina Marqueze, Raul Machado Neto, Debora Botequio Moretti, Jardelina de Souza Todao Bernardino, Loyze Paola Oliveira de Lima, Luiz Aurelio de Campos Crispin. Centro de Genômica Funcional da ESALQ: Luiz Lehmann Coutinho, Ricardo Augusto Brassaloti, Raquel de Lello Rocha Campos Cassano. NGS Soluções Genômicas: Pilar Drummond Sampaio Corrêa Mariani. FZEA-USP Pirassununga: Mirele Daiana Poleti, Jessika Cristina Chagas Lesbon, Elisangela Chicaroni Mattos, Heidge Fukumasu. USP-Botucatu: Rejane Maria Tommasini Grotto, Jayme A. Souza-Neto, Guilherme Targino Valente, Patricia Akemi Assato, Felipe Allan da Silva da Costa, Bianca Cechetto Carlos. Mendelics: Bibiana Santos, João Paulo Kitajima, Erika Freitas, David Schlesinger. Hemocentro Ribeirão Preto: Simone Kashima, Evandra Strazza Rodrigues, Svetoslav Nanev Slavov, Elaine Vieira dos Santos, Rafael dos Santos Bezerra, Luiz Carlos Junior de Alcantara, Marta Giovanetti, Vagner Fonseca, Flavia Aburjaile, Rodrigo Tocantins Calado. FAMERP-SJRP: Cecília Artico Banho, Lívia Sacchetto, Fábio Sossai Possebon, Leila Sabrina Ullmann, Cíntia Bittar, Guilherme Campos, Helena Lage Ferreira, Jorge A. Petroli Marchesi, Maisa C. Pereira Parra, Marília Moraes, Paula Rahal, Paulo Inacio da Costa, João Pessoa Araújo Jr., Maurício Lacerda Nogueira. Prefeitura de Sao Paulo: Melissa Palmieri. |
| EPI_ISL_1966916, EPI_ISL_1966917 | PENITENCIARIA ODON RAMOS MARANHAO IPERO | Instituto Butantan / ESALQ-USP (Piracicaba) | Instituto Butantan: Dimas Tadeu Covas, Sandra Coccuzzo Sampaio, Maria Carolina Elias, José Salvatore Leister Patané, Vincent Louis Viala, Antonio Jorge Martins, Ricardo Haddad, Claudia Renata dos Santos Barros, Elaine Cristina Marqueze, Raul Machado Neto, Debora Botequio Moretti, Jardelina de Souza Todao Bernardino, Loyze Paola Oliveira de Lima, Luiz Aurelio de Campos Crispin. Centro de Genômica Funcional da ESALQ: Luiz Lehmann Coutinho, Ricardo Augusto Brassaloti, Raquel de Lello Rocha Campos Cassano. NGS Soluções Genômicas: Pilar Drummond Sampaio Corrêa Mariani. FZEA-USP Pirassununga: Mirele Daiana Poleti, Jessika Cristina Chagas Lesbon, Elisangela Chicaroni Mattos, Heidge Fukumasu. USP-Botucatu: Rejane Maria Tommasini Grotto, Jayme A. Souza-Neto, Guilherme Targino Valente, Patricia Akemi Assato, Felipe Allan da Silva da Costa, Bianca Cechetto Carlos. Mendelics: Bibiana Santos, João Paulo Kitajima, Erika Freitas, David Schlesinger. Hemocentro Ribeirão Preto: Simone Kashima, Evandra Strazza Rodrigues, Svetoslav Nanev Slavov, Elaine Vieira dos Santos, Rafael dos Santos Bezerra, Luiz Carlos Junior de Alcantara, Marta Giovanetti, Vagner Fonseca, Flavia Aburjaile, Rodrigo Tocantins Calado. FAMERP-SJRP: Cecília Artico Banho, Lívia Sacchetto, Fábio Sossai Possebon, Leila Sabrina Ullmann, Cíntia Bittar, Guilherme Campos, Helena Lage Ferreira, Jorge A. Petroli Marchesi, Maisa C. Pereira Parra, Marília Moraes, Paula Rahal, Paulo Inacio da Costa, João Pessoa Araújo Jr., Maurício Lacerda Nogueira. Prefeitura de Sao Paulo: Melissa Palmieri. |
| EPI_ISL_1966918 | UBS JARDIM VITORINO IPERO | Instituto Butantan / ESALQ-USP (Piracicaba) | Instituto Butantan: Dimas Tadeu Covas, Sandra Coccuzzo Sampaio, Maria Carolina Elias, José Salvatore Leister Patané, Vincent Louis Viala, Antonio Jorge Martins, Ricardo Haddad, Claudia Renata dos Santos Barros, Elaine Cristina Marqueze, Raul Machado Neto, Debora Botequio Moretti, Jardelina de Souza Todao Bernardino, Loyze Paola Oliveira de Lima, Luiz Aurelio de Campos Crispin. Centro de Genômica Funcional da ESALQ: Luiz Lehmann Coutinho, Ricardo Augusto Brassaloti, Raquel de Lello Rocha Campos Cassano. NGS Soluções Genômicas: Pilar Drummond Sampaio Corrêa Mariani. FZEA-USP Pirassununga: Mirele Daiana Poleti, Jessika Cristina Chagas Lesbon, Elisangela Chicaroni Mattos, Heidge Fukumasu. USP-Botucatu: Rejane Maria Tommasini Grotto, Jayme A. Souza-Neto, Guilherme Targino Valente, Patricia Akemi Assato, Felipe Allan da Silva da Costa, Bianca Cechetto Carlos. Mendelics: Bibiana Santos, João Paulo Kitajima, Erika Freitas, David Schlesinger. Hemocentro Ribeirão Preto: Simone Kashima, Evandra Strazza Rodrigues, Svetoslav Nanev Slavov, Elaine Vieira dos Santos, Rafael dos Santos Bezerra, Luiz Carlos Junior de Alcantara, Marta Giovanetti, Vagner Fonseca, Flavia Aburjaile, Rodrigo Tocantins Calado. FAMERP-SJRP: Cecília Artico Banho, Lívia Sacchetto, Fábio Sossai Possebon, Leila Sabrina Ullmann, Cíntia Bittar, Guilherme Campos, Helena Lage Ferreira, Jorge A. Petroli Marchesi, Maisa C. Pereira Parra, Marília Moraes, Paula Rahal, Paulo Inacio da Costa, João Pessoa Araújo Jr., Maurício Lacerda Nogueira. Prefeitura de Sao Paulo: Melissa Palmieri. |
| EPI_ISL_1966919, EPI_ISL_1966920, EPI_ISL_1966921, EPI_ISL_1966922 | HOSPITAL MUNICIPAL DE ITABERA | Instituto Butantan / ESALQ-USP (Piracicaba) | Instituto Butantan: Dimas Tadeu Covas, Sandra Coccuzzo Sampaio, Maria Carolina Elias, José Salvatore Leister Patané, Vincent Louis Viala, Antonio Jorge Martins, Ricardo Haddad, Claudia Renata dos Santos Barros, Elaine Cristina Marqueze, Raul Machado Neto, Debora Botequio Moretti, Jardelina de Souza Todao Bernardino, Loyze Paola Oliveira de Lima, Luiz Aurelio de Campos Crispin. Centro de Genômica Funcional da ESALQ: Luiz Lehmann Coutinho, Ricardo Augusto Brassaloti, Raquel de Lello Rocha Campos Cassano. NGS Soluções Genômicas: Pilar Drummond Sampaio Corrêa Mariani. FZEA-USP Pirassununga: Mirele Daiana Poleti, Jessika Cristina Chagas Lesbon, Elisangela Chicaroni Mattos, Heidge Fukumasu. USP-Botucatu: Rejane Maria Tommasini Grotto, Jayme A. Souza-Neto, Guilherme Targino Valente, Patricia Akemi Assato, Felipe Allan da Silva da Costa, Bianca Cechetto Carlos. Mendelics: Bibiana Santos, João Paulo Kitajima, Erika Freitas, David Schlesinger. Hemocentro Ribeirão Preto: Simone Kashima, Evandra Strazza Rodrigues, Svetoslav Nanev Slavov, Elaine Vieira dos Santos, Rafael dos Santos Bezerra, Luiz Carlos Junior de Alcantara, Marta Giovanetti, Vagner Fonseca, Flavia Aburjaile, Rodrigo Tocantins Calado. FAMERP-SJRP: Cecília Artico Banho, Lívia Sacchetto, Fábio Sossai Possebon, Leila Sabrina Ullmann, Cíntia Bittar, Guilherme Campos, Helena Lage Ferreira, Jorge A. Petroli Marchesi, Maisa C. Pereira Parra, Marília Moraes, Paula Rahal, Paulo Inacio da Costa, João Pessoa Araújo Jr., Maurício Lacerda Nogueira. Prefeitura de Sao Paulo: Melissa Palmieri. |
| EPI_ISL_1966923, EPI_ISL_1966924, EPI_ISL_1966925, EPI_ISL_1966926 | HOSPITAL REGIONAL DE ITAPETININGA | Instituto Butantan / ESALQ-USP (Piracicaba) | Instituto Butantan: Dimas Tadeu Covas, Sandra Coccuzzo Sampaio, Maria Carolina Elias, José Salvatore Leister Patané, Vincent Louis Viala, Antonio Jorge Martins, Ricardo Haddad, Claudia Renata dos Santos Barros, Elaine Cristina Marqueze, Raul Machado Neto, Debora Botequio Moretti, Jardelina de Souza Todao Bernardino, Loyze Paola Oliveira de Lima, Luiz Aurelio de Campos Crispin. Centro de Genômica Funcional da ESALQ: Luiz Lehmann Coutinho, Ricardo Augusto Brassaloti, Raquel de Lello Rocha Campos Cassano. NGS Soluções Genômicas: Pilar Drummond Sampaio Corrêa Mariani. FZEA-USP Pirassununga: Mirele Daiana Poleti, Jessika Cristina Chagas Lesbon, Elisangela Chicaroni Mattos, Heidge Fukumasu. USP-Botucatu: Rejane Maria Tommasini Grotto, Jayme A. Souza-Neto, Guilherme Targino Valente, Patricia Akemi Assato, Felipe Allan da Silva da Costa, Bianca Cechetto Carlos. Mendelics: Bibiana Santos, João Paulo Kitajima, Erika Freitas, David Schlesinger. Hemocentro Ribeirão Preto: Simone Kashima, Evandra Strazza Rodrigues, Svetoslav Nanev Slavov, Elaine Vieira dos Santos, Rafael dos Santos Bezerra, Luiz Carlos Junior de Alcantara, Marta Giovanetti, Vagner Fonseca, Flavia Aburjaile, Rodrigo Tocantins Calado. FAMERP-SJRP: Cecília Artico Banho, Lívia Sacchetto, Fábio Sossai Possebon, Leila Sabrina Ullmann, Cíntia Bittar, Guilherme Campos, Helena Lage Ferreira, Jorge A. Petroli Marchesi, Maisa C. Pereira Parra, Marília Moraes, Paula Rahal, Paulo Inacio da Costa, João Pessoa Araújo Jr., Maurício Lacerda Nogueira. Prefeitura de Sao Paulo: Melissa Palmieri. |
| EPI_ISL_1966927, EPI_ISL_1966928, EPI_ISL_1966929, EPI_ISL_1966930, EPI_ISL_1966931, EPI_ISL_1966932, EPI_ISL_1966933, EPI_ISL_1966934 | POLICLINICA COVID 19 ITAPETININGA | Instituto Butantan / ESALQ-USP (Piracicaba) | Instituto Butantan: Dimas Tadeu Covas, Sandra Coccuzzo Sampaio, Maria Carolina Elias, José Salvatore Leister Patané, Vincent Louis Viala, Antonio Jorge Martins, Ricardo Haddad, Claudia Renata dos Santos Barros, Elaine Cristina Marqueze, Raul Machado Neto, Debora Botequio Moretti, Jardelina de Souza Todao Bernardino, Loyze Paola Oliveira de Lima, Luiz Aurelio de Campos Crispin. Centro de Genômica Funcional da ESALQ: Luiz Lehmann Coutinho, Ricardo Augusto Brassaloti, Raquel de Lello Rocha Campos Cassano. NGS Soluções Genômicas: Pilar Drummond Sampaio Corrêa Mariani. |

| | | | |
|---|---|---|---|
| | | | FZEA-USP Pirassununga: Mirele Daiana Poleti, Jessika Cristina Chagas Lesbon, Elisangela Chicaroni Mattos, Heidge Fukumasu. USP-Botucatu: Rejane Maria Tommasini Grotto, Jayme A. Souza-Neto, Guilherme Targino Valente, Patricia Akemi Assato, Felipe Allan da Silva da Costa, Bianca Cechetto Carlos. Mendelics: Bibiana Santos, João Paulo Kitajima, Erika Freitas, David Schlesinger. Hemocentro Ribeirão Preto: Simone Kashima, Evandra Strazza Rodrigues, Svetoslav Nanev Slavov, Elaine Vieira dos Santos, Rafael dos Santos Bezerra, Luiz Carlos Junior de Alcantara, Marta Giovanetti, Vagner Fonseca, Flavia Aburjaile, Rodrigo Tocantins Calado. FAMERP-SJRP: Cecília Artico Banho, Lívia Sacchetto, Fábio Sossai Possebon, Leila Sabrina Ullmann, Cíntia Bittar, Guilherme Campos, Helena Lage Ferreira, Jorge A. Petroli Marchesi, Maisa C. Pereira Parra, Marília Moraes, Paula Rahal, Paulo Inacio da Costa, João Pessoa Araújo Jr., Maurício Lacerda Nogueira. Prefeitura de Sao Paulo: Melissa Palmieri. |
| EPI_ISL_1966935, EPI_ISL_1966936, EPI_ISL_1966937, EPI_ISL_1966938, EPI_ISL_1966939, EPI_ISL_1966940, EPI_ISL_1966941 | VIGILANCIA SANITARIA E VIG EPIDEMIOLOGICA DE ITAPEVI | Instituto Butantan / ESALQ-USP (Piracicaba) | Instituto Butantan: Dimas Tadeu Covas, Sandra Coccuzzo Sampaio, Maria Carolina Elias, José Salvatore Leister Patané, Vincent Louis Viala, Antonio Jorge Martins, Ricardo Haddad, Claudia Renata dos Santos Barros, Elaine Cristina Marqueze, Raul Machado Neto, Debora Botequio Moretti, Jardelina de Souza Todao Bernardino, Loyze Paola Oliveira de Lima, Luiz Aurelio de Campos Crispin. Centro de Genômica Funcional da ESALQ: Luiz Lehmann Coutinho, Ricardo Augusto Brassaloti, Raquel de Lello Rocha Campos Cassano. NGS Soluções Genômicas: Pilar Drummond Sampaio Corrêa Mariani. FZEA-USP Pirassununga: Mirele Daiana Poleti, Jessika Cristina Chagas Lesbon, Elisangela Chicaroni Mattos, Heidge Fukumasu. USP-Botucatu: Rejane Maria Tommasini Grotto, Jayme A. Souza-Neto, Guilherme Targino Valente, Patricia Akemi Assato, Felipe Allan da Silva da Costa, Bianca Cechetto Carlos. Mendelics: Bibiana Santos, João Paulo Kitajima, Erika Freitas, David Schlesinger. Hemocentro Ribeirão Preto: Simone Kashima, Evandra Strazza Rodrigues, Svetoslav Nanev Slavov, Elaine Vieira dos Santos, Rafael dos Santos Bezerra, Luiz Carlos Junior de Alcantara, Marta Giovanetti, Vagner Fonseca, Flavia Aburjaile, Rodrigo Tocantins Calado. FAMERP-SJRP: Cecília Artico Banho, Lívia Sacchetto, Fábio Sossai Possebon, Leila Sabrina Ullmann, Cíntia Bittar, Guilherme Campos, Helena Lage Ferreira, Jorge A. Petroli Marchesi, Maisa C. Pereira Parra, Marília Moraes, Paula Rahal, Paulo Inacio da Costa, João Pessoa Araújo Jr., Maurício Lacerda Nogueira. Prefeitura de Sao Paulo: Melissa Palmieri. |
| EPI_ISL_1966942, EPI_ISL_1966943, EPI_ISL_1966944, EPI_ISL_1966945, EPI_ISL_1966946, EPI_ISL_1966947, EPI_ISL_1966948, EPI_ISL_1966949 | HOSPITAL MUNICIPAL DE ITAPIRA | Instituto Butantan / ESALQ-USP (Piracicaba) | Instituto Butantan: Dimas Tadeu Covas, Sandra Coccuzzo Sampaio, Maria Carolina Elias, José Salvatore Leister Patané, Vincent Louis Viala, Antonio Jorge Martins, Ricardo Haddad, Claudia Renata dos Santos Barros, Elaine Cristina Marqueze, Raul Machado Neto, Debora Botequio Moretti, Jardelina de Souza Todao Bernardino, Loyze Paola Oliveira de Lima, Luiz Aurelio de Campos Crispin. Centro de Genômica Funcional da ESALQ: Luiz Lehmann Coutinho, Ricardo Augusto Brassaloti, Raquel de Lello Rocha Campos Cassano. NGS Soluções Genômicas: Pilar Drummond Sampaio Corrêa Mariani. FZEA-USP Pirassununga: Mirele Daiana Poleti, Jessika Cristina Chagas Lesbon, Elisangela Chicaroni Mattos, Heidge Fukumasu. USP-Botucatu: Rejane Maria Tommasini Grotto, Jayme A. Souza-Neto, Guilherme Targino Valente, Patricia Akemi Assato, Felipe Allan da Silva da Costa, Bianca Cechetto Carlos. Mendelics: Bibiana Santos, João Paulo Kitajima, Erika Freitas, David Schlesinger. Hemocentro Ribeirão Preto: Simone Kashima, Evandra Strazza Rodrigues, Svetoslav Nanev Slavov, Elaine Vieira dos Santos, Rafael dos Santos Bezerra, Luiz Carlos Junior de Alcantara, Marta Giovanetti, Vagner Fonseca, Flavia Aburjaile, Rodrigo Tocantins Calado. FAMERP-SJRP: Cecília Artico Banho, Lívia Sacchetto, Fábio Sossai Possebon, Leila Sabrina Ullmann, Cíntia Bittar, Guilherme Campos, Helena Lage Ferreira, Jorge A. Petroli Marchesi, Maisa C. Pereira Parra, Marília Moraes, Paula Rahal, Paulo Inacio da Costa, João Pessoa Araújo Jr., Maurício Lacerda Nogueira. Prefeitura de Sao Paulo: Melissa Palmieri. |
| EPI_ISL_1966950, EPI_ISL_1966951 | SECRETARIA MUNICIPAL DA SAUDE DE ITARARE | Instituto Butantan / ESALQ-USP (Piracicaba) | Instituto Butantan: Dimas Tadeu Covas, Sandra Coccuzzo Sampaio, Maria Carolina Elias, José Salvatore Leister Patané, Vincent Louis Viala, Antonio Jorge Martins, Ricardo Haddad, Claudia Renata dos Santos Barros, Elaine Cristina Marqueze, Raul Machado Neto, Debora Botequio Moretti, Jardelina de Souza Todao Bernardino, Loyze Paola Oliveira de Lima, Luiz Aurelio de Campos Crispin. Centro de Genômica Funcional da ESALQ: Luiz Lehmann Coutinho, Ricardo Augusto Brassaloti, Raquel de Lello Rocha Campos Cassano. NGS Soluções Genômicas: Pilar Drummond Sampaio Corrêa Mariani. FZEA-USP Pirassununga: Mirele Daiana Poleti, Jessika Cristina Chagas Lesbon, Elisangela Chicaroni Mattos, Heidge Fukumasu. USP-Botucatu: Rejane Maria Tommasini Grotto, Jayme A. Souza-Neto, Guilherme Targino Valente, Patricia Akemi Assato, Felipe Allan da Silva da Costa, Bianca Cechetto Carlos. Mendelics: Bibiana Santos, João Paulo Kitajima, Erika Freitas, David Schlesinger. Hemocentro Ribeirão Preto: Simone Kashima, Evandra Strazza Rodrigues, Svetoslav Nanev Slavov, Elaine Vieira dos Santos, Rafael dos Santos Bezerra, Luiz Carlos Junior de Alcantara, Marta Giovanetti, Vagner Fonseca, Flavia Aburjaile, Rodrigo Tocantins Calado. FAMERP-SJRP: Cecília Artico Banho, Lívia Sacchetto, Fábio Sossai Possebon, Leila Sabrina Ullmann, Cíntia Bittar, Guilherme Campos, Helena Lage Ferreira, Jorge A. Petroli Marchesi, Maisa C. Pereira Parra, Marília Moraes, Paula Rahal, Paulo Inacio da Costa, João Pessoa Araújo Jr., Maurício Lacerda Nogueira. Prefeitura de Sao Paulo: Melissa Palmieri. |
| EPI_ISL_1966952, EPI_ISL_1966953 | PRONTO ATENDIMENTO MUNICIPAL DE JACUPIRANGA | Instituto Butantan / ESALQ-USP (Piracicaba) | Instituto Butantan: Dimas Tadeu Covas, Sandra Coccuzzo Sampaio, Maria Carolina Elias, José Salvatore Leister Patané, Vincent Louis Viala, Antonio Jorge Martins, Ricardo Haddad, Claudia Renata dos Santos Barros, Elaine Cristina Marqueze, Raul Machado Neto, Debora Botequio Moretti, Jardelina de Souza Todao Bernardino, Loyze Paola Oliveira de Lima, Luiz Aurelio de Campos Crispin. Centro de Genômica Funcional da ESALQ: Luiz Lehmann Coutinho, Ricardo Augusto Brassaloti, Raquel de Lello Rocha Campos Cassano. NGS Soluções Genômicas: Pilar Drummond Sampaio Corrêa Mariani. FZEA-USP Pirassununga: Mirele Daiana Poleti, Jessika Cristina Chagas Lesbon, Elisangela Chicaroni Mattos, Heidge Fukumasu. USP-Botucatu: Rejane Maria Tommasini Grotto, Jayme A. Souza-Neto, Guilherme Targino Valente, Patricia Akemi Assato, Felipe Allan da Silva da Costa, Bianca Cechetto Carlos. Mendelics: Bibiana Santos, João Paulo Kitajima, Erika Freitas, David Schlesinger. Hemocentro Ribeirão Preto: Simone Kashima, Evandra Strazza Rodrigues, Svetoslav Nanev Slavov, Elaine Vieira dos Santos, Rafael dos Santos Bezerra, Luiz Carlos Junior de Alcantara, Marta Giovanetti, Vagner Fonseca, Flavia Aburjaile, Rodrigo Tocantins Calado. FAMERP-SJRP: Cecília Artico Banho, Lívia Sacchetto, Fábio Sossai Possebon, Leila Sabrina Ullmann, Cíntia Bittar, Guilherme Campos, Helena Lage Ferreira, Jorge A. Petroli Marchesi, Maisa C. Pereira Parra, Marília Moraes, Paula Rahal, Paulo Inacio da Costa, João Pessoa Araújo Jr., Maurício Lacerda Nogueira. Prefeitura de Sao Paulo: Melissa Palmieri. |
| EPI_ISL_1966954 | HOSPITAL SANTO ANTONIO DE JUQUIA JUQUIA | Instituto Butantan / ESALQ-USP (Piracicaba) | Instituto Butantan: Dimas Tadeu Covas, Sandra Coccuzzo Sampaio, Maria Carolina Elias, José Salvatore Leister Patané, Vincent Louis Viala, Antonio Jorge Martins, Ricardo Haddad, Claudia Renata dos Santos Barros, Elaine Cristina Marqueze, Raul Machado Neto, Debora Botequio Moretti, Jardelina de Souza Todao Bernardino, Loyze Paola Oliveira de Lima, Luiz Aurelio de Campos Crispin. Centro de Genômica Funcional da ESALQ: Luiz Lehmann Coutinho, Ricardo Augusto Brassaloti, Raquel de Lello Rocha Campos Cassano. NGS Soluções Genômicas: Pilar Drummond Sampaio Corrêa Mariani. FZEA-USP Pirassununga: Mirele Daiana Poleti, Jessika Cristina Chagas Lesbon, Elisangela Chicaroni Mattos, Heidge Fukumasu. USP-Botucatu: Rejane Maria Tommasini Grotto, Jayme A. Souza-Neto, Guilherme Targino Valente, Patricia Akemi Assato, Felipe Allan da Silva da Costa, Bianca Cechetto Carlos. Mendelics: Bibiana Santos, João Paulo Kitajima, Erika Freitas, David Schlesinger. Hemocentro Ribeirão Preto: Simone Kashima, Evandra Strazza Rodrigues, Svetoslav Nanev Slavov, Elaine Vieira dos Santos, Rafael dos Santos Bezerra, Luiz Carlos Junior de Alcantara, Marta Giovanetti, Vagner Fonseca, Flavia Aburjaile, Rodrigo Tocantins Calado. FAMERP-SJRP: Cecília Artico Banho, Lívia Sacchetto, Fábio Sossai Possebon, Leila Sabrina Ullmann, Cíntia Bittar, Guilherme Campos, Helena Lage Ferreira, Jorge A. Petroli Marchesi, Maisa C. Pereira Parra, Marília Moraes, Paula Rahal, Paulo Inacio da Costa, João Pessoa Araújo Jr., Maurício Lacerda Nogueira. Prefeitura de Sao Paulo: Melissa Palmieri. |
| EPI_ISL_1966955, EPI_ISL_1966956 | CENTRO DE SAUDE II MAIRINQUE MAIRINQUE | Instituto Butantan / ESALQ-USP (Piracicaba) | Instituto Butantan: Dimas Tadeu Covas, Sandra Coccuzzo Sampaio, Maria Carolina Elias, José Salvatore Leister Patané, Vincent Louis Viala, Antonio Jorge Martins, Ricardo Haddad, Claudia Renata dos Santos Barros, Elaine Cristina Marqueze, Raul Machado Neto, Debora Botequio Moretti, Jardelina de Souza Todao Bernardino, Loyze Paola Oliveira de Lima, Luiz Aurelio de Campos Crispin. Centro de Genômica Funcional da ESALQ: Luiz Lehmann Coutinho, Ricardo Augusto Brassaloti, Raquel de Lello Rocha Campos Cassano. NGS Soluções Genômicas: Pilar Drummond Sampaio Corrêa Mariani. FZEA-USP Pirassununga: Mirele Daiana Poleti, Jessika Cristina Chagas Lesbon, Elisangela Chicaroni Mattos, Heidge Fukumasu. USP-Botucatu: Rejane Maria Tommasini Grotto, Jayme A. Souza-Neto, Guilherme Targino Valente, Patricia Akemi Assato, Felipe Allan da Silva da Costa, Bianca Cechetto Carlos. Mendelics: Bibiana Santos, João Paulo Kitajima, Erika Freitas, David Schlesinger. Hemocentro Ribeirão Preto: Simone Kashima, Evandra Strazza Rodrigues, Svetoslav Nanev Slavov, Elaine Vieira dos Santos, Rafael dos Santos Bezerra, Luiz Carlos Junior de Alcantara, Marta Giovanetti, Vagner Fonseca, Flavia Aburjaile, Rodrigo Tocantins Calado. FAMERP-SJRP: Cecília Artico Banho, Lívia Sacchetto, Fábio Sossai Possebon, Leila Sabrina Ullmann, Cíntia Bittar, Guilherme Campos, Helena Lage Ferreira, Jorge A. Petroli Marchesi, Maisa C. Pereira Parra, Marília Moraes, Paula Rahal, Paulo Inacio da Costa, João Pessoa Araújo Jr., Maurício Lacerda Nogueira. Prefeitura de Sao Paulo: Melissa Palmieri. |
| EPI_ISL_1966957, EPI_ISL_1966958, EPI_ISL_1966959, EPI_ISL_1966960 | CENTRO DE SAUDE DE MARACAI | Instituto Butantan / ESALQ-USP (Piracicaba) | Instituto Butantan: Dimas Tadeu Covas, Sandra Coccuzzo Sampaio, Maria Carolina Elias, José Salvatore Leister Patané, Vincent Louis Viala, Antonio Jorge Martins, Ricardo Haddad, Claudia Renata dos Santos Barros, Elaine Cristina Marqueze, Raul Machado Neto, Debora Botequio Moretti, Jardelina de Souza Todao Bernardino, Loyze Paola Oliveira de Lima, Luiz Aurelio de Campos Crispin. Centro de Genômica Funcional da ESALQ: Luiz Lehmann Coutinho, Ricardo Augusto Brassaloti, Raquel de Lello Rocha Campos Cassano. NGS Soluções Genômicas: Pilar Drummond Sampaio Corrêa Mariani. FZEA-USP Pirassununga: Mirele Daiana Poleti, Jessika Cristina Chagas Lesbon, Elisangela Chicaroni Mattos, Heidge Fukumasu. USP-Botucatu: Rejane Maria Tommasini Grotto, Jayme A. Souza-Neto, Guilherme Targino Valente, Patricia Akemi Assato, Felipe Allan da Silva da Costa, Bianca Cechetto Carlos. Mendelics: Bibiana Santos, João Paulo Kitajima, Erika Freitas, David Schlesinger. Hemocentro Ribeirão Preto: Simone Kashima, Evandra Strazza Rodrigues, Svetoslav Nanev Slavov, Elaine Vieira dos Santos, Rafael dos Santos Bezerra, Luiz Carlos Junior de Alcantara, Marta Giovanetti, Vagner Fonseca, Flavia Aburjaile, Rodrigo Tocantins Calado. FAMERP-SJRP: Cecília Artico Banho, Lívia Sacchetto, Fábio Sossai Possebon, Leila Sabrina Ullmann, Cíntia Bittar, Guilherme Campos, Helena Lage Ferreira, Jorge A. Petroli Marchesi, Maisa C. Pereira Parra, Marília Moraes, Paula Rahal, Paulo Inacio da Costa, João Pessoa Araújo Jr., Maurício Lacerda Nogueira. Prefeitura de Sao Paulo: Melissa Palmieri. |

| | | | |
|---|---|---|---|
| EPI_ISL_1966961, EPI_ISL_1966962, EPI_ISL_1966963, EPI_ISL_1966964 | CS DE MONTE APRAZIVEL | Instituto Butantan / ESALQ-USP (Piracicaba) | Instituto Butantan: Dimas Tadeu Covas, Sandra Coccuzzo Sampaio, Maria Carolina Elias, José Salvatore Leister Patané, Vincent Louis Viala, Antonio Jorge Martins, Ricardo Haddad, Claudia Renata dos Santos Barros, Elaine Cristina Marqueze, Raul Machado Neto, Debora Botequio Moretti, Jardelina de Souza Todao Bernardino, Loyze Paola Oliveira de Lima, Luiz Aurelio de Campos Crispin. Centro de Genômica Funcional da ESALQ: Luiz Lehmann Coutinho, Ricardo Augusto Brassaloti, Raquel de Lello Rocha Campos Cassano. NGS Soluções Genômicas: Pilar Drummond Sampaio Corrêa Mariani. FZEA-USP Pirassununga: Mirele Daiana Poleti, Jessika Cristina Chagas Lesbon, Elisangela Chicaroni Mattos, Heidge Fukumasu. USP-Botucatu: Rejane Maria Tommasini Grotto, Jayme A. Souza-Neto, Guilherme Targino Valente, Patricia Akemi Assato, Felipe Allan da Silva da Costa, Bianca Cechetto Carlos. Mendelics: Bibiana Santos, João Paulo Kitajima, Erika Freitas, David Schlesinger. Hemocentro Ribeirão Preto: Simone Kashima, Evandra Strazza Rodrigues, Svetoslav Nanev Slavov, Elaine Vieira dos Santos, Rafael dos Santos Bezerra, Luiz Carlos Junior de Alcantara, Marta Giovanetti, Vagner Fonseca, Flavia Aburjaile, Rodrigo Tocantins Calado. FAMERP-SJRP: Cecília Artico Banho, Lívia Sacchetto, Fábio Sossai Possebon, Leila Sabrina Ullmann, Cíntia Bittar, Guilherme Campos, Helena Lage Ferreira, Jorge A. Petroli Marchesi, Maisa C. Pereira Parra, Marília Moraes, Paula Rahal, Paulo Inacio da Costa, João Pessoa Araújo Jr., Maurício Lacerda Nogueira. Prefeitura de Sao Paulo: Melissa Palmieri. |
| EPI_ISL_1966965, EPI_ISL_1966966, EPI_ISL_1966967, EPI_ISL_1966968, EPI_ISL_1966969, EPI_ISL_1966970, EPI_ISL_1966971, EPI_ISL_1966972, EPI_ISL_1966973, EPI_ISL_1966974, EPI_ISL_1966975, EPI_ISL_1966976, EPI_ISL_1966977, EPI_ISL_1966978, EPI_ISL_1966979, EPI_ISL_1966980, EPI_ISL_1966981, EPI_ISL_1966982, EPI_ISL_1966983 | | | |
| see above | CS II EGIDIO BRUNHARA MORRO AGUDO | Instituto Butantan / ESALQ-USP (Piracicaba) | Instituto Butantan: Dimas Tadeu Covas, Sandra Coccuzzo Sampaio, Maria Carolina Elias, José Salvatore Leister Patané, Vincent Louis Viala, Antonio Jorge Martins, Ricardo Haddad, Claudia Renata dos Santos Barros, Elaine Cristina Marqueze, Raul Machado Neto, Debora Botequio Moretti, Jardelina de Souza Todao Bernardino, Loyze Paola Oliveira de Lima, Luiz Aurelio de Campos Crispin. Centro de Genômica Funcional da ESALQ: Luiz Lehmann Coutinho, Ricardo Augusto Brassaloti, Raquel de Lello Rocha Campos Cassano. NGS Soluções Genômicas: Pilar Drummond Sampaio Corrêa Mariani. FZEA-USP Pirassununga: Mirele Daiana Poleti, Jessika Cristina Chagas Lesbon, Elisangela Chicaroni Mattos, Heidge Fukumasu. USP-Botucatu: Rejane Maria Tommasini Grotto, Jayme A. Souza-Neto, Guilherme Targino Valente, Patricia Akemi Assato, Felipe Allan da Silva da Costa, Bianca Cechetto Carlos. Mendelics: Bibiana Santos, João Paulo Kitajima, Erika Freitas, David Schlesinger. Hemocentro Ribeirão Preto: Simone Kashima, Evandra Strazza Rodrigues, Svetoslav Nanev Slavov, Elaine Vieira dos Santos, Rafael dos Santos Bezerra, Luiz Carlos Junior de Alcantara, Marta Giovanetti, Vagner Fonseca, Flavia Aburjaile, Rodrigo Tocantins Calado. FAMERP-SJRP: Cecília Artico Banho, Lívia Sacchetto, Fábio Sossai Possebon, Leila Sabrina Ullmann, Cíntia Bittar, Guilherme Campos, Helena Lage Ferreira, Jorge A. Petroli Marchesi, Maisa C. Pereira Parra, Marília Moraes, Paula Rahal, Paulo Inacio da Costa, João Pessoa Araújo Jr., Maurício Lacerda Nogueira. Prefeitura de Sao Paulo: Melissa Palmieri. |
| EPI_ISL_1966984, EPI_ISL_1966985 | CS DE NEVES PAULISTA | Instituto Butantan / ESALQ-USP (Piracicaba) | Instituto Butantan: Dimas Tadeu Covas, Sandra Coccuzzo Sampaio, Maria Carolina Elias, José Salvatore Leister Patané, Vincent Louis Viala, Antonio Jorge Martins, Ricardo Haddad, Claudia Renata dos Santos Barros, Elaine Cristina Marqueze, Raul Machado Neto, Debora Botequio Moretti, Jardelina de Souza Todao Bernardino, Loyze Paola Oliveira de Lima, Luiz Aurelio de Campos Crispin. Centro de Genômica Funcional da ESALQ: Luiz Lehmann Coutinho, Ricardo Augusto Brassaloti, Raquel de Lello Rocha Campos Cassano. NGS Soluções Genômicas: Pilar Drummond Sampaio Corrêa Mariani. FZEA-USP Pirassununga: Mirele Daiana Poleti, Jessika Cristina Chagas Lesbon, Elisangela Chicaroni Mattos, Heidge Fukumasu. USP-Botucatu: Rejane Maria Tommasini Grotto, Jayme A. Souza-Neto, Guilherme Targino Valente, Patricia Akemi Assato, Felipe Allan da Silva da Costa, Bianca Cechetto Carlos. Mendelics: Bibiana Santos, João Paulo Kitajima, Erika Freitas, David Schlesinger. Hemocentro Ribeirão Preto: Simone Kashima, Evandra Strazza Rodrigues, Svetoslav Nanev Slavov, Elaine Vieira dos Santos, Rafael dos Santos Bezerra, Luiz Carlos Junior de Alcantara, Marta Giovanetti, Vagner Fonseca, Flavia Aburjaile, Rodrigo Tocantins Calado. FAMERP-SJRP: Cecília Artico Banho, Lívia Sacchetto, Fábio Sossai Possebon, Leila Sabrina Ullmann, Cíntia Bittar, Guilherme Campos, Helena Lage Ferreira, Jorge A. Petroli Marchesi, Maisa C. Pereira Parra, Marília Moraes, Paula Rahal, Paulo Inacio da Costa, João Pessoa Araújo Jr., Maurício Lacerda Nogueira. Prefeitura de Sao Paulo: Melissa Palmieri. |
| EPI_ISL_1966986, EPI_ISL_1966987 | UNIDADE MISTA DE SAUDE DE ORIENTE | Instituto Butantan / ESALQ-USP (Piracicaba) | Instituto Butantan: Dimas Tadeu Covas, Sandra Coccuzzo Sampaio, Maria Carolina Elias, José Salvatore Leister Patané, Vincent Louis Viala, Antonio Jorge Martins, Ricardo Haddad, Claudia Renata dos Santos Barros, Elaine Cristina Marqueze, Raul Machado Neto, Debora Botequio Moretti, Jardelina de Souza Todao Bernardino, Loyze Paola Oliveira de Lima, Luiz Aurelio de Campos Crispin. Centro de Genômica Funcional da ESALQ: Luiz Lehmann Coutinho, Ricardo Augusto Brassaloti, Raquel de Lello Rocha Campos Cassano. NGS Soluções Genômicas: Pilar Drummond Sampaio Corrêa Mariani. FZEA-USP Pirassununga: Mirele Daiana Poleti, Jessika Cristina Chagas Lesbon, Elisangela Chicaroni Mattos, Heidge Fukumasu. USP-Botucatu: Rejane Maria Tommasini Grotto, Jayme A. Souza-Neto, Guilherme Targino Valente, Patricia Akemi Assato, Felipe Allan da Silva da Costa, Bianca Cechetto Carlos. Mendelics: Bibiana Santos, João Paulo Kitajima, Erika Freitas, David Schlesinger. Hemocentro Ribeirão Preto: Simone Kashima, Evandra Strazza Rodrigues, Svetoslav Nanev Slavov, Elaine Vieira dos Santos, Rafael dos Santos Bezerra, Luiz Carlos Junior de Alcantara, Marta Giovanetti, Vagner Fonseca, Flavia Aburjaile, Rodrigo Tocantins Calado. FAMERP-SJRP: Cecília Artico Banho, Lívia Sacchetto, Fábio Sossai Possebon, Leila Sabrina Ullmann, Cíntia Bittar, Guilherme Campos, Helena Lage Ferreira, Jorge A. Petroli Marchesi, Maisa C. Pereira Parra, Marília Moraes, Paula Rahal, Paulo Inacio da Costa, João Pessoa Araújo Jr., Maurício Lacerda Nogueira. Prefeitura de Sao Paulo: Melissa Palmieri. |
| EPI_ISL_1966988 | CS DE ORINDIUVA | Instituto Butantan / ESALQ-USP (Piracicaba) | Instituto Butantan: Dimas Tadeu Covas, Sandra Coccuzzo Sampaio, Maria Carolina Elias, José Salvatore Leister Patané, Vincent Louis Viala, Antonio Jorge Martins, Ricardo Haddad, Claudia Renata dos Santos Barros, Elaine Cristina Marqueze, Raul Machado Neto, Debora Botequio Moretti, Jardelina de Souza Todao Bernardino, Loyze Paola Oliveira de Lima, Luiz Aurelio de Campos Crispin. Centro de Genômica Funcional da ESALQ: Luiz Lehmann Coutinho, Ricardo Augusto Brassaloti, Raquel de Lello Rocha Campos Cassano. NGS Soluções Genômicas: Pilar Drummond Sampaio Corrêa Mariani. FZEA-USP Pirassununga: Mirele Daiana Poleti, Jessika Cristina Chagas Lesbon, Elisangela Chicaroni Mattos, Heidge Fukumasu. USP-Botucatu: Rejane Maria Tommasini Grotto, Jayme A. Souza-Neto, Guilherme Targino Valente, Patricia Akemi Assato, Felipe Allan da Silva da Costa, Bianca Cechetto Carlos. Mendelics: Bibiana Santos, João Paulo Kitajima, Erika Freitas, David Schlesinger. Hemocentro Ribeirão Preto: Simone Kashima, Evandra Strazza Rodrigues, Svetoslav Nanev Slavov, Elaine Vieira dos Santos, Rafael dos Santos Bezerra, Luiz Carlos Junior de Alcantara, Marta Giovanetti, Vagner Fonseca, Flavia Aburjaile, Rodrigo Tocantins Calado. FAMERP-SJRP: Cecília Artico Banho, Lívia Sacchetto, Fábio Sossai Possebon, Leila Sabrina Ullmann, Cíntia Bittar, Guilherme Campos, Helena Lage Ferreira, Jorge A. Petroli Marchesi, Maisa C. Pereira Parra, Marília Moraes, Paula Rahal, Paulo Inacio da Costa, João Pessoa Araújo Jr., Maurício Lacerda Nogueira. Prefeitura de Sao Paulo: Melissa Palmieri. |
| EPI_ISL_1966989 | CS II DR MIGUEL VITALIANO ORLANDIA | Instituto Butantan / ESALQ-USP (Piracicaba) | Instituto Butantan: Dimas Tadeu Covas, Sandra Coccuzzo Sampaio, Maria Carolina Elias, José Salvatore Leister Patané, Vincent Louis Viala, Antonio Jorge Martins, Ricardo Haddad, Claudia Renata dos Santos Barros, Elaine Cristina Marqueze, Raul Machado Neto, Debora Botequio Moretti, Jardelina de Souza Todao Bernardino, Loyze Paola Oliveira de Lima, Luiz Aurelio de Campos Crispin. Centro de Genômica Funcional da ESALQ: Luiz Lehmann Coutinho, Ricardo Augusto Brassaloti, Raquel de Lello Rocha Campos Cassano. NGS Soluções Genômicas: Pilar Drummond Sampaio Corrêa Mariani. FZEA-USP Pirassununga: Mirele Daiana Poleti, Jessika Cristina Chagas Lesbon, Elisangela Chicaroni Mattos, Heidge Fukumasu. USP-Botucatu: Rejane Maria Tommasini Grotto, Jayme A. Souza-Neto, Guilherme Targino Valente, Patricia Akemi Assato, Felipe Allan da Silva da Costa, Bianca Cechetto Carlos. Mendelics: Bibiana Santos, João Paulo Kitajima, Erika Freitas, David Schlesinger. Hemocentro Ribeirão Preto: Simone Kashima, Evandra Strazza Rodrigues, Svetoslav Nanev Slavov, Elaine Vieira dos Santos, Rafael dos Santos Bezerra, Luiz Carlos Junior de Alcantara, Marta Giovanetti, Vagner Fonseca, Flavia Aburjaile, Rodrigo Tocantins Calado. FAMERP-SJRP: Cecília Artico Banho, Lívia Sacchetto, Fábio Sossai Possebon, Leila Sabrina Ullmann, Cíntia Bittar, Guilherme Campos, Helena Lage Ferreira, Jorge A. Petroli Marchesi, Maisa C. Pereira Parra, Marília Moraes, Paula Rahal, Paulo Inacio da Costa, João Pessoa Araújo Jr., Maurício Lacerda Nogueira. Prefeitura de Sao Paulo: Melissa Palmieri. |
| EPI_ISL_1966990, EPI_ISL_1966991 | UBS DR HELIO MIGLIARI | Instituto Butantan / ESALQ-USP (Piracicaba) | Instituto Butantan: Dimas Tadeu Covas, Sandra Coccuzzo Sampaio, Maria Carolina Elias, José Salvatore Leister Patané, Vincent Louis Viala, Antonio Jorge Martins, Ricardo Haddad, Claudia Renata dos Santos Barros, Elaine Cristina Marqueze, Raul Machado Neto, Debora Botequio Moretti, Jardelina de Souza Todao Bernardino, Loyze Paola Oliveira de Lima, Luiz Aurelio de Campos Crispin. Centro de Genômica Funcional da ESALQ: Luiz Lehmann Coutinho, Ricardo Augusto Brassaloti, Raquel de Lello Rocha Campos Cassano. NGS Soluções Genômicas: Pilar Drummond Sampaio Corrêa Mariani. FZEA-USP Pirassununga: Mirele Daiana Poleti, Jessika Cristina Chagas Lesbon, Elisangela Chicaroni Mattos, Heidge Fukumasu. USP-Botucatu: Rejane Maria Tommasini Grotto, Jayme A. Souza-Neto, Guilherme Targino Valente, Patricia Akemi Assato, Felipe Allan da Silva da Costa, Bianca Cechetto Carlos. Mendelics: Bibiana Santos, João Paulo Kitajima, Erika Freitas, David Schlesinger. Hemocentro Ribeirão Preto: Simone Kashima, Evandra Strazza Rodrigues, Svetoslav Nanev Slavov, Elaine Vieira dos Santos, Rafael dos Santos Bezerra, Luiz Carlos Junior de Alcantara, Marta Giovanetti, Vagner Fonseca, Flavia Aburjaile, Rodrigo Tocantins Calado. FAMERP-SJRP: Cecília Artico Banho, Lívia Sacchetto, Fábio Sossai Possebon, Leila Sabrina Ullmann, Cíntia Bittar, Guilherme Campos, Helena Lage Ferreira, Jorge A. Petroli Marchesi, Maisa C. Pereira Parra, Marília Moraes, Paula Rahal, Paulo Inacio da Costa, João Pessoa Araújo Jr., Maurício Lacerda Nogueira. Prefeitura de Sao Paulo: Melissa Palmieri. |
| EPI_ISL_1966992, EPI_ISL_1966993 | CS I DE OURINHOS | Instituto Butantan / ESALQ-USP (Piracicaba) | Instituto Butantan: Dimas Tadeu Covas, Sandra Coccuzzo Sampaio, Maria Carolina Elias, José Salvatore Leister Patané, Vincent Louis Viala, Antonio Jorge Martins, Ricardo Haddad, Claudia Renata dos Santos Barros, Elaine Cristina Marqueze, Raul Machado Neto, Debora Botequio Moretti, Jardelina de Souza Todao Bernardino, Loyze Paola Oliveira de Lima, Luiz Aurelio de Campos Crispin. Centro de Genômica Funcional da ESALQ: Luiz Lehmann Coutinho, Ricardo Augusto Brassaloti, Raquel de Lello Rocha Campos Cassano. NGS Soluções Genômicas: Pilar Drummond Sampaio Corrêa Mariani. |

| | | | |
|---|---|---|---|
| | | | FZEA-USP Pirassununga: Mirele Daiana Poleti, Jessika Cristina Chagas Lesbon, Elisangela Chicaroni Mattos, Heidge Fukumasu. USP-Botucatu: Rejane Maria Tommasini Grotto, Jayme A. Souza-Neto, Guilherme Targino Valente, Patricia Akemi Assato, Felipe Allan da Silva da Costa, Bianca Cechetto Carlos. Mendelics: Bibiana Santos, João Paulo Kitajima, Erika Freitas, David Schlesinger. Hemocentro Ribeirão Preto: Simone Kashima, Evandra Strazza Rodrigues, Svetoslav Nanev Slavov, Elaine Vieira dos Santos, Rafael dos Santos Bezerra, Luiz Carlos Junior de Alcantara, Marta Giovanetti, Vagner Fonseca, Flavia Aburjaile, Rodrigo Tocantins Calado. FAMERP-SJRP: Cecília Artico Banho, Lívia Sacchetto, Fábio Sossai Possebon, Leila Sabrina Ullmann, Cíntia Bittar, Guilherme Campos, Helena Lage Ferreira, Jorge A. Petroli Marchesi, Maisa C. Pereira Parra, Marília Moraes, Paula Rahal, Paulo Inacio da Costa, João Pessoa Araújo Jr., Maurício Lacerda Nogueira. Prefeitura de Sao Paulo: Melissa Palmieri. |
| EPI_ISL_1966994 | UBS DR HELIO MIGLIARI | Instituto Butantan / ESALQ-USP (Piracicaba) | Instituto Butantan: Dimas Tadeu Covas, Sandra Coccuzzo Sampaio, Maria Carolina Elias, José Salvatore Leister Patané, Vincent Louis Viala, Antonio Jorge Martins, Ricardo Haddad, Claudia Renata dos Santos Barros, Elaine Cristina Marqueze, Raul Machado Neto, Debora Botequio Moretti, Jardelina de Souza Todao Bernardino, Loyze Paola Oliveira de Lima, Luiz Aurelio de Campos Crispin. Centro de Genômica Funcional da ESALQ: Luiz Lehmann Coutinho, Ricardo Augusto Brassaloti, Raquel de Lello Rocha Campos Cassano. NGS Soluções Genômicas: Pilar Drummond Sampaio Corrêa Mariani. FZEA-USP Pirassununga: Mirele Daiana Poleti, Jessika Cristina Chagas Lesbon, Elisangela Chicaroni Mattos, Heidge Fukumasu. USP-Botucatu: Rejane Maria Tommasini Grotto, Jayme A. Souza-Neto, Guilherme Targino Valente, Patricia Akemi Assato, Felipe Allan da Silva da Costa, Bianca Cechetto Carlos. Mendelics: Bibiana Santos, João Paulo Kitajima, Erika Freitas, David Schlesinger. Hemocentro Ribeirão Preto: Simone Kashima, Evandra Strazza Rodrigues, Svetoslav Nanev Slavov, Elaine Vieira dos Santos, Rafael dos Santos Bezerra, Luiz Carlos Junior de Alcantara, Marta Giovanetti, Vagner Fonseca, Flavia Aburjaile, Rodrigo Tocantins Calado. FAMERP-SJRP: Cecília Artico Banho, Lívia Sacchetto, Fábio Sossai Possebon, Leila Sabrina Ullmann, Cíntia Bittar, Guilherme Campos, Helena Lage Ferreira, Jorge A. Petroli Marchesi, Maisa C. Pereira Parra, Marília Moraes, Paula Rahal, Paulo Inacio da Costa, João Pessoa Araújo Jr., Maurício Lacerda Nogueira. Prefeitura de Sao Paulo: Melissa Palmieri. |
| EPI_ISL_1966995 | UBS III DE PARIQUERA ACU PARIQUERA ACU | Instituto Butantan / ESALQ-USP (Piracicaba) | Instituto Butantan: Dimas Tadeu Covas, Sandra Coccuzzo Sampaio, Maria Carolina Elias, José Salvatore Leister Patané, Vincent Louis Viala, Antonio Jorge Martins, Ricardo Haddad, Claudia Renata dos Santos Barros, Elaine Cristina Marqueze, Raul Machado Neto, Debora Botequio Moretti, Jardelina de Souza Todao Bernardino, Loyze Paola Oliveira de Lima, Luiz Aurelio de Campos Crispin. Centro de Genômica Funcional da ESALQ: Luiz Lehmann Coutinho, Ricardo Augusto Brassaloti, Raquel de Lello Rocha Campos Cassano. NGS Soluções Genômicas: Pilar Drummond Sampaio Corrêa Mariani. FZEA-USP Pirassununga: Mirele Daiana Poleti, Jessika Cristina Chagas Lesbon, Elisangela Chicaroni Mattos, Heidge Fukumasu. USP-Botucatu: Rejane Maria Tommasini Grotto, Jayme A. Souza-Neto, Guilherme Targino Valente, Patricia Akemi Assato, Felipe Allan da Silva da Costa, Bianca Cechetto Carlos. Mendelics: Bibiana Santos, João Paulo Kitajima, Erika Freitas, David Schlesinger. Hemocentro Ribeirão Preto: Simone Kashima, Evandra Strazza Rodrigues, Svetoslav Nanev Slavov, Elaine Vieira dos Santos, Rafael dos Santos Bezerra, Luiz Carlos Junior de Alcantara, Marta Giovanetti, Vagner Fonseca, Flavia Aburjaile, Rodrigo Tocantins Calado. FAMERP-SJRP: Cecília Artico Banho, Lívia Sacchetto, Fábio Sossai Possebon, Leila Sabrina Ullmann, Cíntia Bittar, Guilherme Campos, Helena Lage Ferreira, Jorge A. Petroli Marchesi, Maisa C. Pereira Parra, Marília Moraes, Paula Rahal, Paulo Inacio da Costa, João Pessoa Araújo Jr., Maurício Lacerda Nogueira. Prefeitura de Sao Paulo: Melissa Palmieri. |
| EPI_ISL_1966996, EPI_ISL_1966997, EPI_ISL_1966998, EPI_ISL_1966999, EPI_ISL_1967001, EPI_ISL_1967002, EPI_ISL_1967003 | LABORATORIO MUNICIPAL DE PIRACICABA | Instituto Butantan / ESALQ-USP (Piracicaba) | Instituto Butantan: Dimas Tadeu Covas, Sandra Coccuzzo Sampaio, Maria Carolina Elias, José Salvatore Leister Patané, Vincent Louis Viala, Antonio Jorge Martins, Ricardo Haddad, Claudia Renata dos Santos Barros, Elaine Cristina Marqueze, Raul Machado Neto, Debora Botequio Moretti, Jardelina de Souza Todao Bernardino, Loyze Paola Oliveira de Lima, Luiz Aurelio de Campos Crispin. Centro de Genômica Funcional da ESALQ: Luiz Lehmann Coutinho, Ricardo Augusto Brassaloti, Raquel de Lello Rocha Campos Cassano. NGS Soluções Genômicas: Pilar Drummond Sampaio Corrêa Mariani. FZEA-USP Pirassununga: Mirele Daiana Poleti, Jessika Cristina Chagas Lesbon, Elisangela Chicaroni Mattos, Heidge Fukumasu. USP-Botucatu: Rejane Maria Tommasini Grotto, Jayme A. Souza-Neto, Guilherme Targino Valente, Patricia Akemi Assato, Felipe Allan da Silva da Costa, Bianca Cechetto Carlos. Mendelics: Bibiana Santos, João Paulo Kitajima, Erika Freitas, David Schlesinger. Hemocentro Ribeirão Preto: Simone Kashima, Evandra Strazza Rodrigues, Svetoslav Nanev Slavov, Elaine Vieira dos Santos, Rafael dos Santos Bezerra, Luiz Carlos Junior de Alcantara, Marta Giovanetti, Vagner Fonseca, Flavia Aburjaile, Rodrigo Tocantins Calado. FAMERP-SJRP: Cecília Artico Banho, Lívia Sacchetto, Fábio Sossai Possebon, Leila Sabrina Ullmann, Cíntia Bittar, Guilherme Campos, Helena Lage Ferreira, Jorge A. Petroli Marchesi, Maisa C. Pereira Parra, Marília Moraes, Paula Rahal, Paulo Inacio da Costa, João Pessoa Araújo Jr., Maurício Lacerda Nogueira. Prefeitura de Sao Paulo: Melissa Palmieri. |
| EPI_ISL_1967004 | CSII DR WASHINGTON LUIS M RODRIGUES DA SILVA PITANGUEIRAS | Instituto Butantan / ESALQ-USP (Piracicaba) | Instituto Butantan: Dimas Tadeu Covas, Sandra Coccuzzo Sampaio, Maria Carolina Elias, José Salvatore Leister Patané, Vincent Louis Viala, Antonio Jorge Martins, Ricardo Haddad, Claudia Renata dos Santos Barros, Elaine Cristina Marqueze, Raul Machado Neto, Debora Botequio Moretti, Jardelina de Souza Todao Bernardino, Loyze Paola Oliveira de Lima, Luiz Aurelio de Campos Crispin. Centro de Genômica Funcional da ESALQ: Luiz Lehmann Coutinho, Ricardo Augusto Brassaloti, Raquel de Lello Rocha Campos Cassano. NGS Soluções Genômicas: Pilar Drummond Sampaio Corrêa Mariani. FZEA-USP Pirassununga: Mirele Daiana Poleti, Jessika Cristina Chagas Lesbon, Elisangela Chicaroni Mattos, Heidge Fukumasu. USP-Botucatu: Rejane Maria Tommasini Grotto, Jayme A. Souza-Neto, Guilherme Targino Valente, Patricia Akemi Assato, Felipe Allan da Silva da Costa, Bianca Cechetto Carlos. Mendelics: Bibiana Santos, João Paulo Kitajima, Erika Freitas, David Schlesinger. Hemocentro Ribeirão Preto: Simone Kashima, Evandra Strazza Rodrigues, Svetoslav Nanev Slavov, Elaine Vieira dos Santos, Rafael dos Santos Bezerra, Luiz Carlos Junior de Alcantara, Marta Giovanetti, Vagner Fonseca, Flavia Aburjaile, Rodrigo Tocantins Calado. FAMERP-SJRP: Cecília Artico Banho, Lívia Sacchetto, Fábio Sossai Possebon, Leila Sabrina Ullmann, Cíntia Bittar, Guilherme Campos, Helena Lage Ferreira, Jorge A. Petroli Marchesi, Maisa C. Pereira Parra, Marília Moraes, Paula Rahal, Paulo Inacio da Costa, João Pessoa Araújo Jr., Maurício Lacerda Nogueira. Prefeitura de Sao Paulo: Melissa Palmieri. |
| EPI_ISL_1967005 | PA DE IBITIUVA DR OTAVIO BENETTI PITANGUEIRAS | Instituto Butantan / ESALQ-USP (Piracicaba) | Instituto Butantan: Dimas Tadeu Covas, Sandra Coccuzzo Sampaio, Maria Carolina Elias, José Salvatore Leister Patané, Vincent Louis Viala, Antonio Jorge Martins, Ricardo Haddad, Claudia Renata dos Santos Barros, Elaine Cristina Marqueze, Raul Machado Neto, Debora Botequio Moretti, Jardelina de Souza Todao Bernardino, Loyze Paola Oliveira de Lima, Luiz Aurelio de Campos Crispin. Centro de Genômica Funcional da ESALQ: Luiz Lehmann Coutinho, Ricardo Augusto Brassaloti, Raquel de Lello Rocha Campos Cassano. NGS Soluções Genômicas: Pilar Drummond Sampaio Corrêa Mariani. FZEA-USP Pirassununga: Mirele Daiana Poleti, Jessika Cristina Chagas Lesbon, Elisangela Chicaroni Mattos, Heidge Fukumasu. USP-Botucatu: Rejane Maria Tommasini Grotto, Jayme A. Souza-Neto, Guilherme Targino Valente, Patricia Akemi Assato, Felipe Allan da Silva da Costa, Bianca Cechetto Carlos. Mendelics: Bibiana Santos, João Paulo Kitajima, Erika Freitas, David Schlesinger. Hemocentro Ribeirão Preto: Simone Kashima, Evandra Strazza Rodrigues, Svetoslav Nanev Slavov, Elaine Vieira dos Santos, Rafael dos Santos Bezerra, Luiz Carlos Junior de Alcantara, Marta Giovanetti, Vagner Fonseca, Flavia Aburjaile, Rodrigo Tocantins Calado. FAMERP-SJRP: Cecília Artico Banho, Lívia Sacchetto, Fábio Sossai Possebon, Leila Sabrina Ullmann, Cíntia Bittar, Guilherme Campos, Helena Lage Ferreira, Jorge A. Petroli Marchesi, Maisa C. Pereira Parra, Marília Moraes, Paula Rahal, Paulo Inacio da Costa, João Pessoa Araújo Jr., Maurício Lacerda Nogueira. Prefeitura de Sao Paulo: Melissa Palmieri. |
| EPI_ISL_1967006, EPI_ISL_1967007, EPI_ISL_1967008, EPI_ISL_1967009, EPI_ISL_1967010, EPI_ISL_1967011, EPI_ISL_1967012, EPI_ISL_1967013 | CSII DR WASHINGTON LUIS M RODRIGUES DA SILVA PITANGUEIRAS | Instituto Butantan / ESALQ-USP (Piracicaba) | Instituto Butantan: Dimas Tadeu Covas, Sandra Coccuzzo Sampaio, Maria Carolina Elias, José Salvatore Leister Patané, Vincent Louis Viala, Antonio Jorge Martins, Ricardo Haddad, Claudia Renata dos Santos Barros, Elaine Cristina Marqueze, Raul Machado Neto, Debora Botequio Moretti, Jardelina de Souza Todao Bernardino, Loyze Paola Oliveira de Lima, Luiz Aurelio de Campos Crispin. Centro de Genômica Funcional da ESALQ: Luiz Lehmann Coutinho, Ricardo Augusto Brassaloti, Raquel de Lello Rocha Campos Cassano. NGS Soluções Genômicas: Pilar Drummond Sampaio Corrêa Mariani. FZEA-USP Pirassununga: Mirele Daiana Poleti, Jessika Cristina Chagas Lesbon, Elisangela Chicaroni Mattos, Heidge Fukumasu. USP-Botucatu: Rejane Maria Tommasini Grotto, Jayme A. Souza-Neto, Guilherme Targino Valente, Patricia Akemi Assato, Felipe Allan da Silva da Costa, Bianca Cechetto Carlos. Mendelics: Bibiana Santos, João Paulo Kitajima, Erika Freitas, David Schlesinger. Hemocentro Ribeirão Preto: Simone Kashima, Evandra Strazza Rodrigues, Svetoslav Nanev Slavov, Elaine Vieira dos Santos, Rafael dos Santos Bezerra, Luiz Carlos Junior de Alcantara, Marta Giovanetti, Vagner Fonseca, Flavia Aburjaile, Rodrigo Tocantins Calado. FAMERP-SJRP: Cecília Artico Banho, Lívia Sacchetto, Fábio Sossai Possebon, Leila Sabrina Ullmann, Cíntia Bittar, Guilherme Campos, Helena Lage Ferreira, Jorge A. Petroli Marchesi, Maisa C. Pereira Parra, Marília Moraes, Paula Rahal, Paulo Inacio da Costa, João Pessoa Araújo Jr., Maurício Lacerda Nogueira. Prefeitura de Sao Paulo: Melissa Palmieri. |
| EPI_ISL_1967014, EPI_ISL_1967016, EPI_ISL_1967017, EPI_ISL_1967018, EPI_ISL_1967019 | SANTA CASA DE PORTO FELIZ | Instituto Butantan / ESALQ-USP (Piracicaba) | Instituto Butantan: Dimas Tadeu Covas, Sandra Coccuzzo Sampaio, Maria Carolina Elias, José Salvatore Leister Patané, Vincent Louis Viala, Antonio Jorge Martins, Ricardo Haddad, Claudia Renata dos Santos Barros, Elaine Cristina Marqueze, Raul Machado Neto, Debora Botequio Moretti, Jardelina de Souza Todao Bernardino, Loyze Paola Oliveira de Lima, Luiz Aurelio de Campos Crispin. Centro de Genômica Funcional da ESALQ: Luiz Lehmann Coutinho, Ricardo Augusto Brassaloti, Raquel de Lello Rocha Campos Cassano. NGS Soluções Genômicas: Pilar Drummond Sampaio Corrêa Mariani. FZEA-USP Pirassununga: Mirele Daiana Poleti, Jessika Cristina Chagas Lesbon, Elisangela Chicaroni Mattos, Heidge Fukumasu. USP-Botucatu: Rejane Maria Tommasini Grotto, Jayme A. Souza-Neto, Guilherme Targino Valente, Patricia Akemi Assato, Felipe Allan da Silva da Costa, Bianca Cechetto Carlos. Mendelics: Bibiana Santos, João Paulo Kitajima, Erika Freitas, David Schlesinger. Hemocentro Ribeirão Preto: Simone Kashima, Evandra Strazza Rodrigues, Svetoslav Nanev Slavov, Elaine Vieira dos Santos, Rafael dos Santos Bezerra, Luiz Carlos Junior de Alcantara, Marta Giovanetti, Vagner Fonseca, Flavia Aburjaile, Rodrigo Tocantins Calado. FAMERP-SJRP: Cecília Artico Banho, Lívia Sacchetto, Fábio Sossai Possebon, Leila Sabrina Ullmann, Cíntia Bittar, Guilherme Campos, Helena Lage Ferreira, Jorge A. Petroli Marchesi, Maisa C. Pereira Parra, Marília Moraes, Paula Rahal, Paulo Inacio da Costa, João Pessoa Araújo Jr., Maurício Lacerda Nogueira. Prefeitura de Sao Paulo: Melissa Palmieri. |

| | | | |
|---|---|---|---|
| EPI_ISL_1967020, EPI_ISL_1967021, EPI_ISL_1967022, EPI_ISL_1967023, EPI_ISL_1967024 | PSF DR ANTONIO PIRES DE ALMEIDA PORTO FELIZ | Instituto Butantan / ESALQ-USP (Piracicaba) | Instituto Butantan: Dimas Tadeu Covas, Sandra Coccuzzo Sampaio, Maria Carolina Elias, José Salvatore Leister Patané, Vincent Louis Viala, Antonio Jorge Martins, Ricardo Haddad, Claudia Renata dos Santos Barros, Elaine Cristina Marqueze, Raul Machado Neto, Debora Botequio Moretti, Jardelina de Souza Todao Bernardino, Loyze Paola Oliveira de Lima, Luiz Aurelio de Campos Crispin. Centro de Genômica Funcional da ESALQ: Luiz Lehmann Coutinho, Ricardo Augusto Brassaloti, Raquel de Lello Rocha Campos Cassano. NGS Soluções Genômicas: Pilar Drummond Sampaio Corrêa Mariani. FZEA-USP Pirassununga: Mirele Daiana Poleti, Jessika Cristina Chagas Lesbon, Elisangela Chicaroni Mattos, Heidge Fukumasu. USP-Botucatu: Rejane Maria Tommasini Grotto, Jayme A. Souza-Neto, Guilherme Targino Valente, Patricia Akemi Assato, Felipe Allan da Silva da Costa, Bianca Cechetto Carlos. Mendelics: Bibiana Santos, João Paulo Kitajima, Erika Freitas, David Schlesinger. Hemocentro Ribeirão Preto: Simone Kashima, Evandra Strazza Rodrigues, Svetoslav Nanev Slavov, Elaine Vieira dos Santos, Rafael dos Santos Bezerra, Luiz Carlos Junior de Alcantara, Marta Giovanetti, Vagner Fonseca, Flavia Aburjaile, Rodrigo Tocantins Calado. FAMERP-SJRP: Cecília Artico Banho, Lívia Sacchetto, Fábio Sossai Possebon, Leila Sabrina Ullmann, Cíntia Bittar, Guilherme Campos, Helena Lage Ferreira, Jorge A. Petroli Marchesi, Maisa C. Pereira Parra, Marília Moraes, Paula Rahal, Paulo Inacio da Costa, João Pessoa Araújo Jr., Maurício Lacerda Nogueira. Prefeitura de Sao Paulo: Melissa Palmieri. |
| EPI_ISL_1967025, EPI_ISL_1967026 | HOSPITAL MUNICIPAL MARIA ROSA CARDOSO | Instituto Butantan / ESALQ-USP (Piracicaba) | Instituto Butantan: Dimas Tadeu Covas, Sandra Coccuzzo Sampaio, Maria Carolina Elias, José Salvatore Leister Patané, Vincent Louis Viala, Antonio Jorge Martins, Ricardo Haddad, Claudia Renata dos Santos Barros, Elaine Cristina Marqueze, Raul Machado Neto, Debora Botequio Moretti, Jardelina de Souza Todao Bernardino, Loyze Paola Oliveira de Lima, Luiz Aurelio de Campos Crispin. Centro de Genômica Funcional da ESALQ: Luiz Lehmann Coutinho, Ricardo Augusto Brassaloti, Raquel de Lello Rocha Campos Cassano. NGS Soluções Genômicas: Pilar Drummond Sampaio Corrêa Mariani. FZEA-USP Pirassununga: Mirele Daiana Poleti, Jessika Cristina Chagas Lesbon, Elisangela Chicaroni Mattos, Heidge Fukumasu. USP-Botucatu: Rejane Maria Tommasini Grotto, Jayme A. Souza-Neto, Guilherme Targino Valente, Patricia Akemi Assato, Felipe Allan da Silva da Costa, Bianca Cechetto Carlos. Mendelics: Bibiana Santos, João Paulo Kitajima, Erika Freitas, David Schlesinger. Hemocentro Ribeirão Preto: Simone Kashima, Evandra Strazza Rodrigues, Svetoslav Nanev Slavov, Elaine Vieira dos Santos, Rafael dos Santos Bezerra, Luiz Carlos Junior de Alcantara, Marta Giovanetti, Vagner Fonseca, Flavia Aburjaile, Rodrigo Tocantins Calado. FAMERP-SJRP: Cecília Artico Banho, Lívia Sacchetto, Fábio Sossai Possebon, Leila Sabrina Ullmann, Cíntia Bittar, Guilherme Campos, Helena Lage Ferreira, Jorge A. Petroli Marchesi, Maisa C. Pereira Parra, Marília Moraes, Paula Rahal, Paulo Inacio da Costa, João Pessoa Araújo Jr., Maurício Lacerda Nogueira. Prefeitura de Sao Paulo: Melissa Palmieri. |
| EPI_ISL_1967027 | CENTRO DE SAUDE II | Instituto Butantan / ESALQ-USP (Piracicaba) | Instituto Butantan: Dimas Tadeu Covas, Sandra Coccuzzo Sampaio, Maria Carolina Elias, José Salvatore Leister Patané, Vincent Louis Viala, Antonio Jorge Martins, Ricardo Haddad, Claudia Renata dos Santos Barros, Elaine Cristina Marqueze, Raul Machado Neto, Debora Botequio Moretti, Jardelina de Souza Todao Bernardino, Loyze Paola Oliveira de Lima, Luiz Aurelio de Campos Crispin. Centro de Genômica Funcional da ESALQ: Luiz Lehmann Coutinho, Ricardo Augusto Brassaloti, Raquel de Lello Rocha Campos Cassano. NGS Soluções Genômicas: Pilar Drummond Sampaio Corrêa Mariani. FZEA-USP Pirassununga: Mirele Daiana Poleti, Jessika Cristina Chagas Lesbon, Elisangela Chicaroni Mattos, Heidge Fukumasu. USP-Botucatu: Rejane Maria Tommasini Grotto, Jayme A. Souza-Neto, Guilherme Targino Valente, Patricia Akemi Assato, Felipe Allan da Silva da Costa, Bianca Cechetto Carlos. Mendelics: Bibiana Santos, João Paulo Kitajima, Erika Freitas, David Schlesinger. Hemocentro Ribeirão Preto: Simone Kashima, Evandra Strazza Rodrigues, Svetoslav Nanev Slavov, Elaine Vieira dos Santos, Rafael dos Santos Bezerra, Luiz Carlos Junior de Alcantara, Marta Giovanetti, Vagner Fonseca, Flavia Aburjaile, Rodrigo Tocantins Calado. FAMERP-SJRP: Cecília Artico Banho, Lívia Sacchetto, Fábio Sossai Possebon, Leila Sabrina Ullmann, Cíntia Bittar, Guilherme Campos, Helena Lage Ferreira, Jorge A. Petroli Marchesi, Maisa C. Pereira Parra, Marília Moraes, Paula Rahal, Paulo Inacio da Costa, João Pessoa Araújo Jr., Maurício Lacerda Nogueira. Prefeitura de Sao Paulo: Melissa Palmieri. |
| EPI_ISL_1967028 | AMBULATORIO DE MOLESTIAS INFECCIOSAS DE SALTO | Instituto Butantan / ESALQ-USP (Piracicaba) | Instituto Butantan: Dimas Tadeu Covas, Sandra Coccuzzo Sampaio, Maria Carolina Elias, José Salvatore Leister Patané, Vincent Louis Viala, Antonio Jorge Martins, Ricardo Haddad, Claudia Renata dos Santos Barros, Elaine Cristina Marqueze, Raul Machado Neto, Debora Botequio Moretti, Jardelina de Souza Todao Bernardino, Loyze Paola Oliveira de Lima, Luiz Aurelio de Campos Crispin. Centro de Genômica Funcional da ESALQ: Luiz Lehmann Coutinho, Ricardo Augusto Brassaloti, Raquel de Lello Rocha Campos Cassano. NGS Soluções Genômicas: Pilar Drummond Sampaio Corrêa Mariani. FZEA-USP Pirassununga: Mirele Daiana Poleti, Jessika Cristina Chagas Lesbon, Elisangela Chicaroni Mattos, Heidge Fukumasu. USP-Botucatu: Rejane Maria Tommasini Grotto, Jayme A. Souza-Neto, Guilherme Targino Valente, Patricia Akemi Assato, Felipe Allan da Silva da Costa, Bianca Cechetto Carlos. Mendelics: Bibiana Santos, João Paulo Kitajima, Erika Freitas, David Schlesinger. Hemocentro Ribeirão Preto: Simone Kashima, Evandra Strazza Rodrigues, Svetoslav Nanev Slavov, Elaine Vieira dos Santos, Rafael dos Santos Bezerra, Luiz Carlos Junior de Alcantara, Marta Giovanetti, Vagner Fonseca, Flavia Aburjaile, Rodrigo Tocantins Calado. FAMERP-SJRP: Cecília Artico Banho, Lívia Sacchetto, Fábio Sossai Possebon, Leila Sabrina Ullmann, Cíntia Bittar, Guilherme Campos, Helena Lage Ferreira, Jorge A. Petroli Marchesi, Maisa C. Pereira Parra, Marília Moraes, Paula Rahal, Paulo Inacio da Costa, João Pessoa Araújo Jr., Maurício Lacerda Nogueira. Prefeitura de Sao Paulo: Melissa Palmieri. |
| EPI_ISL_1967029, EPI_ISL_1967030, EPI_ISL_1967031, EPI_ISL_1967032 | HOSP E MATERNIDADE MUNICIPAL N SRA MONTE SERRAT | Instituto Butantan / ESALQ-USP (Piracicaba) | Instituto Butantan: Dimas Tadeu Covas, Sandra Coccuzzo Sampaio, Maria Carolina Elias, José Salvatore Leister Patané, Vincent Louis Viala, Antonio Jorge Martins, Ricardo Haddad, Claudia Renata dos Santos Barros, Elaine Cristina Marqueze, Raul Machado Neto, Debora Botequio Moretti, Jardelina de Souza Todao Bernardino, Loyze Paola Oliveira de Lima, Luiz Aurelio de Campos Crispin. Centro de Genômica Funcional da ESALQ: Luiz Lehmann Coutinho, Ricardo Augusto Brassaloti, Raquel de Lello Rocha Campos Cassano. NGS Soluções Genômicas: Pilar Drummond Sampaio Corrêa Mariani. FZEA-USP Pirassununga: Mirele Daiana Poleti, Jessika Cristina Chagas Lesbon, Elisangela Chicaroni Mattos, Heidge Fukumasu. USP-Botucatu: Rejane Maria Tommasini Grotto, Jayme A. Souza-Neto, Guilherme Targino Valente, Patricia Akemi Assato, Felipe Allan da Silva da Costa, Bianca Cechetto Carlos. Mendelics: Bibiana Santos, João Paulo Kitajima, Erika Freitas, David Schlesinger. Hemocentro Ribeirão Preto: Simone Kashima, Evandra Strazza Rodrigues, Svetoslav Nanev Slavov, Elaine Vieira dos Santos, Rafael dos Santos Bezerra, Luiz Carlos Junior de Alcantara, Marta Giovanetti, Vagner Fonseca, Flavia Aburjaile, Rodrigo Tocantins Calado. FAMERP-SJRP: Cecília Artico Banho, Lívia Sacchetto, Fábio Sossai Possebon, Leila Sabrina Ullmann, Cíntia Bittar, Guilherme Campos, Helena Lage Ferreira, Jorge A. Petroli Marchesi, Maisa C. Pereira Parra, Marília Moraes, Paula Rahal, Paulo Inacio da Costa, João Pessoa Araújo Jr., Maurício Lacerda Nogueira. Prefeitura de Sao Paulo: Melissa Palmieri. |
| EPI_ISL_1967033, EPI_ISL_1967034, EPI_ISL_1967035, EPI_ISL_1967036, EPI_ISL_1967037, EPI_ISL_1967038, EPI_ISL_1967039, EPI_ISL_1967040, EPI_ISL_1967041 | PSF MARIA JOSE SALTO DE PIRAPORA | Instituto Butantan / ESALQ-USP (Piracicaba) | Instituto Butantan: Dimas Tadeu Covas, Sandra Coccuzzo Sampaio, Maria Carolina Elias, José Salvatore Leister Patané, Vincent Louis Viala, Antonio Jorge Martins, Ricardo Haddad, Claudia Renata dos Santos Barros, Elaine Cristina Marqueze, Raul Machado Neto, Debora Botequio Moretti, Jardelina de Souza Todao Bernardino, Loyze Paola Oliveira de Lima, Luiz Aurelio de Campos Crispin. Centro de Genômica Funcional da ESALQ: Luiz Lehmann Coutinho, Ricardo Augusto Brassaloti, Raquel de Lello Rocha Campos Cassano. NGS Soluções Genômicas: Pilar Drummond Sampaio Corrêa Mariani. FZEA-USP Pirassununga: Mirele Daiana Poleti, Jessika Cristina Chagas Lesbon, Elisangela Chicaroni Mattos, Heidge Fukumasu. USP-Botucatu: Rejane Maria Tommasini Grotto, Jayme A. Souza-Neto, Guilherme Targino Valente, Patricia Akemi Assato, Felipe Allan da Silva da Costa, Bianca Cechetto Carlos. Mendelics: Bibiana Santos, João Paulo Kitajima, Erika Freitas, David Schlesinger. Hemocentro Ribeirão Preto: Simone Kashima, Evandra Strazza Rodrigues, Svetoslav Nanev Slavov, Elaine Vieira dos Santos, Rafael dos Santos Bezerra, Luiz Carlos Junior de Alcantara, Marta Giovanetti, Vagner Fonseca, Flavia Aburjaile, Rodrigo Tocantins Calado. FAMERP-SJRP: Cecília Artico Banho, Lívia Sacchetto, Fábio Sossai Possebon, Leila Sabrina Ullmann, Cíntia Bittar, Guilherme Campos, Helena Lage Ferreira, Jorge A. Petroli Marchesi, Maisa C. Pereira Parra, Marília Moraes, Paula Rahal, Paulo Inacio da Costa, João Pessoa Araújo Jr., Maurício Lacerda Nogueira. Prefeitura de Sao Paulo: Melissa Palmieri. |
| EPI_ISL_1967042, EPI_ISL_1967043, EPI_ISL_1967044, EPI_ISL_1967045, EPI_ISL_1967046, EPI_ISL_1967047, EPI_ISL_1967048, EPI_ISL_1967049 | UNIDADE DE PRONTO ATENDIMENTO UPA I SANTA ISABEL | Instituto Butantan / ESALQ-USP (Piracicaba) | Instituto Butantan: Dimas Tadeu Covas, Sandra Coccuzzo Sampaio, Maria Carolina Elias, José Salvatore Leister Patané, Vincent Louis Viala, Antonio Jorge Martins, Ricardo Haddad, Claudia Renata dos Santos Barros, Elaine Cristina Marqueze, Raul Machado Neto, Debora Botequio Moretti, Jardelina de Souza Todao Bernardino, Loyze Paola Oliveira de Lima, Luiz Aurelio de Campos Crispin. Centro de Genômica Funcional da ESALQ: Luiz Lehmann Coutinho, Ricardo Augusto Brassaloti, Raquel de Lello Rocha Campos Cassano. NGS Soluções Genômicas: Pilar Drummond Sampaio Corrêa Mariani. FZEA-USP Pirassununga: Mirele Daiana Poleti, Jessika Cristina Chagas Lesbon, Elisangela Chicaroni Mattos, Heidge Fukumasu. USP-Botucatu: Rejane Maria Tommasini Grotto, Jayme A. Souza-Neto, Guilherme Targino Valente, Patricia Akemi Assato, Felipe Allan da Silva da Costa, Bianca Cechetto Carlos. Mendelics: Bibiana Santos, João Paulo Kitajima, Erika Freitas, David Schlesinger. Hemocentro Ribeirão Preto: Simone Kashima, Evandra Strazza Rodrigues, Svetoslav Nanev Slavov, Elaine Vieira dos Santos, Rafael dos Santos Bezerra, Luiz Carlos Junior de Alcantara, Marta Giovanetti, Vagner Fonseca, Flavia Aburjaile, Rodrigo Tocantins Calado. FAMERP-SJRP: Cecília Artico Banho, Lívia Sacchetto, Fábio Sossai Possebon, Leila Sabrina Ullmann, Cíntia Bittar, Guilherme Campos, Helena Lage Ferreira, Jorge A. Petroli Marchesi, Maisa C. Pereira Parra, Marília Moraes, Paula Rahal, Paulo Inacio da Costa, João Pessoa Araújo Jr., Maurício Lacerda Nogueira. Prefeitura de Sao Paulo: Melissa Palmieri. |
| EPI_ISL_1967050 | CS DE SANTA RITA DOESTE | Instituto Butantan / ESALQ-USP (Piracicaba) | Instituto Butantan: Dimas Tadeu Covas, Sandra Coccuzzo Sampaio, Maria Carolina Elias, José Salvatore Leister Patané, Vincent Louis Viala, Antonio Jorge Martins, Ricardo Haddad, Claudia Renata dos Santos Barros, Elaine Cristina Marqueze, Raul Machado Neto, Debora Botequio Moretti, Jardelina de Souza Todao Bernardino, Loyze Paola Oliveira de Lima, Luiz Aurelio de Campos Crispin. Centro de Genômica Funcional da ESALQ: Luiz Lehmann Coutinho, Ricardo Augusto Brassaloti, Raquel de Lello Rocha Campos Cassano. NGS Soluções Genômicas: Pilar Drummond Sampaio Corrêa Mariani. FZEA-USP Pirassununga: Mirele Daiana Poleti, Jessika Cristina Chagas Lesbon, Elisangela Chicaroni Mattos, Heidge Fukumasu. USP-Botucatu: Rejane Maria Tommasini Grotto, Jayme A. Souza-Neto, Guilherme Targino Valente, Patricia Akemi Assato, Felipe Allan da Silva da Costa, Bianca Cechetto Carlos. Mendelics: Bibiana Santos, João Paulo Kitajima, Erika Freitas, David Schlesinger. Hemocentro Ribeirão Preto: Simone Kashima, Evandra Strazza |

| | | | |
|---|---|---|---|
| | | | Rodrigues, Svetoslav Nanev Slavov, Elaine Vieira dos Santos, Rafael dos Santos Bezerra, Luiz Carlos Junior de Alcantara, Marta Giovanetti, Vagner Fonseca, Flavia Aburjaile, Rodrigo Tocantins Calado. FAMERP-SJRP: Cecília Artico Banho, Lívia Sacchetto, Fábio Sossai Possebon, Leila Sabrina Ullmann, Cíntia Bittar, Guilherme Campos, Helena Lage Ferreira, Jorge A. Petroli Marchesi, Maisa C. Pereira Parra, Marília Moraes, Paula Rahal, Paulo Inacio da Costa, João Pessoa Araújo Jr., Maurício Lacerda Nogueira. Prefeitura de Sao Paulo: Melissa Palmieri. |
| EPI_ISL_1967051 | UBS DE SANTA SALETE | Instituto Butantan / ESALQ-USP (Piracicaba) | Instituto Butantan: Dimas Tadeu Covas, Sandra Coccuzzo Sampaio, Maria Carolina Elias, José Salvatore Leister Patané, Vincent Louis Viala, Antonio Jorge Martins, Ricardo Haddad, Claudia Renata dos Santos Barros, Elaine Cristina Marqueze, Raul Machado Neto, Debora Botequio Moretti, Jardelina de Souza Todao Bernardino, Loyze Paola Oliveira de Lima, Luiz Aurelio de Campos Crispin. Centro de Genômica Funcional da ESALQ: Luiz Lehmann Coutinho, Ricardo Augusto Brassaloti, Raquel de Lello Rocha Campos Cassano. NGS Soluções Genômicas: Pilar Drummond Sampaio Corrêa Mariani. FZEA-USP Pirassununga: Mirele Daiana Poleti, Jessika Cristina Chagas Lesbon, Elisangela Chicaroni Mattos, Heidge Fukumasu. USP-Botucatu: Rejane Maria Tommasini Grotto, Jayme A. Souza-Neto, Guilherme Targino Valente, Patricia Akemi Assato, Felipe Allan da Silva da Costa, Bianca Cechetto Carlos. Mendelics: Bibiana Santos, João Paulo Kitajima, Erika Freitas, David Schlesinger. Hemocentro Ribeirão Preto: Simone Kashima, Evandra Strazza Rodrigues, Svetoslav Nanev Slavov, Elaine Vieira dos Santos, Rafael dos Santos Bezerra, Luiz Carlos Junior de Alcantara, Marta Giovanetti, Vagner Fonseca, Flavia Aburjaile, Rodrigo Tocantins Calado. FAMERP-SJRP: Cecília Artico Banho, Lívia Sacchetto, Fábio Sossai Possebon, Leila Sabrina Ullmann, Cíntia Bittar, Guilherme Campos, Helena Lage Ferreira, Jorge A. Petroli Marchesi, Maisa C. Pereira Parra, Marília Moraes, Paula Rahal, Paulo Inacio da Costa, João Pessoa Araújo Jr., Maurício Lacerda Nogueira. Prefeitura de Sao Paulo: Melissa Palmieri. |
| EPI_ISL_1967052, EPI_ISL_1967053, EPI_ISL_1967054, EPI_ISL_1967055, EPI_ISL_1967056, EPI_ISL_1967057 | UNIDADE DE VIGILANCIA EM SAUDE | Instituto Butantan / ESALQ-USP (Piracicaba) | Instituto Butantan: Dimas Tadeu Covas, Sandra Coccuzzo Sampaio, Maria Carolina Elias, José Salvatore Leister Patané, Vincent Louis Viala, Antonio Jorge Martins, Ricardo Haddad, Claudia Renata dos Santos Barros, Elaine Cristina Marqueze, Raul Machado Neto, Debora Botequio Moretti, Jardelina de Souza Todao Bernardino, Loyze Paola Oliveira de Lima, Luiz Aurelio de Campos Crispin. Centro de Genômica Funcional da ESALQ: Luiz Lehmann Coutinho, Ricardo Augusto Brassaloti, Raquel de Lello Rocha Campos Cassano. NGS Soluções Genômicas: Pilar Drummond Sampaio Corrêa Mariani. FZEA-USP Pirassununga: Mirele Daiana Poleti, Jessika Cristina Chagas Lesbon, Elisangela Chicaroni Mattos, Heidge Fukumasu. USP-Botucatu: Rejane Maria Tommasini Grotto, Jayme A. Souza-Neto, Guilherme Targino Valente, Patricia Akemi Assato, Felipe Allan da Silva da Costa, Bianca Cechetto Carlos. Mendelics: Bibiana Santos, João Paulo Kitajima, Erika Freitas, David Schlesinger. Hemocentro Ribeirão Preto: Simone Kashima, Evandra Strazza Rodrigues, Svetoslav Nanev Slavov, Elaine Vieira dos Santos, Rafael dos Santos Bezerra, Luiz Carlos Junior de Alcantara, Marta Giovanetti, Vagner Fonseca, Flavia Aburjaile, Rodrigo Tocantins Calado. FAMERP-SJRP: Cecília Artico Banho, Lívia Sacchetto, Fábio Sossai Possebon, Leila Sabrina Ullmann, Cíntia Bittar, Guilherme Campos, Helena Lage Ferreira, Jorge A. Petroli Marchesi, Maisa C. Pereira Parra, Marília Moraes, Paula Rahal, Paulo Inacio da Costa, João Pessoa Araújo Jr., Maurício Lacerda Nogueira. Prefeitura de Sao Paulo: Melissa Palmieri. |
| EPI_ISL_1967058, EPI_ISL_1967059, EPI_ISL_1967060 | CENTRO DE SAUDE ESF IV ZONA RURAL DOMINGOS DE S SJRIOPARDO | Instituto Butantan / ESALQ-USP (Piracicaba) | Instituto Butantan: Dimas Tadeu Covas, Sandra Coccuzzo Sampaio, Maria Carolina Elias, José Salvatore Leister Patané, Vincent Louis Viala, Antonio Jorge Martins, Ricardo Haddad, Claudia Renata dos Santos Barros, Elaine Cristina Marqueze, Raul Machado Neto, Debora Botequio Moretti, Jardelina de Souza Todao Bernardino, Loyze Paola Oliveira de Lima, Luiz Aurelio de Campos Crispin. Centro de Genômica Funcional da ESALQ: Luiz Lehmann Coutinho, Ricardo Augusto Brassaloti, Raquel de Lello Rocha Campos Cassano. NGS Soluções Genômicas: Pilar Drummond Sampaio Corrêa Mariani. FZEA-USP Pirassununga: Mirele Daiana Poleti, Jessika Cristina Chagas Lesbon, Elisangela Chicaroni Mattos, Heidge Fukumasu. USP-Botucatu: Rejane Maria Tommasini Grotto, Jayme A. Souza-Neto, Guilherme Targino Valente, Patricia Akemi Assato, Felipe Allan da Silva da Costa, Bianca Cechetto Carlos. Mendelics: Bibiana Santos, João Paulo Kitajima, Erika Freitas, David Schlesinger. Hemocentro Ribeirão Preto: Simone Kashima, Evandra Strazza Rodrigues, Svetoslav Nanev Slavov, Elaine Vieira dos Santos, Rafael dos Santos Bezerra, Luiz Carlos Junior de Alcantara, Marta Giovanetti, Vagner Fonseca, Flavia Aburjaile, Rodrigo Tocantins Calado. FAMERP-SJRP: Cecília Artico Banho, Lívia Sacchetto, Fábio Sossai Possebon, Leila Sabrina Ullmann, Cíntia Bittar, Guilherme Campos, Helena Lage Ferreira, Jorge A. Petroli Marchesi, Maisa C. Pereira Parra, Marília Moraes, Paula Rahal, Paulo Inacio da Costa, João Pessoa Araújo Jr., Maurício Lacerda Nogueira. Prefeitura de Sao Paulo: Melissa Palmieri. |
| EPI_ISL_1967061, EPI_ISL_1967062, EPI_ISL_1967063, EPI_ISL_1967064 | CENTRO DE SAUDE II SAO MIGUEL ARCANJO | Instituto Butantan / ESALQ-USP (Piracicaba) | Instituto Butantan: Dimas Tadeu Covas, Sandra Coccuzzo Sampaio, Maria Carolina Elias, José Salvatore Leister Patané, Vincent Louis Viala, Antonio Jorge Martins, Ricardo Haddad, Claudia Renata dos Santos Barros, Elaine Cristina Marqueze, Raul Machado Neto, Debora Botequio Moretti, Jardelina de Souza Todao Bernardino, Loyze Paola Oliveira de Lima, Luiz Aurelio de Campos Crispin. Centro de Genômica Funcional da ESALQ: Luiz Lehmann Coutinho, Ricardo Augusto Brassaloti, Raquel de Lello Rocha Campos Cassano. NGS Soluções Genômicas: Pilar Drummond Sampaio Corrêa Mariani. FZEA-USP Pirassununga: Mirele Daiana Poleti, Jessika Cristina Chagas Lesbon, Elisangela Chicaroni Mattos, Heidge Fukumasu. USP-Botucatu: Rejane Maria Tommasini Grotto, Jayme A. Souza-Neto, Guilherme Targino Valente, Patricia Akemi Assato, Felipe Allan da Silva da Costa, Bianca Cechetto Carlos. Mendelics: Bibiana Santos, João Paulo Kitajima, Erika Freitas, David Schlesinger. Hemocentro Ribeirão Preto: Simone Kashima, Evandra Strazza Rodrigues, Svetoslav Nanev Slavov, Elaine Vieira dos Santos, Rafael dos Santos Bezerra, Luiz Carlos Junior de Alcantara, Marta Giovanetti, Vagner Fonseca, Flavia Aburjaile, Rodrigo Tocantins Calado. FAMERP-SJRP: Cecília Artico Banho, Lívia Sacchetto, Fábio Sossai Possebon, Leila Sabrina Ullmann, Cíntia Bittar, Guilherme Campos, Helena Lage Ferreira, Jorge A. Petroli Marchesi, Maisa C. Pereira Parra, Marília Moraes, Paula Rahal, Paulo Inacio da Costa, João Pessoa Araújo Jr., Maurício Lacerda Nogueira. Prefeitura de Sao Paulo: Melissa Palmieri. |
| EPI_ISL_1967065, EPI_ISL_1967066 | HOSPITAL SAO MIGUEL ARCANJO | Instituto Butantan / ESALQ-USP (Piracicaba) | Instituto Butantan: Dimas Tadeu Covas, Sandra Coccuzzo Sampaio, Maria Carolina Elias, José Salvatore Leister Patané, Vincent Louis Viala, Antonio Jorge Martins, Ricardo Haddad, Claudia Renata dos Santos Barros, Elaine Cristina Marqueze, Raul Machado Neto, Debora Botequio Moretti, Jardelina de Souza Todao Bernardino, Loyze Paola Oliveira de Lima, Luiz Aurelio de Campos Crispin. Centro de Genômica Funcional da ESALQ: Luiz Lehmann Coutinho, Ricardo Augusto Brassaloti, Raquel de Lello Rocha Campos Cassano. NGS Soluções Genômicas: Pilar Drummond Sampaio Corrêa Mariani. FZEA-USP Pirassununga: Mirele Daiana Poleti, Jessika Cristina Chagas Lesbon, Elisangela Chicaroni Mattos, Heidge Fukumasu. USP-Botucatu: Rejane Maria Tommasini Grotto, Jayme A. Souza-Neto, Guilherme Targino Valente, Patricia Akemi Assato, Felipe Allan da Silva da Costa, Bianca Cechetto Carlos. Mendelics: Bibiana Santos, João Paulo Kitajima, Erika Freitas, David Schlesinger. Hemocentro Ribeirão Preto: Simone Kashima, Evandra Strazza Rodrigues, Svetoslav Nanev Slavov, Elaine Vieira dos Santos, Rafael dos Santos Bezerra, Luiz Carlos Junior de Alcantara, Marta Giovanetti, Vagner Fonseca, Flavia Aburjaile, Rodrigo Tocantins Calado. FAMERP-SJRP: Cecília Artico Banho, Lívia Sacchetto, Fábio Sossai Possebon, Leila Sabrina Ullmann, Cíntia Bittar, Guilherme Campos, Helena Lage Ferreira, Jorge A. Petroli Marchesi, Maisa C. Pereira Parra, Marília Moraes, Paula Rahal, Paulo Inacio da Costa, João Pessoa Araújo Jr., Maurício Lacerda Nogueira. Prefeitura de Sao Paulo: Melissa Palmieri. |
| EPI_ISL_1967067 | HOSP PRONTO SOCORRO ITAQUERA | Instituto Butantan / ESALQ-USP (Piracicaba) | Instituto Butantan: Dimas Tadeu Covas, Sandra Coccuzzo Sampaio, Maria Carolina Elias, José Salvatore Leister Patané, Vincent Louis Viala, Antonio Jorge Martins, Ricardo Haddad, Claudia Renata dos Santos Barros, Elaine Cristina Marqueze, Raul Machado Neto, Debora Botequio Moretti, Jardelina de Souza Todao Bernardino, Loyze Paola Oliveira de Lima, Luiz Aurelio de Campos Crispin. Centro de Genômica Funcional da ESALQ: Luiz Lehmann Coutinho, Ricardo Augusto Brassaloti, Raquel de Lello Rocha Campos Cassano. NGS Soluções Genômicas: Pilar Drummond Sampaio Corrêa Mariani. FZEA-USP Pirassununga: Mirele Daiana Poleti, Jessika Cristina Chagas Lesbon, Elisangela Chicaroni Mattos, Heidge Fukumasu. USP-Botucatu: Rejane Maria Tommasini Grotto, Jayme A. Souza-Neto, Guilherme Targino Valente, Patricia Akemi Assato, Felipe Allan da Silva da Costa, Bianca Cechetto Carlos. Mendelics: Bibiana Santos, João Paulo Kitajima, Erika Freitas, David Schlesinger. Hemocentro Ribeirão Preto: Simone Kashima, Evandra Strazza Rodrigues, Svetoslav Nanev Slavov, Elaine Vieira dos Santos, Rafael dos Santos Bezerra, Luiz Carlos Junior de Alcantara, Marta Giovanetti, Vagner Fonseca, Flavia Aburjaile, Rodrigo Tocantins Calado. FAMERP-SJRP: Cecília Artico Banho, Lívia Sacchetto, Fábio Sossai Possebon, Leila Sabrina Ullmann, Cíntia Bittar, Guilherme Campos, Helena Lage Ferreira, Jorge A. Petroli Marchesi, Maisa C. Pereira Parra, Marília Moraes, Paula Rahal, Paulo Inacio da Costa, João Pessoa Araújo Jr., Maurício Lacerda Nogueira. Prefeitura de Sao Paulo: Melissa Palmieri. |
| EPI_ISL_1967068, EPI_ISL_1967069, EPI_ISL_1967070, EPI_ISL_1967071, EPI_ISL_1967072, EPI_ISL_1967073, EPI_ISL_1967074 | HOSP MUN CIDADE TIRADENTES CARMEN PRUDENTE | Instituto Butantan / ESALQ-USP (Piracicaba) | Instituto Butantan: Dimas Tadeu Covas, Sandra Coccuzzo Sampaio, Maria Carolina Elias, José Salvatore Leister Patané, Vincent Louis Viala, Antonio Jorge Martins, Ricardo Haddad, Claudia Renata dos Santos Barros, Elaine Cristina Marqueze, Raul Machado Neto, Debora Botequio Moretti, Jardelina de Souza Todao Bernardino, Loyze Paola Oliveira de Lima, Luiz Aurelio de Campos Crispin. Centro de Genômica Funcional da ESALQ: Luiz Lehmann Coutinho, Ricardo Augusto Brassaloti, Raquel de Lello Rocha Campos Cassano. NGS Soluções Genômicas: Pilar Drummond Sampaio Corrêa Mariani. FZEA-USP Pirassununga: Mirele Daiana Poleti, Jessika Cristina Chagas Lesbon, Elisangela Chicaroni Mattos, Heidge Fukumasu. USP-Botucatu: Rejane Maria Tommasini Grotto, Jayme A. Souza-Neto, Guilherme Targino Valente, Patricia Akemi Assato, Felipe Allan da Silva da Costa, Bianca Cechetto Carlos. Mendelics: Bibiana Santos, João Paulo Kitajima, Erika Freitas, David Schlesinger. Hemocentro Ribeirão Preto: Simone Kashima, Evandra Strazza Rodrigues, Svetoslav Nanev Slavov, Elaine Vieira dos Santos, Rafael dos Santos Bezerra, Luiz Carlos Junior de Alcantara, Marta Giovanetti, Vagner Fonseca, Flavia Aburjaile, Rodrigo Tocantins Calado. FAMERP-SJRP: Cecília Artico Banho, Lívia Sacchetto, Fábio Sossai Possebon, Leila Sabrina Ullmann, Cíntia Bittar, Guilherme Campos, Helena Lage Ferreira, Jorge A. Petroli Marchesi, Maisa C. Pereira Parra, Marília Moraes, Paula Rahal, Paulo Inacio da Costa, João Pessoa Araújo Jr., Maurício Lacerda Nogueira. Prefeitura de Sao Paulo: Melissa Palmieri. |
| EPI_ISL_1967075 | HOSP MUN JOSANIAS CASTANHA BRAGA | Instituto Butantan / ESALQ-USP (Piracicaba) | Instituto Butantan: Dimas Tadeu Covas, Sandra Coccuzzo Sampaio, Maria Carolina Elias, José Salvatore Leister Patané, Vincent Louis Viala, Antonio Jorge Martins, Ricardo Haddad, Claudia Renata dos Santos Barros, Elaine Cristina Marqueze, Raul Machado Neto, Debora Botequio Moretti, Jardelina de |

| | | | |
|---|---|---|---|
| | | | Souza Todao Bernardino, Loyze Paola Oliveira de Lima, Luiz Aurelio de Campos Crispin. Centro de Genômica Funcional da ESALQ: Luiz Lehmann Coutinho, Ricardo Augusto Brassaloti, Raquel de Lello Rocha Campos Cassano. NGS Soluções Genômicas: Pilar Drummond Sampaio Corrêa Mariani. FZEA-USP Pirassununga: Mirele Daiana Poleti, Jessika Cristina Chagas Lesbon, Elisangela Chicaroni Mattos, Heidge Fukumasu. USP-Botucatu: Rejane Maria Tommasini Grotto, Jayme A. Souza-Neto, Guilherme Targino Valente, Patricia Akemi Assato, Felipe Allan da Silva da Costa, Bianca Cechetto Carlos. Mendelics: Bibiana Santos, João Paulo Kitajima, Erika Freitas, David Schlesinger. Hemocentro Ribeirão Preto: Simone Kashima, Evandra Strazza Rodrigues, Svetoslav Nanev Slavov, Elaine Vieira dos Santos, Rafael dos Santos Bezerra, Luiz Carlos Junior de Alcantara, Marta Giovanetti, Vagner Fonseca, Flavia Aburjaile, Rodrigo Tocantins Calado. FAMERP-SJRP: Cecília Artico Banho, Lívia Sacchetto, Fábio Sossai Possebon, Leila Sabrina Ullmann, Cíntia Bittar, Guilherme Campos, Helena Lage Ferreira, Jorge A. Petroli Marchesi, Maisa C. Pereira Parra, Marília Moraes, Paula Rahal, Paulo Inacio da Costa, João Pessoa Araújo Jr., Maurício Lacerda Nogueira. Prefeitura de Sao Paulo: Melissa Palmieri. |
| EPI_ISL_1967076 | UPA VILA SANTA CATARINA | Instituto Butantan / ESALQ-USP (Piracicaba) | Instituto Butantan: Dimas Tadeu Covas, Sandra Coccuzzo Sampaio, Maria Carolina Elias, José Salvatore Leister Patané, Vincent Louis Viala, Antonio Jorge Martins, Ricardo Haddad, Claudia Renata dos Santos Barros, Elaine Cristina Marqueze, Raul Machado Neto, Debora Botequio Moretti, Jardelina de Souza Todao Bernardino, Loyze Paola Oliveira de Lima, Luiz Aurelio de Campos Crispin. Centro de Genômica Funcional da ESALQ: Luiz Lehmann Coutinho, Ricardo Augusto Brassaloti, Raquel de Lello Rocha Campos Cassano. NGS Soluções Genômicas: Pilar Drummond Sampaio Corrêa Mariani. FZEA-USP Pirassununga: Mirele Daiana Poleti, Jessika Cristina Chagas Lesbon, Elisangela Chicaroni Mattos, Heidge Fukumasu. USP-Botucatu: Rejane Maria Tommasini Grotto, Jayme A. Souza-Neto, Guilherme Targino Valente, Patricia Akemi Assato, Felipe Allan da Silva da Costa, Bianca Cechetto Carlos. Mendelics: Bibiana Santos, João Paulo Kitajima, Erika Freitas, David Schlesinger. Hemocentro Ribeirão Preto: Simone Kashima, Evandra Strazza Rodrigues, Svetoslav Nanev Slavov, Elaine Vieira dos Santos, Rafael dos Santos Bezerra, Luiz Carlos Junior de Alcantara, Marta Giovanetti, Vagner Fonseca, Flavia Aburjaile, Rodrigo Tocantins Calado. FAMERP-SJRP: Cecília Artico Banho, Lívia Sacchetto, Fábio Sossai Possebon, Leila Sabrina Ullmann, Cíntia Bittar, Guilherme Campos, Helena Lage Ferreira, Jorge A. Petroli Marchesi, Maisa C. Pereira Parra, Marília Moraes, Paula Rahal, Paulo Inacio da Costa, João Pessoa Araújo Jr., Maurício Lacerda Nogueira. Prefeitura de Sao Paulo: Melissa Palmieri. |
| EPI_ISL_1967077, EPI_ISL_1967078, EPI_ISL_1967080, EPI_ISL_1967081, EPI_ISL_1967082, EPI_ISL_1967083, EPI_ISL_1967084, EPI_ISL_1967085, EPI_ISL_1967086, EPI_ISL_1967087, EPI_ISL_1967088, EPI_ISL_1967089, EPI_ISL_1967090 | | | |
| see above | HOSPITAL GERAL JESUS TEIXEIRA DA COSTA GUAIANASES SAO PAULO | Instituto Butantan / ESALQ-USP (Piracicaba) | Instituto Butantan: Dimas Tadeu Covas, Sandra Coccuzzo Sampaio, Maria Carolina Elias, José Salvatore Leister Patané, Vincent Louis Viala, Antonio Jorge Martins, Ricardo Haddad, Claudia Renata dos Santos Barros, Elaine Cristina Marqueze, Raul Machado Neto, Debora Botequio Moretti, Jardelina de Souza Todao Bernardino, Loyze Paola Oliveira de Lima, Luiz Aurelio de Campos Crispin. Centro de Genômica Funcional da ESALQ: Luiz Lehmann Coutinho, Ricardo Augusto Brassaloti, Raquel de Lello Rocha Campos Cassano. NGS Soluções Genômicas: Pilar Drummond Sampaio Corrêa Mariani. FZEA-USP Pirassununga: Mirele Daiana Poleti, Jessika Cristina Chagas Lesbon, Elisangela Chicaroni Mattos, Heidge Fukumasu. USP-Botucatu: Rejane Maria Tommasini Grotto, Jayme A. Souza-Neto, Guilherme Targino Valente, Patricia Akemi Assato, Felipe Allan da Silva da Costa, Bianca Cechetto Carlos. Mendelics: Bibiana Santos, João Paulo Kitajima, Erika Freitas, David Schlesinger. Hemocentro Ribeirão Preto: Simone Kashima, Evandra Strazza Rodrigues, Svetoslav Nanev Slavov, Elaine Vieira dos Santos, Rafael dos Santos Bezerra, Luiz Carlos Junior de Alcantara, Marta Giovanetti, Vagner Fonseca, Flavia Aburjaile, Rodrigo Tocantins Calado. FAMERP-SJRP: Cecília Artico Banho, Lívia Sacchetto, Fábio Sossai Possebon, Leila Sabrina Ullmann, Cíntia Bittar, Guilherme Campos, Helena Lage Ferreira, Jorge A. Petroli Marchesi, Maisa C. Pereira Parra, Marília Moraes, Paula Rahal, Paulo Inacio da Costa, João Pessoa Araújo Jr., Maurício Lacerda Nogueira. Prefeitura de Sao Paulo: Melissa Palmieri. |
| EPI_ISL_1967092, EPI_ISL_1967093, EPI_ISL_1967094, EPI_ISL_1967095 | HOSP DO SERV PUB ESTFCO MORATO DE OLIVEIRA SAO PAULO | Instituto Butantan / ESALQ-USP (Piracicaba) | Instituto Butantan: Dimas Tadeu Covas, Sandra Coccuzzo Sampaio, Maria Carolina Elias, José Salvatore Leister Patané, Vincent Louis Viala, Antonio Jorge Martins, Ricardo Haddad, Claudia Renata dos Santos Barros, Elaine Cristina Marqueze, Raul Machado Neto, Debora Botequio Moretti, Jardelina de Souza Todao Bernardino, Loyze Paola Oliveira de Lima, Luiz Aurelio de Campos Crispin. Centro de Genômica Funcional da ESALQ: Luiz Lehmann Coutinho, Ricardo Augusto Brassaloti, Raquel de Lello Rocha Campos Cassano. NGS Soluções Genômicas: Pilar Drummond Sampaio Corrêa Mariani. FZEA-USP Pirassununga: Mirele Daiana Poleti, Jessika Cristina Chagas Lesbon, Elisangela Chicaroni Mattos, Heidge Fukumasu. USP-Botucatu: Rejane Maria Tommasini Grotto, Jayme A. Souza-Neto, Guilherme Targino Valente, Patricia Akemi Assato, Felipe Allan da Silva da Costa, Bianca Cechetto Carlos. Mendelics: Bibiana Santos, João Paulo Kitajima, Erika Freitas, David Schlesinger. Hemocentro Ribeirão Preto: Simone Kashima, Evandra Strazza Rodrigues, Svetoslav Nanev Slavov, Elaine Vieira dos Santos, Rafael dos Santos Bezerra, Luiz Carlos Junior de Alcantara, Marta Giovanetti, Vagner Fonseca, Flavia Aburjaile, Rodrigo Tocantins Calado. FAMERP-SJRP: Cecília Artico Banho, Lívia Sacchetto, Fábio Sossai Possebon, Leila Sabrina Ullmann, Cíntia Bittar, Guilherme Campos, Helena Lage Ferreira, Jorge A. Petroli Marchesi, Maisa C. Pereira Parra, Marília Moraes, Paula Rahal, Paulo Inacio da Costa, João Pessoa Araújo Jr., Maurício Lacerda Nogueira. Prefeitura de Sao Paulo: Melissa Palmieri. |
| EPI_ISL_1967096, EPI_ISL_1967097 | CENTRO DE SAUDE SAO ROQUE DR JOSE CARVALHO BRITO | Instituto Butantan / ESALQ-USP (Piracicaba) | Instituto Butantan: Dimas Tadeu Covas, Sandra Coccuzzo Sampaio, Maria Carolina Elias, José Salvatore Leister Patané, Vincent Louis Viala, Antonio Jorge Martins, Ricardo Haddad, Claudia Renata dos Santos Barros, Elaine Cristina Marqueze, Raul Machado Neto, Debora Botequio Moretti, Jardelina de Souza Todao Bernardino, Loyze Paola Oliveira de Lima, Luiz Aurelio de Campos Crispin. Centro de Genômica Funcional da ESALQ: Luiz Lehmann Coutinho, Ricardo Augusto Brassaloti, Raquel de Lello Rocha Campos Cassano. NGS Soluções Genômicas: Pilar Drummond Sampaio Corrêa Mariani. FZEA-USP Pirassununga: Mirele Daiana Poleti, Jessika Cristina Chagas Lesbon, Elisangela Chicaroni Mattos, Heidge Fukumasu. USP-Botucatu: Rejane Maria Tommasini Grotto, Jayme A. Souza-Neto, Guilherme Targino Valente, Patricia Akemi Assato, Felipe Allan da Silva da Costa, Bianca Cechetto Carlos. Mendelics: Bibiana Santos, João Paulo Kitajima, Erika Freitas, David Schlesinger. Hemocentro Ribeirão Preto: Simone Kashima, Evandra Strazza Rodrigues, Svetoslav Nanev Slavov, Elaine Vieira dos Santos, Rafael dos Santos Bezerra, Luiz Carlos Junior de Alcantara, Marta Giovanetti, Vagner Fonseca, Flavia Aburjaile, Rodrigo Tocantins Calado. FAMERP-SJRP: Cecília Artico Banho, Lívia Sacchetto, Fábio Sossai Possebon, Leila Sabrina Ullmann, Cíntia Bittar, Guilherme Campos, Helena Lage Ferreira, Jorge A. Petroli Marchesi, Maisa C. Pereira Parra, Marília Moraes, Paula Rahal, Paulo Inacio da Costa, João Pessoa Araújo Jr., Maurício Lacerda Nogueira. Prefeitura de Sao Paulo: Melissa Palmieri. |
| EPI_ISL_1967098, EPI_ISL_1967099, EPI_ISL_1967100, EPI_ISL_1967101, EPI_ISL_1967102, EPI_ISL_1967103, EPI_ISL_1967104, EPI_ISL_1967105, EPI_ISL_1967106, EPI_ISL_1967107, EPI_ISL_1967108, EPI_ISL_1967109, EPI_ISL_1967110, EPI_ISL_1967111, EPI_ISL_1967112, EPI_ISL_1967113, EPI_ISL_1967114, EPI_ISL_1967115, EPI_ISL_1967116, EPI_ISL_1967117, EPI_ISL_1967118, EPI_ISL_1967119, EPI_ISL_1967120, EPI_ISL_1967121, EPI_ISL_1967122, EPI_ISL_1967123, EPI_ISL_1967124, EPI_ISL_1967125, EPI_ISL_1967126, EPI_ISL_1967127, EPI_ISL_1967128, EPI_ISL_1967129, EPI_ISL_1967130, EPI_ISL_1967131, EPI_ISL_1967132, EPI_ISL_1967133, EPI_ISL_1967134, EPI_ISL_1967135, EPI_ISL_1967136, EPI_ISL_1967137, EPI_ISL_1967138, EPI_ISL_1967139, EPI_ISL_1967140, EPI_ISL_1967141, EPI_ISL_1967142, EPI_ISL_1967143, EPI_ISL_1967144, EPI_ISL_1967145, EPI_ISL_1967146, EPI_ISL_1967147 | | | |
| see above | SECRETARIA MUNICIPAL DE SAUDE SOROCABA | Instituto Butantan / ESALQ-USP (Piracicaba) | Instituto Butantan: Dimas Tadeu Covas, Sandra Coccuzzo Sampaio, Maria Carolina Elias, José Salvatore Leister Patané, Vincent Louis Viala, Antonio Jorge Martins, Ricardo Haddad, Claudia Renata dos Santos Barros, Elaine Cristina Marqueze, Raul Machado Neto, Debora Botequio Moretti, Jardelina de Souza Todao Bernardino, Loyze Paola Oliveira de Lima, Luiz Aurelio de Campos Crispin. Centro de Genômica Funcional da ESALQ: Luiz Lehmann Coutinho, Ricardo Augusto Brassaloti, Raquel de Lello Rocha Campos Cassano. NGS Soluções Genômicas: Pilar Drummond Sampaio Corrêa Mariani. FZEA-USP Pirassununga: Mirele Daiana Poleti, Jessika Cristina Chagas Lesbon, Elisangela Chicaroni Mattos, Heidge Fukumasu. USP-Botucatu: Rejane Maria Tommasini Grotto, Jayme A. Souza-Neto, Guilherme Targino Valente, Patricia Akemi Assato, Felipe Allan da Silva da Costa, Bianca Cechetto Carlos. Mendelics: Bibiana Santos, João Paulo Kitajima, Erika Freitas, David Schlesinger. Hemocentro Ribeirão Preto: Simone Kashima, Evandra Strazza Rodrigues, Svetoslav Nanev Slavov, Elaine Vieira dos Santos, Rafael dos Santos Bezerra, Luiz Carlos Junior de Alcantara, Marta Giovanetti, Vagner Fonseca, Flavia Aburjaile, Rodrigo Tocantins Calado. FAMERP-SJRP: Cecília Artico Banho, Lívia Sacchetto, Fábio Sossai Possebon, Leila Sabrina Ullmann, Cíntia Bittar, Guilherme Campos, Helena Lage Ferreira, Jorge A. Petroli Marchesi, Maisa C. Pereira Parra, Marília Moraes, Paula Rahal, Paulo Inacio da Costa, João Pessoa Araújo Jr., Maurício Lacerda Nogueira. Prefeitura de Sao Paulo: Melissa Palmieri. |
| EPI_ISL_1967148, EPI_ISL_1967149, EPI_ISL_1967150, EPI_ISL_1967151, EPI_ISL_1967152, EPI_ISL_1967153, EPI_ISL_1967154, EPI_ISL_1967155, EPI_ISL_1967156, EPI_ISL_1967157 | LABORATORIO MUNICIPAL DE SUZANO | Instituto Butantan / ESALQ-USP (Piracicaba) | Instituto Butantan: Dimas Tadeu Covas, Sandra Coccuzzo Sampaio, Maria Carolina Elias, José Salvatore Leister Patané, Vincent Louis Viala, Antonio Jorge Martins, Ricardo Haddad, Claudia Renata dos Santos Barros, Elaine Cristina Marqueze, Raul Machado Neto, Debora Botequio Moretti, Jardelina de Souza Todao Bernardino, Loyze Paola Oliveira de Lima, Luiz Aurelio de Campos Crispin. Centro de Genômica Funcional da ESALQ: Luiz Lehmann Coutinho, Ricardo Augusto Brassaloti, Raquel de Lello Rocha Campos Cassano. NGS Soluções Genômicas: Pilar Drummond Sampaio Corrêa Mariani. FZEA-USP Pirassununga: Mirele Daiana Poleti, Jessika Cristina Chagas Lesbon, Elisangela Chicaroni Mattos, Heidge Fukumasu. USP-Botucatu: Rejane Maria Tommasini Grotto, Jayme A. Souza-Neto, Guilherme Targino Valente, Patricia Akemi Assato, Felipe Allan da Silva da Costa, Bianca Cechetto Carlos. Mendelics: Bibiana Santos, João Paulo Kitajima, Erika Freitas, David Schlesinger. Hemocentro Ribeirão Preto: Simone Kashima, Evandra Strazza Rodrigues, Svetoslav Nanev Slavov, Elaine Vieira dos Santos, Rafael dos Santos Bezerra, Luiz Carlos Junior de Alcantara, Marta Giovanetti, Vagner Fonseca, Flavia Aburjaile, Rodrigo Tocantins Calado. FAMERP-SJRP: Cecília Artico Banho, Lívia Sacchetto, Fábio Sossai Possebon, Leila Sabrina Ullmann, Cíntia Bittar, Guilherme Campos, Helena Lage Ferreira, Jorge A. Petroli Marchesi, Maisa C. Pereira Parra, Marília Moraes, Paula Rahal, Paulo Inacio da Costa, João Pessoa Araújo Jr., Maurício Lacerda Nogueira. Prefeitura de Sao Paulo: Melissa Palmieri. |
| EPI_ISL_1967159, EPI_ISL_1967160, EPI_ISL_1967161, EPI_ISL_1967162 | UBS DE TABAPUA | Instituto Butantan / ESALQ-USP (Piracicaba) | Instituto Butantan: Dimas Tadeu Covas, Sandra Coccuzzo Sampaio, Maria Carolina Elias, José Salvatore Leister Patané, Vincent Louis Viala, Antonio Jorge Martins, Ricardo Haddad, Claudia Renata dos Santos Barros, Elaine Cristina Marqueze, Raul Machado Neto, Debora Botequio Moretti, Jardelina de Souza Todao Bernardino, Loyze Paola Oliveira de Lima, Luiz Aurelio de Campos Crispin. Centro de Genômica Funcional da ESALQ: Luiz Lehmann Coutinho, Ricardo Augusto Brassaloti, Raquel de Lello Rocha Campos Cassano. NGS Soluções Genômicas: Pilar Drummond Sampaio Corrêa Mariani. |

| | | | |
|---|---|---|---|
| | | | FZEA-USP Pirassununga: Mirele Daiana Poleti, Jessika Cristina Chagas Lesbon, Elisangela Chicaroni Mattos, Heidge Fukumasu. USP-Botucatu: Rejane Maria Tommasini Grotto, Jayme A. Souza-Neto, Guilherme Targino Valente, Patricia Akemi Assato, Felipe Allan da Silva da Costa, Bianca Cechetto Carlos. Mendelics: Bibiana Santos, João Paulo Kitajima, Erika Freitas, David Schlesinger. Hemocentro Ribeirão Preto: Simone Kashima, Evandra Strazza Rodrigues, Svetoslav Nanev Slavov, Elaine Vieira dos Santos, Rafael dos Santos Bezerra, Luiz Carlos Junior de Alcantara, Marta Giovanetti, Vagner Fonseca, Flavia Aburjaile, Rodrigo Tocantins Calado. FAMERP-SJRP: Cecília Artico Banho, Lívia Sacchetto, Fábio Sossai Possebon, Leila Sabrina Ullmann, Cíntia Bittar, Guilherme Campos, Helena Lage Ferreira, Jorge A. Petroli Marchesi, Maisa C. Pereira Parra, Marília Moraes, Paula Rahal, Paulo Inacio da Costa, João Pessoa Araújo Jr., Maurício Lacerda Nogueira. Prefeitura de Sao Paulo: Melissa Palmieri. |
| EPI_ISL_1967163, EPI_ISL_1967164 | UBS ENRIQUE GUILHERME ERNESTO MARZEUSKI | Instituto Butantan / ESALQ-USP (Piracicaba) | Instituto Butantan: Dimas Tadeu Covas, Sandra Coccuzzo Sampaio, Maria Carolina Elias, José Salvatore Leister Patané, Vincent Louis Viala, Antonio Jorge Martins, Ricardo Haddad, Claudia Renata dos Santos Barros, Elaine Cristina Marqueze, Raul Machado Neto, Debora Botequio Moretti, Jardelina de Souza Todao Bernardino, Loyze Paola Oliveira de Lima, Luiz Aurelio de Campos Crispin. Centro de Genômica Funcional da ESALQ: Luiz Lehmann Coutinho, Ricardo Augusto Brassaloti, Raquel de Lello Rocha Campos Cassano. NGS Soluções Genômicas: Pilar Drummond Sampaio Corrêa Mariani. FZEA-USP Pirassununga: Mirele Daiana Poleti, Jessika Cristina Chagas Lesbon, Elisangela Chicaroni Mattos, Heidge Fukumasu. USP-Botucatu: Rejane Maria Tommasini Grotto, Jayme A. Souza-Neto, Guilherme Targino Valente, Patricia Akemi Assato, Felipe Allan da Silva da Costa, Bianca Cechetto Carlos. Mendelics: Bibiana Santos, João Paulo Kitajima, Erika Freitas, David Schlesinger. Hemocentro Ribeirão Preto: Simone Kashima, Evandra Strazza Rodrigues, Svetoslav Nanev Slavov, Elaine Vieira dos Santos, Rafael dos Santos Bezerra, Luiz Carlos Junior de Alcantara, Marta Giovanetti, Vagner Fonseca, Flavia Aburjaile, Rodrigo Tocantins Calado. FAMERP-SJRP: Cecília Artico Banho, Lívia Sacchetto, Fábio Sossai Possebon, Leila Sabrina Ullmann, Cíntia Bittar, Guilherme Campos, Helena Lage Ferreira, Jorge A. Petroli Marchesi, Maisa C. Pereira Parra, Marília Moraes, Paula Rahal, Paulo Inacio da Costa, João Pessoa Araújo Jr., Maurício Lacerda Nogueira. Prefeitura de Sao Paulo: Melissa Palmieri. |
| EPI_ISL_1967165, EPI_ISL_1967166, EPI_ISL_1967167, EPI_ISL_1967168, EPI_ISL_1967169, EPI_ISL_1967170, EPI_ISL_1967171, EPI_ISL_1967172, EPI_ISL_1967173, EPI_ISL_1967174, EPI_ISL_1967175, EPI_ISL_1967176, EPI_ISL_1967177, EPI_ISL_1967178, EPI_ISL_1967179, EPI_ISL_1967180, EPI_ISL_1967181, EPI_ISL_1967182, EPI_ISL_1967183, EPI_ISL_1967184, EPI_ISL_1967185, EPI_ISL_1967186, EPI_ISL_1967187, EPI_ISL_1967188, EPI_ISL_1967189, EPI_ISL_1967190, EPI_ISL_1967191 | | | |
| see above | HOSPITAL DE CAMPANHA COVID 19 MUNICIPIO DE TAUBATE | Instituto Butantan / ESALQ-USP (Piracicaba) | Instituto Butantan: Dimas Tadeu Covas, Sandra Coccuzzo Sampaio, Maria Carolina Elias, José Salvatore Leister Patané, Vincent Louis Viala, Antonio Jorge Martins, Ricardo Haddad, Claudia Renata dos Santos Barros, Elaine Cristina Marqueze, Raul Machado Neto, Debora Botequio Moretti, Jardelina de Souza Todao Bernardino, Loyze Paola Oliveira de Lima, Luiz Aurelio de Campos Crispin. Centro de Genômica Funcional da ESALQ: Luiz Lehmann Coutinho, Ricardo Augusto Brassaloti, Raquel de Lello Rocha Campos Cassano. NGS Soluções Genômicas: Pilar Drummond Sampaio Corrêa Mariani. FZEA-USP Pirassununga: Mirele Daiana Poleti, Jessika Cristina Chagas Lesbon, Elisangela Chicaroni Mattos, Heidge Fukumasu. USP-Botucatu: Rejane Maria Tommasini Grotto, Jayme A. Souza-Neto, Guilherme Targino Valente, Patricia Akemi Assato, Felipe Allan da Silva da Costa, Bianca Cechetto Carlos. Mendelics: Bibiana Santos, João Paulo Kitajima, Erika Freitas, David Schlesinger. Hemocentro Ribeirão Preto: Simone Kashima, Evandra Strazza Rodrigues, Svetoslav Nanev Slavov, Elaine Vieira dos Santos, Rafael dos Santos Bezerra, Luiz Carlos Junior de Alcantara, Marta Giovanetti, Vagner Fonseca, Flavia Aburjaile, Rodrigo Tocantins Calado. FAMERP-SJRP: Cecília Artico Banho, Lívia Sacchetto, Fábio Sossai Possebon, Leila Sabrina Ullmann, Cíntia Bittar, Guilherme Campos, Helena Lage Ferreira, Jorge A. Petroli Marchesi, Maisa C. Pereira Parra, Marília Moraes, Paula Rahal, Paulo Inacio da Costa, João Pessoa Araújo Jr., Maurício Lacerda Nogueira. Prefeitura de Sao Paulo: Melissa Palmieri. |
| EPI_ISL_1967192, EPI_ISL_1967193, EPI_ISL_1967194, EPI_ISL_1967195 | SANTA CASA DE MISERICORDIA DE TIETE | Instituto Butantan / ESALQ-USP (Piracicaba) | Instituto Butantan: Dimas Tadeu Covas, Sandra Coccuzzo Sampaio, Maria Carolina Elias, José Salvatore Leister Patané, Vincent Louis Viala, Antonio Jorge Martins, Ricardo Haddad, Claudia Renata dos Santos Barros, Elaine Cristina Marqueze, Raul Machado Neto, Debora Botequio Moretti, Jardelina de Souza Todao Bernardino, Loyze Paola Oliveira de Lima, Luiz Aurelio de Campos Crispin. Centro de Genômica Funcional da ESALQ: Luiz Lehmann Coutinho, Ricardo Augusto Brassaloti, Raquel de Lello Rocha Campos Cassano. NGS Soluções Genômicas: Pilar Drummond Sampaio Corrêa Mariani. FZEA-USP Pirassununga: Mirele Daiana Poleti, Jessika Cristina Chagas Lesbon, Elisangela Chicaroni Mattos, Heidge Fukumasu. USP-Botucatu: Rejane Maria Tommasini Grotto, Jayme A. Souza-Neto, Guilherme Targino Valente, Patricia Akemi Assato, Felipe Allan da Silva da Costa, Bianca Cechetto Carlos. Mendelics: Bibiana Santos, João Paulo Kitajima, Erika Freitas, David Schlesinger. Hemocentro Ribeirão Preto: Simone Kashima, Evandra Strazza Rodrigues, Svetoslav Nanev Slavov, Elaine Vieira dos Santos, Rafael dos Santos Bezerra, Luiz Carlos Junior de Alcantara, Marta Giovanetti, Vagner Fonseca, Flavia Aburjaile, Rodrigo Tocantins Calado. FAMERP-SJRP: Cecília Artico Banho, Lívia Sacchetto, Fábio Sossai Possebon, Leila Sabrina Ullmann, Cíntia Bittar, Guilherme Campos, Helena Lage Ferreira, Jorge A. Petroli Marchesi, Maisa C. Pereira Parra, Marília Moraes, Paula Rahal, Paulo Inacio da Costa, João Pessoa Araújo Jr., Maurício Lacerda Nogueira. Prefeitura de Sao Paulo: Melissa Palmieri. |
| EPI_ISL_1967196, EPI_ISL_1967197 | DEPARTAMENTO DE SAUDE COLETIVA | Instituto Butantan / ESALQ-USP (Piracicaba) | Instituto Butantan: Dimas Tadeu Covas, Sandra Coccuzzo Sampaio, Maria Carolina Elias, José Salvatore Leister Patané, Vincent Louis Viala, Antonio Jorge Martins, Ricardo Haddad, Claudia Renata dos Santos Barros, Elaine Cristina Marqueze, Raul Machado Neto, Debora Botequio Moretti, Jardelina de Souza Todao Bernardino, Loyze Paola Oliveira de Lima, Luiz Aurelio de Campos Crispin. Centro de Genômica Funcional da ESALQ: Luiz Lehmann Coutinho, Ricardo Augusto Brassaloti, Raquel de Lello Rocha Campos Cassano. NGS Soluções Genômicas: Pilar Drummond Sampaio Corrêa Mariani. FZEA-USP Pirassununga: Mirele Daiana Poleti, Jessika Cristina Chagas Lesbon, Elisangela Chicaroni Mattos, Heidge Fukumasu. USP-Botucatu: Rejane Maria Tommasini Grotto, Jayme A. Souza-Neto, Guilherme Targino Valente, Patricia Akemi Assato, Felipe Allan da Silva da Costa, Bianca Cechetto Carlos. Mendelics: Bibiana Santos, João Paulo Kitajima, Erika Freitas, David Schlesinger. Hemocentro Ribeirão Preto: Simone Kashima, Evandra Strazza Rodrigues, Svetoslav Nanev Slavov, Elaine Vieira dos Santos, Rafael dos Santos Bezerra, Luiz Carlos Junior de Alcantara, Marta Giovanetti, Vagner Fonseca, Flavia Aburjaile, Rodrigo Tocantins Calado. FAMERP-SJRP: Cecília Artico Banho, Lívia Sacchetto, Fábio Sossai Possebon, Leila Sabrina Ullmann, Cíntia Bittar, Guilherme Campos, Helena Lage Ferreira, Jorge A. Petroli Marchesi, Maisa C. Pereira Parra, Marília Moraes, Paula Rahal, Paulo Inacio da Costa, João Pessoa Araújo Jr., Maurício Lacerda Nogueira. Prefeitura de Sao Paulo: Melissa Palmieri. |
| EPI_ISL_1967198, EPI_ISL_1967199, EPI_ISL_1967200, EPI_ISL_1967202, EPI_ISL_1967203 | VIGILANCIA EM SAUDE SAE VIGILANCIA EPIDEMIOLOGICA | Instituto Butantan / Mendelics | Instituto Butantan: Dimas Tadeu Covas, Sandra Coccuzzo Sampaio, Maria Carolina Elias, José Salvatore Leister Patané, Vincent Louis Viala, Antonio Jorge Martins, Ricardo Haddad, Claudia Renata dos Santos Barros, Elaine Cristina Marqueze, Raul Machado Neto, Debora Botequio Moretti, Jardelina de Souza Todao Bernardino, Loyze Paola Oliveira de Lima, Luiz Aurelio de Campos Crispin. Centro de Genômica Funcional da ESALQ: Luiz Lehmann Coutinho, Ricardo Augusto Brassaloti, Raquel de Lello Rocha Campos Cassano. NGS Soluções Genômicas: Pilar Drummond Sampaio Corrêa Mariani. FZEA-USP Pirassununga: Mirele Daiana Poleti, Jessika Cristina Chagas Lesbon, Elisangela Chicaroni Mattos, Heidge Fukumasu. USP-Botucatu: Rejane Maria Tommasini Grotto, Jayme A. Souza-Neto, Guilherme Targino Valente, Patricia Akemi Assato, Felipe Allan da Silva da Costa, Bianca Cechetto Carlos. Mendelics: Bibiana Santos, João Paulo Kitajima, Erika Freitas, David Schlesinger. Hemocentro Ribeirão Preto: Simone Kashima, Evandra Strazza Rodrigues, Svetoslav Nanev Slavov, Elaine Vieira dos Santos, Rafael dos Santos Bezerra, Luiz Carlos Junior de Alcantara, Marta Giovanetti, Vagner Fonseca, Flavia Aburjaile, Rodrigo Tocantins Calado. FAMERP-SJRP: Cecília Artico Banho, Lívia Sacchetto, Fábio Sossai Possebon, Leila Sabrina Ullmann, Cíntia Bittar, Guilherme Campos, Helena Lage Ferreira, Jorge A. Petroli Marchesi, Maisa C. Pereira Parra, Marília Moraes, Paula Rahal, Paulo Inacio da Costa, João Pessoa Araújo Jr., Maurício Lacerda Nogueira. Prefeitura de Sao Paulo: Melissa Palmieri. |
| EPI_ISL_1967204, EPI_ISL_1967205, EPI_ISL_1967206, EPI_ISL_1967207, EPI_ISL_1967208, EPI_ISL_1967209, EPI_ISL_1967210, EPI_ISL_1967211 | UBS DR ALFREDO DANTAS DE SOUZA UMUARAMA | Instituto Butantan / Mendelics | Instituto Butantan: Dimas Tadeu Covas, Sandra Coccuzzo Sampaio, Maria Carolina Elias, José Salvatore Leister Patané, Vincent Louis Viala, Antonio Jorge Martins, Ricardo Haddad, Claudia Renata dos Santos Barros, Elaine Cristina Marqueze, Raul Machado Neto, Debora Botequio Moretti, Jardelina de Souza Todao Bernardino, Loyze Paola Oliveira de Lima, Luiz Aurelio de Campos Crispin. Centro de Genômica Funcional da ESALQ: Luiz Lehmann Coutinho, Ricardo Augusto Brassaloti, Raquel de Lello Rocha Campos Cassano. NGS Soluções Genômicas: Pilar Drummond Sampaio Corrêa Mariani. FZEA-USP Pirassununga: Mirele Daiana Poleti, Jessika Cristina Chagas Lesbon, Elisangela Chicaroni Mattos, Heidge Fukumasu. USP-Botucatu: Rejane Maria Tommasini Grotto, Jayme A. Souza-Neto, Guilherme Targino Valente, Patricia Akemi Assato, Felipe Allan da Silva da Costa, Bianca Cechetto Carlos. Mendelics: Bibiana Santos, João Paulo Kitajima, Erika Freitas, David Schlesinger. Hemocentro Ribeirão Preto: Simone Kashima, Evandra Strazza Rodrigues, Svetoslav Nanev Slavov, Elaine Vieira dos Santos, Rafael dos Santos Bezerra, Luiz Carlos Junior de Alcantara, Marta Giovanetti, Vagner Fonseca, Flavia Aburjaile, Rodrigo Tocantins Calado. FAMERP-SJRP: Cecília Artico Banho, Lívia Sacchetto, Fábio Sossai Possebon, Leila Sabrina Ullmann, Cíntia Bittar, Guilherme Campos, Helena Lage Ferreira, Jorge A. Petroli Marchesi, Maisa C. Pereira Parra, Marília Moraes, Paula Rahal, Paulo Inacio da Costa, João Pessoa Araújo Jr., Maurício Lacerda Nogueira. Prefeitura de Sao Paulo: Melissa Palmieri. |
| EPI_ISL_1967212 | CENTRO DE SAUDE DE BARBOSA | Instituto Butantan / Mendelics | Instituto Butantan: Dimas Tadeu Covas, Sandra Coccuzzo Sampaio, Maria Carolina Elias, José Salvatore Leister Patané, Vincent Louis Viala, Antonio Jorge Martins, Ricardo Haddad, Claudia Renata dos Santos Barros, Elaine Cristina Marqueze, Raul Machado Neto, Debora Botequio Moretti, Jardelina de Souza Todao Bernardino, Loyze Paola Oliveira de Lima, Luiz Aurelio de Campos Crispin. Centro de Genômica Funcional da ESALQ: Luiz Lehmann Coutinho, Ricardo Augusto Brassaloti, Raquel de Lello Rocha Campos Cassano. NGS Soluções Genômicas: Pilar Drummond Sampaio Corrêa Mariani. FZEA-USP Pirassununga: Mirele Daiana Poleti, Jessika Cristina Chagas Lesbon, Elisangela Chicaroni Mattos, Heidge Fukumasu. USP-Botucatu: Rejane Maria Tommasini Grotto, Jayme A. Souza-Neto, Guilherme Targino Valente, Patricia Akemi Assato, Felipe Allan da Silva da Costa, Bianca Cechetto Carlos. Mendelics: Bibiana Santos, João Paulo Kitajima, Erika Freitas, David Schlesinger. Hemocentro Ribeirão Preto: Simone Kashima, Evandra Strazza Rodrigues, Svetoslav Nanev Slavov, Elaine Vieira dos Santos, Rafael dos Santos Bezerra, Luiz Carlos Junior de Alcantara, Marta Giovanetti, Vagner |

| | | | |
|---|---|---|---|
| | | | Fonseca, Flavia Aburjaile, Rodrigo Tocantins Calado. FAMERP-SJRP: Cecília Artico Banho, Lívia Sacchetto, Fábio Sossai Possebon, Leila Sabrina Ullmann, Cíntia Bittar, Guilherme Campos, Helena Lage Ferreira, Jorge A. Petroli Marchesi, Maisa C. Pereira Parra, Marília Moraes, Paula Rahal, Paulo Inacio da Costa, João Pessoa Araújo Jr., Maurício Lacerda Nogueira. Prefeitura de Sao Paulo: Melissa Palmieri. |
| EPI_ISL_1967213, EPI_ISL_1967214, EPI_ISL_1967215 | NUCLEO DE SAUDE MARY DOTA DE BAURU | Instituto Butantan / Mendelics | Instituto Butantan: Dimas Tadeu Covas, Sandra Coccuzzo Sampaio, Maria Carolina Elias, José Salvatore Leister Patané, Vincent Louis Viala, Antonio Jorge Martins, Ricardo Haddad, Claudia Renata dos Santos Barros, Elaine Cristina Marqueze, Raul Machado Neto, Debora Botequio Moretti, Jardelina de Souza Todao Bernardino, Loyze Paola Oliveira de Lima, Luiz Aurelio de Campos Crispin. Centro de Genômica Funcional da ESALQ: Luiz Lehmann Coutinho, Ricardo Augusto Brassaloti, Raquel de Lello Rocha Campos Cassano. NGS Soluções Genômicas: Pilar Drummond Sampaio Corrêa Mariani. FZEA-USP Pirassununga: Mirele Daiana Poleti, Jessika Cristina Chagas Lesbon, Elisangela Chicaroni Mattos, Heidge Fukumasu. USP-Botucatu: Rejane Maria Tommasini Grotto, Jayme A. Souza-Neto, Guilherme Targino Valente, Patricia Akemi Assato, Felipe Allan da Silva da Costa, Bianca Cechetto Carlos. Mendelics: Bibiana Santos, João Paulo Kitajima, Erika Freitas, David Schlesinger. Hemocentro Ribeirão Preto: Simone Kashima, Evandra Strazza Rodrigues, Svetoslav Nanev Slavov, Elaine Vieira dos Santos, Rafael dos Santos Bezerra, Luiz Carlos Junior de Alcantara, Marta Giovanetti, Vagner Fonseca, Flavia Aburjaile, Rodrigo Tocantins Calado. FAMERP-SJRP: Cecília Artico Banho, Lívia Sacchetto, Fábio Sossai Possebon, Leila Sabrina Ullmann, Cíntia Bittar, Guilherme Campos, Helena Lage Ferreira, Jorge A. Petroli Marchesi, Maisa C. Pereira Parra, Marília Moraes, Paula Rahal, Paulo Inacio da Costa, João Pessoa Araújo Jr., Maurício Lacerda Nogueira. Prefeitura de Sao Paulo: Melissa Palmieri. |
| EPI_ISL_1967216 | UPA MARY DOTA | Instituto Butantan / Mendelics | Instituto Butantan: Dimas Tadeu Covas, Sandra Coccuzzo Sampaio, Maria Carolina Elias, José Salvatore Leister Patané, Vincent Louis Viala, Antonio Jorge Martins, Ricardo Haddad, Claudia Renata dos Santos Barros, Elaine Cristina Marqueze, Raul Machado Neto, Debora Botequio Moretti, Jardelina de Souza Todao Bernardino, Loyze Paola Oliveira de Lima, Luiz Aurelio de Campos Crispin. Centro de Genômica Funcional da ESALQ: Luiz Lehmann Coutinho, Ricardo Augusto Brassaloti, Raquel de Lello Rocha Campos Cassano. NGS Soluções Genômicas: Pilar Drummond Sampaio Corrêa Mariani. FZEA-USP Pirassununga: Mirele Daiana Poleti, Jessika Cristina Chagas Lesbon, Elisangela Chicaroni Mattos, Heidge Fukumasu. USP-Botucatu: Rejane Maria Tommasini Grotto, Jayme A. Souza-Neto, Guilherme Targino Valente, Patricia Akemi Assato, Felipe Allan da Silva da Costa, Bianca Cechetto Carlos. Mendelics: Bibiana Santos, João Paulo Kitajima, Erika Freitas, David Schlesinger. Hemocentro Ribeirão Preto: Simone Kashima, Evandra Strazza Rodrigues, Svetoslav Nanev Slavov, Elaine Vieira dos Santos, Rafael dos Santos Bezerra, Luiz Carlos Junior de Alcantara, Marta Giovanetti, Vagner Fonseca, Flavia Aburjaile, Rodrigo Tocantins Calado. FAMERP-SJRP: Cecília Artico Banho, Lívia Sacchetto, Fábio Sossai Possebon, Leila Sabrina Ullmann, Cíntia Bittar, Guilherme Campos, Helena Lage Ferreira, Jorge A. Petroli Marchesi, Maisa C. Pereira Parra, Marília Moraes, Paula Rahal, Paulo Inacio da Costa, João Pessoa Araújo Jr., Maurício Lacerda Nogueira. Prefeitura de Sao Paulo: Melissa Palmieri. |
| EPI_ISL_1967217 | NUCLEO DE SAUDE MARY DOTA DE BAURU | Instituto Butantan / Mendelics | Instituto Butantan: Dimas Tadeu Covas, Sandra Coccuzzo Sampaio, Maria Carolina Elias, José Salvatore Leister Patané, Vincent Louis Viala, Antonio Jorge Martins, Ricardo Haddad, Claudia Renata dos Santos Barros, Elaine Cristina Marqueze, Raul Machado Neto, Debora Botequio Moretti, Jardelina de Souza Todao Bernardino, Loyze Paola Oliveira de Lima, Luiz Aurelio de Campos Crispin. Centro de Genômica Funcional da ESALQ: Luiz Lehmann Coutinho, Ricardo Augusto Brassaloti, Raquel de Lello Rocha Campos Cassano. NGS Soluções Genômicas: Pilar Drummond Sampaio Corrêa Mariani. FZEA-USP Pirassununga: Mirele Daiana Poleti, Jessika Cristina Chagas Lesbon, Elisangela Chicaroni Mattos, Heidge Fukumasu. USP-Botucatu: Rejane Maria Tommasini Grotto, Jayme A. Souza-Neto, Guilherme Targino Valente, Patricia Akemi Assato, Felipe Allan da Silva da Costa, Bianca Cechetto Carlos. Mendelics: Bibiana Santos, João Paulo Kitajima, Erika Freitas, David Schlesinger. Hemocentro Ribeirão Preto: Simone Kashima, Evandra Strazza Rodrigues, Svetoslav Nanev Slavov, Elaine Vieira dos Santos, Rafael dos Santos Bezerra, Luiz Carlos Junior de Alcantara, Marta Giovanetti, Vagner Fonseca, Flavia Aburjaile, Rodrigo Tocantins Calado. FAMERP-SJRP: Cecília Artico Banho, Lívia Sacchetto, Fábio Sossai Possebon, Leila Sabrina Ullmann, Cíntia Bittar, Guilherme Campos, Helena Lage Ferreira, Jorge A. Petroli Marchesi, Maisa C. Pereira Parra, Marília Moraes, Paula Rahal, Paulo Inacio da Costa, João Pessoa Araújo Jr., Maurício Lacerda Nogueira. Prefeitura de Sao Paulo: Melissa Palmieri. |
| EPI_ISL_1967218, EPI_ISL_1967219, EPI_ISL_1967220, EPI_ISL_1967221, EPI_ISL_1967222, EPI_ISL_1967223, EPI_ISL_1967224, EPI_ISL_1967225, EPI_ISL_1967226, EPI_ISL_1967227 | UPA DE BEBEDOURO | Instituto Butantan / Mendelics | Instituto Butantan: Dimas Tadeu Covas, Sandra Coccuzzo Sampaio, Maria Carolina Elias, José Salvatore Leister Patané, Vincent Louis Viala, Antonio Jorge Martins, Ricardo Haddad, Claudia Renata dos Santos Barros, Elaine Cristina Marqueze, Raul Machado Neto, Debora Botequio Moretti, Jardelina de Souza Todao Bernardino, Loyze Paola Oliveira de Lima, Luiz Aurelio de Campos Crispin. Centro de Genômica Funcional da ESALQ: Luiz Lehmann Coutinho, Ricardo Augusto Brassaloti, Raquel de Lello Rocha Campos Cassano. NGS Soluções Genômicas: Pilar Drummond Sampaio Corrêa Mariani. FZEA-USP Pirassununga: Mirele Daiana Poleti, Jessika Cristina Chagas Lesbon, Elisangela Chicaroni Mattos, Heidge Fukumasu. USP-Botucatu: Rejane Maria Tommasini Grotto, Jayme A. Souza-Neto, Guilherme Targino Valente, Patricia Akemi Assato, Felipe Allan da Silva da Costa, Bianca Cechetto Carlos. Mendelics: Bibiana Santos, João Paulo Kitajima, Erika Freitas, David Schlesinger. Hemocentro Ribeirão Preto: Simone Kashima, Evandra Strazza Rodrigues, Svetoslav Nanev Slavov, Elaine Vieira dos Santos, Rafael dos Santos Bezerra, Luiz Carlos Junior de Alcantara, Marta Giovanetti, Vagner Fonseca, Flavia Aburjaile, Rodrigo Tocantins Calado. FAMERP-SJRP: Cecília Artico Banho, Lívia Sacchetto, Fábio Sossai Possebon, Leila Sabrina Ullmann, Cíntia Bittar, Guilherme Campos, Helena Lage Ferreira, Jorge A. Petroli Marchesi, Maisa C. Pereira Parra, Marília Moraes, Paula Rahal, Paulo Inacio da Costa, João Pessoa Araújo Jr., Maurício Lacerda Nogueira. Prefeitura de Sao Paulo: Melissa Palmieri. |
| EPI_ISL_1967228 | UBS 09 JOAO CREVELARO BIRIGUI | Instituto Butantan / Mendelics | Instituto Butantan: Dimas Tadeu Covas, Sandra Coccuzzo Sampaio, Maria Carolina Elias, José Salvatore Leister Patané, Vincent Louis Viala, Antonio Jorge Martins, Ricardo Haddad, Claudia Renata dos Santos Barros, Elaine Cristina Marqueze, Raul Machado Neto, Debora Botequio Moretti, Jardelina de Souza Todao Bernardino, Loyze Paola Oliveira de Lima, Luiz Aurelio de Campos Crispin. Centro de Genômica Funcional da ESALQ: Luiz Lehmann Coutinho, Ricardo Augusto Brassaloti, Raquel de Lello Rocha Campos Cassano. NGS Soluções Genômicas: Pilar Drummond Sampaio Corrêa Mariani. FZEA-USP Pirassununga: Mirele Daiana Poleti, Jessika Cristina Chagas Lesbon, Elisangela Chicaroni Mattos, Heidge Fukumasu. USP-Botucatu: Rejane Maria Tommasini Grotto, Jayme A. Souza-Neto, Guilherme Targino Valente, Patricia Akemi Assato, Felipe Allan da Silva da Costa, Bianca Cechetto Carlos. Mendelics: Bibiana Santos, João Paulo Kitajima, Erika Freitas, David Schlesinger. Hemocentro Ribeirão Preto: Simone Kashima, Evandra Strazza Rodrigues, Svetoslav Nanev Slavov, Elaine Vieira dos Santos, Rafael dos Santos Bezerra, Luiz Carlos Junior de Alcantara, Marta Giovanetti, Vagner Fonseca, Flavia Aburjaile, Rodrigo Tocantins Calado. FAMERP-SJRP: Cecília Artico Banho, Lívia Sacchetto, Fábio Sossai Possebon, Leila Sabrina Ullmann, Cíntia Bittar, Guilherme Campos, Helena Lage Ferreira, Jorge A. Petroli Marchesi, Maisa C. Pereira Parra, Marília Moraes, Paula Rahal, Paulo Inacio da Costa, João Pessoa Araújo Jr., Maurício Lacerda Nogueira. Prefeitura de Sao Paulo: Melissa Palmieri. |
| EPI_ISL_1967229 | UBS 01 VILA BANDEIRANTES BIRIGUI | Instituto Butantan / Mendelics | Instituto Butantan: Dimas Tadeu Covas, Sandra Coccuzzo Sampaio, Maria Carolina Elias, José Salvatore Leister Patané, Vincent Louis Viala, Antonio Jorge Martins, Ricardo Haddad, Claudia Renata dos Santos Barros, Elaine Cristina Marqueze, Raul Machado Neto, Debora Botequio Moretti, Jardelina de Souza Todao Bernardino, Loyze Paola Oliveira de Lima, Luiz Aurelio de Campos Crispin. Centro de Genômica Funcional da ESALQ: Luiz Lehmann Coutinho, Ricardo Augusto Brassaloti, Raquel de Lello Rocha Campos Cassano. NGS Soluções Genômicas: Pilar Drummond Sampaio Corrêa Mariani. FZEA-USP Pirassununga: Mirele Daiana Poleti, Jessika Cristina Chagas Lesbon, Elisangela Chicaroni Mattos, Heidge Fukumasu. USP-Botucatu: Rejane Maria Tommasini Grotto, Jayme A. Souza-Neto, Guilherme Targino Valente, Patricia Akemi Assato, Felipe Allan da Silva da Costa, Bianca Cechetto Carlos. Mendelics: Bibiana Santos, João Paulo Kitajima, Erika Freitas, David Schlesinger. Hemocentro Ribeirão Preto: Simone Kashima, Evandra Strazza Rodrigues, Svetoslav Nanev Slavov, Elaine Vieira dos Santos, Rafael dos Santos Bezerra, Luiz Carlos Junior de Alcantara, Marta Giovanetti, Vagner Fonseca, Flavia Aburjaile, Rodrigo Tocantins Calado. FAMERP-SJRP: Cecília Artico Banho, Lívia Sacchetto, Fábio Sossai Possebon, Leila Sabrina Ullmann, Cíntia Bittar, Guilherme Campos, Helena Lage Ferreira, Jorge A. Petroli Marchesi, Maisa C. Pereira Parra, Marília Moraes, Paula Rahal, Paulo Inacio da Costa, João Pessoa Araújo Jr., Maurício Lacerda Nogueira. Prefeitura de Sao Paulo: Melissa Palmieri. |
| EPI_ISL_1967230 | UBS 10 MANOEL SANCHES GRANEIRO COLINAS BIRIGUI | Instituto Butantan / Mendelics | Instituto Butantan: Dimas Tadeu Covas, Sandra Coccuzzo Sampaio, Maria Carolina Elias, José Salvatore Leister Patané, Vincent Louis Viala, Antonio Jorge Martins, Ricardo Haddad, Claudia Renata dos Santos Barros, Elaine Cristina Marqueze, Raul Machado Neto, Debora Botequio Moretti, Jardelina de Souza Todao Bernardino, Loyze Paola Oliveira de Lima, Luiz Aurelio de Campos Crispin. Centro de Genômica Funcional da ESALQ: Luiz Lehmann Coutinho, Ricardo Augusto Brassaloti, Raquel de Lello Rocha Campos Cassano. NGS Soluções Genômicas: Pilar Drummond Sampaio Corrêa Mariani. FZEA-USP Pirassununga: Mirele Daiana Poleti, Jessika Cristina Chagas Lesbon, Elisangela Chicaroni Mattos, Heidge Fukumasu. USP-Botucatu: Rejane Maria Tommasini Grotto, Jayme A. Souza-Neto, Guilherme Targino Valente, Patricia Akemi Assato, Felipe Allan da Silva da Costa, Bianca Cechetto Carlos. Mendelics: Bibiana Santos, João Paulo Kitajima, Erika Freitas, David Schlesinger. Hemocentro Ribeirão Preto: Simone Kashima, Evandra Strazza Rodrigues, Svetoslav Nanev Slavov, Elaine Vieira dos Santos, Rafael dos Santos Bezerra, Luiz Carlos Junior de Alcantara, Marta Giovanetti, Vagner Fonseca, Flavia Aburjaile, Rodrigo Tocantins Calado. FAMERP-SJRP: Cecília Artico Banho, Lívia Sacchetto, Fábio Sossai Possebon, Leila Sabrina Ullmann, Cíntia Bittar, Guilherme Campos, Helena Lage Ferreira, Jorge A. Petroli Marchesi, Maisa C. Pereira Parra, Marília Moraes, Paula Rahal, Paulo Inacio da Costa, João Pessoa Araújo Jr., Maurício Lacerda Nogueira. Prefeitura de Sao Paulo: Melissa Palmieri. |
| EPI_ISL_1967231 | UBS 04 JARDIM COSTA RICA BIRIGUI | Instituto Butantan / Mendelics | Instituto Butantan: Dimas Tadeu Covas, Sandra Coccuzzo Sampaio, Maria Carolina Elias, José Salvatore Leister Patané, Vincent Louis Viala, Antonio Jorge Martins, Ricardo Haddad, Claudia Renata dos Santos Barros, Elaine Cristina Marqueze, Raul Machado Neto, Debora Botequio Moretti, Jardelina de Souza Todao Bernardino, Loyze Paola Oliveira de Lima, Luiz Aurelio de Campos Crispin. Centro de Genômica Funcional da ESALQ: Luiz Lehmann |

| | | | |
|---|---|---|---|
| | | | Coutinho, Ricardo Augusto Brassaloti, Raquel de Lello Rocha Campos Cassano. NGS Soluções Genômicas: Pilar Drummond Sampaio Corrêa Mariani. FZEA-USP Pirassununga: Mirele Daiana Poleti, Jessika Cristina Chagas Lesbon, Elisangela Chicaroni Mattos, Heidge Fukumasu. USP-Botucatu: Rejane Maria Tommasini Grotto, Jayme A. Souza-Neto, Guilherme Targino Valente, Patricia Akemi Assato, Felipe Allan da Silva da Costa, Bianca Cechetto Carlos. Mendelics: Bibiana Santos, João Paulo Kitajima, Erika Freitas, David Schlesinger. Hemocentro Ribeirão Preto: Simone Kashima, Evandra Strazza Rodrigues, Svetoslav Nanev Slavov, Elaine Vieira dos Santos, Rafael dos Santos Bezerra, Luiz Carlos Junior de Alcantara, Marta Giovanetti, Vagner Fonseca, Flavia Aburjaile, Rodrigo Tocantins Calado. FAMERP-SJRP: Cecília Artico Banho, Lívia Sacchetto, Fábio Sossai Possebon, Leila Sabrina Ullmann, Cíntia Bittar, Guilherme Campos, Helena Lage Ferreira, Jorge A. Petroli Marchesi, Maisa C. Pereira Parra, Marília Moraes, Paula Rahal, Paulo Inacio da Costa, João Pessoa Araújo Jr., Maurício Lacerda Nogueira. Prefeitura de Sao Paulo: Melissa Palmieri. |
| EPI_ISL_1967232 | UBS 09 JOAO CREVELARO BIRIGUI | Instituto Butantan / Mendelics | Instituto Butantan: Dimas Tadeu Covas, Sandra Coccuzzo Sampaio, Maria Carolina Elias, José Salvatore Leister Patané, Vincent Louis Viala, Antonio Jorge Martins, Ricardo Haddad, Claudia Renata dos Santos Barros, Elaine Cristina Marqueze, Raul Machado Neto, Debora Botequio Moretti, Jardelina de Souza Todao Bernardino, Loyze Paola Oliveira de Lima, Luiz Aurelio de Campos Crispin. Centro de Genômica Funcional da ESALQ: Luiz Lehmann Coutinho, Ricardo Augusto Brassaloti, Raquel de Lello Rocha Campos Cassano. NGS Soluções Genômicas: Pilar Drummond Sampaio Corrêa Mariani. FZEA-USP Pirassununga: Mirele Daiana Poleti, Jessika Cristina Chagas Lesbon, Elisangela Chicaroni Mattos, Heidge Fukumasu. USP-Botucatu: Rejane Maria Tommasini Grotto, Jayme A. Souza-Neto, Guilherme Targino Valente, Patricia Akemi Assato, Felipe Allan da Silva da Costa, Bianca Cechetto Carlos. Mendelics: Bibiana Santos, João Paulo Kitajima, Erika Freitas, David Schlesinger. Hemocentro Ribeirão Preto: Simone Kashima, Evandra Strazza Rodrigues, Svetoslav Nanev Slavov, Elaine Vieira dos Santos, Rafael dos Santos Bezerra, Luiz Carlos Junior de Alcantara, Marta Giovanetti, Vagner Fonseca, Flavia Aburjaile, Rodrigo Tocantins Calado. FAMERP-SJRP: Cecília Artico Banho, Lívia Sacchetto, Fábio Sossai Possebon, Leila Sabrina Ullmann, Cíntia Bittar, Guilherme Campos, Helena Lage Ferreira, Jorge A. Petroli Marchesi, Maisa C. Pereira Parra, Marília Moraes, Paula Rahal, Paulo Inacio da Costa, João Pessoa Araújo Jr., Maurício Lacerda Nogueira. Prefeitura de Sao Paulo: Melissa Palmieri. |
| EPI_ISL_1967233 | UBS 01 VILA BANDEIRANTES BIRIGUI | Instituto Butantan / Mendelics | Instituto Butantan: Dimas Tadeu Covas, Sandra Coccuzzo Sampaio, Maria Carolina Elias, José Salvatore Leister Patané, Vincent Louis Viala, Antonio Jorge Martins, Ricardo Haddad, Claudia Renata dos Santos Barros, Elaine Cristina Marqueze, Raul Machado Neto, Debora Botequio Moretti, Jardelina de Souza Todao Bernardino, Loyze Paola Oliveira de Lima, Luiz Aurelio de Campos Crispin. Centro de Genômica Funcional da ESALQ: Luiz Lehmann Coutinho, Ricardo Augusto Brassaloti, Raquel de Lello Rocha Campos Cassano. NGS Soluções Genômicas: Pilar Drummond Sampaio Corrêa Mariani. FZEA-USP Pirassununga: Mirele Daiana Poleti, Jessika Cristina Chagas Lesbon, Elisangela Chicaroni Mattos, Heidge Fukumasu. USP-Botucatu: Rejane Maria Tommasini Grotto, Jayme A. Souza-Neto, Guilherme Targino Valente, Patricia Akemi Assato, Felipe Allan da Silva da Costa, Bianca Cechetto Carlos. Mendelics: Bibiana Santos, João Paulo Kitajima, Erika Freitas, David Schlesinger. Hemocentro Ribeirão Preto: Simone Kashima, Evandra Strazza Rodrigues, Svetoslav Nanev Slavov, Elaine Vieira dos Santos, Rafael dos Santos Bezerra, Luiz Carlos Junior de Alcantara, Marta Giovanetti, Vagner Fonseca, Flavia Aburjaile, Rodrigo Tocantins Calado. FAMERP-SJRP: Cecília Artico Banho, Lívia Sacchetto, Fábio Sossai Possebon, Leila Sabrina Ullmann, Cíntia Bittar, Guilherme Campos, Helena Lage Ferreira, Jorge A. Petroli Marchesi, Maisa C. Pereira Parra, Marília Moraes, Paula Rahal, Paulo Inacio da Costa, João Pessoa Araújo Jr., Maurício Lacerda Nogueira. Prefeitura de Sao Paulo: Melissa Palmieri. |
| EPI_ISL_1967234 | UBS 02 JARDIM TOSELAR BIRIGUI | Instituto Butantan / Mendelics | Instituto Butantan: Dimas Tadeu Covas, Sandra Coccuzzo Sampaio, Maria Carolina Elias, José Salvatore Leister Patané, Vincent Louis Viala, Antonio Jorge Martins, Ricardo Haddad, Claudia Renata dos Santos Barros, Elaine Cristina Marqueze, Raul Machado Neto, Debora Botequio Moretti, Jardelina de Souza Todao Bernardino, Loyze Paola Oliveira de Lima, Luiz Aurelio de Campos Crispin. Centro de Genômica Funcional da ESALQ: Luiz Lehmann Coutinho, Ricardo Augusto Brassaloti, Raquel de Lello Rocha Campos Cassano. NGS Soluções Genômicas: Pilar Drummond Sampaio Corrêa Mariani. FZEA-USP Pirassununga: Mirele Daiana Poleti, Jessika Cristina Chagas Lesbon, Elisangela Chicaroni Mattos, Heidge Fukumasu. USP-Botucatu: Rejane Maria Tommasini Grotto, Jayme A. Souza-Neto, Guilherme Targino Valente, Patricia Akemi Assato, Felipe Allan da Silva da Costa, Bianca Cechetto Carlos. Mendelics: Bibiana Santos, João Paulo Kitajima, Erika Freitas, David Schlesinger. Hemocentro Ribeirão Preto: Simone Kashima, Evandra Strazza Rodrigues, Svetoslav Nanev Slavov, Elaine Vieira dos Santos, Rafael dos Santos Bezerra, Luiz Carlos Junior de Alcantara, Marta Giovanetti, Vagner Fonseca, Flavia Aburjaile, Rodrigo Tocantins Calado. FAMERP-SJRP: Cecília Artico Banho, Lívia Sacchetto, Fábio Sossai Possebon, Leila Sabrina Ullmann, Cíntia Bittar, Guilherme Campos, Helena Lage Ferreira, Jorge A. Petroli Marchesi, Maisa C. Pereira Parra, Marília Moraes, Paula Rahal, Paulo Inacio da Costa, João Pessoa Araújo Jr., Maurício Lacerda Nogueira. Prefeitura de Sao Paulo: Melissa Palmieri. |
| EPI_ISL_1967235 | UBS 11 DR SERGIO AUGUSTO CLARK XAVIER SOARES PORTAL II | Instituto Butantan / Mendelics | Instituto Butantan: Dimas Tadeu Covas, Sandra Coccuzzo Sampaio, Maria Carolina Elias, José Salvatore Leister Patané, Vincent Louis Viala, Antonio Jorge Martins, Ricardo Haddad, Claudia Renata dos Santos Barros, Elaine Cristina Marqueze, Raul Machado Neto, Debora Botequio Moretti, Jardelina de Souza Todao Bernardino, Loyze Paola Oliveira de Lima, Luiz Aurelio de Campos Crispin. Centro de Genômica Funcional da ESALQ: Luiz Lehmann Coutinho, Ricardo Augusto Brassaloti, Raquel de Lello Rocha Campos Cassano. NGS Soluções Genômicas: Pilar Drummond Sampaio Corrêa Mariani. FZEA-USP Pirassununga: Mirele Daiana Poleti, Jessika Cristina Chagas Lesbon, Elisangela Chicaroni Mattos, Heidge Fukumasu. USP-Botucatu: Rejane Maria Tommasini Grotto, Jayme A. Souza-Neto, Guilherme Targino Valente, Patricia Akemi Assato, Felipe Allan da Silva da Costa, Bianca Cechetto Carlos. Mendelics: Bibiana Santos, João Paulo Kitajima, Erika Freitas, David Schlesinger. Hemocentro Ribeirão Preto: Simone Kashima, Evandra Strazza Rodrigues, Svetoslav Nanev Slavov, Elaine Vieira dos Santos, Rafael dos Santos Bezerra, Luiz Carlos Junior de Alcantara, Marta Giovanetti, Vagner Fonseca, Flavia Aburjaile, Rodrigo Tocantins Calado. FAMERP-SJRP: Cecília Artico Banho, Lívia Sacchetto, Fábio Sossai Possebon, Leila Sabrina Ullmann, Cíntia Bittar, Guilherme Campos, Helena Lage Ferreira, Jorge A. Petroli Marchesi, Maisa C. Pereira Parra, Marília Moraes, Paula Rahal, Paulo Inacio da Costa, João Pessoa Araújo Jr., Maurício Lacerda Nogueira. Prefeitura de Sao Paulo: Melissa Palmieri. |
| EPI_ISL_1967236, EPI_ISL_1967237, EPI_ISL_1967238, EPI_ISL_1967239, EPI_ISL_1967240, EPI_ISL_1967241 | PRONTO SOCORRO CENTRAL GUIOMAR FERREIRA ROEBBELEN | Instituto Butantan / Mendelics | Instituto Butantan: Dimas Tadeu Covas, Sandra Coccuzzo Sampaio, Maria Carolina Elias, José Salvatore Leister Patané, Vincent Louis Viala, Antonio Jorge Martins, Ricardo Haddad, Claudia Renata dos Santos Barros, Elaine Cristina Marqueze, Raul Machado Neto, Debora Botequio Moretti, Jardelina de Souza Todao Bernardino, Loyze Paola Oliveira de Lima, Luiz Aurelio de Campos Crispin. Centro de Genômica Funcional da ESALQ: Luiz Lehmann Coutinho, Ricardo Augusto Brassaloti, Raquel de Lello Rocha Campos Cassano. NGS Soluções Genômicas: Pilar Drummond Sampaio Corrêa Mariani. FZEA-USP Pirassununga: Mirele Daiana Poleti, Jessika Cristina Chagas Lesbon, Elisangela Chicaroni Mattos, Heidge Fukumasu. USP-Botucatu: Rejane Maria Tommasini Grotto, Jayme A. Souza-Neto, Guilherme Targino Valente, Patricia Akemi Assato, Felipe Allan da Silva da Costa, Bianca Cechetto Carlos. Mendelics: Bibiana Santos, João Paulo Kitajima, Erika Freitas, David Schlesinger. Hemocentro Ribeirão Preto: Simone Kashima, Evandra Strazza Rodrigues, Svetoslav Nanev Slavov, Elaine Vieira dos Santos, Rafael dos Santos Bezerra, Luiz Carlos Junior de Alcantara, Marta Giovanetti, Vagner Fonseca, Flavia Aburjaile, Rodrigo Tocantins Calado. FAMERP-SJRP: Cecília Artico Banho, Lívia Sacchetto, Fábio Sossai Possebon, Leila Sabrina Ullmann, Cíntia Bittar, Guilherme Campos, Helena Lage Ferreira, Jorge A. Petroli Marchesi, Maisa C. Pereira Parra, Marília Moraes, Paula Rahal, Paulo Inacio da Costa, João Pessoa Araújo Jr., Maurício Lacerda Nogueira. Prefeitura de Sao Paulo: Melissa Palmieri. |
| EPI_ISL_1967242, EPI_ISL_1967243 | VIGILANCIA EPIDEMIOLOGICA DE IBATE | Instituto Butantan / Mendelics | Instituto Butantan: Dimas Tadeu Covas, Sandra Coccuzzo Sampaio, Maria Carolina Elias, José Salvatore Leister Patané, Vincent Louis Viala, Antonio Jorge Martins, Ricardo Haddad, Claudia Renata dos Santos Barros, Elaine Cristina Marqueze, Raul Machado Neto, Debora Botequio Moretti, Jardelina de Souza Todao Bernardino, Loyze Paola Oliveira de Lima, Luiz Aurelio de Campos Crispin. Centro de Genômica Funcional da ESALQ: Luiz Lehmann Coutinho, Ricardo Augusto Brassaloti, Raquel de Lello Rocha Campos Cassano. NGS Soluções Genômicas: Pilar Drummond Sampaio Corrêa Mariani. FZEA-USP Pirassununga: Mirele Daiana Poleti, Jessika Cristina Chagas Lesbon, Elisangela Chicaroni Mattos, Heidge Fukumasu. USP-Botucatu: Rejane Maria Tommasini Grotto, Jayme A. Souza-Neto, Guilherme Targino Valente, Patricia Akemi Assato, Felipe Allan da Silva da Costa, Bianca Cechetto Carlos. Mendelics: Bibiana Santos, João Paulo Kitajima, Erika Freitas, David Schlesinger. Hemocentro Ribeirão Preto: Simone Kashima, Evandra Strazza Rodrigues, Svetoslav Nanev Slavov, Elaine Vieira dos Santos, Rafael dos Santos Bezerra, Luiz Carlos Junior de Alcantara, Marta Giovanetti, Vagner Fonseca, Flavia Aburjaile, Rodrigo Tocantins Calado. FAMERP-SJRP: Cecília Artico Banho, Lívia Sacchetto, Fábio Sossai Possebon, Leila Sabrina Ullmann, Cíntia Bittar, Guilherme Campos, Helena Lage Ferreira, Jorge A. Petroli Marchesi, Maisa C. Pereira Parra, Marília Moraes, Paula Rahal, Paulo Inacio da Costa, João Pessoa Araújo Jr., Maurício Lacerda Nogueira. Prefeitura de Sao Paulo: Melissa Palmieri. |
| EPI_ISL_1967244, EPI_ISL_1967245, EPI_ISL_1967246, EPI_ISL_1967247, EPI_ISL_1967248, EPI_ISL_1967249, EPI_ISL_1967250 | LABORATORIO MUNICIPAL DE ANALISES CLINICAS DE ITANHAEM | Instituto Butantan / Mendelics | Instituto Butantan: Dimas Tadeu Covas, Sandra Coccuzzo Sampaio, Maria Carolina Elias, José Salvatore Leister Patané, Vincent Louis Viala, Antonio Jorge Martins, Ricardo Haddad, Claudia Renata dos Santos Barros, Elaine Cristina Marqueze, Raul Machado Neto, Debora Botequio Moretti, Jardelina de Souza Todao Bernardino, Loyze Paola Oliveira de Lima, Luiz Aurelio de Campos Crispin. Centro de Genômica Funcional da ESALQ: Luiz Lehmann Coutinho, Ricardo Augusto Brassaloti, Raquel de Lello Rocha Campos Cassano. NGS Soluções Genômicas: Pilar Drummond Sampaio Corrêa Mariani. FZEA-USP Pirassununga: Mirele Daiana Poleti, Jessika Cristina Chagas Lesbon, Elisangela Chicaroni Mattos, Heidge Fukumasu. USP-Botucatu: Rejane Maria Tommasini Grotto, Jayme A. Souza-Neto, Guilherme Targino Valente, Patricia Akemi Assato, Felipe Allan da Silva da Costa, Bianca Cechetto Carlos. Mendelics: Bibiana Santos, João Paulo Kitajima, Erika Freitas, David Schlesinger. Hemocentro Ribeirão Preto: Simone Kashima, Evandra Strazza Rodrigues, Svetoslav Nanev Slavov, Elaine Vieira dos Santos, Rafael dos Santos Bezerra, Luiz Carlos Junior de Alcantara, Marta Giovanetti, Vagner |

| | | | |
|---|---|---|---|
| | | | Fonseca, Flavia Aburjaile, Rodrigo Tocantins Calado. FAMERP-SJRP: Cecília Artico Banho, Lívia Sacchetto, Fábio Sossai Possebon, Leila Sabrina Ullmann, Cíntia Bittar, Guilherme Campos, Helena Lage Ferreira, Jorge A. Petroli Marchesi, Maisa C. Pereira Parra, Marília Moraes, Paula Rahal, Paulo Inacio da Costa, João Pessoa Araújo Jr., Maurício Lacerda Nogueira. Prefeitura de Sao Paulo: Melissa Palmieri. |
| EPI_ISL_1967251 | CENTRO DE DETENCAO PROVISORIA DE NOVA INDEPENDENCIA | Instituto Butantan / Mendelics | Instituto Butantan: Dimas Tadeu Covas, Sandra Coccuzzo Sampaio, Maria Carolina Elias, José Salvatore Leister Patané, Vincent Louis Viala, Antonio Jorge Martins, Ricardo Haddad, Claudia Renata dos Santos Barros, Elaine Cristina Marqueze, Raul Machado Neto, Debora Botequio Moretti, Jardelina de Souza Todao Bernardino, Loyze Paola Oliveira de Lima, Luiz Aurelio de Campos Crispin. Centro de Genômica Funcional da ESALQ: Luiz Lehmann Coutinho, Ricardo Augusto Brassaloti, Raquel de Lello Rocha Campos Cassano. NGS Soluções Genômicas: Pilar Drummond Sampaio Corrêa Mariani. FZEA-USP Pirassununga: Mirele Daiana Poleti, Jessika Cristina Chagas Lesbon, Elisangela Chicaroni Mattos, Heidge Fukumasu. USP-Botucatu: Rejane Maria Tommasini Grotto, Jayme A. Souza-Neto, Guilherme Targino Valente, Patricia Akemi Assato, Felipe Allan da Silva da Costa, Bianca Cechetto Carlos. Mendelics: Bibiana Santos, João Paulo Kitajima, Erika Freitas, David Schlesinger. Hemocentro Ribeirão Preto: Simone Kashima, Evandra Strazza Rodrigues, Svetoslav Nanev Slavov, Elaine Vieira dos Santos, Rafael dos Santos Bezerra, Luiz Carlos Junior de Alcantara, Marta Giovanetti, Vagner Fonseca, Flavia Aburjaile, Rodrigo Tocantins Calado. FAMERP-SJRP: Cecília Artico Banho, Lívia Sacchetto, Fábio Sossai Possebon, Leila Sabrina Ullmann, Cíntia Bittar, Guilherme Campos, Helena Lage Ferreira, Jorge A. Petroli Marchesi, Maisa C. Pereira Parra, Marília Moraes, Paula Rahal, Paulo Inacio da Costa, João Pessoa Araújo Jr., Maurício Lacerda Nogueira. Prefeitura de Sao Paulo: Melissa Palmieri. |
| EPI_ISL_1967252, EPI_ISL_1967254, EPI_ISL_1967255, EPI_ISL_1967256, EPI_ISL_1967257 | SECRETARIA MUNICIPAL DE SAUDE PORTO FERREIRA | Instituto Butantan / Mendelics | Instituto Butantan: Dimas Tadeu Covas, Sandra Coccuzzo Sampaio, Maria Carolina Elias, José Salvatore Leister Patané, Vincent Louis Viala, Antonio Jorge Martins, Ricardo Haddad, Claudia Renata dos Santos Barros, Elaine Cristina Marqueze, Raul Machado Neto, Debora Botequio Moretti, Jardelina de Souza Todao Bernardino, Loyze Paola Oliveira de Lima, Luiz Aurelio de Campos Crispin. Centro de Genômica Funcional da ESALQ: Luiz Lehmann Coutinho, Ricardo Augusto Brassaloti, Raquel de Lello Rocha Campos Cassano. NGS Soluções Genômicas: Pilar Drummond Sampaio Corrêa Mariani. FZEA-USP Pirassununga: Mirele Daiana Poleti, Jessika Cristina Chagas Lesbon, Elisangela Chicaroni Mattos, Heidge Fukumasu. USP-Botucatu: Rejane Maria Tommasini Grotto, Jayme A. Souza-Neto, Guilherme Targino Valente, Patricia Akemi Assato, Felipe Allan da Silva da Costa, Bianca Cechetto Carlos. Mendelics: Bibiana Santos, João Paulo Kitajima, Erika Freitas, David Schlesinger. Hemocentro Ribeirão Preto: Simone Kashima, Evandra Strazza Rodrigues, Svetoslav Nanev Slavov, Elaine Vieira dos Santos, Rafael dos Santos Bezerra, Luiz Carlos Junior de Alcantara, Marta Giovanetti, Vagner Fonseca, Flavia Aburjaile, Rodrigo Tocantins Calado. FAMERP-SJRP: Cecília Artico Banho, Lívia Sacchetto, Fábio Sossai Possebon, Leila Sabrina Ullmann, Cíntia Bittar, Guilherme Campos, Helena Lage Ferreira, Jorge A. Petroli Marchesi, Maisa C. Pereira Parra, Marília Moraes, Paula Rahal, Paulo Inacio da Costa, João Pessoa Araújo Jr., Maurício Lacerda Nogueira. Prefeitura de Sao Paulo: Melissa Palmieri. |
| EPI_ISL_1967258, EPI_ISL_1967259, EPI_ISL_1967260, EPI_ISL_1967261, EPI_ISL_1967262, EPI_ISL_1967263, EPI_ISL_1967264, EPI_ISL_1967265, EPI_ISL_1967266, EPI_ISL_1967267, EPI_ISL_1967268, EPI_ISL_1967269, EPI_ISL_1967270, EPI_ISL_1967271, EPI_ISL_1967272, EPI_ISL_1967273, EPI_ISL_1967274 | | | |
| see above | SECAO CENTRO DE DIAGNOSTICO SECEDI | Instituto Butantan / Mendelics | Instituto Butantan: Dimas Tadeu Covas, Sandra Coccuzzo Sampaio, Maria Carolina Elias, José Salvatore Leister Patané, Vincent Louis Viala, Antonio Jorge Martins, Ricardo Haddad, Claudia Renata dos Santos Barros, Elaine Cristina Marqueze, Raul Machado Neto, Debora Botequio Moretti, Jardelina de Souza Todao Bernardino, Loyze Paola Oliveira de Lima, Luiz Aurelio de Campos Crispin. Centro de Genômica Funcional da ESALQ: Luiz Lehmann Coutinho, Ricardo Augusto Brassaloti, Raquel de Lello Rocha Campos Cassano. NGS Soluções Genômicas: Pilar Drummond Sampaio Corrêa Mariani. FZEA-USP Pirassununga: Mirele Daiana Poleti, Jessika Cristina Chagas Lesbon, Elisangela Chicaroni Mattos, Heidge Fukumasu. USP-Botucatu: Rejane Maria Tommasini Grotto, Jayme A. Souza-Neto, Guilherme Targino Valente, Patricia Akemi Assato, Felipe Allan da Silva da Costa, Bianca Cechetto Carlos. Mendelics: Bibiana Santos, João Paulo Kitajima, Erika Freitas, David Schlesinger. Hemocentro Ribeirão Preto: Simone Kashima, Evandra Strazza Rodrigues, Svetoslav Nanev Slavov, Elaine Vieira dos Santos, Rafael dos Santos Bezerra, Luiz Carlos Junior de Alcantara, Marta Giovanetti, Vagner Fonseca, Flavia Aburjaile, Rodrigo Tocantins Calado. FAMERP-SJRP: Cecília Artico Banho, Lívia Sacchetto, Fábio Sossai Possebon, Leila Sabrina Ullmann, Cíntia Bittar, Guilherme Campos, Helena Lage Ferreira, Jorge A. Petroli Marchesi, Maisa C. Pereira Parra, Marília Moraes, Paula Rahal, Paulo Inacio da Costa, João Pessoa Araújo Jr., Maurício Lacerda Nogueira. Prefeitura de Sao Paulo: Melissa Palmieri. |
| EPI_ISL_1967275, EPI_ISL_1967276 | PSF PARQUE EMILIA | Instituto Butantan / Mendelics | Instituto Butantan: Dimas Tadeu Covas, Sandra Coccuzzo Sampaio, Maria Carolina Elias, José Salvatore Leister Patané, Vincent Louis Viala, Antonio Jorge Martins, Ricardo Haddad, Claudia Renata dos Santos Barros, Elaine Cristina Marqueze, Raul Machado Neto, Debora Botequio Moretti, Jardelina de Souza Todao Bernardino, Loyze Paola Oliveira de Lima, Luiz Aurelio de Campos Crispin. Centro de Genômica Funcional da ESALQ: Luiz Lehmann Coutinho, Ricardo Augusto Brassaloti, Raquel de Lello Rocha Campos Cassano. NGS Soluções Genômicas: Pilar Drummond Sampaio Corrêa Mariani. FZEA-USP Pirassununga: Mirele Daiana Poleti, Jessika Cristina Chagas Lesbon, Elisangela Chicaroni Mattos, Heidge Fukumasu. USP-Botucatu: Rejane Maria Tommasini Grotto, Jayme A. Souza-Neto, Guilherme Targino Valente, Patricia Akemi Assato, Felipe Allan da Silva da Costa, Bianca Cechetto Carlos. Mendelics: Bibiana Santos, João Paulo Kitajima, Erika Freitas, David Schlesinger. Hemocentro Ribeirão Preto: Simone Kashima, Evandra Strazza Rodrigues, Svetoslav Nanev Slavov, Elaine Vieira dos Santos, Rafael dos Santos Bezerra, Luiz Carlos Junior de Alcantara, Marta Giovanetti, Vagner Fonseca, Flavia Aburjaile, Rodrigo Tocantins Calado. FAMERP-SJRP: Cecília Artico Banho, Lívia Sacchetto, Fábio Sossai Possebon, Leila Sabrina Ullmann, Cíntia Bittar, Guilherme Campos, Helena Lage Ferreira, Jorge A. Petroli Marchesi, Maisa C. Pereira Parra, Marília Moraes, Paula Rahal, Paulo Inacio da Costa, João Pessoa Araújo Jr., Maurício Lacerda Nogueira. Prefeitura de Sao Paulo: Melissa Palmieri. |
| EPI_ISL_1967277, EPI_ISL_1967278, EPI_ISL_1967279 | CENTRO DE SAUDE III TABATINGA | Instituto Butantan / Mendelics | Instituto Butantan: Dimas Tadeu Covas, Sandra Coccuzzo Sampaio, Maria Carolina Elias, José Salvatore Leister Patané, Vincent Louis Viala, Antonio Jorge Martins, Ricardo Haddad, Claudia Renata dos Santos Barros, Elaine Cristina Marqueze, Raul Machado Neto, Debora Botequio Moretti, Jardelina de Souza Todao Bernardino, Loyze Paola Oliveira de Lima, Luiz Aurelio de Campos Crispin. Centro de Genômica Funcional da ESALQ: Luiz Lehmann Coutinho, Ricardo Augusto Brassaloti, Raquel de Lello Rocha Campos Cassano. NGS Soluções Genômicas: Pilar Drummond Sampaio Corrêa Mariani. FZEA-USP Pirassununga: Mirele Daiana Poleti, Jessika Cristina Chagas Lesbon, Elisangela Chicaroni Mattos, Heidge Fukumasu. USP-Botucatu: Rejane Maria Tommasini Grotto, Jayme A. Souza-Neto, Guilherme Targino Valente, Patricia Akemi Assato, Felipe Allan da Silva da Costa, Bianca Cechetto Carlos. Mendelics: Bibiana Santos, João Paulo Kitajima, Erika Freitas, David Schlesinger. Hemocentro Ribeirão Preto: Simone Kashima, Evandra Strazza Rodrigues, Svetoslav Nanev Slavov, Elaine Vieira dos Santos, Rafael dos Santos Bezerra, Luiz Carlos Junior de Alcantara, Marta Giovanetti, Vagner Fonseca, Flavia Aburjaile, Rodrigo Tocantins Calado. FAMERP-SJRP: Cecília Artico Banho, Lívia Sacchetto, Fábio Sossai Possebon, Leila Sabrina Ullmann, Cíntia Bittar, Guilherme Campos, Helena Lage Ferreira, Jorge A. Petroli Marchesi, Maisa C. Pereira Parra, Marília Moraes, Paula Rahal, Paulo Inacio da Costa, João Pessoa Araújo Jr., Maurício Lacerda Nogueira. Prefeitura de Sao Paulo: Melissa Palmieri. |
| EPI_ISL_1967280 | UNIDADE BASICA DE SAUDE DE TAQUARAL CAETANO PITELLI | Instituto Butantan / Mendelics | Instituto Butantan: Dimas Tadeu Covas, Sandra Coccuzzo Sampaio, Maria Carolina Elias, José Salvatore Leister Patané, Vincent Louis Viala, Antonio Jorge Martins, Ricardo Haddad, Claudia Renata dos Santos Barros, Elaine Cristina Marqueze, Raul Machado Neto, Debora Botequio Moretti, Jardelina de Souza Todao Bernardino, Loyze Paola Oliveira de Lima, Luiz Aurelio de Campos Crispin. Centro de Genômica Funcional da ESALQ: Luiz Lehmann Coutinho, Ricardo Augusto Brassaloti, Raquel de Lello Rocha Campos Cassano. NGS Soluções Genômicas: Pilar Drummond Sampaio Corrêa Mariani. FZEA-USP Pirassununga: Mirele Daiana Poleti, Jessika Cristina Chagas Lesbon, Elisangela Chicaroni Mattos, Heidge Fukumasu. USP-Botucatu: Rejane Maria Tommasini Grotto, Jayme A. Souza-Neto, Guilherme Targino Valente, Patricia Akemi Assato, Felipe Allan da Silva da Costa, Bianca Cechetto Carlos. Mendelics: Bibiana Santos, João Paulo Kitajima, Erika Freitas, David Schlesinger. Hemocentro Ribeirão Preto: Simone Kashima, Evandra Strazza Rodrigues, Svetoslav Nanev Slavov, Elaine Vieira dos Santos, Rafael dos Santos Bezerra, Luiz Carlos Junior de Alcantara, Marta Giovanetti, Vagner Fonseca, Flavia Aburjaile, Rodrigo Tocantins Calado. FAMERP-SJRP: Cecília Artico Banho, Lívia Sacchetto, Fábio Sossai Possebon, Leila Sabrina Ullmann, Cíntia Bittar, Guilherme Campos, Helena Lage Ferreira, Jorge A. Petroli Marchesi, Maisa C. Pereira Parra, Marília Moraes, Paula Rahal, Paulo Inacio da Costa, João Pessoa Araújo Jr., Maurício Lacerda Nogueira. Prefeitura de Sao Paulo: Melissa Palmieri. |
| EPI_ISL_1967281 | PRONTO ATENDIMENTO MUNICIPAL | Instituto Butantan / Mendelics | Instituto Butantan: Dimas Tadeu Covas, Sandra Coccuzzo Sampaio, Maria Carolina Elias, José Salvatore Leister Patané, Vincent Louis Viala, Antonio Jorge Martins, Ricardo Haddad, Claudia Renata dos Santos Barros, Elaine Cristina Marqueze, Raul Machado Neto, Debora Botequio Moretti, Jardelina de Souza Todao Bernardino, Loyze Paola Oliveira de Lima, Luiz Aurelio de Campos Crispin. Centro de Genômica Funcional da ESALQ: Luiz Lehmann Coutinho, Ricardo Augusto Brassaloti, Raquel de Lello Rocha Campos Cassano. NGS Soluções Genômicas: Pilar Drummond Sampaio Corrêa Mariani. FZEA-USP Pirassununga: Mirele Daiana Poleti, Jessika Cristina Chagas Lesbon, Elisangela Chicaroni Mattos, Heidge Fukumasu. USP-Botucatu: Rejane Maria Tommasini Grotto, Jayme A. Souza-Neto, Guilherme Targino Valente, Patricia Akemi Assato, Felipe Allan da Silva da Costa, Bianca Cechetto Carlos. Mendelics: Bibiana Santos, João Paulo Kitajima, Erika Freitas, David Schlesinger. Hemocentro Ribeirão Preto: Simone Kashima, Evandra Strazza Rodrigues, Svetoslav Nanev Slavov, Elaine Vieira dos Santos, Rafael dos Santos Bezerra, Luiz Carlos Junior de Alcantara, Marta Giovanetti, Vagner Fonseca, Flavia Aburjaile, Rodrigo Tocantins Calado. FAMERP-SJRP: Cecília Artico Banho, Lívia Sacchetto, Fábio Sossai Possebon, Leila Sabrina Ullmann, Cíntia Bittar, Guilherme Campos, Helena Lage Ferreira, Jorge A. Petroli Marchesi, Maisa C. Pereira Parra, Marília Moraes, Paula Rahal, Paulo Inacio da Costa, João Pessoa Araújo Jr., Maurício Lacerda Nogueira. Prefeitura de Sao Paulo: Melissa Palmieri. |
| EPI_ISL_1967282 | SANTA CASA DE ATIBAIA PRO SAUDE | Instituto Butantan / Mendelics | Instituto Butantan: Dimas Tadeu Covas, Sandra Coccuzzo Sampaio, Maria Carolina Elias, José Salvatore Leister Patané, Vincent Louis Viala, Antonio Jorge Martins, Ricardo Haddad, Claudia Renata dos Santos Barros, Elaine Cristina Marqueze, Raul Machado Neto, Debora Botequio Moretti, Jardelina de |

| | | | |
|---|---|---|---|
| | | | Souza Todao Bernardino, Loyze Paola Oliveira de Lima, Luiz Aurelio de Campos Crispin. Centro de Genômica Funcional da ESALQ: Luiz Lehmann Coutinho, Ricardo Augusto Brassaloti, Raquel de Lello Rocha Campos Cassano. NGS Soluções Genômicas: Pilar Drummond Sampaio Corrêa Mariani. FZEA-USP Pirassununga: Mirele Daiana Poleti, Jessika Cristina Chagas Lesbon, Elisangela Chicaroni Mattos, Heidge Fukumasu. USP-Botucatu: Rejane Maria Tommasini Grotto, Jayme A. Souza-Neto, Guilherme Targino Valente, Patricia Akemi Assato, Felipe Allan da Silva da Costa, Bianca Cechetto Carlos. Mendelics: Bibiana Santos, João Paulo Kitajima, Erika Freitas, David Schlesinger. Hemocentro Ribeirão Preto: Simone Kashima, Evandra Strazza Rodrigues, Svetoslav Nanev Slavov, Elaine Vieira dos Santos, Rafael dos Santos Bezerra, Luiz Carlos Junior de Alcantara, Marta Giovanetti, Vagner Fonseca, Flavia Aburjaile, Rodrigo Tocantins Calado. FAMERP-SJRP: Cecília Artico Banho, Lívia Sacchetto, Fábio Sossai Possebon, Leila Sabrina Ullmann, Cíntia Bittar, Guilherme Campos, Helena Lage Ferreira, Jorge A. Petroli Marchesi, Maisa C. Pereira Parra, Marília Moraes, Paula Rahal, Paulo Inacio da Costa, João Pessoa Araújo Jr., Maurício Lacerda Nogueira. Prefeitura de Sao Paulo: Melissa Palmieri. |
| EPI_ISL_1967283 | VIGILANCIA EM SAUDE DE FCO MORATO | Instituto Butantan / Mendelics | Instituto Butantan: Dimas Tadeu Covas, Sandra Coccuzzo Sampaio, Maria Carolina Elias, José Salvatore Leister Patané, Vincent Louis Viala, Antonio Jorge Martins, Ricardo Haddad, Claudia Renata dos Santos Barros, Elaine Cristina Marqueze, Raul Machado Neto, Debora Botequio Moretti, Jardelina de Souza Todao Bernardino, Loyze Paola Oliveira de Lima, Luiz Aurelio de Campos Crispin. Centro de Genômica Funcional da ESALQ: Luiz Lehmann Coutinho, Ricardo Augusto Brassaloti, Raquel de Lello Rocha Campos Cassano. NGS Soluções Genômicas: Pilar Drummond Sampaio Corrêa Mariani. FZEA-USP Pirassununga: Mirele Daiana Poleti, Jessika Cristina Chagas Lesbon, Elisangela Chicaroni Mattos, Heidge Fukumasu. USP-Botucatu: Rejane Maria Tommasini Grotto, Jayme A. Souza-Neto, Guilherme Targino Valente, Patricia Akemi Assato, Felipe Allan da Silva da Costa, Bianca Cechetto Carlos. Mendelics: Bibiana Santos, João Paulo Kitajima, Erika Freitas, David Schlesinger. Hemocentro Ribeirão Preto: Simone Kashima, Evandra Strazza Rodrigues, Svetoslav Nanev Slavov, Elaine Vieira dos Santos, Rafael dos Santos Bezerra, Luiz Carlos Junior de Alcantara, Marta Giovanetti, Vagner Fonseca, Flavia Aburjaile, Rodrigo Tocantins Calado. FAMERP-SJRP: Cecília Artico Banho, Lívia Sacchetto, Fábio Sossai Possebon, Leila Sabrina Ullmann, Cíntia Bittar, Guilherme Campos, Helena Lage Ferreira, Jorge A. Petroli Marchesi, Maisa C. Pereira Parra, Marília Moraes, Paula Rahal, Paulo Inacio da Costa, João Pessoa Araújo Jr., Maurício Lacerda Nogueira. Prefeitura de Sao Paulo: Melissa Palmieri. |
| EPI_ISL_1967284 | UBS RECREIO SAO JORGE | Instituto Butantan / Mendelics | Instituto Butantan: Dimas Tadeu Covas, Sandra Coccuzzo Sampaio, Maria Carolina Elias, José Salvatore Leister Patané, Vincent Louis Viala, Antonio Jorge Martins, Ricardo Haddad, Claudia Renata dos Santos Barros, Elaine Cristina Marqueze, Raul Machado Neto, Debora Botequio Moretti, Jardelina de Souza Todao Bernardino, Loyze Paola Oliveira de Lima, Luiz Aurelio de Campos Crispin. Centro de Genômica Funcional da ESALQ: Luiz Lehmann Coutinho, Ricardo Augusto Brassaloti, Raquel de Lello Rocha Campos Cassano. NGS Soluções Genômicas: Pilar Drummond Sampaio Corrêa Mariani. FZEA-USP Pirassununga: Mirele Daiana Poleti, Jessika Cristina Chagas Lesbon, Elisangela Chicaroni Mattos, Heidge Fukumasu. USP-Botucatu: Rejane Maria Tommasini Grotto, Jayme A. Souza-Neto, Guilherme Targino Valente, Patricia Akemi Assato, Felipe Allan da Silva da Costa, Bianca Cechetto Carlos. Mendelics: Bibiana Santos, João Paulo Kitajima, Erika Freitas, David Schlesinger. Hemocentro Ribeirão Preto: Simone Kashima, Evandra Strazza Rodrigues, Svetoslav Nanev Slavov, Elaine Vieira dos Santos, Rafael dos Santos Bezerra, Luiz Carlos Junior de Alcantara, Marta Giovanetti, Vagner Fonseca, Flavia Aburjaile, Rodrigo Tocantins Calado. FAMERP-SJRP: Cecília Artico Banho, Lívia Sacchetto, Fábio Sossai Possebon, Leila Sabrina Ullmann, Cíntia Bittar, Guilherme Campos, Helena Lage Ferreira, Jorge A. Petroli Marchesi, Maisa C. Pereira Parra, Marília Moraes, Paula Rahal, Paulo Inacio da Costa, João Pessoa Araújo Jr., Maurício Lacerda Nogueira. Prefeitura de Sao Paulo: Melissa Palmieri. |
| EPI_ISL_1967285, EPI_ISL_1967286 | UBS JARDIM CABUCU | Instituto Butantan / Mendelics | Instituto Butantan: Dimas Tadeu Covas, Sandra Coccuzzo Sampaio, Maria Carolina Elias, José Salvatore Leister Patané, Vincent Louis Viala, Antonio Jorge Martins, Ricardo Haddad, Claudia Renata dos Santos Barros, Elaine Cristina Marqueze, Raul Machado Neto, Debora Botequio Moretti, Jardelina de Souza Todao Bernardino, Loyze Paola Oliveira de Lima, Luiz Aurelio de Campos Crispin. Centro de Genômica Funcional da ESALQ: Luiz Lehmann Coutinho, Ricardo Augusto Brassaloti, Raquel de Lello Rocha Campos Cassano. NGS Soluções Genômicas: Pilar Drummond Sampaio Corrêa Mariani. FZEA-USP Pirassununga: Mirele Daiana Poleti, Jessika Cristina Chagas Lesbon, Elisangela Chicaroni Mattos, Heidge Fukumasu. USP-Botucatu: Rejane Maria Tommasini Grotto, Jayme A. Souza-Neto, Guilherme Targino Valente, Patricia Akemi Assato, Felipe Allan da Silva da Costa, Bianca Cechetto Carlos. Mendelics: Bibiana Santos, João Paulo Kitajima, Erika Freitas, David Schlesinger. Hemocentro Ribeirão Preto: Simone Kashima, Evandra Strazza Rodrigues, Svetoslav Nanev Slavov, Elaine Vieira dos Santos, Rafael dos Santos Bezerra, Luiz Carlos Junior de Alcantara, Marta Giovanetti, Vagner Fonseca, Flavia Aburjaile, Rodrigo Tocantins Calado. FAMERP-SJRP: Cecília Artico Banho, Lívia Sacchetto, Fábio Sossai Possebon, Leila Sabrina Ullmann, Cíntia Bittar, Guilherme Campos, Helena Lage Ferreira, Jorge A. Petroli Marchesi, Maisa C. Pereira Parra, Marília Moraes, Paula Rahal, Paulo Inacio da Costa, João Pessoa Araújo Jr., Maurício Lacerda Nogueira. Prefeitura de Sao Paulo: Melissa Palmieri. |
| EPI_ISL_1967287 | UBS VILA GALVAO | Instituto Butantan / Mendelics | Instituto Butantan: Dimas Tadeu Covas, Sandra Coccuzzo Sampaio, Maria Carolina Elias, José Salvatore Leister Patané, Vincent Louis Viala, Antonio Jorge Martins, Ricardo Haddad, Claudia Renata dos Santos Barros, Elaine Cristina Marqueze, Raul Machado Neto, Debora Botequio Moretti, Jardelina de Souza Todao Bernardino, Loyze Paola Oliveira de Lima, Luiz Aurelio de Campos Crispin. Centro de Genômica Funcional da ESALQ: Luiz Lehmann Coutinho, Ricardo Augusto Brassaloti, Raquel de Lello Rocha Campos Cassano. NGS Soluções Genômicas: Pilar Drummond Sampaio Corrêa Mariani. FZEA-USP Pirassununga: Mirele Daiana Poleti, Jessika Cristina Chagas Lesbon, Elisangela Chicaroni Mattos, Heidge Fukumasu. USP-Botucatu: Rejane Maria Tommasini Grotto, Jayme A. Souza-Neto, Guilherme Targino Valente, Patricia Akemi Assato, Felipe Allan da Silva da Costa, Bianca Cechetto Carlos. Mendelics: Bibiana Santos, João Paulo Kitajima, Erika Freitas, David Schlesinger. Hemocentro Ribeirão Preto: Simone Kashima, Evandra Strazza Rodrigues, Svetoslav Nanev Slavov, Elaine Vieira dos Santos, Rafael dos Santos Bezerra, Luiz Carlos Junior de Alcantara, Marta Giovanetti, Vagner Fonseca, Flavia Aburjaile, Rodrigo Tocantins Calado. FAMERP-SJRP: Cecília Artico Banho, Lívia Sacchetto, Fábio Sossai Possebon, Leila Sabrina Ullmann, Cíntia Bittar, Guilherme Campos, Helena Lage Ferreira, Jorge A. Petroli Marchesi, Maisa C. Pereira Parra, Marília Moraes, Paula Rahal, Paulo Inacio da Costa, João Pessoa Araújo Jr., Maurício Lacerda Nogueira. Prefeitura de Sao Paulo: Melissa Palmieri. |
| EPI_ISL_1967288 | UBS CAVADAS | Instituto Butantan / Mendelics | Instituto Butantan: Dimas Tadeu Covas, Sandra Coccuzzo Sampaio, Maria Carolina Elias, José Salvatore Leister Patané, Vincent Louis Viala, Antonio Jorge Martins, Ricardo Haddad, Claudia Renata dos Santos Barros, Elaine Cristina Marqueze, Raul Machado Neto, Debora Botequio Moretti, Jardelina de Souza Todao Bernardino, Loyze Paola Oliveira de Lima, Luiz Aurelio de Campos Crispin. Centro de Genômica Funcional da ESALQ: Luiz Lehmann Coutinho, Ricardo Augusto Brassaloti, Raquel de Lello Rocha Campos Cassano. NGS Soluções Genômicas: Pilar Drummond Sampaio Corrêa Mariani. FZEA-USP Pirassununga: Mirele Daiana Poleti, Jessika Cristina Chagas Lesbon, Elisangela Chicaroni Mattos, Heidge Fukumasu. USP-Botucatu: Rejane Maria Tommasini Grotto, Jayme A. Souza-Neto, Guilherme Targino Valente, Patricia Akemi Assato, Felipe Allan da Silva da Costa, Bianca Cechetto Carlos. Mendelics: Bibiana Santos, João Paulo Kitajima, Erika Freitas, David Schlesinger. Hemocentro Ribeirão Preto: Simone Kashima, Evandra Strazza Rodrigues, Svetoslav Nanev Slavov, Elaine Vieira dos Santos, Rafael dos Santos Bezerra, Luiz Carlos Junior de Alcantara, Marta Giovanetti, Vagner Fonseca, Flavia Aburjaile, Rodrigo Tocantins Calado. FAMERP-SJRP: Cecília Artico Banho, Lívia Sacchetto, Fábio Sossai Possebon, Leila Sabrina Ullmann, Cíntia Bittar, Guilherme Campos, Helena Lage Ferreira, Jorge A. Petroli Marchesi, Maisa C. Pereira Parra, Marília Moraes, Paula Rahal, Paulo Inacio da Costa, João Pessoa Araújo Jr., Maurício Lacerda Nogueira. Prefeitura de Sao Paulo: Melissa Palmieri. |
| EPI_ISL_1967289, EPI_ISL_1967290, EPI_ISL_1967291, EPI_ISL_1967292, EPI_ISL_1967293 | UNIDADE RESPIRATORIA NOVA HORTOLANDIA | Instituto Butantan / Mendelics | Instituto Butantan: Dimas Tadeu Covas, Sandra Coccuzzo Sampaio, Maria Carolina Elias, José Salvatore Leister Patané, Vincent Louis Viala, Antonio Jorge Martins, Ricardo Haddad, Claudia Renata dos Santos Barros, Elaine Cristina Marqueze, Raul Machado Neto, Debora Botequio Moretti, Jardelina de Souza Todao Bernardino, Loyze Paola Oliveira de Lima, Luiz Aurelio de Campos Crispin. Centro de Genômica Funcional da ESALQ: Luiz Lehmann Coutinho, Ricardo Augusto Brassaloti, Raquel de Lello Rocha Campos Cassano. NGS Soluções Genômicas: Pilar Drummond Sampaio Corrêa Mariani. FZEA-USP Pirassununga: Mirele Daiana Poleti, Jessika Cristina Chagas Lesbon, Elisangela Chicaroni Mattos, Heidge Fukumasu. USP-Botucatu: Rejane Maria Tommasini Grotto, Jayme A. Souza-Neto, Guilherme Targino Valente, Patricia Akemi Assato, Felipe Allan da Silva da Costa, Bianca Cechetto Carlos. Mendelics: Bibiana Santos, João Paulo Kitajima, Erika Freitas, David Schlesinger. Hemocentro Ribeirão Preto: Simone Kashima, Evandra Strazza Rodrigues, Svetoslav Nanev Slavov, Elaine Vieira dos Santos, Rafael dos Santos Bezerra, Luiz Carlos Junior de Alcantara, Marta Giovanetti, Vagner Fonseca, Flavia Aburjaile, Rodrigo Tocantins Calado. FAMERP-SJRP: Cecília Artico Banho, Lívia Sacchetto, Fábio Sossai Possebon, Leila Sabrina Ullmann, Cíntia Bittar, Guilherme Campos, Helena Lage Ferreira, Jorge A. Petroli Marchesi, Maisa C. Pereira Parra, Marília Moraes, Paula Rahal, Paulo Inacio da Costa, João Pessoa Araújo Jr., Maurício Lacerda Nogueira. Prefeitura de Sao Paulo: Melissa Palmieri. |
| EPI_ISL_1967294, EPI_ISL_1967295 | UPA II JUNDIAI | Instituto Butantan / Mendelics | Instituto Butantan: Dimas Tadeu Covas, Sandra Coccuzzo Sampaio, Maria Carolina Elias, José Salvatore Leister Patané, Vincent Louis Viala, Antonio Jorge Martins, Ricardo Haddad, Claudia Renata dos Santos Barros, Elaine Cristina Marqueze, Raul Machado Neto, Debora Botequio Moretti, Jardelina de Souza Todao Bernardino, Loyze Paola Oliveira de Lima, Luiz Aurelio de Campos Crispin. Centro de Genômica Funcional da ESALQ: Luiz Lehmann Coutinho, Ricardo Augusto Brassaloti, Raquel de Lello Rocha Campos Cassano. NGS Soluções Genômicas: Pilar Drummond Sampaio Corrêa Mariani. FZEA-USP Pirassununga: Mirele Daiana Poleti, Jessika Cristina Chagas Lesbon, Elisangela Chicaroni Mattos, Heidge Fukumasu. USP-Botucatu: Rejane Maria Tommasini Grotto, Jayme A. Souza-Neto, Guilherme Targino Valente, Patricia Akemi Assato, Felipe Allan da Silva da Costa, Bianca Cechetto Carlos. Mendelics: Bibiana Santos, João Paulo Kitajima, Erika Freitas, David Schlesinger. Hemocentro Ribeirão Preto: Simone Kashima, Evandra Strazza Rodrigues, Svetoslav Nanev Slavov, Elaine Vieira dos Santos, Rafael dos Santos Bezerra, Luiz Carlos Junior de Alcantara, Marta Giovanetti, Vagner Fonseca, Flavia Aburjaile, Rodrigo Tocantins Calado. FAMERP-SJRP: Cecília Artico Banho, Lívia Sacchetto, Fábio Sossai Possebon, Leila Sabrina |

| | | | |
|---|---|---|---|
| | | | Ullmann, Cíntia Bittar, Guilherme Campos, Helena Lage Ferreira, Jorge A. Petroli Marchesi, Maisa C. Pereira Parra, Marília Moraes, Paula Rahal, Paulo Inacio da Costa, João Pessoa Araújo Jr., Maurício Lacerda Nogueira. Prefeitura de Sao Paulo: Melissa Palmieri. |
| EPI_ISL_1967296, EPI_ISL_1967297 | AMBULATORIO DE ESPECIALIDADE V E MOGI MIRIM | Instituto Butantan / Mendelics | Instituto Butantan: Dimas Tadeu Covas, Sandra Coccuzzo Sampaio, Maria Carolina Elias, José Salvatore Leister Patané, Vincent Louis Viala, Antonio Jorge Martins, Ricardo Haddad, Claudia Renata dos Santos Barros, Elaine Cristina Marqueze, Raul Machado Neto, Debora Botequio Moretti, Jardelina de Souza Todao Bernardino, Loyze Paola Oliveira de Lima, Luiz Aurelio de Campos Crispin. Centro de Genômica Funcional da ESALQ: Luiz Lehmann Coutinho, Ricardo Augusto Brassaloti, Raquel de Lello Rocha Campos Cassano. NGS Soluções Genômicas: Pilar Drummond Sampaio Corrêa Mariani. FZEA-USP Pirassununga: Mirele Daiana Poleti, Jessika Cristina Chagas Lesbon, Elisangela Chicaroni Mattos, Heidge Fukumasu. USP-Botucatu: Rejane Maria Tommasini Grotto, Jayme A. Souza-Neto, Guilherme Targino Valente, Patricia Akemi Assato, Felipe Allan da Silva da Costa, Bianca Cechetto Carlos. Mendelics: Bibiana Santos, João Paulo Kitajima, Erika Freitas, David Schlesinger. Hemocentro Ribeirão Preto: Simone Kashima, Evandra Strazza Rodrigues, Svetoslav Nanev Slavov, Elaine Vieira dos Santos, Rafael dos Santos Bezerra, Luiz Carlos Junior de Alcantara, Marta Giovanetti, Vagner Fonseca, Flavia Aburjaile, Rodrigo Tocantins Calado. FAMERP-SJRP: Cecília Artico Banho, Lívia Sacchetto, Fábio Sossai Possebon, Leila Sabrina Ullmann, Cíntia Bittar, Guilherme Campos, Helena Lage Ferreira, Jorge A. Petroli Marchesi, Maisa C. Pereira Parra, Marília Moraes, Paula Rahal, Paulo Inacio da Costa, João Pessoa Araújo Jr., Maurício Lacerda Nogueira. Prefeitura de Sao Paulo: Melissa Palmieri. |
| EPI_ISL_1967298 | UPA VILA SANTA CATARINA | Instituto Butantan / Mendelics | Instituto Butantan: Dimas Tadeu Covas, Sandra Coccuzzo Sampaio, Maria Carolina Elias, José Salvatore Leister Patané, Vincent Louis Viala, Antonio Jorge Martins, Ricardo Haddad, Claudia Renata dos Santos Barros, Elaine Cristina Marqueze, Raul Machado Neto, Debora Botequio Moretti, Jardelina de Souza Todao Bernardino, Loyze Paola Oliveira de Lima, Luiz Aurelio de Campos Crispin. Centro de Genômica Funcional da ESALQ: Luiz Lehmann Coutinho, Ricardo Augusto Brassaloti, Raquel de Lello Rocha Campos Cassano. NGS Soluções Genômicas: Pilar Drummond Sampaio Corrêa Mariani. FZEA-USP Pirassununga: Mirele Daiana Poleti, Jessika Cristina Chagas Lesbon, Elisangela Chicaroni Mattos, Heidge Fukumasu. USP-Botucatu: Rejane Maria Tommasini Grotto, Jayme A. Souza-Neto, Guilherme Targino Valente, Patricia Akemi Assato, Felipe Allan da Silva da Costa, Bianca Cechetto Carlos. Mendelics: Bibiana Santos, João Paulo Kitajima, Erika Freitas, David Schlesinger. Hemocentro Ribeirão Preto: Simone Kashima, Evandra Strazza Rodrigues, Svetoslav Nanev Slavov, Elaine Vieira dos Santos, Rafael dos Santos Bezerra, Luiz Carlos Junior de Alcantara, Marta Giovanetti, Vagner Fonseca, Flavia Aburjaile, Rodrigo Tocantins Calado. FAMERP-SJRP: Cecília Artico Banho, Lívia Sacchetto, Fábio Sossai Possebon, Leila Sabrina Ullmann, Cíntia Bittar, Guilherme Campos, Helena Lage Ferreira, Jorge A. Petroli Marchesi, Maisa C. Pereira Parra, Marília Moraes, Paula Rahal, Paulo Inacio da Costa, João Pessoa Araújo Jr., Maurício Lacerda Nogueira. Prefeitura de Sao Paulo: Melissa Palmieri. |
| EPI_ISL_1967299 | HOSPITAL E PRONTO SOCORRO PORTINARI | Instituto Butantan / Mendelics | Instituto Butantan: Dimas Tadeu Covas, Sandra Coccuzzo Sampaio, Maria Carolina Elias, José Salvatore Leister Patané, Vincent Louis Viala, Antonio Jorge Martins, Ricardo Haddad, Claudia Renata dos Santos Barros, Elaine Cristina Marqueze, Raul Machado Neto, Debora Botequio Moretti, Jardelina de Souza Todao Bernardino, Loyze Paola Oliveira de Lima, Luiz Aurelio de Campos Crispin. Centro de Genômica Funcional da ESALQ: Luiz Lehmann Coutinho, Ricardo Augusto Brassaloti, Raquel de Lello Rocha Campos Cassano. NGS Soluções Genômicas: Pilar Drummond Sampaio Corrêa Mariani. FZEA-USP Pirassununga: Mirele Daiana Poleti, Jessika Cristina Chagas Lesbon, Elisangela Chicaroni Mattos, Heidge Fukumasu. USP-Botucatu: Rejane Maria Tommasini Grotto, Jayme A. Souza-Neto, Guilherme Targino Valente, Patricia Akemi Assato, Felipe Allan da Silva da Costa, Bianca Cechetto Carlos. Mendelics: Bibiana Santos, João Paulo Kitajima, Erika Freitas, David Schlesinger. Hemocentro Ribeirão Preto: Simone Kashima, Evandra Strazza Rodrigues, Svetoslav Nanev Slavov, Elaine Vieira dos Santos, Rafael dos Santos Bezerra, Luiz Carlos Junior de Alcantara, Marta Giovanetti, Vagner Fonseca, Flavia Aburjaile, Rodrigo Tocantins Calado. FAMERP-SJRP: Cecília Artico Banho, Lívia Sacchetto, Fábio Sossai Possebon, Leila Sabrina Ullmann, Cíntia Bittar, Guilherme Campos, Helena Lage Ferreira, Jorge A. Petroli Marchesi, Maisa C. Pereira Parra, Marília Moraes, Paula Rahal, Paulo Inacio da Costa, João Pessoa Araújo Jr., Maurício Lacerda Nogueira. Prefeitura de Sao Paulo: Melissa Palmieri. |
| EPI_ISL_1967300, EPI_ISL_1967301, EPI_ISL_1967302, EPI_ISL_1967303 | SECRETARIA MUNICIPAL DE SAUDE | Instituto Butantan / Mendelics | Instituto Butantan: Dimas Tadeu Covas, Sandra Coccuzzo Sampaio, Maria Carolina Elias, José Salvatore Leister Patané, Vincent Louis Viala, Antonio Jorge Martins, Ricardo Haddad, Claudia Renata dos Santos Barros, Elaine Cristina Marqueze, Raul Machado Neto, Debora Botequio Moretti, Jardelina de Souza Todao Bernardino, Loyze Paola Oliveira de Lima, Luiz Aurelio de Campos Crispin. Centro de Genômica Funcional da ESALQ: Luiz Lehmann Coutinho, Ricardo Augusto Brassaloti, Raquel de Lello Rocha Campos Cassano. NGS Soluções Genômicas: Pilar Drummond Sampaio Corrêa Mariani. FZEA-USP Pirassununga: Mirele Daiana Poleti, Jessika Cristina Chagas Lesbon, Elisangela Chicaroni Mattos, Heidge Fukumasu. USP-Botucatu: Rejane Maria Tommasini Grotto, Jayme A. Souza-Neto, Guilherme Targino Valente, Patricia Akemi Assato, Felipe Allan da Silva da Costa, Bianca Cechetto Carlos. Mendelics: Bibiana Santos, João Paulo Kitajima, Erika Freitas, David Schlesinger. Hemocentro Ribeirão Preto: Simone Kashima, Evandra Strazza Rodrigues, Svetoslav Nanev Slavov, Elaine Vieira dos Santos, Rafael dos Santos Bezerra, Luiz Carlos Junior de Alcantara, Marta Giovanetti, Vagner Fonseca, Flavia Aburjaile, Rodrigo Tocantins Calado. FAMERP-SJRP: Cecília Artico Banho, Lívia Sacchetto, Fábio Sossai Possebon, Leila Sabrina Ullmann, Cíntia Bittar, Guilherme Campos, Helena Lage Ferreira, Jorge A. Petroli Marchesi, Maisa C. Pereira Parra, Marília Moraes, Paula Rahal, Paulo Inacio da Costa, João Pessoa Araújo Jr., Maurício Lacerda Nogueira. Prefeitura de Sao Paulo: Melissa Palmieri. |
| EPI_ISL_1967304, EPI_ISL_1967305 | PSF PARQUE EMILIA | Instituto Butantan / Mendelics | Instituto Butantan: Dimas Tadeu Covas, Sandra Coccuzzo Sampaio, Maria Carolina Elias, José Salvatore Leister Patané, Vincent Louis Viala, Antonio Jorge Martins, Ricardo Haddad, Claudia Renata dos Santos Barros, Elaine Cristina Marqueze, Raul Machado Neto, Debora Botequio Moretti, Jardelina de Souza Todao Bernardino, Loyze Paola Oliveira de Lima, Luiz Aurelio de Campos Crispin. Centro de Genômica Funcional da ESALQ: Luiz Lehmann Coutinho, Ricardo Augusto Brassaloti, Raquel de Lello Rocha Campos Cassano. NGS Soluções Genômicas: Pilar Drummond Sampaio Corrêa Mariani. FZEA-USP Pirassununga: Mirele Daiana Poleti, Jessika Cristina Chagas Lesbon, Elisangela Chicaroni Mattos, Heidge Fukumasu. USP-Botucatu: Rejane Maria Tommasini Grotto, Jayme A. Souza-Neto, Guilherme Targino Valente, Patricia Akemi Assato, Felipe Allan da Silva da Costa, Bianca Cechetto Carlos. Mendelics: Bibiana Santos, João Paulo Kitajima, Erika Freitas, David Schlesinger. Hemocentro Ribeirão Preto: Simone Kashima, Evandra Strazza Rodrigues, Svetoslav Nanev Slavov, Elaine Vieira dos Santos, Rafael dos Santos Bezerra, Luiz Carlos Junior de Alcantara, Marta Giovanetti, Vagner Fonseca, Flavia Aburjaile, Rodrigo Tocantins Calado. FAMERP-SJRP: Cecília Artico Banho, Lívia Sacchetto, Fábio Sossai Possebon, Leila Sabrina Ullmann, Cíntia Bittar, Guilherme Campos, Helena Lage Ferreira, Jorge A. Petroli Marchesi, Maisa C. Pereira Parra, Marília Moraes, Paula Rahal, Paulo Inacio da Costa, João Pessoa Araújo Jr., Maurício Lacerda Nogueira. Prefeitura de Sao Paulo: Melissa Palmieri. |
| EPI_ISL_1967307, EPI_ISL_1967308, EPI_ISL_1967309 | UPA DR AKIRA TADA | Instituto Butantan / Mendelics | Instituto Butantan: Dimas Tadeu Covas, Sandra Coccuzzo Sampaio, Maria Carolina Elias, José Salvatore Leister Patané, Vincent Louis Viala, Antonio Jorge Martins, Ricardo Haddad, Claudia Renata dos Santos Barros, Elaine Cristina Marqueze, Raul Machado Neto, Debora Botequio Moretti, Jardelina de Souza Todao Bernardino, Loyze Paola Oliveira de Lima, Luiz Aurelio de Campos Crispin. Centro de Genômica Funcional da ESALQ: Luiz Lehmann Coutinho, Ricardo Augusto Brassaloti, Raquel de Lello Rocha Campos Cassano. NGS Soluções Genômicas: Pilar Drummond Sampaio Corrêa Mariani. FZEA-USP Pirassununga: Mirele Daiana Poleti, Jessika Cristina Chagas Lesbon, Elisangela Chicaroni Mattos, Heidge Fukumasu. USP-Botucatu: Rejane Maria Tommasini Grotto, Jayme A. Souza-Neto, Guilherme Targino Valente, Patricia Akemi Assato, Felipe Allan da Silva da Costa, Bianca Cechetto Carlos. Mendelics: Bibiana Santos, João Paulo Kitajima, Erika Freitas, David Schlesinger. Hemocentro Ribeirão Preto: Simone Kashima, Evandra Strazza Rodrigues, Svetoslav Nanev Slavov, Elaine Vieira dos Santos, Rafael dos Santos Bezerra, Luiz Carlos Junior de Alcantara, Marta Giovanetti, Vagner Fonseca, Flavia Aburjaile, Rodrigo Tocantins Calado. FAMERP-SJRP: Cecília Artico Banho, Lívia Sacchetto, Fábio Sossai Possebon, Leila Sabrina Ullmann, Cíntia Bittar, Guilherme Campos, Helena Lage Ferreira, Jorge A. Petroli Marchesi, Maisa C. Pereira Parra, Marília Moraes, Paula Rahal, Paulo Inacio da Costa, João Pessoa Araújo Jr., Maurício Lacerda Nogueira. Prefeitura de Sao Paulo: Melissa Palmieri. |
| EPI_ISL_1967310, EPI_ISL_1967311 | PRONTO SOCORRO MUNICIPAL TAMBAU | Instituto Butantan / Mendelics | Instituto Butantan: Dimas Tadeu Covas, Sandra Coccuzzo Sampaio, Maria Carolina Elias, José Salvatore Leister Patané, Vincent Louis Viala, Antonio Jorge Martins, Ricardo Haddad, Claudia Renata dos Santos Barros, Elaine Cristina Marqueze, Raul Machado Neto, Debora Botequio Moretti, Jardelina de Souza Todao Bernardino, Loyze Paola Oliveira de Lima, Luiz Aurelio de Campos Crispin. Centro de Genômica Funcional da ESALQ: Luiz Lehmann Coutinho, Ricardo Augusto Brassaloti, Raquel de Lello Rocha Campos Cassano. NGS Soluções Genômicas: Pilar Drummond Sampaio Corrêa Mariani. FZEA-USP Pirassununga: Mirele Daiana Poleti, Jessika Cristina Chagas Lesbon, Elisangela Chicaroni Mattos, Heidge Fukumasu. USP-Botucatu: Rejane Maria Tommasini Grotto, Jayme A. Souza-Neto, Guilherme Targino Valente, Patricia Akemi Assato, Felipe Allan da Silva da Costa, Bianca Cechetto Carlos. Mendelics: Bibiana Santos, João Paulo Kitajima, Erika Freitas, David Schlesinger. Hemocentro Ribeirão Preto: Simone Kashima, Evandra Strazza Rodrigues, Svetoslav Nanev Slavov, Elaine Vieira dos Santos, Rafael dos Santos Bezerra, Luiz Carlos Junior de Alcantara, Marta Giovanetti, Vagner Fonseca, Flavia Aburjaile, Rodrigo Tocantins Calado. FAMERP-SJRP: Cecília Artico Banho, Lívia Sacchetto, Fábio Sossai Possebon, Leila Sabrina Ullmann, Cíntia Bittar, Guilherme Campos, Helena Lage Ferreira, Jorge A. Petroli Marchesi, Maisa C. Pereira Parra, Marília Moraes, Paula Rahal, Paulo Inacio da Costa, João Pessoa Araújo Jr., Maurício Lacerda Nogueira. Prefeitura de Sao Paulo: Melissa Palmieri. |
| EPI_ISL_1967312, EPI_ISL_1967313, EPI_ISL_1967314 | UNIDADE DE PRONTO ATENDIMENTO DE VARZEA PAULISTA UPA II | Instituto Butantan / Mendelics | Instituto Butantan: Dimas Tadeu Covas, Sandra Coccuzzo Sampaio, Maria Carolina Elias, José Salvatore Leister Patané, Vincent Louis Viala, Antonio Jorge Martins, Ricardo Haddad, Claudia Renata dos Santos Barros, Elaine Cristina Marqueze, Raul Machado Neto, Debora Botequio Moretti, Jardelina de Souza Todao Bernardino, Loyze Paola Oliveira de Lima, Luiz Aurelio de Campos Crispin. Centro de Genômica Funcional da ESALQ: Luiz Lehmann Coutinho, Ricardo Augusto Brassaloti, Raquel de Lello Rocha Campos Cassano. NGS Soluções Genômicas: Pilar Drummond Sampaio Corrêa Mariani. |

| | | | |
|---|---|---|---|
| | | | FZEA-USP Pirassununga: Mirele Daiana Poleti, Jessika Cristina Chagas Lesbon, Elisangela Chicaroni Mattos, Heidge Fukumasu. USP-Botucatu: Rejane Maria Tommasini Grotto, Jayme A. Souza-Neto, Guilherme Targino Valente, Patricia Akemi Assato, Felipe Allan da Silva da Costa, Bianca Cechetto Carlos. Mendelics: Bibiana Santos, João Paulo Kitajima, Erika Freitas, David Schlesinger. Hemocentro Ribeirão Preto: Simone Kashima, Evandra Strazza Rodrigues, Svetoslav Nanev Slavov, Elaine Vieira dos Santos, Rafael dos Santos Bezerra, Luiz Carlos Junior de Alcantara, Marta Giovanetti, Vagner Fonseca, Flavia Aburjaile, Rodrigo Tocantins Calado. FAMERP-SJRP: Cecília Artico Banho, Lívia Sacchetto, Fábio Sossai Possebon, Leila Sabrina Ullmann, Cíntia Bittar, Guilherme Campos, Helena Lage Ferreira, Jorge A. Petroli Marchesi, Maisa C. Pereira Parra, Marília Moraes, Paula Rahal, Paulo Inacio da Costa, João Pessoa Araújo Jr., Maurício Lacerda Nogueira. Prefeitura de Sao Paulo: Melissa Palmieri. |
| EPI_ISL_2003111 | Pronto Socorro Central de Bauru | Instituto Adolfo Lutz, Interdiciplinary Procedures Center, Strategic Laboratory | Claudio Tavares Sacchi, Claudia Regina Gonçalves, Erica Valessa Ramos Gomes, Karoline Rodrigues Campos, Caio Vinicius Dias Lopes, Leonardo Jose Tadeu de Araujo |
| EPI_ISL_2003112 | CS de Paulo de Faria | Instituto Adolfo Lutz, Interdiciplinary Procedures Center, Strategic Laboratory | Claudio Tavares Sacchi, Claudia Regina Gonçalves, Erica Valessa Ramos Gomes, Karoline Rodrigues Campos, Caio Vinicius Dias Lopes, Leonardo Jose Tadeu de Araujo |
| EPI_ISL_2003113 | Hospital Municipal Dr Mario Gatti Campinas | Instituto Adolfo Lutz, Interdiciplinary Procedures Center, Strategic Laboratory | Claudio Tavares Sacchi, Claudia Regina Gonçalves, Erica Valessa Ramos Gomes, Karoline Rodrigues Campos, Caio Vinicius Dias Lopes, Leonardo Jose Tadeu de Araujo |
| EPI_ISL_2003114, EPI_ISL_2003115, EPI_ISL_2003116, EPI_ISL_2003117, EPI_ISL_2003118, EPI_ISL_2003119, EPI_ISL_2003120, EPI_ISL_2003121, EPI_ISL_2003122, EPI_ISL_2003123, EPI_ISL_2003124, EPI_ISL_2003125, EPI_ISL_2003126, EPI_ISL_2003127, EPI_ISL_2003128, EPI_ISL_2003129, EPI_ISL_2003130, EPI_ISL_2003131, EPI_ISL_2003132, EPI_ISL_2003133 | | | |
| see above | Instituto Adolfo Lutz Central | Instituto Adolfo Lutz, Interdiciplinary Procedures Center, Strategic Laboratory | Claudio Tavares Sacchi, Claudia Regina Gonçalves, Erica Valessa Ramos Gomes, Karoline Rodrigues Campos, Caio Vinicius Dias Lopes, Leonardo Jose Tadeu de Araujo |
| EPI_ISL_2003134, EPI_ISL_2003135, EPI_ISL_2003136, EPI_ISL_2003137, EPI_ISL_2003138, EPI_ISL_2003139, EPI_ISL_2003140, EPI_ISL_2003141, EPI_ISL_2003142, EPI_ISL_2003143, EPI_ISL_2003144, EPI_ISL_2003145, EPI_ISL_2003146, EPI_ISL_2003147, EPI_ISL_2003148, EPI_ISL_2003149 | | | |
| see above | Instituto Adolfo Lutz - Regional de Santos | Instituto Adolfo Lutz, Interdiciplinary Procedures Center, Strategic Laboratory | Claudio Tavares Sacchi, Claudia Regina Gonçalves, Erica Valessa Ramos Gomes, Karoline Rodrigues Campos, Caio Vinicius Dias Lopes, Leonardo Jose Tadeu de Araujo |
| EPI_ISL_2003150, EPI_ISL_2003151 | Instituto Adolfo Lutz - Regional de Bauru | Instituto Adolfo Lutz, Interdiciplinary Procedures Center, Strategic Laboratory | Claudio Tavares Sacchi, Claudia Regina Gonçalves, Erica Valessa Ramos Gomes, Karoline Rodrigues Campos, Caio Vinicius Dias Lopes, Leonardo Jose Tadeu de Araujo |
| EPI_ISL_2003152 | Instituto Adolfo Lutz - Regional de Marilia | Instituto Adolfo Lutz, Interdiciplinary Procedures Center, Strategic Laboratory | Claudio Tavares Sacchi, Claudia Regina Gonçalves, Erica Valessa Ramos Gomes, Karoline Rodrigues Campos, Caio Vinicius Dias Lopes, Leonardo Jose Tadeu de Araujo |
| EPI_ISL_2003153, EPI_ISL_2003154, EPI_ISL_2003155, EPI_ISL_2003156, EPI_ISL_2003157, EPI_ISL_2003158, EPI_ISL_2003159, EPI_ISL_2003160, EPI_ISL_2003161 | Instituto Adolfo Lutz Central | Instituto Adolfo Lutz, Interdiciplinary Procedures Center, Strategic Laboratory | Claudio Tavares Sacchi, Claudia Regina Gonçalves, Erica Valessa Ramos Gomes, Karoline Rodrigues Campos, Caio Vinicius Dias Lopes, Leonardo Jose Tadeu de Araujo |
| EPI_ISL_2003162, EPI_ISL_2003163, EPI_ISL_2003164, EPI_ISL_2003165, EPI_ISL_2003166, EPI_ISL_2003167, EPI_ISL_2003168, EPI_ISL_2003169, EPI_ISL_2003170 | Instituto Adolfo Lutz - Regional de Campinas | Instituto Adolfo Lutz, Interdiciplinary Procedures Center, Strategic Laboratory | Claudio Tavares Sacchi, Claudia Regina Gonçalves, Erica Valessa Ramos Gomes, Karoline Rodrigues Campos, Caio Vinicius Dias Lopes, Leonardo Jose Tadeu de Araujo |
| EPI_ISL_2008906, EPI_ISL_2008907, EPI_ISL_2008908, EPI_ISL_2008909, EPI_ISL_2008910, EPI_ISL_2008911, EPI_ISL_2008912, EPI_ISL_2008913, EPI_ISL_2008914, EPI_ISL_2008915, EPI_ISL_2008916, EPI_ISL_2008917, EPI_ISL_2008918, EPI_ISL_2008919, EPI_ISL_2008920, EPI_ISL_2008921, EPI_ISL_2008922, EPI_ISL_2008923, EPI_ISL_2008924, EPI_ISL_2008925, EPI_ISL_2008926, EPI_ISL_2008927, EPI_ISL_2008928, EPI_ISL_2008929, EPI_ISL_2008930, EPI_ISL_2008931, EPI_ISL_2008933, EPI_ISL_2008934, EPI_ISL_2008935, EPI_ISL_2008936, EPI_ISL_2008937, EPI_ISL_2008938, EPI_ISL_2008939, EPI_ISL_2008940, EPI_ISL_2008941, EPI_ISL_2008942, EPI_ISL_2008943, EPI_ISL_2008944, EPI_ISL_2008945, EPI_ISL_2008946, EPI_ISL_2008947, EPI_ISL_2008948, EPI_ISL_2008949, EPI_ISL_2008950, EPI_ISL_2008951, EPI_ISL_2008952, EPI_ISL_2008953, EPI_ISL_2008954, EPI_ISL_2008955, EPI_ISL_2008956, EPI_ISL_2008957, EPI_ISL_2008958, EPI_ISL_2008959, EPI_ISL_2008960, EPI_ISL_2008961, EPI_ISL_2008962, EPI_ISL_2008963, EPI_ISL_2008964, EPI_ISL_2008965, EPI_ISL_2008966, EPI_ISL_2008967 | | | |
| see above | Laboratorio de Pesquisa em Virologia, FAMERP, SJRP | Laboratorio de Pesquisa em Virologia, FAMERP, SJRP | Cecília Artico Banho; Lívia Sacchetto; Fábio Sossai Possebon; Leila Sabrina Ullmann; Cíntia Bittar; Guilherme Campos; Helena Lage Ferreira; Jorge A. Petroli Marchesi; Maisa C. Pereira Parra; Marília Moraes; Paula Rahal; Paulo Inacio da Costa; João Pessoa Araújo Jr.; Maurício L. Nogueira. |
| EPI_ISL_2017241, EPI_ISL_2017242, EPI_ISL_2017243, EPI_ISL_2017244, EPI_ISL_2017245, EPI_ISL_2017246, EPI_ISL_2017247, EPI_ISL_2017248, EPI_ISL_2017249, EPI_ISL_2017250, EPI_ISL_2017251, EPI_ISL_2017252, EPI_ISL_2017253, EPI_ISL_2017254, EPI_ISL_2017255, EPI_ISL_2017256, EPI_ISL_2017257, EPI_ISL_2017258, EPI_ISL_2017259, EPI_ISL_2017260, EPI_ISL_2017261, EPI_ISL_2017262, EPI_ISL_2017263, EPI_ISL_2017264, EPI_ISL_2017265, EPI_ISL_2017266, EPI_ISL_2017267, EPI_ISL_2017268, EPI_ISL_2017269, EPI_ISL_2017270, EPI_ISL_2017271, EPI_ISL_2017272, EPI_ISL_2017273, EPI_ISL_2017274, EPI_ISL_2017275, EPI_ISL_2017276, EPI_ISL_2017277, EPI_ISL_2017278, EPI_ISL_2017279, EPI_ISL_2017280, EPI_ISL_2017281, EPI_ISL_2017282, EPI_ISL_2017283, EPI_ISL_2017284, EPI_ISL_2017285, EPI_ISL_2017286, EPI_ISL_2017287, EPI_ISL_2017288, EPI_ISL_2017289, EPI_ISL_2017290, EPI_ISL_2017291, EPI_ISL_2017292, EPI_ISL_2017293, EPI_ISL_2017294, EPI_ISL_2017295, EPI_ISL_2017296, EPI_ISL_2017298, EPI_ISL_2017299, EPI_ISL_2017300, EPI_ISL_2017301, EPI_ISL_2017302, EPI_ISL_2017303, EPI_ISL_2017304, EPI_ISL_2017305, EPI_ISL_2017306, EPI_ISL_2017307, EPI_ISL_2017308, EPI_ISL_2017309, EPI_ISL_2017310, EPI_ISL_2017311, EPI_ISL_2017312, EPI_ISL_2017313, EPI_ISL_2017314, EPI_ISL_2017315, EPI_ISL_2017316, EPI_ISL_2017317, EPI_ISL_2017318, EPI_ISL_2017319, EPI_ISL_2017320, EPI_ISL_2017322, EPI_ISL_2017324, EPI_ISL_2017325, EPI_ISL_2017326, EPI_ISL_2017327, EPI_ISL_2017328, EPI_ISL_2017329, EPI_ISL_2017330, EPI_ISL_2017331, EPI_ISL_2017332, EPI_ISL_2017333, EPI_ISL_2017334, EPI_ISL_2017335, EPI_ISL_2017336, EPI_ISL_2017337, EPI_ISL_2017338, EPI_ISL_2017339, EPI_ISL_2017340, EPI_ISL_2017341, EPI_ISL_2017342, EPI_ISL_2017343, EPI_ISL_2017344, EPI_ISL_2017345, EPI_ISL_2017346, EPI_ISL_2017347, EPI_ISL_2017348, EPI_ISL_2017349, EPI_ISL_2017350, EPI_ISL_2017351, EPI_ISL_2017352, EPI_ISL_2017353, EPI_ISL_2017354, EPI_ISL_2017355, EPI_ISL_2017356, EPI_ISL_2017357, EPI_ISL_2017358, EPI_ISL_2017359, EPI_ISL_2017360 | | | |
| see above | HLAGYN - Laboratorio de Imunologia de Transplantes de Goias | HLAGYN - Laboratorio de Imunologia de Transplantes de Goias | Fernando Antonio Vinhal dos Santos, Erika Lopes Rocha Batista, Alessandro Leonardo Alvares Magalhaes, Raphael Bessa Parmigiane, Frederico Rodrigues Vinhal, Sabrina Sara Moreira Duarte, Danielle de Paiva Rezende, Lucas Carlos Gomes Pereira, Paola Cristina Resende Silva |
| EPI_ISL_2017361, EPI_ISL_2017362, EPI_ISL_2017365, EPI_ISL_2017368, EPI_ISL_2017369, EPI_ISL_2017370, EPI_ISL_2017371, EPI_ISL_2017372, EPI_ISL_2017373, EPI_ISL_2017374, EPI_ISL_2017375, EPI_ISL_2017376, EPI_ISL_2017377, EPI_ISL_2017378, EPI_ISL_2017379, EPI_ISL_2017380, EPI_ISL_2017382, EPI_ISL_2017384, EPI_ISL_2017385, EPI_ISL_2017386, EPI_ISL_2017387, EPI_ISL_2017389, EPI_ISL_2017390, EPI_ISL_2017393, EPI_ISL_2017395, EPI_ISL_2017396, EPI_ISL_2017399, EPI_ISL_2017404, EPI_ISL_2017407, EPI_ISL_2017408, EPI_ISL_2017413, EPI_ISL_2017414, EPI_ISL_2017417, EPI_ISL_2017418, EPI_ISL_2017419, EPI_ISL_2017421, EPI_ISL_2017424, EPI_ISL_2017425, EPI_ISL_2017426, EPI_ISL_2017427, EPI_ISL_2017428, EPI_ISL_2017429, EPI_ISL_2017430, EPI_ISL_2017432, EPI_ISL_2017434, EPI_ISL_2017435, EPI_ISL_2017437, EPI_ISL_2017438, EPI_ISL_2017439, EPI_ISL_2017443, EPI_ISL_2017446, EPI_ISL_2017447, EPI_ISL_2017448, EPI_ISL_2017450, EPI_ISL_2017451, EPI_ISL_2017452, EPI_ISL_2017453, EPI_ISL_2017454, EPI_ISL_2017455, EPI_ISL_2017456, EPI_ISL_2017457, EPI_ISL_2017458, EPI_ISL_2017459, EPI_ISL_2017460, EPI_ISL_2017462, EPI_ISL_2017463, EPI_ISL_2017464, EPI_ISL_2017465, EPI_ISL_2017466, EPI_ISL_2017467, EPI_ISL_2017468, EPI_ISL_2017469, EPI_ISL_2017470, EPI_ISL_2017471, EPI_ISL_2017472, EPI_ISL_2017473, EPI_ISL_2017474, EPI_ISL_2017475, EPI_ISL_2017476, EPI_ISL_2017477, EPI_ISL_2017478, EPI_ISL_2017479, EPI_ISL_2017480, EPI_ISL_2017483, EPI_ISL_2017484 | | | |
| see above | HLAGYN - Laboratorio de Imunologia de Transplantes de Goias | HLAGYN - Laboratorio de Imunologia de Transplantes de Goias | Fernando Antonio Vinhal dos Santos, Erika Lopes Rocha Batista, Alessandro Leonardo Alvares Magalhaes, Frederico Rodrigues Vinhal, Sabrina Sara Moreira Duarte, Danielle de Paiva Rezende, Lucas Carlos Gomes Pereira, Paola Cristina Resende Silva |
| EPI_ISL_2038926, EPI_ISL_2038927, EPI_ISL_2038928, EPI_ISL_2038929, EPI_ISL_2038930, EPI_ISL_2038931, EPI_ISL_2038932, EPI_ISL_2038933, EPI_ISL_2038934, EPI_ISL_2038935, EPI_ISL_2038936, EPI_ISL_2038937, EPI_ISL_2038938, EPI_ISL_2038939, EPI_ISL_2038940, EPI_ISL_2038941, EPI_ISL_2038942, EPI_ISL_2038943, EPI_ISL_2038945, EPI_ISL_2038946, EPI_ISL_2038947, EPI_ISL_2038948, EPI_ISL_2038949, EPI_ISL_2038950, EPI_ISL_2038951, EPI_ISL_2038952, EPI_ISL_2038953, EPI_ISL_2038954 | | | |
| see above | Laboratorio Central de Saude Publica do Estado de Santa Catarina (LACEN-SC) | Laboratory of Respiratory Viruses and Measles, Oswaldo Cruz Institute, FIOCRUZ | Paola Resende, Luciana Appolinario, Fernando Motta, Anna Carolina Paixao, Ana Carolina Mendonca, Alice Sampaio Rocha, Taina Venas, Elisa Cavalcante Pereira, Renata Serrano Lopes, Darcita Buerger Rovaris, Sandra Bianchini Fernandes, Marilda Siqueira on behalf of the Fiocruz COVID-19 Genomic Surveillance Network |
| EPI_ISL_2038955 | Laboratorio Central de Saude Publica do Estado do Parana (LACEN-PR) | Laboratory of Respiratory Viruses and Measles, Oswaldo Cruz Institute, FIOCRUZ | Paola Resende, Luciana Appolinario, Fernando Motta, Anna Carolina Paixao, Ana Carolina Mendonca, Alice Sampaio Rocha, Taina Venas, Elisa Cavalcante Pereira, Renata Serrano Lopes, Irina Riediger, Marilda Siqueira on behalf of the Fiocruz COVID-19 Genomic Surveillance Network |
| EPI_ISL_2038956 | Laboratorio Central de Saude Publica do Estado do Rio de Janeiro (LACEN-RJ) | Laboratory of Respiratory Viruses and Measles, Oswaldo Cruz Institute, FIOCRUZ | Paola Resende, Luciana Appolinario, Fernando Motta, Anna Carolina Paixao, Ana Carolina Mendonca, Alice Sampaio Rocha, Taina Venas, Elisa Cavalcante Pereira, Renata Serrano Lopes, Andrea Cony Cavalcanti, Marilda Siqueira on behalf of the Fiocruz COVID-19 Genomic Surveillance Network |
| EPI_ISL_2038957, EPI_ISL_2038958, EPI_ISL_2038959, EPI_ISL_2038960 | Laboratorio Central de Saude Publica do Estado do Rio Grande do Sul (LACEN-RS) | Laboratory of Respiratory Viruses and Measles, Oswaldo Cruz Institute, FIOCRUZ | Paola Resende, Luciana Appolinario, Fernando Motta, Anna Carolina Paixao, Ana Carolina Mendonca, Alice Sampaio Rocha, Taina Venas, Elisa Cavalcante Pereira, Renata Serrano Lopes, Tatiana Schaffer Gregianini, Richard Salvato, Marilda Siqueira on behalf of the Fiocruz COVID-19 Genomic Surveillance Network |
| EPI_ISL_2038961, EPI_ISL_2038962, EPI_ISL_2038963 | Laboratory of Respiratory Viruses and Measles, Oswaldo Cruz Institute, FIOCRUZ | Laboratory of Respiratory Viruses and Measles, Oswaldo Cruz Institute, FIOCRUZ | Paola Resende, Luciana Appolinario, Fernando Motta, Anna Carolina Paixao, Ana Carolina Mendonca, Alice Sampaio Rocha, Taina Venas, Elisa Cavalcante Pereira, Renata Serrano Lopes, Marilda Siqueira on behalf of the Fiocruz COVID-19 Genomic Surveillance Network |
| EPI_ISL_2038964 | Laboratorio Central de Saude Publica do Estado do Alagoas (LACEN-AL) | Laboratory of Respiratory Viruses and Measles, Oswaldo Cruz Institute, FIOCRUZ | Paola Resende, Luciana Appolinario, Fernando Motta, Anna Carolina Paixao, Ana Carolina Mendonca, Alice Sampaio Rocha, Taina Venas, Elisa Cavalcante Pereira, Renata Serrano Lopes, Anderson Brandao Leite, Marilda Siqueira on behalf of the Fiocruz COVID-19 Genomic Surveillance Network |
| EPI_ISL_2038965 | Laboratorio Central de Saude Publica do Estado de Minas Gerais (LACEN-MG) | Laboratory of Respiratory Viruses and Measles, Oswaldo Cruz Institute, FIOCRUZ | Paola Resende, Luciana Appolinario, Fernando Motta, Anna Carolina Paixao, Ana Carolina Mendonca, Alice Sampaio Rocha, Taina Venas, Elisa Cavalcante Pereira, Renata Serrano Lopes, Andre Felipe Leal Bernardes, Marilda Siqueira on behalf of the Fiocruz COVID-19 Genomic Surveillance Network |

| | | | |
|---|---|---|---|
| EPI_ISL_2038966, EPI_ISL_2038967 | Laboratorio Central de Saude Publica do Estado de Santa Catarina (LACEN-SC) | Laboratory of Respiratory Viruses and Measles, Oswaldo Cruz Institute, FIOCRUZ | Paola Resende, Luciana Appolinario, Fernando Motta, Anna Carolina Paixao, Ana Carolina Mendonca, Alice Sampaio Rocha, Taina Venas, Elisa Cavalcante Pereira, Renata Serrano Lopes, Darcita Buerger Rovaris, Sandra Bianchini Fernandes, Marilda Siqueira on behalf of the Fiocruz COVID-19 Genomic Surveillance Network |
| EPI_ISL_2038968 | Laboratorio Central de Saude Publica do Estado do Espitito Santo (LACEN-ES) | Laboratory of Respiratory Viruses and Measles, Oswaldo Cruz Institute, FIOCRUZ | Paola Resende, Luciana Appolinario, Fernando Motta, Anna Carolina Paixao, Ana Carolina Mendonca, Alice Sampaio Rocha, Renata Serrano Lopes, Rodrigo Ribeiro Rodrigues, Marilda Siqueira on behalf of the Fiocruz COVID-19 Genomic Surveillance Network |
| EPI_ISL_2086268 | Laboratory of Molecular Biology of Flavivirus, Oswaldo Cruz Institute, FIOCRUZ | Laboratory of Molecular Biology of Flavivirus, Oswaldo Cruz Institute, FIOCRUZ | Fernando Lopez Tort, Ieda Ribeiro, Lidiane Menezes, Alexandre dos Santos, Marta Pereira, Myrna Bonaldo |
| EPI_ISL_2086585, EPI_ISL_2086586, EPI_ISL_2086587, EPI_ISL_2086588, EPI_ISL_2086589, EPI_ISL_2086590, EPI_ISL_2086591, EPI_ISL_2086592, EPI_ISL_2086593, EPI_ISL_2086594, EPI_ISL_2086595, EPI_ISL_2086596, EPI_ISL_2086597, EPI_ISL_2086598, EPI_ISL_2086599, EPI_ISL_2086600, EPI_ISL_2086601, EPI_ISL_2086602, EPI_ISL_2086603, EPI_ISL_2086604, EPI_ISL_2086605, EPI_ISL_2086606, EPI_ISL_2086607, EPI_ISL_2086608, EPI_ISL_2086609, EPI_ISL_2086610, EPI_ISL_2086611, EPI_ISL_2086612, EPI_ISL_2086613, EPI_ISL_2086614, EPI_ISL_2086615, EPI_ISL_2086616, EPI_ISL_2086617, EPI_ISL_2086618, EPI_ISL_2086619, EPI_ISL_2086620, EPI_ISL_2086621, EPI_ISL_2086622, EPI_ISL_2086763, EPI_ISL_2086764, EPI_ISL_2086765, EPI_ISL_2086766, EPI_ISL_2086767 | | | |
| see above | LABCOVID_HCPA | LABRESIS_HCPA | Wink PL, Martins AF, Volpato F, Monteiro F, Zavascki AP, Barth AL |
| EPI_ISL_2101392 | Unidade de apoio ao diagnóstico da COVID - UNADIG | Bioinformatics Laboratory / LNCC | Luiz G P de Almeida, Alessandra P Lamarca, Ronaldo da Silva F Jr, Liliane Cavalcante, Alexandra L Gerber, Ana Paula de C Guimaraes, Douglas Terra Machado, Cassia Alves, Diana Mariani, Cintia Policarpo, Gleidson da Silva de Oliveira, Mario Sergio Ribeiro, Silvia Carvalho, Flavio Dias da Silva, Marcio Henrique de Oliveira Garcia, Leandro Magalhaes de Souza, Cristiane Gomes da Silva, Caio Luiz Pereira Ribeiro, Andrea Cony Cavalcanti, Claudia Maria Braga de Mello, Amilcar Tanuri, Ana Tereza R Vasconcelos |
| EPI_ISL_2101393 | Laboratorio Central Noel Nutels | Bioinformatics Laboratory / LNCC | Luiz G P de Almeida, Alessandra P Lamarca, Ronaldo da Silva F Jr, Liliane Cavalcante, Alexandra L Gerber, Ana Paula de C Guimaraes, Douglas Terra Machado, Cassia Alves, Diana Mariani, Cintia Policarpo, Gleidson da Silva de Oliveira, Mario Sergio Ribeiro, Silvia Carvalho, Flavio Dias da Silva, Marcio Henrique de Oliveira Garcia, Leandro Magalhaes de Souza, Cristiane Gomes da Silva, Caio Luiz Pereira Ribeiro, Andrea Cony Cavalcanti, Claudia Maria Braga de Mello, Amilcar Tanuri, Ana Tereza R Vasconcelos |
| EPI_ISL_2101394 | Unidade de apoio ao diagnóstico da COVID - UNADIG | Bioinformatics Laboratory / LNCC | Luiz G P de Almeida, Alessandra P Lamarca, Ronaldo da Silva F Jr, Liliane Cavalcante, Alexandra L Gerber, Ana Paula de C Guimaraes, Douglas Terra Machado, Cassia Alves, Diana Mariani, Cintia Policarpo, Gleidson da Silva de Oliveira, Mario Sergio Ribeiro, Silvia Carvalho, Flavio Dias da Silva, Marcio Henrique de Oliveira Garcia, Leandro Magalhaes de Souza, Cristiane Gomes da Silva, Caio Luiz Pereira Ribeiro, Andrea Cony Cavalcanti, Claudia Maria Braga de Mello, Amilcar Tanuri, Ana Tereza R Vasconcelos |
| EPI_ISL_2101395, EPI_ISL_2101396, EPI_ISL_2101397, EPI_ISL_2101398, EPI_ISL_2101399, EPI_ISL_2101400, EPI_ISL_2101401, EPI_ISL_2101402, EPI_ISL_2101403, EPI_ISL_2101404, EPI_ISL_2101405, EPI_ISL_2101406, EPI_ISL_2101409, EPI_ISL_2101410, EPI_ISL_2101411, EPI_ISL_2101412, EPI_ISL_2101413, EPI_ISL_2101414, EPI_ISL_2101415, EPI_ISL_2101416, EPI_ISL_2101417, EPI_ISL_2101418, EPI_ISL_2101419, EPI_ISL_2101420, EPI_ISL_2101421, EPI_ISL_2101422, EPI_ISL_2101423, EPI_ISL_2101424, EPI_ISL_2101425, EPI_ISL_2101426, EPI_ISL_2101427, EPI_ISL_2101428 | | | |
| see above | Laboratorio Central Noel Nutels | Bioinformatics Laboratory / LNCC | Luiz G P de Almeida, Alessandra P Lamarca, Ronaldo da Silva F Jr, Liliane Cavalcante, Alexandra L Gerber, Ana Paula de C Guimaraes, Douglas Terra Machado, Cassia Alves, Diana Mariani, Cintia Policarpo, Gleidson da Silva de Oliveira, Mario Sergio Ribeiro, Silvia Carvalho, Flavio Dias da Silva, Marcio Henrique de Oliveira Garcia, Leandro Magalhaes de Souza, Cristiane Gomes da Silva, Caio Luiz Pereira Ribeiro, Andrea Cony Cavalcanti, Claudia Maria Braga de Mello, Amilcar Tanuri, Ana Tereza R Vasconcelos |
| EPI_ISL_2101429, EPI_ISL_2101430, EPI_ISL_2101431, EPI_ISL_2101432, EPI_ISL_2101433 | Unidade de apoio ao diagnóstico da COVID - UNADIG | Bioinformatics Laboratory / LNCC | Luiz G P de Almeida, Alessandra P Lamarca, Ronaldo da Silva F Jr, Liliane Cavalcante, Alexandra L Gerber, Ana Paula de C Guimaraes, Douglas Terra Machado, Cassia Alves, Diana Mariani, Cintia Policarpo, Gleidson da Silva de Oliveira, Mario Sergio Ribeiro, Silvia Carvalho, Flavio Dias da Silva, Marcio Henrique de Oliveira Garcia, Leandro Magalhaes de Souza, Cristiane Gomes da Silva, Caio Luiz Pereira Ribeiro, Andrea Cony Cavalcanti, Claudia Maria Braga de Mello, Amilcar Tanuri, Ana Tereza R Vasconcelos |
| EPI_ISL_2101434, EPI_ISL_2101435, EPI_ISL_2101436, EPI_ISL_2101437, EPI_ISL_2101438, EPI_ISL_2101439, EPI_ISL_2101440, EPI_ISL_2101441, EPI_ISL_2101442 | Laboratorio Central Noel Nutels | Bioinformatics Laboratory / LNCC | Luiz G P de Almeida, Alessandra P Lamarca, Ronaldo da Silva F Jr, Liliane Cavalcante, Alexandra L Gerber, Ana Paula de C Guimaraes, Douglas Terra Machado, Cassia Alves, Diana Mariani, Cintia Policarpo, Gleidson da Silva de Oliveira, Mario Sergio Ribeiro, Silvia Carvalho, Flavio Dias da Silva, Marcio Henrique de Oliveira Garcia, Leandro Magalhaes de Souza, Cristiane Gomes da Silva, Caio Luiz Pereira Ribeiro, Andrea Cony Cavalcanti, Claudia Maria Braga de Mello, Amilcar Tanuri, Ana Tereza R Vasconcelos |
| EPI_ISL_2101443 | Unidade de apoio ao diagnóstico da COVID - UNADIG | Bioinformatics Laboratory / LNCC | Luiz G P de Almeida, Alessandra P Lamarca, Ronaldo da Silva F Jr, Liliane Cavalcante, Alexandra L Gerber, Ana Paula de C Guimaraes, Douglas Terra Machado, Cassia Alves, Diana Mariani, Cintia Policarpo, Gleidson da Silva de Oliveira, Mario Sergio Ribeiro, Silvia Carvalho, Flavio Dias da Silva, Marcio Henrique de Oliveira Garcia, Leandro Magalhaes de Souza, Cristiane Gomes da Silva, Caio Luiz Pereira Ribeiro, Andrea Cony Cavalcanti, Claudia Maria Braga de Mello, Amilcar Tanuri, Ana Tereza R Vasconcelos |
| EPI_ISL_2101444, EPI_ISL_2101445, EPI_ISL_2101446, EPI_ISL_2101447, EPI_ISL_2101448, EPI_ISL_2101449, EPI_ISL_2101450, EPI_ISL_2101451, EPI_ISL_2101452, EPI_ISL_2101453, EPI_ISL_2101454 | | | |
| see above | Laboratorio Central Noel Nutels | Bioinformatics Laboratory / LNCC | Luiz G P de Almeida, Alessandra P Lamarca, Ronaldo da Silva F Jr, Liliane Cavalcante, Alexandra L Gerber, Ana Paula de C Guimaraes, Douglas Terra Machado, Cassia Alves, Diana Mariani, Cintia Policarpo, Gleidson da Silva de Oliveira, Mario Sergio Ribeiro, Silvia Carvalho, Flavio Dias da Silva, Marcio Henrique de Oliveira Garcia, Leandro Magalhaes de Souza, Cristiane Gomes da Silva, Caio Luiz Pereira Ribeiro, Andrea Cony Cavalcanti, Claudia Maria Braga de Mello, Amilcar Tanuri, Ana Tereza R Vasconcelos |
| EPI_ISL_2101455, EPI_ISL_2101456, EPI_ISL_2101457, EPI_ISL_2101458, EPI_ISL_2101459, EPI_ISL_2101460, EPI_ISL_2101461, EPI_ISL_2101462, EPI_ISL_2101463 | Unidade de apoio ao diagnóstico da COVID - UNADIG | Bioinformatics Laboratory / LNCC | Luiz G P de Almeida, Alessandra P Lamarca, Ronaldo da Silva F Jr, Liliane Cavalcante, Alexandra L Gerber, Ana Paula de C Guimaraes, Douglas Terra Machado, Cassia Alves, Diana Mariani, Cintia Policarpo, Gleidson da Silva de Oliveira, Mario Sergio Ribeiro, Silvia Carvalho, Flavio Dias da Silva, Marcio Henrique de Oliveira Garcia, Leandro Magalhaes de Souza, Cristiane Gomes da Silva, Caio Luiz Pereira Ribeiro, Andrea Cony Cavalcanti, Claudia Maria Braga de Mello, Amilcar Tanuri, Ana Tereza R Vasconcelos |
| EPI_ISL_2101464, EPI_ISL_2101465, EPI_ISL_2101466, EPI_ISL_2101467, EPI_ISL_2101468, EPI_ISL_2101469 | Laboratorio Central Noel Nutels | Bioinformatics Laboratory / LNCC | Luiz G P de Almeida, Alessandra P Lamarca, Ronaldo da Silva F Jr, Liliane Cavalcante, Alexandra L Gerber, Ana Paula de C Guimaraes, Douglas Terra Machado, Cassia Alves, Diana Mariani, Cintia Policarpo, Gleidson da Silva de Oliveira, Mario Sergio Ribeiro, Silvia Carvalho, Flavio Dias da Silva, Marcio Henrique de Oliveira Garcia, Leandro Magalhaes de Souza, Cristiane Gomes da Silva, Caio Luiz Pereira Ribeiro, Andrea Cony Cavalcanti, Claudia Maria Braga de Mello, Amilcar Tanuri, Ana Tereza R Vasconcelos |
| EPI_ISL_2101470, EPI_ISL_2101471, EPI_ISL_2101472, EPI_ISL_2101473 | Unidade de apoio ao diagnóstico da COVID - UNADIG | Bioinformatics Laboratory / LNCC | Luiz G P de Almeida, Alessandra P Lamarca, Ronaldo da Silva F Jr, Liliane Cavalcante, Alexandra L Gerber, Ana Paula de C Guimaraes, Douglas Terra Machado, Cassia Alves, Diana Mariani, Cintia Policarpo, Gleidson da Silva de Oliveira, Mario Sergio Ribeiro, Silvia Carvalho, Flavio Dias da Silva, Marcio Henrique de Oliveira Garcia, Leandro Magalhaes de Souza, Cristiane Gomes da Silva, Caio Luiz Pereira Ribeiro, Andrea Cony Cavalcanti, Claudia Maria Braga de Mello, Amilcar Tanuri, Ana Tereza R Vasconcelos |
| EPI_ISL_2101474 | Laboratorio Central Noel Nutels | Bioinformatics Laboratory / LNCC | Luiz G P de Almeida, Alessandra P Lamarca, Ronaldo da Silva F Jr, Liliane Cavalcante, Alexandra L Gerber, Ana Paula de C Guimaraes, Douglas Terra Machado, Cassia Alves, Diana Mariani, Cintia Policarpo, Gleidson da Silva de Oliveira, Mario Sergio Ribeiro, Silvia Carvalho, Flavio Dias da Silva, Marcio Henrique de Oliveira Garcia, Leandro Magalhaes de Souza, Cristiane Gomes da Silva, Caio Luiz Pereira Ribeiro, Andrea Cony Cavalcanti, Claudia Maria Braga de Mello, Amilcar Tanuri, Ana Tereza R Vasconcelos |
| EPI_ISL_2101475 | Unidade de apoio ao diagnóstico da COVID - UNADIG | Bioinformatics Laboratory / LNCC | Luiz G P de Almeida, Alessandra P Lamarca, Ronaldo da Silva F Jr, Liliane Cavalcante, Alexandra L Gerber, Ana Paula de C Guimaraes, Douglas Terra Machado, Cassia Alves, Diana Mariani, Cintia Policarpo, Gleidson da Silva de Oliveira, Mario Sergio Ribeiro, Silvia Carvalho, Flavio Dias da Silva, Marcio Henrique de Oliveira Garcia, Leandro Magalhaes de Souza, Cristiane Gomes da Silva, Caio Luiz Pereira Ribeiro, Andrea Cony Cavalcanti, Claudia Maria Braga de Mello, Amilcar Tanuri, Ana Tereza R Vasconcelos |
| EPI_ISL_2101476, EPI_ISL_2101477, EPI_ISL_2101478, EPI_ISL_2101479, EPI_ISL_2101480, EPI_ISL_2101481, EPI_ISL_2101482 | Laboratorio Central Noel Nutels | Bioinformatics Laboratory / LNCC | Luiz G P de Almeida, Alessandra P Lamarca, Ronaldo da Silva F Jr, Liliane Cavalcante, Alexandra L Gerber, Ana Paula de C Guimaraes, Douglas Terra Machado, Cassia Alves, Diana Mariani, Cintia Policarpo, Gleidson da Silva de Oliveira, Mario Sergio Ribeiro, Silvia Carvalho, Flavio Dias da Silva, Marcio Henrique de Oliveira Garcia, Leandro Magalhaes de Souza, Cristiane Gomes da Silva, Caio Luiz Pereira Ribeiro, Andrea Cony Cavalcanti, Claudia Maria Braga de Mello, Amilcar Tanuri, Ana Tereza R Vasconcelos |
| EPI_ISL_2101483, EPI_ISL_2101484 | Unidade de apoio ao diagnóstico da COVID - UNADIG | Bioinformatics Laboratory / LNCC | Luiz G P de Almeida, Alessandra P Lamarca, Ronaldo da Silva F Jr, Liliane Cavalcante, Alexandra L Gerber, Ana Paula de C Guimaraes, Douglas Terra Machado, Cassia Alves, Diana Mariani, Cintia Policarpo, Gleidson da Silva de Oliveira, Mario Sergio Ribeiro, Silvia Carvalho, Flavio Dias da Silva, Marcio Henrique de Oliveira Garcia, Leandro Magalhaes de Souza, Cristiane Gomes da Silva, Caio Luiz Pereira Ribeiro, Andrea Cony Cavalcanti, Claudia Maria Braga de Mello, Amilcar Tanuri, Ana Tereza R Vasconcelos |

| | | | |
|---|---|---|---|
| EPI_ISL_2101485, EPI_ISL_2101486, EPI_ISL_2101487, EPI_ISL_2101488, EPI_ISL_2101489, EPI_ISL_2101490, EPI_ISL_2101491, EPI_ISL_2101492, EPI_ISL_2101493 | Laboratorio Central Noel Nutels | Bioinformatics Laboratory / LNCC | Luiz G P de Almeida, Alessandra P Lamarca, Ronaldo da Silva F Jr, Liliane Cavalcante, Alexandra L Gerber, Ana Paula de C Guimaraes, Douglas Terra Machado, Cassia Alves, Diana Mariani, Cintia Policarpo, Gleidson da Silva de Oliveira, Mario Sergio Ribeiro, Silvia Carvalho, Flavio Dias da Silva, Marcio Henrique de Oliveira Garcia, Leandro Magalhaes de Souza, Cristiane Gomes da Silva, Caio Luiz Pereira Ribeiro, Andrea Cony Cavalcanti, Claudia Maria Braga de Mello, Amilcar Tanuri, Ana Tereza R Vasconcelos |
| EPI_ISL_2101494, EPI_ISL_2101495 | Unidade de apoio ao diagnóstico da COVID - UNADIG | Bioinformatics Laboratory / LNCC | Luiz G P de Almeida, Alessandra P Lamarca, Ronaldo da Silva F Jr, Liliane Cavalcante, Alexandra L Gerber, Ana Paula de C Guimaraes, Douglas Terra Machado, Cassia Alves, Diana Mariani, Cintia Policarpo, Gleidson da Silva de Oliveira, Mario Sergio Ribeiro, Silvia Carvalho, Flavio Dias da Silva, Marcio Henrique de Oliveira Garcia, Leandro Magalhaes de Souza, Cristiane Gomes da Silva, Caio Luiz Pereira Ribeiro, Andrea Cony Cavalcanti, Claudia Maria Braga de Mello, Amilcar Tanuri, Ana Tereza R Vasconcelos |
| EPI_ISL_2101496, EPI_ISL_2101497, EPI_ISL_2101498, EPI_ISL_2101499, EPI_ISL_2101500, EPI_ISL_2101501, EPI_ISL_2101502, EPI_ISL_2101503, EPI_ISL_2101504 | Laboratorio Central Noel Nutels | Bioinformatics Laboratory / LNCC | Luiz G P de Almeida, Alessandra P Lamarca, Ronaldo da Silva F Jr, Liliane Cavalcante, Alexandra L Gerber, Ana Paula de C Guimaraes, Douglas Terra Machado, Cassia Alves, Diana Mariani, Cintia Policarpo, Gleidson da Silva de Oliveira, Mario Sergio Ribeiro, Silvia Carvalho, Flavio Dias da Silva, Marcio Henrique de Oliveira Garcia, Leandro Magalhaes de Souza, Cristiane Gomes da Silva, Caio Luiz Pereira Ribeiro, Andrea Cony Cavalcanti, Claudia Maria Braga de Mello, Amilcar Tanuri, Ana Tereza R Vasconcelos |
| EPI_ISL_2101505, EPI_ISL_2101506 | Unidade de apoio ao diagnóstico da COVID - UNADIG | Bioinformatics Laboratory / LNCC | Luiz G P de Almeida, Alessandra P Lamarca, Ronaldo da Silva F Jr, Liliane Cavalcante, Alexandra L Gerber, Ana Paula de C Guimaraes, Douglas Terra Machado, Cassia Alves, Diana Mariani, Cintia Policarpo, Gleidson da Silva de Oliveira, Mario Sergio Ribeiro, Silvia Carvalho, Flavio Dias da Silva, Marcio Henrique de Oliveira Garcia, Leandro Magalhaes de Souza, Cristiane Gomes da Silva, Caio Luiz Pereira Ribeiro, Andrea Cony Cavalcanti, Claudia Maria Braga de Mello, Amilcar Tanuri, Ana Tereza R Vasconcelos |
| EPI_ISL_2101507 | Laboratorio Central Noel Nutels | Bioinformatics Laboratory / LNCC | Luiz G P de Almeida, Alessandra P Lamarca, Ronaldo da Silva F Jr, Liliane Cavalcante, Alexandra L Gerber, Ana Paula de C Guimaraes, Douglas Terra Machado, Cassia Alves, Diana Mariani, Cintia Policarpo, Gleidson da Silva de Oliveira, Mario Sergio Ribeiro, Silvia Carvalho, Flavio Dias da Silva, Marcio Henrique de Oliveira Garcia, Leandro Magalhaes de Souza, Cristiane Gomes da Silva, Caio Luiz Pereira Ribeiro, Andrea Cony Cavalcanti, Claudia Maria Braga de Mello, Amilcar Tanuri, Ana Tereza R Vasconcelos |
| EPI_ISL_2101508, EPI_ISL_2101509, EPI_ISL_2101510, EPI_ISL_2101511, EPI_ISL_2101512, EPI_ISL_2101513 | Unidade de apoio ao diagnóstico da COVID - UNADIG | Bioinformatics Laboratory / LNCC | Luiz G P de Almeida, Alessandra P Lamarca, Ronaldo da Silva F Jr, Liliane Cavalcante, Alexandra L Gerber, Ana Paula de C Guimaraes, Douglas Terra Machado, Cassia Alves, Diana Mariani, Cintia Policarpo, Gleidson da Silva de Oliveira, Mario Sergio Ribeiro, Silvia Carvalho, Flavio Dias da Silva, Marcio Henrique de Oliveira Garcia, Leandro Magalhaes de Souza, Cristiane Gomes da Silva, Caio Luiz Pereira Ribeiro, Andrea Cony Cavalcanti, Claudia Maria Braga de Mello, Amilcar Tanuri, Ana Tereza R Vasconcelos |
| EPI_ISL_2101514, EPI_ISL_2101515, EPI_ISL_2101516, EPI_ISL_2101517, EPI_ISL_2101518, EPI_ISL_2101519 | Laboratorio Central Noel Nutels | Bioinformatics Laboratory / LNCC | Luiz G P de Almeida, Alessandra P Lamarca, Ronaldo da Silva F Jr, Liliane Cavalcante, Alexandra L Gerber, Ana Paula de C Guimaraes, Douglas Terra Machado, Cassia Alves, Diana Mariani, Cintia Policarpo, Gleidson da Silva de Oliveira, Mario Sergio Ribeiro, Silvia Carvalho, Flavio Dias da Silva, Marcio Henrique de Oliveira Garcia, Leandro Magalhaes de Souza, Cristiane Gomes da Silva, Caio Luiz Pereira Ribeiro, Andrea Cony Cavalcanti, Claudia Maria Braga de Mello, Amilcar Tanuri, Ana Tereza R Vasconcelos |
| EPI_ISL_2101520, EPI_ISL_2101521, EPI_ISL_2101522, EPI_ISL_2101523, EPI_ISL_2101524, EPI_ISL_2101525, EPI_ISL_2101526, EPI_ISL_2101527 | Unidade de apoio ao diagnóstico da COVID - UNADIG | Bioinformatics Laboratory / LNCC | Luiz G P de Almeida, Alessandra P Lamarca, Ronaldo da Silva F Jr, Liliane Cavalcante, Alexandra L Gerber, Ana Paula de C Guimaraes, Douglas Terra Machado, Cassia Alves, Diana Mariani, Cintia Policarpo, Gleidson da Silva de Oliveira, Mario Sergio Ribeiro, Silvia Carvalho, Flavio Dias da Silva, Marcio Henrique de Oliveira Garcia, Leandro Magalhaes de Souza, Cristiane Gomes da Silva, Caio Luiz Pereira Ribeiro, Andrea Cony Cavalcanti, Claudia Maria Braga de Mello, Amilcar Tanuri, Ana Tereza R Vasconcelos |
| EPI_ISL_2101528, EPI_ISL_2101529, EPI_ISL_2101530, EPI_ISL_2101531, EPI_ISL_2101532, EPI_ISL_2101533, EPI_ISL_2101534, EPI_ISL_2101535, EPI_ISL_2101536, EPI_ISL_2101537, EPI_ISL_2101538, EPI_ISL_2101539, EPI_ISL_2101540, EPI_ISL_2101541, EPI_ISL_2101542, EPI_ISL_2101543, EPI_ISL_2101544, EPI_ISL_2101545, EPI_ISL_2101546, EPI_ISL_2101547, EPI_ISL_2101548, EPI_ISL_2101549, EPI_ISL_2101550, EPI_ISL_2101551, EPI_ISL_2101552, EPI_ISL_2101553, EPI_ISL_2101554, EPI_ISL_2101555, EPI_ISL_2101556, EPI_ISL_2101557, EPI_ISL_2101558, EPI_ISL_2101559, EPI_ISL_2101560, EPI_ISL_2101561, EPI_ISL_2101562, EPI_ISL_2101563, EPI_ISL_2101564, EPI_ISL_2101565 | | | |
| see above | Laboratorio Central Noel Nutels | Bioinformatics Laboratory / LNCC | Luiz G P de Almeida, Alessandra P Lamarca, Ronaldo da Silva F Jr, Liliane Cavalcante, Alexandra L Gerber, Ana Paula de C Guimaraes, Douglas Terra Machado, Cassia Alves, Diana Mariani, Cintia Policarpo, Gleidson da Silva de Oliveira, Mario Sergio Ribeiro, Silvia Carvalho, Flavio Dias da Silva, Marcio Henrique de Oliveira Garcia, Leandro Magalhaes de Souza, Cristiane Gomes da Silva, Caio Luiz Pereira Ribeiro, Andrea Cony Cavalcanti, Claudia Maria Braga de Mello, Amilcar Tanuri, Ana Tereza R Vasconcelos |
| EPI_ISL_2101566, EPI_ISL_2101567, EPI_ISL_2101568, EPI_ISL_2101569, EPI_ISL_2101570, EPI_ISL_2101571, EPI_ISL_2101572 | Unidade de apoio ao diagnóstico da COVID - UNADIG | Bioinformatics Laboratory / LNCC | Luiz G P de Almeida, Alessandra P Lamarca, Ronaldo da Silva F Jr, Liliane Cavalcante, Alexandra L Gerber, Ana Paula de C Guimaraes, Douglas Terra Machado, Cassia Alves, Diana Mariani, Cintia Policarpo, Gleidson da Silva de Oliveira, Mario Sergio Ribeiro, Silvia Carvalho, Flavio Dias da Silva, Marcio Henrique de Oliveira Garcia, Leandro Magalhaes de Souza, Cristiane Gomes da Silva, Caio Luiz Pereira Ribeiro, Andrea Cony Cavalcanti, Claudia Maria Braga de Mello, Amilcar Tanuri, Ana Tereza R Vasconcelos |
| EPI_ISL_2101573, EPI_ISL_2101574, EPI_ISL_2101575, EPI_ISL_2101576, EPI_ISL_2101577, EPI_ISL_2101578, EPI_ISL_2101579, EPI_ISL_2101580, EPI_ISL_2101581, EPI_ISL_2101582, EPI_ISL_2101583, EPI_ISL_2101584, EPI_ISL_2101585, EPI_ISL_2101586, EPI_ISL_2101587, EPI_ISL_2101588, EPI_ISL_2101589, EPI_ISL_2101590, EPI_ISL_2101591, EPI_ISL_2101592, EPI_ISL_2101593, EPI_ISL_2101594, EPI_ISL_2101595, EPI_ISL_2101596, EPI_ISL_2101597, EPI_ISL_2101598, EPI_ISL_2101599, EPI_ISL_2101600, EPI_ISL_2101601, EPI_ISL_2101602, EPI_ISL_2101603, EPI_ISL_2101604, EPI_ISL_2101605, EPI_ISL_2101606, EPI_ISL_2101607, EPI_ISL_2101608, EPI_ISL_2101609, EPI_ISL_2101610, EPI_ISL_2101611 | | | |
| see above | Laboratorio Central Noel Nutels | Bioinformatics Laboratory / LNCC | Luiz G P de Almeida, Alessandra P Lamarca, Ronaldo da Silva F Jr, Liliane Cavalcante, Alexandra L Gerber, Ana Paula de C Guimaraes, Douglas Terra Machado, Cassia Alves, Diana Mariani, Cintia Policarpo, Gleidson da Silva de Oliveira, Mario Sergio Ribeiro, Silvia Carvalho, Flavio Dias da Silva, Marcio Henrique de Oliveira Garcia, Leandro Magalhaes de Souza, Cristiane Gomes da Silva, Caio Luiz Pereira Ribeiro, Andrea Cony Cavalcanti, Claudia Maria Braga de Mello, Amilcar Tanuri, Ana Tereza R Vasconcelos |
| EPI_ISL_2101612 | Unidade de apoio ao diagnóstico da COVID - UNADIG | Bioinformatics Laboratory / LNCC | Luiz G P de Almeida, Alessandra P Lamarca, Ronaldo da Silva F Jr, Liliane Cavalcante, Alexandra L Gerber, Ana Paula de C Guimaraes, Douglas Terra Machado, Cassia Alves, Diana Mariani, Cintia Policarpo, Gleidson da Silva de Oliveira, Mario Sergio Ribeiro, Silvia Carvalho, Flavio Dias da Silva, Marcio Henrique de Oliveira Garcia, Leandro Magalhaes de Souza, Cristiane Gomes da Silva, Caio Luiz Pereira Ribeiro, Andrea Cony Cavalcanti, Claudia Maria Braga de Mello, Amilcar Tanuri, Ana Tereza R Vasconcelos |
| EPI_ISL_2101613, EPI_ISL_2101615, EPI_ISL_2101616, EPI_ISL_2101617, EPI_ISL_2101618, EPI_ISL_2101619, EPI_ISL_2101620, EPI_ISL_2101621, EPI_ISL_2101622, EPI_ISL_2101623, EPI_ISL_2101624, EPI_ISL_2101625, EPI_ISL_2101626, EPI_ISL_2101627, EPI_ISL_2101628, EPI_ISL_2101629, EPI_ISL_2101630, EPI_ISL_2101631, EPI_ISL_2101632, EPI_ISL_2101633, EPI_ISL_2101634, EPI_ISL_2101635, EPI_ISL_2101636, EPI_ISL_2101637 | | | |
| see above | Laboratorio Central Noel Nutels | Bioinformatics Laboratory / LNCC | Luiz G P de Almeida, Alessandra P Lamarca, Ronaldo da Silva F Jr, Liliane Cavalcante, Alexandra L Gerber, Ana Paula de C Guimaraes, Douglas Terra Machado, Cassia Alves, Diana Mariani, Cintia Policarpo, Gleidson da Silva de Oliveira, Mario Sergio Ribeiro, Silvia Carvalho, Flavio Dias da Silva, Marcio Henrique de Oliveira Garcia, Leandro Magalhaes de Souza, Cristiane Gomes da Silva, Caio Luiz Pereira Ribeiro, Andrea Cony Cavalcanti, Claudia Maria Braga de Mello, Amilcar Tanuri, Ana Tereza R Vasconcelos |
| EPI_ISL_2101638, EPI_ISL_2101639, EPI_ISL_2101640, EPI_ISL_2101641, EPI_ISL_2101642, EPI_ISL_2101643, EPI_ISL_2101644, EPI_ISL_2101645, EPI_ISL_2101646, EPI_ISL_2101647, EPI_ISL_2101648, EPI_ISL_2101649, EPI_ISL_2101650, EPI_ISL_2101651, EPI_ISL_2101652, EPI_ISL_2101653, EPI_ISL_2101654, EPI_ISL_2101655, EPI_ISL_2101656, EPI_ISL_2101657, EPI_ISL_2101658, EPI_ISL_2101659, EPI_ISL_2101660, EPI_ISL_2101661, EPI_ISL_2101662, EPI_ISL_2101663, EPI_ISL_2101664, EPI_ISL_2101665, EPI_ISL_2101666, EPI_ISL_2101667, EPI_ISL_2101668, EPI_ISL_2101669, EPI_ISL_2101670, EPI_ISL_2101671 | | | |
| see above | Unidade de apoio ao diagnóstico da COVID - UNADIG | Bioinformatics Laboratory / LNCC | Luiz G P de Almeida, Alessandra P Lamarca, Ronaldo da Silva F Jr, Liliane Cavalcante, Alexandra L Gerber, Ana Paula de C Guimaraes, Douglas Terra Machado, Cassia Alves, Diana Mariani, Cintia Policarpo, Gleidson da Silva de Oliveira, Mario Sergio Ribeiro, Silvia Carvalho, Flavio Dias da Silva, Marcio Henrique de Oliveira Garcia, Leandro Magalhaes de Souza, Cristiane Gomes da Silva, Caio Luiz Pereira Ribeiro, Andrea Cony Cavalcanti, Claudia Maria Braga de Mello, Amilcar Tanuri, Ana Tereza R Vasconcelos |
| EPI_ISL_2101672, EPI_ISL_2101673, EPI_ISL_2101674, EPI_ISL_2101675 | Laboratorio Central Noel Nutels | Bioinformatics Laboratory / LNCC | Luiz G P de Almeida, Alessandra P Lamarca, Ronaldo da Silva F Jr, Liliane Cavalcante, Alexandra L Gerber, Ana Paula de C Guimaraes, Douglas Terra Machado, Cassia Alves, Diana Mariani, Cintia Policarpo, Gleidson da Silva de Oliveira, Mario Sergio Ribeiro, Silvia Carvalho, Flavio Dias da Silva, Marcio Henrique de Oliveira Garcia, Leandro Magalhaes de Souza, Cristiane Gomes da Silva, Caio Luiz Pereira Ribeiro, Andrea Cony Cavalcanti, Claudia Maria Braga de Mello, Amilcar Tanuri, Ana Tereza R Vasconcelos |
| EPI_ISL_2101676, EPI_ISL_2101677, EPI_ISL_2101678, EPI_ISL_2101679, EPI_ISL_2101680, EPI_ISL_2101681, EPI_ISL_2101682, EPI_ISL_2101683, EPI_ISL_2101684, EPI_ISL_2101685, EPI_ISL_2101686, EPI_ISL_2101687, EPI_ISL_2101688 | | | |
| see above | Unidade de apoio ao diagnóstico da COVID - UNADIG | Bioinformatics Laboratory / LNCC | Luiz G P de Almeida, Alessandra P Lamarca, Ronaldo da Silva F Jr, Liliane Cavalcante, Alexandra L Gerber, Ana Paula de C Guimaraes, Douglas Terra Machado, Cassia Alves, Diana Mariani, Cintia Policarpo, Gleidson da Silva de Oliveira, Mario Sergio Ribeiro, Silvia Carvalho, Flavio Dias da Silva, Marcio Henrique de Oliveira Garcia, Leandro Magalhaes de Souza, Cristiane Gomes da Silva, Caio Luiz Pereira Ribeiro, Andrea Cony Cavalcanti, Claudia Maria Braga de Mello, Amilcar Tanuri, Ana Tereza R Vasconcelos |

| | | | |
|---|---|---|---|
| EPI_ISL_2101689, EPI_ISL_2101690, EPI_ISL_2101691 | Laboratorio Central Noel Nutels | Bioinformatics Laboratory / LNCC | Luiz G P de Almeida, Alessandra P Lamarca, Ronaldo da Silva F Jr, Liliane Cavalcante, Alexandra L Gerber, Ana Paula de C Guimaraes, Douglas Terra Machado, Cassia Alves, Diana Mariani, Cintia Policarpo, Gleidson da Silva de Oliveira, Mario Sergio Ribeiro, Silvia Carvalho, Flavio Dias da Silva, Marcio Henrique de Oliveira Garcia, Leandro Magalhaes de Souza, Cristiane Gomes da Silva, Caio Luiz Pereira Ribeiro, Andrea Cony Cavalcanti, Claudia Maria Braga de Mello, Amilcar Tanuri, Ana Tereza R Vasconcelos |
| EPI_ISL_2101692, EPI_ISL_2101693, EPI_ISL_2101694, EPI_ISL_2101695, EPI_ISL_2101696 | Unidade de apoio ao diagnóstico da COVID - UNADIG | Bioinformatics Laboratory / LNCC | Luiz G P de Almeida, Alessandra P Lamarca, Ronaldo da Silva F Jr, Liliane Cavalcante, Alexandra L Gerber, Ana Paula de C Guimaraes, Douglas Terra Machado, Cassia Alves, Diana Mariani, Cintia Policarpo, Gleidson da Silva de Oliveira, Mario Sergio Ribeiro, Silvia Carvalho, Flavio Dias da Silva, Marcio Henrique de Oliveira Garcia, Leandro Magalhaes de Souza, Cristiane Gomes da Silva, Caio Luiz Pereira Ribeiro, Andrea Cony Cavalcanti, Claudia Maria Braga de Mello, Amilcar Tanuri, Ana Tereza R Vasconcelos |
| EPI_ISL_2101697, EPI_ISL_2101698, EPI_ISL_2101699, EPI_ISL_2101700, EPI_ISL_2101701, EPI_ISL_2101702, EPI_ISL_2101703, EPI_ISL_2101704, EPI_ISL_2101705, EPI_ISL_2101706, EPI_ISL_2101707, EPI_ISL_2101708, EPI_ISL_2101709, EPI_ISL_2101710, EPI_ISL_2101711, EPI_ISL_2101712, EPI_ISL_2101713 | | | |
| see above | Laboratorio Central Noel Nutels | Bioinformatics Laboratory / LNCC | Luiz G P de Almeida, Alessandra P Lamarca, Ronaldo da Silva F Jr, Liliane Cavalcante, Alexandra L Gerber, Ana Paula de C Guimaraes, Douglas Terra Machado, Cassia Alves, Diana Mariani, Cintia Policarpo, Gleidson da Silva de Oliveira, Mario Sergio Ribeiro, Silvia Carvalho, Flavio Dias da Silva, Marcio Henrique de Oliveira Garcia, Leandro Magalhaes de Souza, Cristiane Gomes da Silva, Caio Luiz Pereira Ribeiro, Andrea Cony Cavalcanti, Claudia Maria Braga de Mello, Amilcar Tanuri, Ana Tereza R Vasconcelos |
| EPI_ISL_2101714, EPI_ISL_2101715 | Unidade de apoio ao diagnóstico da COVID - UNADIG | Bioinformatics Laboratory / LNCC | Luiz G P de Almeida, Alessandra P Lamarca, Ronaldo da Silva F Jr, Liliane Cavalcante, Alexandra L Gerber, Ana Paula de C Guimaraes, Douglas Terra Machado, Cassia Alves, Diana Mariani, Cintia Policarpo, Gleidson da Silva de Oliveira, Mario Sergio Ribeiro, Silvia Carvalho, Flavio Dias da Silva, Marcio Henrique de Oliveira Garcia, Leandro Magalhaes de Souza, Cristiane Gomes da Silva, Caio Luiz Pereira Ribeiro, Andrea Cony Cavalcanti, Claudia Maria Braga de Mello, Amilcar Tanuri, Ana Tereza R Vasconcelos |
| EPI_ISL_2101716, EPI_ISL_2101717, EPI_ISL_2101718, EPI_ISL_2101719, EPI_ISL_2101720, EPI_ISL_2101721 | Laboratorio Central Noel Nutels | Bioinformatics Laboratory / LNCC | Luiz G P de Almeida, Alessandra P Lamarca, Ronaldo da Silva F Jr, Liliane Cavalcante, Alexandra L Gerber, Ana Paula de C Guimaraes, Douglas Terra Machado, Cassia Alves, Diana Mariani, Cintia Policarpo, Gleidson da Silva de Oliveira, Mario Sergio Ribeiro, Silvia Carvalho, Flavio Dias da Silva, Marcio Henrique de Oliveira Garcia, Leandro Magalhaes de Souza, Cristiane Gomes da Silva, Caio Luiz Pereira Ribeiro, Andrea Cony Cavalcanti, Claudia Maria Braga de Mello, Amilcar Tanuri, Ana Tereza R Vasconcelos |
| EPI_ISL_2101722 | Unidade de apoio ao diagnóstico da COVID - UNADIG | Bioinformatics Laboratory / LNCC | Luiz G P de Almeida, Alessandra P Lamarca, Ronaldo da Silva F Jr, Liliane Cavalcante, Alexandra L Gerber, Ana Paula de C Guimaraes, Douglas Terra Machado, Cassia Alves, Diana Mariani, Cintia Policarpo, Gleidson da Silva de Oliveira, Mario Sergio Ribeiro, Silvia Carvalho, Flavio Dias da Silva, Marcio Henrique de Oliveira Garcia, Leandro Magalhaes de Souza, Cristiane Gomes da Silva, Caio Luiz Pereira Ribeiro, Andrea Cony Cavalcanti, Claudia Maria Braga de Mello, Amilcar Tanuri, Ana Tereza R Vasconcelos |
| EPI_ISL_2101723 | Laboratorio Central Noel Nutels | Bioinformatics Laboratory / LNCC | Luiz G P de Almeida, Alessandra P Lamarca, Ronaldo da Silva F Jr, Liliane Cavalcante, Alexandra L Gerber, Ana Paula de C Guimaraes, Douglas Terra Machado, Cassia Alves, Diana Mariani, Cintia Policarpo, Gleidson da Silva de Oliveira, Mario Sergio Ribeiro, Silvia Carvalho, Flavio Dias da Silva, Marcio Henrique de Oliveira Garcia, Leandro Magalhaes de Souza, Cristiane Gomes da Silva, Caio Luiz Pereira Ribeiro, Andrea Cony Cavalcanti, Claudia Maria Braga de Mello, Amilcar Tanuri, Ana Tereza R Vasconcelos |
| EPI_ISL_2101724 | Unidade de apoio ao diagnóstico da COVID - UNADIG | Bioinformatics Laboratory / LNCC | Luiz G P de Almeida, Alessandra P Lamarca, Ronaldo da Silva F Jr, Liliane Cavalcante, Alexandra L Gerber, Ana Paula de C Guimaraes, Douglas Terra Machado, Cassia Alves, Diana Mariani, Cintia Policarpo, Gleidson da Silva de Oliveira, Mario Sergio Ribeiro, Silvia Carvalho, Flavio Dias da Silva, Marcio Henrique de Oliveira Garcia, Leandro Magalhaes de Souza, Cristiane Gomes da Silva, Caio Luiz Pereira Ribeiro, Andrea Cony Cavalcanti, Claudia Maria Braga de Mello, Amilcar Tanuri, Ana Tereza R Vasconcelos |
| EPI_ISL_2101725, EPI_ISL_2101730 | Laboratorio Central Noel Nutels | Bioinformatics Laboratory / LNCC | Luiz G P de Almeida, Alessandra P Lamarca, Ronaldo da Silva F Jr, Liliane Cavalcante, Alexandra L Gerber, Ana Paula de C Guimaraes, Douglas Terra Machado, Cassia Alves, Diana Mariani, Cintia Policarpo, Gleidson da Silva de Oliveira, Mario Sergio Ribeiro, Silvia Carvalho, Flavio Dias da Silva, Marcio Henrique de Oliveira Garcia, Leandro Magalhaes de Souza, Cristiane Gomes da Silva, Caio Luiz Pereira Ribeiro, Andrea Cony Cavalcanti, Claudia Maria Braga de Mello, Amilcar Tanuri, Ana Tereza R Vasconcelos |
| EPI_ISL_2101731 | Unidade de apoio ao diagnóstico da COVID - UNADIG | Bioinformatics Laboratory / LNCC | Luiz G P de Almeida, Alessandra P Lamarca, Ronaldo da Silva F Jr, Liliane Cavalcante, Alexandra L Gerber, Ana Paula de C Guimaraes, Douglas Terra Machado, Cassia Alves, Diana Mariani, Cintia Policarpo, Gleidson da Silva de Oliveira, Mario Sergio Ribeiro, Silvia Carvalho, Flavio Dias da Silva, Marcio Henrique de Oliveira Garcia, Leandro Magalhaes de Souza, Cristiane Gomes da Silva, Caio Luiz Pereira Ribeiro, Andrea Cony Cavalcanti, Claudia Maria Braga de Mello, Amilcar Tanuri, Ana Tereza R Vasconcelos |
| EPI_ISL_2101732, EPI_ISL_2101733 | Laboratorio Central Noel Nutels | Bioinformatics Laboratory / LNCC | Luiz G P de Almeida, Alessandra P Lamarca, Ronaldo da Silva F Jr, Liliane Cavalcante, Alexandra L Gerber, Ana Paula de C Guimaraes, Douglas Terra Machado, Cassia Alves, Diana Mariani, Cintia Policarpo, Gleidson da Silva de Oliveira, Mario Sergio Ribeiro, Silvia Carvalho, Flavio Dias da Silva, Marcio Henrique de Oliveira Garcia, Leandro Magalhaes de Souza, Cristiane Gomes da Silva, Caio Luiz Pereira Ribeiro, Andrea Cony Cavalcanti, Claudia Maria Braga de Mello, Amilcar Tanuri, Ana Tereza R Vasconcelos |
| EPI_ISL_2101734 | Unidade de apoio ao diagnóstico da COVID - UNADIG | Bioinformatics Laboratory / LNCC | Luiz G P de Almeida, Alessandra P Lamarca, Ronaldo da Silva F Jr, Liliane Cavalcante, Alexandra L Gerber, Ana Paula de C Guimaraes, Douglas Terra Machado, Cassia Alves, Diana Mariani, Cintia Policarpo, Gleidson da Silva de Oliveira, Mario Sergio Ribeiro, Silvia Carvalho, Flavio Dias da Silva, Marcio Henrique de Oliveira Garcia, Leandro Magalhaes de Souza, Cristiane Gomes da Silva, Caio Luiz Pereira Ribeiro, Andrea Cony Cavalcanti, Claudia Maria Braga de Mello, Amilcar Tanuri, Ana Tereza R Vasconcelos |
| EPI_ISL_2101736, EPI_ISL_2101737, EPI_ISL_2101738, EPI_ISL_2101739 | Laboratorio Central Noel Nutels | Bioinformatics Laboratory / LNCC | Luiz G P de Almeida, Alessandra P Lamarca, Ronaldo da Silva F Jr, Liliane Cavalcante, Alexandra L Gerber, Ana Paula de C Guimaraes, Douglas Terra Machado, Cassia Alves, Diana Mariani, Cintia Policarpo, Gleidson da Silva de Oliveira, Mario Sergio Ribeiro, Silvia Carvalho, Flavio Dias da Silva, Marcio Henrique de Oliveira Garcia, Leandro Magalhaes de Souza, Cristiane Gomes da Silva, Caio Luiz Pereira Ribeiro, Andrea Cony Cavalcanti, Claudia Maria Braga de Mello, Amilcar Tanuri, Ana Tereza R Vasconcelos |
| EPI_ISL_2101740, EPI_ISL_2101741 | Unidade de apoio ao diagnóstico da COVID - UNADIG | Bioinformatics Laboratory / LNCC | Luiz G P de Almeida, Alessandra P Lamarca, Ronaldo da Silva F Jr, Liliane Cavalcante, Alexandra L Gerber, Ana Paula de C Guimaraes, Douglas Terra Machado, Cassia Alves, Diana Mariani, Cintia Policarpo, Gleidson da Silva de Oliveira, Mario Sergio Ribeiro, Silvia Carvalho, Flavio Dias da Silva, Marcio Henrique de Oliveira Garcia, Leandro Magalhaes de Souza, Cristiane Gomes da Silva, Caio Luiz Pereira Ribeiro, Andrea Cony Cavalcanti, Claudia Maria Braga de Mello, Amilcar Tanuri, Ana Tereza R Vasconcelos |
| EPI_ISL_2101742, EPI_ISL_2101743 | Laboratorio Central Noel Nutels | Bioinformatics Laboratory / LNCC | Luiz G P de Almeida, Alessandra P Lamarca, Ronaldo da Silva F Jr, Liliane Cavalcante, Alexandra L Gerber, Ana Paula de C Guimaraes, Douglas Terra Machado, Cassia Alves, Diana Mariani, Cintia Policarpo, Gleidson da Silva de Oliveira, Mario Sergio Ribeiro, Silvia Carvalho, Flavio Dias da Silva, Marcio Henrique de Oliveira Garcia, Leandro Magalhaes de Souza, Cristiane Gomes da Silva, Caio Luiz Pereira Ribeiro, Andrea Cony Cavalcanti, Claudia Maria Braga de Mello, Amilcar Tanuri, Ana Tereza R Vasconcelos |
| EPI_ISL_2101744 | Unidade de apoio ao diagnóstico da COVID - UNADIG | Bioinformatics Laboratory / LNCC | Luiz G P de Almeida, Alessandra P Lamarca, Ronaldo da Silva F Jr, Liliane Cavalcante, Alexandra L Gerber, Ana Paula de C Guimaraes, Douglas Terra Machado, Cassia Alves, Diana Mariani, Cintia Policarpo, Gleidson da Silva de Oliveira, Mario Sergio Ribeiro, Silvia Carvalho, Flavio Dias da Silva, Marcio Henrique de Oliveira Garcia, Leandro Magalhaes de Souza, Cristiane Gomes da Silva, Caio Luiz Pereira Ribeiro, Andrea Cony Cavalcanti, Claudia Maria Braga de Mello, Amilcar Tanuri, Ana Tereza R Vasconcelos |
| EPI_ISL_2101745, EPI_ISL_2101746, EPI_ISL_2101747 | Laboratorio Central Noel Nutels | Bioinformatics Laboratory / LNCC | Luiz G P de Almeida, Alessandra P Lamarca, Ronaldo da Silva F Jr, Liliane Cavalcante, Alexandra L Gerber, Ana Paula de C Guimaraes, Douglas Terra Machado, Cassia Alves, Diana Mariani, Cintia Policarpo, Gleidson da Silva de Oliveira, Mario Sergio Ribeiro, Silvia Carvalho, Flavio Dias da Silva, Marcio Henrique de Oliveira Garcia, Leandro Magalhaes de Souza, Cristiane Gomes da Silva, Caio Luiz Pereira Ribeiro, Andrea Cony Cavalcanti, Claudia Maria Braga de Mello, Amilcar Tanuri, Ana Tereza R Vasconcelos |
| EPI_ISL_2102517, EPI_ISL_2102518, EPI_ISL_2102519, EPI_ISL_2102520, EPI_ISL_2102521 | HLAGYN - Laboratorio de Imunologia de Transplantes de Goias | HLAGYN - Laboratorio de Imunologia de Transplantes de Goias | Fernando Antonio Vinhal dos Santos, Erika Lopes Rocha Batista, Alessandro Leonardo Alvares Magalhaes, Frederico Rodrigues Vinhal, Sabrina Sara Moreira Duarte, Danielle de Paiva Rezende, Lucas Carlos Gomes Pereira, Paola Cristina Resende Silva |
| EPI_ISL_2105677 | Ambulatório IB | Instituto Butantan / Mendelics | Dimas Tadeu Covas, Antonio Jorge Martins, Claudia Renata dos Santos Barros, David Schlesinger, Debora Botequio Moretti, Elaine Cristina Marqueze, Elaine Vieira Santos, Evandra Strazza Rodrigues, Heidge Fukumasu, Jayme Augusto de Souza-Neto, José Salvatore Leister Patané, Luiz Alcantara, Luiz Lehmann Coutinho, Maria Carolina Elias, Maurício Lacerda Nogueira, Rafael dos Santos Bezerra, Raul Machado Neto, Rejane Maria Tommasini Grotto, Ricardo Haddad, Sandra Coccuzzo Sampaio Vessoni, Simone Kashima, Svetoslav Nanev Slavov, Vincent Louis Viala |
| EPI_ISL_2106183 | Biology, UFLA, Universidade Federal de Lavras | Biology, UFLA, Universidade Federal de Lavras | Pylro,V., Barcante,J., Cherem,J., Luciano,P., Fernandes,G. and Melo,D. |

| | | | |
|---|---|---|---|
| EPI_ISL_2106184, EPI_ISL_2106185, EPI_ISL_2106186, EPI_ISL_2106187, EPI_ISL_2106188, EPI_ISL_2106189, EPI_ISL_2106190 | Biology, UFLA, Universidade Federal de Lavras | Biology, UFLA, Universidade Federal de Lavras | Pylro,V., Barcante,J., Cherem,J., Luciano,P., Fernandes,G., Melo,D. |
| EPI_ISL_2107294, EPI_ISL_2107296, EPI_ISL_2107298, EPI_ISL_2107299, EPI_ISL_2107304 | Laboratorio de Pesquisa em Virologia, FAMERP, SJRP | Laboratorio de Pesquisa em Virologia, FAMERP, SJRP | Cecília Artico Banho; Lívia Sacchetto; Fábio Sossai Possebon; Leila Sabrina Ullmann; Cíntia Bittar; Guilherme Campos; Helena Lage Ferreira; Jorge A. Petroli Marchesi; Maisa C. Pereira Parra; Marília Moraes; Paula Rahal; Paulo Inacio da Costa; João Pessoa Araújo Jr.; Maurício L. Nogueira. |
| EPI_ISL_2134584 | Laboratorio de Virologia Molecular / UFRJ | Bioinformatics Laboratory / LNCC | Alice L. Herlinger, Fábio L. Monteiro, Mirela D'arc, Filipe R.R. Moreira, Harrison J. Westgarth, Rafael M. Galliez, Diana Mariani, Luciana J. da Costa, Luiz Almeida, Carolina M. Voloch, Covid19-UFRJ Workgroup, Adriana S.O. Melo, Renato S. Aguiar, André F.A. dos Santos, Terezinha Marta P. P. Castiñeiras, Ana Tereza R. Vasconcelos, Esaú C. João Filho, Claudia C. Escosteguy, Orlando C. Ferreira Jr, Amilcar Tanuri, Luiza M. Higa |
| EPI_ISL_2134822 | HospitalIsraelita Albert Eistein | Virology Laboratory- Institute of Tropical Medicine SP | Fernando Lucas Melo, Camila Malta Romano |
| EPI_ISL_2139494, EPI_ISL_2139495, EPI_ISL_2139496, EPI_ISL_2139497, EPI_ISL_2139498, EPI_ISL_2139499, EPI_ISL_2139500, EPI_ISL_2139501, EPI_ISL_2139502, EPI_ISL_2139504, EPI_ISL_2139505, EPI_ISL_2139506, EPI_ISL_2139507, EPI_ISL_2139508, EPI_ISL_2139509, EPI_ISL_2139510, EPI_ISL_2139511, EPI_ISL_2139512, EPI_ISL_2139513, EPI_ISL_2139515, EPI_ISL_2139516, EPI_ISL_2139518, EPI_ISL_2139519, EPI_ISL_2139520, EPI_ISL_2139521, EPI_ISL_2139522, EPI_ISL_2139523, EPI_ISL_2139525, EPI_ISL_2139526, EPI_ISL_2139527, EPI_ISL_2139528, EPI_ISL_2139529, EPI_ISL_2139530, EPI_ISL_2139531, EPI_ISL_2139532, EPI_ISL_2139534, EPI_ISL_2139535, EPI_ISL_2139536, EPI_ISL_2139538, EPI_ISL_2139539, EPI_ISL_2139540, EPI_ISL_2139541, EPI_ISL_2139542, EPI_ISL_2139543, EPI_ISL_2139545, EPI_ISL_2139546, EPI_ISL_2139547, EPI_ISL_2139548, EPI_ISL_2139549 | | | |
| see above | Laboratorio Exame | Universidade Federal de Ciencias da Saude de Porto Alegre | Vinícius Bonetti Franceschi, Gabriel Dickin Caldana et al. |
| EPI_ISL_2157343 | Lboratorio Central de Saude Publica do Estado do Parana (LACEN/PR) | Laboratory of Respiratory Viruses and Measles, Oswaldo Cruz Institute, FIOCRUZ | Paola Resende, Luciana Appolinario, Fernando Motta, Anna Carolina Paixao, Ana Carolina Mendonca, Alice Sampaio Rocha, Taina Venas, Elisa Cavalcante Pereira, Renata Serrano Lopes, Irina Riediger, Marilda Siqueira on behalf of the Fiocruz COVID-19 Genomic Surveillance Network |
| EPI_ISL_2157375 | Laboratorio Central de Saude Publica do Estado do Espirito Santo (LACEN/ES) | Laboratory of Respiratory Viruses and Measles, Oswaldo Cruz Institute, FIOCRUZ | Paola Resende, Luciana Appolinario, Fernando Motta, Anna Carolina Paixao, Ana Carolina Mendonca, Alice Sampaio Rocha, Renata Serrano Lopes, Rodrigo Ribeiro Rodrigues, Marilda Siqueira on behalf of the Fiocruz COVID-19 Genomic Surveillance Network |
| EPI_ISL_2157376 | Laboratorio Central de Saude Publica do Estado do Espirito Santo (LACEN/ES) | Laboratory of Respiratory Viruses and Measles, Oswaldo Cruz Institute, FIOCRUZ | Paola Resende, Luciana Appolinario, Fernando Motta, Anna Carolina Paixao, Ana Carolina Mendonca, Alice Sampaio Rocha, Taina Venas, Eilisa Cavalcante Pereira, Renata Serrano Lopes, Rodrigo Ribeiro Rodrigues, Marilda Siqueira on behalf of the Fiocruz COVID-19 Genomic Surveillance Network |
| EPI_ISL_2157377 | Laboratório Central de Saude Publica do Estado de Santa Catarina (LACEN/SC) | Laboratory of Respiratory Viruses and Measles, Oswaldo Cruz Institute, FIOCRUZ | Paola Resende, Luciana Appolinario, Fernando Motta, Anna Carolina Paixao, Ana Carolina Mendonca, Alice Sampaio Rocha, Taina Venas, Elisa Cavalcante Pereira, Renata Serrano Lopes, Darcita Buerger Rovaris, Sandra Bianchini Fernandes, Marilda Siqueira on behalf of the Fiocruz COVID-19 Genomic Surveillance Network |
| EPI_ISL_2157379, EPI_ISL_2157380 | Laboratorio Central de Saude Publica do Estado da Paraiba (LACEN-PB) | Laboratory of Respiratory Viruses and Measles, Oswaldo Cruz Institute, FIOCRUZ | Paola Resende, Luciana Appolinario, Fernando Motta, Anna Carolina Paixao, Ana Carolina Mendonca, Alice Sampaio Rocha, Taina Venas, Elisa Cavalcante Pereira, Renata Serrano Lopes, Joao Felipe Bezerra, Dalane Loudal Florentino Teixeira, Marilda Siqueira on behalf of the Fiocruz COVID-19 Genomic Surveillance Network |
| EPI_ISL_2157381, EPI_ISL_2157382 | Laboratorio Central de Saude Publica do Estado de Sergipe (LACEN/SE) | Laboratory of Respiratory Viruses and Measles, Oswaldo Cruz Institute, FIOCRUZ | Paola Resende, Luciana Appolinario, Fernando Motta, Anna Carolina Paixao, Ana Carolina Mendonca, Alice Sampaio Rocha, Tainá Moreira Martins Venas, Elisa Cavalcante Pereira, Renata Serrano Lopes, Cliomar Alves dos Santos, Marilda Siqueira on behalf of the Fiocruz COVID-19 Genomic Surveillance Network |
| EPI_ISL_2157383 | Laboratorio Central de Saude Publica do Estado da Paraiba (LACEN-PB) | Laboratory of Respiratory Viruses and Measles, Oswaldo Cruz Institute, FIOCRUZ | Paola Resende, Luciana Appolinario, Fernando Motta, Anna Carolina Paixao, Ana Carolina Mendonca, Alice Sampaio Rocha, Taina Venas, Elisa Cavalcante Pereira, Renata Serrano Lopes, Joao Felipe Bezerra, Dalane Loudal Florentino Teixeira, Marilda Siqueira on behalf of the Fiocruz COVID-19 Genomic Surveillance Network |
| EPI_ISL_2157384, EPI_ISL_2157385 | Laboratorio Central de Saude Publica do Estado de Sergipe (LACEN/SE) | Laboratory of Respiratory Viruses and Measles, Oswaldo Cruz Institute, FIOCRUZ | Paola Resende, Luciana Appolinario, Fernando Motta, Anna Carolina Paixao, Ana Carolina Mendonca, Alice Sampaio Rocha, Tainá Moreira Martins Venas, Elisa Cavalcante Pereira, Renata Serrano Lopes, Cliomar Alves dos Santos, Marilda Siqueira on behalf of the Fiocruz COVID-19 Genomic Surveillance Network |
| EPI_ISL_2157386, EPI_ISL_2157387, EPI_ISL_2157388 | Lboratorio Central de Saude Publica do Estado do Parana (LACEN/PR) | Laboratory of Respiratory Viruses and Measles, Oswaldo Cruz Institute, FIOCRUZ | Paola Resende, Luciana Appolinario, Fernando Motta, Anna Carolina Paixao, Ana Carolina Mendonca, Alice Sampaio Rocha, Taina Venas, Elisa Cavalcante Pereira, Renata Serrano Lopes, Irina Riediger, Marilda Siqueira on behalf of the Fiocruz COVID-19 Genomic Surveillance Network |
| EPI_ISL_2157389 | Laboratorio Central de Saude Publica do Estado de Sergipe (LACEN/SE) | Laboratory of Respiratory Viruses and Measles, Oswaldo Cruz Institute, FIOCRUZ | Paola Resende, Luciana Appolinario, Fernando Motta, Anna Carolina Paixao, Ana Carolina Mendonca, Alice Sampaio Rocha, Tainá Moreira Martins Venas, Elisa Cavalcante Pereira, Renata Serrano Lopes, Cliomar Alves dos Santos, Marilda Siqueira on behalf of the Fiocruz COVID-19 Genomic Surveillance Network |
| EPI_ISL_2157390, EPI_ISL_2157391, EPI_ISL_2157392, EPI_ISL_2157393 | Laboratório Central de Saude Publica do Estado de Santa Catarina (LACEN/SC) | Laboratory of Respiratory Viruses and Measles, Oswaldo Cruz Institute, FIOCRUZ | Paola Resende, Luciana Appolinario, Fernando Motta, Anna Carolina Paixao, Ana Carolina Mendonca, Alice Sampaio Rocha, Taina Venas, Elisa Cavalcante Pereira, Renata Serrano Lopes, Darcita Buerger Rovaris, Sandra Bianchini Fernandes, Marilda Siqueira on behalf of the Fiocruz COVID-19 Genomic Surveillance Network |
| EPI_ISL_2157394, EPI_ISL_2157395 | Laboratorio Central de Saude Publica do Estado da Paraiba (LACEN-PB) | Laboratory of Respiratory Viruses and Measles, Oswaldo Cruz Institute, FIOCRUZ | Paola Resende, Luciana Appolinario, Fernando Motta, Anna Carolina Paixao, Ana Carolina Mendonca, Alice Sampaio Rocha, Taina Venas, Elisa Cavalcante Pereira, Renata Serrano Lopes, Joao Felipe Bezerra, Dalane Loudal Florentino Teixeira, Marilda Siqueira on behalf of the Fiocruz COVID-19 Genomic Surveillance Network |
| EPI_ISL_2157396 | Laboratório Central de Saude Publica do Estado de Santa Catarina (LACEN/SC) | Laboratory of Respiratory Viruses and Measles, Oswaldo Cruz Institute, FIOCRUZ | Paola Resende, Luciana Appolinario, Fernando Motta, Anna Carolina Paixao, Ana Carolina Mendonca, Alice Sampaio Rocha, Taina Venas, Elisa Cavalcante Pereira, Renata Serrano Lopes, Darcita Buerger Rovaris, Sandra Bianchini Fernandes, Marilda Siqueira on behalf of the Fiocruz COVID-19 Genomic Surveillance Network |
| EPI_ISL_2157397 | Lboratorio Central de Saude Publica do Estado do Parana (LACEN/PR) | Laboratory of Respiratory Viruses and Measles, Oswaldo Cruz Institute, FIOCRUZ | Paola Resende, Luciana Appolinario, Fernando Motta, Anna Carolina Paixao, Ana Carolina Mendonca, Alice Sampaio Rocha, Taina Venas, Elisa Cavalcante Pereira, Renata Serrano Lopes, Irina Riediger, Marilda Siqueira on behalf of the Fiocruz COVID-19 Genomic Surveillance Network |
| EPI_ISL_2157398, EPI_ISL_2157399, EPI_ISL_2157400 | Laboratório Central de Saude Publica do Estado de Santa Catarina (LACEN/SC) | Laboratory of Respiratory Viruses and Measles, Oswaldo Cruz Institute, FIOCRUZ | Paola Resende, Luciana Appolinario, Fernando Motta, Anna Carolina Paixao, Ana Carolina Mendonca, Alice Sampaio Rocha, Taina Venas, Elisa Cavalcante Pereira, Renata Serrano Lopes, Darcita Buerger Rovaris, Sandra Bianchini Fernandes, Marilda Siqueira on behalf of the Fiocruz COVID-19 Genomic Surveillance Network |
| EPI_ISL_2157401 | Laboratorio Central de Saude Publica do Estado da Paraiba (LACEN-PB) | Laboratory of Respiratory Viruses and Measles, Oswaldo Cruz Institute, FIOCRUZ | Paola Resende, Luciana Appolinario, Fernando Motta, Anna Carolina Paixao, Ana Carolina Mendonca, Alice Sampaio Rocha, Taina Venas, Elisa Cavalcante Pereira, Renata Serrano Lopes, Joao Felipe Bezerra, Dalane Loudal Florentino Teixeira, Marilda Siqueira on behalf of the Fiocruz COVID-19 Genomic Surveillance Network |
| EPI_ISL_2157402, EPI_ISL_2157403, EPI_ISL_2157404 | Lboratorio Central de Saude Publica do Estado do Parana (LACEN/PR) | Laboratory of Respiratory Viruses and Measles, Oswaldo Cruz Institute, FIOCRUZ | Paola Resende, Luciana Appolinario, Fernando Motta, Anna Carolina Paixao, Ana Carolina Mendonca, Alice Sampaio Rocha, Taina Venas, Elisa Cavalcante Pereira, Renata Serrano Lopes, Irina Riediger, Marilda Siqueira on behalf of the Fiocruz COVID-19 Genomic Surveillance Network |
| EPI_ISL_2157405, EPI_ISL_2157406, EPI_ISL_2157407 | Laboratório Central de Saude Publica do Estado de Santa Catarina (LACEN/SC) | Laboratory of Respiratory Viruses and Measles, Oswaldo Cruz Institute, FIOCRUZ | Paola Resende, Luciana Appolinario, Fernando Motta, Anna Carolina Paixao, Ana Carolina Mendonca, Alice Sampaio Rocha, Taina Venas, Elisa Cavalcante Pereira, Renata Serrano Lopes, Darcita Buerger Rovaris, Sandra Bianchini Fernandes, Marilda Siqueira on behalf of the Fiocruz COVID-19 Genomic Surveillance Network |
| EPI_ISL_2157408 | Lboratorio Central de Saude Publica do Estado do Parana (LACEN/PR) | Laboratory of Respiratory Viruses and Measles, Oswaldo Cruz Institute, FIOCRUZ | Paola Resende, Luciana Appolinario, Fernando Motta, Anna Carolina Paixao, Ana Carolina Mendonca, Alice Sampaio Rocha, Taina Venas, Elisa Cavalcante Pereira, Renata Serrano Lopes, Irina Riediger, Marilda Siqueira on behalf of the Fiocruz COVID-19 Genomic Surveillance Network |
| EPI_ISL_2157409, EPI_ISL_2157410, EPI_ISL_2157411 | Laboratório Central de Saude Publica do Estado de Santa Catarina (LACEN/SC) | Laboratory of Respiratory Viruses and Measles, Oswaldo Cruz Institute, FIOCRUZ | Paola Resende, Luciana Appolinario, Fernando Motta, Anna Carolina Paixao, Ana Carolina Mendonca, Alice Sampaio Rocha, Taina Venas, Elisa Cavalcante Pereira, Renata Serrano Lopes, Darcita Buerger Rovaris, Sandra Bianchini Fernandes, Marilda Siqueira on behalf of the Fiocruz COVID-19 Genomic Surveillance Network |
| EPI_ISL_2157412 | Laboratorio Central de Saude Publica do Esatado de Alagoas (LACEN/AL) | Laboratory of Respiratory Viruses and Measles, Oswaldo Cruz Institute, FIOCRUZ | Paola Resende, Luciana Appolinario, Fernando Motta, Anna Carolina Paixao, Ana Carolina Mendonca, Alice Sampaio Rocha, Taina Venas, Elisa Cavalcante Pereira, Renata Serrano Lopes, Anderson Brandao Leite, Marilda Siqueira on behalf of the Fiocruz COVID-19 Genomic Surveillance Network |
| EPI_ISL_2157413 | Lboratorio Central de Saude Publica do Estado do Parana (LACEN/PR) | Laboratory of Respiratory Viruses and Measles, Oswaldo Cruz Institute, FIOCRUZ | Paola Resende, Luciana Appolinario, Fernando Motta, Anna Carolina Paixao, Ana Carolina Mendonca, Alice Sampaio Rocha, Taina Venas, Elisa Cavalcante Pereira, Renata Serrano Lopes, Irina Riediger, Marilda Siqueira on behalf of the Fiocruz COVID-19 Genomic Surveillance Network |

| | | | |
|---|---|---|---|
| EPI_ISL_2157414, EPI_ISL_2157415 | Laboratorio Central de Saude Publica do Esatado de Alagoas (LACEN/AL) | Laboratory of Respiratory Viruses and Measles, Oswaldo Cruz Institute, FIOCRUZ | Paola Resende, Luciana Appolinario, Fernando Motta, Anna Carolina Paixao, Ana Carolina Mendonca, Alice Sampaio Rocha, Taina Venas, Elisa Cavalcante Pereira, Renata Serrano Lopes, Anderson Brandao Leite, Marilda Siqueira on behalf of the Fiocruz COVID-19 Genomic Surveillance Network |
| EPI_ISL_2157416, EPI_ISL_2157417 | Lboratorio Central de Saude Publica do Estado do Parana (LACEN/PR) | Laboratory of Respiratory Viruses and Measles, Oswaldo Cruz Institute, FIOCRUZ | Paola Resende, Luciana Appolinario, Fernando Motta, Anna Carolina Paixao, Ana Carolina Mendonca, Alice Sampaio Rocha, Taina Venas, Elisa Cavalcante Pereira, Renata Serrano Lopes, Irina Riediger, Marilda Siqueira on behalf of the Fiocruz COVID-19 Genomic Surveillance Network |
| EPI_ISL_2157418 | Laboratório Central de Saude Publica do Estado de Santa Catarina (LACEN/SC) | Laboratory of Respiratory Viruses and Measles, Oswaldo Cruz Institute, FIOCRUZ | Paola Resende, Luciana Appolinario, Fernando Motta, Anna Carolina Paixao, Ana Carolina Mendonca, Alice Sampaio Rocha, Taina Venas, Elisa Cavalcante Pereira, Renata Serrano Lopes, Darcita Buerger Rovaris, Sandra Bianchini Fernandes, Marilda Siqueira on behalf of the Fiocruz COVID-19 Genomic Surveillance Network |
| EPI_ISL_2157419, EPI_ISL_2157420, EPI_ISL_2157421 | Lboratorio Central de Saude Publica do Estado do Parana (LACEN/PR) | Laboratory of Respiratory Viruses and Measles, Oswaldo Cruz Institute, FIOCRUZ | Paola Resende, Luciana Appolinario, Fernando Motta, Anna Carolina Paixao, Ana Carolina Mendonca, Alice Sampaio Rocha, Taina Venas, Elisa Cavalcante Pereira, Renata Serrano Lopes, Irina Riediger, Marilda Siqueira on behalf of the Fiocruz COVID-19 Genomic Surveillance Network |
| EPI_ISL_2157422 | Laboratorio Central de Saude Publica do Esatado de Alagoas (LACEN/AL) | Laboratory of Respiratory Viruses and Measles, Oswaldo Cruz Institute, FIOCRUZ | Paola Resende, Luciana Appolinario, Fernando Motta, Anna Carolina Paixao, Ana Carolina Mendonca, Alice Sampaio Rocha, Taina Venas, Elisa Cavalcante Pereira, Renata Serrano Lopes, Anderson Brandao Leite, Marilda Siqueira on behalf of the Fiocruz COVID-19 Genomic Surveillance Network |
| EPI_ISL_2157423 | UNIVERSIDADE FEDERAL DE VIÇOSA | Laboratory of Respiratory Viruses and Measles, Oswaldo Cruz Institute, FIOCRUZ | Paola Resende, Luciana Appolinario, Fernando Motta, Anna Carolina Paixao, Ana Carolina Mendonca, Alice Sampaio Rocha, Taina Venas, Elisa Cavalcante Pereira, Renata Serrano Lopes, Rubens Pasa, Marilda Siqueira on behalf of the Fiocruz COVID-19 Genomic Surveillance Network |
| EPI_ISL_2157424 | Lboratorio Central de Saude Publica do Estado do Parana (LACEN/PR) | Laboratory of Respiratory Viruses and Measles, Oswaldo Cruz Institute, FIOCRUZ | Paola Resende, Luciana Appolinario, Fernando Motta, Anna Carolina Paixao, Ana Carolina Mendonca, Alice Sampaio Rocha, Taina Venas, Elisa Cavalcante Pereira, Renata Serrano Lopes, Irina Riediger, Marilda Siqueira on behalf of the Fiocruz COVID-19 Genomic Surveillance Network |
| EPI_ISL_2157425 | Laboratorio Central de Saude Publica do Estado da Paraiba (LACEN-PB) | Laboratory of Respiratory Viruses and Measles, Oswaldo Cruz Institute, FIOCRUZ | Paola Resende, Luciana Appolinario, Fernando Motta, Anna Carolina Paixao, Ana Carolina Mendonca, Alice Sampaio Rocha, Taina Venas, Elisa Cavalcante Pereira, Renata Serrano Lopes, Joao Felipe Bezerra, Dalane Loudal Florentino Teixeira, Marilda Siqueira on behalf of the Fiocruz COVID-19 Genomic Surveillance Network |
| EPI_ISL_2157426 | Laboratório Central de Saude Publica do Estado de Santa Catarina (LACEN/SC) | Laboratory of Respiratory Viruses and Measles, Oswaldo Cruz Institute, FIOCRUZ | Paola Resende, Luciana Appolinario, Fernando Motta, Anna Carolina Paixao, Ana Carolina Mendonca, Alice Sampaio Rocha, Taina Venas, Elisa Cavalcante Pereira, Renata Serrano Lopes, Darcita Buerger Rovaris, Sandra Bianchini Fernandes, Marilda Siqueira on behalf of the Fiocruz COVID-19 Genomic Surveillance Network |
| EPI_ISL_2157427, EPI_ISL_2157428, EPI_ISL_2157429, EPI_ISL_2157430 | Laboratorio Central de Saude Publica do Estado de Sergipe (LACEN/SE) | Laboratory of Respiratory Viruses and Measles, Oswaldo Cruz Institute, FIOCRUZ | Paola Resende, Luciana Appolinario, Fernando Motta, Anna Carolina Paixao, Ana Carolina Mendonca, Alice Sampaio Rocha, Tainá Moreira Martins Venas, Elisa Cavalcante Pereira, Renata Serrano Lopes, Cliomar Alves dos Santos, Marilda Siqueira on behalf of the Fiocruz COVID-19 Genomic Surveillance Network |
| EPI_ISL_2157431 | Lboratorio Central de Saude Publica do Estado do Parana (LACEN/PR) | Laboratory of Respiratory Viruses and Measles, Oswaldo Cruz Institute, FIOCRUZ | Paola Resende, Luciana Appolinario, Fernando Motta, Anna Carolina Paixao, Ana Carolina Mendonca, Alice Sampaio Rocha, Taina Venas, Elisa Cavalcante Pereira, Renata Serrano Lopes, Irina Riediger, Marilda Siqueira on behalf of the Fiocruz COVID-19 Genomic Surveillance Network |
| EPI_ISL_2157432, EPI_ISL_2157433, EPI_ISL_2157434, EPI_ISL_2157435, EPI_ISL_2157436, EPI_ISL_2157437, EPI_ISL_2157438, EPI_ISL_2157439, EPI_ISL_2157440, EPI_ISL_2157441, EPI_ISL_2157442, EPI_ISL_2157443, EPI_ISL_2157444, EPI_ISL_2157445, EPI_ISL_2157446 | | | |
| see above | Laboratorio Central de Saude Publica do Estado de Sergipe (LACEN/SE) | Laboratory of Respiratory Viruses and Measles, Oswaldo Cruz Institute, FIOCRUZ | Paola Resende, Luciana Appolinario, Fernando Motta, Anna Carolina Paixao, Ana Carolina Mendonca, Alice Sampaio Rocha, Tainá Moreira Martins Venas, Elisa Cavalcante Pereira, Renata Serrano Lopes, Cliomar Alves dos Santos, Marilda Siqueira on behalf of the Fiocruz COVID-19 Genomic Surveillance Network |
| EPI_ISL_2157447, EPI_ISL_2157448 | Laboratório Central de Saude Publica do Estado de Santa Catarina (LACEN/SC) | Laboratory of Respiratory Viruses and Measles, Oswaldo Cruz Institute, FIOCRUZ | Paola Resende, Luciana Appolinario, Fernando Motta, Anna Carolina Paixao, Ana Carolina Mendonca, Alice Sampaio Rocha, Taina Venas, Elisa Cavalcante Pereira, Renata Serrano Lopes, Darcita Buerger Rovaris, Sandra Bianchini Fernandes, Marilda Siqueira on behalf of the Fiocruz COVID-19 Genomic Surveillance Network |
| EPI_ISL_2157449 | Laboratorio Central de Saude Publica do Estado de Sergipe (LACEN/SE) | Laboratory of Respiratory Viruses and Measles, Oswaldo Cruz Institute, FIOCRUZ | Paola Resende, Luciana Appolinario, Fernando Motta, Anna Carolina Paixao, Ana Carolina Mendonca, Alice Sampaio Rocha, Tainá Moreira Martins Venas, Elisa Cavalcante Pereira, Renata Serrano Lopes, Cliomar Alves dos Santos, Marilda Siqueira on behalf of the Fiocruz COVID-19 Genomic Surveillance Network |
| EPI_ISL_2157450 | Laboratório Central de Saude Publica do Estado de Santa Catarina (LACEN/SC) | Laboratory of Respiratory Viruses and Measles, Oswaldo Cruz Institute, FIOCRUZ | Paola Resende, Luciana Appolinario, Fernando Motta, Anna Carolina Paixao, Ana Carolina Mendonca, Alice Sampaio Rocha, Taina Venas, Elisa Cavalcante Pereira, Renata Serrano Lopes, Darcita Buerger Rovaris, Sandra Bianchini Fernandes, Marilda Siqueira on behalf of the Fiocruz COVID-19 Genomic Surveillance Network |
| EPI_ISL_2157451 | Laboratorio Central de Saude Publica do Esatado de Alagoas (LACEN/AL) | Laboratory of Respiratory Viruses and Measles, Oswaldo Cruz Institute, FIOCRUZ | Paola Resende, Luciana Appolinario, Fernando Motta, Anna Carolina Paixao, Ana Carolina Mendonca, Alice Sampaio Rocha, Taina Venas, Elisa Cavalcante Pereira, Renata Serrano Lopes, Anderson Brandao Leite, Marilda Siqueira on behalf of the Fiocruz COVID-19 Genomic Surveillance Network |
| EPI_ISL_2157452 | Lboratorio Central de Saude Publica do Estado do Parana (LACEN/PR) | Laboratory of Respiratory Viruses and Measles, Oswaldo Cruz Institute, FIOCRUZ | Paola Resende, Luciana Appolinario, Fernando Motta, Anna Carolina Paixao, Ana Carolina Mendonca, Alice Sampaio Rocha, Taina Venas, Elisa Cavalcante Pereira, Renata Serrano Lopes, Irina Riediger, Marilda Siqueira on behalf of the Fiocruz COVID-19 Genomic Surveillance Network |
| EPI_ISL_2157453, EPI_ISL_2157454, EPI_ISL_2157455, EPI_ISL_2157456, EPI_ISL_2157457, EPI_ISL_2157458, EPI_ISL_2157459, EPI_ISL_2157460 | Laboratorio Central de Saude Publica do Estado de Sergipe (LACEN/SE) | Laboratory of Respiratory Viruses and Measles, Oswaldo Cruz Institute, FIOCRUZ | Paola Resende, Luciana Appolinario, Fernando Motta, Anna Carolina Paixao, Ana Carolina Mendonca, Alice Sampaio Rocha, Tainá Moreira Martins Venas, Elisa Cavalcante Pereira, Renata Serrano Lopes, Cliomar Alves dos Santos, Marilda Siqueira on behalf of the Fiocruz COVID-19 Genomic Surveillance Network |
| EPI_ISL_2157461 | Laboratorio Central de Saude Publica do Esatado de Alagoas (LACEN/AL) | Laboratory of Respiratory Viruses and Measles, Oswaldo Cruz Institute, FIOCRUZ | Paola Resende, Luciana Appolinario, Fernando Motta, Anna Carolina Paixao, Ana Carolina Mendonca, Alice Sampaio Rocha, Taina Venas, Elisa Cavalcante Pereira, Renata Serrano Lopes, Anderson Brandao Leite, Marilda Siqueira on behalf of the Fiocruz COVID-19 Genomic Surveillance Network |
| EPI_ISL_2157462, EPI_ISL_2157463, EPI_ISL_2157464, EPI_ISL_2157465, EPI_ISL_2157466, EPI_ISL_2157467, EPI_ISL_2157468, EPI_ISL_2157469, EPI_ISL_2157470, EPI_ISL_2157471, EPI_ISL_2157472, EPI_ISL_2157473, EPI_ISL_2157474, EPI_ISL_2157475, EPI_ISL_2157476, EPI_ISL_2157477, EPI_ISL_2157478 | | | |
| see above | Laboratorio Central de Saude Publica do Estado de Sergipe (LACEN/SE) | Laboratory of Respiratory Viruses and Measles, Oswaldo Cruz Institute, FIOCRUZ | Paola Resende, Luciana Appolinario, Fernando Motta, Anna Carolina Paixao, Ana Carolina Mendonca, Alice Sampaio Rocha, Tainá Moreira Martins Venas, Elisa Cavalcante Pereira, Renata Serrano Lopes, Cliomar Alves dos Santos, Marilda Siqueira on behalf of the Fiocruz COVID-19 Genomic Surveillance Network |
| EPI_ISL_2157479 | Lboratorio Central de Saude Publica do Estado do Parana (LACEN/PR) | Laboratory of Respiratory Viruses and Measles, Oswaldo Cruz Institute, FIOCRUZ | Paola Resende, Luciana Appolinario, Fernando Motta, Anna Carolina Paixao, Ana Carolina Mendonca, Alice Sampaio Rocha, Taina Venas, Elisa Cavalcante Pereira, Renata Serrano Lopes, Irina Riediger, Marilda Siqueira on behalf of the Fiocruz COVID-19 Genomic Surveillance Network |
| EPI_ISL_2157480, EPI_ISL_2157481 | Laboratorio Central de Saude Publica do Estado de Minas Gerais (LACEN/MG) | Laboratory of Respiratory Viruses and Measles, Oswaldo Cruz Institute, FIOCRUZ | Paola Resende, Luciana Appolinario, Fernando Motta, Anna Carolina Paixao, Ana Carolina Mendonca, Alice Sampaio Rocha, Taina Venas, Elisa Cavalcante Pereira, Renata Serrano Lopes, Andre Felipe Leal Bernardes, Marilda Siqueira on behalf of the Fiocruz COVID-19 Genomic Surveillance Network |
| EPI_ISL_2157482 | Laboratorio Central de Saude Publica do Estado do Parana (LACEN/PR) | Laboratory of Respiratory Viruses and Measles, Oswaldo Cruz Institute, FIOCRUZ | Paola Resende, Luciana Appolinario, Fernando Motta, Anna Carolina Paixao, Ana Carolina Mendonca, Alice Sampaio Rocha, Taina Venas, Elisa Cavalcante Pereira, Renata Serrano Lopes, Irina Riediger, Marilda Siqueira on behalf of the Fiocruz COVID-19 Genomic Surveillance Network |
| EPI_ISL_2157483 | Lboratorio Central de Saude Publica do Estado do Parana (LACEN/PR) | Laboratory of Respiratory Viruses and Measles, Oswaldo Cruz Institute, FIOCRUZ | Paola Resende, Luciana Appolinario, Fernando Motta, Anna Carolina Paixao, Ana Carolina Mendonca, Alice Sampaio Rocha, Taina Venas, Elisa Cavalcante Pereira, Renata Serrano Lopes, Irina Riediger, Marilda Siqueira on behalf of the Fiocruz COVID-19 Genomic Surveillance Network |
| EPI_ISL_2157484 | Laboratório Central de Saude Publica do Estado de Santa Catarina (LACEN/SC) | Laboratory of Respiratory Viruses and Measles, Oswaldo Cruz Institute, FIOCRUZ | Paola Resende, Luciana Appolinario, Fernando Motta, Anna Carolina Paixao, Ana Carolina Mendonca, Alice Sampaio Rocha, Taina Venas, Elisa Cavalcante Pereira, Renata Serrano Lopes, Darcita Buerger Rovaris, Sandra Bianchini Fernandes, Marilda Siqueira on behalf of the Fiocruz COVID-19 Genomic Surveillance Network |
| EPI_ISL_2157485, EPI_ISL_2157486, EPI_ISL_2157487, EPI_ISL_2157488 | Lboratorio Central de Saude Publica do Estado do Parana (LACEN/PR) | Laboratory of Respiratory Viruses and Measles, Oswaldo Cruz Institute, FIOCRUZ | Paola Resende, Luciana Appolinario, Fernando Motta, Anna Carolina Paixao, Ana Carolina Mendonca, Alice Sampaio Rocha, Taina Venas, Elisa Cavalcante Pereira, Renata Serrano Lopes, Irina Riediger, Marilda Siqueira on behalf of the Fiocruz COVID-19 Genomic Surveillance Network |
| EPI_ISL_2157489, EPI_ISL_2157490 | Laboratorio Central de Saude Publica do Esatado de Alagoas (LACEN/AL) | Laboratory of Respiratory Viruses and Measles, Oswaldo Cruz Institute, FIOCRUZ | Paola Resende, Luciana Appolinario, Fernando Motta, Anna Carolina Paixao, Ana Carolina Mendonca, Alice Sampaio Rocha, Taina Venas, Elisa Cavalcante Pereira, Renata Serrano Lopes, Anderson Brandao Leite, Marilda Siqueira on behalf of the Fiocruz COVID-19 Genomic Surveillance Network |
| EPI_ISL_2157491 | Laboratorio Central de Saude Publica do Estado de Minas Gerais (LACEN/MG) | Laboratory of Respiratory Viruses and Measles, Oswaldo Cruz Institute, FIOCRUZ | Paola Resende, Luciana Appolinario, Fernando Motta, Anna Carolina Paixao, Ana Carolina Mendonca, Alice Sampaio Rocha, Taina Venas, Elisa Cavalcante Pereira, Renata Serrano Lopes, Andre Felipe Leal Bernardes, Marilda Siqueira on behalf of the Fiocruz COVID-19 Genomic Surveillance Network |
| EPI_ISL_2157492 | Lboratorio Central de Saude Publica do Estado do Parana | Laboratory of Respiratory Viruses and Measles, Oswaldo Cruz | Paola Resende, Luciana Appolinario, Fernando Motta, Anna Carolina Paixao, Ana Carolina Mendonca, Alice Sampaio Rocha, Taina Venas, Elisa |

| | | | |
|---|---|---|---|
| | (LACEN/PR) | Institute, FIOCRUZ | Cavalcante Pereira, Renata Serrano Lopes, Irina Riediger, Marilda Siqueira on behalf of the Fiocruz COVID-19 Genomic Surveillance Network |
| EPI_ISL_2157493, EPI_ISL_2157494, EPI_ISL_2157495 | Laboratório Central de Saude Publica do Estado de Santa Catarina (LACEN/SC) | Laboratory of Respiratory Viruses and Measles, Oswaldo Cruz Institute, FIOCRUZ | Paola Resende, Luciana Appolinario, Fernando Motta, Anna Carolina Paixao, Ana Carolina Mendonca, Alice Sampaio Rocha, Taina Venas, Elisa Cavalcante Pereira, Renata Serrano Lopes, Darcita Buerger Rovaris, Sandra Bianchini Fernandes, Marilda Siqueira on behalf of the Fiocruz COVID-19 Genomic Surveillance Network |
| EPI_ISL_2157496 | Lboratorio Central de Saude Publica do Estado do Parana (LACEN/PR) | Laboratory of Respiratory Viruses and Measles, Oswaldo Cruz Institute, FIOCRUZ | Paola Resende, Luciana Appolinario, Fernando Motta, Anna Carolina Paixao, Ana Carolina Mendonca, Alice Sampaio Rocha, Taina Venas, Elisa Cavalcante Pereira, Renata Serrano Lopes, Irina Riediger, Marilda Siqueira on behalf of the Fiocruz COVID-19 Genomic Surveillance Network |
| EPI_ISL_2157497, EPI_ISL_2157498, EPI_ISL_2157499, EPI_ISL_2157500, EPI_ISL_2157501, EPI_ISL_2157502, EPI_ISL_2157503 | Laboratório Central de Saude Publica do Estado de Santa Catarina (LACEN/SC) | Laboratory of Respiratory Viruses and Measles, Oswaldo Cruz Institute, FIOCRUZ | Paola Resende, Luciana Appolinario, Fernando Motta, Anna Carolina Paixao, Ana Carolina Mendonca, Alice Sampaio Rocha, Taina Venas, Elisa Cavalcante Pereira, Renata Serrano Lopes, Darcita Buerger Rovaris, Sandra Bianchini Fernandes, Marilda Siqueira on behalf of the Fiocruz COVID-19 Genomic Surveillance Network |
| EPI_ISL_2157504 | Lboratorio Central de Saude Publica do Estado do Parana (LACEN/PR) | Laboratory of Respiratory Viruses and Measles, Oswaldo Cruz Institute, FIOCRUZ | Paola Resende, Luciana Appolinario, Fernando Motta, Anna Carolina Paixao, Ana Carolina Mendonca, Alice Sampaio Rocha, Taina Venas, Elisa Cavalcante Pereira, Renata Serrano Lopes, Irina Riediger, Marilda Siqueira on behalf of the Fiocruz COVID-19 Genomic Surveillance Network |
| EPI_ISL_2157506 | Laboratório Central de Saude Publica do Estado do Paraná (LACEN/PR) | Laboratory of Respiratory Viruses and Measles, Oswaldo Cruz Institute, FIOCRUZ | Paola Resende, Luciana Appolinario, Fernando Motta, Anna Carolina Paixao, Ana Carolina Mendonca, Alice Sampaio Rocha, Renata Serrano Lopes, Maria do Carmo Debur, Taina Venas, Elisa Cavalcante Pereira, Irina Nastassja Riediger, Marilda Siqueira on behalf of the Fiocruz COVID-19 Genomic Surveillance Network |
| EPI_ISL_2157507, EPI_ISL_2157508, EPI_ISL_2157509 | Laboratorio Central de Saude Publica do Esatado de Alagoas (LACEN/AL) | Laboratory of Respiratory Viruses and Measles, Oswaldo Cruz Institute, FIOCRUZ | Paola Resende, Luciana Appolinario, Fernando Motta, Anna Carolina Paixao, Ana Carolina Mendonca, Alice Sampaio Rocha, Taina Venas, Elisa Cavalcante Pereira, Renata Serrano Lopes, Anderson Brandao Leite, Marilda Siqueira on behalf of the Fiocruz COVID-19 Genomic Surveillance Network |
| EPI_ISL_2157510, EPI_ISL_2157511 | Laboratório Central de Saude Publica do Estado de Santa Catarina (LACEN/SC) | Laboratory of Respiratory Viruses and Measles, Oswaldo Cruz Institute, FIOCRUZ | Paola Resende, Luciana Appolinario, Fernando Motta, Anna Carolina Paixao, Ana Carolina Mendonca, Alice Sampaio Rocha, Taina Venas, Elisa Cavalcante Pereira, Renata Serrano Lopes, Darcita Buerger Rovaris, Sandra Bianchini Fernandes, Marilda Siqueira on behalf of the Fiocruz COVID-19 Genomic Surveillance Network |
| EPI_ISL_2157512 | Laboratorio Central de Saude Publica do Esatado de Alagoas (LACEN/AL) | Laboratory of Respiratory Viruses and Measles, Oswaldo Cruz Institute, FIOCRUZ | Paola Resende, Luciana Appolinario, Fernando Motta, Anna Carolina Paixao, Ana Carolina Mendonca, Alice Sampaio Rocha, Taina Venas, Elisa Cavalcante Pereira, Renata Serrano Lopes, Anderson Brandao Leite, Marilda Siqueira on behalf of the Fiocruz COVID-19 Genomic Surveillance Network |
| EPI_ISL_2157513 | Laboratório Central de Saude Publica do Estado de Santa Catarina (LACEN/SC) | Laboratory of Respiratory Viruses and Measles, Oswaldo Cruz Institute, FIOCRUZ | Paola Resende, Luciana Appolinario, Fernando Motta, Anna Carolina Paixao, Ana Carolina Mendonca, Alice Sampaio Rocha, Taina Venas, Elisa Cavalcante Pereira, Renata Serrano Lopes, Darcita Buerger Rovaris, Sandra Bianchini Fernandes, Marilda Siqueira on behalf of the Fiocruz COVID-19 Genomic Surveillance Network |
| EPI_ISL_2157514 | Laboratorio Central de Saude Publica do Esatado de Alagoas (LACEN/AL) | Laboratory of Respiratory Viruses and Measles, Oswaldo Cruz Institute, FIOCRUZ | Paola Resende, Luciana Appolinario, Fernando Motta, Anna Carolina Paixao, Ana Carolina Mendonca, Alice Sampaio Rocha, Taina Venas, Elisa Cavalcante Pereira, Renata Serrano Lopes, Anderson Brandao Leite, Marilda Siqueira on behalf of the Fiocruz COVID-19 Genomic Surveillance Network |
| EPI_ISL_2157515 | Laboratório Central de Saude Publica do Estado de Santa Catarina (LACEN/SC) | Laboratory of Respiratory Viruses and Measles, Oswaldo Cruz Institute, FIOCRUZ | Paola Resende, Luciana Appolinario, Fernando Motta, Anna Carolina Paixao, Ana Carolina Mendonca, Alice Sampaio Rocha, Taina Venas, Elisa Cavalcante Pereira, Renata Serrano Lopes, Darcita Buerger Rovaris, Sandra Bianchini Fernandes, Marilda Siqueira on behalf of the Fiocruz COVID-19 Genomic Surveillance Network |
| EPI_ISL_2157516, EPI_ISL_2157517 | Laboratório Central de Saude Publica do Estado do Paraná (LACEN/PR) | Laboratory of Respiratory Viruses and Measles, Oswaldo Cruz Institute, FIOCRUZ | Paola Resende, Luciana Appolinario, Fernando Motta, Anna Carolina Paixao, Ana Carolina Mendonca, Alice Sampaio Rocha, Renata Serrano Lopes, Maria do Carmo Debur, Taina Venas, Elisa Cavalcante Pereira, Irina Nastassja Riediger, Marilda Siqueira on behalf of the Fiocruz COVID-19 Genomic Surveillance Network |
| EPI_ISL_2157518 | Laboratório Central de Saude Publica do Estado de Santa Catarina (LACEN/SC) | Laboratory of Respiratory Viruses and Measles, Oswaldo Cruz Institute, FIOCRUZ | Paola Resende, Luciana Appolinario, Fernando Motta, Anna Carolina Paixao, Ana Carolina Mendonca, Alice Sampaio Rocha, Taina Venas, Elisa Cavalcante Pereira, Renata Serrano Lopes, Darcita Buerger Rovaris, Sandra Bianchini Fernandes, Marilda Siqueira on behalf of the Fiocruz COVID-19 Genomic Surveillance Network |
| EPI_ISL_2157519, EPI_ISL_2157520, EPI_ISL_2157521, EPI_ISL_2157522, EPI_ISL_2157523 | Laboratorio Central de Saude Publica do Esatado de Alagoas (LACEN/AL) | Laboratory of Respiratory Viruses and Measles, Oswaldo Cruz Institute, FIOCRUZ | Paola Resende, Luciana Appolinario, Fernando Motta, Anna Carolina Paixao, Ana Carolina Mendonca, Alice Sampaio Rocha, Taina Venas, Elisa Cavalcante Pereira, Renata Serrano Lopes, Anderson Brandao Leite, Marilda Siqueira on behalf of the Fiocruz COVID-19 Genomic Surveillance Network |
| EPI_ISL_2157524, EPI_ISL_2157525, EPI_ISL_2157526 | Laboratório Central de Saude Publica do Estado do Rio de Janeiro (LACEN/RJ) | Laboratory of Respiratory Viruses and Measles, Oswaldo Cruz Institute, FIOCRUZ | Paola Resende, Luciana Appolinario, Fernando Motta, Anna Carolina Paixao, Ana Carolina Mendonca, Alice Sampaio Rocha, Taina Venas, Elisa Cavalcante Pereira, Renata Serrano Lopes, Andrea Cony Cavalcanti, Marilda Siqueira on behalf of the Fiocruz COVID-19 Genomic Surveillance Network |
| EPI_ISL_2157527 | Lboratorio Central de Saude Publica do Estado do Parana (LACEN/PR) | Laboratory of Respiratory Viruses and Measles, Oswaldo Cruz Institute, FIOCRUZ | Paola Resende, Luciana Appolinario, Fernando Motta, Anna Carolina Paixao, Ana Carolina Mendonca, Alice Sampaio Rocha, Taina Venas, Elisa Cavalcante Pereira, Renata Serrano Lopes, Irina Riediger, Marilda Siqueira on behalf of the Fiocruz COVID-19 Genomic Surveillance Network |
| EPI_ISL_2157528, EPI_ISL_2157529, EPI_ISL_2157530 | Laboratório Central de Saude Publica do Estado do Rio de Janeiro (LACEN/RJ) | Laboratory of Respiratory Viruses and Measles, Oswaldo Cruz Institute, FIOCRUZ | Paola Resende, Luciana Appolinario, Fernando Motta, Anna Carolina Paixao, Ana Carolina Mendonca, Alice Sampaio Rocha, Taina Venas, Elisa Cavalcante Pereira, Renata Serrano Lopes, Andrea Cony Cavalcanti, Marilda Siqueira on behalf of the Fiocruz COVID-19 Genomic Surveillance Network |
| EPI_ISL_2157531 | Laboratorio Central de Saude Publica do Estado da Bahia (LACEN/BA) | Laboratory of Respiratory Viruses and Measles, Oswaldo Cruz Institute, FIOCRUZ | Paola Resende, Luciana Appolinario, Fernando Motta, Anna Carolina Paixao, Ana Carolina Mendonca, Alice Sampaio Rocha, Taina Venas, Elisa Cavalcante Pereira, Renata Serrano Lopes,Felicidade Pereira, Marilda Siqueira on behalf of the Fiocruz COVID-19 Genomic Surveillance Network |
| EPI_ISL_2157532, EPI_ISL_2157533, EPI_ISL_2157534, EPI_ISL_2157535 | Laboratorio Central de Saude Publica do Estado da Bahia (LACEN/BA) | Laboratory of Respiratory Viruses and Measles, Oswaldo Cruz Institute, FIOCRUZ | Paola Resende, Luciana Appolinario, Fernando Motta, Anna Carolina Paixao, Ana Carolina Mendonca, Alice Sampaio Rocha, Taina Venas, Elisa Cavalcante Pereira, Renata Serrano Lopes,Felicidade Pereira, Marilda Siqueira on behalf of the Fiocruz COVID-19 Genomic Surveillance Network |
| EPI_ISL_2157536, EPI_ISL_2157537, EPI_ISL_2157538, EPI_ISL_2157539 | Laboratório Central de Saude Publica do Estado do Rio de Janeiro (LACEN/RJ) | Laboratory of Respiratory Viruses and Measles, Oswaldo Cruz Institute, FIOCRUZ | Paola Resende, Luciana Appolinario, Fernando Motta, Anna Carolina Paixao, Ana Carolina Mendonca, Alice Sampaio Rocha, Taina Venas, Elisa Cavalcante Pereira, Renata Serrano Lopes, Andrea Cony Cavalcanti, Marilda Siqueira on behalf of the Fiocruz COVID-19 Genomic Surveillance Network |
| EPI_ISL_2157540 | Laboratory of Respiratory Viruses and Measles, Oswaldo Cruz Institute, FIOCRUZ | Laboratory of Respiratory Viruses and Measles, Oswaldo Cruz Institute, FIOCRUZ | Paola Resende, Luciana Appolinario, Fernando Motta, Anna Carolina Paixao, Ana Carolina Mendonca, Alice Sampaio Rocha, Taina Venas, Elisa Cavalcante Pereira, Renata Serrano Lopes, Andrea Cony Cavalcanti, Marilda Siqueira on behalf of the Fiocruz COVID-19 Genomic Surveillance Network |
| EPI_ISL_2157541 | Laboratorio Central de Saude Publica do Esatado de Alagoas (LACEN/AL) | Laboratory of Respiratory Viruses and Measles, Oswaldo Cruz Institute, FIOCRUZ | Paola Resende, Luciana Appolinario, Fernando Motta, Anna Carolina Paixao, Ana Carolina Mendonca, Alice Sampaio Rocha, Taina Venas, Elisa Cavalcante Pereira, Renata Serrano Lopes, Anderson Brandao Leite, Marilda Siqueira on behalf of the Fiocruz COVID-19 Genomic Surveillance Network |
| EPI_ISL_2157542, EPI_ISL_2157543, EPI_ISL_2157544 | Laboratorio Central de Saude Publica do Estado da Bahia (LACEN/BA) | Laboratory of Respiratory Viruses and Measles, Oswaldo Cruz Institute, FIOCRUZ | Paola Resende, Luciana Appolinario, Fernando Motta, Anna Carolina Paixao, Ana Carolina Mendonca, Alice Sampaio Rocha, Taina Venas, Elisa Cavalcante Pereira, Renata Serrano Lopes,Felicidade Pereira, Marilda Siqueira on behalf of the Fiocruz COVID-19 Genomic Surveillance Network |
| EPI_ISL_2157545 | Laboratory of Respiratory Viruses and Measles, Oswaldo Cruz Institute, FIOCRUZ | Laboratory of Respiratory Viruses and Measles, Oswaldo Cruz Institute, FIOCRUZ | Paola Resende, Luciana Appolinario, Fernando Motta, Anna Carolina Paixao, Ana Carolina Mendonca, Alice Sampaio Rocha, Taina Venas, Elisa Cavalcante Pereira, Renata Serrano Lopes, Andrea Cony Cavalcanti, Marilda Siqueira on behalf of the Fiocruz COVID-19 Genomic Surveillance Network |
| EPI_ISL_2157546 | Laboratorio Central de Saude Publica do Estado de Sergipe (LACEN/SE) | Laboratory of Respiratory Viruses and Measles, Oswaldo Cruz Institute, FIOCRUZ | Paola Resende, Luciana Appolinario, Fernando Motta, Anna Carolina Paixao, Ana Carolina Mendonca, Alice Sampaio Rocha, Tainá Moreira Martins Venas, Elisa Cavalcante Pereira, Renata Serrano Lopes, Cliomar Alves dos Santos, Marilda Siqueira on behalf of the Fiocruz COVID-19 Genomic Surveillance Network |
| EPI_ISL_2157547, EPI_ISL_2157548 | Laboratório Central de Saude Publica do Estado de Santa Catarina (LACEN/SC) | Laboratory of Respiratory Viruses and Measles, Oswaldo Cruz Institute, FIOCRUZ | Paola Resende, Luciana Appolinario, Fernando Motta, Anna Carolina Paixao, Ana Carolina Mendonca, Alice Sampaio Rocha, Taina Venas, Elisa Cavalcante Pereira, Renata Serrano Lopes, Darcita Buerger Rovaris, Sandra Bianchini Fernandes, Marilda Siqueira on behalf of the Fiocruz COVID-19 Genomic Surveillance Network |
| EPI_ISL_2157549 | Laboratório Central de Saude Publica do Estado do Rio de Janeiro (LACEN/RJ) | Laboratory of Respiratory Viruses and Measles, Oswaldo Cruz Institute, FIOCRUZ | Paola Resende, Luciana Appolinario, Fernando Motta, Anna Carolina Paixao, Ana Carolina Mendonca, Alice Sampaio Rocha, Taina Venas, Elisa Cavalcante Pereira, Renata Serrano Lopes, Andrea Cony Cavalcanti, Marilda Siqueira on behalf of the Fiocruz COVID-19 Genomic Surveillance Network |
| EPI_ISL_2157550 | Laboratorio Central de Saude Publica do Estado da Paraiba (LACEN-PB) | Laboratory of Respiratory Viruses and Measles, Oswaldo Cruz Institute, FIOCRUZ | Paola Resende, Luciana Appolinario, Fernando Motta, Anna Carolina Paixao, Ana Carolina Mendonca, Alice Sampaio Rocha, Taina Venas, Elisa Cavalcante Pereira, Renata Serrano Lopes, Joao Felipe Bezerra, Dalane Loudal Florentino Teixeira, Marilda Siqueira on behalf of the Fiocruz COVID-19 Genomic Surveillance Network |
| EPI_ISL_2157556 | Laboratorio Central de Saude Publica do Estado de Sergipe (LACEN/SE) | Laboratory of Respiratory Viruses and Measles, Oswaldo Cruz Institute, FIOCRUZ | Paola Resende, Luciana Appolinario, Fernando Motta, Anna Carolina Paixao, Ana Carolina Mendonca, Alice Sampaio Rocha, Tainá Moreira Martins Venas, Elisa Cavalcante Pereira, Renata Serrano Lopes, Cliomar Alves dos Santos, Marilda Siqueira on behalf of the Fiocruz COVID-19 Genomic Surveillance Network |

| | | | |
|---|---|---|---|
| EPI_ISL_2157561 | Laboratorio Central de Saude Publica do Estado do Rio de Janeiro (LACEN/RJ) | Laboratory of Respiratory Viruses and Measles, Oswaldo Cruz Institute, FIOCRUZ | Paola Resende, Luciana Appolinario, Fernando Motta, Anna Carolina Paixao, Ana Carolina Mendonca, Alice Sampaio Rocha, Taina Venas, Elisa Cavalcante Pereira, Renata Serrano Lopes, Andrea Cony Cavalcanti, Marilda Siqueira on behalf of the Fiocruz COVID-19 Genomic Surveillance Network |
| EPI_ISL_2157572 | Laboratório Central de Saude Publica do Estado de Santa Catarina (LACEN/SC) | Laboratory of Respiratory Viruses and Measles, Oswaldo Cruz Institute, FIOCRUZ | Paola Resende, Luciana Appolinario, Fernando Motta, Anna Carolina Paixao, Ana Carolina Mendonca, Alice Sampaio Rocha, Taina Venas, Elisa Cavalcante Pereira, Renata Serrano Lopes, Darcita Buerger Rovaris, Sandra Bianchini Fernandes, Marilda Siqueira on behalf of the Fiocruz COVID-19 Genomic Surveillance Network |
| EPI_ISL_2157573 | Laboratory of Respiratory Viruses and Measles, Oswaldo Cruz Institute, FIOCRUZ | Laboratory of Respiratory Viruses and Measles, Oswaldo Cruz Institute, FIOCRUZ | Paola Resende, Luciana Appolinario, Fernando Motta, Anna Carolina Paixao, Ana Carolina Mendonca, Alice Sampaio Rocha, Taina Venas, Elisa Cavalcante Pereira, Renata Serrano Lopes, Andrea Cony Cavalcanti, Marilda Siqueira on behalf of the Fiocruz COVID-19 Genomic Surveillance Network |
| EPI_ISL_2157574 | Laboratório Central de Saude Publica do Estado de Santa Catarina (LACEN/SC) | Laboratory of Respiratory Viruses and Measles, Oswaldo Cruz Institute, FIOCRUZ | Paola Resende, Luciana Appolinario, Fernando Motta, Anna Carolina Paixao, Ana Carolina Mendonca, Alice Sampaio Rocha, Taina Venas, Elisa Cavalcante Pereira, Renata Serrano Lopes, Darcita Buerger Rovaris, Sandra Bianchini Fernandes, Marilda Siqueira on behalf of the Fiocruz COVID-19 Genomic Surveillance Network |
| EPI_ISL_2157575 | Laboratory of Respiratory Viruses and Measles, Oswaldo Cruz Institute, FIOCRUZ | Laboratory of Respiratory Viruses and Measles, Oswaldo Cruz Institute, FIOCRUZ | Paola Resende, Luciana Appolinario, Fernando Motta, Anna Carolina Paixao, Ana Carolina Mendonca, Alice Sampaio Rocha, Taina Venas, Elisa Cavalcante Pereira, Renata Serrano Lopes, Andrea Cony Cavalcanti, Marilda Siqueira on behalf of the Fiocruz COVID-19 Genomic Surveillance Network |
| EPI_ISL_2157578, EPI_ISL_2157581, EPI_ISL_2157582 | Laboratorio Central de Saude Publica do Estado da Paraiba (LACEN-PB) | Laboratory of Respiratory Viruses and Measles, Oswaldo Cruz Institute, FIOCRUZ | Paola Resende, Luciana Appolinario, Fernando Motta, Anna Carolina Paixao, Ana Carolina Mendonca, Alice Sampaio Rocha, Taina Venas, Elisa Cavalcante Pereira, Renata Serrano Lopes, Joao Felipe Bezerra, Dalane Loudal Florentino Teixeira, Marilda Siqueira on behalf of the Fiocruz COVID-19 Genomic Surveillance Network |
| EPI_ISL_2157586, EPI_ISL_2157587, EPI_ISL_2157588 | Laboratorio Central de Saude Publica do Estado de Sergipe (LACEN/SE) | Laboratory of Respiratory Viruses and Measles, Oswaldo Cruz Institute, FIOCRUZ | Paola Resende, Luciana Appolinario, Fernando Motta, Anna Carolina Paixao, Ana Carolina Mendonca, Alice Sampaio Rocha, Tainá Moreira Martins Venas, Elisa Cavalcante Pereira, Renata Serrano Lopes, Cliomar Alves dos Santos, Marilda Siqueira on behalf of the Fiocruz COVID-19 Genomic Surveillance Network |
| EPI_ISL_2157589 | Laboratorio Central de Saude Publica do Estado da Paraiba (LACEN-PB) | Laboratory of Respiratory Viruses and Measles, Oswaldo Cruz Institute, FIOCRUZ | Paola Resende, Luciana Appolinario, Fernando Motta, Anna Carolina Paixao, Ana Carolina Mendonca, Alice Sampaio Rocha, Taina Venas, Elisa Cavalcante Pereira, Renata Serrano Lopes, Joao Felipe Bezerra, Dalane Loudal Florentino Teixeira, Marilda Siqueira on behalf of the Fiocruz COVID-19 Genomic Surveillance Network |
| EPI_ISL_2157590 | Laboratório Central de Saude Publica do Estado de Santa Catarina (LACEN/SC) | Laboratory of Respiratory Viruses and Measles, Oswaldo Cruz Institute, FIOCRUZ | Paola Resende, Luciana Appolinario, Fernando Motta, Anna Carolina Paixao, Ana Carolina Mendonca, Alice Sampaio Rocha, Taina Venas, Elisa Cavalcante Pereira, Renata Serrano Lopes, Darcita Buerger Rovaris, Sandra Bianchini Fernandes, Marilda Siqueira on behalf of the Fiocruz COVID-19 Genomic Surveillance Network |
| EPI_ISL_2157591 | Laboratorio Central de Saude Publica do Estado da Paraiba (LACEN-PB) | Laboratory of Respiratory Viruses and Measles, Oswaldo Cruz Institute, FIOCRUZ | Paola Resende, Luciana Appolinario, Fernando Motta, Anna Carolina Paixao, Ana Carolina Mendonca, Alice Sampaio Rocha, Taina Venas, Elisa Cavalcante Pereira, Renata Serrano Lopes, Joao Felipe Bezerra, Dalane Loudal Florentino Teixeira, Marilda Siqueira on behalf of the Fiocruz COVID-19 Genomic Surveillance Network |
| EPI_ISL_2157592 | Laboratório Central de Saude Publica do Estado de Santa Catarina (LACEN/SC) | Laboratory of Respiratory Viruses and Measles, Oswaldo Cruz Institute, FIOCRUZ | Paola Resende, Luciana Appolinario, Fernando Motta, Anna Carolina Paixao, Ana Carolina Mendonca, Alice Sampaio Rocha, Taina Venas, Elisa Cavalcante Pereira, Renata Serrano Lopes, Darcita Buerger Rovaris, Sandra Bianchini Fernandes, Marilda Siqueira on behalf of the Fiocruz COVID-19 Genomic Surveillance Network |
| EPI_ISL_2157593 | Lboratorio Central de Saude Publica do Estado do Parana (LACEN/PR) | Laboratory of Respiratory Viruses and Measles, Oswaldo Cruz Institute, FIOCRUZ | Paola Resende, Luciana Appolinario, Fernando Motta, Anna Carolina Paixao, Ana Carolina Mendonca, Alice Sampaio Rocha, Taina Venas, Elisa Cavalcante Pereira, Renata Serrano Lopes, Irina Riediger, Marilda Siqueira on behalf of the Fiocruz COVID-19 Genomic Surveillance Network |
| EPI_ISL_2157594 | Laboratorio Central de Saude Publica do Estado de Sergipe (LACEN/SE) | Laboratory of Respiratory Viruses and Measles, Oswaldo Cruz Institute, FIOCRUZ | Paola Resende, Luciana Appolinario, Fernando Motta, Anna Carolina Paixao, Ana Carolina Mendonca, Alice Sampaio Rocha, Tainá Moreira Martins Venas, Elisa Cavalcante Pereira, Renata Serrano Lopes, Cliomar Alves dos Santos, Marilda Siqueira on behalf of the Fiocruz COVID-19 Genomic Surveillance Network |
| EPI_ISL_2157595 | Lboratorio Central de Saude Publica do Estado do Parana (LACEN/PR) | Laboratory of Respiratory Viruses and Measles, Oswaldo Cruz Institute, FIOCRUZ | Paola Resende, Luciana Appolinario, Fernando Motta, Anna Carolina Paixao, Ana Carolina Mendonca, Alice Sampaio Rocha, Taina Venas, Elisa Cavalcante Pereira, Renata Serrano Lopes, Irina Riediger, Marilda Siqueira on behalf of the Fiocruz COVID-19 Genomic Surveillance Network |
| EPI_ISL_2157596 | Laboratorio Central de Saude Publica do Estado de Sergipe (LACEN/SE) | Laboratory of Respiratory Viruses and Measles, Oswaldo Cruz Institute, FIOCRUZ | Paola Resende, Luciana Appolinario, Fernando Motta, Anna Carolina Paixao, Ana Carolina Mendonca, Alice Sampaio Rocha, Tainá Moreira Martins Venas, Elisa Cavalcante Pereira, Renata Serrano Lopes, Cliomar Alves dos Santos, Marilda Siqueira on behalf of the Fiocruz COVID-19 Genomic Surveillance Network |
| EPI_ISL_2157597 | Lboratorio Central de Saude Publica do Estado do Parana (LACEN/PR) | Laboratory of Respiratory Viruses and Measles, Oswaldo Cruz Institute, FIOCRUZ | Paola Resende, Luciana Appolinario, Fernando Motta, Anna Carolina Paixao, Ana Carolina Mendonca, Alice Sampaio Rocha, Taina Venas, Elisa Cavalcante Pereira, Renata Serrano Lopes, Irina Riediger, Marilda Siqueira on behalf of the Fiocruz COVID-19 Genomic Surveillance Network |
| EPI_ISL_2157598 | Laboratório Central de Saude Publica do Estado da Bahia (LACEN/BA) | Laboratory of Respiratory Viruses and Measles, Oswaldo Cruz Institute, FIOCRUZ | Paola Resende, Luciana Appolinario, Fernando Motta, Anna Carolina Paixao, Ana Carolina Mendonca, Alice Sampaio Rocha, Taina Venas, Elisa Cavalcante Pereira, Renata Serrano Lopes,Felicidade Pereira, Marilda Siqueira on behalf of the Fiocruz COVID-19 Genomic Surveillance Network |
| EPI_ISL_2157599 | Laboratório Central de Saude Publica do Estado do Rio de Janeiro (LACEN/RJ) | Laboratory of Respiratory Viruses and Measles, Oswaldo Cruz Institute, FIOCRUZ | Paola Resende, Luciana Appolinario, Fernando Motta, Anna Carolina Paixao, Ana Carolina Mendonca, Alice Sampaio Rocha, Taina Venas, Elisa Cavalcante Pereira, Renata Serrano Lopes, Andrea Cony Cavalcanti, Marilda Siqueira on behalf of the Fiocruz COVID-19 Genomic Surveillance Network |
| EPI_ISL_2170898 | VIGILANCIA EPIDEMIOLOGICA | Instituto Butantan / Mendelics | Instituto Butantan: Dimas Tadeu Covas, Sandra Coccuzzo Sampaio, Maria Carolina Elias, José Salvatore Leister Patané, Vincent Louis Viala, Antonio Jorge Martins, Ricardo Haddad, Claudia Renata dos Santos Barros, Elaine Cristina Marqueze, Raul Machado Neto, Debora Botequio Moretti, Jardelina de Souza Todao Bernardino, Loyze Paola Oliveira de Lima, Luiz Aurelio de Campos Crispin. Centro de Genômica Funcional da ESALQ: Luiz Lehmann Coutinho, Ricardo Augusto Brassaloti, Raquel de Lello Rocha Campos Cassano. NGS Soluções Genômicas: Pilar Drummond Sampaio Corrêa Mariani. FZEA-USP Pirassununga: Mirele Daiana Poleti, Jessika Cristina Chagas Lesbon, Elisangela Chicaroni Mattos, Heidge Fukumasu. USP-Botucatu: Rejane Maria Tommasini Grotto, Jayme A. Souza-Neto, Guilherme Targino Valente, Patricia Akemi Assato, Felipe Allan da Silva da Costa, Bianca Cechetto Carlos. Mendelics: Bibiana Santos, João Paulo Kitajima, Erika Freitas, David Schlesinger. Hemocentro Ribeirão Preto: Simone Kashima, Evandra Strazza Rodrigues, Svetoslav Nanev Slavov, Elaine Vieira dos Santos, Rafael dos Santos Bezerra, Luiz Carlos Junior de Alcantara, Marta Giovanetti, Vagner Fonseca, Flavia Aburjaile, Rodrigo Tocantins Calado. FAMERP-SJRP: Cecília Artico Banho, Lívia Sacchetto, Fábio Sossai Possebon, Leila Sabrina Ullmann, Cíntia Bittar, Guilherme Campos, Helena Lage Ferreira, Jorge A. Petroli Marchesi, Maisa C. Pereira Parra, Marília Moraes, Paula Rahal, Paulo Inacio da Costa, João Pessoa Araújo Jr., Maurício Lacerda Nogueira. Prefeitura de Sao Paulo: Melissa Palmieri. |
| EPI_ISL_2170899 | SECAO CENTRO DE DIAGNOSTICO SECEDI | Instituto Butantan / Mendelics | Instituto Butantan: Dimas Tadeu Covas, Sandra Coccuzzo Sampaio, Maria Carolina Elias, José Salvatore Leister Patané, Vincent Louis Viala, Antonio Jorge Martins, Ricardo Haddad, Claudia Renata dos Santos Barros, Elaine Cristina Marqueze, Raul Machado Neto, Debora Botequio Moretti, Jardelina de Souza Todao Bernardino, Loyze Paola Oliveira de Lima, Luiz Aurelio de Campos Crispin. Centro de Genômica Funcional da ESALQ: Luiz Lehmann Coutinho, Ricardo Augusto Brassaloti, Raquel de Lello Rocha Campos Cassano. NGS Soluções Genômicas: Pilar Drummond Sampaio Corrêa Mariani. FZEA-USP Pirassununga: Mirele Daiana Poleti, Jessika Cristina Chagas Lesbon, Elisangela Chicaroni Mattos, Heidge Fukumasu. USP-Botucatu: Rejane Maria Tommasini Grotto, Jayme A. Souza-Neto, Guilherme Targino Valente, Patricia Akemi Assato, Felipe Allan da Silva da Costa, Bianca Cechetto Carlos. Mendelics: Bibiana Santos, João Paulo Kitajima, Erika Freitas, David Schlesinger. Hemocentro Ribeirão Preto: Simone Kashima, Evandra Strazza Rodrigues, Svetoslav Nanev Slavov, Elaine Vieira dos Santos, Rafael dos Santos Bezerra, Luiz Carlos Junior de Alcantara, Marta Giovanetti, Vagner Fonseca, Flavia Aburjaile, Rodrigo Tocantins Calado. FAMERP-SJRP: Cecília Artico Banho, Lívia Sacchetto, Fábio Sossai Possebon, Leila Sabrina Ullmann, Cíntia Bittar, Guilherme Campos, Helena Lage Ferreira, Jorge A. Petroli Marchesi, Maisa C. Pereira Parra, Marília Moraes, Paula Rahal, Paulo Inacio da Costa, João Pessoa Araújo Jr., Maurício Lacerda Nogueira. Prefeitura de Sao Paulo: Melissa Palmieri. |
| EPI_ISL_2170900 | AMBULATORIO MEDICO DE ESPECIALIDADES DE PERUIBE | Instituto Butantan / Mendelics | Instituto Butantan: Dimas Tadeu Covas, Sandra Coccuzzo Sampaio, Maria Carolina Elias, José Salvatore Leister Patané, Vincent Louis Viala, Antonio Jorge Martins, Ricardo Haddad, Claudia Renata dos Santos Barros, Elaine Cristina Marqueze, Raul Machado Neto, Debora Botequio Moretti, Jardelina de Souza Todao Bernardino, Loyze Paola Oliveira de Lima, Luiz Aurelio de Campos Crispin. Centro de Genômica Funcional da ESALQ: Luiz Lehmann Coutinho, Ricardo Augusto Brassaloti, Raquel de Lello Rocha Campos Cassano. NGS Soluções Genômicas: Pilar Drummond Sampaio Corrêa Mariani. FZEA-USP Pirassununga: Mirele Daiana Poleti, Jessika Cristina Chagas Lesbon, Elisangela Chicaroni Mattos, Heidge Fukumasu. USP-Botucatu: Rejane Maria Tommasini Grotto, Jayme A. Souza-Neto, Guilherme Targino Valente, Patricia Akemi Assato, Felipe Allan da Silva da Costa, Bianca Cechetto Carlos. Mendelics: Bibiana Santos, João Paulo Kitajima, Erika Freitas, David Schlesinger. Hemocentro Ribeirão Preto: Simone Kashima, Evandra Strazza Rodrigues, Svetoslav Nanev Slavov, Elaine Vieira dos Santos, Rafael dos Santos Bezerra, Luiz Carlos Junior de Alcantara, Marta Giovanetti, Vagner Fonseca, Flavia Aburjaile, Rodrigo Tocantins Calado. FAMERP-SJRP: Cecília Artico Banho, Lívia Sacchetto, Fábio Sossai Possebon, Leila Sabrina |

| | | | |
|---|---|---|---|
| | | | Ullmann, Cíntia Bittar, Guilherme Campos, Helena Lage Ferreira, Jorge A. Petroli Marchesi, Maisa C. Pereira Parra, Marília Moraes, Paula Rahal, Paulo Inacio da Costa, João Pessoa Araújo Jr., Maurício Lacerda Nogueira. Prefeitura de Sao Paulo: Melissa Palmieri. |
| EPI_ISL_2170902 | USF JARDIM SAO DIMAS | Instituto Butantan / Mendelics | Instituto Butantan: Dimas Tadeu Covas, Sandra Coccuzzo Sampaio, Maria Carolina Elias, José Salvatore Leister Patané, Vincent Louis Viala, Antonio Jorge Martins, Ricardo Haddad, Claudia Renata dos Santos Barros, Elaine Cristina Marqueze, Raul Machado Neto, Debora Botequio Moretti, Jardelina de Souza Todao Bernardino, Loyze Paola Oliveira de Lima, Luiz Aurelio de Campos Crispin. Centro de Genômica Funcional da ESALQ: Luiz Lehmann Coutinho, Ricardo Augusto Brassaloti, Raquel de Lello Rocha Campos Cassano. NGS Soluções Genômicas: Pilar Drummond Sampaio Corrêa Mariani. FZEA-USP Pirassununga: Mirele Daiana Poleti, Jessika Cristina Chagas Lesbon, Elisangela Chicaroni Mattos, Heidge Fukumasu. USP-Botucatu: Rejane Maria Tommasini Grotto, Jayme A. Souza-Neto, Guilherme Targino Valente, Patricia Akemi Assato, Felipe Allan da Silva da Costa, Bianca Cechetto Carlos. Mendelics: Bibiana Santos, João Paulo Kitajima, Erika Freitas, David Schlesinger. Hemocentro Ribeirão Preto: Simone Kashima, Evandra Strazza Rodrigues, Svetoslav Nanev Slavov, Elaine Vieira dos Santos, Rafael dos Santos Bezerra, Luiz Carlos Junior de Alcantara, Marta Giovanetti, Vagner Fonseca, Flavia Aburjaile, Rodrigo Tocantins Calado. FAMERP-SJRP: Cecília Artico Banho, Lívia Sacchetto, Fábio Sossai Possebon, Leila Sabrina Ullmann, Cíntia Bittar, Guilherme Campos, Helena Lage Ferreira, Jorge A. Petroli Marchesi, Maisa C. Pereira Parra, Marília Moraes, Paula Rahal, Paulo Inacio da Costa, João Pessoa Araújo Jr., Maurício Lacerda Nogueira. Prefeitura de Sao Paulo: Melissa Palmieri. |
| EPI_ISL_2170906 | SECRETARIA MUNICIPAL DE SAUDE SOROCABA | Instituto Butantan / Mendelics | Instituto Butantan: Dimas Tadeu Covas, Sandra Coccuzzo Sampaio, Maria Carolina Elias, José Salvatore Leister Patané, Vincent Louis Viala, Antonio Jorge Martins, Ricardo Haddad, Claudia Renata dos Santos Barros, Elaine Cristina Marqueze, Raul Machado Neto, Debora Botequio Moretti, Jardelina de Souza Todao Bernardino, Loyze Paola Oliveira de Lima, Luiz Aurelio de Campos Crispin. Centro de Genômica Funcional da ESALQ: Luiz Lehmann Coutinho, Ricardo Augusto Brassaloti, Raquel de Lello Rocha Campos Cassano. NGS Soluções Genômicas: Pilar Drummond Sampaio Corrêa Mariani. FZEA-USP Pirassununga: Mirele Daiana Poleti, Jessika Cristina Chagas Lesbon, Elisangela Chicaroni Mattos, Heidge Fukumasu. USP-Botucatu: Rejane Maria Tommasini Grotto, Jayme A. Souza-Neto, Guilherme Targino Valente, Patricia Akemi Assato, Felipe Allan da Silva da Costa, Bianca Cechetto Carlos. Mendelics: Bibiana Santos, João Paulo Kitajima, Erika Freitas, David Schlesinger. Hemocentro Ribeirão Preto: Simone Kashima, Evandra Strazza Rodrigues, Svetoslav Nanev Slavov, Elaine Vieira dos Santos, Rafael dos Santos Bezerra, Luiz Carlos Junior de Alcantara, Marta Giovanetti, Vagner Fonseca, Flavia Aburjaile, Rodrigo Tocantins Calado. FAMERP-SJRP: Cecília Artico Banho, Lívia Sacchetto, Fábio Sossai Possebon, Leila Sabrina Ullmann, Cíntia Bittar, Guilherme Campos, Helena Lage Ferreira, Jorge A. Petroli Marchesi, Maisa C. Pereira Parra, Marília Moraes, Paula Rahal, Paulo Inacio da Costa, João Pessoa Araújo Jr., Maurício Lacerda Nogueira. Prefeitura de Sao Paulo: Melissa Palmieri. |
| EPI_ISL_2170908 | LABORATORIO DE FRANCA | Instituto Butantan / Mendelics | Instituto Butantan: Dimas Tadeu Covas, Sandra Coccuzzo Sampaio, Maria Carolina Elias, José Salvatore Leister Patané, Vincent Louis Viala, Antonio Jorge Martins, Ricardo Haddad, Claudia Renata dos Santos Barros, Elaine Cristina Marqueze, Raul Machado Neto, Debora Botequio Moretti, Jardelina de Souza Todao Bernardino, Loyze Paola Oliveira de Lima, Luiz Aurelio de Campos Crispin. Centro de Genômica Funcional da ESALQ: Luiz Lehmann Coutinho, Ricardo Augusto Brassaloti, Raquel de Lello Rocha Campos Cassano. NGS Soluções Genômicas: Pilar Drummond Sampaio Corrêa Mariani. FZEA-USP Pirassununga: Mirele Daiana Poleti, Jessika Cristina Chagas Lesbon, Elisangela Chicaroni Mattos, Heidge Fukumasu. USP-Botucatu: Rejane Maria Tommasini Grotto, Jayme A. Souza-Neto, Guilherme Targino Valente, Patricia Akemi Assato, Felipe Allan da Silva da Costa, Bianca Cechetto Carlos. Mendelics: Bibiana Santos, João Paulo Kitajima, Erika Freitas, David Schlesinger. Hemocentro Ribeirão Preto: Simone Kashima, Evandra Strazza Rodrigues, Svetoslav Nanev Slavov, Elaine Vieira dos Santos, Rafael dos Santos Bezerra, Luiz Carlos Junior de Alcantara, Marta Giovanetti, Vagner Fonseca, Flavia Aburjaile, Rodrigo Tocantins Calado. FAMERP-SJRP: Cecília Artico Banho, Lívia Sacchetto, Fábio Sossai Possebon, Leila Sabrina Ullmann, Cíntia Bittar, Guilherme Campos, Helena Lage Ferreira, Jorge A. Petroli Marchesi, Maisa C. Pereira Parra, Marília Moraes, Paula Rahal, Paulo Inacio da Costa, João Pessoa Araújo Jr., Maurício Lacerda Nogueira. Prefeitura de Sao Paulo: Melissa Palmieri. |
| EPI_ISL_2170909 | SECRETARIA MUNICIPAL DE SAUDE SOROCABA | Instituto Butantan / Mendelics | Dimas Tadeu Covas, Antonio Jorge Martins, Claudia Renata dos Santos Barros, David Schlesinger, Debora Botequio Moretti, Elaine Cristina Marqueze, Elaine Vieira Santos, Evandra Strazza Rodrigues, Heidge Fukumasu, Jayme Augusto de Souza-Neto, José Salvatore Leister Patané, Luiz Alcantara, Luiz Lehmann Coutinho, Maria Carolina Elias, Maurício Lacerda Nogueira, Rafael dos Santos Bezerra, Raul Machado Neto, Rejane Maria Tommasini Grotto, Ricardo Haddad, Sandra Coccuzzo Sampaio Vessoni, Simone Kashima, Svetoslav Nanev Slavov, Vincent Louis Viala |
| EPI_ISL_2170910 | SECRETARIA MUNICIPAL DA SAUDE DE GUARIBA | Instituto Butantan / Mendelics | Instituto Butantan: Dimas Tadeu Covas, Sandra Coccuzzo Sampaio, Maria Carolina Elias, José Salvatore Leister Patané, Vincent Louis Viala, Antonio Jorge Martins, Ricardo Haddad, Claudia Renata dos Santos Barros, Elaine Cristina Marqueze, Raul Machado Neto, Debora Botequio Moretti, Jardelina de Souza Todao Bernardino, Loyze Paola Oliveira de Lima, Luiz Aurelio de Campos Crispin. Centro de Genômica Funcional da ESALQ: Luiz Lehmann Coutinho, Ricardo Augusto Brassaloti, Raquel de Lello Rocha Campos Cassano. NGS Soluções Genômicas: Pilar Drummond Sampaio Corrêa Mariani. FZEA-USP Pirassununga: Mirele Daiana Poleti, Jessika Cristina Chagas Lesbon, Elisangela Chicaroni Mattos, Heidge Fukumasu. USP-Botucatu: Rejane Maria Tommasini Grotto, Jayme A. Souza-Neto, Guilherme Targino Valente, Patricia Akemi Assato, Felipe Allan da Silva da Costa, Bianca Cechetto Carlos. Mendelics: Bibiana Santos, João Paulo Kitajima, Erika Freitas, David Schlesinger. Hemocentro Ribeirão Preto: Simone Kashima, Evandra Strazza Rodrigues, Svetoslav Nanev Slavov, Elaine Vieira dos Santos, Rafael dos Santos Bezerra, Luiz Carlos Junior de Alcantara, Marta Giovanetti, Vagner Fonseca, Flavia Aburjaile, Rodrigo Tocantins Calado. FAMERP-SJRP: Cecília Artico Banho, Lívia Sacchetto, Fábio Sossai Possebon, Leila Sabrina Ullmann, Cíntia Bittar, Guilherme Campos, Helena Lage Ferreira, Jorge A. Petroli Marchesi, Maisa C. Pereira Parra, Marília Moraes, Paula Rahal, Paulo Inacio da Costa, João Pessoa Araújo Jr., Maurício Lacerda Nogueira. Prefeitura de Sao Paulo: Melissa Palmieri. |
| EPI_ISL_2170911 | UNIDADE BASICA DE SAUDE DA FAMILIA BILAC | Instituto Butantan / Mendelics | Instituto Butantan: Dimas Tadeu Covas, Sandra Coccuzzo Sampaio, Maria Carolina Elias, José Salvatore Leister Patané, Vincent Louis Viala, Antonio Jorge Martins, Ricardo Haddad, Claudia Renata dos Santos Barros, Elaine Cristina Marqueze, Raul Machado Neto, Debora Botequio Moretti, Jardelina de Souza Todao Bernardino, Loyze Paola Oliveira de Lima, Luiz Aurelio de Campos Crispin. Centro de Genômica Funcional da ESALQ: Luiz Lehmann Coutinho, Ricardo Augusto Brassaloti, Raquel de Lello Rocha Campos Cassano. NGS Soluções Genômicas: Pilar Drummond Sampaio Corrêa Mariani. FZEA-USP Pirassununga: Mirele Daiana Poleti, Jessika Cristina Chagas Lesbon, Elisangela Chicaroni Mattos, Heidge Fukumasu. USP-Botucatu: Rejane Maria Tommasini Grotto, Jayme A. Souza-Neto, Guilherme Targino Valente, Patricia Akemi Assato, Felipe Allan da Silva da Costa, Bianca Cechetto Carlos. Mendelics: Bibiana Santos, João Paulo Kitajima, Erika Freitas, David Schlesinger. Hemocentro Ribeirão Preto: Simone Kashima, Evandra Strazza Rodrigues, Svetoslav Nanev Slavov, Elaine Vieira dos Santos, Rafael dos Santos Bezerra, Luiz Carlos Junior de Alcantara, Marta Giovanetti, Vagner Fonseca, Flavia Aburjaile, Rodrigo Tocantins Calado. FAMERP-SJRP: Cecília Artico Banho, Lívia Sacchetto, Fábio Sossai Possebon, Leila Sabrina Ullmann, Cíntia Bittar, Guilherme Campos, Helena Lage Ferreira, Jorge A. Petroli Marchesi, Maisa C. Pereira Parra, Marília Moraes, Paula Rahal, Paulo Inacio da Costa, João Pessoa Araújo Jr., Maurício Lacerda Nogueira. Prefeitura de Sao Paulo: Melissa Palmieri. |
| EPI_ISL_2170916 | HOSPITAL E MATERNIDADE NOSSA SENHORA DA AJUDA | Instituto Butantan / Mendelics | Instituto Butantan: Dimas Tadeu Covas, Sandra Coccuzzo Sampaio, Maria Carolina Elias, José Salvatore Leister Patané, Vincent Louis Viala, Antonio Jorge Martins, Ricardo Haddad, Claudia Renata dos Santos Barros, Elaine Cristina Marqueze, Raul Machado Neto, Debora Botequio Moretti, Jardelina de Souza Todao Bernardino, Loyze Paola Oliveira de Lima, Luiz Aurelio de Campos Crispin. Centro de Genômica Funcional da ESALQ: Luiz Lehmann Coutinho, Ricardo Augusto Brassaloti, Raquel de Lello Rocha Campos Cassano. NGS Soluções Genômicas: Pilar Drummond Sampaio Corrêa Mariani. FZEA-USP Pirassununga: Mirele Daiana Poleti, Jessika Cristina Chagas Lesbon, Elisangela Chicaroni Mattos, Heidge Fukumasu. USP-Botucatu: Rejane Maria Tommasini Grotto, Jayme A. Souza-Neto, Guilherme Targino Valente, Patricia Akemi Assato, Felipe Allan da Silva da Costa, Bianca Cechetto Carlos. Mendelics: Bibiana Santos, João Paulo Kitajima, Erika Freitas, David Schlesinger. Hemocentro Ribeirão Preto: Simone Kashima, Evandra Strazza Rodrigues, Svetoslav Nanev Slavov, Elaine Vieira dos Santos, Rafael dos Santos Bezerra, Luiz Carlos Junior de Alcantara, Marta Giovanetti, Vagner Fonseca, Flavia Aburjaile, Rodrigo Tocantins Calado. FAMERP-SJRP: Cecília Artico Banho, Lívia Sacchetto, Fábio Sossai Possebon, Leila Sabrina Ullmann, Cíntia Bittar, Guilherme Campos, Helena Lage Ferreira, Jorge A. Petroli Marchesi, Maisa C. Pereira Parra, Marília Moraes, Paula Rahal, Paulo Inacio da Costa, João Pessoa Araújo Jr., Maurício Lacerda Nogueira. Prefeitura de Sao Paulo: Melissa Palmieri. |
| EPI_ISL_2170917, EPI_ISL_2170918 | PRONTO ATENDIMENTO VILA PADRE ANCHIETA | Instituto Butantan / Mendelics | Instituto Butantan: Dimas Tadeu Covas, Sandra Coccuzzo Sampaio, Maria Carolina Elias, José Salvatore Leister Patané, Vincent Louis Viala, Antonio Jorge Martins, Ricardo Haddad, Claudia Renata dos Santos Barros, Elaine Cristina Marqueze, Raul Machado Neto, Debora Botequio Moretti, Jardelina de Souza Todao Bernardino, Loyze Paola Oliveira de Lima, Luiz Aurelio de Campos Crispin. Centro de Genômica Funcional da ESALQ: Luiz Lehmann Coutinho, Ricardo Augusto Brassaloti, Raquel de Lello Rocha Campos Cassano. NGS Soluções Genômicas: Pilar Drummond Sampaio Corrêa Mariani. FZEA-USP Pirassununga: Mirele Daiana Poleti, Jessika Cristina Chagas Lesbon, Elisangela Chicaroni Mattos, Heidge Fukumasu. USP-Botucatu: Rejane Maria Tommasini Grotto, Jayme A. Souza-Neto, Guilherme Targino Valente, Patricia Akemi Assato, Felipe Allan da Silva da Costa, Bianca Cechetto Carlos. Mendelics: Bibiana Santos, João Paulo Kitajima, Erika Freitas, David Schlesinger. Hemocentro Ribeirão Preto: Simone Kashima, Evandra Strazza Rodrigues, Svetoslav Nanev Slavov, Elaine Vieira dos Santos, Rafael dos Santos Bezerra, Luiz Carlos Junior de Alcantara, Marta Giovanetti, Vagner Fonseca, Flavia Aburjaile, Rodrigo Tocantins Calado. FAMERP-SJRP: Cecília Artico Banho, Lívia Sacchetto, Fábio Sossai Possebon, Leila Sabrina Ullmann, Cíntia Bittar, Guilherme Campos, Helena Lage Ferreira, Jorge A. Petroli Marchesi, Maisa C. Pereira Parra, Marília Moraes, Paula Rahal, Paulo Inacio da Costa, João Pessoa Araújo Jr., Maurício Lacerda Nogueira. Prefeitura de Sao Paulo: Melissa Palmieri. |

| | | | |
|---|---|---|---|
| EPI_ISL_2170920 | USAFA MARACANA | Instituto Butantan / Mendelics | Instituto Butantan: Dimas Tadeu Covas, Sandra Coccuzzo Sampaio, Maria Carolina Elias, José Salvatore Leister Patané, Vincent Louis Viala, Antonio Jorge Martins, Ricardo Haddad, Claudia Renata dos Santos Barros, Elaine Cristina Marqueze, Raul Machado Neto, Debora Botequio Moretti, Jardelina de Souza Todao Bernardino, Loyze Paola Oliveira de Lima, Luiz Aurelio de Campos Crispin. Centro de Genômica Funcional da ESALQ: Luiz Lehmann Coutinho, Ricardo Augusto Brassaloti, Raquel de Lello Rocha Campos Cassano. NGS Soluções Genômicas: Pilar Drummond Sampaio Corrêa Mariani. FZEA-USP Pirassununga: Mirele Daiana Poleti, Jessika Cristina Chagas Lesbon, Elisangela Chicaroni Mattos, Heidge Fukumasu. USP-Botucatu: Rejane Maria Tommasini Grotto, Jayme A. Souza-Neto, Guilherme Targino Valente, Patricia Akemi Assato, Felipe Allan da Silva da Costa, Bianca Cechetto Carlos. Mendelics: Bibiana Santos, João Paulo Kitajima, Erika Freitas, David Schlesinger. Hemocentro Ribeirão Preto: Simone Kashima, Evandra Strazza Rodrigues, Svetoslav Nanev Slavov, Elaine Vieira dos Santos, Rafael dos Santos Bezerra, Luiz Carlos Junior de Alcantara, Marta Giovanetti, Vagner Fonseca, Flavia Aburjaile, Rodrigo Tocantins Calado. FAMERP-SJRP: Cecília Artico Banho, Lívia Sacchetto, Fábio Sossai Possebon, Leila Sabrina Ullmann, Cíntia Bittar, Guilherme Campos, Helena Lage Ferreira, Jorge A. Petroli Marchesi, Maisa C. Pereira Parra, Marília Moraes, Paula Rahal, Paulo Inacio da Costa, João Pessoa Araújo Jr., Maurício Lacerda Nogueira. Prefeitura de Sao Paulo: Melissa Palmieri. |
| EPI_ISL_2170922 | SMS SECRETARIA MUNICIPAL DE SAUDE DE BOITUVA | Instituto Butantan / Mendelics | Instituto Butantan: Dimas Tadeu Covas, Sandra Coccuzzo Sampaio, Maria Carolina Elias, José Salvatore Leister Patané, Vincent Louis Viala, Antonio Jorge Martins, Ricardo Haddad, Claudia Renata dos Santos Barros, Elaine Cristina Marqueze, Raul Machado Neto, Debora Botequio Moretti, Jardelina de Souza Todao Bernardino, Loyze Paola Oliveira de Lima, Luiz Aurelio de Campos Crispin. Centro de Genômica Funcional da ESALQ: Luiz Lehmann Coutinho, Ricardo Augusto Brassaloti, Raquel de Lello Rocha Campos Cassano. NGS Soluções Genômicas: Pilar Drummond Sampaio Corrêa Mariani. FZEA-USP Pirassununga: Mirele Daiana Poleti, Jessika Cristina Chagas Lesbon, Elisangela Chicaroni Mattos, Heidge Fukumasu. USP-Botucatu: Rejane Maria Tommasini Grotto, Jayme A. Souza-Neto, Guilherme Targino Valente, Patricia Akemi Assato, Felipe Allan da Silva da Costa, Bianca Cechetto Carlos. Mendelics: Bibiana Santos, João Paulo Kitajima, Erika Freitas, David Schlesinger. Hemocentro Ribeirão Preto: Simone Kashima, Evandra Strazza Rodrigues, Svetoslav Nanev Slavov, Elaine Vieira dos Santos, Rafael dos Santos Bezerra, Luiz Carlos Junior de Alcantara, Marta Giovanetti, Vagner Fonseca, Flavia Aburjaile, Rodrigo Tocantins Calado. FAMERP-SJRP: Cecília Artico Banho, Lívia Sacchetto, Fábio Sossai Possebon, Leila Sabrina Ullmann, Cíntia Bittar, Guilherme Campos, Helena Lage Ferreira, Jorge A. Petroli Marchesi, Maisa C. Pereira Parra, Marília Moraes, Paula Rahal, Paulo Inacio da Costa, João Pessoa Araújo Jr., Maurício Lacerda Nogueira. Prefeitura de Sao Paulo: Melissa Palmieri. |
| EPI_ISL_2170927 | PRONTO SOCORRO DE AGENOR DE CAMPOS MONGAGUA | Instituto Butantan / Mendelics | Instituto Butantan: Dimas Tadeu Covas, Sandra Coccuzzo Sampaio, Maria Carolina Elias, José Salvatore Leister Patané, Vincent Louis Viala, Antonio Jorge Martins, Ricardo Haddad, Claudia Renata dos Santos Barros, Elaine Cristina Marqueze, Raul Machado Neto, Debora Botequio Moretti, Jardelina de Souza Todao Bernardino, Loyze Paola Oliveira de Lima, Luiz Aurelio de Campos Crispin. Centro de Genômica Funcional da ESALQ: Luiz Lehmann Coutinho, Ricardo Augusto Brassaloti, Raquel de Lello Rocha Campos Cassano. NGS Soluções Genômicas: Pilar Drummond Sampaio Corrêa Mariani. FZEA-USP Pirassununga: Mirele Daiana Poleti, Jessika Cristina Chagas Lesbon, Elisangela Chicaroni Mattos, Heidge Fukumasu. USP-Botucatu: Rejane Maria Tommasini Grotto, Jayme A. Souza-Neto, Guilherme Targino Valente, Patricia Akemi Assato, Felipe Allan da Silva da Costa, Bianca Cechetto Carlos. Mendelics: Bibiana Santos, João Paulo Kitajima, Erika Freitas, David Schlesinger. Hemocentro Ribeirão Preto: Simone Kashima, Evandra Strazza Rodrigues, Svetoslav Nanev Slavov, Elaine Vieira dos Santos, Rafael dos Santos Bezerra, Luiz Carlos Junior de Alcantara, Marta Giovanetti, Vagner Fonseca, Flavia Aburjaile, Rodrigo Tocantins Calado. FAMERP-SJRP: Cecília Artico Banho, Lívia Sacchetto, Fábio Sossai Possebon, Leila Sabrina Ullmann, Cíntia Bittar, Guilherme Campos, Helena Lage Ferreira, Jorge A. Petroli Marchesi, Maisa C. Pereira Parra, Marília Moraes, Paula Rahal, Paulo Inacio da Costa, João Pessoa Araújo Jr., Maurício Lacerda Nogueira. Prefeitura de Sao Paulo: Melissa Palmieri. |
| EPI_ISL_2170929 | Ambulatório IB | Instituto Butantan / Mendelics | Instituto Butantan: Dimas Tadeu Covas, Sandra Coccuzzo Sampaio, Maria Carolina Elias, José Salvatore Leister Patané, Vincent Louis Viala, Antonio Jorge Martins, Ricardo Haddad, Claudia Renata dos Santos Barros, Elaine Cristina Marqueze, Raul Machado Neto, Debora Botequio Moretti, Jardelina de Souza Todao Bernardino, Loyze Paola Oliveira de Lima, Luiz Aurelio de Campos Crispin. Centro de Genômica Funcional da ESALQ: Luiz Lehmann Coutinho, Ricardo Augusto Brassaloti, Raquel de Lello Rocha Campos Cassano. NGS Soluções Genômicas: Pilar Drummond Sampaio Corrêa Mariani. FZEA-USP Pirassununga: Mirele Daiana Poleti, Jessika Cristina Chagas Lesbon, Elisangela Chicaroni Mattos, Heidge Fukumasu. USP-Botucatu: Rejane Maria Tommasini Grotto, Jayme A. Souza-Neto, Guilherme Targino Valente, Patricia Akemi Assato, Felipe Allan da Silva da Costa, Bianca Cechetto Carlos. Mendelics: Bibiana Santos, João Paulo Kitajima, Erika Freitas, David Schlesinger. Hemocentro Ribeirão Preto: Simone Kashima, Evandra Strazza Rodrigues, Svetoslav Nanev Slavov, Elaine Vieira dos Santos, Rafael dos Santos Bezerra, Luiz Carlos Junior de Alcantara, Marta Giovanetti, Vagner Fonseca, Flavia Aburjaile, Rodrigo Tocantins Calado. FAMERP-SJRP: Cecília Artico Banho, Lívia Sacchetto, Fábio Sossai Possebon, Leila Sabrina Ullmann, Cíntia Bittar, Guilherme Campos, Helena Lage Ferreira, Jorge A. Petroli Marchesi, Maisa C. Pereira Parra, Marília Moraes, Paula Rahal, Paulo Inacio da Costa, João Pessoa Araújo Jr., Maurício Lacerda Nogueira. Prefeitura de Sao Paulo: Melissa Palmieri. |
| EPI_ISL_2170930 | CENTRO DE SAUDE DE CHAVANTES | Instituto Butantan / Mendelics | Instituto Butantan: Dimas Tadeu Covas, Sandra Coccuzzo Sampaio, Maria Carolina Elias, José Salvatore Leister Patané, Vincent Louis Viala, Antonio Jorge Martins, Ricardo Haddad, Claudia Renata dos Santos Barros, Elaine Cristina Marqueze, Raul Machado Neto, Debora Botequio Moretti, Jardelina de Souza Todao Bernardino, Loyze Paola Oliveira de Lima, Luiz Aurelio de Campos Crispin. Centro de Genômica Funcional da ESALQ: Luiz Lehmann Coutinho, Ricardo Augusto Brassaloti, Raquel de Lello Rocha Campos Cassano. NGS Soluções Genômicas: Pilar Drummond Sampaio Corrêa Mariani. FZEA-USP Pirassununga: Mirele Daiana Poleti, Jessika Cristina Chagas Lesbon, Elisangela Chicaroni Mattos, Heidge Fukumasu. USP-Botucatu: Rejane Maria Tommasini Grotto, Jayme A. Souza-Neto, Guilherme Targino Valente, Patricia Akemi Assato, Felipe Allan da Silva da Costa, Bianca Cechetto Carlos. Mendelics: Bibiana Santos, João Paulo Kitajima, Erika Freitas, David Schlesinger. Hemocentro Ribeirão Preto: Simone Kashima, Evandra Strazza Rodrigues, Svetoslav Nanev Slavov, Elaine Vieira dos Santos, Rafael dos Santos Bezerra, Luiz Carlos Junior de Alcantara, Marta Giovanetti, Vagner Fonseca, Flavia Aburjaile, Rodrigo Tocantins Calado. FAMERP-SJRP: Cecília Artico Banho, Lívia Sacchetto, Fábio Sossai Possebon, Leila Sabrina Ullmann, Cíntia Bittar, Guilherme Campos, Helena Lage Ferreira, Jorge A. Petroli Marchesi, Maisa C. Pereira Parra, Marília Moraes, Paula Rahal, Paulo Inacio da Costa, João Pessoa Araújo Jr., Maurício Lacerda Nogueira. Prefeitura de Sao Paulo: Melissa Palmieri. |
| EPI_ISL_2170931 | UNIDADE RESPIRATORIA NOVA HORTOLANDIA | Instituto Butantan / Mendelics | Instituto Butantan: Dimas Tadeu Covas, Sandra Coccuzzo Sampaio, Maria Carolina Elias, José Salvatore Leister Patané, Vincent Louis Viala, Antonio Jorge Martins, Ricardo Haddad, Claudia Renata dos Santos Barros, Elaine Cristina Marqueze, Raul Machado Neto, Debora Botequio Moretti, Jardelina de Souza Todao Bernardino, Loyze Paola Oliveira de Lima, Luiz Aurelio de Campos Crispin. Centro de Genômica Funcional da ESALQ: Luiz Lehmann Coutinho, Ricardo Augusto Brassaloti, Raquel de Lello Rocha Campos Cassano. NGS Soluções Genômicas: Pilar Drummond Sampaio Corrêa Mariani. FZEA-USP Pirassununga: Mirele Daiana Poleti, Jessika Cristina Chagas Lesbon, Elisangela Chicaroni Mattos, Heidge Fukumasu. USP-Botucatu: Rejane Maria Tommasini Grotto, Jayme A. Souza-Neto, Guilherme Targino Valente, Patricia Akemi Assato, Felipe Allan da Silva da Costa, Bianca Cechetto Carlos. Mendelics: Bibiana Santos, João Paulo Kitajima, Erika Freitas, David Schlesinger. Hemocentro Ribeirão Preto: Simone Kashima, Evandra Strazza Rodrigues, Svetoslav Nanev Slavov, Elaine Vieira dos Santos, Rafael dos Santos Bezerra, Luiz Carlos Junior de Alcantara, Marta Giovanetti, Vagner Fonseca, Flavia Aburjaile, Rodrigo Tocantins Calado. FAMERP-SJRP: Cecília Artico Banho, Lívia Sacchetto, Fábio Sossai Possebon, Leila Sabrina Ullmann, Cíntia Bittar, Guilherme Campos, Helena Lage Ferreira, Jorge A. Petroli Marchesi, Maisa C. Pereira Parra, Marília Moraes, Paula Rahal, Paulo Inacio da Costa, João Pessoa Araújo Jr., Maurício Lacerda Nogueira. Prefeitura de Sao Paulo: Melissa Palmieri. |
| EPI_ISL_2170932, EPI_ISL_2170933 | UNIDADE DE PRONTO ATENDIMENTO UPA DRA ANA OLIVIA BENTIVOGLIO | Instituto Butantan / Mendelics | Instituto Butantan: Dimas Tadeu Covas, Sandra Coccuzzo Sampaio, Maria Carolina Elias, José Salvatore Leister Patané, Vincent Louis Viala, Antonio Jorge Martins, Ricardo Haddad, Claudia Renata dos Santos Barros, Elaine Cristina Marqueze, Raul Machado Neto, Debora Botequio Moretti, Jardelina de Souza Todao Bernardino, Loyze Paola Oliveira de Lima, Luiz Aurelio de Campos Crispin. Centro de Genômica Funcional da ESALQ: Luiz Lehmann Coutinho, Ricardo Augusto Brassaloti, Raquel de Lello Rocha Campos Cassano. NGS Soluções Genômicas: Pilar Drummond Sampaio Corrêa Mariani. FZEA-USP Pirassununga: Mirele Daiana Poleti, Jessika Cristina Chagas Lesbon, Elisangela Chicaroni Mattos, Heidge Fukumasu. USP-Botucatu: Rejane Maria Tommasini Grotto, Jayme A. Souza-Neto, Guilherme Targino Valente, Patricia Akemi Assato, Felipe Allan da Silva da Costa, Bianca Cechetto Carlos. Mendelics: Bibiana Santos, João Paulo Kitajima, Erika Freitas, David Schlesinger. Hemocentro Ribeirão Preto: Simone Kashima, Evandra Strazza Rodrigues, Svetoslav Nanev Slavov, Elaine Vieira dos Santos, Rafael dos Santos Bezerra, Luiz Carlos Junior de Alcantara, Marta Giovanetti, Vagner Fonseca, Flavia Aburjaile, Rodrigo Tocantins Calado. FAMERP-SJRP: Cecília Artico Banho, Lívia Sacchetto, Fábio Sossai Possebon, Leila Sabrina Ullmann, Cíntia Bittar, Guilherme Campos, Helena Lage Ferreira, Jorge A. Petroli Marchesi, Maisa C. Pereira Parra, Marília Moraes, Paula Rahal, Paulo Inacio da Costa, João Pessoa Araújo Jr., Maurício Lacerda Nogueira. Prefeitura de Sao Paulo: Melissa Palmieri. |
| EPI_ISL_2170935 | NUCLEO DE SAUDE VILA FALCAO DE BAURU | Instituto Butantan / Mendelics | Instituto Butantan: Dimas Tadeu Covas, Sandra Coccuzzo Sampaio, Maria Carolina Elias, José Salvatore Leister Patané, Vincent Louis Viala, Antonio Jorge Martins, Ricardo Haddad, Claudia Renata dos Santos Barros, Elaine Cristina Marqueze, Raul Machado Neto, Debora Botequio Moretti, Jardelina de Souza Todao Bernardino, Loyze Paola Oliveira de Lima, Luiz Aurelio de Campos Crispin. Centro de Genômica Funcional da ESALQ: Luiz Lehmann Coutinho, Ricardo Augusto Brassaloti, Raquel de Lello Rocha Campos Cassano. NGS Soluções Genômicas: Pilar Drummond Sampaio Corrêa Mariani. FZEA-USP Pirassununga: Mirele Daiana Poleti, Jessika Cristina Chagas Lesbon, Elisangela Chicaroni Mattos, Heidge Fukumasu. USP-Botucatu: Rejane Maria Tommasini Grotto, Jayme A. Souza-Neto, Guilherme Targino Valente, Patricia Akemi Assato, Felipe Allan da Silva da Costa, Bianca Cechetto Carlos. Mendelics: Bibiana Santos, João Paulo Kitajima, Erika Freitas, David Schlesinger. Hemocentro Ribeirão Preto: Simone Kashima, Evandra Strazza |

| | | | |
|---|---|---|---|
| | | | Rodrigues, Svetoslav Nanev Slavov, Elaine Vieira dos Santos, Rafael dos Santos Bezerra, Luiz Carlos Junior de Alcantara, Marta Giovanetti, Vagner Fonseca, Flavia Aburjaile, Rodrigo Tocantins Calado. FAMERP-SJRP: Cecília Artico Banho, Lívia Sacchetto, Fábio Sossai Possebon, Leila Sabrina Ullmann, Cíntia Bittar, Guilherme Campos, Helena Lage Ferreira, Jorge A. Petroli Marchesi, Maisa C. Pereira Parra, Marília Moraes, Paula Rahal, Paulo Inacio da Costa, João Pessoa Araújo Jr., Maurício Lacerda Nogueira. Prefeitura de Sao Paulo: Melissa Palmieri. |
| EPI_ISL_2170936 | CENTRO DE SAUDE II DR ALCIDES FACUNDO ARROYO | Instituto Butantan / Mendelics | Instituto Butantan: Dimas Tadeu Covas, Sandra Coccuzzo Sampaio, Maria Carolina Elias, José Salvatore Leister Patané, Vincent Louis Viala, Antonio Jorge Martins, Ricardo Haddad, Claudia Renata dos Santos Barros, Elaine Cristina Marqueze, Raul Machado Neto, Debora Botequio Moretti, Jardelina de Souza Todao Bernardino, Loyze Paola Oliveira de Lima, Luiz Aurelio de Campos Crispin. Centro de Genômica Funcional da ESALQ: Luiz Lehmann Coutinho, Ricardo Augusto Brassaloti, Raquel de Lello Rocha Campos Cassano. NGS Soluções Genômicas: Pilar Drummond Sampaio Corrêa Mariani. FZEA-USP Pirassununga: Mirele Daiana Poleti, Jessika Cristina Chagas Lesbon, Elisangela Chicaroni Mattos, Heidge Fukumasu. USP-Botucatu: Rejane Maria Tommasini Grotto, Jayme A. Souza-Neto, Guilherme Targino Valente, Patricia Akemi Assato, Felipe Allan da Silva da Costa, Bianca Cechetto Carlos. Mendelics: Bibiana Santos, João Paulo Kitajima, Erika Freitas, David Schlesinger. Hemocentro Ribeirão Preto: Simone Kashima, Evandra Strazza Rodrigues, Svetoslav Nanev Slavov, Elaine Vieira dos Santos, Rafael dos Santos Bezerra, Luiz Carlos Junior de Alcantara, Marta Giovanetti, Vagner Fonseca, Flavia Aburjaile, Rodrigo Tocantins Calado. FAMERP-SJRP: Cecília Artico Banho, Lívia Sacchetto, Fábio Sossai Possebon, Leila Sabrina Ullmann, Cíntia Bittar, Guilherme Campos, Helena Lage Ferreira, Jorge A. Petroli Marchesi, Maisa C. Pereira Parra, Marília Moraes, Paula Rahal, Paulo Inacio da Costa, João Pessoa Araújo Jr., Maurício Lacerda Nogueira. Prefeitura de Sao Paulo: Melissa Palmieri. |
| EPI_ISL_2170937 | HOSP E MATERNIDADE MUNICIPAL N SRA MONTE SERRAT | Instituto Butantan / Mendelics | Instituto Butantan: Dimas Tadeu Covas, Sandra Coccuzzo Sampaio, Maria Carolina Elias, José Salvatore Leister Patané, Vincent Louis Viala, Antonio Jorge Martins, Ricardo Haddad, Claudia Renata dos Santos Barros, Elaine Cristina Marqueze, Raul Machado Neto, Debora Botequio Moretti, Jardelina de Souza Todao Bernardino, Loyze Paola Oliveira de Lima, Luiz Aurelio de Campos Crispin. Centro de Genômica Funcional da ESALQ: Luiz Lehmann Coutinho, Ricardo Augusto Brassaloti, Raquel de Lello Rocha Campos Cassano. NGS Soluções Genômicas: Pilar Drummond Sampaio Corrêa Mariani. FZEA-USP Pirassununga: Mirele Daiana Poleti, Jessika Cristina Chagas Lesbon, Elisangela Chicaroni Mattos, Heidge Fukumasu. USP-Botucatu: Rejane Maria Tommasini Grotto, Jayme A. Souza-Neto, Guilherme Targino Valente, Patricia Akemi Assato, Felipe Allan da Silva da Costa, Bianca Cechetto Carlos. Mendelics: Bibiana Santos, João Paulo Kitajima, Erika Freitas, David Schlesinger. Hemocentro Ribeirão Preto: Simone Kashima, Evandra Strazza Rodrigues, Svetoslav Nanev Slavov, Elaine Vieira dos Santos, Rafael dos Santos Bezerra, Luiz Carlos Junior de Alcantara, Marta Giovanetti, Vagner Fonseca, Flavia Aburjaile, Rodrigo Tocantins Calado. FAMERP-SJRP: Cecília Artico Banho, Lívia Sacchetto, Fábio Sossai Possebon, Leila Sabrina Ullmann, Cíntia Bittar, Guilherme Campos, Helena Lage Ferreira, Jorge A. Petroli Marchesi, Maisa C. Pereira Parra, Marília Moraes, Paula Rahal, Paulo Inacio da Costa, João Pessoa Araújo Jr., Maurício Lacerda Nogueira. Prefeitura de Sao Paulo: Melissa Palmieri. |
| EPI_ISL_2170940 | NUCLEO DE SAUDE VILA FALCAO DE BAURU | Instituto Butantan / Mendelics | Instituto Butantan: Dimas Tadeu Covas, Sandra Coccuzzo Sampaio, Maria Carolina Elias, José Salvatore Leister Patané, Vincent Louis Viala, Antonio Jorge Martins, Ricardo Haddad, Claudia Renata dos Santos Barros, Elaine Cristina Marqueze, Raul Machado Neto, Debora Botequio Moretti, Jardelina de Souza Todao Bernardino, Loyze Paola Oliveira de Lima, Luiz Aurelio de Campos Crispin. Centro de Genômica Funcional da ESALQ: Luiz Lehmann Coutinho, Ricardo Augusto Brassaloti, Raquel de Lello Rocha Campos Cassano. NGS Soluções Genômicas: Pilar Drummond Sampaio Corrêa Mariani. FZEA-USP Pirassununga: Mirele Daiana Poleti, Jessika Cristina Chagas Lesbon, Elisangela Chicaroni Mattos, Heidge Fukumasu. USP-Botucatu: Rejane Maria Tommasini Grotto, Jayme A. Souza-Neto, Guilherme Targino Valente, Patricia Akemi Assato, Felipe Allan da Silva da Costa, Bianca Cechetto Carlos. Mendelics: Bibiana Santos, João Paulo Kitajima, Erika Freitas, David Schlesinger. Hemocentro Ribeirão Preto: Simone Kashima, Evandra Strazza Rodrigues, Svetoslav Nanev Slavov, Elaine Vieira dos Santos, Rafael dos Santos Bezerra, Luiz Carlos Junior de Alcantara, Marta Giovanetti, Vagner Fonseca, Flavia Aburjaile, Rodrigo Tocantins Calado. FAMERP-SJRP: Cecília Artico Banho, Lívia Sacchetto, Fábio Sossai Possebon, Leila Sabrina Ullmann, Cíntia Bittar, Guilherme Campos, Helena Lage Ferreira, Jorge A. Petroli Marchesi, Maisa C. Pereira Parra, Marília Moraes, Paula Rahal, Paulo Inacio da Costa, João Pessoa Araújo Jr., Maurício Lacerda Nogueira. Prefeitura de Sao Paulo: Melissa Palmieri. |
| EPI_ISL_2170942 | LABORATORIO MUNICIPAL DE ANALISES CLINICAS DE ITANHAEM | Instituto Butantan / Mendelics | Instituto Butantan: Dimas Tadeu Covas, Sandra Coccuzzo Sampaio, Maria Carolina Elias, José Salvatore Leister Patané, Vincent Louis Viala, Antonio Jorge Martins, Ricardo Haddad, Claudia Renata dos Santos Barros, Elaine Cristina Marqueze, Raul Machado Neto, Debora Botequio Moretti, Jardelina de Souza Todao Bernardino, Loyze Paola Oliveira de Lima, Luiz Aurelio de Campos Crispin. Centro de Genômica Funcional da ESALQ: Luiz Lehmann Coutinho, Ricardo Augusto Brassaloti, Raquel de Lello Rocha Campos Cassano. NGS Soluções Genômicas: Pilar Drummond Sampaio Corrêa Mariani. FZEA-USP Pirassununga: Mirele Daiana Poleti, Jessika Cristina Chagas Lesbon, Elisangela Chicaroni Mattos, Heidge Fukumasu. USP-Botucatu: Rejane Maria Tommasini Grotto, Jayme A. Souza-Neto, Guilherme Targino Valente, Patricia Akemi Assato, Felipe Allan da Silva da Costa, Bianca Cechetto Carlos. Mendelics: Bibiana Santos, João Paulo Kitajima, Erika Freitas, David Schlesinger. Hemocentro Ribeirão Preto: Simone Kashima, Evandra Strazza Rodrigues, Svetoslav Nanev Slavov, Elaine Vieira dos Santos, Rafael dos Santos Bezerra, Luiz Carlos Junior de Alcantara, Marta Giovanetti, Vagner Fonseca, Flavia Aburjaile, Rodrigo Tocantins Calado. FAMERP-SJRP: Cecília Artico Banho, Lívia Sacchetto, Fábio Sossai Possebon, Leila Sabrina Ullmann, Cíntia Bittar, Guilherme Campos, Helena Lage Ferreira, Jorge A. Petroli Marchesi, Maisa C. Pereira Parra, Marília Moraes, Paula Rahal, Paulo Inacio da Costa, João Pessoa Araújo Jr., Maurício Lacerda Nogueira. Prefeitura de Sao Paulo: Melissa Palmieri. |
| EPI_ISL_2170944 | UNIDADE DE PRONTO ATENDIMENTO UPA DRA ANA OLIVIA BENTIVOGLIO | Instituto Butantan / Mendelics | Instituto Butantan: Dimas Tadeu Covas, Sandra Coccuzzo Sampaio, Maria Carolina Elias, José Salvatore Leister Patané, Vincent Louis Viala, Antonio Jorge Martins, Ricardo Haddad, Claudia Renata dos Santos Barros, Elaine Cristina Marqueze, Raul Machado Neto, Debora Botequio Moretti, Jardelina de Souza Todao Bernardino, Loyze Paola Oliveira de Lima, Luiz Aurelio de Campos Crispin. Centro de Genômica Funcional da ESALQ: Luiz Lehmann Coutinho, Ricardo Augusto Brassaloti, Raquel de Lello Rocha Campos Cassano. NGS Soluções Genômicas: Pilar Drummond Sampaio Corrêa Mariani. FZEA-USP Pirassununga: Mirele Daiana Poleti, Jessika Cristina Chagas Lesbon, Elisangela Chicaroni Mattos, Heidge Fukumasu. USP-Botucatu: Rejane Maria Tommasini Grotto, Jayme A. Souza-Neto, Guilherme Targino Valente, Patricia Akemi Assato, Felipe Allan da Silva da Costa, Bianca Cechetto Carlos. Mendelics: Bibiana Santos, João Paulo Kitajima, Erika Freitas, David Schlesinger. Hemocentro Ribeirão Preto: Simone Kashima, Evandra Strazza Rodrigues, Svetoslav Nanev Slavov, Elaine Vieira dos Santos, Rafael dos Santos Bezerra, Luiz Carlos Junior de Alcantara, Marta Giovanetti, Vagner Fonseca, Flavia Aburjaile, Rodrigo Tocantins Calado. FAMERP-SJRP: Cecília Artico Banho, Lívia Sacchetto, Fábio Sossai Possebon, Leila Sabrina Ullmann, Cíntia Bittar, Guilherme Campos, Helena Lage Ferreira, Jorge A. Petroli Marchesi, Maisa C. Pereira Parra, Marília Moraes, Paula Rahal, Paulo Inacio da Costa, João Pessoa Araújo Jr., Maurício Lacerda Nogueira. Prefeitura de Sao Paulo: Melissa Palmieri. |
| EPI_ISL_2170945 | POLICLINICA HORTOLANDIA | Instituto Butantan / Mendelics | Instituto Butantan: Dimas Tadeu Covas, Sandra Coccuzzo Sampaio, Maria Carolina Elias, José Salvatore Leister Patané, Vincent Louis Viala, Antonio Jorge Martins, Ricardo Haddad, Claudia Renata dos Santos Barros, Elaine Cristina Marqueze, Raul Machado Neto, Debora Botequio Moretti, Jardelina de Souza Todao Bernardino, Loyze Paola Oliveira de Lima, Luiz Aurelio de Campos Crispin. Centro de Genômica Funcional da ESALQ: Luiz Lehmann Coutinho, Ricardo Augusto Brassaloti, Raquel de Lello Rocha Campos Cassano. NGS Soluções Genômicas: Pilar Drummond Sampaio Corrêa Mariani. FZEA-USP Pirassununga: Mirele Daiana Poleti, Jessika Cristina Chagas Lesbon, Elisangela Chicaroni Mattos, Heidge Fukumasu. USP-Botucatu: Rejane Maria Tommasini Grotto, Jayme A. Souza-Neto, Guilherme Targino Valente, Patricia Akemi Assato, Felipe Allan da Silva da Costa, Bianca Cechetto Carlos. Mendelics: Bibiana Santos, João Paulo Kitajima, Erika Freitas, David Schlesinger. Hemocentro Ribeirão Preto: Simone Kashima, Evandra Strazza Rodrigues, Svetoslav Nanev Slavov, Elaine Vieira dos Santos, Rafael dos Santos Bezerra, Luiz Carlos Junior de Alcantara, Marta Giovanetti, Vagner Fonseca, Flavia Aburjaile, Rodrigo Tocantins Calado. FAMERP-SJRP: Cecília Artico Banho, Lívia Sacchetto, Fábio Sossai Possebon, Leila Sabrina Ullmann, Cíntia Bittar, Guilherme Campos, Helena Lage Ferreira, Jorge A. Petroli Marchesi, Maisa C. Pereira Parra, Marília Moraes, Paula Rahal, Paulo Inacio da Costa, João Pessoa Araújo Jr., Maurício Lacerda Nogueira. Prefeitura de Sao Paulo: Melissa Palmieri. |
| EPI_ISL_2170948, EPI_ISL_2170949, EPI_ISL_2170952, EPI_ISL_2170953 | UPA UNIDADE DE PRONTO ATENDIMENTO 24 HORAS BOM JESUS | Instituto Butantan / Mendelics | Instituto Butantan: Dimas Tadeu Covas, Sandra Coccuzzo Sampaio, Maria Carolina Elias, José Salvatore Leister Patané, Vincent Louis Viala, Antonio Jorge Martins, Ricardo Haddad, Claudia Renata dos Santos Barros, Elaine Cristina Marqueze, Raul Machado Neto, Debora Botequio Moretti, Jardelina de Souza Todao Bernardino, Loyze Paola Oliveira de Lima, Luiz Aurelio de Campos Crispin. Centro de Genômica Funcional da ESALQ: Luiz Lehmann Coutinho, Ricardo Augusto Brassaloti, Raquel de Lello Rocha Campos Cassano. NGS Soluções Genômicas: Pilar Drummond Sampaio Corrêa Mariani. FZEA-USP Pirassununga: Mirele Daiana Poleti, Jessika Cristina Chagas Lesbon, Elisangela Chicaroni Mattos, Heidge Fukumasu. USP-Botucatu: Rejane Maria Tommasini Grotto, Jayme A. Souza-Neto, Guilherme Targino Valente, Patricia Akemi Assato, Felipe Allan da Silva da Costa, Bianca Cechetto Carlos. Mendelics: Bibiana Santos, João Paulo Kitajima, Erika Freitas, David Schlesinger. Hemocentro Ribeirão Preto: Simone Kashima, Evandra Strazza Rodrigues, Svetoslav Nanev Slavov, Elaine Vieira dos Santos, Rafael dos Santos Bezerra, Luiz Carlos Junior de Alcantara, Marta Giovanetti, Vagner Fonseca, Flavia Aburjaile, Rodrigo Tocantins Calado. FAMERP-SJRP: Cecília Artico Banho, Lívia Sacchetto, Fábio Sossai Possebon, Leila Sabrina Ullmann, Cíntia Bittar, Guilherme Campos, Helena Lage Ferreira, Jorge A. Petroli Marchesi, Maisa C. Pereira Parra, Marília Moraes, Paula Rahal, Paulo Inacio da Costa, João Pessoa Araújo Jr., Maurício Lacerda Nogueira. Prefeitura de Sao Paulo: Melissa Palmieri. |
| EPI_ISL_2170954 | LABORATORIO DE FRANCA | Instituto Butantan / Mendelics | Instituto Butantan: Dimas Tadeu Covas, Sandra Coccuzzo Sampaio, Maria Carolina Elias, José Salvatore Leister Patané, Vincent Louis Viala, Antonio Jorge Martins, Ricardo Haddad, Claudia Renata dos Santos Barros, Elaine Cristina Marqueze, Raul Machado Neto, Debora Botequio Moretti, Jardelina de |

| | | | |
|---|---|---|---|
| | | | Souza Todao Bernardino, Loyze Paola Oliveira de Lima, Luiz Aurelio de Campos Crispin. Centro de Genômica Funcional da ESALQ: Luiz Lehmann Coutinho, Ricardo Augusto Brassaloti, Raquel de Lello Rocha Campos Cassano. NGS Soluções Genômicas: Pilar Drummond Sampaio Corrêa Mariani. FZEA-USP Pirassununga: Mirele Daiana Poleti, Jessika Cristina Chagas Lesbon, Elisangela Chicaroni Mattos, Heidge Fukumasu. USP-Botucatu: Rejane Maria Tommasini Grotto, Jayme A. Souza-Neto, Guilherme Targino Valente, Patricia Akemi Assato, Felipe Allan da Silva da Costa, Bianca Cechetto Carlos. Mendelics: Bibiana Santos, João Paulo Kitajima, Erika Freitas, David Schlesinger. Hemocentro Ribeirão Preto: Simone Kashima, Evandra Strazza Rodrigues, Svetoslav Nanev Slavov, Elaine Vieira dos Santos, Rafael dos Santos Bezerra, Luiz Carlos Junior de Alcantara, Marta Giovanetti, Vagner Fonseca, Flavia Aburjaile, Rodrigo Tocantins Calado. FAMERP-SJRP: Cecília Artico Banho, Lívia Sacchetto, Fábio Sossai Possebon, Leila Sabrina Ullmann, Cíntia Bittar, Guilherme Campos, Helena Lage Ferreira, Jorge A. Petroli Marchesi, Maisa C. Pereira Parra, Marília Moraes, Paula Rahal, Paulo Inacio da Costa, João Pessoa Araújo Jr., Maurício Lacerda Nogueira. Prefeitura de Sao Paulo: Melissa Palmieri. |
| EPI_ISL_2170955 | HOSPITAL SANTO ANTONIO DE JUQUIA JUQUIA | Instituto Butantan / Mendelics | Instituto Butantan: Dimas Tadeu Covas, Sandra Coccuzzo Sampaio, Maria Carolina Elias, José Salvatore Leister Patané, Vincent Louis Viala, Antonio Jorge Martins, Ricardo Haddad, Claudia Renata dos Santos Barros, Elaine Cristina Marqueze, Raul Machado Neto, Debora Botequio Moretti, Jardelina de Souza Todao Bernardino, Loyze Paola Oliveira de Lima, Luiz Aurelio de Campos Crispin. Centro de Genômica Funcional da ESALQ: Luiz Lehmann Coutinho, Ricardo Augusto Brassaloti, Raquel de Lello Rocha Campos Cassano. NGS Soluções Genômicas: Pilar Drummond Sampaio Corrêa Mariani. FZEA-USP Pirassununga: Mirele Daiana Poleti, Jessika Cristina Chagas Lesbon, Elisangela Chicaroni Mattos, Heidge Fukumasu. USP-Botucatu: Rejane Maria Tommasini Grotto, Jayme A. Souza-Neto, Guilherme Targino Valente, Patricia Akemi Assato, Felipe Allan da Silva da Costa, Bianca Cechetto Carlos. Mendelics: Bibiana Santos, João Paulo Kitajima, Erika Freitas, David Schlesinger. Hemocentro Ribeirão Preto: Simone Kashima, Evandra Strazza Rodrigues, Svetoslav Nanev Slavov, Elaine Vieira dos Santos, Rafael dos Santos Bezerra, Luiz Carlos Junior de Alcantara, Marta Giovanetti, Vagner Fonseca, Flavia Aburjaile, Rodrigo Tocantins Calado. FAMERP-SJRP: Cecília Artico Banho, Lívia Sacchetto, Fábio Sossai Possebon, Leila Sabrina Ullmann, Cíntia Bittar, Guilherme Campos, Helena Lage Ferreira, Jorge A. Petroli Marchesi, Maisa C. Pereira Parra, Marília Moraes, Paula Rahal, Paulo Inacio da Costa, João Pessoa Araújo Jr., Maurício Lacerda Nogueira. Prefeitura de Sao Paulo: Melissa Palmieri. |
| EPI_ISL_2170956, EPI_ISL_2170957, EPI_ISL_2170958 | AMBULATORIO MEDICO DE ESPECIALIDADES DE PERUIBE | Instituto Butantan / Mendelics | Instituto Butantan: Dimas Tadeu Covas, Sandra Coccuzzo Sampaio, Maria Carolina Elias, José Salvatore Leister Patané, Vincent Louis Viala, Antonio Jorge Martins, Ricardo Haddad, Claudia Renata dos Santos Barros, Elaine Cristina Marqueze, Raul Machado Neto, Debora Botequio Moretti, Jardelina de Souza Todao Bernardino, Loyze Paola Oliveira de Lima, Luiz Aurelio de Campos Crispin. Centro de Genômica Funcional da ESALQ: Luiz Lehmann Coutinho, Ricardo Augusto Brassaloti, Raquel de Lello Rocha Campos Cassano. NGS Soluções Genômicas: Pilar Drummond Sampaio Corrêa Mariani. FZEA-USP Pirassununga: Mirele Daiana Poleti, Jessika Cristina Chagas Lesbon, Elisangela Chicaroni Mattos, Heidge Fukumasu. USP-Botucatu: Rejane Maria Tommasini Grotto, Jayme A. Souza-Neto, Guilherme Targino Valente, Patricia Akemi Assato, Felipe Allan da Silva da Costa, Bianca Cechetto Carlos. Mendelics: Bibiana Santos, João Paulo Kitajima, Erika Freitas, David Schlesinger. Hemocentro Ribeirão Preto: Simone Kashima, Evandra Strazza Rodrigues, Svetoslav Nanev Slavov, Elaine Vieira dos Santos, Rafael dos Santos Bezerra, Luiz Carlos Junior de Alcantara, Marta Giovanetti, Vagner Fonseca, Flavia Aburjaile, Rodrigo Tocantins Calado. FAMERP-SJRP: Cecília Artico Banho, Lívia Sacchetto, Fábio Sossai Possebon, Leila Sabrina Ullmann, Cíntia Bittar, Guilherme Campos, Helena Lage Ferreira, Jorge A. Petroli Marchesi, Maisa C. Pereira Parra, Marília Moraes, Paula Rahal, Paulo Inacio da Costa, João Pessoa Araújo Jr., Maurício Lacerda Nogueira. Prefeitura de Sao Paulo: Melissa Palmieri. |
| EPI_ISL_2170959 | USAFA GUARAMAR | Instituto Butantan / Mendelics | Instituto Butantan: Dimas Tadeu Covas, Sandra Coccuzzo Sampaio, Maria Carolina Elias, José Salvatore Leister Patané, Vincent Louis Viala, Antonio Jorge Martins, Ricardo Haddad, Claudia Renata dos Santos Barros, Elaine Cristina Marqueze, Raul Machado Neto, Debora Botequio Moretti, Jardelina de Souza Todao Bernardino, Loyze Paola Oliveira de Lima, Luiz Aurelio de Campos Crispin. Centro de Genômica Funcional da ESALQ: Luiz Lehmann Coutinho, Ricardo Augusto Brassaloti, Raquel de Lello Rocha Campos Cassano. NGS Soluções Genômicas: Pilar Drummond Sampaio Corrêa Mariani. FZEA-USP Pirassununga: Mirele Daiana Poleti, Jessika Cristina Chagas Lesbon, Elisangela Chicaroni Mattos, Heidge Fukumasu. USP-Botucatu: Rejane Maria Tommasini Grotto, Jayme A. Souza-Neto, Guilherme Targino Valente, Patricia Akemi Assato, Felipe Allan da Silva da Costa, Bianca Cechetto Carlos. Mendelics: Bibiana Santos, João Paulo Kitajima, Erika Freitas, David Schlesinger. Hemocentro Ribeirão Preto: Simone Kashima, Evandra Strazza Rodrigues, Svetoslav Nanev Slavov, Elaine Vieira dos Santos, Rafael dos Santos Bezerra, Luiz Carlos Junior de Alcantara, Marta Giovanetti, Vagner Fonseca, Flavia Aburjaile, Rodrigo Tocantins Calado. FAMERP-SJRP: Cecília Artico Banho, Lívia Sacchetto, Fábio Sossai Possebon, Leila Sabrina Ullmann, Cíntia Bittar, Guilherme Campos, Helena Lage Ferreira, Jorge A. Petroli Marchesi, Maisa C. Pereira Parra, Marília Moraes, Paula Rahal, Paulo Inacio da Costa, João Pessoa Araújo Jr., Maurício Lacerda Nogueira. Prefeitura de Sao Paulo: Melissa Palmieri. |
| EPI_ISL_2170960 | SECRETARIA MUNICIPAL DE SAUDE DE SANTA BARBARA D OESTE | Instituto Butantan / Mendelics | Instituto Butantan: Dimas Tadeu Covas, Sandra Coccuzzo Sampaio, Maria Carolina Elias, José Salvatore Leister Patané, Vincent Louis Viala, Antonio Jorge Martins, Ricardo Haddad, Claudia Renata dos Santos Barros, Elaine Cristina Marqueze, Raul Machado Neto, Debora Botequio Moretti, Jardelina de Souza Todao Bernardino, Loyze Paola Oliveira de Lima, Luiz Aurelio de Campos Crispin. Centro de Genômica Funcional da ESALQ: Luiz Lehmann Coutinho, Ricardo Augusto Brassaloti, Raquel de Lello Rocha Campos Cassano. NGS Soluções Genômicas: Pilar Drummond Sampaio Corrêa Mariani. FZEA-USP Pirassununga: Mirele Daiana Poleti, Jessika Cristina Chagas Lesbon, Elisangela Chicaroni Mattos, Heidge Fukumasu. USP-Botucatu: Rejane Maria Tommasini Grotto, Jayme A. Souza-Neto, Guilherme Targino Valente, Patricia Akemi Assato, Felipe Allan da Silva da Costa, Bianca Cechetto Carlos. Mendelics: Bibiana Santos, João Paulo Kitajima, Erika Freitas, David Schlesinger. Hemocentro Ribeirão Preto: Simone Kashima, Evandra Strazza Rodrigues, Svetoslav Nanev Slavov, Elaine Vieira dos Santos, Rafael dos Santos Bezerra, Luiz Carlos Junior de Alcantara, Marta Giovanetti, Vagner Fonseca, Flavia Aburjaile, Rodrigo Tocantins Calado. FAMERP-SJRP: Cecília Artico Banho, Lívia Sacchetto, Fábio Sossai Possebon, Leila Sabrina Ullmann, Cíntia Bittar, Guilherme Campos, Helena Lage Ferreira, Jorge A. Petroli Marchesi, Maisa C. Pereira Parra, Marília Moraes, Paula Rahal, Paulo Inacio da Costa, João Pessoa Araújo Jr., Maurício Lacerda Nogueira. Prefeitura de Sao Paulo: Melissa Palmieri. |
| EPI_ISL_2170963 | VIGILANCIA EPIDEMIOLOGICA JARDINOPOLIS SP | Instituto Butantan / Mendelics | Instituto Butantan: Dimas Tadeu Covas, Sandra Coccuzzo Sampaio, Maria Carolina Elias, José Salvatore Leister Patané, Vincent Louis Viala, Antonio Jorge Martins, Ricardo Haddad, Claudia Renata dos Santos Barros, Elaine Cristina Marqueze, Raul Machado Neto, Debora Botequio Moretti, Jardelina de Souza Todao Bernardino, Loyze Paola Oliveira de Lima, Luiz Aurelio de Campos Crispin. Centro de Genômica Funcional da ESALQ: Luiz Lehmann Coutinho, Ricardo Augusto Brassaloti, Raquel de Lello Rocha Campos Cassano. NGS Soluções Genômicas: Pilar Drummond Sampaio Corrêa Mariani. FZEA-USP Pirassununga: Mirele Daiana Poleti, Jessika Cristina Chagas Lesbon, Elisangela Chicaroni Mattos, Heidge Fukumasu. USP-Botucatu: Rejane Maria Tommasini Grotto, Jayme A. Souza-Neto, Guilherme Targino Valente, Patricia Akemi Assato, Felipe Allan da Silva da Costa, Bianca Cechetto Carlos. Mendelics: Bibiana Santos, João Paulo Kitajima, Erika Freitas, David Schlesinger. Hemocentro Ribeirão Preto: Simone Kashima, Evandra Strazza Rodrigues, Svetoslav Nanev Slavov, Elaine Vieira dos Santos, Rafael dos Santos Bezerra, Luiz Carlos Junior de Alcantara, Marta Giovanetti, Vagner Fonseca, Flavia Aburjaile, Rodrigo Tocantins Calado. FAMERP-SJRP: Cecília Artico Banho, Lívia Sacchetto, Fábio Sossai Possebon, Leila Sabrina Ullmann, Cíntia Bittar, Guilherme Campos, Helena Lage Ferreira, Jorge A. Petroli Marchesi, Maisa C. Pereira Parra, Marília Moraes, Paula Rahal, Paulo Inacio da Costa, João Pessoa Araújo Jr., Maurício Lacerda Nogueira. Prefeitura de Sao Paulo: Melissa Palmieri. |
| EPI_ISL_2170966 | CS DE MIRASSOLANDIA | Instituto Butantan / Mendelics | Instituto Butantan: Dimas Tadeu Covas, Sandra Coccuzzo Sampaio, Maria Carolina Elias, José Salvatore Leister Patané, Vincent Louis Viala, Antonio Jorge Martins, Ricardo Haddad, Claudia Renata dos Santos Barros, Elaine Cristina Marqueze, Raul Machado Neto, Debora Botequio Moretti, Jardelina de Souza Todao Bernardino, Loyze Paola Oliveira de Lima, Luiz Aurelio de Campos Crispin. Centro de Genômica Funcional da ESALQ: Luiz Lehmann Coutinho, Ricardo Augusto Brassaloti, Raquel de Lello Rocha Campos Cassano. NGS Soluções Genômicas: Pilar Drummond Sampaio Corrêa Mariani. FZEA-USP Pirassununga: Mirele Daiana Poleti, Jessika Cristina Chagas Lesbon, Elisangela Chicaroni Mattos, Heidge Fukumasu. USP-Botucatu: Rejane Maria Tommasini Grotto, Jayme A. Souza-Neto, Guilherme Targino Valente, Patricia Akemi Assato, Felipe Allan da Silva da Costa, Bianca Cechetto Carlos. Mendelics: Bibiana Santos, João Paulo Kitajima, Erika Freitas, David Schlesinger. Hemocentro Ribeirão Preto: Simone Kashima, Evandra Strazza Rodrigues, Svetoslav Nanev Slavov, Elaine Vieira dos Santos, Rafael dos Santos Bezerra, Luiz Carlos Junior de Alcantara, Marta Giovanetti, Vagner Fonseca, Flavia Aburjaile, Rodrigo Tocantins Calado. FAMERP-SJRP: Cecília Artico Banho, Lívia Sacchetto, Fábio Sossai Possebon, Leila Sabrina Ullmann, Cíntia Bittar, Guilherme Campos, Helena Lage Ferreira, Jorge A. Petroli Marchesi, Maisa C. Pereira Parra, Marília Moraes, Paula Rahal, Paulo Inacio da Costa, João Pessoa Araújo Jr., Maurício Lacerda Nogueira. Prefeitura de Sao Paulo: Melissa Palmieri. |
| EPI_ISL_2170967 | CS DE NIPOA | Instituto Butantan / Mendelics | Instituto Butantan: Dimas Tadeu Covas, Sandra Coccuzzo Sampaio, Maria Carolina Elias, José Salvatore Leister Patané, Vincent Louis Viala, Antonio Jorge Martins, Ricardo Haddad, Claudia Renata dos Santos Barros, Elaine Cristina Marqueze, Raul Machado Neto, Debora Botequio Moretti, Jardelina de Souza Todao Bernardino, Loyze Paola Oliveira de Lima, Luiz Aurelio de Campos Crispin. Centro de Genômica Funcional da ESALQ: Luiz Lehmann Coutinho, Ricardo Augusto Brassaloti, Raquel de Lello Rocha Campos Cassano. NGS Soluções Genômicas: Pilar Drummond Sampaio Corrêa Mariani. FZEA-USP Pirassununga: Mirele Daiana Poleti, Jessika Cristina Chagas Lesbon, Elisangela Chicaroni Mattos, Heidge Fukumasu. USP-Botucatu: Rejane Maria Tommasini Grotto, Jayme A. Souza-Neto, Guilherme Targino Valente, Patricia Akemi Assato, Felipe Allan da Silva da Costa, Bianca Cechetto Carlos. Mendelics: Bibiana Santos, João Paulo Kitajima, Erika Freitas, David Schlesinger. Hemocentro Ribeirão Preto: Simone Kashima, Evandra Strazza Rodrigues, Svetoslav Nanev Slavov, Elaine Vieira dos Santos, Rafael dos Santos Bezerra, Luiz Carlos Junior de Alcantara, Marta Giovanetti, Vagner Fonseca, Flavia Aburjaile, Rodrigo Tocantins Calado. FAMERP-SJRP: Cecília Artico Banho, Lívia Sacchetto, Fábio Sossai Possebon, Leila Sabrina |

| | | | |
|---|---|---|---|
| | | | Ullmann, Cíntia Bittar, Guilherme Campos, Helena Lage Ferreira, Jorge A. Petroli Marchesi, Maisa C. Pereira Parra, Marília Moraes, Paula Rahal, Paulo Inacio da Costa, João Pessoa Araújo Jr., Maurício Lacerda Nogueira. Prefeitura de Sao Paulo: Melissa Palmieri. |
| EPI_ISL_2170968 | USF JOAO NASCIMENTO TELLES | Instituto Butantan / Mendelics | Instituto Butantan: Dimas Tadeu Covas, Sandra Coccuzzo Sampaio, Maria Carolina Elias, José Salvatore Leister Patané, Vincent Louis Viala, Antonio Jorge Martins, Ricardo Haddad, Claudia Renata dos Santos Barros, Elaine Cristina Marqueze, Raul Machado Neto, Debora Botequio Moretti, Jardelina de Souza Todao Bernardino, Loyze Paola Oliveira de Lima, Luiz Aurelio de Campos Crispin. Centro de Genômica Funcional da ESALQ: Luiz Lehmann Coutinho, Ricardo Augusto Brassaloti, Raquel de Lello Rocha Campos Cassano. NGS Soluções Genômicas: Pilar Drummond Sampaio Corrêa Mariani. FZEA-USP Pirassununga: Mirele Daiana Poleti, Jessika Cristina Chagas Lesbon, Elisangela Chicaroni Mattos, Heidge Fukumasu. USP-Botucatu: Rejane Maria Tommasini Grotto, Jayme A. Souza-Neto, Guilherme Targino Valente, Patricia Akemi Assato, Felipe Allan da Silva da Costa, Bianca Cechetto Carlos. Mendelics: Bibiana Santos, João Paulo Kitajima, Erika Freitas, David Schlesinger. Hemocentro Ribeirão Preto: Simone Kashima, Evandra Strazza Rodrigues, Svetoslav Nanev Slavov, Elaine Vieira dos Santos, Rafael dos Santos Bezerra, Luiz Carlos Junior de Alcantara, Marta Giovanetti, Vagner Fonseca, Flavia Aburjaile, Rodrigo Tocantins Calado. FAMERP-SJRP: Cecília Artico Banho, Lívia Sacchetto, Fábio Sossai Possebon, Leila Sabrina Ullmann, Cíntia Bittar, Guilherme Campos, Helena Lage Ferreira, Jorge A. Petroli Marchesi, Maisa C. Pereira Parra, Marília Moraes, Paula Rahal, Paulo Inacio da Costa, João Pessoa Araújo Jr., Maurício Lacerda Nogueira. Prefeitura de Sao Paulo: Melissa Palmieri. |
| EPI_ISL_2170974, EPI_ISL_2170975 | PRONTO SOCORRO CENTRAL GUIOMAR FERREIRA ROEBBELEN | Instituto Butantan / Mendelics | Instituto Butantan: Dimas Tadeu Covas, Sandra Coccuzzo Sampaio, Maria Carolina Elias, José Salvatore Leister Patané, Vincent Louis Viala, Antonio Jorge Martins, Ricardo Haddad, Claudia Renata dos Santos Barros, Elaine Cristina Marqueze, Raul Machado Neto, Debora Botequio Moretti, Jardelina de Souza Todao Bernardino, Loyze Paola Oliveira de Lima, Luiz Aurelio de Campos Crispin. Centro de Genômica Funcional da ESALQ: Luiz Lehmann Coutinho, Ricardo Augusto Brassaloti, Raquel de Lello Rocha Campos Cassano. NGS Soluções Genômicas: Pilar Drummond Sampaio Corrêa Mariani. FZEA-USP Pirassununga: Mirele Daiana Poleti, Jessika Cristina Chagas Lesbon, Elisangela Chicaroni Mattos, Heidge Fukumasu. USP-Botucatu: Rejane Maria Tommasini Grotto, Jayme A. Souza-Neto, Guilherme Targino Valente, Patricia Akemi Assato, Felipe Allan da Silva da Costa, Bianca Cechetto Carlos. Mendelics: Bibiana Santos, João Paulo Kitajima, Erika Freitas, David Schlesinger. Hemocentro Ribeirão Preto: Simone Kashima, Evandra Strazza Rodrigues, Svetoslav Nanev Slavov, Elaine Vieira dos Santos, Rafael dos Santos Bezerra, Luiz Carlos Junior de Alcantara, Marta Giovanetti, Vagner Fonseca, Flavia Aburjaile, Rodrigo Tocantins Calado. FAMERP-SJRP: Cecília Artico Banho, Lívia Sacchetto, Fábio Sossai Possebon, Leila Sabrina Ullmann, Cíntia Bittar, Guilherme Campos, Helena Lage Ferreira, Jorge A. Petroli Marchesi, Maisa C. Pereira Parra, Marília Moraes, Paula Rahal, Paulo Inacio da Costa, João Pessoa Araújo Jr., Maurício Lacerda Nogueira. Prefeitura de Sao Paulo: Melissa Palmieri. |
| EPI_ISL_2170978 | SECRETARIA MUNICIPAL DE SAUDE PORTO FERREIRA | Instituto Butantan / Mendelics | Instituto Butantan: Dimas Tadeu Covas, Sandra Coccuzzo Sampaio, Maria Carolina Elias, José Salvatore Leister Patané, Vincent Louis Viala, Antonio Jorge Martins, Ricardo Haddad, Claudia Renata dos Santos Barros, Elaine Cristina Marqueze, Raul Machado Neto, Debora Botequio Moretti, Jardelina de Souza Todao Bernardino, Loyze Paola Oliveira de Lima, Luiz Aurelio de Campos Crispin. Centro de Genômica Funcional da ESALQ: Luiz Lehmann Coutinho, Ricardo Augusto Brassaloti, Raquel de Lello Rocha Campos Cassano. NGS Soluções Genômicas: Pilar Drummond Sampaio Corrêa Mariani. FZEA-USP Pirassununga: Mirele Daiana Poleti, Jessika Cristina Chagas Lesbon, Elisangela Chicaroni Mattos, Heidge Fukumasu. USP-Botucatu: Rejane Maria Tommasini Grotto, Jayme A. Souza-Neto, Guilherme Targino Valente, Patricia Akemi Assato, Felipe Allan da Silva da Costa, Bianca Cechetto Carlos. Mendelics: Bibiana Santos, João Paulo Kitajima, Erika Freitas, David Schlesinger. Hemocentro Ribeirão Preto: Simone Kashima, Evandra Strazza Rodrigues, Svetoslav Nanev Slavov, Elaine Vieira dos Santos, Rafael dos Santos Bezerra, Luiz Carlos Junior de Alcantara, Marta Giovanetti, Vagner Fonseca, Flavia Aburjaile, Rodrigo Tocantins Calado. FAMERP-SJRP: Cecília Artico Banho, Lívia Sacchetto, Fábio Sossai Possebon, Leila Sabrina Ullmann, Cíntia Bittar, Guilherme Campos, Helena Lage Ferreira, Jorge A. Petroli Marchesi, Maisa C. Pereira Parra, Marília Moraes, Paula Rahal, Paulo Inacio da Costa, João Pessoa Araújo Jr., Maurício Lacerda Nogueira. Prefeitura de Sao Paulo: Melissa Palmieri. |
| EPI_ISL_2170980 | SECAO CENTRO DE DIAGNOSTICO SECEDI | Instituto Butantan / Mendelics | Instituto Butantan: Dimas Tadeu Covas, Sandra Coccuzzo Sampaio, Maria Carolina Elias, José Salvatore Leister Patané, Vincent Louis Viala, Antonio Jorge Martins, Ricardo Haddad, Claudia Renata dos Santos Barros, Elaine Cristina Marqueze, Raul Machado Neto, Debora Botequio Moretti, Jardelina de Souza Todao Bernardino, Loyze Paola Oliveira de Lima, Luiz Aurelio de Campos Crispin. Centro de Genômica Funcional da ESALQ: Luiz Lehmann Coutinho, Ricardo Augusto Brassaloti, Raquel de Lello Rocha Campos Cassano. NGS Soluções Genômicas: Pilar Drummond Sampaio Corrêa Mariani. FZEA-USP Pirassununga: Mirele Daiana Poleti, Jessika Cristina Chagas Lesbon, Elisangela Chicaroni Mattos, Heidge Fukumasu. USP-Botucatu: Rejane Maria Tommasini Grotto, Jayme A. Souza-Neto, Guilherme Targino Valente, Patricia Akemi Assato, Felipe Allan da Silva da Costa, Bianca Cechetto Carlos. Mendelics: Bibiana Santos, João Paulo Kitajima, Erika Freitas, David Schlesinger. Hemocentro Ribeirão Preto: Simone Kashima, Evandra Strazza Rodrigues, Svetoslav Nanev Slavov, Elaine Vieira dos Santos, Rafael dos Santos Bezerra, Luiz Carlos Junior de Alcantara, Marta Giovanetti, Vagner Fonseca, Flavia Aburjaile, Rodrigo Tocantins Calado. FAMERP-SJRP: Cecília Artico Banho, Lívia Sacchetto, Fábio Sossai Possebon, Leila Sabrina Ullmann, Cíntia Bittar, Guilherme Campos, Helena Lage Ferreira, Jorge A. Petroli Marchesi, Maisa C. Pereira Parra, Marília Moraes, Paula Rahal, Paulo Inacio da Costa, João Pessoa Araújo Jr., Maurício Lacerda Nogueira. Prefeitura de Sao Paulo: Melissa Palmieri. |
| EPI_ISL_2170982 | SANTA CASA DE ATIBAIA PRO SAUDE | Instituto Butantan / Mendelics | Instituto Butantan: Dimas Tadeu Covas, Sandra Coccuzzo Sampaio, Maria Carolina Elias, José Salvatore Leister Patané, Vincent Louis Viala, Antonio Jorge Martins, Ricardo Haddad, Claudia Renata dos Santos Barros, Elaine Cristina Marqueze, Raul Machado Neto, Debora Botequio Moretti, Jardelina de Souza Todao Bernardino, Loyze Paola Oliveira de Lima, Luiz Aurelio de Campos Crispin. Centro de Genômica Funcional da ESALQ: Luiz Lehmann Coutinho, Ricardo Augusto Brassaloti, Raquel de Lello Rocha Campos Cassano. NGS Soluções Genômicas: Pilar Drummond Sampaio Corrêa Mariani. FZEA-USP Pirassununga: Mirele Daiana Poleti, Jessika Cristina Chagas Lesbon, Elisangela Chicaroni Mattos, Heidge Fukumasu. USP-Botucatu: Rejane Maria Tommasini Grotto, Jayme A. Souza-Neto, Guilherme Targino Valente, Patricia Akemi Assato, Felipe Allan da Silva da Costa, Bianca Cechetto Carlos. Mendelics: Bibiana Santos, João Paulo Kitajima, Erika Freitas, David Schlesinger. Hemocentro Ribeirão Preto: Simone Kashima, Evandra Strazza Rodrigues, Svetoslav Nanev Slavov, Elaine Vieira dos Santos, Rafael dos Santos Bezerra, Luiz Carlos Junior de Alcantara, Marta Giovanetti, Vagner Fonseca, Flavia Aburjaile, Rodrigo Tocantins Calado. FAMERP-SJRP: Cecília Artico Banho, Lívia Sacchetto, Fábio Sossai Possebon, Leila Sabrina Ullmann, Cíntia Bittar, Guilherme Campos, Helena Lage Ferreira, Jorge A. Petroli Marchesi, Maisa C. Pereira Parra, Marília Moraes, Paula Rahal, Paulo Inacio da Costa, João Pessoa Araújo Jr., Maurício Lacerda Nogueira. Prefeitura de Sao Paulo: Melissa Palmieri. |
| EPI_ISL_2170987, EPI_ISL_2170991, EPI_ISL_2170994, EPI_ISL_2170995 | VIGILANCIA EM SAUDE DE FCO MORATO | Instituto Butantan / Mendelics | Instituto Butantan: Dimas Tadeu Covas, Sandra Coccuzzo Sampaio, Maria Carolina Elias, José Salvatore Leister Patané, Vincent Louis Viala, Antonio Jorge Martins, Ricardo Haddad, Claudia Renata dos Santos Barros, Elaine Cristina Marqueze, Raul Machado Neto, Debora Botequio Moretti, Jardelina de Souza Todao Bernardino, Loyze Paola Oliveira de Lima, Luiz Aurelio de Campos Crispin. Centro de Genômica Funcional da ESALQ: Luiz Lehmann Coutinho, Ricardo Augusto Brassaloti, Raquel de Lello Rocha Campos Cassano. NGS Soluções Genômicas: Pilar Drummond Sampaio Corrêa Mariani. FZEA-USP Pirassununga: Mirele Daiana Poleti, Jessika Cristina Chagas Lesbon, Elisangela Chicaroni Mattos, Heidge Fukumasu. USP-Botucatu: Rejane Maria Tommasini Grotto, Jayme A. Souza-Neto, Guilherme Targino Valente, Patricia Akemi Assato, Felipe Allan da Silva da Costa, Bianca Cechetto Carlos. Mendelics: Bibiana Santos, João Paulo Kitajima, Erika Freitas, David Schlesinger. Hemocentro Ribeirão Preto: Simone Kashima, Evandra Strazza Rodrigues, Svetoslav Nanev Slavov, Elaine Vieira dos Santos, Rafael dos Santos Bezerra, Luiz Carlos Junior de Alcantara, Marta Giovanetti, Vagner Fonseca, Flavia Aburjaile, Rodrigo Tocantins Calado. FAMERP-SJRP: Cecília Artico Banho, Lívia Sacchetto, Fábio Sossai Possebon, Leila Sabrina Ullmann, Cíntia Bittar, Guilherme Campos, Helena Lage Ferreira, Jorge A. Petroli Marchesi, Maisa C. Pereira Parra, Marília Moraes, Paula Rahal, Paulo Inacio da Costa, João Pessoa Araújo Jr., Maurício Lacerda Nogueira. Prefeitura de Sao Paulo: Melissa Palmieri. |
| EPI_ISL_2170996 | UPA DR FRANCO DA ROCHA | Instituto Butantan / Mendelics | Instituto Butantan: Dimas Tadeu Covas, Sandra Coccuzzo Sampaio, Maria Carolina Elias, José Salvatore Leister Patané, Vincent Louis Viala, Antonio Jorge Martins, Ricardo Haddad, Claudia Renata dos Santos Barros, Elaine Cristina Marqueze, Raul Machado Neto, Debora Botequio Moretti, Jardelina de Souza Todao Bernardino, Loyze Paola Oliveira de Lima, Luiz Aurelio de Campos Crispin. Centro de Genômica Funcional da ESALQ: Luiz Lehmann Coutinho, Ricardo Augusto Brassaloti, Raquel de Lello Rocha Campos Cassano. NGS Soluções Genômicas: Pilar Drummond Sampaio Corrêa Mariani. FZEA-USP Pirassununga: Mirele Daiana Poleti, Jessika Cristina Chagas Lesbon, Elisangela Chicaroni Mattos, Heidge Fukumasu. USP-Botucatu: Rejane Maria Tommasini Grotto, Jayme A. Souza-Neto, Guilherme Targino Valente, Patricia Akemi Assato, Felipe Allan da Silva da Costa, Bianca Cechetto Carlos. Mendelics: Bibiana Santos, João Paulo Kitajima, Erika Freitas, David Schlesinger. Hemocentro Ribeirão Preto: Simone Kashima, Evandra Strazza Rodrigues, Svetoslav Nanev Slavov, Elaine Vieira dos Santos, Rafael dos Santos Bezerra, Luiz Carlos Junior de Alcantara, Marta Giovanetti, Vagner Fonseca, Flavia Aburjaile, Rodrigo Tocantins Calado. FAMERP-SJRP: Cecília Artico Banho, Lívia Sacchetto, Fábio Sossai Possebon, Leila Sabrina Ullmann, Cíntia Bittar, Guilherme Campos, Helena Lage Ferreira, Jorge A. Petroli Marchesi, Maisa C. Pereira Parra, Marília Moraes, Paula Rahal, Paulo Inacio da Costa, João Pessoa Araújo Jr., Maurício Lacerda Nogueira. Prefeitura de Sao Paulo: Melissa Palmieri. |
| EPI_ISL_2170997 | SANTA CASA DE GUARAREMA | Instituto Butantan / Mendelics | Instituto Butantan: Dimas Tadeu Covas, Sandra Coccuzzo Sampaio, Maria Carolina Elias, José Salvatore Leister Patané, Vincent Louis Viala, Antonio Jorge Martins, Ricardo Haddad, Claudia Renata dos Santos Barros, Elaine Cristina Marqueze, Raul Machado Neto, Debora Botequio Moretti, Jardelina de Souza Todao Bernardino, Loyze Paola Oliveira de Lima, Luiz Aurelio de Campos Crispin. Centro de Genômica Funcional da ESALQ: Luiz Lehmann Coutinho, Ricardo Augusto Brassaloti, Raquel de Lello Rocha Campos Cassano. NGS Soluções Genômicas: Pilar Drummond Sampaio Corrêa Mariani. |

| | | | |
|---|---|---|---|
| | | | FZEA-USP Pirassununga: Mirele Daiana Poleti, Jessika Cristina Chagas Lesbon, Elisangela Chicaroni Mattos, Heidge Fukumasu. USP-Botucatu: Rejane Maria Tommasini Grotto, Jayme A. Souza-Neto, Guilherme Targino Valente, Patricia Akemi Assato, Felipe Allan da Silva da Costa, Bianca Cechetto Carlos. Mendelics: Bibiana Santos, João Paulo Kitajima, Erika Freitas, David Schlesinger. Hemocentro Ribeirão Preto: Simone Kashima, Evandra Strazza Rodrigues, Svetoslav Nanev Slavov, Elaine Vieira dos Santos, Rafael dos Santos Bezerra, Luiz Carlos Junior de Alcantara, Marta Giovanetti, Vagner Fonseca, Flavia Aburjaile, Rodrigo Tocantins Calado. FAMERP-SJRP: Cecília Artico Banho, Lívia Sacchetto, Fábio Sossai Possebon, Leila Sabrina Ullmann, Cíntia Bittar, Guilherme Campos, Helena Lage Ferreira, Jorge A. Petroli Marchesi, Maisa C. Pereira Parra, Marília Moraes, Paula Rahal, Paulo Inacio da Costa, João Pessoa Araújo Jr., Maurício Lacerda Nogueira. Prefeitura de Sao Paulo: Melissa Palmieri. |
| EPI_ISL_2170998 | UBS RECREIO SAO JORGE | Instituto Butantan / Mendelics | Instituto Butantan: Dimas Tadeu Covas, Sandra Coccuzzo Sampaio, Maria Carolina Elias, José Salvatore Leister Patané, Vincent Louis Viala, Antonio Jorge Martins, Ricardo Haddad, Claudia Renata dos Santos Barros, Elaine Cristina Marqueze, Raul Machado Neto, Debora Botequio Moretti, Jardelina de Souza Todao Bernardino, Loyze Paola Oliveira de Lima, Luiz Aurelio de Campos Crispin. Centro de Genômica Funcional da ESALQ: Luiz Lehmann Coutinho, Ricardo Augusto Brassaloti, Raquel de Lello Rocha Campos Cassano. NGS Soluções Genômicas: Pilar Drummond Sampaio Corrêa Mariani. FZEA-USP Pirassununga: Mirele Daiana Poleti, Jessika Cristina Chagas Lesbon, Elisangela Chicaroni Mattos, Heidge Fukumasu. USP-Botucatu: Rejane Maria Tommasini Grotto, Jayme A. Souza-Neto, Guilherme Targino Valente, Patricia Akemi Assato, Felipe Allan da Silva da Costa, Bianca Cechetto Carlos. Mendelics: Bibiana Santos, João Paulo Kitajima, Erika Freitas, David Schlesinger. Hemocentro Ribeirão Preto: Simone Kashima, Evandra Strazza Rodrigues, Svetoslav Nanev Slavov, Elaine Vieira dos Santos, Rafael dos Santos Bezerra, Luiz Carlos Junior de Alcantara, Marta Giovanetti, Vagner Fonseca, Flavia Aburjaile, Rodrigo Tocantins Calado. FAMERP-SJRP: Cecília Artico Banho, Lívia Sacchetto, Fábio Sossai Possebon, Leila Sabrina Ullmann, Cíntia Bittar, Guilherme Campos, Helena Lage Ferreira, Jorge A. Petroli Marchesi, Maisa C. Pereira Parra, Marília Moraes, Paula Rahal, Paulo Inacio da Costa, João Pessoa Araújo Jr., Maurício Lacerda Nogueira. Prefeitura de Sao Paulo: Melissa Palmieri. |
| EPI_ISL_2170999 | UBS JOVAIA | Instituto Butantan / Mendelics | Instituto Butantan: Dimas Tadeu Covas, Sandra Coccuzzo Sampaio, Maria Carolina Elias, José Salvatore Leister Patané, Vincent Louis Viala, Antonio Jorge Martins, Ricardo Haddad, Claudia Renata dos Santos Barros, Elaine Cristina Marqueze, Raul Machado Neto, Debora Botequio Moretti, Jardelina de Souza Todao Bernardino, Loyze Paola Oliveira de Lima, Luiz Aurelio de Campos Crispin. Centro de Genômica Funcional da ESALQ: Luiz Lehmann Coutinho, Ricardo Augusto Brassaloti, Raquel de Lello Rocha Campos Cassano. NGS Soluções Genômicas: Pilar Drummond Sampaio Corrêa Mariani. FZEA-USP Pirassununga: Mirele Daiana Poleti, Jessika Cristina Chagas Lesbon, Elisangela Chicaroni Mattos, Heidge Fukumasu. USP-Botucatu: Rejane Maria Tommasini Grotto, Jayme A. Souza-Neto, Guilherme Targino Valente, Patricia Akemi Assato, Felipe Allan da Silva da Costa, Bianca Cechetto Carlos. Mendelics: Bibiana Santos, João Paulo Kitajima, Erika Freitas, David Schlesinger. Hemocentro Ribeirão Preto: Simone Kashima, Evandra Strazza Rodrigues, Svetoslav Nanev Slavov, Elaine Vieira dos Santos, Rafael dos Santos Bezerra, Luiz Carlos Junior de Alcantara, Marta Giovanetti, Vagner Fonseca, Flavia Aburjaile, Rodrigo Tocantins Calado. FAMERP-SJRP: Cecília Artico Banho, Lívia Sacchetto, Fábio Sossai Possebon, Leila Sabrina Ullmann, Cíntia Bittar, Guilherme Campos, Helena Lage Ferreira, Jorge A. Petroli Marchesi, Maisa C. Pereira Parra, Marília Moraes, Paula Rahal, Paulo Inacio da Costa, João Pessoa Araújo Jr., Maurício Lacerda Nogueira. Prefeitura de Sao Paulo: Melissa Palmieri. |
| EPI_ISL_2171000 | UBS JARDIM MUNHOZ | Instituto Butantan / Mendelics | Instituto Butantan: Dimas Tadeu Covas, Sandra Coccuzzo Sampaio, Maria Carolina Elias, José Salvatore Leister Patané, Vincent Louis Viala, Antonio Jorge Martins, Ricardo Haddad, Claudia Renata dos Santos Barros, Elaine Cristina Marqueze, Raul Machado Neto, Debora Botequio Moretti, Jardelina de Souza Todao Bernardino, Loyze Paola Oliveira de Lima, Luiz Aurelio de Campos Crispin. Centro de Genômica Funcional da ESALQ: Luiz Lehmann Coutinho, Ricardo Augusto Brassaloti, Raquel de Lello Rocha Campos Cassano. NGS Soluções Genômicas: Pilar Drummond Sampaio Corrêa Mariani. FZEA-USP Pirassununga: Mirele Daiana Poleti, Jessika Cristina Chagas Lesbon, Elisangela Chicaroni Mattos, Heidge Fukumasu. USP-Botucatu: Rejane Maria Tommasini Grotto, Jayme A. Souza-Neto, Guilherme Targino Valente, Patricia Akemi Assato, Felipe Allan da Silva da Costa, Bianca Cechetto Carlos. Mendelics: Bibiana Santos, João Paulo Kitajima, Erika Freitas, David Schlesinger. Hemocentro Ribeirão Preto: Simone Kashima, Evandra Strazza Rodrigues, Svetoslav Nanev Slavov, Elaine Vieira dos Santos, Rafael dos Santos Bezerra, Luiz Carlos Junior de Alcantara, Marta Giovanetti, Vagner Fonseca, Flavia Aburjaile, Rodrigo Tocantins Calado. FAMERP-SJRP: Cecília Artico Banho, Lívia Sacchetto, Fábio Sossai Possebon, Leila Sabrina Ullmann, Cíntia Bittar, Guilherme Campos, Helena Lage Ferreira, Jorge A. Petroli Marchesi, Maisa C. Pereira Parra, Marília Moraes, Paula Rahal, Paulo Inacio da Costa, João Pessoa Araújo Jr., Maurício Lacerda Nogueira. Prefeitura de Sao Paulo: Melissa Palmieri. |
| EPI_ISL_2171002, EPI_ISL_2171004 | UNIDADE RESPIRATORIA NOVA HORTOLANDIA | Instituto Butantan / Mendelics | Instituto Butantan: Dimas Tadeu Covas, Sandra Coccuzzo Sampaio, Maria Carolina Elias, José Salvatore Leister Patané, Vincent Louis Viala, Antonio Jorge Martins, Ricardo Haddad, Claudia Renata dos Santos Barros, Elaine Cristina Marqueze, Raul Machado Neto, Debora Botequio Moretti, Jardelina de Souza Todao Bernardino, Loyze Paola Oliveira de Lima, Luiz Aurelio de Campos Crispin. Centro de Genômica Funcional da ESALQ: Luiz Lehmann Coutinho, Ricardo Augusto Brassaloti, Raquel de Lello Rocha Campos Cassano. NGS Soluções Genômicas: Pilar Drummond Sampaio Corrêa Mariani. FZEA-USP Pirassununga: Mirele Daiana Poleti, Jessika Cristina Chagas Lesbon, Elisangela Chicaroni Mattos, Heidge Fukumasu. USP-Botucatu: Rejane Maria Tommasini Grotto, Jayme A. Souza-Neto, Guilherme Targino Valente, Patricia Akemi Assato, Felipe Allan da Silva da Costa, Bianca Cechetto Carlos. Mendelics: Bibiana Santos, João Paulo Kitajima, Erika Freitas, David Schlesinger. Hemocentro Ribeirão Preto: Simone Kashima, Evandra Strazza Rodrigues, Svetoslav Nanev Slavov, Elaine Vieira dos Santos, Rafael dos Santos Bezerra, Luiz Carlos Junior de Alcantara, Marta Giovanetti, Vagner Fonseca, Flavia Aburjaile, Rodrigo Tocantins Calado. FAMERP-SJRP: Cecília Artico Banho, Lívia Sacchetto, Fábio Sossai Possebon, Leila Sabrina Ullmann, Cíntia Bittar, Guilherme Campos, Helena Lage Ferreira, Jorge A. Petroli Marchesi, Maisa C. Pereira Parra, Marília Moraes, Paula Rahal, Paulo Inacio da Costa, João Pessoa Araújo Jr., Maurício Lacerda Nogueira. Prefeitura de Sao Paulo: Melissa Palmieri. |
| EPI_ISL_2171007, EPI_ISL_2171009, EPI_ISL_2171010 | POLICLINICA HORTOLANDIA | Instituto Butantan / Mendelics | Instituto Butantan: Dimas Tadeu Covas, Sandra Coccuzzo Sampaio, Maria Carolina Elias, José Salvatore Leister Patané, Vincent Louis Viala, Antonio Jorge Martins, Ricardo Haddad, Claudia Renata dos Santos Barros, Elaine Cristina Marqueze, Raul Machado Neto, Debora Botequio Moretti, Jardelina de Souza Todao Bernardino, Loyze Paola Oliveira de Lima, Luiz Aurelio de Campos Crispin. Centro de Genômica Funcional da ESALQ: Luiz Lehmann Coutinho, Ricardo Augusto Brassaloti, Raquel de Lello Rocha Campos Cassano. NGS Soluções Genômicas: Pilar Drummond Sampaio Corrêa Mariani. FZEA-USP Pirassununga: Mirele Daiana Poleti, Jessika Cristina Chagas Lesbon, Elisangela Chicaroni Mattos, Heidge Fukumasu. USP-Botucatu: Rejane Maria Tommasini Grotto, Jayme A. Souza-Neto, Guilherme Targino Valente, Patricia Akemi Assato, Felipe Allan da Silva da Costa, Bianca Cechetto Carlos. Mendelics: Bibiana Santos, João Paulo Kitajima, Erika Freitas, David Schlesinger. Hemocentro Ribeirão Preto: Simone Kashima, Evandra Strazza Rodrigues, Svetoslav Nanev Slavov, Elaine Vieira dos Santos, Rafael dos Santos Bezerra, Luiz Carlos Junior de Alcantara, Marta Giovanetti, Vagner Fonseca, Flavia Aburjaile, Rodrigo Tocantins Calado. FAMERP-SJRP: Cecília Artico Banho, Lívia Sacchetto, Fábio Sossai Possebon, Leila Sabrina Ullmann, Cíntia Bittar, Guilherme Campos, Helena Lage Ferreira, Jorge A. Petroli Marchesi, Maisa C. Pereira Parra, Marília Moraes, Paula Rahal, Paulo Inacio da Costa, João Pessoa Araújo Jr., Maurício Lacerda Nogueira. Prefeitura de Sao Paulo: Melissa Palmieri. |
| EPI_ISL_2171011 | UPA II JUNDIAI | Instituto Butantan / Mendelics | Instituto Butantan: Dimas Tadeu Covas, Sandra Coccuzzo Sampaio, Maria Carolina Elias, José Salvatore Leister Patané, Vincent Louis Viala, Antonio Jorge Martins, Ricardo Haddad, Claudia Renata dos Santos Barros, Elaine Cristina Marqueze, Raul Machado Neto, Debora Botequio Moretti, Jardelina de Souza Todao Bernardino, Loyze Paola Oliveira de Lima, Luiz Aurelio de Campos Crispin. Centro de Genômica Funcional da ESALQ: Luiz Lehmann Coutinho, Ricardo Augusto Brassaloti, Raquel de Lello Rocha Campos Cassano. NGS Soluções Genômicas: Pilar Drummond Sampaio Corrêa Mariani. FZEA-USP Pirassununga: Mirele Daiana Poleti, Jessika Cristina Chagas Lesbon, Elisangela Chicaroni Mattos, Heidge Fukumasu. USP-Botucatu: Rejane Maria Tommasini Grotto, Jayme A. Souza-Neto, Guilherme Targino Valente, Patricia Akemi Assato, Felipe Allan da Silva da Costa, Bianca Cechetto Carlos. Mendelics: Bibiana Santos, João Paulo Kitajima, Erika Freitas, David Schlesinger. Hemocentro Ribeirão Preto: Simone Kashima, Evandra Strazza Rodrigues, Svetoslav Nanev Slavov, Elaine Vieira dos Santos, Rafael dos Santos Bezerra, Luiz Carlos Junior de Alcantara, Marta Giovanetti, Vagner Fonseca, Flavia Aburjaile, Rodrigo Tocantins Calado. FAMERP-SJRP: Cecília Artico Banho, Lívia Sacchetto, Fábio Sossai Possebon, Leila Sabrina Ullmann, Cíntia Bittar, Guilherme Campos, Helena Lage Ferreira, Jorge A. Petroli Marchesi, Maisa C. Pereira Parra, Marília Moraes, Paula Rahal, Paulo Inacio da Costa, João Pessoa Araújo Jr., Maurício Lacerda Nogueira. Prefeitura de Sao Paulo: Melissa Palmieri. |
| EPI_ISL_2171013, EPI_ISL_2171014, EPI_ISL_2171015, EPI_ISL_2171016 | POLICLINICA HORTOLANDIA | Instituto Butantan / Mendelics | Instituto Butantan: Dimas Tadeu Covas, Sandra Coccuzzo Sampaio, Maria Carolina Elias, José Salvatore Leister Patané, Vincent Louis Viala, Antonio Jorge Martins, Ricardo Haddad, Claudia Renata dos Santos Barros, Elaine Cristina Marqueze, Raul Machado Neto, Debora Botequio Moretti, Jardelina de Souza Todao Bernardino, Loyze Paola Oliveira de Lima, Luiz Aurelio de Campos Crispin. Centro de Genômica Funcional da ESALQ: Luiz Lehmann Coutinho, Ricardo Augusto Brassaloti, Raquel de Lello Rocha Campos Cassano. NGS Soluções Genômicas: Pilar Drummond Sampaio Corrêa Mariani. FZEA-USP Pirassununga: Mirele Daiana Poleti, Jessika Cristina Chagas Lesbon, Elisangela Chicaroni Mattos, Heidge Fukumasu. USP-Botucatu: Rejane Maria Tommasini Grotto, Jayme A. Souza-Neto, Guilherme Targino Valente, Patricia Akemi Assato, Felipe Allan da Silva da Costa, Bianca Cechetto Carlos. Mendelics: Bibiana Santos, João Paulo Kitajima, Erika Freitas, David Schlesinger. Hemocentro Ribeirão Preto: Simone Kashima, Evandra Strazza Rodrigues, Svetoslav Nanev Slavov, Elaine Vieira dos Santos, Rafael dos Santos Bezerra, Luiz Carlos Junior de Alcantara, Marta Giovanetti, Vagner Fonseca, Flavia Aburjaile, Rodrigo Tocantins Calado. FAMERP-SJRP: Cecília Artico Banho, Lívia Sacchetto, Fábio Sossai Possebon, Leila Sabrina Ullmann, Cíntia Bittar, Guilherme Campos, Helena Lage Ferreira, Jorge A. Petroli Marchesi, Maisa C. Pereira Parra, Marília Moraes, Paula Rahal, Paulo Inacio da Costa, João Pessoa Araújo Jr., Maurício Lacerda Nogueira. Prefeitura de Sao Paulo: Melissa Palmieri. |

| | | | |
|---|---|---|---|
| EPI_ISL_2171017 | HOSPITAL MUNICIPAL DR TABAJARA RAMOS | Instituto Butantan / Mendelics | Instituto Butantan: Dimas Tadeu Covas, Sandra Coccuzzo Sampaio, Maria Carolina Elias, José Salvatore Leister Patané, Vincent Louis Viala, Antonio Jorge Martins, Ricardo Haddad, Claudia Renata dos Santos Barros, Elaine Cristina Marqueze, Raul Machado Neto, Debora Botequio Moretti, Jardelina de Souza Todao Bernardino, Loyze Paola Oliveira de Lima, Luiz Aurelio de Campos Crispin. Centro de Genômica Funcional da ESALQ: Luiz Lehmann Coutinho, Ricardo Augusto Brassaloti, Raquel de Lello Rocha Campos Cassano. NGS Soluções Genômicas: Pilar Drummond Sampaio Corrêa Mariani. FZEA-USP Pirassununga: Mirele Daiana Poleti, Jessika Cristina Chagas Lesbon, Elisangela Chicaroni Mattos, Heidge Fukumasu. USP-Botucatu: Rejane Maria Tommasini Grotto, Jayme A. Souza-Neto, Guilherme Targino Valente, Patricia Akemi Assato, Felipe Allan da Silva da Costa, Bianca Cechetto Carlos. Mendelics: Bibiana Santos, João Paulo Kitajima, Erika Freitas, David Schlesinger. Hemocentro Ribeirão Preto: Simone Kashima, Evandra Strazza Rodrigues, Svetoslav Nanev Slavov, Elaine Vieira dos Santos, Rafael dos Santos Bezerra, Luiz Carlos Junior de Alcantara, Marta Giovanetti, Vagner Fonseca, Flavia Aburjaile, Rodrigo Tocantins Calado. FAMERP-SJRP: Cecília Artico Banho, Lívia Sacchetto, Fábio Sossai Possebon, Leila Sabrina Ullmann, Cíntia Bittar, Guilherme Campos, Helena Lage Ferreira, Jorge A. Petroli Marchesi, Maisa C. Pereira Parra, Marília Moraes, Paula Rahal, Paulo Inacio da Costa, João Pessoa Araújo Jr., Maurício Lacerda Nogueira. Prefeitura de Sao Paulo: Melissa Palmieri. |
| EPI_ISL_2171018 | AMBULATORIO DE ESPECIALIDADE V E MOGI MIRIM | Instituto Butantan / Mendelics | Instituto Butantan: Dimas Tadeu Covas, Sandra Coccuzzo Sampaio, Maria Carolina Elias, José Salvatore Leister Patané, Vincent Louis Viala, Antonio Jorge Martins, Ricardo Haddad, Claudia Renata dos Santos Barros, Elaine Cristina Marqueze, Raul Machado Neto, Debora Botequio Moretti, Jardelina de Souza Todao Bernardino, Loyze Paola Oliveira de Lima, Luiz Aurelio de Campos Crispin. Centro de Genômica Funcional da ESALQ: Luiz Lehmann Coutinho, Ricardo Augusto Brassaloti, Raquel de Lello Rocha Campos Cassano. NGS Soluções Genômicas: Pilar Drummond Sampaio Corrêa Mariani. FZEA-USP Pirassununga: Mirele Daiana Poleti, Jessika Cristina Chagas Lesbon, Elisangela Chicaroni Mattos, Heidge Fukumasu. USP-Botucatu: Rejane Maria Tommasini Grotto, Jayme A. Souza-Neto, Guilherme Targino Valente, Patricia Akemi Assato, Felipe Allan da Silva da Costa, Bianca Cechetto Carlos. Mendelics: Bibiana Santos, João Paulo Kitajima, Erika Freitas, David Schlesinger. Hemocentro Ribeirão Preto: Simone Kashima, Evandra Strazza Rodrigues, Svetoslav Nanev Slavov, Elaine Vieira dos Santos, Rafael dos Santos Bezerra, Luiz Carlos Junior de Alcantara, Marta Giovanetti, Vagner Fonseca, Flavia Aburjaile, Rodrigo Tocantins Calado. FAMERP-SJRP: Cecília Artico Banho, Lívia Sacchetto, Fábio Sossai Possebon, Leila Sabrina Ullmann, Cíntia Bittar, Guilherme Campos, Helena Lage Ferreira, Jorge A. Petroli Marchesi, Maisa C. Pereira Parra, Marília Moraes, Paula Rahal, Paulo Inacio da Costa, João Pessoa Araújo Jr., Maurício Lacerda Nogueira. Prefeitura de Sao Paulo: Melissa Palmieri. |
| EPI_ISL_2171021 | UPA VILA SANTA CATARINA | Instituto Butantan / Mendelics | Instituto Butantan: Dimas Tadeu Covas, Sandra Coccuzzo Sampaio, Maria Carolina Elias, José Salvatore Leister Patané, Vincent Louis Viala, Antonio Jorge Martins, Ricardo Haddad, Claudia Renata dos Santos Barros, Elaine Cristina Marqueze, Raul Machado Neto, Debora Botequio Moretti, Jardelina de Souza Todao Bernardino, Loyze Paola Oliveira de Lima, Luiz Aurelio de Campos Crispin. Centro de Genômica Funcional da ESALQ: Luiz Lehmann Coutinho, Ricardo Augusto Brassaloti, Raquel de Lello Rocha Campos Cassano. NGS Soluções Genômicas: Pilar Drummond Sampaio Corrêa Mariani. FZEA-USP Pirassununga: Mirele Daiana Poleti, Jessika Cristina Chagas Lesbon, Elisangela Chicaroni Mattos, Heidge Fukumasu. USP-Botucatu: Rejane Maria Tommasini Grotto, Jayme A. Souza-Neto, Guilherme Targino Valente, Patricia Akemi Assato, Felipe Allan da Silva da Costa, Bianca Cechetto Carlos. Mendelics: Bibiana Santos, João Paulo Kitajima, Erika Freitas, David Schlesinger. Hemocentro Ribeirão Preto: Simone Kashima, Evandra Strazza Rodrigues, Svetoslav Nanev Slavov, Elaine Vieira dos Santos, Rafael dos Santos Bezerra, Luiz Carlos Junior de Alcantara, Marta Giovanetti, Vagner Fonseca, Flavia Aburjaile, Rodrigo Tocantins Calado. FAMERP-SJRP: Cecília Artico Banho, Lívia Sacchetto, Fábio Sossai Possebon, Leila Sabrina Ullmann, Cíntia Bittar, Guilherme Campos, Helena Lage Ferreira, Jorge A. Petroli Marchesi, Maisa C. Pereira Parra, Marília Moraes, Paula Rahal, Paulo Inacio da Costa, João Pessoa Araújo Jr., Maurício Lacerda Nogueira. Prefeitura de Sao Paulo: Melissa Palmieri. |
| EPI_ISL_2171024 | SECRETARIA MUNICIPAL DE SAUDE | Instituto Butantan / Mendelics | Instituto Butantan: Dimas Tadeu Covas, Sandra Coccuzzo Sampaio, Maria Carolina Elias, José Salvatore Leister Patané, Vincent Louis Viala, Antonio Jorge Martins, Ricardo Haddad, Claudia Renata dos Santos Barros, Elaine Cristina Marqueze, Raul Machado Neto, Debora Botequio Moretti, Jardelina de Souza Todao Bernardino, Loyze Paola Oliveira de Lima, Luiz Aurelio de Campos Crispin. Centro de Genômica Funcional da ESALQ: Luiz Lehmann Coutinho, Ricardo Augusto Brassaloti, Raquel de Lello Rocha Campos Cassano. NGS Soluções Genômicas: Pilar Drummond Sampaio Corrêa Mariani. FZEA-USP Pirassununga: Mirele Daiana Poleti, Jessika Cristina Chagas Lesbon, Elisangela Chicaroni Mattos, Heidge Fukumasu. USP-Botucatu: Rejane Maria Tommasini Grotto, Jayme A. Souza-Neto, Guilherme Targino Valente, Patricia Akemi Assato, Felipe Allan da Silva da Costa, Bianca Cechetto Carlos. Mendelics: Bibiana Santos, João Paulo Kitajima, Erika Freitas, David Schlesinger. Hemocentro Ribeirão Preto: Simone Kashima, Evandra Strazza Rodrigues, Svetoslav Nanev Slavov, Elaine Vieira dos Santos, Rafael dos Santos Bezerra, Luiz Carlos Junior de Alcantara, Marta Giovanetti, Vagner Fonseca, Flavia Aburjaile, Rodrigo Tocantins Calado. FAMERP-SJRP: Cecília Artico Banho, Lívia Sacchetto, Fábio Sossai Possebon, Leila Sabrina Ullmann, Cíntia Bittar, Guilherme Campos, Helena Lage Ferreira, Jorge A. Petroli Marchesi, Maisa C. Pereira Parra, Marília Moraes, Paula Rahal, Paulo Inacio da Costa, João Pessoa Araújo Jr., Maurício Lacerda Nogueira. Prefeitura de Sao Paulo: Melissa Palmieri. |
| EPI_ISL_2171025, EPI_ISL_2171026, EPI_ISL_2171027, EPI_ISL_2171028, EPI_ISL_2171029 | UPA DR AKIRA TADA | Instituto Butantan / Mendelics | Instituto Butantan: Dimas Tadeu Covas, Sandra Coccuzzo Sampaio, Maria Carolina Elias, José Salvatore Leister Patané, Vincent Louis Viala, Antonio Jorge Martins, Ricardo Haddad, Claudia Renata dos Santos Barros, Elaine Cristina Marqueze, Raul Machado Neto, Debora Botequio Moretti, Jardelina de Souza Todao Bernardino, Loyze Paola Oliveira de Lima, Luiz Aurelio de Campos Crispin. Centro de Genômica Funcional da ESALQ: Luiz Lehmann Coutinho, Ricardo Augusto Brassaloti, Raquel de Lello Rocha Campos Cassano. NGS Soluções Genômicas: Pilar Drummond Sampaio Corrêa Mariani. FZEA-USP Pirassununga: Mirele Daiana Poleti, Jessika Cristina Chagas Lesbon, Elisangela Chicaroni Mattos, Heidge Fukumasu. USP-Botucatu: Rejane Maria Tommasini Grotto, Jayme A. Souza-Neto, Guilherme Targino Valente, Patricia Akemi Assato, Felipe Allan da Silva da Costa, Bianca Cechetto Carlos. Mendelics: Bibiana Santos, João Paulo Kitajima, Erika Freitas, David Schlesinger. Hemocentro Ribeirão Preto: Simone Kashima, Evandra Strazza Rodrigues, Svetoslav Nanev Slavov, Elaine Vieira dos Santos, Rafael dos Santos Bezerra, Luiz Carlos Junior de Alcantara, Marta Giovanetti, Vagner Fonseca, Flavia Aburjaile, Rodrigo Tocantins Calado. FAMERP-SJRP: Cecília Artico Banho, Lívia Sacchetto, Fábio Sossai Possebon, Leila Sabrina Ullmann, Cíntia Bittar, Guilherme Campos, Helena Lage Ferreira, Jorge A. Petroli Marchesi, Maisa C. Pereira Parra, Marília Moraes, Paula Rahal, Paulo Inacio da Costa, João Pessoa Araújo Jr., Maurício Lacerda Nogueira. Prefeitura de Sao Paulo: Melissa Palmieri. |
| EPI_ISL_2171032, EPI_ISL_2171033, EPI_ISL_2171035, EPI_ISL_2171036, EPI_ISL_2171037, EPI_ISL_2171038, EPI_ISL_2171039 | UNIDADE DE PRONTO ATENDIMENTO DE VARZEA PAULISTA UPA II | Instituto Butantan / Mendelics | Instituto Butantan: Dimas Tadeu Covas, Sandra Coccuzzo Sampaio, Maria Carolina Elias, José Salvatore Leister Patané, Vincent Louis Viala, Antonio Jorge Martins, Ricardo Haddad, Claudia Renata dos Santos Barros, Elaine Cristina Marqueze, Raul Machado Neto, Debora Botequio Moretti, Jardelina de Souza Todao Bernardino, Loyze Paola Oliveira de Lima, Luiz Aurelio de Campos Crispin. Centro de Genômica Funcional da ESALQ: Luiz Lehmann Coutinho, Ricardo Augusto Brassaloti, Raquel de Lello Rocha Campos Cassano. NGS Soluções Genômicas: Pilar Drummond Sampaio Corrêa Mariani. FZEA-USP Pirassununga: Mirele Daiana Poleti, Jessika Cristina Chagas Lesbon, Elisangela Chicaroni Mattos, Heidge Fukumasu. USP-Botucatu: Rejane Maria Tommasini Grotto, Jayme A. Souza-Neto, Guilherme Targino Valente, Patricia Akemi Assato, Felipe Allan da Silva da Costa, Bianca Cechetto Carlos. Mendelics: Bibiana Santos, João Paulo Kitajima, Erika Freitas, David Schlesinger. Hemocentro Ribeirão Preto: Simone Kashima, Evandra Strazza Rodrigues, Svetoslav Nanev Slavov, Elaine Vieira dos Santos, Rafael dos Santos Bezerra, Luiz Carlos Junior de Alcantara, Marta Giovanetti, Vagner Fonseca, Flavia Aburjaile, Rodrigo Tocantins Calado. FAMERP-SJRP: Cecília Artico Banho, Lívia Sacchetto, Fábio Sossai Possebon, Leila Sabrina Ullmann, Cíntia Bittar, Guilherme Campos, Helena Lage Ferreira, Jorge A. Petroli Marchesi, Maisa C. Pereira Parra, Marília Moraes, Paula Rahal, Paulo Inacio da Costa, João Pessoa Araújo Jr., Maurício Lacerda Nogueira. Prefeitura de Sao Paulo: Melissa Palmieri. |
| EPI_ISL_2172524 | UNIDADE DE PRONTO ATENDIMENTO UPA DRA ANA OLIVIA BENTIVOGLIO | Instituto Butantan / ESALQ-Piracicaba | Instituto Butantan: Dimas Tadeu Covas, Sandra Coccuzzo Sampaio, Maria Carolina Elias, José Salvatore Leister Patané, Vincent Louis Viala, Antonio Jorge Martins, Ricardo Haddad, Claudia Renata dos Santos Barros, Elaine Cristina Marqueze, Raul Machado Neto, Debora Botequio Moretti, Jardelina de Souza Todao Bernardino, Loyze Paola Oliveira de Lima, Luiz Aurelio de Campos Crispin. Centro de Genômica Funcional da ESALQ: Luiz Lehmann Coutinho, Ricardo Augusto Brassaloti, Raquel de Lello Rocha Campos Cassano. NGS Soluções Genômicas: Pilar Drummond Sampaio Corrêa Mariani. FZEA-USP Pirassununga: Mirele Daiana Poleti, Jessika Cristina Chagas Lesbon, Elisangela Chicaroni Mattos, Heidge Fukumasu. USP-Botucatu: Rejane Maria Tommasini Grotto, Jayme A. Souza-Neto, Guilherme Targino Valente, Patricia Akemi Assato, Felipe Allan da Silva da Costa, Bianca Cechetto Carlos. Mendelics: Bibiana Santos, João Paulo Kitajima, Erika Freitas, David Schlesinger. Hemocentro Ribeirão Preto: Simone Kashima, Evandra Strazza Rodrigues, Svetoslav Nanev Slavov, Elaine Vieira dos Santos, Rafael dos Santos Bezerra, Luiz Carlos Junior de Alcantara, Marta Giovanetti, Vagner Fonseca, Flavia Aburjaile, Rodrigo Tocantins Calado. FAMERP-SJRP: Cecília Artico Banho, Lívia Sacchetto, Fábio Sossai Possebon, Leila Sabrina Ullmann, Cíntia Bittar, Guilherme Campos, Helena Lage Ferreira, Jorge A. Petroli Marchesi, Maisa C. Pereira Parra, Marília Moraes, Paula Rahal, Paulo Inacio da Costa, João Pessoa Araújo Jr., Maurício Lacerda Nogueira. Prefeitura de Sao Paulo: Melissa Palmieri. |

EPI_ISL_2187684, EPI_ISL_2187685, EPI_ISL_2187686, EPI_ISL_2187687, EPI_ISL_2187688, EPI_ISL_2187689, EPI_ISL_2187690, EPI_ISL_2187691, EPI_ISL_2187692, EPI_ISL_2187695, EPI_ISL_2187696, EPI_ISL_2187697, EPI_ISL_2187698, EPI_ISL_2187699, EPI_ISL_2187700, EPI_ISL_2187701, EPI_ISL_2187702, EPI_ISL_2187703, EPI_ISL_2187704, EPI_ISL_2187705, EPI_ISL_2187706, EPI_ISL_2187707, EPI_ISL_2187708, EPI_ISL_2187709, EPI_ISL_2187710, EPI_ISL_2187711, EPI_ISL_2187712, EPI_ISL_2187713, EPI_ISL_2187714, EPI_ISL_2187715, EPI_ISL_2187716, EPI_ISL_2187717, EPI_ISL_2187718, EPI_ISL_2187719, EPI_ISL_2187720, EPI_ISL_2187721, EPI_ISL_2187722, EPI_ISL_2187723, EPI_ISL_2187724, EPI_ISL_2187725, EPI_ISL_2187726, EPI_ISL_2187727, EPI_ISL_2187728, EPI_ISL_2187729, EPI_ISL_2187730, EPI_ISL_2187731, EPI_ISL_2187732, EPI_ISL_2187733, EPI_ISL_2187734, EPI_ISL_2187735, EPI_ISL_2187736, EPI_ISL_2187737, EPI_ISL_2187738, EPI_ISL_2187739, EPI_ISL_2187740, EPI_ISL_2187741, EPI_ISL_2187742, EPI_ISL_2187743, EPI_ISL_2187744, EPI_ISL_2187745, EPI_ISL_2187746, EPI_ISL_2187747, EPI_ISL_2187748, EPI_ISL_2187749, EPI_ISL_2187750, EPI_ISL_2187751, EPI_ISL_2187752, EPI_ISL_2187753, EPI_ISL_2187754, EPI_ISL_2187755, EPI_ISL_2187757, EPI_ISL_2187758, EPI_ISL_2187759, EPI_ISL_2187760, EPI_ISL_2187761, EPI_ISL_2187762, EPI_ISL_2187763, EPI_ISL_2187764, EPI_ISL_2187765, EPI_ISL_2187766, EPI_ISL_2187767, EPI_ISL_2187768, EPI_ISL_2187769, EPI_ISL_2187770, EPI_ISL_2187771, EPI_ISL_2187772, EPI_ISL_2187773, EPI_ISL_2187774, EPI_ISL_2187775, EPI_ISL_2187776, EPI_ISL_2187777, EPI_ISL_2187778, EPI_ISL_2187779, EPI_ISL_2187780, EPI_ISL_2187781, EPI_ISL_2187782, EPI_ISL_2187783, EPI_ISL_2187784, EPI_ISL_2187785, EPI_ISL_2187786, EPI_ISL_2187787, EPI_ISL_2187788, EPI_ISL_2187789, EPI_ISL_2187791, EPI_ISL_2187792, EPI_ISL_2187793, EPI_ISL_2187794, EPI_ISL_2187796, EPI_ISL_2187797, EPI_ISL_2187798, EPI_ISL_2187799, EPI_ISL_2187800, EPI_ISL_2187801, EPI_ISL_2187803, EPI_ISL_2187804, EPI_ISL_2187805, EPI_ISL_2187806, EPI_ISL_2187807, EPI_ISL_2187808,

| | | | | |
|---|---|---|---|---|
| EPI_ISL_2187809, EPI_ISL_2187810, EPI_ISL_2187811, EPI_ISL_2187812, EPI_ISL_2187813, EPI_ISL_2187814, EPI_ISL_2187815, EPI_ISL_2187816, EPI_ISL_2187817, EPI_ISL_2187818, EPI_ISL_2187819, EPI_ISL_2187820, EPI_ISL_2187821, EPI_ISL_2187822, EPI_ISL_2187824, EPI_ISL_2187825, EPI_ISL_2187826, EPI_ISL_2187827, EPI_ISL_2187828, EPI_ISL_2187829, EPI_ISL_2187830, EPI_ISL_2187831, EPI_ISL_2187832, EPI_ISL_2187833, EPI_ISL_2187834, EPI_ISL_2187835, EPI_ISL_2187836, EPI_ISL_2187837, EPI_ISL_2187838, EPI_ISL_2187839, EPI_ISL_2187840, EPI_ISL_2187842, EPI_ISL_2187843, EPI_ISL_2187845, EPI_ISL_2187846, EPI_ISL_2187847, EPI_ISL_2187848, EPI_ISL_2187849, EPI_ISL_2187851, EPI_ISL_2187852, EPI_ISL_2187853, EPI_ISL_2187854, EPI_ISL_2187855, EPI_ISL_2187856, EPI_ISL_2187857, EPI_ISL_2187858, EPI_ISL_2187859, EPI_ISL_2187861, EPI_ISL_2187862, EPI_ISL_2187863, EPI_ISL_2187864, EPI_ISL_2187865, EPI_ISL_2187866, EPI_ISL_2187867, EPI_ISL_2187868, EPI_ISL_2187869, EPI_ISL_2187870, EPI_ISL_2187871, EPI_ISL_2187872, EPI_ISL_2187873, EPI_ISL_2187874, EPI_ISL_2187875, EPI_ISL_2187876, EPI_ISL_2187877, EPI_ISL_2187878, EPI_ISL_2187879, EPI_ISL_2187880, EPI_ISL_2187881, EPI_ISL_2187883, EPI_ISL_2187884, EPI_ISL_2187885, EPI_ISL_2187886, EPI_ISL_2187887, EPI_ISL_2187888, EPI_ISL_2187889, EPI_ISL_2187891, EPI_ISL_2187892, EPI_ISL_2187893, EPI_ISL_2187894, EPI_ISL_2187895, EPI_ISL_2187896, EPI_ISL_2187897, EPI_ISL_2187898, EPI_ISL_2187899, EPI_ISL_2187900, EPI_ISL_2187901, EPI_ISL_2187902, EPI_ISL_2187903, EPI_ISL_2187904, EPI_ISL_2187905, EPI_ISL_2187906, EPI_ISL_2187907, EPI_ISL_2187908, EPI_ISL_2187911, EPI_ISL_2187912, EPI_ISL_2187913, EPI_ISL_2187914, EPI_ISL_2187915, EPI_ISL_2187916, EPI_ISL_2187917, EPI_ISL_2187918, EPI_ISL_2187919, EPI_ISL_2187920, EPI_ISL_2187921, EPI_ISL_2187922, EPI_ISL_2187923, EPI_ISL_2187924, EPI_ISL_2187925, EPI_ISL_2187926, EPI_ISL_2187927, EPI_ISL_2187928, EPI_ISL_2187929, EPI_ISL_2187930, EPI_ISL_2187931, EPI_ISL_2187932, EPI_ISL_2187933, EPI_ISL_2187934, EPI_ISL_2187935, EPI_ISL_2187936, EPI_ISL_2187937, EPI_ISL_2187938, EPI_ISL_2187939, EPI_ISL_2187940, EPI_ISL_2187941, EPI_ISL_2187942, EPI_ISL_2187943, EPI_ISL_2187944, EPI_ISL_2187945, EPI_ISL_2187946, EPI_ISL_2187947, EPI_ISL_2187948, EPI_ISL_2187949, EPI_ISL_2187950, EPI_ISL_2187951, EPI_ISL_2187952, EPI_ISL_2187953, EPI_ISL_2187954, EPI_ISL_2187955, EPI_ISL_2187956, EPI_ISL_2187957, EPI_ISL_2187958, EPI_ISL_2187959, EPI_ISL_2187960, EPI_ISL_2187961, EPI_ISL_2187962, EPI_ISL_2187963, EPI_ISL_2187964, EPI_ISL_2187965, EPI_ISL_2187966, EPI_ISL_2187967, EPI_ISL_2187968, EPI_ISL_2187969, EPI_ISL_2187970, EPI_ISL_2187971, EPI_ISL_2187972, EPI_ISL_2187973, EPI_ISL_2187974, EPI_ISL_2187975, EPI_ISL_2187976, EPI_ISL_2187977, EPI_ISL_2187978, EPI_ISL_2187979, EPI_ISL_2187980, EPI_ISL_2187981, EPI_ISL_2187982, EPI_ISL_2187983, EPI_ISL_2187984, EPI_ISL_2187985, EPI_ISL_2187986, EPI_ISL_2187987, EPI_ISL_2187988, EPI_ISL_2187989, EPI_ISL_2187990, EPI_ISL_2187991, EPI_ISL_2187992, EPI_ISL_2187993, EPI_ISL_2187994, EPI_ISL_2187995, EPI_ISL_2187996, EPI_ISL_2187997, EPI_ISL_2187998, EPI_ISL_2187999, EPI_ISL_2188000, EPI_ISL_2188001, EPI_ISL_2188002, EPI_ISL_2188003, EPI_ISL_2188004, EPI_ISL_2188005, EPI_ISL_2188006, EPI_ISL_2188007, EPI_ISL_2188008, EPI_ISL_2188009, EPI_ISL_2188010, EPI_ISL_2188011, EPI_ISL_2188012, EPI_ISL_2188013, EPI_ISL_2188014, EPI_ISL_2188015, EPI_ISL_2188016, EPI_ISL_2188017, EPI_ISL_2188018, EPI_ISL_2188019, EPI_ISL_2188020, EPI_ISL_2188021, EPI_ISL_2188022, EPI_ISL_2188023, EPI_ISL_2188024, EPI_ISL_2188025, EPI_ISL_2188027 | | | | |
| | see above | HLAGYN - Laboratorio de Imunologia de Transplantes de Goias | HLAGYN - Laboratorio de Imunologia de Transplantes de Goias | Fernando Antonio Vinhal dos Santos, Erika Lopes Rocha Batista, Alessandro Leonardo Alvares Magalhaes, Frederico Rodrigues Vinhal, Sabrina Sara Moreira Duarte, Lucas Carlos Gomes Pereira, Daniel Ferreira de Sousa |

We gratefully acknowledge the following Authors from the Originating laboratories responsible for obtaining the specimens, as well as the Submitting laboratories where the genome data were generated and shared via GISAID, on which this research is based.

All Submitters of data may be contacted directly via www.gisaid.org

Authors are sorted alphabetically.

| Accession ID | Originating Laboratory | Submitting Laboratory | Authors |
|---|---|---|---|
| EPI_ISL_2196184 | Laboratorio Central de Saude Publica do Estado do Espitito Santo (LACEN-ES) | Laboratory of Respiratory Viruses and Measles, Oswaldo Cruz Institute, FIOCRUZ | Paola Resende, Luciana Appolinario, Fernando Motta, Anna Carolina Paixao, Ana Carolina Mendonca, Alice Sampaio Rocha, Taina Venas, Eilisa Cavalcante Pereira, Renata Serrano Lopes, Rodrigo Ribeiro Rodrigues, Marilda Siqueira on behalf of the Fiocruz COVID-19 Genomic Surveillance Network |
| EPI_ISL_2196185, EPI_ISL_2196186, EPI_ISL_2196187 | Laboratory of Respiratory Viruses and Measles, Oswaldo Cruz Institute, FIOCRUZ | Laboratory of Respiratory Viruses and Measles, Oswaldo Cruz Institute, FIOCRUZ | Paola Resende, Luciana Appolinario, Fernando Motta, Anna Carolina Paixao, Ana Carolina Mendonca, Alice Sampaio Rocha, Taina Venas, Elisa Cavalcante Pereira, Renata Serrano Lopes, Marilda Siqueira on behalf of the Fiocruz COVID-19 Genomic Surveillance Network |
| EPI_ISL_2196188 | Laboratorio Central de Saude Publica do Estado de Sergipe (LACEN/SE) | Laboratory of Respiratory Viruses and Measles, Oswaldo Cruz Institute, FIOCRUZ | Paola Resende, Luciana Appolinario, Fernando Motta, Anna Carolina Paixao, Ana Carolina Mendonca, Alice Sampaio Rocha, Tainá Moreira Martins Venas, Elisa Cavalcante Pereira, Renata Serrano Lopes, Cliomar Alves dos Santos, Marilda Siqueira on behalf of the Fiocruz COVID-19 Genomic Surveillance Network |
| EPI_ISL_2196189 | Laboratorio Central de Saude Publica do Estado do Rio de Janeiro (LACEN-RJ) | Laboratory of Respiratory Viruses and Measles, Oswaldo Cruz Institute, FIOCRUZ | Paola Resende, Luciana Appolinario, Fernando Motta, Anna Carolina Paixao, Ana Carolina Mendonca, Alice Sampaio Rocha, Taina Venas, Elisa Cavalcante Pereira, Renata Serrano Lopes, Andrea Cony Cavalcanti, Marilda Siqueira on behalf of the Fiocruz COVID-19 Genomic Surveillance Network |
| EPI_ISL_2196190, EPI_ISL_2196191, EPI_ISL_2196192, EPI_ISL_2196193, EPI_ISL_2196194, EPI_ISL_2196195, EPI_ISL_2196196 | Laboratory of Respiratory Viruses and Measles, Oswaldo Cruz Institute, FIOCRUZ | Laboratory of Respiratory Viruses and Measles, Oswaldo Cruz Institute, FIOCRUZ | Paola Resende, Luciana Appolinario, Fernando Motta, Anna Carolina Paixao, Ana Carolina Mendonca, Alice Sampaio Rocha, Taina Venas, Elisa Cavalcante Pereira, Renata Serrano Lopes, Marilda Siqueira on behalf of the Fiocruz COVID-19 Genomic Surveillance Network |
| EPI_ISL_2196197, EPI_ISL_2196198, EPI_ISL_2196199, EPI_ISL_2196200, EPI_ISL_2196201, EPI_ISL_2196202, EPI_ISL_2196203, EPI_ISL_2196204, EPI_ISL_2196205, EPI_ISL_2196206, EPI_ISL_2196207, EPI_ISL_2196208, EPI_ISL_2196209 | | | |
| see above | Laboratorio Central de Saude Publica do Estado do Rio de Janeiro (LACEN-RJ) | Laboratory of Respiratory Viruses and Measles, Oswaldo Cruz Institute, FIOCRUZ | Paola Resende, Luciana Appolinario, Fernando Motta, Anna Carolina Paixao, Ana Carolina Mendonca, Alice Sampaio Rocha, Taina Venas, Elisa Cavalcante Pereira, Renata Serrano Lopes, Andrea Cony Cavalcanti, Marilda Siqueira on behalf of the Fiocruz COVID-19 Genomic Surveillance Network |
| EPI_ISL_2196210, EPI_ISL_2196211, EPI_ISL_2196212, EPI_ISL_2196213, EPI_ISL_2196214, EPI_ISL_2196215, EPI_ISL_2196216, EPI_ISL_2196217, EPI_ISL_2196218, EPI_ISL_2196219, EPI_ISL_2196220, EPI_ISL_2196221, EPI_ISL_2196222, EPI_ISL_2196223, EPI_ISL_2196224, EPI_ISL_2196225, EPI_ISL_2196226, EPI_ISL_2196227, EPI_ISL_2196228, EPI_ISL_2196229, EPI_ISL_2196230, EPI_ISL_2196231, EPI_ISL_2196232, EPI_ISL_2196233, EPI_ISL_2196234, EPI_ISL_2196235 | | | |
| see above | Laboratorio Central de Saude Publica do Estado de Sergipe (LACEN/SE) | Laboratory of Respiratory Viruses and Measles, Oswaldo Cruz Institute, FIOCRUZ | Paola Resende, Luciana Appolinario, Fernando Motta, Anna Carolina Paixao, Ana Carolina Mendonca, Alice Sampaio Rocha, Tainá Moreira Martins Venas, Elisa Cavalcante Pereira, Renata Serrano Lopes, Cliomar Alves dos Santos, Marilda Siqueira on behalf of the Fiocruz COVID-19 Genomic Surveillance Network |
| EPI_ISL_2196236 | Laboratorio Central de Saude Publica do Estado do Rio de Janeiro (LACEN-RJ) | Laboratory of Respiratory Viruses and Measles, Oswaldo Cruz Institute, FIOCRUZ | Paola Resende, Luciana Appolinario, Fernando Motta, Anna Carolina Paixao, Ana Carolina Mendonca, Alice Sampaio Rocha, Taina Venas, Elisa Cavalcante Pereira, Renata Serrano Lopes, Andrea Cony Cavalcanti, Marilda Siqueira on behalf of the Fiocruz COVID-19 Genomic Surveillance Network |
| EPI_ISL_2196237, EPI_ISL_2196238, EPI_ISL_2196239, EPI_ISL_2196240 | Laboratorio Central de Saude Publica do Estado de Sergipe (LACEN/SE) | Laboratory of Respiratory Viruses and Measles, Oswaldo Cruz Institute, FIOCRUZ | Paola Resende, Luciana Appolinario, Fernando Motta, Anna Carolina Paixao, Ana Carolina Mendonca, Alice Sampaio Rocha, Tainá Moreira Martins Venas, Elisa Cavalcante Pereira, Renata Serrano Lopes, Cliomar Alves dos Santos, Marilda Siqueira on behalf of the Fiocruz COVID-19 Genomic Surveillance Network |
| EPI_ISL_2196241 | Laboratorio Central de Saude Publica do Estado de Minas Gerais (LACEN/MG) | Laboratory of Respiratory Viruses and Measles, Oswaldo Cruz Institute, FIOCRUZ | Paola Resende, Luciana Appolinario, Fernando Motta, Anna Carolina Paixao, Ana Carolina Mendonca, Alice Sampaio Rocha, Taina Venas, Elisa Cavalcante Pereira, Renata Serrano Lopes, Andre Felipe Leal Bernardes, Marilda Siqueira on behalf of the Fiocruz COVID-19 Genomic Surveillance Network |
| EPI_ISL_2196242 | Labortorio Central de Saude Publica do Estado do Rio de Janeiro (LACEN/RJ) | Laboratory of Respiratory Viruses and Measles, Oswaldo Cruz Institute, FIOCRUZ | Paola Resende, Luciana Appolinario, Fernando Motta, Anna Carolina Paixao, Ana Carolina Mendonca, Alice Sampaio Rocha, Taina Venas, Elisa Cavalcante Pereira, Renata Serrano Lopes, Andrea Cony Cavalcanti, Marilda Siqueira on behalf of the Fiocruz COVID-19 Genomic Surveillance Network |
| EPI_ISL_2196243 | Laboratorio Central de Saude Publica do Estado do Para (LACEN/PA) | Laboratory of Respiratory Viruses and Measles, Oswaldo Cruz Institute, FIOCRUZ | Paola Resende, Luciana Appolinario, Fernando Motta, Anna Carolina Paixao, Ana Carolina Mendonca, Alice Sampaio Rocha, Taina Venas, Elisa Cavalcante Pereira, Renata Serrano Lopes, Valnete Andrade, Marilda Siqueira on behalf of the Fiocruz COVID-19 Genomic Surveillance Network |
| EPI_ISL_2196244 | Lboratorio Central de Saude Publica do Estado do Parana (LACEN/PR) | Laboratory of Respiratory Viruses and Measles, Oswaldo Cruz Institute, FIOCRUZ | Paola Resende, Luciana Appolinario, Fernando Motta, Anna Carolina Paixao, Ana Carolina Mendonca, Alice Sampaio Rocha, Taina Venas, Elisa Cavalcante Pereira, Renata Serrano Lopes, Irina Riediger, Marilda Siqueira on behalf of the Fiocruz COVID-19 Genomic Surveillance Network |
| EPI_ISL_2196245 | Labortorio Central de Saude Publica do Estado do Rio de Janeiro (LACEN/RJ) | Laboratory of Respiratory Viruses and Measles, Oswaldo Cruz Institute, FIOCRUZ | Paola Resende, Luciana Appolinario, Fernando Motta, Anna Carolina Paixao, Ana Carolina Mendonca, Alice Sampaio Rocha, Taina Venas, Elisa Cavalcante Pereira, Renata Serrano Lopes, Andrea Cony Cavalcanti, Marilda Siqueira on behalf of the Fiocruz COVID-19 Genomic Surveillance Network |
| EPI_ISL_2196246 | Laboratorio Central de Saude Publica do Estado de Minas Gerais (LACEN/MG) | Laboratory of Respiratory Viruses and Measles, Oswaldo Cruz Institute, FIOCRUZ | Paola Resende, Luciana Appolinario, Fernando Motta, Anna Carolina Paixao, Ana Carolina Mendonca, Alice Sampaio Rocha, Taina Venas, Elisa Cavalcante Pereira, Renata Serrano Lopes, Andre Felipe Leal Bernardes, Marilda Siqueira on behalf of the Fiocruz COVID-19 Genomic Surveillance Network |
| EPI_ISL_2196247, EPI_ISL_2196249, EPI_ISL_2196250 | Labortorio Central de Saude Publica do Estado de Santa Catarina (LACEN/SC) | Laboratory of Respiratory Viruses and Measles, Oswaldo Cruz Institute, FIOCRUZ | Paola Resende, Luciana Appolinario, Fernando Motta, Anna Carolina Paixao, Ana Carolina Mendonca, Alice Sampaio Rocha, Taina Venas, Elisa Cavalcante Pereira, Renata Serrano Lopes, Darcita Buerger Rovaris, Sandra Bianchini Fernandes, Marilda Siqueira on behalf of the Fiocruz COVID-19 Genomic Surveillance Network |
| EPI_ISL_2196251 | Laboratory of Respiratory Viruses and Measles, Oswaldo Cruz Institute, FIOCRUZ | Laboratory of Respiratory Viruses and Measles, Oswaldo Cruz Institute, FIOCRUZ | Paola Resende, Luciana Appolinario, Fernando Motta, Anna Carolina Paixao, Ana Carolina Mendonca, Alice Sampaio Rocha, Taina Venas, Elisa Cavalcante Pereira, Renata Serrano Lopes, Marilda Siqueira on behalf of the Fiocruz COVID-19 Genomic Surveillance Network |
| EPI_ISL_2196252 | Lboratorio Central de Saude Publica do Estado do Parana (LACEN/PR) | Laboratory of Respiratory Viruses and Measles, Oswaldo Cruz Institute, FIOCRUZ | Paola Resende, Luciana Appolinario, Fernando Motta, Anna Carolina Paixao, Ana Carolina Mendonca, Alice Sampaio Rocha, Taina Venas, Elisa Cavalcante Pereira, Renata Serrano Lopes, Irina Riediger, Marilda Siqueira on behalf of the Fiocruz COVID-19 Genomic Surveillance Network |
| EPI_ISL_2196253 | Laboratorio Central de Saude Publica do Estado da Bahia (LACEN/BA) | Laboratory of Respiratory Viruses and Measles, Oswaldo Cruz Institute, FIOCRUZ | Paola Resende, Luciana Appolinario, Fernando Motta, Anna Carolina Paixao, Ana Carolina Mendonca, Alice Sampaio Rocha, Taina Venas, Elisa Cavalcante Pereira, Renata Serrano Lopes,Felicidade Pereira, Marilda Siqueira on behalf of the Fiocruz COVID-19 Genomic Surveillance Network |
| EPI_ISL_2196254, EPI_ISL_2196255, EPI_ISL_2196256 | Labortorio Central de Saude Publica do Estado de Santa Catarina (LACEN/SC) | Laboratory of Respiratory Viruses and Measles, Oswaldo Cruz Institute, FIOCRUZ | Paola Resende, Luciana Appolinario, Fernando Motta, Anna Carolina Paixao, Ana Carolina Mendonca, Alice Sampaio Rocha, Taina Venas, Elisa Cavalcante Pereira, Renata Serrano Lopes, Darcita Buerger Rovaris, Sandra Bianchini Fernandes, Marilda Siqueira on behalf of the Fiocruz COVID-19 Genomic Surveillance Network |
| EPI_ISL_2196257, EPI_ISL_2196258 | Laboratorio Central de Saude Publica do Estado de Minas Gerais (LACEN/MG) | Laboratory of Respiratory Viruses and Measles, Oswaldo Cruz Institute, FIOCRUZ | Paola Resende, Luciana Appolinario, Fernando Motta, Anna Carolina Paixao, Ana Carolina Mendonca, Alice Sampaio Rocha, Taina Venas, Elisa Cavalcante Pereira, Renata Serrano Lopes, Andre Felipe Leal Bernardes, Marilda Siqueira on behalf of the Fiocruz COVID-19 Genomic Surveillance Network |
| EPI_ISL_2196259 | Lboratorio Central de Saude Publica do Estado do Parana (LACEN/PR) | Laboratory of Respiratory Viruses and Measles, Oswaldo Cruz Institute, FIOCRUZ | Paola Resende, Luciana Appolinario, Fernando Motta, Anna Carolina Paixao, Ana Carolina Mendonca, Alice Sampaio Rocha, Taina Venas, Elisa Cavalcante Pereira, Renata Serrano Lopes, Irina Riediger, Marilda Siqueira on behalf of the Fiocruz COVID-19 Genomic Surveillance Network |
| EPI_ISL_2196260, EPI_ISL_2196261 | Laboratorio Central de Saude Publica do Estado da Bahia (LACEN/BA) | Laboratory of Respiratory Viruses and Measles, Oswaldo Cruz Institute, FIOCRUZ | Paola Resende, Luciana Appolinario, Fernando Motta, Anna Carolina Paixao, Ana Carolina Mendonca, Alice Sampaio Rocha, Taina Venas, Elisa Cavalcante Pereira, Renata Serrano Lopes,Felicidade Pereira, Marilda Siqueira on behalf of the Fiocruz COVID-19 Genomic Surveillance Network |
| EPI_ISL_2196262 | Laboratorio Central de Saude Publica do Estado do Espirito Santo (LACEN/ES) | Laboratory of Respiratory Viruses and Measles, Oswaldo Cruz Institute, FIOCRUZ | Paola Resende, Luciana Appolinario, Fernando Motta, Anna Carolina Paixao, Ana Carolina Mendonca, Alice Sampaio Rocha, Taina Venas, Elisa Cavalcante Pereira, Renata Serrano Lopes, Rodrigo Ribeiro Rodrigues, Marilda Siqueira on behalf of the Fiocruz COVID-19 Genomic Surveillance Network |

| | | | |
|---|---|---|---|
| EPI_ISL_2196263, EPI_ISL_2196264, EPI_ISL_2196265, EPI_ISL_2196266, EPI_ISL_2196267, EPI_ISL_2196268, EPI_ISL_2196269 | Laboratorio Central de Saude Publica do Estado Maranhao (LACEN-MA) | Laboratory of Respiratory Viruses and Measles, Oswaldo Cruz Institute, FIOCRUZ | Paola Resende, Luciana Appolinario, Fernando Motta, Anna Carolina Paixao, Ana Carolina Mendonca, Alice Sampaio Rocha, Taina Venas, Elisa Cavalcante Pereira, Renata Serrano Lopes, Lidio Gonçalves Lima Neto, Marilda Siqueira on behalf of the Fiocruz COVID-19 Genomic Surveillance Network |
| EPI_ISL_2196270, EPI_ISL_2196271, EPI_ISL_2196272, EPI_ISL_2196273, EPI_ISL_2196274, EPI_ISL_2196275, EPI_ISL_2196276, EPI_ISL_2196277, EPI_ISL_2196278, EPI_ISL_2196279, EPI_ISL_2196280 | | | |
| see above | Laboratorio Central de Saude Publica do Estado de Minas Gerais (LACEN/MG) | Laboratory of Respiratory Viruses and Measles, Oswaldo Cruz Institute, FIOCRUZ | Paola Resende, Luciana Appolinario, Fernando Motta, Anna Carolina Paixao, Ana Carolina Mendonca, Alice Sampaio Rocha, Taina Venas, Elisa Cavalcante Pereira, Renata Serrano Lopes, Andre Felipe Leal Bernardes, Marilda Siqueira on behalf of the Fiocruz COVID-19 Genomic Surveillance Network |
| EPI_ISL_2196281, EPI_ISL_2196282, EPI_ISL_2196283, EPI_ISL_2196284, EPI_ISL_2196285, EPI_ISL_2196286, EPI_ISL_2196287, EPI_ISL_2196288, EPI_ISL_2196289, EPI_ISL_2196290, EPI_ISL_2196291, EPI_ISL_2196292, EPI_ISL_2196293, EPI_ISL_2196294, EPI_ISL_2196295, EPI_ISL_2196296, EPI_ISL_2196297, EPI_ISL_2196298, EPI_ISL_2196299, EPI_ISL_2196300, EPI_ISL_2196301, EPI_ISL_2196302, EPI_ISL_2196303, EPI_ISL_2196304, EPI_ISL_2196305 | | | |
| see above | Laboratorio Central de Saude Publica do Estado do Para (LACEN/PA) | Laboratory of Respiratory Viruses and Measles, Oswaldo Cruz Institute, FIOCRUZ | Paola Resende, Luciana Appolinario, Fernando Motta, Anna Carolina Paixao, Ana Carolina Mendonca, Alice Sampaio Rocha, Taina Venas, Elisa Cavalcante Pereira, Renata Serrano Lopes, Valnete Andrade, Marilda Siqueira on behalf of the Fiocruz COVID-19 Genomic Surveillance Network |
| EPI_ISL_2196306, EPI_ISL_2196307, EPI_ISL_2196308, EPI_ISL_2196309, EPI_ISL_2196310, EPI_ISL_2196311, EPI_ISL_2196312 | Lboratorio Central de Saude Publica do Estado do Parana (LACEN/PR) | Laboratory of Respiratory Viruses and Measles, Oswaldo Cruz Institute, FIOCRUZ | Paola Resende, Luciana Appolinario, Fernando Motta, Anna Carolina Paixao, Ana Carolina Mendonca, Alice Sampaio Rocha, Taina Venas, Elisa Cavalcante Pereira, Renata Serrano Lopes, Irina Riediger, Marilda Siqueira on behalf of the Fiocruz COVID-19 Genomic Surveillance Network |
| EPI_ISL_2196313, EPI_ISL_2196314, EPI_ISL_2196315, EPI_ISL_2196316, EPI_ISL_2196317, EPI_ISL_2196318, EPI_ISL_2196319, EPI_ISL_2196320, EPI_ISL_2196321, EPI_ISL_2196322, EPI_ISL_2196323, EPI_ISL_2196324, EPI_ISL_2196325, EPI_ISL_2196326, EPI_ISL_2196327, EPI_ISL_2196328, EPI_ISL_2196329, EPI_ISL_2196330, EPI_ISL_2196331, EPI_ISL_2196332, EPI_ISL_2196333, EPI_ISL_2196334 | | | |
| see above | Labortorio Central de Saude Publica do Estado do Rio de Janeiro (LACEN/RJ) | Laboratory of Respiratory Viruses and Measles, Oswaldo Cruz Institute, FIOCRUZ | Paola Resende, Luciana Appolinario, Fernando Motta, Anna Carolina Paixao, Ana Carolina Mendonca, Alice Sampaio Rocha, Taina Venas, Elisa Cavalcante Pereira, Renata Serrano Lopes, Andrea Cony Cavalcanti, Marilda Siqueira on behalf of the Fiocruz COVID-19 Genomic Surveillance Network |
| EPI_ISL_2196335, EPI_ISL_2196336, EPI_ISL_2196337, EPI_ISL_2196338, EPI_ISL_2196339, EPI_ISL_2196340, EPI_ISL_2196341, EPI_ISL_2196342, EPI_ISL_2196343, EPI_ISL_2196344, EPI_ISL_2196345, EPI_ISL_2196346, EPI_ISL_2196347, EPI_ISL_2196348, EPI_ISL_2196349, EPI_ISL_2196350 | | | |
| see above | Labortorio Central de Saude Publica do Estado de Santa Catarina (LACEN/SC) | Laboratory of Respiratory Viruses and Measles, Oswaldo Cruz Institute, FIOCRUZ | Paola Resende, Luciana Appolinario, Fernando Motta, Anna Carolina Paixao, Ana Carolina Mendonca, Alice Sampaio Rocha, Taina Venas, Elisa Cavalcante Pereira, Renata Serrano Lopes, Darcita Buerger Rovaris, Sandra Bianchini Fernandes, Marilda Siqueira on behalf of the Fiocruz COVID-19 Genomic Surveillance Network |
| EPI_ISL_2196351, EPI_ISL_2196352 | Laboratorio Central de Saude Publica do Estado de Sergipe (LACEN/SE) | Laboratory of Respiratory Viruses and Measles, Oswaldo Cruz Institute, FIOCRUZ | Paola Resende, Luciana Appolinario, Fernando Motta, Anna Carolina Paixao, Ana Carolina Mendonca, Alice Sampaio Rocha, Tainá Moreira Martins Venas, Elisa Cavalcante Pereira, Renata Serrano Lopes, Cliomar Alves dos Santos, Marilda Siqueira on behalf of the Fiocruz COVID-19 Genomic Surveillance Network |
| EPI_ISL_2196353, EPI_ISL_2196354 | Labortorio Central de Saude Publica do Estado de Santa Catarina (LACEN/SC) | Laboratory of Respiratory Viruses and Measles, Oswaldo Cruz Institute, FIOCRUZ | Paola Resende, Luciana Appolinario, Fernando Motta, Anna Carolina Paixao, Ana Carolina Mendonca, Alice Sampaio Rocha, Taina Venas, Elisa Cavalcante Pereira, Renata Serrano Lopes, Darcita Buerger Rovaris, Sandra Bianchini Fernandes, Marilda Siqueira on behalf of the Fiocruz COVID-19 Genomic Surveillance Network |
| EPI_ISL_2196355 | Laboratorio Central de Saude Publica do Estado Maranhao (LACEN-MA) | Laboratory of Respiratory Viruses and Measles, Oswaldo Cruz Institute, FIOCRUZ | Paola Resende, Luciana Appolinario, Fernando Motta, Anna Carolina Paixao, Ana Carolina Mendonca, Alice Sampaio Rocha, Taina Venas, Elisa Cavalcante Pereira, Renata Serrano Lopes, Lidio Gonçalves Lima Neto, Marilda Siqueira on behalf of the Fiocruz COVID-19 Genomic Surveillance Network |
| EPI_ISL_2196356 | Labortorio Central de Saude Publica do Estado do Rio de Janeiro (LACEN/RJ) | Laboratory of Respiratory Viruses and Measles, Oswaldo Cruz Institute, FIOCRUZ | Paola Resende, Luciana Appolinario, Fernando Motta, Anna Carolina Paixao, Ana Carolina Mendonca, Alice Sampaio Rocha, Taina Venas, Elisa Cavalcante Pereira, Renata Serrano Lopes, Andrea Cony Cavalcanti, Marilda Siqueira on behalf of the Fiocruz COVID-19 Genomic Surveillance Network |
| EPI_ISL_2196357 | Labortorio Central de Saude Publica do Estado de Santa Catarina (LACEN/SC) | Laboratory of Respiratory Viruses and Measles, Oswaldo Cruz Institute, FIOCRUZ | Paola Resende, Luciana Appolinario, Fernando Motta, Anna Carolina Paixao, Ana Carolina Mendonca, Alice Sampaio Rocha, Taina Venas, Elisa Cavalcante Pereira, Renata Serrano Lopes, Darcita Buerger Rovaris, Sandra Bianchini Fernandes, Marilda Siqueira on behalf of the Fiocruz COVID-19 Genomic Surveillance Network |
| EPI_ISL_2196358 | Laboratorio Central de Saude Publica do Estado Maranhao (LACEN-MA) | Laboratory of Respiratory Viruses and Measles, Oswaldo Cruz Institute, FIOCRUZ | Paola Resende, Luciana Appolinario, Fernando Motta, Anna Carolina Paixao, Ana Carolina Mendonca, Alice Sampaio Rocha, Taina Venas, Elisa Cavalcante Pereira, Renata Serrano Lopes, Lidio Gonçalves Lima Neto, Marilda Siqueira on behalf of the Fiocruz COVID-19 Genomic Surveillance Network |
| EPI_ISL_2196359, EPI_ISL_2196360 | Labortorio Central de Saude Publica do Estado de Santa Catarina (LACEN/SC) | Laboratory of Respiratory Viruses and Measles, Oswaldo Cruz Institute, FIOCRUZ | Paola Resende, Luciana Appolinario, Fernando Motta, Anna Carolina Paixao, Ana Carolina Mendonca, Alice Sampaio Rocha, Taina Venas, Elisa Cavalcante Pereira, Renata Serrano Lopes, Darcita Buerger Rovaris, Sandra Bianchini Fernandes, Marilda Siqueira on behalf of the Fiocruz COVID-19 Genomic Surveillance Network |
| EPI_ISL_2196361 | Laboratory of Respiratory Viruses and Measles, Oswaldo Cruz Institute, FIOCRUZ | Laboratory of Respiratory Viruses and Measles, Oswaldo Cruz Institute, FIOCRUZ | Paola Resende, Luciana Appolinario, Fernando Motta, Anna Carolina Paixao, Ana Carolina Mendonca, Alice Sampaio Rocha, Taina Venas, Elisa Cavalcante Pereira, Renata Serrano Lopes, Marilda Siqueira on behalf of the Fiocruz COVID-19 Genomic Surveillance Network |
| EPI_ISL_2196362 | Lboratorio Central de Saude Publica do Estado do Parana (LACEN/PR) | Laboratory of Respiratory Viruses and Measles, Oswaldo Cruz Institute, FIOCRUZ | Paola Resende, Luciana Appolinario, Fernando Motta, Anna Carolina Paixao, Ana Carolina Mendonca, Alice Sampaio Rocha, Taina Venas, Elisa Cavalcante Pereira, Renata Serrano Lopes, Irina Riediger, Marilda Siqueira on behalf of the Fiocruz COVID-19 Genomic Surveillance Network |
| EPI_ISL_2196363, EPI_ISL_2196364, EPI_ISL_2196365 | Laboratorio Central de Saude Publica do Estado de Minas Gerais (LACEN/MG) | Laboratory of Respiratory Viruses and Measles, Oswaldo Cruz Institute, FIOCRUZ | Paola Resende, Luciana Appolinario, Fernando Motta, Anna Carolina Paixao, Ana Carolina Mendonca, Alice Sampaio Rocha, Taina Venas, Elisa Cavalcante Pereira, Renata Serrano Lopes, Andre Felipe Leal Bernardes, Marilda Siqueira on behalf of the Fiocruz COVID-19 Genomic Surveillance Network |
| EPI_ISL_2208862 | SECRETARIA MUNICIPAL DA SAUDE DE JOANOPOLIS | Instituto Butantan / Mendelics | Dimas Tadeu Covas, Antonio Jorge Martins, Claudia Renata dos Santos Barros, David Schlesinger, Debora Botequio Moretti, Elaine Cristina Marqueze, Elaine Vieira Santos, Evandra Strazza Rodrigues, Heidge Fukumasu, Jayme Augusto de Souza-Neto, José Salvatore Leister Patané, Luiz Alcantara, Luiz Lehmann Coutinho, Maria Carolina Elias, Maurício Lacerda Nogueira, Rafael dos Santos Bezerra, Raul Machado Neto, Rejane Maria Tommasini Grotto, Ricardo Haddad, Sandra Coccuzzo Sampaio Vessoni, Simone Kashima, Svetoslav Nanev Slavov, Vincent Louis Viala |
| EPI_ISL_2208870 | HOSPITAL MUNICIPAL DE ITAPIRA | Instituto Butantan / Mendelics | Dimas Tadeu Covas, Antonio Jorge Martins, Claudia Renata dos Santos Barros, David Schlesinger, Debora Botequio Moretti, Elaine Cristina Marqueze, Elaine Vieira Santos, Evandra Strazza Rodrigues, Heidge Fukumasu, Jayme Augusto de Souza-Neto, José Salvatore Leister Patané, Luiz Alcantara, Luiz Lehmann Coutinho, Maria Carolina Elias, Maurício Lacerda Nogueira, Rafael dos Santos Bezerra, Raul Machado Neto, Rejane Maria Tommasini Grotto, Ricardo Haddad, Sandra Coccuzzo Sampaio Vessoni, Simone Kashima, Svetoslav Nanev Slavov, Vincent Louis Viala |
| EPI_ISL_2208874 | AMB MEDICO DR FRANCISCO FELIX DE MENDONCA SAUDADE | Instituto Butantan / ESALQ-Piracicaba | Dimas Tadeu Covas, Antonio Jorge Martins, Claudia Renata dos Santos Barros, David Schlesinger, Debora Botequio Moretti, Elaine Cristina Marqueze, Elaine Vieira Santos, Evandra Strazza Rodrigues, Heidge Fukumasu, Jayme Augusto de Souza-Neto, José Salvatore Leister Patané, Luiz Alcantara, Luiz Lehmann Coutinho, Maria Carolina Elias, Maurício Lacerda Nogueira, Rafael dos Santos Bezerra, Raul Machado Neto, Rejane Maria Tommasini Grotto, Ricardo Haddad, Sandra Coccuzzo Sampaio Vessoni, Simone Kashima, Svetoslav Nanev Slavov, Vincent Louis Viala |
| EPI_ISL_2208881 | CS DE ONDA VERDE | Instituto Butantan / ESALQ-Piracicaba | Dimas Tadeu Covas, Antonio Jorge Martins, Claudia Renata dos Santos Barros, David Schlesinger, Debora Botequio Moretti, Elaine Cristina Marqueze, Elaine Vieira Santos, Evandra Strazza Rodrigues, Heidge Fukumasu, Jayme Augusto de Souza-Neto, José Salvatore Leister Patané, Luiz Alcantara, Luiz Lehmann Coutinho, Maria Carolina Elias, Maurício Lacerda Nogueira, Rafael dos Santos Bezerra, Raul Machado Neto, Rejane Maria Tommasini Grotto, Ricardo Haddad, Sandra Coccuzzo Sampaio Vessoni, Simone Kashima, Svetoslav Nanev Slavov, Vincent Louis Viala |
| EPI_ISL_2208890 | HOSPITAL MUNICIPAL DE IBIUNA IBIUNA SP | Instituto Butantan / Mendelics | Dimas Tadeu Covas, Antonio Jorge Martins, Claudia Renata dos Santos Barros, David Schlesinger, Debora Botequio Moretti, Elaine Cristina Marqueze, Elaine Vieira Santos, Evandra Strazza Rodrigues, Heidge Fukumasu, Jayme Augusto de Souza-Neto, José Salvatore Leister Patané, Luiz Alcantara, Luiz Lehmann Coutinho, Maria Carolina Elias, Maurício Lacerda Nogueira, Rafael dos Santos Bezerra, Raul Machado Neto, Rejane Maria Tommasini Grotto, Ricardo Haddad, Sandra Coccuzzo Sampaio Vessoni, Simone Kashima, Svetoslav Nanev Slavov, Vincent Louis Viala |
| EPI_ISL_2208897 | SERV DE VIG SANITARIA EPIDEMIO E CTRL DE ZOONOZES GUARUJA | Instituto Butantan / Mendelics | Dimas Tadeu Covas, Antonio Jorge Martins, Claudia Renata dos Santos Barros, David Schlesinger, Debora Botequio Moretti, Elaine Cristina Marqueze, Elaine Vieira Santos, Evandra Strazza Rodrigues, Heidge Fukumasu, Jayme Augusto de Souza-Neto, José Salvatore Leister Patané, Luiz Alcantara, Luiz Lehmann Coutinho, Maria Carolina Elias, Maurício Lacerda Nogueira, Rafael dos Santos Bezerra, Raul Machado Neto, Rejane Maria Tommasini Grotto, Ricardo Haddad, Sandra Coccuzzo Sampaio Vessoni, Simone Kashima, Svetoslav Nanev Slavov, Vincent Louis Viala |

| | | | |
|---|---|---|---|
| EPI_ISL_2208904 | UPA DR FRANCO DA ROCHA | Instituto Butantan / Mendelics | Dimas Tadeu Covas, Antonio Jorge Martins, Claudia Renata dos Santos Barros, David Schlesinger, Debora Botequio Moretti, Elaine Cristina Marqueze, Elaine Vieira Santos, Evandra Strazza Rodrigues, Heidge Fukumasu, Jayme Augusto de Souza-Neto, José Salvatore Leister Patané, Luiz Alcantara, Luiz Lehmann Coutinho, Maria Carolina Elias, Maurício Lacerda Nogueira, Rafael dos Santos Bezerra, Raul Machado Neto, Rejane Maria Tommasini Grotto, Ricardo Haddad, Sandra Coccuzzo Sampaio Vessoni, Simone Kashima, Svetoslav Nanev Slavov, Vincent Louis Viala |
| EPI_ISL_2208911 | PENITENCIARIA DE PRESIDENTE PRUDENTE | Instituto Butantan / Mendelics | Dimas Tadeu Covas, Antonio Jorge Martins, Claudia Renata dos Santos Barros, David Schlesinger, Debora Botequio Moretti, Elaine Cristina Marqueze, Elaine Vieira Santos, Evandra Strazza Rodrigues, Heidge Fukumasu, Jayme Augusto de Souza-Neto, José Salvatore Leister Patané, Luiz Alcantara, Luiz Lehmann Coutinho, Maria Carolina Elias, Maurício Lacerda Nogueira, Rafael dos Santos Bezerra, Raul Machado Neto, Rejane Maria Tommasini Grotto, Ricardo Haddad, Sandra Coccuzzo Sampaio Vessoni, Simone Kashima, Svetoslav Nanev Slavov, Vincent Louis Viala |
| EPI_ISL_2208916 | UNIDADE SENTINELA COVID19 | Instituto Butantan / Mendelics | Dimas Tadeu Covas, Antonio Jorge Martins, Claudia Renata dos Santos Barros, David Schlesinger, Debora Botequio Moretti, Elaine Cristina Marqueze, Elaine Vieira Santos, Evandra Strazza Rodrigues, Heidge Fukumasu, Jayme Augusto de Souza-Neto, José Salvatore Leister Patané, Luiz Alcantara, Luiz Lehmann Coutinho, Maria Carolina Elias, Maurício Lacerda Nogueira, Rafael dos Santos Bezerra, Raul Machado Neto, Rejane Maria Tommasini Grotto, Ricardo Haddad, Sandra Coccuzzo Sampaio Vessoni, Simone Kashima, Svetoslav Nanev Slavov, Vincent Louis Viala |
| EPI_ISL_2208922 | UNIDADE RESPIRATORIA NOVA HORTOLANDIA | Instituto Butantan / ESALQ-Piracicaba | Dimas Tadeu Covas, Antonio Jorge Martins, Claudia Renata dos Santos Barros, David Schlesinger, Debora Botequio Moretti, Elaine Cristina Marqueze, Elaine Vieira Santos, Evandra Strazza Rodrigues, Heidge Fukumasu, Jayme Augusto de Souza-Neto, José Salvatore Leister Patané, Luiz Alcantara, Luiz Lehmann Coutinho, Maria Carolina Elias, Maurício Lacerda Nogueira, Rafael dos Santos Bezerra, Raul Machado Neto, Rejane Maria Tommasini Grotto, Ricardo Haddad, Sandra Coccuzzo Sampaio Vessoni, Simone Kashima, Svetoslav Nanev Slavov, Vincent Louis Viala |
| EPI_ISL_2208930 | UBS ALAMBARI | Instituto Butantan / Mendelics | Dimas Tadeu Covas, Antonio Jorge Martins, Claudia Renata dos Santos Barros, David Schlesinger, Debora Botequio Moretti, Elaine Cristina Marqueze, Elaine Vieira Santos, Evandra Strazza Rodrigues, Heidge Fukumasu, Jayme Augusto de Souza-Neto, José Salvatore Leister Patané, Luiz Alcantara, Luiz Lehmann Coutinho, Maria Carolina Elias, Maurício Lacerda Nogueira, Rafael dos Santos Bezerra, Raul Machado Neto, Rejane Maria Tommasini Grotto, Ricardo Haddad, Sandra Coccuzzo Sampaio Vessoni, Simone Kashima, Svetoslav Nanev Slavov, Vincent Louis Viala |
| EPI_ISL_2208938 | CENTRO DE SAUDE III SALES OLIVEIRA | Instituto Butantan / Mendelics | Dimas Tadeu Covas, Antonio Jorge Martins, Claudia Renata dos Santos Barros, David Schlesinger, Debora Botequio Moretti, Elaine Cristina Marqueze, Elaine Vieira Santos, Evandra Strazza Rodrigues, Heidge Fukumasu, Jayme Augusto de Souza-Neto, José Salvatore Leister Patané, Luiz Alcantara, Luiz Lehmann Coutinho, Maria Carolina Elias, Maurício Lacerda Nogueira, Rafael dos Santos Bezerra, Raul Machado Neto, Rejane Maria Tommasini Grotto, Ricardo Haddad, Sandra Coccuzzo Sampaio Vessoni, Simone Kashima, Svetoslav Nanev Slavov, Vincent Louis Viala |
| EPI_ISL_2208944 | UPA DE BEBEDOURO | Instituto Butantan / Mendelics | Dimas Tadeu Covas, Antonio Jorge Martins, Claudia Renata dos Santos Barros, David Schlesinger, Debora Botequio Moretti, Elaine Cristina Marqueze, Elaine Vieira Santos, Evandra Strazza Rodrigues, Heidge Fukumasu, Jayme Augusto de Souza-Neto, José Salvatore Leister Patané, Luiz Alcantara, Luiz Lehmann Coutinho, Maria Carolina Elias, Maurício Lacerda Nogueira, Rafael dos Santos Bezerra, Raul Machado Neto, Rejane Maria Tommasini Grotto, Ricardo Haddad, Sandra Coccuzzo Sampaio Vessoni, Simone Kashima, Svetoslav Nanev Slavov, Vincent Louis Viala |
| EPI_ISL_2208950 | CENTRO DE SAUDE DR RENATO DE CARVALHO RIBEIRO ANGATUBA | Instituto Butantan | Dimas Tadeu Covas, Antonio Jorge Martins, Claudia Renata dos Santos Barros, David Schlesinger, Debora Botequio Moretti, Elaine Cristina Marqueze, Elaine Vieira Santos, Evandra Strazza Rodrigues, Heidge Fukumasu, Jayme Augusto de Souza-Neto, José Salvatore Leister Patané, Luiz Alcantara, Luiz Lehmann Coutinho, Maria Carolina Elias, Maurício Lacerda Nogueira, Rafael dos Santos Bezerra, Raul Machado Neto, Rejane Maria Tommasini Grotto, Ricardo Haddad, Sandra Coccuzzo Sampaio Vessoni, Simone Kashima, Svetoslav Nanev Slavov, Vincent Louis Viala. |
| EPI_ISL_2208958, EPI_ISL_2208968 | VIGILANCIA EPIDEMIOLOGICA JARDINOPOLIS SP | Instituto Butantan / Mendelics | Dimas Tadeu Covas, Antonio Jorge Martins, Claudia Renata dos Santos Barros, David Schlesinger, Debora Botequio Moretti, Elaine Cristina Marqueze, Elaine Vieira Santos, Evandra Strazza Rodrigues, Heidge Fukumasu, Jayme Augusto de Souza-Neto, José Salvatore Leister Patané, Luiz Alcantara, Luiz Lehmann Coutinho, Maria Carolina Elias, Maurício Lacerda Nogueira, Rafael dos Santos Bezerra, Raul Machado Neto, Rejane Maria Tommasini Grotto, Ricardo Haddad, Sandra Coccuzzo Sampaio Vessoni, Simone Kashima, Svetoslav Nanev Slavov, Vincent Louis Viala |
| EPI_ISL_2208974 | CENTRO DE SAUDE II IBITINGA | Instituto Butantan / Mendelics | Dimas Tadeu Covas, Antonio Jorge Martins, Claudia Renata dos Santos Barros, David Schlesinger, Debora Botequio Moretti, Elaine Cristina Marqueze, Elaine Vieira Santos, Evandra Strazza Rodrigues, Heidge Fukumasu, Jayme Augusto de Souza-Neto, José Salvatore Leister Patané, Luiz Alcantara, Luiz Lehmann Coutinho, Maria Carolina Elias, Maurício Lacerda Nogueira, Rafael dos Santos Bezerra, Raul Machado Neto, Rejane Maria Tommasini Grotto, Ricardo Haddad, Sandra Coccuzzo Sampaio Vessoni, Simone Kashima, Svetoslav Nanev Slavov, Vincent Louis Viala |
| EPI_ISL_2208981 | VIGILANCIA EPIDEMIOLOGICA JARDINOPOLIS SP | Instituto Butantan / Mendelics | Dimas Tadeu Covas, Antonio Jorge Martins, Claudia Renata dos Santos Barros, David Schlesinger, Debora Botequio Moretti, Elaine Cristina Marqueze, Elaine Vieira Santos, Evandra Strazza Rodrigues, Heidge Fukumasu, Jayme Augusto de Souza-Neto, José Salvatore Leister Patané, Luiz Alcantara, Luiz Lehmann Coutinho, Maria Carolina Elias, Maurício Lacerda Nogueira, Rafael dos Santos Bezerra, Raul Machado Neto, Rejane Maria Tommasini Grotto, Ricardo Haddad, Sandra Coccuzzo Sampaio Vessoni, Simone Kashima, Svetoslav Nanev Slavov, Vincent Louis Viala |
| EPI_ISL_2208992 | CSII DR WASHINGTON LUIS M RODRIGUES DA SILVA PITANGUEIRAS | Instituto Butantan / Mendelics | Dimas Tadeu Covas, Antonio Jorge Martins, Claudia Renata dos Santos Barros, David Schlesinger, Debora Botequio Moretti, Elaine Cristina Marqueze, Elaine Vieira Santos, Evandra Strazza Rodrigues, Heidge Fukumasu, Jayme Augusto de Souza-Neto, José Salvatore Leister Patané, Luiz Alcantara, Luiz Lehmann Coutinho, Maria Carolina Elias, Maurício Lacerda Nogueira, Rafael dos Santos Bezerra, Raul Machado Neto, Rejane Maria Tommasini Grotto, Ricardo Haddad, Sandra Coccuzzo Sampaio Vessoni, Simone Kashima, Svetoslav Nanev Slavov, Vincent Louis Viala |
| EPI_ISL_2208998 | CENTRO DE SAUDE SAO ROQUE DR JOSE CARVALHO BRITO | Instituto Butantan | Dimas Tadeu Covas, Antonio Jorge Martins, Claudia Renata dos Santos Barros, David Schlesinger, Debora Botequio Moretti, Elaine Cristina Marqueze, Elaine Vieira Santos, Evandra Strazza Rodrigues, Heidge Fukumasu, Jayme Augusto de Souza-Neto, José Salvatore Leister Patané, Luiz Alcantara, Luiz Lehmann Coutinho, Maria Carolina Elias, Maurício Lacerda Nogueira, Rafael dos Santos Bezerra, Raul Machado Neto, Rejane Maria Tommasini Grotto, Ricardo Haddad, Sandra Coccuzzo Sampaio Vessoni, Simone Kashima, Svetoslav Nanev Slavov, Vincent Louis Viala. |
| EPI_ISL_2209004 | HOSPITAL E PRONTO SOCORRO PORTINARI | Instituto Butantan | Dimas Tadeu Covas, Antonio Jorge Martins, Claudia Renata dos Santos Barros, David Schlesinger, Debora Botequio Moretti, Elaine Cristina Marqueze, Elaine Vieira Santos, Evandra Strazza Rodrigues, Heidge Fukumasu, Jayme Augusto de Souza-Neto, José Salvatore Leister Patané, Luiz Alcantara, Luiz Lehmann Coutinho, Maria Carolina Elias, Maurício Lacerda Nogueira, Rafael dos Santos Bezerra, Raul Machado Neto, Rejane Maria Tommasini Grotto, Ricardo Haddad, Sandra Coccuzzo Sampaio Vessoni, Simone Kashima, Svetoslav Nanev Slavov, Vincent Louis Viala |
| EPI_ISL_2209018 | UNIDADE DE PRONTO ATENDIMENTO UPA | Instituto Butantan / ESALQ-Piracicaba | Dimas Tadeu Covas, Antonio Jorge Martins, Claudia Renata dos Santos Barros, David Schlesinger, Debora Botequio Moretti, Elaine Cristina Marqueze, Elaine Vieira Santos, Evandra Strazza Rodrigues, Heidge Fukumasu, Jayme Augusto de Souza-Neto, José Salvatore Leister Patané, Luiz Alcantara, Luiz Lehmann Coutinho, Maria Carolina Elias, Maurício Lacerda Nogueira, Rafael dos Santos Bezerra, Raul Machado Neto, Rejane Maria Tommasini Grotto, Ricardo Haddad, Sandra Coccuzzo Sampaio Vessoni, Simone Kashima, Svetoslav Nanev Slavov, Vincent Louis Viala |
| EPI_ISL_2209024 | HOSP MUN CIDADE TIRADENTES CARMEN PRUDENTE | Instituto Butantan / ESALQ-Piracicaba | Dimas Tadeu Covas, Antonio Jorge Martins, Claudia Renata dos Santos Barros, David Schlesinger, Debora Botequio Moretti, Elaine Cristina Marqueze, Elaine Vieira Santos, Evandra Strazza Rodrigues, Heidge Fukumasu, Jayme Augusto de Souza-Neto, José Salvatore Leister Patané, Luiz Alcantara, Luiz Lehmann Coutinho, Maria Carolina Elias, Maurício Lacerda Nogueira, Rafael dos Santos Bezerra, Raul Machado Neto, Rejane Maria Tommasini Grotto, Ricardo Haddad, Sandra Coccuzzo Sampaio Vessoni, Simone Kashima, Svetoslav Nanev Slavov, Vincent Louis Viala |
| EPI_ISL_2209032 | CENTRO DE SAUDE TEREZINHA DE MORAES ARSILLA | Instituto Butantan / ESALQ-Piracicaba | Dimas Tadeu Covas, Antonio Jorge Martins, Claudia Renata dos Santos Barros, David Schlesinger, Debora Botequio Moretti, Elaine Cristina Marqueze, Elaine Vieira Santos, Evandra Strazza Rodrigues, Heidge Fukumasu, Jayme Augusto de Souza-Neto, José Salvatore Leister Patané, Luiz Alcantara, Luiz Lehmann Coutinho, Maria Carolina Elias, Maurício Lacerda Nogueira, Rafael dos Santos Bezerra, Raul Machado Neto, Rejane Maria Tommasini Grotto, Ricardo Haddad, Sandra Coccuzzo Sampaio Vessoni, Simone Kashima, Svetoslav Nanev Slavov, Vincent Louis Viala |
| EPI_ISL_2209040 | UPA DR AKIRA TADA | Instituto Butantan / Mendelics | Dimas Tadeu Covas, Antonio Jorge Martins, Claudia Renata dos Santos Barros, David Schlesinger, Debora Botequio Moretti, Elaine Cristina Marqueze, Elaine Vieira Santos, Evandra Strazza Rodrigues, Heidge Fukumasu, Jayme Augusto de Souza-Neto, José Salvatore Leister Patané, Luiz Alcantara, Luiz Lehmann Coutinho, Maria Carolina Elias, Maurício Lacerda Nogueira, Rafael dos Santos Bezerra, Raul Machado Neto, Rejane Maria Tommasini Grotto, Ricardo Haddad, Sandra Coccuzzo Sampaio Vessoni, Simone Kashima, Svetoslav Nanev Slavov, Vincent Louis Viala |
| EPI_ISL_2209045 | DEPARTAMENTO DE SAUDE MUNICIPAL SOCORRO SP | Instituto Butantan / Mendelics | Dimas Tadeu Covas, Antonio Jorge Martins, Claudia Renata dos Santos Barros, David Schlesinger, Debora Botequio Moretti, Elaine Cristina Marqueze, Elaine Vieira Santos, Evandra Strazza Rodrigues, Heidge Fukumasu, Jayme Augusto de Souza-Neto, José Salvatore Leister Patané, Luiz Alcantara, Luiz Lehmann Coutinho, Maria Carolina Elias, Maurício Lacerda Nogueira, Rafael dos Santos Bezerra, Raul Machado Neto, Rejane Maria Tommasini Grotto, Ricardo Haddad, Sandra Coccuzzo Sampaio Vessoni, Simone Kashima, Svetoslav Nanev Slavov, Vincent Louis Viala |
| EPI_ISL_2209052 | NUCLEO DE SAUDE MARY DOTA DE BAURU | Instituto Butantan / Mendelics | Dimas Tadeu Covas, Antonio Jorge Martins, Claudia Renata dos Santos Barros, David Schlesinger, Debora Botequio Moretti, Elaine Cristina Marqueze, Elaine Vieira Santos, Evandra Strazza Rodrigues, Heidge Fukumasu, Jayme Augusto de Souza-Neto, José Salvatore Leister Patané, Luiz Alcantara, Luiz |

| | | | |
|---|---|---|---|
| | | | Lehmann Coutinho, Maria Carolina Elias, Maurício Lacerda Nogueira, Rafael dos Santos Bezerra, Raul Machado Neto, Rejane Maria Tommasini Grotto, Ricardo Haddad, Sandra Coccuzzo Sampaio Vessoni, Simone Kashima, Svetoslav Nanev Slavov, Vincent Louis Viala |
| EPI_ISL_2209068 | SECRETARIA MUNICIPAL DE SAUDE SOROCABA | Instituto Butantan / Mendelics | Dimas Tadeu Covas, Antonio Jorge Martins, Claudia Renata dos Santos Barros, David Schlesinger, Debora Botequio Moretti, Elaine Cristina Marqueze, Elaine Vieira Santos, Evandra Strazza Rodrigues, Heidge Fukumasu, Jayme Augusto de Souza-Neto, José Salvatore Leister Patané, Luiz Alcantara, Luiz Lehmann Coutinho, Maria Carolina Elias, Maurício Lacerda Nogueira, Rafael dos Santos Bezerra, Raul Machado Neto, Rejane Maria Tommasini Grotto, Ricardo Haddad, Sandra Coccuzzo Sampaio Vessoni, Simone Kashima, Svetoslav Nanev Slavov, Vincent Louis Viala |
| EPI_ISL_2209074 | CENTRO MUNICIPAL DE EPIDEMIOLOGIA E IMUNIZACOES LIMEIRA | Instituto Butantan | Dimas Tadeu Covas, Antonio Jorge Martins, Claudia Renata dos Santos Barros, David Schlesinger, Debora Botequio Moretti, Elaine Cristina Marqueze, Elaine Vieira Santos, Evandra Strazza Rodrigues, Heidge Fukumasu, Jayme Augusto de Souza-Neto, José Salvatore Leister Patané, Luiz Alcantara, Luiz Lehmann Coutinho, Maria Carolina Elias, Maurício Lacerda Nogueira, Rafael dos Santos Bezerra, Raul Machado Neto, Rejane Maria Tommasini Grotto, Ricardo Haddad, Sandra Coccuzzo Sampaio Vessoni, Simone Kashima, Svetoslav Nanev Slavov, Vincent Louis Viala. |
| EPI_ISL_2209080 | COORDENADORIA MUNICIPAL DE SAUDE DE IRACEMAPOLIS | Instituto Butantan | Dimas Tadeu Covas, Antonio Jorge Martins, Claudia Renata dos Santos Barros, David Schlesinger, Debora Botequio Moretti, Elaine Cristina Marqueze, Elaine Vieira Santos, Evandra Strazza Rodrigues, Heidge Fukumasu, Jayme Augusto de Souza-Neto, José Salvatore Leister Patané, Luiz Alcantara, Luiz Lehmann Coutinho, Maria Carolina Elias, Maurício Lacerda Nogueira, Rafael dos Santos Bezerra, Raul Machado Neto, Rejane Maria Tommasini Grotto, Ricardo Haddad, Sandra Coccuzzo Sampaio Vessoni, Simone Kashima, Svetoslav Nanev Slavov, Vincent Louis Viala. |
| EPI_ISL_2209086 | CENTRO DE APOIO EPIDEMIOLOGICO | Instituto Butantan / Mendelics | Dimas Tadeu Covas, Antonio Jorge Martins, Claudia Renata dos Santos Barros, David Schlesinger, Debora Botequio Moretti, Elaine Cristina Marqueze, Elaine Vieira Santos, Evandra Strazza Rodrigues, Heidge Fukumasu, Jayme Augusto de Souza-Neto, José Salvatore Leister Patané, Luiz Alcantara, Luiz Lehmann Coutinho, Maria Carolina Elias, Maurício Lacerda Nogueira, Rafael dos Santos Bezerra, Raul Machado Neto, Rejane Maria Tommasini Grotto, Ricardo Haddad, Sandra Coccuzzo Sampaio Vessoni, Simone Kashima, Svetoslav Nanev Slavov, Vincent Louis Viala |
| EPI_ISL_2209094 | VIGILANCIA EPIDEMIOLOGICA JARDINOPOLIS SP | Instituto Butantan / Mendelics | Dimas Tadeu Covas, Antonio Jorge Martins, Claudia Renata dos Santos Barros, David Schlesinger, Debora Botequio Moretti, Elaine Cristina Marqueze, Elaine Vieira Santos, Evandra Strazza Rodrigues, Heidge Fukumasu, Jayme Augusto de Souza-Neto, José Salvatore Leister Patané, Luiz Alcantara, Luiz Lehmann Coutinho, Maria Carolina Elias, Maurício Lacerda Nogueira, Rafael dos Santos Bezerra, Raul Machado Neto, Rejane Maria Tommasini Grotto, Ricardo Haddad, Sandra Coccuzzo Sampaio Vessoni, Simone Kashima, Svetoslav Nanev Slavov, Vincent Louis Viala |
| EPI_ISL_2209101 | HOSPITAL DE CAMPANHA COVID 19 MUNICIPIO DE TAUBATE | Instituto Butantan / Mendelics | Dimas Tadeu Covas, Antonio Jorge Martins, Claudia Renata dos Santos Barros, David Schlesinger, Debora Botequio Moretti, Elaine Cristina Marqueze, Elaine Vieira Santos, Evandra Strazza Rodrigues, Heidge Fukumasu, Jayme Augusto de Souza-Neto, José Salvatore Leister Patané, Luiz Alcantara, Luiz Lehmann Coutinho, Maria Carolina Elias, Maurício Lacerda Nogueira, Rafael dos Santos Bezerra, Raul Machado Neto, Rejane Maria Tommasini Grotto, Ricardo Haddad, Sandra Coccuzzo Sampaio Vessoni, Simone Kashima, Svetoslav Nanev Slavov, Vincent Louis Viala |
| EPI_ISL_2209108 | SANTA CASA DE MISERICORDIA SAO JOSE | Instituto Butantan / ESALQ-Piracicaba | Dimas Tadeu Covas, Antonio Jorge Martins, Claudia Renata dos Santos Barros, David Schlesinger, Debora Botequio Moretti, Elaine Cristina Marqueze, Elaine Vieira Santos, Evandra Strazza Rodrigues, Heidge Fukumasu, Jayme Augusto de Souza-Neto, José Salvatore Leister Patané, Luiz Alcantara, Luiz Lehmann Coutinho, Maria Carolina Elias, Maurício Lacerda Nogueira, Rafael dos Santos Bezerra, Raul Machado Neto, Rejane Maria Tommasini Grotto, Ricardo Haddad, Sandra Coccuzzo Sampaio Vessoni, Simone Kashima, Svetoslav Nanev Slavov, Vincent Louis Viala |
| EPI_ISL_2209122 | LABORATORIO MUNICIPAL DE ANALISES CLINICAS DE RIO CLARO | Instituto Butantan / FZEA-USP-Pirassununga | Dimas Tadeu Covas, Antonio Jorge Martins, Claudia Renata dos Santos Barros, David Schlesinger, Debora Botequio Moretti, Elaine Cristina Marqueze, Elaine Vieira Santos, Evandra Strazza Rodrigues, Heidge Fukumasu, Jayme Augusto de Souza-Neto, José Salvatore Leister Patané, Luiz Alcantara, Luiz Lehmann Coutinho, Maria Carolina Elias, Maurício Lacerda Nogueira, Rafael dos Santos Bezerra, Raul Machado Neto, Rejane Maria Tommasini Grotto, Ricardo Haddad, Sandra Coccuzzo Sampaio Vessoni, Simone Kashima, Svetoslav Nanev Slavov, Vincent Louis Viala |
| EPI_ISL_2209129 | UNIDADE DE VIGILANCIA EPIDEMIOLOGICA | Instituto Butantan / FZEA-USP-Pirassununga | Dimas Tadeu Covas, Antonio Jorge Martins, Claudia Renata dos Santos Barros, David Schlesinger, Debora Botequio Moretti, Elaine Cristina Marqueze, Elaine Vieira Santos, Evandra Strazza Rodrigues, Heidge Fukumasu, Jayme Augusto de Souza-Neto, José Salvatore Leister Patané, Luiz Alcantara, Luiz Lehmann Coutinho, Maria Carolina Elias, Maurício Lacerda Nogueira, Rafael dos Santos Bezerra, Raul Machado Neto, Rejane Maria Tommasini Grotto, Ricardo Haddad, Sandra Coccuzzo Sampaio Vessoni, Simone Kashima, Svetoslav Nanev Slavov, Vincent Louis Viala |
| EPI_ISL_2209137, EPI_ISL_2209145 | SECRETARIA MUNICIPAL DE SAUDE DE CORDEIROPOLIS | Instituto Butantan / FZEA-USP-Pirassununga | Dimas Tadeu Covas, Antonio Jorge Martins, Claudia Renata dos Santos Barros, David Schlesinger, Debora Botequio Moretti, Elaine Cristina Marqueze, Elaine Vieira Santos, Evandra Strazza Rodrigues, Heidge Fukumasu, Jayme Augusto de Souza-Neto, José Salvatore Leister Patané, Luiz Alcantara, Luiz Lehmann Coutinho, Maria Carolina Elias, Maurício Lacerda Nogueira, Rafael dos Santos Bezerra, Raul Machado Neto, Rejane Maria Tommasini Grotto, Ricardo Haddad, Sandra Coccuzzo Sampaio Vessoni, Simone Kashima, Svetoslav Nanev Slavov, Vincent Louis Viala |
| EPI_ISL_2209152 | DEPARTAMENTO DE SAUDE COLETIVA | Instituto Butantan | Dimas Tadeu Covas, Antonio Jorge Martins, Claudia Renata dos Santos Barros, David Schlesinger, Debora Botequio Moretti, Elaine Cristina Marqueze, Elaine Vieira Santos, Evandra Strazza Rodrigues, Heidge Fukumasu, Jayme Augusto de Souza-Neto, José Salvatore Leister Patané, Luiz Alcantara, Luiz Lehmann Coutinho, Maria Carolina Elias, Maurício Lacerda Nogueira, Rafael dos Santos Bezerra, Raul Machado Neto, Rejane Maria Tommasini Grotto, Ricardo Haddad, Sandra Coccuzzo Sampaio Vessoni, Simone Kashima, Svetoslav Nanev Slavov, Vincent Louis Viala. |
| EPI_ISL_2209158 | LABORATORIO LOCAL DE ITAPECERICA DA SERRA | Instituto Butantan / Mendelics | Dimas Tadeu Covas, Antonio Jorge Martins, Claudia Renata dos Santos Barros, David Schlesinger, Debora Botequio Moretti, Elaine Cristina Marqueze, Elaine Vieira Santos, Evandra Strazza Rodrigues, Heidge Fukumasu, Jayme Augusto de Souza-Neto, José Salvatore Leister Patané, Luiz Alcantara, Luiz Lehmann Coutinho, Maria Carolina Elias, Maurício Lacerda Nogueira, Rafael dos Santos Bezerra, Raul Machado Neto, Rejane Maria Tommasini Grotto, Ricardo Haddad, Sandra Coccuzzo Sampaio Vessoni, Simone Kashima, Svetoslav Nanev Slavov, Vincent Louis Viala |
| EPI_ISL_2209164 | CS DE PALMARES PAULISTA | Instituto Butantan / ESALQ-Piracicaba | Dimas Tadeu Covas, Antonio Jorge Martins, Claudia Renata dos Santos Barros, David Schlesinger, Debora Botequio Moretti, Elaine Cristina Marqueze, Elaine Vieira Santos, Evandra Strazza Rodrigues, Heidge Fukumasu, Jayme Augusto de Souza-Neto, José Salvatore Leister Patané, Luiz Alcantara, Luiz Lehmann Coutinho, Maria Carolina Elias, Maurício Lacerda Nogueira, Rafael dos Santos Bezerra, Raul Machado Neto, Rejane Maria Tommasini Grotto, Ricardo Haddad, Sandra Coccuzzo Sampaio Vessoni, Simone Kashima, Svetoslav Nanev Slavov, Vincent Louis Viala |
| EPI_ISL_2209177 | CS DE COSMORAMA | Instituto Butantan / ESALQ-Piracicaba | Dimas Tadeu Covas, Antonio Jorge Martins, Claudia Renata dos Santos Barros, David Schlesinger, Debora Botequio Moretti, Elaine Cristina Marqueze, Elaine Vieira Santos, Evandra Strazza Rodrigues, Heidge Fukumasu, Jayme Augusto de Souza-Neto, José Salvatore Leister Patané, Luiz Alcantara, Luiz Lehmann Coutinho, Maria Carolina Elias, Maurício Lacerda Nogueira, Rafael dos Santos Bezerra, Raul Machado Neto, Rejane Maria Tommasini Grotto, Ricardo Haddad, Sandra Coccuzzo Sampaio Vessoni, Simone Kashima, Svetoslav Nanev Slavov, Vincent Louis Viala |
| EPI_ISL_2209183 | UNIDADE BASICA DE SAUDE DR MATHEUS GABRIEL BONASSA | Instituto Butantan / FZEA-USP-Pirassununga | Dimas Tadeu Covas, Antonio Jorge Martins, Claudia Renata dos Santos Barros, David Schlesinger, Debora Botequio Moretti, Elaine Cristina Marqueze, Elaine Vieira Santos, Evandra Strazza Rodrigues, Heidge Fukumasu, Jayme Augusto de Souza-Neto, José Salvatore Leister Patané, Luiz Alcantara, Luiz Lehmann Coutinho, Maria Carolina Elias, Maurício Lacerda Nogueira, Rafael dos Santos Bezerra, Raul Machado Neto, Rejane Maria Tommasini Grotto, Ricardo Haddad, Sandra Coccuzzo Sampaio Vessoni, Simone Kashima, Svetoslav Nanev Slavov, Vincent Louis Viala |
| EPI_ISL_2209189 | CENTRO DE SAUDE II DR GABRIEL MESQUITA VARGEM GDE DO SUL | Instituto Butantan / Mendelics | Dimas Tadeu Covas, Antonio Jorge Martins, Claudia Renata dos Santos Barros, David Schlesinger, Debora Botequio Moretti, Elaine Cristina Marqueze, Elaine Vieira Santos, Evandra Strazza Rodrigues, Heidge Fukumasu, Jayme Augusto de Souza-Neto, José Salvatore Leister Patané, Luiz Alcantara, Luiz Lehmann Coutinho, Maria Carolina Elias, Maurício Lacerda Nogueira, Rafael dos Santos Bezerra, Raul Machado Neto, Rejane Maria Tommasini Grotto, Ricardo Haddad, Sandra Coccuzzo Sampaio Vessoni, Simone Kashima, Svetoslav Nanev Slavov, Vincent Louis Viala |
| EPI_ISL_2209197 | SECRETARIA MUNICIPAL DE SAUDE SOROCABA | Instituto Butantan / Mendelics | Dimas Tadeu Covas, Antonio Jorge Martins, Claudia Renata dos Santos Barros, David Schlesinger, Debora Botequio Moretti, Elaine Cristina Marqueze, Elaine Vieira Santos, Evandra Strazza Rodrigues, Heidge Fukumasu, Jayme Augusto de Souza-Neto, José Salvatore Leister Patané, Luiz Alcantara, Luiz Lehmann Coutinho, Maria Carolina Elias, Maurício Lacerda Nogueira, Rafael dos Santos Bezerra, Raul Machado Neto, Rejane Maria Tommasini Grotto, Ricardo Haddad, Sandra Coccuzzo Sampaio Vessoni, Simone Kashima, Svetoslav Nanev Slavov, Vincent Louis Viala |
| EPI_ISL_2209204 | HOSP E MATERNIDADE MUNICIPAL N SRA MONTE SERRAT | Instituto Butantan | Dimas Tadeu Covas, Antonio Jorge Martins, Claudia Renata dos Santos Barros, David Schlesinger, Debora Botequio Moretti, Elaine Cristina Marqueze, Elaine Vieira Santos, Evandra Strazza Rodrigues, Heidge Fukumasu, Jayme Augusto de Souza-Neto, José Salvatore Leister Patané, Luiz Alcantara, Luiz Lehmann Coutinho, Maria Carolina Elias, Maurício Lacerda Nogueira, Rafael dos Santos Bezerra, Raul Machado Neto, Rejane Maria Tommasini Grotto, Ricardo Haddad, Sandra Coccuzzo Sampaio Vessoni, Simone Kashima, Svetoslav Nanev Slavov, Vincent Louis Viala. |
| EPI_ISL_2209212 | SECRETARIA DE SAUDE DE SAO PEDRO | Instituto Butantan / FZEA-USP-Pirassununga | Dimas Tadeu Covas, Antonio Jorge Martins, Claudia Renata dos Santos Barros, David Schlesinger, Debora Botequio Moretti, Elaine Cristina Marqueze, Elaine Vieira Santos, Evandra Strazza Rodrigues, Heidge Fukumasu, Jayme Augusto de Souza-Neto, José Salvatore Leister Patané, Luiz Alcantara, Luiz Lehmann Coutinho, Maria Carolina Elias, Maurício Lacerda Nogueira, Rafael dos Santos Bezerra, Raul Machado Neto, Rejane Maria Tommasini Grotto, Ricardo Haddad, Sandra Coccuzzo Sampaio Vessoni, Simone Kashima, Svetoslav Nanev Slavov, Vincent Louis Viala |

| | | | |
|---|---|---|---|
| EPI_ISL_2209229, EPI_ISL_2209237, EPI_ISL_2209240 | UNIDADE DE VIGILANCIA EPIDEMIOLOGICA | Instituto Butantan / FZEA-USP-Pirassununga | Dimas Tadeu Covas, Antonio Jorge Martins, Claudia Renata dos Santos Barros, David Schlesinger, Debora Botequio Moretti, Elaine Cristina Marqueze, Elaine Vieira Santos, Evandra Strazza Rodrigues, Heidge Fukumasu, Jayme Augusto de Souza-Neto, José Salvatore Leister Patané, Luiz Alcantara, Luiz Lehmann Coutinho, Maria Carolina Elias, Maurício Lacerda Nogueira, Rafael dos Santos Bezerra, Raul Machado Neto, Rejane Maria Tommasini Grotto, Ricardo Haddad, Sandra Coccuzzo Sampaio Vessoni, Simone Kashima, Svetoslav Nanev Slavov, Vincent Louis Viala |
| EPI_ISL_2209246, EPI_ISL_2209248 | UNIDADE MISTA OSVALDO P CAMARGO ELIAS FAUSTO | Instituto Butantan / FZEA-USP-Pirassununga | Dimas Tadeu Covas, Antonio Jorge Martins, Claudia Renata dos Santos Barros, David Schlesinger, Debora Botequio Moretti, Elaine Cristina Marqueze, Elaine Vieira Santos, Evandra Strazza Rodrigues, Heidge Fukumasu, Jayme Augusto de Souza-Neto, José Salvatore Leister Patané, Luiz Alcantara, Luiz Lehmann Coutinho, Maria Carolina Elias, Maurício Lacerda Nogueira, Rafael dos Santos Bezerra, Raul Machado Neto, Rejane Maria Tommasini Grotto, Ricardo Haddad, Sandra Coccuzzo Sampaio Vessoni, Simone Kashima, Svetoslav Nanev Slavov, Vincent Louis Viala |
| EPI_ISL_2209252 | LABORATORIO MUNICIPAL DE ANALISES CLINICAS DE RIO CLARO | Instituto Butantan / FZEA-USP-Pirassununga | Dimas Tadeu Covas, Antonio Jorge Martins, Claudia Renata dos Santos Barros, David Schlesinger, Debora Botequio Moretti, Elaine Cristina Marqueze, Elaine Vieira Santos, Evandra Strazza Rodrigues, Heidge Fukumasu, Jayme Augusto de Souza-Neto, José Salvatore Leister Patané, Luiz Alcantara, Luiz Lehmann Coutinho, Maria Carolina Elias, Maurício Lacerda Nogueira, Rafael dos Santos Bezerra, Raul Machado Neto, Rejane Maria Tommasini Grotto, Ricardo Haddad, Sandra Coccuzzo Sampaio Vessoni, Simone Kashima, Svetoslav Nanev Slavov, Vincent Louis Viala |
| EPI_ISL_2209255 | SECRETARIA DE SAUDE DE SANTA GERTRUDES | Instituto Butantan / FZEA-USP-Pirassununga | Dimas Tadeu Covas, Antonio Jorge Martins, Claudia Renata dos Santos Barros, David Schlesinger, Debora Botequio Moretti, Elaine Cristina Marqueze, Elaine Vieira Santos, Evandra Strazza Rodrigues, Heidge Fukumasu, Jayme Augusto de Souza-Neto, José Salvatore Leister Patané, Luiz Alcantara, Luiz Lehmann Coutinho, Maria Carolina Elias, Maurício Lacerda Nogueira, Rafael dos Santos Bezerra, Raul Machado Neto, Rejane Maria Tommasini Grotto, Ricardo Haddad, Sandra Coccuzzo Sampaio Vessoni, Simone Kashima, Svetoslav Nanev Slavov, Vincent Louis Viala |
| EPI_ISL_2209259 | SECRETARIA DE SAUDE DE SAO PEDRO | Instituto Butantan / FZEA-USP-Pirassununga | Dimas Tadeu Covas, Antonio Jorge Martins, Claudia Renata dos Santos Barros, David Schlesinger, Debora Botequio Moretti, Elaine Cristina Marqueze, Elaine Vieira Santos, Evandra Strazza Rodrigues, Heidge Fukumasu, Jayme Augusto de Souza-Neto, José Salvatore Leister Patané, Luiz Alcantara, Luiz Lehmann Coutinho, Maria Carolina Elias, Maurício Lacerda Nogueira, Rafael dos Santos Bezerra, Raul Machado Neto, Rejane Maria Tommasini Grotto, Ricardo Haddad, Sandra Coccuzzo Sampaio Vessoni, Simone Kashima, Svetoslav Nanev Slavov, Vincent Louis Viala |
| EPI_ISL_2209261 | UNIDADE DE VIGILANCIA EPIDEMIOLOGICA | Instituto Butantan / FZEA-USP-Pirassununga | Dimas Tadeu Covas, Antonio Jorge Martins, Claudia Renata dos Santos Barros, David Schlesinger, Debora Botequio Moretti, Elaine Cristina Marqueze, Elaine Vieira Santos, Evandra Strazza Rodrigues, Heidge Fukumasu, Jayme Augusto de Souza-Neto, José Salvatore Leister Patané, Luiz Alcantara, Luiz Lehmann Coutinho, Maria Carolina Elias, Maurício Lacerda Nogueira, Rafael dos Santos Bezerra, Raul Machado Neto, Rejane Maria Tommasini Grotto, Ricardo Haddad, Sandra Coccuzzo Sampaio Vessoni, Simone Kashima, Svetoslav Nanev Slavov, Vincent Louis Viala |
| EPI_ISL_2209265 | CENTRO DE SAUDE DR MARIO DIAS DE AGUIAR CAPIVARI | Instituto Butantan / FZEA-USP-Pirassununga | Dimas Tadeu Covas, Antonio Jorge Martins, Claudia Renata dos Santos Barros, David Schlesinger, Debora Botequio Moretti, Elaine Cristina Marqueze, Elaine Vieira Santos, Evandra Strazza Rodrigues, Heidge Fukumasu, Jayme Augusto de Souza-Neto, José Salvatore Leister Patané, Luiz Alcantara, Luiz Lehmann Coutinho, Maria Carolina Elias, Maurício Lacerda Nogueira, Rafael dos Santos Bezerra, Raul Machado Neto, Rejane Maria Tommasini Grotto, Ricardo Haddad, Sandra Coccuzzo Sampaio Vessoni, Simone Kashima, Svetoslav Nanev Slavov, Vincent Louis Viala |
| EPI_ISL_2209269 | HOSPITAL E PRONTO SOCORRO PORTINARI | Instituto Butantan | Dimas Tadeu Covas, Antonio Jorge Martins, Claudia Renata dos Santos Barros, David Schlesinger, Debora Botequio Moretti, Elaine Cristina Marqueze, Elaine Vieira Santos, Evandra Strazza Rodrigues, Heidge Fukumasu, Jayme Augusto de Souza-Neto, José Salvatore Leister Patané, Luiz Alcantara, Luiz Lehmann Coutinho, Maria Carolina Elias, Maurício Lacerda Nogueira, Rafael dos Santos Bezerra, Raul Machado Neto, Rejane Maria Tommasini Grotto, Ricardo Haddad, Sandra Coccuzzo Sampaio Vessoni, Simone Kashima, Svetoslav Nanev Slavov, Vincent Louis Viala. |
| EPI_ISL_2209271 | SECRETARIA MUNICIPAL DE SAUDE DE CORDEIROPOLIS | Instituto Butantan / FZEA-USP-Pirassununga | Dimas Tadeu Covas, Antonio Jorge Martins, Claudia Renata dos Santos Barros, David Schlesinger, Debora Botequio Moretti, Elaine Cristina Marqueze, Elaine Vieira Santos, Evandra Strazza Rodrigues, Heidge Fukumasu, Jayme Augusto de Souza-Neto, José Salvatore Leister Patané, Luiz Alcantara, Luiz Lehmann Coutinho, Maria Carolina Elias, Maurício Lacerda Nogueira, Rafael dos Santos Bezerra, Raul Machado Neto, Rejane Maria Tommasini Grotto, Ricardo Haddad, Sandra Coccuzzo Sampaio Vessoni, Simone Kashima, Svetoslav Nanev Slavov, Vincent Louis Viala |
| EPI_ISL_2209275 | COORDENADORIA MUNICIPAL DE SAUDE DE IRACEMAPOLIS | Instituto Butantan / FZEA-USP-Pirassununga | Dimas Tadeu Covas, Antonio Jorge Martins, Claudia Renata dos Santos Barros, David Schlesinger, Debora Botequio Moretti, Elaine Cristina Marqueze, Elaine Vieira Santos, Evandra Strazza Rodrigues, Heidge Fukumasu, Jayme Augusto de Souza-Neto, José Salvatore Leister Patané, Luiz Alcantara, Luiz Lehmann Coutinho, Maria Carolina Elias, Maurício Lacerda Nogueira, Rafael dos Santos Bezerra, Raul Machado Neto, Rejane Maria Tommasini Grotto, Ricardo Haddad, Sandra Coccuzzo Sampaio Vessoni, Simone Kashima, Svetoslav Nanev Slavov, Vincent Louis Viala |
| EPI_ISL_2209278 | LABORATORIO MUNICIPAL DE ANALISES CLINICAS DE RIO CLARO | Instituto Butantan / FZEA-USP-Pirassununga | Dimas Tadeu Covas, Antonio Jorge Martins, Claudia Renata dos Santos Barros, David Schlesinger, Debora Botequio Moretti, Elaine Cristina Marqueze, Elaine Vieira Santos, Evandra Strazza Rodrigues, Heidge Fukumasu, Jayme Augusto de Souza-Neto, José Salvatore Leister Patané, Luiz Alcantara, Luiz Lehmann Coutinho, Maria Carolina Elias, Maurício Lacerda Nogueira, Rafael dos Santos Bezerra, Raul Machado Neto, Rejane Maria Tommasini Grotto, Ricardo Haddad, Sandra Coccuzzo Sampaio Vessoni, Simone Kashima, Svetoslav Nanev Slavov, Vincent Louis Viala |
| EPI_ISL_2209282 | VIGILANCIA EPIDEMIOLOGICA E CONTROLE DE VETORES PIRASSUNUN | Instituto Butantan / FZEA-USP-Pirassununga | Dimas Tadeu Covas, Antonio Jorge Martins, Claudia Renata dos Santos Barros, David Schlesinger, Debora Botequio Moretti, Elaine Cristina Marqueze, Elaine Vieira Santos, Evandra Strazza Rodrigues, Heidge Fukumasu, Jayme Augusto de Souza-Neto, José Salvatore Leister Patané, Luiz Alcantara, Luiz Lehmann Coutinho, Maria Carolina Elias, Maurício Lacerda Nogueira, Rafael dos Santos Bezerra, Raul Machado Neto, Rejane Maria Tommasini Grotto, Ricardo Haddad, Sandra Coccuzzo Sampaio Vessoni, Simone Kashima, Svetoslav Nanev Slavov, Vincent Louis Viala |
| EPI_ISL_2209285 | SECRETARIA MUNICIPAL DE SAUDE SOROCABA | Instituto Butantan / ESALQ-Piracicaba | Dimas Tadeu Covas, Antonio Jorge Martins, Claudia Renata dos Santos Barros, David Schlesinger, Debora Botequio Moretti, Elaine Cristina Marqueze, Elaine Vieira Santos, Evandra Strazza Rodrigues, Heidge Fukumasu, Jayme Augusto de Souza-Neto, José Salvatore Leister Patané, Luiz Alcantara, Luiz Lehmann Coutinho, Maria Carolina Elias, Maurício Lacerda Nogueira, Rafael dos Santos Bezerra, Raul Machado Neto, Rejane Maria Tommasini Grotto, Ricardo Haddad, Sandra Coccuzzo Sampaio Vessoni, Simone Kashima, Svetoslav Nanev Slavov, Vincent Louis Viala |
| EPI_ISL_2209288 | CENTRO DE SAUDE DR JOAO JOAQUIM FERREIRA CRISTAIS PAULISTA | Instituto Butantan / Mendelics | Dimas Tadeu Covas, Antonio Jorge Martins, Claudia Renata dos Santos Barros, David Schlesinger, Debora Botequio Moretti, Elaine Cristina Marqueze, Elaine Vieira Santos, Evandra Strazza Rodrigues, Heidge Fukumasu, Jayme Augusto de Souza-Neto, José Salvatore Leister Patané, Luiz Alcantara, Luiz Lehmann Coutinho, Maria Carolina Elias, Maurício Lacerda Nogueira, Rafael dos Santos Bezerra, Raul Machado Neto, Rejane Maria Tommasini Grotto, Ricardo Haddad, Sandra Coccuzzo Sampaio Vessoni, Simone Kashima, Svetoslav Nanev Slavov, Vincent Louis Viala |
| EPI_ISL_2209292 | HOSPITAL E MATERNIDADE NOSSA SENHORA DA AJUDA | Instituto Butantan / Mendelics | Dimas Tadeu Covas, Antonio Jorge Martins, Claudia Renata dos Santos Barros, David Schlesinger, Debora Botequio Moretti, Elaine Cristina Marqueze, Elaine Vieira Santos, Evandra Strazza Rodrigues, Heidge Fukumasu, Jayme Augusto de Souza-Neto, José Salvatore Leister Patané, Luiz Alcantara, Luiz Lehmann Coutinho, Maria Carolina Elias, Maurício Lacerda Nogueira, Rafael dos Santos Bezerra, Raul Machado Neto, Rejane Maria Tommasini Grotto, Ricardo Haddad, Sandra Coccuzzo Sampaio Vessoni, Simone Kashima, Svetoslav Nanev Slavov, Vincent Louis Viala |
| EPI_ISL_2209304 | VIGILANCIA EM SAUDE SAE VIGILANCIA EPIDEMIOLOGICA | Instituto Butantan / Mendelics | Dimas Tadeu Covas, Antonio Jorge Martins, Claudia Renata dos Santos Barros, David Schlesinger, Debora Botequio Moretti, Elaine Cristina Marqueze, Elaine Vieira Santos, Evandra Strazza Rodrigues, Heidge Fukumasu, Jayme Augusto de Souza-Neto, José Salvatore Leister Patané, Luiz Alcantara, Luiz Lehmann Coutinho, Maria Carolina Elias, Maurício Lacerda Nogueira, Rafael dos Santos Bezerra, Raul Machado Neto, Rejane Maria Tommasini Grotto, Ricardo Haddad, Sandra Coccuzzo Sampaio Vessoni, Simone Kashima, Svetoslav Nanev Slavov, Vincent Louis Viala |
| EPI_ISL_2209307 | CENTRO DE SAUDE III AFFONSO LUZZI SANTA CRUZ DAS PALMEIRAS | Instituto Butantan / Mendelics | Dimas Tadeu Covas, Antonio Jorge Martins, Claudia Renata dos Santos Barros, David Schlesinger, Debora Botequio Moretti, Elaine Cristina Marqueze, Elaine Vieira Santos, Evandra Strazza Rodrigues, Heidge Fukumasu, Jayme Augusto de Souza-Neto, José Salvatore Leister Patané, Luiz Alcantara, Luiz Lehmann Coutinho, Maria Carolina Elias, Maurício Lacerda Nogueira, Rafael dos Santos Bezerra, Raul Machado Neto, Rejane Maria Tommasini Grotto, Ricardo Haddad, Sandra Coccuzzo Sampaio Vessoni, Simone Kashima, Svetoslav Nanev Slavov, Vincent Louis Viala |
| EPI_ISL_2209309 | VIGILANCIA EM SAUDE SAE VIGILANCIA EPIDEMIOLOGICA | Instituto Butantan / Mendelics | Dimas Tadeu Covas, Antonio Jorge Martins, Claudia Renata dos Santos Barros, David Schlesinger, Debora Botequio Moretti, Elaine Cristina Marqueze, Elaine Vieira Santos, Evandra Strazza Rodrigues, Heidge Fukumasu, Jayme Augusto de Souza-Neto, José Salvatore Leister Patané, Luiz Alcantara, Luiz Lehmann Coutinho, Maria Carolina Elias, Maurício Lacerda Nogueira, Rafael dos Santos Bezerra, Raul Machado Neto, Rejane Maria Tommasini Grotto, Ricardo Haddad, Sandra Coccuzzo Sampaio Vessoni, Simone Kashima, Svetoslav Nanev Slavov, Vincent Louis Viala |
| EPI_ISL_2209312 | CENTRO DE SAUDE III AFFONSO LUZZI SANTA CRUZ DAS PALMEIRAS | Instituto Butantan / Mendelics | Dimas Tadeu Covas, Antonio Jorge Martins, Claudia Renata dos Santos Barros, David Schlesinger, Debora Botequio Moretti, Elaine Cristina Marqueze, Elaine Vieira Santos, Evandra Strazza Rodrigues, Heidge Fukumasu, Jayme Augusto de Souza-Neto, José Salvatore Leister Patané, Luiz Alcantara, Luiz Lehmann Coutinho, Maria Carolina Elias, Maurício Lacerda Nogueira, Rafael dos Santos Bezerra, Raul Machado Neto, Rejane Maria Tommasini Grotto, Ricardo Haddad, Sandra Coccuzzo Sampaio Vessoni, Simone Kashima, Svetoslav Nanev Slavov, Vincent Louis Viala |
| EPI_ISL_2209313 | HOSP E MATERNIDADE MUNICIPAL N SRA MONTE SERRAT | Instituto Butantan / ESALQ-Piracicaba | Dimas Tadeu Covas, Antonio Jorge Martins, Claudia Renata dos Santos Barros, David Schlesinger, Debora Botequio Moretti, Elaine Cristina Marqueze, Elaine Vieira Santos, Evandra Strazza Rodrigues, Heidge Fukumasu, Jayme Augusto de Souza-Neto, José Salvatore Leister Patané, Luiz Alcantara, Luiz |

| | | | |
|---|---|---|---|
| | | | Lehmann Coutinho, Maria Carolina Elias, Maurício Lacerda Nogueira, Rafael dos Santos Bezerra, Raul Machado Neto, Rejane Maria Tommasini Grotto, Ricardo Haddad, Sandra Coccuzzo Sampaio Vessoni, Simone Kashima, Svetoslav Nanev Slavov, Vincent Louis Viala |
| EPI_ISL_2209315 | HOSPITAL E PRONTO SOCORRO PORTINARI | Instituto Butantan | Dimas Tadeu Covas, Antonio Jorge Martins, Claudia Renata dos Santos Barros, David Schlesinger, Debora Botequio Moretti, Elaine Cristina Marqueze, Elaine Vieira Santos, Evandra Strazza Rodrigues, Heidge Fukumasu, Jayme Augusto de Souza-Neto, José Salvatore Leister Patané, Luiz Alcantara, Luiz Lehmann Coutinho, Maria Carolina Elias, Maurício Lacerda Nogueira, Rafael dos Santos Bezerra, Raul Machado Neto, Rejane Maria Tommasini Grotto, Ricardo Haddad, Sandra Coccuzzo Sampaio Vessoni, Simone Kashima, Svetoslav Nanev Slavov, Vincent Louis Viala |
| EPI_ISL_2209319 | HOSPITAL SAO JOSE ITIRAPINA | Instituto Butantan | Dimas Tadeu Covas, Antonio Jorge Martins, Claudia Renata dos Santos Barros, David Schlesinger, Debora Botequio Moretti, Elaine Cristina Marqueze, Elaine Vieira Santos, Evandra Strazza Rodrigues, Heidge Fukumasu, Jayme Augusto de Souza-Neto, José Salvatore Leister Patané, Luiz Alcantara, Luiz Lehmann Coutinho, Maria Carolina Elias, Maurício Lacerda Nogueira, Rafael dos Santos Bezerra, Raul Machado Neto, Rejane Maria Tommasini Grotto, Ricardo Haddad, Sandra Coccuzzo Sampaio Vessoni, Simone Kashima, Svetoslav Nanev Slavov, Vincent Louis Viala. |
| EPI_ISL_2209322 | VIGILANCIA EM SAUDE SAE VIGILANCIA EPIDEMIOLOGICA | Instituto Butantan / Mendelics | Dimas Tadeu Covas, Antonio Jorge Martins, Claudia Renata dos Santos Barros, David Schlesinger, Debora Botequio Moretti, Elaine Cristina Marqueze, Elaine Vieira Santos, Evandra Strazza Rodrigues, Heidge Fukumasu, Jayme Augusto de Souza-Neto, José Salvatore Leister Patané, Luiz Alcantara, Luiz Lehmann Coutinho, Maria Carolina Elias, Maurício Lacerda Nogueira, Rafael dos Santos Bezerra, Raul Machado Neto, Rejane Maria Tommasini Grotto, Ricardo Haddad, Sandra Coccuzzo Sampaio Vessoni, Simone Kashima, Svetoslav Nanev Slavov, Vincent Louis Viala |
| EPI_ISL_2209325 | SECRETARIA MUNICIPAL DE SAUDE DE PIRACAIA | Instituto Butantan / Mendelics | Dimas Tadeu Covas, Antonio Jorge Martins, Claudia Renata dos Santos Barros, David Schlesinger, Debora Botequio Moretti, Elaine Cristina Marqueze, Elaine Vieira Santos, Evandra Strazza Rodrigues, Heidge Fukumasu, Jayme Augusto de Souza-Neto, José Salvatore Leister Patané, Luiz Alcantara, Luiz Lehmann Coutinho, Maria Carolina Elias, Maurício Lacerda Nogueira, Rafael dos Santos Bezerra, Raul Machado Neto, Rejane Maria Tommasini Grotto, Ricardo Haddad, Sandra Coccuzzo Sampaio Vessoni, Simone Kashima, Svetoslav Nanev Slavov, Vincent Louis Viala |
| EPI_ISL_2209329 | SECRETARIA DE SAUDE PUBLICA DE PRAIA GRANDE | Instituto Butantan / Mendelics | Dimas Tadeu Covas, Antonio Jorge Martins, Claudia Renata dos Santos Barros, David Schlesinger, Debora Botequio Moretti, Elaine Cristina Marqueze, Elaine Vieira Santos, Evandra Strazza Rodrigues, Heidge Fukumasu, Jayme Augusto de Souza-Neto, José Salvatore Leister Patané, Luiz Alcantara, Luiz Lehmann Coutinho, Maria Carolina Elias, Maurício Lacerda Nogueira, Rafael dos Santos Bezerra, Raul Machado Neto, Rejane Maria Tommasini Grotto, Ricardo Haddad, Sandra Coccuzzo Sampaio Vessoni, Simone Kashima, Svetoslav Nanev Slavov, Vincent Louis Viala |
| EPI_ISL_2209332 | UPA MARY DOTA | Instituto Butantan / Mendelics | Dimas Tadeu Covas, Antonio Jorge Martins, Claudia Renata dos Santos Barros, David Schlesinger, Debora Botequio Moretti, Elaine Cristina Marqueze, Elaine Vieira Santos, Evandra Strazza Rodrigues, Heidge Fukumasu, Jayme Augusto de Souza-Neto, José Salvatore Leister Patané, Luiz Alcantara, Luiz Lehmann Coutinho, Maria Carolina Elias, Maurício Lacerda Nogueira, Rafael dos Santos Bezerra, Raul Machado Neto, Rejane Maria Tommasini Grotto, Ricardo Haddad, Sandra Coccuzzo Sampaio Vessoni, Simone Kashima, Svetoslav Nanev Slavov, Vincent Louis Viala |
| EPI_ISL_2209335 | SECAO CENTRO DE DIAGNOSTICO SECEDI | Instituto Butantan / Mendelics | Dimas Tadeu Covas, Antonio Jorge Martins, Claudia Renata dos Santos Barros, David Schlesinger, Debora Botequio Moretti, Elaine Cristina Marqueze, Elaine Vieira Santos, Evandra Strazza Rodrigues, Heidge Fukumasu, Jayme Augusto de Souza-Neto, José Salvatore Leister Patané, Luiz Alcantara, Luiz Lehmann Coutinho, Maria Carolina Elias, Maurício Lacerda Nogueira, Rafael dos Santos Bezerra, Raul Machado Neto, Rejane Maria Tommasini Grotto, Ricardo Haddad, Sandra Coccuzzo Sampaio Vessoni, Simone Kashima, Svetoslav Nanev Slavov, Vincent Louis Viala |
| EPI_ISL_2209338 | LABORATORIO LOCAL DE ITAPECERICA DA SERRA | Instituto Butantan | Dimas Tadeu Covas, Antonio Jorge Martins, Claudia Renata dos Santos Barros, David Schlesinger, Debora Botequio Moretti, Elaine Cristina Marqueze, Elaine Vieira Santos, Evandra Strazza Rodrigues, Heidge Fukumasu, Jayme Augusto de Souza-Neto, José Salvatore Leister Patané, Luiz Alcantara, Luiz Lehmann Coutinho, Maria Carolina Elias, Maurício Lacerda Nogueira, Rafael dos Santos Bezerra, Raul Machado Neto, Rejane Maria Tommasini Grotto, Ricardo Haddad, Sandra Coccuzzo Sampaio Vessoni, Simone Kashima, Svetoslav Nanev Slavov, Vincent Louis Viala. |
| EPI_ISL_2209341 | SERV DE VIG SANITARIA EPIDEMIO E CTRL DE ZOONOZES GUARUJA | Instituto Butantan / Mendelics | Dimas Tadeu Covas, Antonio Jorge Martins, Claudia Renata dos Santos Barros, David Schlesinger, Debora Botequio Moretti, Elaine Cristina Marqueze, Elaine Vieira Santos, Evandra Strazza Rodrigues, Heidge Fukumasu, Jayme Augusto de Souza-Neto, José Salvatore Leister Patané, Luiz Alcantara, Luiz Lehmann Coutinho, Maria Carolina Elias, Maurício Lacerda Nogueira, Rafael dos Santos Bezerra, Raul Machado Neto, Rejane Maria Tommasini Grotto, Ricardo Haddad, Sandra Coccuzzo Sampaio Vessoni, Simone Kashima, Svetoslav Nanev Slavov, Vincent Louis Viala |
| EPI_ISL_2209345 | LABORATORIO LOCAL DE ITAPECERICA DA SERRA | Instituto Butantan | Dimas Tadeu Covas, Antonio Jorge Martins, Claudia Renata dos Santos Barros, David Schlesinger, Debora Botequio Moretti, Elaine Cristina Marqueze, Elaine Vieira Santos, Evandra Strazza Rodrigues, Heidge Fukumasu, Jayme Augusto de Souza-Neto, José Salvatore Leister Patané, Luiz Alcantara, Luiz Lehmann Coutinho, Maria Carolina Elias, Maurício Lacerda Nogueira, Rafael dos Santos Bezerra, Raul Machado Neto, Rejane Maria Tommasini Grotto, Ricardo Haddad, Sandra Coccuzzo Sampaio Vessoni, Simone Kashima, Svetoslav Nanev Slavov, Vincent Louis Viala. |
| EPI_ISL_2209348 | PAS JOAO ANTONIO DO NASCIMENTO | Instituto Butantan / Mendelics | Dimas Tadeu Covas, Antonio Jorge Martins, Claudia Renata dos Santos Barros, David Schlesinger, Debora Botequio Moretti, Elaine Cristina Marqueze, Elaine Vieira Santos, Evandra Strazza Rodrigues, Heidge Fukumasu, Jayme Augusto de Souza-Neto, José Salvatore Leister Patané, Luiz Alcantara, Luiz Lehmann Coutinho, Maria Carolina Elias, Maurício Lacerda Nogueira, Rafael dos Santos Bezerra, Raul Machado Neto, Rejane Maria Tommasini Grotto, Ricardo Haddad, Sandra Coccuzzo Sampaio Vessoni, Simone Kashima, Svetoslav Nanev Slavov, Vincent Louis Viala |
| EPI_ISL_2209351 | UNIDADE BASICA DE SAUDE JARDIM DO LAGO | Instituto Butantan / Mendelics | Dimas Tadeu Covas, Antonio Jorge Martins, Claudia Renata dos Santos Barros, David Schlesinger, Debora Botequio Moretti, Elaine Cristina Marqueze, Elaine Vieira Santos, Evandra Strazza Rodrigues, Heidge Fukumasu, Jayme Augusto de Souza-Neto, José Salvatore Leister Patané, Luiz Alcantara, Luiz Lehmann Coutinho, Maria Carolina Elias, Maurício Lacerda Nogueira, Rafael dos Santos Bezerra, Raul Machado Neto, Rejane Maria Tommasini Grotto, Ricardo Haddad, Sandra Coccuzzo Sampaio Vessoni, Simone Kashima, Svetoslav Nanev Slavov, Vincent Louis Viala |
| EPI_ISL_2209353 | CENTRO DE SAUDE II IBITINGA | Instituto Butantan / Mendelics | Dimas Tadeu Covas, Antonio Jorge Martins, Claudia Renata dos Santos Barros, David Schlesinger, Debora Botequio Moretti, Elaine Cristina Marqueze, Elaine Vieira Santos, Evandra Strazza Rodrigues, Heidge Fukumasu, Jayme Augusto de Souza-Neto, José Salvatore Leister Patané, Luiz Alcantara, Luiz Lehmann Coutinho, Maria Carolina Elias, Maurício Lacerda Nogueira, Rafael dos Santos Bezerra, Raul Machado Neto, Rejane Maria Tommasini Grotto, Ricardo Haddad, Sandra Coccuzzo Sampaio Vessoni, Simone Kashima, Svetoslav Nanev Slavov, Vincent Louis Viala |
| EPI_ISL_2209355 | LABORATORIO MUNICIPAL DE SUZANO | Instituto Butantan / Mendelics | Dimas Tadeu Covas, Antonio Jorge Martins, Claudia Renata dos Santos Barros, David Schlesinger, Debora Botequio Moretti, Elaine Cristina Marqueze, Elaine Vieira Santos, Evandra Strazza Rodrigues, Heidge Fukumasu, Jayme Augusto de Souza-Neto, José Salvatore Leister Patané, Luiz Alcantara, Luiz Lehmann Coutinho, Maria Carolina Elias, Maurício Lacerda Nogueira, Rafael dos Santos Bezerra, Raul Machado Neto, Rejane Maria Tommasini Grotto, Ricardo Haddad, Sandra Coccuzzo Sampaio Vessoni, Simone Kashima, Svetoslav Nanev Slavov, Vincent Louis Viala |
| EPI_ISL_2209360 | LABORATORIO MUNICIPAL DE ANALISES CLINICAS DE RIO CLARO | Instituto Butantan | Dimas Tadeu Covas, Antonio Jorge Martins, Claudia Renata dos Santos Barros, David Schlesinger, Debora Botequio Moretti, Elaine Cristina Marqueze, Elaine Vieira Santos, Evandra Strazza Rodrigues, Heidge Fukumasu, Jayme Augusto de Souza-Neto, José Salvatore Leister Patané, Luiz Alcantara, Luiz Lehmann Coutinho, Maria Carolina Elias, Maurício Lacerda Nogueira, Rafael dos Santos Bezerra, Raul Machado Neto, Rejane Maria Tommasini Grotto, Ricardo Haddad, Sandra Coccuzzo Sampaio Vessoni, Simone Kashima, Svetoslav Nanev Slavov, Vincent Louis Viala. |
| EPI_ISL_2209363 | HOSPITAL DE CAMPANHA COVID 19 MUNICIPIO DE TAUBATE | Instituto Butantan / Mendelics | Dimas Tadeu Covas, Antonio Jorge Martins, Claudia Renata dos Santos Barros, David Schlesinger, Debora Botequio Moretti, Elaine Cristina Marqueze, Elaine Vieira Santos, Evandra Strazza Rodrigues, Heidge Fukumasu, Jayme Augusto de Souza-Neto, José Salvatore Leister Patané, Luiz Alcantara, Luiz Lehmann Coutinho, Maria Carolina Elias, Maurício Lacerda Nogueira, Rafael dos Santos Bezerra, Raul Machado Neto, Rejane Maria Tommasini Grotto, Ricardo Haddad, Sandra Coccuzzo Sampaio Vessoni, Simone Kashima, Svetoslav Nanev Slavov, Vincent Louis Viala |
| EPI_ISL_2209366 | UNIDADE BASICA DE SAUDE II ESTRELA DO NORTE | Instituto Butantan / Mendelics | Dimas Tadeu Covas, Antonio Jorge Martins, Claudia Renata dos Santos Barros, David Schlesinger, Debora Botequio Moretti, Elaine Cristina Marqueze, Elaine Vieira Santos, Evandra Strazza Rodrigues, Heidge Fukumasu, Jayme Augusto de Souza-Neto, José Salvatore Leister Patané, Luiz Alcantara, Luiz Lehmann Coutinho, Maria Carolina Elias, Maurício Lacerda Nogueira, Rafael dos Santos Bezerra, Raul Machado Neto, Rejane Maria Tommasini Grotto, Ricardo Haddad, Sandra Coccuzzo Sampaio Vessoni, Simone Kashima, Svetoslav Nanev Slavov, Vincent Louis Viala |
| EPI_ISL_2209368 | CENTRO DE SAUDE II IBITINGA | Instituto Butantan / Mendelics | Dimas Tadeu Covas, Antonio Jorge Martins, Claudia Renata dos Santos Barros, David Schlesinger, Debora Botequio Moretti, Elaine Cristina Marqueze, Elaine Vieira Santos, Evandra Strazza Rodrigues, Heidge Fukumasu, Jayme Augusto de Souza-Neto, José Salvatore Leister Patané, Luiz Alcantara, Luiz Lehmann Coutinho, Maria Carolina Elias, Maurício Lacerda Nogueira, Rafael dos Santos Bezerra, Raul Machado Neto, Rejane Maria Tommasini Grotto, Ricardo Haddad, Sandra Coccuzzo Sampaio Vessoni, Simone Kashima, Svetoslav Nanev Slavov, Vincent Louis Viala |
| EPI_ISL_2209370 | DEPARTAMENTO DE SAUDE MUNICIPAL SOCORRO SP | Instituto Butantan / Mendelics | Dimas Tadeu Covas, Antonio Jorge Martins, Claudia Renata dos Santos Barros, David Schlesinger, Debora Botequio Moretti, Elaine Cristina Marqueze, Elaine Vieira Santos, Evandra Strazza Rodrigues, Heidge Fukumasu, Jayme Augusto de Souza-Neto, José Salvatore Leister Patané, Luiz Alcantara, Luiz Lehmann Coutinho, Maria Carolina Elias, Maurício Lacerda Nogueira, Rafael dos Santos Bezerra, Raul Machado Neto, Rejane Maria Tommasini Grotto, Ricardo Haddad, Sandra Coccuzzo Sampaio Vessoni, Simone Kashima, Svetoslav Nanev Slavov, Vincent Louis Viala |

| | | | |
|---|---|---|---|
| EPI_ISL_2209374 | UPA MARY DOTA | Instituto Butantan / Mendelics | Dimas Tadeu Covas, Antonio Jorge Martins, Claudia Renata dos Santos Barros, David Schlesinger, Debora Botequio Moretti, Elaine Cristina Marqueze, Elaine Vieira Santos, Evandra Strazza Rodrigues, Heidge Fukumasu, Jayme Augusto de Souza-Neto, José Salvatore Leister Patané, Luiz Alcantara, Luiz Lehmann Coutinho, Maria Carolina Elias, Maurício Lacerda Nogueira, Rafael dos Santos Bezerra, Raul Machado Neto, Rejane Maria Tommasini Grotto, Ricardo Haddad, Sandra Coccuzzo Sampaio Vessoni, Simone Kashima, Svetoslav Nanev Slavov, Vincent Louis Viala |
| EPI_ISL_2209377 | HOSP MUN JOSANIAS CASTANHA BRAGA | Instituto Butantan / ESALQ-Piracicaba | Dimas Tadeu Covas, Antonio Jorge Martins, Claudia Renata dos Santos Barros, David Schlesinger, Debora Botequio Moretti, Elaine Cristina Marqueze, Elaine Vieira Santos, Evandra Strazza Rodrigues, Heidge Fukumasu, Jayme Augusto de Souza-Neto, José Salvatore Leister Patané, Luiz Alcantara, Luiz Lehmann Coutinho, Maria Carolina Elias, Maurício Lacerda Nogueira, Rafael dos Santos Bezerra, Raul Machado Neto, Rejane Maria Tommasini Grotto, Ricardo Haddad, Sandra Coccuzzo Sampaio Vessoni, Simone Kashima, Svetoslav Nanev Slavov, Vincent Louis Viala |
| EPI_ISL_2209379 | UPA 24 HORAS DR ALOISIO MUNIZ DE ANDRADE | Instituto Butantan / ESALQ-Piracicaba | Dimas Tadeu Covas, Antonio Jorge Martins, Claudia Renata dos Santos Barros, David Schlesinger, Debora Botequio Moretti, Elaine Cristina Marqueze, Elaine Vieira Santos, Evandra Strazza Rodrigues, Heidge Fukumasu, Jayme Augusto de Souza-Neto, José Salvatore Leister Patané, Luiz Alcantara, Luiz Lehmann Coutinho, Maria Carolina Elias, Maurício Lacerda Nogueira, Rafael dos Santos Bezerra, Raul Machado Neto, Rejane Maria Tommasini Grotto, Ricardo Haddad, Sandra Coccuzzo Sampaio Vessoni, Simone Kashima, Svetoslav Nanev Slavov, Vincent Louis Viala |
| EPI_ISL_2209383, EPI_ISL_2209386 | HOSP MUN DE MOGI DAS CRUZES PREF WALDEMAR COSTA FILHO | Instituto Butantan / ESALQ-Piracicaba | Dimas Tadeu Covas, Antonio Jorge Martins, Claudia Renata dos Santos Barros, David Schlesinger, Debora Botequio Moretti, Elaine Cristina Marqueze, Elaine Vieira Santos, Evandra Strazza Rodrigues, Heidge Fukumasu, Jayme Augusto de Souza-Neto, José Salvatore Leister Patané, Luiz Alcantara, Luiz Lehmann Coutinho, Maria Carolina Elias, Maurício Lacerda Nogueira, Rafael dos Santos Bezerra, Raul Machado Neto, Rejane Maria Tommasini Grotto, Ricardo Haddad, Sandra Coccuzzo Sampaio Vessoni, Simone Kashima, Svetoslav Nanev Slavov, Vincent Louis Viala |
| EPI_ISL_2209389 | UNIDADE DE PRONTO ATENDIMENTO 24HS CIPO | Instituto Butantan / ESALQ-Piracicaba | Dimas Tadeu Covas, Antonio Jorge Martins, Claudia Renata dos Santos Barros, David Schlesinger, Debora Botequio Moretti, Elaine Cristina Marqueze, Elaine Vieira Santos, Evandra Strazza Rodrigues, Heidge Fukumasu, Jayme Augusto de Souza-Neto, José Salvatore Leister Patané, Luiz Alcantara, Luiz Lehmann Coutinho, Maria Carolina Elias, Maurício Lacerda Nogueira, Rafael dos Santos Bezerra, Raul Machado Neto, Rejane Maria Tommasini Grotto, Ricardo Haddad, Sandra Coccuzzo Sampaio Vessoni, Simone Kashima, Svetoslav Nanev Slavov, Vincent Louis Viala |
| EPI_ISL_2209393 | HOSP MUN CIDADE TIRADENTES CARMEN PRUDENTE | Instituto Butantan / ESALQ-Piracicaba | Dimas Tadeu Covas, Antonio Jorge Martins, Claudia Renata dos Santos Barros, David Schlesinger, Debora Botequio Moretti, Elaine Cristina Marqueze, Elaine Vieira Santos, Evandra Strazza Rodrigues, Heidge Fukumasu, Jayme Augusto de Souza-Neto, José Salvatore Leister Patané, Luiz Alcantara, Luiz Lehmann Coutinho, Maria Carolina Elias, Maurício Lacerda Nogueira, Rafael dos Santos Bezerra, Raul Machado Neto, Rejane Maria Tommasini Grotto, Ricardo Haddad, Sandra Coccuzzo Sampaio Vessoni, Simone Kashima, Svetoslav Nanev Slavov, Vincent Louis Viala |
| EPI_ISL_2209396 | CENTRO DE SAUDE DE IBIRAREMA | Instituto Butantan / ESALQ-Piracicaba | Dimas Tadeu Covas, Antonio Jorge Martins, Claudia Renata dos Santos Barros, David Schlesinger, Debora Botequio Moretti, Elaine Cristina Marqueze, Elaine Vieira Santos, Evandra Strazza Rodrigues, Heidge Fukumasu, Jayme Augusto de Souza-Neto, José Salvatore Leister Patané, Luiz Alcantara, Luiz Lehmann Coutinho, Maria Carolina Elias, Maurício Lacerda Nogueira, Rafael dos Santos Bezerra, Raul Machado Neto, Rejane Maria Tommasini Grotto, Ricardo Haddad, Sandra Coccuzzo Sampaio Vessoni, Simone Kashima, Svetoslav Nanev Slavov, Vincent Louis Viala |
| EPI_ISL_2209397 | CS DE COSMORAMA | Instituto Butantan / ESALQ-Piracicaba | Dimas Tadeu Covas, Antonio Jorge Martins, Claudia Renata dos Santos Barros, David Schlesinger, Debora Botequio Moretti, Elaine Cristina Marqueze, Elaine Vieira Santos, Evandra Strazza Rodrigues, Heidge Fukumasu, Jayme Augusto de Souza-Neto, José Salvatore Leister Patané, Luiz Alcantara, Luiz Lehmann Coutinho, Maria Carolina Elias, Maurício Lacerda Nogueira, Rafael dos Santos Bezerra, Raul Machado Neto, Rejane Maria Tommasini Grotto, Ricardo Haddad, Sandra Coccuzzo Sampaio Vessoni, Simone Kashima, Svetoslav Nanev Slavov, Vincent Louis Viala |
| EPI_ISL_2209401, EPI_ISL_2209404 | UPA 24 HORAS ANA JACINTA | Instituto Butantan / ESALQ-Piracicaba | Dimas Tadeu Covas, Antonio Jorge Martins, Claudia Renata dos Santos Barros, David Schlesinger, Debora Botequio Moretti, Elaine Cristina Marqueze, Elaine Vieira Santos, Evandra Strazza Rodrigues, Heidge Fukumasu, Jayme Augusto de Souza-Neto, José Salvatore Leister Patané, Luiz Alcantara, Luiz Lehmann Coutinho, Maria Carolina Elias, Maurício Lacerda Nogueira, Rafael dos Santos Bezerra, Raul Machado Neto, Rejane Maria Tommasini Grotto, Ricardo Haddad, Sandra Coccuzzo Sampaio Vessoni, Simone Kashima, Svetoslav Nanev Slavov, Vincent Louis Viala |
| EPI_ISL_2209407 | SANTA CASA DE MISERICORDIA SAO JOSE | Instituto Butantan / Mendelics | Dimas Tadeu Covas, Antonio Jorge Martins, Claudia Renata dos Santos Barros, David Schlesinger, Debora Botequio Moretti, Elaine Cristina Marqueze, Elaine Vieira Santos, Evandra Strazza Rodrigues, Heidge Fukumasu, Jayme Augusto de Souza-Neto, José Salvatore Leister Patané, Luiz Alcantara, Luiz Lehmann Coutinho, Maria Carolina Elias, Maurício Lacerda Nogueira, Rafael dos Santos Bezerra, Raul Machado Neto, Rejane Maria Tommasini Grotto, Ricardo Haddad, Sandra Coccuzzo Sampaio Vessoni, Simone Kashima, Svetoslav Nanev Slavov, Vincent Louis Viala |
| EPI_ISL_2209408 | PRONTO ATENDIMENTO MUNICIPAL ALUMINIO | Instituto Butantan | Dimas Tadeu Covas, Antonio Jorge Martins, Claudia Renata dos Santos Barros, David Schlesinger, Debora Botequio Moretti, Elaine Cristina Marqueze, Elaine Vieira Santos, Evandra Strazza Rodrigues, Heidge Fukumasu, Jayme Augusto de Souza-Neto, José Salvatore Leister Patané, Luiz Alcantara, Luiz Lehmann Coutinho, Maria Carolina Elias, Maurício Lacerda Nogueira, Rafael dos Santos Bezerra, Raul Machado Neto, Rejane Maria Tommasini Grotto, Ricardo Haddad, Sandra Coccuzzo Sampaio Vessoni, Simone Kashima, Svetoslav Nanev Slavov, Vincent Louis Viala. |
| EPI_ISL_2209413 | PRONTO ATENDIMENTO MUNICIPAL ITALO SANTUCCI | Instituto Butantan | Dimas Tadeu Covas, Antonio Jorge Martins, Claudia Renata dos Santos Barros, David Schlesinger, Debora Botequio Moretti, Elaine Cristina Marqueze, Elaine Vieira Santos, Evandra Strazza Rodrigues, Heidge Fukumasu, Jayme Augusto de Souza-Neto, José Salvatore Leister Patané, Luiz Alcantara, Luiz Lehmann Coutinho, Maria Carolina Elias, Maurício Lacerda Nogueira, Rafael dos Santos Bezerra, Raul Machado Neto, Rejane Maria Tommasini Grotto, Ricardo Haddad, Sandra Coccuzzo Sampaio Vessoni, Simone Kashima, Svetoslav Nanev Slavov, Vincent Louis Viala. |
| EPI_ISL_2209419 | SMS IPERO | Instituto Butantan / Mendelics | Dimas Tadeu Covas, Antonio Jorge Martins, Claudia Renata dos Santos Barros, David Schlesinger, Debora Botequio Moretti, Elaine Cristina Marqueze, Elaine Vieira Santos, Evandra Strazza Rodrigues, Heidge Fukumasu, Jayme Augusto de Souza-Neto, José Salvatore Leister Patané, Luiz Alcantara, Luiz Lehmann Coutinho, Maria Carolina Elias, Maurício Lacerda Nogueira, Rafael dos Santos Bezerra, Raul Machado Neto, Rejane Maria Tommasini Grotto, Ricardo Haddad, Sandra Coccuzzo Sampaio Vessoni, Simone Kashima, Svetoslav Nanev Slavov, Vincent Louis Viala |
| EPI_ISL_2209422 | PRONTO SOCORRO MUNICIPAL DE TAUBATE | Instituto Butantan | Dimas Tadeu Covas, Antonio Jorge Martins, Claudia Renata dos Santos Barros, David Schlesinger, Debora Botequio Moretti, Elaine Cristina Marqueze, Elaine Vieira Santos, Evandra Strazza Rodrigues, Heidge Fukumasu, Jayme Augusto de Souza-Neto, José Salvatore Leister Patané, Luiz Alcantara, Luiz Lehmann Coutinho, Maria Carolina Elias, Maurício Lacerda Nogueira, Rafael dos Santos Bezerra, Raul Machado Neto, Rejane Maria Tommasini Grotto, Ricardo Haddad, Sandra Coccuzzo Sampaio Vessoni, Simone Kashima, Svetoslav Nanev Slavov, Vincent Louis Viala. |
| EPI_ISL_2209425 | PSF DR ANTONIO PIRES DE ALMEIDA PORTO FELIZ | Instituto Butantan | Dimas Tadeu Covas, Antonio Jorge Martins, Claudia Renata dos Santos Barros, David Schlesinger, Debora Botequio Moretti, Elaine Cristina Marqueze, Elaine Vieira Santos, Evandra Strazza Rodrigues, Heidge Fukumasu, Jayme Augusto de Souza-Neto, José Salvatore Leister Patané, Luiz Alcantara, Luiz Lehmann Coutinho, Maria Carolina Elias, Maurício Lacerda Nogueira, Rafael dos Santos Bezerra, Raul Machado Neto, Rejane Maria Tommasini Grotto, Ricardo Haddad, Sandra Coccuzzo Sampaio Vessoni, Simone Kashima, Svetoslav Nanev Slavov, Vincent Louis Viala. |
| EPI_ISL_2209433 | PSF MARIA JOSE SALTO DE PIRAPORA | Instituto Butantan | Dimas Tadeu Covas, Antonio Jorge Martins, Claudia Renata dos Santos Barros, David Schlesinger, Debora Botequio Moretti, Elaine Cristina Marqueze, Elaine Vieira Santos, Evandra Strazza Rodrigues, Heidge Fukumasu, Jayme Augusto de Souza-Neto, José Salvatore Leister Patané, Luiz Alcantara, Luiz Lehmann Coutinho, Maria Carolina Elias, Maurício Lacerda Nogueira, Rafael dos Santos Bezerra, Raul Machado Neto, Rejane Maria Tommasini Grotto, Ricardo Haddad, Sandra Coccuzzo Sampaio Vessoni, Simone Kashima, Svetoslav Nanev Slavov, Vincent Louis Viala. |
| EPI_ISL_2209436 | CENTRO DE SAUDE II MAIRINQUE MAIRINQUE | Instituto Butantan / ESALQ-Piracicaba | Dimas Tadeu Covas, Antonio Jorge Martins, Claudia Renata dos Santos Barros, David Schlesinger, Debora Botequio Moretti, Elaine Cristina Marqueze, Elaine Vieira Santos, Evandra Strazza Rodrigues, Heidge Fukumasu, Jayme Augusto de Souza-Neto, José Salvatore Leister Patané, Luiz Alcantara, Luiz Lehmann Coutinho, Maria Carolina Elias, Maurício Lacerda Nogueira, Rafael dos Santos Bezerra, Raul Machado Neto, Rejane Maria Tommasini Grotto, Ricardo Haddad, Sandra Coccuzzo Sampaio Vessoni, Simone Kashima, Svetoslav Nanev Slavov, Vincent Louis Viala |
| EPI_ISL_2209439 | CENTRO DE ESPECIALIDADES MARIAS DA GLORIA | Instituto Butantan / ESALQ-Piracicaba | Dimas Tadeu Covas, Antonio Jorge Martins, Claudia Renata dos Santos Barros, David Schlesinger, Debora Botequio Moretti, Elaine Cristina Marqueze, Elaine Vieira Santos, Evandra Strazza Rodrigues, Heidge Fukumasu, Jayme Augusto de Souza-Neto, José Salvatore Leister Patané, Luiz Alcantara, Luiz Lehmann Coutinho, Maria Carolina Elias, Maurício Lacerda Nogueira, Rafael dos Santos Bezerra, Raul Machado Neto, Rejane Maria Tommasini Grotto, Ricardo Haddad, Sandra Coccuzzo Sampaio Vessoni, Simone Kashima, Svetoslav Nanev Slavov, Vincent Louis Viala |
| EPI_ISL_2209443 | VIGILANCIA EPIDEMIOLOGICA E CONTROLE DE VETORES PIRASSUNUN | Instituto Butantan / FZEA-USP-Pirassununga | Dimas Tadeu Covas, Antonio Jorge Martins, Claudia Renata dos Santos Barros, David Schlesinger, Debora Botequio Moretti, Elaine Cristina Marqueze, Elaine Vieira Santos, Evandra Strazza Rodrigues, Heidge Fukumasu, Jayme Augusto de Souza-Neto, José Salvatore Leister Patané, Luiz Alcantara, Luiz Lehmann Coutinho, Maria Carolina Elias, Maurício Lacerda Nogueira, Rafael dos Santos Bezerra, Raul Machado Neto, Rejane Maria Tommasini Grotto, Ricardo Haddad, Sandra Coccuzzo Sampaio Vessoni, Simone Kashima, Svetoslav Nanev Slavov, Vincent Louis Viala |
| EPI_ISL_2209445 | CENTRO DE SAUDE DE ANALANDIA | Instituto Butantan / FZEA-USP-Pirassununga | Dimas Tadeu Covas, Antonio Jorge Martins, Claudia Renata dos Santos Barros, David Schlesinger, Debora Botequio Moretti, Elaine Cristina Marqueze, Elaine Vieira Santos, Evandra Strazza Rodrigues, Heidge Fukumasu, Jayme Augusto de Souza-Neto, José Salvatore Leister Patané, Luiz Alcantara, Luiz |

| | | | |
|---|---|---|---|
| | | | Lehmann Coutinho, Maria Carolina Elias, Maurício Lacerda Nogueira, Rafael dos Santos Bezerra, Raul Machado Neto, Rejane Maria Tommasini Grotto, Ricardo Haddad, Sandra Coccuzzo Sampaio Vessoni, Simone Kashima, Svetoslav Nanev Slavov, Vincent Louis Viala |
| EPI_ISL_2209448, EPI_ISL_2209450, EPI_ISL_2209454 | VIGILANCIA EPIDEMIOLOGICA E CONTROLE DE VETORES PIRASSUNUN | Instituto Butantan / FZEA-USP-Pirassununga | Dimas Tadeu Covas, Antonio Jorge Martins, Claudia Renata dos Santos Barros, David Schlesinger, Debora Botequio Moretti, Elaine Cristina Marqueze, Elaine Vieira Santos, Evandra Strazza Rodrigues, Heidge Fukumasu, Jayme Augusto de Souza-Neto, José Salvatore Leister Patané, Luiz Alcantara, Luiz Lehmann Coutinho, Maria Carolina Elias, Maurício Lacerda Nogueira, Rafael dos Santos Bezerra, Raul Machado Neto, Rejane Maria Tommasini Grotto, Ricardo Haddad, Sandra Coccuzzo Sampaio Vessoni, Simone Kashima, Svetoslav Nanev Slavov, Vincent Louis Viala |
| EPI_ISL_2209457, EPI_ISL_2209462 | SECRETARIA MUNICIPAL DE SAUDE DE CORDEIROPOLIS | Instituto Butantan / FZEA-USP-Pirassununga | Dimas Tadeu Covas, Antonio Jorge Martins, Claudia Renata dos Santos Barros, David Schlesinger, Debora Botequio Moretti, Elaine Cristina Marqueze, Elaine Vieira Santos, Evandra Strazza Rodrigues, Heidge Fukumasu, Jayme Augusto de Souza-Neto, José Salvatore Leister Patané, Luiz Alcantara, Luiz Lehmann Coutinho, Maria Carolina Elias, Maurício Lacerda Nogueira, Rafael dos Santos Bezerra, Raul Machado Neto, Rejane Maria Tommasini Grotto, Ricardo Haddad, Sandra Coccuzzo Sampaio Vessoni, Simone Kashima, Svetoslav Nanev Slavov, Vincent Louis Viala |
| EPI_ISL_2209464 | CENTRO DE SAUDE DE MARACAI | Instituto Butantan / Mendelics | Dimas Tadeu Covas, Antonio Jorge Martins, Claudia Renata dos Santos Barros, David Schlesinger, Debora Botequio Moretti, Elaine Cristina Marqueze, Elaine Vieira Santos, Evandra Strazza Rodrigues, Heidge Fukumasu, Jayme Augusto de Souza-Neto, José Salvatore Leister Patané, Luiz Alcantara, Luiz Lehmann Coutinho, Maria Carolina Elias, Maurício Lacerda Nogueira, Rafael dos Santos Bezerra, Raul Machado Neto, Rejane Maria Tommasini Grotto, Ricardo Haddad, Sandra Coccuzzo Sampaio Vessoni, Simone Kashima, Svetoslav Nanev Slavov, Vincent Louis Viala |
| EPI_ISL_2209467 | SANTA CASA DE GUARAREMA | Instituto Butantan | Dimas Tadeu Covas, Antonio Jorge Martins, Claudia Renata dos Santos Barros, David Schlesinger, Debora Botequio Moretti, Elaine Cristina Marqueze, Elaine Vieira Santos, Evandra Strazza Rodrigues, Heidge Fukumasu, Jayme Augusto de Souza-Neto, José Salvatore Leister Patané, Luiz Alcantara, Luiz Lehmann Coutinho, Maria Carolina Elias, Maurício Lacerda Nogueira, Rafael dos Santos Bezerra, Raul Machado Neto, Rejane Maria Tommasini Grotto, Ricardo Haddad, Sandra Coccuzzo Sampaio Vessoni, Simone Kashima, Svetoslav Nanev Slavov, Vincent Louis Viala. |
| EPI_ISL_2209475, EPI_ISL_2209477, EPI_ISL_2209480 | SANTA CASA DE MISERICORDIA DE TIETE | Instituto Butantan | Dimas Tadeu Covas, Antonio Jorge Martins, Claudia Renata dos Santos Barros, David Schlesinger, Debora Botequio Moretti, Elaine Cristina Marqueze, Elaine Vieira Santos, Evandra Strazza Rodrigues, Heidge Fukumasu, Jayme Augusto de Souza-Neto, José Salvatore Leister Patané, Luiz Alcantara, Luiz Lehmann Coutinho, Maria Carolina Elias, Maurício Lacerda Nogueira, Rafael dos Santos Bezerra, Raul Machado Neto, Rejane Maria Tommasini Grotto, Ricardo Haddad, Sandra Coccuzzo Sampaio Vessoni, Simone Kashima, Svetoslav Nanev Slavov, Vincent Louis Viala. |
| EPI_ISL_2209484 | SANTA CASA DE MISERICORDIA DE UBATUBA | Instituto Butantan | Dimas Tadeu Covas, Antonio Jorge Martins, Claudia Renata dos Santos Barros, David Schlesinger, Debora Botequio Moretti, Elaine Cristina Marqueze, Elaine Vieira Santos, Evandra Strazza Rodrigues, Heidge Fukumasu, Jayme Augusto de Souza-Neto, José Salvatore Leister Patané, Luiz Alcantara, Luiz Lehmann Coutinho, Maria Carolina Elias, Maurício Lacerda Nogueira, Rafael dos Santos Bezerra, Raul Machado Neto, Rejane Maria Tommasini Grotto, Ricardo Haddad, Sandra Coccuzzo Sampaio Vessoni, Simone Kashima, Svetoslav Nanev Slavov, Vincent Louis Viala. |
| EPI_ISL_2209487 | SANTA CASA DE PORTO FELIZ | Instituto Butantan | Dimas Tadeu Covas, Antonio Jorge Martins, Claudia Renata dos Santos Barros, David Schlesinger, Debora Botequio Moretti, Elaine Cristina Marqueze, Elaine Vieira Santos, Evandra Strazza Rodrigues, Heidge Fukumasu, Jayme Augusto de Souza-Neto, José Salvatore Leister Patané, Luiz Alcantara, Luiz Lehmann Coutinho, Maria Carolina Elias, Maurício Lacerda Nogueira, Rafael dos Santos Bezerra, Raul Machado Neto, Rejane Maria Tommasini Grotto, Ricardo Haddad, Sandra Coccuzzo Sampaio Vessoni, Simone Kashima, Svetoslav Nanev Slavov, Vincent Louis Viala. |
| EPI_ISL_2209490 | SAUDE COLETIVA CAPAO BONITO | Instituto Butantan | Dimas Tadeu Covas, Antonio Jorge Martins, Claudia Renata dos Santos Barros, David Schlesinger, Debora Botequio Moretti, Elaine Cristina Marqueze, Elaine Vieira Santos, Evandra Strazza Rodrigues, Heidge Fukumasu, Jayme Augusto de Souza-Neto, José Salvatore Leister Patané, Luiz Alcantara, Luiz Lehmann Coutinho, Maria Carolina Elias, Maurício Lacerda Nogueira, Rafael dos Santos Bezerra, Raul Machado Neto, Rejane Maria Tommasini Grotto, Ricardo Haddad, Sandra Coccuzzo Sampaio Vessoni, Simone Kashima, Svetoslav Nanev Slavov, Vincent Louis Viala. |
| EPI_ISL_2209494 | UBS III DE PARIQUERA ACU PARIQUERA ACU | Instituto Butantan / Mendelics | Dimas Tadeu Covas, Antonio Jorge Martins, Claudia Renata dos Santos Barros, David Schlesinger, Debora Botequio Moretti, Elaine Cristina Marqueze, Elaine Vieira Santos, Evandra Strazza Rodrigues, Heidge Fukumasu, Jayme Augusto de Souza-Neto, José Salvatore Leister Patané, Luiz Alcantara, Luiz Lehmann Coutinho, Maria Carolina Elias, Maurício Lacerda Nogueira, Rafael dos Santos Bezerra, Raul Machado Neto, Rejane Maria Tommasini Grotto, Ricardo Haddad, Sandra Coccuzzo Sampaio Vessoni, Simone Kashima, Svetoslav Nanev Slavov, Vincent Louis Viala |
| EPI_ISL_2209496 | SECRETARIA DE SAUDE DE RAFARD | Instituto Butantan | Dimas Tadeu Covas, Antonio Jorge Martins, Claudia Renata dos Santos Barros, David Schlesinger, Debora Botequio Moretti, Elaine Cristina Marqueze, Elaine Vieira Santos, Evandra Strazza Rodrigues, Heidge Fukumasu, Jayme Augusto de Souza-Neto, José Salvatore Leister Patané, Luiz Alcantara, Luiz Lehmann Coutinho, Maria Carolina Elias, Maurício Lacerda Nogueira, Rafael dos Santos Bezerra, Raul Machado Neto, Rejane Maria Tommasini Grotto, Ricardo Haddad, Sandra Coccuzzo Sampaio Vessoni, Simone Kashima, Svetoslav Nanev Slavov, Vincent Louis Viala. |
| EPI_ISL_2209499 | HOSPITAL MUNICIPAL DE ITAPIRA | Instituto Butantan / Mendelics | Dimas Tadeu Covas, Antonio Jorge Martins, Claudia Renata dos Santos Barros, David Schlesinger, Debora Botequio Moretti, Elaine Cristina Marqueze, Elaine Vieira Santos, Evandra Strazza Rodrigues, Heidge Fukumasu, Jayme Augusto de Souza-Neto, José Salvatore Leister Patané, Luiz Alcantara, Luiz Lehmann Coutinho, Maria Carolina Elias, Maurício Lacerda Nogueira, Rafael dos Santos Bezerra, Raul Machado Neto, Rejane Maria Tommasini Grotto, Ricardo Haddad, Sandra Coccuzzo Sampaio Vessoni, Simone Kashima, Svetoslav Nanev Slavov, Vincent Louis Viala |
| EPI_ISL_2209505 | SECRETARIA DE SAUDE DE SANTA GERTRUDES | Instituto Butantan | Dimas Tadeu Covas, Antonio Jorge Martins, Claudia Renata dos Santos Barros, David Schlesinger, Debora Botequio Moretti, Elaine Cristina Marqueze, Elaine Vieira Santos, Evandra Strazza Rodrigues, Heidge Fukumasu, Jayme Augusto de Souza-Neto, José Salvatore Leister Patané, Luiz Alcantara, Luiz Lehmann Coutinho, Maria Carolina Elias, Maurício Lacerda Nogueira, Rafael dos Santos Bezerra, Raul Machado Neto, Rejane Maria Tommasini Grotto, Ricardo Haddad, Sandra Coccuzzo Sampaio Vessoni, Simone Kashima, Svetoslav Nanev Slavov, Vincent Louis Viala. |
| EPI_ISL_2209511 | SECRETARIA MUNICIPAL DE SAUDE SOROCABA | Instituto Butantan / Mendelics | Dimas Tadeu Covas, Antonio Jorge Martins, Claudia Renata dos Santos Barros, David Schlesinger, Debora Botequio Moretti, Elaine Cristina Marqueze, Elaine Vieira Santos, Evandra Strazza Rodrigues, Heidge Fukumasu, Jayme Augusto de Souza-Neto, José Salvatore Leister Patané, Luiz Alcantara, Luiz Lehmann Coutinho, Maria Carolina Elias, Maurício Lacerda Nogueira, Rafael dos Santos Bezerra, Raul Machado Neto, Rejane Maria Tommasini Grotto, Ricardo Haddad, Sandra Coccuzzo Sampaio Vessoni, Simone Kashima, Svetoslav Nanev Slavov, Vincent Louis Viala |
| EPI_ISL_2209516 | SECRETARIA DE SAUDE PUBLICA DE PRAIA GRANDE | Instituto Butantan / Mendelics | Dimas Tadeu Covas, Antonio Jorge Martins, Claudia Renata dos Santos Barros, David Schlesinger, Debora Botequio Moretti, Elaine Cristina Marqueze, Elaine Vieira Santos, Evandra Strazza Rodrigues, Heidge Fukumasu, Jayme Augusto de Souza-Neto, José Salvatore Leister Patané, Luiz Alcantara, Luiz Lehmann Coutinho, Maria Carolina Elias, Maurício Lacerda Nogueira, Rafael dos Santos Bezerra, Raul Machado Neto, Rejane Maria Tommasini Grotto, Ricardo Haddad, Sandra Coccuzzo Sampaio Vessoni, Simone Kashima, Svetoslav Nanev Slavov, Vincent Louis Viala |
| EPI_ISL_2209522 | SECRETARIA DE SAUDE DE SANTA GERTRUDES | Instituto Butantan | Dimas Tadeu Covas, Antonio Jorge Martins, Claudia Renata dos Santos Barros, David Schlesinger, Debora Botequio Moretti, Elaine Cristina Marqueze, Elaine Vieira Santos, Evandra Strazza Rodrigues, Heidge Fukumasu, Jayme Augusto de Souza-Neto, José Salvatore Leister Patané, Luiz Alcantara, Luiz Lehmann Coutinho, Maria Carolina Elias, Maurício Lacerda Nogueira, Rafael dos Santos Bezerra, Raul Machado Neto, Rejane Maria Tommasini Grotto, Ricardo Haddad, Sandra Coccuzzo Sampaio Vessoni, Simone Kashima, Svetoslav Nanev Slavov, Vincent Louis Viala. |
| EPI_ISL_2209527 | CENTRO DE SAUDE II IBITINGA | Instituto Butantan / Mendelics | Dimas Tadeu Covas, Antonio Jorge Martins, Claudia Renata dos Santos Barros, David Schlesinger, Debora Botequio Moretti, Elaine Cristina Marqueze, Elaine Vieira Santos, Evandra Strazza Rodrigues, Heidge Fukumasu, Jayme Augusto de Souza-Neto, José Salvatore Leister Patané, Luiz Alcantara, Luiz Lehmann Coutinho, Maria Carolina Elias, Maurício Lacerda Nogueira, Rafael dos Santos Bezerra, Raul Machado Neto, Rejane Maria Tommasini Grotto, Ricardo Haddad, Sandra Coccuzzo Sampaio Vessoni, Simone Kashima, Svetoslav Nanev Slavov, Vincent Louis Viala |
| EPI_ISL_2209532 | UBS DR WALDOMIRO FERREIRA NEVES SCRPARDO | Instituto Butantan / Mendelics | Dimas Tadeu Covas, Antonio Jorge Martins, Claudia Renata dos Santos Barros, David Schlesinger, Debora Botequio Moretti, Elaine Cristina Marqueze, Elaine Vieira Santos, Evandra Strazza Rodrigues, Heidge Fukumasu, Jayme Augusto de Souza-Neto, José Salvatore Leister Patané, Luiz Alcantara, Luiz Lehmann Coutinho, Maria Carolina Elias, Maurício Lacerda Nogueira, Rafael dos Santos Bezerra, Raul Machado Neto, Rejane Maria Tommasini Grotto, Ricardo Haddad, Sandra Coccuzzo Sampaio Vessoni, Simone Kashima, Svetoslav Nanev Slavov, Vincent Louis Viala |
| EPI_ISL_2209537 | SECRETARIA DE SAUDE DE SAO PEDRO | Instituto Butantan | Dimas Tadeu Covas, Antonio Jorge Martins, Claudia Renata dos Santos Barros, David Schlesinger, Debora Botequio Moretti, Elaine Cristina Marqueze, Elaine Vieira Santos, Evandra Strazza Rodrigues, Heidge Fukumasu, Jayme Augusto de Souza-Neto, José Salvatore Leister Patané, Luiz Alcantara, Luiz Lehmann Coutinho, Maria Carolina Elias, Maurício Lacerda Nogueira, Rafael dos Santos Bezerra, Raul Machado Neto, Rejane Maria Tommasini Grotto, Ricardo Haddad, Sandra Coccuzzo Sampaio Vessoni, Simone Kashima, Svetoslav Nanev Slavov, Vincent Louis Viala. |
| EPI_ISL_2209541 | CENTRO DE SAUDE DR JOAO JOAQUIM FERREIRA CRISTAIS PAULISTA | Instituto Butantan / Mendelics | Dimas Tadeu Covas, Antonio Jorge Martins, Claudia Renata dos Santos Barros, David Schlesinger, Debora Botequio Moretti, Elaine Cristina Marqueze, Elaine Vieira Santos, Evandra Strazza Rodrigues, Heidge Fukumasu, Jayme Augusto de Souza-Neto, José Salvatore Leister Patané, Luiz Alcantara, Luiz Lehmann Coutinho, Maria Carolina Elias, Maurício Lacerda Nogueira, Rafael dos Santos Bezerra, Raul Machado Neto, Rejane Maria Tommasini Grotto, Ricardo Haddad, Sandra Coccuzzo Sampaio Vessoni, Simone Kashima, Svetoslav Nanev Slavov, Vincent Louis Viala |

| | | | |
|---|---|---|---|
| EPI_ISL_2209543 | SANTA CASA DE GUARAREMA | Instituto Butantan | Dimas Tadeu Covas, Antonio Jorge Martins, Claudia Renata dos Santos Barros, David Schlesinger, Debora Botequio Moretti, Elaine Cristina Marqueze, Elaine Vieira Santos, Evandra Strazza Rodrigues, Heidge Fukumasu, Jayme Augusto de Souza-Neto, José Salvatore Leister Patané, Luiz Alcantara, Luiz Lehmann Coutinho, Maria Carolina Elias, Maurício Lacerda Nogueira, Rafael dos Santos Bezerra, Raul Machado Neto, Rejane Maria Tommasini Grotto, Ricardo Haddad, Sandra Coccuzzo Sampaio Vessoni, Simone Kashima, Svetoslav Nanev Slavov, Vincent Louis Viala |
| EPI_ISL_2209551, EPI_ISL_2209557 | SECRETARIA MUNICIPAL DA SAUDE DE GUARATINGUETA | Instituto Butantan | Dimas Tadeu Covas, Antonio Jorge Martins, Claudia Renata dos Santos Barros, David Schlesinger, Debora Botequio Moretti, Elaine Cristina Marqueze, Elaine Vieira Santos, Evandra Strazza Rodrigues, Heidge Fukumasu, Jayme Augusto de Souza-Neto, José Salvatore Leister Patané, Luiz Alcantara, Luiz Lehmann Coutinho, Maria Carolina Elias, Maurício Lacerda Nogueira, Rafael dos Santos Bezerra, Raul Machado Neto, Rejane Maria Tommasini Grotto, Ricardo Haddad, Sandra Coccuzzo Sampaio Vessoni, Simone Kashima, Svetoslav Nanev Slavov, Vincent Louis Viala. |
| EPI_ISL_2209565, EPI_ISL_2209571 | NUCLEO DE SAUDE MARY DOTA DE BAURU | Instituto Butantan / Mendelics | Dimas Tadeu Covas, Antonio Jorge Martins, Claudia Renata dos Santos Barros, David Schlesinger, Debora Botequio Moretti, Elaine Cristina Marqueze, Elaine Vieira Santos, Evandra Strazza Rodrigues, Heidge Fukumasu, Jayme Augusto de Souza-Neto, José Salvatore Leister Patané, Luiz Alcantara, Luiz Lehmann Coutinho, Maria Carolina Elias, Maurício Lacerda Nogueira, Rafael dos Santos Bezerra, Raul Machado Neto, Rejane Maria Tommasini Grotto, Ricardo Haddad, Sandra Coccuzzo Sampaio Vessoni, Simone Kashima, Svetoslav Nanev Slavov, Vincent Louis Viala |
| EPI_ISL_2209579 | UPA DE BEBEDOURO | Instituto Butantan / Mendelics | Dimas Tadeu Covas, Antonio Jorge Martins, Claudia Renata dos Santos Barros, David Schlesinger, Debora Botequio Moretti, Elaine Cristina Marqueze, Elaine Vieira Santos, Evandra Strazza Rodrigues, Heidge Fukumasu, Jayme Augusto de Souza-Neto, José Salvatore Leister Patané, Luiz Alcantara, Luiz Lehmann Coutinho, Maria Carolina Elias, Maurício Lacerda Nogueira, Rafael dos Santos Bezerra, Raul Machado Neto, Rejane Maria Tommasini Grotto, Ricardo Haddad, Sandra Coccuzzo Sampaio Vessoni, Simone Kashima, Svetoslav Nanev Slavov, Vincent Louis Viala |
| EPI_ISL_2209584 | SECRETARIA MUNICIPAL DE SAUDE DE CORDEIROPOLIS | Instituto Butantan | Dimas Tadeu Covas, Antonio Jorge Martins, Claudia Renata dos Santos Barros, David Schlesinger, Debora Botequio Moretti, Elaine Cristina Marqueze, Elaine Vieira Santos, Evandra Strazza Rodrigues, Heidge Fukumasu, Jayme Augusto de Souza-Neto, José Salvatore Leister Patané, Luiz Alcantara, Luiz Lehmann Coutinho, Maria Carolina Elias, Maurício Lacerda Nogueira, Rafael dos Santos Bezerra, Raul Machado Neto, Rejane Maria Tommasini Grotto, Ricardo Haddad, Sandra Coccuzzo Sampaio Vessoni, Simone Kashima, Svetoslav Nanev Slavov, Vincent Louis Viala. |
| EPI_ISL_2209586 | GRIPARIO FALCAO | Instituto Butantan / Mendelics | Dimas Tadeu Covas, Antonio Jorge Martins, Claudia Renata dos Santos Barros, David Schlesinger, Debora Botequio Moretti, Elaine Cristina Marqueze, Elaine Vieira Santos, Evandra Strazza Rodrigues, Heidge Fukumasu, Jayme Augusto de Souza-Neto, José Salvatore Leister Patané, Luiz Alcantara, Luiz Lehmann Coutinho, Maria Carolina Elias, Maurício Lacerda Nogueira, Rafael dos Santos Bezerra, Raul Machado Neto, Rejane Maria Tommasini Grotto, Ricardo Haddad, Sandra Coccuzzo Sampaio Vessoni, Simone Kashima, Svetoslav Nanev Slavov, Vincent Louis Viala |
| EPI_ISL_2209590 | CENTRO DE SAUDE II IBITINGA | Instituto Butantan / Mendelics | Dimas Tadeu Covas, Antonio Jorge Martins, Claudia Renata dos Santos Barros, David Schlesinger, Debora Botequio Moretti, Elaine Cristina Marqueze, Elaine Vieira Santos, Evandra Strazza Rodrigues, Heidge Fukumasu, Jayme Augusto de Souza-Neto, José Salvatore Leister Patané, Luiz Alcantara, Luiz Lehmann Coutinho, Maria Carolina Elias, Maurício Lacerda Nogueira, Rafael dos Santos Bezerra, Raul Machado Neto, Rejane Maria Tommasini Grotto, Ricardo Haddad, Sandra Coccuzzo Sampaio Vessoni, Simone Kashima, Svetoslav Nanev Slavov, Vincent Louis Viala |
| EPI_ISL_2209593 | CENTRO DE SAUDE DE MARACAI | Instituto Butantan / Mendelics | Dimas Tadeu Covas, Antonio Jorge Martins, Claudia Renata dos Santos Barros, David Schlesinger, Debora Botequio Moretti, Elaine Cristina Marqueze, Elaine Vieira Santos, Evandra Strazza Rodrigues, Heidge Fukumasu, Jayme Augusto de Souza-Neto, José Salvatore Leister Patané, Luiz Alcantara, Luiz Lehmann Coutinho, Maria Carolina Elias, Maurício Lacerda Nogueira, Rafael dos Santos Bezerra, Raul Machado Neto, Rejane Maria Tommasini Grotto, Ricardo Haddad, Sandra Coccuzzo Sampaio Vessoni, Simone Kashima, Svetoslav Nanev Slavov, Vincent Louis Viala |
| EPI_ISL_2209600 | UBS II DR JOSE CARLOS MACUCO JANINI | Instituto Butantan / Mendelics | Dimas Tadeu Covas, Antonio Jorge Martins, Claudia Renata dos Santos Barros, David Schlesinger, Debora Botequio Moretti, Elaine Cristina Marqueze, Elaine Vieira Santos, Evandra Strazza Rodrigues, Heidge Fukumasu, Jayme Augusto de Souza-Neto, José Salvatore Leister Patané, Luiz Alcantara, Luiz Lehmann Coutinho, Maria Carolina Elias, Maurício Lacerda Nogueira, Rafael dos Santos Bezerra, Raul Machado Neto, Rejane Maria Tommasini Grotto, Ricardo Haddad, Sandra Coccuzzo Sampaio Vessoni, Simone Kashima, Svetoslav Nanev Slavov, Vincent Louis Viala |
| EPI_ISL_2209603 | HOSPITAL MUNICIPAL DE ITAPIRA | Instituto Butantan / Mendelics | Dimas Tadeu Covas, Antonio Jorge Martins, Claudia Renata dos Santos Barros, David Schlesinger, Debora Botequio Moretti, Elaine Cristina Marqueze, Elaine Vieira Santos, Evandra Strazza Rodrigues, Heidge Fukumasu, Jayme Augusto de Souza-Neto, José Salvatore Leister Patané, Luiz Alcantara, Luiz Lehmann Coutinho, Maria Carolina Elias, Maurício Lacerda Nogueira, Rafael dos Santos Bezerra, Raul Machado Neto, Rejane Maria Tommasini Grotto, Ricardo Haddad, Sandra Coccuzzo Sampaio Vessoni, Simone Kashima, Svetoslav Nanev Slavov, Vincent Louis Viala |
| EPI_ISL_2209607 | NUCLEO DE SAUDE MARY DOTA DE BAURU | Instituto Butantan / Mendelics | Dimas Tadeu Covas, Antonio Jorge Martins, Claudia Renata dos Santos Barros, David Schlesinger, Debora Botequio Moretti, Elaine Cristina Marqueze, Elaine Vieira Santos, Evandra Strazza Rodrigues, Heidge Fukumasu, Jayme Augusto de Souza-Neto, José Salvatore Leister Patané, Luiz Alcantara, Luiz Lehmann Coutinho, Maria Carolina Elias, Maurício Lacerda Nogueira, Rafael dos Santos Bezerra, Raul Machado Neto, Rejane Maria Tommasini Grotto, Ricardo Haddad, Sandra Coccuzzo Sampaio Vessoni, Simone Kashima, Svetoslav Nanev Slavov, Vincent Louis Viala |
| EPI_ISL_2209610 | SECRETARIA MUNICIPAL DE SAUDE DE CORDEIROPOLIS | Instituto Butantan | Dimas Tadeu Covas, Antonio Jorge Martins, Claudia Renata dos Santos Barros, David Schlesinger, Debora Botequio Moretti, Elaine Cristina Marqueze, Elaine Vieira Santos, Evandra Strazza Rodrigues, Heidge Fukumasu, Jayme Augusto de Souza-Neto, José Salvatore Leister Patané, Luiz Alcantara, Luiz Lehmann Coutinho, Maria Carolina Elias, Maurício Lacerda Nogueira, Rafael dos Santos Bezerra, Raul Machado Neto, Rejane Maria Tommasini Grotto, Ricardo Haddad, Sandra Coccuzzo Sampaio Vessoni, Simone Kashima, Svetoslav Nanev Slavov, Vincent Louis Viala. |
| EPI_ISL_2209614 | CENTRO DE SAUDE II IBITINGA | Instituto Butantan / Mendelics | Dimas Tadeu Covas, Antonio Jorge Martins, Claudia Renata dos Santos Barros, David Schlesinger, Debora Botequio Moretti, Elaine Cristina Marqueze, Elaine Vieira Santos, Evandra Strazza Rodrigues, Heidge Fukumasu, Jayme Augusto de Souza-Neto, José Salvatore Leister Patané, Luiz Alcantara, Luiz Lehmann Coutinho, Maria Carolina Elias, Maurício Lacerda Nogueira, Rafael dos Santos Bezerra, Raul Machado Neto, Rejane Maria Tommasini Grotto, Ricardo Haddad, Sandra Coccuzzo Sampaio Vessoni, Simone Kashima, Svetoslav Nanev Slavov, Vincent Louis Viala |
| EPI_ISL_2209617 | SERV DE VIG SANITARIA EPIDEMIO E CTRL DE ZOONOZES GUARUJA | Instituto Butantan / Mendelics | Dimas Tadeu Covas, Antonio Jorge Martins, Claudia Renata dos Santos Barros, David Schlesinger, Debora Botequio Moretti, Elaine Cristina Marqueze, Elaine Vieira Santos, Evandra Strazza Rodrigues, Heidge Fukumasu, Jayme Augusto de Souza-Neto, José Salvatore Leister Patané, Luiz Alcantara, Luiz Lehmann Coutinho, Maria Carolina Elias, Maurício Lacerda Nogueira, Rafael dos Santos Bezerra, Raul Machado Neto, Rejane Maria Tommasini Grotto, Ricardo Haddad, Sandra Coccuzzo Sampaio Vessoni, Simone Kashima, Svetoslav Nanev Slavov, Vincent Louis Viala |
| EPI_ISL_2209618 | UBS II DE REGENTE FEIJO | Instituto Butantan / Mendelics | Dimas Tadeu Covas, Antonio Jorge Martins, Claudia Renata dos Santos Barros, David Schlesinger, Debora Botequio Moretti, Elaine Cristina Marqueze, Elaine Vieira Santos, Evandra Strazza Rodrigues, Heidge Fukumasu, Jayme Augusto de Souza-Neto, José Salvatore Leister Patané, Luiz Alcantara, Luiz Lehmann Coutinho, Maria Carolina Elias, Maurício Lacerda Nogueira, Rafael dos Santos Bezerra, Raul Machado Neto, Rejane Maria Tommasini Grotto, Ricardo Haddad, Sandra Coccuzzo Sampaio Vessoni, Simone Kashima, Svetoslav Nanev Slavov, Vincent Louis Viala |
| EPI_ISL_2209621 | PRONTO SOCORRO CAUCAIA DO ALTO | Instituto Butantan / Mendelics | Dimas Tadeu Covas, Antonio Jorge Martins, Claudia Renata dos Santos Barros, David Schlesinger, Debora Botequio Moretti, Elaine Cristina Marqueze, Elaine Vieira Santos, Evandra Strazza Rodrigues, Heidge Fukumasu, Jayme Augusto de Souza-Neto, José Salvatore Leister Patané, Luiz Alcantara, Luiz Lehmann Coutinho, Maria Carolina Elias, Maurício Lacerda Nogueira, Rafael dos Santos Bezerra, Raul Machado Neto, Rejane Maria Tommasini Grotto, Ricardo Haddad, Sandra Coccuzzo Sampaio Vessoni, Simone Kashima, Svetoslav Nanev Slavov, Vincent Louis Viala |
| EPI_ISL_2209624 | UNIDADE DE PRONTO ATENDIMENTO UPA DRA ANA OLIVIA BENTIVOGLIO | Instituto Butantan / ESALQ-Piracicaba | Dimas Tadeu Covas, Antonio Jorge Martins, Claudia Renata dos Santos Barros, David Schlesinger, Debora Botequio Moretti, Elaine Cristina Marqueze, Elaine Vieira Santos, Evandra Strazza Rodrigues, Heidge Fukumasu, Jayme Augusto de Souza-Neto, José Salvatore Leister Patané, Luiz Alcantara, Luiz Lehmann Coutinho, Maria Carolina Elias, Maurício Lacerda Nogueira, Rafael dos Santos Bezerra, Raul Machado Neto, Rejane Maria Tommasini Grotto, Ricardo Haddad, Sandra Coccuzzo Sampaio Vessoni, Simone Kashima, Svetoslav Nanev Slavov, Vincent Louis Viala |
| EPI_ISL_2209628 | ESF M1 UNI AMERICA DE JALES | Instituto Butantan / ESALQ-Piracicaba | Dimas Tadeu Covas, Antonio Jorge Martins, Claudia Renata dos Santos Barros, David Schlesinger, Debora Botequio Moretti, Elaine Cristina Marqueze, Elaine Vieira Santos, Evandra Strazza Rodrigues, Heidge Fukumasu, Jayme Augusto de Souza-Neto, José Salvatore Leister Patané, Luiz Alcantara, Luiz Lehmann Coutinho, Maria Carolina Elias, Maurício Lacerda Nogueira, Rafael dos Santos Bezerra, Raul Machado Neto, Rejane Maria Tommasini Grotto, Ricardo Haddad, Sandra Coccuzzo Sampaio Vessoni, Simone Kashima, Svetoslav Nanev Slavov, Vincent Louis Viala |
| EPI_ISL_2209630 | SANTA CASA DE JOSE BONIFACIO | Instituto Butantan / ESALQ-Piracicaba | Dimas Tadeu Covas, Antonio Jorge Martins, Claudia Renata dos Santos Barros, David Schlesinger, Debora Botequio Moretti, Elaine Cristina Marqueze, Elaine Vieira Santos, Evandra Strazza Rodrigues, Heidge Fukumasu, Jayme Augusto de Souza-Neto, José Salvatore Leister Patané, Luiz Alcantara, Luiz Lehmann Coutinho, Maria Carolina Elias, Maurício Lacerda Nogueira, Rafael dos Santos Bezerra, Raul Machado Neto, Rejane Maria Tommasini Grotto, Ricardo Haddad, Sandra Coccuzzo Sampaio Vessoni, Simone Kashima, Svetoslav Nanev Slavov, Vincent Louis Viala |
| EPI_ISL_2209632, EPI_ISL_2209641 | SMS IPERO | Instituto Butantan | Dimas Tadeu Covas, Antonio Jorge Martins, Claudia Renata dos Santos Barros, David Schlesinger, Debora Botequio Moretti, Elaine Cristina Marqueze, Elaine Vieira Santos, Evandra Strazza Rodrigues, Heidge Fukumasu, Jayme Augusto de Souza-Neto, José Salvatore Leister Patané, Luiz Alcantara, Luiz |

| | | | |
|---|---|---|---|
| | | | Lehmann Coutinho, Maria Carolina Elias, Maurício Lacerda Nogueira, Rafael dos Santos Bezerra, Raul Machado Neto, Rejane Maria Tommasini Grotto, Ricardo Haddad, Sandra Coccuzzo Sampaio Vessoni, Simone Kashima, Svetoslav Nanev Slavov, Vincent Louis Viala. |
| EPI_ISL_2209653, EPI_ISL_2209661 | SMS SECRETARIA MUNICIPAL DE SAUDE DE BOITUVA | Instituto Butantan | Dimas Tadeu Covas, Antonio Jorge Martins, Claudia Renata dos Santos Barros, David Schlesinger, Debora Botequio Moretti, Elaine Cristina Marqueze, Elaine Vieira Santos, Evandra Strazza Rodrigues, Heidge Fukumasu, Jayme Augusto de Souza-Neto, José Salvatore Leister Patané, Luiz Alcantara, Luiz Lehmann Coutinho, Maria Carolina Elias, Maurício Lacerda Nogueira, Rafael dos Santos Bezerra, Raul Machado Neto, Rejane Maria Tommasini Grotto, Ricardo Haddad, Sandra Coccuzzo Sampaio Vessoni, Simone Kashima, Svetoslav Nanev Slavov, Vincent Louis Viala. |
| EPI_ISL_2209664 | SECRETARIA MUNICIPAL DE SAUDE SOROCABA | Instituto Butantan / Mendelics | Dimas Tadeu Covas, Antonio Jorge Martins, Claudia Renata dos Santos Barros, David Schlesinger, Debora Botequio Moretti, Elaine Cristina Marqueze, Elaine Vieira Santos, Evandra Strazza Rodrigues, Heidge Fukumasu, Jayme Augusto de Souza-Neto, José Salvatore Leister Patané, Luiz Alcantara, Luiz Lehmann Coutinho, Maria Carolina Elias, Maurício Lacerda Nogueira, Rafael dos Santos Bezerra, Raul Machado Neto, Rejane Maria Tommasini Grotto, Ricardo Haddad, Sandra Coccuzzo Sampaio Vessoni, Simone Kashima, Svetoslav Nanev Slavov, Vincent Louis Viala |
| EPI_ISL_2209668 | SECRETARIA DE SAUDE PUBLICA DE PRAIA GRANDE | Instituto Butantan / Mendelics | Dimas Tadeu Covas, Antonio Jorge Martins, Claudia Renata dos Santos Barros, David Schlesinger, Debora Botequio Moretti, Elaine Cristina Marqueze, Elaine Vieira Santos, Evandra Strazza Rodrigues, Heidge Fukumasu, Jayme Augusto de Souza-Neto, José Salvatore Leister Patané, Luiz Alcantara, Luiz Lehmann Coutinho, Maria Carolina Elias, Maurício Lacerda Nogueira, Rafael dos Santos Bezerra, Raul Machado Neto, Rejane Maria Tommasini Grotto, Ricardo Haddad, Sandra Coccuzzo Sampaio Vessoni, Simone Kashima, Svetoslav Nanev Slavov, Vincent Louis Viala |
| EPI_ISL_2209671 | UPA DE BEBEDOURO | Instituto Butantan / Mendelics | Dimas Tadeu Covas, Antonio Jorge Martins, Claudia Renata dos Santos Barros, David Schlesinger, Debora Botequio Moretti, Elaine Cristina Marqueze, Elaine Vieira Santos, Evandra Strazza Rodrigues, Heidge Fukumasu, Jayme Augusto de Souza-Neto, José Salvatore Leister Patané, Luiz Alcantara, Luiz Lehmann Coutinho, Maria Carolina Elias, Maurício Lacerda Nogueira, Rafael dos Santos Bezerra, Raul Machado Neto, Rejane Maria Tommasini Grotto, Ricardo Haddad, Sandra Coccuzzo Sampaio Vessoni, Simone Kashima, Svetoslav Nanev Slavov, Vincent Louis Viala |
| EPI_ISL_2209680 | UBS II DE TANABI MILTON MARTINS PERCHES | Instituto Butantan | Dimas Tadeu Covas, Antonio Jorge Martins, Claudia Renata dos Santos Barros, David Schlesinger, Debora Botequio Moretti, Elaine Cristina Marqueze, Elaine Vieira Santos, Evandra Strazza Rodrigues, Heidge Fukumasu, Jayme Augusto de Souza-Neto, José Salvatore Leister Patané, Luiz Alcantara, Luiz Lehmann Coutinho, Maria Carolina Elias, Maurício Lacerda Nogueira, Rafael dos Santos Bezerra, Raul Machado Neto, Rejane Maria Tommasini Grotto, Ricardo Haddad, Sandra Coccuzzo Sampaio Vessoni, Simone Kashima, Svetoslav Nanev Slavov, Vincent Louis Viala. |
| EPI_ISL_2209692 | LABORATORIO DR PAULO EMILIO DALESSANDRO PINDAMONHANGABA | Instituto Butantan / Mendelics | Dimas Tadeu Covas, Antonio Jorge Martins, Claudia Renata dos Santos Barros, David Schlesinger, Debora Botequio Moretti, Elaine Cristina Marqueze, Elaine Vieira Santos, Evandra Strazza Rodrigues, Heidge Fukumasu, Jayme Augusto de Souza-Neto, José Salvatore Leister Patané, Luiz Alcantara, Luiz Lehmann Coutinho, Maria Carolina Elias, Maurício Lacerda Nogueira, Rafael dos Santos Bezerra, Raul Machado Neto, Rejane Maria Tommasini Grotto, Ricardo Haddad, Sandra Coccuzzo Sampaio Vessoni, Simone Kashima, Svetoslav Nanev Slavov, Vincent Louis Viala |
| EPI_ISL_2209702 | CENTRO DE SAUDE III NELCIDIO DA SILVEIRA BASTOS | Instituto Butantan / Mendelics | Dimas Tadeu Covas, Antonio Jorge Martins, Claudia Renata dos Santos Barros, David Schlesinger, Debora Botequio Moretti, Elaine Cristina Marqueze, Elaine Vieira Santos, Evandra Strazza Rodrigues, Heidge Fukumasu, Jayme Augusto de Souza-Neto, José Salvatore Leister Patané, Luiz Alcantara, Luiz Lehmann Coutinho, Maria Carolina Elias, Maurício Lacerda Nogueira, Rafael dos Santos Bezerra, Raul Machado Neto, Rejane Maria Tommasini Grotto, Ricardo Haddad, Sandra Coccuzzo Sampaio Vessoni, Simone Kashima, Svetoslav Nanev Slavov, Vincent Louis Viala |
| EPI_ISL_2209707 | CENTRO DE SAUDE II IBITINGA | Instituto Butantan / Mendelics | Dimas Tadeu Covas, Antonio Jorge Martins, Claudia Renata dos Santos Barros, David Schlesinger, Debora Botequio Moretti, Elaine Cristina Marqueze, Elaine Vieira Santos, Evandra Strazza Rodrigues, Heidge Fukumasu, Jayme Augusto de Souza-Neto, José Salvatore Leister Patané, Luiz Alcantara, Luiz Lehmann Coutinho, Maria Carolina Elias, Maurício Lacerda Nogueira, Rafael dos Santos Bezerra, Raul Machado Neto, Rejane Maria Tommasini Grotto, Ricardo Haddad, Sandra Coccuzzo Sampaio Vessoni, Simone Kashima, Svetoslav Nanev Slavov, Vincent Louis Viala |
| EPI_ISL_2209716 | PSF DO PARQUE DOS PINHEIROS | Instituto Butantan / Mendelics | Dimas Tadeu Covas, Antonio Jorge Martins, Claudia Renata dos Santos Barros, David Schlesinger, Debora Botequio Moretti, Elaine Cristina Marqueze, Elaine Vieira Santos, Evandra Strazza Rodrigues, Heidge Fukumasu, Jayme Augusto de Souza-Neto, José Salvatore Leister Patané, Luiz Alcantara, Luiz Lehmann Coutinho, Maria Carolina Elias, Maurício Lacerda Nogueira, Rafael dos Santos Bezerra, Raul Machado Neto, Rejane Maria Tommasini Grotto, Ricardo Haddad, Sandra Coccuzzo Sampaio Vessoni, Simone Kashima, Svetoslav Nanev Slavov, Vincent Louis Viala |
| EPI_ISL_2209717 | VIGILANCIA EPIDEMIOLOGICA E CONTROLE DE VETORES PIRASSUNUN | Instituto Butantan / FZEA-USP-Pirassununga | Dimas Tadeu Covas, Antonio Jorge Martins, Claudia Renata dos Santos Barros, David Schlesinger, Debora Botequio Moretti, Elaine Cristina Marqueze, Elaine Vieira Santos, Evandra Strazza Rodrigues, Heidge Fukumasu, Jayme Augusto de Souza-Neto, José Salvatore Leister Patané, Luiz Alcantara, Luiz Lehmann Coutinho, Maria Carolina Elias, Maurício Lacerda Nogueira, Rafael dos Santos Bezerra, Raul Machado Neto, Rejane Maria Tommasini Grotto, Ricardo Haddad, Sandra Coccuzzo Sampaio Vessoni, Simone Kashima, Svetoslav Nanev Slavov, Vincent Louis Viala |
| EPI_ISL_2209721 | CENTRO DE APOIO EPIDEMIOLOGICO | Instituto Butantan / Mendelics | Dimas Tadeu Covas, Antonio Jorge Martins, Claudia Renata dos Santos Barros, David Schlesinger, Debora Botequio Moretti, Elaine Cristina Marqueze, Elaine Vieira Santos, Evandra Strazza Rodrigues, Heidge Fukumasu, Jayme Augusto de Souza-Neto, José Salvatore Leister Patané, Luiz Alcantara, Luiz Lehmann Coutinho, Maria Carolina Elias, Maurício Lacerda Nogueira, Rafael dos Santos Bezerra, Raul Machado Neto, Rejane Maria Tommasini Grotto, Ricardo Haddad, Sandra Coccuzzo Sampaio Vessoni, Simone Kashima, Svetoslav Nanev Slavov, Vincent Louis Viala |
| EPI_ISL_2209724 | UNIDADE BASICA DE SAUDE DE IPEUNA | Instituto Butantan | Dimas Tadeu Covas, Antonio Jorge Martins, Claudia Renata dos Santos Barros, David Schlesinger, Debora Botequio Moretti, Elaine Cristina Marqueze, Elaine Vieira Santos, Evandra Strazza Rodrigues, Heidge Fukumasu, Jayme Augusto de Souza-Neto, José Salvatore Leister Patané, Luiz Alcantara, Luiz Lehmann Coutinho, Maria Carolina Elias, Maurício Lacerda Nogueira, Rafael dos Santos Bezerra, Raul Machado Neto, Rejane Maria Tommasini Grotto, Ricardo Haddad, Sandra Coccuzzo Sampaio Vessoni, Simone Kashima, Svetoslav Nanev Slavov, Vincent Louis Viala. |
| EPI_ISL_2209732 | UNIDADE BASICA DE SAUDE II ESTRELA DO NORTE | Instituto Butantan / Mendelics | Dimas Tadeu Covas, Antonio Jorge Martins, Claudia Renata dos Santos Barros, David Schlesinger, Debora Botequio Moretti, Elaine Cristina Marqueze, Elaine Vieira Santos, Evandra Strazza Rodrigues, Heidge Fukumasu, Jayme Augusto de Souza-Neto, José Salvatore Leister Patané, Luiz Alcantara, Luiz Lehmann Coutinho, Maria Carolina Elias, Maurício Lacerda Nogueira, Rafael dos Santos Bezerra, Raul Machado Neto, Rejane Maria Tommasini Grotto, Ricardo Haddad, Sandra Coccuzzo Sampaio Vessoni, Simone Kashima, Svetoslav Nanev Slavov, Vincent Louis Viala |
| EPI_ISL_2209739 | DEPARTAMENTO DE SAUDE MUNICIPAL SOCORRO SP | Instituto Butantan / Mendelics | Dimas Tadeu Covas, Antonio Jorge Martins, Claudia Renata dos Santos Barros, David Schlesinger, Debora Botequio Moretti, Elaine Cristina Marqueze, Elaine Vieira Santos, Evandra Strazza Rodrigues, Heidge Fukumasu, Jayme Augusto de Souza-Neto, José Salvatore Leister Patané, Luiz Alcantara, Luiz Lehmann Coutinho, Maria Carolina Elias, Maurício Lacerda Nogueira, Rafael dos Santos Bezerra, Raul Machado Neto, Rejane Maria Tommasini Grotto, Ricardo Haddad, Sandra Coccuzzo Sampaio Vessoni, Simone Kashima, Svetoslav Nanev Slavov, Vincent Louis Viala |
| EPI_ISL_2209746 | UPA MARY DOTA | Instituto Butantan / Mendelics | Dimas Tadeu Covas, Antonio Jorge Martins, Claudia Renata dos Santos Barros, David Schlesinger, Debora Botequio Moretti, Elaine Cristina Marqueze, Elaine Vieira Santos, Evandra Strazza Rodrigues, Heidge Fukumasu, Jayme Augusto de Souza-Neto, José Salvatore Leister Patané, Luiz Alcantara, Luiz Lehmann Coutinho, Maria Carolina Elias, Maurício Lacerda Nogueira, Rafael dos Santos Bezerra, Raul Machado Neto, Rejane Maria Tommasini Grotto, Ricardo Haddad, Sandra Coccuzzo Sampaio Vessoni, Simone Kashima, Svetoslav Nanev Slavov, Vincent Louis Viala |
| EPI_ISL_2209754 | CENTRO DE SAUDE DE LUPERCIO | Instituto Butantan / Mendelics | Dimas Tadeu Covas, Antonio Jorge Martins, Claudia Renata dos Santos Barros, David Schlesinger, Debora Botequio Moretti, Elaine Cristina Marqueze, Elaine Vieira Santos, Evandra Strazza Rodrigues, Heidge Fukumasu, Jayme Augusto de Souza-Neto, José Salvatore Leister Patané, Luiz Alcantara, Luiz Lehmann Coutinho, Maria Carolina Elias, Maurício Lacerda Nogueira, Rafael dos Santos Bezerra, Raul Machado Neto, Rejane Maria Tommasini Grotto, Ricardo Haddad, Sandra Coccuzzo Sampaio Vessoni, Simone Kashima, Svetoslav Nanev Slavov, Vincent Louis Viala |
| EPI_ISL_2209769 | UNIDADE BASICA DE SAUDE DR MATHEUS GABRIEL BONASSA | Instituto Butantan | Dimas Tadeu Covas, Antonio Jorge Martins, Claudia Renata dos Santos Barros, David Schlesinger, Debora Botequio Moretti, Elaine Cristina Marqueze, Elaine Vieira Santos, Evandra Strazza Rodrigues, Heidge Fukumasu, Jayme Augusto de Souza-Neto, José Salvatore Leister Patané, Luiz Alcantara, Luiz Lehmann Coutinho, Maria Carolina Elias, Maurício Lacerda Nogueira, Rafael dos Santos Bezerra, Raul Machado Neto, Rejane Maria Tommasini Grotto, Ricardo Haddad, Sandra Coccuzzo Sampaio Vessoni, Simone Kashima, Svetoslav Nanev Slavov, Vincent Louis Viala. |
| EPI_ISL_2209777 | VIGILANCIA EPIDIMIOLOGICA DE ARACARIGUAMA | Instituto Butantan / Mendelics | Dimas Tadeu Covas, Antonio Jorge Martins, Claudia Renata dos Santos Barros, David Schlesinger, Debora Botequio Moretti, Elaine Cristina Marqueze, Elaine Vieira Santos, Evandra Strazza Rodrigues, Heidge Fukumasu, Jayme Augusto de Souza-Neto, José Salvatore Leister Patané, Luiz Alcantara, Luiz Lehmann Coutinho, Maria Carolina Elias, Maurício Lacerda Nogueira, Rafael dos Santos Bezerra, Raul Machado Neto, Rejane Maria Tommasini Grotto, Ricardo Haddad, Sandra Coccuzzo Sampaio Vessoni, Simone Kashima, Svetoslav Nanev Slavov, Vincent Louis Viala |
| EPI_ISL_2209786, EPI_ISL_2209801, EPI_ISL_2209802, EPI_ISL_2209807, EPI_ISL_2209809 | UNIDADE DE PRONTO ATENDIMENTO CENTRAL DE CARAGUATATUBA | Instituto Butantan | Dimas Tadeu Covas, Antonio Jorge Martins, Claudia Renata dos Santos Barros, David Schlesinger, Debora Botequio Moretti, Elaine Cristina Marqueze, Elaine Vieira Santos, Evandra Strazza Rodrigues, Heidge Fukumasu, Jayme Augusto de Souza-Neto, José Salvatore Leister Patané, Luiz Alcantara, Luiz Lehmann Coutinho, Maria Carolina Elias, Maurício Lacerda Nogueira, Rafael dos Santos Bezerra, Raul Machado Neto, Rejane Maria Tommasini Grotto, Ricardo Haddad, Sandra Coccuzzo Sampaio Vessoni, Simone Kashima, Svetoslav Nanev Slavov, Vincent Louis Viala. |

| | | | |
|---|---|---|---|
| EPI_ISL_2209820, EPI_ISL_2209824, EPI_ISL_2209836 | UNIDADE DE PRONTO ATENDIMENTO UPA SUL CARAGUATATUBA | Instituto Butantan | Dimas Tadeu Covas, Antonio Jorge Martins, Claudia Renata dos Santos Barros, David Schlesinger, Debora Botequio Moretti, Elaine Cristina Marqueze, Elaine Vieira Santos, Evandra Strazza Rodrigues, Heidge Fukumasu, Jayme Augusto de Souza-Neto, José Salvatore Leister Patané, Luiz Alcantara, Luiz Lehmann Coutinho, Maria Carolina Elias, Maurício Lacerda Nogueira, Rafael dos Santos Bezerra, Raul Machado Neto, Rejane Maria Tommasini Grotto, Ricardo Haddad, Sandra Coccuzzo Sampaio Vessoni, Simone Kashima, Svetoslav Nanev Slavov, Vincent Louis Viala. |
| EPI_ISL_2209844 | UNIDADE BASICA DE SAUDE JARDIM DO LAGO | Instituto Butantan / Mendelics | Dimas Tadeu Covas, Antonio Jorge Martins, Claudia Renata dos Santos Barros, David Schlesinger, Debora Botequio Moretti, Elaine Cristina Marqueze, Elaine Vieira Santos, Evandra Strazza Rodrigues, Heidge Fukumasu, Jayme Augusto de Souza-Neto, José Salvatore Leister Patané, Luiz Alcantara, Luiz Lehmann Coutinho, Maria Carolina Elias, Maurício Lacerda Nogueira, Rafael dos Santos Bezerra, Raul Machado Neto, Rejane Maria Tommasini Grotto, Ricardo Haddad, Sandra Coccuzzo Sampaio Vessoni, Simone Kashima, Svetoslav Nanev Slavov, Vincent Louis Viala |
| EPI_ISL_2209850 | CS II DR SEBASTIAO RIBEIRO DO VALLE | Instituto Butantan / Mendelics | Dimas Tadeu Covas, Antonio Jorge Martins, Claudia Renata dos Santos Barros, David Schlesinger, Debora Botequio Moretti, Elaine Cristina Marqueze, Elaine Vieira Santos, Evandra Strazza Rodrigues, Heidge Fukumasu, Jayme Augusto de Souza-Neto, José Salvatore Leister Patané, Luiz Alcantara, Luiz Lehmann Coutinho, Maria Carolina Elias, Maurício Lacerda Nogueira, Rafael dos Santos Bezerra, Raul Machado Neto, Rejane Maria Tommasini Grotto, Ricardo Haddad, Sandra Coccuzzo Sampaio Vessoni, Simone Kashima, Svetoslav Nanev Slavov, Vincent Louis Viala |
| EPI_ISL_2209856 | UNIDADE DE VIGILANCIA EPIDEMIOLOGICA | Instituto Butantan | Dimas Tadeu Covas, Antonio Jorge Martins, Claudia Renata dos Santos Barros, David Schlesinger, Debora Botequio Moretti, Elaine Cristina Marqueze, Elaine Vieira Santos, Evandra Strazza Rodrigues, Heidge Fukumasu, Jayme Augusto de Souza-Neto, José Salvatore Leister Patané, Luiz Alcantara, Luiz Lehmann Coutinho, Maria Carolina Elias, Maurício Lacerda Nogueira, Rafael dos Santos Bezerra, Raul Machado Neto, Rejane Maria Tommasini Grotto, Ricardo Haddad, Sandra Coccuzzo Sampaio Vessoni, Simone Kashima, Svetoslav Nanev Slavov, Vincent Louis Viala. |
| EPI_ISL_2209864 | SECRETARIA DE SAUDE PUBLICA DE PRAIA GRANDE | Instituto Butantan / Mendelics | Dimas Tadeu Covas, Antonio Jorge Martins, Claudia Renata dos Santos Barros, David Schlesinger, Debora Botequio Moretti, Elaine Cristina Marqueze, Elaine Vieira Santos, Evandra Strazza Rodrigues, Heidge Fukumasu, Jayme Augusto de Souza-Neto, José Salvatore Leister Patané, Luiz Alcantara, Luiz Lehmann Coutinho, Maria Carolina Elias, Maurício Lacerda Nogueira, Rafael dos Santos Bezerra, Raul Machado Neto, Rejane Maria Tommasini Grotto, Ricardo Haddad, Sandra Coccuzzo Sampaio Vessoni, Simone Kashima, Svetoslav Nanev Slavov, Vincent Louis Viala |
| EPI_ISL_2209872 | SANTA CASA DE MISERICORDIA DE UBATUBA | Instituto Butantan / Mendelics | Dimas Tadeu Covas, Antonio Jorge Martins, Claudia Renata dos Santos Barros, David Schlesinger, Debora Botequio Moretti, Elaine Cristina Marqueze, Elaine Vieira Santos, Evandra Strazza Rodrigues, Heidge Fukumasu, Jayme Augusto de Souza-Neto, José Salvatore Leister Patané, Luiz Alcantara, Luiz Lehmann Coutinho, Maria Carolina Elias, Maurício Lacerda Nogueira, Rafael dos Santos Bezerra, Raul Machado Neto, Rejane Maria Tommasini Grotto, Ricardo Haddad, Sandra Coccuzzo Sampaio Vessoni, Simone Kashima, Svetoslav Nanev Slavov, Vincent Louis Viala |
| EPI_ISL_2209878 | SECRETARIA MUNICIPAL DE SAUDE SOROCABA | Instituto Butantan / Mendelics | Dimas Tadeu Covas, Antonio Jorge Martins, Claudia Renata dos Santos Barros, David Schlesinger, Debora Botequio Moretti, Elaine Cristina Marqueze, Elaine Vieira Santos, Evandra Strazza Rodrigues, Heidge Fukumasu, Jayme Augusto de Souza-Neto, José Salvatore Leister Patané, Luiz Alcantara, Luiz Lehmann Coutinho, Maria Carolina Elias, Maurício Lacerda Nogueira, Rafael dos Santos Bezerra, Raul Machado Neto, Rejane Maria Tommasini Grotto, Ricardo Haddad, Sandra Coccuzzo Sampaio Vessoni, Simone Kashima, Svetoslav Nanev Slavov, Vincent Louis Viala |
| EPI_ISL_2209883 | HOSPITAL DE CAMPANHA COVID 19 MUNICIPIO DE TAUBATE | Instituto Butantan / Mendelics | Dimas Tadeu Covas, Antonio Jorge Martins, Claudia Renata dos Santos Barros, David Schlesinger, Debora Botequio Moretti, Elaine Cristina Marqueze, Elaine Vieira Santos, Evandra Strazza Rodrigues, Heidge Fukumasu, Jayme Augusto de Souza-Neto, José Salvatore Leister Patané, Luiz Alcantara, Luiz Lehmann Coutinho, Maria Carolina Elias, Maurício Lacerda Nogueira, Rafael dos Santos Bezerra, Raul Machado Neto, Rejane Maria Tommasini Grotto, Ricardo Haddad, Sandra Coccuzzo Sampaio Vessoni, Simone Kashima, Svetoslav Nanev Slavov, Vincent Louis Viala |
| EPI_ISL_2209886 | UBS JOSE AUGUSTO BATISTA FILHO MENEGUETI | Instituto Butantan / Mendelics | Dimas Tadeu Covas, Antonio Jorge Martins, Claudia Renata dos Santos Barros, David Schlesinger, Debora Botequio Moretti, Elaine Cristina Marqueze, Elaine Vieira Santos, Evandra Strazza Rodrigues, Heidge Fukumasu, Jayme Augusto de Souza-Neto, José Salvatore Leister Patané, Luiz Alcantara, Luiz Lehmann Coutinho, Maria Carolina Elias, Maurício Lacerda Nogueira, Rafael dos Santos Bezerra, Raul Machado Neto, Rejane Maria Tommasini Grotto, Ricardo Haddad, Sandra Coccuzzo Sampaio Vessoni, Simone Kashima, Svetoslav Nanev Slavov, Vincent Louis Viala |
| EPI_ISL_2209888 | SANTA CASA DE MISERICORDIA SAO JOSE | Instituto Butantan / Mendelics | Dimas Tadeu Covas, Antonio Jorge Martins, Claudia Renata dos Santos Barros, David Schlesinger, Debora Botequio Moretti, Elaine Cristina Marqueze, Elaine Vieira Santos, Evandra Strazza Rodrigues, Heidge Fukumasu, Jayme Augusto de Souza-Neto, José Salvatore Leister Patané, Luiz Alcantara, Luiz Lehmann Coutinho, Maria Carolina Elias, Maurício Lacerda Nogueira, Rafael dos Santos Bezerra, Raul Machado Neto, Rejane Maria Tommasini Grotto, Ricardo Haddad, Sandra Coccuzzo Sampaio Vessoni, Simone Kashima, Svetoslav Nanev Slavov, Vincent Louis Viala |
| EPI_ISL_2209891 | CENTRO DE SAUDE III TABATINGA | Instituto Butantan / Mendelics | Dimas Tadeu Covas, Antonio Jorge Martins, Claudia Renata dos Santos Barros, David Schlesinger, Debora Botequio Moretti, Elaine Cristina Marqueze, Elaine Vieira Santos, Evandra Strazza Rodrigues, Heidge Fukumasu, Jayme Augusto de Souza-Neto, José Salvatore Leister Patané, Luiz Alcantara, Luiz Lehmann Coutinho, Maria Carolina Elias, Maurício Lacerda Nogueira, Rafael dos Santos Bezerra, Raul Machado Neto, Rejane Maria Tommasini Grotto, Ricardo Haddad, Sandra Coccuzzo Sampaio Vessoni, Simone Kashima, Svetoslav Nanev Slavov, Vincent Louis Viala |
| EPI_ISL_2209900 | UNIDADE MISTA DE SAUDE MOMBUCA | Instituto Butantan | Dimas Tadeu Covas, Antonio Jorge Martins, Claudia Renata dos Santos Barros, David Schlesinger, Debora Botequio Moretti, Elaine Cristina Marqueze, Elaine Vieira Santos, Evandra Strazza Rodrigues, Heidge Fukumasu, Jayme Augusto de Souza-Neto, José Salvatore Leister Patané, Luiz Alcantara, Luiz Lehmann Coutinho, Maria Carolina Elias, Maurício Lacerda Nogueira, Rafael dos Santos Bezerra, Raul Machado Neto, Rejane Maria Tommasini Grotto, Ricardo Haddad, Sandra Coccuzzo Sampaio Vessoni, Simone Kashima, Svetoslav Nanev Slavov, Vincent Louis Viala. |
| EPI_ISL_2209904 | VIGILANCIA EPIDEMIOLOGICA JARDINOPOLIS SP | Instituto Butantan / Mendelics | Dimas Tadeu Covas, Antonio Jorge Martins, Claudia Renata dos Santos Barros, David Schlesinger, Debora Botequio Moretti, Elaine Cristina Marqueze, Elaine Vieira Santos, Evandra Strazza Rodrigues, Heidge Fukumasu, Jayme Augusto de Souza-Neto, José Salvatore Leister Patané, Luiz Alcantara, Luiz Lehmann Coutinho, Maria Carolina Elias, Maurício Lacerda Nogueira, Rafael dos Santos Bezerra, Raul Machado Neto, Rejane Maria Tommasini Grotto, Ricardo Haddad, Sandra Coccuzzo Sampaio Vessoni, Simone Kashima, Svetoslav Nanev Slavov, Vincent Louis Viala |
| EPI_ISL_2209905 | SECRETARIA DE SAUDE PUBLICA DE PRAIA GRANDE | Instituto Butantan / Mendelics | Dimas Tadeu Covas, Antonio Jorge Martins, Claudia Renata dos Santos Barros, David Schlesinger, Debora Botequio Moretti, Elaine Cristina Marqueze, Elaine Vieira Santos, Evandra Strazza Rodrigues, Heidge Fukumasu, Jayme Augusto de Souza-Neto, José Salvatore Leister Patané, Luiz Alcantara, Luiz Lehmann Coutinho, Maria Carolina Elias, Maurício Lacerda Nogueira, Rafael dos Santos Bezerra, Raul Machado Neto, Rejane Maria Tommasini Grotto, Ricardo Haddad, Sandra Coccuzzo Sampaio Vessoni, Simone Kashima, Svetoslav Nanev Slavov, Vincent Louis Viala |
| EPI_ISL_2209908 | UNIDADE MISTA OSVALDO P CAMARGO ELIAS FAUSTO | Instituto Butantan | Dimas Tadeu Covas, Antonio Jorge Martins, Claudia Renata dos Santos Barros, David Schlesinger, Debora Botequio Moretti, Elaine Cristina Marqueze, Elaine Vieira Santos, Evandra Strazza Rodrigues, Heidge Fukumasu, Jayme Augusto de Souza-Neto, José Salvatore Leister Patané, Luiz Alcantara, Luiz Lehmann Coutinho, Maria Carolina Elias, Maurício Lacerda Nogueira, Rafael dos Santos Bezerra, Raul Machado Neto, Rejane Maria Tommasini Grotto, Ricardo Haddad, Sandra Coccuzzo Sampaio Vessoni, Simone Kashima, Svetoslav Nanev Slavov, Vincent Louis Viala. |
| EPI_ISL_2209916 | UNIDADE RESPIRATORIA NOVA HORTOLANDIA | Instituto Butantan | Dimas Tadeu Covas, Antonio Jorge Martins, Claudia Renata dos Santos Barros, David Schlesinger, Debora Botequio Moretti, Elaine Cristina Marqueze, Elaine Vieira Santos, Evandra Strazza Rodrigues, Heidge Fukumasu, Jayme Augusto de Souza-Neto, José Salvatore Leister Patané, Luiz Alcantara, Luiz Lehmann Coutinho, Maria Carolina Elias, Maurício Lacerda Nogueira, Rafael dos Santos Bezerra, Raul Machado Neto, Rejane Maria Tommasini Grotto, Ricardo Haddad, Sandra Coccuzzo Sampaio Vessoni, Simone Kashima, Svetoslav Nanev Slavov, Vincent Louis Viala. |
| EPI_ISL_2209927 | CENTRO DE SAUDE III TABATINGA | Instituto Butantan / Mendelics | Dimas Tadeu Covas, Antonio Jorge Martins, Claudia Renata dos Santos Barros, David Schlesinger, Debora Botequio Moretti, Elaine Cristina Marqueze, Elaine Vieira Santos, Evandra Strazza Rodrigues, Heidge Fukumasu, Jayme Augusto de Souza-Neto, José Salvatore Leister Patané, Luiz Alcantara, Luiz Lehmann Coutinho, Maria Carolina Elias, Maurício Lacerda Nogueira, Rafael dos Santos Bezerra, Raul Machado Neto, Rejane Maria Tommasini Grotto, Ricardo Haddad, Sandra Coccuzzo Sampaio Vessoni, Simone Kashima, Svetoslav Nanev Slavov, Vincent Louis Viala |
| EPI_ISL_2209929 | CS ALVINLANDIA | Instituto Butantan / Mendelics | Dimas Tadeu Covas, Antonio Jorge Martins, Claudia Renata dos Santos Barros, David Schlesinger, Debora Botequio Moretti, Elaine Cristina Marqueze, Elaine Vieira Santos, Evandra Strazza Rodrigues, Heidge Fukumasu, Jayme Augusto de Souza-Neto, José Salvatore Leister Patané, Luiz Alcantara, Luiz Lehmann Coutinho, Maria Carolina Elias, Maurício Lacerda Nogueira, Rafael dos Santos Bezerra, Raul Machado Neto, Rejane Maria Tommasini Grotto, Ricardo Haddad, Sandra Coccuzzo Sampaio Vessoni, Simone Kashima, Svetoslav Nanev Slavov, Vincent Louis Viala |
| EPI_ISL_2209934 | UNIDADE SENTINELA COVID19 | Instituto Butantan | Dimas Tadeu Covas, Antonio Jorge Martins, Claudia Renata dos Santos Barros, David Schlesinger, Debora Botequio Moretti, Elaine Cristina Marqueze, Elaine Vieira Santos, Evandra Strazza Rodrigues, Heidge Fukumasu, Jayme Augusto de Souza-Neto, José Salvatore Leister Patané, Luiz Alcantara, Luiz Lehmann Coutinho, Maria Carolina Elias, Maurício Lacerda Nogueira, Rafael dos Santos Bezerra, Raul Machado Neto, Rejane Maria Tommasini Grotto, Ricardo Haddad, Sandra Coccuzzo Sampaio Vessoni, Simone Kashima, Svetoslav Nanev Slavov, Vincent Louis Viala. |
| EPI_ISL_2209937, EPI_ISL_2209947 | SECRETARIA DE SAUDE DE SANTA GERTRUDES | Instituto Butantan / FZEA-USP-Pirassununga | Dimas Tadeu Covas, Antonio Jorge Martins, Claudia Renata dos Santos Barros, David Schlesinger, Debora Botequio Moretti, Elaine Cristina Marqueze, Elaine Vieira Santos, Evandra Strazza Rodrigues, Heidge Fukumasu, Jayme Augusto de Souza-Neto, José Salvatore Leister Patané, Luiz Alcantara, Luiz |

| | | | |
|---|---|---|---|
| | | | Lehmann Coutinho, Maria Carolina Elias, Maurício Lacerda Nogueira, Rafael dos Santos Bezerra, Raul Machado Neto, Rejane Maria Tommasini Grotto, Ricardo Haddad, Sandra Coccuzzo Sampaio Vessoni, Simone Kashima, Svetoslav Nanev Slavov, Vincent Louis Viala |
| EPI_ISL_2209955 | SECRETARIA DE SAUDE DE SAO PEDRO | Instituto Butantan / FZEA-USP-Pirassununga | Dimas Tadeu Covas, Antonio Jorge Martins, Claudia Renata dos Santos Barros, David Schlesinger, Debora Botequio Moretti, Elaine Cristina Marqueze, Elaine Vieira Santos, Evandra Strazza Rodrigues, Heidge Fukumasu, Jayme Augusto de Souza-Neto, José Salvatore Leister Patané, Luiz Alcantara, Luiz Lehmann Coutinho, Maria Carolina Elias, Maurício Lacerda Nogueira, Rafael dos Santos Bezerra, Raul Machado Neto, Rejane Maria Tommasini Grotto, Ricardo Haddad, Sandra Coccuzzo Sampaio Vessoni, Simone Kashima, Svetoslav Nanev Slavov, Vincent Louis Viala |
| EPI_ISL_2209961 | VIGILANCIA EM SAUDE SAE VIGILANCIA EPIDEMIOLOGICA | Instituto Butantan / Mendelics | Dimas Tadeu Covas, Antonio Jorge Martins, Claudia Renata dos Santos Barros, David Schlesinger, Debora Botequio Moretti, Elaine Cristina Marqueze, Elaine Vieira Santos, Evandra Strazza Rodrigues, Heidge Fukumasu, Jayme Augusto de Souza-Neto, José Salvatore Leister Patané, Luiz Alcantara, Luiz Lehmann Coutinho, Maria Carolina Elias, Maurício Lacerda Nogueira, Rafael dos Santos Bezerra, Raul Machado Neto, Rejane Maria Tommasini Grotto, Ricardo Haddad, Sandra Coccuzzo Sampaio Vessoni, Simone Kashima, Svetoslav Nanev Slavov, Vincent Louis Viala |
| EPI_ISL_2209968 | CENTRO DE SAUDE III SALES OLIVEIRA | Instituto Butantan / Mendelics | Dimas Tadeu Covas, Antonio Jorge Martins, Claudia Renata dos Santos Barros, David Schlesinger, Debora Botequio Moretti, Elaine Cristina Marqueze, Elaine Vieira Santos, Evandra Strazza Rodrigues, Heidge Fukumasu, Jayme Augusto de Souza-Neto, José Salvatore Leister Patané, Luiz Alcantara, Luiz Lehmann Coutinho, Maria Carolina Elias, Maurício Lacerda Nogueira, Rafael dos Santos Bezerra, Raul Machado Neto, Rejane Maria Tommasini Grotto, Ricardo Haddad, Sandra Coccuzzo Sampaio Vessoni, Simone Kashima, Svetoslav Nanev Slavov, Vincent Louis Viala |
| EPI_ISL_2209975 | HOSPITAL SAO JOSE ITIRAPINA | Instituto Butantan / FZEA-USP-Pirassununga | Dimas Tadeu Covas, Antonio Jorge Martins, Claudia Renata dos Santos Barros, David Schlesinger, Debora Botequio Moretti, Elaine Cristina Marqueze, Elaine Vieira Santos, Evandra Strazza Rodrigues, Heidge Fukumasu, Jayme Augusto de Souza-Neto, José Salvatore Leister Patané, Luiz Alcantara, Luiz Lehmann Coutinho, Maria Carolina Elias, Maurício Lacerda Nogueira, Rafael dos Santos Bezerra, Raul Machado Neto, Rejane Maria Tommasini Grotto, Ricardo Haddad, Sandra Coccuzzo Sampaio Vessoni, Simone Kashima, Svetoslav Nanev Slavov, Vincent Louis Viala |
| EPI_ISL_2209982 | CENTRO MUNICIPAL DE EPIDEMIOLOGIA E IMUNIZACOES LIMEIRA | Instituto Butantan / FZEA-USP-Pirassununga | Dimas Tadeu Covas, Antonio Jorge Martins, Claudia Renata dos Santos Barros, David Schlesinger, Debora Botequio Moretti, Elaine Cristina Marqueze, Elaine Vieira Santos, Evandra Strazza Rodrigues, Heidge Fukumasu, Jayme Augusto de Souza-Neto, José Salvatore Leister Patané, Luiz Alcantara, Luiz Lehmann Coutinho, Maria Carolina Elias, Maurício Lacerda Nogueira, Rafael dos Santos Bezerra, Raul Machado Neto, Rejane Maria Tommasini Grotto, Ricardo Haddad, Sandra Coccuzzo Sampaio Vessoni, Simone Kashima, Svetoslav Nanev Slavov, Vincent Louis Viala |
| EPI_ISL_2209990 | UNIDADE MISTA DE SAUDE MOMBUCA | Instituto Butantan / FZEA-USP-Pirassununga | Dimas Tadeu Covas, Antonio Jorge Martins, Claudia Renata dos Santos Barros, David Schlesinger, Debora Botequio Moretti, Elaine Cristina Marqueze, Elaine Vieira Santos, Evandra Strazza Rodrigues, Heidge Fukumasu, Jayme Augusto de Souza-Neto, José Salvatore Leister Patané, Luiz Alcantara, Luiz Lehmann Coutinho, Maria Carolina Elias, Maurício Lacerda Nogueira, Rafael dos Santos Bezerra, Raul Machado Neto, Rejane Maria Tommasini Grotto, Ricardo Haddad, Sandra Coccuzzo Sampaio Vessoni, Simone Kashima, Svetoslav Nanev Slavov, Vincent Louis Viala |
| EPI_ISL_2209996 | UNIDADE BASICA DE SAUDE DR MATHEUS GABRIEL BONASSA | Instituto Butantan / FZEA-USP-Pirassununga | Dimas Tadeu Covas, Antonio Jorge Martins, Claudia Renata dos Santos Barros, David Schlesinger, Debora Botequio Moretti, Elaine Cristina Marqueze, Elaine Vieira Santos, Evandra Strazza Rodrigues, Heidge Fukumasu, Jayme Augusto de Souza-Neto, José Salvatore Leister Patané, Luiz Alcantara, Luiz Lehmann Coutinho, Maria Carolina Elias, Maurício Lacerda Nogueira, Rafael dos Santos Bezerra, Raul Machado Neto, Rejane Maria Tommasini Grotto, Ricardo Haddad, Sandra Coccuzzo Sampaio Vessoni, Simone Kashima, Svetoslav Nanev Slavov, Vincent Louis Viala |
| EPI_ISL_2210003 | UPA VILA SANTA CATARINA | Instituto Butantan | Dimas Tadeu Covas, Antonio Jorge Martins, Claudia Renata dos Santos Barros, David Schlesinger, Debora Botequio Moretti, Elaine Cristina Marqueze, Elaine Vieira Santos, Evandra Strazza Rodrigues, Heidge Fukumasu, Jayme Augusto de Souza-Neto, José Salvatore Leister Patané, Luiz Alcantara, Luiz Lehmann Coutinho, Maria Carolina Elias, Maurício Lacerda Nogueira, Rafael dos Santos Bezerra, Raul Machado Neto, Rejane Maria Tommasini Grotto, Ricardo Haddad, Sandra Coccuzzo Sampaio Vessoni, Simone Kashima, Svetoslav Nanev Slavov, Vincent Louis Viala. |
| EPI_ISL_2210010 | USF PAULISTA FERNANDOPOLIS ANTONIO PIVATO | Instituto Butantan | Dimas Tadeu Covas, Antonio Jorge Martins, Claudia Renata dos Santos Barros, David Schlesinger, Debora Botequio Moretti, Elaine Cristina Marqueze, Elaine Vieira Santos, Evandra Strazza Rodrigues, Heidge Fukumasu, Jayme Augusto de Souza-Neto, José Salvatore Leister Patané, Luiz Alcantara, Luiz Lehmann Coutinho, Maria Carolina Elias, Maurício Lacerda Nogueira, Rafael dos Santos Bezerra, Raul Machado Neto, Rejane Maria Tommasini Grotto, Ricardo Haddad, Sandra Coccuzzo Sampaio Vessoni, Simone Kashima, Svetoslav Nanev Slavov, Vincent Louis Viala. |
| EPI_ISL_2210019, EPI_ISL_2210030 | VIGILANCIA EPIDEMIOLOGICA E CONTROLE DE VETORES PIRASSUNUN | Instituto Butantan / FZEA-USP-Pirassununga | Dimas Tadeu Covas, Antonio Jorge Martins, Claudia Renata dos Santos Barros, David Schlesinger, Debora Botequio Moretti, Elaine Cristina Marqueze, Elaine Vieira Santos, Evandra Strazza Rodrigues, Heidge Fukumasu, Jayme Augusto de Souza-Neto, José Salvatore Leister Patané, Luiz Alcantara, Luiz Lehmann Coutinho, Maria Carolina Elias, Maurício Lacerda Nogueira, Rafael dos Santos Bezerra, Raul Machado Neto, Rejane Maria Tommasini Grotto, Ricardo Haddad, Sandra Coccuzzo Sampaio Vessoni, Simone Kashima, Svetoslav Nanev Slavov, Vincent Louis Viala |
| EPI_ISL_2210037 | SECRETARIA MUNICIPAL DE SAUDE DE CORDEIROPOLIS | Instituto Butantan / FZEA-USP-Pirassununga | Dimas Tadeu Covas, Antonio Jorge Martins, Claudia Renata dos Santos Barros, David Schlesinger, Debora Botequio Moretti, Elaine Cristina Marqueze, Elaine Vieira Santos, Evandra Strazza Rodrigues, Heidge Fukumasu, Jayme Augusto de Souza-Neto, José Salvatore Leister Patané, Luiz Alcantara, Luiz Lehmann Coutinho, Maria Carolina Elias, Maurício Lacerda Nogueira, Rafael dos Santos Bezerra, Raul Machado Neto, Rejane Maria Tommasini Grotto, Ricardo Haddad, Sandra Coccuzzo Sampaio Vessoni, Simone Kashima, Svetoslav Nanev Slavov, Vincent Louis Viala |
| EPI_ISL_2210045, EPI_ISL_2210048 | VIGILANCIA EPIDEMIOLOGICA DE LEME | Instituto Butantan / FZEA-USP-Pirassununga | Dimas Tadeu Covas, Antonio Jorge Martins, Claudia Renata dos Santos Barros, David Schlesinger, Debora Botequio Moretti, Elaine Cristina Marqueze, Elaine Vieira Santos, Evandra Strazza Rodrigues, Heidge Fukumasu, Jayme Augusto de Souza-Neto, José Salvatore Leister Patané, Luiz Alcantara, Luiz Lehmann Coutinho, Maria Carolina Elias, Maurício Lacerda Nogueira, Rafael dos Santos Bezerra, Raul Machado Neto, Rejane Maria Tommasini Grotto, Ricardo Haddad, Sandra Coccuzzo Sampaio Vessoni, Simone Kashima, Svetoslav Nanev Slavov, Vincent Louis Viala |
| EPI_ISL_2210056, EPI_ISL_2210063 | UNIDADE BASICA DE SAUDE DE IPEUNA | Instituto Butantan / FZEA-USP-Pirassununga | Dimas Tadeu Covas, Antonio Jorge Martins, Claudia Renata dos Santos Barros, David Schlesinger, Debora Botequio Moretti, Elaine Cristina Marqueze, Elaine Vieira Santos, Evandra Strazza Rodrigues, Heidge Fukumasu, Jayme Augusto de Souza-Neto, José Salvatore Leister Patané, Luiz Alcantara, Luiz Lehmann Coutinho, Maria Carolina Elias, Maurício Lacerda Nogueira, Rafael dos Santos Bezerra, Raul Machado Neto, Rejane Maria Tommasini Grotto, Ricardo Haddad, Sandra Coccuzzo Sampaio Vessoni, Simone Kashima, Svetoslav Nanev Slavov, Vincent Louis Viala |
| EPI_ISL_2210082 | SECRETARIA MUNICIPAL DE SAUDE DE CORDEIROPOLIS | Instituto Butantan / FZEA-USP-Pirassununga | Dimas Tadeu Covas, Antonio Jorge Martins, Claudia Renata dos Santos Barros, David Schlesinger, Debora Botequio Moretti, Elaine Cristina Marqueze, Elaine Vieira Santos, Evandra Strazza Rodrigues, Heidge Fukumasu, Jayme Augusto de Souza-Neto, José Salvatore Leister Patané, Luiz Alcantara, Luiz Lehmann Coutinho, Maria Carolina Elias, Maurício Lacerda Nogueira, Rafael dos Santos Bezerra, Raul Machado Neto, Rejane Maria Tommasini Grotto, Ricardo Haddad, Sandra Coccuzzo Sampaio Vessoni, Simone Kashima, Svetoslav Nanev Slavov, Vincent Louis Viala |
| EPI_ISL_2210083 | VIGILANCIA EM SAUDE DE FCO MORATO | Instituto Butantan | Dimas Tadeu Covas, Antonio Jorge Martins, Claudia Renata dos Santos Barros, David Schlesinger, Debora Botequio Moretti, Elaine Cristina Marqueze, Elaine Vieira Santos, Evandra Strazza Rodrigues, Heidge Fukumasu, Jayme Augusto de Souza-Neto, José Salvatore Leister Patané, Luiz Alcantara, Luiz Lehmann Coutinho, Maria Carolina Elias, Maurício Lacerda Nogueira, Rafael dos Santos Bezerra, Raul Machado Neto, Rejane Maria Tommasini Grotto, Ricardo Haddad, Sandra Coccuzzo Sampaio Vessoni, Simone Kashima, Svetoslav Nanev Slavov, Vincent Louis Viala. |
| EPI_ISL_2210088 | LABORATORIO MUNICIPAL DE ANALISES CLINICAS DE RIO CLARO | Instituto Butantan / FZEA-USP-Pirassununga | Dimas Tadeu Covas, Antonio Jorge Martins, Claudia Renata dos Santos Barros, David Schlesinger, Debora Botequio Moretti, Elaine Cristina Marqueze, Elaine Vieira Santos, Evandra Strazza Rodrigues, Heidge Fukumasu, Jayme Augusto de Souza-Neto, José Salvatore Leister Patané, Luiz Alcantara, Luiz Lehmann Coutinho, Maria Carolina Elias, Maurício Lacerda Nogueira, Rafael dos Santos Bezerra, Raul Machado Neto, Rejane Maria Tommasini Grotto, Ricardo Haddad, Sandra Coccuzzo Sampaio Vessoni, Simone Kashima, Svetoslav Nanev Slavov, Vincent Louis Viala |
| EPI_ISL_2210091 | HOSPITAL SAO JOSE ITIRAPINA | Instituto Butantan / FZEA-USP-Pirassununga | Dimas Tadeu Covas, Antonio Jorge Martins, Claudia Renata dos Santos Barros, David Schlesinger, Debora Botequio Moretti, Elaine Cristina Marqueze, Elaine Vieira Santos, Evandra Strazza Rodrigues, Heidge Fukumasu, Jayme Augusto de Souza-Neto, José Salvatore Leister Patané, Luiz Alcantara, Luiz Lehmann Coutinho, Maria Carolina Elias, Maurício Lacerda Nogueira, Rafael dos Santos Bezerra, Raul Machado Neto, Rejane Maria Tommasini Grotto, Ricardo Haddad, Sandra Coccuzzo Sampaio Vessoni, Simone Kashima, Svetoslav Nanev Slavov, Vincent Louis Viala |
| EPI_ISL_2210101 | HOSPITAL DE CAMPANHA COVID 19 MUNICIPIO DE TAUBATE | Instituto Butantan / Mendelics | Dimas Tadeu Covas, Antonio Jorge Martins, Claudia Renata dos Santos Barros, David Schlesinger, Debora Botequio Moretti, Elaine Cristina Marqueze, Elaine Vieira Santos, Evandra Strazza Rodrigues, Heidge Fukumasu, Jayme Augusto de Souza-Neto, José Salvatore Leister Patané, Luiz Alcantara, Luiz Lehmann Coutinho, Maria Carolina Elias, Maurício Lacerda Nogueira, Rafael dos Santos Bezerra, Raul Machado Neto, Rejane Maria Tommasini Grotto, Ricardo Haddad, Sandra Coccuzzo Sampaio Vessoni, Simone Kashima, Svetoslav Nanev Slavov, Vincent Louis Viala |
| EPI_ISL_2210103 | VIGILANCIA EPIDEMIOLOGICA DE LEME | Instituto Butantan | Dimas Tadeu Covas, Antonio Jorge Martins, Claudia Renata dos Santos Barros, David Schlesinger, Debora Botequio Moretti, Elaine Cristina Marqueze, Elaine Vieira Santos, Evandra Strazza Rodrigues, Heidge Fukumasu, Jayme Augusto de Souza-Neto, José Salvatore Leister Patané, Luiz Alcantara, Luiz Lehmann Coutinho, Maria Carolina Elias, Maurício Lacerda Nogueira, Rafael dos Santos Bezerra, Raul Machado Neto, Rejane Maria Tommasini Grotto, Ricardo Haddad, Sandra Coccuzzo Sampaio Vessoni, Simone Kashima, Svetoslav Nanev Slavov, Vincent Louis Viala. |

| | | | |
|---|---|---|---|
| EPI_ISL_2210110 | VIGILANCIA EM SAUDE SAE VIGILANCIA EPIDEMIOLOGICA | Instituto Butantan / Mendelics | Dimas Tadeu Covas, Antonio Jorge Martins, Claudia Renata dos Santos Barros, David Schlesinger, Debora Botequio Moretti, Elaine Cristina Marqueze, Elaine Vieira Santos, Evandra Strazza Rodrigues, Heidge Fukumasu, Jayme Augusto de Souza-Neto, José Salvatore Leister Patané, Luiz Alcantara, Luiz Lehmann Coutinho, Maria Carolina Elias, Maurício Lacerda Nogueira, Rafael dos Santos Bezerra, Raul Machado Neto, Rejane Maria Tommasini Grotto, Ricardo Haddad, Sandra Coccuzzo Sampaio Vessoni, Simone Kashima, Svetoslav Nanev Slavov, Vincent Louis Viala |
| EPI_ISL_2210119 | CENTRO DE SAUDE III SALES OLIVEIRA | Instituto Butantan / Mendelics | Dimas Tadeu Covas, Antonio Jorge Martins, Claudia Renata dos Santos Barros, David Schlesinger, Debora Botequio Moretti, Elaine Cristina Marqueze, Elaine Vieira Santos, Evandra Strazza Rodrigues, Heidge Fukumasu, Jayme Augusto de Souza-Neto, José Salvatore Leister Patané, Luiz Alcantara, Luiz Lehmann Coutinho, Maria Carolina Elias, Maurício Lacerda Nogueira, Rafael dos Santos Bezerra, Raul Machado Neto, Rejane Maria Tommasini Grotto, Ricardo Haddad, Sandra Coccuzzo Sampaio Vessoni, Simone Kashima, Svetoslav Nanev Slavov, Vincent Louis Viala |
| EPI_ISL_2210126 | UNIDADE BASICA DE SAUDE II ESTRELA DO NORTE | Instituto Butantan / Mendelics | Dimas Tadeu Covas, Antonio Jorge Martins, Claudia Renata dos Santos Barros, David Schlesinger, Debora Botequio Moretti, Elaine Cristina Marqueze, Elaine Vieira Santos, Evandra Strazza Rodrigues, Heidge Fukumasu, Jayme Augusto de Souza-Neto, José Salvatore Leister Patané, Luiz Alcantara, Luiz Lehmann Coutinho, Maria Carolina Elias, Maurício Lacerda Nogueira, Rafael dos Santos Bezerra, Raul Machado Neto, Rejane Maria Tommasini Grotto, Ricardo Haddad, Sandra Coccuzzo Sampaio Vessoni, Simone Kashima, Svetoslav Nanev Slavov, Vincent Louis Viala |
| EPI_ISL_2210133 | SANTA CASA DE MISERICORDIA DE UBATUBA | Instituto Butantan / Mendelics | Dimas Tadeu Covas, Antonio Jorge Martins, Claudia Renata dos Santos Barros, David Schlesinger, Debora Botequio Moretti, Elaine Cristina Marqueze, Elaine Vieira Santos, Evandra Strazza Rodrigues, Heidge Fukumasu, Jayme Augusto de Souza-Neto, José Salvatore Leister Patané, Luiz Alcantara, Luiz Lehmann Coutinho, Maria Carolina Elias, Maurício Lacerda Nogueira, Rafael dos Santos Bezerra, Raul Machado Neto, Rejane Maria Tommasini Grotto, Ricardo Haddad, Sandra Coccuzzo Sampaio Vessoni, Simone Kashima, Svetoslav Nanev Slavov, Vincent Louis Viala |
| EPI_ISL_2210141 | UNIDADE BASICA DE SAUDE JARDIM DO LAGO | Instituto Butantan / Mendelics | Dimas Tadeu Covas, Antonio Jorge Martins, Claudia Renata dos Santos Barros, David Schlesinger, Debora Botequio Moretti, Elaine Cristina Marqueze, Elaine Vieira Santos, Evandra Strazza Rodrigues, Heidge Fukumasu, Jayme Augusto de Souza-Neto, José Salvatore Leister Patané, Luiz Alcantara, Luiz Lehmann Coutinho, Maria Carolina Elias, Maurício Lacerda Nogueira, Rafael dos Santos Bezerra, Raul Machado Neto, Rejane Maria Tommasini Grotto, Ricardo Haddad, Sandra Coccuzzo Sampaio Vessoni, Simone Kashima, Svetoslav Nanev Slavov, Vincent Louis Viala |
| EPI_ISL_2210148 | UBS II DR JOSE CARLOS MACUCO JANINI | Instituto Butantan / Mendelics | Dimas Tadeu Covas, Antonio Jorge Martins, Claudia Renata dos Santos Barros, David Schlesinger, Debora Botequio Moretti, Elaine Cristina Marqueze, Elaine Vieira Santos, Evandra Strazza Rodrigues, Heidge Fukumasu, Jayme Augusto de Souza-Neto, José Salvatore Leister Patané, Luiz Alcantara, Luiz Lehmann Coutinho, Maria Carolina Elias, Maurício Lacerda Nogueira, Rafael dos Santos Bezerra, Raul Machado Neto, Rejane Maria Tommasini Grotto, Ricardo Haddad, Sandra Coccuzzo Sampaio Vessoni, Simone Kashima, Svetoslav Nanev Slavov, Vincent Louis Viala |
| EPI_ISL_2210162 | VIGILANCIA EM SAUDE DE FCO MORATO | Instituto Butantan | Dimas Tadeu Covas, Antonio Jorge Martins, Claudia Renata dos Santos Barros, David Schlesinger, Debora Botequio Moretti, Elaine Cristina Marqueze, Elaine Vieira Santos, Evandra Strazza Rodrigues, Heidge Fukumasu, Jayme Augusto de Souza-Neto, José Salvatore Leister Patané, Luiz Alcantara, Luiz Lehmann Coutinho, Maria Carolina Elias, Maurício Lacerda Nogueira, Rafael dos Santos Bezerra, Raul Machado Neto, Rejane Maria Tommasini Grotto, Ricardo Haddad, Sandra Coccuzzo Sampaio Vessoni, Simone Kashima, Svetoslav Nanev Slavov, Vincent Louis Viala |
| EPI_ISL_2210165 | VIGILANCIA EPIDEMIOLOGICA E CONTROLE DE VETORES PIRASSUNUN | Instituto Butantan | Dimas Tadeu Covas, Antonio Jorge Martins, Claudia Renata dos Santos Barros, David Schlesinger, Debora Botequio Moretti, Elaine Cristina Marqueze, Elaine Vieira Santos, Evandra Strazza Rodrigues, Heidge Fukumasu, Jayme Augusto de Souza-Neto, José Salvatore Leister Patané, Luiz Alcantara, Luiz Lehmann Coutinho, Maria Carolina Elias, Maurício Lacerda Nogueira, Rafael dos Santos Bezerra, Raul Machado Neto, Rejane Maria Tommasini Grotto, Ricardo Haddad, Sandra Coccuzzo Sampaio Vessoni, Simone Kashima, Svetoslav Nanev Slavov, Vincent Louis Viala. |
| EPI_ISL_2210173 | SECRETARIA DE SAUDE PUBLICA DE PRAIA GRANDE | Instituto Butantan / Mendelics | Dimas Tadeu Covas, Antonio Jorge Martins, Claudia Renata dos Santos Barros, David Schlesinger, Debora Botequio Moretti, Elaine Cristina Marqueze, Elaine Vieira Santos, Evandra Strazza Rodrigues, Heidge Fukumasu, Jayme Augusto de Souza-Neto, José Salvatore Leister Patané, Luiz Alcantara, Luiz Lehmann Coutinho, Maria Carolina Elias, Maurício Lacerda Nogueira, Rafael dos Santos Bezerra, Raul Machado Neto, Rejane Maria Tommasini Grotto, Ricardo Haddad, Sandra Coccuzzo Sampaio Vessoni, Simone Kashima, Svetoslav Nanev Slavov, Vincent Louis Viala |
| EPI_ISL_2210183 | SECRETARIA MUNICIPAL DE SAUDE SOROCABA | Instituto Butantan / Mendelics | Dimas Tadeu Covas, Antonio Jorge Martins, Claudia Renata dos Santos Barros, David Schlesinger, Debora Botequio Moretti, Elaine Cristina Marqueze, Elaine Vieira Santos, Evandra Strazza Rodrigues, Heidge Fukumasu, Jayme Augusto de Souza-Neto, José Salvatore Leister Patané, Luiz Alcantara, Luiz Lehmann Coutinho, Maria Carolina Elias, Maurício Lacerda Nogueira, Rafael dos Santos Bezerra, Raul Machado Neto, Rejane Maria Tommasini Grotto, Ricardo Haddad, Sandra Coccuzzo Sampaio Vessoni, Simone Kashima, Svetoslav Nanev Slavov, Vincent Louis Viala |
| EPI_ISL_2210186 | UBS DR WALDOMIRO FERREIRA NEVES SCRPARDO | Instituto Butantan / Mendelics | Dimas Tadeu Covas, Antonio Jorge Martins, Claudia Renata dos Santos Barros, David Schlesinger, Debora Botequio Moretti, Elaine Cristina Marqueze, Elaine Vieira Santos, Evandra Strazza Rodrigues, Heidge Fukumasu, Jayme Augusto de Souza-Neto, José Salvatore Leister Patané, Luiz Alcantara, Luiz Lehmann Coutinho, Maria Carolina Elias, Maurício Lacerda Nogueira, Rafael dos Santos Bezerra, Raul Machado Neto, Rejane Maria Tommasini Grotto, Ricardo Haddad, Sandra Coccuzzo Sampaio Vessoni, Simone Kashima, Svetoslav Nanev Slavov, Vincent Louis Viala |
| EPI_ISL_2210189 | VIGILANCIA EPIDEMIOLOGICA E CONTROLE DE VETORES PIRASSUNUN | Instituto Butantan | Dimas Tadeu Covas, Antonio Jorge Martins, Claudia Renata dos Santos Barros, David Schlesinger, Debora Botequio Moretti, Elaine Cristina Marqueze, Elaine Vieira Santos, Evandra Strazza Rodrigues, Heidge Fukumasu, Jayme Augusto de Souza-Neto, José Salvatore Leister Patané, Luiz Alcantara, Luiz Lehmann Coutinho, Maria Carolina Elias, Maurício Lacerda Nogueira, Rafael dos Santos Bezerra, Raul Machado Neto, Rejane Maria Tommasini Grotto, Ricardo Haddad, Sandra Coccuzzo Sampaio Vessoni, Simone Kashima, Svetoslav Nanev Slavov, Vincent Louis Viala. |
| EPI_ISL_2210198 | SECRETARIA MUNICIPAL DE SAUDE DE TUPI PAULISTA | Instituto Butantan / Mendelics | Dimas Tadeu Covas, Antonio Jorge Martins, Claudia Renata dos Santos Barros, David Schlesinger, Debora Botequio Moretti, Elaine Cristina Marqueze, Elaine Vieira Santos, Evandra Strazza Rodrigues, Heidge Fukumasu, Jayme Augusto de Souza-Neto, José Salvatore Leister Patané, Luiz Alcantara, Luiz Lehmann Coutinho, Maria Carolina Elias, Maurício Lacerda Nogueira, Rafael dos Santos Bezerra, Raul Machado Neto, Rejane Maria Tommasini Grotto, Ricardo Haddad, Sandra Coccuzzo Sampaio Vessoni, Simone Kashima, Svetoslav Nanev Slavov, Vincent Louis Viala |
| EPI_ISL_2210207 | UBS RAIMUNDA CAVALCANTE DE SOUZA | Instituto Butantan / Mendelics | Dimas Tadeu Covas, Antonio Jorge Martins, Claudia Renata dos Santos Barros, David Schlesinger, Debora Botequio Moretti, Elaine Cristina Marqueze, Elaine Vieira Santos, Evandra Strazza Rodrigues, Heidge Fukumasu, Jayme Augusto de Souza-Neto, José Salvatore Leister Patané, Luiz Alcantara, Luiz Lehmann Coutinho, Maria Carolina Elias, Maurício Lacerda Nogueira, Rafael dos Santos Bezerra, Raul Machado Neto, Rejane Maria Tommasini Grotto, Ricardo Haddad, Sandra Coccuzzo Sampaio Vessoni, Simone Kashima, Svetoslav Nanev Slavov, Vincent Louis Viala |
| EPI_ISL_2210219 | CENTRO DE ESPECIALIDADES DE PRIMAVERA | Instituto Butantan / Mendelics | Dimas Tadeu Covas, Antonio Jorge Martins, Claudia Renata dos Santos Barros, David Schlesinger, Debora Botequio Moretti, Elaine Cristina Marqueze, Elaine Vieira Santos, Evandra Strazza Rodrigues, Heidge Fukumasu, Jayme Augusto de Souza-Neto, José Salvatore Leister Patané, Luiz Alcantara, Luiz Lehmann Coutinho, Maria Carolina Elias, Maurício Lacerda Nogueira, Rafael dos Santos Bezerra, Raul Machado Neto, Rejane Maria Tommasini Grotto, Ricardo Haddad, Sandra Coccuzzo Sampaio Vessoni, Simone Kashima, Svetoslav Nanev Slavov, Vincent Louis Viala |
| EPI_ISL_2210223 | SMS SECRETARIA MUNICIPAL DE SAUDE DE BOITUVA | Instituto Butantan / Mendelics | Dimas Tadeu Covas, Antonio Jorge Martins, Claudia Renata dos Santos Barros, David Schlesinger, Debora Botequio Moretti, Elaine Cristina Marqueze, Elaine Vieira Santos, Evandra Strazza Rodrigues, Heidge Fukumasu, Jayme Augusto de Souza-Neto, José Salvatore Leister Patané, Luiz Alcantara, Luiz Lehmann Coutinho, Maria Carolina Elias, Maurício Lacerda Nogueira, Rafael dos Santos Bezerra, Raul Machado Neto, Rejane Maria Tommasini Grotto, Ricardo Haddad, Sandra Coccuzzo Sampaio Vessoni, Simone Kashima, Svetoslav Nanev Slavov, Vincent Louis Viala |
| EPI_ISL_2210225 | UBS JOSE AUGUSTO BATISTA FILHO MENEGUETI | Instituto Butantan / Mendelics | Dimas Tadeu Covas, Antonio Jorge Martins, Claudia Renata dos Santos Barros, David Schlesinger, Debora Botequio Moretti, Elaine Cristina Marqueze, Elaine Vieira Santos, Evandra Strazza Rodrigues, Heidge Fukumasu, Jayme Augusto de Souza-Neto, José Salvatore Leister Patané, Luiz Alcantara, Luiz Lehmann Coutinho, Maria Carolina Elias, Maurício Lacerda Nogueira, Rafael dos Santos Bezerra, Raul Machado Neto, Rejane Maria Tommasini Grotto, Ricardo Haddad, Sandra Coccuzzo Sampaio Vessoni, Simone Kashima, Svetoslav Nanev Slavov, Vincent Louis Viala |
| EPI_ISL_2210228 | DEPARTAMENTO DE SAUDE MUNICIPAL SOCORRO SP | Instituto Butantan / Mendelics | Dimas Tadeu Covas, Antonio Jorge Martins, Claudia Renata dos Santos Barros, David Schlesinger, Debora Botequio Moretti, Elaine Cristina Marqueze, Elaine Vieira Santos, Evandra Strazza Rodrigues, Heidge Fukumasu, Jayme Augusto de Souza-Neto, José Salvatore Leister Patané, Luiz Alcantara, Luiz Lehmann Coutinho, Maria Carolina Elias, Maurício Lacerda Nogueira, Rafael dos Santos Bezerra, Raul Machado Neto, Rejane Maria Tommasini Grotto, Ricardo Haddad, Sandra Coccuzzo Sampaio Vessoni, Simone Kashima, Svetoslav Nanev Slavov, Vincent Louis Viala |
| EPI_ISL_2210231 | VIGILANCIA EPIDEMIOLOGICA E CONTROLE DE VETORES PIRASSUNUN | Instituto Butantan | Dimas Tadeu Covas, Antonio Jorge Martins, Claudia Renata dos Santos Barros, David Schlesinger, Debora Botequio Moretti, Elaine Cristina Marqueze, Elaine Vieira Santos, Evandra Strazza Rodrigues, Heidge Fukumasu, Jayme Augusto de Souza-Neto, José Salvatore Leister Patané, Luiz Alcantara, Luiz Lehmann Coutinho, Maria Carolina Elias, Maurício Lacerda Nogueira, Rafael dos Santos Bezerra, Raul Machado Neto, Rejane Maria Tommasini Grotto, Ricardo Haddad, Sandra Coccuzzo Sampaio Vessoni, Simone Kashima, Svetoslav Nanev Slavov, Vincent Louis Viala. |
| EPI_ISL_2210236 | HOSPITAL DE CAMPANHA COVID 19 MUNICIPIO DE TAUBATE | Instituto Butantan / Mendelics | Dimas Tadeu Covas, Antonio Jorge Martins, Claudia Renata dos Santos Barros, David Schlesinger, Debora Botequio Moretti, Elaine Cristina Marqueze, Elaine Vieira Santos, Evandra Strazza Rodrigues, Heidge Fukumasu, Jayme Augusto de Souza-Neto, José Salvatore Leister Patané, Luiz Alcantara, Luiz |

| | | | |
|---|---|---|---|
| | | | Lehmann Coutinho, Maria Carolina Elias, Maurício Lacerda Nogueira, Rafael dos Santos Bezerra, Raul Machado Neto, Rejane Maria Tommasini Grotto, Ricardo Haddad, Sandra Coccuzzo Sampaio Vessoni, Simone Kashima, Svetoslav Nanev Slavov, Vincent Louis Viala |
| EPI_ISL_2210238 | NUCLEO DE SAUDE MARY DOTA DE BAURU | Instituto Butantan / Mendelics | Dimas Tadeu Covas, Antonio Jorge Martins, Claudia Renata dos Santos Barros, David Schlesinger, Debora Botequio Moretti, Elaine Cristina Marqueze, Elaine Vieira Santos, Evandra Strazza Rodrigues, Heidge Fukumasu, Jayme Augusto de Souza-Neto, José Salvatore Leister Patané, Luiz Alcantara, Luiz Lehmann Coutinho, Maria Carolina Elias, Maurício Lacerda Nogueira, Rafael dos Santos Bezerra, Raul Machado Neto, Rejane Maria Tommasini Grotto, Ricardo Haddad, Sandra Coccuzzo Sampaio Vessoni, Simone Kashima, Svetoslav Nanev Slavov, Vincent Louis Viala |
| EPI_ISL_2210243 | VIGILANCIA EPIDEMIOLOGICA E CONTROLE DE VETORES PIRASSUNUN | Instituto Butantan | Dimas Tadeu Covas, Antonio Jorge Martins, Claudia Renata dos Santos Barros, David Schlesinger, Debora Botequio Moretti, Elaine Cristina Marqueze, Elaine Vieira Santos, Evandra Strazza Rodrigues, Heidge Fukumasu, Jayme Augusto de Souza-Neto, José Salvatore Leister Patané, Luiz Alcantara, Luiz Lehmann Coutinho, Maria Carolina Elias, Maurício Lacerda Nogueira, Rafael dos Santos Bezerra, Raul Machado Neto, Rejane Maria Tommasini Grotto, Ricardo Haddad, Sandra Coccuzzo Sampaio Vessoni, Simone Kashima, Svetoslav Nanev Slavov, Vincent Louis Viala. |
| EPI_ISL_2210246 | VIGILANCIA EPIDEMIOLOGICA JARDINOPOLIS SP | Instituto Butantan / Mendelics | Dimas Tadeu Covas, Antonio Jorge Martins, Claudia Renata dos Santos Barros, David Schlesinger, Debora Botequio Moretti, Elaine Cristina Marqueze, Elaine Vieira Santos, Evandra Strazza Rodrigues, Heidge Fukumasu, Jayme Augusto de Souza-Neto, José Salvatore Leister Patané, Luiz Alcantara, Luiz Lehmann Coutinho, Maria Carolina Elias, Maurício Lacerda Nogueira, Rafael dos Santos Bezerra, Raul Machado Neto, Rejane Maria Tommasini Grotto, Ricardo Haddad, Sandra Coccuzzo Sampaio Vessoni, Simone Kashima, Svetoslav Nanev Slavov, Vincent Louis Viala |
| EPI_ISL_2210250 | SANTA CASA DE MISERICORDIA SAO JOSE | Instituto Butantan / Mendelics | Dimas Tadeu Covas, Antonio Jorge Martins, Claudia Renata dos Santos Barros, David Schlesinger, Debora Botequio Moretti, Elaine Cristina Marqueze, Elaine Vieira Santos, Evandra Strazza Rodrigues, Heidge Fukumasu, Jayme Augusto de Souza-Neto, José Salvatore Leister Patané, Luiz Alcantara, Luiz Lehmann Coutinho, Maria Carolina Elias, Maurício Lacerda Nogueira, Rafael dos Santos Bezerra, Raul Machado Neto, Rejane Maria Tommasini Grotto, Ricardo Haddad, Sandra Coccuzzo Sampaio Vessoni, Simone Kashima, Svetoslav Nanev Slavov, Vincent Louis Viala |
| EPI_ISL_2210253 | HOSPITAL DE CAMPANHA COVID 19 CAIEIRAS | Instituto Butantan / Mendelics | Dimas Tadeu Covas, Antonio Jorge Martins, Claudia Renata dos Santos Barros, David Schlesinger, Debora Botequio Moretti, Elaine Cristina Marqueze, Elaine Vieira Santos, Evandra Strazza Rodrigues, Heidge Fukumasu, Jayme Augusto de Souza-Neto, José Salvatore Leister Patané, Luiz Alcantara, Luiz Lehmann Coutinho, Maria Carolina Elias, Maurício Lacerda Nogueira, Rafael dos Santos Bezerra, Raul Machado Neto, Rejane Maria Tommasini Grotto, Ricardo Haddad, Sandra Coccuzzo Sampaio Vessoni, Simone Kashima, Svetoslav Nanev Slavov, Vincent Louis Viala |
| EPI_ISL_2210257 | VIGILANCIA EPIDEMIOLOGICA E CONTROLE DE VETORES PIRASSUNUN | Instituto Butantan | Dimas Tadeu Covas, Antonio Jorge Martins, Claudia Renata dos Santos Barros, David Schlesinger, Debora Botequio Moretti, Elaine Cristina Marqueze, Elaine Vieira Santos, Evandra Strazza Rodrigues, Heidge Fukumasu, Jayme Augusto de Souza-Neto, José Salvatore Leister Patané, Luiz Alcantara, Luiz Lehmann Coutinho, Maria Carolina Elias, Maurício Lacerda Nogueira, Rafael dos Santos Bezerra, Raul Machado Neto, Rejane Maria Tommasini Grotto, Ricardo Haddad, Sandra Coccuzzo Sampaio Vessoni, Simone Kashima, Svetoslav Nanev Slavov, Vincent Louis Viala. |
| EPI_ISL_2210259 | LABORATORIO MUNICIPAL DE SUZANO | Instituto Butantan / Mendelics | Dimas Tadeu Covas, Antonio Jorge Martins, Claudia Renata dos Santos Barros, David Schlesinger, Debora Botequio Moretti, Elaine Cristina Marqueze, Elaine Vieira Santos, Evandra Strazza Rodrigues, Heidge Fukumasu, Jayme Augusto de Souza-Neto, José Salvatore Leister Patané, Luiz Alcantara, Luiz Lehmann Coutinho, Maria Carolina Elias, Maurício Lacerda Nogueira, Rafael dos Santos Bezerra, Raul Machado Neto, Rejane Maria Tommasini Grotto, Ricardo Haddad, Sandra Coccuzzo Sampaio Vessoni, Simone Kashima, Svetoslav Nanev Slavov, Vincent Louis Viala |
| EPI_ISL_2210269 | UNIDADE DE PRONTO ATENDIMENTO CENTRAL DE CARAGUATATUBA | Instituto Butantan / Mendelics | Dimas Tadeu Covas, Antonio Jorge Martins, Claudia Renata dos Santos Barros, David Schlesinger, Debora Botequio Moretti, Elaine Cristina Marqueze, Elaine Vieira Santos, Evandra Strazza Rodrigues, Heidge Fukumasu, Jayme Augusto de Souza-Neto, José Salvatore Leister Patané, Luiz Alcantara, Luiz Lehmann Coutinho, Maria Carolina Elias, Maurício Lacerda Nogueira, Rafael dos Santos Bezerra, Raul Machado Neto, Rejane Maria Tommasini Grotto, Ricardo Haddad, Sandra Coccuzzo Sampaio Vessoni, Simone Kashima, Svetoslav Nanev Slavov, Vincent Louis Viala |
| EPI_ISL_2210275 | UPA UNIDADE DE PRONTO ATENDIMENTO 24 HORAS BOM JESUS | Instituto Butantan / Mendelics | Dimas Tadeu Covas, Antonio Jorge Martins, Claudia Renata dos Santos Barros, David Schlesinger, Debora Botequio Moretti, Elaine Cristina Marqueze, Elaine Vieira Santos, Evandra Strazza Rodrigues, Heidge Fukumasu, Jayme Augusto de Souza-Neto, José Salvatore Leister Patané, Luiz Alcantara, Luiz Lehmann Coutinho, Maria Carolina Elias, Maurício Lacerda Nogueira, Rafael dos Santos Bezerra, Raul Machado Neto, Rejane Maria Tommasini Grotto, Ricardo Haddad, Sandra Coccuzzo Sampaio Vessoni, Simone Kashima, Svetoslav Nanev Slavov, Vincent Louis Viala |
| EPI_ISL_2210281 | VIGILANCIA EPIDEMIOLOGICA E CONTROLE DE VETORES PIRASSUNUN | Instituto Butantan | Dimas Tadeu Covas, Antonio Jorge Martins, Claudia Renata dos Santos Barros, David Schlesinger, Debora Botequio Moretti, Elaine Cristina Marqueze, Elaine Vieira Santos, Evandra Strazza Rodrigues, Heidge Fukumasu, Jayme Augusto de Souza-Neto, José Salvatore Leister Patané, Luiz Alcantara, Luiz Lehmann Coutinho, Maria Carolina Elias, Maurício Lacerda Nogueira, Rafael dos Santos Bezerra, Raul Machado Neto, Rejane Maria Tommasini Grotto, Ricardo Haddad, Sandra Coccuzzo Sampaio Vessoni, Simone Kashima, Svetoslav Nanev Slavov, Vincent Louis Viala. |
| EPI_ISL_2210285 | VIGILANCIA SANITARIA E VIG EPIDEMIOLOGICA DE ITAPEVI | Instituto Butantan | Dimas Tadeu Covas, Antonio Jorge Martins, Claudia Renata dos Santos Barros, David Schlesinger, Debora Botequio Moretti, Elaine Cristina Marqueze, Elaine Vieira Santos, Evandra Strazza Rodrigues, Heidge Fukumasu, Jayme Augusto de Souza-Neto, José Salvatore Leister Patané, Luiz Alcantara, Luiz Lehmann Coutinho, Maria Carolina Elias, Maurício Lacerda Nogueira, Rafael dos Santos Bezerra, Raul Machado Neto, Rejane Maria Tommasini Grotto, Ricardo Haddad, Sandra Coccuzzo Sampaio Vessoni, Simone Kashima, Svetoslav Nanev Slavov, Vincent Louis Viala. |
| EPI_ISL_2210292 | CENTRO DE SAUDE DE LUPERCIO | Instituto Butantan / Mendelics | Dimas Tadeu Covas, Antonio Jorge Martins, Claudia Renata dos Santos Barros, David Schlesinger, Debora Botequio Moretti, Elaine Cristina Marqueze, Elaine Vieira Santos, Evandra Strazza Rodrigues, Heidge Fukumasu, Jayme Augusto de Souza-Neto, José Salvatore Leister Patané, Luiz Alcantara, Luiz Lehmann Coutinho, Maria Carolina Elias, Maurício Lacerda Nogueira, Rafael dos Santos Bezerra, Raul Machado Neto, Rejane Maria Tommasini Grotto, Ricardo Haddad, Sandra Coccuzzo Sampaio Vessoni, Simone Kashima, Svetoslav Nanev Slavov, Vincent Louis Viala |
| EPI_ISL_2210295 | VIGILANCIA SANITARIA E VIG EPIDEMIOLOGICA DE ITAPEVI | Instituto Butantan | Dimas Tadeu Covas, Antonio Jorge Martins, Claudia Renata dos Santos Barros, David Schlesinger, Debora Botequio Moretti, Elaine Cristina Marqueze, Elaine Vieira Santos, Evandra Strazza Rodrigues, Heidge Fukumasu, Jayme Augusto de Souza-Neto, José Salvatore Leister Patané, Luiz Alcantara, Luiz Lehmann Coutinho, Maria Carolina Elias, Maurício Lacerda Nogueira, Rafael dos Santos Bezerra, Raul Machado Neto, Rejane Maria Tommasini Grotto, Ricardo Haddad, Sandra Coccuzzo Sampaio Vessoni, Simone Kashima, Svetoslav Nanev Slavov, Vincent Louis Viala. |
| EPI_ISL_2210305 | SECRETARIA DE SAUDE PUBLICA DE PRAIA GRANDE | Instituto Butantan / Mendelics | Dimas Tadeu Covas, Antonio Jorge Martins, Claudia Renata dos Santos Barros, David Schlesinger, Debora Botequio Moretti, Elaine Cristina Marqueze, Elaine Vieira Santos, Evandra Strazza Rodrigues, Heidge Fukumasu, Jayme Augusto de Souza-Neto, José Salvatore Leister Patané, Luiz Alcantara, Luiz Lehmann Coutinho, Maria Carolina Elias, Maurício Lacerda Nogueira, Rafael dos Santos Bezerra, Raul Machado Neto, Rejane Maria Tommasini Grotto, Ricardo Haddad, Sandra Coccuzzo Sampaio Vessoni, Simone Kashima, Svetoslav Nanev Slavov, Vincent Louis Viala |
| EPI_ISL_2221767, EPI_ISL_2221771 | Leme Laboratory | Central Public Health Laboratory - LACEN -Bahia, Salvador, Brazil | Stephane Tosta, Luciana Oliveira, Vanessa Nardy,Patrícia Cajado,Marcela Gómez, Breno Dominguez, Jaqueline Gomes, Vagner Fonseca,Marta Giovanetti,Luiz Alcantara, Felicidade Pereira, Arabela Leal |
| EPI_ISL_2221844, EPI_ISL_2221860, EPI_ISL_2221866, EPI_ISL_2221873, EPI_ISL_2221885 | LACEN/PE | WallauLab on behalf of Fiocruz COVID-19 Genomic Surveillance Network | Marcelo Henrique dos Santos Paiva, Duschinka Ribeiro Duarte Guedes, Cássia Docena, Matheus Filgueira Bezerra, Filipe Zimmer Dezordi, Laís Ceschini Machado, Larissa Krokovsky, Elisama Helvecio, Alexandre Freitas da Silva, Antonio Mauro Rezende, Sinval Pinto Brandão Filho, Constância Flávia Junqueira Ayres, Gabriel Luz Wallau |
| EPI_ISL_2222878, EPI_ISL_2222881, EPI_ISL_2222885, EPI_ISL_2222890, EPI_ISL_2222892, EPI_ISL_2222894, EPI_ISL_2222900, EPI_ISL_2222902, EPI_ISL_2222905, EPI_ISL_2222908, EPI_ISL_2222911, EPI_ISL_2222913, EPI_ISL_2222916, EPI_ISL_2222917, EPI_ISL_2222918 | | | |
| see above | HLAGYN - Laboratorio de Imunologia de Transplantes de Goias | HLAGYN - Laboratorio de Imunologia de Transplantes de Goias | Fernando Antonio Vinhal dos Santos, Erika Lopes Rocha Batista, Alessandro Leonardo Alvares Magalhaes, Frederico Rodrigues Vinhal, Sabrina Sara Moreira Duarte, Lucas Carlos Gomes Pereira, Daniel Ferreira de Sousa |
| EPI_ISL_2223407, EPI_ISL_2223414, EPI_ISL_2223416, EPI_ISL_2223419, EPI_ISL_2223422, EPI_ISL_2223425, EPI_ISL_2223427, EPI_ISL_2223430, EPI_ISL_2223431, EPI_ISL_2223433, EPI_ISL_2223435, EPI_ISL_2223438, EPI_ISL_2223443, EPI_ISL_2223446, EPI_ISL_2223452 | | | |
| see above | Instituto de Biotecnologia - UNESP-Botucatu-SP | Instituto de Biotecnologia - UNESP-Botucatu-SP | Fábio Sossai Possebon; Leila Sabrina Ullmann; Cecília Artico Banho; Cíntia Bittar; Guilherme Campos; Helena Lage Ferreira; Jorge A. Petroli Marchesi; Lívia Sacchetto; Maisa C. Pereira Parra; Marília Moraes; Maurício L. Nogueira; Paula Rahal; Paulo Inacio da Costa; João Pessoa Araújo Jr. |
| EPI_ISL_2227557, EPI_ISL_2227558, EPI_ISL_2227559, EPI_ISL_2227560, EPI_ISL_2227561, EPI_ISL_2227562, EPI_ISL_2227563, EPI_ISL_2227564, EPI_ISL_2227565 | HLAGYN - Laboratorio de Imunologia de Transplantes de Goias | HLAGYN - Laboratorio de Imunologia de Transplantes de Goias | Fernando Antonio Vinhal dos Santos, Erika Lopes Rocha Batista, Alessandro Leonardo Alvares Magalhaes, Frederico Rodrigues Vinhal, Sabrina Sara Moreira Duarte, Lucas Carlos Gomes Pereira, Daniel Ferreira de Sousa |
| EPI_ISL_2229838, EPI_ISL_2229839, EPI_ISL_2229840, EPI_ISL_2229841 | Laboratório de Microbiologia Molecular - Universidade FEEVALE | Laboratório de Microbiologia Molecular - Universidade FEEVALE | Alana Witt Hansen, Fágner Henrique Heldt, Fernando Rosado Spilki, Flávio Silveira, Juliana Schons Gularte, Juliane Deise Fleck, Mariana Soares da Silva, Meriane Demoliner, Matheus Nunes Weber, Paula Rodrigues de Almeida, Micheli Filippi. |

| | | | |
|---|---|---|---|
| EPI_ISL_2233906 | Scientific Platform Pasteur USP, University of Sao Paulo | Scientific Platform Pasteur USP, University of Sao Paulo | Cunha,M.P., Muxel,S.M. and Minoprio,P. |
| EPI_ISL_2241317, EPI_ISL_2241318, EPI_ISL_2241319, EPI_ISL_2241320, EPI_ISL_2241322, EPI_ISL_2241323, EPI_ISL_2241324, EPI_ISL_2241325, EPI_ISL_2241326 | Instituto Butantan | Instituto Butantan | Dimas Tadeu Covas, Antonio Jorge Martins, Claudia Renata dos Santos Barros, David Schlesinger, Debora Botequio Moretti, Elaine Cristina Marqueze, Elaine Vieira Santos, Evandra Strazza Rodrigues, Heidge Fukumasu, Jayme Augusto de Souza-Neto, José Salvatore Leister Patané, Luiz Alcantara, Luiz Lehmann Coutinho, Maria Carolina Elias, Maurício Lacerda Nogueira, Rafael dos Santos Bezerra, Raul Machado Neto, Rejane Maria Tommasini Grotto, Ricardo Haddad, Sandra Coccuzzo Sampaio Vessoni, Simone Kashima, Svetoslav Nanev Slavov, Vincent Louis Viala |
| EPI_ISL_2241495, EPI_ISL_2241496, EPI_ISL_2241497, EPI_ISL_2241498, EPI_ISL_2241499 | Laboratório Central de Saúde Pública da Paraiba | Coordenação Geral de Laboratórios de Saúde Pública (CGLAB/DAEVS/SVS/MS) | Vagner Fonseca, et al. |
| EPI_ISL_2241500 | Laboratório Central de Saúde Pública do Piaui | Coordenação Geral de Laboratórios de Saúde Pública (CGLAB/DAEVS/SVS/MS) | Vagner Fonseca, et al. |
| EPI_ISL_2241501, EPI_ISL_2241502, EPI_ISL_2241503, EPI_ISL_2241504, EPI_ISL_2241505, EPI_ISL_2241506, EPI_ISL_2241507, EPI_ISL_2241508, EPI_ISL_2241509, EPI_ISL_2241510, EPI_ISL_2241511, EPI_ISL_2241512, EPI_ISL_2241513 | | | |
| see above | Laboratório Central de Saúde Pública do Rio Grande do Norte | Coordenação Geral de Laboratórios de Saúde Pública (CGLAB/DAEVS/SVS/MS) | Vagner Fonseca, et al. |
| EPI_ISL_2241514, EPI_ISL_2241515, EPI_ISL_2241516, EPI_ISL_2241517, EPI_ISL_2241518, EPI_ISL_2241519, EPI_ISL_2241520, EPI_ISL_2241522, EPI_ISL_2241523, EPI_ISL_2241525, EPI_ISL_2241526, EPI_ISL_2241527 | | | |
| see above | Laboratório Central de Saúde Pública da Paraiba | Coordenação Geral de Laboratórios de Saúde Pública (CGLAB/DAEVS/SVS/MS) | Vagner Fonseca, et al. |
| EPI_ISL_2241528 | Laboratório Central de Saúde Pública da Bahia | Coordenação Geral de Laboratórios de Saúde Pública (CGLAB/DAEVS/SVS/MS) | Vagner Fonseca, et al. |
| EPI_ISL_2241529 | Laboratório Central de Saúde Pública da Paraiba | Coordenação Geral de Laboratórios de Saúde Pública (CGLAB/DAEVS/SVS/MS) | Vagner Fonseca, et al. |
| EPI_ISL_2241532 | Laboratório Central de Saúde Pública de Pernambuco | Coordenação Geral de Laboratórios de Saúde Pública (CGLAB/DAEVS/SVS/MS) | Vagner Fonseca, et al. |
| EPI_ISL_2241534, EPI_ISL_2241539, EPI_ISL_2241540 | Laboratório Central de Saúde Pública do Rio Grande do Norte | Coordenação Geral de Laboratórios de Saúde Pública (CGLAB/DAEVS/SVS/MS) | Vagner Fonseca, et al. |
| EPI_ISL_2241541 | Laboratório Central de Saúde Pública de Sergipe | Coordenação Geral de Laboratórios de Saúde Pública (CGLAB/DAEVS/SVS/MS) | Vagner Fonseca, et al. |
| EPI_ISL_2241544, EPI_ISL_2241545, EPI_ISL_2241546 | Laboratório Central de Saúde Pública do Rio Grande do Norte | Coordenação Geral de Laboratórios de Saúde Pública (CGLAB/DAEVS/SVS/MS) | Vagner Fonseca, et al. |
| EPI_ISL_2241547 | Laboratório Central de Saúde Pública de Pernambuco | Coordenação Geral de Laboratórios de Saúde Pública (CGLAB/DAEVS/SVS/MS) | Vagner Fonseca, et al. |
| EPI_ISL_2241548, EPI_ISL_2241549, EPI_ISL_2241550, EPI_ISL_2241551 | Laboratório Central de Saúde Pública do Rio Grande do Norte | Coordenação Geral de Laboratórios de Saúde Pública (CGLAB/DAEVS/SVS/MS) | Vagner Fonseca, et al. |
| EPI_ISL_2241552 | Laboratório Central de Saúde Pública do Piaui | Coordenação Geral de Laboratórios de Saúde Pública (CGLAB/DAEVS/SVS/MS) | Vagner Fonseca, et al. |
| EPI_ISL_2241553 | Laboratório Central de Saúde Pública do Rio Grande do Norte | Coordenação Geral de Laboratórios de Saúde Pública (CGLAB/DAEVS/SVS/MS) | Vagner Fonseca, et al. |
| EPI_ISL_2241554 | Laboratório Central de Saúde Pública do Piaui | Coordenação Geral de Laboratórios de Saúde Pública (CGLAB/DAEVS/SVS/MS) | Vagner Fonseca, et al. |
| EPI_ISL_2241555 | Laboratório Central de Saúde Pública de Sergipe | Coordenação Geral de Laboratórios de Saúde Pública (CGLAB/DAEVS/SVS/MS) | Vagner Fonseca, et al. |
| EPI_ISL_2241556 | Laboratório Central de Saúde Pública da Paraiba | Coordenação Geral de Laboratórios de Saúde Pública (CGLAB/DAEVS/SVS/MS) | Vagner Fonseca, et al. |
| EPI_ISL_2241557 | Laboratório Central de Saúde Pública de Sergipe | Coordenação Geral de Laboratórios de Saúde Pública (CGLAB/DAEVS/SVS/MS) | Vagner Fonseca, et al. |
| EPI_ISL_2241558, EPI_ISL_2241559, EPI_ISL_2241560, EPI_ISL_2241561, EPI_ISL_2241562, EPI_ISL_2241563, EPI_ISL_2241564 | Laboratório Central de Saúde Pública da Paraiba | Coordenação Geral de Laboratórios de Saúde Pública (CGLAB/DAEVS/SVS/MS) | Vagner Fonseca, et al. |
| EPI_ISL_2241565 | Laboratório Central de Saúde Pública da Bahia | Coordenação Geral de Laboratórios de Saúde Pública (CGLAB/DAEVS/SVS/MS) | Vagner Fonseca, et al. |
| EPI_ISL_2241566, EPI_ISL_2241567, EPI_ISL_2241568, EPI_ISL_2241569, EPI_ISL_2241570, EPI_ISL_2241571, EPI_ISL_2241572, EPI_ISL_2241573 | Laboratório Central de Saúde Pública da Paraiba | Coordenação Geral de Laboratórios de Saúde Pública (CGLAB/DAEVS/SVS/MS) | Vagner Fonseca, et al. |
| EPI_ISL_2241574 | Laboratório Central de Saúde Pública do Rio Grande do Norte | Coordenação Geral de Laboratórios de Saúde Pública (CGLAB/DAEVS/SVS/MS) | Vagner Fonseca, et al. |
| EPI_ISL_2241575 | Laboratório Central de Saúde Pública do Piaui | Coordenação Geral de Laboratórios de Saúde Pública (CGLAB/DAEVS/SVS/MS) | Vagner Fonseca, et al. |
| EPI_ISL_2241576 | Laboratório Central de Saúde Pública da Paraiba | Coordenação Geral de Laboratórios de Saúde Pública (CGLAB/DAEVS/SVS/MS) | Vagner Fonseca, et al. |
| EPI_ISL_2241577 | Laboratório Central de Saúde Pública do Piaui | Coordenação Geral de Laboratórios de Saúde Pública (CGLAB/DAEVS/SVS/MS) | Vagner Fonseca, et al. |
| EPI_ISL_2241578, EPI_ISL_2241579, EPI_ISL_2241580 | Laboratório Central de Saúde Pública de Sergipe | Coordenação Geral de Laboratórios de Saúde Pública (CGLAB/DAEVS/SVS/MS) | Vagner Fonseca, et al. |
| EPI_ISL_2241581 | Laboratório Central de Saúde Pública da Paraiba | Coordenação Geral de Laboratórios de Saúde Pública (CGLAB/DAEVS/SVS/MS) | Vagner Fonseca, et al. |
| EPI_ISL_2241582 | Laboratório Central de Saúde Pública de Sergipe | Coordenação Geral de Laboratórios de Saúde Pública (CGLAB/DAEVS/SVS/MS) | Vagner Fonseca, et al. |
| EPI_ISL_2241583 | Laboratório Central de Saúde Pública da Paraiba | Coordenação Geral de Laboratórios de Saúde Pública (CGLAB/DAEVS/SVS/MS) | Vagner Fonseca, et al. |

| | | | |
|---|---|---|---|
| EPI_ISL_2241584 | Laboratório Central de Saúde Pública de Sergipe | Coordenação Geral de Laboratórios de Saúde Pública (CGLAB/DAEVS/SVS/MS) | Vagner Fonseca, et al. |
| EPI_ISL_2241585, EPI_ISL_2241586 | Laboratório Central de Saúde Pública da Paraiba | Coordenação Geral de Laboratórios de Saúde Pública (CGLAB/DAEVS/SVS/MS) | Vagner Fonseca, et al. |
| EPI_ISL_2241587 | Laboratório Central de Saúde Pública de Sergipe | Coordenação Geral de Laboratórios de Saúde Pública (CGLAB/DAEVS/SVS/MS) | Vagner Fonseca, et al. |
| EPI_ISL_2241588 | Laboratório Central de Saúde Pública da Paraiba | Coordenação Geral de Laboratórios de Saúde Pública (CGLAB/DAEVS/SVS/MS) | Vagner Fonseca, et al. |
| EPI_ISL_2241589, EPI_ISL_2241590 | Laboratório Central de Saúde Pública de Sergipe | Coordenação Geral de Laboratórios de Saúde Pública (CGLAB/DAEVS/SVS/MS) | Vagner Fonseca, et al. |
| EPI_ISL_2241591, EPI_ISL_2241592, EPI_ISL_2241593 | Laboratório Central de Saúde Pública do Piaui | Coordenação Geral de Laboratórios de Saúde Pública (CGLAB/DAEVS/SVS/MS) | Vagner Fonseca, et al. |
| EPI_ISL_2241594 | Laboratório Central de Saúde Pública da Paraiba | Coordenação Geral de Laboratórios de Saúde Pública (CGLAB/DAEVS/SVS/MS) | Vagner Fonseca, et al. |
| EPI_ISL_2241595 | Laboratório Central de Saúde Pública do Piaui | Coordenação Geral de Laboratórios de Saúde Pública (CGLAB/DAEVS/SVS/MS) | Vagner Fonseca, et al. |
| EPI_ISL_2241596 | Laboratório Central de Saúde Pública da Paraiba | Coordenação Geral de Laboratórios de Saúde Pública (CGLAB/DAEVS/SVS/MS) | Vagner Fonseca, et al. |
| EPI_ISL_2241597, EPI_ISL_2241598, EPI_ISL_2241599, EPI_ISL_2241600, EPI_ISL_2241601 | Laboratório Central de Saúde Pública do Piaui | Coordenação Geral de Laboratórios de Saúde Pública (CGLAB/DAEVS/SVS/MS) | Vagner Fonseca, et al. |
| EPI_ISL_2241602 | Laboratório Central de Saúde Pública de Sergipe | Coordenação Geral de Laboratórios de Saúde Pública (CGLAB/DAEVS/SVS/MS) | Vagner Fonseca, et al. |
| EPI_ISL_2241603 | Laboratório Central de Saúde Pública do Piaui | Coordenação Geral de Laboratórios de Saúde Pública (CGLAB/DAEVS/SVS/MS) | Vagner Fonseca, et al. |
| EPI_ISL_2241604 | Laboratório Central de Saúde Pública de Sergipe | Coordenação Geral de Laboratórios de Saúde Pública (CGLAB/DAEVS/SVS/MS) | Vagner Fonseca, et al. |
| EPI_ISL_2241605 | Laboratório Central de Saúde Pública de Pernambuco | Coordenação Geral de Laboratórios de Saúde Pública (CGLAB/DAEVS/SVS/MS) | Vagner Fonseca, et al. |
| EPI_ISL_2241606 | Laboratório Central de Saúde Pública do Rio Grande do Norte | Coordenação Geral de Laboratórios de Saúde Pública (CGLAB/DAEVS/SVS/MS) | Vagner Fonseca, et al. |
| EPI_ISL_2241607 | Laboratório Central de Saúde Pública do Piaui | Coordenação Geral de Laboratórios de Saúde Pública (CGLAB/DAEVS/SVS/MS) | Vagner Fonseca, et al. |
| EPI_ISL_2241608 | Laboratório Central de Saúde Pública do Rio Grande do Norte | Coordenação Geral de Laboratórios de Saúde Pública (CGLAB/DAEVS/SVS/MS) | Vagner Fonseca, et al. |
| EPI_ISL_2241609 | Laboratório Central de Saúde Pública de Sergipe | Coordenação Geral de Laboratórios de Saúde Pública (CGLAB/DAEVS/SVS/MS) | Vagner Fonseca, et al. |
| EPI_ISL_2241610 | Laboratório Central de Saúde Pública do Rio Grande do Norte | Coordenação Geral de Laboratórios de Saúde Pública (CGLAB/DAEVS/SVS/MS) | Vagner Fonseca, et al. |
| EPI_ISL_2245062 | Laboratório Central de Saúde Pública do Acre | Coordenação Geral de Laboratórios de Saúde Pública (CGLAB/DAEVS/SVS/MS) | Vagner Fonseca, et al. |
| EPI_ISL_2245064, EPI_ISL_2245065, EPI_ISL_2245066, EPI_ISL_2245067, EPI_ISL_2245068, EPI_ISL_2245069, EPI_ISL_2245070, EPI_ISL_2245071, EPI_ISL_2245072, EPI_ISL_2245073, EPI_ISL_2245074, EPI_ISL_2245075, EPI_ISL_2245076, EPI_ISL_2245077, EPI_ISL_2245078, EPI_ISL_2245079 | | | |
| see above | Laboratório Central de Saúde Pública do Amapá | Coordenação Geral de Laboratórios de Saúde Pública (CGLAB/DAEVS/SVS/MS) | Vagner Fonseca, et al. |
| EPI_ISL_2245080, EPI_ISL_2245082 | Laboratório Central de Saúde Pública do Ceará | Coordenação Geral de Laboratórios de Saúde Pública (CGLAB/DAEVS/SVS/MS) | Vagner Fonseca, et al. |
| EPI_ISL_2245083, EPI_ISL_2245084, EPI_ISL_2245085, EPI_ISL_2245086 | Laboratório Central de Saúde Pública do Maranhão | Coordenação Geral de Laboratórios de Saúde Pública (CGLAB/DAEVS/SVS/MS) | Vagner Fonseca, et al. |
| EPI_ISL_2245087, EPI_ISL_2245089, EPI_ISL_2245090, EPI_ISL_2245092, EPI_ISL_2245093, EPI_ISL_2245094, EPI_ISL_2245095, EPI_ISL_2245098, EPI_ISL_2245101, EPI_ISL_2245102, EPI_ISL_2245103, EPI_ISL_2245104, EPI_ISL_2245105 | | | |
| see above | Laboratório Central de Saúde Pública do Pará | Coordenação Geral de Laboratórios de Saúde Pública (CGLAB/DAEVS/SVS/MS) | Vagner Fonseca, et al. |
| EPI_ISL_2245107, EPI_ISL_2245108, EPI_ISL_2245109, EPI_ISL_2245110, EPI_ISL_2245111, EPI_ISL_2245112, EPI_ISL_2245113 | Laboratório Central de Saúde Pública de Roraima | Coordenação Geral de Laboratórios de Saúde Pública (CGLAB/DAEVS/SVS/MS) | Vagner Fonseca, et al. |
| EPI_ISL_2245176, EPI_ISL_2245177, EPI_ISL_2245178 | Laboratorio de Saude Publica Dr. Giovanni Cysneiros | Coordenação Geral de Laboratórios de Saúde Pública (CGLAB/DAEVS/SVS/MS) | Vagner Fonseca, et al. |
| EPI_ISL_2245179 | Instituto Adolfo Lutz | Coordenação Geral de Laboratórios de Saúde Pública (CGLAB/DAEVS/SVS/MS) | Vagner Fonseca, et al. |
| EPI_ISL_2245180 | Laboratorio de Saude Publica Dr. Giovanni Cysneiros | Coordenação Geral de Laboratórios de Saúde Pública (CGLAB/DAEVS/SVS/MS) | Vagner Fonseca, et al. |
| EPI_ISL_2245181, EPI_ISL_2245182, EPI_ISL_2245183, EPI_ISL_2245184, EPI_ISL_2245185, EPI_ISL_2245186, EPI_ISL_2245187, EPI_ISL_2245188, EPI_ISL_2245190, EPI_ISL_2245191, EPI_ISL_2245192, EPI_ISL_2245193 | | | |
| see above | Instituto Adolfo Lutz | Coordenação Geral de Laboratórios de Saúde Pública (CGLAB/DAEVS/SVS/MS) | Vagner Fonseca, et al. |
| EPI_ISL_2245194 | Laboratório Central de Saúde Pública do Mato Grosso do Sul | Coordenação Geral de Laboratórios de Saúde Pública (CGLAB/DAEVS/SVS/MS) | Vagner Fonseca, et al. |
| EPI_ISL_2246285 | Laboratório Central de Saúde Pública da Paraiba | Coordenação Geral de Laboratórios de Saúde Pública (CGLAB/DAEVS/SVS/MS) | Vagner Fonseca, et al. |
| EPI_ISL_2246286 | Laboratório Central de Saúde Pública de Sergipe | Coordenação Geral de Laboratórios de Saúde Pública (CGLAB/DAEVS/SVS/MS) | Vagner Fonseca, et al. |

| | | | |
|---|---|---|---|
| EPI_ISL_2246287 | Laboratório Central de Saúde Pública do Piaui | Coordenação Geral de Laboratórios de Saúde Pública (CGLAB/DAEVS/SVS/MS) | Vagner Fonseca, et al. |
| EPI_ISL_2248758, EPI_ISL_2248759, EPI_ISL_2248760, EPI_ISL_2248761, EPI_ISL_2248762, EPI_ISL_2248763, EPI_ISL_2248764, EPI_ISL_2248765, EPI_ISL_2248766, EPI_ISL_2248767 | Laboratório Central de Saúde Pública do Pará | Coordenação Geral de Laboratórios de Saúde Pública (CGLAB/DAEVS/SVS/MS) | Vagner Fonseca, et al. |
| EPI_ISL_2248768, EPI_ISL_2248769, EPI_ISL_2248770, EPI_ISL_2248771, EPI_ISL_2248772, EPI_ISL_2248773, EPI_ISL_2248774, EPI_ISL_2248775, EPI_ISL_2248776, EPI_ISL_2248777, EPI_ISL_2248778, EPI_ISL_2248779 | | | |
| see above | Laboratório Central de Saúde Pública do Maranhão | Coordenação Geral de Laboratórios de Saúde Pública (CGLAB/DAEVS/SVS/MS) | Vagner Fonseca, et al. |
| EPI_ISL_2248780, EPI_ISL_2248781 | Laboratório Central de Saúde Pública do Acre | Coordenação Geral de Laboratórios de Saúde Pública (CGLAB/DAEVS/SVS/MS) | Vagner Fonseca, et al. |
| EPI_ISL_2248782, EPI_ISL_2248783 | Laboratório Central de Saúde Pública do Pará | Coordenação Geral de Laboratórios de Saúde Pública (CGLAB/DAEVS/SVS/MS) | Vagner Fonseca, et al. |
| EPI_ISL_2248784, EPI_ISL_2248785, EPI_ISL_2248786, EPI_ISL_2248787 | Laboratório Central de Saúde Pública do Amapá | Coordenação Geral de Laboratórios de Saúde Pública (CGLAB/DAEVS/SVS/MS) | Vagner Fonseca, et al. |
| EPI_ISL_2248788 | Laboratório Central de Saúde Pública do Ceará | Coordenação Geral de Laboratórios de Saúde Pública (CGLAB/DAEVS/SVS/MS) | Vagner Fonseca, et al. |
| EPI_ISL_2249333, EPI_ISL_2249334, EPI_ISL_2249335, EPI_ISL_2249336, EPI_ISL_2249337, EPI_ISL_2249338, EPI_ISL_2249339, EPI_ISL_2249340, EPI_ISL_2249341, EPI_ISL_2249342, EPI_ISL_2249343, EPI_ISL_2249344 | | | |
| see above | Laboratório Central de Saúde Pública do Espírito Santo | Coordenação Geral de Laboratórios de Saúde Pública (CGLAB/DAEVS/SVS/MS) | Vagner Fonseca, et al. |
| EPI_ISL_2249345, EPI_ISL_2249346, EPI_ISL_2249347, EPI_ISL_2249348, EPI_ISL_2249349, EPI_ISL_2249350, EPI_ISL_2249351, EPI_ISL_2249352, EPI_ISL_2249353, EPI_ISL_2249354, EPI_ISL_2249355, EPI_ISL_2249356, EPI_ISL_2249357, EPI_ISL_2249358, EPI_ISL_2249359, EPI_ISL_2249360, EPI_ISL_2249361, EPI_ISL_2249362 | | | |
| see above | Laboratório Central de Saúde Pública do Rio Grande do Sul | Coordenação Geral de Laboratórios de Saúde Pública (CGLAB/DAEVS/SVS/MS) | Vagner Fonseca, et al. |
| EPI_ISL_2249363, EPI_ISL_2249364 | Fundação Ezequiel Dias | Coordenação Geral de Laboratórios de Saúde Pública (CGLAB/DAEVS/SVS/MS) | Vagner Fonseca, et al. |
| EPI_ISL_2249365, EPI_ISL_2249366, EPI_ISL_2249367, EPI_ISL_2249368, EPI_ISL_2249369, EPI_ISL_2249370, EPI_ISL_2249371, EPI_ISL_2249372, EPI_ISL_2249373, EPI_ISL_2249374, EPI_ISL_2249375, EPI_ISL_2249376, EPI_ISL_2249377, EPI_ISL_2249378, EPI_ISL_2249379, EPI_ISL_2249380, EPI_ISL_2249381, EPI_ISL_2249382, EPI_ISL_2249383, EPI_ISL_2249384, EPI_ISL_2249385, EPI_ISL_2249386 | | | |
| see above | Laboratório Central de Saúde Pública do Rio Grande do Sul | Coordenação Geral de Laboratórios de Saúde Pública (CGLAB/DAEVS/SVS/MS) | Vagner Fonseca, et al. |
| EPI_ISL_2249387, EPI_ISL_2249389, EPI_ISL_2249390, EPI_ISL_2249391, EPI_ISL_2249392, EPI_ISL_2249393, EPI_ISL_2249394, EPI_ISL_2249395, EPI_ISL_2249396 | Laboratório Central de Saúde Pública de Santa Catarina | Coordenação Geral de Laboratórios de Saúde Pública (CGLAB/DAEVS/SVS/MS) | Vagner Fonseca, et al. |
| EPI_ISL_2249397 | Fundação Ezequiel Dias | Coordenação Geral de Laboratórios de Saúde Pública (CGLAB/DAEVS/SVS/MS) | Vagner Fonseca, et al. |
| EPI_ISL_2249398, EPI_ISL_2249399, EPI_ISL_2249400, EPI_ISL_2249401, EPI_ISL_2249402, EPI_ISL_2249403 | Laboratório Central de Saúde Pública de Santa Catarina | Coordenação Geral de Laboratórios de Saúde Pública (CGLAB/DAEVS/SVS/MS) | Vagner Fonseca, et al. |
| EPI_ISL_2249404, EPI_ISL_2249405, EPI_ISL_2249406, EPI_ISL_2249407, EPI_ISL_2249408, EPI_ISL_2249409, EPI_ISL_2249410, EPI_ISL_2249411, EPI_ISL_2249412, EPI_ISL_2249413, EPI_ISL_2249414, EPI_ISL_2249415, EPI_ISL_2249416, EPI_ISL_2249417, EPI_ISL_2249418, EPI_ISL_2249419, EPI_ISL_2249420, EPI_ISL_2249421, EPI_ISL_2249422, EPI_ISL_2249423, EPI_ISL_2249424, EPI_ISL_2249425, EPI_ISL_2249426, EPI_ISL_2249427, EPI_ISL_2249428, EPI_ISL_2249429, EPI_ISL_2249430, EPI_ISL_2249431, EPI_ISL_2249432, EPI_ISL_2249433, EPI_ISL_2249434 | | | |
| see above | Laboratório Central de Saúde Pública do Rio de Janeiro | Coordenação Geral de Laboratórios de Saúde Pública (CGLAB/DAEVS/SVS/MS) | Vagner Fonseca, et al. |
| EPI_ISL_2249435, EPI_ISL_2249436, EPI_ISL_2249437, EPI_ISL_2249438, EPI_ISL_2249439 | Fundação Ezequiel Dias | Coordenação Geral de Laboratórios de Saúde Pública (CGLAB/DAEVS/SVS/MS) | Vagner Fonseca, et al. |
| EPI_ISL_2249440, EPI_ISL_2249441, EPI_ISL_2249442, EPI_ISL_2249443, EPI_ISL_2249444, EPI_ISL_2249445 | Laboratório Central de Saúde Pública do Rio de Janeiro | Coordenação Geral de Laboratórios de Saúde Pública (CGLAB/DAEVS/SVS/MS) | Vagner Fonseca, et al. |
| EPI_ISL_2249446, EPI_ISL_2249447, EPI_ISL_2249448, EPI_ISL_2249449, EPI_ISL_2249450, EPI_ISL_2249451, EPI_ISL_2249452, EPI_ISL_2249453, EPI_ISL_2249454, EPI_ISL_2249455, EPI_ISL_2249456, EPI_ISL_2249457 | | | |
| see above | Laboratório Central de Saúde Pública do Paraá | Coordenação Geral de Laboratórios de Saúde Pública (CGLAB/DAEVS/SVS/MS) | Vagner Fonseca, et al. |
| EPI_ISL_2249458, EPI_ISL_2249459, EPI_ISL_2249460, EPI_ISL_2249461, EPI_ISL_2249462, EPI_ISL_2249463 | Laboratório Central de Saúde Pública de Santa Catarina | Coordenação Geral de Laboratórios de Saúde Pública (CGLAB/DAEVS/SVS/MS) | Vagner Fonseca, et al. |
| EPI_ISL_2258211, EPI_ISL_2258212 | Instituto Adolfo Lutz | Coordenação Geral de Laboratórios de Saúde Pública (CGLAB/DAEVS/SVS/MS) | Vagner Fonseca, et al. |
| EPI_ISL_2274041, EPI_ISL_2274042, EPI_ISL_2274043, EPI_ISL_2274044, EPI_ISL_2274045, EPI_ISL_2274046, EPI_ISL_2274047, EPI_ISL_2274048, EPI_ISL_2274049, EPI_ISL_2274050, EPI_ISL_2274051, EPI_ISL_2274052, EPI_ISL_2274053, EPI_ISL_2274054, EPI_ISL_2274055, EPI_ISL_2274056, EPI_ISL_2274057, EPI_ISL_2274058, EPI_ISL_2274059, EPI_ISL_2274060, EPI_ISL_2274061, EPI_ISL_2274062, EPI_ISL_2274063, EPI_ISL_2274064, EPI_ISL_2274065, EPI_ISL_2274066, EPI_ISL_2274067, EPI_ISL_2274068, EPI_ISL_2274069, EPI_ISL_2274070, EPI_ISL_2274071 | | | |
| see above | Laboratorio Central de Saude Publica do Estado de Alagoas (LACEN/AL) | Laboratory of Respiratory Viruses and Measles, Oswaldo Cruz Institute, FIOCRUZ | Paola Resende, Luciana Appolinario, Fernando Motta, Anna Carolina Paixao, Ana Carolina Mendonca, Alice Sampaio Rocha, Taina Venas, Elisa Cavalcante Pereira, Renata Serrano Lopes, Anderson Brandao Leite, Marilda Siqueira on behalf of the Fiocruz COVID-19 Genomic Surveillance Network |
| EPI_ISL_2274072 | Laboratorio Central de Saude Publica do Estado da Bahia (LACEN/BA) | Laboratory of Respiratory Viruses and Measles, Oswaldo Cruz Institute, FIOCRUZ | Paola Resende, Luciana Appolinario, Fernando Motta, Anna Carolina Paixao, Ana Carolina Mendonca, Alice Sampaio Rocha, Taina Venas, Elisa Cavalcante Pereira, Renata Serrano Lopes,Felicidade Pereira, Marilda Siqueira on behalf of the Fiocruz COVID-19 Genomic Surveillance Network |
| EPI_ISL_2274073, EPI_ISL_2274074 | Laboratorio Central de Saude Publica do Estado da Bahia (LACEN/BA) | Laboratory of Respiratory Viruses and Measles, Oswaldo Cruz Institute, FIOCRUZ | Paola Resende, Luciana Appolinario, Fernando Motta, Anna Carolina Paixao, Ana Carolina Mendonca, Alice Sampaio Rocha, Taina Venas, Elisa Cavalcante Pereira, Renata Serrano Lopes, Anderson Brandao Leite, Marilda Siqueira on behalf of the Fiocruz COVID-19 Genomic Surveillance Network |
| EPI_ISL_2274075 | Laboratorio Central de Saude Publica do Estado da Bahia (LACEN/BA) | Laboratory of Respiratory Viruses and Measles, Oswaldo Cruz Institute, FIOCRUZ | Paola Resende, Luciana Appolinario, Fernando Motta, Anna Carolina Paixao, Ana Carolina Mendonca, Alice Sampaio Rocha, Taina Venas, Elisa Cavalcante Pereira, Renata Serrano Lopes,Felicidade Pereira, Marilda Siqueira on behalf of the Fiocruz COVID-19 Genomic Surveillance Network |
| EPI_ISL_2274076, EPI_ISL_2274077, EPI_ISL_2274078, EPI_ISL_2274079, EPI_ISL_2274080, EPI_ISL_2274081, EPI_ISL_2274082, EPI_ISL_2274083, EPI_ISL_2274084, EPI_ISL_2274085, EPI_ISL_2274086, EPI_ISL_2274087, EPI_ISL_2274088, EPI_ISL_2274089, EPI_ISL_2274090 | | | |
| see above | Laboratorio Central de Saude Publica do Estado Maranhao (LACEN-MA) | Laboratory of Respiratory Viruses and Measles, Oswaldo Cruz Institute, FIOCRUZ | Paola Resende, Luciana Appolinario, Fernando Motta, Anna Carolina Paixao, Ana Carolina Mendonca, Alice Sampaio Rocha, Taina Venas, Elisa Cavalcante Pereira, Renata Serrano Lopes, Lidio Gonçalves Lima Neto, Marilda Siqueira on behalf of the Fiocruz COVID-19 Genomic Surveillance |

| | | | |
|---|---|---|---|
| | | | Network |
| EPI_ISL_2274091 | Laboratorio Central de Saude Publica do Estado do Mato Grosso do Sul (LACEN/MS) | Laboratory of Respiratory Viruses and Measles, Oswaldo Cruz Institute, FIOCRUZ | Paola Resende, Luciana Appolinario, Fernando Motta, Anna Carolina Paixao, Ana Carolina Mendonca, Alice Sampaio Rocha, Taina Venas, Elisa Cavalcante Pereira, Renata Serrano Lopes, Gislene Lichs, Marilda Siqueira on behalf of the Fiocruz COVID-19 Genomic Surveillance Network |
| EPI_ISL_2274092 | Laboratorio Central de Saude Publica do Estado da Paraiba (LACEN-PB) | Laboratory of Respiratory Viruses and Measles, Oswaldo Cruz Institute, FIOCRUZ | Paola Resende, Luciana Appolinario, Fernando Motta, Anna Carolina Paixao, Ana Carolina Mendonca, Alice Sampaio Rocha, Taina Venas, Elisa Cavalcante Pereira, Renata Serrano Lopes, Joao Felipe Bezerra, Dalane Loudal Florentino Teixeira, Marilda Siqueira on behalf of the Fiocruz COVID-19 Genomic Surveillance Network |
| EPI_ISL_2274093 | Laboratorio Central de Saude Publica do Estado da Paraiba (LACEN-PB) | Laboratory of Respiratory Viruses and Measles, Oswaldo Cruz Institute, FIOCRUZ | Paola Resende, Luciana Appolinario, Fernando Motta, Anna Carolina Paixao, Ana Carolina Mendonca, Alice Sampaio Rocha, Taina Venas, Elisa Cavalcante Pereira, Renata Serrano Lopes, Irina Riediger, Marilda Siqueira on behalf of the Fiocruz COVID-19 Genomic Surveillance Network |
| EPI_ISL_2274094, EPI_ISL_2274095, EPI_ISL_2274096 | Lboratorio Central de Saude Publica do Estado do Parana (LACEN/PR) | Laboratory of Respiratory Viruses and Measles, Oswaldo Cruz Institute, FIOCRUZ | Paola Resende, Luciana Appolinario, Fernando Motta, Anna Carolina Paixao, Ana Carolina Mendonca, Alice Sampaio Rocha, Taina Venas, Elisa Cavalcante Pereira, Renata Serrano Lopes, Irina Riediger, Marilda Siqueira on behalf of the Fiocruz COVID-19 Genomic Surveillance Network |
| EPI_ISL_2274097 | Laboratory of Respiratory Viruses and Measles, Oswaldo Cruz Institute, FIOCRUZ | Laboratory of Respiratory Viruses and Measles, Oswaldo Cruz Institute, FIOCRUZ | Paola Resende, Luciana Appolinario, Fernando Motta, Anna Carolina Paixao, Ana Carolina Mendonca, Alice Sampaio Rocha, Taina Venas, Elisa Cavalcante Pereira, Renata Serrano Lopes, Marilda Siqueira on behalf of the Fiocruz COVID-19 Genomic Surveillance Network |
| EPI_ISL_2274098, EPI_ISL_2274099 | Labortorio Central de Saude Publica do Estado do Rio de Janeiro (LACEN/RJ) | Laboratory of Respiratory Viruses and Measles, Oswaldo Cruz Institute, FIOCRUZ | Paola Resende, Luciana Appolinario, Fernando Motta, Anna Carolina Paixao, Ana Carolina Mendonca, Alice Sampaio Rocha, Taina Venas, Elisa Cavalcante Pereira, Renata Serrano Lopes, Andrea Cony Cavalcanti, Marilda Siqueira on behalf of the Fiocruz COVID-19 Genomic Surveillance Network |
| EPI_ISL_2274100, EPI_ISL_2274101, EPI_ISL_2274102, EPI_ISL_2274103, EPI_ISL_2274104, EPI_ISL_2274105, EPI_ISL_2274106, EPI_ISL_2274107 | Laboratory of Respiratory Viruses and Measles, Oswaldo Cruz Institute, FIOCRUZ | Laboratory of Respiratory Viruses and Measles, Oswaldo Cruz Institute, FIOCRUZ | Paola Resende, Luciana Appolinario, Fernando Motta, Anna Carolina Paixao, Ana Carolina Mendonca, Alice Sampaio Rocha, Taina Venas, Elisa Cavalcante Pereira, Renata Serrano Lopes, Marilda Siqueira on behalf of the Fiocruz COVID-19 Genomic Surveillance Network |
| EPI_ISL_2274108, EPI_ISL_2274109, EPI_ISL_2274110, EPI_ISL_2274111, EPI_ISL_2274112, EPI_ISL_2274113, EPI_ISL_2274114, EPI_ISL_2274115, EPI_ISL_2274116, EPI_ISL_2274117, EPI_ISL_2274118, EPI_ISL_2274119, EPI_ISL_2274120, EPI_ISL_2274121, EPI_ISL_2274122, EPI_ISL_2274123, EPI_ISL_2274124, EPI_ISL_2274125, EPI_ISL_2274126, EPI_ISL_2274127, EPI_ISL_2274128, EPI_ISL_2274129, EPI_ISL_2274130, EPI_ISL_2274131, EPI_ISL_2274132 | | | |
| see above | Laboratorio Central de Saude Publica do Estado do Rio Grande do Sul (LACEN-RS) | Laboratory of Respiratory Viruses and Measles, Oswaldo Cruz Institute, FIOCRUZ | Paola Resende, Luciana Appolinario, Fernando Motta, Anna Carolina Paixao, Ana Carolina Mendonca, Alice Sampaio Rocha, Taina Venas, Elisa Cavalcante Pereira, Renata Serrano Lopes, Tatiana Schaffer Gregianini, Richard Salvato, Marilda Siqueira on behalf of the Fiocruz COVID-19 Genomic Surveillance Network |
| EPI_ISL_2274133, EPI_ISL_2274134, EPI_ISL_2274135, EPI_ISL_2274136, EPI_ISL_2274137 | Laboratório Central de Saude Publica do Estado de Santa Catarina (LACEN/SC) | Laboratory of Respiratory Viruses and Measles, Oswaldo Cruz Institute, FIOCRUZ | Paola Resende, Luciana Appolinario, Fernando Motta, Anna Carolina Paixao, Ana Carolina Mendonca, Alice Sampaio Rocha, Taina Venas, Elisa Cavalcante Pereira, Renata Serrano Lopes, Darcita Buerger Rovaris, Sandra Bianchini Fernandes, Marilda Siqueira on behalf of the Fiocruz COVID-19 Genomic Surveillance Network |
| EPI_ISL_2274138 | Laboratorio Central de Saude Publica do Estado do Rio Grande do Sul (LACEN-RS) | Laboratory of Respiratory Viruses and Measles, Oswaldo Cruz Institute, FIOCRUZ | Paola Resende, Luciana Appolinario, Fernando Motta, Anna Carolina Paixao, Ana Carolina Mendonca, Alice Sampaio Rocha, Taina Venas, Elisa Cavalcante Pereira, Renata Serrano Lopes, Tatiana Schaffer Gregianini, Richard Salvato, Marilda Siqueira on behalf of the Fiocruz COVID-19 Genomic Surveillance Network |
| EPI_ISL_2274979 | Aeroporto Internacional de Guarulhos | Instituto Adolfo Lutz, Interdiciplinary Procedures Center, Strategic Laboratory | Claudio Tavares Sacchi, Claudia Regina Gonçalves, Erica Valessa Ramos Gomes, Karoline Rodrigues Campos, Caio Vinicius Dias Lopes, Leonardo Jose Tadeu de Araujo |
| EPI_ISL_2292993, EPI_ISL_2292994, EPI_ISL_2292995, EPI_ISL_2292996, EPI_ISL_2292997, EPI_ISL_2292998, EPI_ISL_2292999, EPI_ISL_2293000, EPI_ISL_2293001, EPI_ISL_2293002, EPI_ISL_2293003, EPI_ISL_2293004, EPI_ISL_2293005, EPI_ISL_2293006, EPI_ISL_2293008, EPI_ISL_2293009 | | | |
| see above | Laboratório Central de Saúde Pública de Santa Catarina | Coordenação Geral de Laboratórios de Saúde Pública (CGLAB/DAEVS/SVS/MS) | Vagner Fonseca, et al. |
| EPI_ISL_2293010, EPI_ISL_2293011, EPI_ISL_2293012, EPI_ISL_2293013, EPI_ISL_2293014, EPI_ISL_2293015, EPI_ISL_2293016 | Laboratório Central de Saúde Pública do Espírito Santo | Coordenação Geral de Laboratórios de Saúde Pública (CGLAB/DAEVS/SVS/MS) | Vagner Fonseca, et al. |
| EPI_ISL_2293017, EPI_ISL_2293018, EPI_ISL_2293019, EPI_ISL_2293020, EPI_ISL_2293021, EPI_ISL_2293022 | Fundação Ezequiel Dias | Coordenação Geral de Laboratórios de Saúde Pública (CGLAB/DAEVS/SVS/MS) | Vagner Fonseca, et al. |
| EPI_ISL_2295387, EPI_ISL_2295388, EPI_ISL_2295389, EPI_ISL_2295396, EPI_ISL_2295399, EPI_ISL_2295403, EPI_ISL_2295404, EPI_ISL_2295405, EPI_ISL_2295406, EPI_ISL_2295408, EPI_ISL_2295409, EPI_ISL_2295410, EPI_ISL_2295413, EPI_ISL_2295414, EPI_ISL_2295416, EPI_ISL_2295417, EPI_ISL_2295418, EPI_ISL_2295419, EPI_ISL_2295420, EPI_ISL_2295423, EPI_ISL_2295424, EPI_ISL_2295425, EPI_ISL_2295427, EPI_ISL_2295428, EPI_ISL_2295429, EPI_ISL_2295431, EPI_ISL_2295432, EPI_ISL_2295433, EPI_ISL_2295434, EPI_ISL_2295435, EPI_ISL_2295436, EPI_ISL_2295437, EPI_ISL_2295438, EPI_ISL_2295439, EPI_ISL_2295440, EPI_ISL_2295441, EPI_ISL_2295442, EPI_ISL_2295443, EPI_ISL_2295444, EPI_ISL_2295445, EPI_ISL_2295447, EPI_ISL_2295448, EPI_ISL_2295449, EPI_ISL_2295450, EPI_ISL_2295451 | | | |
| see above | Instituto de Biotecnologia - UNESP-Botucatu-SP | Instituto de Biotecnologia - UNESP-Botucatu-SP | Fábio Sossai Possebon; Leila Sabrina Ullmann; Cecília Artico Banho; Cíntia Bittar; Guilherme Campos; Helena Lage Ferreira; Jorge A. Petroli Marchesi; Lívia Sacchetto; Maisa C. Pereira Parra; Marília Moraes; Maurício L. Nogueira; Paula Rahal; Paulo Inacio da Costa; João Pessoa Araújo Jr. |
| EPI_ISL_2298733, EPI_ISL_2298735, EPI_ISL_2298737, EPI_ISL_2298739, EPI_ISL_2298741, EPI_ISL_2298743, EPI_ISL_2298745 | Laboratório Central de Saúde Pública do Ceará | Coordenação Geral de Laboratórios de Saúde Pública (CGLAB/DAEVS/SVS/MS) | Vagner Fonseca, et al. |
| EPI_ISL_2298747 | Laboratório Central de Saúde Pública do Amapá | Coordenação Geral de Laboratórios de Saúde Pública (CGLAB/DAEVS/SVS/MS) | Vagner Fonseca, et al. |
| EPI_ISL_2298748 | Laboratório Central de Saúde Pública de Roraima | Coordenação Geral de Laboratórios de Saúde Pública (CGLAB/DAEVS/SVS/MS) | Vagner Fonseca, et al. |
| EPI_ISL_2298750, EPI_ISL_2298752, EPI_ISL_2298754, EPI_ISL_2298755, EPI_ISL_2298757 | Laboratório Central de Saúde Pública do Maranhão | Coordenação Geral de Laboratórios de Saúde Pública (CGLAB/DAEVS/SVS/MS) | Vagner Fonseca, et al. |
| EPI_ISL_2298758 | Laboratório Central de Saúde Pública do Pará | Coordenação Geral de Laboratórios de Saúde Pública (CGLAB/DAEVS/SVS/MS) | Vagner Fonseca, et al. |
| EPI_ISL_2298760 | Laboratório Central de Saúde Pública do Amapá | Coordenação Geral de Laboratórios de Saúde Pública (CGLAB/DAEVS/SVS/MS) | Vagner Fonseca, et al. |
| EPI_ISL_2298762, EPI_ISL_2298764, EPI_ISL_2298765, EPI_ISL_2298767, EPI_ISL_2298769 | Laboratório Central de Saúde Pública de Roraima | Coordenação Geral de Laboratórios de Saúde Pública (CGLAB/DAEVS/SVS/MS) | Vagner Fonseca, et al. |
| EPI_ISL_2298770, EPI_ISL_2298772, EPI_ISL_2298774, EPI_ISL_2298776, EPI_ISL_2298778 | Laboratório Central de Saúde Pública do Maranhão | Coordenação Geral de Laboratórios de Saúde Pública (CGLAB/DAEVS/SVS/MS) | Vagner Fonseca, et al. |
| EPI_ISL_2298779, EPI_ISL_2298781 | Laboratório Central de Saúde Pública do Ceará | Coordenação Geral de Laboratórios de Saúde Pública (CGLAB/DAEVS/SVS/MS) | Vagner Fonseca, et al. |
| EPI_ISL_2298783, EPI_ISL_2298787 | Laboratório Central de Saúde Pública do Pará | Coordenação Geral de Laboratórios de Saúde Pública (CGLAB/DAEVS/SVS/MS) | Vagner Fonseca, et al. |
| EPI_ISL_2298788 | Laboratório Central de Saúde Pública de Roraima | Coordenação Geral de Laboratórios de Saúde Pública (CGLAB/DAEVS/SVS/MS) | Vagner Fonseca, et al. |

EPI_ISL_2298790, EPI_ISL_2298791, EPI_ISL_2298793, EPI_ISL_2298795, EPI_ISL_2298797, EPI_ISL_2298799, EPI_ISL_2298800, EPI_ISL_2298801, EPI_ISL_2298803, EPI_ISL_2298804, EPI_ISL_2298806, EPI_ISL_2298808

| | | | |
|---|---|---|---|
| see above | Laboratório Central de Saúde Pública do Amapá | Coordenação Geral de Laboratórios de Saúde Pública (CGLAB/DAEVS/SVS/MS) | Vagner Fonseca, et al. |
| EPI_ISL_2298810, EPI_ISL_2298812, EPI_ISL_2298814 | Laboratório Central de Saúde Pública do Pará | Coordenação Geral de Laboratórios de Saúde Pública (CGLAB/DAEVS/SVS/MS) | Vagner Fonseca, et al. |
| EPI_ISL_2298816, EPI_ISL_2298817, EPI_ISL_2298819, EPI_ISL_2298821, EPI_ISL_2298823, EPI_ISL_2298824, EPI_ISL_2298826, EPI_ISL_2298828, EPI_ISL_2298830, EPI_ISL_2298832 | Laboratório Central de Saúde Pública do Amapá | Coordenação Geral de Laboratórios de Saúde Pública (CGLAB/DAEVS/SVS/MS) | Vagner Fonseca, et al. |
| EPI_ISL_2298834, EPI_ISL_2298836, EPI_ISL_2298837, EPI_ISL_2298839, EPI_ISL_2298840, EPI_ISL_2298842, EPI_ISL_2298844, EPI_ISL_2298845 | Laboratório Central de Saúde Pública do Maranhão | Coordenação Geral de Laboratórios de Saúde Pública (CGLAB/DAEVS/SVS/MS) | Vagner Fonseca, et al. |
| EPI_ISL_2298847, EPI_ISL_2298849 | Laboratório Central de Saúde Pública de Roraima | Coordenação Geral de Laboratórios de Saúde Pública (CGLAB/DAEVS/SVS/MS) | Vagner Fonseca, et al. |

EPI_ISL_2298850, EPI_ISL_2298852, EPI_ISL_2298854, EPI_ISL_2298856, EPI_ISL_2298859, EPI_ISL_2298861, EPI_ISL_2298863, EPI_ISL_2298865, EPI_ISL_2298867, EPI_ISL_2298869, EPI_ISL_2298871

| | | | |
|---|---|---|---|
| see above | Laboratório Central de Saúde Pública do Amazonas | Coordenação Geral de Laboratórios de Saúde Pública (CGLAB/DAEVS/SVS/MS) | Vagner Fonseca, et al. |
| EPI_ISL_2308399 | Laboratório Central de Saúde Pública do Rio Grande do Norte | Coordenação Geral de Laboratórios de Saúde Pública (CGLAB/DAEVS/SVS/MS) | Vagner Fonseca, et al. |
| EPI_ISL_2308400, EPI_ISL_2308401 | Laboratório Central de Saúde Pública de Alagoas | Coordenação Geral de Laboratórios de Saúde Pública (CGLAB/DAEVS/SVS/MS) | Vagner Fonseca, et al. |
| EPI_ISL_2308402, EPI_ISL_2308403 | Laboratório Central de Saúde Pública de Sergipe | Coordenação Geral de Laboratórios de Saúde Pública (CGLAB/DAEVS/SVS/MS) | Vagner Fonseca, et al. |
| EPI_ISL_2308404 | Laboratório Central de Saúde Pública do Rio Grande do Norte | Coordenação Geral de Laboratórios de Saúde Pública (CGLAB/DAEVS/SVS/MS) | Vagner Fonseca, et al. |
| EPI_ISL_2308405, EPI_ISL_2308406 | Laboratório Central de Saúde Pública de Alagoas | Coordenação Geral de Laboratórios de Saúde Pública (CGLAB/DAEVS/SVS/MS) | Vagner Fonseca, et al. |
| EPI_ISL_2308407 | Laboratório Central de Saúde Pública do Rio Grande do Norte | Coordenação Geral de Laboratórios de Saúde Pública (CGLAB/DAEVS/SVS/MS) | Vagner Fonseca, et al. |
| EPI_ISL_2308408 | Laboratório Central de Saúde Pública de Alagoas | Coordenação Geral de Laboratórios de Saúde Pública (CGLAB/DAEVS/SVS/MS) | Vagner Fonseca, et al. |
| EPI_ISL_2308409 | Laboratório Central de Saúde Pública de Sergipe | Coordenação Geral de Laboratórios de Saúde Pública (CGLAB/DAEVS/SVS/MS) | Vagner Fonseca, et al. |
| EPI_ISL_2308410 | Laboratório Central de Saúde Pública de Alagoas | Coordenação Geral de Laboratórios de Saúde Pública (CGLAB/DAEVS/SVS/MS) | Vagner Fonseca, et al. |
| EPI_ISL_2308411 | Laboratório Central de Saúde Pública de Sergipe | Coordenação Geral de Laboratórios de Saúde Pública (CGLAB/DAEVS/SVS/MS) | Vagner Fonseca, et al. |
| EPI_ISL_2308412 | Laboratório Central de Saúde Pública de Alagoas | Coordenação Geral de Laboratórios de Saúde Pública (CGLAB/DAEVS/SVS/MS) | Vagner Fonseca, et al. |
| EPI_ISL_2308413, EPI_ISL_2308414, EPI_ISL_2308415 | Laboratório Central de Saúde Pública de Sergipe | Coordenação Geral de Laboratórios de Saúde Pública (CGLAB/DAEVS/SVS/MS) | Vagner Fonseca, et al. |
| EPI_ISL_2308416 | Laboratório Central de Saúde Pública de Pernambuco | Coordenação Geral de Laboratórios de Saúde Pública (CGLAB/DAEVS/SVS/MS) | Vagner Fonseca, et al. |
| EPI_ISL_2308417 | Laboratório Central de Saúde Pública de Sergipe | Coordenação Geral de Laboratórios de Saúde Pública (CGLAB/DAEVS/SVS/MS) | Vagner Fonseca, et al. |
| EPI_ISL_2308418, EPI_ISL_2308419, EPI_ISL_2308420, EPI_ISL_2308421, EPI_ISL_2308422, EPI_ISL_2308423, EPI_ISL_2308424, EPI_ISL_2308425, EPI_ISL_2308426 | Laboratório Central de Saúde Pública de Alagoas | Coordenação Geral de Laboratórios de Saúde Pública (CGLAB/DAEVS/SVS/MS) | Vagner Fonseca, et al. |
| EPI_ISL_2308427 | Laboratório Central de Saúde Pública de Sergipe | Coordenação Geral de Laboratórios de Saúde Pública (CGLAB/DAEVS/SVS/MS) | Vagner Fonseca, et al. |
| EPI_ISL_2308428, EPI_ISL_2308429, EPI_ISL_2308430, EPI_ISL_2308431, EPI_ISL_2308432 | Laboratório Central de Saúde Pública de Alagoas | Coordenação Geral de Laboratórios de Saúde Pública (CGLAB/DAEVS/SVS/MS) | Vagner Fonseca, et al. |
| EPI_ISL_2308433 | Laboratório Central de Saúde Pública do Rio Grande do Norte | Coordenação Geral de Laboratórios de Saúde Pública (CGLAB/DAEVS/SVS/MS) | Vagner Fonseca, et al. |
| EPI_ISL_2308434, EPI_ISL_2308435 | Laboratório Central de Saúde Pública de Alagoas | Coordenação Geral de Laboratórios de Saúde Pública (CGLAB/DAEVS/SVS/MS) | Vagner Fonseca, et al. |
| EPI_ISL_2308436, EPI_ISL_2308437 | Laboratório Central de Saúde Pública de Sergipe | Coordenação Geral de Laboratórios de Saúde Pública (CGLAB/DAEVS/SVS/MS) | Vagner Fonseca, et al. |
| EPI_ISL_2308438 | Laboratório Central de Saúde Pública de Pernambuco | Coordenação Geral de Laboratórios de Saúde Pública (CGLAB/DAEVS/SVS/MS) | Vagner Fonseca, et al. |
| EPI_ISL_2308439 | Laboratório Central de Saúde Pública do Rio Grande do Norte | Coordenação Geral de Laboratórios de Saúde Pública (CGLAB/DAEVS/SVS/MS) | Vagner Fonseca, et al. |
| EPI_ISL_2308440, EPI_ISL_2308441, EPI_ISL_2308442, EPI_ISL_2308443, EPI_ISL_2308444, EPI_ISL_2308445 | Laboratório Central de Saúde Pública de Alagoas | Coordenação Geral de Laboratórios de Saúde Pública (CGLAB/DAEVS/SVS/MS) | Vagner Fonseca, et al. |
| EPI_ISL_2308446 | Laboratório Central de Saúde Pública de Pernambuco | Coordenação Geral de Laboratórios de Saúde Pública (CGLAB/DAEVS/SVS/MS) | Vagner Fonseca, et al. |

| | | | |
|---|---|---|---|
| EPI_ISL_2308447, EPI_ISL_2308448, EPI_ISL_2308449, EPI_ISL_2308450, EPI_ISL_2308451, EPI_ISL_2308452 | Laboratório Central de Saúde Pública de Alagoas | Coordenação Geral de Laboratórios de Saúde Pública (CGLAB/DAEVS/SVS/MS) | Vagner Fonseca, et al. |
| EPI_ISL_2308453, EPI_ISL_2308454 | Laboratório Central de Saúde Pública de Sergipe | Coordenação Geral de Laboratórios de Saúde Pública (CGLAB/DAEVS/SVS/MS) | Vagner Fonseca, et al. |
| EPI_ISL_2308455 | Laboratório Central de Saúde Pública de Alagoas | Coordenação Geral de Laboratórios de Saúde Pública (CGLAB/DAEVS/SVS/MS) | Vagner Fonseca, et al. |
| EPI_ISL_2308456, EPI_ISL_2308457, EPI_ISL_2308458 | Laboratório Central de Saúde Pública de Sergipe | Coordenação Geral de Laboratórios de Saúde Pública (CGLAB/DAEVS/SVS/MS) | Vagner Fonseca, et al. |
| EPI_ISL_2308459 | Laboratório Central de Saúde Pública de Alagoas | Coordenação Geral de Laboratórios de Saúde Pública (CGLAB/DAEVS/SVS/MS) | Vagner Fonseca, et al. |
| EPI_ISL_2308460 | Laboratório Central de Saúde Pública do Rio Grande do Norte | Coordenação Geral de Laboratórios de Saúde Pública (CGLAB/DAEVS/SVS/MS) | Vagner Fonseca, et al. |
| EPI_ISL_2308461, EPI_ISL_2308462, EPI_ISL_2308463 | Laboratório Central de Saúde Pública de Alagoas | Coordenação Geral de Laboratórios de Saúde Pública (CGLAB/DAEVS/SVS/MS) | Vagner Fonseca, et al. |
| EPI_ISL_2308464 | Laboratório Central de Saúde Pública do Rio Grande do Norte | Coordenação Geral de Laboratórios de Saúde Pública (CGLAB/DAEVS/SVS/MS) | Vagner Fonseca, et al. |
| EPI_ISL_2308465, EPI_ISL_2308466, EPI_ISL_2308467, EPI_ISL_2308468 | Laboratório Central de Saúde Pública de Sergipe | Coordenação Geral de Laboratórios de Saúde Pública (CGLAB/DAEVS/SVS/MS) | Vagner Fonseca, et al. |
| EPI_ISL_2308469 | Laboratório Central de Saúde Pública de Alagoas | Coordenação Geral de Laboratórios de Saúde Pública (CGLAB/DAEVS/SVS/MS) | Vagner Fonseca, et al. |
| EPI_ISL_2308470, EPI_ISL_2308471, EPI_ISL_2308472, EPI_ISL_2308473, EPI_ISL_2308474, EPI_ISL_2308475, EPI_ISL_2308476, EPI_ISL_2308477, EPI_ISL_2308478 | Laboratório Central de Saúde Pública de Sergipe | Coordenação Geral de Laboratórios de Saúde Pública (CGLAB/DAEVS/SVS/MS) | Vagner Fonseca, et al. |
| EPI_ISL_2308609, EPI_ISL_2308610 | Instituto de Biotecnologia - UNESP-Botucatu-SP | Instituto de Biotecnologia - UNESP-Botucatu-SP | Fábio Sossai Possebon; Leila Sabrina Ullmann; Cecília Artico Banho; Cíntia Bittar; Guilherme Campos; Helena Lage Ferreira; Jorge A. Petroli Marchesi; Lívia Sacchetto; Maisa C. Pereira Parra; Marília Moraes; Maurício L. Nogueira; Paula Rahal; Paulo Inacio da Costa; João Pessoa Araújo Jr. |
| EPI_ISL_2311861 | Fundação Ezequiel Dias | Coordenação Geral de Laboratórios de Saúde Pública (CGLAB/DAEVS/SVS/MS) | Vagner Fonseca, et al. |
| EPI_ISL_2323123, EPI_ISL_2323124, EPI_ISL_2323126, EPI_ISL_2323127, EPI_ISL_2323128, EPI_ISL_2323129, EPI_ISL_2323133, EPI_ISL_2323134, EPI_ISL_2323135, EPI_ISL_2323138 | Instituto de Biotecnologia - UNESP-Botucatu-SP | Instituto de Biotecnologia - UNESP-Botucatu-SP | Fábio Sossai Possebon; Leila Sabrina Ullmann; Cecília Artico Banho; Cíntia Bittar; Guilherme Campos; Helena Lage Ferreira; Jorge A. Petroli Marchesi; Lívia Sacchetto; Maisa C. Pereira Parra; Marília Moraes; Maurício L. Nogueira; Paula Rahal; Paulo Inacio da Costa; João Pessoa Araújo Jr. |
| EPI_ISL_2344420, EPI_ISL_2344421, EPI_ISL_2344422, EPI_ISL_2344423, EPI_ISL_2344424, EPI_ISL_2344425, EPI_ISL_2344426, EPI_ISL_2344427, EPI_ISL_2344428, EPI_ISL_2344429, EPI_ISL_2344430, EPI_ISL_2344431, EPI_ISL_2344432, EPI_ISL_2344433, EPI_ISL_2344434, EPI_ISL_2344435, EPI_ISL_2344436, EPI_ISL_2344437, EPI_ISL_2344438, EPI_ISL_2344439, EPI_ISL_2344440, EPI_ISL_2344441, EPI_ISL_2344442, EPI_ISL_2344443, EPI_ISL_2344444, EPI_ISL_2344445, EPI_ISL_2344446, EPI_ISL_2344447, EPI_ISL_2344448, EPI_ISL_2344449, EPI_ISL_2344450, EPI_ISL_2344451, EPI_ISL_2344452, EPI_ISL_2344453, EPI_ISL_2344454, EPI_ISL_2344455, EPI_ISL_2344456, EPI_ISL_2344457, EPI_ISL_2344458, EPI_ISL_2344459, EPI_ISL_2344460, EPI_ISL_2344461 | | | |
| see above | Instituto Butantan | Instituto de Medicina Tropical de Sao Paulo | Brazil-UK Centre for Arbovirus Discovery Diagnosis Genomics and Epidemiology (CADDE) Genomic Network - Instituto de Medicina Tropical |
| EPI_ISL_2344540 | USF PAULISTA FERNANDOPOLIS ANTONIO PIVATO | Instituto Butantan / UNESP-Botucatu | Dimas Tadeu Covas, Antonio Jorge Martins, Claudia Renata dos Santos Barros, David Schlesinger, Debora Botequio Moretti, Elaine Cristina Marqueze, Elaine Vieira Santos, Evandra Strazza Rodrigues, Heidge Fukumasu, Jayme Augusto de Souza-Neto, José Salvatore Leister Patané, Luiz Alcantara, Luiz Lehmann Coutinho, Maria Carolina Elias, Maurício Lacerda Nogueira, Rafael dos Santos Bezerra, Raul Machado Neto, Rejane Maria Tommasini Grotto, Ricardo Haddad, Sandra Coccuzzo Sampaio Vessoni, Simone Kashima, Svetoslav Nanev Slavov, Vincent Louis Viala |
| EPI_ISL_2344542 | CENTRO DE SAUDE SAO ROQUE DR JOSE CARVALHO BRITO | Instituto Butantan / UNESP-Botucatu | Dimas Tadeu Covas, Antonio Jorge Martins, Claudia Renata dos Santos Barros, David Schlesinger, Debora Botequio Moretti, Elaine Cristina Marqueze, Elaine Vieira Santos, Evandra Strazza Rodrigues, Heidge Fukumasu, Jayme Augusto de Souza-Neto, José Salvatore Leister Patané, Luiz Alcantara, Luiz Lehmann Coutinho, Maria Carolina Elias, Maurício Lacerda Nogueira, Rafael dos Santos Bezerra, Raul Machado Neto, Rejane Maria Tommasini Grotto, Ricardo Haddad, Sandra Coccuzzo Sampaio Vessoni, Simone Kashima, Svetoslav Nanev Slavov, Vincent Louis Viala |
| EPI_ISL_2344544 | ESALQ | Instituto Butantan / ESALQ-Piracicaba | Dimas Tadeu Covas, Antonio Jorge Martins, Claudia Renata dos Santos Barros, David Schlesinger, Debora Botequio Moretti, Elaine Cristina Marqueze, Elaine Vieira Santos, Evandra Strazza Rodrigues, Heidge Fukumasu, Jayme Augusto de Souza-Neto, José Salvatore Leister Patané, Luiz Alcantara, Luiz Lehmann Coutinho, Maria Carolina Elias, Maurício Lacerda Nogueira, Rafael dos Santos Bezerra, Raul Machado Neto, Rejane Maria Tommasini Grotto, Ricardo Haddad, Sandra Coccuzzo Sampaio Vessoni, Simone Kashima, Svetoslav Nanev Slavov, Vincent Louis Viala |
| EPI_ISL_2344545 | DEPARTAMENTO DE SAUDE COLETIVA | Instituto Butantan / UNESP-Botucatu | Dimas Tadeu Covas, Antonio Jorge Martins, Claudia Renata dos Santos Barros, David Schlesinger, Debora Botequio Moretti, Elaine Cristina Marqueze, Elaine Vieira Santos, Evandra Strazza Rodrigues, Heidge Fukumasu, Jayme Augusto de Souza-Neto, José Salvatore Leister Patané, Luiz Alcantara, Luiz Lehmann Coutinho, Maria Carolina Elias, Maurício Lacerda Nogueira, Rafael dos Santos Bezerra, Raul Machado Neto, Rejane Maria Tommasini Grotto, Ricardo Haddad, Sandra Coccuzzo Sampaio Vessoni, Simone Kashima, Svetoslav Nanev Slavov, Vincent Louis Viala |
| EPI_ISL_2344546 | HOSP E MATERNIDADE MUNICIPAL N SRA MONTE SERRAT | Instituto Butantan / UNESP-Botucatu | Dimas Tadeu Covas, Antonio Jorge Martins, Claudia Renata dos Santos Barros, David Schlesinger, Debora Botequio Moretti, Elaine Cristina Marqueze, Elaine Vieira Santos, Evandra Strazza Rodrigues, Heidge Fukumasu, Jayme Augusto de Souza-Neto, José Salvatore Leister Patané, Luiz Alcantara, Luiz Lehmann Coutinho, Maria Carolina Elias, Maurício Lacerda Nogueira, Rafael dos Santos Bezerra, Raul Machado Neto, Rejane Maria Tommasini Grotto, Ricardo Haddad, Sandra Coccuzzo Sampaio Vessoni, Simone Kashima, Svetoslav Nanev Slavov, Vincent Louis Viala |
| EPI_ISL_2344547 | ESALQ | Instituto Butantan / ESALQ-Piracicaba | Dimas Tadeu Covas, Antonio Jorge Martins, Claudia Renata dos Santos Barros, David Schlesinger, Debora Botequio Moretti, Elaine Cristina Marqueze, Elaine Vieira Santos, Evandra Strazza Rodrigues, Heidge Fukumasu, Jayme Augusto de Souza-Neto, José Salvatore Leister Patané, Luiz Alcantara, Luiz Lehmann Coutinho, Maria Carolina Elias, Maurício Lacerda Nogueira, Rafael dos Santos Bezerra, Raul Machado Neto, Rejane Maria Tommasini Grotto, Ricardo Haddad, Sandra Coccuzzo Sampaio Vessoni, Simone Kashima, Svetoslav Nanev Slavov, Vincent Louis Viala |
| EPI_ISL_2344548 | PRONTO ATENDIMENTO MUNICIPAL ALUMINIO | Instituto Butantan / UNESP-Botucatu | Dimas Tadeu Covas, Antonio Jorge Martins, Claudia Renata dos Santos Barros, David Schlesinger, Debora Botequio Moretti, Elaine Cristina Marqueze, Elaine Vieira Santos, Evandra Strazza Rodrigues, Heidge Fukumasu, Jayme Augusto de Souza-Neto, José Salvatore Leister Patané, Luiz Alcantara, Luiz Lehmann Coutinho, Maria Carolina Elias, Maurício Lacerda Nogueira, Rafael dos Santos Bezerra, Raul Machado Neto, Rejane Maria Tommasini Grotto, Ricardo Haddad, Sandra Coccuzzo Sampaio Vessoni, Simone Kashima, Svetoslav Nanev Slavov, Vincent Louis Viala |
| EPI_ISL_2344549 | LABORATORIO MUNICIPAL DE PIRACICABA | Instituto Butantan / ESALQ-Piracicaba | Dimas Tadeu Covas, Antonio Jorge Martins, Claudia Renata dos Santos Barros, David Schlesinger, Debora Botequio Moretti, Elaine Cristina Marqueze, Elaine Vieira Santos, Evandra Strazza Rodrigues, Heidge Fukumasu, Jayme Augusto de Souza-Neto, José Salvatore Leister Patané, Luiz Alcantara, Luiz Lehmann Coutinho, Maria Carolina Elias, Maurício Lacerda Nogueira, Rafael dos Santos Bezerra, Raul Machado Neto, Rejane Maria Tommasini Grotto, Ricardo Haddad, Sandra Coccuzzo Sampaio Vessoni, Simone Kashima, Svetoslav Nanev Slavov, Vincent Louis Viala |
| EPI_ISL_2344550 | HOSPITAL DOS FORNCEDORES | Instituto Butantan / ESALQ-Piracicaba | Dimas Tadeu Covas, Antonio Jorge Martins, Claudia Renata dos Santos Barros, David Schlesinger, Debora Botequio Moretti, Elaine Cristina Marqueze, Elaine Vieira Santos, Evandra Strazza Rodrigues, Heidge Fukumasu, Jayme Augusto de Souza-Neto, José Salvatore Leister Patané, Luiz Alcantara, Luiz Lehmann Coutinho, Maria Carolina Elias, Maurício Lacerda Nogueira, Rafael dos Santos Bezerra, Raul Machado Neto, Rejane Maria Tommasini Grotto, Ricardo Haddad, Sandra Coccuzzo Sampaio Vessoni, Simone Kashima, Svetoslav Nanev Slavov, Vincent Louis Viala |

| | | | |
|---|---|---|---|
| EPI_ISL_2344551 | LABORATORIO MUNICIPAL DE PIRACICABA | Instituto Butantan / ESALQ-Piracicaba | Dimas Tadeu Covas, Antonio Jorge Martins, Claudia Renata dos Santos Barros, David Schlesinger, Debora Botequio Moretti, Elaine Cristina Marqueze, Elaine Vieira Santos, Evandra Strazza Rodrigues, Heidge Fukumasu, Jayme Augusto de Souza-Neto, José Salvatore Leister Patané, Luiz Alcantara, Luiz Lehmann Coutinho, Maria Carolina Elias, Maurício Lacerda Nogueira, Rafael dos Santos Bezerra, Raul Machado Neto, Rejane Maria Tommasini Grotto, Ricardo Haddad, Sandra Coccuzzo Sampaio Vessoni, Simone Kashima, Svetoslav Nanev Slavov, Vincent Louis Viala |
| EPI_ISL_2344552 | HOSPITAL DOS FORNCEDORES | Instituto Butantan / ESALQ-Piracicaba | Dimas Tadeu Covas, Antonio Jorge Martins, Claudia Renata dos Santos Barros, David Schlesinger, Debora Botequio Moretti, Elaine Cristina Marqueze, Elaine Vieira Santos, Evandra Strazza Rodrigues, Heidge Fukumasu, Jayme Augusto de Souza-Neto, José Salvatore Leister Patané, Luiz Alcantara, Luiz Lehmann Coutinho, Maria Carolina Elias, Maurício Lacerda Nogueira, Rafael dos Santos Bezerra, Raul Machado Neto, Rejane Maria Tommasini Grotto, Ricardo Haddad, Sandra Coccuzzo Sampaio Vessoni, Simone Kashima, Svetoslav Nanev Slavov, Vincent Louis Viala |
| EPI_ISL_2344553, EPI_ISL_2344554 | LABORATORIO MUNICIPAL DE PIRACICABA | Instituto Butantan / ESALQ-Piracicaba | Dimas Tadeu Covas, Antonio Jorge Martins, Claudia Renata dos Santos Barros, David Schlesinger, Debora Botequio Moretti, Elaine Cristina Marqueze, Elaine Vieira Santos, Evandra Strazza Rodrigues, Heidge Fukumasu, Jayme Augusto de Souza-Neto, José Salvatore Leister Patané, Luiz Alcantara, Luiz Lehmann Coutinho, Maria Carolina Elias, Maurício Lacerda Nogueira, Rafael dos Santos Bezerra, Raul Machado Neto, Rejane Maria Tommasini Grotto, Ricardo Haddad, Sandra Coccuzzo Sampaio Vessoni, Simone Kashima, Svetoslav Nanev Slavov, Vincent Louis Viala |
| EPI_ISL_2344555 | VIGILANCIA EPIDEMIOLOGICA E CONTROLE DE VETORES PIRASSUNUN | Instituto Butantan / FZEA-USP-Pirassununga | Dimas Tadeu Covas, Antonio Jorge Martins, Claudia Renata dos Santos Barros, David Schlesinger, Debora Botequio Moretti, Elaine Cristina Marqueze, Elaine Vieira Santos, Evandra Strazza Rodrigues, Heidge Fukumasu, Jayme Augusto de Souza-Neto, José Salvatore Leister Patané, Luiz Alcantara, Luiz Lehmann Coutinho, Maria Carolina Elias, Maurício Lacerda Nogueira, Rafael dos Santos Bezerra, Raul Machado Neto, Rejane Maria Tommasini Grotto, Ricardo Haddad, Sandra Coccuzzo Sampaio Vessoni, Simone Kashima, Svetoslav Nanev Slavov, Vincent Louis Viala |
| EPI_ISL_2344556 | HOSPITAL DOS FORNCEDORES | Instituto Butantan / ESALQ-Piracicaba | Dimas Tadeu Covas, Antonio Jorge Martins, Claudia Renata dos Santos Barros, David Schlesinger, Debora Botequio Moretti, Elaine Cristina Marqueze, Elaine Vieira Santos, Evandra Strazza Rodrigues, Heidge Fukumasu, Jayme Augusto de Souza-Neto, José Salvatore Leister Patané, Luiz Alcantara, Luiz Lehmann Coutinho, Maria Carolina Elias, Maurício Lacerda Nogueira, Rafael dos Santos Bezerra, Raul Machado Neto, Rejane Maria Tommasini Grotto, Ricardo Haddad, Sandra Coccuzzo Sampaio Vessoni, Simone Kashima, Svetoslav Nanev Slavov, Vincent Louis Viala |
| EPI_ISL_2344557 | POSTO MEDICO NATAL DIEGUES DE ESTIVA GERBI | Instituto Butantan / FZEA-USP-Pirassununga | Dimas Tadeu Covas, Antonio Jorge Martins, Claudia Renata dos Santos Barros, David Schlesinger, Debora Botequio Moretti, Elaine Cristina Marqueze, Elaine Vieira Santos, Evandra Strazza Rodrigues, Heidge Fukumasu, Jayme Augusto de Souza-Neto, José Salvatore Leister Patané, Luiz Alcantara, Luiz Lehmann Coutinho, Maria Carolina Elias, Maurício Lacerda Nogueira, Rafael dos Santos Bezerra, Raul Machado Neto, Rejane Maria Tommasini Grotto, Ricardo Haddad, Sandra Coccuzzo Sampaio Vessoni, Simone Kashima, Svetoslav Nanev Slavov, Vincent Louis Viala |
| EPI_ISL_2344560, EPI_ISL_2344561 | CENTRO DE SAUDE II DR GABRIEL MESQUITA VARGEM GDE DO SUL | Instituto Butantan / FZEA-USP-Pirassununga | Dimas Tadeu Covas, Antonio Jorge Martins, Claudia Renata dos Santos Barros, David Schlesinger, Debora Botequio Moretti, Elaine Cristina Marqueze, Elaine Vieira Santos, Evandra Strazza Rodrigues, Heidge Fukumasu, Jayme Augusto de Souza-Neto, José Salvatore Leister Patané, Luiz Alcantara, Luiz Lehmann Coutinho, Maria Carolina Elias, Maurício Lacerda Nogueira, Rafael dos Santos Bezerra, Raul Machado Neto, Rejane Maria Tommasini Grotto, Ricardo Haddad, Sandra Coccuzzo Sampaio Vessoni, Simone Kashima, Svetoslav Nanev Slavov, Vincent Louis Viala |
| EPI_ISL_2344562 | UNIDADE BASICA DE SAUDE ALBERTO CARLOS BOINTONI | Instituto Butantan / FZEA-USP-Pirassununga | Dimas Tadeu Covas, Antonio Jorge Martins, Claudia Renata dos Santos Barros, David Schlesinger, Debora Botequio Moretti, Elaine Cristina Marqueze, Elaine Vieira Santos, Evandra Strazza Rodrigues, Heidge Fukumasu, Jayme Augusto de Souza-Neto, José Salvatore Leister Patané, Luiz Alcantara, Luiz Lehmann Coutinho, Maria Carolina Elias, Maurício Lacerda Nogueira, Rafael dos Santos Bezerra, Raul Machado Neto, Rejane Maria Tommasini Grotto, Ricardo Haddad, Sandra Coccuzzo Sampaio Vessoni, Simone Kashima, Svetoslav Nanev Slavov, Vincent Louis Viala |
| EPI_ISL_2344565 | UNIDADE RESPIRATORIA NOVA HORTOLANDIA | Instituto Butantan / FZEA-USP-Pirassununga | Dimas Tadeu Covas, Antonio Jorge Martins, Claudia Renata dos Santos Barros, David Schlesinger, Debora Botequio Moretti, Elaine Cristina Marqueze, Elaine Vieira Santos, Evandra Strazza Rodrigues, Heidge Fukumasu, Jayme Augusto de Souza-Neto, José Salvatore Leister Patané, Luiz Alcantara, Luiz Lehmann Coutinho, Maria Carolina Elias, Maurício Lacerda Nogueira, Rafael dos Santos Bezerra, Raul Machado Neto, Rejane Maria Tommasini Grotto, Ricardo Haddad, Sandra Coccuzzo Sampaio Vessoni, Simone Kashima, Svetoslav Nanev Slavov, Vincent Louis Viala |
| EPI_ISL_2344566, EPI_ISL_2344567 | CENTRO DE SAUDE III AFFONSO LUZZI SANTA CRUZ DAS PALMEIRAS | Instituto Butantan / FZEA-USP-Pirassununga | Dimas Tadeu Covas, Antonio Jorge Martins, Claudia Renata dos Santos Barros, David Schlesinger, Debora Botequio Moretti, Elaine Cristina Marqueze, Elaine Vieira Santos, Evandra Strazza Rodrigues, Heidge Fukumasu, Jayme Augusto de Souza-Neto, José Salvatore Leister Patané, Luiz Alcantara, Luiz Lehmann Coutinho, Maria Carolina Elias, Maurício Lacerda Nogueira, Rafael dos Santos Bezerra, Raul Machado Neto, Rejane Maria Tommasini Grotto, Ricardo Haddad, Sandra Coccuzzo Sampaio Vessoni, Simone Kashima, Svetoslav Nanev Slavov, Vincent Louis Viala |
| EPI_ISL_2344568, EPI_ISL_2344571, EPI_ISL_2344573, EPI_ISL_2344574 | HOSPITAL MUNICIPAL DE ITAPIRA | Instituto Butantan / FZEA-USP-Pirassununga | Dimas Tadeu Covas, Antonio Jorge Martins, Claudia Renata dos Santos Barros, David Schlesinger, Debora Botequio Moretti, Elaine Cristina Marqueze, Elaine Vieira Santos, Evandra Strazza Rodrigues, Heidge Fukumasu, Jayme Augusto de Souza-Neto, José Salvatore Leister Patané, Luiz Alcantara, Luiz Lehmann Coutinho, Maria Carolina Elias, Maurício Lacerda Nogueira, Rafael dos Santos Bezerra, Raul Machado Neto, Rejane Maria Tommasini Grotto, Ricardo Haddad, Sandra Coccuzzo Sampaio Vessoni, Simone Kashima, Svetoslav Nanev Slavov, Vincent Louis Viala |
| EPI_ISL_2344575, EPI_ISL_2344576 | UPA UNIDADE DE PRONTO ATENDIMENTO 24 HORAS BOM JESUS | Instituto Butantan / FZEA-USP-Pirassununga | Dimas Tadeu Covas, Antonio Jorge Martins, Claudia Renata dos Santos Barros, David Schlesinger, Debora Botequio Moretti, Elaine Cristina Marqueze, Elaine Vieira Santos, Evandra Strazza Rodrigues, Heidge Fukumasu, Jayme Augusto de Souza-Neto, José Salvatore Leister Patané, Luiz Alcantara, Luiz Lehmann Coutinho, Maria Carolina Elias, Maurício Lacerda Nogueira, Rafael dos Santos Bezerra, Raul Machado Neto, Rejane Maria Tommasini Grotto, Ricardo Haddad, Sandra Coccuzzo Sampaio Vessoni, Simone Kashima, Svetoslav Nanev Slavov, Vincent Louis Viala |
| EPI_ISL_2344578, EPI_ISL_2344579 | HOSPITAL MUNICIPAL DE ITAPIRA | Instituto Butantan / FZEA-USP-Pirassununga | Dimas Tadeu Covas, Antonio Jorge Martins, Claudia Renata dos Santos Barros, David Schlesinger, Debora Botequio Moretti, Elaine Cristina Marqueze, Elaine Vieira Santos, Evandra Strazza Rodrigues, Heidge Fukumasu, Jayme Augusto de Souza-Neto, José Salvatore Leister Patané, Luiz Alcantara, Luiz Lehmann Coutinho, Maria Carolina Elias, Maurício Lacerda Nogueira, Rafael dos Santos Bezerra, Raul Machado Neto, Rejane Maria Tommasini Grotto, Ricardo Haddad, Sandra Coccuzzo Sampaio Vessoni, Simone Kashima, Svetoslav Nanev Slavov, Vincent Louis Viala |
| EPI_ISL_2344580 | UNIDADE RESPIRATORIA NOVA HORTOLANDIA | Instituto Butantan / FZEA-USP-Pirassununga | Dimas Tadeu Covas, Antonio Jorge Martins, Claudia Renata dos Santos Barros, David Schlesinger, Debora Botequio Moretti, Elaine Cristina Marqueze, Elaine Vieira Santos, Evandra Strazza Rodrigues, Heidge Fukumasu, Jayme Augusto de Souza-Neto, José Salvatore Leister Patané, Luiz Alcantara, Luiz Lehmann Coutinho, Maria Carolina Elias, Maurício Lacerda Nogueira, Rafael dos Santos Bezerra, Raul Machado Neto, Rejane Maria Tommasini Grotto, Ricardo Haddad, Sandra Coccuzzo Sampaio Vessoni, Simone Kashima, Svetoslav Nanev Slavov, Vincent Louis Viala |
| EPI_ISL_2344583 | LABORATORIO DE FRANCA | Instituto Butantan / FZEA-USP-Pirassununga | Dimas Tadeu Covas, Antonio Jorge Martins, Claudia Renata dos Santos Barros, David Schlesinger, Debora Botequio Moretti, Elaine Cristina Marqueze, Elaine Vieira Santos, Evandra Strazza Rodrigues, Heidge Fukumasu, Jayme Augusto de Souza-Neto, José Salvatore Leister Patané, Luiz Alcantara, Luiz Lehmann Coutinho, Maria Carolina Elias, Maurício Lacerda Nogueira, Rafael dos Santos Bezerra, Raul Machado Neto, Rejane Maria Tommasini Grotto, Ricardo Haddad, Sandra Coccuzzo Sampaio Vessoni, Simone Kashima, Svetoslav Nanev Slavov, Vincent Louis Viala |
| EPI_ISL_2344584 | PSF DR ANTONIO PIRES DE ALMEIDA PORTO FELIZ | Instituto Butantan / UNESP-Botucatu | Dimas Tadeu Covas, Antonio Jorge Martins, Claudia Renata dos Santos Barros, David Schlesinger, Debora Botequio Moretti, Elaine Cristina Marqueze, Elaine Vieira Santos, Evandra Strazza Rodrigues, Heidge Fukumasu, Jayme Augusto de Souza-Neto, José Salvatore Leister Patané, Luiz Alcantara, Luiz Lehmann Coutinho, Maria Carolina Elias, Maurício Lacerda Nogueira, Rafael dos Santos Bezerra, Raul Machado Neto, Rejane Maria Tommasini Grotto, Ricardo Haddad, Sandra Coccuzzo Sampaio Vessoni, Simone Kashima, Svetoslav Nanev Slavov, Vincent Louis Viala |
| EPI_ISL_2344585 | VIGILANCIA EPIDEMIOLOGICA E CONTROLE DE VETORES PIRASSUNUN | Instituto Butantan / FZEA-USP-Pirassununga | Dimas Tadeu Covas, Antonio Jorge Martins, Claudia Renata dos Santos Barros, David Schlesinger, Debora Botequio Moretti, Elaine Cristina Marqueze, Elaine Vieira Santos, Evandra Strazza Rodrigues, Heidge Fukumasu, Jayme Augusto de Souza-Neto, José Salvatore Leister Patané, Luiz Alcantara, Luiz Lehmann Coutinho, Maria Carolina Elias, Maurício Lacerda Nogueira, Rafael dos Santos Bezerra, Raul Machado Neto, Rejane Maria Tommasini Grotto, Ricardo Haddad, Sandra Coccuzzo Sampaio Vessoni, Simone Kashima, Svetoslav Nanev Slavov, Vincent Louis Viala |
| EPI_ISL_2344586 | SMS IPERO | Instituto Butantan / UNESP-Botucatu | Dimas Tadeu Covas, Antonio Jorge Martins, Claudia Renata dos Santos Barros, David Schlesinger, Debora Botequio Moretti, Elaine Cristina Marqueze, Elaine Vieira Santos, Evandra Strazza Rodrigues, Heidge Fukumasu, Jayme Augusto de Souza-Neto, José Salvatore Leister Patané, Luiz Alcantara, Luiz Lehmann Coutinho, Maria Carolina Elias, Maurício Lacerda Nogueira, Rafael dos Santos Bezerra, Raul Machado Neto, Rejane Maria Tommasini Grotto, Ricardo Haddad, Sandra Coccuzzo Sampaio Vessoni, Simone Kashima, Svetoslav Nanev Slavov, Vincent Louis Viala |
| EPI_ISL_2344587 | CENTRO DE SAUDE DR RENATO DE CARVALHO RIBEIRO ANGATUBA | Instituto Butantan / UNESP-Botucatu | Dimas Tadeu Covas, Antonio Jorge Martins, Claudia Renata dos Santos Barros, David Schlesinger, Debora Botequio Moretti, Elaine Cristina Marqueze, Elaine Vieira Santos, Evandra Strazza Rodrigues, Heidge Fukumasu, Jayme Augusto de Souza-Neto, José Salvatore Leister Patané, Luiz Alcantara, Luiz Lehmann Coutinho, Maria Carolina Elias, Maurício Lacerda Nogueira, Rafael dos Santos Bezerra, Raul Machado Neto, Rejane Maria Tommasini Grotto, Ricardo Haddad, Sandra Coccuzzo Sampaio Vessoni, Simone Kashima, Svetoslav Nanev Slavov, Vincent Louis Viala |
| EPI_ISL_2344588 | SANTA CASA DE PORTO FELIZ | Instituto Butantan / UNESP-Botucatu | Dimas Tadeu Covas, Antonio Jorge Martins, Claudia Renata dos Santos Barros, David Schlesinger, Debora Botequio Moretti, Elaine Cristina Marqueze, Elaine Vieira Santos, Evandra Strazza Rodrigues, Heidge Fukumasu, Jayme Augusto de Souza-Neto, José Salvatore Leister Patané, Luiz Alcantara, Luiz |

| | | | |
|---|---|---|---|
| | | | Lehmann Coutinho, Maria Carolina Elias, Maurício Lacerda Nogueira, Rafael dos Santos Bezerra, Raul Machado Neto, Rejane Maria Tommasini Grotto, Ricardo Haddad, Sandra Coccuzzo Sampaio Vessoni, Simone Kashima, Svetoslav Nanev Slavov, Vincent Louis Viala |
| EPI_ISL_2344590 | UBS II DE TANABI MILTON MARTINS PERCHES | Instituto Butantan / UNESP-Botucatu | Dimas Tadeu Covas, Antonio Jorge Martins, Claudia Renata dos Santos Barros, David Schlesinger, Debora Botequio Moretti, Elaine Cristina Marqueze, Elaine Vieira Santos, Evandra Strazza Rodrigues, Heidge Fukumasu, Jayme Augusto de Souza-Neto, José Salvatore Leister Patané, Luiz Alcantara, Luiz Lehmann Coutinho, Maria Carolina Elias, Maurício Lacerda Nogueira, Rafael dos Santos Bezerra, Raul Machado Neto, Rejane Maria Tommasini Grotto, Ricardo Haddad, Sandra Coccuzzo Sampaio Vessoni, Simone Kashima, Svetoslav Nanev Slavov, Vincent Louis Viala |
| EPI_ISL_2344591 | SMS SECRETARIA MUNICIPAL DE SAUDE DE BOITUVA | Instituto Butantan / UNESP-Botucatu | Dimas Tadeu Covas, Antonio Jorge Martins, Claudia Renata dos Santos Barros, David Schlesinger, Debora Botequio Moretti, Elaine Cristina Marqueze, Elaine Vieira Santos, Evandra Strazza Rodrigues, Heidge Fukumasu, Jayme Augusto de Souza-Neto, José Salvatore Leister Patané, Luiz Alcantara, Luiz Lehmann Coutinho, Maria Carolina Elias, Maurício Lacerda Nogueira, Rafael dos Santos Bezerra, Raul Machado Neto, Rejane Maria Tommasini Grotto, Ricardo Haddad, Sandra Coccuzzo Sampaio Vessoni, Simone Kashima, Svetoslav Nanev Slavov, Vincent Louis Viala |
| EPI_ISL_2344592 | PRONTO ATENDIMENTO MUNICIPAL ITALO SANTUCCI | Instituto Butantan / UNESP-Botucatu | Dimas Tadeu Covas, Antonio Jorge Martins, Claudia Renata dos Santos Barros, David Schlesinger, Debora Botequio Moretti, Elaine Cristina Marqueze, Elaine Vieira Santos, Evandra Strazza Rodrigues, Heidge Fukumasu, Jayme Augusto de Souza-Neto, José Salvatore Leister Patané, Luiz Alcantara, Luiz Lehmann Coutinho, Maria Carolina Elias, Maurício Lacerda Nogueira, Rafael dos Santos Bezerra, Raul Machado Neto, Rejane Maria Tommasini Grotto, Ricardo Haddad, Sandra Coccuzzo Sampaio Vessoni, Simone Kashima, Svetoslav Nanev Slavov, Vincent Louis Viala |
| EPI_ISL_2344594, EPI_ISL_2344595, EPI_ISL_2344596, EPI_ISL_2344597, EPI_ISL_2344598, EPI_ISL_2344599, EPI_ISL_2344600, EPI_ISL_2344602, EPI_ISL_2344603, EPI_ISL_2344604, EPI_ISL_2344606, EPI_ISL_2344607, EPI_ISL_2344608, EPI_ISL_2344609, EPI_ISL_2344611, EPI_ISL_2344613, EPI_ISL_2344614, EPI_ISL_2344615, EPI_ISL_2344616, EPI_ISL_2344617, EPI_ISL_2344618, EPI_ISL_2344619, EPI_ISL_2344620, EPI_ISL_2344622, EPI_ISL_2344623, EPI_ISL_2344625, EPI_ISL_2344626, EPI_ISL_2344627, EPI_ISL_2344628, EPI_ISL_2344629, EPI_ISL_2344630, EPI_ISL_2344631, EPI_ISL_2344632, EPI_ISL_2344633, EPI_ISL_2344635, EPI_ISL_2344636, EPI_ISL_2344637, EPI_ISL_2344638, EPI_ISL_2344639, EPI_ISL_2344640, EPI_ISL_2344641, EPI_ISL_2344642, EPI_ISL_2344644, EPI_ISL_2344645, EPI_ISL_2344646, EPI_ISL_2344648, EPI_ISL_2344649, EPI_ISL_2344650, EPI_ISL_2344651, EPI_ISL_2344653, EPI_ISL_2344654 | | | |
| see above | Prefeirtura de SP | Instituto Butantan | Dimas Tadeu Covas, Antonio Jorge Martins, Claudia Renata dos Santos Barros, David Schlesinger, Debora Botequio Moretti, Elaine Cristina Marqueze, Elaine Vieira Santos, Evandra Strazza Rodrigues, Heidge Fukumasu, Jayme Augusto de Souza-Neto, José Salvatore Leister Patané, Luiz Alcantara, Luiz Lehmann Coutinho, Maria Carolina Elias, Maurício Lacerda Nogueira, Rafael dos Santos Bezerra, Raul Machado Neto, Rejane Maria Tommasini Grotto, Ricardo Haddad, Sandra Coccuzzo Sampaio Vessoni, Simone Kashima, Svetoslav Nanev Slavov, Vincent Louis Viala |
| EPI_ISL_2344656 | SANTA CASA DE MISERICORDIA DE TIETE | Instituto Butantan / UNESP-Botucatu | Dimas Tadeu Covas, Antonio Jorge Martins, Claudia Renata dos Santos Barros, David Schlesinger, Debora Botequio Moretti, Elaine Cristina Marqueze, Elaine Vieira Santos, Evandra Strazza Rodrigues, Heidge Fukumasu, Jayme Augusto de Souza-Neto, José Salvatore Leister Patané, Luiz Alcantara, Luiz Lehmann Coutinho, Maria Carolina Elias, Maurício Lacerda Nogueira, Rafael dos Santos Bezerra, Raul Machado Neto, Rejane Maria Tommasini Grotto, Ricardo Haddad, Sandra Coccuzzo Sampaio Vessoni, Simone Kashima, Svetoslav Nanev Slavov, Vincent Louis Viala |
| EPI_ISL_2344657 | SAUDE COLETIVA CAPAO BONITO | Instituto Butantan / UNESP-Botucatu | Dimas Tadeu Covas, Antonio Jorge Martins, Claudia Renata dos Santos Barros, David Schlesinger, Debora Botequio Moretti, Elaine Cristina Marqueze, Elaine Vieira Santos, Evandra Strazza Rodrigues, Heidge Fukumasu, Jayme Augusto de Souza-Neto, José Salvatore Leister Patané, Luiz Alcantara, Luiz Lehmann Coutinho, Maria Carolina Elias, Maurício Lacerda Nogueira, Rafael dos Santos Bezerra, Raul Machado Neto, Rejane Maria Tommasini Grotto, Ricardo Haddad, Sandra Coccuzzo Sampaio Vessoni, Simone Kashima, Svetoslav Nanev Slavov, Vincent Louis Viala |
| EPI_ISL_2344658 | UNIDADE SENTINELA COVID19 | Instituto Butantan / UNESP-Botucatu | Dimas Tadeu Covas, Antonio Jorge Martins, Claudia Renata dos Santos Barros, David Schlesinger, Debora Botequio Moretti, Elaine Cristina Marqueze, Elaine Vieira Santos, Evandra Strazza Rodrigues, Heidge Fukumasu, Jayme Augusto de Souza-Neto, José Salvatore Leister Patané, Luiz Alcantara, Luiz Lehmann Coutinho, Maria Carolina Elias, Maurício Lacerda Nogueira, Rafael dos Santos Bezerra, Raul Machado Neto, Rejane Maria Tommasini Grotto, Ricardo Haddad, Sandra Coccuzzo Sampaio Vessoni, Simone Kashima, Svetoslav Nanev Slavov, Vincent Louis Viala |
| EPI_ISL_2344659 | PSF MARIA JOSE SALTO DE PIRAPORA | Instituto Butantan / UNESP-Botucatu | Dimas Tadeu Covas, Antonio Jorge Martins, Claudia Renata dos Santos Barros, David Schlesinger, Debora Botequio Moretti, Elaine Cristina Marqueze, Elaine Vieira Santos, Evandra Strazza Rodrigues, Heidge Fukumasu, Jayme Augusto de Souza-Neto, José Salvatore Leister Patané, Luiz Alcantara, Luiz Lehmann Coutinho, Maria Carolina Elias, Maurício Lacerda Nogueira, Rafael dos Santos Bezerra, Raul Machado Neto, Rejane Maria Tommasini Grotto, Ricardo Haddad, Sandra Coccuzzo Sampaio Vessoni, Simone Kashima, Svetoslav Nanev Slavov, Vincent Louis Viala |
| EPI_ISL_2344660 | COORDENADORIA MUNICIPAL DE SAUDE DE IRACEMAPOLIS | Instituto Butantan / ESALQ-Piracicaba | Dimas Tadeu Covas, Antonio Jorge Martins, Claudia Renata dos Santos Barros, David Schlesinger, Debora Botequio Moretti, Elaine Cristina Marqueze, Elaine Vieira Santos, Evandra Strazza Rodrigues, Heidge Fukumasu, Jayme Augusto de Souza-Neto, José Salvatore Leister Patané, Luiz Alcantara, Luiz Lehmann Coutinho, Maria Carolina Elias, Maurício Lacerda Nogueira, Rafael dos Santos Bezerra, Raul Machado Neto, Rejane Maria Tommasini Grotto, Ricardo Haddad, Sandra Coccuzzo Sampaio Vessoni, Simone Kashima, Svetoslav Nanev Slavov, Vincent Louis Viala |
| EPI_ISL_2344661 | LABORATORIO MUNICIPAL DE ANALISES CLINICAS DE RIO CLARO | Instituto Butantan / ESALQ-Piracicaba | Dimas Tadeu Covas, Antonio Jorge Martins, Claudia Renata dos Santos Barros, David Schlesinger, Debora Botequio Moretti, Elaine Cristina Marqueze, Elaine Vieira Santos, Evandra Strazza Rodrigues, Heidge Fukumasu, Jayme Augusto de Souza-Neto, José Salvatore Leister Patané, Luiz Alcantara, Luiz Lehmann Coutinho, Maria Carolina Elias, Maurício Lacerda Nogueira, Rafael dos Santos Bezerra, Raul Machado Neto, Rejane Maria Tommasini Grotto, Ricardo Haddad, Sandra Coccuzzo Sampaio Vessoni, Simone Kashima, Svetoslav Nanev Slavov, Vincent Louis Viala |
| EPI_ISL_2344662 | SECRETARIA DE SAUDE DE SAO PEDRO | Instituto Butantan / ESALQ-Piracicaba | Dimas Tadeu Covas, Antonio Jorge Martins, Claudia Renata dos Santos Barros, David Schlesinger, Debora Botequio Moretti, Elaine Cristina Marqueze, Elaine Vieira Santos, Evandra Strazza Rodrigues, Heidge Fukumasu, Jayme Augusto de Souza-Neto, José Salvatore Leister Patané, Luiz Alcantara, Luiz Lehmann Coutinho, Maria Carolina Elias, Maurício Lacerda Nogueira, Rafael dos Santos Bezerra, Raul Machado Neto, Rejane Maria Tommasini Grotto, Ricardo Haddad, Sandra Coccuzzo Sampaio Vessoni, Simone Kashima, Svetoslav Nanev Slavov, Vincent Louis Viala |
| EPI_ISL_2344663, EPI_ISL_2344664 | VIGILANCIA EPIDEMIOLOGICA E CONTROLE DE VETORES PIRASSUNUN | Instituto Butantan / FZEA-USP-Pirassununga | Dimas Tadeu Covas, Antonio Jorge Martins, Claudia Renata dos Santos Barros, David Schlesinger, Debora Botequio Moretti, Elaine Cristina Marqueze, Elaine Vieira Santos, Evandra Strazza Rodrigues, Heidge Fukumasu, Jayme Augusto de Souza-Neto, José Salvatore Leister Patané, Luiz Alcantara, Luiz Lehmann Coutinho, Maria Carolina Elias, Maurício Lacerda Nogueira, Rafael dos Santos Bezerra, Raul Machado Neto, Rejane Maria Tommasini Grotto, Ricardo Haddad, Sandra Coccuzzo Sampaio Vessoni, Simone Kashima, Svetoslav Nanev Slavov, Vincent Louis Viala |
| EPI_ISL_2344665, EPI_ISL_2344666 | CENTRO DE ESPECIALIDDS MEDICAS IRMA LEOPOLDINA PIRASSUNUNGA | Instituto Butantan / FZEA-USP-Pirassununga | Dimas Tadeu Covas, Antonio Jorge Martins, Claudia Renata dos Santos Barros, David Schlesinger, Debora Botequio Moretti, Elaine Cristina Marqueze, Elaine Vieira Santos, Evandra Strazza Rodrigues, Heidge Fukumasu, Jayme Augusto de Souza-Neto, José Salvatore Leister Patané, Luiz Alcantara, Luiz Lehmann Coutinho, Maria Carolina Elias, Maurício Lacerda Nogueira, Rafael dos Santos Bezerra, Raul Machado Neto, Rejane Maria Tommasini Grotto, Ricardo Haddad, Sandra Coccuzzo Sampaio Vessoni, Simone Kashima, Svetoslav Nanev Slavov, Vincent Louis Viala |
| EPI_ISL_2344667, EPI_ISL_2344668, EPI_ISL_2344670 | VIGILANCIA EPIDEMIOLOGICA E CONTROLE DE VETORES PIRASSUNUN | Instituto Butantan / FZEA-USP-Pirassununga | Dimas Tadeu Covas, Antonio Jorge Martins, Claudia Renata dos Santos Barros, David Schlesinger, Debora Botequio Moretti, Elaine Cristina Marqueze, Elaine Vieira Santos, Evandra Strazza Rodrigues, Heidge Fukumasu, Jayme Augusto de Souza-Neto, José Salvatore Leister Patané, Luiz Alcantara, Luiz Lehmann Coutinho, Maria Carolina Elias, Maurício Lacerda Nogueira, Rafael dos Santos Bezerra, Raul Machado Neto, Rejane Maria Tommasini Grotto, Ricardo Haddad, Sandra Coccuzzo Sampaio Vessoni, Simone Kashima, Svetoslav Nanev Slavov, Vincent Louis Viala |
| EPI_ISL_2344671 | LABORATORIO MUNICIPAL DE ANALISES CLINICAS DE RIO CLARO | Instituto Butantan / ESALQ-Piracicaba | Dimas Tadeu Covas, Antonio Jorge Martins, Claudia Renata dos Santos Barros, David Schlesinger, Debora Botequio Moretti, Elaine Cristina Marqueze, Elaine Vieira Santos, Evandra Strazza Rodrigues, Heidge Fukumasu, Jayme Augusto de Souza-Neto, José Salvatore Leister Patané, Luiz Alcantara, Luiz Lehmann Coutinho, Maria Carolina Elias, Maurício Lacerda Nogueira, Rafael dos Santos Bezerra, Raul Machado Neto, Rejane Maria Tommasini Grotto, Ricardo Haddad, Sandra Coccuzzo Sampaio Vessoni, Simone Kashima, Svetoslav Nanev Slavov, Vincent Louis Viala |
| EPI_ISL_2344672, EPI_ISL_2344673, EPI_ISL_2344674, EPI_ISL_2344675 | SECRETARIA MUNICIPAL DE SAUDE DE CORDEIROPOLIS | Instituto Butantan / FZEA-USP-Pirassununga | Dimas Tadeu Covas, Antonio Jorge Martins, Claudia Renata dos Santos Barros, David Schlesinger, Debora Botequio Moretti, Elaine Cristina Marqueze, Elaine Vieira Santos, Evandra Strazza Rodrigues, Heidge Fukumasu, Jayme Augusto de Souza-Neto, José Salvatore Leister Patané, Luiz Alcantara, Luiz Lehmann Coutinho, Maria Carolina Elias, Maurício Lacerda Nogueira, Rafael dos Santos Bezerra, Raul Machado Neto, Rejane Maria Tommasini Grotto, Ricardo Haddad, Sandra Coccuzzo Sampaio Vessoni, Simone Kashima, Svetoslav Nanev Slavov, Vincent Louis Viala |
| EPI_ISL_2344676 | SECRETARIA MUNICIPAL DE SAUDE DE CORDEIROPOLIS | Instituto Butantan / ESALQ-Piracicaba | Dimas Tadeu Covas, Antonio Jorge Martins, Claudia Renata dos Santos Barros, David Schlesinger, Debora Botequio Moretti, Elaine Cristina Marqueze, Elaine Vieira Santos, Evandra Strazza Rodrigues, Heidge Fukumasu, Jayme Augusto de Souza-Neto, José Salvatore Leister Patané, Luiz Alcantara, Luiz Lehmann Coutinho, Maria Carolina Elias, Maurício Lacerda Nogueira, Rafael dos Santos Bezerra, Raul Machado Neto, Rejane Maria Tommasini Grotto, Ricardo Haddad, Sandra Coccuzzo Sampaio Vessoni, Simone Kashima, Svetoslav Nanev Slavov, Vincent Louis Viala |
| EPI_ISL_2344677 | SECRETARIA MUNICIPAL DE SAUDE DE CORDEIROPOLIS | Instituto Butantan / FZEA-USP-Pirassununga | Dimas Tadeu Covas, Antonio Jorge Martins, Claudia Renata dos Santos Barros, David Schlesinger, Debora Botequio Moretti, Elaine Cristina Marqueze, Elaine Vieira Santos, Evandra Strazza Rodrigues, Heidge Fukumasu, Jayme Augusto de Souza-Neto, José Salvatore Leister Patané, Luiz Alcantara, Luiz Lehmann Coutinho, Maria Carolina Elias, Maurício Lacerda Nogueira, Rafael dos Santos Bezerra, Raul Machado Neto, Rejane Maria Tommasini Grotto, Ricardo Haddad, Sandra Coccuzzo Sampaio Vessoni, Simone Kashima, Svetoslav Nanev Slavov, Vincent Louis Viala |

| | | | |
|---|---|---|---|
| EPI_ISL_2344678, EPI_ISL_2344679, EPI_ISL_2344680 | CENTRO DE ESPECIALIDDS MEDICAS IRMA LEOPOLDINA PIRASSUNUNGA | Instituto Butantan / FZEA-USP-Pirassununga | Dimas Tadeu Covas, Antonio Jorge Martins, Claudia Renata dos Santos Barros, David Schlesinger, Debora Botequio Moretti, Elaine Cristina Marqueze, Elaine Vieira Santos, Evandra Strazza Rodrigues, Heidge Fukumasu, Jayme Augusto de Souza-Neto, José Salvatore Leister Patané, Luiz Alcantara, Luiz Lehmann Coutinho, Maria Carolina Elias, Maurício Lacerda Nogueira, Rafael dos Santos Bezerra, Raul Machado Neto, Rejane Maria Tommasini Grotto, Ricardo Haddad, Sandra Coccuzzo Sampaio Vessoni, Simone Kashima, Svetoslav Nanev Slavov, Vincent Louis Viala |
| EPI_ISL_2344681 | SANTA CASA DE MISERICORDIA DE TIETE | Instituto Butantan / UNESP-Botucatu | Dimas Tadeu Covas, Antonio Jorge Martins, Claudia Renata dos Santos Barros, David Schlesinger, Debora Botequio Moretti, Elaine Cristina Marqueze, Elaine Vieira Santos, Evandra Strazza Rodrigues, Heidge Fukumasu, Jayme Augusto de Souza-Neto, José Salvatore Leister Patané, Luiz Alcantara, Luiz Lehmann Coutinho, Maria Carolina Elias, Maurício Lacerda Nogueira, Rafael dos Santos Bezerra, Raul Machado Neto, Rejane Maria Tommasini Grotto, Ricardo Haddad, Sandra Coccuzzo Sampaio Vessoni, Simone Kashima, Svetoslav Nanev Slavov, Vincent Louis Viala |
| EPI_ISL_2344683 | CENTRO DE ESPECIALIDDS MEDICAS IRMA LEOPOLDINA PIRASSUNUNGA | Instituto Butantan / ESALQ-Piracicaba | Dimas Tadeu Covas, Antonio Jorge Martins, Claudia Renata dos Santos Barros, David Schlesinger, Debora Botequio Moretti, Elaine Cristina Marqueze, Elaine Vieira Santos, Evandra Strazza Rodrigues, Heidge Fukumasu, Jayme Augusto de Souza-Neto, José Salvatore Leister Patané, Luiz Alcantara, Luiz Lehmann Coutinho, Maria Carolina Elias, Maurício Lacerda Nogueira, Rafael dos Santos Bezerra, Raul Machado Neto, Rejane Maria Tommasini Grotto, Ricardo Haddad, Sandra Coccuzzo Sampaio Vessoni, Simone Kashima, Svetoslav Nanev Slavov, Vincent Louis Viala |
| EPI_ISL_2344684 | UNIDADE DE PRONTO ATENDIMENTO CENTRAL DE CARAGUATATUBA | Instituto Butantan / UNESP-Botucatu | Dimas Tadeu Covas, Antonio Jorge Martins, Claudia Renata dos Santos Barros, David Schlesinger, Debora Botequio Moretti, Elaine Cristina Marqueze, Elaine Vieira Santos, Evandra Strazza Rodrigues, Heidge Fukumasu, Jayme Augusto de Souza-Neto, José Salvatore Leister Patané, Luiz Alcantara, Luiz Lehmann Coutinho, Maria Carolina Elias, Maurício Lacerda Nogueira, Rafael dos Santos Bezerra, Raul Machado Neto, Rejane Maria Tommasini Grotto, Ricardo Haddad, Sandra Coccuzzo Sampaio Vessoni, Simone Kashima, Svetoslav Nanev Slavov, Vincent Louis Viala |
| EPI_ISL_2344685 | COORDENADORIA MUNICIPAL DE SAUDE DE IRACEMAPOLIS | Instituto Butantan / FZEA-USP-Pirassununga | Dimas Tadeu Covas, Antonio Jorge Martins, Claudia Renata dos Santos Barros, David Schlesinger, Debora Botequio Moretti, Elaine Cristina Marqueze, Elaine Vieira Santos, Evandra Strazza Rodrigues, Heidge Fukumasu, Jayme Augusto de Souza-Neto, José Salvatore Leister Patané, Luiz Alcantara, Luiz Lehmann Coutinho, Maria Carolina Elias, Maurício Lacerda Nogueira, Rafael dos Santos Bezerra, Raul Machado Neto, Rejane Maria Tommasini Grotto, Ricardo Haddad, Sandra Coccuzzo Sampaio Vessoni, Simone Kashima, Svetoslav Nanev Slavov, Vincent Louis Viala |
| EPI_ISL_2344686 | PRONTO SOCORRO MUNICIPAL DE TAUBATE | Instituto Butantan / UNESP-Botucatu | Dimas Tadeu Covas, Antonio Jorge Martins, Claudia Renata dos Santos Barros, David Schlesinger, Debora Botequio Moretti, Elaine Cristina Marqueze, Elaine Vieira Santos, Evandra Strazza Rodrigues, Heidge Fukumasu, Jayme Augusto de Souza-Neto, José Salvatore Leister Patané, Luiz Alcantara, Luiz Lehmann Coutinho, Maria Carolina Elias, Maurício Lacerda Nogueira, Rafael dos Santos Bezerra, Raul Machado Neto, Rejane Maria Tommasini Grotto, Ricardo Haddad, Sandra Coccuzzo Sampaio Vessoni, Simone Kashima, Svetoslav Nanev Slavov, Vincent Louis Viala |
| EPI_ISL_2344688 | UNIDADE DE VIGILANCIA EPIDEMIOLOGICA | Instituto Butantan / FZEA-USP-Pirassununga | Dimas Tadeu Covas, Antonio Jorge Martins, Claudia Renata dos Santos Barros, David Schlesinger, Debora Botequio Moretti, Elaine Cristina Marqueze, Elaine Vieira Santos, Evandra Strazza Rodrigues, Heidge Fukumasu, Jayme Augusto de Souza-Neto, José Salvatore Leister Patané, Luiz Alcantara, Luiz Lehmann Coutinho, Maria Carolina Elias, Maurício Lacerda Nogueira, Rafael dos Santos Bezerra, Raul Machado Neto, Rejane Maria Tommasini Grotto, Ricardo Haddad, Sandra Coccuzzo Sampaio Vessoni, Simone Kashima, Svetoslav Nanev Slavov, Vincent Louis Viala |
| EPI_ISL_2344689 | CENTRO DE SAUDE III DE AGUAS DE SAO PEDRO | Instituto Butantan / FZEA-USP-Pirassununga | Dimas Tadeu Covas, Antonio Jorge Martins, Claudia Renata dos Santos Barros, David Schlesinger, Debora Botequio Moretti, Elaine Cristina Marqueze, Elaine Vieira Santos, Evandra Strazza Rodrigues, Heidge Fukumasu, Jayme Augusto de Souza-Neto, José Salvatore Leister Patané, Luiz Alcantara, Luiz Lehmann Coutinho, Maria Carolina Elias, Maurício Lacerda Nogueira, Rafael dos Santos Bezerra, Raul Machado Neto, Rejane Maria Tommasini Grotto, Ricardo Haddad, Sandra Coccuzzo Sampaio Vessoni, Simone Kashima, Svetoslav Nanev Slavov, Vincent Louis Viala |
| EPI_ISL_2344690 | CENTRO MEDICO DR NELSON SALOME DE CONCHAL | Instituto Butantan / FZEA-USP-Pirassununga | Dimas Tadeu Covas, Antonio Jorge Martins, Claudia Renata dos Santos Barros, David Schlesinger, Debora Botequio Moretti, Elaine Cristina Marqueze, Elaine Vieira Santos, Evandra Strazza Rodrigues, Heidge Fukumasu, Jayme Augusto de Souza-Neto, José Salvatore Leister Patané, Luiz Alcantara, Luiz Lehmann Coutinho, Maria Carolina Elias, Maurício Lacerda Nogueira, Rafael dos Santos Bezerra, Raul Machado Neto, Rejane Maria Tommasini Grotto, Ricardo Haddad, Sandra Coccuzzo Sampaio Vessoni, Simone Kashima, Svetoslav Nanev Slavov, Vincent Louis Viala |
| EPI_ISL_2344691 | DIRETORIA MUNICIPAL DE SAUDE DE ENGENHEIRO COELHO | Instituto Butantan / FZEA-USP-Pirassununga | Dimas Tadeu Covas, Antonio Jorge Martins, Claudia Renata dos Santos Barros, David Schlesinger, Debora Botequio Moretti, Elaine Cristina Marqueze, Elaine Vieira Santos, Evandra Strazza Rodrigues, Heidge Fukumasu, Jayme Augusto de Souza-Neto, José Salvatore Leister Patané, Luiz Alcantara, Luiz Lehmann Coutinho, Maria Carolina Elias, Maurício Lacerda Nogueira, Rafael dos Santos Bezerra, Raul Machado Neto, Rejane Maria Tommasini Grotto, Ricardo Haddad, Sandra Coccuzzo Sampaio Vessoni, Simone Kashima, Svetoslav Nanev Slavov, Vincent Louis Viala |
| EPI_ISL_2344692 | VIGILANCIA EPIDEMIOLOGICA E CONTROLE DE VETORES PIRASSUNUN | Instituto Butantan / FZEA-USP-Pirassununga | Dimas Tadeu Covas, Antonio Jorge Martins, Claudia Renata dos Santos Barros, David Schlesinger, Debora Botequio Moretti, Elaine Cristina Marqueze, Elaine Vieira Santos, Evandra Strazza Rodrigues, Heidge Fukumasu, Jayme Augusto de Souza-Neto, José Salvatore Leister Patané, Luiz Alcantara, Luiz Lehmann Coutinho, Maria Carolina Elias, Maurício Lacerda Nogueira, Rafael dos Santos Bezerra, Raul Machado Neto, Rejane Maria Tommasini Grotto, Ricardo Haddad, Sandra Coccuzzo Sampaio Vessoni, Simone Kashima, Svetoslav Nanev Slavov, Vincent Louis Viala |
| EPI_ISL_2344693 | SECRETARIA MUNICIPAL DE SAUDE DE CORDEIROPOLIS | Instituto Butantan / FZEA-USP-Pirassununga | Dimas Tadeu Covas, Antonio Jorge Martins, Claudia Renata dos Santos Barros, David Schlesinger, Debora Botequio Moretti, Elaine Cristina Marqueze, Elaine Vieira Santos, Evandra Strazza Rodrigues, Heidge Fukumasu, Jayme Augusto de Souza-Neto, José Salvatore Leister Patané, Luiz Alcantara, Luiz Lehmann Coutinho, Maria Carolina Elias, Maurício Lacerda Nogueira, Rafael dos Santos Bezerra, Raul Machado Neto, Rejane Maria Tommasini Grotto, Ricardo Haddad, Sandra Coccuzzo Sampaio Vessoni, Simone Kashima, Svetoslav Nanev Slavov, Vincent Louis Viala |
| EPI_ISL_2344694 | VIGILANCIA EPIDEMIOLOGICA DE LEME | Instituto Butantan / FZEA-USP-Pirassununga | Dimas Tadeu Covas, Antonio Jorge Martins, Claudia Renata dos Santos Barros, David Schlesinger, Debora Botequio Moretti, Elaine Cristina Marqueze, Elaine Vieira Santos, Evandra Strazza Rodrigues, Heidge Fukumasu, Jayme Augusto de Souza-Neto, José Salvatore Leister Patané, Luiz Alcantara, Luiz Lehmann Coutinho, Maria Carolina Elias, Maurício Lacerda Nogueira, Rafael dos Santos Bezerra, Raul Machado Neto, Rejane Maria Tommasini Grotto, Ricardo Haddad, Sandra Coccuzzo Sampaio Vessoni, Simone Kashima, Svetoslav Nanev Slavov, Vincent Louis Viala |
| EPI_ISL_2344695 | VIGILANCIA EPIDEMIOLOGICA E CONTROLE DE VETORES PIRASSUNUN | Instituto Butantan / FZEA-USP-Pirassununga | Dimas Tadeu Covas, Antonio Jorge Martins, Claudia Renata dos Santos Barros, David Schlesinger, Debora Botequio Moretti, Elaine Cristina Marqueze, Elaine Vieira Santos, Evandra Strazza Rodrigues, Heidge Fukumasu, Jayme Augusto de Souza-Neto, José Salvatore Leister Patané, Luiz Alcantara, Luiz Lehmann Coutinho, Maria Carolina Elias, Maurício Lacerda Nogueira, Rafael dos Santos Bezerra, Raul Machado Neto, Rejane Maria Tommasini Grotto, Ricardo Haddad, Sandra Coccuzzo Sampaio Vessoni, Simone Kashima, Svetoslav Nanev Slavov, Vincent Louis Viala |
| EPI_ISL_2344696 | UNIDADE DE PRONTO ATENDIMENTO UPA SUL CARAGUATATUBA | Instituto Butantan / UNESP-Botucatu | Dimas Tadeu Covas, Antonio Jorge Martins, Claudia Renata dos Santos Barros, David Schlesinger, Debora Botequio Moretti, Elaine Cristina Marqueze, Elaine Vieira Santos, Evandra Strazza Rodrigues, Heidge Fukumasu, Jayme Augusto de Souza-Neto, José Salvatore Leister Patané, Luiz Alcantara, Luiz Lehmann Coutinho, Maria Carolina Elias, Maurício Lacerda Nogueira, Rafael dos Santos Bezerra, Raul Machado Neto, Rejane Maria Tommasini Grotto, Ricardo Haddad, Sandra Coccuzzo Sampaio Vessoni, Simone Kashima, Svetoslav Nanev Slavov, Vincent Louis Viala |
| EPI_ISL_2344697, EPI_ISL_2344698 | VIGILANCIA EPIDEMIOLOGICA E CONTROLE DE VETORES PIRASSUNUN | Instituto Butantan / FZEA-USP-Pirassununga | Dimas Tadeu Covas, Antonio Jorge Martins, Claudia Renata dos Santos Barros, David Schlesinger, Debora Botequio Moretti, Elaine Cristina Marqueze, Elaine Vieira Santos, Evandra Strazza Rodrigues, Heidge Fukumasu, Jayme Augusto de Souza-Neto, José Salvatore Leister Patané, Luiz Alcantara, Luiz Lehmann Coutinho, Maria Carolina Elias, Maurício Lacerda Nogueira, Rafael dos Santos Bezerra, Raul Machado Neto, Rejane Maria Tommasini Grotto, Ricardo Haddad, Sandra Coccuzzo Sampaio Vessoni, Simone Kashima, Svetoslav Nanev Slavov, Vincent Louis Viala |
| EPI_ISL_2344699, EPI_ISL_2344700 | CENTRO MEDICO DR NELSON SALOME DE CONCHAL | Instituto Butantan / FZEA-USP-Pirassununga | Dimas Tadeu Covas, Antonio Jorge Martins, Claudia Renata dos Santos Barros, David Schlesinger, Debora Botequio Moretti, Elaine Cristina Marqueze, Elaine Vieira Santos, Evandra Strazza Rodrigues, Heidge Fukumasu, Jayme Augusto de Souza-Neto, José Salvatore Leister Patané, Luiz Alcantara, Luiz Lehmann Coutinho, Maria Carolina Elias, Maurício Lacerda Nogueira, Rafael dos Santos Bezerra, Raul Machado Neto, Rejane Maria Tommasini Grotto, Ricardo Haddad, Sandra Coccuzzo Sampaio Vessoni, Simone Kashima, Svetoslav Nanev Slavov, Vincent Louis Viala |
| EPI_ISL_2344701 | UNIDADE MISTA OSVALDO P CAMARGO ELIAS FAUSTO | Instituto Butantan / FZEA-USP-Pirassununga | Dimas Tadeu Covas, Antonio Jorge Martins, Claudia Renata dos Santos Barros, David Schlesinger, Debora Botequio Moretti, Elaine Cristina Marqueze, Elaine Vieira Santos, Evandra Strazza Rodrigues, Heidge Fukumasu, Jayme Augusto de Souza-Neto, José Salvatore Leister Patané, Luiz Alcantara, Luiz Lehmann Coutinho, Maria Carolina Elias, Maurício Lacerda Nogueira, Rafael dos Santos Bezerra, Raul Machado Neto, Rejane Maria Tommasini Grotto, Ricardo Haddad, Sandra Coccuzzo Sampaio Vessoni, Simone Kashima, Svetoslav Nanev Slavov, Vincent Louis Viala |
| EPI_ISL_2344702 | COORDENADORIA MUNICIPAL DE SAUDE DE IRACEMAPOLIS | Instituto Butantan / FZEA-USP-Pirassununga | Dimas Tadeu Covas, Antonio Jorge Martins, Claudia Renata dos Santos Barros, David Schlesinger, Debora Botequio Moretti, Elaine Cristina Marqueze, Elaine Vieira Santos, Evandra Strazza Rodrigues, Heidge Fukumasu, Jayme Augusto de Souza-Neto, José Salvatore Leister Patané, Luiz Alcantara, Luiz Lehmann Coutinho, Maria Carolina Elias, Maurício Lacerda Nogueira, Rafael dos Santos Bezerra, Raul Machado Neto, Rejane Maria Tommasini Grotto, Ricardo Haddad, Sandra Coccuzzo Sampaio Vessoni, Simone Kashima, Svetoslav Nanev Slavov, Vincent Louis Viala |
| EPI_ISL_2344703 | SECRETARIA DE SAUDE DE SANTA GERTRUDES | Instituto Butantan / FZEA-USP-Pirassununga | Dimas Tadeu Covas, Antonio Jorge Martins, Claudia Renata dos Santos Barros, David Schlesinger, Debora Botequio Moretti, Elaine Cristina Marqueze, Elaine Vieira Santos, Evandra Strazza Rodrigues, Heidge Fukumasu, Jayme Augusto de Souza-Neto, José Salvatore Leister Patané, Luiz Alcantara, Luiz |

| | | | |
|---|---|---|---|
| | | | Lehmann Coutinho, Maria Carolina Elias, Maurício Lacerda Nogueira, Rafael dos Santos Bezerra, Raul Machado Neto, Rejane Maria Tommasini Grotto, Ricardo Haddad, Sandra Coccuzzo Sampaio Vessoni, Simone Kashima, Svetoslav Nanev Slavov, Vincent Louis Viala |
| EPI_ISL_2344704 | VIGILANCIA EPIDEMIOLOGICA E CONTROLE DE VETORES PIRASSUNUN | Instituto Butantan / FZEA-USP-Pirassununga | Dimas Tadeu Covas, Antonio Jorge Martins, Claudia Renata dos Santos Barros, David Schlesinger, Debora Botequio Moretti, Elaine Cristina Marqueze, Elaine Vieira Santos, Evandra Strazza Rodrigues, Heidge Fukumasu, Jayme Augusto de Souza-Neto, José Salvatore Leister Patané, Luiz Alcantara, Luiz Lehmann Coutinho, Maria Carolina Elias, Maurício Lacerda Nogueira, Rafael dos Santos Bezerra, Raul Machado Neto, Rejane Maria Tommasini Grotto, Ricardo Haddad, Sandra Coccuzzo Sampaio Vessoni, Simone Kashima, Svetoslav Nanev Slavov, Vincent Louis Viala |
| EPI_ISL_2344705 | VIGILANCIA EPIDEMIOLOGICA DE LEME | Instituto Butantan / FZEA-USP-Pirassununga | Dimas Tadeu Covas, Antonio Jorge Martins, Claudia Renata dos Santos Barros, David Schlesinger, Debora Botequio Moretti, Elaine Cristina Marqueze, Elaine Vieira Santos, Evandra Strazza Rodrigues, Heidge Fukumasu, Jayme Augusto de Souza-Neto, José Salvatore Leister Patané, Luiz Alcantara, Luiz Lehmann Coutinho, Maria Carolina Elias, Maurício Lacerda Nogueira, Rafael dos Santos Bezerra, Raul Machado Neto, Rejane Maria Tommasini Grotto, Ricardo Haddad, Sandra Coccuzzo Sampaio Vessoni, Simone Kashima, Svetoslav Nanev Slavov, Vincent Louis Viala |
| EPI_ISL_2344706 | VIGILANCIA EPIDEMIOLOGICA E CONTROLE DE VETORES PIRASSUNUN | Instituto Butantan / FZEA-USP-Pirassununga | Dimas Tadeu Covas, Antonio Jorge Martins, Claudia Renata dos Santos Barros, David Schlesinger, Debora Botequio Moretti, Elaine Cristina Marqueze, Elaine Vieira Santos, Evandra Strazza Rodrigues, Heidge Fukumasu, Jayme Augusto de Souza-Neto, José Salvatore Leister Patané, Luiz Alcantara, Luiz Lehmann Coutinho, Maria Carolina Elias, Maurício Lacerda Nogueira, Rafael dos Santos Bezerra, Raul Machado Neto, Rejane Maria Tommasini Grotto, Ricardo Haddad, Sandra Coccuzzo Sampaio Vessoni, Simone Kashima, Svetoslav Nanev Slavov, Vincent Louis Viala |
| EPI_ISL_2344707 | UNIDADE DE PRONTO ATENDIMENTO CENTRAL DE CARAGUATATUBA | Instituto Butantan / UNESP-Botucatu | Dimas Tadeu Covas, Antonio Jorge Martins, Claudia Renata dos Santos Barros, David Schlesinger, Debora Botequio Moretti, Elaine Cristina Marqueze, Elaine Vieira Santos, Evandra Strazza Rodrigues, Heidge Fukumasu, Jayme Augusto de Souza-Neto, José Salvatore Leister Patané, Luiz Alcantara, Luiz Lehmann Coutinho, Maria Carolina Elias, Maurício Lacerda Nogueira, Rafael dos Santos Bezerra, Raul Machado Neto, Rejane Maria Tommasini Grotto, Ricardo Haddad, Sandra Coccuzzo Sampaio Vessoni, Simone Kashima, Svetoslav Nanev Slavov, Vincent Louis Viala |
| EPI_ISL_2344709 | UNIDADE DE VIGILANCIA EPIDEMIOLOGICA | Instituto Butantan / FZEA-USP-Pirassununga | Dimas Tadeu Covas, Antonio Jorge Martins, Claudia Renata dos Santos Barros, David Schlesinger, Debora Botequio Moretti, Elaine Cristina Marqueze, Elaine Vieira Santos, Evandra Strazza Rodrigues, Heidge Fukumasu, Jayme Augusto de Souza-Neto, José Salvatore Leister Patané, Luiz Alcantara, Luiz Lehmann Coutinho, Maria Carolina Elias, Maurício Lacerda Nogueira, Rafael dos Santos Bezerra, Raul Machado Neto, Rejane Maria Tommasini Grotto, Ricardo Haddad, Sandra Coccuzzo Sampaio Vessoni, Simone Kashima, Svetoslav Nanev Slavov, Vincent Louis Viala |
| EPI_ISL_2344710 | CENTRO DE SAUDE DR MARIO DIAS DE AGUIAR CAPIVARI | Instituto Butantan / FZEA-USP-Pirassununga | Dimas Tadeu Covas, Antonio Jorge Martins, Claudia Renata dos Santos Barros, David Schlesinger, Debora Botequio Moretti, Elaine Cristina Marqueze, Elaine Vieira Santos, Evandra Strazza Rodrigues, Heidge Fukumasu, Jayme Augusto de Souza-Neto, José Salvatore Leister Patané, Luiz Alcantara, Luiz Lehmann Coutinho, Maria Carolina Elias, Maurício Lacerda Nogueira, Rafael dos Santos Bezerra, Raul Machado Neto, Rejane Maria Tommasini Grotto, Ricardo Haddad, Sandra Coccuzzo Sampaio Vessoni, Simone Kashima, Svetoslav Nanev Slavov, Vincent Louis Viala |
| EPI_ISL_2344712 | CENTRO MEDICO DR NELSON SALOME DE CONCHAL | Instituto Butantan / FZEA-USP-Pirassununga | Dimas Tadeu Covas, Antonio Jorge Martins, Claudia Renata dos Santos Barros, David Schlesinger, Debora Botequio Moretti, Elaine Cristina Marqueze, Elaine Vieira Santos, Evandra Strazza Rodrigues, Heidge Fukumasu, Jayme Augusto de Souza-Neto, José Salvatore Leister Patané, Luiz Alcantara, Luiz Lehmann Coutinho, Maria Carolina Elias, Maurício Lacerda Nogueira, Rafael dos Santos Bezerra, Raul Machado Neto, Rejane Maria Tommasini Grotto, Ricardo Haddad, Sandra Coccuzzo Sampaio Vessoni, Simone Kashima, Svetoslav Nanev Slavov, Vincent Louis Viala |
| EPI_ISL_2344713, EPI_ISL_2344714, EPI_ISL_2344715 | VIGILANCIA EPIDEMIOLOGICA DE LEME | Instituto Butantan / FZEA-USP-Pirassununga | Dimas Tadeu Covas, Antonio Jorge Martins, Claudia Renata dos Santos Barros, David Schlesinger, Debora Botequio Moretti, Elaine Cristina Marqueze, Elaine Vieira Santos, Evandra Strazza Rodrigues, Heidge Fukumasu, Jayme Augusto de Souza-Neto, José Salvatore Leister Patané, Luiz Alcantara, Luiz Lehmann Coutinho, Maria Carolina Elias, Maurício Lacerda Nogueira, Rafael dos Santos Bezerra, Raul Machado Neto, Rejane Maria Tommasini Grotto, Ricardo Haddad, Sandra Coccuzzo Sampaio Vessoni, Simone Kashima, Svetoslav Nanev Slavov, Vincent Louis Viala |
| EPI_ISL_2344717 | UNIDADE DE PRONTO ATENDIMENTO CENTRAL DE CARAGUATATUBA | Instituto Butantan / UNESP-Botucatu | Dimas Tadeu Covas, Antonio Jorge Martins, Claudia Renata dos Santos Barros, David Schlesinger, Debora Botequio Moretti, Elaine Cristina Marqueze, Elaine Vieira Santos, Evandra Strazza Rodrigues, Heidge Fukumasu, Jayme Augusto de Souza-Neto, José Salvatore Leister Patané, Luiz Alcantara, Luiz Lehmann Coutinho, Maria Carolina Elias, Maurício Lacerda Nogueira, Rafael dos Santos Bezerra, Raul Machado Neto, Rejane Maria Tommasini Grotto, Ricardo Haddad, Sandra Coccuzzo Sampaio Vessoni, Simone Kashima, Svetoslav Nanev Slavov, Vincent Louis Viala |
| EPI_ISL_2344718 | UNIDADE DE VIGILANCIA EPIDEMIOLOGICA | Instituto Butantan / FZEA-USP-Pirassununga | Dimas Tadeu Covas, Antonio Jorge Martins, Claudia Renata dos Santos Barros, David Schlesinger, Debora Botequio Moretti, Elaine Cristina Marqueze, Elaine Vieira Santos, Evandra Strazza Rodrigues, Heidge Fukumasu, Jayme Augusto de Souza-Neto, José Salvatore Leister Patané, Luiz Alcantara, Luiz Lehmann Coutinho, Maria Carolina Elias, Maurício Lacerda Nogueira, Rafael dos Santos Bezerra, Raul Machado Neto, Rejane Maria Tommasini Grotto, Ricardo Haddad, Sandra Coccuzzo Sampaio Vessoni, Simone Kashima, Svetoslav Nanev Slavov, Vincent Louis Viala |
| EPI_ISL_2344719 | VIGILANCIA EPIDEMIOLOGICA E CONTROLE DE VETORES PIRASSUNUN | Instituto Butantan / FZEA-USP-Pirassununga | Dimas Tadeu Covas, Antonio Jorge Martins, Claudia Renata dos Santos Barros, David Schlesinger, Debora Botequio Moretti, Elaine Cristina Marqueze, Elaine Vieira Santos, Evandra Strazza Rodrigues, Heidge Fukumasu, Jayme Augusto de Souza-Neto, José Salvatore Leister Patané, Luiz Alcantara, Luiz Lehmann Coutinho, Maria Carolina Elias, Maurício Lacerda Nogueira, Rafael dos Santos Bezerra, Raul Machado Neto, Rejane Maria Tommasini Grotto, Ricardo Haddad, Sandra Coccuzzo Sampaio Vessoni, Simone Kashima, Svetoslav Nanev Slavov, Vincent Louis Viala |
| EPI_ISL_2344720 | VIGILANCIA EPIDEMIOLOGICA DE LEME | Instituto Butantan / FZEA-USP-Pirassununga | Dimas Tadeu Covas, Antonio Jorge Martins, Claudia Renata dos Santos Barros, David Schlesinger, Debora Botequio Moretti, Elaine Cristina Marqueze, Elaine Vieira Santos, Evandra Strazza Rodrigues, Heidge Fukumasu, Jayme Augusto de Souza-Neto, José Salvatore Leister Patané, Luiz Alcantara, Luiz Lehmann Coutinho, Maria Carolina Elias, Maurício Lacerda Nogueira, Rafael dos Santos Bezerra, Raul Machado Neto, Rejane Maria Tommasini Grotto, Ricardo Haddad, Sandra Coccuzzo Sampaio Vessoni, Simone Kashima, Svetoslav Nanev Slavov, Vincent Louis Viala |
| EPI_ISL_2344721, EPI_ISL_2344722 | UNIDADE BASICA DE SAUDE DE IPEUNA | Instituto Butantan / FZEA-USP-Pirassununga | Dimas Tadeu Covas, Antonio Jorge Martins, Claudia Renata dos Santos Barros, David Schlesinger, Debora Botequio Moretti, Elaine Cristina Marqueze, Elaine Vieira Santos, Evandra Strazza Rodrigues, Heidge Fukumasu, Jayme Augusto de Souza-Neto, José Salvatore Leister Patané, Luiz Alcantara, Luiz Lehmann Coutinho, Maria Carolina Elias, Maurício Lacerda Nogueira, Rafael dos Santos Bezerra, Raul Machado Neto, Rejane Maria Tommasini Grotto, Ricardo Haddad, Sandra Coccuzzo Sampaio Vessoni, Simone Kashima, Svetoslav Nanev Slavov, Vincent Louis Viala |
| EPI_ISL_2344725 | UNIDADE MISTA DE SAUDE MOMBUCA | Instituto Butantan / FZEA-USP-Pirassununga | Dimas Tadeu Covas, Antonio Jorge Martins, Claudia Renata dos Santos Barros, David Schlesinger, Debora Botequio Moretti, Elaine Cristina Marqueze, Elaine Vieira Santos, Evandra Strazza Rodrigues, Heidge Fukumasu, Jayme Augusto de Souza-Neto, José Salvatore Leister Patané, Luiz Alcantara, Luiz Lehmann Coutinho, Maria Carolina Elias, Maurício Lacerda Nogueira, Rafael dos Santos Bezerra, Raul Machado Neto, Rejane Maria Tommasini Grotto, Ricardo Haddad, Sandra Coccuzzo Sampaio Vessoni, Simone Kashima, Svetoslav Nanev Slavov, Vincent Louis Viala |
| EPI_ISL_2344726, EPI_ISL_2344727 | SECRETARIA DE SAUDE DE RAFARD | Instituto Butantan / FZEA-USP-Pirassununga | Dimas Tadeu Covas, Antonio Jorge Martins, Claudia Renata dos Santos Barros, David Schlesinger, Debora Botequio Moretti, Elaine Cristina Marqueze, Elaine Vieira Santos, Evandra Strazza Rodrigues, Heidge Fukumasu, Jayme Augusto de Souza-Neto, José Salvatore Leister Patané, Luiz Alcantara, Luiz Lehmann Coutinho, Maria Carolina Elias, Maurício Lacerda Nogueira, Rafael dos Santos Bezerra, Raul Machado Neto, Rejane Maria Tommasini Grotto, Ricardo Haddad, Sandra Coccuzzo Sampaio Vessoni, Simone Kashima, Svetoslav Nanev Slavov, Vincent Louis Viala |
| EPI_ISL_2344728, EPI_ISL_2344729 | LABORATORIO MUNICIPAL DE ANALISES CLINICAS DE RIO CLARO | Instituto Butantan / FZEA-USP-Pirassununga | Dimas Tadeu Covas, Antonio Jorge Martins, Claudia Renata dos Santos Barros, David Schlesinger, Debora Botequio Moretti, Elaine Cristina Marqueze, Elaine Vieira Santos, Evandra Strazza Rodrigues, Heidge Fukumasu, Jayme Augusto de Souza-Neto, José Salvatore Leister Patané, Luiz Alcantara, Luiz Lehmann Coutinho, Maria Carolina Elias, Maurício Lacerda Nogueira, Rafael dos Santos Bezerra, Raul Machado Neto, Rejane Maria Tommasini Grotto, Ricardo Haddad, Sandra Coccuzzo Sampaio Vessoni, Simone Kashima, Svetoslav Nanev Slavov, Vincent Louis Viala |
| EPI_ISL_2344730 | SECRETARIA DE SAUDE DE SANTA GERTRUDES | Instituto Butantan / FZEA-USP-Pirassununga | Dimas Tadeu Covas, Antonio Jorge Martins, Claudia Renata dos Santos Barros, David Schlesinger, Debora Botequio Moretti, Elaine Cristina Marqueze, Elaine Vieira Santos, Evandra Strazza Rodrigues, Heidge Fukumasu, Jayme Augusto de Souza-Neto, José Salvatore Leister Patané, Luiz Alcantara, Luiz Lehmann Coutinho, Maria Carolina Elias, Maurício Lacerda Nogueira, Rafael dos Santos Bezerra, Raul Machado Neto, Rejane Maria Tommasini Grotto, Ricardo Haddad, Sandra Coccuzzo Sampaio Vessoni, Simone Kashima, Svetoslav Nanev Slavov, Vincent Louis Viala |
| EPI_ISL_2344731 | SECRETARIA DE SAUDE DE SAO PEDRO | Instituto Butantan / FZEA-USP-Pirassununga | Dimas Tadeu Covas, Antonio Jorge Martins, Claudia Renata dos Santos Barros, David Schlesinger, Debora Botequio Moretti, Elaine Cristina Marqueze, Elaine Vieira Santos, Evandra Strazza Rodrigues, Heidge Fukumasu, Jayme Augusto de Souza-Neto, José Salvatore Leister Patané, Luiz Alcantara, Luiz Lehmann Coutinho, Maria Carolina Elias, Maurício Lacerda Nogueira, Rafael dos Santos Bezerra, Raul Machado Neto, Rejane Maria Tommasini Grotto, Ricardo Haddad, Sandra Coccuzzo Sampaio Vessoni, Simone Kashima, Svetoslav Nanev Slavov, Vincent Louis Viala |
| EPI_ISL_2344732 | UNIDADE DE PRONTO ATENDIMENTO CENTRAL DE CARAGUATATUBA | Instituto Butantan / UNESP-Botucatu | Dimas Tadeu Covas, Antonio Jorge Martins, Claudia Renata dos Santos Barros, David Schlesinger, Debora Botequio Moretti, Elaine Cristina Marqueze, Elaine Vieira Santos, Evandra Strazza Rodrigues, Heidge Fukumasu, Jayme Augusto de Souza-Neto, José Salvatore Leister Patané, Luiz Alcantara, Luiz Lehmann Coutinho, Maria Carolina Elias, Maurício Lacerda Nogueira, Rafael dos Santos Bezerra, Raul Machado Neto, Rejane Maria Tommasini Grotto, Ricardo Haddad, Sandra Coccuzzo Sampaio Vessoni, Simone Kashima, Svetoslav Nanev Slavov, Vincent Louis Viala |

| | | | |
|---|---|---|---|
| EPI_ISL_2344734 | LABORATORIO MUNICIPAL DE PIRACICABA | Instituto Butantan / ESALQ-Piracicaba | Dimas Tadeu Covas, Antonio Jorge Martins, Claudia Renata dos Santos Barros, David Schlesinger, Debora Botequio Moretti, Elaine Cristina Marqueze, Elaine Vieira Santos, Evandra Strazza Rodrigues, Heidge Fukumasu, Jayme Augusto de Souza-Neto, José Salvatore Leister Patané, Luiz Alcantara, Luiz Lehmann Coutinho, Maria Carolina Elias, Maurício Lacerda Nogueira, Rafael dos Santos Bezerra, Raul Machado Neto, Rejane Maria Tommasini Grotto, Ricardo Haddad, Sandra Coccuzzo Sampaio Vessoni, Simone Kashima, Svetoslav Nanev Slavov, Vincent Louis Viala |
| EPI_ISL_2344735 | CENTRO DE SAUDE PE CESAR CORNEO | Instituto Butantan / ESALQ-Piracicaba | Dimas Tadeu Covas, Antonio Jorge Martins, Claudia Renata dos Santos Barros, David Schlesinger, Debora Botequio Moretti, Elaine Cristina Marqueze, Elaine Vieira Santos, Evandra Strazza Rodrigues, Heidge Fukumasu, Jayme Augusto de Souza-Neto, José Salvatore Leister Patané, Luiz Alcantara, Luiz Lehmann Coutinho, Maria Carolina Elias, Maurício Lacerda Nogueira, Rafael dos Santos Bezerra, Raul Machado Neto, Rejane Maria Tommasini Grotto, Ricardo Haddad, Sandra Coccuzzo Sampaio Vessoni, Simone Kashima, Svetoslav Nanev Slavov, Vincent Louis Viala |
| EPI_ISL_2344736 | UPA 24 HORAS DR ALOISIO MUNIZ DE ANDRADE | Instituto Butantan / ESALQ-Piracicaba | Dimas Tadeu Covas, Antonio Jorge Martins, Claudia Renata dos Santos Barros, David Schlesinger, Debora Botequio Moretti, Elaine Cristina Marqueze, Elaine Vieira Santos, Evandra Strazza Rodrigues, Heidge Fukumasu, Jayme Augusto de Souza-Neto, José Salvatore Leister Patané, Luiz Alcantara, Luiz Lehmann Coutinho, Maria Carolina Elias, Maurício Lacerda Nogueira, Rafael dos Santos Bezerra, Raul Machado Neto, Rejane Maria Tommasini Grotto, Ricardo Haddad, Sandra Coccuzzo Sampaio Vessoni, Simone Kashima, Svetoslav Nanev Slavov, Vincent Louis Viala |
| EPI_ISL_2344737 | SECRETARIA MUNICIPAL DE SAUDE DE CORDEIROPOLIS | Instituto Butantan / ESALQ-Piracicaba | Dimas Tadeu Covas, Antonio Jorge Martins, Claudia Renata dos Santos Barros, David Schlesinger, Debora Botequio Moretti, Elaine Cristina Marqueze, Elaine Vieira Santos, Evandra Strazza Rodrigues, Heidge Fukumasu, Jayme Augusto de Souza-Neto, José Salvatore Leister Patané, Luiz Alcantara, Luiz Lehmann Coutinho, Maria Carolina Elias, Maurício Lacerda Nogueira, Rafael dos Santos Bezerra, Raul Machado Neto, Rejane Maria Tommasini Grotto, Ricardo Haddad, Sandra Coccuzzo Sampaio Vessoni, Simone Kashima, Svetoslav Nanev Slavov, Vincent Louis Viala |
| EPI_ISL_2344739, EPI_ISL_2344740, EPI_ISL_2344741, EPI_ISL_2344742 | SAE CTA SERV ATEND ESPEC TESTAGEM E ACONSELHAMENTO | Instituto Butantan / ESALQ-Piracicaba | Dimas Tadeu Covas, Antonio Jorge Martins, Claudia Renata dos Santos Barros, David Schlesinger, Debora Botequio Moretti, Elaine Cristina Marqueze, Elaine Vieira Santos, Evandra Strazza Rodrigues, Heidge Fukumasu, Jayme Augusto de Souza-Neto, José Salvatore Leister Patané, Luiz Alcantara, Luiz Lehmann Coutinho, Maria Carolina Elias, Maurício Lacerda Nogueira, Rafael dos Santos Bezerra, Raul Machado Neto, Rejane Maria Tommasini Grotto, Ricardo Haddad, Sandra Coccuzzo Sampaio Vessoni, Simone Kashima, Svetoslav Nanev Slavov, Vincent Louis Viala |
| EPI_ISL_2344743 | UBS II DE PIRAPOZINHO C SERVICO DE EACS E ESF | Instituto Butantan / ESALQ-Piracicaba | Dimas Tadeu Covas, Antonio Jorge Martins, Claudia Renata dos Santos Barros, David Schlesinger, Debora Botequio Moretti, Elaine Cristina Marqueze, Elaine Vieira Santos, Evandra Strazza Rodrigues, Heidge Fukumasu, Jayme Augusto de Souza-Neto, José Salvatore Leister Patané, Luiz Alcantara, Luiz Lehmann Coutinho, Maria Carolina Elias, Maurício Lacerda Nogueira, Rafael dos Santos Bezerra, Raul Machado Neto, Rejane Maria Tommasini Grotto, Ricardo Haddad, Sandra Coccuzzo Sampaio Vessoni, Simone Kashima, Svetoslav Nanev Slavov, Vincent Louis Viala |
| EPI_ISL_2344744 | CENTRO DE SAUDE DO INDAIA DO AGUAPEI FLORIDA PAULISTA | Instituto Butantan / ESALQ-Piracicaba | Dimas Tadeu Covas, Antonio Jorge Martins, Claudia Renata dos Santos Barros, David Schlesinger, Debora Botequio Moretti, Elaine Cristina Marqueze, Elaine Vieira Santos, Evandra Strazza Rodrigues, Heidge Fukumasu, Jayme Augusto de Souza-Neto, José Salvatore Leister Patané, Luiz Alcantara, Luiz Lehmann Coutinho, Maria Carolina Elias, Maurício Lacerda Nogueira, Rafael dos Santos Bezerra, Raul Machado Neto, Rejane Maria Tommasini Grotto, Ricardo Haddad, Sandra Coccuzzo Sampaio Vessoni, Simone Kashima, Svetoslav Nanev Slavov, Vincent Louis Viala |
| EPI_ISL_2344745 | SAE CTA SERV ATEND ESPEC TESTAGEM E ACONSELHAMENTO | Instituto Butantan / ESALQ-Piracicaba | Dimas Tadeu Covas, Antonio Jorge Martins, Claudia Renata dos Santos Barros, David Schlesinger, Debora Botequio Moretti, Elaine Cristina Marqueze, Elaine Vieira Santos, Evandra Strazza Rodrigues, Heidge Fukumasu, Jayme Augusto de Souza-Neto, José Salvatore Leister Patané, Luiz Alcantara, Luiz Lehmann Coutinho, Maria Carolina Elias, Maurício Lacerda Nogueira, Rafael dos Santos Bezerra, Raul Machado Neto, Rejane Maria Tommasini Grotto, Ricardo Haddad, Sandra Coccuzzo Sampaio Vessoni, Simone Kashima, Svetoslav Nanev Slavov, Vincent Louis Viala |
| EPI_ISL_2344746 | VIGILANCIA EPIDEMIOLOGICA DE LEME | Instituto Butantan / ESALQ-Piracicaba | Dimas Tadeu Covas, Antonio Jorge Martins, Claudia Renata dos Santos Barros, David Schlesinger, Debora Botequio Moretti, Elaine Cristina Marqueze, Elaine Vieira Santos, Evandra Strazza Rodrigues, Heidge Fukumasu, Jayme Augusto de Souza-Neto, José Salvatore Leister Patané, Luiz Alcantara, Luiz Lehmann Coutinho, Maria Carolina Elias, Maurício Lacerda Nogueira, Rafael dos Santos Bezerra, Raul Machado Neto, Rejane Maria Tommasini Grotto, Ricardo Haddad, Sandra Coccuzzo Sampaio Vessoni, Simone Kashima, Svetoslav Nanev Slavov, Vincent Louis Viala |
| EPI_ISL_2344748 | SECRETARIA MUNICIPAL DE SAUDE | Instituto Butantan / ESALQ-Piracicaba | Dimas Tadeu Covas, Antonio Jorge Martins, Claudia Renata dos Santos Barros, David Schlesinger, Debora Botequio Moretti, Elaine Cristina Marqueze, Elaine Vieira Santos, Evandra Strazza Rodrigues, Heidge Fukumasu, Jayme Augusto de Souza-Neto, José Salvatore Leister Patané, Luiz Alcantara, Luiz Lehmann Coutinho, Maria Carolina Elias, Maurício Lacerda Nogueira, Rafael dos Santos Bezerra, Raul Machado Neto, Rejane Maria Tommasini Grotto, Ricardo Haddad, Sandra Coccuzzo Sampaio Vessoni, Simone Kashima, Svetoslav Nanev Slavov, Vincent Louis Viala |
| EPI_ISL_2344749 | CENTRO DE TRIAGEM COVID 19 | Instituto Butantan / ESALQ-Piracicaba | Dimas Tadeu Covas, Antonio Jorge Martins, Claudia Renata dos Santos Barros, David Schlesinger, Debora Botequio Moretti, Elaine Cristina Marqueze, Elaine Vieira Santos, Evandra Strazza Rodrigues, Heidge Fukumasu, Jayme Augusto de Souza-Neto, José Salvatore Leister Patané, Luiz Alcantara, Luiz Lehmann Coutinho, Maria Carolina Elias, Maurício Lacerda Nogueira, Rafael dos Santos Bezerra, Raul Machado Neto, Rejane Maria Tommasini Grotto, Ricardo Haddad, Sandra Coccuzzo Sampaio Vessoni, Simone Kashima, Svetoslav Nanev Slavov, Vincent Louis Viala |
| EPI_ISL_2344750 | UNIDADE MISTA OSVALDO P CAMARGO ELIAS FAUSTO | Instituto Butantan / ESALQ-Piracicaba | Dimas Tadeu Covas, Antonio Jorge Martins, Claudia Renata dos Santos Barros, David Schlesinger, Debora Botequio Moretti, Elaine Cristina Marqueze, Elaine Vieira Santos, Evandra Strazza Rodrigues, Heidge Fukumasu, Jayme Augusto de Souza-Neto, José Salvatore Leister Patané, Luiz Alcantara, Luiz Lehmann Coutinho, Maria Carolina Elias, Maurício Lacerda Nogueira, Rafael dos Santos Bezerra, Raul Machado Neto, Rejane Maria Tommasini Grotto, Ricardo Haddad, Sandra Coccuzzo Sampaio Vessoni, Simone Kashima, Svetoslav Nanev Slavov, Vincent Louis Viala |
| EPI_ISL_2344751 | UBS II DE TACIBA | Instituto Butantan / ESALQ-Piracicaba | Dimas Tadeu Covas, Antonio Jorge Martins, Claudia Renata dos Santos Barros, David Schlesinger, Debora Botequio Moretti, Elaine Cristina Marqueze, Elaine Vieira Santos, Evandra Strazza Rodrigues, Heidge Fukumasu, Jayme Augusto de Souza-Neto, José Salvatore Leister Patané, Luiz Alcantara, Luiz Lehmann Coutinho, Maria Carolina Elias, Maurício Lacerda Nogueira, Rafael dos Santos Bezerra, Raul Machado Neto, Rejane Maria Tommasini Grotto, Ricardo Haddad, Sandra Coccuzzo Sampaio Vessoni, Simone Kashima, Svetoslav Nanev Slavov, Vincent Louis Viala |
| EPI_ISL_2344752 | CENTRO DE TRIAGEM COVID 19 | Instituto Butantan / ESALQ-Piracicaba | Dimas Tadeu Covas, Antonio Jorge Martins, Claudia Renata dos Santos Barros, David Schlesinger, Debora Botequio Moretti, Elaine Cristina Marqueze, Elaine Vieira Santos, Evandra Strazza Rodrigues, Heidge Fukumasu, Jayme Augusto de Souza-Neto, José Salvatore Leister Patané, Luiz Alcantara, Luiz Lehmann Coutinho, Maria Carolina Elias, Maurício Lacerda Nogueira, Rafael dos Santos Bezerra, Raul Machado Neto, Rejane Maria Tommasini Grotto, Ricardo Haddad, Sandra Coccuzzo Sampaio Vessoni, Simone Kashima, Svetoslav Nanev Slavov, Vincent Louis Viala |
| EPI_ISL_2344753 | UBS II DE PIRAPOZINHO C SERVICO DE EACS E ESF | Instituto Butantan / ESALQ-Piracicaba | Dimas Tadeu Covas, Antonio Jorge Martins, Claudia Renata dos Santos Barros, David Schlesinger, Debora Botequio Moretti, Elaine Cristina Marqueze, Elaine Vieira Santos, Evandra Strazza Rodrigues, Heidge Fukumasu, Jayme Augusto de Souza-Neto, José Salvatore Leister Patané, Luiz Alcantara, Luiz Lehmann Coutinho, Maria Carolina Elias, Maurício Lacerda Nogueira, Rafael dos Santos Bezerra, Raul Machado Neto, Rejane Maria Tommasini Grotto, Ricardo Haddad, Sandra Coccuzzo Sampaio Vessoni, Simone Kashima, Svetoslav Nanev Slavov, Vincent Louis Viala |
| EPI_ISL_2344754 | UPA 24 HORAS ANA JACINTA | Instituto Butantan / ESALQ-Piracicaba | Dimas Tadeu Covas, Antonio Jorge Martins, Claudia Renata dos Santos Barros, David Schlesinger, Debora Botequio Moretti, Elaine Cristina Marqueze, Elaine Vieira Santos, Evandra Strazza Rodrigues, Heidge Fukumasu, Jayme Augusto de Souza-Neto, José Salvatore Leister Patané, Luiz Alcantara, Luiz Lehmann Coutinho, Maria Carolina Elias, Maurício Lacerda Nogueira, Rafael dos Santos Bezerra, Raul Machado Neto, Rejane Maria Tommasini Grotto, Ricardo Haddad, Sandra Coccuzzo Sampaio Vessoni, Simone Kashima, Svetoslav Nanev Slavov, Vincent Louis Viala |
| EPI_ISL_2344755 | UNIDADE BASICA DE SAUDE III DE ALFREDO MARCONDES | Instituto Butantan / ESALQ-Piracicaba | Dimas Tadeu Covas, Antonio Jorge Martins, Claudia Renata dos Santos Barros, David Schlesinger, Debora Botequio Moretti, Elaine Cristina Marqueze, Elaine Vieira Santos, Evandra Strazza Rodrigues, Heidge Fukumasu, Jayme Augusto de Souza-Neto, José Salvatore Leister Patané, Luiz Alcantara, Luiz Lehmann Coutinho, Maria Carolina Elias, Maurício Lacerda Nogueira, Rafael dos Santos Bezerra, Raul Machado Neto, Rejane Maria Tommasini Grotto, Ricardo Haddad, Sandra Coccuzzo Sampaio Vessoni, Simone Kashima, Svetoslav Nanev Slavov, Vincent Louis Viala |
| EPI_ISL_2344756 | CENTRO MEDICO DR NELSON SALOME DE CONCHAL | Instituto Butantan / ESALQ-Piracicaba | Dimas Tadeu Covas, Antonio Jorge Martins, Claudia Renata dos Santos Barros, David Schlesinger, Debora Botequio Moretti, Elaine Cristina Marqueze, Elaine Vieira Santos, Evandra Strazza Rodrigues, Heidge Fukumasu, Jayme Augusto de Souza-Neto, José Salvatore Leister Patané, Luiz Alcantara, Luiz Lehmann Coutinho, Maria Carolina Elias, Maurício Lacerda Nogueira, Rafael dos Santos Bezerra, Raul Machado Neto, Rejane Maria Tommasini Grotto, Ricardo Haddad, Sandra Coccuzzo Sampaio Vessoni, Simone Kashima, Svetoslav Nanev Slavov, Vincent Louis Viala |
| EPI_ISL_2344757 | CENTRO DE TRIAGEM COVID 19 | Instituto Butantan / ESALQ-Piracicaba | Dimas Tadeu Covas, Antonio Jorge Martins, Claudia Renata dos Santos Barros, David Schlesinger, Debora Botequio Moretti, Elaine Cristina Marqueze, Elaine Vieira Santos, Evandra Strazza Rodrigues, Heidge Fukumasu, Jayme Augusto de Souza-Neto, José Salvatore Leister Patané, Luiz Alcantara, Luiz Lehmann Coutinho, Maria Carolina Elias, Maurício Lacerda Nogueira, Rafael dos Santos Bezerra, Raul Machado Neto, Rejane Maria Tommasini Grotto, Ricardo Haddad, Sandra Coccuzzo Sampaio Vessoni, Simone Kashima, Svetoslav Nanev Slavov, Vincent Louis Viala |
| EPI_ISL_2344758 | UPA 24 HORAS DR ALOISIO MUNIZ DE ANDRADE | Instituto Butantan / ESALQ-Piracicaba | Dimas Tadeu Covas, Antonio Jorge Martins, Claudia Renata dos Santos Barros, David Schlesinger, Debora Botequio Moretti, Elaine Cristina Marqueze, Elaine Vieira Santos, Evandra Strazza Rodrigues, Heidge Fukumasu, Jayme Augusto de Souza-Neto, José Salvatore Leister Patané, Luiz Alcantara, Luiz |

| | | | |
|---|---|---|---|
| | | | Lehmann Coutinho, Maria Carolina Elias, Maurício Lacerda Nogueira, Rafael dos Santos Bezerra, Raul Machado Neto, Rejane Maria Tommasini Grotto, Ricardo Haddad, Sandra Coccuzzo Sampaio Vessoni, Simone Kashima, Svetoslav Nanev Slavov, Vincent Louis Viala |
| EPI_ISL_2344759 | CENTRO DE TRIAGEM COVID 19 | Instituto Butantan / ESALQ-Piracicaba | Dimas Tadeu Covas, Antonio Jorge Martins, Claudia Renata dos Santos Barros, David Schlesinger, Debora Botequio Moretti, Elaine Cristina Marqueze, Elaine Vieira Santos, Evandra Strazza Rodrigues, Heidge Fukumasu, Jayme Augusto de Souza-Neto, José Salvatore Leister Patané, Luiz Alcantara, Luiz Lehmann Coutinho, Maria Carolina Elias, Maurício Lacerda Nogueira, Rafael dos Santos Bezerra, Raul Machado Neto, Rejane Maria Tommasini Grotto, Ricardo Haddad, Sandra Coccuzzo Sampaio Vessoni, Simone Kashima, Svetoslav Nanev Slavov, Vincent Louis Viala |
| EPI_ISL_2344760 | UNIDADE BASICA DE SAUDE III DE ALFREDO MARCONDES | Instituto Butantan / ESALQ-Piracicaba | Dimas Tadeu Covas, Antonio Jorge Martins, Claudia Renata dos Santos Barros, David Schlesinger, Debora Botequio Moretti, Elaine Cristina Marqueze, Elaine Vieira Santos, Evandra Strazza Rodrigues, Heidge Fukumasu, Jayme Augusto de Souza-Neto, José Salvatore Leister Patané, Luiz Alcantara, Luiz Lehmann Coutinho, Maria Carolina Elias, Maurício Lacerda Nogueira, Rafael dos Santos Bezerra, Raul Machado Neto, Rejane Maria Tommasini Grotto, Ricardo Haddad, Sandra Coccuzzo Sampaio Vessoni, Simone Kashima, Svetoslav Nanev Slavov, Vincent Louis Viala |
| EPI_ISL_2344761 | UBS II DE PIRAPOZINHO C SERVICO DE EACS E ESF | Instituto Butantan / ESALQ-Piracicaba | Dimas Tadeu Covas, Antonio Jorge Martins, Claudia Renata dos Santos Barros, David Schlesinger, Debora Botequio Moretti, Elaine Cristina Marqueze, Elaine Vieira Santos, Evandra Strazza Rodrigues, Heidge Fukumasu, Jayme Augusto de Souza-Neto, José Salvatore Leister Patané, Luiz Alcantara, Luiz Lehmann Coutinho, Maria Carolina Elias, Maurício Lacerda Nogueira, Rafael dos Santos Bezerra, Raul Machado Neto, Rejane Maria Tommasini Grotto, Ricardo Haddad, Sandra Coccuzzo Sampaio Vessoni, Simone Kashima, Svetoslav Nanev Slavov, Vincent Louis Viala |
| EPI_ISL_2344762 | UPA 24 HORAS DR ALOISIO MUNIZ DE ANDRADE | Instituto Butantan / ESALQ-Piracicaba | Dimas Tadeu Covas, Antonio Jorge Martins, Claudia Renata dos Santos Barros, David Schlesinger, Debora Botequio Moretti, Elaine Cristina Marqueze, Elaine Vieira Santos, Evandra Strazza Rodrigues, Heidge Fukumasu, Jayme Augusto de Souza-Neto, José Salvatore Leister Patané, Luiz Alcantara, Luiz Lehmann Coutinho, Maria Carolina Elias, Maurício Lacerda Nogueira, Rafael dos Santos Bezerra, Raul Machado Neto, Rejane Maria Tommasini Grotto, Ricardo Haddad, Sandra Coccuzzo Sampaio Vessoni, Simone Kashima, Svetoslav Nanev Slavov, Vincent Louis Viala |
| EPI_ISL_2344763 | USF II ALADIA ARAN RODRIGUES | Instituto Butantan / ESALQ-Piracicaba | Dimas Tadeu Covas, Antonio Jorge Martins, Claudia Renata dos Santos Barros, David Schlesinger, Debora Botequio Moretti, Elaine Cristina Marqueze, Elaine Vieira Santos, Evandra Strazza Rodrigues, Heidge Fukumasu, Jayme Augusto de Souza-Neto, José Salvatore Leister Patané, Luiz Alcantara, Luiz Lehmann Coutinho, Maria Carolina Elias, Maurício Lacerda Nogueira, Rafael dos Santos Bezerra, Raul Machado Neto, Rejane Maria Tommasini Grotto, Ricardo Haddad, Sandra Coccuzzo Sampaio Vessoni, Simone Kashima, Svetoslav Nanev Slavov, Vincent Louis Viala |
| EPI_ISL_2344764 | UPA 24 HORAS ANA JACINTA | Instituto Butantan / ESALQ-Piracicaba | Dimas Tadeu Covas, Antonio Jorge Martins, Claudia Renata dos Santos Barros, David Schlesinger, Debora Botequio Moretti, Elaine Cristina Marqueze, Elaine Vieira Santos, Evandra Strazza Rodrigues, Heidge Fukumasu, Jayme Augusto de Souza-Neto, José Salvatore Leister Patané, Luiz Alcantara, Luiz Lehmann Coutinho, Maria Carolina Elias, Maurício Lacerda Nogueira, Rafael dos Santos Bezerra, Raul Machado Neto, Rejane Maria Tommasini Grotto, Ricardo Haddad, Sandra Coccuzzo Sampaio Vessoni, Simone Kashima, Svetoslav Nanev Slavov, Vincent Louis Viala |
| EPI_ISL_2344765, EPI_ISL_2344766 | UPA 24 HORAS DR ALOISIO MUNIZ DE ANDRADE | Instituto Butantan / ESALQ-Piracicaba | Dimas Tadeu Covas, Antonio Jorge Martins, Claudia Renata dos Santos Barros, David Schlesinger, Debora Botequio Moretti, Elaine Cristina Marqueze, Elaine Vieira Santos, Evandra Strazza Rodrigues, Heidge Fukumasu, Jayme Augusto de Souza-Neto, José Salvatore Leister Patané, Luiz Alcantara, Luiz Lehmann Coutinho, Maria Carolina Elias, Maurício Lacerda Nogueira, Rafael dos Santos Bezerra, Raul Machado Neto, Rejane Maria Tommasini Grotto, Ricardo Haddad, Sandra Coccuzzo Sampaio Vessoni, Simone Kashima, Svetoslav Nanev Slavov, Vincent Louis Viala |
| EPI_ISL_2344767 | HOSPITAL MUNICIPAL DE ITAPIRA | Instituto Butantan / ESALQ-Piracicaba | Dimas Tadeu Covas, Antonio Jorge Martins, Claudia Renata dos Santos Barros, David Schlesinger, Debora Botequio Moretti, Elaine Cristina Marqueze, Elaine Vieira Santos, Evandra Strazza Rodrigues, Heidge Fukumasu, Jayme Augusto de Souza-Neto, José Salvatore Leister Patané, Luiz Alcantara, Luiz Lehmann Coutinho, Maria Carolina Elias, Maurício Lacerda Nogueira, Rafael dos Santos Bezerra, Raul Machado Neto, Rejane Maria Tommasini Grotto, Ricardo Haddad, Sandra Coccuzzo Sampaio Vessoni, Simone Kashima, Svetoslav Nanev Slavov, Vincent Louis Viala |
| EPI_ISL_2344768 | UPA 24 HORAS ANA JACINTA | Instituto Butantan / ESALQ-Piracicaba | Dimas Tadeu Covas, Antonio Jorge Martins, Claudia Renata dos Santos Barros, David Schlesinger, Debora Botequio Moretti, Elaine Cristina Marqueze, Elaine Vieira Santos, Evandra Strazza Rodrigues, Heidge Fukumasu, Jayme Augusto de Souza-Neto, José Salvatore Leister Patané, Luiz Alcantara, Luiz Lehmann Coutinho, Maria Carolina Elias, Maurício Lacerda Nogueira, Rafael dos Santos Bezerra, Raul Machado Neto, Rejane Maria Tommasini Grotto, Ricardo Haddad, Sandra Coccuzzo Sampaio Vessoni, Simone Kashima, Svetoslav Nanev Slavov, Vincent Louis Viala |
| EPI_ISL_2344769 | CS DE RINOPOLIS | Instituto Butantan / ESALQ-Piracicaba | Dimas Tadeu Covas, Antonio Jorge Martins, Claudia Renata dos Santos Barros, David Schlesinger, Debora Botequio Moretti, Elaine Cristina Marqueze, Elaine Vieira Santos, Evandra Strazza Rodrigues, Heidge Fukumasu, Jayme Augusto de Souza-Neto, José Salvatore Leister Patané, Luiz Alcantara, Luiz Lehmann Coutinho, Maria Carolina Elias, Maurício Lacerda Nogueira, Rafael dos Santos Bezerra, Raul Machado Neto, Rejane Maria Tommasini Grotto, Ricardo Haddad, Sandra Coccuzzo Sampaio Vessoni, Simone Kashima, Svetoslav Nanev Slavov, Vincent Louis Viala |
| EPI_ISL_2344770 | UPA 24 HORAS ANA JACINTA | Instituto Butantan / ESALQ-Piracicaba | Dimas Tadeu Covas, Antonio Jorge Martins, Claudia Renata dos Santos Barros, David Schlesinger, Debora Botequio Moretti, Elaine Cristina Marqueze, Elaine Vieira Santos, Evandra Strazza Rodrigues, Heidge Fukumasu, Jayme Augusto de Souza-Neto, José Salvatore Leister Patané, Luiz Alcantara, Luiz Lehmann Coutinho, Maria Carolina Elias, Maurício Lacerda Nogueira, Rafael dos Santos Bezerra, Raul Machado Neto, Rejane Maria Tommasini Grotto, Ricardo Haddad, Sandra Coccuzzo Sampaio Vessoni, Simone Kashima, Svetoslav Nanev Slavov, Vincent Louis Viala |
| EPI_ISL_2344771 | VIGILANCIA EPIDEMIOLOGICA E CONTROLE DE VETORES PIRASSUNUN | Instituto Butantan / ESALQ-Piracicaba | Dimas Tadeu Covas, Antonio Jorge Martins, Claudia Renata dos Santos Barros, David Schlesinger, Debora Botequio Moretti, Elaine Cristina Marqueze, Elaine Vieira Santos, Evandra Strazza Rodrigues, Heidge Fukumasu, Jayme Augusto de Souza-Neto, José Salvatore Leister Patané, Luiz Alcantara, Luiz Lehmann Coutinho, Maria Carolina Elias, Maurício Lacerda Nogueira, Rafael dos Santos Bezerra, Raul Machado Neto, Rejane Maria Tommasini Grotto, Ricardo Haddad, Sandra Coccuzzo Sampaio Vessoni, Simone Kashima, Svetoslav Nanev Slavov, Vincent Louis Viala |
| EPI_ISL_2344772 | CS DE RINOPOLIS | Instituto Butantan / ESALQ-Piracicaba | Dimas Tadeu Covas, Antonio Jorge Martins, Claudia Renata dos Santos Barros, David Schlesinger, Debora Botequio Moretti, Elaine Cristina Marqueze, Elaine Vieira Santos, Evandra Strazza Rodrigues, Heidge Fukumasu, Jayme Augusto de Souza-Neto, José Salvatore Leister Patané, Luiz Alcantara, Luiz Lehmann Coutinho, Maria Carolina Elias, Maurício Lacerda Nogueira, Rafael dos Santos Bezerra, Raul Machado Neto, Rejane Maria Tommasini Grotto, Ricardo Haddad, Sandra Coccuzzo Sampaio Vessoni, Simone Kashima, Svetoslav Nanev Slavov, Vincent Louis Viala |
| EPI_ISL_2344773 | UPA 24 HORAS ANA JACINTA | Instituto Butantan / ESALQ-Piracicaba | Dimas Tadeu Covas, Antonio Jorge Martins, Claudia Renata dos Santos Barros, David Schlesinger, Debora Botequio Moretti, Elaine Cristina Marqueze, Elaine Vieira Santos, Evandra Strazza Rodrigues, Heidge Fukumasu, Jayme Augusto de Souza-Neto, José Salvatore Leister Patané, Luiz Alcantara, Luiz Lehmann Coutinho, Maria Carolina Elias, Maurício Lacerda Nogueira, Rafael dos Santos Bezerra, Raul Machado Neto, Rejane Maria Tommasini Grotto, Ricardo Haddad, Sandra Coccuzzo Sampaio Vessoni, Simone Kashima, Svetoslav Nanev Slavov, Vincent Louis Viala |
| EPI_ISL_2344774 | CENTRO DE SAUDE DE CHAVANTES | Instituto Butantan / ESALQ-Piracicaba | Dimas Tadeu Covas, Antonio Jorge Martins, Claudia Renata dos Santos Barros, David Schlesinger, Debora Botequio Moretti, Elaine Cristina Marqueze, Elaine Vieira Santos, Evandra Strazza Rodrigues, Heidge Fukumasu, Jayme Augusto de Souza-Neto, José Salvatore Leister Patané, Luiz Alcantara, Luiz Lehmann Coutinho, Maria Carolina Elias, Maurício Lacerda Nogueira, Rafael dos Santos Bezerra, Raul Machado Neto, Rejane Maria Tommasini Grotto, Ricardo Haddad, Sandra Coccuzzo Sampaio Vessoni, Simone Kashima, Svetoslav Nanev Slavov, Vincent Louis Viala |
| EPI_ISL_2344775 | CENTRO DE SAUDE ESF IV ZONA RURAL DOMINGOS DE S SJRIOPARDO | Instituto Butantan / ESALQ-Piracicaba | Dimas Tadeu Covas, Antonio Jorge Martins, Claudia Renata dos Santos Barros, David Schlesinger, Debora Botequio Moretti, Elaine Cristina Marqueze, Elaine Vieira Santos, Evandra Strazza Rodrigues, Heidge Fukumasu, Jayme Augusto de Souza-Neto, José Salvatore Leister Patané, Luiz Alcantara, Luiz Lehmann Coutinho, Maria Carolina Elias, Maurício Lacerda Nogueira, Rafael dos Santos Bezerra, Raul Machado Neto, Rejane Maria Tommasini Grotto, Ricardo Haddad, Sandra Coccuzzo Sampaio Vessoni, Simone Kashima, Svetoslav Nanev Slavov, Vincent Louis Viala |
| EPI_ISL_2344776 | LABORATORIO MUNICIPAL DE PIRACICABA | Instituto Butantan / ESALQ-Piracicaba | Dimas Tadeu Covas, Antonio Jorge Martins, Claudia Renata dos Santos Barros, David Schlesinger, Debora Botequio Moretti, Elaine Cristina Marqueze, Elaine Vieira Santos, Evandra Strazza Rodrigues, Heidge Fukumasu, Jayme Augusto de Souza-Neto, José Salvatore Leister Patané, Luiz Alcantara, Luiz Lehmann Coutinho, Maria Carolina Elias, Maurício Lacerda Nogueira, Rafael dos Santos Bezerra, Raul Machado Neto, Rejane Maria Tommasini Grotto, Ricardo Haddad, Sandra Coccuzzo Sampaio Vessoni, Simone Kashima, Svetoslav Nanev Slavov, Vincent Louis Viala |
| EPI_ISL_2344777 | CENTRO DE SAUDE DE CHAVANTES | Instituto Butantan / ESALQ-Piracicaba | Dimas Tadeu Covas, Antonio Jorge Martins, Claudia Renata dos Santos Barros, David Schlesinger, Debora Botequio Moretti, Elaine Cristina Marqueze, Elaine Vieira Santos, Evandra Strazza Rodrigues, Heidge Fukumasu, Jayme Augusto de Souza-Neto, José Salvatore Leister Patané, Luiz Alcantara, Luiz Lehmann Coutinho, Maria Carolina Elias, Maurício Lacerda Nogueira, Rafael dos Santos Bezerra, Raul Machado Neto, Rejane Maria Tommasini Grotto, Ricardo Haddad, Sandra Coccuzzo Sampaio Vessoni, Simone Kashima, Svetoslav Nanev Slavov, Vincent Louis Viala |
| EPI_ISL_2344778 | CS IACRI | Instituto Butantan / ESALQ-Piracicaba | Dimas Tadeu Covas, Antonio Jorge Martins, Claudia Renata dos Santos Barros, David Schlesinger, Debora Botequio Moretti, Elaine Cristina Marqueze, Elaine Vieira Santos, Evandra Strazza Rodrigues, Heidge Fukumasu, Jayme Augusto de Souza-Neto, José Salvatore Leister Patané, Luiz Alcantara, Luiz Lehmann Coutinho, Maria Carolina Elias, Maurício Lacerda Nogueira, Rafael dos Santos Bezerra, Raul Machado Neto, Rejane Maria Tommasini Grotto, Ricardo Haddad, Sandra Coccuzzo Sampaio Vessoni, Simone Kashima, Svetoslav Nanev Slavov, Vincent Louis Viala |

| | | | |
|---|---|---|---|
| EPI_ISL_2344779, EPI_ISL_2344780 | CENTRO DE SAUDE ESF IV ZONA RURAL DOMINGOS DE S SJRIOPARDO | Instituto Butantan / ESALQ-Piracicaba | Dimas Tadeu Covas, Antonio Jorge Martins, Claudia Renata dos Santos Barros, David Schlesinger, Debora Botequio Moretti, Elaine Cristina Marqueze, Elaine Vieira Santos, Evandra Strazza Rodrigues, Heidge Fukumasu, Jayme Augusto de Souza-Neto, José Salvatore Leister Patané, Luiz Alcantara, Luiz Lehmann Coutinho, Maria Carolina Elias, Maurício Lacerda Nogueira, Rafael dos Santos Bezerra, Raul Machado Neto, Rejane Maria Tommasini Grotto, Ricardo Haddad, Sandra Coccuzzo Sampaio Vessoni, Simone Kashima, Svetoslav Nanev Slavov, Vincent Louis Viala |
| EPI_ISL_2344781 | CENTRO DE SAUDE DR MARIO DIAS DE AGUIAR CAPIVARI | Instituto Butantan / ESALQ-Piracicaba | Dimas Tadeu Covas, Antonio Jorge Martins, Claudia Renata dos Santos Barros, David Schlesinger, Debora Botequio Moretti, Elaine Cristina Marqueze, Elaine Vieira Santos, Evandra Strazza Rodrigues, Heidge Fukumasu, Jayme Augusto de Souza-Neto, José Salvatore Leister Patané, Luiz Alcantara, Luiz Lehmann Coutinho, Maria Carolina Elias, Maurício Lacerda Nogueira, Rafael dos Santos Bezerra, Raul Machado Neto, Rejane Maria Tommasini Grotto, Ricardo Haddad, Sandra Coccuzzo Sampaio Vessoni, Simone Kashima, Svetoslav Nanev Slavov, Vincent Louis Viala |
| EPI_ISL_2344782 | UNIDADE DE VIGILANCIA EPIDEMIOLOGICA | Instituto Butantan / ESALQ-Piracicaba | Dimas Tadeu Covas, Antonio Jorge Martins, Claudia Renata dos Santos Barros, David Schlesinger, Debora Botequio Moretti, Elaine Cristina Marqueze, Elaine Vieira Santos, Evandra Strazza Rodrigues, Heidge Fukumasu, Jayme Augusto de Souza-Neto, José Salvatore Leister Patané, Luiz Alcantara, Luiz Lehmann Coutinho, Maria Carolina Elias, Maurício Lacerda Nogueira, Rafael dos Santos Bezerra, Raul Machado Neto, Rejane Maria Tommasini Grotto, Ricardo Haddad, Sandra Coccuzzo Sampaio Vessoni, Simone Kashima, Svetoslav Nanev Slavov, Vincent Louis Viala |
| EPI_ISL_2344783, EPI_ISL_2344784 | CENTRO DE SAUDE ESF IV ZONA RURAL DOMINGOS DE S SJRIOPARDO | Instituto Butantan / ESALQ-Piracicaba | Dimas Tadeu Covas, Antonio Jorge Martins, Claudia Renata dos Santos Barros, David Schlesinger, Debora Botequio Moretti, Elaine Cristina Marqueze, Elaine Vieira Santos, Evandra Strazza Rodrigues, Heidge Fukumasu, Jayme Augusto de Souza-Neto, José Salvatore Leister Patané, Luiz Alcantara, Luiz Lehmann Coutinho, Maria Carolina Elias, Maurício Lacerda Nogueira, Rafael dos Santos Bezerra, Raul Machado Neto, Rejane Maria Tommasini Grotto, Ricardo Haddad, Sandra Coccuzzo Sampaio Vessoni, Simone Kashima, Svetoslav Nanev Slavov, Vincent Louis Viala |
| EPI_ISL_2344785 | CENTRO DE SAUDE DE OCAUCU | Instituto Butantan / ESALQ-Piracicaba | Dimas Tadeu Covas, Antonio Jorge Martins, Claudia Renata dos Santos Barros, David Schlesinger, Debora Botequio Moretti, Elaine Cristina Marqueze, Elaine Vieira Santos, Evandra Strazza Rodrigues, Heidge Fukumasu, Jayme Augusto de Souza-Neto, José Salvatore Leister Patané, Luiz Alcantara, Luiz Lehmann Coutinho, Maria Carolina Elias, Maurício Lacerda Nogueira, Rafael dos Santos Bezerra, Raul Machado Neto, Rejane Maria Tommasini Grotto, Ricardo Haddad, Sandra Coccuzzo Sampaio Vessoni, Simone Kashima, Svetoslav Nanev Slavov, Vincent Louis Viala |
| EPI_ISL_2344786 | CENTRO DE ESPECIALIDADES DE AGUAI AGUAI | Instituto Butantan / ESALQ-Piracicaba | Dimas Tadeu Covas, Antonio Jorge Martins, Claudia Renata dos Santos Barros, David Schlesinger, Debora Botequio Moretti, Elaine Cristina Marqueze, Elaine Vieira Santos, Evandra Strazza Rodrigues, Heidge Fukumasu, Jayme Augusto de Souza-Neto, José Salvatore Leister Patané, Luiz Alcantara, Luiz Lehmann Coutinho, Maria Carolina Elias, Maurício Lacerda Nogueira, Rafael dos Santos Bezerra, Raul Machado Neto, Rejane Maria Tommasini Grotto, Ricardo Haddad, Sandra Coccuzzo Sampaio Vessoni, Simone Kashima, Svetoslav Nanev Slavov, Vincent Louis Viala |
| EPI_ISL_2344787 | CENTRO DE SAUDE DE LUTECIA | Instituto Butantan / ESALQ-Piracicaba | Dimas Tadeu Covas, Antonio Jorge Martins, Claudia Renata dos Santos Barros, David Schlesinger, Debora Botequio Moretti, Elaine Cristina Marqueze, Elaine Vieira Santos, Evandra Strazza Rodrigues, Heidge Fukumasu, Jayme Augusto de Souza-Neto, José Salvatore Leister Patané, Luiz Alcantara, Luiz Lehmann Coutinho, Maria Carolina Elias, Maurício Lacerda Nogueira, Rafael dos Santos Bezerra, Raul Machado Neto, Rejane Maria Tommasini Grotto, Ricardo Haddad, Sandra Coccuzzo Sampaio Vessoni, Simone Kashima, Svetoslav Nanev Slavov, Vincent Louis Viala |
| EPI_ISL_2344788 | HOSPITAL MUNICIPAL DE ITAPIRA | Instituto Butantan / ESALQ-Piracicaba | Dimas Tadeu Covas, Antonio Jorge Martins, Claudia Renata dos Santos Barros, David Schlesinger, Debora Botequio Moretti, Elaine Cristina Marqueze, Elaine Vieira Santos, Evandra Strazza Rodrigues, Heidge Fukumasu, Jayme Augusto de Souza-Neto, José Salvatore Leister Patané, Luiz Alcantara, Luiz Lehmann Coutinho, Maria Carolina Elias, Maurício Lacerda Nogueira, Rafael dos Santos Bezerra, Raul Machado Neto, Rejane Maria Tommasini Grotto, Ricardo Haddad, Sandra Coccuzzo Sampaio Vessoni, Simone Kashima, Svetoslav Nanev Slavov, Vincent Louis Viala |
| EPI_ISL_2344789 | PRONTO ATENDIMENTO MUNICIPAL DE JACUPIRANGA | Instituto Butantan / ESALQ-Piracicaba | Dimas Tadeu Covas, Antonio Jorge Martins, Claudia Renata dos Santos Barros, David Schlesinger, Debora Botequio Moretti, Elaine Cristina Marqueze, Elaine Vieira Santos, Evandra Strazza Rodrigues, Heidge Fukumasu, Jayme Augusto de Souza-Neto, José Salvatore Leister Patané, Luiz Alcantara, Luiz Lehmann Coutinho, Maria Carolina Elias, Maurício Lacerda Nogueira, Rafael dos Santos Bezerra, Raul Machado Neto, Rejane Maria Tommasini Grotto, Ricardo Haddad, Sandra Coccuzzo Sampaio Vessoni, Simone Kashima, Svetoslav Nanev Slavov, Vincent Louis Viala |
| EPI_ISL_2344790 | CENTRO DE SAUDE DR PLINIO ALBERS DE GALIA | Instituto Butantan / ESALQ-Piracicaba | Dimas Tadeu Covas, Antonio Jorge Martins, Claudia Renata dos Santos Barros, David Schlesinger, Debora Botequio Moretti, Elaine Cristina Marqueze, Elaine Vieira Santos, Evandra Strazza Rodrigues, Heidge Fukumasu, Jayme Augusto de Souza-Neto, José Salvatore Leister Patané, Luiz Alcantara, Luiz Lehmann Coutinho, Maria Carolina Elias, Maurício Lacerda Nogueira, Rafael dos Santos Bezerra, Raul Machado Neto, Rejane Maria Tommasini Grotto, Ricardo Haddad, Sandra Coccuzzo Sampaio Vessoni, Simone Kashima, Svetoslav Nanev Slavov, Vincent Louis Viala |
| EPI_ISL_2344791 | CENTRO DE SAUDE DE SPEDRO DO TURVO | Instituto Butantan / ESALQ-Piracicaba | Dimas Tadeu Covas, Antonio Jorge Martins, Claudia Renata dos Santos Barros, David Schlesinger, Debora Botequio Moretti, Elaine Cristina Marqueze, Elaine Vieira Santos, Evandra Strazza Rodrigues, Heidge Fukumasu, Jayme Augusto de Souza-Neto, José Salvatore Leister Patané, Luiz Alcantara, Luiz Lehmann Coutinho, Maria Carolina Elias, Maurício Lacerda Nogueira, Rafael dos Santos Bezerra, Raul Machado Neto, Rejane Maria Tommasini Grotto, Ricardo Haddad, Sandra Coccuzzo Sampaio Vessoni, Simone Kashima, Svetoslav Nanev Slavov, Vincent Louis Viala |
| EPI_ISL_2344792, EPI_ISL_2344793 | CENTRO DE ESPECIALIDADES DE AGUAI AGUAI | Instituto Butantan / ESALQ-Piracicaba | Dimas Tadeu Covas, Antonio Jorge Martins, Claudia Renata dos Santos Barros, David Schlesinger, Debora Botequio Moretti, Elaine Cristina Marqueze, Elaine Vieira Santos, Evandra Strazza Rodrigues, Heidge Fukumasu, Jayme Augusto de Souza-Neto, José Salvatore Leister Patané, Luiz Alcantara, Luiz Lehmann Coutinho, Maria Carolina Elias, Maurício Lacerda Nogueira, Rafael dos Santos Bezerra, Raul Machado Neto, Rejane Maria Tommasini Grotto, Ricardo Haddad, Sandra Coccuzzo Sampaio Vessoni, Simone Kashima, Svetoslav Nanev Slavov, Vincent Louis Viala |
| EPI_ISL_2344794, EPI_ISL_2344795, EPI_ISL_2344796 | LABORATORIO DE FRANCA | Instituto Butantan / ESALQ-Piracicaba | Dimas Tadeu Covas, Antonio Jorge Martins, Claudia Renata dos Santos Barros, David Schlesinger, Debora Botequio Moretti, Elaine Cristina Marqueze, Elaine Vieira Santos, Evandra Strazza Rodrigues, Heidge Fukumasu, Jayme Augusto de Souza-Neto, José Salvatore Leister Patané, Luiz Alcantara, Luiz Lehmann Coutinho, Maria Carolina Elias, Maurício Lacerda Nogueira, Rafael dos Santos Bezerra, Raul Machado Neto, Rejane Maria Tommasini Grotto, Ricardo Haddad, Sandra Coccuzzo Sampaio Vessoni, Simone Kashima, Svetoslav Nanev Slavov, Vincent Louis Viala |
| EPI_ISL_2344797, EPI_ISL_2344798, EPI_ISL_2344799 | CENTRO DE SAUDE DR NELSON DA CUNHA BASTOS | Instituto Butantan / ESALQ-Piracicaba | Dimas Tadeu Covas, Antonio Jorge Martins, Claudia Renata dos Santos Barros, David Schlesinger, Debora Botequio Moretti, Elaine Cristina Marqueze, Elaine Vieira Santos, Evandra Strazza Rodrigues, Heidge Fukumasu, Jayme Augusto de Souza-Neto, José Salvatore Leister Patané, Luiz Alcantara, Luiz Lehmann Coutinho, Maria Carolina Elias, Maurício Lacerda Nogueira, Rafael dos Santos Bezerra, Raul Machado Neto, Rejane Maria Tommasini Grotto, Ricardo Haddad, Sandra Coccuzzo Sampaio Vessoni, Simone Kashima, Svetoslav Nanev Slavov, Vincent Louis Viala |
| EPI_ISL_2344800 | HOSPITAL MUNICIPAL DR TABAJARA RAMOS | Instituto Butantan / ESALQ-Piracicaba | Dimas Tadeu Covas, Antonio Jorge Martins, Claudia Renata dos Santos Barros, David Schlesinger, Debora Botequio Moretti, Elaine Cristina Marqueze, Elaine Vieira Santos, Evandra Strazza Rodrigues, Heidge Fukumasu, Jayme Augusto de Souza-Neto, José Salvatore Leister Patané, Luiz Alcantara, Luiz Lehmann Coutinho, Maria Carolina Elias, Maurício Lacerda Nogueira, Rafael dos Santos Bezerra, Raul Machado Neto, Rejane Maria Tommasini Grotto, Ricardo Haddad, Sandra Coccuzzo Sampaio Vessoni, Simone Kashima, Svetoslav Nanev Slavov, Vincent Louis Viala |
| EPI_ISL_2344801 | LABORATORIO MUNICIPAL DE PIRACICABA | Instituto Butantan / ESALQ-Piracicaba | Dimas Tadeu Covas, Antonio Jorge Martins, Claudia Renata dos Santos Barros, David Schlesinger, Debora Botequio Moretti, Elaine Cristina Marqueze, Elaine Vieira Santos, Evandra Strazza Rodrigues, Heidge Fukumasu, Jayme Augusto de Souza-Neto, José Salvatore Leister Patané, Luiz Alcantara, Luiz Lehmann Coutinho, Maria Carolina Elias, Maurício Lacerda Nogueira, Rafael dos Santos Bezerra, Raul Machado Neto, Rejane Maria Tommasini Grotto, Ricardo Haddad, Sandra Coccuzzo Sampaio Vessoni, Simone Kashima, Svetoslav Nanev Slavov, Vincent Louis Viala |
| EPI_ISL_2344802, EPI_ISL_2344803, EPI_ISL_2344804, EPI_ISL_2344805, EPI_ISL_2344806 | SAE CTA SERV ATEND ESPEC TESTAGEM E ACONSELHAMENTO | Instituto Butantan / ESALQ-Piracicaba | Dimas Tadeu Covas, Antonio Jorge Martins, Claudia Renata dos Santos Barros, David Schlesinger, Debora Botequio Moretti, Elaine Cristina Marqueze, Elaine Vieira Santos, Evandra Strazza Rodrigues, Heidge Fukumasu, Jayme Augusto de Souza-Neto, José Salvatore Leister Patané, Luiz Alcantara, Luiz Lehmann Coutinho, Maria Carolina Elias, Maurício Lacerda Nogueira, Rafael dos Santos Bezerra, Raul Machado Neto, Rejane Maria Tommasini Grotto, Ricardo Haddad, Sandra Coccuzzo Sampaio Vessoni, Simone Kashima, Svetoslav Nanev Slavov, Vincent Louis Viala |
| EPI_ISL_2344807 | AMBULATORIO DE ESPECIALIDADE V E MOGI MIRIM | Instituto Butantan / ESALQ-Piracicaba | Dimas Tadeu Covas, Antonio Jorge Martins, Claudia Renata dos Santos Barros, David Schlesinger, Debora Botequio Moretti, Elaine Cristina Marqueze, Elaine Vieira Santos, Evandra Strazza Rodrigues, Heidge Fukumasu, Jayme Augusto de Souza-Neto, José Salvatore Leister Patané, Luiz Alcantara, Luiz Lehmann Coutinho, Maria Carolina Elias, Maurício Lacerda Nogueira, Rafael dos Santos Bezerra, Raul Machado Neto, Rejane Maria Tommasini Grotto, Ricardo Haddad, Sandra Coccuzzo Sampaio Vessoni, Simone Kashima, Svetoslav Nanev Slavov, Vincent Louis Viala |
| EPI_ISL_2344808 | SAE CTA SERV ATEND ESPEC TESTAGEM E ACONSELHAMENTO | Instituto Butantan / ESALQ-Piracicaba | Dimas Tadeu Covas, Antonio Jorge Martins, Claudia Renata dos Santos Barros, David Schlesinger, Debora Botequio Moretti, Elaine Cristina Marqueze, Elaine Vieira Santos, Evandra Strazza Rodrigues, Heidge Fukumasu, Jayme Augusto de Souza-Neto, José Salvatore Leister Patané, Luiz Alcantara, Luiz Lehmann Coutinho, Maria Carolina Elias, Maurício Lacerda Nogueira, Rafael dos Santos Bezerra, Raul Machado Neto, Rejane Maria Tommasini Grotto, Ricardo Haddad, Sandra Coccuzzo Sampaio Vessoni, Simone Kashima, Svetoslav Nanev Slavov, Vincent Louis Viala |
| EPI_ISL_2344809 | VIGILANCIA EPIDEMIOLOGICA DE LEME | Instituto Butantan / ESALQ-Piracicaba | Dimas Tadeu Covas, Antonio Jorge Martins, Claudia Renata dos Santos Barros, David Schlesinger, Debora Botequio Moretti, Elaine Cristina Marqueze, Elaine Vieira Santos, Evandra Strazza Rodrigues, Heidge Fukumasu, Jayme Augusto de Souza-Neto, José Salvatore Leister Patané, Luiz Alcantara, Luiz |

| | | | |
|---|---|---|---|
| | | | Lehmann Coutinho, Maria Carolina Elias, Maurício Lacerda Nogueira, Rafael dos Santos Bezerra, Raul Machado Neto, Rejane Maria Tommasini Grotto, Ricardo Haddad, Sandra Coccuzzo Sampaio Vessoni, Simone Kashima, Svetoslav Nanev Slavov, Vincent Louis Viala |
| EPI_ISL_2344810, EPI_ISL_2344811, EPI_ISL_2344812, EPI_ISL_2344813 | SAE CTA SERV ATEND ESPEC TESTAGEM E ACONSELHAMENTO | Instituto Butantan / ESALQ-Piracicaba | Dimas Tadeu Covas, Antonio Jorge Martins, Claudia Renata dos Santos Barros, David Schlesinger, Debora Botequio Moretti, Elaine Cristina Marqueze, Elaine Vieira Santos, Evandra Strazza Rodrigues, Heidge Fukumasu, Jayme Augusto de Souza-Neto, José Salvatore Leister Patané, Luiz Alcantara, Luiz Lehmann Coutinho, Maria Carolina Elias, Maurício Lacerda Nogueira, Rafael dos Santos Bezerra, Raul Machado Neto, Rejane Maria Tommasini Grotto, Ricardo Haddad, Sandra Coccuzzo Sampaio Vessoni, Simone Kashima, Svetoslav Nanev Slavov, Vincent Louis Viala |
| EPI_ISL_2344814 | ESTRATEGIA DE SAUDE DA FAMILIA DE ARCO IRIS | Instituto Butantan / ESALQ-Piracicaba | Dimas Tadeu Covas, Antonio Jorge Martins, Claudia Renata dos Santos Barros, David Schlesinger, Debora Botequio Moretti, Elaine Cristina Marqueze, Elaine Vieira Santos, Evandra Strazza Rodrigues, Heidge Fukumasu, Jayme Augusto de Souza-Neto, José Salvatore Leister Patané, Luiz Alcantara, Luiz Lehmann Coutinho, Maria Carolina Elias, Maurício Lacerda Nogueira, Rafael dos Santos Bezerra, Raul Machado Neto, Rejane Maria Tommasini Grotto, Ricardo Haddad, Sandra Coccuzzo Sampaio Vessoni, Simone Kashima, Svetoslav Nanev Slavov, Vincent Louis Viala |
| EPI_ISL_2344815, EPI_ISL_2344816, EPI_ISL_2344817 | SAE CTA SERV ATEND ESPEC TESTAGEM E ACONSELHAMENTO | Instituto Butantan / ESALQ-Piracicaba | Dimas Tadeu Covas, Antonio Jorge Martins, Claudia Renata dos Santos Barros, David Schlesinger, Debora Botequio Moretti, Elaine Cristina Marqueze, Elaine Vieira Santos, Evandra Strazza Rodrigues, Heidge Fukumasu, Jayme Augusto de Souza-Neto, José Salvatore Leister Patané, Luiz Alcantara, Luiz Lehmann Coutinho, Maria Carolina Elias, Maurício Lacerda Nogueira, Rafael dos Santos Bezerra, Raul Machado Neto, Rejane Maria Tommasini Grotto, Ricardo Haddad, Sandra Coccuzzo Sampaio Vessoni, Simone Kashima, Svetoslav Nanev Slavov, Vincent Louis Viala |
| EPI_ISL_2344818 | UNIDADE MISTA DE IGUAPE | Instituto Butantan / ESALQ-Piracicaba | Dimas Tadeu Covas, Antonio Jorge Martins, Claudia Renata dos Santos Barros, David Schlesinger, Debora Botequio Moretti, Elaine Cristina Marqueze, Elaine Vieira Santos, Evandra Strazza Rodrigues, Heidge Fukumasu, Jayme Augusto de Souza-Neto, José Salvatore Leister Patané, Luiz Alcantara, Luiz Lehmann Coutinho, Maria Carolina Elias, Maurício Lacerda Nogueira, Rafael dos Santos Bezerra, Raul Machado Neto, Rejane Maria Tommasini Grotto, Ricardo Haddad, Sandra Coccuzzo Sampaio Vessoni, Simone Kashima, Svetoslav Nanev Slavov, Vincent Louis Viala |
| EPI_ISL_2344819 | SAE CTA SERV ATEND ESPEC TESTAGEM E ACONSELHAMENTO | Instituto Butantan / ESALQ-Piracicaba | Dimas Tadeu Covas, Antonio Jorge Martins, Claudia Renata dos Santos Barros, David Schlesinger, Debora Botequio Moretti, Elaine Cristina Marqueze, Elaine Vieira Santos, Evandra Strazza Rodrigues, Heidge Fukumasu, Jayme Augusto de Souza-Neto, José Salvatore Leister Patané, Luiz Alcantara, Luiz Lehmann Coutinho, Maria Carolina Elias, Maurício Lacerda Nogueira, Rafael dos Santos Bezerra, Raul Machado Neto, Rejane Maria Tommasini Grotto, Ricardo Haddad, Sandra Coccuzzo Sampaio Vessoni, Simone Kashima, Svetoslav Nanev Slavov, Vincent Louis Viala |
| EPI_ISL_2344821, EPI_ISL_2344823, EPI_ISL_2344824 | CS DE CATIGUA | Instituto Butantan / ESALQ-Piracicaba | Dimas Tadeu Covas, Antonio Jorge Martins, Claudia Renata dos Santos Barros, David Schlesinger, Debora Botequio Moretti, Elaine Cristina Marqueze, Elaine Vieira Santos, Evandra Strazza Rodrigues, Heidge Fukumasu, Jayme Augusto de Souza-Neto, José Salvatore Leister Patané, Luiz Alcantara, Luiz Lehmann Coutinho, Maria Carolina Elias, Maurício Lacerda Nogueira, Rafael dos Santos Bezerra, Raul Machado Neto, Rejane Maria Tommasini Grotto, Ricardo Haddad, Sandra Coccuzzo Sampaio Vessoni, Simone Kashima, Svetoslav Nanev Slavov, Vincent Louis Viala |
| EPI_ISL_2344825 | UBS JD SALTENSE | Instituto Butantan / ESALQ-Piracicaba | Dimas Tadeu Covas, Antonio Jorge Martins, Claudia Renata dos Santos Barros, David Schlesinger, Debora Botequio Moretti, Elaine Cristina Marqueze, Elaine Vieira Santos, Evandra Strazza Rodrigues, Heidge Fukumasu, Jayme Augusto de Souza-Neto, José Salvatore Leister Patané, Luiz Alcantara, Luiz Lehmann Coutinho, Maria Carolina Elias, Maurício Lacerda Nogueira, Rafael dos Santos Bezerra, Raul Machado Neto, Rejane Maria Tommasini Grotto, Ricardo Haddad, Sandra Coccuzzo Sampaio Vessoni, Simone Kashima, Svetoslav Nanev Slavov, Vincent Louis Viala |
| EPI_ISL_2344826 | LABORATORIO MUNICIPAL DE PIRACICABA | Instituto Butantan / ESALQ-Piracicaba | Dimas Tadeu Covas, Antonio Jorge Martins, Claudia Renata dos Santos Barros, David Schlesinger, Debora Botequio Moretti, Elaine Cristina Marqueze, Elaine Vieira Santos, Evandra Strazza Rodrigues, Heidge Fukumasu, Jayme Augusto de Souza-Neto, José Salvatore Leister Patané, Luiz Alcantara, Luiz Lehmann Coutinho, Maria Carolina Elias, Maurício Lacerda Nogueira, Rafael dos Santos Bezerra, Raul Machado Neto, Rejane Maria Tommasini Grotto, Ricardo Haddad, Sandra Coccuzzo Sampaio Vessoni, Simone Kashima, Svetoslav Nanev Slavov, Vincent Louis Viala |
| EPI_ISL_2344827, EPI_ISL_2344828, EPI_ISL_2344829 | LABORATORIO DE FRANCA | Instituto Butantan / ESALQ-Piracicaba | Dimas Tadeu Covas, Antonio Jorge Martins, Claudia Renata dos Santos Barros, David Schlesinger, Debora Botequio Moretti, Elaine Cristina Marqueze, Elaine Vieira Santos, Evandra Strazza Rodrigues, Heidge Fukumasu, Jayme Augusto de Souza-Neto, José Salvatore Leister Patané, Luiz Alcantara, Luiz Lehmann Coutinho, Maria Carolina Elias, Maurício Lacerda Nogueira, Rafael dos Santos Bezerra, Raul Machado Neto, Rejane Maria Tommasini Grotto, Ricardo Haddad, Sandra Coccuzzo Sampaio Vessoni, Simone Kashima, Svetoslav Nanev Slavov, Vincent Louis Viala |
| EPI_ISL_2344830 | VIGILANCIA EPIDEMIOLOGICA E CONTROLE DE VETORES PIRASSUNUN | Instituto Butantan / ESALQ-Piracicaba | Dimas Tadeu Covas, Antonio Jorge Martins, Claudia Renata dos Santos Barros, David Schlesinger, Debora Botequio Moretti, Elaine Cristina Marqueze, Elaine Vieira Santos, Evandra Strazza Rodrigues, Heidge Fukumasu, Jayme Augusto de Souza-Neto, José Salvatore Leister Patané, Luiz Alcantara, Luiz Lehmann Coutinho, Maria Carolina Elias, Maurício Lacerda Nogueira, Rafael dos Santos Bezerra, Raul Machado Neto, Rejane Maria Tommasini Grotto, Ricardo Haddad, Sandra Coccuzzo Sampaio Vessoni, Simone Kashima, Svetoslav Nanev Slavov, Vincent Louis Viala |
| EPI_ISL_2344831 | CSA COMPLEXO DE SAUDE ANHANGUERA LTDA | Instituto Butantan | Dimas Tadeu Covas, Antonio Jorge Martins, Claudia Renata dos Santos Barros, David Schlesinger, Debora Botequio Moretti, Elaine Cristina Marqueze, Elaine Vieira Santos, Evandra Strazza Rodrigues, Heidge Fukumasu, Jayme Augusto de Souza-Neto, José Salvatore Leister Patané, Luiz Alcantara, Luiz Lehmann Coutinho, Maria Carolina Elias, Maurício Lacerda Nogueira, Rafael dos Santos Bezerra, Raul Machado Neto, Rejane Maria Tommasini Grotto, Ricardo Haddad, Sandra Coccuzzo Sampaio Vessoni, Simone Kashima, Svetoslav Nanev Slavov, Vincent Louis Viala |
| EPI_ISL_2344832 | CENTRO DE SAUDE DR RENATO DE CARVALHO RIBEIRO ANGATUBA | Instituto Butantan / ESALQ-Piracicaba | Dimas Tadeu Covas, Antonio Jorge Martins, Claudia Renata dos Santos Barros, David Schlesinger, Debora Botequio Moretti, Elaine Cristina Marqueze, Elaine Vieira Santos, Evandra Strazza Rodrigues, Heidge Fukumasu, Jayme Augusto de Souza-Neto, José Salvatore Leister Patané, Luiz Alcantara, Luiz Lehmann Coutinho, Maria Carolina Elias, Maurício Lacerda Nogueira, Rafael dos Santos Bezerra, Raul Machado Neto, Rejane Maria Tommasini Grotto, Ricardo Haddad, Sandra Coccuzzo Sampaio Vessoni, Simone Kashima, Svetoslav Nanev Slavov, Vincent Louis Viala |
| EPI_ISL_2344833 | LABORATORIO MUNICIPAL DE PIRACICABA | Instituto Butantan / ESALQ-Piracicaba | Dimas Tadeu Covas, Antonio Jorge Martins, Claudia Renata dos Santos Barros, David Schlesinger, Debora Botequio Moretti, Elaine Cristina Marqueze, Elaine Vieira Santos, Evandra Strazza Rodrigues, Heidge Fukumasu, Jayme Augusto de Souza-Neto, José Salvatore Leister Patané, Luiz Alcantara, Luiz Lehmann Coutinho, Maria Carolina Elias, Maurício Lacerda Nogueira, Rafael dos Santos Bezerra, Raul Machado Neto, Rejane Maria Tommasini Grotto, Ricardo Haddad, Sandra Coccuzzo Sampaio Vessoni, Simone Kashima, Svetoslav Nanev Slavov, Vincent Louis Viala |
| EPI_ISL_2344834, EPI_ISL_2344835 | EMERGENCIA RESPIRATORIA DE NOVA GRANADA | Instituto Butantan / ESALQ-Piracicaba | Dimas Tadeu Covas, Antonio Jorge Martins, Claudia Renata dos Santos Barros, David Schlesinger, Debora Botequio Moretti, Elaine Cristina Marqueze, Elaine Vieira Santos, Evandra Strazza Rodrigues, Heidge Fukumasu, Jayme Augusto de Souza-Neto, José Salvatore Leister Patané, Luiz Alcantara, Luiz Lehmann Coutinho, Maria Carolina Elias, Maurício Lacerda Nogueira, Rafael dos Santos Bezerra, Raul Machado Neto, Rejane Maria Tommasini Grotto, Ricardo Haddad, Sandra Coccuzzo Sampaio Vessoni, Simone Kashima, Svetoslav Nanev Slavov, Vincent Louis Viala |
| EPI_ISL_2344836, EPI_ISL_2344837 | LABORATORIO DE FRANCA | Instituto Butantan / ESALQ-Piracicaba | Dimas Tadeu Covas, Antonio Jorge Martins, Claudia Renata dos Santos Barros, David Schlesinger, Debora Botequio Moretti, Elaine Cristina Marqueze, Elaine Vieira Santos, Evandra Strazza Rodrigues, Heidge Fukumasu, Jayme Augusto de Souza-Neto, José Salvatore Leister Patané, Luiz Alcantara, Luiz Lehmann Coutinho, Maria Carolina Elias, Maurício Lacerda Nogueira, Rafael dos Santos Bezerra, Raul Machado Neto, Rejane Maria Tommasini Grotto, Ricardo Haddad, Sandra Coccuzzo Sampaio Vessoni, Simone Kashima, Svetoslav Nanev Slavov, Vincent Louis Viala |
| EPI_ISL_2344838 | LABORATORIO MUNICIPAL DE PIRACICABA | Instituto Butantan / ESALQ-Piracicaba | Dimas Tadeu Covas, Antonio Jorge Martins, Claudia Renata dos Santos Barros, David Schlesinger, Debora Botequio Moretti, Elaine Cristina Marqueze, Elaine Vieira Santos, Evandra Strazza Rodrigues, Heidge Fukumasu, Jayme Augusto de Souza-Neto, José Salvatore Leister Patané, Luiz Alcantara, Luiz Lehmann Coutinho, Maria Carolina Elias, Maurício Lacerda Nogueira, Rafael dos Santos Bezerra, Raul Machado Neto, Rejane Maria Tommasini Grotto, Ricardo Haddad, Sandra Coccuzzo Sampaio Vessoni, Simone Kashima, Svetoslav Nanev Slavov, Vincent Louis Viala |
| EPI_ISL_2344839, EPI_ISL_2344840, EPI_ISL_2344841, EPI_ISL_2344842, EPI_ISL_2344843, EPI_ISL_2344844 | LABORATORIO DE FRANCA | Instituto Butantan / ESALQ-Piracicaba | Dimas Tadeu Covas, Antonio Jorge Martins, Claudia Renata dos Santos Barros, David Schlesinger, Debora Botequio Moretti, Elaine Cristina Marqueze, Elaine Vieira Santos, Evandra Strazza Rodrigues, Heidge Fukumasu, Jayme Augusto de Souza-Neto, José Salvatore Leister Patané, Luiz Alcantara, Luiz Lehmann Coutinho, Maria Carolina Elias, Maurício Lacerda Nogueira, Rafael dos Santos Bezerra, Raul Machado Neto, Rejane Maria Tommasini Grotto, Ricardo Haddad, Sandra Coccuzzo Sampaio Vessoni, Simone Kashima, Svetoslav Nanev Slavov, Vincent Louis Viala |
| EPI_ISL_2344845 | EMERGENCIA RESPIRATORIA DE NOVA GRANADA | Instituto Butantan / ESALQ-Piracicaba | Dimas Tadeu Covas, Antonio Jorge Martins, Claudia Renata dos Santos Barros, David Schlesinger, Debora Botequio Moretti, Elaine Cristina Marqueze, Elaine Vieira Santos, Evandra Strazza Rodrigues, Heidge Fukumasu, Jayme Augusto de Souza-Neto, José Salvatore Leister Patané, Luiz Alcantara, Luiz Lehmann Coutinho, Maria Carolina Elias, Maurício Lacerda Nogueira, Rafael dos Santos Bezerra, Raul Machado Neto, Rejane Maria Tommasini Grotto, Ricardo Haddad, Sandra Coccuzzo Sampaio Vessoni, Simone Kashima, Svetoslav Nanev Slavov, Vincent Louis Viala |
| EPI_ISL_2344846 | CENTRO DE ESPECIALIDADES MARIAS DA GLORIA | Instituto Butantan / ESALQ-Piracicaba | Dimas Tadeu Covas, Antonio Jorge Martins, Claudia Renata dos Santos Barros, David Schlesinger, Debora Botequio Moretti, Elaine Cristina Marqueze, Elaine Vieira Santos, Evandra Strazza Rodrigues, Heidge Fukumasu, Jayme Augusto de Souza-Neto, José Salvatore Leister Patané, Luiz Alcantara, Luiz Lehmann Coutinho, Maria Carolina Elias, Maurício Lacerda Nogueira, Rafael dos Santos Bezerra, Raul Machado Neto, Rejane Maria Tommasini Grotto, Ricardo Haddad, Sandra Coccuzzo Sampaio Vessoni, Simone Kashima, Svetoslav Nanev Slavov, Vincent Louis Viala |

| | | | |
|---|---|---|---|
| EPI_ISL_2344847 | UBS DE PEDRANOPOLIS | Instituto Butantan / ESALQ-Piracicaba | Dimas Tadeu Covas, Antonio Jorge Martins, Claudia Renata dos Santos Barros, David Schlesinger, Debora Botequio Moretti, Elaine Cristina Marqueze, Elaine Vieira Santos, Evandra Strazza Rodrigues, Heidge Fukumasu, Jayme Augusto de Souza-Neto, José Salvatore Leister Patané, Luiz Alcantara, Luiz Lehmann Coutinho, Maria Carolina Elias, Maurício Lacerda Nogueira, Rafael dos Santos Bezerra, Raul Machado Neto, Rejane Maria Tommasini Grotto, Ricardo Haddad, Sandra Coccuzzo Sampaio Vessoni, Simone Kashima, Svetoslav Nanev Slavov, Vincent Louis Viala |
| EPI_ISL_2344848, EPI_ISL_2344849 | HOSPITAL DE MISERICORDIA ALTINOPOLIS | Instituto Butantan / ESALQ-Piracicaba | Dimas Tadeu Covas, Antonio Jorge Martins, Claudia Renata dos Santos Barros, David Schlesinger, Debora Botequio Moretti, Elaine Cristina Marqueze, Elaine Vieira Santos, Evandra Strazza Rodrigues, Heidge Fukumasu, Jayme Augusto de Souza-Neto, José Salvatore Leister Patané, Luiz Alcantara, Luiz Lehmann Coutinho, Maria Carolina Elias, Maurício Lacerda Nogueira, Rafael dos Santos Bezerra, Raul Machado Neto, Rejane Maria Tommasini Grotto, Ricardo Haddad, Sandra Coccuzzo Sampaio Vessoni, Simone Kashima, Svetoslav Nanev Slavov, Vincent Louis Viala |
| EPI_ISL_2344850 | UBS DE SAO JOAO DAS DUAS PONTES | Instituto Butantan / ESALQ-Piracicaba | Dimas Tadeu Covas, Antonio Jorge Martins, Claudia Renata dos Santos Barros, David Schlesinger, Debora Botequio Moretti, Elaine Cristina Marqueze, Elaine Vieira Santos, Evandra Strazza Rodrigues, Heidge Fukumasu, Jayme Augusto de Souza-Neto, José Salvatore Leister Patané, Luiz Alcantara, Luiz Lehmann Coutinho, Maria Carolina Elias, Maurício Lacerda Nogueira, Rafael dos Santos Bezerra, Raul Machado Neto, Rejane Maria Tommasini Grotto, Ricardo Haddad, Sandra Coccuzzo Sampaio Vessoni, Simone Kashima, Svetoslav Nanev Slavov, Vincent Louis Viala |
| EPI_ISL_2344851 | LABORATORIO MUNICIPAL DE PIRACICABA | Instituto Butantan / ESALQ-Piracicaba | Dimas Tadeu Covas, Antonio Jorge Martins, Claudia Renata dos Santos Barros, David Schlesinger, Debora Botequio Moretti, Elaine Cristina Marqueze, Elaine Vieira Santos, Evandra Strazza Rodrigues, Heidge Fukumasu, Jayme Augusto de Souza-Neto, José Salvatore Leister Patané, Luiz Alcantara, Luiz Lehmann Coutinho, Maria Carolina Elias, Maurício Lacerda Nogueira, Rafael dos Santos Bezerra, Raul Machado Neto, Rejane Maria Tommasini Grotto, Ricardo Haddad, Sandra Coccuzzo Sampaio Vessoni, Simone Kashima, Svetoslav Nanev Slavov, Vincent Louis Viala |
| EPI_ISL_2344852 | CENTRO DE SAUDE DE ANALANDIA | Instituto Butantan / ESALQ-Piracicaba | Dimas Tadeu Covas, Antonio Jorge Martins, Claudia Renata dos Santos Barros, David Schlesinger, Debora Botequio Moretti, Elaine Cristina Marqueze, Elaine Vieira Santos, Evandra Strazza Rodrigues, Heidge Fukumasu, Jayme Augusto de Souza-Neto, José Salvatore Leister Patané, Luiz Alcantara, Luiz Lehmann Coutinho, Maria Carolina Elias, Maurício Lacerda Nogueira, Rafael dos Santos Bezerra, Raul Machado Neto, Rejane Maria Tommasini Grotto, Ricardo Haddad, Sandra Coccuzzo Sampaio Vessoni, Simone Kashima, Svetoslav Nanev Slavov, Vincent Louis Viala |
| EPI_ISL_2344853 | CS DE PALMARES PAULISTA | Instituto Butantan / ESALQ-Piracicaba | Dimas Tadeu Covas, Antonio Jorge Martins, Claudia Renata dos Santos Barros, David Schlesinger, Debora Botequio Moretti, Elaine Cristina Marqueze, Elaine Vieira Santos, Evandra Strazza Rodrigues, Heidge Fukumasu, Jayme Augusto de Souza-Neto, José Salvatore Leister Patané, Luiz Alcantara, Luiz Lehmann Coutinho, Maria Carolina Elias, Maurício Lacerda Nogueira, Rafael dos Santos Bezerra, Raul Machado Neto, Rejane Maria Tommasini Grotto, Ricardo Haddad, Sandra Coccuzzo Sampaio Vessoni, Simone Kashima, Svetoslav Nanev Slavov, Vincent Louis Viala |
| EPI_ISL_2344854, EPI_ISL_2344855, EPI_ISL_2344856 | CS DE SANTA ALBERTINA | Instituto Butantan / ESALQ-Piracicaba | Dimas Tadeu Covas, Antonio Jorge Martins, Claudia Renata dos Santos Barros, David Schlesinger, Debora Botequio Moretti, Elaine Cristina Marqueze, Elaine Vieira Santos, Evandra Strazza Rodrigues, Heidge Fukumasu, Jayme Augusto de Souza-Neto, José Salvatore Leister Patané, Luiz Alcantara, Luiz Lehmann Coutinho, Maria Carolina Elias, Maurício Lacerda Nogueira, Rafael dos Santos Bezerra, Raul Machado Neto, Rejane Maria Tommasini Grotto, Ricardo Haddad, Sandra Coccuzzo Sampaio Vessoni, Simone Kashima, Svetoslav Nanev Slavov, Vincent Louis Viala |
| EPI_ISL_2344857 | CS DE CATIGUA | Instituto Butantan / ESALQ-Piracicaba | Dimas Tadeu Covas, Antonio Jorge Martins, Claudia Renata dos Santos Barros, David Schlesinger, Debora Botequio Moretti, Elaine Cristina Marqueze, Elaine Vieira Santos, Evandra Strazza Rodrigues, Heidge Fukumasu, Jayme Augusto de Souza-Neto, José Salvatore Leister Patané, Luiz Alcantara, Luiz Lehmann Coutinho, Maria Carolina Elias, Maurício Lacerda Nogueira, Rafael dos Santos Bezerra, Raul Machado Neto, Rejane Maria Tommasini Grotto, Ricardo Haddad, Sandra Coccuzzo Sampaio Vessoni, Simone Kashima, Svetoslav Nanev Slavov, Vincent Louis Viala |
| EPI_ISL_2344858 | SECRETARIA MUNICIPAL DE SAUDE DE CORDEIROPOLIS | Instituto Butantan / ESALQ-Piracicaba | Dimas Tadeu Covas, Antonio Jorge Martins, Claudia Renata dos Santos Barros, David Schlesinger, Debora Botequio Moretti, Elaine Cristina Marqueze, Elaine Vieira Santos, Evandra Strazza Rodrigues, Heidge Fukumasu, Jayme Augusto de Souza-Neto, José Salvatore Leister Patané, Luiz Alcantara, Luiz Lehmann Coutinho, Maria Carolina Elias, Maurício Lacerda Nogueira, Rafael dos Santos Bezerra, Raul Machado Neto, Rejane Maria Tommasini Grotto, Ricardo Haddad, Sandra Coccuzzo Sampaio Vessoni, Simone Kashima, Svetoslav Nanev Slavov, Vincent Louis Viala |
| EPI_ISL_2344859 | CS DE PALMARES PAULISTA | Instituto Butantan / ESALQ-Piracicaba | Dimas Tadeu Covas, Antonio Jorge Martins, Claudia Renata dos Santos Barros, David Schlesinger, Debora Botequio Moretti, Elaine Cristina Marqueze, Elaine Vieira Santos, Evandra Strazza Rodrigues, Heidge Fukumasu, Jayme Augusto de Souza-Neto, José Salvatore Leister Patané, Luiz Alcantara, Luiz Lehmann Coutinho, Maria Carolina Elias, Maurício Lacerda Nogueira, Rafael dos Santos Bezerra, Raul Machado Neto, Rejane Maria Tommasini Grotto, Ricardo Haddad, Sandra Coccuzzo Sampaio Vessoni, Simone Kashima, Svetoslav Nanev Slavov, Vincent Louis Viala |
| EPI_ISL_2344860 | CS DE CATIGUA | Instituto Butantan / ESALQ-Piracicaba | Dimas Tadeu Covas, Antonio Jorge Martins, Claudia Renata dos Santos Barros, David Schlesinger, Debora Botequio Moretti, Elaine Cristina Marqueze, Elaine Vieira Santos, Evandra Strazza Rodrigues, Heidge Fukumasu, Jayme Augusto de Souza-Neto, José Salvatore Leister Patané, Luiz Alcantara, Luiz Lehmann Coutinho, Maria Carolina Elias, Maurício Lacerda Nogueira, Rafael dos Santos Bezerra, Raul Machado Neto, Rejane Maria Tommasini Grotto, Ricardo Haddad, Sandra Coccuzzo Sampaio Vessoni, Simone Kashima, Svetoslav Nanev Slavov, Vincent Louis Viala |
| EPI_ISL_2344861 | EMERGENCIA RESPIRATORIA DE NOVA GRANADA | Instituto Butantan / ESALQ-Piracicaba | Dimas Tadeu Covas, Antonio Jorge Martins, Claudia Renata dos Santos Barros, David Schlesinger, Debora Botequio Moretti, Elaine Cristina Marqueze, Elaine Vieira Santos, Evandra Strazza Rodrigues, Heidge Fukumasu, Jayme Augusto de Souza-Neto, José Salvatore Leister Patané, Luiz Alcantara, Luiz Lehmann Coutinho, Maria Carolina Elias, Maurício Lacerda Nogueira, Rafael dos Santos Bezerra, Raul Machado Neto, Rejane Maria Tommasini Grotto, Ricardo Haddad, Sandra Coccuzzo Sampaio Vessoni, Simone Kashima, Svetoslav Nanev Slavov, Vincent Louis Viala |
| EPI_ISL_2344862 | CS DE VALENTIM GENTIL | Instituto Butantan / ESALQ-Piracicaba | Dimas Tadeu Covas, Antonio Jorge Martins, Claudia Renata dos Santos Barros, David Schlesinger, Debora Botequio Moretti, Elaine Cristina Marqueze, Elaine Vieira Santos, Evandra Strazza Rodrigues, Heidge Fukumasu, Jayme Augusto de Souza-Neto, José Salvatore Leister Patané, Luiz Alcantara, Luiz Lehmann Coutinho, Maria Carolina Elias, Maurício Lacerda Nogueira, Rafael dos Santos Bezerra, Raul Machado Neto, Rejane Maria Tommasini Grotto, Ricardo Haddad, Sandra Coccuzzo Sampaio Vessoni, Simone Kashima, Svetoslav Nanev Slavov, Vincent Louis Viala |
| EPI_ISL_2344863 | CS DE BALSAMO | Instituto Butantan / ESALQ-Piracicaba | Dimas Tadeu Covas, Antonio Jorge Martins, Claudia Renata dos Santos Barros, David Schlesinger, Debora Botequio Moretti, Elaine Cristina Marqueze, Elaine Vieira Santos, Evandra Strazza Rodrigues, Heidge Fukumasu, Jayme Augusto de Souza-Neto, José Salvatore Leister Patané, Luiz Alcantara, Luiz Lehmann Coutinho, Maria Carolina Elias, Maurício Lacerda Nogueira, Rafael dos Santos Bezerra, Raul Machado Neto, Rejane Maria Tommasini Grotto, Ricardo Haddad, Sandra Coccuzzo Sampaio Vessoni, Simone Kashima, Svetoslav Nanev Slavov, Vincent Louis Viala |
| EPI_ISL_2344864 | LABORATORIO MUNICIPAL DE PIRACICABA | Instituto Butantan / ESALQ-Piracicaba | Dimas Tadeu Covas, Antonio Jorge Martins, Claudia Renata dos Santos Barros, David Schlesinger, Debora Botequio Moretti, Elaine Cristina Marqueze, Elaine Vieira Santos, Evandra Strazza Rodrigues, Heidge Fukumasu, Jayme Augusto de Souza-Neto, José Salvatore Leister Patané, Luiz Alcantara, Luiz Lehmann Coutinho, Maria Carolina Elias, Maurício Lacerda Nogueira, Rafael dos Santos Bezerra, Raul Machado Neto, Rejane Maria Tommasini Grotto, Ricardo Haddad, Sandra Coccuzzo Sampaio Vessoni, Simone Kashima, Svetoslav Nanev Slavov, Vincent Louis Viala |
| EPI_ISL_2344865 | SANTA CASA DE JOSE BONIFACIO | Instituto Butantan / ESALQ-Piracicaba | Dimas Tadeu Covas, Antonio Jorge Martins, Claudia Renata dos Santos Barros, David Schlesinger, Debora Botequio Moretti, Elaine Cristina Marqueze, Elaine Vieira Santos, Evandra Strazza Rodrigues, Heidge Fukumasu, Jayme Augusto de Souza-Neto, José Salvatore Leister Patané, Luiz Alcantara, Luiz Lehmann Coutinho, Maria Carolina Elias, Maurício Lacerda Nogueira, Rafael dos Santos Bezerra, Raul Machado Neto, Rejane Maria Tommasini Grotto, Ricardo Haddad, Sandra Coccuzzo Sampaio Vessoni, Simone Kashima, Svetoslav Nanev Slavov, Vincent Louis Viala |
| EPI_ISL_2344866 | AMB MEDICO DR FRANCISCO FELIX DE MENDONCA SAUDADE | Instituto Butantan / ESALQ-Piracicaba | Dimas Tadeu Covas, Antonio Jorge Martins, Claudia Renata dos Santos Barros, David Schlesinger, Debora Botequio Moretti, Elaine Cristina Marqueze, Elaine Vieira Santos, Evandra Strazza Rodrigues, Heidge Fukumasu, Jayme Augusto de Souza-Neto, José Salvatore Leister Patané, Luiz Alcantara, Luiz Lehmann Coutinho, Maria Carolina Elias, Maurício Lacerda Nogueira, Rafael dos Santos Bezerra, Raul Machado Neto, Rejane Maria Tommasini Grotto, Ricardo Haddad, Sandra Coccuzzo Sampaio Vessoni, Simone Kashima, Svetoslav Nanev Slavov, Vincent Louis Viala |
| EPI_ISL_2344867 | SECRETARIA MUNICIPAL DE SAUDE SOROCABA | Instituto Butantan / ESALQ-Piracicaba | Dimas Tadeu Covas, Antonio Jorge Martins, Claudia Renata dos Santos Barros, David Schlesinger, Debora Botequio Moretti, Elaine Cristina Marqueze, Elaine Vieira Santos, Evandra Strazza Rodrigues, Heidge Fukumasu, Jayme Augusto de Souza-Neto, José Salvatore Leister Patané, Luiz Alcantara, Luiz Lehmann Coutinho, Maria Carolina Elias, Maurício Lacerda Nogueira, Rafael dos Santos Bezerra, Raul Machado Neto, Rejane Maria Tommasini Grotto, Ricardo Haddad, Sandra Coccuzzo Sampaio Vessoni, Simone Kashima, Svetoslav Nanev Slavov, Vincent Louis Viala |
| EPI_ISL_2344868 | ESF M1 OZIL JOAQUIM REZENDE JALES | Instituto Butantan / ESALQ-Piracicaba | Dimas Tadeu Covas, Antonio Jorge Martins, Claudia Renata dos Santos Barros, David Schlesinger, Debora Botequio Moretti, Elaine Cristina Marqueze, Elaine Vieira Santos, Evandra Strazza Rodrigues, Heidge Fukumasu, Jayme Augusto de Souza-Neto, José Salvatore Leister Patané, Luiz Alcantara, Luiz Lehmann Coutinho, Maria Carolina Elias, Maurício Lacerda Nogueira, Rafael dos Santos Bezerra, Raul Machado Neto, Rejane Maria Tommasini Grotto, Ricardo Haddad, Sandra Coccuzzo Sampaio Vessoni, Simone Kashima, Svetoslav Nanev Slavov, Vincent Louis Viala |
| EPI_ISL_2344869 | E S F DR EDSON GIANETTO JD CARLOS CASSETARI | Instituto Butantan / ESALQ-Piracicaba | Dimas Tadeu Covas, Antonio Jorge Martins, Claudia Renata dos Santos Barros, David Schlesinger, Debora Botequio Moretti, Elaine Cristina Marqueze, Elaine Vieira Santos, Evandra Strazza Rodrigues, Heidge Fukumasu, Jayme Augusto de Souza-Neto, José Salvatore Leister Patané, Luiz Alcantara, Luiz |

| | | | |
|---|---|---|---|
| | | | Lehmann Coutinho, Maria Carolina Elias, Maurício Lacerda Nogueira, Rafael dos Santos Bezerra, Raul Machado Neto, Rejane Maria Tommasini Grotto, Ricardo Haddad, Sandra Coccuzzo Sampaio Vessoni, Simone Kashima, Svetoslav Nanev Slavov, Vincent Louis Viala |
| EPI_ISL_2344870 | ESF RURAL DR JOSE CICERO FONTES XAVIER | Instituto Butantan / ESALQ-Piracicaba | Dimas Tadeu Covas, Antonio Jorge Martins, Claudia Renata dos Santos Barros, David Schlesinger, Debora Botequio Moretti, Elaine Cristina Marqueze, Elaine Vieira Santos, Evandra Strazza Rodrigues, Heidge Fukumasu, Jayme Augusto de Souza-Neto, José Salvatore Leister Patané, Luiz Alcantara, Luiz Lehmann Coutinho, Maria Carolina Elias, Maurício Lacerda Nogueira, Rafael dos Santos Bezerra, Raul Machado Neto, Rejane Maria Tommasini Grotto, Ricardo Haddad, Sandra Coccuzzo Sampaio Vessoni, Simone Kashima, Svetoslav Nanev Slavov, Vincent Louis Viala |
| EPI_ISL_2344871 | CS DE COSMORAMA | Instituto Butantan / ESALQ-Piracicaba | Dimas Tadeu Covas, Antonio Jorge Martins, Claudia Renata dos Santos Barros, David Schlesinger, Debora Botequio Moretti, Elaine Cristina Marqueze, Elaine Vieira Santos, Evandra Strazza Rodrigues, Heidge Fukumasu, Jayme Augusto de Souza-Neto, José Salvatore Leister Patané, Luiz Alcantara, Luiz Lehmann Coutinho, Maria Carolina Elias, Maurício Lacerda Nogueira, Rafael dos Santos Bezerra, Raul Machado Neto, Rejane Maria Tommasini Grotto, Ricardo Haddad, Sandra Coccuzzo Sampaio Vessoni, Simone Kashima, Svetoslav Nanev Slavov, Vincent Louis Viala |
| EPI_ISL_2344872 | CS DE ONDA VERDE | Instituto Butantan / ESALQ-Piracicaba | Dimas Tadeu Covas, Antonio Jorge Martins, Claudia Renata dos Santos Barros, David Schlesinger, Debora Botequio Moretti, Elaine Cristina Marqueze, Elaine Vieira Santos, Evandra Strazza Rodrigues, Heidge Fukumasu, Jayme Augusto de Souza-Neto, José Salvatore Leister Patané, Luiz Alcantara, Luiz Lehmann Coutinho, Maria Carolina Elias, Maurício Lacerda Nogueira, Rafael dos Santos Bezerra, Raul Machado Neto, Rejane Maria Tommasini Grotto, Ricardo Haddad, Sandra Coccuzzo Sampaio Vessoni, Simone Kashima, Svetoslav Nanev Slavov, Vincent Louis Viala |
| EPI_ISL_2344873 | CENTRO DE SAUDE DR RENATO DE CARVALHO RIBEIRO ANGATUBA | Instituto Butantan / ESALQ-Piracicaba | Dimas Tadeu Covas, Antonio Jorge Martins, Claudia Renata dos Santos Barros, David Schlesinger, Debora Botequio Moretti, Elaine Cristina Marqueze, Elaine Vieira Santos, Evandra Strazza Rodrigues, Heidge Fukumasu, Jayme Augusto de Souza-Neto, José Salvatore Leister Patané, Luiz Alcantara, Luiz Lehmann Coutinho, Maria Carolina Elias, Maurício Lacerda Nogueira, Rafael dos Santos Bezerra, Raul Machado Neto, Rejane Maria Tommasini Grotto, Ricardo Haddad, Sandra Coccuzzo Sampaio Vessoni, Simone Kashima, Svetoslav Nanev Slavov, Vincent Louis Viala |
| EPI_ISL_2344874 | CS DE COSMORAMA | Instituto Butantan / ESALQ-Piracicaba | Dimas Tadeu Covas, Antonio Jorge Martins, Claudia Renata dos Santos Barros, David Schlesinger, Debora Botequio Moretti, Elaine Cristina Marqueze, Elaine Vieira Santos, Evandra Strazza Rodrigues, Heidge Fukumasu, Jayme Augusto de Souza-Neto, José Salvatore Leister Patané, Luiz Alcantara, Luiz Lehmann Coutinho, Maria Carolina Elias, Maurício Lacerda Nogueira, Rafael dos Santos Bezerra, Raul Machado Neto, Rejane Maria Tommasini Grotto, Ricardo Haddad, Sandra Coccuzzo Sampaio Vessoni, Simone Kashima, Svetoslav Nanev Slavov, Vincent Louis Viala |
| EPI_ISL_2344875 | CS DE ONDA VERDE | Instituto Butantan / ESALQ-Piracicaba | Dimas Tadeu Covas, Antonio Jorge Martins, Claudia Renata dos Santos Barros, David Schlesinger, Debora Botequio Moretti, Elaine Cristina Marqueze, Elaine Vieira Santos, Evandra Strazza Rodrigues, Heidge Fukumasu, Jayme Augusto de Souza-Neto, José Salvatore Leister Patané, Luiz Alcantara, Luiz Lehmann Coutinho, Maria Carolina Elias, Maurício Lacerda Nogueira, Rafael dos Santos Bezerra, Raul Machado Neto, Rejane Maria Tommasini Grotto, Ricardo Haddad, Sandra Coccuzzo Sampaio Vessoni, Simone Kashima, Svetoslav Nanev Slavov, Vincent Louis Viala |
| EPI_ISL_2344876 | SANTA CASA DE JOSE BONIFACIO | Instituto Butantan / ESALQ-Piracicaba | Dimas Tadeu Covas, Antonio Jorge Martins, Claudia Renata dos Santos Barros, David Schlesinger, Debora Botequio Moretti, Elaine Cristina Marqueze, Elaine Vieira Santos, Evandra Strazza Rodrigues, Heidge Fukumasu, Jayme Augusto de Souza-Neto, José Salvatore Leister Patané, Luiz Alcantara, Luiz Lehmann Coutinho, Maria Carolina Elias, Maurício Lacerda Nogueira, Rafael dos Santos Bezerra, Raul Machado Neto, Rejane Maria Tommasini Grotto, Ricardo Haddad, Sandra Coccuzzo Sampaio Vessoni, Simone Kashima, Svetoslav Nanev Slavov, Vincent Louis Viala |
| EPI_ISL_2344877 | E S F WALDO DSTEFANO JD PRIMAVERA JOSE BONIFACIO | Instituto Butantan / ESALQ-Piracicaba | Dimas Tadeu Covas, Antonio Jorge Martins, Claudia Renata dos Santos Barros, David Schlesinger, Debora Botequio Moretti, Elaine Cristina Marqueze, Elaine Vieira Santos, Evandra Strazza Rodrigues, Heidge Fukumasu, Jayme Augusto de Souza-Neto, José Salvatore Leister Patané, Luiz Alcantara, Luiz Lehmann Coutinho, Maria Carolina Elias, Maurício Lacerda Nogueira, Rafael dos Santos Bezerra, Raul Machado Neto, Rejane Maria Tommasini Grotto, Ricardo Haddad, Sandra Coccuzzo Sampaio Vessoni, Simone Kashima, Svetoslav Nanev Slavov, Vincent Louis Viala |
| EPI_ISL_2344878 | SANTA CASA DE JOSE BONIFACIO | Instituto Butantan / ESALQ-Piracicaba | Dimas Tadeu Covas, Antonio Jorge Martins, Claudia Renata dos Santos Barros, David Schlesinger, Debora Botequio Moretti, Elaine Cristina Marqueze, Elaine Vieira Santos, Evandra Strazza Rodrigues, Heidge Fukumasu, Jayme Augusto de Souza-Neto, José Salvatore Leister Patané, Luiz Alcantara, Luiz Lehmann Coutinho, Maria Carolina Elias, Maurício Lacerda Nogueira, Rafael dos Santos Bezerra, Raul Machado Neto, Rejane Maria Tommasini Grotto, Ricardo Haddad, Sandra Coccuzzo Sampaio Vessoni, Simone Kashima, Svetoslav Nanev Slavov, Vincent Louis Viala |
| EPI_ISL_2344879 | CS DE FLOREAL | Instituto Butantan / ESALQ-Piracicaba | Dimas Tadeu Covas, Antonio Jorge Martins, Claudia Renata dos Santos Barros, David Schlesinger, Debora Botequio Moretti, Elaine Cristina Marqueze, Elaine Vieira Santos, Evandra Strazza Rodrigues, Heidge Fukumasu, Jayme Augusto de Souza-Neto, José Salvatore Leister Patané, Luiz Alcantara, Luiz Lehmann Coutinho, Maria Carolina Elias, Maurício Lacerda Nogueira, Rafael dos Santos Bezerra, Raul Machado Neto, Rejane Maria Tommasini Grotto, Ricardo Haddad, Sandra Coccuzzo Sampaio Vessoni, Simone Kashima, Svetoslav Nanev Slavov, Vincent Louis Viala |
| EPI_ISL_2344880 | SANTA CASA DE JOSE BONIFACIO | Instituto Butantan / ESALQ-Piracicaba | Dimas Tadeu Covas, Antonio Jorge Martins, Claudia Renata dos Santos Barros, David Schlesinger, Debora Botequio Moretti, Elaine Cristina Marqueze, Elaine Vieira Santos, Evandra Strazza Rodrigues, Heidge Fukumasu, Jayme Augusto de Souza-Neto, José Salvatore Leister Patané, Luiz Alcantara, Luiz Lehmann Coutinho, Maria Carolina Elias, Maurício Lacerda Nogueira, Rafael dos Santos Bezerra, Raul Machado Neto, Rejane Maria Tommasini Grotto, Ricardo Haddad, Sandra Coccuzzo Sampaio Vessoni, Simone Kashima, Svetoslav Nanev Slavov, Vincent Louis Viala |
| EPI_ISL_2344881 | CS DE FLOREAL | Instituto Butantan / ESALQ-Piracicaba | Dimas Tadeu Covas, Antonio Jorge Martins, Claudia Renata dos Santos Barros, David Schlesinger, Debora Botequio Moretti, Elaine Cristina Marqueze, Elaine Vieira Santos, Evandra Strazza Rodrigues, Heidge Fukumasu, Jayme Augusto de Souza-Neto, José Salvatore Leister Patané, Luiz Alcantara, Luiz Lehmann Coutinho, Maria Carolina Elias, Maurício Lacerda Nogueira, Rafael dos Santos Bezerra, Raul Machado Neto, Rejane Maria Tommasini Grotto, Ricardo Haddad, Sandra Coccuzzo Sampaio Vessoni, Simone Kashima, Svetoslav Nanev Slavov, Vincent Louis Viala |
| EPI_ISL_2344882 | CS DE MARAPOAMA | Instituto Butantan / ESALQ-Piracicaba | Dimas Tadeu Covas, Antonio Jorge Martins, Claudia Renata dos Santos Barros, David Schlesinger, Debora Botequio Moretti, Elaine Cristina Marqueze, Elaine Vieira Santos, Evandra Strazza Rodrigues, Heidge Fukumasu, Jayme Augusto de Souza-Neto, José Salvatore Leister Patané, Luiz Alcantara, Luiz Lehmann Coutinho, Maria Carolina Elias, Maurício Lacerda Nogueira, Rafael dos Santos Bezerra, Raul Machado Neto, Rejane Maria Tommasini Grotto, Ricardo Haddad, Sandra Coccuzzo Sampaio Vessoni, Simone Kashima, Svetoslav Nanev Slavov, Vincent Louis Viala |
| EPI_ISL_2344883 | SANTA CASA DE JOSE BONIFACIO | Instituto Butantan / ESALQ-Piracicaba | Dimas Tadeu Covas, Antonio Jorge Martins, Claudia Renata dos Santos Barros, David Schlesinger, Debora Botequio Moretti, Elaine Cristina Marqueze, Elaine Vieira Santos, Evandra Strazza Rodrigues, Heidge Fukumasu, Jayme Augusto de Souza-Neto, José Salvatore Leister Patané, Luiz Alcantara, Luiz Lehmann Coutinho, Maria Carolina Elias, Maurício Lacerda Nogueira, Rafael dos Santos Bezerra, Raul Machado Neto, Rejane Maria Tommasini Grotto, Ricardo Haddad, Sandra Coccuzzo Sampaio Vessoni, Simone Kashima, Svetoslav Nanev Slavov, Vincent Louis Viala |
| EPI_ISL_2344884 | LABORATORIO MUNICIPAL DE PIRACICABA | Instituto Butantan / ESALQ-Piracicaba | Dimas Tadeu Covas, Antonio Jorge Martins, Claudia Renata dos Santos Barros, David Schlesinger, Debora Botequio Moretti, Elaine Cristina Marqueze, Elaine Vieira Santos, Evandra Strazza Rodrigues, Heidge Fukumasu, Jayme Augusto de Souza-Neto, José Salvatore Leister Patané, Luiz Alcantara, Luiz Lehmann Coutinho, Maria Carolina Elias, Maurício Lacerda Nogueira, Rafael dos Santos Bezerra, Raul Machado Neto, Rejane Maria Tommasini Grotto, Ricardo Haddad, Sandra Coccuzzo Sampaio Vessoni, Simone Kashima, Svetoslav Nanev Slavov, Vincent Louis Viala |
| EPI_ISL_2344885, EPI_ISL_2344886 | UNIDADE DE PRONTO ATENDIMENTO DE JALES | Instituto Butantan / ESALQ-Piracicaba | Dimas Tadeu Covas, Antonio Jorge Martins, Claudia Renata dos Santos Barros, David Schlesinger, Debora Botequio Moretti, Elaine Cristina Marqueze, Elaine Vieira Santos, Evandra Strazza Rodrigues, Heidge Fukumasu, Jayme Augusto de Souza-Neto, José Salvatore Leister Patané, Luiz Alcantara, Luiz Lehmann Coutinho, Maria Carolina Elias, Maurício Lacerda Nogueira, Rafael dos Santos Bezerra, Raul Machado Neto, Rejane Maria Tommasini Grotto, Ricardo Haddad, Sandra Coccuzzo Sampaio Vessoni, Simone Kashima, Svetoslav Nanev Slavov, Vincent Louis Viala |
| EPI_ISL_2344887 | CS DE ONDA VERDE | Instituto Butantan / ESALQ-Piracicaba | Dimas Tadeu Covas, Antonio Jorge Martins, Claudia Renata dos Santos Barros, David Schlesinger, Debora Botequio Moretti, Elaine Cristina Marqueze, Elaine Vieira Santos, Evandra Strazza Rodrigues, Heidge Fukumasu, Jayme Augusto de Souza-Neto, José Salvatore Leister Patané, Luiz Alcantara, Luiz Lehmann Coutinho, Maria Carolina Elias, Maurício Lacerda Nogueira, Rafael dos Santos Bezerra, Raul Machado Neto, Rejane Maria Tommasini Grotto, Ricardo Haddad, Sandra Coccuzzo Sampaio Vessoni, Simone Kashima, Svetoslav Nanev Slavov, Vincent Louis Viala |
| EPI_ISL_2344888 | SMS SECRETARIA MUNICIPAL DE SAUDE DE BOITUVA | Instituto Butantan / ESALQ-Piracicaba | Dimas Tadeu Covas, Antonio Jorge Martins, Claudia Renata dos Santos Barros, David Schlesinger, Debora Botequio Moretti, Elaine Cristina Marqueze, Elaine Vieira Santos, Evandra Strazza Rodrigues, Heidge Fukumasu, Jayme Augusto de Souza-Neto, José Salvatore Leister Patané, Luiz Alcantara, Luiz Lehmann Coutinho, Maria Carolina Elias, Maurício Lacerda Nogueira, Rafael dos Santos Bezerra, Raul Machado Neto, Rejane Maria Tommasini Grotto, Ricardo Haddad, Sandra Coccuzzo Sampaio Vessoni, Simone Kashima, Svetoslav Nanev Slavov, Vincent Louis Viala |
| EPI_ISL_2344889 | CS DE ONDA VERDE | Instituto Butantan / ESALQ-Piracicaba | Dimas Tadeu Covas, Antonio Jorge Martins, Claudia Renata dos Santos Barros, David Schlesinger, Debora Botequio Moretti, Elaine Cristina Marqueze, Elaine Vieira Santos, Evandra Strazza Rodrigues, Heidge Fukumasu, Jayme Augusto de Souza-Neto, José Salvatore Leister Patané, Luiz Alcantara, Luiz Lehmann Coutinho, Maria Carolina Elias, Maurício Lacerda Nogueira, Rafael dos Santos Bezerra, Raul Machado Neto, Rejane Maria Tommasini Grotto, Ricardo Haddad, Sandra Coccuzzo Sampaio Vessoni, Simone Kashima, Svetoslav Nanev Slavov, Vincent Louis Viala |

| | | | |
|---|---|---|---|
| EPI_ISL_2344890, EPI_ISL_2344891, EPI_ISL_2344892, EPI_ISL_2344894 | SANTA CASA DE PORTO FELIZ | Instituto Butantan / ESALQ-Piracicaba | Dimas Tadeu Covas, Antonio Jorge Martins, Claudia Renata dos Santos Barros, David Schlesinger, Debora Botequio Moretti, Elaine Cristina Marqueze, Elaine Vieira Santos, Evandra Strazza Rodrigues, Heidge Fukumasu, Jayme Augusto de Souza-Neto, José Salvatore Leister Patané, Luiz Alcantara, Luiz Lehmann Coutinho, Maria Carolina Elias, Maurício Lacerda Nogueira, Rafael dos Santos Bezerra, Raul Machado Neto, Rejane Maria Tommasini Grotto, Ricardo Haddad, Sandra Coccuzzo Sampaio Vessoni, Simone Kashima, Svetoslav Nanev Slavov, Vincent Louis Viala |
| EPI_ISL_2344896 | SECRETARIA MUNICIPAL DA SAUDE DE ITAPEVA SP | Instituto Butantan / ESALQ-Piracicaba | Dimas Tadeu Covas, Antonio Jorge Martins, Claudia Renata dos Santos Barros, David Schlesinger, Debora Botequio Moretti, Elaine Cristina Marqueze, Elaine Vieira Santos, Evandra Strazza Rodrigues, Heidge Fukumasu, Jayme Augusto de Souza-Neto, José Salvatore Leister Patané, Luiz Alcantara, Luiz Lehmann Coutinho, Maria Carolina Elias, Maurício Lacerda Nogueira, Rafael dos Santos Bezerra, Raul Machado Neto, Rejane Maria Tommasini Grotto, Ricardo Haddad, Sandra Coccuzzo Sampaio Vessoni, Simone Kashima, Svetoslav Nanev Slavov, Vincent Louis Viala |
| EPI_ISL_2344897 | SECRETARIA MUNICIPAL DE SAUDE SOROCABA | Instituto Butantan / ESALQ-Piracicaba | Dimas Tadeu Covas, Antonio Jorge Martins, Claudia Renata dos Santos Barros, David Schlesinger, Debora Botequio Moretti, Elaine Cristina Marqueze, Elaine Vieira Santos, Evandra Strazza Rodrigues, Heidge Fukumasu, Jayme Augusto de Souza-Neto, José Salvatore Leister Patané, Luiz Alcantara, Luiz Lehmann Coutinho, Maria Carolina Elias, Maurício Lacerda Nogueira, Rafael dos Santos Bezerra, Raul Machado Neto, Rejane Maria Tommasini Grotto, Ricardo Haddad, Sandra Coccuzzo Sampaio Vessoni, Simone Kashima, Svetoslav Nanev Slavov, Vincent Louis Viala |
| EPI_ISL_2344898 | HOSPITAL REGIONAL DE ITAPETININGA | Instituto Butantan / ESALQ-Piracicaba | Dimas Tadeu Covas, Antonio Jorge Martins, Claudia Renata dos Santos Barros, David Schlesinger, Debora Botequio Moretti, Elaine Cristina Marqueze, Elaine Vieira Santos, Evandra Strazza Rodrigues, Heidge Fukumasu, Jayme Augusto de Souza-Neto, José Salvatore Leister Patané, Luiz Alcantara, Luiz Lehmann Coutinho, Maria Carolina Elias, Maurício Lacerda Nogueira, Rafael dos Santos Bezerra, Raul Machado Neto, Rejane Maria Tommasini Grotto, Ricardo Haddad, Sandra Coccuzzo Sampaio Vessoni, Simone Kashima, Svetoslav Nanev Slavov, Vincent Louis Viala |
| EPI_ISL_2344899, EPI_ISL_2344900, EPI_ISL_2344901 | HOSPITAL E PRONTO SOCORRO PORTINARI | Instituto Butantan | Dimas Tadeu Covas, Antonio Jorge Martins, Claudia Renata dos Santos Barros, David Schlesinger, Debora Botequio Moretti, Elaine Cristina Marqueze, Elaine Vieira Santos, Evandra Strazza Rodrigues, Heidge Fukumasu, Jayme Augusto de Souza-Neto, José Salvatore Leister Patané, Luiz Alcantara, Luiz Lehmann Coutinho, Maria Carolina Elias, Maurício Lacerda Nogueira, Rafael dos Santos Bezerra, Raul Machado Neto, Rejane Maria Tommasini Grotto, Ricardo Haddad, Sandra Coccuzzo Sampaio Vessoni, Simone Kashima, Svetoslav Nanev Slavov, Vincent Louis Viala |
| EPI_ISL_2344902 | UBS JARDIM VITORINO IPERO | Instituto Butantan / ESALQ-Piracicaba | Dimas Tadeu Covas, Antonio Jorge Martins, Claudia Renata dos Santos Barros, David Schlesinger, Debora Botequio Moretti, Elaine Cristina Marqueze, Elaine Vieira Santos, Evandra Strazza Rodrigues, Heidge Fukumasu, Jayme Augusto de Souza-Neto, José Salvatore Leister Patané, Luiz Alcantara, Luiz Lehmann Coutinho, Maria Carolina Elias, Maurício Lacerda Nogueira, Rafael dos Santos Bezerra, Raul Machado Neto, Rejane Maria Tommasini Grotto, Ricardo Haddad, Sandra Coccuzzo Sampaio Vessoni, Simone Kashima, Svetoslav Nanev Slavov, Vincent Louis Viala |
| EPI_ISL_2344903 | SECRETARIA MUNICIPAL DA SAUDE DE ITARARE | Instituto Butantan / ESALQ-Piracicaba | Dimas Tadeu Covas, Antonio Jorge Martins, Claudia Renata dos Santos Barros, David Schlesinger, Debora Botequio Moretti, Elaine Cristina Marqueze, Elaine Vieira Santos, Evandra Strazza Rodrigues, Heidge Fukumasu, Jayme Augusto de Souza-Neto, José Salvatore Leister Patané, Luiz Alcantara, Luiz Lehmann Coutinho, Maria Carolina Elias, Maurício Lacerda Nogueira, Rafael dos Santos Bezerra, Raul Machado Neto, Rejane Maria Tommasini Grotto, Ricardo Haddad, Sandra Coccuzzo Sampaio Vessoni, Simone Kashima, Svetoslav Nanev Slavov, Vincent Louis Viala |
| EPI_ISL_2344904, EPI_ISL_2344905 | CENTRO DE SAUDE DR RENATO DE CARVALHO RIBEIRO ANGATUBA | Instituto Butantan / ESALQ-Piracicaba | Dimas Tadeu Covas, Antonio Jorge Martins, Claudia Renata dos Santos Barros, David Schlesinger, Debora Botequio Moretti, Elaine Cristina Marqueze, Elaine Vieira Santos, Evandra Strazza Rodrigues, Heidge Fukumasu, Jayme Augusto de Souza-Neto, José Salvatore Leister Patané, Luiz Alcantara, Luiz Lehmann Coutinho, Maria Carolina Elias, Maurício Lacerda Nogueira, Rafael dos Santos Bezerra, Raul Machado Neto, Rejane Maria Tommasini Grotto, Ricardo Haddad, Sandra Coccuzzo Sampaio Vessoni, Simone Kashima, Svetoslav Nanev Slavov, Vincent Louis Viala |
| EPI_ISL_2344906 | CSA COMPLEXO DE SAUDE ANHANGUERA TUCURUVI | Instituto Butantan | Dimas Tadeu Covas, Antonio Jorge Martins, Claudia Renata dos Santos Barros, David Schlesinger, Debora Botequio Moretti, Elaine Cristina Marqueze, Elaine Vieira Santos, Evandra Strazza Rodrigues, Heidge Fukumasu, Jayme Augusto de Souza-Neto, José Salvatore Leister Patané, Luiz Alcantara, Luiz Lehmann Coutinho, Maria Carolina Elias, Maurício Lacerda Nogueira, Rafael dos Santos Bezerra, Raul Machado Neto, Rejane Maria Tommasini Grotto, Ricardo Haddad, Sandra Coccuzzo Sampaio Vessoni, Simone Kashima, Svetoslav Nanev Slavov, Vincent Louis Viala |
| EPI_ISL_2344907 | CENTRO DE SAUDE DR RENATO DE CARVALHO RIBEIRO ANGATUBA | Instituto Butantan / ESALQ-Piracicaba | Dimas Tadeu Covas, Antonio Jorge Martins, Claudia Renata dos Santos Barros, David Schlesinger, Debora Botequio Moretti, Elaine Cristina Marqueze, Elaine Vieira Santos, Evandra Strazza Rodrigues, Heidge Fukumasu, Jayme Augusto de Souza-Neto, José Salvatore Leister Patané, Luiz Alcantara, Luiz Lehmann Coutinho, Maria Carolina Elias, Maurício Lacerda Nogueira, Rafael dos Santos Bezerra, Raul Machado Neto, Rejane Maria Tommasini Grotto, Ricardo Haddad, Sandra Coccuzzo Sampaio Vessoni, Simone Kashima, Svetoslav Nanev Slavov, Vincent Louis Viala |
| EPI_ISL_2344908 | CENTRO DE SAUDE TEREZINHA DE MORAES ARSILLA | Instituto Butantan / ESALQ-Piracicaba | Dimas Tadeu Covas, Antonio Jorge Martins, Claudia Renata dos Santos Barros, David Schlesinger, Debora Botequio Moretti, Elaine Cristina Marqueze, Elaine Vieira Santos, Evandra Strazza Rodrigues, Heidge Fukumasu, Jayme Augusto de Souza-Neto, José Salvatore Leister Patané, Luiz Alcantara, Luiz Lehmann Coutinho, Maria Carolina Elias, Maurício Lacerda Nogueira, Rafael dos Santos Bezerra, Raul Machado Neto, Rejane Maria Tommasini Grotto, Ricardo Haddad, Sandra Coccuzzo Sampaio Vessoni, Simone Kashima, Svetoslav Nanev Slavov, Vincent Louis Viala |
| EPI_ISL_2344909 | HOSP E MATERNIDADE MUNICIPAL N SRA MONTE SERRAT | Instituto Butantan / ESALQ-Piracicaba | Dimas Tadeu Covas, Antonio Jorge Martins, Claudia Renata dos Santos Barros, David Schlesinger, Debora Botequio Moretti, Elaine Cristina Marqueze, Elaine Vieira Santos, Evandra Strazza Rodrigues, Heidge Fukumasu, Jayme Augusto de Souza-Neto, José Salvatore Leister Patané, Luiz Alcantara, Luiz Lehmann Coutinho, Maria Carolina Elias, Maurício Lacerda Nogueira, Rafael dos Santos Bezerra, Raul Machado Neto, Rejane Maria Tommasini Grotto, Ricardo Haddad, Sandra Coccuzzo Sampaio Vessoni, Simone Kashima, Svetoslav Nanev Slavov, Vincent Louis Viala |
| EPI_ISL_2344910 | PSF MARIA JOSE SALTO DE PIRAPORA | Instituto Butantan / ESALQ-Piracicaba | Dimas Tadeu Covas, Antonio Jorge Martins, Claudia Renata dos Santos Barros, David Schlesinger, Debora Botequio Moretti, Elaine Cristina Marqueze, Elaine Vieira Santos, Evandra Strazza Rodrigues, Heidge Fukumasu, Jayme Augusto de Souza-Neto, José Salvatore Leister Patané, Luiz Alcantara, Luiz Lehmann Coutinho, Maria Carolina Elias, Maurício Lacerda Nogueira, Rafael dos Santos Bezerra, Raul Machado Neto, Rejane Maria Tommasini Grotto, Ricardo Haddad, Sandra Coccuzzo Sampaio Vessoni, Simone Kashima, Svetoslav Nanev Slavov, Vincent Louis Viala |
| EPI_ISL_2344911 | HOSPITAL E PRONTO SOCORRO PORTINARI | Instituto Butantan | Dimas Tadeu Covas, Antonio Jorge Martins, Claudia Renata dos Santos Barros, David Schlesinger, Debora Botequio Moretti, Elaine Cristina Marqueze, Elaine Vieira Santos, Evandra Strazza Rodrigues, Heidge Fukumasu, Jayme Augusto de Souza-Neto, José Salvatore Leister Patané, Luiz Alcantara, Luiz Lehmann Coutinho, Maria Carolina Elias, Maurício Lacerda Nogueira, Rafael dos Santos Bezerra, Raul Machado Neto, Rejane Maria Tommasini Grotto, Ricardo Haddad, Sandra Coccuzzo Sampaio Vessoni, Simone Kashima, Svetoslav Nanev Slavov, Vincent Louis Viala |
| EPI_ISL_2344912 | SECRETARIA MUNICIPAL DA SAUDE DE ITARARE | Instituto Butantan / ESALQ-Piracicaba | Dimas Tadeu Covas, Antonio Jorge Martins, Claudia Renata dos Santos Barros, David Schlesinger, Debora Botequio Moretti, Elaine Cristina Marqueze, Elaine Vieira Santos, Evandra Strazza Rodrigues, Heidge Fukumasu, Jayme Augusto de Souza-Neto, José Salvatore Leister Patané, Luiz Alcantara, Luiz Lehmann Coutinho, Maria Carolina Elias, Maurício Lacerda Nogueira, Rafael dos Santos Bezerra, Raul Machado Neto, Rejane Maria Tommasini Grotto, Ricardo Haddad, Sandra Coccuzzo Sampaio Vessoni, Simone Kashima, Svetoslav Nanev Slavov, Vincent Louis Viala |
| EPI_ISL_2344913 | SMS SECRETARIA MUNICIPAL DE SAUDE DE BOITUVA | Instituto Butantan / ESALQ-Piracicaba | Dimas Tadeu Covas, Antonio Jorge Martins, Claudia Renata dos Santos Barros, David Schlesinger, Debora Botequio Moretti, Elaine Cristina Marqueze, Elaine Vieira Santos, Evandra Strazza Rodrigues, Heidge Fukumasu, Jayme Augusto de Souza-Neto, José Salvatore Leister Patané, Luiz Alcantara, Luiz Lehmann Coutinho, Maria Carolina Elias, Maurício Lacerda Nogueira, Rafael dos Santos Bezerra, Raul Machado Neto, Rejane Maria Tommasini Grotto, Ricardo Haddad, Sandra Coccuzzo Sampaio Vessoni, Simone Kashima, Svetoslav Nanev Slavov, Vincent Louis Viala |
| EPI_ISL_2344914 | HOSP E MATERNIDADE MUNICIPAL N SRA MONTE SERRAT | Instituto Butantan / ESALQ-Piracicaba | Dimas Tadeu Covas, Antonio Jorge Martins, Claudia Renata dos Santos Barros, David Schlesinger, Debora Botequio Moretti, Elaine Cristina Marqueze, Elaine Vieira Santos, Evandra Strazza Rodrigues, Heidge Fukumasu, Jayme Augusto de Souza-Neto, José Salvatore Leister Patané, Luiz Alcantara, Luiz Lehmann Coutinho, Maria Carolina Elias, Maurício Lacerda Nogueira, Rafael dos Santos Bezerra, Raul Machado Neto, Rejane Maria Tommasini Grotto, Ricardo Haddad, Sandra Coccuzzo Sampaio Vessoni, Simone Kashima, Svetoslav Nanev Slavov, Vincent Louis Viala |
| EPI_ISL_2344915 | LABORATORIO MUNICIPAL DE PIRACICABA | Instituto Butantan / ESALQ-Piracicaba | Dimas Tadeu Covas, Antonio Jorge Martins, Claudia Renata dos Santos Barros, David Schlesinger, Debora Botequio Moretti, Elaine Cristina Marqueze, Elaine Vieira Santos, Evandra Strazza Rodrigues, Heidge Fukumasu, Jayme Augusto de Souza-Neto, José Salvatore Leister Patané, Luiz Alcantara, Luiz Lehmann Coutinho, Maria Carolina Elias, Maurício Lacerda Nogueira, Rafael dos Santos Bezerra, Raul Machado Neto, Rejane Maria Tommasini Grotto, Ricardo Haddad, Sandra Coccuzzo Sampaio Vessoni, Simone Kashima, Svetoslav Nanev Slavov, Vincent Louis Viala |
| EPI_ISL_2344916 | UNIDADE PSF PINHEIROS APIAI | Instituto Butantan / ESALQ-Piracicaba | Dimas Tadeu Covas, Antonio Jorge Martins, Claudia Renata dos Santos Barros, David Schlesinger, Debora Botequio Moretti, Elaine Cristina Marqueze, Elaine Vieira Santos, Evandra Strazza Rodrigues, Heidge Fukumasu, Jayme Augusto de Souza-Neto, José Salvatore Leister Patané, Luiz Alcantara, Luiz Lehmann Coutinho, Maria Carolina Elias, Maurício Lacerda Nogueira, Rafael dos Santos Bezerra, Raul Machado Neto, Rejane Maria Tommasini Grotto, Ricardo Haddad, Sandra Coccuzzo Sampaio Vessoni, Simone Kashima, Svetoslav Nanev Slavov, Vincent Louis Viala |
| EPI_ISL_2344917 | LABORATORIO MUNICIPAL DE PIRACICABA | Instituto Butantan / ESALQ-Piracicaba | Dimas Tadeu Covas, Antonio Jorge Martins, Claudia Renata dos Santos Barros, David Schlesinger, Debora Botequio Moretti, Elaine Cristina Marqueze, Elaine Vieira Santos, Evandra Strazza Rodrigues, Heidge Fukumasu, Jayme Augusto de Souza-Neto, José Salvatore Leister Patané, Luiz Alcantara, Luiz |

| | | | |
|---|---|---|---|
| | | | Lehmann Coutinho, Maria Carolina Elias, Maurício Lacerda Nogueira, Rafael dos Santos Bezerra, Raul Machado Neto, Rejane Maria Tommasini Grotto, Ricardo Haddad, Sandra Coccuzzo Sampaio Vessoni, Simone Kashima, Svetoslav Nanev Slavov, Vincent Louis Viala |
| EPI_ISL_2344918 | PSF PALMITALZINHO APIAI | Instituto Butantan / ESALQ-Piracicaba | Dimas Tadeu Covas, Antonio Jorge Martins, Claudia Renata dos Santos Barros, David Schlesinger, Debora Botequio Moretti, Elaine Cristina Marqueze, Elaine Vieira Santos, Evandra Strazza Rodrigues, Heidge Fukumasu, Jayme Augusto de Souza-Neto, José Salvatore Leister Patané, Luiz Alcantara, Luiz Lehmann Coutinho, Maria Carolina Elias, Maurício Lacerda Nogueira, Rafael dos Santos Bezerra, Raul Machado Neto, Rejane Maria Tommasini Grotto, Ricardo Haddad, Sandra Coccuzzo Sampaio Vessoni, Simone Kashima, Svetoslav Nanev Slavov, Vincent Louis Viala |
| EPI_ISL_2344919, EPI_ISL_2344920, EPI_ISL_2344921, EPI_ISL_2344922, EPI_ISL_2344923, EPI_ISL_2344924 | SECRETARIA MUNICIPAL DE SAUDE SOROCABA | Instituto Butantan / ESALQ-Piracicaba | Dimas Tadeu Covas, Antonio Jorge Martins, Claudia Renata dos Santos Barros, David Schlesinger, Debora Botequio Moretti, Elaine Cristina Marqueze, Elaine Vieira Santos, Evandra Strazza Rodrigues, Heidge Fukumasu, Jayme Augusto de Souza-Neto, José Salvatore Leister Patané, Luiz Alcantara, Luiz Lehmann Coutinho, Maria Carolina Elias, Maurício Lacerda Nogueira, Rafael dos Santos Bezerra, Raul Machado Neto, Rejane Maria Tommasini Grotto, Ricardo Haddad, Sandra Coccuzzo Sampaio Vessoni, Simone Kashima, Svetoslav Nanev Slavov, Vincent Louis Viala |
| EPI_ISL_2344925 | CSA COMPLEXO DE SAUDE ANHANGUERA VILA JAGUARA | Instituto Butantan | Dimas Tadeu Covas, Antonio Jorge Martins, Claudia Renata dos Santos Barros, David Schlesinger, Debora Botequio Moretti, Elaine Cristina Marqueze, Elaine Vieira Santos, Evandra Strazza Rodrigues, Heidge Fukumasu, Jayme Augusto de Souza-Neto, José Salvatore Leister Patané, Luiz Alcantara, Luiz Lehmann Coutinho, Maria Carolina Elias, Maurício Lacerda Nogueira, Rafael dos Santos Bezerra, Raul Machado Neto, Rejane Maria Tommasini Grotto, Ricardo Haddad, Sandra Coccuzzo Sampaio Vessoni, Simone Kashima, Svetoslav Nanev Slavov, Vincent Louis Viala |
| EPI_ISL_2344926 | SECRETARIA MUNICIPAL DE SAUDE SOROCABA | Instituto Butantan / ESALQ-Piracicaba | Dimas Tadeu Covas, Antonio Jorge Martins, Claudia Renata dos Santos Barros, David Schlesinger, Debora Botequio Moretti, Elaine Cristina Marqueze, Elaine Vieira Santos, Evandra Strazza Rodrigues, Heidge Fukumasu, Jayme Augusto de Souza-Neto, José Salvatore Leister Patané, Luiz Alcantara, Luiz Lehmann Coutinho, Maria Carolina Elias, Maurício Lacerda Nogueira, Rafael dos Santos Bezerra, Raul Machado Neto, Rejane Maria Tommasini Grotto, Ricardo Haddad, Sandra Coccuzzo Sampaio Vessoni, Simone Kashima, Svetoslav Nanev Slavov, Vincent Louis Viala |
| EPI_ISL_2344927 | POLICLINICA COVID 19 ITAPETININGA | Instituto Butantan / ESALQ-Piracicaba | Dimas Tadeu Covas, Antonio Jorge Martins, Claudia Renata dos Santos Barros, David Schlesinger, Debora Botequio Moretti, Elaine Cristina Marqueze, Elaine Vieira Santos, Evandra Strazza Rodrigues, Heidge Fukumasu, Jayme Augusto de Souza-Neto, José Salvatore Leister Patané, Luiz Alcantara, Luiz Lehmann Coutinho, Maria Carolina Elias, Maurício Lacerda Nogueira, Rafael dos Santos Bezerra, Raul Machado Neto, Rejane Maria Tommasini Grotto, Ricardo Haddad, Sandra Coccuzzo Sampaio Vessoni, Simone Kashima, Svetoslav Nanev Slavov, Vincent Louis Viala |
| EPI_ISL_2344928, EPI_ISL_2344929, EPI_ISL_2344930, EPI_ISL_2344931, EPI_ISL_2344932, EPI_ISL_2344933, EPI_ISL_2344934, EPI_ISL_2344935, EPI_ISL_2344936, EPI_ISL_2344937, EPI_ISL_2344938, EPI_ISL_2344939 | | | |
| see above | SECRETARIA MUNICIPAL DE SAUDE SOROCABA | Instituto Butantan / ESALQ-Piracicaba | Dimas Tadeu Covas, Antonio Jorge Martins, Claudia Renata dos Santos Barros, David Schlesinger, Debora Botequio Moretti, Elaine Cristina Marqueze, Elaine Vieira Santos, Evandra Strazza Rodrigues, Heidge Fukumasu, Jayme Augusto de Souza-Neto, José Salvatore Leister Patané, Luiz Alcantara, Luiz Lehmann Coutinho, Maria Carolina Elias, Maurício Lacerda Nogueira, Rafael dos Santos Bezerra, Raul Machado Neto, Rejane Maria Tommasini Grotto, Ricardo Haddad, Sandra Coccuzzo Sampaio Vessoni, Simone Kashima, Svetoslav Nanev Slavov, Vincent Louis Viala |
| EPI_ISL_2344940 | SMS IPERO | Instituto Butantan / ESALQ-Piracicaba | Dimas Tadeu Covas, Antonio Jorge Martins, Claudia Renata dos Santos Barros, David Schlesinger, Debora Botequio Moretti, Elaine Cristina Marqueze, Elaine Vieira Santos, Evandra Strazza Rodrigues, Heidge Fukumasu, Jayme Augusto de Souza-Neto, José Salvatore Leister Patané, Luiz Alcantara, Luiz Lehmann Coutinho, Maria Carolina Elias, Maurício Lacerda Nogueira, Rafael dos Santos Bezerra, Raul Machado Neto, Rejane Maria Tommasini Grotto, Ricardo Haddad, Sandra Coccuzzo Sampaio Vessoni, Simone Kashima, Svetoslav Nanev Slavov, Vincent Louis Viala |
| EPI_ISL_2344941 | POLICLINICA COVID 19 ITAPETININGA | Instituto Butantan / ESALQ-Piracicaba | Dimas Tadeu Covas, Antonio Jorge Martins, Claudia Renata dos Santos Barros, David Schlesinger, Debora Botequio Moretti, Elaine Cristina Marqueze, Elaine Vieira Santos, Evandra Strazza Rodrigues, Heidge Fukumasu, Jayme Augusto de Souza-Neto, José Salvatore Leister Patané, Luiz Alcantara, Luiz Lehmann Coutinho, Maria Carolina Elias, Maurício Lacerda Nogueira, Rafael dos Santos Bezerra, Raul Machado Neto, Rejane Maria Tommasini Grotto, Ricardo Haddad, Sandra Coccuzzo Sampaio Vessoni, Simone Kashima, Svetoslav Nanev Slavov, Vincent Louis Viala |
| EPI_ISL_2344942 | HOSPITAL E PRONTO SOCORRO PORTINARI | Instituto Butantan | Dimas Tadeu Covas, Antonio Jorge Martins, Claudia Renata dos Santos Barros, David Schlesinger, Debora Botequio Moretti, Elaine Cristina Marqueze, Elaine Vieira Santos, Evandra Strazza Rodrigues, Heidge Fukumasu, Jayme Augusto de Souza-Neto, José Salvatore Leister Patané, Luiz Alcantara, Luiz Lehmann Coutinho, Maria Carolina Elias, Maurício Lacerda Nogueira, Rafael dos Santos Bezerra, Raul Machado Neto, Rejane Maria Tommasini Grotto, Ricardo Haddad, Sandra Coccuzzo Sampaio Vessoni, Simone Kashima, Svetoslav Nanev Slavov, Vincent Louis Viala |
| EPI_ISL_2344943 | CSA COMPLEXO DE SAUDE ANHANGUERA VILA JAGUARA | Instituto Butantan | Dimas Tadeu Covas, Antonio Jorge Martins, Claudia Renata dos Santos Barros, David Schlesinger, Debora Botequio Moretti, Elaine Cristina Marqueze, Elaine Vieira Santos, Evandra Strazza Rodrigues, Heidge Fukumasu, Jayme Augusto de Souza-Neto, José Salvatore Leister Patané, Luiz Alcantara, Luiz Lehmann Coutinho, Maria Carolina Elias, Maurício Lacerda Nogueira, Rafael dos Santos Bezerra, Raul Machado Neto, Rejane Maria Tommasini Grotto, Ricardo Haddad, Sandra Coccuzzo Sampaio Vessoni, Simone Kashima, Svetoslav Nanev Slavov, Vincent Louis Viala |
| EPI_ISL_2344944, EPI_ISL_2344945, EPI_ISL_2344946 | HOSPITAL E PRONTO SOCORRO PORTINARI | Instituto Butantan | Dimas Tadeu Covas, Antonio Jorge Martins, Claudia Renata dos Santos Barros, David Schlesinger, Debora Botequio Moretti, Elaine Cristina Marqueze, Elaine Vieira Santos, Evandra Strazza Rodrigues, Heidge Fukumasu, Jayme Augusto de Souza-Neto, José Salvatore Leister Patané, Luiz Alcantara, Luiz Lehmann Coutinho, Maria Carolina Elias, Maurício Lacerda Nogueira, Rafael dos Santos Bezerra, Raul Machado Neto, Rejane Maria Tommasini Grotto, Ricardo Haddad, Sandra Coccuzzo Sampaio Vessoni, Simone Kashima, Svetoslav Nanev Slavov, Vincent Louis Viala |
| EPI_ISL_2344947, EPI_ISL_2344948 | HOSP E MATERNIDADE MUNICIPAL N SRA MONTE SERRAT | Instituto Butantan / ESALQ-Piracicaba | Dimas Tadeu Covas, Antonio Jorge Martins, Claudia Renata dos Santos Barros, David Schlesinger, Debora Botequio Moretti, Elaine Cristina Marqueze, Elaine Vieira Santos, Evandra Strazza Rodrigues, Heidge Fukumasu, Jayme Augusto de Souza-Neto, José Salvatore Leister Patané, Luiz Alcantara, Luiz Lehmann Coutinho, Maria Carolina Elias, Maurício Lacerda Nogueira, Rafael dos Santos Bezerra, Raul Machado Neto, Rejane Maria Tommasini Grotto, Ricardo Haddad, Sandra Coccuzzo Sampaio Vessoni, Simone Kashima, Svetoslav Nanev Slavov, Vincent Louis Viala |
| EPI_ISL_2344949 | UBS ENRIQUE GUILHERME ERNESTO MARZEUSKI | Instituto Butantan / ESALQ-Piracicaba | Dimas Tadeu Covas, Antonio Jorge Martins, Claudia Renata dos Santos Barros, David Schlesinger, Debora Botequio Moretti, Elaine Cristina Marqueze, Elaine Vieira Santos, Evandra Strazza Rodrigues, Heidge Fukumasu, Jayme Augusto de Souza-Neto, José Salvatore Leister Patané, Luiz Alcantara, Luiz Lehmann Coutinho, Maria Carolina Elias, Maurício Lacerda Nogueira, Rafael dos Santos Bezerra, Raul Machado Neto, Rejane Maria Tommasini Grotto, Ricardo Haddad, Sandra Coccuzzo Sampaio Vessoni, Simone Kashima, Svetoslav Nanev Slavov, Vincent Louis Viala |
| EPI_ISL_2344950 | UNIDADE MISTA DE SAUDE CAPELA DO ALTO | Instituto Butantan / ESALQ-Piracicaba | Dimas Tadeu Covas, Antonio Jorge Martins, Claudia Renata dos Santos Barros, David Schlesinger, Debora Botequio Moretti, Elaine Cristina Marqueze, Elaine Vieira Santos, Evandra Strazza Rodrigues, Heidge Fukumasu, Jayme Augusto de Souza-Neto, José Salvatore Leister Patané, Luiz Alcantara, Luiz Lehmann Coutinho, Maria Carolina Elias, Maurício Lacerda Nogueira, Rafael dos Santos Bezerra, Raul Machado Neto, Rejane Maria Tommasini Grotto, Ricardo Haddad, Sandra Coccuzzo Sampaio Vessoni, Simone Kashima, Svetoslav Nanev Slavov, Vincent Louis Viala |
| EPI_ISL_2344951 | VIGILANCIA EM SAUDE DE FCO MORATO | Instituto Butantan | Dimas Tadeu Covas, Antonio Jorge Martins, Claudia Renata dos Santos Barros, David Schlesinger, Debora Botequio Moretti, Elaine Cristina Marqueze, Elaine Vieira Santos, Evandra Strazza Rodrigues, Heidge Fukumasu, Jayme Augusto de Souza-Neto, José Salvatore Leister Patané, Luiz Alcantara, Luiz Lehmann Coutinho, Maria Carolina Elias, Maurício Lacerda Nogueira, Rafael dos Santos Bezerra, Raul Machado Neto, Rejane Maria Tommasini Grotto, Ricardo Haddad, Sandra Coccuzzo Sampaio Vessoni, Simone Kashima, Svetoslav Nanev Slavov, Vincent Louis Viala |
| EPI_ISL_2344952 | UMS PIRAPORA DO BOM JESUS | Instituto Butantan | Dimas Tadeu Covas, Antonio Jorge Martins, Claudia Renata dos Santos Barros, David Schlesinger, Debora Botequio Moretti, Elaine Cristina Marqueze, Elaine Vieira Santos, Evandra Strazza Rodrigues, Heidge Fukumasu, Jayme Augusto de Souza-Neto, José Salvatore Leister Patané, Luiz Alcantara, Luiz Lehmann Coutinho, Maria Carolina Elias, Maurício Lacerda Nogueira, Rafael dos Santos Bezerra, Raul Machado Neto, Rejane Maria Tommasini Grotto, Ricardo Haddad, Sandra Coccuzzo Sampaio Vessoni, Simone Kashima, Svetoslav Nanev Slavov, Vincent Louis Viala |
| EPI_ISL_2344955, EPI_ISL_2344956, EPI_ISL_2344957 | HOSPITAL DE CAMPANHA COVID 19 MUNICIPIO DE TAUBATE | Instituto Butantan / ESALQ-Piracicaba | Dimas Tadeu Covas, Antonio Jorge Martins, Claudia Renata dos Santos Barros, David Schlesinger, Debora Botequio Moretti, Elaine Cristina Marqueze, Elaine Vieira Santos, Evandra Strazza Rodrigues, Heidge Fukumasu, Jayme Augusto de Souza-Neto, José Salvatore Leister Patané, Luiz Alcantara, Luiz Lehmann Coutinho, Maria Carolina Elias, Maurício Lacerda Nogueira, Rafael dos Santos Bezerra, Raul Machado Neto, Rejane Maria Tommasini Grotto, Ricardo Haddad, Sandra Coccuzzo Sampaio Vessoni, Simone Kashima, Svetoslav Nanev Slavov, Vincent Louis Viala |
| EPI_ISL_2344958, EPI_ISL_2344959 | VIGILANCIA EM SAUDE DE FCO MORATO | Instituto Butantan | Dimas Tadeu Covas, Antonio Jorge Martins, Claudia Renata dos Santos Barros, David Schlesinger, Debora Botequio Moretti, Elaine Cristina Marqueze, Elaine Vieira Santos, Evandra Strazza Rodrigues, Heidge Fukumasu, Jayme Augusto de Souza-Neto, José Salvatore Leister Patané, Luiz Alcantara, Luiz Lehmann Coutinho, Maria Carolina Elias, Maurício Lacerda Nogueira, Rafael dos Santos Bezerra, Raul Machado Neto, Rejane Maria Tommasini Grotto, Ricardo Haddad, Sandra Coccuzzo Sampaio Vessoni, Simone Kashima, Svetoslav Nanev Slavov, Vincent Louis Viala |
| EPI_ISL_2344960 | HOSPITAL MUNICIPAL REYNALDO GUERRA CAJATI | Instituto Butantan / ESALQ-Piracicaba | Dimas Tadeu Covas, Antonio Jorge Martins, Claudia Renata dos Santos Barros, David Schlesinger, Debora Botequio Moretti, Elaine Cristina Marqueze, Elaine Vieira Santos, Evandra Strazza Rodrigues, Heidge Fukumasu, Jayme Augusto de Souza-Neto, José Salvatore Leister Patané, Luiz Alcantara, Luiz |

| | | | |
|---|---|---|---|
| | | | Lehmann Coutinho, Maria Carolina Elias, Maurício Lacerda Nogueira, Rafael dos Santos Bezerra, Raul Machado Neto, Rejane Maria Tommasini Grotto, Ricardo Haddad, Sandra Coccuzzo Sampaio Vessoni, Simone Kashima, Svetoslav Nanev Slavov, Vincent Louis Viala |
| EPI_ISL_2344961 | CENTRO DE APOIO EPIDEMIOLOGICO | Instituto Butantan / ESALQ-Piracicaba | Dimas Tadeu Covas, Antonio Jorge Martins, Claudia Renata dos Santos Barros, David Schlesinger, Debora Botequio Moretti, Elaine Cristina Marqueze, Elaine Vieira Santos, Evandra Strazza Rodrigues, Heidge Fukumasu, Jayme Augusto de Souza-Neto, José Salvatore Leister Patané, Luiz Alcantara, Luiz Lehmann Coutinho, Maria Carolina Elias, Maurício Lacerda Nogueira, Rafael dos Santos Bezerra, Raul Machado Neto, Rejane Maria Tommasini Grotto, Ricardo Haddad, Sandra Coccuzzo Sampaio Vessoni, Simone Kashima, Svetoslav Nanev Slavov, Vincent Louis Viala |
| EPI_ISL_2344962 | VIGILANCIA EM SAUDE DE FCO MORATO | Instituto Butantan | Dimas Tadeu Covas, Antonio Jorge Martins, Claudia Renata dos Santos Barros, David Schlesinger, Debora Botequio Moretti, Elaine Cristina Marqueze, Elaine Vieira Santos, Evandra Strazza Rodrigues, Heidge Fukumasu, Jayme Augusto de Souza-Neto, José Salvatore Leister Patané, Luiz Alcantara, Luiz Lehmann Coutinho, Maria Carolina Elias, Maurício Lacerda Nogueira, Rafael dos Santos Bezerra, Raul Machado Neto, Rejane Maria Tommasini Grotto, Ricardo Haddad, Sandra Coccuzzo Sampaio Vessoni, Simone Kashima, Svetoslav Nanev Slavov, Vincent Louis Viala |
| EPI_ISL_2344963 | CENTRO DE APOIO EPIDEMIOLOGICO | Instituto Butantan / ESALQ-Piracicaba | Dimas Tadeu Covas, Antonio Jorge Martins, Claudia Renata dos Santos Barros, David Schlesinger, Debora Botequio Moretti, Elaine Cristina Marqueze, Elaine Vieira Santos, Evandra Strazza Rodrigues, Heidge Fukumasu, Jayme Augusto de Souza-Neto, José Salvatore Leister Patané, Luiz Alcantara, Luiz Lehmann Coutinho, Maria Carolina Elias, Maurício Lacerda Nogueira, Rafael dos Santos Bezerra, Raul Machado Neto, Rejane Maria Tommasini Grotto, Ricardo Haddad, Sandra Coccuzzo Sampaio Vessoni, Simone Kashima, Svetoslav Nanev Slavov, Vincent Louis Viala |
| EPI_ISL_2344964, EPI_ISL_2344965, EPI_ISL_2344966 | VIGILANCIA EM SAUDE DE FCO MORATO | Instituto Butantan | Dimas Tadeu Covas, Antonio Jorge Martins, Claudia Renata dos Santos Barros, David Schlesinger, Debora Botequio Moretti, Elaine Cristina Marqueze, Elaine Vieira Santos, Evandra Strazza Rodrigues, Heidge Fukumasu, Jayme Augusto de Souza-Neto, José Salvatore Leister Patané, Luiz Alcantara, Luiz Lehmann Coutinho, Maria Carolina Elias, Maurício Lacerda Nogueira, Rafael dos Santos Bezerra, Raul Machado Neto, Rejane Maria Tommasini Grotto, Ricardo Haddad, Sandra Coccuzzo Sampaio Vessoni, Simone Kashima, Svetoslav Nanev Slavov, Vincent Louis Viala |
| EPI_ISL_2344967 | CENTRO DE APOIO EPIDEMIOLOGICO | Instituto Butantan / ESALQ-Piracicaba | Dimas Tadeu Covas, Antonio Jorge Martins, Claudia Renata dos Santos Barros, David Schlesinger, Debora Botequio Moretti, Elaine Cristina Marqueze, Elaine Vieira Santos, Evandra Strazza Rodrigues, Heidge Fukumasu, Jayme Augusto de Souza-Neto, José Salvatore Leister Patané, Luiz Alcantara, Luiz Lehmann Coutinho, Maria Carolina Elias, Maurício Lacerda Nogueira, Rafael dos Santos Bezerra, Raul Machado Neto, Rejane Maria Tommasini Grotto, Ricardo Haddad, Sandra Coccuzzo Sampaio Vessoni, Simone Kashima, Svetoslav Nanev Slavov, Vincent Louis Viala |
| EPI_ISL_2344968 | VIGILANCIA EM SAUDE DE FCO MORATO | Instituto Butantan | Dimas Tadeu Covas, Antonio Jorge Martins, Claudia Renata dos Santos Barros, David Schlesinger, Debora Botequio Moretti, Elaine Cristina Marqueze, Elaine Vieira Santos, Evandra Strazza Rodrigues, Heidge Fukumasu, Jayme Augusto de Souza-Neto, José Salvatore Leister Patané, Luiz Alcantara, Luiz Lehmann Coutinho, Maria Carolina Elias, Maurício Lacerda Nogueira, Rafael dos Santos Bezerra, Raul Machado Neto, Rejane Maria Tommasini Grotto, Ricardo Haddad, Sandra Coccuzzo Sampaio Vessoni, Simone Kashima, Svetoslav Nanev Slavov, Vincent Louis Viala |
| EPI_ISL_2344969 | HOSPITAL MUNICIPAL REYNALDO GUERRA CAJATI | Instituto Butantan / ESALQ-Piracicaba | Dimas Tadeu Covas, Antonio Jorge Martins, Claudia Renata dos Santos Barros, David Schlesinger, Debora Botequio Moretti, Elaine Cristina Marqueze, Elaine Vieira Santos, Evandra Strazza Rodrigues, Heidge Fukumasu, Jayme Augusto de Souza-Neto, José Salvatore Leister Patané, Luiz Alcantara, Luiz Lehmann Coutinho, Maria Carolina Elias, Maurício Lacerda Nogueira, Rafael dos Santos Bezerra, Raul Machado Neto, Rejane Maria Tommasini Grotto, Ricardo Haddad, Sandra Coccuzzo Sampaio Vessoni, Simone Kashima, Svetoslav Nanev Slavov, Vincent Louis Viala |
| EPI_ISL_2344970 | VIGILANCIA EM SAUDE DE FCO MORATO | Instituto Butantan | Dimas Tadeu Covas, Antonio Jorge Martins, Claudia Renata dos Santos Barros, David Schlesinger, Debora Botequio Moretti, Elaine Cristina Marqueze, Elaine Vieira Santos, Evandra Strazza Rodrigues, Heidge Fukumasu, Jayme Augusto de Souza-Neto, José Salvatore Leister Patané, Luiz Alcantara, Luiz Lehmann Coutinho, Maria Carolina Elias, Maurício Lacerda Nogueira, Rafael dos Santos Bezerra, Raul Machado Neto, Rejane Maria Tommasini Grotto, Ricardo Haddad, Sandra Coccuzzo Sampaio Vessoni, Simone Kashima, Svetoslav Nanev Slavov, Vincent Louis Viala |
| EPI_ISL_2344971, EPI_ISL_2344972 | LABORATORIO MUNICIPAL DE PIRACICABA | Instituto Butantan / ESALQ-Piracicaba | Dimas Tadeu Covas, Antonio Jorge Martins, Claudia Renata dos Santos Barros, David Schlesinger, Debora Botequio Moretti, Elaine Cristina Marqueze, Elaine Vieira Santos, Evandra Strazza Rodrigues, Heidge Fukumasu, Jayme Augusto de Souza-Neto, José Salvatore Leister Patané, Luiz Alcantara, Luiz Lehmann Coutinho, Maria Carolina Elias, Maurício Lacerda Nogueira, Rafael dos Santos Bezerra, Raul Machado Neto, Rejane Maria Tommasini Grotto, Ricardo Haddad, Sandra Coccuzzo Sampaio Vessoni, Simone Kashima, Svetoslav Nanev Slavov, Vincent Louis Viala |
| EPI_ISL_2344973 | HOSPITAL DE CAMPANHA COVID 19 MUNICIPIO DE TAUBATE | Instituto Butantan / ESALQ-Piracicaba | Dimas Tadeu Covas, Antonio Jorge Martins, Claudia Renata dos Santos Barros, David Schlesinger, Debora Botequio Moretti, Elaine Cristina Marqueze, Elaine Vieira Santos, Evandra Strazza Rodrigues, Heidge Fukumasu, Jayme Augusto de Souza-Neto, José Salvatore Leister Patané, Luiz Alcantara, Luiz Lehmann Coutinho, Maria Carolina Elias, Maurício Lacerda Nogueira, Rafael dos Santos Bezerra, Raul Machado Neto, Rejane Maria Tommasini Grotto, Ricardo Haddad, Sandra Coccuzzo Sampaio Vessoni, Simone Kashima, Svetoslav Nanev Slavov, Vincent Louis Viala |
| EPI_ISL_2344974 | CENTRO DE APOIO EPIDEMIOLOGICO | Instituto Butantan / ESALQ-Piracicaba | Dimas Tadeu Covas, Antonio Jorge Martins, Claudia Renata dos Santos Barros, David Schlesinger, Debora Botequio Moretti, Elaine Cristina Marqueze, Elaine Vieira Santos, Evandra Strazza Rodrigues, Heidge Fukumasu, Jayme Augusto de Souza-Neto, José Salvatore Leister Patané, Luiz Alcantara, Luiz Lehmann Coutinho, Maria Carolina Elias, Maurício Lacerda Nogueira, Rafael dos Santos Bezerra, Raul Machado Neto, Rejane Maria Tommasini Grotto, Ricardo Haddad, Sandra Coccuzzo Sampaio Vessoni, Simone Kashima, Svetoslav Nanev Slavov, Vincent Louis Viala |
| EPI_ISL_2344975, EPI_ISL_2344976, EPI_ISL_2344977, EPI_ISL_2344978 | HOSPITAL DE CAMPANHA COVID 19 MUNICIPIO DE TAUBATE | Instituto Butantan / ESALQ-Piracicaba | Dimas Tadeu Covas, Antonio Jorge Martins, Claudia Renata dos Santos Barros, David Schlesinger, Debora Botequio Moretti, Elaine Cristina Marqueze, Elaine Vieira Santos, Evandra Strazza Rodrigues, Heidge Fukumasu, Jayme Augusto de Souza-Neto, José Salvatore Leister Patané, Luiz Alcantara, Luiz Lehmann Coutinho, Maria Carolina Elias, Maurício Lacerda Nogueira, Rafael dos Santos Bezerra, Raul Machado Neto, Rejane Maria Tommasini Grotto, Ricardo Haddad, Sandra Coccuzzo Sampaio Vessoni, Simone Kashima, Svetoslav Nanev Slavov, Vincent Louis Viala |
| EPI_ISL_2344979 | LABORATORIO MUNICIPAL DE PIRACICABA | Instituto Butantan / ESALQ-Piracicaba | Dimas Tadeu Covas, Antonio Jorge Martins, Claudia Renata dos Santos Barros, David Schlesinger, Debora Botequio Moretti, Elaine Cristina Marqueze, Elaine Vieira Santos, Evandra Strazza Rodrigues, Heidge Fukumasu, Jayme Augusto de Souza-Neto, José Salvatore Leister Patané, Luiz Alcantara, Luiz Lehmann Coutinho, Maria Carolina Elias, Maurício Lacerda Nogueira, Rafael dos Santos Bezerra, Raul Machado Neto, Rejane Maria Tommasini Grotto, Ricardo Haddad, Sandra Coccuzzo Sampaio Vessoni, Simone Kashima, Svetoslav Nanev Slavov, Vincent Louis Viala |
| EPI_ISL_2344980 | HOSPITAL DE CAMPANHA COVID 19 MUNICIPIO DE TAUBATE | Instituto Butantan / ESALQ-Piracicaba | Dimas Tadeu Covas, Antonio Jorge Martins, Claudia Renata dos Santos Barros, David Schlesinger, Debora Botequio Moretti, Elaine Cristina Marqueze, Elaine Vieira Santos, Evandra Strazza Rodrigues, Heidge Fukumasu, Jayme Augusto de Souza-Neto, José Salvatore Leister Patané, Luiz Alcantara, Luiz Lehmann Coutinho, Maria Carolina Elias, Maurício Lacerda Nogueira, Rafael dos Santos Bezerra, Raul Machado Neto, Rejane Maria Tommasini Grotto, Ricardo Haddad, Sandra Coccuzzo Sampaio Vessoni, Simone Kashima, Svetoslav Nanev Slavov, Vincent Louis Viala |
| EPI_ISL_2344981 | CENTRO DE APOIO EPIDEMIOLOGICO | Instituto Butantan / ESALQ-Piracicaba | Dimas Tadeu Covas, Antonio Jorge Martins, Claudia Renata dos Santos Barros, David Schlesinger, Debora Botequio Moretti, Elaine Cristina Marqueze, Elaine Vieira Santos, Evandra Strazza Rodrigues, Heidge Fukumasu, Jayme Augusto de Souza-Neto, José Salvatore Leister Patané, Luiz Alcantara, Luiz Lehmann Coutinho, Maria Carolina Elias, Maurício Lacerda Nogueira, Rafael dos Santos Bezerra, Raul Machado Neto, Rejane Maria Tommasini Grotto, Ricardo Haddad, Sandra Coccuzzo Sampaio Vessoni, Simone Kashima, Svetoslav Nanev Slavov, Vincent Louis Viala |
| EPI_ISL_2344982 | HOSPITAL E MATERNIDADE NOSSA SENHORA DA AJUDA | Instituto Butantan / ESALQ-Piracicaba | Dimas Tadeu Covas, Antonio Jorge Martins, Claudia Renata dos Santos Barros, David Schlesinger, Debora Botequio Moretti, Elaine Cristina Marqueze, Elaine Vieira Santos, Evandra Strazza Rodrigues, Heidge Fukumasu, Jayme Augusto de Souza-Neto, José Salvatore Leister Patané, Luiz Alcantara, Luiz Lehmann Coutinho, Maria Carolina Elias, Maurício Lacerda Nogueira, Rafael dos Santos Bezerra, Raul Machado Neto, Rejane Maria Tommasini Grotto, Ricardo Haddad, Sandra Coccuzzo Sampaio Vessoni, Simone Kashima, Svetoslav Nanev Slavov, Vincent Louis Viala |
| EPI_ISL_2344983 | LABORATORIO MUNICIPAL DE PIRACICABA | Instituto Butantan / ESALQ-Piracicaba | Dimas Tadeu Covas, Antonio Jorge Martins, Claudia Renata dos Santos Barros, David Schlesinger, Debora Botequio Moretti, Elaine Cristina Marqueze, Elaine Vieira Santos, Evandra Strazza Rodrigues, Heidge Fukumasu, Jayme Augusto de Souza-Neto, José Salvatore Leister Patané, Luiz Alcantara, Luiz Lehmann Coutinho, Maria Carolina Elias, Maurício Lacerda Nogueira, Rafael dos Santos Bezerra, Raul Machado Neto, Rejane Maria Tommasini Grotto, Ricardo Haddad, Sandra Coccuzzo Sampaio Vessoni, Simone Kashima, Svetoslav Nanev Slavov, Vincent Louis Viala |
| EPI_ISL_2344984 | CENTRO DE APOIO EPIDEMIOLOGICO | Instituto Butantan / ESALQ-Piracicaba | Dimas Tadeu Covas, Antonio Jorge Martins, Claudia Renata dos Santos Barros, David Schlesinger, Debora Botequio Moretti, Elaine Cristina Marqueze, Elaine Vieira Santos, Evandra Strazza Rodrigues, Heidge Fukumasu, Jayme Augusto de Souza-Neto, José Salvatore Leister Patané, Luiz Alcantara, Luiz Lehmann Coutinho, Maria Carolina Elias, Maurício Lacerda Nogueira, Rafael dos Santos Bezerra, Raul Machado Neto, Rejane Maria Tommasini Grotto, Ricardo Haddad, Sandra Coccuzzo Sampaio Vessoni, Simone Kashima, Svetoslav Nanev Slavov, Vincent Louis Viala |
| EPI_ISL_2344985 | LABORATORIO MUNICIPAL DE PIRACICABA | Instituto Butantan / ESALQ-Piracicaba | Dimas Tadeu Covas, Antonio Jorge Martins, Claudia Renata dos Santos Barros, David Schlesinger, Debora Botequio Moretti, Elaine Cristina Marqueze, Elaine Vieira Santos, Evandra Strazza Rodrigues, Heidge Fukumasu, Jayme Augusto de Souza-Neto, José Salvatore Leister Patané, Luiz Alcantara, Luiz Lehmann Coutinho, Maria Carolina Elias, Maurício Lacerda Nogueira, Rafael dos Santos Bezerra, Raul Machado Neto, Rejane Maria Tommasini Grotto, Ricardo Haddad, Sandra Coccuzzo Sampaio Vessoni, Simone Kashima, Svetoslav Nanev Slavov, Vincent Louis Viala |

| | | | |
|---|---|---|---|
| EPI_ISL_2344986 | CENTRO DE APOIO EPIDEMIOLOGICO | Instituto Butantan / ESALQ-Piracicaba | Dimas Tadeu Covas, Antonio Jorge Martins, Claudia Renata dos Santos Barros, David Schlesinger, Debora Botequio Moretti, Elaine Cristina Marqueze, Elaine Vieira Santos, Evandra Strazza Rodrigues, Heidge Fukumasu, Jayme Augusto de Souza-Neto, José Salvatore Leister Patané, Luiz Alcantara, Luiz Lehmann Coutinho, Maria Carolina Elias, Maurício Lacerda Nogueira, Rafael dos Santos Bezerra, Raul Machado Neto, Rejane Maria Tommasini Grotto, Ricardo Haddad, Sandra Coccuzzo Sampaio Vessoni, Simone Kashima, Svetoslav Nanev Slavov, Vincent Louis Viala |
| EPI_ISL_2344987 | HOSPITAL DE CAMPANHA COVID 19 MUNICIPIO DE TAUBATE | Instituto Butantan / ESALQ-Piracicaba | Dimas Tadeu Covas, Antonio Jorge Martins, Claudia Renata dos Santos Barros, David Schlesinger, Debora Botequio Moretti, Elaine Cristina Marqueze, Elaine Vieira Santos, Evandra Strazza Rodrigues, Heidge Fukumasu, Jayme Augusto de Souza-Neto, José Salvatore Leister Patané, Luiz Alcantara, Luiz Lehmann Coutinho, Maria Carolina Elias, Maurício Lacerda Nogueira, Rafael dos Santos Bezerra, Raul Machado Neto, Rejane Maria Tommasini Grotto, Ricardo Haddad, Sandra Coccuzzo Sampaio Vessoni, Simone Kashima, Svetoslav Nanev Slavov, Vincent Louis Viala |
| EPI_ISL_2344988 | LABORATORIO MUNICIPAL DE PIRACICABA | Instituto Butantan / ESALQ-Piracicaba | Dimas Tadeu Covas, Antonio Jorge Martins, Claudia Renata dos Santos Barros, David Schlesinger, Debora Botequio Moretti, Elaine Cristina Marqueze, Elaine Vieira Santos, Evandra Strazza Rodrigues, Heidge Fukumasu, Jayme Augusto de Souza-Neto, José Salvatore Leister Patané, Luiz Alcantara, Luiz Lehmann Coutinho, Maria Carolina Elias, Maurício Lacerda Nogueira, Rafael dos Santos Bezerra, Raul Machado Neto, Rejane Maria Tommasini Grotto, Ricardo Haddad, Sandra Coccuzzo Sampaio Vessoni, Simone Kashima, Svetoslav Nanev Slavov, Vincent Louis Viala |
| EPI_ISL_2344989 | HOSPITAL DE CAMPANHA COVID 19 MUNICIPIO DE TAUBATE | Instituto Butantan / ESALQ-Piracicaba | Dimas Tadeu Covas, Antonio Jorge Martins, Claudia Renata dos Santos Barros, David Schlesinger, Debora Botequio Moretti, Elaine Cristina Marqueze, Elaine Vieira Santos, Evandra Strazza Rodrigues, Heidge Fukumasu, Jayme Augusto de Souza-Neto, José Salvatore Leister Patané, Luiz Alcantara, Luiz Lehmann Coutinho, Maria Carolina Elias, Maurício Lacerda Nogueira, Rafael dos Santos Bezerra, Raul Machado Neto, Rejane Maria Tommasini Grotto, Ricardo Haddad, Sandra Coccuzzo Sampaio Vessoni, Simone Kashima, Svetoslav Nanev Slavov, Vincent Louis Viala |
| EPI_ISL_2344991, EPI_ISL_2344992, EPI_ISL_2344993, EPI_ISL_2344994 | CENTRO DE APOIO EPIDEMIOLOGICO | Instituto Butantan / ESALQ-Piracicaba | Dimas Tadeu Covas, Antonio Jorge Martins, Claudia Renata dos Santos Barros, David Schlesinger, Debora Botequio Moretti, Elaine Cristina Marqueze, Elaine Vieira Santos, Evandra Strazza Rodrigues, Heidge Fukumasu, Jayme Augusto de Souza-Neto, José Salvatore Leister Patané, Luiz Alcantara, Luiz Lehmann Coutinho, Maria Carolina Elias, Maurício Lacerda Nogueira, Rafael dos Santos Bezerra, Raul Machado Neto, Rejane Maria Tommasini Grotto, Ricardo Haddad, Sandra Coccuzzo Sampaio Vessoni, Simone Kashima, Svetoslav Nanev Slavov, Vincent Louis Viala |
| EPI_ISL_2344995 | HOSPITAL DE CAMPANHA COVID 19 MUNICIPIO DE TAUBATE | Instituto Butantan / ESALQ-Piracicaba | Dimas Tadeu Covas, Antonio Jorge Martins, Claudia Renata dos Santos Barros, David Schlesinger, Debora Botequio Moretti, Elaine Cristina Marqueze, Elaine Vieira Santos, Evandra Strazza Rodrigues, Heidge Fukumasu, Jayme Augusto de Souza-Neto, José Salvatore Leister Patané, Luiz Alcantara, Luiz Lehmann Coutinho, Maria Carolina Elias, Maurício Lacerda Nogueira, Rafael dos Santos Bezerra, Raul Machado Neto, Rejane Maria Tommasini Grotto, Ricardo Haddad, Sandra Coccuzzo Sampaio Vessoni, Simone Kashima, Svetoslav Nanev Slavov, Vincent Louis Viala |
| EPI_ISL_2344996 | CENTRO DE APOIO EPIDEMIOLOGICO | Instituto Butantan / ESALQ-Piracicaba | Dimas Tadeu Covas, Antonio Jorge Martins, Claudia Renata dos Santos Barros, David Schlesinger, Debora Botequio Moretti, Elaine Cristina Marqueze, Elaine Vieira Santos, Evandra Strazza Rodrigues, Heidge Fukumasu, Jayme Augusto de Souza-Neto, José Salvatore Leister Patané, Luiz Alcantara, Luiz Lehmann Coutinho, Maria Carolina Elias, Maurício Lacerda Nogueira, Rafael dos Santos Bezerra, Raul Machado Neto, Rejane Maria Tommasini Grotto, Ricardo Haddad, Sandra Coccuzzo Sampaio Vessoni, Simone Kashima, Svetoslav Nanev Slavov, Vincent Louis Viala |
| EPI_ISL_2344997 | HOSPITAL DE CAMPANHA COVID 19 MUNICIPIO DE TAUBATE | Instituto Butantan / ESALQ-Piracicaba | Dimas Tadeu Covas, Antonio Jorge Martins, Claudia Renata dos Santos Barros, David Schlesinger, Debora Botequio Moretti, Elaine Cristina Marqueze, Elaine Vieira Santos, Evandra Strazza Rodrigues, Heidge Fukumasu, Jayme Augusto de Souza-Neto, José Salvatore Leister Patané, Luiz Alcantara, Luiz Lehmann Coutinho, Maria Carolina Elias, Maurício Lacerda Nogueira, Rafael dos Santos Bezerra, Raul Machado Neto, Rejane Maria Tommasini Grotto, Ricardo Haddad, Sandra Coccuzzo Sampaio Vessoni, Simone Kashima, Svetoslav Nanev Slavov, Vincent Louis Viala |
| EPI_ISL_2344998 | CENTRO DE APOIO EPIDEMIOLOGICO | Instituto Butantan / ESALQ-Piracicaba | Dimas Tadeu Covas, Antonio Jorge Martins, Claudia Renata dos Santos Barros, David Schlesinger, Debora Botequio Moretti, Elaine Cristina Marqueze, Elaine Vieira Santos, Evandra Strazza Rodrigues, Heidge Fukumasu, Jayme Augusto de Souza-Neto, José Salvatore Leister Patané, Luiz Alcantara, Luiz Lehmann Coutinho, Maria Carolina Elias, Maurício Lacerda Nogueira, Rafael dos Santos Bezerra, Raul Machado Neto, Rejane Maria Tommasini Grotto, Ricardo Haddad, Sandra Coccuzzo Sampaio Vessoni, Simone Kashima, Svetoslav Nanev Slavov, Vincent Louis Viala |
| EPI_ISL_2344999, EPI_ISL_2345000, EPI_ISL_2345001, EPI_ISL_2345002 | UPA VILA SANTA CATARINA | Instituto Butantan | Dimas Tadeu Covas, Antonio Jorge Martins, Claudia Renata dos Santos Barros, David Schlesinger, Debora Botequio Moretti, Elaine Cristina Marqueze, Elaine Vieira Santos, Evandra Strazza Rodrigues, Heidge Fukumasu, Jayme Augusto de Souza-Neto, José Salvatore Leister Patané, Luiz Alcantara, Luiz Lehmann Coutinho, Maria Carolina Elias, Maurício Lacerda Nogueira, Rafael dos Santos Bezerra, Raul Machado Neto, Rejane Maria Tommasini Grotto, Ricardo Haddad, Sandra Coccuzzo Sampaio Vessoni, Simone Kashima, Svetoslav Nanev Slavov, Vincent Louis Viala |
| EPI_ISL_2345004 | SANTA CASA DE MISERICORDIA SAO JOSE | Instituto Butantan / ESALQ-Piracicaba | Dimas Tadeu Covas, Antonio Jorge Martins, Claudia Renata dos Santos Barros, David Schlesinger, Debora Botequio Moretti, Elaine Cristina Marqueze, Elaine Vieira Santos, Evandra Strazza Rodrigues, Heidge Fukumasu, Jayme Augusto de Souza-Neto, José Salvatore Leister Patané, Luiz Alcantara, Luiz Lehmann Coutinho, Maria Carolina Elias, Maurício Lacerda Nogueira, Rafael dos Santos Bezerra, Raul Machado Neto, Rejane Maria Tommasini Grotto, Ricardo Haddad, Sandra Coccuzzo Sampaio Vessoni, Simone Kashima, Svetoslav Nanev Slavov, Vincent Louis Viala |
| EPI_ISL_2345005 | HOSPITAL DE CAMPANHA COVID 19 MUNICIPIO DE TAUBATE | Instituto Butantan / ESALQ-Piracicaba | Dimas Tadeu Covas, Antonio Jorge Martins, Claudia Renata dos Santos Barros, David Schlesinger, Debora Botequio Moretti, Elaine Cristina Marqueze, Elaine Vieira Santos, Evandra Strazza Rodrigues, Heidge Fukumasu, Jayme Augusto de Souza-Neto, José Salvatore Leister Patané, Luiz Alcantara, Luiz Lehmann Coutinho, Maria Carolina Elias, Maurício Lacerda Nogueira, Rafael dos Santos Bezerra, Raul Machado Neto, Rejane Maria Tommasini Grotto, Ricardo Haddad, Sandra Coccuzzo Sampaio Vessoni, Simone Kashima, Svetoslav Nanev Slavov, Vincent Louis Viala |
| EPI_ISL_2345006, EPI_ISL_2345007 | SANTA CASA DE MISERICORDIA SAO JOSE | Instituto Butantan / ESALQ-Piracicaba | Dimas Tadeu Covas, Antonio Jorge Martins, Claudia Renata dos Santos Barros, David Schlesinger, Debora Botequio Moretti, Elaine Cristina Marqueze, Elaine Vieira Santos, Evandra Strazza Rodrigues, Heidge Fukumasu, Jayme Augusto de Souza-Neto, José Salvatore Leister Patané, Luiz Alcantara, Luiz Lehmann Coutinho, Maria Carolina Elias, Maurício Lacerda Nogueira, Rafael dos Santos Bezerra, Raul Machado Neto, Rejane Maria Tommasini Grotto, Ricardo Haddad, Sandra Coccuzzo Sampaio Vessoni, Simone Kashima, Svetoslav Nanev Slavov, Vincent Louis Viala |
| EPI_ISL_2345008 | HOSPITAL DE CAMPANHA COVID 19 MUNICIPIO DE TAUBATE | Instituto Butantan / ESALQ-Piracicaba | Dimas Tadeu Covas, Antonio Jorge Martins, Claudia Renata dos Santos Barros, David Schlesinger, Debora Botequio Moretti, Elaine Cristina Marqueze, Elaine Vieira Santos, Evandra Strazza Rodrigues, Heidge Fukumasu, Jayme Augusto de Souza-Neto, José Salvatore Leister Patané, Luiz Alcantara, Luiz Lehmann Coutinho, Maria Carolina Elias, Maurício Lacerda Nogueira, Rafael dos Santos Bezerra, Raul Machado Neto, Rejane Maria Tommasini Grotto, Ricardo Haddad, Sandra Coccuzzo Sampaio Vessoni, Simone Kashima, Svetoslav Nanev Slavov, Vincent Louis Viala |
| EPI_ISL_2345009, EPI_ISL_2345010 | SANTA CASA DE MISERICORDIA SAO JOSE | Instituto Butantan / ESALQ-Piracicaba | Dimas Tadeu Covas, Antonio Jorge Martins, Claudia Renata dos Santos Barros, David Schlesinger, Debora Botequio Moretti, Elaine Cristina Marqueze, Elaine Vieira Santos, Evandra Strazza Rodrigues, Heidge Fukumasu, Jayme Augusto de Souza-Neto, José Salvatore Leister Patané, Luiz Alcantara, Luiz Lehmann Coutinho, Maria Carolina Elias, Maurício Lacerda Nogueira, Rafael dos Santos Bezerra, Raul Machado Neto, Rejane Maria Tommasini Grotto, Ricardo Haddad, Sandra Coccuzzo Sampaio Vessoni, Simone Kashima, Svetoslav Nanev Slavov, Vincent Louis Viala |
| EPI_ISL_2345011 | HOSPITAL DE CAMPANHA COVID 19 MUNICIPIO DE TAUBATE | Instituto Butantan / ESALQ-Piracicaba | Dimas Tadeu Covas, Antonio Jorge Martins, Claudia Renata dos Santos Barros, David Schlesinger, Debora Botequio Moretti, Elaine Cristina Marqueze, Elaine Vieira Santos, Evandra Strazza Rodrigues, Heidge Fukumasu, Jayme Augusto de Souza-Neto, José Salvatore Leister Patané, Luiz Alcantara, Luiz Lehmann Coutinho, Maria Carolina Elias, Maurício Lacerda Nogueira, Rafael dos Santos Bezerra, Raul Machado Neto, Rejane Maria Tommasini Grotto, Ricardo Haddad, Sandra Coccuzzo Sampaio Vessoni, Simone Kashima, Svetoslav Nanev Slavov, Vincent Louis Viala |
| EPI_ISL_2345012, EPI_ISL_2345013 | SANTA CASA DE MISERICORDIA SAO JOSE | Instituto Butantan / ESALQ-Piracicaba | Dimas Tadeu Covas, Antonio Jorge Martins, Claudia Renata dos Santos Barros, David Schlesinger, Debora Botequio Moretti, Elaine Cristina Marqueze, Elaine Vieira Santos, Evandra Strazza Rodrigues, Heidge Fukumasu, Jayme Augusto de Souza-Neto, José Salvatore Leister Patané, Luiz Alcantara, Luiz Lehmann Coutinho, Maria Carolina Elias, Maurício Lacerda Nogueira, Rafael dos Santos Bezerra, Raul Machado Neto, Rejane Maria Tommasini Grotto, Ricardo Haddad, Sandra Coccuzzo Sampaio Vessoni, Simone Kashima, Svetoslav Nanev Slavov, Vincent Louis Viala |
| EPI_ISL_2345014 | UPA DR FRANCO DA ROCHA | Instituto Butantan | Dimas Tadeu Covas, Antonio Jorge Martins, Claudia Renata dos Santos Barros, David Schlesinger, Debora Botequio Moretti, Elaine Cristina Marqueze, Elaine Vieira Santos, Evandra Strazza Rodrigues, Heidge Fukumasu, Jayme Augusto de Souza-Neto, José Salvatore Leister Patané, Luiz Alcantara, Luiz Lehmann Coutinho, Maria Carolina Elias, Maurício Lacerda Nogueira, Rafael dos Santos Bezerra, Raul Machado Neto, Rejane Maria Tommasini Grotto, Ricardo Haddad, Sandra Coccuzzo Sampaio Vessoni, Simone Kashima, Svetoslav Nanev Slavov, Vincent Louis Viala |
| EPI_ISL_2345015 | HOSPITAL DE CAMPANHA COVID 19 MUNICIPIO DE TAUBATE | Instituto Butantan / ESALQ-Piracicaba | Dimas Tadeu Covas, Antonio Jorge Martins, Claudia Renata dos Santos Barros, David Schlesinger, Debora Botequio Moretti, Elaine Cristina Marqueze, Elaine Vieira Santos, Evandra Strazza Rodrigues, Heidge Fukumasu, Jayme Augusto de Souza-Neto, José Salvatore Leister Patané, Luiz Alcantara, Luiz Lehmann Coutinho, Maria Carolina Elias, Maurício Lacerda Nogueira, Rafael dos Santos Bezerra, Raul Machado Neto, Rejane Maria Tommasini Grotto, Ricardo Haddad, Sandra Coccuzzo Sampaio Vessoni, Simone Kashima, Svetoslav Nanev Slavov, Vincent Louis Viala |
| EPI_ISL_2345016, EPI_ISL_2345017 | UPA DR FRANCO DA ROCHA | Instituto Butantan | Dimas Tadeu Covas, Antonio Jorge Martins, Claudia Renata dos Santos Barros, David Schlesinger, Debora Botequio Moretti, Elaine Cristina Marqueze, Elaine Vieira Santos, Evandra Strazza Rodrigues, Heidge Fukumasu, Jayme Augusto de Souza-Neto, José Salvatore Leister Patané, Luiz Alcantara, Luiz |

| | | | |
|---|---|---|---|
| | | | Lehmann Coutinho, Maria Carolina Elias, Maurício Lacerda Nogueira, Rafael dos Santos Bezerra, Raul Machado Neto, Rejane Maria Tommasini Grotto, Ricardo Haddad, Sandra Coccuzzo Sampaio Vessoni, Simone Kashima, Svetoslav Nanev Slavov, Vincent Louis Viala |
| EPI_ISL_2345018, EPI_ISL_2345021 | HOSPITAL DE CAMPANHA COVID 19 MUNICIPIO DE TAUBATE | Instituto Butantan / ESALQ-Piracicaba | Dimas Tadeu Covas, Antonio Jorge Martins, Claudia Renata dos Santos Barros, David Schlesinger, Debora Botequio Moretti, Elaine Cristina Marqueze, Elaine Vieira Santos, Evandra Strazza Rodrigues, Heidge Fukumasu, Jayme Augusto de Souza-Neto, José Salvatore Leister Patané, Luiz Alcantara, Luiz Lehmann Coutinho, Maria Carolina Elias, Maurício Lacerda Nogueira, Rafael dos Santos Bezerra, Raul Machado Neto, Rejane Maria Tommasini Grotto, Ricardo Haddad, Sandra Coccuzzo Sampaio Vessoni, Simone Kashima, Svetoslav Nanev Slavov, Vincent Louis Viala |
| EPI_ISL_2345022, EPI_ISL_2345023, EPI_ISL_2345026, EPI_ISL_2345027 | HOSP MUN JOSANIAS CASTANHA BRAGA | Instituto Butantan | Dimas Tadeu Covas, Antonio Jorge Martins, Claudia Renata dos Santos Barros, David Schlesinger, Debora Botequio Moretti, Elaine Cristina Marqueze, Elaine Vieira Santos, Evandra Strazza Rodrigues, Heidge Fukumasu, Jayme Augusto de Souza-Neto, José Salvatore Leister Patané, Luiz Alcantara, Luiz Lehmann Coutinho, Maria Carolina Elias, Maurício Lacerda Nogueira, Rafael dos Santos Bezerra, Raul Machado Neto, Rejane Maria Tommasini Grotto, Ricardo Haddad, Sandra Coccuzzo Sampaio Vessoni, Simone Kashima, Svetoslav Nanev Slavov, Vincent Louis Viala |
| EPI_ISL_2345029 | HOSPITAL ESCOLA DA UNIVERSIDADE DE TAUBATE | Instituto Butantan / ESALQ-Piracicaba | Dimas Tadeu Covas, Antonio Jorge Martins, Claudia Renata dos Santos Barros, David Schlesinger, Debora Botequio Moretti, Elaine Cristina Marqueze, Elaine Vieira Santos, Evandra Strazza Rodrigues, Heidge Fukumasu, Jayme Augusto de Souza-Neto, José Salvatore Leister Patané, Luiz Alcantara, Luiz Lehmann Coutinho, Maria Carolina Elias, Maurício Lacerda Nogueira, Rafael dos Santos Bezerra, Raul Machado Neto, Rejane Maria Tommasini Grotto, Ricardo Haddad, Sandra Coccuzzo Sampaio Vessoni, Simone Kashima, Svetoslav Nanev Slavov, Vincent Louis Viala |
| EPI_ISL_2345030 | UPA VILA SANTA CATARINA | Instituto Butantan | Dimas Tadeu Covas, Antonio Jorge Martins, Claudia Renata dos Santos Barros, David Schlesinger, Debora Botequio Moretti, Elaine Cristina Marqueze, Elaine Vieira Santos, Evandra Strazza Rodrigues, Heidge Fukumasu, Jayme Augusto de Souza-Neto, José Salvatore Leister Patané, Luiz Alcantara, Luiz Lehmann Coutinho, Maria Carolina Elias, Maurício Lacerda Nogueira, Rafael dos Santos Bezerra, Raul Machado Neto, Rejane Maria Tommasini Grotto, Ricardo Haddad, Sandra Coccuzzo Sampaio Vessoni, Simone Kashima, Svetoslav Nanev Slavov, Vincent Louis Viala |
| EPI_ISL_2345031, EPI_ISL_2345032 | SANTA CASA DE GUARAREMA | Instituto Butantan | Dimas Tadeu Covas, Antonio Jorge Martins, Claudia Renata dos Santos Barros, David Schlesinger, Debora Botequio Moretti, Elaine Cristina Marqueze, Elaine Vieira Santos, Evandra Strazza Rodrigues, Heidge Fukumasu, Jayme Augusto de Souza-Neto, José Salvatore Leister Patané, Luiz Alcantara, Luiz Lehmann Coutinho, Maria Carolina Elias, Maurício Lacerda Nogueira, Rafael dos Santos Bezerra, Raul Machado Neto, Rejane Maria Tommasini Grotto, Ricardo Haddad, Sandra Coccuzzo Sampaio Vessoni, Simone Kashima, Svetoslav Nanev Slavov, Vincent Louis Viala |
| EPI_ISL_2345033, EPI_ISL_2345034 | LABORATORIO MUNICIPAL DE PIRACICABA | Instituto Butantan / ESALQ-Piracicaba | Dimas Tadeu Covas, Antonio Jorge Martins, Claudia Renata dos Santos Barros, David Schlesinger, Debora Botequio Moretti, Elaine Cristina Marqueze, Elaine Vieira Santos, Evandra Strazza Rodrigues, Heidge Fukumasu, Jayme Augusto de Souza-Neto, José Salvatore Leister Patané, Luiz Alcantara, Luiz Lehmann Coutinho, Maria Carolina Elias, Maurício Lacerda Nogueira, Rafael dos Santos Bezerra, Raul Machado Neto, Rejane Maria Tommasini Grotto, Ricardo Haddad, Sandra Coccuzzo Sampaio Vessoni, Simone Kashima, Svetoslav Nanev Slavov, Vincent Louis Viala |
| EPI_ISL_2345035, EPI_ISL_2345037, EPI_ISL_2345038 | SANTA CASA DE GUARAREMA | Instituto Butantan | Dimas Tadeu Covas, Antonio Jorge Martins, Claudia Renata dos Santos Barros, David Schlesinger, Debora Botequio Moretti, Elaine Cristina Marqueze, Elaine Vieira Santos, Evandra Strazza Rodrigues, Heidge Fukumasu, Jayme Augusto de Souza-Neto, José Salvatore Leister Patané, Luiz Alcantara, Luiz Lehmann Coutinho, Maria Carolina Elias, Maurício Lacerda Nogueira, Rafael dos Santos Bezerra, Raul Machado Neto, Rejane Maria Tommasini Grotto, Ricardo Haddad, Sandra Coccuzzo Sampaio Vessoni, Simone Kashima, Svetoslav Nanev Slavov, Vincent Louis Viala |
| EPI_ISL_2345039, EPI_ISL_2345040, EPI_ISL_2345041 | VIGILANCIA EM SAUDE DE FCO MORATO | Instituto Butantan | Dimas Tadeu Covas, Antonio Jorge Martins, Claudia Renata dos Santos Barros, David Schlesinger, Debora Botequio Moretti, Elaine Cristina Marqueze, Elaine Vieira Santos, Evandra Strazza Rodrigues, Heidge Fukumasu, Jayme Augusto de Souza-Neto, José Salvatore Leister Patané, Luiz Alcantara, Luiz Lehmann Coutinho, Maria Carolina Elias, Maurício Lacerda Nogueira, Rafael dos Santos Bezerra, Raul Machado Neto, Rejane Maria Tommasini Grotto, Ricardo Haddad, Sandra Coccuzzo Sampaio Vessoni, Simone Kashima, Svetoslav Nanev Slavov, Vincent Louis Viala |
| EPI_ISL_2345042 | UNIDADE RESPIRATORIA NOVA HORTOLANDIA | Instituto Butantan / ESALQ-Piracicaba | Dimas Tadeu Covas, Antonio Jorge Martins, Claudia Renata dos Santos Barros, David Schlesinger, Debora Botequio Moretti, Elaine Cristina Marqueze, Elaine Vieira Santos, Evandra Strazza Rodrigues, Heidge Fukumasu, Jayme Augusto de Souza-Neto, José Salvatore Leister Patané, Luiz Alcantara, Luiz Lehmann Coutinho, Maria Carolina Elias, Maurício Lacerda Nogueira, Rafael dos Santos Bezerra, Raul Machado Neto, Rejane Maria Tommasini Grotto, Ricardo Haddad, Sandra Coccuzzo Sampaio Vessoni, Simone Kashima, Svetoslav Nanev Slavov, Vincent Louis Viala |
| EPI_ISL_2345043 | HOSP MUN DE MOGI DAS CRUZES PREF WALDEMAR COSTA FILHO | Instituto Butantan / ESALQ-Piracicaba | Dimas Tadeu Covas, Antonio Jorge Martins, Claudia Renata dos Santos Barros, David Schlesinger, Debora Botequio Moretti, Elaine Cristina Marqueze, Elaine Vieira Santos, Evandra Strazza Rodrigues, Heidge Fukumasu, Jayme Augusto de Souza-Neto, José Salvatore Leister Patané, Luiz Alcantara, Luiz Lehmann Coutinho, Maria Carolina Elias, Maurício Lacerda Nogueira, Rafael dos Santos Bezerra, Raul Machado Neto, Rejane Maria Tommasini Grotto, Ricardo Haddad, Sandra Coccuzzo Sampaio Vessoni, Simone Kashima, Svetoslav Nanev Slavov, Vincent Louis Viala |
| EPI_ISL_2345044, EPI_ISL_2345045 | HOSPITAL DE CAMPANHA COVID 19 CAIEIRAS | Instituto Butantan / ESALQ-Piracicaba | Dimas Tadeu Covas, Antonio Jorge Martins, Claudia Renata dos Santos Barros, David Schlesinger, Debora Botequio Moretti, Elaine Cristina Marqueze, Elaine Vieira Santos, Evandra Strazza Rodrigues, Heidge Fukumasu, Jayme Augusto de Souza-Neto, José Salvatore Leister Patané, Luiz Alcantara, Luiz Lehmann Coutinho, Maria Carolina Elias, Maurício Lacerda Nogueira, Rafael dos Santos Bezerra, Raul Machado Neto, Rejane Maria Tommasini Grotto, Ricardo Haddad, Sandra Coccuzzo Sampaio Vessoni, Simone Kashima, Svetoslav Nanev Slavov, Vincent Louis Viala |
| EPI_ISL_2345046 | UNIDADE DE PRONTO ATENDIMENTO UPA DRA ANA OLIVIA BENTIVOGLIO | Instituto Butantan / ESALQ-Piracicaba | Dimas Tadeu Covas, Antonio Jorge Martins, Claudia Renata dos Santos Barros, David Schlesinger, Debora Botequio Moretti, Elaine Cristina Marqueze, Elaine Vieira Santos, Evandra Strazza Rodrigues, Heidge Fukumasu, Jayme Augusto de Souza-Neto, José Salvatore Leister Patané, Luiz Alcantara, Luiz Lehmann Coutinho, Maria Carolina Elias, Maurício Lacerda Nogueira, Rafael dos Santos Bezerra, Raul Machado Neto, Rejane Maria Tommasini Grotto, Ricardo Haddad, Sandra Coccuzzo Sampaio Vessoni, Simone Kashima, Svetoslav Nanev Slavov, Vincent Louis Viala |
| EPI_ISL_2345047, EPI_ISL_2345049 | UNIDADE RESPIRATORIA NOVA HORTOLANDIA | Instituto Butantan / ESALQ-Piracicaba | Dimas Tadeu Covas, Antonio Jorge Martins, Claudia Renata dos Santos Barros, David Schlesinger, Debora Botequio Moretti, Elaine Cristina Marqueze, Elaine Vieira Santos, Evandra Strazza Rodrigues, Heidge Fukumasu, Jayme Augusto de Souza-Neto, José Salvatore Leister Patané, Luiz Alcantara, Luiz Lehmann Coutinho, Maria Carolina Elias, Maurício Lacerda Nogueira, Rafael dos Santos Bezerra, Raul Machado Neto, Rejane Maria Tommasini Grotto, Ricardo Haddad, Sandra Coccuzzo Sampaio Vessoni, Simone Kashima, Svetoslav Nanev Slavov, Vincent Louis Viala |
| EPI_ISL_2345050, EPI_ISL_2345051 | SECAO CENTRO DE DIAGNOSTICO SECEDI | Instituto Butantan / ESALQ-Piracicaba | Dimas Tadeu Covas, Antonio Jorge Martins, Claudia Renata dos Santos Barros, David Schlesinger, Debora Botequio Moretti, Elaine Cristina Marqueze, Elaine Vieira Santos, Evandra Strazza Rodrigues, Heidge Fukumasu, Jayme Augusto de Souza-Neto, José Salvatore Leister Patané, Luiz Alcantara, Luiz Lehmann Coutinho, Maria Carolina Elias, Maurício Lacerda Nogueira, Rafael dos Santos Bezerra, Raul Machado Neto, Rejane Maria Tommasini Grotto, Ricardo Haddad, Sandra Coccuzzo Sampaio Vessoni, Simone Kashima, Svetoslav Nanev Slavov, Vincent Louis Viala |
| EPI_ISL_2345052, EPI_ISL_2345053 | UNIDADE RESPIRATORIA NOVA HORTOLANDIA | Instituto Butantan / ESALQ-Piracicaba | Dimas Tadeu Covas, Antonio Jorge Martins, Claudia Renata dos Santos Barros, David Schlesinger, Debora Botequio Moretti, Elaine Cristina Marqueze, Elaine Vieira Santos, Evandra Strazza Rodrigues, Heidge Fukumasu, Jayme Augusto de Souza-Neto, José Salvatore Leister Patané, Luiz Alcantara, Luiz Lehmann Coutinho, Maria Carolina Elias, Maurício Lacerda Nogueira, Rafael dos Santos Bezerra, Raul Machado Neto, Rejane Maria Tommasini Grotto, Ricardo Haddad, Sandra Coccuzzo Sampaio Vessoni, Simone Kashima, Svetoslav Nanev Slavov, Vincent Louis Viala |
| EPI_ISL_2345054, EPI_ISL_2345055 | UBS SAO LOURENCO DA SERRA | Instituto Butantan / ESALQ-Piracicaba | Dimas Tadeu Covas, Antonio Jorge Martins, Claudia Renata dos Santos Barros, David Schlesinger, Debora Botequio Moretti, Elaine Cristina Marqueze, Elaine Vieira Santos, Evandra Strazza Rodrigues, Heidge Fukumasu, Jayme Augusto de Souza-Neto, José Salvatore Leister Patané, Luiz Alcantara, Luiz Lehmann Coutinho, Maria Carolina Elias, Maurício Lacerda Nogueira, Rafael dos Santos Bezerra, Raul Machado Neto, Rejane Maria Tommasini Grotto, Ricardo Haddad, Sandra Coccuzzo Sampaio Vessoni, Simone Kashima, Svetoslav Nanev Slavov, Vincent Louis Viala |
| EPI_ISL_2345056, EPI_ISL_2345057, EPI_ISL_2345058 | UPA DR AKIRA TADA | Instituto Butantan / ESALQ-Piracicaba | Dimas Tadeu Covas, Antonio Jorge Martins, Claudia Renata dos Santos Barros, David Schlesinger, Debora Botequio Moretti, Elaine Cristina Marqueze, Elaine Vieira Santos, Evandra Strazza Rodrigues, Heidge Fukumasu, Jayme Augusto de Souza-Neto, José Salvatore Leister Patané, Luiz Alcantara, Luiz Lehmann Coutinho, Maria Carolina Elias, Maurício Lacerda Nogueira, Rafael dos Santos Bezerra, Raul Machado Neto, Rejane Maria Tommasini Grotto, Ricardo Haddad, Sandra Coccuzzo Sampaio Vessoni, Simone Kashima, Svetoslav Nanev Slavov, Vincent Louis Viala |
| EPI_ISL_2345059 | SECAO CENTRO DE DIAGNOSTICO SECEDI | Instituto Butantan / ESALQ-Piracicaba | Dimas Tadeu Covas, Antonio Jorge Martins, Claudia Renata dos Santos Barros, David Schlesinger, Debora Botequio Moretti, Elaine Cristina Marqueze, Elaine Vieira Santos, Evandra Strazza Rodrigues, Heidge Fukumasu, Jayme Augusto de Souza-Neto, José Salvatore Leister Patané, Luiz Alcantara, Luiz Lehmann Coutinho, Maria Carolina Elias, Maurício Lacerda Nogueira, Rafael dos Santos Bezerra, Raul Machado Neto, Rejane Maria Tommasini Grotto, Ricardo Haddad, Sandra Coccuzzo Sampaio Vessoni, Simone Kashima, Svetoslav Nanev Slavov, Vincent Louis Viala |
| EPI_ISL_2345060 | UNIDADE RESPIRATORIA NOVA HORTOLANDIA | Instituto Butantan / ESALQ-Piracicaba | Dimas Tadeu Covas, Antonio Jorge Martins, Claudia Renata dos Santos Barros, David Schlesinger, Debora Botequio Moretti, Elaine Cristina Marqueze, Elaine Vieira Santos, Evandra Strazza Rodrigues, Heidge Fukumasu, Jayme Augusto de Souza-Neto, José Salvatore Leister Patané, Luiz Alcantara, Luiz Lehmann Coutinho, Maria Carolina Elias, Maurício Lacerda Nogueira, Rafael dos Santos Bezerra, Raul Machado Neto, Rejane Maria Tommasini Grotto, Ricardo Haddad, Sandra Coccuzzo Sampaio Vessoni, Simone Kashima, Svetoslav Nanev Slavov, Vincent Louis Viala |

| | | | |
|---|---|---|---|
| EPI_ISL_2345061, EPI_ISL_2345062, EPI_ISL_2345063, EPI_ISL_2345064 | HOSP MUN CIDADE TIRADENTES CARMEN PRUDENTE | Instituto Butantan / ESALQ-Piracicaba | Dimas Tadeu Covas, Antonio Jorge Martins, Claudia Renata dos Santos Barros, David Schlesinger, Debora Botequio Moretti, Elaine Cristina Marqueze, Elaine Vieira Santos, Evandra Strazza Rodrigues, Heidge Fukumasu, Jayme Augusto de Souza-Neto, José Salvatore Leister Patané, Luiz Alcantara, Luiz Lehmann Coutinho, Maria Carolina Elias, Maurício Lacerda Nogueira, Rafael dos Santos Bezerra, Raul Machado Neto, Rejane Maria Tommasini Grotto, Ricardo Haddad, Sandra Coccuzzo Sampaio Vessoni, Simone Kashima, Svetoslav Nanev Slavov, Vincent Louis Viala |
| EPI_ISL_2345065, EPI_ISL_2345066 | HOSPITAL DE CAMPANHA COVID 19 CAIEIRAS | Instituto Butantan / ESALQ-Piracicaba | Dimas Tadeu Covas, Antonio Jorge Martins, Claudia Renata dos Santos Barros, David Schlesinger, Debora Botequio Moretti, Elaine Cristina Marqueze, Elaine Vieira Santos, Evandra Strazza Rodrigues, Heidge Fukumasu, Jayme Augusto de Souza-Neto, José Salvatore Leister Patané, Luiz Alcantara, Luiz Lehmann Coutinho, Maria Carolina Elias, Maurício Lacerda Nogueira, Rafael dos Santos Bezerra, Raul Machado Neto, Rejane Maria Tommasini Grotto, Ricardo Haddad, Sandra Coccuzzo Sampaio Vessoni, Simone Kashima, Svetoslav Nanev Slavov, Vincent Louis Viala |
| EPI_ISL_2345067, EPI_ISL_2345068, EPI_ISL_2345069, EPI_ISL_2345070, EPI_ISL_2345071, EPI_ISL_2345072, EPI_ISL_2345073 | HOSP MUN DE MOGI DAS CRUZES PREF WALDEMAR COSTA FILHO | Instituto Butantan / ESALQ-Piracicaba | Dimas Tadeu Covas, Antonio Jorge Martins, Claudia Renata dos Santos Barros, David Schlesinger, Debora Botequio Moretti, Elaine Cristina Marqueze, Elaine Vieira Santos, Evandra Strazza Rodrigues, Heidge Fukumasu, Jayme Augusto de Souza-Neto, José Salvatore Leister Patané, Luiz Alcantara, Luiz Lehmann Coutinho, Maria Carolina Elias, Maurício Lacerda Nogueira, Rafael dos Santos Bezerra, Raul Machado Neto, Rejane Maria Tommasini Grotto, Ricardo Haddad, Sandra Coccuzzo Sampaio Vessoni, Simone Kashima, Svetoslav Nanev Slavov, Vincent Louis Viala |
| EPI_ISL_2345074, EPI_ISL_2345075 | HOSP MUN CIDADE TIRADENTES CARMEN PRUDENTE | Instituto Butantan / ESALQ-Piracicaba | Dimas Tadeu Covas, Antonio Jorge Martins, Claudia Renata dos Santos Barros, David Schlesinger, Debora Botequio Moretti, Elaine Cristina Marqueze, Elaine Vieira Santos, Evandra Strazza Rodrigues, Heidge Fukumasu, Jayme Augusto de Souza-Neto, José Salvatore Leister Patané, Luiz Alcantara, Luiz Lehmann Coutinho, Maria Carolina Elias, Maurício Lacerda Nogueira, Rafael dos Santos Bezerra, Raul Machado Neto, Rejane Maria Tommasini Grotto, Ricardo Haddad, Sandra Coccuzzo Sampaio Vessoni, Simone Kashima, Svetoslav Nanev Slavov, Vincent Louis Viala |
| EPI_ISL_2345076 | HOSP MUN JOSANIAS CASTANHA BRAGA | Instituto Butantan / ESALQ-Piracicaba | Dimas Tadeu Covas, Antonio Jorge Martins, Claudia Renata dos Santos Barros, David Schlesinger, Debora Botequio Moretti, Elaine Cristina Marqueze, Elaine Vieira Santos, Evandra Strazza Rodrigues, Heidge Fukumasu, Jayme Augusto de Souza-Neto, José Salvatore Leister Patané, Luiz Alcantara, Luiz Lehmann Coutinho, Maria Carolina Elias, Maurício Lacerda Nogueira, Rafael dos Santos Bezerra, Raul Machado Neto, Rejane Maria Tommasini Grotto, Ricardo Haddad, Sandra Coccuzzo Sampaio Vessoni, Simone Kashima, Svetoslav Nanev Slavov, Vincent Louis Viala |
| EPI_ISL_2345078 | CS II DR MIGUEL VITALIANO ORLANDIA | Instituto Butantan / ESALQ-Piracicaba | Dimas Tadeu Covas, Antonio Jorge Martins, Claudia Renata dos Santos Barros, David Schlesinger, Debora Botequio Moretti, Elaine Cristina Marqueze, Elaine Vieira Santos, Evandra Strazza Rodrigues, Heidge Fukumasu, Jayme Augusto de Souza-Neto, José Salvatore Leister Patané, Luiz Alcantara, Luiz Lehmann Coutinho, Maria Carolina Elias, Maurício Lacerda Nogueira, Rafael dos Santos Bezerra, Raul Machado Neto, Rejane Maria Tommasini Grotto, Ricardo Haddad, Sandra Coccuzzo Sampaio Vessoni, Simone Kashima, Svetoslav Nanev Slavov, Vincent Louis Viala |
| EPI_ISL_2345079 | UNIDADE REFERENCIAL SUDOESTE | Instituto Butantan / ESALQ-Piracicaba | Dimas Tadeu Covas, Antonio Jorge Martins, Claudia Renata dos Santos Barros, David Schlesinger, Debora Botequio Moretti, Elaine Cristina Marqueze, Elaine Vieira Santos, Evandra Strazza Rodrigues, Heidge Fukumasu, Jayme Augusto de Souza-Neto, José Salvatore Leister Patané, Luiz Alcantara, Luiz Lehmann Coutinho, Maria Carolina Elias, Maurício Lacerda Nogueira, Rafael dos Santos Bezerra, Raul Machado Neto, Rejane Maria Tommasini Grotto, Ricardo Haddad, Sandra Coccuzzo Sampaio Vessoni, Simone Kashima, Svetoslav Nanev Slavov, Vincent Louis Viala |
| EPI_ISL_2345080 | UNIDADE DE PRONTO ATENDIMENTO UPA DRA ANA OLIVIA BENTIVOGLIO | Instituto Butantan / ESALQ-Piracicaba | Dimas Tadeu Covas, Antonio Jorge Martins, Claudia Renata dos Santos Barros, David Schlesinger, Debora Botequio Moretti, Elaine Cristina Marqueze, Elaine Vieira Santos, Evandra Strazza Rodrigues, Heidge Fukumasu, Jayme Augusto de Souza-Neto, José Salvatore Leister Patané, Luiz Alcantara, Luiz Lehmann Coutinho, Maria Carolina Elias, Maurício Lacerda Nogueira, Rafael dos Santos Bezerra, Raul Machado Neto, Rejane Maria Tommasini Grotto, Ricardo Haddad, Sandra Coccuzzo Sampaio Vessoni, Simone Kashima, Svetoslav Nanev Slavov, Vincent Louis Viala |
| EPI_ISL_2345081 | HOSP MUN JOSANIAS CASTANHA BRAGA | Instituto Butantan / ESALQ-Piracicaba | Dimas Tadeu Covas, Antonio Jorge Martins, Claudia Renata dos Santos Barros, David Schlesinger, Debora Botequio Moretti, Elaine Cristina Marqueze, Elaine Vieira Santos, Evandra Strazza Rodrigues, Heidge Fukumasu, Jayme Augusto de Souza-Neto, José Salvatore Leister Patané, Luiz Alcantara, Luiz Lehmann Coutinho, Maria Carolina Elias, Maurício Lacerda Nogueira, Rafael dos Santos Bezerra, Raul Machado Neto, Rejane Maria Tommasini Grotto, Ricardo Haddad, Sandra Coccuzzo Sampaio Vessoni, Simone Kashima, Svetoslav Nanev Slavov, Vincent Louis Viala |
| EPI_ISL_2345083 | UNIDADE DE PRONTO ATENDIMENTO UPA DRA ANA OLIVIA BENTIVOGLIO | Instituto Butantan / ESALQ-Piracicaba | Dimas Tadeu Covas, Antonio Jorge Martins, Claudia Renata dos Santos Barros, David Schlesinger, Debora Botequio Moretti, Elaine Cristina Marqueze, Elaine Vieira Santos, Evandra Strazza Rodrigues, Heidge Fukumasu, Jayme Augusto de Souza-Neto, José Salvatore Leister Patané, Luiz Alcantara, Luiz Lehmann Coutinho, Maria Carolina Elias, Maurício Lacerda Nogueira, Rafael dos Santos Bezerra, Raul Machado Neto, Rejane Maria Tommasini Grotto, Ricardo Haddad, Sandra Coccuzzo Sampaio Vessoni, Simone Kashima, Svetoslav Nanev Slavov, Vincent Louis Viala |
| EPI_ISL_2345084 | NUCLEO SAUDE III N HABITACIONAL PRESIDENTE GEISEL BAURU | Instituto Butantan / ESALQ-Piracicaba | Dimas Tadeu Covas, Antonio Jorge Martins, Claudia Renata dos Santos Barros, David Schlesinger, Debora Botequio Moretti, Elaine Cristina Marqueze, Elaine Vieira Santos, Evandra Strazza Rodrigues, Heidge Fukumasu, Jayme Augusto de Souza-Neto, José Salvatore Leister Patané, Luiz Alcantara, Luiz Lehmann Coutinho, Maria Carolina Elias, Maurício Lacerda Nogueira, Rafael dos Santos Bezerra, Raul Machado Neto, Rejane Maria Tommasini Grotto, Ricardo Haddad, Sandra Coccuzzo Sampaio Vessoni, Simone Kashima, Svetoslav Nanev Slavov, Vincent Louis Viala |
| EPI_ISL_2345085 | UNIDADE DE PRONTO ATENDIMENTO DE VARZEA PAULISTA UPA II | Instituto Butantan / ESALQ-Piracicaba | Dimas Tadeu Covas, Antonio Jorge Martins, Claudia Renata dos Santos Barros, David Schlesinger, Debora Botequio Moretti, Elaine Cristina Marqueze, Elaine Vieira Santos, Evandra Strazza Rodrigues, Heidge Fukumasu, Jayme Augusto de Souza-Neto, José Salvatore Leister Patané, Luiz Alcantara, Luiz Lehmann Coutinho, Maria Carolina Elias, Maurício Lacerda Nogueira, Rafael dos Santos Bezerra, Raul Machado Neto, Rejane Maria Tommasini Grotto, Ricardo Haddad, Sandra Coccuzzo Sampaio Vessoni, Simone Kashima, Svetoslav Nanev Slavov, Vincent Louis Viala |
| EPI_ISL_2345086 | HOSPITAL GERAL JESUS TEIXEIRA DA COSTA GUAIANASES SAO PAULO | Instituto Butantan / ESALQ-Piracicaba | Dimas Tadeu Covas, Antonio Jorge Martins, Claudia Renata dos Santos Barros, David Schlesinger, Debora Botequio Moretti, Elaine Cristina Marqueze, Elaine Vieira Santos, Evandra Strazza Rodrigues, Heidge Fukumasu, Jayme Augusto de Souza-Neto, José Salvatore Leister Patané, Luiz Alcantara, Luiz Lehmann Coutinho, Maria Carolina Elias, Maurício Lacerda Nogueira, Rafael dos Santos Bezerra, Raul Machado Neto, Rejane Maria Tommasini Grotto, Ricardo Haddad, Sandra Coccuzzo Sampaio Vessoni, Simone Kashima, Svetoslav Nanev Slavov, Vincent Louis Viala |
| EPI_ISL_2345087, EPI_ISL_2345088 | UNIDADE DE PRONTO ATENDIMENTO UPA DRA ANA OLIVIA BENTIVOGLIO | Instituto Butantan / ESALQ-Piracicaba | Dimas Tadeu Covas, Antonio Jorge Martins, Claudia Renata dos Santos Barros, David Schlesinger, Debora Botequio Moretti, Elaine Cristina Marqueze, Elaine Vieira Santos, Evandra Strazza Rodrigues, Heidge Fukumasu, Jayme Augusto de Souza-Neto, José Salvatore Leister Patané, Luiz Alcantara, Luiz Lehmann Coutinho, Maria Carolina Elias, Maurício Lacerda Nogueira, Rafael dos Santos Bezerra, Raul Machado Neto, Rejane Maria Tommasini Grotto, Ricardo Haddad, Sandra Coccuzzo Sampaio Vessoni, Simone Kashima, Svetoslav Nanev Slavov, Vincent Louis Viala |
| EPI_ISL_2345089, EPI_ISL_2345090 | UNIDADE DE PRONTO ATENDIMENTO DE VARZEA PAULISTA UPA II | Instituto Butantan / ESALQ-Piracicaba | Dimas Tadeu Covas, Antonio Jorge Martins, Claudia Renata dos Santos Barros, David Schlesinger, Debora Botequio Moretti, Elaine Cristina Marqueze, Elaine Vieira Santos, Evandra Strazza Rodrigues, Heidge Fukumasu, Jayme Augusto de Souza-Neto, José Salvatore Leister Patané, Luiz Alcantara, Luiz Lehmann Coutinho, Maria Carolina Elias, Maurício Lacerda Nogueira, Rafael dos Santos Bezerra, Raul Machado Neto, Rejane Maria Tommasini Grotto, Ricardo Haddad, Sandra Coccuzzo Sampaio Vessoni, Simone Kashima, Svetoslav Nanev Slavov, Vincent Louis Viala |
| EPI_ISL_2345091 | NUCLEO SAUDE III N HABITACIONAL PRESIDENTE GEISEL BAURU | Instituto Butantan / ESALQ-Piracicaba | Dimas Tadeu Covas, Antonio Jorge Martins, Claudia Renata dos Santos Barros, David Schlesinger, Debora Botequio Moretti, Elaine Cristina Marqueze, Elaine Vieira Santos, Evandra Strazza Rodrigues, Heidge Fukumasu, Jayme Augusto de Souza-Neto, José Salvatore Leister Patané, Luiz Alcantara, Luiz Lehmann Coutinho, Maria Carolina Elias, Maurício Lacerda Nogueira, Rafael dos Santos Bezerra, Raul Machado Neto, Rejane Maria Tommasini Grotto, Ricardo Haddad, Sandra Coccuzzo Sampaio Vessoni, Simone Kashima, Svetoslav Nanev Slavov, Vincent Louis Viala |
| EPI_ISL_2345092 | UNIDADE DE PRONTO ATENDIMENTO DE VARZEA PAULISTA UPA II | Instituto Butantan / ESALQ-Piracicaba | Dimas Tadeu Covas, Antonio Jorge Martins, Claudia Renata dos Santos Barros, David Schlesinger, Debora Botequio Moretti, Elaine Cristina Marqueze, Elaine Vieira Santos, Evandra Strazza Rodrigues, Heidge Fukumasu, Jayme Augusto de Souza-Neto, José Salvatore Leister Patané, Luiz Alcantara, Luiz Lehmann Coutinho, Maria Carolina Elias, Maurício Lacerda Nogueira, Rafael dos Santos Bezerra, Raul Machado Neto, Rejane Maria Tommasini Grotto, Ricardo Haddad, Sandra Coccuzzo Sampaio Vessoni, Simone Kashima, Svetoslav Nanev Slavov, Vincent Louis Viala |
| EPI_ISL_2345093, EPI_ISL_2345094 | UNIDADE DE PRONTO ATENDIMENTO UPA DRA ANA OLIVIA BENTIVOGLIO | Instituto Butantan / ESALQ-Piracicaba | Dimas Tadeu Covas, Antonio Jorge Martins, Claudia Renata dos Santos Barros, David Schlesinger, Debora Botequio Moretti, Elaine Cristina Marqueze, Elaine Vieira Santos, Evandra Strazza Rodrigues, Heidge Fukumasu, Jayme Augusto de Souza-Neto, José Salvatore Leister Patané, Luiz Alcantara, Luiz Lehmann Coutinho, Maria Carolina Elias, Maurício Lacerda Nogueira, Rafael dos Santos Bezerra, Raul Machado Neto, Rejane Maria Tommasini Grotto, Ricardo Haddad, Sandra Coccuzzo Sampaio Vessoni, Simone Kashima, Svetoslav Nanev Slavov, Vincent Louis Viala |
| EPI_ISL_2345095 | UNIDADE DE PRONTO ATENDIMENTO DE VARZEA PAULISTA UPA II | Instituto Butantan / ESALQ-Piracicaba | Dimas Tadeu Covas, Antonio Jorge Martins, Claudia Renata dos Santos Barros, David Schlesinger, Debora Botequio Moretti, Elaine Cristina Marqueze, Elaine Vieira Santos, Evandra Strazza Rodrigues, Heidge Fukumasu, Jayme Augusto de Souza-Neto, José Salvatore Leister Patané, Luiz Alcantara, Luiz Lehmann Coutinho, Maria Carolina Elias, Maurício Lacerda Nogueira, Rafael dos Santos Bezerra, Raul Machado Neto, Rejane Maria Tommasini Grotto, Ricardo Haddad, Sandra Coccuzzo Sampaio Vessoni, Simone Kashima, Svetoslav Nanev Slavov, Vincent Louis Viala |
| EPI_ISL_2345096 | NUCLEO SAUDE III N HABITACIONAL PRESIDENTE GEISEL BAURU | Instituto Butantan / ESALQ-Piracicaba | Dimas Tadeu Covas, Antonio Jorge Martins, Claudia Renata dos Santos Barros, David Schlesinger, Debora Botequio Moretti, Elaine Cristina Marqueze, Elaine Vieira Santos, Evandra Strazza Rodrigues, Heidge Fukumasu, Jayme Augusto de Souza-Neto, José Salvatore Leister Patané, Luiz Alcantara, Luiz |

| | | | |
|---|---|---|---|
| | | | Lehmann Coutinho, Maria Carolina Elias, Maurício Lacerda Nogueira, Rafael dos Santos Bezerra, Raul Machado Neto, Rejane Maria Tommasini Grotto, Ricardo Haddad, Sandra Coccuzzo Sampaio Vessoni, Simone Kashima, Svetoslav Nanev Slavov, Vincent Louis Viala |
| EPI_ISL_2345097 | UNIDADE DE PRONTO ATENDIMENTO UPA DRA ANA OLIVIA BENTIVOGLIO | Instituto Butantan / ESALQ-Piracicaba | Dimas Tadeu Covas, Antonio Jorge Martins, Claudia Renata dos Santos Barros, David Schlesinger, Debora Botequio Moretti, Elaine Cristina Marqueze, Elaine Vieira Santos, Evandra Strazza Rodrigues, Heidge Fukumasu, Jayme Augusto de Souza-Neto, José Salvatore Leister Patané, Luiz Alcantara, Luiz Lehmann Coutinho, Maria Carolina Elias, Maurício Lacerda Nogueira, Rafael dos Santos Bezerra, Raul Machado Neto, Rejane Maria Tommasini Grotto, Ricardo Haddad, Sandra Coccuzzo Sampaio Vessoni, Simone Kashima, Svetoslav Nanev Slavov, Vincent Louis Viala |
| EPI_ISL_2345098 | UNIDADE DE PRONTO ATENDIMENTO DE VARZEA PAULISTA UPA II | Instituto Butantan / ESALQ-Piracicaba | Dimas Tadeu Covas, Antonio Jorge Martins, Claudia Renata dos Santos Barros, David Schlesinger, Debora Botequio Moretti, Elaine Cristina Marqueze, Elaine Vieira Santos, Evandra Strazza Rodrigues, Heidge Fukumasu, Jayme Augusto de Souza-Neto, José Salvatore Leister Patané, Luiz Alcantara, Luiz Lehmann Coutinho, Maria Carolina Elias, Maurício Lacerda Nogueira, Rafael dos Santos Bezerra, Raul Machado Neto, Rejane Maria Tommasini Grotto, Ricardo Haddad, Sandra Coccuzzo Sampaio Vessoni, Simone Kashima, Svetoslav Nanev Slavov, Vincent Louis Viala |
| EPI_ISL_2345099 | UNIDADE DE PRONTO ATENDIMENTO UPA I SANTA ISABEL | Instituto Butantan / ESALQ-Piracicaba | Dimas Tadeu Covas, Antonio Jorge Martins, Claudia Renata dos Santos Barros, David Schlesinger, Debora Botequio Moretti, Elaine Cristina Marqueze, Elaine Vieira Santos, Evandra Strazza Rodrigues, Heidge Fukumasu, Jayme Augusto de Souza-Neto, José Salvatore Leister Patané, Luiz Alcantara, Luiz Lehmann Coutinho, Maria Carolina Elias, Maurício Lacerda Nogueira, Rafael dos Santos Bezerra, Raul Machado Neto, Rejane Maria Tommasini Grotto, Ricardo Haddad, Sandra Coccuzzo Sampaio Vessoni, Simone Kashima, Svetoslav Nanev Slavov, Vincent Louis Viala |
| EPI_ISL_2345100 | UNIDADE DE PRONTO ATENDIMENTO UPA DRA ANA OLIVIA BENTIVOGLIO | Instituto Butantan / ESALQ-Piracicaba | Dimas Tadeu Covas, Antonio Jorge Martins, Claudia Renata dos Santos Barros, David Schlesinger, Debora Botequio Moretti, Elaine Cristina Marqueze, Elaine Vieira Santos, Evandra Strazza Rodrigues, Heidge Fukumasu, Jayme Augusto de Souza-Neto, José Salvatore Leister Patané, Luiz Alcantara, Luiz Lehmann Coutinho, Maria Carolina Elias, Maurício Lacerda Nogueira, Rafael dos Santos Bezerra, Raul Machado Neto, Rejane Maria Tommasini Grotto, Ricardo Haddad, Sandra Coccuzzo Sampaio Vessoni, Simone Kashima, Svetoslav Nanev Slavov, Vincent Louis Viala |
| EPI_ISL_2345101 | NUCLEO SAUDE III N HABITACIONAL PRESIDENTE GEISEL BAURU | Instituto Butantan / ESALQ-Piracicaba | Dimas Tadeu Covas, Antonio Jorge Martins, Claudia Renata dos Santos Barros, David Schlesinger, Debora Botequio Moretti, Elaine Cristina Marqueze, Elaine Vieira Santos, Evandra Strazza Rodrigues, Heidge Fukumasu, Jayme Augusto de Souza-Neto, José Salvatore Leister Patané, Luiz Alcantara, Luiz Lehmann Coutinho, Maria Carolina Elias, Maurício Lacerda Nogueira, Rafael dos Santos Bezerra, Raul Machado Neto, Rejane Maria Tommasini Grotto, Ricardo Haddad, Sandra Coccuzzo Sampaio Vessoni, Simone Kashima, Svetoslav Nanev Slavov, Vincent Louis Viala |
| EPI_ISL_2345102, EPI_ISL_2345103 | UNIDADE DE PRONTO ATENDIMENTO DE VARZEA PAULISTA UPA II | Instituto Butantan / ESALQ-Piracicaba | Dimas Tadeu Covas, Antonio Jorge Martins, Claudia Renata dos Santos Barros, David Schlesinger, Debora Botequio Moretti, Elaine Cristina Marqueze, Elaine Vieira Santos, Evandra Strazza Rodrigues, Heidge Fukumasu, Jayme Augusto de Souza-Neto, José Salvatore Leister Patané, Luiz Alcantara, Luiz Lehmann Coutinho, Maria Carolina Elias, Maurício Lacerda Nogueira, Rafael dos Santos Bezerra, Raul Machado Neto, Rejane Maria Tommasini Grotto, Ricardo Haddad, Sandra Coccuzzo Sampaio Vessoni, Simone Kashima, Svetoslav Nanev Slavov, Vincent Louis Viala |
| EPI_ISL_2345104 | HOSP MUN JOSANIAS CASTANHA BRAGA | Instituto Butantan / ESALQ-Piracicaba | Dimas Tadeu Covas, Antonio Jorge Martins, Claudia Renata dos Santos Barros, David Schlesinger, Debora Botequio Moretti, Elaine Cristina Marqueze, Elaine Vieira Santos, Evandra Strazza Rodrigues, Heidge Fukumasu, Jayme Augusto de Souza-Neto, José Salvatore Leister Patané, Luiz Alcantara, Luiz Lehmann Coutinho, Maria Carolina Elias, Maurício Lacerda Nogueira, Rafael dos Santos Bezerra, Raul Machado Neto, Rejane Maria Tommasini Grotto, Ricardo Haddad, Sandra Coccuzzo Sampaio Vessoni, Simone Kashima, Svetoslav Nanev Slavov, Vincent Louis Viala |
| EPI_ISL_2345105 | UNIDADE DE PRONTO ATENDIMENTO UPA DRA ANA OLIVIA BENTIVOGLIO | Instituto Butantan / ESALQ-Piracicaba | Dimas Tadeu Covas, Antonio Jorge Martins, Claudia Renata dos Santos Barros, David Schlesinger, Debora Botequio Moretti, Elaine Cristina Marqueze, Elaine Vieira Santos, Evandra Strazza Rodrigues, Heidge Fukumasu, Jayme Augusto de Souza-Neto, José Salvatore Leister Patané, Luiz Alcantara, Luiz Lehmann Coutinho, Maria Carolina Elias, Maurício Lacerda Nogueira, Rafael dos Santos Bezerra, Raul Machado Neto, Rejane Maria Tommasini Grotto, Ricardo Haddad, Sandra Coccuzzo Sampaio Vessoni, Simone Kashima, Svetoslav Nanev Slavov, Vincent Louis Viala |
| EPI_ISL_2345106 | UNIDADE DE PRONTO ATENDIMENTO DE VARZEA PAULISTA UPA II | Instituto Butantan / ESALQ-Piracicaba | Dimas Tadeu Covas, Antonio Jorge Martins, Claudia Renata dos Santos Barros, David Schlesinger, Debora Botequio Moretti, Elaine Cristina Marqueze, Elaine Vieira Santos, Evandra Strazza Rodrigues, Heidge Fukumasu, Jayme Augusto de Souza-Neto, José Salvatore Leister Patané, Luiz Alcantara, Luiz Lehmann Coutinho, Maria Carolina Elias, Maurício Lacerda Nogueira, Rafael dos Santos Bezerra, Raul Machado Neto, Rejane Maria Tommasini Grotto, Ricardo Haddad, Sandra Coccuzzo Sampaio Vessoni, Simone Kashima, Svetoslav Nanev Slavov, Vincent Louis Viala |
| EPI_ISL_2345107, EPI_ISL_2345108 | HOSP MUN JOSANIAS CASTANHA BRAGA | Instituto Butantan / ESALQ-Piracicaba | Dimas Tadeu Covas, Antonio Jorge Martins, Claudia Renata dos Santos Barros, David Schlesinger, Debora Botequio Moretti, Elaine Cristina Marqueze, Elaine Vieira Santos, Evandra Strazza Rodrigues, Heidge Fukumasu, Jayme Augusto de Souza-Neto, José Salvatore Leister Patané, Luiz Alcantara, Luiz Lehmann Coutinho, Maria Carolina Elias, Maurício Lacerda Nogueira, Rafael dos Santos Bezerra, Raul Machado Neto, Rejane Maria Tommasini Grotto, Ricardo Haddad, Sandra Coccuzzo Sampaio Vessoni, Simone Kashima, Svetoslav Nanev Slavov, Vincent Louis Viala |
| EPI_ISL_2345109 | UNIDADE REFERENCIAL SUDOESTE | Instituto Butantan / ESALQ-Piracicaba | Dimas Tadeu Covas, Antonio Jorge Martins, Claudia Renata dos Santos Barros, David Schlesinger, Debora Botequio Moretti, Elaine Cristina Marqueze, Elaine Vieira Santos, Evandra Strazza Rodrigues, Heidge Fukumasu, Jayme Augusto de Souza-Neto, José Salvatore Leister Patané, Luiz Alcantara, Luiz Lehmann Coutinho, Maria Carolina Elias, Maurício Lacerda Nogueira, Rafael dos Santos Bezerra, Raul Machado Neto, Rejane Maria Tommasini Grotto, Ricardo Haddad, Sandra Coccuzzo Sampaio Vessoni, Simone Kashima, Svetoslav Nanev Slavov, Vincent Louis Viala |
| EPI_ISL_2345110 | HOSP MAT 8 DE MAIO | Instituto Butantan / ESALQ-Piracicaba | Dimas Tadeu Covas, Antonio Jorge Martins, Claudia Renata dos Santos Barros, David Schlesinger, Debora Botequio Moretti, Elaine Cristina Marqueze, Elaine Vieira Santos, Evandra Strazza Rodrigues, Heidge Fukumasu, Jayme Augusto de Souza-Neto, José Salvatore Leister Patané, Luiz Alcantara, Luiz Lehmann Coutinho, Maria Carolina Elias, Maurício Lacerda Nogueira, Rafael dos Santos Bezerra, Raul Machado Neto, Rejane Maria Tommasini Grotto, Ricardo Haddad, Sandra Coccuzzo Sampaio Vessoni, Simone Kashima, Svetoslav Nanev Slavov, Vincent Louis Viala |
| EPI_ISL_2345111 | COVID 19 CENTRO DE COMBATE AO CORONAVIRUS CCC JANDIRA | Instituto Butantan / ESALQ-Piracicaba | Dimas Tadeu Covas, Antonio Jorge Martins, Claudia Renata dos Santos Barros, David Schlesinger, Debora Botequio Moretti, Elaine Cristina Marqueze, Elaine Vieira Santos, Evandra Strazza Rodrigues, Heidge Fukumasu, Jayme Augusto de Souza-Neto, José Salvatore Leister Patané, Luiz Alcantara, Luiz Lehmann Coutinho, Maria Carolina Elias, Maurício Lacerda Nogueira, Rafael dos Santos Bezerra, Raul Machado Neto, Rejane Maria Tommasini Grotto, Ricardo Haddad, Sandra Coccuzzo Sampaio Vessoni, Simone Kashima, Svetoslav Nanev Slavov, Vincent Louis Viala |
| EPI_ISL_2345112 | NUCLEO SAUDE III N HABITACIONAL PRESIDENTE GEISEL BAURU | Instituto Butantan / ESALQ-Piracicaba | Dimas Tadeu Covas, Antonio Jorge Martins, Claudia Renata dos Santos Barros, David Schlesinger, Debora Botequio Moretti, Elaine Cristina Marqueze, Elaine Vieira Santos, Evandra Strazza Rodrigues, Heidge Fukumasu, Jayme Augusto de Souza-Neto, José Salvatore Leister Patané, Luiz Alcantara, Luiz Lehmann Coutinho, Maria Carolina Elias, Maurício Lacerda Nogueira, Rafael dos Santos Bezerra, Raul Machado Neto, Rejane Maria Tommasini Grotto, Ricardo Haddad, Sandra Coccuzzo Sampaio Vessoni, Simone Kashima, Svetoslav Nanev Slavov, Vincent Louis Viala |
| EPI_ISL_2345113 | UNIDADE DE PRONTO ATENDIMENTO UPA I SANTA ISABEL | Instituto Butantan / ESALQ-Piracicaba | Dimas Tadeu Covas, Antonio Jorge Martins, Claudia Renata dos Santos Barros, David Schlesinger, Debora Botequio Moretti, Elaine Cristina Marqueze, Elaine Vieira Santos, Evandra Strazza Rodrigues, Heidge Fukumasu, Jayme Augusto de Souza-Neto, José Salvatore Leister Patané, Luiz Alcantara, Luiz Lehmann Coutinho, Maria Carolina Elias, Maurício Lacerda Nogueira, Rafael dos Santos Bezerra, Raul Machado Neto, Rejane Maria Tommasini Grotto, Ricardo Haddad, Sandra Coccuzzo Sampaio Vessoni, Simone Kashima, Svetoslav Nanev Slavov, Vincent Louis Viala |
| EPI_ISL_2345114 | UNIDADE DE PRONTO ATENDIMENTO UPA DRA ANA OLIVIA BENTIVOGLIO | Instituto Butantan / ESALQ-Piracicaba | Dimas Tadeu Covas, Antonio Jorge Martins, Claudia Renata dos Santos Barros, David Schlesinger, Debora Botequio Moretti, Elaine Cristina Marqueze, Elaine Vieira Santos, Evandra Strazza Rodrigues, Heidge Fukumasu, Jayme Augusto de Souza-Neto, José Salvatore Leister Patané, Luiz Alcantara, Luiz Lehmann Coutinho, Maria Carolina Elias, Maurício Lacerda Nogueira, Rafael dos Santos Bezerra, Raul Machado Neto, Rejane Maria Tommasini Grotto, Ricardo Haddad, Sandra Coccuzzo Sampaio Vessoni, Simone Kashima, Svetoslav Nanev Slavov, Vincent Louis Viala |
| EPI_ISL_2345115 | SANTA CASA DE GUARAREMA | Instituto Butantan / ESALQ-Piracicaba | Dimas Tadeu Covas, Antonio Jorge Martins, Claudia Renata dos Santos Barros, David Schlesinger, Debora Botequio Moretti, Elaine Cristina Marqueze, Elaine Vieira Santos, Evandra Strazza Rodrigues, Heidge Fukumasu, Jayme Augusto de Souza-Neto, José Salvatore Leister Patané, Luiz Alcantara, Luiz Lehmann Coutinho, Maria Carolina Elias, Maurício Lacerda Nogueira, Rafael dos Santos Bezerra, Raul Machado Neto, Rejane Maria Tommasini Grotto, Ricardo Haddad, Sandra Coccuzzo Sampaio Vessoni, Simone Kashima, Svetoslav Nanev Slavov, Vincent Louis Viala |
| EPI_ISL_2345116, EPI_ISL_2345117 | SECRETARIA MUNICIPAL DE SAUDE DE JARINU | Instituto Butantan / ESALQ-Piracicaba | Dimas Tadeu Covas, Antonio Jorge Martins, Claudia Renata dos Santos Barros, David Schlesinger, Debora Botequio Moretti, Elaine Cristina Marqueze, Elaine Vieira Santos, Evandra Strazza Rodrigues, Heidge Fukumasu, Jayme Augusto de Souza-Neto, José Salvatore Leister Patané, Luiz Alcantara, Luiz Lehmann Coutinho, Maria Carolina Elias, Maurício Lacerda Nogueira, Rafael dos Santos Bezerra, Raul Machado Neto, Rejane Maria Tommasini Grotto, Ricardo Haddad, Sandra Coccuzzo Sampaio Vessoni, Simone Kashima, Svetoslav Nanev Slavov, Vincent Louis Viala |
| EPI_ISL_2345118 | LABORATORIO MUNICIPAL DE PIRACICABA | Instituto Butantan / ESALQ-Piracicaba | Dimas Tadeu Covas, Antonio Jorge Martins, Claudia Renata dos Santos Barros, David Schlesinger, Debora Botequio Moretti, Elaine Cristina Marqueze, Elaine Vieira Santos, Evandra Strazza Rodrigues, Heidge Fukumasu, Jayme Augusto de Souza-Neto, José Salvatore Leister Patané, Luiz Alcantara, Luiz Lehmann Coutinho, Maria Carolina Elias, Maurício Lacerda Nogueira, Rafael dos Santos Bezerra, Raul Machado Neto, Rejane Maria Tommasini Grotto, Ricardo Haddad, Sandra Coccuzzo Sampaio Vessoni, Simone Kashima, Svetoslav Nanev Slavov, Vincent Louis Viala |

| | | | |
|---|---|---|---|
| EPI_ISL_2345119 | UNIDADE DE PRONTO ATENDIMENTO UPA DRA ANA OLIVIA BENTIVOGLIO | Instituto Butantan / ESALQ-Piracicaba | Dimas Tadeu Covas, Antonio Jorge Martins, Claudia Renata dos Santos Barros, David Schlesinger, Debora Botequio Moretti, Elaine Cristina Marqueze, Elaine Vieira Santos, Evandra Strazza Rodrigues, Heidge Fukumasu, Jayme Augusto de Souza-Neto, José Salvatore Leister Patané, Luiz Alcantara, Luiz Lehmann Coutinho, Maria Carolina Elias, Maurício Lacerda Nogueira, Rafael dos Santos Bezerra, Raul Machado Neto, Rejane Maria Tommasini Grotto, Ricardo Haddad, Sandra Coccuzzo Sampaio Vessoni, Simone Kashima, Svetoslav Nanev Slavov, Vincent Louis Viala |
| EPI_ISL_2345120 | SECRETARIA MUNICIPAL DE SAUDE DE JARINU | Instituto Butantan / ESALQ-Piracicaba | Dimas Tadeu Covas, Antonio Jorge Martins, Claudia Renata dos Santos Barros, David Schlesinger, Debora Botequio Moretti, Elaine Cristina Marqueze, Elaine Vieira Santos, Evandra Strazza Rodrigues, Heidge Fukumasu, Jayme Augusto de Souza-Neto, José Salvatore Leister Patané, Luiz Alcantara, Luiz Lehmann Coutinho, Maria Carolina Elias, Maurício Lacerda Nogueira, Rafael dos Santos Bezerra, Raul Machado Neto, Rejane Maria Tommasini Grotto, Ricardo Haddad, Sandra Coccuzzo Sampaio Vessoni, Simone Kashima, Svetoslav Nanev Slavov, Vincent Louis Viala |
| EPI_ISL_2345121 | SANTA CASA DE GUARAREMA | Instituto Butantan / ESALQ-Piracicaba | Dimas Tadeu Covas, Antonio Jorge Martins, Claudia Renata dos Santos Barros, David Schlesinger, Debora Botequio Moretti, Elaine Cristina Marqueze, Elaine Vieira Santos, Evandra Strazza Rodrigues, Heidge Fukumasu, Jayme Augusto de Souza-Neto, José Salvatore Leister Patané, Luiz Alcantara, Luiz Lehmann Coutinho, Maria Carolina Elias, Maurício Lacerda Nogueira, Rafael dos Santos Bezerra, Raul Machado Neto, Rejane Maria Tommasini Grotto, Ricardo Haddad, Sandra Coccuzzo Sampaio Vessoni, Simone Kashima, Svetoslav Nanev Slavov, Vincent Louis Viala |
| EPI_ISL_2345122 | NUCLEO SAUDE III N HABITACIONAL PRESIDENTE GEISEL BAURU | Instituto Butantan / ESALQ-Piracicaba | Dimas Tadeu Covas, Antonio Jorge Martins, Claudia Renata dos Santos Barros, David Schlesinger, Debora Botequio Moretti, Elaine Cristina Marqueze, Elaine Vieira Santos, Evandra Strazza Rodrigues, Heidge Fukumasu, Jayme Augusto de Souza-Neto, José Salvatore Leister Patané, Luiz Alcantara, Luiz Lehmann Coutinho, Maria Carolina Elias, Maurício Lacerda Nogueira, Rafael dos Santos Bezerra, Raul Machado Neto, Rejane Maria Tommasini Grotto, Ricardo Haddad, Sandra Coccuzzo Sampaio Vessoni, Simone Kashima, Svetoslav Nanev Slavov, Vincent Louis Viala |
| EPI_ISL_2345123 | UNIDADE DE PRONTO ATENDIMENTO UPA I SANTA ISABEL | Instituto Butantan / ESALQ-Piracicaba | Dimas Tadeu Covas, Antonio Jorge Martins, Claudia Renata dos Santos Barros, David Schlesinger, Debora Botequio Moretti, Elaine Cristina Marqueze, Elaine Vieira Santos, Evandra Strazza Rodrigues, Heidge Fukumasu, Jayme Augusto de Souza-Neto, José Salvatore Leister Patané, Luiz Alcantara, Luiz Lehmann Coutinho, Maria Carolina Elias, Maurício Lacerda Nogueira, Rafael dos Santos Bezerra, Raul Machado Neto, Rejane Maria Tommasini Grotto, Ricardo Haddad, Sandra Coccuzzo Sampaio Vessoni, Simone Kashima, Svetoslav Nanev Slavov, Vincent Louis Viala |
| EPI_ISL_2345124 | SANTA CASA DE GUARAREMA | Instituto Butantan / ESALQ-Piracicaba | Dimas Tadeu Covas, Antonio Jorge Martins, Claudia Renata dos Santos Barros, David Schlesinger, Debora Botequio Moretti, Elaine Cristina Marqueze, Elaine Vieira Santos, Evandra Strazza Rodrigues, Heidge Fukumasu, Jayme Augusto de Souza-Neto, José Salvatore Leister Patané, Luiz Alcantara, Luiz Lehmann Coutinho, Maria Carolina Elias, Maurício Lacerda Nogueira, Rafael dos Santos Bezerra, Raul Machado Neto, Rejane Maria Tommasini Grotto, Ricardo Haddad, Sandra Coccuzzo Sampaio Vessoni, Simone Kashima, Svetoslav Nanev Slavov, Vincent Louis Viala |
| EPI_ISL_2345125 | SECRETARIA MUNICIPAL DE SAUDE DE JARINU | Instituto Butantan / ESALQ-Piracicaba | Dimas Tadeu Covas, Antonio Jorge Martins, Claudia Renata dos Santos Barros, David Schlesinger, Debora Botequio Moretti, Elaine Cristina Marqueze, Elaine Vieira Santos, Evandra Strazza Rodrigues, Heidge Fukumasu, Jayme Augusto de Souza-Neto, José Salvatore Leister Patané, Luiz Alcantara, Luiz Lehmann Coutinho, Maria Carolina Elias, Maurício Lacerda Nogueira, Rafael dos Santos Bezerra, Raul Machado Neto, Rejane Maria Tommasini Grotto, Ricardo Haddad, Sandra Coccuzzo Sampaio Vessoni, Simone Kashima, Svetoslav Nanev Slavov, Vincent Louis Viala |
| EPI_ISL_2345126, EPI_ISL_2345127 | UNIDADE DE PRONTO ATENDIMENTO UPA I SANTA ISABEL | Instituto Butantan / ESALQ-Piracicaba | Dimas Tadeu Covas, Antonio Jorge Martins, Claudia Renata dos Santos Barros, David Schlesinger, Debora Botequio Moretti, Elaine Cristina Marqueze, Elaine Vieira Santos, Evandra Strazza Rodrigues, Heidge Fukumasu, Jayme Augusto de Souza-Neto, José Salvatore Leister Patané, Luiz Alcantara, Luiz Lehmann Coutinho, Maria Carolina Elias, Maurício Lacerda Nogueira, Rafael dos Santos Bezerra, Raul Machado Neto, Rejane Maria Tommasini Grotto, Ricardo Haddad, Sandra Coccuzzo Sampaio Vessoni, Simone Kashima, Svetoslav Nanev Slavov, Vincent Louis Viala |
| EPI_ISL_2345128 | ESF COHAB IV | Instituto Butantan / ESALQ-Piracicaba | Dimas Tadeu Covas, Antonio Jorge Martins, Claudia Renata dos Santos Barros, David Schlesinger, Debora Botequio Moretti, Elaine Cristina Marqueze, Elaine Vieira Santos, Evandra Strazza Rodrigues, Heidge Fukumasu, Jayme Augusto de Souza-Neto, José Salvatore Leister Patané, Luiz Alcantara, Luiz Lehmann Coutinho, Maria Carolina Elias, Maurício Lacerda Nogueira, Rafael dos Santos Bezerra, Raul Machado Neto, Rejane Maria Tommasini Grotto, Ricardo Haddad, Sandra Coccuzzo Sampaio Vessoni, Simone Kashima, Svetoslav Nanev Slavov, Vincent Louis Viala |
| EPI_ISL_2345129 | CENTRO MEDICO PMESP | Instituto Butantan / ESALQ-Piracicaba | Dimas Tadeu Covas, Antonio Jorge Martins, Claudia Renata dos Santos Barros, David Schlesinger, Debora Botequio Moretti, Elaine Cristina Marqueze, Elaine Vieira Santos, Evandra Strazza Rodrigues, Heidge Fukumasu, Jayme Augusto de Souza-Neto, José Salvatore Leister Patané, Luiz Alcantara, Luiz Lehmann Coutinho, Maria Carolina Elias, Maurício Lacerda Nogueira, Rafael dos Santos Bezerra, Raul Machado Neto, Rejane Maria Tommasini Grotto, Ricardo Haddad, Sandra Coccuzzo Sampaio Vessoni, Simone Kashima, Svetoslav Nanev Slavov, Vincent Louis Viala |
| EPI_ISL_2345130 | HOSPITAL NOSSA SENHORA APARECIDA | Instituto Butantan / ESALQ-Piracicaba | Dimas Tadeu Covas, Antonio Jorge Martins, Claudia Renata dos Santos Barros, David Schlesinger, Debora Botequio Moretti, Elaine Cristina Marqueze, Elaine Vieira Santos, Evandra Strazza Rodrigues, Heidge Fukumasu, Jayme Augusto de Souza-Neto, José Salvatore Leister Patané, Luiz Alcantara, Luiz Lehmann Coutinho, Maria Carolina Elias, Maurício Lacerda Nogueira, Rafael dos Santos Bezerra, Raul Machado Neto, Rejane Maria Tommasini Grotto, Ricardo Haddad, Sandra Coccuzzo Sampaio Vessoni, Simone Kashima, Svetoslav Nanev Slavov, Vincent Louis Viala |
| EPI_ISL_2345131 | HOSPITAL DR OSIRIS FLORINDO COELHO FERRAZ DE VASCONCELOS | Instituto Butantan / ESALQ-Piracicaba | Dimas Tadeu Covas, Antonio Jorge Martins, Claudia Renata dos Santos Barros, David Schlesinger, Debora Botequio Moretti, Elaine Cristina Marqueze, Elaine Vieira Santos, Evandra Strazza Rodrigues, Heidge Fukumasu, Jayme Augusto de Souza-Neto, José Salvatore Leister Patané, Luiz Alcantara, Luiz Lehmann Coutinho, Maria Carolina Elias, Maurício Lacerda Nogueira, Rafael dos Santos Bezerra, Raul Machado Neto, Rejane Maria Tommasini Grotto, Ricardo Haddad, Sandra Coccuzzo Sampaio Vessoni, Simone Kashima, Svetoslav Nanev Slavov, Vincent Louis Viala |
| EPI_ISL_2345132 | NUCLEO SAUDE III N HABITACIONAL PRESIDENTE GEISEL BAURU | Instituto Butantan / ESALQ-Piracicaba | Dimas Tadeu Covas, Antonio Jorge Martins, Claudia Renata dos Santos Barros, David Schlesinger, Debora Botequio Moretti, Elaine Cristina Marqueze, Elaine Vieira Santos, Evandra Strazza Rodrigues, Heidge Fukumasu, Jayme Augusto de Souza-Neto, José Salvatore Leister Patané, Luiz Alcantara, Luiz Lehmann Coutinho, Maria Carolina Elias, Maurício Lacerda Nogueira, Rafael dos Santos Bezerra, Raul Machado Neto, Rejane Maria Tommasini Grotto, Ricardo Haddad, Sandra Coccuzzo Sampaio Vessoni, Simone Kashima, Svetoslav Nanev Slavov, Vincent Louis Viala |
| EPI_ISL_2345133 | UNIDADE DE PRONTO ATENDIMENTO UPA I SANTA ISABEL | Instituto Butantan / ESALQ-Piracicaba | Dimas Tadeu Covas, Antonio Jorge Martins, Claudia Renata dos Santos Barros, David Schlesinger, Debora Botequio Moretti, Elaine Cristina Marqueze, Elaine Vieira Santos, Evandra Strazza Rodrigues, Heidge Fukumasu, Jayme Augusto de Souza-Neto, José Salvatore Leister Patané, Luiz Alcantara, Luiz Lehmann Coutinho, Maria Carolina Elias, Maurício Lacerda Nogueira, Rafael dos Santos Bezerra, Raul Machado Neto, Rejane Maria Tommasini Grotto, Ricardo Haddad, Sandra Coccuzzo Sampaio Vessoni, Simone Kashima, Svetoslav Nanev Slavov, Vincent Louis Viala |
| EPI_ISL_2345134 | UBS VILA RIBEIRO | Instituto Butantan / ESALQ-Piracicaba | Dimas Tadeu Covas, Antonio Jorge Martins, Claudia Renata dos Santos Barros, David Schlesinger, Debora Botequio Moretti, Elaine Cristina Marqueze, Elaine Vieira Santos, Evandra Strazza Rodrigues, Heidge Fukumasu, Jayme Augusto de Souza-Neto, José Salvatore Leister Patané, Luiz Alcantara, Luiz Lehmann Coutinho, Maria Carolina Elias, Maurício Lacerda Nogueira, Rafael dos Santos Bezerra, Raul Machado Neto, Rejane Maria Tommasini Grotto, Ricardo Haddad, Sandra Coccuzzo Sampaio Vessoni, Simone Kashima, Svetoslav Nanev Slavov, Vincent Louis Viala |
| EPI_ISL_2345135 | UNIDADE DE PRONTO ATENDIMENTO UPA I SANTA ISABEL | Instituto Butantan / ESALQ-Piracicaba | Dimas Tadeu Covas, Antonio Jorge Martins, Claudia Renata dos Santos Barros, David Schlesinger, Debora Botequio Moretti, Elaine Cristina Marqueze, Elaine Vieira Santos, Evandra Strazza Rodrigues, Heidge Fukumasu, Jayme Augusto de Souza-Neto, José Salvatore Leister Patané, Luiz Alcantara, Luiz Lehmann Coutinho, Maria Carolina Elias, Maurício Lacerda Nogueira, Rafael dos Santos Bezerra, Raul Machado Neto, Rejane Maria Tommasini Grotto, Ricardo Haddad, Sandra Coccuzzo Sampaio Vessoni, Simone Kashima, Svetoslav Nanev Slavov, Vincent Louis Viala |
| EPI_ISL_2345136 | UNIDADE REFERENCIAL SUDOESTE | Instituto Butantan / ESALQ-Piracicaba | Dimas Tadeu Covas, Antonio Jorge Martins, Claudia Renata dos Santos Barros, David Schlesinger, Debora Botequio Moretti, Elaine Cristina Marqueze, Elaine Vieira Santos, Evandra Strazza Rodrigues, Heidge Fukumasu, Jayme Augusto de Souza-Neto, José Salvatore Leister Patané, Luiz Alcantara, Luiz Lehmann Coutinho, Maria Carolina Elias, Maurício Lacerda Nogueira, Rafael dos Santos Bezerra, Raul Machado Neto, Rejane Maria Tommasini Grotto, Ricardo Haddad, Sandra Coccuzzo Sampaio Vessoni, Simone Kashima, Svetoslav Nanev Slavov, Vincent Louis Viala |
| EPI_ISL_2345137 | UBS JARDIM PARANA | Instituto Butantan / ESALQ-Piracicaba | Dimas Tadeu Covas, Antonio Jorge Martins, Claudia Renata dos Santos Barros, David Schlesinger, Debora Botequio Moretti, Elaine Cristina Marqueze, Elaine Vieira Santos, Evandra Strazza Rodrigues, Heidge Fukumasu, Jayme Augusto de Souza-Neto, José Salvatore Leister Patané, Luiz Alcantara, Luiz Lehmann Coutinho, Maria Carolina Elias, Maurício Lacerda Nogueira, Rafael dos Santos Bezerra, Raul Machado Neto, Rejane Maria Tommasini Grotto, Ricardo Haddad, Sandra Coccuzzo Sampaio Vessoni, Simone Kashima, Svetoslav Nanev Slavov, Vincent Louis Viala |
| EPI_ISL_2345138 | NUCLEO SAUDE III N HABITACIONAL PRESIDENTE GEISEL BAURU | Instituto Butantan / ESALQ-Piracicaba | Dimas Tadeu Covas, Antonio Jorge Martins, Claudia Renata dos Santos Barros, David Schlesinger, Debora Botequio Moretti, Elaine Cristina Marqueze, Elaine Vieira Santos, Evandra Strazza Rodrigues, Heidge Fukumasu, Jayme Augusto de Souza-Neto, José Salvatore Leister Patané, Luiz Alcantara, Luiz Lehmann Coutinho, Maria Carolina Elias, Maurício Lacerda Nogueira, Rafael dos Santos Bezerra, Raul Machado Neto, Rejane Maria Tommasini Grotto, Ricardo Haddad, Sandra Coccuzzo Sampaio Vessoni, Simone Kashima, Svetoslav Nanev Slavov, Vincent Louis Viala |
| EPI_ISL_2345139 | UNIDADE REFERENCIAL SUDOESTE | Instituto Butantan / ESALQ-Piracicaba | Dimas Tadeu Covas, Antonio Jorge Martins, Claudia Renata dos Santos Barros, David Schlesinger, Debora Botequio Moretti, Elaine Cristina Marqueze, Elaine Vieira Santos, Evandra Strazza Rodrigues, Heidge Fukumasu, Jayme Augusto de Souza-Neto, José Salvatore Leister Patané, Luiz Alcantara, Luiz |

| | | | |
|---|---|---|---|
| | | | Lehmann Coutinho, Maria Carolina Elias, Maurício Lacerda Nogueira, Rafael dos Santos Bezerra, Raul Machado Neto, Rejane Maria Tommasini Grotto, Ricardo Haddad, Sandra Coccuzzo Sampaio Vessoni, Simone Kashima, Svetoslav Nanev Slavov, Vincent Louis Viala |
| EPI_ISL_2345140 | UBS JARDIM PARANA | Instituto Butantan / ESALQ-Piracicaba | Dimas Tadeu Covas, Antonio Jorge Martins, Claudia Renata dos Santos Barros, David Schlesinger, Debora Botequio Moretti, Elaine Cristina Marqueze, Elaine Vieira Santos, Evandra Strazza Rodrigues, Heidge Fukumasu, Jayme Augusto de Souza-Neto, José Salvatore Leister Patané, Luiz Alcantara, Luiz Lehmann Coutinho, Maria Carolina Elias, Maurício Lacerda Nogueira, Rafael dos Santos Bezerra, Raul Machado Neto, Rejane Maria Tommasini Grotto, Ricardo Haddad, Sandra Coccuzzo Sampaio Vessoni, Simone Kashima, Svetoslav Nanev Slavov, Vincent Louis Viala |
| EPI_ISL_2345141, EPI_ISL_2345142 | LABORATORIO LOCAL DE ITAPECERICA DA SERRA | Instituto Butantan / ESALQ-Piracicaba | Dimas Tadeu Covas, Antonio Jorge Martins, Claudia Renata dos Santos Barros, David Schlesinger, Debora Botequio Moretti, Elaine Cristina Marqueze, Elaine Vieira Santos, Evandra Strazza Rodrigues, Heidge Fukumasu, Jayme Augusto de Souza-Neto, José Salvatore Leister Patané, Luiz Alcantara, Luiz Lehmann Coutinho, Maria Carolina Elias, Maurício Lacerda Nogueira, Rafael dos Santos Bezerra, Raul Machado Neto, Rejane Maria Tommasini Grotto, Ricardo Haddad, Sandra Coccuzzo Sampaio Vessoni, Simone Kashima, Svetoslav Nanev Slavov, Vincent Louis Viala |
| EPI_ISL_2345143 | HOSPITAL GERAL JESUS TEIXEIRA DA COSTA GUAIANASES SAO PAULO | Instituto Butantan / ESALQ-Piracicaba | Dimas Tadeu Covas, Antonio Jorge Martins, Claudia Renata dos Santos Barros, David Schlesinger, Debora Botequio Moretti, Elaine Cristina Marqueze, Elaine Vieira Santos, Evandra Strazza Rodrigues, Heidge Fukumasu, Jayme Augusto de Souza-Neto, José Salvatore Leister Patané, Luiz Alcantara, Luiz Lehmann Coutinho, Maria Carolina Elias, Maurício Lacerda Nogueira, Rafael dos Santos Bezerra, Raul Machado Neto, Rejane Maria Tommasini Grotto, Ricardo Haddad, Sandra Coccuzzo Sampaio Vessoni, Simone Kashima, Svetoslav Nanev Slavov, Vincent Louis Viala |
| EPI_ISL_2345144, EPI_ISL_2345145, EPI_ISL_2345146, EPI_ISL_2345147, EPI_ISL_2345148 | NUCLEO SAUDE III N HABITACIONAL PRESIDENTE GEISEL BAURU | Instituto Butantan / ESALQ-Piracicaba | Dimas Tadeu Covas, Antonio Jorge Martins, Claudia Renata dos Santos Barros, David Schlesinger, Debora Botequio Moretti, Elaine Cristina Marqueze, Elaine Vieira Santos, Evandra Strazza Rodrigues, Heidge Fukumasu, Jayme Augusto de Souza-Neto, José Salvatore Leister Patané, Luiz Alcantara, Luiz Lehmann Coutinho, Maria Carolina Elias, Maurício Lacerda Nogueira, Rafael dos Santos Bezerra, Raul Machado Neto, Rejane Maria Tommasini Grotto, Ricardo Haddad, Sandra Coccuzzo Sampaio Vessoni, Simone Kashima, Svetoslav Nanev Slavov, Vincent Louis Viala |
| EPI_ISL_2345149, EPI_ISL_2345150 | LABORATORIO LOCAL DE ITAPECERICA DA SERRA | Instituto Butantan / ESALQ-Piracicaba | Dimas Tadeu Covas, Antonio Jorge Martins, Claudia Renata dos Santos Barros, David Schlesinger, Debora Botequio Moretti, Elaine Cristina Marqueze, Elaine Vieira Santos, Evandra Strazza Rodrigues, Heidge Fukumasu, Jayme Augusto de Souza-Neto, José Salvatore Leister Patané, Luiz Alcantara, Luiz Lehmann Coutinho, Maria Carolina Elias, Maurício Lacerda Nogueira, Rafael dos Santos Bezerra, Raul Machado Neto, Rejane Maria Tommasini Grotto, Ricardo Haddad, Sandra Coccuzzo Sampaio Vessoni, Simone Kashima, Svetoslav Nanev Slavov, Vincent Louis Viala |
| EPI_ISL_2345151 | NUCLEO SAUDE III N HABITACIONAL PRESIDENTE GEISEL BAURU | Instituto Butantan / ESALQ-Piracicaba | Dimas Tadeu Covas, Antonio Jorge Martins, Claudia Renata dos Santos Barros, David Schlesinger, Debora Botequio Moretti, Elaine Cristina Marqueze, Elaine Vieira Santos, Evandra Strazza Rodrigues, Heidge Fukumasu, Jayme Augusto de Souza-Neto, José Salvatore Leister Patané, Luiz Alcantara, Luiz Lehmann Coutinho, Maria Carolina Elias, Maurício Lacerda Nogueira, Rafael dos Santos Bezerra, Raul Machado Neto, Rejane Maria Tommasini Grotto, Ricardo Haddad, Sandra Coccuzzo Sampaio Vessoni, Simone Kashima, Svetoslav Nanev Slavov, Vincent Louis Viala |
| EPI_ISL_2345152 | LABORATORIO LOCAL DE ITAPECERICA DA SERRA | Instituto Butantan / ESALQ-Piracicaba | Dimas Tadeu Covas, Antonio Jorge Martins, Claudia Renata dos Santos Barros, David Schlesinger, Debora Botequio Moretti, Elaine Cristina Marqueze, Elaine Vieira Santos, Evandra Strazza Rodrigues, Heidge Fukumasu, Jayme Augusto de Souza-Neto, José Salvatore Leister Patané, Luiz Alcantara, Luiz Lehmann Coutinho, Maria Carolina Elias, Maurício Lacerda Nogueira, Rafael dos Santos Bezerra, Raul Machado Neto, Rejane Maria Tommasini Grotto, Ricardo Haddad, Sandra Coccuzzo Sampaio Vessoni, Simone Kashima, Svetoslav Nanev Slavov, Vincent Louis Viala |
| EPI_ISL_2345154, EPI_ISL_2345155 | UBS BONFIM | Instituto Butantan / ESALQ-Piracicaba | Dimas Tadeu Covas, Antonio Jorge Martins, Claudia Renata dos Santos Barros, David Schlesinger, Debora Botequio Moretti, Elaine Cristina Marqueze, Elaine Vieira Santos, Evandra Strazza Rodrigues, Heidge Fukumasu, Jayme Augusto de Souza-Neto, José Salvatore Leister Patané, Luiz Alcantara, Luiz Lehmann Coutinho, Maria Carolina Elias, Maurício Lacerda Nogueira, Rafael dos Santos Bezerra, Raul Machado Neto, Rejane Maria Tommasini Grotto, Ricardo Haddad, Sandra Coccuzzo Sampaio Vessoni, Simone Kashima, Svetoslav Nanev Slavov, Vincent Louis Viala |
| EPI_ISL_2345156 | UBS DR ALFREDO DANTAS DE SOUZA UMUARAMA | Instituto Butantan / ESALQ-Piracicaba | Dimas Tadeu Covas, Antonio Jorge Martins, Claudia Renata dos Santos Barros, David Schlesinger, Debora Botequio Moretti, Elaine Cristina Marqueze, Elaine Vieira Santos, Evandra Strazza Rodrigues, Heidge Fukumasu, Jayme Augusto de Souza-Neto, José Salvatore Leister Patané, Luiz Alcantara, Luiz Lehmann Coutinho, Maria Carolina Elias, Maurício Lacerda Nogueira, Rafael dos Santos Bezerra, Raul Machado Neto, Rejane Maria Tommasini Grotto, Ricardo Haddad, Sandra Coccuzzo Sampaio Vessoni, Simone Kashima, Svetoslav Nanev Slavov, Vincent Louis Viala |
| EPI_ISL_2345157 | UBS BONFIM | Instituto Butantan / ESALQ-Piracicaba | Dimas Tadeu Covas, Antonio Jorge Martins, Claudia Renata dos Santos Barros, David Schlesinger, Debora Botequio Moretti, Elaine Cristina Marqueze, Elaine Vieira Santos, Evandra Strazza Rodrigues, Heidge Fukumasu, Jayme Augusto de Souza-Neto, José Salvatore Leister Patané, Luiz Alcantara, Luiz Lehmann Coutinho, Maria Carolina Elias, Maurício Lacerda Nogueira, Rafael dos Santos Bezerra, Raul Machado Neto, Rejane Maria Tommasini Grotto, Ricardo Haddad, Sandra Coccuzzo Sampaio Vessoni, Simone Kashima, Svetoslav Nanev Slavov, Vincent Louis Viala |
| EPI_ISL_2345158, EPI_ISL_2345159 | UBS DR ALFREDO DANTAS DE SOUZA UMUARAMA | Instituto Butantan / ESALQ-Piracicaba | Dimas Tadeu Covas, Antonio Jorge Martins, Claudia Renata dos Santos Barros, David Schlesinger, Debora Botequio Moretti, Elaine Cristina Marqueze, Elaine Vieira Santos, Evandra Strazza Rodrigues, Heidge Fukumasu, Jayme Augusto de Souza-Neto, José Salvatore Leister Patané, Luiz Alcantara, Luiz Lehmann Coutinho, Maria Carolina Elias, Maurício Lacerda Nogueira, Rafael dos Santos Bezerra, Raul Machado Neto, Rejane Maria Tommasini Grotto, Ricardo Haddad, Sandra Coccuzzo Sampaio Vessoni, Simone Kashima, Svetoslav Nanev Slavov, Vincent Louis Viala |
| EPI_ISL_2345160 | UNIDADE REFERENCIAL SUDOESTE | Instituto Butantan / ESALQ-Piracicaba | Dimas Tadeu Covas, Antonio Jorge Martins, Claudia Renata dos Santos Barros, David Schlesinger, Debora Botequio Moretti, Elaine Cristina Marqueze, Elaine Vieira Santos, Evandra Strazza Rodrigues, Heidge Fukumasu, Jayme Augusto de Souza-Neto, José Salvatore Leister Patané, Luiz Alcantara, Luiz Lehmann Coutinho, Maria Carolina Elias, Maurício Lacerda Nogueira, Rafael dos Santos Bezerra, Raul Machado Neto, Rejane Maria Tommasini Grotto, Ricardo Haddad, Sandra Coccuzzo Sampaio Vessoni, Simone Kashima, Svetoslav Nanev Slavov, Vincent Louis Viala |
| EPI_ISL_2345161 | LABORATORIO MUNICIPAL DE PIRACICABA | Instituto Butantan / ESALQ-Piracicaba | Dimas Tadeu Covas, Antonio Jorge Martins, Claudia Renata dos Santos Barros, David Schlesinger, Debora Botequio Moretti, Elaine Cristina Marqueze, Elaine Vieira Santos, Evandra Strazza Rodrigues, Heidge Fukumasu, Jayme Augusto de Souza-Neto, José Salvatore Leister Patané, Luiz Alcantara, Luiz Lehmann Coutinho, Maria Carolina Elias, Maurício Lacerda Nogueira, Rafael dos Santos Bezerra, Raul Machado Neto, Rejane Maria Tommasini Grotto, Ricardo Haddad, Sandra Coccuzzo Sampaio Vessoni, Simone Kashima, Svetoslav Nanev Slavov, Vincent Louis Viala |
| EPI_ISL_2345162, EPI_ISL_2345163 | UBS DR ALFREDO DANTAS DE SOUZA UMUARAMA | Instituto Butantan / ESALQ-Piracicaba | Dimas Tadeu Covas, Antonio Jorge Martins, Claudia Renata dos Santos Barros, David Schlesinger, Debora Botequio Moretti, Elaine Cristina Marqueze, Elaine Vieira Santos, Evandra Strazza Rodrigues, Heidge Fukumasu, Jayme Augusto de Souza-Neto, José Salvatore Leister Patané, Luiz Alcantara, Luiz Lehmann Coutinho, Maria Carolina Elias, Maurício Lacerda Nogueira, Rafael dos Santos Bezerra, Raul Machado Neto, Rejane Maria Tommasini Grotto, Ricardo Haddad, Sandra Coccuzzo Sampaio Vessoni, Simone Kashima, Svetoslav Nanev Slavov, Vincent Louis Viala |
| EPI_ISL_2345164 | NUCLEO SAUDE III N HABITACIONAL PRESIDENTE GEISEL BAURU | Instituto Butantan / ESALQ-Piracicaba | Dimas Tadeu Covas, Antonio Jorge Martins, Claudia Renata dos Santos Barros, David Schlesinger, Debora Botequio Moretti, Elaine Cristina Marqueze, Elaine Vieira Santos, Evandra Strazza Rodrigues, Heidge Fukumasu, Jayme Augusto de Souza-Neto, José Salvatore Leister Patané, Luiz Alcantara, Luiz Lehmann Coutinho, Maria Carolina Elias, Maurício Lacerda Nogueira, Rafael dos Santos Bezerra, Raul Machado Neto, Rejane Maria Tommasini Grotto, Ricardo Haddad, Sandra Coccuzzo Sampaio Vessoni, Simone Kashima, Svetoslav Nanev Slavov, Vincent Louis Viala |
| EPI_ISL_2345165 | NUCLEO DE ESPECIALIDADES EM SAUDE | Instituto Butantan / ESALQ-Piracicaba | Dimas Tadeu Covas, Antonio Jorge Martins, Claudia Renata dos Santos Barros, David Schlesinger, Debora Botequio Moretti, Elaine Cristina Marqueze, Elaine Vieira Santos, Evandra Strazza Rodrigues, Heidge Fukumasu, Jayme Augusto de Souza-Neto, José Salvatore Leister Patané, Luiz Alcantara, Luiz Lehmann Coutinho, Maria Carolina Elias, Maurício Lacerda Nogueira, Rafael dos Santos Bezerra, Raul Machado Neto, Rejane Maria Tommasini Grotto, Ricardo Haddad, Sandra Coccuzzo Sampaio Vessoni, Simone Kashima, Svetoslav Nanev Slavov, Vincent Louis Viala |
| EPI_ISL_2345166, EPI_ISL_2345167, EPI_ISL_2345168 | UBS DR ALFREDO DANTAS DE SOUZA UMUARAMA | Instituto Butantan / ESALQ-Piracicaba | Dimas Tadeu Covas, Antonio Jorge Martins, Claudia Renata dos Santos Barros, David Schlesinger, Debora Botequio Moretti, Elaine Cristina Marqueze, Elaine Vieira Santos, Evandra Strazza Rodrigues, Heidge Fukumasu, Jayme Augusto de Souza-Neto, José Salvatore Leister Patané, Luiz Alcantara, Luiz Lehmann Coutinho, Maria Carolina Elias, Maurício Lacerda Nogueira, Rafael dos Santos Bezerra, Raul Machado Neto, Rejane Maria Tommasini Grotto, Ricardo Haddad, Sandra Coccuzzo Sampaio Vessoni, Simone Kashima, Svetoslav Nanev Slavov, Vincent Louis Viala |
| EPI_ISL_2345169, EPI_ISL_2345170 | LABORATORIO MUNICIPAL DE PIRACICABA | Instituto Butantan / ESALQ-Piracicaba | Dimas Tadeu Covas, Antonio Jorge Martins, Claudia Renata dos Santos Barros, David Schlesinger, Debora Botequio Moretti, Elaine Cristina Marqueze, Elaine Vieira Santos, Evandra Strazza Rodrigues, Heidge Fukumasu, Jayme Augusto de Souza-Neto, José Salvatore Leister Patané, Luiz Alcantara, Luiz Lehmann Coutinho, Maria Carolina Elias, Maurício Lacerda Nogueira, Rafael dos Santos Bezerra, Raul Machado Neto, Rejane Maria Tommasini Grotto, Ricardo Haddad, Sandra Coccuzzo Sampaio Vessoni, Simone Kashima, Svetoslav Nanev Slavov, Vincent Louis Viala |
| EPI_ISL_2345171 | CSII DR JOSE FERREIRA TELLES | Instituto Butantan / ESALQ-Piracicaba | Dimas Tadeu Covas, Antonio Jorge Martins, Claudia Renata dos Santos Barros, David Schlesinger, Debora Botequio Moretti, Elaine Cristina Marqueze, Elaine Vieira Santos, Evandra Strazza Rodrigues, Heidge Fukumasu, Jayme Augusto de Souza-Neto, José Salvatore Leister Patané, Luiz Alcantara, Luiz Lehmann Coutinho, Maria Carolina Elias, Maurício Lacerda Nogueira, Rafael dos Santos Bezerra, Raul Machado Neto, Rejane Maria Tommasini Grotto, Ricardo Haddad, Sandra Coccuzzo Sampaio Vessoni, Simone Kashima, Svetoslav Nanev Slavov, Vincent Louis Viala |

| | | | |
|---|---|---|---|
| EPI_ISL_2345172 | NUCLEO SAUDE III N HABITACIONAL PRESIDENTE GEISEL BAURU | Instituto Butantan / ESALQ-Piracicaba | Dimas Tadeu Covas, Antonio Jorge Martins, Claudia Renata dos Santos Barros, David Schlesinger, Debora Botequio Moretti, Elaine Cristina Marqueze, Elaine Vieira Santos, Evandra Strazza Rodrigues, Heidge Fukumasu, Jayme Augusto de Souza-Neto, José Salvatore Leister Patané, Luiz Alcantara, Luiz Lehmann Coutinho, Maria Carolina Elias, Maurício Lacerda Nogueira, Rafael dos Santos Bezerra, Raul Machado Neto, Rejane Maria Tommasini Grotto, Ricardo Haddad, Sandra Coccuzzo Sampaio Vessoni, Simone Kashima, Svetoslav Nanev Slavov, Vincent Louis Viala |
| EPI_ISL_2345173 | UBS DR ALFREDO DANTAS DE SOUZA UMUARAMA | Instituto Butantan / ESALQ-Piracicaba | Dimas Tadeu Covas, Antonio Jorge Martins, Claudia Renata dos Santos Barros, David Schlesinger, Debora Botequio Moretti, Elaine Cristina Marqueze, Elaine Vieira Santos, Evandra Strazza Rodrigues, Heidge Fukumasu, Jayme Augusto de Souza-Neto, José Salvatore Leister Patané, Luiz Alcantara, Luiz Lehmann Coutinho, Maria Carolina Elias, Maurício Lacerda Nogueira, Rafael dos Santos Bezerra, Raul Machado Neto, Rejane Maria Tommasini Grotto, Ricardo Haddad, Sandra Coccuzzo Sampaio Vessoni, Simone Kashima, Svetoslav Nanev Slavov, Vincent Louis Viala |
| EPI_ISL_2345174 | NUCLEO DE ESPECIALIDADES EM SAUDE | Instituto Butantan / ESALQ-Piracicaba | Dimas Tadeu Covas, Antonio Jorge Martins, Claudia Renata dos Santos Barros, David Schlesinger, Debora Botequio Moretti, Elaine Cristina Marqueze, Elaine Vieira Santos, Evandra Strazza Rodrigues, Heidge Fukumasu, Jayme Augusto de Souza-Neto, José Salvatore Leister Patané, Luiz Alcantara, Luiz Lehmann Coutinho, Maria Carolina Elias, Maurício Lacerda Nogueira, Rafael dos Santos Bezerra, Raul Machado Neto, Rejane Maria Tommasini Grotto, Ricardo Haddad, Sandra Coccuzzo Sampaio Vessoni, Simone Kashima, Svetoslav Nanev Slavov, Vincent Louis Viala |
| EPI_ISL_2345175 | UBS DR ALFREDO DANTAS DE SOUZA UMUARAMA | Instituto Butantan / ESALQ-Piracicaba | Dimas Tadeu Covas, Antonio Jorge Martins, Claudia Renata dos Santos Barros, David Schlesinger, Debora Botequio Moretti, Elaine Cristina Marqueze, Elaine Vieira Santos, Evandra Strazza Rodrigues, Heidge Fukumasu, Jayme Augusto de Souza-Neto, José Salvatore Leister Patané, Luiz Alcantara, Luiz Lehmann Coutinho, Maria Carolina Elias, Maurício Lacerda Nogueira, Rafael dos Santos Bezerra, Raul Machado Neto, Rejane Maria Tommasini Grotto, Ricardo Haddad, Sandra Coccuzzo Sampaio Vessoni, Simone Kashima, Svetoslav Nanev Slavov, Vincent Louis Viala |
| EPI_ISL_2345176, EPI_ISL_2345177 | NUCLEO DE ESPECIALIDADES EM SAUDE | Instituto Butantan / ESALQ-Piracicaba | Dimas Tadeu Covas, Antonio Jorge Martins, Claudia Renata dos Santos Barros, David Schlesinger, Debora Botequio Moretti, Elaine Cristina Marqueze, Elaine Vieira Santos, Evandra Strazza Rodrigues, Heidge Fukumasu, Jayme Augusto de Souza-Neto, José Salvatore Leister Patané, Luiz Alcantara, Luiz Lehmann Coutinho, Maria Carolina Elias, Maurício Lacerda Nogueira, Rafael dos Santos Bezerra, Raul Machado Neto, Rejane Maria Tommasini Grotto, Ricardo Haddad, Sandra Coccuzzo Sampaio Vessoni, Simone Kashima, Svetoslav Nanev Slavov, Vincent Louis Viala |
| EPI_ISL_2345179 | UBS DR ALFREDO DANTAS DE SOUZA UMUARAMA | Instituto Butantan / ESALQ-Piracicaba | Dimas Tadeu Covas, Antonio Jorge Martins, Claudia Renata dos Santos Barros, David Schlesinger, Debora Botequio Moretti, Elaine Cristina Marqueze, Elaine Vieira Santos, Evandra Strazza Rodrigues, Heidge Fukumasu, Jayme Augusto de Souza-Neto, José Salvatore Leister Patané, Luiz Alcantara, Luiz Lehmann Coutinho, Maria Carolina Elias, Maurício Lacerda Nogueira, Rafael dos Santos Bezerra, Raul Machado Neto, Rejane Maria Tommasini Grotto, Ricardo Haddad, Sandra Coccuzzo Sampaio Vessoni, Simone Kashima, Svetoslav Nanev Slavov, Vincent Louis Viala |
| EPI_ISL_2345180, EPI_ISL_2345181, EPI_ISL_2345182 | NUCLEO SAUDE III N HABITACIONAL PRESIDENTE GEISEL BAURU | Instituto Butantan / ESALQ-Piracicaba | Dimas Tadeu Covas, Antonio Jorge Martins, Claudia Renata dos Santos Barros, David Schlesinger, Debora Botequio Moretti, Elaine Cristina Marqueze, Elaine Vieira Santos, Evandra Strazza Rodrigues, Heidge Fukumasu, Jayme Augusto de Souza-Neto, José Salvatore Leister Patané, Luiz Alcantara, Luiz Lehmann Coutinho, Maria Carolina Elias, Maurício Lacerda Nogueira, Rafael dos Santos Bezerra, Raul Machado Neto, Rejane Maria Tommasini Grotto, Ricardo Haddad, Sandra Coccuzzo Sampaio Vessoni, Simone Kashima, Svetoslav Nanev Slavov, Vincent Louis Viala |
| EPI_ISL_2345183, EPI_ISL_2345184, EPI_ISL_2345185 | PRONTO SOCORRO MUNICIPAL DE SEVERINIA | Instituto Butantan / ESALQ-Piracicaba | Dimas Tadeu Covas, Antonio Jorge Martins, Claudia Renata dos Santos Barros, David Schlesinger, Debora Botequio Moretti, Elaine Cristina Marqueze, Elaine Vieira Santos, Evandra Strazza Rodrigues, Heidge Fukumasu, Jayme Augusto de Souza-Neto, José Salvatore Leister Patané, Luiz Alcantara, Luiz Lehmann Coutinho, Maria Carolina Elias, Maurício Lacerda Nogueira, Rafael dos Santos Bezerra, Raul Machado Neto, Rejane Maria Tommasini Grotto, Ricardo Haddad, Sandra Coccuzzo Sampaio Vessoni, Simone Kashima, Svetoslav Nanev Slavov, Vincent Louis Viala |
| EPI_ISL_2345186 | LABORATORIO MUNICIPAL DE PIRACICABA | Instituto Butantan / ESALQ-Piracicaba | Dimas Tadeu Covas, Antonio Jorge Martins, Claudia Renata dos Santos Barros, David Schlesinger, Debora Botequio Moretti, Elaine Cristina Marqueze, Elaine Vieira Santos, Evandra Strazza Rodrigues, Heidge Fukumasu, Jayme Augusto de Souza-Neto, José Salvatore Leister Patané, Luiz Alcantara, Luiz Lehmann Coutinho, Maria Carolina Elias, Maurício Lacerda Nogueira, Rafael dos Santos Bezerra, Raul Machado Neto, Rejane Maria Tommasini Grotto, Ricardo Haddad, Sandra Coccuzzo Sampaio Vessoni, Simone Kashima, Svetoslav Nanev Slavov, Vincent Louis Viala |
| EPI_ISL_2345187, EPI_ISL_2345188 | CENTRO DE SAUDE III DE BOCAINA | Instituto Butantan / ESALQ-Piracicaba | Dimas Tadeu Covas, Antonio Jorge Martins, Claudia Renata dos Santos Barros, David Schlesinger, Debora Botequio Moretti, Elaine Cristina Marqueze, Elaine Vieira Santos, Evandra Strazza Rodrigues, Heidge Fukumasu, Jayme Augusto de Souza-Neto, José Salvatore Leister Patané, Luiz Alcantara, Luiz Lehmann Coutinho, Maria Carolina Elias, Maurício Lacerda Nogueira, Rafael dos Santos Bezerra, Raul Machado Neto, Rejane Maria Tommasini Grotto, Ricardo Haddad, Sandra Coccuzzo Sampaio Vessoni, Simone Kashima, Svetoslav Nanev Slavov, Vincent Louis Viala |
| EPI_ISL_2345190, EPI_ISL_2345191 | UBS DR ABELARDO PINHEIRO GUIMARAES SCRPARDO | Instituto Butantan / ESALQ-Piracicaba | Dimas Tadeu Covas, Antonio Jorge Martins, Claudia Renata dos Santos Barros, David Schlesinger, Debora Botequio Moretti, Elaine Cristina Marqueze, Elaine Vieira Santos, Evandra Strazza Rodrigues, Heidge Fukumasu, Jayme Augusto de Souza-Neto, José Salvatore Leister Patané, Luiz Alcantara, Luiz Lehmann Coutinho, Maria Carolina Elias, Maurício Lacerda Nogueira, Rafael dos Santos Bezerra, Raul Machado Neto, Rejane Maria Tommasini Grotto, Ricardo Haddad, Sandra Coccuzzo Sampaio Vessoni, Simone Kashima, Svetoslav Nanev Slavov, Vincent Louis Viala |
| EPI_ISL_2345192 | CENTRO DE SAUDE DE OSVALDO CRUZ | Instituto Butantan / ESALQ-Piracicaba | Dimas Tadeu Covas, Antonio Jorge Martins, Claudia Renata dos Santos Barros, David Schlesinger, Debora Botequio Moretti, Elaine Cristina Marqueze, Elaine Vieira Santos, Evandra Strazza Rodrigues, Heidge Fukumasu, Jayme Augusto de Souza-Neto, José Salvatore Leister Patané, Luiz Alcantara, Luiz Lehmann Coutinho, Maria Carolina Elias, Maurício Lacerda Nogueira, Rafael dos Santos Bezerra, Raul Machado Neto, Rejane Maria Tommasini Grotto, Ricardo Haddad, Sandra Coccuzzo Sampaio Vessoni, Simone Kashima, Svetoslav Nanev Slavov, Vincent Louis Viala |
| EPI_ISL_2345193 | UBS III MURUTINGA DO SUL | Instituto Butantan / ESALQ-Piracicaba | Dimas Tadeu Covas, Antonio Jorge Martins, Claudia Renata dos Santos Barros, David Schlesinger, Debora Botequio Moretti, Elaine Cristina Marqueze, Elaine Vieira Santos, Evandra Strazza Rodrigues, Heidge Fukumasu, Jayme Augusto de Souza-Neto, José Salvatore Leister Patané, Luiz Alcantara, Luiz Lehmann Coutinho, Maria Carolina Elias, Maurício Lacerda Nogueira, Rafael dos Santos Bezerra, Raul Machado Neto, Rejane Maria Tommasini Grotto, Ricardo Haddad, Sandra Coccuzzo Sampaio Vessoni, Simone Kashima, Svetoslav Nanev Slavov, Vincent Louis Viala |
| EPI_ISL_2345194 | CS II DR MIGUEL VITALIANO ORLANDIA | Instituto Butantan / ESALQ-Piracicaba | Dimas Tadeu Covas, Antonio Jorge Martins, Claudia Renata dos Santos Barros, David Schlesinger, Debora Botequio Moretti, Elaine Cristina Marqueze, Elaine Vieira Santos, Evandra Strazza Rodrigues, Heidge Fukumasu, Jayme Augusto de Souza-Neto, José Salvatore Leister Patané, Luiz Alcantara, Luiz Lehmann Coutinho, Maria Carolina Elias, Maurício Lacerda Nogueira, Rafael dos Santos Bezerra, Raul Machado Neto, Rejane Maria Tommasini Grotto, Ricardo Haddad, Sandra Coccuzzo Sampaio Vessoni, Simone Kashima, Svetoslav Nanev Slavov, Vincent Louis Viala |
| EPI_ISL_2345195 | CENTRO DE SAUDE III ALTO ALEGRE | Instituto Butantan / ESALQ-Piracicaba | Dimas Tadeu Covas, Antonio Jorge Martins, Claudia Renata dos Santos Barros, David Schlesinger, Debora Botequio Moretti, Elaine Cristina Marqueze, Elaine Vieira Santos, Evandra Strazza Rodrigues, Heidge Fukumasu, Jayme Augusto de Souza-Neto, José Salvatore Leister Patané, Luiz Alcantara, Luiz Lehmann Coutinho, Maria Carolina Elias, Maurício Lacerda Nogueira, Rafael dos Santos Bezerra, Raul Machado Neto, Rejane Maria Tommasini Grotto, Ricardo Haddad, Sandra Coccuzzo Sampaio Vessoni, Simone Kashima, Svetoslav Nanev Slavov, Vincent Louis Viala |
| EPI_ISL_2345196 | UBS III MURUTINGA DO SUL | Instituto Butantan / ESALQ-Piracicaba | Dimas Tadeu Covas, Antonio Jorge Martins, Claudia Renata dos Santos Barros, David Schlesinger, Debora Botequio Moretti, Elaine Cristina Marqueze, Elaine Vieira Santos, Evandra Strazza Rodrigues, Heidge Fukumasu, Jayme Augusto de Souza-Neto, José Salvatore Leister Patané, Luiz Alcantara, Luiz Lehmann Coutinho, Maria Carolina Elias, Maurício Lacerda Nogueira, Rafael dos Santos Bezerra, Raul Machado Neto, Rejane Maria Tommasini Grotto, Ricardo Haddad, Sandra Coccuzzo Sampaio Vessoni, Simone Kashima, Svetoslav Nanev Slavov, Vincent Louis Viala |
| EPI_ISL_2345197, EPI_ISL_2345198 | CS II DR MIGUEL VITALIANO ORLANDIA | Instituto Butantan / ESALQ-Piracicaba | Dimas Tadeu Covas, Antonio Jorge Martins, Claudia Renata dos Santos Barros, David Schlesinger, Debora Botequio Moretti, Elaine Cristina Marqueze, Elaine Vieira Santos, Evandra Strazza Rodrigues, Heidge Fukumasu, Jayme Augusto de Souza-Neto, José Salvatore Leister Patané, Luiz Alcantara, Luiz Lehmann Coutinho, Maria Carolina Elias, Maurício Lacerda Nogueira, Rafael dos Santos Bezerra, Raul Machado Neto, Rejane Maria Tommasini Grotto, Ricardo Haddad, Sandra Coccuzzo Sampaio Vessoni, Simone Kashima, Svetoslav Nanev Slavov, Vincent Louis Viala |
| EPI_ISL_2345199 | PRONTO SOCORRO MUNICIPAL DE SEVERINIA | Instituto Butantan / ESALQ-Piracicaba | Dimas Tadeu Covas, Antonio Jorge Martins, Claudia Renata dos Santos Barros, David Schlesinger, Debora Botequio Moretti, Elaine Cristina Marqueze, Elaine Vieira Santos, Evandra Strazza Rodrigues, Heidge Fukumasu, Jayme Augusto de Souza-Neto, José Salvatore Leister Patané, Luiz Alcantara, Luiz Lehmann Coutinho, Maria Carolina Elias, Maurício Lacerda Nogueira, Rafael dos Santos Bezerra, Raul Machado Neto, Rejane Maria Tommasini Grotto, Ricardo Haddad, Sandra Coccuzzo Sampaio Vessoni, Simone Kashima, Svetoslav Nanev Slavov, Vincent Louis Viala |
| EPI_ISL_2345200 | CS II DR MIGUEL VITALIANO ORLANDIA | Instituto Butantan / ESALQ-Piracicaba | Dimas Tadeu Covas, Antonio Jorge Martins, Claudia Renata dos Santos Barros, David Schlesinger, Debora Botequio Moretti, Elaine Cristina Marqueze, Elaine Vieira Santos, Evandra Strazza Rodrigues, Heidge Fukumasu, Jayme Augusto de Souza-Neto, José Salvatore Leister Patané, Luiz Alcantara, Luiz Lehmann Coutinho, Maria Carolina Elias, Maurício Lacerda Nogueira, Rafael dos Santos Bezerra, Raul Machado Neto, Rejane Maria Tommasini Grotto, Ricardo Haddad, Sandra Coccuzzo Sampaio Vessoni, Simone Kashima, Svetoslav Nanev Slavov, Vincent Louis Viala |
| EPI_ISL_2345201 | CENTRO DE SAUDE DE OSVALDO CRUZ | Instituto Butantan / ESALQ-Piracicaba | Dimas Tadeu Covas, Antonio Jorge Martins, Claudia Renata dos Santos Barros, David Schlesinger, Debora Botequio Moretti, Elaine Cristina Marqueze, Elaine Vieira Santos, Evandra Strazza Rodrigues, Heidge Fukumasu, Jayme Augusto de Souza-Neto, José Salvatore Leister Patané, Luiz Alcantara, Luiz |

| | | | |
|---|---|---|---|
| | | | Lehmann Coutinho, Maria Carolina Elias, Maurício Lacerda Nogueira, Rafael dos Santos Bezerra, Raul Machado Neto, Rejane Maria Tommasini Grotto, Ricardo Haddad, Sandra Coccuzzo Sampaio Vessoni, Simone Kashima, Svetoslav Nanev Slavov, Vincent Louis Viala |
| EPI_ISL_2345203, EPI_ISL_2345204 | UNIDADE BASICA DE SAUDE DE TARUMA | Instituto Butantan / ESALQ-Piracicaba | Dimas Tadeu Covas, Antonio Jorge Martins, Claudia Renata dos Santos Barros, David Schlesinger, Debora Botequio Moretti, Elaine Cristina Marqueze, Elaine Vieira Santos, Evandra Strazza Rodrigues, Heidge Fukumasu, Jayme Augusto de Souza-Neto, José Salvatore Leister Patané, Luiz Alcantara, Luiz Lehmann Coutinho, Maria Carolina Elias, Maurício Lacerda Nogueira, Rafael dos Santos Bezerra, Raul Machado Neto, Rejane Maria Tommasini Grotto, Ricardo Haddad, Sandra Coccuzzo Sampaio Vessoni, Simone Kashima, Svetoslav Nanev Slavov, Vincent Louis Viala |
| EPI_ISL_2345205 | NUCLEO DE SAUDE VILA FALCAO DE BAURU | Instituto Butantan / ESALQ-Piracicaba | Dimas Tadeu Covas, Antonio Jorge Martins, Claudia Renata dos Santos Barros, David Schlesinger, Debora Botequio Moretti, Elaine Cristina Marqueze, Elaine Vieira Santos, Evandra Strazza Rodrigues, Heidge Fukumasu, Jayme Augusto de Souza-Neto, José Salvatore Leister Patané, Luiz Alcantara, Luiz Lehmann Coutinho, Maria Carolina Elias, Maurício Lacerda Nogueira, Rafael dos Santos Bezerra, Raul Machado Neto, Rejane Maria Tommasini Grotto, Ricardo Haddad, Sandra Coccuzzo Sampaio Vessoni, Simone Kashima, Svetoslav Nanev Slavov, Vincent Louis Viala |
| EPI_ISL_2345206 | CENTRO DE SAUDE III DE IGARACU DO TIETE | Instituto Butantan / ESALQ-Piracicaba | Dimas Tadeu Covas, Antonio Jorge Martins, Claudia Renata dos Santos Barros, David Schlesinger, Debora Botequio Moretti, Elaine Cristina Marqueze, Elaine Vieira Santos, Evandra Strazza Rodrigues, Heidge Fukumasu, Jayme Augusto de Souza-Neto, José Salvatore Leister Patané, Luiz Alcantara, Luiz Lehmann Coutinho, Maria Carolina Elias, Maurício Lacerda Nogueira, Rafael dos Santos Bezerra, Raul Machado Neto, Rejane Maria Tommasini Grotto, Ricardo Haddad, Sandra Coccuzzo Sampaio Vessoni, Simone Kashima, Svetoslav Nanev Slavov, Vincent Louis Viala |
| EPI_ISL_2345207, EPI_ISL_2345208 | NUCLEO DE SAUDE VILA FALCAO DE BAURU | Instituto Butantan / ESALQ-Piracicaba | Dimas Tadeu Covas, Antonio Jorge Martins, Claudia Renata dos Santos Barros, David Schlesinger, Debora Botequio Moretti, Elaine Cristina Marqueze, Elaine Vieira Santos, Evandra Strazza Rodrigues, Heidge Fukumasu, Jayme Augusto de Souza-Neto, José Salvatore Leister Patané, Luiz Alcantara, Luiz Lehmann Coutinho, Maria Carolina Elias, Maurício Lacerda Nogueira, Rafael dos Santos Bezerra, Raul Machado Neto, Rejane Maria Tommasini Grotto, Ricardo Haddad, Sandra Coccuzzo Sampaio Vessoni, Simone Kashima, Svetoslav Nanev Slavov, Vincent Louis Viala |
| EPI_ISL_2345209 | SECRETARIA MUNICIPAL DE SAUDE DE MACATUBA | Instituto Butantan / ESALQ-Piracicaba | Dimas Tadeu Covas, Antonio Jorge Martins, Claudia Renata dos Santos Barros, David Schlesinger, Debora Botequio Moretti, Elaine Cristina Marqueze, Elaine Vieira Santos, Evandra Strazza Rodrigues, Heidge Fukumasu, Jayme Augusto de Souza-Neto, José Salvatore Leister Patané, Luiz Alcantara, Luiz Lehmann Coutinho, Maria Carolina Elias, Maurício Lacerda Nogueira, Rafael dos Santos Bezerra, Raul Machado Neto, Rejane Maria Tommasini Grotto, Ricardo Haddad, Sandra Coccuzzo Sampaio Vessoni, Simone Kashima, Svetoslav Nanev Slavov, Vincent Louis Viala |
| EPI_ISL_2345210 | UNIDADE DE SAUDE DA FAMILIA NOVO HORIZONTE | Instituto Butantan / ESALQ-Piracicaba | Dimas Tadeu Covas, Antonio Jorge Martins, Claudia Renata dos Santos Barros, David Schlesinger, Debora Botequio Moretti, Elaine Cristina Marqueze, Elaine Vieira Santos, Evandra Strazza Rodrigues, Heidge Fukumasu, Jayme Augusto de Souza-Neto, José Salvatore Leister Patané, Luiz Alcantara, Luiz Lehmann Coutinho, Maria Carolina Elias, Maurício Lacerda Nogueira, Rafael dos Santos Bezerra, Raul Machado Neto, Rejane Maria Tommasini Grotto, Ricardo Haddad, Sandra Coccuzzo Sampaio Vessoni, Simone Kashima, Svetoslav Nanev Slavov, Vincent Louis Viala |
| EPI_ISL_2345211 | SECRETARIA MUNICIPAL DE SAUDE DE MACATUBA | Instituto Butantan / ESALQ-Piracicaba | Dimas Tadeu Covas, Antonio Jorge Martins, Claudia Renata dos Santos Barros, David Schlesinger, Debora Botequio Moretti, Elaine Cristina Marqueze, Elaine Vieira Santos, Evandra Strazza Rodrigues, Heidge Fukumasu, Jayme Augusto de Souza-Neto, José Salvatore Leister Patané, Luiz Alcantara, Luiz Lehmann Coutinho, Maria Carolina Elias, Maurício Lacerda Nogueira, Rafael dos Santos Bezerra, Raul Machado Neto, Rejane Maria Tommasini Grotto, Ricardo Haddad, Sandra Coccuzzo Sampaio Vessoni, Simone Kashima, Svetoslav Nanev Slavov, Vincent Louis Viala |
| EPI_ISL_2345212 | UNIDADE DE SAUDE DA FAMILIA NOVO HORIZONTE | Instituto Butantan / ESALQ-Piracicaba | Dimas Tadeu Covas, Antonio Jorge Martins, Claudia Renata dos Santos Barros, David Schlesinger, Debora Botequio Moretti, Elaine Cristina Marqueze, Elaine Vieira Santos, Evandra Strazza Rodrigues, Heidge Fukumasu, Jayme Augusto de Souza-Neto, José Salvatore Leister Patané, Luiz Alcantara, Luiz Lehmann Coutinho, Maria Carolina Elias, Maurício Lacerda Nogueira, Rafael dos Santos Bezerra, Raul Machado Neto, Rejane Maria Tommasini Grotto, Ricardo Haddad, Sandra Coccuzzo Sampaio Vessoni, Simone Kashima, Svetoslav Nanev Slavov, Vincent Louis Viala |
| EPI_ISL_2345215, EPI_ISL_2345216, EPI_ISL_2345217, EPI_ISL_2345218 | NUCLEO DE SAUDE VILA FALCAO DE BAURU | Instituto Butantan / ESALQ-Piracicaba | Dimas Tadeu Covas, Antonio Jorge Martins, Claudia Renata dos Santos Barros, David Schlesinger, Debora Botequio Moretti, Elaine Cristina Marqueze, Elaine Vieira Santos, Evandra Strazza Rodrigues, Heidge Fukumasu, Jayme Augusto de Souza-Neto, José Salvatore Leister Patané, Luiz Alcantara, Luiz Lehmann Coutinho, Maria Carolina Elias, Maurício Lacerda Nogueira, Rafael dos Santos Bezerra, Raul Machado Neto, Rejane Maria Tommasini Grotto, Ricardo Haddad, Sandra Coccuzzo Sampaio Vessoni, Simone Kashima, Svetoslav Nanev Slavov, Vincent Louis Viala |
| EPI_ISL_2345219 | UNIDADE DE SAUDE DA FAMILIA NOVO HORIZONTE | Instituto Butantan / ESALQ-Piracicaba | Dimas Tadeu Covas, Antonio Jorge Martins, Claudia Renata dos Santos Barros, David Schlesinger, Debora Botequio Moretti, Elaine Cristina Marqueze, Elaine Vieira Santos, Evandra Strazza Rodrigues, Heidge Fukumasu, Jayme Augusto de Souza-Neto, José Salvatore Leister Patané, Luiz Alcantara, Luiz Lehmann Coutinho, Maria Carolina Elias, Maurício Lacerda Nogueira, Rafael dos Santos Bezerra, Raul Machado Neto, Rejane Maria Tommasini Grotto, Ricardo Haddad, Sandra Coccuzzo Sampaio Vessoni, Simone Kashima, Svetoslav Nanev Slavov, Vincent Louis Viala |
| EPI_ISL_2345222 | CENTRO DE SAUDE DE AVAI | Instituto Butantan / ESALQ-Piracicaba | Dimas Tadeu Covas, Antonio Jorge Martins, Claudia Renata dos Santos Barros, David Schlesinger, Debora Botequio Moretti, Elaine Cristina Marqueze, Elaine Vieira Santos, Evandra Strazza Rodrigues, Heidge Fukumasu, Jayme Augusto de Souza-Neto, José Salvatore Leister Patané, Luiz Alcantara, Luiz Lehmann Coutinho, Maria Carolina Elias, Maurício Lacerda Nogueira, Rafael dos Santos Bezerra, Raul Machado Neto, Rejane Maria Tommasini Grotto, Ricardo Haddad, Sandra Coccuzzo Sampaio Vessoni, Simone Kashima, Svetoslav Nanev Slavov, Vincent Louis Viala |
| EPI_ISL_2345223, EPI_ISL_2345225 | UBS IV GUARARAPES | Instituto Butantan / ESALQ-Piracicaba | Dimas Tadeu Covas, Antonio Jorge Martins, Claudia Renata dos Santos Barros, David Schlesinger, Debora Botequio Moretti, Elaine Cristina Marqueze, Elaine Vieira Santos, Evandra Strazza Rodrigues, Heidge Fukumasu, Jayme Augusto de Souza-Neto, José Salvatore Leister Patané, Luiz Alcantara, Luiz Lehmann Coutinho, Maria Carolina Elias, Maurício Lacerda Nogueira, Rafael dos Santos Bezerra, Raul Machado Neto, Rejane Maria Tommasini Grotto, Ricardo Haddad, Sandra Coccuzzo Sampaio Vessoni, Simone Kashima, Svetoslav Nanev Slavov, Vincent Louis Viala |
| EPI_ISL_2345226 | NUCLEO DE SAUDE VILA FALCAO DE BAURU | Instituto Butantan / ESALQ-Piracicaba | Dimas Tadeu Covas, Antonio Jorge Martins, Claudia Renata dos Santos Barros, David Schlesinger, Debora Botequio Moretti, Elaine Cristina Marqueze, Elaine Vieira Santos, Evandra Strazza Rodrigues, Heidge Fukumasu, Jayme Augusto de Souza-Neto, José Salvatore Leister Patané, Luiz Alcantara, Luiz Lehmann Coutinho, Maria Carolina Elias, Maurício Lacerda Nogueira, Rafael dos Santos Bezerra, Raul Machado Neto, Rejane Maria Tommasini Grotto, Ricardo Haddad, Sandra Coccuzzo Sampaio Vessoni, Simone Kashima, Svetoslav Nanev Slavov, Vincent Louis Viala |
| EPI_ISL_2345227 | UNIDADE DE SAUDE DA FAMILIA NOVO HORIZONTE | Instituto Butantan / ESALQ-Piracicaba | Dimas Tadeu Covas, Antonio Jorge Martins, Claudia Renata dos Santos Barros, David Schlesinger, Debora Botequio Moretti, Elaine Cristina Marqueze, Elaine Vieira Santos, Evandra Strazza Rodrigues, Heidge Fukumasu, Jayme Augusto de Souza-Neto, José Salvatore Leister Patané, Luiz Alcantara, Luiz Lehmann Coutinho, Maria Carolina Elias, Maurício Lacerda Nogueira, Rafael dos Santos Bezerra, Raul Machado Neto, Rejane Maria Tommasini Grotto, Ricardo Haddad, Sandra Coccuzzo Sampaio Vessoni, Simone Kashima, Svetoslav Nanev Slavov, Vincent Louis Viala |
| EPI_ISL_2345228, EPI_ISL_2345229, EPI_ISL_2345230, EPI_ISL_2345231, EPI_ISL_2345232, EPI_ISL_2345233, EPI_ISL_2345234, EPI_ISL_2345235 | UBS IV GUARARAPES | Instituto Butantan / ESALQ-Piracicaba | Dimas Tadeu Covas, Antonio Jorge Martins, Claudia Renata dos Santos Barros, David Schlesinger, Debora Botequio Moretti, Elaine Cristina Marqueze, Elaine Vieira Santos, Evandra Strazza Rodrigues, Heidge Fukumasu, Jayme Augusto de Souza-Neto, José Salvatore Leister Patané, Luiz Alcantara, Luiz Lehmann Coutinho, Maria Carolina Elias, Maurício Lacerda Nogueira, Rafael dos Santos Bezerra, Raul Machado Neto, Rejane Maria Tommasini Grotto, Ricardo Haddad, Sandra Coccuzzo Sampaio Vessoni, Simone Kashima, Svetoslav Nanev Slavov, Vincent Louis Viala |
| EPI_ISL_2345236 | CENTRO DE SAUDE DE OCAUCU | Instituto Butantan / ESALQ-Piracicaba | Dimas Tadeu Covas, Antonio Jorge Martins, Claudia Renata dos Santos Barros, David Schlesinger, Debora Botequio Moretti, Elaine Cristina Marqueze, Elaine Vieira Santos, Evandra Strazza Rodrigues, Heidge Fukumasu, Jayme Augusto de Souza-Neto, José Salvatore Leister Patané, Luiz Alcantara, Luiz Lehmann Coutinho, Maria Carolina Elias, Maurício Lacerda Nogueira, Rafael dos Santos Bezerra, Raul Machado Neto, Rejane Maria Tommasini Grotto, Ricardo Haddad, Sandra Coccuzzo Sampaio Vessoni, Simone Kashima, Svetoslav Nanev Slavov, Vincent Louis Viala |
| EPI_ISL_2345237, EPI_ISL_2345238 | SECRETARIA MUNICIPAL DE SAUDE DE JARINU | Instituto Butantan / ESALQ-Piracicaba | Dimas Tadeu Covas, Antonio Jorge Martins, Claudia Renata dos Santos Barros, David Schlesinger, Debora Botequio Moretti, Elaine Cristina Marqueze, Elaine Vieira Santos, Evandra Strazza Rodrigues, Heidge Fukumasu, Jayme Augusto de Souza-Neto, José Salvatore Leister Patané, Luiz Alcantara, Luiz Lehmann Coutinho, Maria Carolina Elias, Maurício Lacerda Nogueira, Rafael dos Santos Bezerra, Raul Machado Neto, Rejane Maria Tommasini Grotto, Ricardo Haddad, Sandra Coccuzzo Sampaio Vessoni, Simone Kashima, Svetoslav Nanev Slavov, Vincent Louis Viala |
| EPI_ISL_2345239, EPI_ISL_2345240, EPI_ISL_2345241 | DEPARTAMENTO DE SAUDE MUNICIPAL SOCORRO SP | Instituto Butantan / ESALQ-Piracicaba | Dimas Tadeu Covas, Antonio Jorge Martins, Claudia Renata dos Santos Barros, David Schlesinger, Debora Botequio Moretti, Elaine Cristina Marqueze, Elaine Vieira Santos, Evandra Strazza Rodrigues, Heidge Fukumasu, Jayme Augusto de Souza-Neto, José Salvatore Leister Patané, Luiz Alcantara, Luiz Lehmann Coutinho, Maria Carolina Elias, Maurício Lacerda Nogueira, Rafael dos Santos Bezerra, Raul Machado Neto, Rejane Maria Tommasini Grotto, Ricardo Haddad, Sandra Coccuzzo Sampaio Vessoni, Simone Kashima, Svetoslav Nanev Slavov, Vincent Louis Viala |
| EPI_ISL_2345242 | NUCLEO DE SAUDE VILA FALCAO DE BAURU | Instituto Butantan / ESALQ-Piracicaba | Dimas Tadeu Covas, Antonio Jorge Martins, Claudia Renata dos Santos Barros, David Schlesinger, Debora Botequio Moretti, Elaine Cristina Marqueze, Elaine Vieira Santos, Evandra Strazza Rodrigues, Heidge Fukumasu, Jayme Augusto de Souza-Neto, José Salvatore Leister Patané, Luiz Alcantara, Luiz Lehmann Coutinho, Maria Carolina Elias, Maurício Lacerda Nogueira, Rafael dos Santos Bezerra, Raul Machado Neto, Rejane Maria Tommasini Grotto, Ricardo Haddad, Sandra Coccuzzo Sampaio Vessoni, Simone Kashima, Svetoslav Nanev Slavov, Vincent Louis Viala |

| | | | |
|---|---|---|---|
| EPI_ISL_2345243, EPI_ISL_2345244 | DEPARTAMENTO DE SAUDE MUNICIPAL SOCORRO SP | Instituto Butantan / ESALQ-Piracicaba | Dimas Tadeu Covas, Antonio Jorge Martins, Claudia Renata dos Santos Barros, David Schlesinger, Debora Botequio Moretti, Elaine Cristina Marqueze, Elaine Vieira Santos, Evandra Strazza Rodrigues, Heidge Fukumasu, Jayme Augusto de Souza-Neto, José Salvatore Leister Patané, Luiz Alcantara, Luiz Lehmann Coutinho, Maria Carolina Elias, Maurício Lacerda Nogueira, Rafael dos Santos Bezerra, Raul Machado Neto, Rejane Maria Tommasini Grotto, Ricardo Haddad, Sandra Coccuzzo Sampaio Vessoni, Simone Kashima, Svetoslav Nanev Slavov, Vincent Louis Viala |
| EPI_ISL_2345245 | NUCLEO DE SAUDE VILA FALCAO DE BAURU | Instituto Butantan / ESALQ-Piracicaba | Dimas Tadeu Covas, Antonio Jorge Martins, Claudia Renata dos Santos Barros, David Schlesinger, Debora Botequio Moretti, Elaine Cristina Marqueze, Elaine Vieira Santos, Evandra Strazza Rodrigues, Heidge Fukumasu, Jayme Augusto de Souza-Neto, José Salvatore Leister Patané, Luiz Alcantara, Luiz Lehmann Coutinho, Maria Carolina Elias, Maurício Lacerda Nogueira, Rafael dos Santos Bezerra, Raul Machado Neto, Rejane Maria Tommasini Grotto, Ricardo Haddad, Sandra Coccuzzo Sampaio Vessoni, Simone Kashima, Svetoslav Nanev Slavov, Vincent Louis Viala |
| EPI_ISL_2345246 | CSII DR JOSE FERREIRA TELLES | Instituto Butantan / ESALQ-Piracicaba | Dimas Tadeu Covas, Antonio Jorge Martins, Claudia Renata dos Santos Barros, David Schlesinger, Debora Botequio Moretti, Elaine Cristina Marqueze, Elaine Vieira Santos, Evandra Strazza Rodrigues, Heidge Fukumasu, Jayme Augusto de Souza-Neto, José Salvatore Leister Patané, Luiz Alcantara, Luiz Lehmann Coutinho, Maria Carolina Elias, Maurício Lacerda Nogueira, Rafael dos Santos Bezerra, Raul Machado Neto, Rejane Maria Tommasini Grotto, Ricardo Haddad, Sandra Coccuzzo Sampaio Vessoni, Simone Kashima, Svetoslav Nanev Slavov, Vincent Louis Viala |
| EPI_ISL_2345247 | UNIDADE SAUDE FAMILIA DR PIERRE ANTONIO COPPIETERS AREALVA | Instituto Butantan / ESALQ-Piracicaba | Dimas Tadeu Covas, Antonio Jorge Martins, Claudia Renata dos Santos Barros, David Schlesinger, Debora Botequio Moretti, Elaine Cristina Marqueze, Elaine Vieira Santos, Evandra Strazza Rodrigues, Heidge Fukumasu, Jayme Augusto de Souza-Neto, José Salvatore Leister Patané, Luiz Alcantara, Luiz Lehmann Coutinho, Maria Carolina Elias, Maurício Lacerda Nogueira, Rafael dos Santos Bezerra, Raul Machado Neto, Rejane Maria Tommasini Grotto, Ricardo Haddad, Sandra Coccuzzo Sampaio Vessoni, Simone Kashima, Svetoslav Nanev Slavov, Vincent Louis Viala |
| EPI_ISL_2345248 | LABORATORIO MUNICIPAL DE PIRACICABA | Instituto Butantan / ESALQ-Piracicaba | Dimas Tadeu Covas, Antonio Jorge Martins, Claudia Renata dos Santos Barros, David Schlesinger, Debora Botequio Moretti, Elaine Cristina Marqueze, Elaine Vieira Santos, Evandra Strazza Rodrigues, Heidge Fukumasu, Jayme Augusto de Souza-Neto, José Salvatore Leister Patané, Luiz Alcantara, Luiz Lehmann Coutinho, Maria Carolina Elias, Maurício Lacerda Nogueira, Rafael dos Santos Bezerra, Raul Machado Neto, Rejane Maria Tommasini Grotto, Ricardo Haddad, Sandra Coccuzzo Sampaio Vessoni, Simone Kashima, Svetoslav Nanev Slavov, Vincent Louis Viala |
| EPI_ISL_2345249 | DEPARTAMENTO DE SAUDE MUNICIPAL SOCORRO SP | Instituto Butantan / ESALQ-Piracicaba | Dimas Tadeu Covas, Antonio Jorge Martins, Claudia Renata dos Santos Barros, David Schlesinger, Debora Botequio Moretti, Elaine Cristina Marqueze, Elaine Vieira Santos, Evandra Strazza Rodrigues, Heidge Fukumasu, Jayme Augusto de Souza-Neto, José Salvatore Leister Patané, Luiz Alcantara, Luiz Lehmann Coutinho, Maria Carolina Elias, Maurício Lacerda Nogueira, Rafael dos Santos Bezerra, Raul Machado Neto, Rejane Maria Tommasini Grotto, Ricardo Haddad, Sandra Coccuzzo Sampaio Vessoni, Simone Kashima, Svetoslav Nanev Slavov, Vincent Louis Viala |
| EPI_ISL_2345250, EPI_ISL_2345251 | LABORATORIO MUNICIPAL DE PIRACICABA | Instituto Butantan / ESALQ-Piracicaba | Dimas Tadeu Covas, Antonio Jorge Martins, Claudia Renata dos Santos Barros, David Schlesinger, Debora Botequio Moretti, Elaine Cristina Marqueze, Elaine Vieira Santos, Evandra Strazza Rodrigues, Heidge Fukumasu, Jayme Augusto de Souza-Neto, José Salvatore Leister Patané, Luiz Alcantara, Luiz Lehmann Coutinho, Maria Carolina Elias, Maurício Lacerda Nogueira, Rafael dos Santos Bezerra, Raul Machado Neto, Rejane Maria Tommasini Grotto, Ricardo Haddad, Sandra Coccuzzo Sampaio Vessoni, Simone Kashima, Svetoslav Nanev Slavov, Vincent Louis Viala |
| EPI_ISL_2345252, EPI_ISL_2345253, EPI_ISL_2345254, EPI_ISL_2345255, EPI_ISL_2345256, EPI_ISL_2345257, EPI_ISL_2345258, EPI_ISL_2345259, EPI_ISL_2345260 | DEPARTAMENTO DE SAUDE MUNICIPAL SOCORRO SP | Instituto Butantan / ESALQ-Piracicaba | Dimas Tadeu Covas, Antonio Jorge Martins, Claudia Renata dos Santos Barros, David Schlesinger, Debora Botequio Moretti, Elaine Cristina Marqueze, Elaine Vieira Santos, Evandra Strazza Rodrigues, Heidge Fukumasu, Jayme Augusto de Souza-Neto, José Salvatore Leister Patané, Luiz Alcantara, Luiz Lehmann Coutinho, Maria Carolina Elias, Maurício Lacerda Nogueira, Rafael dos Santos Bezerra, Raul Machado Neto, Rejane Maria Tommasini Grotto, Ricardo Haddad, Sandra Coccuzzo Sampaio Vessoni, Simone Kashima, Svetoslav Nanev Slavov, Vincent Louis Viala |
| EPI_ISL_2345261 | NUCLEO DE SAUDE VILA FALCAO DE BAURU | Instituto Butantan / ESALQ-Piracicaba | Dimas Tadeu Covas, Antonio Jorge Martins, Claudia Renata dos Santos Barros, David Schlesinger, Debora Botequio Moretti, Elaine Cristina Marqueze, Elaine Vieira Santos, Evandra Strazza Rodrigues, Heidge Fukumasu, Jayme Augusto de Souza-Neto, José Salvatore Leister Patané, Luiz Alcantara, Luiz Lehmann Coutinho, Maria Carolina Elias, Maurício Lacerda Nogueira, Rafael dos Santos Bezerra, Raul Machado Neto, Rejane Maria Tommasini Grotto, Ricardo Haddad, Sandra Coccuzzo Sampaio Vessoni, Simone Kashima, Svetoslav Nanev Slavov, Vincent Louis Viala |
| EPI_ISL_2345262 | CS II DR JAHYR DE PAULA RIBEIRO GUARA | Instituto Butantan / ESALQ-Piracicaba | Dimas Tadeu Covas, Antonio Jorge Martins, Claudia Renata dos Santos Barros, David Schlesinger, Debora Botequio Moretti, Elaine Cristina Marqueze, Elaine Vieira Santos, Evandra Strazza Rodrigues, Heidge Fukumasu, Jayme Augusto de Souza-Neto, José Salvatore Leister Patané, Luiz Alcantara, Luiz Lehmann Coutinho, Maria Carolina Elias, Maurício Lacerda Nogueira, Rafael dos Santos Bezerra, Raul Machado Neto, Rejane Maria Tommasini Grotto, Ricardo Haddad, Sandra Coccuzzo Sampaio Vessoni, Simone Kashima, Svetoslav Nanev Slavov, Vincent Louis Viala |
| EPI_ISL_2345263 | NUCLEO DE SAUDE VILA FALCAO DE BAURU | Instituto Butantan / ESALQ-Piracicaba | Dimas Tadeu Covas, Antonio Jorge Martins, Claudia Renata dos Santos Barros, David Schlesinger, Debora Botequio Moretti, Elaine Cristina Marqueze, Elaine Vieira Santos, Evandra Strazza Rodrigues, Heidge Fukumasu, Jayme Augusto de Souza-Neto, José Salvatore Leister Patané, Luiz Alcantara, Luiz Lehmann Coutinho, Maria Carolina Elias, Maurício Lacerda Nogueira, Rafael dos Santos Bezerra, Raul Machado Neto, Rejane Maria Tommasini Grotto, Ricardo Haddad, Sandra Coccuzzo Sampaio Vessoni, Simone Kashima, Svetoslav Nanev Slavov, Vincent Louis Viala |
| EPI_ISL_2345265 | POLICLINICA HORTOLANDIA | Instituto Butantan / ESALQ-Piracicaba | Dimas Tadeu Covas, Antonio Jorge Martins, Claudia Renata dos Santos Barros, David Schlesinger, Debora Botequio Moretti, Elaine Cristina Marqueze, Elaine Vieira Santos, Evandra Strazza Rodrigues, Heidge Fukumasu, Jayme Augusto de Souza-Neto, José Salvatore Leister Patané, Luiz Alcantara, Luiz Lehmann Coutinho, Maria Carolina Elias, Maurício Lacerda Nogueira, Rafael dos Santos Bezerra, Raul Machado Neto, Rejane Maria Tommasini Grotto, Ricardo Haddad, Sandra Coccuzzo Sampaio Vessoni, Simone Kashima, Svetoslav Nanev Slavov, Vincent Louis Viala |
| EPI_ISL_2345266 | LABORATORIO DE FRANCA | Instituto Butantan / ESALQ-Piracicaba | Dimas Tadeu Covas, Antonio Jorge Martins, Claudia Renata dos Santos Barros, David Schlesinger, Debora Botequio Moretti, Elaine Cristina Marqueze, Elaine Vieira Santos, Evandra Strazza Rodrigues, Heidge Fukumasu, Jayme Augusto de Souza-Neto, José Salvatore Leister Patané, Luiz Alcantara, Luiz Lehmann Coutinho, Maria Carolina Elias, Maurício Lacerda Nogueira, Rafael dos Santos Bezerra, Raul Machado Neto, Rejane Maria Tommasini Grotto, Ricardo Haddad, Sandra Coccuzzo Sampaio Vessoni, Simone Kashima, Svetoslav Nanev Slavov, Vincent Louis Viala |
| EPI_ISL_2345267 | LABORATORIO MUNICIPAL DE PIRACICABA | Instituto Butantan / ESALQ-Piracicaba | Dimas Tadeu Covas, Antonio Jorge Martins, Claudia Renata dos Santos Barros, David Schlesinger, Debora Botequio Moretti, Elaine Cristina Marqueze, Elaine Vieira Santos, Evandra Strazza Rodrigues, Heidge Fukumasu, Jayme Augusto de Souza-Neto, José Salvatore Leister Patané, Luiz Alcantara, Luiz Lehmann Coutinho, Maria Carolina Elias, Maurício Lacerda Nogueira, Rafael dos Santos Bezerra, Raul Machado Neto, Rejane Maria Tommasini Grotto, Ricardo Haddad, Sandra Coccuzzo Sampaio Vessoni, Simone Kashima, Svetoslav Nanev Slavov, Vincent Louis Viala |
| EPI_ISL_2345268 | UPA IPIRANGA | Instituto Butantan / ESALQ-Piracicaba | Dimas Tadeu Covas, Antonio Jorge Martins, Claudia Renata dos Santos Barros, David Schlesinger, Debora Botequio Moretti, Elaine Cristina Marqueze, Elaine Vieira Santos, Evandra Strazza Rodrigues, Heidge Fukumasu, Jayme Augusto de Souza-Neto, José Salvatore Leister Patané, Luiz Alcantara, Luiz Lehmann Coutinho, Maria Carolina Elias, Maurício Lacerda Nogueira, Rafael dos Santos Bezerra, Raul Machado Neto, Rejane Maria Tommasini Grotto, Ricardo Haddad, Sandra Coccuzzo Sampaio Vessoni, Simone Kashima, Svetoslav Nanev Slavov, Vincent Louis Viala |
| EPI_ISL_2345269, EPI_ISL_2345270 | UNIDADE DE VIGILANCIA EM SAUDE | Instituto Butantan / ESALQ-Piracicaba | Dimas Tadeu Covas, Antonio Jorge Martins, Claudia Renata dos Santos Barros, David Schlesinger, Debora Botequio Moretti, Elaine Cristina Marqueze, Elaine Vieira Santos, Evandra Strazza Rodrigues, Heidge Fukumasu, Jayme Augusto de Souza-Neto, José Salvatore Leister Patané, Luiz Alcantara, Luiz Lehmann Coutinho, Maria Carolina Elias, Maurício Lacerda Nogueira, Rafael dos Santos Bezerra, Raul Machado Neto, Rejane Maria Tommasini Grotto, Ricardo Haddad, Sandra Coccuzzo Sampaio Vessoni, Simone Kashima, Svetoslav Nanev Slavov, Vincent Louis Viala |
| EPI_ISL_2345271 | LABORATORIO MUNICIPAL DE PIRACICABA | Instituto Butantan / ESALQ-Piracicaba | Dimas Tadeu Covas, Antonio Jorge Martins, Claudia Renata dos Santos Barros, David Schlesinger, Debora Botequio Moretti, Elaine Cristina Marqueze, Elaine Vieira Santos, Evandra Strazza Rodrigues, Heidge Fukumasu, Jayme Augusto de Souza-Neto, José Salvatore Leister Patané, Luiz Alcantara, Luiz Lehmann Coutinho, Maria Carolina Elias, Maurício Lacerda Nogueira, Rafael dos Santos Bezerra, Raul Machado Neto, Rejane Maria Tommasini Grotto, Ricardo Haddad, Sandra Coccuzzo Sampaio Vessoni, Simone Kashima, Svetoslav Nanev Slavov, Vincent Louis Viala |
| EPI_ISL_2345272 | UNIDADE DE VIGILANCIA EM SAUDE | Instituto Butantan / ESALQ-Piracicaba | Dimas Tadeu Covas, Antonio Jorge Martins, Claudia Renata dos Santos Barros, David Schlesinger, Debora Botequio Moretti, Elaine Cristina Marqueze, Elaine Vieira Santos, Evandra Strazza Rodrigues, Heidge Fukumasu, Jayme Augusto de Souza-Neto, José Salvatore Leister Patané, Luiz Alcantara, Luiz Lehmann Coutinho, Maria Carolina Elias, Maurício Lacerda Nogueira, Rafael dos Santos Bezerra, Raul Machado Neto, Rejane Maria Tommasini Grotto, Ricardo Haddad, Sandra Coccuzzo Sampaio Vessoni, Simone Kashima, Svetoslav Nanev Slavov, Vincent Louis Viala |
| EPI_ISL_2345273, EPI_ISL_2345274 | CS IOLANDA FLEURY DE CAMPOS GUEDES NUPORANGA | Instituto Butantan / ESALQ-Piracicaba | Dimas Tadeu Covas, Antonio Jorge Martins, Claudia Renata dos Santos Barros, David Schlesinger, Debora Botequio Moretti, Elaine Cristina Marqueze, Elaine Vieira Santos, Evandra Strazza Rodrigues, Heidge Fukumasu, Jayme Augusto de Souza-Neto, José Salvatore Leister Patané, Luiz Alcantara, Luiz Lehmann Coutinho, Maria Carolina Elias, Maurício Lacerda Nogueira, Rafael dos Santos Bezerra, Raul Machado Neto, Rejane Maria Tommasini Grotto, Ricardo Haddad, Sandra Coccuzzo Sampaio Vessoni, Simone Kashima, Svetoslav Nanev Slavov, Vincent Louis Viala |
| EPI_ISL_2345275 | UNIDADE DE VIGILANCIA EM SAUDE | Instituto Butantan / ESALQ-Piracicaba | Dimas Tadeu Covas, Antonio Jorge Martins, Claudia Renata dos Santos Barros, David Schlesinger, Debora Botequio Moretti, Elaine Cristina Marqueze, |

| | | | |
|---|---|---|---|
| | | | Elaine Vieira Santos, Evandra Strazza Rodrigues, Heidge Fukumasu, Jayme Augusto de Souza-Neto, José Salvatore Leister Patané, Luiz Alcantara, Luiz Lehmann Coutinho, Maria Carolina Elias, Maurício Lacerda Nogueira, Rafael dos Santos Bezerra, Raul Machado Neto, Rejane Maria Tommasini Grotto, Ricardo Haddad, Sandra Coccuzzo Sampaio Vessoni, Simone Kashima, Svetoslav Nanev Slavov, Vincent Louis Viala |
| EPI_ISL_2345276 | NUCLEO DE SAUDE VILA FALCAO DE BAURU | Instituto Butantan / ESALQ-Piracicaba | Dimas Tadeu Covas, Antonio Jorge Martins, Claudia Renata dos Santos Barros, David Schlesinger, Debora Botequio Moretti, Elaine Cristina Marqueze, Elaine Vieira Santos, Evandra Strazza Rodrigues, Heidge Fukumasu, Jayme Augusto de Souza-Neto, José Salvatore Leister Patané, Luiz Alcantara, Luiz Lehmann Coutinho, Maria Carolina Elias, Maurício Lacerda Nogueira, Rafael dos Santos Bezerra, Raul Machado Neto, Rejane Maria Tommasini Grotto, Ricardo Haddad, Sandra Coccuzzo Sampaio Vessoni, Simone Kashima, Svetoslav Nanev Slavov, Vincent Louis Viala |
| EPI_ISL_2345277 | UNIDADE DE VIGILANCIA EM SAUDE | Instituto Butantan / ESALQ-Piracicaba | Dimas Tadeu Covas, Antonio Jorge Martins, Claudia Renata dos Santos Barros, David Schlesinger, Debora Botequio Moretti, Elaine Cristina Marqueze, Elaine Vieira Santos, Evandra Strazza Rodrigues, Heidge Fukumasu, Jayme Augusto de Souza-Neto, José Salvatore Leister Patané, Luiz Alcantara, Luiz Lehmann Coutinho, Maria Carolina Elias, Maurício Lacerda Nogueira, Rafael dos Santos Bezerra, Raul Machado Neto, Rejane Maria Tommasini Grotto, Ricardo Haddad, Sandra Coccuzzo Sampaio Vessoni, Simone Kashima, Svetoslav Nanev Slavov, Vincent Louis Viala |
| EPI_ISL_2345278 | CS IOLANDA FLEURY DE CAMPOS GUEDES NUPORANGA | Instituto Butantan / ESALQ-Piracicaba | Dimas Tadeu Covas, Antonio Jorge Martins, Claudia Renata dos Santos Barros, David Schlesinger, Debora Botequio Moretti, Elaine Cristina Marqueze, Elaine Vieira Santos, Evandra Strazza Rodrigues, Heidge Fukumasu, Jayme Augusto de Souza-Neto, José Salvatore Leister Patané, Luiz Alcantara, Luiz Lehmann Coutinho, Maria Carolina Elias, Maurício Lacerda Nogueira, Rafael dos Santos Bezerra, Raul Machado Neto, Rejane Maria Tommasini Grotto, Ricardo Haddad, Sandra Coccuzzo Sampaio Vessoni, Simone Kashima, Svetoslav Nanev Slavov, Vincent Louis Viala |
| EPI_ISL_2345279 | UNIDADE DE VIGILANCIA EM SAUDE | Instituto Butantan / ESALQ-Piracicaba | Dimas Tadeu Covas, Antonio Jorge Martins, Claudia Renata dos Santos Barros, David Schlesinger, Debora Botequio Moretti, Elaine Cristina Marqueze, Elaine Vieira Santos, Evandra Strazza Rodrigues, Heidge Fukumasu, Jayme Augusto de Souza-Neto, José Salvatore Leister Patané, Luiz Alcantara, Luiz Lehmann Coutinho, Maria Carolina Elias, Maurício Lacerda Nogueira, Rafael dos Santos Bezerra, Raul Machado Neto, Rejane Maria Tommasini Grotto, Ricardo Haddad, Sandra Coccuzzo Sampaio Vessoni, Simone Kashima, Svetoslav Nanev Slavov, Vincent Louis Viala |
| EPI_ISL_2345280 | NUCLEO DE SAUDE VILA FALCAO DE BAURU | Instituto Butantan / ESALQ-Piracicaba | Dimas Tadeu Covas, Antonio Jorge Martins, Claudia Renata dos Santos Barros, David Schlesinger, Debora Botequio Moretti, Elaine Cristina Marqueze, Elaine Vieira Santos, Evandra Strazza Rodrigues, Heidge Fukumasu, Jayme Augusto de Souza-Neto, José Salvatore Leister Patané, Luiz Alcantara, Luiz Lehmann Coutinho, Maria Carolina Elias, Maurício Lacerda Nogueira, Rafael dos Santos Bezerra, Raul Machado Neto, Rejane Maria Tommasini Grotto, Ricardo Haddad, Sandra Coccuzzo Sampaio Vessoni, Simone Kashima, Svetoslav Nanev Slavov, Vincent Louis Viala |
| EPI_ISL_2345281 | UNIDADE DE VIGILANCIA EM SAUDE | Instituto Butantan / ESALQ-Piracicaba | Dimas Tadeu Covas, Antonio Jorge Martins, Claudia Renata dos Santos Barros, David Schlesinger, Debora Botequio Moretti, Elaine Cristina Marqueze, Elaine Vieira Santos, Evandra Strazza Rodrigues, Heidge Fukumasu, Jayme Augusto de Souza-Neto, José Salvatore Leister Patané, Luiz Alcantara, Luiz Lehmann Coutinho, Maria Carolina Elias, Maurício Lacerda Nogueira, Rafael dos Santos Bezerra, Raul Machado Neto, Rejane Maria Tommasini Grotto, Ricardo Haddad, Sandra Coccuzzo Sampaio Vessoni, Simone Kashima, Svetoslav Nanev Slavov, Vincent Louis Viala |
| EPI_ISL_2345282, EPI_ISL_2345283 | NUCLEO DE SAUDE VILA FALCAO DE BAURU | Instituto Butantan / ESALQ-Piracicaba | Dimas Tadeu Covas, Antonio Jorge Martins, Claudia Renata dos Santos Barros, David Schlesinger, Debora Botequio Moretti, Elaine Cristina Marqueze, Elaine Vieira Santos, Evandra Strazza Rodrigues, Heidge Fukumasu, Jayme Augusto de Souza-Neto, José Salvatore Leister Patané, Luiz Alcantara, Luiz Lehmann Coutinho, Maria Carolina Elias, Maurício Lacerda Nogueira, Rafael dos Santos Bezerra, Raul Machado Neto, Rejane Maria Tommasini Grotto, Ricardo Haddad, Sandra Coccuzzo Sampaio Vessoni, Simone Kashima, Svetoslav Nanev Slavov, Vincent Louis Viala |
| EPI_ISL_2345284, EPI_ISL_2345285, EPI_ISL_2345286, EPI_ISL_2345287 | UNIDADE DE VIGILANCIA EM SAUDE | Instituto Butantan / ESALQ-Piracicaba | Dimas Tadeu Covas, Antonio Jorge Martins, Claudia Renata dos Santos Barros, David Schlesinger, Debora Botequio Moretti, Elaine Cristina Marqueze, Elaine Vieira Santos, Evandra Strazza Rodrigues, Heidge Fukumasu, Jayme Augusto de Souza-Neto, José Salvatore Leister Patané, Luiz Alcantara, Luiz Lehmann Coutinho, Maria Carolina Elias, Maurício Lacerda Nogueira, Rafael dos Santos Bezerra, Raul Machado Neto, Rejane Maria Tommasini Grotto, Ricardo Haddad, Sandra Coccuzzo Sampaio Vessoni, Simone Kashima, Svetoslav Nanev Slavov, Vincent Louis Viala |
| EPI_ISL_2345288, EPI_ISL_2345289, EPI_ISL_2345290, EPI_ISL_2345291, EPI_ISL_2345292, EPI_ISL_2345293, EPI_ISL_2345294, EPI_ISL_2345295, EPI_ISL_2345296, EPI_ISL_2345297, EPI_ISL_2345298 | | | |
| see above | LABORATORIO MUNICIPAL DE PIRACICABA | Instituto Butantan / ESALQ-Piracicaba | Dimas Tadeu Covas, Antonio Jorge Martins, Claudia Renata dos Santos Barros, David Schlesinger, Debora Botequio Moretti, Elaine Cristina Marqueze, Elaine Vieira Santos, Evandra Strazza Rodrigues, Heidge Fukumasu, Jayme Augusto de Souza-Neto, José Salvatore Leister Patané, Luiz Alcantara, Luiz Lehmann Coutinho, Maria Carolina Elias, Maurício Lacerda Nogueira, Rafael dos Santos Bezerra, Raul Machado Neto, Rejane Maria Tommasini Grotto, Ricardo Haddad, Sandra Coccuzzo Sampaio Vessoni, Simone Kashima, Svetoslav Nanev Slavov, Vincent Louis Viala |
| EPI_ISL_2345299 | SANTA CASA DE MISERICORDIA DE UBATUBA | Instituto Butantan / UNESP-Botucatu | Dimas Tadeu Covas, Antonio Jorge Martins, Claudia Renata dos Santos Barros, David Schlesinger, Debora Botequio Moretti, Elaine Cristina Marqueze, Elaine Vieira Santos, Evandra Strazza Rodrigues, Heidge Fukumasu, Jayme Augusto de Souza-Neto, José Salvatore Leister Patané, Luiz Alcantara, Luiz Lehmann Coutinho, Maria Carolina Elias, Maurício Lacerda Nogueira, Rafael dos Santos Bezerra, Raul Machado Neto, Rejane Maria Tommasini Grotto, Ricardo Haddad, Sandra Coccuzzo Sampaio Vessoni, Simone Kashima, Svetoslav Nanev Slavov, Vincent Louis Viala |
| EPI_ISL_2345300 | SECRETARIA MUNICIPAL DA SAUDE DE GUARATINGUETA | Instituto Butantan / UNESP-Botucatu | Dimas Tadeu Covas, Antonio Jorge Martins, Claudia Renata dos Santos Barros, David Schlesinger, Debora Botequio Moretti, Elaine Cristina Marqueze, Elaine Vieira Santos, Evandra Strazza Rodrigues, Heidge Fukumasu, Jayme Augusto de Souza-Neto, José Salvatore Leister Patané, Luiz Alcantara, Luiz Lehmann Coutinho, Maria Carolina Elias, Maurício Lacerda Nogueira, Rafael dos Santos Bezerra, Raul Machado Neto, Rejane Maria Tommasini Grotto, Ricardo Haddad, Sandra Coccuzzo Sampaio Vessoni, Simone Kashima, Svetoslav Nanev Slavov, Vincent Louis Viala |
| EPI_ISL_2345301 | UNIDADE DE PRONTO ATENDIMENTO UPA SUL CARAGUATATUBA | Instituto Butantan / UNESP-Botucatu | Dimas Tadeu Covas, Antonio Jorge Martins, Claudia Renata dos Santos Barros, David Schlesinger, Debora Botequio Moretti, Elaine Cristina Marqueze, Elaine Vieira Santos, Evandra Strazza Rodrigues, Heidge Fukumasu, Jayme Augusto de Souza-Neto, José Salvatore Leister Patané, Luiz Alcantara, Luiz Lehmann Coutinho, Maria Carolina Elias, Maurício Lacerda Nogueira, Rafael dos Santos Bezerra, Raul Machado Neto, Rejane Maria Tommasini Grotto, Ricardo Haddad, Sandra Coccuzzo Sampaio Vessoni, Simone Kashima, Svetoslav Nanev Slavov, Vincent Louis Viala |
| EPI_ISL_2345302 | UNIDADE DE PRONTO ATENDIMENTO CENTRAL DE CARAGUATATUBA | Instituto Butantan / UNESP-Botucatu | Dimas Tadeu Covas, Antonio Jorge Martins, Claudia Renata dos Santos Barros, David Schlesinger, Debora Botequio Moretti, Elaine Cristina Marqueze, Elaine Vieira Santos, Evandra Strazza Rodrigues, Heidge Fukumasu, Jayme Augusto de Souza-Neto, José Salvatore Leister Patané, Luiz Alcantara, Luiz Lehmann Coutinho, Maria Carolina Elias, Maurício Lacerda Nogueira, Rafael dos Santos Bezerra, Raul Machado Neto, Rejane Maria Tommasini Grotto, Ricardo Haddad, Sandra Coccuzzo Sampaio Vessoni, Simone Kashima, Svetoslav Nanev Slavov, Vincent Louis Viala |
| EPI_ISL_2345304 | UNIDADE DE PRONTO ATENDIMENTO UPA SUL CARAGUATATUBA | Instituto Butantan / UNESP-Botucatu | Dimas Tadeu Covas, Antonio Jorge Martins, Claudia Renata dos Santos Barros, David Schlesinger, Debora Botequio Moretti, Elaine Cristina Marqueze, Elaine Vieira Santos, Evandra Strazza Rodrigues, Heidge Fukumasu, Jayme Augusto de Souza-Neto, José Salvatore Leister Patané, Luiz Alcantara, Luiz Lehmann Coutinho, Maria Carolina Elias, Maurício Lacerda Nogueira, Rafael dos Santos Bezerra, Raul Machado Neto, Rejane Maria Tommasini Grotto, Ricardo Haddad, Sandra Coccuzzo Sampaio Vessoni, Simone Kashima, Svetoslav Nanev Slavov, Vincent Louis Viala |
| EPI_ISL_2345305 | SECRETARIA MUNICIPAL DA SAUDE DE GUARATINGUETA | Instituto Butantan / UNESP-Botucatu | Dimas Tadeu Covas, Antonio Jorge Martins, Claudia Renata dos Santos Barros, David Schlesinger, Debora Botequio Moretti, Elaine Cristina Marqueze, Elaine Vieira Santos, Evandra Strazza Rodrigues, Heidge Fukumasu, Jayme Augusto de Souza-Neto, José Salvatore Leister Patané, Luiz Alcantara, Luiz Lehmann Coutinho, Maria Carolina Elias, Maurício Lacerda Nogueira, Rafael dos Santos Bezerra, Raul Machado Neto, Rejane Maria Tommasini Grotto, Ricardo Haddad, Sandra Coccuzzo Sampaio Vessoni, Simone Kashima, Svetoslav Nanev Slavov, Vincent Louis Viala |
| EPI_ISL_2345306 | SMS SECRETARIA MUNICIPAL DE SAUDE DE BOITUVA | Instituto Butantan / UNESP-Botucatu | Dimas Tadeu Covas, Antonio Jorge Martins, Claudia Renata dos Santos Barros, David Schlesinger, Debora Botequio Moretti, Elaine Cristina Marqueze, Elaine Vieira Santos, Evandra Strazza Rodrigues, Heidge Fukumasu, Jayme Augusto de Souza-Neto, José Salvatore Leister Patané, Luiz Alcantara, Luiz Lehmann Coutinho, Maria Carolina Elias, Maurício Lacerda Nogueira, Rafael dos Santos Bezerra, Raul Machado Neto, Rejane Maria Tommasini Grotto, Ricardo Haddad, Sandra Coccuzzo Sampaio Vessoni, Simone Kashima, Svetoslav Nanev Slavov, Vincent Louis Viala |
| EPI_ISL_2345308 | SANTA CASA DE MISERICORDIA DE TIETE | Instituto Butantan / UNESP-Botucatu | Dimas Tadeu Covas, Antonio Jorge Martins, Claudia Renata dos Santos Barros, David Schlesinger, Debora Botequio Moretti, Elaine Cristina Marqueze, Elaine Vieira Santos, Evandra Strazza Rodrigues, Heidge Fukumasu, Jayme Augusto de Souza-Neto, José Salvatore Leister Patané, Luiz Alcantara, Luiz Lehmann Coutinho, Maria Carolina Elias, Maurício Lacerda Nogueira, Rafael dos Santos Bezerra, Raul Machado Neto, Rejane Maria Tommasini Grotto, Ricardo Haddad, Sandra Coccuzzo Sampaio Vessoni, Simone Kashima, Svetoslav Nanev Slavov, Vincent Louis Viala |
| EPI_ISL_2345309 | UNIDADE ESF DR LUIZ SPINA | Instituto Butantan / ESALQ-Piracicaba | Dimas Tadeu Covas, Antonio Jorge Martins, Claudia Renata dos Santos Barros, David Schlesinger, Debora Botequio Moretti, Elaine Cristina Marqueze, Elaine Vieira Santos, Evandra Strazza Rodrigues, Heidge Fukumasu, Jayme Augusto de Souza-Neto, José Salvatore Leister Patané, Luiz Alcantara, Luiz Lehmann Coutinho, Maria Carolina Elias, Maurício Lacerda Nogueira, Rafael dos Santos Bezerra, Raul Machado Neto, Rejane Maria Tommasini Grotto, Ricardo Haddad, Sandra Coccuzzo Sampaio Vessoni, Simone Kashima, Svetoslav Nanev Slavov, Vincent Louis Viala |
| EPI_ISL_2345310, EPI_ISL_2345311 | SECRETARIA MUNICIPAL DA SAUDE DE GUARIBA | Instituto Butantan / ESALQ-Piracicaba | Dimas Tadeu Covas, Antonio Jorge Martins, Claudia Renata dos Santos Barros, David Schlesinger, Debora Botequio Moretti, Elaine Cristina Marqueze, |

| | | | |
|---|---|---|---|
| | | | Elaine Vieira Santos, Evandra Strazza Rodrigues, Heidge Fukumasu, Jayme Augusto de Souza-Neto, José Salvatore Leister Patané, Luiz Alcantara, Luiz Lehmann Coutinho, Maria Carolina Elias, Maurício Lacerda Nogueira, Rafael dos Santos Bezerra, Raul Machado Neto, Rejane Maria Tommasini Grotto, Ricardo Haddad, Sandra Coccuzzo Sampaio Vessoni, Simone Kashima, Svetoslav Nanev Slavov, Vincent Louis Viala |
| EPI_ISL_2345312 | AMBULATORIO DE ATENDIMENTO DE DST DE GUARIBA | Instituto Butantan / ESALQ-Piracicaba | Dimas Tadeu Covas, Antonio Jorge Martins, Claudia Renata dos Santos Barros, David Schlesinger, Debora Botequio Moretti, Elaine Cristina Marqueze, Elaine Vieira Santos, Evandra Strazza Rodrigues, Heidge Fukumasu, Jayme Augusto de Souza-Neto, José Salvatore Leister Patané, Luiz Alcantara, Luiz Lehmann Coutinho, Maria Carolina Elias, Maurício Lacerda Nogueira, Rafael dos Santos Bezerra, Raul Machado Neto, Rejane Maria Tommasini Grotto, Ricardo Haddad, Sandra Coccuzzo Sampaio Vessoni, Simone Kashima, Svetoslav Nanev Slavov, Vincent Louis Viala |
| EPI_ISL_2345313, EPI_ISL_2345314 | CENTRO DE SAUDE III BORBOREMA | Instituto Butantan / ESALQ-Piracicaba | Dimas Tadeu Covas, Antonio Jorge Martins, Claudia Renata dos Santos Barros, David Schlesinger, Debora Botequio Moretti, Elaine Cristina Marqueze, Elaine Vieira Santos, Evandra Strazza Rodrigues, Heidge Fukumasu, Jayme Augusto de Souza-Neto, José Salvatore Leister Patané, Luiz Alcantara, Luiz Lehmann Coutinho, Maria Carolina Elias, Maurício Lacerda Nogueira, Rafael dos Santos Bezerra, Raul Machado Neto, Rejane Maria Tommasini Grotto, Ricardo Haddad, Sandra Coccuzzo Sampaio Vessoni, Simone Kashima, Svetoslav Nanev Slavov, Vincent Louis Viala |
| EPI_ISL_2345315, EPI_ISL_2345316, EPI_ISL_2345317, EPI_ISL_2345318, EPI_ISL_2345319 | UNIDADE MISTA DE LUIZ ANTONIO | Instituto Butantan / ESALQ-Piracicaba | Dimas Tadeu Covas, Antonio Jorge Martins, Claudia Renata dos Santos Barros, David Schlesinger, Debora Botequio Moretti, Elaine Cristina Marqueze, Elaine Vieira Santos, Evandra Strazza Rodrigues, Heidge Fukumasu, Jayme Augusto de Souza-Neto, José Salvatore Leister Patané, Luiz Alcantara, Luiz Lehmann Coutinho, Maria Carolina Elias, Maurício Lacerda Nogueira, Rafael dos Santos Bezerra, Raul Machado Neto, Rejane Maria Tommasini Grotto, Ricardo Haddad, Sandra Coccuzzo Sampaio Vessoni, Simone Kashima, Svetoslav Nanev Slavov, Vincent Louis Viala |
| EPI_ISL_2345320, EPI_ISL_2345321 | CENTRO DE SAUDE DE CAJOBI | Instituto Butantan / ESALQ-Piracicaba | Dimas Tadeu Covas, Antonio Jorge Martins, Claudia Renata dos Santos Barros, David Schlesinger, Debora Botequio Moretti, Elaine Cristina Marqueze, Elaine Vieira Santos, Evandra Strazza Rodrigues, Heidge Fukumasu, Jayme Augusto de Souza-Neto, José Salvatore Leister Patané, Luiz Alcantara, Luiz Lehmann Coutinho, Maria Carolina Elias, Maurício Lacerda Nogueira, Rafael dos Santos Bezerra, Raul Machado Neto, Rejane Maria Tommasini Grotto, Ricardo Haddad, Sandra Coccuzzo Sampaio Vessoni, Simone Kashima, Svetoslav Nanev Slavov, Vincent Louis Viala |
| EPI_ISL_2345322, EPI_ISL_2345323, EPI_ISL_2345324, EPI_ISL_2345325, EPI_ISL_2345326 | UNIDADE DE VIGILANCIA EM SAUDE | Instituto Butantan / ESALQ-Piracicaba | Dimas Tadeu Covas, Antonio Jorge Martins, Claudia Renata dos Santos Barros, David Schlesinger, Debora Botequio Moretti, Elaine Cristina Marqueze, Elaine Vieira Santos, Evandra Strazza Rodrigues, Heidge Fukumasu, Jayme Augusto de Souza-Neto, José Salvatore Leister Patané, Luiz Alcantara, Luiz Lehmann Coutinho, Maria Carolina Elias, Maurício Lacerda Nogueira, Rafael dos Santos Bezerra, Raul Machado Neto, Rejane Maria Tommasini Grotto, Ricardo Haddad, Sandra Coccuzzo Sampaio Vessoni, Simone Kashima, Svetoslav Nanev Slavov, Vincent Louis Viala |
| EPI_ISL_2345327 | CENTRO DE SAUDE III CANDIDO RODRIGUES | Instituto Butantan / ESALQ-Piracicaba | Dimas Tadeu Covas, Antonio Jorge Martins, Claudia Renata dos Santos Barros, David Schlesinger, Debora Botequio Moretti, Elaine Cristina Marqueze, Elaine Vieira Santos, Evandra Strazza Rodrigues, Heidge Fukumasu, Jayme Augusto de Souza-Neto, José Salvatore Leister Patané, Luiz Alcantara, Luiz Lehmann Coutinho, Maria Carolina Elias, Maurício Lacerda Nogueira, Rafael dos Santos Bezerra, Raul Machado Neto, Rejane Maria Tommasini Grotto, Ricardo Haddad, Sandra Coccuzzo Sampaio Vessoni, Simone Kashima, Svetoslav Nanev Slavov, Vincent Louis Viala |
| EPI_ISL_2345328 | CENTRO DE SAUDE III SALES OLIVEIRA | Instituto Butantan / ESALQ-Piracicaba | Dimas Tadeu Covas, Antonio Jorge Martins, Claudia Renata dos Santos Barros, David Schlesinger, Debora Botequio Moretti, Elaine Cristina Marqueze, Elaine Vieira Santos, Evandra Strazza Rodrigues, Heidge Fukumasu, Jayme Augusto de Souza-Neto, José Salvatore Leister Patané, Luiz Alcantara, Luiz Lehmann Coutinho, Maria Carolina Elias, Maurício Lacerda Nogueira, Rafael dos Santos Bezerra, Raul Machado Neto, Rejane Maria Tommasini Grotto, Ricardo Haddad, Sandra Coccuzzo Sampaio Vessoni, Simone Kashima, Svetoslav Nanev Slavov, Vincent Louis Viala |
| EPI_ISL_2345329 | CENTRO DE SAUDE III CANDIDO RODRIGUES | Instituto Butantan / ESALQ-Piracicaba | Dimas Tadeu Covas, Antonio Jorge Martins, Claudia Renata dos Santos Barros, David Schlesinger, Debora Botequio Moretti, Elaine Cristina Marqueze, Elaine Vieira Santos, Evandra Strazza Rodrigues, Heidge Fukumasu, Jayme Augusto de Souza-Neto, José Salvatore Leister Patané, Luiz Alcantara, Luiz Lehmann Coutinho, Maria Carolina Elias, Maurício Lacerda Nogueira, Rafael dos Santos Bezerra, Raul Machado Neto, Rejane Maria Tommasini Grotto, Ricardo Haddad, Sandra Coccuzzo Sampaio Vessoni, Simone Kashima, Svetoslav Nanev Slavov, Vincent Louis Viala |
| EPI_ISL_2345330, EPI_ISL_2345331, EPI_ISL_2345332, EPI_ISL_2345333, EPI_ISL_2345334, EPI_ISL_2345335, EPI_ISL_2345336, EPI_ISL_2345337, EPI_ISL_2345338 | UNIDADE DE VIGILANCIA EM SAUDE | Instituto Butantan / ESALQ-Piracicaba | Dimas Tadeu Covas, Antonio Jorge Martins, Claudia Renata dos Santos Barros, David Schlesinger, Debora Botequio Moretti, Elaine Cristina Marqueze, Elaine Vieira Santos, Evandra Strazza Rodrigues, Heidge Fukumasu, Jayme Augusto de Souza-Neto, José Salvatore Leister Patané, Luiz Alcantara, Luiz Lehmann Coutinho, Maria Carolina Elias, Maurício Lacerda Nogueira, Rafael dos Santos Bezerra, Raul Machado Neto, Rejane Maria Tommasini Grotto, Ricardo Haddad, Sandra Coccuzzo Sampaio Vessoni, Simone Kashima, Svetoslav Nanev Slavov, Vincent Louis Viala |
| EPI_ISL_2345339, EPI_ISL_2345340, EPI_ISL_2345341, EPI_ISL_2345342, EPI_ISL_2345343 | UNIDADE MISTA DE VISTA ALEGRE DO ALTO VISTA ALEGRE DO ALTO | Instituto Butantan / ESALQ-Piracicaba | Dimas Tadeu Covas, Antonio Jorge Martins, Claudia Renata dos Santos Barros, David Schlesinger, Debora Botequio Moretti, Elaine Cristina Marqueze, Elaine Vieira Santos, Evandra Strazza Rodrigues, Heidge Fukumasu, Jayme Augusto de Souza-Neto, José Salvatore Leister Patané, Luiz Alcantara, Luiz Lehmann Coutinho, Maria Carolina Elias, Maurício Lacerda Nogueira, Rafael dos Santos Bezerra, Raul Machado Neto, Rejane Maria Tommasini Grotto, Ricardo Haddad, Sandra Coccuzzo Sampaio Vessoni, Simone Kashima, Svetoslav Nanev Slavov, Vincent Louis Viala |
| EPI_ISL_2345344, EPI_ISL_2345345, EPI_ISL_2345346, EPI_ISL_2345348, EPI_ISL_2345349, EPI_ISL_2345350, EPI_ISL_2345351, EPI_ISL_2345352, EPI_ISL_2345353, EPI_ISL_2345354, EPI_ISL_2345355, EPI_ISL_2345356, EPI_ISL_2345357 | | | |
| see above | POLICLINICA HORTOLANDIA | Instituto Butantan / ESALQ-Piracicaba | Dimas Tadeu Covas, Antonio Jorge Martins, Claudia Renata dos Santos Barros, David Schlesinger, Debora Botequio Moretti, Elaine Cristina Marqueze, Elaine Vieira Santos, Evandra Strazza Rodrigues, Heidge Fukumasu, Jayme Augusto de Souza-Neto, José Salvatore Leister Patané, Luiz Alcantara, Luiz Lehmann Coutinho, Maria Carolina Elias, Maurício Lacerda Nogueira, Rafael dos Santos Bezerra, Raul Machado Neto, Rejane Maria Tommasini Grotto, Ricardo Haddad, Sandra Coccuzzo Sampaio Vessoni, Simone Kashima, Svetoslav Nanev Slavov, Vincent Louis Viala |
| EPI_ISL_2345358, EPI_ISL_2345359 | UNIDADE MISTA DE VISTA ALEGRE DO ALTO VISTA ALEGRE DO ALTO | Instituto Butantan / ESALQ-Piracicaba | Dimas Tadeu Covas, Antonio Jorge Martins, Claudia Renata dos Santos Barros, David Schlesinger, Debora Botequio Moretti, Elaine Cristina Marqueze, Elaine Vieira Santos, Evandra Strazza Rodrigues, Heidge Fukumasu, Jayme Augusto de Souza-Neto, José Salvatore Leister Patané, Luiz Alcantara, Luiz Lehmann Coutinho, Maria Carolina Elias, Maurício Lacerda Nogueira, Rafael dos Santos Bezerra, Raul Machado Neto, Rejane Maria Tommasini Grotto, Ricardo Haddad, Sandra Coccuzzo Sampaio Vessoni, Simone Kashima, Svetoslav Nanev Slavov, Vincent Louis Viala |
| EPI_ISL_2345360 | CENTRO DE SAUDE DE CAJOBI | Instituto Butantan / ESALQ-Piracicaba | Dimas Tadeu Covas, Antonio Jorge Martins, Claudia Renata dos Santos Barros, David Schlesinger, Debora Botequio Moretti, Elaine Cristina Marqueze, Elaine Vieira Santos, Evandra Strazza Rodrigues, Heidge Fukumasu, Jayme Augusto de Souza-Neto, José Salvatore Leister Patané, Luiz Alcantara, Luiz Lehmann Coutinho, Maria Carolina Elias, Maurício Lacerda Nogueira, Rafael dos Santos Bezerra, Raul Machado Neto, Rejane Maria Tommasini Grotto, Ricardo Haddad, Sandra Coccuzzo Sampaio Vessoni, Simone Kashima, Svetoslav Nanev Slavov, Vincent Louis Viala |
| EPI_ISL_2345361, EPI_ISL_2345362, EPI_ISL_2345363, EPI_ISL_2345364, EPI_ISL_2345365, EPI_ISL_2345366, EPI_ISL_2345367, EPI_ISL_2345368, EPI_ISL_2345369 | POLICLINICA HORTOLANDIA | Instituto Butantan / ESALQ-Piracicaba | Dimas Tadeu Covas, Antonio Jorge Martins, Claudia Renata dos Santos Barros, David Schlesinger, Debora Botequio Moretti, Elaine Cristina Marqueze, Elaine Vieira Santos, Evandra Strazza Rodrigues, Heidge Fukumasu, Jayme Augusto de Souza-Neto, José Salvatore Leister Patané, Luiz Alcantara, Luiz Lehmann Coutinho, Maria Carolina Elias, Maurício Lacerda Nogueira, Rafael dos Santos Bezerra, Raul Machado Neto, Rejane Maria Tommasini Grotto, Ricardo Haddad, Sandra Coccuzzo Sampaio Vessoni, Simone Kashima, Svetoslav Nanev Slavov, Vincent Louis Viala |
| EPI_ISL_2345370 | UBS CAIC OURINHOS | Instituto Butantan / ESALQ-Piracicaba | Dimas Tadeu Covas, Antonio Jorge Martins, Claudia Renata dos Santos Barros, David Schlesinger, Debora Botequio Moretti, Elaine Cristina Marqueze, Elaine Vieira Santos, Evandra Strazza Rodrigues, Heidge Fukumasu, Jayme Augusto de Souza-Neto, José Salvatore Leister Patané, Luiz Alcantara, Luiz Lehmann Coutinho, Maria Carolina Elias, Maurício Lacerda Nogueira, Rafael dos Santos Bezerra, Raul Machado Neto, Rejane Maria Tommasini Grotto, Ricardo Haddad, Sandra Coccuzzo Sampaio Vessoni, Simone Kashima, Svetoslav Nanev Slavov, Vincent Louis Viala |
| EPI_ISL_2345371 | UBS JARDIM ITAMARATY | Instituto Butantan / ESALQ-Piracicaba | Dimas Tadeu Covas, Antonio Jorge Martins, Claudia Renata dos Santos Barros, David Schlesinger, Debora Botequio Moretti, Elaine Cristina Marqueze, Elaine Vieira Santos, Evandra Strazza Rodrigues, Heidge Fukumasu, Jayme Augusto de Souza-Neto, José Salvatore Leister Patané, Luiz Alcantara, Luiz Lehmann Coutinho, Maria Carolina Elias, Maurício Lacerda Nogueira, Rafael dos Santos Bezerra, Raul Machado Neto, Rejane Maria Tommasini Grotto, Ricardo Haddad, Sandra Coccuzzo Sampaio Vessoni, Simone Kashima, Svetoslav Nanev Slavov, Vincent Louis Viala |
| EPI_ISL_2345372 | CS III VILA ODILON | Instituto Butantan / ESALQ-Piracicaba | Dimas Tadeu Covas, Antonio Jorge Martins, Claudia Renata dos Santos Barros, David Schlesinger, Debora Botequio Moretti, Elaine Cristina Marqueze, Elaine Vieira Santos, Evandra Strazza Rodrigues, Heidge Fukumasu, Jayme Augusto de Souza-Neto, José Salvatore Leister Patané, Luiz Alcantara, Luiz Lehmann Coutinho, Maria Carolina Elias, Maurício Lacerda Nogueira, Rafael dos Santos Bezerra, Raul Machado Neto, Rejane Maria Tommasini Grotto, Ricardo Haddad, Sandra Coccuzzo Sampaio Vessoni, Simone Kashima, Svetoslav Nanev Slavov, Vincent Louis Viala |
| EPI_ISL_2345373, EPI_ISL_2345374, EPI_ISL_2345375 | CENTRO DE SAUDE DR PLINIO ALBERS DE GALIA | Instituto Butantan / ESALQ-Piracicaba | Dimas Tadeu Covas, Antonio Jorge Martins, Claudia Renata dos Santos Barros, David Schlesinger, Debora Botequio Moretti, Elaine Cristina Marqueze, Elaine Vieira Santos, Evandra Strazza Rodrigues, Heidge Fukumasu, Jayme Augusto de Souza-Neto, José Salvatore Leister Patané, Luiz Alcantara, Luiz Lehmann Coutinho, Maria Carolina Elias, Maurício Lacerda Nogueira, Rafael dos Santos Bezerra, Raul Machado Neto, Rejane Maria Tommasini Grotto, Ricardo Haddad, Sandra Coccuzzo Sampaio Vessoni, Simone Kashima, Svetoslav Nanev Slavov, Vincent Louis Viala |

| | | | |
|---|---|---|---|
| EPI_ISL_2345376 | CS DE ALVARO CARVALHO | Instituto Butantan / ESALQ-Piracicaba | Dimas Tadeu Covas, Antonio Jorge Martins, Claudia Renata dos Santos Barros, David Schlesinger, Debora Botequio Moretti, Elaine Cristina Marqueze, Elaine Vieira Santos, Evandra Strazza Rodrigues, Heidge Fukumasu, Jayme Augusto de Souza-Neto, José Salvatore Leister Patané, Luiz Alcantara, Luiz Lehmann Coutinho, Maria Carolina Elias, Maurício Lacerda Nogueira, Rafael dos Santos Bezerra, Raul Machado Neto, Rejane Maria Tommasini Grotto, Ricardo Haddad, Sandra Coccuzzo Sampaio Vessoni, Simone Kashima, Svetoslav Nanev Slavov, Vincent Louis Viala |
| EPI_ISL_2345377 | CENTRO DE SAUDE DR PLINIO ALBERS DE GALIA | Instituto Butantan / ESALQ-Piracicaba | Dimas Tadeu Covas, Antonio Jorge Martins, Claudia Renata dos Santos Barros, David Schlesinger, Debora Botequio Moretti, Elaine Cristina Marqueze, Elaine Vieira Santos, Evandra Strazza Rodrigues, Heidge Fukumasu, Jayme Augusto de Souza-Neto, José Salvatore Leister Patané, Luiz Alcantara, Luiz Lehmann Coutinho, Maria Carolina Elias, Maurício Lacerda Nogueira, Rafael dos Santos Bezerra, Raul Machado Neto, Rejane Maria Tommasini Grotto, Ricardo Haddad, Sandra Coccuzzo Sampaio Vessoni, Simone Kashima, Svetoslav Nanev Slavov, Vincent Louis Viala |
| EPI_ISL_2345378, EPI_ISL_2345379, EPI_ISL_2345380 | CENTRO DE SAUDE DE FLORINEA | Instituto Butantan / ESALQ-Piracicaba | Dimas Tadeu Covas, Antonio Jorge Martins, Claudia Renata dos Santos Barros, David Schlesinger, Debora Botequio Moretti, Elaine Cristina Marqueze, Elaine Vieira Santos, Evandra Strazza Rodrigues, Heidge Fukumasu, Jayme Augusto de Souza-Neto, José Salvatore Leister Patané, Luiz Alcantara, Luiz Lehmann Coutinho, Maria Carolina Elias, Maurício Lacerda Nogueira, Rafael dos Santos Bezerra, Raul Machado Neto, Rejane Maria Tommasini Grotto, Ricardo Haddad, Sandra Coccuzzo Sampaio Vessoni, Simone Kashima, Svetoslav Nanev Slavov, Vincent Louis Viala |
| EPI_ISL_2345381 | CENTRO DE SAUDE DE MARACAI | Instituto Butantan / ESALQ-Piracicaba | Dimas Tadeu Covas, Antonio Jorge Martins, Claudia Renata dos Santos Barros, David Schlesinger, Debora Botequio Moretti, Elaine Cristina Marqueze, Elaine Vieira Santos, Evandra Strazza Rodrigues, Heidge Fukumasu, Jayme Augusto de Souza-Neto, José Salvatore Leister Patané, Luiz Alcantara, Luiz Lehmann Coutinho, Maria Carolina Elias, Maurício Lacerda Nogueira, Rafael dos Santos Bezerra, Raul Machado Neto, Rejane Maria Tommasini Grotto, Ricardo Haddad, Sandra Coccuzzo Sampaio Vessoni, Simone Kashima, Svetoslav Nanev Slavov, Vincent Louis Viala |
| EPI_ISL_2345382, EPI_ISL_2345383 | CS DE OSCAR BRESSANE PSF | Instituto Butantan / ESALQ-Piracicaba | Dimas Tadeu Covas, Antonio Jorge Martins, Claudia Renata dos Santos Barros, David Schlesinger, Debora Botequio Moretti, Elaine Cristina Marqueze, Elaine Vieira Santos, Evandra Strazza Rodrigues, Heidge Fukumasu, Jayme Augusto de Souza-Neto, José Salvatore Leister Patané, Luiz Alcantara, Luiz Lehmann Coutinho, Maria Carolina Elias, Maurício Lacerda Nogueira, Rafael dos Santos Bezerra, Raul Machado Neto, Rejane Maria Tommasini Grotto, Ricardo Haddad, Sandra Coccuzzo Sampaio Vessoni, Simone Kashima, Svetoslav Nanev Slavov, Vincent Louis Viala |
| EPI_ISL_2345384 | CENTRO DE SAUDE DE RIBEIRAO DO SUL | Instituto Butantan / ESALQ-Piracicaba | Dimas Tadeu Covas, Antonio Jorge Martins, Claudia Renata dos Santos Barros, David Schlesinger, Debora Botequio Moretti, Elaine Cristina Marqueze, Elaine Vieira Santos, Evandra Strazza Rodrigues, Heidge Fukumasu, Jayme Augusto de Souza-Neto, José Salvatore Leister Patané, Luiz Alcantara, Luiz Lehmann Coutinho, Maria Carolina Elias, Maurício Lacerda Nogueira, Rafael dos Santos Bezerra, Raul Machado Neto, Rejane Maria Tommasini Grotto, Ricardo Haddad, Sandra Coccuzzo Sampaio Vessoni, Simone Kashima, Svetoslav Nanev Slavov, Vincent Louis Viala |
| EPI_ISL_2345385 | CS DE OSCAR BRESSANE PSF | Instituto Butantan / ESALQ-Piracicaba | Dimas Tadeu Covas, Antonio Jorge Martins, Claudia Renata dos Santos Barros, David Schlesinger, Debora Botequio Moretti, Elaine Cristina Marqueze, Elaine Vieira Santos, Evandra Strazza Rodrigues, Heidge Fukumasu, Jayme Augusto de Souza-Neto, José Salvatore Leister Patané, Luiz Alcantara, Luiz Lehmann Coutinho, Maria Carolina Elias, Maurício Lacerda Nogueira, Rafael dos Santos Bezerra, Raul Machado Neto, Rejane Maria Tommasini Grotto, Ricardo Haddad, Sandra Coccuzzo Sampaio Vessoni, Simone Kashima, Svetoslav Nanev Slavov, Vincent Louis Viala |
| EPI_ISL_2345386, EPI_ISL_2345387 | CENTRO DE SAUDE DE MARACAI | Instituto Butantan / ESALQ-Piracicaba | Dimas Tadeu Covas, Antonio Jorge Martins, Claudia Renata dos Santos Barros, David Schlesinger, Debora Botequio Moretti, Elaine Cristina Marqueze, Elaine Vieira Santos, Evandra Strazza Rodrigues, Heidge Fukumasu, Jayme Augusto de Souza-Neto, José Salvatore Leister Patané, Luiz Alcantara, Luiz Lehmann Coutinho, Maria Carolina Elias, Maurício Lacerda Nogueira, Rafael dos Santos Bezerra, Raul Machado Neto, Rejane Maria Tommasini Grotto, Ricardo Haddad, Sandra Coccuzzo Sampaio Vessoni, Simone Kashima, Svetoslav Nanev Slavov, Vincent Louis Viala |
| EPI_ISL_2345388, EPI_ISL_2345389 | CS DE OSCAR BRESSANE PSF | Instituto Butantan / ESALQ-Piracicaba | Dimas Tadeu Covas, Antonio Jorge Martins, Claudia Renata dos Santos Barros, David Schlesinger, Debora Botequio Moretti, Elaine Cristina Marqueze, Elaine Vieira Santos, Evandra Strazza Rodrigues, Heidge Fukumasu, Jayme Augusto de Souza-Neto, José Salvatore Leister Patané, Luiz Alcantara, Luiz Lehmann Coutinho, Maria Carolina Elias, Maurício Lacerda Nogueira, Rafael dos Santos Bezerra, Raul Machado Neto, Rejane Maria Tommasini Grotto, Ricardo Haddad, Sandra Coccuzzo Sampaio Vessoni, Simone Kashima, Svetoslav Nanev Slavov, Vincent Louis Viala |
| EPI_ISL_2345390 | CENTRO DE SAUDE DE MARACAI | Instituto Butantan / ESALQ-Piracicaba | Dimas Tadeu Covas, Antonio Jorge Martins, Claudia Renata dos Santos Barros, David Schlesinger, Debora Botequio Moretti, Elaine Cristina Marqueze, Elaine Vieira Santos, Evandra Strazza Rodrigues, Heidge Fukumasu, Jayme Augusto de Souza-Neto, José Salvatore Leister Patané, Luiz Alcantara, Luiz Lehmann Coutinho, Maria Carolina Elias, Maurício Lacerda Nogueira, Rafael dos Santos Bezerra, Raul Machado Neto, Rejane Maria Tommasini Grotto, Ricardo Haddad, Sandra Coccuzzo Sampaio Vessoni, Simone Kashima, Svetoslav Nanev Slavov, Vincent Louis Viala |
| EPI_ISL_2345391, EPI_ISL_2345392, EPI_ISL_2345393 | USF SALERNO | Instituto Butantan / ESALQ-Piracicaba | Dimas Tadeu Covas, Antonio Jorge Martins, Claudia Renata dos Santos Barros, David Schlesinger, Debora Botequio Moretti, Elaine Cristina Marqueze, Elaine Vieira Santos, Evandra Strazza Rodrigues, Heidge Fukumasu, Jayme Augusto de Souza-Neto, José Salvatore Leister Patané, Luiz Alcantara, Luiz Lehmann Coutinho, Maria Carolina Elias, Maurício Lacerda Nogueira, Rafael dos Santos Bezerra, Raul Machado Neto, Rejane Maria Tommasini Grotto, Ricardo Haddad, Sandra Coccuzzo Sampaio Vessoni, Simone Kashima, Svetoslav Nanev Slavov, Vincent Louis Viala |
| EPI_ISL_2345394, EPI_ISL_2345395, EPI_ISL_2345396, EPI_ISL_2345397, EPI_ISL_2345398, EPI_ISL_2345399 | POLICLINICA HORTOLANDIA | Instituto Butantan / ESALQ-Piracicaba | Dimas Tadeu Covas, Antonio Jorge Martins, Claudia Renata dos Santos Barros, David Schlesinger, Debora Botequio Moretti, Elaine Cristina Marqueze, Elaine Vieira Santos, Evandra Strazza Rodrigues, Heidge Fukumasu, Jayme Augusto de Souza-Neto, José Salvatore Leister Patané, Luiz Alcantara, Luiz Lehmann Coutinho, Maria Carolina Elias, Maurício Lacerda Nogueira, Rafael dos Santos Bezerra, Raul Machado Neto, Rejane Maria Tommasini Grotto, Ricardo Haddad, Sandra Coccuzzo Sampaio Vessoni, Simone Kashima, Svetoslav Nanev Slavov, Vincent Louis Viala |
| EPI_ISL_2345400 | DEPARTAMENTO DE SAUDE MUNICIPAL SOCORRO SP | Instituto Butantan / ESALQ-Piracicaba | Dimas Tadeu Covas, Antonio Jorge Martins, Claudia Renata dos Santos Barros, David Schlesinger, Debora Botequio Moretti, Elaine Cristina Marqueze, Elaine Vieira Santos, Evandra Strazza Rodrigues, Heidge Fukumasu, Jayme Augusto de Souza-Neto, José Salvatore Leister Patané, Luiz Alcantara, Luiz Lehmann Coutinho, Maria Carolina Elias, Maurício Lacerda Nogueira, Rafael dos Santos Bezerra, Raul Machado Neto, Rejane Maria Tommasini Grotto, Ricardo Haddad, Sandra Coccuzzo Sampaio Vessoni, Simone Kashima, Svetoslav Nanev Slavov, Vincent Louis Viala |
| EPI_ISL_2345401, EPI_ISL_2345402, EPI_ISL_2345403, EPI_ISL_2345404 | POLICLINICA HORTOLANDIA | Instituto Butantan / ESALQ-Piracicaba | Dimas Tadeu Covas, Antonio Jorge Martins, Claudia Renata dos Santos Barros, David Schlesinger, Debora Botequio Moretti, Elaine Cristina Marqueze, Elaine Vieira Santos, Evandra Strazza Rodrigues, Heidge Fukumasu, Jayme Augusto de Souza-Neto, José Salvatore Leister Patané, Luiz Alcantara, Luiz Lehmann Coutinho, Maria Carolina Elias, Maurício Lacerda Nogueira, Rafael dos Santos Bezerra, Raul Machado Neto, Rejane Maria Tommasini Grotto, Ricardo Haddad, Sandra Coccuzzo Sampaio Vessoni, Simone Kashima, Svetoslav Nanev Slavov, Vincent Louis Viala |
| EPI_ISL_2345405, EPI_ISL_2345406, EPI_ISL_2345407 | USF SALERNO | Instituto Butantan / ESALQ-Piracicaba | Dimas Tadeu Covas, Antonio Jorge Martins, Claudia Renata dos Santos Barros, David Schlesinger, Debora Botequio Moretti, Elaine Cristina Marqueze, Elaine Vieira Santos, Evandra Strazza Rodrigues, Heidge Fukumasu, Jayme Augusto de Souza-Neto, José Salvatore Leister Patané, Luiz Alcantara, Luiz Lehmann Coutinho, Maria Carolina Elias, Maurício Lacerda Nogueira, Rafael dos Santos Bezerra, Raul Machado Neto, Rejane Maria Tommasini Grotto, Ricardo Haddad, Sandra Coccuzzo Sampaio Vessoni, Simone Kashima, Svetoslav Nanev Slavov, Vincent Louis Viala |
| EPI_ISL_2345408, EPI_ISL_2345409 | POLICLINICA HORTOLANDIA | Instituto Butantan / ESALQ-Piracicaba | Dimas Tadeu Covas, Antonio Jorge Martins, Claudia Renata dos Santos Barros, David Schlesinger, Debora Botequio Moretti, Elaine Cristina Marqueze, Elaine Vieira Santos, Evandra Strazza Rodrigues, Heidge Fukumasu, Jayme Augusto de Souza-Neto, José Salvatore Leister Patané, Luiz Alcantara, Luiz Lehmann Coutinho, Maria Carolina Elias, Maurício Lacerda Nogueira, Rafael dos Santos Bezerra, Raul Machado Neto, Rejane Maria Tommasini Grotto, Ricardo Haddad, Sandra Coccuzzo Sampaio Vessoni, Simone Kashima, Svetoslav Nanev Slavov, Vincent Louis Viala |
| EPI_ISL_2345410 | USF SALERNO | Instituto Butantan / ESALQ-Piracicaba | Dimas Tadeu Covas, Antonio Jorge Martins, Claudia Renata dos Santos Barros, David Schlesinger, Debora Botequio Moretti, Elaine Cristina Marqueze, Elaine Vieira Santos, Evandra Strazza Rodrigues, Heidge Fukumasu, Jayme Augusto de Souza-Neto, José Salvatore Leister Patané, Luiz Alcantara, Luiz Lehmann Coutinho, Maria Carolina Elias, Maurício Lacerda Nogueira, Rafael dos Santos Bezerra, Raul Machado Neto, Rejane Maria Tommasini Grotto, Ricardo Haddad, Sandra Coccuzzo Sampaio Vessoni, Simone Kashima, Svetoslav Nanev Slavov, Vincent Louis Viala |
| EPI_ISL_2345411, EPI_ISL_2345412 | POLICLINICA HORTOLANDIA | Instituto Butantan / ESALQ-Piracicaba | Dimas Tadeu Covas, Antonio Jorge Martins, Claudia Renata dos Santos Barros, David Schlesinger, Debora Botequio Moretti, Elaine Cristina Marqueze, Elaine Vieira Santos, Evandra Strazza Rodrigues, Heidge Fukumasu, Jayme Augusto de Souza-Neto, José Salvatore Leister Patané, Luiz Alcantara, Luiz Lehmann Coutinho, Maria Carolina Elias, Maurício Lacerda Nogueira, Rafael dos Santos Bezerra, Raul Machado Neto, Rejane Maria Tommasini Grotto, Ricardo Haddad, Sandra Coccuzzo Sampaio Vessoni, Simone Kashima, Svetoslav Nanev Slavov, Vincent Louis Viala |
| EPI_ISL_2345413, EPI_ISL_2345414, EPI_ISL_2345415 | USF SALERNO | Instituto Butantan / ESALQ-Piracicaba | Dimas Tadeu Covas, Antonio Jorge Martins, Claudia Renata dos Santos Barros, David Schlesinger, Debora Botequio Moretti, Elaine Cristina Marqueze, Elaine Vieira Santos, Evandra Strazza Rodrigues, Heidge Fukumasu, Jayme Augusto de Souza-Neto, José Salvatore Leister Patané, Luiz Alcantara, Luiz Lehmann Coutinho, Maria Carolina Elias, Maurício Lacerda Nogueira, Rafael dos Santos Bezerra, Raul Machado Neto, Rejane Maria Tommasini Grotto, Ricardo Haddad, Sandra Coccuzzo Sampaio Vessoni, Simone Kashima, Svetoslav Nanev Slavov, Vincent Louis Viala |

EPI_ISL_2345416, EPI_ISL_2345417, EPI_ISL_2345418, EPI_ISL_2345419, EPI_ISL_2345420, EPI_ISL_2345422, EPI_ISL_2345423, EPI_ISL_2345424, EPI_ISL_2345425, EPI_ISL_2345426, EPI_ISL_2345427, EPI_ISL_2345428, EPI_ISL_2345429, EPI_ISL_2345430

| | | | |
|---|---|---|---|
| see above | AMBULATORIO DE ESPECIALIDADE V E MOGI MIRIM | Instituto Butantan / ESALQ-Piracicaba | Dimas Tadeu Covas, Antonio Jorge Martins, Claudia Renata dos Santos Barros, David Schlesinger, Debora Botequio Moretti, Elaine Cristina Marqueze, Elaine Vieira Santos, Evandra Strazza Rodrigues, Heidge Fukumasu, Jayme Augusto de Souza-Neto, José Salvatore Leister Patané, Luiz Alcantara, Luiz Lehmann Coutinho, Maria Carolina Elias, Maurício Lacerda Nogueira, Rafael dos Santos Bezerra, Raul Machado Neto, Rejane Maria Tommasini Grotto, Ricardo Haddad, Sandra Coccuzzo Sampaio Vessoni, Simone Kashima, Svetoslav Nanev Slavov, Vincent Louis Viala |
| EPI_ISL_2345431, EPI_ISL_2345432, EPI_ISL_2345433, EPI_ISL_2345434, EPI_ISL_2345435, EPI_ISL_2345436, EPI_ISL_2345437 | PRONTO SOCORRO MUNICIPAL TAMBAU | Instituto Butantan / ESALQ-Piracicaba | Dimas Tadeu Covas, Antonio Jorge Martins, Claudia Renata dos Santos Barros, David Schlesinger, Debora Botequio Moretti, Elaine Cristina Marqueze, Elaine Vieira Santos, Evandra Strazza Rodrigues, Heidge Fukumasu, Jayme Augusto de Souza-Neto, José Salvatore Leister Patané, Luiz Alcantara, Luiz Lehmann Coutinho, Maria Carolina Elias, Maurício Lacerda Nogueira, Rafael dos Santos Bezerra, Raul Machado Neto, Rejane Maria Tommasini Grotto, Ricardo Haddad, Sandra Coccuzzo Sampaio Vessoni, Simone Kashima, Svetoslav Nanev Slavov, Vincent Louis Viala |
| EPI_ISL_2345438 | AMBULATORIO DE ESPECIALIDADE V E MOGI MIRIM | Instituto Butantan / ESALQ-Piracicaba | Dimas Tadeu Covas, Antonio Jorge Martins, Claudia Renata dos Santos Barros, David Schlesinger, Debora Botequio Moretti, Elaine Cristina Marqueze, Elaine Vieira Santos, Evandra Strazza Rodrigues, Heidge Fukumasu, Jayme Augusto de Souza-Neto, José Salvatore Leister Patané, Luiz Alcantara, Luiz Lehmann Coutinho, Maria Carolina Elias, Maurício Lacerda Nogueira, Rafael dos Santos Bezerra, Raul Machado Neto, Rejane Maria Tommasini Grotto, Ricardo Haddad, Sandra Coccuzzo Sampaio Vessoni, Simone Kashima, Svetoslav Nanev Slavov, Vincent Louis Viala |
| EPI_ISL_2345439 | HOSPITAL MUNICIPAL DR MARIO GATTI CAMPINAS | Instituto Butantan / ESALQ-Piracicaba | Dimas Tadeu Covas, Antonio Jorge Martins, Claudia Renata dos Santos Barros, David Schlesinger, Debora Botequio Moretti, Elaine Cristina Marqueze, Elaine Vieira Santos, Evandra Strazza Rodrigues, Heidge Fukumasu, Jayme Augusto de Souza-Neto, José Salvatore Leister Patané, Luiz Alcantara, Luiz Lehmann Coutinho, Maria Carolina Elias, Maurício Lacerda Nogueira, Rafael dos Santos Bezerra, Raul Machado Neto, Rejane Maria Tommasini Grotto, Ricardo Haddad, Sandra Coccuzzo Sampaio Vessoni, Simone Kashima, Svetoslav Nanev Slavov, Vincent Louis Viala |
| EPI_ISL_2345440 | CENTRO DE SAUDE II DR ALCIDES FACUNDO ARROYO | Instituto Butantan / ESALQ-Piracicaba | Dimas Tadeu Covas, Antonio Jorge Martins, Claudia Renata dos Santos Barros, David Schlesinger, Debora Botequio Moretti, Elaine Cristina Marqueze, Elaine Vieira Santos, Evandra Strazza Rodrigues, Heidge Fukumasu, Jayme Augusto de Souza-Neto, José Salvatore Leister Patané, Luiz Alcantara, Luiz Lehmann Coutinho, Maria Carolina Elias, Maurício Lacerda Nogueira, Rafael dos Santos Bezerra, Raul Machado Neto, Rejane Maria Tommasini Grotto, Ricardo Haddad, Sandra Coccuzzo Sampaio Vessoni, Simone Kashima, Svetoslav Nanev Slavov, Vincent Louis Viala |
| EPI_ISL_2345441, EPI_ISL_2345442, EPI_ISL_2345443, EPI_ISL_2345444 | PRONTO SOCORRO MUNICIPAL DE SEVERINIA | Instituto Butantan / ESALQ-Piracicaba | Dimas Tadeu Covas, Antonio Jorge Martins, Claudia Renata dos Santos Barros, David Schlesinger, Debora Botequio Moretti, Elaine Cristina Marqueze, Elaine Vieira Santos, Evandra Strazza Rodrigues, Heidge Fukumasu, Jayme Augusto de Souza-Neto, José Salvatore Leister Patané, Luiz Alcantara, Luiz Lehmann Coutinho, Maria Carolina Elias, Maurício Lacerda Nogueira, Rafael dos Santos Bezerra, Raul Machado Neto, Rejane Maria Tommasini Grotto, Ricardo Haddad, Sandra Coccuzzo Sampaio Vessoni, Simone Kashima, Svetoslav Nanev Slavov, Vincent Louis Viala |
| EPI_ISL_2345445, EPI_ISL_2345446 | VIGILANCIA EPIDEMIOLOGICA JARDINOPOLIS SP | Instituto Butantan / ESALQ-Piracicaba | Dimas Tadeu Covas, Antonio Jorge Martins, Claudia Renata dos Santos Barros, David Schlesinger, Debora Botequio Moretti, Elaine Cristina Marqueze, Elaine Vieira Santos, Evandra Strazza Rodrigues, Heidge Fukumasu, Jayme Augusto de Souza-Neto, José Salvatore Leister Patané, Luiz Alcantara, Luiz Lehmann Coutinho, Maria Carolina Elias, Maurício Lacerda Nogueira, Rafael dos Santos Bezerra, Raul Machado Neto, Rejane Maria Tommasini Grotto, Ricardo Haddad, Sandra Coccuzzo Sampaio Vessoni, Simone Kashima, Svetoslav Nanev Slavov, Vincent Louis Viala |
| EPI_ISL_2345447 | VIGILANCIA EPIDIMIOLOGICA DE ARACARIGUAMA | Instituto Butantan / ESALQ-Piracicaba | Dimas Tadeu Covas, Antonio Jorge Martins, Claudia Renata dos Santos Barros, David Schlesinger, Debora Botequio Moretti, Elaine Cristina Marqueze, Elaine Vieira Santos, Evandra Strazza Rodrigues, Heidge Fukumasu, Jayme Augusto de Souza-Neto, José Salvatore Leister Patané, Luiz Alcantara, Luiz Lehmann Coutinho, Maria Carolina Elias, Maurício Lacerda Nogueira, Rafael dos Santos Bezerra, Raul Machado Neto, Rejane Maria Tommasini Grotto, Ricardo Haddad, Sandra Coccuzzo Sampaio Vessoni, Simone Kashima, Svetoslav Nanev Slavov, Vincent Louis Viala |
| EPI_ISL_2345448 | HOSPITAL MUNICIPAL DE IBIUNA IBIUNA SP | Instituto Butantan / ESALQ-Piracicaba | Dimas Tadeu Covas, Antonio Jorge Martins, Claudia Renata dos Santos Barros, David Schlesinger, Debora Botequio Moretti, Elaine Cristina Marqueze, Elaine Vieira Santos, Evandra Strazza Rodrigues, Heidge Fukumasu, Jayme Augusto de Souza-Neto, José Salvatore Leister Patané, Luiz Alcantara, Luiz Lehmann Coutinho, Maria Carolina Elias, Maurício Lacerda Nogueira, Rafael dos Santos Bezerra, Raul Machado Neto, Rejane Maria Tommasini Grotto, Ricardo Haddad, Sandra Coccuzzo Sampaio Vessoni, Simone Kashima, Svetoslav Nanev Slavov, Vincent Louis Viala |
| EPI_ISL_2345449 | PRONTO ATENDIMENTO UNIDADE SAUDE ADALBERTO ROCHA GUAREI | Instituto Butantan / ESALQ-Piracicaba | Dimas Tadeu Covas, Antonio Jorge Martins, Claudia Renata dos Santos Barros, David Schlesinger, Debora Botequio Moretti, Elaine Cristina Marqueze, Elaine Vieira Santos, Evandra Strazza Rodrigues, Heidge Fukumasu, Jayme Augusto de Souza-Neto, José Salvatore Leister Patané, Luiz Alcantara, Luiz Lehmann Coutinho, Maria Carolina Elias, Maurício Lacerda Nogueira, Rafael dos Santos Bezerra, Raul Machado Neto, Rejane Maria Tommasini Grotto, Ricardo Haddad, Sandra Coccuzzo Sampaio Vessoni, Simone Kashima, Svetoslav Nanev Slavov, Vincent Louis Viala |
| EPI_ISL_2345450 | UNIDADE SENTINELA COVID19 | Instituto Butantan / ESALQ-Piracicaba | Dimas Tadeu Covas, Antonio Jorge Martins, Claudia Renata dos Santos Barros, David Schlesinger, Debora Botequio Moretti, Elaine Cristina Marqueze, Elaine Vieira Santos, Evandra Strazza Rodrigues, Heidge Fukumasu, Jayme Augusto de Souza-Neto, José Salvatore Leister Patané, Luiz Alcantara, Luiz Lehmann Coutinho, Maria Carolina Elias, Maurício Lacerda Nogueira, Rafael dos Santos Bezerra, Raul Machado Neto, Rejane Maria Tommasini Grotto, Ricardo Haddad, Sandra Coccuzzo Sampaio Vessoni, Simone Kashima, Svetoslav Nanev Slavov, Vincent Louis Viala |
| EPI_ISL_2345451 | HOSPITAL REGIONAL DE ITAPETININGA | Instituto Butantan / ESALQ-Piracicaba | Dimas Tadeu Covas, Antonio Jorge Martins, Claudia Renata dos Santos Barros, David Schlesinger, Debora Botequio Moretti, Elaine Cristina Marqueze, Elaine Vieira Santos, Evandra Strazza Rodrigues, Heidge Fukumasu, Jayme Augusto de Souza-Neto, José Salvatore Leister Patané, Luiz Alcantara, Luiz Lehmann Coutinho, Maria Carolina Elias, Maurício Lacerda Nogueira, Rafael dos Santos Bezerra, Raul Machado Neto, Rejane Maria Tommasini Grotto, Ricardo Haddad, Sandra Coccuzzo Sampaio Vessoni, Simone Kashima, Svetoslav Nanev Slavov, Vincent Louis Viala |
| EPI_ISL_2345452 | UNIDADE MISTA ARACOIABA DA SERRA | Instituto Butantan / ESALQ-Piracicaba | Dimas Tadeu Covas, Antonio Jorge Martins, Claudia Renata dos Santos Barros, David Schlesinger, Debora Botequio Moretti, Elaine Cristina Marqueze, Elaine Vieira Santos, Evandra Strazza Rodrigues, Heidge Fukumasu, Jayme Augusto de Souza-Neto, José Salvatore Leister Patané, Luiz Alcantara, Luiz Lehmann Coutinho, Maria Carolina Elias, Maurício Lacerda Nogueira, Rafael dos Santos Bezerra, Raul Machado Neto, Rejane Maria Tommasini Grotto, Ricardo Haddad, Sandra Coccuzzo Sampaio Vessoni, Simone Kashima, Svetoslav Nanev Slavov, Vincent Louis Viala |
| EPI_ISL_2345453 | HOSPITAL REGIONAL DE ITAPETININGA | Instituto Butantan / ESALQ-Piracicaba | Dimas Tadeu Covas, Antonio Jorge Martins, Claudia Renata dos Santos Barros, David Schlesinger, Debora Botequio Moretti, Elaine Cristina Marqueze, Elaine Vieira Santos, Evandra Strazza Rodrigues, Heidge Fukumasu, Jayme Augusto de Souza-Neto, José Salvatore Leister Patané, Luiz Alcantara, Luiz Lehmann Coutinho, Maria Carolina Elias, Maurício Lacerda Nogueira, Rafael dos Santos Bezerra, Raul Machado Neto, Rejane Maria Tommasini Grotto, Ricardo Haddad, Sandra Coccuzzo Sampaio Vessoni, Simone Kashima, Svetoslav Nanev Slavov, Vincent Louis Viala |
| EPI_ISL_2345454 | SMS SECRETARIA MUNICIPAL DE SAUDE DE BOITUVA | Instituto Butantan / ESALQ-Piracicaba | Dimas Tadeu Covas, Antonio Jorge Martins, Claudia Renata dos Santos Barros, David Schlesinger, Debora Botequio Moretti, Elaine Cristina Marqueze, Elaine Vieira Santos, Evandra Strazza Rodrigues, Heidge Fukumasu, Jayme Augusto de Souza-Neto, José Salvatore Leister Patané, Luiz Alcantara, Luiz Lehmann Coutinho, Maria Carolina Elias, Maurício Lacerda Nogueira, Rafael dos Santos Bezerra, Raul Machado Neto, Rejane Maria Tommasini Grotto, Ricardo Haddad, Sandra Coccuzzo Sampaio Vessoni, Simone Kashima, Svetoslav Nanev Slavov, Vincent Louis Viala |
| EPI_ISL_2345455 | UNIDADE BASICA DE SAUDE DR APOLONIO MORAES E SOUZA | Instituto Butantan / ESALQ-Piracicaba | Dimas Tadeu Covas, Antonio Jorge Martins, Claudia Renata dos Santos Barros, David Schlesinger, Debora Botequio Moretti, Elaine Cristina Marqueze, Elaine Vieira Santos, Evandra Strazza Rodrigues, Heidge Fukumasu, Jayme Augusto de Souza-Neto, José Salvatore Leister Patané, Luiz Alcantara, Luiz Lehmann Coutinho, Maria Carolina Elias, Maurício Lacerda Nogueira, Rafael dos Santos Bezerra, Raul Machado Neto, Rejane Maria Tommasini Grotto, Ricardo Haddad, Sandra Coccuzzo Sampaio Vessoni, Simone Kashima, Svetoslav Nanev Slavov, Vincent Louis Viala |
| EPI_ISL_2345456 | UNIDADE BASICA DE SAUDE DR WILSON HAYEK SAIHG | Instituto Butantan / ESALQ-Piracicaba | Dimas Tadeu Covas, Antonio Jorge Martins, Claudia Renata dos Santos Barros, David Schlesinger, Debora Botequio Moretti, Elaine Cristina Marqueze, Elaine Vieira Santos, Evandra Strazza Rodrigues, Heidge Fukumasu, Jayme Augusto de Souza-Neto, José Salvatore Leister Patané, Luiz Alcantara, Luiz Lehmann Coutinho, Maria Carolina Elias, Maurício Lacerda Nogueira, Rafael dos Santos Bezerra, Raul Machado Neto, Rejane Maria Tommasini Grotto, Ricardo Haddad, Sandra Coccuzzo Sampaio Vessoni, Simone Kashima, Svetoslav Nanev Slavov, Vincent Louis Viala |
| EPI_ISL_2345457 | UNIDADE BASICA DR JOSE PARASSU CARVALHO | Instituto Butantan / ESALQ-Piracicaba | Dimas Tadeu Covas, Antonio Jorge Martins, Claudia Renata dos Santos Barros, David Schlesinger, Debora Botequio Moretti, Elaine Cristina Marqueze, Elaine Vieira Santos, Evandra Strazza Rodrigues, Heidge Fukumasu, Jayme Augusto de Souza-Neto, José Salvatore Leister Patané, Luiz Alcantara, Luiz Lehmann Coutinho, Maria Carolina Elias, Maurício Lacerda Nogueira, Rafael dos Santos Bezerra, Raul Machado Neto, Rejane Maria Tommasini Grotto, Ricardo Haddad, Sandra Coccuzzo Sampaio Vessoni, Simone Kashima, Svetoslav Nanev Slavov, Vincent Louis Viala |
| EPI_ISL_2345458 | UNIDADE BASICA DE SAUDE DR LOTFALLAH MIZIARA | Instituto Butantan / ESALQ-Piracicaba | Dimas Tadeu Covas, Antonio Jorge Martins, Claudia Renata dos Santos Barros, David Schlesinger, Debora Botequio Moretti, Elaine Cristina Marqueze, Elaine Vieira Santos, Evandra Strazza Rodrigues, Heidge Fukumasu, Jayme Augusto de Souza-Neto, José Salvatore Leister Patané, Luiz Alcantara, Luiz Lehmann Coutinho, Maria Carolina Elias, Maurício Lacerda Nogueira, Rafael dos Santos Bezerra, Raul Machado Neto, Rejane Maria Tommasini Grotto, Ricardo Haddad, Sandra Coccuzzo Sampaio Vessoni, Simone Kashima, Svetoslav Nanev Slavov, Vincent Louis Viala |
| EPI_ISL_2345459, EPI_ISL_2345460 | CENTRO DE REFERENCIA DO IDOSO DR HUMBERTO MENDES DE CARVALHO | Instituto Butantan / ESALQ-Piracicaba | Dimas Tadeu Covas, Antonio Jorge Martins, Claudia Renata dos Santos Barros, David Schlesinger, Debora Botequio Moretti, Elaine Cristina Marqueze, Elaine Vieira Santos, Evandra Strazza Rodrigues, Heidge Fukumasu, Jayme Augusto de Souza-Neto, José Salvatore Leister Patané, Luiz Alcantara, Luiz |

| | | | |
|---|---|---|---|
| | | | Lehmann Coutinho, Maria Carolina Elias, Maurício Lacerda Nogueira, Rafael dos Santos Bezerra, Raul Machado Neto, Rejane Maria Tommasini Grotto, Ricardo Haddad, Sandra Coccuzzo Sampaio Vessoni, Simone Kashima, Svetoslav Nanev Slavov, Vincent Louis Viala |
| EPI_ISL_2345461, EPI_ISL_2345462, EPI_ISL_2345463 | UNIDADE DE SAUDE DA FAMILIA JOSE ADALBERTO LELLIS GARCIA | Instituto Butantan / ESALQ-Piracicaba | Dimas Tadeu Covas, Antonio Jorge Martins, Claudia Renata dos Santos Barros, David Schlesinger, Debora Botequio Moretti, Elaine Cristina Marqueze, Elaine Vieira Santos, Evandra Strazza Rodrigues, Heidge Fukumasu, Jayme Augusto de Souza-Neto, José Salvatore Leister Patané, Luiz Alcantara, Luiz Lehmann Coutinho, Maria Carolina Elias, Maurício Lacerda Nogueira, Rafael dos Santos Bezerra, Raul Machado Neto, Rejane Maria Tommasini Grotto, Ricardo Haddad, Sandra Coccuzzo Sampaio Vessoni, Simone Kashima, Svetoslav Nanev Slavov, Vincent Louis Viala |
| EPI_ISL_2345464, EPI_ISL_2345465 | CS DE URUPES | Instituto Butantan / ESALQ-Piracicaba | Dimas Tadeu Covas, Antonio Jorge Martins, Claudia Renata dos Santos Barros, David Schlesinger, Debora Botequio Moretti, Elaine Cristina Marqueze, Elaine Vieira Santos, Evandra Strazza Rodrigues, Heidge Fukumasu, Jayme Augusto de Souza-Neto, José Salvatore Leister Patané, Luiz Alcantara, Luiz Lehmann Coutinho, Maria Carolina Elias, Maurício Lacerda Nogueira, Rafael dos Santos Bezerra, Raul Machado Neto, Rejane Maria Tommasini Grotto, Ricardo Haddad, Sandra Coccuzzo Sampaio Vessoni, Simone Kashima, Svetoslav Nanev Slavov, Vincent Louis Viala |
| EPI_ISL_2345466, EPI_ISL_2345467, EPI_ISL_2345468, EPI_ISL_2345469, EPI_ISL_2345470 | CS DE BALSAMO | Instituto Butantan / ESALQ-Piracicaba | Dimas Tadeu Covas, Antonio Jorge Martins, Claudia Renata dos Santos Barros, David Schlesinger, Debora Botequio Moretti, Elaine Cristina Marqueze, Elaine Vieira Santos, Evandra Strazza Rodrigues, Heidge Fukumasu, Jayme Augusto de Souza-Neto, José Salvatore Leister Patané, Luiz Alcantara, Luiz Lehmann Coutinho, Maria Carolina Elias, Maurício Lacerda Nogueira, Rafael dos Santos Bezerra, Raul Machado Neto, Rejane Maria Tommasini Grotto, Ricardo Haddad, Sandra Coccuzzo Sampaio Vessoni, Simone Kashima, Svetoslav Nanev Slavov, Vincent Louis Viala |
| EPI_ISL_2345471 | CS DE URUPES | Instituto Butantan / ESALQ-Piracicaba | Dimas Tadeu Covas, Antonio Jorge Martins, Claudia Renata dos Santos Barros, David Schlesinger, Debora Botequio Moretti, Elaine Cristina Marqueze, Elaine Vieira Santos, Evandra Strazza Rodrigues, Heidge Fukumasu, Jayme Augusto de Souza-Neto, José Salvatore Leister Patané, Luiz Alcantara, Luiz Lehmann Coutinho, Maria Carolina Elias, Maurício Lacerda Nogueira, Rafael dos Santos Bezerra, Raul Machado Neto, Rejane Maria Tommasini Grotto, Ricardo Haddad, Sandra Coccuzzo Sampaio Vessoni, Simone Kashima, Svetoslav Nanev Slavov, Vincent Louis Viala |
| EPI_ISL_2345472 | CS DE SEBASTIANOPOLIS DO SUL | Instituto Butantan / ESALQ-Piracicaba | Dimas Tadeu Covas, Antonio Jorge Martins, Claudia Renata dos Santos Barros, David Schlesinger, Debora Botequio Moretti, Elaine Cristina Marqueze, Elaine Vieira Santos, Evandra Strazza Rodrigues, Heidge Fukumasu, Jayme Augusto de Souza-Neto, José Salvatore Leister Patané, Luiz Alcantara, Luiz Lehmann Coutinho, Maria Carolina Elias, Maurício Lacerda Nogueira, Rafael dos Santos Bezerra, Raul Machado Neto, Rejane Maria Tommasini Grotto, Ricardo Haddad, Sandra Coccuzzo Sampaio Vessoni, Simone Kashima, Svetoslav Nanev Slavov, Vincent Louis Viala |
| EPI_ISL_2345473 | CS DE BALSAMO | Instituto Butantan / ESALQ-Piracicaba | Dimas Tadeu Covas, Antonio Jorge Martins, Claudia Renata dos Santos Barros, David Schlesinger, Debora Botequio Moretti, Elaine Cristina Marqueze, Elaine Vieira Santos, Evandra Strazza Rodrigues, Heidge Fukumasu, Jayme Augusto de Souza-Neto, José Salvatore Leister Patané, Luiz Alcantara, Luiz Lehmann Coutinho, Maria Carolina Elias, Maurício Lacerda Nogueira, Rafael dos Santos Bezerra, Raul Machado Neto, Rejane Maria Tommasini Grotto, Ricardo Haddad, Sandra Coccuzzo Sampaio Vessoni, Simone Kashima, Svetoslav Nanev Slavov, Vincent Louis Viala |
| EPI_ISL_2345474 | CS DE URUPES | Instituto Butantan / ESALQ-Piracicaba | Dimas Tadeu Covas, Antonio Jorge Martins, Claudia Renata dos Santos Barros, David Schlesinger, Debora Botequio Moretti, Elaine Cristina Marqueze, Elaine Vieira Santos, Evandra Strazza Rodrigues, Heidge Fukumasu, Jayme Augusto de Souza-Neto, José Salvatore Leister Patané, Luiz Alcantara, Luiz Lehmann Coutinho, Maria Carolina Elias, Maurício Lacerda Nogueira, Rafael dos Santos Bezerra, Raul Machado Neto, Rejane Maria Tommasini Grotto, Ricardo Haddad, Sandra Coccuzzo Sampaio Vessoni, Simone Kashima, Svetoslav Nanev Slavov, Vincent Louis Viala |
| EPI_ISL_2345475 | CS DE BALSAMO | Instituto Butantan / ESALQ-Piracicaba | Dimas Tadeu Covas, Antonio Jorge Martins, Claudia Renata dos Santos Barros, David Schlesinger, Debora Botequio Moretti, Elaine Cristina Marqueze, Elaine Vieira Santos, Evandra Strazza Rodrigues, Heidge Fukumasu, Jayme Augusto de Souza-Neto, José Salvatore Leister Patané, Luiz Alcantara, Luiz Lehmann Coutinho, Maria Carolina Elias, Maurício Lacerda Nogueira, Rafael dos Santos Bezerra, Raul Machado Neto, Rejane Maria Tommasini Grotto, Ricardo Haddad, Sandra Coccuzzo Sampaio Vessoni, Simone Kashima, Svetoslav Nanev Slavov, Vincent Louis Viala |
| EPI_ISL_2345476, EPI_ISL_2345477 | CS DE URUPES | Instituto Butantan / ESALQ-Piracicaba | Dimas Tadeu Covas, Antonio Jorge Martins, Claudia Renata dos Santos Barros, David Schlesinger, Debora Botequio Moretti, Elaine Cristina Marqueze, Elaine Vieira Santos, Evandra Strazza Rodrigues, Heidge Fukumasu, Jayme Augusto de Souza-Neto, José Salvatore Leister Patané, Luiz Alcantara, Luiz Lehmann Coutinho, Maria Carolina Elias, Maurício Lacerda Nogueira, Rafael dos Santos Bezerra, Raul Machado Neto, Rejane Maria Tommasini Grotto, Ricardo Haddad, Sandra Coccuzzo Sampaio Vessoni, Simone Kashima, Svetoslav Nanev Slavov, Vincent Louis Viala |
| EPI_ISL_2345478 | EMERGENCIA RESPIRATORIA DE NOVA GRANADA | Instituto Butantan / ESALQ-Piracicaba | Dimas Tadeu Covas, Antonio Jorge Martins, Claudia Renata dos Santos Barros, David Schlesinger, Debora Botequio Moretti, Elaine Cristina Marqueze, Elaine Vieira Santos, Evandra Strazza Rodrigues, Heidge Fukumasu, Jayme Augusto de Souza-Neto, José Salvatore Leister Patané, Luiz Alcantara, Luiz Lehmann Coutinho, Maria Carolina Elias, Maurício Lacerda Nogueira, Rafael dos Santos Bezerra, Raul Machado Neto, Rejane Maria Tommasini Grotto, Ricardo Haddad, Sandra Coccuzzo Sampaio Vessoni, Simone Kashima, Svetoslav Nanev Slavov, Vincent Louis Viala |
| EPI_ISL_2345479 | UBS DE SANTA SALETE | Instituto Butantan / ESALQ-Piracicaba | Dimas Tadeu Covas, Antonio Jorge Martins, Claudia Renata dos Santos Barros, David Schlesinger, Debora Botequio Moretti, Elaine Cristina Marqueze, Elaine Vieira Santos, Evandra Strazza Rodrigues, Heidge Fukumasu, Jayme Augusto de Souza-Neto, José Salvatore Leister Patané, Luiz Alcantara, Luiz Lehmann Coutinho, Maria Carolina Elias, Maurício Lacerda Nogueira, Rafael dos Santos Bezerra, Raul Machado Neto, Rejane Maria Tommasini Grotto, Ricardo Haddad, Sandra Coccuzzo Sampaio Vessoni, Simone Kashima, Svetoslav Nanev Slavov, Vincent Louis Viala |
| EPI_ISL_2345480, EPI_ISL_2345481 | EMERGENCIA RESPIRATORIA DE NOVA GRANADA | Instituto Butantan / ESALQ-Piracicaba | Dimas Tadeu Covas, Antonio Jorge Martins, Claudia Renata dos Santos Barros, David Schlesinger, Debora Botequio Moretti, Elaine Cristina Marqueze, Elaine Vieira Santos, Evandra Strazza Rodrigues, Heidge Fukumasu, Jayme Augusto de Souza-Neto, José Salvatore Leister Patané, Luiz Alcantara, Luiz Lehmann Coutinho, Maria Carolina Elias, Maurício Lacerda Nogueira, Rafael dos Santos Bezerra, Raul Machado Neto, Rejane Maria Tommasini Grotto, Ricardo Haddad, Sandra Coccuzzo Sampaio Vessoni, Simone Kashima, Svetoslav Nanev Slavov, Vincent Louis Viala |
| EPI_ISL_2345482, EPI_ISL_2345483, EPI_ISL_2345484 | ESF NOVA TANABI II | Instituto Butantan / ESALQ-Piracicaba | Dimas Tadeu Covas, Antonio Jorge Martins, Claudia Renata dos Santos Barros, David Schlesinger, Debora Botequio Moretti, Elaine Cristina Marqueze, Elaine Vieira Santos, Evandra Strazza Rodrigues, Heidge Fukumasu, Jayme Augusto de Souza-Neto, José Salvatore Leister Patané, Luiz Alcantara, Luiz Lehmann Coutinho, Maria Carolina Elias, Maurício Lacerda Nogueira, Rafael dos Santos Bezerra, Raul Machado Neto, Rejane Maria Tommasini Grotto, Ricardo Haddad, Sandra Coccuzzo Sampaio Vessoni, Simone Kashima, Svetoslav Nanev Slavov, Vincent Louis Viala |
| EPI_ISL_2345485, EPI_ISL_2345486, EPI_ISL_2345487, EPI_ISL_2345488, EPI_ISL_2345489, EPI_ISL_2345490, EPI_ISL_2345491 | UBS II DE TANABI MILTON MARTINS PERCHES | Instituto Butantan / ESALQ-Piracicaba | Dimas Tadeu Covas, Antonio Jorge Martins, Claudia Renata dos Santos Barros, David Schlesinger, Debora Botequio Moretti, Elaine Cristina Marqueze, Elaine Vieira Santos, Evandra Strazza Rodrigues, Heidge Fukumasu, Jayme Augusto de Souza-Neto, José Salvatore Leister Patané, Luiz Alcantara, Luiz Lehmann Coutinho, Maria Carolina Elias, Maurício Lacerda Nogueira, Rafael dos Santos Bezerra, Raul Machado Neto, Rejane Maria Tommasini Grotto, Ricardo Haddad, Sandra Coccuzzo Sampaio Vessoni, Simone Kashima, Svetoslav Nanev Slavov, Vincent Louis Viala |
| EPI_ISL_2345492 | CS DE PLANALTO | Instituto Butantan / ESALQ-Piracicaba | Dimas Tadeu Covas, Antonio Jorge Martins, Claudia Renata dos Santos Barros, David Schlesinger, Debora Botequio Moretti, Elaine Cristina Marqueze, Elaine Vieira Santos, Evandra Strazza Rodrigues, Heidge Fukumasu, Jayme Augusto de Souza-Neto, José Salvatore Leister Patané, Luiz Alcantara, Luiz Lehmann Coutinho, Maria Carolina Elias, Maurício Lacerda Nogueira, Rafael dos Santos Bezerra, Raul Machado Neto, Rejane Maria Tommasini Grotto, Ricardo Haddad, Sandra Coccuzzo Sampaio Vessoni, Simone Kashima, Svetoslav Nanev Slavov, Vincent Louis Viala |
| EPI_ISL_2345493, EPI_ISL_2345494, EPI_ISL_2345495, EPI_ISL_2345496, EPI_ISL_2345497 | CS DE ICEM | Instituto Butantan / ESALQ-Piracicaba | Dimas Tadeu Covas, Antonio Jorge Martins, Claudia Renata dos Santos Barros, David Schlesinger, Debora Botequio Moretti, Elaine Cristina Marqueze, Elaine Vieira Santos, Evandra Strazza Rodrigues, Heidge Fukumasu, Jayme Augusto de Souza-Neto, José Salvatore Leister Patané, Luiz Alcantara, Luiz Lehmann Coutinho, Maria Carolina Elias, Maurício Lacerda Nogueira, Rafael dos Santos Bezerra, Raul Machado Neto, Rejane Maria Tommasini Grotto, Ricardo Haddad, Sandra Coccuzzo Sampaio Vessoni, Simone Kashima, Svetoslav Nanev Slavov, Vincent Louis Viala |
| EPI_ISL_2345498 | CS DE PARANAPUA | Instituto Butantan / ESALQ-Piracicaba | Dimas Tadeu Covas, Antonio Jorge Martins, Claudia Renata dos Santos Barros, David Schlesinger, Debora Botequio Moretti, Elaine Cristina Marqueze, Elaine Vieira Santos, Evandra Strazza Rodrigues, Heidge Fukumasu, Jayme Augusto de Souza-Neto, José Salvatore Leister Patané, Luiz Alcantara, Luiz Lehmann Coutinho, Maria Carolina Elias, Maurício Lacerda Nogueira, Rafael dos Santos Bezerra, Raul Machado Neto, Rejane Maria Tommasini Grotto, Ricardo Haddad, Sandra Coccuzzo Sampaio Vessoni, Simone Kashima, Svetoslav Nanev Slavov, Vincent Louis Viala |
| EPI_ISL_2345499 | CS DE ICEM | Instituto Butantan / ESALQ-Piracicaba | Dimas Tadeu Covas, Antonio Jorge Martins, Claudia Renata dos Santos Barros, David Schlesinger, Debora Botequio Moretti, Elaine Cristina Marqueze, Elaine Vieira Santos, Evandra Strazza Rodrigues, Heidge Fukumasu, Jayme Augusto de Souza-Neto, José Salvatore Leister Patané, Luiz Alcantara, Luiz Lehmann Coutinho, Maria Carolina Elias, Maurício Lacerda Nogueira, Rafael dos Santos Bezerra, Raul Machado Neto, Rejane Maria Tommasini Grotto, Ricardo Haddad, Sandra Coccuzzo Sampaio Vessoni, Simone Kashima, Svetoslav Nanev Slavov, Vincent Louis Viala |
| EPI_ISL_2345500, EPI_ISL_2345501, EPI_ISL_2345502 | UBS LUIS FACHIN IPIGUA | Instituto Butantan / ESALQ-Piracicaba | Dimas Tadeu Covas, Antonio Jorge Martins, Claudia Renata dos Santos Barros, David Schlesinger, Debora Botequio Moretti, Elaine Cristina Marqueze, Elaine Vieira Santos, Evandra Strazza Rodrigues, Heidge Fukumasu, Jayme Augusto de Souza-Neto, José Salvatore Leister Patané, Luiz Alcantara, Luiz Lehmann Coutinho, Maria Carolina Elias, Maurício Lacerda Nogueira, Rafael dos Santos Bezerra, Raul Machado Neto, Rejane Maria Tommasini Grotto, Ricardo Haddad, Sandra Coccuzzo Sampaio Vessoni, Simone Kashima, Svetoslav Nanev Slavov, Vincent Louis Viala |

| | | | |
|---|---|---|---|
| EPI_ISL_2345503, EPI_ISL_2345504 | UBS TEREZA GALLO UCHOA | Instituto Butantan / ESALQ-Piracicaba | Dimas Tadeu Covas, Antonio Jorge Martins, Claudia Renata dos Santos Barros, David Schlesinger, Debora Botequio Moretti, Elaine Cristina Marqueze, Elaine Vieira Santos, Evandra Strazza Rodrigues, Heidge Fukumasu, Jayme Augusto de Souza-Neto, José Salvatore Leister Patané, Luiz Alcantara, Luiz Lehmann Coutinho, Maria Carolina Elias, Maurício Lacerda Nogueira, Rafael dos Santos Bezerra, Raul Machado Neto, Rejane Maria Tommasini Grotto, Ricardo Haddad, Sandra Coccuzzo Sampaio Vessoni, Simone Kashima, Svetoslav Nanev Slavov, Vincent Louis Viala |
| EPI_ISL_2345505 | CS DE MARAPOAMA | Instituto Butantan / ESALQ-Piracicaba | Dimas Tadeu Covas, Antonio Jorge Martins, Claudia Renata dos Santos Barros, David Schlesinger, Debora Botequio Moretti, Elaine Cristina Marqueze, Elaine Vieira Santos, Evandra Strazza Rodrigues, Heidge Fukumasu, Jayme Augusto de Souza-Neto, José Salvatore Leister Patané, Luiz Alcantara, Luiz Lehmann Coutinho, Maria Carolina Elias, Maurício Lacerda Nogueira, Rafael dos Santos Bezerra, Raul Machado Neto, Rejane Maria Tommasini Grotto, Ricardo Haddad, Sandra Coccuzzo Sampaio Vessoni, Simone Kashima, Svetoslav Nanev Slavov, Vincent Louis Viala |
| EPI_ISL_2345506 | CS DE PARANAPUA | Instituto Butantan / ESALQ-Piracicaba | Dimas Tadeu Covas, Antonio Jorge Martins, Claudia Renata dos Santos Barros, David Schlesinger, Debora Botequio Moretti, Elaine Cristina Marqueze, Elaine Vieira Santos, Evandra Strazza Rodrigues, Heidge Fukumasu, Jayme Augusto de Souza-Neto, José Salvatore Leister Patané, Luiz Alcantara, Luiz Lehmann Coutinho, Maria Carolina Elias, Maurício Lacerda Nogueira, Rafael dos Santos Bezerra, Raul Machado Neto, Rejane Maria Tommasini Grotto, Ricardo Haddad, Sandra Coccuzzo Sampaio Vessoni, Simone Kashima, Svetoslav Nanev Slavov, Vincent Louis Viala |
| EPI_ISL_2345507 | UBS TEREZA GALLO UCHOA | Instituto Butantan / ESALQ-Piracicaba | Dimas Tadeu Covas, Antonio Jorge Martins, Claudia Renata dos Santos Barros, David Schlesinger, Debora Botequio Moretti, Elaine Cristina Marqueze, Elaine Vieira Santos, Evandra Strazza Rodrigues, Heidge Fukumasu, Jayme Augusto de Souza-Neto, José Salvatore Leister Patané, Luiz Alcantara, Luiz Lehmann Coutinho, Maria Carolina Elias, Maurício Lacerda Nogueira, Rafael dos Santos Bezerra, Raul Machado Neto, Rejane Maria Tommasini Grotto, Ricardo Haddad, Sandra Coccuzzo Sampaio Vessoni, Simone Kashima, Svetoslav Nanev Slavov, Vincent Louis Viala |
| EPI_ISL_2345508 | CS DE PARANAPUA | Instituto Butantan / ESALQ-Piracicaba | Dimas Tadeu Covas, Antonio Jorge Martins, Claudia Renata dos Santos Barros, David Schlesinger, Debora Botequio Moretti, Elaine Cristina Marqueze, Elaine Vieira Santos, Evandra Strazza Rodrigues, Heidge Fukumasu, Jayme Augusto de Souza-Neto, José Salvatore Leister Patané, Luiz Alcantara, Luiz Lehmann Coutinho, Maria Carolina Elias, Maurício Lacerda Nogueira, Rafael dos Santos Bezerra, Raul Machado Neto, Rejane Maria Tommasini Grotto, Ricardo Haddad, Sandra Coccuzzo Sampaio Vessoni, Simone Kashima, Svetoslav Nanev Slavov, Vincent Louis Viala |
| EPI_ISL_2345509 | UBS TEREZA GALLO UCHOA | Instituto Butantan / ESALQ-Piracicaba | Dimas Tadeu Covas, Antonio Jorge Martins, Claudia Renata dos Santos Barros, David Schlesinger, Debora Botequio Moretti, Elaine Cristina Marqueze, Elaine Vieira Santos, Evandra Strazza Rodrigues, Heidge Fukumasu, Jayme Augusto de Souza-Neto, José Salvatore Leister Patané, Luiz Alcantara, Luiz Lehmann Coutinho, Maria Carolina Elias, Maurício Lacerda Nogueira, Rafael dos Santos Bezerra, Raul Machado Neto, Rejane Maria Tommasini Grotto, Ricardo Haddad, Sandra Coccuzzo Sampaio Vessoni, Simone Kashima, Svetoslav Nanev Slavov, Vincent Louis Viala |
| EPI_ISL_2345510, EPI_ISL_2345511, EPI_ISL_2345512, EPI_ISL_2345513 | CENTRO MEDICO PMESP | Instituto Butantan / ESALQ-Piracicaba | Dimas Tadeu Covas, Antonio Jorge Martins, Claudia Renata dos Santos Barros, David Schlesinger, Debora Botequio Moretti, Elaine Cristina Marqueze, Elaine Vieira Santos, Evandra Strazza Rodrigues, Heidge Fukumasu, Jayme Augusto de Souza-Neto, José Salvatore Leister Patané, Luiz Alcantara, Luiz Lehmann Coutinho, Maria Carolina Elias, Maurício Lacerda Nogueira, Rafael dos Santos Bezerra, Raul Machado Neto, Rejane Maria Tommasini Grotto, Ricardo Haddad, Sandra Coccuzzo Sampaio Vessoni, Simone Kashima, Svetoslav Nanev Slavov, Vincent Louis Viala |
| EPI_ISL_2345514, EPI_ISL_2345515, EPI_ISL_2345516, EPI_ISL_2345517, EPI_ISL_2345518, EPI_ISL_2345519, EPI_ISL_2345520, EPI_ISL_2345521, EPI_ISL_2345522, EPI_ISL_2345523, EPI_ISL_2345524 | | | |
| see above | CS DE PALESTINA | Instituto Butantan / ESALQ-Piracicaba | Dimas Tadeu Covas, Antonio Jorge Martins, Claudia Renata dos Santos Barros, David Schlesinger, Debora Botequio Moretti, Elaine Cristina Marqueze, Elaine Vieira Santos, Evandra Strazza Rodrigues, Heidge Fukumasu, Jayme Augusto de Souza-Neto, José Salvatore Leister Patané, Luiz Alcantara, Luiz Lehmann Coutinho, Maria Carolina Elias, Maurício Lacerda Nogueira, Rafael dos Santos Bezerra, Raul Machado Neto, Rejane Maria Tommasini Grotto, Ricardo Haddad, Sandra Coccuzzo Sampaio Vessoni, Simone Kashima, Svetoslav Nanev Slavov, Vincent Louis Viala |
| EPI_ISL_2345525 | UBS ALCIMINIO DE ASSIS LOURENCO BADY BASSITT | Instituto Butantan / ESALQ-Piracicaba | Dimas Tadeu Covas, Antonio Jorge Martins, Claudia Renata dos Santos Barros, David Schlesinger, Debora Botequio Moretti, Elaine Cristina Marqueze, Elaine Vieira Santos, Evandra Strazza Rodrigues, Heidge Fukumasu, Jayme Augusto de Souza-Neto, José Salvatore Leister Patané, Luiz Alcantara, Luiz Lehmann Coutinho, Maria Carolina Elias, Maurício Lacerda Nogueira, Rafael dos Santos Bezerra, Raul Machado Neto, Rejane Maria Tommasini Grotto, Ricardo Haddad, Sandra Coccuzzo Sampaio Vessoni, Simone Kashima, Svetoslav Nanev Slavov, Vincent Louis Viala |
| EPI_ISL_2345526 | CS DE NIPOA | Instituto Butantan / ESALQ-Piracicaba | Dimas Tadeu Covas, Antonio Jorge Martins, Claudia Renata dos Santos Barros, David Schlesinger, Debora Botequio Moretti, Elaine Cristina Marqueze, Elaine Vieira Santos, Evandra Strazza Rodrigues, Heidge Fukumasu, Jayme Augusto de Souza-Neto, José Salvatore Leister Patané, Luiz Alcantara, Luiz Lehmann Coutinho, Maria Carolina Elias, Maurício Lacerda Nogueira, Rafael dos Santos Bezerra, Raul Machado Neto, Rejane Maria Tommasini Grotto, Ricardo Haddad, Sandra Coccuzzo Sampaio Vessoni, Simone Kashima, Svetoslav Nanev Slavov, Vincent Louis Viala |
| EPI_ISL_2345527 | UBS ALCIMINIO DE ASSIS LOURENCO BADY BASSITT | Instituto Butantan / ESALQ-Piracicaba | Dimas Tadeu Covas, Antonio Jorge Martins, Claudia Renata dos Santos Barros, David Schlesinger, Debora Botequio Moretti, Elaine Cristina Marqueze, Elaine Vieira Santos, Evandra Strazza Rodrigues, Heidge Fukumasu, Jayme Augusto de Souza-Neto, José Salvatore Leister Patané, Luiz Alcantara, Luiz Lehmann Coutinho, Maria Carolina Elias, Maurício Lacerda Nogueira, Rafael dos Santos Bezerra, Raul Machado Neto, Rejane Maria Tommasini Grotto, Ricardo Haddad, Sandra Coccuzzo Sampaio Vessoni, Simone Kashima, Svetoslav Nanev Slavov, Vincent Louis Viala |
| EPI_ISL_2345528 | USF ROSA CRUZ | Instituto Butantan / ESALQ-Piracicaba | Dimas Tadeu Covas, Antonio Jorge Martins, Claudia Renata dos Santos Barros, David Schlesinger, Debora Botequio Moretti, Elaine Cristina Marqueze, Elaine Vieira Santos, Evandra Strazza Rodrigues, Heidge Fukumasu, Jayme Augusto de Souza-Neto, José Salvatore Leister Patané, Luiz Alcantara, Luiz Lehmann Coutinho, Maria Carolina Elias, Maurício Lacerda Nogueira, Rafael dos Santos Bezerra, Raul Machado Neto, Rejane Maria Tommasini Grotto, Ricardo Haddad, Sandra Coccuzzo Sampaio Vessoni, Simone Kashima, Svetoslav Nanev Slavov, Vincent Louis Viala |
| EPI_ISL_2345529 | USF EUCALIPTOS | Instituto Butantan / ESALQ-Piracicaba | Dimas Tadeu Covas, Antonio Jorge Martins, Claudia Renata dos Santos Barros, David Schlesinger, Debora Botequio Moretti, Elaine Cristina Marqueze, Elaine Vieira Santos, Evandra Strazza Rodrigues, Heidge Fukumasu, Jayme Augusto de Souza-Neto, José Salvatore Leister Patané, Luiz Alcantara, Luiz Lehmann Coutinho, Maria Carolina Elias, Maurício Lacerda Nogueira, Rafael dos Santos Bezerra, Raul Machado Neto, Rejane Maria Tommasini Grotto, Ricardo Haddad, Sandra Coccuzzo Sampaio Vessoni, Simone Kashima, Svetoslav Nanev Slavov, Vincent Louis Viala |
| EPI_ISL_2345530, EPI_ISL_2345531, EPI_ISL_2345532 | AMBULATORIO DE ESPECIALIDADE V E MOGI MIRIM | Instituto Butantan / ESALQ-Piracicaba | Dimas Tadeu Covas, Antonio Jorge Martins, Claudia Renata dos Santos Barros, David Schlesinger, Debora Botequio Moretti, Elaine Cristina Marqueze, Elaine Vieira Santos, Evandra Strazza Rodrigues, Heidge Fukumasu, Jayme Augusto de Souza-Neto, José Salvatore Leister Patané, Luiz Alcantara, Luiz Lehmann Coutinho, Maria Carolina Elias, Maurício Lacerda Nogueira, Rafael dos Santos Bezerra, Raul Machado Neto, Rejane Maria Tommasini Grotto, Ricardo Haddad, Sandra Coccuzzo Sampaio Vessoni, Simone Kashima, Svetoslav Nanev Slavov, Vincent Louis Viala |
| EPI_ISL_2345533 | CENTRO DE SAUDE III AFFONSO LUZZI SANTA CRUZ DAS PALMEIRAS | Instituto Butantan / ESALQ-Piracicaba | Dimas Tadeu Covas, Antonio Jorge Martins, Claudia Renata dos Santos Barros, David Schlesinger, Debora Botequio Moretti, Elaine Cristina Marqueze, Elaine Vieira Santos, Evandra Strazza Rodrigues, Heidge Fukumasu, Jayme Augusto de Souza-Neto, José Salvatore Leister Patané, Luiz Alcantara, Luiz Lehmann Coutinho, Maria Carolina Elias, Maurício Lacerda Nogueira, Rafael dos Santos Bezerra, Raul Machado Neto, Rejane Maria Tommasini Grotto, Ricardo Haddad, Sandra Coccuzzo Sampaio Vessoni, Simone Kashima, Svetoslav Nanev Slavov, Vincent Louis Viala |
| EPI_ISL_2345534, EPI_ISL_2345535 | UNIDADE DE SAUDE DE ITOBI ALCIBIADES PIRES | Instituto Butantan / ESALQ-Piracicaba | Dimas Tadeu Covas, Antonio Jorge Martins, Claudia Renata dos Santos Barros, David Schlesinger, Debora Botequio Moretti, Elaine Cristina Marqueze, Elaine Vieira Santos, Evandra Strazza Rodrigues, Heidge Fukumasu, Jayme Augusto de Souza-Neto, José Salvatore Leister Patané, Luiz Alcantara, Luiz Lehmann Coutinho, Maria Carolina Elias, Maurício Lacerda Nogueira, Rafael dos Santos Bezerra, Raul Machado Neto, Rejane Maria Tommasini Grotto, Ricardo Haddad, Sandra Coccuzzo Sampaio Vessoni, Simone Kashima, Svetoslav Nanev Slavov, Vincent Louis Viala |
| EPI_ISL_2345536 | USF GUACUANO | Instituto Butantan / ESALQ-Piracicaba | Dimas Tadeu Covas, Antonio Jorge Martins, Claudia Renata dos Santos Barros, David Schlesinger, Debora Botequio Moretti, Elaine Cristina Marqueze, Elaine Vieira Santos, Evandra Strazza Rodrigues, Heidge Fukumasu, Jayme Augusto de Souza-Neto, José Salvatore Leister Patané, Luiz Alcantara, Luiz Lehmann Coutinho, Maria Carolina Elias, Maurício Lacerda Nogueira, Rafael dos Santos Bezerra, Raul Machado Neto, Rejane Maria Tommasini Grotto, Ricardo Haddad, Sandra Coccuzzo Sampaio Vessoni, Simone Kashima, Svetoslav Nanev Slavov, Vincent Louis Viala |
| EPI_ISL_2345537, EPI_ISL_2345538, EPI_ISL_2345539 | CENTRO DE SAUDE III AFFONSO LUZZI SANTA CRUZ DAS PALMEIRAS | Instituto Butantan / ESALQ-Piracicaba | Dimas Tadeu Covas, Antonio Jorge Martins, Claudia Renata dos Santos Barros, David Schlesinger, Debora Botequio Moretti, Elaine Cristina Marqueze, Elaine Vieira Santos, Evandra Strazza Rodrigues, Heidge Fukumasu, Jayme Augusto de Souza-Neto, José Salvatore Leister Patané, Luiz Alcantara, Luiz Lehmann Coutinho, Maria Carolina Elias, Maurício Lacerda Nogueira, Rafael dos Santos Bezerra, Raul Machado Neto, Rejane Maria Tommasini Grotto, Ricardo Haddad, Sandra Coccuzzo Sampaio Vessoni, Simone Kashima, Svetoslav Nanev Slavov, Vincent Louis Viala |
| EPI_ISL_2345540 | UNIDADE DE SAUDE DE ITOBI ALCIBIADES PIRES | Instituto Butantan / ESALQ-Piracicaba | Dimas Tadeu Covas, Antonio Jorge Martins, Claudia Renata dos Santos Barros, David Schlesinger, Debora Botequio Moretti, Elaine Cristina Marqueze, Elaine Vieira Santos, Evandra Strazza Rodrigues, Heidge Fukumasu, Jayme Augusto de Souza-Neto, José Salvatore Leister Patané, Luiz Alcantara, Luiz Lehmann Coutinho, Maria Carolina Elias, Maurício Lacerda Nogueira, Rafael dos Santos Bezerra, Raul Machado Neto, Rejane Maria Tommasini Grotto, Ricardo Haddad, Sandra Coccuzzo Sampaio Vessoni, Simone Kashima, Svetoslav Nanev Slavov, Vincent Louis Viala |

| | | | |
|---|---|---|---|
| EPI_ISL_2345541, EPI_ISL_2345542 | CENTRO DE SAUDE III AFFONSO LUZZI SANTA CRUZ DAS PALMEIRAS | Instituto Butantan / ESALQ-Piracicaba | Dimas Tadeu Covas, Antonio Jorge Martins, Claudia Renata dos Santos Barros, David Schlesinger, Debora Botequio Moretti, Elaine Cristina Marqueze, Elaine Vieira Santos, Evandra Strazza Rodrigues, Heidge Fukumasu, Jayme Augusto de Souza-Neto, José Salvatore Leister Patané, Luiz Alcantara, Luiz Lehmann Coutinho, Maria Carolina Elias, Maurício Lacerda Nogueira, Rafael dos Santos Bezerra, Raul Machado Neto, Rejane Maria Tommasini Grotto, Ricardo Haddad, Sandra Coccuzzo Sampaio Vessoni, Simone Kashima, Svetoslav Nanev Slavov, Vincent Louis Viala |
| EPI_ISL_2345543 | PRONTO SOCORRO MUNICIPAL TAMBAU | Instituto Butantan / ESALQ-Piracicaba | Dimas Tadeu Covas, Antonio Jorge Martins, Claudia Renata dos Santos Barros, David Schlesinger, Debora Botequio Moretti, Elaine Cristina Marqueze, Elaine Vieira Santos, Evandra Strazza Rodrigues, Heidge Fukumasu, Jayme Augusto de Souza-Neto, José Salvatore Leister Patané, Luiz Alcantara, Luiz Lehmann Coutinho, Maria Carolina Elias, Maurício Lacerda Nogueira, Rafael dos Santos Bezerra, Raul Machado Neto, Rejane Maria Tommasini Grotto, Ricardo Haddad, Sandra Coccuzzo Sampaio Vessoni, Simone Kashima, Svetoslav Nanev Slavov, Vincent Louis Viala |
| EPI_ISL_2345544 | CENTRO DE SAUDE III AFFONSO LUZZI SANTA CRUZ DAS PALMEIRAS | Instituto Butantan / ESALQ-Piracicaba | Dimas Tadeu Covas, Antonio Jorge Martins, Claudia Renata dos Santos Barros, David Schlesinger, Debora Botequio Moretti, Elaine Cristina Marqueze, Elaine Vieira Santos, Evandra Strazza Rodrigues, Heidge Fukumasu, Jayme Augusto de Souza-Neto, José Salvatore Leister Patané, Luiz Alcantara, Luiz Lehmann Coutinho, Maria Carolina Elias, Maurício Lacerda Nogueira, Rafael dos Santos Bezerra, Raul Machado Neto, Rejane Maria Tommasini Grotto, Ricardo Haddad, Sandra Coccuzzo Sampaio Vessoni, Simone Kashima, Svetoslav Nanev Slavov, Vincent Louis Viala |
| EPI_ISL_2345545 | PRONTO SOCORRO MUNICIPAL TAMBAU | Instituto Butantan / ESALQ-Piracicaba | Dimas Tadeu Covas, Antonio Jorge Martins, Claudia Renata dos Santos Barros, David Schlesinger, Debora Botequio Moretti, Elaine Cristina Marqueze, Elaine Vieira Santos, Evandra Strazza Rodrigues, Heidge Fukumasu, Jayme Augusto de Souza-Neto, José Salvatore Leister Patané, Luiz Alcantara, Luiz Lehmann Coutinho, Maria Carolina Elias, Maurício Lacerda Nogueira, Rafael dos Santos Bezerra, Raul Machado Neto, Rejane Maria Tommasini Grotto, Ricardo Haddad, Sandra Coccuzzo Sampaio Vessoni, Simone Kashima, Svetoslav Nanev Slavov, Vincent Louis Viala |
| EPI_ISL_2345546 | CENTRO DE SAUDE III AFFONSO LUZZI SANTA CRUZ DAS PALMEIRAS | Instituto Butantan / ESALQ-Piracicaba | Dimas Tadeu Covas, Antonio Jorge Martins, Claudia Renata dos Santos Barros, David Schlesinger, Debora Botequio Moretti, Elaine Cristina Marqueze, Elaine Vieira Santos, Evandra Strazza Rodrigues, Heidge Fukumasu, Jayme Augusto de Souza-Neto, José Salvatore Leister Patané, Luiz Alcantara, Luiz Lehmann Coutinho, Maria Carolina Elias, Maurício Lacerda Nogueira, Rafael dos Santos Bezerra, Raul Machado Neto, Rejane Maria Tommasini Grotto, Ricardo Haddad, Sandra Coccuzzo Sampaio Vessoni, Simone Kashima, Svetoslav Nanev Slavov, Vincent Louis Viala |
| EPI_ISL_2345547, EPI_ISL_2345548 | PRONTO SOCORRO MUNICIPAL TAMBAU | Instituto Butantan / ESALQ-Piracicaba | Dimas Tadeu Covas, Antonio Jorge Martins, Claudia Renata dos Santos Barros, David Schlesinger, Debora Botequio Moretti, Elaine Cristina Marqueze, Elaine Vieira Santos, Evandra Strazza Rodrigues, Heidge Fukumasu, Jayme Augusto de Souza-Neto, José Salvatore Leister Patané, Luiz Alcantara, Luiz Lehmann Coutinho, Maria Carolina Elias, Maurício Lacerda Nogueira, Rafael dos Santos Bezerra, Raul Machado Neto, Rejane Maria Tommasini Grotto, Ricardo Haddad, Sandra Coccuzzo Sampaio Vessoni, Simone Kashima, Svetoslav Nanev Slavov, Vincent Louis Viala |
| EPI_ISL_2345549 | LABORATORIO DE FRANCA | Instituto Butantan / ESALQ-Piracicaba | Dimas Tadeu Covas, Antonio Jorge Martins, Claudia Renata dos Santos Barros, David Schlesinger, Debora Botequio Moretti, Elaine Cristina Marqueze, Elaine Vieira Santos, Evandra Strazza Rodrigues, Heidge Fukumasu, Jayme Augusto de Souza-Neto, José Salvatore Leister Patané, Luiz Alcantara, Luiz Lehmann Coutinho, Maria Carolina Elias, Maurício Lacerda Nogueira, Rafael dos Santos Bezerra, Raul Machado Neto, Rejane Maria Tommasini Grotto, Ricardo Haddad, Sandra Coccuzzo Sampaio Vessoni, Simone Kashima, Svetoslav Nanev Slavov, Vincent Louis Viala |
| EPI_ISL_2345550 | PRONTO SOCORRO MUNICIPAL TAMBAU | Instituto Butantan / ESALQ-Piracicaba | Dimas Tadeu Covas, Antonio Jorge Martins, Claudia Renata dos Santos Barros, David Schlesinger, Debora Botequio Moretti, Elaine Cristina Marqueze, Elaine Vieira Santos, Evandra Strazza Rodrigues, Heidge Fukumasu, Jayme Augusto de Souza-Neto, José Salvatore Leister Patané, Luiz Alcantara, Luiz Lehmann Coutinho, Maria Carolina Elias, Maurício Lacerda Nogueira, Rafael dos Santos Bezerra, Raul Machado Neto, Rejane Maria Tommasini Grotto, Ricardo Haddad, Sandra Coccuzzo Sampaio Vessoni, Simone Kashima, Svetoslav Nanev Slavov, Vincent Louis Viala |
| EPI_ISL_2345551, EPI_ISL_2345552 | LABORATORIO DE FRANCA | Instituto Butantan / ESALQ-Piracicaba | Dimas Tadeu Covas, Antonio Jorge Martins, Claudia Renata dos Santos Barros, David Schlesinger, Debora Botequio Moretti, Elaine Cristina Marqueze, Elaine Vieira Santos, Evandra Strazza Rodrigues, Heidge Fukumasu, Jayme Augusto de Souza-Neto, José Salvatore Leister Patané, Luiz Alcantara, Luiz Lehmann Coutinho, Maria Carolina Elias, Maurício Lacerda Nogueira, Rafael dos Santos Bezerra, Raul Machado Neto, Rejane Maria Tommasini Grotto, Ricardo Haddad, Sandra Coccuzzo Sampaio Vessoni, Simone Kashima, Svetoslav Nanev Slavov, Vincent Louis Viala |
| EPI_ISL_2345553 | UNIDADE DA SAUDE DO ADULTO CASA BRANCA PREFEITURA | Instituto Butantan / ESALQ-Piracicaba | Dimas Tadeu Covas, Antonio Jorge Martins, Claudia Renata dos Santos Barros, David Schlesinger, Debora Botequio Moretti, Elaine Cristina Marqueze, Elaine Vieira Santos, Evandra Strazza Rodrigues, Heidge Fukumasu, Jayme Augusto de Souza-Neto, José Salvatore Leister Patané, Luiz Alcantara, Luiz Lehmann Coutinho, Maria Carolina Elias, Maurício Lacerda Nogueira, Rafael dos Santos Bezerra, Raul Machado Neto, Rejane Maria Tommasini Grotto, Ricardo Haddad, Sandra Coccuzzo Sampaio Vessoni, Simone Kashima, Svetoslav Nanev Slavov, Vincent Louis Viala |

EPI_ISL_2345554, EPI_ISL_2345555, EPI_ISL_2345556, EPI_ISL_2345557, EPI_ISL_2345558, EPI_ISL_2345559, EPI_ISL_2345560, EPI_ISL_2345561, EPI_ISL_2345562, EPI_ISL_2345563, EPI_ISL_2345564, EPI_ISL_2345565, EPI_ISL_2345566, EPI_ISL_2345567, EPI_ISL_2345568, EPI_ISL_2345569, EPI_ISL_2345570, EPI_ISL_2345571, EPI_ISL_2345572, EPI_ISL_2345573, EPI_ISL_2345574, EPI_ISL_2345575, EPI_ISL_2345576, EPI_ISL_2345577, EPI_ISL_2345578, EPI_ISL_2345579, EPI_ISL_2345580, EPI_ISL_2345581, EPI_ISL_2345582, EPI_ISL_2345583, EPI_ISL_2345584

| | | | |
|---|---|---|---|
| see above | LABORATORIO DE FRANCA | Instituto Butantan / ESALQ-Piracicaba | Dimas Tadeu Covas, Antonio Jorge Martins, Claudia Renata dos Santos Barros, David Schlesinger, Debora Botequio Moretti, Elaine Cristina Marqueze, Elaine Vieira Santos, Evandra Strazza Rodrigues, Heidge Fukumasu, Jayme Augusto de Souza-Neto, José Salvatore Leister Patané, Luiz Alcantara, Luiz Lehmann Coutinho, Maria Carolina Elias, Maurício Lacerda Nogueira, Rafael dos Santos Bezerra, Raul Machado Neto, Rejane Maria Tommasini Grotto, Ricardo Haddad, Sandra Coccuzzo Sampaio Vessoni, Simone Kashima, Svetoslav Nanev Slavov, Vincent Louis Viala |
| EPI_ISL_2345585 | UBS II DE NARANDIBA | Instituto Butantan / ESALQ-Piracicaba | Dimas Tadeu Covas, Antonio Jorge Martins, Claudia Renata dos Santos Barros, David Schlesinger, Debora Botequio Moretti, Elaine Cristina Marqueze, Elaine Vieira Santos, Evandra Strazza Rodrigues, Heidge Fukumasu, Jayme Augusto de Souza-Neto, José Salvatore Leister Patané, Luiz Alcantara, Luiz Lehmann Coutinho, Maria Carolina Elias, Maurício Lacerda Nogueira, Rafael dos Santos Bezerra, Raul Machado Neto, Rejane Maria Tommasini Grotto, Ricardo Haddad, Sandra Coccuzzo Sampaio Vessoni, Simone Kashima, Svetoslav Nanev Slavov, Vincent Louis Viala |
| EPI_ISL_2345586 | CENTRO DE ESPECIALIDADES DE PRIMAVERA | Instituto Butantan / ESALQ-Piracicaba | Dimas Tadeu Covas, Antonio Jorge Martins, Claudia Renata dos Santos Barros, David Schlesinger, Debora Botequio Moretti, Elaine Cristina Marqueze, Elaine Vieira Santos, Evandra Strazza Rodrigues, Heidge Fukumasu, Jayme Augusto de Souza-Neto, José Salvatore Leister Patané, Luiz Alcantara, Luiz Lehmann Coutinho, Maria Carolina Elias, Maurício Lacerda Nogueira, Rafael dos Santos Bezerra, Raul Machado Neto, Rejane Maria Tommasini Grotto, Ricardo Haddad, Sandra Coccuzzo Sampaio Vessoni, Simone Kashima, Svetoslav Nanev Slavov, Vincent Louis Viala |
| EPI_ISL_2345587 | UBS II DE REGENTE FEIJO | Instituto Butantan / ESALQ-Piracicaba | Dimas Tadeu Covas, Antonio Jorge Martins, Claudia Renata dos Santos Barros, David Schlesinger, Debora Botequio Moretti, Elaine Cristina Marqueze, Elaine Vieira Santos, Evandra Strazza Rodrigues, Heidge Fukumasu, Jayme Augusto de Souza-Neto, José Salvatore Leister Patané, Luiz Alcantara, Luiz Lehmann Coutinho, Maria Carolina Elias, Maurício Lacerda Nogueira, Rafael dos Santos Bezerra, Raul Machado Neto, Rejane Maria Tommasini Grotto, Ricardo Haddad, Sandra Coccuzzo Sampaio Vessoni, Simone Kashima, Svetoslav Nanev Slavov, Vincent Louis Viala |
| EPI_ISL_2345588 | CENTRO DE SAUDE II JUNQUEIROPOLIS | Instituto Butantan / ESALQ-Piracicaba | Dimas Tadeu Covas, Antonio Jorge Martins, Claudia Renata dos Santos Barros, David Schlesinger, Debora Botequio Moretti, Elaine Cristina Marqueze, Elaine Vieira Santos, Evandra Strazza Rodrigues, Heidge Fukumasu, Jayme Augusto de Souza-Neto, José Salvatore Leister Patané, Luiz Alcantara, Luiz Lehmann Coutinho, Maria Carolina Elias, Maurício Lacerda Nogueira, Rafael dos Santos Bezerra, Raul Machado Neto, Rejane Maria Tommasini Grotto, Ricardo Haddad, Sandra Coccuzzo Sampaio Vessoni, Simone Kashima, Svetoslav Nanev Slavov, Vincent Louis Viala |
| EPI_ISL_2345589 | UBS II DR EXPEDITO SHIZUO KUROCE | Instituto Butantan / ESALQ-Piracicaba | Dimas Tadeu Covas, Antonio Jorge Martins, Claudia Renata dos Santos Barros, David Schlesinger, Debora Botequio Moretti, Elaine Cristina Marqueze, Elaine Vieira Santos, Evandra Strazza Rodrigues, Heidge Fukumasu, Jayme Augusto de Souza-Neto, José Salvatore Leister Patané, Luiz Alcantara, Luiz Lehmann Coutinho, Maria Carolina Elias, Maurício Lacerda Nogueira, Rafael dos Santos Bezerra, Raul Machado Neto, Rejane Maria Tommasini Grotto, Ricardo Haddad, Sandra Coccuzzo Sampaio Vessoni, Simone Kashima, Svetoslav Nanev Slavov, Vincent Louis Viala |
| EPI_ISL_2345590 | CENTRO DE ESPECIALIDADES DE PRIMAVERA | Instituto Butantan / ESALQ-Piracicaba | Dimas Tadeu Covas, Antonio Jorge Martins, Claudia Renata dos Santos Barros, David Schlesinger, Debora Botequio Moretti, Elaine Cristina Marqueze, Elaine Vieira Santos, Evandra Strazza Rodrigues, Heidge Fukumasu, Jayme Augusto de Souza-Neto, José Salvatore Leister Patané, Luiz Alcantara, Luiz Lehmann Coutinho, Maria Carolina Elias, Maurício Lacerda Nogueira, Rafael dos Santos Bezerra, Raul Machado Neto, Rejane Maria Tommasini Grotto, Ricardo Haddad, Sandra Coccuzzo Sampaio Vessoni, Simone Kashima, Svetoslav Nanev Slavov, Vincent Louis Viala |
| EPI_ISL_2345591 | UBS II DR EXPEDITO SHIZUO KUROCE | Instituto Butantan / ESALQ-Piracicaba | Dimas Tadeu Covas, Antonio Jorge Martins, Claudia Renata dos Santos Barros, David Schlesinger, Debora Botequio Moretti, Elaine Cristina Marqueze, Elaine Vieira Santos, Evandra Strazza Rodrigues, Heidge Fukumasu, Jayme Augusto de Souza-Neto, José Salvatore Leister Patané, Luiz Alcantara, Luiz Lehmann Coutinho, Maria Carolina Elias, Maurício Lacerda Nogueira, Rafael dos Santos Bezerra, Raul Machado Neto, Rejane Maria Tommasini Grotto, Ricardo Haddad, Sandra Coccuzzo Sampaio Vessoni, Simone Kashima, Svetoslav Nanev Slavov, Vincent Louis Viala |
| EPI_ISL_2345592 | CENTRO DE SAUDE II JUNQUEIROPOLIS | Instituto Butantan / ESALQ-Piracicaba | Dimas Tadeu Covas, Antonio Jorge Martins, Claudia Renata dos Santos Barros, David Schlesinger, Debora Botequio Moretti, Elaine Cristina Marqueze, Elaine Vieira Santos, Evandra Strazza Rodrigues, Heidge Fukumasu, Jayme Augusto de Souza-Neto, José Salvatore Leister Patané, Luiz Alcantara, Luiz Lehmann Coutinho, Maria Carolina Elias, Maurício Lacerda Nogueira, Rafael dos Santos Bezerra, Raul Machado Neto, Rejane Maria Tommasini Grotto, Ricardo Haddad, Sandra Coccuzzo Sampaio Vessoni, Simone Kashima, Svetoslav Nanev Slavov, Vincent Louis Viala |

| | | | |
|---|---|---|---|
| EPI_ISL_2345593 | UBS III DE RANCHARIA | Instituto Butantan / ESALQ-Piracicaba | Dimas Tadeu Covas, Antonio Jorge Martins, Claudia Renata dos Santos Barros, David Schlesinger, Debora Botequio Moretti, Elaine Cristina Marqueze, Elaine Vieira Santos, Evandra Strazza Rodrigues, Heidge Fukumasu, Jayme Augusto de Souza-Neto, José Salvatore Leister Patané, Luiz Alcantara, Luiz Lehmann Coutinho, Maria Carolina Elias, Maurício Lacerda Nogueira, Rafael dos Santos Bezerra, Raul Machado Neto, Rejane Maria Tommasini Grotto, Ricardo Haddad, Sandra Coccuzzo Sampaio Vessoni, Simone Kashima, Svetoslav Nanev Slavov, Vincent Louis Viala |
| EPI_ISL_2345594 | CENTRO DE ESPECIALIDADES DE PRIMAVERA | Instituto Butantan / ESALQ-Piracicaba | Dimas Tadeu Covas, Antonio Jorge Martins, Claudia Renata dos Santos Barros, David Schlesinger, Debora Botequio Moretti, Elaine Cristina Marqueze, Elaine Vieira Santos, Evandra Strazza Rodrigues, Heidge Fukumasu, Jayme Augusto de Souza-Neto, José Salvatore Leister Patané, Luiz Alcantara, Luiz Lehmann Coutinho, Maria Carolina Elias, Maurício Lacerda Nogueira, Rafael dos Santos Bezerra, Raul Machado Neto, Rejane Maria Tommasini Grotto, Ricardo Haddad, Sandra Coccuzzo Sampaio Vessoni, Simone Kashima, Svetoslav Nanev Slavov, Vincent Louis Viala |
| EPI_ISL_2345595 | UBS III DE RANCHARIA | Instituto Butantan / ESALQ-Piracicaba | Dimas Tadeu Covas, Antonio Jorge Martins, Claudia Renata dos Santos Barros, David Schlesinger, Debora Botequio Moretti, Elaine Cristina Marqueze, Elaine Vieira Santos, Evandra Strazza Rodrigues, Heidge Fukumasu, Jayme Augusto de Souza-Neto, José Salvatore Leister Patané, Luiz Alcantara, Luiz Lehmann Coutinho, Maria Carolina Elias, Maurício Lacerda Nogueira, Rafael dos Santos Bezerra, Raul Machado Neto, Rejane Maria Tommasini Grotto, Ricardo Haddad, Sandra Coccuzzo Sampaio Vessoni, Simone Kashima, Svetoslav Nanev Slavov, Vincent Louis Viala |
| EPI_ISL_2345596 | UBS MARCIA CRISTIANE DA SILVA DE OURO VERDE | Instituto Butantan / ESALQ-Piracicaba | Dimas Tadeu Covas, Antonio Jorge Martins, Claudia Renata dos Santos Barros, David Schlesinger, Debora Botequio Moretti, Elaine Cristina Marqueze, Elaine Vieira Santos, Evandra Strazza Rodrigues, Heidge Fukumasu, Jayme Augusto de Souza-Neto, José Salvatore Leister Patané, Luiz Alcantara, Luiz Lehmann Coutinho, Maria Carolina Elias, Maurício Lacerda Nogueira, Rafael dos Santos Bezerra, Raul Machado Neto, Rejane Maria Tommasini Grotto, Ricardo Haddad, Sandra Coccuzzo Sampaio Vessoni, Simone Kashima, Svetoslav Nanev Slavov, Vincent Louis Viala |
| EPI_ISL_2345597 | UBS III DE RANCHARIA | Instituto Butantan / ESALQ-Piracicaba | Dimas Tadeu Covas, Antonio Jorge Martins, Claudia Renata dos Santos Barros, David Schlesinger, Debora Botequio Moretti, Elaine Cristina Marqueze, Elaine Vieira Santos, Evandra Strazza Rodrigues, Heidge Fukumasu, Jayme Augusto de Souza-Neto, José Salvatore Leister Patané, Luiz Alcantara, Luiz Lehmann Coutinho, Maria Carolina Elias, Maurício Lacerda Nogueira, Rafael dos Santos Bezerra, Raul Machado Neto, Rejane Maria Tommasini Grotto, Ricardo Haddad, Sandra Coccuzzo Sampaio Vessoni, Simone Kashima, Svetoslav Nanev Slavov, Vincent Louis Viala |
| EPI_ISL_2345598 | UBS II DE REGENTE FEIJO | Instituto Butantan / ESALQ-Piracicaba | Dimas Tadeu Covas, Antonio Jorge Martins, Claudia Renata dos Santos Barros, David Schlesinger, Debora Botequio Moretti, Elaine Cristina Marqueze, Elaine Vieira Santos, Evandra Strazza Rodrigues, Heidge Fukumasu, Jayme Augusto de Souza-Neto, José Salvatore Leister Patané, Luiz Alcantara, Luiz Lehmann Coutinho, Maria Carolina Elias, Maurício Lacerda Nogueira, Rafael dos Santos Bezerra, Raul Machado Neto, Rejane Maria Tommasini Grotto, Ricardo Haddad, Sandra Coccuzzo Sampaio Vessoni, Simone Kashima, Svetoslav Nanev Slavov, Vincent Louis Viala |
| EPI_ISL_2345599, EPI_ISL_2345600, EPI_ISL_2345601 | UBS III DE RANCHARIA | Instituto Butantan / ESALQ-Piracicaba | Dimas Tadeu Covas, Antonio Jorge Martins, Claudia Renata dos Santos Barros, David Schlesinger, Debora Botequio Moretti, Elaine Cristina Marqueze, Elaine Vieira Santos, Evandra Strazza Rodrigues, Heidge Fukumasu, Jayme Augusto de Souza-Neto, José Salvatore Leister Patané, Luiz Alcantara, Luiz Lehmann Coutinho, Maria Carolina Elias, Maurício Lacerda Nogueira, Rafael dos Santos Bezerra, Raul Machado Neto, Rejane Maria Tommasini Grotto, Ricardo Haddad, Sandra Coccuzzo Sampaio Vessoni, Simone Kashima, Svetoslav Nanev Slavov, Vincent Louis Viala |
| EPI_ISL_2345602 | PAS JOAO ANTONIO DO NASCIMENTO | Instituto Butantan / ESALQ-Piracicaba | Dimas Tadeu Covas, Antonio Jorge Martins, Claudia Renata dos Santos Barros, David Schlesinger, Debora Botequio Moretti, Elaine Cristina Marqueze, Elaine Vieira Santos, Evandra Strazza Rodrigues, Heidge Fukumasu, Jayme Augusto de Souza-Neto, José Salvatore Leister Patané, Luiz Alcantara, Luiz Lehmann Coutinho, Maria Carolina Elias, Maurício Lacerda Nogueira, Rafael dos Santos Bezerra, Raul Machado Neto, Rejane Maria Tommasini Grotto, Ricardo Haddad, Sandra Coccuzzo Sampaio Vessoni, Simone Kashima, Svetoslav Nanev Slavov, Vincent Louis Viala |
| EPI_ISL_2345603, EPI_ISL_2345604 | HOSPITAL MUNICIPAL REYNALDO GUERRA CAJATI | Instituto Butantan / ESALQ-Piracicaba | Dimas Tadeu Covas, Antonio Jorge Martins, Claudia Renata dos Santos Barros, David Schlesinger, Debora Botequio Moretti, Elaine Cristina Marqueze, Elaine Vieira Santos, Evandra Strazza Rodrigues, Heidge Fukumasu, Jayme Augusto de Souza-Neto, José Salvatore Leister Patané, Luiz Alcantara, Luiz Lehmann Coutinho, Maria Carolina Elias, Maurício Lacerda Nogueira, Rafael dos Santos Bezerra, Raul Machado Neto, Rejane Maria Tommasini Grotto, Ricardo Haddad, Sandra Coccuzzo Sampaio Vessoni, Simone Kashima, Svetoslav Nanev Slavov, Vincent Louis Viala |
| EPI_ISL_2345605, EPI_ISL_2345606 | VIGILANCIA EM SAUDE | Instituto Butantan / ESALQ-Piracicaba | Dimas Tadeu Covas, Antonio Jorge Martins, Claudia Renata dos Santos Barros, David Schlesinger, Debora Botequio Moretti, Elaine Cristina Marqueze, Elaine Vieira Santos, Evandra Strazza Rodrigues, Heidge Fukumasu, Jayme Augusto de Souza-Neto, José Salvatore Leister Patané, Luiz Alcantara, Luiz Lehmann Coutinho, Maria Carolina Elias, Maurício Lacerda Nogueira, Rafael dos Santos Bezerra, Raul Machado Neto, Rejane Maria Tommasini Grotto, Ricardo Haddad, Sandra Coccuzzo Sampaio Vessoni, Simone Kashima, Svetoslav Nanev Slavov, Vincent Louis Viala |
| EPI_ISL_2345607 | HOSPITAL MUNICIPAL REYNALDO GUERRA CAJATI | Instituto Butantan / ESALQ-Piracicaba | Dimas Tadeu Covas, Antonio Jorge Martins, Claudia Renata dos Santos Barros, David Schlesinger, Debora Botequio Moretti, Elaine Cristina Marqueze, Elaine Vieira Santos, Evandra Strazza Rodrigues, Heidge Fukumasu, Jayme Augusto de Souza-Neto, José Salvatore Leister Patané, Luiz Alcantara, Luiz Lehmann Coutinho, Maria Carolina Elias, Maurício Lacerda Nogueira, Rafael dos Santos Bezerra, Raul Machado Neto, Rejane Maria Tommasini Grotto, Ricardo Haddad, Sandra Coccuzzo Sampaio Vessoni, Simone Kashima, Svetoslav Nanev Slavov, Vincent Louis Viala |
| EPI_ISL_2345608, EPI_ISL_2345609 | VIGILANCIA EM SAUDE | Instituto Butantan / ESALQ-Piracicaba | Dimas Tadeu Covas, Antonio Jorge Martins, Claudia Renata dos Santos Barros, David Schlesinger, Debora Botequio Moretti, Elaine Cristina Marqueze, Elaine Vieira Santos, Evandra Strazza Rodrigues, Heidge Fukumasu, Jayme Augusto de Souza-Neto, José Salvatore Leister Patané, Luiz Alcantara, Luiz Lehmann Coutinho, Maria Carolina Elias, Maurício Lacerda Nogueira, Rafael dos Santos Bezerra, Raul Machado Neto, Rejane Maria Tommasini Grotto, Ricardo Haddad, Sandra Coccuzzo Sampaio Vessoni, Simone Kashima, Svetoslav Nanev Slavov, Vincent Louis Viala |
| EPI_ISL_2345610 | HOSPITAL E MATERNIDADE NOSSA SENHORA DA AJUDA | Instituto Butantan / ESALQ-Piracicaba | Dimas Tadeu Covas, Antonio Jorge Martins, Claudia Renata dos Santos Barros, David Schlesinger, Debora Botequio Moretti, Elaine Cristina Marqueze, Elaine Vieira Santos, Evandra Strazza Rodrigues, Heidge Fukumasu, Jayme Augusto de Souza-Neto, José Salvatore Leister Patané, Luiz Alcantara, Luiz Lehmann Coutinho, Maria Carolina Elias, Maurício Lacerda Nogueira, Rafael dos Santos Bezerra, Raul Machado Neto, Rejane Maria Tommasini Grotto, Ricardo Haddad, Sandra Coccuzzo Sampaio Vessoni, Simone Kashima, Svetoslav Nanev Slavov, Vincent Louis Viala |
| EPI_ISL_2345611 | LABORATORIO DR PAULO EMILIO DALESSANDRO PINDAMONHANGABA | Instituto Butantan / ESALQ-Piracicaba | Dimas Tadeu Covas, Antonio Jorge Martins, Claudia Renata dos Santos Barros, David Schlesinger, Debora Botequio Moretti, Elaine Cristina Marqueze, Elaine Vieira Santos, Evandra Strazza Rodrigues, Heidge Fukumasu, Jayme Augusto de Souza-Neto, José Salvatore Leister Patané, Luiz Alcantara, Luiz Lehmann Coutinho, Maria Carolina Elias, Maurício Lacerda Nogueira, Rafael dos Santos Bezerra, Raul Machado Neto, Rejane Maria Tommasini Grotto, Ricardo Haddad, Sandra Coccuzzo Sampaio Vessoni, Simone Kashima, Svetoslav Nanev Slavov, Vincent Louis Viala |
| EPI_ISL_2345612 | HOSPITAL DE CAMPANHA COVID 19 MUNICIPIO DE TAUBATE | Instituto Butantan / ESALQ-Piracicaba | Dimas Tadeu Covas, Antonio Jorge Martins, Claudia Renata dos Santos Barros, David Schlesinger, Debora Botequio Moretti, Elaine Cristina Marqueze, Elaine Vieira Santos, Evandra Strazza Rodrigues, Heidge Fukumasu, Jayme Augusto de Souza-Neto, José Salvatore Leister Patané, Luiz Alcantara, Luiz Lehmann Coutinho, Maria Carolina Elias, Maurício Lacerda Nogueira, Rafael dos Santos Bezerra, Raul Machado Neto, Rejane Maria Tommasini Grotto, Ricardo Haddad, Sandra Coccuzzo Sampaio Vessoni, Simone Kashima, Svetoslav Nanev Slavov, Vincent Louis Viala |
| EPI_ISL_2345613 | LABORATORIO DR PAULO EMILIO DALESSANDRO PINDAMONHANGABA | Instituto Butantan / ESALQ-Piracicaba | Dimas Tadeu Covas, Antonio Jorge Martins, Claudia Renata dos Santos Barros, David Schlesinger, Debora Botequio Moretti, Elaine Cristina Marqueze, Elaine Vieira Santos, Evandra Strazza Rodrigues, Heidge Fukumasu, Jayme Augusto de Souza-Neto, José Salvatore Leister Patané, Luiz Alcantara, Luiz Lehmann Coutinho, Maria Carolina Elias, Maurício Lacerda Nogueira, Rafael dos Santos Bezerra, Raul Machado Neto, Rejane Maria Tommasini Grotto, Ricardo Haddad, Sandra Coccuzzo Sampaio Vessoni, Simone Kashima, Svetoslav Nanev Slavov, Vincent Louis Viala |
| EPI_ISL_2345614 | PRONTO SOCORRO MUNICIPAL DE PARAIBUNA | Instituto Butantan / ESALQ-Piracicaba | Dimas Tadeu Covas, Antonio Jorge Martins, Claudia Renata dos Santos Barros, David Schlesinger, Debora Botequio Moretti, Elaine Cristina Marqueze, Elaine Vieira Santos, Evandra Strazza Rodrigues, Heidge Fukumasu, Jayme Augusto de Souza-Neto, José Salvatore Leister Patané, Luiz Alcantara, Luiz Lehmann Coutinho, Maria Carolina Elias, Maurício Lacerda Nogueira, Rafael dos Santos Bezerra, Raul Machado Neto, Rejane Maria Tommasini Grotto, Ricardo Haddad, Sandra Coccuzzo Sampaio Vessoni, Simone Kashima, Svetoslav Nanev Slavov, Vincent Louis Viala |
| EPI_ISL_2345615, EPI_ISL_2345616, EPI_ISL_2345617, EPI_ISL_2345618, EPI_ISL_2345619, EPI_ISL_2345620, EPI_ISL_2345621, EPI_ISL_2345622 | LABORATORIO DR PAULO EMILIO DALESSANDRO PINDAMONHANGABA | Instituto Butantan / ESALQ-Piracicaba | Dimas Tadeu Covas, Antonio Jorge Martins, Claudia Renata dos Santos Barros, David Schlesinger, Debora Botequio Moretti, Elaine Cristina Marqueze, Elaine Vieira Santos, Evandra Strazza Rodrigues, Heidge Fukumasu, Jayme Augusto de Souza-Neto, José Salvatore Leister Patané, Luiz Alcantara, Luiz Lehmann Coutinho, Maria Carolina Elias, Maurício Lacerda Nogueira, Rafael dos Santos Bezerra, Raul Machado Neto, Rejane Maria Tommasini Grotto, Ricardo Haddad, Sandra Coccuzzo Sampaio Vessoni, Simone Kashima, Svetoslav Nanev Slavov, Vincent Louis Viala |
| EPI_ISL_2345623 | CENTRO DE SAUDE DE NATIVIDADE DA SERRA | Instituto Butantan / ESALQ-Piracicaba | Dimas Tadeu Covas, Antonio Jorge Martins, Claudia Renata dos Santos Barros, David Schlesinger, Debora Botequio Moretti, Elaine Cristina Marqueze, Elaine Vieira Santos, Evandra Strazza Rodrigues, Heidge Fukumasu, Jayme Augusto de Souza-Neto, José Salvatore Leister Patané, Luiz Alcantara, Luiz Lehmann Coutinho, Maria Carolina Elias, Maurício Lacerda Nogueira, Rafael dos Santos Bezerra, Raul Machado Neto, Rejane Maria Tommasini Grotto, Ricardo Haddad, Sandra Coccuzzo Sampaio Vessoni, Simone Kashima, Svetoslav Nanev Slavov, Vincent Louis Viala |
| EPI_ISL_2345624, EPI_ISL_2345625, EPI_ISL_2345626, EPI_ISL_2345627, EPI_ISL_2345628, EPI_ISL_2345629, EPI_ISL_2345631, EPI_ISL_2345632, EPI_ISL_2345633, EPI_ISL_2345634, EPI_ISL_2345635 | | | |

| | | | |
|---|---|---|---|
| see above | LABORATORIO DR PAULO EMILIO DALESSANDRO PINDAMONHANGABA | Instituto Butantan / ESALQ-Piracicaba | Dimas Tadeu Covas, Antonio Jorge Martins, Claudia Renata dos Santos Barros, David Schlesinger, Debora Botequio Moretti, Elaine Cristina Marqueze, Elaine Vieira Santos, Evandra Strazza Rodrigues, Heidge Fukumasu, Jayme Augusto de Souza-Neto, José Salvatore Leister Patané, Luiz Alcantara, Luiz Lehmann Coutinho, Maria Carolina Elias, Maurício Lacerda Nogueira, Rafael dos Santos Bezerra, Raul Machado Neto, Rejane Maria Tommasini Grotto, Ricardo Haddad, Sandra Coccuzzo Sampaio Vessoni, Simone Kashima, Svetoslav Nanev Slavov, Vincent Louis Viala |
| EPI_ISL_2345636 | SANTA CASA DE MISERICORDIA DE UBATUBA | Instituto Butantan / ESALQ-Piracicaba | Dimas Tadeu Covas, Antonio Jorge Martins, Claudia Renata dos Santos Barros, David Schlesinger, Debora Botequio Moretti, Elaine Cristina Marqueze, Elaine Vieira Santos, Evandra Strazza Rodrigues, Heidge Fukumasu, Jayme Augusto de Souza-Neto, José Salvatore Leister Patané, Luiz Alcantara, Luiz Lehmann Coutinho, Maria Carolina Elias, Maurício Lacerda Nogueira, Rafael dos Santos Bezerra, Raul Machado Neto, Rejane Maria Tommasini Grotto, Ricardo Haddad, Sandra Coccuzzo Sampaio Vessoni, Simone Kashima, Svetoslav Nanev Slavov, Vincent Louis Viala |
| EPI_ISL_2345637, EPI_ISL_2345638 | UNIDADE BASICA DE SAUDE DE PIQUETE | Instituto Butantan / ESALQ-Piracicaba | Dimas Tadeu Covas, Antonio Jorge Martins, Claudia Renata dos Santos Barros, David Schlesinger, Debora Botequio Moretti, Elaine Cristina Marqueze, Elaine Vieira Santos, Evandra Strazza Rodrigues, Heidge Fukumasu, Jayme Augusto de Souza-Neto, José Salvatore Leister Patané, Luiz Alcantara, Luiz Lehmann Coutinho, Maria Carolina Elias, Maurício Lacerda Nogueira, Rafael dos Santos Bezerra, Raul Machado Neto, Rejane Maria Tommasini Grotto, Ricardo Haddad, Sandra Coccuzzo Sampaio Vessoni, Simone Kashima, Svetoslav Nanev Slavov, Vincent Louis Viala |
| EPI_ISL_2345639 | SECRETARIA DE SAUDE | Instituto Butantan / ESALQ-Piracicaba | Dimas Tadeu Covas, Antonio Jorge Martins, Claudia Renata dos Santos Barros, David Schlesinger, Debora Botequio Moretti, Elaine Cristina Marqueze, Elaine Vieira Santos, Evandra Strazza Rodrigues, Heidge Fukumasu, Jayme Augusto de Souza-Neto, José Salvatore Leister Patané, Luiz Alcantara, Luiz Lehmann Coutinho, Maria Carolina Elias, Maurício Lacerda Nogueira, Rafael dos Santos Bezerra, Raul Machado Neto, Rejane Maria Tommasini Grotto, Ricardo Haddad, Sandra Coccuzzo Sampaio Vessoni, Simone Kashima, Svetoslav Nanev Slavov, Vincent Louis Viala |
| EPI_ISL_2345640, EPI_ISL_2345641 | LABORATORIO DR PAULO EMILIO DALESSANDRO PINDAMONHANGABA | Instituto Butantan / ESALQ-Piracicaba | Dimas Tadeu Covas, Antonio Jorge Martins, Claudia Renata dos Santos Barros, David Schlesinger, Debora Botequio Moretti, Elaine Cristina Marqueze, Elaine Vieira Santos, Evandra Strazza Rodrigues, Heidge Fukumasu, Jayme Augusto de Souza-Neto, José Salvatore Leister Patané, Luiz Alcantara, Luiz Lehmann Coutinho, Maria Carolina Elias, Maurício Lacerda Nogueira, Rafael dos Santos Bezerra, Raul Machado Neto, Rejane Maria Tommasini Grotto, Ricardo Haddad, Sandra Coccuzzo Sampaio Vessoni, Simone Kashima, Svetoslav Nanev Slavov, Vincent Louis Viala |
| EPI_ISL_2345642 | SMS IPERO | Instituto Butantan / UNESP-Botucatu | Dimas Tadeu Covas, Antonio Jorge Martins, Claudia Renata dos Santos Barros, David Schlesinger, Debora Botequio Moretti, Elaine Cristina Marqueze, Elaine Vieira Santos, Evandra Strazza Rodrigues, Heidge Fukumasu, Jayme Augusto de Souza-Neto, José Salvatore Leister Patané, Luiz Alcantara, Luiz Lehmann Coutinho, Maria Carolina Elias, Maurício Lacerda Nogueira, Rafael dos Santos Bezerra, Raul Machado Neto, Rejane Maria Tommasini Grotto, Ricardo Haddad, Sandra Coccuzzo Sampaio Vessoni, Simone Kashima, Svetoslav Nanev Slavov, Vincent Louis Viala |
| EPI_ISL_2345643 | POLICLINICA COVID 19 ITAPETININGA | Instituto Butantan / Mendelics | Dimas Tadeu Covas, Antonio Jorge Martins, Claudia Renata dos Santos Barros, David Schlesinger, Debora Botequio Moretti, Elaine Cristina Marqueze, Elaine Vieira Santos, Evandra Strazza Rodrigues, Heidge Fukumasu, Jayme Augusto de Souza-Neto, José Salvatore Leister Patané, Luiz Alcantara, Luiz Lehmann Coutinho, Maria Carolina Elias, Maurício Lacerda Nogueira, Rafael dos Santos Bezerra, Raul Machado Neto, Rejane Maria Tommasini Grotto, Ricardo Haddad, Sandra Coccuzzo Sampaio Vessoni, Simone Kashima, Svetoslav Nanev Slavov, Vincent Louis Viala |
| EPI_ISL_2345644 | CENTRO DE SAUDE II IBITINGA | Instituto Butantan / Mendelics | Dimas Tadeu Covas, Antonio Jorge Martins, Claudia Renata dos Santos Barros, David Schlesinger, Debora Botequio Moretti, Elaine Cristina Marqueze, Elaine Vieira Santos, Evandra Strazza Rodrigues, Heidge Fukumasu, Jayme Augusto de Souza-Neto, José Salvatore Leister Patané, Luiz Alcantara, Luiz Lehmann Coutinho, Maria Carolina Elias, Maurício Lacerda Nogueira, Rafael dos Santos Bezerra, Raul Machado Neto, Rejane Maria Tommasini Grotto, Ricardo Haddad, Sandra Coccuzzo Sampaio Vessoni, Simone Kashima, Svetoslav Nanev Slavov, Vincent Louis Viala |
| EPI_ISL_2345645 | VIGILANCIA EPIDEMIOLOGICA JARDINOPOLIS SP | Instituto Butantan / Mendelics | Dimas Tadeu Covas, Antonio Jorge Martins, Claudia Renata dos Santos Barros, David Schlesinger, Debora Botequio Moretti, Elaine Cristina Marqueze, Elaine Vieira Santos, Evandra Strazza Rodrigues, Heidge Fukumasu, Jayme Augusto de Souza-Neto, José Salvatore Leister Patané, Luiz Alcantara, Luiz Lehmann Coutinho, Maria Carolina Elias, Maurício Lacerda Nogueira, Rafael dos Santos Bezerra, Raul Machado Neto, Rejane Maria Tommasini Grotto, Ricardo Haddad, Sandra Coccuzzo Sampaio Vessoni, Simone Kashima, Svetoslav Nanev Slavov, Vincent Louis Viala |
| EPI_ISL_2345646 | UBS DR ALFREDO DANTAS DE SOUZA UMUARAMA | Instituto Butantan / Mendelics | Dimas Tadeu Covas, Antonio Jorge Martins, Claudia Renata dos Santos Barros, David Schlesinger, Debora Botequio Moretti, Elaine Cristina Marqueze, Elaine Vieira Santos, Evandra Strazza Rodrigues, Heidge Fukumasu, Jayme Augusto de Souza-Neto, José Salvatore Leister Patané, Luiz Alcantara, Luiz Lehmann Coutinho, Maria Carolina Elias, Maurício Lacerda Nogueira, Rafael dos Santos Bezerra, Raul Machado Neto, Rejane Maria Tommasini Grotto, Ricardo Haddad, Sandra Coccuzzo Sampaio Vessoni, Simone Kashima, Svetoslav Nanev Slavov, Vincent Louis Viala |
| EPI_ISL_2345648 | SANTA CASA DE MISERICORDIA SAO JOSE | Instituto Butantan / Mendelics | Dimas Tadeu Covas, Antonio Jorge Martins, Claudia Renata dos Santos Barros, David Schlesinger, Debora Botequio Moretti, Elaine Cristina Marqueze, Elaine Vieira Santos, Evandra Strazza Rodrigues, Heidge Fukumasu, Jayme Augusto de Souza-Neto, José Salvatore Leister Patané, Luiz Alcantara, Luiz Lehmann Coutinho, Maria Carolina Elias, Maurício Lacerda Nogueira, Rafael dos Santos Bezerra, Raul Machado Neto, Rejane Maria Tommasini Grotto, Ricardo Haddad, Sandra Coccuzzo Sampaio Vessoni, Simone Kashima, Svetoslav Nanev Slavov, Vincent Louis Viala |
| EPI_ISL_2345649 | CENTRO DE SAUDE II IBITINGA | Instituto Butantan / Mendelics | Dimas Tadeu Covas, Antonio Jorge Martins, Claudia Renata dos Santos Barros, David Schlesinger, Debora Botequio Moretti, Elaine Cristina Marqueze, Elaine Vieira Santos, Evandra Strazza Rodrigues, Heidge Fukumasu, Jayme Augusto de Souza-Neto, José Salvatore Leister Patané, Luiz Alcantara, Luiz Lehmann Coutinho, Maria Carolina Elias, Maurício Lacerda Nogueira, Rafael dos Santos Bezerra, Raul Machado Neto, Rejane Maria Tommasini Grotto, Ricardo Haddad, Sandra Coccuzzo Sampaio Vessoni, Simone Kashima, Svetoslav Nanev Slavov, Vincent Louis Viala |
| EPI_ISL_2345650 | CENTRO DE SAUDE II DR GABRIEL MESQUITA VARGEM GDE DO SUL | Instituto Butantan / Mendelics | Dimas Tadeu Covas, Antonio Jorge Martins, Claudia Renata dos Santos Barros, David Schlesinger, Debora Botequio Moretti, Elaine Cristina Marqueze, Elaine Vieira Santos, Evandra Strazza Rodrigues, Heidge Fukumasu, Jayme Augusto de Souza-Neto, José Salvatore Leister Patané, Luiz Alcantara, Luiz Lehmann Coutinho, Maria Carolina Elias, Maurício Lacerda Nogueira, Rafael dos Santos Bezerra, Raul Machado Neto, Rejane Maria Tommasini Grotto, Ricardo Haddad, Sandra Coccuzzo Sampaio Vessoni, Simone Kashima, Svetoslav Nanev Slavov, Vincent Louis Viala |
| EPI_ISL_2345651 | VIGILANCIA EM SAUDE SAE VIGILANCIA EPIDEMIOLOGICA | Instituto Butantan / Mendelics | Dimas Tadeu Covas, Antonio Jorge Martins, Claudia Renata dos Santos Barros, David Schlesinger, Debora Botequio Moretti, Elaine Cristina Marqueze, Elaine Vieira Santos, Evandra Strazza Rodrigues, Heidge Fukumasu, Jayme Augusto de Souza-Neto, José Salvatore Leister Patané, Luiz Alcantara, Luiz Lehmann Coutinho, Maria Carolina Elias, Maurício Lacerda Nogueira, Rafael dos Santos Bezerra, Raul Machado Neto, Rejane Maria Tommasini Grotto, Ricardo Haddad, Sandra Coccuzzo Sampaio Vessoni, Simone Kashima, Svetoslav Nanev Slavov, Vincent Louis Viala |
| EPI_ISL_2345652 | CENTRO DE SAUDE III SALES OLIVEIRA | Instituto Butantan / Mendelics | Dimas Tadeu Covas, Antonio Jorge Martins, Claudia Renata dos Santos Barros, David Schlesinger, Debora Botequio Moretti, Elaine Cristina Marqueze, Elaine Vieira Santos, Evandra Strazza Rodrigues, Heidge Fukumasu, Jayme Augusto de Souza-Neto, José Salvatore Leister Patané, Luiz Alcantara, Luiz Lehmann Coutinho, Maria Carolina Elias, Maurício Lacerda Nogueira, Rafael dos Santos Bezerra, Raul Machado Neto, Rejane Maria Tommasini Grotto, Ricardo Haddad, Sandra Coccuzzo Sampaio Vessoni, Simone Kashima, Svetoslav Nanev Slavov, Vincent Louis Viala |
| EPI_ISL_2345653 | LABORATORIO DR PAULO EMILIO DALESSANDRO PINDAMONHANGABA | Instituto Butantan / Mendelics | Dimas Tadeu Covas, Antonio Jorge Martins, Claudia Renata dos Santos Barros, David Schlesinger, Debora Botequio Moretti, Elaine Cristina Marqueze, Elaine Vieira Santos, Evandra Strazza Rodrigues, Heidge Fukumasu, Jayme Augusto de Souza-Neto, José Salvatore Leister Patané, Luiz Alcantara, Luiz Lehmann Coutinho, Maria Carolina Elias, Maurício Lacerda Nogueira, Rafael dos Santos Bezerra, Raul Machado Neto, Rejane Maria Tommasini Grotto, Ricardo Haddad, Sandra Coccuzzo Sampaio Vessoni, Simone Kashima, Svetoslav Nanev Slavov, Vincent Louis Viala |
| EPI_ISL_2345654 | VIGILANCIA EM SAUDE SAE VIGILANCIA EPIDEMIOLOGICA | Instituto Butantan / Mendelics | Dimas Tadeu Covas, Antonio Jorge Martins, Claudia Renata dos Santos Barros, David Schlesinger, Debora Botequio Moretti, Elaine Cristina Marqueze, Elaine Vieira Santos, Evandra Strazza Rodrigues, Heidge Fukumasu, Jayme Augusto de Souza-Neto, José Salvatore Leister Patané, Luiz Alcantara, Luiz Lehmann Coutinho, Maria Carolina Elias, Maurício Lacerda Nogueira, Rafael dos Santos Bezerra, Raul Machado Neto, Rejane Maria Tommasini Grotto, Ricardo Haddad, Sandra Coccuzzo Sampaio Vessoni, Simone Kashima, Svetoslav Nanev Slavov, Vincent Louis Viala |
| EPI_ISL_2345655 | SECRETARIA MUNICIPAL DE SAUDE SOROCABA | Instituto Butantan / Mendelics | Dimas Tadeu Covas, Antonio Jorge Martins, Claudia Renata dos Santos Barros, David Schlesinger, Debora Botequio Moretti, Elaine Cristina Marqueze, Elaine Vieira Santos, Evandra Strazza Rodrigues, Heidge Fukumasu, Jayme Augusto de Souza-Neto, José Salvatore Leister Patané, Luiz Alcantara, Luiz Lehmann Coutinho, Maria Carolina Elias, Maurício Lacerda Nogueira, Rafael dos Santos Bezerra, Raul Machado Neto, Rejane Maria Tommasini Grotto, Ricardo Haddad, Sandra Coccuzzo Sampaio Vessoni, Simone Kashima, Svetoslav Nanev Slavov, Vincent Louis Viala |
| EPI_ISL_2345657 | CENTRO DE SAUDE DE MARACAI | Instituto Butantan / Mendelics | Dimas Tadeu Covas, Antonio Jorge Martins, Claudia Renata dos Santos Barros, David Schlesinger, Debora Botequio Moretti, Elaine Cristina Marqueze, Elaine Vieira Santos, Evandra Strazza Rodrigues, Heidge Fukumasu, Jayme Augusto de Souza-Neto, José Salvatore Leister Patané, Luiz Alcantara, Luiz Lehmann Coutinho, Maria Carolina Elias, Maurício Lacerda Nogueira, Rafael dos Santos Bezerra, Raul Machado Neto, Rejane Maria Tommasini Grotto, Ricardo Haddad, Sandra Coccuzzo Sampaio Vessoni, Simone Kashima, Svetoslav Nanev Slavov, Vincent Louis Viala |
| EPI_ISL_2345658 | LABORATORIO DR PAULO EMILIO DALESSANDRO PINDAMONHANGABA | Instituto Butantan / Mendelics | Dimas Tadeu Covas, Antonio Jorge Martins, Claudia Renata dos Santos Barros, David Schlesinger, Debora Botequio Moretti, Elaine Cristina Marqueze, Elaine Vieira Santos, Evandra Strazza Rodrigues, Heidge Fukumasu, Jayme Augusto de Souza-Neto, José Salvatore Leister Patané, Luiz Alcantara, Luiz |

| | | | |
|---|---|---|---|
| | | | Lehmann Coutinho, Maria Carolina Elias, Maurício Lacerda Nogueira, Rafael dos Santos Bezerra, Raul Machado Neto, Rejane Maria Tommasini Grotto, Ricardo Haddad, Sandra Coccuzzo Sampaio Vessoni, Simone Kashima, Svetoslav Nanev Slavov, Vincent Louis Viala |
| EPI_ISL_2345659 | GRIPARIO FALCAO | Instituto Butantan / Mendelics | Dimas Tadeu Covas, Antonio Jorge Martins, Claudia Renata dos Santos Barros, David Schlesinger, Debora Botequio Moretti, Elaine Cristina Marqueze, Elaine Vieira Santos, Evandra Strazza Rodrigues, Heidge Fukumasu, Jayme Augusto de Souza-Neto, José Salvatore Leister Patané, Luiz Alcantara, Luiz Lehmann Coutinho, Maria Carolina Elias, Maurício Lacerda Nogueira, Rafael dos Santos Bezerra, Raul Machado Neto, Rejane Maria Tommasini Grotto, Ricardo Haddad, Sandra Coccuzzo Sampaio Vessoni, Simone Kashima, Svetoslav Nanev Slavov, Vincent Louis Viala |
| EPI_ISL_2345660 | SECRETARIA MUNICIPAL DE SAUDE SOROCABA | Instituto Butantan / Mendelics | Dimas Tadeu Covas, Antonio Jorge Martins, Claudia Renata dos Santos Barros, David Schlesinger, Debora Botequio Moretti, Elaine Cristina Marqueze, Elaine Vieira Santos, Evandra Strazza Rodrigues, Heidge Fukumasu, Jayme Augusto de Souza-Neto, José Salvatore Leister Patané, Luiz Alcantara, Luiz Lehmann Coutinho, Maria Carolina Elias, Maurício Lacerda Nogueira, Rafael dos Santos Bezerra, Raul Machado Neto, Rejane Maria Tommasini Grotto, Ricardo Haddad, Sandra Coccuzzo Sampaio Vessoni, Simone Kashima, Svetoslav Nanev Slavov, Vincent Louis Viala |
| EPI_ISL_2345661 | SECRETARIA DE SAUDE PUBLICA DE PRAIA GRANDE | Instituto Butantan / Mendelics | Dimas Tadeu Covas, Antonio Jorge Martins, Claudia Renata dos Santos Barros, David Schlesinger, Debora Botequio Moretti, Elaine Cristina Marqueze, Elaine Vieira Santos, Evandra Strazza Rodrigues, Heidge Fukumasu, Jayme Augusto de Souza-Neto, José Salvatore Leister Patané, Luiz Alcantara, Luiz Lehmann Coutinho, Maria Carolina Elias, Maurício Lacerda Nogueira, Rafael dos Santos Bezerra, Raul Machado Neto, Rejane Maria Tommasini Grotto, Ricardo Haddad, Sandra Coccuzzo Sampaio Vessoni, Simone Kashima, Svetoslav Nanev Slavov, Vincent Louis Viala |
| EPI_ISL_2345662 | UBS GLENDA RODELLA DUMAS | Instituto Butantan / Mendelics | Dimas Tadeu Covas, Antonio Jorge Martins, Claudia Renata dos Santos Barros, David Schlesinger, Debora Botequio Moretti, Elaine Cristina Marqueze, Elaine Vieira Santos, Evandra Strazza Rodrigues, Heidge Fukumasu, Jayme Augusto de Souza-Neto, José Salvatore Leister Patané, Luiz Alcantara, Luiz Lehmann Coutinho, Maria Carolina Elias, Maurício Lacerda Nogueira, Rafael dos Santos Bezerra, Raul Machado Neto, Rejane Maria Tommasini Grotto, Ricardo Haddad, Sandra Coccuzzo Sampaio Vessoni, Simone Kashima, Svetoslav Nanev Slavov, Vincent Louis Viala |
| EPI_ISL_2345663 | GRIPARIO FALCAO | Instituto Butantan / Mendelics | Dimas Tadeu Covas, Antonio Jorge Martins, Claudia Renata dos Santos Barros, David Schlesinger, Debora Botequio Moretti, Elaine Cristina Marqueze, Elaine Vieira Santos, Evandra Strazza Rodrigues, Heidge Fukumasu, Jayme Augusto de Souza-Neto, José Salvatore Leister Patané, Luiz Alcantara, Luiz Lehmann Coutinho, Maria Carolina Elias, Maurício Lacerda Nogueira, Rafael dos Santos Bezerra, Raul Machado Neto, Rejane Maria Tommasini Grotto, Ricardo Haddad, Sandra Coccuzzo Sampaio Vessoni, Simone Kashima, Svetoslav Nanev Slavov, Vincent Louis Viala |
| EPI_ISL_2345664 | SECRETARIA MUNICIPAL DE SAUDE SOROCABA | Instituto Butantan / Mendelics | Dimas Tadeu Covas, Antonio Jorge Martins, Claudia Renata dos Santos Barros, David Schlesinger, Debora Botequio Moretti, Elaine Cristina Marqueze, Elaine Vieira Santos, Evandra Strazza Rodrigues, Heidge Fukumasu, Jayme Augusto de Souza-Neto, José Salvatore Leister Patané, Luiz Alcantara, Luiz Lehmann Coutinho, Maria Carolina Elias, Maurício Lacerda Nogueira, Rafael dos Santos Bezerra, Raul Machado Neto, Rejane Maria Tommasini Grotto, Ricardo Haddad, Sandra Coccuzzo Sampaio Vessoni, Simone Kashima, Svetoslav Nanev Slavov, Vincent Louis Viala |
| EPI_ISL_2345665 | SECRETARIA DE SAUDE PUBLICA DE PRAIA GRANDE | Instituto Butantan / Mendelics | Dimas Tadeu Covas, Antonio Jorge Martins, Claudia Renata dos Santos Barros, David Schlesinger, Debora Botequio Moretti, Elaine Cristina Marqueze, Elaine Vieira Santos, Evandra Strazza Rodrigues, Heidge Fukumasu, Jayme Augusto de Souza-Neto, José Salvatore Leister Patané, Luiz Alcantara, Luiz Lehmann Coutinho, Maria Carolina Elias, Maurício Lacerda Nogueira, Rafael dos Santos Bezerra, Raul Machado Neto, Rejane Maria Tommasini Grotto, Ricardo Haddad, Sandra Coccuzzo Sampaio Vessoni, Simone Kashima, Svetoslav Nanev Slavov, Vincent Louis Viala |
| EPI_ISL_2345666 | UBS III DE RANCHARIA | Instituto Butantan / Mendelics | Dimas Tadeu Covas, Antonio Jorge Martins, Claudia Renata dos Santos Barros, David Schlesinger, Debora Botequio Moretti, Elaine Cristina Marqueze, Elaine Vieira Santos, Evandra Strazza Rodrigues, Heidge Fukumasu, Jayme Augusto de Souza-Neto, José Salvatore Leister Patané, Luiz Alcantara, Luiz Lehmann Coutinho, Maria Carolina Elias, Maurício Lacerda Nogueira, Rafael dos Santos Bezerra, Raul Machado Neto, Rejane Maria Tommasini Grotto, Ricardo Haddad, Sandra Coccuzzo Sampaio Vessoni, Simone Kashima, Svetoslav Nanev Slavov, Vincent Louis Viala |
| EPI_ISL_2345667 | SECAO CENTRO DE DIAGNOSTICO SECEDI | Instituto Butantan / Mendelics | Dimas Tadeu Covas, Antonio Jorge Martins, Claudia Renata dos Santos Barros, David Schlesinger, Debora Botequio Moretti, Elaine Cristina Marqueze, Elaine Vieira Santos, Evandra Strazza Rodrigues, Heidge Fukumasu, Jayme Augusto de Souza-Neto, José Salvatore Leister Patané, Luiz Alcantara, Luiz Lehmann Coutinho, Maria Carolina Elias, Maurício Lacerda Nogueira, Rafael dos Santos Bezerra, Raul Machado Neto, Rejane Maria Tommasini Grotto, Ricardo Haddad, Sandra Coccuzzo Sampaio Vessoni, Simone Kashima, Svetoslav Nanev Slavov, Vincent Louis Viala |
| EPI_ISL_2345668 | VIGILANCIA EPIDIMIOLOGICA DE ARACARIGUAMA | Instituto Butantan / Mendelics | Dimas Tadeu Covas, Antonio Jorge Martins, Claudia Renata dos Santos Barros, David Schlesinger, Debora Botequio Moretti, Elaine Cristina Marqueze, Elaine Vieira Santos, Evandra Strazza Rodrigues, Heidge Fukumasu, Jayme Augusto de Souza-Neto, José Salvatore Leister Patané, Luiz Alcantara, Luiz Lehmann Coutinho, Maria Carolina Elias, Maurício Lacerda Nogueira, Rafael dos Santos Bezerra, Raul Machado Neto, Rejane Maria Tommasini Grotto, Ricardo Haddad, Sandra Coccuzzo Sampaio Vessoni, Simone Kashima, Svetoslav Nanev Slavov, Vincent Louis Viala |
| EPI_ISL_2345669 | CENTRO DE REFERENCIA DO IDOSO DR HUMBERTO MENDES DE CARVALHO | Instituto Butantan / Mendelics | Dimas Tadeu Covas, Antonio Jorge Martins, Claudia Renata dos Santos Barros, David Schlesinger, Debora Botequio Moretti, Elaine Cristina Marqueze, Elaine Vieira Santos, Evandra Strazza Rodrigues, Heidge Fukumasu, Jayme Augusto de Souza-Neto, José Salvatore Leister Patané, Luiz Alcantara, Luiz Lehmann Coutinho, Maria Carolina Elias, Maurício Lacerda Nogueira, Rafael dos Santos Bezerra, Raul Machado Neto, Rejane Maria Tommasini Grotto, Ricardo Haddad, Sandra Coccuzzo Sampaio Vessoni, Simone Kashima, Svetoslav Nanev Slavov, Vincent Louis Viala |
| EPI_ISL_2345670 | POLICLINICA ALCIA LINEIA VITTI REJANI | Instituto Butantan / Mendelics | Dimas Tadeu Covas, Antonio Jorge Martins, Claudia Renata dos Santos Barros, David Schlesinger, Debora Botequio Moretti, Elaine Cristina Marqueze, Elaine Vieira Santos, Evandra Strazza Rodrigues, Heidge Fukumasu, Jayme Augusto de Souza-Neto, José Salvatore Leister Patané, Luiz Alcantara, Luiz Lehmann Coutinho, Maria Carolina Elias, Maurício Lacerda Nogueira, Rafael dos Santos Bezerra, Raul Machado Neto, Rejane Maria Tommasini Grotto, Ricardo Haddad, Sandra Coccuzzo Sampaio Vessoni, Simone Kashima, Svetoslav Nanev Slavov, Vincent Louis Viala |
| EPI_ISL_2345671 | UBS II DE ALVARES MACHADO | Instituto Butantan / Mendelics | Dimas Tadeu Covas, Antonio Jorge Martins, Claudia Renata dos Santos Barros, David Schlesinger, Debora Botequio Moretti, Elaine Cristina Marqueze, Elaine Vieira Santos, Evandra Strazza Rodrigues, Heidge Fukumasu, Jayme Augusto de Souza-Neto, José Salvatore Leister Patané, Luiz Alcantara, Luiz Lehmann Coutinho, Maria Carolina Elias, Maurício Lacerda Nogueira, Rafael dos Santos Bezerra, Raul Machado Neto, Rejane Maria Tommasini Grotto, Ricardo Haddad, Sandra Coccuzzo Sampaio Vessoni, Simone Kashima, Svetoslav Nanev Slavov, Vincent Louis Viala |
| EPI_ISL_2345672 | PSF MARIA JOSE SALTO DE PIRAPORA | Instituto Butantan / Mendelics | Dimas Tadeu Covas, Antonio Jorge Martins, Claudia Renata dos Santos Barros, David Schlesinger, Debora Botequio Moretti, Elaine Cristina Marqueze, Elaine Vieira Santos, Evandra Strazza Rodrigues, Heidge Fukumasu, Jayme Augusto de Souza-Neto, José Salvatore Leister Patané, Luiz Alcantara, Luiz Lehmann Coutinho, Maria Carolina Elias, Maurício Lacerda Nogueira, Rafael dos Santos Bezerra, Raul Machado Neto, Rejane Maria Tommasini Grotto, Ricardo Haddad, Sandra Coccuzzo Sampaio Vessoni, Simone Kashima, Svetoslav Nanev Slavov, Vincent Louis Viala |
| EPI_ISL_2345673 | UNIDADE BASICA DE SAUDE JARDIM DO LAGO | Instituto Butantan / Mendelics | Dimas Tadeu Covas, Antonio Jorge Martins, Claudia Renata dos Santos Barros, David Schlesinger, Debora Botequio Moretti, Elaine Cristina Marqueze, Elaine Vieira Santos, Evandra Strazza Rodrigues, Heidge Fukumasu, Jayme Augusto de Souza-Neto, José Salvatore Leister Patané, Luiz Alcantara, Luiz Lehmann Coutinho, Maria Carolina Elias, Maurício Lacerda Nogueira, Rafael dos Santos Bezerra, Raul Machado Neto, Rejane Maria Tommasini Grotto, Ricardo Haddad, Sandra Coccuzzo Sampaio Vessoni, Simone Kashima, Svetoslav Nanev Slavov, Vincent Louis Viala |
| EPI_ISL_2345674 | CENTRO DE SAUDE II IBITINGA | Instituto Butantan / Mendelics | Dimas Tadeu Covas, Antonio Jorge Martins, Claudia Renata dos Santos Barros, David Schlesinger, Debora Botequio Moretti, Elaine Cristina Marqueze, Elaine Vieira Santos, Evandra Strazza Rodrigues, Heidge Fukumasu, Jayme Augusto de Souza-Neto, José Salvatore Leister Patané, Luiz Alcantara, Luiz Lehmann Coutinho, Maria Carolina Elias, Maurício Lacerda Nogueira, Rafael dos Santos Bezerra, Raul Machado Neto, Rejane Maria Tommasini Grotto, Ricardo Haddad, Sandra Coccuzzo Sampaio Vessoni, Simone Kashima, Svetoslav Nanev Slavov, Vincent Louis Viala |
| EPI_ISL_2345675 | PAS JOAO ANTONIO DO NASCIMENTO | Instituto Butantan / Mendelics | Dimas Tadeu Covas, Antonio Jorge Martins, Claudia Renata dos Santos Barros, David Schlesinger, Debora Botequio Moretti, Elaine Cristina Marqueze, Elaine Vieira Santos, Evandra Strazza Rodrigues, Heidge Fukumasu, Jayme Augusto de Souza-Neto, José Salvatore Leister Patané, Luiz Alcantara, Luiz Lehmann Coutinho, Maria Carolina Elias, Maurício Lacerda Nogueira, Rafael dos Santos Bezerra, Raul Machado Neto, Rejane Maria Tommasini Grotto, Ricardo Haddad, Sandra Coccuzzo Sampaio Vessoni, Simone Kashima, Svetoslav Nanev Slavov, Vincent Louis Viala |
| EPI_ISL_2345676 | HOSPITAL DE CAMPANHA COVID 19 MUNICIPIO DE TAUBATE | Instituto Butantan / Mendelics | Dimas Tadeu Covas, Antonio Jorge Martins, Claudia Renata dos Santos Barros, David Schlesinger, Debora Botequio Moretti, Elaine Cristina Marqueze, Elaine Vieira Santos, Evandra Strazza Rodrigues, Heidge Fukumasu, Jayme Augusto de Souza-Neto, José Salvatore Leister Patané, Luiz Alcantara, Luiz Lehmann Coutinho, Maria Carolina Elias, Maurício Lacerda Nogueira, Rafael dos Santos Bezerra, Raul Machado Neto, Rejane Maria Tommasini Grotto, Ricardo Haddad, Sandra Coccuzzo Sampaio Vessoni, Simone Kashima, Svetoslav Nanev Slavov, Vincent Louis Viala |
| EPI_ISL_2345677 | UPA UNIDADE DE PRONTO ATENDIMENTO 24 HORAS BOM JESUS | Instituto Butantan / Mendelics | Dimas Tadeu Covas, Antonio Jorge Martins, Claudia Renata dos Santos Barros, David Schlesinger, Debora Botequio Moretti, Elaine Cristina Marqueze, Elaine Vieira Santos, Evandra Strazza Rodrigues, Heidge Fukumasu, Jayme Augusto de Souza-Neto, José Salvatore Leister Patané, Luiz Alcantara, Luiz Lehmann Coutinho, Maria Carolina Elias, Maurício Lacerda Nogueira, Rafael dos Santos Bezerra, Raul Machado Neto, Rejane Maria Tommasini Grotto, Ricardo Haddad, Sandra Coccuzzo Sampaio Vessoni, Simone Kashima, Svetoslav Nanev Slavov, Vincent Louis Viala |

| | | | |
|---|---|---|---|
| EPI_ISL_2345678 | UBS JOSE AUGUSTO BATISTA FILHO MENEGUETI | Instituto Butantan / Mendelics | Dimas Tadeu Covas, Antonio Jorge Martins, Claudia Renata dos Santos Barros, David Schlesinger, Debora Botequio Moretti, Elaine Cristina Marqueze, Elaine Vieira Santos, Evandra Strazza Rodrigues, Heidge Fukumasu, Jayme Augusto de Souza-Neto, José Salvatore Leister Patané, Luiz Alcantara, Luiz Lehmann Coutinho, Maria Carolina Elias, Maurício Lacerda Nogueira, Rafael dos Santos Bezerra, Raul Machado Neto, Rejane Maria Tommasini Grotto, Ricardo Haddad, Sandra Coccuzzo Sampaio Vessoni, Simone Kashima, Svetoslav Nanev Slavov, Vincent Louis Viala |
| EPI_ISL_2345679 | DEPARTAMENTO DE SAUDE MUNICIPAL SOCORRO SP | Instituto Butantan / Mendelics | Dimas Tadeu Covas, Antonio Jorge Martins, Claudia Renata dos Santos Barros, David Schlesinger, Debora Botequio Moretti, Elaine Cristina Marqueze, Elaine Vieira Santos, Evandra Strazza Rodrigues, Heidge Fukumasu, Jayme Augusto de Souza-Neto, José Salvatore Leister Patané, Luiz Alcantara, Luiz Lehmann Coutinho, Maria Carolina Elias, Maurício Lacerda Nogueira, Rafael dos Santos Bezerra, Raul Machado Neto, Rejane Maria Tommasini Grotto, Ricardo Haddad, Sandra Coccuzzo Sampaio Vessoni, Simone Kashima, Svetoslav Nanev Slavov, Vincent Louis Viala |
| EPI_ISL_2345680 | CSII DR WASHINGTON LUIS M RODRIGUES DA SILVA PITANGUEIRAS | Instituto Butantan / Mendelics | Dimas Tadeu Covas, Antonio Jorge Martins, Claudia Renata dos Santos Barros, David Schlesinger, Debora Botequio Moretti, Elaine Cristina Marqueze, Elaine Vieira Santos, Evandra Strazza Rodrigues, Heidge Fukumasu, Jayme Augusto de Souza-Neto, José Salvatore Leister Patané, Luiz Alcantara, Luiz Lehmann Coutinho, Maria Carolina Elias, Maurício Lacerda Nogueira, Rafael dos Santos Bezerra, Raul Machado Neto, Rejane Maria Tommasini Grotto, Ricardo Haddad, Sandra Coccuzzo Sampaio Vessoni, Simone Kashima, Svetoslav Nanev Slavov, Vincent Louis Viala |
| EPI_ISL_2345681 | CENTRO DE SAUDE II MATAO | Instituto Butantan / Mendelics | Dimas Tadeu Covas, Antonio Jorge Martins, Claudia Renata dos Santos Barros, David Schlesinger, Debora Botequio Moretti, Elaine Cristina Marqueze, Elaine Vieira Santos, Evandra Strazza Rodrigues, Heidge Fukumasu, Jayme Augusto de Souza-Neto, José Salvatore Leister Patané, Luiz Alcantara, Luiz Lehmann Coutinho, Maria Carolina Elias, Maurício Lacerda Nogueira, Rafael dos Santos Bezerra, Raul Machado Neto, Rejane Maria Tommasini Grotto, Ricardo Haddad, Sandra Coccuzzo Sampaio Vessoni, Simone Kashima, Svetoslav Nanev Slavov, Vincent Louis Viala |
| EPI_ISL_2345682 | SANTA CASA DE MISERICORDIA SAO JOSE | Instituto Butantan / Mendelics | Dimas Tadeu Covas, Antonio Jorge Martins, Claudia Renata dos Santos Barros, David Schlesinger, Debora Botequio Moretti, Elaine Cristina Marqueze, Elaine Vieira Santos, Evandra Strazza Rodrigues, Heidge Fukumasu, Jayme Augusto de Souza-Neto, José Salvatore Leister Patané, Luiz Alcantara, Luiz Lehmann Coutinho, Maria Carolina Elias, Maurício Lacerda Nogueira, Rafael dos Santos Bezerra, Raul Machado Neto, Rejane Maria Tommasini Grotto, Ricardo Haddad, Sandra Coccuzzo Sampaio Vessoni, Simone Kashima, Svetoslav Nanev Slavov, Vincent Louis Viala |
| EPI_ISL_2345683 | DEPARTAMENTO DE SAUDE MUNICIPAL SOCORRO SP | Instituto Butantan / Mendelics | Dimas Tadeu Covas, Antonio Jorge Martins, Claudia Renata dos Santos Barros, David Schlesinger, Debora Botequio Moretti, Elaine Cristina Marqueze, Elaine Vieira Santos, Evandra Strazza Rodrigues, Heidge Fukumasu, Jayme Augusto de Souza-Neto, José Salvatore Leister Patané, Luiz Alcantara, Luiz Lehmann Coutinho, Maria Carolina Elias, Maurício Lacerda Nogueira, Rafael dos Santos Bezerra, Raul Machado Neto, Rejane Maria Tommasini Grotto, Ricardo Haddad, Sandra Coccuzzo Sampaio Vessoni, Simone Kashima, Svetoslav Nanev Slavov, Vincent Louis Viala |
| EPI_ISL_2345684 | UPA MARY DOTA | Instituto Butantan / Mendelics | Dimas Tadeu Covas, Antonio Jorge Martins, Claudia Renata dos Santos Barros, David Schlesinger, Debora Botequio Moretti, Elaine Cristina Marqueze, Elaine Vieira Santos, Evandra Strazza Rodrigues, Heidge Fukumasu, Jayme Augusto de Souza-Neto, José Salvatore Leister Patané, Luiz Alcantara, Luiz Lehmann Coutinho, Maria Carolina Elias, Maurício Lacerda Nogueira, Rafael dos Santos Bezerra, Raul Machado Neto, Rejane Maria Tommasini Grotto, Ricardo Haddad, Sandra Coccuzzo Sampaio Vessoni, Simone Kashima, Svetoslav Nanev Slavov, Vincent Louis Viala |
| EPI_ISL_2345685 | CENTRO DE SAUDE DR JOAO JOAQUIM FERREIRA CRISTAIS PAULISTA | Instituto Butantan / Mendelics | Dimas Tadeu Covas, Antonio Jorge Martins, Claudia Renata dos Santos Barros, David Schlesinger, Debora Botequio Moretti, Elaine Cristina Marqueze, Elaine Vieira Santos, Evandra Strazza Rodrigues, Heidge Fukumasu, Jayme Augusto de Souza-Neto, José Salvatore Leister Patané, Luiz Alcantara, Luiz Lehmann Coutinho, Maria Carolina Elias, Maurício Lacerda Nogueira, Rafael dos Santos Bezerra, Raul Machado Neto, Rejane Maria Tommasini Grotto, Ricardo Haddad, Sandra Coccuzzo Sampaio Vessoni, Simone Kashima, Svetoslav Nanev Slavov, Vincent Louis Viala |
| EPI_ISL_2345686 | AMBULATORIO CENTRAL DE ESPECIALIDADES ITATIBA | Instituto Butantan / Mendelics | Dimas Tadeu Covas, Antonio Jorge Martins, Claudia Renata dos Santos Barros, David Schlesinger, Debora Botequio Moretti, Elaine Cristina Marqueze, Elaine Vieira Santos, Evandra Strazza Rodrigues, Heidge Fukumasu, Jayme Augusto de Souza-Neto, José Salvatore Leister Patané, Luiz Alcantara, Luiz Lehmann Coutinho, Maria Carolina Elias, Maurício Lacerda Nogueira, Rafael dos Santos Bezerra, Raul Machado Neto, Rejane Maria Tommasini Grotto, Ricardo Haddad, Sandra Coccuzzo Sampaio Vessoni, Simone Kashima, Svetoslav Nanev Slavov, Vincent Louis Viala |
| EPI_ISL_2345687 | NUCLEO DE SAUDE MARY DOTA DE BAURU | Instituto Butantan / Mendelics | Dimas Tadeu Covas, Antonio Jorge Martins, Claudia Renata dos Santos Barros, David Schlesinger, Debora Botequio Moretti, Elaine Cristina Marqueze, Elaine Vieira Santos, Evandra Strazza Rodrigues, Heidge Fukumasu, Jayme Augusto de Souza-Neto, José Salvatore Leister Patané, Luiz Alcantara, Luiz Lehmann Coutinho, Maria Carolina Elias, Maurício Lacerda Nogueira, Rafael dos Santos Bezerra, Raul Machado Neto, Rejane Maria Tommasini Grotto, Ricardo Haddad, Sandra Coccuzzo Sampaio Vessoni, Simone Kashima, Svetoslav Nanev Slavov, Vincent Louis Viala |
| EPI_ISL_2345688 | VIGILANCIA EPIDEMIOLOGICA JARDINOPOLIS SP | Instituto Butantan / Mendelics | Dimas Tadeu Covas, Antonio Jorge Martins, Claudia Renata dos Santos Barros, David Schlesinger, Debora Botequio Moretti, Elaine Cristina Marqueze, Elaine Vieira Santos, Evandra Strazza Rodrigues, Heidge Fukumasu, Jayme Augusto de Souza-Neto, José Salvatore Leister Patané, Luiz Alcantara, Luiz Lehmann Coutinho, Maria Carolina Elias, Maurício Lacerda Nogueira, Rafael dos Santos Bezerra, Raul Machado Neto, Rejane Maria Tommasini Grotto, Ricardo Haddad, Sandra Coccuzzo Sampaio Vessoni, Simone Kashima, Svetoslav Nanev Slavov, Vincent Louis Viala |
| EPI_ISL_2345689 | CENTRO DE APOIO EPIDEMIOLOGICO | Instituto Butantan / Mendelics | Dimas Tadeu Covas, Antonio Jorge Martins, Claudia Renata dos Santos Barros, David Schlesinger, Debora Botequio Moretti, Elaine Cristina Marqueze, Elaine Vieira Santos, Evandra Strazza Rodrigues, Heidge Fukumasu, Jayme Augusto de Souza-Neto, José Salvatore Leister Patané, Luiz Alcantara, Luiz Lehmann Coutinho, Maria Carolina Elias, Maurício Lacerda Nogueira, Rafael dos Santos Bezerra, Raul Machado Neto, Rejane Maria Tommasini Grotto, Ricardo Haddad, Sandra Coccuzzo Sampaio Vessoni, Simone Kashima, Svetoslav Nanev Slavov, Vincent Louis Viala |
| EPI_ISL_2345690 | HOSPITAL MUNICIPAL DE ITAPIRA | Instituto Butantan / Mendelics | Dimas Tadeu Covas, Antonio Jorge Martins, Claudia Renata dos Santos Barros, David Schlesinger, Debora Botequio Moretti, Elaine Cristina Marqueze, Elaine Vieira Santos, Evandra Strazza Rodrigues, Heidge Fukumasu, Jayme Augusto de Souza-Neto, José Salvatore Leister Patané, Luiz Alcantara, Luiz Lehmann Coutinho, Maria Carolina Elias, Maurício Lacerda Nogueira, Rafael dos Santos Bezerra, Raul Machado Neto, Rejane Maria Tommasini Grotto, Ricardo Haddad, Sandra Coccuzzo Sampaio Vessoni, Simone Kashima, Svetoslav Nanev Slavov, Vincent Louis Viala |
| EPI_ISL_2345691 | LABORATORIO MUNICIPAL DE SUZANO | Instituto Butantan / Mendelics | Dimas Tadeu Covas, Antonio Jorge Martins, Claudia Renata dos Santos Barros, David Schlesinger, Debora Botequio Moretti, Elaine Cristina Marqueze, Elaine Vieira Santos, Evandra Strazza Rodrigues, Heidge Fukumasu, Jayme Augusto de Souza-Neto, José Salvatore Leister Patané, Luiz Alcantara, Luiz Lehmann Coutinho, Maria Carolina Elias, Maurício Lacerda Nogueira, Rafael dos Santos Bezerra, Raul Machado Neto, Rejane Maria Tommasini Grotto, Ricardo Haddad, Sandra Coccuzzo Sampaio Vessoni, Simone Kashima, Svetoslav Nanev Slavov, Vincent Louis Viala |
| EPI_ISL_2345692 | UBS DR ALFREDO DANTAS DE SOUZA UMUARAMA | Instituto Butantan / Mendelics | Dimas Tadeu Covas, Antonio Jorge Martins, Claudia Renata dos Santos Barros, David Schlesinger, Debora Botequio Moretti, Elaine Cristina Marqueze, Elaine Vieira Santos, Evandra Strazza Rodrigues, Heidge Fukumasu, Jayme Augusto de Souza-Neto, José Salvatore Leister Patané, Luiz Alcantara, Luiz Lehmann Coutinho, Maria Carolina Elias, Maurício Lacerda Nogueira, Rafael dos Santos Bezerra, Raul Machado Neto, Rejane Maria Tommasini Grotto, Ricardo Haddad, Sandra Coccuzzo Sampaio Vessoni, Simone Kashima, Svetoslav Nanev Slavov, Vincent Louis Viala |
| EPI_ISL_2345694 | CENTRO DE SAUDE II IBITINGA | Instituto Butantan / Mendelics | Dimas Tadeu Covas, Antonio Jorge Martins, Claudia Renata dos Santos Barros, David Schlesinger, Debora Botequio Moretti, Elaine Cristina Marqueze, Elaine Vieira Santos, Evandra Strazza Rodrigues, Heidge Fukumasu, Jayme Augusto de Souza-Neto, José Salvatore Leister Patané, Luiz Alcantara, Luiz Lehmann Coutinho, Maria Carolina Elias, Maurício Lacerda Nogueira, Rafael dos Santos Bezerra, Raul Machado Neto, Rejane Maria Tommasini Grotto, Ricardo Haddad, Sandra Coccuzzo Sampaio Vessoni, Simone Kashima, Svetoslav Nanev Slavov, Vincent Louis Viala |
| EPI_ISL_2345695 | CENTRO DE SAUDE II DR GABRIEL MESQUITA VARGEM GDE DO SUL | Instituto Butantan / Mendelics | Dimas Tadeu Covas, Antonio Jorge Martins, Claudia Renata dos Santos Barros, David Schlesinger, Debora Botequio Moretti, Elaine Cristina Marqueze, Elaine Vieira Santos, Evandra Strazza Rodrigues, Heidge Fukumasu, Jayme Augusto de Souza-Neto, José Salvatore Leister Patané, Luiz Alcantara, Luiz Lehmann Coutinho, Maria Carolina Elias, Maurício Lacerda Nogueira, Rafael dos Santos Bezerra, Raul Machado Neto, Rejane Maria Tommasini Grotto, Ricardo Haddad, Sandra Coccuzzo Sampaio Vessoni, Simone Kashima, Svetoslav Nanev Slavov, Vincent Louis Viala |
| EPI_ISL_2345696 | CENTRO DE SAUDE III SANTA ERNESTINA | Instituto Butantan / Mendelics | Dimas Tadeu Covas, Antonio Jorge Martins, Claudia Renata dos Santos Barros, David Schlesinger, Debora Botequio Moretti, Elaine Cristina Marqueze, Elaine Vieira Santos, Evandra Strazza Rodrigues, Heidge Fukumasu, Jayme Augusto de Souza-Neto, José Salvatore Leister Patané, Luiz Alcantara, Luiz Lehmann Coutinho, Maria Carolina Elias, Maurício Lacerda Nogueira, Rafael dos Santos Bezerra, Raul Machado Neto, Rejane Maria Tommasini Grotto, Ricardo Haddad, Sandra Coccuzzo Sampaio Vessoni, Simone Kashima, Svetoslav Nanev Slavov, Vincent Louis Viala |
| EPI_ISL_2345697 | CENTRO DE SAUDE III SALES OLIVEIRA | Instituto Butantan / Mendelics | Dimas Tadeu Covas, Antonio Jorge Martins, Claudia Renata dos Santos Barros, David Schlesinger, Debora Botequio Moretti, Elaine Cristina Marqueze, Elaine Vieira Santos, Evandra Strazza Rodrigues, Heidge Fukumasu, Jayme Augusto de Souza-Neto, José Salvatore Leister Patané, Luiz Alcantara, Luiz Lehmann Coutinho, Maria Carolina Elias, Maurício Lacerda Nogueira, Rafael dos Santos Bezerra, Raul Machado Neto, Rejane Maria Tommasini Grotto, Ricardo Haddad, Sandra Coccuzzo Sampaio Vessoni, Simone Kashima, Svetoslav Nanev Slavov, Vincent Louis Viala |
| EPI_ISL_2345698 | LABORATORIO DR PAULO EMILIO DALESSANDRO PINDAMONHANGABA | Instituto Butantan / Mendelics | Dimas Tadeu Covas, Antonio Jorge Martins, Claudia Renata dos Santos Barros, David Schlesinger, Debora Botequio Moretti, Elaine Cristina Marqueze, Elaine Vieira Santos, Evandra Strazza Rodrigues, Heidge Fukumasu, Jayme Augusto de Souza-Neto, José Salvatore Leister Patané, Luiz Alcantara, Luiz |

| | | | |
|---|---|---|---|
| | | | Lehmann Coutinho, Maria Carolina Elias, Maurício Lacerda Nogueira, Rafael dos Santos Bezerra, Raul Machado Neto, Rejane Maria Tommasini Grotto, Ricardo Haddad, Sandra Coccuzzo Sampaio Vessoni, Simone Kashima, Svetoslav Nanev Slavov, Vincent Louis Viala |
| EPI_ISL_2345699 | SECRETARIA DE SAUDE PUBLICA DE PRAIA GRANDE | Instituto Butantan / Mendelics | Dimas Tadeu Covas, Antonio Jorge Martins, Claudia Renata dos Santos Barros, David Schlesinger, Debora Botequio Moretti, Elaine Cristina Marqueze, Elaine Vieira Santos, Evandra Strazza Rodrigues, Heidge Fukumasu, Jayme Augusto de Souza-Neto, José Salvatore Leister Patané, Luiz Alcantara, Luiz Lehmann Coutinho, Maria Carolina Elias, Maurício Lacerda Nogueira, Rafael dos Santos Bezerra, Raul Machado Neto, Rejane Maria Tommasini Grotto, Ricardo Haddad, Sandra Coccuzzo Sampaio Vessoni, Simone Kashima, Svetoslav Nanev Slavov, Vincent Louis Viala |
| EPI_ISL_2345700 | SECRETARIA MUNICIPAL DE SAUDE SOROCABA | Instituto Butantan / Mendelics | Dimas Tadeu Covas, Antonio Jorge Martins, Claudia Renata dos Santos Barros, David Schlesinger, Debora Botequio Moretti, Elaine Cristina Marqueze, Elaine Vieira Santos, Evandra Strazza Rodrigues, Heidge Fukumasu, Jayme Augusto de Souza-Neto, José Salvatore Leister Patané, Luiz Alcantara, Luiz Lehmann Coutinho, Maria Carolina Elias, Maurício Lacerda Nogueira, Rafael dos Santos Bezerra, Raul Machado Neto, Rejane Maria Tommasini Grotto, Ricardo Haddad, Sandra Coccuzzo Sampaio Vessoni, Simone Kashima, Svetoslav Nanev Slavov, Vincent Louis Viala |
| EPI_ISL_2345701 | GRIPARIO FALCAO | Instituto Butantan / Mendelics | Dimas Tadeu Covas, Antonio Jorge Martins, Claudia Renata dos Santos Barros, David Schlesinger, Debora Botequio Moretti, Elaine Cristina Marqueze, Elaine Vieira Santos, Evandra Strazza Rodrigues, Heidge Fukumasu, Jayme Augusto de Souza-Neto, José Salvatore Leister Patané, Luiz Alcantara, Luiz Lehmann Coutinho, Maria Carolina Elias, Maurício Lacerda Nogueira, Rafael dos Santos Bezerra, Raul Machado Neto, Rejane Maria Tommasini Grotto, Ricardo Haddad, Sandra Coccuzzo Sampaio Vessoni, Simone Kashima, Svetoslav Nanev Slavov, Vincent Louis Viala |
| EPI_ISL_2345702 | CENTRO DE SAUDE DE MARACAI | Instituto Butantan / Mendelics | Dimas Tadeu Covas, Antonio Jorge Martins, Claudia Renata dos Santos Barros, David Schlesinger, Debora Botequio Moretti, Elaine Cristina Marqueze, Elaine Vieira Santos, Evandra Strazza Rodrigues, Heidge Fukumasu, Jayme Augusto de Souza-Neto, José Salvatore Leister Patané, Luiz Alcantara, Luiz Lehmann Coutinho, Maria Carolina Elias, Maurício Lacerda Nogueira, Rafael dos Santos Bezerra, Raul Machado Neto, Rejane Maria Tommasini Grotto, Ricardo Haddad, Sandra Coccuzzo Sampaio Vessoni, Simone Kashima, Svetoslav Nanev Slavov, Vincent Louis Viala |
| EPI_ISL_2345703, EPI_ISL_2345704 | SECRETARIA DE SAUDE PUBLICA DE PRAIA GRANDE | Instituto Butantan / Mendelics | Dimas Tadeu Covas, Antonio Jorge Martins, Claudia Renata dos Santos Barros, David Schlesinger, Debora Botequio Moretti, Elaine Cristina Marqueze, Elaine Vieira Santos, Evandra Strazza Rodrigues, Heidge Fukumasu, Jayme Augusto de Souza-Neto, José Salvatore Leister Patané, Luiz Alcantara, Luiz Lehmann Coutinho, Maria Carolina Elias, Maurício Lacerda Nogueira, Rafael dos Santos Bezerra, Raul Machado Neto, Rejane Maria Tommasini Grotto, Ricardo Haddad, Sandra Coccuzzo Sampaio Vessoni, Simone Kashima, Svetoslav Nanev Slavov, Vincent Louis Viala |
| EPI_ISL_2345705 | UBS GLENDA RODELLA DUMAS | Instituto Butantan / Mendelics | Dimas Tadeu Covas, Antonio Jorge Martins, Claudia Renata dos Santos Barros, David Schlesinger, Debora Botequio Moretti, Elaine Cristina Marqueze, Elaine Vieira Santos, Evandra Strazza Rodrigues, Heidge Fukumasu, Jayme Augusto de Souza-Neto, José Salvatore Leister Patané, Luiz Alcantara, Luiz Lehmann Coutinho, Maria Carolina Elias, Maurício Lacerda Nogueira, Rafael dos Santos Bezerra, Raul Machado Neto, Rejane Maria Tommasini Grotto, Ricardo Haddad, Sandra Coccuzzo Sampaio Vessoni, Simone Kashima, Svetoslav Nanev Slavov, Vincent Louis Viala |
| EPI_ISL_2345706 | LABORATORIO DR PAULO EMILIO DALESSANDRO PINDAMONHANGABA | Instituto Butantan / Mendelics | Dimas Tadeu Covas, Antonio Jorge Martins, Claudia Renata dos Santos Barros, David Schlesinger, Debora Botequio Moretti, Elaine Cristina Marqueze, Elaine Vieira Santos, Evandra Strazza Rodrigues, Heidge Fukumasu, Jayme Augusto de Souza-Neto, José Salvatore Leister Patané, Luiz Alcantara, Luiz Lehmann Coutinho, Maria Carolina Elias, Maurício Lacerda Nogueira, Rafael dos Santos Bezerra, Raul Machado Neto, Rejane Maria Tommasini Grotto, Ricardo Haddad, Sandra Coccuzzo Sampaio Vessoni, Simone Kashima, Svetoslav Nanev Slavov, Vincent Louis Viala |
| EPI_ISL_2345707 | GRIPARIO FALCAO | Instituto Butantan / Mendelics | Dimas Tadeu Covas, Antonio Jorge Martins, Claudia Renata dos Santos Barros, David Schlesinger, Debora Botequio Moretti, Elaine Cristina Marqueze, Elaine Vieira Santos, Evandra Strazza Rodrigues, Heidge Fukumasu, Jayme Augusto de Souza-Neto, José Salvatore Leister Patané, Luiz Alcantara, Luiz Lehmann Coutinho, Maria Carolina Elias, Maurício Lacerda Nogueira, Rafael dos Santos Bezerra, Raul Machado Neto, Rejane Maria Tommasini Grotto, Ricardo Haddad, Sandra Coccuzzo Sampaio Vessoni, Simone Kashima, Svetoslav Nanev Slavov, Vincent Louis Viala |
| EPI_ISL_2345708 | SMS SECRETARIA MUNICIPAL DE SAUDE DE BOITUVA | Instituto Butantan / Mendelics | Dimas Tadeu Covas, Antonio Jorge Martins, Claudia Renata dos Santos Barros, David Schlesinger, Debora Botequio Moretti, Elaine Cristina Marqueze, Elaine Vieira Santos, Evandra Strazza Rodrigues, Heidge Fukumasu, Jayme Augusto de Souza-Neto, José Salvatore Leister Patané, Luiz Alcantara, Luiz Lehmann Coutinho, Maria Carolina Elias, Maurício Lacerda Nogueira, Rafael dos Santos Bezerra, Raul Machado Neto, Rejane Maria Tommasini Grotto, Ricardo Haddad, Sandra Coccuzzo Sampaio Vessoni, Simone Kashima, Svetoslav Nanev Slavov, Vincent Louis Viala |
| EPI_ISL_2345709 | GRIPARIO FALCAO | Instituto Butantan / Mendelics | Dimas Tadeu Covas, Antonio Jorge Martins, Claudia Renata dos Santos Barros, David Schlesinger, Debora Botequio Moretti, Elaine Cristina Marqueze, Elaine Vieira Santos, Evandra Strazza Rodrigues, Heidge Fukumasu, Jayme Augusto de Souza-Neto, José Salvatore Leister Patané, Luiz Alcantara, Luiz Lehmann Coutinho, Maria Carolina Elias, Maurício Lacerda Nogueira, Rafael dos Santos Bezerra, Raul Machado Neto, Rejane Maria Tommasini Grotto, Ricardo Haddad, Sandra Coccuzzo Sampaio Vessoni, Simone Kashima, Svetoslav Nanev Slavov, Vincent Louis Viala |
| EPI_ISL_2345710 | UBS III DE RANCHARIA | Instituto Butantan / Mendelics | Dimas Tadeu Covas, Antonio Jorge Martins, Claudia Renata dos Santos Barros, David Schlesinger, Debora Botequio Moretti, Elaine Cristina Marqueze, Elaine Vieira Santos, Evandra Strazza Rodrigues, Heidge Fukumasu, Jayme Augusto de Souza-Neto, José Salvatore Leister Patané, Luiz Alcantara, Luiz Lehmann Coutinho, Maria Carolina Elias, Maurício Lacerda Nogueira, Rafael dos Santos Bezerra, Raul Machado Neto, Rejane Maria Tommasini Grotto, Ricardo Haddad, Sandra Coccuzzo Sampaio Vessoni, Simone Kashima, Svetoslav Nanev Slavov, Vincent Louis Viala |
| EPI_ISL_2345711 | UPA DR AKIRA TADA | Instituto Butantan / Mendelics | Dimas Tadeu Covas, Antonio Jorge Martins, Claudia Renata dos Santos Barros, David Schlesinger, Debora Botequio Moretti, Elaine Cristina Marqueze, Elaine Vieira Santos, Evandra Strazza Rodrigues, Heidge Fukumasu, Jayme Augusto de Souza-Neto, José Salvatore Leister Patané, Luiz Alcantara, Luiz Lehmann Coutinho, Maria Carolina Elias, Maurício Lacerda Nogueira, Rafael dos Santos Bezerra, Raul Machado Neto, Rejane Maria Tommasini Grotto, Ricardo Haddad, Sandra Coccuzzo Sampaio Vessoni, Simone Kashima, Svetoslav Nanev Slavov, Vincent Louis Viala |
| EPI_ISL_2345712 | SECAO CENTRO DE DIAGNOSTICO SECEDI | Instituto Butantan / Mendelics | Dimas Tadeu Covas, Antonio Jorge Martins, Claudia Renata dos Santos Barros, David Schlesinger, Debora Botequio Moretti, Elaine Cristina Marqueze, Elaine Vieira Santos, Evandra Strazza Rodrigues, Heidge Fukumasu, Jayme Augusto de Souza-Neto, José Salvatore Leister Patané, Luiz Alcantara, Luiz Lehmann Coutinho, Maria Carolina Elias, Maurício Lacerda Nogueira, Rafael dos Santos Bezerra, Raul Machado Neto, Rejane Maria Tommasini Grotto, Ricardo Haddad, Sandra Coccuzzo Sampaio Vessoni, Simone Kashima, Svetoslav Nanev Slavov, Vincent Louis Viala |
| EPI_ISL_2345713 | SMS SECRETARIA MUNICIPAL DE SAUDE DE BOITUVA | Instituto Butantan / Mendelics | Dimas Tadeu Covas, Antonio Jorge Martins, Claudia Renata dos Santos Barros, David Schlesinger, Debora Botequio Moretti, Elaine Cristina Marqueze, Elaine Vieira Santos, Evandra Strazza Rodrigues, Heidge Fukumasu, Jayme Augusto de Souza-Neto, José Salvatore Leister Patané, Luiz Alcantara, Luiz Lehmann Coutinho, Maria Carolina Elias, Maurício Lacerda Nogueira, Rafael dos Santos Bezerra, Raul Machado Neto, Rejane Maria Tommasini Grotto, Ricardo Haddad, Sandra Coccuzzo Sampaio Vessoni, Simone Kashima, Svetoslav Nanev Slavov, Vincent Louis Viala |
| EPI_ISL_2345714 | PRONTO ATENDIMENTO MUNICIPAL | Instituto Butantan / Mendelics | Dimas Tadeu Covas, Antonio Jorge Martins, Claudia Renata dos Santos Barros, David Schlesinger, Debora Botequio Moretti, Elaine Cristina Marqueze, Elaine Vieira Santos, Evandra Strazza Rodrigues, Heidge Fukumasu, Jayme Augusto de Souza-Neto, José Salvatore Leister Patané, Luiz Alcantara, Luiz Lehmann Coutinho, Maria Carolina Elias, Maurício Lacerda Nogueira, Rafael dos Santos Bezerra, Raul Machado Neto, Rejane Maria Tommasini Grotto, Ricardo Haddad, Sandra Coccuzzo Sampaio Vessoni, Simone Kashima, Svetoslav Nanev Slavov, Vincent Louis Viala |
| EPI_ISL_2345715 | CENTRO DE ESPECIALIDADES DE PRIMAVERA | Instituto Butantan / Mendelics | Dimas Tadeu Covas, Antonio Jorge Martins, Claudia Renata dos Santos Barros, David Schlesinger, Debora Botequio Moretti, Elaine Cristina Marqueze, Elaine Vieira Santos, Evandra Strazza Rodrigues, Heidge Fukumasu, Jayme Augusto de Souza-Neto, José Salvatore Leister Patané, Luiz Alcantara, Luiz Lehmann Coutinho, Maria Carolina Elias, Maurício Lacerda Nogueira, Rafael dos Santos Bezerra, Raul Machado Neto, Rejane Maria Tommasini Grotto, Ricardo Haddad, Sandra Coccuzzo Sampaio Vessoni, Simone Kashima, Svetoslav Nanev Slavov, Vincent Louis Viala |
| EPI_ISL_2345716 | UPA DR FRANCO DA ROCHA | Instituto Butantan / Mendelics | Dimas Tadeu Covas, Antonio Jorge Martins, Claudia Renata dos Santos Barros, David Schlesinger, Debora Botequio Moretti, Elaine Cristina Marqueze, Elaine Vieira Santos, Evandra Strazza Rodrigues, Heidge Fukumasu, Jayme Augusto de Souza-Neto, José Salvatore Leister Patané, Luiz Alcantara, Luiz Lehmann Coutinho, Maria Carolina Elias, Maurício Lacerda Nogueira, Rafael dos Santos Bezerra, Raul Machado Neto, Rejane Maria Tommasini Grotto, Ricardo Haddad, Sandra Coccuzzo Sampaio Vessoni, Simone Kashima, Svetoslav Nanev Slavov, Vincent Louis Viala |
| EPI_ISL_2345717 | UBS II DE ALVARES MACHADO | Instituto Butantan / Mendelics | Dimas Tadeu Covas, Antonio Jorge Martins, Claudia Renata dos Santos Barros, David Schlesinger, Debora Botequio Moretti, Elaine Cristina Marqueze, Elaine Vieira Santos, Evandra Strazza Rodrigues, Heidge Fukumasu, Jayme Augusto de Souza-Neto, José Salvatore Leister Patané, Luiz Alcantara, Luiz Lehmann Coutinho, Maria Carolina Elias, Maurício Lacerda Nogueira, Rafael dos Santos Bezerra, Raul Machado Neto, Rejane Maria Tommasini Grotto, Ricardo Haddad, Sandra Coccuzzo Sampaio Vessoni, Simone Kashima, Svetoslav Nanev Slavov, Vincent Louis Viala |
| EPI_ISL_2345718 | SMS SECRETARIA MUNICIPAL DE SAUDE DE BOITUVA | Instituto Butantan / Mendelics | Dimas Tadeu Covas, Antonio Jorge Martins, Claudia Renata dos Santos Barros, David Schlesinger, Debora Botequio Moretti, Elaine Cristina Marqueze, Elaine Vieira Santos, Evandra Strazza Rodrigues, Heidge Fukumasu, Jayme Augusto de Souza-Neto, José Salvatore Leister Patané, Luiz Alcantara, Luiz Lehmann Coutinho, Maria Carolina Elias, Maurício Lacerda Nogueira, Rafael dos Santos Bezerra, Raul Machado Neto, Rejane Maria Tommasini Grotto, Ricardo Haddad, Sandra Coccuzzo Sampaio Vessoni, Simone Kashima, Svetoslav Nanev Slavov, Vincent Louis Viala |

| | | | |
|---|---|---|---|
| EPI_ISL_2345719 | UNIDADE BASICA DE SAUDE JARDIM DO LAGO | Instituto Butantan / Mendelics | Dimas Tadeu Covas, Antonio Jorge Martins, Claudia Renata dos Santos Barros, David Schlesinger, Debora Botequio Moretti, Elaine Cristina Marqueze, Elaine Vieira Santos, Evandra Strazza Rodrigues, Heidge Fukumasu, Jayme Augusto de Souza-Neto, José Salvatore Leister Patané, Luiz Alcantara, Luiz Lehmann Coutinho, Maria Carolina Elias, Maurício Lacerda Nogueira, Rafael dos Santos Bezerra, Raul Machado Neto, Rejane Maria Tommasini Grotto, Ricardo Haddad, Sandra Coccuzzo Sampaio Vessoni, Simone Kashima, Svetoslav Nanev Slavov, Vincent Louis Viala |
| EPI_ISL_2345720 | CENTRO DE SAUDE II IBITINGA | Instituto Butantan / Mendelics | Dimas Tadeu Covas, Antonio Jorge Martins, Claudia Renata dos Santos Barros, David Schlesinger, Debora Botequio Moretti, Elaine Cristina Marqueze, Elaine Vieira Santos, Evandra Strazza Rodrigues, Heidge Fukumasu, Jayme Augusto de Souza-Neto, José Salvatore Leister Patané, Luiz Alcantara, Luiz Lehmann Coutinho, Maria Carolina Elias, Maurício Lacerda Nogueira, Rafael dos Santos Bezerra, Raul Machado Neto, Rejane Maria Tommasini Grotto, Ricardo Haddad, Sandra Coccuzzo Sampaio Vessoni, Simone Kashima, Svetoslav Nanev Slavov, Vincent Louis Viala |
| EPI_ISL_2345721 | PAS JOAO ANTONIO DO NASCIMENTO | Instituto Butantan / Mendelics | Dimas Tadeu Covas, Antonio Jorge Martins, Claudia Renata dos Santos Barros, David Schlesinger, Debora Botequio Moretti, Elaine Cristina Marqueze, Elaine Vieira Santos, Evandra Strazza Rodrigues, Heidge Fukumasu, Jayme Augusto de Souza-Neto, José Salvatore Leister Patané, Luiz Alcantara, Luiz Lehmann Coutinho, Maria Carolina Elias, Maurício Lacerda Nogueira, Rafael dos Santos Bezerra, Raul Machado Neto, Rejane Maria Tommasini Grotto, Ricardo Haddad, Sandra Coccuzzo Sampaio Vessoni, Simone Kashima, Svetoslav Nanev Slavov, Vincent Louis Viala |
| EPI_ISL_2345722 | HOSPITAL DE CAMPANHA COVID 19 MUNICIPIO DE TAUBATE | Instituto Butantan / Mendelics | Dimas Tadeu Covas, Antonio Jorge Martins, Claudia Renata dos Santos Barros, David Schlesinger, Debora Botequio Moretti, Elaine Cristina Marqueze, Elaine Vieira Santos, Evandra Strazza Rodrigues, Heidge Fukumasu, Jayme Augusto de Souza-Neto, José Salvatore Leister Patané, Luiz Alcantara, Luiz Lehmann Coutinho, Maria Carolina Elias, Maurício Lacerda Nogueira, Rafael dos Santos Bezerra, Raul Machado Neto, Rejane Maria Tommasini Grotto, Ricardo Haddad, Sandra Coccuzzo Sampaio Vessoni, Simone Kashima, Svetoslav Nanev Slavov, Vincent Louis Viala |
| EPI_ISL_2345723 | UNIDADE BASICA DE SAUDE JARDIM DO LAGO | Instituto Butantan / Mendelics | Dimas Tadeu Covas, Antonio Jorge Martins, Claudia Renata dos Santos Barros, David Schlesinger, Debora Botequio Moretti, Elaine Cristina Marqueze, Elaine Vieira Santos, Evandra Strazza Rodrigues, Heidge Fukumasu, Jayme Augusto de Souza-Neto, José Salvatore Leister Patané, Luiz Alcantara, Luiz Lehmann Coutinho, Maria Carolina Elias, Maurício Lacerda Nogueira, Rafael dos Santos Bezerra, Raul Machado Neto, Rejane Maria Tommasini Grotto, Ricardo Haddad, Sandra Coccuzzo Sampaio Vessoni, Simone Kashima, Svetoslav Nanev Slavov, Vincent Louis Viala |
| EPI_ISL_2345724 | CENTRO DE SAUDE II IBITINGA | Instituto Butantan / Mendelics | Dimas Tadeu Covas, Antonio Jorge Martins, Claudia Renata dos Santos Barros, David Schlesinger, Debora Botequio Moretti, Elaine Cristina Marqueze, Elaine Vieira Santos, Evandra Strazza Rodrigues, Heidge Fukumasu, Jayme Augusto de Souza-Neto, José Salvatore Leister Patané, Luiz Alcantara, Luiz Lehmann Coutinho, Maria Carolina Elias, Maurício Lacerda Nogueira, Rafael dos Santos Bezerra, Raul Machado Neto, Rejane Maria Tommasini Grotto, Ricardo Haddad, Sandra Coccuzzo Sampaio Vessoni, Simone Kashima, Svetoslav Nanev Slavov, Vincent Louis Viala |
| EPI_ISL_2345725 | UPA DR AKIRA TADA | Instituto Butantan / Mendelics | Dimas Tadeu Covas, Antonio Jorge Martins, Claudia Renata dos Santos Barros, David Schlesinger, Debora Botequio Moretti, Elaine Cristina Marqueze, Elaine Vieira Santos, Evandra Strazza Rodrigues, Heidge Fukumasu, Jayme Augusto de Souza-Neto, José Salvatore Leister Patané, Luiz Alcantara, Luiz Lehmann Coutinho, Maria Carolina Elias, Maurício Lacerda Nogueira, Rafael dos Santos Bezerra, Raul Machado Neto, Rejane Maria Tommasini Grotto, Ricardo Haddad, Sandra Coccuzzo Sampaio Vessoni, Simone Kashima, Svetoslav Nanev Slavov, Vincent Louis Viala |
| EPI_ISL_2345726 | UBS III DE PARIQUERA ACU PARIQUERA ACU | Instituto Butantan / Mendelics | Dimas Tadeu Covas, Antonio Jorge Martins, Claudia Renata dos Santos Barros, David Schlesinger, Debora Botequio Moretti, Elaine Cristina Marqueze, Elaine Vieira Santos, Evandra Strazza Rodrigues, Heidge Fukumasu, Jayme Augusto de Souza-Neto, José Salvatore Leister Patané, Luiz Alcantara, Luiz Lehmann Coutinho, Maria Carolina Elias, Maurício Lacerda Nogueira, Rafael dos Santos Bezerra, Raul Machado Neto, Rejane Maria Tommasini Grotto, Ricardo Haddad, Sandra Coccuzzo Sampaio Vessoni, Simone Kashima, Svetoslav Nanev Slavov, Vincent Louis Viala |
| EPI_ISL_2345727 | SANTA CASA DE MISERICORDIA SAO JOSE | Instituto Butantan / Mendelics | Dimas Tadeu Covas, Antonio Jorge Martins, Claudia Renata dos Santos Barros, David Schlesinger, Debora Botequio Moretti, Elaine Cristina Marqueze, Elaine Vieira Santos, Evandra Strazza Rodrigues, Heidge Fukumasu, Jayme Augusto de Souza-Neto, José Salvatore Leister Patané, Luiz Alcantara, Luiz Lehmann Coutinho, Maria Carolina Elias, Maurício Lacerda Nogueira, Rafael dos Santos Bezerra, Raul Machado Neto, Rejane Maria Tommasini Grotto, Ricardo Haddad, Sandra Coccuzzo Sampaio Vessoni, Simone Kashima, Svetoslav Nanev Slavov, Vincent Louis Viala |
| EPI_ISL_2345728 | DEPARTAMENTO DE SAUDE MUNICIPAL SOCORRO SP | Instituto Butantan / Mendelics | Dimas Tadeu Covas, Antonio Jorge Martins, Claudia Renata dos Santos Barros, David Schlesinger, Debora Botequio Moretti, Elaine Cristina Marqueze, Elaine Vieira Santos, Evandra Strazza Rodrigues, Heidge Fukumasu, Jayme Augusto de Souza-Neto, José Salvatore Leister Patané, Luiz Alcantara, Luiz Lehmann Coutinho, Maria Carolina Elias, Maurício Lacerda Nogueira, Rafael dos Santos Bezerra, Raul Machado Neto, Rejane Maria Tommasini Grotto, Ricardo Haddad, Sandra Coccuzzo Sampaio Vessoni, Simone Kashima, Svetoslav Nanev Slavov, Vincent Louis Viala |
| EPI_ISL_2345729 | PRONTO SOCORRO MUNICIPAL TAMBAU | Instituto Butantan / Mendelics | Dimas Tadeu Covas, Antonio Jorge Martins, Claudia Renata dos Santos Barros, David Schlesinger, Debora Botequio Moretti, Elaine Cristina Marqueze, Elaine Vieira Santos, Evandra Strazza Rodrigues, Heidge Fukumasu, Jayme Augusto de Souza-Neto, José Salvatore Leister Patané, Luiz Alcantara, Luiz Lehmann Coutinho, Maria Carolina Elias, Maurício Lacerda Nogueira, Rafael dos Santos Bezerra, Raul Machado Neto, Rejane Maria Tommasini Grotto, Ricardo Haddad, Sandra Coccuzzo Sampaio Vessoni, Simone Kashima, Svetoslav Nanev Slavov, Vincent Louis Viala |
| EPI_ISL_2345730 | CENTRO DE SAUDE DR JOAO JOAQUIM FERREIRA CRISTAIS PAULISTA | Instituto Butantan / Mendelics | Dimas Tadeu Covas, Antonio Jorge Martins, Claudia Renata dos Santos Barros, David Schlesinger, Debora Botequio Moretti, Elaine Cristina Marqueze, Elaine Vieira Santos, Evandra Strazza Rodrigues, Heidge Fukumasu, Jayme Augusto de Souza-Neto, José Salvatore Leister Patané, Luiz Alcantara, Luiz Lehmann Coutinho, Maria Carolina Elias, Maurício Lacerda Nogueira, Rafael dos Santos Bezerra, Raul Machado Neto, Rejane Maria Tommasini Grotto, Ricardo Haddad, Sandra Coccuzzo Sampaio Vessoni, Simone Kashima, Svetoslav Nanev Slavov, Vincent Louis Viala |
| EPI_ISL_2345731 | HOSPITAL DE CAMPANHA COVID 19 MUNICIPIO DE TAUBATE | Instituto Butantan / Mendelics | Dimas Tadeu Covas, Antonio Jorge Martins, Claudia Renata dos Santos Barros, David Schlesinger, Debora Botequio Moretti, Elaine Cristina Marqueze, Elaine Vieira Santos, Evandra Strazza Rodrigues, Heidge Fukumasu, Jayme Augusto de Souza-Neto, José Salvatore Leister Patané, Luiz Alcantara, Luiz Lehmann Coutinho, Maria Carolina Elias, Maurício Lacerda Nogueira, Rafael dos Santos Bezerra, Raul Machado Neto, Rejane Maria Tommasini Grotto, Ricardo Haddad, Sandra Coccuzzo Sampaio Vessoni, Simone Kashima, Svetoslav Nanev Slavov, Vincent Louis Viala |
| EPI_ISL_2345732 | DEPARTAMENTO DE SAUDE MUNICIPAL SOCORRO SP | Instituto Butantan / Mendelics | Dimas Tadeu Covas, Antonio Jorge Martins, Claudia Renata dos Santos Barros, David Schlesinger, Debora Botequio Moretti, Elaine Cristina Marqueze, Elaine Vieira Santos, Evandra Strazza Rodrigues, Heidge Fukumasu, Jayme Augusto de Souza-Neto, José Salvatore Leister Patané, Luiz Alcantara, Luiz Lehmann Coutinho, Maria Carolina Elias, Maurício Lacerda Nogueira, Rafael dos Santos Bezerra, Raul Machado Neto, Rejane Maria Tommasini Grotto, Ricardo Haddad, Sandra Coccuzzo Sampaio Vessoni, Simone Kashima, Svetoslav Nanev Slavov, Vincent Louis Viala |
| EPI_ISL_2345733 | NUCLEO DE SAUDE MARY DOTA DE BAURU | Instituto Butantan / Mendelics | Dimas Tadeu Covas, Antonio Jorge Martins, Claudia Renata dos Santos Barros, David Schlesinger, Debora Botequio Moretti, Elaine Cristina Marqueze, Elaine Vieira Santos, Evandra Strazza Rodrigues, Heidge Fukumasu, Jayme Augusto de Souza-Neto, José Salvatore Leister Patané, Luiz Alcantara, Luiz Lehmann Coutinho, Maria Carolina Elias, Maurício Lacerda Nogueira, Rafael dos Santos Bezerra, Raul Machado Neto, Rejane Maria Tommasini Grotto, Ricardo Haddad, Sandra Coccuzzo Sampaio Vessoni, Simone Kashima, Svetoslav Nanev Slavov, Vincent Louis Viala |
| EPI_ISL_2345734 | VIGILANCIA EPIDEMIOLOGICA JARDINOPOLIS SP | Instituto Butantan / Mendelics | Dimas Tadeu Covas, Antonio Jorge Martins, Claudia Renata dos Santos Barros, David Schlesinger, Debora Botequio Moretti, Elaine Cristina Marqueze, Elaine Vieira Santos, Evandra Strazza Rodrigues, Heidge Fukumasu, Jayme Augusto de Souza-Neto, José Salvatore Leister Patané, Luiz Alcantara, Luiz Lehmann Coutinho, Maria Carolina Elias, Maurício Lacerda Nogueira, Rafael dos Santos Bezerra, Raul Machado Neto, Rejane Maria Tommasini Grotto, Ricardo Haddad, Sandra Coccuzzo Sampaio Vessoni, Simone Kashima, Svetoslav Nanev Slavov, Vincent Louis Viala |
| EPI_ISL_2345735 | CENTRO DE APOIO EPIDEMIOLOGICO | Instituto Butantan / Mendelics | Dimas Tadeu Covas, Antonio Jorge Martins, Claudia Renata dos Santos Barros, David Schlesinger, Debora Botequio Moretti, Elaine Cristina Marqueze, Elaine Vieira Santos, Evandra Strazza Rodrigues, Heidge Fukumasu, Jayme Augusto de Souza-Neto, José Salvatore Leister Patané, Luiz Alcantara, Luiz Lehmann Coutinho, Maria Carolina Elias, Maurício Lacerda Nogueira, Rafael dos Santos Bezerra, Raul Machado Neto, Rejane Maria Tommasini Grotto, Ricardo Haddad, Sandra Coccuzzo Sampaio Vessoni, Simone Kashima, Svetoslav Nanev Slavov, Vincent Louis Viala |
| EPI_ISL_2345736 | AMBULATORIO CENTRAL DE ESPECIALIDADES ITATIBA | Instituto Butantan / Mendelics | Dimas Tadeu Covas, Antonio Jorge Martins, Claudia Renata dos Santos Barros, David Schlesinger, Debora Botequio Moretti, Elaine Cristina Marqueze, Elaine Vieira Santos, Evandra Strazza Rodrigues, Heidge Fukumasu, Jayme Augusto de Souza-Neto, José Salvatore Leister Patané, Luiz Alcantara, Luiz Lehmann Coutinho, Maria Carolina Elias, Maurício Lacerda Nogueira, Rafael dos Santos Bezerra, Raul Machado Neto, Rejane Maria Tommasini Grotto, Ricardo Haddad, Sandra Coccuzzo Sampaio Vessoni, Simone Kashima, Svetoslav Nanev Slavov, Vincent Louis Viala |
| EPI_ISL_2345737 | NUCLEO DE SAUDE MARY DOTA DE BAURU | Instituto Butantan / Mendelics | Dimas Tadeu Covas, Antonio Jorge Martins, Claudia Renata dos Santos Barros, David Schlesinger, Debora Botequio Moretti, Elaine Cristina Marqueze, Elaine Vieira Santos, Evandra Strazza Rodrigues, Heidge Fukumasu, Jayme Augusto de Souza-Neto, José Salvatore Leister Patané, Luiz Alcantara, Luiz Lehmann Coutinho, Maria Carolina Elias, Maurício Lacerda Nogueira, Rafael dos Santos Bezerra, Raul Machado Neto, Rejane Maria Tommasini Grotto, Ricardo Haddad, Sandra Coccuzzo Sampaio Vessoni, Simone Kashima, Svetoslav Nanev Slavov, Vincent Louis Viala |
| EPI_ISL_2345738 | VIGILANCIA EPIDEMIOLOGICA JARDINOPOLIS SP | Instituto Butantan / Mendelics | Dimas Tadeu Covas, Antonio Jorge Martins, Claudia Renata dos Santos Barros, David Schlesinger, Debora Botequio Moretti, Elaine Cristina Marqueze, Elaine Vieira Santos, Evandra Strazza Rodrigues, Heidge Fukumasu, Jayme Augusto de Souza-Neto, José Salvatore Leister Patané, Luiz Alcantara, Luiz |

| | | | |
|---|---|---|---|
| | | | Lehmann Coutinho, Maria Carolina Elias, Maurício Lacerda Nogueira, Rafael dos Santos Bezerra, Raul Machado Neto, Rejane Maria Tommasini Grotto, Ricardo Haddad, Sandra Coccuzzo Sampaio Vessoni, Simone Kashima, Svetoslav Nanev Slavov, Vincent Louis Viala |
| EPI_ISL_2345739 | SANTA CASA DE MISERICORDIA SAO JOSE | Instituto Butantan / Mendelics | Dimas Tadeu Covas, Antonio Jorge Martins, Claudia Renata dos Santos Barros, David Schlesinger, Debora Botequio Moretti, Elaine Cristina Marqueze, Elaine Vieira Santos, Evandra Strazza Rodrigues, Heidge Fukumasu, Jayme Augusto de Souza-Neto, José Salvatore Leister Patané, Luiz Alcantara, Luiz Lehmann Coutinho, Maria Carolina Elias, Maurício Lacerda Nogueira, Rafael dos Santos Bezerra, Raul Machado Neto, Rejane Maria Tommasini Grotto, Ricardo Haddad, Sandra Coccuzzo Sampaio Vessoni, Simone Kashima, Svetoslav Nanev Slavov, Vincent Louis Viala |
| EPI_ISL_2345740 | HOSPITAL MUNICIPAL DE ITAPIRA | Instituto Butantan / Mendelics | Dimas Tadeu Covas, Antonio Jorge Martins, Claudia Renata dos Santos Barros, David Schlesinger, Debora Botequio Moretti, Elaine Cristina Marqueze, Elaine Vieira Santos, Evandra Strazza Rodrigues, Heidge Fukumasu, Jayme Augusto de Souza-Neto, José Salvatore Leister Patané, Luiz Alcantara, Luiz Lehmann Coutinho, Maria Carolina Elias, Maurício Lacerda Nogueira, Rafael dos Santos Bezerra, Raul Machado Neto, Rejane Maria Tommasini Grotto, Ricardo Haddad, Sandra Coccuzzo Sampaio Vessoni, Simone Kashima, Svetoslav Nanev Slavov, Vincent Louis Viala |
| EPI_ISL_2345741 | LABORATORIO MUNICIPAL DE SUZANO | Instituto Butantan / Mendelics | Dimas Tadeu Covas, Antonio Jorge Martins, Claudia Renata dos Santos Barros, David Schlesinger, Debora Botequio Moretti, Elaine Cristina Marqueze, Elaine Vieira Santos, Evandra Strazza Rodrigues, Heidge Fukumasu, Jayme Augusto de Souza-Neto, José Salvatore Leister Patané, Luiz Alcantara, Luiz Lehmann Coutinho, Maria Carolina Elias, Maurício Lacerda Nogueira, Rafael dos Santos Bezerra, Raul Machado Neto, Rejane Maria Tommasini Grotto, Ricardo Haddad, Sandra Coccuzzo Sampaio Vessoni, Simone Kashima, Svetoslav Nanev Slavov, Vincent Louis Viala |
| EPI_ISL_2345743 | CENTRO DE SAUDE II IBITINGA | Instituto Butantan / Mendelics | Dimas Tadeu Covas, Antonio Jorge Martins, Claudia Renata dos Santos Barros, David Schlesinger, Debora Botequio Moretti, Elaine Cristina Marqueze, Elaine Vieira Santos, Evandra Strazza Rodrigues, Heidge Fukumasu, Jayme Augusto de Souza-Neto, José Salvatore Leister Patané, Luiz Alcantara, Luiz Lehmann Coutinho, Maria Carolina Elias, Maurício Lacerda Nogueira, Rafael dos Santos Bezerra, Raul Machado Neto, Rejane Maria Tommasini Grotto, Ricardo Haddad, Sandra Coccuzzo Sampaio Vessoni, Simone Kashima, Svetoslav Nanev Slavov, Vincent Louis Viala |
| EPI_ISL_2345744 | VIGILANCIA EPIDEMIOLOGICA JARDINOPOLIS SP | Instituto Butantan / Mendelics | Dimas Tadeu Covas, Antonio Jorge Martins, Claudia Renata dos Santos Barros, David Schlesinger, Debora Botequio Moretti, Elaine Cristina Marqueze, Elaine Vieira Santos, Evandra Strazza Rodrigues, Heidge Fukumasu, Jayme Augusto de Souza-Neto, José Salvatore Leister Patané, Luiz Alcantara, Luiz Lehmann Coutinho, Maria Carolina Elias, Maurício Lacerda Nogueira, Rafael dos Santos Bezerra, Raul Machado Neto, Rejane Maria Tommasini Grotto, Ricardo Haddad, Sandra Coccuzzo Sampaio Vessoni, Simone Kashima, Svetoslav Nanev Slavov, Vincent Louis Viala |
| EPI_ISL_2345745 | UBS DR ALFREDO DANTAS DE SOUZA UMUARAMA | Instituto Butantan / Mendelics | Dimas Tadeu Covas, Antonio Jorge Martins, Claudia Renata dos Santos Barros, David Schlesinger, Debora Botequio Moretti, Elaine Cristina Marqueze, Elaine Vieira Santos, Evandra Strazza Rodrigues, Heidge Fukumasu, Jayme Augusto de Souza-Neto, José Salvatore Leister Patané, Luiz Alcantara, Luiz Lehmann Coutinho, Maria Carolina Elias, Maurício Lacerda Nogueira, Rafael dos Santos Bezerra, Raul Machado Neto, Rejane Maria Tommasini Grotto, Ricardo Haddad, Sandra Coccuzzo Sampaio Vessoni, Simone Kashima, Svetoslav Nanev Slavov, Vincent Louis Viala |
| EPI_ISL_2345746 | USF ARCADAS | Instituto Butantan / Mendelics | Dimas Tadeu Covas, Antonio Jorge Martins, Claudia Renata dos Santos Barros, David Schlesinger, Debora Botequio Moretti, Elaine Cristina Marqueze, Elaine Vieira Santos, Evandra Strazza Rodrigues, Heidge Fukumasu, Jayme Augusto de Souza-Neto, José Salvatore Leister Patané, Luiz Alcantara, Luiz Lehmann Coutinho, Maria Carolina Elias, Maurício Lacerda Nogueira, Rafael dos Santos Bezerra, Raul Machado Neto, Rejane Maria Tommasini Grotto, Ricardo Haddad, Sandra Coccuzzo Sampaio Vessoni, Simone Kashima, Svetoslav Nanev Slavov, Vincent Louis Viala |
| EPI_ISL_2345747 | CENTRO DE SAUDE II IBITINGA | Instituto Butantan / Mendelics | Dimas Tadeu Covas, Antonio Jorge Martins, Claudia Renata dos Santos Barros, David Schlesinger, Debora Botequio Moretti, Elaine Cristina Marqueze, Elaine Vieira Santos, Evandra Strazza Rodrigues, Heidge Fukumasu, Jayme Augusto de Souza-Neto, José Salvatore Leister Patané, Luiz Alcantara, Luiz Lehmann Coutinho, Maria Carolina Elias, Maurício Lacerda Nogueira, Rafael dos Santos Bezerra, Raul Machado Neto, Rejane Maria Tommasini Grotto, Ricardo Haddad, Sandra Coccuzzo Sampaio Vessoni, Simone Kashima, Svetoslav Nanev Slavov, Vincent Louis Viala |
| EPI_ISL_2345748 | CENTRO DE SAUDE III SANTA ERNESTINA | Instituto Butantan / Mendelics | Dimas Tadeu Covas, Antonio Jorge Martins, Claudia Renata dos Santos Barros, David Schlesinger, Debora Botequio Moretti, Elaine Cristina Marqueze, Elaine Vieira Santos, Evandra Strazza Rodrigues, Heidge Fukumasu, Jayme Augusto de Souza-Neto, José Salvatore Leister Patané, Luiz Alcantara, Luiz Lehmann Coutinho, Maria Carolina Elias, Maurício Lacerda Nogueira, Rafael dos Santos Bezerra, Raul Machado Neto, Rejane Maria Tommasini Grotto, Ricardo Haddad, Sandra Coccuzzo Sampaio Vessoni, Simone Kashima, Svetoslav Nanev Slavov, Vincent Louis Viala |
| EPI_ISL_2345749 | UNIDADE BASICA SAUDE NATAL ABADIO LACERDA | Instituto Butantan / Mendelics | Dimas Tadeu Covas, Antonio Jorge Martins, Claudia Renata dos Santos Barros, David Schlesinger, Debora Botequio Moretti, Elaine Cristina Marqueze, Elaine Vieira Santos, Evandra Strazza Rodrigues, Heidge Fukumasu, Jayme Augusto de Souza-Neto, José Salvatore Leister Patané, Luiz Alcantara, Luiz Lehmann Coutinho, Maria Carolina Elias, Maurício Lacerda Nogueira, Rafael dos Santos Bezerra, Raul Machado Neto, Rejane Maria Tommasini Grotto, Ricardo Haddad, Sandra Coccuzzo Sampaio Vessoni, Simone Kashima, Svetoslav Nanev Slavov, Vincent Louis Viala |
| EPI_ISL_2345750 | SECRETARIA DE SAUDE | Instituto Butantan / Mendelics | Dimas Tadeu Covas, Antonio Jorge Martins, Claudia Renata dos Santos Barros, David Schlesinger, Debora Botequio Moretti, Elaine Cristina Marqueze, Elaine Vieira Santos, Evandra Strazza Rodrigues, Heidge Fukumasu, Jayme Augusto de Souza-Neto, José Salvatore Leister Patané, Luiz Alcantara, Luiz Lehmann Coutinho, Maria Carolina Elias, Maurício Lacerda Nogueira, Rafael dos Santos Bezerra, Raul Machado Neto, Rejane Maria Tommasini Grotto, Ricardo Haddad, Sandra Coccuzzo Sampaio Vessoni, Simone Kashima, Svetoslav Nanev Slavov, Vincent Louis Viala |
| EPI_ISL_2345751 | SECRETARIA MUNICIPAL DE SAUDE SOROCABA | Instituto Butantan / Mendelics | Dimas Tadeu Covas, Antonio Jorge Martins, Claudia Renata dos Santos Barros, David Schlesinger, Debora Botequio Moretti, Elaine Cristina Marqueze, Elaine Vieira Santos, Evandra Strazza Rodrigues, Heidge Fukumasu, Jayme Augusto de Souza-Neto, José Salvatore Leister Patané, Luiz Alcantara, Luiz Lehmann Coutinho, Maria Carolina Elias, Maurício Lacerda Nogueira, Rafael dos Santos Bezerra, Raul Machado Neto, Rejane Maria Tommasini Grotto, Ricardo Haddad, Sandra Coccuzzo Sampaio Vessoni, Simone Kashima, Svetoslav Nanev Slavov, Vincent Louis Viala |
| EPI_ISL_2345752 | SECRETARIA DE SAUDE PUBLICA DE PRAIA GRANDE | Instituto Butantan / Mendelics | Dimas Tadeu Covas, Antonio Jorge Martins, Claudia Renata dos Santos Barros, David Schlesinger, Debora Botequio Moretti, Elaine Cristina Marqueze, Elaine Vieira Santos, Evandra Strazza Rodrigues, Heidge Fukumasu, Jayme Augusto de Souza-Neto, José Salvatore Leister Patané, Luiz Alcantara, Luiz Lehmann Coutinho, Maria Carolina Elias, Maurício Lacerda Nogueira, Rafael dos Santos Bezerra, Raul Machado Neto, Rejane Maria Tommasini Grotto, Ricardo Haddad, Sandra Coccuzzo Sampaio Vessoni, Simone Kashima, Svetoslav Nanev Slavov, Vincent Louis Viala |
| EPI_ISL_2345753 | CENTRO DE SAUDE DE MARACAI | Instituto Butantan / Mendelics | Dimas Tadeu Covas, Antonio Jorge Martins, Claudia Renata dos Santos Barros, David Schlesinger, Debora Botequio Moretti, Elaine Cristina Marqueze, Elaine Vieira Santos, Evandra Strazza Rodrigues, Heidge Fukumasu, Jayme Augusto de Souza-Neto, José Salvatore Leister Patané, Luiz Alcantara, Luiz Lehmann Coutinho, Maria Carolina Elias, Maurício Lacerda Nogueira, Rafael dos Santos Bezerra, Raul Machado Neto, Rejane Maria Tommasini Grotto, Ricardo Haddad, Sandra Coccuzzo Sampaio Vessoni, Simone Kashima, Svetoslav Nanev Slavov, Vincent Louis Viala |
| EPI_ISL_2345754 | GRIPARIO FALCAO | Instituto Butantan / Mendelics | Dimas Tadeu Covas, Antonio Jorge Martins, Claudia Renata dos Santos Barros, David Schlesinger, Debora Botequio Moretti, Elaine Cristina Marqueze, Elaine Vieira Santos, Evandra Strazza Rodrigues, Heidge Fukumasu, Jayme Augusto de Souza-Neto, José Salvatore Leister Patané, Luiz Alcantara, Luiz Lehmann Coutinho, Maria Carolina Elias, Maurício Lacerda Nogueira, Rafael dos Santos Bezerra, Raul Machado Neto, Rejane Maria Tommasini Grotto, Ricardo Haddad, Sandra Coccuzzo Sampaio Vessoni, Simone Kashima, Svetoslav Nanev Slavov, Vincent Louis Viala |
| EPI_ISL_2345755 | SECRETARIA MUNICIPAL DE SAUDE SOROCABA | Instituto Butantan / Mendelics | Dimas Tadeu Covas, Antonio Jorge Martins, Claudia Renata dos Santos Barros, David Schlesinger, Debora Botequio Moretti, Elaine Cristina Marqueze, Elaine Vieira Santos, Evandra Strazza Rodrigues, Heidge Fukumasu, Jayme Augusto de Souza-Neto, José Salvatore Leister Patané, Luiz Alcantara, Luiz Lehmann Coutinho, Maria Carolina Elias, Maurício Lacerda Nogueira, Rafael dos Santos Bezerra, Raul Machado Neto, Rejane Maria Tommasini Grotto, Ricardo Haddad, Sandra Coccuzzo Sampaio Vessoni, Simone Kashima, Svetoslav Nanev Slavov, Vincent Louis Viala |
| EPI_ISL_2345756 | GRIPARIO FALCAO | Instituto Butantan / Mendelics | Dimas Tadeu Covas, Antonio Jorge Martins, Claudia Renata dos Santos Barros, David Schlesinger, Debora Botequio Moretti, Elaine Cristina Marqueze, Elaine Vieira Santos, Evandra Strazza Rodrigues, Heidge Fukumasu, Jayme Augusto de Souza-Neto, José Salvatore Leister Patané, Luiz Alcantara, Luiz Lehmann Coutinho, Maria Carolina Elias, Maurício Lacerda Nogueira, Rafael dos Santos Bezerra, Raul Machado Neto, Rejane Maria Tommasini Grotto, Ricardo Haddad, Sandra Coccuzzo Sampaio Vessoni, Simone Kashima, Svetoslav Nanev Slavov, Vincent Louis Viala |
| EPI_ISL_2345757 | UBS DR WALDOMIRO FERREIRA NEVES SCRPARDO | Instituto Butantan / Mendelics | Dimas Tadeu Covas, Antonio Jorge Martins, Claudia Renata dos Santos Barros, David Schlesinger, Debora Botequio Moretti, Elaine Cristina Marqueze, Elaine Vieira Santos, Evandra Strazza Rodrigues, Heidge Fukumasu, Jayme Augusto de Souza-Neto, José Salvatore Leister Patané, Luiz Alcantara, Luiz Lehmann Coutinho, Maria Carolina Elias, Maurício Lacerda Nogueira, Rafael dos Santos Bezerra, Raul Machado Neto, Rejane Maria Tommasini Grotto, Ricardo Haddad, Sandra Coccuzzo Sampaio Vessoni, Simone Kashima, Svetoslav Nanev Slavov, Vincent Louis Viala |
| EPI_ISL_2345758 | SECAO CENTRO DE DIAGNOSTICO SECEDI | Instituto Butantan / Mendelics | Dimas Tadeu Covas, Antonio Jorge Martins, Claudia Renata dos Santos Barros, David Schlesinger, Debora Botequio Moretti, Elaine Cristina Marqueze, Elaine Vieira Santos, Evandra Strazza Rodrigues, Heidge Fukumasu, Jayme Augusto de Souza-Neto, José Salvatore Leister Patané, Luiz Alcantara, Luiz Lehmann Coutinho, Maria Carolina Elias, Maurício Lacerda Nogueira, Rafael dos Santos Bezerra, Raul Machado Neto, Rejane Maria Tommasini Grotto, Ricardo Haddad, Sandra Coccuzzo Sampaio Vessoni, Simone Kashima, Svetoslav Nanev Slavov, Vincent Louis Viala |

| | | | |
|---|---|---|---|
| EPI_ISL_2345759 | PSF DR ANTONIO PIRES DE ALMEIDA PORTO FELIZ | Instituto Butantan / Mendelics | Dimas Tadeu Covas, Antonio Jorge Martins, Claudia Renata dos Santos Barros, David Schlesinger, Debora Botequio Moretti, Elaine Cristina Marqueze, Elaine Vieira Santos, Evandra Strazza Rodrigues, Heidge Fukumasu, Jayme Augusto de Souza-Neto, José Salvatore Leister Patané, Luiz Alcantara, Luiz Lehmann Coutinho, Maria Carolina Elias, Maurício Lacerda Nogueira, Rafael dos Santos Bezerra, Raul Machado Neto, Rejane Maria Tommasini Grotto, Ricardo Haddad, Sandra Coccuzzo Sampaio Vessoni, Simone Kashima, Svetoslav Nanev Slavov, Vincent Louis Viala |
| EPI_ISL_2345760 | SECRETARIA DE SAUDE PUBLICA DE PRAIA GRANDE | Instituto Butantan / Mendelics | Dimas Tadeu Covas, Antonio Jorge Martins, Claudia Renata dos Santos Barros, David Schlesinger, Debora Botequio Moretti, Elaine Cristina Marqueze, Elaine Vieira Santos, Evandra Strazza Rodrigues, Heidge Fukumasu, Jayme Augusto de Souza-Neto, José Salvatore Leister Patané, Luiz Alcantara, Luiz Lehmann Coutinho, Maria Carolina Elias, Maurício Lacerda Nogueira, Rafael dos Santos Bezerra, Raul Machado Neto, Rejane Maria Tommasini Grotto, Ricardo Haddad, Sandra Coccuzzo Sampaio Vessoni, Simone Kashima, Svetoslav Nanev Slavov, Vincent Louis Viala |
| EPI_ISL_2345761 | USF II ALADIA ARAN RODRIGUES | Instituto Butantan / Mendelics | Dimas Tadeu Covas, Antonio Jorge Martins, Claudia Renata dos Santos Barros, David Schlesinger, Debora Botequio Moretti, Elaine Cristina Marqueze, Elaine Vieira Santos, Evandra Strazza Rodrigues, Heidge Fukumasu, Jayme Augusto de Souza-Neto, José Salvatore Leister Patané, Luiz Alcantara, Luiz Lehmann Coutinho, Maria Carolina Elias, Maurício Lacerda Nogueira, Rafael dos Santos Bezerra, Raul Machado Neto, Rejane Maria Tommasini Grotto, Ricardo Haddad, Sandra Coccuzzo Sampaio Vessoni, Simone Kashima, Svetoslav Nanev Slavov, Vincent Louis Viala |
| EPI_ISL_2345762 | UBS MARIA GATTI GIGLIO | Instituto Butantan / Mendelics | Dimas Tadeu Covas, Antonio Jorge Martins, Claudia Renata dos Santos Barros, David Schlesinger, Debora Botequio Moretti, Elaine Cristina Marqueze, Elaine Vieira Santos, Evandra Strazza Rodrigues, Heidge Fukumasu, Jayme Augusto de Souza-Neto, José Salvatore Leister Patané, Luiz Alcantara, Luiz Lehmann Coutinho, Maria Carolina Elias, Maurício Lacerda Nogueira, Rafael dos Santos Bezerra, Raul Machado Neto, Rejane Maria Tommasini Grotto, Ricardo Haddad, Sandra Coccuzzo Sampaio Vessoni, Simone Kashima, Svetoslav Nanev Slavov, Vincent Louis Viala |
| EPI_ISL_2345763 | CENTRO DE SAUDE II JUNQUEIROPOLIS | Instituto Butantan / Mendelics | Dimas Tadeu Covas, Antonio Jorge Martins, Claudia Renata dos Santos Barros, David Schlesinger, Debora Botequio Moretti, Elaine Cristina Marqueze, Elaine Vieira Santos, Evandra Strazza Rodrigues, Heidge Fukumasu, Jayme Augusto de Souza-Neto, José Salvatore Leister Patané, Luiz Alcantara, Luiz Lehmann Coutinho, Maria Carolina Elias, Maurício Lacerda Nogueira, Rafael dos Santos Bezerra, Raul Machado Neto, Rejane Maria Tommasini Grotto, Ricardo Haddad, Sandra Coccuzzo Sampaio Vessoni, Simone Kashima, Svetoslav Nanev Slavov, Vincent Louis Viala |
| EPI_ISL_2345764 | HOSP E MATERNIDADE MUNICIPAL N SRA MONTE SERRAT | Instituto Butantan / Mendelics | Dimas Tadeu Covas, Antonio Jorge Martins, Claudia Renata dos Santos Barros, David Schlesinger, Debora Botequio Moretti, Elaine Cristina Marqueze, Elaine Vieira Santos, Evandra Strazza Rodrigues, Heidge Fukumasu, Jayme Augusto de Souza-Neto, José Salvatore Leister Patané, Luiz Alcantara, Luiz Lehmann Coutinho, Maria Carolina Elias, Maurício Lacerda Nogueira, Rafael dos Santos Bezerra, Raul Machado Neto, Rejane Maria Tommasini Grotto, Ricardo Haddad, Sandra Coccuzzo Sampaio Vessoni, Simone Kashima, Svetoslav Nanev Slavov, Vincent Louis Viala |
| EPI_ISL_2345765 | UPA DE BEBEDOURO | Instituto Butantan / Mendelics | Dimas Tadeu Covas, Antonio Jorge Martins, Claudia Renata dos Santos Barros, David Schlesinger, Debora Botequio Moretti, Elaine Cristina Marqueze, Elaine Vieira Santos, Evandra Strazza Rodrigues, Heidge Fukumasu, Jayme Augusto de Souza-Neto, José Salvatore Leister Patané, Luiz Alcantara, Luiz Lehmann Coutinho, Maria Carolina Elias, Maurício Lacerda Nogueira, Rafael dos Santos Bezerra, Raul Machado Neto, Rejane Maria Tommasini Grotto, Ricardo Haddad, Sandra Coccuzzo Sampaio Vessoni, Simone Kashima, Svetoslav Nanev Slavov, Vincent Louis Viala |
| EPI_ISL_2345766 | CENTRO DE SAUDE II IBITINGA | Instituto Butantan / Mendelics | Dimas Tadeu Covas, Antonio Jorge Martins, Claudia Renata dos Santos Barros, David Schlesinger, Debora Botequio Moretti, Elaine Cristina Marqueze, Elaine Vieira Santos, Evandra Strazza Rodrigues, Heidge Fukumasu, Jayme Augusto de Souza-Neto, José Salvatore Leister Patané, Luiz Alcantara, Luiz Lehmann Coutinho, Maria Carolina Elias, Maurício Lacerda Nogueira, Rafael dos Santos Bezerra, Raul Machado Neto, Rejane Maria Tommasini Grotto, Ricardo Haddad, Sandra Coccuzzo Sampaio Vessoni, Simone Kashima, Svetoslav Nanev Slavov, Vincent Louis Viala |
| EPI_ISL_2345767 | UBS II DE ALVARES MACHADO | Instituto Butantan / Mendelics | Dimas Tadeu Covas, Antonio Jorge Martins, Claudia Renata dos Santos Barros, David Schlesinger, Debora Botequio Moretti, Elaine Cristina Marqueze, Elaine Vieira Santos, Evandra Strazza Rodrigues, Heidge Fukumasu, Jayme Augusto de Souza-Neto, José Salvatore Leister Patané, Luiz Alcantara, Luiz Lehmann Coutinho, Maria Carolina Elias, Maurício Lacerda Nogueira, Rafael dos Santos Bezerra, Raul Machado Neto, Rejane Maria Tommasini Grotto, Ricardo Haddad, Sandra Coccuzzo Sampaio Vessoni, Simone Kashima, Svetoslav Nanev Slavov, Vincent Louis Viala |
| EPI_ISL_2345768 | CENTRO DE SAUDE II IBITINGA | Instituto Butantan / Mendelics | Dimas Tadeu Covas, Antonio Jorge Martins, Claudia Renata dos Santos Barros, David Schlesinger, Debora Botequio Moretti, Elaine Cristina Marqueze, Elaine Vieira Santos, Evandra Strazza Rodrigues, Heidge Fukumasu, Jayme Augusto de Souza-Neto, José Salvatore Leister Patané, Luiz Alcantara, Luiz Lehmann Coutinho, Maria Carolina Elias, Maurício Lacerda Nogueira, Rafael dos Santos Bezerra, Raul Machado Neto, Rejane Maria Tommasini Grotto, Ricardo Haddad, Sandra Coccuzzo Sampaio Vessoni, Simone Kashima, Svetoslav Nanev Slavov, Vincent Louis Viala |
| EPI_ISL_2345769 | HOSPITAL DE CAMPANHA COVID 19 CAIEIRAS | Instituto Butantan / Mendelics | Dimas Tadeu Covas, Antonio Jorge Martins, Claudia Renata dos Santos Barros, David Schlesinger, Debora Botequio Moretti, Elaine Cristina Marqueze, Elaine Vieira Santos, Evandra Strazza Rodrigues, Heidge Fukumasu, Jayme Augusto de Souza-Neto, José Salvatore Leister Patané, Luiz Alcantara, Luiz Lehmann Coutinho, Maria Carolina Elias, Maurício Lacerda Nogueira, Rafael dos Santos Bezerra, Raul Machado Neto, Rejane Maria Tommasini Grotto, Ricardo Haddad, Sandra Coccuzzo Sampaio Vessoni, Simone Kashima, Svetoslav Nanev Slavov, Vincent Louis Viala |
| EPI_ISL_2345770 | UNIDADE BASICA DE SAUDE II ESTRELA DO NORTE | Instituto Butantan / Mendelics | Dimas Tadeu Covas, Antonio Jorge Martins, Claudia Renata dos Santos Barros, David Schlesinger, Debora Botequio Moretti, Elaine Cristina Marqueze, Elaine Vieira Santos, Evandra Strazza Rodrigues, Heidge Fukumasu, Jayme Augusto de Souza-Neto, José Salvatore Leister Patané, Luiz Alcantara, Luiz Lehmann Coutinho, Maria Carolina Elias, Maurício Lacerda Nogueira, Rafael dos Santos Bezerra, Raul Machado Neto, Rejane Maria Tommasini Grotto, Ricardo Haddad, Sandra Coccuzzo Sampaio Vessoni, Simone Kashima, Svetoslav Nanev Slavov, Vincent Louis Viala |
| EPI_ISL_2345771 | CSII DR WASHINGTON LUIS M RODRIGUES DA SILVA PITANGUEIRAS | Instituto Butantan / Mendelics | Dimas Tadeu Covas, Antonio Jorge Martins, Claudia Renata dos Santos Barros, David Schlesinger, Debora Botequio Moretti, Elaine Cristina Marqueze, Elaine Vieira Santos, Evandra Strazza Rodrigues, Heidge Fukumasu, Jayme Augusto de Souza-Neto, José Salvatore Leister Patané, Luiz Alcantara, Luiz Lehmann Coutinho, Maria Carolina Elias, Maurício Lacerda Nogueira, Rafael dos Santos Bezerra, Raul Machado Neto, Rejane Maria Tommasini Grotto, Ricardo Haddad, Sandra Coccuzzo Sampaio Vessoni, Simone Kashima, Svetoslav Nanev Slavov, Vincent Louis Viala |
| EPI_ISL_2345772 | UBS III DE PARIQUERA ACU PARIQUERA ACU | Instituto Butantan / Mendelics | Dimas Tadeu Covas, Antonio Jorge Martins, Claudia Renata dos Santos Barros, David Schlesinger, Debora Botequio Moretti, Elaine Cristina Marqueze, Elaine Vieira Santos, Evandra Strazza Rodrigues, Heidge Fukumasu, Jayme Augusto de Souza-Neto, José Salvatore Leister Patané, Luiz Alcantara, Luiz Lehmann Coutinho, Maria Carolina Elias, Maurício Lacerda Nogueira, Rafael dos Santos Bezerra, Raul Machado Neto, Rejane Maria Tommasini Grotto, Ricardo Haddad, Sandra Coccuzzo Sampaio Vessoni, Simone Kashima, Svetoslav Nanev Slavov, Vincent Louis Viala |
| EPI_ISL_2345773 | DEPARTAMENTO DE SAUDE MUNICIPAL SOCORRO SP | Instituto Butantan / Mendelics | Dimas Tadeu Covas, Antonio Jorge Martins, Claudia Renata dos Santos Barros, David Schlesinger, Debora Botequio Moretti, Elaine Cristina Marqueze, Elaine Vieira Santos, Evandra Strazza Rodrigues, Heidge Fukumasu, Jayme Augusto de Souza-Neto, José Salvatore Leister Patané, Luiz Alcantara, Luiz Lehmann Coutinho, Maria Carolina Elias, Maurício Lacerda Nogueira, Rafael dos Santos Bezerra, Raul Machado Neto, Rejane Maria Tommasini Grotto, Ricardo Haddad, Sandra Coccuzzo Sampaio Vessoni, Simone Kashima, Svetoslav Nanev Slavov, Vincent Louis Viala |
| EPI_ISL_2345774 | HOSPITAL MUNICIPAL DE ITAPIRA | Instituto Butantan / Mendelics | Dimas Tadeu Covas, Antonio Jorge Martins, Claudia Renata dos Santos Barros, David Schlesinger, Debora Botequio Moretti, Elaine Cristina Marqueze, Elaine Vieira Santos, Evandra Strazza Rodrigues, Heidge Fukumasu, Jayme Augusto de Souza-Neto, José Salvatore Leister Patané, Luiz Alcantara, Luiz Lehmann Coutinho, Maria Carolina Elias, Maurício Lacerda Nogueira, Rafael dos Santos Bezerra, Raul Machado Neto, Rejane Maria Tommasini Grotto, Ricardo Haddad, Sandra Coccuzzo Sampaio Vessoni, Simone Kashima, Svetoslav Nanev Slavov, Vincent Louis Viala |
| EPI_ISL_2345775 | CENTRO DE SAUDE DR JOAO JOAQUIM FERREIRA CRISTAIS PAULISTA | Instituto Butantan / Mendelics | Dimas Tadeu Covas, Antonio Jorge Martins, Claudia Renata dos Santos Barros, David Schlesinger, Debora Botequio Moretti, Elaine Cristina Marqueze, Elaine Vieira Santos, Evandra Strazza Rodrigues, Heidge Fukumasu, Jayme Augusto de Souza-Neto, José Salvatore Leister Patané, Luiz Alcantara, Luiz Lehmann Coutinho, Maria Carolina Elias, Maurício Lacerda Nogueira, Rafael dos Santos Bezerra, Raul Machado Neto, Rejane Maria Tommasini Grotto, Ricardo Haddad, Sandra Coccuzzo Sampaio Vessoni, Simone Kashima, Svetoslav Nanev Slavov, Vincent Louis Viala |
| EPI_ISL_2345776 | DEPARTAMENTO DE SAUDE MUNICIPAL SOCORRO SP | Instituto Butantan / Mendelics | Dimas Tadeu Covas, Antonio Jorge Martins, Claudia Renata dos Santos Barros, David Schlesinger, Debora Botequio Moretti, Elaine Cristina Marqueze, Elaine Vieira Santos, Evandra Strazza Rodrigues, Heidge Fukumasu, Jayme Augusto de Souza-Neto, José Salvatore Leister Patané, Luiz Alcantara, Luiz Lehmann Coutinho, Maria Carolina Elias, Maurício Lacerda Nogueira, Rafael dos Santos Bezerra, Raul Machado Neto, Rejane Maria Tommasini Grotto, Ricardo Haddad, Sandra Coccuzzo Sampaio Vessoni, Simone Kashima, Svetoslav Nanev Slavov, Vincent Louis Viala |
| EPI_ISL_2345777 | NUCLEO DE SAUDE MARY DOTA DE BAURU | Instituto Butantan / Mendelics | Dimas Tadeu Covas, Antonio Jorge Martins, Claudia Renata dos Santos Barros, David Schlesinger, Debora Botequio Moretti, Elaine Cristina Marqueze, Elaine Vieira Santos, Evandra Strazza Rodrigues, Heidge Fukumasu, Jayme Augusto de Souza-Neto, José Salvatore Leister Patané, Luiz Alcantara, Luiz Lehmann Coutinho, Maria Carolina Elias, Maurício Lacerda Nogueira, Rafael dos Santos Bezerra, Raul Machado Neto, Rejane Maria Tommasini Grotto, Ricardo Haddad, Sandra Coccuzzo Sampaio Vessoni, Simone Kashima, Svetoslav Nanev Slavov, Vincent Louis Viala |
| EPI_ISL_2345778 | VIGILANCIA EPIDEMIOLOGICA JARDINOPOLIS SP | Instituto Butantan / Mendelics | Dimas Tadeu Covas, Antonio Jorge Martins, Claudia Renata dos Santos Barros, David Schlesinger, Debora Botequio Moretti, Elaine Cristina Marqueze, Elaine Vieira Santos, Evandra Strazza Rodrigues, Heidge Fukumasu, Jayme Augusto de Souza-Neto, José Salvatore Leister Patané, Luiz Alcantara, Luiz |

| | | | |
|---|---|---|---|
| | | | Lehmann Coutinho, Maria Carolina Elias, Maurício Lacerda Nogueira, Rafael dos Santos Bezerra, Raul Machado Neto, Rejane Maria Tommasini Grotto, Ricardo Haddad, Sandra Coccuzzo Sampaio Vessoni, Simone Kashima, Svetoslav Nanev Slavov, Vincent Louis Viala |
| EPI_ISL_2345779 | CENTRO DE APOIO EPIDEMIOLOGICO | Instituto Butantan / Mendelics | Dimas Tadeu Covas, Antonio Jorge Martins, Claudia Renata dos Santos Barros, David Schlesinger, Debora Botequio Moretti, Elaine Cristina Marqueze, Elaine Vieira Santos, Evandra Strazza Rodrigues, Heidge Fukumasu, Jayme Augusto de Souza-Neto, José Salvatore Leister Patané, Luiz Alcantara, Luiz Lehmann Coutinho, Maria Carolina Elias, Maurício Lacerda Nogueira, Rafael dos Santos Bezerra, Raul Machado Neto, Rejane Maria Tommasini Grotto, Ricardo Haddad, Sandra Coccuzzo Sampaio Vessoni, Simone Kashima, Svetoslav Nanev Slavov, Vincent Louis Viala |
| EPI_ISL_2345780 | AMBULATORIO CENTRAL DE ESPECIALIDADES ITATIBA | Instituto Butantan / Mendelics | Dimas Tadeu Covas, Antonio Jorge Martins, Claudia Renata dos Santos Barros, David Schlesinger, Debora Botequio Moretti, Elaine Cristina Marqueze, Elaine Vieira Santos, Evandra Strazza Rodrigues, Heidge Fukumasu, Jayme Augusto de Souza-Neto, José Salvatore Leister Patané, Luiz Alcantara, Luiz Lehmann Coutinho, Maria Carolina Elias, Maurício Lacerda Nogueira, Rafael dos Santos Bezerra, Raul Machado Neto, Rejane Maria Tommasini Grotto, Ricardo Haddad, Sandra Coccuzzo Sampaio Vessoni, Simone Kashima, Svetoslav Nanev Slavov, Vincent Louis Viala |
| EPI_ISL_2345781 | NUCLEO DE SAUDE MARY DOTA DE BAURU | Instituto Butantan / Mendelics | Dimas Tadeu Covas, Antonio Jorge Martins, Claudia Renata dos Santos Barros, David Schlesinger, Debora Botequio Moretti, Elaine Cristina Marqueze, Elaine Vieira Santos, Evandra Strazza Rodrigues, Heidge Fukumasu, Jayme Augusto de Souza-Neto, José Salvatore Leister Patané, Luiz Alcantara, Luiz Lehmann Coutinho, Maria Carolina Elias, Maurício Lacerda Nogueira, Rafael dos Santos Bezerra, Raul Machado Neto, Rejane Maria Tommasini Grotto, Ricardo Haddad, Sandra Coccuzzo Sampaio Vessoni, Simone Kashima, Svetoslav Nanev Slavov, Vincent Louis Viala |
| EPI_ISL_2345782 | VIGILANCIA EPIDEMIOLOGICA JARDINOPOLIS SP | Instituto Butantan / Mendelics | Dimas Tadeu Covas, Antonio Jorge Martins, Claudia Renata dos Santos Barros, David Schlesinger, Debora Botequio Moretti, Elaine Cristina Marqueze, Elaine Vieira Santos, Evandra Strazza Rodrigues, Heidge Fukumasu, Jayme Augusto de Souza-Neto, José Salvatore Leister Patané, Luiz Alcantara, Luiz Lehmann Coutinho, Maria Carolina Elias, Maurício Lacerda Nogueira, Rafael dos Santos Bezerra, Raul Machado Neto, Rejane Maria Tommasini Grotto, Ricardo Haddad, Sandra Coccuzzo Sampaio Vessoni, Simone Kashima, Svetoslav Nanev Slavov, Vincent Louis Viala |
| EPI_ISL_2345783 | SANTA CASA DE MISERICORDIA SAO JOSE | Instituto Butantan / Mendelics | Dimas Tadeu Covas, Antonio Jorge Martins, Claudia Renata dos Santos Barros, David Schlesinger, Debora Botequio Moretti, Elaine Cristina Marqueze, Elaine Vieira Santos, Evandra Strazza Rodrigues, Heidge Fukumasu, Jayme Augusto de Souza-Neto, José Salvatore Leister Patané, Luiz Alcantara, Luiz Lehmann Coutinho, Maria Carolina Elias, Maurício Lacerda Nogueira, Rafael dos Santos Bezerra, Raul Machado Neto, Rejane Maria Tommasini Grotto, Ricardo Haddad, Sandra Coccuzzo Sampaio Vessoni, Simone Kashima, Svetoslav Nanev Slavov, Vincent Louis Viala |
| EPI_ISL_2345784 | HOSPITAL MUNICIPAL DE ITAPIRA | Instituto Butantan / Mendelics | Dimas Tadeu Covas, Antonio Jorge Martins, Claudia Renata dos Santos Barros, David Schlesinger, Debora Botequio Moretti, Elaine Cristina Marqueze, Elaine Vieira Santos, Evandra Strazza Rodrigues, Heidge Fukumasu, Jayme Augusto de Souza-Neto, José Salvatore Leister Patané, Luiz Alcantara, Luiz Lehmann Coutinho, Maria Carolina Elias, Maurício Lacerda Nogueira, Rafael dos Santos Bezerra, Raul Machado Neto, Rejane Maria Tommasini Grotto, Ricardo Haddad, Sandra Coccuzzo Sampaio Vessoni, Simone Kashima, Svetoslav Nanev Slavov, Vincent Louis Viala |
| EPI_ISL_2345785 | LABORATORIO MUNICIPAL DE SUZANO | Instituto Butantan / Mendelics | Dimas Tadeu Covas, Antonio Jorge Martins, Claudia Renata dos Santos Barros, David Schlesinger, Debora Botequio Moretti, Elaine Cristina Marqueze, Elaine Vieira Santos, Evandra Strazza Rodrigues, Heidge Fukumasu, Jayme Augusto de Souza-Neto, José Salvatore Leister Patané, Luiz Alcantara, Luiz Lehmann Coutinho, Maria Carolina Elias, Maurício Lacerda Nogueira, Rafael dos Santos Bezerra, Raul Machado Neto, Rejane Maria Tommasini Grotto, Ricardo Haddad, Sandra Coccuzzo Sampaio Vessoni, Simone Kashima, Svetoslav Nanev Slavov, Vincent Louis Viala |
| EPI_ISL_2345786 | CENTRO DE SAUDE II IBITINGA | Instituto Butantan / Mendelics | Dimas Tadeu Covas, Antonio Jorge Martins, Claudia Renata dos Santos Barros, David Schlesinger, Debora Botequio Moretti, Elaine Cristina Marqueze, Elaine Vieira Santos, Evandra Strazza Rodrigues, Heidge Fukumasu, Jayme Augusto de Souza-Neto, José Salvatore Leister Patané, Luiz Alcantara, Luiz Lehmann Coutinho, Maria Carolina Elias, Maurício Lacerda Nogueira, Rafael dos Santos Bezerra, Raul Machado Neto, Rejane Maria Tommasini Grotto, Ricardo Haddad, Sandra Coccuzzo Sampaio Vessoni, Simone Kashima, Svetoslav Nanev Slavov, Vincent Louis Viala |
| EPI_ISL_2345787 | VIGILANCIA EPIDEMIOLOGICA JARDINOPOLIS SP | Instituto Butantan / Mendelics | Dimas Tadeu Covas, Antonio Jorge Martins, Claudia Renata dos Santos Barros, David Schlesinger, Debora Botequio Moretti, Elaine Cristina Marqueze, Elaine Vieira Santos, Evandra Strazza Rodrigues, Heidge Fukumasu, Jayme Augusto de Souza-Neto, José Salvatore Leister Patané, Luiz Alcantara, Luiz Lehmann Coutinho, Maria Carolina Elias, Maurício Lacerda Nogueira, Rafael dos Santos Bezerra, Raul Machado Neto, Rejane Maria Tommasini Grotto, Ricardo Haddad, Sandra Coccuzzo Sampaio Vessoni, Simone Kashima, Svetoslav Nanev Slavov, Vincent Louis Viala |
| EPI_ISL_2345788 | UBS DR ALFREDO DANTAS DE SOUZA UMUARAMA | Instituto Butantan / Mendelics | Dimas Tadeu Covas, Antonio Jorge Martins, Claudia Renata dos Santos Barros, David Schlesinger, Debora Botequio Moretti, Elaine Cristina Marqueze, Elaine Vieira Santos, Evandra Strazza Rodrigues, Heidge Fukumasu, Jayme Augusto de Souza-Neto, José Salvatore Leister Patané, Luiz Alcantara, Luiz Lehmann Coutinho, Maria Carolina Elias, Maurício Lacerda Nogueira, Rafael dos Santos Bezerra, Raul Machado Neto, Rejane Maria Tommasini Grotto, Ricardo Haddad, Sandra Coccuzzo Sampaio Vessoni, Simone Kashima, Svetoslav Nanev Slavov, Vincent Louis Viala |
| EPI_ISL_2345789 | UNIDADE BASICA SAUDE NATAL ABADIO LACERDA | Instituto Butantan / Mendelics | Dimas Tadeu Covas, Antonio Jorge Martins, Claudia Renata dos Santos Barros, David Schlesinger, Debora Botequio Moretti, Elaine Cristina Marqueze, Elaine Vieira Santos, Evandra Strazza Rodrigues, Heidge Fukumasu, Jayme Augusto de Souza-Neto, José Salvatore Leister Patané, Luiz Alcantara, Luiz Lehmann Coutinho, Maria Carolina Elias, Maurício Lacerda Nogueira, Rafael dos Santos Bezerra, Raul Machado Neto, Rejane Maria Tommasini Grotto, Ricardo Haddad, Sandra Coccuzzo Sampaio Vessoni, Simone Kashima, Svetoslav Nanev Slavov, Vincent Louis Viala |
| EPI_ISL_2345790 | UNIDADE DE PRONTO ATENDIMENTO CENTRAL DE CARAGUATATUBA | Instituto Butantan / Mendelics | Dimas Tadeu Covas, Antonio Jorge Martins, Claudia Renata dos Santos Barros, David Schlesinger, Debora Botequio Moretti, Elaine Cristina Marqueze, Elaine Vieira Santos, Evandra Strazza Rodrigues, Heidge Fukumasu, Jayme Augusto de Souza-Neto, José Salvatore Leister Patané, Luiz Alcantara, Luiz Lehmann Coutinho, Maria Carolina Elias, Maurício Lacerda Nogueira, Rafael dos Santos Bezerra, Raul Machado Neto, Rejane Maria Tommasini Grotto, Ricardo Haddad, Sandra Coccuzzo Sampaio Vessoni, Simone Kashima, Svetoslav Nanev Slavov, Vincent Louis Viala |
| EPI_ISL_2345791 | CENTRO DE SAUDE II IBITINGA | Instituto Butantan / Mendelics | Dimas Tadeu Covas, Antonio Jorge Martins, Claudia Renata dos Santos Barros, David Schlesinger, Debora Botequio Moretti, Elaine Cristina Marqueze, Elaine Vieira Santos, Evandra Strazza Rodrigues, Heidge Fukumasu, Jayme Augusto de Souza-Neto, José Salvatore Leister Patané, Luiz Alcantara, Luiz Lehmann Coutinho, Maria Carolina Elias, Maurício Lacerda Nogueira, Rafael dos Santos Bezerra, Raul Machado Neto, Rejane Maria Tommasini Grotto, Ricardo Haddad, Sandra Coccuzzo Sampaio Vessoni, Simone Kashima, Svetoslav Nanev Slavov, Vincent Louis Viala |
| EPI_ISL_2345792 | CENTRO DE SAUDE III AFFONSO LUZZI SANTA CRUZ DAS PALMEIRAS | Instituto Butantan / Mendelics | Dimas Tadeu Covas, Antonio Jorge Martins, Claudia Renata dos Santos Barros, David Schlesinger, Debora Botequio Moretti, Elaine Cristina Marqueze, Elaine Vieira Santos, Evandra Strazza Rodrigues, Heidge Fukumasu, Jayme Augusto de Souza-Neto, José Salvatore Leister Patané, Luiz Alcantara, Luiz Lehmann Coutinho, Maria Carolina Elias, Maurício Lacerda Nogueira, Rafael dos Santos Bezerra, Raul Machado Neto, Rejane Maria Tommasini Grotto, Ricardo Haddad, Sandra Coccuzzo Sampaio Vessoni, Simone Kashima, Svetoslav Nanev Slavov, Vincent Louis Viala |
| EPI_ISL_2345793 | CENTRO DE SAUDE III TABATINGA | Instituto Butantan / Mendelics | Dimas Tadeu Covas, Antonio Jorge Martins, Claudia Renata dos Santos Barros, David Schlesinger, Debora Botequio Moretti, Elaine Cristina Marqueze, Elaine Vieira Santos, Evandra Strazza Rodrigues, Heidge Fukumasu, Jayme Augusto de Souza-Neto, José Salvatore Leister Patané, Luiz Alcantara, Luiz Lehmann Coutinho, Maria Carolina Elias, Maurício Lacerda Nogueira, Rafael dos Santos Bezerra, Raul Machado Neto, Rejane Maria Tommasini Grotto, Ricardo Haddad, Sandra Coccuzzo Sampaio Vessoni, Simone Kashima, Svetoslav Nanev Slavov, Vincent Louis Viala |
| EPI_ISL_2345794 | CS II DR MIGUEL VITALIANO ORLANDIA | Instituto Butantan / Mendelics | Dimas Tadeu Covas, Antonio Jorge Martins, Claudia Renata dos Santos Barros, David Schlesinger, Debora Botequio Moretti, Elaine Cristina Marqueze, Elaine Vieira Santos, Evandra Strazza Rodrigues, Heidge Fukumasu, Jayme Augusto de Souza-Neto, José Salvatore Leister Patané, Luiz Alcantara, Luiz Lehmann Coutinho, Maria Carolina Elias, Maurício Lacerda Nogueira, Rafael dos Santos Bezerra, Raul Machado Neto, Rejane Maria Tommasini Grotto, Ricardo Haddad, Sandra Coccuzzo Sampaio Vessoni, Simone Kashima, Svetoslav Nanev Slavov, Vincent Louis Viala |
| EPI_ISL_2345795, EPI_ISL_2345797 | GRIPARIO FALCAO | Instituto Butantan / Mendelics | Dimas Tadeu Covas, Antonio Jorge Martins, Claudia Renata dos Santos Barros, David Schlesinger, Debora Botequio Moretti, Elaine Cristina Marqueze, Elaine Vieira Santos, Evandra Strazza Rodrigues, Heidge Fukumasu, Jayme Augusto de Souza-Neto, José Salvatore Leister Patané, Luiz Alcantara, Luiz Lehmann Coutinho, Maria Carolina Elias, Maurício Lacerda Nogueira, Rafael dos Santos Bezerra, Raul Machado Neto, Rejane Maria Tommasini Grotto, Ricardo Haddad, Sandra Coccuzzo Sampaio Vessoni, Simone Kashima, Svetoslav Nanev Slavov, Vincent Louis Viala |
| EPI_ISL_2345798 | SECRETARIA MUNICIPAL DE SAUDE DE PRES BERNARDES SP | Instituto Butantan / Mendelics | Dimas Tadeu Covas, Antonio Jorge Martins, Claudia Renata dos Santos Barros, David Schlesinger, Debora Botequio Moretti, Elaine Cristina Marqueze, Elaine Vieira Santos, Evandra Strazza Rodrigues, Heidge Fukumasu, Jayme Augusto de Souza-Neto, José Salvatore Leister Patané, Luiz Alcantara, Luiz Lehmann Coutinho, Maria Carolina Elias, Maurício Lacerda Nogueira, Rafael dos Santos Bezerra, Raul Machado Neto, Rejane Maria Tommasini Grotto, Ricardo Haddad, Sandra Coccuzzo Sampaio Vessoni, Simone Kashima, Svetoslav Nanev Slavov, Vincent Louis Viala |
| EPI_ISL_2345799 | DEPARTAMENTO DE SAUDE MUNICIPAL SOCORRO SP | Instituto Butantan / Mendelics | Dimas Tadeu Covas, Antonio Jorge Martins, Claudia Renata dos Santos Barros, David Schlesinger, Debora Botequio Moretti, Elaine Cristina Marqueze, Elaine Vieira Santos, Evandra Strazza Rodrigues, Heidge Fukumasu, Jayme Augusto de Souza-Neto, José Salvatore Leister Patané, Luiz Alcantara, Luiz Lehmann Coutinho, Maria Carolina Elias, Maurício Lacerda Nogueira, Rafael dos Santos Bezerra, Raul Machado Neto, Rejane Maria Tommasini Grotto, Ricardo Haddad, Sandra Coccuzzo Sampaio Vessoni, Simone Kashima, Svetoslav Nanev Slavov, Vincent Louis Viala |

| | | | |
|---|---|---|---|
| EPI_ISL_2345800 | NUCLEO DE SAUDE MARY DOTA DE BAURU | Instituto Butantan / Mendelics | Dimas Tadeu Covas, Antonio Jorge Martins, Claudia Renata dos Santos Barros, David Schlesinger, Debora Botequio Moretti, Elaine Cristina Marqueze, Elaine Vieira Santos, Evandra Strazza Rodrigues, Heidge Fukumasu, Jayme Augusto de Souza-Neto, José Salvatore Leister Patané, Luiz Alcantara, Luiz Lehmann Coutinho, Maria Carolina Elias, Maurício Lacerda Nogueira, Rafael dos Santos Bezerra, Raul Machado Neto, Rejane Maria Tommasini Grotto, Ricardo Haddad, Sandra Coccuzzo Sampaio Vessoni, Simone Kashima, Svetoslav Nanev Slavov, Vincent Louis Viala |
| EPI_ISL_2345801 | UBS LAURINDA RODOLFO RUBO | Instituto Butantan / Mendelics | Dimas Tadeu Covas, Antonio Jorge Martins, Claudia Renata dos Santos Barros, David Schlesinger, Debora Botequio Moretti, Elaine Cristina Marqueze, Elaine Vieira Santos, Evandra Strazza Rodrigues, Heidge Fukumasu, Jayme Augusto de Souza-Neto, José Salvatore Leister Patané, Luiz Alcantara, Luiz Lehmann Coutinho, Maria Carolina Elias, Maurício Lacerda Nogueira, Rafael dos Santos Bezerra, Raul Machado Neto, Rejane Maria Tommasini Grotto, Ricardo Haddad, Sandra Coccuzzo Sampaio Vessoni, Simone Kashima, Svetoslav Nanev Slavov, Vincent Louis Viala |
| EPI_ISL_2345802 | SECRETARIA DE SAUDE PUBLICA DE PRAIA GRANDE | Instituto Butantan / Mendelics | Dimas Tadeu Covas, Antonio Jorge Martins, Claudia Renata dos Santos Barros, David Schlesinger, Debora Botequio Moretti, Elaine Cristina Marqueze, Elaine Vieira Santos, Evandra Strazza Rodrigues, Heidge Fukumasu, Jayme Augusto de Souza-Neto, José Salvatore Leister Patané, Luiz Alcantara, Luiz Lehmann Coutinho, Maria Carolina Elias, Maurício Lacerda Nogueira, Rafael dos Santos Bezerra, Raul Machado Neto, Rejane Maria Tommasini Grotto, Ricardo Haddad, Sandra Coccuzzo Sampaio Vessoni, Simone Kashima, Svetoslav Nanev Slavov, Vincent Louis Viala |
| EPI_ISL_2345803 | UBS DR ERNESTO GAION | Instituto Butantan / Mendelics | Dimas Tadeu Covas, Antonio Jorge Martins, Claudia Renata dos Santos Barros, David Schlesinger, Debora Botequio Moretti, Elaine Cristina Marqueze, Elaine Vieira Santos, Evandra Strazza Rodrigues, Heidge Fukumasu, Jayme Augusto de Souza-Neto, José Salvatore Leister Patané, Luiz Alcantara, Luiz Lehmann Coutinho, Maria Carolina Elias, Maurício Lacerda Nogueira, Rafael dos Santos Bezerra, Raul Machado Neto, Rejane Maria Tommasini Grotto, Ricardo Haddad, Sandra Coccuzzo Sampaio Vessoni, Simone Kashima, Svetoslav Nanev Slavov, Vincent Louis Viala |
| EPI_ISL_2345804 | SECRETARIA DE SAUDE PUBLICA DE PRAIA GRANDE | Instituto Butantan / Mendelics | Dimas Tadeu Covas, Antonio Jorge Martins, Claudia Renata dos Santos Barros, David Schlesinger, Debora Botequio Moretti, Elaine Cristina Marqueze, Elaine Vieira Santos, Evandra Strazza Rodrigues, Heidge Fukumasu, Jayme Augusto de Souza-Neto, José Salvatore Leister Patané, Luiz Alcantara, Luiz Lehmann Coutinho, Maria Carolina Elias, Maurício Lacerda Nogueira, Rafael dos Santos Bezerra, Raul Machado Neto, Rejane Maria Tommasini Grotto, Ricardo Haddad, Sandra Coccuzzo Sampaio Vessoni, Simone Kashima, Svetoslav Nanev Slavov, Vincent Louis Viala |
| EPI_ISL_2345805 | SECRETARIA MUNICIPAL DE SAUDE SOROCABA | Instituto Butantan / Mendelics | Dimas Tadeu Covas, Antonio Jorge Martins, Claudia Renata dos Santos Barros, David Schlesinger, Debora Botequio Moretti, Elaine Cristina Marqueze, Elaine Vieira Santos, Evandra Strazza Rodrigues, Heidge Fukumasu, Jayme Augusto de Souza-Neto, José Salvatore Leister Patané, Luiz Alcantara, Luiz Lehmann Coutinho, Maria Carolina Elias, Maurício Lacerda Nogueira, Rafael dos Santos Bezerra, Raul Machado Neto, Rejane Maria Tommasini Grotto, Ricardo Haddad, Sandra Coccuzzo Sampaio Vessoni, Simone Kashima, Svetoslav Nanev Slavov, Vincent Louis Viala |
| EPI_ISL_2345806 | GRIPARIO FALCAO | Instituto Butantan / Mendelics | Dimas Tadeu Covas, Antonio Jorge Martins, Claudia Renata dos Santos Barros, David Schlesinger, Debora Botequio Moretti, Elaine Cristina Marqueze, Elaine Vieira Santos, Evandra Strazza Rodrigues, Heidge Fukumasu, Jayme Augusto de Souza-Neto, José Salvatore Leister Patané, Luiz Alcantara, Luiz Lehmann Coutinho, Maria Carolina Elias, Maurício Lacerda Nogueira, Rafael dos Santos Bezerra, Raul Machado Neto, Rejane Maria Tommasini Grotto, Ricardo Haddad, Sandra Coccuzzo Sampaio Vessoni, Simone Kashima, Svetoslav Nanev Slavov, Vincent Louis Viala |
| EPI_ISL_2345807 | GRUPO TECNICO DE VIGILANCIA SANITARIA E EPIDEMIOLOGICA | Instituto Butantan / Mendelics | Dimas Tadeu Covas, Antonio Jorge Martins, Claudia Renata dos Santos Barros, David Schlesinger, Debora Botequio Moretti, Elaine Cristina Marqueze, Elaine Vieira Santos, Evandra Strazza Rodrigues, Heidge Fukumasu, Jayme Augusto de Souza-Neto, José Salvatore Leister Patané, Luiz Alcantara, Luiz Lehmann Coutinho, Maria Carolina Elias, Maurício Lacerda Nogueira, Rafael dos Santos Bezerra, Raul Machado Neto, Rejane Maria Tommasini Grotto, Ricardo Haddad, Sandra Coccuzzo Sampaio Vessoni, Simone Kashima, Svetoslav Nanev Slavov, Vincent Louis Viala |
| EPI_ISL_2345808 | SECAO CENTRO DE DIAGNOSTICO SECEDI | Instituto Butantan / Mendelics | Dimas Tadeu Covas, Antonio Jorge Martins, Claudia Renata dos Santos Barros, David Schlesinger, Debora Botequio Moretti, Elaine Cristina Marqueze, Elaine Vieira Santos, Evandra Strazza Rodrigues, Heidge Fukumasu, Jayme Augusto de Souza-Neto, José Salvatore Leister Patané, Luiz Alcantara, Luiz Lehmann Coutinho, Maria Carolina Elias, Maurício Lacerda Nogueira, Rafael dos Santos Bezerra, Raul Machado Neto, Rejane Maria Tommasini Grotto, Ricardo Haddad, Sandra Coccuzzo Sampaio Vessoni, Simone Kashima, Svetoslav Nanev Slavov, Vincent Louis Viala |
| EPI_ISL_2345809 | POLICLINICA COVID 19 ITAPETININGA | Instituto Butantan / Mendelics | Dimas Tadeu Covas, Antonio Jorge Martins, Claudia Renata dos Santos Barros, David Schlesinger, Debora Botequio Moretti, Elaine Cristina Marqueze, Elaine Vieira Santos, Evandra Strazza Rodrigues, Heidge Fukumasu, Jayme Augusto de Souza-Neto, José Salvatore Leister Patané, Luiz Alcantara, Luiz Lehmann Coutinho, Maria Carolina Elias, Maurício Lacerda Nogueira, Rafael dos Santos Bezerra, Raul Machado Neto, Rejane Maria Tommasini Grotto, Ricardo Haddad, Sandra Coccuzzo Sampaio Vessoni, Simone Kashima, Svetoslav Nanev Slavov, Vincent Louis Viala |
| EPI_ISL_2345810 | SERV DE VIG SANITARIA EPIDEMIO E CTRL DE ZOONOZES GUARUJA | Instituto Butantan / Mendelics | Dimas Tadeu Covas, Antonio Jorge Martins, Claudia Renata dos Santos Barros, David Schlesinger, Debora Botequio Moretti, Elaine Cristina Marqueze, Elaine Vieira Santos, Evandra Strazza Rodrigues, Heidge Fukumasu, Jayme Augusto de Souza-Neto, José Salvatore Leister Patané, Luiz Alcantara, Luiz Lehmann Coutinho, Maria Carolina Elias, Maurício Lacerda Nogueira, Rafael dos Santos Bezerra, Raul Machado Neto, Rejane Maria Tommasini Grotto, Ricardo Haddad, Sandra Coccuzzo Sampaio Vessoni, Simone Kashima, Svetoslav Nanev Slavov, Vincent Louis Viala |
| EPI_ISL_2345811 | CENTRO DE ESPECIALIDADES DE PRIMAVERA | Instituto Butantan / Mendelics | Dimas Tadeu Covas, Antonio Jorge Martins, Claudia Renata dos Santos Barros, David Schlesinger, Debora Botequio Moretti, Elaine Cristina Marqueze, Elaine Vieira Santos, Evandra Strazza Rodrigues, Heidge Fukumasu, Jayme Augusto de Souza-Neto, José Salvatore Leister Patané, Luiz Alcantara, Luiz Lehmann Coutinho, Maria Carolina Elias, Maurício Lacerda Nogueira, Rafael dos Santos Bezerra, Raul Machado Neto, Rejane Maria Tommasini Grotto, Ricardo Haddad, Sandra Coccuzzo Sampaio Vessoni, Simone Kashima, Svetoslav Nanev Slavov, Vincent Louis Viala |
| EPI_ISL_2345812 | UPA DR FRANCO DA ROCHA | Instituto Butantan / Mendelics | Dimas Tadeu Covas, Antonio Jorge Martins, Claudia Renata dos Santos Barros, David Schlesinger, Debora Botequio Moretti, Elaine Cristina Marqueze, Elaine Vieira Santos, Evandra Strazza Rodrigues, Heidge Fukumasu, Jayme Augusto de Souza-Neto, José Salvatore Leister Patané, Luiz Alcantara, Luiz Lehmann Coutinho, Maria Carolina Elias, Maurício Lacerda Nogueira, Rafael dos Santos Bezerra, Raul Machado Neto, Rejane Maria Tommasini Grotto, Ricardo Haddad, Sandra Coccuzzo Sampaio Vessoni, Simone Kashima, Svetoslav Nanev Slavov, Vincent Louis Viala |
| EPI_ISL_2345813 | UBS II DR JOSE CARLOS MACUCO JANINI | Instituto Butantan / Mendelics | Dimas Tadeu Covas, Antonio Jorge Martins, Claudia Renata dos Santos Barros, David Schlesinger, Debora Botequio Moretti, Elaine Cristina Marqueze, Elaine Vieira Santos, Evandra Strazza Rodrigues, Heidge Fukumasu, Jayme Augusto de Souza-Neto, José Salvatore Leister Patané, Luiz Alcantara, Luiz Lehmann Coutinho, Maria Carolina Elias, Maurício Lacerda Nogueira, Rafael dos Santos Bezerra, Raul Machado Neto, Rejane Maria Tommasini Grotto, Ricardo Haddad, Sandra Coccuzzo Sampaio Vessoni, Simone Kashima, Svetoslav Nanev Slavov, Vincent Louis Viala |
| EPI_ISL_2345814 | POLICLINICA COVID 19 ITAPETININGA | Instituto Butantan / Mendelics | Dimas Tadeu Covas, Antonio Jorge Martins, Claudia Renata dos Santos Barros, David Schlesinger, Debora Botequio Moretti, Elaine Cristina Marqueze, Elaine Vieira Santos, Evandra Strazza Rodrigues, Heidge Fukumasu, Jayme Augusto de Souza-Neto, José Salvatore Leister Patané, Luiz Alcantara, Luiz Lehmann Coutinho, Maria Carolina Elias, Maurício Lacerda Nogueira, Rafael dos Santos Bezerra, Raul Machado Neto, Rejane Maria Tommasini Grotto, Ricardo Haddad, Sandra Coccuzzo Sampaio Vessoni, Simone Kashima, Svetoslav Nanev Slavov, Vincent Louis Viala |
| EPI_ISL_2345815 | CENTRO DE REFERENCIA DO IDOSO DR HUMBERTO MENDES DE CARVALHO | Instituto Butantan / Mendelics | Dimas Tadeu Covas, Antonio Jorge Martins, Claudia Renata dos Santos Barros, David Schlesinger, Debora Botequio Moretti, Elaine Cristina Marqueze, Elaine Vieira Santos, Evandra Strazza Rodrigues, Heidge Fukumasu, Jayme Augusto de Souza-Neto, José Salvatore Leister Patané, Luiz Alcantara, Luiz Lehmann Coutinho, Maria Carolina Elias, Maurício Lacerda Nogueira, Rafael dos Santos Bezerra, Raul Machado Neto, Rejane Maria Tommasini Grotto, Ricardo Haddad, Sandra Coccuzzo Sampaio Vessoni, Simone Kashima, Svetoslav Nanev Slavov, Vincent Louis Viala |
| EPI_ISL_2345816 | CENTRO DE SAUDE II IBITINGA | Instituto Butantan / Mendelics | Dimas Tadeu Covas, Antonio Jorge Martins, Claudia Renata dos Santos Barros, David Schlesinger, Debora Botequio Moretti, Elaine Cristina Marqueze, Elaine Vieira Santos, Evandra Strazza Rodrigues, Heidge Fukumasu, Jayme Augusto de Souza-Neto, José Salvatore Leister Patané, Luiz Alcantara, Luiz Lehmann Coutinho, Maria Carolina Elias, Maurício Lacerda Nogueira, Rafael dos Santos Bezerra, Raul Machado Neto, Rejane Maria Tommasini Grotto, Ricardo Haddad, Sandra Coccuzzo Sampaio Vessoni, Simone Kashima, Svetoslav Nanev Slavov, Vincent Louis Viala |
| EPI_ISL_2345817 | HOSPITAL DE CAMPANHA COVID 19 CAIEIRAS | Instituto Butantan / Mendelics | Dimas Tadeu Covas, Antonio Jorge Martins, Claudia Renata dos Santos Barros, David Schlesinger, Debora Botequio Moretti, Elaine Cristina Marqueze, Elaine Vieira Santos, Evandra Strazza Rodrigues, Heidge Fukumasu, Jayme Augusto de Souza-Neto, José Salvatore Leister Patané, Luiz Alcantara, Luiz Lehmann Coutinho, Maria Carolina Elias, Maurício Lacerda Nogueira, Rafael dos Santos Bezerra, Raul Machado Neto, Rejane Maria Tommasini Grotto, Ricardo Haddad, Sandra Coccuzzo Sampaio Vessoni, Simone Kashima, Svetoslav Nanev Slavov, Vincent Louis Viala |
| EPI_ISL_2345818 | UBS II DR JOSE CARLOS MACUCO JANINI | Instituto Butantan / Mendelics | Dimas Tadeu Covas, Antonio Jorge Martins, Claudia Renata dos Santos Barros, David Schlesinger, Debora Botequio Moretti, Elaine Cristina Marqueze, Elaine Vieira Santos, Evandra Strazza Rodrigues, Heidge Fukumasu, Jayme Augusto de Souza-Neto, José Salvatore Leister Patané, Luiz Alcantara, Luiz Lehmann Coutinho, Maria Carolina Elias, Maurício Lacerda Nogueira, Rafael dos Santos Bezerra, Raul Machado Neto, Rejane Maria Tommasini Grotto, Ricardo Haddad, Sandra Coccuzzo Sampaio Vessoni, Simone Kashima, Svetoslav Nanev Slavov, Vincent Louis Viala |
| EPI_ISL_2345819 | HOSPITAL DE CAMPANHA COVID 19 MUNICIPIO DE TAUBATE | Instituto Butantan / Mendelics | Dimas Tadeu Covas, Antonio Jorge Martins, Claudia Renata dos Santos Barros, David Schlesinger, Debora Botequio Moretti, Elaine Cristina Marqueze, Elaine Vieira Santos, Evandra Strazza Rodrigues, Heidge Fukumasu, Jayme Augusto de Souza-Neto, José Salvatore Leister Patané, Luiz Alcantara, Luiz |

| | | | |
|---|---|---|---|
| | | | Lehmann Coutinho, Maria Carolina Elias, Maurício Lacerda Nogueira, Rafael dos Santos Bezerra, Raul Machado Neto, Rejane Maria Tommasini Grotto, Ricardo Haddad, Sandra Coccuzzo Sampaio Vessoni, Simone Kashima, Svetoslav Nanev Slavov, Vincent Louis Viala |
| EPI_ISL_2345820 | UPA UNIDADE DE PRONTO ATENDIMENTO 24 HORAS BOM JESUS | Instituto Butantan / Mendelics | Dimas Tadeu Covas, Antonio Jorge Martins, Claudia Renata dos Santos Barros, David Schlesinger, Debora Botequio Moretti, Elaine Cristina Marqueze, Elaine Vieira Santos, Evandra Strazza Rodrigues, Heidge Fukumasu, Jayme Augusto de Souza-Neto, José Salvatore Leister Patané, Luiz Alcantara, Luiz Lehmann Coutinho, Maria Carolina Elias, Maurício Lacerda Nogueira, Rafael dos Santos Bezerra, Raul Machado Neto, Rejane Maria Tommasini Grotto, Ricardo Haddad, Sandra Coccuzzo Sampaio Vessoni, Simone Kashima, Svetoslav Nanev Slavov, Vincent Louis Viala |
| EPI_ISL_2345821 | CENTRO DE SAUDE II IBITINGA | Instituto Butantan / Mendelics | Dimas Tadeu Covas, Antonio Jorge Martins, Claudia Renata dos Santos Barros, David Schlesinger, Debora Botequio Moretti, Elaine Cristina Marqueze, Elaine Vieira Santos, Evandra Strazza Rodrigues, Heidge Fukumasu, Jayme Augusto de Souza-Neto, José Salvatore Leister Patané, Luiz Alcantara, Luiz Lehmann Coutinho, Maria Carolina Elias, Maurício Lacerda Nogueira, Rafael dos Santos Bezerra, Raul Machado Neto, Rejane Maria Tommasini Grotto, Ricardo Haddad, Sandra Coccuzzo Sampaio Vessoni, Simone Kashima, Svetoslav Nanev Slavov, Vincent Louis Viala |
| EPI_ISL_2345822 | UBS II DE ALVARES MACHADO | Instituto Butantan / Mendelics | Dimas Tadeu Covas, Antonio Jorge Martins, Claudia Renata dos Santos Barros, David Schlesinger, Debora Botequio Moretti, Elaine Cristina Marqueze, Elaine Vieira Santos, Evandra Strazza Rodrigues, Heidge Fukumasu, Jayme Augusto de Souza-Neto, José Salvatore Leister Patané, Luiz Alcantara, Luiz Lehmann Coutinho, Maria Carolina Elias, Maurício Lacerda Nogueira, Rafael dos Santos Bezerra, Raul Machado Neto, Rejane Maria Tommasini Grotto, Ricardo Haddad, Sandra Coccuzzo Sampaio Vessoni, Simone Kashima, Svetoslav Nanev Slavov, Vincent Louis Viala |
| EPI_ISL_2345823 | SECRETARIA MUNICIPAL DE SAUDE DE PIRACAIA | Instituto Butantan / Mendelics | Dimas Tadeu Covas, Antonio Jorge Martins, Claudia Renata dos Santos Barros, David Schlesinger, Debora Botequio Moretti, Elaine Cristina Marqueze, Elaine Vieira Santos, Evandra Strazza Rodrigues, Heidge Fukumasu, Jayme Augusto de Souza-Neto, José Salvatore Leister Patané, Luiz Alcantara, Luiz Lehmann Coutinho, Maria Carolina Elias, Maurício Lacerda Nogueira, Rafael dos Santos Bezerra, Raul Machado Neto, Rejane Maria Tommasini Grotto, Ricardo Haddad, Sandra Coccuzzo Sampaio Vessoni, Simone Kashima, Svetoslav Nanev Slavov, Vincent Louis Viala |
| EPI_ISL_2345824 | CSII DR WASHINGTON LUIS M RODRIGUES DA SILVA PITANGUEIRAS | Instituto Butantan / Mendelics | Dimas Tadeu Covas, Antonio Jorge Martins, Claudia Renata dos Santos Barros, David Schlesinger, Debora Botequio Moretti, Elaine Cristina Marqueze, Elaine Vieira Santos, Evandra Strazza Rodrigues, Heidge Fukumasu, Jayme Augusto de Souza-Neto, José Salvatore Leister Patané, Luiz Alcantara, Luiz Lehmann Coutinho, Maria Carolina Elias, Maurício Lacerda Nogueira, Rafael dos Santos Bezerra, Raul Machado Neto, Rejane Maria Tommasini Grotto, Ricardo Haddad, Sandra Coccuzzo Sampaio Vessoni, Simone Kashima, Svetoslav Nanev Slavov, Vincent Louis Viala |
| EPI_ISL_2345825 | CENTRO DE SAUDE II MATAO | Instituto Butantan / Mendelics | Dimas Tadeu Covas, Antonio Jorge Martins, Claudia Renata dos Santos Barros, David Schlesinger, Debora Botequio Moretti, Elaine Cristina Marqueze, Elaine Vieira Santos, Evandra Strazza Rodrigues, Heidge Fukumasu, Jayme Augusto de Souza-Neto, José Salvatore Leister Patané, Luiz Alcantara, Luiz Lehmann Coutinho, Maria Carolina Elias, Maurício Lacerda Nogueira, Rafael dos Santos Bezerra, Raul Machado Neto, Rejane Maria Tommasini Grotto, Ricardo Haddad, Sandra Coccuzzo Sampaio Vessoni, Simone Kashima, Svetoslav Nanev Slavov, Vincent Louis Viala |
| EPI_ISL_2345826 | HOSPITAL DE CAMPANHA COVID 19 MUNICIPIO DE TAUBATE | Instituto Butantan / Mendelics | Dimas Tadeu Covas, Antonio Jorge Martins, Claudia Renata dos Santos Barros, David Schlesinger, Debora Botequio Moretti, Elaine Cristina Marqueze, Elaine Vieira Santos, Evandra Strazza Rodrigues, Heidge Fukumasu, Jayme Augusto de Souza-Neto, José Salvatore Leister Patané, Luiz Alcantara, Luiz Lehmann Coutinho, Maria Carolina Elias, Maurício Lacerda Nogueira, Rafael dos Santos Bezerra, Raul Machado Neto, Rejane Maria Tommasini Grotto, Ricardo Haddad, Sandra Coccuzzo Sampaio Vessoni, Simone Kashima, Svetoslav Nanev Slavov, Vincent Louis Viala |
| EPI_ISL_2345827 | DEPARTAMENTO DE SAUDE MUNICIPAL SOCORRO SP | Instituto Butantan / Mendelics | Dimas Tadeu Covas, Antonio Jorge Martins, Claudia Renata dos Santos Barros, David Schlesinger, Debora Botequio Moretti, Elaine Cristina Marqueze, Elaine Vieira Santos, Evandra Strazza Rodrigues, Heidge Fukumasu, Jayme Augusto de Souza-Neto, José Salvatore Leister Patané, Luiz Alcantara, Luiz Lehmann Coutinho, Maria Carolina Elias, Maurício Lacerda Nogueira, Rafael dos Santos Bezerra, Raul Machado Neto, Rejane Maria Tommasini Grotto, Ricardo Haddad, Sandra Coccuzzo Sampaio Vessoni, Simone Kashima, Svetoslav Nanev Slavov, Vincent Louis Viala |
| EPI_ISL_2345828 | UPA MARY DOTA | Instituto Butantan / Mendelics | Dimas Tadeu Covas, Antonio Jorge Martins, Claudia Renata dos Santos Barros, David Schlesinger, Debora Botequio Moretti, Elaine Cristina Marqueze, Elaine Vieira Santos, Evandra Strazza Rodrigues, Heidge Fukumasu, Jayme Augusto de Souza-Neto, José Salvatore Leister Patané, Luiz Alcantara, Luiz Lehmann Coutinho, Maria Carolina Elias, Maurício Lacerda Nogueira, Rafael dos Santos Bezerra, Raul Machado Neto, Rejane Maria Tommasini Grotto, Ricardo Haddad, Sandra Coccuzzo Sampaio Vessoni, Simone Kashima, Svetoslav Nanev Slavov, Vincent Louis Viala |
| EPI_ISL_2345829 | CENTRO DE SAUDE DR JOAO JOAQUIM FERREIRA CRISTAIS PAULISTA | Instituto Butantan / Mendelics | Dimas Tadeu Covas, Antonio Jorge Martins, Claudia Renata dos Santos Barros, David Schlesinger, Debora Botequio Moretti, Elaine Cristina Marqueze, Elaine Vieira Santos, Evandra Strazza Rodrigues, Heidge Fukumasu, Jayme Augusto de Souza-Neto, José Salvatore Leister Patané, Luiz Alcantara, Luiz Lehmann Coutinho, Maria Carolina Elias, Maurício Lacerda Nogueira, Rafael dos Santos Bezerra, Raul Machado Neto, Rejane Maria Tommasini Grotto, Ricardo Haddad, Sandra Coccuzzo Sampaio Vessoni, Simone Kashima, Svetoslav Nanev Slavov, Vincent Louis Viala |
| EPI_ISL_2345830 | CENTRO DE APOIO EPIDEMIOLOGICO | Instituto Butantan / Mendelics | Dimas Tadeu Covas, Antonio Jorge Martins, Claudia Renata dos Santos Barros, David Schlesinger, Debora Botequio Moretti, Elaine Cristina Marqueze, Elaine Vieira Santos, Evandra Strazza Rodrigues, Heidge Fukumasu, Jayme Augusto de Souza-Neto, José Salvatore Leister Patané, Luiz Alcantara, Luiz Lehmann Coutinho, Maria Carolina Elias, Maurício Lacerda Nogueira, Rafael dos Santos Bezerra, Raul Machado Neto, Rejane Maria Tommasini Grotto, Ricardo Haddad, Sandra Coccuzzo Sampaio Vessoni, Simone Kashima, Svetoslav Nanev Slavov, Vincent Louis Viala |
| EPI_ISL_2345831 | SECRETARIA MUNICIPAL DA SAUDE DE JOANOPOLIS | Instituto Butantan / Mendelics | Dimas Tadeu Covas, Antonio Jorge Martins, Claudia Renata dos Santos Barros, David Schlesinger, Debora Botequio Moretti, Elaine Cristina Marqueze, Elaine Vieira Santos, Evandra Strazza Rodrigues, Heidge Fukumasu, Jayme Augusto de Souza-Neto, José Salvatore Leister Patané, Luiz Alcantara, Luiz Lehmann Coutinho, Maria Carolina Elias, Maurício Lacerda Nogueira, Rafael dos Santos Bezerra, Raul Machado Neto, Rejane Maria Tommasini Grotto, Ricardo Haddad, Sandra Coccuzzo Sampaio Vessoni, Simone Kashima, Svetoslav Nanev Slavov, Vincent Louis Viala |
| EPI_ISL_2345832 | CENTRO DE APOIO EPIDEMIOLOGICO | Instituto Butantan / Mendelics | Dimas Tadeu Covas, Antonio Jorge Martins, Claudia Renata dos Santos Barros, David Schlesinger, Debora Botequio Moretti, Elaine Cristina Marqueze, Elaine Vieira Santos, Evandra Strazza Rodrigues, Heidge Fukumasu, Jayme Augusto de Souza-Neto, José Salvatore Leister Patané, Luiz Alcantara, Luiz Lehmann Coutinho, Maria Carolina Elias, Maurício Lacerda Nogueira, Rafael dos Santos Bezerra, Raul Machado Neto, Rejane Maria Tommasini Grotto, Ricardo Haddad, Sandra Coccuzzo Sampaio Vessoni, Simone Kashima, Svetoslav Nanev Slavov, Vincent Louis Viala |
| EPI_ISL_2345833 | HOSPITAL MUNICIPAL DE ITAPIRA | Instituto Butantan / Mendelics | Dimas Tadeu Covas, Antonio Jorge Martins, Claudia Renata dos Santos Barros, David Schlesinger, Debora Botequio Moretti, Elaine Cristina Marqueze, Elaine Vieira Santos, Evandra Strazza Rodrigues, Heidge Fukumasu, Jayme Augusto de Souza-Neto, José Salvatore Leister Patané, Luiz Alcantara, Luiz Lehmann Coutinho, Maria Carolina Elias, Maurício Lacerda Nogueira, Rafael dos Santos Bezerra, Raul Machado Neto, Rejane Maria Tommasini Grotto, Ricardo Haddad, Sandra Coccuzzo Sampaio Vessoni, Simone Kashima, Svetoslav Nanev Slavov, Vincent Louis Viala |
| EPI_ISL_2345834 | USF JARDIM BRASIL | Instituto Butantan / Mendelics | Dimas Tadeu Covas, Antonio Jorge Martins, Claudia Renata dos Santos Barros, David Schlesinger, Debora Botequio Moretti, Elaine Cristina Marqueze, Elaine Vieira Santos, Evandra Strazza Rodrigues, Heidge Fukumasu, Jayme Augusto de Souza-Neto, José Salvatore Leister Patané, Luiz Alcantara, Luiz Lehmann Coutinho, Maria Carolina Elias, Maurício Lacerda Nogueira, Rafael dos Santos Bezerra, Raul Machado Neto, Rejane Maria Tommasini Grotto, Ricardo Haddad, Sandra Coccuzzo Sampaio Vessoni, Simone Kashima, Svetoslav Nanev Slavov, Vincent Louis Viala |
| EPI_ISL_2345835 | GRUPO TECNICO DE VIGILANCIA SANITARIA E EPIDEMIOLOGICA | Instituto Butantan / Mendelics | Dimas Tadeu Covas, Antonio Jorge Martins, Claudia Renata dos Santos Barros, David Schlesinger, Debora Botequio Moretti, Elaine Cristina Marqueze, Elaine Vieira Santos, Evandra Strazza Rodrigues, Heidge Fukumasu, Jayme Augusto de Souza-Neto, José Salvatore Leister Patané, Luiz Alcantara, Luiz Lehmann Coutinho, Maria Carolina Elias, Maurício Lacerda Nogueira, Rafael dos Santos Bezerra, Raul Machado Neto, Rejane Maria Tommasini Grotto, Ricardo Haddad, Sandra Coccuzzo Sampaio Vessoni, Simone Kashima, Svetoslav Nanev Slavov, Vincent Louis Viala |
| EPI_ISL_2345836 | VIGILANCIA EPIDEMIOLOGICA JARDINOPOLIS SP | Instituto Butantan / Mendelics | Dimas Tadeu Covas, Antonio Jorge Martins, Claudia Renata dos Santos Barros, David Schlesinger, Debora Botequio Moretti, Elaine Cristina Marqueze, Elaine Vieira Santos, Evandra Strazza Rodrigues, Heidge Fukumasu, Jayme Augusto de Souza-Neto, José Salvatore Leister Patané, Luiz Alcantara, Luiz Lehmann Coutinho, Maria Carolina Elias, Maurício Lacerda Nogueira, Rafael dos Santos Bezerra, Raul Machado Neto, Rejane Maria Tommasini Grotto, Ricardo Haddad, Sandra Coccuzzo Sampaio Vessoni, Simone Kashima, Svetoslav Nanev Slavov, Vincent Louis Viala |
| EPI_ISL_2345837 | HOSPITAL DE CAMPANHA COVID 19 MUNICIPIO DE TAUBATE | Instituto Butantan / Mendelics | Dimas Tadeu Covas, Antonio Jorge Martins, Claudia Renata dos Santos Barros, David Schlesinger, Debora Botequio Moretti, Elaine Cristina Marqueze, Elaine Vieira Santos, Evandra Strazza Rodrigues, Heidge Fukumasu, Jayme Augusto de Souza-Neto, José Salvatore Leister Patané, Luiz Alcantara, Luiz Lehmann Coutinho, Maria Carolina Elias, Maurício Lacerda Nogueira, Rafael dos Santos Bezerra, Raul Machado Neto, Rejane Maria Tommasini Grotto, Ricardo Haddad, Sandra Coccuzzo Sampaio Vessoni, Simone Kashima, Svetoslav Nanev Slavov, Vincent Louis Viala |
| EPI_ISL_2345838 | HOSPITAL MUNICIPAL DR WALDEMAR TEBALDI | Instituto Butantan / Mendelics | Dimas Tadeu Covas, Antonio Jorge Martins, Claudia Renata dos Santos Barros, David Schlesinger, Debora Botequio Moretti, Elaine Cristina Marqueze, Elaine Vieira Santos, Evandra Strazza Rodrigues, Heidge Fukumasu, Jayme Augusto de Souza-Neto, José Salvatore Leister Patané, Luiz Alcantara, Luiz Lehmann Coutinho, Maria Carolina Elias, Maurício Lacerda Nogueira, Rafael dos Santos Bezerra, Raul Machado Neto, Rejane Maria Tommasini Grotto, Ricardo Haddad, Sandra Coccuzzo Sampaio Vessoni, Simone Kashima, Svetoslav Nanev Slavov, Vincent Louis Viala |

| | | | |
|---|---|---|---|
| EPI_ISL_2345839 | HOSP MUN DE MOGI DAS CRUZES PREF WALDEMAR COSTA FILHO | Instituto Butantan / Mendelics | Dimas Tadeu Covas, Antonio Jorge Martins, Claudia Renata dos Santos Barros, David Schlesinger, Debora Botequio Moretti, Elaine Cristina Marqueze, Elaine Vieira Santos, Evandra Strazza Rodrigues, Heidge Fukumasu, Jayme Augusto de Souza-Neto, José Salvatore Leister Patané, Luiz Alcantara, Luiz Lehmann Coutinho, Maria Carolina Elias, Maurício Lacerda Nogueira, Rafael dos Santos Bezerra, Raul Machado Neto, Rejane Maria Tommasini Grotto, Ricardo Haddad, Sandra Coccuzzo Sampaio Vessoni, Simone Kashima, Svetoslav Nanev Slavov, Vincent Louis Viala |
| EPI_ISL_2345840, EPI_ISL_2345841 | VIGILANCIA EM SAUDE SAE VIGILANCIA EPIDEMIOLOGICA | Instituto Butantan / Mendelics | Dimas Tadeu Covas, Antonio Jorge Martins, Claudia Renata dos Santos Barros, David Schlesinger, Debora Botequio Moretti, Elaine Cristina Marqueze, Elaine Vieira Santos, Evandra Strazza Rodrigues, Heidge Fukumasu, Jayme Augusto de Souza-Neto, José Salvatore Leister Patané, Luiz Alcantara, Luiz Lehmann Coutinho, Maria Carolina Elias, Maurício Lacerda Nogueira, Rafael dos Santos Bezerra, Raul Machado Neto, Rejane Maria Tommasini Grotto, Ricardo Haddad, Sandra Coccuzzo Sampaio Vessoni, Simone Kashima, Svetoslav Nanev Slavov, Vincent Louis Viala |
| EPI_ISL_2345842 | CENTRO DE SAUDE III SALES OLIVEIRA | Instituto Butantan / Mendelics | Dimas Tadeu Covas, Antonio Jorge Martins, Claudia Renata dos Santos Barros, David Schlesinger, Debora Botequio Moretti, Elaine Cristina Marqueze, Elaine Vieira Santos, Evandra Strazza Rodrigues, Heidge Fukumasu, Jayme Augusto de Souza-Neto, José Salvatore Leister Patané, Luiz Alcantara, Luiz Lehmann Coutinho, Maria Carolina Elias, Maurício Lacerda Nogueira, Rafael dos Santos Bezerra, Raul Machado Neto, Rejane Maria Tommasini Grotto, Ricardo Haddad, Sandra Coccuzzo Sampaio Vessoni, Simone Kashima, Svetoslav Nanev Slavov, Vincent Louis Viala |
| EPI_ISL_2345843 | VIGILANCIA EM SAUDE SAE VIGILANCIA EPIDEMIOLOGICA | Instituto Butantan / Mendelics | Dimas Tadeu Covas, Antonio Jorge Martins, Claudia Renata dos Santos Barros, David Schlesinger, Debora Botequio Moretti, Elaine Cristina Marqueze, Elaine Vieira Santos, Evandra Strazza Rodrigues, Heidge Fukumasu, Jayme Augusto de Souza-Neto, José Salvatore Leister Patané, Luiz Alcantara, Luiz Lehmann Coutinho, Maria Carolina Elias, Maurício Lacerda Nogueira, Rafael dos Santos Bezerra, Raul Machado Neto, Rejane Maria Tommasini Grotto, Ricardo Haddad, Sandra Coccuzzo Sampaio Vessoni, Simone Kashima, Svetoslav Nanev Slavov, Vincent Louis Viala |
| EPI_ISL_2345844 | CENTRO DE SAUDE II DR GABRIEL MESQUITA VARGEM GDE DO SUL | Instituto Butantan / Mendelics | Dimas Tadeu Covas, Antonio Jorge Martins, Claudia Renata dos Santos Barros, David Schlesinger, Debora Botequio Moretti, Elaine Cristina Marqueze, Elaine Vieira Santos, Evandra Strazza Rodrigues, Heidge Fukumasu, Jayme Augusto de Souza-Neto, José Salvatore Leister Patané, Luiz Alcantara, Luiz Lehmann Coutinho, Maria Carolina Elias, Maurício Lacerda Nogueira, Rafael dos Santos Bezerra, Raul Machado Neto, Rejane Maria Tommasini Grotto, Ricardo Haddad, Sandra Coccuzzo Sampaio Vessoni, Simone Kashima, Svetoslav Nanev Slavov, Vincent Louis Viala |
| EPI_ISL_2345845 | CENTRO DE ESPECIALIDADES MUNICIPAL CESAR TORREZAN R BONITO | Instituto Butantan / Mendelics | Dimas Tadeu Covas, Antonio Jorge Martins, Claudia Renata dos Santos Barros, David Schlesinger, Debora Botequio Moretti, Elaine Cristina Marqueze, Elaine Vieira Santos, Evandra Strazza Rodrigues, Heidge Fukumasu, Jayme Augusto de Souza-Neto, José Salvatore Leister Patané, Luiz Alcantara, Luiz Lehmann Coutinho, Maria Carolina Elias, Maurício Lacerda Nogueira, Rafael dos Santos Bezerra, Raul Machado Neto, Rejane Maria Tommasini Grotto, Ricardo Haddad, Sandra Coccuzzo Sampaio Vessoni, Simone Kashima, Svetoslav Nanev Slavov, Vincent Louis Viala |
| EPI_ISL_2345846 | CENTRO DE SAUDE DE MARACAI | Instituto Butantan / Mendelics | Dimas Tadeu Covas, Antonio Jorge Martins, Claudia Renata dos Santos Barros, David Schlesinger, Debora Botequio Moretti, Elaine Cristina Marqueze, Elaine Vieira Santos, Evandra Strazza Rodrigues, Heidge Fukumasu, Jayme Augusto de Souza-Neto, José Salvatore Leister Patané, Luiz Alcantara, Luiz Lehmann Coutinho, Maria Carolina Elias, Maurício Lacerda Nogueira, Rafael dos Santos Bezerra, Raul Machado Neto, Rejane Maria Tommasini Grotto, Ricardo Haddad, Sandra Coccuzzo Sampaio Vessoni, Simone Kashima, Svetoslav Nanev Slavov, Vincent Louis Viala |
| EPI_ISL_2345847 | GRIPARIO FALCAO | Instituto Butantan / Mendelics | Dimas Tadeu Covas, Antonio Jorge Martins, Claudia Renata dos Santos Barros, David Schlesinger, Debora Botequio Moretti, Elaine Cristina Marqueze, Elaine Vieira Santos, Evandra Strazza Rodrigues, Heidge Fukumasu, Jayme Augusto de Souza-Neto, José Salvatore Leister Patané, Luiz Alcantara, Luiz Lehmann Coutinho, Maria Carolina Elias, Maurício Lacerda Nogueira, Rafael dos Santos Bezerra, Raul Machado Neto, Rejane Maria Tommasini Grotto, Ricardo Haddad, Sandra Coccuzzo Sampaio Vessoni, Simone Kashima, Svetoslav Nanev Slavov, Vincent Louis Viala |
| EPI_ISL_2345848 | SECRETARIA MUNICIPAL DE SAUDE SOROCABA | Instituto Butantan / Mendelics | Dimas Tadeu Covas, Antonio Jorge Martins, Claudia Renata dos Santos Barros, David Schlesinger, Debora Botequio Moretti, Elaine Cristina Marqueze, Elaine Vieira Santos, Evandra Strazza Rodrigues, Heidge Fukumasu, Jayme Augusto de Souza-Neto, José Salvatore Leister Patané, Luiz Alcantara, Luiz Lehmann Coutinho, Maria Carolina Elias, Maurício Lacerda Nogueira, Rafael dos Santos Bezerra, Raul Machado Neto, Rejane Maria Tommasini Grotto, Ricardo Haddad, Sandra Coccuzzo Sampaio Vessoni, Simone Kashima, Svetoslav Nanev Slavov, Vincent Louis Viala |
| EPI_ISL_2345849 | SECRETARIA DE SAUDE PUBLICA DE PRAIA GRANDE | Instituto Butantan / Mendelics | Dimas Tadeu Covas, Antonio Jorge Martins, Claudia Renata dos Santos Barros, David Schlesinger, Debora Botequio Moretti, Elaine Cristina Marqueze, Elaine Vieira Santos, Evandra Strazza Rodrigues, Heidge Fukumasu, Jayme Augusto de Souza-Neto, José Salvatore Leister Patané, Luiz Alcantara, Luiz Lehmann Coutinho, Maria Carolina Elias, Maurício Lacerda Nogueira, Rafael dos Santos Bezerra, Raul Machado Neto, Rejane Maria Tommasini Grotto, Ricardo Haddad, Sandra Coccuzzo Sampaio Vessoni, Simone Kashima, Svetoslav Nanev Slavov, Vincent Louis Viala |
| EPI_ISL_2345850 | UBS GLENDA RODELLA DUMAS | Instituto Butantan / Mendelics | Dimas Tadeu Covas, Antonio Jorge Martins, Claudia Renata dos Santos Barros, David Schlesinger, Debora Botequio Moretti, Elaine Cristina Marqueze, Elaine Vieira Santos, Evandra Strazza Rodrigues, Heidge Fukumasu, Jayme Augusto de Souza-Neto, José Salvatore Leister Patané, Luiz Alcantara, Luiz Lehmann Coutinho, Maria Carolina Elias, Maurício Lacerda Nogueira, Rafael dos Santos Bezerra, Raul Machado Neto, Rejane Maria Tommasini Grotto, Ricardo Haddad, Sandra Coccuzzo Sampaio Vessoni, Simone Kashima, Svetoslav Nanev Slavov, Vincent Louis Viala |
| EPI_ISL_2345851 | SECAO CENTRO DE DIAGNOSTICO SECEDI | Instituto Butantan / Mendelics | Dimas Tadeu Covas, Antonio Jorge Martins, Claudia Renata dos Santos Barros, David Schlesinger, Debora Botequio Moretti, Elaine Cristina Marqueze, Elaine Vieira Santos, Evandra Strazza Rodrigues, Heidge Fukumasu, Jayme Augusto de Souza-Neto, José Salvatore Leister Patané, Luiz Alcantara, Luiz Lehmann Coutinho, Maria Carolina Elias, Maurício Lacerda Nogueira, Rafael dos Santos Bezerra, Raul Machado Neto, Rejane Maria Tommasini Grotto, Ricardo Haddad, Sandra Coccuzzo Sampaio Vessoni, Simone Kashima, Svetoslav Nanev Slavov, Vincent Louis Viala |
| EPI_ISL_2345852 | SECRETARIA MUNICIPAL DE SAUDE SOROCABA | Instituto Butantan / Mendelics | Dimas Tadeu Covas, Antonio Jorge Martins, Claudia Renata dos Santos Barros, David Schlesinger, Debora Botequio Moretti, Elaine Cristina Marqueze, Elaine Vieira Santos, Evandra Strazza Rodrigues, Heidge Fukumasu, Jayme Augusto de Souza-Neto, José Salvatore Leister Patané, Luiz Alcantara, Luiz Lehmann Coutinho, Maria Carolina Elias, Maurício Lacerda Nogueira, Rafael dos Santos Bezerra, Raul Machado Neto, Rejane Maria Tommasini Grotto, Ricardo Haddad, Sandra Coccuzzo Sampaio Vessoni, Simone Kashima, Svetoslav Nanev Slavov, Vincent Louis Viala |
| EPI_ISL_2345853 | UBS III DE RANCHARIA | Instituto Butantan / Mendelics | Dimas Tadeu Covas, Antonio Jorge Martins, Claudia Renata dos Santos Barros, David Schlesinger, Debora Botequio Moretti, Elaine Cristina Marqueze, Elaine Vieira Santos, Evandra Strazza Rodrigues, Heidge Fukumasu, Jayme Augusto de Souza-Neto, José Salvatore Leister Patané, Luiz Alcantara, Luiz Lehmann Coutinho, Maria Carolina Elias, Maurício Lacerda Nogueira, Rafael dos Santos Bezerra, Raul Machado Neto, Rejane Maria Tommasini Grotto, Ricardo Haddad, Sandra Coccuzzo Sampaio Vessoni, Simone Kashima, Svetoslav Nanev Slavov, Vincent Louis Viala |
| EPI_ISL_2345854 | SECAO CENTRO DE DIAGNOSTICO SECEDI | Instituto Butantan / Mendelics | Dimas Tadeu Covas, Antonio Jorge Martins, Claudia Renata dos Santos Barros, David Schlesinger, Debora Botequio Moretti, Elaine Cristina Marqueze, Elaine Vieira Santos, Evandra Strazza Rodrigues, Heidge Fukumasu, Jayme Augusto de Souza-Neto, José Salvatore Leister Patané, Luiz Alcantara, Luiz Lehmann Coutinho, Maria Carolina Elias, Maurício Lacerda Nogueira, Rafael dos Santos Bezerra, Raul Machado Neto, Rejane Maria Tommasini Grotto, Ricardo Haddad, Sandra Coccuzzo Sampaio Vessoni, Simone Kashima, Svetoslav Nanev Slavov, Vincent Louis Viala |
| EPI_ISL_2345855 | SECRETARIA MUNICIPAL DA SAUDE DE ITARARE | Instituto Butantan / Mendelics | Dimas Tadeu Covas, Antonio Jorge Martins, Claudia Renata dos Santos Barros, David Schlesinger, Debora Botequio Moretti, Elaine Cristina Marqueze, Elaine Vieira Santos, Evandra Strazza Rodrigues, Heidge Fukumasu, Jayme Augusto de Souza-Neto, José Salvatore Leister Patané, Luiz Alcantara, Luiz Lehmann Coutinho, Maria Carolina Elias, Maurício Lacerda Nogueira, Rafael dos Santos Bezerra, Raul Machado Neto, Rejane Maria Tommasini Grotto, Ricardo Haddad, Sandra Coccuzzo Sampaio Vessoni, Simone Kashima, Svetoslav Nanev Slavov, Vincent Louis Viala |
| EPI_ISL_2345856 | SERV DE VIG SANITARIA EPIDEMIO E CTRL DE ZOONOZES GUARUJA | Instituto Butantan / Mendelics | Dimas Tadeu Covas, Antonio Jorge Martins, Claudia Renata dos Santos Barros, David Schlesinger, Debora Botequio Moretti, Elaine Cristina Marqueze, Elaine Vieira Santos, Evandra Strazza Rodrigues, Heidge Fukumasu, Jayme Augusto de Souza-Neto, José Salvatore Leister Patané, Luiz Alcantara, Luiz Lehmann Coutinho, Maria Carolina Elias, Maurício Lacerda Nogueira, Rafael dos Santos Bezerra, Raul Machado Neto, Rejane Maria Tommasini Grotto, Ricardo Haddad, Sandra Coccuzzo Sampaio Vessoni, Simone Kashima, Svetoslav Nanev Slavov, Vincent Louis Viala |
| EPI_ISL_2345857 | CENTRO DE SAUDE DE NOVA GUATAPORANGA | Instituto Butantan / Mendelics | Dimas Tadeu Covas, Antonio Jorge Martins, Claudia Renata dos Santos Barros, David Schlesinger, Debora Botequio Moretti, Elaine Cristina Marqueze, Elaine Vieira Santos, Evandra Strazza Rodrigues, Heidge Fukumasu, Jayme Augusto de Souza-Neto, José Salvatore Leister Patané, Luiz Alcantara, Luiz Lehmann Coutinho, Maria Carolina Elias, Maurício Lacerda Nogueira, Rafael dos Santos Bezerra, Raul Machado Neto, Rejane Maria Tommasini Grotto, Ricardo Haddad, Sandra Coccuzzo Sampaio Vessoni, Simone Kashima, Svetoslav Nanev Slavov, Vincent Louis Viala |
| EPI_ISL_2345858 | UBS II DE ALVARES MACHADO | Instituto Butantan / Mendelics | Dimas Tadeu Covas, Antonio Jorge Martins, Claudia Renata dos Santos Barros, David Schlesinger, Debora Botequio Moretti, Elaine Cristina Marqueze, Elaine Vieira Santos, Evandra Strazza Rodrigues, Heidge Fukumasu, Jayme Augusto de Souza-Neto, José Salvatore Leister Patané, Luiz Alcantara, Luiz Lehmann Coutinho, Maria Carolina Elias, Maurício Lacerda Nogueira, Rafael dos Santos Bezerra, Raul Machado Neto, Rejane Maria Tommasini Grotto, Ricardo Haddad, Sandra Coccuzzo Sampaio Vessoni, Simone Kashima, Svetoslav Nanev Slavov, Vincent Louis Viala |
| EPI_ISL_2345859 | POLICLINICA COVID 19 ITAPETININGA | Instituto Butantan / Mendelics | Dimas Tadeu Covas, Antonio Jorge Martins, Claudia Renata dos Santos Barros, David Schlesinger, Debora Botequio Moretti, Elaine Cristina Marqueze, Elaine Vieira Santos, Evandra Strazza Rodrigues, Heidge Fukumasu, Jayme Augusto de Souza-Neto, José Salvatore Leister Patané, Luiz Alcantara, Luiz |

| | | | |
|---|---|---|---|
| | | | Lehmann Coutinho, Maria Carolina Elias, Maurício Lacerda Nogueira, Rafael dos Santos Bezerra, Raul Machado Neto, Rejane Maria Tommasini Grotto, Ricardo Haddad, Sandra Coccuzzo Sampaio Vessoni, Simone Kashima, Svetoslav Nanev Slavov, Vincent Louis Viala |
| EPI_ISL_2345860 | CENTRO DE REFERENCIA DO IDOSO DR HUMBERTO MENDES DE CARVALHO | Instituto Butantan / Mendelics | Dimas Tadeu Covas, Antonio Jorge Martins, Claudia Renata dos Santos Barros, David Schlesinger, Debora Botequio Moretti, Elaine Cristina Marqueze, Elaine Vieira Santos, Evandra Strazza Rodrigues, Heidge Fukumasu, Jayme Augusto de Souza-Neto, José Salvatore Leister Patané, Luiz Alcantara, Luiz Lehmann Coutinho, Maria Carolina Elias, Maurício Lacerda Nogueira, Rafael dos Santos Bezerra, Raul Machado Neto, Rejane Maria Tommasini Grotto, Ricardo Haddad, Sandra Coccuzzo Sampaio Vessoni, Simone Kashima, Svetoslav Nanev Slavov, Vincent Louis Viala |
| EPI_ISL_2345861 | CENTRO DE SAUDE II IBITINGA | Instituto Butantan / Mendelics | Dimas Tadeu Covas, Antonio Jorge Martins, Claudia Renata dos Santos Barros, David Schlesinger, Debora Botequio Moretti, Elaine Cristina Marqueze, Elaine Vieira Santos, Evandra Strazza Rodrigues, Heidge Fukumasu, Jayme Augusto de Souza-Neto, José Salvatore Leister Patané, Luiz Alcantara, Luiz Lehmann Coutinho, Maria Carolina Elias, Maurício Lacerda Nogueira, Rafael dos Santos Bezerra, Raul Machado Neto, Rejane Maria Tommasini Grotto, Ricardo Haddad, Sandra Coccuzzo Sampaio Vessoni, Simone Kashima, Svetoslav Nanev Slavov, Vincent Louis Viala |
| EPI_ISL_2345862 | UPA DR FRANCO DA ROCHA | Instituto Butantan / Mendelics | Dimas Tadeu Covas, Antonio Jorge Martins, Claudia Renata dos Santos Barros, David Schlesinger, Debora Botequio Moretti, Elaine Cristina Marqueze, Elaine Vieira Santos, Evandra Strazza Rodrigues, Heidge Fukumasu, Jayme Augusto de Souza-Neto, José Salvatore Leister Patané, Luiz Alcantara, Luiz Lehmann Coutinho, Maria Carolina Elias, Maurício Lacerda Nogueira, Rafael dos Santos Bezerra, Raul Machado Neto, Rejane Maria Tommasini Grotto, Ricardo Haddad, Sandra Coccuzzo Sampaio Vessoni, Simone Kashima, Svetoslav Nanev Slavov, Vincent Louis Viala |
| EPI_ISL_2345863 | PAS JOAO ANTONIO DO NASCIMENTO | Instituto Butantan / Mendelics | Dimas Tadeu Covas, Antonio Jorge Martins, Claudia Renata dos Santos Barros, David Schlesinger, Debora Botequio Moretti, Elaine Cristina Marqueze, Elaine Vieira Santos, Evandra Strazza Rodrigues, Heidge Fukumasu, Jayme Augusto de Souza-Neto, José Salvatore Leister Patané, Luiz Alcantara, Luiz Lehmann Coutinho, Maria Carolina Elias, Maurício Lacerda Nogueira, Rafael dos Santos Bezerra, Raul Machado Neto, Rejane Maria Tommasini Grotto, Ricardo Haddad, Sandra Coccuzzo Sampaio Vessoni, Simone Kashima, Svetoslav Nanev Slavov, Vincent Louis Viala |
| EPI_ISL_2345864 | UPA UNIDADE DE PRONTO ATENDIMENTO 24 HORAS BOM JESUS | Instituto Butantan / Mendelics | Dimas Tadeu Covas, Antonio Jorge Martins, Claudia Renata dos Santos Barros, David Schlesinger, Debora Botequio Moretti, Elaine Cristina Marqueze, Elaine Vieira Santos, Evandra Strazza Rodrigues, Heidge Fukumasu, Jayme Augusto de Souza-Neto, José Salvatore Leister Patané, Luiz Alcantara, Luiz Lehmann Coutinho, Maria Carolina Elias, Maurício Lacerda Nogueira, Rafael dos Santos Bezerra, Raul Machado Neto, Rejane Maria Tommasini Grotto, Ricardo Haddad, Sandra Coccuzzo Sampaio Vessoni, Simone Kashima, Svetoslav Nanev Slavov, Vincent Louis Viala |
| EPI_ISL_2345865 | CENTRO DE REFERENCIA DO IDOSO DR HUMBERTO MENDES DE CARVALHO | Instituto Butantan / Mendelics | Dimas Tadeu Covas, Antonio Jorge Martins, Claudia Renata dos Santos Barros, David Schlesinger, Debora Botequio Moretti, Elaine Cristina Marqueze, Elaine Vieira Santos, Evandra Strazza Rodrigues, Heidge Fukumasu, Jayme Augusto de Souza-Neto, José Salvatore Leister Patané, Luiz Alcantara, Luiz Lehmann Coutinho, Maria Carolina Elias, Maurício Lacerda Nogueira, Rafael dos Santos Bezerra, Raul Machado Neto, Rejane Maria Tommasini Grotto, Ricardo Haddad, Sandra Coccuzzo Sampaio Vessoni, Simone Kashima, Svetoslav Nanev Slavov, Vincent Louis Viala |
| EPI_ISL_2345866 | PSF DO PARQUE DOS PINHEIROS | Instituto Butantan / Mendelics | Dimas Tadeu Covas, Antonio Jorge Martins, Claudia Renata dos Santos Barros, David Schlesinger, Debora Botequio Moretti, Elaine Cristina Marqueze, Elaine Vieira Santos, Evandra Strazza Rodrigues, Heidge Fukumasu, Jayme Augusto de Souza-Neto, José Salvatore Leister Patané, Luiz Alcantara, Luiz Lehmann Coutinho, Maria Carolina Elias, Maurício Lacerda Nogueira, Rafael dos Santos Bezerra, Raul Machado Neto, Rejane Maria Tommasini Grotto, Ricardo Haddad, Sandra Coccuzzo Sampaio Vessoni, Simone Kashima, Svetoslav Nanev Slavov, Vincent Louis Viala |
| EPI_ISL_2345867 | SECRETARIA MUNICIPAL DE SAUDE DE PIRACAIA | Instituto Butantan / Mendelics | Dimas Tadeu Covas, Antonio Jorge Martins, Claudia Renata dos Santos Barros, David Schlesinger, Debora Botequio Moretti, Elaine Cristina Marqueze, Elaine Vieira Santos, Evandra Strazza Rodrigues, Heidge Fukumasu, Jayme Augusto de Souza-Neto, José Salvatore Leister Patané, Luiz Alcantara, Luiz Lehmann Coutinho, Maria Carolina Elias, Maurício Lacerda Nogueira, Rafael dos Santos Bezerra, Raul Machado Neto, Rejane Maria Tommasini Grotto, Ricardo Haddad, Sandra Coccuzzo Sampaio Vessoni, Simone Kashima, Svetoslav Nanev Slavov, Vincent Louis Viala |
| EPI_ISL_2345868 | CSII DR WASHINGTON LUIS M RODRIGUES DA SILVA PITANGUEIRAS | Instituto Butantan / Mendelics | Dimas Tadeu Covas, Antonio Jorge Martins, Claudia Renata dos Santos Barros, David Schlesinger, Debora Botequio Moretti, Elaine Cristina Marqueze, Elaine Vieira Santos, Evandra Strazza Rodrigues, Heidge Fukumasu, Jayme Augusto de Souza-Neto, José Salvatore Leister Patané, Luiz Alcantara, Luiz Lehmann Coutinho, Maria Carolina Elias, Maurício Lacerda Nogueira, Rafael dos Santos Bezerra, Raul Machado Neto, Rejane Maria Tommasini Grotto, Ricardo Haddad, Sandra Coccuzzo Sampaio Vessoni, Simone Kashima, Svetoslav Nanev Slavov, Vincent Louis Viala |
| EPI_ISL_2345869 | VIGILANCIA EPIDEMIOLOGICA JARDINOPOLIS SP | Instituto Butantan / Mendelics | Dimas Tadeu Covas, Antonio Jorge Martins, Claudia Renata dos Santos Barros, David Schlesinger, Debora Botequio Moretti, Elaine Cristina Marqueze, Elaine Vieira Santos, Evandra Strazza Rodrigues, Heidge Fukumasu, Jayme Augusto de Souza-Neto, José Salvatore Leister Patané, Luiz Alcantara, Luiz Lehmann Coutinho, Maria Carolina Elias, Maurício Lacerda Nogueira, Rafael dos Santos Bezerra, Raul Machado Neto, Rejane Maria Tommasini Grotto, Ricardo Haddad, Sandra Coccuzzo Sampaio Vessoni, Simone Kashima, Svetoslav Nanev Slavov, Vincent Louis Viala |
| EPI_ISL_2345870 | HOSPITAL DE CAMPANHA COVID 19 MUNICIPIO DE TAUBATE | Instituto Butantan / Mendelics | Dimas Tadeu Covas, Antonio Jorge Martins, Claudia Renata dos Santos Barros, David Schlesinger, Debora Botequio Moretti, Elaine Cristina Marqueze, Elaine Vieira Santos, Evandra Strazza Rodrigues, Heidge Fukumasu, Jayme Augusto de Souza-Neto, José Salvatore Leister Patané, Luiz Alcantara, Luiz Lehmann Coutinho, Maria Carolina Elias, Maurício Lacerda Nogueira, Rafael dos Santos Bezerra, Raul Machado Neto, Rejane Maria Tommasini Grotto, Ricardo Haddad, Sandra Coccuzzo Sampaio Vessoni, Simone Kashima, Svetoslav Nanev Slavov, Vincent Louis Viala |
| EPI_ISL_2345871 | DEPARTAMENTO DE SAUDE MUNICIPAL SOCORRO SP | Instituto Butantan / Mendelics | Dimas Tadeu Covas, Antonio Jorge Martins, Claudia Renata dos Santos Barros, David Schlesinger, Debora Botequio Moretti, Elaine Cristina Marqueze, Elaine Vieira Santos, Evandra Strazza Rodrigues, Heidge Fukumasu, Jayme Augusto de Souza-Neto, José Salvatore Leister Patané, Luiz Alcantara, Luiz Lehmann Coutinho, Maria Carolina Elias, Maurício Lacerda Nogueira, Rafael dos Santos Bezerra, Raul Machado Neto, Rejane Maria Tommasini Grotto, Ricardo Haddad, Sandra Coccuzzo Sampaio Vessoni, Simone Kashima, Svetoslav Nanev Slavov, Vincent Louis Viala |
| EPI_ISL_2345872 | UPA MARY DOTA | Instituto Butantan / Mendelics | Dimas Tadeu Covas, Antonio Jorge Martins, Claudia Renata dos Santos Barros, David Schlesinger, Debora Botequio Moretti, Elaine Cristina Marqueze, Elaine Vieira Santos, Evandra Strazza Rodrigues, Heidge Fukumasu, Jayme Augusto de Souza-Neto, José Salvatore Leister Patané, Luiz Alcantara, Luiz Lehmann Coutinho, Maria Carolina Elias, Maurício Lacerda Nogueira, Rafael dos Santos Bezerra, Raul Machado Neto, Rejane Maria Tommasini Grotto, Ricardo Haddad, Sandra Coccuzzo Sampaio Vessoni, Simone Kashima, Svetoslav Nanev Slavov, Vincent Louis Viala |
| EPI_ISL_2345873 | CENTRO DE SAUDE DR JOAO JOAQUIM FERREIRA CRISTAIS PAULISTA | Instituto Butantan / Mendelics | Dimas Tadeu Covas, Antonio Jorge Martins, Claudia Renata dos Santos Barros, David Schlesinger, Debora Botequio Moretti, Elaine Cristina Marqueze, Elaine Vieira Santos, Evandra Strazza Rodrigues, Heidge Fukumasu, Jayme Augusto de Souza-Neto, José Salvatore Leister Patané, Luiz Alcantara, Luiz Lehmann Coutinho, Maria Carolina Elias, Maurício Lacerda Nogueira, Rafael dos Santos Bezerra, Raul Machado Neto, Rejane Maria Tommasini Grotto, Ricardo Haddad, Sandra Coccuzzo Sampaio Vessoni, Simone Kashima, Svetoslav Nanev Slavov, Vincent Louis Viala |
| EPI_ISL_2345874 | CENTRO DE APOIO EPIDEMIOLOGICO | Instituto Butantan / Mendelics | Dimas Tadeu Covas, Antonio Jorge Martins, Claudia Renata dos Santos Barros, David Schlesinger, Debora Botequio Moretti, Elaine Cristina Marqueze, Elaine Vieira Santos, Evandra Strazza Rodrigues, Heidge Fukumasu, Jayme Augusto de Souza-Neto, José Salvatore Leister Patané, Luiz Alcantara, Luiz Lehmann Coutinho, Maria Carolina Elias, Maurício Lacerda Nogueira, Rafael dos Santos Bezerra, Raul Machado Neto, Rejane Maria Tommasini Grotto, Ricardo Haddad, Sandra Coccuzzo Sampaio Vessoni, Simone Kashima, Svetoslav Nanev Slavov, Vincent Louis Viala |
| EPI_ISL_2345875 | SECRETARIA MUNICIPAL DA SAUDE DE JOANOPOLIS | Instituto Butantan / Mendelics | Dimas Tadeu Covas, Antonio Jorge Martins, Claudia Renata dos Santos Barros, David Schlesinger, Debora Botequio Moretti, Elaine Cristina Marqueze, Elaine Vieira Santos, Evandra Strazza Rodrigues, Heidge Fukumasu, Jayme Augusto de Souza-Neto, José Salvatore Leister Patané, Luiz Alcantara, Luiz Lehmann Coutinho, Maria Carolina Elias, Maurício Lacerda Nogueira, Rafael dos Santos Bezerra, Raul Machado Neto, Rejane Maria Tommasini Grotto, Ricardo Haddad, Sandra Coccuzzo Sampaio Vessoni, Simone Kashima, Svetoslav Nanev Slavov, Vincent Louis Viala |
| EPI_ISL_2345876 | NUCLEO DE SAUDE MARY DOTA DE BAURU | Instituto Butantan / Mendelics | Dimas Tadeu Covas, Antonio Jorge Martins, Claudia Renata dos Santos Barros, David Schlesinger, Debora Botequio Moretti, Elaine Cristina Marqueze, Elaine Vieira Santos, Evandra Strazza Rodrigues, Heidge Fukumasu, Jayme Augusto de Souza-Neto, José Salvatore Leister Patané, Luiz Alcantara, Luiz Lehmann Coutinho, Maria Carolina Elias, Maurício Lacerda Nogueira, Rafael dos Santos Bezerra, Raul Machado Neto, Rejane Maria Tommasini Grotto, Ricardo Haddad, Sandra Coccuzzo Sampaio Vessoni, Simone Kashima, Svetoslav Nanev Slavov, Vincent Louis Viala |
| EPI_ISL_2345877 | VIGILANCIA EPIDEMIOLOGICA JARDINOPOLIS SP | Instituto Butantan / Mendelics | Dimas Tadeu Covas, Antonio Jorge Martins, Claudia Renata dos Santos Barros, David Schlesinger, Debora Botequio Moretti, Elaine Cristina Marqueze, Elaine Vieira Santos, Evandra Strazza Rodrigues, Heidge Fukumasu, Jayme Augusto de Souza-Neto, José Salvatore Leister Patané, Luiz Alcantara, Luiz Lehmann Coutinho, Maria Carolina Elias, Maurício Lacerda Nogueira, Rafael dos Santos Bezerra, Raul Machado Neto, Rejane Maria Tommasini Grotto, Ricardo Haddad, Sandra Coccuzzo Sampaio Vessoni, Simone Kashima, Svetoslav Nanev Slavov, Vincent Louis Viala |
| EPI_ISL_2345878 | CENTRO DE APOIO EPIDEMIOLOGICO | Instituto Butantan / Mendelics | Dimas Tadeu Covas, Antonio Jorge Martins, Claudia Renata dos Santos Barros, David Schlesinger, Debora Botequio Moretti, Elaine Cristina Marqueze, Elaine Vieira Santos, Evandra Strazza Rodrigues, Heidge Fukumasu, Jayme Augusto de Souza-Neto, José Salvatore Leister Patané, Luiz Alcantara, Luiz Lehmann Coutinho, Maria Carolina Elias, Maurício Lacerda Nogueira, Rafael dos Santos Bezerra, Raul Machado Neto, Rejane Maria Tommasini Grotto, Ricardo Haddad, Sandra Coccuzzo Sampaio Vessoni, Simone Kashima, Svetoslav Nanev Slavov, Vincent Louis Viala |

| | | | |
|---|---|---|---|
| EPI_ISL_2345879 | HOSPITAL MUNICIPAL DE ITAPIRA | Instituto Butantan / Mendelics | Dimas Tadeu Covas, Antonio Jorge Martins, Claudia Renata dos Santos Barros, David Schlesinger, Debora Botequio Moretti, Elaine Cristina Marqueze, Elaine Vieira Santos, Evandra Strazza Rodrigues, Heidge Fukumasu, Jayme Augusto de Souza-Neto, José Salvatore Leister Patané, Luiz Alcantara, Luiz Lehmann Coutinho, Maria Carolina Elias, Maurício Lacerda Nogueira, Rafael dos Santos Bezerra, Raul Machado Neto, Rejane Maria Tommasini Grotto, Ricardo Haddad, Sandra Coccuzzo Sampaio Vessoni, Simone Kashima, Svetoslav Nanev Slavov, Vincent Louis Viala |
| EPI_ISL_2345880 | UPA II OROPO | Instituto Butantan / Mendelics | Dimas Tadeu Covas, Antonio Jorge Martins, Claudia Renata dos Santos Barros, David Schlesinger, Debora Botequio Moretti, Elaine Cristina Marqueze, Elaine Vieira Santos, Evandra Strazza Rodrigues, Heidge Fukumasu, Jayme Augusto de Souza-Neto, José Salvatore Leister Patané, Luiz Alcantara, Luiz Lehmann Coutinho, Maria Carolina Elias, Maurício Lacerda Nogueira, Rafael dos Santos Bezerra, Raul Machado Neto, Rejane Maria Tommasini Grotto, Ricardo Haddad, Sandra Coccuzzo Sampaio Vessoni, Simone Kashima, Svetoslav Nanev Slavov, Vincent Louis Viala |
| EPI_ISL_2345881 | VIGILANCIA SANITARIA E VIG EPIDEMIOLOGICA DE ITAPEVI | Instituto Butantan / FZEA-USP-Pirassununga | Dimas Tadeu Covas, Antonio Jorge Martins, Claudia Renata dos Santos Barros, David Schlesinger, Debora Botequio Moretti, Elaine Cristina Marqueze, Elaine Vieira Santos, Evandra Strazza Rodrigues, Heidge Fukumasu, Jayme Augusto de Souza-Neto, José Salvatore Leister Patané, Luiz Alcantara, Luiz Lehmann Coutinho, Maria Carolina Elias, Maurício Lacerda Nogueira, Rafael dos Santos Bezerra, Raul Machado Neto, Rejane Maria Tommasini Grotto, Ricardo Haddad, Sandra Coccuzzo Sampaio Vessoni, Simone Kashima, Svetoslav Nanev Slavov, Vincent Louis Viala |
| EPI_ISL_2345882 | HOSP DO SERV PUB ESTFCO MORATO DE OLIVEIRA SAO PAULO | Instituto Butantan / FZEA-USP-Pirassununga | Dimas Tadeu Covas, Antonio Jorge Martins, Claudia Renata dos Santos Barros, David Schlesinger, Debora Botequio Moretti, Elaine Cristina Marqueze, Elaine Vieira Santos, Evandra Strazza Rodrigues, Heidge Fukumasu, Jayme Augusto de Souza-Neto, José Salvatore Leister Patané, Luiz Alcantara, Luiz Lehmann Coutinho, Maria Carolina Elias, Maurício Lacerda Nogueira, Rafael dos Santos Bezerra, Raul Machado Neto, Rejane Maria Tommasini Grotto, Ricardo Haddad, Sandra Coccuzzo Sampaio Vessoni, Simone Kashima, Svetoslav Nanev Slavov, Vincent Louis Viala |
| EPI_ISL_2345883, EPI_ISL_2345884, EPI_ISL_2345885 | VIGILANCIA SANITARIA E VIG EPIDEMIOLOGICA DE ITAPEVI | Instituto Butantan / FZEA-USP-Pirassununga | Dimas Tadeu Covas, Antonio Jorge Martins, Claudia Renata dos Santos Barros, David Schlesinger, Debora Botequio Moretti, Elaine Cristina Marqueze, Elaine Vieira Santos, Evandra Strazza Rodrigues, Heidge Fukumasu, Jayme Augusto de Souza-Neto, José Salvatore Leister Patané, Luiz Alcantara, Luiz Lehmann Coutinho, Maria Carolina Elias, Maurício Lacerda Nogueira, Rafael dos Santos Bezerra, Raul Machado Neto, Rejane Maria Tommasini Grotto, Ricardo Haddad, Sandra Coccuzzo Sampaio Vessoni, Simone Kashima, Svetoslav Nanev Slavov, Vincent Louis Viala |
| EPI_ISL_2345886 | HOSP DO SERV PUB ESTFCO MORATO DE OLIVEIRA SAO PAULO | Instituto Butantan / FZEA-USP-Pirassununga | Dimas Tadeu Covas, Antonio Jorge Martins, Claudia Renata dos Santos Barros, David Schlesinger, Debora Botequio Moretti, Elaine Cristina Marqueze, Elaine Vieira Santos, Evandra Strazza Rodrigues, Heidge Fukumasu, Jayme Augusto de Souza-Neto, José Salvatore Leister Patané, Luiz Alcantara, Luiz Lehmann Coutinho, Maria Carolina Elias, Maurício Lacerda Nogueira, Rafael dos Santos Bezerra, Raul Machado Neto, Rejane Maria Tommasini Grotto, Ricardo Haddad, Sandra Coccuzzo Sampaio Vessoni, Simone Kashima, Svetoslav Nanev Slavov, Vincent Louis Viala |
| EPI_ISL_2345887 | VIGILANCIA SANITARIA E VIG EPIDEMIOLOGICA DE ITAPEVI | Instituto Butantan / FZEA-USP-Pirassununga | Dimas Tadeu Covas, Antonio Jorge Martins, Claudia Renata dos Santos Barros, David Schlesinger, Debora Botequio Moretti, Elaine Cristina Marqueze, Elaine Vieira Santos, Evandra Strazza Rodrigues, Heidge Fukumasu, Jayme Augusto de Souza-Neto, José Salvatore Leister Patané, Luiz Alcantara, Luiz Lehmann Coutinho, Maria Carolina Elias, Maurício Lacerda Nogueira, Rafael dos Santos Bezerra, Raul Machado Neto, Rejane Maria Tommasini Grotto, Ricardo Haddad, Sandra Coccuzzo Sampaio Vessoni, Simone Kashima, Svetoslav Nanev Slavov, Vincent Louis Viala |
| EPI_ISL_2345889, EPI_ISL_2345890 | SANTA CASA DE GUARAREMA | Instituto Butantan / FZEA-USP-Pirassununga | Dimas Tadeu Covas, Antonio Jorge Martins, Claudia Renata dos Santos Barros, David Schlesinger, Debora Botequio Moretti, Elaine Cristina Marqueze, Elaine Vieira Santos, Evandra Strazza Rodrigues, Heidge Fukumasu, Jayme Augusto de Souza-Neto, José Salvatore Leister Patané, Luiz Alcantara, Luiz Lehmann Coutinho, Maria Carolina Elias, Maurício Lacerda Nogueira, Rafael dos Santos Bezerra, Raul Machado Neto, Rejane Maria Tommasini Grotto, Ricardo Haddad, Sandra Coccuzzo Sampaio Vessoni, Simone Kashima, Svetoslav Nanev Slavov, Vincent Louis Viala |
| EPI_ISL_2345891 | LABORATORIO LOCAL DE ITAPECERICA DA SERRA | Instituto Butantan / FZEA-USP-Pirassununga | Dimas Tadeu Covas, Antonio Jorge Martins, Claudia Renata dos Santos Barros, David Schlesinger, Debora Botequio Moretti, Elaine Cristina Marqueze, Elaine Vieira Santos, Evandra Strazza Rodrigues, Heidge Fukumasu, Jayme Augusto de Souza-Neto, José Salvatore Leister Patané, Luiz Alcantara, Luiz Lehmann Coutinho, Maria Carolina Elias, Maurício Lacerda Nogueira, Rafael dos Santos Bezerra, Raul Machado Neto, Rejane Maria Tommasini Grotto, Ricardo Haddad, Sandra Coccuzzo Sampaio Vessoni, Simone Kashima, Svetoslav Nanev Slavov, Vincent Louis Viala |
| EPI_ISL_2345892, EPI_ISL_2345893 | UBS DO CIPO EMBU GUACU | Instituto Butantan / FZEA-USP-Pirassununga | Dimas Tadeu Covas, Antonio Jorge Martins, Claudia Renata dos Santos Barros, David Schlesinger, Debora Botequio Moretti, Elaine Cristina Marqueze, Elaine Vieira Santos, Evandra Strazza Rodrigues, Heidge Fukumasu, Jayme Augusto de Souza-Neto, José Salvatore Leister Patané, Luiz Alcantara, Luiz Lehmann Coutinho, Maria Carolina Elias, Maurício Lacerda Nogueira, Rafael dos Santos Bezerra, Raul Machado Neto, Rejane Maria Tommasini Grotto, Ricardo Haddad, Sandra Coccuzzo Sampaio Vessoni, Simone Kashima, Svetoslav Nanev Slavov, Vincent Louis Viala |
| EPI_ISL_2345894 | UBS SAO LOURENCO DA SERRA | Instituto Butantan / FZEA-USP-Pirassununga | Dimas Tadeu Covas, Antonio Jorge Martins, Claudia Renata dos Santos Barros, David Schlesinger, Debora Botequio Moretti, Elaine Cristina Marqueze, Elaine Vieira Santos, Evandra Strazza Rodrigues, Heidge Fukumasu, Jayme Augusto de Souza-Neto, José Salvatore Leister Patané, Luiz Alcantara, Luiz Lehmann Coutinho, Maria Carolina Elias, Maurício Lacerda Nogueira, Rafael dos Santos Bezerra, Raul Machado Neto, Rejane Maria Tommasini Grotto, Ricardo Haddad, Sandra Coccuzzo Sampaio Vessoni, Simone Kashima, Svetoslav Nanev Slavov, Vincent Louis Viala |
| EPI_ISL_2345895, EPI_ISL_2345896 | LABORATORIO LOCAL DE ITAPECERICA DA SERRA | Instituto Butantan / FZEA-USP-Pirassununga | Dimas Tadeu Covas, Antonio Jorge Martins, Claudia Renata dos Santos Barros, David Schlesinger, Debora Botequio Moretti, Elaine Cristina Marqueze, Elaine Vieira Santos, Evandra Strazza Rodrigues, Heidge Fukumasu, Jayme Augusto de Souza-Neto, José Salvatore Leister Patané, Luiz Alcantara, Luiz Lehmann Coutinho, Maria Carolina Elias, Maurício Lacerda Nogueira, Rafael dos Santos Bezerra, Raul Machado Neto, Rejane Maria Tommasini Grotto, Ricardo Haddad, Sandra Coccuzzo Sampaio Vessoni, Simone Kashima, Svetoslav Nanev Slavov, Vincent Louis Viala |
| EPI_ISL_2345897 | UBS SAO LOURENCO DA SERRA | Instituto Butantan / FZEA-USP-Pirassununga | Dimas Tadeu Covas, Antonio Jorge Martins, Claudia Renata dos Santos Barros, David Schlesinger, Debora Botequio Moretti, Elaine Cristina Marqueze, Elaine Vieira Santos, Evandra Strazza Rodrigues, Heidge Fukumasu, Jayme Augusto de Souza-Neto, José Salvatore Leister Patané, Luiz Alcantara, Luiz Lehmann Coutinho, Maria Carolina Elias, Maurício Lacerda Nogueira, Rafael dos Santos Bezerra, Raul Machado Neto, Rejane Maria Tommasini Grotto, Ricardo Haddad, Sandra Coccuzzo Sampaio Vessoni, Simone Kashima, Svetoslav Nanev Slavov, Vincent Louis Viala |
| EPI_ISL_2345898, EPI_ISL_2345899 | LABORATORIO LOCAL DE ITAPECERICA DA SERRA | Instituto Butantan / FZEA-USP-Pirassununga | Dimas Tadeu Covas, Antonio Jorge Martins, Claudia Renata dos Santos Barros, David Schlesinger, Debora Botequio Moretti, Elaine Cristina Marqueze, Elaine Vieira Santos, Evandra Strazza Rodrigues, Heidge Fukumasu, Jayme Augusto de Souza-Neto, José Salvatore Leister Patané, Luiz Alcantara, Luiz Lehmann Coutinho, Maria Carolina Elias, Maurício Lacerda Nogueira, Rafael dos Santos Bezerra, Raul Machado Neto, Rejane Maria Tommasini Grotto, Ricardo Haddad, Sandra Coccuzzo Sampaio Vessoni, Simone Kashima, Svetoslav Nanev Slavov, Vincent Louis Viala |
| EPI_ISL_2345900 | UBS DO CIPO EMBU GUACU | Instituto Butantan / FZEA-USP-Pirassununga | Dimas Tadeu Covas, Antonio Jorge Martins, Claudia Renata dos Santos Barros, David Schlesinger, Debora Botequio Moretti, Elaine Cristina Marqueze, Elaine Vieira Santos, Evandra Strazza Rodrigues, Heidge Fukumasu, Jayme Augusto de Souza-Neto, José Salvatore Leister Patané, Luiz Alcantara, Luiz Lehmann Coutinho, Maria Carolina Elias, Maurício Lacerda Nogueira, Rafael dos Santos Bezerra, Raul Machado Neto, Rejane Maria Tommasini Grotto, Ricardo Haddad, Sandra Coccuzzo Sampaio Vessoni, Simone Kashima, Svetoslav Nanev Slavov, Vincent Louis Viala |
| EPI_ISL_2345901, EPI_ISL_2345903 | UBS SAO LOURENCO DA SERRA | Instituto Butantan / FZEA-USP-Pirassununga | Dimas Tadeu Covas, Antonio Jorge Martins, Claudia Renata dos Santos Barros, David Schlesinger, Debora Botequio Moretti, Elaine Cristina Marqueze, Elaine Vieira Santos, Evandra Strazza Rodrigues, Heidge Fukumasu, Jayme Augusto de Souza-Neto, José Salvatore Leister Patané, Luiz Alcantara, Luiz Lehmann Coutinho, Maria Carolina Elias, Maurício Lacerda Nogueira, Rafael dos Santos Bezerra, Raul Machado Neto, Rejane Maria Tommasini Grotto, Ricardo Haddad, Sandra Coccuzzo Sampaio Vessoni, Simone Kashima, Svetoslav Nanev Slavov, Vincent Louis Viala |
| EPI_ISL_2345904 | PRONTO SOCORRO JARDIM JACIRA | Instituto Butantan / FZEA-USP-Pirassununga | Dimas Tadeu Covas, Antonio Jorge Martins, Claudia Renata dos Santos Barros, David Schlesinger, Debora Botequio Moretti, Elaine Cristina Marqueze, Elaine Vieira Santos, Evandra Strazza Rodrigues, Heidge Fukumasu, Jayme Augusto de Souza-Neto, José Salvatore Leister Patané, Luiz Alcantara, Luiz Lehmann Coutinho, Maria Carolina Elias, Maurício Lacerda Nogueira, Rafael dos Santos Bezerra, Raul Machado Neto, Rejane Maria Tommasini Grotto, Ricardo Haddad, Sandra Coccuzzo Sampaio Vessoni, Simone Kashima, Svetoslav Nanev Slavov, Vincent Louis Viala |
| EPI_ISL_2345905 | LABORATORIO LOCAL DE ITAPECERICA DA SERRA | Instituto Butantan / FZEA-USP-Pirassununga | Dimas Tadeu Covas, Antonio Jorge Martins, Claudia Renata dos Santos Barros, David Schlesinger, Debora Botequio Moretti, Elaine Cristina Marqueze, Elaine Vieira Santos, Evandra Strazza Rodrigues, Heidge Fukumasu, Jayme Augusto de Souza-Neto, José Salvatore Leister Patané, Luiz Alcantara, Luiz Lehmann Coutinho, Maria Carolina Elias, Maurício Lacerda Nogueira, Rafael dos Santos Bezerra, Raul Machado Neto, Rejane Maria Tommasini Grotto, Ricardo Haddad, Sandra Coccuzzo Sampaio Vessoni, Simone Kashima, Svetoslav Nanev Slavov, Vincent Louis Viala |
| EPI_ISL_2345906, EPI_ISL_2345907, EPI_ISL_2345908, EPI_ISL_2345909, EPI_ISL_2345910, EPI_ISL_2345911, EPI_ISL_2345912, EPI_ISL_2345913, EPI_ISL_2345914, EPI_ISL_2345915, EPI_ISL_2345916, EPI_ISL_2345917, EPI_ISL_2345918, EPI_ISL_2345920, EPI_ISL_2345921, EPI_ISL_2345922, EPI_ISL_2345923, EPI_ISL_2345924, EPI_ISL_2345925, EPI_ISL_2345926, EPI_ISL_2345927, EPI_ISL_2345928, EPI_ISL_2345930, EPI_ISL_2345931, EPI_ISL_2345932, EPI_ISL_2345933, EPI_ISL_2345934, EPI_ISL_2345935, EPI_ISL_2345936, EPI_ISL_2345937, EPI_ISL_2345938 | | | |
| see above | Prefeirtura de SP | Instituto Butantan | Dimas Tadeu Covas, Antonio Jorge Martins, Claudia Renata dos Santos Barros, David Schlesinger, Debora Botequio Moretti, Elaine Cristina Marqueze, Elaine Vieira Santos, Evandra Strazza Rodrigues, Heidge Fukumasu, Jayme Augusto de Souza-Neto, José Salvatore Leister Patané, Luiz Alcantara, Luiz Lehmann Coutinho, Maria Carolina Elias, Maurício Lacerda Nogueira, Rafael dos Santos Bezerra, Raul Machado Neto, Rejane Maria Tommasini Grotto, Ricardo Haddad, Sandra Coccuzzo Sampaio Vessoni, Simone Kashima, Svetoslav Nanev Slavov, Vincent Louis Viala |

| | | | |
|---|---|---|---|
| EPI_ISL_2345940 | AMBULATORIO MEDICO MUNICIPAL DE AGUDOS | Instituto Butantan / ESALQ-Piracicaba | Dimas Tadeu Covas, Antonio Jorge Martins, Claudia Renata dos Santos Barros, David Schlesinger, Debora Botequio Moretti, Elaine Cristina Marqueze, Elaine Vieira Santos, Evandra Strazza Rodrigues, Heidge Fukumasu, Jayme Augusto de Souza-Neto, José Salvatore Leister Patané, Luiz Alcantara, Luiz Lehmann Coutinho, Maria Carolina Elias, Maurício Lacerda Nogueira, Rafael dos Santos Bezerra, Raul Machado Neto, Rejane Maria Tommasini Grotto, Ricardo Haddad, Sandra Coccuzzo Sampaio Vessoni, Simone Kashima, Svetoslav Nanev Slavov, Vincent Louis Viala |
| EPI_ISL_2345941 | CENTRO INTEGRADO DE SAUDE | Instituto Butantan / ESALQ-Piracicaba | Dimas Tadeu Covas, Antonio Jorge Martins, Claudia Renata dos Santos Barros, David Schlesinger, Debora Botequio Moretti, Elaine Cristina Marqueze, Elaine Vieira Santos, Evandra Strazza Rodrigues, Heidge Fukumasu, Jayme Augusto de Souza-Neto, José Salvatore Leister Patané, Luiz Alcantara, Luiz Lehmann Coutinho, Maria Carolina Elias, Maurício Lacerda Nogueira, Rafael dos Santos Bezerra, Raul Machado Neto, Rejane Maria Tommasini Grotto, Ricardo Haddad, Sandra Coccuzzo Sampaio Vessoni, Simone Kashima, Svetoslav Nanev Slavov, Vincent Louis Viala |
| EPI_ISL_2345942 | AMBULATORIO MEDICO MUNICIPAL DE AGUDOS | Instituto Butantan / ESALQ-Piracicaba | Dimas Tadeu Covas, Antonio Jorge Martins, Claudia Renata dos Santos Barros, David Schlesinger, Debora Botequio Moretti, Elaine Cristina Marqueze, Elaine Vieira Santos, Evandra Strazza Rodrigues, Heidge Fukumasu, Jayme Augusto de Souza-Neto, José Salvatore Leister Patané, Luiz Alcantara, Luiz Lehmann Coutinho, Maria Carolina Elias, Maurício Lacerda Nogueira, Rafael dos Santos Bezerra, Raul Machado Neto, Rejane Maria Tommasini Grotto, Ricardo Haddad, Sandra Coccuzzo Sampaio Vessoni, Simone Kashima, Svetoslav Nanev Slavov, Vincent Louis Viala |
| EPI_ISL_2345943, EPI_ISL_2345944 | CENTRO INTEGRADO DE SAUDE | Instituto Butantan / ESALQ-Piracicaba | Dimas Tadeu Covas, Antonio Jorge Martins, Claudia Renata dos Santos Barros, David Schlesinger, Debora Botequio Moretti, Elaine Cristina Marqueze, Elaine Vieira Santos, Evandra Strazza Rodrigues, Heidge Fukumasu, Jayme Augusto de Souza-Neto, José Salvatore Leister Patané, Luiz Alcantara, Luiz Lehmann Coutinho, Maria Carolina Elias, Maurício Lacerda Nogueira, Rafael dos Santos Bezerra, Raul Machado Neto, Rejane Maria Tommasini Grotto, Ricardo Haddad, Sandra Coccuzzo Sampaio Vessoni, Simone Kashima, Svetoslav Nanev Slavov, Vincent Louis Viala |
| EPI_ISL_2345945, EPI_ISL_2345946 | HOSPITAL SANTA THEREZINHA BROTAS | Instituto Butantan / ESALQ-Piracicaba | Dimas Tadeu Covas, Antonio Jorge Martins, Claudia Renata dos Santos Barros, David Schlesinger, Debora Botequio Moretti, Elaine Cristina Marqueze, Elaine Vieira Santos, Evandra Strazza Rodrigues, Heidge Fukumasu, Jayme Augusto de Souza-Neto, José Salvatore Leister Patané, Luiz Alcantara, Luiz Lehmann Coutinho, Maria Carolina Elias, Maurício Lacerda Nogueira, Rafael dos Santos Bezerra, Raul Machado Neto, Rejane Maria Tommasini Grotto, Ricardo Haddad, Sandra Coccuzzo Sampaio Vessoni, Simone Kashima, Svetoslav Nanev Slavov, Vincent Louis Viala |
| EPI_ISL_2345947 | SECRETARIA MUNICIPAL DE SAUDE DE PEDERNEIRAS | Instituto Butantan / ESALQ-Piracicaba | Dimas Tadeu Covas, Antonio Jorge Martins, Claudia Renata dos Santos Barros, David Schlesinger, Debora Botequio Moretti, Elaine Cristina Marqueze, Elaine Vieira Santos, Evandra Strazza Rodrigues, Heidge Fukumasu, Jayme Augusto de Souza-Neto, José Salvatore Leister Patané, Luiz Alcantara, Luiz Lehmann Coutinho, Maria Carolina Elias, Maurício Lacerda Nogueira, Rafael dos Santos Bezerra, Raul Machado Neto, Rejane Maria Tommasini Grotto, Ricardo Haddad, Sandra Coccuzzo Sampaio Vessoni, Simone Kashima, Svetoslav Nanev Slavov, Vincent Louis Viala |
| EPI_ISL_2345948 | NUCLEO SAUDE III N HABITACIONAL PRESIDENTE GEISEL BAURU | Instituto Butantan / ESALQ-Piracicaba | Dimas Tadeu Covas, Antonio Jorge Martins, Claudia Renata dos Santos Barros, David Schlesinger, Debora Botequio Moretti, Elaine Cristina Marqueze, Elaine Vieira Santos, Evandra Strazza Rodrigues, Heidge Fukumasu, Jayme Augusto de Souza-Neto, José Salvatore Leister Patané, Luiz Alcantara, Luiz Lehmann Coutinho, Maria Carolina Elias, Maurício Lacerda Nogueira, Rafael dos Santos Bezerra, Raul Machado Neto, Rejane Maria Tommasini Grotto, Ricardo Haddad, Sandra Coccuzzo Sampaio Vessoni, Simone Kashima, Svetoslav Nanev Slavov, Vincent Louis Viala |
| EPI_ISL_2345949 | SANTA CASA DE DOIS CORREGOS | Instituto Butantan / ESALQ-Piracicaba | Dimas Tadeu Covas, Antonio Jorge Martins, Claudia Renata dos Santos Barros, David Schlesinger, Debora Botequio Moretti, Elaine Cristina Marqueze, Elaine Vieira Santos, Evandra Strazza Rodrigues, Heidge Fukumasu, Jayme Augusto de Souza-Neto, José Salvatore Leister Patané, Luiz Alcantara, Luiz Lehmann Coutinho, Maria Carolina Elias, Maurício Lacerda Nogueira, Rafael dos Santos Bezerra, Raul Machado Neto, Rejane Maria Tommasini Grotto, Ricardo Haddad, Sandra Coccuzzo Sampaio Vessoni, Simone Kashima, Svetoslav Nanev Slavov, Vincent Louis Viala |
| EPI_ISL_2345950 | NUCLEO SAUDE III N HABITACIONAL PRESIDENTE GEISEL BAURU | Instituto Butantan / ESALQ-Piracicaba | Dimas Tadeu Covas, Antonio Jorge Martins, Claudia Renata dos Santos Barros, David Schlesinger, Debora Botequio Moretti, Elaine Cristina Marqueze, Elaine Vieira Santos, Evandra Strazza Rodrigues, Heidge Fukumasu, Jayme Augusto de Souza-Neto, José Salvatore Leister Patané, Luiz Alcantara, Luiz Lehmann Coutinho, Maria Carolina Elias, Maurício Lacerda Nogueira, Rafael dos Santos Bezerra, Raul Machado Neto, Rejane Maria Tommasini Grotto, Ricardo Haddad, Sandra Coccuzzo Sampaio Vessoni, Simone Kashima, Svetoslav Nanev Slavov, Vincent Louis Viala |
| EPI_ISL_2345951 | SANTA CASA DE DOIS CORREGOS | Instituto Butantan / ESALQ-Piracicaba | Dimas Tadeu Covas, Antonio Jorge Martins, Claudia Renata dos Santos Barros, David Schlesinger, Debora Botequio Moretti, Elaine Cristina Marqueze, Elaine Vieira Santos, Evandra Strazza Rodrigues, Heidge Fukumasu, Jayme Augusto de Souza-Neto, José Salvatore Leister Patané, Luiz Alcantara, Luiz Lehmann Coutinho, Maria Carolina Elias, Maurício Lacerda Nogueira, Rafael dos Santos Bezerra, Raul Machado Neto, Rejane Maria Tommasini Grotto, Ricardo Haddad, Sandra Coccuzzo Sampaio Vessoni, Simone Kashima, Svetoslav Nanev Slavov, Vincent Louis Viala |
| EPI_ISL_2345952, EPI_ISL_2345953, EPI_ISL_2345954 | ESF MINEIROS DO TIETE | Instituto Butantan / ESALQ-Piracicaba | Dimas Tadeu Covas, Antonio Jorge Martins, Claudia Renata dos Santos Barros, David Schlesinger, Debora Botequio Moretti, Elaine Cristina Marqueze, Elaine Vieira Santos, Evandra Strazza Rodrigues, Heidge Fukumasu, Jayme Augusto de Souza-Neto, José Salvatore Leister Patané, Luiz Alcantara, Luiz Lehmann Coutinho, Maria Carolina Elias, Maurício Lacerda Nogueira, Rafael dos Santos Bezerra, Raul Machado Neto, Rejane Maria Tommasini Grotto, Ricardo Haddad, Sandra Coccuzzo Sampaio Vessoni, Simone Kashima, Svetoslav Nanev Slavov, Vincent Louis Viala |
| EPI_ISL_2345955 | MARIA TEREZINHA BOGNAR CHECHETO | Instituto Butantan / ESALQ-Piracicaba | Dimas Tadeu Covas, Antonio Jorge Martins, Claudia Renata dos Santos Barros, David Schlesinger, Debora Botequio Moretti, Elaine Cristina Marqueze, Elaine Vieira Santos, Evandra Strazza Rodrigues, Heidge Fukumasu, Jayme Augusto de Souza-Neto, José Salvatore Leister Patané, Luiz Alcantara, Luiz Lehmann Coutinho, Maria Carolina Elias, Maurício Lacerda Nogueira, Rafael dos Santos Bezerra, Raul Machado Neto, Rejane Maria Tommasini Grotto, Ricardo Haddad, Sandra Coccuzzo Sampaio Vessoni, Simone Kashima, Svetoslav Nanev Slavov, Vincent Louis Viala |
| EPI_ISL_2345956 | NUCLEO DE SAUDE VILA FALCAO DE BAURU | Instituto Butantan / ESALQ-Piracicaba | Dimas Tadeu Covas, Antonio Jorge Martins, Claudia Renata dos Santos Barros, David Schlesinger, Debora Botequio Moretti, Elaine Cristina Marqueze, Elaine Vieira Santos, Evandra Strazza Rodrigues, Heidge Fukumasu, Jayme Augusto de Souza-Neto, José Salvatore Leister Patané, Luiz Alcantara, Luiz Lehmann Coutinho, Maria Carolina Elias, Maurício Lacerda Nogueira, Rafael dos Santos Bezerra, Raul Machado Neto, Rejane Maria Tommasini Grotto, Ricardo Haddad, Sandra Coccuzzo Sampaio Vessoni, Simone Kashima, Svetoslav Nanev Slavov, Vincent Louis Viala |
| EPI_ISL_2345957 | HOSPITAL MATERNIDADE SAO JOSE ITAPUI | Instituto Butantan / ESALQ-Piracicaba | Dimas Tadeu Covas, Antonio Jorge Martins, Claudia Renata dos Santos Barros, David Schlesinger, Debora Botequio Moretti, Elaine Cristina Marqueze, Elaine Vieira Santos, Evandra Strazza Rodrigues, Heidge Fukumasu, Jayme Augusto de Souza-Neto, José Salvatore Leister Patané, Luiz Alcantara, Luiz Lehmann Coutinho, Maria Carolina Elias, Maurício Lacerda Nogueira, Rafael dos Santos Bezerra, Raul Machado Neto, Rejane Maria Tommasini Grotto, Ricardo Haddad, Sandra Coccuzzo Sampaio Vessoni, Simone Kashima, Svetoslav Nanev Slavov, Vincent Louis Viala |
| EPI_ISL_2345958 | SECRETARIA MUNICIPAL DE SAUDE DE MACATUBA | Instituto Butantan / ESALQ-Piracicaba | Dimas Tadeu Covas, Antonio Jorge Martins, Claudia Renata dos Santos Barros, David Schlesinger, Debora Botequio Moretti, Elaine Cristina Marqueze, Elaine Vieira Santos, Evandra Strazza Rodrigues, Heidge Fukumasu, Jayme Augusto de Souza-Neto, José Salvatore Leister Patané, Luiz Alcantara, Luiz Lehmann Coutinho, Maria Carolina Elias, Maurício Lacerda Nogueira, Rafael dos Santos Bezerra, Raul Machado Neto, Rejane Maria Tommasini Grotto, Ricardo Haddad, Sandra Coccuzzo Sampaio Vessoni, Simone Kashima, Svetoslav Nanev Slavov, Vincent Louis Viala |
| EPI_ISL_2345959 | USF JARDIM DO LAGO ARTUR NOGUEIRA | Instituto Butantan | Dimas Tadeu Covas, Antonio Jorge Martins, Claudia Renata dos Santos Barros, David Schlesinger, Debora Botequio Moretti, Elaine Cristina Marqueze, Elaine Vieira Santos, Evandra Strazza Rodrigues, Heidge Fukumasu, Jayme Augusto de Souza-Neto, José Salvatore Leister Patané, Luiz Alcantara, Luiz Lehmann Coutinho, Maria Carolina Elias, Maurício Lacerda Nogueira, Rafael dos Santos Bezerra, Raul Machado Neto, Rejane Maria Tommasini Grotto, Ricardo Haddad, Sandra Coccuzzo Sampaio Vessoni, Simone Kashima, Svetoslav Nanev Slavov, Vincent Louis Viala |
| EPI_ISL_2345960 | PSF PARQUE EMILIA | Instituto Butantan | Dimas Tadeu Covas, Antonio Jorge Martins, Claudia Renata dos Santos Barros, David Schlesinger, Debora Botequio Moretti, Elaine Cristina Marqueze, Elaine Vieira Santos, Evandra Strazza Rodrigues, Heidge Fukumasu, Jayme Augusto de Souza-Neto, José Salvatore Leister Patané, Luiz Alcantara, Luiz Lehmann Coutinho, Maria Carolina Elias, Maurício Lacerda Nogueira, Rafael dos Santos Bezerra, Raul Machado Neto, Rejane Maria Tommasini Grotto, Ricardo Haddad, Sandra Coccuzzo Sampaio Vessoni, Simone Kashima, Svetoslav Nanev Slavov, Vincent Louis Viala |
| EPI_ISL_2345961 | COMPLEXO HOSPITALAR OURO VERDE DE CAMPINAS | Instituto Butantan | Dimas Tadeu Covas, Antonio Jorge Martins, Claudia Renata dos Santos Barros, David Schlesinger, Debora Botequio Moretti, Elaine Cristina Marqueze, Elaine Vieira Santos, Evandra Strazza Rodrigues, Heidge Fukumasu, Jayme Augusto de Souza-Neto, José Salvatore Leister Patané, Luiz Alcantara, Luiz Lehmann Coutinho, Maria Carolina Elias, Maurício Lacerda Nogueira, Rafael dos Santos Bezerra, Raul Machado Neto, Rejane Maria Tommasini Grotto, Ricardo Haddad, Sandra Coccuzzo Sampaio Vessoni, Simone Kashima, Svetoslav Nanev Slavov, Vincent Louis Viala |
| EPI_ISL_2345962, EPI_ISL_2345963 | PRONTO ATENDIMENTO DR SERGIO AROUCA CAMPO GRANDE | Instituto Butantan | Dimas Tadeu Covas, Antonio Jorge Martins, Claudia Renata dos Santos Barros, David Schlesinger, Debora Botequio Moretti, Elaine Cristina Marqueze, Elaine Vieira Santos, Evandra Strazza Rodrigues, Heidge Fukumasu, Jayme Augusto de Souza-Neto, José Salvatore Leister Patané, Luiz Alcantara, Luiz Lehmann Coutinho, Maria Carolina Elias, Maurício Lacerda Nogueira, Rafael dos Santos Bezerra, Raul Machado Neto, Rejane Maria Tommasini Grotto, Ricardo Haddad, Sandra Coccuzzo Sampaio Vessoni, Simone Kashima, Svetoslav Nanev Slavov, Vincent Louis Viala |
| EPI_ISL_2345964 | HOSPITAL MUNICIPAL DR MARIO GATTI CAMPINAS | Instituto Butantan | Dimas Tadeu Covas, Antonio Jorge Martins, Claudia Renata dos Santos Barros, David Schlesinger, Debora Botequio Moretti, Elaine Cristina Marqueze, Elaine Vieira Santos, Evandra Strazza Rodrigues, Heidge Fukumasu, Jayme Augusto de Souza-Neto, José Salvatore Leister Patané, Luiz Alcantara, Luiz |

| | | | |
|---|---|---|---|
| | | | Lehmann Coutinho, Maria Carolina Elias, Maurício Lacerda Nogueira, Rafael dos Santos Bezerra, Raul Machado Neto, Rejane Maria Tommasini Grotto, Ricardo Haddad, Sandra Coccuzzo Sampaio Vessoni, Simone Kashima, Svetoslav Nanev Slavov, Vincent Louis Viala |
| EPI_ISL_2345965, EPI_ISL_2345966 | PA NOVO OSASCO | Instituto Butantan | Dimas Tadeu Covas, Antonio Jorge Martins, Claudia Renata dos Santos Barros, David Schlesinger, Debora Botequio Moretti, Elaine Cristina Marqueze, Elaine Vieira Santos, Evandra Strazza Rodrigues, Heidge Fukumasu, Jayme Augusto de Souza-Neto, José Salvatore Leister Patané, Luiz Alcantara, Luiz Lehmann Coutinho, Maria Carolina Elias, Maurício Lacerda Nogueira, Rafael dos Santos Bezerra, Raul Machado Neto, Rejane Maria Tommasini Grotto, Ricardo Haddad, Sandra Coccuzzo Sampaio Vessoni, Simone Kashima, Svetoslav Nanev Slavov, Vincent Louis Viala |
| EPI_ISL_2345967 | UBS SANTA MARIA | Instituto Butantan | Dimas Tadeu Covas, Antonio Jorge Martins, Claudia Renata dos Santos Barros, David Schlesinger, Debora Botequio Moretti, Elaine Cristina Marqueze, Elaine Vieira Santos, Evandra Strazza Rodrigues, Heidge Fukumasu, Jayme Augusto de Souza-Neto, José Salvatore Leister Patané, Luiz Alcantara, Luiz Lehmann Coutinho, Maria Carolina Elias, Maurício Lacerda Nogueira, Rafael dos Santos Bezerra, Raul Machado Neto, Rejane Maria Tommasini Grotto, Ricardo Haddad, Sandra Coccuzzo Sampaio Vessoni, Simone Kashima, Svetoslav Nanev Slavov, Vincent Louis Viala |
| EPI_ISL_2345968 | SECRETARIA MUNICIPAL DE SAUDE DE MORUNGABA | Instituto Butantan | Dimas Tadeu Covas, Antonio Jorge Martins, Claudia Renata dos Santos Barros, David Schlesinger, Debora Botequio Moretti, Elaine Cristina Marqueze, Elaine Vieira Santos, Evandra Strazza Rodrigues, Heidge Fukumasu, Jayme Augusto de Souza-Neto, José Salvatore Leister Patané, Luiz Alcantara, Luiz Lehmann Coutinho, Maria Carolina Elias, Maurício Lacerda Nogueira, Rafael dos Santos Bezerra, Raul Machado Neto, Rejane Maria Tommasini Grotto, Ricardo Haddad, Sandra Coccuzzo Sampaio Vessoni, Simone Kashima, Svetoslav Nanev Slavov, Vincent Louis Viala |
| EPI_ISL_2345969 | UBS SANTA MARIA | Instituto Butantan | Dimas Tadeu Covas, Antonio Jorge Martins, Claudia Renata dos Santos Barros, David Schlesinger, Debora Botequio Moretti, Elaine Cristina Marqueze, Elaine Vieira Santos, Evandra Strazza Rodrigues, Heidge Fukumasu, Jayme Augusto de Souza-Neto, José Salvatore Leister Patané, Luiz Alcantara, Luiz Lehmann Coutinho, Maria Carolina Elias, Maurício Lacerda Nogueira, Rafael dos Santos Bezerra, Raul Machado Neto, Rejane Maria Tommasini Grotto, Ricardo Haddad, Sandra Coccuzzo Sampaio Vessoni, Simone Kashima, Svetoslav Nanev Slavov, Vincent Louis Viala |
| EPI_ISL_2345970 | UBS IRMA AGUEDA MARIA JAIME | Instituto Butantan | Dimas Tadeu Covas, Antonio Jorge Martins, Claudia Renata dos Santos Barros, David Schlesinger, Debora Botequio Moretti, Elaine Cristina Marqueze, Elaine Vieira Santos, Evandra Strazza Rodrigues, Heidge Fukumasu, Jayme Augusto de Souza-Neto, José Salvatore Leister Patané, Luiz Alcantara, Luiz Lehmann Coutinho, Maria Carolina Elias, Maurício Lacerda Nogueira, Rafael dos Santos Bezerra, Raul Machado Neto, Rejane Maria Tommasini Grotto, Ricardo Haddad, Sandra Coccuzzo Sampaio Vessoni, Simone Kashima, Svetoslav Nanev Slavov, Vincent Louis Viala |
| EPI_ISL_2345971 | UBS SANTA MARIA | Instituto Butantan | Dimas Tadeu Covas, Antonio Jorge Martins, Claudia Renata dos Santos Barros, David Schlesinger, Debora Botequio Moretti, Elaine Cristina Marqueze, Elaine Vieira Santos, Evandra Strazza Rodrigues, Heidge Fukumasu, Jayme Augusto de Souza-Neto, José Salvatore Leister Patané, Luiz Alcantara, Luiz Lehmann Coutinho, Maria Carolina Elias, Maurício Lacerda Nogueira, Rafael dos Santos Bezerra, Raul Machado Neto, Rejane Maria Tommasini Grotto, Ricardo Haddad, Sandra Coccuzzo Sampaio Vessoni, Simone Kashima, Svetoslav Nanev Slavov, Vincent Louis Viala |
| EPI_ISL_2345972 | HOSP DO SERV PUB ESTFCO MORATO DE OLIVEIRA SAO PAULO | Instituto Butantan | Dimas Tadeu Covas, Antonio Jorge Martins, Claudia Renata dos Santos Barros, David Schlesinger, Debora Botequio Moretti, Elaine Cristina Marqueze, Elaine Vieira Santos, Evandra Strazza Rodrigues, Heidge Fukumasu, Jayme Augusto de Souza-Neto, José Salvatore Leister Patané, Luiz Alcantara, Luiz Lehmann Coutinho, Maria Carolina Elias, Maurício Lacerda Nogueira, Rafael dos Santos Bezerra, Raul Machado Neto, Rejane Maria Tommasini Grotto, Ricardo Haddad, Sandra Coccuzzo Sampaio Vessoni, Simone Kashima, Svetoslav Nanev Slavov, Vincent Louis Viala |
| EPI_ISL_2345973, EPI_ISL_2345975 | VIGILANCIA SANITARIA E VIG EPIDEMIOLOGICA DE ITAPEVI | Instituto Butantan | Dimas Tadeu Covas, Antonio Jorge Martins, Claudia Renata dos Santos Barros, David Schlesinger, Debora Botequio Moretti, Elaine Cristina Marqueze, Elaine Vieira Santos, Evandra Strazza Rodrigues, Heidge Fukumasu, Jayme Augusto de Souza-Neto, José Salvatore Leister Patané, Luiz Alcantara, Luiz Lehmann Coutinho, Maria Carolina Elias, Maurício Lacerda Nogueira, Rafael dos Santos Bezerra, Raul Machado Neto, Rejane Maria Tommasini Grotto, Ricardo Haddad, Sandra Coccuzzo Sampaio Vessoni, Simone Kashima, Svetoslav Nanev Slavov, Vincent Louis Viala |
| EPI_ISL_2345976 | UBS JARDIM PRIMAVERA | Instituto Butantan | Dimas Tadeu Covas, Antonio Jorge Martins, Claudia Renata dos Santos Barros, David Schlesinger, Debora Botequio Moretti, Elaine Cristina Marqueze, Elaine Vieira Santos, Evandra Strazza Rodrigues, Heidge Fukumasu, Jayme Augusto de Souza-Neto, José Salvatore Leister Patané, Luiz Alcantara, Luiz Lehmann Coutinho, Maria Carolina Elias, Maurício Lacerda Nogueira, Rafael dos Santos Bezerra, Raul Machado Neto, Rejane Maria Tommasini Grotto, Ricardo Haddad, Sandra Coccuzzo Sampaio Vessoni, Simone Kashima, Svetoslav Nanev Slavov, Vincent Louis Viala |
| EPI_ISL_2345977 | UBS MARINOPOLIS | Instituto Butantan | Dimas Tadeu Covas, Antonio Jorge Martins, Claudia Renata dos Santos Barros, David Schlesinger, Debora Botequio Moretti, Elaine Cristina Marqueze, Elaine Vieira Santos, Evandra Strazza Rodrigues, Heidge Fukumasu, Jayme Augusto de Souza-Neto, José Salvatore Leister Patané, Luiz Alcantara, Luiz Lehmann Coutinho, Maria Carolina Elias, Maurício Lacerda Nogueira, Rafael dos Santos Bezerra, Raul Machado Neto, Rejane Maria Tommasini Grotto, Ricardo Haddad, Sandra Coccuzzo Sampaio Vessoni, Simone Kashima, Svetoslav Nanev Slavov, Vincent Louis Viala |
| EPI_ISL_2345978 | POLICLINICA JARDIM PARAISO | Instituto Butantan | Dimas Tadeu Covas, Antonio Jorge Martins, Claudia Renata dos Santos Barros, David Schlesinger, Debora Botequio Moretti, Elaine Cristina Marqueze, Elaine Vieira Santos, Evandra Strazza Rodrigues, Heidge Fukumasu, Jayme Augusto de Souza-Neto, José Salvatore Leister Patané, Luiz Alcantara, Luiz Lehmann Coutinho, Maria Carolina Elias, Maurício Lacerda Nogueira, Rafael dos Santos Bezerra, Raul Machado Neto, Rejane Maria Tommasini Grotto, Ricardo Haddad, Sandra Coccuzzo Sampaio Vessoni, Simone Kashima, Svetoslav Nanev Slavov, Vincent Louis Viala |
| EPI_ISL_2345979 | USF SACILOTTO ARTUR NOGUEIRA | Instituto Butantan | Dimas Tadeu Covas, Antonio Jorge Martins, Claudia Renata dos Santos Barros, David Schlesinger, Debora Botequio Moretti, Elaine Cristina Marqueze, Elaine Vieira Santos, Evandra Strazza Rodrigues, Heidge Fukumasu, Jayme Augusto de Souza-Neto, José Salvatore Leister Patané, Luiz Alcantara, Luiz Lehmann Coutinho, Maria Carolina Elias, Maurício Lacerda Nogueira, Rafael dos Santos Bezerra, Raul Machado Neto, Rejane Maria Tommasini Grotto, Ricardo Haddad, Sandra Coccuzzo Sampaio Vessoni, Simone Kashima, Svetoslav Nanev Slavov, Vincent Louis Viala |
| EPI_ISL_2345980 | POLICLINICA JARDIM PARAISO | Instituto Butantan | Dimas Tadeu Covas, Antonio Jorge Martins, Claudia Renata dos Santos Barros, David Schlesinger, Debora Botequio Moretti, Elaine Cristina Marqueze, Elaine Vieira Santos, Evandra Strazza Rodrigues, Heidge Fukumasu, Jayme Augusto de Souza-Neto, José Salvatore Leister Patané, Luiz Alcantara, Luiz Lehmann Coutinho, Maria Carolina Elias, Maurício Lacerda Nogueira, Rafael dos Santos Bezerra, Raul Machado Neto, Rejane Maria Tommasini Grotto, Ricardo Haddad, Sandra Coccuzzo Sampaio Vessoni, Simone Kashima, Svetoslav Nanev Slavov, Vincent Louis Viala |
| EPI_ISL_2345981 | UBS JD PRESIDENTE DUTRA | Instituto Butantan | Dimas Tadeu Covas, Antonio Jorge Martins, Claudia Renata dos Santos Barros, David Schlesinger, Debora Botequio Moretti, Elaine Cristina Marqueze, Elaine Vieira Santos, Evandra Strazza Rodrigues, Heidge Fukumasu, Jayme Augusto de Souza-Neto, José Salvatore Leister Patané, Luiz Alcantara, Luiz Lehmann Coutinho, Maria Carolina Elias, Maurício Lacerda Nogueira, Rafael dos Santos Bezerra, Raul Machado Neto, Rejane Maria Tommasini Grotto, Ricardo Haddad, Sandra Coccuzzo Sampaio Vessoni, Simone Kashima, Svetoslav Nanev Slavov, Vincent Louis Viala |
| EPI_ISL_2345982 | UBS JARDIM ACACIO | Instituto Butantan | Dimas Tadeu Covas, Antonio Jorge Martins, Claudia Renata dos Santos Barros, David Schlesinger, Debora Botequio Moretti, Elaine Cristina Marqueze, Elaine Vieira Santos, Evandra Strazza Rodrigues, Heidge Fukumasu, Jayme Augusto de Souza-Neto, José Salvatore Leister Patané, Luiz Alcantara, Luiz Lehmann Coutinho, Maria Carolina Elias, Maurício Lacerda Nogueira, Rafael dos Santos Bezerra, Raul Machado Neto, Rejane Maria Tommasini Grotto, Ricardo Haddad, Sandra Coccuzzo Sampaio Vessoni, Simone Kashima, Svetoslav Nanev Slavov, Vincent Louis Viala |
| EPI_ISL_2345983 | UBS MARIA PIA DE OLIVEIRA | Instituto Butantan | Dimas Tadeu Covas, Antonio Jorge Martins, Claudia Renata dos Santos Barros, David Schlesinger, Debora Botequio Moretti, Elaine Cristina Marqueze, Elaine Vieira Santos, Evandra Strazza Rodrigues, Heidge Fukumasu, Jayme Augusto de Souza-Neto, José Salvatore Leister Patané, Luiz Alcantara, Luiz Lehmann Coutinho, Maria Carolina Elias, Maurício Lacerda Nogueira, Rafael dos Santos Bezerra, Raul Machado Neto, Rejane Maria Tommasini Grotto, Ricardo Haddad, Sandra Coccuzzo Sampaio Vessoni, Simone Kashima, Svetoslav Nanev Slavov, Vincent Louis Viala |
| EPI_ISL_2345984 | USF JARDIM PIRATININGA ITAQUAQUECETUBA | Instituto Butantan | Dimas Tadeu Covas, Antonio Jorge Martins, Claudia Renata dos Santos Barros, David Schlesinger, Debora Botequio Moretti, Elaine Cristina Marqueze, Elaine Vieira Santos, Evandra Strazza Rodrigues, Heidge Fukumasu, Jayme Augusto de Souza-Neto, José Salvatore Leister Patané, Luiz Alcantara, Luiz Lehmann Coutinho, Maria Carolina Elias, Maurício Lacerda Nogueira, Rafael dos Santos Bezerra, Raul Machado Neto, Rejane Maria Tommasini Grotto, Ricardo Haddad, Sandra Coccuzzo Sampaio Vessoni, Simone Kashima, Svetoslav Nanev Slavov, Vincent Louis Viala |
| EPI_ISL_2345985, EPI_ISL_2345986, EPI_ISL_2345987 | UBS JARDIM ODETE | Instituto Butantan | Dimas Tadeu Covas, Antonio Jorge Martins, Claudia Renata dos Santos Barros, David Schlesinger, Debora Botequio Moretti, Elaine Cristina Marqueze, Elaine Vieira Santos, Evandra Strazza Rodrigues, Heidge Fukumasu, Jayme Augusto de Souza-Neto, José Salvatore Leister Patané, Luiz Alcantara, Luiz Lehmann Coutinho, Maria Carolina Elias, Maurício Lacerda Nogueira, Rafael dos Santos Bezerra, Raul Machado Neto, Rejane Maria Tommasini Grotto, Ricardo Haddad, Sandra Coccuzzo Sampaio Vessoni, Simone Kashima, Svetoslav Nanev Slavov, Vincent Louis Viala |
| EPI_ISL_2345988 | UNIDADE DE PRONTO ATENDIMENTO DE VARZEA PAULISTA UPA II | Instituto Butantan | Dimas Tadeu Covas, Antonio Jorge Martins, Claudia Renata dos Santos Barros, David Schlesinger, Debora Botequio Moretti, Elaine Cristina Marqueze, Elaine Vieira Santos, Evandra Strazza Rodrigues, Heidge Fukumasu, Jayme Augusto de Souza-Neto, José Salvatore Leister Patané, Luiz Alcantara, Luiz Lehmann Coutinho, Maria Carolina Elias, Maurício Lacerda Nogueira, Rafael dos Santos Bezerra, Raul Machado Neto, Rejane Maria Tommasini Grotto, Ricardo Haddad, Sandra Coccuzzo Sampaio Vessoni, Simone Kashima, Svetoslav Nanev Slavov, Vincent Louis Viala |

| | | | |
|---|---|---|---|
| EPI_ISL_2345989 | UBS JARDIM ODETE | Instituto Butantan | Dimas Tadeu Covas, Antonio Jorge Martins, Claudia Renata dos Santos Barros, David Schlesinger, Debora Botequio Moretti, Elaine Cristina Marqueze, Elaine Vieira Santos, Evandra Strazza Rodrigues, Heidge Fukumasu, Jayme Augusto de Souza-Neto, José Salvatore Leister Patané, Luiz Alcantara, Luiz Lehmann Coutinho, Maria Carolina Elias, Maurício Lacerda Nogueira, Rafael dos Santos Bezerra, Raul Machado Neto, Rejane Maria Tommasini Grotto, Ricardo Haddad, Sandra Coccuzzo Sampaio Vessoni, Simone Kashima, Svetoslav Nanev Slavov, Vincent Louis Viala |
| EPI_ISL_2345990 | UNIDADE DE SAUDE DA FAMILIA JARDIM AMERICA ITAQUAQUECETUBA | Instituto Butantan | Dimas Tadeu Covas, Antonio Jorge Martins, Claudia Renata dos Santos Barros, David Schlesinger, Debora Botequio Moretti, Elaine Cristina Marqueze, Elaine Vieira Santos, Evandra Strazza Rodrigues, Heidge Fukumasu, Jayme Augusto de Souza-Neto, José Salvatore Leister Patané, Luiz Alcantara, Luiz Lehmann Coutinho, Maria Carolina Elias, Maurício Lacerda Nogueira, Rafael dos Santos Bezerra, Raul Machado Neto, Rejane Maria Tommasini Grotto, Ricardo Haddad, Sandra Coccuzzo Sampaio Vessoni, Simone Kashima, Svetoslav Nanev Slavov, Vincent Louis Viala |
| EPI_ISL_2345991, EPI_ISL_2345992 | PRONTO ATENDIMENTO DE VARGEM GRANDE PAULISTA | Instituto Butantan | Dimas Tadeu Covas, Antonio Jorge Martins, Claudia Renata dos Santos Barros, David Schlesinger, Debora Botequio Moretti, Elaine Cristina Marqueze, Elaine Vieira Santos, Evandra Strazza Rodrigues, Heidge Fukumasu, Jayme Augusto de Souza-Neto, José Salvatore Leister Patané, Luiz Alcantara, Luiz Lehmann Coutinho, Maria Carolina Elias, Maurício Lacerda Nogueira, Rafael dos Santos Bezerra, Raul Machado Neto, Rejane Maria Tommasini Grotto, Ricardo Haddad, Sandra Coccuzzo Sampaio Vessoni, Simone Kashima, Svetoslav Nanev Slavov, Vincent Louis Viala |
| EPI_ISL_2345993 | UBS JARDIM PAINEIRA | Instituto Butantan | Dimas Tadeu Covas, Antonio Jorge Martins, Claudia Renata dos Santos Barros, David Schlesinger, Debora Botequio Moretti, Elaine Cristina Marqueze, Elaine Vieira Santos, Evandra Strazza Rodrigues, Heidge Fukumasu, Jayme Augusto de Souza-Neto, José Salvatore Leister Patané, Luiz Alcantara, Luiz Lehmann Coutinho, Maria Carolina Elias, Maurício Lacerda Nogueira, Rafael dos Santos Bezerra, Raul Machado Neto, Rejane Maria Tommasini Grotto, Ricardo Haddad, Sandra Coccuzzo Sampaio Vessoni, Simone Kashima, Svetoslav Nanev Slavov, Vincent Louis Viala |
| EPI_ISL_2345994 | UBS JARDIM ODETE | Instituto Butantan | Dimas Tadeu Covas, Antonio Jorge Martins, Claudia Renata dos Santos Barros, David Schlesinger, Debora Botequio Moretti, Elaine Cristina Marqueze, Elaine Vieira Santos, Evandra Strazza Rodrigues, Heidge Fukumasu, Jayme Augusto de Souza-Neto, José Salvatore Leister Patané, Luiz Alcantara, Luiz Lehmann Coutinho, Maria Carolina Elias, Maurício Lacerda Nogueira, Rafael dos Santos Bezerra, Raul Machado Neto, Rejane Maria Tommasini Grotto, Ricardo Haddad, Sandra Coccuzzo Sampaio Vessoni, Simone Kashima, Svetoslav Nanev Slavov, Vincent Louis Viala |
| EPI_ISL_2345995, EPI_ISL_2345996 | UPA DR FRANCO DA ROCHA | Instituto Butantan | Dimas Tadeu Covas, Antonio Jorge Martins, Claudia Renata dos Santos Barros, David Schlesinger, Debora Botequio Moretti, Elaine Cristina Marqueze, Elaine Vieira Santos, Evandra Strazza Rodrigues, Heidge Fukumasu, Jayme Augusto de Souza-Neto, José Salvatore Leister Patané, Luiz Alcantara, Luiz Lehmann Coutinho, Maria Carolina Elias, Maurício Lacerda Nogueira, Rafael dos Santos Bezerra, Raul Machado Neto, Rejane Maria Tommasini Grotto, Ricardo Haddad, Sandra Coccuzzo Sampaio Vessoni, Simone Kashima, Svetoslav Nanev Slavov, Vincent Louis Viala |
| EPI_ISL_2345997 | HOSP MUN CIDADE TIRADENTES CARMEN PRUDENTE | Instituto Butantan | Dimas Tadeu Covas, Antonio Jorge Martins, Claudia Renata dos Santos Barros, David Schlesinger, Debora Botequio Moretti, Elaine Cristina Marqueze, Elaine Vieira Santos, Evandra Strazza Rodrigues, Heidge Fukumasu, Jayme Augusto de Souza-Neto, José Salvatore Leister Patané, Luiz Alcantara, Luiz Lehmann Coutinho, Maria Carolina Elias, Maurício Lacerda Nogueira, Rafael dos Santos Bezerra, Raul Machado Neto, Rejane Maria Tommasini Grotto, Ricardo Haddad, Sandra Coccuzzo Sampaio Vessoni, Simone Kashima, Svetoslav Nanev Slavov, Vincent Louis Viala |
| EPI_ISL_2345998 | UPA DR FRANCO DA ROCHA | Instituto Butantan | Dimas Tadeu Covas, Antonio Jorge Martins, Claudia Renata dos Santos Barros, David Schlesinger, Debora Botequio Moretti, Elaine Cristina Marqueze, Elaine Vieira Santos, Evandra Strazza Rodrigues, Heidge Fukumasu, Jayme Augusto de Souza-Neto, José Salvatore Leister Patané, Luiz Alcantara, Luiz Lehmann Coutinho, Maria Carolina Elias, Maurício Lacerda Nogueira, Rafael dos Santos Bezerra, Raul Machado Neto, Rejane Maria Tommasini Grotto, Ricardo Haddad, Sandra Coccuzzo Sampaio Vessoni, Simone Kashima, Svetoslav Nanev Slavov, Vincent Louis Viala |
| EPI_ISL_2345999 | USF PQ DOS TRABALHADORES ARTUR NOGUEIRA | Instituto Butantan | Dimas Tadeu Covas, Antonio Jorge Martins, Claudia Renata dos Santos Barros, David Schlesinger, Debora Botequio Moretti, Elaine Cristina Marqueze, Elaine Vieira Santos, Evandra Strazza Rodrigues, Heidge Fukumasu, Jayme Augusto de Souza-Neto, José Salvatore Leister Patané, Luiz Alcantara, Luiz Lehmann Coutinho, Maria Carolina Elias, Maurício Lacerda Nogueira, Rafael dos Santos Bezerra, Raul Machado Neto, Rejane Maria Tommasini Grotto, Ricardo Haddad, Sandra Coccuzzo Sampaio Vessoni, Simone Kashima, Svetoslav Nanev Slavov, Vincent Louis Viala |
| EPI_ISL_2346000 | UPA DR FRANCO DA ROCHA | Instituto Butantan | Dimas Tadeu Covas, Antonio Jorge Martins, Claudia Renata dos Santos Barros, David Schlesinger, Debora Botequio Moretti, Elaine Cristina Marqueze, Elaine Vieira Santos, Evandra Strazza Rodrigues, Heidge Fukumasu, Jayme Augusto de Souza-Neto, José Salvatore Leister Patané, Luiz Alcantara, Luiz Lehmann Coutinho, Maria Carolina Elias, Maurício Lacerda Nogueira, Rafael dos Santos Bezerra, Raul Machado Neto, Rejane Maria Tommasini Grotto, Ricardo Haddad, Sandra Coccuzzo Sampaio Vessoni, Simone Kashima, Svetoslav Nanev Slavov, Vincent Louis Viala |
| EPI_ISL_2346001 | SANTA CASA DE GUARAREMA | Instituto Butantan | Dimas Tadeu Covas, Antonio Jorge Martins, Claudia Renata dos Santos Barros, David Schlesinger, Debora Botequio Moretti, Elaine Cristina Marqueze, Elaine Vieira Santos, Evandra Strazza Rodrigues, Heidge Fukumasu, Jayme Augusto de Souza-Neto, José Salvatore Leister Patané, Luiz Alcantara, Luiz Lehmann Coutinho, Maria Carolina Elias, Maurício Lacerda Nogueira, Rafael dos Santos Bezerra, Raul Machado Neto, Rejane Maria Tommasini Grotto, Ricardo Haddad, Sandra Coccuzzo Sampaio Vessoni, Simone Kashima, Svetoslav Nanev Slavov, Vincent Louis Viala |
| EPI_ISL_2346002 | HOSPITAL DE CAMPANHA COVID 19 CAIEIRAS | Instituto Butantan | Dimas Tadeu Covas, Antonio Jorge Martins, Claudia Renata dos Santos Barros, David Schlesinger, Debora Botequio Moretti, Elaine Cristina Marqueze, Elaine Vieira Santos, Evandra Strazza Rodrigues, Heidge Fukumasu, Jayme Augusto de Souza-Neto, José Salvatore Leister Patané, Luiz Alcantara, Luiz Lehmann Coutinho, Maria Carolina Elias, Maurício Lacerda Nogueira, Rafael dos Santos Bezerra, Raul Machado Neto, Rejane Maria Tommasini Grotto, Ricardo Haddad, Sandra Coccuzzo Sampaio Vessoni, Simone Kashima, Svetoslav Nanev Slavov, Vincent Louis Viala |
| EPI_ISL_2346003, EPI_ISL_2346004 | HOSP MUN CIDADE TIRADENTES CARMEN PRUDENTE | Instituto Butantan | Dimas Tadeu Covas, Antonio Jorge Martins, Claudia Renata dos Santos Barros, David Schlesinger, Debora Botequio Moretti, Elaine Cristina Marqueze, Elaine Vieira Santos, Evandra Strazza Rodrigues, Heidge Fukumasu, Jayme Augusto de Souza-Neto, José Salvatore Leister Patané, Luiz Alcantara, Luiz Lehmann Coutinho, Maria Carolina Elias, Maurício Lacerda Nogueira, Rafael dos Santos Bezerra, Raul Machado Neto, Rejane Maria Tommasini Grotto, Ricardo Haddad, Sandra Coccuzzo Sampaio Vessoni, Simone Kashima, Svetoslav Nanev Slavov, Vincent Louis Viala |
| EPI_ISL_2346006, EPI_ISL_2346007, EPI_ISL_2346008, EPI_ISL_2346009 | UPA DR FRANCO DA ROCHA | Instituto Butantan | Dimas Tadeu Covas, Antonio Jorge Martins, Claudia Renata dos Santos Barros, David Schlesinger, Debora Botequio Moretti, Elaine Cristina Marqueze, Elaine Vieira Santos, Evandra Strazza Rodrigues, Heidge Fukumasu, Jayme Augusto de Souza-Neto, José Salvatore Leister Patané, Luiz Alcantara, Luiz Lehmann Coutinho, Maria Carolina Elias, Maurício Lacerda Nogueira, Rafael dos Santos Bezerra, Raul Machado Neto, Rejane Maria Tommasini Grotto, Ricardo Haddad, Sandra Coccuzzo Sampaio Vessoni, Simone Kashima, Svetoslav Nanev Slavov, Vincent Louis Viala |
| EPI_ISL_2346010 | PSF PARQUE EMILIA | Instituto Butantan | Dimas Tadeu Covas, Antonio Jorge Martins, Claudia Renata dos Santos Barros, David Schlesinger, Debora Botequio Moretti, Elaine Cristina Marqueze, Elaine Vieira Santos, Evandra Strazza Rodrigues, Heidge Fukumasu, Jayme Augusto de Souza-Neto, José Salvatore Leister Patané, Luiz Alcantara, Luiz Lehmann Coutinho, Maria Carolina Elias, Maurício Lacerda Nogueira, Rafael dos Santos Bezerra, Raul Machado Neto, Rejane Maria Tommasini Grotto, Ricardo Haddad, Sandra Coccuzzo Sampaio Vessoni, Simone Kashima, Svetoslav Nanev Slavov, Vincent Louis Viala |
| EPI_ISL_2346011 | UPA DR FRANCO DA ROCHA | Instituto Butantan | Dimas Tadeu Covas, Antonio Jorge Martins, Claudia Renata dos Santos Barros, David Schlesinger, Debora Botequio Moretti, Elaine Cristina Marqueze, Elaine Vieira Santos, Evandra Strazza Rodrigues, Heidge Fukumasu, Jayme Augusto de Souza-Neto, José Salvatore Leister Patané, Luiz Alcantara, Luiz Lehmann Coutinho, Maria Carolina Elias, Maurício Lacerda Nogueira, Rafael dos Santos Bezerra, Raul Machado Neto, Rejane Maria Tommasini Grotto, Ricardo Haddad, Sandra Coccuzzo Sampaio Vessoni, Simone Kashima, Svetoslav Nanev Slavov, Vincent Louis Viala |
| EPI_ISL_2346012 | VIGILANCIA SANITARIA E VIG EPIDEMIOLOGICA DE ITAPEVI | Instituto Butantan | Dimas Tadeu Covas, Antonio Jorge Martins, Claudia Renata dos Santos Barros, David Schlesinger, Debora Botequio Moretti, Elaine Cristina Marqueze, Elaine Vieira Santos, Evandra Strazza Rodrigues, Heidge Fukumasu, Jayme Augusto de Souza-Neto, José Salvatore Leister Patané, Luiz Alcantara, Luiz Lehmann Coutinho, Maria Carolina Elias, Maurício Lacerda Nogueira, Rafael dos Santos Bezerra, Raul Machado Neto, Rejane Maria Tommasini Grotto, Ricardo Haddad, Sandra Coccuzzo Sampaio Vessoni, Simone Kashima, Svetoslav Nanev Slavov, Vincent Louis Viala |
| EPI_ISL_2346013, EPI_ISL_2346014 | PRONTO ATENDIMENTO DE VARGEM GRANDE PAULISTA | Instituto Butantan | Dimas Tadeu Covas, Antonio Jorge Martins, Claudia Renata dos Santos Barros, David Schlesinger, Debora Botequio Moretti, Elaine Cristina Marqueze, Elaine Vieira Santos, Evandra Strazza Rodrigues, Heidge Fukumasu, Jayme Augusto de Souza-Neto, José Salvatore Leister Patané, Luiz Alcantara, Luiz Lehmann Coutinho, Maria Carolina Elias, Maurício Lacerda Nogueira, Rafael dos Santos Bezerra, Raul Machado Neto, Rejane Maria Tommasini Grotto, Ricardo Haddad, Sandra Coccuzzo Sampaio Vessoni, Simone Kashima, Svetoslav Nanev Slavov, Vincent Louis Viala |
| EPI_ISL_2346015 | UBS RECANTO MONICA | Instituto Butantan | Dimas Tadeu Covas, Antonio Jorge Martins, Claudia Renata dos Santos Barros, David Schlesinger, Debora Botequio Moretti, Elaine Cristina Marqueze, Elaine Vieira Santos, Evandra Strazza Rodrigues, Heidge Fukumasu, Jayme Augusto de Souza-Neto, José Salvatore Leister Patané, Luiz Alcantara, Luiz Lehmann Coutinho, Maria Carolina Elias, Maurício Lacerda Nogueira, Rafael dos Santos Bezerra, Raul Machado Neto, Rejane Maria Tommasini Grotto, Ricardo Haddad, Sandra Coccuzzo Sampaio Vessoni, Simone Kashima, Svetoslav Nanev Slavov, Vincent Louis Viala |
| EPI_ISL_2346016 | UPA DR FRANCO DA ROCHA | Instituto Butantan | Dimas Tadeu Covas, Antonio Jorge Martins, Claudia Renata dos Santos Barros, David Schlesinger, Debora Botequio Moretti, Elaine Cristina Marqueze, Elaine Vieira Santos, Evandra Strazza Rodrigues, Heidge Fukumasu, Jayme Augusto de Souza-Neto, José Salvatore Leister Patané, Luiz Alcantara, Luiz |

| | | | |
|---|---|---|---|
| | | | Lehmann Coutinho, Maria Carolina Elias, Maurício Lacerda Nogueira, Rafael dos Santos Bezerra, Raul Machado Neto, Rejane Maria Tommasini Grotto, Ricardo Haddad, Sandra Coccuzzo Sampaio Vessoni, Simone Kashima, Svetoslav Nanev Slavov, Vincent Louis Viala |
| EPI_ISL_2346017, EPI_ISL_2346018 | UBS JARDIM MARAGOGIPE | Instituto Butantan | Dimas Tadeu Covas, Antonio Jorge Martins, Claudia Renata dos Santos Barros, David Schlesinger, Debora Botequio Moretti, Elaine Cristina Marqueze, Elaine Vieira Santos, Evandra Strazza Rodrigues, Heidge Fukumasu, Jayme Augusto de Souza-Neto, José Salvatore Leister Patané, Luiz Alcantara, Luiz Lehmann Coutinho, Maria Carolina Elias, Maurício Lacerda Nogueira, Rafael dos Santos Bezerra, Raul Machado Neto, Rejane Maria Tommasini Grotto, Ricardo Haddad, Sandra Coccuzzo Sampaio Vessoni, Simone Kashima, Svetoslav Nanev Slavov, Vincent Louis Viala |
| EPI_ISL_2346019, EPI_ISL_2346020 | UPA DR FRANCO DA ROCHA | Instituto Butantan | Dimas Tadeu Covas, Antonio Jorge Martins, Claudia Renata dos Santos Barros, David Schlesinger, Debora Botequio Moretti, Elaine Cristina Marqueze, Elaine Vieira Santos, Evandra Strazza Rodrigues, Heidge Fukumasu, Jayme Augusto de Souza-Neto, José Salvatore Leister Patané, Luiz Alcantara, Luiz Lehmann Coutinho, Maria Carolina Elias, Maurício Lacerda Nogueira, Rafael dos Santos Bezerra, Raul Machado Neto, Rejane Maria Tommasini Grotto, Ricardo Haddad, Sandra Coccuzzo Sampaio Vessoni, Simone Kashima, Svetoslav Nanev Slavov, Vincent Louis Viala |
| EPI_ISL_2346021 | PSF PARQUE EMILIA | Instituto Butantan | Dimas Tadeu Covas, Antonio Jorge Martins, Claudia Renata dos Santos Barros, David Schlesinger, Debora Botequio Moretti, Elaine Cristina Marqueze, Elaine Vieira Santos, Evandra Strazza Rodrigues, Heidge Fukumasu, Jayme Augusto de Souza-Neto, José Salvatore Leister Patané, Luiz Alcantara, Luiz Lehmann Coutinho, Maria Carolina Elias, Maurício Lacerda Nogueira, Rafael dos Santos Bezerra, Raul Machado Neto, Rejane Maria Tommasini Grotto, Ricardo Haddad, Sandra Coccuzzo Sampaio Vessoni, Simone Kashima, Svetoslav Nanev Slavov, Vincent Louis Viala |
| EPI_ISL_2346022 | HOSP MUN JOSANIAS CASTANHA BRAGA | Instituto Butantan | Dimas Tadeu Covas, Antonio Jorge Martins, Claudia Renata dos Santos Barros, David Schlesinger, Debora Botequio Moretti, Elaine Cristina Marqueze, Elaine Vieira Santos, Evandra Strazza Rodrigues, Heidge Fukumasu, Jayme Augusto de Souza-Neto, José Salvatore Leister Patané, Luiz Alcantara, Luiz Lehmann Coutinho, Maria Carolina Elias, Maurício Lacerda Nogueira, Rafael dos Santos Bezerra, Raul Machado Neto, Rejane Maria Tommasini Grotto, Ricardo Haddad, Sandra Coccuzzo Sampaio Vessoni, Simone Kashima, Svetoslav Nanev Slavov, Vincent Louis Viala |
| EPI_ISL_2346023 | UBS JARDIM PAINEIRA | Instituto Butantan | Dimas Tadeu Covas, Antonio Jorge Martins, Claudia Renata dos Santos Barros, David Schlesinger, Debora Botequio Moretti, Elaine Cristina Marqueze, Elaine Vieira Santos, Evandra Strazza Rodrigues, Heidge Fukumasu, Jayme Augusto de Souza-Neto, José Salvatore Leister Patané, Luiz Alcantara, Luiz Lehmann Coutinho, Maria Carolina Elias, Maurício Lacerda Nogueira, Rafael dos Santos Bezerra, Raul Machado Neto, Rejane Maria Tommasini Grotto, Ricardo Haddad, Sandra Coccuzzo Sampaio Vessoni, Simone Kashima, Svetoslav Nanev Slavov, Vincent Louis Viala |
| EPI_ISL_2346024 | HOSP MUN JOSANIAS CASTANHA BRAGA | Instituto Butantan | Dimas Tadeu Covas, Antonio Jorge Martins, Claudia Renata dos Santos Barros, David Schlesinger, Debora Botequio Moretti, Elaine Cristina Marqueze, Elaine Vieira Santos, Evandra Strazza Rodrigues, Heidge Fukumasu, Jayme Augusto de Souza-Neto, José Salvatore Leister Patané, Luiz Alcantara, Luiz Lehmann Coutinho, Maria Carolina Elias, Maurício Lacerda Nogueira, Rafael dos Santos Bezerra, Raul Machado Neto, Rejane Maria Tommasini Grotto, Ricardo Haddad, Sandra Coccuzzo Sampaio Vessoni, Simone Kashima, Svetoslav Nanev Slavov, Vincent Louis Viala |
| EPI_ISL_2346025, EPI_ISL_2346026 | UBS JARDIM CAIUBY | Instituto Butantan | Dimas Tadeu Covas, Antonio Jorge Martins, Claudia Renata dos Santos Barros, David Schlesinger, Debora Botequio Moretti, Elaine Cristina Marqueze, Elaine Vieira Santos, Evandra Strazza Rodrigues, Heidge Fukumasu, Jayme Augusto de Souza-Neto, José Salvatore Leister Patané, Luiz Alcantara, Luiz Lehmann Coutinho, Maria Carolina Elias, Maurício Lacerda Nogueira, Rafael dos Santos Bezerra, Raul Machado Neto, Rejane Maria Tommasini Grotto, Ricardo Haddad, Sandra Coccuzzo Sampaio Vessoni, Simone Kashima, Svetoslav Nanev Slavov, Vincent Louis Viala |
| EPI_ISL_2346027 | UBS PARQUE MARENGO | Instituto Butantan | Dimas Tadeu Covas, Antonio Jorge Martins, Claudia Renata dos Santos Barros, David Schlesinger, Debora Botequio Moretti, Elaine Cristina Marqueze, Elaine Vieira Santos, Evandra Strazza Rodrigues, Heidge Fukumasu, Jayme Augusto de Souza-Neto, José Salvatore Leister Patané, Luiz Alcantara, Luiz Lehmann Coutinho, Maria Carolina Elias, Maurício Lacerda Nogueira, Rafael dos Santos Bezerra, Raul Machado Neto, Rejane Maria Tommasini Grotto, Ricardo Haddad, Sandra Coccuzzo Sampaio Vessoni, Simone Kashima, Svetoslav Nanev Slavov, Vincent Louis Viala |
| EPI_ISL_2346028 | UBS MORRO BRANCO | Instituto Butantan | Dimas Tadeu Covas, Antonio Jorge Martins, Claudia Renata dos Santos Barros, David Schlesinger, Debora Botequio Moretti, Elaine Cristina Marqueze, Elaine Vieira Santos, Evandra Strazza Rodrigues, Heidge Fukumasu, Jayme Augusto de Souza-Neto, José Salvatore Leister Patané, Luiz Alcantara, Luiz Lehmann Coutinho, Maria Carolina Elias, Maurício Lacerda Nogueira, Rafael dos Santos Bezerra, Raul Machado Neto, Rejane Maria Tommasini Grotto, Ricardo Haddad, Sandra Coccuzzo Sampaio Vessoni, Simone Kashima, Svetoslav Nanev Slavov, Vincent Louis Viala |
| EPI_ISL_2346029 | UPA DR FRANCO DA ROCHA | Instituto Butantan | Dimas Tadeu Covas, Antonio Jorge Martins, Claudia Renata dos Santos Barros, David Schlesinger, Debora Botequio Moretti, Elaine Cristina Marqueze, Elaine Vieira Santos, Evandra Strazza Rodrigues, Heidge Fukumasu, Jayme Augusto de Souza-Neto, José Salvatore Leister Patané, Luiz Alcantara, Luiz Lehmann Coutinho, Maria Carolina Elias, Maurício Lacerda Nogueira, Rafael dos Santos Bezerra, Raul Machado Neto, Rejane Maria Tommasini Grotto, Ricardo Haddad, Sandra Coccuzzo Sampaio Vessoni, Simone Kashima, Svetoslav Nanev Slavov, Vincent Louis Viala |
| EPI_ISL_2346030 | UBS JARDIM MARAGOGIPE | Instituto Butantan | Dimas Tadeu Covas, Antonio Jorge Martins, Claudia Renata dos Santos Barros, David Schlesinger, Debora Botequio Moretti, Elaine Cristina Marqueze, Elaine Vieira Santos, Evandra Strazza Rodrigues, Heidge Fukumasu, Jayme Augusto de Souza-Neto, José Salvatore Leister Patané, Luiz Alcantara, Luiz Lehmann Coutinho, Maria Carolina Elias, Maurício Lacerda Nogueira, Rafael dos Santos Bezerra, Raul Machado Neto, Rejane Maria Tommasini Grotto, Ricardo Haddad, Sandra Coccuzzo Sampaio Vessoni, Simone Kashima, Svetoslav Nanev Slavov, Vincent Louis Viala |
| EPI_ISL_2346031 | UBS JARDIM CAIUBY | Instituto Butantan | Dimas Tadeu Covas, Antonio Jorge Martins, Claudia Renata dos Santos Barros, David Schlesinger, Debora Botequio Moretti, Elaine Cristina Marqueze, Elaine Vieira Santos, Evandra Strazza Rodrigues, Heidge Fukumasu, Jayme Augusto de Souza-Neto, José Salvatore Leister Patané, Luiz Alcantara, Luiz Lehmann Coutinho, Maria Carolina Elias, Maurício Lacerda Nogueira, Rafael dos Santos Bezerra, Raul Machado Neto, Rejane Maria Tommasini Grotto, Ricardo Haddad, Sandra Coccuzzo Sampaio Vessoni, Simone Kashima, Svetoslav Nanev Slavov, Vincent Louis Viala |
| EPI_ISL_2346032 | PSF PARQUE EMILIA | Instituto Butantan | Dimas Tadeu Covas, Antonio Jorge Martins, Claudia Renata dos Santos Barros, David Schlesinger, Debora Botequio Moretti, Elaine Cristina Marqueze, Elaine Vieira Santos, Evandra Strazza Rodrigues, Heidge Fukumasu, Jayme Augusto de Souza-Neto, José Salvatore Leister Patané, Luiz Alcantara, Luiz Lehmann Coutinho, Maria Carolina Elias, Maurício Lacerda Nogueira, Rafael dos Santos Bezerra, Raul Machado Neto, Rejane Maria Tommasini Grotto, Ricardo Haddad, Sandra Coccuzzo Sampaio Vessoni, Simone Kashima, Svetoslav Nanev Slavov, Vincent Louis Viala |
| EPI_ISL_2346033 | UBS PARQUE MARENGO | Instituto Butantan | Dimas Tadeu Covas, Antonio Jorge Martins, Claudia Renata dos Santos Barros, David Schlesinger, Debora Botequio Moretti, Elaine Cristina Marqueze, Elaine Vieira Santos, Evandra Strazza Rodrigues, Heidge Fukumasu, Jayme Augusto de Souza-Neto, José Salvatore Leister Patané, Luiz Alcantara, Luiz Lehmann Coutinho, Maria Carolina Elias, Maurício Lacerda Nogueira, Rafael dos Santos Bezerra, Raul Machado Neto, Rejane Maria Tommasini Grotto, Ricardo Haddad, Sandra Coccuzzo Sampaio Vessoni, Simone Kashima, Svetoslav Nanev Slavov, Vincent Louis Viala |
| EPI_ISL_2346034 | UBS JARDIM CAIUBY | Instituto Butantan | Dimas Tadeu Covas, Antonio Jorge Martins, Claudia Renata dos Santos Barros, David Schlesinger, Debora Botequio Moretti, Elaine Cristina Marqueze, Elaine Vieira Santos, Evandra Strazza Rodrigues, Heidge Fukumasu, Jayme Augusto de Souza-Neto, José Salvatore Leister Patané, Luiz Alcantara, Luiz Lehmann Coutinho, Maria Carolina Elias, Maurício Lacerda Nogueira, Rafael dos Santos Bezerra, Raul Machado Neto, Rejane Maria Tommasini Grotto, Ricardo Haddad, Sandra Coccuzzo Sampaio Vessoni, Simone Kashima, Svetoslav Nanev Slavov, Vincent Louis Viala |
| EPI_ISL_2346035, EPI_ISL_2346036 | UPA DR FRANCO DA ROCHA | Instituto Butantan | Dimas Tadeu Covas, Antonio Jorge Martins, Claudia Renata dos Santos Barros, David Schlesinger, Debora Botequio Moretti, Elaine Cristina Marqueze, Elaine Vieira Santos, Evandra Strazza Rodrigues, Heidge Fukumasu, Jayme Augusto de Souza-Neto, José Salvatore Leister Patané, Luiz Alcantara, Luiz Lehmann Coutinho, Maria Carolina Elias, Maurício Lacerda Nogueira, Rafael dos Santos Bezerra, Raul Machado Neto, Rejane Maria Tommasini Grotto, Ricardo Haddad, Sandra Coccuzzo Sampaio Vessoni, Simone Kashima, Svetoslav Nanev Slavov, Vincent Louis Viala |
| EPI_ISL_2346037 | HOSPITAL MUN DR GUIDO GUIDA | Instituto Butantan | Dimas Tadeu Covas, Antonio Jorge Martins, Claudia Renata dos Santos Barros, David Schlesinger, Debora Botequio Moretti, Elaine Cristina Marqueze, Elaine Vieira Santos, Evandra Strazza Rodrigues, Heidge Fukumasu, Jayme Augusto de Souza-Neto, José Salvatore Leister Patané, Luiz Alcantara, Luiz Lehmann Coutinho, Maria Carolina Elias, Maurício Lacerda Nogueira, Rafael dos Santos Bezerra, Raul Machado Neto, Rejane Maria Tommasini Grotto, Ricardo Haddad, Sandra Coccuzzo Sampaio Vessoni, Simone Kashima, Svetoslav Nanev Slavov, Vincent Louis Viala |
| EPI_ISL_2346038 | UPA DR FRANCO DA ROCHA | Instituto Butantan | Dimas Tadeu Covas, Antonio Jorge Martins, Claudia Renata dos Santos Barros, David Schlesinger, Debora Botequio Moretti, Elaine Cristina Marqueze, Elaine Vieira Santos, Evandra Strazza Rodrigues, Heidge Fukumasu, Jayme Augusto de Souza-Neto, José Salvatore Leister Patané, Luiz Alcantara, Luiz Lehmann Coutinho, Maria Carolina Elias, Maurício Lacerda Nogueira, Rafael dos Santos Bezerra, Raul Machado Neto, Rejane Maria Tommasini Grotto, Ricardo Haddad, Sandra Coccuzzo Sampaio Vessoni, Simone Kashima, Svetoslav Nanev Slavov, Vincent Louis Viala |
| EPI_ISL_2346039 | UBS JARDIM CAIUBY | Instituto Butantan | Dimas Tadeu Covas, Antonio Jorge Martins, Claudia Renata dos Santos Barros, David Schlesinger, Debora Botequio Moretti, Elaine Cristina Marqueze, Elaine Vieira Santos, Evandra Strazza Rodrigues, Heidge Fukumasu, Jayme Augusto de Souza-Neto, José Salvatore Leister Patané, Luiz Alcantara, Luiz Lehmann Coutinho, Maria Carolina Elias, Maurício Lacerda Nogueira, Rafael dos Santos Bezerra, Raul Machado Neto, Rejane Maria Tommasini Grotto, Ricardo Haddad, Sandra Coccuzzo Sampaio Vessoni, Simone Kashima, Svetoslav Nanev Slavov, Vincent Louis Viala |

| | | | |
|---|---|---|---|
| EPI_ISL_2346040 | UBS MORRO BRANCO | Instituto Butantan | Dimas Tadeu Covas, Antonio Jorge Martins, Claudia Renata dos Santos Barros, David Schlesinger, Debora Botequio Moretti, Elaine Cristina Marqueze, Elaine Vieira Santos, Evandra Strazza Rodrigues, Heidge Fukumasu, Jayme Augusto de Souza-Neto, José Salvatore Leister Patané, Luiz Alcantara, Luiz Lehmann Coutinho, Maria Carolina Elias, Maurício Lacerda Nogueira, Rafael dos Santos Bezerra, Raul Machado Neto, Rejane Maria Tommasini Grotto, Ricardo Haddad, Sandra Coccuzzo Sampaio Vessoni, Simone Kashima, Svetoslav Nanev Slavov, Vincent Louis Viala |
| EPI_ISL_2346041 | UBS JARDIM MARAGOGIPE | Instituto Butantan | Dimas Tadeu Covas, Antonio Jorge Martins, Claudia Renata dos Santos Barros, David Schlesinger, Debora Botequio Moretti, Elaine Cristina Marqueze, Elaine Vieira Santos, Evandra Strazza Rodrigues, Heidge Fukumasu, Jayme Augusto de Souza-Neto, José Salvatore Leister Patané, Luiz Alcantara, Luiz Lehmann Coutinho, Maria Carolina Elias, Maurício Lacerda Nogueira, Rafael dos Santos Bezerra, Raul Machado Neto, Rejane Maria Tommasini Grotto, Ricardo Haddad, Sandra Coccuzzo Sampaio Vessoni, Simone Kashima, Svetoslav Nanev Slavov, Vincent Louis Viala |
| EPI_ISL_2346042 | UBS JARDIM CAIUBY | Instituto Butantan | Dimas Tadeu Covas, Antonio Jorge Martins, Claudia Renata dos Santos Barros, David Schlesinger, Debora Botequio Moretti, Elaine Cristina Marqueze, Elaine Vieira Santos, Evandra Strazza Rodrigues, Heidge Fukumasu, Jayme Augusto de Souza-Neto, José Salvatore Leister Patané, Luiz Alcantara, Luiz Lehmann Coutinho, Maria Carolina Elias, Maurício Lacerda Nogueira, Rafael dos Santos Bezerra, Raul Machado Neto, Rejane Maria Tommasini Grotto, Ricardo Haddad, Sandra Coccuzzo Sampaio Vessoni, Simone Kashima, Svetoslav Nanev Slavov, Vincent Louis Viala |
| EPI_ISL_2346043 | PSF PARQUE EMILIA | Instituto Butantan | Dimas Tadeu Covas, Antonio Jorge Martins, Claudia Renata dos Santos Barros, David Schlesinger, Debora Botequio Moretti, Elaine Cristina Marqueze, Elaine Vieira Santos, Evandra Strazza Rodrigues, Heidge Fukumasu, Jayme Augusto de Souza-Neto, José Salvatore Leister Patané, Luiz Alcantara, Luiz Lehmann Coutinho, Maria Carolina Elias, Maurício Lacerda Nogueira, Rafael dos Santos Bezerra, Raul Machado Neto, Rejane Maria Tommasini Grotto, Ricardo Haddad, Sandra Coccuzzo Sampaio Vessoni, Simone Kashima, Svetoslav Nanev Slavov, Vincent Louis Viala |
| EPI_ISL_2346044, EPI_ISL_2346045 | UBS JARDIM CAIUBY | Instituto Butantan | Dimas Tadeu Covas, Antonio Jorge Martins, Claudia Renata dos Santos Barros, David Schlesinger, Debora Botequio Moretti, Elaine Cristina Marqueze, Elaine Vieira Santos, Evandra Strazza Rodrigues, Heidge Fukumasu, Jayme Augusto de Souza-Neto, José Salvatore Leister Patané, Luiz Alcantara, Luiz Lehmann Coutinho, Maria Carolina Elias, Maurício Lacerda Nogueira, Rafael dos Santos Bezerra, Raul Machado Neto, Rejane Maria Tommasini Grotto, Ricardo Haddad, Sandra Coccuzzo Sampaio Vessoni, Simone Kashima, Svetoslav Nanev Slavov, Vincent Louis Viala |
| EPI_ISL_2346046 | USF PEQUENO CORACAO | Instituto Butantan | Dimas Tadeu Covas, Antonio Jorge Martins, Claudia Renata dos Santos Barros, David Schlesinger, Debora Botequio Moretti, Elaine Cristina Marqueze, Elaine Vieira Santos, Evandra Strazza Rodrigues, Heidge Fukumasu, Jayme Augusto de Souza-Neto, José Salvatore Leister Patané, Luiz Alcantara, Luiz Lehmann Coutinho, Maria Carolina Elias, Maurício Lacerda Nogueira, Rafael dos Santos Bezerra, Raul Machado Neto, Rejane Maria Tommasini Grotto, Ricardo Haddad, Sandra Coccuzzo Sampaio Vessoni, Simone Kashima, Svetoslav Nanev Slavov, Vincent Louis Viala |
| EPI_ISL_2346047 | UBS JARDIM CAIUBY | Instituto Butantan | Dimas Tadeu Covas, Antonio Jorge Martins, Claudia Renata dos Santos Barros, David Schlesinger, Debora Botequio Moretti, Elaine Cristina Marqueze, Elaine Vieira Santos, Evandra Strazza Rodrigues, Heidge Fukumasu, Jayme Augusto de Souza-Neto, José Salvatore Leister Patané, Luiz Alcantara, Luiz Lehmann Coutinho, Maria Carolina Elias, Maurício Lacerda Nogueira, Rafael dos Santos Bezerra, Raul Machado Neto, Rejane Maria Tommasini Grotto, Ricardo Haddad, Sandra Coccuzzo Sampaio Vessoni, Simone Kashima, Svetoslav Nanev Slavov, Vincent Louis Viala |
| EPI_ISL_2346048 | UBS MARIA PIA DE OLIVEIRA | Instituto Butantan | Dimas Tadeu Covas, Antonio Jorge Martins, Claudia Renata dos Santos Barros, David Schlesinger, Debora Botequio Moretti, Elaine Cristina Marqueze, Elaine Vieira Santos, Evandra Strazza Rodrigues, Heidge Fukumasu, Jayme Augusto de Souza-Neto, José Salvatore Leister Patané, Luiz Alcantara, Luiz Lehmann Coutinho, Maria Carolina Elias, Maurício Lacerda Nogueira, Rafael dos Santos Bezerra, Raul Machado Neto, Rejane Maria Tommasini Grotto, Ricardo Haddad, Sandra Coccuzzo Sampaio Vessoni, Simone Kashima, Svetoslav Nanev Slavov, Vincent Louis Viala |
| EPI_ISL_2346049 | UPA SADAKO SEDOGUTI | Instituto Butantan | Dimas Tadeu Covas, Antonio Jorge Martins, Claudia Renata dos Santos Barros, David Schlesinger, Debora Botequio Moretti, Elaine Cristina Marqueze, Elaine Vieira Santos, Evandra Strazza Rodrigues, Heidge Fukumasu, Jayme Augusto de Souza-Neto, José Salvatore Leister Patané, Luiz Alcantara, Luiz Lehmann Coutinho, Maria Carolina Elias, Maurício Lacerda Nogueira, Rafael dos Santos Bezerra, Raul Machado Neto, Rejane Maria Tommasini Grotto, Ricardo Haddad, Sandra Coccuzzo Sampaio Vessoni, Simone Kashima, Svetoslav Nanev Slavov, Vincent Louis Viala |
| EPI_ISL_2346050, EPI_ISL_2346051, EPI_ISL_2346052, EPI_ISL_2346053, EPI_ISL_2346054, EPI_ISL_2346055, EPI_ISL_2346056, EPI_ISL_2346057, EPI_ISL_2346058, EPI_ISL_2346059, EPI_ISL_2346060, EPI_ISL_2346061, EPI_ISL_2346062, EPI_ISL_2346063, EPI_ISL_2346064, EPI_ISL_2346065, EPI_ISL_2346066, EPI_ISL_2346067, EPI_ISL_2346068, EPI_ISL_2346069, EPI_ISL_2346070, EPI_ISL_2346071, EPI_ISL_2346072, EPI_ISL_2346073, EPI_ISL_2346074, EPI_ISL_2346075, EPI_ISL_2346076, EPI_ISL_2346077, EPI_ISL_2346078, EPI_ISL_2346079, EPI_ISL_2346080, EPI_ISL_2346081, EPI_ISL_2346082 | | | |
| see above | SERRANA | Instituto Butantan / Mendelics | Dimas Tadeu Covas, Antonio Jorge Martins, Claudia Renata dos Santos Barros, David Schlesinger, Debora Botequio Moretti, Elaine Cristina Marqueze, Elaine Vieira Santos, Evandra Strazza Rodrigues, Heidge Fukumasu, Jayme Augusto de Souza-Neto, José Salvatore Leister Patané, Luiz Alcantara, Luiz Lehmann Coutinho, Maria Carolina Elias, Maurício Lacerda Nogueira, Rafael dos Santos Bezerra, Raul Machado Neto, Rejane Maria Tommasini Grotto, Ricardo Haddad, Sandra Coccuzzo Sampaio Vessoni, Simone Kashima, Svetoslav Nanev Slavov, Vincent Louis Viala |
| EPI_ISL_2346083, EPI_ISL_2346085, EPI_ISL_2346086, EPI_ISL_2346087, EPI_ISL_2346088, EPI_ISL_2346089, EPI_ISL_2346090, EPI_ISL_2346091, EPI_ISL_2346092, EPI_ISL_2346093, EPI_ISL_2346094, EPI_ISL_2346095, EPI_ISL_2346096, EPI_ISL_2346097, EPI_ISL_2346098, EPI_ISL_2346099 | | | |
| see above | Prefeirtura de SP | Instituto Butantan | Dimas Tadeu Covas, Antonio Jorge Martins, Claudia Renata dos Santos Barros, David Schlesinger, Debora Botequio Moretti, Elaine Cristina Marqueze, Elaine Vieira Santos, Evandra Strazza Rodrigues, Heidge Fukumasu, Jayme Augusto de Souza-Neto, José Salvatore Leister Patané, Luiz Alcantara, Luiz Lehmann Coutinho, Maria Carolina Elias, Maurício Lacerda Nogueira, Rafael dos Santos Bezerra, Raul Machado Neto, Rejane Maria Tommasini Grotto, Ricardo Haddad, Sandra Coccuzzo Sampaio Vessoni, Simone Kashima, Svetoslav Nanev Slavov, Vincent Louis Viala |
| EPI_ISL_2348593, EPI_ISL_2348594, EPI_ISL_2348595, EPI_ISL_2348596, EPI_ISL_2348597, EPI_ISL_2348598, EPI_ISL_2348599, EPI_ISL_2348600, EPI_ISL_2348601, EPI_ISL_2348602, EPI_ISL_2348603, EPI_ISL_2348604, EPI_ISL_2348605, EPI_ISL_2348606, EPI_ISL_2348607, EPI_ISL_2348608, EPI_ISL_2348609, EPI_ISL_2348610, EPI_ISL_2348611, EPI_ISL_2348612, EPI_ISL_2348613, EPI_ISL_2348614, EPI_ISL_2348615, EPI_ISL_2348616 | | | |
| see above | HLAGYN - Laboratorio de Imunologia de Transplantes de Goias | HLAGYN - Laboratorio de Imunologia de Transplantes de Goias | Fernando Antonio Vinhal dos Santos, Erika Lopes Rocha Batista, Alessandro Leonardo Alvares Magalhaes, Frederico Rodrigues Vinhal, Sabrina Sara Moreira Duarte, Lucas Carlos Gomes Pereira, Daniel Ferreira de Sousa |
| EPI_ISL_2362237 | DIRETORIA MUNICIPAL DE SAUDE DE ENGENHEIRO COELHO | Instituto Butantan / FZEA-USP-Pirassununga | Dimas Tadeu Covas, Antonio Jorge Martins, Claudia Renata dos Santos Barros, David Schlesinger, Debora Botequio Moretti, Elaine Cristina Marqueze, Elaine Vieira Santos, Evandra Strazza Rodrigues, Heidge Fukumasu, Jayme Augusto de Souza-Neto, José Salvatore Leister Patané, Luiz Alcantara, Luiz Lehmann Coutinho, Maria Carolina Elias, Maurício Lacerda Nogueira, Rafael dos Santos Bezerra, Raul Machado Neto, Rejane Maria Tommasini Grotto, Ricardo Haddad, Sandra Coccuzzo Sampaio Vessoni, Simone Kashima, Svetoslav Nanev Slavov, Vincent Louis Viala |
| EPI_ISL_2362239 | UNIDADE MISTA OSVALDO P CAMARGO ELIAS FAUSTO | Instituto Butantan / FZEA-USP-Pirassununga | Dimas Tadeu Covas, Antonio Jorge Martins, Claudia Renata dos Santos Barros, David Schlesinger, Debora Botequio Moretti, Elaine Cristina Marqueze, Elaine Vieira Santos, Evandra Strazza Rodrigues, Heidge Fukumasu, Jayme Augusto de Souza-Neto, José Salvatore Leister Patané, Luiz Alcantara, Luiz Lehmann Coutinho, Maria Carolina Elias, Maurício Lacerda Nogueira, Rafael dos Santos Bezerra, Raul Machado Neto, Rejane Maria Tommasini Grotto, Ricardo Haddad, Sandra Coccuzzo Sampaio Vessoni, Simone Kashima, Svetoslav Nanev Slavov, Vincent Louis Viala |
| EPI_ISL_2362241, EPI_ISL_2362242 | LABORATORIO DR PAULO EMILIO DALESSANDRO PINDAMONHANGABA | Instituto Butantan / Mendelics | Dimas Tadeu Covas, Antonio Jorge Martins, Claudia Renata dos Santos Barros, David Schlesinger, Debora Botequio Moretti, Elaine Cristina Marqueze, Elaine Vieira Santos, Evandra Strazza Rodrigues, Heidge Fukumasu, Jayme Augusto de Souza-Neto, José Salvatore Leister Patané, Luiz Alcantara, Luiz Lehmann Coutinho, Maria Carolina Elias, Maurício Lacerda Nogueira, Rafael dos Santos Bezerra, Raul Machado Neto, Rejane Maria Tommasini Grotto, Ricardo Haddad, Sandra Coccuzzo Sampaio Vessoni, Simone Kashima, Svetoslav Nanev Slavov, Vincent Louis Viala |
| EPI_ISL_2362244 | HOSPITAL DE CAMPANHA COVID 19 MUNICIPIO DE TAUBATE | Instituto Butantan / Mendelics | Dimas Tadeu Covas, Antonio Jorge Martins, Claudia Renata dos Santos Barros, David Schlesinger, Debora Botequio Moretti, Elaine Cristina Marqueze, Elaine Vieira Santos, Evandra Strazza Rodrigues, Heidge Fukumasu, Jayme Augusto de Souza-Neto, José Salvatore Leister Patané, Luiz Alcantara, Luiz Lehmann Coutinho, Maria Carolina Elias, Maurício Lacerda Nogueira, Rafael dos Santos Bezerra, Raul Machado Neto, Rejane Maria Tommasini Grotto, Ricardo Haddad, Sandra Coccuzzo Sampaio Vessoni, Simone Kashima, Svetoslav Nanev Slavov, Vincent Louis Viala |
| EPI_ISL_2362246 | CENTRO DE SAUDE DR JOAO JOAQUIM FERREIRA CRISTAIS PAULISTA | Instituto Butantan / Mendelics | Dimas Tadeu Covas, Antonio Jorge Martins, Claudia Renata dos Santos Barros, David Schlesinger, Debora Botequio Moretti, Elaine Cristina Marqueze, Elaine Vieira Santos, Evandra Strazza Rodrigues, Heidge Fukumasu, Jayme Augusto de Souza-Neto, José Salvatore Leister Patané, Luiz Alcantara, Luiz Lehmann Coutinho, Maria Carolina Elias, Maurício Lacerda Nogueira, Rafael dos Santos Bezerra, Raul Machado Neto, Rejane Maria Tommasini Grotto, Ricardo Haddad, Sandra Coccuzzo Sampaio Vessoni, Simone Kashima, Svetoslav Nanev Slavov, Vincent Louis Viala |
| EPI_ISL_2362247 | GRUPO TECNICO DE VIGILANCIA SANITARIA E EPIDEMIOLOGICA | Instituto Butantan / Mendelics | Dimas Tadeu Covas, Antonio Jorge Martins, Claudia Renata dos Santos Barros, David Schlesinger, Debora Botequio Moretti, Elaine Cristina Marqueze, Elaine Vieira Santos, Evandra Strazza Rodrigues, Heidge Fukumasu, Jayme Augusto de Souza-Neto, José Salvatore Leister Patané, Luiz Alcantara, Luiz Lehmann Coutinho, Maria Carolina Elias, Maurício Lacerda Nogueira, Rafael dos Santos Bezerra, Raul Machado Neto, Rejane Maria Tommasini Grotto, Ricardo Haddad, Sandra Coccuzzo Sampaio Vessoni, Simone Kashima, Svetoslav Nanev Slavov, Vincent Louis Viala |
| EPI_ISL_2362248 | HOSPITAL MUNICIPAL DE ITAPIRA | Instituto Butantan / Mendelics | Dimas Tadeu Covas, Antonio Jorge Martins, Claudia Renata dos Santos Barros, David Schlesinger, Debora Botequio Moretti, Elaine Cristina Marqueze, Elaine Vieira Santos, Evandra Strazza Rodrigues, Heidge Fukumasu, Jayme Augusto de Souza-Neto, José Salvatore Leister Patané, Luiz Alcantara, Luiz |

| | | | |
|---|---|---|---|
| | | | Lehmann Coutinho, Maria Carolina Elias, Maurício Lacerda Nogueira, Rafael dos Santos Bezerra, Raul Machado Neto, Rejane Maria Tommasini Grotto, Ricardo Haddad, Sandra Coccuzzo Sampaio Vessoni, Simone Kashima, Svetoslav Nanev Slavov, Vincent Louis Viala |
| EPI_ISL_2362249 | SANTA CASA DE MISERICORDIA SAO JOSE | Instituto Butantan / Mendelics | Dimas Tadeu Covas, Antonio Jorge Martins, Claudia Renata dos Santos Barros, David Schlesinger, Debora Botequio Moretti, Elaine Cristina Marqueze, Elaine Vieira Santos, Evandra Strazza Rodrigues, Heidge Fukumasu, Jayme Augusto de Souza-Neto, José Salvatore Leister Patané, Luiz Alcantara, Luiz Lehmann Coutinho, Maria Carolina Elias, Maurício Lacerda Nogueira, Rafael dos Santos Bezerra, Raul Machado Neto, Rejane Maria Tommasini Grotto, Ricardo Haddad, Sandra Coccuzzo Sampaio Vessoni, Simone Kashima, Svetoslav Nanev Slavov, Vincent Louis Viala |
| EPI_ISL_2362250 | VIGILANCIA EM SAUDE SAE VIGILANCIA EPIDEMIOLOGICA | Instituto Butantan / Mendelics | Dimas Tadeu Covas, Antonio Jorge Martins, Claudia Renata dos Santos Barros, David Schlesinger, Debora Botequio Moretti, Elaine Cristina Marqueze, Elaine Vieira Santos, Evandra Strazza Rodrigues, Heidge Fukumasu, Jayme Augusto de Souza-Neto, José Salvatore Leister Patané, Luiz Alcantara, Luiz Lehmann Coutinho, Maria Carolina Elias, Maurício Lacerda Nogueira, Rafael dos Santos Bezerra, Raul Machado Neto, Rejane Maria Tommasini Grotto, Ricardo Haddad, Sandra Coccuzzo Sampaio Vessoni, Simone Kashima, Svetoslav Nanev Slavov, Vincent Louis Viala |
| EPI_ISL_2362251 | CENTRO DE SAUDE II IBITINGA | Instituto Butantan / Mendelics | Dimas Tadeu Covas, Antonio Jorge Martins, Claudia Renata dos Santos Barros, David Schlesinger, Debora Botequio Moretti, Elaine Cristina Marqueze, Elaine Vieira Santos, Evandra Strazza Rodrigues, Heidge Fukumasu, Jayme Augusto de Souza-Neto, José Salvatore Leister Patané, Luiz Alcantara, Luiz Lehmann Coutinho, Maria Carolina Elias, Maurício Lacerda Nogueira, Rafael dos Santos Bezerra, Raul Machado Neto, Rejane Maria Tommasini Grotto, Ricardo Haddad, Sandra Coccuzzo Sampaio Vessoni, Simone Kashima, Svetoslav Nanev Slavov, Vincent Louis Viala |
| EPI_ISL_2362252 | SECRETARIA DE SAUDE | Instituto Butantan / Mendelics | Dimas Tadeu Covas, Antonio Jorge Martins, Claudia Renata dos Santos Barros, David Schlesinger, Debora Botequio Moretti, Elaine Cristina Marqueze, Elaine Vieira Santos, Evandra Strazza Rodrigues, Heidge Fukumasu, Jayme Augusto de Souza-Neto, José Salvatore Leister Patané, Luiz Alcantara, Luiz Lehmann Coutinho, Maria Carolina Elias, Maurício Lacerda Nogueira, Rafael dos Santos Bezerra, Raul Machado Neto, Rejane Maria Tommasini Grotto, Ricardo Haddad, Sandra Coccuzzo Sampaio Vessoni, Simone Kashima, Svetoslav Nanev Slavov, Vincent Louis Viala |
| EPI_ISL_2362253 | LABORATORIO DR PAULO EMILIO DALESSANDRO PINDAMONHANGABA | Instituto Butantan / Mendelics | Dimas Tadeu Covas, Antonio Jorge Martins, Claudia Renata dos Santos Barros, David Schlesinger, Debora Botequio Moretti, Elaine Cristina Marqueze, Elaine Vieira Santos, Evandra Strazza Rodrigues, Heidge Fukumasu, Jayme Augusto de Souza-Neto, José Salvatore Leister Patané, Luiz Alcantara, Luiz Lehmann Coutinho, Maria Carolina Elias, Maurício Lacerda Nogueira, Rafael dos Santos Bezerra, Raul Machado Neto, Rejane Maria Tommasini Grotto, Ricardo Haddad, Sandra Coccuzzo Sampaio Vessoni, Simone Kashima, Svetoslav Nanev Slavov, Vincent Louis Viala |
| EPI_ISL_2362254 | HOSP E MATERNIDADE MUNICIPAL N SRA MONTE SERRAT | Instituto Butantan / Mendelics | Dimas Tadeu Covas, Antonio Jorge Martins, Claudia Renata dos Santos Barros, David Schlesinger, Debora Botequio Moretti, Elaine Cristina Marqueze, Elaine Vieira Santos, Evandra Strazza Rodrigues, Heidge Fukumasu, Jayme Augusto de Souza-Neto, José Salvatore Leister Patané, Luiz Alcantara, Luiz Lehmann Coutinho, Maria Carolina Elias, Maurício Lacerda Nogueira, Rafael dos Santos Bezerra, Raul Machado Neto, Rejane Maria Tommasini Grotto, Ricardo Haddad, Sandra Coccuzzo Sampaio Vessoni, Simone Kashima, Svetoslav Nanev Slavov, Vincent Louis Viala |
| EPI_ISL_2362255 | UBS RAIMUNDA CAVALCANTE DE SOUZA | Instituto Butantan / Mendelics | Dimas Tadeu Covas, Antonio Jorge Martins, Claudia Renata dos Santos Barros, David Schlesinger, Debora Botequio Moretti, Elaine Cristina Marqueze, Elaine Vieira Santos, Evandra Strazza Rodrigues, Heidge Fukumasu, Jayme Augusto de Souza-Neto, José Salvatore Leister Patané, Luiz Alcantara, Luiz Lehmann Coutinho, Maria Carolina Elias, Maurício Lacerda Nogueira, Rafael dos Santos Bezerra, Raul Machado Neto, Rejane Maria Tommasini Grotto, Ricardo Haddad, Sandra Coccuzzo Sampaio Vessoni, Simone Kashima, Svetoslav Nanev Slavov, Vincent Louis Viala |
| EPI_ISL_2362256 | UPA DR FRANCO DA ROCHA | Instituto Butantan / Mendelics | Dimas Tadeu Covas, Antonio Jorge Martins, Claudia Renata dos Santos Barros, David Schlesinger, Debora Botequio Moretti, Elaine Cristina Marqueze, Elaine Vieira Santos, Evandra Strazza Rodrigues, Heidge Fukumasu, Jayme Augusto de Souza-Neto, José Salvatore Leister Patané, Luiz Alcantara, Luiz Lehmann Coutinho, Maria Carolina Elias, Maurício Lacerda Nogueira, Rafael dos Santos Bezerra, Raul Machado Neto, Rejane Maria Tommasini Grotto, Ricardo Haddad, Sandra Coccuzzo Sampaio Vessoni, Simone Kashima, Svetoslav Nanev Slavov, Vincent Louis Viala |
| EPI_ISL_2362257 | SANTA CASA DE MISERICORDIA SAO JOSE | Instituto Butantan / Mendelics | Dimas Tadeu Covas, Antonio Jorge Martins, Claudia Renata dos Santos Barros, David Schlesinger, Debora Botequio Moretti, Elaine Cristina Marqueze, Elaine Vieira Santos, Evandra Strazza Rodrigues, Heidge Fukumasu, Jayme Augusto de Souza-Neto, José Salvatore Leister Patané, Luiz Alcantara, Luiz Lehmann Coutinho, Maria Carolina Elias, Maurício Lacerda Nogueira, Rafael dos Santos Bezerra, Raul Machado Neto, Rejane Maria Tommasini Grotto, Ricardo Haddad, Sandra Coccuzzo Sampaio Vessoni, Simone Kashima, Svetoslav Nanev Slavov, Vincent Louis Viala |
| EPI_ISL_2362258 | UBS GETULINO JOSE DIAS | Instituto Butantan / Mendelics | Dimas Tadeu Covas, Antonio Jorge Martins, Claudia Renata dos Santos Barros, David Schlesinger, Debora Botequio Moretti, Elaine Cristina Marqueze, Elaine Vieira Santos, Evandra Strazza Rodrigues, Heidge Fukumasu, Jayme Augusto de Souza-Neto, José Salvatore Leister Patané, Luiz Alcantara, Luiz Lehmann Coutinho, Maria Carolina Elias, Maurício Lacerda Nogueira, Rafael dos Santos Bezerra, Raul Machado Neto, Rejane Maria Tommasini Grotto, Ricardo Haddad, Sandra Coccuzzo Sampaio Vessoni, Simone Kashima, Svetoslav Nanev Slavov, Vincent Louis Viala |
| EPI_ISL_2362259 | HOSPITAL MUNICIPAL DR WALDEMAR TEBALDI | Instituto Butantan / Mendelics | Dimas Tadeu Covas, Antonio Jorge Martins, Claudia Renata dos Santos Barros, David Schlesinger, Debora Botequio Moretti, Elaine Cristina Marqueze, Elaine Vieira Santos, Evandra Strazza Rodrigues, Heidge Fukumasu, Jayme Augusto de Souza-Neto, José Salvatore Leister Patané, Luiz Alcantara, Luiz Lehmann Coutinho, Maria Carolina Elias, Maurício Lacerda Nogueira, Rafael dos Santos Bezerra, Raul Machado Neto, Rejane Maria Tommasini Grotto, Ricardo Haddad, Sandra Coccuzzo Sampaio Vessoni, Simone Kashima, Svetoslav Nanev Slavov, Vincent Louis Viala |
| EPI_ISL_2362260 | CENTRO DE SAUDE III AFFONSO LUZZI SANTA CRUZ DAS PALMEIRAS | Instituto Butantan / Mendelics | Dimas Tadeu Covas, Antonio Jorge Martins, Claudia Renata dos Santos Barros, David Schlesinger, Debora Botequio Moretti, Elaine Cristina Marqueze, Elaine Vieira Santos, Evandra Strazza Rodrigues, Heidge Fukumasu, Jayme Augusto de Souza-Neto, José Salvatore Leister Patané, Luiz Alcantara, Luiz Lehmann Coutinho, Maria Carolina Elias, Maurício Lacerda Nogueira, Rafael dos Santos Bezerra, Raul Machado Neto, Rejane Maria Tommasini Grotto, Ricardo Haddad, Sandra Coccuzzo Sampaio Vessoni, Simone Kashima, Svetoslav Nanev Slavov, Vincent Louis Viala |
| EPI_ISL_2362261 | SANTA CASA DE MISERICORDIA SAO JOSE | Instituto Butantan / Mendelics | Dimas Tadeu Covas, Antonio Jorge Martins, Claudia Renata dos Santos Barros, David Schlesinger, Debora Botequio Moretti, Elaine Cristina Marqueze, Elaine Vieira Santos, Evandra Strazza Rodrigues, Heidge Fukumasu, Jayme Augusto de Souza-Neto, José Salvatore Leister Patané, Luiz Alcantara, Luiz Lehmann Coutinho, Maria Carolina Elias, Maurício Lacerda Nogueira, Rafael dos Santos Bezerra, Raul Machado Neto, Rejane Maria Tommasini Grotto, Ricardo Haddad, Sandra Coccuzzo Sampaio Vessoni, Simone Kashima, Svetoslav Nanev Slavov, Vincent Louis Viala |
| EPI_ISL_2362262 | CENTRO DE SAUDE II DR GABRIEL MESQUITA VARGEM GDE DO SUL | Instituto Butantan / Mendelics | Dimas Tadeu Covas, Antonio Jorge Martins, Claudia Renata dos Santos Barros, David Schlesinger, Debora Botequio Moretti, Elaine Cristina Marqueze, Elaine Vieira Santos, Evandra Strazza Rodrigues, Heidge Fukumasu, Jayme Augusto de Souza-Neto, José Salvatore Leister Patané, Luiz Alcantara, Luiz Lehmann Coutinho, Maria Carolina Elias, Maurício Lacerda Nogueira, Rafael dos Santos Bezerra, Raul Machado Neto, Rejane Maria Tommasini Grotto, Ricardo Haddad, Sandra Coccuzzo Sampaio Vessoni, Simone Kashima, Svetoslav Nanev Slavov, Vincent Louis Viala |
| EPI_ISL_2362263, EPI_ISL_2362264 | LABORATORIO DR PAULO EMILIO DALESSANDRO PINDAMONHANGABA | Instituto Butantan / Mendelics | Dimas Tadeu Covas, Antonio Jorge Martins, Claudia Renata dos Santos Barros, David Schlesinger, Debora Botequio Moretti, Elaine Cristina Marqueze, Elaine Vieira Santos, Evandra Strazza Rodrigues, Heidge Fukumasu, Jayme Augusto de Souza-Neto, José Salvatore Leister Patané, Luiz Alcantara, Luiz Lehmann Coutinho, Maria Carolina Elias, Maurício Lacerda Nogueira, Rafael dos Santos Bezerra, Raul Machado Neto, Rejane Maria Tommasini Grotto, Ricardo Haddad, Sandra Coccuzzo Sampaio Vessoni, Simone Kashima, Svetoslav Nanev Slavov, Vincent Louis Viala |
| EPI_ISL_2362266, EPI_ISL_2362267 | HOSPITAL DE CAMPANHA COVID 19 CAIEIRAS | Instituto Butantan / Mendelics | Dimas Tadeu Covas, Antonio Jorge Martins, Claudia Renata dos Santos Barros, David Schlesinger, Debora Botequio Moretti, Elaine Cristina Marqueze, Elaine Vieira Santos, Evandra Strazza Rodrigues, Heidge Fukumasu, Jayme Augusto de Souza-Neto, José Salvatore Leister Patané, Luiz Alcantara, Luiz Lehmann Coutinho, Maria Carolina Elias, Maurício Lacerda Nogueira, Rafael dos Santos Bezerra, Raul Machado Neto, Rejane Maria Tommasini Grotto, Ricardo Haddad, Sandra Coccuzzo Sampaio Vessoni, Simone Kashima, Svetoslav Nanev Slavov, Vincent Louis Viala |
| EPI_ISL_2362268 | CSII DR WASHINGTON LUIS M RODRIGUES DA SILVA PITANGUEIRAS | Instituto Butantan / Mendelics | Dimas Tadeu Covas, Antonio Jorge Martins, Claudia Renata dos Santos Barros, David Schlesinger, Debora Botequio Moretti, Elaine Cristina Marqueze, Elaine Vieira Santos, Evandra Strazza Rodrigues, Heidge Fukumasu, Jayme Augusto de Souza-Neto, José Salvatore Leister Patané, Luiz Alcantara, Luiz Lehmann Coutinho, Maria Carolina Elias, Maurício Lacerda Nogueira, Rafael dos Santos Bezerra, Raul Machado Neto, Rejane Maria Tommasini Grotto, Ricardo Haddad, Sandra Coccuzzo Sampaio Vessoni, Simone Kashima, Svetoslav Nanev Slavov, Vincent Louis Viala |
| EPI_ISL_2362269 | SANTA CASA DE MISERICORDIA SAO JOSE | Instituto Butantan / Mendelics | Dimas Tadeu Covas, Antonio Jorge Martins, Claudia Renata dos Santos Barros, David Schlesinger, Debora Botequio Moretti, Elaine Cristina Marqueze, Elaine Vieira Santos, Evandra Strazza Rodrigues, Heidge Fukumasu, Jayme Augusto de Souza-Neto, José Salvatore Leister Patané, Luiz Alcantara, Luiz Lehmann Coutinho, Maria Carolina Elias, Maurício Lacerda Nogueira, Rafael dos Santos Bezerra, Raul Machado Neto, Rejane Maria Tommasini Grotto, Ricardo Haddad, Sandra Coccuzzo Sampaio Vessoni, Simone Kashima, Svetoslav Nanev Slavov, Vincent Louis Viala |
| EPI_ISL_2362270 | UBS GUILHERMINA DE NOBREGA ABREU | Instituto Butantan / Mendelics | Dimas Tadeu Covas, Antonio Jorge Martins, Claudia Renata dos Santos Barros, David Schlesinger, Debora Botequio Moretti, Elaine Cristina Marqueze, Elaine Vieira Santos, Evandra Strazza Rodrigues, Heidge Fukumasu, Jayme Augusto de Souza-Neto, José Salvatore Leister Patané, Luiz Alcantara, Luiz Lehmann Coutinho, Maria Carolina Elias, Maurício Lacerda Nogueira, Rafael dos Santos Bezerra, Raul Machado Neto, Rejane Maria Tommasini Grotto, Ricardo Haddad, Sandra Coccuzzo Sampaio Vessoni, Simone Kashima, Svetoslav Nanev Slavov, Vincent Louis Viala |

| | | | |
|---|---|---|---|
| EPI_ISL_2362271 | GRUPO TECNICO DE VIGILANCIA SANITARIA E EPIDEMIOLOGICA | Instituto Butantan / Mendelics | Dimas Tadeu Covas, Antonio Jorge Martins, Claudia Renata dos Santos Barros, David Schlesinger, Debora Botequio Moretti, Elaine Cristina Marqueze, Elaine Vieira Santos, Evandra Strazza Rodrigues, Heidge Fukumasu, Jayme Augusto de Souza-Neto, José Salvatore Leister Patané, Luiz Alcantara, Luiz Lehmann Coutinho, Maria Carolina Elias, Maurício Lacerda Nogueira, Rafael dos Santos Bezerra, Raul Machado Neto, Rejane Maria Tommasini Grotto, Ricardo Haddad, Sandra Coccuzzo Sampaio Vessoni, Simone Kashima, Svetoslav Nanev Slavov, Vincent Louis Viala |
| EPI_ISL_2362272 | HOSPITAL MUNICIPAL DR WALDEMAR TEBALDI | Instituto Butantan / Mendelics | Dimas Tadeu Covas, Antonio Jorge Martins, Claudia Renata dos Santos Barros, David Schlesinger, Debora Botequio Moretti, Elaine Cristina Marqueze, Elaine Vieira Santos, Evandra Strazza Rodrigues, Heidge Fukumasu, Jayme Augusto de Souza-Neto, José Salvatore Leister Patané, Luiz Alcantara, Luiz Lehmann Coutinho, Maria Carolina Elias, Maurício Lacerda Nogueira, Rafael dos Santos Bezerra, Raul Machado Neto, Rejane Maria Tommasini Grotto, Ricardo Haddad, Sandra Coccuzzo Sampaio Vessoni, Simone Kashima, Svetoslav Nanev Slavov, Vincent Louis Viala |
| EPI_ISL_2362274, EPI_ISL_2362275 | LABORATORIO DR PAULO EMILIO DALESSANDRO PINDAMONHANGABA | Instituto Butantan / Mendelics | Dimas Tadeu Covas, Antonio Jorge Martins, Claudia Renata dos Santos Barros, David Schlesinger, Debora Botequio Moretti, Elaine Cristina Marqueze, Elaine Vieira Santos, Evandra Strazza Rodrigues, Heidge Fukumasu, Jayme Augusto de Souza-Neto, José Salvatore Leister Patané, Luiz Alcantara, Luiz Lehmann Coutinho, Maria Carolina Elias, Maurício Lacerda Nogueira, Rafael dos Santos Bezerra, Raul Machado Neto, Rejane Maria Tommasini Grotto, Ricardo Haddad, Sandra Coccuzzo Sampaio Vessoni, Simone Kashima, Svetoslav Nanev Slavov, Vincent Louis Viala |
| EPI_ISL_2375477 | Laboratorio Central de Saude Publica do Estado do Espirito Santo (LACEN/ES) | Laboratory of Respiratory Viruses and Measles, Oswaldo Cruz Institute, FIOCRUZ | Paola Resende, Luciana Appolinario, Fernando Motta, Anna Carolina Paixao, Ana Carolina Mendonca, Alice Sampaio Rocha, Taina Venas, Eilisa Cavalcante Pereira, Renata Serrano Lopes, Rodrigo Ribeiro Rodrigues, Marilda Siqueira on behalf of the Fiocruz COVID-19 Genomic Surveillance Network |
| EPI_ISL_2375797, EPI_ISL_2375798 | Laboratório de Microbiologia Molecular - Universidade FEEVALE | Molecular Microbiology Laboratory | Alana Witt Hansen, Fágner Henrique Heldt, Fernando Rosado Spilki, Flávio Silveira, Juliana Schons Gularte, Juliane Deise Fleck, Mariana Soares da Silva, Meriane Demoliner, Matheus Nunes Weber, Paula Rodrigues de Almeida, Micheli Filippi. |
| EPI_ISL_2375845 | Laboratório de Microbiologia Molecular - Universidade FEEVALE | Molecular Microbiology Laboratory | Alana Witt Hansen, Fágner Henrique Heldt, Fernando Rosado Spilki, Flávio Silveira, Juliana Schons Gularte, Juliane Deise Fleck, Mariana Soares da Silva, Meriane Demoliner, Matheus Nunes Weber, Paula Rodrigues de Almeida, Micheli Filippi. |
| EPI_ISL_2378643, EPI_ISL_2378646, EPI_ISL_2378649, EPI_ISL_2378650, EPI_ISL_2378653, EPI_ISL_2378656, EPI_ISL_2378659, EPI_ISL_2378661, EPI_ISL_2378664, EPI_ISL_2378665, EPI_ISL_2378666, EPI_ISL_2378667, EPI_ISL_2378668, EPI_ISL_2378669, EPI_ISL_2378671, EPI_ISL_2378672, EPI_ISL_2378673, EPI_ISL_2378674, EPI_ISL_2378675, EPI_ISL_2378676, EPI_ISL_2378677, EPI_ISL_2378678, EPI_ISL_2378679, EPI_ISL_2378680, EPI_ISL_2378681, EPI_ISL_2378684, EPI_ISL_2378686, EPI_ISL_2378688, EPI_ISL_2378689, EPI_ISL_2378690, EPI_ISL_2378691, EPI_ISL_2378692, EPI_ISL_2378693, EPI_ISL_2378694 | | | |
| see above | Instituto de Biotecnologia - UNESP-Botucatu-SP | Instituto de Biotecnologia - UNESP-Botucatu-SP | Fábio Sossai Possebon; Leila Sabrina Ullmann; Cecília Artico Banho; Cíntia Bittar; Guilherme Campos; Helena Lage Ferreira; Jorge A. Petroli Marchesi; Lívia Sacchetto; Maisa C. Pereira Parra; Marília Moraes; Maurício L. Nogueira; Paula Rahal; Paulo Inacio da Costa; João Pessoa Araújo Jr. |
| EPI_ISL_2378736 | UPA II JUNDIAI | Instituto Butantan | Dimas Tadeu Covas, Antonio Jorge Martins, Claudia Renata dos Santos Barros, David Schlesinger, Debora Botequio Moretti, Elaine Cristina Marqueze, Elaine Vieira Santos, Evandra Strazza Rodrigues, Heidge Fukumasu, Jayme Augusto de Souza-Neto, José Salvatore Leister Patané, Luiz Alcantara, Luiz Lehmann Coutinho, Maria Carolina Elias, Maurício Lacerda Nogueira, Rafael dos Santos Bezerra, Raul Machado Neto, Rejane Maria Tommasini Grotto, Ricardo Haddad, Sandra Coccuzzo Sampaio Vessoni, Simone Kashima, Svetoslav Nanev Slavov, Vincent Louis Viala |
| EPI_ISL_2378737 | UPA DE BEBEDOURO | Instituto Butantan | Dimas Tadeu Covas, Antonio Jorge Martins, Claudia Renata dos Santos Barros, David Schlesinger, Debora Botequio Moretti, Elaine Cristina Marqueze, Elaine Vieira Santos, Evandra Strazza Rodrigues, Heidge Fukumasu, Jayme Augusto de Souza-Neto, José Salvatore Leister Patané, Luiz Alcantara, Luiz Lehmann Coutinho, Maria Carolina Elias, Maurício Lacerda Nogueira, Rafael dos Santos Bezerra, Raul Machado Neto, Rejane Maria Tommasini Grotto, Ricardo Haddad, Sandra Coccuzzo Sampaio Vessoni, Simone Kashima, Svetoslav Nanev Slavov, Vincent Louis Viala |
| EPI_ISL_2378738 | USAFA FORTE | Instituto Butantan | Dimas Tadeu Covas, Antonio Jorge Martins, Claudia Renata dos Santos Barros, David Schlesinger, Debora Botequio Moretti, Elaine Cristina Marqueze, Elaine Vieira Santos, Evandra Strazza Rodrigues, Heidge Fukumasu, Jayme Augusto de Souza-Neto, José Salvatore Leister Patané, Luiz Alcantara, Luiz Lehmann Coutinho, Maria Carolina Elias, Maurício Lacerda Nogueira, Rafael dos Santos Bezerra, Raul Machado Neto, Rejane Maria Tommasini Grotto, Ricardo Haddad, Sandra Coccuzzo Sampaio Vessoni, Simone Kashima, Svetoslav Nanev Slavov, Vincent Louis Viala |
| EPI_ISL_2378739 | AMBULATORIO AVIACAO | Instituto Butantan | Dimas Tadeu Covas, Antonio Jorge Martins, Claudia Renata dos Santos Barros, David Schlesinger, Debora Botequio Moretti, Elaine Cristina Marqueze, Elaine Vieira Santos, Evandra Strazza Rodrigues, Heidge Fukumasu, Jayme Augusto de Souza-Neto, José Salvatore Leister Patané, Luiz Alcantara, Luiz Lehmann Coutinho, Maria Carolina Elias, Maurício Lacerda Nogueira, Rafael dos Santos Bezerra, Raul Machado Neto, Rejane Maria Tommasini Grotto, Ricardo Haddad, Sandra Coccuzzo Sampaio Vessoni, Simone Kashima, Svetoslav Nanev Slavov, Vincent Louis Viala |
| EPI_ISL_2378740, EPI_ISL_2378741 | UPA DE BEBEDOURO | Instituto Butantan | Dimas Tadeu Covas, Antonio Jorge Martins, Claudia Renata dos Santos Barros, David Schlesinger, Debora Botequio Moretti, Elaine Cristina Marqueze, Elaine Vieira Santos, Evandra Strazza Rodrigues, Heidge Fukumasu, Jayme Augusto de Souza-Neto, José Salvatore Leister Patané, Luiz Alcantara, Luiz Lehmann Coutinho, Maria Carolina Elias, Maurício Lacerda Nogueira, Rafael dos Santos Bezerra, Raul Machado Neto, Rejane Maria Tommasini Grotto, Ricardo Haddad, Sandra Coccuzzo Sampaio Vessoni, Simone Kashima, Svetoslav Nanev Slavov, Vincent Louis Viala |
| EPI_ISL_2378742 | UNIDADE DE SAUDE DR PHEBO DE OLIVEIRA ROGE FERREIRA | Instituto Butantan | Dimas Tadeu Covas, Antonio Jorge Martins, Claudia Renata dos Santos Barros, David Schlesinger, Debora Botequio Moretti, Elaine Cristina Marqueze, Elaine Vieira Santos, Evandra Strazza Rodrigues, Heidge Fukumasu, Jayme Augusto de Souza-Neto, José Salvatore Leister Patané, Luiz Alcantara, Luiz Lehmann Coutinho, Maria Carolina Elias, Maurício Lacerda Nogueira, Rafael dos Santos Bezerra, Raul Machado Neto, Rejane Maria Tommasini Grotto, Ricardo Haddad, Sandra Coccuzzo Sampaio Vessoni, Simone Kashima, Svetoslav Nanev Slavov, Vincent Louis Viala |
| EPI_ISL_2378743 | CSII DR WASHINGTON LUIS M RODRIGUES DA SILVA PITANGUEIRAS | Instituto Butantan | Dimas Tadeu Covas, Antonio Jorge Martins, Claudia Renata dos Santos Barros, David Schlesinger, Debora Botequio Moretti, Elaine Cristina Marqueze, Elaine Vieira Santos, Evandra Strazza Rodrigues, Heidge Fukumasu, Jayme Augusto de Souza-Neto, José Salvatore Leister Patané, Luiz Alcantara, Luiz Lehmann Coutinho, Maria Carolina Elias, Maurício Lacerda Nogueira, Rafael dos Santos Bezerra, Raul Machado Neto, Rejane Maria Tommasini Grotto, Ricardo Haddad, Sandra Coccuzzo Sampaio Vessoni, Simone Kashima, Svetoslav Nanev Slavov, Vincent Louis Viala |
| EPI_ISL_2378744 | CS DE APARECIDA DOESTE | Instituto Butantan | Dimas Tadeu Covas, Antonio Jorge Martins, Claudia Renata dos Santos Barros, David Schlesinger, Debora Botequio Moretti, Elaine Cristina Marqueze, Elaine Vieira Santos, Evandra Strazza Rodrigues, Heidge Fukumasu, Jayme Augusto de Souza-Neto, José Salvatore Leister Patané, Luiz Alcantara, Luiz Lehmann Coutinho, Maria Carolina Elias, Maurício Lacerda Nogueira, Rafael dos Santos Bezerra, Raul Machado Neto, Rejane Maria Tommasini Grotto, Ricardo Haddad, Sandra Coccuzzo Sampaio Vessoni, Simone Kashima, Svetoslav Nanev Slavov, Vincent Louis Viala |
| EPI_ISL_2378745 | HOSPITAL DO RIM E HIPERTENSAO | Instituto Butantan | Dimas Tadeu Covas, Antonio Jorge Martins, Claudia Renata dos Santos Barros, David Schlesinger, Debora Botequio Moretti, Elaine Cristina Marqueze, Elaine Vieira Santos, Evandra Strazza Rodrigues, Heidge Fukumasu, Jayme Augusto de Souza-Neto, José Salvatore Leister Patané, Luiz Alcantara, Luiz Lehmann Coutinho, Maria Carolina Elias, Maurício Lacerda Nogueira, Rafael dos Santos Bezerra, Raul Machado Neto, Rejane Maria Tommasini Grotto, Ricardo Haddad, Sandra Coccuzzo Sampaio Vessoni, Simone Kashima, Svetoslav Nanev Slavov, Vincent Louis Viala |
| EPI_ISL_2378746 | UBS II DE PIRAPOZINHO C SERVICO DE EACS E ESF | Instituto Butantan | Dimas Tadeu Covas, Antonio Jorge Martins, Claudia Renata dos Santos Barros, David Schlesinger, Debora Botequio Moretti, Elaine Cristina Marqueze, Elaine Vieira Santos, Evandra Strazza Rodrigues, Heidge Fukumasu, Jayme Augusto de Souza-Neto, José Salvatore Leister Patané, Luiz Alcantara, Luiz Lehmann Coutinho, Maria Carolina Elias, Maurício Lacerda Nogueira, Rafael dos Santos Bezerra, Raul Machado Neto, Rejane Maria Tommasini Grotto, Ricardo Haddad, Sandra Coccuzzo Sampaio Vessoni, Simone Kashima, Svetoslav Nanev Slavov, Vincent Louis Viala |
| EPI_ISL_2378747 | PROGRAMA SAUDE DA FAMILIA I IEPE | Instituto Butantan | Dimas Tadeu Covas, Antonio Jorge Martins, Claudia Renata dos Santos Barros, David Schlesinger, Debora Botequio Moretti, Elaine Cristina Marqueze, Elaine Vieira Santos, Evandra Strazza Rodrigues, Heidge Fukumasu, Jayme Augusto de Souza-Neto, José Salvatore Leister Patané, Luiz Alcantara, Luiz Lehmann Coutinho, Maria Carolina Elias, Maurício Lacerda Nogueira, Rafael dos Santos Bezerra, Raul Machado Neto, Rejane Maria Tommasini Grotto, Ricardo Haddad, Sandra Coccuzzo Sampaio Vessoni, Simone Kashima, Svetoslav Nanev Slavov, Vincent Louis Viala |
| EPI_ISL_2378748, EPI_ISL_2378749 | PAS JOAO ANTONIO DO NASCIMENTO | Instituto Butantan | Dimas Tadeu Covas, Antonio Jorge Martins, Claudia Renata dos Santos Barros, David Schlesinger, Debora Botequio Moretti, Elaine Cristina Marqueze, Elaine Vieira Santos, Evandra Strazza Rodrigues, Heidge Fukumasu, Jayme Augusto de Souza-Neto, José Salvatore Leister Patané, Luiz Alcantara, Luiz Lehmann Coutinho, Maria Carolina Elias, Maurício Lacerda Nogueira, Rafael dos Santos Bezerra, Raul Machado Neto, Rejane Maria Tommasini Grotto, Ricardo Haddad, Sandra Coccuzzo Sampaio Vessoni, Simone Kashima, Svetoslav Nanev Slavov, Vincent Louis Viala |
| EPI_ISL_2378750 | UNIDADE DE SAUDE DA FAMILIA DE FLORIDA PAULISTA II | Instituto Butantan | Dimas Tadeu Covas, Antonio Jorge Martins, Claudia Renata dos Santos Barros, David Schlesinger, Debora Botequio Moretti, Elaine Cristina Marqueze, Elaine Vieira Santos, Evandra Strazza Rodrigues, Heidge Fukumasu, Jayme Augusto de Souza-Neto, José Salvatore Leister Patané, Luiz Alcantara, Luiz Lehmann Coutinho, Maria Carolina Elias, Maurício Lacerda Nogueira, Rafael dos Santos Bezerra, Raul Machado Neto, Rejane Maria Tommasini Grotto, Ricardo Haddad, Sandra Coccuzzo Sampaio Vessoni, Simone Kashima, Svetoslav Nanev Slavov, Vincent Louis Viala |
| EPI_ISL_2378751 | USF PAULISTA FERNANDOPOLIS ANTONIO PIVATO | Instituto Butantan | Dimas Tadeu Covas, Antonio Jorge Martins, Claudia Renata dos Santos Barros, David Schlesinger, Debora Botequio Moretti, Elaine Cristina Marqueze, Elaine Vieira Santos, Evandra Strazza Rodrigues, Heidge Fukumasu, Jayme Augusto de Souza-Neto, José Salvatore Leister Patané, Luiz Alcantara, Luiz |

| | | | |
|---|---|---|---|
| | | | Lehmann Coutinho, Maria Carolina Elias, Maurício Lacerda Nogueira, Rafael dos Santos Bezerra, Raul Machado Neto, Rejane Maria Tommasini Grotto, Ricardo Haddad, Sandra Coccuzzo Sampaio Vessoni, Simone Kashima, Svetoslav Nanev Slavov, Vincent Louis Viala |
| EPI_ISL_2378752 | UNIDADE REFERENCIAL SUDOESTE | Instituto Butantan | Dimas Tadeu Covas, Antonio Jorge Martins, Claudia Renata dos Santos Barros, David Schlesinger, Debora Botequio Moretti, Elaine Cristina Marqueze, Elaine Vieira Santos, Evandra Strazza Rodrigues, Heidge Fukumasu, Jayme Augusto de Souza-Neto, José Salvatore Leister Patané, Luiz Alcantara, Luiz Lehmann Coutinho, Maria Carolina Elias, Maurício Lacerda Nogueira, Rafael dos Santos Bezerra, Raul Machado Neto, Rejane Maria Tommasini Grotto, Ricardo Haddad, Sandra Coccuzzo Sampaio Vessoni, Simone Kashima, Svetoslav Nanev Slavov, Vincent Louis Viala |
| EPI_ISL_2378753, EPI_ISL_2378754 | CS DE APARECIDA DOESTE | Instituto Butantan | Dimas Tadeu Covas, Antonio Jorge Martins, Claudia Renata dos Santos Barros, David Schlesinger, Debora Botequio Moretti, Elaine Cristina Marqueze, Elaine Vieira Santos, Evandra Strazza Rodrigues, Heidge Fukumasu, Jayme Augusto de Souza-Neto, José Salvatore Leister Patané, Luiz Alcantara, Luiz Lehmann Coutinho, Maria Carolina Elias, Maurício Lacerda Nogueira, Rafael dos Santos Bezerra, Raul Machado Neto, Rejane Maria Tommasini Grotto, Ricardo Haddad, Sandra Coccuzzo Sampaio Vessoni, Simone Kashima, Svetoslav Nanev Slavov, Vincent Louis Viala |
| EPI_ISL_2382535, EPI_ISL_2382536 | Laboratório de Microbiologia Molecular - Universidade FEEVALE | Molecular Microbiology Laboratory | Alana Witt Hansen, Fágner Henrique Heldt, Fernando Rosado Spilki, Flávio Silveira, Juliana Schons Gularte, Juliane Deise Fleck, Mariana Soares da Silva, Meriane Demoliner, Matheus Nunes Weber, Paula Rodrigues de Almeida, Micheli Filippi. |
| EPI_ISL_2383033 | LABORATORIO MUNICIPAL DE PIRACICABA | Instituto Butantan | Dimas Tadeu Covas, Antonio Jorge Martins, Claudia Renata dos Santos Barros, David Schlesinger, Debora Botequio Moretti, Elaine Cristina Marqueze, Elaine Vieira Santos, Evandra Strazza Rodrigues, Heidge Fukumasu, Jayme Augusto de Souza-Neto, José Salvatore Leister Patané, Luiz Alcantara, Luiz Lehmann Coutinho, Maria Carolina Elias, Maurício Lacerda Nogueira, Rafael dos Santos Bezerra, Raul Machado Neto, Rejane Maria Tommasini Grotto, Ricardo Haddad, Sandra Coccuzzo Sampaio Vessoni, Simone Kashima, Svetoslav Nanev Slavov, Vincent Louis Viala |
| EPI_ISL_2383034 | VIGILANCIA EPIDEMIOLOGICA E CONTROLE DE VETORES PIRASSUNUN | Instituto Butantan | Dimas Tadeu Covas, Antonio Jorge Martins, Claudia Renata dos Santos Barros, David Schlesinger, Debora Botequio Moretti, Elaine Cristina Marqueze, Elaine Vieira Santos, Evandra Strazza Rodrigues, Heidge Fukumasu, Jayme Augusto de Souza-Neto, José Salvatore Leister Patané, Luiz Alcantara, Luiz Lehmann Coutinho, Maria Carolina Elias, Maurício Lacerda Nogueira, Rafael dos Santos Bezerra, Raul Machado Neto, Rejane Maria Tommasini Grotto, Ricardo Haddad, Sandra Coccuzzo Sampaio Vessoni, Simone Kashima, Svetoslav Nanev Slavov, Vincent Louis Viala |
| EPI_ISL_2383035 | UNIDADE BASICA DE SAUDE DR MATHEUS GABRIEL BONASSA | Instituto Butantan | Dimas Tadeu Covas, Antonio Jorge Martins, Claudia Renata dos Santos Barros, David Schlesinger, Debora Botequio Moretti, Elaine Cristina Marqueze, Elaine Vieira Santos, Evandra Strazza Rodrigues, Heidge Fukumasu, Jayme Augusto de Souza-Neto, José Salvatore Leister Patané, Luiz Alcantara, Luiz Lehmann Coutinho, Maria Carolina Elias, Maurício Lacerda Nogueira, Rafael dos Santos Bezerra, Raul Machado Neto, Rejane Maria Tommasini Grotto, Ricardo Haddad, Sandra Coccuzzo Sampaio Vessoni, Simone Kashima, Svetoslav Nanev Slavov, Vincent Louis Viala |
| EPI_ISL_2385432, EPI_ISL_2385433, EPI_ISL_2385434, EPI_ISL_2385435, EPI_ISL_2385436, EPI_ISL_2385437, EPI_ISL_2385438, EPI_ISL_2385439, EPI_ISL_2385440, EPI_ISL_2385441, EPI_ISL_2385442, EPI_ISL_2385443, EPI_ISL_2385444, EPI_ISL_2385445, EPI_ISL_2385446, EPI_ISL_2385447, EPI_ISL_2385448, EPI_ISL_2385449, EPI_ISL_2385450, EPI_ISL_2385451, EPI_ISL_2385452, EPI_ISL_2385453, EPI_ISL_2385454, EPI_ISL_2385455, EPI_ISL_2385456, EPI_ISL_2385457, EPI_ISL_2385458, EPI_ISL_2385459, EPI_ISL_2385460, EPI_ISL_2385461, EPI_ISL_2385462, EPI_ISL_2385463, EPI_ISL_2385464, EPI_ISL_2385465, EPI_ISL_2385466, EPI_ISL_2385467, EPI_ISL_2385468, EPI_ISL_2385469, EPI_ISL_2385470, EPI_ISL_2385471, EPI_ISL_2385472, EPI_ISL_2385474, EPI_ISL_2385475, EPI_ISL_2385476, EPI_ISL_2385477, EPI_ISL_2385478, EPI_ISL_2385479, EPI_ISL_2385480, EPI_ISL_2385481, EPI_ISL_2385482, EPI_ISL_2385483, EPI_ISL_2385484, EPI_ISL_2385485, EPI_ISL_2385486, EPI_ISL_2385487, EPI_ISL_2385488, EPI_ISL_2385489, EPI_ISL_2385490, EPI_ISL_2385491, EPI_ISL_2385492, EPI_ISL_2385493, EPI_ISL_2385494, EPI_ISL_2385495, EPI_ISL_2385496, EPI_ISL_2385498, EPI_ISL_2385499, EPI_ISL_2385500, EPI_ISL_2385501, EPI_ISL_2385502, EPI_ISL_2385503, EPI_ISL_2385504, EPI_ISL_2385505, EPI_ISL_2385506, EPI_ISL_2385507, EPI_ISL_2385508, EPI_ISL_2385510, EPI_ISL_2385511, EPI_ISL_2385512, EPI_ISL_2385513, EPI_ISL_2385514, EPI_ISL_2385515, EPI_ISL_2385516, EPI_ISL_2385517, EPI_ISL_2385518, EPI_ISL_2385519, EPI_ISL_2385520, EPI_ISL_2385521, EPI_ISL_2385522, EPI_ISL_2385523, EPI_ISL_2385524, EPI_ISL_2385525, EPI_ISL_2385526, EPI_ISL_2385527, EPI_ISL_2385528, EPI_ISL_2385529, EPI_ISL_2385530, EPI_ISL_2385531, EPI_ISL_2385532, EPI_ISL_2385533, EPI_ISL_2385534, EPI_ISL_2385535, EPI_ISL_2385536, EPI_ISL_2385537, EPI_ISL_2385538, EPI_ISL_2385539, EPI_ISL_2385541, EPI_ISL_2385542, EPI_ISL_2385543, EPI_ISL_2385544, EPI_ISL_2385545, EPI_ISL_2385546, EPI_ISL_2385547, EPI_ISL_2385548, EPI_ISL_2385549, EPI_ISL_2385550, EPI_ISL_2385551, EPI_ISL_2385552, EPI_ISL_2385553, EPI_ISL_2385554, EPI_ISL_2385555, EPI_ISL_2385556, EPI_ISL_2385558, EPI_ISL_2385559, EPI_ISL_2385561, EPI_ISL_2385562, EPI_ISL_2385563, EPI_ISL_2385564, EPI_ISL_2385565, EPI_ISL_2385566, EPI_ISL_2385567, EPI_ISL_2385568, EPI_ISL_2385570, EPI_ISL_2385571, EPI_ISL_2385572, EPI_ISL_2385573, EPI_ISL_2385574, EPI_ISL_2385575, EPI_ISL_2385576, EPI_ISL_2385577, EPI_ISL_2385578, EPI_ISL_2385579, EPI_ISL_2385580, EPI_ISL_2385581, EPI_ISL_2385582, EPI_ISL_2385583, EPI_ISL_2385584, EPI_ISL_2385586, EPI_ISL_2385587, EPI_ISL_2385588, EPI_ISL_2385589, EPI_ISL_2385590, EPI_ISL_2385591, EPI_ISL_2385592, EPI_ISL_2385593, EPI_ISL_2385594, EPI_ISL_2385595, EPI_ISL_2385596, EPI_ISL_2385597, EPI_ISL_2385598, EPI_ISL_2385599, EPI_ISL_2385600, EPI_ISL_2385601, EPI_ISL_2385602, EPI_ISL_2385603, EPI_ISL_2385604, EPI_ISL_2385605, EPI_ISL_2385606, EPI_ISL_2385607, EPI_ISL_2385608, EPI_ISL_2385609, EPI_ISL_2385610, EPI_ISL_2385611, EPI_ISL_2385612, EPI_ISL_2385613, EPI_ISL_2385616, EPI_ISL_2385620, EPI_ISL_2385624, EPI_ISL_2385625, EPI_ISL_2385627, EPI_ISL_2385629, EPI_ISL_2385630, EPI_ISL_2385631, EPI_ISL_2385632, EPI_ISL_2385633, EPI_ISL_2385634, EPI_ISL_2385636, EPI_ISL_2385637, EPI_ISL_2385639, EPI_ISL_2385641, EPI_ISL_2385642, EPI_ISL_2385644, EPI_ISL_2385647, EPI_ISL_2385648, EPI_ISL_2385649, EPI_ISL_2385650, EPI_ISL_2385651, EPI_ISL_2385652, EPI_ISL_2385653, EPI_ISL_2385654, EPI_ISL_2385655, EPI_ISL_2385656, EPI_ISL_2385657, EPI_ISL_2385658, EPI_ISL_2385659, EPI_ISL_2385660, EPI_ISL_2385661, EPI_ISL_2385662, EPI_ISL_2385663, EPI_ISL_2385664, EPI_ISL_2385665, EPI_ISL_2385666, EPI_ISL_2385667, EPI_ISL_2385668 | | | |
| see above | Laboratorio Central Noel Nutels | Bioinformatics Laboratory / LNCC | Luiz G P de Almeida, Alessandra P Lamarca, Ronaldo da Silva F Jr, Liliane Cavalcante, Alexandra L Gerber, Ana Paula de C Guimaraes, Douglas Terra Machado, Cassia Alves, Diana Mariani, Cintia Policarpo, Gleidson da Silva de Oliveira, Mario Sergio Ribeiro, Silvia Carvalho, Flavio Dias da Silva, Marcio Henrique de Oliveira Garcia, Leandro Magalhaes de Souza, Cristiane Gomes da Silva, Caio Luiz Pereira Ribeiro, Andrea Cony Cavalcanti, Claudia Maria Braga de Mello, Amilcar Tanuri, Ana Tereza R Vasconcelos |
| EPI_ISL_2385669, EPI_ISL_2385670, EPI_ISL_2385671, EPI_ISL_2385674, EPI_ISL_2385675, EPI_ISL_2385676, EPI_ISL_2385677, EPI_ISL_2385678, EPI_ISL_2385679, EPI_ISL_2385680, EPI_ISL_2385681, EPI_ISL_2385682, EPI_ISL_2385683, EPI_ISL_2385684, EPI_ISL_2385685, EPI_ISL_2385686, EPI_ISL_2385689, EPI_ISL_2385690, EPI_ISL_2385691, EPI_ISL_2385692, EPI_ISL_2385693, EPI_ISL_2385694, EPI_ISL_2385695, EPI_ISL_2385696, EPI_ISL_2385697, EPI_ISL_2385698, EPI_ISL_2385699, EPI_ISL_2385700, EPI_ISL_2385701, EPI_ISL_2385702, EPI_ISL_2385704, EPI_ISL_2385705, EPI_ISL_2385707, EPI_ISL_2385708, EPI_ISL_2385709, EPI_ISL_2385711, EPI_ISL_2385712, EPI_ISL_2385713, EPI_ISL_2385714, EPI_ISL_2385717, EPI_ISL_2385718, EPI_ISL_2385719, EPI_ISL_2385720, EPI_ISL_2385722, EPI_ISL_2385723, EPI_ISL_2385724, EPI_ISL_2385725, EPI_ISL_2385726, EPI_ISL_2385727, EPI_ISL_2385728, EPI_ISL_2385729, EPI_ISL_2385730, EPI_ISL_2385731, EPI_ISL_2385732, EPI_ISL_2385733, EPI_ISL_2385734, EPI_ISL_2385735, EPI_ISL_2385736, EPI_ISL_2385737, EPI_ISL_2385739, EPI_ISL_2385740, EPI_ISL_2385741, EPI_ISL_2385742, EPI_ISL_2385743, EPI_ISL_2385745, EPI_ISL_2385748, EPI_ISL_2385749, EPI_ISL_2385750, EPI_ISL_2385751, EPI_ISL_2385752, EPI_ISL_2385753, EPI_ISL_2385754, EPI_ISL_2385755, EPI_ISL_2385756, EPI_ISL_2385757, EPI_ISL_2385758, EPI_ISL_2385759, EPI_ISL_2385760, EPI_ISL_2385761, EPI_ISL_2385764, EPI_ISL_2385765, EPI_ISL_2385766, EPI_ISL_2385768, EPI_ISL_2385769, EPI_ISL_2385770, EPI_ISL_2385771, EPI_ISL_2385772, EPI_ISL_2385773, EPI_ISL_2385774, EPI_ISL_2385775, EPI_ISL_2385776, EPI_ISL_2385777, EPI_ISL_2385778, EPI_ISL_2385779, EPI_ISL_2385780, EPI_ISL_2385781, EPI_ISL_2385782, EPI_ISL_2385784, EPI_ISL_2385785, EPI_ISL_2385786 | | | |
| see above | Unidade de apoio ao diagnóstico da COVID - UNADIG | Bioinformatics Laboratory / LNCC | Luiz G P de Almeida, Alessandra P Lamarca, Ronaldo da Silva F Jr, Liliane Cavalcante, Alexandra L Gerber, Ana Paula de C Guimaraes, Douglas Terra Machado, Cassia Alves, Diana Mariani, Cintia Policarpo, Gleidson da Silva de Oliveira, Mario Sergio Ribeiro, Silvia Carvalho, Flavio Dias da Silva, Marcio Henrique de Oliveira Garcia, Leandro Magalhaes de Souza, Cristiane Gomes da Silva, Caio Luiz Pereira Ribeiro, Andrea Cony Cavalcanti, Claudia Maria Braga de Mello, Amilcar Tanuri, Ana Tereza R Vasconcelos |
| EPI_ISL_2399431, EPI_ISL_2399432, EPI_ISL_2399433, EPI_ISL_2399434, EPI_ISL_2399435, EPI_ISL_2399436, EPI_ISL_2399437 | Laboratório de Microbiologia Molecular - Universidade FEEVALE | Molecular Microbiology Laboratory | Alana Witt Hansen, Fágner Henrique Heldt, Fernando Rosado Spilki, Flávio Silveira, Juliana Schons Gularte, Juliane Deise Fleck, Mariana Soares da Silva, Meriane Demoliner, Matheus Nunes Weber, Paula Rodrigues de Almeida, Micheli Filippi. |
| EPI_ISL_2425441, EPI_ISL_2425442, EPI_ISL_2425443 | Instituto Butantan | Instituto de Medicina Tropical de Sao Paulo | Brazil-UK Centre for Arbovirus Discovery Diagnosis Genomics and Epidemiology (CADDE) Genomic Network - Instituto de Medicina Tropical |
| EPI_ISL_2431429, EPI_ISL_2431431, EPI_ISL_2431433, EPI_ISL_2431436, EPI_ISL_2431437, EPI_ISL_2431439, EPI_ISL_2431440, EPI_ISL_2431441, EPI_ISL_2431843, EPI_ISL_2431844, EPI_ISL_2431846, EPI_ISL_2431848, EPI_ISL_2431852, EPI_ISL_2431853, EPI_ISL_2431855 | | | |
| see above | Laboratório de Microbiologia Molecular - Universidade FEEVALE | Molecular Microbiology Laboratory | Alana Witt Hansen, Fágner Henrique Heldt, Fernando Rosado Spilki, Flávio Silveira, Juliana Schons Gularte, Juliane Deise Fleck, Mariana Soares da Silva, Meriane Demoliner, Matheus Nunes Weber, Paula Rodrigues de Almeida, Micheli Filippi. |
