## Supplementary File 2 for "Molecular evolution and structural analyses of the spike glycoprotein from Brazilian SARS-CoV-2 genomes: the impact of the fixation of selected mutations"

**Table S2.** Positively selected sites detected by the HYPHY Evolutionary methods for pervasive site-level selection.

| Lineage | N | Mutation | Site | FUBAR | | | FEL | | | | SLAC | | |
|---|---|---|---|---|---|---|---|---|---|---|---|---|---|
| | | | | α | β | post. p | α | β | LRT | p | dS | dN | p |
| B.1.1<br>B.1.1.28<br>B.1.1.33<br>B.1.1.7<br>P.1<br>P.1.1<br>P.2 | 7 | L→F<br>0.95 | **5** | 0.465 | 15.525 | 1.0000 | 0.000 | 11.735 | 19.399 | 0.0000 | 0.000 | 17.956 | 0.000 |
| B.1.1<br>B.1.1.28<br>B.1.1.33<br>P.1<br>P.1.2<br>P.2 | 6 | S→F<br>0.17 | **12** | - | - | - | 0.000 | 2.736 | 4.545 | 0.0330 | 0.000 | 3.997 | 0.039 |
| B.1.1 | 1 | S→C<br>0.01 | | | | | | | | | | | |
| B.1<br>B.1.1.28<br>P.1<br>P.2 | 4 | R→I<br>0.08 | **21** | 1.150 | 6.195 | 0.9034 | 0.000 | 4.701 | 2.988 | 0.0839 | 0.000 | 4.998 | 0.030 |
| B.1.1.33<br>P.2 | 2 | R→T<br>0.02 | | | | | | | | | | | |
| P.2 | 1 | R→S<br>0.01 | | | | | | | | | | | |
| B.1.1<br>B.1.1.28<br>B.1.1.29<br>B.1.1.33<br>B.1.1.44<br>B.1.1.7<br>P.1<br>P.1.1<br>P.1.2<br>P.2 | 10 | P→S<br>58.45 | **26** | - | - | - | 1.038 | 4.296 | 2.799 | 0.0943 | 1.000 | 6.498 | 0.027 |

| | | | | | | | | | | | | | |
|---|---|---|---|---|---|---|---|---|---|---|---|---|---|
| P.1 | 1 | P→F<br>0.06 | | | | | | | | | | | |
| P.2 | 1 | P→ L<br>0.01 | | | | | | | | | | | |
| P.1<br>P.1.2 | 2 | A→V<br>0.16 | **27** | - | - | - | - | - | - | - | 0.000 | 3.001 | 0.088 |
| B.1.1.28<br>B.1.1.33<br>P.1<br>P.2 | 4 | A→S<br>0.15 | | | | | | | | | | | |
| B.1.1.28<br>P.1<br>P.2 | 3 | T→I<br>0.10 | **29** | - | - | - | 0.000 | 2.265 | 3.610 | 0.0574 | - | - | - |
| P.2 | 1 | T→A<br>0.01 | | | | | | | | | | | |
| B.1.1.28<br>B.1.1.33<br>B.1.195<br>P.1<br>P.2 | 5 | H→Y<br>0.17 | **49** | - | - | - | 0.000 | 3.524 | 3.697 | 0.0545 | - | - | - |
| B.1.1.33<br>P.2 | 2 | T→N<br>0.03 | **63** | - | - | - | 0.000 | 3.180 | 3.885 | 0.0487 | - | - | - |
| B.1.1.28<br>P.2 | 2 | T→E<br>0.02 | | | | | | | | | | | |
| B.1.1<br>B.1.1.28<br>B.1.1.7<br>B.1.525<br>P.1<br>P.2 | 6 | A→V<br>0.12 | **67** | 0.655 | 6.606 | 0.9824 | 0.000 | 4.581 | 7.605 | 0.0058 | 0.000 | 6.500 | 0.005 |
| B.1.1.28<br>B.1.1.7<br>B.1.195<br>P.1<br>P.2 | 5 | A→S<br>0.08 | | | | | | | | | | | |

| | | | | | | | | | | | | | |
|---|---|---|---|---|---|---|---|---|---|---|---|---|---|
| P.1 | 1 | A→P<br>0.01 | | | | | | | | | | | |
| AV.1<br>B.1.1.28<br>B.1.1.7<br>C.37<br>P.1 | 5 | G→V<br>0.11 | **75** | 0.667 | 6.269 | 0.9727 | 0.000 | 4.404 | 7.260 | 0.0070 | 0.000 | 5.999 | 0.008 |
| B.1.1<br>B.1.1.33 | 2 | G→S<br>0.02 | | | | | | | | | | | |
| P.2 | 1 | G→D<br>0.01 | | | | | | | | | | | |
| B.1.1.28<br>C.37<br>P.1<br>P.2<br>P.4 | 5 | T→I<br>0.10 | **76** | - | - | - | 0.000 | 1.810 | 2.888 | 0.0893 | - | - | - |
| P.1 | 1 | T→N<br>0.01 | | | | | | | | | | | |
| B.1.1.33<br>P.1<br>P.2 | 3 | R→S<br>0.10 | **78** | - | - | - | 0.000 | 2.510 | 2.909 | 0.0881 | - | - | - |
| P.1<br>P.2 | 2 | R→M<br>0.04 | | | | | | | | | | | |
| P.1 | 1 | R→T<br>0.01 | | | | | | | | | | | |
| B.1.1.28<br>B.1.1.33<br>B.1.1.7<br>B.1.221<br>P.1<br>P.1.2 | 6 | S→F<br>0.32 | **98** | - | - | - | 0.000 | 2.393 | 3.970 | 0.0463 | 0.00 | 3.496 | 0.059 |
| P.1 | 1 | S→P<br>0.01 | | | | | | | | | | | |

| | | | | | | | | | | | | | |
|---|---|---|---|---|---|---|---|---|---|---|---|---|---|
| B.1.1.28<br>B.1.1.33<br>P.1<br>P.1.1<br>P.1.2<br>P.2 | 6 | D→Y<br>55.04 | **138** | 0.523 | 50.000 | 1.0000 | 0.000 | 38.079 | 38.743 | 0.0000 | 0.000 | 35.868 | 0.000 |
| B.1.1.28<br>B.1.1.33<br>B.1.1.7<br>P.2 | 4 | D→H<br>0.11 | | | | | | | | | | | |
| B.1.1.28<br>B.1.1.33<br>P.2<br>P.4 | 4 | G→V<br>0.77 | **142** | 0.734 | 4.497 | 0.9208 | 0.000 | 3.684 | 6.030 | 0.0141 | 0.000 | 5.498 | 0.012 |
| AV.1<br>B.1.617.2 | 2 | G→D<br>0.03 | | | | | | | | | | | |
| B.1.234<br>P.2 | 2 | G→S<br>0.02 | | | | | | | | | | | |
| B.1.1<br>B.1.1.28<br>B.1.1.33<br>B.1.177.32<br>B.1.177.52<br>B.1.617.2<br>P.1<br>P.2 | 8 | A→V<br>0.49 | **222** | 1.394 | 7.103 | 0.9682 | 1.043 | 5.642 | 4.467 | 0.0345 | 1.000 | 8.000 | 0.010 |
| P.1 | 1 | A→S<br>0.05 | | | | | | | | | | | |
| P.1 | 1 | A→P<br>0.01 | | | | | | | | | | | |
| P.1<br>P.1.2<br>P.2 | 3 | A→S<br>0.50 | **262** | 0.634 | 7.286 | 0.9974 | 0.000 | 6.372 | 10.494 | 0.0012 | 0.000 | 9.001 | 0.001 |
| B.1.1<br>B.1.1.28<br>B.1.351<br>P.2 | 4 | A→D<br>0.04 | | | | | | | | | | | |

| | | | | | | | | | | | | | |
|---|---|---|---|---|---|---|---|---|---|---|---|---|---|
| B.1.1.28<br>B.1.1.33<br>B.1.332 | 3 | A→G<br>0.04 | | | | | | | | | | | |
| B.1.1.33<br>P.1<br>P.2 | 3 | V→F<br>0.04 | **367** | - | - | - | 0.483 | 2.574 | 3.612 | 0.0574 | - | - | - |
| B.1.1.28<br>N.9 | 2 | V→I<br>0.02 | | | | | | | | | | | |
| B.1.1.33 | 1 | V→L<br>0.01 | | | | | | | | | | | |
| B.1.1.28<br>P.1<br>P.1.1<br>P.1.2<br>P.2 | 5 | K→T<br>49.11 | **417** | 0.457 | 50.000 | 1.0000 | 0.000 | 53.895 | 84.598 | 0.0000 | - | - | - |
| B.1.1.28<br>B.1.351 | 2 | K→N<br>0.05 | | | | | | | | | | | |
| P.1.2 | 1 | K→M<br>0.01 | | | | | | | | | | | |
| P.1<br>P.1.2<br>P.2 | 3 | V→F<br>0.04 | **483** | - | - | - | 0.000 | 1.836 | 3.026 | 0.0820 | - | - | - |
| B.1<br>B.1.1<br>B.1.1.28<br>B.1.351<br>B.1.525<br>N.10<br>N.9<br>P.1<br>P.1.1<br>P.1.2<br>P.2 | 11 | E→K<br>71.36 | **484** | 0.842 | 49.941 | 0.9999 | 0.000 | 35.319 | 11.838 | 0.0006 | 0.000 | 23.427 | 0.000 |
| B.1<br>B.1.1.28<br>P.5 | 3 | E→Q<br>0.22 | | | | | | | | | | | |

| | | | | | | | | | | | | | |
|---|---|---|---|---|---|---|---|---|---|---|---|---|---|
| B.1.1.33 | 1 | E→D<br>0.01 | | | | | | | | | | | |
| AV.1<br>B.1.1<br>B.1.1.28<br>B.1.1.7<br>B.1.351<br>B.1.617.1<br>P.1<br>P.1.1<br>P.1.2<br>P.2 | 10 | N→Y<br>61.12 | **501** | 0.873 | 6.320 | 0.9459 | 0.000 | 5.030 | 5.093 | 0.0240 | - | - | - |
| B.1<br>B.1.1.28<br>B.1.1.33<br>P.5 | 4 | N→T<br>0.24 | | | | | | | | | | | |
| B.1.1<br>B.1.1.28<br>B.1.1.33<br>P.1 | 4 | T→I<br>0.13 | **553** | - | - | - | 0.000 | 1.810 | 2.886 | 0.0893 | - | - | - |
| B.1.1<br>B.1.1.7 | 2 | A→D<br>2.61 | **570** | - | - | - | 0.000 | 2.120 | 3.474 | 0.0623 | 0.000 | 2.976 | 0.092 |
| B.1.1.28<br>P.1 | 2 | A→S<br>0.03 | | | | | | | | | | | |
| P.1 | 1 | A→V<br>0.01 | | | | | | | | | | | |
| B.1.1.28<br>B.1.1.7<br>P.1<br>P.1.2<br>P.2 | 5 | T→I<br>0.34 | **572** | 0.743 | 4.142 | 0.9115 | 0.000 | 3.615 | 5.771 | 0.0163 | 0.000 | 3.996 | 0.039 |
| B.1.1.33<br>P.2 | 2 | T→N<br>0.02 | | | | | | | | | | | |
| B.1.1.28<br>P.1<br>P.2 | 3 | L→F<br>0.04 | **585** | - | - | - | 0.000 | 1.667 | 2.781 | 0.0954 | - | - | - |

| | | | | | | | | | | | | | |
|---|---|---|---|---|---|---|---|---|---|---|---|---|---|
| P.1 | 1 | L→V<br>0.02 | | | | | | | | | | | |
| B.1 and derived | 56 | D→G<br>99.14 | **614** | 0.698 | 5.468 | 0.9493 | 0.000 | 4.044 | 6.636 | 0.0100 | 0.000 | 5.984 | 0.008 |
| P.1 | 1 | D→S<br>0.01 | | | | | | | | | | | |
| B.1.1.33<br>N.5<br>P.1<br>P.4 | 4 | A→V<br>0.28 | **653** | - | - | - | 0.000 | 2.810 | 4.662 | 0.0308 | 0.000 | 4.000 | 0.039 |
| P.1 | 1 | A→S<br>0.01 | | | | | | | | | | | |
| B.1.1<br>B.1.1.28<br>B.1.1.33<br>B.1.1.519<br>B.1.1.7<br>B.1.575<br>P.1<br>P.2 | 8 | P→H<br>3.03 | **681** | 0.659 | 6.650 | 0.9833 | 0.000 | 4.507 | 7.461 | 0.0063 | 0.000 | 6.936 | 0.004 |
| P.1<br>P.2<br>P.5 | 3 | P→L<br>0.40 | | | | | | | | | | | |
| B.1.617.2<br>P.1 | 2 | P→R<br>0.03 | | | | | | | | | | | |
| B.1.1.28<br>B.1.1.33<br>P.1<br>P.1.2 | 4 | A→V<br>0.13 | **684** | - | - | - | 0.000 | 3.169 | 2.731 | 0.0984 | 0.000 | 4.500 | 0.026 |
| B.1.1.28<br>P.1 | 2 | A→S<br>0.03 | | | | | | | | | | | |
| B.1.1.28<br>P.1 | 2 | A→T<br>0.03 | | | | | | | | | | | |

| | | | | | | | | | | | | | |
|---|---|---|---|---|---|---|---|---|---|---|---|---|---|
| B.1.1.28<br>B.1.1.7 | 2 | A→S<br>0.04 | **688** | 0.745 | 4.091 | 0.9100 | 0.000 | 3.519 | 5.829 | 0.0158 | 0.000 | 5.000 | 0.017 |
| B.1.1<br>B.1.1.117<br>B.1.1.28<br>N.9<br>P.1<br>P.2 | 6 | A→V<br>0.19 | | | | | | | | | | | |
| P.1 | 1 | A→G<br>0.03 | | | | | | | | | | | |
| B.1.1.28<br>B.1.1.33<br>B.1.1.464<br>P.1<br>P.2 | 5 | S→I<br>1.20 | **689** | - | - | - | 0.000 | 2.830 | 2.970 | 0.0848 | - | - | - |
| B.1.1.33 | 1 | S→G<br>0.01 | | | | | | | | | | | |
| B.1.1.28 | 1 | A→S<br>0.01 | **701** | - | - | - | **-** | - | - | - | 0.000 | 4.000 | 0.039 |
| B.1.1.33<br>B.1.1.7<br>B.1.351<br>P.1<br>P.2 | 5 | A→V<br>0.40 | | | | | | | | | | | |
| B.1.1.28<br>P.1<br>P.2 | 3 | T→I<br>0.03 | **747** | - | - | - | 0.000 | 1.810 | 2.889 | 0.0892 | - | - | - |
| P.1<br>P.2 | 2 | T→A<br>0.06 | | | | | | | | | | | |
| B.1.1.28<br>B.1.1.7<br>N.9<br>P.2 | 4 | G→V<br>0.04 | **769** | - | - | - | - | - | - | - | 0.000 | 3.567 | 0.068 |
| B.1.1.378<br>B.1.195 | 2 | G→R<br>0.02 | | | | | | | | | | | |

| | | | | | | | | | | | | | |
|---|---|---|---|---|---|---|---|---|---|---|---|---|---|
| B.1.1.28<br>B.1.1.7<br>P.1<br>P.2 | 4 | A→S<br>1.27 | **845** | 0.746 | 4.085 | 0.9098 | 0.000 | 3.512 | 5.825 | 0.0158 | 0.000 | 5.000 | 0.017 |
| P.1 | 1 | A→V<br>0.04 | | | | | | | | | | | |
| B.1.1.28<br>P.1<br>P.2 | 3 | A→S<br>0.09 | **846** | - | - | - | 0.000 | 2.120 | 3.496 | 0.0615 | 0.000 | 3.000 | 0.088 |
| B.1.1<br>B.1.1.7<br>P.2 | 3 | A→V<br>0.04 | | | | | | | | | | | |
| B.1.1.28<br>B.1.1.33<br>P.1<br>P.1.1<br>P.1.2<br>P.2 | 6 | T→I<br>58.45 | **1027** | - | - | - | 0.000 | 2.622 | 3.722 | 0.0537 | - | - | - |
| B.1.1.28<br>N.9<br>P.1<br>P.2 | 4 | A→S<br>0.15 | **1078** | - | - | - | 0.000 | 2.110 | 3.496 | 0.0615 | 0.000 | 3.000 | 0.088 |
| N.9 | 1 | A→V<br>0.02 | | | | | | | | | | | |
| B.1.1.33 | 1 | A→T<br>0.01 | | | | | | | | | | | |
| B.1.1<br>B.1.1.28<br>P.1 | 3 | D→Y<br>0.09 | **1084** | - | - | - | 0.000 | 2.640 | 2.731 | 0.0984 | - | - | - |
| P.1 | 1 | D→H<br>0.01 | | | | | | | | | | | |
| B.1.1.33<br>P.1 | 2 | G→V<br>0.04 | **1124** | - | - | - | 0.000 | 1.835 | 3.024 | 0.0820 | - | - | - |

| | | | | | | | | | | | | | |
|---|---|---|---|---|---|---|---|---|---|---|---|---|---|
| B.1.1.7 | 1 | G→A<br>0.01 | | | | | | | | | | | |
| B.1.1.33<br>P.1 | 2 | V→F<br>0.03 | **1133** | - | - | - | 0.000 | 1.102 | 3.416 | 0.0646 | - | - | - |
| B.1.1.33<br>P.1<br><br>P.1 | 2<br><br><br>1 | G→V<br>0.04<br><br>G→A<br>0.03 | **1167** | - | - | - | 0.000 | 1.835 | 3.024 | 0.0820 | - | - | - |
| AV.1<br>B.1<br>B.1.1<br>B.1.1.28<br>B.1.1.332<br>B.1.1.378<br>B.1.464<br>B.1.1.7<br>N.1<br>P.1<br>P.1.1<br>P.1.2<br>P.2<br>P.4<br>P.5 | 15 | V→F<br>82.12 | **1176** | 0.522 | 26.249 | 1.0000 | 0.000 | 16.344 | 19.637 | 0.0000 | 0.000 | 21.365 | 0.000 |
| B.1.1.28<br>B.1.1.33<br>B.1.1.7<br>P.1<br>P.2<br>P.5<br><br>B.1.1.33<br>B.1.1.7<br>P.1<br>P.2 | 6<br><br><br><br><br><br><br>4 | G→C<br>0.28<br><br><br><br><br><br>G→V<br>0.11 | **1219** | 1.334 | 8.108 | 0.9758 | 1.047 | 7.341 | 6.702 | 0.0096 | 1.004 | 9.985 | 0.002 |
| B.1.1.28<br>B.1.1.33<br>B.1.1.7<br>N.9<br>P.1<br>P.1.2 | 7 | V→L<br>0.60 | **1228** | - | - | - | - | - | - | - | 1.000 | 6.012 | 0.038 |

| | | | | | | | | | | | | | |
|---|---|---|---|---|---|---|---|---|---|---|---|---|---|
| P.2 | | | | | | | | | | | | | |
| B.1.1.33 | 1 | V→A<br>0.01 | | | | | | | | | | | |
| P.1<br>P.2 | 2 | D→N<br>0.02 | **1260** | 0.517 | 2.463 | 0.9403 | 0.000 | 2.175 | 5.535 | 0.0186 | - | - | - |
| B.1.1.28<br>P.1<br>P.2 | 3 | D→Y<br>0.04 | | | | | | | | | | | |
| B.1.1.28<br>B.1.1.33<br>B.1.91<br>P.1<br>P.2 | 5 | V→L<br>0.30 | **1264** | 0.900 | 6.706 | 0.9788 | 0.586 | 4.766 | 7.450 | 0.0063 | 1.001 | 6.498 | 0.028 |
| B.1.1.28 | 1 | V→M<br>0.1 | | | | | | | | | | | |

FUBAR inferred 20 sites subject to diversifying positive selection at posterior probability of ≥ 0.9. FEL found 44 sites under pervasive positive diversifying and 239 sites under negative selection at $p \leq 0.1$. SLAC found 29 sites under pervasive positive diversifying and 130 sites under negative selection at p $p \leq 0.1$. N: Number of lineages.
