## Supplementary File 4 for "Molecular evolution and structural analyses of the spike glycoprotein from Brazilian SARS-CoV-2 genomes: the impact of the fixation of selected mutations"

**Table S4.** Coevolutionary analysis: Bayesian Graphical Model (BGM) inference to substitution histories at individual sites.

| Site 1 | Site 2 | P [Site 1 ↔ Site 2] | Subs (1, 2, shared) |
|---|---|---|---|
| 1 | 2 | 0.603 | 3, 2, 1 |
| 1 | 1130 | 0.848 | 3, 1, 1 |
| 9 | 445 | 0.815 | 3, 1, 1 |
| 14 | 132 | 0.668 | 6, 1, 1 |
| 26 | 576 | 0.512 | 14, 1, 1 |
| 62 | 66 | 0.758 | 2, 1, 1 |
| 63 | 66 | 0.573 | 4, 1, 1 |
| 64 | 66 | 0.775 | 2, 1, 1 |
| 65 | 66 | 0.667 | 3, 1, 1 |
| 66 | 68 | 0.563 | 1, 5, 1 |
| 72 | 243 | 0.579 | 2, 4, 1 |
| 88 | 603 | 0.908 | 2, 1, 1 |
| 90 | 91 | 0.801 | 4, 1, 1 |
| 91 | 1153 | 0.720 | 1, 6, 1 |
| 92 | 93 | 0.966 | 1, 1, 1 |
| 147 | 439 | 0.564 | 3, 2, 1 |
| 147 | 1070 | 0.819 | 3, 1, 1 |
| 155 | 670 | 0.918 | 1, 2, 1 |
| 170 | 171 | 0.921 | 2, 1, 1 |
| 180 | 764 | 0.815 | 4, 1, 1 |
| 181 | 849 | 0.839 | 4, 1, 1 |
| 188 | 673 | 0.868 | 3, 1, 1 |
| 210 | 445 | 0.805 | 3, 1, 1 |
| 216 | 333 | 0.765 | 5, 1, 1 |
| 226 | 228 | 0.919 | 1, 2, 1 |
| 232 | 233 | 0.966 | 2, 2, 2 |
| 232 | 234 | 0.521 | 2, 4, 2 |
| 233 | 234 | 0.512 | 2, 4, 2 |
| 244 | 1165 | 0.801 | 1, 4, 1 |
| 262 | 265 | 0.987 | 18, 2, 2 |
| 263 | 265 | 0.665 | 4, 2, 1 |

| | | | |
|---|---|---|---|
| 265 | 266 | 0.932 | 2, 1, 1 |
| 286 | 306 | 0.681 | 1, 6, 1 |
| 303 | 1146 | 0.535 | 3, 3, 1 |
| 310 | 311 | 0.592 | 1, 1, 1 |
| 310 | 316 | 0.596 | 1, 1, 1 |
| 311 | 316 | 0.619 | 1, 1, 1 |
| 312 | 313 | 0.756 | 2, 2, 1 |
| 312 | 314 | 0.996 | 2, 5, 2 |
| 330 | 1042 | 0.922 | 2, 1, 1 |
| 343 | 344 | 0.915 | 1, 2, 1 |
| 355 | 733 | 0.514 | 1, 1, 1 |
| 355 | 814 | 0.516 | 1, 1, 1 |
| 355 | 815 | 0.566 | 1, 1, 1 |
| 367 | 843 | 0.650 | 7, 1, 1 |
| 371 | 490 | 0.778 | 1, 4, 1 |
| 455 | 1020 | 0.586 | 1, 8, 1 |
| 484 | 501 | 0.893 | 51, 8, 3 |
| 528 | 938 | 0.966 | 1, 1, 1 |
| 549 | 575 | 0.804 | 1, 4, 1 |
| 564 | 939 | 0.601 | 1, 10, 1 |
| 574 | 637 | 0.826 | 4, 1, 1 |
| 630 | 632 | 0.874 | 1, 3, 1 |
| 639 | 946 | 0.801 | 2, 2, 1 |
| 681 | 716 | 0.962 | 14, 4, 2 |
| 703 | 1251 | 0.773 | 1, 5, 1 |
| 777 | 778 | 0.719 | 1, 3, 1 |
| 777 | 779 | 0.845 | 1, 2, 1 |
| 777 | 780 | 0.610 | 1, 6, 1 |
| 804 | 1167 | 0.793 | 1, 5, 1 |
| 808 | 1228 | 0.550 | 1, 12, 1 |
| 821 | 843 | 0.613 | 9, 1, 1 |

BGM analysis summary on 574 sites with at least one substitution. Evidence for conditional dependence was reported at posterior probability of 0.5: 62 pairs of conditionally dependent sites identified.
